# TFEB-Mediated Pro-inflammatory Response in Murine Macrophages Induced by Acute Alpha7 Nicotinic Receptor Activation

**DOI:** 10.1101/2024.01.26.577408

**Authors:** Havisha H. Honwad, Mehran Najibi, Balazs Koscso, Milena Bogunovic, Javier E. Irazoqui

## Abstract

Transcription factors TFEB and TFE3 are crucial for regulating autophagy, lysosomal biogenesis, and lipid metabolism, and have significant roles in macrophage function and innate immunity. The alpha7 nicotinic acetylcholine receptor (α7nAChR), a ligand-gated Ca^2+^ channel known for its therapeutic potential in neurological and inflammatory disorders, has been implicated in modulating immune responses by modulating macrophage function. Stimulation of α7nAChR with chemical agonists has been claimed to activate TFEB in pancreatic acinar cells and neurons. However, the impact of α7nAChR activation on TFEB and TFE3 in macrophages remained unknown, posing an important question due to the potential implications for inflammation regulation. This study investigates the effects of acute α7nAChR activation on TFEB-mediated responses in murine macrophages using the specific agonist PNU-282987. We demonstrate that α7nAChR stimulation triggers TFEB nuclear translocation and lysosomal expansion. Surprisingly, PNU-282987 induces a broad pro-inflammatory gene signature without concomitant cytokine secretion, suggesting an uncoupling of gene expression from cytokine release. Mechanistically, TFEB activation requires the lysosomal Ca^2+^ exporter MCOLN1 and the Ca^2+^-dependent phosphatase PPP3/calcineurin. Additionally, PNU-282987 elevates reactive oxygen species (ROS) levels, and ROS are involved in TFEB activation by PNU-282987. Notably, even with α7nAChR deletion, compensatory ROS-mediated TFEB activation persists, suggesting the involvement of additional nicotinic receptors. Our findings reveal a novel α7nAChR-TFEB signaling axis in macrophages, offer new insights into the cholinergic regulation of immune responses, establish a baseline for comparison with disease states, and identify potential therapeutic targets for modulating inflammation.

## Introduction

Transcription factors of the Microphthalmia-TFE (MiT) family, including TFEB, TFE3, MITF, and TFEC, are considered master regulators of autophagy and lysosomal biogenesis [1]. In their resting state during homeostasis, TFEB and TFE3 reside in the cytosol, where they are sequestered by 14-3-3. Under stressful conditions, such as nutrient deprivation, TFEB and TFE3 localize to the nucleus, where they induce broad transcriptional programs that include autophagy and lysosomal biogenesis genes. As a result of TFEB and TFE3 activation, both autophagy and lysosomes are induced, resulting in enhanced cellular clearance of protein aggregates and damaged organelles. More recently, TFEB and TFE3 have also been shown to induce lipid catabolism [2, 3] and to play important roles in the innate immune system [4, 5]. During phagocytosis of bacteria or LPS stimulation, TFEB and TFE3 are activated in macrophages and required for pro-inflammatory macrophage polarization [6, 7], indicating that TFEB can be an important pro-inflammatory transcription factor in these cells. However, the roles and regulation of TFEB and TFE3 in macrophage function are not yet well understood.

Upstream regulation of TFEB and TFE3 involves negative regulation by mTORC1 under nutrient-replete conditions, mediated by phosphorylation [8, 9]. mTORC1 phosphorylation promotes 14-3-3 binding and cytosolic retention of TFEB and TFE3 [8, 10, 11]. Dephosphorylation, mediated by Ca^2+^-dependent protein phosphatase 3 (PPP3, calcineurin), lifts mTORC1-dependent repression of TFEB and TFE3 by enabling their nuclear import [12]. In macrophages, phagocytosis of particles, such as opsonized inert beads or bacterial cells, induces TFEB and TFE3 activation via this mechanism. The Botelho, Kehrl, and our laboratories collectively revealed an upstream pathway in macrophages that activates TFEB and TFE3 during phagocytosis, which involves reactive oxygen species (ROS) production by NADPH oxidase, NAADP-synthesizing enzyme CD38, NAADP-triggered Ca^2+^ release, lysosomal Ca^2+^ exporter mucolipin 1 (MCOLN1), and calcineurin [6, 13, 14]. However, additional stimuli that regulate macrophage function and impinge on TFEB and TFE3 are mostly unknown.

Nicotinic receptors (nAChRs) are pentameric ligand-gated Ca^2+^ channels, for which the ligand is the neurotransmitter acetylcholine (ACh). Their subunit composition varies by subtype, and may include α, β, γ, δ, and ε subunits. The α7nAChR is a homopentameric assembly of type 7 α (α7) subunits, encoded in mice by gene *Chrna7* [15]. Upon acute stimulation, the α7nAChR allows ionotropic Ca^2+^ entry and rapidly desensitizes. However, longer stimulation can lead to metabotropic signaling due to Ca^2+^-induced Ca^2+^ release from intracellular stores [16]. Thus, persistent α7nAChR stimulation is thought to activate downstream Ca^2+^-dependent signaling pathways.

The α7nAChR has been long considered a therapeutic target for neurological disorders including schizophrenia, and more recently attention has turned to its non-neuronal functions. The α7nAChR is known to be expressed in many non-neuronal cell types, including leukocytes, barrier epithelial cells, lung cancer cells, cardiomyocytes, and vascular endothelial cells [17, 18]. Expression in macrophages is less well documented, limited by the nonspecificity of affinity reagents and low levels of expression that border on the limits of detection [18]. Nonetheless, considerable interest in the immunomodulatory functions of the α7nAChR was fueled by the description of a cholinergic cell circuit in the spleen, which represses NF-κB activity, systemic pro-inflammatory cytokine production, and HMGB1 release during sepsis [19]. Several agonistic small molecule compounds, including general agonist carbachol and α7nAChR-specific PNU-282987, were shown to repress the production of pro-inflammatory soluble mediators by phagocytes in *in vitro* and *in vivo* models of sepsis [20]. However, the effects of these small molecules on resting macrophages have not been addressed, leaving a gap in knowledge about the homeostatic functions of α7nAChR that creates uncertainty about the clinical application of α7nAChR agonists as anti-inflammatory drugs.

Recently, the α7nAChR was reported to activate TFEB in pancreatic acinar cells and in a cellular model for amyotrophic lateral sclerosis (ALS) [21, 22]. The first report showed that PNU-282987 induced Ca^2+^ influx and promoted TFEB nuclear localization, lysosomal biogenesis, and autophagic flux in *in vitro* and *in vivo* models of acute pancreatitis, but did not test the effects of α7nAChR stimulation in homeostasis or in α7nAChR-deficient animals, define a pathway linking α7nAChR to TFEB, or define the role of TFEB in the process [21]. The second report showed that PNU-282987 induced TFEB nuclear localization dependent on Ca^2+^ and CAMKK in N2a neuronal cells transfected with SOD1^G85R^, which is causal for familial ALS [22]. In this case, PNU-282987 induced known TFEB target genes, lysosomal biogenesis, and autophagy flux, but again the role of TFEB was not defined and the effect of PNU-282987 on control cells expressing wild type SOD1 was not addressed. Both studies ascribed the effects of PNU-282987 to the α7NAChR, based solely on small molecule inhibitors. Thus, to our knowledge whether α7nAChR stimulation activates TFEB in macrophages, and the effects of such activation, remained an important gap in knowledge.

Here we report the results of our studies of the effects of α7nAChR stimulation in several murine *in vitro* macrophage models. Our results show that stimulation of the α7nAChR with PNU-282987 and other agents acutely triggers TFEB in murine macrophages, which results in lysosomal expansion. The acute transcriptional response to α7nAChR stimulation includes induction of a broad pro-inflammatory gene signature, a large portion of which requires TFEB function. Paradoxically, this pro-inflammatory gene signature is not accompanied by induced secretion of pro-inflammatory cytokines, revealing an uncoupling of secretion from gene expression. Mechanistically, we show that activation of TFEB and TFE3 requires MCOLN1, Ca^2+^, and calcineurin function downstream of α7nAChR. Moreover, we show that ROS accumulate in both PNU-282987-treated and α7nAChR-deleted cells, revealing novel aspects of their mechanisms of function that impact TFEB activation status. These results demonstrate for the first time that the α7nAChR partially mediates the activation of TFEB downstream of PNU-282987 in macrophages, and suggest the existence of additional and unknown parallel pathways for TFEB activation that need to be elucidated.

## Results

### α7nAChR stimulation induces TFEB nuclear localization in macrophages

To examine the effect of α7nAChR stimulation on TFEB, we used immortalized BMDM (iBMDM) transduced to express GFP-tagged TFEB (GFP-TFEB) [7]. DMSO-treated cells exhibited mainly cytosolic GFP-TFEB, es expected in resting cells (**Fig. 1A**). Cells treated with PNU-282987, an α7nAChR agonist [23], showed a marked increase in nuclear GFP-TFEB (**Fig. 1B**), comparable to starved positive controls (**Fig. 1D, G**). Incubation with the cholinergic agent arecoline [24] also induced GFP-TFEB nuclear localization (**Fig. 1C**). Thus, cholinergic stimulation and α7nAChR stimulation both caused GFP-TFEB to localize to the nucleus. In contrast, α7nAChR inhibitor MG624 abrogated these effects [25] (**Fig. 1E-G**). This result suggested that cholinergic stimulation may activate TFEB through the α7nAChR.

**Figure 1.**
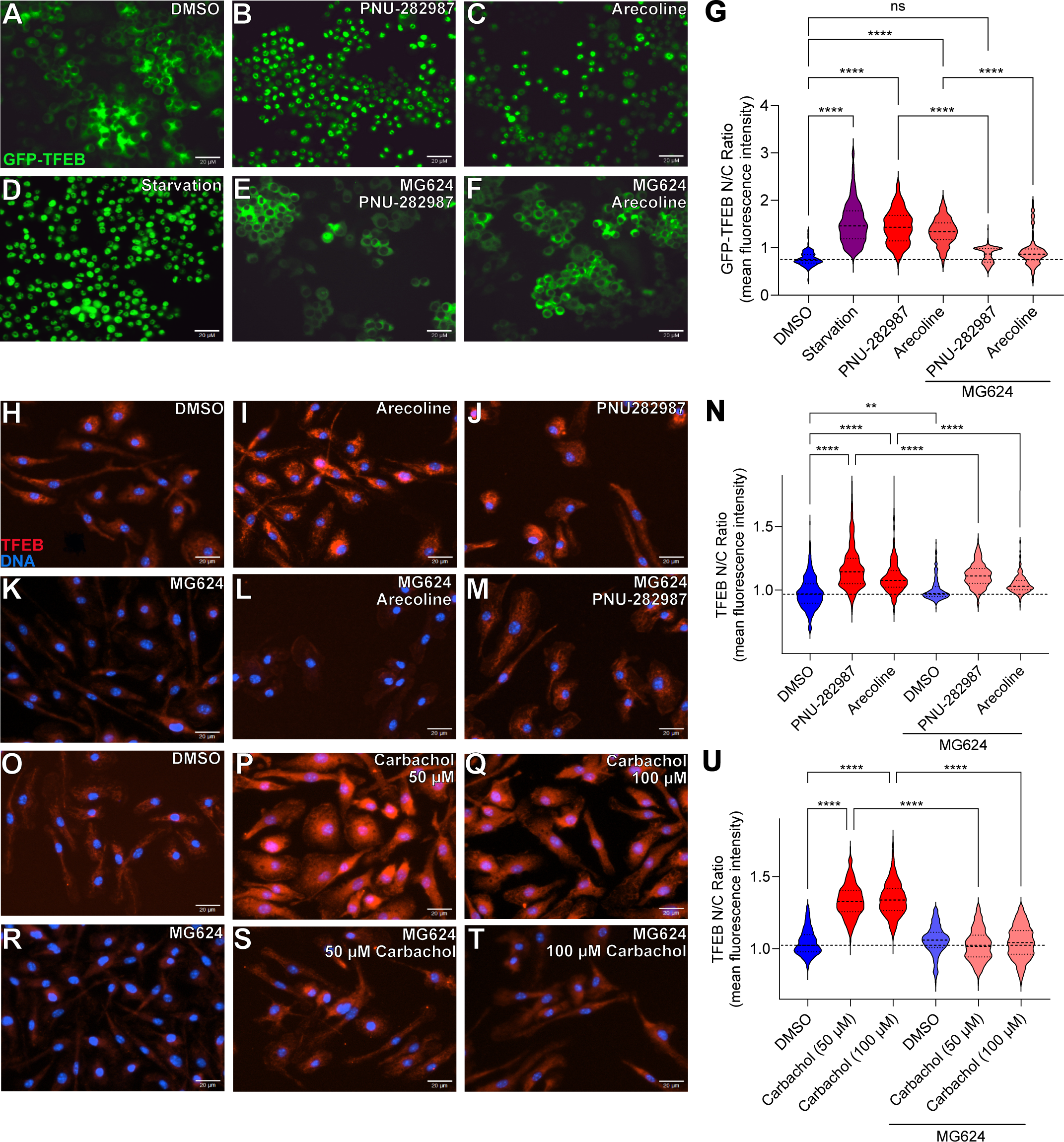
α7nAChR agonism activates TFEB in macrophages. **A-F.** Representative fluorescence micrographs of GFP-TFEB iBMDMs treated with DMSO (negative control) (**A**), PNU-282987 (50 µM, 2 h) (**B**), arecoline (1 µM, 2 h) (**C**), serum starvation (positive control, 2 h) (**D**), MG624 (1 µM, 2 h pre-treatment) followed by addition of PNU-282987 (50 µM, 2 h) (**E**), or MG624 (1 µM, 2 h pre-treatment) followed by addition of arecoline (1 µM, 2 h) (**F**). **G.** Quantification of A-F. Nucleus to cytoplasmic (N/C) ratio of GFP-TFEB (400 cells per condition, 3 biological replicates, \*\*\*\**p* ≤ 0.0001, one-way ANOVA followed by Tukey’s post-hoc test). **H-M.** Representative immunofluorescence micrographs of WT BMDMs stained with anti-TFEB antibody, treated with DMSO (**H**), arecoline (1 µM, 2 h) (**I**), PNU-282987 (50 µM, 2 h) (**J**), MG624 (1 µM, 2 h) (**K**), MG624 (1 µM, 2 h pre-treatment) followed by addition of arecoline (1 µM, 2 h) (**L**), or MG624 (1 µM, 2 h pre-treatment) followed by addition of PNU-282987 (50 µM, 2 h) (**M**). **N.** Quantification of H-M. N/C ratio of TFEB in BMDMs (400 cells per condition, \*\**p* ≤ 0.01, \*\*\*\**p* ≤ 0.0001, one-way ANOVA followed by Tukey’s post-hoc test). **O-T.** Representative immunofluorescence micrographs of WT BMDMs stained with anti-TFEB antibody, treated with DMSO (**O**), carbachol (50 µM, 2 h) (**P**), carbachol (100 µM, 2 h) (**Q**), MG624 (1 µM, 2 h) (**R**), MG624 (1 µM, 2 h pre-treatment) followed by addition of carbachol (50 µM, 2 h) (**S**), or MG624 (1 µM, 2 h pre-treatment) followed by addition of carbachol (100 µM, 2 h) (**T**). **U.** Quantification of O-T. N/C ratio of TFEB in BMDMs (400 cells per condition, \*\**p* ≤ 0.01, \*\*\*\**p* ≤ 0.0001, one-way ANOVA followed by Tukey’s post-hoc test).

To test that conclusion further, we used primary BMDMs, in which we visualized endogenous TFEB by immunofluorescence. In these cells, TFEB signal is lower and more diffuse than that of GFP-TFEB, owing to low TFEB expression and possibly to the existence of several TFEB splice isoforms. Nonetheless, in resting cells TFEB localization was predominantly cytosolic (**Fig. 1H**). As with iBMDM, arecoline and PNU-282987 increased both TFEB nuclear localization and expression, as expected due to TFEB auto-induction [3] (**Fig. 1I, J, N**). MG624 abrogated TFEB activation by arecoline and by PNU-282987, suggesting that they acted through the α7nAChR. Carbachol is another commonly used cholinergic agonist [26]. Carbachol treatment also activated TFEB (**Fig. 1P, Q**), which was inhibited by MG624 (**Fig. 1S-U**). Thus, cholinergic and α7nAChR stimulation activate TFEB in murine macrophages in an MG624-sensitive manner.

### α7nAChR stimulation activates lysosomes in macrophages

When activated in the nucleus, TFEB drives lysosomal biogenesis, which is used as a readout of TFEB activity [27]. To verify the biological significance of TFEB nuclear translocation by cholinergic agents, we measured lysosomes by Lysotracker staining of treated cells. Both carbachol and PNU-282987 caused increased Lysotracker staining compared to vehicle controls (**Fig. 2B, C, I**) but less potently than mTOR inhibitor torin-1 and LPS, which strongly activate TFEB in macrophages (**Fig. 2D, E, J) [6, 28]**. MG624 suppressed lysosomal biogenesis induced by carbachol and PNU-282987 (**Fig. 2G, H, I**). Moreover, MG624 reduced lysosomal staining in the vehicle controls, suggesting that MG624 may repress lysosomes in resting macrophages. Together, these data suggest that α7nAChR stimulation activates TFEB, increasing lysosomal biogenesis.

**Figure 2.**
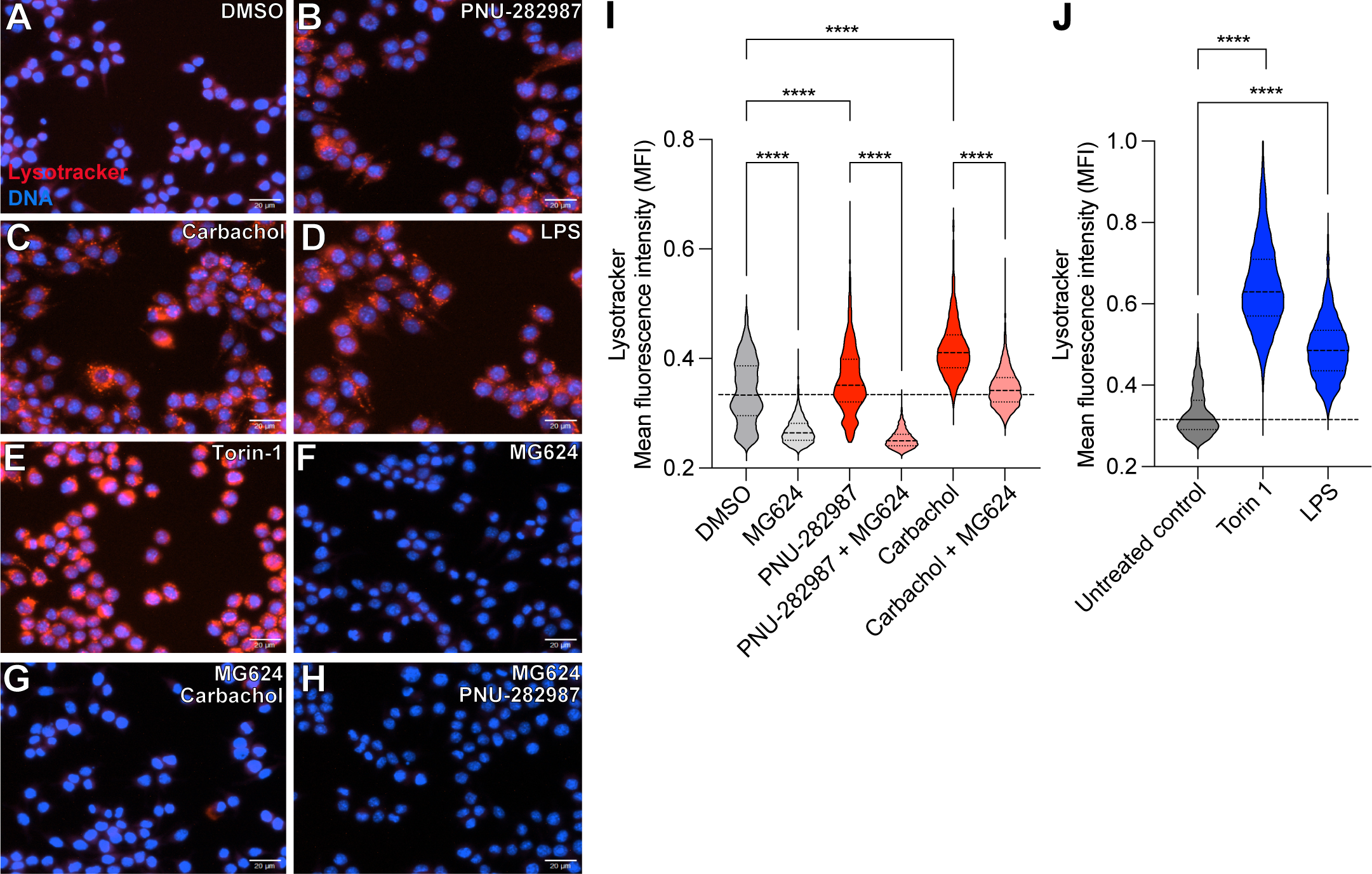
α7nAChR agonism induces lysosomes in macrophages. **A-H.** LysoTracker staining of GFP-TFEB iBMDMs treated with DMSO (**A**), PNU-282987 (50 µM, 2 h) (**B**), Carbachol (50 µM, 2 h) (**C**), LPS (positive control, 100 ng/mL, 2 h) (**D**), Torin-1 (positive control, 1.25 µM, 1 h) (**E**), MG624 (1 µM, 2 h) (**F**), MG624 (1 µM, 2 h pre-treatment) followed by addition of carbachol (50 µM, 2 h) (**G**), or MG624 (1 µM, 2 h pre-treatment) followed by addition of PNU-282987 (50 µM, 2 h). **I.** Quantification of A-C, F-H. LysoTracker mean fluorescence intensity (MFI) per cell. 3 biological replicates (400 cells per condition, \*\*\*\**p* ≤ 0.0001, one-way ANOVA followed by Tukey’s post-hoc test). **J.** Quantification of D and E. LysoTracker MFI per cell. 3 biological replicates (400 cells per condition, \*\*\*\**p* ≤ 0.0001, one-way ANOVA followed by Tukey’s post-hoc test).

### TFEB activation is likely not due to phagocytosis or cytotoxicity

In macrophages, phagocytosis functions upstream of TFEB to drive its nuclear localization [13]. To test phagocytosis as a mechanism for TFEB activation by α7nAChR, we measured uptake and killing of *Escherichia coli* and *Staphylococcus aureus*. In both infection models, carbachol and PNU-282987 had no effect on either measure of macrophage function compared to vehicle controls (**Fig. S1A, B**). Therefore, TFEB activation by carbachol and PNU-282987 is unlikely a result of increased phagocytic activity and does not result in enhanced bacterial killing.

TFEB is known to be activated by diverse cellular stresses [29]]. To test carbachol or PNU-282987 toxicity, we measured cell viability of treated BMDMs. Over a 128-fold concentration range, PNU-282987 and carbachol did not cause significant cell death (**Fig. S1C, D**), confirming and extending previous results using murine peritoneal macrophages [30] and murine macrophage-like cell line RAW264.7 [17]. These data ruled out toxicity as a mechanism of TFEB activation by carbachol and PNU-282987.

### PNU-282987 induces pro-inflammatory genes in macrophages

To characterize the transcriptional effect of acute α7nAChR stimulation with PNU-282987, we performed RNA-seq. Given the rapid kinetics of TFEB activation, we chose to focus on an early treatment time that also yielded lysosomal expansion (2 h, **Fig. 2**) to minimize secondary effects. Compared to vehicle control, PNU-282987 treatment induced 572 induced genes and repressed 119 genes (**Fig. 3A, Table S1**). Surprisingly, the induced genes were enriched for various KEGG pathways that were mostly related to pro-inflammatory signaling (*e.g.* “TNF signaling pathway”, “Cytokine-cytokine receptor interaction”, “NF-κB signaling pathway”), and “Lipid and atherosclerosis” (**Fig. 3B and Table S2**). Complementary Gene Ontology (GO) and Reactome analyses also showed enrichment of pro-inflammatory categories (**Fig. 3B and Table S2**). In contrast, the repressed genes were not significantly enriched for any KEGG pathways or GO categories. These results suggested that PNU-282987 induced a pro-inflammatory gene expression profile in BMDMs.

**Figure 3.**
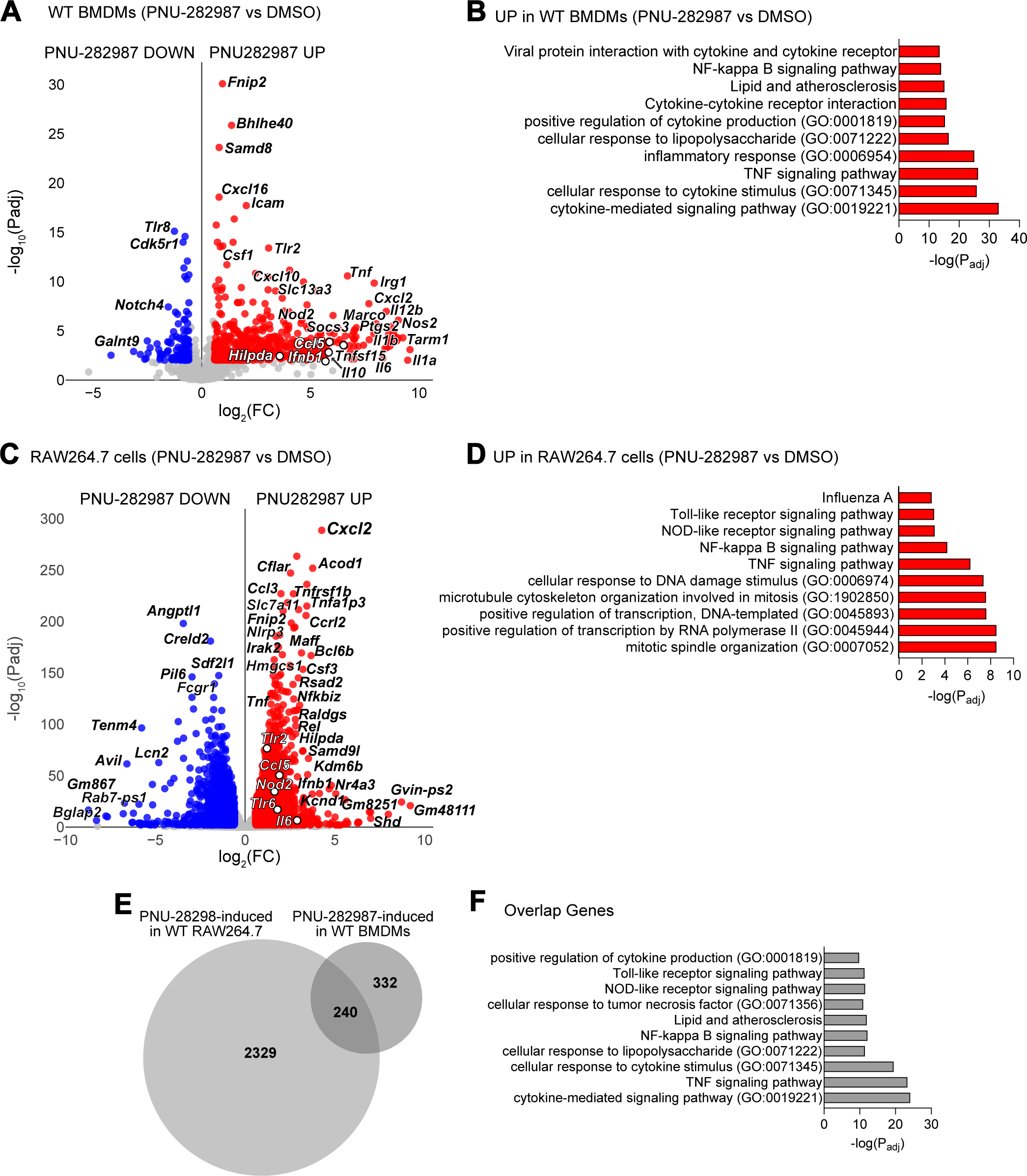
α7nAChR agonism acutely induces pro-inflammatory genes in macrophages. **A.** RNA-seq of wild type (WT) BMDMs treated with PNU-282987 (50 µM, 2 h) v. DMSO controls (fold change (FC) ≥ 1.5; *P*adj. ≤ 0.01, 4 biological replicates). Red indicates upregulated (UP), blue denotes downregulated (DOWN) genes in PNU-282987-treated cells. Notable pro-inflammatory and other genes are highlighted. **B.** Gene ontology (GO) over-representation and KEGG pathway enrichment analysis for PNU-282987-induced genes in A. **C.** RNA-seq of WT RAW264.7 cells treated with PNU-282987 (50 µM, 2 h) v. DMSO controls (FC ≥ 1.5; *P*adj. ≤ 0.01). Red indicates upregulated (UP), blue denotes downregulated (DOWN) genes in PNU-282987-treated cells. Notable pro-inflammatory and other genes are highlighted. **D.** GO over-representation and KEGG pathway enrichment analysis of PNU-282987-induced genes in C. **E.** Overlap between PNU-282987-induced genes in RAW264.7 cells and BMDMs. **F.** GO over-representation and KEGG pathway enrichment analysis of overlap PNU-282987-induced genes in E.

To verify these results in an alternative *in vitro* model, we used RAW264.7 cells. Similar to BMDMs, PNU-282987 induced a clear pro-inflammatory gene signature (**Fig. 3C, D and Table S3**). The induced pro-inflammatory gene categories (*e.g.* “Toll-like receptor signaling pathway”, “NOD-like receptor pathway”, “NF-kappaB signaling pathway”, and “TNF signaling pathway”, **Table S4**) included *Ifnb1, Il1b, Il6,* and *Tnf*, as was the case with BMDMs. 240 genes were induced by PNU-282987 in both cell types; these overlap genes were similarly enriched in pro-inflammatory genes of the same categories as above (**Fig. 3E, F and Table S5**). Thus, in RAW264.7 cells as in BMDMs, PNU-282987 treatment induced an early pro-inflammatory gene expression profile.

To test if the induced RNA expression resulted in cytokine secretion, we performed ELISA of culture supernatants from BMDMs and RAW264.7 cells treated with PNU-282987 for 12 h, to allow for protein production and secretion. PNU-282987 treatment did not induce the secretion of any of the cytokines IL-6, TNF-α, IFN-β1, IL-1β, and IL-10 (**Fig. S2A-E**). PNU-282987 also did not cause measurable lactate dehydrogenase (LDH) release (**Fig. S2F**), ruling out cell death as a reason for the lack of cytokine secretion. Experiments with RAW264.7 cells essentially yielded the same results (**Fig. S2G-K**). Thus, we concluded that PNU-282987 treatment did not increase pro-inflammatory cytokine secretion despite inducing a strong pro-inflammatory gene signature.

In contrast, PNU-282987 had a marked effect on cytokine secretion of LPS-stimulated macrophages. After stimulation with LPS for 12 h, secretion of IL-6, TNF-α, and IFN-β1 was elevated (**Fig. S2A-C**). Subsequent PNU-282987 treatment after LPS stimulation strongly inhibited IL-6 secretion, but not that of TNF-α and IFN-β1 (**Fig. S2A-C**). PNU-282987 treatment for 2 h prior to LPS stimulation only inhibited IL-6 (**Fig. S2A**). These results showed that PNU-282987 inhibits LPS-stimulated IL-6 secretion under both regimens. In parallel experiments with RAW264.7 cells, PNU-282987 treatment prior to LPS stimulation inhibited IL-6, TNF-α, and IFN-β1, but treatment after LPS stimulation did not (**Fig. S2G-K**). Thus, we confirmed and extended prior reports [17, 30] showing that PNU-282987 prophylactically reduced LPS-induced IL-6 in murine macrophages.

### PNU-282987 induces pro-inflammatory genes via the α7nAChR

To verify expression of α7nAChR, we followed several orthogonal approaches. RT-qPCR using previously published [17, 31] and newly designed oligonucleotide primers detected very low expression of nicotinic subunits α3 to α10 and β2 to β4, and of muscarinic receptors M1 to M4 (**Table S6**). α7nAChR was not more highly expressed than any other receptor subunit, except possibly M2. In contrast, *Chrna7* and *Chrnab2* were readily detectable in RNA extracted from murine brains (**Table S6**). We also used immunoblotting to detect α7nAChR in whole cell lysates using an anti-α7nAChR antibody that was previously used for the same application [32, 33]. Whole-cell lysate from WT BMDM did not show the putative α7nAChR band that was clearly visible in mouse brain lysate, and there were no band differences between wild type and *Chrna7* KO BMDMs (**Fig. S3A**). This result confirms the lack of specificity of commercially available antibodies, as previously reported [34, 35]. We also used fluorescently labelled α-bungarotoxin, a standard way to detect α7nAChR surface expression [36], in primary tissue macrophages. Staining of primary murine macrophages from peritoneum, ileum, and colon failed to detect α7nAChR expression (**Fig. S3B-I**). Finally, anti-α7nAChR immunostaining and fluorescence microscopy did not show α7nAChR expression (not shown). Thus, four orthogonal approaches and multiple macrophage cell types did not detect, or barely detected, α7nAChR expression. α7nAChR expression is known to be low and highly variable in leukocytes, with poor correlation of mRNA and protein expression and lack of specific and sensitive tools to reliably detect α7nAChR protein [18].

Therefore, we followed an alternative genetic approach to evaluate the role of α7nAChR. Mice deleted for *Chrna7* were generated previously [37]. We used RNA-seq to measure the PNU-282987-induced response in *Chrna7*^-/-^ (*Chrna7* KO) BMDMs. Compared to vehicle, PNU-282987 induced the expression of just 12 genes in *Chrna7* KO BMDMs (**Fig. 4A and Table S7**), as opposed to 572 genes induced in the wild type (**Fig. 3**). Notably, *Tnf* was still induced, as well as *Rragd* and *Fnip2*, which are mTOR regulators and thus potentially upstream of TFEB. Moreover, only 8 genes were induced in both wild type and *Chrna7* KO backgrounds (**Fig. 4B and Table S8**), supporting the specificity of the PNU-282987 effect in terms of gene induction. Consistently, direct comparison of PNU-282987-treated *Chrna7* KO BMDMs and wild type cells showed a large defect in pro-inflammatory gene expression in the *Chrna7* KO BMDMs (**Fig. 4C, D and Table S9**). The genes that were more highly expressed in PNU-282987-treated *Chrna7* KO BMDMs were enriched in Ras and Rap1 signaling and lysosome categories, suggesting compensatory pathways (**Table S10**). Thus, the induction of pro-inflammatory genes by PNU-282987 is α7nAChR-dependent.

**Figure 4.**
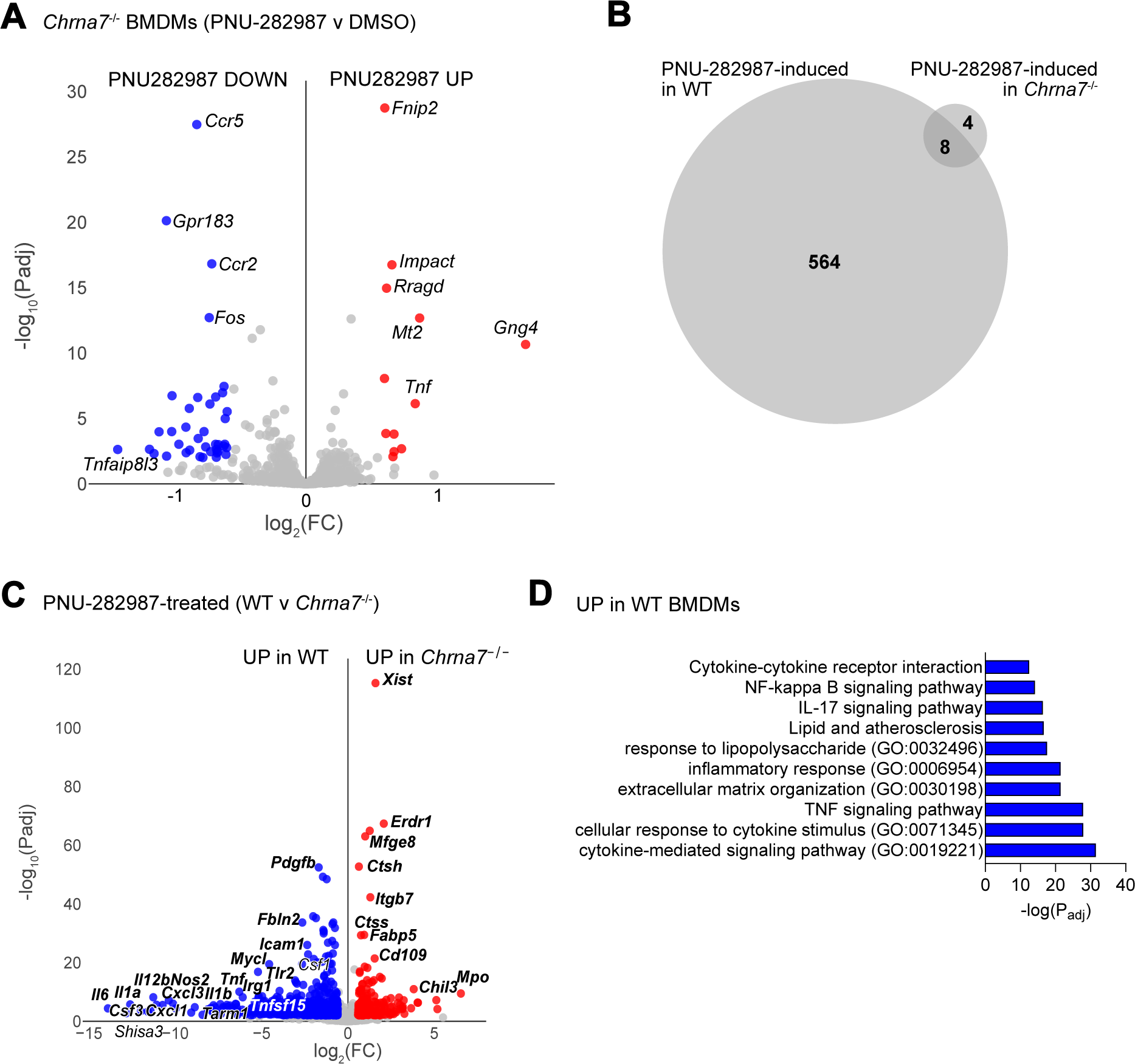
α7nAChR is required for the pro-inflammatory transcriptional response to PNU-282987. **A.** RNA-seq of *Chrna7*^-/-^ BMDMs treated with PNU-282987 (50 µM, 2 h) v. DMSO controls (FC ≥ 1.5; *P*adj. ≤ 0.01, 4 biological replicates). Red indicates upregulated (UP), blue denotes downregulated (DOWN) genes in PNU-282987-treated cells. Notable genes are highlighted. **B.** Overlap of genes induced by PNU-282987 in WT (from Fig. 3A) compared to those induced in *Chrna7*^-/-^ BMDMs. **C.** Differential gene expression of PNU-282987-treated (50 µM, 2 h) WT and *Chrna7*^-/-^ BMDMs (FC ≥ 1.5; *P*adj. ≤ 0.01, 4 biological replicates). Red indicates upregulated (UP), blue denotes downregulated (DOWN) genes in *Chrna7*^-/-^ cells. Notable genes are highlighted. **D.** GO-overrepresentation and KEGG pathway enrichment analysis for genes upregulated in PNU-282987-treated WT BMDMs compared to *Chrna7*^-/-^.

To better understand why *Chrna7* KO BMDMs exhibited such a strongly defective response to PNU-282987, we examined expression of genes encoding macrophage differentiation markers at baseline. Despite a few exceptions (CD11a, CD11c, CD14, and MHC II), macrophage marker genes (CD15, CD16, CD64, CD32, F4/80, CD33, CD68, CD80, and CD86) did not show significant differences between WT and *Chrna7* KO BMDMs (**Fig. S4A, Table S11**) ruling out a broad defect in macrophage *in vitro* differentiation. However, *Chrna7* KO BMDMs showed decreased expression of many immune-related genes at baseline, such as *Il6, Il1a, Il1b, Cxcl2,* and *Nox1* (**Fig. S4B, C** and **Table S12**), but also showed increased expression of other immune-related genes, notably of the inflammatory response (**Fig. S4D** and **Table S13**). This result suggests that deletion of α7nAChR has a non-neutral effect on immune gene expression of *in vitro* differentiated BMDMs.

### TFEB is required for the full transcriptional response to PNU-282987

To define the effect of TFEB activation on the transcriptional response to PNU-282987, we performed RNA-seq of conditionally TFEB-deleted BMDMs [6]. We previously showed that TFEB expression is efficiently abrogated in this line [6]. At baseline, the expression of a set of 23 macrophage differentiation markers was not significantly different between *Tfeb*^fl/fl^ and *Tfeb*^ΔLysM^ BMDMs, showing that *in vitro* macrophage differentiation was not affected by *Tfeb* deletion (**Fig. S5A and Table S14**). TFEB deletion decreased the expression of 665 genes compared to *Tfeb*^fl/fl^ controls (**Fig. S5B and Table S15**). Surprisingly, this gene set was not enriched for autophagy or lysosomal genes, as might have been expected based on data from other cell types [38]. Instead, this TFEB-dependent gene set was enriched in functional categories related to “PI3K-Akt signaling”, “ECM-receptor interaction”, “Proteoglycans in cancer”, and the extracellular matrix (**Fig. S5C and Table S16**). TFEB deletion also resulted in higher expression of 138 genes (**Table S15**), but this gene set did not show significant enrichment of functional categories. These results suggest that TFEB promotes the expression of extracellular matrix genes in resting macrophages.

In PNU-282987-treated cells, deletion of TFEB resulted in decreased expression of TNF-α signaling genes, in addition to the baseline defect in genes related to extracellular matrix and PI3K-AKT compared to *Tfeb*^fl/fl^ controls (**Fig. 5A, B and Tables S17 and S18**). The genes that were more highly expressed in TFEB-deleted cells were not enriched for any informative functional category. Thus, TFEB promotes the expression of TNF signaling genes in response to PNU-282987.

**Figure 5.**
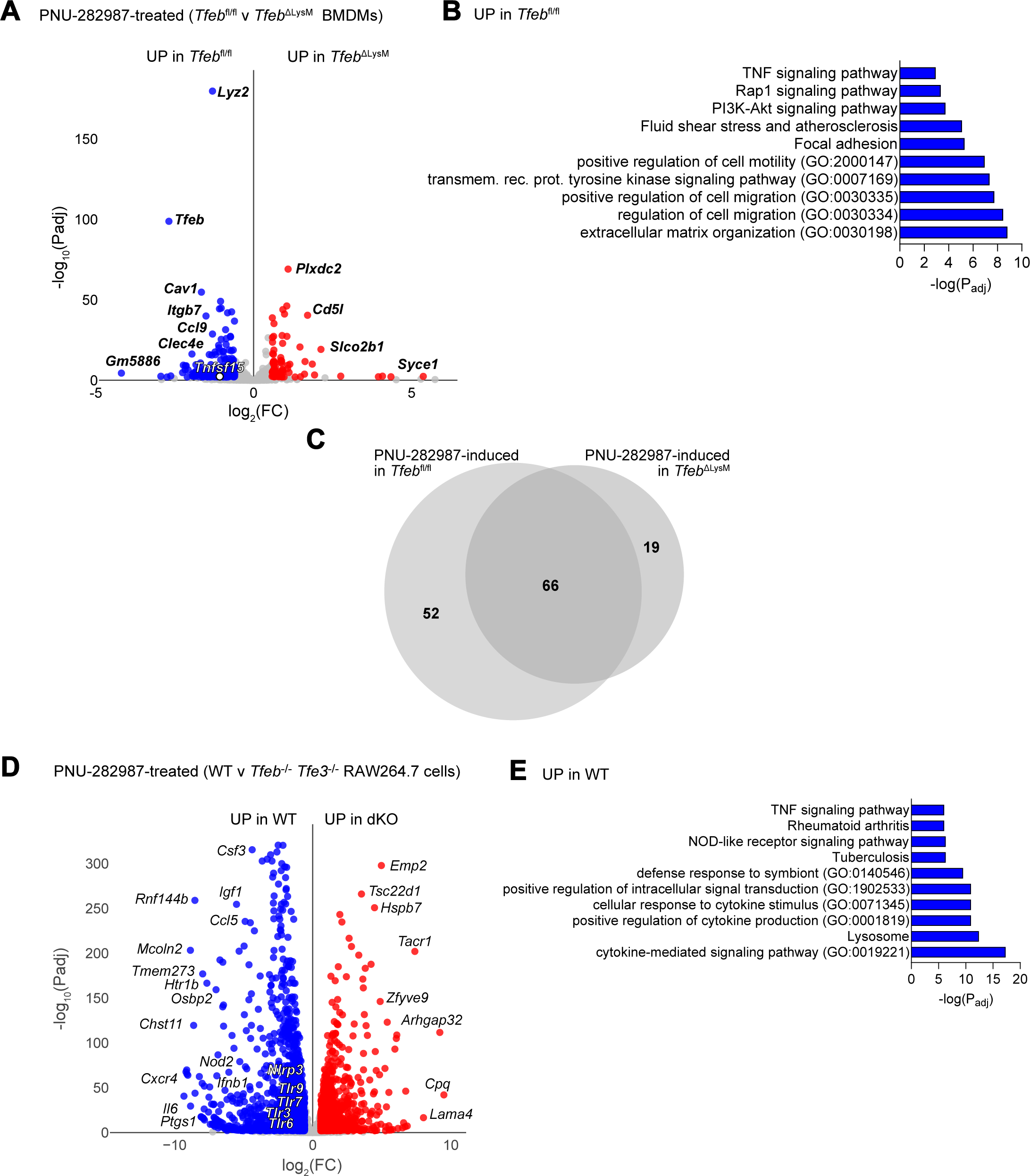
TFEB is required for a large fraction of the transcriptional response to PNU-282987. **A.** Differential gene expression of PNU-282987-treated (50 µM, 2 h) *Tfeb*^f/f^ and *Tfeb*^ΔLysM^ BMDMs (FC ≥ 1.5; *P*adj. ≤ 0.01, 4 biological replicates). Red indicates upregulated (UP), blue denotes downregulated (DOWN) genes in *Chrna7*^-/-^ cells. Notable genes are highlighted. **B.** GO-overrepresentation and KEGG pathway enrichment analysis for genes upregulated in PNU-282987-treated *Tfeb*^f/f^ BMDMs compared to *Tfeb*^ΔLysM^. **C.** Overlap of genes induced by PNU-282987 in *Tfeb*^f/f^ compared to those induced in *Tfeb*^ΔLysM^ BMDMs. **D.** Differential gene expression of PNU-282987-treated (50 µM, 2 h) *Tfeb Tfe3* double knockout (dKO) and WT RAW264.7 cells (FC ≥ 1.5; *P*adj. ≤ 0.01, 4 biological replicates). Red indicates upregulated (UP), blue denotes downregulated (DOWN) genes in dKO cells. Notable genes are highlighted. **E.** GO-overrepresentation and KEGG pathway enrichment analysis for genes upregulated in PNU-282987-treated WT RAW264.7 cells compared to dKO.

Comparing the sets of genes that were induced by PNU-282987 in each genetic background, deletion of TFEB abrogated the induction of 52 out of 118 total that were induced in *Tfeb*^fl/fl^ controls (**Fig. 5C and Table S19**). 19 additional genes were induced only in TFEB-deleted cells. Thus, overall TFEB is required for about 40% of the transcriptional response that is triggered by PNU-282987.

From these analyses in *Tfeb*^fl/fl^, *Tfeb*^ΔLysM^, and wild type BMDMs, gene *Tnfsf15* stood out as a PNU-282987-induced TFEB-dependent gene encoding a soluble signaling mediator. Tumor necrosis factor superfamily-15 (TNFSF15), also called TL1A, is a ligand for death receptor 3 (DR3), which activates signaling pathways involved in immune cell activation, proliferation, apoptosis, and pro-inflammatory responses, and thus could link PNU-282987 to proinflammatory gene expression [39]. To test if PNU-282987-induced TNFSF15 production might cause macrophage activation, we performed ELISA of supernatants from wild type and *Tfeb*^fl/fl^ BMDMs. Treatment with PNU-282987 did not increase soluble TNFSF15 under these conditions (**Fig. S5D**), suggesting that TNFSF15 is not likely to be a mediator of PNU-282987-induced pro-inflammatory gene expression.

To further test these conclusions, we used RAW264.7 cells that were deleted for *Tfeb* and *Tfe3*, which encodes a TFEB paralog that heterodimerizes with TFEB but can also function independently as a homodimer [40, 41]. In this model, PNU-282987-treated *Tfeb Tfe3* double knockout (dKO) cells showed defective expression of 550 genes compared to wild type controls (**Fig. 5D and Table S20**). These genes were enriched in pro-inflammatory pathways, notably TNF and NOD receptor signaling, as well as lysosomal genes (**Fig. 5E and Table S21**), showing that TFEB and TFE3 are important for the pro-inflammatory gene expression program that is triggered by PNU-282987 in RAW264.7 cells. The genes that were more highly expressed in the *Tfeb Tfe3* dKO cells did not show enrichment of any relevant functional categories. These data strongly support the conclusion that PNU-282987 induces a pro-inflammatory gene expression signature in a TFEB and TFE3-dependent manner.

Surprisingly, deletion of *Tfeb* and *Tfe3* abrogated IL-6 and IFN-β1 production induced by LPS (but not that of TNF-α, **Fig. S6A-D**). Moreover, PNU-282987 induced 3-fold higher LDH release in the *Tfeb Tfe3* dKO cells compared to WT (**Fig. S6E**), suggesting that TFEB and TFE3 may mitigate PNU-282987 toxicity. However, at baseline *Tfeb Tfe3* dKO cells showed decreased expression compared to wild type of macrophage differentiation marker genes encoding CD11b, CD15, CD16, CD32, CD64, CD33, and CD86, as well as TLR2 and TLR4 (**Fig. S6F, Tables S22 and 23**), suggesting that the *Tfeb Tfe3* dKO cells may show an impaired macrophage phenotype.

Taken together, these data show that TFEB (and possibly TFE3) is an important mediator of PNU-282987-induced gene transcription, but the upstream mechanism of TFEB activation by PNU-282987 remained unknown.

### The MCOLN1-Ca^2+^-calcineurin pathway is important for α7nAChR stimulation of TFEB

In previous studies, we showed that phagocytosis activates TFEB through a pathway that involves ROS, Ca^2+^ release (potentially through the main lysosomal Ca^2+^ pump MCOLN1), and protein phosphatase 3 (calcineurin), in that sequence [6]. To examine the role of intracellular Ca^2+^ in PNU-282987-induced TFEB translocation, we treated immortalized BMDMs expressing GFP-TFEB [6] with cell-permeable Ca^2+^-chelating agent BAPTA-AM prior to PNU-282987 treatment. BAPTA-AM pretreatment suppressed the PNU-282987-induced nuclear localization of GFP-TFEB (**Fig. 6A-C, E**), suggesting that intracellular Ca^2+^ is required for PNU-282987-induced TFEB nuclear localization. Furthermore, pre-treatment with FK506, which inhibits calcineurin [42], also prevented GFP-TFEB nuclear translocation (**Fig. 6D, E**), suggesting that PNU-282987-induced intracellular Ca^2+^ functions through calcineurin to induce TFEB nuclear translocation. In support of this conclusion, silencing of *Ppp3cb* and *Ppp3r1*, respectively encoding catalytic and regulatory subunits of calcineurin, partially suppressed PNU-282987-induced GFP-TFEB translocation (**Fig. 6F-I, M**). Likewise, silencing of *Trpml1*, which encodes MCOLN1, also prevented GFP-TFEB translocation by PNU-282987 (**Fig. 6K, M**), suggesting that Ca^2+^ export from the lysosome by MCOLN1 is also required. Altogether, these results are consistent with PNU-282987-induced activation of TFEB through the MCOLN1-Ca^2+^-Calcineurin pathway.

**Figure 6.**
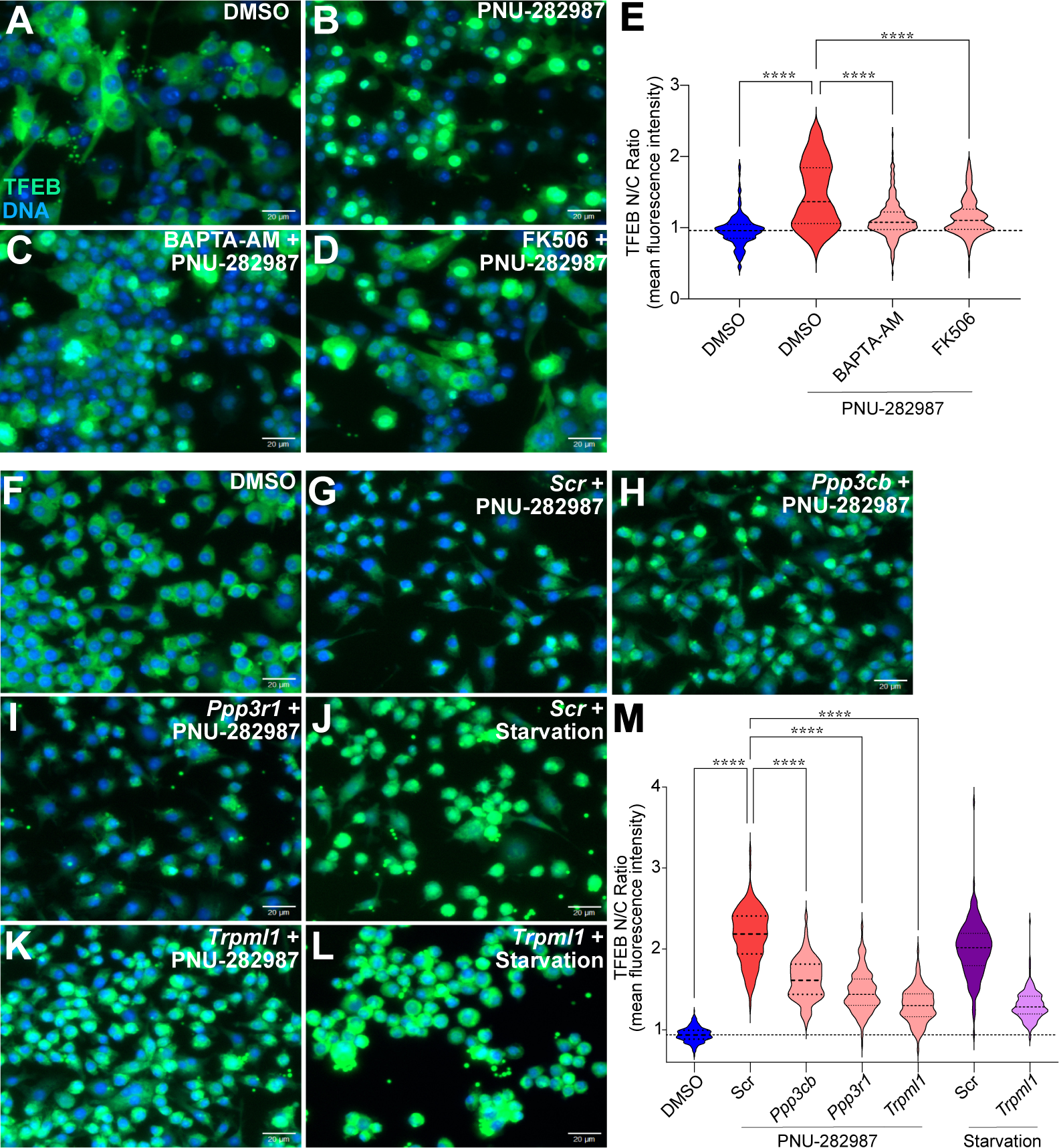
MCOLN1, Ca^2+^, and PPP3/calcineurin are required for TFEB activation by PNU-282987. **A-D.** Representative fluorescence micrographs of GFP-TFEB iBMDMs treated with DMSO (negative control) (**A**), PNU-282987 (50 µM, 2 h) (**B**), BAPTA-AM (10 µM, 3 h pre-treatment) followed by addition of PNU-282987 (50 µM, 2 h) (**C**), or FK506 (5 µM, 6 h pre-treatment) followed by addition of PNU-282987 (50 µM, 2 h) (**D**). **E.** Quantification of A-D. N/C ratio of GFP-TFEB (400 cells per condition, 3 biological replicates, \*\*\*\**p* ≤ 0.0001, one-way ANOVA followed by Tukey’s post-hoc test). **F-L.** Representative fluorescence micrographs of GFP-TFEB iBMDMs treated DMSO (negative control) (**F**), and with siRNA, Scrambled siRNA (Scr) (**G, J**) or directed against *Ppp3cb* (**H**), *Ppp3r1* (**I**), and *Trpml1* (**K, L**) for 48 h prior to treatment with PNU-282987 (50 µM, 2 h) or serum starvation (positive control, **J, L**). **M.** Quantification of F-L. N/C ratio of GFP-TFEB (400 cells per condition, 3 biological replicates, \*\*\*\**p* ≤ 0.0001, one-way ANOVA followed by Tukey’s post-hoc test).

In our previous research, we found that protein kinase D (PKD) 1 is required for TFEB activation during bacterial infection and phagocytosis [7, 43]. To test the role of PKD in PNU-282987-induced activation of TFEB, we used the PKD inhibitor kb-NB142-70 [44]. Pretreatment of BMDMs with kb-NB142-70 suppressed TFEB translocation caused by PNU-282987 (**Fig. S7**), thus implicating PKD in this mechanism of activation as well.

We and others recently showed that oxidative stress and reactive oxygen species can activate TFEB in macrophages and other cell types [6, 45, 46]. To test if PNU-282987 induces ROS production, we used measured ROS levels in PNU-282987-treated BMDMs. PNU-282987 induced the accumulation of both overall ROS and H_2_O_2_ specifically, which were suppressed by MG624 (**Fig**. **S8A, B**). MG624 alone also slightly depressed H_2_O_2_ levels compared to vehicle controls. These results showed that PNU-282987 induces ROS in BMDMs, which could be a mechanism for TFEB activation, further implicating a ROS-MCOLN1-Ca^2+^-calcineurin pathway. Suppression by MG624 is consistent with an α7nAChR-dependent mechanism.

At baseline, *Chrna7* KO BMDMs showed higher total ROS and H_2_O_2_ compared to WT cells (**Fig. S8C, D**). The ROS and H_2_O_2_ levels in *Chrna7* KO BMDMs were similar to those in PNU-282987-treated wild type BMDMs, and treatment with PNU-282987 did not lead to further increases (**Fig. S8C, D**). Thus, both activation and loss of α7nAChR led to increased ROS in macrophages.

To test the functional relevance to TFEB activation, we performed immunofluorescence. In vehicle controls, *Chrna7* KO cells showed slightly less nuclear TFEB than WT (**Fig. S8E**). PNU-282987 induced TFEB translocation in both genotypes, but to a lesser degree in *Chrna7* KO cells, suggesting both α7nAChR-dependent and independent mechanisms of activation (**Fig. S8E**). Pre-incubation with N-acetyl L-cysteine (NAC), a ROS scavenger, reduced TFEB nuclear localization in both genetic backgrounds, as also did FK506, BAPTA-AM, and MCOLN1 inhibitor ML-SI3 [47] (**Fig. S8F, G**). Thus, ROS scavenging and inhibition of Ca^2+^, MCOLN1, and calcineurin all suppressed TFEB nuclear translocation induced by PNU-282987 even in α7nAChR-deleted cells. These results suggest both α7nAChR-dependent and -independent mechanisms of TFEB activation by PNU-282987 and, possibly, other α7nAChR agonists.

## Discussion

For our studies, we focused on resting cell lines and primary macrophages. Resting, or naïve, macrophages are crucial for preserving tissue homeostasis and performing routine clearance of apoptotic cells and debris, ensuring immune surveillance without initiating inflammatory responses [48, 49]. In naive macrophages, we can assess baseline cellular responses to α7nAChR agonists without the confounding effects of prior activation, allowing for a clearer distinction of the direct effects of these agonists on macrophage function [50, 51]. Establishing a baseline with naive macrophages enables comparative studies with activated macrophages, elucidating how α7nAChR might modulate functions in different activation states [52]. Furthermore, naïve macrophages can help us understand how early interventions with α7nAChR agonists might modulate the immune system before inflammation or infection onset, providing insights into designing preventative therapies that maintain immune homeostasis [53]. Understanding the role of naïve macrophages in tissue repair and regeneration offers insights into their potential therapeutic roles in preventing or mitigating chronic inflammatory diseases such as atherosclerosis and fibrosis [54]. Thus, studying naïve macrophages is essential for a comprehensive understanding of the endogenous functions of α7nAChR and of the therapeutic potential of α7nAChR agonists.

Our results show that cholinergic agonists, both nonspecific (arecoline and carbachol) and specific for α7nAChR (PNU-282987), activate TFEB and lysosomal biogenesis in various macrophage models, including RAW264.7, immortalized BMDMs, and primary BMDMs. Therefore, TFEB is a novel mediator of cholinergic signaling in these important cells of the innate immune system. TFEB activation by these agonists is suppressed by α7nAChR-specific antagonist MG624, suggesting that most of the effect is α7nAChR-mediated. However, deletion of α7nAChR reduced but did not abrogate the activation of TFEB by its agonist, suggesting that there may be a second mechanism downstream of PNU-282987 that is also MG624-sensitive. The identity of this second mechanism is currently unknown, but we speculate that it could involve an additional nicotinic receptor that may also be inhibited by MG624. RAW264.7 cells were recently reported to express several nicotinic receptor α subunits (3 to 7, 9, and 10) and β subunits (2 to 4) [55], but more data are required to support this speculation. Nonetheless, an independent study using α7nAChR agonist GTS-21 in peritoneal macrophages reached similar conclusions [56].

Nonetheless, α7nAChR deletion almost completely abrogated the transcriptional response to PNU-282987 stimulation, consistent with the majority of the signaling downstream to be mediated through the α7nAChR. By comparison, TFEB deletion eliminated about 40% of the transcriptional response, showing the biological relevance of TFEB activation and suggesting that transcription factors other than TFEB are involved. Although their identity remains unknown, it is tempting to speculate that an additional Ca^2+^-regulated transcription factor may provide that function. However, our demonstration that modulating α7nAChR activity through agonism and deletion increases ROS and triggers the MCOLN1-Ca^2+^-calcineurin pathway adds a new element to consider for signaling downstream of α7nAChR. This is consistent with prior reports that showed that nicotine induces ROS production in RAW264.7 cells [57]. Thus, it is also possible that a redox-sensitive transcription factor may be involved in parallel to TFEB. Further work to elucidate these scenarios is merited.

Transcriptionally, we find that engaging α7nAChR through PNU-282987 induces a pro-inflammatory response that does not translate into cytokine production by the times that we evaluated. This is surprising for two reasons. First, a large body of literature documents the anti-inflammatory effects of α7nAChR activation with PNU-282987 and other agonists. Indeed, our data recapitulate some of these findings in macrophages that are stimulated with LPS. In their resting state, however, the same macrophages respond by inducing a slew of pro-inflammatory genes in a partially TFEB-dependent manner. This result confirms and extends prior reports of pro-inflammatory responses induced by nicotine in murine RAW264.7 cells and peritoneal macrophages, as well as the aggravation of atherosclerosis *in vivo* [58]. Additionally, many reports document pro-inflammatory activities of nicotinic agonists. Nicotine also induces the production of PGE2, a product of COX-1 and COX-2 induced during inflammation, in primary human monocytes [59, 60], induces the production of CRP by U937 human monocytes [61], induces IL-6 in human monocytic THP-1 cells [62], and at low concentrations potentiates the ATP-induced inflammatory response in rat primary microglial cells [63]. Moreover, treatment of human THP-1 and monocyte-derived macrophages with PNU-282987 induced an activated phenotype and was proposed as an anti-immunosuppression strategy [64]. The exact reasons behind these divergent behaviors are currently unclear. One possibility is that after the initial pro-inflammatory phase, the transcriptome settles to a refractory or less responsive state perhaps through epigenetic mechanisms. Indeed, most reports evaluated PNU-282987 and similar agents on longer time scales than we did, because of our experimental focus on primary transcriptional events that are relevant to the time scale of TFEB activation. Longer stimulation times raise concerns about secondary effects [18].

The second reason for surprise is that the strong pro-inflammatory gene signature does not result in concomitant cytokine production. However, this observation is not without precedent. Cytokine expression is subjected to multiple layers of post-transcriptional control [65, 66]. Moreover, in the case of IL-1β, TLR signaling provides signal 1 to induce its transcription, while the NLRP3 inflammasome provides signal 2 to induce its maturation and secretion through gasdermin-generated membrane pores [67, 68]. Thus, the lack of cytokine production in the supernatants of PNU-282987 cells could be evidence of post-transcriptional control or of necessary but absent co-signals. This “poised” state may help pro-inflammatory responses when signal 2 is presented. Further investigation of this mechanism could offer new insight into the physiological regulation of inflammation.

Although sometimes useful as probes and as proof of concept for therapeutic strategies, small molecules like PNU-282987 fall short of fully recapitulating the endogenous ligand, in this case the neurotransmitter acetylcholine. *In vivo*, the role of TFEB as mediator of acetylcholine signaling is likely to be much more complex. In future research, it will be important to disentangle the cellular sources of acetylcholine, the cellular targets that express receptors capable of engaging TFEB, and the physiological importance of these interactions within the innate immune system and beyond.

## Materials and Methods

### Mice

Mouse colony maintenance and procedures were performed following the Guidelines for the Care and Use of Laboratory Animals, published by the US National Institutes of Health, and were approved by the Institutional Animal Care and Use Committee (IACUC) of the University of Massachusetts Chan Medical School. Wild type mice, strain C57BL/6J (Stock #: 000664) and *Chrna7*^-/-^ mice, strain B6. 129S7-*Chrna7*^tm1Bay/J^ (Stock #: 003232) were from The Jackson Laboratory. *Tfeb*^fl/fl^ mice were a gift from Dr. Andrea Ballabio, TIGEM, Naples, Italy [69]. *Tfeb*^fl/fl^ *LysM-Cre* mice were bred in-house [6]. BMDM genotyping for wild type and *Chrna7*^-/-^ BMDMs was performed by Protocol 28408: Standard PCR Assay - Chrna7<tm1Bay>-Alternate 1 for Jackson lab stock No. 003232.

### Isolation and differentiation of primary murine BMDMs

Bone marrow cells were isolated from separated femurs and tibias from euthanized 8 to 12-week-old mice as previously described [6]. Isolated bone marrow cells from C57BL/6, *Chrna7*^-/-^, *Tfeb*^fl/fl^ and *Tfeb*^ΔLysM^ mice were cultured in BMDM differentiation medium: DMEM containing 4.5 g/L glucose, L-glutamine (Corning, 10-013-CV), sodium pyruvate supplemented with 20% conditioned medium from L929 cells, 10% FBS, 1% AA for 7 days. Cells were seeded in non-tissue culture treated Petri dishes (Fisherbrand, FB0875712) and incubated at 37°C, in a humidified chamber with 5% CO_2_. Cells were seeded for experiments after 7 days of differentiation.

### Cell culture and transfection

G418 sulfate (Life Technologies, 10131) was used for selection. RAW264.7 and *Tfeb*^-/-^ *Tfe3*^-/-^ RAW264.7 cells, a gift from Dr. Rosa Puertollano, NHLBI [5], and iBMDMs [6] were grown in Dulbecco’s Modified Eagle’s Medium (DMEM) containing 4.5 g/L glucose, L-glutamine (Corning, 10-013-CV), sodium pyruvate supplemented with 10% Fetal Bovine Serum (FBS; Gibco, 16000044) and 1% Antibiotic-Antimycotic (Gibco, 15240062). Cells were grown at 37 °C, in a humidified chamber with 5% CO_2_ and passaged 4 to 10 times.

### Drugs and Reagents

PNU-282987 (Sigma-Aldrich, P6499) powder was dissolved in DMSO (Sigma-Aldrich, D2438) - sterile, filtered, cell culture grade, Bio Performance certified, meets EP and USP specifications and thus devoid of endotoxin or pro-inflammatory contaminants; Carbachol (Sigma-Aldrich, 212385); MG624 (Cayman Chemical Company, 24298); Arecoline (Cayman Chemical Company, 13662); LPS from *S. enterica* serotype Typhimurium (Sigma-Aldrich, L6143-1 MG); Torin-1 (Tocris, 427); FK-506 (Cayman Chemical Company, 10,007,965); BAPTA-AM (Cayman Chemical Company, 15,551); kb-NB-142-70 (Cayman Chemical Company, 18,002); ML-SI3 was a gift from Drs. Juan Marugán and Raúl Clavo of the National Center for Advancing Translational Science of the National Institutes of Health (NCATS/NIH); α-bungarotoxin Alexa Fluor 647 conjugate (Thermo Fisher Scientific, B35450).

### Immunofluorescence and fluorescence imaging

Cells were seeded in 96-Well Optical Bottom Plates (Thermo Scientific, 165305) and incubated overnight to attain cell confluency of 0.06 x 10^6^ for experiments. Following experiments, cells were fixed using 4% paraformaldehyde (PFA) (Sigma Aldrich, 158127) and incubated with Hoechst stain (Anaspec, AS-83218) for 20 min at room temperature. For immunofluorescence, cells were washed 3 times with PBS (Gibco Life Technologies,10010) for 5 min, permeabilized with 0.1% Triton X in PBS with shaking at room temperature for 5 min, washed 3 times with PBS and blocked with 5% bovine serum albumin (Sigma Aldrich, A9647) in PBS for 1 h. After washing with PBS (3 times), cells were incubated with primary goat anti-TFEB antibody (Abcam, ab2636) for 2 h. After washing 3 times in PBS, cells were incubated with donkey anti-goat IgG -Alexa 594 secondary antibody (Thermo Fisher Scientific, A11058) along with Hoechst stain (Anaspec, AS-83,218) for 1 h at room temperature. For LysoTracker staining, cells were incubated with 100 nM LysoTracker DND-99 (Thermo Fisher, L7528) added to cell culture media 30 min before fixation step, according to manufacturer’s instructions. Image acquisition for calculating TFEB N:C ratio and for Lysotracker mean fluorescence intensity (MFI) was performed using Lionheart FX automated microscope (Biotek) and Gen5 software (version 3.13).

### Determination of N:C ratio

Nucleus to cytoplasm (N:C) ratio for TFEB was measured using CellPofiler [70], following our previously described protocol [6]. In brief, we used the 3_channels_pipeline.cppipe. algorithm, which uses nuclear stain (Hoechst) as primary object or “seed” region for building image segmentation masks to distinguish individual cells by marking boundaries, define subcellular regions and generate specific fluorescent intensity values. The pipeline calculated N:C ratio as mean fluorescent intensity of nucleus relative to mean fluorescence intensity of the region defined as cytosol.

### RT-qPCR

Cells were lysed using Trizol Reagent (1 mL per 1.5 ml microcentrifuge tube). 100 µL chloroform was added and samples were centrifuged at 12,000 rpm to separate the RNA containing aqueous phase. The aqueous layer was transferred to Purelink RNA mini kit column and total RNA was extracted according to manufacturer’s instructions. cDNA was prepared using iScript gDNA clear cDNA synthesis kit (Bio-Rad, 1725035). Diluted cDNA was used for RT-qPCR, performed using SYBR Green Super mix (Bio-Rad) and ViiA7 Real-Time qPCR system (Applied Bioscience). Ct values were obtained for test genes and relative expression was plotted following normalization to reference gene *Gapdh*. Oligonucleotide sequences are included in **Table S24**.

### RNA sequencing and differential expression analysis

BMDMs differentiated from WT, *Chrna7*^-/-^, *Tfeb*^fl/fl^, *Tfeb*^ΔLysM^ mice were treated with DMSO control or PNU-282987 (50 µL) for 2 h. Total RNA was extracted from samples as described above using Purelink RNA mini kit (Thermo Fisher Scientific, 12183018A). PolyA selection, library preparation and sequencing using Illumina Nova-Seq or HiSeq platform, 2x150bp, ∼350,000,000 paired-end reads was performed from whole cell RNA from 4 replicates per condition by Azenta/Genewiz. Azenta/Genewiz provided raw data in the form of reads in FASTQ format. FASTQ files were analyzed using the RNA-seq pipeline (v1.6.0) on ViaFoundry Inc. software (v1.6.5), which is derived from DolphinNext [71]. Using the pipeline, FASTQ paired end reads were trimmed, quality filtered by FastQC, followed by 3’ adapter sequence removal and aligned to *Mus musculus* reference genome mm10 using STAR, and RSEM was used to estimate gene and isoform expression levels. DEBrowser (v1.28.0) was used for gene count normalization, data analyses, and to generate visual graphs of differentially expressed genes [72]. Differentially expressed gene sets were considered significant for fold change ≥ 1.5; Padj ≤ 0.01 and were used as input for g:Profiler [73]. Comparison of DE genes with public GEO RNA-Seq datasets was performed using Enrichr [74]. Venn diagrams were made using DeepVenn [75].

### ELISA

Secreted TNF-α (DY410-05), IL-1β (DY401-05), IL-6 (DY406-05), IFN-β (DY8234-05), IL-10 (DY417-05), TNFSF15 (DY-1896) were measured using ELISA kits purchased from R&D Systems. Briefly, 96-well plates were coated with capture antibody and incubated at room temperature overnight. Plates were washed with Tris buffered saline (TBS) with 0.5% Tween 20 (Bethyl Laboratories, E106). Wells were blocked for 1 h with Reagent Diluent - TBS containing 1% bovine serum albumin (BSA) (Bethyl Laboratories, E106). After washing, samples or standards diluted in Reagent Diluent were added and plates were incubated for 2 h at room temperature. Detection antibody was added following washing and plates were incubated for 2 h. After washing, Streptavidin-horseradish peroxidase (Strep-HRP) was added for 20 min. TMB One Component HRP Microwell substrate (Bethyl Lab, E102) was added, and reaction was stopped using ELISA Stop Solution (Bethly Laboratories, E115). Optical density (OD) at 450 nm and 540 nm was measured immediately using a SpectraMax Plus 384 Microplate Reader with Softmax Pro software. OD measured at 540 nm was subtracted from 450 nm to correct for optical imperfections as per manufacturer’s guidelines. Concentrations of the samples were interpolated from respective standard curves using GraphPad Prism 10.

### Cell viability assay

Cell viability was determined using CellTiter-Glo 2.0 Assay (Promega, G7570), according to the manufacturer’s instructions. Luminescence was measured using Biotek Synergy H4 Plate Reader Gen5 (version 3.14).

### ROS/Superoxide detection assay

Cellular ROS/Superoxide Detection Assay Kit (Abcam, ab139476) was used according to manufacturer’s instructions. Mean fluorescence intensity (MFI) was visualized on a Lionheart microscope (Biotek) and measured using Gen5 Data Analysis Software (Version 3.13).

### siRNA knockdown

siRNA reagents were purchased from Dharmacon RNAi Technologies: siGENOME NonTargeting siRNA #1 (D-001210-01-05), siGENOME Mouse Ppp3cb (19,056) siRNA (M-063545-00-0005), siGENOME Mouse Ppp3r1 (19,058) siRNA (M-040744-01-0005), siGENOME Mouse Mcoln1 (94178) siRNA (M-044469-00-515 0005). For transfection, we used Lipofectamine RNAiMAX (Thermo Scientific, 13,778,030), according to manufacturer’s instructions.

### Western Blotting

Cell samples were washed 3 times with PBS, harvested, and lysed with 1X SDS sample buffer Blue Loading Pack (Cell Signaling Technology, 7722) at 100 μl per well of 6-well plate. Lysates were heated at 100 °C for 5 min and then centrifuged for 5 min. Supernatants were collected and sonicated, followed by gel electrophoresis using polyacrylamide Novex 4–20% Tris-Glycine Mini Gels (Invitrogen, XP04200BOX), and transfer onto nitrocellulose (Life Technologies, LC2009). After washing with TBS (Life Technologies, 28,358) for 5 min, membranes were soaked in blocking buffer containing 1X TBS with 5% BSA for 1 h at room temperature. After 3 washes with TBS-Tween 20 (Life Technologies, 28,360), membranes were incubated overnight at 4°C with primary antibodies and gentle agitation. Next, membranes were washed thrice with TBS-Tween 20 and incubated with HRP-conjugated secondary antibody (Cell Signaling Technology, 7074 1:2000) for 1 h at room temperature with gentle agitation. Membranes were then washed with TBS-Tween20 and incubated with LumiGLO® (Cell Signaling Technology, 7003) for 1 min and imaged using Gel box imager (Syngene) GeneSys software. Primary antibodies and dilutions were as follows: ACTB/β-actin antibody (Cell Signaling Technology, 4967; 1:1000), α7nAChR antibody (Abcam ab216485, 1:500).

### α-bungarotoxin staining and flow cytometry

α-bungarotoxin, Alexa Fluor 647 Conjugate (Thermo Fisher Scientific, B35450) was used according to manufacturer’s guidelines to stain murine peritoneal macrophages and muscularis macrophages, isolated according to previously described methods [76] by flow cytometry-based cell sorting [77]. Data were analyzed with FlowJo.

### Mouse whole brain lysate for Western Blot

Frozen brain tissue from adult 9-month-old C57Bl/6J mice were homogenized in ProcartaPlex™ Cell Lysis Buffer (ThermoFisher #EPX-99999-000) and centrifuged at 4°C for 20 min at 18000 x g.

### Determination of bactericidal activity

*Escherichia coli* DH5α was purchased from ThermoFisher Scientific (18265017). *Staphylococcus aureus* NCTC8325 is a gift from Fred Ausubel, MGH Research Institute, USA. Bacteria were grown overnight at 37°C in TSB (Sigma-Aldrich, T8907) supplemented with 10 μg/ml nalidixic acid for *S. aureus*, or LB medium (Fisher Scientific, BP1425-500) for *E. coli*. Cultures were diluted 1:50 in the respective media the next day and grown at 37°C to mid-exponential phase. WT and *Tfe3 Tfeb* dKO RAW 264.7 cells were infected with *S. aureus* (MOI = 10) or *E. coli* (MOI = 50) and incubated at 37°C for 30 min as previously described [6]. Lysates were serially diluted, aliquots plated on TSA (Difco, BD, 236,950)-nalidixic acid agar (*S. aureus*) or LB agar (*E. coli*) and incubated overnight at 37°C, and colonies were counted for colony-forming units/ml calculation.

### Statistical analysis

Quantitative data comparisons were analyzed using statistical tests, performed as indicated in each figure legend. *P* ≤ 0.05 were judged to be significant. For normally distributed data and comparisons with a single condition, we used the two-sample two-sided *t*-test. For multiple comparisons, we used one-way ANOVA followed by Tukey’s *post-hoc* test and two-way ANOVA with Šídák’s multiple comparisons test.

## Supporting information

Supplementary Tables_Index

## Acknowledgments

We thank the members of the Irazoqui Lab, Department of Microbiological and Physiological System (MaPS), Department of Pathology, and the Immunology and Microbiology Graduate Program (IMP) for helpful feedback and insights. Dr. Alper Kucukural and Dr. Onur Yukselen provided help with RNA-sequencing data analysis tools. Nicole Briand provided expert technical assistance. Thuyvan Luu and Abigail Hiller from Dr. Paul Greer’s Lab (UMCMS) provided mouse whole brain lysate samples and whole brain lysate cDNA. Amy Parker, Annette Bohigian, Richard Fish, Tracey Rae, Dhruti Desai, and Marie Berardi provided excellent administrative support. This work was supported by the National Institute of General Medical Sciences of the National Institutes of Health under award numbers R01GM101056 and R35GM149284 (JEI), and by the Dr. Marcellette G. Williams Memorial Fund (JEI). The content is solely the responsibility of the authors and does not necessarily represent the official views of the National Institutes of Health.

