## Supplementary Text for "TFEB-Mediated Pro-inflammatory Response in Murine Macrophages Induced by Acute Alpha7 Nicotinic Receptor Activation"

### Supplementary Materials Text

Supplementary Figure Legends and Tables:

**Figure S1. Lack of effect of α7nAChR agonism on phagocytosis, bactericidal killing, and cell viability.**

**A, B.** Quantification of cell-associated or bound bacteria at t = 0 h post infection, internalized bacteria at 1 h post infection, or intracellular bacteria 6 h post infection represented as CFU/mL. RAW264.7 cells were treated with DMSO, PNU282987 (50 µM, 2 h), or carbachol (50 µM, 2 h) prior to infection with *E. coli* DH5α (MOI 50, 30 min) (**A**) or *S. aureus* NTCT8325 (MOI 10, 30 min) (**B**). 4 biological replicates.

**C, D.** Cell viability, measured using CellTiter Glo, normalized to untreated control. WT BMDMs were treated for 24 h with serially diluted concentrations of carbachol (**C**) or PNU-282987 (**D**). 2 biological replicates. *$p<0.05$; ns, not significant, 2-sided 2-sample unpaired *t* test.

**Figure S2. Quantification of cytokine secretion and cell viability during treatment with PNU-282987 and LPS.**

**A-E.** Secreted cytokines measured by ELISA of culture supernatants from WT BMDMs under the following conditions: untreated (unt), DMSO, PNU-282987 (50 µM, 12 h), LPS (100 ng/mL, 12 h), LPS (100 ng/mL, 2 h) followed by addition of PNU282987 (50 µM, 12 h), PNU282987 (50 µM, 2 h) followed by addition of LPS (100 ng/mL, 12 h). Cytokines: IL-6 (**A**), TNF-α (**B**), IFN-β1 (**C**), IL-1β (**D**), and IL-10 (**E**). 3 biological replicates, *$p \leq 0.05$, **$p \leq 0.01$, ***$p \leq 0.001$, ****$p \leq 0.0001$, one-way ANOVA followed by Tukey's post-hoc test.

**F.** Lactate dehydrogenase (LDH) release measured in BMDM culture supernatants. Conditions are same as A-E. 3 biological replicates, *$p \leq 0.05$, **$p \leq 0.01$, ***$p \leq 0.001$, ****$p \leq 0.0001$, one-way ANOVA followed by Tukey's post-hoc test.

**G-J.** Secreted cytokines measured by ELISA of culture supernatants from WT RAW264.7 cells under the following conditions: untreated (unt), DMSO, PNU-282987 (50 µM, 12 h), LPS (100 ng/mL, 12 h), LPS (100 ng/mL, 2 h) followed by addition of PNU282987 (50 µM, 12 h), PNU282987 (50 µM, 2 h) followed by addition of LPS (100 ng/mL, 12 h). Cytokines: IL-6 (**G**), TNF-α (**H**), IFN-β1 (**I**), and IL-1β (**J**). 3 biological replicates, *$p \leq 0.05$, **$p \leq 0.01$, ***$p \leq 0.001$, ****$p \leq 0.0001$, one-way ANOVA followed by Tukey's post-hoc test.

**K.** Lactate dehydrogenase (LDH) release measured in RAW264.7 cell culture supernatants. Conditions are same as G-J. 3 biological replicates.

**Figure S3. α7nAChR protein expression analysis.**

**A.** Anti-α7nAChR immunoblot of whole cell lysates from wild type (WT) and *Chrna7*$^{-/-}$ (KO) BMDMs, including whole brain lysate from wild type mice as positive control. β-actin as loading control. MW, molecular weight standard.

**B, C.** Flow cytometry of mouse peritoneal macrophages stained without (**B**) or with (**C**) Alexa Fluor 647-labelled α-bungarotoxin.

**D, E.** Flow cytometry of mouse small bowel macrophages stained without (**D**) or with (**E**) Alexa Fluor 647-labelled α-bungarotoxin.

**F, G.** Flow cytometry of mouse colon macrophages stained without (**F**) or with (**G**) Alexa Fluor 647-labelled α-bungarotoxin.

**H, I.** Flow cytometry of mouse colonic macrophages from mice infected with *Salmonella enterica* stained (A) without or (B) with Alexa Fluor 647-labelled α-bungarotoxin.

**Figure S4. RNA-seq analysis of baseline WT v. *Chrna7*$^{-/-}$ BMDMs.**

**A.** Expression analysis of macrophage marker genes from RNA-seq analysis. Faint colors represent non-significantly different. Saturated colors and asterisks adjacent to gene names represent significant differences. *$P$adj ≤ 0.05; **$P$adj ≤ 0.01; ***$P$adj ≤ 0.001.

**B.** Differentially expressed genes in *Chrna7*$^{-/-}$ BMDMs compared to WT at baseline. Red, genes upregulated in *Chrna7*$^{-/-}$ BMDMs; blue, genes upregulated in wild type BMDMs (FC ≥ 1.5; $P$adj. ≤ 0.01). 4 biological replicates.

**C.** GO-overrepresentation and KEGG pathway enrichment analysis for genes upregulated in wild type BMDMs compared with *Chrna7*$^{-/-}$.

**D.** GO-overrepresentation and KEGG pathway enrichment analysis for genes upregulated in *Chrna7*$^{-/-}$ BMDMs compared with WT.

**Figure S5. RNA-seq analysis of baseline *Tfeb*$^{f/f}$ v. *Tfeb*$^{\Delta LysM}$ BMDMs.**

**A.** Expression of macrophage marker genes (RNA-seq) in *Tfeb*$^{fl/fl}$ and *Tfeb*$^{\Delta LysM}$ BMDMs. Faint colors represent non-significant differences. Saturated colors and asterisks adjacent to gene names represent significant differences. **** $P$adj ≤ 0.0001.

**B.** Differentially expressed genes in *Tfeb*$^{\Delta LysM}$ BMDMs compared to *Tfeb*$^{fl/fl}$. Red, upregulated genes in *Tfeb*$^{\Delta LysM}$ BMDMs; blue, upregulated genes in *Tfeb*$^{fl/fl}$ BMDMs (FC ≥ 1.5; $P$adj. ≤ 0.01).

**C.** GO-overrepresentation and KEGG pathway enrichment analysis for genes upregulated in *Tfeb*$^{fl/fl}$ BMDMs.

**D.** TNFSF15 in culture supernatants of WT and *Tfeb*$^{fl/fl}$ BMDMs treated with DMSO control or PNU-282987 (50 µM) for 12 h and 24 h, measured by ELISA.

**Figure S6. Quantification of cytokine secretion and cell viability during treatment of WT and *Tfeb Tfe3* dKO RAW264.7 cells with PNU-282987 and LPS.**

**A-D.** ELISA of culture supernatants from RAW264.7 cells treated in the following conditions: untreated (unt), DMSO, PNU-282987 (50 µM, 12 h), LPS (100 ng/mL, 12 h), LPS (100 ng/mL, 2 h) followed by PNU282987 (50 µM, 12 h), PNU282987 (50 µM, 2 h) followed by LPS (100 ng/mL, 12 h). Cytokines: IL-6 (**A**), TNF-α (**B**), IFN-β1 (**C**), and IL-1β (**D**), 3 biological replicates, $^{**}p \le 0.01$, $^{***}p \le 0.001$, one-way ANOVA followed by Tukey's post-hoc test. WT values are included from Fig. S2G-J for comparison.

**E.** LDH release assay using culture supernatants from WT and dKO RAW264.7 cells in the same conditions as A-D. 3 biological replicates, $^{*}p \le 0.05$, $^{**}p \le 0.01$, $^{****}p \le 0.0001$, one-way ANOVA followed by Tukey's post-hoc test. WT values are included from Fig. S2K for comparison.

**F.** Expression of macrophage marker genes from RNA-seq analysis. Faint colors represent non-significantly different. Saturated colors and asterisks adjacent to gene names represent significant differences. ** $P$adj ≤ 0.01; *** $P$adj ≤ 0.001; **** $P$adj ≤ 0.0001.

**Figure S7. Inhibition of protein kinase D prevents TFEB activation by PNU-282987.**

**A-D.** anti-TFEB immunofluorescence micrographs of WT BMDMs treated with DMSO (**A**), PNU282987 (50 µM, 2 h) (**B**), pre-treatment with kb-NB142-70 (10 µM, 3 h) followed by PNU282987 (50 µM, 2 h) (**C**), and kb-NB142-70 (10 µM, 3 h) (**D**).

**E.** Quantification of TFEB N/C ratio from A-D (400 cells per condition, $^{****}p \le 0.0001$, one-way ANOVA followed by Tukey's post-hoc test). 3 biological replicates.

**Figure S8. Elevation of ROS by PNU-282987 and by deletion of *Chrna7.***

**A, B.** Total ROS (**A**) and (**B**) superoxide shown as mean fluorescence intensity (MFI) in WT BMDMs treated with DMSO, PNU-282987 (50 µM, 2 h), or PNU-282987 (50 µM, 2 h) following pre-treatment with MG624 (1 µM, 2 h). Pyocyanin (200 µM, 30 min) and NAC (5 mM, 30 min), positive and negative controls, respectively. 400 cells per condition, $^{*}p \le 0.05$, $^{****}p \le 0.0001$, one-way ANOVA followed by Tukey's post-hoc test, 3 biological replicates.

**C, D.** Total ROS (**C**) and superoxide (**D**) measured in WT and *Chrna7*$^{-/-}$ BMDMs following treatment with PNU-282987 (50 µM, 2 h), NAC (5 mM, 30 min), and Pyocyanin (200 µM, 30 min). 400 cells per condition, 3 biological replicates, $^{****}p \le 0.0001$, one-way ANOVA followed by Tukey's post-hoc test.

**E.** anti-TFEB immunofluorescence quantification of N/C ratio in WT and *Chrna7*$^{-/-}$ BMDMs following treatment with PNU-282987 (50 μM, 2 h). 400 cells per condition, 3 biological replicates, ****$p \leq 0.0001$, one-way ANOVA followed by Tukey's post-hoc test.

**F, G.** anti-TFEB immunofluorescence quantification of N/C ratio in WT and *Chrna7*$^{-/-}$ BMDMs following treatment with PNU-282987 (50 μM, 2 h), pretreatment with NAC (5 mM, 30 min) followed by addition of PNU-282987 (50 μM, 2 h), pretreatment with ML-SI3 (10 μM mM, 1 h) followed by addition of PNU-282987 (50 μM, 2 h), BAPTA-AM (10 μM, 3 h pre-treatment) followed by addition of PNU-282987 (50 μM, 2 h), or FK506 (5 μM, 6 h pre-treatment) followed by addition of PNU-282987 (50 μM, 2 h). 400 cells per condition, 3 biological replicates, ****$p \leq 0.0001$, one-way ANOVA followed by Tukey's post-hoc test. The PNU-282987 controls are included from E for ease of comparison.

**Table S1. Differential gene expression in WT BMDMs (PNU-282987 v. DMSO).**

**Table S2. GO and KEGG Pathway analysis of differentially expressed genes from Table S1.**

**Table S3. Differential gene expression in WT RAW264.7 cells (PNU-282987 v. DMSO).**

**Table S4. GO and KEGG Pathway analysis of differentially expressed genes from Table S3.**

**Table S5. Genes that were induced by PNU-282987 in both WT BMDMs and RAW264.7.**

**Table S6. RT-qPCR analysis of cholinergic receptor subunit expression.**

**Table S7. Differential gene expression in *Chrna7*$^{-/-}$ BMDMs (PNU-282987 v. DMSO).**

**Table S8. Genes that were induced by PNU-282987 in both WT and *Chrna7*$^{-/-}$ BMDMs.**

**Table S9. Differential gene expression in PNU-282987- treated BMDMs (*Chrna7*$^{-/-}$ v. WT).**

**Table S10. GO and KEGG Pathway analysis of differentially expressed genes from Table S9.**

**Table S11. Row-normalized expression of macrophage marker genes in WT and *Chrna7*$^{-/-}$ BMDMs.**

**Table S12. Differential gene expression in baseline BMDMs (*Chrna7*$^{-/-}$ v. WT).**

**Table S13. GO and KEGG Pathway analysis of differentially expressed genes from Table S12.**

**Table S14. Row-normalized expression of macrophage marker genes in *Tfeb*$^{fl/fl}$ and *Tfeb*$^{\Delta LysM}$ BMDMs.**

**Table S15. Differential gene expression in baseline BMDMs (*Tfeb*$^{fl/fl}$ v. *Tfeb*$^{\Delta LysM}$).**

**Table S16. GO and KEGG Pathway analysis of differentially expressed genes from Table S15.**

**Table S17. Differential gene expression in PNU-282987-treated BMDMs (*Tfeb*$^{fl/fl}$ v. *Tfeb*$^{\Delta LysM}$).**

**Table S18. GO and KEGG Pathway analysis of differentially expressed genes from Table S17.**

**Table S19. Genes that were induced by PNU-282987 in *Tfeb*$^{fl/fl}$ but not *Tfeb*$^{\Delta LysM}$ BMDMs.**

**Table S20. Differential gene expression in PNU-282987-treated RAW264.7 cells (WT v. *Tfeb Tfe3* dKO).**

**Table S21. GO and KEGG Pathway analysis of differentially expressed genes from Table S20.**

**Table S22. Differential gene expression in baseline RAW264.7 cells (WT v. *Tfeb Tfe3* dKO).**

**Table S23. Row-normalized expression of macrophage marker genes in WT and *Tfeb Tfe3* dKO RAW264.7 cells.**

**Table S24. Oligonucleotide sequences.**
