## Supplementary Figures for "TFEB-Mediated Pro-inflammatory Response in Murine Macrophages Induced by Acute Alpha7 Nicotinic Receptor Activation"

### FIGURE S1

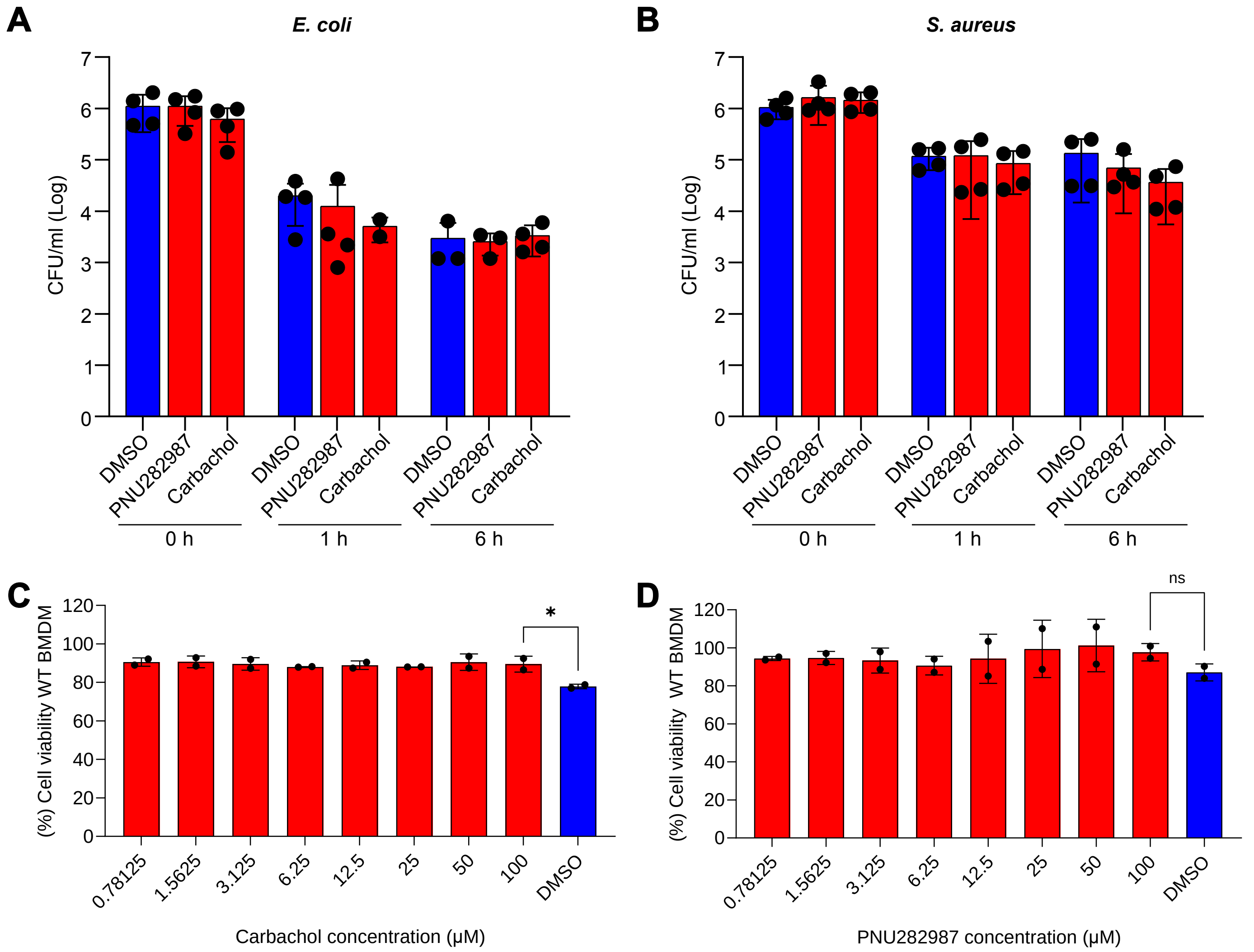

A
E. coli
CFU/ml (Log)
0
1
2
3
4
5
6
7
DMSO
PNU282987
Carbachol
0 h
1 h
6 h
B
S. aureus
C
(%) Cell viability WT BMDM
0
20
40
60
80
100
120
*
0.78125
1.5625
3.125
6.25
12.5
25
50
100
DMSO
Carbachol concentration (μM)
D
ns
PNU282987 concentration (μM)

### FIGURE S2

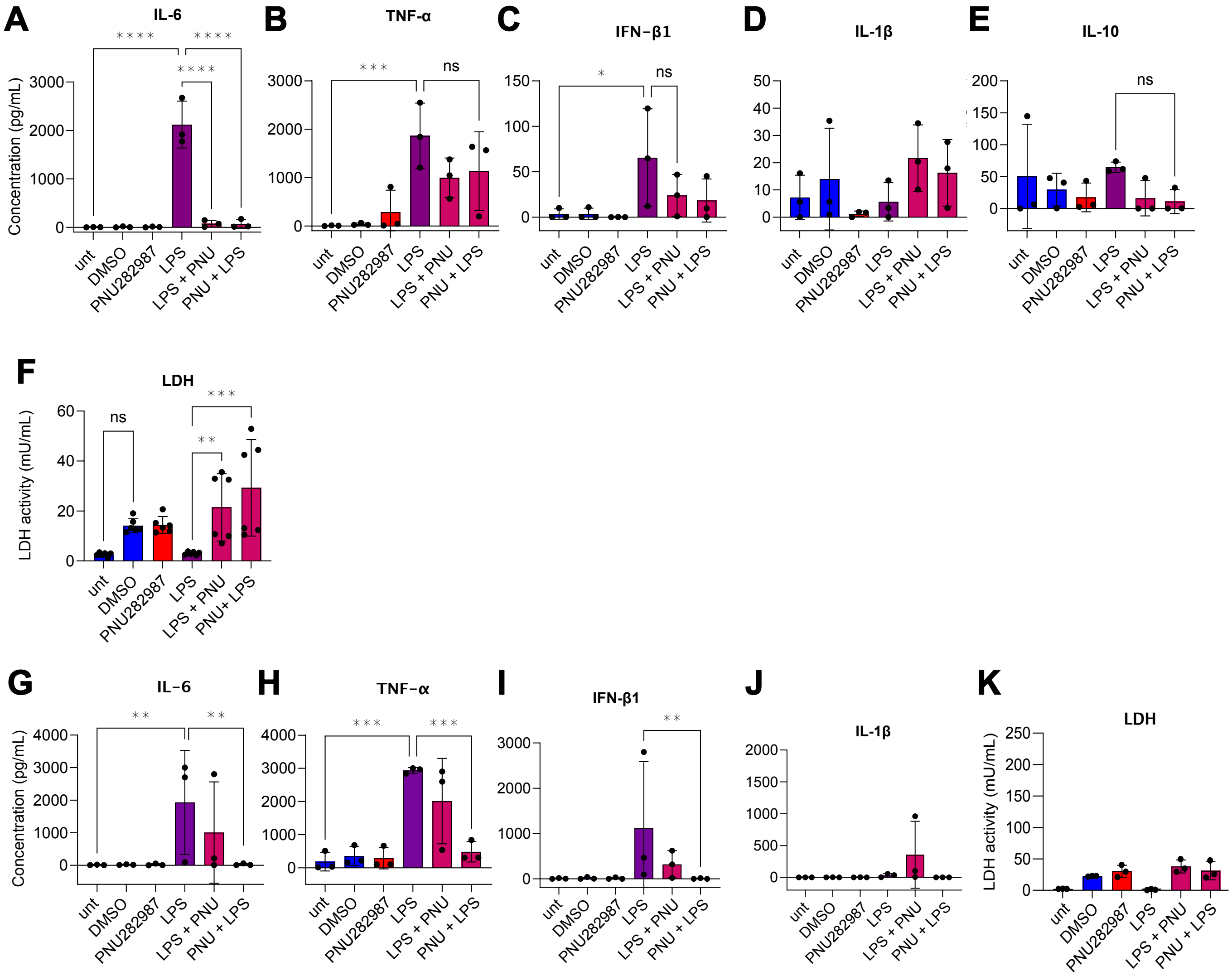

A
IL-6
Concentration (pg/mL)
B
TNF-α
C
IFN-β1
D
IL-1β
E
IL-10
F
LDH
LDH activity (mU/mL)
G
IL-6
Concentration (pg/mL)
H
TNF-α
I
IFN-β1
J
IL-1β
K
LDH
LDH activity (mU/mL)
ns
unt
DMSO
PNU282987
LPS
LPS + PNU
PNU + LPS
PNU+ LPS

### FIGURE S3

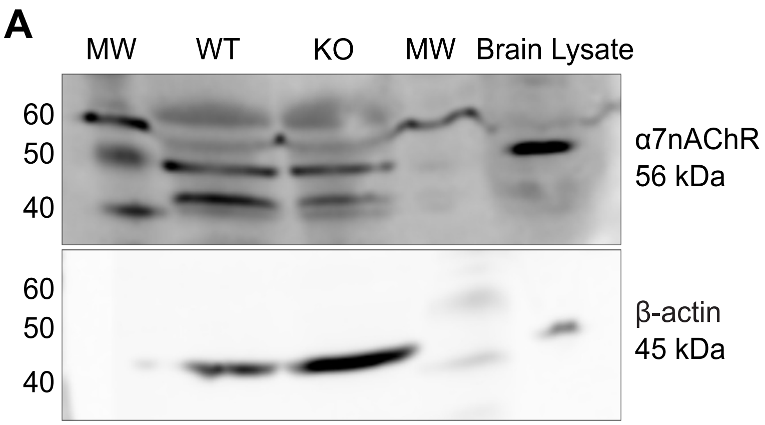
A
MW
WT
KO
MW
Brain Lysate
60
50
40
α7nAChR
56 kDa
60
50
40
β-actin
45 kDa

**B** Steady state peritoneal cells without α-Bungarotoxin

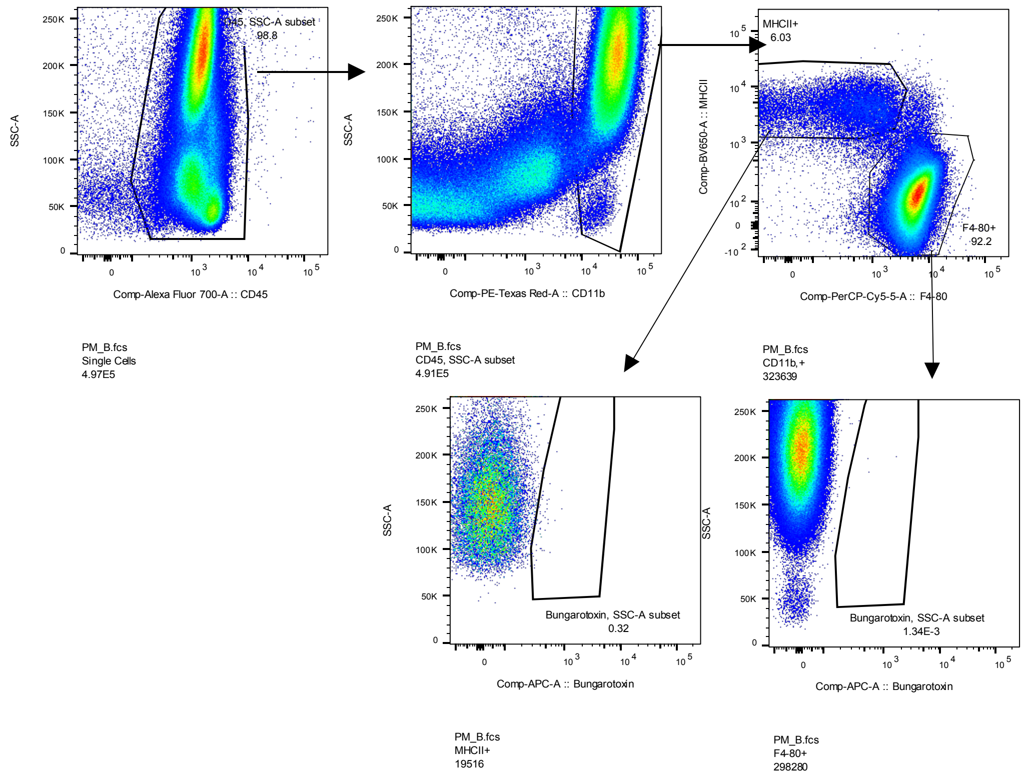


**C** Steady state peritoneal cells with α-Bungarotoxin

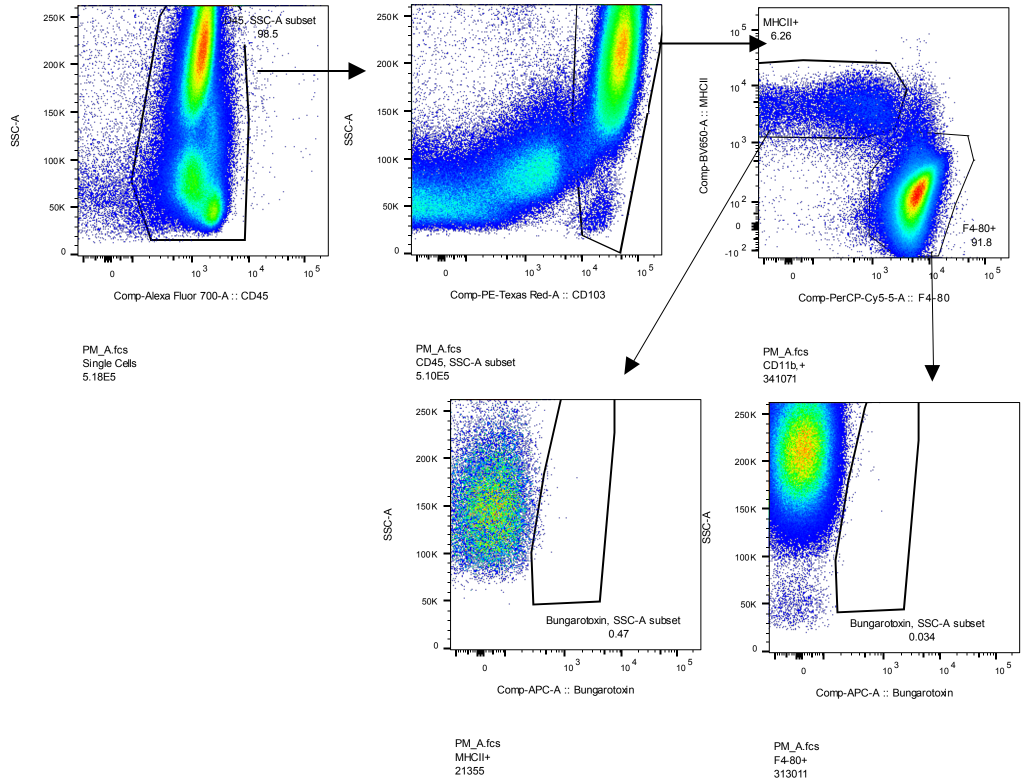

**D** Steady state small bowel macrophages without α-Bungarotoxin

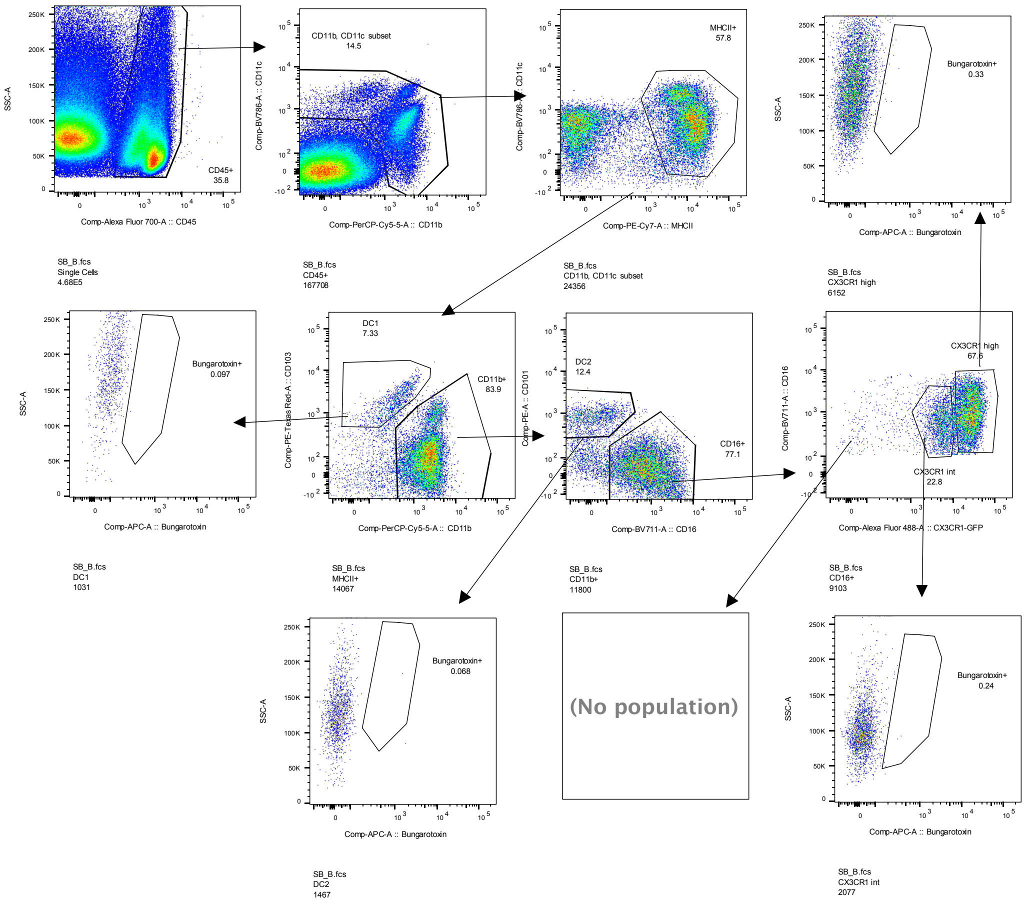


**E** Steady state small bowel macrophages with α-Bungarotoxin

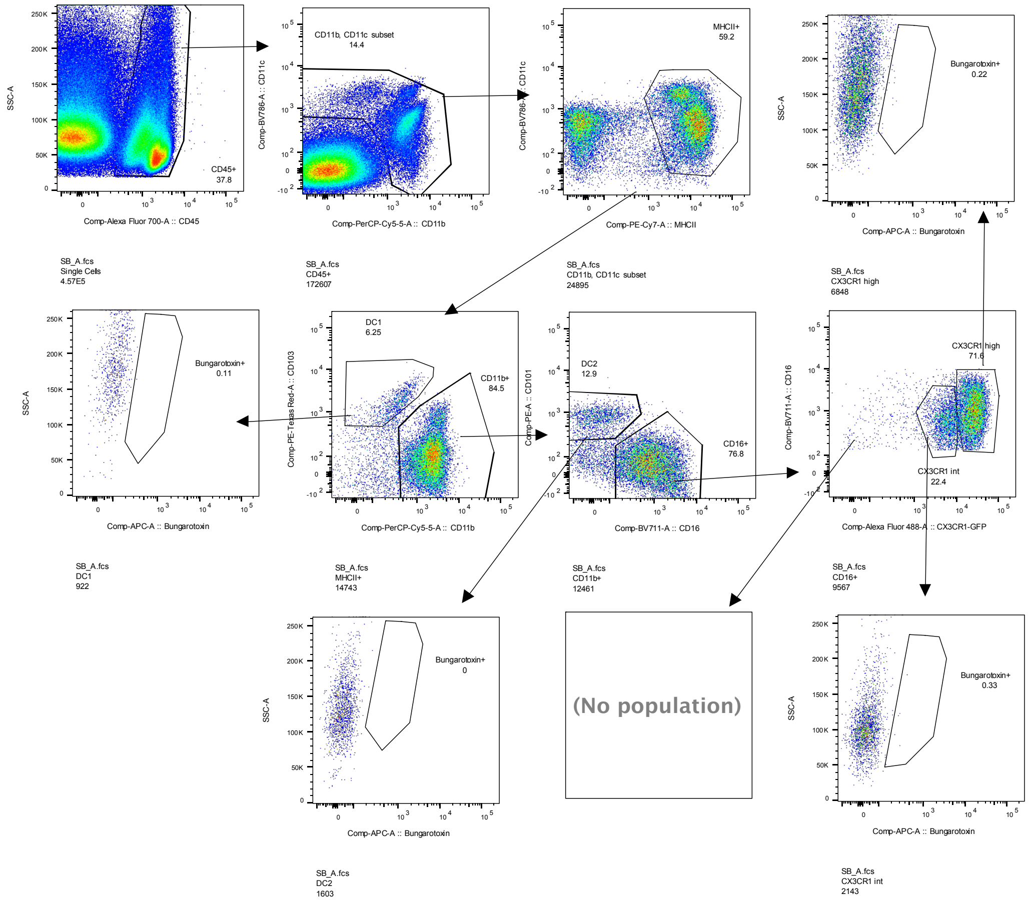

**F** Steady state large bowel macrophages without α-Bungarotoxin

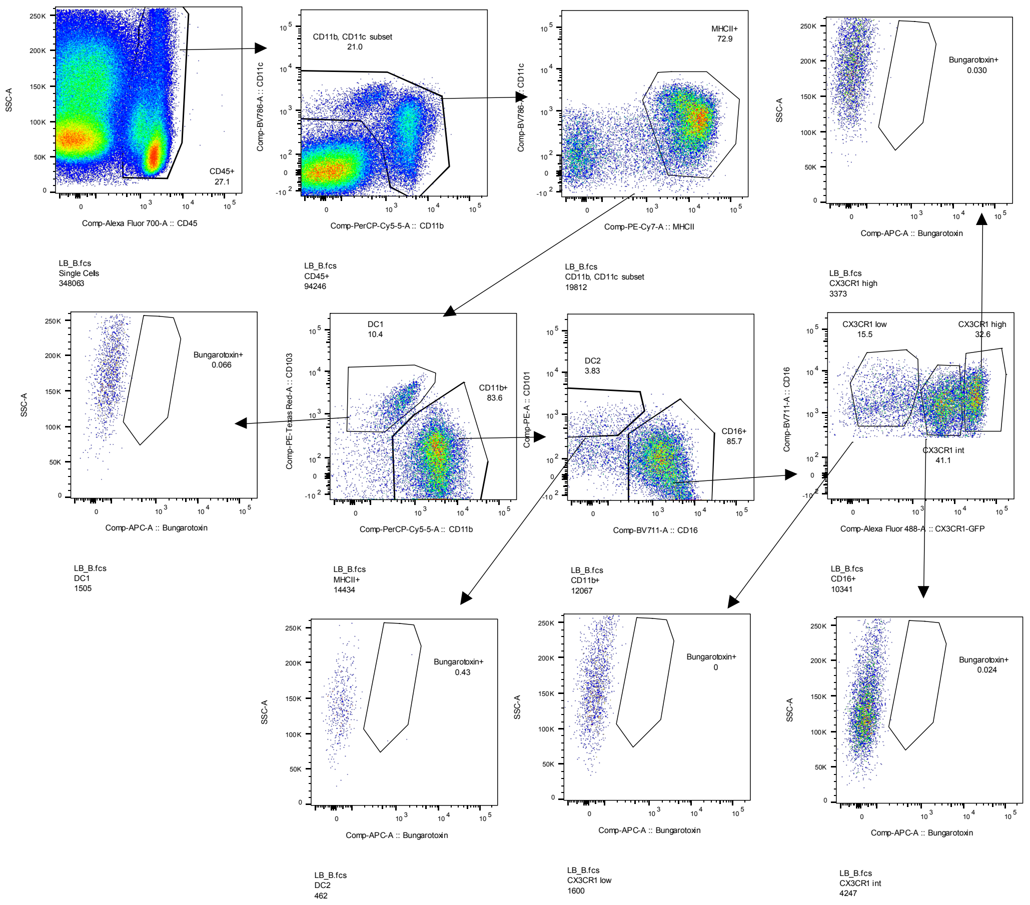


**G** Steady state large bowel macrophages with α-Bungarotoxin

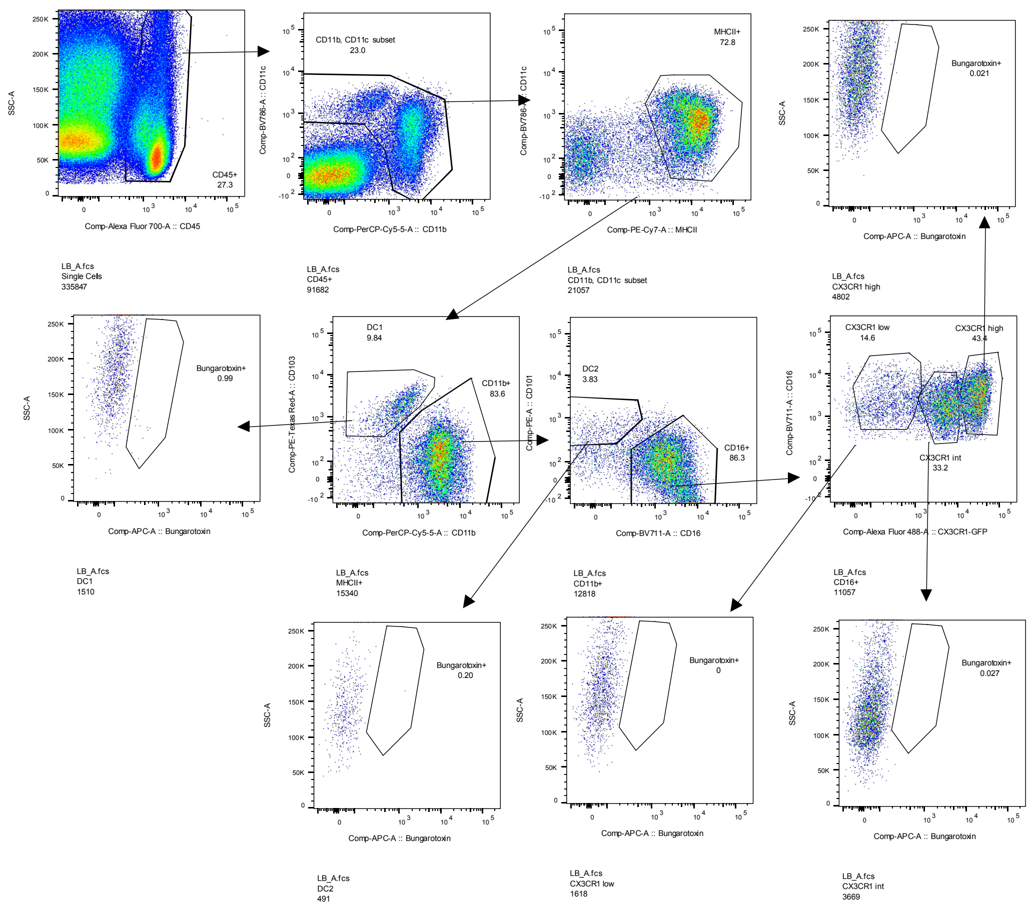

**H** Infected large bowel macrophages without α-Bungarotoxin

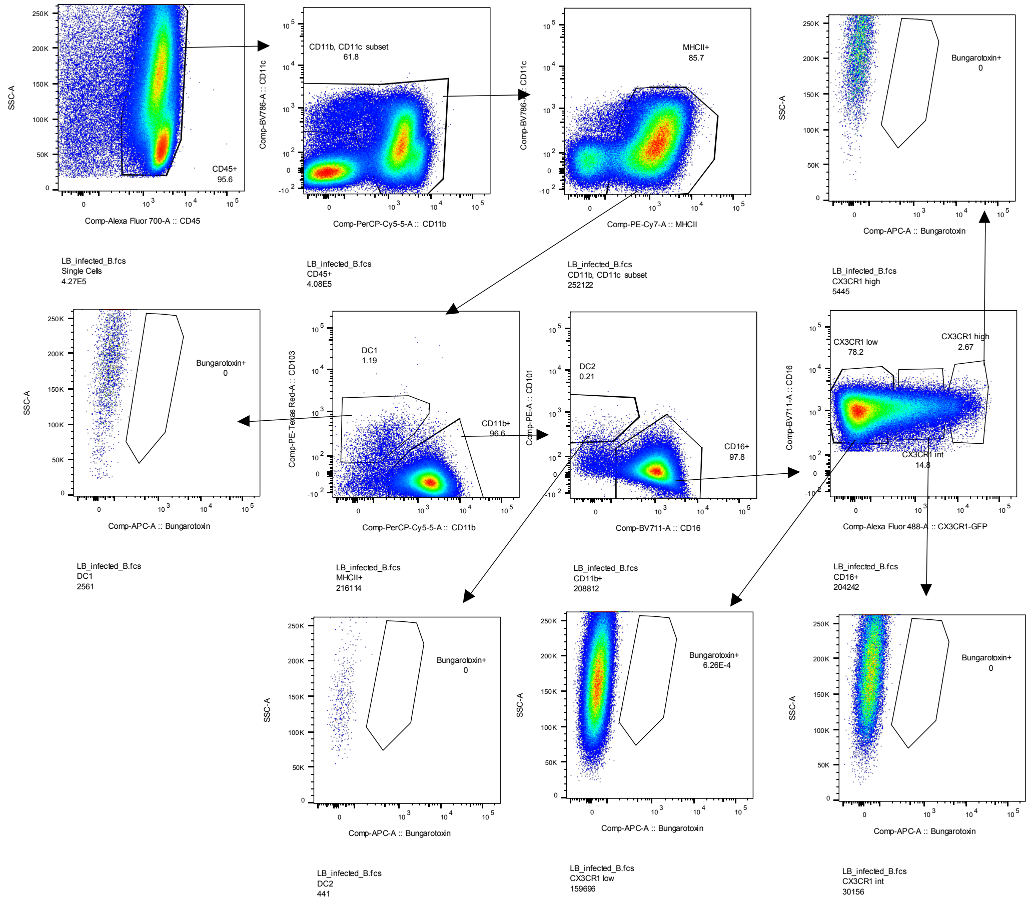


**I** Infected large bowel macrophages with α-Bungarotoxin

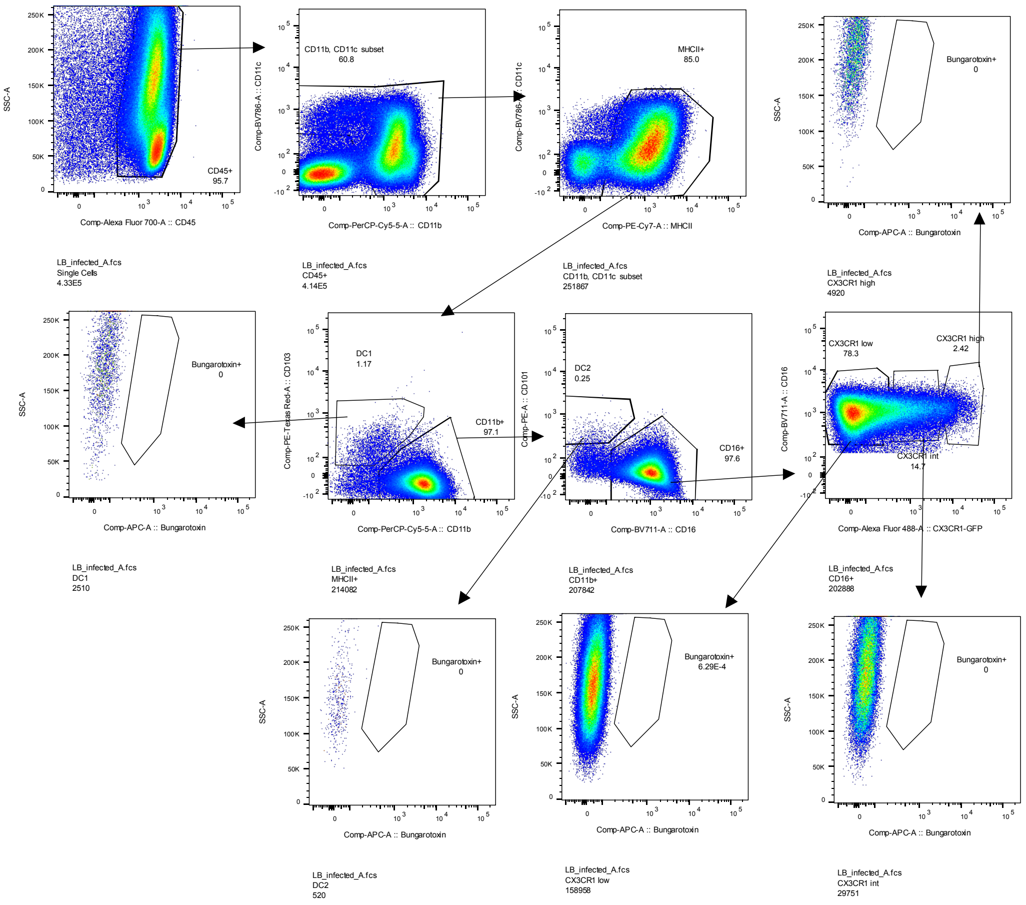

### FIGURE S4

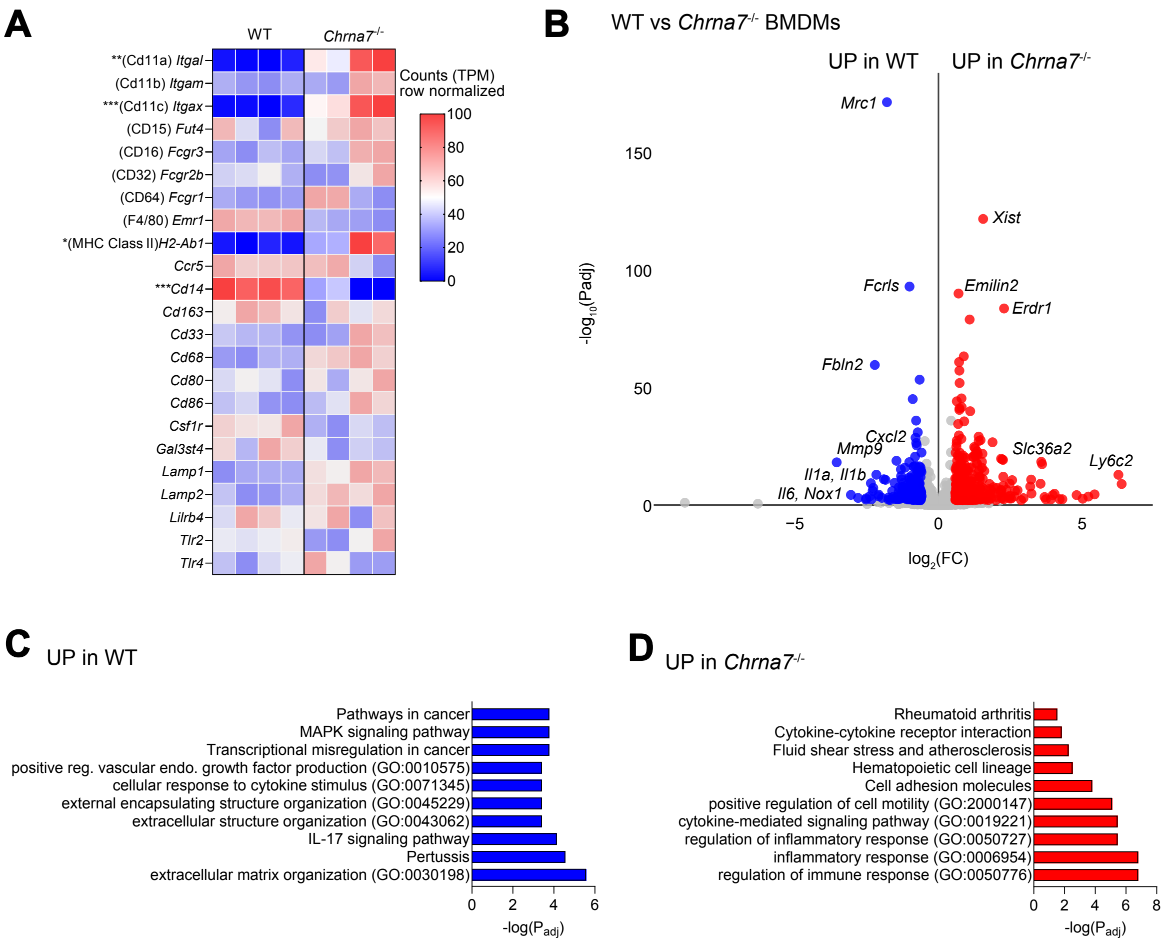
A
WT
Chrna7-/-
**(Cd11a) Itgal
(Cd11b) Itgam
***(Cd11c) Itgax
(CD15) Fut4
(CD16) Fcgr3
(CD32) Fcgr2b
(CD64) Fcgr1
(F4/80) Emr1
*(MHC Class II)H2-Ab1
Ccr5
***Cd14
Cd163
Cd33
Cd68
Cd80
Cd86
Csf1r
Gal3st4
Lamp1
Lamp2
Lilrb4
Tlr2
Tlr4
Counts (TPM)
row normalized
100
80
60
40
20
0
B
WT vs Chrna7-/- BMDMs
UP in WT
UP in Chrna7-/-
Mrc1
Xist
Fcrls
Emilin2
Erdr1
Fbln2
Cxcl2
Mmp9
Slc36a2
Ly6c2
Il1a, Il1b
Il6, Nox1
150
100
50
0
-log10(Padj)
−5
0
5
log2(FC)
C
UP in WT
Pathways in cancer
MAPK signaling pathway
Transcriptional misregulation in cancer
positive reg. vascular endo. growth factor production (GO:0010575)
cellular response to cytokine stimulus (GO:0071345)
external encapsulating structure organization (GO:0045229)
extracellular structure organization (GO:0043062)
IL-17 signaling pathway
Pertussis
extracellular matrix organization (GO:0030198)
0
2
4
6
-log(Padj)
D
UP in Chrna7-/-
Rheumatoid arthritis
Cytokine-cytokine receptor interaction
Fluid shear stress and atherosclerosis
Hematopoietic cell lineage
Cell adhesion molecules
positive regulation of cell motility (GO:2000147)
cytokine-mediated signaling pathway (GO:0019221)
regulation of inflammatory response (GO:0050727)
inflammatory response (GO:0006954)
regulation of immune response (GO:0050776)
0
2
4
6
8
-log(Padj)

### FIGURE S5

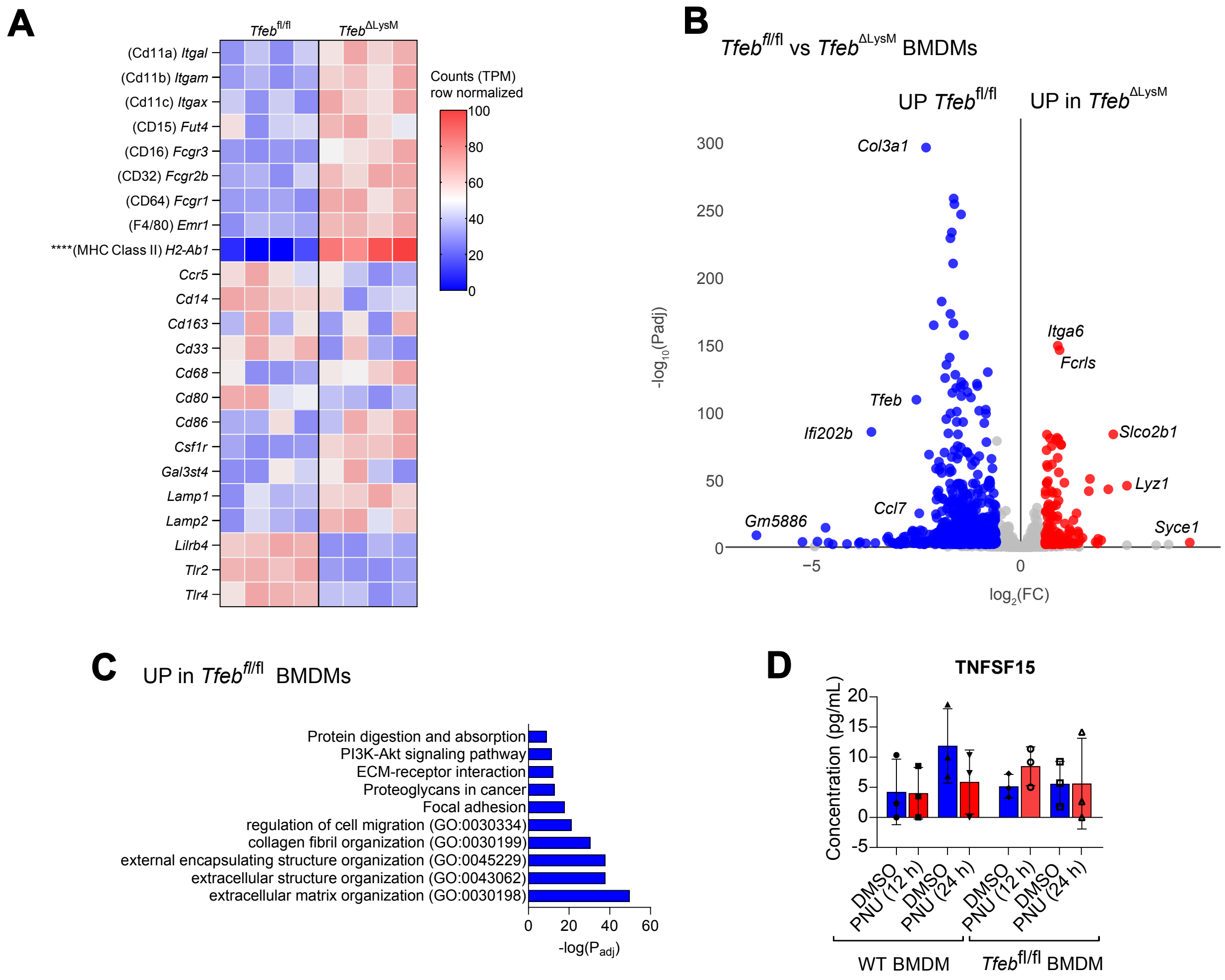
A
Tfeb fl/fl
Tfeb ΔLysM
(Cd11a) Itgal
(Cd11b) Itgam
(Cd11c) Itgax
(CD15) Fut4
(CD16) Fcgr3
(CD32) Fcgr2b
(CD64) Fcgr1
(F4/80) Emr1
****(MHC Class II) H2-Ab1
Ccr5
Cd14
Cd163
Cd33
Cd68
Cd80
Cd86
Csf1r
Gal3st4
Lamp1
Lamp2
Lilrb4
Tlr2
Tlr4
Counts (TPM)
row normalized
100
80
60
40
20
0
B
Tfeb fl/fl vs Tfeb ΔLysM BMDMs
UP Tfeb fl/fl
UP in Tfeb ΔLysM
-log10(Padj)
log2(FC)
Col3a1
Itga6
Fcrls
Tfeb
Ifi202b
Slco2b1
Lyz1
Ccl7
Gm5886
Syce1
300
250
200
150
100
50
0
−5
0
C
UP in Tfeb fl/fl BMDMs
Protein digestion and absorption
PI3K-Akt signaling pathway
ECM-receptor interaction
Proteoglycans in cancer
Focal adhesion
regulation of cell migration (GO:0030334)
collagen fibril organization (GO:0030199)
external encapsulating structure organization (GO:0045229)
extracellular structure organization (GO:0043062)
extracellular matrix organization (GO:0030198)
0
20
40
60
-log(Padj)
D
TNFSF15
Concentration (pg/mL)
20
15
10
5
0
-5
DMSO
PNU (12 h)
DMSO
PNU (24 h)
DMSO
PNU (12 h)
DMSO
PNU (24 h)
WT BMDM
Tfeb fl/fl BMDM

### FIGURE S6

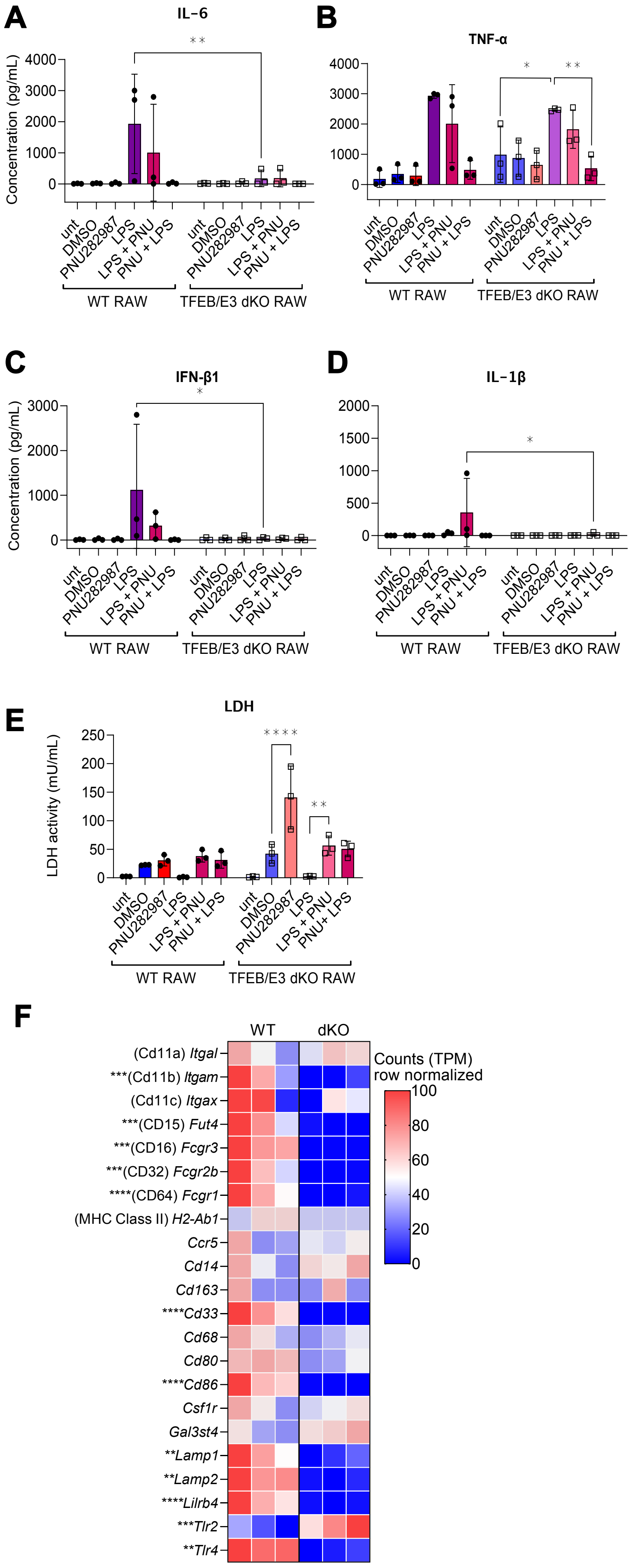
A
IL-6
Concentration (pg/mL)
unt
DMSO
PNU282987
LPS
LPS + PNU
PNU + LPS
WT RAW
TFEB/E3 dKO RAW
B
TNF-α
C
IFN-β1
D
IL-1β
E
LDH
LDH activity (mU/mL)
PNU+ LPS
F
WT
dKO
(Cd11a) Itgal
***(Cd11b) Itgam
(Cd11c) Itgax
***(CD15) Fut4
***(CD16) Fcgr3
***(CD32) Fcgr2b
****(CD64) Fcgr1
(MHC Class II) H2-Ab1
Ccr5
Cd14
Cd163
****Cd33
Cd68
Cd80
****Cd86
Csf1r
Gal3st4
**Lamp1
**Lamp2
****Lilrb4
***Tlr2
**Tlr4
Counts (TPM)
row normalized

### FIGURE S7

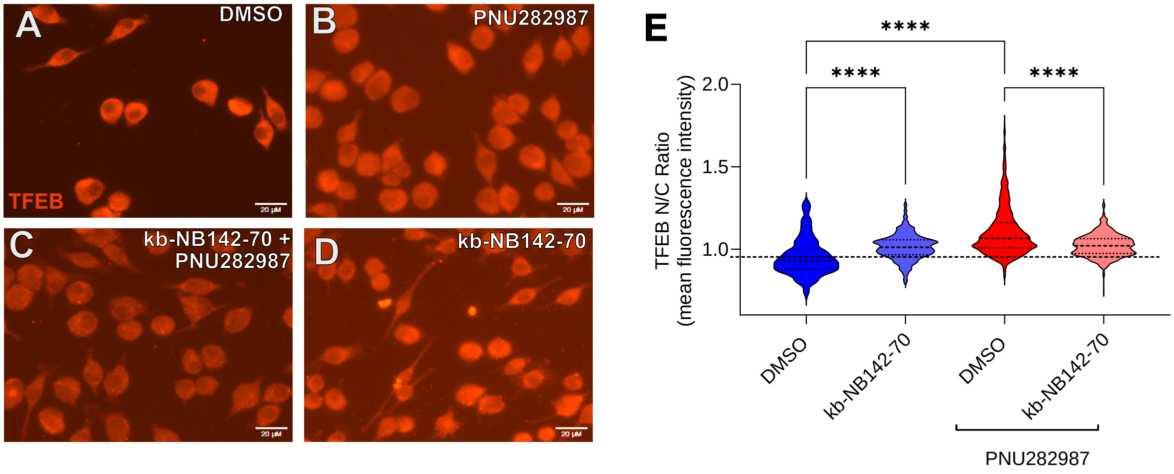
A
DMSO
TFEB
20 µM
B
PNU282987
C
kb-NB142-70 +
PNU282987
D
kb-NB142-70
E
****
****
****
TFEB N/C Ratio
(mean fluorescence intensity)
2.0
1.5
1.0
DMSO
kb-NB142-70
DMSO
kb-NB142-70
PNU282987

#### FIGURE S8

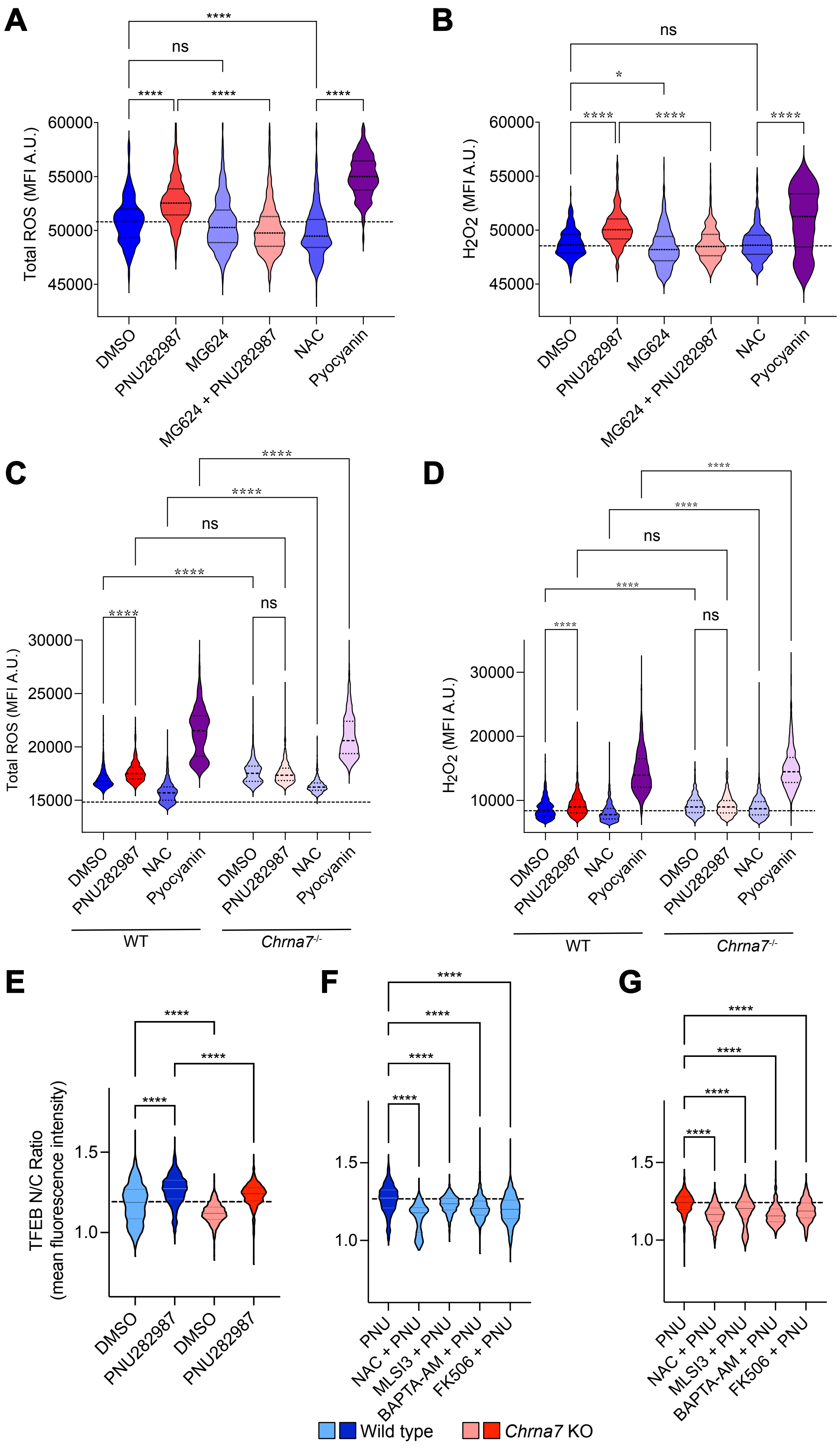
A
Total ROS (MFI A.U.)
60000
55000
50000
45000
****
ns
****
****
****
DMSO
PNU282987
MG624
MG624 + PNU282987
NAC
Pyocyanin
B
H2O2 (MFI A.U.)
ns
*
****
****
****
C
Total ROS (MFI A.U.)
30000
25000
20000
15000
****
****
ns
****
****
ns
WT
Chrna7-/-
D
H2O2 (MFI A.U.)
30000
20000
10000
****
****
ns
****
****
ns
E
TFEB N/C Ratio
(mean fluorescence intensity)
1.5
1.0
****
****
****
F
PNU
NAC + PNU
MLSI3 + PNU
BAPTA-AM + PNU
FK506 + PNU
****
G
Wild type
Chrna7 KO
