## Supplementary material for "TFEB-Mediated Pro-inflammatory Response in Murine Macrophages Induced by Acute Alpha7 Nicotinic Receptor Activation": Table_S1

**Table S1. Differential gene expression in WT BMDMs (PNU-282987 v. DMSO).**

| ID | A1_control_1 | A2_control_2 | A3_control_3 | A4_control_4 | A5_exp_1 | A6_exp_2 | A7_exp_3 | A8_exp_4 | padj | log2FoldChange | pvalue | stat | foldChange | log10padj |
|---|---|---|---|---|---|---|---|---|---|---|---|---|---|---|
| *Tarm1* | 0.891124295 | 1.977122575 | 0 | 0 | 13.95314704 | 646.6473234 | 1508.572273 | 32.88034542 | 0.000816616 | 9.585093563 | 2.79E-05 | 17.55809922 | 768.0697723 | 3.087982111 |
| *Il1f9* | 0 | 0 | 0 | 0.914339478 | 1.073319003 | 242.0542422 | 577.8175374 | 7.109263874 | 0.009701983 | 9.480052768 | 0.000791745 | 11.26048639 | 714.1348771 | 2.013139501 |
| *Bcl2a1a* | 1.782248591 | 0 | 1.036078894 | 0 | 31.1262511 | 589.3494593 | 1045.277131 | 72.86995471 | 5.12E-05 | 9.259035708 | 8.38E-07 | 24.26838914 | 612.6994339 | 4.290954106 |
| *Il12b* | 8.911242953 | 23.72547091 | 15.54118341 | 14.62943166 | 505.5332506 | 12411.41897 | 19577.08283 | 782.0190262 | 8.36E-07 | 9.050215011 | 5.33E-09 | 34.06561341 | 530.1346434 | 6.077743592 |
| *Cxcl3* | 8.020118658 | 6.919929014 | 13.46902562 | 16.45811061 | 65.47245921 | 9084.63482 | 14107.07681 | 389.2321971 | 5.29E-05 | 9.041775406 | 8.70E-07 | 24.19525555 | 527.0424681 | 4.276597524 |
| *Il1a* | 34.75384752 | 37.56532893 | 43.51531355 | 40.23093705 | 122.3583664 | 26560.48337 | 51236.27832 | 1067.278239 | 0.000103499 | 8.983369789 | 2.04E-06 | 22.55726453 | 506.1319677 | 3.985065462 |
| *Nos2* | 4.455621477 | 8.89705159 | 16.5772623 | 6.400376349 | 71.91237323 | 5380.152505 | 11544.89851 | 383.0115912 | 4.00E-05 | 8.904707424 | 6.23E-07 | 24.83959385 | 479.2741848 | 4.397756487 |
| *Cxcl1* | 5.346745772 | 9.885612877 | 9.324710046 | 8.229055306 | 60.10586419 | 5445.635778 | 8937.952357 | 412.3373047 | 1.68E-05 | 8.824896584 | 2.10E-07 | 26.94336435 | 453.480462 | 4.775855363 |
| *Lcn2* | 3.564497181 | 1.977122575 | 5.18039447 | 3.657357914 | 20.39306106 | 1734.137397 | 4123.847325 | 90.6431144 | 0.000120348 | 8.698373634 | 2.47E-06 | 22.18597464 | 415.4046727 | 3.919560781 |
| *Cxcl2* | 37.4272204 | 40.5310128 | 40.40707687 | 33.8305607 | 392.8347552 | 23941.15244 | 36849.14219 | 2022.585572 | 3.56E-06 | 8.698068625 | 3.30E-08 | 30.52050673 | 415.3168587 | 5.448434056 |
| *Il1b* | 105.1526669 | 126.5358448 | 119.1490728 | 129.8362059 | 369.2217372 | 65068.15608 | 128102.6686 | 2670.417243 | 0.000120348 | 8.673128872 | 2.48E-06 | 22.18523886 | 408.1990097 | 3.919560781 |
| *Il6* | 20.49585879 | 6.919929014 | 17.6133412 | 14.62943166 | 26.83297509 | 8359.641438 | 15492.79779 | 279.9272651 | 0.000353542 | 8.661280706 | 9.93E-06 | 19.52576657 | 404.860394 | 3.451559128 |
| *Saa3* | 50.79408483 | 37.56532893 | 34.1906035 | 36.57357914 | 415.3744543 | 17046.69924 | 43497.68777 | 992.6309685 | 2.27E-05 | 8.604680724 | 2.98E-07 | 26.26129892 | 389.2844007 | 4.644391117 |
| *Shisa3* | 4.455621477 | 0 | 1.036078894 | 0.914339478 | 21.46638007 | 894.5482865 | 1419.036583 | 50.65350511 | 0.000489042 | 8.52632784 | 1.48E-05 | 18.75879024 | 368.7061934 | 3.310653753 |
| *H2-M2* | 0.891124295 | 0.988561288 | 0 | 0.914339478 | 40.78612213 | 258.4250605 | 692.3399322 | 30.21437147 | 1.07E-07 | 8.492228586 | 5.10E-10 | 38.63858947 | 360.0937093 | 6.969995654 |
| *Csf3* | 5.346745772 | 2.965683863 | 3.108236682 | 0.914339478 | 6.43991402 | 1341.237758 | 2863.05987 | 64.87203285 | 0.000597694 | 8.43618124 | 1.90E-05 | 18.28725646 | 346.3726577 | 3.223521012 |
| *Slc1a2* | 0 | 0.988561288 | 0 | 0 | 2.146638007 | 119.2731049 | 245.7025925 | 7.109263874 | 0.006065322 | 8.3341257 | 0.000411009 | 12.48146727 | 322.7169849 | 2.217146132 |
| *Marco* | 9.802367249 | 9.885612877 | 3.108236682 | 6.400376349 | 224.3236717 | 2260.342272 | 4965.066371 | 210.6119423 | 1.97E-06 | 8.033253561 | 1.47E-08 | 32.08654293 | 261.9692306 | 5.706541356 |
| *Ppm1n* | 0 | 0 | 0 | 0 | 21.46638007 | 61.97524076 | 70.79566224 | 18.66181767 | 1.44E-10 | 7.939026771 | 3.60E-13 | 52.85125421 | 245.40601 | 9.842032427 |
| *Fpr1* | 0.891124295 | 4.942806439 | 3.108236682 | 4.571697392 | 91.23211529 | 971.7250014 | 2138.445445 | 65.76069084 | 1.91E-05 | 7.920437062 | 2.48E-07 | 26.61372756 | 242.26414 | 4.718266833 |
| *F10* | 2.673372886 | 4.942806439 | 5.18039447 | 1.828678957 | 31.1262511 | 866.4840265 | 2086.389811 | 71.09263874 | 8.00E-05 | 7.711693428 | 1.49E-06 | 23.1587177 | 209.6288439 | 4.096662254 |
| *Irg1* | 202.285215 | 261.9687412 | 199.9632265 | 176.4675193 | 5098.265266 | 53520.88245 | 105667.7314 | 9790.345013 | 1.76E-08 | 7.693973068 | 7.38E-11 | 42.41473664 | 207.0697596 | 7.754479266 |
| *Ptgs2* | 183.5716048 | 241.2089542 | 185.458122 | 163.6667666 | 461.5271715 | 48054.19847 | 84456.10171 | 2052.799944 | 0.000291808 | 7.446860345 | 7.79E-06 | 19.98826107 | 174.4730436 | 3.534902756 |
| *Ccl22* | 13.36686443 | 20.75978704 | 12.43294673 | 13.71509218 | 32.1995701 | 2805.256653 | 5840.642134 | 167.067701 | 0.000391409 | 7.197314185 | 1.14E-05 | 19.25584977 | 146.7599171 | 3.407369274 |
| *Ccl3* | 85.54793235 | 78.09634173 | 84.95846931 | 85.94791098 | 686.9241622 | 17077.10219 | 26715.99248 | 1916.835272 | 4.81E-06 | 7.115633713 | 4.73E-08 | 29.82345807 | 138.6817093 | 5.318004835 |
| *Ptges* | 22.27810738 | 33.61108378 | 36.26276129 | 41.14527653 | 250.0833278 | 4894.874676 | 12587.0523 | 476.3206796 | 4.05E-05 | 7.093971477 | 6.33E-07 | 24.80833537 | 136.6149423 | 4.392958344 |
| *Zfp811* | 0 | 1.977122575 | 2.072157788 | 1.828678957 | 15.02646605 | 232.6994889 | 483.0762835 | 31.10302945 | 8.03E-06 | 7.033814473 | 8.70E-08 | 28.64395828 | 131.0355513 | 5.095491919 |
| *Hcar2* | 11.58461584 | 10.87417417 | 17.6133412 | 15.54377113 | 70.83905423 | 2463.808156 | 4538.210172 | 197.2820725 | 7.34E-05 | 7.030725872 | 1.32E-06 | 23.39440438 | 130.7553233 | 4.134460492 |
| *Edn1* | 5.346745772 | 1.977122575 | 0 | 0.914339478 | 5.366595017 | 419.7945553 | 645.4898616 | 9.775237827 | 0.00719933 | 7.024620817 | 0.000510504 | 12.07689929 | 130.2031748 | 2.142707899 |
| *Cd38* | 1.782248591 | 5.931367726 | 5.18039447 | 2.743018435 | 17.17310405 | 535.5596277 | 1420.077695 | 44.43289922 | 0.000382671 | 7.016275859 | 1.11E-05 | 19.30451494 | 129.4522166 | 3.417174715 |
| *Rasgrp1* | 0 | 0.988561288 | 3.108236682 | 1.828678957 | 15.02646605 | 232.6994889 | 469.5418186 | 30.21437147 | 1.32E-05 | 6.999271754 | 1.59E-07 | 27.47513628 | 127.9354043 | 4.878160416 |
| *Fpr2* | 1.782248591 | 10.87417417 | 13.46902562 | 8.229055306 | 105.1852623 | 1351.761855 | 2783.935306 | 126.1894338 | 6.84E-05 | 6.993525076 | 1.19E-06 | 23.59010148 | 127.4268135 | 4.164899133 |
| *Slc7a11* | 147.0355087 | 133.4557738 | 153.3396763 | 165.4954456 | 1036.826157 | 23735.34787 | 45005.21893 | 1907.948692 | 3.10E-05 | 6.902175169 | 4.38E-07 | 25.51938023 | 119.6084221 | 4.508320525 |
| *Gfi1* | 0 | 0 | 1.036078894 | 0.914339478 | 0 | 99.39425405 | 121.8101836 | 3.554631937 | 0.007031718 | 6.87215519 | 0.000494081 | 12.13787338 | 117.1452941 | 2.15293857 |
| *Vnn3* | 0 | 0 | 1.036078894 | 1.828678957 | 4.293276014 | 101.7329424 | 216.5514374 | 2.665973953 | 0.001516519 | 6.827098582 | 6.15E-05 | 16.05476121 | 113.5432854 | 2.819152216 |
| *Adora2a* | 2.673372886 | 5.931367726 | 2.072157788 | 9.143394785 | 24.68633708 | 728.501415 | 1397.173216 | 58.65142696 | 0.000317693 | 6.798045736 | 8.66E-06 | 19.78577373 | 111.2796314 | 3.497991714 |
| *Tnf* | 103.3704183 | 127.5244061 | 108.7882839 | 96.91998472 | 1276.176295 | 16143.96555 | 24083.01851 | 4144.700839 | 2.68E-11 | 6.70816469 | 5.20E-14 | 56.65442159 | 104.558366 | 10.57131151 |
| *Lad1* | 0.891124295 | 3.954245151 | 1.036078894 | 3.657357914 | 10.73319003 | 380.0368537 | 511.1863258 | 29.32571348 | 3.42E-05 | 6.608515377 | 4.99E-07 | 25.26738126 | 97.58012164 | 4.466228717 |
| *Il1rn* | 184.4627291 | 199.6893801 | 235.1899089 | 213.0410985 | 605.3519179 | 25786.37753 | 51417.43192 | 1393.415719 | 0.000358614 | 6.572170509 | 1.02E-05 | 19.48215976 | 95.15255623 | 3.445372749 |
| *Ccl5* | 38.3183447 | 43.49669666 | 39.37099797 | 30.17320279 | 149.1913415 | 4210.808339 | 9110.777062 | 314.5849264 | 0.000187541 | 6.509122961 | 4.44E-06 | 21.06520025 | 91.08382429 | 3.726904229 |
| *Cfb* | 9.802367249 | 16.80554189 | 20.72157788 | 7.314715828 | 138.4581514 | 982.2490989 | 3201.421491 | 119.0801699 | 8.05E-05 | 6.346769108 | 1.51E-06 | 23.13892339 | 81.38940567 | 4.094067674 |
| *Ccl7* | 8.020118658 | 28.66827734 | 23.82981456 | 16.45811061 | 90.15879629 | 2021.796062 | 3673.045535 | 202.6140204 | 0.000161555 | 6.282664501 | 3.65E-06 | 21.44081329 | 77.85212435 | 3.791678553 |
| *Socs3* | 123.8662771 | 125.5472835 | 99.46357382 | 114.2924348 | 773.8630015 | 11540.25767 | 21503.14129 | 1965.711461 | 5.94E-06 | 6.27153282 | 6.13E-08 | 29.32232031 | 77.25373717 | 5.226569312 |
| *Ccl4* | 52.57633343 | 57.33655469 | 30.04628793 | 60.34640558 | 270.4763889 | 5099.509905 | 7516.833549 | 735.808811 | 1.91E-05 | 6.087315828 | 2.47E-07 | 26.62200485 | 67.99307105 | 4.718266833 |
| *Tnfsf15* | 14.25798873 | 13.83985803 | 9.324710046 | 10.05773426 | 47.22603615 | 1082.812697 | 1892.742852 | 115.525538 | 0.000129584 | 6.045933162 | 2.74E-06 | 21.99131302 | 66.07044526 | 3.887447816 |
| *Cdc42ep2* | 7.128994363 | 11.86273545 | 6.216473364 | 2.743018435 | 54.73926917 | 640.8006026 | 1019.249314 | 125.3007758 | 2.79E-07 | 6.043218258 | 1.60E-09 | 36.41303437 | 65.94622896 | 6.554856546 |
| *Hp* | 4.455621477 | 5.931367726 | 5.18039447 | 2.743018435 | 24.68633708 | 287.6586647 | 742.3133408 | 35.54631937 | 1.51E-05 | 5.898707556 | 1.87E-07 | 27.16481645 | 59.66064029 | 4.81996819 |
| *Bcl2a1d* | 34.75384752 | 30.64539992 | 23.82981456 | 19.20112905 | 112.6984954 | 2286.067843 | 3638.688816 | 244.3809457 | 0.000103499 | 5.856150715 | 2.03E-06 | 22.5655309 | 57.92646475 | 3.985065462 |
| *Robo2* | 0 | 0 | 0 | 1.828678957 | 5.366595017 | 12.86278582 | 79.12456367 | 10.66389581 | 0.001095895 | 5.827128492 | 4.07E-05 | 16.83625071 | 56.77281972 | 2.960230995 |
| *Il10* | 2.673372886 | 3.954245151 | 5.18039447 | 7.314715828 | 12.87982804 | 344.9565288 | 686.0932561 | 23.99376558 | 0.001244678 | 5.80361087 | 4.75E-05 | 16.54561142 | 55.85485837 | 2.904942896 |
| *Slpi* | 1.782248591 | 0.988561288 | 4.144315576 | 8.229055306 | 16.09978505 | 236.2075214 | 559.0775091 | 19.55047565 | 0.000238513 | 5.775312041 | 5.98E-06 | 20.49331254 | 54.76992656 | 3.622487536 |
| *Ccl2* | 76.6366894 | 83.03914817 | 93.24710046 | 83.20489254 | 242.5700948 | 6348.369473 | 10365.31784 | 742.9180749 | 0.000158502 | 5.718576402 | 3.57E-06 | 21.48426264 | 52.65783867 | 3.799965487 |
| *Arg1* | 2.673372886 | 2.965683863 | 3.108236682 | 2.743018435 | 1.073319003 | 121.6117932 | 458.0895792 | 11.5525538 | 0.008750609 | 5.68862401 | 0.000671332 | 11.56698518 | 51.57585793 | 2.057961745 |
| *Ifnb1* | 0 | 0 | 0 | 0 | 2.146638007 | 10.52409749 | 20.8222536 | 2.665973953 | 0.003194992 | 5.680620237 | 0.000172174 | 14.11273229 | 51.29051803 | 2.495530181 |
| *Ccl20* | 0 | 0 | 0 | 0 | 1.073319003 | 16.37081831 | 16.65780282 | 1.777315969 | 0.008983465 | 5.667835881 | 0.000697893 | 11.49485082 | 50.83801801 | 2.04655613 |
| *Il4i1* | 14.25798873 | 18.78266447 | 10.36078894 | 24.68716592 | 16.09978505 | 1045.393684 | 2244.638938 | 110.1935901 | 0.002141599 | 5.648297992 | 0.000100204 | 15.13286 | 50.15417834 | 2.669261902 |
| *Ifi205* | 17.82248591 | 6.919929014 | 3.108236682 | 23.77282644 | 92.30543429 | 735.51748 | 1587.696837 | 79.97921859 | 0.0005659 | 5.591831049 | 1.78E-05 | 18.41084308 | 48.22906877 | 3.247260678 |
| *Socs1* | 14.25798873 | 12.85129674 | 16.5772623 | 15.54377113 | 41.85944113 | 940.1527089 | 1743.863739 | 87.08848246 | 0.000435016 | 5.569658109 | 1.29E-05 | 19.02832548 | 47.49349789 | 3.36149443 |
| *Steap4* | 0 | 1.977122575 | 0 | 0.914339478 | 3.21995701 | 42.09638995 | 88.49457779 | 2.665973953 | 0.004146363 | 5.566237962 | 0.000245517 | 13.44609133 | 47.38104004 | 2.382332638 |
| *Mmp13* | 146.1443844 | 141.3642641 | 168.8808597 | 169.1528035 | 213.5904817 | 8668.348297 | 19634.34403 | 606.9534033 | 0.001919609 | 5.540919071 | 8.47E-05 | 15.45022493 | 46.55677017 | 2.71678717 |
| *Cd274* | 126.5396499 | 97.86756749 | 112.9325994 | 123.4358296 | 450.7939814 | 6460.626513 | 13200.26767 | 760.6912346 | 0.000152709 | 5.501321127 | 3.39E-06 | 21.58336331 | 45.29629442 | 3.816136256 |
| *Arg2* | 9.802367249 | 5.931367726 | 6.216473364 | 8.229055306 | 31.1262511 | 430.3186528 | 848.5068341 | 39.98960929 | 2.71E-05 | 5.480546588 | 3.74E-07 | 25.82616152 | 44.64871 | 4.56731899 |
| *Dnmt3l* | 0.891124295 | 1.977122575 | 2.072157788 | 0 | 12.87982804 | 58.46720827 | 125.9746343 | 16.8845017 | 1.69E-05 | 5.465991304 | 2.12E-07 | 26.91675624 | 44.2005159 | 4.772669418 |
| *2010005H15Rik* | 0 | 0 | 0 | 0 | 2.146638007 | 10.52409749 | 20.8222536 | 2.665973953 | 0.003114788 | 5.454483458 | 0.000166565 | 14.17504108 | 43.84934714 | 2.506571459 |
| *Procr* | 63.26982497 | 81.06202559 | 76.66983815 | 97.8343242 | 108.4052193 | 3771.134933 | 8997.29578 | 273.7066592 | 0.002362065 | 5.366155012 | 0.000116135 | 14.85443343 | 41.24521959 | 2.626708216 |
| *Tnfaip3* | 187.136102 | 149.2727544 | 176.133412 | 181.9535562 | 765.2764494 | 8927.942702 | 17034.68567 | 1588.031818 | 3.73E-05 | 5.349466517 | 5.59E-07 | 25.04879661 | 40.77085531 | 4.428837982 |
| *Met* | 22.27810738 | 12.85129674 | 27.97413014 | 24.68716592 | 72.98569223 | 865.3146823 | 2453.902587 | 111.082248 | 0.000644794 | 5.31848683 | 2.09E-05 | 18.1092538 | 39.90470159 | 3.190579186 |
| *Clec4e* | 644.2828655 | 671.2331144 | 586.420654 | 649.1810297 | 5110.071775 | 30409.96436 | 55589.17043 | 7951.711644 | 8.99E-10 | 5.279101201 | 2.78E-12 | 48.83821227 | 38.83003764 | 9.046280085 |
| *Upp1* | 8.020118658 | 3.954245151 | 4.144315576 | 5.486036871 | 10.73319003 | 251.4089955 | 522.6385653 | 14.21852775 | 0.000523624 | 5.204965234 | 1.63E-05 | 18.58341342 | 36.88507424 | 3.280980051 |
| *Cp* | 19.6047345 | 14.82841932 | 22.79373567 | 27.43018435 | 68.69241622 | 902.7336956 | 2081.184247 | 59.54008495 | 0.001283161 | 5.199111466 | 4.97E-05 | 16.46002319 | 36.75609167 | 2.891718754 |
| *Epha4* | 1.782248591 | 1.977122575 | 5.18039447 | 4.571697392 | 3.21995701 | 150.8453973 | 322.7449308 | 16.8845017 | 0.00109855 | 5.194670817 | 4.09E-05 | 16.82716311 | 36.62281587 | 2.959180249 |
| *Ptx3* | 2.673372886 | 5.931367726 | 0 | 0 | 4.293276014 | 107.5796632 | 171.7835922 | 12.44121178 | 0.006625216 | 5.104647811 | 0.000460708 | 12.26833965 | 34.40741997 | 2.178799954 |
| *Adm* | 3.564497181 | 0.988561288 | 2.072157788 | 0 | 3.21995701 | 60.8058966 | 157.2080147 | 4.443289922 | 0.005408358 | 5.086291637 | 0.00035121 | 12.77538511 | 33.97240943 | 2.266934574 |
| *Mgarp* | 1.782248591 | 0.988561288 | 0 | 6.400376349 | 3.21995701 | 84.1927799 | 214.4692121 | 10.66389581 | 0.005408358 | 5.076081954 | 0.000351538 | 12.77363986 | 33.73284173 | 2.266934574 |
| *Spic* | 2.673372886 | 7.908490302 | 3.108236682 | 2.743018435 | 20.39306106 | 122.7811374 | 361.2660999 | 28.4370555 | 5.83E-05 | 5.023452075 | 9.88E-07 | 23.9522111 | 32.52443466 | 4.234370686 |
| *Tnnt2* | 0 | 0 | 1.036078894 | 0 | 3.21995701 | 14.03212998 | 17.69891556 | 2.665973953 | 0.0030157 | 5.012792635 | 0.000158375 | 14.26992397 | 32.28501148 | 2.520611827 |
| *Gbp5* | 73.96331651 | 69.19929014 | 98.42749493 | 98.74866368 | 270.4763889 | 3346.663001 | 5944.753402 | 553.6339242 | 5.90E-06 | 4.893482796 | 6.05E-08 | 29.34775091 | 29.72248419 | 5.228862426 |
| *Ptgs2os2* | 1.782248591 | 1.977122575 | 4.144315576 | 0.914339478 | 7.513233024 | 66.65261742 | 169.7013668 | 8.886579843 | 0.000763365 | 4.853801369 | 2.57E-05 | 17.70896301 | 28.91610581 | 3.117267857 |
| *Gpr18* | 2.673372886 | 4.942806439 | 4.144315576 | 3.657357914 | 18.24642306 | 144.9986765 | 267.5659587 | 13.32986976 | 0.000117858 | 4.851425468 | 2.41E-06 | 22.24050406 | 28.86852453 | 3.928641774 |
| *Mmp14* | 172.8781133 | 180.9067157 | 192.7106743 | 196.5829879 | 1276.176295 | 5949.623113 | 12640.14905 | 1491.168098 | 2.33E-08 | 4.845061438 | 9.97E-11 | 41.82718597 | 28.74145988 | 7.63226694 |
| *Il27* | 1.782248591 | 2.965683863 | 0 | 0 | 3.21995701 | 45.60442245 | 84.33012707 | 3.554631937 | 0.009293339 | 4.843440211 | 0.000736163 | 11.39514485 | 28.70917983 | 2.031828216 |
| *Ntng2* | 6.237870067 | 8.89705159 | 7.252552258 | 2.743018435 | 15.02646605 | 202.2965406 | 483.0762835 | 14.21852775 | 0.000767549 | 4.83212976 | 2.60E-05 | 17.69023516 | 28.48498537 | 3.114893811 |
| *Tnfrsf9* | 4.455621477 | 7.908490302 | 8.288631152 | 4.571697392 | 37.56616512 | 196.4498198 | 434.1439875 | 47.09887317 | 3.56E-06 | 4.83080787 | 3.28E-08 | 30.53158669 | 28.45889755 | 5.448434056 |
| *Fcrlb* | 3.564497181 | 0.988561288 | 0 | 1.828678957 | 4.293276014 | 53.7898316 | 112.4401694 | 10.66389581 | 0.001724186 | 4.800176862 | 7.41E-05 | 15.70224501 | 27.86103334 | 2.76341576 |
| *Ier3* | 78.41893799 | 116.650232 | 105.6800472 | 103.3203611 | 226.4703097 | 3509.20184 | 7092.059576 | 399.8960929 | 0.000997846 | 4.79636496 | 3.62E-05 | 17.0625284 | 27.78761022 | 3.000936539 |
| *Cish* | 13.36686443 | 6.919929014 | 4.144315576 | 4.571697392 | 24.68633708 | 327.4163663 | 421.6506354 | 32.88034542 | 0.000010584 | 4.792625414 | 2.11E-06 | 22.49298748 | 27.7155823 | 3.975351721 |
| *Mcemp1* | 0 | 1.977122575 | 2.072157788 | 0.914339478 | 9.659871031 | 25.72557164 | 88.49457779 | 7.109263874 | 0.00072581 | 4.750388102 | 2.42E-05 | 17.82783704 | 26.91592501 | 3.139177277 |
| *Ccrl2* | 166.6402432 | 184.8609608 | 174.0612542 | 206.6407221 | 528.0729497 | 6114.50064 | 11902.00016 | 947.3094113 | 0.000296876 | 4.734465092 | 7.95E-06 | 19.94957367 | 26.62048176 | 3.527425573 |
| *Gpr31b* | 1.782248591 | 1.977122575 | 2.072157788 | 0 | 3.21995701 | 46.77376661 | 90.57680315 | 10.66389581 | 0.001614027 | 4.707470431 | 6.72E-05 | 15.8884915 | 26.12701552 | 2.792089216 |
| *Cd69* | 39.209469 | 49.42806439 | 45.58747133 | 53.94602923 | 308.042554 | 1552.889052 | 2388.312488 | 591.8462175 | 1.09E-10 | 4.685218978 | 2.64E-13 | 53.46114814 | 25.72713637 | 9.962634293 |
| *Hdc* | 8.020118658 | 2.965683863 | 7.252552258 | 4.571697392 | 6.43991402 | 139.1519557 | 406.0339452 | 15.10718573 | 0.002011012 | 4.633955585 | 9.10E-05 | 15.31457797 | 24.82902294 | 2.696585234 |
| *Traf1* | 73.07219222 | 100.8332513 | 72.52552258 | 81.37621359 | 331.6555721 | 2510.581923 | 4579.854679 | 596.2895075 | 8.17E-07 | 4.61240004 | 5.14E-09 | 34.13611756 | 24.46080604 | 6.08794671 |
| *Cd40* | 130.9952714 | 159.1583673 | 157.4839919 | 159.0950693 | 491.5801036 | 5418.740862 | 7435.62676 | 1045.06179 | 1.75E-06 | 4.567989525 | 1.23E-08 | 32.43905566 | 23.71929994 | 5.756369097 |
| *Susd2* | 0 | 0 | 2.072157788 | 3.657357914 | 8.586552027 | 30.4029483 | 85.37123975 | 10.66389581 | 0.002013205 | 4.558366403 | 9.13E-05 | 15.30915471 | 23.561613 | 2.696111904 |
| *Pde10a* | 5.346745772 | 0.988561288 | 3.108236682 | 1.828678957 | 9.659871031 | 81.85409157 | 146.7968879 | 12.44121178 | 0.00057501 | 4.46733192 | 1.82E-05 | 18.37066375 | 22.12080404 | 3.240324262 |
| *Col19a1* | 0 | 0.988561288 | 2.072157788 | 0 | 4.293276014 | 30.4029483 | 24.98670432 | 3.554631937 | 0.004297552 | 4.423363029 | 0.000257936 | 13.35353247 | 21.45680001 | 2.36677889 |
| *Mefv* | 11.58461584 | 13.83985803 | 7.252552258 | 5.486036871 | 40.78612213 | 257.2557164 | 464.3362552 | 51.54216309 | 1.86E-05 | 4.41417618 | 2.36E-07 | 26.70944529 | 21.32060069 | 4.73161268 |
| *Atp7b* | 0.891124295 | 0 | 0 | 0 | 2.146638007 | 4.677376661 | 10.4111268 | 5.331947906 | 0.005728443 | 4.285813758 | 0.000383098 | 12.61288044 | 19.50556331 | 2.241963423 |
| *Flrt3* | 19.6047345 | 25.70259348 | 11.39686783 | 25.6015054 | 47.22603615 | 472.4150428 | 999.4681727 | 86.19982448 | 0.000133815 | 4.284613541 | 2.88E-06 | 21.89337141 | 19.48934285 | 3.87349381 |
| *Kdm6b* | 27.62485316 | 38.55389022 | 45.58747133 | 36.57357914 | 48.29935515 | 971.7250014 | 1732.411499 | 131.5213817 | 0.002850515 | 4.281370968 | 0.000147171 | 14.40806289 | 19.44558818 | 2.545076682 |
| *Klf7* | 1.782248591 | 10.87417417 | 5.18039447 | 10.05773426 | 19.31974206 | 163.7081831 | 306.0871279 | 43.54424123 | 0.000224672 | 4.256647008 | 5.56E-06 | 20.63253651 | 19.11518169 | 3.648450906 |
| *Gm6377* | 58.81420349 | 79.08490302 | 81.85023262 | 79.54753463 | 149.1913415 | 2006.594588 | 3320.108336 | 230.1624179 | 0.001617947 | 4.252915555 | 6.75E-05 | 15.88023615 | 19.06580519 | 2.791035651 |
| *Dusp2* | 26.73372886 | 34.59964507 | 38.33491908 | 45.71697392 | 90.15879629 | 975.2330339 | 1465.886653 | 126.1894338 | 7.69E-05 | 4.192104588 | 1.41E-06 | 23.26932798 | 18.27886502 | 4.11405516 |
| *Ak4* | 35.64497181 | 33.61108378 | 33.15452461 | 43.88829497 | 76.20564924 | 695.7597784 | 1780.302683 | 89.75445641 | 0.000358614 | 4.174317008 | 1.01E-05 | 19.4900965 | 18.05488102 | 3.445372749 |
| *Abtb2* | 14.25798873 | 8.89705159 | 13.46902562 | 10.05773426 | 25.75965608 | 242.0542422 | 508.0629878 | 47.98753115 | 0.000171393 | 4.140941304 | 3.93E-06 | 21.30028549 | 17.64198884 | 3.766005716 |
| *Ccr7* | 1.782248591 | 0.988561288 | 2.072157788 | 2.743018435 | 3.21995701 | 39.75770162 | 82.24790171 | 6.22060589 | 0.003662933 | 4.109964429 | 0.000208027 | 13.75715256 | 17.26722604 | 2.43617097 |
| *1700071M16Rik* | 2.673372886 | 0.988561288 | 1.036078894 | 0.914339478 | 6.43991402 | 23.38688331 | 61.42564812 | 5.331947906 | 0.001958651 | 4.088625701 | 8.74E-05 | 15.39155974 | 17.01370809 | 2.708042903 |
| *Lrrc63* | 1.782248591 | 0 | 0 | 0.914339478 | 2.146638007 | 14.03212998 | 27.06892968 | 4.443289922 | 0.009245425 | 4.085844437 | 0.000730923 | 11.40892119 | 16.98094023 | 2.034073099 |
| *Cd200* | 9.802367249 | 7.908490302 | 8.288631152 | 6.400376349 | 20.39306106 | 146.1680207 | 352.9371985 | 22.21644961 | 0.00047919 | 4.062773982 | 1.45E-05 | 18.8026824 | 16.7115539 | 3.319492227 |
| *Gpr84* | 286.9420231 | 273.8314767 | 262.1279602 | 287.1025962 | 1612.125143 | 5177.855964 | 8962.939062 | 2593.992656 | 7.02E-12 | 4.046823852 | 1.19E-14 | 59.55366338 | 16.52781209 | 11.15337467 |
| *Gdf15* | 35.64497181 | 30.64539992 | 46.62355023 | 32.91622123 | 147.0447035 | 838.4197665 | 1113.990568 | 257.7108154 | 1.13E-07 | 4.015129749 | 5.58E-10 | 38.46245902 | 16.16867721 | 6.945280957 |
| *Ralgds* | 138.1242658 | 147.2956319 | 152.3035974 | 133.4935639 | 487.2868275 | 2854.369108 | 5133.726625 | 692.2645698 | 3.00E-06 | 4.004466436 | 2.54E-08 | 31.03215348 | 16.04961111 | 5.523434713 |
| *Nfkbiz* | 348.4295995 | 333.145154 | 335.6895616 | 380.3652231 | 994.9667162 | 6949.412374 | 12583.92896 | 1743.546965 | 5.90E-06 | 3.994130035 | 6.05E-08 | 29.34811213 | 15.93503226 | 5.228862426 |
| *Ccl12* | 4.455621477 | 8.89705159 | 4.144315576 | 7.314715828 | 26.83297509 | 141.490644 | 182.194719 | 28.4370555 | 2.66E-05 | 3.931511431 | 3.61E-07 | 25.89309435 | 15.25818475 | 4.574691639 |

| | | | | | | | | | | | | | | |
|---|---|---|---|---|---|---|---|---|---|---|---|---|---|---|
| *Sod2* | 816.2698545 | 818.5287462 | 850.620772 | 843.0209992 | 2663.977766 | 14904.46073 | 29517.6267 | 3379.566314 | 8.34E-06 | 3.922377778 | 9.22E-08 | 28.53136432 | 15.16189084 | 5.07874154 |
| *Tmc3* | 1.782248591 | 9.885612877 | 20.72157788 | 9.143394785 | 21.46638007 | 160.2001506 | 414.3628466 | 28.4370555 | 0.003194992 | 3.915365451 | 0.000172083 | 14.11373159 | 15.08837415 | 2.495530181 |
| *Plat* | 6.237870067 | 6.919929014 | 6.216473364 | 10.05773426 | 20.39306106 | 164.8775273 | 233.2092403 | 16.8845017 | 0.000510182 | 3.88446577 | 1.57E-05 | 18.64790926 | 14.76864712 | 3.292275151 |
| *Slco3a1* | 122.0840285 | 162.1240512 | 118.1129939 | 152.6946929 | 284.4295359 | 2601.790768 | 4876.571793 | 423.0012005 | 0.000146087 | 3.882465113 | 3.19E-06 | 21.69650229 | 14.7481809 | 3.835389429 |
| *Ppfia3* | 19.6047345 | 22.73690962 | 30.04628793 | 10.97207374 | 35.41952711 | 373.0207887 | 732.9433267 | 84.42250851 | 0.000351927 | 3.879734248 | 9.85E-06 | 19.54001022 | 14.72029061 | 3.453547532 |
| *Nod2* | 47.22958765 | 30.64539992 | 44.55139244 | 32.00188175 | 142.7514275 | 646.6473234 | 1257.664117 | 205.2799944 | 2.54E-06 | 3.866207234 | 2.03E-08 | 31.46850585 | 14.58291508 | 5.595496722 |
| *Rab11fip1* | 373.3810797 | 291.6255799 | 339.8338772 | 352.0206992 | 479.7735945 | 6086.43638 | 12096.68823 | 811.3447397 | 0.004361288 | 3.843195435 | 0.000262465 | 13.32089154 | 14.35215465 | 2.360385276 |
| *Spint2* | 7.128994363 | 3.954245151 | 8.288631152 | 10.97207374 | 18.24642306 | 122.7811374 | 262.3603953 | 29.32571348 | 0.000391409 | 3.831704151 | 1.15E-05 | 19.25085925 | 14.23829163 | 3.407369274 |
| *Dusp16* | 139.0153901 | 133.4557738 | 146.0871241 | 109.7207374 | 197.4906966 | 2564.371755 | 4325.823185 | 346.5766139 | 0.003328178 | 3.814824667 | 0.000181766 | 14.0107709 | 14.07267484 | 2.477793419 |
| *Lix1* | 5.346745772 | 6.919929014 | 2.072157788 | 10.05773426 | 16.09978505 | 113.426384 | 203.0169726 | 7.997921859 | 0.003620619 | 3.797919154 | 0.000203579 | 13.79775372 | 13.90873352 | 2.441217197 |
| *Tnfsf9* | 41.88284188 | 36.57676765 | 39.37099797 | 48.45999236 | 137.3848324 | 698.0984667 | 1134.812821 | 333.2467441 | 9.85E-08 | 3.791477999 | 4.61E-10 | 38.83603679 | 13.84677405 | 7.006502688 |
| *Nfkbie* | 101.5881697 | 125.5472835 | 134.6902562 | 105.14904 | 390.6881172 | 1801.959359 | 3502.303055 | 656.7182504 | 1.11E-06 | 3.765915306 | 7.40E-09 | 33.42587689 | 13.60358793 | 5.956374955 |
| *Egln3* | 26.73372886 | 24.71403219 | 24.86589346 | 30.17320279 | 36.49284612 | 382.3755421 | 973.4403557 | 39.10095131 | 0.002362065 | 3.748285736 | 0.000115774 | 14.86030449 | 13.43836518 | 2.626708216 |
| *Htra4* | 7.128994363 | 4.942806439 | 18.64942009 | 7.314715828 | 21.46638007 | 160.2001506 | 300.8815645 | 13.32986976 | 0.003390154 | 3.707930869 | 0.000187339 | 13.95399521 | 13.06767766 | 2.469780584 |
| *Slamf8* | 15.14911302 | 10.87417417 | 5.18039447 | 10.05773426 | 84.79220127 | 116.9344165 | 266.5248461 | 72.86995471 | 4.86E-09 | 3.704837426 | 1.72E-11 | 45.26233978 | 13.03968783 | 8.313806968 |
| *Ednrb* | 29.40710175 | 36.57676765 | 17.6133412 | 17.37245009 | 76.20564924 | 418.6252112 | 673.5999039 | 129.7440657 | 1.40E-05 | 3.683934868 | 1.71E-07 | 27.34012113 | 12.85212371 | 4.853629927 |
| *Slc7a2* | 600.6177751 | 616.8622435 | 637.1885198 | 749.7583724 | 935.934171 | 9319.672997 | 21659.30819 | 1361.424032 | 0.005816548 | 3.675450689 | 0.000390398 | 12.57760101 | 12.77676494 | 2.235334692 |
| *Siglecg* | 0 | 0.988561288 | 1.036078894 | 0.914339478 | 4.293276014 | 16.37081831 | 13.53446484 | 2.665973953 | 0.005974638 | 3.663897092 | 0.0004039 | 12.51406365 | 12.67485284 | 2.22368843 |
| *Pdpn* | 116.7372827 | 91.93619976 | 112.9325994 | 143.5512981 | 188.9041446 | 1536.518233 | 3841.705789 | 220.3871801 | 0.006453805 | 3.637049941 | 0.000443582 | 12.33904472 | 12.44116723 | 2.190184156 |
| *Adora2b* | 60.59645208 | 69.19929014 | 65.27297032 | 60.34640558 | 136.3115134 | 1006.805326 | 1812.577176 | 211.5006003 | 9.79E-05 | 3.632291149 | 1.90E-06 | 22.6893535 | 12.40019712 | 4.009022918 |
| *Batf* | 24.06035597 | 27.67971606 | 45.58747133 | 36.57357914 | 90.15879629 | 546.0837252 | 947.4125387 | 73.7586127 | 0.000283279 | 3.630213746 | 7.52E-06 | 20.0565766 | 12.38235435 | 3.547785837 |
| *Ets2* | 327.9337407 | 384.5503409 | 401.9986109 | 296.245991 | 800.6959765 | 5402.370044 | 9744.814684 | 1183.692435 | 6.23E-05 | 3.602182737 | 1.07E-06 | 23.79720258 | 12.14409214 | 4.205560409 |
| *Mcmdc2* | 0.891124295 | 0 | 1.036078894 | 0.914339478 | 5.366595017 | 10.52409749 | 14.57557752 | 4.443289922 | 0.001612811 | 3.601091732 | 6.70E-05 | 15.89358945 | 12.13491192 | 2.792416577 |
| *Acpp* | 25.84260457 | 15.8169806 | 13.46902562 | 20.11546853 | 12.87982804 | 211.6512939 | 639.2431855 | 39.98960929 | 0.005646928 | 3.584849473 | 0.000375464 | 12.65050685 | 11.9990599 | 2.248187759 |
| *Phlda1* | 47.22958765 | 45.47381924 | 34.1906035 | 46.6313134 | 83.71888227 | 560.1158552 | 1323.254216 | 99.52969424 | 0.000587245 | 3.57355728 | 1.86E-05 | 18.32569821 | 11.90550803 | 3.231180935 |
| *Fndc9* | 0.891124295 | 0 | 1.036078894 | 0.914339478 | 4.293276014 | 5.846720827 | 15.6166902 | 7.997921859 | 0.002001009 | 3.566569256 | 9.02E-05 | 15.33074569 | 11.84798041 | 2.698750942 |
| *Hilpda* | 49.90296054 | 52.39374825 | 65.27297032 | 69.48980037 | 84.79220127 | 715.6386292 | 1891.701739 | 110.1935901 | 0.001671351 | 3.56328915 | 7.05E-05 | 15.79695182 | 11.82107349 | 2.776932317 |
| *6530402F18Rik* | 63.26982497 | 60.30223855 | 44.55139244 | 37.48791862 | 49.37267416 | 750.7189541 | 1510.654499 | 91.53177238 | 0.0030007 | 3.546208716 | 0.000157346 | 14.28220305 | 11.681946 | 2.522777395 |
| *Fcgr2b* | 586.3597863 | 611.9194371 | 635.116362 | 547.6893476 | 1032.532881 | 8552.583225 | 16031.05305 | 1513.384547 | 0.00031848 | 3.510174428 | 8.71E-06 | 19.775401 | 11.39377904 | 3.496917786 |
| *Pde4b* | 226.345571 | 269.8772316 | 214.4683311 | 215.7841169 | 558.1258818 | 3138.51974 | 5749.024219 | 792.682922 | 4.00E-05 | 3.466062596 | 6.22E-07 | 24.84295365 | 11.05067504 | 4.397756487 |
| *Ppap2b* | 30.29822604 | 23.72547091 | 19.68549899 | 22.85848696 | 54.73926917 | 302.8601388 | 617.3798192 | 87.08848246 | 0.000123336 | 3.457911764 | 2.57E-06 | 22.11571002 | 10.98841779 | 3.908911656 |
| *Zc3h12a* | 193.3739721 | 267.900109 | 251.7671712 | 196.5829879 | 461.5271715 | 3097.592694 | 5598.06288 | 830.8952153 | 7.10E-05 | 3.456942576 | 1.27E-06 | 23.47319165 | 10.98103836 | 4.14844882 |
| *Tma16* | 109.6082883 | 88.9705159 | 101.5357316 | 91.43394785 | 240.4234568 | 1361.116608 | 2438.285896 | 237.2716818 | 0.000147777 | 3.449227366 | 3.27E-06 | 21.6533206 | 10.92247096 | 3.830394239 |
| *Siglece* | 39.209469 | 37.56532893 | 43.51531355 | 34.74490018 | 119.1384094 | 494.6325819 | 969.275905 | 102.1956682 | 7.53E-05 | 3.442316565 | 1.37E-06 | 23.32665495 | 10.87027523 | 4.123156249 |
| *Slc13a3* | 23.16923168 | 10.87417417 | 22.79373567 | 27.43018435 | 194.2707396 | 202.2965406 | 392.4994803 | 99.52969424 | 8.99E-10 | 3.39563858 | 2.89E-12 | 48.76220349 | 10.52419938 | 9.046280085 |
| *Malt1* | 480.3159952 | 566.4456179 | 554.3022083 | 494.6576579 | 787.8161485 | 7059.330726 | 12738.01364 | 1360.535374 | 0.000453476 | 3.388424175 | 1.35E-05 | 18.93866152 | 10.47170297 | 3.343445952 |
| *Cxcl11* | 2.673372886 | 2.965683863 | 2.072157788 | 0.914339478 | 15.02646605 | 10.52409749 | 39.56228184 | 22.21644961 | 3.58E-05 | 3.348884401 | 5.28E-07 | 25.15815845 | 10.18860338 | 4.446406128 |
| *Ifitm1* | 18.7136102 | 23.72547091 | 15.54118341 | 15.54377113 | 30.0529321 | 205.8045731 | 436.2262129 | 44.43289922 | 0.001283161 | 3.284378594 | 4.97E-05 | 16.46065318 | 9.743084608 | 2.891718754 |
| *Gsta3* | 2.673372886 | 2.965683863 | 5.18039447 | 4.571697392 | 21.46638007 | 35.08032496 | 83.28901439 | 6.22060589 | 0.001639454 | 3.248376077 | 6.85E-05 | 15.85158897 | 9.502954202 | 2.785300648 |
| *Mt2* | 255.7526728 | 250.1060058 | 263.1640391 | 252.3576961 | 404.6412643 | 2709.370431 | 5958.287867 | 518.9762628 | 0.000948317 | 3.231175511 | 3.37E-05 | 17.19471843 | 9.390327738 | 3.023046677 |
| *Pim1* | 270.0106615 | 332.1565927 | 246.5867768 | 236.8139249 | 574.2256668 | 3125.656954 | 5553.295035 | 935.7568575 | 5.80E-05 | 3.230462053 | 9.78E-07 | 23.9704593 | 9.385685072 | 4.236414026 |
| *Fzd1* | 28.51597745 | 24.71403219 | 27.97413014 | 29.25886331 | 23.61301808 | 315.7229246 | 639.2431855 | 56.874111 | 0.004006589 | 3.228218182 | 0.000234179 | 13.53481142 | 9.371098557 | 2.39722523 |
| *Vcam1* | 3.564497181 | 12.85129674 | 11.39686783 | 16.45811061 | 42.93276014 | 113.426384 | 200.9347472 | 50.65350511 | 4.34E-05 | 3.205771954 | 6.86E-07 | 24.65531625 | 9.226426308 | 4.362932982 |
| *Gem* | 8.020118658 | 22.73690962 | 25.90197235 | 18.28678957 | 75.13233024 | 182.4176898 | 301.9226772 | 118.1915119 | 3.75E-07 | 3.181843302 | 2.21E-09 | 35.78190648 | 9.074658183 | 6.426262674 |
| *Trem1* | 119.4106556 | 105.7760578 | 122.2573095 | 91.43394785 | 174.9509976 | 1228.980718 | 2302.941248 | 236.3830238 | 0.000846878 | 3.167464319 | 2.91E-05 | 17.4743508 | 8.984662545 | 3.072179012 |
| *Cxcl9* | 4.455621477 | 6.919929014 | 1.036078894 | 5.486036871 | 17.17310405 | 35.08032496 | 98.90570459 | 8.886579843 | 0.004448777 | 3.153746442 | 0.000269166 | 13.27362624 | 8.899636689 | 2.351759407 |
| *Nlrp3* | 759.2378996 | 791.8375915 | 841.2960619 | 734.2146012 | 1330.915564 | 8416.939302 | 15086.76384 | 2723.736722 | 0.000111903 | 3.139830444 | 2.27E-06 | 22.35530422 | 8.814204957 | 3.95115838 |
| *C3* | 1653.035568 | 1632.114686 | 1560.334814 | 1478.486937 | 5171.250958 | 15015.54843 | 30576.4383 | 4948.936315 | 6.19E-06 | 3.139081849 | 6.44E-08 | 29.22590539 | 8.809632573 | 5.208339765 |
| *Dgat2* | 23.16923168 | 21.74834833 | 16.5772623 | 13.71509218 | 19.31974206 | 154.3534298 | 454.9662411 | 31.99168744 | 0.00714332 | 3.13433432 | 0.000504804 | 12.09783574 | 8.780690039 | 2.1460999 |
| *Tnip3* | 413.481673 | 406.2986893 | 403.0346898 | 443.4546471 | 638.624807 | 4936.971066 | 8031.143213 | 941.9774634 | 0.000609422 | 3.126157522 | 1.94E-05 | 18.24541201 | 8.731064274 | 3.215081656 |
| *Il17rd* | 4.455621477 | 4.942806439 | 5.18039447 | 8.229055306 | 4.293276014 | 74.83802658 | 104.111268 | 15.99584372 | 0.007801409 | 3.124029564 | 0.000572707 | 11.86265799 | 8.718195553 | 2.10782698 |
| *Vcan* | 81.98343517 | 83.03914817 | 72.52552258 | 74.06149776 | 89.08547728 | 702.7758433 | 1799.042711 | 122.6348018 | 0.003891117 | 3.122234442 | 0.000226009 | 13.60146814 | 8.707354391 | 2.409925762 |
| *Rsad2* | 867.9550637 | 843.2427784 | 836.1156674 | 837.5349623 | 1779.562908 | 7318.925131 | 18191.36186 | 2111.451371 | 0.000262729 | 3.118704588 | 6.78E-06 | 20.25367687 | 8.686076073 | 3.580491492 |
| *Nfkbia* | 1002.514832 | 977.6871136 | 1009.140843 | 927.1402312 | 2784.189495 | 9534.832324 | 17687.46332 | 3843.445782 | 2.42E-06 | 3.111514015 | 1.92E-08 | 31.57887938 | 8.642891289 | 5.615773229 |
| *Maff* | 73.96331651 | 55.35943211 | 64.23689143 | 35.65923966 | 118.0650904 | 631.4458493 | 963.0292289 | 246.1582617 | 4.57E-05 | 3.094993607 | 7.31E-07 | 24.53110961 | 8.544485401 | 4.339889782 |
| *Gbp6* | 90.00355383 | 66.23360628 | 87.03062709 | 103.3203611 | 238.2768188 | 748.3802658 | 1733.452612 | 239.0489978 | 8.98E-05 | 3.093561358 | 1.72E-06 | 22.8833812 | 8.536006991 | 4.046817541 |
| *Clmp* | 31.18935034 | 24.71403219 | 38.33491908 | 23.77282644 | 32.1995701 | 197.6191639 | 728.778876 | 41.76692526 | 0.008648368 | 3.083779793 | 0.000661395 | 11.59471515 | 8.478328069 | 2.063065835 |
| *Arl5c* | 155.0556274 | 177.9410318 | 159.5561497 | 178.2961983 | 724.4903273 | 1727.121332 | 2229.022248 | 989.9649945 | 3.98E-14 | 3.079312508 | 4.82E-17 | 70.41102203 | 8.452115651 | 13.40021207 |
| *Mybpc3* | 1.782248591 | 4.942806439 | 0 | 5.486036871 | 6.43991402 | 26.8949158 | 37.48005648 | 32.88034542 | 0.002243976 | 3.077681824 | 0.000106623 | 15.01565076 | 8.442567589 | 2.648981835 |
| *Hmox1* | 2635.945666 | 3165.373243 | 2844.036564 | 2779.592015 | 3729.783537 | 25723.23295 | 61697.37853 | 4823.635539 | 0.002345499 | 3.07045732 | 0.000114285 | 14.88471322 | 8.400395886 | 2.629764738 |
| *Slamf7* | 512.3964698 | 547.6629534 | 526.3280781 | 566.8904767 | 1070.099046 | 5407.04742 | 10164.38309 | 1278.778839 | 0.000252838 | 3.056965328 | 6.47E-06 | 20.34512684 | 8.322202122 | 3.597158496 |
| *Cxcl10* | 482.9893681 | 543.7087083 | 546.0135771 | 562.3187793 | 2096.192014 | 3791.013784 | 8760.963202 | 3077.4226 | 6.88E-10 | 3.05352732 | 2.00E-12 | 49.48681546 | 8.302393547 | 9.162622465 |
| *Mycl* | 9.802367249 | 8.89705159 | 5.18039447 | 5.486036871 | 47.22603615 | 43.26573412 | 92.65902851 | 59.54008495 | 3.33E-11 | 3.043030611 | 6.98E-14 | 56.07502308 | 8.242206511 | 10.47809963 |
| *Exoc3l4* | 5.346745772 | 7.908490302 | 4.144315576 | 3.657357914 | 7.513233024 | 67.82196159 | 76.00122563 | 21.32779162 | 0.001787339 | 3.034567756 | 7.74E-05 | 15.62007239 | 8.193999224 | 2.747793005 |
| *Pfkfb3* | 506.1585998 | 594.1253339 | 565.6990761 | 635.4659376 | 792.1094245 | 5453.821187 | 11462.65061 | 1090.383347 | 0.001642865 | 3.03002595 | 6.88E-05 | 15.84400677 | 8.168243927 | 2.784398115 |
| *Vegfc* | 27.62485316 | 21.74834833 | 24.86589346 | 20.11546853 | 103.0386243 | 242.0542422 | 331.0738322 | 88.86579843 | 8.84E-08 | 3.018080713 | 4.07E-10 | 39.08100497 | 8.100891658 | 7.05344664 |
| *Gbgt1* | 0.891124295 | 4.942806439 | 2.072157788 | 0.914339478 | 19.31974206 | 2.338688331 | 24.98670432 | 23.10510759 | 0.007781963 | 3.007574883 | 0.000570652 | 11.86935344 | 8.042114536 | 2.108910818 |
| *Pilra* | 155.9467517 | 114.6731094 | 129.5098617 | 132.5792244 | 287.6494929 | 1282.770549 | 2392.476938 | 307.4756626 | 0.000254359 | 3.002711095 | 6.55E-06 | 20.32161276 | 8.01504764 | 3.594552944 |
| *Rab20* | 59.70532779 | 33.61108378 | 54.91218138 | 66.74678193 | 173.8776786 | 478.2617636 | 786.0400733 | 271.0406852 | 2.43E-07 | 2.989995167 | 1.37E-09 | 36.70717032 | 7.94471335 | 6.61422945 |
| *Sntb1* | 7.128994363 | 8.89705159 | 6.216473364 | 11.88641322 | 11.80650904 | 59.63655243 | 173.8658175 | 23.99376558 | 0.004397921 | 2.977830153 | 0.00026538 | 13.30018789 | 7.878004018 | 2.356752624 |
| *Adamts4* | 19.6047345 | 15.8169806 | 11.39686783 | 11.88641322 | 26.83297509 | 161.3694948 | 239.4559164 | 33.7690034 | 0.001290442 | 2.972094064 | 5.04E-05 | 16.4335649 | 7.846743627 | 2.889261547 |
| *Treml4* | 24.95148027 | 25.70259348 | 19.68549899 | 31.08754227 | 66.54577821 | 182.4176898 | 451.8429031 | 91.53177238 | 7.34E-05 | 2.964275492 | 1.32E-06 | 23.39389744 | 7.804333837 | 4.134460492 |
| *Gbp2* | 396.5503114 | 343.0307668 | 328.4370094 | 362.992773 | 737.3701553 | 2935.053855 | 6313.307291 | 925.9816196 | 0.000197832 | 2.930691981 | 4.77E-06 | 20.92674511 | 7.624760283 | 3.703702394 |
| *AW112010* | 34.75384752 | 56.3479934 | 45.58747133 | 32.91622123 | 77.27896825 | 349.6339054 | 774.5878339 | 83.53385052 | 0.001557851 | 2.92199848 | 6.40E-05 | 15.98139945 | 7.578952578 | 2.807474137 |
| *Psd* | 9.802367249 | 17.79410318 | 9.324710046 | 10.05773426 | 55.81258818 | 105.2409749 | 157.2080147 | 37.32363534 | 4.67E-06 | 2.920104602 | 4.48E-08 | 29.92857634 | 7.569009942 | 5.330737925 |
| *Rnd1* | 34.75384752 | 35.58820636 | 34.1906035 | 42.05961601 | 69.76573522 | 376.5288212 | 571.5708613 | 89.75445641 | 0.00038215 | 2.916895106 | 1.11E-05 | 19.31591999 | 7.552190238 | 3.417766108 |
| *Tnfaip2* | 1867.796523 | 2065.10453 | 1919.854191 | 1771.989909 | 5654.24451 | 16930.93417 | 25862.28008 | 9019.878541 | 1.76E-08 | 2.9139811 | 7.26E-11 | 42.44715482 | 7.536951454 | 7.754479266 |
| *Mmp25* | 3.564497181 | 2.965683863 | 9.324710046 | 2.743018435 | 9.659871031 | 44.43507828 | 65.59009884 | 18.66181767 | 0.002031677 | 2.906212755 | 9.26E-05 | 15.28184499 | 7.496477004 | 2.692145302 |
| *Clec4n* | 85.54793235 | 85.01627075 | 94.28317935 | 84.11923202 | 163.1444885 | 622.0910959 | 1599.149076 | 174.1769649 | 0.001086454 | 2.874139623 | 4.02E-05 | 16.86054011 | 7.331658675 | 2.963988681 |
| *Trem3* | 16.04023732 | 8.89705159 | 8.288631152 | 8.229055306 | 16.09978505 | 71.32999408 | 179.0713809 | 30.21437147 | 0.003089198 | 2.835692972 | 0.00016398 | 14.20447929 | 7.138856351 | 2.510154225 |
| *Ehd1* | 1254.703008 | 1288.095358 | 1314.784116 | 1142.010009 | 2980.606872 | 10202.52784 | 17788.45125 | 4593.473121 | 5.02E-06 | 2.830576731 | 5.03E-08 | 29.70706018 | 7.113584602 | 5.298991345 |
| *Dusp4* | 173.7692376 | 129.5015287 | 181.3138064 | 145.3799771 | 412.1544973 | 1203.255146 | 2133.239881 | 678.9347 | 1.08E-06 | 2.813126321 | 7.04E-09 | 33.52284958 | 7.02805909 | 5.967435347 |
| *Fas* | 56.14083061 | 41.51957408 | 59.05649696 | 51.2030108 | 160.9978505 | 385.8835746 | 698.5866082 | 209.7232843 | 6.18E-07 | 2.806807294 | 3.74E-09 | 34.75417716 | 6.997343398 | 6.208820764 |
| *Slc31a2* | 355.5585938 | 300.5226315 | 365.7358496 | 373.9648467 | 875.8283068 | 2505.904546 | 5152.466653 | 1208.574859 | 1.01E-05 | 2.803200211 | 1.16E-07 | 28.09425179 | 6.979870216 | 4.994705014 |
| *St7* | 75.7455651 | 96.8790062 | 80.81415373 | 91.43394785 | 138.4581514 | 778.7832141 | 1321.171991 | 154.6264893 | 0.00151098 | 2.794564272 | 6.12E-05 | 16.06545779 | 6.938213678 | 2.820741238 |
| *Clec4a1* | 229.9100682 | 274.820038 | 235.1899089 | 169.1528035 | 513.0464836 | 1801.959359 | 3492.933041 | 495.8711552 | 0.000286672 | 2.79377955 | 7.63E-06 | 20.02800546 | 6.934440815 | 3.542615334 |
| *Nox1* | 33.86272322 | 41.51957408 | 42.47923465 | 30.17320279 | 67.61909721 | 280.6425997 | 545.5430443 | 125.3007758 | 0.000196306 | 2.783850467 | 4.70E-06 | 20.95444221 | 6.886879655 | 3.707066711 |
| *Bcl2a1b* | 416.1550459 | 386.5274635 | 449.65824 | 448.9406839 | 622.525022 | 4076.33376 | 5949.958966 | 1006.849496 | 0.00061053 | 2.776296323 | 1.95E-05 | 18.23713663 | 6.850913278 | 3.214292906 |
| *Lcp2* | 367.1432097 | 419.149986 | 431.0088199 | 408.7097469 | 504.4599316 | 3560.652983 | 6159.222614 | 755.3592867 | 0.002186153 | 2.755415918 | 0.000102994 | 15.08101458 | 6.752472775 | 2.660319544 |
| *Src* | 81.98343517 | 76.11921916 | 89.10278488 | 101.4916821 | 173.8776786 | 613.9056868 | 1260.787455 | 277.2612911 | 0.000106199 | 2.737480813 | 2.14E-06 | 22.46658436 | 6.669047908 | 3.973879952 |
| *Fabp3* | 23.16923168 | 36.57676765 | 32.11844571 | 42.05961601 | 53.66595017 | 243.2235864 | 534.0908048 | 56.874111 | 0.003490942 | 2.728747559 | 0.000194317 | 13.88525751 | 6.628799235 | 2.457057351 |
| *Dusp1* | 330.6071136 | 355.8820636 | 340.8699561 | 308.1324043 | 457.2338955 | 2736.265347 | 4929.668539 | 692.2645698 | 0.001399007 | 2.722635693 | 5.55E-05 | 16.24940941 | 6.60077624 | 2.854180104 |
| *Slc2a1* | 610.4201423 | 681.1187272 | 627.8638097 | 640.0376349 | 888.7081348 | 4759.230753 | 10009.2573 | 1117.043086 | 0.002257977 | 2.712331246 | 0.000108381 | 14.98478609 | 6.553798175 | 2.646280408 |
| *Batf2* | 17.82248591 | 10.87417417 | 10.36078894 | 13.71509218 | 22.53969907 | 100.5635982 | 208.222536 | 15.10718573 | 0.009293339 | 2.711555297 | 0.00073696 | 11.39364033 | 6.550274185 | 2.031828216 |
| *Olr1* | 29.40710175 | 41.51957408 | 47.65962912 | 28.34452383 | 40.78612213 | 311.045548 | 500.775199 | 108.4162741 | 0.001939459 | 2.710261149 | 8.61E-05 | 15.42044682 | 6.544400992 | 2.712319433 |
| *Mfsd7a* | 50.79408483 | 44.48525795 | 35.2266824 | 53.94602923 | 82.64556326 | 349.6339054 | 682.969918 | 89.75445641 | 0.001421682 | 2.706722346 | 5.68E-05 | 16.2074323 | 6.528367833 | 2.84719749 |
| *Cd55* | 3.564497181 | 6.919929014 | 4.144315576 | 7.314715828 | 20.39306106 | 37.41901329 | 64.54898616 | 19.55047565 | 9.18E-05 | 2.690121747 | 1.78E-06 | 22.82264481 | 6.453678665 | 4.03733993 |
| *Irak3* | 341.3006051 | 319.3052959 | 345.0142717 | 318.1901385 | 486.2135085 | 2504.735202 | 4952.573018 | 568.74111 | 0.002038604 | 2.684805394 | 9.32E-05 | 15.26895242 | 6.429940529 | 2.690667158 |
| *Zc3h12c* | 150.6000059 | 157.1812447 | 165.772623 | 184.6965747 | 420.7410493 | 1184.545639 | 1926.058458 | 670.0481202 | 4.33E-07 | 2.674047846 | 2.59E-09 | 35.47278908 | 6.382173575 | 6.36325685 |
| *Hivep3* | 385.8568199 | 436.9440892 | 443.4417666 | 338.305607 | 931.640895 | 3175.938753 | 4805.776131 | 1320.545765 | 7.64E-06 | 2.673147229 | 8.20E-08 | 28.75924828 | 6.378190683 | 5.116980791 |
| *Tnfrsf1b* | 5541.901993 | 5672.364669 | 5728.480205 | 5107.500327 | 9201.563816 | 40610.15352 | 74904.93398 | 13009.95289 | 0.00043142 | 2.642939952 | 1.27E-05 | 19.04937457 | 6.246031961 | 3.365099929 |
| *Cflar* | 1141.530222 | 1092.360223 | 1192.526807 | 1117.322843 | 1985.640156 | 8550.244537 | 14551.63193 | 3046.31957 | 0.000178575 | 2.630341525 | 4.18E-06 | 21.18154008 | 6.191725552 | 3.74818018 |
| *Rhou* | 25.84260457 | 16.80554189 | 22.79373567 | 8.229055306 | 33.27288911 | 149.6760532 | 239.4559164 | 29.32571348 | 0.005941822 | 2.615869794 | 0.000400723 | 12.52881861 | 6.129926577 | 2.226080337 |
| *AA467197* | 142.5798873 | 142.3528254 | 136.762414 | 138.0652613 | 191.0507826 | 1089.828762 | 1906.277317 | 232.8283919 | 0.002657033 | 2.610968274 | 0.000135252 | 14.5671449 | 6.109135644 | 2.575603049 |
| *Slfn1* | 16.93136161 | 10.87417417 | 11.39686783 | 10.05773426 | 28.97961309 | 61.97524076 | 179.0713809 | 28.4370555 | 0.002062192 | 2.596501818 | 9.48E-05 | 15.23820914 | 6.04818311 | 2.68567098 |
| *P2ry2* | 115.8461584 | 108.7417417 | 118.1129939 | 101.4916821 | 194.2707396 | 809.1861624 | 1471.092217 | 209.7232843 | 0.001047092 | 2.595150428 | 3.85E-05 | 16.94303476 | 6.042520356 | 2.980015218 |
| *Pstpip2* | 204.9585879 | 212.5406769 | 183.3859642 | 174.6388404 | 306.969235 | 1365.793985 | 2545.520502 | 460.3248359 | 0.000646438 | 2.592743951 | 2.10E-05 | 18.09556443 | 6.032449576 | 3.189473193 |
| *Osm* | 101.5881697 | 97.86756749 | 106.7161261 | 63.08942402 | 123.4316854 | 840.7584549 | 1085.880525 | 145.7399094 | 0.003627832 | 2.57198793 | 0.00020457 | 13.78863118 | 5.946282191 | 2.440352793 |
| *Cd14* | 7758.128115 | 7594.127812 | 7647.298316 | 7457.352787 | 9494.579904 | 54157.00567 | 102690.1492 | 13783.97399 | 0.002596822 | 2.564162092 | 0.000131559 | 14.61931996 | 5.91411416 | 2.585557896 |
| *Ifit1* | 439.3242776 | 516.0289922 | 474.5241334 | 523.9165212 | 872.6083498 | 2773.68436 | 6664.162264 | 1085.051399 | 0.000714096 | 2.54409465 | 2.37E-05 | 17.86803638 | 5.832420145 | 3.146243609 |
| *Wfdc17* | 291.3976446 | 331.1680314 | 325.3287727 | 335.5625886 | 339.1688051 | 2111.835563 | 4491.360101 | 413.2259627 | 0.008612907 | 2.518775342 | 0.000655904 | 11.61022025 | 5.730954103 | 2.064850248 |
| *Peg10* | 2.673372886 | 4.942806439 | 5.18039447 | 3.657357914 | 18.24642306 | 24.55622747 | 36.4389438 | 14.21852775 | 4.57E-05 | 2.513805222 | 7.28E-07 | 24.53954278 | 5.711244793 | 4.339889782 |
| *Tnip1* | 1079.151522 | 1131.902674 | 1117.929127 | 1050.576061 | 2206.743871 | 6980.984667 | 12780.69926 | 2997.443381 | 6.86E-05 | 2.511089428 | 1.20E-06 | 23.57552229 | 5.700503801 | 4.163614561 |
| *E230013L22Rik* | 29.40710175 | 24.71403219 | 27.97413014 | 34.74490018 | 78.35228725 | 182.4176898 | 289.429325 | 111.082248 | 1.21E-06 | 2.498926651 | 8.39E-09 | 33.18159581 | 5.652647177 | 5.917241378 |
| *Lif* | 81.98343517 | 67.22216757 | 54.91218138 | 52.11735027 | 84.79220127 | 522.6968419 | 733.9844393 | 106.6389581 | 0.003740773 | 2.497990707 | 0.000215053 | 13.69476715 | 5.648981231 | 2.427038661 |
| *Igsf6* | 846.5680806 | 896.625088 | 960.4451347 | 928.0545707 | 2238.943441 | 5284.266283 | 10280.98771 | 2594.881314 | 1.16E-05 | 2.489783597 | 1.36E-07 | 27.78568178 | 5.616936903 | 4.936065706 |

| | | | | | | | | | | | | | | |
|---|---|---|---|---|---|---|---|---|---|---|---|---|---|---|
| *Prss50* | 1.782248591 | 6.919929014 | 3.108236682 | 4.571697392 | 17.17310405 | 15.20147415 | 43.72673256 | 15.10718573 | 0.001290442 | 2.483937767 | 5.03E-05 | 16.43647351 | 5.594223011 | 2.889261547 |
| *Tlr2* | 3052.100712 | 3069.482798 | 3086.479025 | 3186.473083 | 9843.40898 | 18458.09765 | 28246.42812 | 12514.08173 | 1.50E-11 | 2.478179029 | 2.67E-14 | 57.96725061 | 5.571937322 | 10.82341445 |
| *Slfn4* | 23.16923168 | 30.64539992 | 36.26276129 | 29.25886331 | 77.27896825 | 157.8614623 | 389.3761423 | 30.21437147 | 0.00563325 | 2.456248538 | 0.00037355 | 12.66005899 | 5.48787849 | 2.249241011 |
| *Mcoln2* | 275.3574073 | 300.5226315 | 301.4989581 | 320.9331569 | 547.3926917 | 2065.061796 | 3208.70928 | 676.2687261 | 0.00026546 | 2.438790253 | 6.87E-06 | 20.22792366 | 5.42186899 | 3.576000288 |
| *Fscn1* | 50.79408483 | 57.33655469 | 58.02041806 | 61.26074506 | 80.49892526 | 376.5288212 | 663.1887771 | 105.7503001 | 0.002579953 | 2.43028035 | 0.000130288 | 14.63761386 | 5.389981609 | 2.588388168 |
| *Ero1l* | 1028.357437 | 1067.646191 | 1142.79502 | 1227.95792 | 1437.174146 | 6545.988637 | 14436.06842 | 1618.246189 | 0.005309785 | 2.427973342 | 0.000339251 | 12.8402044 | 5.381369396 | 2.274923049 |
| *Gbp7* | 259.3171699 | 237.2547091 | 281.8134592 | 252.3576961 | 465.8204475 | 1497.929876 | 3125.420265 | 454.10423 | 0.001298757 | 2.427001412 | 5.10E-05 | 16.4096756 | 5.377745239 | 2.886471943 |
| *Birc3* | 583.6864135 | 523.9374825 | 546.0135771 | 528.4882186 | 1306.229227 | 3334.969559 | 5514.773866 | 1541.821603 | 6.75E-06 | 2.422362996 | 7.13E-08 | 29.02857025 | 5.36048299 | 5.17078644 |
| *Rgl1* | 91.78580242 | 102.8103739 | 83.92239041 | 101.4916821 | 160.9978505 | 475.9230753 | 1216.01961 | 177.7315969 | 0.002171777 | 2.417618445 | 0.000101966 | 15.09994474 | 5.342883075 | 2.66318487 |
| *Jag1* | 185.3538534 | 154.2155609 | 175.0973331 | 172.8101614 | 312.33583 | 997.450573 | 1827.152753 | 530.5288166 | 7.59E-05 | 2.4152339 | 1.38E-06 | 23.3026274 | 5.334059435 | 4.119656888 |
| *Casp4* | 154.1645031 | 164.1011738 | 134.6902562 | 149.037335 | 304.822597 | 831.4037015 | 1611.642429 | 444.3289922 | 5.54E-05 | 2.406540007 | 9.20E-07 | 24.08898162 | 5.302012263 | 4.256868048 |
| *Gadd45b* | 182.6804805 | 186.8380834 | 211.3600944 | 179.2105378 | 216.8104387 | 1322.528251 | 2140.52767 | 345.6879559 | 0.003640285 | 2.404902514 | 0.000206154 | 13.77414805 | 5.295997767 | 2.438864596 |
| *Ass1* | 750.3266567 | 784.9176625 | 698.3171745 | 738.7862986 | 2231.430208 | 3914.964265 | 7806.262874 | 1699.114066 | 1.18E-05 | 2.396605205 | 1.40E-07 | 27.72967368 | 5.265626549 | 4.926477433 |
| *Plaur* | 434.8686561 | 484.395031 | 408.2150842 | 390.4229573 | 511.9731646 | 2707.031743 | 5074.383202 | 702.0398076 | 0.00452789 | 2.388506727 | 0.000275414 | 13.23061283 | 5.236151088 | 2.34410416 |
| *Sdc1* | 523.9810857 | 518.9946761 | 554.3022083 | 536.7172739 | 1037.899476 | 2449.776026 | 6412.212996 | 1119.70906 | 0.000724049 | 2.368424053 | 2.41E-05 | 17.83707228 | 5.163767533 | 3.140232251 |
| *Marcksl1* | 1050.635544 | 1191.216352 | 1023.645947 | 1026.803234 | 2817.462384 | 5823.333943 | 9375.219683 | 3991.851666 | 1.23E-08 | 2.358185861 | 4.58E-11 | 43.34932755 | 5.127252194 | 7.908711717 |
| *Rasip1* | 8.020118658 | 0.988561288 | 5.18039447 | 1.828678957 | 13.95314704 | 32.74163663 | 27.06892968 | 8.886579843 | 0.00991188 | 2.35300989 | 0.000817194 | 11.20176788 | 5.108890058 | 2.003843949 |
| *Gm16712* | 23.16923168 | 16.80554189 | 18.64942009 | 12.8007527 | 32.1995701 | 115.7650724 | 184.2769443 | 27.54839751 | 0.004956919 | 2.330948609 | 0.000311107 | 13.0023092 | 5.031360659 | 2.304788137 |
| *Ppp1r15a* | 127.4307742 | 132.4672126 | 147.1232029 | 126.178848 | 243.6434138 | 765.9204283 | 1309.719751 | 341.244666 | 0.000164456 | 2.318941128 | 3.73E-06 | 21.39986354 | 4.989658669 | 3.783951506 |
| *Pim3* | 123.8662771 | 111.7074255 | 88.06670599 | 132.5792244 | 229.6902667 | 643.1392909 | 1214.978497 | 188.3954927 | 0.001355249 | 2.318270054 | 5.37E-05 | 16.31347683 | 4.987338256 | 2.867980812 |
| *Hip1r* | 8.911242953 | 12.85129674 | 5.18039447 | 8.229055306 | 18.24642306 | 52.62048744 | 83.28901439 | 21.32779162 | 0.002351042 | 2.314697099 | 0.000114935 | 14.87402465 | 4.975001979 | 2.628739563 |
| *Cd300e* | 99.80592108 | 112.6959868 | 112.9325994 | 128.007527 | 198.5640156 | 641.9699468 | 1219.142948 | 185.7295187 | 0.001974884 | 2.307793378 | 8.87E-05 | 15.36234626 | 4.951251997 | 2.704458342 |
| *Slc16a3* | 1347.379935 | 1280.186868 | 1314.784116 | 1351.393749 | 1674.377645 | 7099.088428 | 15342.87756 | 1827.080816 | 0.006605247 | 2.292992358 | 0.000458238 | 12.27837028 | 4.900715368 | 2.180110927 |
| *Nfkb2* | 1037.26868 | 1103.234397 | 1078.558129 | 1120.065861 | 2201.377276 | 6224.418992 | 9969.695023 | 2657.976031 | 5.44E-05 | 2.278555257 | 8.99E-07 | 24.13308734 | 4.8519183 | 4.264701704 |
| *Stard5* | 306.5467576 | 299.5340702 | 340.8699561 | 313.6184411 | 503.3866126 | 1785.58854 | 3213.914843 | 573.1843999 | 0.001478531 | 2.269011111 | 5.94E-05 | 16.12172096 | 4.819926382 | 2.830169678 |
| *Slc28a2* | 147.926633 | 120.6044771 | 163.7004652 | 150.866014 | 200.7106536 | 777.6138699 | 1515.860062 | 234.6057079 | 0.00441847 | 2.226311616 | 0.000266976 | 13.28894288 | 4.679361268 | 2.354728074 |
| *Jak2* | 1039.050928 | 962.8586943 | 965.6255292 | 1098.121714 | 1388.87479 | 5773.052144 | 9734.403557 | 1926.61051 | 0.001974884 | 2.210892764 | 8.87E-05 | 15.36331447 | 4.629616734 | 2.704458342 |
| *2010002M12Rik* | 5.346745772 | 4.942806439 | 7.252552258 | 10.05773426 | 30.0529321 | 33.91098079 | 47.89118328 | 15.99584372 | 0.000133931 | 2.204044536 | 2.90E-06 | 21.88454575 | 4.607692805 | 3.873119931 |
| *Aoah* | 336.8449836 | 353.904941 | 307.7154315 | 262.4154303 | 701.9506282 | 1575.106591 | 2722.509658 | 716.2583354 | 6.45E-05 | 2.180455476 | 1.11E-06 | 23.72073343 | 4.532966428 | 4.190339865 |
| *Ankrd33b* | 203.1763393 | 183.8723995 | 207.2157788 | 155.4377113 | 290.8694499 | 1044.22434 | 1681.396978 | 374.1250114 | 0.00118093 | 2.177070156 | 4.46E-05 | 16.66548054 | 4.522342176 | 2.9277759 |
| *Lpar1* | 16.04023732 | 19.77122575 | 38.33491908 | 20.11546853 | 52.59263117 | 128.6278582 | 163.4546907 | 76.42458665 | 8.34E-05 | 2.16380126 | 1.58E-06 | 23.04580833 | 4.480939537 | 4.078615473 |
| *Arrdc4* | 242.3858083 | 212.5406769 | 202.0353843 | 252.3576961 | 392.8347552 | 1162.3281 | 2084.307585 | 433.6650963 | 0.000951739 | 2.163023468 | 3.39E-05 | 17.18357179 | 4.478524406 | 3.021482053 |
| *Ccl9* | 6159.451129 | 6241.775971 | 6325.261648 | 5663.41873 | 9367.928262 | 32788.4104 | 54904.11829 | 12002.21474 | 0.001007022 | 2.160798803 | 3.67E-05 | 17.03384621 | 4.471623751 | 2.996961081 |
| *Slc39a14* | 449.1266149 | 459.6809988 | 466.2355023 | 436.1399312 | 551.6859678 | 2683.644859 | 4136.340677 | 718.9243093 | 0.004556481 | 2.159259884 | 0.00027752 | 13.21633034 | 4.466856427 | 2.341370399 |
| *Plek* | 4758.603737 | 4770.796775 | 4796.0092 | 4760.051325 | 6193.05065 | 27091.36562 | 43028.14595 | 8664.415347 | 0.002426946 | 2.154593028 | 0.000120418 | 14.78612219 | 4.452430302 | 2.614939948 |
| *Ampd3* | 450.0177691 | 538.7659018 | 463.1272656 | 414.1957838 | 630.038255 | 2635.701749 | 4145.710691 | 885.9920104 | 0.00209248 | 2.152604149 | 9.71E-05 | 15.19301799 | 4.446296474 | 2.67933869 |
| *Foxp4* | 159.5112489 | 163.1126125 | 133.6541773 | 150.866014 | 293.0160879 | 726.1627267 | 1181.662892 | 492.3165233 | 8.03E-06 | 2.148897565 | 8.74E-08 | 28.63452291 | 4.434887674 | 5.095491919 |
| *Prdx5* | 1418.669878 | 1338.511984 | 1254.691541 | 1346.822052 | 2073.652315 | 6667.600431 | 12616.20346 | 2306.067469 | 0.002016212 | 2.142682401 | 9.16E-05 | 15.30264295 | 4.415823171 | 2.695463706 |
| *Gch1* | 301.2000118 | 282.7285283 | 302.535037 | 269.7301462 | 502.3132936 | 1416.075784 | 2526.780474 | 660.2728823 | 0.000306048 | 2.142577249 | 8.25E-06 | 19.87991587 | 4.41550133 | 3.514210458 |
| *Dancr* | 1.782248591 | 1.977122575 | 3.108236682 | 3.657357914 | 8.586552027 | 10.52409749 | 19.78114092 | 7.109263874 | 0.008776736 | 2.125578694 | 0.00067546 | 11.55558546 | 4.36378098 | 2.056666941 |
| *Slc11a2* | 564.081679 | 580.2854759 | 586.420654 | 581.5199083 | 1246.123363 | 2790.055178 | 4543.415735 | 1419.186801 | 1.63E-05 | 2.112366289 | 2.02E-07 | 27.01375849 | 4.323999298 | 4.788861057 |
| *Pilrb2* | 20.49585879 | 18.78266447 | 26.93805124 | 21.94414748 | 51.51931216 | 109.9183515 | 176.9891556 | 41.76692526 | 0.000676447 | 2.108053651 | 2.22E-05 | 17.99442233 | 4.3110929 | 3.169766479 |
| *Ebi3* | 462.4935093 | 461.6581214 | 523.2198415 | 435.2255918 | 563.4924768 | 2535.13815 | 4286.260903 | 729.5882051 | 0.005757961 | 2.107745208 | 0.000385536 | 12.60102029 | 4.310171299 | 2.239731311 |
| *Slc15a3* | 3464.69126 | 3447.11321 | 3420.096429 | 3350.139849 | 3918.687681 | 17274.72135 | 31264.61378 | 5790.495426 | 0.005275545 | 2.089931618 | 0.000336212 | 12.85704133 | 4.257278935 | 2.277732643 |
| *Dst* | 1254.703008 | 1129.925552 | 1215.320543 | 1390.710347 | 2201.377276 | 5384.829881 | 10675.56942 | 2911.243545 | 0.00028135 | 2.084891424 | 7.44E-06 | 20.07546516 | 4.242431673 | 3.550753195 |
| *Aqp9* | 56.14083061 | 76.11921916 | 67.34512811 | 76.80451619 | 107.3319003 | 368.3434121 | 604.886467 | 91.53177238 | 0.006118114 | 2.083880425 | 0.00041508 | 12.4630564 | 4.23945974 | 2.213382414 |
| *Ikbke* | 571.2106733 | 586.2168436 | 572.9516284 | 600.7210374 | 1330.915564 | 2468.485533 | 4570.484665 | 1468.951648 | 9.39E-06 | 2.077420814 | 1.04E-07 | 28.28900702 | 4.220520155 | 5.027532408 |
| *Oasl1* | 264.6639157 | 236.2661478 | 290.1020903 | 200.2403458 | 373.5150132 | 868.8227148 | 2533.02715 | 379.4569593 | 0.007591857 | 2.067444694 | 0.000548322 | 11.94370992 | 4.19143627 | 2.119651955 |
| *Gbp3* | 315.4580006 | 305.4654379 | 347.0864295 | 341.0486255 | 551.6859678 | 1257.044978 | 3003.610082 | 617.6172991 | 0.001764643 | 2.05236631 | 7.60E-05 | 15.65483426 | 4.147857438 | 2.75334317 |
| *Icam1* | 1346.48881 | 1490.750422 | 1398.706507 | 1355.051107 | 3753.396555 | 6063.049497 | 7552.23138 | 5762.947028 | 1.93E-18 | 2.04873348 | 7.80E-22 | 92.20941072 | 4.137425917 | 17.71374365 |
| *Tfec* | 144.3621358 | 139.3871416 | 151.2675185 | 160.0094087 | 246.8633708 | 722.6546942 | 1117.113906 | 349.2425878 | 0.000252838 | 2.03315877 | 6.46E-06 | 20.34699296 | 4.093000294 | 3.597158496 |
| *Stx11* | 172.8781133 | 153.2269996 | 146.0871241 | 137.1509218 | 284.4295359 | 666.5261742 | 1100.456103 | 410.5599887 | 4.00E-05 | 2.014185406 | 6.18E-07 | 24.85591886 | 4.039524291 | 4.397756487 |
| *Pilrb1* | 33.86272322 | 36.57676765 | 25.90197235 | 31.08754227 | 76.20564924 | 142.6599882 | 251.9492685 | 44.43289922 | 0.002256556 | 2.013783152 | 0.000108131 | 14.9891483 | 4.038398142 | 2.646553839 |
| *Clec2d* | 127.4307742 | 172.9982254 | 133.6541773 | 151.7803534 | 301.60264 | 602.2122451 | 1113.990568 | 339.46735 | 0.000105149 | 2.008370357 | 2.09E-06 | 22.51352255 | 4.023275012 | 3.97817461 |
| *Cebpb* | 1227.969279 | 1262.392764 | 1186.310334 | 1212.414148 | 2588.845436 | 5819.825911 | 8957.733498 | 2164.77085 | 0.000196306 | 1.998099324 | 4.70E-06 | 20.95587292 | 3.994733676 | 3.707066711 |
| *Ptgir* | 44.55621477 | 72.164974 | 56.98433917 | 59.4320661 | 127.7249614 | 243.2235864 | 428.9384241 | 129.7440657 | 0.000167449 | 1.995640631 | 3.82E-06 | 21.35551705 | 3.987931505 | 3.776118068 |
| *Bcam* | 4.455621477 | 7.908490302 | 4.144315576 | 6.400376349 | 9.659871031 | 26.8949158 | 33.31560576 | 21.32779162 | 0.001969432 | 1.992124357 | 8.82E-05 | 15.37436657 | 3.978223573 | 2.705659039 |
| *Cd83* | 557.8438089 | 544.6972695 | 568.8073128 | 536.7172739 | 1034.679519 | 2480.178975 | 3291.998294 | 1951.492934 | 1.89E-07 | 1.987864085 | 9.91E-10 | 37.34202318 | 3.966493225 | 6.723446448 |
| *Acsl1* | 1034.595307 | 1003.389707 | 1072.341655 | 1047.833042 | 2067.212401 | 4038.914747 | 8163.364523 | 2211.869723 | 0.000178635 | 1.986794327 | 4.19E-06 | 21.17429029 | 3.963553162 | 3.748032333 |
| *Isg15* | 613.9846395 | 633.6677854 | 649.6214665 | 621.7508454 | 993.8933972 | 2419.373078 | 5527.267218 | 1004.183522 | 0.003543709 | 1.981037687 | 0.000197826 | 13.85162465 | 3.947769308 | 2.450541669 |
| *Flnb* | 1000.732584 | 1054.794894 | 1173.877387 | 1056.976437 | 1451.127293 | 4849.270264 | 8748.469849 | 1787.091206 | 0.003628039 | 1.973701284 | 0.000205167 | 13.78315649 | 3.927745041 | 2.440328049 |
| *Slfn2* | 2756.247446 | 2916.255799 | 2915.526008 | 2870.111623 | 5483.586788 | 12251.21882 | 20434.95968 | 6604.506139 | 5.55E-05 | 1.966288323 | 9.26E-07 | 24.07563783 | 3.907614976 | 4.255961669 |
| *Lrrc25* | 434.8686561 | 538.7659018 | 471.4158968 | 403.22371 | 504.4599316 | 2101.311465 | 3872.939169 | 715.3696774 | 0.009413239 | 1.960613891 | 0.000752543 | 11.35477408 | 3.892275666 | 2.026260912 |
| *Sdc4* | 1367.875793 | 1403.757029 | 1222.573095 | 1230.700938 | 2138.051455 | 5879.462463 | 9269.026189 | 3031.212384 | 0.000202126 | 1.959231322 | 4.89E-06 | 20.87922565 | 3.888547393 | 3.694378828 |
| *St3gal1* | 998.0592108 | 1083.463171 | 1050.583998 | 977.4289025 | 1949.14731 | 4498.467004 | 6868.220349 | 2492.685646 | 2.60E-05 | 1.943635157 | 3.50E-07 | 25.94945089 | 3.846736895 | 4.584777719 |
| *Ppfibp2* | 164.8579946 | 184.8609608 | 184.4220431 | 183.7822352 | 294.0894069 | 777.6138699 | 1323.254216 | 359.0178257 | 0.000750234 | 1.939494874 | 2.52E-05 | 17.74652742 | 3.835713257 | 3.124803427 |
| *Rbpms* | 56.14083061 | 42.50813537 | 27.97413014 | 36.57357914 | 88.01215828 | 164.8775273 | 286.305987 | 87.08848246 | 0.000568774 | 1.937424946 | 1.79E-05 | 18.39630899 | 3.830213857 | 3.245060247 |
| *Cav1* | 189.8094749 | 178.9295931 | 222.7569622 | 237.7282644 | 302.675959 | 698.0984667 | 1782.384908 | 389.2321791 | 0.004242098 | 1.935665748 | 0.000252897 | 13.39053749 | 3.825546205 | 2.372419351 |
| *H2-Ab1* | 319.913622 | 270.8657928 | 352.266824 | 317.275799 | 501.2399746 | 1257.044978 | 2552.808291 | 478.0979956 | 0.003891117 | 1.925908928 | 0.000225979 | 13.60171321 | 3.799761658 | 2.409925762 |
| *H2-Q7* | 108.717164 | 127.5244061 | 128.4737829 | 96.00564524 | 191.0507826 | 509.8340561 | 788.1222987 | 260.3767894 | 0.000363658 | 1.92498231 | 1.04E-05 | 19.43383891 | 3.79732192 | 3.439306894 |
| *Ch25h* | 80.20118658 | 84.02770946 | 74.59768037 | 64.00376349 | 119.1384094 | 343.7871846 | 450.8017904 | 221.2758381 | 7.38E-05 | 1.905886682 | 1.33E-06 | 23.37305651 | 3.747391433 | 4.131698382 |
| *Gstt1* | 14.25798873 | 9.885612877 | 13.46902562 | 15.54377113 | 19.31974206 | 64.31392909 | 92.65902851 | 23.10510759 | 0.00948846 | 1.903033086 | 0.000760853 | 11.33437928 | 3.739986561 | 2.022804279 |
| *Mx2* | 106.9349154 | 78.09634173 | 85.9945482 | 103.3203611 | 128.7982804 | 339.1098079 | 727.7377653 | 184.8408607 | 0.004039279 | 1.882107312 | 0.000237221 | 13.51058717 | 3.686130921 | 2.393696125 |
| *Stk40* | 412.5905487 | 404.3215667 | 374.0244807 | 373.9648467 | 529.1462687 | 1768.048378 | 2851.60763 | 615.8399831 | 0.004199011 | 1.881042322 | 0.000249312 | 13.41731682 | 3.683410841 | 2.376852972 |
| *Zhx2* | 109.6082883 | 124.5587223 | 118.1129939 | 103.3203611 | 174.9509976 | 546.0837252 | 719.4088618 | 223.0531541 | 0.000909036 | 1.868143275 | 3.19E-05 | 17.30112175 | 3.650624483 | 3.041419069 |
| *Bcl3* | 293.1798932 | 328.2023475 | 299.4268004 | 213.955438 | 530.2195877 | 1130.755808 | 1770.932669 | 693.1532278 | 5.90E-05 | 1.862066472 | 1.00E-06 | 23.91973476 | 3.635279958 | 4.229107704 |
| *Cmpk2* | 884.8864253 | 901.5678944 | 872.3784287 | 1014.002482 | 1541.286089 | 3027.432044 | 7074.36066 | 1565.815368 | 0.002437623 | 1.846504391 | 0.000121331 | 14.77189286 | 3.596277606 | 2.613033391 |
| *Usp18* | 498.1384811 | 511.0861858 | 523.2198415 | 538.5459528 | 889.7814538 | 1888.490827 | 3743.841197 | 878.8827465 | 0.00155242 | 1.837335809 | 6.34E-05 | 15.99730352 | 3.573495097 | 2.80899079 |
| *Rapgef2* | 494.5739839 | 397.4016377 | 472.4519757 | 435.2255918 | 610.7185129 | 1807.80608 | 3293.039407 | 714.4810194 | 0.004956919 | 1.83611847 | 0.000311051 | 13.0026456 | 3.570481072 | 2.304788137 |
| *Arhgef3* | 367.1432097 | 343.0307668 | 385.4213486 | 319.104478 | 473.3336805 | 1572.767902 | 2332.092403 | 632.7244848 | 0.002579953 | 1.824494129 | 0.000130112 | 14.64016245 | 3.54182794 | 2.588388168 |
| *Relb* | 351.9940967 | 335.1222765 | 339.8338772 | 363.9071124 | 799.6226575 | 1275.754484 | 1904.195092 | 941.0888054 | 1.27E-08 | 1.822668322 | 4.81E-11 | 43.25400732 | 3.537348406 | 7.896895098 |
| *Ptafr* | 729.8307979 | 686.0615337 | 763.5901449 | 647.3523508 | 1662.571136 | 2650.903223 | 3686.58 | 1992.371201 | 4.06E-10 | 1.821637454 | 1.08E-12 | 50.69150701 | 3.53482172 | 9.39146186 |
| *Dcbld2* | 237.0390626 | 241.2089542 | 260.0558024 | 245.0429802 | 347.7553571 | 1014.990706 | 1624.135781 | 478.0979956 | 0.00155242 | 1.816966023 | 6.33E-05 | 16.00374305 | 3.523394515 | 2.80899079 |
| *Sh3d21* | 4.455621477 | 5.931367726 | 5.18039447 | 2.743018435 | 9.659871031 | 22.21753914 | 22.90447896 | 9.775237827 | 0.009571717 | 1.816266824 | 0.000773809 | 11.30302514 | 3.521687323 | 2.019010159 |
| *Tlr1* | 310.1112548 | 393.4473925 | 335.6895616 | 320.0188175 | 539.8794587 | 1349.423167 | 2191.542109 | 610.5080352 | 0.001290442 | 1.787117326 | 5.02E-05 | 16.43992745 | 3.451246038 | 2.889261547 |
| *Rilpl2* | 375.1633283 | 352.9163797 | 352.266824 | 325.5048543 | 712.6838183 | 1343.576446 | 2077.019796 | 710.9263874 | 4.57E-05 | 1.784645523 | 7.34E-07 | 24.52380047 | 3.445338 | 4.339889782 |
| *Fgr* | 102.479294 | 119.6159158 | 122.2573095 | 135.3222428 | 286.5761739 | 349.6539054 | 724.6144252 | 269.2633692 | 2.44E-05 | 1.764770322 | 3.27E-07 | 26.08493282 | 3.398199099 | 4.61264589 |
| *Rtn2* | 8.911242953 | 13.83985803 | 11.39686783 | 9.143394785 | 25.75965608 | 38.58835746 | 63.50787348 | 18.66181767 | 0.003340842 | 1.759249964 | 0.000182997 | 13.99808343 | 3.385220864 | 2.47614407 |
| *Clec4d* | 2128.895942 | 2275.668084 | 2145.719389 | 1989.602705 | 3369.148352 | 7869.686232 | 14060.22674 | 3543.079383 | 0.001958651 | 1.755875545 | 8.72E-05 | 15.39498797 | 3.377312191 | 2.708042903 |
| *Col27a1* | 28.51597745 | 24.71403219 | 54.91218138 | 28.34452383 | 47.22603615 | 157.8614623 | 171.7835922 | 82.64519254 | 0.003556386 | 1.753033006 | 0.000199045 | 13.84007674 | 3.370664432 | 2.448991061 |
| *Denr* | 757.455651 | 730.5467916 | 741.8324881 | 691.2406457 | 866.1684358 | 3151.382525 | 4741.227144 | 1007.738154 | 0.009522054 | 1.741282802 | 0.000765851 | 11.32222011 | 3.343323141 | 2.021269348 |
| *Vasp* | 2074.53736 | 2119.475401 | 2064.905236 | 2099.323443 | 3951.960571 | 7467.43184 | 11557.39186 | 4571.256671 | 2.00E-05 | 1.720653458 | 2.62E-07 | 26.51427468 | 3.295856566 | 4.698599158 |
| *Gsap* | 563.1905547 | 596.1024565 | 646.5132298 | 647.3523508 | 1145.231377 | 2065.061796 | 3678.251098 | 1194.356331 | 0.000206989 | 1.720191017 | 5.04E-06 | 20.82096152 | 3.294800282 | 3.684053531 |
| *Rassf4* | 4060.853414 | 4152.94597 | 4134.990866 | 4008.464274 | 6640.624674 | 15403.77069 | 23921.64605 | 7926.82922 | 0.000380223 | 1.720153656 | 1.10E-05 | 19.33270036 | 3.294714957 | 3.419961206 |
| *Il20rb* | 24.06035597 | 23.72547091 | 33.15452461 | 12.8007527 | 41.85944113 | 90.03950073 | 132.2213104 | 42.65558325 | 0.005568649 | 1.712068627 | 0.000368024 | 12.68792475 | 3.276302637 | 2.25425014 |
| *Phospho1* | 40.99171759 | 36.57676765 | 30.04628793 | 25.6015054 | 65.47245921 | 76.00737074 | 230.0859023 | 64.87203285 | 0.005693093 | 1.711348762 | 0.000379356 | 12.63122848 | 3.27466826 | 2.244651756 |
| *Sema4a* | 188.0272263 | 221.4377285 | 218.6126466 | 197.4973274 | 310.2117284 | 703.9451875 | 1309.719751 | 375.9023274 | 0.002113777 | 1.709332351 | 9.86E-05 | 15.16407012 | 3.270094552 | 2.674940734 |
| *Hck* | 2044.239134 | 2060.161724 | 1992.379713 | 1972.230255 | 2911.914456 | 7100.257772 | 12993.08625 | 3297.80978 | 0.003194992 | 1.704747909 | 0.000172455 | 14.10967246 | 3.259719688 | 2.495530181 |
| *St3gal3* | 171.0958647 | 173.9867866 | 176.133412 | 177.3818588 | 189.9774636 | 717.9773175 | 1091.086089 | 277.2612911 | 0.009608187 | 1.703922863 | 0.000779172 | 11.29020288 | 3.257856059 | 2.017358537 |
| *Cers6* | 422.392916 | 405.310128 | 490.0653169 | 475.4565288 | 667.6044201 | 1730.629365 | 2544.47939 | 828.2292414 | 0.001141256 | 1.686160859 | 4.27E-05 | 16.74850171 | 3.217992267 | 2.942617037 |
| *Txnrd1* | 4312.150465 | 4753.991233 | 4538.025556 | 4244.363859 | 5833.488784 | 16555.57469 | 27804.99634 | 7077.272187 | 0.00491144 | 1.682001952 | 0.000306573 | 13.02980357 | 3.208729006 | 2.3087912 |
| *Fam20c* | 2518.317259 | 2510.945671 | 2489.697582 | 2609.524872 | 5813.095722 | 8054.442611 | 12487.10548 | 6046.428925 | 1.40E-08 | 1.677595784 | 5.53E-11 | 42.98096343 | 3.198944103 | 7.854384925 |
| *Mnda* | 938.353883 | 1015.252443 | 959.4090558 | 954.5704155 | 1344.868711 | 3296.381202 | 6395.555193 | 1326.766371 | 0.00819222 | 1.676555584 | 0.000612753 | 11.73683016 | 3.19663846 | 2.086598375 |
| *Gng4* | 14.25798873 | 10.87417417 | 14.50510452 | 5.486036871 | 19.31974206 | 46.77376661 | 39.56228184 | 38.21229333 | 0.001290442 | 1.676035541 | 5.04E-05 | 16.43419475 | 3.195486387 | 2.889261547 |
| *Pvr* | 313.675752 | 360.82487 | 341.906035 | 304.4750463 | 444.3540674 | 1257.044978 | 1998.936345 | 502.0917611 | 0.005408358 | 1.669640129 | 0.000350931 | 12.77687251 | 3.181352267 | 2.266934574 |
| *Slc25a37* | 653.1941085 | 667.2788692 | 756.3375926 | 710.4417748 | 1190.310775 | 2307.116038 | 3976.009325 | 1345.428188 | 0.000360494 | 1.66172022 | 1.03E-05 | 19.46132218 | 3.163935567 | 3.443102472 |
| *Rffl* | 378.7278255 | 341.0536443 | 340.8699561 | 284.3595778 | 433.6208774 | 1329.544316 | 1959.374064 | 502.0917611 | 0.006732633 | 1.6510481 | 0.000469264 | 12.23400389 | 3.140617182 | 2.171815032 |
| *Emr4* | 49.90296054 | 36.57676765 | 25.90197235 | 24.68716592 | 91.23211529 | 111.0876957 | 188.4413951 | 39.98960929 | 0.008067138 | 1.648712765 | 0.000597766 | 11.78292095 | 3.135537481 | 2.093280504 |
| *Fabp4* | 44.55621477 | 47.45094181 | 62.16473364 | 41.14527653 | 120.2117284 | 139.1519557 | 212.3869867 | 135.9646716 | 1.34E-08 | 1.641350552 | 5.18E-11 | 43.10695346 | 3.119577286 | 7.873397475 |
| *Sqstm1* | 8760.642948 | 9245.025163 | 8585.985794 | 8918.467273 | 16060.07225 | 31503.30116 | 43689.2525 | 19512.26336 | 1.04E-05 | 1.64119132 | 1.20E-07 | 28.02221796 | 3.119232993 | 4.983124618 |
| *Ifrd1* | 656.7586057 | 618.8393661 | 611.2865474 | 630.8942422 | 1126.984954 | 2212.399161 | 3309.697209 | 1177.471829 | 0.000153912 | 1.63609289 | 3.45E-06 | 21.54750377 | 3.108229197 | 3.812728085 |
| *Sh3bp4* | 137.2331415 | 120.6044771 | 129.5098617 | 107.8920585 | 174.9509976 | 470.0763545 | 653.818763 | 235.4943658 | 0.001587953 | 1.631002801 | 6.56E-05 | 15.9349268 | 3.097282125 | 2.799162302 |
| *Cybb* | 10972.41345 | 11079.79491 | 11730.48524 | 11901.95699 | 16463.64019 | 35380.84641 | 71017.41924 | 18412.10478 | 0.004142987 | 1.628712808 | 0.000244919 | 13.45066405 | 3.092369702 | 2.382686387 |
| *Cx3cl1* | 114.9550341 | 122.5815997 | 132.6180984 | 90.51960837 | 118.0650904 | 478.2617636 | 599.6809036 | 221.2758381 | 0.008156412 | 1.621254334 | 0.00060798 | 11.75138324 | 3.07642396 | 2.088500844 |
| *Car13* | 461.602385 | 516.0289922 | 452.7664767 | 414.1957838 | 715.9037753 | 1680.347566 | 2601.740587 | 654.9409344 | 0.003764705 | 1.61559424 | 0.000217148 | 13.67656179 | 3.064377938 | 2.424268997 |
| *Hspa1a* | 53.46745772 | 33.61108378 | 33.15452461 | 38.4022581 | 74.05901124 | 116.9344165 | 231.1270149 | 64.87203285 | 0.004571629 | 1.615395507 | 0.000279181 | 13.20515169 | 3.063955846 | 2.339929004 |

| | | | | | | | | | | | | | | |
|---|---|---|---|---|---|---|---|---|---|---|---|---|---|---|
| *Sema6d* | 10.69349154 | 11.86273545 | 15.54118341 | 13.71509218 | 19.31974206 | 40.92704579 | 70.79566224 | 27.54839751 | 0.00715535 | 1.614848979 | 0.000506231 | 12.09257114 | 3.062795365 | 2.145369141 |
| *Hif1a* | 1907.897116 | 1934.61444 | 1970.622056 | 2064.578542 | 2559.865823 | 6869.896971 | 11740.62769 | 2804.604598 | 0.007883687 | 1.605657689 | 0.000580656 | 11.83699182 | 3.043344577 | 2.103270638 |
| *Fam219aos* | 14.25798873 | 10.87417417 | 10.36078894 | 4.571697392 | 24.68633708 | 30.4029483 | 34.35671844 | 31.99168744 | 0.000205026 | 1.601662068 | 4.98E-06 | 20.84556155 | 3.034927532 | 3.688191146 |
| *Klra2* | 40.10059329 | 47.45094181 | 41.44315576 | 34.74490018 | 69.76573522 | 126.2891699 | 242.5792544 | 55.09679503 | 0.009508636 | 1.591806382 | 0.000764005 | 11.32670224 | 3.014265259 | 2.0218818 |
| *Syk* | 4581.270002 | 4495.976737 | 4449.95885 | 4364.142331 | 5813.095722 | 15107.92662 | 25965.35024 | 6904.872538 | 0.005792409 | 1.588099338 | 0.00038831 | 12.58762104 | 3.006529967 | 2.237140781 |
| *Map3k8* | 207.6319608 | 200.6779414 | 195.818911 | 195.6686484 | 276.9163029 | 715.6386292 | 1101.497215 | 283.481897 | 0.006478947 | 1.571434112 | 0.000445833 | 12.32959676 | 2.971999991 | 2.188495541 |
| *Osmr* | 16.04023732 | 12.85129674 | 21.75765677 | 20.11546853 | 39.71280313 | 57.2978641 | 84.33012707 | 29.32571348 | 0.002087162 | 1.571363241 | 9.65E-05 | 15.20374063 | 2.971853998 | 2.680443769 |
| *Cacng8* | 13.36686443 | 17.79410318 | 15.54118341 | 11.88641322 | 32.1995701 | 33.91098079 | 77.04233831 | 30.21437147 | 0.002955427 | 1.567870712 | 0.000153541 | 14.3282778 | 2.964668328 | 2.529379754 |
| *Itgal* | 3085.07231 | 3059.597186 | 2988.05153 | 3228.532699 | 4351.23524 | 10374.42143 | 17395.95177 | 4456.619791 | 0.005405571 | 1.565140125 | 0.000347551 | 12.79497936 | 2.959062409 | 2.267158439 |
| *Pik3r5* | 2948.730293 | 2927.129973 | 2972.510347 | 2824.394649 | 5245.30997 | 8882.33828 | 13788.49633 | 6537.856791 | 8.04E-06 | 1.561513965 | 8.82E-08 | 28.61639983 | 2.951634257 | 5.094632461 |
| *Fmnl2* | 263.7727914 | 260.98018 | 280.7773803 | 272.4731646 | 399.2746693 | 879.3468123 | 1449.22885 | 453.215572 | 0.002186557 | 1.56098337 | 0.000103189 | 15.0774357 | 2.950548902 | 2.66023915 |
| *Nfkbib* | 343.0828537 | 326.225225 | 284.9216958 | 349.2776808 | 474.4069995 | 1057.087125 | 1694.931443 | 616.7286411 | 0.001250228 | 1.559615736 | 4.79E-05 | 16.53083348 | 2.947753192 | 2.903010723 |
| *Plagl2* | 926.7692672 | 1063.691946 | 919.001979 | 896.0526889 | 1193.530732 | 3380.573982 | 5275.317949 | 1340.984898 | 0.00904258 | 1.556051109 | 0.000703944 | 11.47880447 | 2.940478841 | 2.043707647 |
| *Lacc1* | 621.1136339 | 669.2559918 | 659.9822555 | 637.2946165 | 996.0400352 | 2109.496874 | 3230.572646 | 1172.139881 | 0.000714096 | 1.53677149 | 2.36E-05 | 17.8703031 | 2.901444816 | 3.146243609 |
| *Rnf149* | 2661.78827 | 2597.939064 | 2697.94944 | 2583.923366 | 3966.987037 | 8118.75654 | 14762.9778 | 3701.260505 | 0.005252021 | 1.53505969 | 0.000333441 | 12.87252472 | 2.898004208 | 2.279673561 |
| *Agpat4* | 172.8781133 | 214.5177994 | 211.3600944 | 233.156567 | 339.1688051 | 699.2678109 | 957.8236655 | 413.2259627 | 0.000215532 | 1.534040691 | 5.32E-06 | 20.71858717 | 2.895958024 | 3.66648919 |
| *Rcl1* | 216.5432038 | 204.6321866 | 227.9373567 | 234.0709065 | 323.06902 | 723.8240383 | 1203.526258 | 291.4798189 | 0.007923137 | 1.524845262 | 0.00058484 | 11.82362299 | 2.877558508 | 2.101102827 |
| *Mocs1* | 369.8165826 | 314.3624895 | 347.0864295 | 300.8176884 | 488.3601466 | 953.0154947 | 1786.549359 | 595.4008495 | 0.00240202 | 1.521023107 | 0.000118783 | 14.81190409 | 2.869945036 | 2.619423412 |
| *Txlnb* | 22.27810738 | 33.61108378 | 13.46902562 | 34.74490018 | 48.29935515 | 76.00737074 | 116.6046202 | 55.09679503 | 0.003698384 | 1.503786639 | 0.000211532 | 13.72576766 | 2.835860642 | 2.431987996 |
| *Pfkl* | 1507.782308 | 1425.505377 | 1445.330057 | 1585.464656 | 2000.666622 | 4635.280271 | 8183.145664 | 2014.58765 | 0.009530726 | 1.496934071 | 0.000767416 | 11.31842991 | 2.822422703 | 2.020873999 |
| *Gadd45a* | 287.8331474 | 259.9916187 | 236.2259878 | 204.8120432 | 587.1054949 | 686.405025 | 839.13682 | 676.2687261 | 4.46E-17 | 1.495607251 | 2.16E-20 | 85.64003208 | 2.819828167 | 16.35069572 |
| *Trim13* | 47.22958765 | 81.06202559 | 87.03062709 | 84.11923202 | 120.2117284 | 201.1271964 | 391.4583677 | 126.1894338 | 0.006924189 | 1.48690144 | 0.000485967 | 12.16875709 | 2.802863409 | 2.159631101 |
| *Mllt6* | 982.0189735 | 1092.360223 | 970.8059237 | 954.5704155 | 1757.023209 | 2945.577952 | 4196.725213 | 2270.52115 | 2.69E-06 | 1.481599377 | 2.21E-08 | 31.298967 | 2.792581484 | 5.57053877 |
| *3110043O21Rik* | 584.5775377 | 588.1939662 | 528.4002359 | 598.8923584 | 1084.052193 | 1709.58117 | 2392.476938 | 1236.123256 | 1.15E-06 | 1.481191099 | 7.88E-09 | 33.30326981 | 2.791791304 | 5.939335945 |
| *Jdp2* | 427.7396618 | 420.1385473 | 458.98295 | 432.4825733 | 672.9710151 | 1388.011524 | 1896.907303 | 887.7693263 | 0.000123914 | 1.478043485 | 2.59E-06 | 22.099295 | 2.785706926 | 3.906878077 |
| *Ablim2* | 8.911242953 | 5.931367726 | 14.50510452 | 14.62943166 | 32.1995701 | 25.72557164 | 22.90447896 | 40.87826728 | 0.002121271 | 1.477663751 | 9.91E-05 | 15.15412411 | 2.784973792 | 2.673403748 |
| *Pirb* | 2750.9007 | 2803.559812 | 2808.809882 | 2607.696193 | 3690.070734 | 8164.360962 | 14826.48567 | 3843.445782 | 0.008632628 | 1.476256184 | 0.000659495 | 11.6000658 | 2.782257953 | 2.063856996 |
| *Dram1* | 597.0532779 | 620.8164887 | 604.0339952 | 585.1772662 | 826.4556326 | 1860.426567 | 3167.064772 | 836.2271632 | 0.00773678 | 1.474674887 | 0.000565467 | 11.88635569 | 2.779210071 | 2.111439733 |
| *Cd82* | 923.20477 | 872.8996171 | 876.5227443 | 921.6541943 | 1600.318634 | 2564.371755 | 3962.47486 | 1836.856054 | 1.91E-05 | 1.470931789 | 2.48E-07 | 26.61593184 | 2.772008706 | 4.718266833 |
| *B4galt5* | 1511.346805 | 1493.716106 | 1465.015556 | 1228.872259 | 2052.185935 | 4665.68322 | 6699.560095 | 2261.63457 | 0.002416784 | 1.460031418 | 0.000119708 | 14.79727388 | 2.751143547 | 2.616762075 |
| *Slc2a6* | 1163.80833 | 1189.239229 | 1221.537016 | 1262.70282 | 2099.411971 | 3562.991672 | 5015.039779 | 2562.000969 | 6.32E-06 | 1.452523358 | 6.63E-08 | 29.16929843 | 2.736863264 | 5.199005219 |
| *Gdpd1* | 215.6520795 | 220.4491672 | 234.15383 | 254.186375 | 433.6208774 | 631.4458493 | 973.4403557 | 488.7618914 | 5.90E-06 | 1.450705189 | 6.00E-08 | 29.36445433 | 2.73341628 | 5.228862426 |
| *Nr1h3* | 194.2650964 | 200.6779414 | 173.0251753 | 178.2961983 | 446.5007054 | 454.8748803 | 662.1476644 | 464.7681258 | 1.03E-14 | 1.44233646 | 9.22E-18 | 73.6738587 | 2.717606285 | 13.98855453 |
| *Slfn5* | 2275.040326 | 2174.834833 | 2402.666955 | 2381.854341 | 3314.409083 | 6443.086351 | 12137.29162 | 3037.43299 | 0.009413239 | 1.432900068 | 0.000751724 | 11.35679662 | 2.699888948 | 2.026260912 |
| *Rhbdf2* | 957.0674932 | 1024.149494 | 963.5533714 | 930.7975891 | 1441.467422 | 2765.498951 | 4598.594707 | 1564.92671 | 0.002145229 | 1.419954266 | 0.000100547 | 15.12640979 | 2.675770286 | 2.668526263 |
| *Mapkapk2* | 2970.117276 | 2968.649547 | 2805.701645 | 2906.685202 | 3978.793546 | 8584.155517 | 13722.90623 | 4740.990346 | 0.003075804 | 1.41302095 | 0.000163021 | 14.21551808 | 2.662941896 | 2.512041356 |
| *Orai2* | 1230.642652 | 1176.387932 | 1199.779359 | 1060.633795 | 1634.664842 | 3087.068596 | 5793.792064 | 1907.060034 | 0.004142987 | 1.412231368 | 0.000244976 | 13.45022751 | 2.661484875 | 2.382686387 |
| *Gpr132* | 120.3017799 | 139.3871416 | 153.3396763 | 112.4637559 | 221.1037147 | 362.4966912 | 480.9940581 | 326.1374802 | 3.52E-06 | 1.405224829 | 3.18E-08 | 30.59211312 | 2.648590544 | 5.452983364 |
| *Pla2g7* | 430.4130347 | 425.0813537 | 397.8542953 | 392.2516363 | 643.991402 | 1161.158756 | 2055.15643 | 494.9824973 | 0.009608187 | 1.40397925 | 0.000779754 | 11.28881471 | 2.646304819 | 2.017358537 |
| *Helz2* | 2568.220219 | 2525.77409 | 2655.470205 | 2599.467137 | 3588.105428 | 6926.025491 | 12907.71501 | 3628.39055 | 0.008287171 | 1.386140936 | 0.000622407 | 11.70773928 | 2.613785835 | 2.081593716 |
| *Bhlhe40* | 502.5941026 | 625.7592951 | 608.1783108 | 549.5180266 | 1318.035736 | 1382.164803 | 1663.698063 | 1553.374157 | 1.37E-26 | 1.373245209 | 2.21E-30 | 131.2211935 | 2.590526261 | 25.86237932 |
| *Rusc2* | 517.7432156 | 665.3017466 | 626.8277309 | 562.3187793 | 820.0157186 | 1469.865616 | 2721.468545 | 1124.15235 | 0.00228606 | 1.371018335 | 0.000110467 | 14.94882107 | 2.586530736 | 2.640912438 |
| *Plcxd2* | 69.50769504 | 70.18785143 | 71.48944368 | 71.31847932 | 130.9449184 | 219.8367031 | 240.4970291 | 138.6306456 | 3.26E-06 | 1.367557696 | 2.87E-08 | 30.79551425 | 2.580333777 | 5.486717492 |
| *N4bp1* | 949.0473745 | 978.6756749 | 967.697687 | 902.4530653 | 1363.115134 | 2443.929305 | 4511.141242 | 1404.079615 | 0.005568649 | 1.356057726 | 0.00036832 | 12.68642593 | 2.559847252 | 2.25425014 |
| *Nlrc4* | 97.13254819 | 96.8790062 | 82.88631152 | 109.7207374 | 140.6047894 | 256.0863722 | 437.2673256 | 150.1831993 | 0.0057222 | 1.346914748 | 0.000382219 | 12.6171749 | 2.543675697 | 2.242436998 |
| *5730508B09Rik* | 96.2414239 | 76.11921916 | 122.2573095 | 106.977719 | 140.6047894 | 263.1024372 | 436.2262129 | 181.2862288 | 0.004266492 | 1.346353501 | 0.00025504 | 13.37471377 | 2.542686332 | 2.369929109 |
| *Osgin2* | 285.1597745 | 277.7857219 | 241.4063823 | 272.4731646 | 384.2482032 | 729.6707591 | 1210.814047 | 393.675487 | 0.005625327 | 1.335617587 | 0.000372571 | 12.66496608 | 2.523834986 | 2.249852206 |
| *Rbpj* | 2165.432038 | 2357.718671 | 2393.342245 | 2071.893258 | 2754.136563 | 6050.186711 | 10131.06749 | 3586.623625 | 0.007361457 | 1.325194233 | 0.000525404 | 12.02327226 | 2.505666198 | 2.133036204 |
| *Icosl* | 550.7148145 | 553.5943211 | 561.5547605 | 524.8308607 | 882.2682208 | 1478.051025 | 1933.346247 | 1194.356331 | 2.64E-06 | 1.324852314 | 2.15E-08 | 31.35006031 | 2.505072424 | 5.577689267 |
| *Gm21188* | 220.9988252 | 236.2661478 | 208.2518577 | 206.6407221 | 412.1544973 | 554.2691344 | 821.4379045 | 375.9023274 | 3.82E-05 | 1.310446464 | 5.79E-07 | 24.98065954 | 2.480182811 | 4.418135521 |
| *Tmem176a* | 254.8615485 | 246.1517606 | 223.7930411 | 223.0988328 | 311.262511 | 616.2443751 | 1053.606032 | 361.6837996 | 0.007708897 | 1.305120593 | 0.000561563 | 11.89925865 | 2.471043817 | 2.113007776 |
| *Tmem132a* | 725.3751764 | 703.8556369 | 666.1987288 | 603.4640558 | 860.8018407 | 1932.925905 | 2888.046574 | 987.2990206 | 0.00773678 | 1.305009191 | 0.000564497 | 11.88955028 | 2.470853015 | 2.111439733 |
| *Ifit2* | 1582.636749 | 1540.178486 | 1554.118341 | 1606.494464 | 2453.607242 | 3448.395943 | 7185.759717 | 2332.727209 | 0.00491144 | 1.295192401 | 0.000306668 | 13.02922269 | 2.454097216 | 2.3087912 |
| *Ntn1* | 11.58461584 | 14.82841932 | 12.43294673 | 10.05773426 | 21.46638007 | 36.24966912 | 42.68561988 | 19.55047565 | 0.009245425 | 1.29060211 | 0.000727031 | 11.41883887 | 2.446301309 | 2.034073099 |
| *Cd302* | 409.0260516 | 416.1843021 | 388.5295852 | 384.022581 | 536.6595017 | 1060.595158 | 1700.137006 | 582.9596377 | 0.006605247 | 1.279953317 | 0.000458252 | 12.27831399 | 2.428311191 | 2.180110927 |
| *Junb* | 2904.174079 | 2754.131748 | 2763.22241 | 2838.109741 | 4366.261706 | 6835.98599 | 10518.36141 | 5556.778376 | 6.78E-05 | 1.276522244 | 1.18E-06 | 23.61575208 | 2.42254295 | 4.168673064 |
| *Impact* | 605.0733965 | 534.8116567 | 589.5288907 | 551.3467055 | 1002.479949 | 1602.001506 | 1863.591697 | 1049.505079 | 2.12E-06 | 1.274226015 | 1.63E-08 | 31.89515739 | 2.418690237 | 5.672994891 |
| *1700047I17Rik2* | 971.3254819 | 957.9158878 | 957.336898 | 957.313434 | 1190.310775 | 2801.74862 | 3901.049212 | 1399.636325 | 0.007572965 | 1.273446967 | 0.000544941 | 11.95523451 | 2.417384511 | 2.120734054 |
| *Ccdc86* | 371.5988312 | 379.6075345 | 354.3389817 | 307.2180648 | 480.8469135 | 895.7176306 | 1430.488822 | 545.6360024 | 0.004936942 | 1.246654428 | 0.000308658 | 13.017106 | 2.37290514 | 2.306541967 |
| *Irs2* | 102.479294 | 103.7989352 | 93.24710046 | 91.43394785 | 187.8308256 | 285.3199763 | 298.7993391 | 153.7378313 | 3.69E-05 | 1.241250473 | 5.51E-07 | 25.07616466 | 2.364033489 | 4.432660819 |
| *Slc39a4* | 127.4307742 | 146.3070706 | 126.4016251 | 148.1229955 | 281.2095789 | 316.8922688 | 488.2818469 | 206.1686524 | 0.000192405 | 1.23635701 | 4.58E-06 | 21.00584197 | 2.356028531 | 3.715783974 |
| *Hspa1b* | 61.48757638 | 73.15353529 | 60.09257585 | 86.86225046 | 118.0650904 | 154.3534298 | 255.0726066 | 127.9667497 | 0.000952345 | 1.217671473 | 3.40E-05 | 17.17807041 | 2.325710417 | 3.021205776 |
| *Parp14* | 4963.562325 | 4828.133329 | 5106.832868 | 4999.608268 | 7104.298484 | 11723.8446 | 19616.64512 | 7628.240137 | 0.003041528 | 1.211276916 | 0.000160954 | 14.23952933 | 2.315424822 | 2.516908205 |
| *Akap6* | 16.04023732 | 16.80554189 | 18.64942009 | 31.08754227 | 36.49284612 | 37.41901329 | 71.83677492 | 44.43289922 | 0.009027956 | 1.202492587 | 0.000702077 | 11.48373945 | 2.301369421 | 2.044410548 |
| *Psme2* | 1703.829653 | 1574.778131 | 1515.783422 | 1468.429202 | 2112.291799 | 3805.045914 | 6321.636193 | 2117.671977 | 0.00970187 | 1.196775934 | 0.000789512 | 11.26572711 | 2.29226833 | 2.01314453 |
| *Ptrh1* | 51.68520913 | 57.33655469 | 60.09257585 | 63.08942402 | 83.71888227 | 162.538839 | 190.5236204 | 95.97506231 | 0.001794966 | 1.196538917 | 7.79E-05 | 15.60850256 | 2.291891769 | 2.745943745 |
| *Fam49a* | 528.4367071 | 460.6695601 | 577.0959439 | 546.7750081 | 723.4170083 | 1279.262517 | 1932.305134 | 901.9878541 | 0.001818023 | 1.194724825 | 7.92E-05 | 15.57735361 | 2.289011681 | 2.740400742 |
| *Zmynd15* | 152.3822545 | 163.1126125 | 137.7984929 | 174.6388404 | 228.6169477 | 363.6660354 | 563.2419598 | 280.815923 | 0.001010348 | 1.193090251 | 3.70E-05 | 17.01920165 | 2.286419698 | 2.995529043 |
| *Hvcn1* | 1407.976387 | 1246.575784 | 1187.346412 | 1356.879786 | 1916.94774 | 3102.270071 | 4697.500412 | 2061.686524 | 0.001339291 | 1.179782414 | 5.29E-05 | 16.33978783 | 2.265426075 | 2.873125106 |
| *Susd6* | 1640.559828 | 1624.206196 | 1690.880755 | 1632.095969 | 2087.605462 | 4089.196546 | 6402.842982 | 2330.949893 | 0.009093821 | 1.178439294 | 0.000710867 | 11.46061572 | 2.263317991 | 2.041253593 |
| *Ugt1a6a* | 21.38698309 | 35.58820636 | 34.1906035 | 36.57357914 | 52.59263117 | 68.99130575 | 114.5223948 | 50.65350511 | 0.009439562 | 1.167422628 | 0.00075617 | 11.34584367 | 2.246100728 | 2.025048178 |
| *Polr3d* | 109.6082883 | 146.3070706 | 165.772623 | 138.9796007 | 211.4438437 | 360.1580029 | 470.5829313 | 216.8325482 | 0.002301075 | 1.167220915 | 0.000111564 | 14.93018054 | 2.245786708 | 2.638069319 |
| *Tnfsf10* | 57.9230792 | 60.30223855 | 48.69570802 | 71.31847932 | 83.71888227 | 139.1519557 | 205.0991979 | 106.6389581 | 0.0030007 | 1.163979639 | 0.000157218 | 14.28372604 | 2.240746804 | 2.522777395 |
| *Dnajb2* | 140.7976387 | 140.3757029 | 145.0510452 | 138.0652613 | 166.3644455 | 362.4966912 | 507.0218751 | 226.607786 | 0.007819126 | 1.16132764 | 0.000574639 | 11.8563881 | 2.236631589 | 2.106841808 |
| *Tank* | 652.3029842 | 594.1253339 | 707.6418846 | 598.8923584 | 890.8547728 | 1492.083155 | 2188.418853 | 1120.597718 | 0.000818979 | 1.156716706 | 2.80E-05 | 17.54811618 | 2.229494601 | 3.086727497 |
| *Csf1* | 433.9775318 | 405.310128 | 375.0605596 | 384.9369204 | 681.5575672 | 909.7497606 | 968.2347923 | 1004.183522 | 1.97E-12 | 1.156095406 | 2.86E-15 | 62.35938212 | 2.228534671 | 11.7053746 |
| *Cpeb4* | 1680.660421 | 1633.103247 | 1792.416487 | 1791.191038 | 2295.829348 | 4525.36192 | 5882.286642 | 2649.978109 | 0.002452944 | 1.154385231 | 0.000122291 | 14.75702558 | 2.225894523 | 2.610312389 |
| *Cacnb1* | 12.47574013 | 15.8169806 | 16.5772623 | 16.45811061 | 36.49284612 | 30.4029483 | 39.56228184 | 29.32571348 | 0.001028507 | 1.148811855 | 3.78E-05 | 16.98120233 | 2.217312104 | 2.987792875 |
| *Nupr1* | 168.4224918 | 169.0439802 | 163.7004652 | 191.096951 | 257.5965608 | 370.6821004 | 554.9130584 | 338.578692 | 0.000117793 | 1.136014252 | 2.39E-06 | 22.24916974 | 2.197730147 | 3.928881813 |
| *Spryd7* | 415.2639216 | 401.3558828 | 457.9468711 | 462.6557761 | 590.3254519 | 984.5877872 | 1593.943513 | 646.9430126 | 0.005493473 | 1.135056558 | 0.00035985 | 12.72993041 | 2.196271727 | 2.260153035 |
| *Cmklr1* | 112.2816612 | 97.86756749 | 73.56160147 | 116.1211138 | 167.4377645 | 231.5301447 | 348.7727478 | 128.8554077 | 0.008944004 | 1.130275689 | 0.000692662 | 11.50883315 | 2.189005666 | 2.048468021 |
| *Clcn7* | 2082.557478 | 2185.709007 | 2203.739807 | 2213.615877 | 3122.284981 | 5028.179911 | 7609.492578 | 3219.607877 | 0.002290683 | 1.127709418 | 0.000110875 | 14.94186233 | 2.185115316 | 2.640035101 |
| *Slc3a2* | 4654.342195 | 4716.425904 | 4745.241334 | 4953.891294 | 6494.65329 | 11080.70531 | 17544.83088 | 6546.74337 | 0.007086879 | 1.127586902 | 0.000499672 | 12.11688935 | 2.184929761 | 2.149544975 |
| *Irak2* | 1211.037917 | 1212.9647 | 1120.001284 | 1192.29868 | 1571.339021 | 2807.595341 | 4131.135114 | 1806.641682 | 0.003501765 | 1.123058317 | 0.000195202 | 13.87671806 | 2.178082073 | 2.455712982 |
| *Ninj1* | 1924.828478 | 1868.380834 | 1811.065907 | 1611.066161 | 2829.268893 | 3861.174434 | 5638.666274 | 3260.486144 | 4.00E-05 | 1.11142127 | 6.22E-07 | 24.84284663 | 2.160583922 | 4.397756487 |
| *Nfe2l2* | 2997.74213 | 3057.620063 | 2920.706402 | 2832.623704 | 4612.051758 | 6911.993361 | 8587.097384 | 5359.496303 | 3.52E-06 | 1.108949864 | 3.18E-08 | 30.59595569 | 2.156885907 | 5.452983364 |
| *Psmd10* | 484.7716167 | 454.7381924 | 466.2355023 | 460.8270972 | 698.7306712 | 989.2651638 | 1639.752471 | 686.9326219 | 0.002676787 | 1.104698022 | 0.000137338 | 14.53831809 | 2.150538593 | 2.572386244 |
| *Pde2a* | 191.5917235 | 190.7923285 | 212.3961733 | 168.238464 | 276.9163029 | 358.9886587 | 680.8876927 | 307.4756626 | 0.004864273 | 1.090170249 | 0.000302153 | 13.05699934 | 2.128991587 | 2.312982074 |
| *Fam219a* | 343.973978 | 349.9506959 | 367.8080074 | 279.7878804 | 464.7471285 | 809.1861624 | 1135.853934 | 444.3289922 | 0.008619543 | 1.088998012 | 0.000657105 | 11.60681862 | 2.127262415 | 2.06451574 |
| *Plxna1* | 3689.254583 | 3626.042803 | 3852.141328 | 3298.936838 | 5690.737356 | 7902.427869 | 11308.56593 | 5794.050058 | 0.000176234 | 1.085322113 | 4.11E-06 | 21.21344297 | 2.121849178 | 3.753909117 |
| *H2-Q8* | 109.6082883 | 105.7760578 | 113.9686783 | 93.26262681 | 216.8104387 | 185.9257223 | 216.5514374 | 271.0406852 | 3.81E-09 | 1.080145357 | 1.29E-11 | 45.82907726 | 2.114249089 | 8.419263729 |
| *Gm12250* | 356.4497181 | 362.8019926 | 323.2566149 | 326.4191938 | 494.8000606 | 739.0255125 | 1127.525032 | 532.3061326 | 0.001686772 | 1.079560678 | 7.21E-05 | 15.75442211 | 2.113392424 | 2.772943647 |
| *Ptpn23* | 1072.022527 | 1144.753971 | 1216.356622 | 1018.574179 | 1517.673071 | 2453.284059 | 3540.824224 | 1778.204627 | 0.001676436 | 1.061312875 | 7.10E-05 | 15.78398261 | 2.086829706 | 2.77561294 |
| *Rnd3* | 66.83432215 | 74.14209658 | 72.52552258 | 98.74866368 | 114.8451334 | 180.0790015 | 241.5381417 | 114.63688 | 0.005535915 | 1.058326584 | 0.000363744 | 12.70980267 | 2.082514563 | 2.256810566 |
| *Zfp263* | 690.6213289 | 778.9862947 | 763.5901449 | 770.7881804 | 1100.151978 | 1591.477409 | 2320.640164 | 1209.463517 | 0.00051967 | 1.050380865 | 1.61E-05 | 18.60779417 | 2.07107653 | 3.284272406 |
| *Mrpl52* | 192.4828478 | 202.655064 | 200.9993054 | 237.7282644 | 270.4763889 | 429.1493087 | 677.7643546 | 341.244666 | 0.004559173 | 1.043061703 | 0.000278052 | 13.21274289 | 2.060596035 | 2.34111397 |
| *Clec5a* | 929.44264 | 957.9158878 | 1015.357316 | 1065.205492 | 1536.992813 | 2038.16688 | 2859.936532 | 1704.446014 | 3.31E-05 | 1.036504027 | 4.76E-07 | 25.35887355 | 2.051250986 | 4.479569904 |
| *Odc1* | 1505.108935 | 1622.229073 | 1589.345023 | 1554.377113 | 2203.523914 | 3799.199193 | 4477.825636 | 2306.067469 | 0.000967221 | 1.027763577 | 3.47E-05 | 17.14007287 | 2.038861219 | 3.014474501 |
| *Myd88* | 941.0272559 | 905.5221396 | 990.4914226 | 991.1439947 | 1336.282159 | 1973.852951 | 3080.65242 | 1380.08585 | 0.003714252 | 1.021325824 | 0.00021274 | 13.7150793 | 2.029783455 | 2.430128629 |
| *Il13ra1* | 232.5834411 | 201.6665027 | 219.6487255 | 230.4135486 | 333.8022101 | 444.3507828 | 738.1488901 | 279.0386071 | 0.009722841 | 1.021022889 | 0.000796836 | 11.24858749 | 2.029357289 | 2.012206818 |
| *Pnrc1* | 1103.211878 | 1090.3831 | 1059.908709 | 949.9987182 | 1391.021428 | 2363.244558 | 3189.969251 | 1537.378313 | 0.003390154 | 1.01264811 | 0.000187258 | 13.95481308 | 2.017611089 | 2.469780584 |
| *Slc25a33* | 133.6686443 | 134.4443351 | 129.5098617 | 116.1211138 | 238.2768188 | 270.1185022 | 345.6494097 | 183.0635448 | 9.09E-05 | 1.012024609 | 1.75E-06 | 22.84929359 | 2.01673931 | 4.041548238 |
| *Usb1* | 422.392916 | 471.5437342 | 433.0809777 | 420.5961601 | 616.085108 | 859.4679615 | 1401.337667 | 616.7286411 | 0.005400015 | 0.999210014 | 0.000346758 | 12.79925178 | 1.998905147 | 2.267605003 |
| *Irf1* | 775.278137 | 705.8327594 | 630.9720464 | 703.127059 | 1040.046114 | 1358.77792 | 1891.701739 | 1328.543687 | 1.71E-05 | 0.996905996 | 2.17E-07 | 26.87430511 | 1.995715396 | 4.765905313 |
| *Ly96* | 62.37870067 | 56.3479934 | 56.98433917 | 60.34640558 | 88.01215828 | 122.7811374 | 173.8658175 | 86.19982448 | 0.003671703 | 0.993555 | 0.000208821 | 13.74999446 | 1.99108526 | 2.435132472 |
| *Ciita* | 33.86272322 | 28.66827734 | 45.58747133 | 30.17320279 | 64.3991402 | 59.63655243 | 56.22008472 | 91.53177238 | 0.002746152 | 0.985119854 | 0.00014134 | 14.48420708 | 1.979477754 | 2.561275381 |
| *Mina* | 407.243803 | 407.2872505 | 413.3954787 | 440.7116286 | 553.8326058 | 957.6928714 | 1158.758413 | 622.949247 | 0.001774735 | 0.980437437 | 7.66E-05 | 15.64051138 | 1.973063567 | 2.7508665 |
| *Snx20* | 723.5929278 | 729.5582303 | 757.3736715 | 739.7006381 | 902.6612819 | 1573.937246 | 2287.324558 | 1052.171053 | 0.008944004 | 0.979115248 | 0.000691719 | 11.5113679 | 1.971256139 | 2.048468021 |
| *Ctsz* | 9657.113989 | 9825.310639 | 9253.220602 | 9355.521544 | 13872.64812 | 18675.59566 | 28710.76437 | 13727.98854 | 0.00181221 | 0.977172728 | 7.88E-05 | 15.5869127 | 1.968603723 | 2.741791415 |
| *Mtdh* | 1968.493568 | 1923.740266 | 1954.044794 | 1939.314034 | 2512.639787 | 4310.202593 | 5930.177825 | 2545.116467 | 0.008983465 | 0.974407806 | 0.000697685 | 11.49540365 | 1.96483451 | 2.04655613 |
| *Tmem176b* | 861.7171936 | 864.0025655 | 811.249774 | 779.9315752 | 1198.897327 | 1775.064443 | 2389.3536 | 1140.148194 | 0.002087162 | 0.971183172 | 9.66E-05 | 15.20111001 | 1.960447723 | 2.680443769 |
| *Lilrb4* | 6482.929249 | 6879.398001 | 6657.842973 | 6566.786135 | 8804.435785 | 14396.96536 | 18890.98958 | 9974.297216 | 0.001671123 | 0.969626412 | 7.04E-05 | 15.80083085 | 1.958333417 | 2.776991513 |
| *Arl8a* | 1055.091166 | 1179.353616 | 1153.155809 | 1045.090024 | 1607.831867 | 2325.825545 | 3173.311448 | 1572.924632 | 0.001473764 | 0.969428712 | 5.91E-05 | 16.13164406 | 1.958065074 | 2.831572157 |
| *Ust* | 158.6201246 | 132.4672126 | 135.7263351 | 145.3799771 | 267.2564318 | 289.997353 | 302.9637899 | 261.2654474 | 2.44E-14 | 0.966698009 | 2.55E-17 | 71.6619179 | 1.954362395 | 13.61341586 |
| *Gcnt2* | 342.1917294 | 316.3396121 | 370.916244 | 341.962965 | 490.5067846 | 681.7276484 | 1051.523807 | 446.9949661 | 0.007673286 | 0.961314601 | 0.000557112 | 11.91408229 | 1.947083292 | 2.11501861 |
| *Nfkbid* | 188.9183506 | 174.9753479 | 163.7004652 | 152.6946929 | 232.9102237 | 341.4484963 | 467.4595933 | 282.593239 | 0.000931299 | 0.960554111 | 3.30E-05 | 17.23775596 | 1.946057194 | 3.030910978 |

| | | | | | | | | | | | | | | |
|---|---|---|---|---|---|---|---|---|---|---|---|---|---|---|
| *Fnip2* | 2886.351593 | 2716.566419 | 2940.391901 | 2540.949411 | 5675.71089 | 5328.701361 | 4959.860807 | 5505.236213 | 8.24E-31 | 0.95392608 | 6.65E-35 | 151.904994 | 1.937137122 | 30.08412812 |
| *Mvp* | 2016.61428 | 2153.086485 | 2102.204076 | 2014.289871 | 2645.731343 | 4001.495734 | 6479.88532 | 2921.018794 | 0.009245425 | 0.95360026 | 0.000728543 | 11.41498024 | 1.936699686 | 2.034073099 |
| *Rrs1* | 292.2887689 | 306.4539992 | 323.2566149 | 352.9350387 | 412.1544973 | 714.469285 | 894.3157921 | 447.8836241 | 0.005408358 | 0.952962129 | 0.000350852 | 12.77729582 | 1.935843237 | 2.266934574 |
| *Cln5* | 1447.185856 | 1522.384383 | 1471.232029 | 1492.202029 | 2105.851885 | 3164.245311 | 4042.640536 | 2163.882192 | 0.000703651 | 0.951759886 | 2.32E-05 | 17.9100186 | 1.93423071 | 3.152642427 |
| *Rnf19a* | 626.4603796 | 573.3655469 | 617.5030208 | 616.2648085 | 797.4760195 | 1361.116608 | 1748.02819 | 796.2375539 | 0.007378614 | 0.950166852 | 0.000527979 | 12.01415846 | 1.932096097 | 2.13202522 |
| *Nfe2l1* | 3267.752791 | 3154.499069 | 3345.498749 | 3209.331569 | 4415.63438 | 6534.295196 | 9463.714261 | 4642.34931 | 0.00283183 | 0.949158242 | 0.000145978 | 14.42339266 | 1.930745812 | 2.547932833 |
| *Bhlhe41* | 516.8520913 | 427.0584763 | 460.0190289 | 403.22371 | 889.7814538 | 827.895669 | 750.6422422 | 1007.738154 | 4.17E-10 | 0.94424999 | 1.14E-12 | 50.58044848 | 1.924188304 | 9.379851705 |
| *Eif1a* | 962.414239 | 939.1332233 | 913.8215845 | 931.7119286 | 1313.74246 | 2113.004907 | 2429.956995 | 1352.537452 | 0.00066754 | 0.943830776 | 2.18E-05 | 18.02905637 | 1.92362926 | 3.175523006 |
| *Ppp1r12b* | 78.41893799 | 65.24504499 | 63.20081253 | 58.51772662 | 115.9184524 | 99.39425405 | 183.2358317 | 109.3049321 | 0.002426946 | 0.935758586 | 0.000120603 | 14.78322956 | 1.912896194 | 2.614939948 |
| *Gm14446* | 64.16094926 | 50.41662567 | 42.47923465 | 54.86036871 | 89.08547728 | 80.68474741 | 145.7557752 | 88.86579843 | 0.004701492 | 0.929563337 | 0.00028787 | 13.14771778 | 1.904699411 | 2.327764312 |
| *Tnfrsf14* | 42.77396618 | 49.42806439 | 55.94826027 | 42.97395549 | 68.69241622 | 94.71687739 | 128.0568596 | 71.98129673 | 0.005253494 | 0.928075685 | 0.000334383 | 12.86725004 | 1.902736369 | 2.27955175 |
| *Pgs1* | 604.1822722 | 650.4733273 | 554.3022083 | 622.6651849 | 835.0421847 | 1142.44925 | 1733.452612 | 893.1012742 | 0.002984995 | 0.920788628 | 0.0001558 | 14.30078866 | 1.893149874 | 2.525056402 |
| *Mxd1* | 304.764509 | 308.4311218 | 289.0660114 | 281.6165594 | 447.5740244 | 529.7129069 | 850.5890595 | 404.3393829 | 0.003454114 | 0.91461021 | 0.000191988 | 13.90791647 | 1.885059695 | 2.461663277 |
| *Rnf19b* | 1657.491189 | 1668.691454 | 1699.169386 | 1758.274817 | 2782.042857 | 2781.869769 | 4139.464015 | 3026.769095 | 2.24E-07 | 0.908184343 | 1.25E-09 | 36.89322789 | 1.876682175 | 6.649427397 |
| *Gp49a* | 2152.956298 | 2230.194265 | 2168.513125 | 2080.122314 | 2713.350441 | 4489.112251 | 5885.40998 | 3081.86589 | 0.004250194 | 0.90551323 | 0.000253722 | 13.38442462 | 1.87321076 | 2.371591216 |
| *Ldlr* | 867.9550637 | 894.6479654 | 900.3525589 | 1053.319079 | 1600.318634 | 1689.702319 | 2082.22536 | 1556.040131 | 7.44E-10 | 0.898382821 | 2.22E-12 | 49.27816656 | 1.863975401 | 9.128330545 |
| *Myo10* | 702.2059447 | 652.4504499 | 655.8379399 | 768.0451619 | 908.0278769 | 1196.239081 | 2034.334177 | 1037.952526 | 0.007618395 | 0.897524019 | 0.000551897 | 11.93160109 | 1.862866152 | 2.118136538 |
| *Nrp2* | 8029.029901 | 7819.519786 | 8070.018505 | 8399.122449 | 10868.42823 | 15103.24924 | 22021.61541 | 12069.75274 | 0.001692251 | 0.894140697 | 7.25E-05 | 15.7435799 | 1.858502587 | 2.771535265 |
| *Nadk* | 2384.648614 | 2366.615723 | 2397.486561 | 2527.234319 | 3474.333614 | 4309.033249 | 6417.418559 | 3708.369769 | 0.000460261 | 0.888179162 | 1.38E-05 | 18.89488141 | 1.85083869 | 3.336996062 |
| *Ifit3* | 908.055657 | 918.3734363 | 940.7596357 | 904.2817442 | 1490.840096 | 1400.87431 | 2571.548319 | 1315.213817 | 0.002222754 | 0.884594388 | 0.000105256 | 15.04000385 | 1.846245479 | 2.653108656 |
| *Itga5* | 4682.858172 | 4957.634858 | 5157.600734 | 4815.826033 | 6182.31746 | 8899.878442 | 13479.28587 | 7643.347323 | 0.003257425 | 0.884307341 | 0.00017764 | 14.05395854 | 1.845878176 | 2.487125525 |
| *Slc7a7* | 186.2449777 | 186.8380834 | 157.4839919 | 191.096951 | 244.7167328 | 356.6499704 | 487.2407342 | 243.4922877 | 0.006790567 | 0.883102957 | 0.000474282 | 12.21415903 | 1.844337851 | 2.168093986 |
| *Rnf14* | 771.7136398 | 702.8670756 | 598.8536007 | 734.2146012 | 899.4413249 | 1245.351536 | 1997.895233 | 1033.509236 | 0.0088288 | 0.882298916 | 0.000680179 | 11.54264126 | 1.843310253 | 2.05409833 |
| *Cep85l* | 40.99171759 | 37.56532893 | 33.15452461 | 37.48791862 | 62.2525022 | 65.48327326 | 93.70014119 | 54.20813704 | 0.003693441 | 0.880457036 | 0.000210865 | 13.7316992 | 1.840958412 | 2.432568862 |
| *Daam1* | 402.7881815 | 407.2872505 | 456.9107922 | 417.8531417 | 573.1523478 | 827.895669 | 1144.182835 | 548.3019763 | 0.006597502 | 0.876412899 | 0.000455587 | 12.28919821 | 1.835805097 | 2.180620476 |
| *Plekhm3* | 1139.747974 | 1189.239229 | 1191.490728 | 1082.577943 | 1511.233157 | 2210.060472 | 3210.791505 | 1517.827837 | 0.00819222 | 0.876270401 | 0.000613229 | 11.73538434 | 1.835623779 | 2.086598375 |
| *Acot7* | 1129.054482 | 1026.126617 | 1009.140843 | 1034.11795 | 1487.620139 | 1855.74919 | 2731.879672 | 1516.939179 | 0.001313211 | 0.854502693 | 5.18E-05 | 16.38092953 | 1.808135371 | 2.881665416 |
| *Srfbp1* | 153.2733788 | 164.1011738 | 152.3035974 | 131.6648849 | 187.8308256 | 316.8922688 | 350.8549731 | 230.1624179 | 0.002665597 | 0.851806616 | 0.000136333 | 14.5521444 | 1.804759523 | 2.57420543 |
| *Dnajb6* | 2007.703037 | 1872.335079 | 1940.575768 | 2031.662321 | 2443.947371 | 3896.254759 | 5102.493244 | 2719.293432 | 0.005535915 | 0.850750699 | 0.00036397 | 12.70864064 | 1.803439091 | 2.256810566 |
| *Mvd* | 219.2165767 | 231.3233413 | 191.6745954 | 228.5848696 | 362.7818232 | 389.391607 | 408.1161705 | 410.5599887 | 3.00E-14 | 0.85070956 | 3.39E-17 | 71.10342937 | 1.803387667 | 13.52267193 |
| *Siah2* | 155.9467517 | 173.9867866 | 137.7984929 | 149.037335 | 210.3705247 | 278.3039113 | 410.1983959 | 209.7232843 | 0.008590051 | 0.845100187 | 0.000650005 | 11.62702183 | 1.796389491 | 2.066004264 |
| *Ube2f* | 452.691142 | 456.7153149 | 389.5656641 | 465.3987946 | 559.1992008 | 834.911734 | 1146.265061 | 628.2811949 | 0.005646928 | 0.844283302 | 0.000375824 | 12.64871406 | 1.795372624 | 2.248187759 |
| *Col18a1* | 1210.146793 | 1162.548074 | 1183.202097 | 1134.695293 | 1588.512125 | 2070.908517 | 3141.036955 | 1603.139004 | 0.00475234 | 0.841146084 | 0.000292517 | 13.11771514 | 1.791472732 | 2.323092503 |
| *Sema4b* | 209.4142094 | 220.4491672 | 189.6024376 | 174.6388404 | 374.5883322 | 304.029483 | 403.9517198 | 335.0240601 | 1.67E-07 | 0.836637777 | 8.64E-10 | 37.61087149 | 1.785883259 | 6.776576708 |
| *Ppp1r13l* | 81.09231088 | 107.7531804 | 95.31925825 | 83.20489254 | 113.7718144 | 180.0790015 | 220.7158881 | 137.7419876 | 0.008156412 | 0.828563719 | 0.000607598 | 11.75255546 | 1.77591646 | 2.088500844 |
| *Ripk2* | 226.345571 | 222.4262897 | 246.5867768 | 240.4712828 | 272.6230269 | 452.536192 | 570.5297486 | 362.5724576 | 0.005149107 | 0.825112235 | 0.000325246 | 12.9191006 | 1.771672861 | 2.288268096 |
| *Plekho2* | 1405.303014 | 1543.14417 | 1506.458712 | 1425.455247 | 2190.644086 | 2318.80948 | 3537.700886 | 2366.496212 | 8.30E-05 | 0.824592675 | 1.57E-06 | 23.06352059 | 1.771034941 | 4.080764406 |
| *Eef1e1* | 204.9585879 | 226.3805349 | 251.7671712 | 226.7561907 | 316.629106 | 450.1975036 | 522.6385653 | 317.2509004 | 0.001193155 | 0.820020399 | 4.51E-05 | 16.64189611 | 1.765430955 | 2.923303138 |
| *Cry1* | 186.2449777 | 200.6779414 | 182.3498853 | 206.6407221 | 242.5700948 | 371.8514446 | 443.5140016 | 310.1416365 | 0.001083054 | 0.817539001 | 4.00E-05 | 16.87062805 | 1.762397069 | 2.965349776 |
| *Acot9* | 487.4449896 | 466.6009278 | 547.049656 | 438.8829497 | 588.1788139 | 924.9512348 | 1202.485145 | 666.4934882 | 0.009785466 | 0.801780587 | 0.000805193 | 11.2292235 | 1.743251335 | 2.009418472 |
| *1700112E06Rik* | 35.64497181 | 48.4395031 | 44.55139244 | 40.23093705 | 64.3991402 | 73.66868241 | 81.20678903 | 73.7586127 | 0.000632331 | 0.800658782 | 2.04E-05 | 18.15117414 | 1.741896352 | 3.199055482 |
| *Samd8* | 847.4592049 | 896.625088 | 850.620772 | 803.7044016 | 1401.754618 | 1609.017571 | 1456.516639 | 1452.067146 | 2.42E-24 | 0.800315561 | 5.86E-28 | 120.1511758 | 1.741482 | 23.61580225 |
| *Il2rg* | 1780.466342 | 1918.797459 | 1958.18911 | 1857.93782 | 2695.104018 | 3315.090709 | 4427.852228 | 2649.978109 | 0.000360249 | 0.800287522 | 1.02E-05 | 19.4680351 | 1.741448155 | 3.443396779 |
| *Mafk* | 449.1266449 | 459.6809988 | 449.65824 | 416.0244627 | 591.3987709 | 805.6781299 | 1042.153793 | 647.8316706 | 0.00091489 | 0.798616716 | 3.23E-05 | 17.27586467 | 1.739432526 | 3.038631271 |
| *Zdhhc18* | 604.1822722 | 697.9242691 | 616.4669419 | 601.6353768 | 859.7285217 | 1068.780567 | 1540.846766 | 909.9857759 | 0.001924555 | 0.797128149 | 8.52E-05 | 15.44025377 | 1.737638711 | 2.715669764 |
| *Irgm1* | 1094.300635 | 1096.314468 | 1122.073442 | 1130.123595 | 1620.711695 | 1865.103944 | 2772.483067 | 1458.287752 | 0.002527338 | 0.796367101 | 0.000126906 | 14.68719107 | 1.736722317 | 2.597336658 |
| *Parp16* | 64.16094926 | 51.40518696 | 44.55139244 | 45.71697392 | 78.35228725 | 92.37818906 | 117.6457328 | 70.20398076 | 0.008944004 | 0.795267205 | 0.000691419 | 11.51217224 | 1.735398762 | 2.048468021 |
| *Ralgapa2* | 1164.699454 | 1258.438519 | 1285.773907 | 1278.246591 | 1583.14553 | 2326.994889 | 2815.168687 | 1929.276484 | 0.000491427 | 0.795209172 | 1.50E-05 | 18.73939567 | 1.735328957 | 3.308540887 |
| *Traf3* | 808.2497359 | 849.1741462 | 863.0537187 | 945.4270208 | 1457.567207 | 1411.398408 | 1529.394527 | 1612.025584 | 2.69E-19 | 0.794992945 | 8.69E-23 | 96.55265939 | 1.735068889 | 18.56975116 |
| *Eif6* | 1333.121946 | 1206.044771 | 1209.104069 | 1221.557543 | 1583.14553 | 2252.156862 | 3087.940209 | 1691.116144 | 0.005490052 | 0.793395764 | 0.000359183 | 12.73339994 | 1.73314909 | 2.26042353 |
| *Klhl12* | 105.1526669 | 100.8332513 | 103.6078894 | 100.5773426 | 140.6047894 | 196.4498198 | 223.8392262 | 148.4058834 | 0.000767549 | 0.788034439 | 2.60E-05 | 17.68948505 | 1.726720336 | 3.114893811 |
| *Herpud1* | 683.4923345 | 617.8508048 | 653.7657821 | 704.9557379 | 1162.404481 | 1062.933846 | 1040.071567 | 1324.989055 | 6.93E-11 | 0.78774151 | 1.62E-13 | 54.41735781 | 1.726369773 | 10.15926785 |
| *Atf5* | 340.4094808 | 394.4359538 | 309.7875893 | 361.164094 | 566.7124338 | 601.042901 | 701.7099463 | 558.0772141 | 2.60E-08 | 0.7873995 | 1.13E-10 | 41.57932084 | 1.725960563 | 7.585326527 |
| *Gas2l3* | 1722.543263 | 1710.211028 | 1726.107437 | 1639.410685 | 2518.006382 | 2935.053855 | 3515.83752 | 2751.285119 | 4.78E-09 | 0.78574012 | 1.66E-11 | 45.33987896 | 1.723976513 | 8.321017 |
| *Zfp503* | 106.0437911 | 99.84469006 | 118.1129939 | 96.91998472 | 122.3583664 | 208.1432614 | 232.1681276 | 159.9584372 | 0.007361457 | 0.779301422 | 0.000525564 | 12.02270448 | 1.716299609 | 2.133036204 |
| *Tgif1* | 1280.545612 | 1284.141113 | 1267.124487 | 1130.123595 | 1645.398032 | 2131.714413 | 2961.965574 | 1771.095363 | 0.001969432 | 0.778236195 | 8.81E-05 | 15.37600619 | 1.715032832 | 2.705659039 |
| *Enpp4* | 256.6437971 | 215.5063607 | 174.0612542 | 234.0709065 | 394.9813933 | 304.029483 | 422.6917481 | 385.6775652 | 0.000124906 | 0.77513273 | 2.62E-06 | 22.07660271 | 1.71134749 | 3.90341772 |
| *1810022K09Rik* | 133.6686443 | 149.2727544 | 137.7984929 | 139.8939402 | 187.8308256 | 226.8527681 | 338.361621 | 193.7274406 | 0.006836376 | 0.755609065 | 0.000478701 | 12.1968581 | 1.688344213 | 2.165174083 |
| *Anpep* | 5355.657015 | 5958.058881 | 5963.670114 | 4819.483391 | 7411.267719 | 9345.398569 | 13137.80091 | 7409.630273 | 0.004939941 | 0.755494534 | 0.000309244 | 13.01355393 | 1.688210186 | 2.306278198 |
| *Atp6v0c* | 7499.70207 | 7508.12298 | 7154.124763 | 7098.017372 | 10573.2655 | 12023.19671 | 16305.90679 | 10420.40352 | 0.000123067 | 0.753304222 | 2.55E-06 | 22.12735443 | 1.685649075 | 3.909859373 |
| *Uchl1* | 735.1775437 | 790.8490302 | 742.868567 | 851.2500545 | 1144.158058 | 1254.706289 | 1839.646105 | 1009.51547 | 0.003102607 | 0.749932519 | 0.000164942 | 14.19347164 | 1.681714167 | 2.508273174 |
| *Slc30a1* | 503.4852269 | 484.395031 | 515.9672892 | 470.8848314 | 669.7510581 | 879.3468123 | 1117.113906 | 652.2749605 | 0.002095385 | 0.748515129 | 9.75E-05 | 15.1838418 | 1.680062763 | 2.678736224 |
| *Myc* | 63.26982497 | 85.01627075 | 82.88631152 | 68.57546089 | 109.4785383 | 102.9022865 | 139.5090991 | 144.8512514 | 0.002062192 | 0.73690247 | 9.48E-05 | 15.23704672 | 1.666593744 | 2.68567098 |
| *Eif1* | 2349.003643 | 2596.950503 | 2405.775192 | 2350.766799 | 3289.722745 | 4092.704579 | 5580.363964 | 3096.084417 | 0.004138582 | 0.726886863 | 0.000243721 | 13.4598677 | 1.655063835 | 2.383148443 |
| *Mtmr14* | 875.084058 | 919.3619976 | 878.5949021 | 1036.860969 | 1378.1416 | 1516.639382 | 1884.413951 | 1340.984898 | 1.34E-05 | 0.721878368 | 1.62E-07 | 27.44396984 | 1.649328039 | 4.874066584 |
| *Asns* | 930.3337643 | 951.9845201 | 938.6874779 | 1047.833042 | 1528.406261 | 1547.042331 | 1955.209613 | 1349.871478 | 4.41E-06 | 0.72143443 | 4.18E-08 | 30.06370811 | 1.648820595 | 5.355643828 |
| *Gnl3* | 348.4295995 | 321.2824185 | 322.220536 | 293.5029726 | 516.2664406 | 553.0997902 | 526.803016 | 522.5308948 | 1.03E-14 | 0.71991517 | 9.94E-18 | 73.52458313 | 1.647085184 | 13.98855453 |
| *Naa25* | 471.4047522 | 451.7725085 | 498.353948 | 555.9184029 | 627.891617 | 836.0810782 | 1113.990568 | 678.046042 | 0.00545083 | 0.719131623 | 0.000354858 | 12.7560578 | 1.646190873 | 2.263537355 |
| *Slc4a7* | 1234.207149 | 1268.324132 | 1305.459406 | 1270.017536 | 1483.326863 | 2337.518986 | 2764.154165 | 1758.654151 | 0.005551806 | 0.716310559 | 0.000365463 | 12.70098744 | 1.642975035 | 2.25556572 |
| *Rasgef1b* | 782.4071313 | 804.6888882 | 765.6623026 | 768.9595014 | 1162.404481 | 1207.932523 | 1242.047427 | 1500.943335 | 8.99E-10 | 0.712984196 | 2.90E-12 | 48.75467029 | 1.639191259 | 9.046280085 |
| *Hsd17b7* | 270.0106615 | 254.0602509 | 292.1742481 | 276.1305225 | 356.3419091 | 478.2617636 | 583.0231008 | 369.6817215 | 0.002266679 | 0.709718472 | 0.000108982 | 14.97435933 | 1.635484937 | 2.644609949 |
| *Nat6* | 158.6201246 | 157.1812447 | 174.0612542 | 124.3501691 | 214.6638007 | 232.6994889 | 320.6627054 | 233.7170499 | 0.001958651 | 0.707106238 | 8.73E-05 | 15.39235261 | 1.632526304 | 2.708042903 |
| *Larp1* | 2039.783512 | 2070.047337 | 2033.822869 | 1924.684602 | 2642.511386 | 3243.760715 | 4639.198102 | 2598.435946 | 0.006496988 | 0.701796939 | 0.000447598 | 12.32221924 | 1.626529444 | 2.187287942 |
| *Tor3a* | 1674.422551 | 1744.810673 | 1806.921591 | 1850.623104 | 2725.15695 | 2480.178975 | 3596.003196 | 2700.631614 | 1.15E-05 | 0.700825268 | 1.34E-07 | 27.8070572 | 1.625434327 | 4.937858508 |
| *Wsb2* | 1175.392946 | 1196.159158 | 1185.274255 | 1108.179448 | 1439.320784 | 1851.071814 | 2521.574911 | 1715.998568 | 0.002087162 | 0.690395026 | 9.64E-05 | 15.20529061 | 1.613725314 | 2.680443769 |
| *Acp2* | 2190.383518 | 2271.713839 | 2260.724147 | 2143.211738 | 2991.340063 | 3234.405961 | 5043.149822 | 2993.000091 | 0.004788103 | 0.685804863 | 0.000295104 | 13.1012161 | 1.60859915 | 2.319836484 |
| *Mapk6* | 1878.490015 | 1739.867866 | 1891.88006 | 1769.246891 | 2262.556459 | 2997.029096 | 3929.159254 | 2509.570148 | 0.003114788 | 0.684328639 | 0.000166033 | 14.18106968 | 1.606954009 | 2.506571459 |
| *Dtx3l* | 1634.321958 | 1651.885912 | 1756.153725 | 1737.245009 | 2130.538222 | 2718.725184 | 3820.883535 | 2179.878036 | 0.009245425 | 0.678362437 | 0.000726841 | 11.41932588 | 1.600322242 | 2.034073099 |
| *H2-Eb1* | 272.6840344 | 270.8657928 | 254.8754079 | 247.7859987 | 374.5883322 | 380.0368537 | 576.7764247 | 337.690034 | 0.005568649 | 0.673450707 | 0.000368368 | 12.68617919 | 1.594883126 | 2.25425014 |
| *Rrp9* | 227.2366953 | 244.1746381 | 221.7208833 | 227.6705301 | 291.9427689 | 383.5448862 | 437.2673256 | 352.7972198 | 0.000143397 | 0.670242884 | 3.12E-06 | 21.73924634 | 1.591340854 | 3.84346055 |
| *Pofut2* | 749.4355324 | 778.9862947 | 768.7705393 | 802.7900621 | 910.1745149 | 1335.391037 | 1658.492499 | 1029.065946 | 0.008737784 | 0.670025239 | 0.000669643 | 11.57166864 | 1.591100802 | 2.058598711 |
| *Zbtb21* | 421.5017917 | 377.6304119 | 406.1429264 | 375.7935257 | 542.0260967 | 641.9699468 | 824.5612425 | 507.423709 | 0.002255413 | 0.669603888 | 0.000107811 | 14.99474316 | 1.590636177 | 2.646773936 |
| *Cxcl16* | 4805.833325 | 4699.620362 | 4775.287622 | 4963.034689 | 7134.351416 | 7227.716286 | 8468.410539 | 7742.877017 | 1.82E-16 | 0.667947639 | 1.03E-19 | 82.55283863 | 1.588811135 | 15.73947383 |
| *Slc1a4* | 183.5716048 | 153.2269996 | 148.1592818 | 206.6407221 | 356.3419091 | 253.7476839 | 242.5792544 | 246.1582617 | 0.005941822 | 0.665684295 | 0.000400634 | 12.52923473 | 1.586320514 | 2.226080337 |
| *Adam17* | 6115.786039 | 5858.214191 | 6180.210603 | 6631.704237 | 8374.034865 | 9815.474924 | 12434.00874 | 8554.221757 | 0.000229233 | 0.660474808 | 5.70E-06 | 20.58783785 | 1.580602733 | 3.639723503 |
| *H2-Aa* | 310.1112548 | 287.6713347 | 260.0558024 | 298.07467 | 360.6351851 | 549.5917577 | 562.2008472 | 355.4631937 | 0.009722841 | 0.659518514 | 0.0007965 | 11.24937092 | 1.579555374 | 2.012206818 |
| *Atf4* | 3192.007226 | 2967.660986 | 2997.37624 | 3196.530817 | 4481.106839 | 4566.288966 | 5963.493431 | 4483.279531 | 3.00E-06 | 0.658041673 | 2.53E-08 | 31.03960881 | 1.57793926 | 5.523434713 |
| *Adck3* | 199.6118422 | 188.815206 | 230.0095145 | 227.6705301 | 315.555787 | 354.3112821 | 408.1161705 | 257.7108154 | 0.002339804 | 0.657256881 | 0.000113819 | 14.89242471 | 1.577081132 | 2.630820493 |
| *Nufip1* | 194.2650964 | 213.5292382 | 196.8549899 | 213.955438 | 268.3297509 | 343.7871846 | 406.0339452 | 265.7087373 | 0.002663773 | 0.647961733 | 0.000135907 | 14.55804991 | 1.56695281 | 2.574502803 |
| *Slc38a7* | 992.712465 | 944.0760298 | 959.4090558 | 1000.287389 | 1382.434876 | 1579.783967 | 1855.262796 | 1284.110787 | 4.90E-05 | 0.646533758 | 7.94E-07 | 24.37224152 | 1.565402613 | 4.310068586 |
| *Mdm2* | 2086.121975 | 2057.19604 | 2052.472289 | 2123.096269 | 2564.159099 | 3174.769409 | 4235.246382 | 3032.9897 | 0.00127024 | 0.644824483 | 4.89E-05 | 16.49109775 | 1.563549054 | 2.896114175 |
| *Gtf2h3* | 125.6485256 | 149.2727544 | 102.5718105 | 128.007527 | 194.2707396 | 174.2322806 | 211.345874 | 209.7232843 | 0.000423117 | 0.644499286 | 1.25E-05 | 19.09168263 | 1.563196656 | 3.373539751 |
| *Plxnd1* | 3527.961085 | 3919.645506 | 3758.894227 | 3342.825133 | 5190.5707 | 5546.199376 | 6275.827235 | 5658.085386 | 2.90E-10 | 0.640055925 | 7.47E-13 | 51.41569872 | 1.558389568 | 9.538324098 |
| *Rab12* | 741.4154137 | 681.1187272 | 646.5132298 | 636.380277 | 941.300766 | 1009.144015 | 1319.089765 | 941.9774634 | 0.000209947 | 0.63814324 | 5.13E-06 | 20.78745447 | 1.556324867 | 3.677891069 |
| *Mex3d* | 210.3053337 | 173.9867866 | 204.1075421 | 220.3558143 | 250.0833278 | 315.7229246 | 397.7050437 | 295.0344508 | 0.003556386 | 0.637204469 | 0.000199349 | 13.83720558 | 1.555312486 | 2.448991061 |
| *Ppif* | 101.5881697 | 95.89044491 | 73.56160147 | 94.17696628 | 138.4581514 | 132.1358907 | 171.7835922 | 126.1894338 | 0.003257425 | 0.634748521 | 0.000177605 | 14.05432867 | 1.552667078 | 2.487125525 |
| *Tollip* | 1584.418997 | 1616.297705 | 1597.633655 | 1676.898604 | 2041.452744 | 2635.701749 | 3267.01159 | 2079.459683 | 0.003242671 | 0.630244003 | 0.000176312 | 14.0680687 | 1.547826756 | 2.489097072 |
| *Optn* | 312.7846277 | 377.6304119 | 366.7719285 | 328.2478728 | 472.2603615 | 558.946511 | 585.1053261 | 524.3082107 | 2.16E-06 | 0.628765869 | 1.67E-08 | 31.83900088 | 1.546241719 | 5.664986557 |
| *Tbcel* | 648.738487 | 704.8441982 | 670.3430444 | 642.7806534 | 796.4027005 | 1116.723678 | 1337.829794 | 859.3322708 | 0.007651105 | 0.623948321 | 0.000554884 | 11.92154614 | 1.541087012 | 2.11627586 |
| *Tcirg1* | 5672.00614 | 5635.787901 | 5422.836931 | 5294.93992 | 7153.671158 | 8533.873718 | 11100.34339 | 7137.70093 | 0.00208346 | 0.6231576 | 9.60E-05 | 15.2143587 | 1.540242595 | 2.681214858 |
| *Zfand5* | 1524.713669 | 1525.350067 | 1620.42739 | 1524.203911 | 2033.939511 | 2414.695701 | 3212.87373 | 1853.740555 | 0.008455463 | 0.619094185 | 0.000637093 | 11.66434767 | 1.535910537 | 2.072862623 |
| *Lrp12* | 891.1242953 | 816.5516237 | 783.2756438 | 817.4194938 | 1203.190603 | 1231.319406 | 1552.299006 | 1069.055555 | 0.000172198 | 0.611276659 | 4.00E-06 | 21.26450117 | 1.527610413 | 3.763972507 |
| *Abhd17c* | 470.5136279 | 506.1433793 | 440.3335299 | 446.1976655 | 630.038255 | 722.6546942 | 904.7269189 | 587.4029276 | 0.002206004 | 0.610003528 | 0.000104285 | 15.05749826 | 1.526262941 | 2.656393614 |
| *Fzd5* | 200.5029665 | 208.5864317 | 211.3600944 | 247.7859987 | 305.895916 | 319.2309571 | 301.9226772 | 391.8981711 | 0.000268718 | 0.605169558 | 7.05E-06 | 20.1809029 | 1.521157522 | 3.570703619 |
| *Mapk7* | 342.1917294 | 312.3853669 | 318.0762205 | 330.0765517 | 413.2278163 | 491.1245494 | 644.4487489 | 430.9991224 | 0.003688388 | 0.603113207 | 0.000210068 | 13.7388158 | 1.518990879 | 2.433163451 |
| *Mob1b* | 297.6355146 | 312.3853669 | 349.1585873 | 297.1603305 | 390.6881172 | 530.882251 | 550.7486077 | 435.4424123 | 0.000823888 | 0.602610094 | 2.82E-05 | 17.53227173 | 1.518461251 | 3.084131614 |
| *Sgms1* | 449.1266449 | 414.2071796 | 421.6841098 | 416.9388022 | 534.5128637 | 664.1874859 | 843.3012707 | 523.4195528 | 0.008703098 | 0.591333557 | 0.000666283 | 11.5810233 | 1.506638768 | 2.060326132 |
| *Cstb* | 2867.637982 | 3196.018643 | 2971.474268 | 3091.381777 | 3722.270304 | 4653.989778 | 6056.152459 | 3827.449938 | 0.007911438 | 0.590463763 | 0.000583338 | 11.82841046 | 1.505730696 | 2.101744577 |
| *P2rx4* | 1493.524319 | 1423.528254 | 1419.428085 | 1448.313734 | 1807.469202 | 2015.949341 | 2916.156616 | 1955.047565 | 0.005408358 | 0.587813337 | 0.000350324 | 12.78011104 | 1.502967004 | 2.266934574 |
| *Ccdc93* | 938.353883 | 837.3114107 | 938.6874779 | 959.1421129 | 1183.870861 | 1501.437908 | 1636.629133 | 1195.244989 | 0.000556685 | 0.58627949 | 1.74E-05 | 18.45192626 | 1.501369926 | 3.254390784 |
| *Tcf4* | 1541.645031 | 1567.858202 | 1642.185047 | 1526.946929 | 1048.632666 | 1068.780567 | 926.5902851 | 1120.597718 | 2.08E-11 | -0.59155294 | 3.87E-14 | 57.23512483 | 0.663628183 | 10.68106891 |
| *Fmnl3* | 1972.058066 | 2028.527762 | 2195.451176 | 2174.29928 | 1472.593673 | 1281.601205 | 1270.15747 | 1518.716495 | 1.41E-08 | -0.593760992 | 5.68E-11 | 42.92735701 | 0.662613273 | 7.851258723 |
| *Fam134b* | 2073.646235 | 2233.159949 | 2231.713938 | 2130.410985 | 1605.685229 | 1605.509539 | 1116.072793 | 1413.854853 | 0.000265835 | -0.594768941 | 6.91E-06 | 20.21927886 | 0.662150495 | 3.575388595 |
| *Zbtb12* | 91.78580242 | 118.6273545 | 103.6078894 | 109.7207374 | 68.69241622 | 68.99130575 | 62.4667608 | 79.0905606 | 0.003768627 | -0.595837907 | 0.000217704 | 13.67175591 | 0.661660056 | 2.423816901 |
| *Tmc6* | 449.1266449 | 438.9212118 | 456.9107922 | 404.1380495 | 346.6820381 | 254.917028 | 213.4280994 | 340.356008 | 0.008944004 | -0.596607202 | 0.000692277 | 11.509869 | 0.66130733 | 2.048468021 |
| *Dmwd* | 201.3940907 | 203.6436253 | 175.0973331 | 199.3260063 | 137.3848324 | 153.1840857 | 111.3990568 | 114.63688 | 0.000731686 | -0.600148109 | 2.45E-05 | 17.80330089 | 0.659686227 | 3.135675064 |

| | | | | | | | | | | | | | | |
|---|---|---|---|---|---|---|---|---|---|---|---|---|---|---|
| *Zfp516* | 885.7775496 | 921.3391202 | 1022.609868 | 887.8236336 | 672.9710151 | 584.6720827 | 468.500706 | 722.4789412 | 0.001653375 | -0.601348447 | 6.94E-05 | 15.82829781 | 0.65913759 | 2.78162873 |
| *Tifab* | 729.8307979 | 691.9929014 | 722.1469891 | 693.9836642 | 384.2482032 | 486.4471728 | 537.2141428 | 454.992888 | 1.00E-05 | -0.607527356 | 1.13E-07 | 28.13029321 | 0.65632061 | 4.998169683 |
| *Zc3h12d* | 513.2875941 | 522.9489212 | 485.9210013 | 548.6036871 | 341.3154431 | 342.6178404 | 295.6760011 | 375.9023274 | 2.01E-07 | -0.610179251 | 1.07E-09 | 37.18780797 | 0.6551153 | 6.695733779 |
| *Ehd4* | 4689.096042 | 4563.198904 | 4610.551078 | 4288.252154 | 3531.219521 | 2836.828945 | 2069.732008 | 3406.226054 | 0.006605247 | -0.615803849 | 0.000458254 | 12.2783079 | 0.652566192 | 2.180110927 |
| *Naa16* | 333.2804865 | 383.5617796 | 350.1946662 | 343.7916439 | 243.6434138 | 249.0703072 | 182.194719 | 244.3809457 | 7.98E-05 | -0.617685676 | 1.47E-06 | 23.18374745 | 0.65171555 | 4.098250991 |
| *Card6* | 163.9668703 | 222.4262897 | 177.1694909 | 161.8380877 | 126.6516424 | 104.0716307 | 99.94681727 | 137.7419876 | 0.00546286 | -0.62598626 | 0.000356963 | 12.74499403 | 0.647976654 | 2.262579906 |
| *Epb4.1* | 2108.400083 | 1901.991918 | 2114.637023 | 2202.643804 | 1561.67915 | 1295.633335 | 1055.688267 | 1479.615544 | 0.000266961 | -0.626761089 | 6.98E-06 | 20.19934656 | 0.647628738 | 3.573552603 |
| *Zfp958* | 173.7692376 | 185.8495221 | 167.8447808 | 200.2403458 | 101.9653053 | 122.7811374 | 112.4401694 | 131.5213817 | 0.00010584 | -0.632553314 | 2.12E-06 | 22.48539409 | 0.645033809 | 3.975351721 |
| *Zranb3* | 662.9964757 | 684.0844111 | 607.1422319 | 736.0432802 | 470.1137235 | 470.0763545 | 302.9637899 | 475.4320216 | 0.00240202 | -0.646688173 | 0.0001187 | 14.81323316 | 0.638744924 | 2.619423412 |
| *Cttnbp2nl* | 5397.539857 | 5186.981077 | 5422.836931 | 5133.101832 | 3502.239908 | 3451.903976 | 2698.564066 | 3771.464485 | 1.79E-06 | -0.655049081 | 1.29E-08 | 32.34346463 | 0.635053886 | 5.747488594 |
| *Rassf2* | 2962.097158 | 2903.404502 | 3151.751995 | 2793.307107 | 2132.68486 | 2001.917211 | 1196.238469 | 2152.329638 | 0.009530726 | -0.658282465 | 0.000768485 | 11.31584413 | 0.63363219 | 2.020873999 |
| *Plxdc2* | 1276.981115 | 1399.802783 | 1408.031217 | 1236.186975 | 846.8486937 | 881.6855006 | 644.4487489 | 994.4082844 | 0.000232213 | -0.6594503 | 5.79E-06 | 20.55689987 | 0.633119484 | 3.634112577 |
| *Ctla2a* | 493.6828596 | 516.0289922 | 539.7971038 | 509.2870895 | 324.142339 | 355.4806263 | 308.1693533 | 314.5849264 | 8.30E-13 | -0.662345086 | 1.14E-15 | 64.17464442 | 0.631850396 | 12.08077804 |
| *Chd9* | 1286.783482 | 1267.335571 | 1361.407667 | 1249.902067 | 977.7936121 | 817.3715715 | 528.8852414 | 931.3135676 | 0.008246513 | -0.665843649 | 0.000618023 | 11.72089341 | 0.630320003 | 2.083769269 |
| *Pwwp2a* | 496.3562325 | 507.1319406 | 522.1837626 | 505.6297316 | 383.1748842 | 316.8922688 | 244.6614798 | 335.0240601 | 0.000131366 | -0.666267107 | 2.81E-06 | 21.94325156 | 0.630135019 | 3.88151666 |
| *Ctla2b* | 1482.830827 | 1484.819054 | 1472.268108 | 1595.52299 | 1051.852623 | 1016.16008 | 659.0243264 | 1073.498845 | 0.001556869 | -0.667182116 | 6.38E-05 | 15.98631579 | 0.629735491 | 2.807747906 |
| *Fzd7* | 1113.014245 | 1100.268713 | 1018.465553 | 1129.209256 | 711.6104993 | 729.6707591 | 583.0231008 | 679.823358 | 5.62E-11 | -0.690470469 | 1.27E-13 | 54.89723647 | 0.619651746 | 10.25007461 |
| *Map3k14* | 666.5609729 | 664.3131854 | 668.2708866 | 586.0916057 | 444.3540674 | 395.2383279 | 318.5804801 | 442.5516762 | 1.01E-05 | -0.690597074 | 1.16E-07 | 28.08740136 | 0.61959737 | 4.994705014 |
| *Mta3* | 712.8994363 | 665.3017466 | 656.8740188 | 726.8998854 | 510.8998456 | 406.9317695 | 308.1693533 | 481.6526275 | 0.000991761 | -0.693271652 | 3.58E-05 | 17.08396863 | 0.618449778 | 3.003593004 |
| *Ctnnd2* | 233.4745654 | 270.8657928 | 271.4526702 | 240.4712828 | 197.4906966 | 136.8132673 | 112.4401694 | 179.5089128 | 0.005301221 | -0.695973671 | 0.000338276 | 12.84558884 | 0.617292569 | 2.275624059 |
| *Il16* | 491.0094867 | 446.8297021 | 435.1531355 | 461.7414366 | 327.362296 | 278.3039113 | 242.5792544 | 281.704581 | 1.09E-07 | -0.700348362 | 5.27E-10 | 38.57356314 | 0.615423585 | 6.962825726 |
| *Tbxas1* | 1249.356262 | 1287.106797 | 1243.294673 | 1394.367705 | 947.74068 | 810.3555066 | 490.3640722 | 914.4290659 | 0.008954609 | -0.709944264 | 0.000694206 | 11.50469508 | 0.611343757 | 2.047953349 |
| *Stbd1* | 106.0437911 | 101.8218126 | 77.70591705 | 102.4060216 | 67.61909721 | 63.14458493 | 51.01452132 | 55.09679503 | 0.001658602 | -0.718966374 | 6.97E-05 | 15.81868902 | 0.607532556 | 2.780257706 |
| *Bcl2* | 288.7242717 | 261.9687412 | 297.3546426 | 290.7599542 | 194.2707396 | 190.6030989 | 124.9335216 | 181.2862288 | 0.000183862 | -0.72137913 | 4.33E-06 | 21.11245094 | 0.606517371 | 3.735508208 |
| *Mxd4* | 392.09469 | 307.4425605 | 318.0762205 | 395.9089942 | 256.5232418 | 228.0221122 | 152.0024513 | 221.2758381 | 0.002663773 | -0.721737387 | 0.000136025 | 14.55640874 | 0.606366776 | 2.574502803 |
| *Rundc3b* | 55.24970631 | 48.4395031 | 61.12865474 | 57.60338714 | 26.83297509 | 30.4029483 | 36.4389438 | 39.98960929 | 0.009245425 | -0.723974883 | 0.000729846 | 11.41165944 | 0.605427082 | 2.034073099 |
| *Adap2* | 651.4118599 | 658.3818176 | 585.3845751 | 604.3783953 | 427.1809634 | 406.9317695 | 228.0036769 | 440.7743602 | 0.007686924 | -0.733739211 | 0.000559342 | 11.90663939 | 0.601343317 | 2.114247388 |
| *Vamp1* | 153.2733788 | 129.5015287 | 159.5561497 | 155.4377113 | 99.81866732 | 90.03950073 | 62.4667608 | 105.7503001 | 0.001904103 | -0.736785744 | 8.37E-05 | 15.47247686 | 0.600074802 | 2.720309474 |
| *Pdlim5* | 4057.288917 | 3990.821919 | 4272.789359 | 4660.388322 | 3207.077182 | 2286.067843 | 1679.314753 | 3000.998013 | 0.006374848 | -0.739032893 | 0.00043507 | 12.37522007 | 0.59914085 | 2.195530135 |
| *Zbtb4* | 1215.493539 | 1301.935216 | 1212.212306 | 1197.784717 | 794.2560625 | 722.6546942 | 550.7486077 | 881.5487204 | 6.18E-05 | -0.739807025 | 1.06E-06 | 23.82213042 | 0.598819445 | 4.209141031 |
| *Rab39* | 133.6686443 | 108.7417417 | 117.076915 | 149.037335 | 84.79220127 | 85.36212407 | 60.38453544 | 74.64727068 | 0.001010348 | -0.7417855 | 3.70E-05 | 17.02003611 | 0.597998802 | 2.995529043 |
| *Sfxn5* | 284.2686502 | 294.5912637 | 270.4165913 | 269.7301462 | 186.7575066 | 146.1680207 | 110.3579441 | 220.3871801 | 0.006605247 | -0.750719491 | 0.000456945 | 12.28364378 | 0.594307095 | 2.180110927 |
| *Rps6ka5* | 452.691142 | 454.7381924 | 479.7045279 | 424.253518 | 308.0452554 | 261.933093 | 189.4825077 | 313.6962685 | 0.000367315 | -0.753871427 | 1.05E-05 | 19.40935857 | 0.593010097 | 3.434961816 |
| *Bin2* | 1127.272234 | 1122.017062 | 1103.424022 | 1109.093787 | 709.4638613 | 654.8327326 | 564.2830725 | 691.3759118 | 2.63E-15 | -0.767701631 | 1.91E-18 | 76.78384854 | 0.587352446 | 14.58053944 |
| *Ssh2* | 2238.50423 | 2107.612665 | 2418.208139 | 2348.023781 | 1433.954189 | 1420.753161 | 1035.907117 | 1454.73312 | 7.69E-07 | -0.769464423 | 4.77E-09 | 34.2795218 | 0.586635213 | 6.11434621 |
| *Hdgfrp3* | 166.6402432 | 191.7808898 | 157.4839919 | 156.3520508 | 97.67202931 | 121.6117932 | 76.00122563 | 97.75237827 | 0.000172198 | -0.777766642 | 4.00E-06 | 21.26577494 | 0.583269022 | 3.763972507 |
| *Evi2b* | 2465.740925 | 2513.911355 | 2491.76974 | 2676.271654 | 1812.835797 | 1438.293323 | 981.7692572 | 1674.231642 | 0.001209985 | -0.78051657 | 4.60E-05 | 16.6072576 | 0.582158309 | 2.917219916 |
| *Lbh* | 373.3810797 | 368.7333603 | 345.0142717 | 369.3931493 | 233.9835427 | 229.1914564 | 194.6880171 | 191.9501246 | 3.07E-11 | -0.782126957 | 6.20E-14 | 56.30803413 | 0.581508846 | 10.51253492 |
| *Pkib* | 232.5834411 | 219.4606059 | 242.4424612 | 243.2143013 | 195.3440586 | 129.7972023 | 91.61791583 | 128.8554077 | 0.006642373 | -0.782137912 | 0.000462437 | 12.26134967 | 0.58150443 | 2.177676725 |
| *Hdac9* | 605.9645208 | 529.8688502 | 564.6629972 | 562.3187793 | 344.5354001 | 333.2630871 | 236.3325783 | 398.118777 | 0.000103499 | -0.785032056 | 2.05E-06 | 22.55173956 | 0.580339062 | 3.985065462 |
| *Dock8* | 4698.898409 | 4853.835923 | 5178.322312 | 5230.936156 | 3610.645127 | 2788.885834 | 1691.808105 | 3487.09393 | 0.009949114 | -0.785743569 | 0.000821067 | 11.19299568 | 0.580052919 | 2.002215578 |
| *Erf* | 928.5515158 | 926.2819266 | 955.2647402 | 866.7938256 | 581.7388998 | 523.8661861 | 344.6082971 | 672.7140941 | 0.00188771 | -0.79169687 | 8.28E-05 | 15.49229194 | 0.577668219 | 2.724064741 |
| *Mamdc2* | 93.56805101 | 120.6044771 | 96.35533714 | 94.17696628 | 60.10586419 | 47.94311078 | 45.80895792 | 77.31324463 | 0.003793662 | -0.79769304 | 0.000219736 | 13.65431161 | 0.575268333 | 2.420941326 |
| *F630028O10Rik* | 257.5349214 | 247.1403219 | 234.15383 | 228.5848696 | 113.7718144 | 154.3534298 | 180.1124936 | 108.4162741 | 0.00052825 | -0.800717717 | 1.65E-05 | 18.5591375 | 0.574063519 | 3.277160848 |
| *4632428N05Rik* | 4381.65816 | 4407.994782 | 4303.871726 | 4165.730064 | 2671.490999 | 2560.863722 | 1855.262796 | 2764.614989 | 7.06E-17 | -0.808754816 | 4.34E-09 | 34.46600139 | 0.570874365 | 6.150280407 |
| *Vav3* | 1002.514832 | 980.6527974 | 1079.594208 | 892.395331 | 655.7979111 | 592.8574918 | 357.1016492 | 646.0543546 | 0.001578633 | -0.812382065 | 6.49E-05 | 15.95259711 | 0.569440865 | 2.801718759 |
| *Bcl6* | 1603.132607 | 1818.952769 | 1819.354538 | 1627.524272 | 1045.412709 | 831.4037015 | 954.7003275 | 1053.059711 | 4.19E-12 | -0.821077195 | 6.42E-15 | 60.76766864 | 0.566019164 | 11.37777405 |
| *Tnfrsf11a* | 1031.921934 | 958.9044491 | 982.2027915 | 949.9987182 | 659.0178681 | 485.2778286 | 425.8150861 | 648.7203285 | 2.87E-05 | -0.82128803 | 4.01E-07 | 25.68855034 | 0.565936452 | 4.541403277 |
| *Cetn4* | 49.01183624 | 56.3479934 | 43.51531355 | 60.34640558 | 32.1995701 | 37.41901329 | 20.8222536 | 28.4370555 | 0.00908324 | -0.824927476 | 0.000709223 | 11.46491925 | 0.56451058 | 2.041759198 |
| *Herc3* | 749.4355324 | 726.5925465 | 778.0952494 | 720.499509 | 471.1870425 | 390.5609512 | 288.3882123 | 524.3082107 | 0.000351927 | -0.827868067 | 9.85E-06 | 19.54096471 | 0.563361131 | 3.453547532 |
| *Tlr13* | 4422.649878 | 4922.046652 | 4815.694699 | 5018.809397 | 3255.376537 | 2496.549793 | 2059.320881 | 2961.008404 | 7.39E-06 | -0.832103391 | 7.87E-08 | 28.83840801 | 0.561709695 | 5.131453612 |
| *Golm1* | 1334.904194 | 1404.74559 | 1449.474373 | 1562.606169 | 1031.459562 | 801.0007532 | 458.0895792 | 917.0950398 | 0.006809731 | -0.84237229 | 0.000476286 | 12.2062936 | 0.557725721 | 2.166870036 |
| *D3Ertd751e* | 110.4994126 | 159.1583673 | 149.1953607 | 129.8362059 | 85.86552027 | 87.7008124 | 46.8500706 | 84.42250851 | 0.005225381 | -0.846994918 | 0.000330907 | 12.88680371 | 0.555941537 | 2.281882033 |
| *Plekhg5* | 76.6366894 | 71.17641272 | 68.381207 | 58.51772662 | 41.85944113 | 32.74163663 | 30.19226772 | 46.21021518 | 0.001400107 | -0.854141814 | 5.57E-05 | 16.24407303 | 0.553194296 | 2.85383869 |
| *Tlr8* | 1466.79059 | 1391.894293 | 1519.927737 | 1506.831461 | 860.8018407 | 901.5643514 | 692.3399322 | 788.2396321 | 1.01E-14 | -0.860768358 | 8.13E-18 | 73.92063665 | 0.550659207 | 13.99651208 |
| *Fmnl1* | 4424.432126 | 4655.135104 | 4642.669524 | 4347.68422 | 3107.258515 | 2426.389143 | 1355.528709 | 3048.985544 | 0.008603052 | -0.862460014 | 0.000653071 | 11.61826962 | 0.550013901 | 2.065347451 |
| *Gatm* | 2460.394179 | 2437.792136 | 2632.67647 | 2912.171239 | 1778.489589 | 1376.318083 | 800.6156509 | 1762.208783 | 0.007604155 | -0.868920473 | 0.000550252 | 11.93716231 | 0.547556418 | 2.118949035 |
| *P2ry13* | 108.717164 | 85.01627075 | 97.39141603 | 87.77658993 | 50.44599316 | 44.43507828 | 44.76784524 | 62.2060589 | 7.98E-05 | -0.900021436 | 1.47E-06 | 23.18247403 | 0.535878769 | 4.098250991 |
| *Lrig1* | 46.33846336 | 49.42806439 | 59.05649696 | 52.11735027 | 27.90629409 | 23.38688331 | 23.94559164 | 33.7690034 | 0.001457543 | -0.909768885 | 5.83E-05 | 16.15642029 | 0.532270352 | 2.836378506 |
| *Ftx* | 121.1929042 | 82.05058688 | 75.63375926 | 120.6928112 | 74.05901124 | 50.28179911 | 45.80895792 | 41.76692526 | 0.008630383 | -0.921241492 | 0.000658627 | 11.60251435 | 0.528054414 | 2.063969945 |
| *Dab2* | 10654.28208 | 12253.21716 | 12796.61042 | 10725.20208 | 7318.962284 | 6634.858794 | 3072.323518 | 7418.516853 | 0.009335506 | -0.92549514 | 0.00074181 | 11.38145536 | 0.52649979 | 2.029862134 |
| *Pced1b* | 94.45917531 | 83.03914817 | 81.85023262 | 72.2328188 | 51.51931216 | 35.08032496 | 31.2333804 | 55.09679503 | 0.001250228 | -0.930709663 | 4.79E-05 | 16.5291741 | 0.524600227 | 2.903010723 |
| *Arrdc3* | 1568.37876 | 1343.45479 | 1461.907319 | 1392.539026 | 940.227447 | 730.8401033 | 385.2116916 | 932.2022255 | 0.007045087 | -0.948025272 | 0.000495589 | 12.13219125 | 0.518341471 | 2.152113666 |
| *Grasp* | 56.14083061 | 70.18785143 | 65.27297032 | 61.26074506 | 21.46638034 | 38.58835746 | 37.48005648 | 32.88034542 | 0.000630995 | -0.95503837 | 2.03E-05 | 18.15996804 | 0.515827871 | 3.199973867 |
| *Il1rl1* | 48.12071195 | 42.50813537 | 47.65962912 | 31.08754227 | 18.24642306 | 18.70950664 | 17.69891556 | 30.21437147 | 0.009363592 | -0.979889551 | 0.000744797 | 11.37399061 | 0.507018554 | 2.028557496 |
| *Zfp467* | 193.3739721 | 194.7465737 | 181.3138064 | 174.6388404 | 128.7982804 | 81.85409157 | 52.055634 | 113.748222 | 0.004167356 | -0.980977471 | 0.000247096 | 13.43406278 | 0.506636362 | 2.380135355 |
| *Ica1* | 112.2816612 | 91.93619976 | 95.31925825 | 74.06149776 | 52.59263117 | 50.28179911 | 24.98670432 | 57.76276898 | 0.00452789 | -1.006929838 | 0.000275228 | 13.2318751 | 0.49760406 | 2.34410416 |
| *B3galnt1* | 362.6875882 | 305.4654379 | 294.2464059 | 310.8754227 | 180.3175926 | 170.7242481 | 79.12456367 | 196.3934145 | 0.003194992 | -1.022540622 | 0.000172688 | 14.10712571 | 0.492248726 | 2.495530181 |
| *Epha2* | 106.9349154 | 113.6845481 | 105.6800472 | 85.94791098 | 61.17918319 | 42.09638995 | 37.48005648 | 59.54008495 | 5.65E-05 | -1.034128199 | 9.48E-07 | 24.03145675 | 0.488310871 | 4.248089821 |
| *Ssbp2* | 37.4272204 | 49.42806439 | 59.05649696 | 61.26074506 | 22.53969907 | 35.08032496 | 18.74002824 | 23.10510759 | 0.003346306 | -1.065928039 | 0.000183566 | 13.99224348 | 0.477665293 | 2.475434292 |
| *Lgalsl* | 72.18106792 | 76.11921916 | 73.56160147 | 77.71885567 | 34.34620811 | 31.57229246 | 31.2333804 | 43.54424123 | 1.95E-06 | -1.078235397 | 1.45E-08 | 32.12366192 | 0.473607753 | 5.710143939 |
| *Xk* | 95.3502996 | 79.08490302 | 100.4996527 | 94.17696628 | 57.95922618 | 43.26573412 | 24.98670432 | 47.98753115 | 0.000460261 | -1.082537371 | 1.38E-05 | 18.90086428 | 0.472197605 | 3.336996062 |
| *Scn3b* | 56.14083061 | 50.41662567 | 56.98433917 | 55.77470819 | 22.53969907 | 31.57229246 | 11.45223948 | 36.43497736 | 0.007781963 | -1.098992886 | 0.000569815 | 11.87208678 | 0.466842274 | 2.108910818 |
| *Tox2* | 40.99171759 | 35.58820636 | 34.1906035 | 29.25886331 | 20.39306106 | 19.87885081 | 10.4111268 | 15.10718573 | 0.003210714 | -1.100589092 | 0.000173797 | 14.09508725 | 0.466326043 | 2.493398382 |
| *Kitl* | 401.8970572 | 392.4588312 | 397.8542953 | 438.8829497 | 242.5700948 | 180.0790015 | 86.41235243 | 246.1582617 | 0.004717409 | -1.110002398 | 0.000289225 | 13.13891816 | 0.463293261 | 2.326296484 |
| *Zfp69* | 48.12071195 | 46.46238052 | 48.69570802 | 33.8305607 | 16.09978505 | 31.57229246 | 12.49335216 | 22.21644961 | 0.006375917 | -1.110019114 | 0.000435768 | 12.37222668 | 0.463287893 | 2.195457324 |
| *Cytip* | 799.3384929 | 784.9176625 | 843.3682197 | 747.0153539 | 419.6677303 | 423.3025878 | 244.6614798 | 379.4569593 | 2.08E-07 | -1.114111468 | 1.14E-09 | 37.0647192 | 0.461975591 | 6.681284831 |
| *Col14a1* | 112.2816612 | 83.03914817 | 116.0408361 | 112.4637559 | 69.76573522 | 51.45114327 | 23.94559164 | 45.3215572 | 0.002993877 | -1.15663732 | 0.000156505 | 14.29228843 | 0.448556829 | 2.523765985 |
| *Zfp36l1* | 1548.774025 | 1653.863034 | 1555.15442 | 1466.600523 | 932.714214 | 606.8896218 | 366.4716633 | 867.3301927 | 0.000895476 | -1.165954565 | 3.11E-05 | 17.35165247 | 0.445669283 | 3.047946202 |
| *Cxcl14* | 973.1077305 | 1061.714823 | 1041.259288 | 1190.470001 | 618.2311746 | 450.1975036 | 284.2237616 | 536.7494225 | 3.80E-05 | -1.174901151 | 5.74E-07 | 24.99893606 | 0.442914106 | 4.419936065 |
| *4930577N17Rik* | 30.29822604 | 23.72547091 | 30.04628793 | 19.20112905 | 13.95314704 | 7.016064992 | 14.57557752 | 9.775237827 | 0.009249783 | -1.183607809 | 0.000732013 | 11.40615109 | 0.440249171 | 2.033868465 |
| *Fgf11* | 109.6082883 | 87.98195461 | 112.9325994 | 93.26262681 | 41.85944113 | 52.62048744 | 33.31560576 | 47.09887317 | 1.28E-07 | -1.208708907 | 6.43E-10 | 38.18738934 | 0.432655633 | 6.891126027 |
| *Cdk5r1* | 192.4828478 | 173.9867866 | 174.0612542 | 155.4377113 | 74.05901124 | 79.51540324 | 71.83677492 | 66.64934882 | 7.83E-16 | -1.258383567 | 5.06E-19 | 79.40633277 | 0.418012049 | 15.10596891 |
| *Gpr155* | 463.3846336 | 414.2071796 | 544.9774982 | 560.4901003 | 258.6698798 | 191.7724431 | 120.7690709 | 235.4943658 | 1.25E-05 | -1.296892355 | 1.49E-07 | 27.60635719 | 0.407001959 | 4.901756112 |
| *Gpr65* | 1687.789415 | 1564.892518 | 1733.35999 | 1563.520508 | 894.0747298 | 596.3655243 | 241.5381417 | 885.1033524 | 0.005312963 | -1.323201978 | 0.000339882 | 12.83672371 | 0.39964696 | 2.274663186 |
| *Phtf1os* | 47.22958765 | 47.45094181 | 61.12865474 | 60.34640558 | 34.34620811 | 17.54016248 | 8.328901439 | 24.88242356 | 0.005451312 | -1.341043663 | 0.000355769 | 12.75126276 | 0.394734996 | 2.263498982 |
| *Bbs12* | 104.2615426 | 116.650232 | 121.2212306 | 98.74866368 | 50.44599316 | 51.45114327 | 20.8222536 | 50.65350511 | 4.84E-05 | -1.345742538 | 7.81E-07 | 24.40473185 | 0.393451431 | 4.315227058 |
| *Pde3b* | 924.0958943 | 955.9387652 | 1026.754184 | 948.1700392 | 474.4069995 | 450.1975036 | 113.4812821 | 465.6567838 | 0.000608856 | -1.358117693 | 0.000654901 | 11.6130665 | 0.390090916 | 2.065054571 |
| *Ramp1* | 177.3337348 | 149.2727544 | 130.5459406 | 129.8362059 | 81.57224426 | 50.28179911 | 20.8222536 | 75.53592867 | 0.005408358 | -1.360983131 | 0.00035139 | 12.77443063 | 0.389316898 | 2.266934574 |
| *Psma8* | 24.06035597 | 17.79410318 | 34.1906035 | 31.08754227 | 10.73319003 | 4.677376661 | 9.370014119 | 15.10718573 | 0.007591857 | -1.397708678 | 0.000548029 | 11.94470448 | 0.379531444 | 2.119651555 |
| *Tal1* | 684.3834588 | 682.1072885 | 696.2450168 | 673.8681956 | 320.922382 | 232.6994889 | 119.7279582 | 352.7972198 | 0.000187541 | -1.414173394 | 4.45E-06 | 21.06140825 | 0.375224674 | 3.726904229 |
| *Ccr2* | 641.6094926 | 563.479934 | 675.5234389 | 593.4063215 | 286.5761739 | 238.5462097 | 90.57680315 | 253.2675255 | 0.000152981 | -1.509415363 | 3.42E-06 | 21.56604644 | 0.351253532 | 3.815362048 |
| *Notch4* | 212.9787066 | 210.5635543 | 253.839329 | 267.9014672 | 114.8451334 | 73.66868241 | 52.055634 | 84.42250851 | 3.78E-08 | -1.539899374 | 1.68E-10 | 40.80886508 | 0.343909441 | 7.422108453 |
| *Fabp7* | 32.08047463 | 36.57676765 | 26.93805124 | 27.43018435 | 18.24642306 | 4.677376661 | 8.328901439 | 10.66389581 | 0.001053888 | -1.544571741 | 3.89E-05 | 16.92659559 | 0.342797444 | 2.977205714 |
| *Lifr* | 345.7562266 | 332.1565927 | 361.591534 | 328.2478728 | 155.6312555 | 111.0876957 | 46.8500706 | 147.5172254 | 0.000131822 | -1.567578678 | 2.83E-06 | 21.92937002 | 0.337374145 | 3.880011027 |
| *Pcyt1b* | 125.6485256 | 116.650232 | 133.6541773 | 123.4358296 | 59.03254519 | 51.45114327 | 12.49335216 | 44.43289922 | 0.001220145 | -1.57858003 | 4.65E-05 | 16.58737725 | 0.334811264 | 2.913588591 |
| *C5ar2* | 258.4260457 | 274.820038 | 242.4424612 | 257.8437329 | 154.5579365 | 76.00737074 | 30.19226772 | 74.64727068 | 0.002038604 | -1.624478738 | 9.32E-05 | 15.26872773 | 0.32432705 | 2.690667158 |
| *Gprc5c* | 47.22958765 | 38.55389022 | 35.2266824 | 31.08754227 | 13.95314704 | 16.37081831 | 1.04111268 | 15.10718573 | 0.009785466 | -1.712138298 | 0.000803858 | 11.23230279 | 0.305207369 | 2.009418472 |
| *Gpr183* | 331.4982379 | 363.7905539 | 379.2048752 | 360.2497545 | 144.8980655 | 101.7329424 | 22.90447896 | 142.1852775 | 0.002647058 | -1.800205531 | 0.000134531 | 14.57722075 | 0.28713368 | 2.577356553 |
| *Hes1* | 108.717164 | 131.4786513 | 118.1129939 | 118.8641322 | 32.1995701 | 36.24966912 | 6.24667608 | 55.09679503 | 0.002588561 | -1.873688571 | 0.000130932 | 14.62832806 | 0.272874867 | 2.586941623 |
| *Dppa3* | 20.49585879 | 24.71403219 | 26.93805124 | 30.17320279 | 9.659871031 | 3.508032496 | 1.04111268 | 12.44121178 | 0.005158552 | -1.911145858 | 0.000326259 | 12.91328122 | 0.265881286 | 2.287472186 |
| *Igsf11* | 17.82248591 | 16.80554189 | 25.90197235 | 16.45811061 | 5.366595017 | 4.677376661 | 1.04111268 | 8.886579843 | 0.002038604 | -1.913162031 | 9.34E-05 | 15.26541341 | 0.265509975 | 2.690667158 |
| *Cecr6* | 13.36686443 | 15.8169806 | 27.97413014 | 13.71509218 | 3.21995701 | 8.185409157 | 1.04111268 | 6.22060589 | 0.008283091 | -1.923578799 | 0.000621432 | 11.71065544 | 0.263599804 | 2.081807576 |
| *S1pr4* | 31.18935034 | 33.61108378 | 23.82981456 | 29.25886331 | 16.09978505 | 3.508032496 | 1.04111268 | 7.109263874 | 0.00081276 | -1.979692795 | 2.77E-05 | 17.57159931 | 0.253543853 | 3.090037491 |
| *D5Ertd605e* | 16.04023732 | 25.70259348 | 20.72157788 | 19.20112905 | 6.43991402 | 5.846720827 | 1.04111268 | 6.22060589 | 0.000212183 | -2.055809738 | 5.20E-06 | 20.76084917 | 0.24051358 | 3.67328859 |
| *A430078G23Rik* | 17.82248591 | 14.82841932 | 11.39686763 | 14.62943166 | 5.366595017 | 2.338688331 | 0 | 3.554631937 | 0.00052825 | -2.376308699 | 1.65E-05 | 18.5567731 | 0.192601561 | 3.277160848 |
| *Skint3* | 10.69349154 | 10.87417417 | 10.36078894 | 8.229055306 | 0 | 2.338688331 | 3.12333804 | 1.777315969 | 0.001821987 | -2.47194364 | 7.96E-05 | 15.56838074 | 0.18024815 | 2.739454662 |
| *Kcne3* | 29.40710175 | 19.77122575 | 14.50510452 | 23.77282644 | 5.366595017 | 4.677376661 | 0 | 5.331947906 | 0.000106199 | -2.506780041 | 2.14E-06 | 22.46345605 | 0.175947871 | 3.973879952 |
| *Scel* | 7.128994363 | 14.82841932 | 5.18039447 | 9.143394785 | 1.073319003 | 1.169344165 | 1.04111268 | 2.665973953 | 0.006383404 | -2.56365846 | 0.000437198 | 12.36610687 | 0.169146067 | 2.194947664 |
| *Dlx2* | 11.58461584 | 9.885612877 | 4.144315576 | 12.8007527 | 2.146638007 | 1.169344165 | 3.12333804 | 0 | 0.007781963 | -2.622464084 | 0.000570343 | 11.87036164 | 0.162390137 | 2.108910818 |
| *Mrvi1* | 12.47574013 | 6.919929014 | 14.50510452 | 9.143394785 | 2.146638007 | 0 | 0 | 3.554631937 | 0.002524297 | -2.824966978 | 0.000126256 | 14.69687482 | 0.141123781 | 2.597859526 |
| *Galnt9* | 7.128994363 | 7.908490302 | 15.54118341 | 14.62943166 | 3.21995701 | 0 | 0 | 1.777315969 | 0.001263123 | -3.137360569 | 4.85E-05 | 16.50573053 | 0.113647624 | 2.898554473 |
| *4930447K03Rik* | 4.455621477 | 3.954245151 | 5.18039447 | 9.143394785 | 1.073319003 | 0 | 0 | 0 | 0.003041528 | -4.177842126 | 0.000160959 | 14.23947369 | 0.055251517 | 2.516908205 |
