## Supplementary material for "TFEB-Mediated Pro-inflammatory Response in Murine Macrophages Induced by Acute Alpha7 Nicotinic Receptor Activation": Table_S2

**Table S2. GO and KEGG Pathway analysis of differentially expressed genes from Table S1.**

| source | term_name | term_id | highlighted | adjusted_p_value | negative_log10_of_adjusted_p_value | term_size | query_size | intersection_size | effective_domain_size | intersections |
|---|---|---|---|---|---|---|---|---|---|---|
| GO:MF | binding | GO:0005488 | TRUE | 7.59E-139 | 138.1197484 | 16204 | 2300 | 2007 | 24945 | [illegible] |
| GO:MF | protein binding | GO:0005515 | FALSE | 3.98E-92 | 91.40058648 | 10378 | 2300 | 1426 | 24945 | [illegible] |
| GO:MF | ion binding | GO:0043167 | FALSE | 1.14E-87 | 86.94302577 | 5976 | 2300 | 967 | 24945 | [illegible] |
| GO:MF | DNA binding | GO:0003677 | FALSE | 8.34E-86 | 85.07883038 | 2342 | 2300 | 518 | 24945 | [illegible] |
| GO:MF | heterocyclic compound binding | GO:1901363 | FALSE | 3.32E-65 | 64.47853111 | 6779 | 2300 | 994 | 24945 | [illegible] |
| GO:MF | organic cyclic compound binding | GO:0097159 | FALSE | 3.97E-63 | 62.40164789 | 6865 | 2300 | 997 | 24945 | [illegible] |
| GO:MF | metal ion binding | GO:0046872 | FALSE | 5.34E-63 | 62.27248169 | 4138 | 2300 | 695 | 24945 | [illegible] |
| GO:MF | cation binding | GO:0043169 | FALSE | 2.95E-61 | 60.52965796 | 4237 | 2300 | 702 | 24945 | [illegible] |
| GO:MF | nucleic acid binding | GO:0003676 | FALSE | 9.10E-55 | 54.0408129 | 4663 | 2300 | 733 | 24945 | [illegible] |
| GO:MF | double-stranded DNA binding | GO:0003690 | FALSE | 1.43E-52 | 51.84470728 | 1485 | 2300 | 331 | 24945 | [illegible] |
| GO:MF | sequence-specific DNA binding | GO:0043565 | FALSE | 2.02E-52 | 51.6951187 | 1501 | 2300 | 333 | 24945 | [illegible] |
| GO:MF | transcription regulator activity | GO:0140110 | TRUE | 3.11E-48 | 47.50706361 | 1605 | 2300 | 339 | 24945 | [illegible] |
| GO:MF | transcription regulatory region nucleic acid binding | GO:0001067 | FALSE | 4.71E-48 | 47.32680527 | 1313 | 2300 | 297 | 24945 | [illegible] |
| GO:MF | transcription cis-regulatory region binding | GO:0000976 | FALSE | 6.39E-47 | 46.19437403 | 1308 | 2300 | 294 | 24945 | [illegible] |
| GO:MF | sequence-specific double-stranded DNA binding | GO:1990837 | FALSE | 1.19E-46 | 45.92442473 | 1388 | 2300 | 305 | 24945 | [illegible] |
| GO:MF | enzyme binding | GO:0019899 | FALSE | 3.84E-45 | 44.41605592 | 2297 | 2300 | 424 | 24945 | [illegible] |
| GO:MF | cis-regulatory region sequence-specific DNA binding | GO:0000987 | FALSE | 9.94E-39 | 38.00275487 | 1086 | 2300 | 245 | 24945 | [illegible] |
| GO:MF | RNA polymerase II transcription regulatory region sequence-specific DNA binding | GO:0000977 | FALSE | 3.83E-38 | 37.41646495 | 1205 | 2300 | 261 | 24945 | [illegible] |
| GO:MF | RNA polymerase II cis-regulatory region sequence-specific DNA binding | GO:0000978 | FALSE | 1.94E-37 | 36.71305201 | 1063 | 2300 | 239 | 24945 | [illegible] |
| GO:MF | DNA-binding transcription factor activity | GO:0003700 | FALSE | 5.76E-32 | 31.23992839 | 1159 | 2300 | 241 | 24945 | [illegible] |
| GO:MF | catalytic activity, acting on a protein | GO:0140096 | TRUE | 1.15E-29 | 28.93990617 | 2322 | 2300 | 385 | 24945 | [illegible] |
| GO:MF | DNA-binding transcription factor activity, RNA polymerase II-specific | GO:0000981 | FALSE | 1.62E-29 | 28.79049147 | 1077 | 2300 | 224 | 24945 | [illegible] |
| GO:MF | catalytic activity | GO:0003824 | FALSE | 5.61E-28 | 27.25111335 | 5721 | 2300 | 757 | 24945 | [illegible] |
| GO:MF | purine ribonucleoside triphosphate binding | GO:0035639 | FALSE | 1.31E-27 | 26.88207017 | 1835 | 2300 | 319 | 24945 | [illegible] |
| GO:MF | purine ribonucleotide binding | GO:0032555 | FALSE | 3.08E-26 | 25.51111237 | 1915 | 2300 | 325 | 24945 | [illegible] |
| GO:MF | ribonucleotide binding | GO:0032553 | FALSE | 8.78E-26 | 25.05629149 | 1934 | 2300 | 326 | 24945 | [illegible] |
| GO:MF | transferase activity | GO:0016740 | FALSE | 1.18E-25 | 24.92986487 | 2300 | 2300 | 370 | 24945 | [illegible] |
| GO:MF | carbohydrate derivative binding | GO:0097367 | FALSE | 6.74E-25 | 24.17147622 | 2270 | 2300 | 364 | 24945 | [illegible] |
| GO:MF | purine nucleotide binding | GO:0017076 | FALSE | 4.54E-24 | 23.34268533 | 2034 | 2300 | 333 | 24945 | [illegible] |
| GO:MF | nucleotide binding | GO:0000166 | FALSE | 9.39E-23 | 22.02726393 | 2194 | 2300 | 348 | 24945 | [illegible] |
| GO:MF | nucleoside phosphate binding | GO:1901265 | FALSE | 9.39E-23 | 22.02726393 | 2194 | 2300 | 348 | 24945 | [illegible] |
| GO:MF | ATP binding | GO:0005524 | FALSE | 1.04E-21 | 20.9826633 | 1484 | 2300 | 258 | 24945 | [illegible] |
| GO:MF | protein-containing complex binding | GO:0044877 | FALSE | 1.08E-21 | 20.96798606 | 2047 | 2300 | 327 | 24945 | [illegible] |
| GO:MF | adenyl ribonucleotide binding | GO:0032559 | FALSE | 9.11E-21 | 20.04066918 | 1554 | 2300 | 264 | 24945 | [illegible] |
| GO:MF | kinase binding | GO:0019900 | FALSE | 1.03E-20 | 19.98617471 | 843 | 2300 | 171 | 24945 | [illegible] |
| GO:MF | anion binding | GO:0043168 | FALSE | 1.33E-20 | 19.87504811 | 2470 | 2300 | 373 | 24945 | [illegible] |
| GO:MF | chromatin binding | GO:0003682 | FALSE | 3.33E-20 | 19.47725936 | 613 | 2300 | 137 | 24945 | [illegible] |
| GO:MF | protein kinase binding | GO:0019901 | FALSE | 8.02E-19 | 18.09604625 | 753 | 2300 | 154 | 24945 | [illegible] |
| GO:MF | adenyl nucleotide binding | GO:0030554 | FALSE | 2.33E-18 | 17.63205833 | 1672 | 2300 | 271 | 24945 | [illegible] |
| GO:MF | transcription factor binding | GO:0008134 | FALSE | 3.37E-18 | 17.4724985 | 643 | 2300 | 137 | 24945 | [illegible] |
| GO:MF | ubiquitin-like protein transferase activity | GO:0019787 | FALSE | 3.42E-18 | 17.46554022 | 448 | 2300 | 108 | 24945 | [illegible] |
| GO:MF | ubiquitin-protein transferase activity | GO:0004842 | FALSE | 1.29E-17 | 16.89084621 | 423 | 2300 | 103 | 24945 | [illegible] |
| GO:MF | aminoacyltransferase activity | GO:0016755 | FALSE | 1.87E-17 | 16.72765085 | 464 | 2300 | 109 | 24945 | [illegible] |
| GO:MF | ubiquitin-like protein ligase activity | GO:0061659 | FALSE | 1.41E-16 | 15.84983794 | 347 | 2300 | 89 | 24945 | [illegible] |
| GO:MF | acyltransferase activity | GO:0016746 | FALSE | 3.56E-16 | 15.44875338 | 705 | 2300 | 141 | 24945 | [illegible] |
| GO:MF | DNA-binding transcription repressor activity | GO:0001217 | FALSE | 4.41E-16 | 15.3559057 | 334 | 2300 | 86 | 24945 | [illegible] |
| GO:MF | DNA-binding transcription repressor activity, RNA polymerase II-specific | GO:0001227 | FALSE | 5.61E-16 | 15.25099591 | 329 | 2300 | 85 | 24945 | [illegible] |
| GO:MF | ubiquitin protein ligase activity | GO:0061630 | FALSE | 8.45E-16 | 15.07327301 | 331 | 2300 | 85 | 24945 | [illegible] |
| GO:MF | DNA-binding transcription factor binding | GO:0140297 | FALSE | 1.63E-15 | 14.78739719 | 525 | 2300 | 114 | 24945 | [illegible] |
| GO:MF | transcription coregulator activity | GO:0003712 | FALSE | 1.75E-15 | 14.75713101 | 471 | 2300 | 106 | 24945 | [illegible] |
| GO:MF | transferase activity, transferring phosphorus-containing groups | GO:0016772 | FALSE | 4.24E-15 | 14.37254264 | 943 | 2300 | 170 | 24945 | [illegible] |
| GO:MF | small molecule binding | GO:0036094 | FALSE | 9.54E-15 | 14.02057489 | 2625 | 2300 | 368 | 24945 | [illegible] |
| GO:MF | RNA polymerase II-specific DNA-binding transcription factor binding | GO:0061629 | FALSE | 4.92E-14 | 13.30769767 | 378 | 2300 | 89 | 24945 | [illegible] |
| GO:MF | identical protein binding | GO:0042802 | FALSE | 5.61E-14 | 13.2507566 | 2350 | 2300 | 334 | 24945 | [illegible] |
| GO:MF | enzyme regulator activity | GO:0030234 | TRUE | 9.53E-14 | 13.02075931 | 1286 | 2300 | 209 | 24945 | [illegible] |
| GO:MF | zinc ion binding | GO:0008270 | FALSE | 4.30E-13 | 12.36621478 | 741 | 2300 | 138 | 24945 | [illegible] |
| GO:MF | ribonucleoside triphosphate phosphatase activity | GO:0017111 | TRUE | 1.01E-12 | 11.99489692 | 666 | 2300 | 127 | 24945 | [illegible] |
| GO:MF | DNA-binding transcription activator activity, RNA polymerase II-specific | GO:0001228 | FALSE | 1.27E-12 | 11.89639327 | 508 | 2300 | 105 | 24945 | [illegible] |
| GO:MF | ATP-dependent activity | GO:0140657 | FALSE | 1.55E-12 | 11.80898617 | 574 | 2300 | 114 | 24945 | [illegible] |
| GO:MF | DNA-binding transcription activator activity | GO:0001216 | FALSE | 2.89E-12 | 11.53905733 | 514 | 2300 | 105 | 24945 | [illegible] |
| GO:MF | small GTPase binding | GO:0031267 | FALSE | 4.66E-12 | 11.33187179 | 287 | 2300 | 71 | 24945 | [illegible] |
| GO:MF | enzyme activator activity | GO:0008047 | FALSE | 7.34E-12 | 11.13417014 | 579 | 2300 | 113 | 24945 | [illegible] |
| GO:MF | GTPase binding | GO:0051020 | FALSE | 1.87E-11 | 10.7277102 | 327 | 2300 | 76 | 24945 | [illegible] |
| GO:MF | kinase regulator activity | GO:0019207 | FALSE | 1.88E-11 | 10.72534717 | 269 | 2300 | 67 | 24945 | [illegible] |
| GO:MF | tubulin binding | GO:0015631 | FALSE | 1.95E-11 | 10.7103178 | 374 | 2300 | 83 | 24945 | [illegible] |
| GO:MF | histone modifying activity | GO:0140993 | FALSE | 2.45E-11 | 10.61007927 | 203 | 2300 | 56 | 24945 | [illegible] |
| GO:MF | hydrolase activity, acting on acid anhydrides, in phosphorus-containing anhydrides | GO:0016818 | FALSE | 2.59E-11 | 10.58606127 | 725 | 2300 | 131 | 24945 | [illegible] |
| GO:MF | hydrolase activity, acting on acid anhydrides | GO:0016817 | FALSE | 2.59E-11 | 10.58606127 | 725 | 2300 | 131 | 24945 | [illegible] |
| GO:MF | histone binding | GO:0042393 | FALSE | 4.53E-11 | 10.34432152 | 280 | 2300 | 68 | 24945 | [illegible] |
| GO:MF | pyrophosphatase activity | GO:0016462 | FALSE | 5.26E-11 | 10.27868162 | 724 | 2300 | 130 | 24945 | [illegible] |
| GO:MF | protein serine/threonine kinase activity | GO:0004674 | FALSE | 5.35E-11 | 10.27197276 | 436 | 2300 | 91 | 24945 | [illegible] |
| GO:MF | protein domain specific binding | GO:0019904 | FALSE | 8.70E-11 | 10.06026461 | 791 | 2300 | 138 | 24945 | [illegible] |
| GO:MF | transcription coactivator activity | GO:0003713 | FALSE | 1.05E-10 | 9.979330389 | 259 | 2300 | 64 | 24945 | [illegible] |
| GO:MF | ubiquitin-like protein ligase binding | GO:0044389 | FALSE | 4.82E-10 | 9.316871726 | 347 | 2300 | 76 | 24945 | [illegible] |
| GO:MF | molecular function regulator activity | GO:0098772 | FALSE | 5.11E-10 | 9.291649763 | 2145 | 2300 | 294 | 24945 | [illegible] |
| GO:MF | catalytic activity, acting on a nucleic acid | GO:0140640 | FALSE | 7.20E-10 | 9.142433972 | 626 | 2300 | 114 | 24945 | [illegible] |
| GO:MF | kinase activity | GO:0016301 | FALSE | 8.41E-10 | 9.075319188 | 775 | 2300 | 133 | 24945 | [illegible] |
| GO:MF | protein serine kinase activity | GO:0106310 | FALSE | 1.30E-09 | 8.884471584 | 333 | 2300 | 73 | 24945 | [illegible] |
| GO:MF | protein kinase activity | GO:0004672 | FALSE | 1.39E-09 | 8.857587348 | 594 | 2300 | 109 | 24945 | [illegible] |
| GO:MF | ATP hydrolysis activity | GO:0016887 | FALSE | 2.89E-09 | 8.538853199 | 359 | 2300 | 76 | 24945 | [illegible] |
| GO:MF | double-stranded RNA binding | GO:0003725 | TRUE | 4.41E-09 | 8.355455013 | 80 | 2300 | 30 | 24945 | [illegible] |
| GO:MF | microtubule binding | GO:0008017 | FALSE | 1.10E-08 | 7.957020946 | 259 | 2300 | 60 | 24945 | [illegible] |
| GO:MF | protein kinase regulator activity | GO:0019887 | FALSE | 1.21E-08 | 7.91818263 | 227 | 2300 | 55 | 24945 | [illegible] |
| GO:MF | molecular adaptor activity | GO:0060090 | TRUE | 1.28E-08 | 7.891195287 | 615 | 2300 | 109 | 24945 | [illegible] |
| GO:MF | ubiquitin protein ligase binding | GO:0031625 | FALSE | 1.37E-08 | 7.863644097 | 328 | 2300 | 70 | 24945 | [illegible] |
| GO:MF | mRNA binding | GO:0003729 | TRUE | 1.72E-08 | 7.764117689 | 386 | 2300 | 78 | 24945 | [illegible] |
| GO:MF | nucleoside-triphosphatase regulator activity | GO:0060589 | FALSE | 2.19E-08 | 7.658666668 | 446 | 2300 | 86 | 24945 | [illegible] |
| GO:MF | GTPase regulator activity | GO:0030695 | FALSE | 3.30E-08 | 7.482124366 | 442 | 2300 | 85 | 24945 | [illegible] |
| GO:MF | transition metal ion binding | GO:0046914 | FALSE | 3.60E-08 | 7.443327175 | 1082 | 2300 | 165 | 24945 | [illegible] |
| GO:MF | phosphotransferase activity, alcohol group as acceptor | GO:0016773 | FALSE | 4.28E-08 | 7.368608343 | 698 | 2300 | 118 | 24945 | [illegible] |
| GO:MF | pattern recognition receptor activity | GO:0038187 | TRUE | 8.51E-08 | 7.070154869 | 38 | 2300 | 19 | 24945 | [illegible] |
| GO:MF | ATP-dependent chromatin remodeler activity | GO:0140658 | FALSE | 8.51E-08 | 7.070154869 | 38 | 2300 | 19 | 24945 | [illegible] |
| GO:MF | deubiquitinase activity | GO:0101005 | TRUE | 1.12E-07 | 6.95188367 | 100 | 2300 | 32 | 24945 | [illegible] |
| GO:MF | ATP-dependent activity, acting on DNA | GO:0008094 | FALSE | 2.16E-07 | 6.664585388 | 119 | 2300 | 35 | 24945 | [illegible] |
| GO:MF | ubiquitin-like protein peptidase activity | GO:0019783 | FALSE | 2.56E-07 | 6.592360342 | 114 | 2300 | 34 | 24945 | [illegible] |
| GO:MF | cysteine-type deubiquitinase activity | GO:0004843 | FALSE | 3.31E-07 | 6.479745004 | 93 | 2300 | 30 | 24945 | [illegible] |
| GO:MF | helicase activity | GO:0004386 | FALSE | 6.11E-07 | 6.213830002 | 153 | 2300 | 40 | 24945 | [illegible] |
| GO:MF | cytoskeletal protein binding | GO:0008092 | FALSE | 7.07E-07 | 6.150564805 | 1005 | 2300 | 151 | 24945 | [illegible] |
| GO:MF | nuclear receptor binding | GO:0016922 | FALSE | 1.13466E-06 | 5.945136067 | 156 | 2300 | 40 | 24945 | [illegible] |
| GO:MF | kinase activator activity | GO:0019209 | FALSE | 1.76863E-06 | 5.752363085 | 152 | 2300 | 39 | 24945 | [illegible] |
| GO:MF | regulatory RNA binding | GO:0061980 | TRUE | 4.84938E-06 | 5.314313779 | 46 | 2300 | 19 | 24945 | [illegible] |
| GO:MF | molecular function activator activity | GO:0140677 | FALSE | 2.06187E-05 | 4.685738532 | 1197 | 2300 | 167 | 24945 | [illegible] |
| GO:MF | RNA polymerase binding | GO:0070063 | FALSE | 2.59534E-05 | 4.585806131 | 60 | 2300 | 21 | 24945 | [illegible] |
| GO:MF | miRNA binding | GO:0035198 | FALSE | 3.96781E-05 | 4.40144963 | 33 | 2300 | 15 | 24945 | [illegible] |
| GO:MF | catalytic activity, acting on DNA | GO:0140097 | FALSE | 4.4866E-05 | 4.348082184 | 238 | 2300 | 49 | 24945 | [illegible] |
| GO:MF | ribonucleoprotein complex binding | GO:0043021 | FALSE | 5.63878E-05 | 4.248814584 | 191 | 2300 | 42 | 24945 | [illegible] |
| GO:MF | protein-macromolecule adaptor activity | GO:0030674 | FALSE | 6.31889E-05 | 4.199359416 | 397 | 2300 | 70 | 24945 | [illegible] |
| GO:MF | hydrolase activity | GO:0016787 | FALSE | 7.74088E-05 | 4.111209824 | 2446 | 2300 | 299 | 24945 | [illegible] |
| GO:MF | nucleotidyltransferase activity | GO:0016779 | TRUE | 8.56828E-05 | 4.067106593 | 147 | 2300 | 35 | 24945 | [illegible] |
| GO:MF | modification-dependent protein binding | GO:0140030 | FALSE | 8.97182E-05 | 4.047119609 | 236 | 2300 | 48 | 24945 | [illegible] |
| GO:MF | protein kinase activator activity | GO:0030295 | FALSE | 0.000120759 | 3.918082126 | 136 | 2300 | 33 | 24945 | [illegible] |
| GO:MF | peptide-lysine-N-acetyltransferase activity | GO:0061733 | FALSE | 0.000206672 | 3.684718313 | 46 | 2300 | 17 | 24945 | [illegible] |
| GO:MF | transcription coregulator binding | GO:0001221 | FALSE | 0.000208221 | 3.681475994 | 139 | 2300 | 33 | 24945 | [illegible] |
| GO:MF | guanyl-nucleotide exchange factor activity | GO:0005085 | FALSE | 0.000275793 | 3.559416342 | 223 | 2300 | 45 | 24945 | [illegible] |
| GO:MF | mRNA 3'-UTR binding | GO:0003730 | FALSE | 0.000276527 | 3.558263214 | 115 | 2300 | 29 | 24945 | [illegible] |
| GO:MF | peptide N-acetyltransferase activity | GO:0034212 | FALSE | 0.000291887 | 3.534785954 | 52 | 2300 | 18 | 24945 | [illegible] |
| GO:MF | single-stranded RNA binding | GO:0003727 | TRUE | 0.000302724 | 3.518953846 | 97 | 2300 | 26 | 24945 | [illegible] |
| GO:MF | GTP binding | GO:0005525 | FALSE | 0.000415219 | 3.381722335 | 385 | 2300 | 66 | 24945 | [illegible] |
| GO:MF | basal RNA polymerase II transcription machinery binding | GO:0001099 | FALSE | 0.000675273 | 3.170520337 | 60 | 2300 | 19 | 24945 | [illegible] |
| GO:MF | basal transcription machinery binding | GO:0001098 | FALSE | 0.000675273 | 3.170520337 | 60 | 2300 | 19 | 24945 | [illegible] |
| GO:MF | single-stranded DNA binding | GO:0003697 | FALSE | 0.000717756 | 3.144023457 | 120 | 2300 | 29 | 24945 | [illegible] |
| GO:MF | GTPase activator activity | GO:0005096 | FALSE | 0.001876986 | 2.726538864 | 231 | 2300 | 44 | 24945 | [illegible] |
| GO:MF | mitogen-activated protein kinase binding | GO:0051019 | FALSE | 0.00222163 | 2.653328255 | 33 | 2300 | 13 | 24945 | [illegible] |
| GO:MF | histone acetyltransferase activity | GO:0004402 | FALSE | 0.002379344 | 2.623542713 | 43 | 2300 | 15 | 24945 | [illegible] |
| GO:MF | protein homodimerization activity | GO:0042803 | FALSE | 0.002628081 | 2.580361185 | 778 | 2300 | 110 | 24945 | [illegible] |
| GO:MF | catalytic activity, acting on RNA | GO:0140098 | FALSE | 0.002976783 | 2.526252806 | 407 | 2300 | 66 | 24945 | [illegible] |
| GO:MF | guanyl nucleotide binding | GO:0019001 | FALSE | 0.003519039 | 2.453575949 | 409 | 2300 | 66 | 24945 | [illegible] |
| GO:MF | guanyl ribonucleotide binding | GO:0032561 | FALSE | 0.003519039 | 2.453575949 | 409 | 2300 | 66 | 24945 | [illegible] |
| GO:MF | phosphatidylinositol binding | GO:0035091 | TRUE | 0.004395388 | 2.357002739 | 292 | 2300 | 51 | 24945 | [illegible] |
| GO:MF | RNA polymerase II complex binding | GO:0000993 | FALSE | 0.004798125 | 2.318928458 | 35 | 2300 | 13 | 24945 | [illegible] |
| GO:MF | RNA polymerase core enzyme binding | GO:0043175 | FALSE | 0.004802295 | 2.318551196 | 40 | 2300 | 14 | 24945 | [illegible] |
| GO:MF | growth factor receptor binding | GO:0070851 | TRUE | 0.005689424 | 2.244931694 | 146 | 2300 | 31 | 24945 | [illegible] |
| GO:MF | ribosome binding | GO:0043022 | FALSE | 0.00782277 | 2.106639465 | 114 | 2300 | 26 | 24945 | [illegible] |
| GO:MF | ubiquitin-like protein binding | GO:0032182 | FALSE | 0.008211897 | 2.085556498 | 121 | 2300 | 27 | 24945 | [illegible] |
| GO:MF | GTPase activity | GO:0003924 | FALSE | 0.011529452 | 1.93819132 | 310 | 2300 | 52 | 24945 | [illegible] |
| GO:MF | K48-linked deubiquitinase activity | GO:1990380 | FALSE | 0.012548671 | 1.901402284 | 19 | 2300 | 9 | 24945 | [illegible] |
| GO:MF | transcription corepressor activity | GO:0003714 | FALSE | 0.012858339 | 1.890815121 | 188 | 2300 | 36 | 24945 | [illegible] |
| GO:MF | N-acetyltransferase activity | GO:0008080 | FALSE | 0.016357469 | 1.786283887 | 73 | 2300 | 19 | 24945 | [illegible] |
| GO:MF | histone H4 acetyltransferase activity | GO:0010485 | FALSE | 0.017165574 | 1.765341679 | 24 | 2300 | 10 | 24945 | [illegible] |
| GO:MF | cytokine receptor binding | GO:0005126 | TRUE | 0.018037807 | 1.743816261 | 315 | 2300 | 52 | 24945 | [illegible] |
| GO:MF | histone acetyltransferase binding | GO:0035035 | FALSE | 0.019422616 | 1.711692268 | 29 | 2300 | 11 | 24945 | [illegible] |
| GO:MF | NF-kappaB binding | GO:0051059 | FALSE | 0.019674629 | 1.706093437 | 34 | 2300 | 12 | 24945 | [illegible] |
| GO:MF | RNA exonuclease activity | GO:0004532 | TRUE | 0.019674629 | 1.706093437 | 34 | 2300 | 12 | 24945 | [illegible] |
| GO:MF | cysteine-type peptidase activity | GO:0008234 | FALSE | 0.020208898 | 1.694457372 | 177 | 2300 | 34 | 24945 | [illegible] |
| GO:MF | acetyltransferase activity | GO:0016407 | FALSE | 0.025113827 | 1.600087107 | 101 | 2300 | 23 | 24945 | [illegible] |
| GO:MF | nuclear estrogen receptor binding | GO:0030331 | FALSE | 0.038117932 | 1.41887069 | 47 | 2300 | 14 | 24945 | [illegible] |
| GO:MF | promoter-specific chromatin binding | GO:1990841 | FALSE | 0.038764064 | 1.411570701 | 71 | 2300 | 18 | 24945 | [illegible] |
| GO:MF | molecular condensate scaffold activity | GO:0140693 | FALSE | 0.039133008 | 1.407456774 | 26 | 2300 | 10 | 24945 | [illegible] |
| GO:MF | poly(A)-specific ribonuclease activity | GO:0004535 | TRUE | 0.039453817 | 1.403910971 | 13 | 2300 | 7 | 24945 | [illegible] |
| GO:MF | protein dimerization activity | GO:0046983 | FALSE | 0.040518087 | 1.392351072 | 1150 | 2300 | 145 | 24945 | [illegible] |
| GO:MF | N-acyltransferase activity | GO:0016410 | FALSE | 0.046206247 | 1.335299303 | 98 | 2300 | 22 | 24945 | [illegible] |
| GO:BP | regulation of nitrogen compound metabolic process | GO:0051171 | TRUE | 2.18E-175 | 174.6614015 | 5660 | 2278 | 1059 | 26856 | [illegible] |
| GO:BP | regulation of primary metabolic process | GO:0080090 | FALSE | 7.32E-170 | 169.135553 | 5856 | 2278 | 1071 | 26856 | [illegible] |
| GO:BP | regulation of nucleobase-containing compound metabolic process | GO:0019219 | FALSE | 2.14E-165 | 164.6705996 | 4033 | 2278 | 851 | 26856 | [illegible] |
| GO:BP | regulation of macromolecule metabolic process | GO:0060255 | FALSE | 2.16E-159 | 158.6664713 | 6707 | 2278 | 1143 | 26856 | [illegible] |
| GO:BP | regulation of cellular metabolic process | GO:0031323 | FALSE | 1.10E-157 | 156.9573553 | 6594 | 2278 | 1128 | 26856 | [illegible] |
| GO:BP | regulation of metabolic process | GO:0019222 | FALSE | 5.59E-157 | 156.2526954 | 7198 | 2278 | 1189 | 26856 | [illegible] |
| GO:BP | regulation of RNA metabolic process | GO:0051252 | FALSE | 3.67E-154 | 153.4354218 | 3743 | 2278 | 796 | 26856 | [illegible] |
| GO:BP | regulation of cellular process | GO:0050794 | FALSE | 2.30E-144 | 143.6378585 | 12664 | 2278 | 1657 | 26856 | [illegible] |
| GO:BP | regulation of biological process | GO:0050789 | FALSE | 2.26E-143 | 142.6462394 | 13256 | 2278 | 1702 | 26856 | [illegible] |
| GO:BP | regulation of biosynthetic process | GO:0009889 | FALSE | 6.13E-143 | 142.2123945 | 5714 | 2278 | 1006 | 26856 | [illegible] |
| GO:BP | regulation of macromolecule biosynthetic process | GO:0010556 | FALSE | 1.64E-142 | 141.7845955 | 5521 | 2278 | 984 | 26856 | [illegible] |
| GO:BP | regulation of cellular biosynthetic process | GO:0031326 | FALSE | 2.58E-142 | 141.5887292 | 5670 | 2278 | 1000 | 26856 | [illegible] |
| GO:BP | biological regulation | GO:0065007 | FALSE | 2.75E-141 | 140.5605918 | 13668 | 2278 | 1730 | 26856 | [illegible] |
| GO:BP | regulation of gene expression | GO:0010468 | FALSE | 4.54E-137 | 136.3425244 | 5389 | 2278 | 959 | 26856 | [illegible] |
| GO:BP | RNA biosynthetic process | GO:0032774 | FALSE | 2.78E-134 | 133.5566783 | 3577 | 2278 | 741 | 26856 | [illegible] |
| GO:BP | regulation of RNA biosynthetic process | GO:2001141 | FALSE | 1.43E-133 | 132.8457457 | 3444 | 2278 | 723 | 26856 | [illegible] |
| GO:BP | DNA-templated transcription | GO:0006351 | FALSE | 2.65E-132 | 131.5772594 | 3549 | 2278 | 734 | 26856 | [illegible] |
| GO:BP | regulation of DNA-templated transcription | GO:0006355 | FALSE | 8.44E-132 | 131.0737458 | 3429 | 2278 | 718 | 26856 | [illegible] |
| GO:BP | nucleobase-containing compound biosynthetic process | GO:0034654 | FALSE | 3.34E-130 | 129.4763192 | 3978 | 2278 | 783 | 26856 | [illegible] |
| GO:BP | organic cyclic compound biosynthetic process | GO:1901362 | FALSE | 6.71E-130 | 129.172977 | 4208 | 2278 | 810 | 26856 | [illegible] |
| GO:BP | aromatic compound biosynthetic process | GO:0019438 | FALSE | 7.43E-130 | 129.1291788 | 4058 | 2278 | 792 | 26856 | [illegible] |
| GO:BP | heterocycle biosynthetic process | GO:0018130 | FALSE | 6.99E-129 | 128.1557825 | 4048 | 2278 | 789 | 26856 | [illegible] |
| GO:BP | cellular nitrogen compound biosynthetic process | GO:0044271 | FALSE | 1.54E-125 | 124.8119759 | 4822 | 2278 | 873 | 26856 | [illegible] |
| GO:BP | positive regulation of biological process | GO:0048518 | FALSE | 2.47E-114 | 113.6072023 | 6614 | 2278 | 1043 | 26856 | [illegible] |
| GO:BP | positive regulation of cellular process | GO:0048522 | FALSE | 7.13E-110 | 109.1467634 | 6091 | 2278 | 979 | 26856 | [illegible] |
| GO:BP | positive regulation of metabolic process | GO:0009893 | FALSE | 1.49E-100 | 99.82809398 | 3905 | 2278 | 717 | 26856 | [illegible] |
| GO:BP | positive regulation of macromolecule metabolic process | GO:0010604 | FALSE | 2.09E-97 | 96.67971448 | 3580 | 2278 | 672 | 26856 | [illegible] |
| GO:BP | positive regulation of cellular metabolic process | GO:0031325 | FALSE | 2.16E-97 | 96.66518351 | 3522 | 2278 | 665 | 26856 | [illegible] |
| GO:BP | transcription by RNA polymerase II | GO:0006366 | FALSE | 1.18E-91 | 90.92688764 | 2425 | 2278 | 516 | 26856 | [illegible] |
| GO:BP | regulation of transcription by RNA polymerase II | GO:0006357 | FALSE | 1.08E-89 | 88.96605351 | 2354 | 2278 | 503 | 26856 | [illegible] |
| GO:BP | positive regulation of nitrogen compound metabolic process | GO:0051173 | FALSE | 1.94E-89 | 88.71221152 | 3152 | 2278 | 604 | 26856 | [illegible] |
| GO:BP | negative regulation of cellular process | GO:0048523 | FALSE | 2.74E-84 | 83.56260661 | 5584 | 2278 | 867 | 26856 | [illegible] |
| GO:BP | positive regulation of nucleobase-containing compound metabolic process | GO:0045935 | FALSE | 5.06E-82 | 81.29618164 | 2078 | 2278 | 452 | 26856 | [illegible] |
| GO:BP | positive regulation of biosynthetic process | GO:0009891 | FALSE | 6.51E-82 | 81.18652236 | 2798 | 2278 | 545 | 26856 | [illegible] |
| GO:BP | positive regulation of cellular biosynthetic process | GO:0031328 | FALSE | 4.15E-81 | 80.38168158 | 2771 | 2278 | 540 | 26856 | [illegible] |
| GO:BP | positive regulation of macromolecule biosynthetic process | GO:0010557 | FALSE | 5.21E-81 | 80.28292622 | 2669 | 2278 | 527 | 26856 | [illegible] |
| GO:BP | cellular process | GO:0009987 | FALSE | 1.90E-79 | 78.7211377 | 20940 | 2278 | 2105 | 26856 | [illegible] |
| GO:BP | nitrogen compound metabolic process | GO:0006807 | FALSE | 1.90E-79 | 78.72074175 | 12481 | 2278 | 1496 | 26856 | [illegible] |
| GO:BP | negative regulation of biological process | GO:0048519 | FALSE | 7.67E-79 | 78.11547414 | 6031 | 2278 | 900 | 26856 | [illegible] |
| GO:BP | macromolecule metabolic process | GO:0043170 | FALSE | 1.38E-78 | 77.85896194 | 12617 | 2278 | 1505 | 26856 | [illegible] |
| GO:BP | primary metabolic process | GO:0044238 | FALSE | 2.41E-78 | 77.61871653 | 13086 | 2278 | 1543 | 26856 | [illegible] |
| GO:BP | macromolecule modification | GO:0043412 | FALSE | 4.66E-78 | 77.33192977 | 3597 | 2278 | 633 | 26856 | [illegible] |
| GO:BP | positive regulation of RNA metabolic process | GO:0051254 | FALSE | 1.49E-76 | 75.82799926 | 1872 | 2278 | 414 | 26856 | [illegible] |
| GO:BP | protein modification process | GO:0036211 | FALSE | 1.94E-74 | 73.7118235 | 3401 | 2278 | 602 | 26856 | [illegible] |
| GO:BP | metabolic process | GO:0008152 | FALSE | 6.59E-72 | 71.18095638 | 14599 | 2278 | 1647 | 26856 | [illegible] |
| GO:BP | negative regulation of nitrogen compound metabolic process | GO:0051172 | FALSE | 2.94E-71 | 70.53095424 | 2493 | 2278 | 485 | 26856 | [illegible] |
| GO:BP | organic substance metabolic process | GO:0071704 | FALSE | 2.43E-69 | 68.61414709 | 14220 | 2278 | 1610 | 26856 | [illegible] |
| GO:BP | positive regulation of RNA biosynthetic process | GO:1902680 | FALSE | 1.41E-68 | 67.8507522 | 1731 | 2278 | 380 | 26856 | [illegible] |
| GO:BP | positive regulation of DNA-templated transcription | GO:0045893 | FALSE | 5.12E-68 | 67.29095145 | 1724 | 2278 | 378 | 26856 | [illegible] |
| GO:BP | cellular metabolic process | GO:0044237 | FALSE | 4.94E-66 | 65.30600236 | 13077 | 2278 | 1508 | 26856 | [illegible] |
| GO:BP | negative regulation of cellular metabolic process | GO:0031324 | FALSE | 2.53E-64 | 63.59743239 | 3180 | 2278 | 553 | 26856 | [illegible] |
| GO:BP | response to stress | GO:0006950 | FALSE | 2.94E-63 | 62.53190982 | 4038 | 2278 | 648 | 26856 | [illegible] |
| GO:BP | protein metabolic process | GO:0019538 | FALSE | 7.48E-63 | 62.12628249 | 5451 | 2278 | 798 | 26856 | [illegible] |
| GO:BP | negative regulation of metabolic process | GO:0009892 | FALSE | 3.22E-62 | 61.49263358 | 3553 | 2278 | 591 | 26856 | [illegible] |
| GO:BP | negative regulation of macromolecule metabolic process | GO:0010605 | FALSE | 8.34E-62 | 61.07897568 | 3326 | 2278 | 564 | 26856 | [illegible] |
| GO:BP | nucleic acid metabolic process | GO:0090304 | FALSE | 5.65E-60 | 59.24779609 | 7754 | 2278 | 1018 | 26856 | [illegible] |
| GO:BP | regulation of response to stimulus | GO:0048583 | FALSE | 1.14E-59 | 58.94290363 | 3958 | 2278 | 630 | 26856 | [illegible] |
| GO:BP | nucleobase-containing compound metabolic process | GO:0006139 | FALSE | 1.36E-57 | 56.86627467 | 8265 | 2278 | 1059 | 26856 | [illegible] |
| GO:BP | heterocycle metabolic process | GO:0046483 | FALSE | 6.85E-57 | 56.16460165 | 8406 | 2278 | 1070 | 26856 | [illegible] |
| GO:BP | negative regulation of nucleobase-containing compound metabolic process | GO:0045934 | FALSE | 2.02E-56 | 55.69545827 | 1637 | 2278 | 344 | 26856 | [illegible] |
| GO:BP | cellular aromatic compound metabolic process | GO:0006725 | FALSE | 2.13E-56 | 55.67090626 | 8487 | 2278 | 1076 | 26856 | [illegible] |
| GO:BP | intracellular signal transduction | GO:0035556 | FALSE | 7.96E-56 | 55.09915271 | 2555 | 2278 | 459 | 26856 | [illegible] |
| GO:BP | organic cyclic compound metabolic process | GO:1901360 | FALSE | 1.17E-55 | 54.93103423 | 8717 | 2278 | 1095 | 26856 | [illegible] |
| GO:BP | negative regulation of biosynthetic process | GO:0009890 | FALSE | 1.52E-55 | 54.81888622 | 2778 | 2278 | 485 | 26856 | [illegible] |
| GO:BP | negative regulation of cellular biosynthetic process | GO:0031327 | FALSE | 5.85E-55 | 54.23260207 | 2756 | 2278 | 481 | 26856 | [illegible] |
| GO:BP | negative regulation of macromolecule biosynthetic process | GO:0010558 | FALSE | 1.73E-54 | 53.76235393 | 2698 | 2278 | 473 | 26856 | [illegible] |
| GO:BP | cellular nitrogen compound metabolic process | GO:0034641 | FALSE | 1.11E-53 | 52.95502154 | 8916 | 2278 | 1107 | 26856 | [illegible] |
| GO:BP | organonitrogen compound metabolic process | GO:1901564 | FALSE | 2.92E-53 | 52.53409621 | 6389 | 2278 | 865 | 26856 | [illegible] |
| GO:BP | positive regulation of transcription by RNA polymerase II | GO:0045944 | FALSE | 4.91E-52 | 51.3088917 | 1311 | 2278 | 291 | 26856 | [illegible] |
| GO:BP | negative regulation of RNA metabolic process | GO:0051253 | FALSE | 4.97E-52 | 51.30391011 | 1523 | 2278 | 320 | 26856 | [illegible] |
| GO:BP | cellular component organization or biogenesis | GO:0071840 | FALSE | 1.29E-50 | 49.88829012 | 7758 | 2278 | 989 | 26856 | [illegible] |
| GO:BP | cell cycle | GO:0007049 | FALSE | 2.90E-50 | 49.53691121 | 1802 | 2278 | 353 | 26856 | [illegible] |
| GO:BP | regulation of signal transduction | GO:0009966 | FALSE | 5.28E-50 | 49.27721941 | 2989 | 2278 | 496 | 26856 | [illegible] |
| GO:BP | cellular component organization | GO:0016043 | FALSE | 6.32E-49 | 48.1992818 | 7553 | 2278 | 964 | 26856 | [illegible] |
| GO:BP | regulation of response to stress | GO:0080134 | FALSE | 1.47E-48 | 47.83302831 | 1463 | 2278 | 305 | 26856 | [illegible] |
| GO:BP | RNA metabolic process | GO:0016070 | FALSE | 5.69E-47 | 46.24521989 | 7202 | 2278 | 924 | 26856 | [illegible] |
| GO:BP | biosynthetic process | GO:0009058 | FALSE | 5.81E-47 | 46.2359709 | 10877 | 2278 | 1260 | 26856 | [illegible] |
| GO:BP | organic substance biosynthetic process | GO:1901576 | FALSE | 6.57E-47 | 46.1825545 | 10822 | 2278 | 1255 | 26856 | [illegible] |
| GO:BP | cellular biosynthetic process | GO:0044249 | FALSE | 1.62E-46 | 45.79116816 | 10735 | 2278 | 1246 | 26856 | [illegible] |
| GO:BP | regulation of signaling | GO:0023051 | FALSE | 3.82E-46 | 45.41787981 | 3588 | 2278 | 553 | 26856 | [illegible] |
| GO:BP | regulation of cell communication | GO:0010646 | FALSE | 1.14E-45 | 44.94278091 | 3591 | 2278 | 552 | 26856 | [illegible] |
| GO:BP | negative regulation of RNA biosynthetic process | GO:1902679 | FALSE | 1.89E-45 | 44.72302206 | 1413 | 2278 | 292 | 26856 | [illegible] |
| GO:BP | negative regulation of DNA-templated transcription | GO:0045892 | FALSE | 5.40E-45 | 44.26744666 | 1405 | 2278 | 290 | 26856 | [illegible] |
| GO:BP | cellular response to stress | GO:0033554 | FALSE | 1.21E-44 | 43.91834787 | 2005 | 2278 | 366 | 26856 | [illegible] |
| GO:BP | organelle organization | GO:0006996 | FALSE | 1.98E-44 | 43.70402763 | 3468 | 2278 | 535 | 26856 | [illegible] |
| GO:BP | macromolecule biosynthetic process | GO:0009059 | FALSE | 2.77E-44 | 43.55676084 | 9910 | 2278 | 1165 | 26856 | [illegible] |
| GO:BP | regulation of protein metabolic process | GO:0051246 | FALSE | 3.81E-44 | 43.41890247 | 2455 | 2278 | 419 | 26856 | [illegible] |
| GO:BP | gene expression | GO:0010467 | FALSE | 1.39E-43 | 42.85760728 | 9415 | 2278 | 1118 | 26856 | [illegible] |
| GO:BP | regulation of cellular component organization | GO:0051128 | FALSE | 1.93E-43 | 42.71504594 | 2564 | 2278 | 430 | 26856 | [illegible] |
| GO:BP | post-translational protein modification | GO:0043687 | FALSE | 5.47E-43 | 42.26231719 | 1213 | 2278 | 260 | 26856 | [illegible] |
| GO:BP | protein-DNA complex organization | GO:0071824 | FALSE | 9.79E-42 | 41.00921969 | 830 | 2278 | 203 | 26856 | [illegible] |
| GO:BP | positive regulation of response to stimulus | GO:0048584 | FALSE | 2.01E-41 | 40.69736602 | 2293 | 2278 | 393 | 26856 | [illegible] |
| GO:BP | cell cycle process | GO:0022402 | FALSE | 3.75E-41 | 40.42570406 | 1240 | 2278 | 260 | 26856 | [illegible] |
| GO:BP | chromatin organization | GO:0006325 | FALSE | 1.74E-39 | 38.75973212 | 743 | 2278 | 186 | 26856 | [illegible] |
| GO:BP | cellular localization | GO:0051641 | FALSE | 2.26E-39 | 38.64502496 | 3510 | 2278 | 525 | 26856 | [illegible] |
| GO:BP | immune system process | GO:0002376 | FALSE | 4.16E-39 | 38.38038731 | 2663 | 2278 | 430 | 26856 | [illegible] |
| GO:BP | regulation of immune system process | GO:0002682 | FALSE | 2.31E-38 | 37.63607384 | 1545 | 2278 | 294 | 26856 | [illegible] |
| GO:BP | regulation of cell cycle | GO:0051726 | FALSE | 2.69E-37 | 36.57088843 | 1055 | 2278 | 227 | 26856 | [illegible] |
| GO:BP | negative regulation of transcription by RNA polymerase II | GO:0000122 | FALSE | 3.22E-37 | 36.49240338 | 1020 | 2278 | 222 | 26856 | [illegible] |
| GO:BP | regulation of cytokine production | GO:0001817 | FALSE | 6.60E-37 | 36.18013296 | 800 | 2278 | 190 | 26856 | [illegible] |
| GO:BP | cytokine production | GO:0001816 | FALSE | 2.03E-36 | 35.6930983 | 806 | 2278 | 190 | 26856 | [illegible] |
| GO:BP | regulation of multicellular organismal process | GO:0051239 | FALSE | 2.88E-36 | 35.54028124 | 3173 | 2278 | 479 | 26856 | [illegible] |
| GO:BP | biological process involved in interspecies interaction between organisms | GO:0044419 | FALSE | 5.69E-36 | 35.24505292 | 1881 | 2278 | 330 | 26856 | [illegible] |
| GO:BP | regulation of response to external stimulus | GO:0032101 | FALSE | 1.14E-35 | 34.94308935 | 1101 | 2278 | 230 | 26856 | [illegible] |
| GO:BP | phosphorylation | GO:0016310 | FALSE | 1.64E-35 | 34.7858178 | 1808 | 2278 | 320 | 26856 | [illegible] |
| GO:BP | protein phosphorylation | GO:0006468 | FALSE | 3.36E-35 | 34.4738399 | 1579 | 2278 | 291 | 26856 | [illegible] |
| GO:BP | regulation of molecular function | GO:0065009 | FALSE | 2.95E-34 | 33.53005284 | 2535 | 2278 | 402 | 26856 | [illegible] |
| GO:BP | positive regulation of response to external stimulus | GO:0032103 | FALSE | 3.11E-34 | 33.50783719 | 612 | 2278 | 157 | 26856 | [illegible] |
| GO:BP | regulation of intracellular signal transduction | GO:1902531 | FALSE | 4.55E-34 | 33.34191705 | 1641 | 2278 | 296 | 26856 | [illegible] |
| GO:BP | DNA damage response | GO:0006974 | FALSE | 7.56E-34 | 33.12134807 | 853 | 2278 | 192 | 26856 | [illegible] |
| GO:BP | innate immune response | GO:0045087 | FALSE | 1.25E-33 | 32.90470134 | 927 | 2278 | 202 | 26856 | [illegible] |
| GO:BP | chromatin remodeling | GO:0006338 | FALSE | 1.50E-33 | 32.82312086 | 626 | 2278 | 158 | 26856 | [illegible] |
| GO:BP | multicellular organism development | GO:0007275 | FALSE | 1.56E-33 | 32.80757021 | 4828 | 2278 | 644 | 26856 | [illegible] |
| GO:BP | regulation of response to biotic stimulus | GO:0002831 | FALSE | 3.32E-33 | 32.47903138 | 515 | 2278 | 140 | 26856 | [illegible] |
| GO:BP | defense response to other organism | GO:0098542 | FALSE | 1.55E-32 | 31.80919015 | 1219 | 2278 | 239 | 26856 | [illegible] |
| GO:BP | regulation of defense response | GO:0031347 | FALSE | 2.41E-32 | 31.61836869 | 789 | 2278 | 180 | 26856 | [illegible] |
| GO:BP | response to external stimulus | GO:0009605 | FALSE | 4.35E-32 | 31.36149777 | 2987 | 2278 | 446 | 26856 | [illegible] |
| GO:BP | cell population proliferation | GO:0008283 | FALSE | 4.66E-32 | 31.33119475 | 3413 | 2278 | 492 | 26856 | [illegible] |
| GO:BP | defense response | GO:0006952 | FALSE | 5.51E-32 | 31.25858264 | 1782 | 2278 | 308 | 26856 | [illegible] |
| GO:BP | anatomical structure development | GO:0048856 | FALSE | 5.74E-32 | 31.2411308 | 6222 | 2278 | 776 | 26856 | [illegible] |
| GO:BP | cellular response to organic substance | GO:0071310 | FALSE | 1.11E-31 | 30.9534394 | 2441 | 2278 | 384 | 26856 | [illegible] |
| GO:BP | response to biotic stimulus | GO:0009607 | FALSE | 1.17E-31 | 30.93286328 | 1764 | 2278 | 305 | 26856 | [illegible] |
| GO:BP | regulation of innate immune response | GO:0045088 | FALSE | 1.44E-31 | 30.84265914 | 421 | 2278 | 122 | 26856 | [illegible] |
| GO:BP | protein modification by small protein conjugation or removal | GO:0070647 | FALSE | 1.60E-31 | 30.79592791 | 965 | 2278 | 203 | 26856 | [illegible] |
| GO:BP | regulation of protein modification process | GO:0031399 | FALSE | 1.84E-31 | 30.73423985 | 1434 | 2278 | 264 | 26856 | [illegible] |
| GO:BP | regulation of immune response | GO:0050776 | FALSE | 3.36E-31 | 30.47324639 | 897 | 2278 | 193 | 26856 | [illegible] |
| GO:BP | response to other organism | GO:0051707 | FALSE | 7.44E-31 | 30.12813998 | 1723 | 2278 | 298 | 26856 | [illegible] |
| GO:BP | response to external biotic stimulus | GO:0043207 | FALSE | 1.15E-30 | 29.94045911 | 1727 | 2278 | 298 | 26856 | [illegible] |
| GO:BP | developmental process | GO:0032502 | FALSE | 1.19E-30 | 29.92558673 | 6808 | 2278 | 827 | 26856 | [illegible] |
| GO:BP | cellular response to stimulus | GO:0051716 | FALSE | 1.34E-30 | 29.87436354 | 8758 | 2278 | 1008 | 26856 | [illegible] |
| GO:BP | cellular macromolecule localization | GO:0070727 | FALSE | 1.39E-30 | 29.85617928 | 2594 | 2278 | 398 | 26856 | [illegible] |
| GO:BP | macromolecule catabolic process | GO:0009057 | FALSE | 2.05E-30 | 29.68903262 | 1406 | 2278 | 258 | 26856 | [illegible] |
| GO:BP | protein localization | GO:0008104 | FALSE | 3.27E-30 | 29.48536214 | 2586 | 2278 | 396 | 26856 | [illegible] |
| GO:BP | localization | GO:0051179 | FALSE | 4.81E-30 | 29.31795321 | 5386 | 2278 | 687 | 26856 | [illegible] |
| GO:BP | chromosome organization | GO:0051276 | FALSE | 8.94E-30 | 29.04864869 | 590 | 2278 | 146 | 26856 | [illegible] |
| GO:BP | phosphate-containing compound metabolic process | GO:0006796 | FALSE | 9.61E-30 | 29.01709598 | 2734 | 2278 | 411 | 26856 | [illegible] |
| GO:BP | regulation of organelle organization | GO:0033043 | FALSE | 1.01E-29 | 28.99535664 | 1175 | 2278 | 227 | 26856 | [illegible] |
| GO:BP | phosphorus metabolic process | GO:0006793 | FALSE | 1.32E-29 | 28.87907968 | 2756 | 2278 | 413 | 26856 | [illegible] |
| GO:BP | response to stimulus | GO:0050896 | FALSE | 2.20E-29 | 28.65839047 | 10392 | 2278 | 1149 | 26856 | [illegible] |
| GO:BP | activation of innate immune response | GO:0002218 | FALSE | 4.49E-29 | 28.34748931 | 266 | 2278 | 91 | 26856 | [illegible] |
| GO:BP | cellular response to chemical stimulus | GO:0070887 | FALSE | 1.02E-28 | 27.99251308 | 3150 | 2278 | 453 | 26856 | [illegible] |
| GO:BP | positive regulation of gene expression | GO:0010628 | FALSE | 1.09E-28 | 27.96173133 | 1256 | 2278 | 235 | 26856 | [illegible] |
| GO:BP | DNA metabolic process | GO:0006259 | FALSE | 1.28E-28 | 27.89285348 | 1018 | 2278 | 204 | 26856 | [illegible] |
| GO:BP | positive regulation of immune system process | GO:0002684 | FALSE | 1.39E-28 | 27.85588285 | 1087 | 2278 | 213 | 26856 | [illegible] |
| GO:BP | immune response | GO:0006955 | FALSE | 1.54E-28 | 27.81303314 | 1799 | 2278 | 301 | 26856 | [illegible] |
| GO:BP | macromolecule localization | GO:0033036 | FALSE | 2.24E-28 | 27.64879911 | 3086 | 2278 | 445 | 26856 | [illegible] |
| GO:BP | response to virus | GO:0009615 | FALSE | 2.53E-28 | 27.59671039 | 427 | 2278 | 118 | 26856 | [illegible] |
| GO:BP | positive regulation of cytokine production | GO:0001819 | FALSE | 3.75E-28 | 27.42636623 | 504 | 2278 | 130 | 26856 | [illegible] |
| GO:BP | defense response to virus | GO:0051607 | FALSE | 6.23E-28 | 27.20534293 | 347 | 2278 | 104 | 26856 | [illegible] |
| GO:BP | defense response to symbiont | GO:0140546 | FALSE | 8.17E-28 | 27.08784855 | 348 | 2278 | 104 | 26856 | [illegible] |
| GO:BP | negative regulation of response to stimulus | GO:0048585 | FALSE | 9.24E-28 | 27.03415536 | 1707 | 2278 | 288 | 26856 | [illegible] |
| GO:BP | positive regulation of defense response | GO:0031349 | FALSE | 1.16E-27 | 26.93459898 | 471 | 2278 | 124 | 26856 | [illegible] |
| GO:BP | positive regulation of innate immune response | GO:0045089 | FALSE | 1.26E-27 | 26.90064851 | 338 | 2278 | 102 | 26856 | [illegible] |
| GO:BP | canonical NF-kappaB signal transduction | GO:0007249 | FALSE | 2.58E-27 | 26.5888833 | 257 | 2278 | 87 | 26856 | [illegible] |
| GO:BP | chromosome segregation | GO:0007059 | TRUE | 5.40E-27 | 26.26738451 | 397 | 2278 | 111 | 26856 | [illegible] |
| GO:BP | positive regulation of response to biotic stimulus | GO:0002833 | FALSE | 5.48E-27 | 26.26141147 | 367 | 2278 | 106 | 26856 | [illegible] |
| GO:BP | programmed cell death | GO:0012501 | FALSE | 9.61E-27 | 26.01746622 | 2127 | 2278 | 334 | 26856 | [illegible] |
| GO:BP | response to cytokine | GO:0034097 | FALSE | 1.59E-26 | 25.79783007 | 1178 | 2278 | 220 | 26856 | [illegible] |
| GO:BP | cell death | GO:0008219 | FALSE | 1.64E-26 | 25.78520353 | 2133 | 2278 | 334 | 26856 | [illegible] |
| GO:BP | positive regulation of signal transduction | GO:0009967 | FALSE | 2.23E-26 | 25.65176876 | 1562 | 2278 | 267 | 26856 | [illegible] |
| GO:BP | negative regulation of protein metabolic process | GO:0051248 | FALSE | 2.36E-26 | 25.62744554 | 1041 | 2278 | 202 | 26856 | [illegible] |
| GO:BP | cell division | GO:0051301 | FALSE | 2.69E-26 | 25.56995579 | 605 | 2278 | 142 | 26856 | [illegible] |
| GO:BP | positive regulation of molecular function | GO:0044093 | FALSE | 2.86E-26 | 25.54381938 | 1498 | 2278 | 259 | 26856 | [illegible] |
| GO:BP | cytoskeleton organization | GO:0007010 | FALSE | 2.92E-26 | 25.53487081 | 1416 | 2278 | 249 | 26856 | [illegible] |
| GO:BP | positive regulation of protein metabolic process | GO:0051247 | FALSE | 3.19E-26 | 25.49552954 | 1425 | 2278 | 250 | 26856 | [illegible] |
| GO:BP | mitotic cell cycle | GO:0000278 | FALSE | 3.41E-26 | 25.46777483 | 870 | 2278 | 179 | 26856 | [illegible] |
| GO:BP | positive regulation of immune response | GO:0050778 | FALSE | 3.55E-26 | 25.45024255 | 725 | 2278 | 159 | 26856 | [illegible] |
| GO:BP | regulation of catalytic activity | GO:0050790 | FALSE | 4.86E-26 | 25.31333673 | 1772 | 2278 | 291 | 26856 | [illegible] |
| GO:BP | activation of immune response | GO:0002253 | FALSE | 5.51E-26 | 25.25904046 | 502 | 2278 | 126 | 26856 | [illegible] |
| GO:BP | apoptotic process | GO:0006915 | FALSE | 7.19E-26 | 25.14298964 | 2053 | 2278 | 323 | 26856 | [illegible] |
| GO:BP | mitotic cell cycle process | GO:1903047 | FALSE | 8.00E-26 | 25.09702953 | 730 | 2278 | 159 | 26856 | [illegible] |
| GO:BP | regulation of developmental process | GO:0050793 | FALSE | 1.78E-25 | 24.74914677 | 2702 | 2278 | 394 | 26856 | [illegible] |
| GO:BP | response to organic substance | GO:0010033 | FALSE | 2.13E-25 | 24.67204837 | 3551 | 2278 | 484 | 26856 | [illegible] |
| GO:BP | cell surface receptor signaling pathway | GO:0007166 | FALSE | 2.60E-25 | 24.58493856 | 2753 | 2278 | 399 | 26856 | [illegible] |
| GO:BP | regulation of catabolic process | GO:0009894 | FALSE | 3.66E-25 | 24.43680845 | 962 | 2278 | 189 | 26856 | [illegible] |
| GO:BP | positive regulation of cellular component organization | GO:0051130 | FALSE | 4.19E-25 | 24.37828137 | 1245 | 2278 | 225 | 26856 | [illegible] |
| GO:BP | regulation of cell population proliferation | GO:0042127 | FALSE | 5.85E-25 | 24.23259393 | 2344 | 2278 | 353 | 26856 | [illegible] |
| GO:BP | hemopoiesis | GO:0030097 | FALSE | 6.92E-25 | 24.16012608 | 1036 | 2278 | 198 | 26856 | [illegible] |
| GO:BP | protein modification by small protein conjugation | GO:0032446 | FALSE | 7.00E-25 | 24.1546515 | 824 | 2278 | 170 | 26856 | [illegible] |
| GO:BP | pattern recognition receptor signaling pathway | GO:0002221 | FALSE | 9.17E-25 | 24.0374072 | 227 | 2278 | 78 | 26856 | [illegible] |
| GO:BP | regulation of cell cycle process | GO:0010564 | FALSE | 1.63E-24 | 23.78745447 | 713 | 2278 | 154 | 26856 | [illegible] |
| GO:BP | positive regulation of multicellular organismal process | GO:0051240 | FALSE | 3.25E-24 | 23.4886782 | 1799 | 2278 | 289 | 26856 | [illegible] |
| GO:BP | system development | GO:0048731 | FALSE | 4.61E-24 | 23.33591904 | 4076 | 2278 | 533 | 26856 | [illegible] |
| GO:BP | immune response-activating signaling pathway | GO:0002757 | FALSE | 5.24E-24 | 23.28102106 | 427 | 2278 | 111 | 26856 | [illegible] |
| GO:BP | negative regulation of signal transduction | GO:0009968 | FALSE | 7.62E-24 | 23.1178369 | 1303 | 2278 | 229 | 26856 | [illegible] |
| GO:BP | nuclear chromosome segregation | GO:0098813 | FALSE | 2.48E-23 | 22.60469586 | 305 | 2278 | 90 | 26856 | [illegible] |
| GO:BP | establishment of localization in cell | GO:0051649 | FALSE | 2.90E-23 | 22.53767822 | 2203 | 2278 | 332 | 26856 | [illegible] |
| GO:BP | innate immune response-activating signaling pathway | GO:0002758 | FALSE | 3.20E-23 | 22.49501092 | 238 | 2278 | 78 | 26856 | [illegible] |
| GO:BP | intracellular receptor signaling pathway | GO:0030522 | FALSE | 4.44E-23 | 22.35261695 | 337 | 2278 | 95 | 26856 | [illegible] |
| GO:BP | mitotic sister chromatid segregation | GO:0000070 | FALSE | 5.62E-23 | 22.25041219 | 177 | 2278 | 66 | 26856 | [illegible] |
| GO:BP | regulation of phosphorylation | GO:0042325 | FALSE | 7.13E-23 | 22.14672419 | 1177 | 2278 | 211 | 26856 | [illegible] |
| GO:BP | cellular response to cytokine stimulus | GO:0071345 | FALSE | 7.74E-23 | 22.11112502 | 1066 | 2278 | 197 | 26856 | [illegible] |
| GO:BP | regulation of programmed cell death | GO:0043067 | FALSE | 7.89E-23 | 22.10276284 | 1704 | 2278 | 274 | 26856 | [illegible] |
| GO:BP | regulation of cellular component biogenesis | GO:0044087 | FALSE | 9.96E-23 | 22.00163946 | 1013 | 2278 | 190 | 26856 | [illegible] |
| GO:BP | immune response-regulating signaling pathway | GO:0002764 | FALSE | 9.98E-23 | 22.0009328 | 441 | 2278 | 111 | 26856 | [illegible] |
| GO:BP | protein localization to organelle | GO:0033365 | FALSE | 1.05E-22 | 21.97806464 | 990 | 2278 | 187 | 26856 | [illegible] |
| GO:BP | protein ubiquitination | GO:0016567 | FALSE | 1.13E-22 | 21.94628478 | 770 | 2278 | 158 | 26856 | [illegible] |
| GO:BP | positive regulation of catabolic process | GO:0009896 | FALSE | 1.85E-22 | 21.73288925 | 524 | 2278 | 123 | 26856 | [illegible] |
| GO:BP | microtubule cytoskeleton organization | GO:0000226 | FALSE | 4.28E-22 | 21.3683522 | 662 | 2278 | 142 | 26856 | [illegible] |
| GO:BP | regulation of apoptotic process | GO:0042981 | FALSE | 5.48E-22 | 21.26149759 | 1656 | 2278 | 266 | 26856 | [illegible] |
| GO:BP | cell development | GO:0048468 | FALSE | 6.30E-22 | 21.2004629 | 3164 | 2278 | 431 | 26856 | [illegible] |
| GO:BP | regulation of phosphorus metabolic process | GO:0051174 | FALSE | 6.50E-22 | 21.18702029 | 1328 | 2278 | 227 | 26856 | [illegible] |
| GO:BP | negative regulation of signaling | GO:0023057 | FALSE | 6.56E-22 | 21.18320912 | 1445 | 2278 | 241 | 26856 | [illegible] |
| GO:BP | regulation of protein phosphorylation | GO:0001932 | FALSE | 6.56E-22 | 21.18294406 | 1076 | 2278 | 196 | 26856 | [illegible] |
| GO:BP | sister chromatid segregation | GO:0000819 | FALSE | 8.76E-22 | 21.05753153 | 216 | 2278 | 72 | 26856 | [illegible] |
| GO:BP | viral process | GO:0016032 | FALSE | 1.18E-21 | 20.92634415 | 357 | 2278 | 96 | 26856 | [illegible] |
| GO:BP | regulation of phosphate metabolic process | GO:0019220 | FALSE | 1.39E-21 | 20.85780844 | 1327 | 2278 | 226 | 26856 | [illegible] |
| GO:BP | catabolic process | GO:0009056 | FALSE | 1.52E-21 | 20.81847335 | 2537 | 2278 | 363 | 26856 | [illegible] |
| GO:BP | signaling | GO:0023052 | FALSE | 1.90E-21 | 20.72111796 | 7418 | 2278 | 844 | 26856 | [illegible] |
| GO:BP | regulation of cellular response to stress | GO:0080135 | FALSE | 1.93E-21 | 20.71515742 | 650 | 2278 | 139 | 26856 | [illegible] |
| GO:BP | positive regulation of intracellular signal transduction | GO:1902533 | FALSE | 2.73E-21 | 20.56373109 | 985 | 2278 | 183 | 26856 | [illegible] |
| GO:BP | negative regulation of cell communication | GO:0010648 | FALSE | 4.64E-21 | 20.33375175 | 1448 | 2278 | 239 | 26856 | [illegible] |
| GO:BP | regulation of viral process | GO:0050792 | FALSE | 6.47E-21 | 20.18906037 | 185 | 2278 | 65 | 26856 | [illegible] |
| GO:BP | cell communication | GO:0007154 | FALSE | 7.56E-21 | 20.12127082 | 7536 | 2278 | 852 | 26856 | [illegible] |
| GO:BP | regulation of canonical NF-kappaB signal transduction | GO:0043122 | FALSE | 9.00E-21 | 20.04591921 | 218 | 2278 | 71 | 26856 | [illegible] |
| GO:BP | signal transduction | GO:0007165 | FALSE | 9.43E-21 | 20.02559663 | 6862 | 2278 | 789 | 26856 | [illegible] |
| GO:BP | non-membrane-bounded organelle assembly | GO:0140694 | FALSE | 1.33E-20 | 19.87776584 | 387 | 2278 | 99 | 26856 | [illegible] |
| GO:BP | peptidyl-amino acid modification | GO:0018193 | FALSE | 2.12E-20 | 19.67419976 | 1058 | 2278 | 190 | 26856 | [illegible] |
| GO:BP | positive regulation of cellular catabolic process | GO:0031331 | FALSE | 2.46E-20 | 19.60863171 | 333 | 2278 | 90 | 26856 | [illegible] |
| GO:BP | nuclear division | GO:0000280 | FALSE | 3.52E-20 | 19.45396392 | 431 | 2278 | 105 | 26856 | [illegible] |
| GO:BP | positive regulation of signaling | GO:0023056 | FALSE | 1.37E-19 | 18.8641515 | 1940 | 2278 | 291 | 26856 | [illegible] |
| GO:BP | positive regulation of organelle organization | GO:0010638 | FALSE | 1.58E-19 | 18.80029737 | 528 | 2278 | 118 | 26856 | [illegible] |
| GO:BP | anatomical structure morphogenesis | GO:0009653 | FALSE | 1.66E-19 | 18.78102866 | 2773 | 2278 | 381 | 26856 | [illegible] |
| GO:BP | cell differentiation | GO:0030154 | FALSE | 1.80E-19 | 18.74360548 | 4737 | 2278 | 580 | 26856 | [illegible] |
| GO:BP | cellular developmental process | GO:0048869 | FALSE | 1.90E-19 | 18.72160196 | 4738 | 2278 | 580 | 26856 | [illegible] |
| GO:BP | organelle assembly | GO:0070925 | FALSE | 2.07E-19 | 18.6841007 | 950 | 2278 | 174 | 26856 | [illegible] |
| GO:BP | organelle fission | GO:0048285 | FALSE | 2.11E-19 | 18.6765753 | 488 | 2278 | 112 | 26856 | [illegible] |
| GO:BP | positive regulation of cell communication | GO:0010647 | FALSE | 3.02E-19 | 18.51945854 | 1941 | 2278 | 290 | 26856 | [illegible] |
| GO:BP | positive regulation of catalytic activity | GO:0043085 | FALSE | 3.14E-19 | 18.5034296 | 1074 | 2278 | 189 | 26856 | [illegible] |
| GO:BP | mitotic nuclear division | GO:0140014 | FALSE | 3.34E-19 | 18.47657487 | 265 | 2278 | 77 | 26856 | [illegible] |
| GO:BP | regulation of cellular catabolic process | GO:0031329 | FALSE | 3.66E-19 | 18.43687519 | 590 | 2278 | 126 | 26856 | [illegible] |
| GO:BP | regulation of DNA metabolic process | GO:0051052 | FALSE | 8.29E-19 | 18.0817029 | 538 | 2278 | 118 | 26856 | [illegible] |
| GO:BP | regulation of mRNA metabolic process | GO:1903311 | FALSE | 8.93E-19 | 18.04904301 | 293 | 2278 | 81 | 26856 | [illegible] |
| GO:BP | regulation of transferase activity | GO:0051338 | FALSE | 1.86E-18 | 17.73081142 | 819 | 2278 | 155 | 26856 | [illegible] |
| GO:BP | regulation of protein kinase activity | GO:0045859 | FALSE | 1.97E-18 | 17.70561615 | 601 | 2278 | 126 | 26856 | [illegible] |
| GO:BP | establishment of localization | GO:0051234 | FALSE | 3.27E-18 | 17.48486105 | 4723 | 2278 | 573 | 26856 | [illegible] |
| GO:BP | regulation of multicellular organismal development | GO:2000026 | FALSE | 9.66E-18 | 17.01501701 | 1540 | 2278 | 240 | 26856 | [illegible] |
| GO:BP | regulation of cell development | GO:0060284 | FALSE | 1.31E-17 | 16.88176409 | 930 | 2278 | 167 | 26856 | [illegible] |
| GO:BP | microtubule-based process | GO:0007017 | FALSE | 1.82E-17 | 16.74009393 | 957 | 2278 | 170 | 26856 | [illegible] |
| GO:BP | positive regulation of protein modification process | GO:0031401 | FALSE | 1.83E-17 | 16.737463 | 949 | 2278 | 169 | 26856 | [illegible] |
| GO:BP | cytosolic pattern recognition receptor signaling pathway | GO:0002753 | FALSE | 1.94E-17 | 16.7117179 | 146 | 2278 | 53 | 26856 | [illegible] |
| GO:BP | leukocyte differentiation | GO:0002521 | FALSE | 2.01E-17 | 16.69688297 | 669 | 2278 | 133 | 26856 | [illegible] |
| GO:BP | cell activation | GO:0001775 | FALSE | 2.76E-17 | 16.55944249 | 1133 | 2278 | 191 | 26856 | [illegible] |
| GO:BP | RNA catabolic process | GO:0006401 | FALSE | 3.28E-17 | 16.48458923 | 290 | 2278 | 78 | 26856 | [illegible] |
| GO:BP | negative regulation of gene expression | GO:0010629 | FALSE | 3.77E-17 | 16.42315184 | 1486 | 2278 | 232 | 26856 | [illegible] |
| GO:BP | cell migration | GO:0016477 | FALSE | 5.33E-17 | 16.27310905 | 1525 | 2278 | 236 | 26856 | [illegible] |
| GO:BP | regulation of cell differentiation | GO:0045595 | FALSE | 6.50E-17 | 16.18718101 | 1749 | 2278 | 261 | 26856 | [illegible] |
| GO:BP | intracellular transport | GO:0046907 | FALSE | 1.01E-16 | 15.99474885 | 1559 | 2278 | 239 | 26856 | [illegible] |
| GO:BP | regulation of kinase activity | GO:0043549 | FALSE | 1.07E-16 | 15.97181031 | 689 | 2278 | 134 | 26856 | [illegible] |
| GO:BP | cellular response to biotic stimulus | GO:0071216 | FALSE | 1.54E-16 | 15.81265168 | 355 | 2278 | 87 | 26856 | [illegible] |
| GO:BP | mRNA catabolic process | GO:0006402 | FALSE | 1.68E-16 | 15.77403367 | 242 | 2278 | 69 | 26856 | [illegible] |
| GO:BP | regulation of cellular localization | GO:0060341 | FALSE | 2.03E-16 | 15.69162305 | 1078 | 2278 | 182 | 26856 | [illegible] |
| GO:BP | regulation of viral life cycle | GO:1903900 | FALSE | 2.08E-16 | 15.68263047 | 158 | 2278 | 54 | 26856 | [illegible] |
| GO:BP | regulation of localization | GO:0032879 | FALSE | 2.48E-16 | 15.6048978 | 2290 | 2278 | 318 | 26856 | [illegible] |
| GO:BP | negative regulation of viral process | GO:0048525 | FALSE | 4.09E-16 | 15.38786489 | 105 | 2278 | 43 | 26856 | [illegible] |
| GO:BP | positive regulation of cell population proliferation | GO:0008284 | FALSE | 4.26E-16 | 15.37024337 | 1427 | 2278 | 222 | 26856 | [illegible] |
| GO:BP | regulation of chromosome organization | GO:0033044 | FALSE | 5.80E-16 | 15.23642524 | 241 | 2278 | 68 | 26856 | [illegible] |
| GO:BP | modification-dependent macromolecule catabolic process | GO:0043632 | FALSE | 6.82E-16 | 15.16648299 | 665 | 2278 | 129 | 26856 | [illegible] |
| GO:BP | protein catabolic process | GO:0030163 | FALSE | 8.69E-16 | 15.06088625 | 1026 | 2278 | 174 | 26856 | [illegible] |
| GO:BP | nervous system development | GO:0007399 | FALSE | 8.77E-16 | 15.0571956 | 2517 | 2278 | 340 | 26856 | [illegible] |
| GO:BP | cellular response to molecule of bacterial origin | GO:0071219 | FALSE | 1.20E-15 | 14.92009389 | 326 | 2278 | 81 | 26856 | [illegible] |
| GO:BP | leukocyte activation | GO:0045321 | FALSE | 1.98E-15 | 14.70304054 | 1001 | 2278 | 170 | 26856 | [illegible] |
| GO:BP | regulation of amide metabolic process | GO:0034248 | FALSE | 2.52E-15 | 14.59824981 | 460 | 2278 | 100 | 26856 | [illegible] |
| GO:BP | DNA repair | GO:0006281 | FALSE | 3.27E-15 | 14.48577628 | 571 | 2278 | 115 | 26856 | [illegible] |
| GO:BP | cytokine-mediated signaling pathway | GO:0019221 | FALSE | 4.06E-15 | 14.39094407 | 463 | 2278 | 100 | 26856 | [illegible] |
| GO:BP | modification-dependent protein catabolic process | GO:0019941 | FALSE | 4.16E-15 | 14.38057831 | 656 | 2278 | 126 | 26856 | [illegible] |
| GO:BP | apoptotic signaling pathway | GO:0097190 | FALSE | 4.33E-15 | 14.36329989 | 641 | 2278 | 124 | 26856 | [illegible] |
| GO:BP | establishment of chromosome localization | GO:0051303 | FALSE | 4.69E-15 | 14.32893667 | 106 | 2278 | 42 | 26856 | [illegible] |
| GO:BP | positive regulation of programmed cell death | GO:0043068 | FALSE | 5.36E-15 | 14.27081513 | 635 | 2278 | 123 | 26856 | [illegible] |
| GO:BP | proteolysis involved in protein catabolic process | GO:0051603 | FALSE | 5.94E-15 | 14.22594333 | 776 | 2278 | 141 | 26856 | [illegible] |
| GO:BP | ubiquitin-dependent protein catabolic process | GO:0006511 | FALSE | 6.39E-15 | 14.19439172 | 644 | 2278 | 124 | 26856 | [illegible] |
| GO:BP | growth | GO:0040007 | FALSE | 6.62E-15 | 14.1794665 | 1038 | 2278 | 173 | 26856 | [illegible] |
| GO:BP | chromosome localization | GO:0050000 | FALSE | 7.59E-15 | 14.11983788 | 112 | 2278 | 43 | 26856 | [illegible] |
| GO:BP | negative regulation of translation | GO:0017148 | FALSE | 7.93E-15 | 14.10060994 | 198 | 2278 | 59 | 26856 | [illegible] |
| GO:BP | myeloid cell differentiation | GO:0030099 | FALSE | 9.14E-15 | 14.03904377 | 461 | 2278 | 99 | 26856 | [illegible] |
| GO:BP | regulation of translation | GO:0006417 | FALSE | 9.81E-15 | 14.00831116 | 398 | 2278 | 90 | 26856 | [illegible] |
| GO:BP | regulation of DNA-binding transcription factor activity | GO:0051090 | FALSE | 1.17E-14 | 13.93315467 | 427 | 2278 | 94 | 26856 | [illegible] |
| GO:BP | regulation of protein localization | GO:0032880 | FALSE | 1.45E-14 | 13.83756745 | 971 | 2278 | 164 | 26856 | [illegible] |
| GO:BP | negative regulation of cellular component organization | GO:0051129 | FALSE | 1.84E-14 | 13.73622176 | 754 | 2278 | 137 | 26856 | [illegible] |
| GO:BP | metaphase chromosome alignment | GO:0051310 | FALSE | 2.32E-14 | 13.6341279 | 91 | 2278 | 38 | 26856 | [illegible] |
| GO:BP | cell motility | GO:0048870 | FALSE | 2.64E-14 | 13.57920105 | 1720 | 2278 | 249 | 26856 | [illegible] |
| GO:BP | cellular response to lipid | GO:0071396 | FALSE | 2.93E-14 | 13.53300022 | 774 | 2278 | 139 | 26856 | [illegible] |
| GO:BP | response to type II interferon | GO:0034341 | TRUE | 3.51E-14 | 13.45431514 | 158 | 2278 | 51 | 26856 | [illegible] |
| GO:BP | negative regulation of immune system process | GO:0002683 | FALSE | 3.52E-14 | 13.45363369 | 521 | 2278 | 106 | 26856 | [illegible] |
| GO:BP | tumor necrosis factor superfamily cytokine production | GO:0071706 | FALSE | 4.40E-14 | 13.35652683 | 187 | 2278 | 56 | 26856 | [illegible] |
| GO:BP | regulation of tumor necrosis factor superfamily cytokine production | GO:1903555 | FALSE | 4.40E-14 | 13.35652683 | 187 | 2278 | 56 | 26856 | [illegible] |
| GO:BP | positive regulation of apoptotic process | GO:0043065 | FALSE | 4.99E-14 | 13.30158157 | 614 | 2278 | 118 | 26856 | [illegible] |
| GO:BP | organic substance catabolic process | GO:1901575 | FALSE | 5.28E-14 | 13.27734658 | 2156 | 2278 | 295 | 26856 | [illegible] |
| GO:BP | establishment of protein localization | GO:0045184 | FALSE | 5.48E-14 | 13.26094589 | 1721 | 2278 | 248 | 26856 | [illegible] |
| GO:BP | regulation of cell migration | GO:0030334 | FALSE | 6.03E-14 | 13.21943879 | 977 | 2278 | 163 | 26856 | [illegible] |
| GO:BP | mononuclear cell differentiation | GO:1903131 | FALSE | 6.04E-14 | 13.21882447 | 510 | 2278 | 104 | 26856 | [illegible] |
| GO:BP | neurogenesis | GO:0022008 | FALSE | 7.75E-14 | 13.11042778 | 1882 | 2278 | 265 | 26856 | [illegible] |
| GO:BP | regulation of tumor necrosis factor production | GO:0032680 | FALSE | 9.32E-14 | 13.03068533 | 184 | 2278 | 55 | 26856 | [illegible] |
| GO:BP | tumor necrosis factor production | GO:0032640 | FALSE | 9.32E-14 | 13.03068533 | 184 | 2278 | 55 | 26856 | [illegible] |
| GO:BP | regulation of RNA stability | GO:0043487 | FALSE | 1.06E-13 | 12.97309232 | 173 | 2278 | 53 | 26856 | [illegible] |
| GO:BP | regulation of mRNA catabolic process | GO:0061013 | FALSE | 1.15E-13 | 12.93997117 | 179 | 2278 | 54 | 26856 | [illegible] |
| GO:BP | positive regulation of mRNA catabolic process | GO:0061014 | FALSE | 1.33E-13 | 12.87504081 | 95 | 2278 | 38 | 26856 | [illegible] |
| GO:BP | protein localization to nucleus | GO:0034504 | FALSE | 1.43E-13 | 12.84572481 | 331 | 2278 | 78 | 26856 | [illegible] |
| GO:BP | cell projection organization | GO:0030030 | FALSE | 1.46E-13 | 12.83643017 | 1689 | 2278 | 243 | 26856 | [illegible] |
| GO:BP | regulation of hemopoiesis | GO:1903706 | FALSE | 1.59E-13 | 12.79824714 | 422 | 2278 | 91 | 26856 | [illegible] |
| GO:BP | positive regulation of cell differentiation | GO:0045597 | FALSE | 1.61E-13 | 12.79221118 | 962 | 2278 | 160 | 26856 | [illegible] |
| GO:BP | negative regulation of amide metabolic process | GO:0034249 | FALSE | 1.64E-13 | 12.78446389 | 222 | 2278 | 61 | 26856 | [illegible] |
| GO:BP | MAPK cascade | GO:0000165 | FALSE | 1.70E-13 | 12.77001203 | 806 | 2278 | 141 | 26856 | [illegible] |
| GO:BP | regulation of mRNA stability | GO:0043488 | FALSE | 1.83E-13 | 12.73650982 | 158 | 2278 | 50 | 26856 | [illegible] |
| GO:BP | cellular response to lipopolysaccharide | GO:0071222 | FALSE | 1.87E-13 | 12.72876391 | 319 | 2278 | 76 | 26856 | [illegible] |
| GO:BP | spindle organization | GO:0007051 | FALSE | 2.07E-13 | 12.68467689 | 187 | 2278 | 55 | 26856 | [illegible] |
| GO:BP | regulation of type I interferon production | GO:0032479 | FALSE | 2.19E-13 | 12.65951339 | 121 | 2278 | 43 | 26856 | [illegible] |
| GO:BP | type I interferon production | GO:0032606 | FALSE | 2.19E-13 | 12.65951339 | 121 | 2278 | 43 | 26856 | [illegible] |
| GO:BP | response to bacterium | GO:0009617 | FALSE | 2.55E-13 | 12.59262491 | 950 | 2278 | 158 | 26856 | [illegible] |
| GO:BP | response to abiotic stimulus | GO:0009628 | FALSE | 2.77E-13 | 12.55698632 | 1303 | 2278 | 199 | 26856 | [illegible] |
| GO:BP | protein polyubiquitination | GO:0000209 | FALSE | 2.80E-13 | 12.55315357 | 262 | 2278 | 67 | 26856 | [illegible] |
| GO:BP | regulation of pattern recognition receptor signaling pathway | GO:0062207 | FALSE | 2.89E-13 | 12.53873489 | 143 | 2278 | 47 | 26856 | [illegible] |
| GO:BP | regulation of response to cytokine stimulus | GO:0060759 | FALSE | 2.91E-13 | 12.53631155 | 154 | 2278 | 49 | 26856 | [illegible] |
| GO:BP | multicellular organismal-level homeostasis | GO:0048871 | FALSE | 3.19E-13 | 12.49684191 | 894 | 2278 | 151 | 26856 | [illegible] |
| GO:BP | response to molecule of bacterial origin | GO:0002237 | FALSE | 3.19E-13 | 12.49598549 | 470 | 2278 | 97 | 26856 | [illegible] |
| GO:BP | centrosome cycle | GO:0007098 | FALSE | 3.29E-13 | 12.48343166 | 138 | 2278 | 46 | 26856 | [illegible] |
| GO:BP | establishment of organelle localization | GO:0051656 | FALSE | 3.35E-13 | 12.47557647 | 463 | 2278 | 96 | 26856 | [illegible] |
| GO:BP | animal organ development | GO:0048513 | FALSE | 3.68E-13 | 12.43393443 | 3278 | 2278 | 407 | 26856 | [illegible] |
| GO:BP | organelle localization | GO:0051640 | FALSE | 4.20E-13 | 12.37688249 | 600 | 2278 | 114 | 26856 | [illegible] |
| GO:BP | negative regulation of protein modification process | GO:0031400 | FALSE | 4.27E-13 | 12.36965113 | 472 | 2278 | 97 | 26856 | [illegible] |
| GO:BP | regulation of locomotion | GO:0040012 | FALSE | 4.94E-13 | 12.30635302 | 1084 | 2278 | 173 | 26856 | [illegible] |
| GO:BP | protein-containing complex organization | GO:0043933 | FALSE | 5.18E-13 | 12.28603492 | 3445 | 2278 | 423 | 26856 | [illegible] |
| GO:BP | positive regulation of cellular component biogenesis | GO:0044089 | FALSE | 5.20E-13 | 12.28372331 | 533 | 2278 | 105 | 26856 | [illegible] |
| GO:BP | cellular response to type II interferon | GO:0071346 | FALSE | 5.65E-13 | 12.24785925 | 129 | 2278 | 44 | 26856 | [illegible] |
| GO:BP | RNA destabilization | GO:0050779 | FALSE | 6.40E-13 | 12.1941421 | 94 | 2278 | 37 | 26856 | [illegible] |
| GO:BP | peptidyl-serine phosphorylation | GO:0018105 | FALSE | 6.47E-13 | 12.18908574 | 319 | 2278 | 75 | 26856 | [illegible] |
| GO:BP | regulation of cell motility | GO:2000145 | FALSE | 6.61E-13 | 12.17955415 | 1036 | 2278 | 167 | 26856 | [illegible] |
| GO:BP | mRNA destabilization | GO:0061157 | FALSE | 8.77E-13 | 12.05690182 | 90 | 2278 | 36 | 26856 | [illegible] |
| GO:BP | positive regulation of developmental process | GO:0051094 | FALSE | 8.82E-13 | 12.05437005 | 1469 | 2278 | 216 | 26856 | [illegible] |
| GO:BP | response to lipid | GO:0033993 | FALSE | 9.12E-13 | 12.03994784 | 1134 | 2278 | 178 | 26856 | [illegible] |
| GO:BP | locomotion | GO:0040011 | FALSE | 1.09E-12 | 11.96331336 | 1267 | 2278 | 193 | 26856 | [illegible] |
| GO:BP | transport | GO:0006810 | FALSE | 1.19E-12 | 11.92575681 | 4547 | 2278 | 529 | 26856 | [illegible] |
| GO:BP | negative regulation of molecular function | GO:0044092 | FALSE | 1.32E-12 | 11.8806885 | 1001 | 2278 | 162 | 26856 | [illegible] |
| GO:BP | positive regulation of canonical NF-kappaB signal transduction | GO:0043123 | FALSE | 1.32E-12 | 11.87940305 | 148 | 2278 | 47 | 26856 | [illegible] |
| GO:BP | nitrogen compound transport | GO:0071705 | FALSE | 1.37E-12 | 11.86216257 | 2104 | 2278 | 284 | 26856 | [illegible] |

| | | | | | | | | | | |
|---|---|---|---|---|---|---|---|---|---|---|
| GO:BP | positive regulation of cell cycle | GO:0045787 | FALSE | 1.43E-12 | 11.84403906 | 344 | 2278 | 78 | 26856 | [illegible] |
| GO:BP | positive regulation of phosphorus metabolic process | GO:0010562 | FALSE | 1.49E-12 | 11.82647032 | 868 | 2278 | 146 | 26856 | [illegible] |
| GO:BP | positive regulation of phosphate metabolic process | GO:0045937 | FALSE | 1.49E-12 | 11.82647032 | 868 | 2278 | 146 | 26856 | [illegible] |
| GO:BP | plasma membrane bounded cell projection organization | GO:0120036 | FALSE | 1.75E-12 | 11.7564241 | 1651 | 2278 | 235 | 26856 | [illegible] |
| GO:BP | inflammatory response | GO:0006954 | FALSE | 2.19E-12 | 11.65974699 | 814 | 2278 | 139 | 26856 | [illegible] |
| GO:BP | spindle assembly | GO:0051225 | FALSE | 2.80E-12 | 11.55231681 | 118 | 2278 | 41 | 26856 | [illegible] |
| GO:BP | viral life cycle | GO:0019058 | FALSE | 2.81E-12 | 11.55087182 | 280 | 2278 | 68 | 26856 | [illegible] |
| GO:BP | negative regulation of intracellular signal transduction | GO:1902532 | FALSE | 2.87E-12 | 11.54186661 | 546 | 2278 | 105 | 26856 | [illegible] |
| GO:BP | negative regulation of phosphorus metabolic process | GO:0010563 | FALSE | 3.23E-12 | 11.49014341 | 420 | 2278 | 88 | 26856 | [illegible] |
| GO:BP | negative regulation of phosphate metabolic process | GO:0045936 | FALSE | 3.23E-12 | 11.49014341 | 420 | 2278 | 88 | 26856 | [illegible] |
| GO:BP | negative regulation of phosphorylation | GO:0042326 | FALSE | 3.34E-12 | 11.47652412 | 363 | 2278 | 80 | 26856 | [illegible] |
| GO:BP | positive regulation of phosphorylation | GO:0042327 | FALSE | 3.60E-12 | 11.44409945 | 786 | 2278 | 135 | 26856 | [illegible] |
| GO:BP | embryo development | GO:0009790 | FALSE | 3.67E-12 | 11.43555335 | 1238 | 2278 | 188 | 26856 | [illegible] |
| GO:BP | negative regulation of cell cycle | GO:0045786 | FALSE | 3.93E-12 | 11.40565104 | 364 | 2278 | 80 | 26856 | [illegible] |
| GO:BP | microtubule organizing center organization | GO:0031023 | FALSE | 4.20E-12 | 11.37645178 | 152 | 2278 | 47 | 26856 | [illegible] |
| GO:BP | viral genome replication | GO:0019079 | FALSE | 4.38E-12 | 11.3584181 | 130 | 2278 | 43 | 26856 | [illegible] |
| GO:BP | negative regulation of immune response | GO:0050777 | FALSE | 5.07E-12 | 11.29508833 | 206 | 2278 | 56 | 26856 | [illegible] |
| GO:BP | positive regulation of DNA metabolic process | GO:0051054 | FALSE | 5.31E-12 | 11.27460182 | 324 | 2278 | 74 | 26856 | [illegible] |
| GO:BP | peptidyl-serine modification | GO:0018209 | FALSE | 5.47E-12 | 11.26189646 | 338 | 2278 | 76 | 26856 | [illegible] |
| GO:BP | protein transport | GO:0015031 | FALSE | 6.20E-12 | 11.20748957 | 1605 | 2278 | 228 | 26856 | [illegible] |
| GO:BP | response to lipopolysaccharide | GO:0032496 | FALSE | 6.37E-12 | 11.19567239 | 454 | 2278 | 92 | 26856 | [illegible] |
| GO:BP | homeostatic process | GO:0042592 | FALSE | 6.59E-12 | 11.18106675 | 1828 | 2278 | 252 | 26856 | [illegible] |
| GO:BP | lymphocyte activation | GO:0046649 | FALSE | 6.66E-12 | 11.17648361 | 825 | 2278 | 139 | 26856 | [illegible] |
| GO:BP | positive regulation of protein phosphorylation | GO:0001934 | FALSE | 7.67E-12 | 11.11497025 | 728 | 2278 | 127 | 26856 | [illegible] |
| GO:BP | response to interferon-beta | GO:0035456 | FALSE | 8.35E-12 | 11.07849027 | 72 | 2278 | 31 | 26856 | [illegible] |
| GO:BP | nucleocytoplasmic transport | GO:0006913 | FALSE | 9.01E-12 | 11.04546269 | 355 | 2278 | 78 | 26856 | [illegible] |
| GO:BP | nuclear transport | GO:0051169 | FALSE | 9.01E-12 | 11.04546269 | 355 | 2278 | 78 | 26856 | [illegible] |
| GO:BP | cellular catabolic process | GO:0044248 | FALSE | 9.52E-12 | 11.0212862 | 1438 | 2278 | 209 | 26856 | [illegible] |
| GO:BP | developmental growth | GO:0048589 | FALSE | 1.05E-11 | 10.97877028 | 780 | 2278 | 133 | 26856 | [illegible] |
| GO:BP | regulation of viral genome replication | GO:0045069 | FALSE | 1.43E-11 | 10.8452003 | 97 | 2278 | 36 | 26856 | [illegible] |
| GO:BP | negative regulation of protein phosphorylation | GO:0001933 | FALSE | 2.07E-11 | 10.68311615 | 332 | 2278 | 74 | 26856 | [illegible] |
| GO:BP | intrinsic apoptotic signaling pathway | GO:0097193 | FALSE | 2.87E-11 | 10.54238606 | 320 | 2278 | 72 | 26856 | [illegible] |
| GO:BP | negative regulation of cell cycle process | GO:0010948 | FALSE | 3.02E-11 | 10.52018126 | 286 | 2278 | 67 | 26856 | [illegible] |
| GO:BP | negative regulation of multicellular organismal process | GO:0051241 | FALSE | 3.14E-11 | 10.50328917 | 1212 | 2278 | 182 | 26856 | [illegible] |
| GO:BP | lipopolysaccharide-mediated signaling pathway | GO:0031663 | FALSE | 3.36E-11 | 10.47320323 | 66 | 2278 | 29 | 26856 | [illegible] |
| GO:BP | regulation of cytokine-mediated signaling pathway | GO:0001959 | FALSE | 4.03E-11 | 10.39472961 | 143 | 2278 | 44 | 26856 | [illegible] |
| GO:BP | response to oxygen-containing compound | GO:1901700 | FALSE | 4.09E-11 | 10.38781649 | 2161 | 2278 | 284 | 26856 | [illegible] |
| GO:BP | attachment of spindle microtubules to kinetochore | GO:0008608 | FALSE | 4.43E-11 | 10.35372501 | 42 | 2278 | 23 | 26856 | [illegible] |
| GO:BP | positive regulation of cell cycle process | GO:0090068 | FALSE | 4.79E-11 | 10.31981234 | 255 | 2278 | 62 | 26856 | [illegible] |
| GO:BP | cellular response to oxygen-containing compound | GO:1901701 | FALSE | 4.95E-11 | 10.3055932 | 1571 | 2278 | 221 | 26856 | [illegible] |
| GO:BP | in utero embryonic development | GO:0001701 | FALSE | 5.16E-11 | 10.28774646 | 515 | 2278 | 98 | 26856 | [illegible] |
| GO:BP | positive regulation of DNA-binding transcription factor activity | GO:0051091 | FALSE | 6.20E-11 | 10.20766561 | 263 | 2278 | 63 | 26856 | [illegible] |
| GO:BP | mitotic spindle organization | GO:0007052 | FALSE | 7.92E-11 | 10.10134482 | 123 | 2278 | 40 | 26856 | [illegible] |
| GO:BP | homeostasis of number of cells | GO:0048872 | FALSE | 8.15E-11 | 10.08889853 | 369 | 2278 | 78 | 26856 | [illegible] |
| GO:BP | positive regulation of mRNA metabolic process | GO:1903313 | FALSE | 8.56E-11 | 10.06766903 | 140 | 2278 | 43 | 26856 | [illegible] |
| GO:BP | cell cycle phase transition | GO:0044770 | FALSE | 9.04E-11 | 10.04400756 | 535 | 2278 | 100 | 26856 | [illegible] |
| GO:BP | lymphocyte differentiation | GO:0030098 | FALSE | 9.25E-11 | 10.03400521 | 451 | 2278 | 89 | 26856 | [illegible] |
| GO:BP | regulation of cytoskeleton organization | GO:0051493 | FALSE | 9.62E-11 | 10.016756 | 520 | 2278 | 98 | 26856 | [illegible] |
| GO:BP | cellular response to interferon-beta | GO:0035458 | FALSE | 1.09E-10 | 9.960793886 | 64 | 2278 | 28 | 26856 | [illegible] |
| GO:BP | positive regulation of cell development | GO:0010720 | FALSE | 1.12E-10 | 9.950191545 | 529 | 2278 | 99 | 26856 | [illegible] |
| GO:BP | positive regulation of type I interferon production | GO:0032481 | FALSE | 1.20E-10 | 9.919629802 | 78 | 2278 | 31 | 26856 | [illegible] |
| GO:BP | microtubule cytoskeleton organization involved in mitosis | GO:1902850 | FALSE | 1.31E-10 | 9.881925605 | 159 | 2278 | 46 | 26856 | [illegible] |
| GO:BP | nuclear-transcribed mRNA catabolic process | GO:0000956 | FALSE | 1.69E-10 | 9.772418591 | 120 | 2278 | 39 | 26856 | [illegible] |
| GO:BP | heterocycle catabolic process | GO:0046700 | FALSE | 1.70E-10 | 9.77013729 | 448 | 2278 | 88 | 26856 | [illegible] |
| GO:BP | regulation of leukocyte activation | GO:0002694 | FALSE | 2.02E-10 | 9.694650921 | 597 | 2278 | 107 | 26856 | [illegible] |
| GO:BP | positive regulation of tumor necrosis factor production | GO:0032760 | FALSE | 2.19E-10 | 9.660460829 | 110 | 2278 | 37 | 26856 | [illegible] |
| GO:BP | organonitrogen compound catabolic process | GO:1901565 | FALSE | 2.60E-10 | 9.585179437 | 1348 | 2278 | 194 | 26856 | [illegible] |
| GO:BP | regulation of viral-induced cytoplasmic pattern recognition receptor signaling pathway | GO:0039531 | FALSE | 2.78E-10 | 9.556336003 | 95 | 2278 | 34 | 26856 | [illegible] |
| GO:BP | regulation of leukocyte differentiation | GO:1902105 | FALSE | 3.30E-10 | 9.481343693 | 335 | 2278 | 72 | 26856 | [illegible] |
| GO:BP | positive regulation of autophagy | GO:0010508 | FALSE | 3.56E-10 | 9.449069206 | 157 | 2278 | 45 | 26856 | [illegible] |
| GO:BP | negative regulation of viral genome replication | GO:0045071 | FALSE | 3.56E-10 | 9.448707951 | 62 | 2278 | 27 | 26856 | [illegible] |
| GO:BP | nucleobase-containing compound catabolic process | GO:0034655 | FALSE | 3.59E-10 | 9.444551785 | 416 | 2278 | 83 | 26856 | [illegible] |
| GO:BP | autophagy | GO:0006914 | FALSE | 3.93E-10 | 9.405305975 | 485 | 2278 | 92 | 26856 | [illegible] |
| GO:BP | process utilizing autophagic mechanism | GO:0061919 | FALSE | 3.93E-10 | 9.405305975 | 485 | 2278 | 92 | 26856 | [illegible] |
| GO:BP | positive regulation of tumor necrosis factor superfamily cytokine production | GO:1903557 | FALSE | 4.13E-10 | 9.383715064 | 112 | 2278 | 37 | 26856 | [illegible] |
| GO:BP | epigenetic regulation of gene expression | GO:0040029 | FALSE | 4.36E-10 | 9.360868423 | 240 | 2278 | 58 | 26856 | [illegible] |
| GO:BP | response to ionizing radiation | GO:0010212 | FALSE | 4.57E-10 | 9.340329392 | 158 | 2278 | 45 | 26856 | [illegible] |
| GO:BP | regulation of RNA splicing | GO:0043484 | FALSE | 4.99E-10 | 9.301576565 | 189 | 2278 | 50 | 26856 | [illegible] |
| GO:BP | positive regulation of smooth muscle cell proliferation | GO:0048661 | FALSE | 5.65E-10 | 9.248198397 | 113 | 2278 | 37 | 26856 | [illegible] |
| GO:BP | cellular nitrogen compound catabolic process | GO:0044270 | FALSE | 5.98E-10 | 9.223061361 | 450 | 2278 | 87 | 26856 | [illegible] |
| GO:BP | regulation of transport | GO:0051049 | FALSE | 7.36E-10 | 9.133172673 | 1924 | 2278 | 254 | 26856 | [illegible] |
| GO:BP | negative regulation of chromosome organization | GO:2001251 | FALSE | 8.41E-10 | 9.074994759 | 93 | 2278 | 33 | 26856 | [illegible] |
| GO:BP | sister chromatid cohesion | GO:0007062 | FALSE | 8.54E-10 | 9.068769141 | 55 | 2278 | 25 | 26856 | [illegible] |
| GO:BP | regulation of cell activation | GO:0050865 | FALSE | 8.83E-10 | 9.054217021 | 651 | 2278 | 112 | 26856 | [illegible] |
| GO:BP | head development | GO:0060322 | FALSE | 1.20E-09 | 8.922288285 | 787 | 2278 | 128 | 26856 | [illegible] |
| GO:BP | peptidyl-tyrosine phosphorylation | GO:0018108 | FALSE | 1.26E-09 | 8.900082095 | 315 | 2278 | 68 | 26856 | [illegible] |
| GO:BP | proteolysis | GO:0006508 | FALSE | 1.30E-09 | 8.885107428 | 1829 | 2278 | 243 | 26856 | [illegible] |
| GO:BP | regulation of organelle assembly | GO:1902115 | FALSE | 1.34E-09 | 8.871340271 | 226 | 2278 | 55 | 26856 | [illegible] |
| GO:BP | cell cycle checkpoint signaling | GO:0000075 | FALSE | 1.36E-09 | 8.865747029 | 181 | 2278 | 48 | 26856 | [illegible] |
| GO:BP | positive regulation of response to cytokine stimulus | GO:0060760 | FALSE | 1.43E-09 | 8.844113013 | 65 | 2278 | 27 | 26856 | [illegible] |
| GO:BP | positive regulation of hydrolase activity | GO:0051345 | FALSE | 1.48E-09 | 8.831166698 | 543 | 2278 | 98 | 26856 | [illegible] |
| GO:BP | positive regulation of NF-kappaB transcription factor activity | GO:0051092 | FALSE | 1.57E-09 | 8.804900139 | 151 | 2278 | 43 | 26856 | [illegible] |
| GO:BP | regulation of MAPK cascade | GO:0043408 | FALSE | 1.61E-09 | 8.794414998 | 723 | 2278 | 120 | 26856 | [illegible] |
| GO:BP | regulation of post-translational protein modification | GO:1901873 | FALSE | 1.63E-09 | 8.78760847 | 331 | 2278 | 70 | 26856 | [illegible] |
| GO:BP | peptidyl-tyrosine modification | GO:0018212 | FALSE | 1.72E-09 | 8.763828302 | 317 | 2278 | 68 | 26856 | [illegible] |
| GO:BP | DNA recombination | GO:0006310 | FALSE | 1.96E-09 | 8.708022574 | 325 | 2278 | 69 | 26856 | [illegible] |
| GO:BP | regulation of microtubule cytoskeleton organization | GO:0070507 | FALSE | 1.99E-09 | 8.700655453 | 158 | 2278 | 44 | 26856 | [illegible] |
| GO:BP | T cell differentiation | GO:0030217 | FALSE | 2.06E-09 | 8.686527631 | 311 | 2278 | 67 | 26856 | [illegible] |
| GO:BP | proteasomal protein catabolic process | GO:0010498 | FALSE | 2.12E-09 | 8.673951152 | 491 | 2278 | 91 | 26856 | [illegible] |
| GO:BP | regulation of cell cycle phase transition | GO:1901987 | FALSE | 2.51E-09 | 8.600627425 | 423 | 2278 | 82 | 26856 | [illegible] |
| GO:BP | regulation of cell adhesion | GO:0030155 | FALSE | 2.60E-09 | 8.584992605 | 787 | 2278 | 127 | 26856 | [illegible] |
| GO:BP | histone modification | GO:0016570 | FALSE | 2.97E-09 | 8.527267278 | 278 | 2278 | 62 | 26856 | [illegible] |
| GO:BP | regulation of apoptotic signaling pathway | GO:2001233 | FALSE | 3.09E-09 | 8.510304273 | 417 | 2278 | 81 | 26856 | [illegible] |
| GO:BP | cellular component biogenesis | GO:0044085 | FALSE | 3.21E-09 | 8.493056925 | 4347 | 2278 | 491 | 26856 | [illegible] |
| GO:BP | tissue development | GO:0009888 | FALSE | 3.29E-09 | 8.48280146 | 2105 | 2278 | 270 | 26856 | [illegible] |
| GO:BP | extrinsic apoptotic signaling pathway | GO:0097191 | FALSE | 3.47E-09 | 8.459332432 | 231 | 2278 | 55 | 26856 | [illegible] |
| GO:BP | regulation of hydrolase activity | GO:0051336 | FALSE | 3.53E-09 | 8.451735245 | 963 | 2278 | 147 | 26856 | [illegible] |
| GO:BP | small GTPase mediated signal transduction | GO:0007264 | FALSE | 4.01E-09 | 8.397304021 | 419 | 2278 | 81 | 26856 | [illegible] |
| GO:BP | negative regulation of response to biotic stimulus | GO:0002832 | FALSE | 4.05E-09 | 8.39211459 | 161 | 2278 | 44 | 26856 | [illegible] |
| GO:BP | regulation of microtubule-based process | GO:0032886 | FALSE | 4.09E-09 | 8.3879556 | 266 | 2278 | 60 | 26856 | [illegible] |
| GO:BP | regulation of mitotic cell cycle | GO:0007346 | FALSE | 4.24E-09 | 8.373091059 | 489 | 2278 | 90 | 26856 | [illegible] |
| GO:BP | negative regulation of response to external stimulus | GO:0032102 | FALSE | 4.32E-09 | 8.364217387 | 458 | 2278 | 86 | 26856 | [illegible] |
| GO:BP | negative regulation of organelle organization | GO:0010639 | FALSE | 5.01E-09 | 8.300392782 | 368 | 2278 | 74 | 26856 | [illegible] |
| GO:BP | regulation of growth | GO:0040008 | FALSE | 5.06E-09 | 8.296013284 | 660 | 2278 | 111 | 26856 | [illegible] |
| GO:BP | myeloid cell homeostasis | GO:0002262 | FALSE | 5.11E-09 | 8.29127634 | 200 | 2278 | 50 | 26856 | [illegible] |
| GO:BP | central nervous system development | GO:0007417 | FALSE | 6.15E-09 | 8.211297881 | 1014 | 2278 | 152 | 26856 | [illegible] |
| GO:BP | negative regulation of defense response | GO:0031348 | FALSE | 6.42E-09 | 8.192444106 | 304 | 2278 | 65 | 26856 | [illegible] |
| GO:BP | multicellular organismal process | GO:0032501 | FALSE | 6.54E-09 | 8.184712883 | 9064 | 2278 | 920 | 26856 | [illegible] |
| GO:BP | leukocyte cell-cell adhesion | GO:0007159 | FALSE | 6.56E-09 | 8.183383027 | 400 | 2278 | 78 | 26856 | [illegible] |
| GO:BP | negative regulation of cell population proliferation | GO:0008285 | FALSE | 7.52E-09 | 8.123967875 | 885 | 2278 | 137 | 26856 | [illegible] |
| GO:BP | regulation of lymphocyte activation | GO:0051249 | FALSE | 7.60E-09 | 8.118962043 | 494 | 2278 | 90 | 26856 | [illegible] |
| GO:BP | aromatic compound catabolic process | GO:0019439 | FALSE | 8.12E-09 | 8.090457284 | 471 | 2278 | 87 | 26856 | [illegible] |
| GO:BP | non-canonical NF-kappaB signal transduction | GO:0038061 | FALSE | 8.30E-09 | 8.080697299 | 128 | 2278 | 38 | 26856 | [illegible] |
| GO:BP | regulation of smooth muscle cell proliferation | GO:0048660 | FALSE | 9.32E-09 | 8.030393631 | 190 | 2278 | 48 | 26856 | [illegible] |
| GO:BP | circulatory system development | GO:0072359 | FALSE | 1.07E-08 | 7.970966646 | 1210 | 2278 | 173 | 26856 | [illegible] |
| GO:BP | myeloid leukocyte differentiation | GO:0002573 | FALSE | 1.11E-08 | 7.95290434 | 265 | 2278 | 59 | 26856 | [illegible] |
| GO:BP | regulation of immune effector process | GO:0002697 | FALSE | 1.15E-08 | 7.93949912 | 435 | 2278 | 82 | 26856 | [illegible] |
| GO:BP | interferon-mediated signaling pathway | GO:0140888 | FALSE | 1.21E-08 | 7.91834529 | 80 | 2278 | 29 | 26856 | [illegible] |
| GO:BP | response to peptide | GO:1901652 | FALSE | 1.34E-08 | 7.872172549 | 555 | 2278 | 97 | 26856 | [illegible] |
| GO:BP | T cell activation | GO:0042110 | FALSE | 1.42E-08 | 7.848951401 | 588 | 2278 | 101 | 26856 | [illegible] |
| GO:BP | negative regulation of cell cycle phase transition | GO:1901988 | FALSE | 1.51E-08 | 7.819923723 | 246 | 2278 | 56 | 26856 | [illegible] |
| GO:BP | negative regulation of gene expression, epigenetic | GO:0045814 | FALSE | 1.52E-08 | 7.817279633 | 113 | 2278 | 35 | 26856 | [illegible] |
| GO:BP | regulation of interferon-beta production | GO:0032648 | FALSE | 1.65E-08 | 7.78153828 | 66 | 2278 | 26 | 26856 | [illegible] |
| GO:BP | toll-like receptor signaling pathway | GO:0002224 | FALSE | 1.65E-08 | 7.78153828 | 66 | 2278 | 26 | 26856 | [illegible] |
| GO:BP | interferon-beta production | GO:0032608 | FALSE | 1.65E-08 | 7.78153828 | 66 | 2278 | 26 | 26856 | [illegible] |
| GO:BP | negative regulation of apoptotic process | GO:0043066 | FALSE | 1.68E-08 | 7.774102867 | 1045 | 2278 | 154 | 26856 | [illegible] |
| GO:BP | immune effector process | GO:0002252 | FALSE | 1.79E-08 | 7.747742149 | 673 | 2278 | 111 | 26856 | [illegible] |
| GO:BP | regulation of protein-containing complex assembly | GO:0043254 | FALSE | 1.88E-08 | 7.72663532 | 439 | 2278 | 82 | 26856 | [illegible] |
| GO:BP | positive regulation of transferase activity | GO:0051347 | FALSE | 1.90E-08 | 7.721335421 | 526 | 2278 | 93 | 26856 | [illegible] |
| GO:BP | response to endogenous stimulus | GO:0009719 | FALSE | 1.90E-08 | 7.720161606 | 1944 | 2278 | 250 | 26856 | [illegible] |
| GO:BP | negative regulation of programmed cell death | GO:0043069 | FALSE | 1.97E-08 | 7.70531614 | 1074 | 2278 | 157 | 26856 | [illegible] |
| GO:BP | organic cyclic compound catabolic process | GO:1901361 | FALSE | 2.11E-08 | 7.675914858 | 495 | 2278 | 89 | 26856 | [illegible] |
| GO:BP | myeloid leukocyte activation | GO:0002274 | FALSE | 2.15E-08 | 7.666731113 | 255 | 2278 | 57 | 26856 | [illegible] |
| GO:BP | double-strand break repair | GO:0006302 | FALSE | 2.39E-08 | 7.621753326 | 291 | 2278 | 62 | 26856 | [illegible] |
| GO:BP | positive regulation of protein kinase activity | GO:0045860 | FALSE | 2.39E-08 | 7.621483135 | 372 | 2278 | 73 | 26856 | [illegible] |
| GO:BP | organic substance transport | GO:0071702 | FALSE | 2.40E-08 | 7.620271879 | 2860 | 2278 | 342 | 26856 | [illegible] |
| GO:BP | positive regulation of interleukin-6 production | GO:0032755 | FALSE | 2.43E-08 | 7.613597771 | 109 | 2278 | 34 | 26856 | [illegible] |
| GO:BP | positive regulation of transport | GO:0051050 | FALSE | 2.45E-08 | 7.61069656 | 1068 | 2278 | 156 | 26856 | [illegible] |
| GO:BP | anatomical structure formation involved in morphogenesis | GO:0048646 | FALSE | 2.47E-08 | 7.608032405 | 1213 | 2278 | 172 | 26856 | [illegible] |
| GO:BP | positive regulation of protein localization | GO:1903829 | FALSE | 2.65E-08 | 7.576821548 | 513 | 2278 | 91 | 26856 | [illegible] |
| GO:BP | regulation of binding | GO:0051098 | FALSE | 2.69E-08 | 7.570298533 | 411 | 2278 | 78 | 26856 | [illegible] |
| GO:BP | regulation of neurogenesis | GO:0050767 | FALSE | 2.86E-08 | 7.544379626 | 466 | 2278 | 85 | 26856 | [illegible] |
| GO:BP | immune system development | GO:0002520 | FALSE | 3.12E-08 | 7.506171397 | 216 | 2278 | 51 | 26856 | [illegible] |
| GO:BP | mitotic cell cycle phase transition | GO:0044772 | FALSE | 3.77E-08 | 7.423698759 | 437 | 2278 | 81 | 26856 | [illegible] |
| GO:BP | negative regulation of innate immune response | GO:0045824 | FALSE | 3.88E-08 | 7.410975147 | 105 | 2278 | 33 | 26856 | [illegible] |
| GO:BP | response to hormone | GO:0009725 | FALSE | 3.94E-08 | 7.404685052 | 985 | 2278 | 146 | 26856 | [illegible] |
| GO:BP | negative regulation of protein kinase activity | GO:0006469 | FALSE | 4.57E-08 | 7.340288081 | 198 | 2278 | 48 | 26856 | [illegible] |
| GO:BP | smooth muscle cell proliferation | GO:0048659 | FALSE | 4.57E-08 | 7.340288081 | 198 | 2278 | 48 | 26856 | [illegible] |
| GO:BP | brain development | GO:0007420 | FALSE | 4.61E-08 | 7.335970484 | 734 | 2278 | 117 | 26856 | [illegible] |
| GO:BP | cytokinesis | GO:0000910 | FALSE | 4.62E-08 | 7.335381203 | 172 | 2278 | 44 | 26856 | [illegible] |
| GO:BP | circadian rhythm | GO:0007623 | FALSE | 4.87E-08 | 7.312487315 | 239 | 2278 | 54 | 26856 | [illegible] |
| GO:BP | signal transduction in response to DNA damage | GO:0042770 | FALSE | 5.19E-08 | 7.28485605 | 179 | 2278 | 45 | 26856 | [illegible] |
| GO:BP | heterochromatin organization | GO:0070828 | FALSE | 6.17E-08 | 7.209529434 | 101 | 2278 | 32 | 26856 | [illegible] |
| GO:BP | embryo development ending in birth or egg hatching | GO:0009792 | FALSE | 7.19E-08 | 7.143138008 | 808 | 2278 | 125 | 26856 | [illegible] |
| GO:BP | regulation of peptidyl-tyrosine phosphorylation | GO:0050730 | FALSE | 7.94E-08 | 7.10005108 | 263 | 2278 | 57 | 26856 | [illegible] |
| GO:BP | mitotic spindle assembly | GO:0090307 | FALSE | 7.98E-08 | 7.097730084 | 70 | 2278 | 26 | 26856 | [illegible] |
| GO:BP | generation of neurons | GO:0048699 | FALSE | 8.10E-08 | 7.091768787 | 1634 | 2278 | 215 | 26856 | [illegible] |
| GO:BP | heterochromatin formation | GO:0031507 | FALSE | 8.35E-08 | 7.07816333 | 91 | 2278 | 30 | 26856 | [illegible] |
| GO:BP | regulation of cell projection organization | GO:0031344 | FALSE | 8.40E-08 | 7.075886574 | 758 | 2278 | 119 | 26856 | [illegible] |
| GO:BP | intrinsic apoptotic signaling pathway in response to DNA damage | GO:0008630 | FALSE | 9.11E-08 | 7.040388338 | 108 | 2278 | 33 | 26856 | [illegible] |
| GO:BP | positive regulation of chromosome organization | GO:2001252 | FALSE | 9.11E-08 | 7.040388338 | 108 | 2278 | 33 | 26856 | [illegible] |
| GO:BP | regulation of interleukin-1 beta production | GO:0032651 | FALSE | 1.10E-07 | 6.957670712 | 103 | 2278 | 32 | 26856 | [illegible] |
| GO:BP | interleukin-1 beta production | GO:0032611 | FALSE | 1.10E-07 | 6.957670712 | 103 | 2278 | 32 | 26856 | [illegible] |
| GO:BP | circadian regulation of gene expression | GO:0032922 | FALSE | 1.15E-07 | 6.937915334 | 66 | 2278 | 25 | 26856 | [illegible] |
| GO:BP | positive regulation of kinase activity | GO:0033674 | FALSE | 1.18E-07 | 6.927160839 | 431 | 2278 | 79 | 26856 | [illegible] |
| GO:BP | proteasome-mediated ubiquitin-dependent protein catabolic process | GO:0043161 | FALSE | 1.28E-07 | 6.893445708 | 416 | 2278 | 77 | 26856 | [illegible] |
| GO:BP | regulation of biological quality | GO:0065008 | FALSE | 1.33E-07 | 6.877579342 | 3186 | 2278 | 370 | 26856 | [illegible] |
| GO:BP | biological process involved in symbiotic interaction | GO:0044403 | FALSE | 1.35E-07 | 6.870513918 | 288 | 2278 | 60 | 26856 | [illegible] |
| GO:BP | response to radiation | GO:0009314 | FALSE | 1.36E-07 | 6.865168953 | 448 | 2278 | 81 | 26856 | [illegible] |
| GO:BP | regulation of T cell activation | GO:0050863 | FALSE | 1.44E-07 | 6.840138273 | 363 | 2278 | 70 | 26856 | [illegible] |
| GO:BP | negative regulation of cytokine production | GO:0001818 | FALSE | 1.49E-07 | 6.825503862 | 296 | 2278 | 61 | 26856 | [illegible] |
| GO:BP | nuclear-transcribed mRNA catabolic process, deadenylation-dependent decay | GO:0000288 | FALSE | 1.50E-07 | 6.823774917 | 57 | 2278 | 23 | 26856 | [illegible] |
| GO:BP | chordate embryonic development | GO:0043009 | FALSE | 1.52E-07 | 6.817877487 | 791 | 2278 | 122 | 26856 | [illegible] |
| GO:BP | regulation of autophagy | GO:0010506 | FALSE | 1.56E-07 | 6.806346049 | 289 | 2278 | 60 | 26856 | [illegible] |
| GO:BP | positive regulation of interferon-beta production | GO:0032728 | FALSE | 1.61E-07 | 6.791926934 | 48 | 2278 | 21 | 26856 | [illegible] |
| GO:BP | positive regulation of organelle assembly | GO:1902117 | FALSE | 1.78E-07 | 6.748460012 | 88 | 2278 | 29 | 26856 | [illegible] |
| GO:BP | regulation of protein modification by small protein conjugation or removal | GO:1903320 | FALSE | 1.81E-07 | 6.741787119 | 254 | 2278 | 55 | 26856 | [illegible] |
| GO:BP | negative regulation of catalytic activity | GO:0043086 | FALSE | 2.00E-07 | 6.699731371 | 640 | 2278 | 104 | 26856 | [illegible] |
| GO:BP | cellular component assembly | GO:0022607 | FALSE | 2.16E-07 | 6.665991172 | 4073 | 2278 | 454 | 26856 | [illegible] |
| GO:BP | positive regulation of cell migration | GO:0030335 | FALSE | 2.17E-07 | 6.664259572 | 574 | 2278 | 96 | 26856 | [illegible] |
| GO:BP | centrosome duplication | GO:0051298 | FALSE | 2.22E-07 | 6.652850521 | 78 | 2278 | 27 | 26856 | [illegible] |
| GO:BP | regulation of protein serine/threonine kinase activity | GO:0071900 | FALSE | 2.23E-07 | 6.652342139 | 336 | 2278 | 66 | 26856 | [illegible] |
| GO:BP | muscle cell proliferation | GO:0033002 | FALSE | 2.27E-07 | 6.643127437 | 277 | 2278 | 58 | 26856 | [illegible] |
| GO:BP | cellular response to endogenous stimulus | GO:0071495 | FALSE | 2.29E-07 | 6.640968682 | 1643 | 2278 | 214 | 26856 | [illegible] |
| GO:BP | negative regulation of kinase activity | GO:0033673 | FALSE | 2.54E-07 | 6.595879669 | 221 | 2278 | 50 | 26856 | [illegible] |
| GO:BP | positive regulation of cell projection organization | GO:0031346 | FALSE | 2.97E-07 | 6.52711162 | 439 | 2278 | 79 | 26856 | [illegible] |
| GO:BP | positive regulation of hemopoiesis | GO:1903708 | FALSE | 3.33E-07 | 6.477254614 | 202 | 2278 | 47 | 26856 | [illegible] |
| GO:BP | positive regulation of leukocyte differentiation | GO:1902107 | FALSE | 3.33E-07 | 6.477254614 | 202 | 2278 | 47 | 26856 | [illegible] |
| GO:BP | regulation of leukocyte cell-cell adhesion | GO:1903037 | FALSE | 3.37E-07 | 6.472284588 | 362 | 2278 | 69 | 26856 | [illegible] |
| GO:BP | regulation of nervous system development | GO:0051960 | FALSE | 3.39E-07 | 6.470092147 | 562 | 2278 | 94 | 26856 | [illegible] |
| GO:BP | negative regulation of transferase activity | GO:0051348 | FALSE | 3.39E-07 | 6.470011013 | 258 | 2278 | 55 | 26856 | [illegible] |
| GO:BP | regulation of anatomical structure morphogenesis | GO:0022603 | FALSE | 3.49E-07 | 6.457511297 | 925 | 2278 | 136 | 26856 | [illegible] |
| GO:BP | cell morphogenesis | GO:0000902 | FALSE | 3.49E-07 | 6.457203105 | 1033 | 2278 | 148 | 26856 | [illegible] |
| GO:BP | regulation of chromosome segregation | GO:0051983 | FALSE | 3.56E-07 | 6.449054669 | 125 | 2278 | 35 | 26856 | [illegible] |
| GO:BP | negative regulation of mitotic cell cycle | GO:0045930 | FALSE | 3.56E-07 | 6.44847554 | 223 | 2278 | 50 | 26856 | [illegible] |
| GO:BP | adaptive immune response based on somatic recombination of immune receptors built from i | GO:0002460 | FALSE | 3.56E-07 | 6.44810961 | 332 | 2278 | 65 | 26856 | [illegible] |
| GO:BP | Ras protein signal transduction | GO:0007265 | FALSE | 3.56E-07 | 6.44810961 | 332 | 2278 | 65 | 26856 | [illegible] |
| GO:BP | response to organic cyclic compound | GO:0014070 | FALSE | 4.21E-07 | 6.375524803 | 1127 | 2278 | 158 | 26856 | [illegible] |
| GO:BP | signal transduction by p53 class mediator | GO:0072331 | FALSE | 4.27E-07 | 6.369329987 | 157 | 2278 | 40 | 26856 | [illegible] |
| GO:BP | regulation of plasma membrane bounded cell projection organization | GO:0120035 | FALSE | 4.75E-07 | 6.323676264 | 744 | 2278 | 115 | 26856 | [illegible] |
| GO:BP | regulation of myeloid cell differentiation | GO:0045637 | FALSE | 5.87E-07 | 6.231546446 | 226 | 2278 | 50 | 26856 | [illegible] |
| GO:BP | mitotic sister chromatid cohesion | GO:0007064 | FALSE | 6.04E-07 | 6.218685581 | 26 | 2278 | 15 | 26856 | [illegible] |
| GO:BP | positive regulation of interferon-alpha production | GO:0032727 | FALSE | 6.04E-07 | 6.218685581 | 26 | 2278 | 15 | 26856 | [illegible] |
| GO:BP | response to nitrogen compound | GO:1901698 | FALSE | 6.14E-07 | 6.212181157 | 1328 | 2278 | 179 | 26856 | [illegible] |
| GO:BP | cellular response to virus | GO:0098586 | FALSE | 6.45E-07 | 6.190383638 | 51 | 2278 | 21 | 26856 | [illegible] |
| GO:BP | regulation of inflammatory response | GO:0050727 | FALSE | 6.48E-07 | 6.188408663 | 375 | 2278 | 70 | 26856 | [illegible] |
| GO:BP | leukocyte activation involved in immune response | GO:0002366 | FALSE | 6.57E-07 | 6.182297648 | 314 | 2278 | 62 | 26856 | [illegible] |
| GO:BP | response to peptide hormone | GO:0043434 | FALSE | 7.03E-07 | 6.153149637 | 479 | 2278 | 83 | 26856 | [illegible] |
| GO:BP | positive regulation of locomotion | GO:0040017 | FALSE | 7.04E-07 | 6.152641355 | 620 | 2278 | 100 | 26856 | [illegible] |
| GO:BP | mitotic metaphase chromosome alignment | GO:0007080 | FALSE | 7.24E-07 | 6.14034354 | 56 | 2278 | 22 | 26856 | [illegible] |
| GO:BP | regulation of adaptive immune response | GO:0002819 | FALSE | 8.13E-07 | 6.08964627 | 228 | 2278 | 50 | 26856 | [illegible] |
| GO:BP | positive regulation of leukocyte activation | GO:0002696 | FALSE | 8.25E-07 | 6.083568935 | 377 | 2278 | 70 | 26856 | [illegible] |
| GO:BP | neuron differentiation | GO:0030182 | FALSE | 8.56E-07 | 6.067771917 | 1552 | 2278 | 202 | 26856 | [illegible] |
| GO:BP | regulation of cell-cell adhesion | GO:0022407 | FALSE | 8.66E-07 | 6.062290354 | 481 | 2278 | 83 | 26856 | [illegible] |
| GO:BP | vasculature development | GO:0001944 | FALSE | 9.12E-07 | 6.039811264 | 787 | 2278 | 119 | 26856 | [illegible] |
| GO:BP | cell activation involved in immune response | GO:0002263 | FALSE | 9.81E-07 | 6.008417846 | 317 | 2278 | 62 | 26856 | [illegible] |
| GO:BP | alpha-beta T cell activation | GO:0046631 | FALSE | 9.96E-07 | 6.001721303 | 181 | 2278 | 43 | 26856 | [illegible] |
| GO:BP | blood vessel development | GO:0001568 | FALSE | 1.26003E-06 | 5.899619086 | 756 | 2278 | 115 | 26856 | [illegible] |
| GO:BP | regulation of mRNA processing | GO:0050684 | FALSE | 1.36314E-06 | 5.865458185 | 137 | 2278 | 36 | 26856 | [illegible] |
| GO:BP | positive regulation of cell motility | GO:2000147 | FALSE | 1.37388E-06 | 5.862052542 | 602 | 2278 | 97 | 26856 | [illegible] |
| GO:BP | regulation of establishment of protein localization | GO:0070201 | FALSE | 1.37388E-06 | 5.862052542 | 602 | 2278 | 97 | 26856 | [illegible] |
| GO:BP | cellular response to hormone stimulus | GO:0032870 | FALSE | 1.41268E-06 | 5.849957696 | 662 | 2278 | 104 | 26856 | [illegible] |
| GO:BP | regulation of interferon-alpha production | GO:0032647 | FALSE | 1.42095E-06 | 5.847421438 | 31 | 2278 | 16 | 26856 | [illegible] |
| GO:BP | interferon-alpha production | GO:0032607 | FALSE | 1.42095E-06 | 5.847421438 | 31 | 2278 | 16 | 26856 | [illegible] |
| GO:BP | positive regulation of cell activation | GO:0050867 | FALSE | 1.53564E-06 | 5.813710636 | 398 | 2278 | 72 | 26856 | [illegible] |
| GO:BP | chemokine production | GO:0032602 | FALSE | 1.55205E-06 | 5.809094404 | 107 | 2278 | 31 | 26856 | [illegible] |
| GO:BP | regulation of chemokine production | GO:0032642 | FALSE | 1.55205E-06 | 5.809094404 | 107 | 2278 | 31 | 26856 | [illegible] |
| GO:BP | positive regulation of cytokine-mediated signaling pathway | GO:0001961 | FALSE | 1.60756E-06 | 5.793833532 | 58 | 2278 | 22 | 26856 | [illegible] |
| GO:BP | neuron projection development | GO:0031175 | FALSE | 1.72239E-06 | 5.763868185 | 1121 | 2278 | 155 | 26856 | [illegible] |
| GO:BP | regulation of mitotic cell cycle phase transition | GO:1901990 | FALSE | 1.73646E-06 | 5.760334848 | 329 | 2278 | 63 | 26856 | [illegible] |
| GO:BP | mitotic cytokinesis | GO:0000281 | FALSE | 1.74372E-06 | 5.758523929 | 79 | 2278 | 26 | 26856 | [illegible] |
| GO:BP | chromosome separation | GO:0051304 | FALSE | 1.74372E-06 | 5.758523929 | 79 | 2278 | 26 | 26856 | [illegible] |
| GO:BP | protein acetylation | GO:0006473 | FALSE | 1.74593E-06 | 5.757972095 | 164 | 2278 | 40 | 26856 | [illegible] |
| GO:BP | production of molecular mediator of immune response | GO:0002440 | FALSE | 1.8026E-06 | 5.744100048 | 233 | 2278 | 50 | 26856 | [illegible] |
| GO:BP | intracellular protein transport | GO:0006886 | FALSE | 1.8217E-06 | 5.739523499 | 867 | 2278 | 127 | 26856 | [illegible] |
| GO:BP | viral gene expression | GO:0019080 | FALSE | 1.97307E-06 | 5.70485782 | 74 | 2278 | 25 | 26856 | [illegible] |
| GO:BP | interleukin-6 production | GO:0032635 | FALSE | 2.11778E-06 | 5.67411899 | 165 | 2278 | 40 | 26856 | [illegible] |
| GO:BP | regulation of interleukin-6 production | GO:0032675 | FALSE | 2.11778E-06 | 5.67411899 | 165 | 2278 | 40 | 26856 | [illegible] |
| GO:BP | regulation of adaptive immune response based on somatic recombination of immune receptor | GO:0002822 | FALSE | 2.19707E-06 | 5.658155461 | 213 | 2278 | 47 | 26856 | [illegible] |
| GO:BP | regulation of chromosome separation | GO:1905818 | FALSE | 3.03949E-06 | 5.517199909 | 70 | 2278 | 24 | 26856 | [illegible] |
| GO:BP | regulation of lymphocyte differentiation | GO:0045619 | FALSE | 3.05801E-06 | 5.514561248 | 208 | 2278 | 46 | 26856 | [illegible] |
| GO:BP | response to tumor necrosis factor | GO:0034612 | FALSE | 3.09386E-06 | 5.509499251 | 251 | 2278 | 52 | 26856 | [illegible] |
| GO:BP | cellular response to ionizing radiation | GO:0071479 | FALSE | 3.21756E-06 | 5.492473576 | 81 | 2278 | 26 | 26856 | [illegible] |
| GO:BP | regulation of defense response to virus | GO:0050688 | FALSE | 3.29081E-06 | 5.482697453 | 110 | 2278 | 31 | 26856 | [illegible] |
| GO:BP | nucleotide-binding oligomerization domain containing 2 signaling pathway | GO:0070431 | FALSE | 3.67204E-06 | 5.435092868 | 18 | 2278 | 12 | 26856 | [illegible] |
| GO:BP | neuroinflammatory response | GO:0150076 | FALSE | 3.73074E-06 | 5.428205585 | 76 | 2278 | 25 | 26856 | [illegible] |
| GO:BP | cell projection assembly | GO:0030031 | FALSE | 3.83664E-06 | 5.41604936 | 588 | 2278 | 94 | 26856 | [illegible] |
| GO:BP | positive regulation of lymphocyte activation | GO:0051251 | FALSE | 3.86275E-06 | 5.413102862 | 320 | 2278 | 61 | 26856 | [illegible] |
| GO:BP | regulation of interleukin-1 production | GO:0032652 | FALSE | 3.93997E-06 | 5.404507075 | 123 | 2278 | 33 | 26856 | [illegible] |
| GO:BP | interleukin-1 production | GO:0032612 | FALSE | 3.93997E-06 | 5.404507075 | 123 | 2278 | 33 | 26856 | [illegible] |
| GO:BP | DNA replication | GO:0006260 | FALSE | 4.0463E-06 | 5.392942245 | 275 | 2278 | 55 | 26856 | [illegible] |
| GO:BP | immune response-regulating cell surface receptor signaling pathway | GO:0002768 | FALSE | 4.34884E-06 | 5.361626361 | 283 | 2278 | 56 | 26856 | [illegible] |
| GO:BP | positive regulation of interleukin-1 beta production | GO:0032731 | FALSE | 4.64487E-06 | 5.333026262 | 66 | 2278 | 23 | 26856 | [illegible] |
| GO:BP | cellular response to external stimulus | GO:0071496 | FALSE | 4.68589E-06 | 5.329207518 | 408 | 2278 | 72 | 26856 | [illegible] |
| GO:BP | immune response-activating cell surface receptor signaling pathway | GO:0002429 | FALSE | 4.93762E-06 | 5.306482072 | 269 | 2278 | 54 | 26856 | [illegible] |
| GO:BP | supramolecular fiber organization | GO:0097435 | FALSE | 5.11331E-06 | 5.29129794 | 774 | 2278 | 115 | 26856 | [illegible] |
| GO:BP | cytoskeleton-dependent cytokinesis | GO:0061640 | FALSE | 5.39805E-06 | 5.267762895 | 106 | 2278 | 30 | 26856 | [illegible] |
| GO:BP | forebrain development | GO:0030900 | FALSE | 5.50273E-06 | 5.25942174 | 450 | 2278 | 77 | 26856 | [illegible] |
| GO:BP | rhythmic process | GO:0048511 | FALSE | 5.56229E-06 | 5.254746181 | 354 | 2278 | 65 | 26856 | [illegible] |
| GO:BP | regulation of T cell differentiation | GO:0045580 | FALSE | 5.62529E-06 | 5.249855172 | 177 | 2278 | 41 | 26856 | [illegible] |
| GO:BP | regulation of DNA binding | GO:0051101 | FALSE | 5.75893E-06 | 5.2396582 | 131 | 2278 | 34 | 26856 | [illegible] |
| GO:BP | protein K63-linked ubiquitination | GO:0070534 | FALSE | 5.82739E-06 | 5.234526284 | 72 | 2278 | 24 | 26856 | [illegible] |
| GO:BP | cellular response to organic cyclic compound | GO:0071407 | FALSE | 6.85593E-06 | 5.163933764 | 681 | 2278 | 104 | 26856 | [illegible] |
| GO:BP | positive regulation of leukocyte cell-cell adhesion | GO:1903039 | FALSE | 6.91881E-06 | 5.1599687 | 264 | 2278 | 53 | 26856 | [illegible] |
| GO:BP | cellular response to growth factor stimulus | GO:0071363 | FALSE | 6.95394E-06 | 5.157769123 | 725 | 2278 | 109 | 26856 | [illegible] |
| GO:BP | positive regulation of post-translational protein modification | GO:1901875 | FALSE | 6.98235E-06 | 5.155998183 | 199 | 2278 | 44 | 26856 | [illegible] |
| GO:BP | plasma membrane bounded cell projection assembly | GO:0120031 | FALSE | 7.14283E-06 | 5.146129626 | 578 | 2278 | 92 | 26856 | [illegible] |
| GO:BP | cellular component disassembly | GO:0022411 | FALSE | 8.25911E-06 | 5.083066719 | 454 | 2278 | 77 | 26856 | [illegible] |
| GO:BP | adaptive immune response | GO:0002250 | FALSE | 9.34232E-06 | 5.029545321 | 480 | 2278 | 80 | 26856 | [illegible] |
| GO:BP | stress-activated protein kinase signaling cascade | GO:0031098 | FALSE | 9.38516E-06 | 5.027558533 | 244 | 2278 | 50 | 26856 | [illegible] |
| GO:BP | alpha-beta T cell differentiation | GO:0046632 | FALSE | 9.53334E-06 | 5.020754905 | 127 | 2278 | 33 | 26856 | [illegible] |
| GO:BP | regulation of protein catabolic process | GO:0042176 | FALSE | 9.93623E-06 | 5.002778316 | 359 | 2278 | 65 | 26856 | [illegible] |
| GO:BP | positive regulation of cell adhesion | GO:0045785 | FALSE | 9.98803E-06 | 5.000520308 | 489 | 2278 | 81 | 26856 | [illegible] |
| GO:BP | cell growth | GO:0016049 | FALSE | 1.16205E-05 | 4.934774873 | 541 | 2278 | 87 | 26856 | [illegible] |
| GO:BP | neuron development | GO:0048666 | FALSE | 1.18824E-05 | 4.925094879 | 1275 | 2278 | 168 | 26856 | [illegible] |
| GO:BP | leukocyte mediated immunity | GO:0002443 | FALSE | 1.20585E-05 | 4.918708247 | 425 | 2278 | 73 | 26856 | [illegible] |
| GO:BP | response to type I interferon | GO:0034340 | FALSE | 1.24824E-05 | 4.903703419 | 69 | 2278 | 23 | 26856 | [illegible] |
| GO:BP | positive regulation of GTPase activity | GO:0043547 | FALSE | 1.2492E-05 | 4.903367399 | 246 | 2278 | 50 | 26856 | [illegible] |
| GO:BP | ncRNA metabolic process | GO:0034660 | FALSE | 1.26542E-05 | 4.897765633 | 619 | 2278 | 96 | 26856 | [illegible] |
| GO:BP | nuclear-transcribed mRNA poly(A) tail shortening | GO:0000289 | FALSE | 1.37582E-05 | 4.861438033 | 23 | 2278 | 13 | 26856 | [illegible] |
| GO:BP | positive regulation of establishment of protein localization | GO:1904951 | FALSE | 1.38476E-05 | 4.858625147 | 354 | 2278 | 64 | 26856 | [illegible] |
| GO:BP | regulation of extrinsic apoptotic signaling pathway | GO:2001236 | FALSE | 1.47582E-05 | 4.830967448 | 162 | 2278 | 38 | 26856 | [illegible] |
| GO:BP | vesicle-mediated transport | GO:0016192 | FALSE | 1.50106E-05 | 4.823600613 | 1529 | 2278 | 194 | 26856 | [illegible] |
| GO:BP | regulation of GTPase activity | GO:0043087 | FALSE | 1.51166E-05 | 4.820545242 | 339 | 2278 | 62 | 26856 | [illegible] |
| GO:BP | response to growth factor | GO:0070848 | FALSE | 1.52156E-05 | 4.817709524 | 753 | 2278 | 111 | 26856 | [illegible] |
| GO:BP | viral transcription | GO:0019083 | FALSE | 1.55316E-05 | 4.80878418 | 54 | 2278 | 20 | 26856 | [illegible] |
| GO:BP | protein localization to microtubule cytoskeleton | GO:0072698 | TRUE | 1.55316E-05 | 4.80878418 | 54 | 2278 | 20 | 26856 | [illegible] |
| GO:BP | cytokine production involved in immune response | GO:0002367 | FALSE | 1.57836E-05 | 4.801792623 | 123 | 2278 | 32 | 26856 | [illegible] |
| GO:BP | regulation of cytokine production involved in immune response | GO:0002718 | FALSE | 1.57836E-05 | 4.801792623 | 123 | 2278 | 32 | 26856 | [illegible] |
| GO:BP | enzyme-linked receptor protein signaling pathway | GO:0007167 | FALSE | 1.71156E-05 | 4.766608622 | 963 | 2278 | 134 | 26856 | [illegible] |
| GO:BP | gliogenesis | GO:0042063 | FALSE | 1.75045E-05 | 4.756851152 | 380 | 2278 | 67 | 26856 | [illegible] |
| GO:BP | regulation of proteolysis | GO:0030162 | FALSE | 1.7647E-05 | 4.753329598 | 755 | 2278 | 111 | 26856 | [illegible] |
| GO:BP | regulation of non-canonical NF-kappaB signal transduction | GO:1901222 | FALSE | 1.8592E-05 | 4.730673755 | 99 | 2278 | 28 | 26856 | [illegible] |
| GO:BP | response to chemical | GO:0042221 | FALSE | 1.91013E-05 | 4.718937784 | 5560 | 2278 | 579 | 26856 | [illegible] |
| GO:BP | response to organonitrogen compound | GO:0010243 | FALSE | 1.91543E-05 | 4.717734642 | 1236 | 2278 | 163 | 26856 | [illegible] |
| GO:BP | telomere organization | GO:0032200 | FALSE | 1.92255E-05 | 4.716122456 | 150 | 2278 | 36 | 26856 | [illegible] |
| GO:BP | positive regulation of pattern recognition receptor signaling pathway | GO:0062208 | FALSE | 1.93843E-05 | 4.712549843 | 65 | 2278 | 22 | 26856 | [illegible] |
| GO:BP | regulation of developmental growth | GO:0048638 | FALSE | 1.9554E-05 | 4.708763863 | 373 | 2278 | 66 | 26856 | [illegible] |
| GO:BP | protein deubiquitination | GO:0016579 | FALSE | 1.99661E-05 | 4.699705867 | 137 | 2278 | 34 | 26856 | [illegible] |
| GO:BP | mitotic cell cycle checkpoint signaling | GO:0007093 | FALSE | 1.99661E-05 | 4.699705867 | 137 | 2278 | 34 | 26856 | [illegible] |
| GO:BP | nucleotide-binding oligomerization domain containing signaling pathway | GO:0070423 | FALSE | 2.11968E-05 | 4.673729726 | 20 | 2278 | 12 | 26856 | [illegible] |
| GO:BP | protein acylation | GO:0043543 | FALSE | 2.13569E-05 | 4.670461127 | 206 | 2278 | 44 | 26856 | [illegible] |
| GO:BP | positive regulation of amide metabolic process | GO:0034250 | FALSE | 2.17428E-05 | 4.662685146 | 171 | 2278 | 39 | 26856 | [illegible] |
| GO:BP | leukocyte homeostasis | GO:0001776 | FALSE | 2.20805E-05 | 4.655991549 | 131 | 2278 | 33 | 26856 | [illegible] |
| GO:BP | regulation of protein transport | GO:0051223 | FALSE | 2.22685E-05 | 4.652309896 | 574 | 2278 | 90 | 26856 | [illegible] |
| GO:BP | intracellular steroid hormone receptor signaling pathway | GO:0030518 | FALSE | 2.22769E-05 | 4.652144933 | 112 | 2278 | 30 | 26856 | [illegible] |
| GO:BP | regulation of tyrosine phosphorylation of STAT protein | GO:0042509 | FALSE | 2.32749E-05 | 4.633111389 | 71 | 2278 | 23 | 26856 | [illegible] |
| GO:BP | protein autophosphorylation | GO:0046777 | FALSE | 2.45195E-05 | 4.610488477 | 236 | 2278 | 48 | 26856 | [illegible] |
| GO:BP | regulation of production of molecular mediator of immune response | GO:0002700 | FALSE | 2.55269E-05 | 4.593002445 | 200 | 2278 | 43 | 26856 | [illegible] |
| GO:BP | regulation of type II interferon production | GO:0032649 | FALSE | 2.61359E-05 | 4.582763104 | 119 | 2278 | 31 | 26856 | [illegible] |
| GO:BP | positive regulation of interleukin-1 production | GO:0032732 | FALSE | 2.64264E-05 | 4.577961317 | 77 | 2278 | 24 | 26856 | [illegible] |
| GO:BP | peptide biosynthetic process | GO:0043043 | FALSE | 2.75129E-05 | 4.560464323 | 779 | 2278 | 113 | 26856 | [illegible] |
| GO:BP | regulation of cell projection assembly | GO:0060491 | FALSE | 2.90442E-05 | 4.536940469 | 208 | 2278 | 44 | 26856 | [illegible] |
| GO:BP | regulation of sister chromatid cohesion | GO:0007063 | FALSE | 2.90879E-05 | 4.536286903 | 17 | 2278 | 11 | 26856 | [illegible] |
| GO:BP | protein modification by small protein removal | GO:0070646 | FALSE | 2.92093E-05 | 4.534478685 | 159 | 2278 | 37 | 26856 | [illegible] |
| GO:BP | cellular response to peptide | GO:1901653 | FALSE | 2.96337E-05 | 4.52821340 | 393 | 2278 | 68 | 26856 | [illegible] |
| GO:BP | positive regulation of cytoskeleton organization | GO:0051495 | FALSE | 3.04358E-05 | 4.516615972 | 194 | 2278 | 42 | 26856 | [illegible] |
| GO:BP | regulation of protein ubiquitination | GO:0031396 | FALSE | 3.12397E-05 | 4.505293383 | 223 | 2278 | 46 | 26856 | [illegible] |
| GO:BP | regulation of mitotic sister chromatid segregation | GO:0033047 | FALSE | 3.21618E-05 | 4.492659435 | 56 | 2278 | 20 | 26856 | [illegible] |
| GO:BP | type II interferon production | GO:0032609 | FALSE | 3.23986E-05 | 4.489473804 | 120 | 2278 | 31 | 26856 | [illegible] |
| GO:BP | sexual reproduction | GO:0019953 | FALSE | 3.26277E-05 | 4.486413182 | 1236 | 2278 | 162 | 26856 | [illegible] |
| GO:BP | regulation of DNA replication | GO:0006275 | FALSE | 3.30896E-05 | 4.480308961 | 133 | 2278 | 33 | 26856 | [illegible] |
| GO:BP | stress-activated MAPK cascade | GO:0051403 | FALSE | 3.43529E-05 | 4.464036609 | 231 | 2278 | 47 | 26856 | [illegible] |
| GO:BP | positive regulation of chemokine production | GO:0032722 | FALSE | 3.51148E-05 | 4.454509893 | 78 | 2278 | 24 | 26856 | [illegible] |
| GO:BP | intrinsic apoptotic signaling pathway by p53 class mediator | GO:0072332 | FALSE | 3.88432E-05 | 4.410684647 | 90 | 2278 | 26 | 26856 | [illegible] |
| GO:BP | regulation of protein stability | GO:0031647 | FALSE | 3.94579E-05 | 4.403866216 | 308 | 2278 | 57 | 26856 | [illegible] |
| GO:BP | epithelial cell proliferation | GO:0050673 | FALSE | 3.9826E-05 | 4.399833627 | 487 | 2278 | 79 | 26856 | [illegible] |
| GO:BP | positive regulation of T cell activation | GO:0050870 | FALSE | 4.29316E-05 | 4.367223373 | 240 | 2278 | 48 | 26856 | [illegible] |
| GO:BP | regulation of mitotic sister chromatid separation | GO:0010965 | FALSE | 4.56035E-05 | 4.341001392 | 57 | 2278 | 20 | 26856 | [illegible] |
| GO:BP | nucleotide-binding domain, leucine rich repeat containing receptor signaling pathway | GO:0035872 | FALSE | 4.56204E-05 | 4.340841363 | 21 | 2278 | 12 | 26856 | [illegible] |
| GO:BP | negative regulation of chromosome segregation | GO:0051985 | FALSE | 4.79695E-05 | 4.319034775 | 52 | 2278 | 19 | 26856 | [illegible] |
| GO:BP | epithelial cell apoptotic process | GO:1904019 | FALSE | 4.82383E-05 | 4.316607984 | 155 | 2278 | 36 | 26856 | [illegible] |
| GO:BP | cellular response to radiation | GO:0071478 | FALSE | 5.07111E-05 | 4.294897355 | 176 | 2278 | 39 | 26856 | [illegible] |
| GO:BP | blood vessel morphogenesis | GO:0048514 | FALSE | 5.16701E-05 | 4.286760586 | 654 | 2278 | 98 | 26856 | [illegible] |
| GO:BP | tube morphogenesis | GO:0035239 | FALSE | 5.19822E-05 | 4.28414457 | 925 | 2278 | 128 | 26856 | [illegible] |
| GO:BP | regulation of nuclear-transcribed mRNA catabolic process, deadenylation-dependent decay | GO:1900151 | FALSE | 5.31614E-05 | 4.274403201 | 25 | 2278 | 13 | 26856 | [illegible] |
| GO:BP | regulation of epithelial cell proliferation | GO:0050678 | FALSE | 5.39701E-05 | 4.26784708 | 407 | 2278 | 69 | 26856 | [illegible] |
| GO:BP | steroid hormone mediated signaling pathway | GO:0043401 | FALSE | 5.48506E-05 | 4.260818303 | 129 | 2278 | 32 | 26856 | [illegible] |
| GO:BP | regulation of plasma membrane bounded cell projection assembly | GO:0120032 | FALSE | 5.49915E-05 | 4.259704102 | 205 | 2278 | 43 | 26856 | [illegible] |
| GO:BP | positive regulation of cell-cell adhesion | GO:0022409 | FALSE | 5.62766E-05 | 4.249671781 | 311 | 2278 | 57 | 26856 | [illegible] |
| GO:BP | tyrosine phosphorylation of STAT protein | GO:0007260 | FALSE | 5.64224E-05 | 4.248548258 | 74 | 2278 | 23 | 26856 | [illegible] |
| GO:BP | negative regulation of cytokine-mediated signaling pathway | GO:0001960 | FALSE | 5.64224E-05 | 4.248548258 | 74 | 2278 | 23 | 26856 | [illegible] |
| GO:BP | protein-containing complex disassembly | GO:0032984 | FALSE | 5.64548E-05 | 4.248299294 | 242 | 2278 | 48 | 26856 | [illegible] |
| GO:BP | cellular response to type I interferon | GO:0071357 | FALSE | 5.73834E-05 | 4.241213957 | 63 | 2278 | 21 | 26856 | [illegible] |
| GO:BP | type I interferon-mediated signaling pathway | GO:0060337 | FALSE | 5.73834E-05 | 4.241213957 | 63 | 2278 | 21 | 26856 | [illegible] |
| GO:BP | positive regulation of peptidyl-tyrosine phosphorylation | GO:0050731 | FALSE | 5.82418E-05 | 4.234765259 | 191 | 2278 | 41 | 26856 | [illegible] |
| GO:BP | import into nucleus | GO:0051170 | FALSE | 5.92608E-05 | 4.227232732 | 184 | 2278 | 40 | 26856 | [illegible] |
| GO:BP | T cell differentiation involved in immune response | GO:0002292 | FALSE | 6.09785E-05 | 4.21482341 | 80 | 2278 | 24 | 26856 | [illegible] |
| GO:BP | protein localization to cytoskeleton | GO:0044380 | FALSE | 6.40673E-05 | 4.193363563 | 58 | 2278 | 20 | 26856 | [illegible] |
| GO:BP | translation | GO:0006412 | FALSE | 6.44775E-05 | 4.190591843 | 755 | 2278 | 109 | 26856 | [illegible] |
| GO:BP | lymphocyte activation involved in immune response | GO:0002285 | FALSE | 6.74386E-05 | 4.171091569 | 221 | 2278 | 45 | 26856 | [illegible] |
| GO:BP | glial cell activation | GO:0061900 | FALSE | 6.88383E-05 | 4.162169712 | 53 | 2278 | 19 | 26856 | [illegible] |
| GO:BP | muscle structure development | GO:0061061 | FALSE | 6.89952E-05 | 4.161181124 | 729 | 2278 | 106 | 26856 | [illegible] |
| GO:BP | protein import into nucleus | GO:0006606 | FALSE | 7.03536E-05 | 4.15271394 | 178 | 2278 | 39 | 26856 | [illegible] |
| GO:BP | regulation of leukocyte apoptotic process | GO:2000106 | FALSE | 7.17025E-05 | 4.144465985 | 111 | 2278 | 29 | 26856 | [illegible] |
| GO:BP | positive regulation of protein-containing complex assembly | GO:0031334 | FALSE | 7.4075E-05 | 4.130328052 | 207 | 2278 | 43 | 26856 | [illegible] |
| GO:BP | amide biosynthetic process | GO:0043604 | FALSE | 7.46431E-05 | 4.127010115 | 903 | 2278 | 125 | 26856 | [illegible] |
| GO:BP | B cell activation | GO:0042113 | FALSE | 7.69509E-05 | 4.113786474 | 298 | 2278 | 55 | 26856 | [illegible] |
| GO:BP | ameboidal-type cell migration | GO:0001667 | FALSE | 7.87331E-05 | 4.103842491 | 444 | 2278 | 73 | 26856 | [illegible] |
| GO:BP | negative regulation of cysteine-type endopeptidase activity | GO:2000117 | FALSE | 7.9847E-05 | 4.097741394 | 99 | 2278 | 27 | 26856 | [illegible] |
| GO:BP | regulation of leukocyte mediated immunity | GO:0002703 | FALSE | 8.22375E-05 | 4.084930002 | 283 | 2278 | 53 | 26856 | [illegible] |
| GO:BP | RNA localization | GO:0006403 | FALSE | 8.26718E-05 | 4.082642799 | 179 | 2278 | 39 | 26856 | [illegible] |
| GO:BP | negative regulation of mitotic cell cycle phase transition | GO:1901991 | FALSE | 8.32562E-05 | 4.079583192 | 172 | 2278 | 38 | 26856 | [illegible] |
| GO:BP | reproduction | GO:0000003 | FALSE | 8.70749E-05 | 4.060107031 | 1812 | 2278 | 219 | 26856 | [illegible] |
| GO:BP | regulation of alpha-beta T cell activation | GO:0046634 | FALSE | 8.89716E-05 | 4.050748517 | 112 | 2278 | 29 | 26856 | [illegible] |
| GO:BP | positive regulation of non-canonical NF-kappaB signal transduction | GO:1901224 | FALSE | 9.03802E-05 | 4.043926611 | 70 | 2278 | 22 | 26856 | [illegible] |
| GO:BP | AIM2 inflammasome complex assembly | GO:0140970 | FALSE | 9.08412E-05 | 4.041717187 | 15 | 2278 | 10 | 26856 | [illegible] |
| GO:BP | reproductive process | GO:0022414 | FALSE | 9.08567E-05 | 4.041642923 | 1803 | 2278 | 218 | 26856 | [illegible] |
| GO:BP | regulation of intracellular transport | GO:0032386 | FALSE | 9.09071E-05 | 4.041402298 | 339 | 2278 | 60 | 26856 | [illegible] |
| GO:BP | telomere maintenance | GO:0000723 | FALSE | 9.18051E-05 | 4.037132985 | 145 | 2278 | 34 | 26856 | [illegible] |
| GO:BP | negative regulation of MAPK cascade | GO:0043409 | FALSE | 9.69973E-05 | 4.013240479 | 180 | 2278 | 39 | 26856 | [illegible] |
| GO:BP | negative regulation of canonical NF-kappaB signal transduction | GO:0043124 | FALSE | 9.77763E-05 | 4.009766547 | 54 | 2278 | 19 | 26856 | [illegible] |
| GO:BP | negative regulation of sister chromatid segregation | GO:0033046 | FALSE | 0.000102466 | 3.98941832 | 49 | 2278 | 18 | 26856 | [illegible] |
| GO:BP | negative regulation of mitotic sister chromatid separation | GO:2000816 | FALSE | 0.000102466 | 3.98941832 | 49 | 2278 | 18 | 26856 | [illegible] |
| GO:BP | negative regulation of mitotic metaphase/anaphase transition | GO:0045841 | FALSE | 0.000102466 | 3.98941832 | 49 | 2278 | 18 | 26856 | [illegible] |
| GO:BP | negative regulation of mitotic sister chromatid segregation | GO:0033048 | FALSE | 0.000102466 | 3.98941832 | 49 | 2278 | 18 | 26856 | [illegible] |
| GO:BP | peptide metabolic process | GO:0006518 | FALSE | 0.000104094 | 3.982575504 | 899 | 2278 | 124 | 26856 | [illegible] |
| GO:BP | cellular response to steroid hormone stimulus | GO:0071383 | FALSE | 0.000108919 | 3.962898119 | 217 | 2278 | 44 | 26856 | [illegible] |
| GO:BP | blastocyst development | GO:0001824 | FALSE | 0.000110014 | 3.958550337 | 146 | 2278 | 34 | 26856 | [illegible] |
| GO:BP | positive regulation of neurogenesis | GO:0050769 | FALSE | 0.000111163 | 3.954039784 | 309 | 2278 | 56 | 26856 | [illegible] |
| GO:BP | T-helper cell differentiation | GO:0042093 | FALSE | 0.000120509 | 3.9189809 | 71 | 2278 | 22 | 26856 | [illegible] |
| GO:BP | positive regulation of immune effector process | GO:0002699 | FALSE | 0.000123055 | 3.909902393 | 302 | 2278 | 55 | 26856 | [illegible] |
| GO:BP | mitotic sister chromatid separation | GO:0051306 | FALSE | 0.000123176 | 3.909472159 | 60 | 2278 | 20 | 26856 | [illegible] |
| GO:BP | negative regulation of catabolic process | GO:0009895 | FALSE | 0.000126322 | 3.898519304 | 334 | 2278 | 59 | 26856 | [illegible] |
| GO:BP | cellular component morphogenesis | GO:0032989 | FALSE | 0.000128289 | 3.89180975 | 847 | 2278 | 118 | 26856 | [illegible] |
| GO:BP | positive regulation of proteolysis | GO:0045862 | FALSE | 0.000133589 | 3.874229272 | 375 | 2278 | 64 | 26856 | [illegible] |
| GO:BP | modulation by host of symbiont process | GO:0051851 | FALSE | 0.000134192 | 3.872272895 | 83 | 2278 | 24 | 26856 | [illegible] |
| GO:BP | hematopoietic progenitor cell differentiation | GO:0002244 | FALSE | 0.000135383 | 3.868435923 | 175 | 2278 | 38 | 26856 | [illegible] |
| GO:BP | cellular response to tumor necrosis factor | GO:0071356 | FALSE | 0.000135594 | 3.867759229 | 226 | 2278 | 45 | 26856 | [illegible] |
| GO:BP | negative regulation of peptidyl-tyrosine phosphorylation | GO:0050732 | FALSE | 0.000137525 | 3.861619481 | 55 | 2278 | 19 | 26856 | [illegible] |
| GO:BP | negative regulation of nuclear division | GO:0051784 | FALSE | 0.000144249 | 3.840887529 | 66 | 2278 | 21 | 26856 | [illegible] |
| GO:BP | positive regulation of lymphocyte differentiation | GO:0045621 | FALSE | 0.000144415 | 3.840386796 | 134 | 2278 | 32 | 26856 | [illegible] |
| GO:BP | transmembrane receptor protein tyrosine kinase signaling pathway | GO:0007169 | FALSE | 0.00014593 | 3.835855762 | 597 | 2278 | 90 | 26856 | [illegible] |
| GO:BP | kinetochore organization | GO:0051383 | FALSE | 0.0001513 | 3.820161371 | 19 | 2278 | 11 | 26856 | [illegible] |
| GO:BP | P-body assembly | GO:0033962 | FALSE | 0.0001513 | 3.820161371 | 19 | 2278 | 11 | 26856 | [illegible] |
| GO:BP | CD4-positive, alpha-beta T cell differentiation involved in immune response | GO:0002294 | FALSE | 0.000159689 | 3.79672581 | 72 | 2278 | 22 | 26856 | [illegible] |
| GO:BP | DNA damage checkpoint signaling | GO:0000077 | FALSE | 0.000166916 | 3.777502783 | 115 | 2278 | 29 | 26856 | [illegible] |
| GO:BP | positive regulation of plasma membrane bounded cell projection assembly | GO:0120034 | FALSE | 0.000166916 | 3.777502783 | 115 | 2278 | 29 | 26856 | [illegible] |
| GO:BP | negative regulation of response to cytokine stimulus | GO:0060761 | FALSE | 0.000169118 | 3.771811077 | 78 | 2278 | 23 | 26856 | [illegible] |
| GO:BP | negative regulation of RNA catabolic process | GO:1902369 | FALSE | 0.00017284 | 3.762355634 | 84 | 2278 | 24 | 26856 | [illegible] |
| GO:BP | regulation of supramolecular fiber organization | GO:1902903 | FALSE | 0.000176954 | 3.752138707 | 386 | 2278 | 65 | 26856 | [illegible] |
| GO:BP | erythrocyte homeostasis | GO:0034101 | FALSE | 0.000189821 | 3.721655496 | 156 | 2278 | 35 | 26856 | [illegible] |
| GO:BP | cellular response to extracellular stimulus | GO:0031668 | FALSE | 0.00019449 | 3.711101859 | 306 | 2278 | 55 | 26856 | [illegible] |
| GO:BP | positive regulation of protein transport | GO:0051222 | FALSE | 0.000194536 | 3.710999466 | 338 | 2278 | 59 | 26856 | [illegible] |
| GO:BP | regulation of secretion by cell | GO:1903530 | FALSE | 0.000198569 | 3.702089644 | 672 | 2278 | 98 | 26856 | [illegible] |
| GO:BP | negative regulation of metaphase/anaphase transition of cell cycle | GO:1902100 | FALSE | 0.000209802 | 3.678191218 | 51 | 2278 | 18 | 26856 | [illegible] |
| GO:BP | negative regulation of chromosome separation | GO:1905819 | FALSE | 0.000209802 | 3.678191218 | 51 | 2278 | 18 | 26856 | [illegible] |
| GO:BP | alpha-beta T cell differentiation involved in immune response | GO:0002293 | FALSE | 0.000210342 | 3.677073736 | 73 | 2278 | 22 | 26856 | [illegible] |
| GO:BP | response to steroid hormone | GO:0048545 | FALSE | 0.000211417 | 3.67485932 | 355 | 2278 | 61 | 26856 | [illegible] |
| GO:BP | cellular response to peptide hormone stimulus | GO:0071375 | FALSE | 0.000217866 | 3.661810966 | 331 | 2278 | 58 | 26856 | [illegible] |
| GO:BP | alternative mRNA splicing, via spliceosome | GO:0000380 | FALSE | 0.00021952 | 3.658526253 | 79 | 2278 | 23 | 26856 | [illegible] |
| GO:BP | regulation of cytokinesis | GO:0032465 | FALSE | 0.000221579 | 3.654470678 | 85 | 2278 | 24 | 26856 | [illegible] |
| GO:BP | developmental cell growth | GO:0048588 | FALSE | 0.00023065 | 3.637045603 | 276 | 2278 | 51 | 26856 | [illegible] |
| GO:BP | response to extracellular stimulus | GO:0009991 | FALSE | 0.000253486 | 3.596045941 | 613 | 2278 | 91 | 26856 | [illegible] |
| GO:BP | leukocyte apoptotic process | GO:0071887 | FALSE | 0.000259152 | 3.586445918 | 144 | 2278 | 33 | 26856 | [illegible] |
| GO:BP | maintenance of cell number | GO:0098727 | FALSE | 0.000260058 | 3.584930553 | 201 | 2278 | 41 | 26856 | [illegible] |
| GO:BP | negative regulation of developmental process | GO:0051093 | FALSE | 0.000264479 | 3.577608233 | 1074 | 2278 | 141 | 26856 | [illegible] |
| GO:BP | regulation of centrosome cycle | GO:0046605 | FALSE | 0.000264643 | 3.577339523 | 57 | 2278 | 19 | 26856 | [illegible] |
| GO:BP | negative regulation of mitotic nuclear division | GO:0045839 | FALSE | 0.000264643 | 3.577339523 | 57 | 2278 | 19 | 26856 | [illegible] |
| GO:BP | CD4-positive, alpha-beta T cell differentiation | GO:0043367 | FALSE | 0.000274914 | 3.560802781 | 92 | 2278 | 25 | 26856 | [illegible] |
| GO:BP | alpha-beta T cell activation involved in immune response | GO:0002287 | FALSE | 0.000275462 | 3.559938298 | 74 | 2278 | 22 | 26856 | [illegible] |
| GO:BP | regulation of AIM2 inflammasome complex assembly | GO:0140971 | FALSE | 0.000278047 | 3.55881838 | 13 | 2278 | 9 | 26856 | [illegible] |
| GO:BP | regulation of lipopolysaccharide-mediated signaling pathway | GO:0031664 | FALSE | 0.000300083 | 3.522758825 | 28 | 2278 | 13 | 26856 | [illegible] |
| GO:BP | macrophage activation | GO:0042116 | FALSE | 0.000302832 | 3.518798301 | 105 | 2278 | 27 | 26856 | [illegible] |
| GO:BP | erythrocyte differentiation | GO:0030218 | FALSE | 0.000308459 | 3.510802297 | 145 | 2278 | 33 | 26856 | [illegible] |
| GO:BP | mitotic spindle assembly checkpoint signaling | GO:0007094 | FALSE | 0.000315931 | 3.500407108 | 47 | 2278 | 17 | 26856 | [illegible] |
| GO:BP | spindle assembly checkpoint signaling | GO:0071173 | FALSE | 0.000315931 | 3.500407108 | 47 | 2278 | 17 | 26856 | [illegible] |
| GO:BP | mitotic spindle checkpoint signaling | GO:0071174 | FALSE | 0.000315931 | 3.500407108 | 47 | 2278 | 17 | 26856 | [illegible] |
| GO:BP | cellular response to nutrient levels | GO:0031669 | FALSE | 0.000322117 | 3.49198606 | 271 | 2278 | 50 | 26856 | [illegible] |
| GO:BP | inflammatory cell apoptotic process | GO:0006925 | FALSE | 0.000329454 | 3.482205113 | 24 | 2278 | 12 | 26856 | [illegible] |
| GO:BP | hormone-mediated signaling pathway | GO:0009755 | FALSE | 0.000343324 | 3.464295447 | 181 | 2278 | 38 | 26856 | [illegible] |
| GO:BP | microtubule polymerization or depolymerization | GO:0031109 | FALSE | 0.000346213 | 3.460657099 | 132 | 2278 | 31 | 26856 | [illegible] |
| GO:BP | organonitrogen compound biosynthetic process | GO:1901566 | FALSE | 0.000347404 | 3.45916525 | 1774 | 2278 | 212 | 26856 | [illegible] |
| GO:BP | cellular response to environmental stimulus | GO:0104004 | FALSE | 0.000360564 | 3.443017286 | 385 | 2278 | 64 | 26856 | [illegible] |
| GO:BP | cellular response to abiotic stimulus | GO:0071214 | FALSE | 0.000360564 | 3.443017286 | 385 | 2278 | 64 | 26856 | [illegible] |
| GO:BP | positive regulation of apoptotic signaling pathway | GO:2001235 | FALSE | 0.000362625 | 3.440542655 | 167 | 2278 | 36 | 26856 | [illegible] |
| GO:BP | protein autoubiquitination | GO:0051865 | FALSE | 0.0003643 | 3.438541025 | 81 | 2278 | 23 | 26856 | [illegible] |
| GO:BP | stem cell population maintenance | GO:0019827 | FALSE | 0.000364495 | 3.438308905 | 196 | 2278 | 40 | 26856 | [illegible] |
| GO:BP | protein localization to centrosome | GO:0071539 | FALSE | 0.000397147 | 3.40104844 | 33 | 2278 | 14 | 26856 | [illegible] |
| GO:BP | regulation of cell morphogenesis | GO:0022604 | FALSE | 0.000407903 | 3.389443424 | 273 | 2278 | 50 | 26856 | [illegible] |
| GO:BP | establishment of protein localization to organelle | GO:0072594 | FALSE | 0.000421106 | 3.375608531 | 454 | 2278 | 72 | 26856 | [illegible] |
| GO:BP | positive regulation of inflammatory response | GO:0050729 | FALSE | 0.000427714 | 3.368846251 | 140 | 2278 | 32 | 26856 | [illegible] |
| GO:BP | spindle checkpoint signaling | GO:0031577 | FALSE | 0.000450576 | 3.346231501 | 48 | 2278 | 17 | 26856 | [illegible] |
| GO:BP | regulation of intrinsic apoptotic signaling pathway | GO:2001242 | FALSE | 0.000462686 | 3.334713264 | 183 | 2278 | 38 | 26856 | [illegible] |
| GO:BP | response to decreased oxygen levels | GO:0036293 | FALSE | 0.000462821 | 3.334586493 | 371 | 2278 | 62 | 26856 | [illegible] |
| GO:BP | regulation of cysteine-type endopeptidase activity | GO:2000116 | FALSE | 0.000464593 | 3.33292691 | 243 | 2278 | 46 | 26856 | [illegible] |
| GO:BP | peptidyl-lysine modification | GO:0018205 | FALSE | 0.000464831 | 3.33270498 | 282 | 2278 | 51 | 26856 | [illegible] |
| GO:BP | type 2 immune response | GO:0042092 | FALSE | 0.000467967 | 3.329785203 | 43 | 2278 | 16 | 26856 | [illegible] |
| GO:BP | cell part morphogenesis | GO:0032990 | FALSE | 0.000478101 | 3.320480705 | 748 | 2278 | 105 | 26856 | [illegible] |
| GO:BP | nuclear export | GO:0051168 | FALSE | 0.000480658 | 3.318163929 | 176 | 2278 | 37 | 26856 | [illegible] |
| GO:BP | T cell proliferation | GO:0042098 | FALSE | 0.000482572 | 3.316437808 | 228 | 2278 | 44 | 26856 | [illegible] |
| GO:BP | negative regulation of mRNA metabolic process | GO:1903312 | FALSE | 0.000541229 | 3.266619324 | 95 | 2278 | 25 | 26856 | [illegible] |
| GO:BP | CD4-positive, alpha-beta T cell activation | GO:0035710 | FALSE | 0.000544162 | 3.264271903 | 121 | 2278 | 29 | 26856 | [illegible] |
| GO:BP | regulation of chromatin organization | GO:1902275 | FALSE | 0.000569164 | 3.244762701 | 54 | 2278 | 18 | 26856 | [illegible] |
| GO:BP | RNA stabilization | GO:0043489 | FALSE | 0.000585474 | 3.232492586 | 71 | 2278 | 21 | 26856 | [illegible] |
| GO:BP | positive regulation of translation | GO:0045727 | FALSE | 0.000603822 | 3.219091064 | 142 | 2278 | 32 | 26856 | [illegible] |
| GO:BP | regulation of sister chromatid segregation | GO:0033045 | FALSE | 0.000620493 | 3.207263462 | 102 | 2278 | 26 | 26856 | [illegible] |
| GO:BP | centrosome localization | GO:0051642 | TRUE | 0.000622386 | 3.20594003 | 34 | 2278 | 14 | 26856 | [illegible] |
| GO:BP | protein localization to microtubule organizing center | GO:1905508 | FALSE | 0.000622386 | 3.20594003 | 34 | 2278 | 14 | 26856 | [illegible] |
| GO:BP | regulation of centrosome duplication | GO:0010824 | FALSE | 0.00063551 | 3.196877676 | 49 | 2278 | 17 | 26856 | [illegible] |
| GO:BP | regulation of leukocyte proliferation | GO:0070663 | FALSE | 0.000639531 | 3.194138595 | 269 | 2278 | 49 | 26856 | [illegible] |
| GO:BP | multicellular organism growth | GO:0035264 | FALSE | 0.000639868 | 3.193909445 | 200 | 2278 | 40 | 26856 | [illegible] |
| GO:BP | DNA integrity checkpoint signaling | GO:0031570 | FALSE | 0.000656496 | 3.182768165 | 122 | 2278 | 29 | 26856 | [illegible] |
| GO:BP | positive regulation of MAPK cascade | GO:0043410 | FALSE | 0.000670579 | 3.173550096 | 520 | 2278 | 79 | 26856 | [illegible] |
| GO:BP | endolysosomal toll-like receptor signaling pathway | GO:0140894 | FALSE | 0.00067752 | 3.169077562 | 44 | 2278 | 16 | 26856 | [illegible] |
| GO:BP | protein-DNA complex assembly | GO:0065004 | FALSE | 0.000678848 | 3.168227166 | 193 | 2278 | 39 | 26856 | [illegible] |
| GO:BP | tube development | GO:0035295 | FALSE | 0.00068458 | 3.164576054 | 1178 | 2278 | 150 | 26856 | [illegible] |
| GO:BP | positive regulation of nervous system development | GO:0051962 | FALSE | 0.000701908 | 3.153719554 | 367 | 2278 | 61 | 26856 | [illegible] |
| GO:BP | regulation of stress-activated protein kinase signaling cascade | GO:0070302 | FALSE | 0.000734231 | 3.134167195 | 201 | 2278 | 40 | 26856 | [illegible] |
| GO:BP | leukocyte proliferation | GO:0070661 | FALSE | 0.000749515 | 3.125219716 | 376 | 2278 | 62 | 26856 | [illegible] |
| GO:BP | regulation of DNA-templated transcription initiation | GO:2000142 | FALSE | 0.000751487 | 3.124078437 | 84 | 2278 | 23 | 26856 | [illegible] |
| GO:BP | extrinsic apoptotic signaling pathway via death domain receptors | GO:0008625 | FALSE | 0.000767375 | 3.114992112 | 78 | 2278 | 22 | 26856 | [illegible] |
| GO:BP | cell surface toll-like receptor signaling pathway | GO:0140895 | FALSE | 0.00077915 | 3.10837866 | 55 | 2278 | 18 | 26856 | [illegible] |
| GO:BP | positive regulation of leukocyte apoptotic process | GO:2000108 | FALSE | 0.000816242 | 3.088181326 | 30 | 2278 | 13 | 26856 | [illegible] |
| GO:BP | regulation of small GTPase mediated signal transduction | GO:0051056 | FALSE | 0.000873644 | 3.058665653 | 225 | 2278 | 43 | 26856 | [illegible] |
| GO:BP | positive regulation of cytoplasmic mRNA processing body assembly | GO:0010606 | FALSE | 0.000881466 | 3.05479437 | 6 | 2278 | 6 | 26856 | CNOT6,PAN2,PAN3,CNOT6L,CNOT2,CNOT1 |
| GO:BP | positive regulation of smooth muscle cell migration | GO:0014911 | FALSE | 0.000886718 | 3.052214274 | 61 | 2278 | 19 | 26856 | [illegible] |
| GO:BP | phosphatidylinositol phosphate biosynthetic process | GO:0046854 | TRUE | 0.000886718 | 3.052214274 | 61 | 2278 | 19 | 26856 | [illegible] |
| GO:BP | cell surface pattern recognition receptor signaling pathway | GO:0002752 | FALSE | 0.000886718 | 3.052214274 | 61 | 2278 | 19 | 26856 | [illegible] |
| GO:BP | metaphase/anaphase transition of mitotic cell cycle | GO:0007091 | FALSE | 0.000897795 | 3.046822774 | 91 | 2278 | 24 | 26856 | [illegible] |
| GO:BP | cellular response to decreased oxygen levels | GO:0036294 | FALSE | 0.000951944 | 3.021388776 | 188 | 2278 | 38 | 26856 | [illegible] |
| GO:BP | microtubule organizing center localization | GO:0061842 | FALSE | 0.000956277 | 3.019416089 | 35 | 2278 | 14 | 26856 | [illegible] |
| GO:BP | microglial cell activation | GO:0001774 | FALSE | 0.000968641 | 3.013837138 | 45 | 2278 | 16 | 26856 | [illegible] |
| GO:BP | regulation of type I interferon-mediated signaling pathway | GO:0060338 | FALSE | 0.000968641 | 3.013837138 | 45 | 2278 | 16 | 26856 | [illegible] |
| GO:BP | macroautophagy | GO:0016236 | FALSE | 0.000986182 | 3.006042737 | 257 | 2278 | 47 | 26856 | [illegible] |
| GO:BP | meiotic cell cycle process | GO:1903046 | FALSE | 0.001023362 | 2.989970686 | 211 | 2278 | 41 | 26856 | [illegible] |
| GO:BP | miRNA metabolic process | GO:0010586 | FALSE | 0.001030622 | 2.986900462 | 98 | 2278 | 25 | 26856 | [illegible] |
| GO:BP | response to gamma radiation | GO:0010332 | FALSE | 0.00105745 | 2.975740279 | 56 | 2278 | 18 | 26856 | [illegible] |

| | | | | | | | | | | |
|---|---|---|---|---|---|---|---|---|---|---|
| GO:BP | response to nutrient levels | GO:0031667 | FALSE | 0.001067207 | 2.971751178 | 579 | 2278 | 85 | 26856 | [illegible] |
| GO:BP | regulation of T cell proliferation | GO:0042129 | FALSE | 0.001095248 | 2.960487633 | 189 | 2278 | 38 | 26856 | [illegible] |
| GO:BP | osteoclast differentiation | GO:0030316 | FALSE | 0.001135684 | 2.94474233 | 125 | 2278 | 29 | 26856 | [illegible] |
| GO:BP | regulation of cell division | GO:0051302 | FALSE | 0.001218457 | 2.914189842 | 175 | 2278 | 36 | 26856 | [illegible] |
| GO:BP | epithelial cell migration | GO:0010631 | FALSE | 0.001223906 | 2.912252094 | 315 | 2278 | 54 | 26856 | [illegible] |
| GO:BP | peptidyl-threonine phosphorylation | GO:0018107 | FALSE | 0.001238164 | 2.907221743 | 112 | 2278 | 27 | 26856 | [illegible] |
| GO:BP | biological process involved in interaction with symbiont | GO:0051702 | FALSE | 0.001238164 | 2.907221743 | 112 | 2278 | 27 | 26856 | [illegible] |
| GO:BP | positive regulation of viral process | GO:0048524 | FALSE | 0.001244386 | 2.905044879 | 68 | 2278 | 20 | 26856 | [illegible] |
| GO:BP | cell projection morphogenesis | GO:0048858 | FALSE | 0.001258601 | 2.90011189 | 717 | 2278 | 100 | 26856 | [illegible] |
| GO:BP | localization within membrane | GO:0051668 | FALSE | 0.001258601 | 2.90011189 | 717 | 2278 | 100 | 26856 | [illegible] |
| GO:BP | B cell differentiation | GO:0030183 | FALSE | 0.001272702 | 2.895273362 | 168 | 2278 | 35 | 26856 | [illegible] |
| GO:BP | centriole replication | GO:0007099 | FALSE | 0.00136848 | 2.863761611 | 46 | 2278 | 16 | 26856 | [illegible] |
| GO:BP | positive regulation of growth | GO:0045927 | FALSE | 0.001377131 | 2.861024691 | 308 | 2278 | 53 | 26856 | [illegible] |
| GO:BP | mononuclear cell proliferation | GO:0032943 | FALSE | 0.001404482 | 2.852483799 | 341 | 2278 | 57 | 26856 | [illegible] |
| GO:BP | cellular response to oxygen levels | GO:0071453 | FALSE | 0.001434494 | 2.843301297 | 206 | 2278 | 40 | 26856 | [illegible] |
| GO:BP | regulation of MAP kinase activity | GO:0043405 | FALSE | 0.001444132 | 2.840393002 | 191 | 2278 | 38 | 26856 | [illegible] |
| GO:BP | positive regulation of vascular associated smooth muscle cell proliferation | GO:1904707 | FALSE | 0.001454091 | 2.837408558 | 41 | 2278 | 15 | 26856 | [illegible] |
| GO:BP | regulation of mitotic nuclear division | GO:0007088 | FALSE | 0.001496515 | 2.824919015 | 113 | 2278 | 27 | 26856 | [illegible] |
| GO:BP | angiogenesis | GO:0001525 | FALSE | 0.001509291 | 2.821226876 | 557 | 2278 | 82 | 26856 | [illegible] |
| GO:BP | regulation of protein localization to nucleus | GO:1900180 | FALSE | 0.001535285 | 2.813811095 | 162 | 2278 | 34 | 26856 | [illegible] |
| GO:BP | regulation of alpha-beta T cell differentiation | GO:0046637 | FALSE | 0.001613246 | 2.792299327 | 69 | 2278 | 20 | 26856 | [illegible] |
| GO:BP | epithelium migration | GO:0090132 | FALSE | 0.001662358 | 2.779275439 | 318 | 2278 | 54 | 26856 | [illegible] |
| GO:BP | response to macrophage colony-stimulating factor | GO:0036005 | FALSE | 0.001662502 | 2.779237709 | 15 | 2278 | 9 | 26856 | [illegible] |
| GO:BP | regulation of mononuclear cell proliferation | GO:0032944 | FALSE | 0.001708987 | 2.767261264 | 246 | 2278 | 45 | 26856 | [illegible] |
| GO:BP | metaphase/anaphase transition of cell cycle | GO:0044784 | FALSE | 0.001712668 | 2.766326741 | 94 | 2278 | 24 | 26856 | [illegible] |
| GO:BP | heart development | GO:0007507 | FALSE | 0.001753468 | 2.756102067 | 649 | 2278 | 92 | 26856 | [illegible] |
| GO:BP | meiotic cell cycle | GO:0051321 | FALSE | 0.001821713 | 2.739520054 | 352 | 2278 | 58 | 26856 | [illegible] |
| GO:BP | regulation of mitotic metaphase/anaphase transition | GO:0030071 | FALSE | 0.001856178 | 2.731380286 | 88 | 2278 | 23 | 26856 | [illegible] |
| GO:BP | regulation of vesicle-mediated transport | GO:0060627 | FALSE | 0.001905253 | 2.720047243 | 587 | 2278 | 85 | 26856 | [illegible] |
| GO:BP | regulation of protein-containing complex disassembly | GO:0043244 | FALSE | 0.001922951 | 2.716031734 | 128 | 2278 | 29 | 26856 | [illegible] |
| GO:BP | positive regulation of DNA repair | GO:0045739 | FALSE | 0.001922951 | 2.716031734 | 128 | 2278 | 29 | 26856 | [illegible] |
| GO:BP | type II interferon-mediated signaling pathway | GO:0060333 | FALSE | 0.001964541 | 2.706738916 | 23 | 2278 | 11 | 26856 | [illegible] |
| GO:BP | regulation of spindle assembly | GO:0090169 | FALSE | 0.002013733 | 2.69599812 | 32 | 2278 | 13 | 26856 | [illegible] |
| GO:BP | smooth muscle cell migration | GO:0014909 | FALSE | 0.002055202 | 2.687145524 | 108 | 2278 | 26 | 26856 | [illegible] |
| GO:BP | monocyte differentiation | GO:0030224 | FALSE | 0.002084729 | 2.680950324 | 42 | 2278 | 15 | 26856 | [illegible] |
| GO:BP | regulation of T cell cytokine production | GO:0002724 | FALSE | 0.002139616 | 2.669664223 | 37 | 2278 | 14 | 26856 | [illegible] |
| GO:BP | T cell cytokine production | GO:0002369 | FALSE | 0.002139616 | 2.669664223 | 37 | 2278 | 14 | 26856 | [illegible] |
| GO:BP | establishment of RNA localization | GO:0051236 | FALSE | 0.002150975 | 2.667364613 | 157 | 2278 | 33 | 26856 | [illegible] |
| GO:BP | regulation of mRNA splicing, via spliceosome | GO:0048024 | FALSE | 0.002168429 | 2.663854712 | 115 | 2278 | 27 | 26856 | [illegible] |
| GO:BP | peptidyl-threonine modification | GO:0018210 | FALSE | 0.002243068 | 2.649157563 | 122 | 2278 | 28 | 26856 | [illegible] |
| GO:BP | response to interleukin-1 | GO:0070555 | FALSE | 0.002281507 | 2.641778104 | 129 | 2278 | 29 | 26856 | [illegible] |
| GO:BP | DNA-templated transcription initiation | GO:0006352 | FALSE | 0.002287263 | 2.640683837 | 136 | 2278 | 30 | 26856 | [illegible] |
| GO:BP | positive regulation of protein catabolic process | GO:0045732 | FALSE | 0.002290854 | 2.640002625 | 202 | 2278 | 39 | 26856 | [illegible] |
| GO:BP | positive regulation of cytokine production involved in immune response | GO:0002720 | FALSE | 0.002473318 | 2.606719979 | 83 | 2278 | 22 | 26856 | [illegible] |
| GO:BP | positive regulation of myeloid cell differentiation | GO:0045639 | FALSE | 0.002483035 | 2.605017168 | 109 | 2278 | 26 | 26856 | [illegible] |
| GO:BP | regulation of intracellular protein transport | GO:0033157 | FALSE | 0.002571874 | 2.589750317 | 226 | 2278 | 42 | 26856 | [illegible] |
| GO:BP | positive regulation of type II interferon production | GO:0032729 | FALSE | 0.002600706 | 2.584908753 | 77 | 2278 | 21 | 26856 | [illegible] |
| GO:BP | plasma membrane bounded cell projection morphogenesis | GO:0120039 | FALSE | 0.002614451 | 2.582619566 | 710 | 2278 | 98 | 26856 | [illegible] |
| GO:BP | leukocyte activation involved in inflammatory response | GO:0002269 | FALSE | 0.002642308 | 2.57801663 | 48 | 2278 | 16 | 26856 | [illegible] |
| GO:BP | negative regulation of mRNA catabolic process | GO:1902373 | FALSE | 0.002665205 | 2.574269382 | 71 | 2278 | 20 | 26856 | [illegible] |
| GO:BP | negative regulation of DNA-binding transcription factor activity | GO:0043433 | FALSE | 0.002770223 | 2.557485277 | 166 | 2278 | 34 | 26856 | [illegible] |
| GO:BP | cellular response to interleukin-1 | GO:0071347 | FALSE | 0.002817571 | 2.55012514 | 103 | 2278 | 25 | 26856 | [illegible] |
| GO:BP | regulation of secretion | GO:0051046 | FALSE | 0.00285424 | 2.544509569 | 767 | 2278 | 104 | 26856 | [illegible] |
| GO:BP | androgen receptor signaling pathway | GO:0030521 | FALSE | 0.002950852 | 2.530052535 | 43 | 2278 | 15 | 26856 | [illegible] |
| GO:BP | tissue migration | GO:0090130 | FALSE | 0.003014811 | 2.520739959 | 324 | 2278 | 54 | 26856 | [illegible] |
| GO:BP | regulation of type 2 immune response | GO:0002828 | FALSE | 0.003064514 | 2.513638435 | 33 | 2278 | 13 | 26856 | [illegible] |
| GO:BP | regulation of nuclear division | GO:0051783 | FALSE | 0.003104085 | 2.508066422 | 145 | 2278 | 31 | 26856 | [illegible] |
| GO:BP | regulation of T-helper cell differentiation | GO:0045622 | FALSE | 0.003123428 | 2.505368559 | 38 | 2278 | 14 | 26856 | [illegible] |
| GO:BP | regulation of smooth muscle cell migration | GO:0014910 | FALSE | 0.003163332 | 2.499855191 | 97 | 2278 | 24 | 26856 | [illegible] |
| GO:BP | protein stabilization | GO:0050821 | FALSE | 0.003211472 | 2.493295795 | 197 | 2278 | 38 | 26856 | [illegible] |
| GO:BP | regulation of stress-activated MAPK cascade | GO:0032872 | FALSE | 0.003211472 | 2.493295795 | 197 | 2278 | 38 | 26856 | [illegible] |
| GO:BP | response to hypoxia | GO:0001666 | FALSE | 0.003224377 | 2.491554132 | 333 | 2278 | 55 | 26856 | [illegible] |
| GO:BP | negative regulation of protein serine/threonine kinase activity | GO:0071901 | FALSE | 0.003412844 | 2.466883602 | 104 | 2278 | 25 | 26856 | [illegible] |
| GO:BP | negative regulation of metaphase/anaphase transition of cell cycle | GO:1902099 | FALSE | 0.003507119 | 2.45504948 | 91 | 2278 | 23 | 26856 | [illegible] |
| GO:BP | negative regulation of cellular catabolic process | GO:0031330 | FALSE | 0.003727998 | 2.428524296 | 183 | 2278 | 36 | 26856 | [illegible] |
| GO:BP | attachment of mitotic spindle microtubules to kinetochore | GO:0051315 | FALSE | 0.003755497 | 2.425332543 | 20 | 2278 | 10 | 26856 | [illegible] |
| GO:BP | macromolecule methylation | GO:0043414 | FALSE | 0.003777338 | 2.42281418 | 245 | 2278 | 44 | 26856 | [illegible] |
| GO:BP | regulation of DNA repair | GO:0006282 | FALSE | 0.003813215 | 2.418708741 | 206 | 2278 | 39 | 26856 | [illegible] |
| GO:BP | regulation of miRNA metabolic process | GO:2000628 | FALSE | 0.003830274 | 2.416770157 | 85 | 2278 | 22 | 26856 | [illegible] |
| GO:BP | B cell activation involved in immune response | GO:0002312 | FALSE | 0.003855022 | 2.413973144 | 98 | 2278 | 24 | 26856 | [illegible] |
| GO:BP | antiviral innate immune response | GO:0140374 | FALSE | 0.004078 | 2.389552829 | 55 | 2278 | 17 | 26856 | [illegible] |
| GO:BP | inflammasome-mediated signaling pathway | GO:0141084 | FALSE | 0.004078 | 2.389552829 | 55 | 2278 | 17 | 26856 | [illegible] |
| GO:BP | positive regulation of protein modification by small protein conjugation or removal | GO:1903322 | FALSE | 0.004223613 | 2.374315844 | 147 | 2278 | 31 | 26856 | [illegible] |
| GO:BP | regulation of telomere maintenance | GO:0032204 | FALSE | 0.004303614 | 2.366166716 | 92 | 2278 | 23 | 26856 | [illegible] |
| GO:BP | mRNA stabilization | GO:0048255 | FALSE | 0.004331179 | 2.363393907 | 61 | 2278 | 18 | 26856 | [illegible] |
| GO:BP | platelet-derived growth factor receptor signaling pathway | GO:0048008 | FALSE | 0.004331179 | 2.363393907 | 61 | 2278 | 18 | 26856 | [illegible] |
| GO:BP | positive regulation of T cell differentiation | GO:0045582 | FALSE | 0.004413236 | 2.355242834 | 119 | 2278 | 27 | 26856 | [illegible] |
| GO:BP | post-embryonic development | GO:0009791 | FALSE | 0.004449475 | 2.351691246 | 126 | 2278 | 28 | 26856 | [illegible] |
| GO:BP | double-strand break repair via homologous recombination | GO:0000724 | FALSE | 0.004488015 | 2.34794569 | 162 | 2278 | 33 | 26856 | [illegible] |
| GO:BP | regulation of cell growth | GO:0001558 | FALSE | 0.004558106 | 2.341215618 | 448 | 2278 | 68 | 26856 | [illegible] |
| GO:BP | epithelium development | GO:0060429 | FALSE | 0.004867741 | 2.312672516 | 1307 | 2278 | 159 | 26856 | [illegible] |
| GO:BP | muscle tissue development | GO:0060537 | FALSE | 0.004873724 | 2.312139035 | 484 | 2278 | 72 | 26856 | [illegible] |
| GO:BP | JNK cascade | GO:0007254 | FALSE | 0.004877487 | 2.31180388 | 170 | 2278 | 34 | 26856 | [illegible] |
| GO:BP | actin filament-based process | GO:0030029 | FALSE | 0.004895944 | 2.310163514 | 748 | 2278 | 101 | 26856 | [illegible] |
| GO:BP | regulation of inflammasome-mediated signaling pathway | GO:0141085 | FALSE | 0.004899231 | 2.309872067 | 50 | 2278 | 16 | 26856 | [illegible] |
| GO:BP | centriole assembly | GO:0098534 | FALSE | 0.004899231 | 2.309872067 | 50 | 2278 | 16 | 26856 | [illegible] |
| GO:BP | regulation of intracellular steroid hormone receptor signaling pathway | GO:0033143 | FALSE | 0.005127802 | 2.290068795 | 80 | 2278 | 21 | 26856 | [illegible] |
| GO:BP | regulation of nucleocytoplasmic transport | GO:0046822 | FALSE | 0.005192639 | 2.284611849 | 134 | 2278 | 29 | 26856 | [illegible] |
| GO:BP | positive regulation of tyrosine phosphorylation of STAT protein | GO:0042531 | FALSE | 0.005394649 | 2.268036772 | 56 | 2278 | 17 | 26856 | [illegible] |
| GO:BP | maintenance of sister chromatid cohesion | GO:0034086 | FALSE | 0.005498345 | 2.259768016 | 13 | 2278 | 8 | 26856 | [illegible] |
| GO:BP | regulation of nuclear-transcribed mRNA poly(A) tail shortening | GO:0060211 | FALSE | 0.005498345 | 2.259768016 | 13 | 2278 | 8 | 26856 | [illegible] |
| GO:BP | cellular response to macrophage colony-stimulating factor stimulus | GO:0036006 | FALSE | 0.005498345 | 2.259768016 | 13 | 2278 | 8 | 26856 | [illegible] |
| GO:BP | endocytosis | GO:0006897 | FALSE | 0.005505322 | 2.259217259 | 722 | 2278 | 98 | 26856 | [illegible] |
| GO:BP | negative regulation of type I interferon-mediated signaling pathway | GO:0060339 | FALSE | 0.005519462 | 2.25810323 | 25 | 2278 | 11 | 26856 | [illegible] |
| GO:BP | female gamete generation | GO:0007292 | TRUE | 0.005597833 | 2.251980083 | 171 | 2278 | 34 | 26856 | [illegible] |
| GO:BP | regulation of spindle organization | GO:0090224 | FALSE | 0.00570575 | 2.243687281 | 45 | 2278 | 15 | 26856 | [illegible] |
| GO:BP | neuron projection morphogenesis | GO:0048812 | FALSE | 0.005798919 | 2.236652948 | 695 | 2278 | 95 | 26856 | [illegible] |
| GO:BP | meiotic nuclear division | GO:0140013 | FALSE | 0.005817857 | 2.235236932 | 194 | 2278 | 37 | 26856 | [illegible] |
| GO:BP | fat cell differentiation | GO:0045444 | FALSE | 0.005879173 | 2.230683776 | 257 | 2278 | 45 | 26856 | [illegible] |
| GO:BP | regulation of myeloid leukocyte differentiation | GO:0002761 | FALSE | 0.005919669 | 2.227702584 | 142 | 2278 | 30 | 26856 | [illegible] |
| GO:BP | regulation of gliogenesis | GO:0014013 | FALSE | 0.00608294 | 2.215886479 | 135 | 2278 | 29 | 26856 | [illegible] |
| GO:BP | negative regulation of cysteine-type endopeptidase activity involved in apoptotic process | GO:0043154 | FALSE | 0.006373008 | 2.195655558 | 81 | 2278 | 21 | 26856 | [illegible] |
| GO:BP | protein K48-linked ubiquitination | GO:0070936 | FALSE | 0.006373008 | 2.195655558 | 81 | 2278 | 21 | 26856 | [illegible] |
| GO:BP | RNA 3'-end processing | GO:0031123 | TRUE | 0.006412095 | 2.193000053 | 94 | 2278 | 23 | 26856 | [illegible] |
| GO:BP | cellular response to nitrogen compound | GO:1901699 | FALSE | 0.006476667 | 2.188648464 | 828 | 2278 | 109 | 26856 | [illegible] |
| GO:BP | regulation of CD4-positive, alpha-beta T cell differentiation | GO:0043370 | FALSE | 0.006578642 | 2.181863757 | 51 | 2278 | 16 | 26856 | [illegible] |
| GO:BP | regulation of lymphocyte proliferation | GO:0050670 | FALSE | 0.006580881 | 2.18171599 | 242 | 2278 | 43 | 26856 | [illegible] |
| GO:BP | liver morphogenesis | GO:0072576 | TRUE | 0.006715415 | 2.172927133 | 35 | 2278 | 13 | 26856 | [illegible] |
| GO:BP | protein K63-linked deubiquitination | GO:0070536 | FALSE | 0.006715415 | 2.172927133 | 35 | 2278 | 13 | 26856 | [illegible] |
| GO:BP | amide metabolic process | GO:0043603 | FALSE | 0.00673862 | 2.171429055 | 1186 | 2278 | 146 | 26856 | [illegible] |
| GO:BP | regulation of transcription initiation by RNA polymerase II | GO:0060260 | FALSE | 0.00682634 | 2.165812082 | 75 | 2278 | 20 | 26856 | [illegible] |
| GO:BP | recombinational repair | GO:0000725 | FALSE | 0.006847136 | 2.164491054 | 165 | 2278 | 33 | 26856 | [illegible] |
| GO:BP | response to oxygen levels | GO:0070482 | FALSE | 0.007041273 | 2.152348847 | 436 | 2278 | 66 | 26856 | [illegible] |
| GO:BP | positive regulation of RIG-I signaling pathway | GO:1900246 | FALSE | 0.00710882 | 2.148202498 | 10 | 2278 | 7 | 26856 | [illegible] |
| GO:BP | muscle cell migration | GO:0014812 | FALSE | 0.007329359 | 2.134934018 | 122 | 2278 | 27 | 26856 | [illegible] |
| GO:BP | negative regulation of viral-induced cytoplasmic pattern recognition receptor signaling pathway | GO:0039532 | FALSE | 0.007804429 | 2.10765888 | 46 | 2278 | 15 | 26856 | [illegible] |
| GO:BP | telencephalon development | GO:0021537 | FALSE | 0.007842366 | 2.105552911 | 309 | 2278 | 51 | 26856 | [illegible] |
| GO:BP | response to transforming growth factor beta | GO:0071559 | FALSE | 0.007988239 | 2.097548926 | 268 | 2278 | 46 | 26856 | [illegible] |
| GO:BP | regulation of DNA biosynthetic process | GO:2000278 | FALSE | 0.008516204 | 2.069753942 | 130 | 2278 | 28 | 26856 | [illegible] |
| GO:BP | maintenance of location | GO:0051235 | FALSE | 0.008790798 | 2.055971718 | 378 | 2278 | 59 | 26856 | [illegible] |
| GO:BP | negative regulation of inflammasome-mediated signaling pathway | GO:0141086 | FALSE | 0.008832665 | 2.053908218 | 26 | 2278 | 11 | 26856 | [illegible] |
| GO:BP | regulation of hematopoietic progenitor cell differentiation | GO:1901532 | FALSE | 0.008923428 | 2.049468285 | 41 | 2278 | 14 | 26856 | [illegible] |
| GO:BP | negative regulation of apoptotic signaling pathway | GO:2001234 | FALSE | 0.00914159 | 2.038978265 | 245 | 2278 | 43 | 26856 | [illegible] |
| GO:BP | regulation of epithelial cell migration | GO:0010632 | FALSE | 0.009158158 | 2.038191851 | 237 | 2278 | 42 | 26856 | [illegible] |
| GO:BP | regulation of peptidyl-serine phosphorylation | GO:0033135 | FALSE | 0.009614582 | 2.017069585 | 160 | 2278 | 32 | 26856 | [illegible] |
| GO:BP | negative regulation of microtubule depolymerization | GO:0007026 | FALSE | 0.009787934 | 2.009308984 | 31 | 2278 | 12 | 26856 | [illegible] |
| GO:BP | positive regulation of developmental growth | GO:0048639 | FALSE | 0.009830995 | 2.007402512 | 206 | 2278 | 38 | 26856 | [illegible] |
| GO:BP | import into cell | GO:0098657 | FALSE | 0.010259283 | 1.988882978 | 932 | 2278 | 119 | 26856 | [illegible] |
| GO:BP | protein localization to cell periphery | GO:1990778 | FALSE | 0.010404735 | 1.982768982 | 380 | 2278 | 59 | 26856 | [illegible] |
| GO:BP | regulation of proteasomal protein catabolic process | GO:0061136 | FALSE | 0.010480495 | 1.97961821 | 191 | 2278 | 36 | 26856 | [illegible] |
| GO:BP | positive regulation of macroautophagy | GO:0016239 | FALSE | 0.010611132 | 1.97423828 | 77 | 2278 | 20 | 26856 | [illegible] |
| GO:BP | regulation of microtubule polymerization or depolymerization | GO:0031110 | FALSE | 0.01067216 | 1.971747668 | 90 | 2278 | 22 | 26856 | [illegible] |
| GO:BP | negative regulation of binding | GO:0051100 | FALSE | 0.010913749 | 1.962026043 | 176 | 2278 | 34 | 26856 | [illegible] |
| GO:BP | cellular response to transforming growth factor beta stimulus | GO:0071560 | FALSE | 0.011057146 | 1.956356949 | 263 | 2278 | 45 | 26856 | [illegible] |
| GO:BP | methylation | GO:0032259 | FALSE | 0.011116222 | 1.954042807 | 296 | 2278 | 49 | 26856 | [illegible] |
| GO:BP | neuron apoptotic process | GO:0051402 | FALSE | 0.011117686 | 1.953985592 | 338 | 2278 | 54 | 26856 | [illegible] |
| GO:BP | suppression of viral release by host | GO:0044790 | FALSE | 0.011216461 | 1.950144159 | 22 | 2278 | 10 | 26856 | [illegible] |
| GO:BP | protein dephosphorylation | GO:0006470 | FALSE | 0.011399264 | 1.943123194 | 239 | 2278 | 42 | 26856 | [illegible] |
| GO:BP | positive regulation of epithelial cell proliferation | GO:0050679 | FALSE | 0.011412021 | 1.942637428 | 231 | 2278 | 41 | 26856 | [illegible] |
| GO:BP | cellular response to hypoxia | GO:0071456 | FALSE | 0.011746242 | 1.930101058 | 154 | 2278 | 31 | 26856 | [illegible] |
| GO:BP | RNA transport | GO:0050658 | FALSE | 0.011746242 | 1.930101058 | 154 | 2278 | 31 | 26856 | [illegible] |
| GO:BP | nucleic acid transport | GO:0050657 | FALSE | 0.011746242 | 1.930101058 | 154 | 2278 | 31 | 26856 | [illegible] |
| GO:BP | regulation of type II interferon-mediated signaling pathway | GO:0060334 | FALSE | 0.011870131 | 1.9255445 | 14 | 2278 | 8 | 26856 | [illegible] |
| GO:BP | regulation of response to type II interferon | GO:0060330 | FALSE | 0.011870131 | 1.9255445 | 14 | 2278 | 8 | 26856 | [illegible] |
| GO:BP | negative regulation of cell differentiation | GO:0045596 | FALSE | 0.011873424 | 1.925424027 | 801 | 2278 | 105 | 26856 | [illegible] |
| GO:BP | regulation of T cell mediated immunity | GO:0002709 | FALSE | 0.012048826 | 1.919055274 | 111 | 2278 | 25 | 26856 | [illegible] |
| GO:BP | response to insulin | GO:0032868 | FALSE | 0.012234247 | 1.912422747 | 297 | 2278 | 49 | 26856 | [illegible] |
| GO:BP | positive regulation of interleukin-10 production | GO:0032733 | FALSE | 0.012342777 | 1.908587115 | 42 | 2278 | 14 | 26856 | [illegible] |
| GO:BP | myeloid cell apoptotic process | GO:0033028 | FALSE | 0.012342777 | 1.908587115 | 42 | 2278 | 14 | 26856 | [illegible] |
| GO:BP | negative regulation of growth | GO:0045926 | FALSE | 0.01244895 | 1.904867267 | 256 | 2278 | 44 | 26856 | [illegible] |
| GO:BP | positive regulation of production of molecular mediator of immune response | GO:0002702 | FALSE | 0.013032265 | 1.884980096 | 140 | 2278 | 29 | 26856 | [illegible] |
| GO:BP | regulation of protein tyrosine kinase activity | GO:0061097 | FALSE | 0.013140438 | 1.881390143 | 78 | 2278 | 20 | 26856 | [illegible] |
| GO:BP | secretion by cell | GO:0032940 | FALSE | 0.013175729 | 1.880225354 | 908 | 2278 | 116 | 26856 | [illegible] |
| GO:BP | neuron migration | GO:0001764 | FALSE | 0.013405174 | 1.872727559 | 193 | 2278 | 36 | 26856 | [illegible] |
| GO:BP | peptidyl-lysine acetylation | GO:0018394 | FALSE | 0.013556835 | 1.867841698 | 133 | 2278 | 28 | 26856 | [illegible] |
| GO:BP | wound healing | GO:0042060 | TRUE | 0.01370955 | 1.862976788 | 392 | 2278 | 60 | 26856 | [illegible] |
| GO:BP | positive regulation of binding | GO:0051099 | FALSE | 0.013746663 | 1.861802723 | 201 | 2278 | 37 | 26856 | [illegible] |
| GO:BP | activation of protein kinase activity | GO:0032147 | FALSE | 0.014106167 | 1.850590985 | 105 | 2278 | 24 | 26856 | [illegible] |
| GO:BP | regulation of extrinsic apoptotic signaling pathway via death domain receptors | GO:1902041 | FALSE | 0.014168596 | 1.848673176 | 48 | 2278 | 15 | 26856 | [illegible] |
| GO:BP | somatic diversification of immune receptors | GO:0002200 | FALSE | 0.014591664 | 1.835895167 | 85 | 2278 | 21 | 26856 | [illegible] |
| GO:BP | positive regulation of protein localization to nucleus | GO:1900182 | FALSE | 0.016398006 | 1.785208957 | 99 | 2278 | 23 | 26856 | [illegible] |
| GO:BP | regulation of fibroblast proliferation | GO:0048145 | FALSE | 0.01686682 | 1.772966792 | 113 | 2278 | 25 | 26856 | [illegible] |
| GO:BP | regulation of defense response to virus by host | GO:0050691 | FALSE | 0.01687681 | 1.772709646 | 43 | 2278 | 14 | 26856 | [illegible] |
| GO:BP | intrinsic apoptotic signaling pathway in response to DNA damage by p53 class mediator | GO:0042771 | FALSE | 0.01687681 | 1.772709646 | 43 | 2278 | 14 | 26856 | [illegible] |
| GO:BP | cell-cell adhesion | GO:0098609 | FALSE | 0.017015344 | 1.769159261 | 913 | 2278 | 116 | 26856 | [illegible] |
| GO:BP | neuron projection extension | GO:1990138 | FALSE | 0.017397361 | 1.759516615 | 203 | 2278 | 37 | 26856 | [illegible] |
| GO:BP | positive regulation of DNA-templated transcription initiation | GO:2000144 | FALSE | 0.017681448 | 1.752482176 | 73 | 2278 | 19 | 26856 | [illegible] |
| GO:BP | innate immune response activating cell surface receptor signaling pathway | GO:0002220 | FALSE | 0.017681448 | 1.752482176 | 73 | 2278 | 19 | 26856 | [illegible] |
| GO:BP | regulation of vascular associated smooth muscle cell proliferation | GO:1904705 | FALSE | 0.017681448 | 1.752482176 | 73 | 2278 | 19 | 26856 | [illegible] |
| GO:BP | vacuolar transport | GO:0007034 | FALSE | 0.017764648 | 1.750443403 | 157 | 2278 | 31 | 26856 | [illegible] |
| GO:BP | regulation of osteoclast differentiation | GO:0045670 | FALSE | 0.017775304 | 1.75018296 | 86 | 2278 | 21 | 26856 | [illegible] |
| GO:BP | lymphocyte proliferation | GO:0046651 | FALSE | 0.018078208 | 1.742844617 | 335 | 2278 | 53 | 26856 | [illegible] |
| GO:BP | negative regulation of AIM2 inflammasome complex assembly | GO:0140972 | FALSE | 0.018107832 | 1.742133539 | 11 | 2278 | 7 | 26856 | [illegible] |
| GO:BP | cerebral cortex development | GO:0021987 | FALSE | 0.018300194 | 1.73754431 | 135 | 2278 | 28 | 26856 | [illegible] |
| GO:BP | response to endoplasmic reticulum stress | GO:0034976 | FALSE | 0.018742571 | 1.727170843 | 260 | 2278 | 44 | 26856 | [illegible] |
| GO:BP | positive regulation of protein ubiquitination | GO:0031398 | FALSE | 0.019022675 | 1.720728417 | 128 | 2278 | 27 | 26856 | [illegible] |
| GO:BP | DNA biosynthetic process | GO:0071897 | FALSE | 0.019222397 | 1.716192468 | 196 | 2278 | 36 | 26856 | [illegible] |
| GO:BP | regulation of alternative mRNA splicing, via spliceosome | GO:0000381 | FALSE | 0.019621848 | 1.707260093 | 61 | 2278 | 17 | 26856 | [illegible] |
| GO:BP | regulation of nuclease activity | GO:0032069 | FALSE | 0.020928685 | 1.679258067 | 28 | 2278 | 11 | 26856 | [illegible] |
| GO:BP | positive regulation of type II interferon-mediated signaling pathway | GO:0060335 | FALSE | 0.0212331 | 1.672982293 | 8 | 2278 | 6 | 26856 | [illegible] |
| GO:BP | regulation of cytoplasmic mRNA processing body assembly | GO:0010603 | FALSE | 0.0212331 | 1.672982293 | 8 | 2278 | 6 | 26856 | [illegible] |
| GO:BP | positive regulation of response to type II interferon | GO:0060332 | FALSE | 0.0212331 | 1.672982293 | 8 | 2278 | 6 | 26856 | [illegible] |
| GO:BP | actin cytoskeleton organization | GO:0030036 | FALSE | 0.021776095 | 1.662020007 | 661 | 2278 | 89 | 26856 | [illegible] |
| GO:BP | regulation of chemotaxis | GO:0050920 | FALSE | 0.021817136 | 1.661202262 | 237 | 2278 | 41 | 26856 | [illegible] |
| GO:BP | regulation of CD4-positive, alpha-beta T cell activation | GO:2000514 | FALSE | 0.021911439 | 1.659329102 | 74 | 2278 | 19 | 26856 | [illegible] |
| GO:BP | regulation of attachment of spindle microtubules to kinetochore | GO:0051988 | FALSE | 0.02240716 | 1.649613188 | 19 | 2278 | 9 | 26856 | [illegible] |
| GO:BP | regulation of circadian rhythm | GO:0042752 | FALSE | 0.022881492 | 1.640515663 | 122 | 2278 | 26 | 26856 | [illegible] |
| GO:BP | protein localization to condensed chromosome | GO:1903083 | FALSE | 0.023535992 | 1.628267501 | 15 | 2278 | 8 | 26856 | [illegible] |
| GO:BP | dendritic cell cytokine production | GO:0002371 | FALSE | 0.023535992 | 1.628267501 | 15 | 2278 | 8 | 26856 | [illegible] |
| GO:BP | regulation of dendritic cell cytokine production | GO:0002730 | FALSE | 0.023535992 | 1.628267501 | 15 | 2278 | 8 | 26856 | [illegible] |
| GO:BP | protein localization to kinetochore | GO:0034501 | FALSE | 0.023535992 | 1.628267501 | 15 | 2278 | 8 | 26856 | [illegible] |
| GO:BP | positive regulation of intracellular transport | GO:0032388 | FALSE | 0.023874948 | 1.622057565 | 190 | 2278 | 35 | 26856 | [illegible] |
| GO:BP | positive regulation of leukocyte proliferation | GO:0070665 | FALSE | 0.024458393 | 1.611572089 | 167 | 2278 | 32 | 26856 | [illegible] |
| GO:BP | phospholipid biosynthetic process | GO:0008654 | FALSE | 0.02449739 | 1.610880175 | 230 | 2278 | 40 | 26856 | [illegible] |
| GO:BP | nucleus organization | GO:0006997 | FALSE | 0.024515412 | 1.610560806 | 137 | 2278 | 28 | 26856 | [illegible] |
| GO:BP | protein localization to chromosome | GO:0034502 | FALSE | 0.024515412 | 1.610560806 | 137 | 2278 | 28 | 26856 | [illegible] |
| GO:BP | regulation of cysteine-type endopeptidase activity involved in apoptotic process | GO:0043281 | FALSE | 0.024569695 | 1.609600238 | 206 | 2278 | 37 | 26856 | [illegible] |
| GO:BP | G1/S transition of mitotic cell cycle | GO:0000082 | FALSE | 0.024652237 | 1.608143658 | 222 | 2278 | 39 | 26856 | [illegible] |
| GO:BP | p38MAPK cascade | GO:0038066 | FALSE | 0.024792418 | 1.605681112 | 50 | 2278 | 15 | 26856 | [illegible] |
| GO:BP | regulation of cytokine production involved in inflammatory response | GO:1900015 | FALSE | 0.024910896 | 1.603610652 | 62 | 2278 | 17 | 26856 | [illegible] |
| GO:BP | cytokine production involved in inflammatory response | GO:0002534 | FALSE | 0.024910896 | 1.603610652 | 62 | 2278 | 17 | 26856 | [illegible] |
| GO:BP | regulation of interleukin-10 production | GO:0032653 | FALSE | 0.024910896 | 1.603610652 | 62 | 2278 | 17 | 26856 | [illegible] |
| GO:BP | interleukin-10 production | GO:0032613 | FALSE | 0.024910896 | 1.603610652 | 62 | 2278 | 17 | 26856 | [illegible] |
| GO:BP | regulation of neuron apoptotic process | GO:0043523 | FALSE | 0.025346257 | 1.596086165 | 288 | 2278 | 47 | 26856 | [illegible] |
| GO:BP | interleukin-12 production | GO:0032615 | FALSE | 0.025391547 | 1.595310846 | 56 | 2278 | 16 | 26856 | [illegible] |
| GO:BP | regulation of interleukin-12 production | GO:0032655 | FALSE | 0.025391547 | 1.595310846 | 56 | 2278 | 16 | 26856 | [illegible] |
| GO:BP | negative regulation of DNA binding | GO:0043392 | FALSE | 0.025391547 | 1.595310846 | 56 | 2278 | 16 | 26856 | [illegible] |
| GO:BP | positive regulation of gliogenesis | GO:0014015 | FALSE | 0.02609059 | 1.58351607 | 88 | 2278 | 21 | 26856 | [illegible] |
| GO:BP | positive regulation of interleukin-12 production | GO:0032735 | FALSE | 0.026755346 | 1.572589435 | 39 | 2278 | 13 | 26856 | [illegible] |
| GO:BP | response to protozoan | GO:0001562 | FALSE | 0.026755346 | 1.572589435 | 39 | 2278 | 13 | 26856 | [illegible] |
| GO:BP | cell adhesion | GO:0007155 | FALSE | 0.026943661 | 1.569543393 | 1648 | 2278 | 169 | 26856 | [illegible] |
| GO:BP | positive regulation of alpha-beta T cell activation | GO:0046635 | FALSE | 0.027029597 | 1.568160422 | 75 | 2278 | 19 | 26856 | [illegible] |
| GO:BP | chemotaxis | GO:0006935 | FALSE | 0.027449297 | 1.561467189 | 454 | 2278 | 66 | 26856 | [illegible] |
| GO:BP | receptor signaling pathway via JAK-STAT | GO:0007259 | FALSE | 0.027507782 | 1.56054442 | 207 | 2278 | 37 | 26856 | [illegible] |
| GO:BP | hepatocyte apoptotic process | GO:0097284 | FALSE | 0.029037069 | 1.537047218 | 24 | 2278 | 10 | 26856 | [illegible] |
| GO:BP | cellular response to platelet-derived growth factor stimulus | GO:0036120 | FALSE | 0.029037069 | 1.537047218 | 24 | 2278 | 10 | 26856 | [illegible] |
| GO:BP | DNA damage response, signal transduction by p53 class mediator | GO:0030330 | FALSE | 0.029488595 | 1.530345921 | 69 | 2278 | 18 | 26856 | [illegible] |
| GO:BP | negative regulation of NF-kappaB transcription factor activity | GO:0032088 | FALSE | 0.029630004 | 1.528268294 | 82 | 2278 | 20 | 26856 | [illegible] |
| GO:BP | glycerolipid biosynthetic process | GO:0045017 | FALSE | 0.030578009 | 1.514590796 | 224 | 2278 | 39 | 26856 | [illegible] |
| GO:BP | neural precursor cell proliferation | GO:0061351 | FALSE | 0.030765326 | 1.511938477 | 208 | 2278 | 37 | 26856 | [illegible] |
| GO:BP | response to oxidative stress | GO:0006979 | FALSE | 0.030934512 | 1.509556725 | 420 | 2278 | 62 | 26856 | [illegible] |
| GO:BP | neutrophil homeostasis | GO:0001780 | FALSE | 0.031135248 | 1.506747673 | 29 | 2278 | 11 | 26856 | [illegible] |
| GO:BP | positive regulation of cytokine production involved in inflammatory response | GO:1900017 | FALSE | 0.031135248 | 1.506747673 | 29 | 2278 | 11 | 26856 | [illegible] |
| GO:BP | regulation of epidermis development | GO:0045682 | FALSE | 0.031438059 | 1.502544273 | 63 | 2278 | 17 | 26856 | [illegible] |
| GO:BP | taxis | GO:0042330 | FALSE | 0.031687118 | 1.499117264 | 456 | 2278 | 66 | 26856 | [illegible] |
| GO:BP | regulation of viral entry into host cell | GO:0046596 | FALSE | 0.032381492 | 1.48970315 | 51 | 2278 | 15 | 26856 | [illegible] |
| GO:BP | lung alveolus development | GO:0048286 | FALSE | 0.032536561 | 1.487628353 | 57 | 2278 | 16 | 26856 | [illegible] |
| GO:BP | mRNA transcription | GO:0009299 | FALSE | 0.032536561 | 1.487628353 | 57 | 2278 | 16 | 26856 | [illegible] |
| GO:BP | response to unfolded protein | GO:0006986 | FALSE | 0.032740294 | 1.484917427 | 110 | 2278 | 24 | 26856 | [illegible] |
| GO:BP | phosphatidylinositol biosynthetic process | GO:0006661 | FALSE | 0.032740294 | 1.484917427 | 110 | 2278 | 24 | 26856 | [illegible] |
| GO:BP | regulation of interleukin-8 production | GO:0032677 | FALSE | 0.033195963 | 1.478914733 | 76 | 2278 | 19 | 26856 | [illegible] |
| GO:BP | interleukin-8 production | GO:0032637 | FALSE | 0.033195963 | 1.478914733 | 76 | 2278 | 19 | 26856 | [illegible] |
| GO:BP | regulation of DNA recombination | GO:0000018 | FALSE | 0.034461359 | 1.462667603 | 132 | 2278 | 27 | 26856 | [illegible] |
| GO:BP | protein polymerization | GO:0051258 | TRUE | 0.034751896 | 1.459021502 | 283 | 2278 | 46 | 26856 | [illegible] |
| GO:BP | regulation of signal transduction by p53 class mediator | GO:1901796 | FALSE | 0.035950386 | 1.444296439 | 83 | 2278 | 20 | 26856 | [illegible] |
| GO:BP | negative regulation of cell development | GO:0010721 | FALSE | 0.036758382 | 1.434643618 | 309 | 2278 | 49 | 26856 | [illegible] |
| GO:BP | negative regulation of DNA metabolic process | GO:0051053 | FALSE | 0.037491967 | 1.42606717 | 140 | 2278 | 28 | 26856 | [illegible] |
| GO:BP | T-helper 2 cell differentiation | GO:0045064 | FALSE | 0.037667611 | 1.424031923 | 20 | 2278 | 9 | 26856 | [illegible] |
| GO:BP | regulation of proteolysis involved in protein catabolic process | GO:1903050 | FALSE | 0.037787762 | 1.422648828 | 226 | 2278 | 39 | 26856 | [illegible] |
| GO:BP | phosphatidylinositol 3-kinase/protein kinase B signal transduction | GO:0043491 | FALSE | 0.038170606 | 1.418270941 | 218 | 2278 | 38 | 26856 | [illegible] |
| GO:BP | lipid biosynthetic process | GO:0008610 | FALSE | 0.03936263 | 1.404915896 | 690 | 2278 | 91 | 26856 | [illegible] |
| GO:BP | response to topologically incorrect protein | GO:0035966 | FALSE | 0.039786096 | 1.400268674 | 133 | 2278 | 27 | 26856 | [illegible] |
| GO:BP | maintenance of mitotic sister chromatid cohesion | GO:0034088 | FALSE | 0.040258602 | 1.395141313 | 12 | 2278 | 7 | 26856 | [illegible] |
| GO:BP | MyD88-dependent toll-like receptor signaling pathway | GO:0002755 | FALSE | 0.040258602 | 1.395141313 | 12 | 2278 | 7 | 26856 | [illegible] |
| GO:BP | positive regulation of nuclear-transcribed mRNA poly(A) tail shortening | GO:0060213 | FALSE | 0.040258602 | 1.395141313 | 12 | 2278 | 7 | 26856 | [illegible] |
| GO:BP | vascular associated smooth muscle cell proliferation | GO:1990874 | FALSE | 0.040594442 | 1.391533428 | 77 | 2278 | 19 | 26856 | [illegible] |
| GO:BP | somatic diversification of immune receptors via germline recombination within a single locus | GO:0002562 | FALSE | 0.040594442 | 1.391533428 | 77 | 2278 | 19 | 26856 | [illegible] |
| GO:BP | somatic cell DNA recombination | GO:0016444 | FALSE | 0.040594442 | 1.391533428 | 77 | 2278 | 19 | 26856 | [illegible] |
| GO:BP | regulation of anatomical structure size | GO:0090066 | FALSE | 0.04111624 | 1.385986604 | 560 | 2278 | 77 | 26856 | [illegible] |
| GO:BP | regeneration | GO:0031099 | FALSE | 0.041949622 | 1.377271948 | 227 | 2278 | 39 | 26856 | [illegible] |
| GO:BP | defense response to protozoan | GO:0042832 | FALSE | 0.041990714 | 1.376846742 | 35 | 2278 | 12 | 26856 | [illegible] |
| GO:BP | regulation of microtubule depolymerization | GO:0031114 | FALSE | 0.041990714 | 1.376846742 | 35 | 2278 | 12 | 26856 | [illegible] |
| GO:BP | positive regulation of epithelial cell migration | GO:0010634 | FALSE | 0.042389054 | 1.372746278 | 156 | 2278 | 30 | 26856 | [illegible] |
| GO:BP | T cell mediated immunity | GO:0002456 | FALSE | 0.043042342 | 1.366104108 | 141 | 2278 | 28 | 26856 | [illegible] |
| GO:BP | positive regulation of double-strand break repair | GO:2000781 | FALSE | 0.04345583 | 1.361951955 | 84 | 2278 | 20 | 26856 | [illegible] |
| GO:BP | positive regulation of T cell cytokine production | GO:0002726 | FALSE | 0.044715474 | 1.34954216 | 25 | 2278 | 10 | 26856 | [illegible] |
| GO:BP | histone acetylation | GO:0016573 | FALSE | 0.045041352 | 1.346388578 | 112 | 2278 | 24 | 26856 | [illegible] |
| GO:BP | maintenance of protein location in cell | GO:0032507 | TRUE | 0.045137708 | 1.345460499 | 71 | 2278 | 18 | 26856 | [illegible] |
| GO:BP | regulation of cilium assembly | GO:1902017 | FALSE | 0.045137708 | 1.345460499 | 71 | 2278 | 18 | 26856 | [illegible] |
| GO:BP | stress granule assembly | GO:0034063 | FALSE | 0.04539724 | 1.342970554 | 30 | 2278 | 11 | 26856 | [illegible] |
| GO:BP | protein K48-linked deubiquitination | GO:0071108 | FALSE | 0.04539724 | 1.342970554 | 30 | 2278 | 11 | 26856 | [illegible] |
| GO:BP | lymphocyte mediated immunity | GO:0002449 | FALSE | 0.045429481 | 1.342662228 | 320 | 2278 | 50 | 26856 | [illegible] |
| GO:BP | cell cycle G2/M phase transition | GO:0044839 | FALSE | 0.045853868 | 1.338624022 | 149 | 2278 | 29 | 26856 | [illegible] |
| GO:BP | protein localization to plasma membrane | GO:0072659 | FALSE | 0.045871538 | 1.338456686 | 303 | 2278 | 48 | 26856 | [illegible] |
| GO:BP | regulation of actin filament-based process | GO:0032970 | FALSE | 0.046497045 | 1.332574646 | 390 | 2278 | 58 | 26856 | [illegible] |
| GO:BP | glycerophospholipid biosynthetic process | GO:0046474 | FALSE | 0.046736052 | 1.33034798 | 180 | 2278 | 33 | 26856 | [illegible] |
| GO:BP | embryonic morphogenesis | GO:0048598 | FALSE | 0.046946288 | 1.328398741 | 646 | 2278 | 86 | 26856 | [illegible] |
| GO:BP | T cell activation involved in immune response | GO:0002286 | FALSE | 0.048461599 | 1.314602256 | 127 | 2278 | 26 | 26856 | [illegible] |
| GO:BP | regulation of mitotic spindle organization | GO:0060236 | FALSE | 0.049404598 | 1.306232632 | 41 | 2278 | 13 | 26856 | [illegible] |
| GO:BP | multicellular organismal reproductive process | GO:0048609 | FALSE | 0.049694089 | 1.303695263 | 1132 | 2278 | 136 | 26856 | [illegible] |
| GO:BP | gland development | GO:0048732 | FALSE | 0.049701927 | 1.303626775 | 517 | 2278 | 72 | 26856 | [illegible] |
| GO:CC | intracellular anatomical structure | GO:0005622 | TRUE | 8.70E-153 | 152.0606455 | 17319 | 2205 | 1945 | 26959 | [illegible] |
| GO:CC | nucleoplasm | GO:0005654 | FALSE | 6.69E-134 | 133.1744348 | 3826 | 2205 | 752 | 26959 | [illegible] |
| GO:CC | intracellular organelle | GO:0043229 | FALSE | 5.13E-133 | 132.2902287 | 15733 | 2205 | 1810 | 26959 | [illegible] |
| GO:CC | cytoplasm | GO:0005737 | FALSE | 1.93E-130 | 129.7140464 | 11583 | 2205 | 1495 | 26959 | [illegible] |
| GO:CC | organelle | GO:0043226 | FALSE | 7.20E-130 | 129.1426369 | 16054 | 2205 | 1826 | 26959 | [illegible] |
| GO:CC | membrane-bounded organelle | GO:0043227 | FALSE | 3.21E-114 | 113.4938335 | 14825 | 2205 | 1710 | 26959 | [illegible] |
| GO:CC | intracellular membrane-bounded organelle | GO:0043231 | FALSE | 4.35E-114 | 113.3610826 | 14349 | 2205 | 1675 | 26959 | [illegible] |
| GO:CC | nucleus | GO:0005634 | FALSE | 6.09E-97 | 96.21550592 | 10010 | 2205 | 1288 | 26959 | [illegible] |
| GO:CC | cellular anatomical entity | GO:0110165 | FALSE | 3.15E-82 | 81.50139921 | 22588 | 2205 | 2126 | 26959 | [illegible] |
| GO:CC | cytosol | GO:0005829 | FALSE | 7.32E-71 | 70.13546168 | 4169 | 2205 | 662 | 26959 | [illegible] |
| GO:CC | nuclear lumen | GO:0031981 | FALSE | 4.45E-63 | 62.35212091 | 5855 | 2205 | 815 | 26959 | [illegible] |
| GO:CC | intracellular organelle lumen | GO:0070013 | FALSE | 2.49E-58 | 57.60437213 | 6395 | 2205 | 854 | 26959 | [illegible] |
| GO:CC | organelle lumen | GO:0043233 | FALSE | 2.68E-58 | 57.5715586 | 6396 | 2205 | 854 | 26959 | [illegible] |
| GO:CC | membrane-enclosed lumen | GO:0031974 | FALSE | 2.68E-58 | 57.5715586 | 6396 | 2205 | 854 | 26959 | [illegible] |
| GO:CC | chromosome | GO:0005694 | FALSE | 3.51E-44 | 43.45507043 | 1420 | 2205 | 282 | 26959 | [illegible] |
| GO:CC | microtubule cytoskeleton | GO:0015630 | FALSE | 1.33E-39 | 38.87513715 | 1362 | 2205 | 265 | 26959 | [illegible] |
| GO:CC | nuclear body | GO:0016604 | FALSE | 2.25E-38 | 37.64818934 | 927 | 2205 | 205 | 26959 | [illegible] |
| GO:CC | intracellular non-membrane-bounded organelle | GO:0043232 | FALSE | 1.45E-36 | 35.8397796 | 6529 | 2205 | 796 | 26959 | [illegible] |
| GO:CC | non-membrane-bounded organelle | GO:0043228 | FALSE | 1.53E-36 | 35.81424783 | 6530 | 2205 | 796 | 26959 | [illegible] |
| GO:CC | protein-containing complex | GO:0032991 | FALSE | 2.74E-32 | 31.56272046 | 7392 | 2205 | 859 | 26959 | [illegible] |
| GO:CC | cytoskeleton | GO:0005856 | FALSE | 1.28E-31 | 30.89136044 | 2364 | 2205 | 363 | 26959 | [illegible] |
| GO:CC | microtubule organizing center | GO:0005815 | FALSE | 8.74E-31 | 30.05825117 | 836 | 2205 | 178 | 26959 | [illegible] |
| GO:CC | centrosome | GO:0005813 | FALSE | 2.50E-28 | 27.60251753 | 646 | 2205 | 147 | 26959 | [illegible] |
| GO:CC | supramolecular complex | GO:0099080 | FALSE | 3.96E-26 | 25.40257479 | 1404 | 2205 | 239 | 26959 | [illegible] |
| GO:CC | catalytic complex | GO:1902494 | FALSE | 2.08E-25 | 24.68270239 | 1794 | 2205 | 282 | 26959 | [illegible] |
| GO:CC | protein-DNA complex | GO:0032993 | FALSE | 3.21E-25 | 24.49291266 | 854 | 2205 | 169 | 26959 | [illegible] |
| GO:CC | chromosomal region | GO:0098687 | FALSE | 3.46E-25 | 24.46083116 | 380 | 2205 | 102 | 26959 | [illegible] |
| GO:CC | chromatin | GO:0000785 | FALSE | 1.17E-24 | 23.93010412 | 764 | 2205 | 156 | 26959 | [illegible] |
| GO:CC | spindle | GO:0005819 | FALSE | 7.81E-24 | 23.10736441 | 420 | 2205 | 106 | 26959 | [illegible] |
| GO:CC | ribonucleoprotein granule | GO:0035770 | FALSE | 1.33E-21 | 20.87693138 | 266 | 2205 | 78 | 26959 | [illegible] |
| GO:CC | cytoplasmic ribonucleoprotein granule | GO:0036464 | FALSE | 1.60E-21 | 20.79492466 | 249 | 2205 | 75 | 26959 | [illegible] |
| GO:CC | transferase complex | GO:1990234 | FALSE | 1.31E-20 | 19.88146506 | 880 | 2205 | 162 | 26959 | [illegible] |
| GO:CC | chromosome, centromeric region | GO:0000775 | FALSE | 2.28E-20 | 19.64192789 | 271 | 2205 | 77 | 26959 | [illegible] |
| GO:CC | nuclear speck | GO:0016607 | FALSE | 5.99E-18 | 17.2223844 | 458 | 2205 | 101 | 26959 | [illegible] |
| GO:CC | condensed chromosome, centromeric region | GO:0000779 | FALSE | 4.41E-16 | 15.3558001 | 172 | 2205 | 54 | 26959 | [illegible] |
| GO:CC | intracellular protein-containing complex | GO:0140535 | FALSE | 2.42E-15 | 14.61585897 | 948 | 2205 | 157 | 26959 | [illegible] |
| GO:CC | endomembrane system | GO:0012505 | FALSE | 2.80E-15 | 14.55313963 | 4293 | 2205 | 498 | 26959 | [illegible] |
| GO:CC | kinetochore | GO:0000776 | FALSE | 1.22E-13 | 12.91291922 | 163 | 2205 | 49 | 26959 | [illegible] |
| GO:CC | spindle pole | GO:0000922 | FALSE | 1.01E-12 | 11.99411781 | 171 | 2205 | 49 | 26959 | [illegible] |
| GO:CC | condensed chromosome | GO:0000793 | FALSE | 3.87E-12 | 11.41205051 | 360 | 2205 | 76 | 26959 | [illegible] |
| GO:CC | intracellular vesicle | GO:0097708 | FALSE | 3.89E-12 | 11.41019955 | 2066 | 2205 | 267 | 26959 | [illegible] |
| GO:CC | midbody | GO:0030496 | FALSE | 8.39E-12 | 11.07598324 | 218 | 2205 | 55 | 26959 | [illegible] |
| GO:CC | cytoplasmic vesicle | GO:0031410 | FALSE | 9.62E-12 | 11.01689297 | 2062 | 2205 | 265 | 26959 | [illegible] |
| GO:CC | vesicle | GO:0031982 | FALSE | 3.41E-11 | 10.46730982 | 2214 | 2205 | 278 | 26959 | [illegible] |
| GO:CC | P-body | GO:0000932 | FALSE | 8.37E-11 | 10.07702692 | 95 | 2205 | 33 | 26959 | [illegible] |
| GO:CC | mitotic spindle | GO:0072686 | FALSE | 1.77E-10 | 9.75145085 | 174 | 2205 | 46 | 26959 | [illegible] |
| GO:CC | perinuclear region of cytoplasm | GO:0048471 | FALSE | 2.85E-10 | 9.544616436 | 804 | 2205 | 126 | 26959 | [illegible] |
| GO:CC | PML body | GO:0016605 | FALSE | 3.69E-10 | 9.432855285 | 105 | 2205 | 34 | 26959 | [illegible] |
| GO:CC | microtubule | GO:0005874 | FALSE | 4.06E-10 | 9.391637301 | 463 | 2205 | 85 | 26959 | [illegible] |
| GO:CC | centriole | GO:0005814 | FALSE | 9.30E-10 | 9.031383762 | 156 | 2205 | 42 | 26959 | [illegible] |
| GO:CC | nuclear periphery | GO:0034399 | FALSE | 1.98E-09 | 8.702295702 | 153 | 2205 | 41 | 26959 | [illegible] |
| GO:CC | cell projection | GO:0042995 | FALSE | 6.07E-09 | 8.21676087 | 2574 | 2205 | 303 | 26959 | [illegible] |
| GO:CC | centriolar satellite | GO:0034451 | FALSE | 6.55E-08 | 7.18347402 | 118 | 2205 | 33 | 26959 | [illegible] |
| GO:CC | nuclear envelope | GO:0005635 | FALSE | 7.83E-08 | 7.106058144 | 494 | 2205 | 83 | 26959 | [illegible] |
| GO:CC | endosome | GO:0005768 | FALSE | 9.53E-08 | 7.02082248 | 959 | 2205 | 135 | 26959 | [illegible] |
| GO:CC | cytoplasmic stress granule | GO:0010494 | FALSE | 1.02E-07 | 6.991490817 | 90 | 2205 | 28 | 26959 | [illegible] |
| GO:CC | nuclear matrix | GO:0016363 | FALSE | 2.70E-07 | 6.568804621 | 124 | 2205 | 33 | 26959 | [illegible] |
| GO:CC | plasma membrane bounded cell projection | GO:0120025 | FALSE | 4.40E-07 | 6.35683313 | 2476 | 2205 | 284 | 26959 | [illegible] |
| GO:CC | spliceosomal complex | GO:0005681 | FALSE | 6.53E-07 | 6.185329351 | 218 | 2205 | 46 | 26959 | [illegible] |
| GO:CC | transcription regulator complex | GO:0005667 | FALSE | 9.15E-07 | 6.038420273 | 528 | 2205 | 84 | 26959 | [illegible] |
| GO:CC | organelle membrane | GO:0031090 | FALSE | 1.06092E-06 | 5.974318909 | 3319 | 2205 | 361 | 26959 | [illegible] |
| GO:CC | endosome membrane | GO:0010008 | FALSE | 3.38881E-06 | 5.469952252 | 456 | 2205 | 74 | 26959 | [illegible] |
| GO:CC | Golgi apparatus | GO:0005794 | FALSE | 6.14656E-06 | 5.211367924 | 1527 | 2205 | 186 | 26959 | [illegible] |
| GO:CC | methyltransferase complex | GO:0034708 | FALSE | 9.34741E-06 | 5.029308809 | 108 | 2205 | 28 | 26959 | [illegible] |
| GO:CC | mitotic spindle pole | GO:0097431 | FALSE | 1.16862E-05 | 4.932326214 | 40 | 2205 | 16 | 26959 | [illegible] |
| GO:CC | cell junction | GO:0030054 | FALSE | 1.17618E-05 | 4.929526143 | 2275 | 2205 | 257 | 26959 | [illegible] |
| GO:CC | chromosome, telomeric region | GO:0000781 | FALSE | 1.37279E-05 | 4.86239715 | 123 | 2205 | 30 | 26959 | [illegible] |
| GO:CC | early endosome | GO:0005769 | FALSE | 1.45277E-05 | 4.837803673 | 420 | 2205 | 68 | 26959 | [illegible] |
| GO:CC | fibrillar center | GO:0001650 | FALSE | 2.51586E-05 | 4.599313526 | 154 | 2205 | 34 | 26959 | [illegible] |
| GO:CC | polymeric cytoskeletal fiber | GO:0099513 | FALSE | 2.67694E-05 | 4.572361067 | 799 | 2205 | 109 | 26959 | [illegible] |
| GO:CC | bounding membrane of organelle | GO:0098588 | FALSE | 3.47425E-05 | 4.45913865 | 1718 | 2205 | 201 | 26959 | [illegible] |
| GO:CC | vesicle membrane | GO:0012506 | FALSE | 3.9978E-05 | 4.398178796 | 920 | 2205 | 121 | 26959 | [illegible] |
| GO:CC | supramolecular polymer | GO:0099081 | FALSE | 5.18069E-05 | 4.285612457 | 1051 | 2205 | 134 | 26959 | [illegible] |
| GO:CC | serine/threonine protein kinase complex | GO:1902554 | FALSE | 5.88003E-05 | 4.230620652 | 117 | 2205 | 28 | 26959 | [illegible] |
| GO:CC | nuclear membrane | GO:0031965 | FALSE | 5.91808E-05 | 4.227818997 | 316 | 2205 | 54 | 26959 | [illegible] |
| GO:CC | cytoplasmic vesicle membrane | GO:0030659 | FALSE | 6.01613E-05 | 4.220682961 | 898 | 2205 | 118 | 26959 | [illegible] |
| GO:CC | supramolecular fiber | GO:0099512 | FALSE | 6.03804E-05 | 4.219103866 | 1044 | 2205 | 133 | 26959 | [illegible] |
| GO:CC | acetyltransferase complex | GO:1902493 | FALSE | 6.34141E-05 | 4.197814054 | 104 | 2205 | 26 | 26959 | [illegible] |
| GO:CC | transferase complex, transferring phosphorus-containing groups | GO:0061695 | FALSE | 6.40188E-05 | 4.193692213 | 292 | 2205 | 51 | 26959 | [illegible] |
| GO:CC | ruffle | GO:0001726 | FALSE | 6.48973E-05 | 4.18777336 | 182 | 2205 | 37 | 26959 | [illegible] |
| GO:CC | anchoring junction | GO:0070161 | FALSE | 7.03756E-05 | 4.152578167 | 729 | 2205 | 100 | 26959 | [illegible] |
| GO:CC | omegasome | GO:1990462 | FALSE | 8.5197E-05 | 4.069575605 | 11 | 2205 | 8 | 26959 | [illegible] |
| GO:CC | protein acetyltransferase complex | GO:0031248 | FALSE | 8.79657E-05 | 4.055686662 | 99 | 2205 | 25 | 26959 | [illegible] |
| GO:CC | cytoplasmic side of plasma membrane | GO:0009898 | TRUE | 9.32046E-05 | 4.030565654 | 177 | 2205 | 36 | 26959 | [illegible] |
| GO:CC | transcription repressor complex | GO:0017053 | FALSE | 0.000155289 | 3.808859496 | 76 | 2205 | 21 | 26959 | [illegible] |
| GO:CC | somatodendritic compartment | GO:0036477 | FALSE | 0.000238849 | 3.621877388 | 1100 | 2205 | 136 | 26959 | [illegible] |
| GO:CC | heterochromatin | GO:0000792 | FALSE | 0.000240332 | 3.619188349 | 104 | 2205 | 25 | 26959 | [illegible] |
| GO:CC | site of DNA damage | GO:0090734 | FALSE | 0.000252146 | 3.598347602 | 118 | 2205 | 27 | 26959 | [illegible] |
| GO:CC | sex chromosome | GO:0000803 | FALSE | 0.000399104 | 3.398914406 | 39 | 2205 | 14 | 26959 | [illegible] |
| GO:CC | transcription elongation factor complex | GO:0008023 | FALSE | 0.000416041 | 3.380864008 | 50 | 2205 | 16 | 26959 | [illegible] |
| GO:CC | cell leading edge | GO:0031252 | FALSE | 0.000492515 | 3.307580341 | 452 | 2205 | 67 | 26959 | [illegible] |
| GO:CC | cytoplasmic side of membrane | GO:0098562 | FALSE | 0.000508316 | 3.293866462 | 213 | 2205 | 39 | 26959 | [illegible] |
| GO:CC | histone methyltransferase complex | GO:0035097 | FALSE | 0.000534861 | 3.27175887 | 75 | 2205 | 20 | 26959 | [illegible] |
| GO:CC | nuclear inclusion body | GO:0042405 | FALSE | 0.00057202 | 3.242588605 | 13 | 2205 | 8 | 26959 | [illegible] |
| GO:CC | RNA polymerase II transcription regulator complex | GO:0090575 | FALSE | 0.00078216 | 3.106704318 | 241 | 2205 | 42 | 26959 | [illegible] |
| GO:CC | postsynaptic density | GO:0014069 | FALSE | 0.000788568 | 3.103160716 | 431 | 2205 | 64 | 26959 | [illegible] |
| GO:CC | SWI/SNF superfamily-type complex | GO:0070603 | FALSE | 0.00080955 | 3.091756208 | 90 | 2205 | 22 | 26959 | [illegible] |
| GO:CC | organelle envelope | GO:0031967 | FALSE | 0.001008535 | 2.996309098 | 1303 | 2205 | 153 | 26959 | [illegible] |
| GO:CC | envelope | GO:0031975 | FALSE | 0.001008535 | 2.996309098 | 1303 | 2205 | 153 | 26959 | [illegible] |
| GO:CC | phagocytic vesicle | GO:0045335 | FALSE | 0.001030126 | 2.987109852 | 112 | 2205 | 25 | 26959 | [illegible] |
| GO:CC | neuron projection | GO:0043005 | FALSE | 0.001113891 | 2.953157338 | 1585 | 2205 | 180 | 26959 | [illegible] |
| GO:CC | symbiont-containing vacuole | GO:0020003 | FALSE | 0.001191222 | 2.924007178 | 5 | 2205 | 5 | 26959 | [illegible] |
| GO:CC | host intracellular region | GO:0043656 | FALSE | 0.001191222 | 2.924007178 | 5 | 2205 | 5 | 26959 | [illegible] |
| GO:CC | symbiont-containing vacuole membrane | GO:0020005 | FALSE | 0.001191222 | 2.924007178 | 5 | 2205 | 5 | 26959 | [illegible] |
| GO:CC | host cell cytoplasm | GO:0030430 | FALSE | 0.001191222 | 2.924007178 | 5 | 2205 | 5 | 26959 | [illegible] |
| GO:CC | host cell part | GO:0033643 | FALSE | 0.001191222 | 2.924007178 | 5 | 2205 | 5 | 26959 | [illegible] |
| GO:CC | host intracellular part | GO:0033646 | TRUE | 0.001191222 | 2.924007178 | 5 | 2205 | 5 | 26959 | [illegible] |
| GO:CC | host cell cytoplasm part | GO:0033655 | FALSE | 0.001191222 | 2.924007178 | 5 | 2205 | 5 | 26959 | [illegible] |
| GO:CC | endocytic vesicle | GO:0030139 | FALSE | 0.001482711 | 2.828943634 | 206 | 2205 | 37 | 26959 | [illegible] |
| GO:CC | asymmetric synapse | GO:0032279 | FALSE | 0.001567301 | 2.804847624 | 449 | 2205 | 65 | 26959 | [illegible] |
| GO:CC | histone acetyltransferase complex | GO:0000123 | FALSE | 0.002045306 | 2.689241762 | 88 | 2205 | 21 | 26959 | [illegible] |
| GO:CC | postsynaptic specialization | GO:0099572 | FALSE | 0.002051261 | 2.687979072 | 471 | 2205 | 67 | 26959 | [illegible] |
| GO:CC | cyclin/CDK positive transcription elongation factor complex | GO:0008024 | FALSE | 0.002355215 | 2.627969489 | 8 | 2205 | 6 | 26959 | [illegible] |
| GO:CC | neuron to neuron synapse | GO:0098984 | FALSE | 0.002868582 | 2.542332702 | 485 | 2205 | 68 | 26959 | [illegible] |
| GO:CC | bicellular tight junction | GO:0005923 | FALSE | 0.003020182 | 2.519966849 | 141 | 2205 | 28 | 26959 | [illegible] |
| GO:CC | tight junction | GO:0070160 | FALSE | 0.0031714 | 2.498749004 | 149 | 2205 | 29 | 26959 | [illegible] |
| GO:CC | spindle microtubule | GO:0005876 | FALSE | 0.00332715 | 2.47792764 | 77 | 2205 | 19 | 26959 | [illegible] |
| GO:CC | apical junction complex | GO:0043296 | FALSE | 0.003819772 | 2.417962583 | 166 | 2205 | 31 | 26959 | [illegible] |
| GO:CC | catalytic step 2 spliceosome | GO:0071013 | FALSE | 0.004241343 | 2.372496608 | 99 | 2205 | 22 | 26959 | [illegible] |
| GO:CC | early endosome membrane | GO:0031901 | FALSE | 0.004358737 | 2.360639307 | 175 | 2205 | 32 | 26959 | [illegible] |
| GO:CC | CCR4-NOT complex | GO:0030014 | FALSE | 0.004570921 | 2.339996286 | 16 | 2205 | 8 | 26959 | [illegible] |
| GO:CC | cyclin-dependent protein kinase holoenzyme complex | GO:0000307 | FALSE | 0.004807257 | 2.318102628 | 53 | 2205 | 15 | 26959 | [illegible] |
| GO:CC | actin cytoskeleton | GO:0015629 | FALSE | 0.004941827 | 2.306112486 | 493 | 2205 | 68 | 26959 | [illegible] |
| GO:CC | protein kinase complex | GO:1902911 | FALSE | 0.005962557 | 2.22456745 | 146 | 2205 | 28 | 26959 | [illegible] |
| GO:CC | Flemming body | GO:0090543 | FALSE | 0.006336975 | 2.198118033 | 31 | 2205 | 11 | 26959 | [illegible] |
| GO:CC | nuclear chromosome | GO:0000228 | FALSE | 0.006940994 | 2.158578343 | 315 | 2205 | 48 | 26959 | [illegible] |
| GO:CC | cell body | GO:0044297 | FALSE | 0.007055459 | 2.151474747 | 829 | 2205 | 102 | 26959 | [illegible] |
| GO:CC | neuronal cell body | GO:0043025 | FALSE | 0.008360039 | 2.077791691 | 743 | 2205 | 93 | 26959 | [illegible] |
| GO:CC | ubiquitin ligase complex | GO:0000151 | FALSE | 0.011398904 | 1.943136901 | 339 | 2205 | 50 | 26959 | [illegible] |
| GO:CC | euchromatin | GO:0000791 | FALSE | 0.011973363 | 1.921783854 | 70 | 2205 | 17 | 26959 | [illegible] |
| GO:CC | outer kinetochore | GO:0000940 | FALSE | 0.013388084 | 1.873281558 | 18 | 2205 | 8 | 26959 | [illegible] |
| GO:CC | inclusion body | GO:0016234 | FALSE | 0.014880494 | 1.827382641 | 78 | 2205 | 18 | 26959 | [illegible] |
| GO:CC | cohesin complex | GO:0008278 | FALSE | 0.015288863 | 1.815624797 | 10 | 2205 | 6 | 26959 | [illegible] |
| GO:CC | postsynapse | GO:0098794 | FALSE | 0.015820686 | 1.800774677 | 866 | 2205 | 104 | 26959 | [illegible] |
| GO:CC | nuclear pore | GO:0005643 | FALSE | 0.017812569 | 1.749273447 | 79 | 2205 | 18 | 26959 | [illegible] |
| GO:CC | lytic vacuole | GO:0000323 | FALSE | 0.019629325 | 1.707094644 | 534 | 2205 | 70 | 26959 | [illegible] |
| GO:CC | lysosome | GO:0005764 | FALSE | 0.019629325 | 1.707094644 | 534 | 2205 | 70 | 26959 | [illegible] |
| GO:CC | synapse | GO:0045202 | FALSE | 0.023153408 | 1.635385083 | 1660 | 2205 | 179 | 26959 | [illegible] |
| GO:CC | cell-cell junction | GO:0005911 | FALSE | 0.023390244 | 1.630965252 | 537 | 2205 | 70 | 26959 | [illegible] |
| GO:CC | vacuole | GO:0005773 | FALSE | 0.024047357 | 1.618932651 | 635 | 2205 | 80 | 26959 | [illegible] |
| GO:CC | extrinsic component of membrane | GO:0019898 | FALSE | 0.024407506 | 1.612476598 | 268 | 2205 | 41 | 26959 | [illegible] |
| GO:CC | kinesin complex | GO:0005871 | TRUE | 0.039572466 | 1.402606888 | 43 | 2205 | 12 | 26959 | [illegible] |
| GO:CC | pericentric heterochromatin | GO:0005721 | FALSE | 0.040441709 | 1.393170505 | 37 | 2205 | 11 | 26959 | [illegible] |
| KEGG | Herpes simplex virus 1 infection | KEGG:05168 | FALSE | 7.22E-38 | 37.1412999 | 444 | 1023 | 153 | 8947 | [illegible] |
| KEGG | TNF signaling pathway | KEGG:04668 | FALSE | 1.84E-10 | 9.734304114 | 115 | 1023 | 42 | 8947 | [illegible] |
| KEGG | NF-kappa B signaling pathway | KEGG:04064 | FALSE | 5.80E-08 | 7.236625403 | 105 | 1023 | 36 | 8947 | [illegible] |
| KEGG | NOD-like receptor signaling pathway | KEGG:04621 | FALSE | 8.63803E-06 | 5.063585389 | 208 | 1023 | 51 | 8947 | [illegible] |
| KEGG | Cell cycle | KEGG:04110 | FALSE | 1.87787E-05 | 4.726333676 | 155 | 1023 | 41 | 8947 | [illegible] |
| KEGG | Influenza A | KEGG:05164 | FALSE | 5.21619E-05 | 4.282646662 | 172 | 1023 | 43 | 8947 | [illegible] |
| KEGG | Hepatitis B | KEGG:05161 | FALSE | 8.13265E-05 | 4.089767873 | 163 | 1023 | 41 | 8947 | [illegible] |
| KEGG | Toll-like receptor signaling pathway | KEGG:04620 | FALSE | 0.000130492 | 3.884417397 | 99 | 1023 | 29 | 8947 | [illegible] |
| KEGG | Lipid and atherosclerosis | KEGG:05417 | FALSE | 0.00018846 | 3.724779666 | 216 | 1023 | 49 | 8947 | [illegible] |
| KEGG | Measles | KEGG:05162 | FALSE | 0.000196076 | 3.707574915 | 145 | 1023 | 37 | 8947 | [illegible] |
| KEGG | Yersinia infection | KEGG:05135 | FALSE | 0.000252101 | 3.598425802 | 135 | 1023 | 35 | 8947 | [illegible] |
| KEGG | Hepatitis C | KEGG:05160 | FALSE | 0.000258689 | 3.587222599 | 164 | 1023 | 40 | 8947 | [illegible] |
| KEGG | Chronic myeloid leukemia | KEGG:05220 | FALSE | 0.000924244 | 3.034213199 | 76 | 1023 | 23 | 8947 | [illegible] |
| KEGG | Transcriptional misregulation in cancer | KEGG:05202 | FALSE | 0.001381651 | 2.859601563 | 212 | 1023 | 46 | 8947 | [illegible] |
| KEGG | FoxO signaling pathway | KEGG:04068 | FALSE | 0.002585951 | 2.587379732 | 131 | 1023 | 32 | 8947 | [illegible] |
| KEGG | Steroid biosynthesis | KEGG:00100 | FALSE | 0.002865195 | 2.542845748 | 20 | 1023 | 10 | 8947 | [illegible] |
| KEGG | RNA degradation | KEGG:03018 | FALSE | 0.003657859 | 2.436772989 | 82 | 1023 | 23 | 8947 | [illegible] |
| KEGG | Small cell lung cancer | KEGG:05222 | FALSE | 0.003699632 | 2.431841469 | 93 | 1023 | 25 | 8947 | [illegible] |
| KEGG | Viral life cycle - HIV-1 | KEGG:03250 | FALSE | 0.007961649 | 2.098996949 | 59 | 1023 | 18 | 8947 | [illegible] |
| KEGG | Pathways in cancer | KEGG:05200 | FALSE | 0.010499044 | 1.978850233 | 543 | 1023 | 91 | 8947 | [illegible] |
| KEGG | RIG-I-like receptor signaling pathway | KEGG:04622 | FALSE | 0.011186568 | 1.951303151 | 71 | 1023 | 20 | 8947 | [illegible] |
| KEGG | Prostate cancer | KEGG:05215 | FALSE | 0.011389619 | 1.943490798 | 99 | 1023 | 25 | 8947 | [illegible] |
| KEGG | Epstein-Barr virus infection | KEGG:05169 | FALSE | 0.012530377 | 1.90203586 | 224 | 1023 | 45 | 8947 | [illegible] |
| KEGG | C-type lectin receptor signaling pathway | KEGG:04625 | FALSE | 0.014267948 | 1.845638479 | 112 | 1023 | 27 | 8947 | [illegible] |
| KEGG | Focal adhesion | KEGG:04510 | FALSE | 0.019948887 | 1.700081329 | 202 | 1023 | 41 | 8947 | [illegible] |
| KEGG | Human papillomavirus infection | KEGG:05165 | FALSE | 0.024936336 | 1.603167366 | 354 | 1023 | 63 | 8947 | [illegible] |
| KEGG | Circadian rhythm | KEGG:04710 | FALSE | 0.028982686 | 1.53786137 | 34 | 1023 | 12 | 8947 | [illegible] |
| KEGG | Chemokine signaling pathway | KEGG:04062 | FALSE | 0.041759062 | 1.379249269 | 189 | 1023 | 38 | 8947 | [illegible] |
| REAC | Gene expression (Transcription) | REAC:R-MMU-74160 | FALSE | 2.25E-12 | 11.64780377 | 1002 | 1115 | 216 | 8412 | [illegible] |
| REAC | Mitotic Prometaphase | REAC:R-MMU-68877 | FALSE | 4.05E-11 | 10.3925962 | 178 | 1115 | 62 | 8412 | [illegible] |
| REAC | RNA Polymerase II Transcription | REAC:R-MMU-73857 | FALSE | 3.41E-10 | 9.466955954 | 907 | 1115 | 193 | 8412 | [illegible] |
| REAC | Signaling by Rho GTPases | REAC:R-MMU-194315 | FALSE | 5.40E-08 | 7.267718932 | 552 | 1115 | 126 | 8412 | [illegible] |
| REAC | Resolution of Sister Chromatid Cohesion | REAC:R-MMU-2500257 | FALSE | 1.58E-07 | 6.801124728 | 113 | 1115 | 41 | 8412 | [illegible] |
| REAC | Signaling by Rho GTPases, Miro GTPases and RHOBTB3 | REAC:R-MMU-9716542 | FALSE | 1.68E-07 | 6.77404563 | 567 | 1115 | 127 | 8412 | [illegible] |
| REAC | Generic Transcription Pathway | REAC:R-MMU-212436 | FALSE | 4.33E-07 | 6.363669468 | 792 | 1115 | 163 | 8412 | [illegible] |
| REAC | Amplification of signal from unattached kinetochores via a MAD2 inhibitory signal | REAC:R-MMU-141444 | FALSE | 6.65837E-06 | 5.176632262 | 86 | 1115 | 32 | 8412 | [illegible] |
| REAC | Amplification of signal from the kinetochores | REAC:R-MMU-141424 | FALSE | 6.65837E-06 | 5.176632262 | 86 | 1115 | 32 | 8412 | [illegible] |
| REAC | Cell Cycle | REAC:R-MMU-1640170 | FALSE | 9.62209E-06 | 5.016730571 | 567 | 1115 | 121 | 8412 | [illegible] |
| REAC | Mitotic Telophase/Cytokinesis | REAC:R-MMU-68884 | FALSE | 3.38935E-05 | 4.469883652 | 12 | 1115 | 10 | 8412 | [illegible] |
| REAC | Establishment of Sister Chromatid Cohesion | REAC:R-MMU-2468052 | FALSE | 4.43645E-05 | 4.352964835 | 10 | 1115 | 9 | 8412 | [illegible] |
| REAC | RHO GTPase Effectors | REAC:R-MMU-195258 | FALSE | 0.000132115 | 3.879048611 | 232 | 1115 | 59 | 8412 | [illegible] |
| REAC | M Phase | REAC:R-MMU-68886 | FALSE | 0.000132406 | 3.878091616 | 372 | 1115 | 84 | 8412 | [illegible] |
| REAC | Mitotic Spindle Checkpoint | REAC:R-MMU-69618 | FALSE | 0.00018793 | 3.726003388 | 102 | 1115 | 33 | 8412 | [illegible] |
| REAC | RHO GTPases Activate Formins | REAC:R-MMU-5663220 | FALSE | 0.000245886 | 3.609266569 | 127 | 1115 | 38 | 8412 | [illegible] |
| REAC | Cohesin Loading onto Chromatin | REAC:R-MMU-2470946 | FALSE | 0.000303592 | 3.5177092 | 9 | 1115 | 8 | 8412 | [illegible] |
| REAC | Cell Cycle, Mitotic | REAC:R-MMU-69278 | FALSE | 0.000376917 | 3.423753823 | 488 | 1115 | 102 | 8412 | [illegible] |
| REAC | EML4 and NUDC in mitotic spindle formation | REAC:R-MMU-9648025 | FALSE | 0.000472647 | 3.325462979 | 101 | 1115 | 32 | 8412 | [illegible] |
| REAC | TP53 regulates transcription of additional cell cycle genes whose exact role in the p53 pathway | REAC:R-MMU-6804115 | FALSE | 0.000500011 | 3.301020671 | 17 | 1115 | 11 | 8412 | [illegible] |
| REAC | SUMO E3 ligases SUMOylate target proteins | REAC:R-MMU-3108232 | FALSE | 0.001214575 | 2.915575777 | 150 | 1115 | 41 | 8412 | [illegible] |
| REAC | Cholesterol biosynthesis | REAC:R-MMU-191273 | FALSE | 0.001281635 | 2.892235561 | 28 | 1115 | 14 | 8412 | [illegible] |
| REAC | Mitotic Anaphase | REAC:R-MMU-68882 | FALSE | 0.002795134 | 2.553597337 | 214 | 1115 | 52 | 8412 | [illegible] |
| REAC | Mitotic Metaphase and Anaphase | REAC:R-MMU-2555396 | FALSE | 0.003224732 | 2.4915064 | 215 | 1115 | 52 | 8412 | [illegible] |
| REAC | Separation of Sister Chromatids | REAC:R-MMU-2467813 | FALSE | 0.003339256 | 2.476350266 | 177 | 1115 | 45 | 8412 | [illegible] |
| REAC | SUMOylation | REAC:R-MMU-2990846 | FALSE | 0.003516231 | 2.45392256 | 156 | 1115 | 41 | 8412 | [illegible] |
| REAC | RHO GTPase cycle | REAC:R-MMU-9012999 | FALSE | 0.003662082 | 2.436271953 | 378 | 1115 | 80 | 8412 | [illegible] |
| REAC | TP53 Regulates Transcription of Cell Cycle Genes | REAC:R-MMU-6791312 | FALSE | 0.004027196 | 2.394997286 | 34 | 1115 | 15 | 8412 | [illegible] |
| REAC | Cytokine Signaling in Immune system | REAC:R-MMU-1280215 | FALSE | 0.004978464 | 2.302904658 | 381 | 1115 | 80 | 8412 | [illegible] |
| REAC | Deubiquitination | REAC:R-MMU-5688426 | FALSE | 0.009552039 | 2.019903911 | 240 | 1115 | 55 | 8412 | [illegible] |
| REAC | Death Receptor Signaling | REAC:R-MMU-73887 | FALSE | 0.022590955 | 1.646065405 | 130 | 1115 | 34 | 8412 | [illegible] |
| REAC | TRIF(TICAM1)-mediated TLR4 signaling | REAC:R-MMU-937061 | FALSE | 0.034263372 | 1.465169894 | 96 | 1115 | 27 | 8412 | [illegible] |
| REAC | MyD88-independent TLR4 cascade | REAC:R-MMU-166166 | FALSE | 0.034263372 | 1.465169894 | 96 | 1115 | 27 | 8412 | [illegible] |
| REAC | Toll-like Receptor Cascades | REAC:R-MMU-168898 | FALSE | 0.037048805 | 1.431225792 | 133 | 1115 | 34 | 8412 | [illegible] |
| REAC | Toll Like Receptor 4 (TLR4) Cascade | REAC:R-MMU-166016 | FALSE | 0.039455801 | 1.403889136 | 107 | 1115 | 29 | 8412 | [illegible] |
| REAC | DDX58/IFIH1-mediated induction of interferon-alpha/beta | REAC:R-MMU-168928 | FALSE | 0.04494828 | 1.347286925 | 32 | 1115 | 13 | 8412 | [illegible] |
| REAC | RHOB GTPase cycle | REAC:R-MMU-9013026 | FALSE | 0.045988803 | 1.337347894 | 63 | 1115 | 20 | 8412 | [illegible] |
| REAC | HDR through MMEJ (alt-NHEJ) | REAC:R-MMU-5685939 | FALSE | 0.048400755 | 1.315147868 | 8 | 1115 | 6 | 8412 | [illegible] |
| WP | mRNA processing | WP:WP310 | FALSE | 1.26E-07 | 6.899001002 | 446 | 612 | 104 | 4529 | [illegible] |
| WP | Cholesterol biosynthesis | WP:WP103 | FALSE | 1.32534E-05 | 4.87767177 | 15 | 612 | 11 | 4529 | [illegible] |
| WP | Toll-like receptor signaling | WP:WP88 | FALSE | 0.000304315 | 3.516676147 | 32 | 612 | 15 | 4529 | [illegible] |
| WP | Type II interferon signaling (IFNG) | WP:WP1253 | FALSE | 0.001694936 | 2.770846706 | 32 | 612 | 14 | 4529 | [illegible] |
| WP | Fibrin complement receptor 3 signaling pathway | WP:WP5128 | FALSE | 0.001698781 | 2.769862671 | 44 | 612 | 17 | 4529 | [illegible] |
| WP | Apoptosis | WP:WP1254 | FALSE | 0.002738131 | 2.562545826 | 77 | 612 | 24 | 4529 | [illegible] |
| WP | IL-7 signaling pathway | WP:WP297 | FALSE | 0.005024049 | 2.298946137 | 43 | 612 | 16 | 4529 | [illegible] |
| WP | Cholesterol metabolism with Bloch and Kandutsch-Russell pathways | WP:WP4346 | FALSE | 0.00547776 | 2.261396985 | 52 | 612 | 18 | 4529 | [illegible] |

| | | | | | | | | | | |
|---|---|---|---|---|---|---|---|---|---|---|
| WP | Mechanisms associated with pluripotency | WP:WP1763 | FALSE | 0.00589655 | 2.229401996 | 290 | 612 | 62 | 4529 | [illegible] |
| WP | Kit receptor signaling pathway | WP:WP407 | FALSE | 0.022638579 | 1.645150835 | 67 | 612 | 20 | 4529 | [illegible] |
| WP | Adipogenesis genes | WP:WP447 | FALSE | 0.029224158 | 1.534257991 | 131 | 612 | 32 | 4529 | [illegible] |
| TF | Factor: FOXN4; motif: NNWANNCGWMCGCGTCNNNNMT; match class: 1 | TF:M04662_1 | FALSE | 2.59E-97 | 96.58632329 | 12782 | 2370 | 1855 | 21787 | [illegible] |
| TF | Factor: ZF5; motif: GSGCGCGR; match class: 1 | TF:M00716_1 | FALSE | 3.18E-80 | 79.49760592 | 12083 | 2370 | 1746 | 21787 | [illegible] |
| TF | Factor: FOXN4; motif: NNWANNCGWMCGCGTCNNNNMT | TF:M04662 | FALSE | 2.53E-76 | 75.59608846 | 15998 | 2370 | 2096 | 21787 | [illegible] |
| TF | Factor: E2F-1; motif: GNGGGCGGGRMN | TF:M10209 | FALSE | 1.57E-75 | 74.80287033 | 14255 | 2370 | 1942 | 21787 | [illegible] |
| TF | Factor: E2F-1; motif: GNGGGCGGGRMN; match class: 1 | TF:M10209_1 | FALSE | 1.48E-74 | 73.82940503 | 10651 | 2370 | 1584 | 21787 | [illegible] |
| TF | Factor: ZF5; motif: GSGCGCGR | TF:M00716 | FALSE | 1.13E-71 | 70.94647135 | 14843 | 2370 | 1986 | 21787 | [illegible] |
| TF | Factor: Kaiso; motif: SARNYCTCGCGAGAN | TF:M10276 | FALSE | 4.41E-70 | 69.35525998 | 13501 | 2370 | 1858 | 21787 | [illegible] |
| TF | Factor: Kaiso; motif: SARNYCTCGCGAGAN; match class: 1 | TF:M10276_1 | FALSE | 1.94E-68 | 67.71209036 | 9980 | 2370 | 1495 | 21787 | [illegible] |
| TF | Factor: E2F; motif: GGCGSG | TF:M00803 | FALSE | 6.08E-67 | 66.21632903 | 10440 | 2370 | 1540 | 21787 | [illegible] |
| TF | Factor: ZF5; motif: NRNGNGCGCGCWN; match class: 1 | TF:M00333_1 | FALSE | 1.74E-65 | 64.76017277 | 11212 | 2370 | 1617 | 21787 | [illegible] |
| TF | Factor: E2F-4; motif: NTTTCSCGCC | TF:M07380 | FALSE | 6.27E-61 | 60.20286467 | 10581 | 2370 | 1537 | 21787 | [illegible] |
| TF | Factor: ZF5; motif: GYCGCGCARNGCNN | TF:M02933 | FALSE | 5.47E-59 | 58.26217868 | 9518 | 2370 | 1416 | 21787 | [illegible] |
| TF | Factor: ZF5; motif: NRNGNGCGCGCWN | TF:M00333 | FALSE | 5.90E-59 | 58.22907888 | 14484 | 2370 | 1921 | 21787 | [illegible] |
| TF | Factor: RNF96; motif: BCCCGCRGCC | TF:M01199 | FALSE | 8.88E-59 | 58.05152723 | 6364 | 2370 | 1051 | 21787 | [illegible] |
| TF | Factor: E2F-3; motif: GGCGGGN | TF:M02089 | FALSE | 2.20E-58 | 57.65686082 | 11292 | 2370 | 1604 | 21787 | [illegible] |
| TF | Factor: ELK-1; motif: ACCGGAWRTN | TF:M01981 | FALSE | 2.38E-55 | 54.62262697 | 14698 | 2370 | 1931 | 21787 | [illegible] |
| TF | Factor: E2F; motif: GGCGSG; match class: 1 | TF:M00803_1 | FALSE | 2.78E-54 | 53.55576522 | 7057 | 2370 | 1120 | 21787 | [illegible] |
| TF | Factor: ELK-1; motif: ACCGGAWRTN; match class: 1 | TF:M01981_1 | FALSE | 4.18E-53 | 52.37831199 | 9022 | 2370 | 1342 | 21787 | [illegible] |
| TF | Factor: ZAC; motif: KGGGCCR | TF:M05547 | FALSE | 1.52E-52 | 51.81814553 | 6693 | 2370 | 1071 | 21787 | [illegible] |
| TF | Factor: BEN; motif: CAGCGRNV | TF:M01240 | FALSE | 5.44E-52 | 51.26458411 | 16242 | 2370 | 2061 | 21787 | [illegible] |
| TF | Factor: IRF6; motif: NNNACTCYCGGKNNN; match class: 1 | TF:M02874_1 | FALSE | 1.98E-51 | 50.70415602 | 7153 | 2370 | 1122 | 21787 | [illegible] |
| TF | Factor: Foxn1; motif: RGAMGC | TF:M12470 | FALSE | 5.18E-51 | 50.28552334 | 9728 | 2370 | 1413 | 21787 | [illegible] |
| TF | Factor: E2F-1; motif: NNNSSCGCSAANN | TF:M07250 | FALSE | 8.66E-51 | 50.06226668 | 9038 | 2370 | 1336 | 21787 | [illegible] |
| TF | Factor: BEN; motif: CAGCGRNV; match class: 1 | TF:M01240_1 | FALSE | 9.13E-51 | 50.03975663 | 12010 | 2370 | 1654 | 21787 | [illegible] |
| TF | Factor: E2F-7; motif: GRGGCGGGAANNN | TF:M10212 | FALSE | 5.22E-50 | 49.2819394 | 8115 | 2370 | 1229 | 21787 | [illegible] |
| TF | Factor: TEL1; motif: CNCGGAANNN | TF:M01993 | FALSE | 2.82E-49 | 48.54974191 | 10419 | 2370 | 1482 | 21787 | [illegible] |
| TF | Factor: RNF96; motif: BCCCGCRGCC; match class: 1 | TF:M01199_1 | FALSE | 1.46E-47 | 46.83598973 | 2733 | 2370 | 544 | 21787 | [illegible] |
| TF | Factor: Egr-2; motif: GCGTGGGCGG | TF:M05494 | FALSE | 2.75E-47 | 46.56031075 | 7247 | 2370 | 1119 | 21787 | [illegible] |
| TF | Factor: E2F-3; motif: GGCGGGN; match class: 1 | TF:M02089_1 | FALSE | 4.47E-47 | 46.34980356 | 6914 | 2370 | 1079 | 21787 | [illegible] |
| TF | Factor: MOVO-B; motif: GNGGGGG | TF:M01104 | FALSE | 6.05E-46 | 45.21835459 | 7656 | 2370 | 1162 | 21787 | [illegible] |
| TF | Factor: IRF4; motif: NNNAYTCTCGGWNNN; match class: 1 | TF:M02872_1 | FALSE | 8.75E-46 | 45.05816715 | 8130 | 2370 | 1216 | 21787 | [illegible] |
| TF | Factor: IRF6; motif: NNNACTCYCGGKNNN | TF:M02874 | FALSE | 2.24E-45 | 44.64943635 | 13418 | 2370 | 1778 | 21787 | [illegible] |
| TF | Factor: Elk-1; motif: NNNNCCGGAARTNN | TF:M00025 | FALSE | 7.41E-45 | 44.13026417 | 9833 | 2370 | 1403 | 21787 | [illegible] |
| TF | Factor: SP100; motif: NNCGTCGNNTAAWNN | TF:M02913 | FALSE | 8.44E-45 | 44.0736601 | 8495 | 2370 | 1254 | 21787 | [illegible] |
| TF | Factor: Sp1; motif: NGGGGGCGGGGYN | TF:M00196 | FALSE | 2.20E-44 | 43.65716632 | 8810 | 2370 | 1288 | 21787 | [illegible] |
| TF | Factor: AP-2; motif: MKCCCSCNGGCG | TF:M00189 | FALSE | 5.45E-44 | 43.26376546 | 8064 | 2370 | 1202 | 21787 | [illegible] |
| TF | Factor: E2F-4; motif: NTTTCSCGCC; match class: 1 | TF:M07380_1 | FALSE | 1.14E-43 | 42.94320337 | 5386 | 2370 | 882 | 21787 | [illegible] |
| TF | Factor: ZF5; motif: GYCGCGCARNGCNN; match class: 1 | TF:M02933_1 | FALSE | 1.78E-43 | 42.75055238 | 5846 | 2370 | 938 | 21787 | [illegible] |
| TF | Factor: E2F-4; motif: NGGGGGCGGGRMNN | TF:M10211 | FALSE | 4.67E-43 | 42.33085074 | 7000 | 2370 | 1075 | 21787 | [illegible] |
| TF | Factor: Sp1; motif: NNGGGGCGGGGNN | TF:M00932 | FALSE | 8.47E-43 | 42.07224446 | 8960 | 2370 | 1299 | 21787 | [illegible] |
| TF | Factor: Sp1; motif: NGGGGCGGGGN | TF:M07395 | FALSE | 4.34E-42 | 41.36288933 | 9145 | 2370 | 1317 | 21787 | [illegible] |
| TF | Factor: PEA3; motif: RCCGGAAGYN | TF:M01991 | FALSE | 4.82E-42 | 41.31721754 | 9964 | 2370 | 1407 | 21787 | [illegible] |
| TF | Factor: GABP-alpha; motif: CTTCCK | TF:M01660 | FALSE | 5.73E-42 | 41.24166716 | 10764 | 2370 | 1493 | 21787 | [illegible] |
| TF | Factor: Sp1; motif: GGGGCGGGGC | TF:M00931 | FALSE | 6.81E-42 | 41.16674881 | 8678 | 2370 | 1264 | 21787 | [illegible] |
| TF | Factor: BCL6B; motif: NNNNCCGCCCCWNNNN | TF:M02844 | FALSE | 3.36E-41 | 40.47329242 | 13692 | 2370 | 1791 | 21787 | [illegible] |
| TF | Factor: AP-2; motif: MKCCCSCNGGCG; match class: 1 | TF:M00189_1 | FALSE | 5.69E-41 | 40.24451207 | 3676 | 2370 | 654 | 21787 | [illegible] |
| TF | Factor: SP2; motif: GNNGGGGGCGGGGSN | TF:M03807 | FALSE | 2.41E-40 | 39.61767705 | 7110 | 2370 | 1078 | 21787 | [illegible] |
| TF | Factor: Sp1; motif: CCCCGCCCCN | TF:M00933 | FALSE | 6.95E-40 | 39.1580908 | 8781 | 2370 | 1268 | 21787 | [illegible] |
| TF | Factor: MOVO-B; motif: GNGGGGG; match class: 1 | TF:M01104_1 | FALSE | 9.91E-40 | 39.00401168 | 3279 | 2370 | 597 | 21787 | [illegible] |
| TF | Factor: E2F-4; motif: GCGGGAAANA | TF:M02090 | FALSE | 2.43E-39 | 38.61473052 | 10353 | 2370 | 1439 | 21787 | [illegible] |
| TF | Factor: ZBED6; motif: NRRGCTCGCCNN | TF:M01598 | FALSE | 9.92E-39 | 38.00334826 | 5567 | 2370 | 887 | 21787 | [illegible] |
| TF | Factor: AP-2; motif: GSCCSCRGGCNRNRNN | TF:M00800 | FALSE | 1.02E-38 | 37.99036831 | 6746 | 2370 | 1029 | 21787 | [illegible] |
| TF | Factor: WT1; motif: NGCGGGGGGGTSMMCYN | TF:M05599 | FALSE | 4.36E-38 | 37.36011594 | 5326 | 2370 | 855 | 21787 | [illegible] |
| TF | Factor: IRF4; motif: NNNAYTCTCGGWNNN | TF:M02872 | FALSE | 2.95E-37 | 36.52992623 | 14528 | 2370 | 1859 | 21787 | [illegible] |
| TF | Factor: E2F-4; motif: GNNGGCGGGAAN | TF:M09682 | FALSE | 3.06E-37 | 36.51402589 | 5932 | 2370 | 926 | 21787 | [illegible] |
| TF | Factor: ZAC; motif: KGGGCCR; match class: 1 | TF:M05547_1 | FALSE | 6.68E-37 | 36.17530874 | 2520 | 2370 | 484 | 21787 | [illegible] |
| TF | Factor: Sp1; motif: GGNGGGGGNGGGGGMGGGGCNGGG; match class: 1 | TF:M10375_1 | FALSE | 1.13E-36 | 35.94698132 | 7410 | 2370 | 1099 | 21787 | [illegible] |
| TF | Factor: WT1; motif: CGCCCCCNCN | TF:M02036 | FALSE | 4.06E-36 | 35.39189769 | 7541 | 2370 | 1112 | 21787 | [illegible] |
| TF | Factor: Sp1; motif: GGNGGGGGNGGGGGMGGGGCNGGG | TF:M10375 | FALSE | 4.83E-36 | 35.3161746 | 10761 | 2370 | 1470 | 21787 | [illegible] |
| TF | Factor: KLF3; motif: NNNNNNGGGCGGGGCNNGN | TF:M10278 | FALSE | 1.42E-35 | 34.8484353 | 7344 | 2370 | 1087 | 21787 | [illegible] |
| TF | Factor: Klf17; motif: NGGGCGG | TF:M05407 | FALSE | 3.70E-35 | 34.43231549 | 6566 | 2370 | 994 | 21787 | [illegible] |
| TF | Factor: Sp2; motif: NYSGCCCCGCCCCCY | TF:M03567 | FALSE | 4.44E-35 | 34.35258252 | 6434 | 2370 | 978 | 21787 | [illegible] |
| TF | Factor: WT1; motif: GNGGGGGCGGGG | TF:M03893 | FALSE | 6.97E-35 | 34.15698703 | 6668 | 2370 | 1005 | 21787 | [illegible] |
| TF | Factor: Pax-5; motif: RRMSWGANWYCTNRAGCGKRACSRYNSM; match class: 1 | TF:M00144_1 | FALSE | 1.16E-34 | 33.93431491 | 5527 | 2370 | 867 | 21787 | [illegible] |
| TF | Factor: Egr-1; motif: GCGGGGGCGG | TF:M07354 | FALSE | 2.83E-34 | 33.54837645 | 6109 | 2370 | 936 | 21787 | [illegible] |
| TF | Factor: BCL6B; motif: NNNNCCGCCCCWNNNN; match class: 1 | TF:M02844_1 | FALSE | 7.32E-34 | 33.13571183 | 8549 | 2370 | 1218 | 21787 | [illegible] |
| TF | Factor: KROX; motif: CCCGCCCCCRCCCC | TF:M00982 | FALSE | 1.55E-33 | 32.80925544 | 6000 | 2370 | 920 | 21787 | [illegible] |
| TF | Factor: SP1:SP3; motif: CCSCCCCCYCC | TF:M01219 | FALSE | 1.84E-33 | 32.73561475 | 6699 | 2370 | 1003 | 21787 | [illegible] |
| TF | Factor: ELF4; motif: CCCGGAARTN | TF:M01979 | FALSE | 2.24E-33 | 32.64980366 | 7188 | 2370 | 1060 | 21787 | [illegible] |
| TF | Factor: LKLF; motif: NGGGCGG | TF:M05391 | FALSE | 2.78E-33 | 32.55604024 | 7260 | 2370 | 1068 | 21787 | [illegible] |
| TF | Factor: SP6; motif: WGGGCGG | TF:M05455 | FALSE | 2.78E-33 | 32.55604024 | 7260 | 2370 | 1068 | 21787 | [illegible] |
| TF | Factor: CPBP; motif: NGGGCGG | TF:M05301 | FALSE | 2.78E-33 | 32.55604024 | 7260 | 2370 | 1068 | 21787 | [illegible] |
| TF | Factor: Sp3; motif: AGGGCGG | TF:M05665 | FALSE | 2.78E-33 | 32.55604024 | 7260 | 2370 | 1068 | 21787 | [illegible] |
| TF | Factor: Sp1; motif: NGGGGCGGGGN; match class: 1 | TF:M07395_1 | FALSE | 2.78E-33 | 32.55587431 | 4545 | 2370 | 740 | 21787 | [illegible] |
| TF | Factor: TEL1; motif: CNCGGAANNN; match class: 1 | TF:M01993_1 | FALSE | 3.63E-33 | 32.43993645 | 4398 | 2370 | 721 | 21787 | [illegible] |
| TF | Factor: GABP-alpha; motif: CTTCCK; match class: 1 | TF:M01660_1 | FALSE | 5.08E-33 | 32.29373773 | 4688 | 2370 | 757 | 21787 | [illegible] |
| TF | Factor: Pax-3; motif: NNNNNNCGTCACGSTYNNNNN | TF:M00327 | FALSE | 5.79E-33 | 32.23746223 | 12888 | 2370 | 1680 | 21787 | [illegible] |
| TF | Factor: ERG; motif: ACCGGAART | TF:M01752 | FALSE | 6.83E-33 | 32.16528388 | 9536 | 2370 | 1324 | 21787 | [illegible] |
| TF | Factor: E2F-7; motif: GRGGCGGGAANNN; match class: 1 | TF:M10212_1 | FALSE | 1.11E-32 | 31.95598872 | 3082 | 2370 | 548 | 21787 | [illegible] |
| TF | Factor: CNOT3; motif: GGCCGCGSSS | TF:M01253 | FALSE | 1.65E-32 | 31.78328098 | 2471 | 2370 | 464 | 21787 | [illegible] |
| TF | Factor: Hes1; motif: NNCACGYGNN | TF:M07991 | FALSE | 2.11E-32 | 31.6760273 | 15237 | 2370 | 1910 | 21787 | [illegible] |
| TF | Factor: Zfx; motif: SNSCAGGCCKCGSCSS | TF:M01593 | FALSE | 5.34E-32 | 31.27209947 | 2678 | 2370 | 491 | 21787 | [illegible] |
| TF | Factor: Zfp536; motif: NCGGAKG | TF:M05541 | FALSE | 6.49E-32 | 31.1875331 | 6330 | 2370 | 953 | 21787 | [illegible] |
| TF | Factor: Ets2; motif: ACCGGAWRYN | TF:M01989 | FALSE | 1.16E-31 | 30.9359642 | 7908 | 2370 | 1136 | 21787 | [illegible] |
| TF | Factor: E2F-1; motif: NTTSGCGG | TF:M00430 | FALSE | 1.33E-31 | 30.87771996 | 6985 | 2370 | 1029 | 21787 | [illegible] |
| TF | Factor: Kaiso; motif: GCMGGGRGCRGS; match class: 1 | TF:M03876_1 | FALSE | 1.72E-31 | 30.76413603 | 5860 | 2370 | 895 | 21787 | [illegible] |
| TF | Factor: SAP-1a; motif: NRRCCGGAAGYRN | TF:M10220 | FALSE | 2.04E-31 | 30.68965416 | 9105 | 2370 | 1270 | 21787 | [illegible] |
| TF | Factor: Foxn1; motif: RGAMGC; match class: 1 | TF:M12470_1 | FALSE | 4.02E-31 | 30.39569677 | 4363 | 2370 | 709 | 21787 | [illegible] |
| TF | Factor: CTCF; motif: NAGGGGGCGCNNKNNNN; match class: 1 | TF:M08995_1 | FALSE | 5.85E-31 | 30.23288578 | 9366 | 2370 | 1297 | 21787 | [illegible] |
| TF | Factor: Egr-1; motif: GCGGGGGCGG | TF:M01873 | FALSE | 7.01E-31 | 30.15399755 | 5019 | 2370 | 790 | 21787 | [illegible] |
| TF | Factor: Sp1; motif: NNGGGGCGGGGNN; match class: 1 | TF:M00932_1 | FALSE | 8.11E-31 | 30.09094833 | 4380 | 2370 | 710 | 21787 | [illegible] |
| TF | Factor: Atf-1; motif: RTGACGTA | TF:M07980 | FALSE | 9.31E-31 | 30.03118962 | 8808 | 2370 | 1234 | 21787 | [illegible] |
| TF | Factor: E2F-1; motif: NKTSSCGC | TF:M00428 | FALSE | 1.21E-30 | 29.91705967 | 7911 | 2370 | 1132 | 21787 | [illegible] |
| TF | Factor: PEA3; motif: RCCGGAAGYN; match class: 1 | TF:M01991_1 | FALSE | 1.40E-30 | 29.85343189 | 4166 | 2370 | 682 | 21787 | [illegible] |
| TF | Factor: Tcf65; motif: NNCNCGNGNN; match class: 1 | TF:M08005_1 | FALSE | 1.87E-30 | 29.7287382 | 6330 | 2370 | 947 | 21787 | [illegible] |
| TF | Factor: Tcf65; motif: NNCNCGNGNN | TF:M08005 | FALSE | 1.87E-30 | 29.7287382 | 6330 | 2370 | 947 | 21787 | [illegible] |
| TF | Factor: E2F-4; motif: GCGGGAAANA; match class: 1 | TF:M02090_1 | FALSE | 2.34E-30 | 29.6308472 | 4713 | 2370 | 750 | 21787 | [illegible] |
| TF | Factor: BTEB2; motif: RGGGNGKGGN | TF:M07277 | FALSE | 4.55E-30 | 29.34221687 | 7610 | 2370 | 1095 | 21787 | [illegible] |
| TF | Factor: Elk-1; motif: NNNNCCGGAARTNN; match class: 1 | TF:M00025_1 | FALSE | 4.59E-30 | 29.33787596 | 4079 | 2370 | 669 | 21787 | [illegible] |
| TF | Factor: CTCF; motif: NAGGGGGCGCNNKNNNN | TF:M08995 | FALSE | 7.15E-30 | 29.14575656 | 14346 | 2370 | 1814 | 21787 | [illegible] |
| TF | Factor: E2F-1; motif: NNNSSCGCSAANN; match class: 1 | TF:M07250_1 | FALSE | 1.25E-29 | 28.90284114 | 4139 | 2370 | 675 | 21787 | [illegible] |
| TF | Factor: c-ets-1; motif: ACCGGAWRYN | TF:M01986 | FALSE | 1.45E-29 | 28.83723054 | 8401 | 2370 | 1183 | 21787 | [illegible] |
| TF | Factor: Zic1; motif: NNCCCCCCGGGGGGG | TF:M02835 | FALSE | 1.84E-29 | 28.73412205 | 9543 | 2370 | 1310 | 21787 | [illegible] |
| TF | Factor: SP2; motif: GGGCGGGAC | TF:M01783 | FALSE | 2.53E-29 | 28.59676048 | 8128 | 2370 | 1151 | 21787 | [illegible] |
| TF | Factor: Egr-2; motif: GCGTGGGCGG; match class: 1 | TF:M05494_1 | FALSE | 5.37E-29 | 28.269817 | 2809 | 2370 | 499 | 21787 | [illegible] |
| TF | Factor: Sp1; motif: GGGGCGGGGC; match class: 1 | TF:M00931_1 | FALSE | 1.10E-28 | 27.9569577 | 4199 | 2370 | 679 | 21787 | [illegible] |
| TF | Factor: Sp1; motif: CCCCGCCCCN; match class: 1 | TF:M00933_1 | FALSE | 1.49E-28 | 27.82555482 | 4006 | 2370 | 654 | 21787 | [illegible] |
| TF | Factor: Elk-1; motif: NCCGGAAGTGN; match class: 1 | TF:M10219_1 | FALSE | 3.43E-28 | 27.46472632 | 7699 | 2370 | 1097 | 21787 | [illegible] |
| TF | Factor: Kaiso; motif: GCMGGGRGCRGS | TF:M03876 | FALSE | 4.06E-28 | 27.39176137 | 11148 | 2370 | 1478 | 21787 | [illegible] |
| TF | Factor: AP2; motif: GCCYGSGGSN | TF:M08867 | FALSE | 4.66E-28 | 27.33127673 | 7279 | 2370 | 1048 | 21787 | [illegible] |
| TF | Factor: Elk-1; motif: NCCGGAAGTGN | TF:M10219 | FALSE | 4.91E-28 | 27.30909028 | 14552 | 2370 | 1827 | 21787 | [illegible] |
| TF | Factor: SP1; motif: GGGGYGGGGNS | TF:M01303 | FALSE | 6.21E-28 | 27.20722002 | 7163 | 2370 | 1034 | 21787 | [illegible] |
| TF | Factor: AP-2alpha; motif: NGCCYSNNGSN | TF:M01857 | FALSE | 6.97E-28 | 27.15680624 | 6186 | 2370 | 919 | 21787 | [illegible] |
| TF | Factor: CREB,; motif: NTGACGTNA | TF:M00981 | FALSE | 1.18E-27 | 26.92938413 | 8123 | 2370 | 1143 | 21787 | [illegible] |
| TF | Factor: SP100; motif: NNCGTCGNNTAAWNN; match class: 1 | TF:M02913_1 | FALSE | 2.22E-27 | 26.65396783 | 3068 | 2370 | 528 | 21787 | [illegible] |
| TF | Factor: Pax-5; motif: RRMSWGANWYCTNRAGCGKRACSRYNSM | TF:M00144 | FALSE | 3.61E-27 | 26.44236991 | 12002 | 2370 | 1564 | 21787 | [illegible] |
| TF | Factor: Pax-3; motif: NNNNNNCGTCACGSTYNNNNN; match class: 1 | TF:M00327_1 | FALSE | 5.76E-27 | 26.23975567 | 6169 | 2370 | 913 | 21787 | [illegible] |
| TF | Factor: AP-2; motif: GSCCSCRGGCNRNRNN; match class: 1 | TF:M00800_1 | FALSE | 6.64E-27 | 26.17776583 | 2368 | 2370 | 432 | 21787 | [illegible] |
| TF | Factor: CREM; motif: TGACGTCASYN | TF:M01820 | FALSE | 1.28E-26 | 25.89197714 | 5763 | 2370 | 863 | 21787 | [illegible] |
| TF | Factor: EGR; motif: CGCCCCCGCNN | TF:M08878 | FALSE | 1.87E-26 | 25.72823998 | 5844 | 2370 | 872 | 21787 | [illegible] |
| TF | Factor: GABPalpha; motif: ACCGGAARYN | TF:M02039 | FALSE | 2.95E-26 | 25.53054188 | 6170 | 2370 | 910 | 21787 | [illegible] |
| TF | Factor: Zic1; motif: NNCCCCCCGGGGGGG; match class: 1 | TF:M02835_1 | FALSE | 3.05E-26 | 25.51511203 | 6611 | 2370 | 962 | 21787 | [illegible] |
| TF | Factor: WT1; motif: GNGGGGGCGGGG; match class: 1 | TF:M03893_1 | FALSE | 6.05E-26 | 25.21800356 | 2560 | 2370 | 455 | 21787 | [illegible] |
| TF | Factor: c-Ets-1; motif: NNNRCCGGAWRYNNNN | TF:M01078 | FALSE | 2.05E-25 | 24.6882598 | 7296 | 2370 | 1038 | 21787 | [illegible] |
| TF | Factor: Sp1; motif: NGGGGGCGGGGYN; match class: 1 | TF:M00196_1 | FALSE | 2.35E-25 | 24.62938327 | 4256 | 2370 | 673 | 21787 | [illegible] |
| TF | Factor: Fli-1; motif: GNNRCCGGAAGYGS | TF:M10234 | FALSE | 2.82E-25 | 24.54916524 | 6461 | 2370 | 940 | 21787 | [illegible] |
| TF | Factor: MAZ; motif: NKGGGAGGGGRGGR | TF:M02023 | FALSE | 3.04E-25 | 24.51693884 | 7957 | 2370 | 1113 | 21787 | [illegible] |
| TF | Factor: E2F-4; motif: GNNGGCGGGAAN; match class: 1 | TF:M09682_1 | FALSE | 4.19E-25 | 24.37755106 | 1635 | 2370 | 323 | 21787 | [illegible] |
| TF | Factor: TAFII250; motif: NAARAWGGCGGMGG | TF:M10388 | FALSE | 4.94E-25 | 24.3058781 | 6115 | 2370 | 898 | 21787 | [illegible] |
| TF | Factor: CREB; motif: NSTGACGTAANN | TF:M00177 | FALSE | 7.70E-25 | 24.1134863 | 5050 | 2370 | 769 | 21787 | [illegible] |
| TF | Factor: ERF; motif: ACCGGAARTN | TF:M01984 | FALSE | 1.05E-24 | 23.97908626 | 6018 | 2370 | 885 | 21787 | [illegible] |
| TF | Factor: CPBP; motif: SNCCCNN; match class: 1 | TF:M01822_1 | FALSE | 1.85E-24 | 23.73355263 | 19901 | 2370 | 2291 | 21787 | [illegible] |
| TF | Factor: WT1; motif: CGCCCCCNCN; match class: 1 | TF:M02036_1 | FALSE | 2.50E-24 | 23.60186616 | 3153 | 2370 | 528 | 21787 | [illegible] |
| TF | Factor: Sp1; motif: GGGGCGGGGT | TF:M00008 | FALSE | 2.80E-24 | 23.55357808 | 8368 | 2370 | 1155 | 21787 | [illegible] |
| TF | Factor: Egr-3; motif: NTGCGTGGGCGK | TF:M00245 | FALSE | 5.09E-24 | 23.29296135 | 6078 | 2370 | 889 | 21787 | [illegible] |
| TF | Factor: SP2; motif: GNNGGGGGCGGGGSN; match class: 1 | TF:M03807_1 | FALSE | 7.71E-24 | 23.11282032 | 2961 | 2370 | 501 | 21787 | [illegible] |
| TF | Factor: Hes1; motif: NNCACGYGNN; match class: 1 | TF:M07991_1 | FALSE | 9.10E-24 | 23.04118888 | 11150 | 2370 | 1458 | 21787 | [illegible] |
| TF | Factor: Ets2; motif: ACCGGAWRYN; match class: 1 | TF:M01989_1 | FALSE | 2.17E-23 | 22.66446758 | 2709 | 2370 | 466 | 21787 | [illegible] |
| TF | Factor: Egr-1; motif: WTGCGTGGGCGK | TF:M00243 | FALSE | 2.72E-23 | 22.56539087 | 4357 | 2370 | 677 | 21787 | [illegible] |
| TF | Factor: BTEB2; motif: RGGGNGKGGN; match class: 1 | TF:M07277_1 | FALSE | 2.97E-23 | 22.52752996 | 3123 | 2370 | 520 | 21787 | [illegible] |
| TF | Factor: E2F-1; motif: TTTSGCGS | TF:M00431 | FALSE | 4.03E-23 | 22.39511539 | 5995 | 2370 | 875 | 21787 | [illegible] |
| TF | Factor: Tfap2a; motif: TGCCCYNRGGGCA | TF:M04157 | FALSE | 5.46E-23 | 22.26277209 | 8673 | 2370 | 1183 | 21787 | [illegible] |
| TF | Factor: ERG; motif: ACCGGAART; match class: 1 | TF:M01752_1 | FALSE | 8.18E-23 | 22.08717495 | 3657 | 2370 | 587 | 21787 | [illegible] |
| TF | Factor: CREB,; motif: NTGACGTNA; match class: 1 | TF:M00981_1 | FALSE | 3.74E-22 | 21.42673909 | 2751 | 2370 | 467 | 21787 | [illegible] |
| TF | Factor: CREB; motif: NSTGACGTMANN | TF:M00178 | FALSE | 3.94E-22 | 21.40489696 | 5182 | 2370 | 773 | 21787 | [illegible] |
| TF | Factor: GABP-alpha; motif: GGNRCCGGAAGTGN | TF:M10247 | FALSE | 4.48E-22 | 21.34891054 | 3998 | 2370 | 627 | 21787 | [illegible] |
| TF | Factor: E2F1; motif: NNNNNGCGSSAAAN | TF:M08874 | FALSE | 5.34E-22 | 21.27274498 | 5871 | 2370 | 855 | 21787 | [illegible] |
| TF | Factor: CREB; motif: NNTKACGTCANNNS | TF:M00916 | FALSE | 9.90E-22 | 21.00424214 | 5907 | 2370 | 858 | 21787 | [illegible] |
| TF | Factor: ATF-1; motif: TNACGTCAN | TF:M01861 | FALSE | 1.03E-21 | 20.98679105 | 6280 | 2370 | 902 | 21787 | [illegible] |
| TF | Factor: AP-2; motif: SNNNCCNCAGGCN | TF:M00915 | FALSE | 1.38E-21 | 20.8601375 | 7126 | 2370 | 1000 | 21787 | [illegible] |
| TF | Factor: Sp2; motif: NYSGCCCCGCCCCCY; match class: 1 | TF:M03567_1 | FALSE | 1.83E-21 | 20.73722871 | 2492 | 2370 | 430 | 21787 | [illegible] |
| TF | Factor: CREB; motif: NNGNTGACGTNN | TF:M00113 | FALSE | 2.39E-21 | 20.6220325 | 4604 | 2370 | 699 | 21787 | [illegible] |
| TF | Factor: CREB1; motif: NRRTGACGTMA | TF:M10200 | FALSE | 2.53E-21 | 20.5964831 | 4662 | 2370 | 706 | 21787 | [illegible] |
| TF | Factor: NRF-1; motif: GCGCMTGCGCN | TF:M07052 | FALSE | 3.14E-21 | 20.50308741 | 1918 | 2370 | 351 | 21787 | [illegible] |
| TF | Factor: E2F; motif: TTTSGCGSG | TF:M00939 | FALSE | 4.47E-21 | 20.34935022 | 5472 | 2370 | 803 | 21787 | [illegible] |
| TF | Factor: E2F-1; motif: TTTSGCGCGMNR | TF:M00516 | FALSE | 6.59E-21 | 20.1811793 | 6008 | 2370 | 866 | 21787 | [illegible] |
| TF | Factor: AP2; motif: GCCYGSGGSN; match class: 1 | TF:M08867_1 | FALSE | 7.01E-21 | 20.15433662 | 2689 | 2370 | 454 | 21787 | [illegible] |
| TF | Factor: ATF2; motif: VGTGACGTMACN | TF:M00179 | FALSE | 7.56E-21 | 20.12160822 | 6002 | 2370 | 865 | 21787 | [illegible] |
| TF | Factor: KLF3; motif: NNNNNNGGGCGGGGCNNGN; match class: 1 | TF:M10278_1 | FALSE | 8.67E-21 | 20.06205725 | 3136 | 2370 | 512 | 21787 | [illegible] |
| TF | Factor: Egr-1; motif: NNGCGKGGGCGGGG | TF:M10213 | FALSE | 1.04E-20 | 19.98192853 | 4408 | 2370 | 672 | 21787 | [illegible] |
| TF | Factor: E2F; motif: NCSCGCSAAAN | TF:M00919 | FALSE | 1.05E-20 | 19.98020911 | 5212 | 2370 | 770 | 21787 | [illegible] |
| TF | Factor: Atf-1; motif: RTGACGTA; match class: 1 | TF:M07980_1 | FALSE | 1.42E-20 | 19.84752303 | 3104 | 2370 | 507 | 21787 | [illegible] |
| TF | Factor: CREB1; motif: NNNNTGACGTNANNN | TF:M07248 | FALSE | 1.84E-20 | 19.73461835 | 4336 | 2370 | 662 | 21787 | [illegible] |
| TF | Factor: ATF-2; motif: NNTGACGTCAN | TF:M07312 | FALSE | 2.04E-20 | 19.68955116 | 4160 | 2370 | 640 | 21787 | [illegible] |
| TF | Factor: Tfap2a; motif: TGCCCYNRGGGCA; match class: 1 | TF:M04157_1 | FALSE | 2.45E-20 | 19.61006253 | 4748 | 2370 | 712 | 21787 | [illegible] |
| TF | Factor: CREB; motif: CNNTGACGTMA | TF:M00917 | FALSE | 2.72E-20 | 19.56515809 | 5629 | 2370 | 818 | 21787 | [illegible] |
| TF | Factor: ESE-1; motif: NSMGGAARYN | TF:M01977 | FALSE | 5.40E-20 | 19.26798949 | 7667 | 2370 | 1054 | 21787 | [illegible] |
| TF | Factor: AP-2gamma; motif: GCCYNCRGSN | TF:M03811 | FALSE | 6.16E-20 | 19.21073203 | 7103 | 2370 | 989 | 21787 | [illegible] |
| TF | Factor: E2F-1; motif: TTGGCGCGRAANNGNM | TF:M00938 | FALSE | 2.15E-19 | 18.66820037 | 5231 | 2370 | 766 | 21787 | [illegible] |
| TF | Factor: AP-2; motif: SNNNCCNCAGGCN; match class: 1 | TF:M00915_1 | FALSE | 2.65E-19 | 18.57610342 | 2425 | 2370 | 413 | 21787 | [illegible] |
| TF | Factor: GKLF; motif: NNNRGGNGNGGSN | TF:M07289 | FALSE | 2.70E-19 | 18.56852194 | 15310 | 2370 | 1864 | 21787 | [illegible] |
| TF | Factor: AP-2beta; motif: GCNNNGGSCNGVGGGN | TF:M01858 | FALSE | 2.91E-19 | 18.53583331 | 5120 | 2370 | 752 | 21787 | [illegible] |
| TF | Factor: EGR-1; motif: NGCGKGGGCGKNN | TF:M08996 | FALSE | 3.01E-19 | 18.52147991 | 4001 | 2370 | 615 | 21787 | [illegible] |
| TF | Factor: Osx; motif: CCNCCCCCNNN | TF:M07329 | FALSE | 3.46E-19 | 18.46043144 | 6174 | 2370 | 877 | 21787 | [illegible] |
| TF | Factor: AP-2alpha; motif: GCCNNNRGS | TF:M00469 | FALSE | 4.24E-19 | 18.37248323 | 3276 | 2370 | 523 | 21787 | [illegible] |
| TF | Factor: E2F-3; motif: GGGCGGGARN | TF:M10210 | FALSE | 4.48E-19 | 18.3490278 | 2522 | 2370 | 425 | 21787 | [illegible] |
| TF | Factor: YY1; motif: CAARATGGCGGC | TF:M10408 | FALSE | 4.57E-19 | 18.33997429 | 5103 | 2370 | 749 | 21787 | [illegible] |
| TF | Factor: Erm; motif: NCCGGAWGTN | TF:M01992 | FALSE | 9.30E-19 | 18.03151001 | 5257 | 2370 | 766 | 21787 | [illegible] |
| TF | Factor: CREB1; motif: NNNNACGTCANN | TF:M03544 | FALSE | 9.51E-19 | 18.02195285 | 4205 | 2370 | 638 | 21787 | [illegible] |
| TF | Factor: WT1; motif: NNGGGNGGGSGN | TF:M07436 | FALSE | 9.81E-19 | 18.00825914 | 7743 | 2370 | 1056 | 21787 | [illegible] |
| TF | Factor: Zic3; motif: NMCCCCCGGGGGGGN | TF:M02837 | FALSE | 1.00E-18 | 17.99946479 | 3312 | 2370 | 526 | 21787 | [illegible] |
| TF | Factor: E2F; motif: TTTSGCGSG | TF:M00918 | FALSE | 1.08E-18 | 17.9648214 | 4812 | 2370 | 712 | 21787 | [illegible] |
| TF | Factor: Pax-3; motif: TCGTCACRCTTHM; match class: 1 | TF:M00360_1 | FALSE | 1.38E-18 | 17.86036059 | 5448 | 2370 | 788 | 21787 | [illegible] |
| TF | Factor: Egr-1; motif: GCGGGGGCGG; match class: 1 | TF:M07354_1 | FALSE | 1.86E-18 | 17.73155124 | 2113 | 2370 | 368 | 21787 | [illegible] |
| TF | Factor: GKLF; motif: GCCMCRCCCNNN | TF:M01588 | FALSE | 2.26E-18 | 17.6455116 | 7857 | 2370 | 1067 | 21787 | [illegible] |
| TF | Factor: AP-2gamma; motif: GCCYNNGGS | TF:M00470 | FALSE | 3.04E-18 | 17.51678277 | 7024 | 2370 | 971 | 21787 | [illegible] |
| TF | Factor: GKLF; motif: NNRRGRRNGNSNNN | TF:M07040 | FALSE | 3.39E-18 | 17.46989233 | 12956 | 2370 | 1620 | 21787 | [illegible] |
| TF | Factor: EGR-1; motif: TGCGTGGGCGK | TF:M01972 | FALSE | 3.58E-18 | 17.4466317 | 4170 | 2370 | 631 | 21787 | [illegible] |
| TF | Factor: E2F-4; motif: NGGGGGCGGGRMNN; match class: 1 | TF:M10211_1 | FALSE | 6.17E-18 | 17.20961734 | 2427 | 2370 | 408 | 21787 | [illegible] |
| TF | Factor: XBP-1; motif: WNNGMCACGTC | TF:M01770 | FALSE | 7.93E-18 | 17.10069576 | 10526 | 2370 | 1360 | 21787 | [illegible] |
| TF | Factor: ATF-1; motif: NNNTGACGTNNN | TF:M07034 | FALSE | 8.62E-18 | 17.06473956 | 4331 | 2370 | 649 | 21787 | [illegible] |
| TF | Factor: Sp4; motif: NNNNMCGCCCCCTNNNN | TF:M02810 | FALSE | 8.84E-18 | 17.05347273 | 4015 | 2370 | 610 | 21787 | [illegible] |
| TF | Factor: CREB; motif: TGACGTMA | TF:M00039 | FALSE | 1.09E-17 | 16.96285657 | 5419 | 2370 | 780 | 21787 | [illegible] |
| TF | Factor: E2F; motif: TTTSGCGC | TF:M00050 | FALSE | 1.65E-17 | 16.78364636 | 6163 | 2370 | 867 | 21787 | [illegible] |
| TF | Factor: Tax/CREB; motif: GGGGGTTGACGYANA | TF:M00114 | FALSE | 1.69E-17 | 16.77145464 | 5335 | 2370 | 769 | 21787 | [illegible] |
| TF | Factor: CREM; motif: TGACGTCASYN; match class: 1 | TF:M01820_1 | FALSE | 1.76E-17 | 16.75523819 | 1816 | 2370 | 324 | 21787 | [illegible] |
| TF | Factor: IRF-9; motif: NNRAAWCGAAACTNA | TF:M02771 | FALSE | 2.08E-17 | 16.68183989 | 406 | 2370 | 111 | 21787 | [illegible] |
| TF | Factor: WT1; motif: SMCNCCNSC | TF:M01118 | FALSE | 3.32E-17 | 16.47947147 | 8117 | 2370 | 1090 | 21787 | [illegible] |
| TF | Factor: ZBP89; motif: CCCCKCCCCCNN | TF:M07397 | FALSE | 3.43E-17 | 16.4651515 | 8100 | 2370 | 1088 | 21787 | [illegible] |
| TF | Factor: AP-2alpha; motif: NSCCNCRGGSN; match class: 1 | TF:M07348_1 | FALSE | 3.58E-17 | 16.44636589 | 1679 | 2370 | 304 | 21787 | [illegible] |
| TF | Factor: WT1; motif: NGCGGGGGGGTSMMCYN; match class: 1 | TF:M05599_1 | FALSE | 3.85E-17 | 16.41485472 | 1687 | 2370 | 305 | 21787 | [illegible] |
| TF | Factor: Egr1; motif: NCCGCCCCCGCANN | TF:M02744 | FALSE | 3.88E-17 | 16.41114237 | 3462 | 2370 | 538 | 21787 | [illegible] |
| TF | Factor: MAZ; motif: NKGGGAGGGGRGGR; match class: 1 | TF:M02023_1 | FALSE | 4.87E-17 | 16.31252304 | 4165 | 2370 | 625 | 21787 | [illegible] |
| TF | Factor: Egr1; motif: NNMCGCCCMCTCAMWN | TF:M04506 | FALSE | 5.26E-17 | 16.27923287 | 4362 | 2370 | 649 | 21787 | [illegible] |
| TF | Factor: SP6; motif: WGGGCGG; match class: 1 | TF:M05455_1 | FALSE | 1.18E-16 | 15.92733633 | 2367 | 2370 | 395 | 21787 | [illegible] |
| TF | Factor: LKLF; motif: NGGGCGG; match class: 1 | TF:M05391_1 | FALSE | 1.18E-16 | 15.92733633 | 2367 | 2370 | 395 | 21787 | [illegible] |
| TF | Factor: CPBP; motif: NGGGCGG; match class: 1 | TF:M05301_1 | FALSE | 1.18E-16 | 15.92733633 | 2367 | 2370 | 395 | 21787 | [illegible] |
| TF | Factor: Sp3; motif: AGGGCGG; match class: 1 | TF:M05665_1 | FALSE | 1.18E-16 | 15.92733633 | 2367 | 2370 | 395 | 21787 | [illegible] |
| TF | Factor: ATF-1; motif: TNACGTCAN; match class: 1 | TF:M01861_1 | FALSE | 1.35E-16 | 15.86866902 | 1898 | 2370 | 332 | 21787 | [illegible] |
| TF | Factor: CREB; motif: NSTGACGTAANN; match class: 1 | TF:M00177_1 | FALSE | 2.14E-16 | 15.67053662 | 1356 | 2370 | 256 | 21787 | [illegible] |
| TF | Factor: Pet-1; motif: GCNGGAAGYG | TF:M10233 | FALSE | 2.20E-16 | 15.65804147 | 13301 | 2370 | 1646 | 21787 | [illegible] |
| TF | Factor: E2F-2; motif: NNNANGGCGCGCNNN | TF:M02742 | FALSE | 2.45E-16 | 15.61059477 | 3539 | 2370 | 544 | 21787 | [illegible] |
| TF | Factor: AP-2gamma; motif: GCCYNNGGS; match class: 1 | TF:M00470_1 | FALSE | 2.69E-16 | 15.56998654 | 3413 | 2370 | 528 | 21787 | [illegible] |
| TF | Factor: Zfp536; motif: NCGGAKG; match class: 1 | TF:M05541_1 | FALSE | 3.57E-16 | 15.44711482 | 1719 | 2370 | 306 | 21787 | [illegible] |
| TF | Factor: BEN; motif: CWGCGAYA | TF:M01241 | FALSE | 3.76E-16 | 15.42451241 | 6578 | 2370 | 908 | 21787 | [illegible] |
| TF | Factor: AhR:Arnt; motif: GRGKATYGCGTGMCWNSCC | TF:M00237 | FALSE | 6.06E-16 | 15.21748074 | 5860 | 2370 | 823 | 21787 | [illegible] |
| TF | Factor: GKLF; motif: WGGGYGKGGCCN | TF:M09692 | FALSE | 6.32E-16 | 15.19904646 | 8986 | 2370 | 1180 | 21787 | [illegible] |
| TF | Factor: Klf7; motif: NNNRCCMCGCCCNNNN | TF:M02773 | FALSE | 7.00E-16 | 15.15472209 | 4203 | 2370 | 624 | 21787 | [illegible] |
| TF | Factor: ZBP89; motif: CCCCKCCCCCNN; match class: 1 | TF:M07397_1 | FALSE | 8.91E-16 | 15.0500806 | 3882 | 2370 | 584 | 21787 | [illegible] |
| TF | Factor: E2F; motif: NNTTTCGCGCN | TF:M08875 | FALSE | 9.30E-16 | 15.03163562 | 5168 | 2370 | 740 | 21787 | [illegible] |
| TF | Factor: SP1:SP3; motif: CCSCCCCCYCC; match class: 1 | TF:M01219_1 | FALSE | 9.88E-16 | 15.00517703 | 3394 | 2370 | 523 | 21787 | [illegible] |
| TF | Factor: SAP-1a; motif: NRRCCGGAAGYRN; match class: 1 | TF:M10220_1 | FALSE | 1.01E-15 | 14.99475723 | 3299 | 2370 | 511 | 21787 | [illegible] |
| TF | Factor: Zic2; motif: NMCCCCCGGGGGGGN | TF:M02836 | FALSE | 1.12E-15 | 14.9504978 | 3797 | 2370 | 573 | 21787 | [illegible] |
| TF | Factor: TAFII250; motif: NAARAWGGCGGMGG; match class: 1 | TF:M10388_1 | FALSE | 1.36E-15 | 14.86606915 | 1172 | 2370 | 227 | 21787 | [illegible] |
| TF | Factor: ZBED6; motif: NRRGCTCGCCNN; match class: 1 | TF:M01598_1 | FALSE | 1.56E-15 | 14.80673959 | 1369 | 2370 | 255 | 21787 | [illegible] |
| TF | Factor: Elk-1; motif: RACCGGAAGTR | TF:M01165 | FALSE | 1.94E-15 | 14.71272643 | 8877 | 2370 | 1165 | 21787 | [illegible] |
| TF | Factor: AhR,; motif: NRCGTGNGN | TF:M00976 | FALSE | 2.02E-15 | 14.69465654 | 4026 | 2370 | 600 | 21787 | [illegible] |
| TF | Factor: E2F; motif: NKCGCGCSAAAN | TF:M00920 | FALSE | 2.73E-15 | 14.56452256 | 4863 | 2370 | 701 | 21787 | [illegible] |
| TF | Factor: USF; motif: GYCACGTGNC | TF:M00187 | FALSE | 3.18E-15 | 14.49692459 | 3533 | 2370 | 538 | 21787 | [illegible] |
| TF | Factor: Pet-1; motif: GCNGGAAGYG; match class: 1 | TF:M10233_1 | FALSE | 5.24E-15 | 14.28099648 | 6092 | 2370 | 845 | 21787 | [illegible] |
| TF | Factor: EGR; motif: CGCCCCCGCNN; match class: 1 | TF:M08878_1 | FALSE | 5.36E-15 | 14.27109889 | 1861 | 2370 | 321 | 21787 | [illegible] |
| TF | Factor: Spic; motif: NNVGGAAGTN | TF:M02042 | FALSE | 5.81E-15 | 14.23547867 | 6197 | 2370 | 857 | 21787 | [illegible] |
| TF | Factor: Fli-1; motif: GNNRCCGGAAGYGS; match class: 1 | TF:M10234_1 | FALSE | 7.29E-15 | 14.13723634 | 1637 | 2370 | 290 | 21787 | [illegible] |
| TF | Factor: SP2; motif: GGGCGGGAC; match class: 1 | TF:M01783_1 | FALSE | 7.75E-15 | 14.1106487 | 3300 | 2370 | 507 | 21787 | [illegible] |
| TF | Factor: Osx; motif: CCNCCCCCNNN; match class: 1 | TF:M07329_1 | FALSE | 8.02E-15 | 14.0958951 | 2015 | 2370 | 341 | 21787 | [illegible] |
| TF | Factor: ATF-3; motif: SNTGACGYNATN | TF:M07313 | FALSE | 8.73E-15 | 14.05902999 | 2891 | 2370 | 455 | 21787 | [illegible] |
| TF | Factor: Sp1; motif: GGGGCGGGGT; match class: 1 | TF:M00008_1 | FALSE | 1.20E-14 | 13.92024994 | 3395 | 2370 | 518 | 21787 | [illegible] |
| TF | Factor: KROX; motif: CCCGCCCCCRCCCC; match class: 1 | TF:M00982_1 | FALSE | 1.33E-14 | 13.87767196 | 2044 | 2370 | 344 | 21787 | [illegible] |
| TF | Factor: E2F1; motif: NNNNNGCGSSAAAN; match class: 1 | TF:M08874_1 | FALSE | 1.34E-14 | 13.87254571 | 1499 | 2370 | 270 | 21787 | [illegible] |
| TF | Factor: ATF2; motif: VGTGACGTMACN; match class: 1 | TF:M00179_1 | FALSE | 1.82E-14 | 13.73997796 | 1802 | 2370 | 311 | 21787 | [illegible] |
| TF | Factor: ER81; motif: RCCGGAWRYN | TF:M01987 | FALSE | 1.86E-14 | 13.72979406 | 4033 | 2370 | 596 | 21787 | [illegible] |
| TF | Factor: Mitf; motif: NRTCACGNGN | TF:M07996 | FALSE | 1.89E-14 | 13.7245879 | 2848 | 2370 | 448 | 21787 | [illegible] |
| TF | Factor: ZNF151; motif: GGSGGNGGGGGAGGGGMGG | TF:M10412 | FALSE | 2.17E-14 | 13.66312658 | 3816 | 2370 | 569 | 21787 | [illegible] |
| TF | Factor: c-ets-1; motif: ACCGGAWRYN; match class: 1 | TF:M01986_1 | FALSE | 2.23E-14 | 13.65087006 | 2882 | 2370 | 452 | 21787 | [illegible] |
| TF | Factor: GKLF; motif: NNNRGGNGNGGSN; match class: 1 | TF:M07289_1 | FALSE | 2.24E-14 | 13.64921435 | 10990 | 2370 | 1390 | 21787 | [illegible] |
| TF | Factor: ctcf; motif: CCNCNAGRKGGCRSTN | TF:M07249 | FALSE | 2.56E-14 | 13.59103945 | 5580 | 2370 | 781 | 21787 | [illegible] |
| TF | Factor: ZFP740; motif: NNNTWCCCCCCGGNRNN | TF:M02938 | FALSE | 6.15E-14 | 13.21102249 | 3151 | 2370 | 484 | 21787 | [illegible] |
| TF | Factor: SP1; motif: GGGGYGGGGNS; match class: 1 | TF:M01303_1 | FALSE | 8.28E-14 | 13.08415397 | 2769 | 2370 | 435 | 21787 | [illegible] |
| TF | Factor: Elf-1; motif: NNANCCGGAAGTGN | TF:M10216 | FALSE | 9.12E-14 | 13.03994984 | 3086 | 2370 | 475 | 21787 | [illegible] |
| TF | Factor: Sp4; motif: SCCCCKCCCCCSN | TF:M07617 | FALSE | 9.12E-14 | 13.03976826 | 2708 | 2370 | 427 | 21787 | [illegible] |
| TF | Factor: MECP2; motif: CCGGNNTTWA | TF:M01299 | FALSE | 1.02E-13 | 12.98988062 | 5018 | 2370 | 711 | 21787 | [illegible] |
| TF | Factor: Elf-1; motif: AWCCCGGAAGTN | TF:M07415 | FALSE | 1.05E-13 | 12.98078388 | 8345 | 2370 | 1095 | 21787 | [illegible] |
| TF | Factor: Pax-3; motif: TCGTCACRCTTHM | TF:M00360 | FALSE | 1.08E-13 | 12.96583183 | 12893 | 2370 | 1588 | 21787 | [illegible] |
| TF | Factor: Irf-4; motif: NNNANCGAAACYNNA | TF:M02768 | FALSE | 1.28E-13 | 12.89167878 | 5031 | 2370 | 712 | 21787 | [illegible] |
| TF | Factor: CREB; motif: NNTKACGTCANNNS; match class: 1 | TF:M00916_1 | FALSE | 1.29E-13 | 12.88987027 | 1767 | 2370 | 303 | 21787 | [illegible] |
| TF | Factor: CKROX; motif: SCCCTCCCC | TF:M01175 | FALSE | 1.43E-13 | 12.84381542 | 6344 | 2370 | 866 | 21787 | [illegible] |
| TF | Factor: NRF-1; motif: YGCGCMTGCGC | TF:M02102 | FALSE | 1.60E-13 | 12.79721332 | 2062 | 2370 | 342 | 21787 | [illegible] |
| TF | Factor: Ehf; motif: NNNANCCGGAAGTNN | TF:M02745 | FALSE | 1.75E-13 | 12.75713107 | 3821 | 2370 | 565 | 21787 | [illegible] |
| TF | Factor: YY1; motif: GCCGCCATTTTG | TF:M02044 | FALSE | 1.84E-13 | 12.73504002 | 6410 | 2370 | 873 | 21787 | [illegible] |
| TF | Factor: Tfap2a; motif: TGCCCYNRGGGCA | TF:M04155 | FALSE | 1.95E-13 | 12.70923439 | 6541 | 2370 | 888 | 21787 | [illegible] |
| TF | Factor: ATF-3; motif: SNTGACGYNATN; match class: 1 | TF:M07313_1 | FALSE | 1.96E-13 | 12.70869981 | 546 | 2370 | 126 | 21787 | [illegible] |
| TF | Factor: NGFI-C; motif: WTGCGTGGGYGG | TF:M00244 | FALSE | 2.28E-13 | 12.64190413 | 5026 | 2370 | 710 | 21787 | [illegible] |
| TF | Factor: Egr-1; motif: WTGCGTGGGCGK; match class: 1 | TF:M00243_1 | FALSE | 2.76E-13 | 12.55983639 | 859 | 2370 | 174 | 21787 | [illegible] |
| TF | Factor: MAZ; motif: GGGGAGGG | TF:M00649 | FALSE | 3.64E-13 | 12.43866733 | 8420 | 2370 | 1100 | 21787 | [illegible] |
| TF | Factor: Egr-2; motif: NTGCGTRGGCGK | TF:M00246 | FALSE | 4.17E-13 | 12.38004963 | 5376 | 2370 | 750 | 21787 | [illegible] |
| TF | Factor: c-Ets-1; motif: NNNRCCGGAWRYNNNN; match class: 1 | TF:M01078_1 | FALSE | 4.72E-13 | 12.32597489 | 2337 | 2370 | 376 | 21787 | [illegible] |
| TF | Factor: Egr-1; motif: GCGGGGGCGG; match class: 1 | TF:M01873_1 | FALSE | 4.74E-13 | 12.32410512 | 1518 | 2370 | 267 | 21787 | [illegible] |
| TF | Factor: Egr-1; motif: GCGTGGGCGT | TF:M07597 | FALSE | 4.95E-13 | 12.30500854 | 4732 | 2370 | 673 | 21787 | [illegible] |
| TF | Factor: AP-2alpha; motif: GCCNNNRGS; match class: 1 | TF:M00469_1 | FALSE | 6.41E-13 | 12.19322896 | 2028 | 2370 | 335 | 21787 | [illegible] |
| TF | Factor: AP-2alpha; motif: NSCCNCRGGSN | TF:M07348 | FALSE | 1.01E-12 | 11.99404236 | 6233 | 2370 | 848 | 21787 | [illegible] |
| TF | Factor: E2F-1; motif: NTTSGCGG; match class: 1 | TF:M00430_1 | FALSE | 1.40E-12 | 11.85346379 | 1820 | 2370 | 306 | 21787 | [illegible] |
| TF | Factor: CREB; motif: CNNTGACGTMA; match class: 1 | TF:M00917_1 | FALSE | 1.45E-12 | 11.83764261 | 1664 | 2370 | 285 | 21787 | [illegible] |
| TF | Factor: Zfx; motif: SNSCAGGCCKCGSCSS; match class: 1 | TF:M01593_1 | FALSE | 1.57E-12 | 11.80423401 | 504 | 2370 | 117 | 21787 | [illegible] |
| TF | Factor: CREB; motif: NSTGACGTMANN; match class: 1 | TF:M00178_1 | FALSE | 1.62E-12 | 11.79165922 | 1562 | 2370 | 271 | 21787 | [illegible] |
| TF | Factor: ATF-2; motif: NNNTGACGTNAN | TF:M01862 | FALSE | 1.64E-12 | 11.7854158 | 3311 | 2370 | 497 | 21787 | [illegible] |
| TF | Factor: CNOT3; motif: GGCCGCGSSS; match class: 1 | TF:M01253_1 | FALSE | 4.23E-12 | 11.37328497 | 523 | 2370 | 119 | 21787 | [illegible] |
| TF | Factor: FKLF; motif: BGGGNGGVMD | TF:M01837 | FALSE | 4.85E-12 | 11.3139229 | 4981 | 2370 | 697 | 21787 | [illegible] |
| TF | Factor: BTEB3; motif: BNRNGGGAGGNGT | TF:M01865 | FALSE | 4.91E-12 | 11.30883782 | 8952 | 2370 | 1152 | 21787 | [illegible] |
| TF | Factor: Sp5; motif: RNGGRGGNGGRGNNGGGGGAGGRG | TF:M10376 | FALSE | 4.98E-12 | 11.30262239 | 4847 | 2370 | 681 | 21787 | [illegible] |
| TF | Factor: AP-2alpha; motif: NGCCYSNNGSN; match class: 1 | TF:M01857_1 | FALSE | 5.55E-12 | 11.2560114 | 2204 | 2370 | 354 | 21787 | [illegible] |
| TF | Factor: ER71; motif: ACCGGAARYN | TF:M01988 | FALSE | 6.20E-12 | 11.20760964 | 4309 | 2370 | 616 | 21787 | [illegible] |
| TF | Factor: HIC1; motif: NSNNNNTGCCCSSNN | TF:M01072 | FALSE | 6.69E-12 | 11.17428756 | 2864 | 2370 | 438 | 21787 | [illegible] |
| TF | Factor: ETV3; motif: ANCGGAARYN | TF:M01990 | FALSE | 8.20E-12 | 11.08636656 | 3203 | 2370 | 480 | 21787 | [illegible] |
| TF | Factor: E2F; motif: TTTSGCGC; match class: 1 | TF:M00050_1 | FALSE | 8.39E-12 | 11.07604095 | 1450 | 2370 | 253 | 21787 | [illegible] |
| TF | Factor: ELF4; motif: CCCGGAARTN; match class: 1 | TF:M01979_1 | FALSE | 1.06E-11 | 10.9756651 | 2260 | 2370 | 360 | 21787 | [illegible] |
| TF | Factor: AP-2beta; motif: GCNNNGGSCNGVGGGN; match class: 1 | TF:M01858_1 | FALSE | 1.72E-11 | 10.76407291 | 1525 | 2370 | 262 | 21787 | [illegible] |
| TF | Factor: Elf5; motif: CSMGGAARTN | TF:M01980 | FALSE | 1.91E-11 | 10.7192724 | 6198 | 2370 | 836 | 21787 | [illegible] |
| TF | Factor: GKLF; motif: NNRRGRRNGNSNNN; match class: 1 | TF:M07040_1 | FALSE | 2.15E-11 | 10.66751873 | 8461 | 2370 | 1093 | 21787 | [illegible] |
| TF | Factor: USF2; motif: NGTCACGTGRNSN | TF:M09722 | FALSE | 3.64E-11 | 10.43948192 | 1872 | 2370 | 307 | 21787 | [illegible] |
| TF | Factor: ATF-3; motif: RTKRCGTCANN | TF:M01863 | FALSE | 5.27E-11 | 10.27842952 | 2053 | 2370 | 330 | 21787 | [illegible] |
| TF | Factor: GKLF; motif: WGGGYGKGGCCN; match class: 1 | TF:M09692_1 | FALSE | 5.36E-11 | 10.27105333 | 3611 | 2370 | 526 | 21787 | [illegible] |
| TF | Factor: ATF-1; motif: NNNTGACGTNNN; match class: 1 | TF:M07034_1 | FALSE | 5.74E-11 | 10.24123282 | 1269 | 2370 | 225 | 21787 | [illegible] |
| TF | Factor: CREB1; motif: NNNNTGACGTNANNN; match class: 1 | TF:M07248_1 | FALSE | 5.97E-11 | 10.22407812 | 1313 | 2370 | 231 | 21787 | [illegible] |
| TF | Factor: CREB1; motif: NRRTGACGTMA; match class: 1 | TF:M10200_1 | FALSE | 6.85E-11 | 10.1642463 | 1395 | 2370 | 242 | 21787 | [illegible] |
| TF | Factor: YY1; motif: GCCGCCATTTTG; match class: 1 | TF:M02044_1 | FALSE | 1.06E-10 | 9.973306336 | 1125 | 2370 | 204 | 21787 | [illegible] |
| TF | Factor: GABP; motif: VCCGGAAGNGCR | TF:M00341 | FALSE | 1.23E-10 | 9.909510428 | 7830 | 2370 | 1017 | 21787 | [illegible] |
| TF | Factor: YY1; motif: NNNNAARATGGNNNN | TF:M07370 | FALSE | 1.23E-10 | 9.908264477 | 6330 | 2370 | 846 | 21787 | [illegible] |
| TF | Factor: WT1; motif: SMCNCCNSC; match class: 1 | TF:M01118_1 | FALSE | 1.26E-10 | 9.899433048 | 4598 | 2370 | 643 | 21787 | [illegible] |
| TF | Factor: DEAF1; motif: NCGNNYTCGGGNRTTTCCGDARNNN | TF:M01001 | FALSE | 1.34E-10 | 9.873795406 | 1845 | 2370 | 301 | 21787 | [illegible] |
| TF | Factor: Pet-1; motif: ACCGGAARYN | TF:M02037 | FALSE | 1.63E-10 | 9.787414616 | 2976 | 2370 | 445 | 21787 | [illegible] |
| TF | Factor: CREB; motif: TGACGTMA; match class: 1 | TF:M00039_1 | FALSE | 1.78E-10 | 9.749420733 | 1585 | 2370 | 266 | 21787 | [illegible] |
| TF | Factor: Kid3; motif: CCACN; match class: 1 | TF:M01160_1 | FALSE | 2.08E-10 | 9.681443248 | 21113 | 2370 | 2348 | 21787 | [illegible] |
| TF | Factor: ZFP740; motif: NNNTWCCCCCCGGNRNN; match class: 1 | TF:M02938_1 | FALSE | 2.83E-10 | 9.548335129 | 483 | 2370 | 108 | 21787 | [illegible] |
| TF | Factor: Elk-1; motif: AACCGGAAGTR | TF:M01163 | FALSE | 2.94E-10 | 9.531308838 | 6066 | 2370 | 813 | 21787 | [illegible] |
| TF | Factor: Fli-1; motif: NNNRCCGGAAGTGGN | TF:M09685 | FALSE | 3.90E-10 | 9.408449242 | 4640 | 2370 | 645 | 21787 | [illegible] |
| TF | Factor: GABPalpha_GABPbeta; motif: CTTCCKGY | TF:M03791 | FALSE | 4.15E-10 | 9.382209919 | 2953 | 2370 | 440 | 21787 | [illegible] |
| TF | Factor: EHF; motif: CSCGGAARTN | TF:M01974 | FALSE | 4.37E-10 | 9.35968826 | 6885 | 2370 | 906 | 21787 | [illegible] |
| TF | Factor: ATF-1; motif: CYYTGACGTCA | TF:M00691 | FALSE | 5.39E-10 | 9.268696452 | 5036 | 2370 | 691 | 21787 | [illegible] |
| TF | Factor: EGR-1; motif: NGCGKGGGCGKNN; match class: 1 | TF:M08996_1 | FALSE | 5.88E-10 | 9.230769673 | 924 | 2370 | 173 | 21787 | [illegible] |
| TF | Factor: ATF-2; motif: NNTGACGTCAN; match class: 1 | TF:M07312_1 | FALSE | 6.21E-10 | 9.207110844 | 1261 | 2370 | 220 | 21787 | [illegible] |
| TF | Factor: ctcf; motif: YGGCCASCAGRGGGCGCYNN | TF:M10203 | FALSE | 6.35E-10 | 9.197040458 | 6650 | 2370 | 878 | 21787 | [illegible] |
| TF | Factor: GKLF; motif: GCCMCRCCCNNN; match class: 1 | TF:M01588_1 | FALSE | 6.67E-10 | 9.175608954 | 2986 | 2370 | 443 | 21787 | [illegible] |
| TF | Factor: ERG; motif: ACCGGAARYN | TF:M01985 | FALSE | 7.48E-10 | 9.126149897 | 2628 | 2370 | 398 | 21787 | [illegible] |
| TF | Factor: Zic2; motif: NMCCCCCGGGGGGGN; match class: 1 | TF:M02836_1 | FALSE | 8.19E-10 | 9.086953331 | 2479 | 2370 | 379 | 21787 | [illegible] |
| TF | Factor: Klf17; motif: NGGGCGG; match class: 1 | TF:M05407_1 | FALSE | 8.58E-10 | 9.066454725 | 1735 | 2370 | 283 | 21787 | [illegible] |
| TF | Factor: BTEB2; motif: NGGGSWGGGNGGGGC | TF:M10280 | FALSE | 8.95E-10 | 9.048191772 | 3048 | 2370 | 450 | 21787 | [illegible] |
| TF | Factor: Tax/CREB; motif: GGGGGTTGACGYANA; match class: 1 | TF:M00114_1 | FALSE | 9.05E-10 | 9.043488037 | 999 | 2370 | 183 | 21787 | [illegible] |
| TF | Factor: E2F-2; motif: NNNWYGGCGCCAANNNN | TF:M02846 | FALSE | 1.51E-09 | 8.821528839 | 1981 | 2370 | 314 | 21787 | [illegible] |
| TF | Factor: E2F-1; motif: NKTSSCGC; match class: 1 | TF:M00428_1 | FALSE | 1.69E-09 | 8.77315959 | 2931 | 2370 | 434 | 21787 | [illegible] |
| TF | Factor: AP-2gamma; motif: GCCYNCRGSN; match class: 1 | TF:M03811_1 | FALSE | 2.15E-09 | 8.666882938 | 2330 | 2370 | 358 | 21787 | [illegible] |
| TF | Factor: Egr-1; motif: NNGCGKGGGCGGGG; match class: 1 | TF:M10213_1 | FALSE | 2.19E-09 | 8.659800075 | 1167 | 2370 | 205 | 21787 | [illegible] |
| TF | Factor: YY1; motif: CAARATGGCGGC; match class: 1 | TF:M10408_1 | FALSE | 2.59E-09 | 8.586675334 | 738 | 2370 | 144 | 21787 | [illegible] |
| TF | Factor: ESE-1; motif: NSMGGAARTN | TF:M01214 | FALSE | 2.60E-09 | 8.585395808 | 7851 | 2370 | 1010 | 21787 | [illegible] |
| TF | Factor: CKROX; motif: SCCCTCCCC; match class: 1 | TF:M01175_1 | FALSE | 2.75E-09 | 8.560234763 | 1975 | 2370 | 312 | 21787 | [illegible] |
| TF | Factor: ATF1; motif: NNATGACGANYANN | TF:M02842 | FALSE | 2.83E-09 | 8.548230371 | 6541 | 2370 | 861 | 21787 | [illegible] |
| TF | Factor: USF; motif: GYCACGTGNC; match class: 1 | TF:M00187_1 | FALSE | 3.61E-09 | 8.442450962 | 1342 | 2370 | 228 | 21787 | [illegible] |
| TF | Factor: CPBP; motif: SNCCCNN | TF:M01822 | FALSE | 4.53E-09 | 8.344267282 | 21076 | 2370 | 2343 | 21787 | [illegible] |
| TF | Factor: YY1; motif: NYNKCCATNTT | TF:M01035 | FALSE | 4.58E-09 | 8.338870264 | 6519 | 2370 | 857 | 21787 | [illegible] |
| TF | Factor: E2F-1; motif: TWSGCGCGAAAAYKR | TF:M00024 | FALSE | 5.10E-09 | 8.292775168 | 8585 | 2370 | 1090 | 21787 | [illegible] |
| TF | Factor: MAZ; motif: GGGGAGGG; match class: 1 | TF:M00649_1 | FALSE | 5.28E-09 | 8.276984618 | 3042 | 2370 | 445 | 21787 | [illegible] |
| TF | Factor: E2F-1; motif: TWSGCGCGAAAAYKR; match class: 1 | TF:M00024_1 | FALSE | 5.57E-09 | 8.253880929 | 2244 | 2370 | 345 | 21787 | [illegible] |
| TF | Factor: MAX; motif: NNGNCACGCGACNN | TF:M02881 | FALSE | 6.50E-09 | 8.187196969 | 5438 | 2370 | 731 | 21787 | [illegible] |
| TF | Factor: SP4; motif: SCCCCGCCCCS | TF:M01273 | FALSE | 6.87E-09 | 8.162908438 | 2358 | 2370 | 359 | 21787 | [illegible] |
| TF | Factor: MAX; motif: NNGNCACGCGACNN; match class: 1 | TF:M02881_1 | FALSE | 6.92E-09 | 8.160100599 | 1568 | 2370 | 257 | 21787 | [illegible] |
| TF | Factor: ELF1; motif: NCCGGAARTN | TF:M01975 | FALSE | 7.31E-09 | 8.135878691 | 3016 | 2370 | 441 | 21787 | [illegible] |
| TF | Factor: Klf15; motif: RGGGMGGRGNNGGGGNGG | TF:M10277 | FALSE | 7.59E-09 | 8.119927202 | 3163 | 2370 | 459 | 21787 | [illegible] |
| TF | Factor: ATF-4; motif: KACGTCAKS; match class: 1 | TF:M01864_1 | FALSE | 7.96E-09 | 8.0990591 | 1442 | 2370 | 240 | 21787 | [illegible] |
| TF | Factor: CREB; motif: NNGNTGACGTNN; match class: 1 | TF:M00113_1 | FALSE | 8.72E-09 | 8.059397888 | 798 | 2370 | 151 | 21787 | [illegible] |
| TF | Factor: Arnt; motif: NGTCACGTGN | TF:M07979 | FALSE | 9.74E-09 | 8.011332972 | 855 | 2370 | 159 | 21787 | [illegible] |
| TF | Factor: CREB1; motif: NNNNACGTCANN; match class: 1 | TF:M03544_1 | FALSE | 1.06E-08 | 7.975671101 | 1282 | 2370 | 218 | 21787 | [illegible] |
| TF | Factor: ESE-1; motif: NSMGGAARYN; match class: 1 | TF:M01977_1 | FALSE | 1.14E-08 | 7.942198186 | 2367 | 2370 | 359 | 21787 | [illegible] |
| TF | Factor: NGFI-C; motif: WTGCGTGGGYGG; match class: 1 | TF:M00244_1 | FALSE | 1.49E-08 | 7.825502088 | 1154 | 2370 | 200 | 21787 | [illegible] |
| TF | Factor: EGR-1; motif: TGCGTGGGCGK; match class: 1 | TF:M01972_1 | FALSE | 1.57E-08 | 7.805389139 | 860 | 2370 | 159 | 21787 | [illegible] |
| TF | Factor: Tcfap2b; motif: NYGCCYNARGGCAN | TF:M02820 | FALSE | 1.90E-08 | 7.722264814 | 7908 | 2370 | 1010 | 21787 | [illegible] |
| TF | Factor: nerf; motif: NCCGGAARTN | TF:M01976 | FALSE | 1.99E-08 | 7.702179213 | 2785 | 2370 | 410 | 21787 | [illegible] |
| TF | Factor: SREBP-1; motif: ATCACGTGAY | TF:M04189 | FALSE | 2.12E-08 | 7.673837577 | 261 | 2370 | 67 | 21787 | [illegible] |
| TF | Factor: SREBP-1; motif: ATCACGTGAY; match class: 1 | TF:M04189_1 | FALSE | 2.12E-08 | 7.673837577 | 261 | 2370 | 67 | 21787 | [illegible] |
| TF | Factor: Wt1; motif: NNCCTCCCCCACNN | TF:M12438 | FALSE | 2.18E-08 | 7.662160838 | 5987 | 2370 | 791 | 21787 | [illegible] |
| TF | Factor: Egr-3; motif: NTGCGTGGGCGK; match class: 1 | TF:M00245_1 | FALSE | 2.19E-08 | 7.660345315 | 1661 | 2370 | 267 | 21787 | [illegible] |
| TF | Factor: GMEB1; motif: NGNNCRACGTYGTYNN | TF:M02865 | FALSE | 2.57E-08 | 7.589949201 | 4199 | 2370 | 581 | 21787 | [illegible] |
| TF | Factor: USF; motif: NNRYCACGTGRYNN; match class: 1 | TF:M00121_1 | FALSE | 3.01E-08 | 7.521861294 | 2584 | 2370 | 384 | 21787 | [illegible] |
| TF | Factor: USF; motif: NNRYCACGTGRYNN | TF:M00121 | FALSE | 3.01E-08 | 7.521861294 | 2584 | 2370 | 384 | 21787 | [illegible] |
| TF | Factor: c-Myc:Max; motif: GCCAYGYGSN | TF:M00322 | FALSE | 3.26E-08 | 7.486319122 | 7347 | 2370 | 945 | 21787 | [illegible] |
| TF | Factor: Zic3; motif: NMCCCCCGGGGGGGN; match class: 1 | TF:M02837_1 | FALSE | 3.71E-08 | 7.430702902 | 2041 | 2370 | 315 | 21787 | [illegible] |
| TF | Factor: GKLF; motif: NNNRNWGGGTGKGGC | TF:M10279 | FALSE | 4.66E-08 | 7.331699562 | 9301 | 2370 | 1162 | 21787 | [illegible] |
| TF | Factor: REX1; motif: AANATGGCKGCN | TF:M01744 | FALSE | 6.17E-08 | 7.20968735 | 564 | 2370 | 114 | 21787 | [illegible] |
| TF | Factor: ATF-4; motif: NNTKRCRTCANS; match class: 1 | TF:M07246_1 | FALSE | 6.99E-08 | 7.155763916 | 701 | 2370 | 134 | 21787 | [illegible] |
| TF | Factor: HES-1; motif: GSCACGMGMC | TF:M02011 | FALSE | 7.08E-08 | 7.149938235 | 3597 | 2370 | 506 | 21787 | [illegible] |
| TF | Factor: NRF-1; motif: CNSTGCGCATGCGCNNS | TF:M10328 | FALSE | 7.17E-08 | 7.144701316 | 1013 | 2370 | 178 | 21787 | [illegible] |
| TF | Factor: WT1; motif: NNGGGNGGGSGN; match class: 1 | TF:M07436_1 | FALSE | 7.25E-08 | 7.139836993 | 4014 | 2370 | 556 | 21787 | [illegible] |
| TF | Factor: ATF-2; motif: NNNTGACGTNAN; match class: 1 | TF:M01862_1 | FALSE | 8.12E-08 | 7.090351997 | 1102 | 2370 | 190 | 21787 | [illegible] |
| TF | Factor: ATF4; motif: CVTGACGYMABG; match class: 1 | TF:M00514_1 | FALSE | 9.07E-08 | 7.042272514 | 1281 | 2370 | 214 | 21787 | [illegible] |
| TF | Factor: CREM; motif: NNTGACGT | TF:M08803 | FALSE | 1.05E-07 | 6.978320476 | 1216 | 2370 | 205 | 21787 | [illegible] |
| TF | Factor: Stra13; motif: NYSTCACGTGABNN; match class: 1 | TF:M00985_1 | FALSE | 1.30E-07 | 6.887123148 | 584 | 2370 | 116 | 21787 | [illegible] |
| TF | Factor: REX-1; motif: GGCMGCCATTTT | TF:M10415 | FALSE | 1.37E-07 | 6.864470466 | 571 | 2370 | 114 | 21787 | [illegible] |
| TF | Factor: Elk-1; motif: RACCGGAAGTR; match class: 1 | TF:M01165_1 | FALSE | 1.45E-07 | 6.838140913 | 2510 | 2370 | 371 | 21787 | [illegible] |
| TF | Factor: Zfp281; motif: NNCCCCCCCCCCMYC | TF:M02831 | FALSE | 1.52E-07 | 6.818917505 | 3973 | 2370 | 549 | 21787 | [illegible] |
| TF | Factor: HIC1; motif: NNNGGKTGCCCSNNNNNN | TF:M01073 | FALSE | 1.64E-07 | 6.784538084 | 2868 | 2370 | 415 | 21787 | [illegible] |
| TF | Factor: usf1; motif: NNGTCACGTGRCS | TF:M10409 | FALSE | 1.66E-07 | 6.778759968 | 2257 | 2370 | 339 | 21787 | [illegible] |
| TF | Factor: BTEB2; motif: GNAGGGGGNGGGSSNN | TF:M03814 | FALSE | 1.76E-07 | 6.754791027 | 4984 | 2370 | 668 | 21787 | [illegible] |
| TF | Factor: Egr1; motif: NCCGCCCCCGCANN; match class: 1 | TF:M02744_1 | FALSE | 2.01E-07 | 6.697510604 | 670 | 2370 | 128 | 21787 | [illegible] |
| TF | Factor: Mitf; motif: NRTCACGNGN; match class: 1 | TF:M07996_1 | FALSE | 2.04E-07 | 6.690916472 | 521 | 2370 | 106 | 21787 | [illegible] |
| TF | Factor: GABPalpha; motif: ACCGGAARYN; match class: 1 | TF:M02039_1 | FALSE | 2.06E-07 | 6.687154744 | 1619 | 2370 | 257 | 21787 | [illegible] |
| TF | Factor: E2F; motif: NNTTTCGCGCN; match class: 1 | TF:M08875_1 | FALSE | 2.26E-07 | 6.646554783 | 1204 | 2370 | 202 | 21787 | [illegible] |
| TF | Factor: USF; motif: NRCCACGTGASN; match class: 1 | TF:M00796_1 | FALSE | 2.26E-07 | 6.645021389 | 502 | 2370 | 103 | 21787 | [illegible] |
| TF | Factor: Fli-1; motif: ACCGGAARYN | TF:M02038 | FALSE | 2.81E-07 | 6.550558015 | 2116 | 2370 | 320 | 21787 | [illegible] |
| TF | Factor: AHR; motif: CACGCN | TF:M01855 | FALSE | 3.08E-07 | 6.511291659 | 8485 | 2370 | 1065 | 21787 | [illegible] |
| TF | Factor: ATF6; motif: TGACGTGG | TF:M00483 | FALSE | 3.34E-07 | 6.476785865 | 2607 | 2370 | 381 | 21787 | [illegible] |
| TF | Factor: Arnt; motif: NGTCACGTGN; match class: 1 | TF:M07979_1 | FALSE | 3.49E-07 | 6.457297629 | 179 | 2370 | 50 | 21787 | [illegible] |
| TF | Factor: Zfp740; motif: NCCCCCCCAC | TF:M03944 | FALSE | 3.91E-07 | 6.407454686 | 7315 | 2370 | 933 | 21787 | [illegible] |
| TF | Factor: BEN; motif: CWGCGAYA; match class: 1 | TF:M01241_1 | FALSE | 4.42E-07 | 6.354559788 | 1585 | 2370 | 251 | 21787 | [illegible] |
| TF | Factor: Tfap2a; motif: TGCCCYNRGGGCA; match class: 1 | TF:M04155_1 | FALSE | 5.37E-07 | 6.270131063 | 3180 | 2370 | 450 | 21787 | [illegible] |
| TF | Factor: USF2; motif: NGTCACGTGRNSN; match class: 1 | TF:M09722_1 | FALSE | 6.03E-07 | 6.220025516 | 571 | 2370 | 112 | 21787 | [illegible] |
| TF | Factor: Elf-1; motif: AWCCCGGAAGTN; match class: 1 | TF:M07415_1 | FALSE | 6.05E-07 | 6.21836094 | 2498 | 2370 | 366 | 21787 | [illegible] |
| TF | Factor: BTEB3; motif: BNRNGGGAGGNGT; match class: 1 | TF:M01865_1 | FALSE | 7.51E-07 | 6.124507968 | 3097 | 2370 | 439 | 21787 | [illegible] |
| TF | Factor: Stra13; motif: NYSTCACGTGABNN | TF:M00985 | FALSE | 8.78E-07 | 6.056644103 | 727 | 2370 | 134 | 21787 | [illegible] |
| TF | Factor: HIF-1alpha; motif: GNACGTGM | TF:M10257 | FALSE | 1.01189E-06 | 5.994867506 | 6612 | 2370 | 850 | 21787 | [illegible] |
| TF | Factor: TFEA; motif: RGTCACGTGA; match class: 1 | TF:M10403_1 | FALSE | 1.13089E-06 | 5.946579703 | 625 | 2370 | 119 | 21787 | [illegible] |
| TF | Factor: Klf7; motif: NNNRCCMCGCCCNNNN; match class: 1 | TF:M02773_1 | FALSE | 1.20294E-06 | 5.919754453 | 997 | 2370 | 171 | 21787 | [illegible] |
| TF | Factor: YY1; motif: GCCATNTTN | TF:M00793 | FALSE | 1.37436E-06 | 5.861900923 | 5057 | 2370 | 670 | 21787 | [illegible] |
| TF | Factor: XBP-1; motif: WNNGMCACGTC; match class: 1 | TF:M01770_1 | FALSE | 1.55453E-06 | 5.808401212 | 3854 | 2370 | 528 | 21787 | [illegible] |
| TF | Factor: USF2; motif: NNNCCACGTGACN | TF:M03860 | FALSE | 1.89426E-06 | 5.722561083 | 4061 | 2370 | 552 | 21787 | [illegible] |
| TF | Factor: ATF-2; motif: TTACGTAA; match class: 1 | TF:M00040_1 | FALSE | 1.99744E-06 | 5.699526839 | 7065 | 2370 | 899 | 21787 | [illegible] |
| TF | Factor: ERF; motif: ACCGGAARTN; match class: 1 | TF:M01984_1 | FALSE | 2.17139E-06 | 5.663262438 | 1611 | 2370 | 251 | 21787 | [illegible] |
| TF | Factor: SREBP-2; motif: NNGYCACNNSMN | TF:M01177 | FALSE | 2.24636E-06 | 5.648520695 | 7699 | 2370 | 970 | 21787 | [illegible] |
| TF | Factor: TFEA; motif: RGTCACGTGA | TF:M10403 | FALSE | 2.27077E-06 | 5.643827171 | 2590 | 2370 | 374 | 21787 | [illegible] |
| TF | Factor: E2F-3; motif: YTCCCGCCCNCNSNN | TF:M07083 | FALSE | 2.49757E-06 | 5.602483037 | 633 | 2370 | 119 | 21787 | [illegible] |
| TF | Factor: SREBP-1; motif: NATCACGTGAY; match class: 1 | TF:M00220_1 | FALSE | 2.51402E-06 | 5.599630647 | 647 | 2370 | 121 | 21787 | [illegible] |
| TF | Factor: E2F; motif: TTTSGCGSG; match class: 1 | TF:M00939_1 | FALSE | 2.86953E-06 | 5.542189737 | 1293 | 2370 | 209 | 21787 | [illegible] |
| TF | Factor: MAZ; motif: CCCTCCCYCYN | TF:M07297 | FALSE | 3.07239E-06 | 5.512523716 | 3245 | 2370 | 453 | 21787 | [illegible] |
| TF | Factor: Zfp740; motif: NCCCCCCCAC; match class: 1 | TF:M03944_1 | FALSE | 3.11603E-06 | 5.506397913 | 4464 | 2370 | 598 | 21787 | [illegible] |
| TF | Factor: USF1; motif: NNNNGTCACGTGGN | TF:M07067 | FALSE | 3.37545E-06 | 5.471668256 | 3297 | 2370 | 459 | 21787 | [illegible] |
| TF | Factor: Tcfap2b; motif: NYGCCYNARGGCAN; match class: 1 | TF:M02820_1 | FALSE | 4.10969E-06 | 5.386190528 | 5226 | 2370 | 686 | 21787 | [illegible] |
| TF | Factor: c-Rel; motif: SGGRNTTTCC; match class: 1 | TF:M00053_1 | FALSE | 4.36467E-06 | 5.360048513 | 337 | 2370 | 74 | 21787 | [illegible] |
| TF | Factor: AhR; motif: CCYCNRRSTNGCGTGASA | TF:M00139 | FALSE | 5.05831E-06 | 5.295994139 | 3490 | 2370 | 481 | 21787 | [illegible] |
| TF | Factor: Sp5; motif: RNGGRGGNGGRGNNGGGGGAGGRG; match class: 1 | TF:M10376_1 | FALSE | 5.12093E-06 | 5.290650951 | 1948 | 2370 | 292 | 21787 | [illegible] |
| TF | Factor: MECP2; motif: CCGGNNTTWA; match class: 1 | TF:M01299_1 | FALSE | 5.28252E-06 | 5.277159001 | 855 | 2370 | 149 | 21787 | [illegible] |
| TF | Factor: AhR; motif: NTNGCGTGNNN | TF:M00778 | FALSE | 5.62641E-06 | 5.249768976 | 3501 | 2370 | 482 | 21787 | [illegible] |
| TF | Factor: YY1; motif: NNNNAARATGGNNNN; match class: 1 | TF:M07370_1 | FALSE | 5.63467E-06 | 5.249131853 | 1062 | 2370 | 177 | 21787 | [illegible] |
| TF | Factor: NRF-1; motif: YGCGCMTGCGC; match class: 1 | TF:M02102_1 | FALSE | 5.67067E-06 | 5.246365952 | 572 | 2370 | 109 | 21787 | [illegible] |
| TF | Factor: USF; motif: NRCCACGTGASN | TF:M00796 | FALSE | 5.81825E-06 | 5.235207636 | 2651 | 2370 | 379 | 21787 | [illegible] |
| TF | Factor: NRF-1; motif: CGCATGCGCR | TF:M00652 | FALSE | 7.05797E-06 | 5.151320484 | 1499 | 2370 | 234 | 21787 | [illegible] |
| TF | Factor: ZNF151; motif: GGSGGNGGGGGAGGGGMGG; match class: 1 | TF:M10412_1 | FALSE | 7.17753E-06 | 5.144024691 | 1178 | 2370 | 192 | 21787 | [illegible] |
| TF | Factor: E2F-1; motif: TTTSGCGS; match class: 1 | TF:M00431_1 | FALSE | 7.31368E-06 | 5.135864131 | 1585 | 2370 | 245 | 21787 | [illegible] |
| TF | Factor: ZN451; motif: CKGGGGGG | TF:M06457 | FALSE | 7.85787E-06 | 5.104695124 | 1980 | 2370 | 295 | 21787 | [illegible] |
| TF | Factor: Erm; motif: NCCGGAWGTN; match class: 1 | TF:M01992_1 | FALSE | 7.98683E-06 | 5.09762543 | 1225 | 2370 | 198 | 21787 | [illegible] |
| TF | Factor: Pax-4; motif: NNNNNYCACCCB | TF:M00378 | FALSE | 8.25527E-06 | 5.083268681 | 18695 | 2370 | 2123 | 21787 | [illegible] |
| TF | Factor: SAP-1a; motif: NACCGGAAGTV | TF:M01167 | FALSE | 1.18116E-05 | 4.927692302 | 2172 | 2370 | 318 | 21787 | [illegible] |
| TF | Factor: ATF-2; motif: TTACGTAA | TF:M00040 | FALSE | 1.42447E-05 | 4.846347943 | 7734 | 2370 | 967 | 21787 | [illegible] |
| TF | Factor: Bhlhe40; motif: NKCACGTGMN | TF:M04167 | FALSE | 1.51627E-05 | 4.819222143 | 147 | 2370 | 41 | 21787 | [illegible] |
| TF | Factor: C-Myc; motif: NGCCACGTGNN; match class: 1 | TF:M07601_1 | FALSE | 1.57768E-05 | 4.801980023 | 4296 | 2370 | 573 | 21787 | [illegible] |
| TF | Factor: E2F-1; motif: TTTSGCGCGMNR; match class: 1 | TF:M00516_1 | FALSE | 1.59626E-05 | 4.796895518 | 1677 | 2370 | 255 | 21787 | [illegible] |
| TF | Factor: E2F; motif: NCSCGCSAAAN; match class: 1 | TF:M00919_1 | FALSE | 1.72606E-05 | 4.762945221 | 1176 | 2370 | 190 | 21787 | [illegible] |
| TF | Factor: YY1; motif: NYNKCCATNTT; match class: 1 | TF:M01035_1 | FALSE | 1.88107E-05 | 4.72559443 | 1147 | 2370 | 186 | 21787 | [illegible] |
| TF | Factor: HES-1; motif: NNCKYGTGNNN | TF:M07042 | FALSE | 2.17187E-05 | 4.663167099 | 4050 | 2370 | 543 | 21787 | [illegible] |
| TF | Factor: GMEB1; motif: NGNNCRACGTYGTYNN; match class: 1 | TF:M02865_1 | FALSE | 2.39083E-05 | 4.621451879 | 1598 | 2370 | 244 | 21787 | [illegible] |
| TF | Factor: ATF-3; motif: RTKRCGTCANN; match class: 1 | TF:M01863_1 | FALSE | 2.71656E-05 | 4.565980696 | 498 | 2370 | 96 | 21787 | [illegible] |
| TF | Factor: NF-Y; motif: RGCCAATCRGN | TF:M07302 | FALSE | 2.74926E-05 | 4.560783732 | 1877 | 2370 | 279 | 21787 | [illegible] |
| TF | Factor: E2F-3; motif: NNNWYGGCGCCAMNNNN | TF:M02847 | FALSE | 2.77512E-05 | 4.556718285 | 1229 | 2370 | 196 | 21787 | [illegible] |
| TF | Factor: MYF6; motif: NNNRACAGNCNCNCC | TF:M02885 | FALSE | 3.07798E-05 | 4.511734257 | 14271 | 2370 | 1672 | 21787 | [illegible] |
| TF | Factor: GABP; motif: VCCGGAAGNGCR; match class: 1 | TF:M00341_1 | FALSE | 3.24082E-05 | 4.489345419 | 1992 | 2370 | 293 | 21787 | [illegible] |
| TF | Factor: NF-YA; motif: CRGCCAATCAGNRN | TF:M02106 | FALSE | 3.39682E-05 | 4.468927948 | 2025 | 2370 | 297 | 21787 | [illegible] |
| TF | Factor: TCFAP2B; motif: NTWGCCKSMGGCRNN | TF:M02924 | FALSE | 3.88376E-05 | 4.410747779 | 5912 | 2370 | 757 | 21787 | [illegible] |
| TF | Factor: EHF; motif: CSCGGAARTN; match class: 1 | TF:M01974_1 | FALSE | 4.15175E-05 | 4.381768615 | 1608 | 2370 | 244 | 21787 | [illegible] |
| TF | Factor: RelA-p65; motif: KGGRNTTTCCC; match class: 1 | TF:M10399_1 | FALSE | 4.3445E-05 | 4.362059775 | 482 | 2370 | 93 | 21787 | [illegible] |
| TF | Factor: ctcf; motif: CCNCNAGRKGGCRSTN; match class: 1 | TF:M07249_1 | FALSE | 4.7071E-05 | 4.32724643 | 1322 | 2370 | 207 | 21787 | [illegible] |
| TF | Factor: NF-kappaB2; motif: GGGRAANYCCC | TF:M10312 | FALSE | 4.83931E-05 | 4.315216185 | 832 | 2370 | 142 | 21787 | [illegible] |
| TF | Factor: Zfp740; motif: NNNCCCCCCCCMNNNN | TF:M02834 | FALSE | 4.87701E-05 | 4.311846121 | 5197 | 2370 | 674 | 21787 | [illegible] |
| TF | Factor: Egr-2; motif: NTGCGTRGGCGK; match class: 1 | TF:M00246_1 | FALSE | 5.63148E-05 | 4.249377481 | 1202 | 2370 | 191 | 21787 | [illegible] |
| TF | Factor: usf1; motif: NNGTCACGTGRCS; match class: 1 | TF:M10409_1 | FALSE | 5.91747E-05 | 4.227863601 | 377 | 2370 | 77 | 21787 | [illegible] |
| TF | Factor: Sp4; motif: NNNNMCGCCCCCTNNNN; match class: 1 | TF:M02810_1 | FALSE | 8.2656E-05 | 4.082725525 | 773 | 2370 | 133 | 21787 | [illegible] |
| TF | Factor: c-Ets-1(p54); motif: NCMGGAWGYN | TF:M00032 | FALSE | 9.63487E-05 | 4.01615402 | 8493 | 2370 | 1044 | 21787 | [illegible] |
| TF | Factor: Pax-4; motif: NNNNNYCACCCB; match class: 1 | TF:M00378_1 | FALSE | 9.67626E-05 | 4.01429262 | 14437 | 2370 | 1685 | 21787 | [illegible] |
| TF | Factor: ESE-1; motif: NSMGGAARTN; match class: 1 | TF:M01214_1 | FALSE | 9.84254E-05 | 4.006892921 | 2282 | 2370 | 326 | 21787 | [illegible] |
| TF | Factor: PLAG1; motif: CCCCCKWNNNGGSCCC | TF:M01973 | FALSE | 0.000112482 | 3.948917055 | 4276 | 2370 | 564 | 21787 | [illegible] |
| TF | Factor: SP4; motif: NNKGGGCGKGNCN | TF:M09005 | FALSE | 0.000115995 | 3.935559083 | 1946 | 2370 | 284 | 21787 | [illegible] |
| TF | Factor: NF-Y; motif: NNNNRRCCAATSR | TF:M00775 | FALSE | 0.000116884 | 3.932245554 | 1882 | 2370 | 276 | 21787 | [illegible] |
| TF | Factor: Barhl1; motif: NNTAAAYGNN; match class: 1 | TF:M04293_1 | FALSE | 0.000141715 | 3.848583622 | 1271 | 2370 | 198 | 21787 | [illegible] |
| TF | Factor: VDR; motif: GGGKNARNRRGGWSA; match class: 1 | TF:M00444_1 | FALSE | 0.000144289 | 3.840767516 | 3798 | 2370 | 507 | 21787 | [illegible] |
| TF | Factor: NRF-1; motif: GCGCMTGCGCN; match class: 1 | TF:M07052_1 | FALSE | 0.000182124 | 3.739631671 | 645 | 2370 | 114 | 21787 | [illegible] |
| TF | Factor: MITF; motif: NCNYGTGACY | TF:M10295 | FALSE | 0.000184972 | 3.732893007 | 5205 | 2370 | 670 | 21787 | [illegible] |
| TF | Factor: ZN451; motif: CKGGGGGG; match class: 1 | TF:M06457_1 | FALSE | 0.000211548 | 3.674591157 | 220 | 2370 | 51 | 21787 | [illegible] |
| TF | Factor: Egr1; motif: NNMCGCCCMCTCAMWN; match class: 1 | TF:M04506_1 | FALSE | 0.000214387 | 3.668800995 | 815 | 2370 | 137 | 21787 | [illegible] |
| TF | Factor: GABP-beta; motif: ASMGGAAGKGN | TF:M01876 | FALSE | 0.00021733 | 3.662880082 | 2546 | 2370 | 356 | 21787 | [illegible] |
| TF | Factor: AhR,; motif: NRCGTGNGN; match class: 1 | TF:M00976_1 | FALSE | 0.000266447 | 3.574388677 | 885 | 2370 | 146 | 21787 | [illegible] |
| TF | Factor: IRF-3; motif: NARAANSGAAACYN | TF:M02767 | FALSE | 0.000292529 | 3.533831557 | 2611 | 2370 | 363 | 21787 | [illegible] |
| TF | Factor: DEC2; motif: SNTCACGTGS | TF:M01843 | FALSE | 0.000320813 | 3.493747818 | 4265 | 2370 | 559 | 21787 | [illegible] |
| TF | Factor: Egr-1; motif: GCGTGGGCGT; match class: 1 | TF:M07597_1 | FALSE | 0.00036361 | 3.439363647 | 882 | 2370 | 145 | 21787 | [illegible] |
| TF | Factor: Wt1; motif: NNCCTCCCCCACNN; match class: 1 | TF:M12438_1 | FALSE | 0.000382173 | 3.417740253 | 1579 | 2370 | 235 | 21787 | [illegible] |
| TF | Factor: E2F-1; motif: TTGGCGCGRAANNGNM; match class: 1 | TF:M00938_1 | FALSE | 0.000382486 | 3.417384314 | 1492 | 2370 | 224 | 21787 | [illegible] |
| TF | Factor: E2F; motif: TTTSGCGSG; match class: 1 | TF:M00918_1 | FALSE | 0.000389455 | 3.409542349 | 981 | 2370 | 158 | 21787 | [illegible] |
| TF | Factor: MAZR; motif: NSGGGGGGGGMCN | TF:M00491 | FALSE | 0.000409976 | 3.387242044 | 3854 | 2370 | 510 | 21787 | [illegible] |
| TF | Factor: NF-Y; motif: NNNRRCCAATSRGNNN | TF:M00287 | FALSE | 0.000423713 | 3.372927818 | 1998 | 2370 | 287 | 21787 | [illegible] |
| TF | Factor: ICSBP; motif: RAARTGAAACTG; match class: 1 | TF:M00699_1 | FALSE | 0.000502568 | 3.298805013 | 72 | 2370 | 24 | 21787 | [illegible] |
| TF | Factor: N-Myc; motif: GSSCACGYGS | TF:M03834 | FALSE | 0.000503877 | 3.297675894 | 1387 | 2370 | 210 | 21787 | [illegible] |
| TF | Factor: Spic; motif: NNVGGAAGTN; match class: 1 | TF:M02042_1 | FALSE | 0.000533017 | 3.273258769 | 1341 | 2370 | 204 | 21787 | [illegible] |
| TF | Factor: FKLF; motif: BGGGNGGVMD; match class: 1 | TF:M01837_1 | FALSE | 0.000537798 | 3.269381201 | 986 | 2370 | 158 | 21787 | [illegible] |
| TF | Factor: Klf7; motif: NGGGCGGGGG | TF:M05437 | FALSE | 0.000546242 | 3.26261453 | 508 | 2370 | 93 | 21787 | [illegible] |
| TF | Factor: ATF-4; motif: NNTKRCRTCANS | TF:M07246 | FALSE | 0.000555652 | 3.255196761 | 3515 | 2370 | 469 | 21787 | [illegible] |
| TF | Factor: E2F-3; motif: GGGCGGGARN; match class: 1 | TF:M10210_1 | FALSE | 0.00056234 | 3.250000969 | 297 | 2370 | 62 | 21787 | [illegible] |
| TF | Factor: NF-kappaB; motif: NGGGANTTYCCMNNNN | TF:M00774 | FALSE | 0.000640876 | 3.193225874 | 1661 | 2370 | 244 | 21787 | [illegible] |
| TF | Factor: HIF-1alpha; motif: NCACGT | TF:M02012 | FALSE | 0.00066687 | 3.175958764 | 6353 | 2370 | 796 | 21787 | [illegible] |
| TF | Factor: IRF; motif: RRAANTGAAASYGNV; match class: 1 | TF:M00772_1 | FALSE | 0.000670784 | 3.17341734 | 78 | 2370 | 25 | 21787 | [illegible] |
| TF | Factor: ATF-2; motif: RRTGANGTCAY; match class: 1 | TF:M10179_1 | FALSE | 0.000673615 | 3.17158812 | 846 | 2370 | 139 | 21787 | [illegible] |
| TF | Factor: Atf-1; motif: NNNRTGACGTCAYNRN; match class: 1 | TF:M02738_1 | FALSE | 0.000701925 | 3.153709352 | 5015 | 2370 | 643 | 21787 | [illegible] |
| TF | Factor: USF; motif: NNRNCACGTGNYNN; match class: 1 | TF:M00122_1 | FALSE | 0.000758826 | 3.119857706 | 2544 | 2370 | 352 | 21787 | [illegible] |
| TF | Factor: USF; motif: NNRNCACGTGNYNN | TF:M00122 | FALSE | 0.000758826 | 3.119857706 | 2544 | 2370 | 352 | 21787 | [illegible] |
| TF | Factor: AhR; motif: NNNKNGCGTGNSNNNNN | TF:M07347 | FALSE | 0.000835848 | 3.077872648 | 1232 | 2370 | 189 | 21787 | [illegible] |
| TF | Factor: NF-kappaB; motif: GGGAMTTYCC | TF:M00054 | FALSE | 0.000857683 | 3.066673439 | 2308 | 2370 | 323 | 21787 | [illegible] |
| TF | Factor: E2F-3; motif: NNNANGGCGCGCSNW | TF:M02743 | FALSE | 0.000963104 | 3.016326873 | 1806 | 2370 | 261 | 21787 | [illegible] |
| TF | Factor: IRF-2; motif: NNRAAAGTGAAASNNA | TF:M01882 | FALSE | 0.00098093 | 3.008361885 | 295 | 2370 | 61 | 21787 | [illegible] |
| TF | Factor: YY1; motif: GCCATNTTN; match class: 1 | TF:M00793_1 | FALSE | 0.001010221 | 2.995583451 | 681 | 2370 | 116 | 21787 | [illegible] |
| TF | Factor: DEC2; motif: SNTCACGTGS; match class: 1 | TF:M01843_1 | FALSE | 0.00101065 | 2.995399266 | 996 | 2370 | 158 | 21787 | [illegible] |
| TF | Factor: USF1; motif: NNNNGTCACGTGGN; match class: 1 | TF:M07067_1 | FALSE | 0.001124761 | 2.948939924 | 816 | 2370 | 134 | 21787 | [illegible] |
| TF | Factor: E2F; motif: NKCGCGCSAAAN; match class: 1 | TF:M00920_1 | FALSE | 0.001209862 | 2.917264084 | 1068 | 2370 | 167 | 21787 | [illegible] |

| | | | | | | | | | | |
|---|---|---|---|---|---|---|---|---|---|---|
| TF | Factor: Tfap2a; motif: NGCCYSAGGCN | TF:M04156 | FALSE | 0.001223863 | 2.912267022 | 7827 | 2370 | 959 | 21787 | SHD,IFITM1,ADAMTS14,TSPYL3,MSS51,BCL6B,CENP [illegible] |
| TF | Factor: BTEB2; motif: GNAGGGGGNGGGSSNN; match class: 1 | TF:M03814_1 | FALSE | 0.001259848 | 2.899681911 | 1115 | 2370 | 173 | 21787 | GM37240,GDF15,PIM1,RORA,CSF3,IGLON5,HBEGF [illegible] |
| TF | Factor: Elf5; motif: CSMGGAARTN; match class: 1 | TF:M01980_1 | FALSE | 0.001411051 | 2.850457251 | 1312 | 2370 | 198 | 21787 | KCND1,B930094E09RIK,RORA,NPPA,ZFP994,ZFP947 [illegible] |
| TF | Factor: IRF-1; motif: NNRAAANNGAAASN | TF:M07045 | FALSE | 0.001512208 | 2.820388394 | 679 | 2370 | 115 | 21787 | KLHL38,GDF15,PLEK2,CCRL2,GBP5,CXCL10,RSAD2 [illegible] |
| TF | Factor: NF-YC; motif: NYARCCAATCAGAR | TF:M10314 | FALSE | 0.001581348 | 2.800972437 | 791 | 2370 | 130 | 21787 | CXCL10,REC114,ZFP966,GAS2L3,KLF9,CREBRF,SLC [illegible] |
| TF | Factor: Crem; motif: NATGACGTMA | TF:M07985 | FALSE | 0.001804027 | 2.743756886 | 1129 | 2370 | 174 | 21787 | B930094E09RIK,DIO2,CENPE,OR8D1B,REL,TRAF1 [illegible] |
| TF | Factor: AhR:Arnt; motif: GRGKATYGCGTGMCWNSCC; match class: 1 | TF:M00237_1 | FALSE | 0.001910338 | 2.718889738 | 1302 | 2370 | 196 | 21787 | CD83,TNFAIP3,MAFF,ZHX2,REL,BCL2L11,GNG3,KLF9 [illegible] |
| TF | Factor: Irf-4; motif: NNNANCGAAACYNNA; match class: 1 | TF:M02768_1 | FALSE | 0.001939645 | 2.712277787 | 809 | 2370 | 132 | 21787 | SHD,CCDC18,ZHX2,KLHL24,MKI67,ERBIN,LIPG,KNL [illegible] |
| TF | Factor: Zfp281; motif: NNCCCCCCCCCCMYC; match class: 1 | TF:M02831_1 | FALSE | 0.001966979 | 2.706200189 | 2494 | 2370 | 343 | 21787 | CXCL2,FAM167A,TNFAIP3,RORA,IGLON5,PRKCZ,MKI [illegible] |
| TF | Factor: SP4; motif: NNWAGGCGTGNCNNN | TF:M02914 | FALSE | 0.002525191 | 2.597705702 | 4929 | 2370 | 628 | 21787 | GM37240,GM8251,FAM167A,PIM1,IL4I1,CENPE,GA [illegible] |
| TF | Factor: ATF-1; motif: CYYTGACGTCA; match class: 1 | TF:M00691_1 | FALSE | 0.002576667 | 2.588941737 | 950 | 2370 | 150 | 21787 | BCL6B,RORA,MAFF,LIPG,EGR2,EGR1,ZFC3H1,ARL5B [illegible] |
| TF | Factor: NF-YA; motif: NYRRCCAATCAGAR | TF:M10313 | FALSE | 0.003009062 | 2.521568926 | 1374 | 2370 | 204 | 21787 | CENPF,CXCL10,REC114,ZFP966,GAS2L3,KLF9,LIPG [illegible] |
| TF | Factor: Atf-1; motif: NNNRTGACGTCAYNRN | TF:M02738 | FALSE | 0.003079072 | 2.511580208 | 6137 | 2370 | 765 | 21787 | SHD,4930503B20RIK,NR4A3,CD83,FAM167A,ADAMT [illegible] |
| TF | Factor: Zfp740; motif: NNNCCCCCCCCMNNNN; match class: 1 | TF:M02834_1 | FALSE | 0.003210696 | 2.493400751 | 3824 | 2370 | 499 | 21787 | OR7C70,TMEM236,CXCL2,FAM167A,CISH,TNFAIP3 [illegible] |
| TF | Factor: c-Myc:Max; motif: NNACCACGTGGTNN; match class: 1 | TF:M00118_1 | FALSE | 0.003270366 | 2.485403695 | 7921 | 2370 | 965 | 21787 | IFITM1,4930503B20RIK,GM8251,OR7C70,NR4A3,CD [illegible] |
| TF | Factor: c-Myc:Max; motif: NNACCACGTGGTNN | TF:M00118 | FALSE | 0.003270366 | 2.485403695 | 7921 | 2370 | 965 | 21787 | IFITM1,4930503B20RIK,GM8251,OR7C70,NR4A3,CD [illegible] |
| TF | Factor: VDR; motif: GGGKNARNRRGGWSA | TF:M00444 | FALSE | 0.004552963 | 2.341705895 | 10039 | 2370 | 1197 | 21787 | SHD,NR4A3,CD83,TMEM236,GDF15,KCND1,CXCL2 [illegible] |
| TF | Factor: Arnt; motif: NNNNNRTCACGTGAYNNNNN; match class: 1 | TF:M00539_1 | FALSE | 0.004713875 | 2.326621969 | 7059 | 2370 | 867 | 21787 | 4930503B20RIK,OR7C70,NR4A3,CD83,GDF15,KCND [illegible] |
| TF | Factor: Arnt; motif: NNNNNRTCACGTGAYNNNNN | TF:M00539 | FALSE | 0.004713875 | 2.326621969 | 7059 | 2370 | 867 | 21787 | 4930503B20RIK,OR7C70,NR4A3,CD83,GDF15,KCND [illegible] |
| TF | Factor: Spi-B; motif: ASNGGAASTN | TF:M02041 | FALSE | 0.005133934 | 2.289549692 | 9962 | 2370 | 1188 | 21787 | GM37240,IFITM1,4930503B20RIK,CD83,TMEM236 [illegible] |
| TF | Factor: ATF4; motif: CVTGACGYMABG | TF:M00514 | FALSE | 0.005286787 | 2.27680822 | 5375 | 2370 | 676 | 21787 | FAP,IFITM1,NR4A3,XIRP1,FAM167A,BCL6B,DIO2,NEO [illegible] |
| TF | Factor: Net; motif: CACTTCCKGK | TF:M03832 | FALSE | 0.005739216 | 2.241147446 | 1002 | 2370 | 155 | 21787 | B930094E09RIK,CENPF,CHD2,ZFP433,TASOR2,GM4 [illegible] |
| TF | Factor: ICSBP; motif: RAARTGAAACTG | TF:M00699 | FALSE | 0.005926549 | 2.227198115 | 1340 | 2370 | 198 | 21787 | KLHL38,GDF15,XIRP1,PLEK2,MSS51,CCRL2,GBP5 [illegible] |
| TF | Factor: USF2; motif: CASGYG; match class: 1 | TF:M00726_1 | FALSE | 0.006106082 | 2.214237348 | 6270 | 2370 | 777 | 21787 | 4930503B20RIK,OR7C70,NR4A3,CD83,GDF15,KCND [illegible] |
| TF | Factor: USF2; motif: CASGYG | TF:M00726 | FALSE | 0.006106082 | 2.214237348 | 6270 | 2370 | 777 | 21787 | 4930503B20RIK,OR7C70,NR4A3,CD83,GDF15,KCND [illegible] |
| TF | Factor: BTEB2; motif: NGGGSWGGGNGGGGC; match class: 1 | TF:M10280_1 | FALSE | 0.006267165 | 2.202928845 | 595 | 2370 | 101 | 21787 | KANSL1L,HIVEP3,ABCG4,IRKL,EEA1,9930021J03RIK [illegible] |
| TF | Factor: PDEF; motif: NCCGGATRYN | TF:M02040 | FALSE | 0.006478129 | 2.188550421 | 2643 | 2370 | 357 | 21787 | FAP,TSPYL3,TNFRSF8,PIM1,CSF3,ZFP811,FBN1,GBP7 [illegible] |
| TF | Factor: KLF; motif: GGGNGGGG | TF:M07461 | FALSE | 0.007302675 | 2.136517997 | 8496 | 2370 | 1025 | 21787 | NR4A3,GDF15,KCND1,XIRP1,FAM167A,ADAMTS14 [illegible] |
| TF | Factor: CREB; motif: CGTCAN | TF:M00801 | FALSE | 0.00747501 | 2.126388216 | 4222 | 2370 | 542 | 21787 | GM8251,NR4A3,IFNB1,B930094E09RIK,DIO2,PIM1 [illegible] |
| TF | Factor: IRF-5; motif: NNNANCGAGARTNNN | TF:M02873 | FALSE | 0.007731042 | 2.111761954 | 1764 | 2370 | 250 | 21787 | XIRP1,BCL6B,RORA,CCDC18,ZFP966,IL0R1,MALT1 [illegible] |
| TF | Factor: Sp4; motif: SCCCCKCCCCCSN; match class: 1 | TF:M07617_1 | FALSE | 0.007820842 | 2.106746488 | 460 | 2370 | 82 | 21787 | GM37240,MAFF,MALT1,KANSL1L,ABCG4,IRKL,ARID [illegible] |
| TF | Factor: ATF-4; motif: KACGTCAKS | TF:M01864 | FALSE | 0.009009649 | 2.045292129 | 5334 | 2370 | 669 | 21787 | GM37240,OR7C70,NR4A3,REG2,B930094E09RIK [illegible] |
| TF | Factor: IRF-7; motif: TNSGAAWNCGAAANTNNN | TF:M00453 | FALSE | 0.009242835 | 2.034194797 | 3380 | 2370 | 443 | 21787 | 4930503B20RIK,KLHL38,GDF15,PLEK2,KDM6B,CCR [illegible] |
| TF | Factor: STAT1; motif: NTTTCASTTYCNNTTTCYN; match class: 1 | TF:M09713_1 | FALSE | 0.009807671 | 2.008434095 | 5425 | 2370 | 679 | 21787 | GM8251,OR7C70,KLHL38,ZFP976,REG2,IFNB1,CISH [illegible] |
| TF | Factor: C-Myc; motif: NGCCACGTGNN | TF:M07601 | FALSE | 0.009854864 | 2.00634936 | 6192 | 2370 | 766 | 21787 | 4930503B20RIK,OR7C70,NR4A3,CD83,GDF15,KCND [illegible] |
| TF | Factor: ctcf; motif: YGGCCASCAGRGGGCGCYNN; match class: 1 | TF:M10203_1 | FALSE | 0.010382587 | 1.98369442 | 1641 | 2370 | 234 | 21787 | TNFRSF8,PIM1,TNFAIP3,ACSS3,HBEGF,DLL1,FBN1 [illegible] |
| TF | Factor: AP-2alpha; motif: NNNSCCTGRGGC | TF:M10175 | FALSE | 0.010789815 | 1.966985983 | 4956 | 2370 | 625 | 21787 | SHD,NR4A3,CD83,ZFP976,TSPYL3,CENPF,LIF,NEO1 [illegible] |
| TF | Factor: Pax-5; motif: BCNNNRNGCANBGNTGNRTAGCSGCHNB | TF:M00143 | FALSE | 0.011711484 | 1.931388056 | 10862 | 2370 | 1282 | 21787 | SHD,GM37240,4930503B20RIK,NR4A3,CD83,GDF1 [illegible] |
| TF | Factor: IRF; motif: NNGAAANTGAAANN | TF:M08887 | FALSE | 0.012023651 | 1.919963654 | 1507 | 2370 | 217 | 21787 | KLHL38,GDF15,ZFP976,PLEK2,CCRL2,GBP5,CXCL10 [illegible] |
| TF | Factor: Ehf; motif: NNNANCCGGAAGTNN; match class: 1 | TF:M02745_1 | FALSE | 0.012827403 | 1.89186126 | 732 | 2370 | 118 | 21787 | B930094E09RIK,TRIM5,PARP14,ZFP677,MTERF1B [illegible] |
| TF | Factor: IRF-1; motif: NRAANNGAAASTGAAASYNN | TF:M10267 | FALSE | 0.013320222 | 1.875488529 | 1285 | 2370 | 189 | 21787 | KLHL38,GDF15,ZFP976,CCRL2,GBP5,CXCL10,A1CF [illegible] |
| TF | Factor: SAP1A; motif: ACCGGAWRTN | TF:M01983 | FALSE | 0.013711676 | 1.862909458 | 532 | 2370 | 91 | 21787 | B930094E09RIK,CE [illegible] |
| TF | Factor: Barhl1; motif: NNTAAAYGNN | TF:M04293 | FALSE | 0.014467983 | 1.839591998 | 6665 | 2370 | 795 | 21787 | SHD,FAP,GM8251,B [illegible] |
| TF | Factor: LKLF; motif: GGGGTGGKSN | TF:M07261 | FALSE | 0.014975159 | 1.824628559 | 9199 | 2370 | 1099 | 21787 | SHD,FAP,OR7C70,N [illegible] |
| TF | Factor: Blimp-1; motif: NRGRAAGKGAAAGKRN | TF:M01066 | FALSE | 0.015811952 | 1.801014523 | 454 | 2370 | 80 | 21787 | LIF,TNFAIP3,CXCL10 [illegible] |
| TF | Factor: Elk-1; motif: AACCGGAAGTR; match class: 1 | TF:M01163_1 | FALSE | 0.015907651 | 1.798393937 | 1233 | 2370 | 182 | 21787 | B930094E09RIK,CE [illegible] |
| TF | Factor: GABP-alpha; motif: GGNRCCGGAAGTGN; match class: 1 | TF:M10247_1 | FALSE | 0.016990665 | 1.769789633 | 874 | 2370 | 136 | 21787 | B930094E09RIK,IRK [illegible] |
| TF | Factor: SREBP-2; motif: NTCACCYNNNN | TF:M03852 | FALSE | 0.017200352 | 1.764462664 | 7795 | 2370 | 943 | 21787 | GM37240,4930503 [illegible] |
| TF | Factor: HAIRYLIKE; motif: NNNNCANGTG | TF:M08885 | FALSE | 0.017675182 | 1.752636111 | 5730 | 2370 | 711 | 21787 | KLHL38,CD83,GDF1 [illegible] |
| TF | Factor: FOXO1; motif: NWAAACAAN | TF:M01216 | FALSE | 0.017843546 | 1.748518838 | 5089 | 2370 | 638 | 21787 | IFITM1,4930503B20 [illegible] |
| TF | Factor: Max; motif: NNANCACGTGNTNN; match class: 1 | TF:M00119_1 | FALSE | 0.018561671 | 1.731382922 | 7305 | 2370 | 888 | 21787 | 4930503B20RIK,GM [illegible] |
| TF | Factor: Max; motif: NNANCACGTGNTNN | TF:M00119 | FALSE | 0.018561671 | 1.731382922 | 7305 | 2370 | 888 | 21787 | 4930503B20RIK,GM [illegible] |
| TF | Factor: RBP-Jkappa; motif: TTCCCACK | TF:M01111 | FALSE | 0.018780259 | 1.726298428 | 1094 | 2370 | 164 | 21787 | FAP,PLEK2,CCDC18 [illegible] |
| TF | Factor: MYF6; motif: NNNRACAGNCNCNCC; match class: 1 | TF:M02885_1 | FALSE | 0.019037518 | 1.720389668 | 7020 | 2370 | 856 | 21787 | GM37240,IFITM1,NR [illegible] |
| TF | Factor: E2F-2; motif: NNNWYGGCGCCAANNNN; match class: 1 | TF:M02846_1 | FALSE | 0.020607658 | 1.68597136 | 450 | 2370 | 79 | 21787 | PIM1,CREBRF,ZC3H [illegible] |
| TF | Factor: c-Myb; motif: NNNNNAACCGTTANNNN | TF:M02779 | FALSE | 0.021183763 | 1.673996884 | 4720 | 2370 | 595 | 21787 | ZFP976,SRMS,TNFRS [illegible] |
| TF | Factor: SREBP-2; motif: NNGYCACNNSMN; match class: 1 | TF:M01177_1 | FALSE | 0.023886738 | 1.621843151 | 2146 | 2370 | 293 | 21787 | SHD,KLHL38,ADAMT [illegible] |
| TF | Factor: GKLF; motif: NNNRNWGGGTGKGGC; match class: 1 | TF:M10279_1 | FALSE | 0.024584096 | 1.609345757 | 3455 | 2370 | 448 | 21787 | GM37240,PIM1,CSF [illegible] |
| TF | Factor: STAT1; motif: NNNSANTTCCGGGAANTGNSN | TF:M00224 | FALSE | 0.026210839 | 1.581519081 | 5976 | 2370 | 737 | 21787 | CD83,TMEM236,KCN [illegible] |
| TF | Factor: Klf15; motif: RGGGMGGRGNNGGGGGNGG; match class: 1 | TF:M10277_1 | FALSE | 0.026462017 | 1.577377055 | 1022 | 2370 | 154 | 21787 | FAM167A,CSF3,IGLO [illegible] |
| TF | Factor: FOXP3; motif: NNNWAAACANWD; match class: 1 | TF:M07419_1 | FALSE | 0.027543538 | 1.559980271 | 3811 | 2370 | 489 | 21787 | IFITM1,OR7C70,ZFP9 [illegible] |
| TF | Factor: USF2; motif: NNNCCACGTGACN; match class: 1 | TF:M03860_1 | FALSE | 0.029826957 | 1.525391051 | 1040 | 2370 | 156 | 21787 | KCND1,TNFAIP3,GU [illegible] |
| TF | Factor: E2F-2; motif: NNNANGGCGCGCNNN; match class: 1 | TF:M02742_1 | FALSE | 0.031399536 | 1.503076764 | 1112 | 2370 | 165 | 21787 | PIM1,RORA,CHD2,IG [illegible] |
| TF | Factor: Egr-2; motif: NNNMCRCCCACRCNN | TF:M07086 | FALSE | 0.031438928 | 1.502532277 | 3532 | 2370 | 456 | 21787 | OR7C70,XIRP1,ADAM [illegible] |
| TF | Factor: c-Myc:Max; motif: NNNNNNNCACGTGNNNNNNN; match class: 1 | TF:M00615_1 | FALSE | 0.034173015 | 1.466316704 | 7207 | 2370 | 874 | 21787 | 4930503B20RIK,OR7 [illegible] |
| TF | Factor: c-Myc:Max; motif: NNNNNNNCACGTGNNNNNNN | TF:M00615 | FALSE | 0.034173015 | 1.466316704 | 7207 | 2370 | 874 | 21787 | 4930503B20RIK,OR7 [illegible] |
| TF | Factor: Arnt; motif: NDDNNCACGTGNNNNN | TF:M00236 | FALSE | 0.036053538 | 1.443052115 | 2366 | 2370 | 318 | 21787 | OR7C70,KCND1,AD [illegible] |
| [illegible] | [illegible] | [illegible] | [illegible] | [illegible] | [illegible] | [illegible] | [illegible] | [illegible] | [illegible] | [illegible] |
