## Supplementary material for "TFEB-Mediated Pro-inflammatory Response in Murine Macrophages Induced by Acute Alpha7 Nicotinic Receptor Activation": Table_S3

**Table S3. Differential gene expression in WT RAW264.7 cells (PNU-282987 v. DMSO).**

| ID | Gene | counts_WT_CO1 | counts_WT_CO2 | counts_WT_CO3 | counts_WT_PN1 | counts_WT_PN2 | counts_WT_PN3 | padj | log2FoldChange | pvalue | stat | foldChange | log10padj |
|---|---|---|---|---|---|---|---|---|---|---|---|---|---|
| ENSMUSG00000113388 | *Gm48111 predicted gene* | 0 | 0 | 0 | 86.87012143 | 184.575906 | 62.81701342 | 1.48E-21 | 9.201900852 | 1.22E-22 | 95.88658374 | 588.9089759 | 20.83073226 |
| ENSMUSG00000099241 | *Gvin-ps2* | 0 | 0 | 0 | 99.52110999 | 77.08758427 | 60.88418224 | 4.13E-25 | 8.713543067 | 2.80E-26 | 112.4817082 | 419.795551 | 24.38415414 |
| ENSMUSG00000072693 | *None* | 0 | 0 | 0 | 25.30197712 | 70.57314053 | 47.35436397 | 3.80E-13 | 7.977013408 | 5.36E-14 | 56.59353282 | 251.9534513 | 12.41995249 |
| ENSMUSG00000039154 | *Shd* | 0 | 0 | 0 | 27.83217483 | 24.97203434 | 19.32831182 | 3.39E-09 | 6.993460552 | 6.53E-10 | 38.15496097 | 127.4211146 | 8.469665537 |
| ENSMUSG00000000392 | *Fap* | 0 | 0 | 1.341985292 | 46.38695805 | 57.54425305 | 36.72379246 | 3.42E-15 | 6.992159235 | 4.18E-16 | 66.14965508 | 127.306232 | 14.46603693 |
| ENSMUSG00000113921 | *Gm48054* | 0.798863467 | 0 | 0 | 41.32656262 | 41.2581437 | 51.22002633 | 1.83E-15 | 6.921975357 | 2.19E-16 | 67.42025784 | 121.2612977 | 14.7370628 |
| ENSMUSG00000111327 | *Gm48401* | 0 | 0 | 0 | 6.747193898 | 22.80055309 | 16.42906505 | 2.39E-05 | 6.335712569 | 6.85E-06 | 20.23539225 | 80.76803685 | 4.621377154 |
| ENSMUSG00000114956 | *Gm47893* | 0 | 0 | 0 | 21.92838017 | 9.771665612 | 12.56340268 | 1.40E-05 | 6.294045114 | 3.88E-06 | 21.32168422 | 78.46868475 | 4.854729573 |
| ENSMUSG00000102805 | *Gm37240* | 0 | 0 | 0 | 3.373596949 | 16.28610935 | 24.16038978 | 0.000172964 | 6.268742226 | 5.54E-05 | 16.25309286 | 77.10445029 | 3.762044794 |
| ENSMUSG00000025491 | *Ifitm1* | 0 | 0 | 0 | 7.590593135 | 9.771665612 | 12.56340268 | 0.000311393 | 5.72120793 | 0.000102787 | 15.08481509 | 52.75397612 | 3.506691531 |
| ENSMUSG00000090036 | *Gm15900* | 0 | 1.009272028 | 0 | 23.61517864 | 17.37184998 | 14.49623387 | 2.23E-06 | 5.655083243 | 5.68E-07 | 25.01658461 | 50.39061795 | 5.65252222 |
| ENSMUSG00000090202 | *4930503B20Rik* | 0 | 0 | 0 | 5.060395423 | 10.85740624 | 11.59698709 | 0.00082745 | 5.596919388 | 0.000291053 | 13.12711339 | 48.39947146 | 3.082258449 |
| ENSMUSG00000091844 | *Gm8251* | 2.396590402 | 3.027816084 | 0 | 96.99091228 | 83.60202801 | 104.3728838 | 2.73E-27 | 5.578135164 | 1.65E-28 | 122.6688538 | 47.77338388 | 26.56311103 |
| ENSMUSG00000085738 | *Gm12335* | 0 | 0 | 0 | 9.27739161 | 9.771665612 | 7.731324729 | 0.000626856 | 5.562474858 | 0.000216536 | 13.68185796 | 47.25761311 | 3.202831945 |
| ENSMUSG00000111529 | *Gm48141* | 0 | 0 | 0 | 12.65098856 | 1.085740624 | 8.69774032 | 0.008882601 | 5.315952473 | 0.003684308 | 8.433202904 | 39.83466322 | 2.051459865 |
| ENSMUSG00000051190 | *Or7c70* | 0 | 0 | 0 | 13.4943878 | 2.171481247 | 4.832077956 | 0.009335509 | 5.187623445 | 0.003890347 | 8.334299968 | 36.44435455 | 2.029861976 |
| ENSMUSG00000090551 | *None* | 0 | 0 | 0 | 9.27739161 | 2.171481247 | 8.69774032 | 0.007654697 | 5.159541024 | 0.003131718 | 8.729095409 | 35.74181578 | 2.116072 |
| ENSMUSG00000022357 | *Klhl38* | 0 | 0 | 0 | 5.060395423 | 9.771665612 | 4.832077956 | 0.005709093 | 5.111532497 | 0.00228512 | 9.305138899 | 34.57200804 | 2.243432866 |
| ENSMUSG00000028341 | *Nr4a3* | 3.994317336 | 3.027816084 | 4.025955875 | 125.6664863 | 132.4603561 | 108.2385462 | 4.03E-33 | 5.051961627 | 1.88E-34 | 149.8363825 | 33.17355287 | 32.39443779 |
| ENSMUSG00000110820 | *None* | 0 | 0 | 0 | 9.27739161 | 6.514443741 | 2.899246773 | 0.008653816 | 5.047570542 | 0.003581242 | 8.484809047 | 33.07273709 | 2.062792336 |
| ENSMUSG00000015396 | *Cd83* | 207.7045015 | 209.9285819 | 213.3756614 | 6702.493738 | 6856.452038 | 6862.517113 | 0 | 5.018341077 | 0 | 3499.701229 | 32.40941505 | Inf |
| ENSMUSG00000109195 | *Gm45038* | 0 | 4.037088113 | 0 | 52.29075271 | 43.42962494 | 34.79096128 | 1.98E-10 | 5.012678283 | 3.43E-11 | 43.9125251 | 32.28245259 | 9.702959549 |
| ENSMUSG00000061531 | *Tmem236* | 1.597726935 | 0 | 0 | 26.14537635 | 16.28610935 | 13.52981828 | 1.65E-05 | 4.856899955 | 4.62E-06 | 20.98695357 | 28.97827785 | 4.783279095 |
| ENSMUSG00000038508 | *Gdf15* | 8.78749814 | 3.027816084 | 8.051911749 | 190.6082276 | 160.6896123 | 206.8129365 | 2.37E-41 | 4.803413747 | 7.36E-43 | 188.3303733 | 27.92361359 | 40.62601192 |
| ENSMUSG00000093385 | *A330044P14Rik* | 0 | 1.009272028 | 0 | 7.590593135 | 9.771665612 | 12.56340268 | 0.001212686 | 4.759142611 | 0.000436435 | 12.36936747 | 27.07975179 | 2.916251468 |
| ENSMUSG00000074158 | *Zfp976* | 0 | 0 | 1.341985292 | 10.12079085 | 11.94314686 | 6.764909138 | 0.001930962 | 4.705748633 | 0.000716019 | 11.44719646 | 26.0958526 | 2.714226289 |
| ENSMUSG00000009731 | *Kcnd1* | 8.78749814 | 4.037088113 | 4.025955875 | 147.5948665 | 161.7753529 | 150.7608322 | 6.31E-38 | 4.68768932 | 2.31E-39 | 172.315791 | 25.77122696 | 37.19998159 |
| ENSMUSG00000079243 | *Xirp1* | 2.396590402 | 2.018544056 | 1.341985292 | 50.60395423 | 55.3727718 | 41.55587042 | 1.35E-13 | 4.623424777 | 1.83E-14 | 58.70806382 | 24.64844582 | 12.87127562 |
| ENSMUSG00000023140 | *Reg2* | 0 | 2.018544056 | 0 | 6.747193898 | 17.37184998 | 22.2275586 | 0.000535802 | 4.510199251 | 0.000183122 | 13.99679515 | 22.78795016 | 3.270995919 |
| ENSMUSG00000058427 | *Cxcl2* | 1050.505459 | 1270.673483 | 951.4675717 | 21183.65864 | 21093.76883 | 20894.8715 | 1.95E-289 | 4.268983348 | 5.74E-293 | 1338.166909 | 19.27933451 | 288.7108453 |
| ENSMUSG00000114805 | *Gm9626* | 1.597726935 | 0 | 0 | 13.4943878 | 11.94314686 | 8.69774032 | 0.001417163 | 4.155795429 | 0.000514792 | 12.06130657 | 17.82457075 | 2.84858033 |
| ENSMUSG00000048806 | *Ifnb1* | 10.38522507 | 9.083448253 | 4.025955875 | 147.5948665 | 145.4892436 | 143.9959231 | 1.95E-33 | 4.142029909 | 8.91E-35 | 151.3223578 | 17.65530585 | 32.71091381 |
| ENSMUSG00000111824 | *Gm47102* | 12.78181548 | 22.20398462 | 0 | 173.7402429 | 141.1462811 | 290.8910929 | 0.003743788 | 4.099871045 | 0.001453634 | 10.13648064 | 17.14684266 | 2.426688775 |
| ENSMUSG00000111581 | *Gm47364* | 1.597726935 | 0 | 0 | 15.18118627 | 11.94314686 | 4.832077956 | 0.0044218 | 4.074749412 | 0.001734377 | 9.811354848 | 16.85084942 | 2.354400917 |
| ENSMUSG00000035095 | *Fam167a* | 0.798863467 | 1.009272028 | 1.341985292 | 13.4943878 | 9.771665612 | 27.05963655 | 0.000200476 | 4.047898059 | 6.47E-05 | 15.95894418 | 16.54012301 | 3.697937104 |
| ENSMUSG00000059901 | *Adamts14* | 0 | 6.055632169 | 2.683970583 | 46.38695805 | 42.34388432 | 46.38794837 | 8.57E-09 | 4.027520178 | 1.72E-09 | 36.26822689 | 16.30813815 | 8.06718437 |
| ENSMUSG00000092124 | *B930094E09Rik* | 1.597726935 | 2.018544056 | 0 | 13.4943878 | 9.771665612 | 39.62303924 | 0.000277938 | 4.014395187 | 9.12E-05 | 15.31095454 | 16.16044687 | 3.556051663 |
| ENSMUSG00000027579 | *Srms* | 2.396590402 | 1.009272028 | 0 | 21.08498093 | 21.71481247 | 19.32831182 | 9.71E-06 | 4.001501663 | 2.66E-06 | 22.0490557 | 16.01666264 | 5.012808196 |
| ENSMUSG00000021118 | *Plek2* | 3.994317336 | 2.018544056 | 0 | 31.20577178 | 44.51536557 | 29.95888332 | 5.56E-08 | 3.963955738 | 1.21E-08 | 32.47775534 | 15.6052086 | 7.255139469 |
| ENSMUSG00000074671 | *Tspyl3* | 0.798863467 | 5.046360141 | 1.341985292 | 28.67557407 | 33.65795933 | 35.75737687 | 2.86E-07 | 3.785013211 | 6.65E-08 | 29.16346739 | 13.78486478 | 6.542965187 |
| ENSMUSG00000028602 | *Tnfrsf8* | 4.793180804 | 10.09272028 | 8.051911749 | 128.1966841 | 96.6309155 | 80.21249406 | 1.40E-17 | 3.769136169 | 1.46E-18 | 77.30847583 | 13.63399231 | 16.8538997 |
| ENSMUSG00000022126 | *Acod1* | 2273.565428 | 2268.843519 | 1787.524408 | 29031.48854 | 28818.81337 | 28331.43947 | 1.45E-252 | 3.765065924 | 6.01E-256 | 1167.821751 | 13.59558122 | 251.8374725 |
| ENSMUSG00000032578 | *Cish* | 12.78181548 | 14.12980839 | 9.393897041 | 144.2212696 | 176.9757216 | 173.9548064 | 8.09E-33 | 3.742779685 | 3.81E-34 | 148.4334529 | 13.38717536 | 32.09214957 |
| ENSMUSG00000079652 | *Garin1a* | 3.994317336 | 7.064904197 | 0 | 53.97755118 | 49.94406868 | 53.15285751 | 1.25E-10 | 3.735237683 | 2.14E-11 | 44.84119316 | 13.3173736 | 9.902719175 |
| ENSMUSG00000021815 | *Mss51* | 8.78749814 | 6.055632169 | 4.025955875 | 92.7739161 | 105.3168405 | 57.01851988 | 5.32E-14 | 3.698129502 | 7.02E-15 | 60.59128329 | 12.9791995 | 13.27423262 |
| ENSMUSG00000112149 | *Gm47611* | 0 | 0 | 5.367941166 | 24.45857788 | 15.20036873 | 18.36189623 | 0.001754679 | 3.680221031 | 0.000646406 | 11.63735051 | 12.81908187 | 2.755802386 |
| ENSMUSG00000000317 | *Bcl6b* | 108.6454316 | 87.80666645 | 83.20308807 | 1211.964704 | 1191.057464 | 1221.549307 | 1.44E-167 | 3.676410076 | 2.30E-170 | 774.1066444 | 12.78526427 | 166.8404894 |
| ENSMUSG00000073902 | *Gvin3* | 23.96590402 | 6.055632169 | 2.683970583 | 151.8118627 | 149.8322061 | 140.1302607 | 1.06E-11 | 3.66467934 | 1.66E-12 | 49.84734687 | 12.68172717 | 10.97347433 |
| ENSMUSG00000024810 | *Il33* | 0.798863467 | 1.009272028 | 0 | 9.27739161 | 5.428703118 | 9.664155911 | 0.009715813 | 3.636977129 | 0.004063738 | 8.255113104 | 12.44053936 | 2.012520837 |
| ENSMUSG00000007682 | *Dio2* | 2.396590402 | 1.009272028 | 4.025955875 | 24.45857788 | 39.08666245 | 22.2275586 | 3.11E-06 | 3.584028857 | 8.07E-07 | 24.34072277 | 11.99223668 | 5.506694661 |
| ENSMUSG00000026605 | *Cenpf* | 345.9078813 | 206.9007658 | 99.30691157 | 2735.143726 | 2518.918247 | 2479.822407 | 5.50E-11 | 3.563916477 | 9.10E-12 | 46.51255664 | 11.82621477 | 10.25971695 |
| ENSMUSG00000063286 | *Gvin-ps7* | 637.4930469 | 438.0240602 | 232.1634554 | 5423.057095 | 5101.89519 | 4894.894969 | 9.00E-16 | 3.557185757 | 1.06E-16 | 68.8618708 | 11.77116949 | 15.04592612 |
| ENSMUSG00000018476 | *Kdm6b* | 238.8601767 | 223.0491182 | 123.4626468 | 2275.491142 | 2315.88475 | 2236.285678 | 1.66E-67 | 3.529256384 | 1.98E-69 | 310.1997678 | 11.54548108 | 66.78062877 |
| ENSMUSG00000113664 | *Gm47184* | 3.994317336 | 1.009272028 | 0 | 29.5189733 | 9.771665612 | 23.19397419 | 0.000642718 | 3.485293496 | 0.000222205 | 13.63333458 | 11.19896496 | 3.191979835 |
| ENSMUSG00000034394 | *Lif* | 167.7613281 | 304.8001525 | 155.6702938 | 2370.795256 | 2447.259366 | 2214.058119 | 1.00E-51 | 3.481936166 | 1.99E-53 | 236.7718559 | 11.17293388 | 50.99871882 |
| ENSMUSG00000032340 | *Neo1* | 2.396590402 | 1.009272028 | 5.367941166 | 26.98877559 | 43.42962494 | 20.29472741 | 1.06E-05 | 3.451330508 | 2.91E-06 | 21.87660356 | 10.93840522 | 4.975466243 |
| ENSMUSG00000024014 | *Pim1* | 363.4828776 | 442.0611483 | 355.6261023 | 4103.137289 | 4300.61861 | 4230.967458 | 7.26E-237 | 3.441998227 | 3.86E-240 | 1095.08818 | 10.86787692 | 236.1387837 |
| ENSMUSG00000019850 | *Tnfaip3* | 166.9624647 | 162.4927965 | 151.6443379 | 1777.885592 | 1804.500916 | 1664.167648 | 9.96E-216 | 3.440299397 | 7.64E-219 | 997.1061496 | 10.8550871 | 215.0016418 |
| ENSMUSG00000043953 | *Ccrl2* | 160.5715569 | 144.3259 | 159.6962497 | 1676.677684 | 1565.637979 | 1648.704998 | 1.47E-206 | 3.396512084 | 1.39E-209 | 954.5081914 | 10.53057336 | 205.8323278 |
| ENSMUSG00000082674 | *Gm11914* | 30.35681176 | 15.13908042 | 5.367941166 | 168.6798474 | 191.0903497 | 194.2495338 | 8.05E-16 | 3.371481861 | 9.43E-17 | 69.08595901 | 10.3494476 | 15.09417951 |
| ENSMUSG00000113795 | *Gm48119* | 5.592044271 | 3.027816084 | 6.709926458 | 64.09834203 | 53.20129055 | 35.75737687 | 3.86E-09 | 3.340921448 | 7.50E-10 | 37.88710567 | 10.13252232 | 8.413883663 |
| ENSMUSG00000026628 | *Atf3* | 1795.845074 | 1871.19034 | 1818.39007 | 18182.00076 | 18419.58968 | 18265.25467 | 0 | 3.322473216 | 0 | 3100.119635 | 10.00377921 | Inf |
| ENSMUSG00000074141 | *Il4i1* | 16.77613281 | 21.19471259 | 16.1038235 | 154.3420604 | 167.204056 | 214.5442612 | 1.88E-27 | 3.305090607 | 1.13E-28 | 123.4224937 | 9.88396983 | 26.72470076 |
| ENSMUSG00000083822 | *Hmgb1-ps5* | 0.798863467 | 1.009272028 | 1.341985292 | 5.060395423 | 9.771665612 | 15.46264946 | 0.008706086 | 3.304890323 | 0.003605442 | 8.472557395 | 9.882597768 | 2.060177038 |
| ENSMUSG00000082765 | *Gm14411* | 0 | 9.083448253 | 0 | 32.04917101 | 30.40073746 | 28.02605214 | 0.002506775 | 3.304098096 | 0.000943147 | 10.93597695 | 9.877172427 | 2.600884676 |
| ENSMUSG00000112049 | *Gm48269* | 10.38522507 | 4.037088113 | 0 | 41.32656262 | 64.05869679 | 50.25361074 | 1.10E-06 | 3.3009711 | 2.70E-07 | 26.44992876 | 9.855787157 | 5.95734999 |
| ENSMUSG00000032238 | *Rora* | 3.195453869 | 5.046360141 | 6.709926458 | 39.63976415 | 48.85832806 | 50.25361074 | 2.21E-09 | 3.272512734 | 4.20E-10 | 39.01726735 | 9.663278451 | 8.65524475 |
| ENSMUSG00000111928 | *Gm48082* | 11.98295201 | 4.037088113 | 6.709926458 | 74.21913288 | 76.00184365 | 77.31324729 | 2.94E-13 | 3.271591771 | 4.11E-14 | 57.1170332 | 9.657111741 | 12.53209446 |
| ENSMUSG00000105504 | *Gbp5* | 16.77613281 | 15.13908042 | 20.12977937 | 151.8118627 | 159.6038717 | 181.6861311 | 2.24E-29 | 3.260071663 | 1.23E-30 | 132.3922112 | 9.580305509 | 28.64981493 |
| ENSMUSG00000056531 | *Ccdc18* | 19.17272321 | 12.11126434 | 6.709926458 | 121.4494902 | 136.8033186 | 117.9027021 | 5.04E-19 | 3.235875395 | 4.82E-20 | 84.05126273 | 9.420968597 | 18.29735259 |
| ENSMUSG00000055523 | *Gucy2g* | 14.37954241 | 13.12053637 | 9.393897041 | 126.5098856 | 114.0027655 | 115.0034553 | 2.04E-22 | 3.235818934 | 1.61E-23 | 99.89247973 | 9.420599908 | 21.69121117 |
| ENSMUSG00000047735 | *Samd9l* | 969.0213858 | 716.58314 | 566.317793 | 7271.788223 | 6953.082953 | 6879.912593 | 6.03E-75 | 3.223632287 | 5.69E-77 | 344.8193303 | 9.341357932 | 74.21977647 |
| ENSMUSG00000038067 | *Csf3* | 145.393151 | 188.7338693 | 174.4580879 | 1564.505585 | 1560.209276 | 1589.753647 | 3.24E-154 | 3.221888437 | 5.92E-157 | 712.4315955 | 9.330073442 | 153.4899401 |
| ENSMUSG00000034855 | *Cxcl10* | 359.4885603 | 364.3472022 | 218.7436025 | 2987.320098 | 2930.413943 | 2841.261838 | 1.56E-74 | 3.206559391 | 1.49E-76 | 342.8940117 | 9.231463558 | 73.805885 |
| ENSMUSG00000082419 | *Gm11425* | 47.13294457 | 7.064904197 | 21.47176466 | 245.429178 | 208.4621997 | 257.0665472 | 9.04E-11 | 3.205119419 | 1.53E-11 | 45.49824405 | 9.222254113 | 10.04405431 |
| ENSMUSG00000114217 | *Gm7054* | 0 | 3.027816084 | 8.051911749 | 46.38695805 | 7.600184365 | 40.58945483 | 0.004675507 | 3.199903545 | 0.001840788 | 9.701888053 | 9.18897247 | 2.33017126 |
| ENSMUSG00000074269 | *Rec114* | 3.994317336 | 0 | 2.683970583 | 18.55478322 | 10.85740624 | 32.8581301 | 0.001400777 | 3.189634311 | 0.000508509 | 12.08419987 | 9.12379677 | 2.853630973 |
| ENSMUSG00000045328 | *Cenpe* | 540.8305674 | 399.6717231 | 195.9298526 | 3608.905336 | 3515.628139 | 3259.719789 | 2.95E-12 | 3.18917769 | 4.45E-13 | 52.4349544 | 9.120909495 | 11.52976126 |
| ENSMUSG00000090939 | *Gm17147* | 2.396590402 | 2.018544056 | 1.341985292 | 19.39818246 | 16.28610935 | 18.36189623 | 0.00016583 | 3.185865207 | 5.30E-05 | 16.33814733 | 9.099991559 | 3.780336961 |
| ENSMUSG00000112557 | *Gm47626* | 23.96590402 | 6.055632169 | 2.683970583 | 100.3645092 | 128.1173936 | 85.04457202 | 1.59E-07 | 3.182863996 | 3.61E-08 | 30.34974813 | 9.081080699 | 6.797538496 |
| ENSMUSG00000086513 | *Gvin-ps1* | 44.73635417 | 21.19471259 | 21.47176466 | 242.8989803 | 307.2645965 | 266.7307032 | 2.80E-25 | 3.181898146 | 1.89E-26 | 113.2589888 | 9.075003161 | 24.55213665 |
| ENSMUSG00000020178 | *Adora2a* | 1.597726935 | 0 | 2.683970583 | 19.39818246 | 8.685924988 | 8.69774032 | 0.00727852 | 3.175763761 | 0.002966564 | 8.827919596 | 9.036497884 | 2.137956927 |
| ENSMUSG00000035948 | *Acss3* | 3.994317336 | 1.009272028 | 1.341985292 | 21.08498093 | 17.37184998 | 22.2275586 | 0.000160982 | 3.162910137 | 5.14E-05 | 16.39718837 | 8.956345214 | 3.793223588 |
| ENSMUSG00000042622 | *Maff* | 168.5601916 | 180.659693 | 197.2718379 | 1551.011197 | 1563.466498 | 1681.563129 | 3.86E-170 | 3.143272173 | 5.70E-173 | 786.0933252 | 8.835257444 | 169.4132685 |
| ENSMUSG00000114367 | *Gm48216* | 12.78181548 | 19.17616853 | 0 | 113.0154978 | 100.973878 | 70.54833815 | 7.33E-07 | 3.101247007 | 1.77E-07 | 27.27030794 | 8.581602086 | 6.135174025 |
| ENSMUSG00000013367 | *Iglon5* | 0.798863467 | 4.037088113 | 1.341985292 | 16.86798474 | 21.71481247 | 13.52981828 | 0.000878639 | 3.089855597 | 0.000309992 | 13.0090341 | 8.51410922 | 3.056189765 |
| ENSMUSG00000071757 | *Zhx2* | 5.592044271 | 11.10199231 | 5.367941166 | 55.66434966 | 61.88721554 | 70.54833815 | 5.41E-11 | 3.085917732 | 8.94E-12 | 46.54817083 | 8.490901475 | 10.26714547 |
| ENSMUSG00000112667 | *Gm48270* | 11.98295201 | 0 | 6.709926458 | 77.59272983 | 52.11554993 | 30.92529892 | 0.000154577 | 3.07416487 | 4.92E-05 | 16.47942052 | 8.422011643 | 3.810855913 |
| ENSMUSG00000042607 | *Asb4* | 5.592044271 | 1.009272028 | 2.683970583 | 29.5189733 | 22.80055309 | 28.02605214 | 3.59E-05 | 3.048526029 | 1.05E-05 | 19.41360648 | 8.273662046 | 4.444837579 |
| ENSMUSG00000111009 | *None* | 3.994317336 | 0 | 1.341985292 | 21.92838017 | 15.20036873 | 9.664155911 | 0.004792557 | 3.040660359 | 0.001894225 | 9.649311814 | 8.228676228 | 2.319432709 |
| ENSMUSG00000028525 | *Pde4b* | 133.410199 | 158.4557084 | 128.830588 | 1217.025099 | 1160.656727 | 1085.284709 | 2.87E-119 | 3.03869686 | 9.82E-122 | 550.4969145 | 8.217484671 | 118.5425209 |
| ENSMUSG00000055202 | *Zfp811* | 5.592044271 | 10.09272028 | 5.367941166 | 59.03794661 | 54.28703118 | 57.01851988 | 1.70E-10 | 3.012601035 | 2.93E-11 | 44.22426715 | 8.070181021 | 9.769307066 |
| ENSMUSG00000052595 | *A1cf* | 3.994317336 | 4.037088113 | 9.393897041 | 53.13415195 | 38.00092182 | 41.55587042 | 1.07E-07 | 3.006419049 | 2.39E-08 | 31.14746733 | 8.035674068 | 6.968768053 |
| ENSMUSG00000095649 | *Gvin-ps3* | 13.58067894 | 5.046360141 | 10.73588233 | 80.96632677 | 53.20129055 | 102.4400527 | 1.31E-09 | 2.99759666 | 2.45E-10 | 40.07143313 | 7.986684147 | 8.881507623 |
| ENSMUSG00000058515 | *Or8d1b* | 6.390907738 | 2.018544056 | 1.341985292 | 27.83217483 | 34.74369995 | 21.261143 | 6.28E-05 | 2.995555966 | 1.89E-05 | 18.29258575 | 7.975394957 | 4.20209623 |
| ENSMUSG00000089756 | *Zfp966* | 35.94885603 | 18.16689651 | 13.41985292 | 176.2704406 | 178.0614623 | 199.0816118 | 1.81E-18 | 2.980322037 | 1.79E-19 | 81.46290194 | 7.891623 | 17.74212149 |
| ENSMUSG00000015312 | *Gadd45b* | 496.8930767 | 412.7922595 | 476.4047785 | 3581.916561 | 3628.545164 | 3713.935117 | 2.88E-212 | 2.977280382 | 2.38E-215 | 981.0366347 | 7.875002505 | 211.5408547 |
| ENSMUSG00000020641 | *Rsad2* | 1803.034846 | 1669.335935 | 1356.74713 | 13124.97893 | 12557.67605 | 12185.53419 | 7.97E-146 | 2.968490616 | 1.74E-148 | 673.4891167 | 7.827169112 | 145.098357 |
| ENSMUSG00000078671 | *Chd2* | 303.5681176 | 336.0875854 | 240.2153672 | 2230.790982 | 2363.657337 | 2160.905262 | 7.38E-115 | 2.932095649 | 2.74E-117 | 530.058133 | 7.632182392 | 114.1320229 |
| ENSMUSG00000041616 | *Nppa* | 2.396590402 | 2.018544056 | 1.341985292 | 10.96419008 | 19.54333122 | 14.49623387 | 0.001712295 | 2.916306092 | 0.000630273 | 11.68436936 | 7.549107506 | 2.766421409 |
| ENSMUSG00000024486 | *Hbegf* | 40.74203683 | 39.3616091 | 40.25955875 | 304.4671246 | 302.921634 | 301.5216644 | 1.11E-49 | 2.915948782 | 2.39E-51 | 227.236226 | 7.547238063 | 48.9539943 |
| ENSMUSG00000014773 | *Dll1* | 2.396590402 | 8.074176225 | 0 | 30.36237254 | 26.05777497 | 25.12680537 | 0.000343295 | 2.910831728 | 0.000113925 | 14.89066998 | 7.520516395 | 3.464332908 |
| ENSMUSG00000096463 | *None* | 1.597726935 | 9.083448253 | 4.025955875 | 24.45857788 | 43.42962494 | 39.62303924 | 4.83E-05 | 2.89283554 | 1.44E-05 | 18.81874283 | 7.427288099 | 4.315986431 |
| ENSMUSG00000074802 | *Gas2l3* | 760.5180209 | 863.9368561 | 838.7408072 | 6244.527952 | 5975.916392 | 5910.597755 | 3.39E-264 | 2.882678589 | 1.20E-267 | 1221.654852 | 7.375181677 | 263.4700047 |
| ENSMUSG00000029053 | *Prkcz* | 1.597726935 | 1.009272028 | 4.025955875 | 19.39818246 | 10.85740624 | 15.46264946 | 0.002549896 | 2.881029828 | 0.000960425 | 10.90234771 | 7.366757888 | 2.593477594 |
| ENSMUSG00000060441 | *Trim5* | 9.586361607 | 6.055632169 | 4.025955875 | 57.35114813 | 42.34388432 | 50.25361074 | 2.03E-08 | 2.871848569 | 4.23E-09 | 34.51319681 | 7.320024966 | 7.692037485 |
| ENSMUSG00000027536 | *Chmp4c* | 3.994317336 | 3.027816084 | 4.025955875 | 32.89257025 | 24.97203434 | 22.2275586 | 2.35E-05 | 2.862865895 | 6.71E-06 | 20.27416853 | 7.274589779 | 4.629462888 |
| ENSMUSG00000022900 | *Ildr1* | 1.597726935 | 3.027816084 | 0 | 10.96419008 | 13.02888748 | 11.59698709 | 0.007464588 | 2.858658255 | 0.003047247 | 8.778961057 | 7.253404243 | 2.12699416 |
| ENSMUSG00000072621 | *Slfn10-ps* | 272.4124423 | 264.4292714 | 199.9558084 | 1782.102588 | 1762.157032 | 1773.37261 | 3.95E-114 | 2.838599639 | 1.52E-116 | 526.6449298 | 7.153253846 | 113.4030305 |
| ENSMUSG00000032688 | *Malt1* | 877.9509505 | 918.4375456 | 849.4766895 | 6373.568036 | 6224.550995 | 6208.253757 | 0 | 2.828037188 | 0 | 1606.981014 | 7.101073737 | Inf |
| ENSMUSG00000020275 | *Rel* | 524.853298 | 473.3485812 | 362.3360287 | 3271.545641 | 3197.506136 | 3193.037113 | 4.14E-99 | 2.819818266 | 2.12E-101 | 457.0353816 | 7.060734484 | 98.3834554 |
| ENSMUSG00000062901 | *Klhl24* | 166.9624647 | 154.4186203 | 213.3756614 | 1297.991426 | 1180.200058 | 1247.642528 | 1.61E-91 | 2.81414903 | 9.91E-94 | 421.7983661 | 7.033042959 | 90.7920048 |
| ENSMUSG00000031004 | *Mki67* | 2641.042623 | 1849.995628 | 1027.960733 | 13564.38993 | 13066.8884 | 12094.69112 | 6.17E-12 | 2.810188865 | 9.49E-13 | 50.94642814 | 7.013763893 | 11.20942679 |
| ENSMUSG00000024795 | *Kif20b* | 428.1908185 | 259.3829112 | 161.038235 | 2060.424337 | 1967.36201 | 1907.704377 | 2.19E-10 | 2.802519089 | 3.81E-11 | 43.7099595 | 6.976575673 | 9.660009613 |

| | | | | | | | | | | | | | |
|---|---|---|---|---|---|---|---|---|---|---|---|---|---|
| ENSMUSG00000096433 | *Zfp994* | 170.1579185 | 106.982835 | 85.88705866 | 837.4954426 | 869.6782395 | 860.1098761 | 5.54E-36 | 2.802137684 | 2.23E-37 | 163.2249941 | 6.974731518 | 35.25633629 |
| ENSMUSG00000026821 | *Ralgds* | 447.3635417 | 513.7194623 | 359.6520581 | 3086.841208 | 3134.53318 | 3019.082307 | 1.71E-105 | 2.80084914 | 7.69E-108 | 486.635125 | 6.968504816 | 104.765822 |
| ENSMUSG00000034647 | *Ankrd12* | 234.8658594 | 216.993486 | 118.0947057 | 1419.440916 | 1333.289486 | 1250.541775 | 1.43E-37 | 2.799037293 | 5.32E-39 | 170.6555115 | 6.959758724 | 36.84432681 |
| ENSMUSG00000113393 | *Gm48415* | 40.74203683 | 8.074176225 | 6.709926458 | 150.9684635 | 156.3466498 | 87.94381879 | 1.01E-05 | 2.78655097 | 2.78E-06 | 21.96594917 | 6.899782907 | 4.994938841 |
| ENSMUSG00000073490 | *Ifi207* | 1061.689548 | 1051.661453 | 676.3605869 | 6539.717686 | 6526.386888 | 6219.850745 | 7.03E-65 | 2.785764279 | 9.05E-67 | 297.9858146 | 6.896021533 | 64.15284117 |
| ENSMUSG00000027204 | *Fbn1* | 2.396590402 | 1.009272028 | 2.683970583 | 11.80758932 | 15.20036873 | 14.49623387 | 0.003119212 | 2.779594143 | 0.0011929 | 10.50130969 | 6.866591518 | 2.505955165 |
| ENSMUSG00000041695 | *Kcnj2* | 480.1169438 | 537.941991 | 507.2704402 | 3688.184864 | 3442.883517 | 3316.738309 | 2.69E-195 | 2.777958539 | 3.02E-198 | 902.3549617 | 6.858811178 | 194.5698566 |
| ENSMUSG00000029161 | *Cgref1* | 3.195453869 | 4.037088113 | 8.051911749 | 36.2661672 | 26.05777497 | 36.72379246 | 1.06E-05 | 2.772633215 | 2.91E-06 | 21.87423951 | 6.833540375 | 4.975059024 |
| ENSMUSG00000033581 | *Igf2bp2* | 117.4329297 | 252.318007 | 122.1206615 | 918.4617693 | 1225.801164 | 1213.817982 | 1.55E-22 | 2.769814196 | 1.21E-23 | 100.4493681 | 6.820200706 | 21.80908512 |
| ENSMUSG00000096795 | *Zfp433* | 23.16704055 | 41.38015315 | 4.025955875 | 163.619452 | 152.0036873 | 155.5929102 | 1.33E-07 | 2.751951399 | 3.00E-08 | 30.70535474 | 6.736276697 | 6.875327646 |
| ENSMUSG00000095193 | *Gm20939* | 79.88634673 | 50.46360141 | 50.99544108 | 425.9166148 | 414.7529182 | 402.0288859 | 4.71E-39 | 2.751604919 | 1.64E-40 | 177.5736973 | 6.734659093 | 38.32663596 |
| ENSMUSG00000040253 | *Gbp7* | 83.0818006 | 55.50996155 | 41.60154404 | 434.3506072 | 465.7827275 | 336.3126257 | 3.16E-25 | 2.751604094 | 2.14E-26 | 113.0159023 | 6.734655242 | 24.49965103 |
| ENSMUSG00000045502 | *Hcar2* | 149.3874684 | 104.9642909 | 130.1725733 | 915.9315716 | 838.1917614 | 839.8151487 | 5.48E-71 | 2.748015111 | 5.82E-73 | 326.4079841 | 6.717922306 | 70.26111055 |
| ENSMUSG00000039431 | *Mtmr7* | 37.54658296 | 56.51923358 | 57.70536754 | 330.612501 | 279.0353403 | 388.4990676 | 7.15E-29 | 2.738385316 | 4.02E-30 | 130.039375 | 6.673230405 | 28.14568769 |
| ENSMUSG00000040829 | *Zmynd15* | 28.75908482 | 16.14835245 | 14.76183821 | 142.5344711 | 145.4892436 | 119.8355333 | 2.68E-16 | 2.729970459 | 3.04E-17 | 71.31814769 | 6.634420517 | 15.57205183 |
| ENSMUSG00000026875 | *Traf1* | 1313.33154 | 1239.386051 | 1076.272204 | 7902.650853 | 8001.908396 | 7954.566731 | 2.29E-194 | 2.71313984 | 2.70E-197 | 897.9754666 | 6.557472445 | 193.6308505 |
| ENSMUSG00000027381 | *Bcl2l11* | 424.9953646 | 349.2081217 | 265.7130877 | 2238.381576 | 2334.342341 | 2284.606457 | 1.79E-66 | 2.711141112 | 2.21E-68 | 305.3835565 | 6.54839393 | 65.74829351 |
| ENSMUSG00000074903 | *Gm2058* | 2.396590402 | 0 | 10.73588233 | 23.61517864 | 20.62907185 | 35.75737687 | 0.005497107 | 2.710176928 | 0.002195728 | 9.378268829 | 6.544018959 | 2.259865843 |
| ENSMUSG00000035356 | *Nfkbiz* | 490.5021689 | 499.5896539 | 382.4658081 | 3059.852433 | 3000.987084 | 2962.063787 | 2.39E-125 | 2.709314292 | 7.76E-128 | 578.5491564 | 6.540107238 | 124.6216773 |
| ENSMUSG00000026004 | *Kansl1l* | 185.3363244 | 219.0120301 | 218.7436025 | 1402.572932 | 1375.63337 | 1271.802918 | 9.47E-114 | 2.708566467 | 3.69E-116 | 524.8702261 | 6.536718035 | 113.0235589 |
| ENSMUSG00000063383 | *Zfp947* | 29.55794829 | 10.09272028 | 6.709926458 | 117.234294 | 97.71665612 | 99.54080589 | 4.79E-08 | 2.701608694 | 1.03E-08 | 32.78036302 | 6.505268906 | 7.319673252 |
| ENSMUSG00000067928 | *Zfp760* | 39.94317336 | 20.18544056 | 29.52367641 | 205.7894139 | 168.2897967 | 215.5106768 | 5.57E-20 | 2.700660353 | 5.03E-21 | 88.52003105 | 6.500994137 | 19.25447882 |
| ENSMUSG00000028599 | *Tnfrsf1b* | 1341.291762 | 1438.21264 | 1213.154704 | 8537.730479 | 8690.267951 | 8561.475722 | 1.13E-227 | 2.689098976 | 7.33E-231 | 1052.398029 | 6.449105079 | 226.9472668 |
| ENSMUSG00000028634 | *Hivep3* | 630.3032757 | 631.8042896 | 299.26272 | 3339.860979 | 3540.600173 | 3123.455191 | 4.38E-13 | 2.677590541 | 6.20E-14 | 56.30706853 | 6.397864958 | 12.3583153 |
| ENSMUSG00000021709 | *Erbin* | 1098.437268 | 1035.513101 | 852.1606601 | 6491.643929 | 6297.295617 | 6265.272277 | 1.74E-139 | 2.669496439 | 4.20E-142 | 644.1406956 | 6.362070855 | 138.7603647 |
| ENSMUSG00000024330 | *Col11a2* | 42.33976377 | 39.36161091 | 28.18169112 | 226.8743948 | 270.3494153 | 210.6785989 | 5.00E-27 | 2.661515333 | 3.05E-28 | 121.4461053 | 6.326972533 | 26.30136807 |
| ENSMUSG00000071658 | *Gng3* | 4.793180804 | 2.018544056 | 2.683970583 | 21.92838017 | 19.54333122 | 20.29472741 | 0.00042577 | 2.654670324 | 0.000143155 | 14.46017346 | 6.297024692 | 3.370825194 |
| ENSMUSG00000028268 | *Gbp3* | 104.6511142 | 66.61195386 | 52.33742637 | 472.3035729 | 471.2114306 | 486.1070423 | 2.34E-29 | 2.649375026 | 1.28E-30 | 132.3021008 | 6.273954323 | 28.63056925 |
| ENSMUSG00000050533 | *Gm9845* | 43.9374907 | 30.27816084 | 71.12522045 | 296.8765315 | 295.3214496 | 300.5552488 | 6.60E-22 | 2.647185764 | 5.37E-23 | 97.50754185 | 6.264440939 | 21.18033588 |
| ENSMUSG00000019971 | *Cep290* | 159.7726935 | 106.982835 | 71.12522045 | 727.8535417 | 728.5319584 | 682.2894073 | 9.33E-27 | 2.641617224 | 5.83E-28 | 120.1626039 | 6.240307942 | 26.03044447 |
| ENSMUSG00000043421 | *Hilpda* | 339.5169736 | 316.9114168 | 444.1971315 | 2287.298731 | 2226.854019 | 2290.404951 | 1.30E-88 | 2.63697191 | 8.82E-91 | 408.2485188 | 6.220247174 | 87.88630098 |
| ENSMUSG00000067297 | *Ifit1bl2* | 6.390907738 | 18.16689651 | 5.367941166 | 53.97755118 | 82.51628739 | 51.22002633 | 1.80E-06 | 2.635668123 | 4.54E-07 | 25.44853315 | 6.214628374 | 5.7439264 |
| ENSMUSG00000090675 | *Or5v1b* | 33.55226563 | 12.11126434 | 17.44580879 | 127.3532848 | 147.6607248 | 126.6004424 | 4.28E-12 | 2.632633107 | 6.49E-13 | 51.69228551 | 6.201568325 | 11.36889442 |
| ENSMUSG00000108312 | *Gm44521* | 22.36817708 | 18.16689651 | 10.73588233 | 132.4136802 | 92.287953 | 101.4736371 | 9.09E-13 | 2.630408882 | 1.31E-13 | 54.83402471 | 6.192014639 | 12.0416706 |
| ENSMUSG00000032131 | *Abcg4* | 8.78749814 | 7.064904197 | 8.051911749 | 42.16996186 | 53.20129055 | 52.18644192 | 3.34E-08 | 2.616903392 | 7.09E-09 | 33.51004186 | 6.134319851 | 7.476417314 |
| ENSMUSG00000033863 | *Klf9* | 115.8352028 | 102.9457469 | 84.54507337 | 623.2720363 | 604.7575273 | 637.8342901 | 1.45E-64 | 2.602554904 | 1.88E-66 | 296.5315703 | 6.073612657 | 63.83798836 |
| ENSMUSG00000032712 | *Resf1* | 433.7828627 | 324.9855931 | 216.0596319 | 2001.38639 | 2067.250147 | 1852.618688 | 5.62E-34 | 2.593779595 | 2.50E-35 | 153.8507986 | 6.036781546 | 33.25044365 |
| ENSMUSG00000112122 | *Gm47628* | 2.396590402 | 5.046360141 | 0 | 19.39818246 | 10.85740624 | 16.42906505 | 0.008019884 | 2.585694338 | 0.003294287 | 8.636860328 | 6.003044397 | 2.095831905 |
| ENSMUSG00000053846 | *Lipg* | 11.98295201 | 4.037088113 | 2.683970583 | 48.91715576 | 33.65795933 | 35.75737687 | 5.70E-05 | 2.581150848 | 1.71E-05 | 18.48503414 | 5.984168706 | 4.244001864 |
| ENSMUSG00000034422 | *Parp14* | 1590.537163 | 1394.813943 | 1437.266247 | 9190.521488 | 8821.642566 | 8357.562032 | 3.23E-199 | 2.574338393 | 3.24E-202 | 920.6163364 | 5.955977869 | 198.4913002 |
| ENSMUSG00000108857 | *Gm44578* | 4.793180804 | 5.046360141 | 0 | 16.86798474 | 26.05777497 | 19.32831182 | 0.002119671 | 2.561937021 | 0.000789748 | 11.26517416 | 5.90499984 | 2.673731475 |
| ENSMUSG00000048249 | *Crebrf* | 119.0306566 | 122.1219154 | 162.3802203 | 790.2650853 | 787.1619521 | 774.0988885 | 2.00E-67 | 2.560613795 | 2.40E-69 | 309.8084744 | 5.899536315 | 66.69966694 |
| ENSMUSG00000045624 | *Esf1* | 462.5419476 | 344.1617616 | 346.2322052 | 2366.57826 | 2299.598641 | 2168.636586 | 4.96E-80 | 2.560366354 | 4.15E-82 | 368.4100631 | 5.898574548 | 79.30489777 |
| ENSMUSG00000062743 | *Zfp677* | 16.77613281 | 13.12053637 | 10.73588233 | 86.0267222 | 81.43054677 | 77.31324729 | 2.70E-12 | 2.559283119 | 4.04E-13 | 52.62562908 | 5.894147317 | 11.56919502 |
| ENSMUSG00000050064 | *Zfp697* | 4.793180804 | 3.027816084 | 0 | 13.4943878 | 14.11462811 | 22.225586 | 0.006380642 | 2.554816104 | 0.002574249 | 9.087056291 | 5.875925499 | 2.195135591 |
| ENSMUSG00000030156 | *Cd69* | 37.54658296 | 41.38015315 | 16.1038235 | 176.2704406 | 209.5479403 | 186.5182091 | 1.70E-16 | 2.55334742 | 1.91E-17 | 72.24006541 | 5.869946766 | 15.76973428 |
| ENSMUSG00000069892 | *9930111J21Rik2* | 320.3442504 | 213.96567 | 174.4580879 | 1498.720445 | 1396.262442 | 1297.896139 | 1.88E-35 | 2.553336116 | 7.76E-37 | 160.7496518 | 5.869900773 | 34.72684131 |
| ENSMUSG00000031245 | *Hmgn5* | 210.8999554 | 158.4557084 | 108.7008086 | 1013.765883 | 943.5086019 | 879.4381879 | 3.99E-33 | 2.553079956 | 1.86E-34 | 149.8618875 | 5.868858629 | 32.39946323 |
| ENSMUSG00000078606 | *Gvin2* | 1572.163304 | 1205.070802 | 813.2430867 | 7285.282611 | 6987.826653 | 6607.383397 | 1.62E-36 | 2.537215976 | 6.28E-38 | 165.7472815 | 5.804677746 | 35.79179932 |
| ENSMUSG00000079138 | *Gm8818* | 23.16704055 | 15.13908042 | 0 | 59.03794661 | 85.77350926 | 86.01098761 | 0.000376752 | 2.536702426 | 0.00012574 | 14.70459523 | 5.802611846 | 3.423944678 |
| ENSMUSG00000026031 | *Cflar* | 1673.618964 | 1842.930723 | 1830.467938 | 10515.50169 | 10502.36905 | 9889.330744 | 7.53E-248 | 2.532648897 | 3.56E-251 | 1145.862474 | 5.786331176 | 247.1229533 |
| ENSMUSG00000037868 | *Egr2* | 15.97726935 | 19.17616853 | 0 | 69.15873745 | 73.8303624 | 66.68267579 | 0.000138297 | 2.525070673 | 4.37E-05 | 16.70354084 | 5.756016282 | 3.859187342 |
| ENSMUSG00000020593 | *Lpin1* | 261.2283538 | 262.4107273 | 201.2977937 | 1394.982338 | 1414.720032 | 1370.377308 | 1.22E-93 | 2.516138692 | 7.20E-96 | 431.6237759 | 5.720489851 | 92.91350917 |
| ENSMUSG00000039967 | *Zfp292* | 413.0124126 | 305.8094245 | 261.6871318 | 1940.661645 | 1853.359244 | 1844.887363 | 1.67E-54 | 2.514168845 | 2.99E-56 | 249.7199823 | 5.712684461 | 53.7769405 |
| ENSMUSG00000084372 | *Gm13988* | 7.189771206 | 5.046360141 | 1.341985292 | 21.08498093 | 29.31499684 | 31.89171451 | 0.000294702 | 2.512349244 | 9.71E-05 | 15.19254201 | 5.70548387 | 3.530616834 |
| ENSMUSG00000042268 | *Slc26a9* | 11.98295201 | 7.064904197 | 2.683970583 | 48.91715576 | 41.2581437 | 40.58945483 | 7.05E-06 | 2.511311201 | 1.90E-06 | 22.68994606 | 5.701380156 | 5.151891443 |
| ENSMUSG00000093930 | *Hmgcs1* | 1235.841784 | 1074.87471 | 1019.908822 | 6469.715549 | 6162.663779 | 6357.081758 | 4.53E-158 | 2.507820804 | 7.75E-161 | 730.2900332 | 5.687603147 | 157.3442482 |
| ENSMUSG00000051160 | *Or7g33* | 7.988634673 | 2.018544056 | 1.341985292 | 24.45857788 | 20.62907185 | 24.16038978 | 0.001661189 | 2.507299498 | 0.000609907 | 11.74549564 | 5.685548349 | 2.779580894 |
| ENSMUSG00000021367 | *Edn1* | 31.95453869 | 16.14835245 | 16.1038235 | 134.943878 | 131.3746155 | 105.3392994 | 2.96E-12 | 2.494376322 | 4.45E-13 | 52.43180253 | 5.634846544 | 11.52923438 |
| ENSMUSG00000024290 | *Rock1* | 607.1362351 | 472.3393092 | 246.9252936 | 2520.076921 | 2488.517509 | 2466.292589 | 3.57E-10 | 2.491976073 | 6.34E-11 | 42.71408306 | 5.625479499 | 9.447628605 |
| ENSMUSG00000021775 | *Nr1d2* | 179.7442801 | 118.0848273 | 122.1206615 | 810.506667 | 754.5897334 | 817.5875901 | 8.89E-44 | 2.490149511 | 2.48E-45 | 199.6612649 | 5.61836172 | 43.05117427 |
| ENSMUSG00000069727 | *Zfp975* | 41.5409003 | 33.30597693 | 42.94352933 | 240.3687826 | 198.6905341 | 216.4770924 | 2.14E-26 | 2.481586068 | 1.35E-27 | 118.4986659 | 5.585111443 | 25.67053355 |
| ENSMUSG00000000204 | *Slfn4* | 3090.003891 | 2310.223672 | 1469.473894 | 13423.54226 | 12451.27347 | 12327.59728 | 2.28E-14 | 2.474716653 | 2.93E-15 | 62.31304313 | 5.55858106 | 13.64260522 |
| ENSMUSG00000079362 | *Gm43302* | 53.52385231 | 10.09272028 | 28.18169112 | 172.8968436 | 191.0903497 | 151.7272478 | 2.64E-07 | 2.470624538 | 6.11E-08 | 29.32843553 | 5.542836827 | 6.578503078 |
| ENSMUSG00000083899 | *Gm12346* | 471.3294457 | 288.6518 | 480.4307344 | 1659.809699 | 2385.37215 | 2797.773136 | 1.57E-25 | 2.464302149 | 1.04E-26 | 114.4412281 | 5.518599348 | 24.80418596 |
| ENSMUSG00000033799 | *Tasor2* | 683.827128 | 595.4704966 | 381.1238228 | 3132.384767 | 3123.675774 | 2915.675838 | 2.23E-39 | 2.459926558 | 7.63E-41 | 179.0983469 | 5.501887185 | 38.6521216 |
| ENSMUSG00000027326 | *Knl1* | 501.6862575 | 275.5312637 | 150.3023526 | 1700.292862 | 1674.212042 | 1727.951077 | 4.15E-06 | 2.456934113 | 1.09E-06 | 23.7586593 | 5.490486971 | 5.381597369 |
| ENSMUSG00000037876 | *Jmjd1c* | 1382.832662 | 1315.081453 | 1038.696616 | 7195.882292 | 6731.591866 | 6560.029933 | 8.29E-95 | 2.451821443 | 4.70E-97 | 437.0713125 | 5.471064039 | 94.08140902 |
| ENSMUSG00000042851 | *Zc3h6* | 19.97158668 | 14.12980839 | 5.367941166 | 88.55691991 | 70.57314053 | 65.7162602 | 8.45E-08 | 2.451255049 | 1.86E-08 | 31.63379756 | 5.468916555 | 7.072919879 |
| ENSMUSG00000047414 | *Flrt2* | 711.7873494 | 611.6188491 | 554.2399254 | 3530.469207 | 3331.052233 | 3414.346283 | 1.52E-112 | 2.446362242 | 6.18E-115 | 519.2457188 | 5.450400491 | 111.8191872 |
| ENSMUSG00000073427 | *Gm4924* | 45.53521764 | 29.26888882 | 28.18169112 | 192.2950261 | 201.947756 | 177.8204688 | 3.03E-20 | 2.444098872 | 2.69E-21 | 89.76098403 | 5.441856344 | 19.51918478 |
| ENSMUSG00000046311 | *Zfp62* | 353.0976525 | 298.7445203 | 225.453529 | 1667.400292 | 1572.152423 | 1552.063439 | 1.19E-54 | 2.438492975 | 2.13E-56 | 250.3953462 | 5.420751886 | 53.92274009 |
| ENSMUSG00000106631 | *Gm42669* | 23.16704055 | 19.17616853 | 18.78779408 | 113.0154978 | 121.6029498 | 99.54080589 | 4.43E-15 | 2.435085246 | 5.45E-16 | 65.62764914 | 5.40796287 | 14.35373067 |
| ENSMUSG00000045799 | *Gm9800* | 8.78749814 | 27.25034476 | 8.051911749 | 70.84553593 | 122.6886905 | 45.42153278 | 7.19E-05 | 2.429509641 | 2.19E-05 | 18.01754105 | 5.38710297 | 4.143345835 |
| ENSMUSG00000079083 | *Jrkl* | 55.12157924 | 42.38942518 | 30.8656617 | 242.0555811 | 221.4910872 | 235.8054042 | 3.99E-23 | 2.414015609 | 3.00E-24 | 103.2164568 | 5.32955696 | 22.39904821 |
| ENSMUSG00000029061 | *Mmp23* | 1.597726935 | 3.027816084 | 6.709926458 | 14.33778703 | 26.05777497 | 16.42906505 | 0.004709359 | 2.411581999 | 0.001855505 | 9.687255084 | 5.32057438 | 2.327038208 |
| ENSMUSG00000033022 | *Cdo1* | 7.988634673 | 2.018544056 | 9.393897041 | 26.98877559 | 41.2581437 | 33.82454569 | 0.000174117 | 2.409291829 | 5.58E-05 | 16.24015707 | 5.312135067 | 3.759159579 |
| ENSMUSG00000112101 | *Gm47924* | 45.53521764 | 13.12053637 | 30.8656617 | 175.4270413 | 133.5460967 | 170.089144 | 2.53E-09 | 2.406210373 | 4.81E-10 | 38.75150458 | 5.300800971 | 8.597735473 |
| ENSMUSG00000112516 | *Gm47968* | 3.994317336 | 9.083448253 | 8.051911749 | 35.42276796 | 51.02980931 | 23.19397419 | 0.000142759 | 2.404900191 | 4.52E-05 | 16.63940388 | 5.295989239 | 3.845395018 |
| ENSMUSG00000095332 | *Gm9821* | 8.78749814 | 3.027816084 | 1.341985292 | 18.55478322 | 19.54333122 | 35.75737687 | 0.003358469 | 2.399066202 | 0.001291536 | 10.35458347 | 5.274616491 | 2.473858602 |
| ENSMUSG00000032740 | *Ccdc88a* | 378.6612835 | 272.5034476 | 144.9344115 | 1448.11649 | 1417.977254 | 1319.157282 | 1.14E-08 | 2.390229791 | 2.31E-09 | 35.69059065 | 5.242408553 | 7.944568156 |
| ENSMUSG00000024401 | *Tnf* | 6064.971444 | 6269.597839 | 5029.760873 | 30660.09247 | 30212.90433 | 30062.28979 | 4.08E-132 | 2.388001363 | 1.13E-134 | 609.9752313 | 5.234317231 | 131.3889456 |
| ENSMUSG00000044033 | *Ccdc141* | 4.793180804 | 0 | 8.051911749 | 18.55478322 | 28.22925621 | 18.36189623 | 0.008880349 | 2.386899496 | 0.00368285 | 8.43392271 | 5.230321016 | 2.051569979 |
| ENSMUSG00000114025 | *Gm49331* | 4.793180804 | 9.083448253 | 14.76183821 | 40.48316339 | 46.68684681 | 56.05210429 | 7.80E-06 | 2.386762859 | 2.12E-06 | 22.48487681 | 5.229825677 | 5.107806543 |
| ENSMUSG00000058114 | *Or14j6* | 3.195453869 | 3.027816084 | 1.341985292 | 15.18118627 | 15.20036873 | 10.6305715 | 0.008497037 | 2.385049614 | 0.003507334 | 8.522753419 | 5.223618783 | 2.070732509 |
| ENSMUSG00000106170 | *Gm43393* | 5.592044271 | 2.018544056 | 5.367941166 | 24.45857788 | 23.88629372 | 19.32831182 | 0.000805901 | 2.382224033 | 0.00028295 | 13.1800178 | 5.213398113 | 3.093718222 |
| ENSMUSG00000016494 | *Cd34* | 1.597726935 | 10.09272028 | 5.367941166 | 29.5189733 | 29.31499684 | 27.05963655 | 0.000996647 | 2.366785495 | 0.000354391 | 12.75852201 | 5.157906051 | 3.001458652 |
| ENSMUSG00000041153 | *Osgin2* | 784.4839249 | 769.0652854 | 1048.090513 | 4500.37833 | 4410.278413 | 4413.620005 | 5.75E-76 | 2.361032532 | 5.26E-78 | 349.5697079 | 5.13737908 | 75.24047378 |
| ENSMUSG00000027737 | *Slc7a11* | 1751.907584 | 1888.347965 | 1987.480217 | 9748.008384 | 9670.691734 | 9428.350507 | 4.93E-219 | 2.36002375 | 3.50E-222 | 1012.471403 | 5.133788104 | 218.3067163 |
| ENSMUSG00000022141 | *Nipbl* | 762.1157478 | 609.600305 | 405.279558 | 3081.780813 | 3072.645965 | 2978.492852 | 1.03E-33 | 2.356264191 | 4.64E-35 | 152.6200284 | 5.120427239 | 32.98889703 |
| ENSMUSG00000114123 | *Gm48416* | 65.50680432 | 45.41724127 | 9.393897041 | 262.2971628 | 209.5479403 | 154.6264946 | 5.40E-06 | 2.354655979 | 1.44E-06 | 23.23091866 | 5.114722536 | 5.267990559 |
| ENSMUSG00000102443 | *Gm37102* | 5.592044271 | 5.046360141 | 2.683970583 | 17.71138398 | 31.48647808 | 21.261143 | 0.000914564 | 2.35199141 | 0.000323854 | 12.92712835 | 5.105284676 | 3.038785748 |
| ENSMUSG00000021125 | *Arg2* | 36.7477195 | 17.15762448 | 14.76183821 | 125.6664863 | 114.0027655 | 120.8019489 | 5.00E-10 | 2.348125244 | 8.97E-11 | 42.03369664 | 5.091621737 | 9.301282787 |
| ENSMUSG00000041064 | *Pif1* | 203.7101842 | 199.8358616 | 111.3847792 | 863.6408189 | 930.4797144 | 834.9830707 | 4.21E-29 | 2.338185593 | 2.33E-30 | 131.1201786 | 5.056662859 | 28.3757179 |
| ENSMUSG00000031538 | *Plat* | 7.988634673 | 14.12980839 | 1.341985292 | 45.54355881 | 38.00092182 | 38.65662365 | 0.00020484 | 2.332993815 | 6.63E-05 | 15.91438367 | 5.038498318 | 3.688585494 |
| ENSMUSG00000055945 | *Prr18* | 24.76476749 | 19.17616853 | 13.41985292 | 102.0513077 | 99.88813737 | 93.74231234 | 4.24E-12 | 2.327712234 | 6.44E-13 | 51.70849585 | 5.020086538 | 11.37231123 |
| ENSMUSG00000038167 | *Plekhg6* | 3.994317336 | 3.027816084 | 9.393897041 | 26.14537635 | 27.14351559 | 25.12680537 | 0.000532807 | 2.326409321 | 0.000182036 | 14.00798411 | 5.015554891 | 3.273430267 |
| ENSMUSG00000109771 | *Gm35315* | 31.95453869 | 44.40796924 | 16.1038235 | 188.9214291 | 158.518131 | 122.7347801 | 2.25E-10 | 2.323677213 | 3.92E-11 | 43.65561198 | 5.006065657 | 9.648310196 |
| ENSMUSG00000083287 | *Idi1-ps1* | 161.3704204 | 177.631877 | 230.8214701 | 954.7279365 | 832.7630583 | 1035.997514 | 8.74E-44 | 2.321879816 | 2.43E-45 | 199.6986337 | 4.999832682 | 43.05840796 |
| ENSMUSG00000049038 | *Mterf2* | 7.988634673 | 3.027816084 | 1.341985292 | 24.45857788 | 24.97203434 | 16.42906505 | 0.00353695 | 2.321157263 | 0.001366017 | 10.25111869 | 4.997329215 | 2.451371127 |
| ENSMUSG00000048620 | *Or6z3* | 18.37385975 | 16.14835245 | 4.025955875 | 70.84553593 | 60.80147492 | 68.61550697 | 1.18E-06 | 2.318204314 | 2.91E-07 | 26.30579907 | 4.987110998 | 5.927028546 |
| ENSMUSG00000044468 | *Tent5c* | 210.1010919 | 253.3272791 | 289.868823 | 1215.338301 | 1232.315608 | 1273.735749 | 5.04E-67 | 2.316476441 | 6.16E-69 | 307.9329901 | 4.981141661 | 66.29744072 |
| ENSMUSG00000040264 | *Gbp2b* | 24.76476749 | 31.28743287 | 25.49772054 | 165.3062505 | 127.031653 | 111.137793 | 1.07E-13 | 2.30989045 | 1.44E-14 | 59.1748834 | 4.958454269 | 12.97090383 |
| ENSMUSG00000112105 | *Rps7-ps2* | 27.16135789 | 5.046360141 | 5.367941166 | 59.88134584 | 69.48739991 | 62.81701342 | 0.000404243 | 2.297801067 | 0.000135535 | 14.56321145 | 4.917077408 | 3.393357487 |
| ENSMUSG00000112038 | *Gm47056* | 11.98295201 | 8.074176225 | 6.709926458 | 40.48316339 | 51.02980931 | 43.4887016 | 2.61E-06 | 2.292613805 | 6.72E-07 | 24.69505564 | 4.899429625 | 5.582586572 |
| ENSMUSG00000036499 | *Eea1* | 1611.307614 | 1339.303981 | 1274.886027 | 7141.904741 | 6881.424072 | 6709.823449 | 3.33E-105 | 2.291985494 | 1.51E-107 | 485.2846567 | 4.897296329 | 104.4776476 |
| ENSMUSG00000073130 | *Gm1141* | 2.396590402 | 3.027816084 | 12.07786762 | 32.04917101 | 19.54333122 | 28.02605214 | 0.003686811 | 2.284228945 | 0.001429771 | 10.16699556 | 4.871037012 | 2.433349114 |
| ENSMUSG00000033952 | *Aspm* | 359.4885603 | 267.4570875 | 217.4016172 | 1426.18811 | 1385.405036 | 1304.661048 | 1.10E-38 | 2.274933797 | 3.91E-40 | 175.8464538 | 4.839754262 | 37.96037401 |
| ENSMUSG00000090641 | *Zfp712* | 21.56931362 | 14.12980839 | 10.73588233 | 69.15873745 | 83.60202801 | 78.27966288 | 5.36E-09 | 2.270310538 | 1.05E-09 | 37.2248669 | 4.824268947 | 8.270892003 |
| ENSMUSG00000078153 | *Psme2b* | 31.95453869 | 36.33379301 | 16.1038235 | 131.570281 | 119.4314686 | 163.3242349 | 1.22E-11 | 2.266241804 | 1.92E-12 | 49.56216794 | 4.810683226 | 10.91410285 |
| ENSMUSG00000027670 | *Ocstamp* | 21.56931362 | 22.20398462 | 14.76183821 | 95.30411381 | 110.7455436 | 80.21249406 | 5.52E-11 | 2.265677558 | 9.15E-12 | 46.50158792 | 4.808802109 | 10.25775015 |
| ENSMUSG00000079065 | *None* | 59.11589658 | 57.5285056 | 40.25955875 | 262.2971628 | 258.4062684 | 242.5703134 | 3.82E-26 | 2.26014063 | 2.46E-27 | 117.3031446 | 4.79038175 | 25.41799974 |
| ENSMUSG00000021701 | *Plk2* | 438.5760435 | 456.1909567 | 407.9635286 | 2154.041652 | 2003.19145 | 2077.793521 | 1.83E-132 | 2.255909785 | 4.96E-135 | 611.6245004 | 4.776354063 | 131.7383242 |
| ENSMUSG00000053178 | *Mterf1b* | 15.17840588 | 19.17616853 | 32.207647 | 107.9551024 | 97.71665612 | 101.4736371 | 1.89E-10 | 2.253368491 | 3.26E-11 | 44.01138043 | 4.767947965 | 9.724005531 |
| ENSMUSG00000032841 | *Prr5l* | 3.195453869 | 5.046360141 | 9.393897041 | 39.63976415 | 27.14351559 | 13.52981828 | 0.003558048 | 2.249459428 | 0.001375216 | 10.23873906 | 4.755046429 | 2.448788247 |
| ENSMUSG00000018930 | *Ccl4* | 12616.45074 | 11213.01223 | 9489.177996 | 53676.45766 | 52000.46142 | 52593.30288 | 6.24E-93 | 2.247566181 | 3.72E-95 | 428.3488945 | 4.74881048 | 92.20505284 |

| | | | | | | | | | | | | | |
|---|---|---|---|---|---|---|---|---|---|---|---|---|---|
| ENSMUSG00000090222 | *Ifi203-ps* | 49.52953497 | 60.55632169 | 52.33742637 | 266.514159 | 255.1490465 | 243.536729 | 1.33E-28 | 2.239249875 | 7.53E-30 | 128.7922906 | 4.72151507 | 27.8764597 |
| ENSMUSG00000108314 | *Prkcz2* | 7.988634673 | 11.10199231 | 9.393897041 | 37.10956644 | 42.34388432 | 54.1192731 | 4.92E-06 | 2.236876672 | 1.30E-06 | 23.4162977 | 4.71375466 | 5.307724743 |
| ENSMUSG00000015882 | *Lcorl* | 214.8942727 | 145.3351721 | 123.4626468 | 753.1555188 | 760.0184365 | 762.5019014 | 4.88E-29 | 2.218779733 | 2.71E-30 | 130.8193204 | 4.65499537 | 28.31128268 |
| ENSMUSG00000070327 | *Rnf213* | 10266.19442 | 9187.403272 | 4935.821902 | 37973.20726 | 39157.23559 | 35596.95188 | 2.76E-12 | 2.208330881 | 4.14E-13 | 52.57340399 | 4.621402932 | 11.55833087 |
| ENSMUSG00000062488 | *Ifit3b* | 66.30566778 | 43.39869721 | 44.28551462 | 259.7669651 | 213.8909028 | 245.4695601 | 1.33E-19 | 2.203951045 | 1.22E-20 | 86.76411786 | 4.607394223 | 18.87763068 |
| ENSMUSG00000061755 | *Bod1l* | 520.0601172 | 482.4320294 | 299.26272 | 2092.473508 | 2032.506447 | 1873.879831 | 6.65E-32 | 2.197527668 | 3.26E-33 | 144.1697021 | 4.586926113 | 31.17709691 |
| ENSMUSG00000034349 | *Smc4* | 1313.33154 | 942.6600743 | 625.3651458 | 4557.729478 | 4426.564522 | 4233.866705 | 1.19E-23 | 2.194574886 | 8.67E-25 | 105.678703 | 4.577547593 | 22.92596444 |
| ENSMUSG00000084408 | *Gm11870* | 3.994317336 | 13.12053637 | 9.393897041 | 39.63976415 | 39.08666245 | 39.62303924 | 9.72E-05 | 2.192699655 | 3.01E-05 | 17.41164607 | 4.571601511 | 4.012362953 |
| ENSMUSG00000081137 | *BC022960* | 14.37954241 | 6.055632169 | 10.73588233 | 59.03794661 | 42.34388432 | 42.52228601 | 1.40E-05 | 2.191975242 | 3.88E-06 | 21.32389924 | 4.569306573 | 4.855139138 |
| ENSMUSG00000055027 | *Smyd1* | 29.55794829 | 25.2318007 | 13.41985292 | 105.4249047 | 103.1453592 | 112.1042086 | 7.29E-11 | 2.191790615 | 1.22E-11 | 45.93511414 | 4.568721858 | 10.13741178 |
| ENSMUSG00000056260 | *Lrif1* | 353.0976525 | 310.8557847 | 226.7955143 | 1402.572932 | 1357.175779 | 1335.586347 | 1.83E-44 | 2.190319468 | 4.90E-46 | 202.8861819 | 4.56406541 | 43.73705789 |
| ENSMUSG00000045932 | *Ifit2* | 782.0873345 | 604.5539449 | 607.9193371 | 3100.335596 | 3066.131521 | 2908.910929 | 8.03E-73 | 2.18126425 | 7.96E-75 | 334.9661115 | 4.535508319 | 72.09510764 |
| ENSMUSG00000020689 | *Itgb3* | 27.16135789 | 29.26888882 | 18.78779408 | 130.7268818 | 92.287953 | 120.8019489 | 1.99E-11 | 2.175348088 | 3.18E-12 | 48.57404232 | 4.51694732 | 10.70145631 |
| ENSMUSG00000110277 | *Gm45871* | 95.06475261 | 68.63049791 | 93.93897041 | 382.9032537 | 421.2673619 | 358.5401843 | 1.55E-30 | 2.173931838 | 8.01E-32 | 137.813407 | 4.512515346 | 29.81058652 |
| ENSMUSG00000044906 | *4930503L19Rik* | 201.3135938 | 130.1960916 | 140.9084556 | 708.4553593 | 741.5608459 | 697.7520568 | 1.71E-33 | 2.172569232 | 7.80E-35 | 151.5871971 | 4.508255348 | 32.76711556 |
| ENSMUSG00000042677 | *Zc3h12a* | 627.9066853 | 589.4148644 | 458.9589697 | 2495.618343 | 2558.004909 | 2518.47903 | 2.59E-69 | 2.168687268 | 2.87E-71 | 318.6355633 | 4.496140963 | 68.58703485 |
| ENSMUSG00000052727 | *Map1b* | 52.72498884 | 47.43578532 | 29.52367641 | 205.7894139 | 191.0903497 | 198.1151962 | 6.57E-18 | 2.168514529 | 6.74E-19 | 78.83988656 | 4.495602658 | 17.18217448 |
| ENSMUSG00000047534 | *Mis18bp1* | 317.1487965 | 251.308735 | 221.4275731 | 1180.758932 | 1180.200058 | 1206.066658 | 1.10E-51 | 2.163675228 | 2.21E-53 | 236.5662726 | 4.480548106 | 50.95775032 |
| ENSMUSG00000073555 | *Gm4951* | 6.390907738 | 2.018544056 | 2.683970583 | 16.02458551 | 18.4575906 | 17.39548064 | 0.007769335 | 2.16224369 | 0.003184362 | 8.698705013 | 4.476104414 | 2.109616153 |
| ENSMUSG00000095307 | *Gm12573* | 7.988634673 | 12.11126434 | 4.025955875 | 22.77177941 | 48.85832806 | 38.65662365 | 0.000540648 | 2.157198158 | 0.000184915 | 13.97847768 | 4.460477491 | 3.267085477 |
| ENSMUSG00000033943 | *Mga* | 842.0020945 | 766.0374694 | 613.2872782 | 3467.214264 | 3280.022424 | 3194.969944 | 1.68E-67 | 2.157100366 | 2.02E-69 | 310.1576859 | 4.460175152 | 66.77360618 |
| ENSMUSG00000037475 | *Thoc2* | 784.4839249 | 589.4148644 | 523.3742637 | 2866.714007 | 2870.698209 | 2729.157629 | 9.94E-48 | 2.152391793 | 2.33E-49 | 218.1160109 | 4.445642054 | 47.00243705 |
| ENSMUSG00000016356 | *Col20a1* | 23.16704055 | 30.27816084 | 29.52367641 | 122.2928894 | 125.9459123 | 115.9698709 | 3.27E-14 | 2.148575339 | 4.27E-15 | 61.57237554 | 4.433897253 | 13.48558117 |
| ENSMUSG00000067942 | *Zfp160* | 79.88634673 | 66.61195386 | 34.89161758 | 292.6595353 | 266.0064528 | 260.9322096 | 6.03E-15 | 2.14838927 | 7.47E-16 | 65.0040071 | 4.433325437 | 14.22002927 |
| ENSMUSG00000022346 | *Myc* | 141.3988337 | 141.2980839 | 85.88705866 | 542.3057095 | 541.7845712 | 568.2523676 | 5.43E-29 | 2.146950867 | 3.03E-30 | 130.5997546 | 4.428907503 | 28.26508738 |
| ENSMUSG00000038418 | *Egr1* | 435.3805897 | 378.4770106 | 193.245882 | 1513.058232 | 1470.092804 | 1488.28001 | 1.63E-18 | 2.143176863 | 1.60E-19 | 81.67523623 | 4.417336886 | 17.78748002 |
| ENSMUSG00000054203 | *Ifi205* | 12.78181548 | 8.074176225 | 13.41985292 | 49.760555 | 44.51536557 | 56.05210429 | 2.07E-06 | 2.139472094 | 5.26E-07 | 25.16747276 | 4.406007936 | 5.684795496 |
| ENSMUSG00000061331 | *Gm17132* | 15.17840588 | 20.18544056 | 16.1038235 | 85.18332296 | 99.88813737 | 41.55587042 | 5.79E-06 | 2.138774818 | 1.55E-06 | 23.09010834 | 4.403878963 | 5.237630241 |
| ENSMUSG00000102049 | *None* | 39.1443099 | 26.24107273 | 55.02139695 | 120.6060909 | 207.3764591 | 197.1487806 | 1.47E-09 | 2.138675382 | 2.74E-10 | 39.85184511 | 4.40357544 | 8.834057889 |
| ENSMUSG00000034640 | *Tiparp* | 687.0225819 | 632.8135616 | 786.4033808 | 3053.948638 | 3156.247093 | 3032.612125 | 3.35E-102 | 2.137634589 | 1.60E-104 | 471.3816983 | 4.400399748 | 101.4743671 |
| ENSMUSG00000113684 | *Gm48418* | 15.17840588 | 10.09272028 | 6.709926458 | 50.60395423 | 43.42962494 | 51.22002633 | 6.52E-06 | 2.136098481 | 1.75E-06 | 22.84747904 | 4.39571692 | 5.185684977 |
| ENSMUSG00000113069 | *Gm48541* | 15.97726935 | 19.17616853 | 9.393897041 | 52.29075271 | 77.08758427 | 70.54833815 | 4.65E-07 | 2.134956526 | 1.10E-07 | 28.19148401 | 4.392238899 | 6.332314867 |
| ENSMUSG00000033730 | *Egr3* | 63.11021392 | 52.48214546 | 12.07786762 | 199.02422 | 165.0325748 | 206.8129365 | 2.82E-06 | 2.134325583 | 7.26E-07 | 24.54610654 | 4.390318432 | 5.550214656 |
| ENSMUSG00000044716 | *Dok7* | 31.95453869 | 23.21325665 | 25.49772054 | 102.0513077 | 122.6886905 | 132.398936 | 1.96E-12 | 2.130881681 | 2.91E-13 | 53.2713177 | 4.379850666 | 11.70765835 |
| ENSMUSG00000032691 | *Nlrp3* | 2191.282491 | 2146.721604 | 1813.022129 | 9075.819192 | 8996.446807 | 8781.818477 | 1.53E-119 | 2.124097336 | 5.13E-122 | 551.7920406 | 4.359302554 | 118.8167093 |
| ENSMUSG00000061175 | *Fnip2* | 2433.338121 | 2628.144361 | 2447.781172 | 10882.38036 | 11199.41453 | 10562.92241 | 5.76E-211 | 2.11999256 | 1.10E-214 | 974.9129799 | 4.346917032 | 210.2397263 |
| ENSMUSG00000034163 | *Zfc3h1* | 421.0010473 | 345.1710336 | 185.1939702 | 1424.501312 | 1404.948367 | 1313.358788 | 6.06E-18 | 2.114681471 | 6.20E-19 | 79.00414568 | 4.330943859 | 17.2175329 |
| ENSMUSG00000066406 | *Akap13* | 1801.437119 | 1805.587658 | 1111.163821 | 6794.424255 | 7031.256278 | 6567.760357 | 2.00E-34 | 2.109767366 | 8.70E-36 | 155.9468286 | 4.3162169 | 33.69916493 |
| ENSMUSG00000113486 | *Gm48676* | 14.37954241 | 21.19471259 | 9.393897041 | 65.7851405 | 74.91610303 | 56.05210429 | 5.96E-07 | 2.106161209 | 1.42E-07 | 27.6900023 | 4.305441568 | 6.224939015 |
| ENSMUSG00000029385 | *Ccng2* | 93.46702567 | 55.50996155 | 53.67941166 | 320.4910701 | 281.2068215 | 278.3276902 | 7.77E-17 | 2.097301631 | 8.52E-18 | 73.82883099 | 4.279082907 | 16.10936713 |
| ENSMUSG00000098188 | *Sowahc* | 207.7045015 | 198.8265895 | 157.0122791 | 734.6007356 | 877.2784238 | 812.7555121 | 4.15E-44 | 2.093347703 | 1.12E-45 | 201.2359411 | 4.267371476 | 43.38172361 |
| ENSMUSG00000079415 | *Cntf* | 16.77613281 | 11.10199231 | 6.709926458 | 59.03794661 | 57.54425305 | 35.75737687 | 4.19E-05 | 2.093269293 | 1.24E-05 | 19.10577279 | 4.267139552 | 4.37794931 |
| ENSMUSG00000075081 | *Or4a74* | 9.586361607 | 7.064904197 | 2.683970583 | 26.98877559 | 38.00092182 | 21.261143 | 0.001594257 | 2.09273719 | 0.000583921 | 11.82654921 | 4.265566012 | 2.797441656 |
| ENSMUSG00000028248 | *Pnisr* | 409.0180953 | 334.0690413 | 303.2886759 | 1623.543532 | 1469.007064 | 1388.739204 | 1.79E-51 | 2.091045959 | 3.63E-53 | 235.5735963 | 4.260568538 | 50.7463823 |
| ENSMUSG00000001627 | *Ifrd1* | 3189.861825 | 2610.986737 | 2501.460583 | 11881.80845 | 11654.33985 | 11872.41554 | 2.67E-88 | 2.09101278 | 1.84E-90 | 406.7761049 | 4.260470554 | 87.5732776 |
| ENSMUSG00000008307 | *1700109H08Rik* | 31.95453869 | 19.17616853 | 21.47176466 | 113.0154978 | 100.973878 | 99.54080589 | 1.32E-10 | 2.087465983 | 2.26E-11 | 44.73538157 | 4.250009256 | 9.880299377 |
| ENSMUSG00000112226 | *Gm48786* | 30.35681176 | 27.25034476 | 6.709926458 | 105.4249047 | 93.37369363 | 82.14532525 | 6.31E-06 | 2.084516002 | 1.69E-06 | 22.91349057 | 4.241327839 | 5.199646033 |
| ENSMUSG00000027646 | *Src* | 41.5409003 | 67.62122589 | 41.60154404 | 213.380007 | 195.4333122 | 230.9733263 | 6.14E-16 | 2.08384903 | 7.14E-17 | 69.63258017 | 4.239367484 | 15.21190726 |
| ENSMUSG00000026193 | *Fn1* | 6.390907738 | 7.064904197 | 2.683970583 | 26.14537635 | 13.02888748 | 30.92529892 | 0.004967107 | 2.083212129 | 0.001967724 | 9.579403004 | 4.23749636 | 2.303896448 |
| ENSMUSG00000060477 | *Irak2* | 794.0702865 | 835.6772393 | 864.2385277 | 3490.829443 | 3548.200358 | 3459.767816 | 2.19E-168 | 2.076614438 | 3.36E-171 | 777.948429 | 4.218161809 | 167.6594042 |
| ENSMUSG00000111777 | *Gm47147* | 12.78181548 | 14.12980839 | 22.81374996 | 52.29075271 | 70.57314053 | 82.14532525 | 1.09E-06 | 2.076269451 | 2.66E-07 | 26.48149778 | 4.217153253 | 5.964029091 |
| ENSMUSG00000020948 | *Klhl28* | 170.956782 | 196.8080455 | 139.5664703 | 754.8423173 | 706.8171459 | 688.0879009 | 2.11E-43 | 2.074726622 | 5.93E-45 | 197.9225126 | 4.212645808 | 42.67630473 |
| ENSMUSG00000080059 | *Rps19-ps3* | 7.988634673 | 2.018544056 | 8.051911749 | 23.61517864 | 26.05777497 | 26.09322096 | 0.002398665 | 2.07251805 | 0.00090049 | 11.02174964 | 4.20620175 | 2.620030386 |
| ENSMUSG00000047193 | *Dync2h1* | 250.8431287 | 252.318007 | 194.5878673 | 1023.043275 | 971.7378581 | 958.6842664 | 1.12E-56 | 2.071528443 | 1.87E-58 | 259.8286293 | 4.203317523 | 55.9524315 |
| ENSMUSG00000054304 | *D130007C19Rik* | 18.37385975 | 11.10199231 | 6.709926458 | 58.19454737 | 56.45851243 | 42.52228601 | 2.39E-05 | 2.067351694 | 6.85E-06 | 20.23379946 | 4.191166088 | 4.621105069 |
| ENSMUSG00000069053 | *Uba1y* | 9.586361607 | 7.064904197 | 2.683970583 | 28.67557407 | 24.97203434 | 30.92529892 | 0.00102303 | 2.065236971 | 0.000364436 | 12.70624529 | 4.185027118 | 2.990111519 |
| ENSMUSG00000002617 | *Zfp40* | 51.12726191 | 23.21325665 | 6.709926458 | 131.570281 | 100.973878 | 115.0034553 | 0.000224225 | 2.064023194 | 7.28E-05 | 15.73570575 | 4.181507626 | 3.649315005 |
| ENSMUSG00000035164 | *Zc3h12c* | 631.1021392 | 589.4148644 | 587.7895577 | 2545.378898 | 2593.83435 | 2431.501627 | 2.13E-133 | 2.063033215 | 5.65E-136 | 615.9622906 | 4.178639253 | 132.6722477 |
| ENSMUSG00000020300 | *Cpeb4* | 389.0465086 | 367.3750182 | 305.9726465 | 1441.369296 | 1533.06576 | 1436.093568 | 1.53E-69 | 2.044863267 | 1.68E-71 | 319.7086307 | 4.126341619 | 68.81612912 |
| ENSMUSG00000042595 | *Fam199x* | 135.007926 | 107.992107 | 116.7527204 | 545.6793065 | 459.2682838 | 482.24138 | 1.94E-35 | 2.042066451 | 8.04E-37 | 160.679317 | 4.118350022 | 34.7120935 |
| ENSMUSG00000056290 | *Ms4a4b* | 7.189771206 | 2.018544056 | 5.367941166 | 18.55478322 | 23.88629372 | 18.36189623 | 0.006444265 | 2.037966783 | 0.002601438 | 9.067842414 | 4.106663628 | 2.190826618 |
| ENSMUSG00000031098 | *Syt8* | 7.189771206 | 13.12053637 | 4.025955875 | 32.04917101 | 36.9151812 | 32.8581301 | 0.000432308 | 2.036440136 | 0.000145455 | 14.43014852 | 4.102320292 | 3.364206577 |
| ENSMUSG00000040943 | *Tet2* | 243.6533575 | 272.5034476 | 203.9817643 | 972.4393205 | 999.9671143 | 995.4080589 | 8.27E-57 | 2.035257435 | 1.37E-58 | 260.445472 | 4.098958645 | 56.08227618 |
| ENSMUSG00000003882 | *Il7r* | 337.1203582 | 361.3193861 | 479.0887491 | 1648.845509 | 1495.064839 | 1653.537076 | 1.49E-44 | 2.035154285 | 3.95E-46 | 203.3121654 | 4.098665589 | 43.82615824 |
| ENSMUSG00000051413 | *Plagl2* | 1286.969046 | 1365.545054 | 778.3514691 | 4610.86363 | 4932.519653 | 4517.992889 | 1.25E-27 | 2.032616315 | 7.37E-29 | 124.2659406 | 4.091461609 | 26.90278396 |
| ENSMUSG00000060509 | *Xcr1* | 9.586361607 | 8.074176225 | 4.025955875 | 43.85676034 | 30.40073746 | 16.42906505 | 0.003058587 | 2.028220806 | 0.001168631 | 10.53929257 | 4.079014981 | 2.514479192 |
| ENSMUSG00000020696 | *Rffl* | 309.1601618 | 296.7259763 | 264.3711024 | 1154.613556 | 1293.117083 | 1111.37793 | 3.84E-63 | 2.025181991 | 5.05E-65 | 289.9691061 | 4.070432207 | 62.4160011 |
| ENSMUSG00000033031 | *Cip2a* | 643.0850912 | 534.9141749 | 465.6688962 | 2305.010115 | 2217.082353 | 2186.998483 | 2.67E-57 | 2.022711025 | 4.32E-59 | 262.7474617 | 4.06346657 | 56.57308765 |
| ENSMUSG00000048621 | *Gm6377* | 9.586361607 | 7.064904197 | 6.709926458 | 41.32656262 | 21.71481247 | 32.8581301 | 0.000608736 | 2.022499044 | 0.000209828 | 13.74095874 | 4.062869551 | 3.215570992 |
| ENSMUSG00000025907 | *Rb1cc1* | 402.6271875 | 365.3564742 | 269.7390436 | 1425.344711 | 1411.462811 | 1385.839958 | 1.25E-42 | 2.015352725 | 3.67E-44 | 194.295829 | 4.042794089 | 41.90351544 |
| ENSMUSG00000039219 | *Arid4b* | 238.0613133 | 208.9193098 | 136.8824997 | 826.5312525 | 783.9047281 | 770.2332261 | 3.65E-26 | 2.014355903 | 2.35E-27 | 117.400793 | 4.040001709 | 25.43739029 |
| ENSMUSG00000074220 | *Zfp382* | 95.06475261 | 80.74176225 | 49.65345579 | 333.9860979 | 312.6932996 | 276.3948591 | 1.77E-16 | 2.01090284 | 1.99E-17 | 72.15440241 | 4.030343603 | 15.75179518 |
| ENSMUSG00000031504 | *Rab20* | 331.5283389 | 348.1988497 | 410.6474992 | 1416.067319 | 1501.579282 | 1448.656971 | 5.49E-72 | 2.009603194 | 5.57E-74 | 331.0880233 | 4.026714519 | 71.2606969 |
| ENSMUSG00000054051 | *Ercc6* | 279.6022136 | 311.8650567 | 221.4275731 | 1170.638141 | 1041.225258 | 1076.586969 | 4.58E-45 | 2.009295981 | 1.20E-46 | 205.6845864 | 4.025857148 | 44.33895466 |
| ENSMUSG00000026872 | *Zeb2* | 1134.386124 | 1284.803292 | 1394.322718 | 5073.889811 | 5122.524262 | 5019.56258 | 5.18E-101 | 1.999745144 | 2.54E-103 | 465.8692448 | 3.999293451 | 100.2854025 |
| ENSMUSG00000053541 | *Gvin-ps6* | 6.390907738 | 10.09272028 | 1.341985292 | 27.83217483 | 19.54333122 | 26.09322096 | 0.00606038 | 1.996316879 | 0.002436456 | 9.187709646 | 3.989801243 | 2.217500172 |
| ENSMUSG00000114216 | *Gm48592* | 18.37385975 | 14.12980839 | 9.393897041 | 59.03794661 | 58.62999367 | 54.1192731 | 1.81E-06 | 1.99622868 | 4.57E-07 | 25.43864453 | 3.989557335 | 5.741903746 |
| ENSMUSG00000000982 | *Ccl3* | 14191.01063 | 15066.41284 | 14531.01674 | 60618.47678 | 56774.46295 | 57292.9819 | 9.20E-228 | 1.996224312 | 5.43E-231 | 1052.99851 | 3.989545256 | 227.0363906 |
| ENSMUSG00000035173 | *Ccdc186* | 358.6896968 | 268.4663595 | 318.0505141 | 1335.944392 | 1315.917636 | 1291.941332 | 3.03E-44 | 1.995296336 | 8.16E-46 | 201.8695824 | 3.986979911 | 43.51809249 |
| ENSMUSG00000041235 | *Chd7* | 904.313445 | 861.918312 | 594.4994841 | 3068.286425 | 3217.049468 | 3155.346905 | 1.26E-39 | 1.995201373 | 4.25E-41 | 180.2601379 | 3.986717484 | 38.90127407 |
| ENSMUSG00000056919 | *Cep162* | 250.0442653 | 311.8650567 | 309.9986023 | 1200.157115 | 1149.79932 | 1093.982449 | 2.78E-56 | 1.989275724 | 4.73E-58 | 257.9786961 | 3.970376235 | 55.55525793 |
| ENSMUSG00000100131 | *Gm28439* | 3127.550474 | 2656.403978 | 3122.799773 | 11711.44181 | 11195.07157 | 12446.4664 | 4.52E-101 | 1.989051818 | 2.19E-103 | 466.1670601 | 3.969760081 | 100.3449463 |
| ENSMUSG00000017418 | *Arl5b* | 355.4942429 | 399.6717231 | 472.3788226 | 1620.169935 | 1602.55316 | 1611.01479 | 4.11E-61 | 1.986032598 | 5.82E-63 | 280.5066693 | 3.961460996 | 60.38601083 |
| ENSMUSG00000074580 | *4931440P22Rik* | 34.35112909 | 32.2967049 | 13.41985292 | 107.9551024 | 92.287953 | 124.6676113 | 1.05E-07 | 1.98573751 | 2.33E-08 | 31.19603348 | 3.960650804 | 6.979287876 |
| ENSMUSG00000020826 | *Nos2* | 451.357859 | 567.2108798 | 393.2016904 | 1870.659508 | 1871.816835 | 1858.417182 | 1.04E-47 | 1.984803288 | 2.43E-49 | 218.0284071 | 3.958086901 | 46.98442023 |
| ENSMUSG00000030103 | *Bhlhe40* | 772.5009729 | 784.2043659 | 720.6461015 | 2998.284288 | 3027.044858 | 3005.552488 | 2.77E-149 | 1.984715554 | 5.23E-152 | 689.6874665 | 3.957846207 | 148.557878 |
| ENSMUSG00000044098 | *Rsbn1* | 952.245253 | 903.2984652 | 762.2476456 | 3467.214264 | 3445.054999 | 3445.271582 | 5.88E-88 | 1.979602175 | 4.17E-90 | 405.1502602 | 3.943843149 | 87.23035988 |
| ENSMUSG00000042133 | *Ppig* | 696.6089435 | 659.0546344 | 618.6552194 | 2628.875422 | 2666.578971 | 2499.150719 | 3.92E-115 | 1.977012262 | 1.41E-117 | 531.3832249 | 3.936769547 | 114.4062924 |
| ENSMUSG00000031779 | *Ccl22* | 7.988634673 | 13.12053637 | 4.025955875 | 32.04917101 | 28.22925621 | 40.58945483 | 0.000806523 | 1.975538244 | 0.000283264 | 13.17794253 | 3.932749359 | 3.093383254 |
| ENSMUSG00000096910 | *Zfp955b* | 67.90339472 | 50.46360141 | 38.91757345 | 242.8989803 | 199.7762747 | 185.5517935 | 1.21E-13 | 1.975349435 | 1.63E-14 | 58.9324366 | 3.932234705 | 12.91891077 |
| ENSMUSG00000074867 | *Zfp808* | 16.77613281 | 25.2318007 | 6.709926458 | 63.25494279 | 60.80147492 | 71.51475374 | 2.14E-05 | 1.97250462 | 6.09E-06 | 20.46056953 | 3.924488469 | 4.669610249 |
| ENSMUSG00000110444 | *Gm10033* | 108.6454316 | 48.44505735 | 45.62749991 | 278.3217483 | 268.177934 | 259.965794 | 3.05E-09 | 1.97047177 | 5.87E-10 | 38.3641176 | 3.918962506 | 8.51501642 |
| ENSMUSG00000078234 | *Klhdc7a* | 23.96590402 | 14.12980839 | 18.78779408 | 88.55691991 | 67.31591866 | 68.61550697 | 2.37E-07 | 1.969458248 | 5.45E-08 | 29.54836879 | 3.916210323 | 6.625456552 |
| ENSMUSG00000001156 | *Mxd1* | 103.8522507 | 114.0477392 | 106.016838 | 413.2656262 | 433.2105088 | 419.4243665 | 5.38E-39 | 1.966807093 | 1.88E-40 | 177.3049003 | 3.909020344 | 38.26941178 |
| ENSMUSG00000090894 | *Or5v1* | 8.78749814 | 7.064904197 | 9.393897041 | 41.32656262 | 31.48647808 | 25.12680537 | 0.000447844 | 1.965000259 | 0.000151052 | 14.35904623 | 3.904127743 | 3.348873278 |
| ENSMUSG00000039982 | *Dtx4* | 1679.211008 | 1718.790264 | 1388.954777 | 6084.282097 | 6294.038395 | 6319.39155 | 3.16E-89 | 1.963205972 | 2.09E-91 | 411.1202304 | 3.899275178 | 88.49992019 |
| ENSMUSG00000033740 | *St18* | 103.0533873 | 104.9642909 | 122.1206615 | 421.6996186 | 435.38199 | 419.4243665 | 2.77E-37 | 1.961158032 | 1.05E-38 | 169.3025667 | 3.893743991 | 36.55770037 |
| ENSMUSG00000022322 | *Shcbp1* | 341.913564 | 274.5219917 | 257.661176 | 1191.723122 | 1079.22618 | 1137.471151 | 2.06E-47 | 1.954698604 | 4.89E-49 | 216.638523 | 3.876349369 | 46.68665652 |
| ENSMUSG00000027959 | *Sass6* | 798.8634673 | 748.8798449 | 687.0964693 | 2959.487923 | 2817.496918 | 2865.422228 | 1.08E-108 | 1.946965497 | 4.52E-111 | 501.4856619 | 3.855627019 | 107.9675212 |
| ENSMUSG00000040865 | *Ino80d* | 545.6237482 | 561.1552476 | 351.6001464 | 1755.113813 | 2072.67885 | 1811.062818 | 2.33E-28 | 1.945323066 | 1.33E-29 | 127.6588612 | 3.851240094 | 27.63205215 |
| ENSMUSG00000007033 | *Hspa1l* | 18.37385975 | 12.11126434 | 16.1038235 | 61.56814432 | 59.7157343 | 58.95135106 | 7.01E-07 | 1.944143888 | 1.69E-07 | 27.36260759 | 3.848093592 | 6.15409646 |
| ENSMUSG00000040407 | *Akap9* | 742.1441611 | 619.6930253 | 479.0887491 | 2409.591621 | 2420.11585 | 2275.908717 | 1.65E-37 | 1.942764902 | 6.20E-39 | 170.3517824 | 3.844417185 | 36.78142362 |
| ENSMUSG00000037820 | *Tgm2* | 107.8465681 | 113.0384672 | 112.7267645 | 409.8920293 | 409.3242151 | 458.0809902 | 3.83E-36 | 1.939248537 | 1.53E-37 | 163.9750336 | 3.835058374 | 35.41635698 |
| ENSMUSG00000073929 | *Trim30e-ps1* | 15.17840588 | 6.055632169 | 8.051911749 | 43.85676034 | 38.00092182 | 32.8581301 | 0.000434787 | 1.933336991 | 0.000146443 | 14.41740127 | 3.819376099 | 3.361723795 |
| ENSMUSG00000062515 | *Fabp4* | 127.0192913 | 114.0477392 | 119.4366909 | 494.231953 | 431.0390276 | 453.2489122 | 9.53E-37 | 1.933251808 | 3.67E-38 | 166.8177517 | 3.819150592 | 36.02099276 |
| ENSMUSG00000052593 | *Adam17* | 1729.539407 | 1647.13195 | 1418.478453 | 6417.424796 | 5820.655483 | 6099.048796 | 1.21E-87 | 1.932766814 | 8.68E-90 | 403.6852885 | 3.817919844 | 86.91864115 |
| ENSMUSG00000035107 | *Dcbld2* | 662.2578144 | 619.6930253 | 512.6383814 | 2290.672328 | 2372.343262 | 2210.192457 | 6.39E-66 | 1.931187748 | 8.03E-68 | 302.8132391 | 3.813690452 | 65.19449298 |
| ENSMUSG00000035133 | *Arhgap5* | 740.5464342 | 580.3314162 | 323.4184553 | 2148.981256 | 2118.279957 | 2022.707832 | 1.38E-15 | 1.930760482 | 1.64E-16 | 67.99524326 | 3.812561163 | 14.85984255 |
| ENSMUSG00000054519 | *Zfp867* | 44.73635417 | 48.44505735 | 57.70536754 | 193.9818246 | 200.8620154 | 173.9548064 | 2.40E-17 | 1.929700681 | 2.55E-18 | 76.21165575 | 3.809761491 | 16.62000066 |
| ENSMUSG00000096916 | *Zfp850* | 167.7613281 | 121.1126434 | 162.3802203 | 608.09085 | 538.5273493 | 569.2187832 | 8.70E-32 | 1.927362401 | 4.28E-33 | 143.6288999 | 3.803591736 | 31.06042311 |
| ENSMUSG00000039232 | *Stx11* | 149.3874684 | 188.7338693 | 198.6138231 | 692.4307738 | 687.2738147 | 644.5991993 | 5.82E-39 | 1.926663626 | 2.04E-40 | 177.1404706 | 3.801749898 | 38.23497182 |
| ENSMUSG00000030921 | *Trim30a* | 1687.998506 | 1674.382295 | 1697.611394 | 6641.768993 | 6362.440054 | 6216.951498 | 1.46E-187 | 1.925804918 | 1.81E-190 | 866.5744123 | 3.799487729 | 186.8349087 |
| ENSMUSG00000053347 | *Zfp943* | 50.32839844 | 51.47287344 | 26.83970583 | 150.9684635 | 169.3755373 | 176.8540532 | 1.10E-11 | 1.922667338 | 1.72E-12 | 49.78025054 | 3.791233563 | 10.95944174 |
| ENSMUSG00000022124 | *Fbxl3* | 422.5987742 | 465.274405 | 464.3269109 | 1697.762665 | 1653.58297 | 1735.682402 | 1.37E-95 | 1.915426935 | 7.65E-98 | 440.6919431 | 3.772254297 | 94.86486466 |
| ENSMUSG00000026896 | *Ifih1* | 870.7611793 | 916.4190015 | 832.0308807 | 3290.943824 | 3350.595564 | 3246.189971 | 8.40E-139 | 1.914683697 | 2.13E-141 | 640.8970782 | 3.770311432 | 138.0756139 |
| ENSMUSG00000003198 | *Zfp959* | 86.27725447 | 75.69540211 | 83.20308807 | 327.238904 | 282.2925621 | 313.1186515 | 3.19E-26 | 1.911819198 | 2.04E-27 | 117.6738339 | 3.762832833 | 25.49557694 |

| | | | | | | | | | | | | | |
|---|---|---|---|---|---|---|---|---|---|---|---|---|---|
| ENSMUSG00000031770 | *Herpud1* | 1251.02019 | 1039.550189 | 1038.696616 | 4055.063533 | 4260.446207 | 4222.269718 | 3.68E-83 | 1.909856245 | 2.85E-85 | 382.9428479 | 3.757716547 | 82.43399193 |
| ENSMUSG00000039158 | *Akna* | 1263.802005 | 1064.78199 | 523.3742637 | 3544.806994 | 3662.203123 | 3500.357271 | 1.67E-06 | 1.907426332 | 4.19E-07 | 25.60699022 | 3.7513928 | 5.777962685 |
| ENSMUSG00000052688 | *Rab7b* | 682.2294011 | 679.2400749 | 821.2949984 | 2678.635977 | 2813.153956 | 2666.340616 | 7.53E-79 | 1.906791564 | 6.49E-81 | 362.9275276 | 3.749742596 | 78.1232066 |
| ENSMUSG00000003992 | *Ssbp2* | 10.38522507 | 15.13908042 | 1.341985292 | 37.10956644 | 36.9151812 | 29.95888332 | 0.00474715 | 1.904105925 | 0.001873197 | 9.669820029 | 3.742768782 | 2.323567065 |
| ENSMUSG00000035385 | *Ccl2* | 1684.004189 | 1541.158387 | 1707.005291 | 6301.035701 | 6092.090639 | 6038.164613 | 6.06E-148 | 1.902478673 | 1.25E-150 | 683.3435083 | 3.738549599 | 147.2172223 |
| ENSMUSG00000037921 | *Ddx60* | 153.3817857 | 154.4186203 | 118.0947057 | 540.6189111 | 538.5273493 | 524.763666 | 7.32E-37 | 1.900325501 | 2.80E-38 | 167.3516952 | 3.732974106 | 36.13562007 |
| ENSMUSG00000033671 | *Cep350* | 524.853298 | 612.6281211 | 442.8551462 | 2029.218565 | 1981.476638 | 1894.174559 | 1.24E-50 | 1.897782664 | 2.63E-52 | 231.6289715 | 3.726400308 | 49.90598211 |
| ENSMUSG00000056529 | *Ptafr* | 499.2896671 | 638.8691938 | 790.4293367 | 2488.02775 | 2320.227713 | 2305.8676 | 1.33E-32 | 1.888657969 | 6.35E-34 | 147.4215803 | 3.702906109 | 31.87528506 |
| ENSMUSG00000043336 | *Filip1l* | 116.6340662 | 79.73249022 | 63.0733087 | 334.8294972 | 289.8927465 | 348.8760284 | 8.56E-15 | 1.888202831 | 1.07E-15 | 64.28982231 | 3.701738109 | 14.06734234 |
| ENSMUSG00000020838 | *Slc6a4* | 11.18408854 | 21.19471259 | 8.051911749 | 53.13415195 | 47.77258744 | 50.25361074 | 7.69E-05 | 1.887480033 | 2.35E-05 | 17.88502122 | 3.699883983 | 4.114286005 |
| ENSMUSG00000034765 | *Dusp5* | 516.8646633 | 512.7101903 | 453.5910285 | 1825.115949 | 1880.50276 | 1798.499415 | 1.90E-91 | 1.886511607 | 1.18E-93 | 421.4491055 | 3.697401226 | 90.72014076 |
| ENSMUSG00000028312 | *Smc2* | 1488.28264 | 1119.282679 | 1139.345512 | 4761.832093 | 4645.884128 | 4460.974369 | 2.07E-52 | 1.8855141 | 4.01E-54 | 239.9592963 | 3.694845657 | 51.68459919 |
| ENSMUSG00000049658 | *Bdp1* | 853.9850465 | 799.3434463 | 656.2308076 | 2892.859384 | 2923.899499 | 2730.124045 | 1.66E-62 | 1.88283552 | 2.23E-64 | 287.0123784 | 3.687991983 | 61.77946646 |
| ENSMUSG00000046404 | *Yod1* | 148.5886049 | 157.4464364 | 92.59698511 | 504.3527439 | 502.6979087 | 476.4428864 | 2.47E-21 | 1.881049883 | 2.06E-22 | 94.84362852 | 3.683430145 | 20.6071867 |
| ENSMUSG00000031229 | *Atrx* | 903.5145815 | 649.9711861 | 449.5650727 | 2539.475103 | 2485.260287 | 2374.483107 | 6.22E-19 | 1.880727545 | 5.97E-20 | 83.62894882 | 3.682607256 | 18.20618468 |
| ENSMUSG00000058402 | *Zfp420* | 55.92044271 | 65.60268183 | 69.78323516 | 243.7423796 | 217.1481247 | 236.7718198 | 3.18E-20 | 1.880461421 | 2.84E-21 | 89.65313764 | 3.681928015 | 19.49752599 |
| ENSMUSG00000047648 | *Fbxo30* | 300.3726637 | 309.8465126 | 307.3146318 | 1152.926757 | 1088.997845 | 1131.672657 | 1.24E-78 | 1.880037681 | 1.07E-80 | 361.9219991 | 3.680846739 | 77.90722416 |
| ENSMUSG00000044071 | *Tafa2* | 156.5772396 | 138.2702679 | 152.9863232 | 563.3906905 | 542.8703118 | 540.2263154 | 6.13E-44 | 1.877866353 | 1.67E-45 | 200.4466801 | 3.675311049 | 43.21232815 |
| ENSMUSG00000108218 | *Or2y7* | 198.1181399 | 123.1311874 | 101.9908822 | 541.4623103 | 507.0408712 | 518.9651724 | 2.26E-15 | 1.87387039 | 2.72E-16 | 66.9982468 | 3.665145292 | 14.64623381 |
| ENSMUSG00000062232 | *Rapgef2* | 1000.177061 | 1069.82835 | 916.5759541 | 3739.632218 | 3651.345717 | 3570.905609 | 8.58E-108 | 1.873628753 | 3.70E-110 | 497.2894078 | 3.664531468 | 107.0665633 |
| ENSMUSG00000020205 | *Phlda1* | 524.853298 | 542.9883511 | 292.5527936 | 1621.856733 | 1651.411488 | 1729.883908 | 1.25E-19 | 1.873237464 | 1.15E-20 | 86.87916614 | 3.663537705 | 18.90150304 |
| ENSMUSG00000039115 | *Itga9* | 10.38522507 | 11.10199231 | 9.393897041 | 43.0133611 | 32.57221871 | 37.69020805 | 0.000133874 | 1.872011797 | 4.23E-05 | 16.76733952 | 3.660426603 | 3.873303974 |
| ENSMUSG00000052749 | *Trim30b* | 7.988634673 | 8.074176225 | 8.051911749 | 28.67557407 | 32.57221871 | 27.05963655 | 0.000676405 | 1.871330947 | 0.00023481 | 13.52976409 | 3.658699549 | 3.169793301 |
| ENSMUSG00000057110 | *Cntrl* | 665.4532683 | 529.8678148 | 462.9849256 | 2007.290185 | 2106.33681 | 1977.286299 | 1.35E-41 | 1.870592771 | 4.16E-43 | 189.4628915 | 3.656828 | 40.86825071 |
| ENSMUSG00000030717 | *Nupr1* | 301.1715272 | 239.1974707 | 256.3191907 | 1029.790469 | 953.2802675 | 938.389539 | 8.25E-48 | 1.869242686 | 1.92E-49 | 218.497927 | 3.653407514 | 47.08355215 |
| ENSMUSG00000006724 | *Cyp27b1* | 17.57499628 | 10.09272028 | 8.051911749 | 37.10956644 | 59.7157343 | 37.69020805 | 0.000382895 | 1.868656912 | 0.00012797 | 14.67144304 | 3.65192443 | 3.416920775 |
| ENSMUSG00000113889 | *Gm48501* | 29.55794829 | 12.11126434 | 13.41985292 | 75.06253211 | 76.00184365 | 55.08568869 | 5.29E-05 | 1.867436394 | 1.58E-05 | 18.63443991 | 3.648836215 | 4.276245818 |
| ENSMUSG00000040165 | *Cd209c* | 79.88634673 | 84.77885036 | 71.12522045 | 291.8161361 | 293.1499684 | 278.3276902 | 1.55E-25 | 1.865166544 | 1.03E-26 | 114.4695405 | 3.643099867 | 24.81000086 |
| ENSMUSG00000022676 | *Snai2* | 23.96590402 | 26.24107273 | 30.8656617 | 107.9551024 | 86.85924988 | 96.64155911 | 1.75E-09 | 1.864421479 | 3.29E-10 | 39.49683777 | 3.641218912 | 8.758118959 |
| ENSMUSG00000017291 | *Taok1* | 1084.057725 | 1047.624365 | 810.5591161 | 3636.737511 | 3741.462189 | 3361.193426 | 2.10E-51 | 1.864198603 | 4.25E-53 | 235.2573967 | 3.640656438 | 50.67869551 |
| ENSMUSG00000054072 | *Iigp1* | 28.75908482 | 19.17616853 | 18.78779408 | 89.40031915 | 73.8303624 | 83.11174084 | 9.66E-08 | 1.861618644 | 2.14E-08 | 31.36248164 | 3.634151703 | 7.015137607 |
| ENSMUSG00000040423 | *Rc3h1* | 1472.30537 | 1534.093483 | 1529.863232 | 5600.170935 | 5513.390886 | 5358.774453 | 7.10E-176 | 1.861481213 | 1.01E-178 | 812.5559252 | 3.633805529 | 175.148992 |
| ENSMUSG00000031433 | *Rbm41* | 162.1692839 | 152.4000762 | 112.7267645 | 534.7151164 | 508.1266118 | 525.7300816 | 4.76E-30 | 1.86129617 | 2.52E-31 | 135.5344326 | 3.63333948 | 29.32242049 |
| ENSMUSG00000028381 | *Ugcg* | 627.1078218 | 574.275784 | 558.2658813 | 2261.153355 | 2063.992925 | 2075.86069 | 8.36E-86 | 1.859787945 | 6.22E-88 | 395.1649767 | 3.629543093 | 85.07788033 |
| ENSMUSG00000056267 | *Cep70* | 156.5772396 | 139.2795399 | 122.1206615 | 521.2207286 | 501.6121681 | 506.4017698 | 2.92E-36 | 1.859085204 | 1.16E-37 | 164.5357268 | 3.627775561 | 35.53432858 |
| ENSMUSG00000069184 | *Zfp72* | 45.53521764 | 44.40796924 | 46.9694852 | 151.8118627 | 172.6327591 | 172.0219752 | 1.11E-15 | 1.858972295 | 1.31E-16 | 68.44041512 | 3.627491655 | 14.95528938 |
| ENSMUSG00000038668 | *Lpar1* | 5.592044271 | 25.2318007 | 16.1038235 | 67.47193898 | 47.77258744 | 51.22002633 | 0.001225568 | 1.851223585 | 0.000441505 | 12.34780442 | 3.608060637 | 2.911662553 |
| ENSMUSG00000032253 | *Phip* | 818.835054 | 672.1751707 | 456.2749991 | 2475.376761 | 2318.056231 | 2249.815496 | 4.61E-23 | 1.850001236 | 3.49E-24 | 102.9185525 | 3.605004938 | 22.33645745 |
| ENSMUSG00000069910 | *Spdl1* | 566.3941983 | 563.1737917 | 692.4644104 | 2224.887188 | 2170.395506 | 2142.543366 | 5.60E-68 | 1.849100791 | 6.51E-70 | 312.4129987 | 3.602755609 | 67.25187464 |
| ENSMUSG00000060044 | *Tmem26* | 414.6101395 | 449.1260525 | 318.0505141 | 1491.129851 | 1388.662258 | 1352.981828 | 6.11E-40 | 1.834506668 | 2.05E-41 | 181.7161051 | 3.5664943 | 39.21383257 |
| ENSMUSG00000071267 | *Zfp942* | 162.1692839 | 93.86229862 | 87.22904395 | 414.9524247 | 420.1816213 | 402.9953015 | 6.20E-14 | 1.834315837 | 8.21E-15 | 60.28410534 | 3.566022577 | 13.20781619 |
| ENSMUSG00000024293 | *Esco1* | 535.2385231 | 433.9869721 | 492.508602 | 1752.583615 | 1828.38721 | 1638.074427 | 4.39E-60 | 1.833944174 | 6.50E-62 | 275.6973401 | 3.565104028 | 59.35741196 |
| ENSMUSG00000006005 | *Tpr* | 2333.480188 | 2146.721604 | 1415.794483 | 7065.15541 | 7055.142572 | 6925.334126 | 2.45E-28 | 1.833932368 | 1.40E-29 | 127.5581093 | 3.565074853 | 27.61090166 |
| ENSMUSG00000074909 | *Ranbp6* | 124.6227009 | 126.1590035 | 115.4107351 | 415.795824 | 464.6969869 | 430.0549381 | 2.44E-34 | 1.833901822 | 1.07E-35 | 155.5428885 | 3.564999371 | 33.61324498 |
| ENSMUSG00000093942 | *Or13a18* | 8.78749814 | 9.083448253 | 22.81374996 | 40.48316339 | 51.02980931 | 48.32077956 | 0.000473028 | 1.833855359 | 0.0001603 | 14.24718654 | 3.56488456 | 3.325113013 |
| ENSMUSG00000028979 | *Masp2* | 34.35112909 | 36.33379301 | 16.1038235 | 123.1362886 | 95.54517487 | 96.64155911 | 2.92E-07 | 1.833429868 | 6.80E-08 | 29.12229448 | 3.563833331 | 6.534177204 |
| ENSMUSG00000045210 | *Vcpip1* | 1018.550921 | 969.910419 | 860.2125719 | 3325.523192 | 3524.314064 | 3307.074153 | 2.60E-89 | 1.8300676 | 1.70E-91 | 411.5300449 | 3.555537322 | 88.58523055 |
| ENSMUSG00000023892 | *Zfp51* | 123.024974 | 90.83448253 | 92.59698511 | 370.2522651 | 375.6662558 | 352.7416908 | 5.58E-24 | 1.828376786 | 4.02E-25 | 107.1999002 | 3.551372734 | 23.25298026 |
| ENSMUSG00000024423 | *Impact* | 755.7248401 | 831.6401512 | 1022.592792 | 3277.449436 | 2991.215418 | 2967.86228 | 3.03E-48 | 1.827878221 | 6.91E-50 | 220.5365702 | 3.550145665 | 47.51822679 |
| ENSMUSG00000033792 | *Atp7a* | 330.7294755 | 390.5882749 | 319.3924994 | 1288.714034 | 1241.001533 | 1162.597956 | 6.07E-54 | 1.825354885 | 1.12E-55 | 247.0852408 | 3.543941736 | 53.21663028 |
| ENSMUSG00000035992 | *Fnip1* | 647.878272 | 611.6188491 | 845.4507337 | 2602.730046 | 2427.716034 | 2402.50916 | 5.20E-43 | 1.824978179 | 1.49E-44 | 196.0888997 | 3.543016488 | 42.28425756 |
| ENSMUSG00000026358 | *Rgs1* | 3100.389117 | 2738.155012 | 2798.039333 | 10361.15963 | 9911.726152 | 10343.54607 | 8.39E-127 | 1.824773865 | 2.67E-129 | 585.2732393 | 3.542514763 | 126.0763251 |
| ENSMUSG00000049164 | *Zfp518a* | 274.0101693 | 193.7802294 | 151.6443379 | 720.2629486 | 766.5328802 | 725.7781089 | 4.33E-19 | 1.824160685 | 4.13E-20 | 84.35554292 | 3.541009428 | 18.36311961 |
| ENSMUSG00000078546 | *Zfp995* | 47.13294457 | 47.43578532 | 48.31147049 | 180.4874368 | 165.0325748 | 159.4585725 | 1.04E-15 | 1.824152796 | 1.22E-16 | 68.57738942 | 3.540990066 | 14.98436765 |
| ENSMUSG00000094568 | *Smarce1-ps1* | 39.1443099 | 70.64904197 | 68.44124987 | 185.5478322 | 219.319606 | 218.4099236 | 2.57E-11 | 1.823650711 | 4.15E-12 | 48.05074143 | 3.539757948 | 10.59049923 |
| ENSMUSG00000041133 | *Smc1a* | 1533.018994 | 1419.036472 | 1346.011247 | 5184.375111 | 5095.380746 | 4935.484424 | 6.35E-118 | 1.821213624 | 2.21E-120 | 544.2800831 | 3.533783424 | 117.1975199 |
| ENSMUSG00000035614 | *Togaram1* | 483.3123977 | 474.3578532 | 307.3146318 | 1465.827874 | 1556.952054 | 1463.153205 | 2.55E-27 | 1.819213612 | 1.53E-28 | 122.8170619 | 3.528887925 | 26.59427412 |
| ENSMUSG00000078903 | *Gm14391* | 87.07611793 | 98.90865876 | 41.60154404 | 241.2121818 | 293.1499684 | 279.2941058 | 6.10E-10 | 1.819201395 | 1.10E-10 | 41.62624841 | 3.528858041 | 9.214902776 |
| ENSMUSG00000034158 | *Lrrc58* | 878.749814 | 876.0481204 | 595.8414694 | 2712.371947 | 2959.72894 | 2641.213811 | 1.16E-32 | 1.818216137 | 5.52E-34 | 147.6995232 | 3.526448905 | 31.93496558 |
| ENSMUSG00000006678 | *Pola1* | 597.5498735 | 494.5432938 | 535.4521313 | 2001.38639 | 1868.559613 | 1874.846247 | 1.25E-68 | 1.816919879 | 1.43E-70 | 315.4388147 | 3.523281824 | 67.90209322 |
| ENSMUSG00000040549 | *Ckap5* | 1649.65306 | 1565.380916 | 1492.287644 | 5711.499635 | 5490.590333 | 5403.22957 | 1.68E-134 | 1.816690276 | 4.36E-137 | 621.0756996 | 3.522721142 | 133.77464 |
| ENSMUSG00000091350 | *Vmn2r92* | 7.189771206 | 10.09272028 | 4.025955875 | 24.45857788 | 24.97203434 | 27.05963655 | 0.003366275 | 1.813836212 | 0.001295928 | 10.34831616 | 3.515759079 | 2.472850414 |
| ENSMUSG00000025402 | *Nab2* | 1250.221326 | 1101.115783 | 869.6064689 | 3737.10202 | 3763.177001 | 3826.039325 | 2.10E-43 | 1.81039939 | 5.89E-45 | 197.937945 | 3.507393724 | 42.67876117 |
| ENSMUSG00000046138 | *9930021J03Rik* | 243.6533575 | 243.2345588 | 186.5359555 | 770.0235036 | 793.6763958 | 813.7219277 | 3.43E-40 | 1.809070571 | 1.14E-41 | 182.8821861 | 3.504164666 | 39.46457125 |
| ENSMUSG00000094472 | *None* | 47.93180804 | 14.12980839 | 9.393897041 | 102.0513077 | 85.77350926 | 68.61550697 | 0.002205081 | 1.807445762 | 0.000823652 | 11.18716254 | 3.500220387 | 2.656575376 |
| ENSMUSG00000032657 | *Entrep3* | 17.57499628 | 22.20398462 | 10.73588233 | 59.03794661 | 60.80147492 | 59.91776665 | 7.66E-06 | 1.80545139 | 2.08E-06 | 22.52241381 | 3.495385047 | 5.115537139 |
| ENSMUSG00000034295 | *Fhod3* | 51.12726191 | 45.41724127 | 53.67941166 | 177.9572391 | 149.8322061 | 195.2159494 | 6.14E-14 | 1.804762124 | 8.13E-15 | 60.30420964 | 3.493715483 | 13.21205848 |
| ENSMUSG00000020422 | *Tns3* | 4194.832067 | 4488.232709 | 3285.179994 | 14140.43161 | 14053.82663 | 13638.05682 | 3.72E-52 | 1.804519545 | 7.27E-54 | 238.7766206 | 3.493128086 | 51.42935243 |
| ENSMUSG00000039286 | *Fndc3b* | 1844.575746 | 1987.256623 | 1285.621909 | 5932.470235 | 6139.863226 | 5824.586768 | 9.56E-33 | 1.804368128 | 4.52E-34 | 148.0943437 | 3.492761487 | 32.01965036 |
| ENSMUSG00000021431 | *Snrnp48* | 405.0237779 | 412.7922595 | 300.6047053 | 1348.59538 | 1271.40227 | 1299.82897 | 1.78E-40 | 1.801825183 | 5.84E-42 | 184.2083634 | 3.486610447 | 39.74865598 |
| ENSMUSG00000037434 | *Slc30a1* | 907.5088988 | 953.7620666 | 848.1347042 | 3086.841208 | 3214.877986 | 3156.313321 | 7.86E-113 | 1.801130404 | 3.16E-115 | 520.5860703 | 3.484931755 | 112.1044583 |
| ENSMUSG00000005371 | *Fbxo11* | 396.2362798 | 426.9220679 | 222.7695584 | 1244.013875 | 1234.487089 | 1179.993437 | 2.70E-17 | 1.799863973 | 2.88E-18 | 75.96827206 | 3.481873942 | 16.56839907 |
| ENSMUSG00000046667 | *Rbm12b1* | 159.7726935 | 210.9378539 | 151.6443379 | 665.4419982 | 627.5580804 | 527.6629128 | 4.65E-24 | 1.799127384 | 3.33E-25 | 107.5725474 | 3.480096676 | 23.33214272 |
| ENSMUSG00000064061 | *Dzip3* | 223.6817708 | 164.5113406 | 111.3847792 | 607.2474508 | 603.6717867 | 543.1255622 | 7.31E-15 | 1.797983108 | 9.09E-16 | 64.61787284 | 3.477337525 | 14.13636857 |
| ENSMUSG00000066152 | *Slc31a2* | 555.2101098 | 620.7022973 | 656.2308076 | 2123.679279 | 2137.823288 | 2068.129365 | 2.02E-80 | 1.793722778 | 1.68E-82 | 370.210208 | 3.467083971 | 79.69377782 |
| ENSMUSG00000041147 | *Brca2* | 181.3420071 | 124.1404595 | 83.20308807 | 474.8337706 | 445.1536557 | 441.6519251 | 2.03E-12 | 1.792465151 | 3.01E-13 | 53.20066035 | 3.464062959 | 11.6927283 |
| ENSMUSG00000050714 | *Zbtb26* | 116.6340662 | 60.55632169 | 49.65345579 | 241.2121818 | 285.549784 | 270.5963655 | 1.38E-08 | 1.792085049 | 2.85E-09 | 35.28501219 | 3.463150415 | 7.85874785 |
| ENSMUSG00000073664 | *Nbeal1* | 270.0158519 | 252.318007 | 197.2718379 | 861.1106212 | 820.8199114 | 820.4868369 | 1.65E-37 | 1.787162338 | 6.17E-39 | 170.3600825 | 3.451353717 | 36.78246834 |
| ENSMUSG00000042063 | *Zfp386* | 205.3079111 | 202.8636777 | 198.6138231 | 695.8043707 | 680.759371 | 718.0467842 | 8.10E-52 | 1.785816721 | 1.60E-53 | 237.2092003 | 3.448136103 | 51.09149155 |
| ENSMUSG00000057396 | *Zfp759* | 35.14999256 | 35.32452098 | 38.91757345 | 139.1608741 | 111.8312842 | 123.7011957 | 3.31E-11 | 1.785211189 | 5.40E-12 | 47.5370581 | 3.446689145 | 10.47970923 |
| ENSMUSG00000020982 | *Nemf* | 650.2748624 | 490.5062057 | 535.4521313 | 1912.82947 | 2041.192372 | 1828.458298 | 3.67E-45 | 1.782540346 | 9.50E-47 | 206.1497777 | 3.440314237 | 44.43553714 |
| ENSMUSG00000110185 | *Igip* | 19.97158668 | 7.064904197 | 8.051911749 | 42.16996186 | 45.60110619 | 36.72379246 | 0.001413198 | 1.780438312 | 0.000513268 | 12.06683251 | 3.435305284 | 2.849797084 |
| ENSMUSG00000035042 | *Ccl5* | 371.4715123 | 321.957777 | 198.6138231 | 1033.164066 | 1022.767667 | 1026.333358 | 2.30E-18 | 1.779680574 | 2.29E-19 | 80.96993941 | 3.433501453 | 17.63869888 |
| ENSMUSG00000029254 | *Stap1* | 4198.027521 | 2887.527273 | 2815.485142 | 11383.3595 | 11348.161 | 11263.57371 | 3.05E-32 | 1.77863807 | 1.48E-33 | 145.7426198 | 3.431021271 | 31.51518932 |
| ENSMUSG00000062101 | *Zfp119b* | 15.17840588 | 25.2318007 | 28.18169112 | 63.25494279 | 83.60202801 | 84.07815643 | 3.70E-06 | 1.777725902 | 9.69E-07 | 23.98962829 | 3.428852636 | 5.431359055 |
| ENSMUSG00000047473 | *Zfp30* | 10.38522507 | 25.2318007 | 17.44580879 | 59.03794661 | 51.02980931 | 69.58192256 | 9.09E-05 | 1.77757232 | 2.81E-05 | 17.54550487 | 3.428487637 | 4.041532528 |
| ENSMUSG00000026535 | *Ifi202b* | 1259.008824 | 1060.744902 | 1363.457056 | 4241.454764 | 4145.357701 | 4212.605562 | 1.54E-64 | 1.775923475 | 2.00E-66 | 296.4029123 | 3.424571483 | 63.81193559 |
| ENSMUSG00000021712 | *Trim23* | 201.3135938 | 189.7431413 | 212.0336761 | 739.6611311 | 670.9877054 | 647.4984461 | 2.26E-42 | 1.774654282 | 6.71E-44 | 193.0959957 | 3.421560084 | 41.64684457 |
| ENSMUSG00000028175 | *Depdc1a* | 255.6363095 | 114.0477392 | 107.3588233 | 576.8850783 | 548.2990149 | 517.9987568 | 1.16E-08 | 1.773188158 | 2.37E-09 | 35.64684249 | 3.418084725 | 7.936046636 |
| ENSMUSG00000022303 | *Dcstamp* | 412.2135491 | 282.5961679 | 366.3619846 | 1203.530712 | 1200.82913 | 1228.314216 | 1.20E-33 | 1.773155461 | 5.47E-35 | 152.2936059 | 3.41800726 | 32.91981821 |
| ENSMUSG00000020841 | *Cpd* | 3760.250341 | 3523.36865 | 2457.175069 | 11181.78709 | 11350.33248 | 10764.90327 | 1.04E-32 | 1.772120344 | 4.94E-34 | 147.9197472 | 3.415555758 | 31.98202558 |
| ENSMUSG00000090963 | *Gm17655* | 47.93180804 | 29.26888882 | 46.9694852 | 161.0892543 | 132.4603561 | 128.5332736 | 1.07E-09 | 1.767733977 | 1.98E-10 | 40.48761622 | 3.405186881 | 8.969879272 |
| ENSMUSG00000040651 | *Tasor* | 211.6988188 | 155.4278923 | 127.4886027 | 608.9342493 | 554.8134586 | 536.3606531 | 1.95E-19 | 1.767702441 | 1.82E-20 | 85.98011392 | 3.405112446 | 18.71098046 |
| ENSMUSG00000091952 | *Gm17709* | 48.7306715 | 98.90865876 | 13.41985292 | 143.3778703 | 200.8620154 | 207.7793521 | 0.001304908 | 1.765725334 | 0.000471551 | 12.22493428 | 3.400449188 | 2.884420063 |
| ENSMUSG00000002983 | *Relb* | 413.0124126 | 352.2359378 | 261.6871318 | 1084.611419 | 1182.371539 | 1244.743281 | 5.53E-26 | 1.765025496 | 3.59E-27 | 116.5557698 | 3.398800062 | 25.25751137 |
| ENSMUSG00000024778 | *Fas* | 82.28293713 | 55.50996155 | 76.49316162 | 240.3687826 | 269.2636746 | 220.3427548 | 3.42E-15 | 1.764944581 | 4.18E-16 | 66.15089678 | 3.398609442 | 14.46610077 |
| ENSMUSG00000054702 | *Ap1s3* | 550.416929 | 545.0068952 | 453.5910285 | 1734.872231 | 1756.728329 | 1787.868844 | 2.35E-65 | 1.762860488 | 2.98E-67 | 300.2013926 | 3.393703412 | 64.62952776 |
| ENSMUSG00000016496 | *Cd274* | 139.0022433 | 181.6689651 | 182.5099996 | 566.7642874 | 581.9569742 | 546.024809 | 2.05E-30 | 1.762232836 | 1.07E-31 | 137.2384191 | 3.392227285 | 29.68877802 |
| ENSMUSG00000024190 | *Dusp1* | 382.6556008 | 368.3842903 | 330.1283817 | 1262.568658 | 1216.029498 | 1204.153827 | 2.38E-65 | 1.762194794 | 3.04E-67 | 300.1611796 | 3.392137837 | 64.622782 |
| ENSMUSG00000050310 | *Rictor* | 434.5817262 | 390.5882749 | 382.4658081 | 1362.933167 | 1357.175779 | 1381.974295 | 8.12E-72 | 1.758950294 | 8.38E-74 | 330.2723638 | 3.384517774 | 71.09055586 |
| ENSMUSG00000041308 | *Sntb2* | 783.6850614 | 985.0494995 | 797.1392632 | 2910.570768 | 2898.927465 | 2858.657319 | 7.90E-59 | 1.756051236 | 1.21E-60 | 269.8687424 | 3.377723503 | 58.10244111 |
| ENSMUSG00000005871 | *Apc* | 569.5896522 | 614.6466651 | 379.7818375 | 1702.82306 | 1819.701285 | 1779.171103 | 1.54E-25 | 1.755927137 | 1.02E-26 | 114.4801782 | 3.377432968 | 24.81164532 |
| ENSMUSG00000057894 | *Zfp329* | 50.32839844 | 55.50996155 | 52.33742637 | 204.1026154 | 167.204056 | 161.3914037 | 7.72E-14 | 1.755844763 | 1.03E-14 | 59.83660999 | 3.377240133 | 13.11254397 |
| ENSMUSG00000033713 | *Foxn3* | 366.6783315 | 349.2081217 | 264.3711024 | 1120.034187 | 1150.885061 | 1041.796007 | 2.60E-35 | 1.747870015 | 1.08E-36 | 160.0858616 | 3.35862335 | 34.58428453 |
| ENSMUSG00000024079 | *Eif2ak2* | 1473.104234 | 1448.30536 | 1753.974776 | 5299.077408 | 5212.640734 | 5150.028685 | 1.60E-76 | 1.746740395 | 1.45E-78 | 352.1447295 | 3.355994601 | 75.7955771 |
| ENSMUSG00000017485 | *Top2b* | 762.9146113 | 693.3698833 | 590.4735283 | 2311.757309 | 2294.169938 | 2277.841548 | 1.33E-57 | 1.744240897 | 2.13E-59 | 264.1573616 | 3.350185307 | 56.8756157 |
| ENSMUSG00000039146 | *Ifi44l* | 1801.437119 | 1386.739767 | 1097.743968 | 4797.254861 | 4772.915781 | 4804.051903 | 3.57E-27 | 1.743250521 | 2.16E-28 | 122.1290162 | 3.347886274 | 26.44791147 |
| ENSMUSG00000058258 | *Idi1* | 1449.937193 | 1297.923828 | 1551.334997 | 5003.044275 | 4718.62875 | 4641.694084 | 1.18E-81 | 1.741780276 | 9.36E-84 | 375.9743808 | 3.344476193 | 80.92667492 |
| ENSMUSG00000113450 | *Zfp935* | 40.74203683 | 73.67685805 | 40.25955875 | 134.943878 | 221.4910872 | 162.3578193 | 2.69E-07 | 1.740830337 | 6.23E-08 | 29.29198077 | 3.342274756 | 6.570664912 |
| ENSMUSG00000028053 | *Ash1l* | 953.8429799 | 915.4097295 | 547.5299989 | 2802.615665 | 2782.753218 | 2494.318641 | 6.92E-20 | 1.737633846 | 6.29E-21 | 88.07905497 | 3.334877679 | 19.16020478 |
| ENSMUSG00000048279 | *Sacs* | 786.8805153 | 691.3513393 | 526.0582343 | 2225.730587 | 2276.798088 | 2202.461132 | 9.77E-34 | 1.736679973 | 4.40E-35 | 152.7253701 | 3.332673474 | 33.01021539 |
| ENSMUSG00000019823 | *Mical1* | 7.189771206 | 17.15762448 | 8.051911749 | 29.5189733 | 38.00092182 | 40.58945483 | 0.002068196 | 1.736508242 | 0.00076947 | 11.31346513 | 3.332276793 | 2.684408341 |
| ENSMUSG00000028270 | *Gbp2* | 43.9374907 | 54.50068952 | 34.89161758 | 146.7514673 | 158.518131 | 143.0295075 | 1.37E-11 | 1.736203366 | 2.17E-12 | 49.32246192 | 3.331572678 | 10.86330696 |
| ENSMUSG00000065947 | *mt-Nd4l* | 4708.501276 | 4676.966578 | 4339.980433 | 14882.62294 | 15900.67143 | 14912.75899 | 2.51E-139 | 1.734398632 | 6.21E-142 | 643.3591528 | 3.327407664 | 138.6008574 |
| ENSMUSG00000046798 | *Cldn12* | 148.5886049 | 158.4557084 | 128.830588 | 447.844995 | 504.86939 | 505.4353542 | 3.22E-30 | 1.734026069 | 1.69E-31 | 136.3289508 | 3.326548504 | 29.49182455 |
| ENSMUSG00000039735 | *Fnbp1l* | 1183.116795 | 1005.23494 | 1080.29816 | 3699.992454 | 3648.088495 | 3530.316154 | 3.22E-86 | 1.733044963 | 2.37E-88 | 397.0843954 | 3.32428705 | 85.49226321 |

| | | | | | | | | | | | | | |
|---|---|---|---|---|---|---|---|---|---|---|---|---|---|
| ENSMUSG00000110519 | *Or7g23* | 33.55226563 | 31.28743287 | 37.57558816 | 114.7022963 | 107.4883217 | 115.9698709 | 2.01E-10 | 1.73192382 | 3.49E-11 | 43.87942962 | 3.321704692 | 9.696059502 |
| ENSMUSG00000000600 | *Krit1* | 912.3020796 | 887.1501127 | 1101.769924 | 3279.136234 | 3225.735393 | 3098.328385 | 2.49E-62 | 1.730642619 | 3.37E-64 | 286.1861351 | 3.318756127 | 61.6032379 |
| ENSMUSG00000049606 | *Zfp644* | 556.8078367 | 546.0161672 | 495.1925726 | 1814.151759 | 1773.014438 | 1729.883908 | 3.34E-81 | 1.729983342 | 2.68E-83 | 373.8758707 | 3.31723988 | 80.47621434 |
| ENSMUSG00000078879 | *Zfp973* | 37.54658296 | 8.074176225 | 9.393897041 | 46.38695805 | 77.08758427 | 63.78342901 | 0.007624941 | 1.727534414 | 0.003118108 | 8.737037147 | 3.311613749 | 2.117763522 |
| ENSMUSG00000003847 | *Nfat5* | 2980.559596 | 2937.990874 | 1768.736614 | 8422.184783 | 8943.245516 | 8099.529069 | 8.49E-22 | 1.726731507 | 6.94E-23 | 96.99843631 | 3.309771242 | 21.07118637 |
| ENSMUSG00000032515 | *Csrnp1* | 658.263497 | 599.5075847 | 507.2704402 | 1815.838558 | 2024.906263 | 2021.741417 | 5.28E-47 | 1.725909616 | 1.28E-48 | 214.7171974 | 3.307886229 | 46.27712124 |
| ENSMUSG00000052137 | *Rbm12b2* | 220.486317 | 165.5206126 | 234.847426 | 603.0304546 | 708.9886272 | 738.3415116 | 3.75E-24 | 1.725828776 | 2.67E-25 | 108.0143296 | 3.30770088 | 23.42607728 |
| ENSMUSG00000039473 | *Ubn1* | 71.09884859 | 98.90865876 | 260.3451466 | 575.1982798 | 472.2971712 | 370.1371714 | 0.00941839 | 1.725141396 | 0.00392822 | 8.316700863 | 3.306125285 | 2.026023315 |
| ENSMUSG00000022489 | *Pde1b* | 241.2567671 | 221.0305742 | 154.3283085 | 705.0817623 | 681.8451116 | 668.7595891 | 6.83E-24 | 1.724684902 | 4.95E-25 | 106.7911907 | 3.305079333 | 23.16554437 |
| ENSMUSG00000060989 | *Gm11847* | 11.18408854 | 40.37088113 | 17.44580879 | 43.85676034 | 110.7455436 | 73.44758493 | 0.003587883 | 1.724676175 | 0.001388441 | 10.22108546 | 3.305059342 | 2.445161771 |
| ENSMUSG00000111331 | *None* | 7.189771206 | 14.12980839 | 4.025955875 | 27.83217483 | 28.22925621 | 28.99246773 | 0.00587114 | 1.722252974 | 0.002354832 | 9.250095294 | 3.299512708 | 2.231277552 |
| ENSMUSG00000026601 | *Axdnd1* | 8.78749814 | 4.037088113 | 8.051911749 | 22.77177941 | 26.05777497 | 20.29472741 | 0.008207819 | 1.721798695 | 0.003376328 | 8.592054961 | 3.298473914 | 2.08577222 |
| ENSMUSG00000027939 | *Nup210l* | 54.32271578 | 30.27816084 | 29.52367641 | 113.0154978 | 132.4603561 | 138.1974295 | 1.19E-07 | 1.721794818 | 2.65E-08 | 30.94613109 | 3.298465048 | 6.926016307 |
| ENSMUSG00000034297 | *Med13* | 1285.371319 | 1066.800534 | 705.8842633 | 3427.5745 | 3519.971102 | 3155.346905 | 1.06E-19 | 1.720941614 | 9.70E-21 | 87.22220892 | 3.296514926 | 18.9757145 |
| ENSMUSG00000046733 | *Gprc5a* | 109.444295 | 129.1868196 | 155.6702938 | 445.3147973 | 401.7240307 | 436.8198472 | 4.95E-23 | 1.719049326 | 3.76E-24 | 102.7727602 | 3.292193939 | 22.30517289 |
| ENSMUSG00000021377 | *Dek* | 1691.19396 | 1290.858924 | 1143.371468 | 4569.537067 | 4473.251369 | 4564.380837 | 2.56E-34 | 1.719039396 | 1.12E-35 | 155.4402379 | 3.292171279 | 33.59139882 |
| ENSMUSG00000060519 | *Tor3a* | 3876.085543 | 3744.399224 | 3168.427273 | 11822.77051 | 11921.43205 | 11786.40455 | 1.75E-78 | 1.71820228 | 1.54E-80 | 361.2092774 | 3.29026157 | 77.75790184 |
| ENSMUSG00000056383 | *AI987944* | 207.7045015 | 168.5484287 | 179.8260291 | 624.1154356 | 596.0716023 | 616.5731471 | 1.75E-36 | 1.717739828 | 6.82E-38 | 165.5827044 | 3.289207054 | 35.75650719 |
| ENSMUSG00000002699 | *Lcp2* | 3058.848216 | 2886.518 | 2571.243819 | 9338.959754 | 9313.483069 | 9313.347052 | 1.78E-94 | 1.713633579 | 1.03E-96 | 435.5080022 | 3.279858509 | 93.75012079 |
| ENSMUSG00000037416 | *Dmxl1* | 1312.532677 | 1295.905284 | 1034.67066 | 3956.385822 | 4106.271038 | 3911.083897 | 3.63E-57 | 1.7133434 | 5.92E-59 | 262.1208003 | 3.279198875 | 56.43965374 |
| ENSMUSG00000020463 | *Ppp4r3b* | 1142.374758 | 1058.726358 | 991.7271304 | 3608.061937 | 3499.34203 | 3378.588907 | 5.30E-90 | 1.71243758 | 3.35E-92 | 414.7748928 | 3.277140621 | 89.27559842 |
| ENSMUSG00000045382 | *Cxcr4* | 45.53521764 | 61.56559372 | 77.83514691 | 206.6328131 | 218.2338653 | 172.0219752 | 5.40E-11 | 1.712131771 | 8.92E-12 | 46.55136337 | 3.276446037 | 10.26769801 |
| ENSMUSG00000017009 | *Sdc4* | 4059.824141 | 3925.058917 | 3816.606169 | 12969.79347 | 12984.37212 | 12713.1971 | 1.98E-186 | 1.711315298 | 2.57E-189 | 861.2744633 | 3.274592304 | 185.7029137 |
| ENSMUSG00000109179 | *Gm35339* | 9.586361607 | 9.083448253 | 4.025955875 | 22.77177941 | 33.65795933 | 20.29472741 | 0.008228838 | 1.711011576 | 0.003386431 | 8.586613986 | 3.273902996 | 2.084661487 |
| ENSMUSG00000030047 | *Arhgap25* | 802.0589212 | 974.9567792 | 917.9179394 | 2920.691559 | 2962.986162 | 2923.407163 | 3.33E-71 | 1.7108437 | 3.52E-73 | 327.4105132 | 3.273522056 | 70.47704941 |
| ENSMUSG00000042608 | *Stk40* | 1559.381488 | 1531.065667 | 1124.583674 | 4533.2709 | 4684.970791 | 4588.541227 | 4.39E-40 | 1.709031429 | 1.46E-41 | 182.3841798 | 3.269412536 | 39.35738753 |
| ENSMUSG00000063894 | *Zkscan8* | 166.1636012 | 130.1960916 | 136.8824997 | 495.9187515 | 450.5823588 | 477.409302 | 1.07E-27 | 1.708777922 | 6.26E-29 | 124.5883617 | 3.268838095 | 26.97204331 |
| ENSMUSG00000026034 | *Clk1* | 603.1419178 | 537.941991 | 602.5513959 | 1936.444649 | 1858.787948 | 1900.939468 | 2.02E-80 | 1.708520699 | 1.68E-82 | 370.2117816 | 3.268255333 | 79.69377782 |
| ENSMUSG00000059149 | *Mfsd4a* | 147.7897414 | 207.9100378 | 120.7786762 | 527.1245233 | 555.8991993 | 479.3421332 | 1.09E-16 | 1.707038959 | 1.21E-17 | 73.13472903 | 3.264900349 | 15.96062783 |
| ENSMUSG00000029313 | *Aff1* | 2106.602963 | 2054.877849 | 1362.115071 | 6072.474508 | 6193.064517 | 5789.795806 | 1.46E-27 | 1.706871346 | 8.65E-29 | 123.9469421 | 3.264521055 | 26.8351352 |
| ENSMUSG00000064289 | *Tank* | 843.5998215 | 855.8626799 | 950.1255864 | 3048.888243 | 2821.839881 | 2757.183681 | 2.82E-79 | 1.706657884 | 2.41E-81 | 364.8999667 | 3.264038069 | 78.54970203 |
| ENSMUSG00000028214 | *Gem* | 15.17840588 | 9.083448253 | 14.76183821 | 54.82095042 | 30.40073746 | 41.55587042 | 0.000829374 | 1.706099347 | 0.000291779 | 13.1224466 | 3.262774647 | 3.081249518 |
| ENSMUSG00000025815 | *Dhtkd1* | 26.36249442 | 29.26888882 | 12.07786762 | 86.0267222 | 85.77350926 | 53.15285751 | 9.48E-05 | 1.704593312 | 2.93E-05 | 17.46223557 | 3.259370402 | 4.023258297 |
| ENSMUSG00000053007 | *Creb5* | 110.2431585 | 131.2053637 | 89.91301453 | 345.7936873 | 394.1238464 | 345.010366 | 8.04E-20 | 1.701036453 | 7.33E-21 | 87.77663985 | 3.25134456 | 19.09465007 |
| ENSMUSG00000023341 | *Mx2* | 392.2419624 | 360.310114 | 351.6001464 | 1264.255457 | 1184.54302 | 1146.168891 | 2.23E-58 | 1.699216402 | 3.47E-60 | 267.7733651 | 3.247245369 | 57.65238604 |
| ENSMUSG00000034560 | *Washc4* | 1200.691791 | 1164.69992 | 1331.249409 | 4066.871122 | 3995.525495 | 3917.848806 | 1.43E-98 | 1.699024708 | 7.50E-101 | 454.516368 | 3.246813929 | 97.84513435 |
| ENSMUSG00000034391 | *Fbxo15* | 33.55226563 | 35.32452098 | 21.47176466 | 84.33992372 | 95.54517487 | 117.9027021 | 3.21E-07 | 1.698565195 | 7.49E-08 | 28.93433182 | 3.245779951 | 6.493909298 |
| ENSMUSG00000047022 | *Mipol1* | 16.77613281 | 21.19471259 | 30.8656617 | 75.06253211 | 83.60202801 | 59.91776665 | 1.18E-05 | 1.698077872 | 3.26E-06 | 21.65771521 | 3.244683754 | 4.927500068 |
| ENSMUSG00000055240 | *Zfp101* | 111.8408854 | 89.8252105 | 107.3588233 | 331.4559002 | 337.6653339 | 334.3797945 | 6.34E-24 | 1.697496167 | 4.59E-25 | 106.9396559 | 3.243375739 | 23.19772537 |
| ENSMUSG00000021843 | *Ktn1* | 459.3464937 | 391.5975469 | 366.3619846 | 1379.801152 | 1317.003376 | 1266.004424 | 3.85E-47 | 1.696226951 | 9.32E-49 | 215.3551591 | 3.240523623 | 46.41417788 |
| ENSMUSG00000020994 | *Pnn* | 985.7975186 | 810.4454386 | 701.8583075 | 2820.327049 | 2676.350637 | 2621.885499 | 8.96E-39 | 1.696049053 | 3.17E-40 | 176.2636082 | 3.240124061 | 38.04786266 |
| ENSMUSG00000095990 | *Zfp97* | 126.2204278 | 106.982835 | 116.7527204 | 393.8674438 | 389.7808839 | 352.7416908 | 7.36E-26 | 1.69564184 | 4.80E-27 | 115.9789335 | 3.239209637 | 25.13316121 |
| ENSMUSG00000009145 | *Dqx1* | 19.17272321 | 13.12053637 | 17.44580879 | 62.41154356 | 56.45851243 | 42.52228601 | 6.76E-05 | 1.694965657 | 2.05E-05 | 18.14063513 | 3.237691793 | 4.169989671 |
| ENSMUSG00000004508 | *Gab2* | 970.6191128 | 955.7806106 | 782.377425 | 2872.617802 | 3028.130599 | 2892.481864 | 6.11E-60 | 1.694496982 | 9.13E-62 | 275.0226633 | 3.236640165 | 59.21382632 |
| ENSMUSG00000046230 | *Vps13a* | 621.5157776 | 580.3314162 | 475.0627932 | 1912.82947 | 1806.672398 | 1720.219752 | 1.54E-44 | 1.692022286 | 4.11E-46 | 203.2368497 | 3.231093023 | 43.81165416 |
| ENSMUSG00000038608 | *Dock10* | 2330.284734 | 2358.66873 | 2606.135436 | 8127.838449 | 7845.561746 | 7543.840104 | 1.34E-106 | 1.690329609 | 5.87E-109 | 491.7708197 | 3.227304287 | 105.8717125 |
| ENSMUSG00000027550 | *Lrrcc1* | 371.4715123 | 299.7537924 | 248.2672789 | 1039.91126 | 948.937305 | 997.34089 | 2.55E-27 | 1.690211584 | 1.53E-28 | 122.8122728 | 3.227040276 | 26.59365225 |
| ENSMUSG00000112148 | *Lilrb4a* | 9654.265002 | 9633.501509 | 8486.714984 | 31314.57028 | 28979.50298 | 29314.28413 | 3.65E-99 | 1.68946884 | 1.85E-101 | 457.3068991 | 3.225379324 | 98.43752227 |
| ENSMUSG00000022041 | *Chrna2* | 10.38522507 | 10.09272028 | 13.41985292 | 48.07375652 | 31.48647808 | 28.02605214 | 0.001553072 | 1.687320129 | 0.000567737 | 11.87889342 | 3.220579107 | 2.808808346 |
| ENSMUSG00000043259 | *Fam13c* | 119.0306566 | 135.2424518 | 115.4107351 | 380.373056 | 432.1247682 | 380.7677429 | 7.55E-26 | 1.687183359 | 4.93E-27 | 115.9273626 | 3.220273805 | 25.12226024 |
| ENSMUSG00000089857 | *Zfp882* | 23.96590402 | 24.22252868 | 25.49772054 | 71.68893516 | 89.03073113 | 76.3468317 | 2.96E-07 | 1.686823465 | 6.89E-08 | 29.09628924 | 3.219470576 | 6.528568183 |
| ENSMUSG00000027981 | *Rnpc3* | 374.6669662 | 273.5127196 | 334.1543376 | 1080.394423 | 1018.424705 | 1066.922813 | 4.84E-35 | 1.685771786 | 2.04E-36 | 158.830488 | 3.21712454 | 34.31550396 |
| ENSMUSG00000072620 | *Slfn2* | 4368.984303 | 3980.568879 | 4101.107051 | 13449.68764 | 13054.94526 | 13540.44885 | 1.69E-143 | 1.68486845 | 3.79E-146 | 662.7375338 | 3.215110784 | 142.7717782 |
| ENSMUSG00000103051 | *Gm7871* | 13.58067894 | 12.11126434 | 16.1038235 | 42.16996186 | 35.82944058 | 55.08568869 | 0.000274997 | 1.683830424 | 9.02E-05 | 15.33207348 | 3.21279833 | 3.56067223 |
| ENSMUSG00000019726 | *Lyst* | 909.1066258 | 1003.216396 | 983.6752187 | 3123.107375 | 3143.219105 | 3004.586073 | 8.84E-100 | 1.680648678 | 4.43E-102 | 460.1611479 | 3.205720572 | 99.05358028 |
| ENSMUSG00000055994 | *Nod2* | 177.3476897 | 179.650421 | 134.1985292 | 527.9679225 | 518.9840181 | 537.3270687 | 2.52E-28 | 1.677648047 | 1.44E-29 | 127.5015951 | 3.199059991 | 27.59898248 |
| ENSMUSG00000003226 | *Ranbp2* | 3375.198149 | 3058.094245 | 2845.008818 | 10181.51559 | 9802.066349 | 9712.476691 | 1.94E-88 | 1.6769286 | 1.33E-90 | 407.4339106 | 3.197465072 | 87.71273959 |
| ENSMUSG00000020638 | *Cmpk2* | 1644.859879 | 1559.325283 | 1057.48441 | 4502.908528 | 4719.714491 | 4420.384914 | 6.52E-27 | 1.676182063 | 4.01E-28 | 120.9020833 | 3.195810939 | 26.18603592 |
| ENSMUSG00000060183 | *Cxcl11* | 15.97726935 | 19.17616853 | 18.78779408 | 49.760555 | 72.74462178 | 49.28719515 | 5.07E-05 | 1.674897926 | 1.51E-05 | 18.72424523 | 3.192967628 | 4.29541359 |
| ENSMUSG00000040296 | *Rigi* | 1620.095112 | 1514.917314 | 1539.257129 | 5127.023963 | 4880.404103 | 4924.853852 | 3.79E-128 | 1.674711473 | 1.12E-130 | 591.6085592 | 3.192554998 | 127.4209286 |
| ENSMUSG00000017929 | *B4galt5* | 3369.606105 | 3891.75294 | 3112.063891 | 10981.90147 | 11307.98859 | 10842.21652 | 8.76E-68 | 1.6747063 | 1.03E-69 | 311.4996549 | 3.192543549 | 67.05729967 |
| ENSMUSG00000024736 | *Tmem132a* | 472.9271726 | 458.2095008 | 303.2886759 | 1353.655776 | 1321.346339 | 1275.66858 | 5.83E-24 | 1.671163758 | 4.21E-25 | 107.1090688 | 3.184713865 | 23.23414145 |
| ENSMUSG00000025898 | *Cwf19l2* | 221.2851804 | 175.6133329 | 186.5359555 | 630.8626295 | 623.2151179 | 612.7074848 | 9.89E-34 | 1.670750452 | 4.46E-35 | 152.6990001 | 3.183801633 | 33.00502101 |
| ENSMUSG00000049470 | *Aff4* | 1493.07582 | 1485.648425 | 1331.249409 | 4664.841181 | 4664.341719 | 4398.157355 | 1.69E-96 | 1.668979043 | 9.38E-99 | 444.8817571 | 3.179894811 | 95.77211103 |
| ENSMUSG00000028071 | *Sh2d2a* | 7.988634673 | 15.13908042 | 12.07786762 | 43.85676034 | 31.48647808 | 34.79096128 | 0.001332689 | 1.667290036 | 0.000482061 | 12.18380769 | 3.176174191 | 2.875271305 |
| ENSMUSG00000021549 | *Rasa1* | 1640.066698 | 1366.554326 | 1010.514925 | 4165.548833 | 4404.84971 | 4192.310834 | 1.67E-25 | 1.664930448 | 1.11E-26 | 114.3136992 | 3.170983671 | 24.77702606 |
| ENSMUSG00000032244 | *Fem1b* | 2147.345 | 2124.517619 | 1998.216099 | 6591.165039 | 6773.93575 | 6512.674669 | 1.50E-139 | 1.663056088 | 3.53E-142 | 644.4884626 | 3.166866579 | 138.8252743 |
| ENSMUSG00000001542 | *Ell2* | 1686.400779 | 1770.263137 | 2049.21154 | 5895.360668 | 5678.423461 | 5813.956196 | 1.54E-72 | 1.66155642 | 1.56E-74 | 333.6268337 | 3.163576363 | 71.81110917 |
| ENSMUSG00000021990 | *Spata13* | 611.9294159 | 664.1009945 | 513.9803667 | 1939.818246 | 1918.503682 | 1819.760558 | 6.20E-49 | 1.66116543 | 1.37E-50 | 223.7540335 | 3.162719105 | 48.20745146 |
| ENSMUSG00000095403 | *Gm21092* | 12.78181548 | 22.20398462 | 28.18169112 | 88.55691991 | 33.65795933 | 73.44758493 | 0.001810378 | 1.66068016 | 0.000667994 | 11.57625408 | 3.161655463 | 2.74223066 |
| ENSMUSG00000024056 | *Ndc80* | 404.2249144 | 342.1432175 | 399.9116169 | 1260.03846 | 1203.000611 | 1159.698709 | 9.53E-51 | 1.660603008 | 2.01E-52 | 232.1680743 | 3.161486388 | 50.02112111 |
| ENSMUSG00000007812 | *Zfp655* | 683.0282645 | 639.8784658 | 805.1911749 | 2261.996754 | 2255.083275 | 2174.43508 | 3.27E-55 | 1.657093483 | 5.77E-57 | 252.9969372 | 3.153805048 | 54.48555965 |
| ENSMUSG00000052512 | *Nav2* | 866.766862 | 771.0838295 | 464.3269109 | 2185.247424 | 2315.88475 | 2146.409028 | 1.09E-16 | 1.656694894 | 1.21E-17 | 73.1400321 | 3.152933833 | 15.96156291 |
| ENSMUSG00000027677 | *Ttc14* | 460.9442206 | 383.5233707 | 311.3405876 | 1277.749844 | 1226.886905 | 1158.732294 | 2.79E-28 | 1.656323722 | 1.60E-29 | 127.2957501 | 3.15212276 | 27.55483186 |
| ENSMUSG00000058006 | *Mdn1* | 2368.630181 | 2281.964056 | 1403.716615 | 6448.630568 | 6486.214485 | 6168.630718 | 7.51E-21 | 1.656203714 | 6.43E-22 | 92.59063347 | 3.151860567 | 20.12436564 |
| ENSMUSG00000021366 | *Hivep1* | 379.460147 | 339.1154015 | 202.639779 | 966.5355259 | 1023.853408 | 928.7253831 | 2.42E-15 | 1.655345822 | 2.93E-16 | 66.85256018 | 3.149986885 | 14.61626357 |
| ENSMUSG00000072763 | *5430403G16Rik* | 31.95453869 | 30.27816084 | 29.52367641 | 110.4853001 | 109.659803 | 69.58192256 | 1.05E-06 | 1.655191503 | 2.56E-07 | 26.55332975 | 3.149649962 | 5.979444525 |
| ENSMUSG00000054404 | *Slfn5* | 1231.847467 | 1296.914556 | 1390.296762 | 4212.77919 | 4082.384745 | 4006.759041 | 9.43E-99 | 1.653155855 | 4.90E-101 | 455.3679943 | 3.145208929 | 98.02556003 |
| ENSMUSG00000000489 | *Pdgfb* | 4724.478546 | 4468.047269 | 2942.973744 | 12749.66627 | 12646.70678 | 12687.10388 | 1.00E-24 | 1.649051814 | 6.92E-26 | 110.6893437 | 3.136274453 | 23.99974875 |
| ENSMUSG00000022272 | *Myo10* | 1597.726935 | 1563.362372 | 960.8614687 | 4173.139426 | 4351.648419 | 4405.88868 | 8.11E-21 | 1.647085999 | 6.98E-22 | 92.429137 | 3.132003878 | 20.09071797 |
| ENSMUSG00000113691 | *Gm38699* | 12.78181548 | 8.074176225 | 9.393897041 | 37.95296568 | 41.2581437 | 16.42906505 | 0.009247472 | 1.645387585 | 0.00384711 | 8.354605559 | 3.128318894 | 2.033976997 |
| ENSMUSG00000034303 | *Ccdc15* | 31.15567522 | 16.14835245 | 24.15573525 | 74.21913288 | 74.91610303 | 76.3468317 | 7.15E-06 | 1.644321383 | 1.93E-06 | 22.66288256 | 3.12600781 | 5.14596429 |
| ENSMUSG00000023284 | *Zfp605* | 39.94317336 | 34.31524896 | 32.207647 | 114.7022963 | 110.7455436 | 110.1713774 | 1.39E-09 | 1.644108296 | 2.60E-10 | 39.95296298 | 3.125546132 | 8.855992291 |
| ENSMUSG00000078616 | *Trim30c* | 14.37954241 | 6.055632169 | 9.393897041 | 35.42276796 | 27.14351559 | 31.89171451 | 0.004011629 | 1.64124543 | 0.001564261 | 10.00134337 | 3.119349986 | 2.396679211 |
| ENSMUSG00000046079 | *Lrrc8d* | 2105.005236 | 2093.230186 | 1885.489335 | 6437.666378 | 6347.239685 | 6186.992614 | 2.93E-108 | 1.639006541 | 1.25E-110 | 499.4580741 | 3.114512885 | 107.532434 |
| ENSMUSG00000018846 | *Pank3* | 2047.487067 | 2043.775857 | 1988.822202 | 6349.952857 | 6390.66931 | 6163.79864 | 5.51E-156 | 1.635835758 | 9.75E-159 | 720.6330935 | 3.107675265 | 155.2591109 |
| ENSMUSG00000027306 | *Nusap1* | 551.2157924 | 380.4955546 | 377.0978669 | 1379.801152 | 1327.860783 | 1371.343724 | 1.71E-24 | 1.63383073 | 1.20E-25 | 109.6036724 | 3.103359282 | 23.7659522 |
| ENSMUSG00000051285 | *Pcmtd1* | 146.1920145 | 136.2517238 | 178.4840438 | 505.1961431 | 450.5823588 | 463.8794837 | 4.35E-25 | 1.633802747 | 2.96E-26 | 112.3740624 | 3.103299088 | 24.36132941 |
| ENSMUSG00000028037 | *Ifi44* | 4349.012716 | 3303.347348 | 3163.059332 | 11643.12647 | 11134.27009 | 10790.03007 | 9.95E-38 | 1.632954533 | 3.68E-39 | 171.391113 | 3.101475079 | 37.00221778 |
| ENSMUSG00000057143 | *Trim12c* | 431.3862723 | 442.0611483 | 518.0063225 | 1450.646688 | 1399.519664 | 1442.858478 | 2.04E-54 | 1.632157898 | 3.65E-56 | 249.3200084 | 3.099762964 | 53.69117288 |
| ENSMUSG00000035569 | *Ankrd11* | 1981.980262 | 1863.116164 | 1172.895145 | 5132.084359 | 5461.275337 | 4968.342554 | 2.64E-20 | 1.63085545 | 2.34E-21 | 90.03518348 | 3.096965798 | 19.57850822 |
| ENSMUSG00000027580 | *Helz2* | 2827.177811 | 2767.423901 | 1752.632791 | 7552.640169 | 7767.388421 | 7439.467221 | 1.38E-22 | 1.629823299 | 1.08E-23 | 100.6902681 | 3.094750919 | 21.86026546 |
| ENSMUSG00000026536 | *Ifi211* | 242.8544941 | 270.4849035 | 285.8428671 | 850.9898304 | 817.5626895 | 791.4943691 | 8.88E-43 | 1.629441866 | 2.59E-44 | 194.9900967 | 3.093932809 | 42.05153814 |
| ENSMUSG00000063364 | *3300002I08Rik* | 12.78181548 | 11.10199231 | 8.051911749 | 37.10956644 | 31.48647808 | 31.89171451 | 0.001435701 | 1.629219966 | 0.000522204 | 12.03465827 | 3.093456969 | 2.842935971 |
| ENSMUSG00000037126 | *Psd* | 166.1636012 | 136.2517238 | 112.7267645 | 400.6146377 | 452.75384 | 443.5847563 | 1.34E-19 | 1.628116691 | 1.24E-20 | 86.73510308 | 3.091092208 | 18.87153693 |
| ENSMUSG00000024974 | *Smc3* | 971.4179762 | 908.3448253 | 742.1178662 | 2821.170449 | 2615.549162 | 2684.702512 | 1.01E-45 | 1.626997479 | 2.58E-47 | 208.7480885 | 3.088695135 | 44.99647074 |
| ENSMUSG00000039634 | *Zfp189* | 73.49543899 | 91.84375456 | 100.6488969 | 267.3575582 | 277.9495996 | 266.7307032 | 2.74E-17 | 1.625053714 | 2.93E-18 | 75.93419279 | 3.084536491 | 16.56162385 |
| ENSMUSG00000012443 | *Kif11* | 1012.160013 | 790.259998 | 790.4293367 | 2644.056609 | 2815.325437 | 2549.404329 | 7.43E-41 | 1.623395807 | 2.38E-42 | 185.9979884 | 3.080993859 | 40.12905177 |
| ENSMUSG00000020189 | *Osbpl8* | 3281.731124 | 3139.84528 | 3295.915876 | 10059.2227 | 10074.58725 | 9775.293704 | 1.63E-163 | 1.622089351 | 2.70E-166 | 755.389201 | 3.078205077 | 162.7865413 |
| ENSMUSG00000002028 | *Kmt2a* | 949.0497991 | 999.1793079 | 477.7467638 | 2518.390122 | 2654.635825 | 2304.901185 | 4.44E-12 | 1.621168051 | 6.76E-13 | 51.61196383 | 3.076239975 | 11.35230745 |
| ENSMUSG00000062949 | *Atp11c* | 432.1851358 | 360.310114 | 244.2413231 | 1126.781381 | 1033.625074 | 1045.66167 | 2.45E-17 | 1.620737789 | 2.60E-18 | 76.17114869 | 3.075322669 | 16.61157435 |
| ENSMUSG00000076617 | *Ighm* | 142.9965606 | 111.0199231 | 61.73132341 | 311.2143185 | 336.5795933 | 334.3797945 | 4.36E-09 | 1.618829634 | 8.50E-10 | 37.64232919 | 3.071257838 | 8.360706326 |
| ENSMUSG00000059142 | *Zfp945* | 64.70794085 | 46.42651329 | 30.8656617 | 149.281665 | 128.1173936 | 166.2234817 | 2.25E-07 | 1.617675388 | 5.17E-08 | 29.65037023 | 3.068801623 | 6.646972526 |
| ENSMUSG00000071855 | *Ccdc112* | 43.9374907 | 39.3616091 | 34.89161758 | 122.2928894 | 120.5172092 | 122.7347801 | 4.64E-10 | 1.615576987 | 8.30E-11 | 42.18483243 | 3.064341293 | 9.333847108 |
| ENSMUSG00000069045 | *Ddx3y* | 612.7282794 | 492.5247497 | 519.3483078 | 1712.943851 | 1599.295939 | 1676.731051 | 3.44E-47 | 1.614885538 | 8.32E-49 | 215.5830965 | 3.062872979 | 46.46284092 |
| ENSMUSG00000035247 | *Hectd1* | 2357.446092 | 2281.964056 | 2065.315364 | 7173.110513 | 6809.765191 | 6573.558851 | 1.01E-86 | 1.614771837 | 7.37E-89 | 399.4185407 | 3.062631599 | 85.99689449 |
| ENSMUSG00000032409 | *Atr* | 444.1680878 | 450.1353245 | 457.6169844 | 1350.282179 | 1409.291329 | 1371.343724 | 9.41E-73 | 1.612419854 | 9.39E-75 | 334.6388631 | 3.057642749 | 72.02641297 |
| ENSMUSG00000083068 | *Gm11800* | 35.94885603 | 31.28743287 | 13.41985292 | 92.7739161 | 81.43054677 | 78.27966288 | 4.28E-05 | 1.611673316 | 1.27E-05 | 19.06250056 | 3.056060947 | 4.368796049 |
| ENSMUSG00000089736 | *Tgfbr3l* | 19.17272321 | 16.14835245 | 6.709926458 | 47.23035729 | 45.60110619 | 39.62303924 | 0.001152576 | 1.611333208 | 0.000413714 | 12.46921506 | 3.055340582 | 2.938330357 |
| ENSMUSG00000073489 | *Ifi204* | 972.2168397 | 795.3063582 | 790.4293367 | 2674.418981 | 2601.434534 | 2559.068485 | 6.49E-52 | 1.611234775 | 1.27E-53 | 237.6632031 | 3.055132127 | 51.18788039 |
| ENSMUSG00000090015 | *Gm15446* | 71.09884859 | 54.50068952 | 60.38933812 | 176.2704406 | 191.0903497 | 203.9136897 | 3.79E-13 | 1.609707484 | 5.34E-14 | 56.60084063 | 3.051899563 | 12.42138458 |
| ENSMUSG00000080950 | *Gm7278* | 218.0897266 | 149.3722602 | 186.5359555 | 549.0529034 | 525.4984618 | 620.4388095 | 3.24E-20 | 1.60918168 | 2.89E-21 | 89.61472091 | 3.050787471 | 19.48966689 |
| ENSMUSG00000027365 | *Trpm7* | 1554.588307 | 1425.092104 | 1398.348674 | 4544.23509 | 4493.880441 | 4328.575433 | 3.71E-98 | 1.608180545 | 1.99E-100 | 452.5660649 | 3.048671161 | 97.43035311 |
| ENSMUSG00000059474 | *Mbtd1* | 315.5510696 | 348.1988497 | 318.0505141 | 1026.416872 | 972.8235987 | 992.5088121 | 5.81E-53 | 1.607928347 | 1.10E-54 | 242.528975 | 3.048138267 | 52.23555206 |
| ENSMUSG00000025779 | *Ly96* | 125.4215644 | 85.78812239 | 101.9908822 | 323.8653071 | 313.7790402 | 321.8163918 | 3.81E-17 | 1.606673505 | 4.10E-18 | 75.27496151 | 3.04548818 | 16.41949546 |
| ENSMUSG00000084071 | *Gm13378* | 43.9374907 | 24.22252868 | 30.8656617 | 105.4249047 | 103.1453592 | 96.64155911 | 6.44E-07 | 1.606519534 | 1.54E-07 | 27.53430613 | 3.045163169 | 6.191159429 |

| | | | | | | | | | | | | | |
|---|---|---|---|---|---|---|---|---|---|---|---|---|---|
| ENSMUSG00000027470 | *Mylk2* | 31.95453869 | 22.20398462 | 38.91757345 | 90.24371838 | 86.85924988 | 103.4064683 | 8.52E-07 | 1.605384702 | 2.07E-07 | 26.96622252 | 3.042768768 | 6.069400739 |
| ENSMUSG00000098934 | *Gvin-ps4* | 23.16704055 | 8.074176225 | 10.73588233 | 42.16996186 | 43.42962494 | 45.42153278 | 0.002592428 | 1.604311926 | 0.000977669 | 10.86938746 | 3.040507032 | 2.586293246 |
| ENSMUSG00000098090 | *2700099C18Rik* | 47.93180804 | 48.44505735 | 33.54963229 | 141.6910719 | 119.4314686 | 138.1974295 | 1.95E-09 | 1.603623369 | 3.68E-10 | 39.27491499 | 3.039056231 | 8.710662771 |
| ENSMUSG00000040321 | *Zfp770* | 317.94766 | 209.9285819 | 236.1894113 | 813.0368647 | 751.3325115 | 767.3339794 | 4.34E-21 | 1.602858939 | 3.68E-22 | 93.69817228 | 3.037446376 | 20.36227573 |
| ENSMUSG00000082593 | *Gm11331* | 36.7477195 | 21.19471259 | 28.18169112 | 91.08711762 | 71.65888115 | 100.5072215 | 6.06E-06 | 1.601459856 | 1.62E-06 | 22.99675624 | 3.034502178 | 5.217309804 |
| ENSMUSG00000048852 | *Gm12185* | 8.78749814 | 8.074176225 | 8.051911749 | 23.61517864 | 22.80055309 | 28.99246773 | 0.006455261 | 1.59285331 | 0.00260702 | 9.063923103 | 3.016453432 | 2.190086187 |
| ENSMUSG00000053774 | *Ubxn7* | 131.0136086 | 117.0755553 | 99.30691157 | 349.1672842 | 362.6373683 | 345.010366 | 2.85E-21 | 1.591235694 | 2.38E-22 | 94.55566484 | 3.013073141 | 20.54554321 |
| ENSMUSG00000041483 | *Zfp281* | 477.7203534 | 444.0796924 | 411.9894845 | 1418.597517 | 1325.689301 | 1282.433489 | 1.17E-51 | 1.589434147 | 2.35E-53 | 236.4373021 | 3.009312952 | 50.93090651 |
| ENSMUSG00000093625 | *Gm18953* | 37.54658296 | 26.24107273 | 18.78779408 | 68.31533822 | 100.973878 | 84.07815643 | 2.97E-05 | 1.58457379 | 8.62E-06 | 19.79572317 | 2.999191808 | 4.52738986 |
| ENSMUSG00000082838 | *Gm14519* | 137.4045164 | 161.4835245 | 166.4061761 | 473.1469721 | 492.9262431 | 420.3907821 | 5.12E-24 | 1.582843377 | 3.68E-25 | 107.3775776 | 2.995596641 | 23.29084725 |
| ENSMUSG00000056342 | *Usp34* | 1679.211008 | 1422.064288 | 1211.812718 | 4361.217456 | 4445.022113 | 4133.359483 | 1.75E-37 | 1.582178378 | 6.58E-39 | 170.2326743 | 2.994216163 | 36.75609288 |
| ENSMUSG00000031246 | *Sh3bgrl* | 1154.35771 | 871.0017603 | 980.9912481 | 3137.445163 | 2914.127834 | 2960.130956 | 2.06E-41 | 1.581961744 | 6.37E-43 | 188.6164283 | 2.993766587 | 40.68597483 |
| ENSMUSG00000020134 | *Peli1* | 739.7475707 | 674.1937148 | 687.0964693 | 2150.668055 | 2011.877375 | 2133.845625 | 2.00E-74 | 1.581570242 | 1.92E-76 | 342.3927952 | 2.992954282 | 73.69940399 |
| ENSMUSG00000044501 | *Zfp758* | 73.49543899 | 54.50068952 | 45.62749991 | 194.8252238 | 156.3466498 | 174.921222 | 9.52E-10 | 1.580724758 | 1.75E-10 | 40.72960425 | 2.991200792 | 9.02143609 |
| ENSMUSG00000062627 | *Mysm1* | 331.5283389 | 306.8186966 | 287.1848524 | 904.9673816 | 947.8515644 | 927.7589675 | 4.32E-47 | 1.580362204 | 1.05E-48 | 215.1218084 | 2.990449187 | 46.36433109 |
| ENSMUSG00000038056 | *Kmt2c* | 655.0680432 | 607.5817609 | 327.4444111 | 1669.08709 | 1712.212963 | 1388.739204 | 7.07E-12 | 1.580294915 | 1.09E-12 | 50.66747165 | 2.990309712 | 11.15036208 |
| ENSMUSG00000078899 | *Gm4631* | 562.399881 | 465.274405 | 433.4612492 | 1519.805425 | 1412.548551 | 1454.455465 | 1.44E-38 | 1.58007533 | 5.17E-40 | 175.2928253 | 2.989854609 | 37.8423399 |
| ENSMUSG00000018920 | *Cxcl16* | 895.5259468 | 772.0931015 | 750.169778 | 2403.687826 | 2340.856784 | 2497.217887 | 6.51E-58 | 1.578808387 | 1.03E-59 | 265.6068547 | 2.987230136 | 57.1867141 |
| ENSMUSG00000049985 | *Ankrd55* | 107.8465681 | 82.76030631 | 123.4626468 | 310.3709193 | 311.607559 | 309.2529892 | 1.24E-16 | 1.575938859 | 1.38E-17 | 72.87823724 | 2.981294425 | 15.9055763 |
| ENSMUSG00000025019 | *Lcor* | 154.9795127 | 145.3351721 | 151.6443379 | 443.6279988 | 499.4406868 | 404.9281327 | 4.59E-24 | 1.574654467 | 3.28E-25 | 107.6066692 | 2.978641442 | 23.33854528 |
| ENSMUSG00000040703 | *Cyp2s1* | 18.37385975 | 17.15762448 | 21.47176466 | 66.62853974 | 54.28703118 | 47.35436397 | 7.95E-05 | 1.57439361 | 2.43E-05 | 17.81594 | 2.978102914 | 4.09977742 |
| ENSMUSG00000030657 | *Xylt1* | 712.5862128 | 714.5645959 | 440.1711756 | 1814.995159 | 1994.505525 | 1767.574116 | 1.95E-18 | 1.573899464 | 1.93E-19 | 81.31355306 | 2.97708304 | 17.71085818 |
| ENSMUSG00000091183 | *Gm5141* | 31.95453869 | 33.30597693 | 20.12977937 | 93.61731533 | 78.1733249 | 86.01098761 | 1.99E-06 | 1.573249753 | 5.03E-07 | 25.25141038 | 2.975742627 | 5.70218279 |
| ENSMUSG00000026039 | *Sgo2a* | 373.8681027 | 288.6518 | 261.6871318 | 930.2693587 | 928.3082331 | 901.6657465 | 9.11E-26 | 1.569415499 | 5.98E-27 | 115.5439928 | 2.96784449 | 25.04065274 |
| ENSMUSG00000090790 | *Gm17377* | 22.36817708 | 31.28743287 | 53.67941166 | 102.0513077 | 95.54517487 | 113.0706242 | 2.53E-05 | 1.568314666 | 7.28E-06 | 20.11729023 | 2.965580771 | 4.596617089 |
| ENSMUSG00000042406 | *Atf4* | 5062.397792 | 4690.087115 | 4499.676682 | 14180.07138 | 14115.71385 | 13995.63059 | 1.79E-113 | 1.568236861 | 7.06E-116 | 523.5740127 | 2.965420842 | 112.7480857 |
| ENSMUSG00000032724 | *Abtb2* | 155.7783761 | 138.2702679 | 72.46720574 | 364.3484705 | 347.4369995 | 385.5998209 | 1.98E-09 | 1.56657163 | 3.74E-10 | 39.24477503 | 2.961999979 | 8.704229844 |
| ENSMUSG00000020611 | *Gna13* | 2158.529089 | 2105.341451 | 2082.761172 | 6343.205663 | 6277.752285 | 6184.093368 | 5.08E-148 | 1.566302117 | 1.02E-150 | 683.7561756 | 2.961446691 | 147.2943734 |
| ENSMUSG00000032265 | *Tent5a* | 217.2908631 | 221.0305742 | 206.6657349 | 646.887215 | 627.5580804 | 638.8007057 | 8.32E-40 | 1.566274313 | 2.80E-41 | 181.0931814 | 2.961389619 | 39.08012054 |
| ENSMUSG00000002250 | *Ppard* | 543.2271578 | 453.1631406 | 383.8077934 | 1331.727396 | 1425.577439 | 1350.082581 | 5.39E-30 | 1.565958689 | 2.87E-31 | 135.2802816 | 2.960741815 | 29.268279 |
| ENSMUSG00000027620 | *Rbm39* | 3413.543596 | 2952.120682 | 3420.720508 | 9686.44024 | 9719.550062 | 9522.092819 | 4.13E-82 | 1.563809547 | 3.22E-84 | 378.1061031 | 2.956334564 | 81.38426296 |
| ENSMUSG00000020074 | *Ccar1* | 1209.479289 | 1032.485285 | 860.2125719 | 3241.183269 | 3033.559302 | 2917.60867 | 5.09E-33 | 1.563657183 | 2.38E-34 | 149.3719227 | 2.956022361 | 32.2934649 |
| ENSMUSG00000050350 | *Gpr18* | 15.97726935 | 13.12053637 | 9.393897041 | 35.42276796 | 44.2581437 | 39.62303924 | 0.001009595 | 1.563504286 | 0.000359358 | 12.73248813 | 2.955709099 | 2.995852736 |
| ENSMUSG00000037225 | *Fgf2* | 11.98295201 | 9.083448253 | 9.393897041 | 31.20577178 | 33.65795933 | 26.09322096 | 0.003422465 | 1.562628288 | 0.001318772 | 10.31606466 | 2.953914949 | 2.465661022 |
| ENSMUSG00000040429 | *Mterf1a* | 56.71930618 | 45.41724127 | 60.38933812 | 171.2100452 | 176.9757216 | 130.4661048 | 1.65E-09 | 1.562187591 | 3.10E-10 | 39.61361173 | 2.953012761 | 8.783543309 |
| ENSMUSG00000023947 | *Nfkbie* | 255.6363095 | 198.8265895 | 199.9558084 | 632.5494279 | 634.0725242 | 677.4573294 | 9.51E-27 | 1.56195232 | 5.95E-28 | 120.1220922 | 2.95253123 | 26.02188531 |
| ENSMUSG00000031712 | *Il15* | 59.11589658 | 41.38015315 | 44.28551462 | 154.3420604 | 146.5749842 | 130.4661048 | 2.18E-09 | 1.561217746 | 4.13E-10 | 39.04962502 | 2.95102828 | 8.662037852 |
| ENSMUSG00000021188 | *Trip11* | 581.5726042 | 456.1909567 | 348.9161758 | 1388.235144 | 1436.434845 | 1286.299152 | 7.71E-19 | 1.561159556 | 7.45E-20 | 83.19162889 | 2.950908435 | 18.11277191 |
| ENSMUSG00000087370 | *Tmem170b* | 227.6760882 | 219.0120301 | 142.2504409 | 608.09085 | 577.6140117 | 566.3195364 | 3.19E-17 | 1.561000266 | 3.42E-18 | 75.63007152 | 2.950583459 | 16.49664918 |
| ENSMUSG00000003452 | *Bicd1* | 22.36817708 | 26.24107273 | 13.41985292 | 59.03794661 | 67.31591866 | 59.91776665 | 6.40E-05 | 1.560180047 | 1.94E-05 | 18.25202188 | 2.948906433 | 4.193696825 |
| ENSMUSG00000029422 | *Rsrc2* | 881.9452679 | 854.8534078 | 1030.644704 | 2720.805939 | 2706.751375 | 2710.795733 | 1.25E-60 | 1.559990937 | 1.80E-62 | 278.2529098 | 2.948519913 | 59.9038286 |
| ENSMUSG00000034751 | *Mast4* | 28.75908482 | 27.25034476 | 29.52367641 | 68.31533822 | 92.287953 | 91.80948116 | 1.94E-06 | 1.559464214 | 4.90E-07 | 25.30289781 | 2.947443614 | 5.713067496 |
| ENSMUSG00000112328 | *Gm47325* | 15.17840588 | 23.21325665 | 6.709926458 | 48.07375652 | 38.00092182 | 49.28719515 | 0.002787831 | 1.559259896 | 0.00105827 | 10.72273719 | 2.947026218 | 2.554733582 |
| ENSMUSG00000006740 | *Kif5b* | 2183.293856 | 2055.887121 | 1803.628232 | 6146.693641 | 5927.058064 | 5750.172767 | 5.86E-65 | 1.558459922 | 7.51E-67 | 298.3580299 | 2.945392546 | 64.23193922 |
| ENSMUSG00000075376 | *Rc3h2* | 627.9066853 | 612.6281211 | 573.0277195 | 1837.766938 | 1838.158876 | 1668.999726 | 1.10E-61 | 1.556368295 | 1.53E-63 | 283.1673084 | 2.941125396 | 60.9584977 |
| ENSMUSG00000017548 | *Suz12* | 1622.491702 | 1455.370265 | 1091.034042 | 4297.962513 | 4017.240307 | 3960.371092 | 6.62E-28 | 1.555429394 | 3.86E-29 | 125.5467274 | 2.939211943 | 27.17916137 |
| ENSMUSG00000032036 | *Kirrel3* | 11.98295201 | 13.12053637 | 9.393897041 | 34.57936873 | 36.9151812 | 30.92529892 | 0.001738659 | 1.554625434 | 0.000640299 | 11.65500751 | 2.937574487 | 2.759785513 |
| ENSMUSG00000051341 | *Zfp52* | 144.5942876 | 129.1868196 | 114.0687498 | 366.035269 | 397.3810682 | 385.5998209 | 3.37E-22 | 1.554590155 | 2.70E-23 | 98.87042766 | 2.937502654 | 21.4728843 |
| ENSMUSG00000067931 | *Zfp948* | 77.48975633 | 67.62122589 | 83.20308807 | 210.8498093 | 231.2627528 | 226.1412483 | 2.96E-15 | 1.553061062 | 3.60E-16 | 66.44436076 | 2.934390883 | 14.52844584 |
| ENSMUSG00000043241 | *Upf2* | 482.5135342 | 441.0518763 | 515.3223519 | 1489.443053 | 1383.233554 | 1338.485594 | 3.67E-49 | 1.552372211 | 7.99E-51 | 224.8306931 | 2.93299012 | 48.43527347 |
| ENSMUSG00000098549 | *Gm5269* | 138.2033798 | 94.87157064 | 88.57102924 | 310.3709193 | 298.5786715 | 344.0439504 | 6.20E-13 | 1.551748724 | 8.86E-14 | 55.6048641 | 2.931722848 | 12.20734549 |
| ENSMUSG00000083736 | *Gm6039* | 42.33976377 | 17.15762448 | 120.7786762 | 190.6082276 | 130.2888748 | 201.014443 | 0.00889988 | 1.549942622 | 0.003692001 | 8.429410313 | 2.928054937 | 2.050615842 |
| ENSMUSG00000044791 | *Setd2* | 850.7895927 | 795.3063582 | 585.1055871 | 2210.549401 | 2275.712347 | 2060.39804 | 3.13E-27 | 1.548298886 | 1.90E-28 | 122.3909864 | 2.924720754 | 26.50398629 |
| ENSMUSG00000078651 | *Aoc2* | 123.8238374 | 116.0662832 | 83.20308807 | 328.0823033 | 327.8936683 | 298.6224177 | 1.33E-15 | 1.546905315 | 1.57E-16 | 68.07664419 | 2.921896985 | 14.87712285 |
| ENSMUSG00000091618 | *H60c* | 37.54658296 | 34.31524896 | 33.54963229 | 97.83431152 | 105.3168405 | 106.305715 | 2.29E-08 | 1.546074194 | 4.80E-09 | 34.26930602 | 2.920214196 | 7.640192051 |
| ENSMUSG00000015880 | *Ncapg* | 919.4918508 | 823.565975 | 789.0873514 | 2412.965218 | 2615.549162 | 2385.113679 | 1.25E-58 | 1.54603739 | 1.92E-60 | 268.9518969 | 2.920139701 | 57.9042842 |
| ENSMUSG00000093593 | *Gm20683* | 24.76476749 | 43.39869721 | 26.83970583 | 52.29075271 | 117.2599873 | 108.2385462 | 0.000540134 | 1.544009491 | 0.000184667 | 13.98100362 | 2.916037942 | 3.267498257 |
| ENSMUSG00000079429 | *Mroh2a* | 112.6397489 | 119.0940993 | 67.09926458 | 272.4179536 | 318.1220027 | 290.8910929 | 3.81E-11 | 1.543697022 | 6.22E-12 | 47.25928897 | 2.915406434 | 10.41958252 |
| ENSMUSG00000094520 | *Or51q1* | 24.76476749 | 14.12980839 | 18.78779408 | 53.97755118 | 62.97295617 | 53.15285751 | 0.00014284 | 1.543259618 | 4.52E-05 | 16.63798519 | 2.914522659 | 3.845151132 |
| ENSMUSG00000026565 | *Pou2f1* | 412.2135491 | 391.5975469 | 234.847426 | 964.8487274 | 1033.625074 | 1044.695254 | 5.66E-15 | 1.543131129 | 7.00E-16 | 65.1324897 | 2.914263099 | 14.24689674 |
| ENSMUSG00000106666 | *Gm43587* | 55.92044271 | 61.56559372 | 37.57558816 | 163.619452 | 156.3466498 | 137.2310139 | 4.51E-09 | 1.542779612 | 8.80E-10 | 37.57315443 | 2.913553115 | 8.345854931 |
| ENSMUSG00000072774 | *Zfp951* | 31.95453869 | 17.15762448 | 42.94352933 | 104.5815054 | 82.51628739 | 77.31324729 | 0.000100419 | 1.540773167 | 3.12E-05 | 17.34597464 | 2.909503875 | 3.998183853 |
| ENSMUSG00000083777 | *Gm13361* | 19.17272321 | 13.12053637 | 12.07786762 | 49.760555 | 35.82944058 | 45.42153278 | 0.000851806 | 1.540578003 | 0.000300073 | 13.0699404 | 2.90911031 | 3.069659177 |
| ENSMUSG00000055480 | *Zfp458* | 34.35112909 | 35.32452098 | 28.18169112 | 99.52110999 | 80.34480614 | 106.305715 | 4.91E-07 | 1.539774968 | 1.16E-07 | 28.08178115 | 2.907491487 | 6.308782003 |
| ENSMUSG00000053641 | *Dennd4a* | 535.2385231 | 607.5817609 | 552.8979401 | 1709.570254 | 1710.041482 | 1502.776244 | 7.55E-48 | 1.537987129 | 1.76E-49 | 218.6781898 | 2.903890652 | 47.12177232 |
| ENSMUSG00000022034 | *Esco2* | 226.8772247 | 173.5947888 | 212.0336761 | 573.5114813 | 584.1284555 | 623.3380563 | 6.38E-26 | 1.537752437 | 4.15E-27 | 116.2678358 | 2.903418298 | 25.19524725 |
| ENSMUSG00000039934 | *Gsap* | 674.2407664 | 514.7287344 | 564.9758077 | 1749.210018 | 1687.240929 | 1665.134064 | 1.20E-37 | 1.536934112 | 4.42E-39 | 171.0229493 | 2.901771888 | 36.9225036 |
| ENSMUSG00000022329 | *Stk3* | 358.6896968 | 389.5790029 | 609.2613223 | 1451.490087 | 1351.747076 | 1110.411514 | 1.65E-13 | 1.533524659 | 2.25E-14 | 58.29619336 | 2.894922365 | 12.78355125 |
| ENSMUSG00000058624 | *Gda* | 49.52953497 | 42.38942518 | 42.94352933 | 140.8476726 | 130.2888748 | 120.8019489 | 1.25E-09 | 1.532337632 | 2.34E-10 | 40.16326127 | 2.892541447 | 8.901785198 |
| ENSMUSG00000079173 | *Zan* | 31.95453869 | 30.27816084 | 24.15573525 | 90.24371838 | 102.0596186 | 59.91776665 | 2.65E-05 | 1.531825119 | 7.65E-06 | 20.02460214 | 2.891514064 | 4.576504509 |
| ENSMUSG00000020901 | *Pik3r5* | 2812.798268 | 2969.278307 | 2380.681907 | 7802.286344 | 8068.138574 | 7746.787378 | 1.32E-56 | 1.531338313 | 2.22E-58 | 259.485492 | 2.890538549 | 55.87916189 |
| ENSMUSG00000045392 | *Or5m3b* | 63.11021392 | 65.60268183 | 52.33742637 | 165.3062505 | 187.8331279 | 173.9548064 | 4.43E-12 | 1.529863634 | 6.73E-13 | 51.62088564 | 2.887585439 | 11.35377552 |
| ENSMUSG00000058291 | *Zfp68* | 349.9021987 | 314.8928728 | 371.7299258 | 1050.03205 | 974.99508 | 960.6170976 | 1.81E-41 | 1.529769371 | 5.58E-43 | 188.8802048 | 2.887396776 | 40.74272389 |
| ENSMUSG00000032000 | *Birc3* | 631.9010026 | 592.4426805 | 583.7636018 | 1764.391204 | 1816.444063 | 1646.772167 | 6.59E-62 | 1.528916506 | 9.06E-64 | 284.213988 | 2.885690363 | 61.18102081 |
| ENSMUSG00000022906 | *Parp9* | 1568.168986 | 1549.232563 | 1826.441982 | 4774.483082 | 4767.487078 | 4693.880526 | 3.64E-68 | 1.528292567 | 4.21E-70 | 313.2827976 | 2.884442626 | 67.43913885 |
| ENSMUSG00000021670 | *Hmgcr* | 6197.582779 | 6390.710482 | 5210.928887 | 17400.16966 | 17176.41666 | 16696.76217 | 1.34E-60 | 1.525714967 | 1.95E-62 | 278.0986188 | 2.87929372 | 59.87197415 |
| ENSMUSG00000020863 | *Luc7l3* | 1292.56109 | 1093.041606 | 1503.023526 | 3845.900522 | 3651.345717 | 3683.009818 | 1.94E-36 | 1.525657095 | 7.57E-38 | 165.3776124 | 2.879178223 | 35.71302011 |
| ENSMUSG00000035293 | *G2e3* | 317.94766 | 267.4570875 | 318.0505141 | 925.2089632 | 863.1637957 | 809.8562654 | 1.59E-33 | 1.525136934 | 7.24E-35 | 151.7358685 | 2.878140327 | 32.7990471 |
| ENSMUSG00000072872 | *Rybp* | 2498.046062 | 2415.187963 | 2592.715583 | 7235.522056 | 7226.68959 | 7107.986673 | 6.73E-131 | 1.523753283 | 1.91E-133 | 604.3381058 | 2.8753813 | 130.1719932 |
| ENSMUSG00000043940 | *Wdfy3* | 1985.97458 | 1983.219535 | 972.9393363 | 4777.856679 | 4868.460956 | 4569.212915 | 1.44E-05 | 1.523567381 | 4.01E-06 | 21.25824096 | 2.87501081 | 4.841369265 |
| ENSMUSG00000064370 | *mt-Cytb* | 33522.70768 | 33076.87218 | 43857.42131 | 102533.7321 | 107946.5043 | 106959.012 | 3.09E-38 | 1.523109438 | 1.12E-39 | 173.7564213 | 2.874098364 | 37.51037474 |
| ENSMUSG00000068959 | *Zfp619* | 83.88066407 | 123.1311874 | 111.3847792 | 323.8653071 | 294.235709 | 289.9246773 | 2.70E-14 | 1.522398746 | 3.49E-15 | 61.96877403 | 2.872682892 | 13.56906509 |
| ENSMUSG00000005893 | *Nr2c2* | 729.3623456 | 760.9911092 | 583.7636018 | 2035.965759 | 2010.791635 | 1927.032689 | 2.71E-39 | 1.521724399 | 9.32E-41 | 178.6986668 | 2.871340451 | 38.56634732 |
| ENSMUSG00000031530 | *Dusp4* | 3613.259463 | 3843.307883 | 3244.920435 | 10116.57385 | 10243.96278 | 10370.60571 | 2.20E-77 | 1.520926982 | 1.95E-79 | 356.1441317 | 2.869753821 | 76.65787472 |
| ENSMUSG00000038774 | *Ascc3* | 1545.800809 | 1333.248349 | 1701.63735 | 4546.765288 | 4324.504904 | 4252.228601 | 4.19E-48 | 1.520060969 | 9.56E-50 | 219.8897574 | 2.868031697 | 47.37826469 |
| ENSMUSG00000066613 | *Zfp932* | 141.3988337 | 109.001379 | 152.9863232 | 342.4200903 | 407.1527338 | 403.9617171 | 2.07E-16 | 1.518881049 | 2.33E-17 | 71.84249168 | 2.865687013 | 15.68405981 |
| ENSMUSG00000048118 | *Arid4a* | 231.6704055 | 199.8358016 | 131.5145586 | 584.4756714 | 514.6410556 | 527.6629128 | 1.81E-13 | 1.518822077 | 2.49E-14 | 58.09908485 | 2.865569876 | 12.74153192 |
| ENSMUSG00000038305 | *Spats2l* | 47.93180804 | 21.19471259 | 45.62749991 | 126.5098856 | 103.1453592 | 98.5743903 | 3.18E-05 | 1.517984781 | 9.26E-06 | 19.65914039 | 2.863907271 | 4.497938253 |
| ENSMUSG00000107092 | *Gm7993* | 2788.033501 | 2728.062292 | 2282.716981 | 7436.251075 | 7525.268262 | 7398.877766 | 1.23E-59 | 1.517891362 | 1.85E-61 | 273.6164218 | 2.863721829 | 58.90907286 |
| ENSMUSG00000089809 | *Rasgef1b* | 1062.488411 | 1019.364748 | 1049.432498 | 3091.058204 | 2902.184687 | 2943.701891 | 4.38E-92 | 1.513127271 | 2.64E-94 | 424.4403607 | 2.854280799 | 91.35862304 |
| ENSMUSG00000015340 | *Cybb* | 11762.46569 | 10088.68319 | 9997.790422 | 30999.13896 | 29983.81306 | 29919.26029 | 1.30E-71 | 1.5126892 | 1.36E-73 | 329.3015117 | 2.853414233 | 70.88648454 |
| ENSMUSG00000024402 | *Lta* | 15.17840588 | 13.12053637 | 9.393897041 | 35.42276796 | 42.34388432 | 31.89171451 | 0.002146395 | 1.511964679 | 0.000800464 | 11.24015594 | 2.85198161 | 2.668290443 |
| ENSMUSG00000094410 | *Zbed6* | 221.2851804 | 181.6689651 | 87.22904395 | 556.6434966 | 397.3810682 | 453.2489122 | 8.51E-07 | 1.511430878 | 2.07E-07 | 26.96939228 | 2.850926564 | 6.070007427 |
| ENSMUSG00000038538 | *Ubn2* | 561.6010175 | 494.5432938 | 335.4963229 | 1326.667 | 1359.347261 | 1296.929723 | 9.37E-18 | 1.510398735 | 9.69E-19 | 78.1209531 | 2.848887664 | 17.02810295 |
| ENSMUSG00000034480 | *Diaph2* | 386.6499182 | 381.5048266 | 328.7863964 | 1083.76802 | 1008.653039 | 1043.728838 | 1.26E-42 | 1.509852118 | 3.72E-44 | 194.2699552 | 2.847808465 | 41.8983959 |
| ENSMUSG00000057835 | *Zfp119a* | 48.7306715 | 52.48214546 | 49.65345579 | 123.9796879 | 157.4323904 | 148.828001 | 1.06E-09 | 1.509746738 | 1.96E-10 | 40.50142777 | 2.847600456 | 8.972671431 |
| ENSMUSG00000099342 | *Gm18180* | 51.92612537 | 32.2967049 | 45.62749991 | 177.1138398 | 87.94499051 | 104.3728838 | 7.19E-05 | 1.509503866 | 2.19E-05 | 18.01558098 | 2.847121116 | 4.142982856 |
| ENSMUSG00000094747 | *Or4f14b* | 61.51248698 | 42.38942518 | 29.52367641 | 124.8230871 | 129.2031342 | 133.3653516 | 1.24E-06 | 1.508880774 | 3.08E-07 | 26.20149798 | 2.845891724 | 5.90544182 |
| ENSMUSG00000050106 | *Tmc8* | 146.1920145 | 138.2702679 | 135.5405144 | 381.2164552 | 432.1247682 | 385.5998209 | 2.71E-23 | 1.508173063 | 2.02E-24 | 104.0006225 | 2.84449602 | 22.56683724 |
| ENSMUSG00000068130 | *Zfp442* | 16.77613281 | 16.14835245 | 6.709926458 | 37.10956644 | 40.17240307 | 38.65662365 | 0.002882046 | 1.507020992 | 0.001096756 | 10.65665433 | 2.842225441 | 2.54029906 |
| ENSMUSG00000041685 | *Fcho2* | 452.955586 | 443.0704204 | 331.470367 | 1232.206286 | 1105.283955 | 1164.530787 | 3.53E-27 | 1.505862408 | 2.14E-28 | 122.1523876 | 2.839943855 | 26.45260472 |
| ENSMUSG00000028874 | *Fgr* | 302.7692541 | 262.4107273 | 273.7649995 | 774.2404998 | 838.1917614 | 774.0988885 | 1.28E-36 | 1.503605354 | 4.95E-38 | 166.2206121 | 2.83550432 | 35.89254963 |
| ENSMUSG00000079555 | *Haus3* | 341.913564 | 301.7723364 | 295.2367641 | 879.6654044 | 898.9932363 | 894.9008374 | 2.44E-41 | 1.503419348 | 7.60E-43 | 188.2657841 | 2.835138763 | 40.61273615 |
| ENSMUSG00000097449 | *None* | 55.12157924 | 62.57486574 | 21.47176466 | 141.6910719 | 132.4603561 | 126.6004424 | 4.67E-05 | 1.503201702 | 1.39E-05 | 18.88930571 | 2.834711084 | 4.331103696 |
| ENSMUSG00000061371 | *Zfp873* | 35.94885603 | 32.2967049 | 46.9694852 | 108.7985016 | 119.4314686 | 94.70872793 | 3.22E-07 | 1.501768473 | 7.53E-08 | 28.92438203 | 2.83189637 | 6.491788257 |
| ENSMUSG00000024943 | *Smc5* | 929.8770759 | 783.1950938 | 931.3377923 | 2651.647202 | 2373.429003 | 2452.76277 | 1.98E-47 | 1.500488357 | 4.70E-49 | 216.7180695 | 2.829384719 | 46.70292981 |
| ENSMUSG00000011179 | *Odc1* | 27602.33052 | 24084.25841 | 22829.85378 | 70916.38146 | 69065.04681 | 70727.12504 | 3.87E-67 | 1.499429666 | 4.70E-69 | 308.4701178 | 2.827309197 | 66.41234942 |
| ENSMUSG00000038708 | *Golga4* | 1269.39405 | 1138.458848 | 1108.479851 | 3360.102561 | 3408.139817 | 3188.205035 | 1.36E-67 | 1.498796693 | 1.61E-69 | 310.6049813 | 2.826069005 | 66.86674611 |
| ENSMUSG00000020889 | *Nr1d1* | 77.48975633 | 53.49141749 | 40.25955875 | 148.4382658 | 201.947756 | 141.0966763 | 1.57E-06 | 1.498172251 | 3.94E-07 | 25.72564306 | 2.824846063 | 5.803540721 |
| ENSMUSG00000029561 | *Oasl2* | 1134.386124 | 859.899768 | 587.7895577 | 2530.197712 | 2439.659181 | 2326.162328 | 1.55E-13 | 1.495059201 | 2.12E-14 | 58.41908167 | 2.818757176 | 12.80910947 |
| ENSMUSG00000031138 | *F9* | 71.09884859 | 51.47287344 | 83.20308807 | 197.3554215 | 179.1472029 | 199.0816118 | 3.22E-10 | 1.494249613 | 5.70E-11 | 42.92018393 | 2.817175835 | 9.492517295 |
| ENSMUSG00000089901 | *Gm8113* | 193.3249591 | 260.3921833 | 261.6871318 | 711.8289562 | 613.4434523 | 677.4573294 | 9.30E-21 | 1.493963462 | 8.04E-22 | 92.14898709 | 2.816617118 | 20.03161666 |
| ENSMUSG00000048108 | *Tmem72* | 282.7976674 | 253.3272791 | 220.0855878 | 714.3591539 | 782.8189896 | 644.5991993 | 3.24E-24 | 1.493428801 | 2.29E-25 | 108.3154445 | 2.815573477 | 23.48916688 |
| ENSMUSG00000041328 | *Pcf11* | 1052.103186 | 935.5951701 | 845.4507337 | 2702.251156 | 2677.436378 | 2617.053421 | 3.01E-50 | 1.492893244 | 6.42E-52 | 229.852979 | 2.814528475 | 49.5210858 |
| ENSMUSG00000039981 | *Zc3h12d* | 40.74203683 | 28.25961679 | 38.91757345 | 110.4853001 | 90.11647176 | 102.4400527 | 6.91E-07 | 1.490613644 | 1.66E-07 | 27.39330921 | 2.810084755 | 6.160458338 |
| ENSMUSG00000035112 | *Wnk4* | 9.586361607 | 21.19471259 | 13.41985292 | 41.32656262 | 42.34388432 | 39.62303924 | 0.002474833 | 1.490446886 | 0.000930107 | 10.96177388 | 2.809759961 | 2.606454031 |

| | | | | | | | | | | | | | |
|---|---|---|---|---|---|---|---|---|---|---|---|---|---|
| ENSMUSG00000022974 | *Paxbp1* | 575.1816964 | 522.8029106 | 425.4093374 | 1442.212696 | 1407.119848 | 1443.824893 | 8.18E-33 | 1.488043848 | 3.86E-34 | 148.4089897 | 2.805083755 | 32.08734534 |
| ENSMUSG00000035597 | *Prpf39* | 726.9657552 | 575.285056 | 539.4780872 | 1732.342033 | 1767.585735 | 1683.49596 | 3.10E-31 | 1.487439518 | 1.55E-32 | 141.070149 | 2.803908981 | 30.50919996 |
| ENSMUSG00000096856 | *Gm10778* | 107.8465681 | 93.86229862 | 26.83970583 | 256.3933681 | 200.8620154 | 190.3838715 | 0.0005611 | 1.487314571 | 0.000192398 | 13.90390893 | 2.803666153 | 3.250959825 |
| ENSMUSG00000041842 | *Fhdc1* | 34.35112909 | 29.26888882 | 25.49772054 | 70.84553593 | 93.37369363 | 88.91023438 | 5.31E-06 | 1.487197455 | 1.41E-06 | 23.26288966 | 2.803438566 | 5.274632273 |
| ENSMUSG00000030739 | *Myh14* | 219.6874535 | 183.6875091 | 97.96492628 | 472.3035729 | 483.1545775 | 461.9466526 | 1.16E-08 | 1.486746069 | 2.38E-09 | 35.63601602 | 2.802561571 | 7.933758542 |
| ENSMUSG00000047945 | *Marcksl1* | 3975.943477 | 3905.882749 | 3192.583009 | 10319.83307 | 10235.27686 | 10504.93748 | 1.51E-51 | 1.486635161 | 3.06E-53 | 235.9154578 | 2.802346132 | 50.81966085 |
| ENSMUSG00000024521 | *Pmaip1* | 297.9760733 | 296.7259763 | 307.3146318 | 929.4259594 | 778.4760271 | 813.7219277 | 1.83E-33 | 1.485715814 | 8.34E-35 | 151.4533597 | 2.800560926 | 32.73842525 |
| ENSMUSG00000028339 | *Col15a1* | 9.586361607 | 15.13908042 | 18.78779408 | 34.57936873 | 31.48647808 | 53.15285751 | 0.00482016 | 1.485351155 | 0.001905989 | 9.637940029 | 2.799853138 | 2.316938526 |
| ENSMUSG00000053693 | *Mast1* | 32.75340216 | 46.42651329 | 32.207647 | 103.7381062 | 115.0885061 | 93.74231234 | 7.21E-07 | 1.485083518 | 1.74E-07 | 27.30308994 | 2.799333781 | 6.142006107 |
| ENSMUSG00000059834 | *Sclt1* | 198.1181399 | 210.9378539 | 165.0641909 | 533.0283179 | 546.1275337 | 536.3606531 | 2.62E-26 | 1.484150466 | 1.66E-27 | 118.0871086 | 2.797523918 | 25.58244648 |
| ENSMUSG00000008496 | *Pou2f2* | 726.9657552 | 630.7950176 | 328.7863964 | 1603.30195 | 1543.923167 | 1584.921569 | 7.49E-10 | 1.483970378 | 1.36E-10 | 41.2145703 | 2.797174733 | 9.125700049 |
| ENSMUSG00000040274 | *Cdk6* | 2148.143864 | 1842.930723 | 1388.954777 | 4944.006329 | 5306.014427 | 4814.682475 | 4.03E-23 | 1.483350999 | 3.04E-24 | 103.1926829 | 2.795974104 | 22.39429511 |
| ENSMUSG00000029787 | *Avl9* | 688.6203088 | 811.4547106 | 834.7148513 | 2246.815568 | 2161.709582 | 2088.424092 | 7.21E-47 | 1.480576856 | 1.77E-48 | 214.0842522 | 2.790602921 | 46.14219461 |
| ENSMUSG00000039633 | *Lonrf1* | 123.8238374 | 106.982835 | 73.80919103 | 267.3575582 | 314.8647808 | 278.3276902 | 2.89E-11 | 1.479755661 | 4.69E-12 | 47.81040057 | 2.789014938 | 10.53852803 |
| ENSMUSG00000047036 | *Zfp445* | 1032.930463 | 981.0124113 | 956.8355129 | 2881.051794 | 2778.410256 | 2629.616823 | 6.75E-73 | 1.478577233 | 6.61E-75 | 335.3383695 | 2.786737732 | 72.17098101 |
| ENSMUSG00000032745 | *Gpbp1* | 1674.417827 | 1524.000762 | 1988.822202 | 4815.809645 | 4764.229856 | 4826.279462 | 8.61E-43 | 1.475634428 | 2.50E-44 | 195.0565958 | 2.781059148 | 42.06517151 |
| ENSMUSG00000022533 | *Atp13a3* | 2259.984749 | 2251.685895 | 1877.437423 | 6053.076326 | 5933.572508 | 5782.064482 | 2.51E-56 | 1.473749949 | 4.25E-58 | 258.1945594 | 2.777428839 | 55.59953536 |
| ENSMUSG00000095362 | *Gm14325* | 91.86929874 | 88.81593848 | 110.0427939 | 229.4045925 | 287.7212652 | 286.059015 | 2.24E-13 | 1.47274899 | 3.11E-14 | 57.66365994 | 2.775502494 | 12.64948231 |
| ENSMUSG00000061544 | *Zfp229* | 43.9374907 | 35.32452098 | 65.75727928 | 121.4494902 | 137.8890592 | 137.2310139 | 6.31E-07 | 1.471254593 | 1.51E-07 | 27.57322231 | 2.772629015 | 6.199683686 |
| ENSMUSG00000068284 | *Usf3* | 452.955586 | 421.8757078 | 253.6352201 | 1022.199876 | 1099.855252 | 1019.568449 | 1.98E-13 | 1.469936133 | 2.73E-14 | 57.91724363 | 2.770096303 | 12.70343294 |
| ENSMUSG00000035623 | *Rsf1* | 556.0089732 | 442.0611483 | 444.1971315 | 1350.282179 | 1344.146892 | 1314.325204 | 1.91E-34 | 1.469252157 | 8.28E-36 | 156.043934 | 2.768783323 | 33.71920444 |
| ENSMUSG00000033964 | *Zbtb41* | 544.8248847 | 471.3300371 | 395.885661 | 1391.608741 | 1223.629683 | 1310.459542 | 4.88E-27 | 1.469097675 | 2.98E-28 | 121.4945472 | 2.768486862 | 26.3115525 |
| ENSMUSG00000031604 | *Msmo1* | 3379.192467 | 3138.836008 | 3636.78014 | 9495.832012 | 9170.165307 | 9429.316923 | 8.88E-81 | 1.469068854 | 7.23E-83 | 371.8966214 | 2.768431556 | 80.05161949 |
| ENSMUSG00000098905 | *Zfp953* | 63.90907738 | 59.54704966 | 76.49316162 | 170.3666459 | 169.3755373 | 209.7121833 | 1.79E-10 | 1.468147671 | 3.10E-11 | 44.11404425 | 2.766664433 | 9.74607345 |
| ENSMUSG00000028300 | *C9orf72* | 941.8600279 | 971.9289631 | 1148.73941 | 2842.255429 | 2712.180078 | 2895.381111 | 2.65E-48 | 1.468114444 | 6.01E-50 | 220.8130982 | 2.766600713 | 47.5762877 |
| ENSMUSG00000111594 | *Gm3365* | 25.56363095 | 20.18544056 | 12.07786762 | 59.88134584 | 45.60110619 | 57.98493547 | 0.000670991 | 1.46787221 | 0.000232679 | 13.54687313 | 2.76613623 | 3.173283026 |
| ENSMUSG00000026131 | *Dst* | 3179.4766 | 3031.853173 | 1826.441982 | 7264.19763 | 7772.817124 | 7154.374621 | 1.78E-15 | 1.463934363 | 2.13E-16 | 67.47816207 | 2.758596335 | 14.74960371 |
| ENSMUSG00000026365 | *Cfh* | 150.9851953 | 158.4557084 | 177.1420585 | 476.520569 | 424.5245838 | 431.0213536 | 6.24E-23 | 1.461782648 | 4.75E-24 | 102.3092372 | 2.754485079 | 22.20490559 |
| ENSMUSG00000036478 | *Btg1* | 1680.808735 | 1834.856547 | 1869.385511 | 4910.270359 | 4974.863537 | 4924.853852 | 8.83E-90 | 1.461412613 | 5.68E-92 | 413.7199526 | 2.753778676 | 89.05401343 |
| ENSMUSG00000025225 | *Nfkb2* | 1828.598477 | 1682.456471 | 1371.508968 | 4437.966786 | 4487.365997 | 4538.287616 | 3.30E-38 | 1.460720002 | 1.20E-39 | 173.622525 | 2.752456954 | 37.48184075 |
| ENSMUSG00000023961 | *Enpp4* | 268.418125 | 335.0783133 | 326.1024258 | 866.1710166 | 834.9345395 | 842.7143955 | 4.03E-31 | 1.458717788 | 2.03E-32 | 140.5336694 | 2.748639666 | 30.39494371 |
| ENSMUSG00000026103 | *Gls* | 2191.282491 | 2274.899151 | 2015.661908 | 5897.047467 | 6104.033786 | 5806.224871 | 8.00E-84 | 1.456526767 | 6.14E-86 | 386.0028241 | 2.744468474 | 83.09684875 |
| ENSMUSG00000086414 | *Gm15645* | 279.6022136 | 273.5127196 | 249.6092642 | 763.2763097 | 730.7034397 | 715.1475374 | 2.28E-36 | 1.455716353 | 8.96E-38 | 165.0417758 | 2.742927239 | 35.64227413 |
| ENSMUSG00000091736 | *Yy2* | 51.92612537 | 57.5285056 | 38.91757345 | 135.7872772 | 149.8322061 | 125.6340268 | 2.10E-08 | 1.455571321 | 4.39E-09 | 34.44305224 | 2.742651511 | 7.676878388 |
| ENSMUSG00000034252 | *Senp6* | 901.1179911 | 901.2799211 | 736.749925 | 2366.57826 | 2406.001222 | 2209.226041 | 1.91E-42 | 1.455305943 | 5.66E-44 | 193.4337376 | 2.742147058 | 41.7196974 |
| ENSMUSG00000050555 | *Hyls1* | 154.1806492 | 145.3351721 | 143.5924262 | 410.7354285 | 393.0381057 | 413.625873 | 2.18E-24 | 1.454904822 | 1.53E-25 | 109.1116234 | 2.741384749 | 23.66143668 |
| ENSMUSG00000054409 | *Tmem74* | 28.75908482 | 43.39869721 | 42.94352933 | 91.93051686 | 112.9170249 | 107.2721306 | 2.00E-06 | 1.45437514 | 5.07E-07 | 25.23741235 | 2.740378441 | 5.699334051 |
| ENSMUSG00000051335 | *Gfod1* | 718.1782571 | 835.6772393 | 654.8888223 | 1953.312633 | 2010.791635 | 2095.189002 | 1.63E-39 | 1.453578657 | 5.57E-41 | 179.7243578 | 2.738865949 | 38.78805818 |
| ENSMUSG00000022876 | *Samsn1* | 240.4579037 | 253.3272791 | 293.8947788 | 708.4553593 | 712.2458491 | 719.0131998 | 1.47E-31 | 1.450399852 | 7.31E-33 | 142.5669398 | 2.732837831 | 30.83237773 |
| ENSMUSG00000066263 | *Or51k1* | 46.3340811 | 26.24107273 | 26.83970583 | 83.49652449 | 100.973878 | 91.80948116 | 3.08E-05 | 1.448359515 | 8.97E-06 | 19.71879142 | 2.728975638 | 4.510878616 |
| ENSMUSG00000020859 | *Spag9* | 1707.970093 | 1727.873712 | 1739.212938 | 4737.373516 | 4747.943747 | 4632.029928 | 2.23E-120 | 1.448281001 | 7.36E-123 | 555.667914 | 2.728827125 | 119.6521722 |
| ENSMUSG00000021108 | *Prkch* | 1118.408854 | 1058.726358 | 858.8705866 | 2730.92673 | 2771.895812 | 2803.57163 | 9.72E-39 | 1.448093142 | 3.46E-40 | 176.0917384 | 2.728471817 | 38.01249817 |
| ENSMUSG00000047959 | *Kcna3* | 16.77613281 | 13.12053637 | 8.051911749 | 39.63976415 | 31.48647808 | 34.79096128 | 0.004765453 | 1.447003842 | 0.001881544 | 9.661651642 | 2.726412475 | 2.321895836 |
| ENSMUSG00000021025 | *Nfkbia* | 2146.546137 | 2112.406355 | 2195.487937 | 5910.541855 | 5815.22678 | 5861.31056 | 4.36E-128 | 1.446813254 | 1.31E-130 | 591.2914491 | 2.726052325 | 127.3605531 |
| ENSMUSG00000029253 | *Cenpc1* | 438.5760435 | 397.6531791 | 485.7986755 | 1234.736483 | 1141.113395 | 1215.750814 | 8.59E-39 | 1.446311038 | 3.04E-40 | 176.351059 | 2.725103526 | 38.06623444 |
| ENSMUSG00000090215 | *Trim34b* | 69.50112165 | 91.84375456 | 81.86110278 | 238.6819841 | 228.0055309 | 192.3167026 | 2.29E-11 | 1.444742568 | 3.69E-12 | 48.28235555 | 2.722142455 | 10.63989283 |
| ENSMUSG00000026981 | *Il1rn* | 1543.404219 | 1268.654939 | 1285.621909 | 3763.247397 | 3707.804229 | 3696.539636 | 1.16E-48 | 1.444039362 | 2.60E-50 | 222.4838792 | 2.720815937 | 47.93502358 |
| ENSMUSG00000046753 | *Ccdc66* | 237.2624498 | 185.7060532 | 167.7481614 | 576.8850783 | 545.041793 | 494.8047827 | 1.86E-17 | 1.442282617 | 1.96E-18 | 76.73496595 | 2.717504863 | 16.73120778 |
| ENSMUSG00000095253 | *Zfp799* | 111.042022 | 101.9364748 | 80.51911749 | 291.8161361 | 270.3494153 | 242.5703134 | 4.68E-13 | 1.442243198 | 6.64E-14 | 56.17071829 | 2.717430614 | 12.32946442 |
| ENSMUSG00000059208 | *Hnrnpm* | 3628.437868 | 3579.887884 | 2475.962863 | 8601.828821 | 8740.21202 | 8928.713646 | 5.84E-24 | 1.438594307 | 4.22E-25 | 107.1063144 | 2.710566321 | 23.23389252 |
| ENSMUSG00000064363 | *mt-Nd4* | 19511.44133 | 21061.48868 | 20163.32901 | 54453.22835 | 55949.30007 | 53983.97492 | 1.15E-130 | 1.436512623 | 3.34E-133 | 603.2210146 | 2.706658029 | 129.9379743 |
| ENSMUSG00000018143 | *Mafk* | 1131.19067 | 1223.237698 | 1081.640145 | 2994.067292 | 3133.44744 | 3173.708801 | 8.17E-69 | 1.435179349 | 9.25E-71 | 316.3024115 | 2.704157813 | 68.08795479 |
| ENSMUSG00000021112 | *Pals1* | 371.4715123 | 395.634635 | 497.8765432 | 1171.481541 | 1040.139517 | 1188.691177 | 7.32E-25 | 1.434017619 | 5.03E-26 | 111.3219014 | 2.701981168 | 24.13572127 |
| ENSMUSG00000058922 | *Gm10052* | 1882.122329 | 1627.955781 | 1229.258527 | 4340.975874 | 4213.75936 | 4262.859172 | 6.86E-23 | 1.432966926 | 5.26E-24 | 102.1080765 | 2.70001407 | 22.16382414 |
| ENSMUSG00000062310 | *Glrp1* | 181.3420071 | 148.3629881 | 155.6702938 | 475.6771698 | 419.0958807 | 419.4243665 | 5.74E-20 | 1.431991708 | 5.20E-21 | 88.45656302 | 2.69818956 | 19.24082789 |
| ENSMUSG00000050944 | *Efcab5* | 9.586361607 | 12.11126434 | 12.07786762 | 26.98877559 | 32.57221871 | 30.92529892 | 0.006269835 | 1.431696131 | 0.002526214 | 9.121510018 | 2.697636816 | 2.2027439 |
| ENSMUSG00000039384 | *Dusp10* | 61.51248698 | 57.5285056 | 44.28551462 | 180.4874368 | 134.6318373 | 128.5332736 | 1.84E-07 | 1.430850366 | 4.19E-08 | 30.06127836 | 2.696055818 | 6.735969876 |
| ENSMUSG00000074896 | *Ifit3* | 696.6089435 | 589.4148644 | 589.131543 | 1765.234604 | 1683.983707 | 1617.7797 | 1.44E-41 | 1.430271106 | 4.44E-43 | 189.3345049 | 2.694973536 | 40.84105901 |
| ENSMUSG00000031762 | *Mt2* | 7749.774496 | 8308.327336 | 10435.27763 | 23600.84086 | 22566.03312 | 25175.12615 | 7.32E-33 | 1.42962077 | 3.45E-34 | 148.6352471 | 2.693758974 | 32.1357159 |
| ENSMUSG00000037270 | *Bltp1* | 969.8202493 | 810.4454386 | 687.0964693 | 2268.743948 | 2230.111241 | 2160.905262 | 8.81E-28 | 1.428011872 | 5.16E-29 | 124.973014 | 2.69075656 | 27.05491715 |
| ENSMUSG00000020290 | *Xpo1* | 3974.34575 | 3651.546198 | 3231.500582 | 9816.323722 | 9803.15209 | 9595.540404 | 8.54E-56 | 1.426712437 | 1.48E-57 | 255.7041702 | 2.688334087 | 55.06839741 |
| ENSMUSG00000028793 | *Rnf19b* | 1947.629133 | 1917.616853 | 1707.005291 | 4845.328618 | 5131.210187 | 5025.361074 | 3.64E-70 | 1.426594281 | 3.96E-72 | 322.588162 | 2.688113923 | 69.43865138 |
| ENSMUSG00000054598 | *9130230L23Rik* | 27.96022136 | 33.30597693 | 30.8656617 | 80.12292754 | 83.60202801 | 83.11174084 | 3.09E-06 | 1.426172699 | 7.99E-07 | 24.35933473 | 2.68732852 | 5.510692908 |
| ENSMUSG00000025862 | *Stag2* | 1861.351879 | 1582.53854 | 1927.090879 | 4854.60601 | 4936.862615 | 4632.996344 | 2.36E-50 | 1.425916256 | 5.02E-52 | 230.3403557 | 2.686850883 | 49.62616965 |
| ENSMUSG00000071551 | *Akr1c19* | 58.31703311 | 71.658314 | 40.25955875 | 151.8118627 | 148.7464654 | 161.3914037 | 6.65E-08 | 1.42499977 | 1.45E-08 | 32.11622693 | 2.685144578 | 7.177278799 |
| ENSMUSG00000025612 | *Bach1* | 507.2783017 | 568.2201518 | 610.6033076 | 1459.92408 | 1596.038717 | 1457.354711 | 4.82E-40 | 1.424989968 | 1.61E-41 | 182.1911457 | 2.685126334 | 39.31678054 |
| ENSMUSG00000029714 | *Gigyf1* | 468.9328553 | 467.292949 | 289.868823 | 1068.586834 | 1186.714502 | 1044.695254 | 1.46E-14 | 1.421787094 | 1.85E-15 | 63.21398115 | 2.679171796 | 13.83665994 |
| ENSMUSG00000005045 | *Chd5* | 86.27725447 | 76.70467414 | 32.207647 | 163.619452 | 175.889981 | 191.350287 | 3.34E-05 | 1.421703285 | 9.77E-06 | 19.55641229 | 2.679016163 | 4.475905361 |
| ENSMUSG00000038174 | *Hycc2* | 321.1431138 | 314.8928728 | 354.284117 | 869.5446136 | 932.6511956 | 842.7143955 | 2.00E-36 | 1.420953653 | 7.84E-38 | 165.3078374 | 2.677624493 | 35.69843365 |
| ENSMUSG00000027379 | *Bub1* | 1232.64633 | 1071.846894 | 1133.977571 | 3155.999946 | 2964.071902 | 3094.462723 | 1.61E-61 | 1.420819578 | 2.25E-63 | 282.4025759 | 2.677375664 | 60.79369133 |
| ENSMUSG00000030016 | *Zfp638* | 1653.647377 | 1395.823215 | 1415.794483 | 4031.448354 | 4048.726785 | 3892.722001 | 4.93E-54 | 1.420779421 | 9.01E-56 | 247.5212408 | 2.677301139 | 53.30750363 |
| ENSMUSG00000030203 | *Dusp16* | 840.4043676 | 719.6109561 | 770.2995573 | 2174.283234 | 2132.394585 | 1935.730429 | 4.19E-45 | 1.419923478 | 1.09E-46 | 205.8777116 | 2.675713184 | 44.37814875 |
| ENSMUSG00000030779 | *Rbbp6* | 1563.375805 | 1612.816701 | 1119.215733 | 3950.482027 | 3814.206811 | 3746.793247 | 1.37E-24 | 1.419897841 | 9.52E-26 | 110.0562888 | 2.675665635 | 23.86438104 |
| ENSMUSG00000035969 | *Rusc2* | 1859.754152 | 2049.831489 | 1737.870953 | 4889.185378 | 5185.497218 | 5037.924477 | 6.69E-62 | 1.418830254 | 9.27E-64 | 284.1688574 | 2.673686389 | 61.17489853 |
| ENSMUSG00000032254 | *Kif23* | 938.6645741 | 868.9832162 | 611.9452929 | 2203.802207 | 2096.565144 | 2186.032067 | 2.57E-20 | 1.418519904 | 2.27E-21 | 90.09537715 | 2.673111292 | 19.59085288 |
| ENSMUSG00000063663 | *Brwd3* | 274.0101693 | 289.6610721 | 297.9207347 | 844.2426365 | 687.2738147 | 764.4347326 | 2.24E-27 | 1.418394006 | 1.34E-28 | 123.0746444 | 2.672878031 | 26.6498012 |
| ENSMUSG00000081739 | *Mdm4-ps* | 17.57499628 | 26.24107273 | 10.73588233 | 50.60395423 | 49.94406868 | 47.35436397 | 0.001702283 | 1.418186785 | 0.0006263 | 11.69613583 | 2.67249414 | 2.768968258 |
| ENSMUSG00000026482 | *Rgl1* | 1044.114552 | 966.882603 | 896.4461747 | 2568.150677 | 2538.461578 | 2679.870434 | 7.46E-58 | 1.417711882 | 1.18E-59 | 265.3257486 | 2.671614559 | 57.12706192 |
| ENSMUSG00000040481 | *Bptf* | 1256.612234 | 1349.396702 | 815.9270572 | 3066.599627 | 3090.017815 | 2998.787579 | 9.43E-17 | 1.417297475 | 1.04E-17 | 73.43976647 | 2.670847263 | 16.0256558 |
| ENSMUSG00000058503 | *Fam133b* | 474.5248996 | 485.4598455 | 566.317793 | 1393.29554 | 1305.06023 | 1358.780321 | 1.11E-40 | 1.416583079 | 3.60E-42 | 185.1717509 | 2.669525036 | 39.95346125 |
| ENSMUSG00000029634 | *Rnf6* | 870.7611793 | 793.2878141 | 740.7758809 | 2166.69264 | 2168.224025 | 2094.222586 | 8.88E-54 | 1.414508964 | 1.65E-55 | 246.3208401 | 2.665689904 | 53.05136062 |
| ENSMUSG00000091549 | *Gm6548* | 217.2908631 | 227.0862063 | 271.0810289 | 613.9946447 | 664.4732616 | 617.5395627 | 1.84E-25 | 1.414483619 | 1.22E-26 | 114.1233081 | 2.665643075 | 24.73571212 |
| ENSMUSG00000048327 | *Ckap2l* | 637.4930469 | 497.5711099 | 599.8674253 | 1627.760528 | 1508.093726 | 1488.28001 | 2.12E-32 | 1.413759857 | 1.02E-33 | 146.476336 | 2.664306128 | 31.67346608 |
| ENSMUSG00000049516 | *Spty2d1* | 599.1476005 | 678.2308029 | 587.7895577 | 1672.460687 | 1634.039638 | 1663.201232 | 3.83E-51 | 1.412962939 | 7.92E-53 | 234.0210005 | 2.662834821 | 50.41659042 |
| ENSMUSG00000039997 | *Ifi203* | 1415.586064 | 1248.469499 | 1316.487571 | 3597.941146 | 3534.08573 | 3470.398388 | 8.74E-71 | 1.412066695 | 9.33E-73 | 325.4672725 | 2.661181104 | 70.05862076 |
| ENSMUSG00000035671 | *Zswim4* | 664.6544048 | 620.7022973 | 531.4261754 | 1578.843372 | 1678.555004 | 1596.518557 | 1.50E-38 | 1.411915762 | 5.39E-40 | 175.2103746 | 2.660902709 | 37.82504746 |
| ENSMUSG00000021392 | *Nol8* | 887.5373122 | 765.0281973 | 860.2125719 | 2309.227112 | 2271.369384 | 2102.920326 | 6.29E-49 | 1.411009198 | 1.40E-50 | 223.7215112 | 2.65923117 | 48.2015146 |
| ENSMUSG00000024235 | *Map3k8* | 206.905638 | 154.4186203 | 159.6962497 | 426.760014 | 478.811615 | 489.0062891 | 8.12E-17 | 1.410732485 | 8.91E-18 | 73.74049908 | 2.658721171 | 16.09042437 |
| ENSMUSG00000063317 | *Usp31* | 988.194109 | 927.5209939 | 591.8155136 | 2318.504503 | 2263.7692 | 2097.121833 | 2.37E-15 | 1.409829121 | 2.86E-16 | 66.89613296 | 2.657056896 | 14.62458923 |
| ENSMUSG00000020580 | *Rock2* | 1118.408854 | 975.9660512 | 762.2476456 | 2576.58467 | 2491.774731 | 2541.673005 | 1.67E-24 | 1.409692228 | 1.17E-25 | 109.6539693 | 2.656804789 | 23.77660466 |
| ENSMUSG00000033355 | *Rtp4* | 1129.592943 | 961.8362428 | 1160.817277 | 3005.874881 | 2831.611546 | 2788.10898 | 3.85E-46 | 1.408256953 | 9.73E-48 | 210.6867832 | 2.654162962 | 45.4143975 |
| ENSMUSG00000004837 | *Grap* | 1472.30537 | 1521.982218 | 1753.974776 | 4090.486301 | 4086.727707 | 4402.989433 | 3.86E-51 | 1.408173039 | 8.00E-53 | 234.0002751 | 2.654008587 | 50.41330996 |
| ENSMUSG00000035208 | *Slfn8* | 603.9407813 | 504.6360141 | 590.4735283 | 1556.071593 | 1548.266129 | 1405.16827 | 1.64E-37 | 1.408148656 | 6.10E-39 | 170.3833061 | 2.653963734 | 36.78625199 |
| ENSMUSG00000037007 | *Zfp113* | 84.67952753 | 132.2146357 | 108.7008086 | 291.8161361 | 281.2068215 | 286.059015 | 1.16E-11 | 1.407190993 | 1.82E-12 | 49.66474361 | 2.652202612 | 10.93550709 |
| ENSMUSG00000024399 | *Ltb* | 1585.743983 | 1412.980839 | 979.6492628 | 3455.406675 | 3586.20128 | 3528.383323 | 2.90E-18 | 1.407031874 | 2.91E-19 | 80.49553221 | 2.651910109 | 17.53827414 |
| ENSMUSG00000027115 | *Kif18a* | 265.2226711 | 153.4093483 | 170.432132 | 528.8113217 | 511.3838337 | 530.5621595 | 6.70E-11 | 1.406026602 | 1.12E-11 | 46.11025835 | 2.650062898 | 10.17424658 |
| ENSMUSG00000044701 | *Il27* | 31.95453869 | 45.41724127 | 24.15573525 | 94.46071457 | 87.94499051 | 88.91023438 | 2.02E-05 | 1.40590439 | 5.74E-06 | 20.57303656 | 2.649838418 | 4.693864361 |
| ENSMUSG00000040721 | *Zfhx2* | 166.9624647 | 178.641149 | 89.91301453 | 350.0106835 | 433.2105088 | 380.7677429 | 3.02E-08 | 1.40585705 | 6.38E-09 | 33.7139793 | 2.64975147 | 7.519286886 |
| ENSMUSG00000025314 | *Ptprj* | 1358.067894 | 1368.57287 | 1085.666101 | 3460.46707 | 3460.255367 | 3195.93636 | 2.58E-37 | 1.405213339 | 9.76E-39 | 169.4491779 | 2.64856945 | 36.58836944 |
| ENSMUSG00000027104 | *Atf2* | 526.4510249 | 464.2651329 | 462.9849256 | 1328.353799 | 1259.459123 | 1266.97084 | 5.89E-44 | 1.403012295 | 1.60E-45 | 200.5312892 | 2.644531748 | 43.22984843 |
| ENSMUSG00000028034 | *Fubp1* | 1876.530285 | 1947.895014 | 1913.671026 | 5010.634868 | 5010.692978 | 5142.29736 | 6.71E-112 | 1.402350953 | 2.77E-114 | 516.2470445 | 2.643319752 | 111.1730294 |
| ENSMUSG00000034459 | *Ifit1* | 1399.608795 | 1185.894633 | 1500.339556 | 3661.196089 | 3488.484623 | 3636.621869 | 3.63E-42 | 1.401895195 | 1.10E-43 | 192.1110524 | 2.64248484 | 41.44040022 |
| ENSMUSG00000021720 | *Rnf180* | 47.93180804 | 68.63049791 | 50.99544108 | 145.908068 | 125.9459123 | 170.089144 | 1.98E-07 | 1.401472832 | 4.53E-08 | 29.9074565 | 2.641711339 | 6.70288566 |
| ENSMUSG00000113175 | *Gm9973* | 81.48407366 | 72.66758603 | 79.1771322 | 208.3196116 | 230.1770122 | 178.7868844 | 3.87E-11 | 1.401423474 | 6.33E-12 | 47.22596202 | 2.641620962 | 10.41266864 |
| ENSMUSG00000105096 | *Gbp10* | 22.36817708 | 22.20398462 | 34.89161758 | 55.66434966 | 56.45851243 | 94.70872793 | 0.000939169 | 1.40018775 | 0.000333066 | 12.87463463 | 2.63935928 | 3.027256293 |
| ENSMUSG00000015568 | *Lpl* | 6664.917908 | 5828.545962 | 4867.380652 | 15769.87894 | 14987.56357 | 15041.29226 | 3.68E-33 | 1.398721692 | 1.71E-34 | 150.0254244 | 2.636678542 | 32.43465849 |
| ENSMUSG00000059897 | *Zfp930* | 325.1374312 | 206.9007658 | 228.1374996 | 681.4665837 | 654.701596 | 678.423745 | 2.41E-14 | 1.398703648 | 3.11E-15 | 62.19346806 | 2.636645565 | 13.61762745 |
| ENSMUSG00000053835 | *H2-T24* | 1640.865562 | 1416.008655 | 814.585072 | 3306.12501 | 3423.340186 | 3466.532725 | 6.20E-11 | 1.394774186 | 1.03E-11 | 46.26528937 | 2.629473917 | 10.20768934 |
| ENSMUSG00000066760 | *Psg16* | 251.6419922 | 261.4014553 | 268.3970583 | 669.6589944 | 731.7891803 | 650.3976928 | 6.66E-31 | 1.394644234 | 3.40E-32 | 139.5160187 | 2.629237076 | 30.17644726 |
| ENSMUSG00000080717 | *B230307C23Rik* | 23.96590402 | 52.48214546 | 28.18169112 | 75.90593135 | 106.4025811 | 92.77589675 | 0.000429274 | 1.393626161 | 0.000144409 | 14.44374592 | 2.627382345 | 3.367265904 |
| ENSMUSG00000041992 | *Rapgef5* | 754.1271131 | 747.8705728 | 703.2002928 | 2001.38639 | 1989.076822 | 1811.062818 | 2.56E-54 | 1.393455898 | 4.62E-56 | 248.8542699 | 2.627072286 | 53.59106246 |
| ENSMUSG00000039879 | *Heca* | 184.5374609 | 216.993486 | 173.1161026 | 521.2207286 | 501.6121681 | 488.0398735 | 2.11E-22 | 1.391677843 | 1.67E-23 | 99.81842417 | 2.623836535 | 21.67529646 |
| ENSMUSG00000033306 | *Lpp* | 335.5226563 | 306.8186966 | 152.9863232 | 737.9743326 | 744.8180678 | 614.640316 | 1.10E-07 | 1.391168125 | 2.45E-08 | 31.10214629 | 2.622909673 | 6.959306024 |
| ENSMUSG00000020694 | *Tlk2* | 744.5407515 | 648.9619141 | 726.0140427 | 1874.033105 | 1819.701285 | 1856.484351 | 5.69E-53 | 1.388605368 | 1.08E-54 | 242.5782798 | 2.618254556 | 52.24495141 |
| ENSMUSG00000049421 | *Zfp260* | 313.9533426 | 352.2359378 | 434.8032345 | 994.3677007 | 912.0221238 | 958.6842664 | 9.74E-23 | 1.388553673 | 7.53E-24 | 101.3951947 | 2.618160739 | 22.01152453 |
| ENSMUSG00000068457 | *Uty* | 383.4544643 | 252.318007 | 301.9466906 | 827.3746517 | 791.5049146 | 843.6808111 | 4.02E-17 | 1.388247975 | 4.33E-18 | 75.16457579 | 2.617606026 | 16.395691 |

| | | | | | | | | | | | | | |
|---|---|---|---|---|---|---|---|---|---|---|---|---|---|
| ENSMUSG00000000078 | *Klf6* | 1770.281444 | 1829.810187 | 2242.457422 | 5113.529575 | 5164.868146 | 4990.570113 | 7.33E-40 | 1.388169847 | 2.46E-41 | 181.3501206 | 2.617464275 | 39.13469223 |
| ENSMUSG00000063273 | *Naa15* | 2103.407509 | 1911.561221 | 1780.814482 | 5161.603332 | 5070.408712 | 4948.047827 | 3.65E-60 | 1.386957404 | 5.39E-62 | 276.0731412 | 2.61526548 | 59.43757612 |
| ENSMUSG00000051095 | *Or6x1* | 54.32271578 | 68.63049791 | 55.02139695 | 185.5478322 | 136.8033186 | 142.0630919 | 8.35E-08 | 1.386463105 | 1.84E-08 | 31.65942628 | 2.614369585 | 7.07830192 |
| ENSMUSG00000074994 | *Qser1* | 202.1124572 | 164.5113406 | 146.2763968 | 565.0774889 | 456.0110619 | 323.749223 | 9.89E-09 | 1.385497835 | 2.00E-09 | 35.97735034 | 2.612620964 | 8.004669229 |
| ENSMUSG00000048874 | *Phf3* | 1086.454316 | 1140.477392 | 1217.180659 | 3062.38263 | 2980.358012 | 2934.037735 | 7.50E-69 | 1.385050928 | 8.46E-71 | 316.4821679 | 2.61181177 | 68.12484313 |
| ENSMUSG00000015757 | *Ppil4* | 325.9362947 | 327.0041371 | 304.6306612 | 827.3746517 | 788.2476927 | 889.1023438 | 1.91E-34 | 1.384731902 | 8.30E-36 | 156.0403565 | 2.611234279 | 33.71901315 |
| ENSMUSG00000028233 | *Tgs1* | 833.2145964 | 854.8534078 | 850.8186748 | 2304.166716 | 2195.367541 | 2121.282223 | 1.74E-66 | 1.383746191 | 2.15E-68 | 305.4409587 | 2.609450782 | 65.75872577 |
| ENSMUSG00000036368 | *Rmdn2* | 190.1295052 | 201.8544056 | 234.847426 | 535.5585156 | 535.2701274 | 553.7561337 | 3.91E-24 | 1.383433411 | 2.79E-25 | 107.9278137 | 2.608885107 | 23.40747964 |
| ENSMUSG00000020687 | *Cdc27* | 1290.1645 | 1270.673483 | 1578.174703 | 3705.052849 | 3551.45758 | 3517.752752 | 3.16E-40 | 1.383140624 | 1.04E-41 | 183.0522207 | 2.608355704 | 39.49997448 |
| ENSMUSG00000007617 | *Homer1* | 399.4317336 | 436.0055162 | 399.9116169 | 1093.888811 | 1047.739702 | 1079.486215 | 1.80E-44 | 1.382736864 | 4.81E-46 | 202.9215565 | 2.607625817 | 43.74381711 |
| ENSMUSG00000018199 | *Ro60* | 441.7714974 | 433.9869721 | 457.6169844 | 1219.555297 | 1135.684692 | 1116.210008 | 4.72E-45 | 1.382620929 | 1.24E-46 | 205.6214847 | 2.607416276 | 44.32616337 |
| ENSMUSG00000037253 | *Mex3c* | 850.7895927 | 950.7342505 | 624.0231606 | 2082.352717 | 2201.881985 | 2053.633131 | 3.76E-20 | 1.382281695 | 3.38E-21 | 89.31023025 | 2.606803243 | 19.42482752 |
| ENSMUSG00000041408 | *Wapl* | 2162.523406 | 2222.417006 | 2270.639113 | 5836.322722 | 5884.71418 | 5608.109675 | 9.48E-103 | 1.381485363 | 4.48E-105 | 473.9286542 | 2.60536475 | 102.0232017 |
| ENSMUSG00000066798 | *Zbtb6* | 443.3692243 | 400.6809952 | 429.4352933 | 1149.55316 | 1086.826364 | 1082.385462 | 2.92E-43 | 1.381386632 | 8.25E-45 | 197.265779 | 2.605186457 | 42.53389176 |
| ENSMUSG00000045005 | *Fzd5* | 184.5374609 | 159.4649804 | 112.7267645 | 402.3014362 | 414.7529182 | 384.6334053 | 2.12E-12 | 1.380998957 | 3.15E-13 | 53.11524445 | 2.604486497 | 11.67453378 |
| ENSMUSG00000037617 | *Spag1* | 39.1443099 | 36.33379301 | 36.23360287 | 80.12292754 | 108.5740624 | 103.4064683 | 4.02E-06 | 1.379190887 | 1.06E-06 | 23.82463498 | 2.601224446 | 5.395703008 |
| ENSMUSG00000020716 | *Nf1* | 512.0714825 | 472.3393092 | 332.8123523 | 1148.709761 | 1169.342652 | 1124.907748 | 5.88E-18 | 1.379071679 | 6.00E-19 | 79.06670441 | 2.601009518 | 17.23053267 |
| ENSMUSG00000101523 | *Csnk2a1-ps3* | 510.4737556 | 649.9711861 | 625.3651458 | 1685.111676 | 1478.778729 | 1466.052452 | 1.86E-27 | 1.378293271 | 1.11E-28 | 123.4477141 | 2.599606519 | 26.72974605 |
| ENSMUSG00000035158 | *Mitf* | 316.349933 | 332.0504973 | 365.0199993 | 881.3522029 | 893.5645332 | 848.512889 | 6.76E-36 | 1.378130066 | 2.74E-37 | 162.8155766 | 2.599312455 | 35.17006128 |
| ENSMUSG00000039531 | *Zup1* | 194.1238226 | 197.8173175 | 263.0291171 | 590.3794661 | 541.7845712 | 556.6553805 | 1.91E-18 | 1.378046009 | 1.88E-19 | 81.35667605 | 2.599161013 | 17.71981642 |
| ENSMUSG00000036391 | *Sec24a* | 651.0737258 | 657.0360903 | 362.3360287 | 1448.95989 | 1548.266129 | 1357.813906 | 4.30E-11 | 1.37786368 | 7.05E-12 | 47.01326658 | 2.59883255 | 10.36679156 |
| ENSMUSG00000049717 | *Lig4* | 125.4215644 | 132.2146357 | 124.8046321 | 366.035269 | 290.9784871 | 335.3462101 | 1.35E-15 | 1.37747415 | 1.60E-16 | 68.04867905 | 2.598130956 | 14.87117976 |
| ENSMUSG00000042842 | *Serpinb6b* | 123.8238374 | 143.316628 | 134.1985292 | 337.3596949 | 380.0092182 | 323.749223 | 6.38E-17 | 1.377116287 | 6.95E-18 | 74.2317382 | 2.597486565 | 16.19521778 |
| ENSMUSG00000053062 | *Jam2* | 18.37385975 | 24.22252868 | 18.78779408 | 56.50774889 | 45.60110619 | 57.01851988 | 0.000503204 | 1.376944342 | 0.000171239 | 14.12297991 | 2.597177007 | 3.298255968 |
| ENSMUSG00000078495 | *Zfp984* | 139.8011068 | 116.0662832 | 123.4626468 | 340.7332918 | 349.6084808 | 298.6224177 | 9.94E-16 | 1.376903438 | 1.17E-16 | 68.66264562 | 2.597103371 | 15.00270743 |
| ENSMUSG00000091474 | *2610021A01Rik* | 197.3192764 | 153.4093483 | 126.1466174 | 413.2656262 | 423.4388432 | 413.625873 | 1.78E-13 | 1.376466392 | 2.45E-14 | 58.13099315 | 2.596316731 | 12.74838964 |
| ENSMUSG00000084353 | *Gm15452* | 247.6476749 | 219.0120301 | 198.6138231 | 540.6189111 | 579.785493 | 617.5395627 | 3.13E-22 | 1.376457609 | 2.51E-23 | 99.01556087 | 2.596300925 | 21.50410168 |
| ENSMUSG00000004535 | *Tax1bp1* | 1936.445045 | 1606.761069 | 2066.657349 | 5077.263408 | 4814.173925 | 4661.022396 | 5.60E-37 | 1.376166998 | 2.14E-38 | 167.8862136 | 2.595777989 | 36.251704 |
| ENSMUSG00000035236 | *Scai* | 153.3817857 | 83.76957834 | 122.1206615 | 326.3955048 | 327.8936683 | 281.226937 | 1.14E-08 | 1.375089373 | 2.31E-09 | 35.69061255 | 2.59383979 | 7.944568156 |
| ENSMUSG00000111585 | *Gm47355* | 24.76476749 | 9.083448253 | 32.207647 | 56.50774889 | 51.02980931 | 61.85059783 | 0.006316594 | 1.374938302 | 0.002545799 | 9.107382439 | 2.593568191 | 2.199517033 |
| ENSMUSG00000066440 | *Zfyve26* | 781.288471 | 765.0281973 | 624.0231606 | 1822.585752 | 1936.961272 | 1883.543987 | 3.81E-36 | 1.373574846 | 1.52E-37 | 163.990613 | 2.591118231 | 35.41911842 |
| ENSMUSG00000021270 | *Hsp90aa1* | 15331.78766 | 13431.39215 | 17683.34019 | 41119.92981 | 40086.62956 | 39085.71217 | 3.86E-38 | 1.373108303 | 1.41E-39 | 173.3012607 | 2.590280443 | 37.41309148 |
| ENSMUSG00000025255 | *Zfhx4* | 464.938538 | 368.3842903 | 229.4794848 | 961.4751304 | 926.1367519 | 878.4717723 | 1.03E-09 | 1.372730172 | 1.90E-10 | 40.57187756 | 2.589601617 | 8.987634924 |
| ENSMUSG00000021519 | *Mterf3* | 436.1794531 | 454.1724127 | 406.6215433 | 1095.575609 | 1130.255989 | 1140.370398 | 8.13E-45 | 1.372640281 | 2.14E-46 | 204.5301726 | 2.589440271 | 44.0899909 |
| ENSMUSG00000029720 | *Gm20605* | 206.1067746 | 207.9100378 | 76.49316162 | 452.9053904 | 442.9821744 | 382.7005741 | 3.01E-05 | 1.372560256 | 8.74E-06 | 19.76968002 | 2.58929664 | 4.521560574 |
| ENSMUSG00000040225 | *Prrc2c* | 5099.944375 | 4992.868723 | 2955.051612 | 11476.97682 | 11537.07987 | 10772.63459 | 7.62E-14 | 1.371921647 | 1.02E-14 | 59.86483579 | 2.588150743 | 13.11831196 |
| ENSMUSG00000049409 | *Prokr1* | 36.7477195 | 26.24107273 | 17.44580879 | 60.72474508 | 74.91610303 | 77.31324729 | 0.000433454 | 1.371281018 | 0.000145943 | 14.42384106 | 2.587001729 | 3.363056653 |
| ENSMUSG00000027162 | *Lin7c* | 902.715718 | 806.4083505 | 948.7836011 | 2333.685689 | 2268.112163 | 2259.479652 | 7.70E-52 | 1.369994261 | 1.51E-53 | 237.3160567 | 2.584695379 | 51.11348988 |
| ENSMUSG00000106948 | *Gm42785* | 62.31135045 | 58.53777763 | 36.23360287 | 120.6060909 | 171.5470185 | 119.8355333 | 1.06E-05 | 1.369797936 | 2.92E-06 | 21.865998 | 2.584343673 | 4.973345673 |
| ENSMUSG00000028256 | *Odf2l* | 176.5488263 | 203.8729497 | 173.1161026 | 479.894166 | 489.6690212 | 461.9466526 | 7.67E-23 | 1.369078569 | 5.90E-24 | 101.8808779 | 2.583055369 | 22.11526834 |
| ENSMUSG00000032690 | *Oas2* | 2351.055184 | 2549.421143 | 2011.635952 | 5917.289048 | 6133.348782 | 5809.124118 | 3.19E-42 | 1.368082912 | 9.58E-44 | 192.3874521 | 2.581273324 | 41.49648335 |
| ENSMUSG00000003382 | *Etv3* | 1735.131451 | 1876.2367 | 1528.521247 | 4441.340383 | 4516.680994 | 4313.112783 | 2.44E-48 | 1.366714618 | 5.52E-50 | 220.9815444 | 2.578826328 | 47.61189698 |
| ENSMUSG00000020120 | *Plek* | 10717.55228 | 9678.91875 | 9948.136966 | 26648.8857 | 26040.40312 | 25564.59163 | 5.08E-94 | 1.366456657 | 2.97E-96 | 433.3923881 | 2.578365263 | 93.29407752 |
| ENSMUSG00000040435 | *Ppp1r15a* | 1210.278153 | 1133.412488 | 1042.722572 | 2951.89733 | 2893.498762 | 2904.078851 | 1.63E-59 | 1.366192788 | 2.47E-61 | 273.0363273 | 2.577893722 | 58.7877488 |
| ENSMUSG00000095865 | *Gm13237* | 802.0589212 | 593.4519525 | 726.0140427 | 1848.731128 | 1913.074979 | 1711.522012 | 2.64E-26 | 1.366003183 | 1.68E-27 | 118.0648612 | 2.577554949 | 25.57838251 |
| ENSMUSG00000052825 | *Gm9892* | 1095.241814 | 789.250726 | 1376.876909 | 2682.852974 | 2742.580815 | 2968.828696 | 3.21E-14 | 1.365685909 | 4.18E-15 | 61.61105905 | 2.576988161 | 13.49352406 |
| ENSMUSG00000043279 | *Trim56* | 867.5657255 | 933.576626 | 666.9666899 | 2092.473508 | 2232.282722 | 2050.733884 | 7.06E-25 | 1.36546226 | 4.84E-26 | 111.3965726 | 2.576588702 | 24.1513328 |
| ENSMUSG00000024491 | *Rbm27* | 964.228205 | 875.0388484 | 832.0308807 | 2327.781895 | 2345.199747 | 2226.621522 | 2.19E-52 | 1.365347948 | 4.27E-54 | 239.8372916 | 2.576384554 | 51.65931474 |
| ENSMUSG00000021796 | *Bmpr1a* | 1955.617768 | 1807.606202 | 2226.353599 | 5253.533849 | 5161.610924 | 4996.368606 | 2.66E-47 | 1.365150269 | 6.37E-49 | 216.112847 | 2.576031562 | 46.57520561 |
| ENSMUSG00000070738 | *Dgkd* | 5126.30687 | 5145.268799 | 4109.158963 | 12100.24886 | 12706.42252 | 12228.05647 | 1.34E-40 | 1.363832863 | 4.36E-42 | 184.7896233 | 2.573680316 | 39.87241048 |
| ENSMUSG00000057335 | *Cep170* | 1310.136086 | 1398.851031 | 1407.742571 | 3562.518378 | 3585.115539 | 3427.876102 | 1.27E-78 | 1.362784849 | 1.11E-80 | 361.8524091 | 2.571811401 | 77.89501574 |
| ENSMUSG00000042508 | *Dmtf1* | 543.2271578 | 604.5539449 | 559.6078666 | 1540.890406 | 1451.635214 | 1391.638451 | 3.82E-44 | 1.361526923 | 1.03E-45 | 201.4057052 | 2.569569945 | 43.41781945 |
| ENSMUSG00000029919 | *Hpgds* | 1163.944072 | 1264.617851 | 1697.611394 | 3659.50929 | 3415.740002 | 3504.222933 | 9.18E-22 | 1.361309541 | 7.51E-23 | 96.84193898 | 2.569182796 | 21.03717312 |
| ENSMUSG00000038039 | *Gcc2* | 428.9896819 | 341.1339455 | 249.6092642 | 905.8107808 | 859.9065739 | 866.8747852 | 3.26E-13 | 1.360059663 | 4.57E-14 | 56.90545721 | 2.56695795 | 12.48628718 |
| ENSMUSG00000028649 | *Macf1* | 4845.106929 | 4845.515007 | 3516.001464 | 11316.73097 | 11849.77317 | 10754.2727 | 1.87E-25 | 1.360042318 | 1.25E-26 | 114.0887354 | 2.566927088 | 24.72852455 |
| ENSMUSG00000031592 | *Pcm1* | 794.0702865 | 716.58314 | 528.7422049 | 1806.561166 | 1794.729251 | 1650.63783 | 1.88E-19 | 1.35981705 | 1.76E-20 | 86.04735219 | 2.56652631 | 18.72547264 |
| ENSMUSG00000046434 | *Hnrnpa1* | 3873.688953 | 2949.092866 | 2292.110878 | 8377.484623 | 7636.013805 | 7381.482285 | 1.49E-15 | 1.358783215 | 1.78E-16 | 67.83610019 | 2.564687796 | 14.82629932 |
| ENSMUSG00000006262 | *Mob1b* | 710.9884859 | 629.7857456 | 548.8719842 | 1695.232467 | 1578.666867 | 1586.854401 | 2.20E-31 | 1.357937431 | 1.10E-32 | 141.7613179 | 2.563184683 | 30.6582836 |
| ENSMUSG00000023827 | *Agpat4* | 826.0248252 | 892.1964729 | 904.4980865 | 2258.623157 | 2174.738469 | 2275.908717 | 2.67E-62 | 1.357910308 | 3.63E-64 | 286.038953 | 2.563136494 | 61.57305913 |
| ENSMUSG00000025531 | *Chm* | 490.5021689 | 450.1353245 | 489.8246314 | 1238.953479 | 1224.715423 | 1193.523255 | 6.48E-47 | 1.354629962 | 1.58E-48 | 214.2998898 | 2.55731515 | 46.18818858 |
| ENSMUSG00000018849 | *Wwc1* | 96.66247954 | 100.9272028 | 123.4626468 | 274.9481513 | 280.1210809 | 258.9993784 | 9.07E-14 | 1.35455342 | 1.22E-14 | 59.51098527 | 2.557179477 | 13.04260217 |
| ENSMUSG00000031137 | *Fgf13* | 53.52385231 | 60.55632169 | 34.89161758 | 138.3174749 | 128.1173936 | 118.8691177 | 1.49E-06 | 1.353952964 | 3.72E-07 | 25.83301159 | 2.556115389 | 5.826367472 |
| ENSMUSG00000035840 | *Lysmd3* | 390.6442355 | 330.0319532 | 354.284117 | 951.3543396 | 915.2793457 | 885.2366815 | 1.45E-32 | 1.35317386 | 6.94E-34 | 147.2429075 | 2.554735373 | 31.83783834 |
| ENSMUSG00000062248 | *Cks2* | 305.964708 | 272.5034476 | 222.7695584 | 767.4933059 | 641.6727085 | 647.4984461 | 1.82E-17 | 1.352663716 | 1.92E-18 | 76.77176468 | 2.553832165 | 16.73905658 |
| ENSMUSG00000074733 | *Zfp950* | 86.27725447 | 90.83448253 | 106.016838 | 267.3575582 | 229.0912716 | 220.3427548 | 1.05E-11 | 1.351721622 | 1.65E-12 | 49.86724746 | 2.552165033 | 10.97755118 |
| ENSMUSG00000068882 | *Ssb* | 2225.63362 | 1944.867198 | 2241.115437 | 5407.875909 | 5459.103855 | 5466.046583 | 3.07E-62 | 1.349339738 | 4.19E-64 | 285.7527701 | 2.547954894 | 61.51258357 |
| ENSMUSG00000027351 | *Spred1* | 4095.772997 | 4006.809952 | 3114.747862 | 9570.894544 | 9664.17729 | 9362.634247 | 5.14E-33 | 1.349036084 | 2.40E-34 | 149.3504123 | 2.547418664 | 32.28931111 |
| ENSMUSG00000032487 | *Ptgs2* | 5794.156728 | 5430.892783 | 4898.246314 | 14205.37335 | 13725.93296 | 13158.71469 | 2.31E-57 | 1.348835685 | 3.72E-59 | 263.045854 | 2.547064838 | 56.63653943 |
| ENSMUSG00000018425 | *Dhx40* | 362.6840141 | 340.1246735 | 309.9986023 | 875.4484082 | 796.9336177 | 915.1955648 | 5.87E-29 | 1.348439934 | 3.29E-30 | 130.4385776 | 2.546366239 | 28.23120057 |
| ENSMUSG00000020330 | *Hmmr* | 1198.295201 | 902.2891932 | 774.3255132 | 2459.352176 | 2484.174547 | 2392.845004 | 2.26E-18 | 1.347904911 | 2.25E-19 | 81.00486985 | 2.545422095 | 17.64586115 |
| ENSMUSG00000037826 | *Ppm1k* | 111.042022 | 85.78812239 | 96.62294099 | 279.1651475 | 222.5768278 | 246.4359757 | 3.43E-11 | 1.346918458 | 5.59E-12 | 47.46883681 | 2.543682239 | 10.46506704 |
| ENSMUSG00000060336 | *Zfp937* | 23.16704055 | 27.25034476 | 46.9694852 | 85.18332296 | 81.43054677 | 75.38041611 | 0.000279122 | 1.346373721 | 9.16E-05 | 15.30258479 | 2.54272197 | 3.554205576 |
| ENSMUSG00000020561 | *Polr1f* | 699.0055339 | 580.3314162 | 670.9926458 | 1638.724718 | 1642.725563 | 1679.630297 | 1.55E-40 | 1.346263472 | 5.04E-42 | 184.5040939 | 2.542527664 | 39.81086378 |
| ENSMUSG00000025925 | *Terf1* | 292.384029 | 213.96567 | 220.0855878 | 645.2004165 | 604.7575273 | 606.9089912 | 1.33E-17 | 1.346153723 | 1.38E-18 | 77.41613698 | 2.542334257 | 16.87683614 |
| ENSMUSG00000002413 | *Braf* | 457.7487668 | 419.8571637 | 314.0245582 | 1013.765883 | 1027.11063 | 1005.072215 | 7.59E-20 | 1.346068978 | 6.92E-21 | 87.89130826 | 2.542184923 | 19.119546 |
| ENSMUSG00000040661 | *Rad54l2* | 994.5850168 | 975.9660512 | 633.4170576 | 2128.739675 | 2324.570675 | 2179.267158 | 6.90E-16 | 1.345179623 | 8.05E-17 | 69.39684885 | 2.540618265 | 15.16132166 |
| ENSMUSG00000024843 | *Chka* | 385.0521912 | 360.310114 | 356.9680875 | 990.1507045 | 898.9932363 | 913.2627336 | 6.58E-36 | 1.34406657 | 2.66E-37 | 162.8757046 | 2.53865891 | 35.18193281 |
| ENSMUSG00000026436 | *Elk4* | 599.9464639 | 674.1937148 | 491.1666167 | 1565.348984 | 1504.836504 | 1422.56375 | 5.13E-25 | 1.343915947 | 3.49E-26 | 112.0443787 | 2.538393878 | 24.2902419 |
| ENSMUSG00000027132 | *Katnbl1* | 631.1021392 | 608.591033 | 699.1743369 | 1660.653098 | 1586.267051 | 1655.469908 | 2.38E-47 | 1.342158019 | 5.67E-49 | 216.3454338 | 2.535302722 | 46.62379799 |
| ENSMUSG00000036676 | *Tmtc3* | 612.7282794 | 587.3963204 | 560.9498519 | 1540.890406 | 1492.893357 | 1438.992815 | 6.70E-50 | 1.341844026 | 1.44E-51 | 228.2495421 | 2.534750991 | 49.17379622 |
| ENSMUSG00000074519 | *Zfp971* | 69.50112165 | 46.42651329 | 89.91301453 | 182.1742352 | 168.2897967 | 166.2234817 | 1.01E-06 | 1.340913848 | 2.45E-07 | 26.63803164 | 2.533117235 | 5.997645197 |
| ENSMUSG00000027829 | *Ccnl1* | 834.8123233 | 872.0110323 | 732.7239692 | 2097.533903 | 2124.7944 | 1964.722897 | 8.03E-43 | 1.339394275 | 2.33E-44 | 195.1987037 | 2.530450539 | 42.09530459 |
| ENSMUSG00000020493 | *Prr11* | 516.0657999 | 441.0518763 | 322.07647 | 1117.503989 | 1112.884139 | 1017.635617 | 3.69E-15 | 1.337306377 | 4.51E-16 | 65.99804629 | 2.526791067 | 14.4330505 |
| ENSMUSG00000034832 | *Tet3* | 1456.328101 | 1563.362372 | 708.5682339 | 3040.45425 | 3232.249836 | 3147.61558 | 0.000370709 | 1.336444502 | 0.000123515 | 14.73824969 | 2.525281997 | 3.430966773 |
| ENSMUSG00000041935 | *Rimoc1* | 388.2476451 | 437.0147882 | 428.093308 | 1071.117031 | 1056.425627 | 1022.467695 | 7.80E-39 | 1.333129493 | 2.75E-40 | 176.5492328 | 2.519486093 | 38.10792416 |
| ENSMUSG00000112808 | *Gm4739* | 2710.543745 | 1873.208884 | 2042.501614 | 5716.56003 | 5447.160708 | 5528.863597 | 3.44E-22 | 1.33172968 | 2.76E-23 | 98.82583229 | 2.51704268 | 21.46374407 |
| ENSMUSG00000031488 | *Rab11fip1* | 548.0203386 | 528.8585427 | 448.2230874 | 1299.678225 | 1341.975411 | 1211.885151 | 5.23E-32 | 1.3315935 | 2.55E-33 | 144.6605496 | 2.516805102 | 31.28179029 |
| ENSMUSG00000033487 | *Fndc3a* | 1708.768957 | 1879.264516 | 1869.385511 | 4805.688854 | 4563.367841 | 4346.937329 | 7.23E-61 | 1.330992573 | 1.03E-62 | 279.3657125 | 2.515756992 | 60.14098051 |
| ENSMUSG00000027122 | *Arl14ep* | 667.8498587 | 553.0810714 | 607.9193371 | 1615.109539 | 1562.380757 | 1428.362244 | 2.64E-33 | 1.330745262 | 1.22E-34 | 150.7029249 | 2.515325772 | 32.57775926 |
| ENSMUSG00000025245 | *Lztfl1* | 263.6249442 | 215.984214 | 253.6352201 | 610.6210478 | 603.6717867 | 631.069381 | 2.15E-24 | 1.330649421 | 1.51E-25 | 109.1370891 | 2.515158678 | 23.66665178 |
| ENSMUSG00000034243 | *Golgb1* | 1197.496337 | 1178.829729 | 1023.934777 | 2895.389581 | 2879.384134 | 2793.907474 | 6.02E-51 | 1.330209221 | 1.26E-52 | 233.0917586 | 2.514391362 | 50.22011169 |
| ENSMUSG00000001774 | *Chordc1* | 1642.463289 | 1665.298846 | 2200.855878 | 4697.733751 | 4654.570053 | 4476.437018 | 8.10E-27 | 1.330125416 | 5.03E-28 | 120.4539042 | 2.514245308 | 26.09164932 |
| ENSMUSG00000024298 | *Zfp871* | 365.879468 | 342.1432175 | 355.6261023 | 879.6654044 | 928.3082331 | 866.8747852 | 3.76E-37 | 1.328867647 | 1.43E-38 | 168.6874096 | 2.512054296 | 36.4246881 |
| ENSMUSG00000032212 | *Sltm* | 1096.040677 | 941.6508023 | 1053.458454 | 2711.528548 | 2576.4625 | 2474.990329 | 3.35E-46 | 1.328167591 | 8.44E-48 | 210.9694365 | 2.510835638 | 45.47504744 |
| ENSMUSG00000029267 | *Mtf2* | 575.9805599 | 493.5340218 | 559.6078666 | 1456.550483 | 1293.117083 | 1338.485594 | 1.16E-34 | 1.327645173 | 4.98E-36 | 157.0536734 | 2.509926597 | 33.93628999 |
| ENSMUSG00000024991 | *Eif3a* | 6935.732623 | 6608.71324 | 6393.217929 | 17109.19693 | 16439.19878 | 16511.21037 | 3.66E-100 | 1.327624591 | 1.81E-102 | 461.9448581 | 2.50989079 | 99.43660487 |
| ENSMUSG00000021733 | *Slc4a7* | 1080.063408 | 1218.191338 | 955.4935276 | 2812.736456 | 2711.094337 | 2651.844382 | 2.75E-36 | 1.327226758 | 1.09E-37 | 164.6590401 | 2.509198765 | 35.55996814 |
| ENSMUSG00000016239 | *Lonrf3* | 850.7895927 | 851.8255917 | 779.6934544 | 2009.820382 | 2134.566066 | 2095.189002 | 1.04E-56 | 1.326664678 | 1.73E-58 | 259.9835584 | 2.508221361 | 55.98467048 |
| ENSMUSG00000039478 | *Micu3* | 67.10453125 | 76.70467414 | 56.36338224 | 148.4382658 | 167.204056 | 189.4174559 | 2.55E-08 | 1.325739828 | 5.36E-09 | 34.05395175 | 2.506613964 | 7.593351289 |
| ENSMUSG00000082292 | *Gm12250* | 75.89202939 | 64.5934098 | 71.12522045 | 178.8006383 | 162.8610935 | 189.4174559 | 5.81E-10 | 1.325653699 | 1.05E-10 | 41.72613987 | 2.506464324 | 9.235980736 |
| ENSMUSG00000052336 | *Cx3cr1* | 1327.711083 | 1164.69992 | 650.8628664 | 2709.841749 | 2609.034718 | 2562.934148 | 1.58E-09 | 1.323616506 | 2.96E-10 | 39.70309941 | 2.502927506 | 8.802624578 |
| ENSMUSG00000030142 | *Clec4e* | 8629.323174 | 8447.606876 | 11978.56071 | 24133.86917 | 24469.33643 | 24031.8565 | 1.27E-22 | 1.322269151 | 9.89E-24 | 100.8565876 | 2.500591075 | 21.89541583 |
| ENSMUSG00000083594 | *Gm13722* | 63.90907738 | 66.61195386 | 42.94352933 | 154.3420604 | 142.2320217 | 142.0630919 | 1.80E-07 | 1.321718218 | 4.09E-08 | 30.10525653 | 2.499636338 | 6.745501458 |
| ENSMUSG00000027313 | *Chac1* | 234.8658594 | 197.8173175 | 147.6183821 | 484.1111622 | 503.7836493 | 475.4764708 | 5.12E-13 | 1.321603669 | 7.28E-14 | 55.99108392 | 2.499437875 | 12.29086719 |
| ENSMUSG00000054387 | *Mdm4* | 693.4134896 | 693.3698833 | 642.8109546 | 1719.691045 | 1717.641666 | 1641.940089 | 4.45E-54 | 1.320726638 | 8.12E-56 | 247.7301704 | 2.4979189 | 53.35165068 |
| ENSMUSG00000015305 | *Sash1* | 1252.617917 | 1384.721223 | 1037.35463 | 3021.899467 | 3157.333733 | 3007.48532 | 2.09E-30 | 1.319494652 | 1.09E-31 | 137.1978241 | 2.49578672 | 29.68087994 |
| ENSMUSG00000008976 | *Gabpa* | 991.3895629 | 919.4468176 | 1042.722572 | 2520.076921 | 2478.745844 | 2363.852536 | 2.92E-54 | 1.319455839 | 5.27E-56 | 248.5907657 | 2.495719576 | 53.53503662 |
| ENSMUSG00000045664 | *Cdc42ep2* | 272.4124423 | 323.976321 | 236.1894113 | 671.3457928 | 670.9877054 | 739.3079272 | 2.68E-19 | 1.317361496 | 2.52E-20 | 85.33106886 | 2.492099198 | 18.57252891 |
| ENSMUSG00000028701 | *Lurap1* | 22.36817708 | 22.20398462 | 14.76183821 | 44.70015957 | 64.05869679 | 41.55587042 | 0.002674313 | 1.317356483 | 0.001010759 | 10.8077534 | 2.492090539 | 2.572787828 |
| ENSMUSG00000028224 | *Nbn* | 541.6294308 | 514.7287344 | 554.2399254 | 1341.004787 | 1288.77412 | 1378.108633 | 6.27E-47 | 1.316683498 | 1.53E-48 | 214.3720412 | 2.490928304 | 46.20287819 |
| ENSMUSG00000017421 | *Zfp207* | 3398.36519 | 3087.363134 | 3230.158597 | 8077.234495 | 7989.965249 | 8123.689459 | 5.05E-88 | 1.315520799 | 3.55E-90 | 405.4703413 | 2.488921621 | 87.29640097 |
| ENSMUSG00000049502 | *Dtx3l* | 1147.167939 | 1211.126434 | 1060.16838 | 2800.928867 | 2936.928387 | 2777.478409 | 8.12E-56 | 1.31453168 | 1.40E-57 | 255.8110547 | 2.487215788 | 55.09021744 |
| ENSMUSG00000029798 | *Herc6* | 338.7181101 | 339.1154015 | 292.5527936 | 833.2784464 | 788.2476927 | 798.2592783 | 1.88E-30 | 1.312821915 | 9.80E-32 | 137.4110254 | 2.484269889 | 29.72554554 |
| ENSMUSG00000031939 | *Taf1d* | 1559.381488 | 1113.227047 | 1265.49213 | 3194.796311 | 3235.507058 | 3326.402465 | 2.13E-24 | 1.307175379 | 1.49E-25 | 109.1660211 | 2.474565756 | 23.67226092 |
| ENSMUSG00000001168 | *Oas1h* | 25.56363095 | 19.17616853 | 13.41985292 | 45.54355881 | 54.28703118 | 47.35436397 | 0.002408953 | 1.306258009 | 0.000904495 | 11.01352525 | 2.472992748 | 2.618171743 |
| ENSMUSG00000040128 | *Pnrc1* | 238.0613133 | 240.2067427 | 264.3711024 | 618.2116409 | 625.3865992 | 585.6478482 | 8.06E-27 | 1.305421687 | 5.00E-28 | 120.46532 | 2.471559583 | 26.09373585 |

| | | | | | | | | | | | | | |
|---|---|---|---|---|---|---|---|---|---|---|---|---|---|
| ENSMUSG00000044641 | *Pard6b* | 207.7045015 | 201.8544056 | 136.8824997 | 434.3506072 | 483.1545775 | 444.5511719 | 1.41E-12 | 1.30521608 | 2.07E-13 | 53.94118462 | 2.471207372 | 11.84946721 |
| ENSMUSG00000027201 | *Myef2* | 1131.989533 | 996.1514918 | 1198.392865 | 2821.170449 | 2599.263053 | 2786.176149 | 3.19E-41 | 1.304440075 | 1.00E-42 | 187.7187474 | 2.469878503 | 40.49578391 |
| ENSMUSG00000050721 | *Plekho2* | 5864.456713 | 5725.600216 | 4297.036903 | 13094.61656 | 13070.14563 | 13064.97238 | 7.97E-27 | 1.303313427 | 4.94E-28 | 120.4891072 | 2.467950447 | 26.09853033 |
| ENSMUSG00000107653 | *Gm31520* | 18.37385975 | 30.27816084 | 22.81374996 | 50.60395423 | 55.3727718 | 69.58192256 | 0.001201634 | 1.302938623 | 0.000432316 | 12.38707531 | 2.46730937 | 2.920227834 |
| ENSMUSG00000104377 | *Gm37515* | 79.08748326 | 68.63049791 | 68.44124987 | 175.4270413 | 203.0334966 | 157.5257414 | 6.86E-09 | 1.301751913 | 1.36E-09 | 36.7239605 | 2.465280683 | 8.163605999 |
| ENSMUSG00000022228 | *Zscan26* | 476.1226265 | 472.3393092 | 434.8032345 | 1129.311579 | 1128.084508 | 1158.732294 | 1.29E-42 | 1.300554175 | 3.82E-44 | 194.2165286 | 2.463234834 | 41.88794848 |
| ENSMUSG00000037533 | *Rapgef6* | 933.0725298 | 1102.125055 | 1019.908822 | 2626.345225 | 2484.174547 | 2405.408406 | 1.56E-42 | 1.3003386 | 4.61E-44 | 193.8404782 | 2.462866793 | 41.80674128 |
| ENSMUSG00000078185 | *Chml* | 278.8033501 | 259.3829112 | 166.4061761 | 620.7418386 | 566.7566055 | 558.5882117 | 3.96E-11 | 1.299472067 | 6.48E-12 | 47.17947947 | 2.461387954 | 10.4026827 |
| ENSMUSG00000091144 | *Phf11c* | 57.51816964 | 44.40796924 | 44.28551462 | 116.3890947 | 127.031653 | 119.8355333 | 5.89E-07 | 1.298438182 | 1.41E-07 | 27.71258446 | 2.459624669 | 6.22968693 |
| ENSMUSG00000021770 | *Samd8* | 575.1816964 | 661.0731784 | 605.2353665 | 1587.277364 | 1461.406879 | 1474.750192 | 2.79E-39 | 1.29812665 | 9.63E-41 | 178.6350793 | 2.459093602 | 38.55395289 |
| ENSMUSG00000056144 | *Trim34a* | 112.6397489 | 96.8901147 | 118.0947057 | 269.0443567 | 260.5777497 | 273.4956123 | 1.13E-13 | 1.297151211 | 1.53E-14 | 59.0575996 | 2.457431515 | 12.94558821 |
| ENSMUSG00000052748 | *Swt1* | 428.1908185 | 332.0504973 | 386.491764 | 934.4863549 | 971.7378581 | 916.1619804 | 6.62E-25 | 1.296594848 | 4.54E-26 | 111.5259314 | 2.45648401 | 24.1789229 |
| ENSMUSG00000056537 | *Rlim* | 2002.750712 | 2060.933481 | 2144.492496 | 5330.283179 | 4970.520575 | 4933.551593 | 6.53E-72 | 1.296491967 | 6.67E-74 | 330.7282099 | 2.45630884 | 71.18484736 |
| ENSMUSG00000020949 | *Fkbp3* | 1247.025872 | 969.910419 | 1253.414262 | 2843.942228 | 2723.037484 | 2948.533969 | 3.66E-28 | 1.295555255 | 2.12E-29 | 126.7399246 | 2.454714528 | 27.43599444 |
| ENSMUSG00000033883 | *Zfp267* | 272.4124423 | 306.8186966 | 303.2886759 | 705.0817623 | 686.1880741 | 768.3003949 | 1.71E-26 | 1.295141814 | 1.08E-27 | 118.9470176 | 2.454011167 | 25.76747931 |
| ENSMUSG00000028385 | *Snx30* | 1461.920145 | 1640.067046 | 1532.547203 | 3789.392773 | 3895.637357 | 3683.009818 | 6.67E-62 | 1.295038223 | 9.21E-64 | 284.1831308 | 2.453834966 | 61.17615678 |
| ENSMUSG00000032410 | *Xrn1* | 685.4248549 | 657.0360903 | 633.4170576 | 1604.988748 | 1666.611857 | 1584.921569 | 1.37E-51 | 1.294370018 | 2.75E-53 | 236.1264906 | 2.4526987 | 50.86440685 |
| ENSMUSG00000038344 | *Txlng* | 178.1465532 | 145.3351721 | 127.4886027 | 413.2656262 | 368.0660714 | 332.4469633 | 5.45E-12 | 1.29423579 | 8.34E-13 | 51.20014061 | 2.452470512 | 11.26355464 |
| ENSMUSG00000022119 | *Rbm26* | 914.69867 | 851.8255917 | 625.3651458 | 2030.061964 | 1953.247382 | 1899.973052 | 1.87E-19 | 1.294216324 | 1.74E-20 | 86.06325214 | 2.452437422 | 18.7286899 |
| ENSMUSG00000022237 | *Ankrd33b* | 343.5112909 | 425.9127959 | 383.8077934 | 911.7145754 | 1014.081742 | 898.7664997 | 8.53E-25 | 1.294206473 | 5.89E-26 | 111.0098157 | 2.452420676 | 24.06921353 |
| ENSMUSG00000069495 | *Epc2* | 352.2987891 | 353.2452098 | 318.0505141 | 848.4596327 | 859.9065739 | 807.9234342 | 1.06E-32 | 1.293162821 | 5.01E-34 | 147.8919255 | 2.450647226 | 31.97648437 |
| ENSMUSG00000001151 | *Pcnt* | 811.6452828 | 739.7963966 | 428.093308 | 1601.615152 | 1688.32667 | 1572.358167 | 5.86E-10 | 1.292231267 | 1.06E-10 | 41.70703375 | 2.449065344 | 9.232140303 |
| ENSMUSG00000036501 | *Fam13b* | 448.1624051 | 535.9234469 | 461.6429403 | 1114.130392 | 1175.857095 | 1250.541775 | 1.90E-29 | 1.292097752 | 1.04E-30 | 132.7310456 | 2.448838704 | 28.7215951 |
| ENSMUSG00000071281 | *Zfp65* | 142.1976972 | 175.6133329 | 166.4061761 | 357.6012766 | 403.895512 | 420.3907821 | 2.70E-15 | 1.29178699 | 3.27E-16 | 66.63329589 | 2.448311271 | 14.56901598 |
| ENSMUSG00000112023 | *Lilrb4b* | 8595.770908 | 7854.154923 | 8505.502778 | 21403.78584 | 20142.66005 | 19528.35985 | 8.88E-72 | 1.291280438 | 9.22E-74 | 330.0824568 | 2.447451784 | 71.05166856 |
| ENSMUSG00000021281 | *Tnfaip2* | 3575.71288 | 3936.16091 | 3796.47639 | 9123.049549 | 9183.194194 | 9359.735 | 5.69E-85 | 1.291266329 | 4.26E-87 | 391.3242512 | 2.447427849 | 84.24520037 |
| ENSMUSG00000016756 | *Cmah* | 469.7317188 | 385.5419147 | 331.470367 | 1050.03205 | 961.9661925 | 903.5985777 | 1.18E-17 | 1.290142919 | 1.22E-18 | 77.65832285 | 2.445522807 | 16.92959609 |
| ENSMUSG00000057561 | *Eif1a* | 3068.434578 | 3021.760452 | 3691.801537 | 8199.527384 | 7912.877665 | 7773.847015 | 1.04E-40 | 1.289320184 | 3.36E-42 | 185.3080912 | 2.444128579 | 39.98243206 |
| ENSMUSG00000028163 | *Nfkb1* | 3734.68671 | 3847.344971 | 3012.756979 | 8440.739566 | 8770.612757 | 8686.143333 | 5.67E-35 | 1.288310673 | 2.39E-36 | 158.5111229 | 2.442418924 | 34.24633081 |
| ENSMUSG00000053580 | *Tanc2* | 85.478391 | 89.8252105 | 65.75727928 | 179.6440375 | 197.6047935 | 216.4770924 | 3.73E-09 | 1.287730433 | 7.23E-10 | 37.95776527 | 2.4414368 | 8.428556801 |
| ENSMUSG00000046186 | *Cd109* | 344.3101544 | 301.7723364 | 328.7863964 | 786.8914883 | 814.3054677 | 781.8302132 | 6.87E-31 | 1.287616888 | 3.51E-32 | 139.4527641 | 2.441244659 | 30.16311696 |
| ENSMUSG00000048154 | *Kmt2d* | 1747.114403 | 1492.71333 | 615.9712488 | 3082.624212 | 3390.767967 | 2943.701891 | 0.008643475 | 1.287558997 | 0.003574922 | 8.488022395 | 2.441146702 | 2.063311597 |
| ENSMUSG00000020142 | *Slc1a4* | 448.9612686 | 464.2651329 | 382.4658081 | 1034.007465 | 1109.626917 | 1031.165436 | 7.05E-30 | 1.287254074 | 3.76E-31 | 134.7393231 | 2.440630803 | 29.15187751 |
| ENSMUSG00000032531 | *Amotl2* | 102.2545238 | 56.51923358 | 38.91757345 | 172.0534444 | 170.4612779 | 146.8951699 | 0.000332655 | 1.286318875 | 0.000110257 | 14.9524164 | 2.439049225 | 3.478005899 |
| ENSMUSG00000060301 | *2610008E11Rik* | 46.3340811 | 42.38942518 | 37.57558816 | 111.3286993 | 103.1453592 | 95.67514352 | 2.80E-06 | 1.286107212 | 7.20E-07 | 24.55992499 | 2.43869141 | 5.553030477 |
| ENSMUSG00000031617 | *Tmem184c* | 309.1601618 | 265.4385434 | 316.7085288 | 721.1063478 | 730.7034397 | 716.113953 | 9.10E-27 | 1.283827302 | 5.68E-28 | 120.2136943 | 2.434840558 | 26.04111916 |
| ENSMUSG00000032661 | *Oas3* | 4251.551373 | 3707.056159 | 2057.263452 | 8009.762556 | 8422.090017 | 7942.969743 | 3.46E-05 | 1.282646777 | 1.01E-05 | 19.48503417 | 2.432848997 | 4.460466477 |
| ENSMUSG00000049659 | *Aftph* | 1186.312249 | 1079.92107 | 1311.11963 | 2955.270927 | 2903.270427 | 2830.631266 | 1.13E-42 | 1.2825398 | 3.31E-44 | 194.5032591 | 2.432668605 | 41.94704003 |
| ENSMUSG00000022148 | *Fyb* | 2263.180203 | 2020.5626 | 1791.550364 | 4922.077948 | 5048.6939 | 4827.245878 | 7.62E-38 | 1.282163001 | 2.80E-39 | 171.9331174 | 2.432033331 | 37.11780645 |
| ENSMUSG00000024122 | *Pdpk1* | 1131.989533 | 1089.004518 | 966.2294099 | 2502.365537 | 2734.980631 | 2531.042433 | 2.38E-42 | 1.281865881 | 7.11E-44 | 192.9795777 | 2.431532511 | 41.62315509 |
| ENSMUSG00000026153 | *Fam135a* | 233.2681324 | 260.3921833 | 234.847426 | 650.2608119 | 567.8423461 | 550.8568869 | 8.62E-21 | 1.280954933 | 7.44E-22 | 92.3019896 | 2.429997675 | 20.06430332 |
| ENSMUSG00000015659 | *Serac1* | 162.9681473 | 175.6133329 | 171.7741173 | 449.5317934 | 395.209587 | 391.3983144 | 5.94E-18 | 1.279860181 | 6.07E-19 | 79.04613503 | 2.428154432 | 17.22651272 |
| ENSMUSG00000060771 | *Tsga10* | 61.51248698 | 58.53777763 | 83.20308807 | 159.4024558 | 157.4323904 | 171.0555596 | 5.21E-08 | 1.279359261 | 1.13E-08 | 32.60878595 | 2.427311496 | 7.282762612 |
| ENSMUSG00000079339 | *Ifit1bl1* | 52.72498884 | 35.32452098 | 55.02139695 | 111.3286993 | 109.659803 | 124.6676113 | 6.84E-06 | 1.276551397 | 1.85E-06 | 22.74864779 | 2.422591904 | 5.164683571 |
| ENSMUSG00000054717 | *Hmgb2* | 2915.851656 | 2170.944133 | 2521.590363 | 6041.268736 | 6069.290086 | 6325.190044 | 5.03E-29 | 1.276114768 | 2.80E-30 | 130.7561333 | 2.421858821 | 28.29884079 |
| ENSMUSG00000043415 | *Otud1* | 48.7306715 | 64.5934098 | 52.33742637 | 128.1966841 | 153.0894279 | 119.8355333 | 1.06E-06 | 1.275741406 | 2.59E-07 | 26.53467422 | 2.421232138 | 5.975460297 |
| ENSMUSG00000028842 | *Ago3* | 341.1147005 | 403.7088113 | 346.2322052 | 885.5691991 | 959.7947112 | 797.2928627 | 1.71E-22 | 1.27563124 | 1.35E-23 | 100.2448254 | 2.421047257 | 21.76682957 |
| ENSMUSG00000036768 | *Kif15* | 747.7362054 | 586.3870483 | 566.317793 | 1523.179022 | 1594.952976 | 1498.910582 | 1.89E-23 | 1.275630772 | 1.40E-24 | 104.7359874 | 2.421046472 | 22.72398181 |
| ENSMUSG00000004891 | *Nes* | 103.8522507 | 87.80666645 | 57.70536754 | 215.9102047 | 206.2907185 | 189.4174559 | 3.61E-07 | 1.275439284 | 8.46E-08 | 28.69698333 | 2.420725149 | 6.441893849 |
| ENSMUSG00000068246 | *Apol9b* | 124.6227009 | 64.5934098 | 50.99544108 | 189.7648284 | 158.518131 | 239.6710666 | 0.000452786 | 1.275333935 | 0.000153093 | 14.33377688 | 2.420548388 | 3.344106898 |
| ENSMUSG00000044763 | *Trmt10c* | 686.2237184 | 583.3592323 | 611.9452929 | 1612.579342 | 1473.350026 | 1474.750192 | 3.02E-34 | 1.275101625 | 1.33E-35 | 155.1071608 | 2.420158651 | 33.52036441 |
| ENSMUSG00000061533 | *Cep128* | 316.349933 | 284.6147119 | 287.1848524 | 723.6365455 | 705.7314053 | 725.7781089 | 1.97E-29 | 1.275088934 | 1.08E-30 | 132.6507273 | 2.420137361 | 28.70449443 |
| ENSMUSG00000079509 | *Zfx* | 706.1953051 | 637.8599218 | 717.962131 | 1708.726855 | 1694.841113 | 1573.324582 | 2.12E-41 | 1.272087843 | 6.57E-43 | 188.5558251 | 2.41510823 | 40.67357397 |
| ENSMUSG00000027068 | *Dhrs9* | 1827.799613 | 1789.439306 | 2422.283451 | 5044.370838 | 4778.344484 | 4743.167721 | 8.25E-24 | 1.272064518 | 6.00E-25 | 106.4070888 | 2.415069184 | 23.08348783 |
| ENSMUSG00000028133 | *Rwdd3* | 46.3340811 | 38.35233707 | 38.91757345 | 100.3645092 | 105.3168405 | 94.70872793 | 4.29E-06 | 1.271189689 | 1.13E-06 | 23.69107911 | 2.413605165 | 5.367321368 |
| ENSMUSG00000034154 | *Ino80* | 813.2430097 | 827.6030631 | 677.7025722 | 1896.804885 | 1910.903497 | 1795.600168 | 5.74E-34 | 1.26900644 | 2.56E-35 | 153.8030972 | 2.409955388 | 33.2411709 |
| ENSMUSG00000069049 | *Eif2s3y* | 614.3260063 | 546.0161672 | 477.7467638 | 1416.910719 | 1309.403192 | 1235.079125 | 1.75E-24 | 1.268994607 | 1.23E-25 | 109.5544904 | 2.409935621 | 23.75591032 |
| ENSMUSG00000021690 | *Jmy* | 218.0897266 | 215.984214 | 213.3756614 | 506.8829416 | 512.4695743 | 542.1591466 | 4.39E-24 | 1.268890288 | 3.13E-25 | 107.696777 | 2.409761369 | 23.35793169 |
| ENSMUSG00000104713 | *Gbp6* | 59.91476005 | 82.76030631 | 53.67941166 | 177.9572391 | 112.9170249 | 182.6525467 | 3.19E-05 | 1.268492847 | 9.30E-06 | 19.65073659 | 2.409097607 | 4.496185249 |
| ENSMUSG00000019947 | *Arid5b* | 161.3704204 | 213.96567 | 163.7222056 | 408.2052308 | 450.5823588 | 440.6855096 | 2.04E-14 | 1.26803768 | 2.61E-15 | 62.53942042 | 2.408337662 | 13.69032951 |
| ENSMUSG00000042323 | *Pbrm1* | 1360.464485 | 1270.673483 | 1124.583674 | 3084.311011 | 3085.674852 | 2890.549033 | 3.30E-40 | 1.267439103 | 1.09E-41 | 182.965669 | 2.407338646 | 39.48202393 |
| ENSMUSG00000039968 | *Rsbn1l* | 617.5214602 | 524.8214546 | 531.4261754 | 1341.004787 | 1345.232633 | 1354.914659 | 4.67E-35 | 1.2673513 | 1.96E-36 | 158.9032895 | 2.407192138 | 34.33080221 |
| ENSMUSG00000021176 | *Efcab11* | 84.67952753 | 64.5934098 | 93.93897041 | 169.5232467 | 201.947756 | 211.6450145 | 5.14E-08 | 1.265789954 | 1.11E-08 | 32.63560568 | 2.404588382 | 7.288636789 |
| ENSMUSG00000040433 | *Zbtb38* | 721.373711 | 729.7036763 | 789.0873514 | 1751.740216 | 1898.960351 | 1725.05183 | 7.13E-44 | 1.265048331 | 1.97E-45 | 200.1199266 | 2.403352611 | 43.14663968 |
| ENSMUSG00000030208 | *Emp1* | 2297.531332 | 2461.614477 | 2484.014775 | 5821.141535 | 5874.942514 | 5697.01991 | 6.14E-80 | 1.26503126 | 5.18E-82 | 367.9676193 | 2.403324173 | 79.21161047 |
| ENSMUSG00000020024 | *Cep83* | 873.9566332 | 724.6573162 | 774.3255132 | 1897.648284 | 1942.389976 | 1868.081338 | 3.15E-37 | 1.263672608 | 1.20E-38 | 169.0440174 | 2.401061919 | 36.50190393 |
| ENSMUSG00000038535 | *Zfp280d* | 329.930612 | 254.3365511 | 250.9512495 | 703.3949639 | 653.6158554 | 658.1290176 | 1.12E-17 | 1.263263085 | 1.17E-18 | 77.75185253 | 2.40038045 | 16.9489288 |
| ENSMUSG00000051344 | *Plekhm3* | 2562.754003 | 2770.451717 | 2428.993378 | 6350.796256 | 6358.097092 | 5931.858898 | 9.82E-57 | 1.263174346 | 1.63E-58 | 260.0975208 | 2.400232809 | 56.00797421 |
| ENSMUSG00000024259 | *Slc25a46* | 689.4191723 | 660.0639064 | 789.0873514 | 1683.424878 | 1720.898888 | 1712.488427 | 2.39E-38 | 1.262488827 | 8.63E-40 | 174.272811 | 2.399092572 | 37.62101738 |
| ENSMUSG00000026384 | *Ptpn4* | 739.7475707 | 676.2122589 | 469.694852 | 1421.971114 | 1598.210198 | 1517.272478 | 2.75E-14 | 1.261633807 | 3.57E-15 | 61.92598696 | 2.39767116 | 13.56022248 |
| ENSMUSG00000071266 | *Zfp946* | 138.2033798 | 100.9272028 | 162.3802203 | 333.1426987 | 320.293484 | 302.48808 | 7.25E-10 | 1.258961865 | 1.32E-10 | 41.27918581 | 2.393234665 | 9.13977505 |
| ENSMUSG00000037818 | *Abhd18* | 31.95453869 | 30.27816084 | 41.60154404 | 75.06253211 | 85.77350926 | 85.04457202 | 6.12E-05 | 1.257376577 | 1.84E-05 | 18.34536797 | 2.390606332 | 4.213363399 |
| ENSMUSG00000058755 | *Osm* | 354.6953795 | 445.0889644 | 469.694852 | 1014.609282 | 1032.539333 | 974.1469159 | 1.47E-21 | 1.257357339 | 1.21E-22 | 95.89157692 | 2.390574454 | 20.83151639 |
| ENSMUSG00000024989 | *Cep55* | 663.8555413 | 487.4783896 | 630.733087 | 1490.286452 | 1419.062995 | 1351.048996 | 1.08E-20 | 1.257247005 | 9.40E-22 | 91.83999699 | 2.390391636 | 19.96499019 |
| ENSMUSG00000069170 | *Adgrv1* | 38.34544643 | 17.15762448 | 22.81374996 | 64.94174127 | 73.8303624 | 51.22002633 | 0.004287685 | 1.256634443 | 0.001679242 | 9.870781074 | 2.389376901 | 2.367777132 |
| ENSMUSG00000025746 | *Il6* | 24.76476749 | 32.2967049 | 32.207647 | 70.84553593 | 74.91610303 | 65.7162602 | 0.000170488 | 1.256495697 | 5.46E-05 | 16.28285583 | 2.389147122 | 3.768306441 |
| ENSMUSG00000040552 | *C3ar1* | 3276.139079 | 3562.730259 | 3432.798376 | 8415.437589 | 8181.055599 | 7915.910107 | 7.23E-77 | 1.255353016 | 6.44E-79 | 353.7570472 | 2.387255557 | 76.14095936 |
| ENSMUSG00000038459 | *Abhd17c* | 774.8975633 | 783.1950938 | 543.5040431 | 1680.051281 | 1649.240007 | 1701.857856 | 6.87E-18 | 1.254952872 | 7.05E-19 | 78.74871815 | 2.386593523 | 17.16288213 |
| ENSMUSG00000037234 | *Hook3* | 544.0260212 | 594.4612246 | 504.5864696 | 1340.161388 | 1343.061151 | 1240.877619 | 9.62E-34 | 1.253392567 | 4.33E-35 | 152.7572509 | 2.384013767 | 33.0166134 |
| ENSMUSG00000078866 | *Zfp970* | 111.042022 | 75.69540211 | 100.6488969 | 224.3441971 | 222.5768278 | 238.704651 | 2.75E-09 | 1.251235806 | 5.26E-10 | 38.57729676 | 2.380452442 | 8.560315902 |
| ENSMUSG00000052144 | *Ppp4r2* | 1537.812175 | 1491.704058 | 1452.028085 | 3779.271982 | 3475.455736 | 3416.279115 | 4.26E-57 | 1.250863697 | 6.97E-59 | 261.795273 | 2.379838539 | 56.37026925 |
| ENSMUSG00000049076 | *Acap2* | 280.401077 | 309.8465126 | 224.1115437 | 937.8599518 | 351.779962 | 647.4984461 | 8.63E-05 | 1.249738404 | 2.65E-05 | 17.65293215 | 2.377983004 | 4.064151652 |
| ENSMUSG00000021930 | *Spryd7* | 1308.538359 | 1211.126434 | 1437.266247 | 3199.856706 | 2966.243384 | 3229.760906 | 3.31E-41 | 1.249719283 | 1.04E-42 | 187.6421953 | 2.377951487 | 40.4798903 |
| ENSMUSG00000096054 | *Syne1* | 1170.33498 | 1016.336932 | 637.4430135 | 2239.224975 | 2302.855863 | 2186.998483 | 1.28E-10 | 1.249335102 | 2.20E-11 | 44.78681553 | 2.377318338 | 9.891258217 |
| ENSMUSG00000055024 | *Ep300* | 1495.472411 | 1392.795399 | 837.3988219 | 2905.510372 | 3078.074668 | 2885.716955 | 6.72E-11 | 1.248843993 | 1.12E-11 | 46.10171761 | 2.37650921 | 10.17266027 |
| ENSMUSG00000091775 | *Gm16494* | 91.86929874 | 62.57486574 | 111.3847792 | 193.9818246 | 192.1760904 | 241.6038978 | 1.72E-06 | 1.248145762 | 4.33E-07 | 25.54095013 | 2.375359313 | 5.764220445 |
| ENSMUSG00000044348 | *Slc25a53* | 55.12157924 | 39.3616091 | 52.33742637 | 100.3645092 | 132.4603561 | 116.9362865 | 1.06E-05 | 1.247823933 | 2.91E-06 | 21.87398019 | 2.37482949 | 4.975059024 |
| ENSMUSG00000009207 | *Lnpk* | 421.7999107 | 453.1631406 | 509.9544108 | 1121.720986 | 1131.34173 | 1022.467695 | 1.58E-27 | 1.247568785 | 9.36E-29 | 123.7912156 | 2.374409526 | 26.80191599 |
| ENSMUSG00000019943 | *Atp2b1* | 2208.857487 | 2170.944133 | 2375.313966 | 5324.379385 | 5459.103855 | 5234.106842 | 9.21E-72 | 1.246867217 | 9.62E-74 | 329.9984597 | 2.373255155 | 71.03583448 |
| ENSMUSG00000043190 | *Rfesd* | 95.06475261 | 75.69540211 | 79.1771322 | 197.3554215 | 193.261831 | 205.8465209 | 8.23E-10 | 1.246316611 | 1.50E-10 | 41.02396689 | 2.372349572 | 9.084472386 |
| ENSMUSG00000042029 | *Ncapg2* | 1338.895171 | 1268.654939 | 1001.121027 | 2928.282152 | 2895.670243 | 2754.284435 | 1.96E-25 | 1.246045392 | 1.31E-26 | 113.9880174 | 2.371903626 | 24.70723336 |
| ENSMUSG00000048280 | *Zfp738* | 226.8772247 | 183.6875091 | 195.9298526 | 512.7867362 | 492.9262431 | 436.8198472 | 4.24E-16 | 1.24531539 | 4.88E-17 | 70.38519059 | 2.370703749 | 15.37273426 |
| ENSMUSG00000021669 | *Cert1* | 845.1975484 | 865.9554001 | 771.6415426 | 1949.095637 | 2041.192372 | 1903.838715 | 7.15E-46 | 1.244357264 | 1.82E-47 | 209.4376681 | 2.369129834 | 45.14591635 |
| ENSMUSG00000056592 | *Zfp658* | 67.90339472 | 47.43578532 | 79.1771322 | 161.0892543 | 158.518131 | 138.1974295 | 2.28E-06 | 1.243680798 | 5.84E-07 | 24.96453872 | 2.368019233 | 5.641200179 |
| ENSMUSG00000024542 | *Cep192* | 917.8941239 | 891.1872008 | 704.542278 | 2072.231926 | 1966.276269 | 1929.931935 | 8.74E-26 | 1.243563129 | 5.73E-27 | 115.6287222 | 2.3678261 | 25.05842692 |
| ENSMUSG00000036792 | *Mbd5* | 173.3533724 | 179.650421 | 115.4107351 | 356.7578773 | 386.523662 | 374.9692494 | 1.46E-10 | 1.242750255 | 2.51E-11 | 44.52493467 | 2.366492345 | 9.83466561 |
| ENSMUSG00000069833 | *Ahnak* | 12723.49844 | 13374.87292 | 11272.67645 | 29698.61734 | 31129.26942 | 27589.2323 | 3.79E-43 | 1.242135458 | 1.08E-44 | 196.7296119 | 2.365484091 | 42.42139862 |
| ENSMUSG00000034653 | *Ythdc2* | 230.871542 | 225.0676623 | 213.3756614 | 559.1736943 | 526.5842024 | 499.6368606 | 9.76E-22 | 1.2412325 | 8.00E-23 | 96.71741015 | 2.364004039 | 21.01048435 |
| ENSMUSG00000109901 | *Chmp1b* | 349.1033352 | 317.9206889 | 461.6429403 | 900.7503854 | 870.7639801 | 883.3038503 | 4.08E-16 | 1.241126752 | 4.69E-17 | 70.4608521 | 2.363830765 | 15.38916828 |
| ENSMUSG00000074221 | *Zfp568* | 604.7396447 | 565.1923358 | 471.0368373 | 1343.534985 | 1275.745233 | 1273.735749 | 3.16E-26 | 1.240636134 | 2.02E-27 | 117.6993639 | 2.363027034 | 25.50076622 |
| ENSMUSG00000114827 | *Gm46432* | 236.4635863 | 180.659693 | 206.6657349 | 537.2453141 | 446.2393963 | 492.8719515 | 1.56E-14 | 1.240286025 | 1.99E-15 | 63.07593097 | 2.362453651 | 13.80662445 |
| ENSMUSG00000003929 | *Zfp81* | 172.5545089 | 172.5855168 | 177.1420585 | 387.9636491 | 429.9532869 | 415.5587042 | 1.08E-18 | 1.239906269 | 1.05E-19 | 82.51281018 | 2.361831872 | 17.96627627 |
| ENSMUSG00000042271 | *Nxt2* | 734.1555264 | 657.0360903 | 734.0659545 | 1695.232467 | 1670.95482 | 1649.671414 | 2.32E-44 | 1.239327684 | 6.22E-46 | 202.409283 | 2.360884862 | 43.63395216 |
| ENSMUSG00000099974 | *Bcl2a1d* | 1672.021237 | 1333.248349 | 1413.110512 | 3541.433397 | 3421.168705 | 3471.364803 | 2.35E-33 | 1.237906333 | 1.08E-34 | 150.947563 | 2.358560049 | 32.62955587 |
| ENSMUSG00000032621 | *Srek1* | 930.6759394 | 855.8626799 | 921.9438953 | 2149.824656 | 2112.851253 | 2120.315807 | 3.13E-55 | 1.236833002 | 5.50E-57 | 253.0906234 | 2.35680599 | 54.50452821 |
| ENSMUSG00000031095 | *Cul4b* | 939.4634375 | 799.3434463 | 990.3851451 | 2162.475644 | 2150.852175 | 2111.618067 | 6.24E-34 | 1.236755082 | 2.79E-35 | 153.6308639 | 2.356678701 | 33.20467962 |
| ENSMUSG00000038002 | *Cramp1* | 365.879468 | 414.8108036 | 241.5573525 | 786.0480891 | 849.0491676 | 784.72946 | 9.41E-11 | 1.23667301 | 1.60E-11 | 45.41375982 | 2.356544639 | 10.02638108 |
| ENSMUSG00000035898 | *Uba6* | 1035.327054 | 928.5302659 | 999.7790422 | 2356.457469 | 2370.171781 | 2260.446068 | 6.19E-51 | 1.236616971 | 1.30E-52 | 233.0321778 | 2.356453105 | 50.2083405 |
| ENSMUSG00000020817 | *Rabep1* | 1243.031555 | 1226.265514 | 1442.634188 | 2994.910691 | 3230.078355 | 2977.526436 | 7.80E-39 | 1.236491939 | 2.75E-40 | 176.5486288 | 2.35624889 | 38.10792416 |
| ENSMUSG00000026499 | *Acbd3* | 1012.160013 | 971.9289631 | 921.9438953 | 2317.661104 | 2137.823288 | 2399.609913 | 4.92E-44 | 1.236328118 | 1.33E-45 | 200.8923321 | 2.355981349 | 43.30768957 |
| ENSMUSG00000027995 | *Tlr2* | 3465.469721 | 3764.584665 | 4212.49183 | 9185.461093 | 8807.527938 | 8935.478556 | 7.08E-44 | 1.236025458 | 1.95E-45 | 200.1403519 | 2.355487144 | 43.15016595 |
| ENSMUSG00000036941 | *Elac1* | 137.4045164 | 126.1590035 | 124.8046321 | 312.0577178 | 331.1508902 | 274.4620279 | 4.44E-13 | 1.235668525 | 6.28E-14 | 56.28225596 | 2.354904452 | 12.35301588 |
| ENSMUSG00000028576 | *Ift74* | 152.5829223 | 153.4093483 | 193.245882 | 403.1448354 | 363.7231089 | 399.1296391 | 2.41E-14 | 1.235015316 | 3.11E-15 | 62.19398937 | 2.353838463 | 13.61762745 |
| ENSMUSG00000029178 | *Klf3* | 158.97383 | 144.3259 | 115.4107351 | 344.1068888 | 315.9505215 | 333.4133789 | 1.75E-12 | 1.234631535 | 2.58E-13 | 53.50142815 | 2.353212387 | 11.75627572 |
| ENSMUSG00000082588 | *Gm15443* | 22.36817708 | 22.20398462 | 21.47176466 | 59.03794661 | 49.94406868 | 46.38794837 | 0.001421145 | 1.234534019 | 0.000516323 | 12.05577214 | 2.353053332 | 2.847361552 |
| ENSMUSG00000029009 | *Mthfr* | 1004.970242 | 1092.032334 | 897.78816 | 2373.325454 | 2368.0003 | 2317.464588 | 1.16E-38 | 1.234471466 | 4.16E-40 | 175.7236178 | 2.352951308 | 37.93498378 |

| | | | | | | | | | | | | | |
|---|---|---|---|---|---|---|---|---|---|---|---|---|---|
| ENSMUSG00000026596 | *Abl2* | 1913.278004 | 2012.488424 | 1395.664703 | 4055.906932 | 4339.705272 | 4140.124392 | 2.57E-19 | 1.234232806 | 2.42E-20 | 85.41202069 | 2.3525621 | 18.58949263 |
| ENSMUSG00000041313 | *Slc7a1* | 5525.738603 | 5375.382822 | 3291.88992 | 11147.20772 | 11438.27747 | 10800.66065 | 2.15E-12 | 1.2333743 | 3.20E-13 | 53.08013076 | 2.351162575 | 11.66694267 |
| ENSMUSG00000004151 | *Etv1* | 672.6430395 | 708.5089638 | 779.6934544 | 1665.713494 | 1704.612779 | 1694.126531 | 2.99E-41 | 1.233071331 | 9.36E-43 | 187.8524731 | 2.350668878 | 40.52415647 |
| ENSMUSG00000022914 | *Brwd1* | 971.4179762 | 987.0680435 | 798.4812485 | 2203.802207 | 2155.195138 | 2133.845625 | 1.47E-32 | 1.232132775 | 7.05E-34 | 147.213737 | 2.34914013 | 31.83199743 |
| ENSMUSG00000024383 | *Map3k2* | 748.5350688 | 786.2229099 | 680.3865428 | 1766.078003 | 1740.44222 | 1704.757103 | 8.21E-44 | 1.231347632 | 8.21E-44 | 192.6933223 | 2.34786203 | 41.56238961 |
| ENSMUSG00000029605 | *Oas1b* | 257.2340365 | 228.0954784 | 226.7955143 | 519.5339301 | 570.0138274 | 588.547095 | 3.10E-20 | 1.230804309 | 2.76E-21 | 89.70790031 | 2.346977985 | 19.50810881 |
| ENSMUSG00000081953 | *Gm12435* | 653.4703162 | 526.8399987 | 552.8979401 | 1227.14589 | 1505.922245 | 1345.250503 | 5.02E-21 | 1.230788124 | 4.27E-22 | 93.40292181 | 2.346951655 | 20.29900314 |
| ENSMUSG00000021510 | *Zfp729a* | 320.3442504 | 253.3272791 | 241.5573525 | 638.4532226 | 615.6149336 | 667.7931735 | 6.69E-17 | 1.229079508 | 7.29E-18 | 74.13648377 | 2.344173752 | 16.1747323 |
| ENSMUSG00000040483 | *Xaf1* | 1164.742935 | 1034.503829 | 979.6492628 | 2528.510913 | 2398.401037 | 2538.773758 | 5.01E-39 | 1.228672233 | 1.75E-40 | 177.448992 | 2.343512082 | 38.30014115 |
| ENSMUSG00000022507 | *Hapstr1* | 706.9941685 | 725.6665882 | 735.4079398 | 1698.606064 | 1702.441298 | 1673.831804 | 5.35E-54 | 1.228388068 | 9.86E-56 | 247.3424263 | 2.343050531 | 53.27131177 |
| ENSMUSG00000021189 | *Atxn3* | 502.4851209 | 529.8678148 | 638.7849988 | 1319.919806 | 1295.288564 | 1281.467074 | 3.32E-26 | 1.227664142 | 2.12E-27 | 117.5975183 | 2.341875112 | 25.47926712 |
| ENSMUSG00000052656 | *Rnf103* | 248.4465383 | 275.5312637 | 245.5833083 | 610.6210478 | 603.6717867 | 588.547095 | 2.93E-24 | 1.227355231 | 2.07E-25 | 108.5209721 | 2.341373723 | 23.53347537 |
| ENSMUSG00000054321 | *Taf4b* | 279.6022136 | 330.0319532 | 250.9512495 | 647.7306142 | 737.2178834 | 635.901459 | 4.90E-17 | 1.227075974 | 5.31E-18 | 74.76364887 | 2.340920555 | 16.30963503 |
| ENSMUSG00000029817 | *Tra2a* | 1409.195156 | 1367.563598 | 1665.403747 | 3571.79577 | 3392.939449 | 3410.480621 | 3.03E-36 | 1.226438631 | 1.20E-37 | 164.4611962 | 2.339886629 | 35.51869496 |
| ENSMUSG00000022021 | *Diaph3* | 127.0192913 | 140.2888119 | 193.245882 | 348.323885 | 375.6662558 | 343.0775349 | 3.18E-10 | 1.225803026 | 5.63E-11 | 42.94509834 | 2.338855978 | 9.497903476 |
| ENSMUSG00000048271 | *Rbm33* | 1446.741739 | 1498.768962 | 1218.522645 | 3269.858843 | 3351.681305 | 3123.455191 | 1.32E-34 | 1.224195584 | 5.69E-36 | 156.7884122 | 2.33625149 | 33.87892207 |
| ENSMUSG00000040455 | *Usp45* | 565.5953348 | 575.285056 | 520.6902931 | 1321.606605 | 1306.14597 | 1262.138762 | 4.09E-40 | 1.224153541 | 1.36E-41 | 182.5297659 | 2.336183409 | 39.38840187 |
| ENSMUSG00000055204 | *Ankrd17* | 3389.577692 | 3295.273172 | 2497.434628 | 7379.743326 | 7032.342019 | 7019.076438 | 1.18E-23 | 1.2215162 | 8.61E-25 | 105.6919871 | 2.331916612 | 22.92852518 |
| ENSMUSG00000027368 | *Dusp2* | 762.1157478 | 733.7407645 | 633.4170576 | 1642.941714 | 1688.32667 | 1648.704998 | 7.97E-35 | 1.220872797 | 3.40E-36 | 157.8134275 | 2.330876873 | 34.09871429 |
| ENSMUSG00000041135 | *Ripk2* | 704.5975781 | 599.5075847 | 740.7758809 | 1578.843372 | 1574.323904 | 1607.149128 | 3.76E-31 | 1.220832313 | 1.90E-32 | 140.6746012 | 2.330811466 | 30.42525368 |
| ENSMUSG00000038070 | *Cntln* | 216.4919996 | 194.7895014 | 159.6962497 | 461.3393828 | 470.12569 | 408.793795 | 9.65E-14 | 1.220521212 | 1.30E-14 | 59.38432282 | 2.330308907 | 13.01541309 |
| ENSMUSG00000058690 | *Ccser2* | 334.7237928 | 373.4306504 | 426.7513227 | 930.2693587 | 851.2206489 | 845.6136422 | 1.86E-21 | 1.218607514 | 1.54E-22 | 95.42497258 | 2.327219862 | 20.73071024 |
| ENSMUSG00000045312 | *Lhfpl2* | 1691.992824 | 2023.590416 | 1888.173305 | 4436.279988 | 4332.105088 | 4258.99351 | 6.07E-43 | 1.218261923 | 1.75E-44 | 195.768229 | 2.326662454 | 42.21694552 |
| ENSMUSG00000027660 | *Skil* | 2082.637059 | 2284.991872 | 2302.84676 | 5327.752982 | 5171.38259 | 5004.099931 | 4.35E-59 | 1.218170976 | 6.65E-61 | 271.0661757 | 2.326515785 | 58.36174126 |
| ENSMUSG00000024477 | *Pggt1b* | 292.384029 | 296.7259763 | 303.2886759 | 737.1309333 | 688.3595553 | 644.5991993 | 1.35E-24 | 1.216153843 | 9.42E-26 | 110.0776897 | 2.323265195 | 23.86870177 |
| ENSMUSG00000004609 | *Cd33* | 1332.504263 | 1323.155629 | 1312.461615 | 3182.988721 | 3036.816524 | 2999.753995 | 7.22E-67 | 1.216121993 | 8.86E-69 | 307.2086855 | 2.323213905 | 66.14174489 |
| ENSMUSG00000035678 | *Tnfsf9* | 1620.893975 | 1729.892256 | 1524.495291 | 3745.536013 | 3867.408101 | 3723.599273 | 2.39E-55 | 1.215977376 | 4.18E-57 | 253.6377168 | 2.322981034 | 54.62086874 |
| ENSMUSG00000068134 | *Zfp120* | 375.4658296 | 305.8094245 | 363.678014 | 844.2426365 | 787.1619521 | 795.3600315 | 7.67E-23 | 1.215661121 | 5.90E-24 | 101.8805091 | 2.322471852 | 22.11526834 |
| ENSMUSG00000027708 | *Dcun1d1* | 723.7703014 | 710.5275078 | 870.9484542 | 1788.849782 | 1748.042404 | 1797.532999 | 7.18E-31 | 1.214921335 | 3.67E-32 | 139.3635432 | 2.321281255 | 30.14410763 |
| ENSMUSG00000053477 | *Tcf4* | 632.6998661 | 652.9990022 | 558.2658813 | 1450.646688 | 1435.349104 | 1405.16827 | 2.64E-36 | 1.214644245 | 1.04E-37 | 164.7467156 | 2.320835462 | 35.57847024 |
| ENSMUSG00000033499 | *Larp4b* | 1821.408705 | 1816.689651 | 1681.50757 | 4260.009547 | 4120.385666 | 3972.934495 | 1.56E-60 | 1.214093342 | 2.28E-62 | 277.789071 | 2.319949404 | 59.80627842 |
| ENSMUSG00000002602 | *Axl* | 313.1544792 | 286.633256 | 158.3542644 | 585.3190706 | 592.8143805 | 591.4463418 | 1.95E-07 | 1.213794017 | 4.46E-08 | 29.93709886 | 2.319468119 | 6.709300833 |
| ENSMUSG00000030521 | *Mphosph10* | 1092.04636 | 1037.531645 | 1391.638747 | 2709.841749 | 2712.180078 | 2726.258383 | 1.69E-23 | 1.213478473 | 1.25E-24 | 104.9557389 | 2.318960865 | 22.77100232 |
| ENSMUSG00000027878 | *Notch2* | 1845.374609 | 1956.978463 | 1184.973012 | 3785.175777 | 3981.410867 | 3812.509507 | 1.34E-12 | 1.213377317 | 1.95E-13 | 54.04955181 | 2.318798274 | 11.87306874 |
| ENSMUSG00000002881 | *Nab1* | 1253.41678 | 1110.199231 | 1074.930219 | 2748.638114 | 2655.721565 | 2581.296044 | 6.65E-41 | 1.212874701 | 2.12E-42 | 186.2282854 | 2.317990576 | 40.17691549 |
| ENSMUSG00000066232 | *Ipo7* | 6056.183946 | 6122.244123 | 7244.036604 | 15252.03181 | 15072.25134 | 14620.90148 | 7.53E-42 | 1.211172362 | 2.30E-43 | 190.6413649 | 2.315257026 | 41.12297766 |
| ENSMUSG00000041219 | *Arhgap11a* | 816.4384636 | 786.2229099 | 630.733087 | 1676.677684 | 1780.614623 | 1730.850324 | 3.20E-25 | 1.211051387 | 2.16E-26 | 112.9950077 | 2.315062891 | 24.49545314 |
| ENSMUSG00000038379 | *Ttk* | 411.4146857 | 387.5604588 | 432.1192639 | 958.9449327 | 947.8515644 | 935.4902922 | 2.74E-33 | 1.209610585 | 1.27E-34 | 150.6240782 | 2.312752022 | 32.56219291 |
| ENSMUSG00000057093 | *Zfp607b* | 19.17272321 | 18.16689651 | 21.47176466 | 46.38695805 | 43.42962494 | 45.42153278 | 0.002971786 | 1.209442334 | 0.001133011 | 10.59651396 | 2.312482318 | 2.52698251 |
| ENSMUSG00000017667 | *Zfp334* | 18.37385975 | 18.16689651 | 18.78779408 | 34.57936873 | 41.2581437 | 52.18644192 | 0.006510588 | 1.20930565 | 0.002631729 | 9.046675852 | 2.312263238 | 2.186379766 |
| ENSMUSG00000035476 | *Tab3* | 289.1885752 | 257.3643672 | 276.4489701 | 661.225002 | 611.2719711 | 631.069381 | 9.42E-24 | 1.208956993 | 6.86E-25 | 106.1418594 | 2.311704499 | 23.02606569 |
| ENSMUSG00000079470 | *Utp14b* | 336.3215197 | 310.8557847 | 316.7085288 | 754.8423173 | 738.303624 | 738.3415116 | 4.92E-29 | 1.208808876 | 2.74E-30 | 130.8011158 | 2.311467177 | 28.30776111 |
| ENSMUSG00000021338 | *Carmil1* | 36.7477195 | 40.37088113 | 48.31147049 | 104.5815054 | 89.03073113 | 92.77589675 | 2.82E-05 | 1.207777181 | 8.17E-06 | 19.89832372 | 2.3098148 | 4.549373266 |
| ENSMUSG00000020387 | *Jade2* | 1596.928071 | 1446.286816 | 1214.496689 | 3312.872204 | 3297.394274 | 3244.257139 | 8.13E-28 | 1.207757468 | 4.76E-29 | 125.1344952 | 2.309783237 | 27.08982041 |
| ENSMUSG00000023030 | *Slc11a2* | 2283.15179 | 2365.733634 | 1996.874114 | 5205.460092 | 5089.952043 | 5063.051282 | 2.35E-45 | 1.206938284 | 6.07E-47 | 207.0419376 | 2.308472081 | 44.62821297 |
| ENSMUSG00000067430 | *Zfp763* | 90.2715718 | 69.63976994 | 88.57102924 | 188.9214291 | 205.2049779 | 179.7532999 | 1.33E-08 | 1.206646415 | 2.74E-09 | 35.36292234 | 2.308005104 | 7.875622227 |
| ENSMUSG00000096292 | *None* | 47.93180804 | 26.24107273 | 17.44580879 | 80.12292754 | 66.23017804 | 69.58192256 | 0.006179108 | 1.206299904 | 0.002486376 | 9.150591978 | 2.307450827 | 2.209074224 |
| ENSMUSG00000042129 | *Rassf4* | 2295.134742 | 2343.529649 | 2258.561246 | 5353.054959 | 5291.899799 | 5264.065725 | 1.36E-89 | 1.205394882 | 8.83E-92 | 412.8397653 | 2.306003786 | 88.86638911 |
| ENSMUSG00000083405 | *Gm15725* | 96.66247954 | 83.76957834 | 49.65345579 | 215.0668055 | 148.7464654 | 172.0219752 | 6.40E-05 | 1.20478978 | 1.93E-05 | 18.25248088 | 2.305036795 | 4.193716679 |
| ENSMUSG00000059423 | *Zfp933* | 131.0136086 | 142.307356 | 116.7527204 | 285.0689422 | 304.0073746 | 314.0850671 | 5.18E-13 | 1.204782743 | 7.38E-14 | 55.96322178 | 2.305025551 | 12.28525275 |
| ENSMUSG00000024073 | *Birc6* | 3202.64364 | 3343.718229 | 2211.59176 | 6798.641251 | 6957.425916 | 6446.958408 | 6.74E-16 | 1.204770387 | 7.85E-17 | 69.44659339 | 2.30500581 | 15.17161577 |
| ENSMUSG00000017831 | *Rab5a* | 852.3873196 | 850.8163197 | 892.4202189 | 1956.68623 | 2017.306079 | 1999.513858 | 2.60E-55 | 1.203902296 | 4.56E-57 | 253.4651358 | 2.303619271 | 54.58471133 |
| ENSMUSG00000090093 | *Gm14399* | 87.8749814 | 109.001379 | 80.51911749 | 217.5970032 | 195.4333122 | 227.1076639 | 7.06E-09 | 1.203005899 | 1.40E-09 | 36.66259026 | 2.302188396 | 8.150937167 |
| ENSMUSG00000033752 | *Mnd1* | 89.47270834 | 78.72321819 | 83.20308807 | 221.8139994 | 169.3755373 | 187.4846247 | 2.96E-08 | 1.202847182 | 6.23E-09 | 33.76009963 | 2.301935137 | 7.529218705 |
| ENSMUSG00000021068 | *Nin* | 1169.536116 | 1237.367506 | 903.1561012 | 2589.235658 | 2713.265818 | 2326.162328 | 2.62E-19 | 1.201911062 | 2.47E-20 | 85.37578819 | 2.300441967 | 18.58207898 |
| ENSMUSG00000055560 | *Zfp459* | 15.97726935 | 16.14835245 | 26.83970583 | 45.54355881 | 47.77258744 | 39.62303924 | 0.007185097 | 1.201302171 | 0.002923853 | 8.854386961 | 2.299471267 | 2.143567341 |
| ENSMUSG00000057133 | *Chd6* | 357.8908333 | 340.1246735 | 272.4230142 | 785.2046899 | 771.9615834 | 684.2222385 | 3.84E-17 | 1.20038255 | 4.13E-18 | 75.25691315 | 2.298005977 | 16.4157637 |
| ENSMUSG00000091876 | *Gm3851* | 339.5169736 | 250.299463 | 326.1024258 | 688.2137776 | 668.8162241 | 748.0056675 | 1.27E-15 | 1.200365074 | 1.51E-16 | 68.16105931 | 2.29797814 | 14.89484909 |
| ENSMUSG00000071337 | *Tia1* | 417.8055934 | 409.7644434 | 348.9161758 | 942.076948 | 826.2486145 | 943.2216169 | 3.53E-22 | 1.200115076 | 2.84E-23 | 98.76808757 | 2.297579968 | 21.45203814 |
| ENSMUSG00000022952 | *Runx1* | 722.9714379 | 843.7514155 | 582.4216165 | 1632.820923 | 1632.953898 | 1682.529544 | 3.09E-18 | 1.199973526 | 3.12E-19 | 80.36245654 | 2.297354552 | 17.51004395 |
| ENSMUSG00000068250 | *Amn1* | 111.8408854 | 104.9642909 | 132.8565439 | 255.5499689 | 295.3214496 | 248.3688069 | 1.76E-10 | 1.199423829 | 3.03E-11 | 44.15693842 | 2.296479379 | 9.755261874 |
| ENSMUSG00000021356 | *Irf4* | 65.50680432 | 65.60268183 | 73.80919103 | 155.1854597 | 169.3755373 | 143.9959231 | 7.08E-08 | 1.198468034 | 1.55E-08 | 31.99059429 | 2.294958451 | 7.150130974 |
| ENSMUSG00000020220 | *Vps13d* | 1351.676987 | 1355.452334 | 1152.765365 | 2875.148 | 3031.387821 | 2964.963034 | 4.30E-39 | 1.197636865 | 1.50E-40 | 177.7590578 | 2.293636654 | 38.36637383 |
| ENSMUSG00000035851 | *Ythdc1* | 1059.292958 | 1010.2813 | 958.1774981 | 2311.757309 | 2343.028266 | 2300.069107 | 1.06E-52 | 1.196858891 | 2.05E-54 | 241.3002936 | 2.292400143 | 51.97301505 |
| ENSMUSG00000010608 | *Rbm25* | 1854.162108 | 1881.28306 | 1571.464776 | 4018.797365 | 4277.818057 | 3882.09143 | 3.63E-35 | 1.196322033 | 1.52E-36 | 159.4116068 | 2.29154725 | 34.44003581 |
| ENSMUSG00000026088 | *Mitd1* | 356.2931064 | 317.9206889 | 399.9116169 | 821.4708571 | 786.0762115 | 844.6472266 | 2.67E-22 | 1.195784369 | 2.12E-23 | 99.34538967 | 2.290693394 | 21.57382937 |
| ENSMUSG00000028967 | *Errfi1* | 289.1885752 | 298.7445203 | 283.1588965 | 678.936386 | 643.8441898 | 670.6924202 | 2.03E-26 | 1.193872536 | 1.28E-27 | 118.5978456 | 2.28765982 | 25.6918428 |
| ENSMUSG00000071042 | *Rasgrp3* | 1936.445045 | 1997.349344 | 2290.768893 | 4874.847591 | 4669.770422 | 4670.686552 | 5.07E-42 | 1.193583433 | 1.54E-43 | 191.4412492 | 2.28720144 | 41.29504771 |
| ENSMUSG00000027469 | *Tpx2* | 2338.273369 | 1920.64467 | 1890.857276 | 4879.064587 | 4581.825431 | 4619.466526 | 6.62E-31 | 1.193418562 | 3.37E-32 | 139.5293814 | 2.286940073 | 30.17886649 |
| ENSMUSG00000073761 | *4933427I04Rik* | 39.94317336 | 26.24107273 | 17.44580879 | 68.31533822 | 53.20129055 | 73.44758493 | 0.00483915 | 1.192810913 | 0.001914069 | 9.630169833 | 2.28597704 | 2.315230871 |
| ENSMUSG00000001833 | *Septin7* | 2553.966505 | 2306.186584 | 2484.014775 | 5686.197657 | 5569.849399 | 5520.165857 | 3.45E-64 | 1.191553675 | 4.50E-66 | 294.7882165 | 2.283985791 | 63.46210015 |
| ENSMUSG00000022816 | *Fstl1* | 16.77613281 | 24.22252868 | 16.1038235 | 41.32656262 | 39.08666245 | 50.25361074 | 0.007002634 | 1.191377632 | 0.002845853 | 8.903751854 | 2.283707106 | 2.154738566 |
| ENSMUSG00000096006 | *Gm21596* | 890.732766 | 887.1501127 | 866.9224983 | 2105.967895 | 2002.10571 | 1925.099858 | 6.55E-48 | 1.189285962 | 1.52E-49 | 218.9727848 | 2.280398507 | 47.18382236 |
| ENSMUSG00000049323 | *Smcr8* | 3916.028717 | 4329.777001 | 3717.299258 | 9215.823465 | 9283.082331 | 8786.650555 | 3.89E-49 | 1.189043297 | 8.49E-51 | 224.710446 | 2.280014971 | 48.41022236 |
| ENSMUSG00000070883 | *Ccdc173* | 31.15567522 | 15.13908042 | 25.49772054 | 60.72474508 | 42.34388432 | 60.88418224 | 0.007674885 | 1.187158358 | 0.003141702 | 8.723292064 | 2.277037984 | 2.1149281 |
| ENSMUSG00000094786 | *Gm14403* | 17.57499628 | 23.21325665 | 36.23360287 | 67.47193898 | 57.54425305 | 46.38794837 | 0.006696185 | 1.186847411 | 0.002713804 | 8.990542453 | 2.276547262 | 2.174172578 |
| ENSMUSG00000024294 | *Mib1* | 1209.479289 | 1230.302602 | 837.3988219 | 2476.220161 | 2517.832506 | 2481.755238 | 6.78E-16 | 1.186753567 | 7.90E-17 | 69.43332957 | 2.276399184 | 15.16891505 |
| ENSMUSG00000021738 | *Atxn7* | 409.8169587 | 457.2002287 | 401.2536022 | 992.6809022 | 955.4517487 | 940.3223702 | 1.88E-29 | 1.186589638 | 1.03E-30 | 132.7492055 | 2.276140538 | 28.72509749 |
| ENSMUSG00000059975 | *Zfp74* | 91.86929874 | 70.64904197 | 72.46720574 | 207.4762124 | 181.3186841 | 148.828001 | 1.26E-06 | 1.186501066 | 3.11E-07 | 26.18106181 | 2.276000802 | 5.900980267 |
| ENSMUSG00000037860 | *Aim2* | 826.8236886 | 812.4639827 | 1202.418821 | 2227.417386 | 2238.797166 | 1983.084793 | 1.61E-13 | 1.185784387 | 2.20E-14 | 58.34447329 | 2.274870447 | 12.79327259 |
| ENSMUSG00000030660 | *Pik3c2a* | 1135.184987 | 1102.125055 | 1123.241689 | 2563.933681 | 2616.634903 | 2463.393342 | 1.53E-58 | 1.185261792 | 2.37E-60 | 268.5353716 | 2.274046558 | 57.81516355 |
| ENSMUSG00000066829 | *Zfp810* | 40.74203683 | 42.38942518 | 36.23360287 | 90.24371838 | 95.54517487 | 86.9774032 | 2.88E-05 | 1.184904574 | 8.35E-06 | 19.85601686 | 2.273483563 | 4.540470119 |
| ENSMUSG00000024242 | *Map4k3* | 252.4408557 | 341.1339455 | 344.8902199 | 700.0213669 | 685.1023335 | 737.375096 | 8.11E-15 | 1.184597728 | 1.02E-15 | 64.39974417 | 2.273000069 | 14.09095825 |
| ENSMUSG00000031347 | *Cetn2* | 689.4191723 | 539.9605351 | 693.8063957 | 1433.778703 | 1454.892436 | 1478.615854 | 8.46E-23 | 1.18388921 | 6.52E-24 | 101.6810451 | 2.271884056 | 22.07286888 |
| ENSMUSG00000051316 | *Taf7* | 188.5317783 | 183.6875091 | 272.4230142 | 541.4623103 | 427.7818057 | 484.1742112 | 1.87E-09 | 1.18358974 | 3.53E-10 | 39.35688035 | 2.271412515 | 8.728079436 |
| ENSMUSG00000036202 | *Rif1* | 1266.997459 | 1129.375399 | 819.9530131 | 2590.079058 | 2425.544553 | 2302.968354 | 3.24E-14 | 1.183074321 | 4.22E-15 | 61.59215569 | 2.270601172 | 13.48955117 |
| ENSMUSG00000032413 | *Rasa2* | 684.6259915 | 616.6652092 | 734.0659545 | 1599.084954 | 1470.092804 | 1543.365699 | 1.59E-31 | 1.183048742 | 7.90E-33 | 142.4132934 | 2.270560915 | 30.79799834 |
| ENSMUSG00000041225 | *Arhgap12* | 418.6044569 | 361.3193861 | 373.071911 | 921.8353663 | 851.2206489 | 850.4457202 | 3.77E-25 | 1.183008217 | 2.55E-26 | 112.6655481 | 2.270497136 | 24.42404523 |
| ENSMUSG00000004980 | *Hnrnpa2b1* | 9953.838802 | 9588.084267 | 8928.228144 | 21728.49455 | 21462.92065 | 21462.15745 | 1.36E-72 | 1.182782772 | 1.36E-74 | 333.895773 | 2.270142361 | 71.86713533 |
| ENSMUSG00000037857 | *Nufip2* | 2014.733665 | 1974.136087 | 2076.051246 | 4712.914938 | 4599.197281 | 4441.646057 | 4.52E-70 | 1.182067326 | 4.93E-72 | 322.1477663 | 2.269016855 | 69.34507974 |
| ENSMUSG00000027804 | *Ppid* | 2188.8859 | 2047.812945 | 1858.649629 | 4765.20569 | 4389.649341 | 4690.981279 | 5.47E-41 | 1.181941299 | 1.73E-42 | 186.629728 | 2.268818653 | 40.26239216 |
| ENSMUSG00000031715 | *Smarca5* | 2090.625694 | 1984.228807 | 2029.081761 | 4574.597463 | 4763.144116 | 4516.060057 | 1.50E-70 | 1.181783955 | 1.61E-72 | 324.3825826 | 2.268571224 | 69.82475564 |
| ENSMUSG00000028187 | *Rpf1* | 730.1612091 | 672.1751707 | 726.0140427 | 1674.990885 | 1554.780573 | 1595.552141 | 1.83E-40 | 1.181520192 | 6.01E-42 | 184.1528541 | 2.268156506 | 39.73731935 |
| ENSMUSG00000029826 | *Zc3hav1* | 3578.10947 | 3568.785891 | 2883.926391 | 7656.378276 | 7741.330646 | 7366.986051 | 9.69E-32 | 1.18109979 | 4.79E-33 | 143.4057853 | 2.267495661 | 31.01371933 |
| ENSMUSG00000020380 | *Rad50* | 671.0453125 | 713.5553239 | 573.0277195 | 1470.88827 | 1537.408723 | 1441.892062 | 1.37E-27 | 1.180623706 | 8.06E-29 | 124.0881819 | 2.266747519 | 26.86474758 |
| ENSMUSG00000022521 | *Crebbp* | 658.263497 | 702.4533316 | 428.093308 | 1355.342574 | 1421.234476 | 1290.164814 | 3.21E-11 | 1.180436154 | 5.23E-12 | 47.5977629 | 2.266452858 | 10.49300027 |
| ENSMUSG00000022263 | *Trio* | 1020.947511 | 1118.273407 | 748.8277927 | 2188.621021 | 2199.710503 | 2168.636586 | 4.04E-16 | 1.179934408 | 4.65E-17 | 70.48004911 | 2.26566476 | 15.39317156 |
| ENSMUSG00000061665 | *Cd2ap* | 830.0191425 | 750.8983889 | 772.9835279 | 1870.659508 | 1813.186841 | 1653.537076 | 6.92E-35 | 1.179655037 | 2.94E-36 | 158.1045824 | 2.265226068 | 34.15992372 |
| ENSMUSG00000036572 | *Upf3b* | 254.8374461 | 223.0491182 | 269.7390436 | 560.0170935 | 570.0138274 | 559.5546273 | 5.96E-20 | 1.179192585 | 5.41E-21 | 88.37797221 | 2.264500071 | 19.2244223 |
| ENSMUSG00000024483 | *Ankhd1* | 1958.014358 | 2018.544056 | 1775.446541 | 4448.087577 | 4348.391197 | 4232.900289 | 4.01E-51 | 1.178071751 | 8.34E-53 | 233.916852 | 2.262741458 | 50.39635362 |
| ENSMUSG00000049878 | *Rlf* | 624.7112314 | 587.3963204 | 597.1834547 | 1360.40297 | 1369.118926 | 1367.478061 | 7.95E-43 | 1.177490249 | 2.30E-44 | 195.222908 | 2.261829607 | 42.09970352 |
| ENSMUSG00000083902 | *Tent2-ps1* | 127.8181548 | 128.1775476 | 122.1206615 | 307.8407216 | 237.7771966 | 309.2529892 | 3.91E-10 | 1.177356242 | 6.95E-11 | 42.53201424 | 2.261619522 | 9.408213592 |
| ENSMUSG00000054737 | *Zfp182* | 71.89771206 | 96.8901147 | 59.04735283 | 164.4628513 | 178.0614623 | 175.8876376 | 1.57E-06 | 1.177088579 | 3.94E-07 | 25.72255292 | 2.261199964 | 5.80305005 |
| ENSMUSG00000028630 | *Dyrk2* | 434.5817262 | 424.9035238 | 303.2886759 | 894.8465907 | 892.4787926 | 854.3113826 | 1.01E-14 | 1.176552442 | 1.28E-15 | 63.94895717 | 2.260359809 | 13.99424246 |
| ENSMUSG00000017830 | *Dhx58* | 1365.257666 | 1035.513101 | 803.8491896 | 2455.13518 | 2376.686225 | 2423.770303 | 1.24E-11 | 1.175816299 | 1.96E-12 | 49.52044258 | 2.259206742 | 10.90551581 |
| ENSMUSG00000032905 | *Atg12* | 671.844176 | 613.6373931 | 672.334631 | 1493.660049 | 1460.321139 | 1467.018867 | 8.72E-41 | 1.175658045 | 2.80E-42 | 185.674521 | 2.258958936 | 40.05923825 |
| ENSMUSG00000071302 | *2610044O15Rik8* | 151.7840588 | 142.307356 | 123.4626468 | 328.0823033 | 298.5786715 | 321.8163918 | 9.14E-13 | 1.175438641 | 1.32E-13 | 54.82218484 | 2.25861542 | 12.0392318 |
| ENSMUSG00000069014 | *Gm5641* | 7568.432489 | 7298.046035 | 7138.019766 | 17073.77416 | 15824.66959 | 16795.33656 | 2.01E-75 | 1.175002611 | 1.88E-77 | 347.0321327 | 2.257932895 | 74.69619863 |
| ENSMUSG00000038095 | *Sbno1* | 1947.629133 | 1951.932102 | 2211.59176 | 4665.68458 | 4613.31191 | 4494.798914 | 7.18E-49 | 1.174399849 | 1.60E-50 | 223.4515484 | 2.256989723 | 48.14378654 |
| ENSMUSG00000074656 | *Eif2s2* | 2971.772098 | 2737.14574 | 3365.699111 | 6775.026073 | 6860.795 | 6825.79332 | 8.01E-38 | 1.174118852 | 2.95E-39 | 171.8286078 | 2.256550166 | 37.09637616 |
| ENSMUSG00000028382 | *Ptbp3* | 3461.475404 | 3370.968574 | 3509.291537 | 8032.534335 | 7831.447118 | 7466.526857 | 5.25E-75 | 1.174049879 | 4.93E-77 | 345.1074319 | 2.256442286 | 74.27979439 |
| ENSMUSG00000021027 | *Ralgapa1* | 578.3771503 | 561.1552476 | 438.8291903 | 1157.143753 | 1244.258755 | 1169.362865 | 4.92E-20 | 1.171584932 | 4.44E-21 | 88.76871217 | 2.252590286 | 19.30793998 |
| ENSMUSG00000033014 | *Trim33* | 1135.184987 | 1155.616472 | 970.2553658 | 2510.799529 | 2476.574362 | 2370.617445 | 7.93E-35 | 1.171007641 | 3.38E-36 | 157.8248196 | 2.251689098 | 34.100481 |
| ENSMUSG00000074862 | *BC025920* | 19.97158668 | 20.18454056 | 29.52367641 | 51.44735347 | 47.77258744 | 55.08568869 | 0.003288139 | 1.170211308 | 0.001262743 | 10.39620865 | 2.250446563 | 2.483049832 |
| ENSMUSG00000004591 | *Pkn2* | 1016.15433 | 981.0124113 | 1029.302719 | 2251.032564 | 2284.398272 | 2272.043055 | 8.17E-58 | 1.169932861 | 1.30E-59 | 265.1378205 | 2.250012257 | 57.08771263 |
| ENSMUSG00000034317 | *Trim59* | 559.2044271 | 431.968428 | 360.9940434 | 1067.743434 | 972.8235987 | 1013.769955 | 1.49E-12 | 1.16927413 | 2.18E-13 | 53.83405714 | 2.248985143 | 11.82683936 |

| | | | | | | | | | | | | | |
|---|---|---|---|---|---|---|---|---|---|---|---|---|---|
| ENSMUSG00000034610 | *Tut4* | 437.7771801 | 386.5511868 | 195.9298526 | 709.2987585 | 811.0482458 | 783.7630444 | 6.82E-06 | 1.168530455 | 1.84E-06 | 22.7573839 | 2.247826142 | 5.166467811 |
| ENSMUSG00000105175 | *Gm43097* | 213.2965458 | 225.0676623 | 147.6183821 | 441.9412003 | 428.8675463 | 455.1817434 | 1.05E-10 | 1.168072684 | 1.79E-11 | 45.1851682 | 2.247113014 | 9.977049853 |
| ENSMUSG00000011831 | *Evi5* | 627.9066853 | 623.7301134 | 615.9712488 | 1411.006924 | 1433.177623 | 1350.082581 | 5.07E-42 | 1.16637169 | 1.54E-43 | 191.4360335 | 2.244465141 | 41.29475177 |
| ENSMUSG00000034413 | *Neurl1b* | 111.042022 | 107.992107 | 61.73132341 | 222.6573986 | 201.947756 | 212.61143 | 4.12E-06 | 1.166179836 | 1.08E-06 | 23.77785554 | 2.244166686 | 5.385441105 |
| ENSMUSG00000028086 | *Fbxw7* | 430.5874089 | 480.4134854 | 318.0505141 | 943.7637465 | 914.193605 | 910.3634868 | 9.94E-14 | 1.1659064 | 1.34E-14 | 59.32254465 | 2.243741386 | 13.00254141 |
| ENSMUSG00000037262 | *Kin* | 182.1408705 | 205.8914937 | 226.7955143 | 461.3393828 | 458.1825431 | 450.3496655 | 2.95E-16 | 1.164946329 | 3.36E-17 | 71.12198054 | 2.242248739 | 15.53045032 |
| ENSMUSG00000096696 | *Zfp960* | 47.13294457 | 63.58413777 | 59.04735283 | 122.2928894 | 110.7455436 | 144.9623387 | 1.52E-05 | 1.164473103 | 4.23E-06 | 21.15711998 | 2.241513368 | 4.819283321 |
| ENSMUSG00000090186 | *Gm52951* | 72.69657552 | 59.54704966 | 89.91301453 | 196.5120223 | 153.0894279 | 143.9959231 | 9.29E-06 | 1.163786918 | 2.54E-06 | 22.13726659 | 2.240447496 | 5.032202798 |
| ENSMUSG00000020608 | *Smc6* | 1425.172426 | 1126.347583 | 1230.600512 | 2791.651475 | 2853.326359 | 2836.42976 | 7.89E-29 | 1.163242877 | 4.44E-30 | 129.8404469 | 2.239602781 | 28.10307492 |
| ENSMUSG00000022378 | *Cyrib* | 1113.615673 | 920.4560897 | 938.0477188 | 2261.996754 | 2215.996613 | 2189.897729 | 1.16E-30 | 1.162863233 | 5.96E-32 | 138.4003436 | 2.239013509 | 29.93696277 |
| ENSMUSG00000032293 | *Ireb2* | 1517.041724 | 1550.241835 | 1415.794483 | 3364.319557 | 3413.56852 | 3264.551867 | 4.47E-56 | 1.161837454 | 7.67E-58 | 257.0168394 | 2.237422101 | 55.35008895 |
| ENSMUSG00000035649 | *Zcchc7* | 190.1295052 | 198.8265895 | 157.0122791 | 419.1694209 | 401.7240307 | 406.8609639 | 9.42E-15 | 1.161447023 | 1.19E-15 | 64.09577779 | 2.236816678 | 14.02579324 |
| ENSMUSG00000024691 | *Fam111a* | 2919.047109 | 2784.581526 | 3444.876243 | 7065.99881 | 6841.251669 | 6533.935812 | 5.62E-32 | 1.161140917 | 2.74E-33 | 144.515009 | 2.236342129 | 31.25049636 |
| ENSMUSG00000000001 | *Gnai3* | 3418.336777 | 3122.687655 | 3612.624405 | 7681.680253 | 7481.838637 | 7517.746883 | 2.89E-51 | 1.160237915 | 5.93E-53 | 234.5972269 | 2.234942811 | 50.53975715 |
| ENSMUSG00000047141 | *Zfp654* | 193.3249591 | 197.8173175 | 159.6962497 | 412.422227 | 407.1527338 | 418.457951 | 3.10E-15 | 1.16019352 | 3.78E-16 | 66.34734817 | 2.234874037 | 14.50812479 |
| ENSMUSG00000044162 | *Tnip3* | 1425.971289 | 1159.65356 | 1549.993012 | 3165.277337 | 2992.301159 | 3074.167995 | 3.76E-23 | 1.159852202 | 2.82E-24 | 103.3397307 | 2.234345366 | 22.42538995 |
| ENSMUSG00000069135 | *Cep43* | 515.2669364 | 600.5168567 | 564.9758077 | 1320.763206 | 1167.17117 | 1257.306684 | 9.62E-27 | 1.158801157 | 6.03E-28 | 120.0961264 | 2.232718172 | 26.01661025 |
| ENSMUSG00000025326 | *Ube3a* | 1891.708691 | 1692.549191 | 1694.927423 | 3912.529062 | 3964.039017 | 3924.613716 | 4.22E-51 | 1.158678478 | 8.79E-53 | 233.8115794 | 2.232528323 | 50.37462907 |
| ENSMUSG00000019841 | *Rev3l* | 345.1090179 | 332.0504973 | 202.639779 | 684.8401806 | 656.8730773 | 633.0022122 | 7.06E-09 | 1.158232986 | 1.40E-09 | 36.66439069 | 2.231839044 | 8.151109681 |
| ENSMUSG00000070336 | *Fbxo47* | 17.57499628 | 23.21325665 | 28.18169112 | 50.60395423 | 45.60110619 | 55.08568869 | 0.00447203 | 1.158082304 | 0.001756718 | 9.787817704 | 2.231605952 | 2.349495254 |
| ENSMUSG00000111640 | *None* | 398.6328702 | 346.1803057 | 340.864264 | 863.6408189 | 743.7323271 | 819.5204213 | 3.11E-20 | 1.156883151 | 2.77E-21 | 89.69931504 | 2.229751835 | 19.50698452 |
| ENSMUSG00000031010 | *Usp9x* | 2739.302829 | 2665.487426 | 2274.665069 | 5779.814973 | 5974.830651 | 5384.867674 | 2.69E-33 | 1.156681091 | 1.24E-34 | 150.663278 | 2.229439564 | 32.56965008 |
| ENSMUSG00000104882 | *Gm43096* | 160.5715569 | 110.0106511 | 92.59698511 | 282.5387445 | 289.8927465 | 245.4695601 | 1.50E-06 | 1.156538221 | 3.74E-07 | 25.82301497 | 2.229218793 | 5.824323544 |
| ENSMUSG00000000282 | *Mnt* | 642.2862277 | 631.8042896 | 405.279558 | 1209.434506 | 1246.430236 | 1300.795386 | 3.17E-11 | 1.15636118 | 5.16E-12 | 47.62614858 | 2.228945251 | 10.49913122 |
| ENSMUSG00000040021 | *Lats1* | 1274.986094 | 1111.208503 | 974.2813216 | 2565.62048 | 2529.775653 | 2409.274069 | 1.76E-24 | 1.155694445 | 1.23E-25 | 109.5499925 | 2.227915391 | 23.75529119 |
| ENSMUSG00000057596 | *Trim30d* | 67.10453125 | 43.39869721 | 57.70536754 | 118.9192924 | 128.1173936 | 129.4996892 | 1.65E-05 | 1.155580779 | 4.63E-06 | 20.98356189 | 2.227739866 | 4.782601536 |
| ENSMUSG00000027680 | *Fxr1* | 1520.237178 | 1457.388809 | 1881.463379 | 3685.654667 | 3516.71388 | 3603.763739 | 2.60E-25 | 1.155500041 | 1.75E-26 | 113.4167938 | 2.227615198 | 24.58479849 |
| ENSMUSG00000104058 | *None* | 43.13862723 | 50.46360141 | 53.67941166 | 108.7985016 | 107.4883217 | 109.2049618 | 9.53E-06 | 1.155416523 | 2.61E-06 | 22.08643032 | 2.227486245 | 5.020983172 |
| ENSMUSG00000039145 | *Camk1d* | 536.83625 | 585.3777763 | 518.0063225 | 1250.761069 | 1226.886905 | 1177.09419 | 1.34E-32 | 1.154516294 | 6.40E-34 | 147.4065659 | 2.226096747 | 31.87254071 |
| ENSMUSG00000022500 | *Litaf* | 4567.102442 | 4993.877995 | 4135.998668 | 10037.29432 | 10182.07557 | 10250.77018 | 2.34E-41 | 1.152872877 | 7.27E-43 | 188.355165 | 2.223562378 | 40.6305988 |
| ENSMUSG00000045427 | *Hnrnph2* | 1382.033798 | 1452.342448 | 1725.793085 | 3481.552051 | 3331.052233 | 3306.107737 | 2.71E-29 | 1.152742785 | 1.49E-30 | 132.0040889 | 2.223361881 | 28.56677293 |
| ENSMUSG00000038024 | *Dennd4c* | 965.825932 | 1053.679997 | 1127.267645 | 2355.61407 | 2358.228634 | 2265.278146 | 1.41E-38 | 1.15252319 | 5.06E-40 | 175.3344065 | 2.223023485 | 37.85070634 |
| ENSMUSG00000021895 | *Arhgef3* | 183.7385975 | 158.4557084 | 202.639779 | 415.795824 | 365.8945901 | 424.2564445 | 7.63E-13 | 1.152064986 | 1.10E-13 | 55.1853013 | 2.22231756 | 12.11733214 |
| ENSMUSG00000025949 | *Pikfyve* | 1171.932707 | 1229.29333 | 1117.873748 | 2671.045384 | 2629.66379 | 2523.311108 | 5.24E-49 | 1.151452486 | 1.15E-50 | 224.1122367 | 2.221374268 | 48.28092124 |
| ENSMUSG00000033319 | *Fem1c* | 712.5862128 | 662.0824505 | 614.6292635 | 1481.85246 | 1451.635214 | 1494.078504 | 1.45E-34 | 1.149982843 | 6.26E-36 | 156.6007097 | 2.219112554 | 33.8396872 |
| ENSMUSG00000113961 | *Gm48498* | 55.12157924 | 50.46360141 | 59.04735283 | 113.858897 | 122.6886905 | 127.566858 | 3.10E-06 | 1.148332146 | 8.02E-07 | 24.35244449 | 2.21657495 | 5.509238438 |
| ENSMUSG00000026360 | *Rgs2* | 2085.832513 | 1750.077697 | 1839.861835 | 4243.141563 | 4227.873988 | 4120.796081 | 2.99E-38 | 1.148178332 | 1.08E-39 | 173.8212238 | 2.216338641 | 37.5238186 |
| ENSMUSG00000082127 | *Gm13577* | 536.0373866 | 486.4691176 | 483.1147049 | 1117.503989 | 1087.912105 | 1138.437566 | 8.03E-32 | 1.147776987 | 3.95E-33 | 143.7898685 | 2.215722162 | 31.09509591 |
| ENSMUSG00000101932 | *Gm10074* | 119.8295201 | 50.46360141 | 97.96492628 | 166.993049 | 194.3475716 | 233.8725731 | 0.00067726 | 1.145868941 | 0.000235186 | 13.52675436 | 2.21279368 | 3.169244459 |
| ENSMUSG00000011008 | *Mcoln2* | 218.88859 | 295.7167042 | 332.8123523 | 634.2362264 | 619.9578961 | 608.8418224 | 2.02E-11 | 1.145670847 | 3.23E-12 | 48.54196638 | 2.212489867 | 10.69483355 |
| ENSMUSG00000003282 | *Plag1* | 23.16704055 | 15.13908042 | 22.81374996 | 48.07375652 | 43.42962494 | 43.4887016 | 0.007041419 | 1.145330474 | 0.002862446 | 8.893136185 | 2.211967938 | 2.152339827 |
| ENSMUSG00000039156 | *Stim2* | 596.7510101 | 673.1844428 | 586.4475724 | 1325.823601 | 1383.233554 | 1396.470529 | 8.81E-32 | 1.14384025 | 4.35E-33 | 143.5970981 | 2.209684277 | 31.05502942 |
| ENSMUSG00000071796 | *6820431F20Rik* | 3590.092422 | 3096.446582 | 3075.830288 | 7462.396451 | 7124.629972 | 6985.251893 | 2.73E-39 | 1.142824842 | 9.39E-41 | 178.6841053 | 2.208129589 | 38.56391339 |
| ENSMUSG00000000838 | *Fmr1* | 658.263497 | 668.1380826 | 524.716249 | 1382.33135 | 1265.973567 | 1448.656971 | 6.22E-20 | 1.142091643 | 5.65E-21 | 88.29219747 | 2.20700767 | 19.20615534 |
| ENSMUSG00000072812 | *None* | 418.6044569 | 517.7565504 | 336.8383082 | 874.605009 | 1004.310077 | 937.4231234 | 5.05E-12 | 1.141028549 | 7.71E-13 | 51.35457319 | 2.205381968 | 11.29688166 |
| ENSMUSG00000029505 | *Ep400* | 1606.514433 | 1611.807429 | 832.0308807 | 2946.836935 | 3041.159487 | 2956.265293 | 1.78E-07 | 1.140956195 | 4.06E-08 | 30.12290398 | 2.205271366 | 6.749340938 |
| ENSMUSG00000023852 | *Chd1* | 1933.249591 | 1786.41149 | 1811.680144 | 4182.416817 | 4094.327891 | 3928.479378 | 2.11E-55 | 1.140778007 | 3.68E-57 | 253.8933824 | 2.204999009 | 54.6751332 |
| ENSMUSG00000026779 | *Mastl* | 227.6760882 | 218.0027581 | 201.2977937 | 470.6167744 | 471.2114306 | 489.9727047 | 2.49E-18 | 1.140373556 | 2.49E-19 | 80.80470028 | 2.204380936 | 17.60341111 |
| ENSMUSG00000033066 | *Gas7* | 3856.91282 | 3866.52114 | 2968.471465 | 8040.968328 | 7953.050068 | 7575.731819 | 1.90E-23 | 1.139317471 | 1.40E-24 | 104.7258724 | 2.202767871 | 22.72235511 |
| ENSMUSG00000020385 | *Clk4* | 547.2214751 | 462.2465889 | 526.0582343 | 1211.964704 | 1087.912105 | 1082.385462 | 1.41E-23 | 1.139288567 | 1.04E-24 | 105.3256154 | 2.202723739 | 22.84962916 |
| ENSMUSG00000007613 | *Tgfbr1* | 691.0168992 | 670.1566267 | 685.754484 | 1544.264003 | 1444.035029 | 1519.205309 | 5.96E-41 | 1.13926247 | 1.89E-42 | 186.4521608 | 2.202683895 | 40.22498108 |
| ENSMUSG00000109925 | *Gm45315* | 651.0737258 | 580.3314162 | 661.5987487 | 1471.731669 | 1340.88967 | 1350.082581 | 1.76E-29 | 1.138298918 | 9.57E-31 | 132.8876472 | 2.20121325 | 28.75491198 |
| ENSMUSG00000099703 | *None* | 57.51816964 | 80.74176225 | 96.62294099 | 197.3554215 | 147.6607248 | 165.2570661 | 2.95E-05 | 1.138126991 | 8.55E-06 | 19.81148555 | 2.200950945 | 4.530719372 |
| ENSMUSG00000038664 | *Herc1* | 1366.855393 | 1332.239077 | 897.78816 | 2632.249019 | 2720.866003 | 2565.833394 | 3.39E-13 | 1.135751446 | 4.75E-14 | 56.83202782 | 2.197329837 | 12.47043662 |
| ENSMUSG00000035696 | *Rnf38* | 423.3976377 | 380.4955546 | 297.9207347 | 786.0480891 | 838.1917614 | 810.822681 | 1.09E-14 | 1.135729805 | 1.37E-15 | 63.80453569 | 2.197296876 | 13.96362386 |
| ENSMUSG00000037958 | *Nsrp1* | 322.7408408 | 346.1803057 | 386.491764 | 876.2918075 | 740.4751053 | 690.9871477 | 2.96E-15 | 1.134388606 | 3.59E-16 | 66.44873133 | 2.195255112 | 14.52900136 |
| ENSMUSG00000114147 | *Gm18939* | 100.6567969 | 99.91793079 | 71.12522045 | 204.9460146 | 212.8051622 | 184.5853779 | 1.97E-07 | 1.134235916 | 4.51E-08 | 29.91746984 | 2.195022786 | 6.705016398 |
| ENSMUSG00000063234 | *Gpr84* | 2497.247199 | 2532.263519 | 2231.72154 | 5237.509263 | 5271.270727 | 5431.255622 | 5.98E-49 | 1.132822958 | 1.31E-50 | 223.8432667 | 2.19287406 | 48.22342569 |
| ENSMUSG00000075470 | *Alg10b* | 1243.031555 | 1216.172794 | 746.1438221 | 2255.24956 | 2704.579893 | 2080.692768 | 6.88E-09 | 1.132589611 | 1.36E-09 | 36.71883098 | 2.192519405 | 8.162592465 |
| ENSMUSG00000041671 | *Pyroxd1* | 179.7442801 | 191.7616853 | 179.8260291 | 411.5788278 | 370.2375526 | 426.1892757 | 6.08E-15 | 1.13254671 | 7.55E-16 | 64.9853036 | 2.192454207 | 14.21611372 |
| ENSMUSG00000002297 | *Dbf4* | 1025.740692 | 1009.272028 | 1036.012645 | 2320.191302 | 2205.139206 | 2201.494717 | 2.57E-51 | 1.131952571 | 5.27E-53 | 234.8309696 | 2.191551484 | 50.58947839 |
| ENSMUSG00000017677 | *Wsb1* | 2438.131302 | 2318.297849 | 2215.617716 | 5161.603332 | 5104.066671 | 5029.226736 | 7.29E-61 | 1.131865498 | 1.04E-62 | 279.3417307 | 2.191419218 | 60.13754532 |
| ENSMUSG00000036197 | *Gxylt1* | 490.5021689 | 430.959156 | 397.2276463 | 1049.188651 | 969.5663768 | 879.4381879 | 2.79E-18 | 1.131430321 | 2.80E-19 | 80.57645497 | 2.190758294 | 17.55478392 |
| ENSMUSG00000031529 | *Tnks* | 734.1555264 | 749.8891169 | 471.0368373 | 1429.561707 | 1461.406879 | 1402.269023 | 5.51E-11 | 1.130444394 | 9.13E-12 | 46.50644304 | 2.189261658 | 10.25867152 |
| ENSMUSG00000029208 | *Guf1* | 545.6237482 | 413.8015315 | 434.8032345 | 1033.164066 | 912.0221238 | 1113.310761 | 1.53E-14 | 1.129803826 | 1.95E-15 | 63.11595695 | 2.188289824 | 13.81524847 |
| ENSMUSG00000027665 | *Pik3ca* | 961.8316146 | 1018.355476 | 1092.376027 | 2293.202526 | 2213.825131 | 2200.528301 | 5.32E-41 | 1.129713574 | 1.68E-42 | 186.6891629 | 2.188152934 | 40.27428991 |
| ENSMUSG00000059839 | *Zfp874b* | 76.69089286 | 73.67685805 | 104.6748527 | 188.0780299 | 193.261831 | 171.0555596 | 5.61E-07 | 1.129382482 | 1.34E-07 | 27.81139372 | 2.187650821 | 6.250901395 |
| ENSMUSG00000063450 | *Syne2* | 19.17272321 | 31.28743287 | 24.15573525 | 53.13415195 | 51.02980931 | 57.98493547 | 0.004268137 | 1.128579928 | 0.001670965 | 9.879872263 | 2.186434195 | 2.369761598 |
| ENSMUSG00000057113 | *Npm1* | 25089.90492 | 22844.87236 | 26696.1134 | 54235.63135 | 54690.92669 | 54192.72069 | 4.68E-50 | 1.12819095 | 9.99E-52 | 228.9712743 | 2.18584477 | 49.32999998 |
| ENSMUSG00000073678 | *Pgap1* | 313.1544792 | 315.9021448 | 375.7558816 | 746.408325 | 730.7034397 | 708.3826283 | 2.12E-20 | 1.128165182 | 1.87E-21 | 90.48288862 | 2.185805729 | 19.67271361 |
| ENSMUSG00000031278 | *Acsl4* | 2653.824438 | 2416.197235 | 2581.979701 | 5705.59584 | 5580.706805 | 5440.919778 | 2.67E-58 | 1.128102092 | 4.18E-60 | 267.4041902 | 2.185710144 | 57.57356529 |
| ENSMUSG00000042390 | *Gatad2b* | 1271.79064 | 1225.256242 | 885.7102924 | 2480.437157 | 2554.747687 | 2374.483107 | 1.09E-15 | 1.128057165 | 1.28E-16 | 68.47769826 | 2.18564208 | 14.96306561 |
| ENSMUSG00000021500 | *Ddx46* | 1860.553015 | 1800.541298 | 1771.420585 | 4003.616179 | 3959.696054 | 3915.915975 | 3.11E-68 | 1.127680547 | 3.58E-70 | 313.6063055 | 2.185071589 | 67.50738895 |
| ENSMUSG00000035367 | *Rmi1* | 397.8340067 | 476.3763973 | 477.7467638 | 1041.598058 | 915.2793457 | 984.7774874 | 3.11E-20 | 1.126893246 | 2.77E-21 | 89.69884412 | 2.183879487 | 19.50698452 |
| ENSMUSG00000022708 | *Zbtb20* | 193.3249591 | 191.7616853 | 157.0122791 | 349.1672842 | 432.1247682 | 409.7602106 | 2.00E-11 | 1.126474198 | 3.20E-12 | 48.56313561 | 2.183245246 | 10.69936141 |
| ENSMUSG00000038679 | *Trps1* | 292.384029 | 329.0226812 | 249.6092642 | 707.61196 | 627.5580804 | 571.1516144 | 3.61E-13 | 1.126359549 | 5.07E-14 | 56.70078146 | 2.183071752 | 12.4429109 |
| ENSMUSG00000061589 | *Dot1l* | 4744.450132 | 4597.234088 | 3099.986023 | 8898.705352 | 9212.509191 | 9064.011829 | 1.17E-13 | 1.126227886 | 1.59E-14 | 58.98523923 | 2.18287253 | 12.92999677 |
| ENSMUSG00000020397 | *Med7* | 297.9760733 | 366.3657462 | 370.3879405 | 795.3254807 | 694.8739991 | 757.6698234 | 1.14E-16 | 1.125900518 | 1.26E-17 | 73.0547154 | 2.182377262 | 15.94325317 |
| ENSMUSG00000048379 | *Socs4* | 520.0601172 | 434.9962441 | 441.5131609 | 1059.309442 | 1003.224336 | 993.4752277 | 6.44E-23 | 1.125552421 | 4.91E-24 | 102.2421255 | 2.181850757 | 22.19107688 |
| ENSMUSG00000024737 | *Slc15a3* | 3534.970843 | 3581.906428 | 2916.134038 | 7098.047981 | 7454.695121 | 7347.657739 | 3.18E-31 | 1.124926018 | 1.60E-32 | 141.0120875 | 2.180903625 | 30.49701453 |
| ENSMUSG00000028282 | *Casp8ap2* | 549.6180655 | 487.4783896 | 367.7039699 | 1002.801693 | 1047.739702 | 1025.366942 | 2.18E-13 | 1.123729417 | 3.01E-14 | 57.72473537 | 2.179095489 | 12.66204256 |
| ENSMUSG00000030067 | *Foxp1* | 925.0838951 | 883.1130246 | 713.9361751 | 1799.813972 | 1862.045169 | 1850.685857 | 1.02E-22 | 1.123584731 | 7.88E-24 | 101.3068735 | 2.17887696 | 21.99249131 |
| ENSMUSG00000024378 | *Stard4* | 1179.921341 | 1083.958158 | 1191.682939 | 2709.841749 | 2401.658259 | 2409.274069 | 3.29E-34 | 1.12288071 | 1.46E-35 | 154.9230745 | 2.17781395 | 33.48304093 |
| ENSMUSG00000017631 | *Abr* | 7685.865419 | 7733.04228 | 6116.768959 | 15360.83031 | 15766.03959 | 15755.47338 | 1.78E-27 | 1.121709116 | 1.06E-28 | 123.5392623 | 2.176046092 | 26.74892482 |
| ENSMUSG00000059588 | *Calcrl* | 1508.254226 | 1477.574249 | 1410.426541 | 3409.019717 | 3044.416708 | 3113.791035 | 2.58E-42 | 1.121048969 | 7.73E-44 | 192.8143909 | 2.175050608 | 41.58795858 |
| ENSMUSG00000030275 | *Etnk1* | 2438.131302 | 2531.254247 | 2368.60404 | 5291.486814 | 5490.590333 | 5177.088322 | 8.02E-63 | 1.120210537 | 1.07E-64 | 288.4810693 | 2.17378693 | 62.09564607 |
| ENSMUSG00000038729 | *Pakap* | 23.96590402 | 44.40796924 | 25.49772054 | 66.62853974 | 62.97295617 | 74.41400052 | 0.003922492 | 1.120027431 | 0.001526494 | 10.04635863 | 2.173511051 | 2.406437958 |
| ENSMUSG00000041390 | *Mdfic* | 1378.039481 | 1503.815322 | 1784.840438 | 3446.129283 | 3303.908717 | 3371.823997 | 8.03E-25 | 1.119855163 | 5.54E-26 | 111.1316866 | 2.173251534 | 24.0951686 |
| ENSMUSG00000043535 | *Setx* | 1104.828175 | 1090.01379 | 838.7408072 | 2170.066237 | 2348.456969 | 2089.390508 | 5.42E-19 | 1.119676521 | 5.19E-20 | 83.90424758 | 2.172982447 | 18.26586057 |
| ENSMUSG00000109916 | *2410039M03Rik* | 41.5409003 | 41.38015315 | 45.62749991 | 103.7381062 | 89.03073113 | 85.04457202 | 8.82E-05 | 1.119225135 | 2.71E-05 | 17.6084266 | 2.172302678 | 4.054738717 |
| ENSMUSG00000032400 | *Zwilch* | 484.1112612 | 412.7922595 | 441.5131609 | 957.2581343 | 938.0798988 | 1015.702786 | 3.40E-24 | 1.118452704 | 2.41E-25 | 108.2150921 | 2.17113992 | 23.46790234 |
| ENSMUSG00000022895 | *Ets2* | 6779.954247 | 7084.080365 | 5439.066387 | 14148.8656 | 14071.19848 | 13682.51194 | 2.40E-25 | 1.117609672 | 1.61E-26 | 113.5855509 | 2.169871595 | 24.62061564 |
| ENSMUSG00000023025 | *Larp4* | 1726.343953 | 1282.784748 | 1635.88007 | 3548.180591 | 3240.935761 | 3283.880179 | 1.60E-19 | 1.116716607 | 1.49E-20 | 86.3750062 | 2.168528805 | 18.79495122 |
| ENSMUSG00000035004 | *Igsf6* | 295.5794829 | 303.7908805 | 319.3924994 | 709.2987585 | 652.5301148 | 624.3044719 | 1.71E-20 | 1.116156499 | 1.50E-21 | 90.91915647 | 2.167687064 | 19.76612937 |
| ENSMUSG00000056018 | *Ccdc7b* | 49.52953497 | 35.32452098 | 20.12977937 | 70.00213669 | 80.34480614 | 82.14532525 | 0.004796313 | 1.116142582 | 0.001895993 | 9.647598455 | 2.167666155 | 2.31909247 |
| ENSMUSG00000045225 | *Or5w19* | 51.92612537 | 26.24107273 | 29.52367641 | 74.21913288 | 92.287953 | 70.54833815 | 0.004212031 | 1.115492228 | 0.001647127 | 9.90631395 | 2.16668921 | 2.375508436 |
| ENSMUSG00000035954 | *Dock4* | 377.86242 | 386.5511868 | 304.6306612 | 787.7348876 | 788.2476927 | 750.9049143 | 2.18E-18 | 1.115386785 | 2.17E-19 | 81.08056967 | 2.166530858 | 17.66172472 |
| ENSMUSG00000026395 | *Ptprc* | 3750.663979 | 3541.535547 | 4001.800139 | 8547.851269 | 7972.593399 | 7918.809354 | 2.47E-47 | 1.114434174 | 5.90E-49 | 216.2663858 | 2.165100771 | 46.60762797 |
| ENSMUSG00000070923 | *Klhl9* | 599.1476005 | 659.0546344 | 594.4994841 | 1442.212696 | 1267.059308 | 1299.82897 | 5.46E-27 | 1.113979365 | 3.35E-28 | 121.2624968 | 2.164418331 | 26.26284984 |
| ENSMUSG00000038215 | *Cep44* | 254.8374461 | 208.9193098 | 185.1939702 | 496.7621507 | 442.9821744 | 473.5436397 | 6.86E-12 | 1.113839065 | 1.06E-12 | 50.72862454 | 2.164207855 | 11.16372802 |
| ENSMUSG00000056763 | *Cspp1* | 198.1181399 | 205.8914937 | 159.6962497 | 432.6638087 | 438.6392119 | 354.6745219 | 1.12E-10 | 1.113783018 | 1.90E-11 | 45.06910418 | 2.164123779 | 9.951762762 |
| ENSMUSG00000036390 | *Gadd45a* | 74.29430246 | 87.80666645 | 81.86110278 | 153.4986612 | 205.2049779 | 169.1227284 | 1.15E-06 | 1.113656613 | 2.83E-07 | 26.36199652 | 2.163934173 | 5.938831977 |
| ENSMUSG00000030530 | *Furin* | 2532.397191 | 2689.709955 | 2046.52757 | 5191.122305 | 5279.956652 | 5257.300816 | 4.97E-24 | 1.112132276 | 3.57E-25 | 107.437714 | 2.16164899 | 23.3033114 |
| ENSMUSG00000033446 | *Lpar6* | 127.8181548 | 98.90865876 | 84.54507337 | 243.7423796 | 226.9197903 | 208.7457677 | 4.08E-07 | 1.112071884 | 9.62E-08 | 28.44943648 | 2.161558503 | 6.389039072 |
| ENSMUSG00000048720 | *Tbc1d12* | 102.2545238 | 87.80666645 | 80.51911749 | 180.4874368 | 206.2907185 | 202.9472741 | 6.73E-08 | 1.111602651 | 1.47E-08 | 32.08981423 | 2.160855577 | 7.1717263 |
| ENSMUSG00000032086 | *Bace1* | 82.28293713 | 79.73249022 | 46.9694852 | 140.0042734 | 166.1183154 | 151.7272478 | 5.20E-05 | 1.111390083 | 1.55E-05 | 18.66897776 | 2.160537218 | 4.283770142 |
| ENSMUSG00000040133 | *Gpr176* | 34.35112909 | 48.44505735 | 48.31147049 | 114.7022963 | 90.11647176 | 75.38041611 | 0.000826515 | 1.110714558 | 0.000290675 | 13.12954497 | 2.159525808 | 3.082749222 |
| ENSMUSG00000018501 | *Ncor1* | 2568.346047 | 2646.311258 | 2147.176466 | 5293.173613 | 5386.359233 | 5234.106842 | 5.05E-31 | 1.110524717 | 2.56E-32 | 140.0769625 | 2.159241659 | 30.29709866 |
| ENSMUSG00000025758 | *Plk4* | 934.6702567 | 747.8705728 | 805.1911749 | 1784.632786 | 1773.014438 | 1823.62622 | 3.68E-25 | 1.110497482 | 2.49E-26 | 112.71397 | 2.159200898 | 24.43427349 |
| ENSMUSG00000090272 | *Mndal* | 1226.255422 | 1062.763446 | 983.6752187 | 2411.278419 | 2381.029187 | 2290.404951 | 7.23E-27 | 1.11044001 | 4.46E-28 | 120.6930115 | 2.159114884 | 26.14109787 |
| ENSMUSG00000033972 | *Zfp944* | 123.024974 | 143.316628 | 110.0427939 | 266.514159 | 286.6355246 | 262.8650408 | 5.23E-10 | 1.110224909 | 9.42E-11 | 41.93778654 | 2.158792992 | 9.281121985 |
| ENSMUSG00000022035 | *Ccdc25* | 560.0032906 | 526.8399987 | 613.2872782 | 1250.761069 | 1222.543942 | 1184.825515 | 7.16E-29 | 1.108961423 | 4.03E-30 | 130.0331741 | 2.156903188 | 28.14478694 |
| ENSMUSG00000030655 | *Smg1* | 2792.826682 | 2781.55371 | 2124.362716 | 5436.551483 | 5849.97048 | 5329.781985 | 1.97E-20 | 1.108362563 | 1.72E-21 | 90.63962278 | 2.156008047 | 19.70623946 |
| ENSMUSG00000058331 | *Zfp85* | 31.15567522 | 24.22252868 | 40.25955875 | 74.21913288 | 60.80147492 | 68.61550697 | 0.001919109 | 1.107841749 | 0.000711285 | 11.45952532 | 2.155229868 | 2.716900257 |
| ENSMUSG00000030102 | *Itpr1* | 618.3203237 | 614.6466651 | 496.5345579 | 1211.121305 | 1279.002455 | 1250.541775 | 5.34E-22 | 1.107350456 | 4.33E-23 | 97.93235409 | 2.154496053 | 21.27256107 |
| ENSMUSG00000031093 | *Dock11* | 965.0270685 | 869.9924883 | 959.5194834 | 2082.352717 | 1992.334044 | 1941.528923 | 2.22E-37 | 1.106664443 | 8.37E-39 | 169.7556032 | 2.153471817 | 36.65393952 |

| | | | | | | | | | | | | | |
|---|---|---|---|---|---|---|---|---|---|---|---|---|---|
| ENSMUSG00000080186 | *Gm14448* | 108.6454316 | 110.0106511 | 97.96492628 | 257.2367674 | 216.0623841 | 209.7121833 | 1.22E-08 | 1.106587283 | 2.49E-09 | 35.54479682 | 2.153356646 | 7.914674974 |
| ENSMUSG00000000386 | *Mx1* | 517.6635268 | 494.5432938 | 393.2016904 | 1022.199876 | 1045.56822 | 971.2476691 | 9.96E-18 | 1.106079505 | 1.03E-18 | 77.99790317 | 2.152598874 | 17.00154444 |
| ENSMUSG00000027177 | *Hipk3* | 1418.781518 | 1456.379537 | 1217.180659 | 2987.320098 | 2895.670243 | 2940.802644 | 2.64E-33 | 1.106025949 | 1.21E-34 | 150.7088485 | 2.152518965 | 32.57849684 |
| ENSMUSG00000081603 | *Gm14681* | 5472.214751 | 4880.839528 | 5787.982562 | 11316.73097 | 11813.94372 | 11582.49086 | 1.03E-40 | 1.105108051 | 3.32E-42 | 185.3353472 | 2.151149886 | 39.98758755 |
| ENSMUSG00000036990 | *Otud4* | 1970.796174 | 1892.385053 | 1541.9411 | 3933.614042 | 3952.09587 | 3754.524572 | 6.05E-25 | 1.104440889 | 4.13E-26 | 111.7106223 | 2.150155335 | 24.21825749 |
| ENSMUSG00000085028 | *Slc2a4rg-ps* | 50.32839844 | 44.40796924 | 44.28551462 | 91.93051686 | 105.3168405 | 103.4064683 | 4.64E-05 | 1.104346738 | 1.38E-05 | 18.89959251 | 2.150015019 | 4.333359253 |
| ENSMUSG00000038766 | *Gabpb2* | 2236.018845 | 2244.620991 | 1869.385511 | 4639.539204 | 4651.312831 | 4378.829043 | 2.84E-31 | 1.104316859 | 1.43E-32 | 141.2408868 | 2.149970491 | 30.54602303 |
| ENSMUSG00000020572 | *Nampt* | 3078.020939 | 2715.951028 | 3438.166317 | 6737.073107 | 6562.216329 | 6534.902227 | 3.22E-29 | 1.104197273 | 1.78E-30 | 131.6568254 | 2.149792286 | 28.49219621 |
| ENSMUSG00000100967 | *Gm29666* | 26.36249442 | 30.27816084 | 36.23360287 | 61.56814432 | 57.54425305 | 78.27966288 | 0.002113958 | 1.103923143 | 0.000787369 | 11.27077278 | 2.149383837 | 2.674903679 |
| ENSMUSG00000030386 | *Zfp606* | 95.86361607 | 91.84375456 | 67.09926458 | 187.2346307 | 192.1760904 | 173.9548064 | 5.66E-07 | 1.103662128 | 1.35E-07 | 27.79488342 | 2.148995002 | 6.247410274 |
| ENSMUSG00000027171 | *Prrg4* | 184.5374609 | 229.1047504 | 234.847426 | 479.894166 | 444.067915 | 460.980237 | 2.56E-13 | 1.103515849 | 3.57E-14 | 57.39461102 | 2.148777121 | 12.59210337 |
| ENSMUSG00000066026 | *Dhrs3* | 2473.281295 | 2465.651565 | 2481.330804 | 5411.249506 | 5164.868146 | 5359.740868 | 7.68E-73 | 1.103015162 | 7.56E-75 | 335.0689261 | 2.148031517 | 72.11490697 |
| ENSMUSG00000020464 | *Pnpt1* | 1245.428146 | 1207.089346 | 1454.712056 | 2840.568631 | 2816.411178 | 2718.527058 | 8.80E-31 | 1.102786242 | 4.51E-32 | 138.9515606 | 2.147690705 | 30.05551468 |
| ENSMUSG00000036943 | *Rab8b* | 1634.474654 | 1724.845896 | 1952.588599 | 3909.998864 | 3827.235698 | 3652.084519 | 1.86E-33 | 1.102769621 | 8.51E-35 | 151.4130729 | 2.147665962 | 32.73018079 |
| ENSMUSG00000022353 | *Mtss1* | 722.1725744 | 717.592412 | 548.8719842 | 1443.899494 | 1455.978176 | 1384.873542 | 1.84E-18 | 1.102512642 | 1.82E-19 | 81.42855138 | 2.147283444 | 17.73483272 |
| ENSMUSG00000040297 | *Suco* | 466.5362649 | 568.2201518 | 567.6597783 | 1193.409921 | 1151.970802 | 1084.318293 | 1.08E-20 | 1.102404951 | 9.34E-22 | 91.85173675 | 2.147123165 | 19.96727089 |
| ENSMUSG00000094915 | *ENSMUSG00000094915* | 121.427247 | 98.90865876 | 91.25499982 | 227.7177941 | 222.5768278 | 224.2084171 | 1.58E-08 | 1.101971252 | 3.27E-09 | 35.01651602 | 2.146477801 | 7.800361595 |
| ENSMUSG00000019917 | *Septin10* | 297.1772098 | 266.4478154 | 257.661176 | 613.9946447 | 568.9280867 | 585.6478482 | 1.27E-18 | 1.101800111 | 1.23E-19 | 82.19212276 | 2.146223188 | 17.89713092 |
| ENSMUSG00000035284 | *Vps13c* | 3091.601618 | 2928.907426 | 2271.981099 | 5993.19498 | 6106.205267 | 5713.448975 | 9.73E-20 | 1.101551019 | 8.89E-21 | 87.3942459 | 2.145852659 | 19.01209085 |
| ENSMUSG00000054178 | *None* | 56.71930618 | 38.35233707 | 42.94352933 | 109.6419008 | 108.5740624 | 80.21249406 | 0.000475173 | 1.101488016 | 0.000161095 | 14.23787579 | 2.14575895 | 3.323148156 |
| ENSMUSG00000044149 | *Nkrf* | 364.2817411 | 327.0041371 | 260.3451466 | 756.5291158 | 664.4732616 | 630.1029654 | 5.63E-12 | 1.101080057 | 8.63E-13 | 51.13325575 | 2.145152267 | 11.24925923 |
| ENSMUSG00000027309 | *Dnaaf9* | 838.8066407 | 782.1858218 | 644.1529399 | 1609.205745 | 1711.127223 | 1553.996271 | 3.28E-20 | 1.100826979 | 2.93E-21 | 89.58866264 | 2.144775997 | 19.48423335 |
| ENSMUSG00000009569 | *Mrtfb* | 733.356663 | 757.9632931 | 526.0582343 | 1414.380521 | 1563.466498 | 1359.746737 | 4.17E-13 | 1.100023815 | 5.90E-14 | 56.40484322 | 2.14358231 | 12.3795487 |
| ENSMUSG00000036840 | *Siah1a* | 154.9795127 | 135.2424518 | 181.1680144 | 317.9615124 | 318.1220027 | 369.1707558 | 5.57E-10 | 1.09941094 | 1.01E-10 | 41.81100435 | 2.142671882 | 9.254099967 |
| ENSMUSG00000020640 | *Itsn2* | 2400.584719 | 2486.846277 | 2620.897274 | 5403.658913 | 5504.704961 | 5161.625672 | 3.71E-54 | 1.099048348 | 6.71E-56 | 248.1077768 | 2.142133434 | 53.43115487 |
| ENSMUSG00000040123 | *Zmym5* | 326.7351581 | 319.9392329 | 339.5222788 | 709.2987585 | 701.3884428 | 698.7184724 | 1.24E-24 | 1.098923661 | 8.59E-26 | 110.2611256 | 2.141948304 | 23.9077841 |
| ENSMUSG00000023832 | *Acat2* | 1850.16779 | 1973.126815 | 2118.994775 | 4127.595867 | 4260.446207 | 4322.776939 | 8.29E-44 | 1.098886538 | 2.29E-45 | 199.8124354 | 2.141893189 | 43.08139412 |
| ENSMUSG00000025912 | *Mybl1* | 184.5374609 | 135.2424518 | 107.3588233 | 318.8049177 | 306.1788558 | 298.6224177 | 7.01E-07 | 1.098625323 | 1.69E-07 | 27.36175276 | 2.141505412 | 6.154011038 |
| ENSMUSG00000064354 | *mt-Co2* | 12048.45881 | 20652.73351 | 15749.53938 | 36625.45528 | 39441.69963 | 27674.27687 | 0.000153038 | 1.098425822 | 4.86E-05 | 16.50015127 | 2.141209298 | 3.815201206 |
| ENSMUSG00000036057 | *Ptpn23* | 2025.11889 | 2001.386432 | 1693.585438 | 4101.450491 | 4174.672698 | 3982.598651 | 9.14E-33 | 1.097535124 | 4.32E-34 | 148.1851788 | 2.139887755 | 32.03896451 |
| ENSMUSG00000091058 | *Gm17538* | 49.52953497 | 44.40796924 | 63.0733087 | 121.4494902 | 109.659803 | 101.4736371 | 6.55E-05 | 1.097254404 | 1.98E-05 | 18.20456334 | 2.139471415 | 4.183468757 |
| ENSMUSG00000029684 | *Wasl* | 276.4067597 | 268.4663595 | 170.432132 | 479.0507667 | 528.7556837 | 532.4949907 | 2.67E-08 | 1.096592209 | 5.61E-09 | 33.96535272 | 2.138489626 | 7.573941496 |
| ENSMUSG00000028886 | *Eya3* | 2558.759686 | 2660.441066 | 2281.374996 | 5443.298677 | 5377.673308 | 5233.140426 | 3.41E-41 | 1.096516872 | 1.07E-42 | 187.5766253 | 2.138377958 | 40.46712984 |
| ENSMUSG00000020962 | *Gtf2a1* | 825.2259617 | 844.7606876 | 789.0873514 | 1828.489546 | 1725.241851 | 1701.857856 | 1.11E-38 | 1.094698799 | 3.99E-40 | 175.8091669 | 2.135684889 | 37.95294899 |
| ENSMUSG00000030059 | *Tmf1* | 491.3010324 | 514.7287344 | 430.7772786 | 999.4280961 | 1071.625995 | 1006.03863 | 1.65E-23 | 1.093923619 | 1.22E-24 | 105.0096283 | 2.134537664 | 22.78176929 |
| ENSMUSG00000027699 | *Ect2* | 885.1407218 | 856.8719519 | 841.4247778 | 1907.769075 | 1822.958507 | 1787.868844 | 1.17E-41 | 1.09382254 | 3.59E-43 | 189.756875 | 2.134388118 | 40.93158725 |
| ENSMUSG00000024054 | *Smchd1* | 1366.855393 | 1333.248349 | 1167.527204 | 2784.904281 | 2799.039328 | 2686.635343 | 4.26E-35 | 1.093615082 | 1.79E-36 | 159.0872628 | 2.134081218 | 34.37039016 |
| ENSMUSG00000028191 | *Bcl10* | 889.9339026 | 972.9382351 | 1021.250807 | 2166.69264 | 1942.389976 | 2025.607079 | 1.08E-30 | 1.092228109 | 5.53E-32 | 138.5487429 | 2.132030547 | 29.96842065 |
| ENSMUSG00000024528 | *Srfbp1* | 533.6407961 | 500.598926 | 640.1269841 | 1232.206286 | 1125.913027 | 1198.355333 | 2.10E-19 | 1.092002072 | 1.97E-20 | 85.82511089 | 2.131696534 | 18.67748714 |
| ENSMUSG00000046879 | *Irgm1* | 2631.456261 | 2534.282063 | 3062.410435 | 5906.324858 | 5916.200658 | 5698.952741 | 5.28E-33 | 1.091913928 | 2.47E-34 | 149.294846 | 2.131566298 | 32.27771259 |
| ENSMUSG00000052551 | *Adarb2* | 61.51248698 | 58.53777763 | 72.46720574 | 140.8476726 | 130.2888748 | 136.2645983 | 2.99E-06 | 1.091749234 | 7.73E-07 | 24.4233908 | 2.131322979 | 5.524341331 |
| ENSMUSG00000031153 | *Gripap1* | 2422.952896 | 2196.175933 | 1972.718379 | 4694.360154 | 4734.914859 | 4632.996344 | 3.68E-32 | 1.091013981 | 1.79E-33 | 145.3638501 | 2.13023705 | 31.43396894 |
| ENSMUSG00000034485 | *Uaca* | 336.3215197 | 332.0504973 | 183.8519849 | 598.8134584 | 640.5869679 | 584.6814326 | 1.13E-06 | 1.089803286 | 2.76E-07 | 26.40830481 | 2.128450127 | 5.948409471 |
| ENSMUSG00000070544 | *Top1* | 1741.522359 | 1804.578386 | 1587.5686 | 3800.356963 | 3677.403492 | 3455.902154 | 7.56E-38 | 1.089326327 | 2.77E-39 | 171.9530071 | 2.127746572 | 37.12145151 |
| ENSMUSG00000063972 | *Nr6a1* | 158.97383 | 163.5020686 | 127.4886027 | 328.0823033 | 336.5795893 | 298.6224177 | 1.59E-10 | 1.088945439 | 2.73E-11 | 44.35952327 | 2.127184897 | 9.798717697 |
| ENSMUSG00000026675 | *Hsd17b7* | 1148.765666 | 1192.959537 | 1187.656983 | 2460.195575 | 2541.7188 | 2499.150719 | 8.35E-53 | 1.088348347 | 1.59E-54 | 241.8024402 | 2.126304696 | 52.07848747 |
| ENSMUSG00000027882 | *Stxbp3* | 492.0998958 | 541.9790791 | 743.4598515 | 1299.678225 | 1196.486167 | 1266.004424 | 1.29E-11 | 1.087512359 | 2.05E-12 | 49.4392876 | 2.125072939 | 10.8883609 |
| ENSMUSG00000079625 | *Tm4sf19* | 605.5385082 | 655.0175463 | 622.6811753 | 1324.980202 | 1399.519664 | 1275.66858 | 2.50E-31 | 1.0870628 | 1.25E-32 | 141.4959999 | 2.124410846 | 30.60129479 |
| ENSMUSG00000068245 | *Phf11d* | 2489.258564 | 1734.938616 | 1643.931982 | 4200.128201 | 4163.815291 | 4110.165509 | 1.94E-12 | 1.086252933 | 2.87E-13 | 53.29355538 | 2.123218628 | 11.71240172 |
| ENSMUSG00000048490 | *Nrip1* | 258.8317634 | 342.1432175 | 198.6138231 | 549.8963027 | 573.2710492 | 579.8493547 | 3.77E-08 | 1.085434479 | 8.04E-09 | 33.26660763 | 2.122014448 | 7.423983112 |
| ENSMUSG00000036898 | *Zfp157* | 122.2261105 | 123.1311874 | 123.4626468 | 273.2613529 | 245.3773809 | 262.8650408 | 9.92E-11 | 1.085274736 | 1.68E-11 | 45.30675509 | 2.1217795 | 10.00340891 |
| ENSMUSG00000021929 | *Kpna3* | 2315.905192 | 2198.194477 | 2983.233303 | 5419.683498 | 5292.98554 | 5177.088322 | 7.70E-19 | 1.085189648 | 7.43E-20 | 83.19600078 | 2.121654364 | 18.11346686 |
| ENSMUSG00000028180 | *Zranb2* | 1425.971289 | 1254.525131 | 1525.876276 | 2972.138912 | 2961.900421 | 2977.526436 | 2.13E-31 | 1.084526227 | 1.06E-32 | 141.8244447 | 2.120678949 | 30.67157313 |
| ENSMUSG00000022280 | *Rnf19a* | 265.2226711 | 313.8836008 | 394.5436757 | 745.5649257 | 717.6745522 | 590.4799262 | 1.84E-09 | 1.084343902 | 3.48E-10 | 39.38634561 | 2.120410959 | 8.734225282 |
| ENSMUSG00000079316 | *Rab9* | 290.7863021 | 281.5868959 | 322.07647 | 655.3212073 | 632.9867835 | 601.1104977 | 3.94E-19 | 1.083735181 | 3.75E-20 | 84.55032309 | 2.119516476 | 18.40401988 |
| ENSMUSG00000040033 | *Stat2* | 1362.062212 | 1301.960916 | 1033.328674 | 2574.897871 | 2634.006753 | 2645.079473 | 7.95E-21 | 1.083704932 | 6.83E-22 | 92.47231979 | 2.119472037 | 20.09989616 |
| ENSMUSG00000019907 | *Ppp1r12a* | 1080.862271 | 1126.347583 | 1100.427939 | 2435.736997 | 2255.083275 | 2314.565341 | 1.29E-44 | 1.083543809 | 3.41E-46 | 203.6075682 | 2.119235342 | 43.88964807 |
| ENSMUSG00000025959 | *Klf7* | 347.5056083 | 359.300842 | 279.1329406 | 728.696941 | 680.759371 | 689.0543165 | 4.58E-16 | 1.083525363 | 5.28E-17 | 70.22925615 | 2.119208246 | 15.33929468 |
| ENSMUSG00000031639 | *Tlr3* | 329.1317485 | 341.1339455 | 389.1757345 | 756.5291158 | 742.6465865 | 735.4422648 | 8.21E-21 | 1.083394837 | 7.07E-22 | 92.40419778 | 2.119016523 | 20.08584083 |
| ENSMUSG00000041488 | *Stx3* | 1453.132647 | 1545.195475 | 2059.947422 | 3587.820355 | 3513.456658 | 3592.166752 | 1.10E-15 | 1.082426056 | 1.29E-16 | 68.46007936 | 2.117594064 | 14.95940278 |
| ENSMUSG00000063334 | *Krr1* | 1021.746375 | 896.233561 | 1125.92566 | 2290.672328 | 2071.59311 | 2067.162949 | 1.18E-22 | 1.0810178 | 9.13E-24 | 101.0142226 | 2.115528028 | 21.92931855 |
| ENSMUSG00000043384 | *Gprasp1* | 283.5965309 | 310.8557847 | 269.7390436 | 607.2474508 | 644.9299304 | 578.8829391 | 5.22E-18 | 1.080352997 | 5.32E-19 | 79.30730832 | 2.114553404 | 17.28266947 |
| ENSMUSG00000000555 | *Itga5* | 5169.445497 | 5138.203895 | 4101.107051 | 10070.18689 | 10284.13519 | 10128.0354 | 9.04E-26 | 1.080114406 | 5.94E-27 | 115.5594288 | 2.114203731 | 25.04364276 |
| ENSMUSG00000035021 | *Baz1a* | 2211.254077 | 2327.381297 | 2066.657349 | 4713.758337 | 4801.145037 | 4452.276628 | 3.23E-43 | 1.079240895 | 9.14E-45 | 197.0613656 | 2.112924028 | 42.4910944 |
| ENSMUSG00000052812 | *Atad2b* | 444.1680878 | 430.959156 | 366.3619846 | 857.7370243 | 934.8226769 | 841.7479799 | 3.31E-19 | 1.078959564 | 3.14E-20 | 84.89833895 | 2.112512041 | 18.47965211 |
| ENSMUSG00000029823 | *Luc7l2* | 1754.304174 | 1628.965053 | 1325.881468 | 3299.377816 | 3284.365386 | 3375.68966 | 1.79E-21 | 1.077966449 | 1.48E-22 | 95.49912007 | 2.11105834 | 20.74666614 |
| ENSMUSG00000025044 | *Msr1* | 5651.160168 | 4914.145505 | 5723.567268 | 11623.72829 | 11402.44803 | 11350.55112 | 1.22E-40 | 1.077748482 | 3.97E-42 | 184.9781951 | 2.11073942 | 39.91199822 |
| ENSMUSG00000078901 | *Gm14440* | 77.48975633 | 63.58413777 | 60.38933812 | 172.8968436 | 158.5740624 | 144.9623387 | 0.000134944 | 1.077185796 | 4.26E-05 | 16.75152233 | 2.10991634 | 3.869846025 |
| ENSMUSG00000039908 | *Slc26a11* | 100.6567969 | 89.8252105 | 72.46720574 | 208.3196116 | 174.8042404 | 175.8876376 | 1.44E-06 | 1.076807565 | 3.59E-07 | 25.90211559 | 2.109363256 | 5.841208089 |
| ENSMUSG00000018899 | *Irf1* | 727.7646187 | 699.4255155 | 606.5773518 | 1490.286452 | 1344.146892 | 1462.186789 | 1.30E-23 | 1.075347337 | 9.49E-25 | 105.4993787 | 2.107229338 | 22.88701293 |
| ENSMUSG00000094441 | *Zfp955a* | 150.1863319 | 119.0940993 | 151.6443379 | 279.1651475 | 283.3783027 | 323.749223 | 2.87E-09 | 1.075308565 | 5.50E-10 | 38.49004703 | 2.10717284 | 8.542105611 |
| ENSMUSG00000027160 | *Ccdc34* | 324.3385677 | 296.7259763 | 263.0291171 | 628.3324317 | 606.9290086 | 635.901459 | 3.19E-17 | 1.074376418 | 3.43E-18 | 75.62455023 | 2.105811672 | 16.49567348 |
| ENSMUSG00000027009 | *Itga4* | 7051.327958 | 7226.387721 | 6405.295796 | 15176.12587 | 14940.87672 | 14404.42439 | 2.81E-40 | 1.074351389 | 9.26E-42 | 183.2928058 | 2.105775139 | 39.55111589 |
| ENSMUSG00000032998 | *Foxj3* | 867.5657255 | 765.0281973 | 626.7071311 | 1696.075866 | 1495.064839 | 1579.123076 | 1.58E-15 | 1.074141962 | 1.89E-16 | 67.71322792 | 2.105469478 | 14.80030983 |
| ENSMUSG00000039841 | *Zfp800* | 623.1135045 | 508.6731022 | 442.8551462 | 1142.805966 | 1085.740624 | 1099.780943 | 1.61E-14 | 1.073878888 | 2.05E-15 | 63.01198869 | 2.105085584 | 13.79272691 |
| ENSMUSG00000023830 | *Igf2r* | 897.9225372 | 923.4839057 | 628.0491164 | 1703.666459 | 1828.38721 | 1634.208765 | 2.35E-12 | 1.072899524 | 3.51E-13 | 52.89960656 | 2.103657044 | 11.62823287 |
| ENSMUSG00000031231 | *Cox7b* | 1612.90534 | 1456.379537 | 2167.306246 | 3615.65253 | 3600.315908 | 3781.584208 | 1.48E-13 | 1.072369435 | 2.02E-14 | 58.50868764 | 2.102884241 | 12.82839573 |
| ENSMUSG00000018076 | *Med13l* | 745.339615 | 805.3990785 | 469.694852 | 1411.850323 | 1489.636136 | 1357.813906 | 1.14E-08 | 1.072119388 | 2.31E-09 | 35.6900163 | 2.102519802 | 7.944568156 |
| ENSMUSG00000103847 | *Gm20056* | 299.5738002 | 356.2730259 | 360.9940434 | 705.9251616 | 752.4182521 | 671.6588358 | 8.94E-17 | 1.071773588 | 9.82E-18 | 73.54812951 | 2.102015909 | 16.04879937 |
| ENSMUSG00000030107 | *Usp18* | 4009.495742 | 3869.548598 | 4017.903963 | 8436.52257 | 8415.575573 | 8150.749096 | 1.16E-75 | 1.071604178 | 1.07E-77 | 348.1462217 | 2.101769091 | 74.93605459 |
| ENSMUSG00000044617 | *Zbtb39* | 327.5340216 | 349.2081217 | 248.2672789 | 621.5852378 | 689.445296 | 642.6663681 | 3.52E-12 | 1.071021511 | 5.31E-13 | 52.08572967 | 2.100920413 | 11.45405677 |
| ENSMUSG00000082938 | *Gm2810* | 84.67952763 | 38.35233707 | 79.1771322 | 141.6910719 | 157.4323904 | 125.6340268 | 0.001623895 | 1.069920269 | 0.000595735 | 11.78925752 | 2.099317345 | 2.789442017 |
| ENSMUSG00000068917 | *Clk2* | 591.9578293 | 641.8970099 | 485.7986755 | 1231.362866 | 1193.228945 | 1194.489671 | 2.05E-18 | 1.069240986 | 2.04E-19 | 81.20152404 | 2.098329128 | 17.68753189 |
| ENSMUSG00000031015 | *Swap70* | 2292.738151 | 2209.29647 | 1936.484776 | 4546.765288 | 4536.224325 | 4441.646057 | 1.08E-35 | 1.06877112 | 4.42E-37 | 161.8668523 | 2.097645843 | 34.96593937 |
| ENSMUSG00000022105 | *Rb1* | 322.7408408 | 246.2623749 | 195.9298526 | 520.3773294 | 549.3847555 | 546.024809 | 2.18E-08 | 1.068609363 | 4.56E-09 | 34.369066 | 2.097410665 | 7.661596568 |
| ENSMUSG00000036403 | *Cep135* | 331.5283389 | 288.6518 | 318.0505141 | 673.0325913 | 690.5310366 | 605.9425756 | 2.29E-17 | 1.068495972 | 2.42E-18 | 76.31209315 | 2.097245822 | 16.64063923 |
| ENSMUSG00000028522 | *Mier1* | 807.6509654 | 825.584519 | 889.7362483 | 1866.442512 | 1711.127223 | 1701.857856 | 7.82E-32 | 1.068407426 | 3.84E-33 | 143.8451788 | 2.097117106 | 31.10666709 |
| ENSMUSG00000046962 | *Zbtb21* | 271.6135789 | 235.1603826 | 309.9986023 | 539.7755118 | 554.8134586 | 611.7410692 | 4.14E-13 | 1.06769928 | 5.85E-14 | 56.42177997 | 2.09608799 | 12.38292672 |
| ENSMUSG00000021635 | *Rad17* | 588.7623754 | 524.8214546 | 743.4598515 | 1272.689449 | 1341.975411 | 1265.038009 | 2.40E-14 | 1.06713034 | 3.09E-15 | 62.20767918 | 2.095261541 | 13.62036415 |
| ENSMUSG00000027438 | *Napb* | 217.2908631 | 189.7431413 | 209.3497055 | 431.8204095 | 467.9542088 | 393.3311456 | 6.70E-13 | 1.067057075 | 9.60E-14 | 55.44699441 | 2.095155139 | 12.17390092 |
| ENSMUSG00000020362 | *Cnot6* | 1815.816661 | 1767.235321 | 1443.976174 | 3485.769047 | 3610.087573 | 3453.969323 | 1.13E-24 | 1.06694427 | 7.87E-26 | 110.4333383 | 2.094991326 | 23.9451435 |
| ENSMUSG00000021713 | *Ppwd1* | 325.9362947 | 328.0134091 | 386.491764 | 741.3479295 | 727.4462178 | 698.7184724 | 8.75E-19 | 1.065476982 | 8.47E-20 | 82.93692619 | 2.092861705 | 18.05787376 |
| ENSMUSG00000031644 | *Nek1* | 300.3726637 | 253.3272791 | 208.0077202 | 509.4131393 | 541.7845712 | 553.7561337 | 5.87E-11 | 1.064967906 | 9.76E-12 | 46.37552548 | 2.092123338 | 10.23104513 |
| ENSMUSG00000044583 | *Tlr7* | 2356.647228 | 2378.85417 | 3270.418155 | 5688.727855 | 5574.192361 | 5467.979415 | 1.69E-15 | 1.064893576 | 2.02E-16 | 67.57922413 | 2.092015551 | 14.77143655 |
| ENSMUSG00000026977 | *Marchf7* | 730.9600726 | 669.1473547 | 744.8018368 | 1491.129851 | 1482.035951 | 1510.507569 | 7.12E-34 | 1.064754617 | 3.19E-35 | 153.3659031 | 2.09181406 | 33.14734733 |
| ENSMUSG00000039585 | *Myo9a* | 577.5782868 | 548.0347113 | 523.3742637 | 1145.336164 | 1166.08543 | 1144.23606 | 1.85E-30 | 1.063643279 | 9.62E-32 | 137.4490103 | 2.090203313 | 29.73336092 |
| ENSMUSG00000027881 | *Prpf38b* | 1558.582625 | 1525.010035 | 1374.192939 | 3140.818759 | 3108.475405 | 3076.100827 | 7.03E-41 | 1.062393722 | 2.24E-42 | 186.1138581 | 2.088393715 | 40.15274083 |
| ENSMUSG00000001467 | *Cyp51* | 3866.499182 | 3690.907807 | 4126.604771 | 8187.719795 | 7862.933596 | 8334.368058 | 1.78E-49 | 1.062262191 | 3.85E-51 | 226.2874798 | 2.088203324 | 48.74945571 |
| ENSMUSG00000021754 | *Map3k1* | 480.9158073 | 466.283677 | 374.4138963 | 911.7145754 | 916.3650863 | 944.1880325 | 1.43E-17 | 1.062099028 | 1.49E-18 | 77.27217012 | 2.08796717 | 16.84616179 |
| ENSMUSG00000074829 | *2010315B03Rik* | 105.4499777 | 89.8252105 | 88.57102924 | 210.8498093 | 190.0046091 | 194.2495338 | 8.11E-08 | 1.061342908 | 1.78E-08 | 31.71882857 | 2.086873147 | 7.091119408 |
| ENSMUSG00000038550 | *Ciart* | 83.88066407 | 84.77885036 | 80.51911749 | 188.0780299 | 161.7753529 | 170.089144 | 2.59E-07 | 1.061152015 | 6.00E-08 | 29.36465436 | 2.086597038 | 6.58639862 |
| ENSMUSG00000038212 | *Mfsd14b* | 1064.885002 | 1023.401837 | 944.7576452 | 2057.05074 | 2191.024578 | 2090.356924 | 2.48E-35 | 1.060066398 | 1.03E-36 | 160.1822391 | 2.085027481 | 34.60472529 |
| ENSMUSG00000026385 | *Dbi* | 1251.02019 | 1153.597928 | 1226.574556 | 2502.365537 | 2400.572519 | 2666.340616 | 1.34E-36 | 1.059662381 | 5.20E-38 | 166.1235279 | 2.084443664 | 35.87332727 |
| ENSMUSG00000024975 | *Pdcd4* | 56.71930618 | 36.33379301 | 41.60154404 | 104.5815054 | 66.23017804 | 111.137793 | 0.003230838 | 1.059238388 | 0.00123845 | 10.43208318 | 2.083831157 | 2.490684814 |
| ENSMUSG00000112825 | *Gm9118* | 266.8203981 | 207.9100378 | 318.0505141 | 504.3527439 | 572.1853086 | 570.1851988 | 2.42E-09 | 1.059215699 | 4.61E-10 | 38.83643103 | 2.083798386 | 8.616226995 |
| ENSMUSG00000028134 | *Ptbp2* | 142.1976972 | 167.5391567 | 130.1725933 | 331.4559002 | 281.2068215 | 305.3873268 | 2.02E-09 | 1.059046981 | 3.82E-10 | 39.20030811 | 2.083554707 | 8.694609834 |
| ENSMUSG00000038388 | *Pals2* | 1643.262152 | 1484.639153 | 1605.014409 | 3335.643983 | 3253.964649 | 3269.383945 | 1.17E-45 | 1.058558173 | 2.99E-47 | 208.4546539 | 2.082848886 | 44.93345146 |
| ENSMUSG00000031609 | *Sap30* | 1157.553164 | 1115.245591 | 913.8919835 | 2227.417386 | 2174.738469 | 2250.781912 | 9.46E-23 | 1.058421728 | 7.31E-24 | 101.454273 | 2.082651906 | 22.02391674 |
| ENSMUSG00000024287 | *Thoc1* | 1149.564529 | 1092.032334 | 1308.435659 | 2493.088145 | 2369.086041 | 2514.613368 | 8.16E-29 | 1.057936855 | 4.60E-30 | 129.7715305 | 2.081952069 | 28.08845135 |
| ENSMUSG00000042349 | *Ikbke* | 693.4134896 | 606.5724889 | 495.1925726 | 1250.761069 | 1241.001533 | 1259.239515 | 3.13E-15 | 1.05740569 | 3.81E-16 | 66.33217987 | 2.081185687 | 14.50499315 |
| ENSMUSG00000031349 | *Nsdhl* | 1135.98385 | 1025.420381 | 1147.397424 | 2316.817705 | 2256.169016 | 2304.901185 | 5.04E-38 | 1.056361354 | 1.84E-39 | 172.7708558 | 2.079679706 | 37.2979607 |
| ENSMUSG00000037646 | *Vps13b* | 1188.708839 | 1261.590035 | 1132.635586 | 2561.403483 | 2485.260287 | 2410.240484 | 3.19E-39 | 1.056294553 | 1.10E-40 | 178.3616588 | 2.079583412 | 38.49573614 |
| ENSMUSG00000034801 | *Sos2* | 730.9600726 | 724.6573162 | 610.6033076 | 1397.512536 | 1490.721876 | 1421.597335 | 5.62E-24 | 1.05586543 | 4.05E-25 | 107.1856357 | 2.078964942 | 23.25020994 |
| ENSMUSG00000031314 | *Taf1* | 1154.35771 | 1217.182066 | 1174.23713 | 2501.522138 | 2473.31714 | 2390.912172 | 8.71E-47 | 1.055139757 | 2.14E-48 | 213.7033829 | 2.077919489 | 46.060152 |
| ENSMUSG00000036097 | *Slf2* | 913.8998066 | 887.1501127 | 1001.121027 | 1974.397614 | 1969.533491 | 1867.114922 | 8.85E-34 | 1.054859915 | 3.97E-35 | 152.928927 | 2.077516471 | 33.05299182 |
| ENSMUSG00000005267 | *Zfp287* | 191.7272321 | 184.6967811 | 212.0336761 | 398.08444 | 434.2962494 | 386.5662365 | 3.17E-13 | 1.054380166 | 4.43E-14 | 56.96629125 | 2.076825735 | 12.49953865 |

| | | | | | | | | | | | | | |
|---|---|---|---|---|---|---|---|---|---|---|---|---|---|
| ENSMUSG00000024811 | *Tnks2* | 2242.409753 | 2439.410492 | 2171.332202 | 4841.111622 | 4760.972634 | 4632.029928 | 7.16E-45 | 1.0539344 | 1.88E-46 | 204.7877844 | 2.076184134 | 44.14522866 |
| ENSMUSG00000038784 | *Cnot4* | 455.3521764 | 536.932719 | 385.1497787 | 953.8845373 | 982.5952643 | 930.6582143 | 2.20E-14 | 1.053589666 | 2.83E-15 | 62.38137879 | 2.075688086 | 13.65687743 |
| ENSMUSG00000027109 | *Sp3* | 1030.533873 | 1000.18858 | 947.4416158 | 2086.569713 | 2081.364775 | 2022.707832 | 3.41E-41 | 1.053243093 | 1.07E-42 | 187.5762742 | 2.075189511 | 40.46712984 |
| ENSMUSG00000018189 | *Uchl5* | 1419.580381 | 1299.942372 | 1405.0586 | 2897.07638 | 2828.354324 | 2827.73202 | 4.45E-45 | 1.052328106 | 1.16E-46 | 205.745013 | 2.073873801 | 44.35116007 |
| ENSMUSG00000055116 | *Arntl* | 542.4282943 | 519.7750945 | 599.8674253 | 1163.047548 | 1191.057464 | 1084.318293 | 4.76E-24 | 1.051986593 | 3.41E-25 | 107.5284972 | 2.073382934 | 23.32284758 |
| ENSMUSG00000026970 | *Rbms1* | 1203.088382 | 1162.681376 | 966.2294099 | 2348.023476 | 2274.626606 | 2297.16986 | 2.84E-24 | 1.051000502 | 2.00E-25 | 108.5848878 | 2.071966248 | 23.54638936 |
| ENSMUSG00000037236 | *Matr3* | 3316.082253 | 3059.103517 | 3529.421317 | 6929.368133 | 6835.822966 | 6743.647995 | 5.86E-43 | 1.050876199 | 1.69E-44 | 195.8414365 | 2.071787735 | 42.23203368 |
| ENSMUSG00000030766 | *Arhgap17* | 1175.12816 | 1161.672104 | 899.1301453 | 2269.587347 | 2252.911794 | 2196.662639 | 6.59E-19 | 1.050744067 | 6.34E-20 | 83.51060476 | 2.071597995 | 18.18114096 |
| ENSMUSG00000027534 | *Snx16* | 206.1067746 | 220.0213021 | 273.7649995 | 500.9791469 | 466.8684681 | 470.6443929 | 8.17E-12 | 1.049876178 | 1.27E-12 | 50.37973572 | 2.070352148 | 11.08768211 |
| ENSMUSG00000029238 | *Clock* | 1305.342906 | 1269.664211 | 1512.417424 | 2967.078517 | 2768.63859 | 2706.930071 | 5.27E-27 | 1.049121753 | 3.23E-28 | 121.3360268 | 2.069269788 | 26.27810944 |
| ENSMUSG00000029415 | *Sdad1* | 1679.211008 | 1623.918693 | 1736.528967 | 3600.471344 | 3276.765202 | 3541.913141 | 3.32E-42 | 1.048976609 | 1.00E-43 | 192.295678 | 2.069061616 | 41.47900448 |
| ENSMUSG00000026305 | *Lrrfip1* | 1938.042772 | 2034.692409 | 2174.016172 | 4305.553106 | 4315.818979 | 4076.340963 | 2.28E-42 | 1.048438612 | 6.80E-44 | 193.0675201 | 2.068290184 | 41.64149032 |
| ENSMUSG00000072762 | *4930522L14Rik* | 59.91476005 | 48.44505735 | 71.12522045 | 130.7268818 | 122.6886905 | 115.0034553 | 5.92E-05 | 1.048008246 | 1.78E-05 | 18.41091307 | 2.067673291 | 4.227792282 |
| ENSMUSG00000034663 | *Bmp2k* | 856.3816369 | 780.1672778 | 708.5682339 | 1668.243691 | 1622.096492 | 1572.358167 | 2.63E-25 | 1.047974136 | 1.77E-26 | 113.3902994 | 2.067624405 | 24.58013782 |
| ENSMUSG00000021870 | *Slmap* | 941.0611645 | 952.7527946 | 736.749925 | 1892.587888 | 1779.528882 | 1780.137519 | 1.43E-18 | 1.047953487 | 1.40E-19 | 81.94474842 | 2.067594811 | 17.8451358 |
| ENSMUSG00000041238 | *Rbbp8* | 587.1646485 | 526.8399987 | 614.6292635 | 1159.673951 | 1277.916714 | 1127.806995 | 1.10E-21 | 1.045434892 | 9.00E-23 | 96.48346841 | 2.063988443 | 20.95979437 |
| ENSMUSG00000053470 | *Kdm3a* | 760.5180209 | 727.6851323 | 813.2430867 | 1579.686771 | 1577.581126 | 1584.921569 | 2.74E-34 | 1.045431627 | 1.20E-35 | 155.3045555 | 2.063983771 | 33.56291753 |
| ENSMUSG00000073647 | *Gm10557* | 432.1851358 | 375.4491945 | 413.3314698 | 883.0390014 | 838.1917614 | 799.2256939 | 2.83E-20 | 1.044989133 | 2.51E-21 | 89.89640962 | 2.063350817 | 19.54862387 |
| ENSMUSG00000072969 | *Armcx5* | 86.27725447 | 85.78812239 | 50.99544108 | 172.0534444 | 152.0036873 | 141.0966763 | 0.000124496 | 1.044500824 | 3.92E-05 | 16.91193247 | 2.062652552 | 3.904846171 |
| ENSMUSG00000004364 | *Cul3* | 1266.198596 | 1237.367506 | 1186.314998 | 2555.499689 | 2576.4625 | 2483.688069 | 2.34E-46 | 1.043567508 | 5.88E-48 | 211.6886985 | 2.061318601 | 45.63094714 |
| ENSMUSG00000024639 | *Gnaq* | 1327.711083 | 1322.146357 | 1009.172939 | 2379.229248 | 2707.837115 | 2466.292589 | 2.31E-16 | 1.042674422 | 2.61E-17 | 71.62086973 | 2.060042958 | 15.63687225 |
| ENSMUSG00000069011 | *Gm10254* | 303.5681176 | 272.5034476 | 169.0901467 | 462.182782 | 548.2990149 | 534.4278219 | 2.31E-06 | 1.04223079 | 5.90E-07 | 24.94576898 | 2.059409587 | 5.637072359 |
| ENSMUSG00000034574 | *Daam1* | 347.5056083 | 400.6809952 | 440.1711756 | 818.0972601 | 826.2486145 | 790.5279535 | 4.50E-17 | 1.041709263 | 4.86E-18 | 74.93556937 | 2.058665256 | 16.34650509 |
| ENSMUSG00000027423 | *Snx5* | 6357.355473 | 5549.986883 | 5904.735283 | 12259.65131 | 12170.06665 | 12247.38479 | 2.30E-46 | 1.041661323 | 5.76E-48 | 211.7283837 | 2.058596848 | 45.63858434 |
| ENSMUSG00000052155 | *Acvr2a* | 154.1806492 | 134.2331797 | 159.6962497 | 307.8407216 | 326.8079277 | 286.059015 | 3.08E-10 | 1.040392681 | 5.44E-11 | 43.01171041 | 2.056787405 | 9.511386591 |
| ENSMUSG00000096361 | *Gm5814* | 71.89771206 | 38.35233707 | 40.25955875 | 99.52110999 | 121.6029498 | 92.77589675 | 0.003227261 | 1.040082267 | 0.001236507 | 10.43498218 | 2.056344909 | 2.491165845 |
| ENSMUSG00000044150 | *Bclaf3* | 196.520413 | 198.8265895 | 93.93897041 | 318.8049117 | 337.6653339 | 356.6073531 | 0.000178795 | 1.039042663 | 5.74E-05 | 16.18818548 | 2.054863644 | 3.74764498 |
| ENSMUSG00000027522 | *Stx16* | 650.2748624 | 590.4241365 | 468.3528667 | 1239.796879 | 1117.227102 | 1167.430034 | 5.61E-14 | 1.039028283 | 7.41E-15 | 60.48607847 | 2.054843162 | 13.2514098 |
| ENSMUSG00000030031 | *Kbtbd8* | 243.6533575 | 221.0305742 | 283.1588965 | 522.9075271 | 542.8703118 | 464.8458993 | 4.97E-12 | 1.038715686 | 7.59E-13 | 51.38571923 | 2.054397976 | 11.30360406 |
| ENSMUSG00000026683 | *Nuf2* | 853.9850465 | 681.258619 | 736.749925 | 1545.107403 | 1574.323904 | 1556.895517 | 2.05E-21 | 1.038569486 | 1.70E-22 | 95.22817613 | 2.054189798 | 20.68908538 |
| ENSMUSG00000012640 | *Zfp715* | 523.2555711 | 450.1353245 | 492.508602 | 1021.356476 | 978.2523018 | 1012.80354 | 2.78E-23 | 1.037833254 | 2.08E-24 | 103.9489338 | 2.053141776 | 22.55585066 |
| ENSMUSG00000001280 | *Sp1* | 2211.254077 | 2201.222293 | 1916.354996 | 4368.808049 | 4355.991382 | 4257.060679 | 5.45E-36 | 1.034588529 | 2.20E-37 | 163.259715 | 2.048529302 | 35.26328514 |
| ENSMUSG00000043090 | *Zfp866* | 174.9510993 | 165.5206126 | 122.1206615 | 350.8540827 | 300.7501527 | 302.48808 | 7.25E-08 | 1.034359254 | 1.59E-08 | 31.94295248 | 2.048203773 | 7.139947827 |
| ENSMUSG00000039068 | *Zzz3* | 1143.972485 | 1278.74766 | 1384.928821 | 2652.490601 | 2605.777497 | 2522.344693 | 1.59E-27 | 1.034097199 | 9.47E-29 | 123.7686655 | 2.047831765 | 26.79784376 |
| ENSMUSG00000057137 | *Tmem140* | 198.9170034 | 180.659693 | 213.3756614 | 392.1806453 | 424.5245838 | 395.2639768 | 6.35E-13 | 1.034082327 | 9.09E-14 | 55.55478595 | 2.047810655 | 12.19699924 |
| ENSMUSG00000024472 | *Dcp2* | 1120.006581 | 1058.726358 | 1084.324116 | 2229.947583 | 2297.427159 | 2152.207521 | 7.28E-41 | 1.032561248 | 2.33E-42 | 186.0413832 | 2.045652721 | 40.13772292 |
| ENSMUSG00000026065 | *Slc9a4* | 775.6964267 | 712.5460519 | 853.5026454 | 1667.400292 | 1521.122614 | 1589.753647 | 4.64E-24 | 1.032001246 | 3.32E-25 | 107.5801976 | 2.044858827 | 23.33346123 |
| ENSMUSG00000024789 | *Jak2* | 949.8486626 | 987.0680435 | 1056.142424 | 2167.53604 | 2057.478482 | 1879.678325 | 5.41E-27 | 1.030738435 | 3.31E-28 | 121.2840589 | 2.043069717 | 26.26715181 |
| ENSMUSG00000022747 | *St3gal6* | 2820.786903 | 2251.685895 | 2133.756614 | 4930.511941 | 4878.232622 | 4929.68593 | 3.26E-19 | 1.030634578 | 3.09E-20 | 84.930896 | 2.042922646 | 18.48626201 |
| ENSMUSG00000051444 | *Bbs12* | 142.9965606 | 124.1404595 | 104.6748527 | 248.802775 | 260.5777497 | 257.0665472 | 3.17E-08 | 1.030428092 | 6.72E-09 | 33.61390341 | 2.042630273 | 7.498277656 |
| ENSMUSG00000037172 | *Dennd11* | 517.6635268 | 546.0161672 | 560.9498519 | 1177.385335 | 1128.084508 | 1007.005046 | 1.89E-21 | 1.030355119 | 1.56E-22 | 95.38925896 | 2.042526956 | 20.72318556 |
| ENSMUSG00000051498 | *Tlr6* | 331.5283389 | 286.633256 | 365.0199993 | 698.3345684 | 652.5301148 | 649.4312772 | 5.35E-15 | 1.029135296 | 6.61E-16 | 65.24591547 | 2.040800696 | 14.27127152 |
| ENSMUSG00000043263 | *Ifi209* | 130.2147452 | 114.0477392 | 139.5664703 | 286.7557407 | 243.2058997 | 250.3016381 | 1.77E-08 | 1.029120014 | 3.67E-09 | 34.79346411 | 2.040779078 | 7.752343593 |
| ENSMUSG00000036257 | *Pnpla8* | 464.1396745 | 449.1260525 | 585.1055871 | 1042.441457 | 1019.510446 | 981.8782406 | 1.58E-15 | 1.028442004 | 1.89E-16 | 67.71759897 | 2.039820218 | 14.80105789 |
| ENSMUSG00000037110 | *Ralgapa2* | 380.2590104 | 408.7551714 | 371.7299258 | 774.2404998 | 853.3921301 | 743.1735896 | 6.59E-19 | 1.028408658 | 6.34E-20 | 83.51009431 | 2.039773071 | 18.18114096 |
| ENSMUSG00000109488 | *Gm6828* | 75.09316593 | 69.63976994 | 93.93897041 | 167.8364482 | 185.6616466 | 130.4661048 | 5.48E-05 | 1.028304318 | 1.64E-05 | 18.56730019 | 2.039625554 | 4.261549975 |
| ENSMUSG00000026491 | *Ahctf1* | 1823.805296 | 1763.198233 | 1788.866394 | 3621.556325 | 3698.032564 | 3649.185272 | 1.88E-58 | 1.028301397 | 2.91E-60 | 268.1202459 | 2.039621423 | 57.72634112 |
| ENSMUSG00000027206 | *Cops2* | 1858.156425 | 1591.621988 | 2088.129114 | 3761.560598 | 3683.917936 | 3836.669897 | 6.17E-21 | 1.027989077 | 5.26E-22 | 92.98759592 | 2.039179927 | 20.20967533 |
| ENSMUSG00000066551 | *Hmgb1* | 2884.69598 | 2457.577389 | 2690.680509 | 5636.437102 | 5283.213874 | 5455.416012 | 5.71E-34 | 1.027100332 | 2.54E-35 | 153.8160017 | 2.037924115 | 33.24341547 |
| ENSMUSG00000021306 | *Gpr137b* | 1473.104234 | 1578.501452 | 1469.473894 | 3123.107375 | 3065.04578 | 3024.8808 | 1.91E-46 | 1.026867129 | 4.79E-48 | 212.0984131 | 2.037594724 | 45.7182878 |
| ENSMUSG00000027091 | *Zc3h15* | 2035.504115 | 1872.199612 | 2524.274333 | 4424.472399 | 4321.247682 | 4337.273173 | 3.19E-18 | 1.026043024 | 3.21E-19 | 80.30133783 | 2.036431129 | 17.4968649 |
| ENSMUSG00000027806 | *Tsc22d2* | 602.3430543 | 578.3128721 | 342.2062493 | 1066.900035 | 994.5384112 | 1050.493748 | 3.64E-07 | 1.025989786 | 8.52E-08 | 28.68388366 | 2.036355983 | 6.439284394 |
| ENSMUSG00000040938 | *Slc16a11* | 25.56363095 | 36.33379301 | 40.25955875 | 66.62853974 | 64.05869679 | 74.41400052 | 0.003514201 | 1.025846504 | 0.001356817 | 10.26358639 | 2.03615375 | 2.454173447 |
| ENSMUSG00000013662 | *Atad1* | 1606.514433 | 1572.44582 | 2191.461981 | 3590.350553 | 3612.259055 | 3711.03587 | 3.19E-14 | 1.025123751 | 4.15E-15 | 61.62771492 | 2.035133945 | 13.49680438 |
| ENSMUSG00000018377 | *Vezf1* | 909.9054892 | 952.7527946 | 870.9484542 | 1930.540854 | 1819.701285 | 1812.029233 | 3.18E-34 | 1.023896358 | 1.40E-35 | 154.998699 | 2.033403263 | 33.49782657 |
| ENSMUSG00000027245 | *Hypk* | 150.1863319 | 186.7153252 | 127.4886027 | 374.4692613 | 339.8368152 | 231.9397419 | 3.59E-05 | 1.023644027 | 1.05E-05 | 19.41423843 | 2.033047646 | 4.44489384 |
| ENSMUSG00000064367 | *mt-Nd5* | 21550.93976 | 21206.82386 | 14363.26858 | 39170.83417 | 38440.64678 | 38518.42622 | 3.32E-12 | 1.023452958 | 5.01E-13 | 52.20071588 | 2.03277841 | 11.4789787 |
| ENSMUSG00000035726 | *Supt16* | 2954.995965 | 2828.989495 | 2987.259259 | 6225.12977 | 5946.601395 | 5647.732715 | 9.42E-47 | 1.022957937 | 2.32E-48 | 213.5410406 | 2.032081038 | 46.02577902 |
| ENSMUSG00000070709 | *Zfp974* | 85.478391 | 83.76957834 | 69.78323516 | 163.619452 | 181.3186841 | 143.9959231 | 6.29E-06 | 1.022213398 | 1.69E-06 | 22.92244651 | 2.031032602 | 5.201382853 |
| ENSMUSG00000067851 | *Arfgef1* | 1714.361001 | 1650.159766 | 1315.145586 | 3149.252752 | 3276.765202 | 3095.429138 | 2.30E-19 | 1.022164742 | 2.16E-20 | 85.64210194 | 2.030964105 | 18.63811519 |
| ENSMUSG00000059729 | *Or2y1* | 60.71362351 | 109.001379 | 104.6748527 | 154.3420604 | 165.0325748 | 233.8725731 | 0.00110235 | 1.02161106 | 0.000394189 | 12.55953823 | 2.030184806 | 2.957680354 |
| ENSMUSG00000057135 | *Scimp* | 434.5817262 | 445.0889644 | 421.3833815 | 878.8220052 | 895.7360144 | 869.774032 | 1.54E-25 | 1.021607759 | 1.02E-26 | 114.4788125 | 2.03018016 | 24.81164532 |
| ENSMUSG00000044934 | *Zfp367* | 540.0317039 | 525.8307267 | 467.0108814 | 1056.779244 | 1021.681927 | 1042.762423 | 1.13E-23 | 1.021294293 | 8.22E-25 | 105.7845676 | 2.029739094 | 22.94846452 |
| ENSMUSG00000029638 | *Glcci1* | 38.34544643 | 64.5934098 | 63.0733087 | 96.14751304 | 96.6309155 | 141.0966763 | 0.002819975 | 1.021229819 | 0.001070971 | 10.70066345 | 2.029648387 | 2.549754809 |
| ENSMUSG00000042167 | *Tent2* | 639.0907738 | 601.5261288 | 544.8460284 | 1207.747708 | 1216.029498 | 1211.885151 | 4.26E-25 | 1.021002499 | 2.89E-26 | 112.4188073 | 2.029328608 | 24.37075346 |
| ENSMUSG00000026083 | *Eif5b* | 1952.422314 | 2051.850033 | 2065.315364 | 4123.378871 | 4084.556226 | 4096.635691 | 3.67E-56 | 1.020703372 | 6.27E-58 | 257.4169067 | 2.028907893 | 55.43580264 |
| ENSMUSG00000094574 | *Gm15027* | 36.7477195 | 29.26888882 | 60.38933812 | 83.49652449 | 100.973878 | 68.61550697 | 0.008465041 | 1.020189282 | 0.003493128 | 8.530140059 | 2.02818504 | 2.072370955 |
| ENSMUSG00000009535 | *Rnmt* | 1119.207718 | 1063.772718 | 1448.00213 | 2412.965218 | 2401.658259 | 2530.076018 | 1.67E-15 | 1.0192307 | 2.00E-16 | 67.60132691 | 2.026837883 | 14.77609074 |
| ENSMUSG00000030061 | *Uba3* | 675.0396299 | 575.285056 | 735.4079398 | 1372.210559 | 1319.174858 | 1324.955775 | 4.23E-19 | 1.018880743 | 4.03E-20 | 84.40643758 | 2.026346291 | 18.37327378 |
| ENSMUSG00000022704 | *Qtrt2* | 368.2760584 | 407.7458994 | 347.5741905 | 720.2629486 | 779.5617677 | 781.8302132 | 2.92E-18 | 1.018868443 | 2.94E-19 | 80.47694901 | 2.026329015 | 17.53444453 |
| ENSMUSG00000041431 | *Ccnb1* | 3059.64708 | 2469.688653 | 2204.881834 | 5369.922943 | 5101.89519 | 5211.879283 | 2.33E-16 | 1.018423687 | 2.64E-17 | 71.59795397 | 2.025704432 | 15.63205521 |
| ENSMUSG00000024186 | *Rgs11* | 35.94885603 | 52.48214546 | 29.52367641 | 70.00213669 | 94.45943425 | 76.3468317 | 0.004328374 | 1.018103492 | 0.001695944 | 9.852573077 | 2.025254892 | 2.36367526 |
| ENSMUSG00000078894 | *2210418O10Rik* | 296.3783464 | 290.6703441 | 389.1757345 | 658.6948043 | 617.7864148 | 689.0543165 | 1.46E-11 | 1.017704072 | 2.32E-12 | 49.18925951 | 2.024694263 | 10.83494727 |
| ENSMUSG00000047702 | *Or2y16* | 189.3306417 | 150.3815322 | 118.0947057 | 320.4917101 | 295.3214496 | 318.9171451 | 7.26E-07 | 1.017027029 | 1.75E-07 | 27.2903404 | 2.023744317 | 6.139354977 |
| ENSMUSG00000038069 | *Cdkn2aip* | 280.401077 | 212.9563979 | 206.6657349 | 500.9791469 | 447.3251369 | 475.4764708 | 9.63E-10 | 1.016528967 | 1.77E-10 | 40.70563428 | 2.023045779 | 9.016387888 |
| ENSMUSG00000054893 | *Zfp667* | 27.96022136 | 33.30597693 | 32.207647 | 69.15873745 | 60.80147492 | 57.98493547 | 0.003156561 | 1.016327502 | 0.001208115 | 10.47789236 | 2.022763292 | 2.50078576 |
| ENSMUSG00000024384 | *Iws1* | 822.8293713 | 926.5117218 | 916.5759541 | 1841.140535 | 1797.986473 | 1743.413726 | 4.25E-30 | 1.016308491 | 2.24E-31 | 135.7684643 | 2.022736636 | 29.37167032 |
| ENSMUSG00000026234 | *Ncl* | 20989.33874 | 20249.0247 | 22169.59702 | 44069.29694 | 42410.1145 | 41740.4558 | 7.01E-57 | 1.016046181 | 1.15E-58 | 260.7900861 | 2.022368897 | 56.15429103 |
| ENSMUSG00000040250 | *Ints13* | 1061.689548 | 997.1607638 | 1131.293601 | 2125.366078 | 2157.366619 | 2158.972431 | 6.34E-35 | 1.015625609 | 2.68E-36 | 158.2863247 | 2.021779426 | 34.19781909 |
| ENSMUSG00000048285 | *Frmd6* | 87.07611793 | 130.1960916 | 101.9908822 | 226.8743948 | 200.8620154 | 215.5106768 | 4.01E-06 | 1.015621244 | 1.05E-06 | 23.82965657 | 2.021773309 | 5.396738307 |
| ENSMUSG00000015127 | *Unkl* | 176.5488263 | 168.5484287 | 146.2763968 | 305.3105239 | 387.6094026 | 306.3537424 | 2.19E-08 | 1.015513011 | 4.59E-09 | 34.3580284 | 2.021621638 | 7.659256132 |
| ENSMUSG00000005667 | *Mthfd2* | 4348.213852 | 4144.070948 | 3721.325213 | 8518.332296 | 7949.792846 | 8220.331018 | 8.10E-35 | 1.014378793 | 3.46E-36 | 157.7785459 | 2.020032904 | 34.0916927 |
| ENSMUSG00000026889 | *Rbm18* | 507.2783017 | 525.8307267 | 489.8246314 | 1059.309442 | 1062.94007 | 956.7514352 | 3.50E-23 | 1.014272175 | 2.63E-24 | 103.4811548 | 2.019883625 | 22.45570897 |
| ENSMUSG00000029729 | *Zkscan1* | 631.1021392 | 680.249347 | 696.4903663 | 1375.584156 | 1353.918558 | 1316.258035 | 2.21E-29 | 1.013987934 | 1.21E-30 | 132.4168079 | 2.019485705 | 28.65472804 |
| ENSMUSG00000021149 | *Gtpbp4* | 3267.351581 | 3129.752559 | 3813.922198 | 6952.983312 | 6758.735382 | 6868.315606 | 3.32E-28 | 1.012334879 | 1.91E-29 | 126.9417947 | 2.017173082 | 27.47915768 |
| ENSMUSG00000027630 | *Tbl1xr1* | 683.0282645 | 609.600305 | 624.0231606 | 1317.389609 | 1280.088195 | 1274.702165 | 1.88E-27 | 1.012114343 | 1.13E-28 | 123.4207461 | 2.016864753 | 26.72470076 |
| ENSMUSG00000032582 | *Rbm6* | 1171.133843 | 1068.819078 | 1264.150145 | 2367.421659 | 2356.057153 | 2331.960821 | 4.98E-30 | 1.011605477 | 2.64E-31 | 135.4424456 | 2.016153492 | 29.30278381 |
| ENSMUSG00000106190 | *Gm20768* | 263.6249442 | 176.6226049 | 159.6962497 | 409.0486301 | 350.6942214 | 457.1145746 | 8.89E-06 | 1.011564176 | 2.43E-06 | 22.22457594 | 2.016095775 | 5.051201979 |
| ENSMUSG00000021028 | *Mbip* | 227.6760882 | 166.5298846 | 234.847426 | 376.1560598 | 399.5525495 | 490.9391203 | 2.31E-07 | 1.011091872 | 5.31E-08 | 29.59905881 | 2.015435862 | 6.636257213 |
| ENSMUSG00000021822 | *Plau* | 18715.77331 | 17639.04724 | 14642.40152 | 35291.19768 | 33807.79154 | 33696.01242 | 1.69E-23 | 1.011014387 | 1.24E-24 | 104.9668005 | 2.015327618 | 22.77307892 |
| ENSMUSG00000052837 | *Junb* | 5452.243164 | 5220.964202 | 4376.214036 | 9787.648148 | 10073.50151 | 10485.60916 | 6.88E-25 | 1.010922116 | 4.72E-26 | 111.4498022 | 2.015198727 | 24.16261983 |
| ENSMUSG00000058355 | *Abce1* | 4567.102442 | 4288.396848 | 5731.61918 | 9971.509182 | 9709.778396 | 9686.38347 | 1.69E-18 | 1.010352664 | 1.67E-19 | 81.59676829 | 2.014403457 | 17.77127851 |
| ENSMUSG00000039840 | *Epg5* | 483.3123977 | 501.608198 | 281.8169112 | 840.8690395 | 889.2215707 | 831.1174084 | 1.17E-06 | 1.009962802 | 2.89E-07 | 26.32347301 | 2.013859175 | 5.93058643 |
| ENSMUSG00000064128 | *Cenpj* | 178.9454167 | 214.974942 | 224.1115437 | 427.6034133 | 411.4956963 | 398.1632235 | 2.34E-11 | 1.00993962 | 3.77E-12 | 48.24068524 | 2.013826815 | 10.63145879 |
| ENSMUSG00000034858 | *Atosa* | 64.70794085 | 63.58413777 | 61.73132341 | 127.3532848 | 130.2888748 | 125.6340268 | 1.20E-05 | 1.009378963 | 3.31E-06 | 21.62818233 | 2.013044358 | 4.920999194 |
| ENSMUSG00000089774 | *Slc5a3* | 1300.549725 | 1366.554326 | 888.394263 | 2342.119682 | 2542.80454 | 2284.606457 | 2.68E-10 | 1.009237003 | 4.71E-11 | 43.29601325 | 2.012846286 | 9.57158125 |
| ENSMUSG00000010290 | *AI597479* | 503.2839844 | 467.292949 | 509.9544108 | 1038.224461 | 990.1954487 | 947.0872793 | 3.18E-23 | 1.008226139 | 2.38E-24 | 103.6756546 | 2.011436424 | 22.49731883 |
| ENSMUSG00000019861 | *Gopc* | 666.2521317 | 635.8413777 | 560.9498519 | 1228.832689 | 1237.744311 | 1293.064061 | 6.49E-23 | 1.0081163 | 4.95E-24 | 102.2255405 | 2.01128329 | 22.18790125 |
| ENSMUSG00000049764 | *Zfp280b* | 730.1612091 | 759.9818372 | 653.546837 | 1533.299813 | 1381.062073 | 1400.336192 | 7.34E-23 | 1.006948892 | 5.63E-24 | 101.9707373 | 2.009656447 | 22.1340487 |
| ENSMUSG00000021714 | *Cenpk* | 121.427247 | 95.88084267 | 108.7008086 | 202.4158169 | 230.1770122 | 225.1748327 | 2.44E-07 | 1.006784001 | 5.63E-08 | 29.48646564 | 2.009426769 | 6.612586497 |
| ENSMUSG00000032407 | *U2surp* | 1535.415584 | 1526.019307 | 1540.599115 | 3159.373543 | 3155.162252 | 2931.138488 | 8.63E-44 | 1.006617768 | 2.39E-45 | 199.7292046 | 2.009195247 | 43.06415621 |
| ENSMUSG00000054280 | *Prr14l* | 1326.113356 | 1406.925207 | 1007.830954 | 2610.320639 | 2543.890281 | 2371.583861 | 4.14E-14 | 1.006073849 | 5.43E-15 | 61.09685081 | 2.008437893 | 13.3830393 |
| ENSMUSG00000053289 | *Ddx10* | 802.0589212 | 746.8613008 | 986.3591893 | 1681.738079 | 1748.042404 | 1649.671414 | 1.06E-16 | 1.006049463 | 1.18E-17 | 73.19358964 | 2.008403944 | 15.97265148 |
| ENSMUSG00000005233 | *Spc25* | 466.5362649 | 397.6531791 | 487.1406608 | 888.942796 | 921.7937894 | 899.7329153 | 2.61E-18 | 1.005835129 | 2.62E-19 | 80.70826829 | 2.008105587 | 17.58298637 |
| ENSMUSG00000102976 | *Zc3h11a* | 2518.816512 | 2700.811947 | 2761.80573 | 5467.757255 | 5301.671465 | 5240.871751 | 3.24E-47 | 1.005645647 | 7.81E-49 | 215.7086984 | 2.007841862 | 46.48917722 |
| ENSMUSG00000092260 | *Zfp963* | 45.53521764 | 42.38942518 | 38.91757345 | 96.99091228 | 98.80239674 | 59.91776665 | 0.003648112 | 1.005275541 | 0.001413797 | 10.1877116 | 2.007326841 | 2.437931778 |
| ENSMUSG00000026484 | *Rnf2* | 614.3260063 | 569.2294239 | 766.2736015 | 1349.43878 | 1287.68838 | 1262.138762 | 9.82E-15 | 1.004499438 | 1.24E-15 | 64.01097236 | 2.006247282 | 14.00770979 |
| ENSMUSG00000038068 | *Rnf144b* | 371.4715123 | 422.8849798 | 508.6124255 | 874.605009 | 817.5626895 | 908.4306557 | 1.39E-12 | 1.004299074 | 2.04E-13 | 53.96944315 | 2.00596867 | 11.85553782 |
| ENSMUSG00000005682 | *Pan2* | 244.452221 | 276.5405357 | 190.5619114 | 472.3035729 | 508.1266118 | 454.2153278 | 2.19E-09 | 1.004095025 | 4.16E-10 | 39.03549715 | 2.005684975 | 8.6590301 |
| ENSMUSG00000028578 | *Caap1* | 244.452221 | 150.3815322 | 205.3237496 | 411.5788278 | 386.523662 | 407.8273795 | 5.13E-07 | 1.003492815 | 1.22E-07 | 27.9916602 | 2.004847936 | 6.289854222 |
| ENSMUSG00000028367 | *Txn1* | 5391.529541 | 5177.565504 | 5584.000798 | 10932.98431 | 10450.2535 | 10976.54828 | 2.51E-56 | 1.002856215 | 4.26E-58 | 258.18875 | 2.003963476 | 55.59953536 |
| ENSMUSG00000056211 | *R3hdm1* | 996.9816072 | 847.7885036 | 630.733087 | 1541.733806 | 1724.15611 | 1708.622765 | 9.68E-10 | 1.002653928 | 1.78E-10 | 40.69553911 | 2.003682512 | 9.014283701 |
| ENSMUSG00000066571 | *Garre1* | 822.8293713 | 909.3540974 | 805.1911749 | 1691.015471 | 1753.471107 | 1642.906505 | 1.27E-28 | 1.002475916 | 7.20E-30 | 128.881071 | 2.003435295 | 27.89543288 |
| ENSMUSG00000058881 | *Zfp516* | 393.8396894 | 346.1803057 | 268.3970583 | 682.3099829 | 701.3884428 | 648.4648616 | 2.72E-10 | 1.002466224 | 4.78E-11 | 43.26451001 | 2.003421836 | 9.56488099 |
| ENSMUSG00000022674 | *Ube2v2* | 592.7566927 | 583.3592323 | 594.4994841 | 1253.291267 | 1140.027655 | 1150.034553 | 2.66E-26 | 1.001873334 | 1.69E-27 | 118.0492094 | 2.002598679 | 25.57535851 |
| ENSMUSG00000030471 | *Zdhhc13* | 713.3850763 | 706.4904197 | 980.9912481 | 1622.700132 | 1575.409645 | 1594.585725 | 1.60E-12 | 1.001663714 | 2.35E-13 | 53.68623503 | 2.002307727 | 11.79521252 |

| | | | | | | | | | | | | | |
|---|---|---|---|---|---|---|---|---|---|---|---|---|---|
| ENSMUSG00000022867 | *Usp25* | 1525.030359 | 1588.594172 | 1493.629629 | 3028.646661 | 3099.78948 | 3094.462723 | 6.15E-48 | 1.00052031 | 1.42E-49 | 219.1040626 | 2.000721434 | 47.21134866 |
| ENSMUSG00000069083 | *Gm10259* | 1711.96441 | 1114.236319 | 1273.544042 | 2831.291239 | 2896.755984 | 2482.721654 | 5.24E-10 | 1.0003895 | 9.43E-11 | 41.93657034 | 2.000540034 | 9.280994232 |
| ENSMUSG00000031608 | *Galnt7* | 3616.454916 | 3777.705201 | 3530.763302 | 7636.136694 | 7165.888115 | 7055.800231 | 7.01E-48 | 1.000356364 | 1.63E-49 | 218.831083 | 2.000494086 | 47.15401939 |
| ENSMUSG00000037415 | *Ranbp10* | 1170.33498 | 1175.801913 | 954.1515423 | 2232.477781 | 2157.366619 | 2223.722275 | 9.66E-22 | 0.999713981 | 7.91E-23 | 96.73854622 | 1.999603533 | 21.0148075 |
| ENSMUSG00000033004 | *Mycbp2* | 1999.555259 | 1923.672486 | 1472.157865 | 3544.806994 | 3843.521807 | 3416.279115 | 5.48E-15 | 0.999703883 | 6.77E-16 | 65.19869127 | 1.999589536 | 14.26107173 |
| ENSMUSG00000048047 | *Zbtb33* | 414.6101395 | 403.7088113 | 403.9375727 | 860.267222 | 758.9326959 | 822.4196681 | 4.27E-20 | 0.998598769 | 3.85E-21 | 89.05014845 | 1.998058424 | 19.36944148 |
| ENSMUSG00000047213 | *Ythdf3* | 1231.847467 | 1147.542296 | 1096.401983 | 2311.757309 | 2325.656416 | 2315.531756 | 1.36E-35 | 0.997478787 | 5.58E-37 | 161.406273 | 1.996507908 | 34.86719052 |
| ENSMUSG00000046675 | *Tmem251* | 714.1839398 | 726.6758603 | 717.962131 | 1399.199335 | 1404.948367 | 1505.675491 | 5.60E-31 | 0.99745692 | 2.85E-32 | 139.8640388 | 1.996477648 | 30.25154591 |
| ENSMUSG00000001998 | *Ap4e1* | 348.3044717 | 296.7259763 | 362.3360287 | 676.4061883 | 667.7304835 | 662.9610955 | 6.57E-16 | 0.996726614 | 7.65E-17 | 69.4973509 | 1.995467268 | 15.182352 |
| ENSMUSG00000097352 | *C920009B18Rik* | 317.1487965 | 325.9948651 | 314.0245582 | 691.5873745 | 634.0725242 | 583.715017 | 1.41E-15 | 0.996676327 | 1.68E-16 | 67.95143222 | 1.995397715 | 14.8504089 |
| ENSMUSG00000039285 | *Azi2* | 1119.207718 | 1172.774097 | 1272.202056 | 2467.786168 | 2370.171781 | 2259.479652 | 8.40E-30 | 0.996548228 | 4.51E-31 | 134.382416 | 1.995220548 | 29.07572372 |
| ENSMUSG00000038170 | *Pde4dip* | 1687.199643 | 1710.716088 | 1661.377791 | 3478.178454 | 3346.252602 | 3262.619036 | 1.82E-47 | 0.995318939 | 4.30E-49 | 216.8968033 | 1.993521188 | 46.7408371 |
| ENSMUSG00000033439 | *Trmt13* | 214.8942727 | 226.0769343 | 248.2672789 | 473.1469721 | 419.0958807 | 474.5100552 | 1.65E-12 | 0.994683024 | 2.43E-13 | 53.62517785 | 1.992642672 | 11.78223904 |
| ENSMUSG00000111877 | *Gm6477* | 242.8544941 | 232.1325665 | 260.3451466 | 483.2677629 | 491.8405025 | 485.1406267 | 3.36E-15 | 0.993015537 | 4.10E-16 | 66.18887359 | 1.990340879 | 14.47404863 |
| ENSMUSG00000025104 | *Hdgfl3* | 54.32271578 | 39.3616091 | 33.54963229 | 83.49652449 | 81.43054677 | 91.80948116 | 0.001973607 | 0.99239175 | 0.000732531 | 11.40483667 | 1.989480488 | 2.704739403 |
| ENSMUSG00000032018 | *Sc5d* | 1616.899658 | 1499.778234 | 1890.857276 | 3380.344143 | 3214.877986 | 3349.596439 | 5.53E-22 | 0.991948364 | 4.49E-23 | 97.86176199 | 1.988869151 | 21.25739462 |
| ENSMUSG00000051037 | *Zfp455* | 27.96022136 | 34.31524896 | 46.9694852 | 75.06253211 | 62.97295617 | 75.38041611 | 0.005174596 | 0.991573757 | 0.00205572 | 9.499105943 | 1.988352793 | 2.286123568 |
| ENSMUSG00000020130 | *Tbc1d15* | 1415.586064 | 1366.554326 | 1584.884629 | 2924.065155 | 2870.698209 | 2872.187137 | 6.36E-32 | 0.991225726 | 3.12E-33 | 144.2607076 | 1.987873187 | 31.19647077 |
| ENSMUSG00000022701 | *Ccdc191* | 91.07043527 | 91.84375456 | 106.016838 | 204.1026154 | 187.8331279 | 178.7868844 | 6.76E-07 | 0.990805998 | 1.62E-07 | 27.43732469 | 1.987294934 | 6.170129607 |
| ENSMUSG00000039943 | *Plcb4* | 922.6873047 | 947.7064344 | 986.3591893 | 1958.373029 | 1852.273504 | 1854.551519 | 9.55E-35 | 0.990113291 | 4.10E-36 | 157.4432127 | 1.986340967 | 34.02021555 |
| ENSMUSG00000022314 | *Rad21* | 2841.557353 | 2751.275549 | 2602.10948 | 5525.951802 | 5313.614612 | 5440.919778 | 2.16E-48 | 0.989172522 | 4.85E-50 | 221.2400079 | 1.985046113 | 47.6659994 |
| ENSMUSG00000026020 | *Nop58* | 3558.137883 | 3142.873096 | 4310.456756 | 7119.132961 | 7208.232 | 7511.94839 | 1.19E-16 | 0.988789123 | 1.32E-17 | 72.97019079 | 1.984518653 | 15.92511696 |
| ENSMUSG00000028577 | *Plaa* | 1858.156425 | 1809.624746 | 1788.866394 | 3788.549374 | 3566.657948 | 3467.499141 | 5.50E-42 | 0.98756489 | 1.68E-43 | 191.2701524 | 1.982835357 | 41.25938565 |
| ENSMUSG00000029469 | *Ift81* | 71.09884859 | 81.75103428 | 107.3588233 | 161.9326535 | 176.9757216 | 171.0555596 | 2.73E-05 | 0.986675828 | 7.90E-06 | 19.96283153 | 1.981613811 | 4.563228485 |
| ENSMUSG00000061244 | *Exoc5* | 1230.24974 | 1162.681376 | 1223.890586 | 2458.508777 | 2458.116772 | 2247.882665 | 1.34E-33 | 0.986180762 | 6.10E-35 | 152.0753139 | 1.980933929 | 32.87267286 |
| ENSMUSG00000110669 | *Gm10060* | 93.46702567 | 70.64904197 | 71.12522045 | 139.1608741 | 170.4612779 | 160.4249881 | 4.00E-05 | 0.985386238 | 1.18E-05 | 19.19743961 | 1.979843285 | 4.397763252 |
| ENSMUSG00000042699 | *Dhx9* | 4160.480938 | 4314.63792 | 4682.186682 | 8791.593649 | 8655.524251 | 8562.442137 | 6.15E-44 | 0.984251717 | 1.68E-45 | 200.4362699 | 1.978286968 | 43.21099748 |
| ENSMUSG00000018736 | *Ndel1* | 1420.379245 | 1470.509345 | 1719.083158 | 3015.152273 | 3040.073746 | 3043.242696 | 2.69E-25 | 0.983597365 | 1.82E-26 | 113.3400704 | 1.977389895 | 24.56951627 |
| ENSMUSG00000078779 | *Zfp59* | 129.4158817 | 171.5762448 | 187.8779408 | 317.9615124 | 334.4081121 | 306.3537424 | 1.99E-07 | 0.982580217 | 4.56E-08 | 29.89679754 | 1.975996261 | 6.700610256 |
| ENSMUSG00000029014 | *Dnajc2* | 1711.165547 | 1515.926586 | 1892.199261 | 3453.719876 | 3326.709271 | 3319.637556 | 2.59E-23 | 0.981910554 | 1.93E-24 | 104.0928966 | 1.975079266 | 22.58671961 |
| ENSMUSG00000040738 | *Ints8* | 1107.224766 | 1168.737009 | 1300.383747 | 2404.531225 | 2348.456969 | 2294.270613 | 9.39E-28 | 0.981851405 | 5.50E-29 | 124.8448107 | 1.974998291 | 27.02729631 |
| ENSMUSG00000045962 | *Wnk1* | 4511.182 | 4459.973092 | 2946.9997 | 7767.706975 | 8051.852464 | 7728.425482 | 2.51E-10 | 0.98155014 | 4.40E-11 | 43.42721823 | 1.974585913 | 9.599824436 |
| ENSMUSG00000026222 | *Sp100* | 1827.799613 | 1731.9108 | 1317.829556 | 3267.328645 | 3319.109086 | 3055.806099 | 4.32E-14 | 0.98088548 | 5.68E-15 | 61.01037781 | 1.973676416 | 13.36435408 |
| ENSMUSG00000041852 | *Tcf20* | 1330.107673 | 1410.962295 | 1328.565439 | 2678.635977 | 2742.580815 | 2605.456434 | 1.56E-40 | 0.979558601 | 5.09E-42 | 184.4813331 | 1.971862017 | 39.80668088 |
| ENSMUSG00000079184 | *Mphosph8* | 564.7964714 | 580.3314162 | 554.2399254 | 1162.204149 | 1078.140439 | 1109.445099 | 5.21E-26 | 0.97900197 | 3.38E-27 | 116.6787209 | 1.971101366 | 25.28325006 |
| ENSMUSG00000014075 | *Dynlt2b* | 501.6862575 | 415.8200756 | 497.8765432 | 910.8711762 | 908.7649019 | 969.3148379 | 5.63E-17 | 0.978872834 | 6.12E-18 | 74.48340293 | 1.970924939 | 16.24940392 |
| ENSMUSG00000031540 | *Kat6a* | 1333.303127 | 1334.257621 | 1034.67066 | 2448.387986 | 2447.259366 | 2416.038978 | 1.93E-17 | 0.978829202 | 2.04E-18 | 76.65416782 | 1.970865334 | 16.71368291 |
| ENSMUSG00000069793 | *Slfn9* | 916.296397 | 799.3434463 | 802.5072043 | 1675.834284 | 1644.897045 | 1651.604245 | 7.08E-26 | 0.978536568 | 4.62E-27 | 116.057838 | 1.970465607 | 25.15004751 |
| ENSMUSG00000027176 | *Cstf3* | 832.4157329 | 878.0666645 | 952.809557 | 1764.391204 | 1758.89981 | 1711.522012 | 1.58E-28 | 0.978526604 | 8.96E-30 | 128.4461304 | 1.970451997 | 27.80207135 |
| ENSMUSG00000075040 | *Zfp408* | 228.4749516 | 213.96567 | 175.8000732 | 414.1090255 | 438.6392119 | 373.0364182 | 1.62E-09 | 0.978249244 | 3.04E-10 | 39.64994112 | 1.970073212 | 8.791076623 |
| ENSMUSG00000029249 | *Rest* | 681.4305376 | 758.9725652 | 697.8323516 | 1391.608741 | 1437.520586 | 1381.00788 | 2.47E-27 | 0.977565531 | 1.48E-28 | 122.879196 | 1.969139788 | 26.60744765 |
| ENSMUSG00000023845 | *Lnpep* | 1343.688352 | 1315.081453 | 1202.418821 | 2636.466016 | 2591.662868 | 2381.248017 | 8.78E-29 | 0.976721702 | 4.95E-30 | 129.6242198 | 1.96798838 | 28.05667479 |
| ENSMUSG00000114293 | *Gm48078* | 56.71930618 | 53.49141749 | 37.57558816 | 104.5815054 | 112.9170249 | 76.3468317 | 0.002520964 | 0.975606884 | 0.000948783 | 10.92493877 | 1.966468238 | 2.598433402 |
| ENSMUSG00000066107 | *Gm12666* | 41.5409003 | 34.31524896 | 42.94352933 | 77.59272983 | 79.25906552 | 76.3468317 | 0.001321744 | 0.975362924 | 0.000477869 | 12.20010356 | 1.966135735 | 2.878852527 |
| ENSMUSG00000029004 | *Kmt2e* | 552.0146559 | 616.6652092 | 342.2062493 | 992.6809022 | 1002.138596 | 984.7774874 | 1.23E-06 | 0.97517825 | 3.05E-07 | 26.21900315 | 1.965884073 | 5.909171398 |
| ENSMUSG00000020530 | *Ggnbp2* | 930.6759394 | 973.9475072 | 1002.463013 | 1906.925675 | 1873.988316 | 1921.234195 | 9.20E-36 | 0.974256552 | 3.75E-37 | 162.1939993 | 1.964628526 | 35.03615643 |
| ENSMUSG00000042726 | *Trafd1* | 1821.408705 | 1875.227428 | 1570.122791 | 3516.13142 | 3369.053155 | 3471.364803 | 7.77E-27 | 0.973672876 | 4.81E-28 | 120.5437214 | 1.96383385 | 26.10965926 |
| ENSMUSG00000111469 | *None* | 30.35681176 | 31.28743287 | 41.60154404 | 68.31533822 | 66.23017804 | 65.7162602 | 0.003791156 | 0.973263177 | 0.001473145 | 10.11190647 | 1.963276237 | 2.421228396 |
| ENSMUSG00000050890 | *Pdik1l* | 584.7680581 | 552.0717994 | 527.4002196 | 1123.407784 | 1102.026733 | 1047.594501 | 1.89E-23 | 0.972534617 | 1.40E-24 | 104.7348748 | 1.962285033 | 22.72398181 |
| ENSMUSG00000020077 | *Srgn* | 2820.786903 | 2364.724362 | 3023.492862 | 5341.247369 | 5323.386277 | 5428.356375 | 2.99E-21 | 0.971840125 | 2.51E-22 | 94.45551898 | 1.961340646 | 20.52418344 |
| ENSMUSG00000090100 | *Ttbk2* | 218.0897266 | 232.1325665 | 234.847426 | 470.6167744 | 445.1536557 | 424.2564445 | 3.41E-13 | 0.971708499 | 4.78E-14 | 56.81720607 | 1.961161709 | 12.46734606 |
| ENSMUSG00000028484 | *Psip1* | 1431.563333 | 1150.570112 | 1121.899704 | 2431.520001 | 2482.003065 | 2364.818951 | 1.49E-17 | 0.971663071 | 1.55E-18 | 77.18799275 | 1.961099957 | 16.82789715 |
| ENSMUSG00000018372 | *Cep95* | 175.7499628 | 160.4742525 | 144.9344115 | 337.3596949 | 317.0362621 | 293.7903397 | 3.49E-09 | 0.971595947 | 6.75E-10 | 38.09281765 | 1.961008715 | 8.4566312 |
| ENSMUSG00000033105 | *Lss* | 3153.114105 | 3089.381678 | 2626.265215 | 5889.456874 | 5743.567899 | 5758.870508 | 2.99E-27 | 0.970169944 | 1.81E-28 | 122.4878856 | 1.959071353 | 26.52434862 |
| ENSMUSG00000040389 | *Wdr47* | 377.0635566 | 378.4770106 | 397.2276463 | 769.1801044 | 750.2467709 | 734.4758493 | 1.35E-20 | 0.969885302 | 1.18E-21 | 91.39540542 | 1.958684869 | 19.86948101 |
| ENSMUSG00000052557 | *Gan* | 818.0361905 | 739.7963966 | 528.7422049 | 1432.091905 | 1461.406879 | 1204.153827 | 1.61E-08 | 0.969515469 | 3.34E-09 | 34.9771692 | 1.958182826 | 7.792082032 |
| ENSMUSG00000021244 | *Ylpm1* | 1020.148648 | 990.0958596 | 566.317793 | 1631.134125 | 1756.728329 | 1660.301986 | 2.11E-06 | 0.967037513 | 5.37E-07 | 25.12440334 | 1.954822361 | 5.675800917 |
| ENSMUSG00000029202 | *Pds5a* | 3117.964113 | 3013.686276 | 2894.662274 | 5933.313634 | 5907.514733 | 5812.023365 | 1.75E-52 | 0.966584793 | 3.38E-54 | 240.2989963 | 1.954209031 | 51.75602065 |
| ENSMUSG00000020128 | *Vps54* | 682.2294011 | 630.7950176 | 497.8765432 | 1248.230871 | 1159.570986 | 1140.370398 | 1.01E-12 | 0.965363657 | 1.46E-13 | 54.61684886 | 1.952555636 | 11.99491976 |
| ENSMUSG00000064138 | *Fam172a* | 273.2113058 | 252.318007 | 292.5527936 | 533.8717172 | 532.0129055 | 527.6629128 | 7.67E-15 | 0.965358155 | 9.58E-16 | 64.51539631 | 1.952548189 | 14.11522044 |
| ENSMUSG00000032560 | *Dnajc13* | 2037.900705 | 2182.046125 | 2172.674187 | 4195.911205 | 4185.530104 | 4087.93795 | 1.46E-46 | 0.965026846 | 3.60E-48 | 212.6629618 | 1.952099846 | 45.83526032 |
| ENSMUSG00000059005 | *Hnrnpa3* | 4620.626295 | 4171.321292 | 4468.811021 | 9146.664728 | 8801.013495 | 7930.406341 | 5.40E-30 | 0.964492538 | 2.87E-31 | 135.2748271 | 1.951377012 | 29.2675677 |
| ENSMUSG00000020420 | *Zfp607a* | 107.8465681 | 82.76030631 | 75.15117632 | 196.5120223 | 146.5749842 | 178.7868844 | 0.000103069 | 0.963905066 | 3.20E-05 | 17.2939576 | 1.950582564 | 3.986873949 |
| ENSMUSG00000093954 | *Gm16867* | 53.52385231 | 64.5934098 | 85.88705866 | 129.8834825 | 112.9170249 | 149.7944166 | 0.000705959 | 0.963166455 | 0.000245819 | 13.44378595 | 1.949584188 | 3.151220538 |
| ENSMUSG00000027714 | *Exosc9* | 1007.366832 | 890.1779288 | 951.4675717 | 1834.393341 | 1865.302391 | 1857.450766 | 1.28E-29 | 0.962523425 | 6.92E-31 | 133.529481 | 1.948715423 | 28.89295395 |
| ENSMUSG00000036693 | *Nop14* | 1553.789444 | 1432.157008 | 1580.858673 | 3036.237254 | 2972.757827 | 2877.019215 | 1.42E-35 | 0.961038457 | 5.85E-37 | 161.3111456 | 1.94671064 | 34.84703094 |
| ENSMUSG00000042496 | *Prdm10* | 211.6988188 | 271.4941756 | 209.3497055 | 457.9657858 | 427.7818057 | 461.9466526 | 9.22E-10 | 0.959854296 | 1.69E-10 | 40.79252882 | 1.94511344 | 9.035280899 |
| ENSMUSG00000021785 | *Ngly1* | 267.6192615 | 277.5498077 | 316.7085288 | 581.9454737 | 542.8703118 | 543.1255622 | 9.80E-14 | 0.95982661 | 1.32E-14 | 59.35335275 | 1.945076112 | 13.00895957 |
| ENSMUSG00000044950 | *Pwwp2a* | 273.2113058 | 254.3365511 | 186.5359555 | 485.7979606 | 494.0119837 | 417.4915354 | 1.10E-07 | 0.959296493 | 2.44E-08 | 31.10655682 | 1.944361527 | 6.960075408 |
| ENSMUSG00000030469 | *Zfp719* | 300.3726637 | 307.8279686 | 369.0459552 | 646.887215 | 624.3008585 | 618.5059783 | 9.83E-14 | 0.959036849 | 1.32E-14 | 59.34650018 | 1.944011629 | 13.00763779 |
| ENSMUSG00000069539 | *Scyl2* | 838.0077772 | 775.1209176 | 913.8919835 | 1613.422741 | 1611.239085 | 1678.663882 | 3.69E-24 | 0.958839212 | 2.62E-25 | 108.0478183 | 1.943745335 | 23.43299594 |
| ENSMUSG00000028901 | *Gmeb1* | 484.9101246 | 488.4876616 | 360.9940434 | 872.0748113 | 895.7360144 | 836.9159019 | 1.22E-11 | 0.958609371 | 1.93E-12 | 49.55381686 | 1.943435693 | 10.91257915 |
| ENSMUSG00000038612 | *Mcl1* | 8165.183499 | 8010.592087 | 7656.026088 | 15650.11625 | 15122.1954 | 15552.52611 | 2.12E-60 | 0.95850083 | 3.11E-62 | 277.1678996 | 1.943289485 | 59.67266484 |
| ENSMUSG00000037552 | *Plekhg2* | 107.0477046 | 85.78812239 | 77.83514691 | 175.4270413 | 198.6905341 | 156.5593258 | 3.74E-05 | 0.957936848 | 1.10E-05 | 19.33069941 | 1.942529958 | 4.426681794 |
| ENSMUSG00000029647 | *Pan3* | 532.8419327 | 605.5632169 | 401.2536022 | 983.4035106 | 1030.367852 | 985.743903 | 8.01E-10 | 0.957472371 | 1.46E-10 | 41.07937314 | 1.94190466 | 9.096362814 |
| ENSMUSG00000036533 | *Cdc42ep3* | 482.5135342 | 515.7380064 | 518.0063225 | 959.788332 | 960.8804519 | 1017.635617 | 1.61E-22 | 0.956868868 | 1.26E-23 | 100.3718278 | 1.941092498 | 21.79241034 |
| ENSMUSG00000002107 | *Celf2* | 1374.844027 | 1309.02582 | 1176.921101 | 2604.416845 | 2500.460656 | 2401.542744 | 6.68E-26 | 0.956699753 | 4.36E-27 | 116.17316 | 1.940864973 | 25.17490825 |
| ENSMUSG00000025915 | *Sgk3* | 936.2679837 | 829.6216071 | 1076.272204 | 1889.214291 | 1796.900732 | 1815.894896 | 2.71E-17 | 0.955838548 | 2.89E-18 | 75.96172449 | 1.939706736 | 16.56742585 |
| ENSMUSG00000026069 | *Il1rl1* | 421.7999107 | 406.7366273 | 398.5696316 | 784.3612906 | 822.9913927 | 776.9981353 | 1.78E-20 | 0.955574548 | 1.56E-21 | 90.83719471 | 1.93935182 | 19.74872634 |
| ENSMUSG00000028403 | *Zdhhc21* | 940.262301 | 908.3448253 | 888.394263 | 1822.585752 | 1824.044248 | 1666.100479 | 1.43E-28 | 0.955384119 | 8.12E-30 | 128.6431113 | 1.939095852 | 27.84472177 |
| ENSMUSG00000057842 | *Zfp595* | 48.7306715 | 75.69540211 | 63.0733087 | 102.0513077 | 135.7175779 | 124.6676113 | 0.001001927 | 0.955347766 | 0.000356505 | 12.74739708 | 1.93904699 | 2.999163779 |
| ENSMUSG00000020273 | *Papolg* | 321.1431138 | 299.7537924 | 378.4398522 | 664.5985989 | 617.7864148 | 646.5320305 | 3.76E-13 | 0.955310676 | 5.29E-14 | 56.61801651 | 1.93899714 | 12.42481398 |
| ENSMUSG00000045294 | *Insig1* | 3218.62091 | 3097.455854 | 2869.164553 | 5978.013794 | 5840.198814 | 6006.272899 | 1.24E-40 | 0.955162638 | 4.04E-42 | 184.9420279 | 1.938798187 | 39.90489293 |
| ENSMUSG00000053830 | *Gm9923* | 107.8465681 | 80.74176225 | 120.7786762 | 229.4045925 | 219.319606 | 148.828001 | 0.000413997 | 0.95447196 | 0.000139025 | 14.51530714 | 1.937870225 | 3.383002514 |
| ENSMUSG00000020806 | *Rhbdf2* | 3368.807242 | 3538.507731 | 2650.420951 | 6133.199253 | 6181.12137 | 6221.783576 | 9.33E-17 | 0.954440713 | 1.03E-17 | 73.46053891 | 1.937828254 | 16.02999339 |
| ENSMUSG00000056708 | *Ier5* | 1742.321222 | 1894.403597 | 1490.945659 | 3232.749276 | 3366.881674 | 3348.630023 | 2.23E-20 | 0.954281711 | 1.96E-21 | 90.38262256 | 1.937614694 | 19.65128874 |
| ENSMUSG00000044211 | *Gm7887* | 401.828324 | 412.7922595 | 598.52544 | 950.5109404 | 866.4210176 | 908.4306557 | 2.08E-08 | 0.954210863 | 4.33E-09 | 34.46846077 | 1.937519544 | 7.682178907 |
| ENSMUSG00000040648 | *Ppip5k2* | 893.1293564 | 948.7157064 | 1029.302719 | 1936.444649 | 1843.587579 | 1765.641285 | 2.91E-24 | 0.953148249 | 2.05E-25 | 108.5333563 | 1.936092993 | 23.53582563 |
| ENSMUSG00000026113 | *Inpp4a* | 154.9795127 | 187.7245972 | 155.6702938 | 305.3105239 | 358.2944058 | 302.48808 | 3.58E-08 | 0.952718605 | 7.62E-09 | 33.37066106 | 1.935516498 | 7.446619244 |
| ENSMUSG00000066415 | *Msl2* | 1117.609991 | 1141.486664 | 785.0613955 | 1980.301409 | 1976.047935 | 1946.361001 | 7.52E-11 | 0.952180051 | 1.26E-11 | 45.87012937 | 1.934794111 | 10.12376863 |
| ENSMUSG00000055660 | *Mettl4* | 363.4828776 | 339.1154015 | 346.2322052 | 682.3099829 | 686.1880741 | 662.9610955 | 2.05E-18 | 0.951793633 | 2.03E-19 | 81.21135773 | 1.934275956 | 17.68917622 |
| ENSMUSG00000031429 | *Psmd10* | 806.0532385 | 677.2215309 | 931.3377923 | 1568.722581 | 1534.151501 | 1554.962686 | 7.73E-14 | 0.950678683 | 1.03E-14 | 59.8334594 | 1.932781677 | 13.11204062 |
| ENSMUSG00000022911 | *Arl13b* | 182.1408705 | 180.659693 | 189.2199261 | 379.5296568 | 364.8088495 | 320.8499763 | 4.84E-10 | 0.950547855 | 8.69E-11 | 42.09640228 | 1.932606414 | 9.315066227 |
| ENSMUSG00000033454 | *Zbtb1* | 512.0714825 | 471.3300371 | 369.0459552 | 856.893625 | 897.9074957 | 871.7068632 | 1.30E-11 | 0.950516333 | 2.07E-12 | 49.42086469 | 1.932564189 | 10.88444443 |
| ENSMUSG00000031200 | *Mtcp1* | 45.53521764 | 53.49141749 | 26.83970583 | 79.2795283 | 78.1733249 | 88.91023438 | 0.006578779 | 0.950370875 | 0.002661952 | 9.025801483 | 1.93236935 | 2.181854674 |
| ENSMUSG00000087403 | *Kantr* | 35.14999256 | 50.46360141 | 56.36338224 | 84.33992372 | 98.80239674 | 87.94381879 | 0.002596063 | 0.949835544 | 0.000979193 | 10.86650287 | 1.931652452 | 2.585684763 |
| ENSMUSG00000074305 | *Peak1* | 4589.47062 | 4461.991636 | 3330.807494 | 8053.619316 | 8071.395796 | 7796.074574 | 3.71E-14 | 0.94903361 | 4.86E-15 | 61.31808607 | 1.930579026 | 13.43066779 |
| ENSMUSG00000058799 | *Nap1l1* | 8240.276665 | 7407.047414 | 7307.109912 | 14722.37709 | 14630.3549 | 14971.71034 | 9.74E-40 | 0.948787089 | 3.30E-41 | 180.767234 | 1.930249165 | 39.01123474 |
| ENSMUSG00000023106 | *Denr* | 1802.235982 | 1847.977084 | 2223.669628 | 3860.238309 | 3737.119226 | 3712.968701 | 3.86E-21 | 0.947514108 | 3.26E-22 | 93.93694834 | 1.928546735 | 20.41314983 |
| ENSMUSG00000035235 | *Trim13* | 150.1863319 | 145.3351721 | 177.1420585 | 312.0577178 | 300.7501527 | 292.8239241 | 6.17E-09 | 0.946402915 | 1.22E-09 | 36.93687051 | 1.927061902 | 8.209610007 |
| ENSMUSG00000021273 | *Fdft1* | 2140.155229 | 2115.434171 | 2266.613157 | 4228.803775 | 4135.586035 | 4191.344419 | 1.68E-46 | 0.946230213 | 4.17E-48 | 212.3715845 | 1.926831231 | 45.77479936 |
| ENSMUSG00000041836 | *Ptpre* | 305.1658445 | 343.1524896 | 300.6047053 | 593.753063 | 639.5012273 | 597.2448353 | 8.11E-15 | 0.946079222 | 1.01E-15 | 64.4013447 | 1.926629582 | 14.09110633 |
| ENSMUSG00000039630 | *Hnrnpu* | 10150.35922 | 9580.010091 | 7497.671824 | 17609.33267 | 17488.02422 | 17361.65609 | 3.72E-16 | 0.945613627 | 4.27E-17 | 70.64774176 | 1.926007909 | 15.42919522 |
| ENSMUSG00000051034 | *Zfp11* | 65.50680432 | 76.70467414 | 63.0733087 | 139.1608741 | 124.8601717 | 131.4325204 | 5.85E-05 | 0.944971204 | 1.76E-05 | 18.43336147 | 1.925150459 | 4.232652761 |
| ENSMUSG00000021481 | *Zfp346* | 691.8157627 | 792.2785421 | 607.9193371 | 1278.593244 | 1402.776886 | 1352.981828 | 5.34E-15 | 0.944341777 | 6.59E-16 | 65.2524916 | 1.924310727 | 14.27251312 |
| ENSMUSG00000056602 | *Fry* | 110.2431585 | 90.83448253 | 85.88705866 | 185.5478322 | 185.6616466 | 184.5853779 | 4.13E-06 | 0.942620714 | 1.09E-06 | 23.77079564 | 1.92201649 | 5.38404327 |
| ENSMUSG00000038844 | *Kif16b* | 553.6123828 | 631.8042896 | 479.0887491 | 1046.658453 | 1120.484324 | 1039.863176 | 7.19E-14 | 0.942399608 | 9.56E-15 | 59.9844007 | 1.921721947 | 13.14323208 |
| ENSMUSG00000024750 | *Zfand5* | 1338.096308 | 1440.231184 | 1248.046321 | 2598.51305 | 2567.776575 | 2579.363213 | 4.53E-30 | 0.942368213 | 2.40E-31 | 135.6364419 | 1.921680128 | 29.34424868 |
| ENSMUSG00000025241 | *Fyco1* | 218.88859 | 207.9100378 | 150.3023526 | 398.08444 | 361.5516276 | 356.6073531 | 2.59E-07 | 0.941860316 | 5.98E-08 | 29.37001199 | 1.921003724 | 6.5874885 |
| ENSMUSG00000064341 | *mt-Nd1* | 34578.00632 | 32418.82682 | 44473.39256 | 69752.49052 | 72612.16142 | 71728.33159 | 7.12E-15 | 0.941682462 | 8.86E-16 | 64.66987635 | 1.920766919 | 14.14762467 |
| ENSMUSG00000041702 | *Btbd7* | 681.4305376 | 703.4626036 | 652.2048517 | 1258.351662 | 1364.775964 | 1289.198399 | 5.70E-25 | 0.939285238 | 3.89E-26 | 111.8307053 | 1.917577967 | 24.24418728 |
| ENSMUSG00000042408 | *Zmym6* | 142.9965606 | 151.3908042 | 132.8565439 | 312.901117 | 254.0633059 | 252.2344693 | 2.98E-07 | 0.938418579 | 6.94E-08 | 29.08269263 | 1.916426381 | 6.525630416 |
| ENSMUSG00000027361 | *Gabpb1* | 574.382833 | 579.3221442 | 538.1361019 | 1130.998377 | 1035.796555 | 1078.5198 | 3.00E-22 | 0.938281612 | 2.39E-23 | 99.10691092 | 1.916244448 | 21.52346271 |
| ENSMUSG00000034724 | *Cnot6l* | 696.6089435 | 652.9990022 | 542.1620578 | 1265.098856 | 1195.400427 | 1176.127774 | 4.35E-15 | 0.938024991 | 5.34E-16 | 65.66574639 | 1.915903624 | 14.36157782 |
| ENSMUSG00000024878 | *Cbwd1* | 359.4885603 | 367.3750182 | 444.1971315 | 718.5761501 | 787.1619521 | 727.7109401 | 3.01E-13 | 0.937239911 | 4.21E-14 | 57.06845956 | 1.914861319 | 12.52173455 |
| ENSMUSG00000056608 | *Chd9* | 901.9168546 | 923.4839057 | 746.1438221 | 1614.26614 | 1749.128145 | 1572.358167 | 8.48E-17 | 0.936918393 | 9.31E-18 | 73.6534978 | 1.914434622 | 16.07174945 |

| | | | | | | | | | | | | | |
|---|---|---|---|---|---|---|---|---|---|---|---|---|---|
| ENSMUSG00000023919 | *Cenpq* | 484.9101246 | 371.4121064 | 378.4398522 | 766.6499066 | 813.219727 | 793.4272003 | 1.73E-11 | 0.936707578 | 2.76E-12 | 48.85398667 | 1.914154894 | 10.76200198 |
| ENSMUSG00000032462 | *Pik3cb* | 1379.637208 | 1316.090725 | 1248.046321 | 2431.520001 | 2577.54584 | 2550.370745 | 3.30E-32 | 0.936181113 | 1.60E-33 | 145.5831316 | 1.913456512 | 31.48138026 |
| ENSMUSG00000025323 | *Sp4* | 166.9624647 | 137.2609958 | 150.3023526 | 271.5745544 | 305.0931152 | 295.7231709 | 3.07E-08 | 0.935241727 | 6.49E-09 | 33.68050139 | 1.912211003 | 7.512298265 |
| ENSMUSG00000025757 | *Hspa4l* | 700.6032608 | 717.592412 | 781.0354397 | 1409.320125 | 1353.918558 | 1431.26149 | 6.61E-25 | 0.935161986 | 4.53E-26 | 111.5315057 | 1.912105314 | 24.17976978 |
| ENSMUSG00000046269 | *Usp27x* | 43.9374907 | 67.62122589 | 48.31147049 | 104.5815054 | 102.0596186 | 98.5743903 | 0.001397345 | 0.935133739 | 0.000507181 | 12.08907664 | 1.912067877 | 2.854696323 |
| ENSMUSG00000039458 | *Mtmr12* | 512.0714825 | 548.0347113 | 454.9330138 | 969.9091228 | 998.8813737 | 935.4902922 | 1.99E-17 | 0.934868678 | 2.10E-18 | 76.59313124 | 1.911716613 | 16.70147506 |
| ENSMUSG00000039783 | *Kmo* | 72.69657552 | 51.47287344 | 48.31147049 | 120.6060909 | 114.0027655 | 98.5743903 | 0.001260328 | 0.93447607 | 0.000454697 | 12.29284976 | 1.911196438 | 2.899516438 |
| ENSMUSG00000079737 | *Bfar* | 93.46702567 | 131.2053637 | 130.1725733 | 255.5499689 | 205.2049779 | 211.6450145 | 3.11E-05 | 0.933650235 | 9.05E-06 | 19.70204297 | 1.910102734 | 4.507248605 |
| ENSMUSG00000036810 | *Cnep1r1* | 460.9442206 | 444.0796924 | 574.3697048 | 928.5825602 | 979.3380425 | 907.4642401 | 5.69E-13 | 0.933623327 | 8.11E-14 | 55.7778922 | 1.910067109 | 12.24503171 |
| ENSMUSG00000036698 | *Ago2* | 3387.979965 | 3282.152636 | 2340.422348 | 5624.629513 | 5937.91547 | 5662.228948 | 2.28E-11 | 0.933512152 | 3.67E-12 | 48.29345897 | 1.909919923 | 10.64192183 |
| ENSMUSG00000043257 | *Pigv* | 271.6135789 | 208.9193098 | 224.1115437 | 442.7845995 | 480.9830962 | 428.1221069 | 2.65E-09 | 0.933472109 | 5.07E-10 | 38.64954404 | 1.909866912 | 8.575989439 |
| ENSMUSG00000030201 | *Lrp6* | 1066.482729 | 974.9567792 | 813.2430867 | 1822.585752 | 1844.673319 | 1799.465831 | 5.36E-16 | 0.933280685 | 6.20E-17 | 69.91098508 | 1.909613519 | 15.27121875 |
| ENSMUSG00000014850 | *Msh3* | 458.5476302 | 442.0611483 | 424.0673521 | 858.5804235 | 875.1069426 | 801.158525 | 5.06E-19 | 0.933248487 | 4.84E-20 | 84.04259795 | 1.909570901 | 18.29571718 |
| ENSMUSG00000048371 | *Pdp2* | 771.7021094 | 741.8149407 | 644.1529399 | 1407.633327 | 1423.405957 | 1300.795386 | 1.47E-18 | 0.933131252 | 1.44E-19 | 81.8873472 | 1.909415733 | 17.83278417 |
| ENSMUSG00000107023 | *Gm42715* | 83.0818006 | 97.89938673 | 48.31147049 | 185.5478322 | 122.6886905 | 132.398936 | 0.005435551 | 0.933020417 | 0.002167291 | 9.402167109 | 1.909269048 | 2.264756425 |
| ENSMUSG00000015597 | *Zfp318* | 605.5385082 | 644.924826 | 391.8597051 | 1050.03205 | 1092.255067 | 1003.139384 | 3.48E-07 | 0.932829746 | 8.13E-08 | 28.77540831 | 1.90901673 | 6.459042639 |
| ENSMUSG00000022698 | *Naa50* | 4806.761483 | 4512.455238 | 3919.939037 | 8675.204554 | 8290.715402 | 8323.737486 | 3.65E-24 | 0.932740878 | 2.59E-25 | 108.073355 | 1.908899141 | 23.43792934 |
| ENSMUSG00000034311 | *Kif4* | 738.9487072 | 609.600305 | 508.6124255 | 1244.857274 | 1201.91487 | 1110.411514 | 4.04E-10 | 0.932639209 | 7.20E-11 | 42.46519085 | 1.908764623 | 9.393807866 |
| ENSMUSG00000046111 | *Cep295* | 750.9316593 | 755.9447491 | 822.6369837 | 1508.841235 | 1548.266129 | 1381.00788 | 1.94E-22 | 0.932225638 | 1.53E-23 | 99.98922178 | 1.908217523 | 21.71210016 |
| ENSMUSG00000015749 | *Anp32e* | 4563.906989 | 4170.31202 | 4621.797344 | 8748.580288 | 8162.598008 | 8568.240631 | 2.19E-37 | 0.932182684 | 8.23E-39 | 169.787445 | 1.90816071 | 36.66021395 |
| ENSMUSG00000032883 | *Acsl3* | 826.8236886 | 772.0931015 | 876.3163954 | 1593.181159 | 1571.066682 | 1551.097024 | 2.97E-26 | 0.931896013 | 1.89E-27 | 117.8257609 | 1.907781587 | 25.52763949 |
| ENSMUSG00000032030 | *Cul5* | 921.8884413 | 904.3077372 | 964.8874246 | 1731.498634 | 1815.358323 | 1771.439779 | 1.27E-31 | 0.931282655 | 6.31E-33 | 142.8585821 | 1.906970671 | 30.89459465 |
| ENSMUSG00000027706 | *Sec62* | 1007.366832 | 964.8640589 | 1257.440218 | 2147.294458 | 1960.847566 | 2034.304819 | 1.28E-14 | 0.930784781 | 1.62E-15 | 63.47911219 | 1.906312689 | 13.89350087 |
| ENSMUSG00000020124 | *Usp15* | 2164.121133 | 1898.440685 | 2685.312568 | 4466.64236 | 4206.159176 | 4173.948938 | 8.00E-13 | 0.930340859 | 1.15E-13 | 55.089262 | 1.9057262 | 12.09664659 |
| ENSMUSG00000049550 | *Clip1* | 1802.235982 | 1673.373023 | 1535.231173 | 3169.494334 | 3258.307611 | 3133.119346 | 1.81E-27 | 0.929576809 | 1.08E-28 | 123.5016697 | 1.904717196 | 26.74112641 |
| ENSMUSG00000048756 | *Foxo3* | 838.0077772 | 854.8534078 | 559.6078666 | 1416.910719 | 1438.606326 | 1445.757724 | 1.09E-08 | 0.929287795 | 2.21E-09 | 35.78058012 | 1.904335664 | 7.962836514 |
| ENSMUSG00000020152 | *Actr2* | 8711.606111 | 8286.123351 | 9609.956673 | 17447.40002 | 16431.5986 | 16760.5456 | 2.34E-32 | 0.928875312 | 1.13E-33 | 146.2768288 | 1.90379127 | 31.6303835 |
| ENSMUSG00000025409 | *Mbd6* | 680.6316741 | 678.2308029 | 532.7681607 | 1151.239959 | 1259.459123 | 1202.220995 | 7.66E-14 | 0.928336999 | 1.02E-14 | 59.8513704 | 1.903081042 | 13.1156091 |
| ENSMUSG00000096887 | *Gm20594* | 420.2021838 | 428.940612 | 564.9758077 | 937.8599518 | 801.2765802 | 939.3559546 | 1.11E-09 | 0.927907423 | 2.05E-10 | 40.41652523 | 1.902514465 | 8.955878518 |
| ENSMUSG00000068742 | *Cry2* | 246.8488114 | 213.96567 | 139.5664703 | 376.999459 | 370.2375526 | 402.9953015 | 2.29E-05 | 0.926927689 | 6.53E-06 | 20.32474427 | 1.901222907 | 4.640673739 |
| ENSMUSG00000029516 | *Cit* | 357.8908333 | 302.7816084 | 157.0122791 | 490.0149568 | 558.0706805 | 515.0995101 | 0.000713654 | 0.926818549 | 0.000248625 | 13.42249537 | 1.901079085 | 3.146512482 |
| ENSMUSG00000039701 | *Usp53* | 172.5545089 | 173.5947888 | 171.7741173 | 356.7578773 | 301.8358933 | 324.7156386 | 2.59E-09 | 0.926743578 | 4.94E-10 | 38.70259605 | 1.900980296 | 8.587390994 |
| ENSMUSG00000056019 | *Zfp709* | 119.8295201 | 77.71394617 | 134.1985292 | 199.04222 | 209.5479403 | 219.3763392 | 0.000163225 | 0.925506015 | 5.21E-05 | 16.36850752 | 1.899350308 | 3.787213851 |
| ENSMUSG00000026577 | *Blzf1* | 830.818006 | 750.8983889 | 880.3423512 | 1575.469775 | 1596.038717 | 1498.910582 | 6.33E-22 | 0.925419465 | 5.14E-23 | 97.59365965 | 1.899236367 | 21.19890857 |
| ENSMUSG00000021706 | *Zfyve16* | 459.3464937 | 424.9035238 | 509.9544108 | 864.4842182 | 907.6791613 | 868.8076164 | 9.91E-17 | 0.925053085 | 1.09E-17 | 73.33822765 | 1.898754107 | 16.00377965 |
| ENSMUSG00000021062 | *Rab15* | 57.51816964 | 76.70467414 | 42.94352933 | 111.3286993 | 109.659803 | 117.9027021 | 0.001676582 | 0.92483869 | 0.000616152 | 11.72653535 | 1.898471959 | 2.775575101 |
| ENSMUSG00000026021 | *Sumo1* | 1686.400779 | 1358.48015 | 2113.626834 | 3278.292835 | 3173.619843 | 3329.301711 | 1.11E-09 | 0.92456913 | 2.05E-10 | 40.41991807 | 1.898117273 | 8.956494396 |
| ENSMUSG00000040078 | *Ptges3-ps* | 1183.915659 | 1061.754174 | 1142.029483 | 2261.153355 | 2181.252913 | 1988.833287 | 3.44E-23 | 0.924505055 | 2.58E-24 | 103.5173186 | 1.898032973 | 22.46329477 |
| ENSMUSG00000043262 | *Uevld* | 365.879468 | 385.5419147 | 410.6474992 | 743.034728 | 714.4173303 | 740.2743428 | 1.98E-17 | 0.924206369 | 2.09E-18 | 76.60232962 | 1.897640058 | 16.70325521 |
| ENSMUSG00000071793 | *2610005L07Rik* | 626.3089584 | 705.4811477 | 707.2262486 | 1274.376247 | 1288.77412 | 1295.963308 | 1.51E-22 | 0.923634241 | 1.18E-23 | 100.4987723 | 1.896887661 | 21.81959019 |
| ENSMUSG00000025854 | *Fam20c* | 5149.47391 | 5325.928492 | 4284.959036 | 9074.132393 | 9489.37305 | 9445.745988 | 2.26E-21 | 0.92333278 | 1.87E-22 | 95.03030154 | 1.896491335 | 20.64629258 |
| ENSMUSG00000033545 | *Znrf1* | 1155.955437 | 1280.766204 | 1148.73941 | 2227.417386 | 2331.085119 | 2243.050587 | 1.06E-28 | 0.923026662 | 5.97E-30 | 129.2514496 | 1.89608897 | 27.97556675 |
| ENSMUSG00000033904 | *Ccp110* | 364.2817411 | 362.3286581 | 378.4398522 | 738.8177318 | 665.5590022 | 686.1550697 | 1.55E-16 | 0.922464131 | 1.73E-17 | 72.4320909 | 1.895349798 | 15.81039061 |
| ENSMUSG00000021945 | *Zmym2* | 777.2941537 | 748.8798449 | 822.6369837 | 1527.396019 | 1478.778729 | 1438.0264 | 4.23E-26 | 0.922211332 | 2.73E-27 | 117.0978471 | 1.89501771 | 25.37383976 |
| ENSMUSG00000034379 | *Wdr5b* | 36.7477195 | 44.40796924 | 37.57558816 | 83.49652449 | 69.48739991 | 71.51475374 | 0.003484258 | 0.922043696 | 0.001344433 | 10.28050252 | 1.894797529 | 2.457889726 |
| ENSMUSG00000004113 | *Cacna1b* | 58.31703311 | 54.50068952 | 57.70536754 | 126.5098856 | 82.51628739 | 113.0706242 | 0.001595093 | 0.922006437 | 0.000584415 | 11.82497345 | 1.894748595 | 2.797214056 |
| ENSMUSG00000064368 | *mt-Nd6* | 6202.37596 | 6050.585809 | 3683.749625 | 10121.63425 | 10133.21724 | 9941.517186 | 2.13E-07 | 0.921409354 | 4.89E-08 | 29.76145084 | 1.893964585 | 6.671057527 |
| ENSMUSG00000030641 | *Ddias* | 327.5340216 | 298.7445203 | 264.3711024 | 595.4398615 | 554.8134586 | 544.0919778 | 6.75E-12 | 0.921328587 | 1.04E-12 | 50.76380074 | 1.893858557 | 11.17085441 |
| ENSMUSG00000024317 | *Rnf138* | 309.1601618 | 288.6518 | 250.9512495 | 526.281124 | 560.2421618 | 529.5957439 | 4.61E-12 | 0.921078489 | 7.02E-13 | 51.53841171 | 1.893530276 | 11.33671019 |
| ENSMUSG00000039801 | *Cplane1* | 297.9760733 | 336.0875854 | 210.6916908 | 564.2340897 | 503.7836493 | 538.2934843 | 7.13E-07 | 0.920881636 | 1.72E-07 | 27.32792833 | 1.893271925 | 6.146840637 |
| ENSMUSG00000039086 | *Ss18l1* | 66.30566778 | 69.63976994 | 56.36338224 | 134.1004787 | 129.2031342 | 102.4400527 | 0.00042278 | 0.920819655 | 0.0001421 | 14.47410601 | 1.893190588 | 3.373885145 |
| ENSMUSG00000031706 | *Rfx1* | 425.7942281 | 439.0333322 | 299.26272 | 716.0459524 | 782.8189896 | 714.1811218 | 2.12E-08 | 0.920019744 | 4.43E-09 | 34.4228511 | 1.892141188 | 7.672689538 |
| ENSMUSG00000062783 | *Csprs* | 2099.413192 | 2073.044746 | 2245.141393 | 4276.034133 | 3958.610313 | 3889.822754 | 2.70E-32 | 0.919165692 | 1.30E-33 | 145.9908671 | 1.891021403 | 31.56839964 |
| ENSMUSG00000028035 | *Dnajb4* | 218.0897266 | 267.4570875 | 320.7344847 | 530.4981202 | 470.12569 | 512.2002633 | 1.03E-07 | 0.917854266 | 2.29E-08 | 31.22802054 | 1.889303224 | 6.985982643 |
| ENSMUSG00000041025 | *Iffo2* | 1261.405415 | 1228.284058 | 924.6278659 | 2095.847105 | 2250.740313 | 2118.382976 | 1.99E-12 | 0.917774312 | 2.95E-13 | 53.24151812 | 1.889198522 | 11.70141591 |
| ENSMUSG00000093384 | *Gm20689* | 121.427247 | 118.0848273 | 142.2504409 | 244.5857788 | 242.1201591 | 230.0069107 | 2.42E-07 | 0.916693798 | 5.58E-08 | 29.50572158 | 1.887784126 | 6.616679132 |
| ENSMUSG00000046210 | *Or4q3* | 54.32271578 | 58.53777763 | 81.86110278 | 112.1720986 | 149.8322061 | 102.4400527 | 0.002563767 | 0.916414818 | 0.000966255 | 10.89113781 | 1.887419114 | 2.591121422 |
| ENSMUSG00000031274 | *Col4a5* | 973.0157032 | 1042.578005 | 692.4644104 | 1781.259189 | 1711.127223 | 1629.376687 | 2.92E-09 | 0.916362014 | 5.60E-10 | 38.45632252 | 1.887350034 | 8.535135242 |
| ENSMUSG00000078995 | *Zfp456* | 68.70225819 | 57.5285056 | 61.73132341 | 121.4494902 | 130.2888748 | 104.3728858 | 0.000324812 | 0.916350624 | 0.000107523 | 14.99979054 | 1.887335133 | 3.488367678 |
| ENSMUSG00000020914 | *Top2a* | 6091.333938 | 5464.19876 | 5590.710724 | 11049.37341 | 10967.06604 | 10353.21023 | 4.37E-36 | 0.916278546 | 1.75E-37 | 163.7070058 | 1.887240843 | 35.35972471 |
| ENSMUSG00000032267 | *Usp28* | 368.2760584 | 374.4399224 | 312.6825729 | 662.0684012 | 635.1582648 | 701.6177192 | 2.55E-13 | 0.916263559 | 3.55E-14 | 57.40606795 | 1.887221238 | 12.59426527 |
| ENSMUSG00000033669 | *Zfp7* | 226.8772247 | 168.5484287 | 148.9603674 | 310.3709193 | 381.0949589 | 344.0439504 | 8.86E-06 | 0.915955955 | 2.42E-06 | 22.23019035 | 1.886818897 | 5.052377962 |
| ENSMUSG00000026434 | *Nucks1* | 2899.874386 | 2842.110031 | 2732.282054 | 5419.683498 | 5355.958496 | 5219.610608 | 1.68E-47 | 0.915558028 | 3.97E-49 | 217.0572334 | 1.886298542 | 46.77474876 |
| ENSMUSG00000032425 | *Zfp949* | 56.71930618 | 59.54704966 | 71.12522045 | 129.0400833 | 95.54517487 | 125.6340268 | 0.000784533 | 0.915543308 | 0.000274892 | 13.23416706 | 1.886279297 | 3.105389 |
| ENSMUSG00000038323 | *1700066M21Rik* | 168.5601916 | 128.1775476 | 134.1985292 | 275.7915506 | 251.8918247 | 288.9582617 | 1.08E-06 | 0.915029898 | 2.64E-07 | 26.49658161 | 1.885608148 | 5.967315554 |
| ENSMUSG00000020954 | *Strn3* | 925.8827586 | 899.2613771 | 864.2385277 | 1699.449463 | 1714.384445 | 1664.167648 | 1.62E-30 | 0.914773384 | 8.39E-32 | 137.7194235 | 1.885272913 | 29.79052714 |
| ENSMUSG00000026987 | *Baz2b* | 496.0942132 | 472.3393092 | 501.902499 | 934.4863549 | 976.0808206 | 860.1098761 | 4.39E-18 | 0.914283241 | 4.47E-19 | 79.65021847 | 1.884632517 | 17.35729105 |
| ENSMUSG00000029207 | *Apbb2* | 557.6067002 | 599.5075847 | 433.4612492 | 947.9807426 | 1133.513211 | 924.8597207 | 3.78E-09 | 0.913851374 | 7.35E-10 | 37.92520462 | 1.884068441 | 8.421969049 |
| ENSMUSG00000018707 | *Dync1h1* | 8146.809639 | 7712.856839 | 5118.331902 | 12959.67268 | 13814.96369 | 12750.88731 | 1.43E-08 | 0.9133854 | 2.95E-09 | 35.21600043 | 1.883460007 | 7.84385633 |
| ENSMUSG00000045098 | *Kmt5b* | 748.5350688 | 677.2215309 | 730.0399986 | 1362.089768 | 1372.376148 | 1326.888607 | 1.06E-24 | 0.913170773 | 7.35E-26 | 110.5700132 | 1.883179829 | 23.97397669 |
| ENSMUSG00000069208 | *Zfp825* | 190.9283687 | 135.2424518 | 195.9298526 | 292.6595353 | 350.6942214 | 339.2118725 | 4.59E-06 | 0.912736717 | 1.21E-06 | 23.55638044 | 1.882613332 | 5.338087081 |
| ENSMUSG00000062127 | *Cttnbp2nl* | 2843.953944 | 2737.14574 | 2577.953745 | 5233.292267 | 5072.580193 | 5062.084866 | 2.63E-39 | 0.912178362 | 9.03E-41 | 178.7630964 | 1.881884861 | 38.57966859 |
| ENSMUSG00000090112 | *Shprh* | 441.7714974 | 412.7922595 | 538.1361019 | 915.9315716 | 878.3641645 | 816.6211745 | 9.31E-12 | 0.912083799 | 1.45E-12 | 50.11498407 | 1.881761514 | 11.03090416 |
| ENSMUSG00000063108 | *Zfp26* | 451.357859 | 454.1724127 | 582.4216165 | 997.7412976 | 884.8786082 | 904.5649933 | 3.15E-11 | 0.911798843 | 5.13E-12 | 47.63715392 | 1.881389871 | 10.5010966 |
| ENSMUSG00000036890 | *Gtdc1* | 205.3079111 | 198.8265895 | 205.3237496 | 391.3372461 | 373.4947745 | 380.7677429 | 1.97E-11 | 0.91128418 | 3.16E-12 | 48.58965721 | 1.88071883 | 10.70459359 |
| ENSMUSG00000041598 | *Cdc42ep4* | 745.339615 | 659.0546344 | 496.5345579 | 1150.39656 | 1194.314686 | 1242.81045 | 3.44E-09 | 0.911029803 | 6.63E-10 | 38.12518084 | 1.880387249 | 8.463436082 |
| ENSMUSG00000031226 | *Pbdc1* | 1133.58726 | 1071.846894 | 1379.56088 | 2199.585211 | 2224.682538 | 2302.968354 | 5.91E-16 | 0.910851544 | 6.87E-17 | 69.71014897 | 1.880154924 | 15.22832519 |
| ENSMUSG00000022419 | *Deptor* | 54.32271578 | 55.50996155 | 59.04735283 | 116.3890947 | 103.1453592 | 96.64155911 | 0.000492257 | 0.910809077 | 0.000167194 | 14.16795069 | 1.88009958 | 3.307808051 |
| ENSMUSG00000020883 | *Fbxl20* | 260.4294903 | 281.5868959 | 300.6047053 | 543.1491088 | 517.8982774 | 516.0659257 | 4.82E-13 | 0.910584072 | 6.84E-14 | 56.11343093 | 1.87980638 | 12.31717148 |
| ENSMUSG00000028514 | *Usp24* | 2128.172277 | 2161.860684 | 1707.005291 | 3866.985503 | 3816.378292 | 3603.763739 | 5.48E-17 | 0.910543537 | 5.94E-18 | 74.54007117 | 1.879753564 | 16.26116159 |
| ENSMUSG00000084067 | *Gm14269* | 242.0556306 | 204.8822217 | 222.7695584 | 419.1694209 | 438.6392119 | 403.9617171 | 1.14E-10 | 0.910361572 | 1.95E-11 | 45.02479382 | 1.879516489 | 9.942387395 |
| ENSMUSG00000057181 | *5730455P16Rik* | 351.4999256 | 374.4399224 | 379.7818375 | 709.2987585 | 687.2738147 | 676.4909138 | 5.58E-17 | 0.910150938 | 6.06E-18 | 74.50265386 | 1.879242099 | 16.25340281 |
| ENSMUSG00000029135 | *Fosl2* | 964.228205 | 1012.299844 | 654.8888223 | 1648.845509 | 1695.926854 | 1609.081959 | 1.94E-08 | 0.909362966 | 4.03E-09 | 34.60688171 | 1.878215974 | 7.711701531 |
| ENSMUSG00000005615 | *Pcyt1a* | 1727.94168 | 1838.893635 | 1947.220658 | 3534.686203 | 3372.310377 | 3431.741764 | 8.80E-32 | 0.909065027 | 4.34E-33 | 143.6023052 | 1.877828133 | 31.05564872 |
| ENSMUSG00000036282 | *Naa30* | 493.6976228 | 517.7565504 | 355.6261023 | 873.7616098 | 867.5067582 | 835.9494863 | 4.25E-09 | 0.90901425 | 8.28E-10 | 37.69382172 | 1.877762041 | 8.371907659 |
| ENSMUSG00000032826 | *Ank2* | 393.8396894 | 311.8650567 | 309.9986023 | 670.5023936 | 643.8441898 | 600.1440821 | 2.57E-10 | 0.908622689 | 4.50E-11 | 43.38342929 | 1.877252469 | 9.590398469 |
| ENSMUSG00000068036 | *Afdn* | 2149.74159 | 2280.954784 | 1884.147349 | 3973.253807 | 3986.83957 | 3907.218235 | 8.53E-24 | 0.908601902 | 6.21E-25 | 106.3393962 | 1.877225421 | 23.06900494 |
| ENSMUSG00000034248 | *Slc25a37* | 1563.375805 | 1730.901528 | 1374.192939 | 2985.6333 | 2991.215418 | 2796.806721 | 1.94E-18 | 0.908596708 | 1.92E-19 | 81.32133564 | 1.877218662 | 17.71230939 |
| ENSMUSG00000000275 | *Trim25* | 7936.708548 | 7525.132242 | 5864.475724 | 13394.86669 | 13492.49873 | 13159.6811 | 8.04E-15 | 0.908445803 | 1.01E-15 | 64.41843304 | 1.877022316 | 14.09466815 |
| ENSMUSG00000069874 | *Irgm2* | 670.2464491 | 735.7593085 | 719.3041863 | 1350.282179 | 1333.289486 | 1299.82897 | 1.91E-24 | 0.908438263 | 1.34E-25 | 109.3819473 | 1.877012507 | 23.71920486 |
| ENSMUSG00000054237 | *Fra10ac1* | 174.9510993 | 161.4835245 | 148.9603674 | 330.612501 | 288.8070059 | 294.7567553 | 3.10E-08 | 0.908391095 | 6.56E-09 | 33.66089318 | 1.87695114 | 7.50828431 |
| ENSMUSG00000006599 | *Gtf2h1* | 2176.902948 | 2246.639535 | 3251.630361 | 4919.547751 | 4739.257822 | 4726.738656 | 3.05E-09 | 0.907865831 | 5.86E-10 | 38.3661323 | 1.876267894 | 8.515331378 |
| ENSMUSG00000007850 | *Hnrnph1* | 2260.783612 | 2031.664593 | 1864.01757 | 3969.88021 | 3725.176079 | 3869.528027 | 2.16E-22 | 0.907690654 | 1.71E-23 | 99.76930055 | 1.876040085 | 21.66517227 |
| ENSMUSG00000021466 | *Ptch1* | 714.1839398 | 730.7129484 | 449.5650727 | 1179.915533 | 1208.429314 | 1174.194943 | 5.57E-07 | 0.906743716 | 1.33E-07 | 27.82795967 | 1.874809117 | 6.254512426 |
| ENSMUSG00000058093 | *Zfp729b* | 231.6704055 | 237.1789266 | 254.9772054 | 441.9412003 | 458.1825431 | 453.2489122 | 1.80E-12 | 0.906729615 | 2.65E-13 | 53.45106448 | 1.874790794 | 11.74583677 |
| ENSMUSG00000037266 | *Rsrp1* | 2469.286977 | 2162.869956 | 1485.577178 | 3702.522651 | 3809.863848 | 3965.20317 | 8.58E-08 | 0.906022902 | 1.89E-08 | 31.60472506 | 1.87387264 | 7.066650945 |
| ENSMUSG00000018160 | *Med1* | 1282.974728 | 1297.923828 | 1025.276763 | 2340.432883 | 2215.996613 | 2212.125288 | 5.14E-16 | 0.905655677 | 5.95E-17 | 69.99400941 | 1.873395723 | 15.28905538 |
| ENSMUSG00000063047 | *Zfp780b* | 157.3761031 | 149.3722602 | 171.7741173 | 302.7803262 | 326.8079277 | 264.797872 | 1.33E-07 | 0.90533886 | 3.00E-08 | 30.70671721 | 1.87298437 | 6.875518859 |
| ENSMUSG00000025613 | *Cct8* | 8037.365344 | 7417.140135 | 8270.655352 | 15261.3092 | 14530.46677 | 14618.00223 | 9.02E-39 | 0.904731547 | 3.20E-40 | 176.2456575 | 1.872196089 | 38.04466607 |
| ENSMUSG00000029290 | *Zfp326* | 288.3897117 | 298.7445203 | 248.2672789 | 543.992508 | 520.0697587 | 505.4353542 | 6.82E-12 | 0.904558915 | 1.05E-12 | 50.74179845 | 1.871972077 | 11.16647726 |
| ENSMUSG00000008333 | *Snrpb2* | 1334.10199 | 1189.931721 | 1603.672423 | 2595.982852 | 2473.31714 | 2644.113057 | 6.99E-14 | 0.904291102 | 9.29E-15 | 60.04124296 | 1.871624608 | 13.15538885 |
| ENSMUSG00000028114 | *Mettl14* | 307.5624349 | 308.8372406 | 291.2108083 | 541.4623103 | 575.4425305 | 584.6814326 | 2.01E-14 | 0.903640629 | 2.57E-15 | 62.56975783 | 1.870780932 | 13.69661874 |
| ENSMUSG00000038975 | *Rabggtb* | 1314.929267 | 1126.347583 | 1191.682939 | 2273.804344 | 2336.513822 | 2193.763392 | 2.99E-23 | 0.903501127 | 2.24E-24 | 103.7981123 | 1.870600045 | 22.52381943 |
| ENSMUSG00000013646 | *Sh3bp5l* | 929.8770759 | 841.7328715 | 759.563675 | 1593.181159 | 1598.210198 | 1556.895517 | 7.39E-19 | 0.903372426 | 7.12E-20 | 83.28013802 | 1.870433178 | 18.13141997 |
| ENSMUSG00000032475 | *Nck1* | 443.3692243 | 319.9392329 | 457.6169844 | 824.844454 | 728.5319584 | 726.7445245 | 3.23E-08 | 0.903066854 | 6.84E-09 | 33.5798261 | 1.87003705 | 7.491274059 |
| ENSMUSG00000041075 | *Fzd7* | 262.0272173 | 230.1140224 | 158.3542644 | 393.8674438 | 427.7818057 | 403.9617171 | 7.48E-06 | 0.90272421 | 2.03E-06 | 22.5700971 | 1.869592965 | 5.125843717 |
| ENSMUSG00000027589 | *Pcmtd2* | 234.8658594 | 264.4292714 | 264.3711024 | 538.9321126 | 449.4966182 | 433.9206004 | 3.54E-09 | 0.902319608 | 6.83E-10 | 38.06778998 | 1.869068713 | 8.451325923 |
| ENSMUSG00000020594 | *Pum2* | 3945.586665 | 4030.023208 | 4290.326977 | 7652.161279 | 7637.099546 | 7611.489196 | 6.84E-46 | 0.901726589 | 1.74E-47 | 209.5309183 | 1.868300591 | 45.16525465 |
| ENSMUSG00000019920 | *Lims1* | 1845.374609 | 1686.493559 | 2092.155069 | 3667.943283 | 3434.197592 | 3392.118725 | 2.76E-19 | 0.901625137 | 2.60E-20 | 85.2708266 | 1.868169215 | 18.55984073 |
| ENSMUSG00000038446 | *Cdc40* | 850.7895927 | 878.0666645 | 975.6233069 | 1771.138398 | 1634.039638 | 1633.242349 | 3.31E-21 | 0.901312646 | 2.79E-22 | 94.24650148 | 1.867764609 | 20.4801557 |
| ENSMUSG00000054293 | *P2ry10b* | 344.3101544 | 291.6796161 | 441.5131609 | 719.4195494 | 611.2719711 | 671.6588358 | 2.28E-07 | 0.900259824 | 5.25E-08 | 29.6218707 | 1.866402085 | 6.641145286 |
| ENSMUSG00000032119 | *Hinfp* | 746.9373419 | 657.0360903 | 605.2353665 | 1274.376247 | 1268.145048 | 1219.616476 | 1.62E-16 | 0.900191826 | 1.81E-17 | 72.34129556 | 1.866314119 | 15.79109705 |
| ENSMUSG00000096356 | *Or8b40* | 113.4386124 | 123.1311874 | 112.7267645 | 232.7781895 | 200.8620154 | 217.443508 | 1.04E-06 | 0.899757979 | 2.54E-07 | 26.57456611 | 1.865752965 | 5.984113577 |
| ENSMUSG00000068798 | *Rap1a* | 1928.45641 | 1614.835245 | 2108.258893 | 3689.871663 | 3380.996302 | 3444.305167 | 2.01E-15 | 0.896933215 | 2.41E-16 | 67.2364724 | 1.862103437 | 14.69786035 |
| ENSMUSG00000024515 | *Smad4* | 1191.10543 | 1156.625744 | 854.8446307 | 2038.495956 | 2054.22126 | 1879.678325 | 7.68E-11 | 0.895885824 | 1.29E-11 | 45.82687735 | 1.860752048 | 10.1144848 |
| ENSMUSG00000020664 | *Dld* | 2817.591449 | 2683.654323 | 3617.992346 | 5671.85987 | 5558.991993 | 5706.684066 | 1.19E-13 | 0.894535399 | 1.62E-14 | 58.95111819 | 1.859011119 | 12.9226553 |

| | | | | | | | | | | | | | |
|---|---|---|---|---|---|---|---|---|---|---|---|---|---|
| ENSMUSG00000020064 | *Herc4* | 732.5577995 | 774.1116456 | 777.0094838 | 1471.731669 | 1401.691145 | 1365.54523 | 1.74E-24 | 0.894477755 | 1.22E-25 | 109.56925 | 1.858936842 | 23.75877742 |
| ENSMUSG00000024276 | *Zfp397* | 423.3976377 | 449.1260525 | 386.491764 | 792.795283 | 839.277502 | 713.2147063 | 3.54E-13 | 0.894164415 | 4.97E-14 | 56.74324175 | 1.858533142 | 12.4513765 |
| ENSMUSG00000053870 | *Fpgt* | 302.7692541 | 369.3935623 | 471.0368373 | 673.8759905 | 711.1601084 | 728.6773557 | 2.58E-07 | 0.894127412 | 5.96E-08 | 29.37573155 | 1.858485474 | 6.588548596 |
| ENSMUSG00000004880 | *Lbr* | 2462.89607 | 2376.835626 | 1912.32904 | 4247.358559 | 4248.50306 | 4064.743976 | 2.69E-16 | 0.893673897 | 3.06E-17 | 71.30581445 | 1.857901346 | 15.56978858 |
| ENSMUSG00000035725 | *Prkx* | 816.4384636 | 885.1315687 | 823.978969 | 1489.443053 | 1560.209276 | 1642.906505 | 4.60E-23 | 0.893447353 | 3.48E-24 | 102.9249355 | 1.857609626 | 22.33751836 |
| ENSMUSG00000027782 | *Kpna4* | 3100.389117 | 3127.734015 | 3391.196832 | 6074.161307 | 5885.79992 | 5883.538119 | 8.28E-39 | 0.892669432 | 2.92E-40 | 176.4257757 | 1.856608246 | 38.08182431 |
| ENSMUSG00000037400 | *Atp11b* | 1174.329297 | 1195.987353 | 1253.414262 | 2283.925134 | 2269.197903 | 2157.039599 | 4.30E-31 | 0.890619791 | 2.18E-32 | 140.3999901 | 1.853972431 | 30.36672532 |
| ENSMUSG00000020170 | *Frs2* | 405.8226414 | 475.3671253 | 383.8077934 | 774.2404998 | 820.8199114 | 752.8377455 | 8.07E-13 | 0.889718505 | 1.16E-13 | 55.07219461 | 1.852814572 | 12.09323154 |
| ENSMUSG00000064127 | *Med14* | 873.1577697 | 916.4190015 | 699.1743369 | 1541.733806 | 1593.867235 | 1487.313595 | 1.85E-13 | 0.889673831 | 2.54E-14 | 58.06200744 | 1.8527572 | 12.73390551 |
| ENSMUSG00000025969 | *Nrp2* | 11630.65322 | 11975.01261 | 9554.935276 | 20174.95315 | 21090.51161 | 20171.99263 | 2.76E-19 | 0.889260714 | 2.61E-20 | 85.26568589 | 1.852226737 | 18.55898298 |
| ENSMUSG00000023977 | *Ubr2* | 2422.154033 | 2465.651565 | 1810.338158 | 4224.586779 | 4187.701585 | 4007.725456 | 1.85E-12 | 0.889224075 | 2.74E-13 | 53.38616682 | 1.852179697 | 11.73218344 |
| ENSMUSG00000098134 | *Rnf113a2* | 197.3192764 | 209.9285819 | 221.4275731 | 386.2768507 | 378.9234776 | 394.2975612 | 1.27E-10 | 0.888439199 | 2.17E-11 | 44.80825527 | 1.851172321 | 9.895713706 |
| ENSMUSG00000066036 | *Ubr4* | 5982.688507 | 5415.753703 | 3314.70367 | 8966.177291 | 9447.029166 | 8827.240009 | 2.76E-06 | 0.887968725 | 7.10E-07 | 24.58915189 | 1.850568739 | 5.559318728 |
| ENSMUSG00000026478 | *Lamc1* | 3570.919699 | 3412.348727 | 2065.315364 | 5700.535444 | 5739.224936 | 5314.319336 | 1.69E-06 | 0.887706132 | 4.23E-07 | 25.58429679 | 1.850231937 | 5.773365351 |
| ENSMUSG00000014226 | *Cacybp* | 2509.230151 | 2079.100378 | 2789.987421 | 4556.04268 | 4352.73416 | 4732.53715 | 3.37E-14 | 0.8875297 | 4.40E-15 | 61.51080457 | 1.850005679 | 13.47258911 |
| ENSMUSG00000033237 | *Arid2* | 929.8770759 | 905.3170092 | 684.4124987 | 1491.129851 | 1651.411488 | 1531.768712 | 4.42E-11 | 0.88735485 | 7.26E-12 | 46.95668986 | 1.849781478 | 10.35456703 |
| ENSMUSG00000037965 | *Zc3h7a* | 1194.300884 | 1177.820457 | 1107.137866 | 2305.853515 | 2133.480325 | 1999.513858 | 3.47E-22 | 0.886954365 | 2.78E-23 | 98.80626649 | 1.84926806 | 21.46009191 |
| ENSMUSG00000021224 | *Numb* | 1320.521311 | 1311.044365 | 1188.998968 | 2407.904822 | 2323.484934 | 2340.658562 | 1.61E-28 | 0.886146757 | 9.15E-30 | 128.4047919 | 1.848233146 | 27.79392699 |
| ENSMUSG00000037674 | *Rfx7* | 1204.686109 | 1260.580763 | 970.2553658 | 2116.088686 | 2186.681616 | 2059.431625 | 1.43E-14 | 0.886086441 | 1.82E-15 | 63.25507867 | 1.848155876 | 13.84531826 |
| ENSMUSG00000042272 | *Sestd1* | 742.9430246 | 817.5103428 | 673.6766163 | 1402.572932 | 1437.520586 | 1295.963308 | 2.41E-16 | 0.886058435 | 2.72E-17 | 71.53624795 | 1.848119999 | 15.6187002 |
| ENSMUSG00000061136 | *Prpf40a* | 2992.542548 | 2851.193479 | 3376.434993 | 5761.26019 | 5696.881052 | 5563.654558 | 1.67E-25 | 0.885738854 | 1.11E-26 | 114.3146867 | 1.847710656 | 24.77702606 |
| ENSMUSG00000026775 | *Yme1l1* | 2362.239273 | 2052.859305 | 2938.947788 | 4663.154383 | 4341.876754 | 4567.280084 | 2.33E-11 | 0.885372567 | 3.76E-12 | 48.24518122 | 1.847241599 | 10.63229568 |
| ENSMUSG00000035798 | *Zdhhc17* | 280.401077 | 349.2081217 | 347.5741905 | 597.12666 | 589.5571586 | 610.7746536 | 6.26E-11 | 0.885364095 | 1.04E-11 | 46.24582247 | 1.84723075 | 10.20368192 |
| ENSMUSG00000108484 | *Gm18537* | 99.85793341 | 87.80666645 | 87.22904395 | 172.8968436 | 199.7762747 | 137.2310139 | 0.000228182 | 0.885022519 | 7.42E-05 | 15.70021965 | 1.846793448 | 3.641719551 |
| ENSMUSG00000040195 | *Nemp1* | 459.3464937 | 400.6809952 | 300.6047053 | 743.8781272 | 719.8460334 | 690.0207321 | 1.17E-07 | 0.884683726 | 2.60E-08 | 30.98328759 | 1.84635981 | 6.933528037 |
| ENSMUSG00000000732 | *Icosl* | 441.7714974 | 432.9777001 | 272.4230142 | 676.4061883 | 774.1330646 | 676.4909138 | 6.18E-06 | 0.883937556 | 1.66E-06 | 22.95678503 | 1.845405109 | 5.208854002 |
| ENSMUSG00000022351 | *Sqle* | 3243.385677 | 3069.196238 | 2897.346244 | 5665.956076 | 5573.106621 | 5770.467495 | 3.60E-35 | 0.883796339 | 1.51E-36 | 159.4328496 | 1.845224481 | 34.44406509 |
| ENSMUSG00000001630 | *Stk38l* | 694.2123531 | 721.6295001 | 782.377425 | 1401.729532 | 1325.689301 | 1313.358788 | 4.92E-21 | 0.882187852 | 4.17E-22 | 93.44707123 | 1.843168354 | 20.30838868 |
| ENSMUSG00000029920 | *Smarcad1* | 897.1236738 | 844.7606876 | 938.0477188 | 1725.594839 | 1623.182232 | 1583.955154 | 6.35E-23 | 0.88202702 | 4.84E-24 | 102.2721012 | 1.842962889 | 22.19710077 |
| ENSMUSG00000040464 | *Gtpbp10* | 261.2283538 | 196.8080455 | 315.3665435 | 499.2923484 | 454.9253213 | 464.8458993 | 3.33E-06 | 0.881533964 | 8.67E-07 | 24.20250637 | 1.842333145 | 5.477011079 |
| ENSMUSG00000060657 | *Marf1* | 1048.108869 | 1080.930342 | 924.6278659 | 1928.854056 | 1979.305157 | 1726.018246 | 1.97E-17 | 0.881289408 | 2.08E-18 | 76.60939041 | 1.842020871 | 16.70456514 |
| ENSMUSG00000044201 | *Cdc25c* | 364.2817411 | 299.7537924 | 191.9038967 | 522.9075271 | 504.86939 | 557.6217961 | 9.93E-05 | 0.880932558 | 3.08E-05 | 17.36881059 | 1.841565304 | 4.002988934 |
| ENSMUSG00000046785 | *Epm2aip1* | 296.3783464 | 258.3736392 | 222.7695584 | 506.0395423 | 479.8973556 | 453.2489122 | 8.94E-09 | 0.880216509 | 1.80E-09 | 36.18053722 | 1.840651512 | 8.048426717 |
| ENSMUSG00000019883 | *Echdc1* | 670.2464491 | 631.8042896 | 935.3637482 | 1440.525897 | 1310.488933 | 1354.914659 | 3.01E-08 | 0.880191013 | 6.36E-09 | 33.72179717 | 1.840618983 | 7.520910779 |
| ENSMUSG00000034525 | *Ice1* | 1379.637208 | 1392.795399 | 1049.432498 | 2261.996754 | 2517.832506 | 2268.177392 | 7.08E-12 | 0.880127906 | 1.10E-12 | 50.6654365 | 1.840538472 | 11.15007741 |
| ENSMUSG00000044786 | *Zfp36* | 1374.045164 | 1259.571491 | 750.169778 | 1981.144808 | 2171.481247 | 2085.524846 | 1.34E-05 | 0.879809221 | 3.71E-06 | 21.40723162 | 1.84013195 | 4.87300033 |
| ENSMUSG00000030795 | *Fus* | 4581.481985 | 4109.755699 | 2608.819407 | 7007.804262 | 6903.138885 | 6894.408827 | 1.28E-06 | 0.879738185 | 3.16E-07 | 26.14641513 | 1.840041347 | 5.893674557 |
| ENSMUSG00000045322 | *Tlr9* | 467.3351284 | 355.2637539 | 246.9252936 | 670.5023936 | 690.5310366 | 616.5731471 | 6.82E-05 | 0.879529672 | 2.07E-05 | 18.12163126 | 1.839775425 | 4.166161976 |
| ENSMUSG00000062007 | *Hsh2d* | 118.2317932 | 121.1126434 | 111.3847792 | 247.9593757 | 180.2329435 | 216.4770924 | 2.38E-05 | 0.879196987 | 6.81E-06 | 20.24552242 | 1.839351222 | 4.623497761 |
| ENSMUSG00000028068 | *Iqgap3* | 1502.662182 | 1303.97946 | 708.5682339 | 2143.077462 | 2172.566988 | 2153.173937 | 0.000166274 | 0.877568889 | 5.31E-05 | 16.3326279 | 1.837276664 | 3.779175728 |
| ENSMUSG00000021704 | *Mtx3* | 331.5283389 | 352.2359378 | 314.0245582 | 670.5023936 | 598.2430836 | 565.3531208 | 1.42E-11 | 0.877359848 | 2.26E-12 | 49.24286825 | 1.837010468 | 10.84633195 |
| ENSMUSG00000038369 | *Ncoa6* | 960.2338877 | 923.4839057 | 589.131543 | 1515.588429 | 1573.238164 | 1465.086036 | 9.20E-07 | 0.877253844 | 2.24E-07 | 26.81367474 | 1.836875496 | 6.036385941 |
| ENSMUSG00000020423 | *Btg2* | 354.6953795 | 361.3193861 | 312.6825729 | 640.1400211 | 611.2719711 | 642.6663681 | 1.26E-13 | 0.876338571 | 1.71E-14 | 58.84197099 | 1.835710518 | 12.89951088 |
| ENSMUSG00000005506 | *Celf1* | 2712.141471 | 2839.082215 | 2298.820804 | 4803.158656 | 4980.29224 | 4639.761253 | 4.03E-19 | 0.876219468 | 3.83E-20 | 84.50555105 | 1.835558975 | 18.39445515 |
| ENSMUSG00000026812 | *Tsc1* | 827.6225521 | 835.6772393 | 712.5941898 | 1442.212696 | 1498.322061 | 1432.227906 | 5.03E-19 | 0.876147806 | 4.80E-20 | 84.05860915 | 1.835467801 | 18.29869823 |
| ENSMUSG00000022801 | *Lrch3* | 719.775984 | 842.7421435 | 621.33919 | 1322.450004 | 1317.003376 | 1375.209386 | 4.93E-12 | 0.875736343 | 7.52E-13 | 51.4024719 | 1.834944393 | 11.30714222 |
| ENSMUSG00000039501 | *Znfx1* | 3374.399286 | 3264.995011 | 2263.929187 | 5460.166662 | 5594.821433 | 5294.024608 | 2.89E-09 | 0.875548733 | 5.54E-10 | 38.4769333 | 1.834705789 | 8.539321078 |
| ENSMUSG00000038943 | *Prc1* | 1735.930314 | 1434.175552 | 1380.902865 | 2749.481513 | 2864.183765 | 2746.55311 | 3.56E-16 | 0.874820456 | 4.07E-17 | 70.74291922 | 1.833779858 | 15.44902889 |
| ENSMUSG00000002996 | *Hbp1* | 361.0862872 | 488.4876616 | 550.2139695 | 909.1843777 | 847.963427 | 798.2592783 | 2.57E-07 | 0.874442239 | 5.94E-08 | 29.38332171 | 1.833299177 | 6.590138695 |
| ENSMUSG00000025616 | *Usp16* | 972.2168397 | 939.6322582 | 1254.756248 | 2056.20734 | 1935.875532 | 1798.499415 | 4.67E-11 | 0.873970196 | 7.69E-12 | 46.84307753 | 1.832699428 | 10.33079347 |
| ENSMUSG00000042599 | *Kdm7a* | 512.0714825 | 534.9141749 | 543.5040431 | 986.7771076 | 974.99508 | 947.0872793 | 2.29E-20 | 0.873279099 | 2.02E-21 | 90.32773147 | 1.831821717 | 19.64011369 |
| ENSMUSG00000031631 | *Cfap97* | 417.8055934 | 359.300842 | 452.2490432 | 772.5537013 | 741.5608459 | 732.5430181 | 2.17E-12 | 0.873250715 | 3.23E-13 | 53.06341396 | 1.831785678 | 11.66341893 |
| ENSMUSG00000000804 | *Usp32* | 543.2271578 | 726.6758603 | 758.2216897 | 1282.81024 | 1283.345417 | 1138.437566 | 3.59E-09 | 0.873169157 | 6.94E-10 | 38.03683969 | 1.831682126 | 8.444702085 |
| ENSMUSG00000074165 | *Zfp788* | 173.3533724 | 131.2053637 | 130.1725733 | 269.0443567 | 238.8629372 | 292.8239241 | 1.35E-05 | 0.873080681 | 3.74E-06 | 21.39542013 | 1.831569798 | 4.870629298 |
| ENSMUSG00000079297 | *Gm2223* | 754.1271131 | 744.8427568 | 861.5545572 | 1354.499175 | 1508.093726 | 1451.556218 | 1.66E-17 | 0.872592403 | 1.74E-18 | 76.96519125 | 1.830950012 | 16.78037208 |
| ENSMUSG00000019774 | *Mtrf1l* | 131.8124721 | 168.5484287 | 104.6748527 | 247.9593757 | 251.8918247 | 245.4695601 | 4.73E-05 | 0.872245755 | 1.40E-05 | 18.8635188 | 1.830510127 | 4.325491712 |
| ENSMUSG00000020063 | *Sirt1* | 513.6692095 | 484.4505735 | 362.3360287 | 850.1464311 | 813.219727 | 837.8823175 | 4.03E-09 | 0.872201239 | 7.85E-10 | 37.79745139 | 1.830453645 | 8.394320543 |
| ENSMUSG00000041037 | *Irgq* | 637.4930469 | 688.3235232 | 480.4307344 | 1030.633868 | 1128.084508 | 1157.765878 | 4.89E-09 | 0.872167269 | 9.57E-10 | 37.4109679 | 1.830410545 | 8.310896872 |
| ENSMUSG00000024072 | *Yipf4* | 864.3702716 | 760.9911092 | 764.9316162 | 1411.006924 | 1452.720954 | 1520.171725 | 1.83E-19 | 0.871978193 | 1.71E-20 | 86.10146856 | 1.830170672 | 18.73680799 |
| ENSMUSG00000013663 | *Pten* | 1816.615525 | 1899.449957 | 1492.287644 | 3282.509831 | 3235.507058 | 3022.947969 | 1.45E-15 | 0.871442584 | 1.72E-16 | 67.89486814 | 1.829491337 | 14.83902822 |
| ENSMUSG00000019189 | *Rnf145* | 1175.12816 | 1278.74766 | 1155.449336 | 2143.920861 | 2335.428081 | 2128.047132 | 2.51E-23 | 0.871339212 | 1.87E-24 | 104.1606744 | 1.829360255 | 22.60019851 |
| ENSMUSG00000031352 | *Hccs* | 731.758936 | 765.0281973 | 885.7102924 | 1488.599654 | 1424.491698 | 1430.295075 | 4.85E-17 | 0.87095198 | 5.25E-18 | 74.78502665 | 1.828869305 | 16.31386381 |
| ENSMUSG00000026504 | *Sdccag8* | 207.7045015 | 183.6875091 | 217.4016172 | 418.3260217 | 349.6084808 | 342.1111193 | 1.20E-07 | 0.870526507 | 2.69E-08 | 30.91606108 | 1.828330022 | 6.919745431 |
| ENSMUSG00000074746 | *Pdzd8* | 1155.955437 | 1099.097239 | 889.7362483 | 1938.131447 | 1990.162563 | 1833.290376 | 1.38E-13 | 0.869966792 | 1.88E-14 | 58.6498633 | 1.827620831 | 12.85880577 |
| ENSMUSG00000074735 | *None* | 36.7477195 | 47.43578532 | 45.62749991 | 67.47193898 | 73.8303624 | 94.70872793 | 0.008653513 | 0.869524089 | 0.003580605 | 8.485132299 | 1.827060097 | 2.062807562 |
| ENSMUSG00000017119 | *Nbr1* | 1362.062212 | 1543.176931 | 1694.927423 | 2929.125551 | 2704.579893 | 2753.318019 | 3.10E-17 | 0.869238283 | 3.31E-18 | 75.69314517 | 1.826698182 | 16.50932656 |
| ENSMUSG00000058163 | *Gm5431* | 151.7840588 | 170.5669728 | 189.2199261 | 325.5521056 | 294.235709 | 308.2865736 | 1.29E-07 | 0.869227458 | 2.91E-08 | 30.76536235 | 1.826684476 | 6.888189187 |
| ENSMUSG00000041498 | *Kif14* | 402.6271875 | 377.4677385 | 248.2672789 | 631.7060287 | 617.7864148 | 638.8007057 | 3.51E-06 | 0.869203377 | 9.16E-07 | 24.09761913 | 1.826653986 | 5.454341891 |
| ENSMUSG00000021192 | *Golga5* | 714.1839398 | 839.7143274 | 815.9270572 | 1443.899494 | 1461.406879 | 1410.000347 | 1.62E-18 | 0.867408021 | 1.59E-19 | 81.69730148 | 1.824382228 | 17.79180706 |
| ENSMUSG00000057522 | *Spop* | 1746.315539 | 1725.855168 | 1611.724335 | 3204.917101 | 3088.932074 | 2989.123423 | 1.17E-30 | 0.867275954 | 6.03E-32 | 138.3755718 | 1.824215228 | 29.93204251 |
| ENSMUSG00000042197 | *Zfp451* | 863.5714081 | 853.8441358 | 746.1438221 | 1548.481 | 1500.493542 | 1453.489049 | 2.29E-19 | 0.866605513 | 2.15E-20 | 85.65244301 | 1.823367685 | 18.64011279 |
| ENSMUSG00000082491 | *Gm5909* | 98.26020648 | 85.78812239 | 127.4886027 | 182.1742352 | 172.6327591 | 208.7457677 | 0.000179594 | 0.866466272 | 5.76E-05 | 16.17904279 | 1.823191713 | 3.745708642 |
| ENSMUSG00000021147 | *Wdr37* | 494.4964863 | 469.3114931 | 547.5299989 | 941.2335488 | 939.1656394 | 867.8412008 | 2.59E-15 | 0.866220206 | 3.14E-16 | 66.71380528 | 1.822880776 | 14.58654175 |
| ENSMUSG00000059810 | *Rgs3* | 1039.321371 | 1083.958158 | 1065.536321 | 1883.310497 | 1981.476638 | 1945.394585 | 7.05E-30 | 0.86576896 | 3.77E-31 | 134.7362556 | 1.822310704 | 29.15168616 |
| ENSMUSG00000032497 | *Lrrfip2* | 1366.056529 | 1352.424518 | 1454.712056 | 2588.392259 | 2548.233243 | 2456.628433 | 9.73E-31 | 0.865257595 | 5.00E-32 | 138.7492732 | 1.821664899 | 30.01177604 |
| ENSMUSG00000008429 | *Herpud2* | 579.9748773 | 758.9725652 | 524.716249 | 1163.047548 | 1142.199136 | 1093.016034 | 7.60E-09 | 0.86488058 | 1.52E-09 | 36.51236302 | 1.821188911 | 8.118912281 |
| ENSMUSG00000037808 | *Fam76b* | 473.7260361 | 406.7366273 | 414.6734551 | 823.1576555 | 746.989549 | 793.4272003 | 1.66E-13 | 0.864821811 | 2.28E-14 | 58.27316678 | 1.821114725 | 12.7786551 |
| ENSMUSG00000031765 | *Mt1* | 10133.58308 | 11706.54625 | 12677.73505 | 20833.64796 | 20009.11395 | 21968.55922 | 1.63E-18 | 0.864076409 | 1.60E-19 | 81.68135751 | 1.820174047 | 17.78856438 |
| ENSMUSG00000020283 | *Pex13* | 436.1794531 | 459.2187728 | 523.3742637 | 845.9294349 | 872.9354613 | 853.344967 | 1.41E-14 | 0.863882994 | 1.79E-15 | 63.28472694 | 1.819930042 | 13.85145182 |
| ENSMUSG00000063656 | *Gm10135* | 99.85793341 | 81.75103428 | 69.78323516 | 136.6306764 | 168.2897967 | 157.5257414 | 0.000401916 | 0.863535451 | 0.000134707 | 14.57475065 | 1.819491677 | 3.395864189 |
| ENSMUSG00000034173 | *Zbed5* | 58.31703311 | 63.58413777 | 83.20308807 | 121.4494902 | 94.45943425 | 153.660079 | 0.005089934 | 0.863412272 | 0.002020773 | 9.530570104 | 1.819336332 | 2.293287873 |
| ENSMUSG00000032525 | *Nktr* | 1979.583672 | 2020.5626 | 2014.319923 | 3619.026127 | 3746.890892 | 3573.804856 | 1.74E-40 | 0.863208453 | 5.68E-42 | 184.2633853 | 1.819079321 | 39.75988478 |
| ENSMUSG00000051495 | *Irf2bp2* | 842.800958 | 782.1858218 | 520.6902931 | 1241.483677 | 1306.14597 | 1366.511646 | 1.00E-06 | 0.862949738 | 2.45E-07 | 26.6404648 | 1.818753139 | 5.998087076 |
| ENSMUSG00000039789 | *Zfp597* | 243.6533575 | 245.2531028 | 248.2672789 | 444.471398 | 462.5255056 | 432.9541848 | 1.12E-11 | 0.862424156 | 1.75E-12 | 49.74517193 | 1.818090678 | 10.95200421 |
| ENSMUSG00000020166 | *Cnot2* | 1244.629282 | 1189.931721 | 1323.197497 | 2423.929408 | 2199.710503 | 2196.662639 | 1.62E-22 | 0.861828254 | 1.27E-23 | 100.3576218 | 1.817339875 | 21.78962256 |
| ENSMUSG00000025899 | *Alkbh8* | 529.6464788 | 519.7750945 | 583.7636018 | 1004.488492 | 989.1097081 | 966.4155911 | 3.28E-18 | 0.861641394 | 3.32E-19 | 80.24005632 | 1.817104505 | 17.4836498 |
| ENSMUSG00000047126 | *Cltc* | 11962.18156 | 12617.9189 | 10863.37093 | 21613.79225 | 21659.4397 | 21108.44934 | 6.61E-30 | 0.860825648 | 3.52E-31 | 134.8700989 | 1.816077346 | 29.1800026 |
| ENSMUSG00000035310 | *None* | 681.4305376 | 631.8042896 | 830.6888954 | 1353.655776 | 1244.258755 | 1283.399905 | 6.54E-12 | 0.860594268 | 1.01E-12 | 50.82653624 | 1.815786106 | 11.18456332 |
| ENSMUSG00000034893 | *Cog3* | 699.8043973 | 661.0731784 | 515.3223519 | 1136.902172 | 1361.518742 | 916.1619804 | 2.00E-06 | 0.86042714 | 5.07E-07 | 25.23524624 | 1.815575771 | 5.698947328 |
| ENSMUSG00000111619 | *Gm48348* | 173.3533724 | 154.4186203 | 83.20308807 | 267.3575582 | 223.6625685 | 260.9322096 | 0.002497562 | 0.859773992 | 0.000939534 | 10.94308952 | 1.814753995 | 2.602483739 |
| ENSMUSG00000075284 | *Wipf1* | 2209.656351 | 2004.414248 | 1611.724335 | 3514.444622 | 3605.744611 | 3463.633479 | 1.68E-12 | 0.859170636 | 2.47E-13 | 53.58740009 | 1.813995197 | 11.7742366 |
| ENSMUSG00000026219 | *Trip12* | 5943.544197 | 6440.164812 | 6343.564473 | 11365.64812 | 11445.87765 | 11147.60384 | 3.80E-43 | 0.859087596 | 1.09E-44 | 196.7201783 | 1.813890788 | 42.42023807 |
| ENSMUSG00000002307 | *Daxx* | 1632.876927 | 1571.436548 | 1293.673821 | 2766.349498 | 2755.609703 | 2650.877966 | 5.41E-16 | 0.858917596 | 6.27E-17 | 69.889279 | 1.813677062 | 15.26666089 |
| ENSMUSG00000032218 | *Ccnb2* | 1374.045164 | 1090.01379 | 920.60191 | 2076.448922 | 1993.419785 | 2081.659183 | 4.68E-09 | 0.858826865 | 9.15E-10 | 37.49771825 | 1.813563002 | 8.329558437 |
| ENSMUSG00000021621 | *Zcchc9* | 407.4203683 | 401.6902672 | 456.2749991 | 808.8198685 | 723.1032553 | 754.7705767 | 1.01E-13 | 0.85871047 | 1.35E-14 | 59.2991583 | 1.813416692 | 12.9975985 |
| ENSMUSG00000096401 | *Gm21811* | 142.9965606 | 135.2424518 | 108.7008086 | 236.9951857 | 241.0344184 | 229.0404951 | 3.99E-06 | 0.858452809 | 1.05E-06 | 23.84187645 | 1.81309285 | 5.399104418 |
| ENSMUSG00000032178 | *Ilf3* | 2359.842682 | 2260.769343 | 2054.579481 | 4145.307251 | 3918.43791 | 4051.214158 | 2.64E-26 | 0.858301225 | 1.68E-27 | 118.0652132 | 1.812902359 | 25.57838251 |
| ENSMUSG00000049511 | *Htr1b* | 377.86242 | 291.6796161 | 225.453529 | 552.4265004 | 510.2980931 | 568.2523676 | 1.46E-05 | 0.857603007 | 4.08E-06 | 21.2284088 | 1.812025185 | 4.834793945 |
| ENSMUSG00000021597 | *Slf1* | 381.0578739 | 407.7458994 | 430.7772786 | 711.8289562 | 726.3604772 | 765.4011482 | 1.30E-14 | 0.857592732 | 1.65E-15 | 63.44116842 | 1.812012279 | 13.88553945 |
| ENSMUSG00000024283 | *Wac* | 1840.581429 | 1797.513482 | 1779.472497 | 3347.451573 | 3302.822977 | 3169.843139 | 1.24E-36 | 0.857463834 | 4.79E-38 | 166.2864114 | 1.811850392 | 35.90625914 |
| ENSMUSG00000082500 | *Gm15195* | 59.11589658 | 54.50068952 | 61.73132341 | 112.1720986 | 102.0596186 | 102.4400527 | 0.000789369 | 0.856898 | 0.000276773 | 13.22138476 | 1.811139912 | 3.102719798 |
| ENSMUSG00000043131 | *Mob1a* | 7403.067751 | 7463.566648 | 8233.079763 | 14140.43161 | 14108.11366 | 13564.60924 | 3.26E-34 | 0.85665526 | 1.44E-35 | 154.9407416 | 1.810835205 | 33.48632229 |
| ENSMUSG00000020069 | *Hnrnph3* | 189.3306417 | 174.6040609 | 126.1466174 | 339.8898926 | 271.4351559 | 281.226937 | 5.20E-05 | 0.856115227 | 1.55E-05 | 18.66969664 | 1.810157496 | 4.283848116 |
| ENSMUSG00000024070 | *Prkd3* | 1820.609842 | 1840.912179 | 1874.753452 | 3358.415763 | 3364.710192 | 3293.544335 | 9.43E-41 | 0.856111494 | 3.03E-42 | 185.5128514 | 1.810152812 | 40.02554164 |
| ENSMUSG00000027968 | *Larp7* | 794.0702865 | 879.0759365 | 931.3377923 | 1584.747167 | 1541.751685 | 1575.257414 | 1.11E-19 | 0.856082601 | 1.02E-20 | 87.11541664 | 1.81011656 | 18.95282318 |
| ENSMUSG00000037369 | *Kdm6a* | 402.6271875 | 432.9777001 | 385.1497787 | 813.0368647 | 689.445296 | 706.4497971 | 6.36E-12 | 0.855399596 | 9.80E-13 | 50.88429893 | 1.809259813 | 11.19667929 |
| ENSMUSG00000026492 | *Tfb2m* | 431.3862723 | 410.7737155 | 536.7941166 | 893.1597922 | 737.2178834 | 854.3113826 | 6.42E-09 | 0.855263035 | 1.27E-09 | 36.85662157 | 1.809088562 | 8.192382289 |
| ENSMUSG00000037287 | *Tbcel* | 689.4191723 | 611.6188491 | 688.4384545 | 1179.915533 | 1169.342652 | 1248.608944 | 3.55E-18 | 0.85514003 | 3.59E-19 | 80.08332786 | 1.808934324 | 17.44996289 |
| ENSMUSG00000092416 | *Zfp141* | 183.7385975 | 154.4186203 | 183.8519849 | 326.3955048 | 319.2077433 | 297.6560021 | 1.03E-07 | 0.854560522 | 2.29E-08 | 31.23390874 | 1.808207849 | 6.987184514 |
| ENSMUSG00000052707 | *Tnrc6a* | 1405.200839 | 1474.546433 | 1307.093674 | 2648.273605 | 2468.974178 | 2457.594848 | 5.27E-24 | 0.854216361 | 3.79E-25 | 107.317354 | 1.807776546 | 23.27800662 |
| ENSMUSG00000036333 | *Kidins220* | 1699.182595 | 1692.549191 | 1678.8236 | 3198.169908 | 3043.330968 | 2921.474332 | 1.41E-31 | 0.853706614 | 6.97E-33 | 142.6605235 | 1.807137917 | 30.85180691 |
| ENSMUSG00000062822 | *4833420G17Rik* | 481.7146708 | 415.8200756 | 440.1711756 | 804.6028723 | 841.4489833 | 776.0317197 | 2.88E-14 | 0.853446636 | 3.74E-15 | 61.8329436 | 1.806812294 | 13.54049235 |
| ENSMUSG00000026782 | *Abi2* | 377.86242 | 374.4399224 | 377.0978669 | 685.6835799 | 730.7034397 | 624.3044719 | 4.98E-13 | 0.852833345 | 7.08E-14 | 56.04639517 | 1.80604438 | 12.30272467 |
| ENSMUSG00000096592 | *Gm15801* | 262.0272173 | 312.8743287 | 338.1802935 | 544.8359073 | 572.1853086 | 523.7972504 | 2.43E-09 | 0.85257372 | 4.62E-10 | 38.83137101 | 1.805719397 | 8.61523579 |
| ENSMUSG00000002083 | *Bbc3* | 865.9679985 | 827.6030631 | 668.3086752 | 1330.883996 | 1415.805773 | 1529.835881 | 1.68E-11 | 0.852333838 | 2.67E-12 | 48.91610897 | 1.805419179 | 10.7752747 |
| ENSMUSG00000051977 | *Prdm9* | 43.13862723 | 39.3616091 | 52.33742637 | 84.33992372 | 86.85924988 | 70.54833815 | 0.006055136 | 0.852131314 | 0.002432993 | 9.190312894 | 1.805165754 | 2.217876079 |

| | | | | | | | | | | | | | |
|---|---|---|---|---|---|---|---|---|---|---|---|---|---|
| ENSMUSG00000097715 | *Gpr137b-ps* | 2066.65979 | 2220.398462 | 1968.692423 | 3834.092932 | 3814.206811 | 3648.218857 | 3.31E-28 | 0.8517624 | 1.91E-29 | 126.9495172 | 1.804704211 | 27.48040286 |
| ENSMUSG00000025060 | *Slk* | 2352.652911 | 2122.499075 | 1807.654188 | 3817.224948 | 3866.32236 | 3670.446415 | 4.87E-15 | 0.851658125 | 6.01E-16 | 65.43406308 | 1.804573774 | 14.31211147 |
| ENSMUSG00000030347 | *D6Wsu163e* | 1075.270227 | 987.0680435 | 1328.565439 | 2055.363941 | 1953.247382 | 2097.121833 | 8.10E-12 | 0.851392267 | 1.26E-12 | 50.39743165 | 1.804241261 | 11.09143287 |
| ENSMUSG00000042743 | *Sgtb* | 79.88634673 | 79.73249022 | 106.016838 | 150.1250642 | 160.6896123 | 164.2906505 | 0.000174258 | 0.850986623 | 5.59E-05 | 16.23827029 | 1.803734033 | 3.758807046 |
| ENSMUSG00000027014 | *Cwc22* | 820.4327809 | 826.593791 | 899.1301453 | 1548.481 | 1556.952054 | 1476.683023 | 1.36E-22 | 0.850386612 | 1.06E-23 | 100.7204259 | 1.802984022 | 21.86654904 |
| ENSMUSG00000047368 | *Abhd17b* | 279.6022136 | 255.3458231 | 242.8993378 | 473.9903713 | 441.8964338 | 490.9391203 | 2.50E-10 | 0.850321766 | 4.37E-11 | 43.43999282 | 1.802902984 | 9.602220274 |
| ENSMUSG00000067336 | *Bmpr2* | 530.4453423 | 568.2201518 | 377.0978669 | 900.7503854 | 884.8786082 | 884.2702659 | 1.25E-07 | 0.850192674 | 2.80E-08 | 30.84303451 | 1.802741668 | 6.904317802 |
| ENSMUSG00000041763 | *Tpp2* | 2823.183493 | 2669.524514 | 2893.320289 | 5228.231872 | 5095.380746 | 4776.025851 | 2.08E-30 | 0.848956009 | 1.09E-31 | 137.2080665 | 1.801197036 | 29.68263015 |
| ENSMUSG00000020893 | *Per1* | 680.6316741 | 701.4440596 | 499.2185284 | 1037.381062 | 1188.885983 | 1170.329281 | 1.83E-08 | 0.84762773 | 3.79E-09 | 34.72975891 | 1.79953945 | 7.738379641 |
| ENSMUSG00000036721 | *Zscan12* | 156.5772396 | 117.0755553 | 191.9038967 | 316.274714 | 257.3205278 | 258.9993784 | 0.000213981 | 0.846809695 | 6.94E-05 | 15.82761923 | 1.798519367 | 3.669623923 |
| ENSMUSG00000034998 | *Foxn2* | 913.8998066 | 965.8733309 | 1044.064557 | 1845.357531 | 1797.986473 | 1605.216297 | 6.38E-17 | 0.846728828 | 6.95E-18 | 74.22998513 | 1.798418559 | 16.19506743 |
| ENSMUSG00000032812 | *Arap1* | 4983.310309 | 5043.332325 | 4149.418521 | 8255.191734 | 8643.581104 | 8595.300267 | 2.34E-19 | 0.845774084 | 2.19E-20 | 85.6079087 | 1.797228798 | 18.63087882 |
| ENSMUSG00000033282 | *Rpgrip1l* | 154.9795127 | 157.4464364 | 187.8779408 | 303.6237254 | 306.1788558 | 284.1261838 | 2.56E-07 | 0.845687728 | 5.90E-08 | 29.39493643 | 1.797121223 | 6.592519716 |
| ENSMUSG00000034327 | *Kctd9* | 528.0487519 | 566.2016078 | 515.3223519 | 958.1015335 | 985.8524862 | 950.9529417 | 2.52E-18 | 0.845414693 | 2.52E-19 | 80.78286199 | 1.796781144 | 17.59886796 |
| ENSMUSG00000022241 | *Tars* | 3924.816215 | 3357.848038 | 3436.824332 | 6323.807481 | 6338.55376 | 6572.592435 | 1.46E-23 | 0.842563222 | 1.07E-24 | 105.2601952 | 1.793233333 | 22.83564041 |
| ENSMUSG00000038437 | *Mllt6* | 1846.972336 | 1854.032716 | 1162.159262 | 2922.378357 | 3070.474483 | 2733.989707 | 1.41E-06 | 0.841653238 | 3.51E-07 | 25.94809882 | 1.792102603 | 5.851232224 |
| ENSMUSG00000079575 | *Rbpj-ps3* | 3272.943625 | 3186.271793 | 2479.988819 | 5283.896221 | 5450.41793 | 5299.823102 | 2.76E-13 | 0.841508729 | 3.86E-14 | 57.24073478 | 1.791923104 | 12.55867663 |
| ENSMUSG00000030232 | *Aebp2* | 1522.633769 | 1436.194096 | 1133.977571 | 2503.208936 | 2375.600484 | 2466.292589 | 4.48E-12 | 0.840996624 | 6.82E-13 | 51.59459542 | 1.791287148 | 11.34863382 |
| ENSMUSG00000094627 | *Gm10182* | 925.0838951 | 934.5858981 | 822.6369837 | 1641.254916 | 1578.666867 | 1592.652894 | 8.95E-21 | 0.840543142 | 7.73E-22 | 92.22688541 | 1.790724183 | 20.04841523 |
| ENSMUSG00000046139 | *Patl1* | 1139.978168 | 1272.692027 | 968.9133805 | 2090.786709 | 2006.448672 | 1966.655728 | 3.27E-13 | 0.840435869 | 4.58E-14 | 56.90332007 | 1.790591036 | 12.48599811 |
| ENSMUSG00000022789 | *Dnm1l* | 1104.029312 | 975.9660512 | 1154.107351 | 1993.795797 | 1849.016282 | 1940.562507 | 2.36E-18 | 0.840121859 | 2.35E-19 | 80.91763552 | 1.790201347 | 17.62771785 |
| ENSMUSG00000090210 | *Itga10* | 46.3340811 | 60.55632169 | 45.62749991 | 93.61731533 | 107.4883217 | 72.48116933 | 0.008717033 | 0.839708808 | 0.003611004 | 8.469753325 | 1.789688877 | 2.059631335 |
| ENSMUSG00000034023 | *Fancd2* | 556.8078367 | 499.5896539 | 405.279558 | 883.8824006 | 852.3063895 | 890.0687594 | 1.09E-09 | 0.839152589 | 2.02E-10 | 40.44799183 | 1.78899901 | 8.962042528 |
| ENSMUSG00000033382 | *Trappc8* | 1216.669061 | 1164.69992 | 1115.189777 | 2044.399751 | 2158.45236 | 2062.330871 | 5.33E-26 | 0.838927041 | 3.46E-27 | 116.6297061 | 1.788719343 | 25.27291224 |
| ENSMUSG00000021470 | *Ercc6l2* | 560.0032906 | 533.9049029 | 512.6383814 | 989.3073053 | 994.5384112 | 894.9008374 | 8.41E-16 | 0.838731657 | 9.86E-17 | 68.99735753 | 1.788477114 | 15.0751064 |
| ENSMUSG00000026718 | *Stam* | 513.6692095 | 553.0810714 | 564.9758077 | 961.4751304 | 944.5943425 | 1004.105799 | 2.23E-17 | 0.837957644 | 2.36E-18 | 76.36237201 | 1.787517845 | 16.65121221 |
| ENSMUSG00000029482 | *Aacs* | 3029.290268 | 2989.463747 | 2384.707863 | 4911.957158 | 5104.066671 | 5019.56258 | 4.35E-15 | 0.837791911 | 5.34E-16 | 65.66515181 | 1.787312511 | 14.36157782 |
| ENSMUSG00000002428 | *Hltf* | 644.6828181 | 607.5817609 | 747.4858074 | 1141.962567 | 1214.943758 | 1208.019489 | 4.16E-14 | 0.83753374 | 5.46E-15 | 61.08704172 | 1.786992699 | 13.38107071 |
| ENSMUSG00000022858 | *Tra2b* | 3696.341263 | 3619.249493 | 4313.140727 | 7273.475022 | 6586.102623 | 6904.072983 | 4.23E-19 | 0.837485324 | 4.03E-20 | 84.4054428 | 1.78693273 | 18.37327378 |
| ENSMUSG00000038518 | *Jarid2* | 576.7794234 | 533.9049029 | 386.491764 | 901.5937846 | 883.7829466 | 900.6993309 | 1.21E-07 | 0.83723468 | 2.71E-08 | 30.90676681 | 1.786622307 | 6.9178944 |
| ENSMUSG00000072568 | *Lratd2* | 792.4725595 | 843.7514155 | 735.4079398 | 1461.610878 | 1394.090961 | 1384.873542 | 2.94E-19 | 0.836532976 | 2.78E-20 | 85.13891432 | 1.785753534 | 18.53141015 |
| ENSMUSG00000102644 | *Thap6* | 61.51248698 | 76.70467414 | 63.0733087 | 127.3532848 | 121.6029498 | 110.1713774 | 0.000848839 | 0.836125936 | 0.000298977 | 13.0767895 | 1.785249775 | 3.071174434 |
| ENSMUSG00000040302 | *Rbm48* | 115.0363393 | 111.0199231 | 132.8565439 | 231.9347902 | 201.947756 | 202.9472741 | 1.32E-05 | 0.835853633 | 3.65E-06 | 21.44228969 | 1.784912848 | 4.88066363 |
| ENSMUSG00000044519 | *Zfp488* | 100.6567969 | 85.78812239 | 55.02139695 | 135.7872772 | 168.2897967 | 132.398936 | 0.004454892 | 0.835461978 | 0.001749197 | 9.795707353 | 1.784428355 | 2.351162825 |
| ENSMUSG00000025982 | *Sf3b1* | 5522.543149 | 5439.976232 | 6153.002562 | 10588.87742 | 10171.21816 | 9762.730302 | 1.40E-26 | 0.835296533 | 8.79E-28 | 119.3469819 | 1.784223733 | 25.85423282 |
| ENSMUSG00000102070 | *Gm28661* | 42319.79218 | 31280.36797 | 54915.38011 | 71060.60273 | 72720.73548 | 85510.38433 | 4.88E-06 | 0.835291885 | 1.29E-06 | 23.43503415 | 1.784217984 | 5.311664334 |
| ENSMUSG00000094870 | *Zfp131* | 1664.831466 | 1516.935858 | 1473.49985 | 2875.991399 | 2599.263053 | 2837.396176 | 4.54E-21 | 0.835036328 | 3.84E-22 | 93.61015311 | 1.783901959 | 20.34326541 |
| ENSMUSG00000026159 | *Agfg1* | 2124.17796 | 2040.748041 | 1845.229776 | 3625.773321 | 3569.91517 | 3537.081064 | 2.62E-25 | 0.834557958 | 1.76E-26 | 113.4002984 | 1.783310549 | 24.58194515 |
| ENSMUSG00000025261 | *Huwe1* | 6719.240623 | 6562.286727 | 4337.296462 | 10469.11473 | 10975.75196 | 9987.905134 | 1.48E-07 | 0.834546981 | 3.36E-08 | 30.48966355 | 1.78329698 | 6.828296027 |
| ENSMUSG00000028164 | *Manba* | 1922.065502 | 1871.19034 | 1732.503011 | 3409.019717 | 3221.39243 | 3231.693737 | 2.76E-27 | 0.834203733 | 1.66E-28 | 122.648874 | 1.782872746 | 26.5591628 |
| ENSMUSG00000046062 | *Ppp1r15b* | 3498.223123 | 3567.776619 | 2811.459186 | 5875.119087 | 6050.832495 | 5697.01991 | 4.20E-15 | 0.834029149 | 5.15E-16 | 65.73970768 | 1.782657009 | 14.37696261 |
| ENSMUSG00000100838 | *Gm29094* | 201.3135938 | 158.4557084 | 288.5268377 | 422.5430179 | 337.6653339 | 388.4990676 | 0.000499657 | 0.833884252 | 0.000169973 | 14.13693824 | 1.782477977 | 3.301327738 |
| ENSMUSG00000023952 | *Gtpbp2* | 1159.150891 | 1117.264135 | 901.8141159 | 1880.780299 | 1927.189607 | 1870.014169 | 1.95E-13 | 0.833541728 | 2.69E-14 | 57.94949471 | 1.782054833 | 12.71018185 |
| ENSMUSG00000030960 | *Eef1akmt2* | 468.9328553 | 465.274405 | 556.923896 | 893.1597922 | 892.4787926 | 860.1098761 | 3.07E-13 | 0.832618868 | 4.30E-14 | 57.02500184 | 1.780915256 | 12.51232088 |
| ENSMUSG00000046876 | *Atxn1* | 297.9760733 | 346.1803057 | 205.3237496 | 525.4377248 | 505.9551306 | 488.0398735 | 2.64E-05 | 0.832511385 | 7.62E-06 | 20.03155478 | 1.78078258 | 4.577948751 |
| ENSMUSG00000022231 | *Sema5a* | 553.6123828 | 489.4969336 | 382.4658081 | 847.6162334 | 956.5374894 | 744.1400052 | 1.01E-06 | 0.832301547 | 2.47E-07 | 26.62819144 | 1.780523586 | 5.995643331 |
| ENSMUSG00000039396 | *Neil3* | 148.5886049 | 165.5206126 | 140.9084556 | 256.3933681 | 270.3494153 | 285.0925994 | 6.66E-07 | 0.831857897 | 1.60E-07 | 27.46727095 | 1.779976133 | 6.176742633 |
| ENSMUSG00000111375 | *Btbd8* | 54.32271578 | 88.81593848 | 87.22904395 | 145.908068 | 129.2031342 | 130.4661048 | 0.003276698 | 0.830817453 | 0.001258156 | 10.4029286 | 1.778692911 | 2.484563568 |
| ENSMUSG00000102525 | *Gm37784* | 89.47270834 | 73.67685805 | 83.20308807 | 147.5948665 | 145.4892436 | 145.9287543 | 0.000148858 | 0.830785269 | 4.72E-05 | 16.55582604 | 1.778653232 | 3.827226783 |
| ENSMUSG00000035093 | *Secisbp2l* | 1130.391806 | 1152.588656 | 1140.687498 | 1999.699591 | 2056.392741 | 2033.338404 | 1.12E-30 | 0.830745411 | 5.78E-32 | 138.4616723 | 1.778604093 | 29.94987725 |
| ENSMUSG00000071669 | *Snx29* | 301.9703906 | 323.976321 | 289.868823 | 525.4377248 | 527.6699431 | 577.9165235 | 2.83E-11 | 0.830738673 | 4.59E-12 | 47.85451897 | 1.778595786 | 10.54782668 |
| ENSMUSG00000006932 | *Ctnnb1* | 12034.07927 | 13111.45292 | 13337.99181 | 22872.14391 | 23194.67694 | 22360.92395 | 1.26E-35 | 0.830650877 | 5.17E-37 | 161.5554622 | 1.778487552 | 34.89916033 |
| ENSMUSG00000024360 | *Etf1* | 8141.217595 | 7273.823507 | 8879.916674 | 14293.93027 | 14145.02884 | 14753.30041 | 8.78E-21 | 0.830465598 | 7.58E-22 | 92.26572141 | 1.778259163 | 20.05664115 |
| ENSMUSG00000043019 | *Edem3* | 791.6736961 | 880.0852085 | 960.8614687 | 1584.747167 | 1520.036873 | 1561.727595 | 2.47E-16 | 0.830060191 | 2.79E-17 | 71.4868311 | 1.777759531 | 15.60804983 |
| ENSMUSG00000099083 | *Atf7* | 880.347541 | 841.7328715 | 606.5773518 | 1458.237281 | 1353.918558 | 1337.519178 | 3.48E-08 | 0.829613709 | 7.40E-09 | 33.42775001 | 1.777209439 | 7.458765543 |
| ENSMUSG00000038507 | *Parp12* | 3302.501574 | 3157.002904 | 2560.507936 | 5381.730533 | 5386.359233 | 5275.662712 | 8.82E-15 | 0.829364116 | 1.11E-15 | 64.22812577 | 1.776902 | 14.05435534 |
| ENSMUSG00000032570 | *Atp2c1* | 1509.05309 | 1649.150494 | 1638.564041 | 2824.544045 | 2832.697287 | 2854.791656 | 4.98E-29 | 0.828986075 | 2.77E-30 | 130.7777028 | 1.776436445 | 28.30309953 |
| ENSMUSG00000066324 | *Bpnt2* | 1227.853149 | 1337.285437 | 1244.020365 | 2284.768534 | 2237.711425 | 2243.050587 | 2.67E-27 | 0.82895406 | 1.61E-28 | 122.7187188 | 1.776397025 | 26.57360036 |
| ENSMUSG00000035161 | *Ints6* | 593.5555562 | 612.6281211 | 438.8291903 | 1052.562248 | 973.9093393 | 902.6321621 | 5.11E-08 | 0.828048273 | 1.10E-08 | 32.65131334 | 1.775282075 | 7.291909239 |
| ENSMUSG00000036707 | *Cab39* | 3743.474208 | 3868.539684 | 4023.271904 | 7162.146323 | 6756.5639 | 6721.420436 | 6.92E-35 | 0.827885316 | 2.93E-36 | 158.106418 | 1.775081563 | 34.15992372 |
| ENSMUSG00000072704 | *Smim10l1* | 335.5226563 | 325.9948651 | 229.4794848 | 541.4623103 | 523.3269806 | 524.763666 | 1.05E-06 | 0.826951602 | 2.58E-07 | 26.54324254 | 1.7739331 | 5.977281711 |
| ENSMUSG00000020400 | *Tnip1* | 3823.360554 | 3923.040373 | 3605.914478 | 6496.704324 | 6816.279635 | 6831.591814 | 1.35E-34 | 0.826565099 | 5.84E-36 | 156.7377798 | 1.773457921 | 33.86845263 |
| ENSMUSG00000015839 | *Nfe2l2* | 2784.039183 | 2926.888882 | 2992.6272 | 5350.524761 | 5170.296849 | 4903.592709 | 1.78E-30 | 0.826524318 | 9.26E-32 | 137.5240017 | 1.77340779 | 29.74877596 |
| ENSMUSG00000103318 | *Gm38356* | 80.6852102 | 67.62122589 | 73.80919103 | 120.6060909 | 133.5460967 | 141.0966763 | 0.000463894 | 0.82628965 | 0.000157068 | 14.28552441 | 1.773119352 | 3.333580842 |
| ENSMUSG00000029911 | *Ssbp1* | 971.4179762 | 835.6772393 | 1076.272204 | 1703.666459 | 1611.239085 | 1789.801675 | 1.47E-12 | 0.826242739 | 2.15E-13 | 53.86203955 | 1.773061697 | 11.83274274 |
| ENSMUSG00000031555 | *Adam9* | 1701.579185 | 1756.133329 | 1529.863232 | 2988.163498 | 2920.642277 | 2944.668306 | 3.69E-24 | 0.826101663 | 2.62E-25 | 108.0475482 | 1.772888325 | 23.43299594 |
| ENSMUSG00000053113 | *Socs3* | 945.0554818 | 966.882603 | 707.2262486 | 1502.094042 | 1601.46742 | 1552.063439 | 7.20E-10 | 0.826049157 | 1.31E-10 | 41.29186923 | 1.772823803 | 9.142452235 |
| ENSMUSG00000025058 | *Tasl* | 507.2783017 | 414.8108036 | 552.8979401 | 852.6766288 | 906.5934207 | 851.4121358 | 5.90E-10 | 0.825983753 | 1.07E-10 | 41.69378019 | 1.772743434 | 9.229479488 |
| ENSMUSG00000027329 | *Spef1* | 63.11021392 | 61.56559372 | 59.04735283 | 91.93051686 | 128.1173936 | 107.2721306 | 0.002532554 | 0.825970975 | 0.000953295 | 10.91615079 | 1.772727733 | 2.596441353 |
| ENSMUSG00000042688 | *Mapk6* | 3288.920895 | 3275.087731 | 3420.720508 | 5949.338219 | 6026.946201 | 5715.381806 | 1.29E-39 | 0.825829785 | 4.38E-41 | 180.2011949 | 1.772554253 | 38.88916046 |
| ENSMUSG00000066880 | *Zfp617* | 254.8374461 | 286.633256 | 394.5436757 | 520.3773294 | 548.2990149 | 580.8157703 | 8.22E-06 | 0.825784932 | 2.23E-06 | 22.38282314 | 1.772499146 | 5.085297209 |
| ENSMUSG00000051451 | *Crebzf* | 423.3976377 | 430.959156 | 287.1848524 | 684.8401806 | 664.4732616 | 683.2558229 | 1.32E-06 | 0.825759354 | 3.27E-07 | 26.08260604 | 1.772467721 | 5.880458989 |
| ENSMUSG00000030180 | *Kdm5a* | 1469.109916 | 1480.602065 | 1566.096835 | 2713.215346 | 2679.607859 | 2601.590771 | 2.48E-30 | 0.825651123 | 1.30E-31 | 136.8509587 | 1.772334756 | 29.60550649 |
| ENSMUSG00000022636 | *Alcam* | 5860.462396 | 5748.813472 | 5598.762636 | 10275.13291 | 10318.87889 | 9904.793393 | 4.15E-46 | 0.825215587 | 1.05E-47 | 210.5337776 | 1.771799784 | 45.38202999 |
| ENSMUSG00000037355 | *Uvssa* | 99.85793341 | 120.1033713 | 68.44124987 | 173.7402429 | 174.8042404 | 166.2234817 | 0.001198858 | 0.824193797 | 0.000431247 | 12.39169938 | 1.77054535 | 2.921232083 |
| ENSMUSG00000028945 | *Rheb* | 1934.847318 | 1835.865819 | 2055.921467 | 3390.464934 | 3297.394274 | 3619.226389 | 7.72E-24 | 0.824048263 | 5.61E-25 | 106.5425202 | 1.770366753 | 23.11210965 |
| ENSMUSG00000004085 | *Map3k20* | 1473.903097 | 1352.424518 | 1218.522645 | 2394.410435 | 2455.94529 | 2322.296665 | 7.03E-18 | 0.823438826 | 7.22E-19 | 78.70262793 | 1.769619056 | 17.15299935 |
| ENSMUSG00000049504 | *Proser1* | 671.844176 | 674.1937148 | 366.3619846 | 1028.10367 | 1054.254145 | 956.7514352 | 0.000202591 | 0.823349066 | 6.55E-05 | 15.93700805 | 1.769508959 | 3.693380298 |
| ENSMUSG00000067367 | *Lyar* | 1182.317932 | 1003.216396 | 1286.963895 | 2093.316907 | 2045.535335 | 1998.547442 | 7.22E-14 | 0.823327971 | 9.61E-15 | 59.9749481 | 1.769483086 | 13.14153166 |
| ENSMUSG00000039756 | *Dnttip2* | 2720.130106 | 2584.745664 | 3414.010582 | 5229.91867 | 5196.354624 | 4986.70445 | 1.21E-12 | 0.823253208 | 1.75E-13 | 54.26395757 | 1.769391389 | 11.91887348 |
| ENSMUSG00000026893 | *Gca* | 128.6170182 | 122.1219154 | 106.016838 | 222.6573986 | 222.5768278 | 189.4174559 | 2.48E-05 | 0.823175321 | 7.12E-06 | 20.16075999 | 1.769295868 | 4.60550615 |
| ENSMUSG00000063281 | *Zfp35* | 666.2521317 | 704.4718756 | 889.7362483 | 1417.754118 | 1282.259676 | 1284.366321 | 7.70E-10 | 0.822583339 | 1.40E-10 | 41.15915958 | 1.768570021 | 9.113669619 |
| ENSMUSG00000027763 | *Mbnl1* | 7536.47795 | 6911.494849 | 6652.22109 | 13017.86723 | 12155.95202 | 12142.04549 | 5.62E-27 | 0.822051706 | 3.45E-28 | 121.1999616 | 1.767918422 | 26.25041243 |
| ENSMUSG00000058706 | *0610030E20Rik* | 619.1191871 | 627.7672015 | 501.902499 | 1034.850864 | 1074.883217 | 990.5759809 | 9.05E-12 | 0.821092008 | 1.41E-12 | 50.17396115 | 1.766742773 | 11.0434628 |
| ENSMUSG00000037029 | *Zfp146* | 945.0554818 | 831.6401512 | 1025.276763 | 1738.245828 | 1596.038717 | 1607.149128 | 2.12E-14 | 0.820766213 | 2.72E-15 | 62.45948511 | 1.766343846 | 13.67330257 |
| ENSMUSG00000061898 | *Rbak* | 127.8181548 | 120.1033713 | 162.3802203 | 245.429178 | 238.8629372 | 234.8389886 | 1.60E-05 | 0.820306764 | 4.49E-06 | 21.04436864 | 1.765781415 | 4.795471085 |
| ENSMUSG00000053175 | *Bcl3* | 556.8078367 | 471.3300371 | 330.1283817 | 784.3612906 | 893.4136595 | 770.2332261 | 1.66E-05 | 0.819586829 | 4.66E-06 | 20.97332552 | 1.764900473 | 4.780463584 |
| ENSMUSG00000028252 | *Ccnc* | 802.0589212 | 648.9619141 | 793.1133073 | 1276.063046 | 1299.631526 | 1382.940711 | 1.32E-12 | 0.818988381 | 1.92E-13 | 54.08563868 | 1.764168523 | 11.88069355 |
| ENSMUSG00000024535 | *Snx24* | 256.435173 | 216.993486 | 271.0810289 | 436.8808049 | 432.1247682 | 441.6519251 | 9.19E-09 | 0.81888201 | 1.85E-09 | 36.12590412 | 1.764038454 | 8.036633443 |
| ENSMUSG00000035248 | *Tut7* | 1311.733813 | 1343.341069 | 1387.612791 | 2395.253834 | 2403.829741 | 2324.229497 | 6.46E-30 | 0.818674379 | 3.44E-31 | 134.9157058 | 1.763784594 | 29.18949771 |
| ENSMUSG00000037628 | *Cdkn3* | 114.2374758 | 107.992107 | 85.88705866 | 194.8252238 | 158.518131 | 193.2831182 | 0.000223491 | 0.818657944 | 7.26E-05 | 15.74259994 | 1.763764501 | 3.650739933 |
| ENSMUSG00000029298 | *Gbp9* | 237.2624498 | 210.9378539 | 193.245882 | 382.9032537 | 400.6382901 | 353.7081064 | 9.61E-08 | 0.818284389 | 2.13E-08 | 31.37224443 | 1.763307871 | 7.017085617 |
| ENSMUSG00000027312 | *Atrn* | 1922.864366 | 1956.978463 | 1421.162424 | 3223.471885 | 3144.304846 | 2985.257761 | 5.85E-10 | 0.817257907 | 1.06E-10 | 41.71029907 | 1.762053718 | 9.232723954 |
| ENSMUSG00000034206 | *Polq* | 366.6783315 | 329.0226812 | 365.0199993 | 620.7418386 | 681.8451116 | 565.3531208 | 5.12E-10 | 0.815772047 | 9.20E-11 | 41.98385084 | 1.760239878 | 9.290355012 |
| ENSMUSG00000028412 | *Slc44a1* | 1659.239422 | 1684.475015 | 1575.490732 | 2944.306737 | 2841.383212 | 2878.952046 | 4.09E-31 | 0.815688826 | 2.07E-32 | 140.498947 | 1.760138344 | 30.38785798 |
| ENSMUSG00000041827 | *Oasl1* | 5982.688507 | 5447.041136 | 4066.215433 | 9171.123306 | 9131.078644 | 8985.732166 | 2.59E-09 | 0.815587679 | 4.95E-10 | 38.69530222 | 1.760014945 | 8.585902386 |
| ENSMUSG00000053581 | *Zfand2a* | 1753.505311 | 1931.746662 | 1747.26485 | 3190.579314 | 3214.877986 | 3157.279736 | 3.37E-27 | 0.81548004 | 2.04E-28 | 122.2418899 | 1.759883636 | 26.47177356 |
| ENSMUSG00000023169 | *Slc38a1* | 6560.266793 | 6372.543586 | 5202.876975 | 10564.41885 | 10919.29345 | 10444.05329 | 9.45E-16 | 0.81521738 | 1.11E-16 | 68.76231589 | 1.759563256 | 15.02443889 |
| ENSMUSG00000025220 | *Oga* | 2676.991479 | 2571.625128 | 2121.678746 | 4265.913342 | 4417.878597 | 4300.549381 | 2.16E-15 | 0.815167921 | 2.60E-16 | 67.08429141 | 1.759502935 | 14.66497606 |
| ENSMUSG00000002222 | *Rmnd5a* | 659.861224 | 715.5738679 | 555.5819107 | 1136.058773 | 1149.79932 | 1120.07567 | 4.05E-12 | 0.814791265 | 6.15E-13 | 51.79955814 | 1.759043627 | 11.39211575 |
| ENSMUSG00000034593 | *Myo5a* | 3248.178858 | 3391.154015 | 3363.015141 | 6023.557352 | 5795.683449 | 5763.702585 | 9.35E-40 | 0.814441584 | 3.15E-41 | 180.8557655 | 1.758617322 | 39.02904578 |
| ENSMUSG00000108686 | *Gm45080* | 406.6215048 | 345.1710336 | 367.7039699 | 676.4061883 | 669.9019647 | 626.2373031 | 2.08E-11 | 0.814282045 | 3.33E-12 | 48.48488246 | 1.758422858 | 10.68267198 |
| ENSMUSG00000026349 | *Ccnt2* | 718.1782571 | 646.94337 | 673.6766163 | 1213.651502 | 1204.086352 | 1171.295696 | 9.25E-19 | 0.814163519 | 8.96E-20 | 82.8255705 | 1.758278398 | 18.03393739 |
| ENSMUSG00000023882 | *Zfp54* | 75.89202939 | 67.62122589 | 84.54507337 | 138.3174749 | 125.9459123 | 134.3317672 | 0.000467708 | 0.812827083 | 0.000158442 | 14.26912821 | 1.756650377 | 3.330024742 |
| ENSMUSG00000058835 | *Abi1* | 2248.001797 | 2123.508347 | 2789.987421 | 4154.584643 | 4120.385666 | 4288.952393 | 1.36E-12 | 0.812531167 | 1.69E-13 | 54.33913729 | 1.756290102 | 11.93443018 |
| ENSMUSG00000041143 | *Tmco4* | 643.8839546 | 658.0453623 | 646.8369105 | 1140.275769 | 1120.484324 | 1158.732294 | 7.16E-21 | 0.811494881 | 6.13E-22 | 92.68667222 | 1.755029014 | 20.14484166 |
| ENSMUSG00000026761 | *Orc4* | 780.4896075 | 659.0546344 | 794.4552926 | 1306.425418 | 1263.802086 | 1342.351256 | 4.13E-14 | 0.809547814 | 5.42E-15 | 61.10271317 | 1.752662017 | 13.38413718 |
| ENSMUSG00000001020 | *S100a4* | 16602.77944 | 15079.53337 | 19088.39879 | 29186.674 | 30232.44766 | 29543.32462 | 7.54E-17 | 0.809397854 | 8.26E-18 | 73.89048328 | 1.752479847 | 16.12248755 |
| ENSMUSG00000063550 | *Nup98* | 2450.913118 | 2483.818461 | 2140.46654 | 4051.689936 | 4282.161019 | 4078.273795 | 2.89E-21 | 0.809191954 | 2.42E-22 | 94.52542637 | 1.752229753 | 20.53921511 |
| ENSMUSG00000037845 | *Fdxacb1* | 127.8181548 | 132.2146357 | 128.830588 | 246.2725773 | 207.3764591 | 226.1412483 | 7.18E-06 | 0.808375977 | 1.94E-06 | 22.65266977 | 1.751238986 | 5.143845601 |
| ENSMUSG00000056216 | *Cebpg* | 908.3077623 | 831.6401512 | 1002.463013 | 1599.084954 | 1561.295017 | 1631.309518 | 2.89E-16 | 0.807715936 | 3.29E-17 | 71.16148068 | 1.750437968 | 15.53891998 |
| ENSMUSG00000032316 | *Clk3* | 930.6759394 | 906.3262813 | 858.8705866 | 1521.492224 | 1576.495385 | 1628.410271 | 2.02E-21 | 0.807446457 | 1.67E-22 | 95.25525562 | 1.750111036 | 20.69471691 |
| ENSMUSG00000025278 | *Flnb* | 2171.310904 | 2097.267274 | 1281.595953 | 3370.223352 | 3266.993536 | 3081.89932 | 1.29E-05 | 0.806666006 | 3.58E-06 | 21.47921306 | 1.749164539 | 4.88884128 |
| ENSMUSG00000048661 | *Lemd3* | 370.6726488 | 345.1710336 | 281.8169112 | 593.753063 | 550.4704961 | 608.8418224 | 2.02E-08 | 0.80641686 | 4.20E-09 | 34.52923746 | 1.748862494 | 7.695370297 |
| ENSMUSG00000022973 | *Synj1* | 653.4703162 | 713.5553239 | 538.1361019 | 1156.300354 | 1153.056542 | 1030.19902 | 9.13E-10 | 0.806217423 | 1.67E-10 | 40.81404253 | 1.748620749 | 9.03950325 |
| ENSMUSG00000048911 | *Rnf24* | 329.1317485 | 450.1353245 | 291.2108083 | 622.4286371 | 632.9867835 | 619.4723939 | 5.37E-06 | 0.805730527 | 1.43E-06 | 23.23916103 | 1.748030706 | 5.269755602 |

| | | | | | | | | | | | | | |
|---|---|---|---|---|---|---|---|---|---|---|---|---|---|
| ENSMUSG00000034928 | *Rnf44* | 1259.807688 | 1443.259 | 877.6583807 | 2033.435561 | 2211.65365 | 2023.674248 | 2.87E-06 | 0.805719769 | 7.40E-07 | 24.50817618 | 1.748017671 | 5.542262101 |
| ENSMUSG00000037071 | *Scd1* | 4985.706899 | 4332.804817 | 3208.686832 | 7263.354231 | 7529.611224 | 7107.986673 | 7.94E-08 | 0.804981358 | 1.74E-08 | 31.7605493 | 1.747123216 | 7.10033208 |
| ENSMUSG00000020661 | *Dnmt3a* | 5019.259165 | 5160.40788 | 4118.55286 | 8347.96565 | 8346.088173 | 8277.349538 | 5.81E-16 | 0.803590743 | 6.75E-17 | 69.74428795 | 1.745439974 | 15.23562122 |
| ENSMUSG00000096544 | *Gm4617* | 6161.633923 | 5824.508874 | 6476.421017 | 10628.51719 | 10857.40624 | 10727.21306 | 9.22E-34 | 0.803441246 | 4.14E-35 | 152.8443104 | 1.745259115 | 33.03506996 |
| ENSMUSG00000020590 | *Snx13* | 495.2953497 | 541.9790791 | 558.2658813 | 929.4259594 | 921.7937894 | 924.8597207 | 1.21E-15 | 0.802972213 | 1.43E-16 | 68.26658034 | 1.744691808 | 14.91787263 |
| ENSMUSG00000094483 | *Purb* | 2108.999554 | 1888.347965 | 1513.759409 | 3269.858843 | 3192.077433 | 3165.011061 | 1.82E-10 | 0.80256727 | 3.14E-11 | 44.08549168 | 1.744202167 | 9.740004069 |
| ENSMUSG00000071054 | *Safb* | 1665.630329 | 1661.261758 | 1074.930219 | 2542.8487 | 2533.032875 | 2612.221343 | 1.17E-06 | 0.802360513 | 2.89E-07 | 26.32420259 | 1.743952219 | 5.930646325 |
| ENSMUSG00000019838 | *Slc16a10* | 2430.142667 | 2310.223672 | 1614.408306 | 3667.099883 | 3627.459423 | 3796.080442 | 9.10E-08 | 0.801744904 | 2.01E-08 | 31.48512168 | 1.743208219 | 7.04106335 |
| ENSMUSG00000031292 | *Cdkl5* | 66.30566778 | 67.62122589 | 89.91301453 | 117.232494 | 131.3746155 | 138.1974295 | 0.001594453 | 0.801361906 | 0.000584087 | 11.82602021 | 1.742745504 | 2.797388244 |
| ENSMUSG00000052485 | *Tmem171* | 768.5066555 | 868.9832162 | 887.0522777 | 1486.912855 | 1457.063917 | 1444.791309 | 2.36E-17 | 0.801242444 | 2.51E-18 | 76.2447185 | 1.742601202 | 16.62654758 |
| ENSMUSG00000046731 | *Kctd11* | 192.5260956 | 148.3629881 | 138.224485 | 265.6707597 | 262.7492309 | 312.1522359 | 6.30E-05 | 0.801239013 | 1.90E-05 | 18.28477621 | 1.742597059 | 4.200485686 |
| ENSMUSG00000037625 | *Cldn11* | 421.0010473 | 357.282298 | 284.5008818 | 621.5852378 | 629.7295617 | 610.7746536 | 7.42E-07 | 0.801007695 | 1.79E-07 | 27.24576818 | 1.742317678 | 6.129846751 |
| ENSMUSG00000009907 | *Vps4b* | 2230.426801 | 2140.665972 | 2462.54301 | 4025.544559 | 4050.898267 | 3813.475923 | 3.12E-22 | 0.800375911 | 2.49E-23 | 99.02517143 | 1.74155485 | 21.50585835 |
| ENSMUSG00000029016 | *Clcn6* | 452.1567225 | 488.4876616 | 311.3405876 | 712.6723555 | 771.9615834 | 704.5169659 | 6.68E-06 | 0.800275476 | 1.80E-06 | 22.79880533 | 1.741433613 | 5.175257715 |
| ENSMUSG00000014907 | *Naf1* | 588.7623754 | 591.4334085 | 407.9635286 | 962.3185297 | 884.8786082 | 927.7589675 | 5.51E-07 | 0.800175086 | 1.31E-07 | 27.84875515 | 1.74131244 | 6.258857481 |
| ENSMUSG00000043542 | *Zc2hc1a* | 139.8011068 | 104.9642909 | 127.4886027 | 231.091391 | 198.6905341 | 218.4099236 | 7.55E-05 | 0.79897341 | 2.30E-05 | 17.92067846 | 1.739862638 | 4.122086876 |
| ENSMUSG00000048546 | *Tob2* | 1106.425902 | 1188.922449 | 921.9438953 | 1857.16512 | 1957.590344 | 1793.667337 | 3.76E-12 | 0.7989576 | 5.70E-13 | 51.94743707 | 1.739843572 | 11.42431699 |
| ENSMUSG00000097891 | *Gm3650* | 55.12157924 | 66.61195386 | 50.99544108 | 113.0154978 | 110.7455436 | 77.31324729 | 0.009477401 | 0.7984008 | 0.003956748 | 8.303558407 | 1.739172218 | 2.023310757 |
| ENSMUSG00000006010 | *Odr4* | 681.4305376 | 617.6744812 | 772.9835279 | 1206.904308 | 1241.001533 | 1147.135307 | 2.19E-12 | 0.797963826 | 3.26E-13 | 53.04426417 | 1.738645526 | 11.65935717 |
| ENSMUSG00000035239 | *Neu3* | 98.26020648 | 80.74176225 | 50.99544108 | 134.1004787 | 145.4892436 | 125.6340268 | 0.007660318 | 0.797824022 | 0.003134835 | 8.727281685 | 1.73847705 | 2.115753183 |
| ENSMUSG00000020078 | *Vps26a* | 2726.521014 | 2832.017311 | 3485.135802 | 5331.126579 | 5175.725553 | 5198.349465 | 1.08E-13 | 0.797784938 | 1.46E-14 | 59.15188874 | 1.738429954 | 12.96601879 |
| ENSMUSG00000028274 | *Rngtt* | 490.5021689 | 496.5618378 | 630.733087 | 1007.862088 | 898.9932363 | 893.9344218 | 5.60E-09 | 0.797666479 | 1.10E-09 | 37.13515731 | 1.738287218 | 8.251695147 |
| ENSMUSG00000049232 | *Tigd2* | 277.2056231 | 348.1988497 | 434.8032345 | 634.2362264 | 585.2141961 | 612.7074848 | 8.70E-06 | 0.797475815 | 2.37E-06 | 22.2670532 | 1.738057504 | 5.060433697 |
| ENSMUSG00000047454 | *Gphn* | 540.0317039 | 577.3036001 | 536.7941166 | 958.9449327 | 983.6810049 | 931.6246298 | 1.35E-16 | 0.796396193 | 1.50E-17 | 72.7065199 | 1.736757338 | 15.86848177 |
| ENSMUSG00000086699 | *Gm16510* | 189.3306417 | 159.4649804 | 167.7481614 | 295.189733 | 269.2636746 | 334.3797945 | 5.56E-06 | 0.796003233 | 1.48E-06 | 23.17218723 | 1.736284347 | 5.255304167 |
| ENSMUSG00000038013 | *Wipf2* | 664.6544048 | 671.1658987 | 453.5910285 | 1100.636005 | 1065.111552 | 949.9865261 | 2.11E-06 | 0.79573514 | 5.38E-07 | 25.12247898 | 1.735961726 | 5.675468122 |
| ENSMUSG00000041268 | *Dmxl2* | 448.9612686 | 450.1353245 | 328.7863964 | 719.4195494 | 732.8749209 | 688.0879009 | 1.07E-07 | 0.795106086 | 2.38E-08 | 31.15519818 | 1.735204964 | 6.970267275 |
| ENSMUSG00000042202 | *Slc35e2* | 313.9533426 | 339.1154015 | 242.8993378 | 486.6413599 | 563.4993836 | 512.2002633 | 1.09E-06 | 0.794886539 | 2.68E-07 | 26.47028941 | 1.734940924 | 5.96161384 |
| ENSMUSG00000036103 | *Colec12* | 3368.807242 | 3501.164666 | 3758.900802 | 6183.803207 | 6191.978776 | 6046.862354 | 8.98E-30 | 0.794696835 | 4.83E-31 | 134.2456315 | 1.734712807 | 29.04675788 |
| ENSMUSG00000028249 | *Sdcbp* | 14075.97429 | 13043.83169 | 19253.46298 | 27171.79323 | 26244.52235 | 27005.51728 | 1.97E-08 | 0.794539888 | 4.09E-09 | 34.58074744 | 1.734524103 | 7.706246824 |
| ENSMUSG00000024817 | *Uhrf2* | 1284.572455 | 1291.868196 | 1472.157865 | 2371.638655 | 2318.056231 | 2315.531756 | 5.36E-20 | 0.793822231 | 4.84E-21 | 88.5962715 | 1.733661492 | 19.27064983 |
| ENSMUSG00000051705 | *Senp8* | 122.2261105 | 140.2888119 | 132.8565439 | 220.9706002 | 241.0344184 | 222.275586 | 7.78E-06 | 0.793369687 | 2.11E-06 | 22.49003621 | 1.733117763 | 5.108878615 |
| ENSMUSG00000041936 | *Agrn* | 552.8135194 | 453.1631406 | 350.2581611 | 749.7819219 | 814.3054677 | 796.3264471 | 4.95E-06 | 0.792620561 | 1.31E-06 | 23.40382943 | 1.732218068 | 5.305103298 |
| ENSMUSG00000091562 | *None* | 1105.627039 | 1139.46812 | 955.4935276 | 1896.804885 | 1864.216651 | 1792.700922 | 3.68E-16 | 0.792384597 | 4.22E-17 | 70.67033416 | 1.731934772 | 15.43394522 |
| ENSMUSG00000033790 | *Tubgcp5* | 556.0089732 | 532.8956309 | 606.5773518 | 1059.309442 | 908.7649019 | 959.650682 | 2.99E-12 | 0.79175678 | 4.51E-13 | 52.40856273 | 1.731181252 | 11.52443511 |
| ENSMUSG00000041459 | *Tardbp* | 4194.832067 | 3777.705201 | 4271.539183 | 7205.159684 | 6900.967403 | 7083.826283 | 4.60E-26 | 0.791673081 | 2.98E-27 | 116.9274408 | 1.731080818 | 25.33692189 |
| ENSMUSG00000026771 | *Spopl* | 373.8681027 | 309.8465126 | 352.9421317 | 602.1870554 | 611.2719711 | 582.7486015 | 3.16E-10 | 0.791397997 | 5.60E-11 | 42.95533434 | 1.730750778 | 9.499886345 |
| ENSMUSG00000020053 | *Igf1* | 683.827128 | 515.7380064 | 548.8719842 | 1045.815054 | 981.5095237 | 1005.072215 | 3.55E-09 | 0.790895771 | 6.86E-10 | 38.06015326 | 1.73014838 | 8.449758768 |
| ENSMUSG00000058046 | *4933430I17Rik* | 79.88634673 | 76.70467414 | 120.7786762 | 178.8006383 | 137.8890592 | 157.5257414 | 0.003639345 | 0.790396167 | 0.00140986 | 10.19285374 | 1.729549336 | 2.438976768 |
| ENSMUSG00000035401 | *Emsy* | 730.1612091 | 662.0824505 | 625.3651458 | 1223.772293 | 1141.113395 | 1131.672657 | 2.51E-14 | 0.78993312 | 3.24E-15 | 62.11586599 | 1.728994308 | 13.60091034 |
| ENSMUSG00000027018 | *Hat1* | 2089.027967 | 1851.0049 | 2463.884995 | 3673.847077 | 3731.690523 | 3653.050934 | 3.69E-12 | 0.789549093 | 5.58E-13 | 51.98800838 | 1.728534134 | 11.43295186 |
| ENSMUSG00000038733 | *Wdr26* | 4858.687608 | 4469.056541 | 3721.325213 | 7572.881751 | 7507.896412 | 7475.224597 | 1.35E-13 | 0.788520513 | 1.84E-14 | 58.69388485 | 1.727302202 | 12.86833435 |
| ENSMUSG00000053950 | *Adnp2* | 850.7895927 | 904.3077372 | 688.4384545 | 1416.067319 | 1379.976333 | 1434.160737 | 5.98E-11 | 0.788325743 | 9.96E-12 | 46.33721721 | 1.727069025 | 10.22301659 |
| ENSMUSG00000033991 | *Skic3* | 840.4043676 | 846.7792316 | 991.7271304 | 1609.205745 | 1534.151501 | 1470.88453 | 2.88E-14 | 0.788224985 | 3.73E-15 | 61.83528678 | 1.72694841 | 13.54081161 |
| ENSMUSG00000039087 | *Rreb1* | 859.5770908 | 888.1593848 | 496.5345579 | 1335.944392 | 1328.946523 | 1217.683645 | 0.000153757 | 0.787421959 | 4.89E-05 | 16.48984943 | 1.725987431 | 3.813164245 |
| ENSMUSG00000034235 | *Usp54* | 260.4294903 | 273.5127196 | 208.0077202 | 437.7242041 | 433.2105088 | 415.5587042 | 2.39E-07 | 0.786772262 | 5.51E-08 | 29.52966839 | 1.725210333 | 6.621711463 |
| ENSMUSG00000028954 | *Nub1* | 2525.20742 | 2312.242216 | 2545.746098 | 4335.915479 | 4198.558991 | 4196.176497 | 1.46E-27 | 0.786409891 | 8.65E-29 | 123.9482208 | 1.724777055 | 26.8351352 |
| ENSMUSG00000003778 | *Brd8* | 842.800958 | 995.1422197 | 885.7102924 | 1572.096178 | 1634.039638 | 1490.212842 | 9.06E-15 | 0.786369492 | 1.14E-15 | 64.17420307 | 1.724728758 | 14.04267312 |
| ENSMUSG00000033326 | *Kdm4a* | 1500.265592 | 1448.30536 | 1221.206615 | 2380.916047 | 2451.602328 | 2370.617445 | 4.20E-15 | 0.785715254 | 5.16E-16 | 65.73572531 | 1.7239468 | 14.37629412 |
| ENSMUSG00000036777 | *Anln* | 1772.678034 | 1561.343828 | 1448.00213 | 2840.568631 | 2731.723409 | 2680.83685 | 1.32E-15 | 0.785000865 | 1.57E-16 | 68.082586 | 1.723093353 | 14.87799875 |
| ENSMUSG00000032103 | *Pus3* | 374.6669662 | 370.4028343 | 473.7208079 | 655.3212073 | 735.0464022 | 701.6177192 | 2.14E-08 | 0.784854437 | 4.46E-09 | 34.41011874 | 1.722918474 | 7.670021138 |
| ENSMUSG00000046982 | *Tshz1* | 455.3521764 | 448.1167805 | 296.5787494 | 659.5382035 | 731.7891803 | 685.1886541 | 1.35E-05 | 0.784229789 | 3.73E-06 | 21.39846274 | 1.722172658 | 4.871179283 |
| ENSMUSG00000021451 | *Sema4d* | 5802.145363 | 5879.009564 | 4288.984992 | 9065.698401 | 9410.113984 | 9038.885024 | 5.38E-10 | 0.784086299 | 9.69E-11 | 41.88311962 | 1.722001381 | 9.269264799 |
| ENSMUSG00000070713 | *Hmgn2-ps* | 569.5896522 | 505.6452861 | 497.8765432 | 949.6675411 | 990.1954487 | 774.0988885 | 1.05E-08 | 0.784034756 | 2.13E-09 | 35.85315946 | 1.72193986 | 7.978762302 |
| ENSMUSG00000057367 | *Birc2* | 377.86242 | 334.0690413 | 362.3360287 | 620.7418386 | 601.5003055 | 627.2037186 | 1.33E-11 | 0.782841004 | 2.10E-12 | 49.38615471 | 1.720515636 | 10.87708406 |
| ENSMUSG00000019843 | *Fyn* | 1569.766713 | 1594.649804 | 1206.444777 | 2454.29178 | 2535.204356 | 2541.673005 | 9.80E-11 | 0.782432665 | 1.66E-11 | 45.3317087 | 1.720028733 | 10.0086403 |
| ENSMUSG00000066568 | *Lsm14a* | 2654.623302 | 2651.357618 | 2792.671392 | 4702.794147 | 4628.512278 | 4587.574811 | 1.54E-35 | 0.782304017 | 6.35E-37 | 161.1474463 | 1.719875362 | 34.81251066 |
| ENSMUSG00000016520 | *Lnx2* | 378.6612835 | 325.9948651 | 277.7909553 | 587.0058691 | 556.9849399 | 553.7561337 | 1.44E-07 | 0.781599348 | 3.26E-08 | 30.54496819 | 1.719035511 | 6.840449599 |
| ENSMUSG00000032534 | *Cep63* | 476.1226265 | 426.9220679 | 467.0108814 | 730.3837394 | 821.905652 | 805.0241874 | 6.17E-12 | 0.781541893 | 9.48E-13 | 50.94982144 | 1.718967053 | 11.20988526 |
| ENSMUSG00000069237 | *Fam8a1* | 315.5510696 | 318.9299609 | 273.7649995 | 557.4868958 | 488.5832806 | 518.9651724 | 5.38E-09 | 0.781448906 | 1.06E-09 | 37.21619919 | 1.718856262 | 8.269092153 |
| ENSMUSG00000024241 | *Sos1* | 562.399881 | 630.7950176 | 403.9375727 | 925.2089632 | 915.2793457 | 912.296318 | 2.92E-06 | 0.78100101 | 7.53E-07 | 24.47323973 | 1.718322713 | 5.534883875 |
| ENSMUSG00000044827 | *Tlr1* | 425.7942281 | 418.8478917 | 471.0368373 | 805.4462716 | 715.5030709 | 732.5430181 | 1.56E-11 | 0.780920978 | 2.49E-12 | 49.05528342 | 1.718227393 | 10.80576798 |
| ENSMUSG00000054469 | *Lclat1* | 418.6044569 | 428.940612 | 326.1024258 | 644.3570172 | 708.9886272 | 672.6252514 | 3.40E-08 | 0.780854685 | 7.22E-09 | 33.47560751 | 1.718148441 | 7.469090386 |
| ENSMUSG00000035834 | *Polr3g* | 294.7806194 | 243.2345588 | 252.2932348 | 452.9053904 | 462.5255056 | 447.4504187 | 2.12E-08 | 0.780614766 | 4.44E-09 | 34.42262767 | 1.717862739 | 7.672689538 |
| ENSMUSG00000048949 | *Gm6206* | 147.7897414 | 153.4093483 | 146.2763968 | 239.5253834 | 271.4351559 | 258.9993784 | 2.63E-06 | 0.780601792 | 6.77E-07 | 24.68075046 | 1.71784729 | 5.57956254 |
| ENSMUSG00000042444 | *Mindy2* | 788.4782422 | 816.5010708 | 798.4812485 | 1448.11649 | 1371.290408 | 1306.593879 | 2.27E-18 | 0.780371675 | 2.26E-19 | 80.99559279 | 1.717573306 | 17.64407957 |
| ENSMUSG00000025860 | *Xiap* | 1446.741739 | 1392.795399 | 1642.589997 | 2678.635977 | 2558.004909 | 2448.897108 | 1.51E-16 | 0.780195008 | 1.69E-17 | 72.47893915 | 1.717362991 | 15.82001154 |
| ENSMUSG00000022048 | *Dpysl2* | 381.8567374 | 405.7273553 | 268.3970583 | 646.0438157 | 589.5571586 | 584.6814326 | 8.24E-06 | 0.779675324 | 2.24E-06 | 22.37624894 | 1.716744479 | 5.083904857 |
| ENSMUSG00000047804 | *Akap10* | 1255.014507 | 1210.117162 | 1206.444777 | 2122.83588 | 2109.594032 | 2073.927859 | 4.39E-27 | 0.779406223 | 2.68E-28 | 121.7049882 | 1.71642429 | 26.35719883 |
| ENSMUSG00000042524 | *Sun2* | 1829.39734 | 1758.151873 | 1272.202056 | 2657.550997 | 2804.468031 | 2892.481864 | 4.33E-08 | 0.779397389 | 9.28E-09 | 32.98688139 | 1.71641378 | 7.363901398 |
| ENSMUSG00000007827 | *Ankrd26* | 339.5169736 | 397.6531791 | 301.9466906 | 637.6098233 | 536.355868 | 611.7410692 | 2.60E-07 | 0.779074124 | 6.01E-08 | 29.36131267 | 1.716029227 | 6.585760497 |
| ENSMUSG00000026104 | *Stat1* | 2603.49604 | 2501.985358 | 2237.089481 | 4269.286939 | 4123.642888 | 4219.370471 | 2.83E-21 | 0.778832894 | 2.37E-22 | 94.56993207 | 1.715742316 | 20.54836702 |
| ENSMUSG00000032307 | *Ube2q2* | 941.0611645 | 850.8163197 | 649.5208811 | 1442.212696 | 1387.576517 | 1370.377308 | 6.47E-08 | 0.778616442 | 1.41E-08 | 32.17022649 | 1.715484918 | 7.188880019 |
| ENSMUSG00000028106 | *Rprd2* | 961.0327511 | 969.910419 | 739.4338956 | 1554.384794 | 1564.552239 | 1472.817361 | 5.54E-10 | 0.778329716 | 9.98E-11 | 41.82458817 | 1.71514401 | 9.256690897 |
| ENSMUSG00000006850 | *Tmco6* | 286.7919848 | 294.7074322 | 260.3451466 | 464.7129797 | 514.6410556 | 468.7115617 | 3.19E-09 | 0.777514818 | 6.14E-10 | 38.27731125 | 1.714175495 | 8.495992038 |
| ENSMUSG00000024397 | *Aif1* | 88.67384487 | 83.76957834 | 81.86110278 | 140.0042734 | 144.4035029 | 152.6936634 | 0.000291685 | 0.777130081 | 9.60E-05 | 15.21332798 | 1.71371842 | 3.53508585 |
| ENSMUSG00000004319 | *Clcn3* | 1566.571259 | 1605.751797 | 1515.101394 | 2692.973764 | 2681.77934 | 2660.542122 | 1.35E-29 | 0.776663475 | 7.34E-31 | 133.4133595 | 1.713164247 | 28.86849848 |
| ENSMUSG00000038702 | *Dsel* | 226.8772247 | 196.8080455 | 206.6657349 | 331.4559002 | 384.3521807 | 367.2379246 | 3.84E-07 | 0.775916959 | 9.02E-08 | 28.57469135 | 1.712278008 | 6.416105868 |
| ENSMUSG00000030498 | *Gas2* | 88.67384487 | 126.1590035 | 103.3328674 | 192.2950261 | 181.3186841 | 169.1227284 | 0.000544755 | 0.775214792 | 0.000186472 | 13.96271948 | 1.711444836 | 3.263799058 |
| ENSMUSG00000068966 | *Zbtb34* | 355.4942429 | 343.1524896 | 261.6871318 | 595.4398615 | 528.7556837 | 526.6964972 | 9.35E-07 | 0.775194977 | 2.28E-07 | 26.78092385 | 1.71142133 | 6.029236022 |
| ENSMUSG00000074643 | *Cpne1* | 569.5896522 | 611.6188491 | 488.4826461 | 860.267222 | 1079.22618 | 925.8261363 | 2.97E-08 | 0.775074651 | 6.26E-09 | 33.75299284 | 1.711278598 | 7.527753677 |
| ENSMUSG00000020491 | *2810021J22Rik* | 109.444295 | 102.9457469 | 100.6488969 | 206.6328131 | 172.6327591 | 156.5593258 | 0.000352105 | 0.77456123 | 0.000116911 | 14.84186366 | 1.710669702 | 3.453327226 |
| ENSMUSG00000030166 | *Rad52* | 319.5453869 | 255.3458231 | 259.0031613 | 501.8225462 | 477.7258744 | 451.3160811 | 2.28E-07 | 0.773272912 | 5.24E-08 | 29.62644605 | 1.709142767 | 6.641835676 |
| ENSMUSG00000022812 | *Gsk3b* | 1194.300884 | 1209.10789 | 899.1301453 | 1802.34417 | 1976.047935 | 1878.711909 | 3.71E-09 | 0.773267868 | 7.18E-10 | 37.97119333 | 1.709136791 | 8.431016892 |
| ENSMUSG00000013736 | *Trnt1* | 1213.473607 | 1054.689269 | 1392.980733 | 2059.580937 | 2122.622919 | 2064.263703 | 2.27E-11 | 0.772714915 | 3.66E-12 | 48.30113711 | 1.708481843 | 10.64341544 |
| ENSMUSG00000052821 | *Cysltr1* | 253.2397191 | 186.7153252 | 259.0031613 | 381.2164552 | 432.1247682 | 379.8013273 | 1.42E-05 | 0.772068684 | 3.94E-06 | 21.29388322 | 1.707716728 | 4.848892109 |
| ENSMUSG00000025981 | *Coq10b* | 359.4885603 | 316.9114168 | 348.9161758 | 599.6568577 | 599.3288242 | 551.8233025 | 2.75E-10 | 0.771276884 | 4.83E-11 | 43.24340535 | 1.706779732 | 9.560342589 |
| ENSMUSG00000021911 | *Parg* | 888.3361756 | 766.0374694 | 775.6674985 | 1392.452141 | 1328.946523 | 1433.194322 | 2.21E-14 | 0.771121252 | 2.84E-15 | 62.37236144 | 1.706595622 | 13.65508842 |
| ENSMUSG00000023940 | *Sgo1* | 439.374907 | 380.4955546 | 383.8077934 | 680.6231844 | 710.0743678 | 668.7595891 | 5.80E-11 | 0.770132841 | 9.62E-12 | 46.40356276 | 1.70542681 | 10.23679654 |
| ENSMUSG00000068823 | *Csde1* | 8281.018702 | 7914.711245 | 10772.11594 | 15674.57482 | 15457.68926 | 14842.21065 | 3.88E-10 | 0.770041666 | 6.90E-11 | 42.54711564 | 1.705319033 | 9.411422626 |
| ENSMUSG00000053012 | *Krcc1* | 884.3418583 | 802.3712624 | 968.9133805 | 1620.169935 | 1441.863548 | 1458.321127 | 1.52E-12 | 0.770030459 | 2.23E-13 | 53.79487199 | 1.705305787 | 11.8183527 |
| ENSMUSG00000022770 | *Dlg1* | 818.0361905 | 861.918312 | 801.165219 | 1513.901631 | 1377.804851 | 1338.485594 | 7.25E-16 | 0.769746221 | 8.47E-17 | 69.2978328 | 1.704969843 | 15.13995781 |
| ENSMUSG00000034021 | *Pds5b* | 895.5259468 | 902.2891932 | 825.3209543 | 1539.203608 | 1510.265207 | 1428.362244 | 3.70E-18 | 0.769398532 | 3.75E-19 | 79.99540239 | 1.704558995 | 17.43190225 |
| ENSMUSG00000034484 | *Snx2* | 2581.127863 | 2306.186584 | 2787.30345 | 4590.622048 | 4221.359544 | 4258.027095 | 3.84E-16 | 0.769135375 | 4.42E-17 | 70.58201861 | 1.704248101 | 15.41517344 |
| ENSMUSG00000028483 | *Snapc3* | 293.981756 | 243.2345588 | 307.3146318 | 463.0261812 | 497.2692056 | 477.409302 | 7.77E-08 | 0.7690907 | 1.70E-08 | 31.80508151 | 1.704195327 | 7.109822845 |
| ENSMUSG00000068686 | *Cd59b* | 102.2545238 | 67.62122589 | 84.54507337 | 166.1496497 | 116.1742467 | 151.7272478 | 0.006116285 | 0.768396766 | 0.002460014 | 9.170097619 | 1.703375809 | 2.213512255 |
| ENSMUSG00000032434 | *Cmtm6* | 1177.524751 | 1243.423139 | 1057.48441 | 1923.79366 | 2037.93515 | 1973.420637 | 7.02E-17 | 0.768327406 | 7.67E-18 | 74.03735493 | 1.703293918 | 16.15386308 |
| ENSMUSG00000029649 | *Pomp* | 2299.927922 | 2305.177312 | 2977.865362 | 4347.723068 | 4120.385666 | 4429.082654 | 8.91E-11 | 0.767889313 | 1.51E-11 | 45.52549138 | 1.70277677 | 10.04994421 |
| ENSMUSG00000039318 | *Rab3gap2* | 3441.503817 | 3504.192482 | 2772.541612 | 5486.312038 | 5507.962183 | 5567.52022 | 1.63E-13 | 0.767850765 | 2.22E-14 | 58.32380434 | 1.702731273 | 12.78908476 |
| ENSMUSG00000031601 | *Cnot7* | 1139.978168 | 993.1236757 | 1131.293601 | 1927.167257 | 1790.386288 | 1838.122454 | 2.11E-16 | 0.767720368 | 2.38E-17 | 71.80365794 | 1.702577379 | 15.67574038 |
| ENSMUSG00000022897 | *Dyrk1a* | 1318.124721 | 1261.590035 | 1077.614189 | 2170.909637 | 2059.649963 | 2007.245183 | 8.83E-14 | 0.767485918 | 1.18E-14 | 59.56445643 | 1.702300719 | 13.05421114 |
| ENSMUSG00000051510 | *Mafg* | 3800.193514 | 3996.717231 | 3199.292935 | 6148.380439 | 6453.642266 | 6126.108432 | 9.69E-15 | 0.767147868 | 1.22E-15 | 64.03920707 | 1.701901885 | 14.0137303 |
| ENSMUSG00000071637 | *Cebpd* | 1551.392853 | 1466.472257 | 1505.707497 | 2436.580396 | 2584.062684 | 2679.870434 | 1.31E-22 | 0.766659928 | 1.02E-23 | 100.8046601 | 1.701326375 | 21.88435985 |
| ENSMUSG00000062078 | *Qki* | 2079.441605 | 2171.953405 | 1654.667864 | 3570.108971 | 3347.338342 | 3139.884256 | 2.31E-10 | 0.766490971 | 4.04E-11 | 43.59604474 | 1.701127141 | 9.635679168 |
| ENSMUSG00000074151 | *Nlrc5* | 1223.059968 | 1238.376779 | 900.4721306 | 1847.887729 | 1964.104788 | 1918.334948 | 1.41E-08 | 0.766218784 | 2.90E-09 | 35.2482553 | 1.700806227 | 7.850924774 |
| ENSMUSG00000031931 | *Ankrd49* | 550.416929 | 535.9234469 | 527.4002196 | 942.9203472 | 930.4797144 | 873.6396944 | 4.29E-15 | 0.766159115 | 5.27E-16 | 65.69219733 | 1.700735885 | 14.36712017 |
| ENSMUSG00000029414 | *Kntc1* | 1120.006581 | 980.0031393 | 764.9316162 | 1566.192384 | 1677.469263 | 1640.007258 | 1.19E-07 | 0.765562045 | 2.67E-08 | 30.9301427 | 1.700032167 | 6.922782095 |
| ENSMUSG00000027035 | *Cers6* | 1891.708691 | 2024.599688 | 1814.364114 | 3239.49647 | 3395.11093 | 3111.858203 | 2.44E-21 | 0.765196775 | 2.03E-22 | 94.86977721 | 1.699601799 | 20.61261601 |
| ENSMUSG00000020015 | *Cdk17* | 657.4646336 | 658.0453623 | 649.5208811 | 1211.964704 | 1052.082664 | 1073.687722 | 7.62E-14 | 0.765110245 | 1.02E-14 | 59.86448947 | 1.699499863 | 13.11831196 |
| ENSMUSG00000037608 | *Bclaf1* | 3518.19471 | 3288.208268 | 4125.262786 | 6155.971033 | 6262.551917 | 6141.571082 | 8.04E-15 | 0.764692899 | 1.01E-15 | 64.42035184 | 1.699008299 | 14.09488619 |
| ENSMUSG00000027715 | *Ccna2* | 2640.243759 | 2073.044746 | 1771.420585 | 3779.271982 | 3622.03072 | 3626.957714 | 9.71E-08 | 0.764377813 | 2.15E-08 | 31.35235898 | 1.698637274 | 7.012988901 |
| ENSMUSG00000032423 | *Syncrip* | 4400.938841 | 4283.350487 | 4601.667565 | 7708.669028 | 7468.809749 | 7375.683791 | 1.17E-33 | 0.764069331 | 5.30E-35 | 152.3538763 | 1.698274104 | 32.93242622 |
| ENSMUSG00000039449 | *Prpf18* | 1009.763423 | 998.1700358 | 1129.951615 | 1806.561166 | 1759.985551 | 1751.145051 | 2.33E-18 | 0.764022541 | 2.33E-19 | 80.93783316 | 1.698219025 | 17.63189989 |
| ENSMUSG00000028414 | *Fktn* | 574.382833 | 648.9619141 | 727.356028 | 1119.190788 | 1162.828208 | 1020.534864 | 7.14E-10 | 0.763900269 | 1.30E-10 | 41.31109068 | 1.698075104 | 9.146441055 |
| ENSMUSG00000036432 | *Siah2* | 647.878272 | 656.0268183 | 495.1925726 | 999.4280961 | 991.2811893 | 1073.687722 | 1.50E-08 | 0.763586569 | 3.10E-09 | 35.11903413 | 1.697705914 | 7.822604806 |
| ENSMUSG00000029169 | *Dhx15* | 5401.115902 | 5265.372171 | 6197.288076 | 9625.715495 | 9514.345084 | 9467.973546 | 3.85E-20 | 0.763275915 | 3.46E-21 | 89.25941067 | 1.697340388 | 19.41452697 |

| | | | | | | | | | | | | | |
|---|---|---|---|---|---|---|---|---|---|---|---|---|---|
| ENSMUSG00000022508 | *Bcl6* | 917.8941239 | 1051.661453 | 628.0491164 | 1466.671274 | 1441.863548 | 1508.574738 | 1.92E-05 | 0.763158273 | 5.43E-06 | 20.67818371 | 1.697201987 | 4.716086316 |
| ENSMUSG00000037286 | *Stag1* | 1412.39061 | 1413.990111 | 1744.580879 | 2608.633841 | 2602.520275 | 2531.042433 | 7.01E-13 | 0.762854626 | 1.01E-13 | 55.35621791 | 1.696844811 | 12.15420266 |
| ENSMUSG00000059263 | *Usp47* | 1723.148499 | 1759.161145 | 2196.829922 | 3281.666432 | 3243.107243 | 3096.395554 | 1.17E-11 | 0.762722344 | 1.83E-12 | 49.65450284 | 1.696689233 | 10.9334032 |
| ENSMUSG00000021413 | *Prpf4b* | 1367.654256 | 1387.749039 | 1489.603674 | 2536.101506 | 2415.772887 | 2241.117756 | 2.85E-18 | 0.762626127 | 2.87E-19 | 80.5261476 | 1.69657608 | 17.54449309 |
| ENSMUSG00000027244 | *Atg13* | 1965.20413 | 1928.718846 | 1576.832718 | 3082.624212 | 3141.047624 | 3072.235164 | 1.43E-13 | 0.762588573 | 1.95E-14 | 58.57983853 | 1.696531919 | 12.84372535 |
| ENSMUSG00000032754 | *Slc8b1* | 1422.775835 | 1598.686893 | 1368.824997 | 2438.267195 | 2618.806384 | 2393.811419 | 3.67E-16 | 0.761993004 | 4.21E-17 | 70.67770269 | 1.695831706 | 15.43512006 |
| ENSMUSG00000028309 | *Rnf20* | 1194.300884 | 1249.478771 | 1296.357792 | 2193.681416 | 2070.507369 | 2068.129365 | 1.11E-21 | 0.76192245 | 9.13E-23 | 96.45514205 | 1.695748774 | 20.95389321 |
| ENSMUSG00000032384 | *Csnk1g1* | 774.8975633 | 765.0281973 | 692.4644104 | 1260.88186 | 1337.632448 | 1194.489671 | 1.89E-14 | 0.761432872 | 2.42E-15 | 62.69347035 | 1.69517342 | 13.72369832 |
| ENSMUSG00000051235 | *Gen1* | 360.2874237 | 366.3657462 | 352.9421317 | 658.6948043 | 584.1284555 | 586.6142638 | 1.65E-10 | 0.761432238 | 2.83E-11 | 44.28803787 | 1.695172676 | 9.783007953 |
| ENSMUSG00000045362 | *Tnfrsf26* | 796.4668769 | 896.233561 | 897.78816 | 1525.70922 | 1457.063917 | 1398.40336 | 2.58E-15 | 0.760981718 | 3.13E-16 | 66.72071617 | 1.694643395 | 14.58785271 |
| ENSMUSG00000033777 | *Tlr13* | 1345.286079 | 1269.664211 | 1717.741173 | 2391.036838 | 2526.518431 | 2412.173315 | 1.87E-09 | 0.760770768 | 3.53E-10 | 39.35815013 | 1.694395623 | 8.728226029 |
| ENSMUSG00000044864 | *Ankrd50* | 687.8214453 | 785.2136379 | 707.2262486 | 1240.640278 | 1231.229867 | 1220.582892 | 1.36E-15 | 0.760505485 | 1.61E-16 | 68.02441017 | 1.694084087 | 14.86605062 |
| ENSMUSG00000045767 | *B230219D22Rik* | 623.1135045 | 565.1923358 | 528.7422049 | 1031.477267 | 966.309155 | 918.0948116 | 1.46E-11 | 0.76037362 | 2.32E-12 | 49.19414074 | 1.693929251 | 10.8357053 |
| ENSMUSG00000021109 | *Hif1a* | 2265.576793 | 2099.285819 | 2500.118598 | 3965.663213 | 3815.292551 | 3835.703481 | 3.42E-18 | 0.760277653 | 3.45E-19 | 80.15931937 | 1.693816576 | 17.46641189 |
| ENSMUSG00000050737 | *Ptges* | 109.444295 | 110.0106511 | 76.49316162 | 158.5590566 | 157.4323904 | 189.4174559 | 0.001561512 | 0.759994695 | 0.000571006 | 11.86819815 | 1.693484397 | 2.806454566 |
| ENSMUSG00000014444 | *Piezo1* | 7580.415441 | 7363.648717 | 5335.733519 | 11071.30179 | 11931.20371 | 11329.28997 | 1.93E-08 | 0.75889895 | 4.00E-09 | 34.62183506 | 1.692198664 | 7.714673352 |
| ENSMUSG00000039191 | *Rbpj* | 10910.87724 | 10788.10871 | 8611.519616 | 17621.14026 | 17090.64316 | 16586.59079 | 5.16E-13 | 0.758684004 | 7.34E-14 | 55.97427151 | 1.691946562 | 12.28751361 |
| ENSMUSG00000032860 | *P2ry2* | 551.2157924 | 533.9049029 | 483.1147049 | 858.5804235 | 874.021202 | 926.7925519 | 1.01E-12 | 0.757666513 | 1.47E-13 | 54.61424337 | 1.690753702 | 11.99452099 |
| ENSMUSG00000024539 | *Ptpn2* | 624.7112314 | 617.6744812 | 730.0399986 | 1126.781381 | 1046.653961 | 1151.000969 | 6.47E-12 | 0.757632376 | 9.97E-13 | 50.84961362 | 1.690713696 | 11.18933726 |
| ENSMUSG00000074166 | *AW146154* | 120.6283836 | 136.2517238 | 162.3802203 | 235.3083872 | 211.7194216 | 256.1001316 | 0.000157797 | 0.757543167 | 5.03E-05 | 16.43716978 | 1.690609154 | 3.801901439 |
| ENSMUSG00000034157 | *Cipc* | 1042.516825 | 1073.865438 | 924.6278659 | 1723.064642 | 1777.357401 | 1650.63783 | 8.90E-16 | 0.75752017 | 1.04E-16 | 68.88335083 | 1.690582205 | 15.05043793 |
| ENSMUSG00000056673 | *Kdm5d* | 567.1930618 | 529.8678148 | 625.3651458 | 953.8845373 | 1039.053777 | 912.296318 | 3.39E-11 | 0.756454358 | 5.53E-12 | 47.49083644 | 1.689333725 | 10.46978376 |
| ENSMUSG00000030539 | *Sema4b* | 337.9192467 | 317.9206889 | 287.1848524 | 533.0283179 | 555.8991993 | 510.2674321 | 3.47E-09 | 0.755759854 | 6.70E-10 | 38.10551651 | 1.688520686 | 8.459192098 |
| ENSMUSG00000103588 | *Gm18445* | 117.4329297 | 88.81593848 | 143.5924262 | 190.6082276 | 170.4612779 | 226.1412483 | 0.003029267 | 0.755246281 | 0.001156356 | 10.55880996 | 1.687919711 | 2.51866244 |
| ENSMUSG00000078853 | *Igtp* | 885.1407218 | 836.6865113 | 920.60191 | 1533.299813 | 1488.550395 | 1431.26149 | 6.80E-18 | 0.754516975 | 6.98E-19 | 78.76939188 | 1.687066656 | 17.1671772 |
| ENSMUSG00000002052 | *Supt6* | 3459.078813 | 3556.674627 | 2816.827127 | 5594.267141 | 5552.477549 | 5452.516765 | 2.41E-13 | 0.754295984 | 3.35E-14 | 57.51763444 | 1.686808251 | 12.6179786 |
| ENSMUSG00000049969 | *Plekhf2* | 775.6964267 | 739.7963966 | 911.2080129 | 1366.306764 | 1377.804851 | 1338.485594 | 2.30E-12 | 0.754145441 | 3.42E-13 | 52.94895324 | 1.686632245 | 11.63897192 |
| ENSMUSG00000030451 | *Herc2* | 1651.250787 | 1744.022065 | 1108.479851 | 2477.06356 | 2636.178234 | 2488.520147 | 4.74E-06 | 0.753147224 | 1.25E-06 | 23.49119249 | 1.685465649 | 5.324051221 |
| ENSMUSG00000021133 | *Susd6* | 1999.555259 | 2163.879228 | 1702.979335 | 3295.16082 | 3369.053155 | 3233.626568 | 3.82E-13 | 0.752993404 | 5.39E-14 | 56.58272669 | 1.685285954 | 12.41774767 |
| ENSMUSG00000022253 | *Nadk2* | 242.0556306 | 178.641149 | 197.2718379 | 393.8674438 | 335.4938527 | 314.0850671 | 6.80E-05 | 0.751930954 | 2.07E-05 | 18.12802483 | 1.684045308 | 4.167535792 |
| ENSMUSG00000023027 | *Atf1* | 1101.632721 | 1190.940993 | 1142.029483 | 1908.612474 | 1928.275347 | 1941.528923 | 1.78E-22 | 0.751204454 | 1.41E-23 | 100.1580039 | 1.683197485 | 21.74846033 |
| ENSMUSG00000038495 | *Otud7b* | 1668.02692 | 1806.59693 | 1539.257129 | 2951.89733 | 2812.068215 | 2680.83685 | 2.86E-16 | 0.75108042 | 3.25E-17 | 71.18598827 | 1.68305278 | 15.54408934 |
| ENSMUSG00000031600 | *Vps37a* | 432.1851358 | 469.3114931 | 362.3360287 | 710.1421578 | 707.9028866 | 716.113953 | 5.79E-09 | 0.750950326 | 1.14E-09 | 37.07074857 | 1.682901019 | 8.237610476 |
| ENSMUSG00000003541 | *Ier3* | 2058.671155 | 2302.149496 | 2200.855878 | 3721.077435 | 3511.285177 | 3799.946104 | 3.84E-21 | 0.750550982 | 3.24E-22 | 93.94797489 | 1.68243525 | 20.41526517 |
| ENSMUSG00000016409 | *Nkap* | 399.4317336 | 366.3657462 | 383.8077934 | 622.4286371 | 613.4434523 | 698.7184724 | 4.50E-10 | 0.749882014 | 8.06E-11 | 42.24393732 | 1.681655297 | 9.346543162 |
| ENSMUSG00000071072 | *Ptges3* | 3677.967403 | 3617.230949 | 3427.430435 | 6236.937359 | 5870.599552 | 5930.892483 | 6.01E-29 | 0.749773791 | 3.37E-30 | 130.3908886 | 1.681529153 | 28.22122468 |
| ENSMUSG00000032555 | *Topbp1* | 2647.433531 | 2703.839763 | 2698.732421 | 4419.412003 | 4659.998756 | 4455.175875 | 1.91E-32 | 0.749633166 | 9.18E-34 | 146.688237 | 1.681365256 | 31.71872382 |
| ENSMUSG00000020115 | *Tbk1* | 1121.604308 | 1185.894633 | 1317.829556 | 2065.484732 | 1968.447751 | 2046.868222 | 3.38E-16 | 0.74944775 | 3.86E-17 | 70.84537256 | 1.681149179 | 15.47068546 |
| ENSMUSG00000025234 | *Arih1* | 2614.680128 | 2674.570875 | 2470.594922 | 4574.597463 | 4361.420085 | 4112.09834 | 3.13E-22 | 0.749152953 | 2.51E-23 | 99.01423791 | 1.680805692 | 21.50410168 |
| ENSMUSG00000032217 | *Rnf111* | 732.5577995 | 755.9447491 | 505.9284549 | 1130.154978 | 1149.79932 | 1080.452631 | 3.41E-06 | 0.748364932 | 8.87E-07 | 24.15991998 | 1.679887863 | 5.467864091 |
| ENSMUSG00000011960 | *Ccnt1* | 1371.648573 | 1411.971567 | 1332.591394 | 2278.864739 | 2406.001222 | 2233.386431 | 1.73E-22 | 0.747970081 | 1.36E-23 | 100.2182474 | 1.679428157 | 21.76134495 |
| ENSMUSG00000020888 | *Dvl2* | 1254.215644 | 1205.070802 | 1040.038601 | 1928.010656 | 1916.332201 | 2044.935391 | 6.66E-14 | 0.747963181 | 8.85E-15 | 60.13687895 | 1.679420124 | 13.17629792 |
| ENSMUSG00000032624 | *Eml4* | 1239.836101 | 1172.774097 | 994.411101 | 2003.073188 | 1881.588501 | 1845.853779 | 4.84E-12 | 0.747231085 | 7.38E-13 | 51.44109126 | 1.678568119 | 11.31551761 |
| ENSMUSG00000027778 | *Ift80* | 198.9170034 | 167.5391567 | 208.0077202 | 330.612501 | 320.293484 | 311.1858203 | 2.79E-06 | 0.746894216 | 7.17E-07 | 24.56928043 | 1.678176219 | 5.555039423 |
| ENSMUSG00000025235 | *Bbs4* | 174.1522359 | 192.7709574 | 222.7695584 | 335.6728964 | 331.1508902 | 316.0178983 | 3.97E-06 | 0.746399708 | 1.04E-06 | 23.85361306 | 1.677601095 | 5.401556768 |
| ENSMUSG00000034343 | *Ube2f* | 1568.168986 | 1457.388809 | 1576.832718 | 2593.452654 | 2540.633059 | 2583.228875 | 3.20E-24 | 0.746170797 | 2.26E-25 | 108.3457748 | 1.677334932 | 23.49545007 |
| ENSMUSG00000066491 | *Cox6c2* | 627.1078218 | 540.9698071 | 801.165219 | 1046.658453 | 1104.198214 | 1143.269644 | 1.50E-06 | 0.745862889 | 3.73E-07 | 25.82698899 | 1.676976984 | 5.825115056 |
| ENSMUSG00000027199 | *Gatm* | 1338.096308 | 1261.590035 | 1468.131909 | 2273.804344 | 2301.770122 | 2236.285678 | 7.15E-18 | 0.745730735 | 7.35E-19 | 78.66777737 | 1.676823377 | 17.14558729 |
| ENSMUSG00000018217 | *Pmp22* | 3897.654857 | 3909.919837 | 3572.364846 | 6410.677602 | 6351.582648 | 6318.425135 | 1.74E-28 | 0.744648314 | 9.90E-30 | 128.2483665 | 1.675565767 | 27.76014873 |
| ENSMUSG00000037913 | *Tmem156* | 227.6760882 | 265.4385434 | 358.3100728 | 462.182782 | 467.9542088 | 487.0734579 | 8.84E-05 | 0.74438262 | 2.72E-05 | 17.60245863 | 1.675257215 | 4.053625544 |
| ENSMUSG00000037447 | *Arid5a* | 278.8033501 | 286.633256 | 197.2718379 | 434.3506072 | 424.5245838 | 426.1892757 | 2.09E-05 | 0.744287115 | 5.94E-06 | 20.50840551 | 1.675146318 | 4.679923373 |
| ENSMUSG00000060068 | *Gm12222* | 137.4045164 | 110.0106511 | 139.5664703 | 237.8385849 | 178.0614623 | 230.0069107 | 0.000681478 | 0.743054844 | 0.000236732 | 13.51446484 | 1.673716111 | 3.166548023 |
| ENSMUSG00000020680 | *Taf15* | 2395.791538 | 2297.103136 | 1896.225217 | 3780.115381 | 3636.145348 | 3623.092051 | 1.18E-12 | 0.742679832 | 1.71E-13 | 54.31519192 | 1.673281104 | 11.92931469 |
| ENSMUSG00000014353 | *Tmem87b* | 460.1453572 | 546.0161672 | 634.7590429 | 938.703351 | 866.4210176 | 928.7253831 | 1.43E-07 | 0.742560858 | 3.23E-08 | 30.56408839 | 1.67314312 | 6.844275627 |
| ENSMUSG00000032743 | *Katnip* | 443.3692243 | 451.1445966 | 426.7513227 | 676.4061883 | 775.2188052 | 762.5019014 | 8.26E-11 | 0.742177902 | 1.39E-11 | 45.67695846 | 1.672699051 | 10.08276865 |
| ENSMUSG00000058446 | *Znrf2* | 1387.625843 | 1336.276165 | 1108.479851 | 2174.283234 | 2160.623841 | 2086.491261 | 2.86E-12 | 0.741664223 | 4.30E-13 | 52.50326791 | 1.672103584 | 11.54401647 |
| ENSMUSG00000041552 | *Ptchd1* | 365.0806046 | 348.1988497 | 280.4749259 | 518.6905309 | 570.0138274 | 581.7821859 | 4.08E-07 | 0.7415398 | 9.62E-08 | 28.4491651 | 1.671959382 | 6.389039072 |
| ENSMUSG00000019777 | *Hdac2* | 1436.356514 | 1307.007276 | 1556.702938 | 2381.759446 | 2368.0003 | 2431.501627 | 2.54E-16 | 0.74152025 | 2.88E-17 | 71.42178381 | 1.671936726 | 15.59441125 |
| ENSMUSG00000049130 | *C5ar1* | 1588.939436 | 1550.241835 | 1214.496689 | 2430.676602 | 2424.458812 | 2435.36729 | 2.40E-10 | 0.741140711 | 4.19E-11 | 43.52126408 | 1.671496937 | 9.619817365 |
| ENSMUSG00000030929 | *Eri2* | 359.4885603 | 285.623984 | 426.7513227 | 598.8134584 | 591.7286398 | 595.3120041 | 7.15E-06 | 0.741130518 | 1.93E-06 | 22.66046982 | 1.671485127 | 5.145513802 |
| ENSMUSG00000033088 | *Triobp* | 635.0964565 | 592.4426805 | 424.0673521 | 901.5937846 | 909.8506425 | 959.650682 | 4.45E-06 | 0.741071021 | 1.18E-06 | 23.61730343 | 1.671416196 | 5.351449252 |
| ENSMUSG00000029782 | *Tmem209* | 1290.1645 | 1255.534403 | 1188.998968 | 2092.473508 | 2111.765513 | 2045.901806 | 3.46E-22 | 0.740773241 | 2.77E-23 | 98.81415637 | 1.671071242 | 21.46150296 |
| ENSMUSG00000034083 | *Ccdc174* | 272.4124423 | 314.8928728 | 406.6215433 | 522.9075271 | 570.0138274 | 557.6217961 | 2.00E-05 | 0.739546954 | 5.66E-06 | 20.59917191 | 1.66965144 | 4.699431744 |
| ENSMUSG00000042719 | *Naa25* | 2161.724542 | 2123.508347 | 2250.509334 | 3823.972142 | 3585.115539 | 3494.558778 | 1.53E-23 | 0.739395925 | 1.12E-24 | 105.1636706 | 1.669476661 | 22.81483402 |
| ENSMUSG00000034300 | *Fam53c* | 1437.155378 | 1582.53854 | 1242.67838 | 2333.685689 | 2465.716956 | 2325.195912 | 6.21E-12 | 0.739106836 | 9.56E-13 | 50.93328968 | 1.669142163 | 11.20701967 |
| ENSMUSG00000031922 | *Cep57* | 553.6123828 | 506.6545581 | 570.3437489 | 915.0881724 | 907.6791613 | 895.867253 | 1.62E-13 | 0.738938675 | 2.21E-14 | 58.33187263 | 1.668947619 | 12.79067845 |
| ENSMUSG00000056600 | *Or2h2* | 74.29430246 | 71.658314 | 76.49316162 | 133.2570795 | 128.1173936 | 109.2049618 | 0.002221166 | 0.738564157 | 0.000830184 | 11.17250535 | 1.668514423 | 2.653418947 |
| ENSMUSG00000042082 | *Arsb* | 2180.098402 | 2150.758692 | 1533.889188 | 3282.509831 | 3321.280567 | 3194.003529 | 1.24E-07 | 0.738532731 | 2.77E-08 | 30.85915863 | 1.668478078 | 6.907583487 |
| ENSMUSG00000020415 | *Pttg1* | 954.6418434 | 810.4454386 | 728.6980133 | 1432.935304 | 1382.147814 | 1356.84749 | 1.07E-09 | 0.73835902 | 1.98E-10 | 40.48462591 | 1.668277193 | 8.969492197 |
| ENSMUSG00000035517 | *Tdrd7* | 1211.87588 | 1185.894633 | 1241.336395 | 2138.017066 | 1933.704051 | 1990.816118 | 2.59E-19 | 0.737642623 | 2.44E-20 | 85.39825565 | 1.667448984 | 18.58674157 |
| ENSMUSG00000029502 | *Golga3* | 1541.007628 | 1478.583521 | 1313.8036 | 2411.278419 | 2449.430847 | 2376.415939 | 7.07E-17 | 0.73716786 | 7.74E-18 | 74.01932107 | 1.66690035 | 16.15036453 |
| ENSMUSG00000024077 | *Strn* | 683.827128 | 704.4718756 | 680.3865428 | 1163.047548 | 1119.398583 | 1164.530787 | 2.45E-17 | 0.736727334 | 2.61E-18 | 76.1656184 | 1.66639144 | 16.61059902 |
| ENSMUSG00000028329 | *Xpa* | 138.2033798 | 87.80666645 | 159.6962497 | 221.8139994 | 208.4621997 | 209.7121833 | 0.004182577 | 0.736516488 | 0.001634868 | 9.920062525 | 1.666147919 | 2.378556045 |
| ENSMUSG00000037926 | *Ssh2* | 1382.033798 | 1555.288195 | 1227.916542 | 2402.844427 | 2256.169016 | 2282.673626 | 5.30E-12 | 0.735466126 | 8.11E-13 | 51.25584411 | 1.664935313 | 11.27571067 |
| ENSMUSG00000083798 | *Gm14584* | 141.3988337 | 165.5206126 | 208.0077202 | 259.7669651 | 280.1210809 | 311.1858203 | 0.000315827 | 0.735388184 | 0.000104381 | 15.05576386 | 1.664845367 | 3.500550576 |
| ENSMUSG00000018412 | *Kansl1* | 1034.52819 | 1157.635016 | 1048.090513 | 1784.632786 | 1833.815913 | 1776.271856 | 7.95E-18 | 0.735250084 | 8.19E-19 | 78.4540058 | 1.66468601 | 17.09983614 |
| ENSMUSG00000027770 | *Dhx36* | 864.3702716 | 805.3990785 | 877.6583807 | 1464.141076 | 1337.632448 | 1434.160737 | 7.09E-16 | 0.734969665 | 8.29E-17 | 69.34011959 | 1.664362473 | 15.14904977 |
| ENSMUSG00000015501 | *Hivep2* | 695.0112165 | 729.7036763 | 709.9102192 | 1254.978065 | 1184.54302 | 1110.411514 | 2.34E-14 | 0.734783908 | 3.01E-15 | 62.25996389 | 1.664148188 | 13.63129793 |
| ENSMUSG00000020719 | *Ddx5* | 15553.87171 | 14658.66694 | 13288.33836 | 24656.7767 | 23776.63392 | 23936.18136 | 6.01E-21 | 0.734016513 | 5.11E-22 | 93.04387293 | 1.663263234 | 20.22142291 |
| ENSMUSG00000003119 | *Cdk12* | 1227.054286 | 1229.29333 | 1061.510366 | 1991.265599 | 1953.247382 | 1914.469286 | 1.13E-15 | 0.733219134 | 1.33E-16 | 68.40551559 | 1.662344201 | 14.9480376 |
| ENSMUSG00000055436 | *Srsf11* | 2568.346047 | 2608.968193 | 2435.703304 | 4219.526384 | 4374.448972 | 4066.676807 | 2.62E-25 | 0.732811718 | 1.76E-26 | 113.400288 | 1.661874823 | 24.58194515 |
| ENSMUSG00000084230 | *Gm14388* | 112.6397489 | 82.76030631 | 126.1466174 | 160.2458551 | 180.2329435 | 192.3167026 | 0.002238528 | 0.732737737 | 0.000837334 | 11.15659637 | 1.661789604 | 2.650037535 |
| ENSMUSG00000034973 | *Dop1a* | 576.7794234 | 510.6916462 | 505.9284549 | 871.2314121 | 925.0510113 | 859.1434605 | 1.41E-11 | 0.73257512 | 2.24E-12 | 49.26032539 | 1.661602302 | 10.85003536 |
| ENSMUSG00000029068 | *Ccnl2* | 1238.238374 | 1294.896012 | 1091.034042 | 2007.290185 | 1993.419785 | 2028.506326 | 2.01E-15 | 0.731895041 | 2.41E-16 | 67.23703662 | 1.660819216 | 14.69786035 |
| ENSMUSG00000004843 | *Chmp2b* | 1052.90205 | 972.9382351 | 980.9912481 | 1676.677684 | 1593.867235 | 1726.018246 | 1.68E-17 | 0.731403068 | 1.76E-18 | 76.94292176 | 1.660252957 | 16.77571921 |
| ENSMUSG00000025437 | *Usp33* | 914.69867 | 958.8084267 | 1060.16838 | 1673.304087 | 1571.066682 | 1612.947622 | 5.87E-15 | 0.731164585 | 7.27E-16 | 65.05788203 | 1.659978533 | 14.23169615 |
| ENSMUSG00000038047 | *Haus6* | 657.4646336 | 680.249347 | 650.8628664 | 1079.551024 | 1135.684692 | 1085.284709 | 3.33E-16 | 0.729951319 | 3.80E-17 | 70.8793089 | 1.658583125 | 15.47770677 |
| ENSMUSG00000093752 | *Gm20716* | 699.8043973 | 662.0824505 | 569.0017636 | 850.1464311 | 1302.888748 | 1056.292241 | 3.56E-05 | 0.72969137 | 1.04E-05 | 19.4288704 | 1.658284304 | 4.447958647 |
| ENSMUSG00000042606 | *Hirip3* | 851.5884561 | 845.7699596 | 535.4521313 | 1238.953479 | 1259.459123 | 1212.851567 | 4.74E-05 | 0.729154815 | 1.41E-05 | 18.85661468 | 1.657667683 | 4.324092311 |
| ENSMUSG00000040565 | *Btaf1* | 2086.631377 | 1933.765206 | 2383.365878 | 3554.927785 | 3480.884439 | 3562.207869 | 3.04E-14 | 0.728408212 | 3.95E-15 | 61.72524144 | 1.656810053 | 13.51693538 |
| ENSMUSG00000040520 | *Manea* | 353.0976525 | 362.3286581 | 281.8169112 | 577.7284775 | 565.6708649 | 515.0995101 | 5.43E-07 | 0.727860718 | 1.29E-07 | 27.87930177 | 1.656181423 | 6.265390918 |
| ENSMUSG00000040760 | *Appl1* | 875.5543601 | 768.0560134 | 912.5499982 | 1436.308901 | 1460.321139 | 1332.6871 | 2.85E-12 | 0.72760781 | 4.29E-13 | 52.50633037 | 1.655891116 | 11.54448973 |
| ENSMUSG00000039456 | *Morc3* | 1409.99402 | 1315.081453 | 1484.235732 | 2357.300868 | 2312.627528 | 2293.304198 | 1.39E-19 | 0.727604905 | 1.29E-20 | 86.66377921 | 1.655887781 | 18.85698372 |
| ENSMUSG00000041731 | *Pgm5* | 100.6567969 | 76.70467414 | 88.57102924 | 141.6910719 | 134.6318373 | 165.2570661 | 0.002283293 | 0.727576238 | 0.000854753 | 11.11840486 | 1.655854878 | 2.641438305 |
| ENSMUSG00000027427 | *Polr3f* | 636.6941834 | 681.258619 | 672.334631 | 1078.707624 | 1102.026733 | 1111.37793 | 5.92E-16 | 0.727473896 | 6.88E-17 | 69.70664613 | 1.655737419 | 15.22777444 |
| ENSMUSG00000021495 | *Fam193b* | 456.9499033 | 462.2465889 | 344.8902199 | 642.6702188 | 706.8171459 | 751.8713299 | 1.94E-06 | 0.727091715 | 4.91E-07 | 25.29976934 | 1.655298859 | 5.712464371 |
| ENSMUSG00000065954 | *Tacc1* | 1932.450727 | 1969.089727 | 1582.200659 | 3076.720417 | 2992.301159 | 3018.115891 | 1.81E-12 | 0.726693249 | 2.67E-13 | 53.43746941 | 1.654841736 | 11.74300586 |
| ENSMUSG00000016487 | *Ppfibp1* | 1823.006432 | 1972.117543 | 1859.991614 | 3265.641847 | 3008.587268 | 3079.966489 | 3.21E-21 | 0.726634502 | 2.70E-22 | 94.30841416 | 1.654774352 | 20.49282496 |
| ENSMUSG00000028184 | *Adgrl2* | 991.3895629 | 1001.197852 | 786.4033808 | 1551.011197 | 1550.43761 | 1506.641907 | 4.49E-10 | 0.726004944 | 8.02E-11 | 42.25191105 | 1.654052406 | 9.348170613 |
| ENSMUSG00000001767 | *Crnkl1* | 838.8066407 | 826.593791 | 1176.921101 | 1594.024558 | 1553.694832 | 1541.432868 | 1.39E-06 | 0.725914586 | 3.47E-07 | 25.96919228 | 1.653948813 | 5.85587458 |
| ENSMUSG00000045639 | *Zfp629* | 408.2192318 | 397.6531791 | 275.1069848 | 626.6456333 | 591.7286398 | 577.9165235 | 2.73E-05 | 0.725892341 | 7.89E-06 | 19.96359725 | 1.653923311 | 4.563234782 |
| ENSMUSG00000112508 | *None* | 4312.264996 | 4284.359759 | 5181.405211 | 7873.975279 | 7528.525484 | 7367.952467 | 5.06E-14 | 0.725778348 | 6.67E-15 | 60.6917345 | 1.653792633 | 13.29619994 |
| ENSMUSG00000021520 | *Uqcrb* | 989.791836 | 1046.615093 | 1376.876909 | 1867.285911 | 1896.788869 | 1866.148506 | 1.04E-07 | 0.725340092 | 2.30E-08 | 31.22333125 | 1.653290327 | 6.985048948 |
| ENSMUSG00000030177 | *Ccdc77* | 310.7578888 | 248.2809189 | 296.5787494 | 457.9657858 | 497.2692056 | 460.980237 | 5.10E-07 | 0.725289126 | 1.21E-07 | 28.00235594 | 1.653231922 | 6.292038745 |
| ENSMUSG00000000791 | *Il12rb1* | 123.8238374 | 119.0940993 | 97.96492628 | 171.2100452 | 194.3475716 | 201.9808585 | 0.000461697 | 0.724624365 | 0.000156231 | 14.29558733 | 1.652470326 | 3.335642951 |
| ENSMUSG00000042423 | *Fbrs* | 1323.716765 | 1238.376779 | 861.5545572 | 1866.442512 | 1898.960351 | 1900.939468 | 5.02E-06 | 0.723882931 | 1.33E-06 | 23.37683909 | 1.651621302 | 5.299203064 |
| ENSMUSG00000033054 | *Npat* | 575.9805599 | 591.4334085 | 564.9758077 | 1011.235685 | 939.1656394 | 910.3634868 | 3.79E-13 | 0.723662461 | 5.34E-14 | 56.6013565 | 1.651368924 | 12.42138458 |
| ENSMUSG00000060090 | *Rp2* | 472.9271726 | 342.1432175 | 462.9849256 | 744.7215265 | 670.9877054 | 692.9199788 | 3.63E-06 | 0.723191377 | 9.47E-07 | 24.03206303 | 1.650829789 | 5.440342133 |
| ENSMUSG00000014956 | *Ppp1cb* | 2370.227907 | 2310.223672 | 2654.446907 | 4229.647175 | 3964.039017 | 3900.453326 | 3.14E-17 | 0.723008767 | 3.37E-18 | 75.66172017 | 1.650620848 | 16.50313242 |
| ENSMUSG00000035365 | *Parpbp* | 307.5624349 | 266.4478154 | 257.661176 | 460.4959835 | 467.9542088 | 449.3832499 | 1.11E-07 | 0.721905922 | 2.48E-08 | 31.07718537 | 1.649359539 | 6.95384881 |
| ENSMUSG00000022014 | *Epsti1* | 1198.295201 | 1010.2813 | 938.0477188 | 1645.471912 | 1711.127223 | 1842.954532 | 1.24E-09 | 0.721300292 | 2.31E-10 | 40.18877715 | 1.648667298 | 8.907182457 |
| ENSMUSG00000027901 | *Dennd2d* | 1012.160013 | 954.7713386 | 879.0003659 | 1691.015471 | 1515.69391 | 1491.179257 | 1.79E-12 | 0.721059888 | 2.64E-13 | 53.45709052 | 1.648392596 | 11.74699534 |
| ENSMUSG00000009470 | *Tnpo1* | 6556.272476 | 6550.175463 | 7033.344913 | 11326.85176 | 10958.38011 | 10896.33579 | 1.80E-30 | 0.720940831 | 9.36E-32 | 137.503685 | 1.648256569 | 29.74482584 |
| ENSMUSG00000022788 | *Fgd4* | 518.4623903 | 418.8478917 | 597.1834547 | 892.316393 | 811.0482458 | 818.5540057 | 1.77E-06 | 0.720435837 | 4.45E-07 | 25.4883135 | 1.647679722 | 5.752575934 |
| ENSMUSG00000034247 | *Plekhm1* | 2613.881265 | 2957.167042 | 2380.681907 | 4308.083304 | 4535.138585 | 4264.792004 | 2.33E-13 | 0.720154686 | 3.23E-14 | 57.58807604 | 1.647358655 | 12.63316307 |
| ENSMUSG00000041649 | *Klf8* | 163.7670108 | 169.5577007 | 154.3283085 | 302.7803262 | 254.0633059 | 246.4359757 | 4.64E-05 | 0.719823424 | 1.38E-05 | 18.90172847 | 1.646980443 | 4.33375917 |

| | | | | | | | | | | | | | |
|---|---|---|---|---|---|---|---|---|---|---|---|---|---|
| ENSMUSG00000089791 | *Gm3555* | 72.69657552 | 95.88084267 | 89.91301453 | 144.2212696 | 128.1173936 | 150.7608322 | 0.002660782 | 0.719785085 | 0.001005331 | 10.81772212 | 1.646936676 | 2.574990643 |
| ENSMUSG00000021111 | *Papola* | 4953.752361 | 4837.440831 | 5503.481681 | 8556.285262 | 8456.833717 | 8162.346083 | 3.14E-21 | 0.719776809 | 2.64E-22 | 94.35480239 | 1.646927228 | 20.50269737 |
| ENSMUSG00000031583 | *Wrn* | 317.94766 | 278.5590798 | 311.3405876 | 509.4131393 | 466.8684681 | 517.9987568 | 3.91E-08 | 0.719522585 | 8.34E-09 | 33.19383477 | 1.646637041 | 7.40809057 |
| ENSMUSG00000028709 | *Mob3c* | 163.7670108 | 160.4742525 | 114.0687498 | 240.3687826 | 223.6625685 | 262.8650408 | 0.000561035 | 0.719449741 | 0.000192343 | 13.90444911 | 1.646553902 | 3.251009885 |
| ENSMUSG00000030342 | *Cd9* | 6144.058927 | 5891.120828 | 6503.260723 | 10359.47283 | 9896.525784 | 10248.83734 | 8.38E-27 | 0.71906602 | 5.22E-28 | 120.3819801 | 1.646116017 | 26.07672766 |
| ENSMUSG00000021391 | *Cenpp* | 161.3704204 | 133.2239077 | 139.5664703 | 209.1630108 | 251.8918247 | 257.0665472 | 0.000199473 | 0.719025717 | 6.44E-05 | 15.9691371 | 1.646070033 | 3.700116773 |
| ENSMUSG00000036568 | *Bicral* | 407.4203683 | 390.5882749 | 268.3970583 | 596.2832607 | 572.1853086 | 595.3120041 | 4.53E-05 | 0.718778122 | 1.34E-05 | 18.94993312 | 1.645787559 | 4.344214189 |
| ENSMUSG00000003038 | *Hmgn2* | 3586.098105 | 3165.07708 | 3109.37992 | 5395.224921 | 5304.928687 | 5540.460584 | 2.01E-18 | 0.718726789 | 1.99E-19 | 81.25307267 | 1.645729 | 17.6978261 |
| ENSMUSG00000046185 | *Zfp84* | 284.3953944 | 238.1881986 | 220.0855878 | 430.133611 | 399.5525495 | 398.1632235 | 3.80E-06 | 0.718089019 | 9.94E-07 | 23.93892848 | 1.645001636 | 5.420314337 |
| ENSMUSG00000070697 | *Utp3* | 1051.304323 | 1014.318388 | 1117.873748 | 1756.800611 | 1741.52796 | 1728.917493 | 6.59E-19 | 0.717365481 | 6.33E-20 | 83.51275131 | 1.644176845 | 18.18119147 |
| ENSMUSG00000051329 | *Nup160* | 1628.88261 | 1423.07356 | 1180.947057 | 2399.47083 | 2343.028266 | 2227.587938 | 1.84E-08 | 0.716856678 | 3.82E-09 | 34.71127008 | 1.643597086 | 7.734378646 |
| ENSMUSG00000036769 | *Wdr44* | 131.8124721 | 161.4835245 | 148.9603674 | 232.7781895 | 239.9486778 | 252.2344693 | 5.52E-05 | 0.716803394 | 1.66E-05 | 18.54968388 | 1.643536383 | 4.257707478 |
| ENSMUSG00000041777 | *Cir1* | 131.8124721 | 169.5577007 | 241.5573525 | 304.4671246 | 293.1499684 | 287.0254306 | 0.003462854 | 0.71672114 | 0.001335357 | 10.29300158 | 1.643442682 | 2.460565817 |
| ENSMUSG00000033589 | *Reep4* | 704.5975781 | 609.600305 | 467.0108814 | 1009.548887 | 986.9382268 | 940.3223702 | 6.84E-06 | 0.716143804 | 1.84E-06 | 22.75050524 | 1.642785142 | 5.165008325 |
| ENSMUSG00000021832 | *Psmc6* | 1110.42022 | 897.242833 | 1370.166983 | 1851.261326 | 1780.614623 | 1906.737961 | 3.18E-06 | 0.715586741 | 8.25E-07 | 24.29926969 | 1.642150941 | 5.497645233 |
| ENSMUSG00000027787 | *Nmd3* | 1468.311053 | 1382.702679 | 1846.571761 | 2489.714548 | 2530.861394 | 2680.83685 | 7.40E-09 | 0.715351931 | 1.47E-09 | 36.56911634 | 1.641883689 | 8.131042765 |
| ENSMUSG00000028121 | *Bcar3* | 3900.051447 | 4350.971713 | 3681.065655 | 6506.825115 | 6604.560213 | 6469.185967 | 7.80E-18 | 0.714025405 | 8.03E-19 | 78.4929199 | 1.640374708 | 17.10764448 |
| ENSMUSG00000048550 | *Thnsl1* | 142.1976972 | 161.4835245 | 158.3542644 | 247.1159765 | 245.3773809 | 262.8650408 | 2.11E-05 | 0.713483274 | 5.99E-06 | 20.4903522 | 1.63975841 | 4.676007554 |
| ENSMUSG00000027304 | *Rtf1* | 909.1066258 | 843.7514155 | 815.9270572 | 1472.575068 | 1373.461889 | 1372.310139 | 1.03E-14 | 0.713454401 | 1.30E-15 | 63.91319231 | 1.639725593 | 13.98717118 |
| ENSMUSG00000042446 | *Zmym4* | 626.3089584 | 581.3406882 | 628.0491164 | 1028.947069 | 994.5384112 | 983.8110718 | 4.72E-14 | 0.712855143 | 6.22E-15 | 60.83156521 | 1.639044635 | 13.32627101 |
| ENSMUSG00000008730 | *Hipk1* | 4074.203683 | 4587.141368 | 3483.793817 | 6617.310415 | 6898.795922 | 6398.637629 | 1.23E-10 | 0.712721281 | 2.11E-11 | 44.8695345 | 1.638892562 | 9.908704855 |
| ENSMUSG00000026017 | *Carf* | 85.478391 | 93.86229862 | 71.12522045 | 156.0288589 | 140.0605404 | 115.9698709 | 0.00521224 | 0.712610448 | 0.002072714 | 9.484001201 | 1.63876666 | 2.282975632 |
| ENSMUSG00000024660 | *Incenp* | 2973.369825 | 3113.604207 | 2144.492496 | 4545.921889 | 4448.279335 | 4506.395901 | 3.65E-07 | 0.712588412 | 8.57E-08 | 28.67249487 | 1.638741631 | 6.437167643 |
| ENSMUSG00000021271 | *Zfp839* | 153.3817857 | 172.5855168 | 158.3542644 | 273.2613529 | 259.492009 | 259.965794 | 1.12E-05 | 0.712534701 | 3.08E-06 | 21.7656048 | 1.638680621 | 4.950994651 |
| ENSMUSG00000008348 | *Ubc* | 34384.68136 | 30669.75839 | 19772.81129 | 45884.2921 | 47492.46636 | 45632.21138 | 6.58E-05 | 0.712452032 | 1.99E-05 | 18.19460507 | 1.638586725 | 4.18145193 |
| ENSMUSG00000030452 | *Nipa2* | 1240.634965 | 1385.730495 | 1454.712056 | 2298.262921 | 2209.482169 | 2164.770924 | 7.61E-15 | 0.712019196 | 9.50E-16 | 64.53198649 | 1.638095191 | 14.1184665 |
| ENSMUSG00000030604 | *Zfp626* | 269.2169885 | 232.1325665 | 289.868823 | 466.3997782 | 418.0101401 | 407.8273795 | 2.41E-06 | 0.711937604 | 6.18E-07 | 24.85544308 | 1.638002551 | 5.617325081 |
| ENSMUSG00000040268 | *Plekha1* | 1406.798566 | 1413.990111 | 1673.455659 | 2484.654153 | 2452.688069 | 2409.274069 | 1.20E-13 | 0.711668342 | 1.62E-14 | 58.94177863 | 1.637696866 | 12.92078335 |
| ENSMUSG00000002897 | *Il17ra* | 1369.251983 | 1390.776855 | 999.7790422 | 2000.542991 | 2147.594953 | 2020.775001 | 2.71E-07 | 0.711568376 | 6.27E-08 | 29.27706898 | 1.637583392 | 6.567544036 |
| ENSMUSG00000015766 | *Eps8* | 1965.20413 | 1953.950646 | 2430.335363 | 3494.20304 | 3364.710192 | 3522.58483 | 2.34E-11 | 0.711299939 | 3.78E-12 | 48.2331832 | 1.637278721 | 10.62995607 |
| ENSMUSG00000025878 | *Uimc1* | 703.7987147 | 723.6480442 | 810.5591161 | 1253.291267 | 1185.628761 | 1213.817982 | 4.26E-13 | 0.711035986 | 6.03E-14 | 56.36272058 | 1.636979194 | 12.37042626 |
| ENSMUSG00000013698 | *Pea15a* | 4305.075225 | 4443.82474 | 4773.441682 | 7410.949098 | 7506.810671 | 7193.99766 | 1.30E-24 | 0.710367619 | 9.04E-26 | 110.158879 | 1.636220996 | 23.88612093 |
| ENSMUSG00000095115 | *Itpripl2* | 1994.762078 | 2213.333558 | 1501.681541 | 3062.38263 | 3257.221871 | 3026.813631 | 7.13E-07 | 0.709375382 | 1.72E-07 | 27.32745583 | 1.635096045 | 6.146840637 |
| ENSMUSG00000042744 | *Hectd4* | 750.9316593 | 775.1209176 | 391.8597051 | 1004.488492 | 1047.739702 | 1091.083202 | 0.003021419 | 0.709094513 | 0.001153182 | 10.56389089 | 1.634777751 | 2.519789044 |
| ENSMUSG00000026276 | *Septin2* | 519.2612537 | 573.266512 | 735.4079398 | 882.1956021 | 1049.911183 | 1047.594501 | 9.02E-06 | 0.709069284 | 2.46E-06 | 22.19355684 | 1.634749162 | 5.044561128 |
| ENSMUSG00000030966 | *Trim21* | 545.6237482 | 543.9976232 | 597.1834547 | 862.7974197 | 970.6521175 | 920.0276428 | 2.49E-11 | 0.708745082 | 4.02E-12 | 48.11444275 | 1.634381843 | 10.60429159 |
| ENSMUSG00000026781 | *Acbd5* | 361.0862872 | 354.2544819 | 334.1543376 | 580.2586752 | 547.2132743 | 590.4799262 | 1.12E-09 | 0.708668332 | 2.08E-10 | 40.39074695 | 1.634294898 | 8.950597973 |
| ENSMUSG00000048000 | *Gigyf2* | 1572.962167 | 1730.901528 | 1712.373232 | 2754.541909 | 2787.096181 | 2647.012304 | 1.01E-19 | 0.708364256 | 9.23E-21 | 87.32006954 | 1.633950475 | 18.99636365 |
| ENSMUSG00000064357 | *mt-Atp6* | 13367.3824 | 16021.18417 | 21875.70224 | 27762.17269 | 31835.00082 | 24143.96071 | 8.16E-05 | 0.70816218 | 2.50E-05 | 17.76169566 | 1.633721626 | 4.088315427 |
| ENSMUSG00000027936 | *Crtc2* | 1227.054286 | 1336.276165 | 905.8400718 | 1897.648284 | 1897.87461 | 1881.611156 | 1.23E-06 | 0.708059164 | 3.04E-07 | 26.22186415 | 1.633604973 | 5.909711151 |
| ENSMUSG00000030276 | *Ttll3* | 205.3079111 | 222.0398462 | 150.3023526 | 295.189733 | 326.8079277 | 327.6148854 | 0.000227934 | 0.707680857 | 7.41E-05 | 15.7026148 | 1.633176662 | 3.642190683 |
| ENSMUSG00000039361 | *Picalm* | 4883.452376 | 4796.060678 | 5220.322784 | 8116.03086 | 8037.737836 | 8166.211745 | 1.07E-28 | 0.707569003 | 6.07E-30 | 129.2215312 | 1.633050045 | 27.96947409 |
| ENSMUSG00000030528 | *Blm* | 761.3168843 | 631.8042896 | 708.5682339 | 1174.011738 | 1136.770433 | 1123.941332 | 2.98E-11 | 0.707517044 | 4.84E-12 | 47.75102866 | 1.632991232 | 10.52569187 |
| ENSMUSG00000018848 | *Rars* | 6582.63497 | 5460.161672 | 6444.21337 | 10162.11741 | 9905.211709 | 10120.30407 | 3.78E-15 | 0.707482147 | 4.63E-16 | 65.94670475 | 1.632951732 | 14.42215641 |
| ENSMUSG00000032249 | *Anp32a* | 1982.779126 | 1855.041988 | 2327.002495 | 3333.113786 | 3371.224636 | 3349.596439 | 2.87E-12 | 0.707320449 | 4.32E-13 | 52.4936393 | 1.63276872 | 11.54205768 |
| ENSMUSG00000007659 | *Bcl2l1* | 4826.733069 | 4950.479298 | 4581.537785 | 7680.836853 | 7884.648408 | 7885.951224 | 1.33E-29 | 0.707090785 | 7.18E-31 | 133.4573141 | 1.632508818 | 28.87764077 |
| ENSMUSG00000021277 | *Traf3* | 1409.99402 | 1499.778234 | 999.7790422 | 2140.547264 | 2181.252913 | 2069.095781 | 3.53E-06 | 0.706674315 | 9.21E-07 | 24.08705097 | 1.632037622 | 5.452253295 |
| ENSMUSG00000034621 | *Gpatch8* | 1026.539555 | 983.0309554 | 760.9056603 | 1533.299813 | 1495.064839 | 1503.74266 | 3.69E-08 | 0.706300633 | 7.87E-09 | 33.30615525 | 1.631614952 | 7.432574776 |
| ENSMUSG00000037725 | *Ckap2* | 561.6010175 | 479.4042134 | 410.6474992 | 777.6140967 | 782.8189896 | 817.5875901 | 4.79E-07 | 0.705749102 | 1.13E-07 | 28.13378311 | 1.630991317 | 6.320018103 |
| ENSMUSG00000031662 | *Snx20* | 1310.136086 | 1299.942372 | 1269.518086 | 2075.605523 | 2083.536257 | 2167.670171 | 3.92E-22 | 0.704682564 | 3.15E-23 | 98.55939141 | 1.629786024 | 21.40658865 |
| ENSMUSG00000039218 | *Srrm2* | 9284.391217 | 8826.083886 | 5613.524474 | 12884.61015 | 13477.29836 | 12309.23538 | 5.09E-05 | 0.704472859 | 1.52E-05 | 18.71574893 | 1.629549141 | 4.293650419 |
| ENSMUSG00000042628 | *Zfyve1* | 380.2590104 | 503.626742 | 445.5391168 | 762.4329105 | 687.2738147 | 711.2818751 | 5.98E-07 | 0.704269801 | 1.43E-07 | 27.68229184 | 1.6293198 | 6.223422386 |
| ENSMUSG00000014313 | *Cox6c* | 1723.947362 | 1426.101376 | 1975.402349 | 2682.009574 | 2759.952665 | 2898.280358 | 3.24E-08 | 0.70365538 | 6.87E-09 | 33.56992271 | 1.628626045 | 7.489304604 |
| ENSMUSG00000017776 | *Crk* | 2215.248395 | 2446.475396 | 2883.926391 | 4225.430179 | 4086.727707 | 3959.404677 | 2.05E-10 | 0.703393864 | 3.56E-11 | 43.84005506 | 1.628330853 | 9.687470246 |
| ENSMUSG00000032235 | *Ice2* | 275.6078962 | 255.3458231 | 352.9421317 | 536.4019149 | 442.9821744 | 452.2824966 | 6.17E-05 | 0.703361209 | 1.86E-05 | 18.32662297 | 1.628293997 | 4.209430676 |
| ENSMUSG00000017057 | *Il13ra1* | 567.1930618 | 514.7287344 | 629.3911017 | 964.8487274 | 922.87953 | 891.035175 | 9.01E-10 | 0.70286062 | 1.65E-10 | 40.84109159 | 1.627729106 | 9.045234783 |
| ENSMUSG00000039990 | *Edrf1* | 506.4794383 | 515.7380064 | 618.6552194 | 913.4013739 | 836.0202801 | 910.3634868 | 8.00E-09 | 0.702521311 | 1.60E-09 | 36.40798707 | 1.627346323 | 8.097066968 |
| ENSMUSG00000041297 | *Cdk13* | 1357.269031 | 1310.035093 | 911.2080129 | 1953.312633 | 1931.532569 | 1946.361001 | 4.69E-06 | 0.701623303 | 1.24E-06 | 23.51101265 | 1.626333695 | 5.32842836 |
| ENSMUSG00000025940 | *Tmem70* | 656.6657701 | 520.7843665 | 598.52544 | 959.788332 | 957.62323 | 974.1469159 | 3.10E-09 | 0.701503051 | 5.95E-10 | 38.33619028 | 1.626198142 | 8.508934652 |
| ENSMUSG00000032782 | *Cntrob* | 221.2851804 | 229.1047504 | 136.8824997 | 333.9860979 | 306.1788558 | 320.8499763 | 0.001670797 | 0.701381145 | 0.00061373 | 11.73386485 | 1.626060736 | 2.777076239 |
| ENSMUSG00000041220 | *Elovl6* | 857.9793639 | 797.3249022 | 823.978969 | 1386.548346 | 1313.746155 | 1332.6871 | 4.49E-16 | 0.701379072 | 5.18E-17 | 70.26823233 | 1.626058399 | 15.34765321 |
| ENSMUSG00000024597 | *Slc12a2* | 488.904442 | 494.5432938 | 365.0199993 | 760.746112 | 739.3893646 | 700.6513036 | 1.93E-06 | 0.701032048 | 4.87E-07 | 25.31275715 | 1.625667316 | 5.71508476 |
| ENSMUSG00000003779 | *Kif20a* | 2180.098402 | 1880.273788 | 1439.950218 | 3041.297649 | 2914.127834 | 2995.888332 | 1.98E-06 | 0.700571999 | 5.02E-07 | 25.25661348 | 1.625149004 | 5.703253247 |
| ENSMUSG00000047123 | *Ticam1* | 1000.975925 | 972.9382351 | 774.3255132 | 1429.561707 | 1567.80946 | 1480.548686 | 1.10E-08 | 0.700298692 | 2.23E-09 | 35.76543836 | 1.624841161 | 7.959840279 |
| ENSMUSG00000050029 | *Rap2c* | 865.9679985 | 850.8163197 | 993.0691157 | 1525.70922 | 1501.579282 | 1366.511646 | 2.20E-11 | 0.700266759 | 3.54E-12 | 48.36216802 | 1.624805196 | 10.65677261 |
| ENSMUSG00000022247 | *Brix1* | 1393.217887 | 1285.812564 | 1274.886027 | 2151.511454 | 2096.565144 | 2183.13282 | 8.50E-19 | 0.700073916 | 8.22E-20 | 82.99613853 | 1.624588026 | 18.07061811 |
| ENSMUSG00000039929 | *Urb1* | 1375.642891 | 1303.97946 | 964.8874246 | 1895.961485 | 2067.250147 | 1969.554975 | 8.30E-07 | 0.699909074 | 2.01E-07 | 27.01997279 | 1.624402411 | 6.080843994 |
| ENSMUSG00000031652 | *N4bp1* | 1234.244057 | 1275.719844 | 1089.692057 | 1902.708679 | 1993.419785 | 1961.82365 | 2.22E-14 | 0.699827019 | 2.86E-15 | 62.36103761 | 1.624310024 | 13.65279068 |
| ENSMUSG00000034342 | *Cbl* | 1823.805296 | 1658.233942 | 1105.79588 | 2458.508777 | 2599.263053 | 2402.50916 | 6.45E-05 | 0.699198148 | 1.95E-05 | 18.23827878 | 1.623602142 | 4.19064812 |
| ENSMUSG00000011427 | *Zfp790* | 268.418125 | 271.4941756 | 327.4444111 | 488.3281584 | 428.8675463 | 482.24138 | 2.09E-06 | 0.697915337 | 5.33E-07 | 25.13890287 | 1.622159114 | 5.678965703 |
| ENSMUSG00000027303 | *Ptpra* | 3327.266341 | 3172.141984 | 2952.367641 | 5080.637005 | 5097.552228 | 5161.625672 | 7.24E-22 | 0.69725085 | 5.89E-23 | 97.32220082 | 1.62141214 | 21.14031484 |
| ENSMUSG00000032328 | *Tmem30a* | 1982.779126 | 1782.374402 | 2100.206981 | 3205.760501 | 3172.534102 | 3123.455191 | 5.74E-16 | 0.697028864 | 6.66E-17 | 69.77118114 | 1.621162674 | 15.24110089 |
| ENSMUSG00000038481 | *Cdk19* | 411.4146857 | 387.5604588 | 414.6734551 | 662.0684012 | 634.0725242 | 669.7260047 | 1.44E-10 | 0.696866982 | 2.48E-11 | 44.5505082 | 1.620980776 | 9.84018909 |
| ENSMUSG00000006191 | *Cdkal1* | 274.8090327 | 306.8186966 | 407.9635286 | 528.8113217 | 514.6410556 | 550.8568869 | 6.54E-05 | 0.696553899 | 1.98E-05 | 18.20802865 | 1.620629041 | 4.184174256 |
| ENSMUSG00000036550 | *Cnot1* | 4318.655904 | 4594.206272 | 4365.478153 | 7292.029805 | 7089.886272 | 7135.046309 | 3.71E-30 | 0.696524099 | 1.95E-31 | 136.0435847 | 1.620595567 | 29.43087264 |
| ENSMUSG00000030269 | *Mtmr14* | 1419.580381 | 1365.545054 | 1230.600512 | 2159.102047 | 2081.364775 | 2276.875133 | 2.49E-14 | 0.69635765 | 3.22E-15 | 62.12645087 | 1.620408604 | 13.60304626 |
| ENSMUSG00000036822 | *Topors* | 1146.369076 | 1122.310495 | 1278.911983 | 1976.927812 | 1860.959429 | 1897.073805 | 4.82E-15 | 0.695742681 | 5.94E-16 | 65.45565505 | 1.619718028 | 14.31666177 |
| ENSMUSG00000029209 | *Gnpda2* | 376.2646931 | 333.0597693 | 556.923896 | 710.985557 | 664.4732616 | 666.8267579 | 0.000527574 | 0.695660252 | 0.000180155 | 14.02751801 | 1.619625488 | 3.277716706 |
| ENSMUSG00000047248 | *C2cd3* | 649.4759989 | 685.2957071 | 497.8765432 | 1019.669678 | 1002.138596 | 953.8521884 | 7.01E-07 | 0.694954036 | 1.69E-07 | 27.36442839 | 1.618832856 | 6.1543988 |
| ENSMUSG00000026667 | *Uhmk1* | 2686.57784 | 2844.128575 | 2768.515656 | 4592.308847 | 4491.70896 | 4345.970913 | 6.27E-26 | 0.69488617 | 4.08E-27 | 116.3032672 | 1.618756707 | 25.20261222 |
| ENSMUSG00000041341 | *Atg2b* | 976.211157 | 1070.837622 | 777.0094838 | 1522.335623 | 1521.122614 | 1536.60079 | 8.62E-08 | 0.694650533 | 1.90E-08 | 31.59330122 | 1.618492335 | 7.064328727 |
| ENSMUSG00000020076 | *Ddx50* | 920.2907143 | 904.3077372 | 1277.569998 | 1704.509858 | 1729.584813 | 1574.290998 | 3.84E-06 | 0.69424831 | 1.01E-06 | 23.918422 | 1.618041163 | 5.415786266 |
| ENSMUSG00000106924 | *Gm42857* | 124.6227009 | 145.3351721 | 178.4840438 | 215.9102047 | 246.4631215 | 258.0329628 | 0.000907477 | 0.693971567 | 0.000321077 | 12.94325116 | 1.617730814 | 3.042164175 |
| ENSMUSG00000041775 | *Mapk1ip1* | 91.07043527 | 84.77885036 | 59.04735283 | 119.7626917 | 128.1173936 | 136.2645983 | 0.00911742 | 0.693386304 | 0.003787625 | 8.382924317 | 1.617074677 | 2.040128028 |
| ENSMUSG00000028760 | *Eif4g3* | 3007.720954 | 3011.667732 | 2047.869555 | 4403.387418 | 4478.680072 | 4168.150445 | 4.40E-06 | 0.692635355 | 1.16E-06 | 23.64210023 | 1.616233178 | 5.356657007 |
| ENSMUSG00000023089 | *Ndufa5* | 504.8817113 | 507.6638302 | 723.3300721 | 916.7749709 | 912.0221238 | 965.4491755 | 2.01E-05 | 0.692148394 | 5.71E-06 | 20.58402513 | 1.615687734 | 4.696176468 |
| ENSMUSG00000059820 | *Nkapd1* | 794.86915 | 726.6758603 | 938.0477188 | 1351.968977 | 1311.574673 | 1297.896139 | 4.13E-09 | 0.691107649 | 8.05E-10 | 37.74915809 | 1.614522615 | 8.383964216 |
| ENSMUSG00000022387 | *Brd1* | 1440.350832 | 1531.065667 | 1072.246248 | 2058.737538 | 2321.313453 | 2158.006015 | 1.44E-06 | 0.690771127 | 3.59E-07 | 25.90171288 | 1.614146056 | 5.841208089 |
| ENSMUSG00000035329 | *Fbxo33* | 654.2691797 | 670.1566267 | 618.6552194 | 1102.322803 | 1032.539333 | 1004.105799 | 1.30E-12 | 0.690766001 | 1.90E-13 | 54.11051167 | 1.614140322 | 11.88601531 |
| ENSMUSG00000057278 | *Snrpg* | 831.6168694 | 743.8334847 | 670.9926458 | 1197.626917 | 1191.057464 | 1246.676113 | 3.72E-10 | 0.690063041 | 6.62E-11 | 42.62760046 | 1.613354015 | 9.429004546 |
| ENSMUSG00000062931 | *Zfp938* | 76.69089286 | 93.86229862 | 108.7008086 | 153.4986612 | 159.6038717 | 133.3653516 | 0.00472276 | 0.689324722 | 0.0018619 | 9.68093334 | 1.612528571 | 2.32580411 |
| ENSMUSG00000022369 | *Mtbp* | 1012.160013 | 993.1236757 | 1068.220292 | 1718.847645 | 1697.012595 | 1534.667959 | 3.51E-14 | 0.689125292 | 4.60E-15 | 61.42600612 | 1.612305679 | 13.45427679 |
| ENSMUSG00000046873 | *Mbtps2* | 1046.511142 | 1089.004518 | 1250.730292 | 1821.742352 | 1750.213885 | 1874.846247 | 4.28E-12 | 0.689115173 | 6.50E-13 | 51.69081579 | 1.61229437 | 11.36873814 |
| ENSMUSG00000002477 | *Snrpd1* | 1600.123525 | 1428.11992 | 1839.861835 | 2621.284829 | 2597.091572 | 2619.952668 | 1.65E-10 | 0.688987643 | 2.84E-11 | 44.28287985 | 1.612151855 | 9.782161615 |
| ENSMUSG00000020390 | *Ube2b* | 1667.228056 | 1534.093483 | 2018.345878 | 2880.208395 | 2668.750453 | 2851.892409 | 4.67E-09 | 0.688607028 | 9.13E-10 | 37.5018124 | 1.611726589 | 8.330338907 |
| ENSMUSG00000042228 | *Lyn* | 4992.896671 | 4813.218302 | 5158.591461 | 8203.744381 | 7945.449883 | 7958.432393 | 2.99E-29 | 0.688421827 | 1.65E-30 | 131.8057518 | 1.611519702 | 28.52431281 |
| ENSMUSG00000056153 | *Socs6* | 553.6123828 | 578.3128721 | 679.0445575 | 981.7167121 | 950.0230456 | 975.1133314 | 1.62E-09 | 0.688135365 | 3.05E-10 | 39.64103757 | 1.61119975 | 8.789369555 |
| ENSMUSG00000054823 | *Nsd3* | 1278.980411 | 1278.74766 | 1103.11191 | 1992.952398 | 1960.847566 | 1955.058741 | 4.32E-14 | 0.687968955 | 5.68E-15 | 61.00915442 | 1.611013915 | 13.36427926 |
| ENSMUSG00000043467 | *Zbtb37* | 209.3022284 | 236.1696546 | 238.8733819 | 352.5408812 | 363.7231089 | 382.7005741 | 2.00E-06 | 0.68782559 | 5.06E-07 | 25.23941865 | 1.610853832 | 5.699684762 |
| ENSMUSG00000059273 | *Zc3h4* | 1644.061016 | 1646.122678 | 1205.102792 | 2446.701187 | 2517.832506 | 2283.640042 | 4.28E-07 | 0.686888129 | 1.01E-07 | 28.35854775 | 1.609807441 | 6.36881966 |
| ENSMUSG00000037058 | *Paip2* | 1402.804249 | 1089.004518 | 1410.426541 | 2089.94331 | 2047.706816 | 2137.711288 | 1.10E-08 | 0.685866238 | 2.23E-09 | 35.76373365 | 1.608667585 | 7.959586572 |
| ENSMUSG00000020184 | *Mdm2* | 4709.30014 | 4634.577153 | 5231.058666 | 7873.13188 | 7755.445274 | 7799.940236 | 6.92E-21 | 0.68566431 | 5.92E-22 | 92.75555737 | 1.608442442 | 20.15965808 |
| ENSMUSG00000041124 | *Msantd4* | 802.8577846 | 852.8348638 | 1093.718013 | 1435.465502 | 1500.493542 | 1472.817361 | 2.89E-07 | 0.685118055 | 6.70E-08 | 29.14850107 | 1.607833544 | 6.539720767 |
| ENSMUSG00000029632 | *Ndufa4* | 2623.467627 | 2588.782752 | 3287.863964 | 4442.183783 | 4484.108775 | 4723.839409 | 7.15E-10 | 0.684768378 | 1.30E-10 | 41.30764096 | 1.607443888 | 9.145815502 |
| ENSMUSG00000021113 | *Snapc1* | 339.5169736 | 309.8465126 | 360.9940434 | 567.6076867 | 555.8991993 | 496.7376138 | 2.25E-07 | 0.684050525 | 5.17E-08 | 29.6508954 | 1.606644259 | 6.646978499 |
| ENSMUSG00000091228 | *Gm20390* | 1208.680426 | 1815.680379 | 1606.356394 | 2266.21375 | 2820.75414 | 2347.423471 | 3.34E-05 | 0.683748161 | 9.76E-06 | 19.55719999 | 1.606307569 | 4.4759967 |
| ENSMUSG00000068479 | *Mfap1a* | 298.7749368 | 198.8265895 | 210.6916908 | 406.5184323 | 401.7240307 | 334.3797945 | 0.001112162 | 0.683246484 | 0.000398157 | 12.54082151 | 1.605749095 | 2.953831873 |
| ENSMUSG00000061778 | *Mospd2* | 563.9976079 | 630.7950176 | 787.7453661 | 1067.743434 | 1033.625074 | 1069.822059 | 1.29E-06 | 0.683059804 | 3.20E-07 | 26.1272189 | 1.605541331 | 5.889667273 |
| ENSMUSG00000007817 | *Zmiz1* | 3322.47316 | 3317.477156 | 1870.727496 | 4577.97106 | 4683.88505 | 4411.687174 | 0.000673413 | 0.68300751 | 0.000233691 | 13.53872267 | 1.605483135 | 3.171718674 |
| ENSMUSG00000020790 | *Ankfy1* | 4610.24107 | 4806.153398 | 4314.482712 | 7469.143645 | 7309.205878 | 7274.210154 | 2.55E-23 | 0.682921732 | 1.90E-24 | 104.1255283 | 1.60538768 | 22.59318405 |
| ENSMUSG00000036291 | *Ap5m1* | 305.964708 | 173.5947888 | 254.9772054 | 407.3618316 | 349.6084808 | 423.2900289 | 0.003379896 | 0.682459423 | 0.001301372 | 10.34057935 | 1.60487332 | 2.471096687 |
| ENSMUSG00000063659 | *Zbtb18* | 600.7453274 | 745.8520288 | 645.4949252 | 1043.284856 | 1088.997845 | 1063.05715 | 2.82E-09 | 0.682380722 | 5.40E-10 | 38.52624128 | 1.604785774 | 8.549892331 |
| ENSMUSG00000028273 | *Pdlim5* | 1067.281592 | 1027.438925 | 972.9393363 | 1635.351121 | 1591.695754 | 1701.857856 | 8.90E-16 | 0.682001206 | 1.04E-16 | 68.88535469 | 1.604363674 | 15.0506607 |
| ENSMUSG00000022781 | *Pak2* | 3498.223123 | 3398.218919 | 3077.172273 | 5353.054959 | 5329.900721 | 5323.983492 | 1.02E-19 | 0.681196904 | 9.34E-21 | 87.29773909 | 1.60346949 | 18.99174025 |
| ENSMUSG00000079685 | *Ulbp1* | 986.5963821 | 1057.717085 | 959.5194834 | 1665.713494 | 1562.380757 | 1587.820816 | 2.96E-15 | 0.68058243 | 3.59E-16 | 66.44783975 | 1.602786685 | 14.52900136 |
| ENSMUSG00000003031 | *Cdkn1b* | 384.2533278 | 389.5790029 | 350.2581611 | 644.3570172 | 578.6997524 | 580.8157703 | 2.15E-08 | 0.680271277 | 4.49E-09 | 34.40023817 | 1.602441042 | 7.668184524 |

| | | | | | | | | | | | | | |
|---|---|---|---|---|---|---|---|---|---|---|---|---|---|
| ENSMUSG00000044224 | *Dnajc21* | 603.1419178 | 575.285056 | 491.1666167 | 868.7012143 | 909.8506425 | 906.4978245 | 4.05E-09 | 0.679875074 | 7.88E-10 | 37.78958459 | 1.602001028 | 8.392700919 |
| ENSMUSG00000038914 | *Dido1* | 1526.628086 | 1571.436548 | 1309.776445 | 2358.987667 | 2328.913638 | 2382.214432 | 6.37E-13 | 0.679383605 | 9.11E-14 | 55.5501877 | 1.601455383 | 12.19616249 |
| ENSMUSG00000062593 | *Gm49339* | 5563.285186 | 3739.352864 | 6822.653222 | 8719.904714 | 9280.91085 | 7794.141742 | 0.000510018 | 0.678060058 | 0.000173739 | 14.09572148 | 1.599986861 | 3.292414248 |
| ENSMUSG00000040249 | *Lrp1* | 11156.12832 | 10423.76151 | 5753.090945 | 14234.89233 | 15084.19448 | 14410.22288 | 0.001473735 | 0.677622487 | 0.000536821 | 11.98320842 | 1.599501657 | 2.831580659 |
| ENSMUSG00000042460 | *C1galt1* | 874.7554967 | 958.8084267 | 836.0568366 | 1428.718308 | 1414.720032 | 1429.328659 | 1.01E-13 | 0.677181889 | 1.36E-14 | 59.28631322 | 1.599013245 | 12.99511615 |
| ENSMUSG00000002055 | *Spag5* | 1413.988337 | 1158.644288 | 920.60191 | 1945.72204 | 1864.216651 | 1786.902428 | 1.46E-05 | 0.677126823 | 4.06E-06 | 21.23546123 | 1.598952213 | 4.836299919 |
| ENSMUSG00000023460 | *Rab12* | 948.2509357 | 945.6878904 | 915.2339688 | 1497.877045 | 1473.350026 | 1522.104556 | 1.06E-17 | 0.67665468 | 1.10E-18 | 77.87053284 | 1.598429019 | 16.97428218 |
| ENSMUSG00000051378 | *Kif18b* | 838.0077772 | 838.7050554 | 527.4002196 | 1151.239959 | 1256.201901 | 1123.941332 | 0.000255625 | 0.6761021 | 8.36E-05 | 15.47487719 | 1.597816907 | 3.592396652 |
| ENSMUSG00000059060 | *Rad51b* | 127.8181548 | 97.89938673 | 100.6488969 | 182.1742352 | 187.8331279 | 154.6264946 | 0.002795755 | 0.676084466 | 0.001061608 | 10.71691032 | 1.597797377 | 2.553500951 |
| ENSMUSG00000031333 | *Abcb7* | 1053.700913 | 1046.615093 | 1132.635586 | 1863.068915 | 1635.125379 | 1656.436323 | 1.47E-12 | 0.675301398 | 2.15E-13 | 53.86155622 | 1.596930358 | 11.83274274 |
| ENSMUSG00000028676 | *Srsf10* | 2065.062063 | 1860.088348 | 2372.629995 | 3495.046439 | 3316.937605 | 3231.693737 | 3.09E-10 | 0.674876555 | 5.46E-11 | 43.00374207 | 1.596460165 | 9.509762817 |
| ENSMUSG00000047044 | *D030056L22Rik* | 543.2271578 | 489.4969336 | 327.4444111 | 716.8893516 | 737.2178834 | 726.7445245 | 0.000391661 | 0.674859179 | 0.000131085 | 14.62611438 | 1.596440937 | 3.407089239 |
| ENSMUSG00000038187 | *Btbd10* | 643.8839546 | 703.4626036 | 730.0399986 | 1055.092446 | 1110.712658 | 1143.269644 | 1.51E-11 | 0.674505934 | 2.40E-12 | 49.12768896 | 1.596050096 | 10.82163792 |
| ENSMUSG00000092428 | *Gm20545* | 369.0749219 | 360.310114 | 452.2490432 | 633.3928272 | 613.4434523 | 630.1029654 | 3.62E-07 | 0.674132564 | 8.47E-08 | 28.69578144 | 1.59563709 | 6.441733775 |
| ENSMUSG00000021504 | *B4galt7* | 1140.777031 | 1190.940993 | 1069.562277 | 1744.149623 | 1752.385366 | 1934.764013 | 2.25E-13 | 0.673768911 | 3.13E-14 | 57.65245182 | 1.595234936 | 12.64719232 |
| ENSMUSG00000025591 | *Tma16* | 861.9736812 | 784.2043659 | 1117.873748 | 1502.937441 | 1402.776886 | 1492.145673 | 3.06E-06 | 0.6734497 | 7.91E-07 | 24.37848467 | 1.594882013 | 5.514713253 |
| ENSMUSG00000048495 | *Tyw5* | 336.3215197 | 260.3921833 | 381.1238228 | 511.0999378 | 504.86939 | 539.2598998 | 6.09E-05 | 0.673314797 | 1.83E-05 | 18.35586532 | 1.594732887 | 4.21567112 |
| ENSMUSG00000057886 | *Cbx3-ps6* | 542.4282943 | 529.8678148 | 613.2872782 | 913.4013739 | 863.1637957 | 902.6321621 | 2.99E-10 | 0.673010059 | 5.27E-11 | 43.07384722 | 1.59439607 | 9.524453666 |
| ENSMUSG00000027652 | *Ralgapb* | 1617.698521 | 1685.484287 | 1689.559482 | 2658.394396 | 2679.607859 | 2612.221343 | 9.46E-23 | 0.672067316 | 7.32E-24 | 101.4532276 | 1.593354536 | 22.02391674 |
| ENSMUSG00000111559 | *None* | 218.0897266 | 248.2809189 | 136.8824997 | 307.8407216 | 314.8647808 | 344.0439504 | 0.004775481 | 0.67198005 | 0.001886067 | 9.657240843 | 1.59325816 | 2.320982906 |
| ENSMUSG00000031370 | *Zrsr2* | 530.4453423 | 510.6916462 | 493.8505873 | 790.2650853 | 832.7630583 | 827.251746 | 2.12E-11 | 0.671609134 | 3.40E-12 | 48.444065 | 1.592848588 | 10.67395226 |
| ENSMUSG00000052031 | *Tagap1* | 551.2157924 | 565.1923358 | 590.4735283 | 926.8957617 | 891.3930519 | 894.9008374 | 2.29E-12 | 0.671518517 | 3.42E-13 | 52.95300639 | 1.592748543 | 11.63969613 |
| ENSMUSG00000111228 | *Gm48789* | 315.5510696 | 300.7630644 | 283.1588965 | 527.9679225 | 435.38199 | 470.6443929 | 1.97E-06 | 0.67140313 | 4.98E-07 | 25.27147127 | 1.592621159 | 5.706396411 |
| ENSMUSG00000029670 | *Ing3* | 405.8226414 | 392.6068189 | 468.3528667 | 640.9834203 | 678.5878897 | 690.9871477 | 3.61E-08 | 0.670701266 | 7.69E-09 | 33.35142123 | 1.591846546 | 7.442563321 |
| ENSMUSG00000024251 | *Thada* | 737.3509803 | 636.8506498 | 460.300955 | 990.9941037 | 978.2523018 | 960.6170976 | 0.000113135 | 0.670397989 | 3.53E-05 | 17.10721238 | 1.591511949 | 3.946404108 |
| ENSMUSG00000028232 | *Tmem68* | 1009.763423 | 972.9382351 | 1375.534924 | 1809.934763 | 1652.497229 | 1869.047753 | 7.42E-06 | 0.669907552 | 2.01E-06 | 22.58694092 | 1.590971015 | 5.129367252 |
| ENSMUSG00000033902 | *Mapkbp1* | 496.8930767 | 440.0426043 | 375.7558816 | 617.3682417 | 707.9028866 | 771.1996417 | 1.59E-05 | 0.669489644 | 4.45E-06 | 21.06154499 | 1.590510222 | 4.799181249 |
| ENSMUSG00000026064 | *Ptp4a1* | 2533.994918 | 2635.209265 | 2938.947788 | 4416.038406 | 4331.019347 | 4134.325899 | 2.47E-15 | 0.669434152 | 2.98E-16 | 66.81405528 | 1.590449046 | 14.60799263 |
| ENSMUSG00000045482 | *Trrap* | 2911.857338 | 2837.063671 | 1685.533526 | 3929.397046 | 4036.783638 | 3864.695949 | 0.000461996 | 0.669007721 | 0.000156371 | 14.29389885 | 1.589979011 | 3.33536158 |
| ENSMUSG00000024052 | *Lpin2* | 1013.75774 | 1003.216396 | 1003.804998 | 1581.37357 | 1639.468398 | 1583.955154 | 3.61E-18 | 0.668929951 | 3.65E-19 | 80.0480546 | 1.589893304 | 17.44271645 |
| ENSMUSG00000040054 | *Baz2a* | 1151.96112 | 1177.820457 | 774.3255132 | 1628.603927 | 1694.841113 | 1621.645362 | 4.13E-05 | 0.668814791 | 1.22E-05 | 19.13225188 | 1.5897664 | 4.383540258 |
| ENSMUSG00000028771 | *Ptpn12* | 1342.889489 | 1358.48015 | 1651.983894 | 2266.21375 | 2359.314375 | 2282.673626 | 2.33E-10 | 0.66875195 | 4.06E-11 | 43.58408814 | 1.589697154 | 9.633319466 |
| ENSMUSG00000021715 | *Cwc27* | 468.1339918 | 494.5432938 | 625.3651458 | 830.7482487 | 856.649352 | 826.2853304 | 1.28E-06 | 0.668407535 | 3.16E-07 | 26.1473667 | 1.589317689 | 5.893785114 |
| ENSMUSG00000000682 | *Cd52* | 2699.359656 | 2460.605205 | 2859.770656 | 4182.416817 | 4265.87491 | 4288.952393 | 1.40E-16 | 0.668192111 | 1.55E-17 | 72.64184703 | 1.58908039 | 15.85471101 |
| ENSMUSG00000020770 | *Unk* | 337.9192467 | 331.0412252 | 287.1848524 | 501.8225462 | 490.7547618 | 532.4949907 | 3.08E-07 | 0.668141129 | 7.17E-08 | 29.01890197 | 1.589024236 | 6.51210042 |
| ENSMUSG00000070047 | *Fat1* | 583.9691946 | 645.934098 | 383.8077934 | 840.0256403 | 850.1349082 | 879.4381879 | 0.000496735 | 0.666782145 | 0.000168862 | 14.14928022 | 1.587528118 | 3.30387539 |
| ENSMUSG00000038866 | *Zcchc2* | 1368.453119 | 1397.841759 | 998.4370569 | 1954.156033 | 1976.047935 | 2056.532378 | 1.63E-06 | 0.666572306 | 4.09E-07 | 25.65334618 | 1.58729723 | 5.787884677 |
| ENSMUSG00000100622 | *Gm20379* | 397.8340067 | 339.1154015 | 366.3619846 | 598.8134584 | 565.6708649 | 588.547095 | 5.41E-08 | 0.666155269 | 1.17E-08 | 32.53190993 | 1.58683846 | 7.266413532 |
| ENSMUSG00000028060 | *Khdc4* | 1706.372366 | 1832.838003 | 1869.385511 | 2814.423255 | 2876.126912 | 2881.851293 | 1.66E-19 | 0.665975552 | 1.54E-20 | 86.30762974 | 1.586640799 | 18.78098236 |
| ENSMUSG00000046351 | *Zfp322a* | 163.7670108 | 193.7802294 | 218.7436025 | 328.9257025 | 300.7501527 | 278.3276902 | 0.000257944 | 0.66575729 | 8.44E-05 | 15.45697247 | 1.586400777 | 3.588474416 |
| ENSMUSG00000025980 | *Hspd1* | 16334.36132 | 15650.78134 | 16829.83754 | 26079.59121 | 25785.25407 | 25563.62522 | 3.26E-31 | 0.665689441 | 1.64E-32 | 140.9601189 | 1.586326172 | 30.48667021 |
| ENSMUSG00000028890 | *Mtf1* | 917.8941239 | 900.2706491 | 811.9011014 | 1356.185973 | 1471.178545 | 1353.948243 | 4.42E-12 | 0.665386399 | 6.71E-13 | 51.62683485 | 1.585992995 | 11.35475419 |
| ENSMUSG00000024999 | *Noc3l* | 831.6168694 | 922.4746337 | 1034.67066 | 1475.105266 | 1439.692067 | 1495.044919 | 5.80E-10 | 0.665169344 | 1.05E-10 | 41.72827489 | 1.585754399 | 9.2362916 |
| ENSMUSG00000026708 | *Cenpl* | 518.4623903 | 456.1909567 | 421.3833815 | 734.6007356 | 708.9886272 | 776.9981353 | 9.02E-08 | 0.66449193 | 1.99E-08 | 31.50182735 | 1.585009986 | 7.044567643 |
| ENSMUSG00000028126 | *Pip5k1a* | 1906.088233 | 2091.211642 | 1831.809923 | 3172.024531 | 3036.816564 | 3032.612125 | 4.72E-16 | 0.664264723 | 5.45E-17 | 70.16712637 | 1.584760386 | 15.32628257 |
| ENSMUSG00000061119 | *Prcp* | 620.7169141 | 589.4148644 | 499.2185284 | 938.703351 | 896.8217551 | 880.4046035 | 2.27E-08 | 0.663173066 | 4.76E-09 | 34.28325805 | 1.583561685 | 7.643183231 |
| ENSMUSG00000026510 | *Trp53bp2* | 242.8544941 | 315.9021448 | 226.7955143 | 403.9882346 | 438.6392119 | 403.9617171 | 0.000114191 | 0.66306763 | 3.57E-05 | 17.08740291 | 1.583445958 | 3.942368704 |
| ENSMUSG00000033396 | *Spg11* | 864.3702716 | 919.4468176 | 747.4858074 | 1299.678225 | 1339.803929 | 1377.142217 | 6.19E-10 | 0.662885323 | 1.12E-10 | 41.59361925 | 1.583245878 | 9.208078555 |
| ENSMUSG00000036636 | *Clcn7* | 1790.25303 | 1897.431413 | 1613.06632 | 2773.096692 | 2965.157643 | 2661.508538 | 2.11E-12 | 0.662508614 | 3.13E-13 | 53.12303612 | 1.582832523 | 11.67608403 |
| ENSMUSG00000080904 | *Gm11966* | 101.4556603 | 117.0755553 | 108.7008086 | 181.330836 | 154.1751685 | 180.7197155 | 0.001440312 | 0.662199471 | 0.000524171 | 12.02765053 | 1.582493387 | 2.841543414 |
| ENSMUSG00000029512 | *Ulk1* | 402.6271875 | 424.9035238 | 273.7649995 | 565.0774889 | 575.4425305 | 609.808238 | 0.000299159 | 0.662050018 | 9.86E-05 | 15.16318409 | 1.58232946 | 3.524097904 |
| ENSMUSG00000050954 | *Zfp169* | 95.06475261 | 81.75103428 | 96.62294099 | 140.8476726 | 134.6318373 | 156.5593258 | 0.003252659 | 0.661863934 | 0.001247772 | 10.41823404 | 1.582125379 | 2.487761479 |
| ENSMUSG00000074238 | *Ap1ar* | 839.6055041 | 837.6957834 | 798.4812485 | 1298.834825 | 1256.201901 | 1364.578815 | 5.38E-14 | 0.661607563 | 7.11E-15 | 60.5679759 | 1.581844256 | 13.2692845 |
| ENSMUSG00000041846 | *Ppp4r3a* | 1833.391657 | 1987.256623 | 1618.434262 | 2803.459065 | 2938.014127 | 2872.187137 | 8.30E-12 | 0.661548608 | 1.29E-12 | 50.34780974 | 1.581779616 | 11.08111248 |
| ENSMUSG00000020492 | *Ska2* | 405.8226414 | 422.8849798 | 410.6474992 | 662.0684012 | 621.0436367 | 675.5244982 | 1.54E-09 | 0.661300594 | 2.89E-10 | 39.74667662 | 1.581507716 | 8.812041515 |
| ENSMUSG00000020288 | *Ahsa2* | 614.3260063 | 677.2215309 | 665.6247046 | 1029.790469 | 1060.768589 | 1000.240137 | 7.34E-12 | 0.661128309 | 1.14E-12 | 50.59300312 | 1.581318865 | 11.1343804 |
| ENSMUSG00000026946 | *Nmi* | 978.6077474 | 708.5089638 | 730.0399986 | 1294.617829 | 1273.573751 | 1262.138762 | 4.59E-06 | 0.660843948 | 1.21E-06 | 23.55726809 | 1.581007211 | 5.338190418 |
| ENSMUSG00000046057 | *Gm15428* | 496.8930767 | 394.625363 | 500.5605137 | 741.3479295 | 702.4741834 | 752.8377455 | 8.06E-07 | 0.659253622 | 1.95E-07 | 27.0791445 | 1.57926538 | 6.093715146 |
| ENSMUSG00000057346 | *Apol9a* | 1122.403172 | 1120.291951 | 1050.774483 | 1692.702269 | 1774.100179 | 1741.480895 | 1.21E-16 | 0.659004098 | 1.34E-17 | 72.93519961 | 1.578992259 | 15.91767524 |
| ENSMUSG00000069895 | *Atxn1l* | 1357.269031 | 1387.749039 | 1056.142424 | 2012.35058 | 1963.019047 | 2035.271235 | 5.37E-08 | 0.658443437 | 1.16E-08 | 32.54764156 | 1.578378749 | 7.269692155 |
| ENSMUSG00000058192 | *Zfp846* | 114.2374758 | 96.8901147 | 124.8046321 | 187.2346307 | 157.4323904 | 182.6525467 | 0.002281755 | 0.657060373 | 0.000854402 | 11.11994706 | 1.576866335 | 2.641730941 |
| ENSMUSG00000031691 | *Tnpo2* | 3098.79139 | 3115.622751 | 2352.500216 | 4471.702756 | 4564.453581 | 4482.235512 | 4.58E-08 | 0.656693691 | 9.85E-09 | 32.87034341 | 1.576465602 | 7.339061273 |
| ENSMUSG00000030822 | *Prr14* | 648.6771354 | 635.8413777 | 566.317793 | 963.1619289 | 986.9382268 | 976.079747 | 6.99E-11 | 0.656531755 | 1.17E-11 | 46.02208405 | 1.57628866 | 10.15577377 |
| ENSMUSG00000026618 | *Iars2* | 1432.362197 | 1298.9331 | 1269.518086 | 2126.209477 | 2065.078666 | 2119.349391 | 5.73E-15 | 0.6553424 | 7.09E-16 | 65.1070492 | 1.574989706 | 14.2417044 |
| ENSMUSG00000026819 | *Slc25a25* | 1249.422463 | 1252.506587 | 1044.064557 | 1856.321721 | 1935.875532 | 1802.365077 | 1.83E-10 | 0.654804851 | 3.15E-11 | 44.07831143 | 1.574402975 | 9.738559326 |
| ENSMUSG00000034271 | *Jdp2* | 1072.873637 | 1177.820457 | 1231.942498 | 1804.030968 | 1814.227572 | 1851.652273 | 9.74E-14 | 0.654123022 | 1.31E-14 | 59.36594952 | 1.573659074 | 13.01154897 |
| ENSMUSG00000020272 | *Stk10* | 2780.044866 | 2871.37892 | 2002.242055 | 3920.963054 | 4063.927154 | 4063.777561 | 2.94E-06 | 0.653259743 | 7.59E-07 | 24.45991413 | 1.572717711 | 5.532078607 |
| ENSMUSG00000038122 | *Tbc1d32* | 206.905638 | 190.7524133 | 242.8993378 | 331.4559002 | 368.0660714 | 304.4209112 | 0.000141617 | 0.653049981 | 4.48E-05 | 16.65570757 | 1.57248906 | 3.848884964 |
| ENSMUSG00000020108 | *Ddit4* | 202.1124572 | 252.318007 | 217.4016172 | 312.0577178 | 372.4090339 | 372.0700026 | 0.000125555 | 0.652933933 | 3.95E-05 | 16.89513322 | 1.572362578 | 3.901165563 |
| ENSMUSG00000046603 | *Tcaim* | 137.4045164 | 115.0570112 | 162.3802203 | 232.7781895 | 211.7194216 | 203.9136897 | 0.001618488 | 0.652526322 | 0.00059356 | 11.79606617 | 1.571918393 | 2.79089061 |
| ENSMUSG00000048410 | *Zfp407* | 595.9521466 | 668.1380826 | 562.2918371 | 999.4280961 | 929.3939738 | 944.1880325 | 2.67E-09 | 0.652311775 | 5.11E-10 | 38.63538005 | 1.571684647 | 8.572971988 |
| ENSMUSG00000022364 | *Tbc1d31* | 670.2464491 | 593.4519525 | 530.0841901 | 1022.199876 | 906.5934207 | 895.867253 | 4.39E-07 | 0.651285717 | 1.04E-07 | 28.30433714 | 1.570567248 | 6.357092495 |
| ENSMUSG00000037331 | *Larp1* | 6291.848668 | 5916.352629 | 3732.061096 | 8300.735293 | 8616.437589 | 8128.521537 | 0.000265519 | 0.651243135 | 8.70E-05 | 15.40038353 | 1.570520893 | 3.575904373 |
| ENSMUSG00000031197 | *Vbp1* | 1030.533873 | 943.6693463 | 1261.466174 | 1706.196657 | 1685.069448 | 1679.630297 | 1.28E-07 | 0.651101361 | 2.88E-08 | 30.78835904 | 1.570366565 | 6.89299422 |
| ENSMUSG00000045757 | *Zfp764* | 288.3897117 | 306.8186966 | 297.9207347 | 470.6167744 | 474.4686525 | 456.148159 | 1.09E-07 | 0.650796342 | 2.42E-08 | 31.12322811 | 1.570034588 | 6.963690323 |
| ENSMUSG00000030965 | *Abraxas2* | 992.9872898 | 1055.698541 | 1249.388306 | 1724.75144 | 1676.383523 | 1763.708454 | 3.66E-09 | 0.650720208 | 7.08E-10 | 37.99934981 | 1.569951736 | 8.436887259 |
| ENSMUSG00000043881 | *Kbtbd7* | 134.2090625 | 119.0940993 | 96.62294099 | 196.5120223 | 176.9757216 | 179.7532999 | 0.002497562 | 0.650199952 | 0.000939429 | 10.94329626 | 1.569385692 | 2.602483739 |
| ENSMUSG00000040859 | *Bsdc1* | 800.4611942 | 925.5024498 | 968.9133805 | 1418.597517 | 1416.891514 | 1380.041464 | 4.15E-10 | 0.649089404 | 7.40E-11 | 42.40926065 | 1.568178086 | 9.381820062 |
| ENSMUSG00000039988 | *Ankrd13c* | 1099.236131 | 1239.386051 | 1070.904263 | 1817.525356 | 1790.386288 | 1740.51448 | 1.64E-12 | 0.648819715 | 2.41E-13 | 53.63709323 | 1.567884967 | 11.78469857 |
| ENSMUSG00000035764 | *Fbxo45* | 574.382833 | 546.0161672 | 458.9589697 | 836.6520433 | 876.1926832 | 771.1996417 | 4.33E-07 | 0.648031348 | 1.02E-07 | 28.33658078 | 1.567028424 | 6.364000266 |
| ENSMUSG00000034667 | *Xpot* | 6299.837303 | 6223.171325 | 5561.187048 | 9476.43383 | 9462.229534 | 9413.854273 | 1.07E-18 | 0.647989276 | 1.03E-19 | 82.54151011 | 1.566982726 | 17.97231824 |
| ENSMUSG00000034575 | *Tent4a* | 1435.557651 | 1486.657697 | 1097.743968 | 2120.305682 | 2066.164407 | 2123.215054 | 3.63E-07 | 0.647954073 | 8.51E-08 | 28.68643101 | 1.566944491 | 6.439746263 |
| ENSMUSG00000028028 | *Alpk1* | 579.1760138 | 525.8307267 | 450.9070579 | 875.4484082 | 758.9326959 | 810.822681 | 1.39E-06 | 0.647937423 | 3.46E-07 | 25.97680827 | 1.566926407 | 5.857381913 |
| ENSMUSG00000021583 | *Erap1* | 3308.093618 | 3101.492942 | 3451.58617 | 5200.399697 | 5167.039628 | 5076.5811 | 2.75E-20 | 0.647880897 | 2.44E-21 | 89.9508701 | 1.566865015 | 19.56028969 |
| ENSMUSG00000019852 | *Arfgef3* | 1069.678183 | 1222.228426 | 815.9270572 | 1680.89468 | 1640.554082 | 1555.929102 | 1.93E-05 | 0.647860733 | 5.47E-06 | 20.66508537 | 1.566843115 | 4.713387381 |
| ENSMUSG00000036580 | *Spg20* | 690.2180357 | 682.267891 | 680.3865428 | 1066.056636 | 1015.167483 | 1134.571904 | 1.01E-11 | 0.647550275 | 1.57E-12 | 49.96012059 | 1.566505978 | 10.9976151 |
| ENSMUSG00000098022 | *Zfp82* | 87.07611793 | 88.81593848 | 96.62294099 | 148.4382658 | 144.4035029 | 132.398936 | 0.003156561 | 0.647364057 | 0.001207983 | 10.47809377 | 1.566303792 | 2.50078576 |
| ENSMUSG00000078862 | *Gm14326* | 146.1920145 | 139.2795399 | 162.3802203 | 274.9481513 | 199.7762747 | 223.2420016 | 0.001815483 | 0.647202684 | 0.000670092 | 11.57042326 | 1.566128602 | 2.741007738 |
| ENSMUSG00000024645 | *Timm21* | 409.8169587 | 323.976321 | 437.487205 | 608.9342493 | 594.9858617 | 626.2373031 | 1.05E-05 | 0.646407742 | 2.87E-06 | 21.89912118 | 1.565265884 | 4.980375817 |
| ENSMUSG00000020522 | *Mfap3* | 1108.822493 | 1171.764825 | 980.9912481 | 1771.981797 | 1694.841113 | 1645.805752 | 6.49E-11 | 0.646389215 | 1.08E-11 | 46.17387251 | 1.565245783 | 10.1880407 |
| ENSMUSG00000030330 | *Ing4* | 419.4033203 | 361.3193861 | 366.3619846 | 602.1870554 | 573.2710492 | 624.3044719 | 2.03E-07 | 0.646200313 | 4.65E-08 | 29.85745521 | 1.565040848 | 6.692245857 |
| ENSMUSG00000040325 | *Dcaf1* | 703.7987147 | 793.2878141 | 873.6324248 | 1215.338301 | 1235.57283 | 1247.642528 | 3.73E-09 | 0.645869113 | 7.23E-10 | 37.95629843 | 1.564681603 | 8.428362427 |
| ENSMUSG00000021707 | *Dhfr* | 1634.474654 | 1302.970188 | 1666.745732 | 2418.869012 | 2396.229556 | 2386.080094 | 1.14E-08 | 0.645832986 | 2.31E-09 | 35.6895777 | 1.564642422 | 7.944568156 |
| ENSMUSG00000079614 | *Seh1l* | 1674.417827 | 1512.89877 | 1830.467938 | 2629.718822 | 2533.032875 | 2678.904019 | 1.06E-11 | 0.64567699 | 1.66E-12 | 49.84583629 | 1.564473249 | 10.97330389 |
| ENSMUSG00000036112 | *Metap2* | 3773.032156 | 3572.82298 | 4509.070579 | 6317.903686 | 6063.861383 | 6147.359575 | 2.95E-10 | 0.645347268 | 5.19E-11 | 43.10289899 | 1.564115736 | 9.530320692 |
| ENSMUSG00000064345 | *mt-Nd2* | 20295.12639 | 19529.41374 | 24748.89275 | 33281.3773 | 34218.20149 | 33479.53532 | 2.68E-10 | 0.645237975 | 4.71E-11 | 43.29615053 | 1.563997249 | 9.57158125 |
| ENSMUSG00000050549 | *Fam241a* | 174.1522359 | 162.4927965 | 166.4061761 | 232.7781895 | 262.7492309 | 292.8239241 | 0.0002837 | 0.645172678 | 9.32E-05 | 15.2691421 | 1.563926463 | 3.547141011 |
| ENSMUSG00000042396 | *Rbm7* | 1131.989533 | 1110.199231 | 1508.391468 | 2080.665918 | 1835.987394 | 1935.730429 | 3.99E-06 | 0.644895231 | 1.05E-06 | 23.84008287 | 1.563625731 | 5.398895256 |
| ENSMUSG00000028885 | *Smpdl3b* | 1572.163304 | 1381.693407 | 1151.42338 | 2193.681416 | 2199.710503 | 2036.237651 | 3.92E-07 | 0.64447695 | 9.23E-08 | 28.52857462 | 1.563172455 | 6.40619816 |
| ENSMUSG00000019948 | *Actr6* | 260.4294903 | 284.6147119 | 374.4138963 | 430.133611 | 482.0688369 | 517.9987568 | 0.000400729 | 0.644463989 | 0.000134286 | 14.58065876 | 1.563158411 | 3.397149402 |
| ENSMUSG00000029836 | *Cbx3* | 1737.528041 | 1758.151873 | 1970.034408 | 2991.537094 | 2740.409334 | 2798.739552 | 1.84E-13 | 0.644450915 | 2.53E-14 | 58.07060743 | 1.563144245 | 12.73543148 |
| ENSMUSG00000044573 | *Acp1* | 1518.639451 | 1396.832487 | 1696.269408 | 2424.772807 | 2413.601406 | 2360.953289 | 1.31E-11 | 0.644410411 | 2.07E-12 | 49.41764613 | 1.563100361 | 10.8838938 |
| ENSMUSG00000074182 | *Znhit6* | 548.819202 | 560.1459756 | 591.8155136 | 919.3051686 | 875.1069426 | 858.1770449 | 7.98E-11 | 0.644324422 | 1.35E-11 | 45.74691371 | 1.563007198 | 10.09782243 |
| ENSMUSG00000041303 | *Gtf3c3* | 730.1612091 | 750.8983889 | 907.1820571 | 1252.447867 | 1256.201901 | 1211.885151 | 1.65E-08 | 0.64389989 | 3.41E-09 | 34.93519307 | 1.562547329 | 7.783121761 |
| ENSMUSG00000020650 | *Bcap29* | 988.9929725 | 901.2799211 | 1103.11191 | 1539.203608 | 1539.580204 | 1589.753647 | 3.17E-10 | 0.643615918 | 5.61E-11 | 42.95157834 | 1.562239797 | 9.499197287 |
| ENSMUSG00000098950 | *Gm28036* | 675.8384933 | 659.0546344 | 518.0063225 | 982.5601114 | 959.7947112 | 961.5835132 | 5.62E-07 | 0.643283427 | 1.34E-07 | 27.80810672 | 1.561881962 | 6.250271086 |
| ENSMUSG00000041360 | *Pum3* | 1966.002993 | 1934.774478 | 2484.014775 | 3448.659481 | 3213.792246 | 3292.577919 | 3.07E-08 | 0.642745375 | 6.47E-09 | 33.68653064 | 1.561297404 | 7.513523016 |
| ENSMUSG00000054611 | *Kdm2a* | 2262.381339 | 2578.690032 | 1850.597717 | 3482.395451 | 3480.884439 | 3487.793868 | 3.00E-07 | 0.641971949 | 6.99E-08 | 29.06722978 | 1.560460619 | 6.522494448 |
| ENSMUSG00000022255 | *Mtdh* | 4702.909232 | 4722.38382 | 4781.493594 | 7560.230763 | 7246.232922 | 7346.691324 | 1.38E-29 | 0.641296547 | 7.49E-31 | 133.3740642 | 1.559730256 | 28.8603749 |
| ENSMUSG00000025404 | *R3hdm2* | 1672.820101 | 1653.187582 | 1139.345512 | 2275.491142 | 2403.829741 | 2296.203445 | 2.65E-05 | 0.641184244 | 7.65E-06 | 20.02434965 | 1.559608848 | 4.576504509 |
| ENSMUSG00000040725 | *Hnrnpul1* | 4223.591152 | 4019.930488 | 3016.782935 | 5757.886593 | 5905.343252 | 5905.765677 | 8.04E-07 | 0.640688992 | 1.95E-07 | 27.08391952 | 1.559073553 | 6.094682121 |
| ENSMUSG00000032913 | *Lrig2* | 306.7635714 | 281.5868959 | 309.9986023 | 506.0395423 | 442.9821744 | 449.3832499 | 2.03E-06 | 0.640678955 | 5.15E-07 | 25.20862304 | 1.559062706 | 5.693356213 |
| ENSMUSG00000023015 | *Racgap1* | 1846.972336 | 1510.880226 | 1171.553159 | 2289.828929 | 2441.830662 | 2341.624977 | 6.28E-05 | 0.640579557 | 1.89E-05 | 18.29355407 | 1.558955294 | 4.202177949 |
| ENSMUSG00000034826 | *Nup54* | 1429.965606 | 1361.507966 | 1536.573159 | 2309.227112 | 2199.710503 | 2227.587938 | 7.43E-15 | 0.640196359 | 9.26E-16 | 64.58160033 | 1.558541272 | 14.12899101 |
| ENSMUSG00000048732 | *Klhl11* | 278.8033501 | 288.6518 | 275.1069848 | 460.4959835 | 437.5534713 | 414.5922886 | 8.36E-07 | 0.63965733 | 2.03E-07 | 27.00631141 | 1.557959067 | 6.077985864 |
| ENSMUSG00000020409 | *Slu7* | 1155.955437 | 1109.189959 | 1468.131909 | 1947.408839 | 1923.932385 | 1930.898351 | 2.35E-07 | 0.63905413 | 5.40E-08 | 29.56622489 | 1.557307811 | 6.629346131 |

| | | | | | | | | | | | | |
|---|---|---|---|---|---|---|---|---|---|---|---|---|
| ENSMUSG00000036087 | *Slain2* | 2182.494993 | 2183.055397 | 1829.125952 | 3324.679793 | 3189.905952 | 3142.783502 | 2.67E-11 | 0.638873725 | 4.33E-12 | 47.96863971 | 1.557113087 | 10.5729471 |
| ENSMUSG00000048799 | *Cep120* | 691.8157627 | 634.8321057 | 740.7758809 | 1015.452682 | 1042.310999 | 1156.799463 | 9.34E-09 | 0.638507771 | 1.88E-09 | 36.0933006 | 1.556718159 | 8.029876356 |
| ENSMUSG00000031668 | *Eif2ak3* | 360.2874237 | 370.4028343 | 276.4489701 | 553.2698996 | 512.4695743 | 508.3346009 | 3.23E-05 | 0.638255967 | 9.42E-06 | 19.62639818 | 1.556446478 | 4.491111776 |
| ENSMUSG00000056429 | *Tgoln1* | 3991.920746 | 4585.122824 | 5724.909254 | 7602.400724 | 7444.923456 | 7197.863323 | 1.45E-06 | 0.638198956 | 3.61E-07 | 25.89411251 | 1.556384973 | 5.839600962 |
| ENSMUSG00000015755 | *Tab2* | 2335.077915 | 2225.444822 | 2065.315364 | 3521.191815 | 3530.828508 | 3268.417529 | 1.02E-14 | 0.637761769 | 1.29E-15 | 63.92841842 | 1.555913405 | 13.99032463 |
| ENSMUSG00000040782 | *Cop1* | 897.9225372 | 868.9832162 | 767.6155867 | 1318.233008 | 1348.489854 | 1286.299152 | 7.80E-11 | 0.63740667 | 1.31E-11 | 45.79560539 | 1.555530486 | 10.10785657 |
| ENSMUSG00000036591 | *Arhgap21* | 1326.912219 | 1421.055016 | 1010.514925 | 1940.661645 | 2004.277191 | 1910.603624 | 2.69E-06 | 0.637141326 | 6.91E-07 | 24.64063133 | 1.555244415 | 5.570820767 |
| ENSMUSG00000036461 | *Elf1* | 1134.386124 | 1188.922449 | 821.2949984 | 1665.713494 | 1728.499073 | 1501.809829 | 3.45E-05 | 0.635955804 | 1.01E-05 | 19.49052022 | 1.553966932 | 4.461626234 |
| ENSMUSG00000027164 | *Traf6* | 1092.845223 | 1174.792641 | 1248.046321 | 1809.934763 | 1846.844801 | 1795.600168 | 1.86E-13 | 0.635778685 | 2.56E-14 | 58.04940602 | 1.553776164 | 12.73149581 |
| ENSMUSG00000027810 | *Eif2a* | 1173.530433 | 1310.035093 | 1631.854114 | 2198.741811 | 2032.506447 | 2146.409028 | 1.99E-06 | 0.635001896 | 5.04E-07 | 25.2487092 | 1.55293979 | 5.701675667 |
| ENSMUSG00000022983 | *Scaf4* | 1356.470167 | 1153.597928 | 727.356028 | 1749.210018 | 1679.640745 | 1606.182712 | 0.002321957 | 0.634462586 | 0.000870323 | 11.08492612 | 1.552359377 | 2.634145757 |
| ENSMUSG00000024002 | *Brd4* | 2106.602963 | 2156.814324 | 1229.258527 | 2805.145863 | 2888.070059 | 2841.261838 | 0.001363581 | 0.634221615 | 0.000494202 | 12.13741757 | 1.552100111 | 2.865318991 |
| ENSMUSG00000029416 | *Slc15a4* | 1217.467924 | 1275.719844 | 1073.588233 | 1820.055554 | 1867.473873 | 1857.450766 | 1.46E-11 | 0.633988035 | 2.32E-12 | 49.1932558 | 1.551848838 | 10.83567074 |
| ENSMUSG00000032599 | *Ip6k2* | 398.6328702 | 427.9313399 | 479.0887491 | 662.9118005 | 692.7025178 | 662.9610955 | 1.24E-07 | 0.633840101 | 2.78E-08 | 30.85752562 | 1.551689719 | 6.907332311 |
| ENSMUSG00000036023 | *Parp2* | 453.7544494 | 375.4491945 | 352.9421317 | 748.9385227 | 606.9290086 | 481.2749644 | 0.001604517 | 0.633748973 | 0.000588342 | 11.81250562 | 1.55159171 | 2.794655549 |
| ENSMUSG00000090035 | *Galnt4* | 214.8942727 | 196.8080455 | 197.2718379 | 313.7445163 | 318.1220027 | 315.0514827 | 1.61E-05 | 0.633129197 | 4.50E-06 | 21.03863049 | 1.550925296 | 4.794261925 |
| ENSMUSG00000026116 | *Tmem131* | 1414.787201 | 1368.57287 | 799.8232337 | 1903.552078 | 1862.045169 | 1800.432246 | 0.001831503 | 0.633056901 | 0.000676545 | 11.55260277 | 1.550847579 | 2.737192442 |
| ENSMUSG00000024867 | *Pip5k1b* | 986.5963821 | 1065.791262 | 1078.956174 | 1663.183296 | 1574.323904 | 1610.048375 | 7.97E-14 | 0.633009128 | 1.07E-14 | 59.77055262 | 1.550796224 | 13.09854272 |
| ENSMUSG00000030256 | *Bhlhe41* | 611.9294159 | 568.2201518 | 509.9544108 | 885.5691991 | 897.9074957 | 845.6136422 | 1.59E-08 | 0.632487813 | 3.28E-09 | 35.00775088 | 1.550235949 | 7.798654881 |
| ENSMUSG00000014599 | *Csf1* | 358.6896968 | 342.1432175 | 216.0596319 | 437.7242041 | 482.0688369 | 509.3010165 | 0.002995066 | 0.632413346 | 0.00114277 | 10.58065721 | 1.550155933 | 2.523593653 |
| ENSMUSG00000042632 | *Pla2g6* | 374.6669662 | 384.5326427 | 305.9726465 | 513.6301355 | 542.8703118 | 601.1104977 | 1.54E-05 | 0.632255998 | 4.29E-06 | 21.13012912 | 1.549986873 | 4.813533866 |
| ENSMUSG00000022672 | *Prkdc* | 502.4851209 | 509.6823742 | 598.52544 | 821.4708571 | 838.1917614 | 828.2181616 | 2.56E-08 | 0.632213109 | 5.38E-09 | 34.04628384 | 1.549940795 | 7.591761786 |
| ENSMUSG00000089929 | *Bcl2a1b* | 1997.957532 | 1692.549191 | 1724.4511 | 2827.074243 | 2653.550084 | 2917.60867 | 9.33E-11 | 0.631782803 | 1.58E-11 | 45.43418244 | 1.549478571 | 10.03030072 |
| ENSMUSG00000023206 | *Il15ra* | 161.3704204 | 141.2980839 | 128.830588 | 201.5724177 | 261.6634903 | 209.7121833 | 0.002551086 | 0.631184558 | 0.000961023 | 10.90119304 | 1.548836179 | 2.593274923 |
| ENSMUSG00000059486 | *Kbtbd2* | 1056.097504 | 1082.948886 | 1087.008086 | 1725.594839 | 1681.812226 | 1585.887985 | 1.20E-14 | 0.631154211 | 1.52E-15 | 63.60994107 | 1.5488036 | 13.9217373 |
| ENSMUSG00000038485 | *Socs7* | 1321.320175 | 1293.88674 | 874.9744101 | 1812.464961 | 1814.272582 | 1786.902428 | 8.56E-05 | 0.630504369 | 2.63E-05 | 17.66865895 | 1.54810612 | 4.067325395 |
| ENSMUSG00000021693 | *Kif2a* | 653.4703162 | 652.9990022 | 657.5727928 | 1021.356476 | 1027.11063 | 990.5759809 | 1.43E-12 | 0.629969598 | 2.08E-13 | 53.92435545 | 1.547532382 | 11.84592279 |
| ENSMUSG00000042997 | *Nhlrc3* | 1029.735009 | 1113.227047 | 1017.224851 | 1651.375707 | 1648.154267 | 1592.652894 | 4.05E-14 | 0.629939582 | 5.31E-15 | 61.1422471 | 1.547500185 | 13.39246715 |
| ENSMUSG00000047888 | *Tnrc6b* | 222.8829074 | 190.7524133 | 127.4886027 | 269.0443567 | 301.8358933 | 273.4956123 | 0.007736949 | 0.629881437 | 0.003168935 | 8.707557819 | 1.547437818 | 2.111430254 |
| ENSMUSG00000037652 | *Phc3* | 563.9976079 | 613.6373931 | 497.8765432 | 846.7728342 | 885.9643488 | 864.9419541 | 4.90E-08 | 0.628797913 | 1.06E-08 | 32.73388416 | 1.546276064 | 7.309883352 |
| ENSMUSG00000073791 | *Efcab7* | 169.3590551 | 166.5298846 | 194.5878673 | 281.6953452 | 290.9784871 | 244.5031446 | 0.000306499 | 0.628557855 | 0.000101117 | 15.11573035 | 1.546018791 | 3.513570839 |
| ENSMUSG00000021697 | *Depdc1b* | 99.85793341 | 88.81593848 | 77.83514691 | 131.570281 | 133.5460967 | 149.7944166 | 0.007358844 | 0.628124552 | 0.003000605 | 8.80710034 | 1.545554526 | 2.133190388 |
| ENSMUSG00000049723 | *Mmp12* | 199.7158668 | 219.0120301 | 252.2932348 | 363.5050712 | 334.4081121 | 332.4469633 | 8.69E-05 | 0.627845365 | 2.67E-05 | 17.63737122 | 1.545255462 | 4.061098558 |
| ENSMUSG00000001305 | *Rrp15* | 1067.281592 | 1012.299844 | 962.203454 | 1608.362345 | 1568.895201 | 1530.802296 | 2.11E-13 | 0.627842545 | 2.91E-14 | 57.792156 | 1.545252442 | 12.67637242 |
| ENSMUSG00000006169 | *Clint1* | 2541.983553 | 2550.430415 | 2289.426907 | 3771.681389 | 3791.406257 | 3852.132546 | 6.07E-17 | 0.627390487 | 6.60E-18 | 74.33237357 | 1.544768324 | 16.21688619 |
| ENSMUSG00000033450 | *Tagap* | 252.4408557 | 308.8372406 | 234.847426 | 425.9166148 | 422.3531026 | 383.6669897 | 9.38E-05 | 0.62727699 | 2.90E-05 | 17.4824035 | 1.544646802 | 4.027699746 |
| ENSMUSG00000041642 | *Kif21b* | 1056.097504 | 1056.707813 | 650.8628664 | 1441.369296 | 1427.74892 | 1408.067516 | 0.000745224 | 0.626985671 | 0.000260371 | 13.33591389 | 1.544334927 | 3.127713025 |
| ENSMUSG00000029177 | *Cenpa* | 2030.710934 | 1711.72536 | 1501.681541 | 2638.152814 | 2669.836193 | 2801.638799 | 2.99E-07 | 0.626644054 | 6.97E-08 | 29.07293679 | 1.543969287 | 6.523663712 |
| ENSMUSG00000027951 | *Adar* | 5658.349939 | 5650.914086 | 4580.1958 | 8274.589916 | 8138.711714 | 8130.454368 | 3.15E-10 | 0.626488375 | 5.58E-11 | 42.96394979 | 1.543802688 | 9.501509264 |
| ENSMUSG00000028059 | *Arhgef2* | 5355.580685 | 5148.296616 | 3719.983228 | 7333.356368 | 7364.57865 | 7264.545999 | 5.66E-06 | 0.625902363 | 1.51E-06 | 23.13530658 | 1.543175735 | 5.24716687 |
| ENSMUSG00000029657 | *Hsph1* | 6065.770307 | 6415.942283 | 6356.984326 | 9904.880642 | 9589.261187 | 9553.984534 | 8.02E-26 | 0.625183321 | 5.25E-27 | 115.8009596 | 1.542406804 | 25.09575458 |
| ENSMUSG00000020458 | *Rtn4* | 7624.352932 | 7694.689943 | 6259.0194 | 10891.65775 | 11131.01287 | 11269.37221 | 1.16E-10 | 0.624997832 | 1.98E-11 | 44.99083936 | 1.542208508 | 9.935157798 |
| ENSMUSG00000026516 | *Nvl* | 1894.105281 | 1825.773099 | 2051.895511 | 3037.080653 | 2852.240618 | 2995.888332 | 7.63E-15 | 0.624083255 | 9.53E-16 | 64.52576148 | 1.541231156 | 14.1172998 |
| ENSMUSG00000022200 | *Golph3* | 1195.099747 | 1301.960916 | 1305.751689 | 2105.124496 | 1822.958507 | 1921.234195 | 6.27E-11 | 0.62330209 | 1.05E-11 | 46.24133671 | 1.540396862 | 10.20284132 |
| ENSMUSG00000024772 | *Ehd1* | 8012.600577 | 8150.880899 | 7779.488735 | 11961.08798 | 12272.12627 | 12655.21217 | 3.33E-26 | 0.623283616 | 2.13E-27 | 117.5901267 | 1.540377138 | 25.4780487 |
| ENSMUSG00000037313 | *Tacc3* | 1777.471215 | 1576.482908 | 1256.098233 | 2371.638655 | 2368.0003 | 2373.516692 | 2.57E-06 | 0.623150232 | 6.59E-07 | 24.73204332 | 1.540234729 | 5.590222087 |
| ENSMUSG00000027366 | *Sppl2a* | 2591.513088 | 2438.40122 | 2870.506539 | 4237.237768 | 3936.895501 | 3982.598651 | 1.89E-12 | 0.623097145 | 2.79E-13 | 53.34757329 | 1.540178054 | 11.72399779 |
| ENSMUSG00000042772 | *Smg7* | 1985.97458 | 1948.904286 | 1735.186982 | 2960.331323 | 2901.098946 | 2881.851293 | 4.09E-14 | 0.62286006 | 5.36E-15 | 61.12220882 | 1.53992497 | 13.38824236 |
| ENSMUSG00000032193 | *Ldlr* | 6978.87125 | 6631.926497 | 4774.783667 | 9465.469639 | 9778.180056 | 9080.440894 | 1.15E-05 | 0.622811928 | 3.18E-06 | 21.70393315 | 1.539873595 | 4.937498903 |
| ENSMUSG00000074634 | *Tmem267* | 186.9340513 | 188.7338693 | 154.3283085 | 258.9235658 | 293.1499684 | 268.6635343 | 0.000314326 | 0.622243841 | 0.000103848 | 15.06543209 | 1.539267361 | 3.502620001 |
| ENSMUSG00000039217 | *Il18* | 308.3612984 | 246.2623749 | 224.1115437 | 355.0710789 | 433.2105088 | 417.4915354 | 0.000643079 | 0.621938167 | 0.000222406 | 13.63163946 | 1.53894126 | 3.19173598 |
| ENSMUSG00000036779 | *Tent4b* | 1019.349784 | 1030.466741 | 821.2949984 | 1491.973251 | 1477.692989 | 1458.321127 | 4.17E-08 | 0.621603943 | 8.93E-09 | 33.06167468 | 1.53858478 | 7.3803692 |
| ENSMUSG00000040167 | *Ikzf5* | 349.9021987 | 361.3193861 | 477.7467638 | 599.6568577 | 605.8432679 | 614.640316 | 5.72E-05 | 0.621474036 | 1.72E-05 | 18.4799047 | 1.538446246 | 4.24291832 |
| ENSMUSG00000031174 | *Rpgr* | 87.8749814 | 104.9642909 | 110.0427939 | 162.7760528 | 140.0605404 | 159.4585725 | 0.005655721 | 0.620705908 | 0.002262422 | 9.323429129 | 1.537627356 | 2.247512019 |
| ENSMUSG00000026622 | *Nek2* | 1579.353075 | 1395.823215 | 1054.800439 | 2084.882914 | 2060.735704 | 2062.330871 | 2.90E-05 | 0.620626946 | 8.39E-06 | 19.84591431 | 1.5375432 | 4.538263342 |
| ENSMUSG00000026238 | *Ptma* | 24577.03457 | 22169.66937 | 26898.75318 | 37985.85825 | 36847.86528 | 38322.24385 | 9.36E-13 | 0.619776496 | 1.35E-13 | 54.77356171 | 1.536637105 | 12.02884202 |
| ENSMUSG00000091625 | *Lsm5* | 321.1431138 | 288.6518 | 394.5436757 | 513.6301355 | 478.811615 | 544.0919778 | 9.75E-05 | 0.619700019 | 3.02E-05 | 17.40619489 | 1.536555651 | 4.01120021 |
| ENSMUSG00000038563 | *Efl1* | 734.9543899 | 678.2308029 | 648.1788958 | 1084.611419 | 1098.769511 | 990.5759809 | 2.89E-09 | 0.619474245 | 5.55E-10 | 38.47509193 | 1.536315206 | 8.539044927 |
| ENSMUSG00000049775 | *Tmsb4x* | 13152.48813 | 11768.11185 | 15512.00798 | 20580.62819 | 20377.18002 | 21146.13955 | 1.14E-08 | 0.619424829 | 2.33E-09 | 35.67982632 | 1.536262585 | 7.942897361 |
| ENSMUSG00000040446 | *Rprd1a* | 532.0430692 | 514.7287344 | 599.8674253 | 875.4484082 | 824.0771333 | 822.4196681 | 1.85E-08 | 0.619391825 | 3.83E-09 | 34.70610565 | 1.536227441 | 7.73335007 |
| ENSMUSG00000046519 | *Golph3l* | 941.8600279 | 962.8455148 | 1273.544042 | 1622.700132 | 1679.640745 | 1566.559673 | 5.69E-06 | 0.618654532 | 1.52E-06 | 23.12259487 | 1.535442548 | 5.244583969 |
| ENSMUSG00000089936 | *Gm16199* | 83.0818006 | 93.86229862 | 87.22904395 | 136.6306764 | 127.031653 | 141.0966763 | 0.005935267 | 0.61839647 | 0.002382654 | 9.22858756 | 1.535167921 | 2.226559746 |
| ENSMUSG00000040613 | *Apobec1* | 4692.524007 | 4558.881751 | 4915.692123 | 7425.286885 | 7087.714791 | 7196.896907 | 7.82E-22 | 0.616472507 | 6.37E-23 | 97.16632393 | 1.533122001 | 21.10706913 |
| ENSMUSG00000028678 | *Kif2c* | 1180.720205 | 962.8455148 | 876.3163954 | 1565.348984 | 1586.267051 | 1489.246426 | 1.84E-06 | 0.616396938 | 4.64E-07 | 25.40989296 | 1.533041698 | 5.735837738 |
| ENSMUSG00000058325 | *Dock1* | 643.0850912 | 695.3884274 | 595.8414694 | 1025.573472 | 995.6241518 | 948.0536949 | 4.90E-09 | 0.616261874 | 9.60E-10 | 37.40392753 | 1.532898182 | 8.309460057 |
| ENSMUSG00000002227 | *Mov10* | 2436.533575 | 2066.989114 | 1478.867791 | 3077.563817 | 3105.218183 | 2993.955501 | 0.000286409 | 0.615468565 | 9.42E-05 | 15.24915548 | 1.532055504 | 3.543013103 |
| ENSMUSG00000039005 | *Tlr4* | 877.1520871 | 1019.364748 | 1371.508968 | 1682.581478 | 1706.78426 | 1605.216297 | 0.000205659 | 0.615175441 | 6.66E-05 | 15.90613847 | 1.531744256 | 3.686851983 |
| ENSMUSG00000030145 | *Zfp248* | 87.07611793 | 84.77885036 | 89.91301453 | 134.943878 | 140.0605404 | 125.6340268 | 0.00590311 | 0.615120268 | 0.002368351 | 9.239612132 | 1.531685679 | 2.228919124 |
| ENSMUSG00000060427 | *Zfp868* | 299.5738002 | 279.5683518 | 326.1024258 | 483.2677629 | 475.5543931 | 424.2564445 | 8.95E-06 | 0.615026295 | 2.44E-06 | 22.21067896 | 1.531585912 | 5.048340265 |
| ENSMUSG00000041417 | *Pik3r1* | 663.8555413 | 674.1937148 | 562.2918371 | 941.2335488 | 1027.11063 | 950.9529417 | 7.80E-08 | 0.614776194 | 1.71E-08 | 31.79554313 | 1.531320425 | 7.107923539 |
| ENSMUSG00000024498 | *Tcerg1* | 1768.683717 | 1652.17831 | 1235.968453 | 2310.070511 | 2517.832506 | 2310.699678 | 1.57E-05 | 0.613815415 | 4.39E-06 | 21.08579933 | 1.530300963 | 4.804403848 |
| ENSMUSG00000090553 | *Snrpe* | 1406.798566 | 1164.69992 | 1657.351835 | 2108.498093 | 2180.167172 | 2175.401496 | 5.19E-06 | 0.613789364 | 1.38E-06 | 23.30865295 | 1.530273331 | 5.28448364 |
| ENSMUSG00000049124 | *Gm8186* | 1131.989533 | 1059.73563 | 1248.046321 | 1709.570254 | 1702.441298 | 1841.988117 | 3.02E-10 | 0.613155061 | 5.32E-11 | 43.05408026 | 1.52960067 | 9.520356414 |
| ENSMUSG00000026788 | *Zbtb43* | 618.3203237 | 676.2122589 | 668.3086752 | 1001.114895 | 984.7667456 | 1010.870708 | 8.87E-11 | 0.612871677 | 1.50E-11 | 45.53527066 | 1.529300245 | 10.05196088 |
| ENSMUSG00000022043 | *Trim35* | 4831.52625 | 4760.736157 | 3801.844331 | 6777.55627 | 6945.482769 | 6769.741216 | 9.00E-09 | 0.612532311 | 1.81E-09 | 36.16818017 | 1.52894055 | 8.045800319 |
| ENSMUSG00000027952 | *Pmvk* | 1080.862271 | 1052.670725 | 920.60191 | 1570.40938 | 1559.123535 | 1550.130608 | 1.09E-10 | 0.612322362 | 1.85E-11 | 45.12463474 | 1.528718066 | 9.963927121 |
| ENSMUSG00000041912 | *Tdrkh* | 930.6759394 | 837.6957834 | 1013.198895 | 1495.346848 | 1401.691145 | 1347.183334 | 1.19E-08 | 0.611916698 | 2.44E-09 | 35.58507274 | 1.528288275 | 7.922877192 |
| ENSMUSG00000024006 | *Stk38* | 1151.96112 | 1234.33969 | 1293.673821 | 1881.623698 | 1834.901654 | 1896.10739 | 2.22E-13 | 0.611666228 | 3.08E-14 | 57.68027616 | 1.528022967 | 12.65296627 |
| ENSMUSG00000035692 | *Isg15* | 2688.974431 | 2204.250109 | 2151.202422 | 3568.422173 | 3544.943136 | 3661.748675 | 7.68E-09 | 0.611551975 | 1.53E-09 | 36.49082662 | 1.527901961 | 8.114755779 |
| ENSMUSG00000042447 | *Mios* | 426.5930915 | 361.3193861 | 373.071911 | 599.6568577 | 619.9578961 | 558.5882117 | 2.00E-06 | 0.610967637 | 5.08E-07 | 25.2339111 | 1.527283237 | 5.69877761 |
| ENSMUSG00000036712 | *Cyld* | 849.9907292 | 813.4732547 | 813.2430867 | 1245.700673 | 1277.916714 | 1263.105178 | 2.64E-13 | 0.610925243 | 3.68E-14 | 57.33368609 | 1.527238358 | 12.57883431 |
| ENSMUSG00000057234 | *Mettl15* | 114.2374758 | 103.9550189 | 124.8046321 | 169.5232467 | 168.2897967 | 183.6189623 | 0.002568936 | 0.608859424 | 0.000968354 | 10.88711719 | 1.525053045 | 2.590246702 |
| ENSMUSG00000034761 | *Map4k5* | 737.3509803 | 703.4626036 | 692.4644104 | 1093.045411 | 1121.570064 | 1042.762423 | 4.21E-11 | 0.608519884 | 6.89E-12 | 47.05707991 | 1.524694164 | 10.37618741 |
| ENSMUSG00000049739 | *Zfp646* | 1589.7383 | 1520.972946 | 1284.279924 | 2191.151218 | 2295.255678 | 2227.587938 | 3.72E-09 | 0.608453025 | 7.21E-10 | 37.96341627 | 1.524623507 | 8.429682526 |
| ENSMUSG00000021886 | *Gpr65* | 194.922686 | 149.3722602 | 218.7436025 | 271.5745544 | 296.4071902 | 287.9918462 | 0.001840719 | 0.608147474 | 0.000680493 | 11.54178548 | 1.524300638 | 2.735012426 |
| ENSMUSG00000025958 | *Creb1* | 1215.071334 | 1186.903905 | 1500.339556 | 2116.088686 | 1935.875532 | 1884.510403 | 3.76E-07 | 0.608102289 | 8.82E-08 | 28.61706616 | 1.524252898 | 6.425064213 |
| ENSMUSG00000021661 | *Ankra2* | 230.0726786 | 265.4385434 | 312.6825729 | 409.8920293 | 400.6382901 | 413.625873 | 0.00015683 | 0.608090392 | 5.00E-05 | 16.44987676 | 1.524240329 | 3.804571167 |
| ENSMUSG00000066892 | *Fbxl12* | 112.6397489 | 111.0199231 | 96.62294099 | 178.8006383 | 175.889981 | 135.2981828 | 0.008715553 | 0.608085645 | 0.003609877 | 8.470321269 | 1.524235313 | 2.059705054 |
| ENSMUSG00000028228 | *Cpne3* | 823.6282348 | 828.6123351 | 1053.458454 | 1450.646688 | 1416.891514 | 1246.676113 | 5.70E-06 | 0.608075983 | 1.52E-06 | 23.1207923 | 1.524225105 | 5.244272844 |
| ENSMUSG00000066878 | *Gm10184* | 3880.079861 | 3542.544819 | 3631.412199 | 5586.676547 | 5471.047002 | 5792.695053 | 5.91E-18 | 0.607655761 | 6.04E-19 | 79.05437958 | 1.5237812 | 17.22807429 |
| ENSMUSG00000035437 | *Rabgap1* | 1078.465681 | 1222.228426 | 1488.261688 | 1929.697455 | 1976.047935 | 1855.517935 | 3.53E-06 | 0.607595487 | 9.20E-07 | 24.08910805 | 1.52371754 | 5.45261897 |
| ENSMUSG00000034042 | *Gpbp1l1* | 1127.196352 | 1202.042986 | 1439.950218 | 1939.818246 | 1901.131832 | 1887.409649 | 6.33E-08 | 0.607147406 | 1.38E-08 | 32.21955344 | 1.523244368 | 7.198847359 |
| ENSMUSG00000040451 | *Sgms1* | 500.887394 | 533.9049029 | 538.1361019 | 803.7594731 | 793.6763958 | 793.4272003 | 1.09E-09 | 0.606657979 | 2.01E-10 | 40.45718572 | 1.522727702 | 8.963808974 |
| ENSMUSG00000027739 | *Rab33b* | 307.5624349 | 261.4014553 | 289.868823 | 454.5921889 | 440.8106932 | 412.6594574 | 1.26E-05 | 0.605697737 | 3.49E-06 | 21.52656574 | 1.521714529 | 4.898731485 |
| ENSMUSG00000052040 | *Klf13* | 805.254375 | 916.4190015 | 621.33919 | 1208.591107 | 1168.256911 | 1195.456086 | 5.65E-05 | 0.605505886 | 1.70E-05 | 18.50182819 | 1.521512183 | 4.247658007 |
| ENSMUSG00000032637 | *Atxn2l* | 4846.704656 | 4461.991636 | 3024.834847 | 6247.901549 | 6461.242451 | 6059.425756 | 0.000243155 | 0.604845324 | 7.93E-05 | 15.57454191 | 1.520815693 | 3.614116228 |
| ENSMUSG00000031959 | *Wdr59* | 400.2305971 | 371.4121064 | 366.3619846 | 557.4868958 | 623.2151179 | 554.7225493 | 8.36E-07 | 0.604717732 | 2.03E-07 | 27.00691886 | 1.520681198 | 6.078016716 |
| ENSMUSG00000054381 | *Zfp747* | 246.8488114 | 236.1696546 | 267.055073 | 397.2410407 | 353.9514433 | 385.5998209 | 2.27E-05 | 0.604652759 | 6.49E-06 | 20.33939422 | 1.520612714 | 4.64363984 |
| ENSMUSG00000048222 | *Mfap1b* | 472.1283092 | 516.7472784 | 582.4216165 | 779.3008952 | 718.7602928 | 883.3038503 | 1.26E-05 | 0.604507459 | 3.49E-06 | 21.52527848 | 1.520459575 | 4.898532698 |
| ENSMUSG00000022064 | *Pibf1* | 159.7726935 | 188.7338693 | 210.6916908 | 256.3933681 | 300.7501527 | 288.9582617 | 0.00105341 | 0.604491905 | 0.000375632 | 12.64967087 | 1.520443182 | 2.977402424 |
| ENSMUSG00000025340 | *Rabgef1* | 704.5975781 | 764.0189253 | 802.5072043 | 1158.830552 | 1171.514133 | 1113.310761 | 4.17E-10 | 0.603849309 | 7.43E-11 | 42.40142421 | 1.519766107 | 9.38022366 |
| ENSMUSG00000009741 | *Ubp1* | 1246.227009 | 1202.042986 | 1030.644704 | 1709.570254 | 1837.073135 | 1750.178636 | 4.08E-09 | 0.603096449 | 7.95E-10 | 37.77194479 | 1.518973235 | 8.388905532 |
| ENSMUSG00000025429 | *Pstpip2* | 312.3556157 | 348.1988497 | 326.1024258 | 485.7979606 | 490.7547618 | 520.8980036 | 1.28E-06 | 0.602959368 | 3.17E-07 | 26.14251106 | 1.518828913 | 5.892899919 |
| ENSMUSG00000026648 | *Dclre1c* | 341.1147005 | 302.7816084 | 314.0245582 | 490.8583561 | 521.1554993 | 445.5175875 | 9.42E-06 | 0.602432346 | 2.58E-06 | 22.10792336 | 1.51827418 | 5.025752009 |
| ENSMUSG00000052681 | *Rap1b* | 6744.804254 | 6243.356766 | 6150.318591 | 9821.384118 | 9519.773787 | 9723.107262 | 1.01E-19 | 0.602231865 | 9.20E-21 | 87.32690921 | 1.518063212 | 18.99758541 |
| ENSMUSG00000035762 | *Tmem161b* | 321.1431138 | 268.4663595 | 366.3619846 | 498.4489492 | 490.7547618 | 457.1145746 | 0.000120749 | 0.601822005 | 3.79E-05 | 16.97385917 | 1.517632001 | 3.918115708 |
| ENSMUSG00000038855 | *Itpkb* | 1604.916706 | 1663.280302 | 1380.902865 | 2380.072647 | 2319.141972 | 2365.785367 | 2.76E-10 | 0.601678123 | 4.86E-11 | 43.23473016 | 1.517480653 | 9.558562951 |
| ENSMUSG00000038005 | *Hpf1* | 1311.733813 | 1172.774097 | 1217.180659 | 1959.216428 | 1840.330357 | 1820.726974 | 6.45E-12 | 0.601518567 | 9.95E-13 | 50.85491268 | 1.517312836 | 11.19034351 |
| ENSMUSG00000113611 | *Gm47750* | 494.4964863 | 452.1538686 | 468.3528667 | 694.9609715 | 712.2458491 | 742.207174 | 2.02E-08 | 0.601166937 | 4.21E-09 | 34.52456873 | 1.516943063 | 7.694451711 |
| ENSMUSG00000064120 | *Mocs1* | 313.1544792 | 301.7723364 | 242.8993378 | 444.471398 | 436.4677307 | 427.1556913 | 5.59E-05 | 0.601166442 | 1.68E-05 | 18.52376375 | 1.516942543 | 4.252314605 |
| ENSMUSG00000026627 | *Pacc1* | 869.9623159 | 910.3633694 | 860.2125719 | 1322.450004 | 1305.06023 | 1379.075049 | 1.33E-12 | 0.601165025 | 1.95E-13 | 54.05834107 | 1.516941053 | 11.87483577 |
| ENSMUSG00000002486 | *Tchp* | 184.5374609 | 165.5206126 | 155.6702938 | 253.0197712 | 272.5208965 | 245.4695601 | 0.000440188 | 0.600761269 | 0.00014834 | 14.39315975 | 1.516516577 | 3.356361371 |
| ENSMUSG00000097221 | *1810049J17Rik* | 272.4124423 | 207.9100378 | 228.1374996 | 375.3126606 | 312.6932996 | 388.4990676 | 0.000911299 | 0.600699407 | 0.00032258 | 12.93450534 | 1.516451552 | 3.04033926 |
| ENSMUSG00000058013 | *Septin11* | 4199.625248 | 4363.082978 | 4103.791021 | 6528.753495 | 6524.215407 | 6158.966562 | 3.97E-20 | 0.600681373 | 3.58E-21 | 89.19628721 | 1.516432596 | 19.4009544 |
| ENSMUSG00000041997 | *Tlk1* | 1110.42022 | 1191.950265 | 1269.518086 | 1832.706543 | 1781.700363 | 1790.76809 | 4.52E-12 | 0.600652697 | 6.89E-13 | 51.57567385 | 1.516402455 | 11.34478476 |
| ENSMUSG00000062190 | *Lancl2* | 615.1248698 | 516.7472784 | 507.2704402 | 846.7728342 | 792.5906552 | 852.3785514 | 8.38E-07 | 0.600258365 | 2.03E-07 | 27.00136306 | 1.515988033 | 6.076979668 |

| | | | | | | | | | | | | | |
|---|---|---|---|---|---|---|---|---|---|---|---|---|---|
| ENSMUSG00000021891 | *Mettl6* | 499.2896671 | 490.5062057 | 677.7025722 | 849.3030319 | 832.7630583 | 834.9830707 | 7.86E-05 | 0.59960433 | 2.41E-05 | 17.83771011 | 1.515300927 | 4.104327444 |
| ENSMUSG00000061143 | *Maml3* | 312.3556157 | 311.8650567 | 216.0596319 | 411.5788278 | 451.6680994 | 417.4915354 | 0.001024844 | 0.599448856 | 0.000365143 | 12.70262358 | 1.515137637 | 2.989342347 |
| ENSMUSG00000038342 | *Mlxip* | 965.825932 | 1043.587277 | 650.8628664 | 1321.606605 | 1458.149657 | 1259.239515 | 0.000918453 | 0.599349793 | 0.00032534 | 12.91856147 | 1.515033604 | 3.036942861 |
| ENSMUSG00000020659 | *Cbll1* | 368.2760584 | 415.8200756 | 283.1588965 | 517.8471317 | 517.8982774 | 586.6142638 | 0.00053196 | 0.598996373 | 0.000181684 | 14.0116242 | 1.514662508 | 3.274120942 |
| ENSMUSG00000027531 | *Impa1* | 1479.495141 | 1488.676242 | 1810.338158 | 2539.475103 | 2294.169938 | 2388.012926 | 4.48E-08 | 0.598378493 | 9.62E-09 | 32.91623707 | 1.514013944 | 7.34895612 |
| ENSMUSG00000057409 | *Zfp53* | 142.9965606 | 117.0755553 | 161.038235 | 219.2838017 | 209.5479403 | 205.8465209 | 0.002881754 | 0.597958239 | 0.001096305 | 10.65741569 | 1.51357298 | 2.540343103 |
| ENSMUSG00000024474 | *Ik* | 5286.079563 | 5223.992018 | 5533.005357 | 8262.782327 | 8097.453571 | 7907.212367 | 8.23E-23 | 0.597618369 | 6.34E-24 | 101.7370229 | 1.513216454 | 22.08480917 |
| ENSMUSG00000022961 | *Son* | 7986.238082 | 8090.324578 | 6602.567634 | 11729.99659 | 11502.33617 | 11098.31665 | 6.65E-10 | 0.597596311 | 1.21E-10 | 41.4502447 | 1.513193318 | 9.176933587 |
| ENSMUSG00000015189 | *Casd1* | 566.3941983 | 597.4890407 | 625.3651458 | 871.2314121 | 959.7947112 | 872.6732788 | 1.23E-08 | 0.597502795 | 2.53E-09 | 35.51565683 | 1.513095235 | 7.908553369 |
| ENSMUSG00000017652 | *Cd40* | 970.6191128 | 945.6878904 | 1218.522645 | 1635.351121 | 1538.494464 | 1557.861933 | 1.01E-06 | 0.597481303 | 2.47E-07 | 26.62411981 | 1.513072695 | 5.994938011 |
| ENSMUSG00000040599 | *Mis12* | 992.1884264 | 976.9753232 | 1164.843233 | 1603.30195 | 1617.753529 | 1509.541153 | 8.47E-09 | 0.59703206 | 1.70E-09 | 36.29462478 | 1.51260161 | 8.072190494 |
| ENSMUSG00000056268 | *Dennd1b* | 605.5385082 | 690.3420672 | 680.3865428 | 1036.537663 | 1002.138596 | 944.1880325 | 1.94E-08 | 0.596914879 | 4.02E-09 | 34.61206282 | 1.512478756 | 7.712740052 |
| ENSMUSG00000021051 | *Ppp2r5e* | 905.9111719 | 1014.318388 | 944.7576452 | 1460.767479 | 1364.775964 | 1500.843413 | 2.44E-10 | 0.595760137 | 4.27E-11 | 43.48435078 | 1.511268644 | 9.61177177 |
| ENSMUSG00000095562 | *Gm55594* | 270.8147154 | 334.0690413 | 324.7604405 | 465.556379 | 427.7818057 | 506.4017698 | 9.51E-05 | 0.59553263 | 2.94E-05 | 17.45594994 | 1.511030341 | 4.021988084 |
| ENSMUSG00000021767 | *Kat6b* | 135.8067894 | 163.5020686 | 162.3802203 | 236.1517864 | 241.0344184 | 217.443508 | 0.001138146 | 0.595422529 | 0.000408266 | 12.49397988 | 1.510915029 | 2.943802066 |
| ENSMUSG00000031176 | *Dynlt3* | 720.5748475 | 579.3221442 | 927.3118364 | 1129.311579 | 1133.513211 | 1094.948865 | 0.000576221 | 0.59532249 | 0.000197957 | 13.85037768 | 1.510810264 | 3.239410911 |
| ENSMUSG00000006705 | *Pknox1* | 919.4918508 | 918.4375456 | 860.2125719 | 1325.823601 | 1341.975411 | 1413.86601 | 6.75E-12 | 0.595169843 | 1.04E-12 | 50.76307961 | 1.510650418 | 11.17085441 |
| ENSMUSG00000041057 | *Wdr43* | 3371.203832 | 3199.392329 | 3809.896243 | 5471.130852 | 5096.466487 | 5098.808659 | 5.67E-11 | 0.594924171 | 9.40E-12 | 46.44964798 | 1.510393196 | 10.24670207 |
| ENSMUSG00000042138 | *Msantd2* | 262.8260807 | 270.4849035 | 224.1115437 | 358.4446758 | 388.6951432 | 402.0288859 | 7.18E-05 | 0.594904851 | 2.19E-05 | 18.01930292 | 1.510372969 | 4.14366358 |
| ENSMUSG00000038205 | *Prkab2* | 241.2567671 | 202.8636777 | 216.0596319 | 325.5521056 | 344.1797777 | 330.5141322 | 9.06E-05 | 0.594825539 | 2.80E-05 | 17.55162031 | 1.51028994 | 4.042846248 |
| ENSMUSG00000035024 | *Ncapd3* | 841.2032311 | 848.7977757 | 719.3041163 | 1205.21751 | 1254.03042 | 1187.724761 | 1.00E-08 | 0.594238671 | 2.02E-09 | 35.95587481 | 1.5096757 | 8.000009595 |
| ENSMUSG00000061024 | *Rrs1* | 2519.615376 | 2437.391948 | 2081.419187 | 3482.395451 | 3478.712958 | 3676.244909 | 6.12E-10 | 0.594064832 | 1.11E-10 | 41.61638424 | 1.509493801 | 9.212994298 |
| ENSMUSG00000034160 | *Ogt* | 1386.028116 | 1628.965053 | 1838.519849 | 2542.005301 | 2461.373994 | 2308.766847 | 9.13E-07 | 0.593708985 | 2.22E-07 | 26.82842248 | 1.509121524 | 6.039489722 |
| ENSMUSG00000027883 | *Gpsm2* | 794.86915 | 707.4996917 | 704.542278 | 1164.734347 | 1029.282111 | 1138.437566 | 3.48E-08 | 0.592828771 | 7.41E-09 | 33.42437359 | 1.508201063 | 7.458132065 |
| ENSMUSG00000052299 | *Ltn1* | 1230.24974 | 1348.38743 | 1293.673821 | 1989.578801 | 1973.876454 | 1872.913416 | 4.46E-13 | 0.592828415 | 6.32E-14 | 56.26924035 | 1.50820069 | 12.35050306 |
| ENSMUSG00000035011 | *Zbtb7a* | 2533.196055 | 2613.005281 | 2093.497055 | 3653.605496 | 3627.459423 | 3644.353194 | 5.99E-09 | 0.592140434 | 1.18E-09 | 36.99958593 | 1.507481643 | 8.222280863 |
| ENSMUSG00000028820 | *Sfpq* | 6131.277111 | 5998.103663 | 4315.824698 | 8179.285803 | 8444.89057 | 8153.648342 | 1.60E-05 | 0.590660116 | 4.48E-06 | 21.04854226 | 1.505935643 | 4.796325599 |
| ENSMUSG00000030213 | *Atf7ip* | 855.5827735 | 935.5951701 | 717.962131 | 1279.436643 | 1298.545786 | 1208.019489 | 1.29E-06 | 0.590457115 | 3.21E-07 | 26.12063727 | 1.505723757 | 5.888393645 |
| ENSMUSG00000039046 | *Usp6nl* | 388.2476451 | 375.4491945 | 308.656617 | 508.5697401 | 578.6997524 | 535.3942375 | 3.77E-05 | 0.590317176 | 1.11E-05 | 19.3154473 | 1.505577712 | 4.4234746 |
| ENSMUSG00000032349 | *Elovl5* | 2317.502919 | 2294.07532 | 2382.023892 | 3543.963595 | 3510.199436 | 3463.633479 | 1.08E-20 | 0.589396787 | 9.41E-22 | 91.83722243 | 1.504617512 | 19.96467682 |
| ENSMUSG00000038876 | *Rnf146* | 468.1339918 | 458.2095008 | 515.3223519 | 752.3121196 | 692.7025178 | 717.0803686 | 1.17E-07 | 0.588937023 | 2.61E-08 | 30.97617565 | 1.50413809 | 6.932271544 |
| ENSMUSG00000066595 | *Flvcr1* | 895.5259468 | 1107.171415 | 903.1561012 | 1456.550483 | 1441.863548 | 1471.850945 | 2.00E-07 | 0.588495013 | 4.58E-08 | 29.88836591 | 1.503677326 | 6.698833707 |
| ENSMUSG00000055319 | *Sec23ip* | 1516.242861 | 1499.778234 | 1414.452497 | 2244.28537 | 2263.7692 | 2158.006015 | 6.75E-15 | 0.587732881 | 8.39E-16 | 64.77638965 | 1.502883189 | 14.17048231 |
| ENSMUSG00000018666 | *Cbx1* | 1172.73157 | 1166.718465 | 1265.49213 | 1779.572391 | 1824.044248 | 1805.264324 | 1.29E-13 | 0.587186905 | 1.75E-14 | 58.78932154 | 1.502314543 | 12.88845676 |
| ENSMUSG00000040711 | *Sh3pxd2b* | 4071.807093 | 4274.267039 | 3432.798376 | 6066.570713 | 5906.428992 | 5728.911624 | 3.48E-09 | 0.586822256 | 6.73E-10 | 38.09875111 | 1.501934873 | 8.457819085 |
| ENSMUSG00000060743 | *H3f3a* | 6794.333789 | 6066.734161 | 7767.410867 | 10344.29164 | 10228.76241 | 10388.00119 | 7.06E-09 | 0.586297896 | 1.40E-09 | 36.66245873 | 1.501389081 | 8.150937167 |
| ENSMUSG00000010755 | *Cars* | 2630.657398 | 2648.329802 | 2367.262054 | 3803.73056 | 3879.351248 | 3807.677429 | 9.42E-15 | 0.586128153 | 1.19E-15 | 64.0946854 | 1.501212442 | 14.02575619 |
| ENSMUSG00000033417 | *Cacul1* | 2091.424557 | 2192.138845 | 2438.387275 | 3344.921375 | 3377.73908 | 3353.462101 | 2.82E-12 | 0.585892252 | 4.24E-13 | 52.52943274 | 1.500966993 | 11.54911992 |
| ENSMUSG00000030967 | *Zranb1* | 667.8498587 | 712.5460519 | 772.9835279 | 1114.973792 | 1037.968036 | 1068.855644 | 9.35E-09 | 0.585625643 | 1.88E-09 | 36.08932733 | 1.500689642 | 8.029118027 |
| ENSMUSG00000023951 | *Vegfa* | 358.6896968 | 399.6717231 | 366.3619846 | 546.5227057 | 536.355868 | 604.0097445 | 2.89E-06 | 0.58528274 | 7.45E-07 | 24.49618582 | 1.500332996 | 5.539658827 |
| ENSMUSG00000042625 | *Safb2* | 1515.443997 | 1486.657697 | 1044.064557 | 1987.892002 | 2098.736625 | 1994.68178 | 0.000101815 | 0.585219127 | 3.16E-05 | 17.3179404 | 1.500266844 | 3.992190069 |
| ENSMUSG00000007041 | *Clic1* | 18870.75282 | 18764.38555 | 21341.59209 | 13028.83142 | 13019.11582 | 13260.18833 | 7.34E-15 | -0.585078934 | 9.15E-16 | 64.60647278 | 0.666612865 | 14.13406174 |
| ENSMUSG00000091478 | *Gm10039* | 3859.30941 | 4186.460373 | 4691.580579 | 2892.015984 | 2777.324515 | 2814.202201 | 4.06E-10 | -0.585114746 | 7.23E-11 | 42.45450633 | 0.666596318 | 9.391723038 |
| ENSMUSG00000042694 | *Stn1* | 820.4327809 | 836.6865113 | 1011.85691 | 606.4040516 | 589.5571586 | 577.9165235 | 2.07E-06 | -0.585611763 | 5.28E-07 | 25.16014872 | 0.666366711 | 5.683246893 |
| ENSMUSG00000059439 | *Bcas3* | 521.6578441 | 434.9962441 | 445.5391168 | 331.4559002 | 311.607559 | 292.8239241 | 5.06E-05 | -0.585626016 | 1.51E-05 | 18.72529473 | 0.666360128 | 4.295566656 |
| ENSMUSG00000026641 | *Usf1* | 984.1997917 | 1150.570112 | 1198.392865 | 666.2853974 | 719.8460334 | 832.083824 | 1.27E-05 | -0.586075487 | 3.52E-06 | 21.5113199 | 0.666152556 | 4.895742275 |
| ENSMUSG00000033813 | *Tcea1* | 6063.373717 | 5679.173702 | 8293.469102 | 4421.942201 | 4353.8199 | 4565.347252 | 5.34E-05 | -0.586157472 | 1.60E-05 | 18.6180885 | 0.666114702 | 4.272606662 |
| ENSMUSG00000056228 | *Cars2* | 1034.52819 | 972.9382351 | 993.0691157 | 624.9588348 | 691.6167772 | 684.2222385 | 2.28E-09 | -0.586761872 | 4.34E-10 | 38.95421904 | 0.665835699 | 8.641623555 |
| ENSMUSG00000000552 | *Zfp385a* | 969.0213858 | 959.8176988 | 888.394263 | 557.4868958 | 701.3884428 | 621.4052251 | 1.72E-06 | -0.586809428 | 4.32E-07 | 25.54381743 | 0.665813751 | 5.764763969 |
| ENSMUSG00000024792 | *Zfpl1* | 790.8748326 | 878.0666645 | 774.3255132 | 517.8471317 | 521.1554993 | 588.547095 | 4.78E-07 | -0.586874709 | 1.13E-07 | 28.13486527 | 0.665783624 | 6.320152563 |
| ENSMUSG00000025381 | *Cnpy2* | 2123.379096 | 1883.301605 | 2383.365878 | 1408.476726 | 1426.663179 | 1413.86601 | 4.81E-08 | -0.587568756 | 1.04E-08 | 32.77297259 | 0.665463408 | 7.318259995 |
| ENSMUSG00000002332 | *Dhrs1* | 1008.165696 | 1055.698541 | 1066.878307 | 693.274173 | 733.9606615 | 655.2297708 | 1.25E-09 | -0.587829041 | 2.33E-10 | 40.16428858 | 0.665343359 | 8.901875996 |
| ENSMUSG00000015714 | *Cers2* | 14102.33679 | 15031.08832 | 14937.63828 | 9681.379844 | 9728.235987 | 9898.028484 | 8.09E-24 | -0.588368855 | 5.88E-25 | 106.4475904 | 0.665094454 | 23.09201143 |
| ENSMUSG00000020457 | *Drg1* | 5616.010175 | 5180.59332 | 5902.051312 | 3662.882887 | 3590.544242 | 3849.233299 | 1.23E-13 | -0.588385881 | 1.67E-14 | 58.8864112 | 0.665086605 | 12.90913063 |
| ENSMUSG00000028803 | *Nipal3* | 746.9373419 | 875.0388484 | 683.0705134 | 500.1357477 | 518.9840181 | 516.0659257 | 6.75E-06 | -0.588502719 | 1.82E-06 | 22.7758852 | 0.665032744 | 5.170458745 |
| ENSMUSG00000020974 | *Pole2* | 680.6316741 | 690.3420672 | 767.6155867 | 461.3393828 | 459.2682838 | 498.670445 | 2.47E-07 | -0.588621394 | 5.69E-08 | 29.46540954 | 0.664978041 | 6.607979244 |
| ENSMUSG00000018841 | *Rad51d* | 685.4248549 | 765.0281973 | 794.4552926 | 504.3527439 | 513.5553149 | 471.6108085 | 3.64E-07 | -0.589101328 | 8.54E-08 | 28.68024491 | 0.664756863 | 6.438796412 |
| ENSMUSG00000000902 | *Smarcb1* | 2288.743834 | 2458.586661 | 2301.504775 | 1515.588429 | 1567.80946 | 1603.283466 | 1.29E-14 | -0.589190552 | 1.63E-15 | 63.46583679 | 0.664715752 | 13.89077611 |
| ENSMUSG00000018661 | *Cog1* | 1416.384928 | 1427.110648 | 1162.159262 | 877.978606 | 843.6204645 | 944.1880325 | 6.56E-07 | -0.589443336 | 1.57E-07 | 27.4978501 | 0.664599293 | 6.183080215 |
| ENSMUSG00000040048 | *Ndufb10* | 1654.446241 | 1670.345207 | 1957.95654 | 1136.902172 | 1147.627839 | 1221.549307 | 1.03E-08 | -0.589520798 | 2.08E-09 | 35.89363336 | 0.66456361 | 7.987151034 |
| ENSMUSG00000039316 | *Rftn1* | 2088.229103 | 2191.129573 | 2108.258893 | 1417.754118 | 1414.720032 | 1410.966763 | 1.65E-16 | -0.589946426 | 1.85E-17 | 72.29551102 | 0.664367578 | 15.78147903 |
| ENSMUSG00000083844 | *Ube2d-ps* | 429.7885454 | 433.9869721 | 579.7376459 | 311.2143185 | 313.7790402 | 330.5141322 | 0.000442739 | -0.59004332 | 0.000149226 | 14.38194935 | 0.664322959 | 3.353851874 |
| ENSMUSG00000004070 | *Hmox2* | 2124.976823 | 2194.157389 | 2544.404113 | 1532.456414 | 1527.637057 | 1492.145673 | 1.36E-09 | -0.590578839 | 2.54E-10 | 40.00201371 | 0.664076412 | 8.86648606 |
| ENSMUSG00000030979 | *Uros* | 464.1396745 | 424.9035238 | 570.3437489 | 315.4313147 | 331.1508902 | 319.8835607 | 0.000172222 | -0.590790967 | 5.52E-05 | 16.26228144 | 0.663978776 | 3.763910921 |
| ENSMUSG00000056665 | *Them6* | 1752.706447 | 1999.367888 | 1787.524408 | 1182.445731 | 1160.656727 | 1334.619931 | 9.08E-09 | -0.590846466 | 1.83E-09 | 36.15120873 | 0.663953234 | 8.0421455 |
| ENSMUSG00000032020 | *Ubash3b* | 1891.708691 | 1975.145359 | 1705.663306 | 1235.579883 | 1242.087273 | 1225.41497 | 4.53E-11 | -0.591149938 | 7.45E-12 | 46.90507657 | 0.663813586 | 10.34359785 |
| ENSMUSG00000047798 | *Cd300lf* | 1702.378049 | 1858.069804 | 1865.359555 | 1197.626917 | 1183.45728 | 1216.717229 | 7.54E-13 | -0.591227828 | 1.08E-13 | 55.21002022 | 0.663777748 | 12.1226154 |
| ENSMUSG00000055943 | *Emc7* | 2915.052792 | 2861.2862 | 3592.494625 | 2177.656831 | 1969.533491 | 2063.297287 | 1.94E-07 | -0.591644881 | 4.44E-08 | 29.9474004 | 0.663585892 | 6.711383794 |
| ENSMUSG00000044709 | *Gemin7* | 1257.411098 | 1256.543675 | 1480.209777 | 861.1106212 | 896.8217551 | 888.1359282 | 1.71E-08 | -0.59193327 | 3.55E-09 | 34.85470495 | 0.663453257 | 7.765757038 |
| ENSMUSG00000026520 | *Pycr2* | 3143.527744 | 3278.115547 | 2984.575288 | 2076.448922 | 2077.021813 | 2086.491261 | 1.71E-16 | -0.592804724 | 1.91E-17 | 72.23234477 | 0.663052622 | 15.76826368 |
| ENSMUSG00000046806 | *Cyren* | 337.1203832 | 394.625363 | 395.885661 | 269.8877559 | 269.2636746 | 206.8129365 | 0.000743722 | -0.593238683 | 0.000259802 | 13.34001474 | 0.662853207 | 3.128589395 |
| ENSMUSG00000043008 | *Klhl6* | 657.4646336 | 683.277163 | 693.8063957 | 479.0507667 | 419.0958807 | 445.5175875 | 1.29E-07 | -0.595255578 | 2.90E-08 | 30.77210189 | 0.661927182 | 6.889469743 |
| ENSMUSG00000040385 | *Ppp1ca* | 13086.18246 | 14013.74211 | 15157.72387 | 9111.24196 | 9422.057131 | 9432.216169 | 1.02E-14 | -0.595268809 | 1.29E-15 | 63.93026044 | 0.661921112 | 13.99052751 |
| ENSMUSG00000042570 | *Mier2* | 1372.447437 | 1411.971567 | 1215.838674 | 835.8086441 | 948.937305 | 867.8412008 | 2.14E-08 | -0.595539904 | 4.48E-09 | 34.40280639 | 0.661796743 | 7.668634924 |
| ENSMUSG00000053128 | *Rnf26* | 1386.028116 | 1340.313253 | 980.9912481 | 797.0122792 | 841.4489833 | 819.5204213 | 6.80E-05 | -0.595796314 | 2.06E-05 | 18.12858683 | 0.661679132 | 4.167579581 |
| ENSMUSG00000038543 | *BC028528* | 617.5214602 | 703.4626036 | 1025.276763 | 525.4377248 | 500.5264275 | 521.8644192 | 0.003049415 | -0.596091125 | 0.001164587 | 10.5457001 | 0.661543933 | 2.515783516 |
| ENSMUSG00000054484 | *Tmem62* | 492.0998958 | 457.2002287 | 555.5819107 | 302.7803262 | 340.9225558 | 350.8088596 | 3.15E-05 | -0.59618656 | 9.18E-06 | 19.67480438 | 0.661500173 | 4.501410327 |
| ENSMUSG00000031505 | *Naxd* | 930.6759394 | 1051.661453 | 1171.553159 | 684.8401806 | 663.387521 | 732.5430181 | 1.14E-06 | -0.596769097 | 2.79E-07 | 26.38977124 | 0.661233124 | 5.944659917 |
| ENSMUSG00000065979 | *Cpped1* | 1240.634965 | 1482.620609 | 1818.39007 | 940.3901495 | 1041.225258 | 1016.669202 | 9.03E-05 | -0.597167306 | 2.79E-05 | 17.55779277 | 0.661050638 | 4.044089878 |
| ENSMUSG00000091498 | *Mpc1-ps* | 460.1453572 | 427.9313399 | 551.5559548 | 338.2030941 | 308.3503371 | 301.5216644 | 9.97E-05 | -0.597465885 | 3.09E-05 | 17.3609342 | 0.660913841 | 4.001354453 |
| ENSMUSG00000042097 | *Zfp239* | 346.7067448 | 402.6995392 | 389.1757345 | 272.4179536 | 252.9775653 | 225.1748327 | 0.000118678 | -0.597578184 | 3.72E-05 | 17.00920316 | 0.660862398 | 3.925628215 |
| ENSMUSG00000028064 | *Sema4a* | 1830.995067 | 1889.357237 | 1470.81588 | 1171.481541 | 1150.885061 | 1112.344345 | 5.20E-07 | -0.597656147 | 1.23E-07 | 27.9658577 | 0.660826686 | 6.28427909 |
| ENSMUSG00000030844 | *Rgs10* | 1281.377002 | 1283.79402 | 1574.148747 | 881.3522029 | 913.1078644 | 936.4567078 | 2.68E-07 | -0.597774604 | 6.20E-08 | 29.29968066 | 0.660772429 | 6.572279974 |
| ENSMUSG00000003617 | *Cp* | 488.1055785 | 547.0254393 | 713.9361751 | 394.710843 | 364.8088495 | 391.3983144 | 0.000603875 | -0.59828174 | 0.000207992 | 13.75747007 | 0.660540195 | 3.219052627 |
| ENSMUSG00000044080 | *S100a1* | 1485.087186 | 1659.243214 | 2160.596319 | 1138.58897 | 1147.627839 | 1212.851567 | 6.54E-05 | -0.598299693 | 1.98E-05 | 18.2106221 | 0.660531975 | 4.184596647 |
| ENSMUSG00000056666 | *Retsat* | 2784.838047 | 2867.341832 | 3105.353965 | 1889.214291 | 1888.102944 | 2001.446689 | 8.27E-14 | -0.598571266 | 1.11E-14 | 59.69519238 | 0.660407649 | 13.08248757 |
| ENSMUSG00000027339 | *Rassf2* | 1837.385975 | 1764.207505 | 1596.962497 | 1125.937982 | 1225.801164 | 1084.318293 | 1.47E-09 | -0.599695612 | 2.75E-10 | 39.84378286 | 0.659893169 | 8.832676298 |
| ENSMUSG00000031584 | *Gsr* | 3245.782268 | 3721.185968 | 4208.465874 | 2548.752495 | 2354.971413 | 2463.393342 | 8.19E-08 | -0.59980679 | 1.80E-08 | 31.69779175 | 0.659842317 | 7.086648347 |
| ENSMUSG00000001785 | *Pwp1* | 3425.526548 | 3544.563363 | 3993.748228 | 2448.387986 | 2407.086962 | 2368.684614 | 2.77E-12 | -0.600576902 | 4.15E-13 | 52.56879976 | 0.659490187 | 11.55748366 |
| ENSMUSG00000037787 | *Coa8* | 1007.366832 | 953.7620666 | 1011.85691 | 695.8043707 | 622.1293773 | 639.7671213 | 1.47E-09 | -0.601007406 | 2.74E-10 | 39.84966561 | 0.659293423 | 8.83371034 |
| ENSMUSG00000045594 | *Glb1* | 1972.393901 | 2326.372025 | 2261.245216 | 1483.539258 | 1492.893357 | 1345.250503 | 3.20E-09 | -0.601070247 | 6.15E-10 | 38.271898 | 0.659264706 | 8.495026059 |
| ENSMUSG00000108352 | *Usf2-ps1* | 478.5192169 | 509.6823742 | 547.5299989 | 340.7332918 | 336.5795933 | 332.4469633 | 1.31E-06 | -0.601095639 | 3.26E-07 | 26.08751208 | 0.659253103 | 5.881356092 |
| ENSMUSG00000034473 | *Sec22a* | 802.0589212 | 760.9911092 | 879.0003659 | 567.6076867 | 534.1843868 | 504.4689386 | 1.51E-07 | -0.601525916 | 3.42E-08 | 30.45603776 | 0.659056513 | 6.820881671 |
| ENSMUSG00000071454 | *Dtnb* | 511.2726191 | 581.3406882 | 509.9544108 | 363.5050712 | 347.4369995 | 345.010366 | 1.72E-06 | -0.601556561 | 4.34E-07 | 25.53845067 | 0.659042514 | 5.763759691 |
| ENSMUSG00000019986 | *Ahi1* | 573.5839695 | 603.5446728 | 683.0705134 | 387.9636491 | 444.067915 | 392.36473 | 5.40E-06 | -0.601557049 | 1.44E-06 | 23.22980199 | 0.659042291 | 5.267834632 |
| ENSMUSG00000025142 | *Aspscr1* | 2308.71542 | 2452.531028 | 2563.191907 | 1594.024558 | 1586.267051 | 1641.940089 | 1.76E-14 | -0.60161964 | 2.25E-15 | 62.83622495 | 0.659013699 | 13.7543736 |
| ENSMUSG00000026553 | *Copa* | 15925.34322 | 16403.69827 | 16262.17776 | 10879.85016 | 10640.25811 | 10492.37407 | 8.65E-30 | -0.601921351 | 4.64E-31 | 134.3229817 | 0.658875894 | 29.06320056 |
| ENSMUSG00000022265 | *Ank* | 8461.561846 | 9100.605878 | 7466.806162 | 5437.394882 | 5566.592177 | 5483.442064 | 2.38E-11 | -0.602683616 | 3.84E-12 | 48.20583377 | 0.658527861 | 10.62405746 |
| ENSMUSG00000034201 | *Gas2l1* | 1950.824587 | 2123.508347 | 1851.939702 | 1265.098856 | 1361.518742 | 1279.534243 | 3.11E-11 | -0.602766257 | 5.06E-12 | 47.66543208 | 0.65849014 | 10.5072034 |
| ENSMUSG00000041439 | *Mfsd6* | 1483.489459 | 1648.141222 | 1591.594556 | 1055.092446 | 1019.510446 | 1032.131851 | 2.61E-12 | -0.602969034 | 3.89E-13 | 52.69574394 | 0.658397593 | 11.58418527 |
| ENSMUSG00000006715 | *Gmnn* | 1426.770153 | 1302.970188 | 1516.443379 | 942.076948 | 916.3650863 | 933.557461 | 2.64E-10 | -0.603358371 | 4.62E-11 | 43.32988589 | 0.658219937 | 9.578660897 |
| ENSMUSG00000016481 | *Cr1l* | 4142.905941 | 4174.349108 | 4972.055505 | 2887.798988 | 2944.528571 | 2908.910929 | 2.33E-10 | -0.603430117 | 4.06E-11 | 43.58437629 | 0.658187204 | 9.633319466 |
| ENSMUSG00000029816 | *Gpnmb* | 52327.15483 | 58383.35901 | 59342.58959 | 37061.49268 | 37933.60591 | 36920.94124 | 5.11E-17 | -0.603501509 | 5.54E-18 | 74.67765756 | 0.658154634 | 16.29119198 |
| ENSMUSG00000097530 | *Kansl2-ps* | 591.9578293 | 474.3578532 | 633.4170576 | 365.1918697 | 351.779962 | 400.0960547 | 0.000108387 | -0.603633613 | 3.38E-05 | 17.19367573 | 0.658094371 | 3.965022431 |
| ENSMUSG00000050022 | *Amz1* | 367.477195 | 552.0717994 | 591.8155136 | 326.3955048 | 337.6653339 | 327.6148854 | 0.003361259 | -0.60368782 | 0.001293402 | 10.35191807 | 0.658069645 | 2.47349805 |
| ENSMUSG00000032109 | *Nlrx1* | 822.0305078 | 869.9924883 | 779.6934544 | 526.281124 | 574.3567899 | 528.6293283 | 1.54E-08 | -0.603843659 | 3.18E-09 | 35.06856227 | 0.657998564 | 7.811596961 |
| ENSMUSG00000040415 | *Dtx3* | 909.1066258 | 954.7713386 | 805.1911749 | 607.2474508 | 535.2701274 | 614.640316 | 5.10E-07 | -0.603929917 | 1.21E-07 | 28.00380652 | 0.657959224 | 6.292256301 |
| ENSMUSG00000025508 | *Rplp2* | 24471.58459 | 23644.2158 | 26626.33017 | 16065.91207 | 15688.95201 | 17414.80895 | 2.09E-14 | -0.604011289 | 2.68E-15 | 62.49123798 | 0.657922114 | 13.68010471 |
| ENSMUSG00000073889 | *Il11ra1* | 227.6760882 | 250.299463 | 233.5054407 | 161.9326535 | 161.7753529 | 143.9959231 | 0.00045212 | -0.60464188 | 0.000152788 | 14.3375343 | 0.657634604 | 3.34474619 |
| ENSMUSG00000018677 | *Slc25a39* | 7040.383737 | 7495.863353 | 7074.946457 | 4619.297622 | 4894.518731 | 4695.813357 | 1.21E-20 | -0.60518046 | 1.05E-21 | 91.61627646 | 0.657389145 | 19.91736694 |
| ENSMUSG00000030744 | *Rps3* | 63659.03312 | 62633.40352 | 80199.72499 | 44219.42201 | 44262.388 | 47244.19259 | 3.95E-08 | -0.60532748 | 8.43E-09 | 33.17220772 | 0.657322157 | 7.403500334 |
| ENSMUSG00000041881 | *Ndufa7* | 1741.522359 | 1853.023444 | 2261.245216 | 1322.450004 | 1213.858017 | 1305.627464 | 6.96E-07 | -0.605682954 | 1.67E-07 | 27.38007195 | 0.657160215 | 6.157592188 |
| ENSMUSG00000038517 | *Tbkbp1* | 3858.510547 | 3837.252251 | 3141.587567 | 2325.251697 | 2459.202512 | 2338.725731 | 4.88E-09 | -0.606704895 | 9.55E-10 | 37.41455843 | 0.656694877 | 8.311565446 |
| ENSMUSG00000031832 | *Taf1c* | 941.0611645 | 895.224289 | 852.1606601 | 553.2698996 | 618.8721554 | 597.2448353 | 7.75E-09 | -0.607033396 | 1.55E-09 | 36.47115603 | 0.656545364 | 8.110757688 |
| ENSMUSG00000024142 | *Mlst8* | 1067.281592 | 1134.42176 | 1198.392865 | 720.2629486 | 803.4480614 | 707.4162127 | 6.23E-09 | -0.607075841 | 1.23E-09 | 36.91856745 | 0.656526049 | 8.205662707 |
| ENSMUSG00000039105 | *Atp6v1g1* | 4707.702413 | 5131.138991 | 7010.531163 | 3729.511427 | 3645.917014 | 3676.244909 | 4.28E-05 | -0.607681999 | 1.27E-05 | 19.0613714 | 0.656250263 | 4.368625742 |

| | | | | | | | | | | | | | |
|---|---|---|---|---|---|---|---|---|---|---|---|---|---|
| ENSMUSG00000026170 | *Cyp27a1* | 508.0771652 | 467.292949 | 461.6429403 | 344.950288 | 262.7492309 | 334.3797945 | 8.96E-05 | -0.607773169 | 2.76E-05 | 17.5739123 | 0.656208793 | 4.047605215 |
| ENSMUSG00000004929 | *Thop1* | 2561.156276 | 2704.849035 | 2384.707863 | 1518.962026 | 1761.071291 | 1745.346558 | 6.53E-10 | -0.607809375 | 1.19E-10 | 41.48703636 | 0.656192325 | 9.184966044 |
| ENSMUSG00000024781 | *Lipa* | 12360.01557 | 14158.06801 | 16168.23879 | 9322.935168 | 9392.742134 | 9281.455337 | 1.02E-08 | -0.608165634 | 2.07E-09 | 35.90731196 | 0.656030305 | 7.989692911 |
| ENSMUSG00000005054 | *Cstb* | 8777.112915 | 9364.025877 | 14097.55549 | 7056.721418 | 6858.623519 | 7226.85579 | 0.000400674 | -0.60830387 | 0.000134244 | 14.58124661 | 0.655967449 | 3.397208393 |
| ENSMUSG00000003037 | *Rab8a* | 3603.673101 | 3633.379301 | 4397.6858 | 2561.403483 | 2444.002144 | 2619.952668 | 3.65E-09 | -0.608350728 | 7.05E-10 | 38.00554007 | 0.655946143 | 8.438132706 |
| ENSMUSG00000024177 | *Nme4* | 315.5510696 | 306.8186966 | 369.0459552 | 218.4404024 | 245.3773809 | 185.5517935 | 0.000687827 | -0.608398895 | 0.00023914 | 13.49547101 | 0.655924244 | 3.162520972 |
| ENSMUSG00000032436 | *Cmtm7* | 2528.402874 | 2669.524514 | 2478.646833 | 1676.677684 | 1678.555004 | 1680.596713 | 2.35E-17 | -0.608660489 | 2.49E-18 | 76.25375769 | 0.655805321 | 16.62829377 |
| ENSMUSG00000031821 | *Gins2* | 1906.088233 | 1979.182447 | 2131.072643 | 1323.293403 | 1322.432079 | 1294.996892 | 1.31E-13 | -0.60872954 | 1.78E-14 | 58.75656513 | 0.655773933 | 12.88160406 |
| ENSMUSG00000023967 | *Mrps18a* | 2116.988188 | 2196.175933 | 2645.05301 | 1487.756254 | 1543.923167 | 1525.003803 | 2.16E-08 | -0.609155079 | 4.51E-09 | 34.39056302 | 0.655580533 | 7.666148262 |
| ENSMUSG00000040913 | *Fbxw4* | 906.7100354 | 902.2891932 | 841.4247778 | 566.7642874 | 603.6717867 | 570.1851988 | 7.73E-10 | -0.609378588 | 1.41E-10 | 41.14963457 | 0.655478976 | 9.111693793 |
| ENSMUSG00000025503 | *Taldo1* | 10743.91477 | 11434.04281 | 14164.65475 | 7873.975279 | 7719.615834 | 8218.398187 | 9.78E-08 | -0.609582171 | 2.17E-08 | 31.33758763 | 0.655386486 | 7.009800271 |
| ENSMUSG00000079057 | *Cyp4v3* | 1033.729327 | 1086.985974 | 1257.440218 | 800.3858761 | 682.9308522 | 724.8116933 | 6.66E-07 | -0.609628379 | 1.60E-07 | 27.46724835 | 0.655365495 | 6.176742633 |
| ENSMUSG00000022545 | *Ercc4* | 805.254375 | 857.8812239 | 759.563675 | 513.6301355 | 521.1554993 | 554.7225493 | 1.44E-08 | -0.609747022 | 2.97E-09 | 35.20107053 | 0.655311601 | 7.840651296 |
| ENSMUSG00000070283 | *Ndufaf3* | 904.313445 | 813.4732547 | 939.3897041 | 592.0662645 | 561.3279024 | 585.6478482 | 1.27E-08 | -0.609822426 | 2.61E-09 | 35.45562155 | 0.655277352 | 7.89566768 |
| ENSMUSG00000039849 | *Pcif1* | 1480.294005 | 1418.0272 | 1131.293601 | 850.9898304 | 856.649352 | 937.4231234 | 2.77E-06 | -0.609842583 | 7.13E-07 | 24.58098529 | 0.655268197 | 5.557577822 |
| ENSMUSG00000058886 | *Deaf1* | 655.8669066 | 685.2957071 | 539.4780872 | 446.1581965 | 397.3810682 | 390.4318988 | 2.16E-05 | -0.609945201 | 6.14E-06 | 20.44385184 | 0.655221589 | 4.666176995 |
| ENSMUSG00000021748 | *Pdhb* | 5707.879474 | 5892.1301 | 7384.945059 | 4061.810727 | 4209.416398 | 4161.385535 | 6.40E-08 | -0.610085716 | 1.40E-08 | 32.19311878 | 0.655157775 | 7.193644551 |
| ENSMUSG00000068267 | *Cenpb* | 8148.407366 | 8297.225343 | 7092.392266 | 5033.406648 | 5235.441287 | 5155.827179 | 3.55E-13 | -0.610370482 | 4.98E-14 | 56.7361364 | 0.65502847 | 12.44998979 |
| ENSMUSG00000000301 | *Pemt* | 236.4635863 | 229.1047504 | 273.7649995 | 156.8722581 | 160.6896123 | 165.2570661 | 0.000448665 | -0.610380645 | 0.000151435 | 14.35428148 | 0.655023856 | 3.348078081 |
| ENSMUSG00000031545 | *Gpat4* | 7150.626896 | 7041.69094 | 6500.576752 | 4481.823547 | 4569.882285 | 4506.395901 | 1.96E-19 | -0.610665671 | 1.83E-20 | 85.96196271 | 0.654894459 | 18.7072684 |
| ENSMUSG00000083380 | *Ndufb4c* | 1326.912219 | 1267.645667 | 1817.048085 | 995.2110999 | 916.3650863 | 972.2140847 | 8.02E-05 | -0.610842906 | 2.46E-05 | 17.79620567 | 0.65481401 | 4.095691402 |
| ENSMUSG00000032198 | *Dock6* | 452.955586 | 370.4028343 | 330.1283817 | 211.6932085 | 264.9207121 | 282.1933526 | 0.001925673 | -0.610861416 | 0.000713945 | 11.45258912 | 0.654805609 | 2.715417508 |
| ENSMUSG00000058064 | *Gm10036* | 257.2340365 | 248.2809189 | 288.5268377 | 192.2950261 | 153.0894279 | 172.0219752 | 0.000664113 | -0.610866314 | 0.000230151 | 13.56737814 | 0.654803385 | 3.177757931 |
| ENSMUSG00000031708 | *Tecr* | 4188.441159 | 4603.28972 | 4608.377491 | 2941.776539 | 2910.870612 | 2914.709423 | 2.39E-17 | -0.611232994 | 2.54E-18 | 76.21972781 | 0.65463698 | 16.62153457 |
| ENSMUSG00000025410 | *Dctn2* | 5068.7887 | 5441.994776 | 5555.819107 | 3388.778135 | 3622.03072 | 3504.222933 | 2.28E-17 | -0.611309096 | 2.41E-18 | 76.31917143 | 0.654602448 | 16.64195375 |
| ENSMUSG00000048779 | *P2ry6* | 3254.569766 | 3318.486429 | 3061.06845 | 2111.87169 | 2111.765513 | 2086.491261 | 2.62E-18 | -0.61131366 | 2.62E-19 | 80.70490599 | 0.654600378 | 17.58250326 |
| ENSMUSG00000042737 | *Dpm3* | 662.2578144 | 670.1566267 | 777.0094838 | 471.4601736 | 457.0968025 | 448.4168343 | 2.99E-07 | -0.611354801 | 6.96E-08 | 29.0749468 | 0.654581711 | 6.524004174 |
| ENSMUSG00000051184 | *Zfp524* | 245.2510845 | 271.4941756 | 241.5573525 | 173.7402429 | 176.9757216 | 145.9287543 | 0.000562862 | -0.61154438 | 0.000193102 | 13.89704495 | 0.6544957 | 3.249597983 |
| ENSMUSG00000079557 | *Marchf2* | 639.0907738 | 644.924826 | 727.356028 | 429.2902118 | 427.7818057 | 456.148159 | 1.51E-07 | -0.611893294 | 3.42E-08 | 30.45533986 | 0.654337431 | 6.820838779 |
| ENSMUSG00000024725 | *Ostf1* | 4513.57859 | 4975.711099 | 5728.935209 | 3279.979634 | 3282.193905 | 3387.286647 | 2.80E-09 | -0.61220566 | 5.35E-10 | 38.54515251 | 0.654195772 | 8.553565154 |
| ENSMUSG00000029993 | *Nfu1* | 1024.142965 | 977.9845953 | 1160.817277 | 673.8759905 | 690.5310366 | 701.6177192 | 1.12E-08 | -0.612264888 | 2.28E-09 | 35.71489705 | 0.654168915 | 7.949456828 |
| ENSMUSG00000001642 | *Akr1b3* | 9646.276368 | 10468.16948 | 12515.35483 | 6861.052795 | 7280.976622 | 7193.99766 | 1.59E-08 | -0.612534481 | 3.28E-09 | 35.01118211 | 0.654046684 | 7.799296073 |
| ENSMUSG00000010721 | *Lmbr1* | 200.5147303 | 163.5020686 | 217.4016172 | 127.3532848 | 131.3746155 | 120.8019489 | 0.003272407 | -0.612794523 | 0.001256315 | 10.4056322 | 0.653928804 | 2.485132659 |
| ENSMUSG00000038712 | *Mindy1* | 2699.359656 | 3129.752559 | 3140.245582 | 1938.131447 | 1904.389054 | 2017.875754 | 2.33E-11 | -0.612828962 | 3.76E-12 | 48.24664756 | 0.653913194 | 10.6324615 |
| ENSMUSG00000012117 | *Dhdds* | 2501.241516 | 2439.410492 | 2282.716981 | 1524.865821 | 1634.039638 | 1570.425336 | 1.12E-14 | -0.612845271 | 1.41E-15 | 63.74918895 | 0.653905802 | 13.95182802 |
| ENSMUSG00000051790 | *Nlgn2* | 422.5987742 | 492.5247497 | 315.3665435 | 305.3105239 | 254.0633059 | 246.4359757 | 0.003496343 | -0.61286005 | 0.001349509 | 10.27354964 | 0.653899103 | 2.456385968 |
| ENSMUSG00000024213 | *Nudt3* | 2919.047109 | 3188.290337 | 3159.033376 | 1990.4222 | 2035.763669 | 2028.506326 | 7.77E-17 | -0.613288735 | 8.52E-18 | 73.82872129 | 0.653704832 | 16.10936713 |
| ENSMUSG00000067825 | *Pex26* | 426.5930915 | 563.1737917 | 473.7208079 | 319.6483109 | 327.8936683 | 308.2865736 | 7.30E-05 | -0.613645666 | 2.22E-05 | 17.98727346 | 0.653543121 | 4.136777212 |
| ENSMUSG00000028398 | *Dmac1* | 610.331689 | 571.2479679 | 752.8537485 | 431.8204095 | 424.5245838 | 403.9617171 | 2.42E-05 | -0.614177189 | 6.95E-06 | 20.20767164 | 0.653302385 | 4.615531345 |
| ENSMUSG00000030271 | *Ogg1* | 729.3623456 | 781.1765498 | 797.1392632 | 481.5809645 | 500.5264275 | 523.7972504 | 7.70E-09 | -0.614396874 | 1.54E-09 | 36.48433407 | 0.653202911 | 8.113437457 |
| ENSMUSG00000033068 | *Entpd6* | 809.2486924 | 903.2984652 | 862.8965424 | 565.0774889 | 512.4695743 | 602.0769133 | 7.16E-08 | -0.614633144 | 1.57E-08 | 31.96702133 | 0.653095945 | 7.144977905 |
| ENSMUSG00000073402 | *Gm8909* | 1688.79737 | 1689.521375 | 1647.957938 | 1138.58897 | 1105.283955 | 1038.89676 | 4.72E-14 | -0.614664634 | 6.22E-15 | 60.83023509 | 0.65308169 | 13.32617202 |
| ENSMUSG00000021607 | *Mrpl36* | 858.7782273 | 847.7885036 | 879.0003659 | 583.6322722 | 558.0706805 | 545.0583934 | 2.00E-10 | -0.614805066 | 3.47E-11 | 43.8941438 | 0.653018122 | 9.699028818 |
| ENSMUSG00000075227 | *Znhit2* | 979.4066109 | 969.910419 | 1025.276763 | 663.7551997 | 605.8432679 | 668.7595891 | 2.66E-10 | -0.615554571 | 4.66E-11 | 43.31321415 | 0.652678956 | 9.575106786 |
| ENSMUSG00000031534 | *Smim19* | 484.1112612 | 439.0333322 | 421.3833815 | 303.6237254 | 268.177934 | 306.3537424 | 9.89E-06 | -0.616011221 | 2.71E-06 | 22.01165533 | 0.652472399 | 5.004720239 |
| ENSMUSG00000037722 | *Gnpnat1* | 863.5714081 | 753.926205 | 997.0950716 | 581.9454737 | 567.8423461 | 552.7897181 | 4.22E-06 | -0.616242519 | 1.11E-06 | 23.72486608 | 0.652367801 | 5.374556491 |
| ENSMUSG00000029467 | *Atp2a2* | 20577.12519 | 21450.05841 | 17833.64254 | 13100.52035 | 13167.86228 | 12777.94695 | 1.28E-12 | -0.616630677 | 1.86E-13 | 54.1488395 | 0.652192304 | 11.89431076 |
| ENSMUSG00000066233 | *Tmem42* | 198.9170034 | 229.1047504 | 256.3191907 | 150.9684635 | 153.0894279 | 140.1302607 | 0.001258543 | -0.617010189 | 0.00045383 | 12.2964113 | 0.652020763 | 2.900131999 |
| ENSMUSG00000038236 | *Hoxa7* | 170.956782 | 158.4557084 | 165.0641909 | 102.0513077 | 109.659803 | 111.137793 | 0.002032675 | -0.617599113 | 0.000755415 | 11.34769906 | 0.651754655 | 2.691932129 |
| ENSMUSG00000028542 | *Slc6a9* | 1340.492898 | 1264.617851 | 994.411101 | 707.61196 | 814.3054677 | 829.1845772 | 1.93E-05 | -0.617717178 | 5.46E-06 | 20.66773604 | 0.65170132 | 4.713897953 |
| ENSMUSG00000073705 | *Cenps* | 389.0465086 | 455.1816847 | 530.0841901 | 317.1181132 | 282.2925621 | 291.8575085 | 0.000188137 | -0.61814039 | 6.05E-05 | 16.08684643 | 0.651510173 | 3.725525261 |
| ENSMUSG00000056131 | *Pgm3* | 707.793032 | 730.7129484 | 746.1438221 | 468.0865767 | 482.0688369 | 471.6108085 | 1.10E-09 | -0.618800831 | 2.03E-10 | 40.43442444 | 0.651211991 | 8.959303754 |
| ENSMUSG00000049922 | *Slc35c1* | 2018.727982 | 1965.052639 | 1677.481614 | 1211.121305 | 1255.116161 | 1225.41497 | 8.05E-10 | -0.619084966 | 1.47E-10 | 41.06753315 | 0.651083749 | 9.094012752 |
| ENSMUSG00000035397 | *Klf16* | 1818.213252 | 1814.671107 | 1591.594556 | 1119.190788 | 1136.770433 | 1149.068138 | 1.21E-11 | -0.619661417 | 1.91E-12 | 49.57102873 | 0.65082365 | 10.91573915 |
| ENSMUSG00000032673 | *Prorsd1* | 322.7408408 | 324.9855931 | 295.2367641 | 206.6328131 | 203.0334966 | 204.8801053 | 2.53E-05 | -0.620246188 | 7.28E-06 | 20.11798385 | 0.650559904 | 4.596685491 |
| ENSMUSG00000029310 | *Nudt9* | 2207.25976 | 2179.018309 | 2419.599481 | 1436.308901 | 1409.291329 | 1575.257414 | 5.14E-12 | -0.621235144 | 7.86E-13 | 51.31654227 | 0.650114103 | 11.28897136 |
| ENSMUSG00000024369 | *Nelfe* | 2361.440409 | 2253.704439 | 2596.741539 | 1583.060368 | 1510.265207 | 1589.753647 | 1.18E-12 | -0.621389852 | 1.71E-13 | 54.30723846 | 0.650044391 | 11.92790998 |
| ENSMUSG00000048920 | *Fkrp* | 1805.431436 | 1989.275167 | 1694.927423 | 1181.602331 | 1166.08543 | 1222.515723 | 3.84E-11 | -0.621566087 | 6.29E-12 | 47.23825864 | 0.649964989 | 10.41523648 |
| ENSMUSG00000079109 | *Pms2* | 643.8839546 | 601.5261288 | 763.5896309 | 447.844995 | 444.067915 | 410.7266262 | 5.22E-06 | -0.621586378 | 1.39E-06 | 23.29723692 | 0.649955847 | 5.282002398 |
| ENSMUSG00000024847 | *Aip* | 1972.393901 | 2074.054018 | 2237.089481 | 1370.523761 | 1315.917636 | 1391.638451 | 4.74E-13 | -0.621699997 | 6.73E-14 | 56.14643967 | 0.649904662 | 12.32428221 |
| ENSMUSG00000079283 | *2310009B15Rik* | 206.1067746 | 246.2623749 | 263.0291171 | 175.4270413 | 136.8033186 | 149.7944166 | 0.00219124 | -0.621757244 | 0.000818223 | 11.19943301 | 0.649878874 | 2.659310052 |
| ENSMUSG00000060568 | *Fam78b* | 184.5374609 | 205.8914937 | 171.7741173 | 113.858897 | 138.9747998 | 114.0370398 | 0.00272614 | -0.621762515 | 0.001031956 | 10.76933453 | 0.6498765 | 2.564451807 |
| ENSMUSG00000001794 | *Capns1* | 7242.496194 | 7617.985268 | 7971.392632 | 4880.751386 | 4926.005209 | 5019.56258 | 8.46E-21 | -0.622375203 | 7.29E-22 | 92.3427385 | 0.649600567 | 20.07265118 |
| ENSMUSG00000002731 | *Prkra* | 528.8476153 | 545.0068952 | 589.131543 | 367.7220676 | 370.2375526 | 340.1782881 | 2.85E-07 | -0.622446879 | 6.62E-08 | 29.17270302 | 0.649568294 | 6.544814582 |
| ENSMUSG00000029003 | *Mad2l2* | 652.6714528 | 656.0268183 | 605.2353665 | 383.7466529 | 402.8097713 | 458.0809902 | 5.81E-07 | -0.623157826 | 1.39E-07 | 27.7401696 | 0.649248272 | 6.235557613 |
| ENSMUSG00000040414 | *Slc25a28* | 846.7952753 | 815.4917987 | 802.5072043 | 488.3281584 | 561.3279024 | 553.7561337 | 1.14E-08 | -0.623284751 | 2.31E-09 | 35.69176367 | 0.649191155 | 7.944568156 |
| ENSMUSG00000081094 | *Rpl19-ps11* | 19418.77316 | 19276.08647 | 24633.48201 | 13872.23065 | 13261.23598 | 13966.63812 | 1.60E-08 | -0.623477745 | 3.31E-09 | 34.99118106 | 0.649104316 | 7.79508323 |
| ENSMUSG00000001366 | *Fbxo9* | 731.758936 | 783.1950938 | 803.8491896 | 510.2565385 | 496.183465 | 495.7711582 | 1.73E-09 | -0.623591987 | 3.26E-10 | 39.51094515 | 0.649052918 | 8.761120215 |
| ENSMUSG00000022516 | *Nudt16l1* | 496.8930767 | 574.275784 | 656.2308076 | 411.5788278 | 327.8936683 | 376.9020805 | 0.000124211 | -0.623802216 | 3.91E-05 | 16.91663971 | 0.648958345 | 3.905841623 |
| ENSMUSG00000019039 | *Dalrd3* | 1453.93151 | 1497.75969 | 1453.370071 | 931.9561571 | 947.8515644 | 978.9789938 | 6.30E-15 | -0.624089901 | 7.82E-16 | 64.91455157 | 0.648828951 | 14.20072681 |
| ENSMUSG00000026873 | *Phf19* | 973.8145666 | 952.7527946 | 695.148381 | 581.9454737 | 559.1564211 | 562.453874 | 5.16E-05 | -0.625084499 | 1.54E-05 | 18.68580862 | 0.6483818 | 4.287432192 |
| ENSMUSG00000028743 | *Akr7a5* | 840.4043676 | 864.9461281 | 807.8751455 | 610.6210478 | 514.6410556 | 502.5361074 | 2.16E-07 | -0.625395038 | 4.96E-08 | 29.7322977 | 0.648242252 | 6.664880245 |
| ENSMUSG00000031818 | *Cox4i1* | 14238.94244 | 14775.74249 | 16945.24828 | 9921.748627 | 9832.467087 | 10029.461 | 1.85E-13 | -0.625516881 | 2.54E-14 | 58.0606067 | 0.648187507 | 12.73378246 |
| ENSMUSG00000038023 | *Atp6v0a2* | 3851.320776 | 4110.764971 | 3770.978669 | 2515.016525 | 2581.891203 | 2510.747706 | 4.30E-19 | -0.625549022 | 4.10E-20 | 84.37008286 | 0.648173066 | 18.36604467 |
| ENSMUSG00000022706 | *Mrpl40* | 1620.893975 | 1615.844517 | 2007.609996 | 1163.890947 | 1111.798399 | 1117.176423 | 5.28E-08 | -0.625900591 | 1.14E-08 | 32.58436961 | 0.648015133 | 7.277662376 |
| ENSMUSG00000025076 | *Casp7* | 790.0759691 | 785.2136379 | 868.2644836 | 520.3773294 | 505.9551306 | 553.7561337 | 4.96E-09 | -0.626370172 | 9.72E-10 | 37.3808931 | 0.647804245 | 8.304592479 |
| ENSMUSG00000032939 | *Nup93* | 5569.676094 | 5394.55899 | 5296.815946 | 3508.540827 | 3488.484623 | 3537.081064 | 1.23E-25 | -0.626807656 | 8.11E-27 | 114.9414352 | 0.647607835 | 24.91063917 |
| ENSMUSG00000021213 | *Akr1c13* | 297.9760733 | 258.3736392 | 373.071011 | 180.4874368 | 207.3764591 | 212.61143 | 0.001736268 | -0.62691334 | 0.000639116 | 11.6578662 | 0.647560396 | 2.760373277 |
| ENSMUSG00000042870 | *Tom1* | 3744.273071 | 3967.448343 | 3978.986389 | 2495.618343 | 2529.775653 | 2540.706589 | 1.73E-21 | -0.627193104 | 1.43E-22 | 95.56975605 | 0.647434835 | 20.7615406 |
| ENSMUSG00000035342 | *Lzts2* | 1055.29864 | 1169.746281 | 966.2294099 | 683.1533822 | 689.445296 | 695.8192256 | 8.57E-09 | -0.627360143 | 1.72E-09 | 36.26814432 | 0.647359877 | 8.06718437 |
| ENSMUSG00000010047 | *Hyal2* | 1096.839541 | 1200.024441 | 1124.583674 | 710.985557 | 743.7323271 | 760.5690702 | 1.99E-11 | -0.627415251 | 3.18E-12 | 48.57531619 | 0.64733515 | 10.70157813 |
| ENSMUSG00000024742 | *Fen1* | 4759.628538 | 4712.291099 | 4464.785065 | 2983.103102 | 3025.959118 | 3017.149476 | 8.59E-23 | -0.627476492 | 6.63E-24 | 101.6490394 | 0.647307672 | 22.066184 |
| ENSMUSG00000043015 | *Nemp2* | 188.5317783 | 157.4464364 | 134.1985292 | 97.83431152 | 99.88813737 | 115.0034553 | 0.008648614 | -0.627482687 | 0.003578068 | 8.486422052 | 0.647304892 | 2.063053499 |
| ENSMUSG00000032583 | *Mon1a* | 524.0544345 | 515.7380064 | 571.6857342 | 328.9257025 | 387.6094026 | 326.6484698 | 2.89E-06 | -0.6276398 | 7.46E-07 | 24.49126629 | 0.647234403 | 5.538649294 |
| ENSMUSG00000041957 | *Pkp2* | 1037.723644 | 1094.050879 | 1138.003527 | 663.7551997 | 724.1889959 | 726.7445245 | 1.41E-10 | -0.628074235 | 2.42E-11 | 44.60063379 | 0.647039532 | 9.851009117 |
| ENSMUSG00000082241 | *Gng5-ps* | 285.9931213 | 310.8557847 | 367.7039699 | 214.2234063 | 182.4044248 | 224.2084171 | 0.000491806 | -0.628138911 | 0.000167012 | 14.17000406 | 0.647010527 | 3.308206554 |
| ENSMUSG00000073409 | *H2-Q6* | 4173.262769 | 4613.382441 | 4432.577418 | 2793.338274 | 2876.126829 | 2880.884877 | 1.64E-18 | -0.62832749 | 1.61E-19 | 81.66425299 | 0.646925959 | 17.78532712 |
| ENSMUSG00000037408 | *Cnnm4* | 638.2919104 | 619.6930253 | 429.4352933 | 343.2634896 | 372.4090339 | 379.8013273 | 0.000615637 | -0.628332173 | 0.000212443 | 13.71770005 | 0.646923859 | 3.210675006 |
| ENSMUSG00000029148 | *Nrbp1* | 3486.240171 | 3477.951409 | 3827.342051 | 2337.902686 | 2322.399194 | 2315.531756 | 1.53E-17 | -0.628395916 | 1.60E-18 | 77.13247681 | 0.646895277 | 16.81642423 |
| ENSMUSG00000020483 | *Dynll2* | 3389.577692 | 3388.126198 | 2815.485142 | 2033.435561 | 2101.993847 | 2075.86069 | 1.65E-10 | -0.628499669 | 2.84E-11 | 44.28487665 | 0.646848756 | 9.782455647 |
| ENSMUSG00000060600 | *Eno3* | 568.7907887 | 627.7672015 | 689.7804398 | 390.4938468 | 421.2673619 | 405.8945483 | 8.07E-07 | -0.628542766 | 1.96E-07 | 27.07664914 | 0.646829433 | 6.093260293 |
| ENSMUSG00000054342 | *Kcnn4* | 11875.9043 | 11610.66541 | 11090.16645 | 7292.873204 | 7531.782706 | 7545.772936 | 1.87E-25 | -0.628630489 | 1.25E-26 | 114.0813749 | 0.646790104 | 24.72729663 |
| ENSMUSG00000071477 | *Zfp777* | 978.6077474 | 970.9196911 | 864.2385277 | 630.8626295 | 567.8423461 | 622.3716407 | 6.90E-09 | -0.628840159 | 1.37E-09 | 36.71161523 | 0.646696111 | 8.161243246 |
| ENSMUSG00000041346 | *Wrap53* | 690.2180357 | 703.4626036 | 633.4170576 | 420.8562194 | 444.067915 | 448.4168343 | 1.56E-08 | -0.629266162 | 3.22E-09 | 35.04584109 | 0.646505182 | 7.806777912 |
| ENSMUSG00000026172 | *Bcs1l* | 448.9612686 | 472.3393092 | 519.3483078 | 307.8407216 | 302.921634 | 317.9507295 | 1.21E-06 | -0.629274301 | 2.99E-07 | 26.25787516 | 0.646501534 | 5.91749842 |
| ENSMUSG00000017756 | *Slc12a7* | 1654.446241 | 1667.31739 | 1206.444777 | 981.7167121 | 993.4526706 | 956.7514352 | 1.06E-05 | -0.629288842 | 2.91E-06 | 21.87733308 | 0.646495018 | 4.975538065 |
| ENSMUSG00000055725 | *Paqr3* | 400.2305971 | 430.959156 | 448.2230874 | 256.3933681 | 289.8927465 | 280.2605214 | 4.22E-06 | -0.629448924 | 1.11E-06 | 23.72354989 | 0.646423287 | 5.37435683 |
| ENSMUSG00000030432 | *Rpl28* | 9599.942286 | 9820.216834 | 10505.06086 | 6498.391123 | 6332.039317 | 6494.312772 | 2.91E-22 | -0.630399868 | 2.32E-23 | 99.16931061 | 0.645997341 | 21.53650423 |
| ENSMUSG00000051373 | *Plpp7* | 264.4238077 | 303.7908805 | 387.8337492 | 188.9214291 | 217.1481247 | 208.7457677 | 0.00167566 | -0.631634007 | 0.000615714 | 11.72785835 | 0.645444965 | 2.775814092 |
| ENSMUSG00000030678 | *Maz* | 6608.997465 | 7011.412779 | 6241.573591 | 4125.065669 | 4477.594332 | 4214.538393 | 2.28E-16 | -0.632545714 | 2.58E-17 | 71.64245207 | 0.645037207 | 15.64139566 |
| ENSMUSG00000080268 | *Brms1* | 2031.509797 | 2238.565358 | 2646.394995 | 1431.248506 | 1497.23632 | 1527.90305 | 6.38E-08 | -0.632601256 | 1.39E-08 | 32.20027209 | 0.645012375 | 7.194890654 |
| ENSMUSG00000062963 | *Ufc1* | 2258.387022 | 2181.036853 | 2718.862201 | 1586.433905 | 1498.322061 | 1525.003803 | 6.02E-09 | -0.633138265 | 1.19E-09 | 36.9888878 | 0.644772329 | 8.220287917 |
| ENSMUSG00000025144 | *Cenpx* | 1101.632721 | 1279.756932 | 1545.967056 | 835.8086441 | 893.5645332 | 798.2592783 | 1.17E-05 | -0.633492 | 3.22E-06 | 21.68024163 | 0.644614256 | 4.932414255 |
| ENSMUSG00000030263 | *Irag2* | 4015.087787 | 3952.309262 | 4503.702638 | 2585.862061 | 2775.153034 | 2673.105525 | 3.10E-14 | -0.633831386 | 4.03E-15 | 61.68354816 | 0.644462632 | 13.50813412 |
| ENSMUSG00000026836 | *Acvr1* | 942.6588914 | 950.7342505 | 815.9270572 | 637.6098233 | 564.5851243 | 545.0583934 | 1.89E-07 | -0.633855494 | 4.32E-08 | 29.9992986 | 0.644451863 | 6.722558519 |
| ENSMUSG00000040532 | *Abhd11* | 1036.125917 | 1097.078695 | 1087.008086 | 697.4911692 | 659.0445585 | 716.113953 | 9.18E-12 | -0.633971684 | 1.43E-12 | 50.14576335 | 0.644399963 | 11.03738688 |
| ENSMUSG00000047446 | *Arl4a* | 301.1715272 | 264.4292714 | 411.9894845 | 227.7177941 | 203.0334966 | 196.182365 | 0.003395136 | -0.634186669 | 0.00130764 | 10.33170961 | 0.644303944 | 2.469142835 |
| ENSMUSG00000026239 | *Pde6d* | 1056.896367 | 1077.902526 | 1359.4311 | 769.1801044 | 728.5319584 | 748.0056675 | 8.88E-07 | -0.634484568 | 2.16E-07 | 26.88446247 | 0.644170917 | 6.051767116 |
| ENSMUSG00000037824 | *Tspan14* | 2369.429044 | 2725.034476 | 2678.602642 | 1615.109539 | 1719.813148 | 1669.966141 | 1.08E-12 | -0.634503521 | 1.56E-13 | 54.49072217 | 0.644162454 | 11.96757739 |
| ENSMUSG00000037103 | *Dcaf15* | 1040.919098 | 957.7991547 | 909.8660276 | 635.0796256 | 623.2151179 | 617.5395627 | 4.45E-10 | -0.635203685 | 7.96E-11 | 42.26736079 | 0.643849907 | 9.351294543 |
| ENSMUSG00000039754 | *Alkbh4* | 317.94766 | 292.6888882 | 271.0810289 | 173.7402429 | 216.0623841 | 180.7197155 | 0.000262063 | -0.6353489 | 8.58E-05 | 15.42581983 | 0.643785103 | 3.581593605 |
| ENSMUSG00000000149 | *Gna12* | 2066.65979 | 2385.919075 | 1890.857276 | 1326.667 | 1355.004298 | 1403.235438 | 5.78E-09 | -0.635956492 | 1.14E-09 | 37.07399529 | 0.64351403 | 8.238203397 |
| ENSMUSG00000020788 | *Atp2a3* | 2895.880069 | 3143.882368 | 2545.746098 | 1831.019744 | 1906.560535 | 1791.734506 | 2.84E-10 | -0.635972938 | 4.99E-11 | 43.18034489 | 0.643506695 | 9.547074553 |
| ENSMUSG00000001751 | *Naglu* | 1782.264396 | 1749.068425 | 1646.615953 | 1125.937982 | 1092.255067 | 1116.210008 | 1.85E-15 | -0.636053959 | 2.21E-16 | 67.40405941 | 0.643470556 | 14.73370862 |
| ENSMUSG00000058603 | *Rpl28-ps1* | 14327.61629 | 14542.60065 | 16370.87857 | 9658.608065 | 9393.827875 | 10051.68856 | 8.35E-16 | -0.636075105 | 9.78E-17 | 69.01242827 | 0.643461125 | 15.07820638 |
| ENSMUSG00000038697 | *Taf5l* | 1390.022433 | 1565.380916 | 1508.391468 | 953.0411381 | 961.9661925 | 953.8521884 | 9.03E-13 | -0.636716047 | 1.30E-13 | 54.84577503 | 0.64317532 | 12.04408952 |

| | | | | | | | | | | | | | |
|---|---|---|---|---|---|---|---|---|---|---|---|---|---|
| ENSMUSG00000025579 | *Gaa* | 4046.243462 | 4577.048648 | 3930.674919 | 2608.633841 | 2769.724331 | 2697.265915 | 2.17E-13 | -0.637050078 | 3.00E-14 | 57.73472851 | 0.643026421 | 12.66406377 |
| ENSMUSG00000039616 | *Mocos* | 730.1612091 | 796.3156302 | 951.4675717 | 542.3057095 | 515.7267962 | 530.5621595 | 1.96E-06 | -0.637056252 | 4.98E-07 | 25.27308351 | 0.643023669 | 5.706658234 |
| ENSMUSG00000038648 | *Creb3l2* | 403.426051 | 467.292949 | 347.5741905 | 249.6461742 | 284.4640434 | 251.2680537 | 0.000149267 | -0.638095689 | 4.74E-05 | 16.55028062 | 0.642560548 | 3.826037532 |
| ENSMUSG00000037946 | *Fgd3* | 4046.243462 | 4258.118687 | 3879.679478 | 2628.875422 | 2620.977865 | 2581.296044 | 5.00E-20 | -0.63817939 | 4.51E-21 | 88.73462505 | 0.64252327 | 19.30074057 |
| ENSMUSG00000056938 | *Acbd4* | 671.844176 | 637.8599218 | 609.2613223 | 377.8428583 | 435.38199 | 423.2900289 | 1.13E-07 | -0.638254671 | 2.53E-08 | 31.03647219 | 0.642489743 | 6.945198886 |
| ENSMUSG00000002870 | *Mcm2* | 8048.549433 | 8063.074233 | 6832.047119 | 4911.113758 | 5022.636125 | 4809.850397 | 5.49E-13 | -0.638618468 | 7.82E-14 | 55.85098077 | 0.64232775 | 12.26063828 |
| ENSMUSG00000049086 | *Bmyc* | 364.2817411 | 369.3935623 | 267.055073 | 199.8856192 | 234.5199747 | 211.6450145 | 0.000818947 | -0.638801846 | 0.00028782 | 13.14803848 | 0.642246111 | 3.086744261 |
| ENSMUSG00000025371 | *Chmp6* | 1532.22013 | 1516.935858 | 1729.819041 | 1028.947069 | 1037.968036 | 997.34089 | 5.92E-12 | -0.639293914 | 9.08E-13 | 51.0325686 | 0.642027094 | 11.22781827 |
| ENSMUSG00000025156 | *Gps1* | 5715.069245 | 5927.454621 | 5971.834547 | 3764.090796 | 3739.290708 | 3801.878935 | 1.71E-27 | -0.639319386 | 1.02E-28 | 123.6266257 | 0.642015758 | 26.76761643 |
| ENSMUSG00000031521 | *Aga* | 711.7873494 | 803.3805344 | 927.3118364 | 540.6189111 | 551.5562368 | 471.6108085 | 7.06E-06 | -0.640067957 | 1.91E-06 | 22.68536307 | 0.641682722 | 5.150950505 |
| ENSMUSG00000022472 | *Desi1* | 1473.104234 | 1594.649804 | 1535.231173 | 937.8599518 | 968.4806362 | 1046.628085 | 1.53E-12 | -0.640232752 | 2.24E-13 | 53.78024766 | 0.641609429 | 11.81546986 |
| ENSMUSG00000055652 | *Klhl25* | 1197.496337 | 1268.654939 | 1203.760806 | 791.1084845 | 733.9606615 | 828.2181616 | 1.15E-11 | -0.640245078 | 1.80E-12 | 49.68737513 | 0.641603947 | 10.94002742 |
| ENSMUSG00000042558 | *Adprs* | 1515.443997 | 1722.827352 | 1810.338158 | 1025.573472 | 1033.625074 | 1176.127774 | 8.56E-09 | -0.640382966 | 1.72E-09 | 36.27008543 | 0.641542627 | 8.067361465 |
| ENSMUSG00000090124 | *Ugt1a7c* | 2672.198298 | 2654.385434 | 2889.294333 | 1779.572391 | 1758.89981 | 1725.05183 | 8.67E-18 | -0.641176265 | 8.95E-19 | 78.27797467 | 0.641189957 | 17.06188099 |
| ENSMUSG00000000823 | *Zfp512b* | 558.4055636 | 632.8135616 | 477.7467638 | 365.1918697 | 363.7231089 | 343.0775349 | 1.36E-05 | -0.64156271 | 3.77E-06 | 21.38023828 | 0.641018229 | 4.867440322 |
| ENSMUSG00000029156 | *Sgcb* | 755.7248401 | 848.7977757 | 879.0003659 | 517.8471317 | 546.1275337 | 524.763666 | 8.01E-09 | -0.642314863 | 1.60E-09 | 36.40521979 | 0.64068412 | 8.09657841 |
| ENSMUSG00000047921 | *Trappc9* | 847.5941388 | 842.7421435 | 712.5941898 | 538.9321126 | 520.0697587 | 483.2077956 | 1.43E-07 | -0.642416428 | 3.24E-08 | 30.55984919 | 0.640639017 | 6.843440242 |
| ENSMUSG00000025511 | *Tspan4* | 4084.588908 | 4499.334701 | 4470.153006 | 2702.251156 | 2723.037484 | 2934.037735 | 3.94E-16 | -0.642495468 | 4.53E-17 | 70.5316318 | 0.64060392 | 15.40430424 |
| ENSMUSG00000038214 | *Bend3* | 1070.477046 | 968.901147 | 701.8583075 | 608.9342493 | 597.157343 | 553.7561337 | 0.000179798 | -0.642536064 | 5.77E-05 | 16.17654335 | 0.640585895 | 3.745215461 |
| ENSMUSG00000035206 | *Sppl2b* | 1293.359954 | 1348.38743 | 1127.267645 | 747.2517242 | 822.9913927 | 848.512889 | 1.20E-08 | -0.642598341 | 2.47E-09 | 35.56573975 | 0.640558243 | 7.919093843 |
| ENSMUSG00000021257 | *Angel1* | 279.6022136 | 274.5219917 | 221.4275731 | 150.1250642 | 155.2609092 | 193.2831182 | 0.001333734 | -0.642685332 | 0.000482518 | 12.18204103 | 0.64051962 | 2.874930842 |
| ENSMUSG00000096472 | *Cdkn2d* | 635.89532 | 620.7022973 | 652.2048517 | 414.1090255 | 408.2384745 | 398.1632235 | 1.83E-09 | -0.644063119 | 3.45E-10 | 39.39954096 | 0.63990821 | 8.736888114 |
| ENSMUSG00000022681 | *Ntan1* | 1159.150891 | 1224.24697 | 1638.564041 | 856.0502258 | 836.0202801 | 876.5389412 | 1.63E-05 | -0.644178623 | 4.57E-06 | 21.00969646 | 0.63985698 | 4.788160481 |
| ENSMUSG00000020265 | *Sumo3* | 5176.635268 | 5428.874239 | 6473.737046 | 3673.003678 | 3620.94498 | 3625.024882 | 1.74E-10 | -0.644556685 | 3.00E-11 | 44.17503334 | 0.639689326 | 9.758979207 |
| ENSMUSG00000050552 | *Lamtor4* | 1343.688352 | 1182.866817 | 1551.334997 | 815.5670624 | 880.5356457 | 909.3970713 | 4.03E-07 | -0.645069312 | 9.50E-08 | 28.47379429 | 0.639462068 | 6.394237303 |
| ENSMUSG00000028063 | *Lmna* | 27895.51341 | 28357.51617 | 25939.2337 | 16810.6336 | 17621.57032 | 18121.25875 | 2.14E-21 | -0.645387272 | 1.77E-22 | 95.14032675 | 0.63932115 | 20.67012172 |
| ENSMUSG00000038871 | *Bpgm* | 404.2249144 | 525.8307267 | 527.4002196 | 316.274714 | 285.549784 | 326.6484698 | 7.88E-05 | -0.645540705 | 2.41E-05 | 17.83236015 | 0.639253161 | 4.103357704 |
| ENSMUSG00000028933 | *Xrcc2* | 574.382833 | 579.3221442 | 536.7941166 | 372.7824629 | 353.9514433 | 354.6745219 | 2.22E-08 | -0.645567033 | 4.65E-09 | 34.32910797 | 0.639241496 | 7.653170016 |
| ENSMUSG00000068396 | *Rpl34-ps1* | 3344.841338 | 3232.698306 | 4486.25683 | 2547.909096 | 1926.103866 | 2592.893031 | 0.000102749 | -0.645626077 | 3.19E-05 | 17.30022143 | 0.639215334 | 3.98822304 |
| ENSMUSG00000021556 | *Golm1* | 1460.322418 | 1642.08559 | 1889.51529 | 1027.260271 | 1118.312842 | 1039.863176 | 7.30E-08 | -0.64638044 | 1.60E-08 | 31.92680969 | 0.638881186 | 7.13650998 |
| ENSMUSG00000029461 | *Fam168a* | 1042.516825 | 1048.633637 | 821.2949984 | 599.6568577 | 638.4154867 | 627.2037186 | 7.16E-07 | -0.646400289 | 1.73E-07 | 27.31862784 | 0.638872396 | 6.145283357 |
| ENSMUSG00000017747 | *Ghdc* | 486.5078516 | 491.5154777 | 495.1925726 | 285.9123414 | 337.6653339 | 318.9171451 | 5.38E-07 | -0.646699655 | 1.28E-07 | 27.89696583 | 0.638739841 | 6.269247958 |
| ENSMUSG00000079190 | *ENSMUSG00000079190* | 2036.302978 | 1980.191719 | 2321.634554 | 1439.682498 | 1325.689301 | 1275.66858 | 1.79E-10 | -0.647427116 | 3.10E-11 | 44.11351615 | 0.638417845 | 9.74607345 |
| ENSMUSG00000004100 | *Ppan* | 3844.131005 | 4100.67225 | 3705.22139 | 2460.195575 | 2416.858628 | 2560.034901 | 4.11E-18 | -0.64784934 | 4.18E-19 | 79.78175072 | 0.638231031 | 17.38595075 |
| ENSMUSG00000015668 | *Pdzd11* | 1657.641695 | 1653.187582 | 2006.268011 | 1104.853001 | 1158.485245 | 1125.874164 | 2.14E-09 | -0.647951269 | 4.06E-10 | 39.08172499 | 0.638185941 | 8.668908304 |
| ENSMUSG00000025169 | *Ogfod3* | 359.4885603 | 318.9299609 | 293.8947788 | 198.1988207 | 217.1481247 | 207.7793521 | 5.10E-05 | -0.64854088 | 1.52E-05 | 18.71120496 | 0.637925175 | 4.292728274 |
| ENSMUSG00000024414 | *Mrpl27* | 1880.524602 | 1812.652563 | 2285.400951 | 1292.087631 | 1185.628761 | 1330.754269 | 4.21E-08 | -0.64856195 | 9.03E-09 | 33.03870234 | 0.637915859 | 7.375357439 |
| ENSMUSG00000022575 | *Gsdmd* | 2086.631377 | 2237.556086 | 2518.906392 | 1353.655776 | 1501.579282 | 1506.641907 | 5.20E-10 | -0.648701438 | 9.35E-11 | 41.95242675 | 0.637854184 | 9.283946227 |
| ENSMUSG00000032892 | *Rangrf* | 371.4715123 | 356.2730259 | 354.284117 | 270.7311552 | 231.2627528 | 187.4846247 | 0.000318942 | -0.648815049 | 0.000105486 | 15.03589354 | 0.637803956 | 3.496288494 |
| ENSMUSG00000049288 | *Lix1l* | 908.3077623 | 839.7143274 | 876.3163954 | 505.1961431 | 573.2710492 | 597.2448353 | 8.47E-09 | -0.64922714 | 1.70E-09 | 36.29387514 | 0.6376218 | 8.072151387 |
| ENSMUSG00000044496 | *2510039O18Rik* | 5399.518175 | 5978.927495 | 4962.661608 | 3301.064615 | 3557.972023 | 3562.207869 | 2.93E-12 | -0.649618297 | 4.42E-13 | 52.44878313 | 0.637448945 | 11.53264917 |
| ENSMUSG00000051256 | *Jagn1* | 1657.641695 | 1801.55057 | 2002.242055 | 1166.421145 | 1199.743389 | 1110.411514 | 2.19E-10 | -0.649789003 | 3.80E-11 | 43.71216002 | 0.637373524 | 9.660270063 |
| ENSMUSG00000066900 | *Suds3* | 3555.741293 | 3495.109033 | 3956.172639 | 2202.115408 | 2366.914559 | 2444.06503 | 8.56E-14 | -0.649803077 | 1.15E-14 | 59.62643021 | 0.637367306 | 13.0675049 |
| ENSMUSG00000041429 | *Nthl1* | 270.0158519 | 319.9392329 | 271.0810289 | 185.5478322 | 184.575906 | 178.7868844 | 7.86E-05 | -0.649823159 | 2.40E-05 | 17.83994961 | 0.637358434 | 4.104754767 |
| ENSMUSG00000033059 | *Pygb* | 3236.195906 | 3228.661218 | 2967.12948 | 2036.809158 | 2029.249225 | 1949.260247 | 3.77E-18 | -0.649899497 | 3.83E-19 | 79.95483453 | 0.63732471 | 17.42323807 |
| ENSMUSG00000043162 | *Pyurf* | 283.5965309 | 256.3550951 | 335.4963229 | 204.9460146 | 182.4044248 | 168.1563129 | 0.000487931 | -0.650103006 | 0.000165639 | 14.18553966 | 0.637234815 | 3.31164149 |
| ENSMUSG00000051355 | *Commd1* | 2082.637059 | 2147.730876 | 2940.289774 | 1512.214832 | 1439.692067 | 1611.01479 | 1.11E-05 | -0.650511658 | 3.06E-06 | 21.77677959 | 0.63705434 | 4.953431084 |
| ENSMUSG00000022885 | *St6gal1* | 289.1885752 | 292.6888882 | 358.3100728 | 224.3441971 | 187.8331279 | 183.6189623 | 0.000317626 | -0.65054171 | 0.000104994 | 15.04470968 | 0.63704107 | 3.498084582 |
| ENSMUSG00000027935 | *Rab13* | 147.7897414 | 182.6782371 | 190.5619114 | 112.1720986 | 111.8312842 | 106.305715 | 0.002463721 | -0.650709273 | 0.000925786 | 10.97040426 | 0.636967084 | 2.608408476 |
| ENSMUSG00000078619 | *Smarcd2* | 2383.009723 | 2491.892637 | 2348.47426 | 1528.239418 | 1539.580204 | 1532.735128 | 3.65E-20 | -0.65128902 | 3.27E-21 | 89.3717577 | 0.636711171 | 19.43747696 |
| ENSMUSG00000067567 | *Hdac8* | 429.7885454 | 401.6902672 | 430.7772786 | 262.2971628 | 277.9495996 | 263.8314564 | 5.23E-07 | -0.651395052 | 1.24E-07 | 27.9539045 | 0.636664377 | 6.281704201 |
| ENSMUSG00000015605 | *Srf* | 3561.333337 | 3378.033478 | 2571.243819 | 2004.759987 | 2032.506447 | 2021.741417 | 5.89E-07 | -0.651724202 | 1.41E-07 | 27.71386533 | 0.636519138 | 6.229867159 |
| ENSMUSG00000064105 | *Cnnm2* | 603.1419178 | 599.5075847 | 543.5040431 | 361.8182728 | 371.3232933 | 380.7677429 | 2.47E-08 | -0.651788467 | 5.18E-09 | 34.11881516 | 0.636490785 | 7.607462084 |
| ENSMUSG00000075033 | *Nxpe3* | 711.7873494 | 587.3963204 | 501.902499 | 376.999459 | 404.9812526 | 368.2043402 | 6.45E-05 | -0.652188789 | 1.95E-05 | 18.23654238 | 0.636314195 | 4.190336972 |
| ENSMUSG00000032350 | *Gclc* | 3077.222076 | 3734.306504 | 3945.436757 | 2325.251697 | 2237.711425 | 2274.942302 | 2.59E-09 | -0.652432501 | 4.94E-10 | 38.70268474 | 0.636206713 | 8.587390994 |
| ENSMUSG00000028958 | *Tmub1* | 607.1362351 | 594.4612246 | 653.546837 | 381.2164552 | 401.7240307 | 396.2303924 | 7.58E-09 | -0.65250579 | 1.51E-09 | 36.5204503 | 0.636174394 | 8.120457116 |
| ENSMUSG00000015013 | *Trappc2l* | 1556.984898 | 1615.844517 | 1987.480217 | 1050.87545 | 1015.167483 | 1211.885151 | 7.28E-07 | -0.652722821 | 1.76E-07 | 27.28305998 | 0.636078699 | 6.137932111 |
| ENSMUSG00000006315 | *Tmem147* | 2044.291613 | 2115.434171 | 2337.738378 | 1374.740757 | 1387.576517 | 1365.54523 | 3.08E-14 | -0.652815776 | 3.99E-15 | 61.7024426 | 0.636037717 | 13.51210423 |
| ENSMUSG00000005505 | *Kbtbd4* | 1173.530433 | 1342.331797 | 1335.275365 | 864.4842182 | 834.9345395 | 746.0728364 | 2.83E-09 | -0.653287452 | 5.43E-10 | 38.51776216 | 0.635829804 | 8.548139258 |
| ENSMUSG00000032177 | *Pde4a* | 809.2486924 | 834.6679673 | 578.3956606 | 442.7845995 | 500.5264275 | 473.5436397 | 0.00010705 | -0.653588562 | 3.33E-05 | 17.21869908 | 0.635697111 | 3.970413596 |
| ENSMUSG00000036860 | *Mrpl55* | 531.2442057 | 483.4413015 | 575.7116901 | 346.6370865 | 304.0073746 | 357.5737687 | 2.47E-06 | -0.653847406 | 6.33E-07 | 24.80922091 | 0.635583067 | 5.607411586 |
| ENSMUSG00000034595 | *Ppp1r18* | 9011.978774 | 8861.408407 | 7815.722338 | 5403.658913 | 5438.474783 | 5491.173389 | 1.07E-16 | -0.653896714 | 1.18E-17 | 73.18556956 | 0.635561344 | 15.97111894 |
| ENSMUSG00000052139 | *Babam2* | 1854.162108 | 1873.208884 | 1762.026688 | 1148.709761 | 1178.028577 | 1164.530787 | 1.17E-17 | -0.654220216 | 1.21E-18 | 77.67687327 | 0.635418846 | 16.93342874 |
| ENSMUSG00000000378 | *Ccm2* | 1803.833709 | 1886.329421 | 2081.419187 | 1243.170476 | 1184.54302 | 1233.146294 | 5.15E-13 | -0.654506 | 7.34E-14 | 55.97628144 | 0.635292988 | 12.28777751 |
| ENSMUSG00000018572 | *Phf23* | 3225.011817 | 3304.35662 | 3707.90536 | 2137.173667 | 2163.881063 | 2196.662639 | 4.21E-15 | -0.654667788 | 5.17E-16 | 65.73238829 | 0.635221748 | 14.37576773 |
| ENSMUSG00000053730 | *Tmem39b* | 814.8407366 | 814.4825267 | 760.9056603 | 524.5943256 | 492.9262431 | 501.5696918 | 1.32E-10 | -0.654695088 | 2.25E-11 | 44.73766272 | 0.635209728 | 9.880655617 |
| ENSMUSG00000002778 | *Kdelr1* | 2827.177811 | 3179.206889 | 3164.401317 | 1980.301409 | 1964.104788 | 1876.779078 | 2.78E-15 | -0.654713199 | 3.37E-16 | 66.57570996 | 0.635201754 | 14.55654025 |
| ENSMUSG00000039244 | *Ints15* | 2867.120984 | 2822.933863 | 2753.753818 | 1728.968436 | 1793.64351 | 1843.920948 | 1.09E-20 | -0.65483457 | 9.47E-22 | 91.82488123 | 0.635148318 | 19.96226372 |
| ENSMUSG00000059689 | *Zfp637* | 468.9328553 | 471.3300371 | 508.6124255 | 327.238904 | 296.4071902 | 293.7903397 | 3.11E-07 | -0.655216719 | 7.26E-08 | 28.99390386 | 0.634980099 | 6.506716015 |
| ENSMUSG00000028882 | *Ppp1r8* | 2743.297147 | 3013.686276 | 3018.124921 | 1906.082276 | 1829.472951 | 1831.357545 | 4.02E-17 | -0.655251203 | 4.33E-18 | 75.16328262 | 0.634964921 | 16.39564445 |
| ENSMUSG00000033653 | *Vps8* | 1421.976972 | 1613.825973 | 1558.044923 | 937.0165526 | 972.8235987 | 1003.139384 | 1.77E-12 | -0.656408902 | 2.61E-13 | 53.47884789 | 0.634455594 | 11.75145778 |
| ENSMUSG00000054676 | *1600014C10Rik* | 828.4214156 | 816.5010708 | 883.0263218 | 539.7755118 | 535.2701274 | 526.6964972 | 2.89E-11 | -0.656433018 | 4.68E-12 | 47.81637045 | 0.634444989 | 10.53969244 |
| ENSMUSG00000005043 | *Sgsh* | 902.715718 | 804.3898064 | 738.0919103 | 551.5831012 | 484.2403181 | 517.0323413 | 1.74E-07 | -0.656815984 | 3.95E-08 | 30.17247819 | 0.634276597 | 6.760104571 |
| ENSMUSG00000013822 | *Elof1* | 1952.422314 | 2017.534784 | 2320.292569 | 1342.691586 | 1265.973567 | 1374.242971 | 4.57E-11 | -0.657165146 | 7.52E-12 | 46.88721867 | 0.634123107 | 10.33979648 |
| ENSMUSG00000068744 | *Psrc1* | 670.2464491 | 728.6944043 | 734.0659545 | 407.3618316 | 460.3540244 | 484.1742112 | 6.07E-08 | -0.657805528 | 1.32E-08 | 32.30043828 | 0.633841696 | 7.21645498 |
| ENSMUSG00000009863 | *Sdhb* | 7672.28474 | 7670.467414 | 9189.915276 | 5097.50499 | 5169.211109 | 5274.696296 | 1.78E-12 | -0.658061762 | 2.62E-13 | 53.473212 | 0.633729131 | 11.75038562 |
| ENSMUSG00000018648 | *Dusp14* | 186.1351879 | 180.659693 | 142.2504409 | 107.1117031 | 112.9170249 | 104.3728838 | 0.002491227 | -0.658457517 | 0.000936856 | 10.94837689 | 0.633555312 | 2.603586766 |
| ENSMUSG00000021916 | *Glt8d1* | 321.1431138 | 381.5048266 | 450.9070579 | 229.4045925 | 219.319606 | 279.2941058 | 0.000664986 | -0.658503069 | 0.000230492 | 13.56459439 | 0.633535308 | 3.177187792 |
| ENSMUSG00000025358 | *Cdk2* | 2411.768808 | 2628.144361 | 2351.158231 | 1604.145349 | 1502.665023 | 1575.257414 | 1.91E-15 | -0.658569812 | 2.29E-16 | 67.3380798 | 0.633506 | 14.71960222 |
| ENSMUSG00000028672 | *Hmgcl* | 2329.485871 | 2305.177312 | 2332.370437 | 1440.525897 | 1448.377992 | 1522.104556 | 1.72E-20 | -0.659466398 | 1.50E-21 | 90.91454957 | 0.63311242 | 19.76541156 |
| ENSMUSG00000009563 | *Tor2a* | 2069.855244 | 2100.295091 | 2137.782569 | 1324.980202 | 1334.375226 | 1332.6871 | 7.36E-21 | -0.659513228 | 6.30E-22 | 92.63242647 | 0.633091869 | 20.13323745 |
| ENSMUSG00000031217 | *Efnb1* | 196.520413 | 219.0120301 | 170.432132 | 115.5456955 | 132.4603561 | 124.6676113 | 0.00117891 | -0.659565116 | 0.000423793 | 12.42425953 | 0.6330691 | 2.928519354 |
| ENSMUSG00000022841 | *Ap2m1* | 13009.49156 | 13897.67583 | 14679.9771 | 8767.97847 | 8691.353692 | 8856.232477 | 2.32E-21 | -0.659857641 | 1.93E-22 | 94.97150137 | 0.63294075 | 20.63431759 |
| ENSMUSG00000023051 | *Tarbp2* | 1646.457606 | 1760.170417 | 1778.130511 | 1014.609282 | 1159.570986 | 1107.512267 | 8.71E-13 | -0.660069824 | 1.26E-13 | 54.91903099 | 0.632847667 | 12.05992159 |
| ENSMUSG00000057788 | *Ddx49* | 2657.818756 | 2635.209265 | 2679.944627 | 1648.845509 | 1683.983707 | 1712.488427 | 5.35E-23 | -0.66024668 | 4.07E-24 | 102.6158058 | 0.632770093 | 22.27143986 |
| ENSMUSG00000053559 | *Smagp* | 146.1920145 | 207.9100378 | 155.6702938 | 107.1117031 | 103.1453592 | 112.1042086 | 0.005475933 | -0.660247466 | 0.002186947 | 9.385614377 | 0.632769749 | 2.261541877 |
| ENSMUSG00000030761 | *Myo7a* | 5955.527149 | 5998.103663 | 4764.047785 | 3480.708652 | 3594.887205 | 3502.290102 | 5.23E-10 | -0.661121961 | 9.41E-11 | 41.93967937 | 0.632386309 | 9.281257513 |
| ENSMUSG00000032040 | *Dcps* | 2033.906388 | 1942.848654 | 2147.176466 | 1327.510399 | 1279.002455 | 1261.172346 | 3.74E-16 | -0.66165082 | 4.29E-17 | 70.6384759 | 0.632154533 | 15.42737878 |
| ENSMUSG00000029771 | *Irf5* | 7323.181405 | 7339.426189 | 5880.579547 | 4244.828361 | 4400.506747 | 4345.004498 | 1.91E-10 | -0.661896958 | 3.31E-11 | 43.98358011 | 0.632046691 | 9.718429678 |
| ENSMUSG00000059070 | *Rpl18* | 44568.59284 | 44916.64234 | 53161.40534 | 29255.83274 | 29609.23255 | 31281.90627 | 2.79E-13 | -0.662002157 | 3.89E-14 | 57.22288793 | 0.632000604 | 12.5549193 |
| ENSMUSG00000031781 | *Ciapin1* | 3799.39465 | 4129.941139 | 4131.972713 | 2581.645065 | 2491.774731 | 2542.63942 | 1.89E-20 | -0.662305123 | 1.66E-21 | 90.71848297 | 0.631867898 | 19.72296333 |
| ENSMUSG00000040322 | *Slc25a24* | 3470.262902 | 3312.430796 | 3742.796978 | 2259.466557 | 2149.766435 | 2235.319262 | 4.58E-17 | -0.662536373 | 4.95E-18 | 74.90017548 | 0.631766624 | 16.33895665 |
| ENSMUSG00000047205 | *Dusp18* | 247.6476749 | 329.0226812 | 245.5833083 | 162.7760528 | 173.7184998 | 183.6189623 | 0.000742564 | -0.66257987 | 0.000259354 | 13.34325247 | 0.631747577 | 3.129265846 |
| ENSMUSG00000007610 | *Gtpbp3* | 1416.384928 | 1361.507966 | 1252.072277 | 818.0972601 | 895.7360144 | 836.9159019 | 6.77E-12 | -0.662818819 | 1.05E-12 | 50.75484021 | 0.631642951 | 11.16919718 |
| ENSMUSG00000079426 | *Arpc4* | 9734.151349 | 9977.66327 | 10878.13277 | 6437.666378 | 6405.869679 | 6471.118798 | 1.06E-21 | -0.662885432 | 8.67E-23 | 96.55659573 | 0.631613787 | 20.97552249 |
| ENSMUSG00000031029 | *Eif3f* | 10036.12174 | 9725.345263 | 9770.994908 | 6267.299732 | 6226.722476 | 6157.033731 | 8.26E-35 | -0.663176563 | 3.53E-36 | 157.7372759 | 0.631486342 | 34.08327437 |
| ENSMUSG00000031657 | *Heatr3* | 2939.81756 | 3138.836008 | 3232.842567 | 1992.952398 | 1887.017204 | 1993.715364 | 3.62E-18 | -0.663348934 | 3.67E-19 | 80.03934331 | 0.631410898 | 17.44105476 |
| ENSMUSG00000034088 | *Hdlbp* | 23151.06328 | 23393.91634 | 23668.59459 | 15084.19536 | 14568.46769 | 14670.18867 | 2.89E-38 | -0.663554535 | 1.04E-39 | 173.8953245 | 0.631320921 | 37.53929183 |
| ENSMUSG00000038178 | *Slc43a2* | 1924.462093 | 1825.773099 | 1399.690659 | 1053.405647 | 1093.340808 | 1109.445099 | 9.15E-07 | -0.663615979 | 2.23E-07 | 26.82300964 | 0.631294034 | 6.03837855 |
| ENSMUSG00000025651 | *Uqcrc1* | 10589.73412 | 10365.22373 | 10790.90373 | 6599.599031 | 6516.615223 | 6916.636386 | 6.41E-29 | -0.664039027 | 3.59E-30 | 130.2605176 | 0.631108943 | 28.19315837 |
| ENSMUSG00000021033 | *Gstz1* | 931.4748029 | 919.4468176 | 1133.977571 | 647.7306142 | 610.1862304 | 620.4388095 | 5.11E-08 | -0.66461709 | 1.10E-08 | 32.64857416 | 0.63085612 | 7.291415868 |
| ENSMUSG00000030881 | *Arfip2* | 2469.286977 | 2597.8662 | 2698.732421 | 1704.509858 | 1558.037795 | 1630.343102 | 1.32E-16 | -0.664702947 | 1.46E-17 | 72.76117343 | 0.630818577 | 15.88027789 |
| ENSMUSG00000018900 | *Slc22a5* | 343.5112909 | 404.7180833 | 338.1802935 | 221.8139994 | 229.0912716 | 234.8389886 | 1.22E-05 | -0.665268582 | 3.38E-06 | 21.58867893 | 0.630571302 | 4.912333065 |
| ENSMUSG00000000743 | *Chmp1a* | 4002.305971 | 4251.053783 | 4560.066021 | 2671.045384 | 2740.409334 | 2662.474954 | 1.04E-17 | -0.665445441 | 1.08E-18 | 77.91060811 | 0.630494005 | 16.98284619 |
| ENSMUSG00000023939 | *Mrpl14* | 1100.034994 | 1137.449576 | 1254.756248 | 705.0817623 | 728.5319584 | 764.4347326 | 3.45E-11 | -0.665953884 | 5.63E-12 | 47.45517532 | 0.630271842 | 10.46235463 |
| ENSMUSG00000024965 | *Fermt3* | 12282.52581 | 11782.24166 | 10130.64697 | 6894.788764 | 7380.864759 | 7283.87431 | 5.66E-13 | -0.665970422 | 8.08E-14 | 55.78704362 | 0.630264617 | 12.24687364 |
| ENSMUSG00000025464 | *Paox* | 190.9283687 | 174.6040609 | 199.9558084 | 111.3286993 | 117.2599873 | 127.566858 | 0.000604841 | -0.666201927 | 0.000208396 | 13.75382531 | 0.630163489 | 3.21835877 |
| ENSMUSG00000029390 | *Tmed2* | 13714.88801 | 14535.53575 | 16749.31842 | 9430.890271 | 9439.428981 | 9479.570533 | 1.55E-13 | -0.666238289 | 2.12E-14 | 58.4187649 | 0.630147606 | 12.80910947 |
| ENSMUSG00000109946 | *Brd3os* | 143.7954241 | 145.3351721 | 152.9863232 | 109.6419008 | 79.25906552 | 87.94381879 | 0.004513396 | -0.667496053 | 0.001774566 | 9.769231932 | 0.629598473 | 2.345496598 |
| ENSMUSG00000020477 | *Mrps24* | 1541.007628 | 1661.261758 | 1913.671026 | 1111.600195 | 977.1665612 | 1120.07567 | 1.19E-08 | -0.670281946 | 2.44E-09 | 35.58495747 | 0.62838387 | 7.922877192 |
| ENSMUSG00000044134 | *Pheta1* | 664.6544048 | 709.5182358 | 641.4689693 | 403.1448354 | 408.2384745 | 456.148159 | 1.11E-08 | -0.670439284 | 2.26E-09 | 35.73719034 | 0.628315343 | 7.954048197 |
| ENSMUSG00000055413 | *H2-Q5* | 1823.805296 | 1863.116164 | 1892.199261 | 1088.828415 | 1137.856173 | 1278.567827 | 7.09E-13 | -0.670449151 | 1.02E-13 | 55.3339143 | 0.628311046 | 12.14963219 |
| ENSMUSG00000036748 | *Cuedc2* | 5343.597733 | 5661.006806 | 6444.21337 | 3527.09561 | 3704.547008 | 3726.498519 | 2.89E-13 | -0.670479959 | 4.04E-14 | 57.14736833 | 0.628297629 | 12.53842643 |
| ENSMUSG00000050705 | *2310061I04Rik* | 664.6544048 | 736.7685805 | 636.1010282 | 443.6279988 | 442.9821744 | 394.2975612 | 4.67E-08 | -0.670747353 | 1.00E-08 | 32.83200426 | 0.628181189 | 7.330853498 |
| ENSMUSG00000020444 | *Guk1* | 1016.15433 | 1086.985974 | 1202.418821 | 724.4799448 | 695.9597397 | 651.3641084 | 1.08E-09 | -0.670941181 | 2.00E-10 | 40.47122329 | 0.628096798 | 8.966651865 |
| ENSMUSG00000026999 | *Nup35* | 1816.615525 | 1718.790264 | 1893.541246 | 1117.503989 | 1178.028577 | 1113.310761 | 9.36E-16 | -0.671124799 | 1.10E-16 | 68.78199191 | 0.628016862 | 15.02855374 |

| | | | | | | | | | | | | | |
|---|---|---|---|---|---|---|---|---|---|---|---|---|---|
| ENSMUSG00000024855 | *Pacs1* | 809.2486924 | 854.8534078 | 660.2567634 | 481.5809645 | 514.6410556 | 466.7787305 | 8.70E-07 | -0.671582943 | 2.12E-07 | 26.92439615 | 0.62781746 | 6.060529768 |
| ENSMUSG00000053460 | *Ggcx* | 460.1453572 | 482.4320294 | 458.9589697 | 335.6728964 | 279.0353403 | 262.8650408 | 4.96E-06 | -0.672868963 | 1.32E-06 | 23.39909271 | 0.627258072 | 5.304130587 |
| ENSMUSG00000024818 | *Slc25a45* | 1493.07582 | 1540.149115 | 1579.516688 | 896.5333892 | 1001.052855 | 994.4416433 | 5.62E-14 | -0.673834948 | 7.43E-15 | 60.48024171 | 0.62683822 | 13.25050964 |
| ENSMUSG00000003531 | *Dgcr6* | 729.3623456 | 675.2029868 | 877.6583807 | 460.4959835 | 504.86939 | 462.9130682 | 1.14E-06 | -0.67395672 | 2.80E-07 | 26.3839763 | 0.626785313 | 5.94356559 |
| ENSMUSG00000000693 | *Loxl3* | 190.9283687 | 194.7895014 | 170.432132 | 110.4853001 | 105.3168405 | 133.3653516 | 0.001094143 | -0.674236207 | 0.000390867 | 12.57535515 | 0.626663901 | 2.96092577 |
| ENSMUSG00000003299 | *Mrpl4* | 3662.788998 | 3659.620374 | 3907.861169 | 2274.647743 | 2397.315297 | 2363.852536 | 4.94E-22 | -0.67423995 | 3.99E-23 | 98.09181157 | 0.626662275 | 21.30658396 |
| ENSMUSG00000031442 | *Mcf2l* | 332.3272024 | 392.6068189 | 304.6306612 | 227.7177941 | 192.1760904 | 225.1748327 | 0.000101029 | -0.674482046 | 3.14E-05 | 17.33373834 | 0.626557125 | 3.995552749 |
| ENSMUSG00000036845 | *Lin37* | 509.6748921 | 426.9220679 | 501.902499 | 285.9123414 | 307.2645965 | 308.2865736 | 1.20E-06 | -0.675224914 | 2.95E-07 | 26.28310459 | 0.626234583 | 5.922549033 |
| ENSMUSG00000030062 | *Rpn1* | 22019.87261 | 21763.94201 | 25296.42275 | 14343.69083 | 14430.57863 | 14476.90556 | 3.15E-17 | -0.675335456 | 3.38E-18 | 75.65374549 | 0.626186601 | 16.50161737 |
| ENSMUSG00000107470 | *Gm3375* | 2588.317634 | 2577.68076 | 3176.479185 | 1671.617288 | 1690.498151 | 1856.484351 | 1.89E-09 | -0.675563729 | 3.57E-10 | 39.33428606 | 0.62608753 | 8.723325625 |
| ENSMUSG00000022354 | *Ndufb9* | 3021.301633 | 3039.927349 | 3992.406242 | 2083.196116 | 2014.048857 | 2189.897729 | 1.14E-07 | -0.676001057 | 2.55E-08 | 31.02087246 | 0.625897771 | 6.941823206 |
| ENSMUSG00000044066 | *Cep68* | 641.4873642 | 606.5724889 | 587.7895577 | 394.710843 | 375.6662558 | 379.8013273 | 1.90E-09 | -0.676006749 | 3.60E-10 | 39.31940974 | 0.625895301 | 8.720152556 |
| ENSMUSG00000070003 | *Ssbp4* | 4257.143417 | 4559.891023 | 4558.724035 | 2616.224434 | 2744.752296 | 3007.48532 | 5.04E-16 | -0.676400508 | 5.82E-17 | 70.03605229 | 0.625724497 | 15.29786863 |
| ENSMUSG00000068697 | *Myoz1* | 266.0215346 | 244.2438308 | 257.661176 | 171.2100452 | 141.1462811 | 167.1898973 | 9.06E-05 | -0.676593057 | 2.79E-05 | 17.55274574 | 0.625640991 | 4.043020233 |
| ENSMUSG00000033416 | *Gucd1* | 729.3623456 | 895.224289 | 881.6843365 | 526.281124 | 536.355868 | 502.5361074 | 6.69E-08 | -0.676764625 | 1.46E-08 | 32.10419726 | 0.625566592 | 7.174706987 |
| ENSMUSG00000042389 | *Tsen2* | 480.1169438 | 527.8492707 | 571.6857342 | 366.8786682 | 279.0353403 | 338.2454569 | 1.47E-05 | -0.676827611 | 4.10E-06 | 21.21757115 | 0.625539282 | 4.832614114 |
| ENSMUSG00000033502 | *Cdc14a* | 436.9783166 | 433.9869721 | 420.0413962 | 275.7915506 | 274.6923778 | 257.0665472 | 1.05E-07 | -0.677963663 | 2.32E-08 | 31.20355505 | 0.625046894 | 6.980855324 |
| ENSMUSG00000038963 | *Slco4a1* | 19777.46286 | 21006.98799 | 17482.04239 | 11936.6294 | 12410.01533 | 12077.29564 | 2.39E-15 | -0.678004409 | 2.88E-16 | 66.88186933 | 0.625029241 | 14.62187186 |
| ENSMUSG00000063065 | *Mapk3* | 3709.123079 | 3847.344971 | 4032.665801 | 2413.808617 | 2423.373072 | 2401.542744 | 2.97E-23 | -0.678075947 | 2.22E-24 | 103.8130394 | 0.624998249 | 22.52674882 |
| ENSMUSG00000001105 | *Ift20* | 1223.059968 | 1277.738388 | 1753.974776 | 916.7749709 | 854.4778707 | 882.3374347 | 1.15E-05 | -0.678401782 | 3.18E-06 | 21.70764617 | 0.624857108 | 4.93824645 |
| ENSMUSG00000024038 | *Ndufv3* | 1421.976972 | 1528.037851 | 1729.819041 | 1034.007465 | 892.4787926 | 991.5423965 | 2.99E-09 | -0.678468993 | 5.74E-10 | 38.40837558 | 0.624827999 | 8.524598825 |
| ENSMUSG00000031482 | *Slc25a15* | 477.7203534 | 531.8863588 | 597.1834547 | 347.4804857 | 308.3503371 | 344.0439504 | 2.06E-06 | -0.678558103 | 5.25E-07 | 25.17081572 | 0.624789406 | 5.685447541 |
| ENSMUSG00000031672 | *Got2* | 15052.18545 | 15465.07529 | 15240.92696 | 9618.124901 | 9566.460634 | 9403.223702 | 1.53E-39 | -0.678566758 | 5.21E-41 | 179.8547587 | 0.624785658 | 38.81427408 |
| ENSMUSG00000043740 | *B430306N03Rik* | 542.4282943 | 522.8029106 | 567.6597783 | 324.7087063 | 398.4668088 | 297.6560021 | 6.34E-06 | -0.679204743 | 1.70E-06 | 22.90528845 | 0.624509428 | 5.197888373 |
| ENSMUSG00000041607 | *Mbp* | 2678.589206 | 2661.450338 | 2082.761172 | 1587.277364 | 1513.522429 | 1538.533621 | 8.53E-09 | -0.679351448 | 1.71E-09 | 36.27749335 | 0.624445926 | 8.068884467 |
| ENSMUSG00000025212 | *Sfxn3* | 882.7441314 | 850.8163197 | 640.1269841 | 446.1581965 | 504.86939 | 535.3942375 | 2.44E-05 | -0.679424284 | 7.01E-06 | 20.19131773 | 0.624414401 | 4.612176042 |
| ENSMUSG00000027248 | *Pdia3* | 36482.49682 | 40549.52227 | 47568.01064 | 25697.53136 | 26186.9781 | 25862.24763 | 7.87E-11 | -0.680337159 | 1.33E-11 | 45.77700778 | 0.624019424 | 10.10419019 |
| ENSMUSG00000031556 | *Tm2d2* | 1235.841784 | 1209.10789 | 1344.669262 | 780.9876937 | 802.3623208 | 778.9309664 | 8.77E-14 | -0.68041238 | 1.18E-14 | 59.57686166 | 0.623986888 | 13.05675752 |
| ENSMUSG00000029998 | *Pcyox1* | 1737.528041 | 1949.913558 | 2084.103158 | 1195.940118 | 1185.628761 | 1214.784398 | 2.92E-12 | -0.680418069 | 4.40E-13 | 52.45718049 | 0.623984428 | 11.53433591 |
| ENSMUSG00000027502 | *Rtf2* | 4240.367284 | 4364.09225 | 4738.550064 | 2770.566494 | 2689.379525 | 2857.690903 | 4.53E-19 | -0.680812068 | 4.32E-20 | 84.26689233 | 0.623814042 | 18.34391484 |
| ENSMUSG00000032806 | *Slc10a3* | 930.6759394 | 989.0865876 | 870.9484542 | 528.8113217 | 614.5291929 | 601.1104977 | 2.31E-09 | -0.680840613 | 4.40E-10 | 38.9284419 | 0.623801699 | 8.636023772 |
| ENSMUSG00000039263 | *Npepl1* | 4984.109172 | 4818.264662 | 5010.973079 | 2998.284288 | 3068.303002 | 3173.708801 | 7.42E-27 | -0.680944411 | 4.59E-28 | 120.6363192 | 0.62375682 | 26.12951651 |
| ENSMUSG00000034974 | *Dapk3* | 1505.857636 | 1515.926586 | 1505.707497 | 917.6183701 | 853.3921301 | 1051.460163 | 8.57E-11 | -0.681230474 | 1.45E-11 | 45.60362724 | 0.623633151 | 10.06681362 |
| ENSMUSG00000064254 | *Ethe1* | 698.2066704 | 825.584519 | 728.6980133 | 460.4959835 | 411.4956963 | 531.5285751 | 1.65E-06 | -0.681248942 | 4.13E-07 | 25.63397387 | 0.623625168 | 5.783729246 |
| ENSMUSG00000025578 | *Cbx8* | 300.3726637 | 353.2452098 | 378.4398522 | 221.8139994 | 205.2049779 | 213.5778456 | 2.60E-05 | -0.681520392 | 7.48E-06 | 20.06663024 | 0.623507841 | 4.585647478 |
| ENSMUSG00000024654 | *Asrgl1* | 587.1646485 | 605.5632169 | 597.1834547 | 368.5654667 | 389.7808839 | 357.5737687 | 1.12E-09 | -0.682251484 | 2.07E-10 | 40.39696856 | 0.623191956 | 8.95180784 |
| ENSMUSG00000030839 | *Sergef* | 1077.666817 | 1143.505208 | 1346.011247 | 741.3479295 | 730.7034397 | 746.0728364 | 3.88E-09 | -0.682342378 | 7.55E-10 | 37.87178999 | 0.623152694 | 8.410737836 |
| ENSMUSG00000022442 | *Ttll1* | 389.0465086 | 399.6717231 | 363.678014 | 241.2121818 | 261.6634903 | 216.4770924 | 3.25E-06 | -0.683080786 | 8.44E-07 | 24.25470613 | 0.62283383 | 5.488288513 |
| ENSMUSG00000025648 | *Pfkfb4* | 990.5906994 | 1032.485285 | 1091.034042 | 663.7551997 | 661.2160397 | 611.7410692 | 4.01E-12 | -0.683159393 | 6.08E-13 | 51.82046288 | 0.622799895 | 11.39657059 |
| ENSMUSG00000055612 | *Cdca7* | 3439.107227 | 3284.17118 | 3163.059332 | 2060.424337 | 2062.907185 | 2038.170482 | 1.87E-22 | -0.683164954 | 1.48E-23 | 100.0643599 | 0.622797494 | 21.7282515 |
| ENSMUSG00000048997 | *Atxn7l2* | 365.879468 | 316.9114168 | 276.4489701 | 173.7402429 | 207.3764591 | 219.3763392 | 0.000282306 | -0.683253344 | 9.27E-05 | 15.27912646 | 0.622759339 | 3.549280637 |
| ENSMUSG00000052456 | *Get3* | 6148.852108 | 7125.460519 | 8280.049249 | 4508.812322 | 4452.622297 | 4451.310213 | 2.64E-09 | -0.683725614 | 5.03E-10 | 38.66365006 | 0.62255551 | 8.578993921 |
| ENSMUSG00000063146 | *Clip2* | 2131.367731 | 1997.349344 | 1528.521247 | 1102.322803 | 1225.801164 | 1199.321749 | 1.04E-06 | -0.683727059 | 2.53E-07 | 26.57873129 | 0.622554887 | 5.984945024 |
| ENSMUSG00000050299 | *Gm9843* | 15029.01841 | 14669.76893 | 17158.62394 | 9743.791388 | 9457.886572 | 9960.845498 | 1.43E-16 | -0.683850936 | 1.59E-17 | 72.59964711 | 0.622501433 | 15.84565483 |
| ENSMUSG00000073405 | *H2-T-ps* | 206.905638 | 241.2160147 | 232.1634554 | 155.1854597 | 109.659803 | 156.5593258 | 0.001504088 | -0.684325562 | 0.000548676 | 11.94250626 | 0.622296673 | 2.822726669 |
| ENSMUSG00000044948 | *Cfap43* | 616.7225967 | 634.8321057 | 448.2230874 | 344.1068888 | 348.5227402 | 368.2043402 | 4.77E-05 | -0.684492832 | 1.42E-05 | 18.84243739 | 0.622224526 | 4.321036827 |
| ENSMUSG00000002372 | *Ranbp3* | 4198.027521 | 4414.555851 | 4247.383448 | 2618.754632 | 2636.178234 | 2746.55311 | 9.62E-26 | -0.684544021 | 6.33E-27 | 115.4329874 | 0.622202449 | 25.016733 |
| ENSMUSG00000043964 | *Orai3* | 953.8429799 | 1002.207124 | 908.5240424 | 613.9946447 | 606.9290086 | 562.453874 | 1.21E-11 | -0.684685819 | 1.90E-12 | 49.58417083 | 0.622141298 | 10.91848568 |
| ENSMUSG00000041707 | *Tmem273* | 846.7952753 | 859.899768 | 1129.951615 | 631.7060287 | 553.727718 | 574.0508611 | 4.41E-06 | -0.684964196 | 1.16E-06 | 23.63854746 | 0.622021263 | 5.355952377 |
| ENSMUSG00000022554 | *Hgh1* | 1016.15433 | 955.7806106 | 962.203454 | 557.4868958 | 641.6727085 | 629.1365498 | 7.12E-11 | -0.6851533 | 1.19E-11 | 45.98305076 | 0.621939736 | 10.14742697 |
| ENSMUSG00000026939 | *Tmem141* | 840.4043676 | 943.6693463 | 1231.942498 | 603.0304546 | 612.3577117 | 656.1961864 | 2.02E-05 | -0.685220834 | 5.74E-06 | 20.5736766 | 0.621910623 | 4.69391919 |
| ENSMUSG00000024948 | *Map4k2* | 602.3430543 | 685.2957071 | 453.5910285 | 384.5900522 | 336.5795933 | 363.3722623 | 0.000142615 | -0.685291622 | 4.51E-05 | 16.64168099 | 0.621880109 | 3.845835434 |
| ENSMUSG00000016503 | *Gtf3a* | 1723.947362 | 1716.77172 | 1826.441982 | 1110.756795 | 988.0239674 | 1171.295696 | 1.53E-12 | -0.686022955 | 2.24E-13 | 53.78148784 | 0.621564944 | 11.81556912 |
| ENSMUSG00000060098 | *Prmt7* | 3277.736806 | 3203.429417 | 3344.227346 | 2008.976983 | 2019.47756 | 2076.827105 | 1.20E-25 | -0.686171015 | 7.94E-27 | 114.9819924 | 0.621501158 | 24.91913275 |
| ENSMUSG00000024953 | *Prdx5* | 3893.66054 | 4131.959683 | 5044.522711 | 2659.237795 | 2650.292862 | 2807.437292 | 1.21E-09 | -0.686261686 | 2.25E-10 | 40.23367359 | 0.621462099 | 8.916887866 |
| ENSMUSG00000025486 | *Sirt3* | 255.6363095 | 237.1789266 | 268.3970583 | 145.908068 | 176.9757216 | 150.7608322 | 9.92E-05 | -0.686312681 | 3.07E-05 | 17.37228215 | 0.621440133 | 4.003699507 |
| ENSMUSG00000070691 | *Runx3* | 1288.566773 | 1096.069423 | 705.8842633 | 651.1042111 | 642.7584491 | 630.1029654 | 0.001102722 | -0.686470209 | 0.000394452 | 12.55829159 | 0.621372281 | 2.957533965 |
| ENSMUSG00000034126 | *Pomt2* | 1143.173622 | 1193.968809 | 1036.012645 | 728.696941 | 653.6158554 | 714.1811218 | 7.41E-11 | -0.686681501 | 1.24E-11 | 45.90059017 | 0.621281284 | 10.13021637 |
| ENSMUSG00000027490 | *E2f1* | 1519.438315 | 1491.704058 | 1350.037203 | 875.4484082 | 981.5095237 | 857.2106293 | 1.69E-11 | -0.686870388 | 2.69E-12 | 48.89989479 | 0.621199947 | 10.77184546 |
| ENSMUSG00000038976 | *Ppp1r9b* | 6776.758793 | 6700.556995 | 5151.881534 | 3732.041625 | 3923.866614 | 3921.714469 | 6.37E-09 | -0.686952357 | 1.26E-09 | 36.87199154 | 0.621164654 | 8.195676553 |
| ENSMUSG00000028643 | *Svbp* | 529.6464788 | 587.3963204 | 716.6201457 | 393.0240446 | 373.4947745 | 368.2043402 | 5.95E-06 | -0.687156482 | 1.59E-06 | 23.03388914 | 0.621076772 | 5.225506524 |
| ENSMUSG00000001473 | *Tubb6* | 8459.165255 | 8453.662508 | 7273.56028 | 4913.643956 | 4973.777797 | 5136.498867 | 4.36E-16 | -0.687498396 | 5.02E-17 | 70.32934612 | 0.620929596 | 15.36066268 |
| ENSMUSG00000031834 | *Pik3r2* | 2166.517723 | 2272.880607 | 1747.26485 | 1185.819328 | 1372.376148 | 1288.231983 | 4.79E-08 | -0.68766927 | 1.03E-08 | 32.77939735 | 0.620856057 | 7.319576472 |
| ENSMUSG00000068264 | *Ap5s1* | 484.9101246 | 497.5711099 | 492.508602 | 296.0331323 | 326.8079277 | 293.7903397 | 2.44E-08 | -0.687716051 | 5.13E-09 | 34.13917247 | 0.620835925 | 7.611883394 |
| ENSMUSG00000074754 | *Smim26* | 270.8147154 | 219.0120301 | 395.885661 | 196.5120223 | 186.7473873 | 164.2906505 | 0.007605492 | -0.687819101 | 0.003109257 | 8.742220477 | 0.620791582 | 2.118872664 |
| ENSMUSG00000051238 | *Swsap1* | 230.0726786 | 218.0027581 | 209.3497055 | 150.9684635 | 118.345728 | 138.1974295 | 0.000272551 | -0.688236021 | 8.93E-05 | 15.35065534 | 0.620612207 | 3.5645527 |
| ENSMUSG00000002804 | *Nudt14* | 633.4987296 | 547.0254393 | 662.940734 | 375.3126606 | 371.3232933 | 396.2303924 | 8.99E-08 | -0.688255586 | 1.98E-08 | 31.50995162 | 0.62060379 | 7.046268601 |
| ENSMUSG00000030220 | *Arhgdib* | 17237.07703 | 17559.31475 | 20741.72467 | 11697.94742 | 11315.58878 | 11445.25985 | 2.34E-14 | -0.688313614 | 3.01E-15 | 62.25727112 | 0.620578829 | 13.63110197 |
| ENSMUSG00000003559 | *As3mt* | 885.1407218 | 923.4839057 | 936.7057335 | 582.7888729 | 564.5851243 | 553.7561337 | 4.98E-13 | -0.688612597 | 7.09E-14 | 56.04394218 | 0.620450234 | 12.30236313 |
| ENSMUSG00000035765 | *Dym* | 1112.81681 | 1120.291951 | 994.411101 | 680.6231844 | 653.6158554 | 669.7260047 | 2.70E-12 | -0.689312862 | 4.05E-13 | 52.6201476 | 0.620149149 | 11.56815412 |
| ENSMUSG00000083240 | *Gm13453* | 1466.713326 | 1579.510724 | 2171.332202 | 1052.562248 | 1093.340808 | 1085.284709 | 1.08E-05 | -0.689367927 | 2.97E-06 | 21.83439728 | 0.620125479 | 4.966286568 |
| ENSMUSG00000028636 | *Ppcs* | 656.6657701 | 631.8042896 | 665.6247046 | 402.3014362 | 432.1247682 | 377.8684961 | 1.47E-09 | -0.689434168 | 2.76E-10 | 39.83817521 | 0.620097007 | 8.831703249 |
| ENSMUSG00000001211 | *Agpat3* | 3356.82429 | 3614.203133 | 3561.628964 | 2057.05074 | 2197.539022 | 2273.00947 | 5.68E-19 | -0.690185807 | 5.45E-20 | 83.81053043 | 0.619774023 | 18.24554045 |
| ENSMUSG00000032418 | *Me1* | 1515.443997 | 1878.255244 | 2137.782569 | 1089.671815 | 1196.486167 | 1138.437566 | 7.16E-07 | -0.690193905 | 1.72E-07 | 27.31939233 | 0.619770544 | 6.145348731 |
| ENSMUSG00000052934 | *Fbxo31* | 981.8032013 | 1063.772718 | 874.9744101 | 591.2228653 | 616.7006742 | 604.97616 | 8.62E-10 | -0.690510759 | 1.58E-10 | 40.92995281 | 0.619634441 | 9.064282113 |
| ENSMUSG00000036992 | *Nxt1* | 1717.556455 | 1643.094862 | 1952.588599 | 1051.718849 | 1092.255067 | 1144.23606 | 8.18E-12 | -0.690949988 | 1.27E-12 | 50.37610817 | 0.619445822 | 11.08704461 |
| ENSMUSG00000024900 | *Cpt1a* | 1455.529237 | 1640.067046 | 1387.612791 | 936.1731533 | 956.5374894 | 886.2030971 | 1.15E-11 | -0.691033365 | 1.81E-12 | 49.67910009 | 0.619410024 | 10.93835885 |
| ENSMUSG00000002489 | *Tiam1* | 139.0022433 | 155.4278923 | 138.224485 | 95.30411381 | 80.34480614 | 91.80948116 | 0.001855215 | -0.691055362 | 0.00068618 | 11.52631267 | 0.61940058 | 2.731605813 |
| ENSMUSG00000024782 | *Ak3* | 1814.218934 | 1913.579765 | 2076.051246 | 1270.159251 | 1173.685614 | 1144.23606 | 2.39E-13 | -0.691549202 | 3.32E-14 | 57.53463237 | 0.619188593 | 12.62154731 |
| ENSMUSG00000017404 | *Rpl19* | 36229.2571 | 35586.93171 | 45843.55954 | 23585.65967 | 23923.2089 | 25321.0549 | 5.20E-10 | -0.691913051 | 9.35E-11 | 41.95377513 | 0.619032452 | 9.283960548 |
| ENSMUSG00000019464 | *Ptger1* | 259.6306269 | 221.0305742 | 154.3283085 | 144.2212696 | 127.031653 | 123.7011957 | 0.006316512 | -0.691947085 | 0.002545393 | 9.10767412 | 0.619017849 | 2.199522686 |
| ENSMUSG00000022827 | *Rabl3* | 419.4033203 | 487.4783896 | 436.1452197 | 276.6349498 | 279.0353403 | 275.4284435 | 2.52E-07 | -0.691967687 | 5.82E-08 | 29.4229888 | 0.619009009 | 6.598584215 |
| ENSMUSG00000025217 | *Btrc* | 595.1532831 | 661.0731784 | 594.4994841 | 382.0598545 | 404.9812526 | 359.5065999 | 1.03E-08 | -0.69198617 | 2.07E-09 | 35.90263551 | 0.619001079 | 7.988826158 |
| ENSMUSG00000000631 | *Myo18a* | 3046.066401 | 3029.834628 | 2398.127716 | 1742.462824 | 1726.327591 | 1781.103934 | 9.17E-10 | -0.692083989 | 1.68E-10 | 40.80448549 | 0.61895911 | 9.037798481 |
| ENSMUSG00000053929 | *Cyhr1* | 1731.137134 | 1697.595551 | 1578.174703 | 1040.754659 | 960.8804519 | 1096.881696 | 1.31E-13 | -0.692932757 | 1.78E-14 | 58.75664635 | 0.618595071 | 12.88160406 |
| ENSMUSG00000020526 | *Znhit3* | 878.749814 | 886.1408407 | 1107.137866 | 560.0170935 | 603.6717867 | 609.808238 | 1.22E-07 | -0.692938929 | 2.72E-08 | 30.89468261 | 0.618592424 | 6.915418899 |
| ENSMUSG00000029082 | *Bst1* | 931.4748029 | 1000.18858 | 1210.470733 | 622.4286371 | 665.5590022 | 651.3641084 | 8.61E-08 | -0.69348735 | 1.90E-08 | 31.5977364 | 0.61835732 | 7.065204324 |
| ENSMUSG00000026189 | *Pecr* | 197.3192764 | 251.308735 | 280.4749259 | 161.9326535 | 148.7464654 | 137.2310139 | 0.000989424 | -0.694706421 | 0.000351647 | 12.77305846 | 0.617835031 | 3.004617384 |
| ENSMUSG00000024937 | *Ehbp1l1* | 7454.993877 | 7253.638066 | 5806.770356 | 4138.560057 | 4203.987694 | 4334.373926 | 1.75E-10 | -0.695154503 | 3.02E-11 | 44.16428363 | 0.61764317 | 9.756742903 |
| ENSMUSG00000078681 | *Tm2d3* | 981.0043378 | 1043.587277 | 1348.695218 | 672.1891921 | 712.2458491 | 694.85281 | 1.32E-06 | -0.695376439 | 3.27E-07 | 26.08498523 | 0.617548162 | 5.880890946 |
| ENSMUSG00000002280 | *Ciao3* | 1269.39405 | 1472.527889 | 1567.43882 | 798.6990777 | 897.9074957 | 961.5835132 | 2.17E-08 | -0.695907137 | 4.54E-09 | 34.37847999 | 0.617321038 | 7.663574656 |
| ENSMUSG00000008855 | *Hdac5* | 2811.200541 | 2364.724362 | 1767.394629 | 1410.163525 | 1438.606326 | 1440.925646 | 1.92E-05 | -0.696339494 | 5.41E-06 | 20.68653493 | 0.617136062 | 4.717799052 |
| ENSMUSG00000006941 | *Eif1b* | 1723.947362 | 1643.094862 | 2015.661908 | 1090.515214 | 1126.998767 | 1099.780943 | 3.81E-11 | -0.696740306 | 6.22E-12 | 47.25838877 | 0.616964632 | 10.41953988 |
| ENSMUSG00000025436 | *Atp23* | 430.5874089 | 440.0426043 | 401.2536022 | 241.2121818 | 262.7492309 | 282.1933526 | 4.80E-07 | -0.697501878 | 1.14E-07 | 28.1269523 | 0.616639034 | 6.318701958 |
| ENSMUSG00000022677 | *Cep20* | 865.1691351 | 982.0216834 | 1064.194336 | 625.802234 | 567.8423461 | 596.2784197 | 5.27E-09 | -0.697829367 | 1.03E-09 | 37.26081191 | 0.616499074 | 8.278505289 |
| ENSMUSG00000022503 | *Nubp1* | 2409.372217 | 2262.787887 | 2230.379555 | 1463.297677 | 1413.634292 | 1379.075049 | 2.36E-19 | -0.698333377 | 2.21E-20 | 85.59111398 | 0.616283736 | 18.62746346 |
| ENSMUSG00000040701 | *Ap1g2* | 1804.632573 | 1944.867198 | 1564.75485 | 1113.286993 | 1098.769511 | 1065.956397 | 5.94E-11 | -0.698492204 | 9.88E-12 | 46.35175348 | 0.616215892 | 10.22593032 |
| ENSMUSG00000021868 | *Ppif* | 1656.043968 | 1689.521375 | 1543.283085 | 960.6317312 | 932.6511956 | 1120.07567 | 3.97E-11 | -0.698779471 | 6.50E-12 | 47.17343493 | 0.616093205 | 10.40149993 |
| ENSMUSG00000001228 | *Uhrf1* | 8072.515337 | 8143.815995 | 6746.16006 | 4739.903713 | 4795.716334 | 4616.567279 | 7.58E-14 | -0.698828928 | 1.01E-14 | 59.87521315 | 0.616072085 | 13.12029393 |
| ENSMUSG00000009894 | *Snap47* | 1631.2792 | 1738.975704 | 1649.299923 | 1094.73221 | 950.0230456 | 1043.728838 | 3.97E-14 | -0.699272972 | 5.20E-15 | 61.1844033 | 0.615882495 | 13.4015708 |
| ENSMUSG00000050493 | *Fam167b* | 1658.440558 | 1677.410111 | 1745.922864 | 1029.790469 | 1037.968036 | 1059.191488 | 1.44E-19 | -0.699639121 | 1.33E-20 | 86.59255141 | 0.615726207 | 18.84189576 |
| ENSMUSG00000022822 | *Abcc5* | 1184.714522 | 1474.546433 | 1461.421982 | 798.6990777 | 865.335277 | 870.7404476 | 6.77E-09 | -0.699697636 | 1.34E-09 | 36.75024674 | 0.615701233 | 8.169203355 |
| ENSMUSG00000030811 | *Fbxl19* | 867.5657255 | 1021.383292 | 818.6110278 | 542.3057095 | 553.727718 | 572.1180299 | 1.20E-08 | -0.700164254 | 2.46E-09 | 35.57120422 | 0.615502126 | 7.920061611 |
| ENSMUSG00000071203 | *Naip5* | 614.3260063 | 630.7950176 | 614.6292635 | 399.7712384 | 363.7231089 | 379.8013273 | 2.98E-10 | -0.700298659 | 5.25E-11 | 43.08269791 | 0.615444787 | 9.526272948 |
| ENSMUSG00000056698 | *Elmod3* | 289.9874386 | 356.2730259 | 328.7863964 | 205.7894139 | 210.633681 | 182.6525467 | 2.44E-05 | -0.700599451 | 6.99E-06 | 20.19690605 | 0.615316485 | 4.613355417 |
| ENSMUSG00000058267 | *Mrps14* | 2418.958579 | 2556.486047 | 3427.430435 | 1752.583615 | 1666.611857 | 1745.346558 | 7.19E-07 | -0.700725898 | 1.73E-07 | 27.30813931 | 0.615262557 | 6.143033931 |
| ENSMUSG00000022842 | *Ece2* | 1448.339466 | 1468.490801 | 1564.75485 | 960.6317312 | 868.5924988 | 922.9268895 | 4.56E-15 | -0.700945356 | 5.61E-16 | 65.57025028 | 0.615168972 | 14.34129041 |
| ENSMUSG00000015133 | *Lrrk1* | 1939.640499 | 1956.978463 | 1316.487571 | 995.2110999 | 1154.142283 | 1062.090735 | 2.40E-05 | -0.701060951 | 6.88E-06 | 20.22681445 | 0.615119684 | 4.619608843 |
| ENSMUSG00000057672 | *Pkn1* | 6117.696432 | 6118.207035 | 5345.127416 | 3518.661618 | 3632.888126 | 3667.547168 | 8.22E-18 | -0.701261398 | 8.47E-19 | 78.38635496 | 0.615034226 | 17.08521179 |
| ENSMUSG00000003865 | *Gys1* | 1802.235982 | 1709.706816 | 1528.521247 | 1046.658453 | 993.4526706 | 1063.05715 | 8.25E-13 | -0.701436947 | 1.19E-13 | 55.02806973 | 0.614959392 | 12.08365966 |
| ENSMUSG00000041084 | *Ostc* | 6313.417982 | 5958.742054 | 8293.469102 | 4173.139426 | 4049.812526 | 4417.485667 | 1.17E-07 | -0.701618584 | 2.60E-08 | 30.98179492 | 0.614881973 | 6.933308787 |
| ENSMUSG00000025792 | *Slc25a10* | 1618.497385 | 1663.280302 | 1387.612791 | 878.8220052 | 998.8813737 | 998.3073056 | 4.45E-10 | -0.701728221 | 7.96E-11 | 42.26740897 | 0.614835247 | 9.351294543 |
| ENSMUSG00000038695 | *Josd2* | 1443.546285 | 1567.39946 | 1627.828159 | 899.9069861 | 968.4806362 | 980.911825 | 7.21E-14 | -0.702165014 | 9.59E-15 | 59.97778423 | 0.614649126 | 13.14196484 |
| ENSMUSG00000020869 | *Lrrc59* | 19400.3993 | 19555.65482 | 18705.93298 | 11781.44394 | 11990.91945 | 11673.33393 | 5.78E-39 | -0.702190338 | 2.02E-40 | 177.1587677 | 0.614638338 | 38.23823541 |
| ENSMUSG00000037012 | *Hk1* | 7421.441611 | 7674.504502 | 6613.303517 | 4450.617775 | 4502.566366 | 4395.258108 | 1.32E-18 | -0.702201173 | 1.29E-19 | 82.1064571 | 0.614633722 | 17.8788334 |
| ENSMUSG00000005575 | *Ube2m* | 5024.851209 | 5053.425045 | 5264.608299 | 3075.877018 | 3157.333733 | 3194.969944 | 1.12E-29 | -0.702214879 | 6.03E-31 | 133.803007 | 0.614627882 | 28.95136417 |

| | | | | | | | | | | | | | |
|---|---|---|---|---|---|---|---|---|---|---|---|---|---|
| ENSMUSG00000030815 | *Phkg2* | 1219.065651 | 1241.404595 | 1331.249409 | 713.5157547 | 801.2765802 | 814.6883433 | 2.57E-12 | -0.702632245 | 3.85E-13 | 52.71942781 | 0.614450099 | 11.58925085 |
| ENSMUSG00000026000 | *Lancl1* | 1008.165696 | 1064.78199 | 1138.003527 | 647.7306142 | 712.2458491 | 611.7410692 | 1.01E-10 | -0.702634802 | 1.72E-11 | 45.27048814 | 0.61444901 | 9.995668501 |
| ENSMUSG00000009292 | *Trpm2* | 155.7783761 | 194.7895014 | 185.1939702 | 117.232494 | 103.1453592 | 107.2721306 | 0.000841346 | -0.702783806 | 0.000296239 | 13.09402779 | 0.614385552 | 3.075025504 |
| ENSMUSG00000025158 | *Rfng* | 677.4362203 | 681.258619 | 579.7376459 | 398.08444 | 385.4379214 | 409.7602106 | 1.03E-08 | -0.702918115 | 2.07E-09 | 35.90228563 | 0.614328358 | 7.988826158 |
| ENSMUSG00000055053 | *Nfic* | 2681.78466 | 2580.708576 | 1944.536687 | 1428.718308 | 1530.894279 | 1472.817361 | 1.63E-07 | -0.703053198 | 3.70E-08 | 30.30210263 | 0.614270839 | 6.787323474 |
| ENSMUSG00000053898 | *Ech1* | 2331.083598 | 2331.418385 | 2674.576686 | 1502.937441 | 1472.264286 | 1525.970218 | 3.20E-15 | -0.703234098 | 3.90E-16 | 66.28676156 | 0.61419382 | 14.49519639 |
| ENSMUSG00000001576 | *Ergic1* | 4313.06386 | 4696.142747 | 5019.02499 | 2918.161361 | 2858.755062 | 2832.564098 | 7.91E-18 | -0.703300665 | 8.15E-19 | 78.46397431 | 0.614165482 | 17.10177889 |
| ENSMUSG00000004788 | *Eif2b2* | 2428.544941 | 2486.846277 | 3010.073009 | 1657.279501 | 1518.951132 | 1681.563129 | 2.51E-10 | -0.704401338 | 4.39E-11 | 43.4307648 | 0.613697096 | 9.600318713 |
| ENSMUSG00000002835 | *Chaf1a* | 2819.189176 | 2924.870338 | 2139.124555 | 1636.19452 | 1635.125379 | 1570.425336 | 6.44E-08 | -0.704531914 | 1.40E-08 | 32.1823728 | 0.613641554 | 7.191360076 |
| ENSMUSG00000038268 | *Ovca2* | 897.1236738 | 1025.420381 | 998.4370569 | 562.5472912 | 553.727718 | 673.591667 | 1.65E-08 | -0.705245367 | 3.42E-09 | 34.9271824 | 0.613338166 | 7.781552263 |
| ENSMUSG00000046229 | *Scand1* | 1980.382535 | 1863.116164 | 2306.872716 | 1202.687312 | 1189.971723 | 1375.209386 | 1.49E-09 | -0.705596534 | 2.80E-10 | 39.81276678 | 0.613188891 | 8.826463978 |
| ENSMUSG00000032046 | *Abhd12* | 2966.180054 | 2970.287579 | 3032.886759 | 1803.187569 | 1806.672398 | 1888.376065 | 5.35E-26 | -0.705720469 | 3.48E-27 | 116.6208136 | 0.613136218 | 25.27135997 |
| ENSMUSG00000003873 | *Bax* | 10012.9547 | 11457.25606 | 13823.79049 | 6882.981175 | 7195.203112 | 7551.571429 | 1.02E-08 | -0.706071198 | 2.07E-09 | 35.90819262 | 0.612987178 | 7.98976235 |
| ENSMUSG00000021773 | *Comtd1* | 1746.315539 | 1763.198233 | 1960.640511 | 1119.190788 | 1095.512289 | 1133.605488 | 4.95E-16 | -0.706155179 | 5.72E-17 | 70.07001985 | 0.612951496 | 15.30512532 |
| ENSMUSG00000037706 | *Cd81* | 2851.143715 | 2925.87961 | 2937.605803 | 1747.52322 | 1780.614623 | 1812.029233 | 2.16E-26 | -0.706323701 | 1.36E-27 | 118.4761131 | 0.612879901 | 25.66600131 |
| ENSMUSG00000055817 | *Mta3* | 789.2771057 | 815.4917987 | 971.5973511 | 506.0395423 | 487.49754 | 581.7821859 | 1.83E-07 | -0.706488055 | 4.16E-08 | 30.07227097 | 0.612810085 | 6.738226899 |
| ENSMUSG00000034639 | *Setmar* | 105.4499777 | 127.1682755 | 142.2504409 | 72.5323344 | 84.68776864 | 71.51475374 | 0.006055136 | -0.707378634 | 0.002433276 | 9.190100226 | 0.612431913 | 2.217876079 |
| ENSMUSG00000053040 | *Aph1c* | 476.1226265 | 464.2651329 | 585.1055871 | 319.6483109 | 301.8358933 | 309.2529892 | 1.24E-06 | -0.707586343 | 3.07E-07 | 26.20626786 | 0.612343745 | 5.906410938 |
| ENSMUSG00000024165 | *Jpt2* | 4412.12293 | 4650.725506 | 5185.431166 | 2940.089741 | 2882.641356 | 2893.44828 | 4.90E-17 | -0.707967148 | 5.30E-18 | 74.76649933 | 0.612182136 | 16.3100252 |
| ENSMUSG00000019763 | *Rmnd1* | 351.4999256 | 355.2637539 | 390.5177198 | 226.0309956 | 209.5479403 | 233.8725731 | 8.26E-07 | -0.708429573 | 2.00E-07 | 27.02979296 | 0.611985946 | 6.082944661 |
| ENSMUSG00000030339 | *Ltbr* | 3183.470917 | 3402.256007 | 3369.725067 | 1965.963622 | 2046.621075 | 2078.759937 | 2.59E-23 | -0.708461607 | 1.93E-24 | 104.0963022 | 0.611972357 | 22.58712211 |
| ENSMUSG00000029119 | *Man2b2* | 4330.638856 | 4779.912325 | 4462.101094 | 2740.204122 | 2809.896734 | 2755.25085 | 1.93E-23 | -0.708592976 | 1.43E-24 | 104.6881216 | 0.611916635 | 22.71442696 |
| ENSMUSG00000057789 | *Bak1* | 8496.711838 | 8812.96335 | 8673.250939 | 5245.943256 | 5378.759049 | 5273.729881 | 2.71E-37 | -0.708686796 | 1.03E-38 | 169.348842 | 0.611876843 | 36.56713163 |
| ENSMUSG00000026123 | *Plekhb2* | 5431.472714 | 5907.269181 | 6026.855944 | 3578.542964 | 3486.313142 | 3554.476544 | 1.87E-23 | -0.708759797 | 1.38E-24 | 104.76146 | 0.611845882 | 22.72911506 |
| ENSMUSG00000041625 | *Ggact* | 360.2874237 | 380.4955546 | 448.2230874 | 268.2009574 | 263.8349715 | 193.2831182 | 0.000178718 | -0.709299729 | 5.73E-05 | 16.18935246 | 0.61161694 | 3.747832612 |
| ENSMUSG00000034165 | *Ccnd3* | 2923.84029 | 3030.8439 | 2516.222422 | 1691.85887 | 1723.07037 | 1770.473363 | 1.54E-13 | -0.70937473 | 2.10E-14 | 58.43816593 | 0.611585145 | 12.81301648 |
| ENSMUSG00000059323 | *Tonsl* | 3164.298194 | 3137.826735 | 3138.903597 | 1875.719904 | 1938.047013 | 1960.857234 | 1.01E-27 | -0.709663304 | 5.90E-29 | 124.705179 | 0.611462825 | 26.99717354 |
| ENSMUSG00000079884 | *Tmed2b* | 2590.714224 | 2596.856928 | 3234.184553 | 1759.330809 | 1543.923167 | 1840.055286 | 9.23E-09 | -0.709849194 | 1.86E-09 | 36.1175405 | 0.611384044 | 8.034896829 |
| ENSMUSG00000001143 | *Lman2l* | 881.9452679 | 920.4560897 | 971.5973511 | 605.5606523 | 515.7267962 | 570.1851988 | 1.34E-10 | -0.709967606 | 2.29E-11 | 44.70563737 | 0.611333865 | 9.874001221 |
| ENSMUSG00000082121 | *Gm8199* | 322.7408408 | 369.3935623 | 399.9116169 | 206.6328131 | 219.319606 | 239.6710666 | 1.01E-05 | -0.710430422 | 2.77E-06 | 21.96906711 | 0.611137781 | 4.99545748 |
| ENSMUSG00000030493 | *Faap24* | 1080.862271 | 1034.503829 | 1030.644704 | 630.0192302 | 646.015671 | 648.4648616 | 2.41E-15 | -0.710532321 | 2.91E-16 | 66.86549503 | 0.611094617 | 14.61868906 |
| ENSMUSG00000031749 | *St3gal2* | 739.7475707 | 789.250726 | 661.5987487 | 420.8562194 | 465.7827275 | 455.1817434 | 4.92E-09 | -0.710765121 | 9.63E-10 | 37.39761037 | 0.610996017 | 8.308184237 |
| ENSMUSG00000068335 | *Dok1* | 1822.207569 | 1918.626125 | 2043.843599 | 1174.011738 | 1193.228945 | 1162.597956 | 1.57E-17 | -0.711081057 | 1.64E-18 | 77.07792687 | 0.610862229 | 16.80467359 |
| ENSMUSG00000044894 | *Uqcrq* | 6033.815768 | 5826.527418 | 8026.414029 | 3837.466529 | 3971.639201 | 4334.373926 | 1.33E-07 | -0.711117261 | 2.99E-08 | 30.71166509 | 0.6108469 | 6.876512523 |
| ENSMUSG00000041765 | *Ubac2* | 1075.270227 | 1282.784748 | 1264.150145 | 710.1421578 | 718.7602928 | 780.8637976 | 3.15E-10 | -0.711207027 | 5.57E-11 | 42.96450962 | 0.610808894 | 9.501509264 |
| ENSMUSG00000031513 | *Leprotl1* | 2876.707346 | 2868.351104 | 2941.631759 | 1842.827333 | 1697.012595 | 1761.775623 | 2.33E-23 | -0.711288794 | 1.73E-24 | 104.3080707 | 0.610774276 | 22.63216157 |
| ENSMUSG00000027523 | *Gnas* | 5834.099902 | 5860.842667 | 5547.767195 | 3383.71774 | 3685.003676 | 3466.532725 | 7.75E-25 | -0.711572261 | 5.34E-26 | 111.2048895 | 0.61065428 | 24.11046065 |
| ENSMUSG00000029762 | *Akr1b8* | 1743.120086 | 1645.113406 | 2074.709261 | 1091.358613 | 1129.170249 | 1111.37793 | 1.88E-10 | -0.711751642 | 3.25E-11 | 44.02140022 | 0.610578357 | 9.726080456 |
| ENSMUSG00000074916 | *Chst14* | 495.2953497 | 458.2095008 | 452.2490432 | 286.7557407 | 289.8927465 | 283.1597682 | 1.28E-08 | -0.711944127 | 2.62E-09 | 35.44675673 | 0.6104969 | 7.893815999 |
| ENSMUSG00000030987 | *Stim1* | 1782.264396 | 1840.912179 | 1665.403747 | 1091.358613 | 1110.712658 | 1028.266189 | 1.63E-16 | -0.712496218 | 1.82E-17 | 72.32740198 | 0.610263319 | 15.78826847 |
| ENSMUSG00000018171 | *Vmp1* | 4506.388819 | 4763.763973 | 6000.016238 | 3128.167771 | 3103.046702 | 3078.033658 | 1.76E-09 | -0.71309963 | 3.31E-10 | 39.48036203 | 0.610008128 | 8.754726992 |
| ENSMUSG00000010609 | *Psen2* | 2205.662033 | 2264.806431 | 2090.813084 | 1254.978065 | 1432.091882 | 1320.123697 | 5.75E-16 | -0.713162783 | 6.67E-17 | 69.7680884 | 0.609981425 | 15.24064079 |
| ENSMUSG00000080981 | *Gm12161* | 154.9795127 | 135.2424518 | 116.7527204 | 84.33992372 | 76.00184365 | 88.91023438 | 0.003298329 | -0.713488909 | 0.001266851 | 10.39021136 | 0.609843553 | 2.481705991 |
| ENSMUSG00000026126 | *Ptpn18* | 1583.347392 | 1443.259 | 1712.373232 | 923.5221648 | 891.3930519 | 1072.721306 | 8.76E-10 | -0.713547122 | 1.60E-10 | 40.89787255 | 0.609818946 | 9.057293238 |
| ENSMUSG00000028538 | *St3gal3* | 878.749814 | 904.3077372 | 1064.194336 | 538.0887134 | 591.7286398 | 604.0097445 | 5.12E-09 | -0.713594755 | 1.00E-09 | 37.31684826 | 0.609798812 | 8.290722913 |
| ENSMUSG00000056629 | *Fkbp2* | 1858.156425 | 1925.69103 | 2230.379555 | 1267.629054 | 1187.800242 | 1206.086658 | 1.92E-12 | -0.713605224 | 2.84E-13 | 53.31484248 | 0.609794387 | 11.71693483 |
| ENSMUSG00000041571 | *Selenow* | 2171.310904 | 2164.8885 | 1864.01757 | 1211.121305 | 1279.002455 | 1295.963308 | 1.21E-13 | -0.713653798 | 1.64E-14 | 58.92230527 | 0.609773856 | 12.9168639 |
| ENSMUSG00000093989 | *Rnasek* | 1914.076868 | 2164.8885 | 2469.252936 | 1255.821464 | 1332.203745 | 1398.40336 | 1.07E-09 | -0.714564719 | 1.98E-10 | 40.48693616 | 0.609388965 | 8.969866932 |
| ENSMUSG00000029456 | *Acad10* | 400.2305971 | 321.957777 | 283.1588965 | 213.380007 | 197.6047935 | 203.9136897 | 0.00011926 | -0.714619692 | 3.74E-05 | 16.99920574 | 0.609365745 | 3.923505012 |
| ENSMUSG00000024889 | *Rce1* | 1133.58726 | 1325.174173 | 1328.565439 | 775.083899 | 789.3334333 | 740.2743428 | 1.71E-11 | -0.714723363 | 2.72E-12 | 48.87857572 | 0.609321958 | 10.7672858 |
| ENSMUSG00000099693 | *Gm29284* | 919.4918508 | 863.9368561 | 1136.661542 | 582.7888729 | 583.0427149 | 609.808238 | 9.43E-08 | -0.714785171 | 2.09E-08 | 31.41238972 | 0.609295854 | 7.025375149 |
| ENSMUSG00000044600 | *Smim7* | 2045.090476 | 2219.38919 | 2759.121759 | 1434.622103 | 1426.663179 | 1412.899594 | 1.18E-08 | -0.714798218 | 2.40E-09 | 35.61629028 | 0.609290344 | 7.929486379 |
| ENSMUSG00000027495 | *Fam210b* | 468.1339918 | 457.2002287 | 528.7422049 | 296.8765315 | 270.3494153 | 316.0178983 | 2.40E-07 | -0.714844575 | 5.52E-08 | 29.52405786 | 0.609270767 | 6.620565504 |
| ENSMUSG00000001472 | *Tcf25* | 11642.63617 | 12061.81001 | 12480.46321 | 7340.103562 | 7400.40809 | 7297.404129 | 2.01E-34 | -0.715103671 | 8.75E-36 | 155.9333892 | 0.609161356 | 33.69681698 |
| ENSMUSG00000004661 | *Arid3b* | 145.393151 | 155.4278923 | 115.4107351 | 74.21913288 | 76.00184365 | 104.3728838 | 0.007965304 | -0.715118539 | 0.003270927 | 8.64982543 | 0.609155079 | 2.098797637 |
| ENSMUSG00000038286 | *Bphl* | 448.1624051 | 432.9777001 | 440.1711756 | 286.7557407 | 256.2347872 | 260.9322096 | 3.52E-08 | -0.715122522 | 7.49E-09 | 33.40197995 | 0.609153397 | 7.453467663 |
| ENSMUSG00000004567 | *Mcoln1* | 1930.054137 | 2057.905665 | 2139.124555 | 1190.036324 | 1290.945601 | 1249.575359 | 3.65E-17 | -0.715213786 | 3.92E-18 | 75.35976543 | 0.609114863 | 16.4379105 |
| ENSMUSG00000020778 | *Ten1* | 1172.73157 | 1279.756932 | 1610.38235 | 808.8198685 | 825.1628739 | 834.9830707 | 1.77E-07 | -0.716009257 | 4.03E-08 | 30.13582934 | 0.608779103 | 6.752122863 |
| ENSMUSG00000022102 | *Dok2* | 5549.704507 | 5229.038378 | 4619.113373 | 3079.250615 | 3062.874299 | 3235.559399 | 3.04E-15 | -0.716271079 | 3.70E-16 | 66.38801846 | 0.608668631 | 14.51666453 |
| ENSMUSG00000060152 | *Pop5* | 1565.772396 | 1551.251107 | 1751.290805 | 979.1865144 | 957.62323 | 1021.50128 | 7.44E-15 | -0.716657062 | 9.28E-16 | 64.57766449 | 0.608505808 | 14.12832908 |
| ENSMUSG00000089832 | *Shkbp1* | 1514.645134 | 1668.326663 | 1536.573159 | 857.7370243 | 1002.138596 | 1013.769955 | 9.99E-12 | -0.716980243 | 1.56E-12 | 49.97361665 | 0.60836951 | 11.00043795 |
| ENSMUSG00000006641 | *Slc5a6* | 791.6736961 | 778.1487337 | 693.8063957 | 458.8091851 | 480.9830962 | 440.6855096 | 2.48E-10 | -0.716982282 | 4.33E-11 | 43.45853718 | 0.608368651 | 9.606189418 |
| ENSMUSG00000060550 | *H2-Q7* | 1339.694035 | 1412.980839 | 1332.591394 | 818.9406594 | 822.9913927 | 843.6808111 | 1.00E-17 | -0.717163924 | 1.04E-18 | 77.98947706 | 0.608292059 | 16.99993937 |
| ENSMUSG00000089917 | *Uckl1* | 965.825932 | 1019.364748 | 1073.588233 | 610.6210478 | 617.7864148 | 628.1701342 | 8.23E-14 | -0.718218046 | 1.10E-14 | 59.70605568 | 0.607847766 | 13.08469335 |
| ENSMUSG00000025155 | *Dus1l* | 2319.899509 | 2271.871335 | 2141.808525 | 1386.548346 | 1377.804851 | 1330.754269 | 5.83E-21 | -0.718621261 | 4.96E-22 | 93.10308378 | 0.607677904 | 20.23411464 |
| ENSMUSG00000082120 | *Gm15720* | 272.4124423 | 234.1511105 | 276.4489701 | 188.9214291 | 152.0036873 | 133.3653516 | 0.00041108 | -0.7192482 | 0.000137997 | 14.52929221 | 0.607413888 | 3.386073739 |
| ENSMUSG00000027663 | *Zmat3* | 1803.833709 | 2097.267274 | 2022.371834 | 1256.664863 | 1246.430236 | 1092.049618 | 6.27E-12 | -0.719415357 | 9.66E-13 | 50.91180545 | 0.607343515 | 11.20259903 |
| ENSMUSG00000058392 | *Rrp1b* | 2576.334682 | 2547.402599 | 2140.46654 | 1448.95989 | 1512.436689 | 1454.455465 | 5.86E-13 | -0.719929779 | 8.37E-14 | 55.71669554 | 0.607126993 | 12.23187174 |
| ENSMUSG00000010142 | *Tnfrsf13b* | 123.024974 | 100.9272028 | 140.9084556 | 75.90593135 | 79.25906552 | 65.7162602 | 0.006454999 | -0.719960489 | 0.002606533 | 9.06426471 | 0.607114069 | 2.190103852 |
| ENSMUSG00000042831 | *Alkbh6* | 658.263497 | 668.1380826 | 685.754484 | 403.9882346 | 401.7240307 | 413.625873 | 5.94E-12 | -0.721398291 | 9.12E-13 | 51.02478431 | 0.606509315 | 11.22626294 |
| ENSMUSG00000078931 | *Pdf* | 750.1327958 | 591.4334085 | 621.33919 | 406.5184323 | 375.6662558 | 409.7602106 | 2.56E-07 | -0.721688517 | 5.91E-08 | 29.39235969 | 0.606387317 | 6.592053211 |
| ENSMUSG00000066839 | *Ecsit* | 1051.304323 | 1030.466741 | 1014.54088 | 629.175831 | 601.5003055 | 646.5320305 | 3.01E-15 | -0.722114452 | 3.66E-16 | 66.41280273 | 0.606208316 | 14.52191448 |
| ENSMUSG00000015747 | *Vps45* | 1604.916706 | 1531.065667 | 1893.541246 | 989.3073053 | 1001.052855 | 1054.35941 | 5.61E-11 | -0.722322538 | 9.30E-12 | 46.4698204 | 0.606120886 | 10.25086395 |
| ENSMUSG00000028898 | *Trnau1ap* | 1116.012264 | 1117.264135 | 1341.985292 | 739.6611311 | 749.1610303 | 674.5580826 | 7.49E-10 | -0.722352471 | 1.36E-10 | 41.21349988 | 0.60610831 | 9.125602877 |
| ENSMUSG00000041355 | *Ssr2* | 13118.93586 | 13309.27024 | 15326.81401 | 8335.314662 | 8373.231689 | 8592.401021 | 2.22E-18 | -0.722420457 | 2.21E-19 | 81.04310644 | 0.606079749 | 17.65374933 |
| ENSMUSG00000021241 | *Isca2* | 557.6067002 | 596.4797686 | 712.5941898 | 390.4938468 | 373.4947745 | 363.3722623 | 2.29E-07 | -0.722553349 | 5.26E-08 | 29.61982835 | 0.606023923 | 6.64079921 |
| ENSMUSG00000037638 | *Zbtb42* | 474.5248996 | 457.2002287 | 405.279558 | 276.6349498 | 259.492009 | 275.4284435 | 1.10E-07 | -0.723210919 | 2.45E-08 | 31.10208994 | 0.605747765 | 6.959306024 |
| ENSMUSG00000031158 | *Timm17b* | 957.0384338 | 1027.438925 | 1198.392865 | 644.3570172 | 652.5301148 | 626.2373031 | 1.10E-09 | -0.723802819 | 2.04E-10 | 40.42567966 | 0.605499293 | 8.95749838 |
| ENSMUSG00000029344 | *Tpst2* | 2085.03365 | 2164.8885 | 2031.765731 | 1249.07427 | 1226.886905 | 1327.855022 | 5.51E-20 | -0.724022674 | 4.97E-21 | 88.54279432 | 0.605407027 | 19.25919286 |
| ENSMUSG00000022814 | *Umps* | 2919.845973 | 2902.666353 | 3210.028817 | 1857.16512 | 1792.557769 | 1812.995649 | 8.48E-21 | -0.724108865 | 7.31E-22 | 92.33700315 | 0.605370859 | 20.07168991 |
| ENSMUSG00000022214 | *Dcaf11* | 2166.517723 | 2236.546814 | 2364.578084 | 1299.678225 | 1402.776886 | 1391.638451 | 5.27E-19 | -0.724631824 | 5.04E-20 | 83.96251687 | 0.605151459 | 18.27839327 |
| ENSMUSG00000038811 | *Gngt2* | 662.2578144 | 665.1102665 | 850.8186748 | 441.0978011 | 412.581437 | 460.0138214 | 5.14E-07 | -0.72508846 | 1.22E-07 | 27.98858363 | 0.60495995 | 6.28927166 |
| ENSMUSG00000053291 | *Rab4b* | 1233.445193 | 1255.534403 | 1272.202056 | 785.2046899 | 715.5030709 | 771.1996417 | 2.77E-16 | -0.725388057 | 3.15E-17 | 71.24553163 | 0.604834334 | 15.55674505 |
| ENSMUSG00000015806 | *Qdpr* | 2647.433531 | 2724.025204 | 3553.577052 | 1766.921402 | 1763.242773 | 1862.282844 | 1.64E-08 | -0.725584682 | 3.40E-09 | 34.94141216 | 0.604751907 | 7.784354536 |
| ENSMUSG00000036835 | *Psenen* | 1112.017946 | 1238.376779 | 1321.855512 | 743.8781272 | 673.1591866 | 799.2256939 | 5.23E-10 | -0.725684473 | 9.42E-11 | 41.93792655 | 0.604710078 | 9.281121985 |
| ENSMUSG00000107002 | *Ncbp2as2* | 881.9452679 | 895.224289 | 1054.800439 | 556.6434966 | 562.413643 | 589.5135106 | 3.47E-10 | -0.726022238 | 6.15E-11 | 42.77061593 | 0.604568519 | 9.459681114 |
| ENSMUSG00000029403 | *Cdkl2* | 219.6874535 | 308.8372406 | 374.4138963 | 197.3554215 | 152.0036873 | 193.2831182 | 0.003509272 | -0.726329611 | 0.001354707 | 10.26645754 | 0.604439726 | 2.454782931 |
| ENSMUSG00000027187 | *Cat* | 3501.418577 | 4060.301369 | 4668.766829 | 2474.533362 | 2516.746765 | 2392.845004 | 3.51E-10 | -0.726976422 | 6.23E-11 | 42.74630841 | 0.604168795 | 9.454638764 |
| ENSMUSG00000032606 | *Nicn1* | 267.6192615 | 261.4014553 | 350.2581611 | 186.3912314 | 176.9757216 | 165.2570661 | 0.000126424 | -0.727035186 | 3.98E-05 | 16.88022178 | 0.604144187 | 3.898171737 |
| ENSMUSG00000024906 | *Mus81* | 827.6225521 | 783.1950938 | 734.0659545 | 452.9053904 | 453.8395806 | 512.2002633 | 2.83E-10 | -0.727127907 | 4.98E-11 | 43.18678149 | 0.60410536 | 9.548357546 |
| ENSMUSG00000017664 | *Slc35c2* | 4937.775091 | 5011.03562 | 4781.493594 | 2978.886106 | 3049.845412 | 2871.220721 | 1.63E-29 | -0.727416069 | 8.87E-31 | 133.0372145 | 0.603984709 | 28.78715955 |
| ENSMUSG00000041949 | *Tango6* | 196.520413 | 206.9007658 | 161.038235 | 121.4494902 | 116.1742467 | 104.3728838 | 0.000499482 | -0.727489606 | 0.000169884 | 14.13792368 | 0.603953924 | 3.301479879 |
| ENSMUSG00000051065 | *Mb21d2* | 486.5078516 | 543.9976232 | 468.3528667 | 313.7445163 | 310.5218183 | 281.226937 | 5.42E-08 | -0.728080368 | 1.17E-08 | 32.52949633 | 0.603706664 | 7.265992532 |
| ENSMUSG00000005057 | *Sh2b2* | 1071.27591 | 991.1051316 | 755.5377191 | 549.8963027 | 578.6997524 | 576.9501079 | 1.42E-06 | -0.728093454 | 3.53E-07 | 25.93497725 | 0.603701188 | 5.848383181 |
| ENSMUSG00000021996 | *Esd* | 11621.06686 | 12607.82618 | 17667.23636 | 8629.660995 | 8270.08633 | 8383.655253 | 1.47E-06 | -0.728314936 | 3.67E-07 | 25.86039888 | 0.603608515 | 5.832426839 |
| ENSMUSG00000024767 | *Otub1* | 2556.363095 | 2618.051641 | 2679.944627 | 1561.131988 | 1581.924089 | 1595.552141 | 2.20E-26 | -0.728426075 | 1.39E-27 | 118.4381925 | 0.603562018 | 25.65850882 |
| ENSMUSG00000068923 | *Syt11* | 471.3294457 | 511.7009183 | 422.7253668 | 294.3463338 | 263.8349715 | 290.8910929 | 2.39E-07 | -0.728457107 | 5.50E-08 | 29.53358251 | 0.603549036 | 6.622477271 |
| ENSMUSG00000037991 | *Rmi2* | 148.5886049 | 134.2331797 | 116.7527204 | 73.37573364 | 79.25906552 | 89.87664997 | 0.002722665 | -0.728846021 | 0.00103048 | 10.77198431 | 0.603386356 | 2.565005838 |
| ENSMUSG00000025026 | *Add3* | 1140.777031 | 1055.698541 | 955.4935276 | 675.562789 | 614.5291929 | 613.6739004 | 1.03E-10 | -0.729046399 | 1.75E-11 | 45.22997457 | 0.603302556 | 9.986835269 |
| ENSMUSG00000031903 | *Pla2g15* | 1744.717813 | 1820.726739 | 1733.844997 | 1071.960431 | 1087.912105 | 1036.963929 | 6.04E-21 | -0.729553752 | 5.15E-22 | 93.02990665 | 0.60309043 | 20.2186591 |
| ENSMUSG00000026473 | *Glul* | 925.8827586 | 902.2891932 | 790.4293367 | 530.4981202 | 525.4984618 | 525.7300816 | 2.84E-11 | -0.730280727 | 4.60E-12 | 47.85077346 | 0.602786609 | 10.54715513 |
| ENSMUSG00000020647 | *Ncoa1* | 670.2464491 | 783.1950938 | 611.9452929 | 430.133611 | 437.5534713 | 378.8349117 | 2.64E-07 | -0.730319849 | 6.12E-08 | 29.32556624 | 0.602770264 | 6.577970742 |
| ENSMUSG00000023992 | *Trem2* | 10546.5955 | 10270.35216 | 10491.64101 | 6279.107321 | 6365.697276 | 6211.153004 | 1.22E-42 | -0.731612875 | 3.57E-44 | 194.3519669 | 0.602230268 | 41.91489478 |
| ENSMUSG00000022400 | *Rbx1* | 431.3862723 | 445.0889644 | 430.7772786 | 251.3329727 | 261.6634903 | 274.4620279 | 1.01E-08 | -0.732225624 | 2.04E-09 | 35.93317475 | 0.60197454 | 7.995203848 |
| ENSMUSG00000024891 | *Slc29a2* | 316.349933 | 314.8928728 | 265.7130877 | 207.4762124 | 156.3466498 | 175.8876376 | 7.09E-05 | -0.732510581 | 2.16E-05 | 18.04564637 | 0.601855651 | 4.149336184 |
| ENSMUSG00000021054 | *Sgpp1* | 844.3986849 | 811.4547106 | 909.8660276 | 523.7509263 | 525.4984618 | 492.8719515 | 2.73E-12 | -0.732606088 | 4.09E-13 | 52.59831012 | 0.60181581 | 11.56349633 |
| ENSMUSG00000050627 | *Gpd1l* | 3144.326607 | 3244.80957 | 3404.616685 | 2000.542991 | 1911.989238 | 1974.387053 | 1.10E-24 | -0.733023529 | 7.60E-26 | 110.5030409 | 0.601641701 | 23.9600444 |
| ENSMUSG00000043419 | *Rnf227* | 155.7783761 | 168.5484287 | 213.3756614 | 100.3645092 | 105.3168405 | 115.9698709 | 0.001135587 | -0.733914607 | 0.000407146 | 12.49910751 | 0.601270213 | 2.944779662 |
| ENSMUSG00000028692 | *Akr1a1* | 30380.77766 | 34160.83034 | 38741.77338 | 20685.20969 | 20692.0448 | 20706.42046 | 7.31E-14 | -0.734177475 | 9.73E-15 | 59.94970003 | 0.601160667 | 13.13634551 |
| ENSMUSG00000052419 | *Smim30* | 1672.820101 | 1739.984977 | 2171.332202 | 1188.349525 | 1081.397661 | 1081.419046 | 5.04E-09 | -0.734245827 | 9.89E-10 | 37.34629233 | 0.601132186 | 8.297149023 |
| ENSMUSG00000036114 | *Rpp25l* | 1120.006581 | 977.9845953 | 1006.488969 | 580.2586752 | 642.7584491 | 646.5320305 | 1.75E-11 | -0.734447696 | 2.80E-12 | 48.82363482 | 0.601048079 | 10.75576335 |
| ENSMUSG00000026701 | *Prdx6* | 9058.312856 | 9459.90672 | 10527.87461 | 5804.273551 | 5792.426227 | 5846.814326 | 7.88E-21 | -0.735138367 | 6.77E-22 | 92.48935534 | 0.600760404 | 20.10333618 |
| ENSMUSG00000047423 | *AI837181* | 1286.170182 | 1259.571491 | 1190.340954 | 743.8781272 | 753.5039928 | 749.9384987 | 2.71E-17 | -0.735189549 | 2.89E-18 | 75.96166387 | 0.600739091 | 16.56742585 |
| ENSMUSG00000041362 | *Shtn1* | 2365.434727 | 2465.651565 | 2996.653156 | 1579.686771 | 1531.98002 | 1584.921569 | 3.16E-11 | -0.735259251 | 5.14E-12 | 47.63175818 | 0.600710068 | 10.50005888 |
| ENSMUSG00000086784 | *Isoc2a* | 932.2736663 | 1000.18858 | 1160.817277 | 585.3190706 | 635.1582648 | 633.9686278 | 1.22E-09 | -0.735849787 | 2.27E-10 | 40.21557746 | 0.600464231 | 8.913002712 |
| ENSMUSG00000055301 | *Adh7* | 622.314641 | 627.7672015 | 806.5331602 | 420.0128201 | 410.4099557 | 400.0960547 | 3.53E-07 | -0.736610226 | 8.25E-08 | 28.7453878 | 0.600147812 | 6.452529784 |

| | | | | | | | | | | | | | |
|---|---|---|---|---|---|---|---|---|---|---|---|---|---|
| ENSMUSG00000112693 | *Gm5512* | 298.7749368 | 292.6888882 | 292.5527936 | 181.330836 | 200.8620154 | 148.828001 | 2.48E-05 | -0.737443232 | 7.12E-06 | 20.16019701 | 0.599801389 | 4.605467607 |
| ENSMUSG00000049357 | *Brd8dc* | 158.1749665 | 160.4742525 | 187.8779408 | 96.99091228 | 107.4883217 | 98.5743903 | 0.000393558 | -0.73750363 | 0.000131813 | 14.61568266 | 0.599776279 | 3.404991543 |
| ENSMUSG00000035769 | *Xylb* | 527.2498884 | 499.5896539 | 469.694852 | 301.0935277 | 310.5218183 | 287.9918462 | 4.37E-09 | -0.737816581 | 8.52E-10 | 37.63681821 | 0.599646189 | 8.359610764 |
| ENSMUSG00000060166 | *Zdhhc8* | 450.5589956 | 455.1816847 | 365.0199993 | 253.8631704 | 257.3205278 | 253.2008849 | 8.42E-07 | -0.738256828 | 2.04E-07 | 26.99032748 | 0.599463231 | 6.074605743 |
| ENSMUSG00000092224 | *Gm8815* | 718.9771206 | 776.1301896 | 567.6597783 | 359.2880751 | 463.6112463 | 417.4915354 | 1.12E-05 | -0.738384276 | 3.09E-06 | 21.75784208 | 0.599410277 | 4.94933054 |
| ENSMUSG00000047635 | *Mtrfr* | 393.0408259 | 444.0796924 | 406.6215433 | 241.2121818 | 239.9486778 | 263.8314564 | 9.60E-08 | -0.738815877 | 2.13E-08 | 31.37599036 | 0.599230982 | 7.017812296 |
| ENSMUSG00000039680 | *Mrps6* | 3570.120835 | 3742.38068 | 4851.276829 | 2403.687826 | 2449.430847 | 2427.635965 | 4.96E-09 | -0.739440296 | 9.73E-10 | 37.37895094 | 0.598971683 | 8.304290788 |
| ENSMUSG00000015092 | *Edf1* | 3634.029913 | 3786.78865 | 4674.13477 | 2293.202526 | 2354.971413 | 2589.993784 | 3.69E-10 | -0.739812959 | 6.55E-11 | 42.64796897 | 0.598816982 | 9.433382706 |
| ENSMUSG00000090083 | *Rnf8* | 655.0680432 | 624.7393854 | 644.1529399 | 374.4692613 | 377.837737 | 400.0960547 | 1.22E-11 | -0.739970274 | 1.92E-12 | 49.56470572 | 0.598751689 | 10.91450207 |
| ENSMUSG00000024349 | *Sting1* | 2109.798417 | 1981.200991 | 1862.675585 | 1231.362886 | 1135.684692 | 1198.355333 | 7.05E-17 | -0.740566971 | 7.70E-18 | 74.027981 | 0.598504097 | 16.15203533 |
| ENSMUSG00000024885 | *Aldh3b1* | 2461.298343 | 2569.606584 | 2942.973744 | 1594.867958 | 1576.495385 | 1595.552141 | 3.51E-15 | -0.740570076 | 4.29E-16 | 66.09837696 | 0.598502809 | 14.45494803 |
| ENSMUSG00000049932 | *H2ax* | 1736.729178 | 1542.167659 | 1180.947057 | 873.7616098 | 857.7350926 | 941.2887858 | 1.12E-06 | -0.740854696 | 2.75E-07 | 26.41777727 | 0.598384746 | 5.950330305 |
| ENSMUSG00000058625 | *Gm17383* | 2317.502919 | 2024.599688 | 1908.303085 | 1242.327076 | 1188.885983 | 1312.392373 | 1.64E-12 | -0.740916811 | 2.41E-13 | 53.63978561 | 0.598358983 | 11.78511922 |
| ENSMUSG00000009828 | *Cilk1* | 446.5646782 | 459.2187728 | 468.3528667 | 264.8273605 | 289.8927465 | 267.6971187 | 4.50E-09 | -0.740959804 | 8.79E-10 | 37.57645552 | 0.598341152 | 8.34643382 |
| ENSMUSG00000049600 | *Zbtb45* | 931.4748029 | 990.0958596 | 789.0873514 | 504.3527439 | 562.413643 | 558.5882117 | 5.86E-09 | -0.741408515 | 1.15E-09 | 37.0441051 | 0.598155083 | 8.231806433 |
| ENSMUSG00000032554 | *Trf* | 233.2681324 | 240.2067427 | 171.7741173 | 151.8118627 | 121.6029498 | 113.0706242 | 0.001460942 | -0.742116913 | 0.000531902 | 12.00036216 | 0.597861447 | 2.835367098 |
| ENSMUSG00000020099 | *Unc5b* | 299.5738002 | 283.6054399 | 202.639779 | 156.0288589 | 156.3466498 | 159.4585725 | 0.000364406 | -0.743296236 | 0.000121318 | 14.77208546 | 0.597372928 | 3.438414311 |
| ENSMUSG00000023067 | *Cdkn1a* | 17509.48948 | 19010.64792 | 18744.85055 | 11039.25262 | 10950.77993 | 11019.07057 | 3.03E-34 | -0.743316866 | 1.33E-35 | 155.0950351 | 0.597364386 | 33.51829714 |
| ENSMUSG00000024030 | *Abcg1* | 1880.524602 | 2137.638156 | 1645.273967 | 1121.720986 | 1150.885061 | 1112.344345 | 1.16E-10 | -0.743904582 | 1.97E-11 | 44.99575526 | 0.597121085 | 9.936097693 |
| ENSMUSG00000005107 | *Slc2a9* | 1036.924781 | 1130.384672 | 1140.687498 | 672.1891921 | 663.387521 | 636.8678746 | 7.00E-15 | -0.743974972 | 8.71E-16 | 64.70306695 | 0.597091952 | 14.15473406 |
| ENSMUSG00000024608 | *Rps14* | 33402.87816 | 33931.72559 | 42497.99021 | 21266.31177 | 21740.87025 | 22560.97197 | 3.51E-12 | -0.744122021 | 5.31E-13 | 52.0866016 | 0.597031095 | 11.45408 |
| ENSMUSG00000024525 | *Impa2* | 655.0680432 | 769.0652854 | 628.0491164 | 407.3618316 | 371.3232933 | 446.4840031 | 8.70E-08 | -0.744241142 | 1.92E-08 | 31.57585926 | 0.596981802 | 7.060660442 |
| ENSMUSG00000047037 | *Nipa1* | 307.5624349 | 357.282298 | 387.8337492 | 242.0555811 | 187.8331279 | 195.2159494 | 4.16E-05 | -0.744308899 | 1.23E-05 | 19.12171617 | 0.596953765 | 4.381229832 |
| ENSMUSG00000022791 | *Tnk2* | 1533.018994 | 1630.983597 | 1305.751689 | 838.3388418 | 917.4508269 | 916.1619804 | 9.41E-11 | -0.744713607 | 1.60E-11 | 45.41385031 | 0.596786329 | 10.02638108 |
| ENSMUSG00000029270 | *Dipk1a* | 289.9874386 | 273.5127196 | 288.5268377 | 183.0176345 | 165.0325748 | 159.4585725 | 2.83E-06 | -0.745263168 | 7.30E-07 | 24.53407155 | 0.596559041 | 5.547601605 |
| ENSMUSG00000026632 | *Tatdn3* | 205.3079111 | 190.7524133 | 218.7436025 | 115.5456955 | 105.3168405 | 144.9623387 | 0.000370709 | -0.745814023 | 0.000123526 | 14.73808115 | 0.596331304 | 3.430966773 |
| ENSMUSG00000070337 | *Gpr179* | 626.3089584 | 726.6758603 | 539.4780872 | 367.7220674 | 416.9243994 | 345.9767816 | 2.04E-06 | -0.746600824 | 5.20E-07 | 25.18929118 | 0.596006172 | 5.689406457 |
| ENSMUSG00000051343 | *Rab11fip5* | 1442.747422 | 1529.047123 | 1164.843233 | 802.9160738 | 807.7910239 | 858.1770449 | 3.35E-09 | -0.746708585 | 6.46E-10 | 38.17826715 | 0.595961656 | 8.474719792 |
| ENSMUSG00000030872 | *Gga2* | 2827.976674 | 3044.973709 | 3371.067052 | 1814.995159 | 1811.01536 | 1877.745494 | 2.49E-16 | -0.746714157 | 2.82E-17 | 71.4633428 | 0.595959354 | 15.60333255 |
| ENSMUSG00000032376 | *Usp3* | 1533.817857 | 1354.443062 | 1555.360953 | 842.555838 | 869.6782395 | 935.4902922 | 1.47E-13 | -0.746873929 | 2.00E-14 | 58.53155734 | 0.595893357 | 12.83325595 |
| ENSMUSG00000010307 | *Tmem86a* | 240.4579037 | 319.9392329 | 375.7558816 | 166.1496497 | 176.9757216 | 212.61143 | 0.000902906 | -0.746961111 | 0.000319246 | 12.95395458 | 0.595857349 | 3.044357352 |
| ENSMUSG00000023723 | *Mrps23* | 2384.60745 | 2388.946891 | 2584.663671 | 1492.81665 | 1434.263364 | 1450.589802 | 1.66E-22 | -0.747612723 | 1.31E-23 | 100.3035256 | 0.595588283 | 21.77874084 |
| ENSMUSG00000059555 | *Tor4a* | 4321.851358 | 4595.215544 | 4330.586536 | 2642.36981 | 2594.92009 | 2651.844382 | 2.00E-30 | -0.747731092 | 1.04E-31 | 137.2864961 | 0.595539419 | 29.6988018 |
| ENSMUSG00000023094 | *Msrb2* | 409.0180953 | 369.3935623 | 414.6734551 | 221.8139994 | 236.6914559 | 252.2344693 | 1.44E-07 | -0.747782791 | 3.25E-08 | 30.55250685 | 0.595518078 | 6.842023678 |
| ENSMUSG00000033918 | *Parl* | 2273.565428 | 2275.908423 | 2286.742937 | 1370.523761 | 1377.804851 | 1322.056529 | 1.14E-26 | -0.74797981 | 7.15E-28 | 119.7558106 | 0.595436757 | 25.94333152 |
| ENSMUSG00000031155 | *Pim2* | 510.4737556 | 497.5711099 | 465.6688962 | 285.9123414 | 302.921634 | 290.8910929 | 1.39E-09 | -0.74798849 | 2.61E-10 | 39.94926769 | 0.595433175 | 8.855445117 |
| ENSMUSG00000061104 | *Sap18b* | 725.3680283 | 723.6480442 | 637.4430135 | 492.5451545 | 340.9225558 | 406.8609639 | 4.15E-06 | -0.748879171 | 1.09E-06 | 23.75869438 | 0.595065684 | 5.381597369 |
| ENSMUSG00000025260 | *Hsd17b10* | 1597.726935 | 1491.704058 | 2067.999334 | 975.8129175 | 1010.824521 | 1078.5198 | 1.04E-07 | -0.749036893 | 2.30E-08 | 31.22171789 | 0.595000632 | 6.98480335 |
| ENSMUSG00000029066 | *Mrpl20* | 2956.593692 | 3016.714092 | 3807.212272 | 1833.549942 | 1997.762747 | 1984.051209 | 3.12E-10 | -0.749111263 | 5.51E-11 | 42.98804203 | 0.594969961 | 9.506422657 |
| ENSMUSG00000029446 | *Psph* | 5073.581881 | 4698.161291 | 5119.673887 | 2996.59749 | 2941.271349 | 2914.709423 | 1.17E-27 | -0.750027689 | 6.91E-29 | 124.3943519 | 0.594592146 | 26.9300177 |
| ENSMUSG00000040430 | *Pitpnc1* | 146.990878 | 222.0398462 | 182.5099996 | 94.46071457 | 119.4314686 | 114.0370398 | 0.002631428 | -0.750187839 | 0.000993464 | 10.83970868 | 0.594526145 | 2.579808442 |
| ENSMUSG00000056413 | *Adap1* | 4893.038737 | 5104.897918 | 4808.3333 | 2946.836935 | 2850.069137 | 3005.552488 | 1.74E-30 | -0.750238241 | 9.03E-32 | 137.574475 | 0.594505375 | 29.75932087 |
| ENSMUSG00000030207 | *Fam234b* | 514.4680729 | 517.7565504 | 369.0459552 | 256.3933681 | 311.607559 | 268.6635343 | 4.95E-05 | -0.750296463 | 1.48E-05 | 18.76914682 | 0.594481383 | 4.305209352 |
| ENSMUSG00000022750 | *Klhl22* | 1553.789444 | 1635.020686 | 1599.646467 | 932.7995564 | 996.7098924 | 917.128396 | 2.53E-19 | -0.750696378 | 2.38E-20 | 85.44669466 | 0.594316616 | 18.59683581 |
| ENSMUSG00000100774 | *Gm7329* | 107.8465681 | 113.0384672 | 148.9603674 | 65.7851405 | 71.65888115 | 81.17890965 | 0.005184415 | -0.750860529 | 0.002060468 | 9.494873042 | 0.594248998 | 2.285300256 |
| ENSMUSG00000021725 | *Parp8* | 187.7329148 | 173.5947888 | 148.9603674 | 110.4853001 | 86.85924988 | 106.305715 | 0.000759746 | -0.750861655 | 0.000265759 | 13.29751064 | 0.594248534 | 3.119331437 |
| ENSMUSG00000040751 | *Lat2* | 13467.24033 | 15006.86579 | 20219.69239 | 9715.115814 | 9331.940659 | 9883.53225 | 2.28E-07 | -0.750867614 | 5.25E-08 | 29.62276052 | 0.59424608 | 6.641233107 |
| ENSMUSG00000032126 | *Hmbs* | 2154.534771 | 2131.582523 | 2206.223819 | 1284.497038 | 1271.40227 | 1298.862554 | 2.38E-26 | -0.751485536 | 1.51E-27 | 118.2722243 | 0.593991612 | 25.62257428 |
| ENSMUSG00000090877 | *Hspa1b* | 249.2454018 | 304.8001525 | 236.1894113 | 139.1608741 | 156.3466498 | 174.921222 | 0.000122696 | -0.751695235 | 3.85E-05 | 16.9417144 | 0.593905281 | 3.911170407 |
| ENSMUSG00000001424 | *Snd1* | 13842.70616 | 14840.3359 | 13433.27277 | 8438.209369 | 8331.973545 | 8244.491408 | 1.32E-30 | -0.75170723 | 6.85E-32 | 138.1238762 | 0.593900343 | 29.87799037 |
| ENSMUSG00000031819 | *Emc8* | 2379.814269 | 2359.678002 | 2657.130877 | 1450.646688 | 1422.320217 | 1515.339647 | 1.66E-18 | -0.751733498 | 1.63E-19 | 81.63797337 | 0.593889529 | 17.77981274 |
| ENSMUSG00000001942 | *Siae* | 938.6645741 | 1014.318388 | 1091.034042 | 597.12666 | 632.9867835 | 575.0172767 | 2.84E-12 | -0.751978054 | 4.26E-13 | 52.5184511 | 0.593788866 | 11.54686221 |
| ENSMUSG00000038372 | *Gmds* | 1325.314492 | 1537.121299 | 1791.550364 | 910.8711762 | 966.309155 | 881.3710191 | 8.86E-09 | -0.752568342 | 1.78E-09 | 36.20012399 | 0.593545963 | 8.052664219 |
| ENSMUSG00000047731 | *Wbp1l* | 3070.032305 | 3232.698306 | 2755.095803 | 1796.440375 | 1784.957585 | 1798.499415 | 3.95E-18 | -0.752577132 | 4.01E-19 | 79.86276028 | 0.593542347 | 17.40325252 |
| ENSMUSG00000035027 | *Map2k2* | 5140.686412 | 5434.929872 | 5231.058666 | 2936.716144 | 3228.992614 | 3218.163918 | 5.50E-25 | -0.752683959 | 3.75E-26 | 111.9032303 | 0.593498398 | 24.25969874 |
| ENSMUSG00000006335 | *Tfpt* | 583.1703311 | 620.7022973 | 638.7849988 | 355.0710789 | 381.0949589 | 356.6073531 | 5.82E-11 | -0.752754918 | 9.67E-12 | 46.39512351 | 0.593469207 | 10.23508039 |
| ENSMUSG00000025868 | *Higd2a* | 813.2430097 | 864.9461281 | 1001.121027 | 508.5697401 | 511.3838337 | 566.3195364 | 1.62E-09 | -0.753113943 | 3.04E-10 | 39.64714819 | 0.593321537 | 8.790592002 |
| ENSMUSG00000017801 | *Mlx* | 2663.4108 | 2771.460989 | 3137.561612 | 1707.040056 | 1640.554082 | 1732.783155 | 1.91E-16 | -0.753114676 | 2.15E-17 | 71.99983677 | 0.593321235 | 15.71800616 |
| ENSMUSG00000026179 | *Pnkd* | 719.775984 | 709.5182358 | 858.8705866 | 453.7487896 | 439.7249525 | 460.0138214 | 1.08E-09 | -0.75368049 | 2.00E-10 | 40.47017988 | 0.593088585 | 8.966558626 |
| ENSMUSG00000046380 | *Jrk* | 312.3556157 | 344.1617616 | 276.4489701 | 182.1742352 | 187.8331279 | 184.5853779 | 3.33E-06 | -0.754410503 | 8.65E-07 | 24.20789745 | 0.592788554 | 5.478128188 |
| ENSMUSG00000031378 | *Abcd1* | 1586.542846 | 1661.261758 | 1329.907424 | 904.9673816 | 915.2793457 | 896.8336686 | 1.07E-11 | -0.754639287 | 1.68E-12 | 49.83110555 | 0.592694557 | 10.97020717 |
| ENSMUSG00000003644 | *Rps6ka1* | 4849.101246 | 4735.504356 | 4192.362051 | 2733.456928 | 2688.293784 | 2748.485941 | 3.83E-20 | -0.754676763 | 3.44E-21 | 89.27010166 | 0.592679161 | 19.41658888 |
| ENSMUSG00000064267 | *Hvcn1* | 153.3817857 | 261.4014553 | 246.9252936 | 107.9551024 | 154.1751685 | 129.4996892 | 0.00766149 | -0.754945181 | 0.003135767 | 8.726740021 | 0.592568901 | 2.115686788 |
| ENSMUSG00000015474 | *Ppt2* | 1966.002993 | 2080.10965 | 1902.935143 | 1098.949206 | 1198.657648 | 1230.247048 | 1.12E-17 | -0.755195835 | 1.17E-18 | 77.75372763 | 0.592465957 | 16.94909425 |
| ENSMUSG00000031887 | *Tradd* | 1185.513385 | 1238.376779 | 1363.457056 | 728.696941 | 745.9038084 | 765.4011482 | 1.11E-14 | -0.755557304 | 1.40E-15 | 63.76781612 | 0.592317533 | 13.95573175 |
| ENSMUSG00000031730 | *Dhodh* | 576.7794234 | 629.7857456 | 587.7895577 | 333.1426987 | 343.094037 | 386.5662365 | 1.11E-09 | -0.755893453 | 2.06E-10 | 40.4109313 | 0.592179538 | 8.954773329 |
| ENSMUSG00000020346 | *Mgat1* | 2104.206373 | 2203.240837 | 2050.553525 | 1253.291267 | 1297.460045 | 1215.750814 | 2.97E-22 | -0.756223709 | 2.37E-23 | 99.12246655 | 0.592043995 | 21.52655289 |
| ENSMUSG00000029001 | *Fbxo44* | 178.9454167 | 197.8173175 | 218.7436025 | 107.9551024 | 123.7744311 | 119.8355333 | 0.000126459 | -0.757201146 | 3.98E-05 | 16.87902713 | 0.591643016 | 3.898050305 |
| ENSMUSG00000059248 | *Septin9* | 6129.679385 | 6533.017838 | 6147.63462 | 3650.231899 | 3794.663479 | 3681.076987 | 2.88E-32 | -0.757851537 | 1.39E-33 | 145.8583954 | 0.591376354 | 31.53996986 |
| ENSMUSG00000028378 | *Ptgr1* | 791.6736961 | 873.0203043 | 1089.692057 | 524.5943256 | 567.8423461 | 532.4949907 | 2.19E-07 | -0.758431536 | 5.02E-08 | 29.70963069 | 0.591138653 | 6.659923789 |
| ENSMUSG00000033644 | *Piwil2* | 511.2726191 | 636.8506498 | 569.0017636 | 344.1068888 | 340.9225558 | 328.581301 | 2.31E-08 | -0.758773503 | 4.84E-09 | 34.25289335 | 0.590998551 | 7.636651683 |
| ENSMUSG00000033147 | *Slc22a15* | 209.3022284 | 209.9285819 | 152.9863232 | 100.3645092 | 116.1742467 | 123.7011957 | 0.000880469 | -0.759195473 | 0.000310741 | 13.00451127 | 0.590825716 | 3.055286073 |
| ENSMUSG00000054836 | *Elp6* | 616.7225967 | 634.8321057 | 574.3697048 | 339.8898926 | 391.9523651 | 349.842444 | 6.11E-10 | -0.759304912 | 1.11E-10 | 41.62213541 | 0.590780899 | 9.214130512 |
| ENSMUSG00000082286 | *Pisd-ps1* | 254.0385826 | 269.4756315 | 246.9252936 | 151.8118627 | 163.9468342 | 140.1302607 | 6.10E-06 | -0.759571873 | 1.63E-06 | 22.98554224 | 0.59067159 | 5.214967961 |
| ENSMUSG00000022422 | *Dscc1* | 201.3135938 | 176.6226049 | 234.847426 | 121.4494902 | 129.2031342 | 110.1713774 | 0.000247182 | -0.760383793 | 8.07E-05 | 15.54109863 | 0.590339265 | 3.606983591 |
| ENSMUSG00000051671 | *Coa6* | 255.6363095 | 216.993486 | 299.26272 | 150.9684635 | 146.5749842 | 156.5593258 | 8.00E-05 | -0.760436493 | 2.45E-05 | 17.80281641 | 0.590317701 | 4.096949367 |
| ENSMUSG00000031813 | *Mvb12a* | 3175.482282 | 3238.753938 | 3507.949552 | 1915.359668 | 1944.561457 | 1992.748949 | 5.79E-24 | -0.760586879 | 4.18E-25 | 107.1250544 | 0.590256169 | 23.23728954 |
| ENSMUSG00000025422 | *Agap2* | 847.5941388 | 850.8163197 | 683.0705134 | 468.0865767 | 498.3549462 | 442.6183407 | 1.11E-08 | -0.760985297 | 2.25E-09 | 35.74720463 | 0.590093185 | 7.956028182 |
| ENSMUSG00000037366 | *Pafah2* | 722.1725744 | 848.7977757 | 811.9011014 | 456.2789873 | 488.5832806 | 459.0474058 | 5.26E-11 | -0.762250484 | 8.68E-12 | 46.60541737 | 0.589575924 | 10.27936766 |
| ENSMUSG00000025545 | *Clybl* | 178.1465532 | 233.1418385 | 289.868823 | 153.4986612 | 132.4603561 | 124.6676113 | 0.001832042 | -0.762400358 | 0.000676852 | 11.55175872 | 0.589514679 | 2.737064667 |
| ENSMUSG00000029136 | *Rbks* | 197.3192764 | 177.631877 | 240.2153672 | 122.2928894 | 112.9170249 | 125.6340268 | 0.000245065 | -0.762485919 | 8.00E-05 | 15.55906555 | 0.589479718 | 3.610718348 |
| ENSMUSG00000009687 | *Fxyd5* | 20615.47064 | 20982.76546 | 25458.80297 | 13339.20234 | 13101.6321 | 13075.60295 | 3.83E-15 | -0.762708617 | 4.69E-16 | 65.92272502 | 0.589388731 | 14.41708178 |
| ENSMUSG00000039168 | *Dap* | 4155.687757 | 4718.346732 | 4844.566902 | 2773.940091 | 2687.208043 | 2617.053421 | 9.11E-19 | -0.763058331 | 8.82E-20 | 82.85707696 | 0.589245879 | 18.04059506 |
| ENSMUSG00000030223 | *Ptpro* | 823.6282348 | 863.9368561 | 952.809557 | 547.366105 | 512.4695743 | 491.9055359 | 1.12E-11 | -0.763200144 | 1.75E-12 | 49.74100228 | 0.589187961 | 10.95124473 |
| ENSMUSG00000042489 | *Clspn* | 1190.306566 | 1190.940993 | 1254.756248 | 700.8647661 | 710.0743678 | 729.6437713 | 1.51E-18 | -0.763323461 | 1.48E-19 | 81.83824213 | 0.589137601 | 17.82225528 |
| ENSMUSG00000037419 | *Endod1* | 2355.049502 | 2555.476775 | 2135.098599 | 1432.091905 | 1323.51782 | 1396.470529 | 3.38E-15 | -0.763385472 | 4.12E-16 | 66.1755558 | 0.589112279 | 14.4713242 |
| ENSMUSG00000035530 | *Eif1* | 15933.33186 | 15543.79851 | 15612.65688 | 9349.923944 | 9122.392719 | 9260.194194 | 1.21E-49 | -0.763836202 | 2.61E-51 | 227.0558958 | 0.588928256 | 48.91585668 |
| ENSMUSG00000042766 | *Trim46* | 462.5419476 | 470.3207651 | 371.7299258 | 230.2479918 | 272.5208965 | 268.6635343 | 2.44E-06 | -0.764253292 | 6.25E-07 | 24.83369628 | 0.588758018 | 5.612525747 |
| ENSMUSG00000096100 | *ENSMUSG00000096100* | 357.8908333 | 331.0412252 | 393.2016904 | 210.0064101 | 232.3484934 | 194.2495338 | 1.14E-06 | -0.764386019 | 2.81E-07 | 26.3753705 | 0.588703855 | 5.941734877 |
| ENSMUSG00000035203 | *Epn1* | 4339.426354 | 4445.843284 | 3944.094772 | 2449.231385 | 2596.005831 | 2453.729186 | 4.41E-22 | -0.76446972 | 3.56E-23 | 98.32043153 | 0.588669702 | 21.3551371 |
| ENSMUSG00000039183 | *Nubp2* | 2662.611936 | 2779.535165 | 2989.943229 | 1705.353258 | 1612.324826 | 1640.007258 | 2.75E-21 | -0.764505929 | 2.30E-22 | 94.62895917 | 0.588654927 | 20.56070395 |
| ENSMUSG00000050592 | *Fam78a* | 1165.541799 | 1148.551568 | 883.0263218 | 608.09085 | 690.5310366 | 587.5806794 | 1.18E-07 | -0.764746038 | 2.65E-08 | 30.95114368 | 0.588556965 | 6.926908794 |
| ENSMUSG00000030589 | *Rasgrp4* | 257.2340365 | 233.1418385 | 185.1939702 | 126.5098856 | 140.0605404 | 133.3653516 | 0.000185429 | -0.765454378 | 5.96E-05 | 16.11535143 | 0.588268064 | 3.731822984 |
| ENSMUSG00000025188 | *Hps1* | 1715.958728 | 1650.159766 | 1453.370071 | 911.7145754 | 991.2811893 | 936.4567078 | 3.42E-14 | -0.766055473 | 4.47E-15 | 61.47927473 | 0.588023015 | 13.46583054 |
| ENSMUSG00000028757 | *Ddost* | 11495.64529 | 12183.93192 | 11860.46601 | 6991.779677 | 7089.886272 | 6814.196333 | 3.26E-39 | -0.76615926 | 1.13E-40 | 178.3168781 | 0.587980714 | 38.4866987 |
| ENSMUSG00000027575 | *Arfgap1* | 1839.782565 | 1952.941374 | 1842.545805 | 1077.020826 | 1126.998767 | 1110.411514 | 1.07E-22 | -0.766384221 | 8.26E-24 | 101.2126587 | 0.587889037 | 21.97216548 |
| ENSMUSG00000062785 | *Kcnc3* | 460.1453572 | 454.1724127 | 355.6261023 | 242.0555811 | 230.1770122 | 276.3948591 | 5.31E-06 | -0.766477164 | 1.41E-06 | 23.26586924 | 0.587851164 | 5.275164663 |
| ENSMUSG00000023050 | *Map3k12* | 234.8658594 | 218.0027581 | 245.5833083 | 119.7626917 | 131.3746155 | 159.4585725 | 8.02E-05 | -0.766611483 | 2.46E-05 | 17.79742777 | 0.587796436 | 4.095803058 |
| ENSMUSG00000026177 | *Slc11a1* | 3177.878873 | 3435.561984 | 3498.555655 | 1927.167257 | 2005.362932 | 2008.211598 | 1.90E-24 | -0.766712625 | 1.33E-25 | 109.3949141 | 0.58775523 | 23.72168015 |
| ENSMUSG00000029580 | *Actb* | 94579.84248 | 96088.75271 | 83978.75557 | 54508.0493 | 54130.68453 | 52718.93691 | 4.91E-25 | -0.767364471 | 3.35E-26 | 112.1296549 | 0.587489727 | 24.30854561 |
| ENSMUSG00000028199 | *Cryz* | 844.3986849 | 879.0759365 | 1205.102792 | 569.2944851 | 592.8143805 | 554.7225493 | 1.53E-06 | -0.767432827 | 3.82E-07 | 25.78373393 | 0.587461892 | 5.815792746 |
| ENSMUSG00000056412 | *Psenen-ps* | 458.5476302 | 555.0996155 | 730.0399986 | 357.6012766 | 324.6364464 | 338.2454569 | 6.63E-05 | -0.767446062 | 2.01E-05 | 18.18240043 | 0.587456503 | 4.178753456 |
| ENSMUSG00000024381 | *Bin1* | 4217.200244 | 4405.472403 | 4684.870653 | 2605.260244 | 2581.891203 | 2617.053421 | 5.57E-27 | -0.768946118 | 3.42E-28 | 121.2196313 | 0.586846006 | 26.25388396 |
| ENSMUSG00000001665 | *Gstt3* | 335.5226563 | 329.0226812 | 348.9161758 | 168.6798474 | 221.4910872 | 205.8465209 | 3.43E-06 | -0.769115503 | 8.93E-07 | 24.14710019 | 0.58677711 | 5.465304314 |
| ENSMUSG00000022428 | *Cby1* | 397.8340067 | 424.9035238 | 519.3483078 | 279.1651475 | 245.3773809 | 258.9993784 | 1.68E-06 | -0.769564515 | 4.21E-07 | 25.59472155 | 0.586594514 | 5.775405578 |
| ENSMUSG00000068328 | *Aup1* | 8412.032311 | 8617.164576 | 9051.690791 | 4997.98388 | 5178.982774 | 5116.204139 | 4.16E-35 | -0.769825048 | 1.75E-36 | 159.137126 | 0.586488592 | 34.38067466 |
| ENSMUSG00000021482 | *Prxl2c* | 938.6645741 | 904.3077372 | 1392.980733 | 602.1870554 | 653.6158554 | 638.8007057 | 2.07E-05 | -0.769937653 | 5.87E-06 | 20.53064754 | 0.586442817 | 4.684479959 |
| ENSMUSG00000009566 | *Fpgs* | 1487.483776 | 1512.89877 | 1425.18838 | 842.555838 | 883.7928676 | 871.7068632 | 9.91E-21 | -0.770033258 | 8.57E-22 | 92.02168257 | 0.586403956 | 20.00397518 |
| ENSMUSG00000042105 | *Inpp5f* | 377.86242 | 411.7829875 | 465.6688962 | 258.9235658 | 212.8051622 | 260.9322096 | 1.88E-06 | -0.770172593 | 4.75E-07 | 25.36330874 | 0.586347324 | 5.725857828 |
| ENSMUSG00000017188 | *Coa3* | 1414.787201 | 1420.045744 | 1558.044923 | 858.5804235 | 864.2495364 | 848.512889 | 1.29E-18 | -0.770874522 | 1.26E-19 | 82.15675332 | 0.586062112 | 17.8896222 |
| ENSMUSG00000042265 | *Trem1* | 464.1396745 | 460.2280448 | 583.7636018 | 314.5879155 | 273.6066371 | 291.8575085 | 6.78E-07 | -0.770962739 | 1.63E-07 | 27.42973572 | 0.586026277 | 6.168532104 |
| ENSMUSG00000044816 | *D630023F18Rik* | 131.8124721 | 144.3259 | 182.5099996 | 104.5815054 | 85.77350926 | 76.3468317 | 0.002773161 | -0.771009112 | 0.001051556 | 10.73451505 | 0.586007441 | 2.557024963 |
| ENSMUSG00000009013 | *Dynll1* | 8184.356222 | 7501.918985 | 9431.472629 | 4860.509804 | 4730.571897 | 5118.136971 | 2.03E-14 | -0.771544481 | 2.60E-15 | 62.54631598 | 0.58579002 | 13.69164982 |
| ENSMUSG00000046410 | *Kcnk6* | 1352.47585 | 1267.645667 | 991.7271304 | 668.8155951 | 731.7891803 | 719.9796154 | 2.95E-08 | -0.771562181 | 6.21E-09 | 33.76673902 | 0.585782833 | 7.530457746 |
| ENSMUSG00000015149 | *Sirt2* | 2879.9028 | 3199.392329 | 3314.70367 | 1789.693181 | 1889.188685 | 1820.726974 | 1.36E-19 | -0.771590959 | 1.26E-20 | 86.70492544 | 0.585771148 | 18.86546503 |
| ENSMUSG00000095463 | *Entpd4* | 1368.453119 | 1457.388809 | 1336.61735 | 762.4329105 | 836.0202801 | 841.7479799 | 1.29E-16 | -0.772059531 | 1.43E-17 | 72.80215421 | 0.585580927 | 15.88906505 |

| | | | | | | | | | | | | | |
|---|---|---|---|---|---|---|---|---|---|---|---|---|---|
| ENSMUSG00000002043 | *Trappc6a* | 1275.784957 | 1399.860303 | 1629.170144 | 793.6386822 | 818.6484302 | 904.5649933 | 4.69E-10 | -0.772235877 | 8.40E-11 | 42.16186982 | 0.585509354 | 9.329033744 |
| ENSMUSG00000044434 | *Gm9791* | 274.8090327 | 254.3365511 | 267.055073 | 147.5948665 | 158.518131 | 160.4249881 | 1.58E-06 | -0.77334835 | 3.96E-07 | 25.71142252 | 0.585058037 | 5.800749936 |
| ENSMUSG00000002396 | *Ocel1* | 1024.142965 | 1017.346204 | 1074.930219 | 590.3794661 | 615.6149336 | 615.6067315 | 2.94E-17 | -0.774142587 | 3.15E-18 | 75.79524169 | 0.584736037 | 16.53154249 |
| ENSMUSG00000043510 | *Hscb* | 540.0317039 | 605.5632169 | 707.2262486 | 360.1314743 | 381.0949589 | 339.2118725 | 1.19E-07 | -0.774403287 | 2.67E-08 | 30.93046997 | 0.584630383 | 6.922782095 |
| ENSMUSG00000073491 | *Ifi213* | 643.0850912 | 555.0996155 | 770.2995573 | 377.8428583 | 402.8097713 | 367.2379246 | 5.23E-07 | -0.775724858 | 1.24E-07 | 27.9529221 | 0.584095082 | 6.281591545 |
| ENSMUSG00000028670 | *Lypla2* | 3205.839094 | 3368.95003 | 3180.505141 | 1822.585752 | 2005.362932 | 1869.047753 | 1.24E-24 | -0.776876787 | 8.64E-26 | 110.2493323 | 0.583628894 | 23.90556924 |
| ENSMUSG00000010538 | *Tsacc* | 129.4158817 | 145.3351721 | 220.0855878 | 81.80972601 | 102.0596186 | 103.4064683 | 0.007452128 | -0.777462883 | 0.003041281 | 8.782536021 | 0.583391842 | 2.127719712 |
| ENSMUSG00000033917 | *Gde1* | 1989.170034 | 2093.230186 | 2202.197863 | 1200.157115 | 1225.801164 | 1237.011957 | 3.38E-22 | -0.777537898 | 2.71E-23 | 98.86105606 | 0.583361509 | 21.47114901 |
| ENSMUSG00000073436 | *Eme2* | 981.8032013 | 999.1793079 | 819.9530131 | 545.6793065 | 577.6140117 | 514.1330945 | 2.35E-10 | -0.777764456 | 4.11E-11 | 43.56009505 | 0.583269906 | 9.628288347 |
| ENSMUSG00000005374 | *Tbl2* | 2180.098402 | 2037.720225 | 1668.087717 | 1180.758932 | 1122.655805 | 1132.639073 | 6.04E-11 | -0.778179858 | 1.01E-11 | 46.31812006 | 0.583101986 | 10.2190917 |
| ENSMUSG00000005580 | *Adcy9* | 294.7806194 | 305.8094245 | 191.9038967 | 130.7268818 | 172.6327591 | 161.3914037 | 0.001492521 | -0.77853035 | 0.000544192 | 11.95779511 | 0.582960343 | 2.826079569 |
| ENSMUSG00000040964 | *Arhgef10l* | 3423.928821 | 3373.99639 | 2607.477421 | 1868.12931 | 1833.815913 | 1780.137519 | 9.33E-11 | -0.779918169 | 1.58E-11 | 45.43416652 | 0.582399827 | 10.03030072 |
| ENSMUSG00000071078 | *Nr2c2ap* | 1485.886049 | 1645.113406 | 1873.411467 | 956.414735 | 957.62323 | 995.4080589 | 2.42E-12 | -0.780183161 | 3.61E-13 | 52.84600168 | 0.582292862 | 11.61672378 |
| ENSMUSG00000017734 | *Dbndd2* | 345.9078813 | 366.3657462 | 311.3405876 | 172.8968436 | 158.518131 | 264.797872 | 0.000734802 | -0.780630356 | 0.000256426 | 13.36454584 | 0.582112395 | 3.133829876 |
| ENSMUSG00000003234 | *Abcf3* | 1158.352028 | 1176.811185 | 1370.166983 | 646.887215 | 767.6186209 | 741.2407584 | 7.82E-11 | -0.780757985 | 1.32E-11 | 45.79099114 | 0.582060901 | 10.10698589 |
| ENSMUSG00000031828 | *Klhl36* | 391.443099 | 410.7737155 | 409.3055139 | 232.7781895 | 222.5768278 | 248.3688069 | 6.09E-09 | -0.781761228 | 1.20E-09 | 36.96516906 | 0.581656279 | 8.215394407 |
| ENSMUSG00000018567 | *Gabarap* | 9508.871851 | 10659.93116 | 14539.06865 | 6624.901008 | 6706.619832 | 6848.987294 | 1.68E-07 | -0.781952406 | 3.82E-08 | 30.24079817 | 0.581579207 | 6.774613732 |
| ENSMUSG00000037979 | *Ccdc92* | 285.9931213 | 309.8465126 | 272.4230142 | 173.7402429 | 153.0894279 | 177.8204688 | 1.67E-06 | -0.782210781 | 4.19E-07 | 25.60385762 | 0.58147506 | 5.777359743 |
| ENSMUSG00000059316 | *Slc27a4* | 1846.173473 | 1902.477773 | 1873.411467 | 1112.443594 | 1069.454514 | 1085.284709 | 7.78E-26 | -0.782292606 | 5.09E-27 | 115.8648737 | 0.581442081 | 25.10896795 |
| ENSMUSG00000039640 | *Mrpl12* | 6648.940638 | 6491.637685 | 7845.246014 | 3978.314202 | 4058.498451 | 4160.41912 | 5.91E-17 | -0.782311457 | 6.42E-18 | 74.38725292 | 0.581434484 | 16.22872359 |
| ENSMUSG00000018169 | *Mfng* | 1973.192764 | 2084.146738 | 1897.567202 | 1173.168339 | 1123.741545 | 1165.497203 | 2.70E-22 | -0.782568626 | 2.15E-23 | 99.31922206 | 0.581330849 | 21.5687354 |
| ENSMUSG00000113149 | *Gm49383* | 409.8169587 | 406.7366273 | 401.2536022 | 241.2121818 | 272.5208965 | 195.2159494 | 2.08E-06 | -0.78263731 | 5.29E-07 | 25.15663492 | 0.581303174 | 5.682556318 |
| ENSMUSG00000002781 | *Tmem143* | 535.2385231 | 576.2943281 | 461.6429403 | 302.7803262 | 318.1220027 | 295.7231709 | 1.79E-08 | -0.783213064 | 3.72E-09 | 34.76619062 | 0.581071232 | 7.74638321 |
| ENSMUSG00000017774 | *Myo1c* | 8607.75386 | 8436.504883 | 6947.457854 | 4550.982284 | 4707.771344 | 4680.350708 | 2.14E-15 | -0.784015164 | 2.58E-16 | 67.10418756 | 0.580748262 | 14.66914591 |
| ENSMUSG00000001482 | *Def8* | 1589.7383 | 1758.151873 | 1556.702938 | 942.9203472 | 921.7937894 | 981.8782406 | 9.14E-18 | -0.785186252 | 9.45E-19 | 78.17125389 | 0.580277039 | 17.03891408 |
| ENSMUSG00000022193 | *Psmb5* | 2971.772098 | 3230.679762 | 3685.091611 | 1952.469234 | 1781.700363 | 1996.614611 | 1.43E-13 | -0.785346597 | 1.95E-14 | 58.5814471 | 0.580212549 | 12.84389221 |
| ENSMUSG00000040441 | *Slc26a10* | 123.024974 | 138.2702679 | 104.6748527 | 67.47193898 | 66.23017804 | 79.24607847 | 0.002111666 | -0.785939592 | 0.000786391 | 11.27308032 | 0.579974111 | 2.675374702 |
| ENSMUSG00000031934 | *Panx1* | 838.8066407 | 931.558082 | 925.9698511 | 533.8717172 | 539.6130899 | 487.0734579 | 2.91E-13 | -0.787296116 | 4.07E-14 | 57.13596836 | 0.579429035 | 12.53609247 |
| ENSMUSG00000027994 | *Mcub* | 1171.932707 | 1083.958158 | 1070.904263 | 626.6456333 | 667.7304835 | 635.901459 | 1.43E-16 | -0.787952374 | 1.60E-17 | 72.58808948 | 0.579165522 | 15.84334165 |
| ENSMUSG00000011837 | *Snapc2* | 859.5770908 | 850.8163197 | 929.995807 | 482.4243637 | 495.0977243 | 549.8904714 | 4.91E-13 | -0.788375888 | 6.98E-14 | 56.0738566 | 0.578995528 | 12.30861033 |
| ENSMUSG00000072235 | *Tuba1a* | 12011.71109 | 11662.13829 | 10060.86373 | 6516.102507 | 6647.989838 | 6372.544408 | 1.36E-19 | -0.788500971 | 1.26E-20 | 86.70808718 | 0.578945331 | 18.86588193 |
| ENSMUSG00000064264 | *Zfp428* | 142.1976972 | 142.307356 | 124.8046321 | 70.84553593 | 78.1733249 | 88.91023438 | 0.000736977 | -0.788760843 | 0.000257228 | 13.35868591 | 0.578841056 | 3.132546321 |
| ENSMUSG00000019428 | *Fkbp8* | 11403.776 | 11914.45629 | 12293.92726 | 6727.795715 | 6745.706494 | 7120.550075 | 1.41E-34 | -0.789862382 | 6.11E-36 | 156.6491899 | 0.578399263 | 33.84968802 |
| ENSMUSG00000095500 | *ENSMUSG00000095500* | 98.26020648 | 81.75103428 | 97.96492628 | 54.82095042 | 45.60110619 | 59.91776665 | 0.006084409 | -0.789903303 | 0.002446475 | 9.18019816 | 0.578382857 | 2.215781603 |
| ENSMUSG00000084883 | *Ccdc85c* | 640.6885008 | 557.1181595 | 393.2016904 | 312.901117 | 305.0931152 | 304.4209112 | 6.02E-05 | -0.791153528 | 1.81E-05 | 18.37856055 | 0.577881853 | 4.220503598 |
| ENSMUSG00000004207 | *Psap* | 29608.27669 | 35057.0639 | 34730.57934 | 18998.41122 | 19248.00977 | 19147.59211 | 6.35E-21 | -0.792178278 | 5.42E-22 | 92.92842476 | 0.577471528 | 20.19699153 |
| ENSMUSG00000037752 | *Xkr8* | 351.4999256 | 426.9220679 | 409.3055139 | 249.6461742 | 216.0623841 | 217.443508 | 4.55E-07 | -0.792797202 | 1.07E-07 | 28.23628783 | 0.577223843 | 6.342042072 |
| ENSMUSG00000113949 | *Scamp4* | 3129.148201 | 3255.911563 | 3054.358523 | 1779.572391 | 1892.445907 | 1778.204688 | 5.88E-28 | -0.793284657 | 3.43E-29 | 125.7842365 | 0.577028845 | 27.23070059 |
| ENSMUSG00000028048 | *Gba* | 6859.04173 | 7078.024733 | 7686.89175 | 4229.647175 | 4167.072513 | 4072.475301 | 3.92E-28 | -0.79358834 | 2.27E-29 | 126.6001282 | 0.576907394 | 27.4066063 |
| ENSMUSG00000036106 | *Prr5* | 174.1522359 | 183.6875091 | 136.8824997 | 92.7739161 | 94.45943425 | 99.54080589 | 0.000375405 | -0.793681533 | 0.000125201 | 14.71268267 | 0.576870129 | 3.425500484 |
| ENSMUSG00000028463 | *Car9* | 170.956782 | 167.5391567 | 179.8260291 | 94.46071457 | 96.6309155 | 107.2721306 | 6.34E-05 | -0.79553913 | 1.91E-05 | 18.27225145 | 0.576127836 | 4.197969772 |
| ENSMUSG00000039345 | *Mettl22* | 464.1396745 | 414.8108036 | 425.4093374 | 285.0689422 | 248.6346028 | 217.443508 | 3.90E-07 | -0.795641244 | 9.17E-08 | 28.54197337 | 0.576087059 | 6.408985519 |
| ENSMUSG00000003549 | *Ercc1* | 1402.804249 | 1387.749039 | 1190.340954 | 793.6386822 | 767.6186209 | 733.5094337 | 4.83E-14 | -0.79667886 | 6.38E-15 | 60.78124387 | 0.575672875 | 13.31575272 |
| ENSMUSG00000045752 | *Tssc4* | 1417.982654 | 1311.044365 | 1154.107351 | 735.4441349 | 758.9326959 | 745.1064208 | 2.74E-13 | -0.797041326 | 3.82E-14 | 57.25839668 | 0.57552826 | 12.56239302 |
| ENSMUSG00000057191 | *AB124611* | 2847.948261 | 2843.119303 | 3099.986023 | 1725.594839 | 1720.898888 | 1608.115544 | 7.82E-24 | -0.797418419 | 5.68E-25 | 106.5174131 | 0.575377847 | 23.10696041 |
| ENSMUSG00000044629 | *Cnrip1* | 129.4158817 | 141.2980839 | 150.3023526 | 73.37573364 | 95.54517487 | 73.44758493 | 0.0008718 | -0.797741 | 0.000307528 | 13.02397963 | 0.575249209 | 3.059583223 |
| ENSMUSG00000046949 | *Nqo2* | 1605.715569 | 1660.252486 | 2085.445143 | 1020.513077 | 1026.024889 | 1027.299773 | 6.76E-11 | -0.79776465 | 1.13E-11 | 46.08860342 | 0.575239779 | 10.16990658 |
| ENSMUSG00000001323 | *Srr* | 277.2056231 | 354.2544819 | 430.7772786 | 253.8631704 | 184.575906 | 169.1227284 | 0.000990099 | -0.798672925 | 0.000351946 | 12.77147257 | 0.574877741 | 3.004321299 |
| ENSMUSG00000039660 | *Spout1* | 1525.829223 | 1604.742525 | 1517.785365 | 894.8465907 | 852.3063895 | 922.9268895 | 2.36E-21 | -0.799452824 | 1.97E-22 | 94.93408435 | 0.574567054 | 20.62641663 |
| ENSMUSG00000027194 | *Ttc17* | 2113.792734 | 2486.846277 | 2478.646833 | 1327.510399 | 1371.290408 | 1364.578815 | 6.88E-17 | -0.799831546 | 7.51E-18 | 74.07882802 | 0.574416245 | 16.16251787 |
| ENSMUSG00000072572 | *Slc39a2* | 151.7840588 | 155.4278923 | 152.9863232 | 94.46071457 | 79.25906552 | 89.87664997 | 0.000158028 | -0.799980922 | 5.04E-05 | 16.43404742 | 0.574356773 | 3.801266531 |
| ENSMUSG00000051682 | *Treml4* | 102.2545238 | 132.2146357 | 158.3542644 | 76.74933059 | 71.65888115 | 75.38041611 | 0.003156293 | -0.800294321 | 0.00120764 | 10.4786191 | 0.574232018 | 2.500822638 |
| ENSMUSG00000081051 | *Gm15427* | 111.8408854 | 122.1219154 | 124.8046321 | 89.40031915 | 53.20129055 | 61.85059783 | 0.006266439 | -0.800368366 | 0.002524475 | 9.122769208 | 0.574202547 | 2.202979211 |
| ENSMUSG00000051483 | *Cbr1* | 480.9158073 | 537.941991 | 569.0017636 | 316.274714 | 299.6644121 | 292.8239241 | 1.20E-09 | -0.800490971 | 2.22E-10 | 40.25970807 | 0.574153751 | 8.922261988 |
| ENSMUSG00000062825 | *Actg1* | 55960.38588 | 54644.00615 | 46001.91381 | 30376.71033 | 30219.41878 | 29289.15732 | 1.15E-18 | -0.80107428 | 1.12E-19 | 82.38064255 | 0.573921657 | 17.93750007 |
| ENSMUSG00000034255 | *Arhgap27* | 2430.142667 | 2198.194477 | 1486.919703 | 1150.39656 | 1198.657648 | 1164.530787 | 4.16E-06 | -0.801179074 | 1.09E-06 | 23.7570315 | 0.57387997 | 5.38132754 |
| ENSMUSG00000042066 | *Tmcc2* | 742.9430246 | 796.3156302 | 695.148381 | 445.3147973 | 428.8675463 | 408.793795 | 1.58E-12 | -0.801421103 | 2.32E-13 | 53.71571081 | 0.573783703 | 11.80137883 |
| ENSMUSG00000003199 | *Mpnd* | 3398.36519 | 3584.934244 | 4098.42308 | 2068.858329 | 2180.167172 | 2098.088248 | 1.65E-17 | -0.803072697 | 1.73E-18 | 76.97719338 | 0.573127212 | 16.78276701 |
| ENSMUSG00000040548 | *Tex2* | 925.0838951 | 1052.670725 | 1002.463013 | 547.366105 | 568.9280867 | 590.4799262 | 1.14E-14 | -0.803422572 | 1.44E-15 | 63.71615753 | 0.572988237 | 13.94474869 |
| ENSMUSG00000032477 | *Cdc25a* | 2672.997162 | 2581.717848 | 2293.452863 | 1456.550483 | 1448.377992 | 1423.530166 | 1.09E-19 | -0.803788218 | 9.96E-21 | 87.16932107 | 0.572843034 | 18.96438058 |
| ENSMUSG00000030329 | *Pianp* | 1855.759835 | 1946.885742 | 1508.391468 | 991.837503 | 1046.653961 | 1007.971462 | 1.08E-11 | -0.80394004 | 1.68E-12 | 49.82284294 | 0.572782754 | 10.96854207 |
| ENSMUSG00000072381 | *Gm10355* | 115.8352028 | 98.90865876 | 152.9863232 | 60.72474508 | 60.80147492 | 87.94381879 | 0.009345113 | -0.805286732 | 0.0038949 | 8.332174815 | 0.572248336 | 2.02941544 |
| ENSMUSG00000028795 | *Ccdc28b* | 177.3476897 | 215.984214 | 183.8519849 | 111.3286993 | 121.6029498 | 97.6079747 | 0.000108474 | -0.805475849 | 3.38E-05 | 17.19142761 | 0.572173328 | 3.964673048 |
| ENSMUSG00000061981 | *Flot2* | 1039.321371 | 1098.087967 | 1167.527204 | 604.7172531 | 615.6149336 | 667.7931735 | 2.27E-15 | -0.805845304 | 2.74E-16 | 66.98280555 | 0.57202682 | 14.64325791 |
| ENSMUSG00000030798 | *Cd37* | 3612.460599 | 3616.221677 | 3334.833449 | 1995.482595 | 2055.307 | 1994.68178 | 3.22E-29 | -0.806303703 | 1.78E-30 | 131.6523113 | 0.571845094 | 28.49167237 |
| ENSMUSG00000045165 | *AI467606* | 1473.903097 | 1609.788885 | 1456.054041 | 871.2314121 | 857.7350926 | 866.8747852 | 2.37E-20 | -0.806629794 | 2.09E-21 | 90.26111295 | 0.571715855 | 19.6257816 |
| ENSMUSG00000024663 | *Rab3il1* | 2202.466579 | 2344.538921 | 2400.811687 | 1298.834825 | 1355.004298 | 1316.258035 | 2.18E-25 | -0.806642352 | 1.46E-26 | 113.7760842 | 0.571710879 | 24.66158197 |
| ENSMUSG00000004415 | *Col26a1* | 147.7897414 | 191.7616853 | 181.1680144 | 106.2683039 | 103.1453592 | 86.9774032 | 0.000333145 | -0.807912134 | 0.000110478 | 14.94863214 | 0.571207911 | 3.477366786 |
| ENSMUSG00000030281 | *Il17rc* | 984.9986552 | 1072.856166 | 962.203454 | 582.7888729 | 564.5851243 | 577.9165235 | 2.18E-16 | -0.807991892 | 2.46E-17 | 71.73617783 | 0.571176333 | 15.66111586 |
| ENSMUSG00000054708 | *Ankrd24* | 164.5658743 | 187.7245972 | 244.2413231 | 129.0400833 | 108.5740624 | 100.5072215 | 0.000826294 | -0.8085307 | 0.000290549 | 13.13036109 | 0.570963054 | 3.082865513 |
| ENSMUSG00000006519 | *Cyba* | 15447.62287 | 16315.89161 | 19509.78217 | 9317.031374 | 9848.753196 | 10092.27802 | 3.36E-15 | -0.809150329 | 4.09E-16 | 66.1905898 | 0.570717881 | 14.47421672 |
| ENSMUSG00000007867 | *Ift43* | 350.7010621 | 383.5233707 | 481.7727197 | 225.1875963 | 243.2058997 | 223.2420016 | 2.93E-06 | -0.809171211 | 7.57E-07 | 24.46324463 | 0.570709621 | 5.532730117 |
| ENSMUSG00000049686 | *Orai1* | 1300.549725 | 1266.636395 | 1260.124189 | 700.0213669 | 722.0175147 | 763.468317 | 3.11E-20 | -0.809453133 | 2.77E-21 | 89.69924546 | 0.570598107 | 19.50698452 |
| ENSMUSG00000025993 | *Slc40a1* | 2668.203981 | 3012.677004 | 2929.553891 | 1674.990885 | 1659.011673 | 1575.257414 | 2.68E-22 | -0.809702346 | 2.13E-23 | 99.3347349 | 0.57049955 | 21.57181517 |
| ENSMUSG00000026792 | *Lrsam1* | 740.5464342 | 766.0374694 | 770.2995573 | 430.133611 | 432.1247682 | 435.8534316 | 7.53E-16 | -0.809733503 | 8.80E-17 | 69.22071281 | 0.570487229 | 15.12319544 |
| ENSMUSG00000038828 | *Tmem214* | 4777.203534 | 4870.746808 | 4385.607933 | 2775.62689 | 2638.349715 | 2592.893031 | 2.17E-26 | -0.809949631 | 1.37E-27 | 118.4639075 | 0.570401772 | 25.66373419 |
| ENSMUSG00000027111 | *Itga6* | 631.9010026 | 601.5261288 | 540.8200725 | 360.9748735 | 321.3792246 | 329.5477166 | 2.35E-10 | -0.811674136 | 4.11E-11 | 43.56296709 | 0.569720358 | 9.628778982 |
| ENSMUSG00000020937 | *Plcd3* | 642.2862277 | 595.4704966 | 534.110146 | 338.2030941 | 336.5795933 | 337.2790413 | 1.39E-10 | -0.811907457 | 2.39E-11 | 44.62266564 | 0.569628227 | 9.855746645 |
| ENSMUSG00000054612 | *Mgmt* | 301.9703906 | 327.0041371 | 407.9635286 | 203.2592162 | 172.6327591 | 211.6450145 | 1.33E-05 | -0.811925483 | 3.69E-06 | 21.41875915 | 0.569621109 | 4.875518938 |
| ENSMUSG00000041168 | *Lonp1* | 11383.80441 | 10638.73645 | 8641.043292 | 5719.933627 | 5934.658248 | 5790.762222 | 1.64E-13 | -0.814168099 | 2.24E-14 | 58.30890324 | 0.568736342 | 12.78598261 |
| ENSMUSG00000052298 | *Cdc42se2* | 4468.842236 | 4727.43018 | 4478.204918 | 2647.430206 | 2538.461578 | 2588.060953 | 1.57E-35 | -0.814506762 | 6.47E-37 | 161.1111804 | 0.568602851 | 34.8052082 |
| ENSMUSG00000002059 | *Rab34* | 1381.234935 | 1444.268272 | 1646.615953 | 753.9989181 | 880.5356457 | 906.4978245 | 8.29E-12 | -0.814871904 | 1.29E-12 | 50.35040508 | 0.568458958 | 11.08152168 |
| ENSMUSG00000030748 | *Il4ra* | 319.5453869 | 305.8094245 | 319.3924994 | 176.2704406 | 190.0046091 | 171.0555596 | 4.67E-08 | -0.815533701 | 1.00E-08 | 32.83339923 | 0.568198252 | 7.331046047 |
| ENSMUSG00000048752 | *Prss50* | 133.410199 | 128.1775476 | 162.3802203 | 70.84553593 | 82.51628739 | 86.9774032 | 0.000677067 | -0.815637268 | 0.000235079 | 13.52760974 | 0.568157464 | 3.169368588 |
| ENSMUSG00000028343 | *Erp44* | 3997.51279 | 4192.516005 | 4829.805064 | 2431.520001 | 2379.943447 | 2578.396797 | 2.33E-17 | -0.816043076 | 2.47E-18 | 76.27274461 | 0.567997673 | 16.63222758 |
| ENSMUSG00000024963 | *Dnajc4* | 352.2987891 | 358.29157 | 413.3314698 | 201.5724177 | 228.0055309 | 207.7793521 | 7.24E-08 | -0.816304338 | 1.59E-08 | 31.94603735 | 0.567894822 | 7.140520541 |
| ENSMUSG00000033379 | *Atp6v0b* | 11082.63288 | 12396.88832 | 15258.37276 | 7168.893516 | 7206.060519 | 7612.455611 | 1.74E-11 | -0.8167257 | 2.78E-12 | 48.83875098 | 0.567728983 | 10.75894985 |
| ENSMUSG00000019689 | *Fmc1* | 459.3464937 | 490.5062057 | 613.2872782 | 249.6461742 | 302.921634 | 333.4133789 | 5.48E-06 | -0.81695881 | 1.46E-06 | 23.20016157 | 0.567637257 | 5.261525726 |
| ENSMUSG00000005881 | *Ergic3* | 3072.428895 | 3335.644053 | 3628.728228 | 1918.733265 | 1916.332201 | 1855.517935 | 1.65E-20 | -0.817373942 | 1.44E-21 | 90.99519506 | 0.567473944 | 19.7822314 |
| ENSMUSG00000053411 | *Cbx7* | 222.8829074 | 293.6981602 | 269.7390436 | 151.8118627 | 130.2888748 | 162.3578193 | 3.51E-05 | -0.817607644 | 1.03E-05 | 19.4568824 | 0.567382027 | 4.454240647 |
| ENSMUSG00000030879 | *Mrpl17* | 3053.256172 | 2955.148498 | 3565.65492 | 1738.245828 | 1805.586657 | 1884.510403 | 1.61E-16 | -0.817691652 | 1.81E-17 | 72.34636717 | 0.567348989 | 15.79198405 |
| ENSMUSG00000044475 | *Ascc1* | 830.0191425 | 892.1964729 | 998.4370569 | 476.520569 | 528.7556837 | 536.3606531 | 1.08E-11 | -0.81772978 | 1.68E-12 | 49.82033474 | 0.567333995 | 10.96815062 |
| ENSMUSG00000060019 | *Gm10073* | 120.6283836 | 92.85302659 | 127.4886027 | 64.09834203 | 53.20129055 | 75.38041611 | 0.004229728 | -0.818307716 | 0.001654297 | 9.89832033 | 0.56710677 | 2.373687555 |
| ENSMUSG00000018858 | *Mrpl58* | 1685.601916 | 1700.623367 | 1955.27257 | 1060.99624 | 939.1656394 | 1023.434111 | 1.95E-15 | -0.818326263 | 2.34E-16 | 67.29306812 | 0.567099479 | 14.70990056 |
| ENSMUSG00000030584 | *Dpf1* | 147.7897414 | 118.0848273 | 130.1725733 | 64.09834203 | 80.34480614 | 81.17890965 | 0.001009595 | -0.819194773 | 0.000359412 | 12.7322075 | 0.566758185 | 2.995852736 |
| ENSMUSG00000028405 | *Aco1* | 3195.453869 | 3496.118305 | 3369.725067 | 1882.467098 | 1935.875532 | 1881.611156 | 9.79E-30 | -0.81954276 | 5.28E-31 | 134.0689007 | 0.566621496 | 29.00905174 |
| ENSMUSG00000030532 | *Hddc3* | 1552.99058 | 1585.566356 | 1842.545805 | 965.6921266 | 898.9932363 | 951.9193573 | 1.88E-15 | -0.81982986 | 2.25E-16 | 67.36635802 | 0.566508748 | 14.72561762 |
| ENSMUSG00000001930 | *Vwf* | 135.8067894 | 180.659693 | 221.4275731 | 108.7985016 | 81.43054677 | 112.1042086 | 0.00285376 | -0.819850438 | 0.001084813 | 10.67690727 | 0.566500668 | 2.544582558 |
| ENSMUSG00000003585 | *Sec14l2* | 100.6567969 | 115.0570112 | 126.1466174 | 66.62853974 | 62.97295617 | 62.81701342 | 0.001175432 | -0.820877069 | 0.000422334 | 12.43069751 | 0.566097686 | 2.929802512 |
| ENSMUSG00000006526 | *Stimate* | 421.0010473 | 378.4770106 | 417.3574257 | 228.5611933 | 224.7483091 | 234.8389886 | 1.53E-09 | -0.821979188 | 2.86E-10 | 39.76551827 | 0.565665392 | 8.816094537 |
| ENSMUSG00000042670 | *Immp1l* | 627.9066853 | 618.6837532 | 887.0522777 | 414.9524247 | 411.4956963 | 376.9020805 | 2.06E-06 | -0.822339928 | 5.23E-07 | 25.17587781 | 0.565523967 | 5.686486698 |
| ENSMUSG00000002395 | *Use1* | 2561.95514 | 2369.770722 | 2794.013377 | 1422.814513 | 1489.636136 | 1453.489049 | 9.35E-20 | -0.822705859 | 8.53E-21 | 87.475723 | 0.565380543 | 19.02929206 |
| ENSMUSG00000001039 | *B9d1* | 245.2510845 | 227.0862063 | 296.5787494 | 144.2212696 | 127.031653 | 161.3914037 | 3.29E-05 | -0.823110221 | 9.60E-06 | 19.5889815 | 0.565222099 | 4.482870257 |
| ENSMUSG00000022553 | *Maf1* | 2917.449383 | 2827.980223 | 2973.839406 | 1617.639737 | 1609.067604 | 1699.925025 | 2.04E-31 | -0.82323012 | 1.02E-32 | 141.9125349 | 0.565175127 | 30.69032039 |
| ENSMUSG00000003546 | *Klc4* | 1109.621356 | 1336.276165 | 1515.101394 | 729.5403402 | 740.4751053 | 763.468317 | 7.00E-10 | -0.824040561 | 1.27E-10 | 41.35114726 | 0.564857726 | 9.155199613 |
| ENSMUSG00000066270 | *Gm10157* | 1981.980262 | 2042.766585 | 1983.454261 | 1045.815054 | 1096.59803 | 1251.508191 | 1.26E-17 | -0.824402705 | 1.31E-18 | 77.51843054 | 0.564715954 | 16.89908283 |
| ENSMUSG00000030045 | *Mrpl19* | 2429.343804 | 2316.279305 | 3016.782935 | 1445.586293 | 1503.750764 | 1430.295075 | 3.21E-12 | -0.824483773 | 4.84E-13 | 52.26815259 | 0.564684222 | 11.4937231 |
| ENSMUSG00000025521 | *Tmem192* | 1731.935997 | 1932.755934 | 2369.946025 | 1163.047548 | 1112.884139 | 1125.874164 | 1.89E-10 | -0.824814045 | 3.27E-11 | 44.00905729 | 0.564554965 | 9.723644607 |
| ENSMUSG00000018401 | *Mtmr4* | 1240.634965 | 1325.174173 | 982.3332334 | 635.9230249 | 659.0445585 | 710.3154595 | 2.05E-09 | -0.825707585 | 3.89E-10 | 39.16635791 | 0.564205414 | 8.687193418 |
| ENSMUSG00000054894 | *Dmac2l* | 153.3817857 | 145.3351721 | 194.5878673 | 84.33992372 | 94.45943425 | 98.5743903 | 0.00034166 | -0.825723634 | 0.000113362 | 14.90001139 | 0.564199138 | 3.466406245 |
| ENSMUSG00000001911 | *Nfix* | 549.6180655 | 576.2943281 | 360.9940434 | 281.6953452 | 281.2068215 | 278.3276902 | 3.68E-05 | -0.8261336 | 1.08E-05 | 19.36777621 | 0.564038834 | 4.43459012 |
| ENSMUSG00000046822 | *Slc39a3* | 1047.310006 | 1114.236319 | 1248.046321 | 613.1512455 | 657.9588179 | 649.4312772 | 2.75E-14 | -0.826145642 | 3.56E-15 | 61.93176323 | 0.564034126 | 13.56129852 |
| ENSMUSG00000029534 | *St7* | 814.0418732 | 837.6957834 | 971.5973511 | 498.4489492 | 510.2980931 | 467.7451461 | 2.76E-12 | -0.826350767 | 4.13E-13 | 52.58005003 | 0.563953936 | 11.55962951 |
| ENSMUSG00000028073 | *Pear1* | 301.1715272 | 363.3379301 | 288.5268377 | 184.704433 | 165.0325748 | 187.4846247 | 1.55E-06 | -0.826760612 | 3.88E-07 | 25.75527921 | 0.563793749 | 5.809799974 |
| ENSMUSG00000033114 | *Slc35d2* | 173.3533724 | 197.8173175 | 254.9772054 | 135.7872772 | 94.45943425 | 119.8355333 | 0.000883293 | -0.827383535 | 0.000311895 | 12.99757469 | 0.563550368 | 3.053895077 |
| ENSMUSG00000057069 | *Ero1b* | 2402.98131 | 2360.687274 | 2512.196466 | 1427.031509 | 1303.974489 | 1364.578815 | 3.45E-26 | -0.827485791 | 2.21E-27 | 117.5168273 | 0.563510426 | 25.46239847 |

| | | | | | | | | | | | | | |
|---|---|---|---|---|---|---|---|---|---|---|---|---|---|
| ENSMUSG00000024208 | *Uqcc2* | 1461.121282 | 1593.640532 | 2065.315364 | 945.4505449 | 898.9932363 | 1035.997514 | 2.09E-08 | -0.827660719 | 4.35E-09 | 34.45989831 | 0.563442104 | 7.680391375 |
| ENSMUSG00000007589 | *Tinf2* | 408.2192318 | 332.0504973 | 454.9330138 | 208.3196116 | 263.8349715 | 201.014443 | 1.55E-05 | -0.828660046 | 4.32E-06 | 21.11514075 | 0.563051954 | 4.810595635 |
| ENSMUSG00000044026 | *Slc35g1* | 786.8805153 | 812.4639827 | 825.3209543 | 430.133611 | 498.3549462 | 437.7862628 | 7.18E-14 | -0.828779915 | 9.54E-15 | 59.98848595 | 0.563005174 | 13.14394071 |
| ENSMUSG00000025277 | *Abhd6* | 623.912368 | 632.8135616 | 751.5117632 | 373.6258621 | 393.0381057 | 361.4394311 | 1.32E-10 | -0.828979815 | 2.27E-11 | 44.72650677 | 0.562927169 | 9.878480574 |
| ENSMUSG00000046814 | *Gchfr* | 132.6113356 | 152.4000762 | 183.8519849 | 91.08711762 | 89.03073113 | 82.14532525 | 0.000432807 | -0.828998178 | 0.000145674 | 14.42731426 | 0.562920004 | 3.363705316 |
| ENSMUSG00000050708 | *Ftl1* | 233495.8085 | 250158.1649 | 279484.5408 | 142061.3241 | 140493.7509 | 147021.7703 | 2.79E-23 | -0.829007574 | 2.08E-24 | 103.9431182 | 0.562916338 | 22.55491907 |
| ENSMUSG00000022389 | *Tef* | 1004.970242 | 1068.819078 | 1101.769924 | 599.6568577 | 604.7575273 | 580.8157703 | 2.26E-18 | -0.829106778 | 2.25E-19 | 81.01046134 | 0.562877631 | 17.6468326 |
| ENSMUSG00000020156 | *Pwwp3a* | 1587.34171 | 1728.882984 | 1272.202056 | 880.5088037 | 824.0771333 | 880.4046035 | 3.13E-10 | -0.829235157 | 5.54E-11 | 42.97666223 | 0.562827546 | 9.504041465 |
| ENSMUSG00000020319 | *Wdpcp* | 387.4487816 | 404.7180833 | 340.864264 | 237.8385849 | 203.0334966 | 196.182365 | 1.87E-07 | -0.831047634 | 4.25E-08 | 30.02969835 | 0.562120902 | 6.729254648 |
| ENSMUSG00000012519 | *Mlkl* | 1334.900854 | 1348.38743 | 1653.325879 | 764.9631082 | 825.1628739 | 844.6472266 | 3.61E-12 | -0.831052255 | 5.46E-13 | 52.03197036 | 0.562119101 | 11.44250575 |
| ENSMUSG00000023973 | *Cnpy3* | 2614.680128 | 2780.544438 | 2887.952347 | 1504.624239 | 1604.724642 | 1536.60079 | 3.82E-26 | -0.833647545 | 2.46E-27 | 117.308822 | 0.561108804 | 25.41844749 |
| ENSMUSG00000019710 | *Mrpl24* | 4861.084198 | 4833.403743 | 6108.717047 | 2955.270927 | 2945.614312 | 2961.097371 | 1.86E-14 | -0.833699986 | 2.38E-15 | 62.72548858 | 0.561088409 | 13.73055802 |
| ENSMUSG00000054452 | *Tle5* | 5834.099902 | 6162.615004 | 6586.463811 | 3359.259162 | 3491.741845 | 3571.872025 | 3.66E-28 | -0.833738673 | 2.12E-29 | 126.7384675 | 0.561073363 | 27.43599444 |
| ENSMUSG00000041632 | *Mrps27* | 1741.522359 | 1731.9108 | 1859.991614 | 985.0903091 | 993.4526706 | 1010.870708 | 1.30E-25 | -0.834054789 | 8.58E-27 | 114.8293487 | 0.560950437 | 24.88647987 |
| ENSMUSG00000036299 | *BC031181* | 1637.670108 | 1644.104134 | 2210.249775 | 1022.199876 | 989.1097081 | 1064.023566 | 1.63E-09 | -0.834519028 | 3.06E-10 | 39.63570349 | 0.56076996 | 8.788319747 |
| ENSMUSG00000031851 | *Ntpcr* | 317.1487965 | 295.7167042 | 292.5527936 | 155.1854597 | 166.1183154 | 187.4846247 | 2.55E-07 | -0.834942452 | 5.89E-08 | 29.39922692 | 0.560605401 | 6.593730244 |
| ENSMUSG00000020441 | *2310033P09Rik* | 598.348737 | 724.6573162 | 634.7590429 | 361.8182728 | 364.8088495 | 370.1371714 | 3.97E-11 | -0.835383622 | 6.50E-12 | 47.17259307 | 0.560433996 | 10.40146987 |
| ENSMUSG00000049744 | *Arhgap15* | 369.0749219 | 299.7537924 | 454.9330138 | 232.7781895 | 182.4044248 | 211.6450145 | 5.10E-05 | -0.836141086 | 1.52E-05 | 18.71094576 | 0.560139826 | 4.292728274 |
| ENSMUSG00000023938 | *Aars2* | 524.0544345 | 522.8029106 | 522.0322784 | 305.3105239 | 284.4640434 | 287.9918462 | 1.45E-12 | -0.836508272 | 2.13E-13 | 53.88309659 | 0.559997281 | 11.83715365 |
| ENSMUSG00000074272 | *Ceacam1* | 699.8043973 | 749.8891169 | 614.6292635 | 387.9636491 | 401.7240307 | 368.2043402 | 3.49E-11 | -0.837244626 | 5.69E-12 | 47.43312436 | 0.55971153 | 10.4576263 |
| ENSMUSG00000041096 | *Tspyl2* | 635.89532 | 656.0268183 | 518.0063225 | 338.2030941 | 322.4649652 | 353.7081064 | 2.53E-09 | -0.838426425 | 4.83E-10 | 38.74646242 | 0.559253225 | 8.596746806 |
| ENSMUSG00000001285 | *Myg1* | 1834.190521 | 1768.244593 | 1788.866394 | 993.5243015 | 1015.167483 | 1007.005046 | 7.09E-29 | -0.838885002 | 3.98E-30 | 130.0585355 | 0.559075488 | 28.14942343 |
| ENSMUSG00000040714 | *Klc3* | 183.7385975 | 257.3643672 | 203.9817643 | 118.0758932 | 119.4314686 | 122.7347801 | 7.81E-05 | -0.839397376 | 2.39E-05 | 17.85216211 | 0.558876967 | 4.107458182 |
| ENSMUSG00000032527 | *Pccb* | 943.4577549 | 965.8733309 | 1031.986689 | 553.2698996 | 548.2990149 | 538.2934843 | 2.73E-18 | -0.840656405 | 2.74E-19 | 80.61589259 | 0.558389452 | 17.56319591 |
| ENSMUSG00000040390 | *Map3k10* | 957.0384338 | 980.0031393 | 980.9912481 | 549.8963027 | 548.2990149 | 529.5957439 | 5.86E-20 | -0.841432232 | 5.31E-21 | 88.41389071 | 0.558089252 | 19.23174239 |
| ENSMUSG00000026421 | *Csrp1* | 3949.580982 | 3986.624511 | 4360.110212 | 2278.864739 | 2213.825151 | 2363.852536 | 3.12E-28 | -0.841635015 | 1.80E-29 | 127.0664712 | 0.558010813 | 27.50555082 |
| ENSMUSG00000037754 | *Ppp1r16b* | 292.384029 | 288.6518 | 264.3711024 | 145.908068 | 178.0614623 | 149.7944166 | 7.23E-07 | -0.841938204 | 1.74E-07 | 27.29680506 | 0.557893557 | 6.14070075 |
| ENSMUSG00000060063 | *Alox5ap* | 6438.839546 | 6296.848184 | 7627.844397 | 3812.164552 | 3671.974789 | 3869.528027 | 7.14E-19 | -0.842125328 | 6.88E-20 | 83.34817186 | 0.557821201 | 18.1461006 |
| ENSMUSG00000038074 | *Fkbp14* | 70.29998512 | 71.658314 | 85.88705866 | 36.2661672 | 40.17240307 | 50.25361074 | 0.009387242 | -0.842431936 | 0.003914675 | 8.322975205 | 0.557702663 | 2.027461989 |
| ENSMUSG00000074622 | *Mafb* | 734.9543899 | 696.3976994 | 509.9544108 | 339.8898926 | 337.6653339 | 407.8273795 | 2.14E-06 | -0.842505277 | 5.45E-07 | 25.0971761 | 0.557674312 | 5.670071195 |
| ENSMUSG00000036985 | *Zdhhc9* | 1064.086138 | 1107.171415 | 980.9912481 | 565.9208882 | 588.471418 | 605.9425756 | 4.04E-17 | -0.842792694 | 4.36E-18 | 75.15152396 | 0.557563222 | 16.39329574 |
| ENSMUSG00000042380 | *Smim12* | 1024.142965 | 1080.930342 | 1033.328674 | 595.4398615 | 602.5860461 | 551.8233025 | 1.05E-18 | -0.842891491 | 1.02E-19 | 82.57255195 | 0.557525041 | 17.97887458 |
| ENSMUSG00000046417 | *Lrrc75a* | 1056.896367 | 1200.024441 | 1019.908822 | 607.2474508 | 606.9290086 | 612.7074848 | 1.41E-15 | -0.843785197 | 1.68E-16 | 67.94808759 | 0.557179778 | 14.84988802 |
| ENSMUSG00000082536 | *Gm13456* | 232.469269 | 182.6782371 | 217.4016172 | 97.83431152 | 117.2599873 | 138.1974295 | 0.000153038 | -0.843938136 | 4.86E-05 | 16.50014977 | 0.557120715 | 3.815201206 |
| ENSMUSG00000048388 | *Fam171b* | 259.6306269 | 199.8358616 | 230.8214701 | 126.5098856 | 138.9747998 | 119.8355333 | 1.72E-05 | -0.84525594 | 4.83E-06 | 20.9041326 | 0.556612055 | 4.765141985 |
| ENSMUSG00000025730 | *Rab40c* | 1080.862271 | 1160.662832 | 1062.852351 | 580.2586752 | 614.5291929 | 645.5656149 | 1.34E-17 | -0.845686951 | 1.40E-18 | 77.39810308 | 0.55644579 | 16.87311648 |
| ENSMUSG00000032456 | *Nmnat3* | 213.2965458 | 300.7630644 | 260.3451466 | 148.4382658 | 152.0036873 | 129.4996892 | 3.97E-05 | -0.846230659 | 1.17E-05 | 19.21194419 | 0.556236122 | 4.400714882 |
| ENSMUSG00000021368 | *Tbc1d7* | 386.6499182 | 359.300842 | 433.4612492 | 254.7065696 | 186.7473873 | 211.6450145 | 1.37E-06 | -0.846261976 | 3.40E-07 | 26.00788511 | 0.556224047 | 5.864063697 |
| ENSMUSG00000060032 | *H2aj* | 1040.120234 | 1145.523752 | 1431.898306 | 644.3570172 | 634.0725242 | 728.6773557 | 6.92E-09 | -0.84715119 | 1.37E-09 | 36.70494138 | 0.555881321 | 8.159885544 |
| ENSMUSG00000029098 | *Acox3* | 1090.448633 | 1108.180687 | 1041.380586 | 610.6210478 | 599.3288242 | 589.5135106 | 7.55E-21 | -0.849247372 | 6.47E-22 | 92.57818804 | 0.555074233 | 20.12193376 |
| ENSMUSG00000060992 | *Copz1* | 6357.355473 | 6914.522665 | 7529.879471 | 3958.07262 | 3821.806995 | 3757.423818 | 1.61E-23 | -0.849629036 | 1.18E-24 | 105.0600148 | 0.554927408 | 22.79246416 |
| ENSMUSG00000037474 | *Dtl* | 1326.912219 | 1298.9331 | 1328.565439 | 771.7103021 | 732.8749209 | 688.0879009 | 4.60E-21 | -0.849931527 | 3.90E-22 | 93.57897643 | 0.554811068 | 20.33702949 |
| ENSMUSG00000071649 | *B3gat3* | 1807.828026 | 1851.0049 | 1966.008452 | 986.7771076 | 1073.797477 | 1059.191488 | 1.67E-23 | -0.850028486 | 1.23E-24 | 104.9848648 | 0.554773782 | 22.7766901 |
| ENSMUSG00000027459 | *Fam110a* | 642.2862277 | 689.3327952 | 617.3132341 | 371.0956644 | 365.8945901 | 344.0439504 | 1.61E-13 | -0.851156331 | 2.20E-14 | 58.34755376 | 0.55434025 | 12.79376505 |
| ENSMUSG00000031805 | *Jak3* | 907.5088988 | 862.9275841 | 691.1224251 | 473.1469721 | 445.1536557 | 448.4168343 | 5.78E-10 | -0.851840286 | 1.04E-10 | 41.73683733 | 0.55407751 | 9.23805169 |
| ENSMUSG00000032322 | *Pstpip1* | 3057.250489 | 3285.180452 | 3316.045655 | 1788.849782 | 1777.357401 | 1780.137519 | 1.37E-32 | -0.852280248 | 6.56E-34 | 147.3559783 | 0.553908565 | 31.86201936 |
| ENSMUSG00000035227 | *Spcs2* | 7606.777936 | 7924.803965 | 9589.826893 | 4766.04909 | 4525.465558 | 4615.600863 | 1.47E-16 | -0.852465554 | 1.64E-17 | 72.53968558 | 0.553837423 | 15.83291986 |
| ENSMUSG00000038009 | *Dnajc22* | 237.2624498 | 260.3921833 | 312.6825729 | 142.5344711 | 162.8610935 | 142.0630919 | 6.65E-06 | -0.852533539 | 1.79E-06 | 22.80736439 | 0.553811325 | 5.177001453 |
| ENSMUSG00000028673 | *Fuca1* | 2639.444896 | 2654.385434 | 3124.141759 | 1532.456414 | 1584.09557 | 1541.432868 | 4.19E-19 | -0.852541157 | 3.99E-20 | 84.42764276 | 0.553808401 | 18.37761144 |
| ENSMUSG00000038831 | *Ralgps1* | 313.9533426 | 301.7723364 | 220.0855878 | 130.7268818 | 159.6038717 | 174.921222 | 9.72E-05 | -0.853175909 | 3.01E-05 | 17.41290644 | 0.553564791 | 4.012485351 |
| ENSMUSG00000040212 | *Emp3* | 7761.757448 | 8045.916608 | 8579.311969 | 4390.736429 | 4570.968025 | 4527.657044 | 1.03E-35 | -0.853928436 | 4.20E-37 | 161.970559 | 0.553276121 | 34.98796927 |
| ENSMUSG00000030060 | *Hmces* | 974.6134301 | 994.1329477 | 1108.479851 | 562.5472912 | 555.8991993 | 580.8157703 | 6.99E-17 | -0.853998753 | 7.63E-18 | 74.04677628 | 0.553249155 | 16.15570115 |
| ENSMUSG00000030934 | *Oat* | 7043.579191 | 7226.387721 | 8182.084322 | 4153.741243 | 4201.816213 | 4057.979067 | 1.31E-25 | -0.854345809 | 8.68E-27 | 114.8050039 | 0.553116081 | 24.88153611 |
| ENSMUSG00000074093 | *Svip* | 80.6852102 | 90.83448253 | 93.93897041 | 51.44735347 | 56.45851243 | 38.65662365 | 0.00472336 | -0.854586709 | 0.001862415 | 9.680425072 | 0.55302373 | 2.325748981 |
| ENSMUSG00000043683 | *Fem1a* | 3512.602666 | 3514.285202 | 3008.731024 | 1829.332946 | 1926.103866 | 1799.465831 | 2.09E-20 | -0.854684234 | 1.83E-21 | 90.52084396 | 0.552986347 | 19.6807527 |
| ENSMUSG00000034278 | *Dnajc17* | 211.6988188 | 188.7338693 | 224.1115437 | 122.2928894 | 137.8890592 | 85.04457202 | 0.000347728 | -0.855529486 | 0.000115437 | 14.86579402 | 0.552662456 | 3.458760069 |
| ENSMUSG00000058569 | *Tmed9* | 9226.074184 | 9866.643347 | 11102.24432 | 5567.278365 | 5523.162552 | 5582.016454 | 3.59E-23 | -0.856330657 | 2.70E-24 | 103.4292449 | 0.552355631 | 22.44467124 |
| ENSMUSG00000026869 | *Psmd5* | 3730.692392 | 3779.723745 | 5159.933446 | 2339.589484 | 2264.854941 | 2387.04651 | 2.97E-10 | -0.856790093 | 5.24E-11 | 43.08578133 | 0.552179758 | 9.526811974 |
| ENSMUSG00000031157 | *Pqbp1* | 1684.004189 | 1607.770341 | 1798.260291 | 923.5221648 | 918.5365675 | 965.4491755 | 7.82E-23 | -0.857137826 | 6.02E-24 | 101.8399463 | 0.552046682 | 22.10704149 |
| ENSMUSG00000040997 | *Abhd4* | 1050.505459 | 1130.384672 | 1265.49213 | 640.9834203 | 631.9010429 | 625.2708875 | 2.11E-15 | -0.857446998 | 2.54E-16 | 67.1341548 | 0.55192839 | 14.67553421 |
| ENSMUSG00000039043 | *Arpin* | 965.0270685 | 1063.772718 | 1022.592792 | 542.3057095 | 566.7566055 | 574.0508611 | 1.89E-18 | -0.858013854 | 1.87E-19 | 81.37059204 | 0.551711572 | 17.72261516 |
| ENSMUSG00000023186 | *Vwa5a* | 4985.706899 | 5307.761596 | 6717.978369 | 3110.456387 | 3195.334655 | 3074.167995 | 1.20E-12 | -0.858135666 | 1.74E-13 | 54.27836904 | 0.551664991 | 11.92188231 |
| ENSMUSG00000007837 | *Prrg2* | 260.4294903 | 257.3643672 | 253.6352201 | 140.8476726 | 136.8033186 | 147.8615854 | 1.11E-07 | -0.85860286 | 2.48E-08 | 31.07393856 | 0.551486372 | 6.953237292 |
| ENSMUSG00000028799 | *Zfp362* | 421.0010473 | 439.0333322 | 370.3879405 | 215.0668055 | 224.7483091 | 240.6374822 | 3.75E-09 | -0.858644681 | 7.27E-10 | 37.94555737 | 0.551470385 | 8.426235625 |
| ENSMUSG00000032431 | *Crtap* | 1035.327054 | 1092.032334 | 1021.250807 | 597.12666 | 511.3838337 | 626.2373031 | 3.17E-14 | -0.85866893 | 4.12E-15 | 61.64024935 | 0.551461117 | 13.4993722 |
| ENSMUSG00000030082 | *Sec61a1* | 19660.02993 | 20674.9375 | 18913.9407 | 10832.6198 | 10735.80329 | 11105.08156 | 5.27E-43 | -0.858770002 | 1.51E-44 | 196.0587314 | 0.551422484 | 42.2785662 |
| ENSMUSG00000050410 | *Tcf19* | 1187.909976 | 1177.820457 | 1095.059998 | 612.3078462 | 681.8451116 | 617.5395627 | 2.81E-18 | -0.859204992 | 2.82E-19 | 80.55994042 | 0.551256248 | 17.55140975 |
| ENSMUSG00000024777 | *Ppp2r5b* | 997.7804706 | 1013.309116 | 1049.432498 | 574.3548806 | 542.8703118 | 567.285952 | 2.00E-20 | -0.859320648 | 1.76E-21 | 90.60339537 | 0.551212058 | 19.69858015 |
| ENSMUSG00000031974 | *Abcb10* | 336.3215197 | 363.3379301 | 340.864264 | 165.3062505 | 209.5479403 | 200.0480274 | 9.35E-08 | -0.859791128 | 2.07E-08 | 31.43087053 | 0.55103233 | 7.029392961 |
| ENSMUSG00000029602 | *Rasal1* | 526.4510249 | 554.0903434 | 421.3833815 | 262.2971628 | 300.7501527 | 267.6971187 | 5.47E-08 | -0.859861769 | 1.19E-08 | 32.51114708 | 0.55100535 | 7.262128711 |
| ENSMUSG00000074129 | *Rpl13a* | 52292.80371 | 52937.32715 | 69215.57538 | 31048.89952 | 31230.2433 | 33743.36678 | 6.94E-13 | -0.861261177 | 9.95E-14 | 55.37627921 | 0.550471136 | 12.15845622 |
| ENSMUSG00000056724 | *Nbeal2* | 1948.427997 | 1803.569114 | 1197.05088 | 935.3297541 | 929.3939738 | 863.0091229 | 1.39E-06 | -0.861290756 | 3.46E-07 | 25.97628124 | 0.55045985 | 5.85736635 |
| ENSMUSG00000036875 | *Dna2* | 1857.357561 | 1841.921451 | 1885.489335 | 1087.141617 | 945.6800831 | 1036.963929 | 6.81E-23 | -0.861665891 | 5.22E-24 | 102.1234556 | 0.550316736 | 22.1668613 |
| ENSMUSG00000025791 | *Pgm1* | 2633.852852 | 3040.936621 | 3514.659478 | 1699.449463 | 1660.097413 | 1692.1937 | 1.89E-13 | -0.861739843 | 2.60E-14 | 58.01308242 | 0.550288528 | 12.72366268 |
| ENSMUSG00000033166 | *Dis3* | 295.5794829 | 265.4385434 | 233.5054407 | 123.9796879 | 140.0605404 | 174.921222 | 3.29E-05 | -0.862079551 | 9.61E-06 | 19.58833277 | 0.550158968 | 4.482810589 |
| ENSMUSG00000070394 | *Tmem256* | 1252.617917 | 1304.988732 | 1756.658747 | 767.4933059 | 750.2467709 | 851.4121358 | 1.29E-08 | -0.862358858 | 2.65E-09 | 35.42716803 | 0.550052467 | 7.889697976 |
| ENSMUSG00000040146 | *Rgl3* | 114.2374758 | 125.1497315 | 142.2504409 | 66.62853974 | 76.00184365 | 66.68267579 | 0.000391888 | -0.862592749 | 0.000131184 | 14.62469351 | 0.549963299 | 3.406838391 |
| ENSMUSG00000032239 | *Rp9* | 1040.120234 | 1021.383292 | 1037.35463 | 562.5472912 | 575.4425305 | 566.3195364 | 4.03E-22 | -0.862967402 | 3.25E-23 | 98.50144353 | 0.549820498 | 21.3941987 |
| ENSMUSG00000003363 | *Pld3* | 4120.537764 | 4616.410257 | 4513.096535 | 2387.663241 | 2370.171781 | 2521.378277 | 2.95E-28 | -0.863536386 | 1.70E-29 | 127.1794982 | 0.549603697 | 27.5298388 |
| ENSMUSG00000040026 | *Saa3* | 208.503365 | 176.6226049 | 254.9772054 | 119.7626917 | 108.5740624 | 121.7683645 | 7.06E-05 | -0.863575007 | 2.15E-05 | 18.05374594 | 0.549588984 | 4.151099524 |
| ENSMUSG00000020868 | *Xylt2* | 1409.99402 | 1413.990111 | 1198.392865 | 704.2383631 | 743.7323271 | 765.4011482 | 7.65E-16 | -0.864534397 | 8.95E-17 | 69.18803075 | 0.54922363 | 15.11621802 |
| ENSMUSG00000068290 | *Ddrgk1* | 1955.617768 | 1996.340072 | 2547.088083 | 1163.890947 | 1173.685614 | 1227.347801 | 2.55E-12 | -0.8646401 | 3.80E-13 | 52.74292234 | 0.549183391 | 11.59427445 |
| ENSMUSG00000025232 | *Hexa* | 13409.72216 | 14693.99146 | 16405.77019 | 8050.245719 | 8203.856152 | 8167.178161 | 2.38E-22 | -0.865661773 | 1.89E-23 | 99.57446854 | 0.548794614 | 21.62277219 |
| ENSMUSG00000024236 | *Svil* | 289.9874386 | 342.1432175 | 279.1329406 | 152.6552619 | 184.575906 | 164.2906505 | 7.81E-07 | -0.866056193 | 1.89E-07 | 27.14313539 | 0.548644599 | 6.107349838 |
| ENSMUSG00000048481 | *Mypop* | 309.1601618 | 319.9392329 | 264.3711024 | 155.1854597 | 161.7753529 | 174.921222 | 2.28E-07 | -0.866063417 | 5.25E-08 | 29.62371982 | 0.548641852 | 6.641336466 |
| ENSMUSG00000043372 | *Hexim2* | 209.3022284 | 227.0862063 | 213.3756614 | 135.7872772 | 104.2310999 | 115.0034553 | 7.74E-06 | -0.866611441 | 2.10E-06 | 22.50156604 | 0.548433483 | 5.111201977 |
| ENSMUSG00000025533 | *Asl* | 2273.565428 | 2278.93624 | 2592.715583 | 1328.353799 | 1285.516898 | 1295.963308 | 6.61E-23 | -0.868068274 | 5.06E-24 | 102.1857556 | 0.547879955 | 22.17951461 |
| ENSMUSG00000014778 | *Fhod1* | 866.766862 | 693.3698833 | 422.7253668 | 328.9257025 | 369.151812 | 390.4318988 | 0.000519689 | -0.869248108 | 0.000177309 | 14.05746079 | 0.547432082 | 3.284256185 |
| ENSMUSG00000028884 | *Rpa2* | 3172.286829 | 2948.083594 | 3310.677714 | 1765.234604 | 1708.955741 | 1685.428791 | 2.69E-28 | -0.869322321 | 1.55E-29 | 127.3650346 | 0.547403923 | 27.5695466 |
| ENSMUSG00000113996 | *Gm48455* | 92.66816221 | 91.84375456 | 79.1771322 | 57.35114813 | 41.2581437 | 45.42153278 | 0.003664826 | -0.870149204 | 0.001420812 | 10.17858435 | 0.547090267 | 2.435946626 |
| ENSMUSG00000029616 | *Erp29* | 12431.11441 | 12703.70702 | 14549.80453 | 7278.535417 | 7229.946812 | 7193.99766 | 1.53E-26 | -0.870336874 | 9.61E-28 | 119.1715137 | 0.547019105 | 25.81622365 |
| ENSMUSG00000053436 | *Mapk14* | 4087.784362 | 4253.072327 | 3756.216831 | 2151.511454 | 2311.541788 | 2158.972431 | 1.02E-26 | -0.870413869 | 6.37E-28 | 119.9858336 | 0.546989912 | 25.99927328 |
| ENSMUSG00000061887 | *Ssbp3* | 1142.374758 | 1189.931721 | 1005.146983 | 594.5964622 | 584.1284555 | 648.4648616 | 4.61E-15 | -0.870964674 | 5.68E-16 | 65.54587108 | 0.546781117 | 14.33612632 |
| ENSMUSG00000063952 | *Brpf3* | 682.2294011 | 680.249347 | 460.300955 | 329.7691018 | 343.094037 | 326.6484698 | 1.36E-06 | -0.871269377 | 3.39E-07 | 26.0149281 | 0.546665647 | 5.865441807 |
| ENSMUSG00000005823 | *Gpr108* | 2335.077915 | 2433.35486 | 2712.152274 | 1395.825738 | 1359.347261 | 1326.888607 | 1.71E-22 | -0.872089528 | 1.35E-23 | 100.2447444 | 0.546354964 | 21.76682957 |
| ENSMUSG00000089847 | *Timm10b* | 847.5941388 | 767.0467414 | 786.4033808 | 415.795824 | 427.7818057 | 469.6779773 | 7.37E-15 | -0.872366172 | 9.18E-16 | 64.59869682 | 0.546250208 | 14.13255361 |
| ENSMUSG00000107068 | *Gm42742* | 1111.219083 | 1024.411109 | 1166.185218 | 607.2474508 | 624.4112212 | 668.7595891 | 2.29E-12 | -0.872939671 | 3.41E-13 | 52.9539009 | 0.546033106 | 11.63972188 |
| ENSMUSG00000006241 | *Ccdc159* | 93.46702567 | 78.72321819 | 83.20308807 | 51.44735347 | 44.51536557 | 43.4887016 | 0.002777273 | -0.872997644 | 0.001053279 | 10.73148521 | 0.546011165 | 2.556381462 |
| ENSMUSG00000028102 | *Pex11b* | 1641.664425 | 1661.261758 | 1583.542644 | 876.2918075 | 869.6782395 | 922.9268895 | 4.32E-27 | -0.873204009 | 2.63E-28 | 121.743624 | 0.545933068 | 26.36481467 |
| ENSMUSG00000046179 | *E2f8* | 977.0100205 | 949.7249785 | 593.1574989 | 455.4355881 | 460.3540244 | 461.9466526 | 8.42E-06 | -0.874245855 | 2.29E-06 | 22.33375724 | 0.545538963 | 5.074672682 |
| ENSMUSG00000060376 | *Bckdha* | 1133.58726 | 1069.82835 | 1093.718013 | 567.6076867 | 622.1293773 | 610.7746536 | 2.53E-20 | -0.874837386 | 2.23E-21 | 90.12638663 | 0.545315328 | 19.59736983 |
| ENSMUSG00000034361 | *Cpne2* | 1207.881563 | 1452.342448 | 1372.850953 | 705.0817623 | 775.2188052 | 718.0467842 | 1.65E-15 | -0.875044532 | 1.98E-16 | 67.62689091 | 0.545237036 | 14.78150721 |
| ENSMUSG00000020471 | *Pold2* | 3381.589057 | 3322.523517 | 3271.760141 | 1763.547805 | 1849.016282 | 1829.424714 | 9.63E-40 | -0.87523526 | 3.25E-41 | 180.7945058 | 0.545164959 | 39.01643054 |
| ENSMUSG00000032850 | *Rnft2* | 100.6567969 | 120.1033713 | 99.30691157 | 59.03794661 | 56.45851243 | 58.95135106 | 0.000737299 | -0.87565372 | 0.000257384 | 13.35754922 | 0.545006854 | 3.132356521 |
| ENSMUSG00000024664 | *Fads3* | 178.1465532 | 277.5498077 | 264.3711024 | 121.4494902 | 133.5460967 | 136.2645983 | 0.000197262 | -0.875715879 | 6.36E-05 | 15.99231124 | 0.544983373 | 3.704956389 |
| ENSMUSG00000004285 | *Atp6v1f* | 4446.474059 | 4514.473782 | 6194.604106 | 2637.309415 | 2778.410256 | 2839.329007 | 2.00E-10 | -0.875811613 | 3.47E-11 | 43.89215088 | 0.54494721 | 9.69873445 |
| ENSMUSG00000030956 | *Fam53b* | 1550.59399 | 1603.733253 | 1258.782203 | 780.1442944 | 825.1628739 | 801.158525 | 1.74E-13 | -0.877384599 | 2.38E-14 | 58.18895304 | 0.544353372 | 12.76043815 |
| ENSMUSG00000008892 | *Vdac3* | 7601.984755 | 7558.438219 | 9242.252703 | 4397.483623 | 4474.33711 | 4404.922264 | 2.52E-19 | -0.877625037 | 2.37E-20 | 85.45228163 | 0.544262659 | 18.59779034 |
| ENSMUSG00000039740 | *Alg2* | 1147.167939 | 1261.590035 | 966.2294099 | 640.1400211 | 589.5571586 | 608.8418224 | 5.77E-12 | -0.87766265 | 8.84E-13 | 51.08519061 | 0.544248469 | 11.23878298 |
| ENSMUSG00000021508 | *Cxcl14* | 639.0907738 | 605.5632169 | 564.9758077 | 344.950288 | 325.7221871 | 315.0514827 | 3.58E-13 | -0.878288991 | 5.03E-14 | 56.71788277 | 0.544012237 | 12.44650555 |
| ENSMUSG00000004626 | *Stxbp2* | 3973.546886 | 4175.35838 | 4408.421683 | 2209.706002 | 2301.770122 | 2315.531756 | 1.50E-32 | -0.878587654 | 7.17E-34 | 147.1789671 | 0.543899629 | 31.82493181 |
| ENSMUSG00000029192 | *Tbc1d14* | 2538.788099 | 2691.728499 | 2738.99198 | 1484.382658 | 1444.035029 | 1402.269023 | 2.36E-31 | -0.878656134 | 1.18E-32 | 141.6186721 | 0.543873812 | 30.62760588 |
| ENSMUSG00000029406 | *Pitpnm2* | 891.5316295 | 900.2706491 | 735.4079398 | 430.133611 | 494.0119837 | 454.2153278 | 1.27E-11 | -0.878959463 | 2.01E-12 | 49.47056495 | 0.543759474 | 10.89496086 |
| ENSMUSG00000032123 | *Dpagt1* | 1874.133694 | 1844.949267 | 1868.043526 | 962.3185297 | 1034.710814 | 1042.762423 | 1.37E-28 | -0.879090718 | 7.74E-30 | 128.736669 | 0.543710005 | 27.86423728 |
| ENSMUSG00000026860 | *Sh3glb2* | 1693.590551 | 1810.634018 | 1952.588599 | 996.8978984 | 982.5952643 | 981.8782406 | 1.48E-22 | -0.879703144 | 1.16E-23 | 100.5428043 | 0.543479249 | 21.82858891 |
| ENSMUSG00000036442 | *Thap11* | 1460.322418 | 1491.704058 | 1384.928821 | 742.1913288 | 771.9615834 | 844.6472266 | 1.27E-20 | -0.880012528 | 1.11E-21 | 91.51639063 | 0.543362713 | 19.89573958 |

| | | | | | | | | | | | | | |
|---|---|---|---|---|---|---|---|---|---|---|---|---|---|
| ENSMUSG00000019853 | *Hebp2* | 242.8544941 | 268.4663595 | 284.5008818 | 144.2212696 | 123.7744311 | 162.3578193 | 1.97E-06 | -0.880900096 | 4.99E-07 | 25.26695012 | 0.543028531 | 5.705479578 |
| ENSMUSG00000022721 | *Trmt2a* | 2995.738002 | 3212.512866 | 3407.300655 | 1721.377843 | 1656.840192 | 1838.122454 | 4.02E-24 | -0.881066389 | 2.87E-25 | 107.8709603 | 0.542965942 | 23.39538136 |
| ENSMUSG00000030409 | *Dmpk* | 1302.147452 | 1215.163522 | 687.0964693 | 554.1132989 | 631.9010429 | 556.6553805 | 0.000108449 | -0.881488347 | 3.38E-05 | 17.1922244 | 0.542807159 | 3.964772929 |
| ENSMUSG00000031827 | *Cotl1* | 7696.250644 | 7789.561513 | 7937.842999 | 4182.416817 | 4189.873066 | 4329.541848 | 5.29E-54 | -0.88270516 | 9.72E-56 | 247.3709723 | 0.542349532 | 53.27614112 |
| ENSMUSG00000040213 | *Kyat3* | 422.5987742 | 475.3671253 | 617.3132341 | 270.7311552 | 261.6634903 | 286.059015 | 7.08E-07 | -0.883050558 | 1.70E-07 | 27.34379486 | 0.542219702 | 6.15019092 |
| ENSMUSG00000048807 | *Slc35e4* | 4910.613733 | 5049.387957 | 5112.963961 | 2682.009574 | 2731.723409 | 2757.183681 | 6.11E-48 | -0.88313371 | 1.41E-49 | 219.1221444 | 0.542188452 | 47.21418068 |
| ENSMUSG00000071657 | *Bscl2* | 1749.510993 | 1919.635398 | 1970.034408 | 980.0299137 | 1009.73878 | 1064.989981 | 1.66E-22 | -0.883271633 | 1.30E-23 | 100.3133133 | 0.54213662 | 21.78056059 |
| ENSMUSG00000048440 | *Cyp4f16* | 795.6680134 | 826.593791 | 810.5591161 | 432.6638087 | 410.4099557 | 474.5100552 | 3.37E-16 | -0.883674363 | 3.84E-17 | 70.85628707 | 0.541985303 | 15.47286358 |
| ENSMUSG00000028969 | *Cdk5* | 1184.714522 | 1200.024441 | 1379.56088 | 700.8647661 | 673.1591866 | 661.9946799 | 1.76E-17 | -0.883715161 | 1.85E-18 | 76.84925591 | 0.541969977 | 16.75560987 |
| ENSMUSG00000036555 | *Iqce* | 695.0112165 | 673.1844428 | 585.1055871 | 367.7220674 | 347.4369995 | 345.010366 | 1.26E-12 | -0.884284233 | 1.83E-13 | 54.17979874 | 0.541756239 | 11.90062537 |
| ENSMUSG00000035776 | *Cd99l2* | 1370.050846 | 1536.112027 | 1684.191541 | 850.9898304 | 826.2486145 | 805.0241874 | 2.25E-16 | -0.884367993 | 2.54E-17 | 71.67670126 | 0.541724786 | 15.64870689 |
| ENSMUSG00000011832 | *Evi5l* | 352.2987891 | 348.1988497 | 312.6825729 | 193.1384253 | 182.4044248 | 173.9548064 | 4.56E-09 | -0.884852639 | 8.92E-10 | 37.54832444 | 0.541542835 | 8.340564096 |
| ENSMUSG00000058099 | *Nfam1* | 1862.949606 | 1872.199612 | 1672.113673 | 974.126119 | 1001.052855 | 956.7514352 | 7.69E-24 | -0.884957675 | 5.58E-25 | 106.5523602 | 0.541503409 | 23.11391285 |
| ENSMUSG00000000530 | *Acvrl1* | 717.3793936 | 702.4533316 | 750.169778 | 412.422227 | 378.9234776 | 380.7677429 | 3.43E-16 | -0.885839257 | 3.92E-17 | 70.8159484 | 0.541172616 | 15.46488107 |
| ENSMUSG00000040495 | *Chrm4* | 245.2510845 | 248.2809189 | 217.4016172 | 133.2570795 | 131.3746155 | 120.8019489 | 4.75E-07 | -0.886524498 | 1.12E-07 | 28.15067514 | 0.540915634 | 6.323483363 |
| ENSMUSG00000025736 | *Jmjd8* | 961.8316146 | 1047.624365 | 1061.510366 | 537.2453141 | 576.5282711 | 546.024809 | 6.56E-19 | -0.886878954 | 6.30E-20 | 83.5230594 | 0.540782753 | 18.18318919 |
| ENSMUSG00000006445 | *Epha2* | 635.0964565 | 547.0254393 | 424.0673521 | 257.2367674 | 304.0073746 | 310.2194048 | 1.90E-06 | -0.888037234 | 4.81E-07 | 25.33994672 | 0.540348755 | 5.720902372 |
| ENSMUSG00000074657 | *Kif5a* | 1159.949755 | 1228.284058 | 1037.35463 | 587.0058691 | 685.1023335 | 582.7486015 | 9.26E-14 | -0.888077719 | 1.24E-14 | 59.4677537 | 0.540333592 | 13.03344362 |
| ENSMUSG00000033970 | *Rfc3* | 1328.509946 | 1307.007276 | 1394.322718 | 737.9743326 | 647.1014116 | 788.5951224 | 5.78E-17 | -0.888618111 | 6.28E-18 | 74.43164618 | 0.540131236 | 16.23825391 |
| ENSMUSG00000030688 | *Stard10* | 82.28293713 | 80.74176225 | 76.49316162 | 32.04917101 | 49.94406868 | 48.32077956 | 0.006376599 | -0.888664858 | 0.002571488 | 9.089018479 | 0.540113735 | 2.195410908 |
| ENSMUSG00000028843 | *Sh3bgrl3* | 11708.14298 | 12570.48311 | 12856.21909 | 6681.408757 | 6560.044848 | 6810.330671 | 9.71E-42 | -0.888668662 | 2.98E-43 | 190.1320707 | 0.540112311 | 41.0126485 |
| ENSMUSG00000025161 | *Slc16a3* | 9964.224027 | 10064.46066 | 8998.01138 | 5090.757796 | 5297.328502 | 5293.058193 | 1.69E-34 | -0.888883183 | 7.32E-36 | 156.2880372 | 0.540032005 | 33.7719547 |
| ENSMUSG00000037613 | *Tnfrsf23* | 1477.098551 | 1552.260379 | 1139.345512 | 710.985557 | 796.9336177 | 747.0392519 | 1.56E-10 | -0.889197422 | 2.69E-11 | 44.39099438 | 0.539914391 | 9.805550182 |
| ENSMUSG00000023010 | *Tmbim6* | 12829.74728 | 13899.69437 | 13499.03005 | 7152.868931 | 7225.60385 | 7329.295843 | 5.24E-48 | -0.889876714 | 1.20E-49 | 219.4305897 | 0.539660233 | 47.28034646 |
| ENSMUSG00000057469 | *E2f6* | 936.2679837 | 976.9753232 | 971.5973511 | 500.9791469 | 497.2692056 | 557.6217961 | 1.66E-18 | -0.890020136 | 1.64E-19 | 81.63252373 | 0.539606587 | 17.7788755 |
| ENSMUSG00000000958 | *Slc7a7* | 321.1431138 | 301.7723364 | 248.2672789 | 145.0646688 | 89.03073113 | 235.8054042 | 0.009013926 | -0.890512657 | 0.003742503 | 8.40470895 | 0.539422402 | 2.045085992 |
| ENSMUSG00000018442 | *Derl2* | 3088.406165 | 3004.602828 | 3893.099331 | 1799.813972 | 1744.785182 | 1835.223208 | 2.53E-14 | -0.891109283 | 3.28E-15 | 62.09310185 | 0.53919937 | 13.5960883 |
| ENSMUSG00000048534 | *Jaml* | 107.0477046 | 146.3444441 | 136.8824997 | 72.5323344 | 81.43054677 | 56.05210429 | 0.001279418 | -0.89122198 | 0.000461962 | 12.26326848 | 0.539157252 | 2.892987592 |
| ENSMUSG00000036686 | *Cc2d1a* | 2073.849561 | 2049.831489 | 1843.887791 | 1014.609282 | 1092.255067 | 1112.344345 | 1.49E-22 | -0.892574919 | 1.17E-23 | 100.5280656 | 0.538651875 | 21.82568535 |
| ENSMUSG00000034708 | *Grn* | 50441.03819 | 52827.3165 | 44378.11161 | 26426.2283 | 26901.39543 | 26149.27306 | 1.65E-27 | -0.893628806 | 9.81E-29 | 123.6987307 | 0.538258534 | 26.78296785 |
| ENSMUSG00000036775 | *Decr2* | 528.8476153 | 517.7565504 | 585.1055871 | 306.9973224 | 295.3214496 | 273.4956123 | 2.78E-12 | -0.894329513 | 4.17E-13 | 52.56132624 | 0.537997169 | 11.55600189 |
| ENSMUSG00000025920 | *Stau2* | 1464.316736 | 1463.444441 | 1830.467938 | 811.3500662 | 847.963427 | 895.867253 | 1.85E-13 | -0.895183185 | 2.54E-14 | 58.06305834 | 0.537678919 | 12.73395124 |
| ENSMUSG00000019461 | *Plscr3* | 1470.707643 | 1495.741146 | 1321.855512 | 791.9518838 | 777.3902865 | 737.375096 | 5.54E-21 | -0.896063399 | 4.71E-22 | 93.20624506 | 0.537350972 | 20.25645011 |
| ENSMUSG00000036854 | *Hspb6* | 84.67952753 | 101.9364748 | 81.86110278 | 45.54355881 | 46.68684681 | 52.18644192 | 0.001849218 | -0.896892778 | 0.000683853 | 11.53262865 | 0.537042148 | 2.733012004 |
| ENSMUSG00000041736 | *Tspo* | 4754.036494 | 4934.330946 | 5355.863298 | 2649.117004 | 2666.578971 | 2758.150097 | 1.94E-32 | -0.897083266 | 9.34E-34 | 146.6533872 | 0.536971243 | 31.71163856 |
| ENSMUSG00000020085 | *Aifm2* | 901.1179911 | 1020.37402 | 989.0431598 | 518.6905309 | 526.5842024 | 516.0659257 | 2.04E-18 | -0.897102652 | 2.02E-19 | 81.22040055 | 0.536964028 | 17.69090487 |
| ENSMUSG00000053398 | *Phgdh* | 25888.76838 | 27526.8853 | 29926.272 | 14684.42412 | 14905.04728 | 15136.00099 | 2.47E-33 | -0.897777854 | 1.13E-34 | 150.8438109 | 0.53671278 | 32.60743758 |
| ENSMUSG00000031983 | *2310022B05Rik* | 379.460147 | 396.6439071 | 311.3405876 | 205.7894139 | 184.575906 | 194.2495338 | 4.15E-08 | -0.898058046 | 8.89E-09 | 33.06986808 | 0.536608553 | 7.382079742 |
| ENSMUSG00000032553 | *Srprb* | 5087.16256 | 5399.605351 | 5672.571827 | 2828.761042 | 2995.55838 | 2842.228254 | 1.17E-33 | -0.898432309 | 5.29E-35 | 152.3595847 | 0.536469364 | 32.93310868 |
| ENSMUSG00000020585 | *Laptm4a* | 7295.221183 | 7549.35477 | 9305.326011 | 4268.44354 | 4168.158254 | 4513.160811 | 1.70E-16 | -0.89851192 | 1.90E-17 | 72.24164928 | 0.536439762 | 15.76985422 |
| ENSMUSG00000022564 | *Grina* | 2871.115301 | 3264.995011 | 3146.955509 | 1778.728991 | 1616.667788 | 1579.123076 | 7.41E-23 | -0.898848647 | 5.69E-24 | 101.9519117 | 0.53631457 | 22.13025539 |
| ENSMUSG00000024883 | *Rin1* | 198.9170034 | 196.8080455 | 131.5145586 | 103.7381062 | 79.25906552 | 100.5072215 | 0.000550974 | -0.899020947 | 0.000188731 | 13.94008205 | 0.536250522 | 3.25886877 |
| ENSMUSG00000040699 | *Limd2* | 4615.833114 | 4446.852556 | 4597.641609 | 2455.978579 | 2420.11585 | 2435.36729 | 2.15E-49 | -0.901618761 | 4.66E-51 | 225.9053344 | 0.535285782 | 48.66729502 |
| ENSMUSG00000075701 | *Selenos* | 4142.905941 | 4042.134473 | 4806.991314 | 2327.781895 | 2257.254756 | 2362.88612 | 6.66E-23 | -0.901936073 | 5.09E-24 | 102.1719815 | 0.535168062 | 22.1768301 |
| ENSMUSG00000038046 | *Mrm3* | 502.4851209 | 529.8678148 | 591.8155136 | 298.56333 | 262.7492309 | 304.4209112 | 4.28E-11 | -0.90236847 | 7.02E-12 | 47.02299657 | 0.535007689 | 10.36879135 |
| ENSMUSG00000053158 | *Fes* | 4627.017203 | 4438.77838 | 3796.47639 | 2303.323317 | 2336.513822 | 2242.084171 | 3.06E-21 | -0.903328187 | 2.57E-22 | 94.40763047 | 0.534651907 | 20.51398242 |
| ENSMUSG00000026458 | *Ppfia4* | 995.3838802 | 867.9739442 | 756.8797044 | 478.2073675 | 444.067915 | 481.2749644 | 2.47E-11 | -0.903354366 | 3.99E-12 | 48.12784851 | 0.534642205 | 10.60710223 |
| ENSMUSG00000020307 | *Cdc34* | 5126.30687 | 5205.825121 | 5327.681607 | 2693.817164 | 2708.922856 | 2967.86228 | 2.11E-36 | -0.903421505 | 8.28E-38 | 165.1972977 | 0.534617325 | 35.67559482 |
| ENSMUSG00000024327 | *Slc39a7* | 10146.3649 | 10297.6025 | 9816.622407 | 5307.5114 | 5325.557759 | 5545.292662 | 3.66E-53 | -0.903597132 | 6.89E-55 | 243.4701921 | 0.534552247 | 52.43670387 |
| ENSMUSG00000027953 | *Slc50a1* | 1616.899658 | 1638.048502 | 1813.022129 | 859.4238227 | 914.193605 | 932.5910454 | 1.24E-22 | -0.904108743 | 9.67E-24 | 100.9013372 | 0.534362717 | 21.90489772 |
| ENSMUSG00000110126 | *Gm9347* | 5901.204433 | 6239.319678 | 6355.642341 | 3220.098288 | 3291.965571 | 3366.025504 | 3.72E-43 | -0.904534444 | 1.06E-44 | 196.7712322 | 0.534205063 | 42.42958174 |
| ENSMUSG00000041115 | *Iqsec2* | 412.2135491 | 441.0518763 | 338.1802935 | 207.4762124 | 220.4053466 | 210.6785989 | 1.65E-08 | -0.904985909 | 3.41E-09 | 34.9347742 | 0.53403792 | 7.783121761 |
| ENSMUSG00000069307 | *H2bc23* | 81.48407366 | 123.1311874 | 91.25499982 | 52.29075271 | 42.34388432 | 62.81701342 | 0.005236544 | -0.905816944 | 0.002083615 | 9.474377948 | 0.533730387 | 2.280955276 |
| ENSMUSG00000059108 | *Ifitm6* | 1012.160013 | 982.0216834 | 1238.652424 | 552.4265004 | 583.0427149 | 586.6142638 | 4.15E-13 | -0.9064036 | 5.86E-14 | 56.4176281 | 0.533513396 | 12.38219114 |
| ENSMUSG00000018537 | *Pcgf2* | 571.1873791 | 590.4241365 | 445.5391168 | 272.4179536 | 321.3792246 | 266.7307032 | 3.83E-08 | -0.906697477 | 8.18E-09 | 33.23262919 | 0.53340473 | 7.416569374 |
| ENSMUSG00000064284 | *Cdpf1* | 438.5760435 | 460.2280448 | 527.4002196 | 247.1159765 | 254.0633059 | 257.0665472 | 5.85E-11 | -0.907503458 | 9.72E-12 | 46.38498256 | 0.533106819 | 10.232987 |
| ENSMUSG00000032802 | *Srxn1* | 8384.072089 | 9724.335991 | 10604.36777 | 5154.856138 | 5108.409634 | 5033.092399 | 3.24E-20 | -0.908008073 | 2.89E-21 | 89.61555536 | 0.532920386 | 19.48966689 |
| ENSMUSG00000061474 | *Mrps36* | 681.4305376 | 805.3990785 | 946.0996305 | 443.6279988 | 425.6103244 | 423.2900289 | 8.38E-10 | -0.908306492 | 1.53E-10 | 40.98834842 | 0.532810164 | 9.07697832 |
| ENSMUSG00000021476 | *Habp4* | 696.6089435 | 825.584519 | 939.3897041 | 402.3014362 | 465.7827275 | 440.6855096 | 6.68E-10 | -0.909293875 | 1.22E-10 | 41.44054644 | 0.532445632 | 9.174920052 |
| ENSMUSG00000043998 | *Mgat2* | 5826.91013 | 6153.531556 | 6417.373664 | 3235.279474 | 3252.878908 | 3301.275659 | 5.47E-41 | -0.909611579 | 1.73E-42 | 186.6272373 | 0.532328392 | 40.26239216 |
| ENSMUSG00000053801 | *Grwd1* | 3263.357264 | 3477.951409 | 3309.335729 | 1722.221242 | 1820.787026 | 1808.163571 | 2.40E-37 | -0.909798553 | 9.08E-39 | 169.5938783 | 0.532259407 | 36.6192959 |
| ENSMUSG00000019558 | *Slc6a8* | 3713.916259 | 4553.835391 | 4362.794183 | 2197.898412 | 2293.084197 | 2227.587938 | 1.21E-20 | -0.910192018 | 1.05E-21 | 91.62272984 | 0.532114264 | 19.91848851 |
| ENSMUSG00000002820 | *Atg4d* | 1690.395097 | 1668.326663 | 1799.602276 | 918.4617693 | 900.0789769 | 921.9604739 | 2.37E-29 | -0.910969893 | 1.30E-30 | 132.2771851 | 0.531827435 | 28.62558529 |
| ENSMUSG00000033565 | *Rbfox2* | 457.7487668 | 415.8200756 | 338.1802935 | 218.4404024 | 198.6905341 | 228.0740795 | 8.83E-08 | -0.913401234 | 1.95E-08 | 31.54424416 | 0.530931913 | 7.053822148 |
| ENSMUSG00000021646 | *Mccc2* | 437.7771801 | 463.2558609 | 524.716249 | 253.0197712 | 255.1490465 | 246.4359757 | 3.16E-11 | -0.913688161 | 5.15E-12 | 47.62935212 | 0.53082633 | 10.49968341 |
| ENSMUSG00000017493 | *Igfbp4* | 1410.792883 | 1576.482908 | 1327.223453 | 751.4687204 | 763.2756584 | 776.9981353 | 1.54E-19 | -0.913969896 | 1.43E-20 | 86.46073626 | 0.530722679 | 18.81322543 |
| ENSMUSG00000034171 | *Faah* | 120.6283836 | 131.2053637 | 108.7008086 | 71.68893516 | 57.54425305 | 61.85059783 | 0.000272866 | -0.914538068 | 8.94E-05 | 15.34812758 | 0.530513707 | 3.56404988 |
| ENSMUSG00000001054 | *Rmnd5b* | 577.5782868 | 547.0254393 | 554.2399254 | 285.9123414 | 295.3214496 | 310.2194048 | 8.57E-15 | -0.914910689 | 1.08E-15 | 64.28665739 | 0.530376703 | 14.06684903 |
| ENSMUSG00000063439 | *B9d2* | 763.7134747 | 714.5645959 | 811.9011014 | 384.5900522 | 428.8675463 | 400.0960547 | 1.11E-15 | -0.916374576 | 1.31E-16 | 68.43625783 | 0.529838809 | 14.954591 |
| ENSMUSG00000020334 | *Slc22a4* | 59.91476005 | 71.658314 | 55.02139695 | 27.83217483 | 34.74369995 | 36.72379246 | 0.009832081 | -0.917988091 | 0.004115849 | 8.231984841 | 0.529246566 | 2.007354558 |
| ENSMUSG00000032491 | *Nradd* | 519.2612537 | 501.608198 | 532.7681607 | 270.7311552 | 274.6923778 | 276.3948591 | 1.39E-14 | -0.91817882 | 1.76E-15 | 63.31632735 | 0.529176603 | 13.85821722 |
| ENSMUSG00000026791 | *Slc2a8* | 345.1090179 | 328.0134091 | 260.3451466 | 149.281665 | 190.0046091 | 157.5257414 | 1.89E-06 | -0.918523421 | 4.77E-07 | 25.35366602 | 0.529050219 | 5.723788329 |
| ENSMUSG00000029723 | *Spacdr* | 3600.477647 | 3692.926351 | 3136.219626 | 1813.30836 | 1873.988316 | 1833.290376 | 6.37E-25 | -0.919024012 | 4.36E-26 | 111.6064199 | 0.528866679 | 24.19580593 |
| ENSMUSG00000024194 | *Cuta* | 2291.939288 | 2478.772101 | 2755.095803 | 1298.834825 | 1326.775042 | 1349.116165 | 5.08E-22 | -0.919516089 | 4.12E-23 | 98.03101566 | 0.528686323 | 21.29388336 |
| ENSMUSG00000026389 | *Steap3* | 649.4759989 | 685.2957071 | 703.2002928 | 357.6012766 | 361.5516276 | 356.6073531 | 2.63E-17 | -0.919985518 | 2.81E-18 | 76.0211369 | 0.528514326 | 16.57954516 |
| ENSMUSG00000037089 | *Slc35b2* | 2854.339169 | 2811.83187 | 2820.853083 | 1404.25973 | 1510.265207 | 1568.492504 | 3.08E-33 | -0.921594822 | 1.43E-34 | 150.3873565 | 0.527925105 | 32.51156018 |
| ENSMUSG00000006575 | *Rundc3a* | 139.0022433 | 118.0848273 | 144.9344115 | 77.59272983 | 54.28703118 | 79.24607847 | 0.00044889 | -0.921702463 | 0.000151537 | 14.35300921 | 0.527885717 | 3.347860542 |
| ENSMUSG00000020253 | *Ppm1m* | 1871.737104 | 2015.51624 | 2117.65279 | 1023.886674 | 1042.310999 | 1100.747358 | 2.51E-25 | -0.921702588 | 1.69E-26 | 113.4867831 | 0.527885671 | 24.59974671 |
| ENSMUSG00000083166 | *Gm8648* | 78.28861979 | 78.72321819 | 63.0733087 | 37.10956644 | 49.94406868 | 29.95888332 | 0.008520877 | -0.923338528 | 0.003518181 | 8.517134199 | 0.527287416 | 2.069515703 |
| ENSMUSG00000043889 | *Gm8399* | 57.51816964 | 56.51923358 | 60.38933812 | 28.67557407 | 30.40073746 | 32.8581301 | 0.007770872 | -0.923578001 | 0.00318626 | 8.69761907 | 0.527199899 | 2.109530258 |
| ENSMUSG00000017861 | *Mybl2* | 3765.043521 | 3550.618995 | 2708.126318 | 1719.691045 | 1850.102023 | 1718.286921 | 2.67E-12 | -0.92394956 | 4.00E-13 | 52.643162 | 0.527064139 | 11.57290078 |
| ENSMUSG00000019055 | *Plod1* | 4503.992229 | 4740.550716 | 4458.075138 | 2289.828929 | 2430.973256 | 2503.982797 | 1.80E-37 | -0.92396764 | 6.78E-39 | 170.1736633 | 0.527057534 | 36.74388668 |
| ENSMUSG00000038545 | *Cul7* | 93.46702567 | 123.1311874 | 130.1725733 | 64.94174127 | 56.45851243 | 59.91776665 | 0.000691401 | -0.924477829 | 0.000240464 | 13.48510942 | 0.52687118 | 3.160270223 |
| ENSMUSG00000028758 | *Kif17* | 223.6817708 | 219.0120301 | 202.639779 | 108.7985016 | 118.345728 | 114.0370398 | 2.89E-07 | -0.924509522 | 6.71E-08 | 29.14598779 | 0.526859606 | 6.539267724 |
| ENSMUSG00000055041 | *Commd5* | 990.5906994 | 1035.513101 | 1235.968453 | 583.6322722 | 540.6988305 | 589.5135106 | 4.78E-14 | -0.924907866 | 6.30E-15 | 60.80528117 | 0.526714154 | 13.32086132 |
| ENSMUSG00000049823 | *Zbtb12* | 348.3044717 | 316.9114168 | 303.2886759 | 158.5590566 | 172.6327591 | 180.7197155 | 4.51E-09 | -0.925119546 | 8.81E-10 | 37.57267627 | 0.526636877 | 8.345854931 |
| ENSMUSG00000071661 | *Zbtb3* | 123.8238374 | 130.1960916 | 195.9298526 | 77.59272983 | 78.1733249 | 79.24607847 | 0.000692839 | -0.926863407 | 0.000241087 | 13.48025474 | 0.526000688 | 3.159367616 |
| ENSMUSG00000068874 | *Selenbp1* | 222.0840439 | 172.5855168 | 174.4580879 | 102.8947069 | 98.80239674 | 98.5743903 | 1.26E-05 | -0.927675759 | 3.48E-06 | 21.53068446 | 0.525704592 | 4.899478699 |
| ENSMUSG00000025986 | *Slc39a10* | 1457.925828 | 1554.278923 | 1912.32904 | 880.5088037 | 838.1917614 | 864.9419541 | 8.11E-14 | -0.927977216 | 1.08E-14 | 59.73620498 | 0.525594755 | 13.09115488 |
| ENSMUSG00000038534 | *Osbpl7* | 472.1283092 | 475.3671253 | 421.3833815 | 227.7177941 | 232.3484934 | 260.9322096 | 2.51E-11 | -0.928144745 | 4.05E-12 | 48.09919464 | 0.525533725 | 10.60107281 |
| ENSMUSG00000028825 | *Rhd* | 344.3101544 | 356.2730259 | 440.1711756 | 205.7894139 | 199.7762747 | 191.350287 | 1.38E-08 | -0.928381024 | 2.84E-09 | 35.29162518 | 0.525447663 | 7.859973264 |
| ENSMUSG00000023277 | *Twf2* | 3934.402576 | 4267.202135 | 4341.322418 | 2151.511454 | 2180.167172 | 2251.748327 | 3.95E-36 | -0.929265409 | 1.58E-37 | 163.9119219 | 0.525125657 | 35.40321088 |
| ENSMUSG00000028700 | *Pomgnt1* | 2302.324513 | 2326.372025 | 2663.840804 | 1157.987153 | 1310.488933 | 1358.780321 | 8.46E-19 | -0.929674274 | 8.18E-20 | 83.00604826 | 0.524976856 | 18.07253026 |
| ENSMUSG00000020321 | *Mdh1* | 11478.86916 | 13223.48211 | 18476.45349 | 7537.458983 | 7538.297149 | 7587.328806 | 6.01E-09 | -0.929702086 | 1.19E-09 | 36.99253154 | 0.524966735 | 8.220839609 |
| ENSMUSG00000054716 | *Zfp771* | 655.8669066 | 738.7871246 | 724.6720574 | 342.4200903 | 397.3810682 | 373.0364182 | 2.24E-14 | -0.929845245 | 2.88E-15 | 62.34675029 | 0.524914645 | 13.6498393 |
| ENSMUSG00000024118 | *Tedc2* | 495.2953497 | 501.608198 | 508.6124255 | 284.2255429 | 281.2068215 | 224.2084171 | 9.40E-11 | -0.930171284 | 1.59E-11 | 45.41663047 | 0.524796032 | 10.02671508 |
| ENSMUSG00000041199 | *Rpusd1* | 587.1646485 | 669.1473547 | 569.0017636 | 307.8407216 | 310.5218183 | 340.1782881 | 6.29E-13 | -0.930387647 | 8.99E-14 | 55.57554887 | 0.524717333 | 12.20104848 |
| ENSMUSG00000025538 | *Sumf2* | 350.7010621 | 400.6809952 | 452.2490432 | 224.3441971 | 192.1760904 | 211.6450145 | 1.38E-08 | -0.93092239 | 2.84E-09 | 35.28789971 | 0.52452288 | 7.859267148 |
| ENSMUSG00000038740 | *Mvb12b* | 1596.129208 | 1585.566356 | 1297.699777 | 775.9272983 | 814.3054677 | 763.468317 | 2.68E-16 | -0.930973034 | 3.05E-17 | 71.3132585 | 0.524504467 | 15.57120145 |
| ENSMUSG00000029557 | *Mrm2* | 413.0124126 | 436.0055162 | 456.2749991 | 226.8743948 | 238.8629372 | 217.443508 | 3.56E-12 | -0.932361834 | 5.38E-13 | 52.06143998 | 0.5239998 | 11.44885415 |
| ENSMUSG00000071537 | *Klrg2* | 1150.363393 | 1174.792641 | 1082.98213 | 604.7172531 | 579.785493 | 602.0769133 | 5.10E-24 | -0.93249648 | 3.67E-25 | 107.3849019 | 0.523950898 | 23.29209554 |
| ENSMUSG00000002963 | *Pnkp* | 1696.786005 | 1616.853789 | 1511.075438 | 845.9294349 | 850.1349082 | 834.9830707 | 2.01E-26 | -0.932645848 | 1.27E-27 | 118.621499 | 0.523896654 | 25.69661522 |
| ENSMUSG00000017307 | *Acot8* | 421.0010473 | 431.968428 | 438.8291903 | 195.668623 | 244.2916403 | 237.7382354 | 2.31E-10 | -0.933448553 | 4.03E-11 | 43.60045469 | 0.523605243 | 9.636510889 |
| ENSMUSG00000020133 | *2310011J03Rik* | 2366.23359 | 2445.466124 | 2396.785731 | 1206.060909 | 1276.830973 | 1291.13123 | 1.31E-36 | -0.934131618 | 5.07E-38 | 166.1714121 | 0.523357393 | 35.88246499 |
| ENSMUSG00000039637 | *Coro7* | 2914.253929 | 3005.6121 | 2513.538451 | 1453.176886 | 1482.035951 | 1482.481517 | 3.40E-23 | -0.934189676 | 2.55E-24 | 103.5422595 | 0.523336332 | 22.46842013 |
| ENSMUSG00000023990 | *Tfeb* | 873.1577697 | 898.252105 | 666.9666899 | 393.8674438 | 399.5525495 | 485.1406267 | 6.86E-09 | -0.934212453 | 1.36E-09 | 36.72538566 | 0.523328069 | 8.163794253 |
| ENSMUSG00000036752 | *Tubb4b* | 13452.06193 | 12311.1002 | 12087.26152 | 6462.124956 | 6770.678529 | 6576.458098 | 2.28E-40 | -0.93457333 | 7.49E-42 | 183.7140446 | 0.52319718 | 39.6423009 |
| ENSMUSG00000027984 | *Hadh* | 1859.754152 | 1936.793022 | 2321.634554 | 1055.935845 | 1039.053777 | 1098.814527 | 2.60E-17 | -0.935832213 | 2.77E-18 | 76.04897133 | 0.522740842 | 16.58542632 |
| ENSMUSG00000097394 | *None* | 1082.459998 | 1085.976702 | 1246.704336 | 561.703892 | 579.785493 | 639.7671213 | 5.50E-17 | -0.937246543 | 5.96E-18 | 74.53346933 | 0.52222863 | 16.25994562 |
| ENSMUSG00000025466 | *Fuom* | 519.2612537 | 491.5154777 | 666.9666899 | 296.8765315 | 276.863859 | 298.6224177 | 4.50E-09 | -0.938545799 | 8.78E-10 | 37.57909256 | 0.521758535 | 8.34688962 |
| ENSMUSG00000024952 | *None* | 2618.674446 | 2526.207886 | 2322.97654 | 1254.978065 | 1308.317451 | 1336.552763 | 5.74E-29 | -0.939019959 | 3.21E-30 | 130.4857439 | 0.521587081 | 28.24106158 |
| ENSMUSG00000033809 | *Alg3* | 361.8851507 | 409.7644434 | 416.0154404 | 258.0801666 | 142.2320217 | 216.4770924 | 4.24E-05 | -0.939281326 | 1.25E-05 | 19.07862574 | 0.521492595 | 4.372205572 |
| ENSMUSG00000001270 | *Ckb* | 10275.78078 | 10225.94419 | 9513.333732 | 5170.880723 | 5078.008896 | 5396.464661 | 3.86E-45 | -0.940241523 | 1.00E-46 | 206.0436018 | 0.521145627 | 44.41335868 |
| ENSMUSG00000029033 | *Acap3* | 1116.012264 | 1124.329039 | 997.0950716 | 565.0774889 | 533.0986462 | 589.5135106 | 1.66E-19 | -0.941102608 | 1.55E-20 | 86.29974433 | 0.52083467 | 18.77980145 |
| ENSMUSG00000075486 | *Commd6* | 1084.856589 | 1283.79402 | 1913.671026 | 732.0705379 | 743.7323271 | 750.9049143 | 2.27E-06 | -0.941311815 | 5.81E-07 | 24.97400491 | 0.520759149 | 5.643232211 |
| ENSMUSG00000024955 | *Esrra* | 2679.388069 | 2789.627886 | 2365.920069 | 1300.521624 | 1406.034108 | 1378.108633 | 1.72E-23 | -0.941385963 | 1.27E-24 | 104.9261663 | 0.520732385 | 22.7648682 |

| | | | | | | | | | | | | | |
|---|---|---|---|---|---|---|---|---|---|---|---|---|---|
| ENSMUSG00000016552 | *Foxred2* | 1798.241665 | 2047.812945 | 1474.841835 | 909.1843777 | 933.7369363 | 930.6582143 | 9.88E-13 | -0.941460057 | 1.43E-13 | 54.66441131 | 0.520705642 | 12.00507624 |
| ENSMUSG00000048878 | *Hexim1* | 2068.257517 | 2048.822217 | 2022.371834 | 1063.526438 | 1056.425627 | 1077.553384 | 6.12E-39 | -0.941497381 | 2.15E-40 | 177.0361544 | 0.520692171 | 38.21292386 |
| ENSMUSG00000026820 | *Ptges2* | 1600.922388 | 1598.686893 | 1720.425144 | 808.8198685 | 878.3641645 | 872.6732788 | 5.60E-27 | -0.942397192 | 3.44E-28 | 121.2083402 | 0.520367516 | 26.25182961 |
| ENSMUSG00000024570 | *Rbfa* | 1743.918949 | 1681.447199 | 1800.944261 | 924.365564 | 926.1367519 | 867.8412008 | 4.11E-30 | -0.942405336 | 2.17E-31 | 135.8359051 | 0.520364578 | 29.38593514 |
| ENSMUSG00000029804 | *Herc3* | 796.4668769 | 957.7991547 | 1040.038601 | 474.8337706 | 494.0119837 | 482.24138 | 8.81E-13 | -0.942523581 | 1.27E-13 | 54.89682956 | 0.52032193 | 12.05519349 |
| ENSMUSG00000101431 | *Gm7901* | 4985.706899 | 5209.862209 | 5932.916974 | 2841.41203 | 2777.324515 | 2762.982175 | 8.26E-27 | -0.94334361 | 5.14E-28 | 120.4136684 | 0.520026263 | 26.08325308 |
| ENSMUSG00000019433 | *Gipc1* | 3213.028865 | 3243.800298 | 3485.135802 | 1665.713494 | 1756.728329 | 1741.480895 | 4.48E-37 | -0.944596587 | 1.71E-38 | 168.3345756 | 0.519574817 | 36.34829455 |
| ENSMUSG00000033429 | *Mcee* | 449.7601321 | 432.9777001 | 670.9926458 | 260.6103643 | 258.4062684 | 285.0925994 | 2.84E-06 | -0.945593525 | 7.31E-07 | 24.53099894 | 0.519215902 | 5.547108009 |
| ENSMUSG00000113786 | *Gm49384* | 307.5624349 | 297.7352483 | 299.26272 | 161.0892543 | 148.7464654 | 159.4585725 | 5.20E-10 | -0.945773218 | 9.35E-11 | 41.95283847 | 0.519151236 | 9.283946227 |
| ENSMUSG00000044469 | *Tnfaip8l1* | 127.8181548 | 151.3908042 | 162.3802203 | 69.15873745 | 79.25906552 | 80.21249406 | 5.11E-05 | -0.946047185 | 1.52E-05 | 18.7062996 | 0.519052659 | 4.291755954 |
| ENSMUSG00000027611 | *Procr* | 1163.145208 | 1431.147736 | 1698.953379 | 766.6499066 | 731.7891803 | 725.7781089 | 1.90E-10 | -0.946104796 | 3.29E-11 | 43.99651099 | 0.519031932 | 9.721150747 |
| ENSMUSG00000026974 | *Zmynd19* | 3122.757294 | 3312.430796 | 3516.001464 | 1702.82306 | 1754.556848 | 1703.790687 | 8.27E-34 | -0.946250483 | 3.71E-35 | 153.0660268 | 0.518979521 | 33.08238296 |
| ENSMUSG00000026435 | *Slc45a3* | 87.07611793 | 129.1868196 | 83.20308807 | 51.44735347 | 51.02980931 | 53.15285751 | 0.002332897 | -0.94632896 | 0.000874699 | 11.07562603 | 0.518951292 | 2.632104363 |
| ENSMUSG00000081600 | *Gm12286* | 75.89202939 | 76.70467414 | 88.57102924 | 26.98877559 | 54.28703118 | 44.45511719 | 0.009835944 | -0.947102028 | 0.004118047 | 8.231016017 | 0.518673286 | 2.007183948 |
| ENSMUSG00000038880 | *Mrps34* | 1193.50202 | 1272.692027 | 1207.786762 | 621.5852378 | 616.7006742 | 666.8267579 | 7.23E-25 | -0.947302137 | 4.97E-26 | 111.3464995 | 0.518601349 | 24.14073688 |
| ENSMUSG00000089715 | *Cbx6* | 2633.053988 | 2730.080836 | 1937.826761 | 1254.978065 | 1234.487089 | 1298.862554 | 8.49E-12 | -0.947803688 | 1.32E-12 | 50.30231744 | 0.518421089 | 11.07120988 |
| ENSMUSG00000020412 | *Ascc2* | 3211.431138 | 3268.022827 | 3012.756979 | 1681.738079 | 1642.725563 | 1593.61931 | 7.74E-38 | -0.949211281 | 2.85E-39 | 171.8995412 | 0.517915528 | 37.1111712 |
| ENSMUSG00000042286 | *Stab1* | 452.955586 | 357.282298 | 272.4230142 | 194.8252238 | 193.261831 | 174.921222 | 1.40E-05 | -0.94938197 | 3.89E-06 | 21.31697967 | 0.517854256 | 4.85375599 |
| ENSMUSG00000037251 | *Pomk* | 968.2225224 | 1157.635016 | 1309.777645 | 573.5114813 | 600.4145648 | 601.1104977 | 1.71E-12 | -0.950194726 | 2.53E-13 | 53.54609444 | 0.5175626 | 11.76580203 |
| ENSMUSG00000005986 | *Ankrd13d* | 75.89202939 | 88.81593848 | 65.75727928 | 41.32656262 | 31.48647808 | 46.38794837 | 0.004374515 | -0.950658688 | 0.001714539 | 9.832512972 | 0.517396182 | 2.359070053 |
| ENSMUSG00000076435 | *Acsf2* | 261.2283538 | 259.3829112 | 268.3970583 | 129.8834825 | 129.2031342 | 148.828001 | 1.19E-08 | -0.950919307 | 2.44E-09 | 35.58499012 | 0.517302724 | 7.922877192 |
| ENSMUSG00000035521 | *Gnptg* | 893.9282199 | 1004.225668 | 1198.392865 | 545.6793065 | 523.3269806 | 528.6293283 | 1.81E-12 | -0.951088854 | 2.67E-13 | 53.43668928 | 0.517241934 | 11.74300586 |
| ENSMUSG00000002885 | *Adgre5* | 97.46134301 | 113.0384672 | 96.62294099 | 54.82095042 | 60.80147492 | 43.4887016 | 0.000672646 | -0.951671794 | 0.000233306 | 13.54181876 | 0.517032977 | 3.172213475 |
| ENSMUSG00000025650 | *Col7a1* | 71.89771206 | 89.8252105 | 92.59698511 | 47.23035729 | 48.85832806 | 34.79096128 | 0.002682304 | -0.952445824 | 0.001013938 | 10.80194025 | 0.516755655 | 2.571491922 |
| ENSMUSG00000028179 | *Cth* | 2609.088084 | 2326.372025 | 2321.634554 | 1226.302491 | 1263.802086 | 1262.138762 | 5.71E-29 | -0.953158545 | 3.19E-30 | 130.4975226 | 0.51650043 | 28.24317977 |
| ENSMUSG00000028863 | *Meaf6* | 881.1464044 | 894.2150169 | 935.3637482 | 500.9791469 | 451.6680994 | 443.5847563 | 1.23E-19 | -0.954178847 | 1.13E-20 | 86.91612314 | 0.51613528 | 18.90933965 |
| ENSMUSG00000042476 | *Abcb4* | 1021.746375 | 1142.495936 | 1113.847792 | 553.2698996 | 584.1284555 | 553.7561337 | 7.33E-22 | -0.954210276 | 5.97E-23 | 97.29706781 | 0.516124036 | 21.13511711 |
| ENSMUSG00000039062 | *Anpep* | 2017.130255 | 2499.966814 | 2164.622275 | 1146.179563 | 1158.485245 | 1143.269644 | 6.90E-20 | -0.954226364 | 6.27E-21 | 88.08371878 | 0.516118281 | 19.16094684 |
| ENSMUSG00000064109 | *Hcst* | 234.0669959 | 253.3272791 | 283.1588965 | 128.1966841 | 137.8890592 | 130.4661048 | 5.50E-08 | -0.954570201 | 1.19E-08 | 32.49863344 | 0.515995289 | 7.259568784 |
| ENSMUSG00000010154 | *Spire2* | 382.6556008 | 365.3564742 | 389.1757345 | 187.2346307 | 183.4901654 | 215.5106768 | 6.61E-11 | -0.955045646 | 1.10E-11 | 46.13606979 | 0.515825269 | 10.17981467 |
| ENSMUSG00000030008 | *Pradc1* | 285.9931213 | 314.8928728 | 355.6261023 | 172.8968436 | 159.6038717 | 158.4921569 | 9.85E-09 | -0.955234027 | 1.99E-09 | 35.98716584 | 0.515757919 | 8.00672992 |
| ENSMUSG00000031824 | *6430548M08Rik* | 2601.898313 | 2361.696546 | 1960.640511 | 1147.022963 | 1175.857095 | 1252.474606 | 4.37E-15 | -0.955335773 | 5.38E-16 | 65.65315069 | 0.515721547 | 14.35914179 |
| ENSMUSG00000062995 | *Ica1* | 1106.425902 | 981.0124113 | 1021.250807 | 522.9075271 | 533.0986462 | 548.9240558 | 3.23E-21 | -0.955478845 | 2.72E-22 | 94.29513423 | 0.515670405 | 20.49021639 |
| ENSMUSG00000032171 | *Pin1* | 4126.129809 | 4345.925353 | 4923.744035 | 2240.068374 | 2285.484013 | 2376.415939 | 7.87E-26 | -0.955813663 | 5.15E-27 | 115.8413863 | 0.515550743 | 25.10421623 |
| ENSMUSG00000105835 | *Gm43552* | 160.5715569 | 171.5762448 | 177.1420585 | 83.49652449 | 72.74462178 | 105.3392994 | 2.83E-05 | -0.956735156 | 8.18E-06 | 19.89423213 | 0.515221551 | 4.54853235 |
| ENSMUSG00000026162 | *Nhej1* | 157.3761031 | 188.7338693 | 205.3237496 | 95.30411381 | 94.45943425 | 92.77589675 | 6.19E-06 | -0.957663974 | 1.66E-06 | 22.95307195 | 0.514889954 | 5.208110767 |
| ENSMUSG00000037190 | *Cyb561d2* | 688.6203088 | 790.259998 | 782.377425 | 382.9032537 | 388.6951432 | 389.4654832 | 6.16E-17 | -0.959528343 | 6.71E-18 | 74.30140258 | 0.514225 | 16.21030829 |
| ENSMUSG00000037278 | *Tmem97* | 5127.105733 | 5246.196002 | 6169.106385 | 2886.955589 | 2733.89489 | 2877.019215 | 2.89E-24 | -0.960020723 | 2.04E-25 | 108.5505486 | 0.514049529 | 23.5392291 |
| ENSMUSG00000038175 | *Mylip* | 204.5090476 | 262.4107273 | 225.453529 | 98.67771076 | 130.2888748 | 127.566858 | 6.07E-06 | -0.960492206 | 1.63E-06 | 22.9930402 | 0.513881562 | 5.21656608 |
| ENSMUSG00000043822 | *Adamtsl5* | 406.6215048 | 484.4505735 | 477.7467638 | 218.4404024 | 237.7771966 | 245.4695601 | 4.69E-11 | -0.962327558 | 7.73E-12 | 46.83197303 | 0.513228233 | 10.32864377 |
| ENSMUSG00000036599 | *Chst12* | 868.3645889 | 881.0944806 | 805.1911749 | 427.6034133 | 449.4966182 | 435.8534316 | 8.12E-21 | -0.962994359 | 6.99E-22 | 92.42604117 | 0.512991079 | 20.09033657 |
| ENSMUSG00000052151 | *Plpp2* | 1815.017798 | 1950.92283 | 2137.782569 | 974.9695182 | 993.4526706 | 1056.292241 | 4.03E-23 | -0.96322539 | 3.04E-24 | 103.1916713 | 0.512908936 | 22.39429511 |
| ENSMUSG00000002580 | *Mien1* | 1787.85644 | 1936.793022 | 2486.698745 | 1006.17529 | 1050.996924 | 1124.907748 | 4.44E-12 | -0.963355478 | 6.75E-13 | 51.61660925 | 0.512862688 | 11.35299812 |
| ENSMUSG00000005413 | *Hmox1* | 12469.45986 | 14926.12402 | 16713.08482 | 7626.859302 | 7536.125668 | 7446.23213 | 5.23E-18 | -0.96383323 | 5.33E-19 | 79.30166944 | 0.512692881 | 17.28168133 |
| ENSMUSG00000000409 | *Lck* | 391.443099 | 367.3750182 | 383.8077934 | 172.0534444 | 218.2338653 | 197.1487806 | 3.75E-10 | -0.964386533 | 6.66E-11 | 42.61521712 | 0.51249629 | 9.426399228 |
| ENSMUSG00000028070 | *Naxe* | 2412.567671 | 2478.772101 | 2610.161392 | 1291.244232 | 1252.94468 | 1294.996892 | 8.28E-38 | -0.964910789 | 3.05E-39 | 171.759416 | 0.51231009 | 37.0819603 |
| ENSMUSG00000063904 | *Dpp3* | 3985.529838 | 4138.015315 | 3795.134404 | 1896.804885 | 2121.537178 | 2092.289755 | 1.20E-32 | -0.964948509 | 5.72E-34 | 147.6279264 | 0.512296695 | 31.91985378 |
| ENSMUSG00000070348 | *Ccnd1* | 39598.86321 | 40449.60434 | 37156.88875 | 19719.51757 | 20156.77468 | 20116.90694 | 1.39E-57 | -0.966287354 | 2.23E-59 | 264.0639858 | 0.511821496 | 56.8568628 |
| ENSMUSG00000049709 | *Nlrp10* | 444.1680878 | 437.0147882 | 446.8811021 | 220.1272009 | 232.3484934 | 227.1076639 | 3.21E-14 | -0.967532676 | 4.18E-15 | 61.61282316 | 0.511379887 | 13.49371651 |
| ENSMUSG00000074364 | *Ehd2* | 925.8827586 | 980.0031393 | 649.5208811 | 421.6996186 | 425.6103244 | 461.9466526 | 1.99E-08 | -0.967764115 | 4.14E-09 | 34.55878034 | 0.511297857 | 7.701715434 |
| ENSMUSG00000024446 | *Rpp21* | 474.5248996 | 489.4969336 | 524.716249 | 239.5253834 | 231.2627528 | 288.9582617 | 2.33E-11 | -0.9680159 | 3.76E-12 | 48.24833352 | 0.511208631 | 10.63267591 |
| ENSMUSG00000023349 | *Clec4n* | 127.0192913 | 150.3815322 | 191.9038967 | 81.80972601 | 76.00184365 | 80.21249406 | 0.000119296 | -0.968568776 | 3.74E-05 | 16.99827982 | 0.511012761 | 3.923374956 |
| ENSMUSG00000020873 | *Slc35b1* | 1112.017946 | 1277.738388 | 1225.232571 | 702.5515646 | 656.8730773 | 486.1070423 | 3.49E-10 | -0.968903277 | 6.19E-11 | 42.75964854 | 0.510894292 | 9.457456036 |
| ENSMUSG00000089753 | *Cd300ld2* | 139.0022433 | 131.2053637 | 143.5924262 | 53.13415195 | 96.6309155 | 62.81701342 | 0.000941112 | -0.969258184 | 0.00033381 | 12.87045472 | 0.510768626 | 3.026358549 |
| ENSMUSG00000017176 | *Nt5c3b* | 1423.574699 | 1455.370265 | 1690.901467 | 770.0235036 | 780.6475083 | 779.897382 | 3.75E-21 | -0.969297659 | 3.16E-22 | 93.99820147 | 0.510754651 | 20.42598121 |
| ENSMUSG00000028717 | *Tal1* | 1849.368927 | 1988.265895 | 1850.597717 | 997.7412976 | 999.9671143 | 906.4978245 | 1.78E-28 | -0.969957856 | 1.01E-29 | 128.2020295 | 0.510520976 | 27.75045883 |
| ENSMUSG00000030108 | *Slc6a13* | 1116.811127 | 1079.92107 | 1315.145586 | 561.703892 | 606.9290086 | 621.4052251 | 1.37E-16 | -0.970718737 | 1.53E-17 | 72.67886796 | 0.510251797 | 15.86262734 |
| ENSMUSG00000068580 | *Zfyve19* | 880.347541 | 903.2984652 | 1131.293601 | 508.5697401 | 444.067915 | 530.5621595 | 9.89E-12 | -0.970842231 | 1.54E-12 | 49.99447646 | 0.510208122 | 11.00472616 |
| ENSMUSG00000098112 | *Bin2* | 1962.807539 | 2006.432792 | 1704.32132 | 999.4280961 | 936.9941581 | 959.650682 | 4.35E-23 | -0.971201825 | 3.29E-24 | 103.0356787 | 0.510080967 | 22.36145648 |
| ENSMUSG00000039835 | *Nhsl1* | 268.418125 | 288.6518 | 249.6092642 | 121.4494902 | 158.518131 | 133.3653516 | 1.26E-07 | -0.971478343 | 2.83E-08 | 30.82183557 | 0.509983211 | 6.899916073 |
| ENSMUSG00000037295 | *Ldlrap1* | 4988.902353 | 5240.14037 | 5099.544108 | 2568.150677 | 2574.291018 | 2672.139109 | 1.90E-53 | -0.971852612 | 3.57E-55 | 244.7776074 | 0.509850926 | 52.72053553 |
| ENSMUSG00000032898 | *Fbxo21* | 1611.307614 | 1661.261758 | 1470.81588 | 807.13307 | 855.5636114 | 757.6698234 | 2.00E-23 | -0.972626718 | 1.48E-24 | 104.6141768 | 0.509577429 | 22.69856532 |
| ENSMUSG00000031840 | *Rab3a* | 112.6397489 | 105.973563 | 85.88705866 | 45.54355881 | 45.60110619 | 64.74984461 | 0.00119816 | -0.972915747 | 0.000430925 | 12.39309348 | 0.509475351 | 2.921485068 |
| ENSMUSG00000011267 | *Zfp296* | 286.7919848 | 233.1418385 | 177.1420585 | 106.2683039 | 142.2320217 | 109.2049618 | 0.000116304 | -0.973001757 | 3.64E-05 | 17.05008429 | 0.509444978 | 3.934405553 |
| ENSMUSG00000022671 | *Mzt2* | 595.9521466 | 509.6823742 | 676.3605869 | 292.6595353 | 309.4360777 | 304.4209112 | 3.80E-11 | -0.97317921 | 6.21E-12 | 47.26117317 | 0.50938232 | 10.41984312 |
| ENSMUSG00000030590 | *Fam98c* | 647.878272 | 691.3513393 | 666.9666899 | 362.661672 | 320.293984 | 336.3126257 | 2.56E-17 | -0.974170238 | 2.72E-18 | 76.07953906 | 0.509032531 | 16.59190845 |
| ENSMUSG00000042213 | *Zfand4* | 146.990878 | 204.8822217 | 195.9298526 | 104.5815054 | 90.11647176 | 82.14532525 | 4.48E-05 | -0.976714624 | 1.33E-05 | 18.97102097 | 0.508135575 | 4.348584129 |
| ENSMUSG00000020520 | *Galnt10* | 778.0930171 | 788.241454 | 752.8537485 | 403.1448354 | 385.4379214 | 389.4654832 | 1.43E-21 | -0.97725901 | 1.18E-22 | 95.95490573 | 0.507943871 | 20.84509715 |
| ENSMUSG00000024205 | *Rpl36-ps2* | 1733.533724 | 1333.248349 | 1807.654188 | 693.274173 | 723.1032553 | 1059.191488 | 3.28E-07 | -0.977307025 | 7.67E-08 | 28.88660821 | 0.507926966 | 6.483538357 |
| ENSMUSG00000003948 | *Mmd* | 1397.212204 | 1339.303981 | 1481.551762 | 706.7685608 | 720.931774 | 713.2147063 | 2.36E-28 | -0.977378748 | 1.35E-29 | 127.6371004 | 0.507901716 | 27.6277388 |
| ENSMUSG00000029047 | *Pex10* | 659.0623605 | 697.4069714 | 701.8583075 | 332.2992995 | 344.1797777 | 368.2043402 | 3.63E-18 | -0.977669022 | 3.68E-19 | 80.03613209 | 0.507799535 | 17.44060181 |
| ENSMUSG00000060260 | *Pwwp2b* | 1137.581577 | 1184.885361 | 1174.23713 | 587.0058691 | 593.9001211 | 593.379173 | 4.62E-29 | -0.97831588 | 2.56E-30 | 130.9330688 | 0.507571905 | 28.33524476 |
| ENSMUSG00000110949 | *Nudt8* | 227.6760882 | 290.6703441 | 285.8428671 | 129.8834825 | 140.0605404 | 137.2310139 | 1.40E-07 | -0.978526423 | 3.15E-08 | 30.61315157 | 0.507497836 | 6.855144529 |
| ENSMUSG00000024790 | *Sac3d1* | 1034.52819 | 1089.004518 | 1081.640145 | 546.5227057 | 535.2701274 | 543.1255622 | 7.12E-27 | -0.978738146 | 4.39E-28 | 120.7232387 | 0.507423364 | 26.14729976 |
| ENSMUSG00000021631 | *Mrps36-ps1* | 202.1124572 | 178.641149 | 236.1894113 | 129.0400833 | 72.74462178 | 109.2049618 | 0.000235311 | -0.978776027 | 7.66E-05 | 15.63962911 | 0.507410041 | 3.628357423 |
| ENSMUSG00000047721 | *Bola2* | 2708.147154 | 2694.756315 | 3424.746464 | 1418.597517 | 1458.149657 | 1597.484972 | 4.66E-16 | -0.979284169 | 5.38E-17 | 70.19378121 | 0.507231354 | 15.3319289 |
| ENSMUSG00000057963 | *Itpk1* | 1285.371319 | 1250.488043 | 1072.246248 | 601.3436561 | 618.8721554 | 613.6739004 | 1.66E-19 | -0.979297455 | 1.54E-20 | 86.30182164 | 0.507226683 | 18.77998234 |
| ENSMUSG00000024187 | *Fam234a* | 1388.424706 | 1349.396702 | 1410.426541 | 657.0080058 | 741.5608459 | 707.4162127 | 1.03E-26 | -0.979318431 | 6.46E-28 | 119.9579184 | 0.507219308 | 25.9871706 |
| ENSMUSG00000067370 | *B3galt4* | 168.5601916 | 186.7153252 | 214.7176466 | 90.24371838 | 92.287953 | 105.3392994 | 3.41E-06 | -0.979625058 | 8.87E-07 | 24.15976014 | 0.507111516 | 5.467864091 |
| ENSMUSG00000018659 | *Pnpo* | 790.0759691 | 807.4176225 | 736.749925 | 370.2522651 | 428.8675463 | 387.532652 | 1.35E-17 | -0.979870943 | 1.41E-18 | 77.37851845 | 0.507025094 | 16.86905569 |
| ENSMUSG00000036632 | *Alg5* | 1795.046211 | 1643.094862 | 2070.683305 | 928.5825602 | 893.5645332 | 965.4491755 | 2.15E-18 | -0.98096879 | 2.14E-19 | 81.10629871 | 0.50663941 | 17.66712119 |
| ENSMUSG00000031399 | *Fam3a* | 602.3430543 | 665.1102665 | 645.4949252 | 350.0106835 | 310.5218183 | 306.3537424 | 1.41E-15 | -0.98101189 | 1.68E-16 | 67.94438214 | 0.506624275 | 14.84928757 |
| ENSMUSG00000002768 | *Mea1* | 3048.462991 | 3088.372406 | 3978.986389 | 1751.740216 | 1657.925932 | 1704.757103 | 4.62E-16 | -0.982594836 | 5.33E-17 | 70.21119494 | 0.506068704 | 15.33554058 |
| ENSMUSG00000038991 | *Txndc5* | 12561.32916 | 13758.39629 | 14414.26402 | 6734.542909 | 6779.364454 | 7087.691945 | 5.14E-38 | -0.983165383 | 1.88E-39 | 172.7267781 | 0.505868607 | 37.28903673 |
| ENSMUSG00000015542 | *Nat9* | 587.9635119 | 606.5724889 | 552.8979401 | 289.2859384 | 279.0353403 | 316.0178983 | 1.13E-15 | -0.98359763 | 1.34E-16 | 68.39776248 | 0.505717066 | 14.94654708 |
| ENSMUSG00000039748 | *Exo1* | 1123.202035 | 1106.162143 | 1033.328674 | 528.8113217 | 605.8432679 | 517.9987568 | 6.08E-20 | -0.984425269 | 5.52E-21 | 88.3374818 | 0.505427032 | 19.21581519 |
| ENSMUSG00000023904 | *Hcfc1r1* | 516.0657999 | 456.1909567 | 504.5864696 | 245.429178 | 254.0633059 | 247.4023913 | 2.85E-14 | -0.984469175 | 3.69E-15 | 61.85825698 | 0.50541165 | 13.54568012 |
| ENSMUSG00000078881 | *Gm14434* | 95.86361607 | 111.0199231 | 134.1985292 | 57.35114813 | 40.17240307 | 73.44758493 | 0.002288593 | -0.985258038 | 0.000856879 | 11.1137959 | 0.505135367 | 2.640431397 |
| ENSMUSG00000004665 | *Cnn2* | 225.2794978 | 244.2438308 | 218.7436025 | 122.2928894 | 115.0885061 | 110.1713774 | 3.07E-08 | -0.985457698 | 6.47E-09 | 33.68669953 | 0.505065465 | 7.513523016 |
| ENSMUSG00000020758 | *Itgb4* | 55.12157924 | 81.75103428 | 53.67941166 | 32.89257025 | 31.48647808 | 31.89171451 | 0.007917515 | -0.986064383 | 0.003250368 | 8.661314086 | 0.504853118 | 2.101411086 |
| ENSMUSG00000070960 | *Gm19680* | 150.9851953 | 173.5947888 | 157.0122791 | 84.33992372 | 91.20221238 | 67.64909138 | 1.41E-05 | -0.986612888 | 3.91E-06 | 21.30844509 | 0.504661212 | 4.852006863 |
| ENSMUSG00000015776 | *Med22* | 2331.083598 | 2458.586661 | 2227.695584 | 1165.577746 | 1184.54302 | 1192.556839 | 2.79E-36 | -0.987020696 | 1.10E-37 | 164.6326501 | 0.504518579 | 35.55485289 |
| ENSMUSG00000055681 | *Cope* | 5967.510101 | 6334.191249 | 7258.798442 | 3241.183269 | 3324.537789 | 3297.409997 | 5.04E-27 | -0.987176112 | 3.08E-28 | 121.4259816 | 0.504464232 | 26.29738225 |
| ENSMUSG00000034108 | *Ccs* | 658.263497 | 687.3142512 | 758.2216897 | 353.3842804 | 347.4369995 | 357.5737687 | 1.68E-17 | -0.98764062 | 1.76E-18 | 76.94054767 | 0.504301835 | 16.7754412 |
| ENSMUSG00000016526 | *Dyrk3* | 829.220279 | 867.9739442 | 994.411101 | 452.9053904 | 449.4966182 | 451.3160811 | 9.87E-18 | -0.988166525 | 1.02E-18 | 78.01861469 | 0.504118036 | 17.00585048 |
| ENSMUSG00000068220 | *Lgals1* | 15581.03307 | 15171.37713 | 21357.69591 | 8708.940524 | 8791.241829 | 8764.422996 | 1.28E-13 | -0.988216948 | 1.74E-14 | 58.80760206 | 0.504100417 | 12.89230285 |
| ENSMUSG00000029610 | *Aimp2* | 2522.011966 | 2477.762829 | 2878.55845 | 1281.123441 | 1288.77412 | 1396.470529 | 1.99E-25 | -0.988889116 | 1.33E-26 | 113.956206 | 0.503865605 | 24.70064949 |
| ENSMUSG00000024193 | *Phf1* | 421.7999107 | 560.1459756 | 437.487205 | 204.1026154 | 254.0633059 | 258.0329628 | 4.04E-08 | -0.989292278 | 8.64E-09 | 33.12489644 | 0.503724819 | 7.393413019 |
| ENSMUSG00000040563 | *Plppr2* | 131.0136086 | 137.2609958 | 114.0687498 | 68.31533822 | 59.7157343 | 64.74984461 | 3.75E-05 | -0.989320648 | 1.10E-05 | 19.32507829 | 0.503714914 | 4.425490604 |
| ENSMUSG00000006395 | *Hyi* | 602.3430543 | 563.1737917 | 666.9666899 | 312.901117 | 245.3773809 | 362.4058467 | 7.14E-09 | -0.989556409 | 1.42E-09 | 36.6417153 | 0.503632605 | 8.146573574 |
| ENSMUSG00000026938 | *Fcna* | 365.879468 | 379.4862826 | 453.5910285 | 205.7894139 | 205.2049779 | 190.3838715 | 2.16E-10 | -0.990360976 | 3.76E-11 | 43.73406337 | 0.503351816 | 9.664835874 |
| ENSMUSG00000049562 | *Ap5b1* | 1716.757591 | 1839.902907 | 1529.863232 | 829.0614502 | 871.8497207 | 862.0427073 | 1.33E-22 | -0.990811774 | 1.03E-23 | 100.7684407 | 0.503194558 | 21.87674773 |
| ENSMUSG00000041654 | *Slc39a11* | 2789.631228 | 2865.323288 | 3188.557053 | 1475.105266 | 1488.550395 | 1481.515101 | 1.58E-32 | -0.991097055 | 7.59E-34 | 147.0674153 | 0.503095066 | 31.80108068 |
| ENSMUSG00000075419 | *Dolk* | 782.0873345 | 803.3805344 | 732.7239692 | 402.3014362 | 388.6951432 | 375.9356649 | 2.62E-20 | -0.991258837 | 2.32E-21 | 90.04840187 | 0.503038658 | 19.58112052 |
| ENSMUSG00000068129 | *Cst7* | 415.409003 | 402.6995392 | 432.1192639 | 225.1875963 | 199.7762747 | 201.9808585 | 6.33E-13 | -0.99206891 | 9.06E-14 | 55.56142666 | 0.502756276 | 12.19828714 |
| ENSMUSG00000023048 | *Prr13* | 1407.597429 | 1575.473636 | 1630.512129 | 796.1688799 | 748.0752896 | 770.2332261 | 3.54E-24 | -0.992595438 | 2.51E-25 | 108.1333107 | 0.502572822 | 23.45070569 |
| ENSMUSG00000021094 | *Dhrs7* | 1269.39405 | 1297.923828 | 1303.067718 | 681.4665837 | 612.3577117 | 648.4648616 | 4.54E-28 | -0.992621653 | 2.64E-29 | 126.3036547 | 0.50256369 | 27.34261015 |
| ENSMUSG00000027890 | *Gstm4* | 63.90907738 | 47.43578532 | 67.09926458 | 30.36237254 | 26.05777497 | 32.8581301 | 0.007411835 | -0.993447538 | 0.003023962 | 8.792953971 | 0.502276075 | 2.130074286 |
| ENSMUSG00000082809 | *Gm14150* | 58.31703311 | 110.0106511 | 119.4366909 | 42.16996186 | 54.28703118 | 47.35436397 | 0.009325019 | -0.993720438 | 0.003883223 | 8.337629504 | 0.502181074 | 2.030350273 |
| ENSMUSG00000075307 | *Klhl41* | 541.6294308 | 550.0532553 | 731.3819839 | 303.6237254 | 342.0082964 | 267.6971187 | 1.21E-08 | -0.994560304 | 2.49E-09 | 35.5502759 | 0.501888814 | 7.915771374 |
| ENSMUSG00000034792 | *Gna15* | 752.5293862 | 793.2878141 | 828.0049249 | 387.9636491 | 386.523662 | 414.5922886 | 2.50E-20 | -0.995215429 | 2.21E-21 | 90.15072778 | 0.501660959 | 19.60242284 |
| ENSMUSG00000028671 | *Gale* | 1333.303127 | 1397.841759 | 1391.638747 | 640.9834203 | 710.0743678 | 718.0467842 | 8.76E-27 | -0.995337889 | 5.47E-28 | 120.2897334 | 0.501618378 | 26.05735527 |
| ENSMUSG00000000278 | *Scpep1* | 7752.96995 | 8943.159441 | 9615.324614 | 4505.438725 | 4325.590644 | 4351.769407 | 1.13E-25 | -0.996434289 | 7.45E-27 | 115.1076574 | 0.50123731 | 24.9458755 |
| ENSMUSG00000004668 | *Abca13* | 119.0306566 | 121.1126434 | 157.0122791 | 69.15873745 | 51.02980931 | 77.31324729 | 0.000325413 | -0.996461905 | 0.00010776 | 14.99562939 | 0.501227715 | 3.487564863 |
| ENSMUSG00000029452 | *Tmem116* | 103.0533873 | 135.2424518 | 128.830588 | 53.13415195 | 61.88721554 | 68.61550697 | 0.000179009 | -0.997366541 | 5.74E-05 | 16.18556831 | 0.500913521 | 3.747124852 |
| ENSMUSG00000047557 | *Lxn* | 536.0373866 | 526.8399987 | 571.6857342 | 259.7669651 | 276.863859 | 281.226937 | 2.81E-16 | -0.998420946 | 3.20E-17 | 71.21763738 | 0.500547558 | 15.55083053 |

| | | | | | | | | | | | | | |
|---|---|---|---|---|---|---|---|---|---|---|---|---|---|
| ENSMUSG00000042215 | *Bag2* | 1021.746375 | 1168.737009 | 1235.968453 | 565.0774889 | 549.3847555 | 597.2448353 | 1.38E-18 | -0.99846677 | 1.35E-19 | 82.01576278 | 0.500531659 | 17.86021658 |
| ENSMUSG00000021238 | *Aldh6a1* | 104.6511142 | 99.91793079 | 144.9344115 | 61.56814432 | 64.05869679 | 48.32077956 | 0.000619544 | -0.998643497 | 0.000213937 | 13.70453589 | 0.500470349 | 3.207927933 |
| ENSMUSG00000007038 | *Neu1* | 4139.710487 | 4642.651329 | 5476.641975 | 2337.902686 | 2291.998456 | 2498.184303 | 5.71E-18 | -0.9993473 | 5.83E-19 | 79.12452248 | 0.50022626 | 17.24299155 |
| ENSMUSG00000047415 | *Gpr68* | 456.1510398 | 439.0333322 | 344.8902199 | 206.6328131 | 234.5199747 | 181.6861311 | 2.88E-08 | -0.999522942 | 6.06E-09 | 33.81574776 | 0.500165363 | 7.541155136 |
| ENSMUSG00000027808 | *Serp1* | 13439.28011 | 13972.36196 | 18232.21217 | 7467.456846 | 7490.524562 | 7861.790834 | 3.68E-16 | -0.999852016 | 4.22E-17 | 70.67356926 | 0.50005129 | 15.43443382 |
| ENSMUSG00000066357 | *Wdr6* | 1815.816661 | 2029.646049 | 1541.9411 | 863.6408189 | 947.8515644 | 885.2366815 | 3.60E-16 | -1.000296742 | 4.12E-17 | 70.72001748 | 0.499897168 | 15.44421132 |
| ENSMUSG00000048897 | *Zfp710* | 3650.007182 | 3763.575393 | 2651.762936 | 1586.433965 | 1698.098335 | 1747.279389 | 1.16E-12 | -1.001397926 | 1.68E-13 | 54.34830397 | 0.49951575 | 11.93627937 |
| ENSMUSG00000081911 | *Mrps36-ps2* | 77.48975633 | 90.83448253 | 130.1725733 | 32.89257025 | 60.80147492 | 55.08568869 | 0.0077932 | -1.003037904 | 0.003197033 | 8.691466312 | 0.49894825 | 2.108284193 |
| ENSMUSG00000024646 | *Cyb5a* | 3575.71288 | 4023.967576 | 6312.698811 | 2317.661104 | 2293.084197 | 2327.128743 | 1.61E-07 | -1.003044641 | 3.64E-08 | 30.33364787 | 0.498945921 | 6.794046954 |
| ENSMUSG00000002393 | *Nr2f6* | 1377.240618 | 1393.804671 | 1464.105953 | 659.5382035 | 745.9038084 | 706.4497971 | 3.12E-27 | -1.004462919 | 1.89E-28 | 122.3992601 | 0.49845566 | 26.50537345 |
| ENSMUSG00000022175 | *Lrp10* | 3481.44699 | 3570.804436 | 3206.002861 | 1620.169935 | 1817.529804 | 1680.596713 | 6.04E-32 | -1.00453033 | 2.95E-33 | 144.36787 | 0.49843237 | 31.21885212 |
| ENSMUSG00000025223 | *Ldb1* | 2682.583523 | 2771.460989 | 2908.082127 | 1347.751981 | 1420.148736 | 1398.40336 | 9.32E-41 | -1.004615344 | 2.99E-42 | 185.5405923 | 0.498402999 | 40.03080004 |
| ENSMUSG00000005873 | *Reep5* | 5390.730677 | 5914.334085 | 6857.54484 | 3059.009033 | 2870.698209 | 3112.824619 | 1.17E-21 | -1.005287618 | 9.60E-23 | 96.35637261 | 0.498170806 | 20.93253972 |
| ENSMUSG00000028032 | *Papss1* | 2065.062063 | 2248.658079 | 2416.91551 | 1145.336164 | 1100.940992 | 1101.713774 | 4.39E-28 | -1.005435827 | 2.55E-29 | 126.3718351 | 0.498119631 | 27.35708916 |
| ENSMUSG00000020834 | *Dhrs13* | 882.7441314 | 858.8904959 | 739.4338956 | 381.2164552 | 452.75384 | 405.8945483 | 5.53E-15 | -1.005566936 | 6.84E-16 | 65.17943208 | 0.498074365 | 14.25703477 |
| ENSMUSG00000024352 | *Spata24* | 120.6283836 | 120.1033713 | 116.7527204 | 58.19454737 | 66.23017804 | 54.1192731 | 3.74E-05 | -1.005736736 | 1.10E-05 | 19.33324838 | 0.498015747 | 4.426999566 |
| ENSMUSG00000063458 | *Lrmda* | 329.930612 | 308.8372406 | 378.4398522 | 154.3420604 | 173.7184998 | 177.8204688 | 1.10E-09 | -1.00619239 | 2.05E-10 | 40.42306771 | 0.49785848 | 8.95705617 |
| ENSMUSG00000003402 | *Prkcsh* | 7761.757448 | 8483.940669 | 8426.325645 | 4069.40132 | 4066.098635 | 4137.225146 | 2.36E-51 | -1.007027294 | 4.81E-53 | 235.0149315 | 0.497570447 | 50.62708346 |
| ENSMUSG00000038650 | *Rnh1* | 24658.51864 | 26236.02637 | 29664.58487 | 13049.073 | 13430.61151 | 13564.60924 | 5.11E-32 | -1.008288463 | 2.49E-33 | 144.7076938 | 0.497135673 | 31.29157123 |
| ENSMUSG00000036882 | *Arhgap33* | 65.50680432 | 46.42651329 | 60.38933812 | 22.77177941 | 30.40073746 | 32.8581301 | 0.00928693 | -1.009655978 | 0.003864622 | 8.34635355 | 0.496664667 | 2.032127806 |
| ENSMUSG00000032417 | *Rwdd2a* | 73.49543899 | 55.50996155 | 48.31147049 | 28.67557407 | 27.14351559 | 32.8581301 | 0.007741434 | -1.010383741 | 0.003171685 | 8.70597611 | 0.496414189 | 2.111178568 |
| ENSMUSG00000027811 | *4930579G24Rik* | 790.8748326 | 753.926205 | 731.3819839 | 382.0598545 | 386.523662 | 362.4058467 | 3.24E-21 | -1.011042778 | 2.73E-22 | 94.28816458 | 0.496187475 | 20.4889921 |
| ENSMUSG00000038055 | *Dexi* | 289.1885752 | 361.3193861 | 395.885661 | 175.4270413 | 182.4044248 | 159.4585725 | 3.12E-08 | -1.011441351 | 6.61E-09 | 33.64661603 | 0.496050412 | 7.505217939 |
| ENSMUSG00000020903 | *Stx8* | 1009.763423 | 1050.652181 | 1401.032644 | 594.5964622 | 551.5562368 | 565.3531208 | 1.14E-11 | -1.012753049 | 1.79E-12 | 49.69955288 | 0.495599608 | 10.9423965 |
| ENSMUSG00000018750 | *Zbtb4* | 699.0055339 | 855.8626799 | 677.7025722 | 363.5050712 | 412.581437 | 331.4805478 | 1.22E-11 | -1.013366078 | 1.93E-12 | 49.55384356 | 0.495389063 | 10.91257915 |
| ENSMUSG00000069607 | *Cd300ld3* | 535.2385231 | 571.2479679 | 539.4780872 | 283.3821437 | 268.177934 | 262.8650408 | 4.24E-17 | -1.013901457 | 4.57E-18 | 75.05683999 | 0.49520526 | 16.37294363 |
| ENSMUSG00000022365 | *Derl1* | 10116.80695 | 10777.00672 | 11392.11314 | 5436.551483 | 5139.896112 | 5402.263154 | 6.28E-44 | -1.014190434 | 1.72E-45 | 200.3910485 | 0.495106078 | 43.20206828 |
| ENSMUSG00000033762 | *Recql4* | 1078.465681 | 1012.299844 | 885.7102924 | 535.5585156 | 469.0399494 | 469.6779773 | 2.61E-16 | -1.014722198 | 2.96E-17 | 71.3725336 | 0.49492362 | 15.58379687 |
| ENSMUSG00000024944 | *Arl2* | 1104.029312 | 1284.803292 | 1419.820438 | 637.6098233 | 628.643821 | 613.6739004 | 5.09E-17 | -1.015791973 | 5.52E-18 | 74.68721037 | 0.494556765 | 16.29305681 |
| ENSMUSG00000023909 | *Paqr4* | 387.4487816 | 427.9313399 | 378.4398522 | 173.7402429 | 201.947756 | 215.5106768 | 5.28E-11 | -1.01724262 | 8.73E-12 | 46.59475022 | 0.494059732 | 10.27715868 |
| ENSMUSG00000081251 | *Gm12164* | 204.5090476 | 202.8636777 | 233.5054407 | 112.1720986 | 84.68776864 | 117.9027021 | 1.30E-06 | -1.017827814 | 3.23E-07 | 26.10522248 | 0.493859369 | 5.885133125 |
| ENSMUSG00000038618 | *Rassf7* | 224.4806343 | 258.3736392 | 216.0596319 | 111.3286993 | 125.9459123 | 108.2385462 | 5.42E-08 | -1.020331435 | 1.17E-08 | 32.5281764 | 0.49300308 | 7.265815895 |
| ENSMUSG00000058672 | *Tubb2a* | 1236.640647 | 1234.33969 | 1421.162424 | 651.1042111 | 621.0436367 | 642.6663681 | 1.40E-23 | -1.020508709 | 1.03E-24 | 105.3436082 | 0.492942505 | 22.85322284 |
| ENSMUSG00000003161 | *Sri* | 3425.526548 | 3698.981983 | 4731.840138 | 1928.010656 | 1900.046091 | 2009.178014 | 2.30E-15 | -1.021221393 | 2.78E-16 | 66.95525185 | 0.492699054 | 14.63740058 |
| ENSMUSG00000046671 | *Mtfr1l* | 2334.279051 | 2609.977465 | 2661.156833 | 1225.459092 | 1285.516898 | 1230.247048 | 8.75E-32 | -1.022668419 | 4.31E-33 | 143.6148544 | 0.492205124 | 31.05787264 |
| ENSMUSG00000100228 | *Gm29257* | 47.13294457 | 67.62122589 | 67.09926458 | 24.45857788 | 38.00092182 | 27.05963655 | 0.009711624 | -1.023490122 | 0.004061412 | 8.25615219 | 0.491924862 | 2.012708141 |
| ENSMUSG00000034786 | *Gpsm3* | 1508.254226 | 1629.974325 | 1470.81588 | 727.8535417 | 718.7602928 | 820.4868369 | 8.32E-25 | -1.024003025 | 5.74E-26 | 111.0611902 | 0.491750006 | 24.08009676 |
| ENSMUSG00000040848 | *Sft2d2* | 4388.157026 | 4718.346732 | 4722.446241 | 2377.54245 | 2219.253835 | 2196.662639 | 3.18E-43 | -1.024484915 | 9.00E-45 | 197.0929083 | 0.491585779 | 42.49707248 |
| ENSMUSG00000060675 | *Plaat3* | 84.67952753 | 91.84375456 | 107.3588233 | 48.07375652 | 52.11554993 | 38.65662365 | 0.000470138 | -1.026508336 | 0.000159293 | 14.25904744 | 0.490896799 | 3.327774318 |
| ENSMUSG00000030407 | *Qpctl* | 723.7703014 | 808.4268945 | 681.7285281 | 360.1314743 | 361.5516276 | 366.271509 | 5.06E-18 | -1.026534116 | 5.15E-19 | 79.36880948 | 0.490888027 | 17.29593706 |
| ENSMUSG00000043091 | *Tuba1c* | 15901.37732 | 15898.05299 | 15703.91188 | 7830.118518 | 7645.785471 | 7828.932704 | 8.50E-90 | -1.027365175 | 5.42E-92 | 413.8136806 | 0.490605334 | 89.0704124 |
| ENSMUSG00000042747 | *Krtcap2* | 7474.1666 | 7578.623659 | 8124.378955 | 3834.092932 | 3616.602017 | 3913.983144 | 2.87E-47 | -1.027462413 | 6.89E-49 | 215.9586111 | 0.490572268 | 46.54262846 |
| ENSMUSG00000029545 | *Acads* | 1874.932558 | 2034.692409 | 2324.318525 | 994.3677007 | 989.1097081 | 1069.822059 | 1.98E-21 | -1.027938312 | 1.64E-22 | 95.29834817 | 0.490410471 | 20.70386115 |
| ENSMUSG00000021453 | *Gadd45g* | 243.6533575 | 249.2901909 | 213.3756614 | 97.83431152 | 131.3746155 | 118.8691177 | 2.14E-07 | -1.02947775 | 4.91E-08 | 29.75139849 | 0.489887454 | 6.668935651 |
| ENSMUSG00000024787 | *Snx15* | 989.791836 | 1157.635016 | 1131.293601 | 530.4981202 | 522.2412399 | 550.8568869 | 1.75E-21 | -1.029761713 | 1.45E-22 | 95.54318393 | 0.48979104 | 20.75602212 |
| ENSMUSG00000024388 | *Myo7b* | 59.91476005 | 62.57486574 | 56.36338224 | 23.61517864 | 33.65795933 | 30.92529892 | 0.004131845 | -1.030209005 | 0.001614063 | 9.943637416 | 0.489639209 | 2.383855962 |
| ENSMUSG00000025499 | *Hras* | 2658.617619 | 2871.37892 | 3250.288376 | 1379.801152 | 1399.519664 | 1510.507569 | 5.58E-24 | -1.03209899 | 4.02E-25 | 107.2003219 | 0.488998183 | 23.25298026 |
| ENSMUSG00000098369 | *None* | 114.2374758 | 236.1696546 | 217.4016172 | 60.72474508 | 129.2031342 | 87.94381879 | 0.007903047 | -1.032171246 | 0.003243962 | 8.664909034 | 0.488973693 | 2.102205438 |
| ENSMUSG00000037580 | *Gch1* | 3687.553765 | 3694.944895 | 4160.154404 | 1776.198794 | 1819.701285 | 2041.069728 | 5.13E-28 | -1.033354532 | 2.98E-29 | 126.0604886 | 0.488572805 | 27.29027704 |
| ENSMUSG00000002014 | *Ssr4* | 6363.74638 | 7140.599599 | 9958.872848 | 3817.224948 | 3688.260898 | 3953.606183 | 4.37E-11 | -1.033376341 | 7.17E-12 | 46.98036381 | 0.488565419 | 10.35965685 |
| ENSMUSG00000006442 | *Srm* | 9050.324221 | 9059.225725 | 8567.234101 | 4298.805912 | 4335.36231 | 4401.056602 | 3.19E-67 | -1.0335723 | 3.86E-69 | 308.8648513 | 0.488499063 | 66.49622728 |
| ENSMUSG00000035914 | *Cd276* | 1761.493945 | 1757.142601 | 1319.171542 | 770.8669028 | 804.5338021 | 790.5279535 | 8.78E-15 | -1.034258746 | 1.10E-15 | 64.23963278 | 0.488266687 | 14.0566877 |
| ENSMUSG00000030592 | *Ryr1* | 493.6976228 | 425.9127959 | 516.6643372 | 233.6215887 | 226.9197903 | 239.6710666 | 2.44E-13 | -1.034499374 | 3.39E-14 | 57.49201725 | 0.488185255 | 12.61250665 |
| ENSMUSG00000035637 | *Grhpr* | 660.6600874 | 695.3884274 | 740.7758809 | 328.9257025 | 328.9794089 | 363.3722623 | 1.26E-18 | -1.03541408 | 1.23E-19 | 82.20242924 | 0.487875831 | 17.89913242 |
| ENSMUSG00000020877 | *Scrn2* | 230.0726786 | 295.7167042 | 273.7649995 | 112.1720986 | 134.6318373 | 143.0295075 | 1.64E-07 | -1.036315862 | 3.71E-08 | 30.2964464 | 0.487570971 | 6.786170133 |
| ENSMUSG00000028944 | *Prkag2* | 810.0475558 | 824.575247 | 666.9666899 | 367.7220674 | 402.8097713 | 354.6745219 | 8.51E-15 | -1.036575411 | 1.07E-15 | 64.30199486 | 0.487483262 | 14.06982108 |
| ENSMUSG00000051319 | *Mtln* | 1221.462241 | 1340.313253 | 1940.510732 | 638.4532226 | 737.2178834 | 816.6211745 | 3.97E-08 | -1.036755565 | 8.49E-09 | 33.15939463 | 0.487422392 | 7.400878486 |
| ENSMUSG00000017754 | *Pltp* | 278.8033501 | 337.0968574 | 236.1894113 | 129.0400833 | 147.6607248 | 140.1302607 | 1.92E-07 | -1.037114652 | 4.39E-08 | 29.96858063 | 0.487301088 | 6.716015447 |
| ENSMUSG00000035585 | *Tsen34* | 1370.84971 | 1350.405974 | 1261.466174 | 653.6344089 | 594.9858617 | 691.9535632 | 4.80E-25 | -1.037464057 | 3.27E-26 | 112.1786073 | 0.487183083 | 24.31889213 |
| ENSMUSG00000070490 | *Gm10293* | 50.32839844 | 68.63049791 | 71.12522045 | 24.45857788 | 27.14351559 | 40.58945483 | 0.008640938 | -1.038228736 | 0.003573362 | 8.488816161 | 0.486924927 | 2.063439102 |
| ENSMUSG00000038503 | *Mesd* | 3622.046961 | 3721.185968 | 4467.469035 | 1906.925675 | 1927.189607 | 1909.637208 | 9.80E-25 | -1.038967041 | 6.77E-26 | 110.7332236 | 0.486675805 | 24.00899099 |
| ENSMUSG00000072582 | *Ptrh2* | 963.4293415 | 912.3819134 | 1123.241689 | 554.9566981 | 456.0110619 | 444.5511719 | 1.12E-13 | -1.039281731 | 1.52E-14 | 59.07768295 | 0.48656966 | 12.949831 |
| ENSMUSG00000031373 | *Car5b* | 516.8646633 | 585.3777763 | 554.2399254 | 292.6595353 | 243.2058997 | 267.6971187 | 1.68E-14 | -1.039430916 | 2.14E-15 | 62.93092262 | 0.486519348 | 13.77505336 |
| ENSMUSG00000036661 | *Dennd3* | 500.887394 | 519.7750945 | 367.7039699 | 227.7177941 | 210.633681 | 238.704651 | 5.22E-09 | -1.0395996 | 1.02E-09 | 37.277857 | 0.486462466 | 8.282170533 |
| ENSMUSG00000095041 | *ENSMUSG00000095041* | 3649.208319 | 4261.146503 | 4423.183521 | 2052.833743 | 2007.534413 | 1934.764013 | 3.84E-27 | -1.039686504 | 2.34E-28 | 121.9772504 | 0.486433163 | 26.4155343 |
| ENSMUSG00000042293 | *Gm5617* | 149.3874684 | 167.5391567 | 194.5878673 | 85.18332296 | 73.8303624 | 87.94381879 | 3.62E-06 | -1.04089043 | 9.44E-07 | 24.0389065 | 0.486027405 | 5.441787529 |
| ENSMUSG00000058818 | *Pirb* | 355.4942429 | 408.7551714 | 383.8077934 | 220.9706002 | 178.0614623 | 156.5593258 | 5.92E-09 | -1.042681413 | 1.17E-09 | 37.0259144 | 0.485424418 | 8.227884946 |
| ENSMUSG00000020308 | *Tpgs1* | 976.211157 | 990.0958596 | 1179.605071 | 506.8829416 | 483.1545775 | 533.4614063 | 2.93E-18 | -1.042800273 | 2.95E-19 | 80.47295852 | 0.485384427 | 17.53382199 |
| ENSMUSG00000009621 | *Vav2* | 1010.562286 | 1061.754174 | 943.4156599 | 452.0619912 | 489.6690212 | 522.8308348 | 2.29E-21 | -1.044439253 | 1.90E-22 | 95.00018634 | 0.484833317 | 20.6403026 |
| ENSMUSG00000036073 | *Galt* | 530.4453423 | 545.0068952 | 705.8842633 | 263.9839613 | 301.8358933 | 294.7567553 | 7.41E-11 | -1.046870493 | 1.24E-11 | 45.90080772 | 0.484016961 | 10.13021637 |
| ENSMUSG00000081309 | *Gm12397* | 281.998804 | 286.633256 | 253.6352201 | 129.8834825 | 157.4323904 | 112.1042086 | 4.42E-08 | -1.047500606 | 9.49E-09 | 32.94234741 | 0.483805607 | 7.354669645 |
| ENSMUSG00000028407 | *Smim27* | 77.48975633 | 74.68613008 | 81.86110278 | 37.95296568 | 29.31499684 | 45.42153278 | 0.001136077 | -1.047720687 | 0.000407389 | 12.49799413 | 0.483731809 | 2.944592321 |
| ENSMUSG00000028041 | *Adam15* | 16589.99763 | 17817.68838 | 15525.42784 | 7825.058123 | 8274.429292 | 8056.040368 | 6.92E-44 | -1.047909094 | 1.90E-45 | 200.1891573 | 0.48366864 | 43.15988338 |
| ENSMUSG00000003762 | *Coq8b* | 939.4634375 | 881.0944806 | 915.2339688 | 449.5317934 | 419.0958807 | 453.2489122 | 2.33E-25 | -1.048626636 | 1.56E-26 | 113.6437745 | 0.483428142 | 24.63298568 |
| ENSMUSG00000097989 | *Gm4335* | 77.48975633 | 76.70467414 | 100.6488969 | 48.07375652 | 29.31499684 | 44.45511719 | 0.001901943 | -1.048811657 | 0.000704473 | 11.47740794 | 0.483366147 | 2.720802553 |
| ENSMUSG00000048707 | *Tprn* | 1400.407658 | 1432.157008 | 1150.081395 | 649.4174127 | 640.5869679 | 636.8678746 | 1.40E-19 | -1.049631991 | 1.29E-20 | 86.65076263 | 0.483091378 | 18.85440219 |
| ENSMUSG00000030431 | *Tmem238* | 417.0067299 | 513.7194623 | 464.3269109 | 228.5611933 | 207.3764591 | 235.8054042 | 2.02E-12 | -1.051169588 | 3.00E-13 | 53.20933839 | 0.482576782 | 11.6944742 |
| ENSMUSG00000027931 | *Npr1* | 49.52953497 | 59.54704966 | 56.36338224 | 31.20577178 | 23.88629372 | 24.16038978 | 0.005012446 | -1.051577497 | 0.001987939 | 9.560639138 | 0.482440358 | 2.299950316 |
| ENSMUSG00000025372 | *Baiap2* | 801.2600577 | 821.5474309 | 862.8965424 | 425.9166148 | 375.6662558 | 394.2975612 | 4.40E-22 | -1.051851371 | 3.55E-23 | 98.32655836 | 0.482348782 | 21.35616274 |
| ENSMUSG00000019478 | *Rab4a* | 102.2545238 | 105.973563 | 91.25499982 | 57.35114813 | 41.2581437 | 45.42153278 | 0.000205786 | -1.052106402 | 6.66E-05 | 15.90462875 | 0.482263523 | 3.686584746 |
| ENSMUSG00000021485 | *Mxd3* | 109.444295 | 89.8252105 | 80.51911749 | 41.32656262 | 40.17240307 | 54.1192731 | 0.000604394 | -1.052401043 | 0.000208206 | 13.75553682 | 0.482165041 | 3.218680073 |
| ENSMUSG00000002274 | *Metrn* | 1594.531481 | 1646.122678 | 1740.554923 | 786.8914883 | 753.5039928 | 858.1770449 | 2.82E-29 | -1.05263393 | 1.55E-30 | 131.9242102 | 0.482087213 | 28.54976322 |
| ENSMUSG00000009108 | *Gnat2* | 115.8352028 | 66.61195386 | 95.2809557 | 49.760555 | 44.51536557 | 39.62303924 | 0.00196248 | -1.054073867 | 0.000728286 | 11.41563539 | 0.481606288 | 2.707194717 |
| ENSMUSG00000013033 | *Adgrl1* | 254.8374461 | 272.5034476 | 181.1680144 | 121.4494902 | 109.659803 | 111.137793 | 2.11E-06 | -1.054437084 | 5.38E-07 | 25.12202147 | 0.481485053 | 5.675465652 |
| ENSMUSG00000023008 | *Fmnl3* | 3463.871994 | 3535.479915 | 3303.967788 | 1651.375707 | 1705.69852 | 1605.216297 | 2.79E-49 | -1.054927975 | 6.05E-51 | 225.3846723 | 0.48132125 | 48.55491974 |
| ENSMUSG00000114547 | *Gm3226* | 225.2794978 | 190.7524133 | 179.8260291 | 59.88134584 | 97.71665612 | 130.4661048 | 0.000645206 | -1.055267419 | 0.000223217 | 13.62479831 | 0.481208016 | 3.190301703 |
| ENSMUSG00000042642 | *Flad1* | 1481.891732 | 1538.130571 | 1543.283085 | 740.5045303 | 699.2169616 | 752.8377455 | 6.68E-36 | -1.055907771 | 2.71E-37 | 162.8411791 | 0.480994475 | 35.17502284 |
| ENSMUSG00000039959 | *Hip1* | 2832.769855 | 2822.933863 | 2100.206981 | 1271.002651 | 1247.515976 | 1212.851567 | 5.72E-16 | -1.056881837 | 6.63E-17 | 69.77923618 | 0.480669831 | 15.24265345 |
| ENSMUSG00000031445 | *Proz* | 46.3340811 | 69.63976994 | 52.33742637 | 21.92838017 | 28.22925621 | 30.92529892 | 0.008313346 | -1.056997421 | 0.003427587 | 8.564618923 | 0.480631323 | 2.080224137 |
| ENSMUSG00000004558 | *Ndrg2* | 2549.972188 | 2847.156391 | 3160.375362 | 1347.751981 | 1359.347261 | 1401.302607 | 1.54E-25 | -1.057213616 | 1.02E-26 | 114.4888513 | 0.480559304 | 24.81307049 |
| ENSMUSG00000039195 | *Bbln* | 750.1327958 | 925.5024498 | 997.0950716 | 426.760014 | 395.209587 | 457.1145746 | 2.13E-13 | -1.059547428 | 2.94E-14 | 57.7718253 | 0.479782543 | 12.67206902 |
| ENSMUSG00000070476 | *Fam217b* | 310.7578888 | 298.7445203 | 287.1848524 | 145.0646688 | 153.0894279 | 132.398936 | 3.05E-11 | -1.061740357 | 4.96E-12 | 47.70239188 | 0.479053817 | 10.51507551 |
| ENSMUSG00000021610 | *Clptm1l* | 9875.550183 | 10588.27285 | 11039.17101 | 5132.084359 | 4906.461878 | 5035.02523 | 1.05E-51 | -1.062908552 | 2.09E-53 | 236.6746279 | 0.478666069 | 50.9788082 |
| ENSMUSG00000015980 | *Lrrc27* | 340.3158371 | 377.4677385 | 411.9894845 | 188.9214291 | 169.3755373 | 179.7532999 | 7.27E-12 | -1.063728234 | 1.13E-12 | 50.61121977 | 0.478394187 | 11.1382459 |
| ENSMUSG00000019578 | *Ubxn6* | 3176.281146 | 3319.495701 | 3923.964992 | 1581.37357 | 1717.641666 | 1682.529544 | 2.13E-24 | -1.063867217 | 1.50E-25 | 109.1617187 | 0.478348103 | 23.67168338 |
| ENSMUSG00000025268 | *Maged2* | 85.478391 | 83.76957834 | 146.2763968 | 48.07375652 | 56.45851243 | 45.42153278 | 0.002034434 | -1.064176691 | 0.000756189 | 11.34579712 | 0.478245503 | 2.691556325 |
| ENSMUSG00000050002 | *Idnk* | 728.5634822 | 735.7593085 | 865.580513 | 346.6370865 | 377.837737 | 387.532652 | 3.11E-17 | -1.0651924 | 3.33E-18 | 75.68476669 | 0.47790892 | 16.5077231 |
| ENSMUSG00000005447 | *Pafah1b3* | 786.0816518 | 752.916933 | 873.6324248 | 389.6504476 | 382.1806995 | 378.8349117 | 4.15E-21 | -1.065384679 | 3.51E-22 | 93.79066109 | 0.47784523 | 20.38196318 |
| ENSMUSG00000017697 | *Ada* | 454.5533129 | 451.1445966 | 579.7376459 | 222.6573986 | 245.3773809 | 239.6710666 | 1.61E-11 | -1.066463708 | 2.57E-12 | 48.99458918 | 0.47748797 | 10.79249061 |
| ENSMUSG00000014686 | *Ceacam16* | 125.4215644 | 150.3815322 | 174.4580879 | 70.00213669 | 79.25906552 | 64.74984461 | 1.66E-05 | -1.067529796 | 4.65E-06 | 20.97637308 | 0.477135258 | 4.781063187 |
| ENSMUSG00000027488 | *Snta1* | 486.5078516 | 548.0347113 | 566.317793 | 238.6819841 | 256.2347872 | 267.6971187 | 2.26E-15 | -1.067951428 | 2.72E-16 | 66.99397444 | 0.476995834 | 14.64550556 |
| ENSMUSG00000031398 | *Plxna3* | 127.0192913 | 110.0106511 | 84.54507337 | 43.0133611 | 46.68684681 | 64.74984461 | 0.000728804 | -1.067964892 | 0.000254118 | 13.38150462 | 0.476991383 | 3.137389477 |
| ENSMUSG00000002661 | *Alkbh7* | 441.7714974 | 476.3763973 | 489.8246314 | 221.8139994 | 206.2907185 | 241.6038978 | 7.90E-15 | -1.068413749 | 9.88E-16 | 64.45511922 | 0.476843002 | 14.10234461 |
| ENSMUSG00000032012 | *Nectin1* | 1360.464485 | 1310.035093 | 919.2599247 | 558.330295 | 562.413643 | 591.4463418 | 3.33E-11 | -1.07062301 | 5.42E-12 | 47.52759438 | 0.476113351 | 10.47776996 |
| ENSMUSG00000039450 | *Dcxr* | 520.0601172 | 539.9605351 | 560.9498519 | 248.802775 | 256.2347872 | 265.7642876 | 1.35E-18 | -1.07107529 | 1.32E-19 | 82.05923466 | 0.475964114 | 17.86871978 |
| ENSMUSG00000110234 | *Gm45799* | 517.6635268 | 566.2016078 | 668.3086752 | 277.478349 | 238.8629372 | 314.0850671 | 7.43E-11 | -1.072234753 | 1.25E-11 | 45.89320789 | 0.475581746 | 10.12873241 |
| ENSMUSG00000029475 | *Kdm2b* | 2215.248395 | 2193.148117 | 1643.931982 | 962.3185297 | 956.5374894 | 960.6170976 | 3.14E-16 | -1.073384852 | 3.58E-17 | 70.99789153 | 0.475202769 | 15.50336049 |
| ENSMUSG00000039063 | *Echdc3* | 150.9851953 | 133.2239077 | 106.016838 | 64.94174127 | 57.54425305 | 63.78342901 | 3.57E-05 | -1.073405273 | 1.05E-05 | 19.42337321 | 0.475196043 | 4.446796119 |
| ENSMUSG00000022856 | *Tmem41a* | 365.879468 | 357.282298 | 326.1024258 | 193.1384253 | 163.9468342 | 141.0966763 | 8.04E-10 | -1.07398363 | 1.47E-10 | 41.07141889 | 0.475005581 | 9.094735791 |
| ENSMUSG00000038119 | *Cdon* | 83.88066407 | 63.58413777 | 55.02139695 | 28.67557407 | 33.65795933 | 34.79096128 | 0.002826324 | -1.074496638 | 0.00107355 | 10.69621476 | 0.474836704 | 2.548777981 |
| ENSMUSG00000042155 | *Klhl23* | 211.6988188 | 192.7709574 | 205.3237496 | 96.99091228 | 99.88813737 | 92.77589675 | 9.27E-09 | -1.075340226 | 1.87E-09 | 36.1083858 | 0.474559134 | 8.03311113 |
| ENSMUSG00000026853 | *Crat* | 1026.539555 | 999.1793079 | 944.7576452 | 461.3393828 | 465.7827275 | 483.2077956 | 3.86E-28 | -1.076669511 | 2.24E-29 | 126.632413 | 0.474122081 | 27.4132289 |
| ENSMUSG00000069792 | *Wfdc17* | 315.5510696 | 361.3193861 | 546.1880136 | 183.0176345 | 197.6047935 | 197.1487806 | 3.29E-06 | -1.076794688 | 8.55E-07 | 24.22916182 | 0.474080945 | 5.482824548 |
| ENSMUSG00000017718 | *Afmid* | 99.05906994 | 80.74176225 | 128.830588 | 49.760555 | 42.34388432 | 53.15285751 | 0.000518788 | -1.076828906 | 0.000176971 | 14.06105164 | 0.474069701 | 3.285010267 |
| ENSMUSG00000039648 | *Kyat1* | 83.0818006 | 84.77885036 | 68.44124987 | 38.79636491 | 35.82944058 | 37.69020805 | 0.00042823 | -1.077396135 | 0.000144032 | 14.44865856 | 0.473883346 | 3.368322504 |
| ENSMUSG00000035559 | *Mpv17l2* | 1161.547481 | 1247.460227 | 1260.124189 | 601.3436561 | 583.0427149 | 551.8233025 | 1.17E-29 | -1.077574619 | 6.34E-31 | 133.7042787 | 0.473824723 | 28.9307169 |

| | | | | | | | | | | | | | |
|---|---|---|---|---|---|---|---|---|---|---|---|---|---|
| ENSMUSG00000037822 | *Smim14* | 1596.928071 | 1598.686893 | 1626.486173 | 739.6611311 | 811.0482458 | 734.4758493 | 3.96E-36 | -1.078334802 | 1.59E-37 | 163.9041297 | 0.473575121 | 35.4021487 |
| ENSMUSG00000025138 | *Sirt7* | 1852.564381 | 1964.043367 | 1963.324481 | 900.7503854 | 935.9084175 | 898.7664997 | 3.26E-42 | -1.078924129 | 9.80E-44 | 192.3411717 | 0.47338171 | 41.48723437 |
| ENSMUSG00000035228 | *Ccdc106* | 674.2407664 | 714.5645959 | 783.7194102 | 337.3596949 | 350.6942214 | 338.2454569 | 3.45E-20 | -1.079376057 | 3.09E-21 | 89.48505732 | 0.473233445 | 19.46177637 |
| ENSMUSG00000050957 | *Insl6* | 151.7840588 | 98.90865876 | 182.5099996 | 64.94174127 | 62.97295617 | 76.3468317 | 0.000406331 | -1.079692462 | 0.000136259 | 14.55317177 | 0.473129669 | 3.391119833 |
| ENSMUSG00000046245 | *Pilra* | 167.7613281 | 160.4742525 | 224.1115437 | 89.40031915 | 91.20221238 | 79.24607847 | 3.41E-06 | -1.080872569 | 8.87E-07 | 24.15838485 | 0.472742813 | 5.467652372 |
| ENSMUSG00000107476 | *Zfp862-ps* | 1086.454316 | 1099.097239 | 987.7011746 | 479.0507667 | 499.4406868 | 522.8308348 | 3.82E-26 | -1.082235698 | 2.46E-27 | 117.3042018 | 0.472296354 | 25.41799974 |
| ENSMUSG00000040713 | *Creg1* | 7935.909684 | 9768.74396 | 12930.02828 | 4695.203554 | 4890.175769 | 4874.600242 | 1.27E-11 | -1.082746351 | 2.01E-12 | 49.47530617 | 0.47212921 | 10.8958483 |
| ENSMUSG00000045948 | *Mrps12* | 766.9089286 | 722.6387721 | 833.372866 | 338.2030941 | 373.4947745 | 384.6334053 | 1.63E-19 | -1.082960736 | 1.51E-20 | 86.34127921 | 0.472059057 | 18.78782055 |
| ENSMUSG00000025494 | *Sigirr* | 523.2555711 | 526.8399987 | 645.4949252 | 277.478349 | 252.9775653 | 266.7307032 | 4.63E-14 | -1.083317697 | 6.10E-15 | 60.86903102 | 0.471942271 | 13.3343417 |
| ENSMUSG00000050382 | *Kif7* | 189.3306417 | 213.96567 | 144.9344115 | 80.12292754 | 90.11647176 | 89.87664997 | 4.13E-06 | -1.08360648 | 1.08E-06 | 23.77177776 | 0.471847813 | 5.384167313 |
| ENSMUSG00000021280 | *Exoc3l4* | 129.4158817 | 125.1497315 | 182.5099996 | 55.66434966 | 70.57314053 | 79.24607847 | 0.000110917 | -1.084827771 | 3.46E-05 | 17.14732431 | 0.471448546 | 3.955000924 |
| ENSMUSG00000021484 | *Lman2* | 14303.65038 | 14528.47085 | 16176.2907 | 6911.656749 | 7089.886272 | 7213.325972 | 1.17E-48 | -1.084864899 | 2.63E-50 | 222.4580902 | 0.471436414 | 47.93054298 |
| ENSMUSG00000053687 | *Dpep2* | 2186.48931 | 2327.381297 | 2591.373598 | 1090.515214 | 1167.17117 | 1089.150371 | 2.15E-29 | -1.085052969 | 1.18E-30 | 132.4732675 | 0.471374961 | 28.66661107 |
| ENSMUSG00000033751 | *Gadd45gip1* | 1550.59399 | 1481.611337 | 1556.702938 | 706.7685608 | 689.445296 | 765.4011482 | 2.58E-34 | -1.085574221 | 1.13E-35 | 155.4216511 | 0.471204682 | 33.58792201 |
| ENSMUSG00000062661 | *Ncs1* | 1282.974728 | 1486.657697 | 1284.279924 | 629.175831 | 690.5310366 | 591.4463418 | 7.90E-22 | -1.086049625 | 6.45E-23 | 97.14157448 | 0.471049434 | 21.10226866 |
| ENSMUSG00000034880 | *Mrpl34* | 745.339615 | 954.7713386 | 1088.350071 | 419.1694209 | 456.0110619 | 435.8534316 | 8.64E-12 | -1.086243837 | 1.34E-12 | 50.26558187 | 0.470986027 | 11.0635749 |
| ENSMUSG00000020262 | *Adarb1* | 305.1658445 | 304.8001525 | 228.1374996 | 138.3174749 | 129.2031342 | 128.5332736 | 1.63E-08 | -1.086256883 | 3.36E-09 | 34.96184045 | 0.470981768 | 7.788787044 |
| ENSMUSG00000020571 | *Pdia6* | 31920.18756 | 34417.18543 | 43309.89131 | 17223.05582 | 17150.35889 | 17194.4662 | 1.06E-20 | -1.088195456 | 9.14E-22 | 91.89448623 | 0.470349327 | 19.97635683 |
| ENSMUSG00000035845 | *Alg12* | 1034.52819 | 1119.282679 | 932.6797776 | 460.4959835 | 483.1545775 | 510.2674321 | 2.82E-21 | -1.088382821 | 2.35E-22 | 94.58029658 | 0.470288246 | 20.55033436 |
| ENSMUSG00000025875 | *Tspan17* | 262.8260807 | 258.3736392 | 292.5527936 | 123.9796879 | 131.3746155 | 126.6004424 | 6.97E-11 | -1.089114518 | 1.17E-11 | 46.02805942 | 0.470049788 | 10.15694522 |
| ENSMUSG00000040093 | *Bmf* | 74.29430246 | 98.90865876 | 104.6748527 | 38.79636491 | 48.85832806 | 42.52228601 | 0.000406589 | -1.090443854 | 0.000136369 | 14.5516462 | 0.469616872 | 3.390844617 |
| ENSMUSG00000062619 | *2310039H08Rik* | 237.2624498 | 231.1232944 | 261.6871318 | 114.7022963 | 103.1453592 | 123.7011957 | 1.66E-09 | -1.090898968 | 3.12E-10 | 39.60043529 | 0.469468749 | 8.780749391 |
| ENSMUSG00000030035 | *Wbp1* | 746.9373419 | 776.1301896 | 885.7102924 | 387.9636491 | 386.523662 | 353.7081064 | 3.07E-19 | -1.091081172 | 2.90E-20 | 85.05521446 | 0.469409462 | 18.51329713 |
| ENSMUSG00000032182 | *Yipf2* | 911.5032162 | 981.0124113 | 1005.146983 | 414.9524247 | 477.7258744 | 466.7787305 | 3.90E-23 | -1.092276903 | 2.93E-24 | 103.2639453 | 0.469020568 | 22.40911775 |
| ENSMUSG00000039745 | *Htatip2* | 2483.66652 | 2675.580147 | 3452.928155 | 1360.40297 | 1276.830973 | 1397.436945 | 5.95E-16 | -1.092363961 | 6.93E-17 | 69.69364298 | 0.468992267 | 15.22513182 |
| ENSMUSG00000024610 | *Cd74* | 15078.54795 | 15735.56019 | 14553.83049 | 7119.132961 | 7038.856463 | 7108.953088 | 9.78E-75 | -1.093178346 | 9.30E-77 | 343.8413969 | 0.4687276 | 74.00951364 |
| ENSMUSG00000114380 | *Gm47942* | 151.7840588 | 165.5206126 | 242.8993378 | 81.80972601 | 94.45943425 | 85.04457202 | 2.73E-05 | -1.09334111 | 7.89E-06 | 19.96402653 | 0.468674722 | 4.563234782 |
| ENSMUSG00000069808 | *Tlcd3a* | 677.4362203 | 737.7778526 | 783.7194102 | 339.0464934 | 335.4938527 | 353.7081064 | 7.67E-21 | -1.093480431 | 6.58E-22 | 92.54514854 | 0.468629464 | 20.1149824 |
| ENSMUSG00000026814 | *Eng* | 664.6544048 | 666.1195386 | 543.5040431 | 276.6349498 | 301.8358933 | 302.48808 | 2.40E-15 | -1.094033536 | 2.90E-16 | 66.86862881 | 0.468449834 | 14.61916724 |
| ENSMUSG00000095687 | *Rnaset2a* | 1281.377002 | 1433.16628 | 1673.455659 | 716.0459524 | 667.7304835 | 666.8267579 | 1.36E-18 | -1.094515513 | 1.32E-19 | 82.05402212 | 0.46829336 | 17.86783691 |
| ENSMUSG00000031803 | *B3gnt3* | 301.9703906 | 302.7816084 | 249.6092642 | 133.2570795 | 122.6886905 | 144.9623387 | 8.25E-10 | -1.094747639 | 1.51E-10 | 41.02024989 | 0.468218019 | 9.083786733 |
| ENSMUSG00000047617 | *Paxx* | 214.8942727 | 224.0583902 | 284.5008818 | 109.6419008 | 106.4025811 | 120.8019489 | 5.64E-08 | -1.095660215 | 1.22E-08 | 32.44839004 | 0.467921941 | 7.248813242 |
| ENSMUSG00000064179 | *Tnnt1* | 64.70794085 | 46.42651329 | 85.88705866 | 27.83217483 | 39.08666245 | 25.12680537 | 0.008198047 | -1.096153096 | 0.003371824 | 8.594485776 | 0.467762108 | 2.086289611 |
| ENSMUSG00000037904 | *Ankrd9* | 202.1124572 | 199.8358616 | 233.5054407 | 100.3645092 | 112.9170249 | 83.11174084 | 1.28E-07 | -1.098783293 | 2.87E-08 | 30.79113552 | 0.466910102 | 6.893501628 |
| ENSMUSG00000025577 | *Cbx2* | 614.3260063 | 541.9790791 | 471.0368373 | 265.6707597 | 254.0633059 | 241.6038978 | 2.51E-13 | -1.099679727 | 3.49E-14 | 57.43439035 | 0.466620072 | 12.60015087 |
| ENSMUSG00000026566 | *Mpzl1* | 697.4078069 | 770.0745575 | 783.7194102 | 363.5050712 | 343.094037 | 341.1447037 | 4.88E-22 | -1.100020026 | 3.94E-23 | 98.11710402 | 0.46651002 | 21.31181415 |
| ENSMUSG00000026202 | *Tuba4a* | 6257.497539 | 6402.821747 | 6334.170576 | 2998.284288 | 2986.872455 | 2874.119968 | 9.41E-77 | -1.100197536 | 8.44E-79 | 353.2174096 | 0.466452624 | 76.02631553 |
| ENSMUSG00000024942 | *Capn1* | 2379.015406 | 2558.504591 | 2411.547569 | 1171.481541 | 1088.997845 | 1164.530787 | 1.60E-43 | -1.100594881 | 4.47E-45 | 198.4872157 | 0.466324172 | 42.79680098 |
| ENSMUSG00000062861 | *Zfp28* | 169.3590551 | 233.1418385 | 236.1894113 | 96.99091228 | 100.973878 | 98.5743903 | 6.08E-07 | -1.101314999 | 1.46E-07 | 27.64809448 | 0.466091465 | 6.216066131 |
| ENSMUSG00000018102 | *H2bc4* | 121.427247 | 130.1960916 | 182.5099996 | 70.00213669 | 61.88721554 | 68.61550697 | 3.64E-05 | -1.102696873 | 1.07E-05 | 19.38537902 | 0.465645237 | 4.438505901 |
| ENSMUSG00000025366 | *Esyt1* | 9790.071792 | 9424.582199 | 7888.189544 | 4087.956103 | 4328.847866 | 4207.773484 | 2.57E-29 | -1.102823822 | 1.41E-30 | 132.1113508 | 0.465604265 | 28.58977263 |
| ENSMUSG00000073471 | *Rsph3a* | 202.1124572 | 261.4014553 | 327.4444111 | 150.1250642 | 59.7157343 | 156.5593258 | 0.002981055 | -1.103017675 | 0.001136897 | 10.59018362 | 0.465541706 | 2.525629947 |
| ENSMUSG00000026956 | *Uap1l1* | 8107.66533 | 8876.547487 | 10319.86689 | 4217.839585 | 4247.417319 | 4238.698783 | 5.42E-28 | -1.103324989 | 3.16E-29 | 125.9481091 | 0.46544255 | 27.26612366 |
| ENSMUSG00000007950 | *Abhd8* | 905.1123084 | 996.1514918 | 1085.666101 | 458.8091851 | 415.8386588 | 512.2002633 | 8.28E-18 | -1.103631522 | 8.54E-19 | 78.37060149 | 0.465343667 | 17.08199689 |
| ENSMUSG00000058135 | *Gstm1* | 384.2533278 | 558.1274316 | 453.5910285 | 221.8139994 | 224.7483091 | 201.9808585 | 1.19E-09 | -1.104626575 | 2.21E-10 | 40.27122118 | 0.465022821 | 8.924683466 |
| ENSMUSG00000025507 | *Pidd1* | 633.4987296 | 648.9619141 | 424.0673521 | 260.6103643 | 256.2347872 | 278.3276902 | 6.13E-09 | -1.105593843 | 1.21E-09 | 36.95029188 | 0.464711147 | 8.212340164 |
| ENSMUSG00000047180 | *Neurl3* | 1447.540603 | 1459.407353 | 1171.553159 | 630.0192302 | 648.1871523 | 620.4388095 | 7.87E-21 | -1.105643704 | 6.76E-22 | 92.49278393 | 0.464695087 | 20.10378999 |
| ENSMUSG00000019738 | *Polr2i* | 906.7100354 | 916.4190015 | 1107.137866 | 440.2544018 | 456.0110619 | 462.9130682 | 1.31E-19 | -1.105674225 | 1.21E-20 | 86.78654482 | 0.464685256 | 18.88227758 |
| ENSMUSG00000045917 | *Tmem268* | 947.4520722 | 1074.87471 | 1014.54088 | 477.3639683 | 479.8973556 | 452.2824966 | 2.76E-26 | -1.106396037 | 1.76E-27 | 117.96952 | 0.464452822 | 25.5583126 |
| ENSMUSG00000067071 | *Hes6* | 561.6010175 | 665.1102665 | 719.3041163 | 271.5745544 | 296.4071902 | 334.3797945 | 4.06E-13 | -1.106511517 | 5.73E-14 | 56.46341725 | 0.464415646 | 12.39194157 |
| ENSMUSG00000009772 | *Nuak2* | 667.0509952 | 769.0652854 | 617.3132341 | 317.1181132 | 313.7790402 | 323.749223 | 3.59E-17 | -1.10653257 | 3.86E-18 | 75.39381747 | 0.464408869 | 16.44516203 |
| ENSMUSG00000090639 | *None* | 628.7055488 | 583.3592323 | 622.6811753 | 232.7781895 | 279.0353403 | 340.1782881 | 1.20E-11 | -1.108576044 | 1.89E-12 | 49.59577143 | 0.463751533 | 10.9208909 |
| ENSMUSG00000022571 | *Pycrl* | 1894.105281 | 2100.295091 | 2500.118598 | 1014.609282 | 979.3380425 | 1012.80354 | 2.28E-20 | -1.108971974 | 2.01E-21 | 90.33692946 | 0.46362428 | 19.64184167 |
| ENSMUSG00000003228 | *Grk5* | 421.7999107 | 502.61747 | 436.1452197 | 208.3196116 | 235.6057153 | 187.4846247 | 1.70E-12 | -1.109309544 | 2.50E-13 | 53.56640656 | 0.463515811 | 11.76978814 |
| ENSMUSG00000057337 | *Chst3* | 573.5839695 | 567.2108798 | 289.868823 | 225.1875963 | 212.8051622 | 226.1412483 | 2.28E-05 | -1.110931228 | 6.51E-06 | 20.33279229 | 0.462995081 | 4.642320949 |
| ENSMUSG00000019528 | *Gyg* | 2788.033501 | 2900.647809 | 3403.274699 | 1387.391745 | 1460.321139 | 1358.780321 | 1.38E-27 | -1.110936473 | 8.15E-29 | 124.0658341 | 0.462993398 | 26.86029038 |
| ENSMUSG00000026411 | *Tmem9* | 685.4248549 | 831.6401512 | 916.5759541 | 399.7712384 | 340.9225558 | 380.7677429 | 7.51E-14 | -1.112936105 | 1.00E-14 | 59.89455805 | 0.462352115 | 13.12437031 |
| ENSMUSG00000029145 | *Eif2b4* | 2121.781369 | 2112.406355 | 2371.28801 | 953.8845373 | 1054.254145 | 1042.762423 | 3.17E-33 | -1.114083665 | 1.47E-34 | 150.3229642 | 0.461984494 | 32.49859085 |
| ENSMUSG00000049411 | *Tmem241* | 444.9669513 | 411.7829875 | 472.3788226 | 204.9460146 | 204.1192372 | 203.9136897 | 3.15E-16 | -1.11457991 | 3.59E-17 | 70.99050569 | 0.461825612 | 15.50195953 |
| ENSMUSG00000030680 | *Pagr1a* | 524.0544345 | 444.0796924 | 560.9498519 | 209.1630108 | 281.2068215 | 216.4770924 | 1.71E-10 | -1.115453221 | 2.94E-11 | 44.21808874 | 0.461546138 | 9.768085199 |
| ENSMUSG00000008822 | *Acyp1* | 203.7101842 | 252.318007 | 402.5955875 | 135.7872772 | 136.8033186 | 121.7683645 | 6.72E-05 | -1.115503843 | 2.04E-05 | 18.15465442 | 0.461529944 | 4.172595258 |
| ENSMUSG00000037922 | *Bank1* | 664.6544048 | 613.6373931 | 870.9484542 | 338.2030941 | 321.3792246 | 328.581301 | 5.09E-12 | -1.11699966 | 7.78E-13 | 51.3363146 | 0.461051668 | 11.29301017 |
| ENSMUSG00000067714 | *Lpar5* | 400.2305971 | 472.3393092 | 366.3619846 | 172.8968436 | 208.4621997 | 191.350287 | 2.59E-11 | -1.118075903 | 4.19E-12 | 48.03551057 | 0.460707854 | 10.58735856 |
| ENSMUSG00000029073 | *Cptp* | 770.9032459 | 859.899768 | 806.5331602 | 391.3372461 | 359.3801464 | 370.1371714 | 4.07E-24 | -1.118518097 | 2.90E-25 | 107.8469671 | 0.460566666 | 23.39048323 |
| ENSMUSG00000018909 | *Arrb1* | 6172.818012 | 6547.147647 | 5465.906092 | 2862.497011 | 2803.38229 | 2710.795733 | 3.54E-36 | -1.118829048 | 1.41E-37 | 164.145319 | 0.460467408 | 35.45098311 |
| ENSMUSG00000030727 | *Rabep2* | 1013.75774 | 1129.375399 | 998.4370569 | 471.4601736 | 469.0399494 | 505.4353542 | 2.90E-26 | -1.119957673 | 1.85E-27 | 117.8740254 | 0.460107324 | 25.53780555 |
| ENSMUSG00000019158 | *Tmem160* | 1938.841635 | 2021.571872 | 2615.529333 | 1003.645092 | 972.8235987 | 1040.829592 | 1.49E-17 | -1.122074561 | 1.56E-18 | 77.17828848 | 0.459432697 | 16.82600827 |
| ENSMUSG00000030257 | *Srgap3* | 742.1441611 | 798.3341743 | 487.1406608 | 323.8653071 | 309.4360777 | 299.5888332 | 1.26E-08 | -1.122932285 | 2.57E-09 | 35.48172961 | 0.459159632 | 7.901238978 |
| ENSMUSG00000032415 | *Ube2cbp* | 93.46702567 | 84.77885036 | 108.7008086 | 44.70015957 | 36.9151812 | 49.28719515 | 0.000123575 | -1.123423274 | 3.88E-05 | 16.92745089 | 0.459003394 | 3.908070427 |
| ENSMUSG00000029499 | *Pxmp2* | 34.35112909 | 45.41724127 | 45.62749991 | 20.24158169 | 19.54333122 | 17.39548064 | 0.009934279 | -1.123731003 | 0.004163321 | 8.211173726 | 0.458905498 | 2.002863652 |
| ENSMUSG00000001435 | *Col18a1* | 225.2794978 | 265.4385434 | 222.7695584 | 118.0758932 | 99.88813737 | 108.2385462 | 2.90E-09 | -1.126450921 | 5.56E-10 | 38.47093846 | 0.458041137 | 8.538254222 |
| ENSMUSG00000042210 | *Abhd14a* | 147.7897414 | 160.4742525 | 178.4840438 | 70.84553593 | 90.11647176 | 61.85059783 | 2.98E-06 | -1.127698446 | 7.69E-07 | 24.43349482 | 0.457645232 | 5.526420762 |
| ENSMUSG00000028479 | *Gne* | 1302.946315 | 1399.860303 | 1384.928821 | 641.8268195 | 614.5291929 | 611.7410692 | 1.02E-36 | -1.127980779 | 3.94E-38 | 166.6740951 | 0.45755568 | 35.9902788 |
| ENSMUSG00000035711 | *Dok3* | 2943.811877 | 3390.144743 | 3722.667199 | 1471.731669 | 1539.580204 | 1586.854401 | 3.80E-26 | -1.128041158 | 2.44E-27 | 117.3202524 | 0.457536531 | 25.42055213 |
| ENSMUSG00000053907 | *Mat2a* | 15680.09214 | 15714.36548 | 17479.35842 | 7663.125469 | 7294.005509 | 7375.683791 | 6.02E-55 | -1.129520531 | 1.07E-56 | 251.7679461 | 0.457067603 | 54.22053374 |
| ENSMUSG00000032204 | *Aqp9* | 1266.997459 | 1342.331797 | 1076.272204 | 563.3906905 | 539.6130899 | 581.7821859 | 7.08E-22 | -1.13096336 | 5.76E-23 | 97.36750705 | 0.456610721 | 21.149937 |
| ENSMUSG00000020268 | *Lyrm7* | 79.08748326 | 50.46360141 | 68.44124987 | 28.67557407 | 39.08666245 | 23.19397419 | 0.004209765 | -1.131315813 | 0.001645992 | 9.907581943 | 0.456499184 | 2.375742112 |
| ENSMUSG00000033857 | *Engase* | 402.6271875 | 382.5140987 | 299.26272 | 168.6798474 | 160.6896123 | 167.1898973 | 9.93E-11 | -1.13173529 | 1.69E-11 | 45.30355896 | 0.456366472 | 10.0028511 |
| ENSMUSG00000110206 | *Flt3l* | 270.8147154 | 253.3272791 | 224.1115437 | 116.3890947 | 109.659803 | 115.9698709 | 2.88E-10 | -1.133273862 | 5.07E-11 | 43.14866402 | 0.455880036 | 9.540188077 |
| ENSMUSG00000028470 | *Hint2* | 643.0850912 | 645.934098 | 857.5286013 | 325.5521056 | 324.6364464 | 325.6820542 | 1.34E-13 | -1.133282182 | 1.82E-14 | 58.71878051 | 0.455877407 | 12.87345257 |
| ENSMUSG00000006219 | *Fblim1* | 587.1646485 | 732.7314924 | 709.9102192 | 309.5275201 | 299.6644121 | 314.0850671 | 9.05E-17 | -1.133329046 | 9.95E-18 | 73.52307372 | 0.455862599 | 16.04351965 |
| ENSMUSG00000020284 | *Cfap410* | 445.7658147 | 478.3949413 | 546.1880136 | 198.1988207 | 234.5199747 | 236.7718198 | 1.60E-13 | -1.133793513 | 2.18E-14 | 58.3616062 | 0.45571586 | 12.79667949 |
| ENSMUSG00000031168 | *Ebp* | 2317.502919 | 2383.90053 | 2717.520215 | 1193.409921 | 1096.59803 | 1085.284709 | 7.99E-31 | -1.134118592 | 4.09E-32 | 139.1464672 | 0.455613187 | 30.09763821 |
| ENSMUSG00000053897 | *Slc39a8* | 117.4329297 | 122.1219154 | 162.3802203 | 48.07375652 | 77.08758427 | 57.98493547 | 0.000109613 | -1.134939853 | 3.42E-05 | 17.17087141 | 0.4553539 | 3.960137968 |
| ENSMUSG00000017466 | *Timp2* | 73.49543899 | 52.48214546 | 56.36338224 | 22.77177941 | 24.97203434 | 35.75737687 | 0.003812867 | -1.135343782 | 0.001481806 | 10.10110248 | 0.455226427 | 2.418748342 |
| ENSMUSG00000031451 | *Gas6* | 333.9249293 | 493.5340218 | 535.4521313 | 185.5478322 | 222.5768278 | 210.6785989 | 9.45E-08 | -1.136433236 | 2.09E-08 | 31.40687327 | 0.454882792 | 7.024372928 |
| ENSMUSG00000027954 | *Efna1* | 55.92044271 | 52.48214546 | 44.28551462 | 22.77177941 | 23.88629372 | 23.19397419 | 0.002991421 | -1.136967913 | 0.001141026 | 10.58348042 | 0.454714239 | 2.5241225 |
| ENSMUSG00000020941 | *Map3k14* | 961.0327511 | 968.901147 | 806.5331602 | 387.9636491 | 419.0958807 | 436.8198472 | 3.14E-20 | -1.141110339 | 2.80E-21 | 89.68195522 | 0.453410486 | 19.50356464 |
| ENSMUSG00000049037 | *Clec4a1* | 94.26588914 | 116.0662832 | 138.224485 | 56.50774889 | 49.94406868 | 50.25361074 | 4.70E-05 | -1.141202006 | 1.40E-05 | 18.87492182 | 0.453381678 | 4.328001587 |
| ENSMUSG00000022236 | *Ropn1l* | 80.6852102 | 94.87157064 | 88.57102924 | 33.73596949 | 46.68684681 | 39.62303924 | 0.000145877 | -1.142965795 | 4.62E-05 | 16.59595296 | 0.452827727 | 3.836012011 |
| ENSMUSG00000016933 | *Plcg1* | 629.5044122 | 728.6944043 | 578.3956606 | 315.4313147 | 290.9784871 | 270.5963655 | 1.32E-15 | -1.143300533 | 1.57E-16 | 68.08091056 | 0.452722673 | 14.87784618 |
| ENSMUSG00000078532 | *Nkain1* | 87.8749814 | 125.1497315 | 110.0427939 | 44.70015957 | 52.11554993 | 49.28719515 | 6.56E-05 | -1.143477673 | 1.99E-05 | 18.2031466 | 0.45266709 | 4.183287851 |
| ENSMUSG00000020649 | *Rrm2* | 4815.548981 | 4619.438073 | 3620.676317 | 1972.710816 | 2007.534413 | 1932.831182 | 4.83E-22 | -1.143679002 | 3.90E-23 | 98.13940959 | 0.452603924 | 21.31638902 |
| ENSMUSG00000003500 | *Impdh1* | 5011.27053 | 5273.446347 | 4682.186682 | 2302.479918 | 2246.39735 | 2223.722275 | 2.06E-52 | -1.144372793 | 3.98E-54 | 239.9751552 | 0.452386319 | 51.68673509 |
| ENSMUSG00000025140 | *Pycr1* | 831.6168694 | 816.5010708 | 845.4507337 | 379.5296568 | 363.7231089 | 382.7005741 | 9.62E-30 | -1.145753699 | 5.18E-31 | 134.1071099 | 0.451953516 | 29.01693379 |
| ENSMUSG00000059498 | *Fcgr3* | 1930.853 | 1887.338693 | 2403.495657 | 965.6921266 | 894.6502738 | 946.1208637 | 7.49E-22 | -1.146398561 | 6.10E-23 | 97.25305763 | 0.451751545 | 21.1257787 |
| ENSMUSG00000038776 | *Ephx1* | 95.86361607 | 117.0755553 | 157.0122791 | 48.91715576 | 55.3727718 | 61.85059783 | 0.000134744 | -1.146649755 | 4.25E-05 | 16.7546876 | 0.451672895 | 3.870489339 |
| ENSMUSG00000059895 | *Ptp4a3* | 109.444295 | 161.4835245 | 218.7436025 | 75.06253211 | 58.62999367 | 85.04457202 | 0.000436203 | -1.151346946 | 0.000146946 | 14.4109446 | 0.45020471 | 3.360310878 |
| ENSMUSG00000040857 | *Erf* | 1621.692839 | 1697.595551 | 1319.171542 | 663.7551997 | 667.7304835 | 757.6698234 | 2.28E-19 | -1.15249993 | 2.14E-20 | 85.65979272 | 0.449845055 | 18.64145324 |
| ENSMUSG00000039824 | *Myl6b* | 133.410199 | 152.4000762 | 135.5405144 | 57.35114813 | 59.7157343 | 72.48116933 | 1.29E-06 | -1.153495794 | 3.20E-07 | 26.12489133 | 0.449534643 | 5.889247126 |
| ENSMUSG00000028101 | *Pias3* | 743.741888 | 858.8904959 | 752.8537485 | 346.6370865 | 352.8657027 | 358.5401843 | 4.20E-23 | -1.15504816 | 3.17E-24 | 103.1122732 | 0.449051196 | 22.37722968 |
| ENSMUSG00000068114 | *Ccdc134* | 622.314641 | 634.8321057 | 536.7941166 | 337.3596949 | 200.8620154 | 265.7642876 | 7.55E-09 | -1.155131428 | 1.50E-09 | 36.52903017 | 0.449025279 | 8.122240164 |
| ENSMUSG00000038181 | *Chpf2* | 3719.508304 | 3991.670871 | 3816.606169 | 1728.968436 | 1681.812226 | 1763.708454 | 3.14E-61 | -1.155189204 | 4.43E-63 | 281.0517287 | 0.449007297 | 60.50297422 |
| ENSMUSG00000024854 | *Pold4* | 523.2555711 | 536.932719 | 560.9498519 | 237.8385849 | 249.7203434 | 239.6710666 | 2.28E-21 | -1.155696744 | 1.89E-22 | 95.01333399 | 0.448849364 | 20.64287868 |
| ENSMUSG00000036528 | *Ppfibp2* | 291.5851656 | 358.29157 | 386.491764 | 151.8118627 | 163.9468342 | 147.8615854 | 7.24E-11 | -1.156534852 | 1.21E-11 | 45.94850253 | 0.448588689 | 10.14007409 |
| ENSMUSG00000028042 | *Zbtb7b* | 4217.200244 | 4496.306885 | 3744.138963 | 1837.766938 | 1831.644432 | 1912.536455 | 9.32E-37 | -1.158921293 | 3.58E-38 | 166.8650149 | 0.447847267 | 36.03065002 |
| ENSMUSG00000032202 | *Rab27a* | 341.913564 | 302.7816084 | 299.26272 | 146.7514673 | 144.4035029 | 132.398936 | 8.51E-13 | -1.159084362 | 1.23E-13 | 54.96466843 | 0.447796649 | 12.06982816 |
| ENSMUSG00000037649 | *H2-DMa* | 225.2794978 | 303.7908805 | 267.055073 | 104.5815054 | 117.2599873 | 134.3317672 | 1.98E-08 | -1.159793912 | 4.12E-09 | 34.56710008 | 0.447576467 | 7.703448464 |
| ENSMUSG00000041754 | *Trem3* | 985.7975186 | 991.1051316 | 1158.133307 | 475.6771698 | 447.3251369 | 472.5772241 | 7.79E-25 | -1.164117756 | 5.37E-26 | 111.1928148 | 0.446237061 | 24.10818778 |
| ENSMUSG00000024055 | *Cyp4f13* | 302.7692541 | 350.2173938 | 264.3711024 | 140.0042734 | 145.4892436 | 124.6676113 | 2.69E-10 | -1.165041691 | 4.72E-11 | 43.28865095 | 0.445951372 | 9.570093211 |
| ENSMUSG00000019188 | *H13* | 8841.820856 | 9055.188636 | 9184.547335 | 4102.29389 | 4056.32697 | 3914.94956 | 7.87E-88 | -1.16515226 | 5.62E-90 | 404.5528777 | 0.445917196 | 87.10392518 |
| ENSMUSG00000113308 | *Gm48445* | 279.6022136 | 222.0398462 | 206.6657349 | 102.0513077 | 100.973878 | 114.0370398 | 1.95E-08 | -1.165802425 | 4.05E-09 | 34.59734196 | 0.445716284 | 7.709825866 |
| ENSMUSG00000072772 | *Grcc10* | 1767.884853 | 1912.570493 | 2329.686466 | 877.978606 | 849.0491676 | 948.0536949 | 2.43E-20 | -1.165806022 | 2.14E-21 | 90.21028574 | 0.445715173 | 19.61520587 |
| ENSMUSG00000049577 | *Zfpm1* | 1335.699717 | 1283.79402 | 1030.644704 | 522.0641278 | 539.6130899 | 566.3195364 | 1.38E-19 | -1.167780759 | 1.28E-20 | 86.67361084 | 0.445105502 | 18.85886567 |
| ENSMUSG00000038351 | *Sgsm2* | 385.8510547 | 485.4598455 | 351.6001464 | 171.2100452 | 184.575906 | 189.4174559 | 4.47E-11 | -1.168531852 | 7.34E-12 | 46.93511081 | 0.444873833 | 10.35009735 |
| ENSMUSG00000067924 | *Rtl8b* | 231.6704055 | 259.3829112 | 236.1894113 | 83.49652449 | 119.4314686 | 121.7683645 | 4.10E-08 | -1.169000675 | 8.78E-09 | 33.0945591 | 0.444729289 | 7.38735554 |

| | | | | | | | | | | | | | |
|---|---|---|---|---|---|---|---|---|---|---|---|---|---|
| ENSMUSG00000052087 | *Rgs14* | 1174.329297 | 1142.495936 | 1014.54088 | 516.1603332 | 460.3540244 | 505.4353542 | 3.04E-26 | -1.169361998 | 1.94E-27 | 117.7729295 | 0.44461792 | 25.51647307 |
| ENSMUSG00000005718 | *Tfap4* | 496.0942132 | 556.1088875 | 536.7941166 | 242.0555811 | 241.0344184 | 222.275586 | 1.65E-19 | -1.169909918 | 1.53E-20 | 86.31684054 | 0.444449091 | 18.78272947 |
| ENSMUSG00000035561 | *Aldh1b1* | 316.349933 | 313.8836008 | 324.7604405 | 145.908068 | 150.9179467 | 127.566858 | 1.27E-13 | -1.169962927 | 1.72E-14 | 58.82638834 | 0.444432761 | 12.89626047 |
| ENSMUSG00000004748 | *Mtfp1* | 587.1646485 | 657.0360903 | 670.9926458 | 288.4425391 | 251.8918247 | 308.2865736 | 4.47E-18 | -1.170093413 | 4.55E-19 | 79.61348378 | 0.444392566 | 17.34946825 |
| ENSMUSG00000051950 | *B3glct* | 506.4794383 | 617.6744812 | 683.0705134 | 280.0085468 | 255.1490465 | 263.8314564 | 4.43E-14 | -1.172179015 | 5.83E-15 | 60.95815909 | 0.443750603 | 13.35322467 |
| ENSMUSG00000011263 | *Exoc3l2* | 61.51248698 | 87.80666645 | 57.70536754 | 26.98877559 | 29.31499684 | 35.75737687 | 0.001440312 | -1.173530346 | 0.000524221 | 12.02747221 | 0.443335149 | 2.841543414 |
| ENSMUSG00000052673 | *Gm9887* | 77.48975633 | 89.8252105 | 79.1771322 | 32.04917101 | 35.82944058 | 41.55587042 | 0.000126424 | -1.174063281 | 3.98E-05 | 16.87991345 | 0.44317141 | 3.898171737 |
| ENSMUSG00000028100 | *Nudt17* | 47.93180804 | 54.50068952 | 45.62749991 | 25.30197712 | 22.80055309 | 17.39548064 | 0.003464304 | -1.175466261 | 0.00133612 | 10.29194693 | 0.442740648 | 2.460384059 |
| ENSMUSG00000017400 | *Stac2* | 189.3306417 | 205.8914937 | 214.7176466 | 89.40031915 | 96.6309155 | 83.11174084 | 1.47E-09 | -1.178549658 | 2.75E-10 | 39.8460396 | 0.441795412 | 8.83304111 |
| ENSMUSG00000015363 | *Trabd* | 4308.270679 | 4360.055162 | 4043.401683 | 1849.574527 | 1889.188685 | 1878.711909 | 4.02E-66 | -1.179039024 | 5.03E-68 | 303.7466242 | 0.441645579 | 65.39579519 |
| ENSMUSG00000045503 | *Sys1* | 1862.949606 | 1941.839382 | 1866.70154 | 768.3367051 | 839.277502 | 896.8336686 | 1.65E-37 | -1.180284605 | 6.17E-39 | 170.3596997 | 0.44126444 | 36.78246834 |
| ENSMUSG00000013160 | *Atp6v0d1* | 7634.738157 | 7932.878141 | 9397.922996 | 3661.196089 | 3519.971102 | 3812.509507 | 5.81E-33 | -1.182639767 | 2.73E-34 | 149.0996537 | 0.440544675 | 32.23613929 |
| ENSMUSG00000027074 | *Slc43a3* | 1879.725739 | 1898.440685 | 1940.510732 | 860.267222 | 839.277502 | 817.5875901 | 9.68E-53 | -1.182908749 | 1.85E-54 | 241.5005604 | 0.440462546 | 52.01400852 |
| ENSMUSG00000023982 | *Guca1a* | 482.5135342 | 559.1367036 | 680.3865428 | 256.3933681 | 264.9207121 | 234.8389886 | 1.66E-12 | -1.183188793 | 2.44E-13 | 53.61184675 | 0.440377055 | 11.77946647 |
| ENSMUSG00000026856 | *Dolpp1* | 756.5237035 | 817.5103428 | 654.8888223 | 335.6728964 | 344.1797777 | 302.48808 | 1.36E-18 | -1.18475203 | 1.33E-19 | 82.04597252 | 0.439900141 | 17.86633048 |
| ENSMUSG00000067925 | *Rtl8a* | 85.478391 | 107.992107 | 60.38933812 | 42.16996186 | 38.00092127 | 31.89171451 | 0.000883749 | -1.184968261 | 0.000312108 | 12.99629693 | 0.439834214 | 3.0536713 |
| ENSMUSG00000042073 | *Abhd14b* | 39.94317336 | 54.50068952 | 41.60154404 | 21.92838017 | 17.37184998 | 20.29472741 | 0.005223441 | -1.185169534 | 0.002078094 | 9.479246022 | 0.439772856 | 2.282043288 |
| ENSMUSG00000006179 | *Prss16* | 408.2192318 | 349.2081217 | 336.8383082 | 172.0534444 | 155.2609092 | 154.6264946 | 2.52E-13 | -1.185178752 | 3.50E-14 | 57.42931559 | 0.439770046 | 12.59921447 |
| ENSMUSG00000040462 | *Os9* | 5681.516979 | 6694.501363 | 6994.427339 | 2865.027209 | 2837.040249 | 2808.403708 | 3.30E-35 | -1.185866851 | 1.38E-36 | 159.6077108 | 0.439560346 | 34.48165789 |
| ENSMUSG00000040904 | *Gm21988* | 127.0192913 | 147.3537161 | 163.7222056 | 59.03794661 | 66.23017804 | 66.68267579 | 6.58E-07 | -1.186295768 | 1.58E-07 | 27.48985968 | 0.439429683 | 6.181499712 |
| ENSMUSG00000031144 | *Syp* | 267.6192615 | 346.1803057 | 331.4707467 | 145.908068 | 143.3177623 | 124.6676113 | 1.27E-10 | -1.187364431 | 2.18E-11 | 44.80241814 | 0.4391043 | 9.894568978 |
| ENSMUSG00000036880 | *Acaa2* | 1381.234935 | 1481.611337 | 1877.437423 | 704.2383631 | 660.1302991 | 713.2147063 | 2.84E-18 | -1.187394928 | 2.85E-19 | 80.53783647 | 0.439095018 | 17.54680699 |
| ENSMUSG00000000884 | *Gnb1l* | 434.5817262 | 366.3657462 | 386.491764 | 134.1004787 | 200.8620154 | 188.4510403 | 1.10E-09 | -1.188268204 | 2.03E-10 | 40.43762882 | 0.438829311 | 8.959877557 |
| ENSMUSG00000019822 | *Smpd2* | 398.6328702 | 430.959156 | 430.7772786 | 192.2950261 | 182.4044248 | 176.8540532 | 8.80E-18 | -1.189123204 | 9.09E-19 | 78.2476118 | 0.43856932 | 17.05545379 |
| ENSMUSG00000045980 | *Tmem104* | 660.6600874 | 786.2229099 | 711.2522045 | 302.7803262 | 332.2366308 | 311.1858203 | 2.60E-21 | -1.190093799 | 2.17E-22 | 94.73994253 | 0.438274365 | 20.58474606 |
| ENSMUSG00000013539 | *Tango2* | 567.9919252 | 583.3592323 | 770.2995573 | 312.901117 | 279.0353403 | 246.4359757 | 6.53E-12 | -1.191812393 | 1.01E-12 | 50.8299028 | 0.437752586 | 11.18514212 |
| ENSMUSG00000003575 | *Crtc1* | 790.8748326 | 854.8534078 | 676.3605869 | 343.2634896 | 325.7221871 | 346.9431972 | 9.39E-20 | -1.194509406 | 8.58E-21 | 87.465579 | 0.436935004 | 19.02747351 |
| ENSMUSG00000006782 | *Cnp* | 1315.728131 | 1338.294709 | 1535.231173 | 630.0192302 | 611.2719711 | 583.715017 | 3.59E-30 | -1.196205422 | 1.89E-31 | 136.1081529 | 0.43642165 | 29.44450787 |
| ENSMUSG00000024158 | *Hagh* | 1405.999702 | 1536.112027 | 1715.057203 | 717.7327509 | 621.0436367 | 688.0879009 | 2.92E-25 | -1.196917896 | 1.98E-26 | 113.1748253 | 0.436206177 | 24.53408154 |
| ENSMUSG00000035936 | *Aldh5a1* | 92.66816221 | 73.67685805 | 46.9694852 | 24.45857788 | 38.00092127 | 31.89171451 | 0.003645208 | -1.196920203 | 0.001412346 | 10.18960462 | 0.436205479 | 2.438277698 |
| ENSMUSG00000042647 | *Acad12* | 189.3306417 | 220.0213021 | 307.3146318 | 116.3890947 | 103.1453592 | 90.84306557 | 1.78E-06 | -1.198075218 | 4.47E-07 | 25.47943441 | 0.435856395 | 5.750679021 |
| ENSMUSG00000056367 | *Actr3b* | 34.35112909 | 46.42651329 | 64.41529399 | 16.86798474 | 22.80055309 | 23.19397419 | 0.009447809 | -1.19831677 | 0.003941765 | 8.310448734 | 0.435783425 | 2.024668885 |
| ENSMUSG00000090231 | *Cfb* | 234.0669959 | 242.2252868 | 230.8214701 | 113.858897 | 100.973878 | 92.77589675 | 6.23E-11 | -1.198353625 | 1.04E-11 | 46.25510272 | 0.435772293 | 10.2055852 |
| ENSMUSG00000020635 | *Fkbp1b* | 127.0192913 | 140.2888119 | 175.8000732 | 74.21913288 | 67.31591866 | 50.25361074 | 9.75E-06 | -1.198952385 | 2.67E-06 | 22.04055385 | 0.435591472 | 5.01097821 |
| ENSMUSG00000032763 | *Ilvbl* | 1201.490655 | 1298.9331 | 1237.310439 | 589.5360668 | 511.3838337 | 523.7972504 | 1.47E-31 | -1.199612567 | 7.31E-33 | 142.5670618 | 0.43539219 | 30.83237773 |
| ENSMUSG00000028334 | *Nans* | 6550.680432 | 6909.476305 | 8055.937705 | 3100.335596 | 3158.419474 | 3105.093294 | 1.34E-36 | -1.199638308 | 5.20E-38 | 166.1246347 | 0.435384421 | 35.87332727 |
| ENSMUSG00000027999 | *Pla2g12a* | 1107.224766 | 1156.625744 | 1245.362351 | 500.9791469 | 522.2412399 | 502.5361074 | 4.19E-34 | -1.200109322 | 1.86E-35 | 154.4377333 | 0.435242299 | 33.37755629 |
| ENSMUSG00000023047 | *Amhr2* | 352.2987891 | 395.634635 | 343.5482346 | 141.6910719 | 166.1183154 | 168.1563129 | 6.52E-14 | -1.201351815 | 8.65E-15 | 60.18049909 | 0.434867617 | 13.18553631 |
| ENSMUSG00000042978 | *Sbk1* | 334.7237928 | 409.7644434 | 322.07647 | 169.5232467 | 161.7753529 | 132.398936 | 6.72E-11 | -1.20195819 | 1.12E-11 | 46.10319381 | 0.434684877 | 10.17283406 |
| ENSMUSG00000063160 | *Numbl* | 83.0818006 | 104.9642909 | 59.04735283 | 38.79636491 | 27.14351559 | 41.55587042 | 0.00122721 | -1.2021121 | 0.000442169 | 12.34499977 | 0.434638507 | 2.911081237 |
| ENSMUSG00000031387 | *Renbp* | 3630.035595 | 4161.228572 | 5308.893813 | 1912.82947 | 1883.759982 | 1893.208143 | 4.07E-18 | -1.202159409 | 4.13E-19 | 79.80522022 | 0.434624254 | 17.39085732 |
| ENSMUSG00000050335 | *Lgals3* | 60072.93501 | 65981.15884 | 81442.40337 | 29498.73172 | 30287.82043 | 30329.0205 | 1.20E-25 | -1.203175938 | 7.92E-27 | 114.9876548 | 0.434318124 | 24.91998386 |
| ENSMUSG00000063382 | *Bcl9l* | 453.7544494 | 492.5247497 | 330.1283817 | 191.4516269 | 185.6616466 | 178.7868844 | 2.19E-10 | -1.203190396 | 3.81E-11 | 43.70965906 | 0.434313771 | 9.660009613 |
| ENSMUSG00000095098 | *Ccdc85b* | 473.7260361 | 441.0518763 | 441.5131609 | 187.2346307 | 205.2049779 | 198.1151962 | 3.30E-19 | -1.203277203 | 3.13E-20 | 84.9068225 | 0.434287639 | 18.481245 |
| ENSMUSG00000033948 | *Zswim5* | 43.13862723 | 38.35233707 | 45.62749991 | 21.08498093 | 17.37184998 | 16.42906505 | 0.0048503 | -1.203875098 | 0.001919051 | 9.625395254 | 0.434107695 | 2.314241443 |
| ENSMUSG00000032281 | *Acsbg1* | 1406.798566 | 1501.796778 | 1650.641909 | 621.5852378 | 668.8162241 | 687.1214853 | 4.90E-31 | -1.203927662 | 2.48E-32 | 140.1369024 | 0.434091879 | 30.30970076 |
| ENSMUSG00000051615 | *Rap2a* | 589.5612389 | 699.4255155 | 579.7376459 | 286.7557407 | 252.9775653 | 270.5963655 | 3.06E-18 | -1.204208431 | 3.08E-19 | 80.38318974 | 0.434007407 | 17.51434661 |
| ENSMUSG00000027002 | *Nckap1* | 138.2033798 | 119.0940993 | 138.224485 | 54.82095042 | 60.80147492 | 56.05210429 | 4.65E-07 | -1.206125125 | 1.10E-07 | 28.1930759 | 0.433431189 | 6.332563523 |
| ENSMUSG00000057375 | *Yipf1* | 1149.564529 | 1332.239077 | 1727.13507 | 620.7418386 | 624.3008585 | 575.0172767 | 3.87E-14 | -1.207086385 | 5.07E-15 | 61.23200325 | 0.433142492 | 13.41187508 |
| ENSMUSG00000035504 | *Reep6* | 417.8055934 | 390.5882749 | 351.6001464 | 158.5590566 | 174.8042404 | 171.0555596 | 2.41E-15 | -1.207483181 | 2.91E-16 | 66.85993684 | 0.433023378 | 14.61767668 |
| ENSMUSG00000049515 | *Espnl* | 149.3874684 | 152.4000762 | 144.9344115 | 59.03794661 | 62.97295617 | 71.51475374 | 6.56E-08 | -1.209542081 | 1.43E-08 | 32.14318345 | 0.432405842 | 7.183069872 |
| ENSMUSG00000028641 | *P3h1* | 1023.344102 | 1057.717085 | 966.2294099 | 444.471398 | 439.7249525 | 433.9206004 | 6.40E-35 | -1.210183702 | 2.71E-36 | 158.2667761 | 0.432213577 | 34.19415381 |
| ENSMUSG00000003573 | *Homer3* | 966.6247954 | 1116.254863 | 1288.30588 | 514.4735347 | 427.7818057 | 510.2674321 | 6.81E-17 | -1.210740424 | 7.43E-18 | 74.09973223 | 0.432046823 | 16.16688193 |
| ENSMUSG00000028393 | *Alad* | 2485.264247 | 2647.32053 | 2590.031613 | 1092.202012 | 1109.626917 | 1131.672657 | 2.41E-60 | -1.211765827 | 3.54E-62 | 276.9092686 | 0.431739852 | 59.6180502 |
| ENSMUSG00000028497 | *Hacd4* | 192.5260956 | 237.1789266 | 210.6916908 | 100.3645092 | 91.20221238 | 84.07815643 | 2.14E-09 | -1.212282839 | 4.05E-10 | 39.08812626 | 0.431585159 | 8.670197073 |
| ENSMUSG00000075705 | *Msrb1* | 2111.396144 | 2319.307121 | 2628.949186 | 1015.452682 | 1003.224336 | 1016.669202 | 3.25E-31 | -1.215915893 | 1.63E-32 | 140.9715712 | 0.430499691 | 30.488665 |
| ENSMUSG00000027510 | *Rbm38* | 2353.451775 | 2446.475396 | 1991.506173 | 935.3297541 | 1020.596186 | 965.4491755 | 5.39E-30 | -1.219076828 | 2.87E-31 | 135.2808605 | 0.429557502 | 29.268279 |
| ENSMUSG00000001761 | *Smo* | 90.2715718 | 88.81593848 | 99.30691157 | 37.10956644 | 49.94406868 | 32.8581301 | 5.70E-05 | -1.219388669 | 1.71E-05 | 18.48728249 | 0.429464662 | 4.244428855 |
| ENSMUSG00000002803 | *Btbd6* | 674.2407664 | 725.6665882 | 641.4689693 | 323.0219079 | 269.2636746 | 283.1597682 | 4.12E-21 | -1.219531239 | 3.48E-22 | 93.80722453 | 0.429422224 | 20.38529407 |
| ENSMUSG00000022436 | *Sh3bp1* | 2308.71542 | 2410.141603 | 2025.055805 | 958.9449327 | 998.8813737 | 940.3223702 | 7.35E-36 | -1.220131372 | 2.99E-37 | 162.64716 | 0.429243629 | 35.13389881 |
| ENSMUSG00000014867 | *Surf4* | 17989.60642 | 20060.29083 | 20611.55209 | 8418.811186 | 8354.774098 | 8402.017149 | 9.50E-61 | -1.220141707 | 1.37E-62 | 278.8038829 | 0.429240554 | 60.02212116 |
| ENSMUSG00000021384 | *Susd3* | 1416.384928 | 1371.600686 | 1297.699777 | 540.6189111 | 604.7575273 | 611.7410692 | 7.29E-35 | -1.220269816 | 3.10E-36 | 157.9980879 | 0.42920244 | 34.13725782 |
| ENSMUSG00000090996 | *Gm20458* | 758.9202939 | 769.0652854 | 1013.198895 | 399.7712384 | 374.5805151 | 312.1522359 | 1.64E-13 | -1.222341739 | 2.24E-14 | 58.30570415 | 0.428586485 | 12.78546364 |
| ENSMUSG00000037583 | *Nr0b2* | 189.3306417 | 181.6689651 | 147.6183821 | 70.00213669 | 77.08758427 | 76.3468317 | 3.26E-08 | -1.223433386 | 6.92E-09 | 33.55738894 | 0.428262308 | 7.486808443 |
| ENSMUSG00000110040 | *Gm49369* | 20132.15824 | 25978.662 | 30461.72413 | 11034.19222 | 11262.38749 | 10456.6167 | 3.30E-18 | -1.225042754 | 3.33E-19 | 80.22974444 | 0.427784835 | 17.48163711 |
| ENSMUSG00000037239 | *Spred3* | 253.2397191 | 247.2716469 | 173.1161026 | 91.93051686 | 102.0596186 | 95.67514352 | 5.55E-08 | -1.226415743 | 1.20E-08 | 32.47970906 | 0.427377913 | 7.255457915 |
| ENSMUSG00000055675 | *Kbtbd11* | 2059.470019 | 1997.349344 | 1430.556321 | 730.3837394 | 770.8758427 | 845.6136422 | 1.71E-15 | -1.227311815 | 2.04E-16 | 67.56216774 | 0.427112547 | 14.76789381 |
| ENSMUSG00000042724 | *Map3k9* | 328.3328851 | 283.6054399 | 253.6352201 | 126.5098856 | 120.5172092 | 123.7011957 | 1.26E-11 | -1.228117143 | 1.99E-12 | 49.49616568 | 0.426874195 | 10.90030385 |
| ENSMUSG00000072825 | *Cep170b* | 712.5862128 | 784.2043659 | 555.5819107 | 278.3217483 | 294.235709 | 305.3873268 | 1.81E-14 | -1.228749775 | 2.32E-15 | 62.77672694 | 0.426686989 | 13.74145497 |
| ENSMUSG00000029455 | *Aldh2* | 8933.690155 | 10599.37484 | 11542.41549 | 4251.575555 | 4524.281178 | 4478.369849 | 3.39E-31 | -1.229015086 | 1.71E-32 | 140.8793499 | 0.426608588 | 30.46951753 |
| ENSMUSG00000022415 | *Syngr1* | 965.0270685 | 1135.431032 | 1425.18838 | 505.1961431 | 491.8405025 | 503.502523 | 3.09E-15 | -1.229365844 | 3.76E-16 | 66.35823736 | 0.426504881 | 14.51031352 |
| ENSMUSG00000052160 | *Pld4* | 11365.43055 | 11990.15169 | 12558.29836 | 4996.297081 | 5177.897034 | 5118.136971 | 4.75E-76 | -1.231503503 | 4.31E-78 | 349.9648263 | 0.425873391 | 75.32370561 |
| ENSMUSG00000037960 | *Card19* | 3662.788998 | 3964.420527 | 4447.339256 | 1673.304087 | 1713.298704 | 1752.111467 | 9.80E-39 | -1.231533884 | 3.49E-40 | 176.0723944 | 0.425864423 | 38.00899372 |
| ENSMUSG00000076441 | *Ass1* | 1129.592943 | 1169.746281 | 1186.314998 | 495.0753523 | 525.4984618 | 463.8794837 | 2.04E-36 | -1.23158819 | 8.01E-38 | 165.2632361 | 0.425848393 | 35.68934523 |
| ENSMUSG00000071648 | *Rom1* | 54.32271578 | 59.54704966 | 77.83514691 | 21.92838017 | 32.57221871 | 27.05963655 | 0.001114536 | -1.231659497 | 0.000399073 | 12.536529 | 0.425827345 | 2.95290579 |
| ENSMUSG00000026817 | *Ak1* | 174.1522359 | 193.7802294 | 177.1420585 | 84.33992372 | 70.57314053 | 76.3468317 | 2.55E-09 | -1.232526148 | 4.86E-10 | 38.7341089 | 0.42557162 | 8.594133863 |
| ENSMUSG00000038502 | *Ptov1* | 1517.840588 | 1843.939995 | 1894.883232 | 725.323344 | 717.6745522 | 790.5279535 | 1.32E-25 | -1.233066993 | 8.72E-27 | 114.7959742 | 0.42541211 | 24.87994613 |
| ENSMUSG00000022945 | *Chaf1b* | 2113.792734 | 2025.60896 | 2324.318525 | 909.1843477 | 893.5645332 | 943.2216169 | 1.63E-42 | -1.233617052 | 4.83E-44 | 193.7507577 | 0.425249443 | 41.78802519 |
| ENSMUSG00000038375 | *Trp53inp2* | 1009.763423 | 1254.525131 | 1001.121027 | 477.3639683 | 456.0110619 | 454.2153278 | 4.63E-22 | -1.234616147 | 3.73E-23 | 98.22568295 | 0.424955551 | 21.33467522 |
| ENSMUSG00000059089 | *Fcgr4* | 309.9590253 | 298.7445203 | 414.6734551 | 143.3778703 | 144.4035029 | 144.9623387 | 8.25E-11 | -1.236114284 | 1.39E-11 | 45.68111436 | 0.424514494 | 10.08353813 |
| ENSMUSG00000043445 | *Pgp* | 2330.284734 | 2246.639535 | 2461.201025 | 974.9695182 | 961.9661925 | 1048.560916 | 6.10E-49 | -1.236314687 | 1.35E-50 | 223.7919992 | 0.424455529 | 48.21457245 |
| ENSMUSG00000006736 | *Tspan31* | 5018.460302 | 5301.705964 | 5847.029915 | 2223.200389 | 2368.0003 | 2258.513236 | 2.87E-48 | -1.238478578 | 6.53E-50 | 220.6499804 | 0.423819368 | 47.54183756 |
| ENSMUSG00000040883 | *Tmem205* | 1345.286879 | 1506.843138 | 1811.680144 | 627.4890325 | 702.4741834 | 644.5991993 | 2.23E-20 | -1.238561551 | 1.96E-21 | 90.38835149 | 0.423794993 | 19.65225467 |
| ENSMUSG00000036820 | *Amdhd2* | 3851.320776 | 3785.779378 | 4152.102492 | 1652.219106 | 1634.039638 | 1705.723518 | 9.82E-63 | -1.238918062 | 1.31E-64 | 288.0686252 | 0.42369028 | 62.00770232 |
| ENSMUSG00000082926 | *Gm8172* | 154.9795127 | 117.0755553 | 134.1985292 | 80.12292754 | 46.68684681 | 44.45511719 | 0.000100235 | -1.239328293 | 3.11E-05 | 17.35017572 | 0.423569821 | 3.998978627 |
| ENSMUSG00000025150 | *Cbr2* | 86.27725447 | 79.73249022 | 101.9908822 | 37.10956644 | 40.17240307 | 35.75737687 | 2.77E-05 | -1.241560821 | 8.02E-06 | 19.93228717 | 0.422914866 | 4.556822596 |
| ENSMUSG00000020282 | *Rhbdf1* | 315.5510696 | 373.4306504 | 309.9986023 | 139.1608741 | 131.3746155 | 151.7272478 | 1.99E-13 | -1.24217863 | 2.75E-14 | 57.90718301 | 0.422733799 | 12.70134734 |
| ENSMUSG00000029101 | *Rgs12* | 75.89202939 | 90.83448253 | 56.36338224 | 32.04917101 | 35.82944058 | 27.05963655 | 0.00049807 | -1.242346842 | 0.000169374 | 14.14357821 | 0.422684513 | 3.30270987 |
| ENSMUSG00000038312 | *Edem2* | 3254.569766 | 3564.748803 | 3569.680875 | 1381.487951 | 1517.865392 | 1488.28001 | 3.11E-51 | -1.243602294 | 6.40E-53 | 234.4429945 | 0.422316847 | 50.50737057 |
| ENSMUSG00000034993 | *Vat1* | 17313.76793 | 20774.85543 | 21890.46408 | 8310.012684 | 8503.520564 | 8509.28928 | 3.58E-36 | -1.24382636 | 1.42E-37 | 164.1215047 | 0.422251262 | 35.4464256 |
| ENSMUSG00000007891 | *Ctsd* | 109842.9279 | 135231.3498 | 167469.0285 | 57290.42339 | 57699.51396 | 59150.43267 | 7.04E-19 | -1.244264343 | 6.77E-20 | 83.37879235 | 0.422123092 | 18.15256151 |
| ENSMUSG00000096755 | *Ftl1-ps2* | 63.90907738 | 44.40796924 | 88.57102924 | 26.14537635 | 19.54333122 | 36.72379246 | 0.005422112 | -1.244642157 | 0.002161293 | 9.407248737 | 0.42201256 | 2.265831498 |
| ENSMUSG00000038816 | *Ctnnal1* | 237.2624498 | 225.0676623 | 199.9558084 | 102.0513077 | 81.43054677 | 95.67514352 | 3.60E-10 | -1.245954675 | 6.39E-11 | 42.69606783 | 0.421628801 | 9.443773212 |
| ENSMUSG00000029674 | *Limk1* | 691.8157627 | 666.1195386 | 513.9803667 | 274.1047521 | 282.2925621 | 234.8389886 | 3.16E-14 | -1.246319797 | 4.12E-15 | 61.64286008 | 0.421522107 | 13.49975112 |
| ENSMUSG00000024939 | *Fam89b* | 1819.012115 | 1845.958539 | 2062.631393 | 790.2650853 | 807.7910239 | 810.822681 | 3.65E-43 | -1.248107989 | 1.03E-44 | 196.8149477 | 0.420999963 | 42.43822025 |
| ENSMUSG00000095762 | *Gm16378* | 575.9805599 | 667.1288106 | 513.9803667 | 225.1875963 | 228.0055309 | 287.0254306 | 6.40E-14 | -1.249093927 | 8.49E-15 | 60.21875848 | 0.42071235 | 13.19359144 |
| ENSMUSG00000019139 | *Isyna1* | 10423.57052 | 10971.79622 | 11212.28711 | 4562.789873 | 4472.165628 | 4663.921643 | 9.49E-89 | -1.250741471 | 6.39E-91 | 408.8925625 | 0.420232175 | 88.02270094 |
| ENSMUSG00000033065 | *Pfkm* | 570.3885156 | 605.5632169 | 559.6078666 | 284.2255429 | 168.2897967 | 274.4620279 | 8.26E-10 | -1.251619911 | 1.51E-10 | 41.01681878 | 0.419976378 | 9.083164559 |
| ENSMUSG00000025701 | *Alox5* | 2584.323317 | 2578.690032 | 2518.906392 | 1106.539799 | 1055.339886 | 1062.090735 | 6.18E-68 | -1.252354631 | 7.22E-70 | 312.2058879 | 0.419762551 | 67.20895617 |
| ENSMUSG00000024875 | *Yif1a* | 2152.138181 | 2343.529649 | 2788.645436 | 1043.284856 | 978.2523018 | 1031.165436 | 6.07E-27 | -1.252682528 | 3.73E-28 | 121.0457509 | 0.419667158 | 26.21707053 |
| ENSMUSG00000028980 | *H6pd* | 1915.674595 | 2001.386432 | 1672.113673 | 750.6253211 | 791.5049146 | 805.0241874 | 2.23E-34 | -1.253750227 | 9.73E-36 | 155.7229946 | 0.419356689 | 33.6520162 |
| ENSMUSG00000027230 | *Creb3l1* | 156.5772396 | 140.2888119 | 146.2763968 | 58.19454737 | 51.02980931 | 76.3468317 | 4.91E-07 | -1.255123001 | 1.16E-07 | 28.08004264 | 0.418957847 | 6.308500102 |
| ENSMUSG00000038451 | *Spsb2* | 637.4930469 | 688.3235232 | 764.9316162 | 290.1293376 | 280.1210809 | 302.48808 | 4.64E-23 | -1.256400204 | 3.52E-24 | 102.9026024 | 0.418587112 | 22.33329901 |
| ENSMUSG00000034758 | *Tle6* | 35.94885603 | 47.43578532 | 55.02139695 | 17.71138398 | 19.54333122 | 20.29472741 | 0.003544958 | -1.256519284 | 0.001369529 | 10.24638299 | 0.418552563 | 2.450388923 |
| ENSMUSG00000004500 | *Zfp324* | 602.3430543 | 649.9711861 | 503.2444843 | 238.6819841 | 235.6057153 | 261.8986252 | 2.64E-17 | -1.256721255 | 2.81E-18 | 76.01634443 | 0.418493971 | 16.57873155 |
| ENSMUSG00000044328 | *Trp53i13* | 487.306715 | 459.2187728 | 416.0154404 | 198.1988207 | 193.261831 | 179.7532999 | 1.37E-18 | -1.257241922 | 1.34E-19 | 82.02880074 | 0.418342965 | 17.86281944 |
| ENSMUSG00000034382 | *AI661453* | 60.71362351 | 64.5934098 | 42.94352933 | 23.61517864 | 24.97203434 | 22.2275586 | 0.001174546 | -1.259427838 | 0.000421946 | 12.43241295 | 0.417709587 | 2.930130078 |
| ENSMUSG00000018599 | *Mief2* | 659.861224 | 628.7764735 | 630.733087 | 251.3329727 | 251.8918247 | 298.6224177 | 4.33E-23 | -1.260240124 | 3.27E-24 | 103.04587 | 0.417474469 | 22.36335168 |
| ENSMUSG00000081665 | *Pira1* | 160.5715569 | 142.307356 | 284.5008818 | 86.0267222 | 80.34480614 | 77.31324729 | 6.14E-05 | -1.26052635 | 1.85E-05 | 18.33708653 | 0.417391652 | 4.211560725 |
| ENSMUSG00000028773 | *Fabp3* | 92.66816221 | 104.9642909 | 124.8046321 | 39.63976415 | 49.94406868 | 44.45511719 | 8.91E-06 | -1.263554132 | 2.43E-06 | 22.21993412 | 0.416516591 | 5.050339847 |
| ENSMUSG00000114515 | *Aldoa* | 64.70794085 | 67.62122589 | 52.33742637 | 27.83217483 | 27.14351559 | 22.2275586 | 0.000448343 | -1.26375022 | 0.000151247 | 14.35661942 | 0.416459983 | 3.348389487 |
| ENSMUSG00000039886 | *Tmem120a* | 3582.902651 | 3975.522519 | 4709.026388 | 1718.004246 | 1603.638901 | 1773.37261 | 2.09E-27 | -1.266430317 | 1.25E-28 | 123.2111951 | 0.415687043 | 26.67926217 |
| ENSMUSG00000049097 | *Ankrd34a* | 127.0192913 | 141.2980839 | 79.1771322 | 53.13415195 | 41.2581437 | 50.25361074 | 9.11E-05 | -1.268720031 | 2.81E-05 | 17.54022051 | 0.415027825 | 4.040408591 |
| ENSMUSG00000083429 | *Gm15198* | 304.366981 | 452.1538686 | 217.4016172 | 153.4986612 | 158.518134 | 92.77589675 | 7.57E-05 | -1.26910253 | 2.31E-05 | 17.91601074 | 0.414917804 | 4.121105776 |
| ENSMUSG00000038422 | *Hdhd3* | 273.2113058 | 285.623984 | 348.9161758 | 122.2928894 | 98.80239674 | 153.660079 | 6.09E-09 | -1.269946509 | 1.20E-09 | 36.96696399 | 0.414675148 | 8.215664397 |
| ENSMUSG00000018669 | *Cdk5rap3* | 2722.526697 | 2413.169419 | 2726.914112 | 1047.501853 | 1123.741545 | 1090.116787 | 1.83E-46 | -1.270031438 | 4.55E-48 | 212.2006736 | 0.414650737 | 45.73854368 |

| | | | | | | | | | | | | | |
|---|---|---|---|---|---|---|---|---|---|---|---|---|---|
| ENSMUSG00000071176 | *Arhgef10* | 975.4122936 | 1031.476013 | 782.377425 | 367.7220674 | 408.2384745 | 382.7005741 | 8.68E-20 | -1.271169905 | 7.91E-21 | 87.62461048 | 0.414323655 | 19.06155021 |
| ENSMUSG00000052713 | *Zfp608* | 165.3647377 | 172.5855168 | 116.7527204 | 68.31533822 | 60.80147492 | 59.91776665 | 9.61E-07 | -1.271401212 | 2.34E-07 | 26.72644519 | 0.414257232 | 6.017097251 |
| ENSMUSG00000015599 | *Ttbk1* | 178.9454167 | 178.641149 | 108.7008086 | 66.62853974 | 67.31591866 | 59.91776665 | 6.45E-06 | -1.274827516 | 1.73E-06 | 22.86956138 | 0.413274566 | 5.190388593 |
| ENSMUSG00000049604 | *Hoxb13* | 117.4329297 | 111.0199231 | 132.8565439 | 46.38695805 | 57.54425305 | 45.42153278 | 9.77E-07 | -1.275775761 | 2.38E-07 | 26.69409068 | 0.413003021 | 6.009930415 |
| ENSMUSG00000021250 | *Fos* | 515.2669364 | 561.1552476 | 332.8123523 | 162.7760528 | 245.3773809 | 175.8876376 | 2.95E-07 | -1.27698642 | 6.86E-08 | 29.10445668 | 0.412656589 | 6.530288844 |
| ENSMUSG00000032997 | *Chpf* | 1726.343953 | 1758.151873 | 1564.75485 | 685.6835799 | 690.5310366 | 701.6177192 | 1.26E-45 | -1.282641489 | 3.25E-47 | 208.288517 | 0.411042225 | 44.8982033 |
| ENSMUSG00000048232 | *Fbxo10* | 43.9374907 | 63.58413777 | 71.12522045 | 22.77177941 | 26.05777497 | 24.16038978 | 0.001135381 | -1.282674853 | 0.000407006 | 12.49975412 | 0.411032719 | 2.94485848 |
| ENSMUSG00000074419 | *Pira13* | 49.52953497 | 72.66758603 | 41.60154404 | 16.02458551 | 21.71481247 | 29.95888332 | 0.005063826 | -1.284260362 | 0.002009512 | 9.54082723 | 0.410581247 | 2.295521258 |
| ENSMUSG00000033021 | *Gmppa* | 2814.395995 | 3012.677004 | 3097.302053 | 1178.228734 | 1247.515976 | 1236.045541 | 2.26E-59 | -1.284812471 | 3.45E-61 | 272.3753072 | 0.41042415 | 58.64537411 |
| ENSMUSG00000032478 | *Nme6* | 2183.293856 | 2081.118922 | 2407.521613 | 904.1239823 | 905.50768 | 925.8261363 | 6.10E-46 | -1.285080312 | 1.55E-47 | 209.760931 | 0.41034796 | 45.21442849 |
| ENSMUSG00000009739 | *Pou6f1* | 119.8295201 | 101.9364748 | 79.1771322 | 48.07375652 | 39.08666245 | 36.72379246 | 3.74E-05 | -1.285350723 | 1.10E-05 | 19.33119341 | 0.410271054 | 4.426681794 |
| ENSMUSG00000072680 | *None* | 601.5441909 | 512.7101903 | 642.8109546 | 223.5007979 | 252.9775653 | 244.5031446 | 7.61E-19 | -1.285493527 | 7.34E-20 | 83.2210809 | 0.410230446 | 18.1187112 |
| ENSMUSG00000001288 | *Rarg* | 539.2328404 | 547.0254393 | 442.8551462 | 212.5366078 | 208.4621997 | 207.7793521 | 1.20E-18 | -1.285648707 | 1.17E-19 | 82.2995658 | 0.410186323 | 17.92021262 |
| ENSMUSG00000022558 | *Mroh1* | 7148.230305 | 7832.96021 | 7504.38175 | 3100.335596 | 3016.187452 | 3099.294801 | 8.98E-82 | -1.286433752 | 7.05E-84 | 376.5396313 | 0.40996318 | 81.04648885 |
| ENSMUSG00000033735 | *Spr* | 2732.113058 | 2682.645051 | 3098.644038 | 1094.73221 | 1141.113395 | 1251.508191 | 9.14E-39 | -1.286899379 | 3.25E-40 | 176.2173059 | 0.409830887 | 38.03919706 |
| ENSMUSG00000038990 | *Cables2* | 1000.975925 | 1108.180687 | 927.3118364 | 391.3372461 | 428.8675463 | 425.2228601 | 1.59E-28 | -1.288433239 | 9.06E-30 | 128.4253311 | 0.40939539 | 27.79797095 |
| ENSMUSG00000008318 | *Relt* | 826.8236886 | 841.7328715 | 770.2995573 | 333.1426987 | 326.8079277 | 339.2118725 | 2.70E-33 | -1.288754714 | 1.25E-34 | 150.6559368 | 0.409304175 | 32.56860123 |
| ENSMUSG00000020658 | *Efr3b* | 357.8908333 | 309.8465126 | 175.8000732 | 118.0758932 | 104.2310999 | 123.7011957 | 5.66E-06 | -1.290858308 | 1.51E-06 | 23.13379968 | 0.408707803 | 5.246922599 |
| ENSMUSG00000028337 | *Coro2a* | 295.5794829 | 279.5683518 | 226.7955143 | 102.0513077 | 99.88813737 | 126.6004424 | 1.92E-10 | -1.292418876 | 3.32E-11 | 43.9781074 | 0.408265941 | 9.717363378 |
| ENSMUSG00000026932 | *Nacc2* | 79.88634673 | 116.0662832 | 97.96492628 | 34.57936873 | 36.9151812 | 48.32077956 | 4.80E-05 | -1.292528424 | 1.43E-05 | 18.83248347 | 0.408234942 | 4.318942805 |
| ENSMUSG00000026864 | *Hspa5* | 144525.5853 | 138854.6364 | 165685.53 | 61675.25602 | 61184.74136 | 60376.81406 | 3.91E-54 | -1.293186361 | 7.11E-56 | 247.9926925 | 0.40804881 | 53.40747703 |
| ENSMUSG00000050014 | *Apol10b* | 79.08748326 | 76.70467414 | 87.22904395 | 37.10956644 | 32.57221871 | 28.99246773 | 2.83E-05 | -1.293275729 | 8.21E-06 | 19.88903417 | 0.408023534 | 4.547528607 |
| ENSMUSG00000046668 | *Cxxc5* | 48.7306715 | 59.54704966 | 52.33742637 | 24.45857788 | 18.4575906 | 22.2275586 | 0.000776624 | -1.293349284 | 0.000271983 | 13.25410621 | 0.408002732 | 3.109789146 |
| ENSMUSG00000010607 | *Pigyl* | 408.2192318 | 387.5604588 | 504.5864696 | 156.0288589 | 184.575906 | 189.4174559 | 9.62E-14 | -1.293375577 | 1.29E-14 | 59.39246499 | 0.407995296 | 13.01701921 |
| ENSMUSG00000083044 | *Gm12416* | 69.50112165 | 42.38942518 | 50.99544108 | 29.5189733 | 17.37184998 | 19.32831182 | 0.003519504 | -1.294089429 | 0.001359072 | 10.26052258 | 0.407793468 | 2.453518586 |
| ENSMUSG00000020377 | *Ltc4s* | 123.024974 | 114.0477392 | 130.1725733 | 60.72474508 | 44.51536557 | 43.4887016 | 1.31E-06 | -1.294683725 | 3.26E-07 | 26.09079108 | 0.407625518 | 5.881990391 |
| ENSMUSG00000020964 | *Sel1l* | 11649.02708 | 12105.20871 | 11579.99108 | 4933.042139 | 4842.403181 | 4620.432941 | 1.67E-98 | -1.295371509 | 8.88E-101 | 454.1801541 | 0.407431235 | 97.77681895 |
| ENSMUSG00000018398 | *Septin8* | 2094.620011 | 2247.648807 | 2137.782569 | 893.1597922 | 880.5356457 | 862.0427073 | 4.23E-61 | -1.297393638 | 6.02E-63 | 280.4396965 | 0.406860566 | 60.37322218 |
| ENSMUSG00000000811 | *Txnrd3* | 235.6647228 | 227.0862063 | 216.0596319 | 80.12292754 | 118.345728 | 79.24607847 | 6.49E-09 | -1.298322686 | 1.29E-09 | 36.83497906 | 0.406598645 | 8.187820082 |
| ENSMUSG00000019066 | *Rab3d* | 868.3645889 | 1039.550189 | 1021.250807 | 382.0598545 | 404.9812526 | 402.0288859 | 1.23E-27 | -1.299346478 | 7.24E-29 | 124.3018794 | 0.40631021 | 26.91021439 |
| ENSMUSG00000032383 | *Ppib* | 17150.79978 | 19616.21114 | 26100.27193 | 8563.875855 | 8483.977232 | 8436.808111 | 1.55E-19 | -1.302473963 | 1.44E-20 | 86.4464742 | 0.405430361 | 18.81036978 |
| ENSMUSG00000025877 | *Hk3* | 17949.66325 | 18518.12317 | 17467.28055 | 7167.206718 | 7429.723087 | 7272.277323 | 3.84E-114 | -1.302557267 | 1.45E-116 | 526.7324603 | 0.405406952 | 113.4153401 |
| ENSMUSG00000039234 | *Sec24d* | 3599.678784 | 4014.884128 | 3312.019699 | 1501.250642 | 1458.149657 | 1466.052452 | 2.04E-43 | -1.304374154 | 5.72E-45 | 197.9964415 | 0.404896716 | 42.69061371 |
| ENSMUSG00000032038 | *St3gal4* | 1486.684913 | 1668.326663 | 1662.719776 | 634.2362264 | 664.4732616 | 642.6663681 | 6.85E-44 | -1.310458286 | 1.88E-45 | 200.2129758 | 0.403192781 | 43.1641461 |
| ENSMUSG00000041895 | *Wipi1* | 159.7726935 | 150.3815322 | 148.9603674 | 54.82095042 | 66.23017804 | 64.74984461 | 5.43E-09 | -1.311754318 | 1.07E-09 | 37.19744169 | 0.402830739 | 8.2651756 |
| ENSMUSG00000024909 | *Efemp2* | 305.1658445 | 281.5868959 | 265.7130877 | 134.1004787 | 93.37369363 | 115.0034553 | 1.92E-11 | -1.311825483 | 3.07E-12 | 48.64487394 | 0.402810869 | 10.71650027 |
| ENSMUSG00000037348 | *Paqr7* | 2194.477945 | 2497.94827 | 2673.234701 | 931.9561571 | 981.5095237 | 1050.493748 | 8.51E-35 | -1.31230235 | 3.65E-36 | 157.6747871 | 0.402677746 | 34.07021873 |
| ENSMUSG00000050288 | *Fzd2* | 75.09316593 | 91.84375456 | 65.75727928 | 36.2661672 | 28.22925621 | 28.99246773 | 8.95E-05 | -1.313513076 | 2.76E-05 | 17.57750642 | 0.402339956 | 4.048342928 |
| ENSMUSG00000024959 | *Bad* | 1481.092868 | 1606.761069 | 1996.874114 | 646.0438157 | 691.6167772 | 705.4833815 | 6.80E-23 | -1.313735835 | 5.20E-24 | 102.128144 | 0.402277838 | 22.16755445 |
| ENSMUSG00000045092 | *S1pr1* | 393.0408259 | 385.5419147 | 315.3665435 | 142.5344711 | 143.3177623 | 155.5929102 | 2.90E-15 | -1.314125127 | 3.52E-16 | 66.48704088 | 0.402169303 | 14.537216 |
| ENSMUSG00000025353 | *Ormdl2* | 3411.945869 | 3437.580528 | 4800.281388 | 1572.096178 | 1531.98002 | 1571.391751 | 1.69E-20 | -1.316072453 | 1.47E-21 | 90.94810815 | 0.401626828 | 19.7721903 |
| ENSMUSG00000013707 | *Tnfaip8l2* | 2645.835804 | 2882.480912 | 3483.793817 | 1173.168339 | 1216.029498 | 1223.482138 | 4.65E-29 | -1.317599424 | 2.58E-30 | 130.9174723 | 0.401201965 | 28.3322948 |
| ENSMUSG00000036764 | *Dnajc12* | 174.1522359 | 245.2531028 | 257.661176 | 86.87012143 | 94.45943425 | 88.91023438 | 1.07E-08 | -1.320013466 | 2.17E-09 | 35.81047076 | 0.4005312 | 7.969373111 |
| ENSMUSG00000048458 | *Inka2* | 116.6340662 | 117.0755553 | 97.96492628 | 47.23035729 | 41.2581437 | 44.45511719 | 6.61E-07 | -1.320789742 | 1.59E-07 | 27.48158629 | 0.400315743 | 6.179748723 |
| ENSMUSG00000021867 | *None* | 127.0192913 | 253.3272791 | 127.4886027 | 79.2795283 | 41.2581437 | 82.14532525 | 0.000907059 | -1.321173778 | 0.000320875 | 12.94442736 | 0.400209196 | 3.042364594 |
| ENSMUSG00000003380 | *Rabac1* | 3090.003891 | 3758.529033 | 4612.403447 | 1496.190247 | 1508.093726 | 1573.324582 | 3.07E-20 | -1.323060226 | 2.73E-21 | 89.73301201 | 0.39968623 | 19.51333296 |
| ENSMUSG00000031385 | *Plxnb3* | 56.71930618 | 44.40796924 | 49.65345579 | 27.83217483 | 19.54333122 | 12.56340268 | 0.00354498 | -1.324466884 | 0.001369747 | 10.24608962 | 0.399296717 | 2.450386194 |
| ENSMUSG00000032192 | *Gnb5* | 338.7181101 | 319.9392329 | 327.4444111 | 113.858897 | 131.3746155 | 148.828001 | 5.13E-15 | -1.3269264 | 6.33E-16 | 65.3307909 | 0.398616573 | 14.28956083 |
| ENSMUSG00000041420 | *Meis3* | 442.5703609 | 480.4134854 | 534.110146 | 172.0534444 | 231.2627528 | 176.8540532 | 7.87E-15 | -1.330094071 | 9.83E-16 | 64.46488776 | 0.397742306 | 14.10429268 |
| ENSMUSG00000025512 | *Chid1* | 3903.246901 | 3822.113171 | 4466.12705 | 1612.579342 | 1604.724642 | 1627.443855 | 8.98E-52 | -1.330393052 | 1.78E-53 | 236.9974515 | 0.397659888 | 51.04661689 |
| ENSMUSG00000046312 | *Myorg* | 88.67384487 | 77.71394617 | 110.0427939 | 40.48316339 | 35.82944058 | 32.8581301 | 1.83E-05 | -1.33040384 | 5.15E-06 | 20.77883006 | 0.397656914 | 4.737640708 |
| ENSMUSG00000051503 | *Ccdc121rt3* | 38.34544643 | 38.35233707 | 36.23360287 | 16.86798474 | 18.4575906 | 9.664155911 | 0.005934773 | -1.331509088 | 0.002382105 | 9.229009236 | 0.397352386 | 2.226595912 |
| ENSMUSG00000074738 | *Fndc10* | 73.49543899 | 92.85302659 | 72.46720574 | 31.20577178 | 33.65795933 | 29.95888332 | 2.83E-05 | -1.335635475 | 8.20E-06 | 19.89063481 | 0.396217505 | 4.547803694 |
| ENSMUSG00000021061 | *Sptb* | 39.1443099 | 40.37088113 | 26.83970583 | 11.80758932 | 17.37184998 | 13.52981828 | 0.007452128 | -1.336398894 | 0.003041028 | 8.782687455 | 0.396007897 | 2.127719712 |
| ENSMUSG00000044456 | *Rin3* | 1542.605355 | 1585.566356 | 1289.647865 | 612.3078462 | 576.5282711 | 561.4874584 | 5.38E-31 | -1.336987625 | 2.73E-32 | 139.9486111 | 0.395846328 | 30.26953588 |
| ENSMUSG00000026922 | *Agpat2* | 1496.271274 | 1618.872333 | 1782.156467 | 646.887215 | 599.3288242 | 680.3565762 | 2.36E-35 | -1.343292022 | 9.79E-37 | 160.2880277 | 0.394120303 | 34.62722128 |
| ENSMUSG00000047654 | *Tssk6* | 169.3590551 | 145.3351721 | 197.2718379 | 58.19454737 | 77.08758427 | 66.68267579 | 3.19E-08 | -1.343445764 | 6.77E-09 | 33.60121102 | 0.394078306 | 7.495564867 |
| ENSMUSG00000032359 | *Ctsh* | 167.7613281 | 174.6040609 | 225.453529 | 72.5323344 | 77.08758427 | 72.48116933 | 2.32E-09 | -1.348086086 | 4.42E-10 | 38.91989702 | 0.392812818 | 8.634257557 |
| ENSMUSG00000050538 | *B230217C12Rik* | 206.1067746 | 195.7987735 | 229.4794848 | 83.49652449 | 84.68776864 | 79.24607847 | 4.64E-12 | -1.348356243 | 7.08E-13 | 51.52231107 | 0.392739268 | 11.33331664 |
| ENSMUSG00000037049 | *Smpd1* | 1182.317932 | 1184.885361 | 1103.11191 | 467.2431774 | 475.5543931 | 421.3571977 | 4.48E-40 | -1.348565054 | 1.49E-41 | 182.3417067 | 0.392682428 | 39.34888392 |
| ENSMUSG00000048240 | *Gng7* | 78.28861979 | 95.88084267 | 85.88705866 | 39.63976415 | 38.00092182 | 24.16038978 | 4.61E-05 | -1.349919057 | 1.37E-05 | 18.91179689 | 0.392314059 | 4.335878527 |
| ENSMUSG00000044339 | *Alkbh2* | 722.1725744 | 731.7222204 | 978.3072775 | 315.4313147 | 311.607559 | 323.749223 | 2.86E-18 | -1.351364735 | 2.88E-19 | 80.52010438 | 0.391921131 | 17.54341985 |
| ENSMUSG00000080115 | *Eef1akmt3* | 143.7954241 | 171.5762448 | 111.3847792 | 49.760555 | 56.45851243 | 61.85059783 | 7.89E-07 | -1.35237118 | 1.91E-07 | 27.12161897 | 0.391647816 | 6.102833976 |
| ENSMUSG00000015943 | *Bola1* | 1279.779275 | 1207.089346 | 1454.712056 | 489.1715576 | 477.7258744 | 574.0508611 | 1.56E-27 | -1.353265755 | 9.25E-29 | 123.8153282 | 0.391405042 | 26.80676126 |
| ENSMUSG00000024150 | *Mcfd2* | 3353.628836 | 4072.412634 | 4931.795946 | 1620.169935 | 1678.555004 | 1533.701543 | 7.89E-22 | -1.35376088 | 6.44E-23 | 97.14704249 | 0.391270737 | 21.1031541 |
| ENSMUSG00000022474 | *Pmm1* | 712.5862128 | 805.3990785 | 828.0049249 | 311.2143185 | 282.2925621 | 319.8835607 | 7.68E-28 | -1.356865534 | 4.49E-29 | 125.2503191 | 0.390429634 | 27.11473062 |
| ENSMUSG00000026839 | *Upp2* | 49.52953497 | 47.43578532 | 64.41529399 | 13.4943878 | 32.57221871 | 17.39548064 | 0.004290289 | -1.357569542 | 0.001680515 | 9.869386963 | 0.390239159 | 2.3675135 |
| ENSMUSG00000034041 | *Lyl1* | 3717.111713 | 3847.344971 | 3674.355728 | 1400.042734 | 1509.179467 | 1476.683023 | 4.57E-81 | -1.358535326 | 3.70E-83 | 373.2344819 | 0.389978008 | 80.33974933 |
| ENSMUSG00000092486 | *None* | 357.0919699 | 341.1339455 | 432.1192639 | 137.4740757 | 135.7175779 | 166.2234817 | 2.18E-14 | -1.359101491 | 2.80E-15 | 62.4027424 | 0.389824997 | 13.66118927 |
| ENSMUSG00000084333 | *Gm11599* | 51.92612537 | 44.40796924 | 57.70536754 | 21.08498093 | 15.20036873 | 23.19397419 | 0.000834229 | -1.361796617 | 0.000293634 | 13.11056975 | 0.389097437 | 3.078714801 |
| ENSMUSG00000020453 | *Patz1* | 1870.93824 | 1776.31877 | 1503.023526 | 651.9476104 | 685.1023335 | 668.7595891 | 4.35E-34 | -1.363031224 | 1.93E-35 | 154.3637177 | 0.388764604 | 33.36195912 |
| ENSMUSG00000071073 | *Lrrc73* | 71.89771206 | 78.72321819 | 83.20308807 | 30.36237254 | 26.05777497 | 33.82454569 | 1.74E-05 | -1.365680485 | 4.90E-06 | 20.87546027 | 0.38805136 | 4.758823505 |
| ENSMUSG00000020712 | *Tcam1* | 126.2204278 | 128.1775476 | 126.1466174 | 48.07375652 | 51.02980931 | 48.32077956 | 1.15E-08 | -1.369760806 | 2.34E-09 | 35.66566042 | 0.386955399 | 7.939990642 |
| ENSMUSG00000068699 | *Flnc* | 366.6783315 | 422.8849798 | 320.7344847 | 127.3532848 | 152.0036873 | 151.7272478 | 2.66E-14 | -1.37061609 | 3.44E-15 | 61.99857168 | 0.386726065 | 13.57543866 |
| ENSMUSG00000028124 | *Gclm* | 4694.920597 | 5889.102284 | 7073.604472 | 2266.21375 | 2158.45236 | 2378.34877 | 1.97E-21 | -1.375304746 | 1.63E-22 | 95.31088625 | 0.385471273 | 20.7063021 |
| ENSMUSG00000042675 | *Ypel3* | 187.7329148 | 183.6875091 | 193.245882 | 72.5323344 | 80.34480614 | 64.74984461 | 2.06E-11 | -1.377120581 | 3.30E-12 | 48.49893646 | 0.384986409 | 10.68562441 |
| ENSMUSG00000038515 | *Grtp1* | 153.3817857 | 158.4557084 | 177.1420585 | 64.09834203 | 46.68684681 | 76.3468317 | 5.82E-08 | -1.377163808 | 1.26E-08 | 32.38550248 | 0.384974874 | 7.235112912 |
| ENSMUSG00000070371 | *Prss36* | 104.6511142 | 84.77885036 | 130.1725733 | 44.70015957 | 35.82944058 | 41.55587042 | 6.45E-06 | -1.377302719 | 1.73E-06 | 22.87067564 | 0.384937808 | 5.190544986 |
| ENSMUSG00000044551 | *9930012K11Rik* | 164.5658743 | 196.8080455 | 202.639779 | 75.90593135 | 70.57314053 | 69.58192256 | 1.21E-10 | -1.377416505 | 2.06E-11 | 44.90874759 | 0.384907449 | 9.917251723 |
| ENSMUSG00000025477 | *Inpp5a* | 384.2533278 | 414.8108036 | 343.5482346 | 143.3778703 | 141.1462811 | 155.5929102 | 2.40E-18 | -1.378920555 | 2.40E-19 | 80.87950932 | 0.384506381 | 17.61959549 |
| ENSMUSG00000022844 | *Pdia5* | 1437.155378 | 1676.400839 | 1984.796246 | 626.6456333 | 624.3008585 | 705.4833815 | 5.30E-23 | -1.379810004 | 4.02E-24 | 102.6380731 | 0.384269398 | 22.27598406 |
| ENSMUSG00000033594 | *Spata2l* | 97.46134301 | 112.0291951 | 115.4107351 | 56.50774889 | 28.22925621 | 38.65662365 | 2.95E-05 | -1.380867754 | 8.55E-06 | 19.81115437 | 0.383987764 | 4.530719372 |
| ENSMUSG00000038604 | *Ripor1* | 1386.028116 | 1281.775476 | 950.1255864 | 453.7487896 | 456.0110619 | 481.2749644 | 8.11E-19 | -1.381652543 | 7.83E-20 | 83.09177117 | 0.383778942 | 18.09109858 |
| ENSMUSG00000056917 | *Sipa1* | 4710.897867 | 5082.693934 | 3944.094772 | 1688.485273 | 1715.470185 | 1856.484351 | 3.05E-35 | -1.385611198 | 1.27E-36 | 159.7681086 | 0.382727323 | 34.51608944 |
| ENSMUSG00000004460 | *Dnajb11* | 12248.97354 | 13280.00135 | 16035.38225 | 5336.186974 | 5241.955731 | 5318.184998 | 1.33E-37 | -1.386286021 | 4.94E-39 | 170.8020346 | 0.382548343 | 36.8749472 |
| ENSMUSG00000026198 | *Abcb6* | 2445.321073 | 2639.246354 | 2810.1172 | 980.0299137 | 1007.567299 | 1030.19902 | 4.77E-56 | -1.386310355 | 8.22E-58 | 256.8790032 | 0.382541891 | 55.32153247 |
| ENSMUSG00000037196 | *Pacrg* | 44.73635417 | 57.5285056 | 99.30691157 | 26.98877559 | 26.05777497 | 23.19397419 | 0.00177184 | -1.386445032 | 0.000652937 | 11.6186515 | 0.382506182 | 2.751575516 |
| ENSMUSG00000026456 | *Cyb5r1* | 8119.648282 | 8027.749712 | 9144.287776 | 3188.049117 | 3199.677618 | 3276.148854 | 1.77E-70 | -1.387503524 | 1.91E-72 | 324.0422449 | 0.382225643 | 69.7530054 |
| ENSMUSG00000024856 | *Cdk2ap2* | 2637.847169 | 2554.467503 | 2607.477421 | 981.7167121 | 1004.310077 | 995.4080589 | 6.47E-85 | -1.387911766 | 4.88E-87 | 391.0520595 | 0.3821175 | 84.18935061 |
| ENSMUSG00000030588 | *Yif1b* | 3507.809485 | 3935.151638 | 4546.646168 | 1491.973251 | 1462.49262 | 1616.813284 | 3.18E-34 | -1.390027595 | 1.41E-35 | 154.9920736 | 0.381557504 | 33.49695971 |
| ENSMUSG00000074785 | *Plxnc1* | 3800.193514 | 4374.18497 | 3999.116169 | 1615.109539 | 1528.722798 | 1493.112088 | 3.31E-58 | -1.391877098 | 5.19E-60 | 266.970941 | 0.381068669 | 57.48077267 |
| ENSMUSG00000023393 | *Slc17a9* | 738.1498438 | 774.1116456 | 716.6201457 | 269.0443567 | 301.8358933 | 280.2605214 | 1.01E-33 | -1.391987728 | 4.56E-35 | 152.6536427 | 0.381039449 | 32.99567616 |
| ENSMUSG00000046207 | *Pik3r6* | 694.2123531 | 672.1751707 | 742.1178662 | 259.7669651 | 259.492009 | 283.1597682 | 4.87E-32 | -1.39219119 | 2.37E-33 | 144.8038984 | 0.380985715 | 31.31207745 |
| ENSMUSG00000060671 | *Atp8b2* | 5842.088536 | 6202.985885 | 4676.818741 | 2070.545127 | 2188.853097 | 2113.550898 | 3.57E-33 | -1.392488621 | 1.66E-34 | 150.0881541 | 0.380907178 | 32.44781889 |
| ENSMUSG00000007783 | *Cpt1c* | 32.75340216 | 26.24107273 | 33.54963229 | 12.65098856 | 10.85740624 | 11.59698709 | 0.006664662 | -1.392636296 | 0.002699849 | 8.999964308 | 0.38086819 | 2.17622189 |
| ENSMUSG00000006356 | *Crip2* | 30.35681176 | 44.40796924 | 38.91757345 | 14.33778703 | 16.28610935 | 12.56340268 | 0.003481585 | -1.393816215 | 0.001343196 | 10.28220077 | 0.380556821 | 2.458223023 |
| ENSMUSG00000028988 | *Ctnnbip1* | 858.7782273 | 968.901147 | 929.995807 | 342.4200903 | 340.9225558 | 364.3386779 | 7.84E-37 | -1.394943993 | 3.01E-38 | 167.2107253 | 0.38025945 | 36.10549663 |
| ENSMUSG00000022131 | *Gpr180* | 1070.477046 | 1181.857545 | 1277.569998 | 451.2185919 | 421.2673619 | 465.8123149 | 7.93E-35 | -1.395926717 | 3.38E-36 | 157.8242638 | 0.380000516 | 34.100481 |
| ENSMUSG00000110841 | *Gpx4-ps2* | 4122.135491 | 4219.76635 | 5610.840504 | 1757.64401 | 1748.042404 | 1791.734506 | 3.69E-27 | -1.396299635 | 2.24E-28 | 122.0607868 | 0.379902303 | 26.43339846 |
| ENSMUSG00000040219 | *Ttc12* | 804.4555116 | 906.3262813 | 1180.947057 | 389.6504476 | 373.4947745 | 332.4469633 | 1.09E-16 | -1.39678658 | 1.21E-17 | 73.14313821 | 0.379774099 | 15.96201444 |
| ENSMUSG00000003420 | *Fcgrt* | 106.2488411 | 115.0570112 | 135.5405144 | 59.88134584 | 41.2581437 | 32.8581301 | 8.90E-06 | -1.400157855 | 2.43E-06 | 22.22225275 | 0.378887683 | 5.050770408 |
| ENSMUSG00000040855 | *Reps2* | 40.74203683 | 35.32452098 | 69.78323516 | 20.24158169 | 11.94314686 | 22.2275586 | 0.005558279 | -1.400821859 | 0.002220818 | 9.357442285 | 0.378713339 | 2.255059677 |
| ENSMUSG00000106964 | *None* | 35.14999256 | 48.44505735 | 79.1771322 | 15.18118627 | 22.80055309 | 23.19397419 | 0.004465026 | -1.403106618 | 0.00175344 | 9.791252622 | 0.378114055 | 2.350176002 |
| ENSMUSG00000064307 | *Lrrc51* | 76.69089286 | 115.0570112 | 130.1725733 | 42.16996186 | 36.9151812 | 41.55587042 | 1.54E-05 | -1.403490968 | 4.31E-06 | 21.11989098 | 0.378013334 | 4.811580508 |
| ENSMUSG00000059674 | *Cdh24* | 59.11589658 | 53.49141749 | 53.67941166 | 21.92838017 | 20.62907185 | 20.29472741 | 0.000147085 | -1.404799365 | 4.67E-05 | 16.57925972 | 0.377670665 | 3.83243177 |
| ENSMUSG00000044345 | *Marveld1* | 758.9202939 | 806.4083505 | 653.546837 | 288.4425391 | 272.5208965 | 275.4284435 | 4.34E-27 | -1.409268001 | 2.64E-28 | 121.7305032 | 0.376502669 | 26.36236352 |
| ENSMUSG00000004562 | *Arhgef40* | 1913.278004 | 1873.208884 | 1470.81588 | 683.9967814 | 585.2141961 | 708.3826283 | 1.09E-24 | -1.411205158 | 7.52E-26 | 110.5239362 | 0.375997465 | 23.96425234 |
| ENSMUSG00000030605 | *Mfge8* | 5313.240921 | 6171.698452 | 6398.58587 | 2220.670192 | 2236.625685 | 2260.446068 | 1.71E-53 | -1.411927613 | 3.20E-55 | 244.9946546 | 0.375809225 | 52.7663498 |
| ENSMUSG00000035824 | *Tk2* | 657.4646336 | 749.8891169 | 893.7622041 | 309.5275201 | 258.4062684 | 292.8239241 | 2.72E-19 | -1.413165476 | 2.57E-20 | 85.29648157 | 0.375486911 | 18.56520398 |
| ENSMUSG00000047786 | *Lix1* | 248.4465383 | 231.1232944 | 319.3924994 | 128.1966841 | 68.40165928 | 100.5072215 | 1.34E-07 | -1.416643907 | 3.03E-08 | 30.69035861 | 0.374582679 | 6.872084897 |
| ENSMUSG00000025732 | *Mcrip2* | 640.6885008 | 634.8321057 | 734.0659545 | 258.9235658 | 259.492009 | 230.9733263 | 1.58E-27 | -1.420307041 | 9.37E-29 | 123.7890388 | 0.373632786 | 26.80187146 |
| ENSMUSG00000029860 | *Zyx* | 2962.9846 | 2849.174935 | 2285.400951 | 937.8599518 | 1094.426549 | 997.34089 | 8.20E-30 | -1.420421424 | 4.39E-31 | 134.4330332 | 0.373603163 | 29.08631768 |
| ENSMUSG00000025130 | *P4hb* | 68373.12644 | 76571.45023 | 87740.34034 | 28918.47305 | 28816.64189 | 29177.05311 | 1.89E-46 | -1.420681956 | 4.71E-48 | 212.1302562 | 0.373535702 | 45.72420923 |
| ENSMUSG00000020829 | *Slc46a1* | 294.7806194 | 319.9392329 | 279.1329406 | 120.6060909 | 110.7455436 | 101.4736371 | 1.58E-16 | -1.424716583 | 1.77E-17 | 72.39112555 | 0.372492535 | 15.80160479 |
| ENSMUSG00000044813 | *Shb* | 508.8760287 | 610.609577 | 500.5605137 | 200.7290185 | 212.8051622 | 190.3838715 | 5.08E-22 | -1.425357208 | 4.11E-23 | 98.03400796 | 0.372327167 | 21.29422321 |
| ENSMUSG00000010021 | *Kif19a* | 51.92612537 | 47.43578532 | 41.60154404 | 23.61517864 | 11.94314686 | 16.42906505 | 0.0014689 | -1.428722982 | 0.000534886 | 11.98993605 | 0.371459549 | 2.833007785 |

| | | | | | | | | | | | | | |
|---|---|---|---|---|---|---|---|---|---|---|---|---|---|
| ENSMUSG00000023262 | *Acy1* | 709.390759 | 721.6295001 | 966.2294099 | 307.8407216 | 281.2068215 | 298.6224177 | 2.43E-19 | -1.428764372 | 2.28E-20 | 85.52783204 | 0.371448892 | 18.61383796 |
| ENSMUSG00000022761 | *Lztr1* | 55.12157924 | 38.35233707 | 24.15573525 | 15.18118627 | 18.4575906 | 10.6305715 | 0.00922686 | -1.430432252 | 0.00383799 | 8.358917925 | 0.371019713 | 2.034946062 |
| ENSMUSG00000031788 | *Kifc3* | 1482.690595 | 1529.047123 | 1152.765365 | 514.4735347 | 500.5264275 | 529.5957439 | 7.25E-27 | -1.43280763 | 4.48E-28 | 120.6838767 | 0.370409337 | 26.13951311 |
| ENSMUSG00000090607 | *Gm17052* | 85.478391 | 92.85302659 | 99.30691157 | 32.89257025 | 38.00092182 | 31.89171451 | 9.07E-07 | -1.432980373 | 2.21E-07 | 26.84167334 | 0.370364988 | 6.042256957 |
| ENSMUSG00000072898 | *Gm10434* | 59.91476005 | 53.49141749 | 61.73132341 | 36.2661672 | 15.20036873 | 12.56340268 | 0.004268895 | -1.434176884 | 0.001671631 | 9.879139231 | 0.37005795 | 2.369684514 |
| ENSMUSG00000031586 | *Rbpms* | 421.0010473 | 392.6068189 | 356.9680875 | 145.908068 | 159.6038717 | 128.5332736 | 1.16E-18 | -1.436528149 | 1.13E-19 | 82.36954263 | 0.369455331 | 17.93532477 |
| ENSMUSG00000040722 | *Scamp5* | 716.5805302 | 733.7407645 | 606.5773518 | 245.429178 | 261.6634903 | 253.2008849 | 1.28E-27 | -1.440028434 | 7.52E-29 | 124.2251255 | 0.36856004 | 26.89428578 |
| ENSMUSG00000019437 | *Tlcd1* | 266.8203981 | 244.2438308 | 248.2672789 | 120.6060909 | 72.74462178 | 85.04457202 | 3.91E-10 | -1.440112087 | 6.97E-11 | 42.52647031 | 0.36853867 | 9.407270533 |
| ENSMUSG00000040466 | *Blvrb* | 1748.71213 | 1978.173175 | 2590.031613 | 746.408325 | 760.0184365 | 818.5540057 | 4.34E-21 | -1.440390792 | 3.67E-22 | 93.70010297 | 0.368467482 | 20.36239639 |
| ENSMUSG00000051339 | *2900026A02Rik* | 1560.180352 | 1440.231184 | 932.6797776 | 444.471398 | 517.8982774 | 486.1070423 | 6.60E-14 | -1.443975239 | 8.76E-15 | 60.1577489 | 0.367553142 | 13.18070974 |
| ENSMUSG00000024066 | *Xdh* | 993.7861533 | 957.7991547 | 760.9056603 | 355.0710789 | 325.7221871 | 316.0178983 | 3.46E-24 | -1.44616554 | 2.46E-25 | 108.1787996 | 0.366995546 | 23.46031173 |
| ENSMUSG00000074063 | *Osgin1* | 787.6793787 | 904.3077372 | 814.585072 | 306.1539231 | 312.6932996 | 300.5552488 | 5.63E-36 | -1.447011772 | 2.27E-37 | 163.1907703 | 0.366780343 | 35.2494951 |
| ENSMUSG00000075706 | *Gpx4* | 7548.460902 | 8086.287489 | 10279.60733 | 3123.950775 | 2991.215418 | 3385.353816 | 3.50E-30 | -1.44713147 | 1.84E-31 | 136.1601986 | 0.366749913 | 29.45540358 |
| ENSMUSG00000035547 | *Capn5* | 576.7794234 | 645.934098 | 559.6078666 | 219.2838017 | 205.2049779 | 227.1076639 | 4.87E-28 | -1.451375781 | 2.83E-29 | 126.1653292 | 0.365672546 | 27.31277995 |
| ENSMUSG00000028108 | *Ecm1* | 327.5340216 | 309.8465126 | 256.3191907 | 93.61731533 | 124.8601717 | 110.1713774 | 2.18E-13 | -1.453132224 | 3.02E-14 | 57.72363516 | 0.36522762 | 12.66198473 |
| ENSMUSG00000034266 | *Batf* | 49.52953497 | 62.57486574 | 108.7008086 | 22.77177941 | 28.22925621 | 28.99246773 | 0.000831504 | -1.454040355 | 0.000292626 | 13.11701389 | 0.364997793 | 3.080135839 |
| ENSMUSG00000053395 | *Cacng8* | 468.9328553 | 475.3671253 | 426.7513227 | 152.6552619 | 154.1751685 | 194.2495338 | 3.66E-20 | -1.454376376 | 3.28E-21 | 89.36780157 | 0.36491279 | 19.43689449 |
| ENSMUSG00000028807 | *Zbtb8a* | 346.7067448 | 325.9948651 | 347.5741905 | 123.1362886 | 145.4892436 | 104.3728838 | 1.92E-16 | -1.454949086 | 2.16E-17 | 71.99720898 | 0.364767959 | 15.71765574 |
| ENSMUSG00000002058 | *Unc119* | 2607.490357 | 3031.853173 | 3105.353965 | 1040.754659 | 1087.912105 | 1059.191488 | 9.99E-51 | -1.454996679 | 2.11E-52 | 232.0671338 | 0.364755926 | 50.00032333 |
| ENSMUSG00000072437 | *Nanos1* | 214.8942727 | 173.5947888 | 163.7222056 | 70.00213669 | 62.97295617 | 68.61550697 | 1.32E-10 | -1.45775021 | 2.25E-11 | 44.73872684 | 0.364060416 | 9.880741861 |
| ENSMUSG00000069633 | *Pex11g* | 84.67952753 | 105.973563 | 154.3283085 | 44.70015957 | 39.08666245 | 40.58945483 | 1.13E-05 | -1.457952726 | 3.11E-06 | 21.74677367 | 0.364009315 | 4.946918066 |
| ENSMUSG00000081669 | *Npm3-ps1* | 162.1692839 | 88.81593848 | 92.59698511 | 37.95296568 | 43.42962494 | 44.45511719 | 3.30E-05 | -1.460293592 | 9.63E-06 | 19.58399542 | 0.363419165 | 4.481939412 |
| ENSMUSG00000084289 | *Fth1-ps* | 47.13294457 | 44.40796924 | 48.31147049 | 16.86798474 | 19.54333122 | 14.49623387 | 0.000406875 | -1.461027006 | 0.000136514 | 14.54965534 | 0.363234463 | 3.390538535 |
| ENSMUSG00000056904 | *Gm5620* | 62.31135045 | 80.74176225 | 91.25499982 | 34.57936873 | 29.31499684 | 20.29472741 | 7.17E-05 | -1.46411949 | 2.18E-05 | 18.02278672 | 0.362456687 | 4.144374129 |
| ENSMUSG00000025171 | *Ubtd1* | 1731.137134 | 1933.765206 | 2231.72154 | 694.1175722 | 688.3595553 | 750.9049143 | 2.23E-34 | -1.464837359 | 9.71E-36 | 155.7278243 | 0.362276377 | 33.65248432 |
| ENSMUSG00000003809 | *Gcdh* | 666.2521317 | 679.2400749 | 782.377425 | 266.514159 | 248.6346028 | 253.2008849 | 1.31E-30 | -1.465085468 | 6.77E-32 | 138.1462646 | 0.36221408 | 29.8823906 |
| ENSMUSG00000021266 | *Wars1* | 14085.56066 | 14551.6841 | 14539.06865 | 5139.674952 | 5349.444052 | 5142.29736 | 3.58E-148 | -1.465866871 | 6.97E-151 | 684.5138431 | 0.362017948 | 147.4461736 |
| ENSMUSG00000022546 | *Gpt* | 98.26020648 | 82.76030631 | 190.5619114 | 35.42276796 | 21.71481247 | 76.3468317 | 0.00570501 | -1.469245295 | 0.002283149 | 9.306720015 | 0.361171186 | 2.243743626 |
| ENSMUSG00000053746 | *Ptrh1* | 1019.349784 | 1091.023062 | 1541.9411 | 424.2298163 | 471.2114306 | 420.3907821 | 2.41E-17 | -1.470989509 | 2.57E-18 | 76.19759032 | 0.360734795 | 16.61714851 |
| ENSMUSG00000047649 | *Polr1g* | 1792.649621 | 1725.855168 | 1902.935143 | 652.7910096 | 648.1871523 | 647.4984461 | 1.23E-65 | -1.47513545 | 1.55E-67 | 301.5011196 | 0.359699622 | 64.91066718 |
| ENSMUSG00000041762 | *Gpr155* | 173.3533724 | 143.316628 | 147.6183821 | 61.56814432 | 59.7157343 | 45.42153278 | 2.10E-09 | -1.480340931 | 3.98E-10 | 39.12347217 | 0.358404106 | 8.677789155 |
| ENSMUSG00000055782 | *Abcd2* | 83.0818006 | 73.67685805 | 110.0427939 | 33.73596949 | 34.74369995 | 26.09322096 | 8.30E-06 | -1.488807873 | 2.26E-06 | 22.3617709 | 0.356306851 | 5.080819361 |
| ENSMUSG00000050373 | *Snx21* | 332.3272024 | 368.3842903 | 318.0505141 | 105.4249047 | 122.6886905 | 134.3317672 | 7.44E-18 | -1.495381428 | 7.65E-19 | 78.58726636 | 0.354687052 | 17.12813691 |
| ENSMUSG00000079602 | *9230102O04Rik* | 34.35112909 | 57.5285056 | 64.41529399 | 20.24158169 | 14.11462811 | 20.29472741 | 0.0012208 | -1.496376872 | 0.000439572 | 12.3559993 | 0.354442407 | 2.913355417 |
| ENSMUSG00000062300 | *Nectin2* | 158.1749665 | 188.7338693 | 179.8260291 | 48.91715576 | 70.57314053 | 67.64909138 | 5.57E-10 | -1.498234453 | 1.01E-10 | 41.81102543 | 0.353986328 | 9.254099967 |
| ENSMUSG00000028931 | *Kcnab2* | 14139.88337 | 14883.7346 | 13380.93534 | 4843.641819 | 5188.75444 | 4971.241801 | 3.74E-98 | -1.499313886 | 2.03E-100 | 452.5291032 | 0.353721573 | 97.42705574 |
| ENSMUSG00000035165 | *Kcne3* | 106.2488411 | 81.75103428 | 85.88705866 | 24.45857788 | 31.48647808 | 41.55587042 | 8.72E-06 | -1.499810648 | 2.38E-06 | 22.26274635 | 0.353599797 | 5.059553592 |
| ENSMUSG00000032353 | *Tmed3* | 3229.804998 | 3361.885126 | 4337.296462 | 1341.848186 | 1314.831895 | 1203.187411 | 7.52E-31 | -1.500400315 | 3.85E-32 | 139.2689683 | 0.353455301 | 30.12392679 |
| ENSMUSG00000022799 | *Arhgap31* | 376.2646931 | 423.8942518 | 254.9772054 | 127.3532848 | 132.4603561 | 114.0370398 | 4.84E-11 | -1.501875276 | 8.00E-12 | 46.76593078 | 0.353094126 | 10.3147851 |
| ENSMUSG00000025351 | *Cd63* | 7405.464342 | 9477.064344 | 13172.92762 | 3586.976956 | 3404.882595 | 3617.293558 | 1.04E-16 | -1.501916284 | 1.15E-17 | 73.23500848 | 0.353084089 | 15.98130043 |
| ENSMUSG00000049625 | *Tifab* | 541.6294308 | 502.61747 | 462.9849256 | 148.4382658 | 184.575906 | 201.014443 | 5.91E-20 | -1.50284962 | 5.35E-21 | 88.39721055 | 0.352855739 | 19.22836326 |
| ENSMUSG00000033033 | *Calhm2* | 953.8429799 | 960.8269708 | 815.9270572 | 320.4917101 | 324.6364464 | 319.8835607 | 4.77E-37 | -1.503494097 | 1.82E-38 | 168.2084422 | 0.352698148 | 36.3214159 |
| ENSMUSG00000064281 | *Rpl19-ps1* | 39.94317336 | 54.50068952 | 59.04735283 | 24.45857788 | 18.4575906 | 10.6305715 | 0.001477244 | -1.503793794 | 0.000538274 | 11.97817237 | 0.352624888 | 2.830547675 |
| ENSMUSG00000034853 | *Acot11* | 549.6180655 | 506.6545581 | 453.5910285 | 156.8722581 | 180.2329435 | 197.1487806 | 1.29E-21 | -1.504540121 | 1.06E-22 | 96.16353429 | 0.352442517 | 20.89055088 |
| ENSMUSG00000030742 | *Lat* | 215.6931362 | 240.2067427 | 305.9726465 | 80.96632677 | 100.973878 | 86.01098761 | 1.15E-11 | -1.505165606 | 1.80E-12 | 49.69052581 | 0.352289747 | 10.9405616 |
| ENSMUSG00000029570 | *Lfng* | 2486.06311 | 2277.926968 | 1842.545805 | 775.083899 | 784.9904708 | 765.4011482 | 7.21E-33 | -1.50817368 | 3.39E-34 | 148.6673595 | 0.351555975 | 32.14219075 |
| ENSMUSG00000025348 | *Itga7* | 940.262301 | 961.8362428 | 1085.666101 | 324.7087063 | 340.9225558 | 380.7677429 | 8.86E-36 | -1.512224329 | 3.61E-37 | 162.271784 | 0.350570297 | 35.05252109 |
| ENSMUSG00000089762 | *Ier5l* | 122.2261105 | 171.5762448 | 151.6443379 | 43.0133611 | 55.3727718 | 57.01851988 | 1.54E-08 | -1.520851117 | 3.18E-09 | 35.07241114 | 0.34848027 | 7.812331087 |
| ENSMUSG00000028583 | *Pdpn* | 37.54658296 | 40.37088113 | 50.99544108 | 16.02458551 | 11.94314686 | 16.42906505 | 0.000728492 | -1.521313746 | 0.000253966 | 13.38262399 | 0.348368541 | 3.137575154 |
| ENSMUSG00000028461 | *Ccdc107* | 1320.521311 | 1344.350341 | 1607.698379 | 495.0753523 | 489.6690212 | 500.6032762 | 3.16E-40 | -1.521818364 | 1.05E-41 | 183.051439 | 0.348246712 | 39.49997448 |
| ENSMUSG00000036918 | *Ttc7* | 5091.95574 | 4887.904432 | 4064.873448 | 1599.928353 | 1613.410567 | 1676.731051 | 2.31E-48 | -1.523042527 | 5.22E-50 | 221.0955567 | 0.347951341 | 47.63563004 |
| ENSMUSG00000030284 | *Creld1* | 2109.798417 | 2133.601067 | 2353.842201 | 770.8669028 | 760.0184365 | 759.6026546 | 2.49E-69 | -1.524548752 | 2.74E-71 | 318.7269182 | 0.347588257 | 68.60461794 |
| ENSMUSG00000026930 | *Gpsm1* | 451.357859 | 493.5340218 | 421.3833815 | 160.2458551 | 157.4323904 | 157.5257414 | 3.74E-26 | -1.525076835 | 2.41E-27 | 117.3501455 | 0.347461049 | 25.42669914 |
| ENSMUSG00000032122 | *Slc37a2* | 2626.66308 | 2723.015932 | 2513.538451 | 969.9091228 | 883.7928676 | 876.5389412 | 6.96E-72 | -1.525475455 | 7.15E-74 | 330.58956 | 0.347365058 | 71.15715256 |
| ENSMUSG00000028167 | *Bdh2* | 55.12157924 | 51.47287344 | 116.7527204 | 15.18118627 | 29.31499684 | 32.8581301 | 0.002734427 | -1.526319761 | 0.001035416 | 10.76314004 | 0.34716183 | 2.563133685 |
| ENSMUSG00000050232 | *Cxcr3* | 198.9170034 | 172.5855168 | 130.1725733 | 54.82095042 | 58.62999367 | 61.85059783 | 3.40E-09 | -1.526638346 | 6.56E-10 | 38.14776324 | 0.347085176 | 8.468196506 |
| ENSMUSG00000031101 | *Sash3* | 142.1976972 | 155.4278923 | 142.2504409 | 48.07375652 | 60.80147492 | 44.45511719 | 4.34E-10 | -1.526764734 | 7.74E-11 | 42.32196421 | 0.347054771 | 9.36300989 |
| ENSMUSG00000037349 | *Nudt22* | 766.1100651 | 795.3063582 | 829.3469102 | 269.0443567 | 280.1210809 | 278.3276902 | 7.20E-44 | -1.529778605 | 1.99E-45 | 200.0965443 | 0.346330511 | 43.1424661 |
| ENSMUSG00000030659 | *Nucb2* | 2359.842682 | 2637.22781 | 3996.432198 | 1055.092446 | 1065.111552 | 990.5759809 | 1.53E-16 | -1.530497126 | 1.70E-17 | 72.46020308 | 0.346158067 | 15.81611809 |
| ENSMUSG00000038523 | *1700003F12Rik* | 45.53521764 | 57.5285056 | 95.2809557 | 28.67557407 | 18.4575906 | 20.29472741 | 0.000734288 | -1.534197421 | 0.000256203 | 13.36617457 | 0.345271361 | 3.134133584 |
| ENSMUSG00000036957 | *Lrfn3* | 39.94317336 | 45.41724127 | 21.47176466 | 13.4943878 | 13.02888748 | 10.6305715 | 0.004714511 | -1.536121907 | 0.001857813 | 9.684970891 | 0.344811093 | 2.326563364 |
| ENSMUSG00000043155 | *Hpdl* | 123.024974 | 106.982835 | 122.1206615 | 46.38695805 | 31.48647808 | 42.52228601 | 3.22E-08 | -1.539066979 | 6.82E-09 | 33.58577908 | 0.344107924 | 7.49248229 |
| ENSMUSG00000037572 | *Wdhd1* | 1578.554211 | 1715.762448 | 1826.441982 | 602.1870554 | 554.8134586 | 599.1776665 | 1.71E-53 | -1.541090396 | 3.20E-55 | 244.9958622 | 0.343625642 | 52.7663498 |
| ENSMUSG00000040813 | *Tex264* | 3383.985647 | 3864.502596 | 4233.963595 | 1312.329213 | 1256.201901 | 1367.478061 | 1.10E-48 | -1.543342189 | 2.46E-50 | 222.5939355 | 0.343089722 | 47.95788102 |
| ENSMUSG00000022901 | *Cd86* | 230.0726786 | 198.8265895 | 241.5573525 | 71.68893516 | 80.34480614 | 77.31324729 | 1.06E-14 | -1.548643829 | 1.34E-15 | 63.85582722 | 0.341831243 | 13.97472819 |
| ENSMUSG00000015337 | *Endog* | 623.1135045 | 576.2943281 | 693.8063957 | 217.5970032 | 187.8331279 | 239.6710666 | 2.44E-25 | -1.549083801 | 1.64E-26 | 113.5499825 | 0.341727013 | 24.61320734 |
| ENSMUSG00000000202 | *Btbd17* | 45.53521764 | 54.50068952 | 48.31147049 | 23.61517864 | 10.85740624 | 15.46264946 | 0.000608792 | -1.5515816 | 0.000209901 | 13.74030818 | 0.341135878 | 3.215530987 |
| ENSMUSG00000081534 | *Slc48a1* | 2684.980114 | 3143.882368 | 3330.807494 | 984.2469099 | 1119.398583 | 1019.568449 | 2.95E-44 | -1.552011329 | 7.93E-46 | 201.9265036 | 0.341034281 | 43.52955995 |
| ENSMUSG00000046841 | *Ckap4* | 3263.357264 | 3313.440068 | 2804.749259 | 999.4280961 | 1120.484324 | 1079.486215 | 1.28E-53 | -1.553908828 | 2.37E-55 | 245.5915083 | 0.340586032 | 52.89373036 |
| ENSMUSG00000004105 | *Angptl2* | 1409.195156 | 1408.943751 | 1307.093674 | 452.0619912 | 500.5264275 | 455.1817434 | 7.17E-57 | -1.554079229 | 1.19E-58 | 260.7368058 | 0.340545806 | 56.14423117 |
| ENSMUSG00000031066 | *Usp11* | 69.50112165 | 68.63049791 | 57.70536754 | 18.55478322 | 31.48647808 | 17.39548064 | 7.41E-05 | -1.555918534 | 2.26E-05 | 17.9578378 | 0.340111918 | 4.130397629 |
| ENSMUSG00000071324 | *Armc2* | 219.6874535 | 179.650421 | 218.7436025 | 65.7851405 | 79.25906552 | 65.7162602 | 5.83E-13 | -1.556168911 | 8.32E-14 | 55.72900585 | 0.340052897 | 12.2344117 |
| ENSMUSG00000073002 | *Vamp5* | 94.26588914 | 128.1775476 | 112.7267645 | 37.95296568 | 36.9151812 | 38.65662365 | 4.05E-08 | -1.557598499 | 8.67E-09 | 33.11904616 | 0.339716101 | 7.392465787 |
| ENSMUSG00000047409 | *Ctdspl* | 154.1806492 | 172.5855168 | 116.7527204 | 57.35114813 | 51.02980931 | 42.52228601 | 1.39E-08 | -1.558289272 | 2.87E-09 | 35.27229069 | 0.339553481 | 7.856035587 |
| ENSMUSG00000042659 | *Arrdc4* | 64.70794085 | 64.5934098 | 107.3588233 | 23.61517864 | 26.05777497 | 29.95888332 | 4.17E-05 | -1.561514262 | 1.23E-05 | 19.11568139 | 0.338795293 | 4.379943512 |
| ENSMUSG00000075702 | *Selenom* | 1292.56109 | 1351.415246 | 1837.177864 | 508.5697401 | 476.6401337 | 529.5957439 | 7.13E-24 | -1.562103778 | 5.17E-25 | 106.7042839 | 0.338656883 | 23.14685299 |
| ENSMUSG00000030826 | *Bcat2* | 3491.832216 | 3794.862826 | 4403.053741 | 1281.123441 | 1292.031342 | 1381.974295 | 8.79E-48 | -1.56243343 | 2.05E-49 | 218.3668916 | 0.338579509 | 47.05606715 |
| ENSMUSG00000022453 | *Naga* | 158.97383 | 199.8358616 | 114.0687498 | 48.07375652 | 58.62999367 | 54.1192731 | 8.55E-08 | -1.565130179 | 1.88E-08 | 31.61035433 | 0.337947213 | 7.067793515 |
| ENSMUSG00000041263 | *Rusc1* | 1139.978168 | 1224.24697 | 1155.449336 | 382.9032537 | 425.6103244 | 381.7341585 | 5.69E-53 | -1.566051543 | 1.08E-54 | 242.5816107 | 0.337731455 | 52.24495141 |
| ENSMUSG00000018809 | *Smyd4* | 1191.904293 | 1325.174173 | 1402.37463 | 414.9524247 | 445.1536557 | 460.0138214 | 1.32E-45 | -1.568954461 | 3.41E-47 | 208.1915396 | 0.337052572 | 44.87804199 |
| ENSMUSG00000031791 | *Tmem38a* | 67.10453125 | 72.66758603 | 81.86110278 | 21.08498093 | 32.57221871 | 21.261143 | 1.02E-05 | -1.570978467 | 2.79E-06 | 21.95865128 | 0.336580041 | 4.993380827 |
| ENSMUSG00000031129 | *Slc9a9* | 202.1124572 | 209.9285819 | 205.3237496 | 76.74933059 | 67.31591866 | 62.81701342 | 5.75E-15 | -1.57321045 | 7.12E-16 | 65.09944691 | 0.336059723 | 14.24023601 |
| ENSMUSG00000044927 | *H1f10* | 80.6852102 | 56.51923358 | 79.1771322 | 28.67557407 | 17.37184998 | 26.09322096 | 3.21E-05 | -1.573502578 | 9.37E-06 | 19.63545368 | 0.335991682 | 4.493082171 |
| ENSMUSG00000002289 | *Angptl4* | 401.828324 | 397.6531791 | 342.2062493 | 118.9192924 | 119.4314686 | 145.9287543 | 1.95E-20 | -1.574463232 | 1.71E-21 | 90.65919798 | 0.335768028 | 19.71024352 |
| ENSMUSG00000028212 | *Ccne2* | 1892.507554 | 1511.889498 | 1878.779408 | 588.6926676 | 608.0147492 | 576.9501079 | 7.87E-40 | -1.57481135 | 2.64E-41 | 181.2061983 | 0.335687018 | 39.10403328 |
| ENSMUSG00000006651 | *Aplp1* | 241.2567671 | 267.4570875 | 288.5268377 | 70.00213669 | 119.4314686 | 79.24607847 | 7.71E-11 | -1.575260124 | 1.30E-11 | 45.81961317 | 0.335582613 | 10.11302675 |
| ENSMUSG00000001281 | *Itgb7* | 1929.255274 | 2111.397083 | 2198.171908 | 729.5403402 | 682.9308522 | 676.4909138 | 4.16E-63 | -1.576188588 | 5.49E-65 | 289.8010836 | 0.335366715 | 62.38133362 |
| ENSMUSG00000018821 | *Avpi1* | 700.6032608 | 734.7500365 | 917.9179394 | 241.2121818 | 247.5488622 | 295.7231709 | 2.54E-23 | -1.582137481 | 1.89E-24 | 104.1385784 | 0.333986691 | 22.59570002 |
| ENSMUSG00000032679 | *Cd59a* | 234.0669959 | 219.0120301 | 344.8902199 | 82.65312525 | 81.43054677 | 100.5072215 | 1.53E-10 | -1.586352127 | 2.62E-11 | 44.44044261 | 0.333012416 | 9.816221269 |
| ENSMUSG00000028909 | *Ptpru* | 55.12157924 | 77.71394617 | 45.62749991 | 14.33778703 | 14.11462811 | 30.92529892 | 0.001073582 | -1.59171524 | 0.000383268 | 12.61204922 | 0.331776765 | 2.969164976 |
| ENSMUSG00000020092 | *Pald1* | 982.6020648 | 1171.764825 | 895.1041894 | 309.5275201 | 355.0371839 | 345.9767816 | 1.12E-29 | -1.596293245 | 6.04E-31 | 133.8007635 | 0.330725629 | 28.95134832 |
| ENSMUSG00000003814 | *Calr* | 88735.35736 | 100018.858 | 112338.9307 | 33471.14213 | 33566.75712 | 32468.66462 | 1.95E-61 | -1.597297425 | 2.74E-63 | 282.0121082 | 0.33049551 | 60.71043261 |
| ENSMUSG00000003849 | *Nqo1* | 37.54658296 | 26.24107273 | 49.65345579 | 12.65098856 | 10.85740624 | 13.52981828 | 0.001901452 | -1.601845487 | 0.000704179 | 11.4781842 | 0.329455271 | 2.720914745 |
| ENSMUSG00000025357 | *Dgka* | 194.922686 | 202.8636777 | 226.7955143 | 59.88134584 | 79.25906552 | 66.68267579 | 6.49E-14 | -1.604535762 | 8.61E-15 | 60.18994051 | 0.328841489 | 13.18742627 |
| ENSMUSG00000047821 | *Trim16* | 307.5624349 | 304.8001525 | 327.4444111 | 105.4249047 | 114.0027655 | 88.91023438 | 2.58E-20 | -1.608039147 | 2.28E-21 | 90.08332489 | 0.328043912 | 19.58849695 |
| ENSMUSG00000091742 | *Gm5093* | 35.14999256 | 43.39869721 | 45.62749991 | 21.92838017 | 4.342962494 | 13.52981828 | 0.007779251 | -1.613444318 | 0.003190393 | 8.695256204 | 0.326817169 | 2.109062225 |
| ENSMUSG00000082160 | *Gm11578* | 34.35112909 | 45.41724127 | 64.41529399 | 18.55478322 | 17.37184998 | 10.6305715 | 0.001016758 | -1.614920376 | 0.000362122 | 12.71815896 | 0.326482965 | 2.992782566 |
| ENSMUSG00000035863 | *Palm* | 116.6340662 | 118.0848273 | 79.1771322 | 30.36237254 | 38.00092182 | 34.79096128 | 2.73E-07 | -1.619138666 | 6.33E-08 | 29.26061776 | 0.325529757 | 6.564077966 |
| ENSMUSG00000049588 | *Ccdc69* | 307.5624349 | 315.9021448 | 468.3528667 | 118.0758932 | 109.659803 | 124.6676113 | 1.94E-13 | -1.624704446 | 2.68E-14 | 57.957269 | 0.324276314 | 12.71171242 |
| ENSMUSG00000002279 | *Lmf1* | 1750.309857 | 1883.301605 | 2121.678746 | 649.4174127 | 613.4434523 | 600.1440821 | 1.06E-51 | -1.62474677 | 2.11E-53 | 236.6519783 | 0.324266801 | 50.97515608 |
| ENSMUSG00000023191 | *P3h3* | 778.8918806 | 921.4653617 | 915.2339688 | 259.7669651 | 286.6355246 | 296.6895865 | 1.78E-36 | -1.632895635 | 6.94E-38 | 165.5487792 | 0.322440387 | 35.74975397 |
| ENSMUSG00000027602 | *Map1lc3a* | 305.1658445 | 343.1524896 | 448.2230874 | 120.6060909 | 118.345728 | 112.1042086 | 5.27E-16 | -1.636899092 | 6.10E-17 | 69.94500452 | 0.32154686 | 15.27848749 |
| ENSMUSG00000034265 | *Zdhhc14* | 390.6442355 | 430.959156 | 323.4184553 | 112.1720986 | 130.2888748 | 126.6004424 | 1.26E-19 | -1.63982546 | 1.16E-20 | 86.87443803 | 0.320895294 | 18.90074345 |
| ENSMUSG00000028756 | *Pink1* | 833.2145964 | 905.3170092 | 903.1561012 | 273.2613529 | 255.1490465 | 316.0178983 | 1.58E-39 | -1.643242233 | 5.38E-41 | 179.7915269 | 0.320136209 | 38.80197331 |
| ENSMUSG00000049858 | *Suox* | 364.2817411 | 447.1075085 | 399.9116169 | 123.9796879 | 143.3177623 | 120.8019489 | 1.13E-22 | -1.64380105 | 8.73E-24 | 101.1029453 | 0.320012231 | 21.94844108 |
| ENSMUSG00000053198 | *Prx* | 32.75340216 | 24.22252868 | 25.49772054 | 9.27739161 | 5.428703118 | 11.59698709 | 0.005443663 | -1.644091109 | 0.002171489 | 9.398618981 | 0.319947897 | 2.264108777 |
| ENSMUSG00000025586 | *Cpeb1* | 136.6056529 | 157.4464364 | 174.4580879 | 57.35114813 | 49.94406868 | 41.55587042 | 9.35E-11 | -1.644878367 | 1.58E-11 | 45.42751875 | 0.319773354 | 10.02897807 |
| ENSMUSG00000057244 | *Gm6139* | 34.35112909 | 48.44505735 | 29.52367641 | 6.747193898 | 14.11462811 | 15.46264946 | 0.002860897 | -1.647332541 | 0.001087695 | 10.67199986 | 0.319229849 | 2.543497814 |
| ENSMUSG00000073910 | *Mob3b* | 128.6170182 | 151.3908042 | 108.7008086 | 48.07375652 | 38.00092182 | 37.69020805 | 3.69E-09 | -1.649328019 | 7.14E-10 | 37.98292834 | 0.318788608 | 8.433364341 |
| ENSMUSG00000072494 | *Ppp1r3e* | 29.55794829 | 35.32452098 | 32.207647 | 15.18118627 | 6.514443741 | 8.69774032 | 0.003078655 | -1.651610977 | 0.001176663 | 10.52663505 | 0.318284548 | 2.511638941 |
| ENSMUSG00000029591 | *Ung* | 1305.342906 | 1272.692027 | 1013.198895 | 385.4334514 | 386.523662 | 373.0364182 | 3.18E-36 | -1.651746475 | 1.26E-37 | 164.3618747 | 0.318254656 | 35.49764469 |
| ENSMUSG00000031497 | *Tnfsf13b* | 221.2851804 | 274.5219917 | 260.3451466 | 82.65312525 | 85.77350926 | 71.51475374 | 1.73E-16 | -1.653662096 | 1.95E-17 | 72.19635372 | 0.317832356 | 15.76079957 |
| ENSMUSG00000022440 | *C1qtnf6* | 120.6283836 | 172.5855168 | 166.4061761 | 10.96419008 | 64.05869679 | 71.51475374 | 0.003694751 | -1.654750396 | 0.001433068 | 10.16274898 | 0.317592688 | 2.432414874 |
| ENSMUSG00000019872 | *Smpdl3a* | 776.4952902 | 873.0203043 | 1227.916542 | 283.3821437 | 302.921634 | 325.6820542 | 1.52E-18 | -1.655212377 | 1.49E-19 | 81.81738394 | 0.317491005 | 17.81793314 |
| ENSMUSG00000034957 | *Cebpa* | 2305.519967 | 2381.881986 | 2313.582643 | 757.372515 | 692.7025178 | 766.3675638 | 4.06E-90 | -1.657992447 | 2.54E-92 | 415.3267796 | 0.31687979 | 89.39164857 |
| ENSMUSG00000028751 | *Pla2g2e* | 340.3158371 | 388.5697308 | 481.7727197 | 114.7022963 | 138.9747998 | 128.5332736 | 1.13E-17 | -1.661394248 | 1.17E-18 | 77.74458734 | 0.316133483 | 16.94757807 |

| | | | | | | | | | | | | | |
|---|---|---|---|---|---|---|---|---|---|---|---|---|---|
| ENSMUSG00000026209 | *Dnpep* | 4289.097956 | 4384.27769 | 4859.328741 | 1379.801152 | 1416.891514 | 1477.649439 | 3.97E-88 | -1.662050181 | 2.77E-90 | 405.9678511 | 0.315989784 | 87.40103387 |
| ENSMUSG00000028755 | *Cda* | 74.29430246 | 103.9550189 | 119.4366909 | 28.67557407 | 38.00092182 | 27.05963655 | 1.13E-06 | -1.664411909 | 2.77E-07 | 26.40153011 | 0.315472923 | 5.947095083 |
| ENSMUSG00000046694 | *Tent5b* | 59.91476005 | 63.58413777 | 59.04735283 | 10.12079085 | 24.97203434 | 23.19397419 | 0.000155192 | -1.664873802 | 4.94E-05 | 16.47154245 | 0.315371938 | 3.809131827 |
| ENSMUSG00000049580 | *Tsku* | 88.67384487 | 56.51923358 | 69.78323516 | 21.08498093 | 22.80055309 | 24.16038978 | 6.98E-06 | -1.667094417 | 1.88E-06 | 22.710547 | 0.314886887 | 5.15635766 |
| ENSMUSG00000060224 | *Pyroxd2* | 91.86929874 | 141.2980839 | 108.7008086 | 31.20577178 | 35.82944058 | 40.58945483 | 9.55E-08 | -1.668353514 | 2.11E-08 | 31.38704537 | 0.314612192 | 7.020169382 |
| ENSMUSG00000021322 | *Aoah* | 440.9726339 | 545.0068952 | 708.5682339 | 203.2592162 | 185.6616466 | 142.0630919 | 7.30E-14 | -1.669482826 | 9.72E-15 | 59.95219914 | 0.314366017 | 13.1367045 |
| ENSMUSG00000079003 | *Samd1* | 2537.190372 | 2644.292714 | 2290.768893 | 796.1688799 | 755.675474 | 797.2928627 | 1.96E-73 | -1.669884355 | 1.90E-75 | 337.8195778 | 0.314278535 | 72.70873582 |
| ENSMUSG00000022136 | *Dnajc3* | 8570.207277 | 9282.274843 | 12430.80976 | 3127.324372 | 3209.449283 | 3173.708801 | 1.10E-33 | -1.670552755 | 5.00E-35 | 152.4721268 | 0.314132964 | 32.95713935 |
| ENSMUSG00000020331 | *Hcn2* | 134.2090625 | 168.5484287 | 166.4061761 | 37.95296568 | 59.7157343 | 50.25361074 | 5.58E-10 | -1.671145729 | 1.01E-10 | 41.80796115 | 0.314003876 | 9.253566012 |
| ENSMUSG00000050824 | *Sstr5* | 31.95453869 | 20.18544056 | 38.91757345 | 10.12079085 | 6.514443741 | 11.59698709 | 0.004739905 | -1.673042443 | 0.001869499 | 9.673450585 | 0.313591325 | 2.324230365 |
| ENSMUSG00000029596 | *Sdsl* | 71.09884859 | 60.55632169 | 100.6488969 | 24.45857788 | 20.62907185 | 27.05963655 | 7.63E-06 | -1.674051734 | 2.07E-06 | 22.53143181 | 0.313372018 | 5.117481307 |
| ENSMUSG00000031169 | *Porcn* | 56.71930618 | 48.44505735 | 45.62749991 | 12.65098856 | 20.62907185 | 14.49623387 | 8.75E-05 | -1.677482605 | 2.69E-05 | 17.62417065 | 0.312627673 | 4.058167482 |
| ENSMUSG00000023019 | *Gpd1* | 37.54658296 | 25.2318007 | 55.02139695 | 9.27739161 | 10.85740624 | 16.42906505 | 0.002624996 | -1.680581686 | 0.00099088 | 10.84452994 | 0.311956833 | 2.580871319 |
| ENSMUSG00000041000 | *Trim62* | 75.09316593 | 71.658314 | 76.49316162 | 18.55478322 | 30.40073746 | 21.261143 | 1.79E-06 | -1.681523152 | 4.51E-07 | 25.46148927 | 0.311753324 | 5.746741221 |
| ENSMUSG00000031214 | *Ophn1* | 158.97383 | 189.7431413 | 198.6138231 | 58.19454737 | 65.14443741 | 46.38794837 | 2.11E-12 | -1.687224505 | 3.13E-13 | 53.12373807 | 0.310523744 | 11.67608403 |
| ENSMUSG00000061848 | *Gm5805* | 31.15567522 | 44.40796924 | 53.67941166 | 12.65098856 | 21.71481247 | 5.798493547 | 0.004089557 | -1.689853055 | 0.001595854 | 9.964524313 | 0.309958494 | 2.388323735 |
| ENSMUSG00000084013 | *Gm14270* | 189.3306417 | 81.75103428 | 67.09926458 | 32.04917101 | 44.51536557 | 28.99246773 | 0.000362797 | -1.693076395 | 0.00012074 | 14.78110025 | 0.309266743 | 3.440336552 |
| ENSMUSG00000028555 | *Ttc39a* | 277.2056231 | 252.318007 | 316.7085288 | 83.49652449 | 93.37369363 | 84.07815643 | 3.60E-19 | -1.696405604 | 3.41E-20 | 84.73364933 | 0.308553892 | 18.44374831 |
| ENSMUSG00000036634 | *Mag* | 150.1863319 | 113.0384672 | 167.7481614 | 51.44735347 | 38.00092182 | 42.52228601 | 2.17E-09 | -1.697402782 | 4.12E-10 | 39.0574079 | 0.308340696 | 8.663634076 |
| ENSMUSG00000005803 | *Sqor* | 1279.779275 | 1206.080074 | 1362.115071 | 379.5296568 | 418.0101401 | 387.532652 | 1.29E-59 | -1.699689715 | 1.95E-61 | 273.5135996 | 0.307852307 | 58.88837074 |
| ENSMUSG00000002732 | *Fkbp7* | 55.92044271 | 73.67685805 | 87.22904395 | 26.98877559 | 19.54333122 | 19.32831182 | 8.55E-06 | -1.700671161 | 2.33E-06 | 22.30236106 | 0.30764295 | 5.067854773 |
| ENSMUSG00000021256 | *Vash1* | 24.76476749 | 27.25034476 | 20.12977937 | 5.060395423 | 9.771665612 | 7.731324729 | 0.006582355 | -1.700987899 | 0.002663953 | 9.02442805 | 0.307575416 | 2.181618699 |
| ENSMUSG00000025510 | *Cd151* | 2540.385826 | 2763.386813 | 3091.934112 | 813.0368647 | 887.0500894 | 878.4717723 | 4.40E-60 | -1.702508619 | 6.55E-62 | 275.6848555 | 0.307251377 | 59.35641802 |
| ENSMUSG00000015947 | *Fcgr1* | 6208.766868 | 6064.715617 | 6242.915576 | 1984.518405 | 1830.558691 | 1855.517935 | 4.86E-140 | -1.706397857 | 1.12E-142 | 646.7831018 | 0.3064242 | 139.3133229 |
| ENSMUSG00000059714 | *Flot1* | 1251.02019 | 1325.174173 | 1472.157865 | 382.0598545 | 434.2962494 | 422.3236133 | 4.69E-51 | -1.708395078 | 9.81E-53 | 233.5938856 | 0.306000289 | 50.32838546 |
| ENSMUSG00000052435 | *Cebpe* | 623.912368 | 571.2479679 | 605.2353665 | 169.5232467 | 180.2329435 | 200.0480274 | 1.40E-37 | -1.713234791 | 5.20E-39 | 170.7011072 | 0.30497549 | 36.85359523 |
| ENSMUSG00000029361 | *Nos1* | 283.5965309 | 313.8836008 | 296.5787494 | 75.90593135 | 102.0596186 | 95.67514352 | 6.75E-20 | -1.71416929 | 6.13E-21 | 88.12823665 | 0.304778008 | 19.17043903 |
| ENSMUSG00000006095 | *Tbcb* | 4039.053691 | 4468.047269 | 5671.229842 | 1425.344711 | 1344.146892 | 1544.332115 | 5.86E-36 | -1.715658781 | 2.37E-37 | 163.1084584 | 0.304463506 | 35.23214752 |
| ENSMUSG00000020865 | *Abcc3* | 738.1498438 | 796.3156302 | 717.962131 | 220.1272009 | 245.3773809 | 219.3763392 | 4.45E-45 | -1.720672832 | 1.16E-46 | 205.745756 | 0.303407188 | 44.35116007 |
| ENSMUSG00000021236 | *Entpd5* | 1984.376853 | 2180.027581 | 2545.746098 | 656.1646066 | 698.1312209 | 678.423745 | 6.06E-49 | -1.721765463 | 1.34E-50 | 223.8092471 | 0.303177488 | 48.2171716 |
| ENSMUSG00000030089 | *Slc41a3* | 393.8396894 | 415.8200756 | 399.9116169 | 123.9796879 | 119.4314686 | 119.8355333 | 3.28E-32 | -1.734177284 | 1.59E-33 | 145.5990592 | 0.300580374 | 31.48433367 |
| ENSMUSG00000037031 | *Tspan15* | 27.96022136 | 29.26888882 | 36.23360287 | 8.433992372 | 11.94314686 | 7.731324729 | 0.00133839 | -1.736004042 | 0.000484598 | 12.17401917 | 0.300200016 | 2.873417232 |
| ENSMUSG00000050947 | *Amigo1* | 419.4033203 | 464.2651329 | 330.1283817 | 119.7626917 | 117.2599873 | 127.566858 | 1.14E-20 | -1.738713371 | 9.92E-22 | 91.73275475 | 0.29963678 | 19.94234039 |
| ENSMUSG00000031410 | *Nxf7* | 167.7613281 | 144.3259 | 167.7481614 | 44.70015957 | 48.85832806 | 50.25361074 | 1.04E-13 | -1.740417724 | 1.40E-14 | 59.23292672 | 0.299283008 | 12.983524 |
| ENSMUSG00000022090 | *Pdlim2* | 2409.372217 | 2461.614477 | 2602.10948 | 743.034728 | 685.1023335 | 794.3936159 | 1.49E-81 | -1.747739305 | 1.19E-83 | 375.499415 | 0.297768014 | 80.82649331 |
| ENSMUSG00000037493 | *Cib2* | 32.75340216 | 28.25961679 | 38.91757345 | 10.96419008 | 11.94314686 | 6.764909138 | 0.001054831 | -1.748161624 | 0.000376263 | 12.64653303 | 0.297680861 | 2.976817231 |
| ENSMUSG00000032575 | *Manf* | 23284.47348 | 22687.42592 | 26084.16811 | 7251.546642 | 7081.200347 | 7110.88592 | 6.43E-115 | -1.748341409 | 2.35E-117 | 530.3639048 | 0.297643768 | 114.1915963 |
| ENSMUSG00000032115 | *Hyou1* | 51060.15738 | 53809.33818 | 46899.70197 | 15042.0254 | 15225.34076 | 14840.27782 | 8.13E-127 | -1.750518248 | 2.54E-129 | 585.3728311 | 0.297195001 | 126.0898701 |
| ENSMUSG00000040364 | *Sec1* | 47.93180804 | 75.69540211 | 61.73132341 | 16.86798474 | 20.62907185 | 17.39548064 | 1.30E-05 | -1.75644539 | 3.60E-06 | 21.46859943 | 0.295976515 | 4.886530019 |
| ENSMUSG00000026307 | *Scly* | 2843.953944 | 3122.687655 | 3520.02742 | 883.0390014 | 947.8515644 | 953.8521884 | 1.75E-62 | -1.767673688 | 2.36E-64 | 286.8970915 | 0.293681911 | 61.75625456 |
| ENSMUSG00000036995 | *Asap3* | 99.85793341 | 106.982835 | 92.59698511 | 28.67557407 | 31.48647808 | 28.02605214 | 1.04E-09 | -1.768748372 | 1.92E-10 | 40.54865745 | 0.293463225 | 8.98275213 |
| ENSMUSG00000039713 | *Plekhg5* | 349.1033352 | 293.6981602 | 248.2672789 | 90.24371838 | 89.03073113 | 83.11174084 | 6.24E-18 | -1.769483607 | 6.39E-19 | 78.94467609 | 0.293313706 | 17.20471015 |
| ENSMUSG00000024084 | *Qpct* | 939.4634375 | 1188.922449 | 1658.69382 | 420.8562194 | 362.6373683 | 323.749223 | 8.14E-17 | -1.771136899 | 8.94E-18 | 73.73429923 | 0.292977769 | 16.08929386 |
| ENSMUSG00000034930 | *Rtkn* | 70.29998512 | 75.69540211 | 107.3588233 | 32.89257025 | 23.88629372 | 16.42906505 | 6.70E-06 | -1.774468199 | 1.81E-06 | 22.79088668 | 0.29230204 | 5.17375371 |
| ENSMUSG00000081467 | *Gm13337* | 23.16704055 | 21.19471259 | 36.23360287 | 10.96419008 | 3.257221871 | 8.69774032 | 0.008158529 | -1.778348747 | 0.003354608 | 8.603808393 | 0.291516865 | 2.088388115 |
| ENSMUSG00000037235 | *Mxd4* | 801.2600577 | 751.907661 | 693.8063957 | 220.1272009 | 203.0334966 | 230.0069107 | 1.50E-44 | -1.783460588 | 3.98E-46 | 203.3003877 | 0.290485773 | 43.82455446 |
| ENSMUSG00000031570 | *Plpp5* | 2138.557502 | 2364.724362 | 2548.430069 | 667.9721959 | 642.7584491 | 719.9796154 | 2.27E-66 | -1.794198585 | 2.82E-68 | 304.8977788 | 0.28833171 | 65.64452793 |
| ENSMUSG00000035273 | *Hpse* | 36.7477195 | 27.25034476 | 33.54963229 | 7.590593135 | 13.02888748 | 7.731324729 | 0.000966182 | -1.800724203 | 0.000343102 | 12.81908252 | 0.287030469 | 3.014940908 |
| ENSMUSG00000020256 | *Aldh1l2* | 6362.947517 | 7496.872625 | 8547.104322 | 2139.703865 | 2255.083275 | 2026.573495 | 2.50E-51 | -1.802593751 | 5.10E-53 | 234.8946343 | 0.286658755 | 50.60210802 |
| ENSMUSG00000017386 | *Traf4* | 808.4498289 | 793.2878141 | 838.7408072 | 234.464988 | 243.2058997 | 221.3091704 | 3.16E-57 | -1.803778677 | 5.13E-59 | 262.4063235 | 0.286423411 | 56.50031256 |
| ENSMUSG00000028838 | *Extl1* | 219.6874535 | 253.3272791 | 226.7955143 | 72.5323344 | 53.20129055 | 73.44758493 | 3.11E-17 | -1.805666098 | 3.34E-18 | 75.6794385 | 0.28604894 | 16.50679041 |
| ENSMUSG00000047250 | *Ptgs1* | 203.7101842 | 139.2795399 | 136.8824997 | 47.23035729 | 38.00092182 | 52.18644192 | 3.64E-10 | -1.806823376 | 6.46E-11 | 42.67458428 | 0.285819574 | 9.439147695 |
| ENSMUSG00000062082 | *Cd200r4* | 587.1646485 | 733.7407645 | 1066.878307 | 226.0309956 | 224.7483091 | 227.1076639 | 2.02E-16 | -1.813113588 | 2.28E-17 | 71.88845568 | 0.284576101 | 15.69394835 |
| ENSMUSG00000013584 | *Aldh1a2* | 55.92044271 | 39.3616091 | 48.31147049 | 16.86798474 | 16.28610935 | 7.731324729 | 0.000166274 | -1.813361691 | 5.31E-05 | 16.33238013 | 0.284527166 | 3.779175728 |
| ENSMUSG00000017607 | *Tns4* | 70.29998512 | 62.57486574 | 65.75727928 | 16.86798474 | 19.54333122 | 20.29472741 | 3.98E-07 | -1.814731617 | 9.37E-08 | 28.49981007 | 0.284257118 | 6.39996402 |
| ENSMUSG00000040616 | *Tmem51* | 362.6840141 | 439.0333322 | 497.8765432 | 118.0758932 | 118.345728 | 131.4325204 | 2.21E-23 | -1.816204305 | 1.64E-24 | 104.413414 | 0.2839671 | 22.65490639 |
| ENSMUSG00000020592 | *Sdc1* | 1165.541799 | 1232.321146 | 1062.852351 | 336.5162957 | 297.4929309 | 343.0775349 | 5.29E-55 | -1.823880356 | 9.36E-57 | 252.0322127 | 0.282460227 | 54.2766993 |
| ENSMUSG00000015846 | *Rxra* | 211.6988188 | 198.8265895 | 187.8779408 | 64.94174127 | 55.3727718 | 47.35436397 | 1.72E-16 | -1.833520663 | 1.93E-17 | 72.21526189 | 0.280579077 | 15.76473254 |
| ENSMUSG00000038400 | *Pmepa1* | 718.9771206 | 751.907661 | 625.3651458 | 212.5366078 | 191.0903497 | 184.5853779 | 1.58E-39 | -1.833555516 | 5.37E-41 | 179.7959755 | 0.280572299 | 38.80219259 |
| ENSMUSG00000026674 | *Ddr2* | 14.37954241 | 20.18544056 | 28.18169112 | 4.216996186 | 6.514443741 | 6.764909138 | 0.009925761 | -1.836473919 | 0.004159165 | 8.212985863 | 0.280005308 | 2.003236204 |
| ENSMUSG00000050821 | *Fam131a* | 85.478391 | 84.77885036 | 59.04735283 | 24.45857788 | 19.54333122 | 20.29472741 | 3.80E-07 | -1.836988456 | 8.91E-08 | 28.5969747 | 0.279905462 | 6.420776413 |
| ENSMUSG00000002799 | *Jag2* | 304.366981 | 358.29157 | 217.4016172 | 89.40031915 | 76.00184365 | 81.17890965 | 2.29E-14 | -1.837213996 | 2.95E-15 | 62.29974038 | 0.279861707 | 13.63987075 |
| ENSMUSG00000040502 | *Marchf9* | 570.3885156 | 600.5168567 | 480.4307344 | 172.8968436 | 142.2320217 | 145.9287543 | 4.44E-30 | -1.839516213 | 2.35E-31 | 135.6767241 | 0.279415467 | 29.35257499 |
| ENSMUSG00000070570 | *Slc17a7* | 25.56363095 | 52.48214546 | 50.99544108 | 16.86798474 | 10.85740624 | 7.731324729 | 0.001167008 | -1.841977743 | 0.00041917 | 12.44474443 | 0.278939134 | 2.932926185 |
| ENSMUSG00000036587 | *Fut7* | 295.5794829 | 351.2266658 | 355.6261023 | 101.2079085 | 91.20221238 | 85.04457202 | 3.27E-24 | -1.846948576 | 2.31E-25 | 108.2980176 | 0.277979698 | 23.48571073 |
| ENSMUSG00000022861 | *Dgkg* | 278.8033501 | 250.299463 | 260.3451466 | 83.49652449 | 47.77258744 | 86.9774032 | 5.75E-14 | -1.847134127 | 7.61E-15 | 60.43415592 | 0.277943948 | 13.24053545 |
| ENSMUSG00000024968 | *Rcor2* | 373.0692392 | 327.0041371 | 291.2108083 | 100.3645092 | 92.287953 | 83.11174084 | 1.42E-22 | -1.848401356 | 1.10E-23 | 100.6375538 | 0.277699916 | 21.84903577 |
| ENSMUSG00000023046 | *Igfbp6* | 92.66816221 | 92.85302659 | 122.1206615 | 31.20577178 | 32.57221871 | 21.261143 | 1.20E-08 | -1.849064316 | 2.46E-09 | 35.5733114 | 0.277572334 | 7.920406086 |
| ENSMUSG00000034449 | *Dhrs11* | 646.280545 | 700.4347875 | 787.7453661 | 200.7290185 | 187.8331279 | 198.1151962 | 3.37E-42 | -1.858527531 | 1.02E-43 | 192.2613399 | 0.275757584 | 41.47235707 |
| ENSMUSG00000070284 | *Gmppb* | 2354.250638 | 2306.186584 | 2306.872716 | 654.4778081 | 593.9001211 | 669.7260047 | 6.80E-106 | -1.859628987 | 3.01E-108 | 488.506198 | 0.275547131 | 105.1671979 |
| ENSMUSG00000030824 | *Nucb1* | 9910.700175 | 11879.13177 | 12700.5488 | 3105.395991 | 3097.617999 | 3275.182438 | 3.21E-69 | -1.863139046 | 3.58E-71 | 318.1962411 | 0.274877543 | 68.49364346 |
| ENSMUSG00000032024 | *Clmp* | 45.53521764 | 44.40796924 | 45.62749991 | 5.903794661 | 17.37184998 | 14.49623387 | 0.000259923 | -1.866717699 | 8.51E-05 | 15.44166249 | 0.274196546 | 3.585156028 |
| ENSMUSG00000112597 | *Gm4777* | 24.76476749 | 20.18544056 | 17.44580879 | 5.060395423 | 5.428703118 | 6.764909138 | 0.005278315 | -1.867028577 | 0.002101482 | 9.458715318 | 0.274137467 | 2.277504715 |
| ENSMUSG00000002068 | *Ccne1* | 2521.213103 | 2499.966814 | 2203.539849 | 672.1891921 | 652.5301148 | 657.162602 | 4.25E-89 | -1.86705625 | 2.84E-91 | 410.5121207 | 0.274132209 | 88.37141139 |
| ENSMUSG00000040950 | *Mgl2* | 37.54658296 | 55.50996155 | 67.09926458 | 10.96419008 | 17.37184998 | 15.46264946 | 5.85E-05 | -1.869801215 | 1.76E-05 | 18.43408561 | 0.273611123 | 4.23273253 |
| ENSMUSG00000041845 | *Rhod* | 492.0998958 | 526.8399987 | 555.5819107 | 157.7156574 | 124.8601717 | 145.9287543 | 2.98E-35 | -1.870111468 | 1.24E-36 | 159.8185175 | 0.273552289 | 34.52648854 |
| ENSMUSG00000089704 | *Galnt2* | 407.4203683 | 380.4955546 | 385.1497787 | 107.1117031 | 80.34480614 | 132.398936 | 6.59E-21 | -1.87116551 | 5.63E-22 | 92.85333232 | 0.273352503 | 20.1808137 |
| ENSMUSG00000074796 | *Slc4a11* | 31.95453869 | 48.44505735 | 33.54963229 | 13.4943878 | 7.600184365 | 9.664155911 | 0.000353724 | -1.873640765 | 0.000117511 | 14.83220898 | 0.27288391 | 3.451335596 |
| ENSMUSG00000028862 | *Map3k6* | 204.5090476 | 219.0120301 | 123.4626468 | 54.82095042 | 53.20129055 | 41.55587042 | 5.57E-10 | -1.876412118 | 1.00E-10 | 41.81213446 | 0.272360215 | 9.254099967 |
| ENSMUSG00000033720 | *Sfxn5* | 321.1431138 | 347.1895777 | 322.07647 | 101.2079085 | 80.34480614 | 86.9774032 | 7.80E-27 | -1.876965966 | 4.84E-28 | 120.5326155 | 0.272255676 | 26.10764176 |
| ENSMUSG00000050711 | *Scg2* | 19.97158668 | 15.13908042 | 34.89161758 | 8.433992372 | 4.342962494 | 5.798493547 | 0.008674121 | -1.880550468 | 0.003590668 | 8.480026706 | 0.271580073 | 2.06177453 |
| ENSMUSG00000110545 | *Gm7730* | 77.48975633 | 81.75103428 | 103.3328674 | 22.77177941 | 38.00092182 | 10.6305715 | 4.18E-05 | -1.885170205 | 1.23E-05 | 19.11096521 | 0.270711822 | 4.379044036 |
| ENSMUSG00000056069 | *Otulinl* | 96.66247954 | 205.8914937 | 318.0505141 | 63.25494279 | 60.80147492 | 42.52228601 | 1.17E-05 | -1.890656016 | 3.22E-06 | 21.68174381 | 0.269684402 | 4.932661362 |
| ENSMUSG00000018340 | *Anxa6* | 3334.456112 | 3857.437692 | 4062.189477 | 996.0544992 | 996.7098924 | 1033.098267 | 7.61E-80 | -1.894012677 | 6.46E-82 | 367.5278172 | 0.269057667 | 79.11887681 |
| ENSMUSG00000020844 | *Nxn* | 185.3363244 | 174.6040609 | 174.4580879 | 44.70015957 | 52.11554993 | 47.35436397 | 1.15E-17 | -1.896285913 | 1.19E-18 | 77.70736885 | 0.26863405 | 16.93964126 |
| ENSMUSG00000027298 | *Tyro3* | 156.5772396 | 171.5762448 | 127.4886027 | 36.2661672 | 40.17240307 | 46.38794837 | 5.56E-13 | -1.898786094 | 7.93E-14 | 55.82374254 | 0.268168912 | 12.25480101 |
| ENSMUSG00000026544 | *Dusp23* | 71.09884859 | 90.83448253 | 73.80919103 | 22.77177941 | 24.97203434 | 15.46264946 | 1.24E-07 | -1.90124305 | 2.78E-08 | 30.85216005 | 0.267712601 | 6.906245798 |
| ENSMUSG00000009378 | *Slc16a12* | 489.7033054 | 682.267891 | 791.771322 | 150.9684635 | 212.8051622 | 161.3914037 | 1.15E-17 | -1.903202912 | 1.20E-18 | 77.70282904 | 0.267349167 | 16.93888949 |
| ENSMUSG00000047793 | *Sned1* | 22.36817708 | 24.22252868 | 33.54963229 | 6.747193898 | 9.771665612 | 4.832077956 | 0.002119671 | -1.908503113 | 0.000789664 | 11.26537081 | 0.266368777 | 2.673731475 |
| ENSMUSG00000038022 | *Mindy4* | 53.52385231 | 61.56559372 | 84.54507337 | 16.02458551 | 15.20036873 | 21.261143 | 2.22E-06 | -1.915506974 | 5.67E-07 | 25.02026037 | 0.265078768 | 5.653149371 |
| ENSMUSG00000082360 | *Gm8814* | 15.97726935 | 27.25034476 | 17.44580879 | 5.903794661 | 3.257221871 | 6.764909138 | 0.008270426 | -1.915598065 | 0.003407939 | 8.575086006 | 0.265062032 | 2.082472141 |
| ENSMUSG00000043687 | *1190005I06Rik* | 30.35681176 | 56.51923358 | 42.94352933 | 2.530197712 | 20.62907185 | 11.59698709 | 0.007397652 | -1.923587257 | 0.003017739 | 8.796712198 | 0.263598258 | 2.130906124 |
| ENSMUSG00000061451 | *Tmem151a* | 42.33976377 | 64.5934098 | 48.31147049 | 16.86798474 | 14.11462811 | 9.664155911 | 2.60E-05 | -1.925660795 | 7.47E-06 | 20.06808101 | 0.263219669 | 4.585798944 |
| ENSMUSG00000022769 | *Sdf2l1* | 11866.31794 | 12506.89897 | 13097.77645 | 3281.666432 | 3278.936683 | 3297.409997 | 1.14E-181 | -1.926047467 | 1.55E-184 | 839.2864193 | 0.26314913 | 180.9419732 |
| ENSMUSG00000080774 | *Gm9143* | 13.58067894 | 22.20398462 | 18.78779408 | 5.903794661 | 4.342962494 | 3.865662365 | 0.009290348 | -1.933637528 | 0.003866592 | 8.345427381 | 0.261768331 | 2.031968012 |
| ENSMUSG00000035064 | *Eef2k* | 903.5145815 | 1015.32766 | 781.0354397 | 245.429178 | 231.2627528 | 227.1076639 | 5.19E-43 | -1.940577995 | 1.48E-44 | 196.0966447 | 0.260512049 | 42.28505332 |
| ENSMUSG00000024679 | *Ms4a6d* | 3955.173027 | 4341.888265 | 6693.822634 | 1303.051822 | 1303.974489 | 1291.13123 | 3.84E-26 | -1.942518135 | 2.48E-27 | 117.2889758 | 0.260161947 | 25.41529439 |
| ENSMUSG00000002908 | *Kcnn1* | 27.96022136 | 45.41724127 | 36.23360287 | 12.65098856 | 8.685924988 | 6.764909138 | 0.000359927 | -1.947011435 | 0.0001197 | 14.79741253 | 0.259352928 | 3.443785179 |
| ENSMUSG00000038967 | *Pdk2* | 38.34544643 | 39.3616091 | 34.89161758 | 6.747193898 | 13.02888748 | 9.664155911 | 0.000117675 | -1.957417147 | 3.69E-05 | 17.02640605 | 0.257489026 | 3.929317512 |
| ENSMUSG00000056185 | *Snx32* | 230.871542 | 210.9378539 | 346.2322052 | 71.68893516 | 66.23017804 | 63.78342901 | 3.30E-14 | -1.958350521 | 4.31E-15 | 61.55279829 | 0.257322494 | 13.48145934 |
| ENSMUSG00000034438 | *Gbp8* | 186.9340513 | 185.7060532 | 213.3756614 | 58.19454737 | 40.17240307 | 50.25361074 | 4.15E-17 | -1.96653513 | 4.48E-18 | 75.09982018 | 0.2558668 | 16.38216034 |
| ENSMUSG00000032591 | *Mst1* | 727.7646187 | 802.3712624 | 889.7362483 | 243.7423796 | 180.2329435 | 189.4174559 | 1.07E-34 | -1.973232817 | 4.58E-36 | 157.2227337 | 0.254681696 | 33.97263558 |
| ENSMUSG00000012819 | *Cdh23* | 47.13294457 | 49.45432938 | 57.70536754 | 13.4943878 | 11.94314686 | 13.52981828 | 2.35E-06 | -1.975586684 | 6.00E-07 | 24.91258442 | 0.254266502 | 5.629797379 |
| ENSMUSG00000075602 | *Ly6a* | 19.97158668 | 16.14835245 | 28.18169112 | 5.903794661 | 5.428703118 | 4.832077956 | 0.004007716 | -1.977303883 | 0.001562025 | 10.00397729 | 0.253964036 | 2.397103111 |
| ENSMUSG00000074547 | *Rps8-ps4* | 28.75908482 | 44.40796924 | 45.62749991 | 8.433992372 | 13.02888748 | 8.69774032 | 0.000143904 | -1.980639691 | 4.56E-05 | 16.62355601 | 0.253377497 | 3.841928224 |
| ENSMUSG00000030745 | *Il21r* | 71.89771206 | 55.50996155 | 40.25955875 | 14.33778703 | 11.94314686 | 16.42906505 | 9.70E-06 | -1.981174003 | 2.65E-06 | 22.052286 | 0.253283675 | 5.013445421 |
| ENSMUSG00000042745 | *Id1* | 1263.802005 | 1142.495936 | 1097.743968 | 286.7557407 | 270.3494153 | 325.6820542 | 1.63E-59 | -1.989715417 | 2.46E-61 | 273.0474424 | 0.25178855 | 58.78847802 |
| ENSMUSG00000048200 | *Cracr2b* | 70.29998512 | 52.48214546 | 73.80919103 | 10.12079085 | 23.88629372 | 15.46264946 | 5.31E-06 | -2.009476838 | 1.41E-06 | 23.2656742 | 0.248363171 | 5.275164663 |
| ENSMUSG00000046908 | *Ltb4r1* | 26.36249442 | 25.2318007 | 33.54963229 | 6.747193898 | 6.514443741 | 7.731324729 | 0.000479678 | -2.011344015 | 0.000162724 | 14.21894896 | 0.24804194 | 3.319050166 |
| ENSMUSG00000020577 | *Tspan13* | 277.2056231 | 304.8001525 | 319.3924994 | 91.93051686 | 52.11554993 | 77.31324729 | 2.05E-18 | -2.013296885 | 2.04E-19 | 81.20220958 | 0.247706411 | 17.68753189 |
| ENSMUSG00000026223 | *Itm2c* | 10717.55228 | 12239.44189 | 13419.85292 | 2984.789901 | 3019.444674 | 2991.056255 | 8.96E-95 | -2.015426431 | 5.13E-97 | 436.8964129 | 0.247341044 | 94.04784403 |
| ENSMUSG00000113061 | *Rps18-ps5* | 15.97726935 | 35.32452098 | 33.54963229 | 7.590593135 | 4.342962494 | 8.69774032 | 0.002608028 | -2.015972403 | 0.000984167 | 10.85711807 | 0.247247459 | 2.583687742 |
| ENSMUSG00000067001 | *Serpinb7* | 51.12726191 | 27.25034476 | 60.38933812 | 12.65098856 | 7.600184365 | 13.52981828 | 0.00018415 | -2.022662493 | 5.92E-05 | 16.12950421 | 0.246103573 | 3.734829052 |
| ENSMUSG00000103735 | *Gm38317* | 280.401077 | 269.4756315 | 283.1588965 | 74.21913288 | 70.57314053 | 57.98493547 | 8.50E-27 | -2.036285849 | 5.29E-28 | 120.3529608 | 0.243790557 | 26.07078612 |
| ENSMUSG00000022817 | *Itgb5* | 3240.190223 | 3155.993632 | 2744.359921 | 781.8310929 | 646.015671 | 788.5951224 | 1.30E-68 | -2.043381347 | 1.49E-70 | 315.3490134 | 0.242594484 | 67.8847761 |
| ENSMUSG00000032194 | *Kank2* | 996.9816072 | 1029.457469 | 801.165219 | 218.4404024 | 239.9486778 | 229.0404951 | 1.67E-46 | -2.044430467 | 4.13E-48 | 212.3915731 | 0.242418135 | 45.77711398 |

| | | | | | | | | | | | | | |
|---|---|---|---|---|---|---|---|---|---|---|---|---|---|
| ENSMUSG00000000730 | *Dnmt3l* | 672.6430395 | 711.5367798 | 991.7271304 | 175.4270413 | 198.6905341 | 201.014443 | 3.54E-28 | -2.044678159 | 2.04E-29 | 126.8095229 | 0.242376518 | 27.45065657 |
| ENSMUSG00000054723 | *Vmac* | 191.7272321 | 185.7060532 | 167.7481614 | 45.54355881 | 36.9151812 | 49.28719515 | 2.10E-18 | -2.046828999 | 2.09E-19 | 81.15517034 | 0.242015441 | 17.67760332 |
| ENSMUSG00000055452 | *Gm7353* | 422.5987742 | 349.2081217 | 285.8428671 | 83.49652449 | 91.20221238 | 82.14532525 | 2.42E-22 | -2.049204429 | 1.92E-23 | 99.54416106 | 0.241617285 | 21.61677244 |
| ENSMUSG00000007594 | *Hapln4* | 387.4487816 | 416.8293476 | 362.3360287 | 96.99091228 | 87.94499051 | 96.64155911 | 4.39E-36 | -2.049865052 | 1.77E-37 | 163.6919499 | 0.241508178 | 35.35707382 |
| ENSMUSG00000025068 | *Gsto1* | 3451.090179 | 4330.786273 | 6028.197929 | 1076.177427 | 1111.798399 | 1114.277177 | 2.10E-29 | -2.06351022 | 1.15E-30 | 132.5225011 | 0.239233243 | 28.67691278 |
| ENSMUSG00000018983 | *E2f2* | 1918.870048 | 2087.174554 | 1728.477055 | 428.4468125 | 464.6969869 | 480.3085488 | 6.28E-76 | -2.064053104 | 5.78E-78 | 349.3799377 | 0.239143237 | 75.20194137 |
| ENSMUSG00000037703 | *Lzts3* | 341.913564 | 373.4306504 | 267.055073 | 84.33992372 | 69.48739991 | 80.21249406 | 4.25E-23 | -2.069489199 | 3.21E-24 | 103.0868758 | 0.238243837 | 22.37200163 |
| ENSMUSG00000032470 | *Mras* | 96.66247954 | 147.3537161 | 189.2199261 | 23.61517864 | 36.9151812 | 42.52228601 | 2.33E-08 | -2.071225173 | 4.89E-09 | 34.23469877 | 0.237957334 | 7.632713578 |
| ENSMUSG00000036181 | *H1f2* | 1004.970242 | 1147.542296 | 1480.209777 | 301.9369269 | 268.177934 | 291.8575085 | 1.53E-35 | -2.07154561 | 6.29E-37 | 161.1680199 | 0.237904488 | 34.81638381 |
| ENSMUSG00000029442 | *Wdr66* | 377.86242 | 448.1167805 | 590.4735283 | 99.52110999 | 132.4603561 | 104.3728838 | 1.49E-20 | -2.07412396 | 1.29E-21 | 91.20552344 | 0.23747969 | 19.82810058 |
| ENSMUSG00000079110 | *Capn3* | 23.96590402 | 12.11126434 | 21.47176466 | 3.373596949 | 7.600184365 | 2.899246773 | 0.009129826 | -2.079405889 | 0.003793318 | 8.380194864 | 0.23661183 | 2.03953752 |
| ENSMUSG00000007872 | *Id3* | 1610.50875 | 1862.106892 | 1660.035806 | 376.999459 | 406.0669932 | 429.0885225 | 9.15E-81 | -2.083220664 | 7.51E-83 | 371.8224403 | 0.235987007 | 80.03860388 |
| ENSMUSG00000000562 | *Adora3* | 83.0818006 | 51.47287344 | 85.88705866 | 16.02458551 | 15.20036873 | 20.29472741 | 2.41E-07 | -2.094345558 | 5.56E-08 | 29.50952779 | 0.234174265 | 6.61742087 |
| ENSMUSG00000063954 | *H2ac19* | 119.0306566 | 97.89938673 | 112.7267645 | 31.20577178 | 29.31499684 | 16.42906505 | 2.80E-10 | -2.097148161 | 4.92E-11 | 43.20950857 | 0.233719796 | 9.55325639 |
| ENSMUSG00000059900 | *Tmem40* | 29.55794829 | 31.28743287 | 25.49772054 | 8.433992372 | 9.771665612 | 1.932831182 | 0.001601962 | -2.10738006 | 0.000587311 | 11.81577246 | 0.23206807 | 2.795347658 |
| ENSMUSG00000022947 | *Cbr3* | 661.4589509 | 758.9725652 | 994.411101 | 201.5724177 | 171.5470185 | 184.5853779 | 4.31E-30 | -2.108915532 | 2.28E-31 | 135.7370118 | 0.231821209 | 29.36527642 |
| ENSMUSG00000042106 | *Inka1* | 591.9578293 | 534.9141749 | 603.8933812 | 132.4136802 | 134.6318373 | 128.5332736 | 1.01E-52 | -2.129070454 | 1.93E-54 | 241.4160724 | 0.228605108 | 51.99692518 |
| ENSMUSG00000074006 | *Omp* | 25.56363095 | 11.10199231 | 22.81374996 | 2.530197712 | 5.428703118 | 5.798493547 | 0.007298617 | -2.132190595 | 0.002975616 | 8.822359551 | 0.228111234 | 2.136759432 |
| ENSMUSG00000027803 | *Wwtr1* | 15.17840588 | 18.16689651 | 21.47176466 | 5.060395423 | 3.257221871 | 3.865662365 | 0.003933777 | -2.14506062 | 0.001531582 | 10.04022838 | 0.226085344 | 2.405190249 |
| ENSMUSG00000030410 | *Dmwd* | 2376.618815 | 2217.370646 | 1562.070879 | 472.3035729 | 450.5823588 | 466.7787305 | 2.85E-39 | -2.148529315 | 9.85E-41 | 178.5900653 | 0.225542417 | 38.54486712 |
| ENSMUSG00000041801 | *Phlda3* | 516.0657999 | 618.6837532 | 681.7285281 | 133.2570795 | 114.0027655 | 153.660079 | 5.91E-34 | -2.174186243 | 2.64E-35 | 153.7423464 | 0.22156682 | 33.22846917 |
| ENSMUSG00000029490 | *Mfsd7a* | 193.3249591 | 243.2345588 | 254.9772054 | 46.38695805 | 57.54425305 | 48.32077956 | 5.18E-21 | -2.184090544 | 4.40E-22 | 93.34081411 | 0.220050942 | 20.28567682 |
| ENSMUSG00000091383 | *H2af-ps2* | 86.27725447 | 109.001379 | 134.1985292 | 22.77177941 | 22.80055309 | 26.09322096 | 3.07E-11 | -2.191915404 | 4.99E-12 | 47.6919862 | 0.218860665 | 10.51292816 |
| ENSMUSG00000017499 | *Cdc6* | 3026.094814 | 2703.839763 | 2650.420951 | 611.464447 | 567.8423461 | 653.2969396 | 9.54E-105 | -2.193194246 | 4.39E-107 | 483.1569372 | 0.218666748 | 104.0202728 |
| ENSMUSG00000049422 | *Chchd10* | 259.6306269 | 344.1617616 | 479.0887491 | 72.5323344 | 62.97295617 | 98.5743903 | 1.26E-14 | -2.203625296 | 1.59E-15 | 63.51288963 | 0.217091434 | 13.90054331 |
| ENSMUSG00000030148 | *Clec4a2* | 202.1124572 | 263.4199993 | 428.093308 | 72.5323344 | 58.62999367 | 60.88418224 | 1.70E-12 | -2.20918753 | 2.50E-13 | 53.56337356 | 0.216256061 | 11.76944777 |
| ENSMUSG00000024617 | *Camk2a* | 17.57499628 | 38.35233707 | 33.54963229 | 5.903794661 | 6.514443741 | 6.764909138 | 0.000544223 | -2.213767541 | 0.000186258 | 13.96487897 | 0.215570618 | 3.264223033 |
| ENSMUSG00000029650 | *Slc46a3* | 386.6499182 | 330.0319532 | 312.6825729 | 72.5323344 | 77.08758427 | 67.64909138 | 5.33E-33 | -2.249863424 | 2.50E-34 | 149.2738846 | 0.210244006 | 32.27367742 |
| ENSMUSG00000042770 | *Hebp1* | 408.2192318 | 480.4134854 | 739.4338956 | 123.1362886 | 110.7455436 | 106.305715 | 7.51E-20 | -2.253145092 | 6.83E-21 | 87.91534034 | 0.209766312 | 19.12454104 |
| ENSMUSG00000028480 | *Glipr2* | 498.4908036 | 512.7101903 | 504.5864696 | 100.3645092 | 106.4025811 | 107.2721306 | 4.27E-57 | -2.272677135 | 7.00E-59 | 261.78662 | 0.206945512 | 56.36994817 |
| ENSMUSG00000048120 | *Entpd1* | 11.98295201 | 26.24107273 | 36.23360287 | 4.216996186 | 1.085740624 | 9.664155911 | 0.009573214 | -2.288086853 | 0.003999574 | 8.284008403 | 0.204746848 | 2.018942236 |
| ENSMUSG00000063851 | *Rnf183* | 635.89532 | 630.7950176 | 817.2690425 | 145.908068 | 138.9747998 | 140.1302607 | 3.05E-45 | -2.28925403 | 7.90E-47 | 206.5188583 | 0.20458127 | 44.51507631 |
| ENSMUSG00000064202 | *Spata6l* | 24.76476749 | 17.15762448 | 12.07786762 | 6.747193898 | 2.171481247 | 1.932831182 | 0.008266656 | -2.292564631 | 0.003404434 | 8.576959588 | 0.204112348 | 2.082670147 |
| ENSMUSG00000060803 | *Gstp1* | 1254.215644 | 1541.158387 | 1948.562643 | 320.4917101 | 310.5218183 | 329.5477166 | 2.97E-41 | -2.301791992 | 9.28E-43 | 187.8681792 | 0.202811028 | 40.52676477 |
| ENSMUSG00000108840 | *Gm5320* | 23.96590402 | 18.16689651 | 29.52367641 | 4.216996186 | 5.428703118 | 4.832077956 | 0.000544068 | -2.308416896 | 0.000186172 | 13.96573957 | 0.201881848 | 3.264346942 |
| ENSMUSG00000027636 | *Sla2* | 19.17272321 | 21.19471259 | 30.8656617 | 3.373596949 | 1.085740624 | 9.664155911 | 0.004781223 | -2.314848698 | 0.001889181 | 9.654209889 | 0.200983825 | 2.320460997 |
| ENSMUSG00000020773 | *Trim47* | 39.94317336 | 44.40796924 | 36.23360287 | 4.216996186 | 8.685924988 | 11.59698709 | 1.87E-05 | -2.319616692 | 5.29E-06 | 20.72965151 | 0.200320685 | 4.727033273 |
| ENSMUSG00000018381 | *Abi3* | 618.3203237 | 498.5803819 | 458.9589697 | 104.5815054 | 106.4025811 | 102.4400527 | 3.82E-39 | -2.334353819 | 1.33E-40 | 177.9966027 | 0.198284826 | 38.4175049 |
| ENSMUSG00000025823 | *Pdia4* | 24863.82656 | 28824.80912 | 36001.43942 | 5914.758851 | 5908.600473 | 5925.093989 | 3.37E-69 | -2.337105301 | 3.78E-71 | 318.0889173 | 0.197907021 | 68.47255738 |
| ENSMUSG00000018459 | *Slc13a3* | 51.12726191 | 89.8252105 | 83.20308807 | 10.96419008 | 17.37184998 | 15.46264946 | 4.71E-08 | -2.360485705 | 1.01E-08 | 32.81531945 | 0.194725577 | 7.327245005 |
| ENSMUSG00000071303 | *Rps8-ps1* | 42.33976377 | 26.24107273 | 69.78323516 | 10.12079085 | 8.685924988 | 7.731324729 | 4.25E-05 | -2.366262548 | 1.26E-05 | 19.0765432 | 0.193947415 | 4.371818368 |
| ENSMUSG00000039481 | *Nrtn* | 19.97158668 | 38.35233707 | 29.52367641 | 8.433992372 | 5.428703118 | 2.899246773 | 0.00045235 | -2.367490796 | 0.000152892 | 14.33624924 | 0.193782367 | 3.344525589 |
| ENSMUSG00000036186 | *Dipk1b* | 48.7306715 | 34.31524896 | 45.62749991 | 8.433992372 | 8.685924988 | 6.764909138 | 1.21E-06 | -2.431134496 | 2.98E-07 | 26.26118473 | 0.18541958 | 5.918035135 |
| ENSMUSG00000030825 | *Hsd17b14* | 51.92612537 | 94.87157064 | 95.2809557 | 10.12079085 | 19.54333122 | 15.46264946 | 7.07E-08 | -2.431452093 | 1.55E-08 | 31.99250118 | 0.185378766 | 7.150440178 |
| ENSMUSG00000040841 | *Six5* | 107.0477046 | 110.0106511 | 75.15117632 | 17.71138398 | 15.20036873 | 21.261143 | 6.21E-12 | -2.434860254 | 9.56E-13 | 50.93244201 | 0.184941352 | 11.20699861 |
| ENSMUSG00000019590 | *Cyb561* | 263.6249442 | 233.1418385 | 254.9772054 | 46.38695805 | 49.94406868 | 41.55587042 | 6.06E-32 | -2.449532764 | 2.96E-33 | 144.3603246 | 0.183069992 | 31.21772665 |
| ENSMUSG00000020017 | *Hal* | 694.2123531 | 817.5103428 | 991.7271304 | 168.6798474 | 134.6318373 | 143.9959231 | 1.07E-42 | -2.477865941 | 3.13E-44 | 194.6149299 | 0.179509744 | 41.97053595 |
| ENSMUSG00000025355 | *Mmp19* | 388.2476451 | 393.616091 | 532.7681607 | 69.15873745 | 95.54517487 | 71.51475374 | 8.17E-30 | -2.479271931 | 4.37E-31 | 134.4422468 | 0.179334886 | 29.08785448 |
| ENSMUSG00000018672 | *Copz2* | 29.55794829 | 31.28743287 | 57.70536754 | 8.433992372 | 7.600184365 | 4.832077956 | 2.65E-05 | -2.48706489 | 7.65E-06 | 20.02402125 | 0.17836879 | 4.576504509 |
| ENSMUSG00000030747 | *Dgat2* | 97.46134301 | 106.982835 | 110.0427939 | 17.71138398 | 21.71481247 | 16.42906505 | 2.57E-15 | -2.497398136 | 3.11E-16 | 66.73509154 | 0.177095795 | 14.59080806 |
| ENSMUSG00000026271 | *Gpr35* | 109.444295 | 87.80666645 | 81.86110278 | 16.86798474 | 13.02888748 | 19.32831182 | 1.44E-12 | -2.50267948 | 2.10E-13 | 53.90878972 | 0.176448677 | 11.84265754 |
| ENSMUSG00000034586 | *Hid1* | 185.3363244 | 201.8544056 | 144.9344115 | 27.83217483 | 35.82944058 | 30.92529892 | 2.40E-20 | -2.504414322 | 2.12E-21 | 90.23188085 | 0.176236625 | 19.61965567 |
| ENSMUSG00000023036 | *Pcdhgc4* | 27.16135789 | 60.55632169 | 29.52367641 | 12.65098856 | 7.600184365 | 0 | 0.004921829 | -2.520509073 | 0.001949386 | 9.596594499 | 0.17428145 | 2.307873517 |
| ENSMUSG00000044092 | *C130050O18Rik* | 1426.770153 | 1351.415246 | 1289.647865 | 230.2479918 | 279.0353403 | 199.0816118 | 7.53E-74 | -2.525966375 | 7.29E-76 | 339.7344799 | 0.173623439 | 73.12313169 |
| ENSMUSG00000000154 | *Slc22a18* | 33.55226563 | 54.50068952 | 41.60154404 | 4.216996186 | 11.94314686 | 6.764909138 | 8.70E-06 | -2.527442776 | 2.37E-06 | 22.26783376 | 0.17344585 | 5.060516226 |
| ENSMUSG00000027546 | *Atp9a* | 229.2738151 | 316.9114168 | 366.3619846 | 54.82095042 | 41.2581437 | 58.95135106 | 2.42E-22 | -2.546885413 | 1.92E-23 | 99.54517796 | 0.171124068 | 21.61677244 |
| ENSMUSG00000100230 | *Gm28650* | 21.56931362 | 26.24107273 | 30.8656617 | 4.216996186 | 3.257221871 | 5.798493547 | 9.64E-05 | -2.551547087 | 2.98E-05 | 17.42886365 | 0.17057202 | 4.015965559 |
| ENSMUSG00000022469 | *Rapgef3* | 28.75908482 | 47.43578532 | 28.18169112 | 9.27739161 | 3.257221871 | 4.832077956 | 5.54E-05 | -2.55736537 | 1.66E-05 | 18.5422646 | 0.1698855 | 4.256102588 |
| ENSMUSG00000013419 | *Zfp651* | 869.1634524 | 839.7143274 | 669.6506605 | 137.4740757 | 119.4314686 | 143.0295075 | 3.12E-56 | -2.57238349 | 5.32E-58 | 257.7442832 | 0.168126205 | 55.50566527 |
| ENSMUSG00000055148 | *Klf2* | 418.6044569 | 441.0518763 | 373.071911 | 65.7851405 | 77.08758427 | 64.74984461 | 1.07E-47 | -2.57702487 | 2.53E-49 | 217.9538832 | 0.167586185 | 46.96925382 |
| ENSMUSG00000020432 | *Tcn2* | 565.5953348 | 699.4255155 | 943.4156599 | 113.858897 | 118.345728 | 131.4325204 | 3.14E-33 | -2.599580216 | 1.45E-34 | 150.348765 | 0.164986488 | 32.50367802 |
| ENSMUSG00000000318 | *Clec10a* | 1097.638404 | 1190.940993 | 1713.715217 | 207.4762124 | 234.5199747 | 213.5778456 | 7.60E-43 | -2.60907921 | 2.20E-44 | 195.3165863 | 0.163903753 | 42.11926324 |
| ENSMUSG00000081058 | *H3c15* | 37.54658296 | 31.28743287 | 24.15573525 | 5.060395423 | 3.257221871 | 6.764909138 | 2.23E-05 | -2.623084982 | 6.36E-06 | 20.37626932 | 0.162320264 | 4.651559786 |
| ENSMUSG00000079484 | *Phyhd1* | 774.0986998 | 950.7342505 | 1290.98985 | 163.619452 | 180.2329435 | 144.9623387 | 1.22E-35 | -2.623680262 | 5.00E-37 | 161.6229626 | 0.162253301 | 34.91328214 |
| ENSMUSG00000109032 | *Gm7972* | 15.97726935 | 13.12053637 | 25.49772054 | 0.843399237 | 2.171481247 | 5.798493547 | 0.004162449 | -2.636425742 | 0.001626755 | 9.929219541 | 0.160826189 | 2.380651095 |
| ENSMUSG00000035202 | *Lars2* | 14939.5457 | 38597.59017 | 31917.77817 | 4825.930435 | 4552.510435 | 4207.773484 | 2.95E-11 | -2.652985001 | 4.78E-12 | 47.77345347 | 0.158990779 | 10.53050154 |
| ENSMUSG00000043629 | *1700019D03Rik* | 28.75908482 | 20.18544056 | 22.81374996 | 3.373596949 | 0 | 7.731324729 | 0.00189414 | -2.672276425 | 0.000701359 | 11.48564105 | 0.156878937 | 2.722587999 |
| ENSMUSG00000023495 | *Pcbp4* | 252.4408557 | 282.5961679 | 254.9772054 | 43.85676034 | 44.51536557 | 34.79096128 | 3.71E-36 | -2.681491498 | 1.48E-37 | 164.0448098 | 0.155880082 | 35.43031508 |
| ENSMUSG00000026548 | *Slamf9* | 363.4828776 | 359.300842 | 426.7513227 | 70.84553593 | 44.51536557 | 60.88418224 | 1.52E-37 | -2.690974344 | 5.67E-39 | 170.5304584 | 0.154858841 | 36.8177033 |
| ENSMUSG00000049551 | *Fzd9* | 9.586361607 | 17.15762448 | 10.73588233 | 1.686798474 | 3.257221871 | 0.966415591 | 0.008044011 | -2.69145156 | 0.003305147 | 8.630864613 | 0.154807625 | 2.094527339 |
| ENSMUSG00000055538 | *Zcchc24* | 154.9795127 | 162.4927965 | 178.4840438 | 22.77177941 | 32.57221871 | 21.261143 | 5.89E-23 | -2.704068513 | 4.48E-24 | 102.4255487 | 0.153459673 | 22.23006764 |
| ENSMUSG00000035498 | *Cdcp1* | 63.11021392 | 97.89938673 | 143.5924262 | 12.65098856 | 15.20036873 | 18.36189623 | 7.97E-10 | -2.714590271 | 1.45E-10 | 41.08910418 | 0.152344543 | 9.098384592 |
| ENSMUSG00000039543 | *Cfap70* | 47.93180804 | 31.28743287 | 9.393897041 | 1.686798474 | 4.342962494 | 7.731324729 | 0.002230916 | -2.716925293 | 0.00083396 | 11.16408685 | 0.152098171 | 2.65151686 |
| ENSMUSG00000036867 | *Smad6* | 802.0589212 | 928.5302659 | 860.2125719 | 128.1966841 | 137.8890592 | 125.6340268 | 1.16E-91 | -2.726707037 | 7.07E-94 | 422.471062 | 0.151070405 | 90.93422717 |
| ENSMUSG00000006675 | *P4htm* | 18.37385975 | 17.15762448 | 8.051911749 | 3.373596949 | 2.171481247 | 0.966415591 | 0.0047513 | -2.738362037 | 0.001875395 | 9.667665196 | 0.149854879 | 2.323187574 |
| ENSMUSG00000025176 | *Hoga1* | 67.10453125 | 87.80666645 | 97.96492628 | 13.4943878 | 7.600184365 | 15.46264946 | 1.02E-11 | -2.768319159 | 1.60E-12 | 49.92647243 | 0.146775273 | 10.99049593 |
| ENSMUSG00000044734 | *Serpinb1a* | 123.024974 | 146.3444441 | 183.8519849 | 22.77177941 | 19.54333122 | 23.19397419 | 1.67E-19 | -2.778354151 | 1.55E-20 | 86.29175719 | 0.145757886 | 18.77832254 |
| ENSMUSG00000026807 | *Ak8* | 12.78181548 | 16.14835245 | 26.83970583 | 0 | 3.257221871 | 4.832077956 | 0.004829307 | -2.811104171 | 0.001909891 | 9.634183717 | 0.14248637 | 2.316115169 |
| ENSMUSG00000074607 | *Tox2* | 26.36249442 | 40.37088113 | 29.52367641 | 1.686798474 | 6.514443741 | 5.798493547 | 2.41E-05 | -2.816618612 | 6.91E-06 | 20.2186252 | 0.141942782 | 4.617928632 |
| ENSMUSG00000006435 | *Neurl1a* | 18.37385975 | 23.21325665 | 20.12977937 | 2.530197712 | 1.085740624 | 4.832077956 | 0.000311393 | -2.851055727 | 0.000102775 | 15.08502556 | 0.138594727 | 3.506691531 |
| ENSMUSG00000052631 | *Sh2d6* | 1445.942876 | 1422.064288 | 1238.652424 | 181.330836 | 204.1192372 | 176.8540532 | 1.36E-115 | -2.873159697 | 4.81E-118 | 533.5356927 | 0.136487459 | 114.8673908 |
| ENSMUSG00000027296 | *Itpka* | 21.56931362 | 15.13908042 | 12.07786762 | 1.686798474 | 2.171481247 | 2.899246773 | 0.001092269 | -2.875078681 | 0.000390068 | 12.57917784 | 0.136306033 | 2.961670566 |
| ENSMUSG00000027805 | *Pfn2* | 37.54658296 | 54.50068952 | 36.23360287 | 4.216996186 | 9.771665612 | 3.865662365 | 7.42E-07 | -2.877577497 | 1.79E-07 | 27.24564573 | 0.136070148 | 6.129846751 |
| ENSMUSG00000020101 | *Vsir* | 3869.694636 | 3934.142366 | 3906.519184 | 517.8471317 | 533.0986462 | 500.6032762 | 0 | -2.916572654 | 0 | 1488.303063 | 0.132441517 | Inf |
| ENSMUSG00000034685 | *Fam171a2* | 60.71362351 | 62.57486574 | 67.09926458 | 4.216996186 | 6.514443741 | 14.49623387 | 4.23E-09 | -2.924891618 | 8.24E-10 | 37.70157794 | 0.131680022 | 8.373502929 |
| ENSMUSG00000024424 | *Ttc39c* | 17.57499628 | 14.12980839 | 20.12977937 | 0.843399237 | 3.257221871 | 2.899246773 | 0.000785586 | -2.924986766 | 0.000275354 | 13.23102071 | 0.131671337 | 3.10480627 |
| ENSMUSG00000037709 | *Fam13a* | 22.36817708 | 15.13908042 | 25.49772054 | 5.903794661 | 0 | 1.932831182 | 0.001543221 | -2.934237992 | 0.000563862 | 11.89164727 | 0.130829701 | 2.811571787 |
| ENSMUSG00000055963 | *Triqk* | 18.37385975 | 26.24107273 | 45.62749991 | 3.373596949 | 4.342962494 | 3.865662365 | 3.80E-05 | -2.952495881 | 1.12E-05 | 19.30051667 | 0.129184431 | 4.420165843 |
| ENSMUSG00000023272 | *Creld2* | 23392.32005 | 25737.44599 | 30916.65715 | 3472.27466 | 3413.56852 | 3448.170829 | 8.51E-147 | -2.953221912 | 1.81E-149 | 678.0110905 | 0.129119436 | 146.0698407 |
| ENSMUSG00000024011 | *Pi16* | 1594.531481 | 1787.420762 | 1913.671026 | 208.3196116 | 238.8629372 | 234.8389886 | 6.56E-127 | -2.958256912 | 2.01E-129 | 585.8383517 | 0.128669595 | 126.1828561 |
| ENSMUSG00000027221 | *Chst1* | 74.29430246 | 98.90865876 | 119.4366909 | 12.65098856 | 14.11462811 | 10.6305715 | 2.17E-14 | -2.963010606 | 2.79E-15 | 62.41269626 | 0.128246325 | 13.66318452 |
| ENSMUSG00000000686 | *Abhd15* | 218.0897266 | 238.1881986 | 224.1115437 | 24.45857788 | 38.00092182 | 25.12680537 | 3.06E-33 | -2.973042941 | 1.41E-34 | 150.4038165 | 0.127357609 | 32.5140429 |
| ENSMUSG00000041954 | *Tnfrsf18* | 11.18408854 | 18.16689651 | 9.393897041 | 0.843399237 | 3.257221871 | 0.966415591 | 0.004715444 | -2.992496536 | 0.001858459 | 9.684332099 | 0.125651819 | 2.326477415 |
| ENSMUSG00000031844 | *Hsd17b2* | 8.78749814 | 22.20398462 | 16.10238235 | 1.686798474 | 3.257221871 | 0.966415591 | 0.00216399 | -3.013714617 | 0.000807665 | 11.22353537 | 0.123817349 | 2.664744717 |
| ENSMUSG00000050777 | *Tmem37* | 159.7726935 | 169.5577007 | 224.1115437 | 22.77177941 | 19.54333122 | 24.16038978 | 5.14E-26 | -3.046407934 | 3.33E-27 | 116.7062051 | 0.121043043 | 25.28887276 |
| ENSMUSG00000104864 | *Gm43655* | 32.75340216 | 29.26888882 | 25.49772054 | 10.12079085 | 0 | 0 | 0.008414162 | -3.0561623 | 0.003470643 | 8.541893861 | 0.120227406 | 2.074989108 |
| ENSMUSG00000043832 | *Clec4a3* | 927.4804855 | 1042.578005 | 1401.032644 | 123.1362886 | 133.5460967 | 144.9623387 | 3.56E-58 | -3.068048204 | 5.61E-60 | 266.8165896 | 0.11924096 | 57.44876065 |
| ENSMUSG00000003355 | *Fkbp11* | 1635.273518 | 2103.322907 | 2991.285215 | 278.3217483 | 247.5488622 | 274.4620279 | 1.67E-46 | -3.069705196 | 4.14E-48 | 212.3869283 | 0.119104085 | 45.77711398 |
| ENSMUSG00000002504 | *Nherf2* | 178.1465532 | 309.8465126 | 316.7085288 | 26.98877559 | 38.00092182 | 28.99246773 | 1.71E-22 | -3.101686814 | 1.34E-23 | 100.2489186 | 0.11649284 | 21.76709324 |
| ENSMUSG00000013083 | *2200002J24Rik* | 9.586361607 | 16.14835245 | 24.15573525 | 2.530197712 | 1.085740624 | 1.932831182 | 0.001030276 | -3.120106181 | 0.0003672 | 12.69211947 | 0.115014991 | 2.987046623 |
| ENSMUSG00000083365 | *Gm12895* | 119.8295201 | 381.5048266 | 291.2108083 | 33.73596949 | 28.22925621 | 28.02605214 | 3.56E-13 | -3.133011641 | 5.00E-14 | 56.73065875 | 0.113990726 | 12.44914485 |
| ENSMUSG00000083558 | *Gm12896* | 119.8295201 | 381.5048266 | 291.2108083 | 33.73596949 | 28.22925621 | 28.02605214 | 3.56E-13 | -3.133011641 | 5.00E-14 | 56.73065875 | 0.113990726 | 12.44914485 |
| ENSMUSG00000027412 | *Lpin3* | 36.7477195 | 53.49141749 | 64.41529399 | 8.433992372 | 3.257221871 | 3.865662365 | 1.09E-08 | -3.267838601 | 2.22E-09 | 35.77134492 | 0.103820367 | 7.961030619 |
| ENSMUSG00000089722 | *Cd300ld5* | 19.17272321 | 27.25034876 | 75.15117632 | 0 | 7.600184365 | 4.832077956 | 0.000589671 | -3.321024517 | 0.000202821 | 13.80476573 | 0.100062651 | 3.229390457 |
| ENSMUSG00000022475 | *Hdac7* | 16.77613281 | 7.064904197 | 22.81374996 | 3.373596949 | 1.085740624 | 0 | 0.003574653 | -3.323630724 | 0.001382056 | 10.22958773 | 0.099882052 | 2.44676617 |
| ENSMUSG00000028039 | *Efna3* | 15.17840588 | 13.12053637 | 20.12977937 | 1.686798474 | 2.171481247 | 0.966415591 | 0.000325298 | -3.329995586 | 0.000107703 | 14.99663667 | 0.099442365 | 3.487719272 |
| ENSMUSG00000029814 | *Igf2bp3* | 23.96590402 | 36.33379301 | 29.52367641 | 0.843399237 | 4.342962494 | 3.865662365 | 3.42E-06 | -3.355231366 | 8.92E-07 | 24.14813817 | 0.097718033 | 5.465440015 |
| ENSMUSG00000046456 | *Tmem150b* | 31.95453869 | 40.37088113 | 33.54963229 | 5.903794661 | 1.085740624 | 2.899246773 | 3.09E-07 | -3.363073684 | 7.19E-08 | 29.01233339 | 0.097188291 | 6.510737792 |
| ENSMUSG00000010830 | *Kdelr3* | 43.9374907 | 53.49141749 | 81.86110278 | 6.747193898 | 5.428703118 | 4.832077956 | 1.72E-10 | -3.376906255 | 2.97E-11 | 44.19604173 | 0.096260902 | 9.763342684 |
| ENSMUSG00000092574 | *2810047C21Rik1* | 23.16704055 | 11.10199231 | 46.9694852 | 0 | 0 | 7.731324729 | 0.008610306 | -3.389222075 | 0.00355917 | 8.49605661 | 0.095442651 | 2.064981432 |
| ENSMUSG00000017446 | *C1qtnf1* | 20.77045015 | 11.10199231 | 9.393897041 | 0.843399237 | 3.257221871 | 0 | 0.003349372 | -3.409580619 | 0.001287445 | 10.36044072 | 0.094105273 | 2.475036568 |
| ENSMUSG00000053338 | *Tarm1* | 755.7248401 | 847.7885036 | 976.9652922 | 75.06253211 | 72.74462178 | 91.80948116 | 3.21E-87 | -3.425687611 | 2.33E-89 | 401.7151127 | 0.093060477 | 86.49331103 |
| ENSMUSG00000030785 | *Cox6a2* | 1068.080456 | 1211.126434 | 1586.226615 | 128.1966841 | 97.71665612 | 131.4325204 | 1.32E-67 | -3.428872302 | 1.56E-69 | 310.6760207 | 0.092855276 | 66.88005542 |
| ENSMUSG00000021186 | *Fbln5* | 35.14999256 | 34.31524896 | 61.73132341 | 5.903794661 | 0 | 5.798493547 | 1.19E-06 | -3.429818791 | 2.94E-07 | 26.29129061 | 0.092794378 | 5.924078049 |
| ENSMUSG00000038742 | *Angptl6* | 1525.030359 | 1487.666969 | 1622.460217 | 150.1250642 | 142.2320217 | 133.3653516 | 8.36E-199 | -3.44169184 | 8.88E-202 | 918.5993851 | 0.092033835 | 198.0776731 |
| ENSMUSG00000032268 | *Tmprss5* | 10.38522507 | 13.12053637 | 28.18169112 | 2.530197712 | 0 | 1.932831182 | 0.000978193 | -3.452800274 | 0.000347483 | 12.79534697 | 0.091327916 | 3.009575424 |

| | | | | | | | | | | | | | |
|---|---|---|---|---|---|---|---|---|---|---|---|---|---|
| ENSMUSG00000061619 | *Vma21-ps* | 19.17272321 | 20.18544056 | 60.38933812 | 6.747193898 | 1.085740624 | 0 | 0.000505247 | -3.60025153 | 0.000171964 | 14.11503263 | 0.082454867 | 3.296496488 |
| ENSMUSG00000079547 | *H2-DMb1* | 15.17840588 | 13.12053637 | 8.051911749 | 0 | 3.257221871 | 0 | 0.006614344 | -3.605118378 | 0.002678684 | 9.014347864 | 0.082177179 | 2.179513213 |
| ENSMUSG00000027624 | *Epb41l1* | 530.4453423 | 572.25724 | 601.2094106 | 43.85676034 | 43.42962494 | 41.55587042 | 2.41E-103 | -3.722427368 | 1.12E-105 | 476.6866957 | 0.075759606 | 102.6178947 |
| ENSMUSG00000029005 | *Draxin* | 4.793180804 | 20.18544056 | 13.41985292 | 0.843399237 | 0 | 1.932831182 | 0.003452166 | -3.74843381 | 0.001331032 | 10.29898834 | 0.074406176 | 2.46190835 |
| ENSMUSG00000022505 | *Emp2* | 504.0828479 | 535.9234469 | 626.7071311 | 41.32656262 | 45.60110619 | 35.75737687 | 3.46E-84 | -3.764240052 | 2.63E-86 | 387.6901185 | 0.073595428 | 83.46082872 |
| ENSMUSG00000020787 | *P2rx1* | 7.988634673 | 9.083448253 | 9.393897041 | 0.843399237 | 0 | 0.966415591 | 0.004580745 | -3.806163877 | 0.001801587 | 9.741450584 | 0.071487565 | 2.33906388 |
| ENSMUSG00000018924 | *Alox15* | 55.12157924 | 61.56559372 | 85.88705866 | 6.747193898 | 6.514443741 | 0.966415591 | 5.10E-12 | -3.819486992 | 7.80E-13 | 51.33176499 | 0.070830425 | 11.29217139 |
| ENSMUSG00000026969 | *Fam166a* | 20.77045015 | 17.15762448 | 14.76183821 | 3.373596949 | 0 | 0 | 0.000375953 | -3.849770788 | 0.000125429 | 14.70926386 | 0.069359111 | 3.424866576 |
| ENSMUSG00000026809 | *Spaca9* | 27.16135789 | 28.25961679 | 59.04735283 | 0.843399237 | 2.171481247 | 4.832077956 | 5.11E-07 | -3.876827674 | 1.21E-07 | 27.99812932 | 0.068070444 | 6.291198141 |
| ENSMUSG00000031502 | *Col4a1* | 209.3022284 | 209.9285819 | 205.3237496 | 15.18118627 | 13.02888748 | 11.59698709 | 4.51E-48 | -3.964151432 | 1.03E-49 | 219.7360047 | 0.064072477 | 47.34584794 |
| ENSMUSG00000082280 | *Gm5847* | 11.18408854 | 8.074176225 | 12.07786762 | 0.843399237 | 1.085740624 | 0 | 0.001658618 | -4.031352146 | 0.000608865 | 11.74867806 | 0.061156424 | 2.780253664 |
| ENSMUSG00000054545 | *Ugt1a6a* | 727.7646187 | 889.1686568 | 1211.812718 | 48.07375652 | 78.1733249 | 33.82454569 | 1.20E-43 | -4.151482434 | 3.34E-45 | 199.0659258 | 0.056270304 | 42.92217508 |
| ENSMUSG00000048065 | *Cyb5r2* | 11.18408854 | 16.14835245 | 25.49772054 | 0.843399237 | 2.171481247 | 0 | 0.000108067 | -4.171566503 | 3.36E-05 | 17.19965356 | 0.05549238 | 3.966306728 |
| ENSMUSG00000059430 | *Actg2* | 16.77613281 | 17.15762448 | 21.47176466 | 0.843399237 | 1.085740624 | 0.966415591 | 8.62E-06 | -4.264121467 | 2.35E-06 | 22.28682986 | 0.052044104 | 5.064530401 |
| ENSMUSG00000084148 | *Gm12380* | 14.37954241 | 10.09272028 | 13.41985292 | 0 | 0 | 1.932831182 | 0.000646573 | -4.304533765 | 0.000223729 | 13.62050391 | 0.05060649 | 3.189382393 |
| ENSMUSG00000035860 | *Cdhr3* | 63.11021392 | 79.73249022 | 104.6748527 | 5.903794661 | 6.514443741 | 0 | 3.33E-14 | -4.31260046 | 4.36E-15 | 61.53246418 | 0.050324318 | 13.47717046 |
| ENSMUSG00000108442 | *Rpl15-ps5* | 21.56931362 | 21.19471259 | 18.78779408 | 1.686798474 | 0 | 0.966415591 | 2.37E-06 | -4.448411175 | 6.06E-07 | 24.89303375 | 0.045803093 | 5.625493265 |
| ENSMUSG00000054675 | *Tmem119* | 138.2033798 | 152.4000762 | 154.3283085 | 7.590593135 | 5.428703118 | 6.764909138 | 3.84E-38 | -4.473151898 | 1.40E-39 | 173.3155817 | 0.045024314 | 37.41551453 |
| ENSMUSG00000082804 | *Rpl17-ps1* | 6.390907738 | 8.074176225 | 9.393897041 | 0 | 1.085740624 | 0 | 0.004656073 | -4.522882569 | 0.001832312 | 9.710369218 | 0.04349874 | 2.331980227 |
| ENSMUSG00000026822 | *Lcn2* | 849.9907292 | 1156.625744 | 1678.8236 | 52.29075271 | 39.08666245 | 39.62303924 | 1.96E-63 | -4.805321725 | 2.57E-65 | 291.3190233 | 0.035764654 | 62.70818175 |
| ENSMUSG00000009596 | *Taf7l* | 21.56931362 | 25.2318007 | 65.75727928 | 0 | 0 | 3.865662365 | 2.09E-06 | -4.86059665 | 5.33E-07 | 25.13952148 | 0.034420297 | 5.678998405 |
| ENSMUSG00000022818 | *Cyp2ab1* | 9.586361607 | 9.083448253 | 13.41985292 | 0 | 1.085740624 | 0 | 0.000577857 | -4.950500219 | 0.000198655 | 13.84376071 | 0.032340814 | 3.238179964 |
| ENSMUSG00000026580 | *Selp* | 7.988634673 | 4.037088113 | 5.367941166 | 0 | 0 | 0 | 0.008531717 | -5.044057591 | 0.00352316 | 8.514560531 | 0.0303101 | 2.068963568 |
| ENSMUSG00000052131 | *Akr1b7* | 139.0022433 | 188.7338693 | 382.4658081 | 5.903794661 | 3.257221871 | 10.6305715 | 2.44E-22 | -5.148057277 | 1.94E-23 | 99.52011168 | 0.028202028 | 21.61182149 |
| ENSMUSG00000039518 | *Cdsn* | 194.922686 | 280.5776238 | 348.9161758 | 5.903794661 | 6.514443741 | 10.6305715 | 3.30E-42 | -5.160847026 | 9.95E-44 | 192.3111376 | 0.027953117 | 41.48152971 |
| ENSMUSG00000005611 | *Irag1* | 9.586361607 | 14.12980839 | 13.41985292 | 0 | 1.085740624 | 0 | 0.000156627 | -5.163483789 | 4.99E-05 | 16.4526782 | 0.027902075 | 3.805132464 |
| ENSMUSG00000063975 | *Slco1a5* | 1.597726935 | 12.11126434 | 6.709926458 | 0 | 0 | 0 | 0.009218076 | -5.260079501 | 0.003833249 | 8.361164378 | 0.026095059 | 2.035359704 |
| ENSMUSG00000066810 | *None* | 8.78749814 | 5.046360141 | 6.709926458 | 0 | 0 | 0 | 0.003414596 | -5.281755079 | 0.001315539 | 10.32059527 | 0.025705928 | 2.466660637 |
| ENSMUSG00000022871 | *Fetub* | 3.994317336 | 11.10199231 | 6.709926458 | 0 | 0 | 0 | 0.003316585 | -5.358530998 | 0.00127445 | 10.37916942 | 0.024373699 | 2.479308901 |
| ENSMUSG00000095159 | *Tubb4b-ps1* | 11.18408854 | 6.055632169 | 5.367941166 | 0 | 0 | 0 | 0.002251105 | -5.427925856 | 0.000842171 | 11.14591012 | 0.023229053 | 2.6476043 |
| ENSMUSG00000044405 | *Adig* | 19.97158668 | 50.46360141 | 56.36338224 | 0.843399237 | 1.085740624 | 0.966415591 | 5.50E-10 | -5.454278338 | 9.92E-11 | 41.83808098 | 0.022808599 | 9.259545676 |
| ENSMUSG00000113963 | *Gm6554* | 5.592044271 | 10.09272028 | 9.393897041 | 0 | 0 | 0 | 0.001053982 | -5.55727495 | 0.000375898 | 12.64834677 | 0.021237019 | 2.977166707 |
| ENSMUSG00000017943 | *Gdap1l1* | 10.38522507 | 7.064904197 | 8.051911749 | 0 | 0 | 0 | 0.000796018 | -5.593551523 | 0.000279198 | 13.20503306 | 0.020709671 | 3.099076989 |
| ENSMUSG00000048078 | *Tenm4* | 468.1339918 | 651.9897302 | 548.8719842 | 6.747193898 | 14.11462811 | 10.6305715 | 3.38E-97 | -5.761484543 | 1.86E-99 | 448.1129 | 0.018434032 | 96.47066435 |
| ENSMUSG00000072214 | *Septin5* | 241.2567671 | 447.1075085 | 562.2918371 | 0 | 14.11462811 | 8.69774032 | 0.000153135 | -5.789943466 | 4.87E-05 | 16.49859179 | 0.018073961 | 3.814924969 |
| ENSMUSG00000046110 | *Serinc4* | 18.37385975 | 11.10199231 | 0 | 0 | 0 | 0 | 0.009992923 | -5.811934215 | 0.004190257 | 8.199472445 | 0.017800552 | 2.000307459 |
| ENSMUSG00000032936 | *Camkv* | 11.98295201 | 9.083448253 | 10.73588233 | 0 | 0 | 0 | 0.000127029 | -5.909273998 | 4.00E-05 | 16.87013517 | 0.016639155 | 3.896097065 |
| ENSMUSG00000028972 | *Car6* | 616.7225967 | 843.7514155 | 1503.023526 | 16.02458551 | 10.85740624 | 22.2275586 | 6.44E-23 | -5.909325461 | 4.91E-24 | 102.2415573 | 0.016638562 | 22.19107688 |
| ENSMUSG00000106956 | *Gm19617* | 14.37954241 | 7.064904197 | 10.73588233 | 0 | 0 | 0 | 0.000169333 | -5.929419884 | 5.41E-05 | 16.29713694 | 0.01640842 | 3.771259089 |
| ENSMUSG00000029797 | *Sspo* | 35.14999256 | 62.57486574 | 44.28551462 | 0.843399237 | 0 | 0.966415591 | 3.16E-14 | -6.220266429 | 4.11E-15 | 61.64592367 | 0.013412608 | 13.50003274 |
| ENSMUSG00000039007 | *Cpq* | 15.17840588 | 8.074176225 | 17.44580879 | 0 | 0 | 0 | 2.23E-05 | -6.25919607 | 6.36E-06 | 20.37793197 | 0.013055522 | 4.651847234 |
| ENSMUSG00000028445 | *Enho* | 14.37954241 | 6.055632169 | 21.47176466 | 0 | 0 | 0 | 4.48E-05 | -6.29646822 | 1.33E-05 | 18.97065098 | 0.012722551 | 4.348584129 |
| ENSMUSG00000025432 | *Avil* | 4464.049055 | 6666.241746 | 8701.43263 | 69.15873745 | 78.1733249 | 59.91776665 | 3.39E-62 | -6.581441623 | 4.64E-64 | 285.5473289 | 0.010442119 | 61.46969349 |
| ENSMUSG00000033207 | *Mamdc2* | 79.08748326 | 81.75103428 | 138.224485 | 2.530197712 | 0 | 0 | 4.60E-24 | -6.713732309 | 3.29E-25 | 107.6001055 | 0.009527198 | 23.33746544 |
| ENSMUSG00000038319 | *Kcnh2* | 38.34544643 | 40.37088113 | 34.89161758 | 0.843399237 | 0 | 0 | 1.01E-13 | -6.784716377 | 1.35E-14 | 59.29842285 | 0.009069782 | 12.9975985 |
| ENSMUSG00000111686 | *Rab7-ps1* | 18.37385975 | 35.32452098 | 55.02139695 | 0 | 0 | 0 | 1.06E-10 | -7.668395184 | 1.80E-11 | 45.17902994 | 0.004915674 | 9.975839345 |
| ENSMUSG00000030205 | *Gprc5d* | 27.16135789 | 26.24107273 | 61.73132341 | 0 | 0 | 0 | 1.73E-11 | -7.751382955 | 2.76E-12 | 48.84973266 | 0.004640889 | 10.76122079 |
| ENSMUSG00000074486 | *Bglap2* | 150.9851953 | 333.0597693 | 626.7071311 | 3.373596949 | 0 | 0 | 2.86E-07 | -8.273087942 | 6.65E-08 | 29.16387756 | 0.003232603 | 6.542965187 |
| ENSMUSG00000050157 | *Gm867* | 38.34544643 | 67.62122589 | 119.4366909 | 0 | 0 | 0 | 2.84E-17 | -8.723188035 | 3.04E-18 | 75.86542407 | 0.00236624 | 16.54673878 |
