## Supplementary material for "TFEB-Mediated Pro-inflammatory Response in Murine Macrophages Induced by Acute Alpha7 Nicotinic Receptor Activation": Table_S4

**Table S4. GO and KEGG Pathway analysis of differentially expressed genes from Table S3.**

| Term | Library | p-value | q-value | z-score | combined score | converted -Log(Padj) |
|---|---|---|---|---|---|---|
| mitotic spindle organization (GO:0007052) | GO_Biological_Process_2021 | 7.76E-13 | 3.09E-09 | 3.717 | 103.6 | 8.510660374 |
| positive regulation of transcription by RNA polymerase II (GO:0045944) | GO_Biological_Process_2021 | 1.26E-12 | 3.09E-09 | 1.872 | 51.3 | 8.510660374 |
| positive regulation of transcription, DNA-templated (GO:0045893) | GO_Biological_Process_2021 | 1.48E-11 | 2.41E-08 | 1.71 | 42.65 | 7.617892864 |
| microtubule cytoskeleton organization involved in mitosis (GO:1902850) | GO_Biological_Process_2021 | 2.08E-11 | 2.54E-08 | 3.857 | 94.88 | 7.594824454 |
| cellular response to DNA damage stimulus (GO:0006974) | GO_Biological_Process_2021 | 4.29E-11 | 4.20E-08 | 2.398 | 57.24 | 7.376864468 |
| TNF signaling pathway | KEGG_2021_Human | 1.97E-09 | 5.90E-07 | 3.665 | 73.48 | 6.229449891 |
| NF-kappa B signaling pathway | KEGG_2021_Human | 4.10E-07 | 0.000061526 | 3.182 | 46.79 | 4.210941319 |
| NOD-like receptor signaling pathway | KEGG_2021_Human | 7.7281E-06 | 0.0007728 | 2.267 | 26.69 | 3.111932887 |
| Toll-like receptor signaling pathway | KEGG_2021_Human | 0.000011748 | 0.0008811 | 2.771 | 31.46 | 3.054974799 |
| Influenza A | KEGG_2021_Human | 0.000024311 | 0.001429 | 2.212 | 23.5 | 2.844967771 |
