## Supplementary material for "TFEB-Mediated Pro-inflammatory Response in Murine Macrophages Induced by Acute Alpha7 Nicotinic Receptor Activation": Table_S5

**Table S5. Genes that were induced by PNU-282987 in both WT BMDMs and RAW264.7.**

*Abhd17c*
*Abtb2*
*Adam17*
*Adora2a*
*Agpat4*
*Ankrd33b*
*Arg2*
*Arhgef3*
*Atf4*
*B4galt5*
*Bcl2a1b*
*Bcl2a1d*
*Bcl3*
*Bhlhe40*
*Bhlhe41*
*Birc3*
*Ccl2*
*Ccl22*
*Ccl3*
*Ccl4*
*Ccl5*
*Ccrl2*
*Cd274*
*Cd40*
*Cd69*
*Cd83*
*Cdc42ep2*
*Cers6*
*Cflar*
*Cish*
*Clcn7*
*Clec4e*
*Cmpk2*
*Cpeb4*
*Csf1*
*Csf3*
*Cxcl10*
*Cxcl11*
*Cxcl16*
*Cxcl2*
*Cybb*
*Daam1*
*Dcbld2*
*Denr*
*Dst*
*Dtx3l*
*Dusp1*
*Dusp16*
*Dusp2*
*Dusp4*
*Edn1*
*Ehd1*
*Eif1a*
*Enpp4*
*Ets2*
*Fabp4*
*Fam20c*
*Fas*
*Fgr*
*Flnb*
*Fnip2*
*Fzd5*
*Gadd45a*
*Gadd45b*
*Gas2l3*
*Gbp2*
*Gbp3*
*Gbp5*
*Gbp6*
*Gbp7*
*Gdf15*
*Gem*
*Gm12250*
*Gm6377*
*Gpr18*
*Gpr84*
*Gsap*
*Hcar2*
*Helz2*
*Herpud1*

*Hif1a*
*Hilpda*
*Hivep3*
*Hsd17b7*
*Icosl*
*Ier3*
*Ifi205*
*Ifit1*
*Ifit2*
*Ifit3*
*Ifitm1*
*Ifnb1*
*Ifrd1*
*Igsf6*
*Ikbke*
*Il13ra1*
*Il1rn*
*Il27*
*Il4i1*
*Il6*
*Impact*
*Irak2*
*Irf1*
*Irgm1*
*Isg15*
*Itga5*
*Jak2*
*Jdp2*
*Junb*
*Kdm6b*
*Klf7*
*Larp1*
*Lcp2*
*Ldlr*
*Lif*
*Lpar1*
*Ly96*
*Maff*
*Mafk*
*Malt1*
*Map3k8*
*Mapk6*
*Marcksl1*
*Mcoln2*
*Mdm2*
*Mllt6*
*Mob1b*
*Mocs1*
*Mt2*
*Mtdh*
*Mtmr14*
*Mx2*
*Mxd1*
*Myc*
*Myo10*
*N4bp1*
*Naa25*
*Nfe2l2*
*Nfkb2*
*Nfkbia*
*Nfkbie*
*Nfkbiz*
*Nlrp3*
*Nod2*
*Nos2*
*Nrp2*
*Nupr1*
*Oasl1*
*Odc1*
*Osgin2*
*Osm*
*P2ry2*
*Parp14*
*Pde4b*
*Phlda1*
*Pik3r5*
*Pim1*
*Plagl2*
*Plat*
*Plek*
*Plekhm3*

*Plekho2*
*Pnrc1*
*Ppp1r15a*
*Psd*
*Psmd10*
*Pstpip2*
*Ptafr*
*Ptges*
*Ptgs2*
*Ptpn23*
*Rab11fip1*
*Rab12*
*Rab20*
*Ralgapa2*
*Ralgds*
*Rapgef2*
*Rasgef1b*
*Rassf4*
*Rbpj*
*Relb*
*Rffl*
*Rgl1*
*Rhbdf2*
*Ripk2*
*Rnf19a*
*Rnf19b*
*Rrs1*
*Rsad2*
*Rusc2*
*Samd8*
*Sdc4*
*Sema4b*
*Sgms1*
*Siah2*
*Slc11a2*
*Slc15a3*
*Slc1a4*
*Slc25a37*
*Slc30a1*
*Slc31a2*
*Slc4a7*
*Slc7a11*
*Slfn2*
*Slfn4*
*Slfn5*
*Snx20*
*Socs3*
*Spryd7*
*Src*
*Srfbp1*
*Stk40*
*Stx11*
*Susd6*
*Tank*
*Tbcel*
*Tlr1*
*Tlr2*
*Tma16*
*Tmem132a*
*Tnf*
*Tnfaip2*
*Tnfaip3*
*Tnfrsf1b*
*Tnfsf9*
*Tnip1*
*Tnip3*
*Tor3a*
*Traf1*
*Traf3*
*Trim13*
*Ube2f*
*Usp18*
*Zbtb21*
*Zc3h12a*
*Zc3h12c*
*Zfand5*
*Zfp811*
*Zhx2*
*Zmynd15*
