## Supplementary material for "TFEB-Mediated Pro-inflammatory Response in Murine Macrophages Induced by Acute Alpha7 Nicotinic Receptor Activation": Table_S6

**Table S6. RT-qPCR analysis of cholinergic receptor subunit expression.**
**Each column represents a the average of two technical replicates in a biological replicate/run**

**Raw Ct Values**

| Macrophages | Wild type BMDMs | | | | | | | | Chrna7-/- BMDMs | | | |
|---|---|---|---|---|---|---|---|---|---|---|---|---|
| α3 | 35.0 | 32.6 | 35.6 | 33.9 | 34.3 | 33.8 | 34.7 | 38.1 | Undetermined* | 39.14 | 35.40 | 37.19 |
| α4 | 36.0 | 33.0 | 33.3 | 33.2 | 34.7 | 33.4 | 33.0 | 33.3 | 33.2 | 34.1 | 33.4 | 33.4 |
| α5 | 33.7 | Undetermined* | 34.0 | 33.5 | 34.0 | 33.7 | 31.9 | 31.9 | 34.5 | 35.1 | 36.1 | 36.7 |
| α6 | 32.5 | 32.6 | 32.3 | 32.6 | 32.9 | 32.9 | 35.9 | 35.6 | 36.5 | 37.5 | 32.6 | 32.7 |
| α7 | **Oligo pair 1** | **Oligo pair 1** | **Oligo pair 1** | **Oligo pair 1** | **Oligo pair 3** | **Oligo pair 3** | **Oligo pair 4** | **Oligo pair 4** | **Oligo pair 1** | **Oligo pair 1** | **Oligo pair 1** | **Oligo pair 1** |
| | 36.2 | 37.3 | 35.1 | 34.8 | 38.2 | 39.0 | 37.2 | 37.6 | 34.1 | 32.6 | 32.3 | 33.1 |
| α9 | 32.8 | 32.8 | 33.7 | 32.7 | 33.9 | 32.3 | 33.5 | 33.5 | 33.2 | 34.1 | 32.5 | 33.5 |
| α10 | 33.6 | 32.7 | 32.6 | 33.1 | 32.7 | 34.2 | 26.6 | 26.5 | 26.3 | 26.7 | 33.0 | 32.8 |
| β2 | 29.4 | 29.8 | 29.2 | 29.4 | 30.4 | 29.0 | 30.0 | 30.6 | 30.6 | 30.8 | 28.9 | 29.0 |
| β3 | 33.1 | 33.0 | 31.4 | 31.5 | 31.5 | 33.4 | 33.0 | 33.0 | 32.8 | 33.1 | 31.6 | 32.3 |
| β4 | 36.2 | 32.2 | 32.8 | 34.6 | 36.4 | 33.4 | 32.3 | 32.8 | 32.1 | 33.0 | 33.0 | 33.9 |
| M1 | 32.3 | 33.3 | 36.5 | 36.5 | 38.3 | 31.3 | 36.9 | 36.9 | 37.9 | 37.5 | 36.8 | 37.8 |
| M2 | Undetermined* | Undetermined* | 39.0 | Undetermined* | 35.5 | Undetermined* | Undetermined* | Undetermined* | Undetermined* | Undetermined* | 35.3 | Undetermined* |
| M3 | 33.5 | 33.8 | 32.2 | 33.4 | 31.3 | 33.5 | 32.1 | 33.6 | 34.1 | 33.3 | 33.9 | 33.7 |
| M4 | Undetermined* | Undetermined* | 30.9 | 30.6 | 32.8 | 33.0 | 34.2 | 35.4 | 36.1 | Undetermined* | 31.9 | 33.6 |
| M5 | Undetermined* | Undetermined* | Undetermined* | Undetermined* | Undetermined* | 36.7 | Undetermined* | Undetermined* | Undetermined* | Undetermined* | Undetermined* | Undetermined* |
| *Gapdh* | 24.6 | 23.8 | 18.2 | 18.1 | 21.9 | 24.3 | 17.6 | 18.2 | 17.9 | 19.0 | 18.5 | 18.3 |
| *B actin* | 18.2 | 17.7 | 14.4 | 14.4 | 16.2 | 17.9 | 16.0 | 16.5 | 16.5 | 16.7 | 14.6 | 14.6 |

| Macrophages | Tfebfl/fl BMDMs | | TfebΔLysMCre BMDMs | | Wild type RAW264.7 | | Tfeb-/- Tfe3-/- RAW 264.7 | |
|---|---|---|---|---|---|---|---|---|
| α3 | 36.3 | 36.5 | 36.1 | 35.2 | 31.6 | 29.9 | 29.8 | 29.7 |
| α4 | 32.7 | 34.2 | 33.3 | 33.7 | 32.4 | 33.0 | 31.2 | 32.2 |
| α5 | 34.9 | 34.1 | 34.9 | 34.1 | 34.4 | 32.5 | 36.9 | 36.5 |
| α6 | 32.5 | 32.3 | 33.4 | 32.8 | 36.7 | 31.3 | 36.5 | 33.9 |
| α7 | **Oligo pair 1** | **Oligo pair 1** | **Oligo pair 1** | **Oligo pair 1** | **Oligo pair 1** | **Oligo pair 1** | **Oligo pair 1** | **Oligo pair 1** |
| | 33.6 | 32.7 | 33.4 | 32.9 | 36.2 | 39.9 | Undetermined* | 38.9 |
| α9 | 35.9 | 33.5 | Undetermined* | Undetermined* | 32.3 | 33.0 | 33.2 | 34.5 |
| α10 | 32.8 | 32.8 | 33.3 | 32.7 | 27.9 | 27.9 | 27.6 | 27.8 |
| β2 | 29.3 | 28.7 | 28.8 | 28.7 | 27.5 | 30.5 | 29.7 | 30.2 |
| β3 | 31.4 | 31.8 | 31.7 | 31.4 | 34.5 | 33.8 | 34.5 | 33.8 |
| β4 | 32.6 | 33.9 | 33.6 | 32.8 | 33.4 | 32.8 | 32.2 | 32.6 |
| M1 | 36.6 | 37.3 | 34.7 | 36.3 | 35.8 | 33.3 | 33.7 | 33.1 |
| M2 | Undetermined* | Undetermined* | Undetermined* | Undetermined* | Undetermined* | Undetermined* | Undetermined* | 34.4 |
| M3 | 35.0 | Undetermined* | 32.4 | 32.4 | 36.2 | Undetermined* | 39.1 | Undetermined* |
| M4 | 30.5 | 32.7 | 33.7 | 31.3 | 33.6 | 34.1 | 32.6 | 33.2 |
| M5 | Undetermined* | Undetermined* | Undetermined* | Undetermined* | Undetermined* | 38.5 | Undetermined* | 37.7 |
| *Gapdh* | 18.2 | 19.6 | 18.8 | 19.8 | 18.6 | 19.3 | 18.3 | 20.5 |
| *B actin* | 14.2 | 14.1 | 14.4 | 14.2 | 18.2 | 18.4 | 17.7 | 18.0 |

| Brain Lysate | PBS | PBS | PBS | LPS | LPS | LPS |
|---|---|---|---|---|---|---|
| α7 (pair 1) | 25.8 | 25.2 | 25.9 | 25.6 | 26.2 | 26.3 |
| β2 | 25.0 | 24.4 | 25.1 | 24.3 | 25.5 | 25.7 |
| *Gapdh* | 19.0 | 18.7 | 18.9 | 18.6 | 19.5 | 19.6 |

**ΔCt (gene - *Gapdh*)**

| Macrophages | Wild type BMDMs | | | | | | | | Chrna7-/- BMDMs | | | |
|---|---|---|---|---|---|---|---|---|---|---|---|---|
| α3 | 10.4 | 8.8 | 17.4 | 15.8 | 12.4 | 9.5 | 17.1 | 19.9 | | 20.2 | 16.9 | 18.9 |
| α4 | 11.4 | 9.2 | 15.1 | 15.1 | 12.7 | 9.2 | 15.4 | 15.1 | 15.4 | 15.2 | 14.9 | 15.1 |
| α5 | 9.1 | | 15.8 | 15.4 | 12.1 | 9.4 | 14.3 | 13.6 | 16.6 | 16.1 | 17.6 | 18.3 |
| α6 | 7.9 | 8.8 | 14.1 | 14.5 | 11.0 | 8.6 | 18.3 | 17.4 | 18.6 | 18.5 | 14.1 | 14.4 |
| α7 | **Oligo pair 1** | **Oligo pair 1** | **Oligo pair 1** | **Oligo pair 1** | **Oligo pair 3** | **Oligo pair 3** | **Oligo pair 4** | **Oligo pair 4** | **Oligo pair 1** | **Oligo pair 1** | **Oligo pair 1** | **Oligo pair 1** |
| | 11.6 | 13.5 | 16.9 | 16.7 | 16.3 | 14.8 | 19.6 | 19.4 | 16.2 | 13.7 | 13.8 | 14.8 |
| α9 | 8.2 | 9.0 | 15.5 | 14.6 | 11.9 | 8.1 | 16.0 | 15.2 | 15.3 | 15.1 | 14.0 | 15.2 |
| α10 | 9.1 | 8.9 | 14.4 | 15.0 | 10.8 | 9.9 | 9.0 | 8.3 | 8.5 | 7.7 | 14.5 | 14.5 |
| β2 | 4.8 | 6.0 | 11.0 | 11.3 | 8.5 | 4.8 | 12.4 | 12.3 | 12.7 | 11.9 | 10.5 | 10.7 |
| β3 | 8.5 | 9.2 | 13.2 | 13.4 | 9.5 | 9.2 | 15.4 | 14.8 | 15.0 | 14.2 | 13.1 | 14.0 |
| β4 | 11.7 | 8.4 | 14.6 | 16.6 | 14.5 | 9.2 | 14.7 | 14.5 | 14.3 | 14.0 | 14.5 | 15.6 |
| M1 | 7.8 | 9.5 | 18.3 | 18.4 | 16.3 | 7.0 | 19.3 | 18.7 | 20.1 | 18.6 | 18.4 | 19.5 |
| M2 | | | 20.8 | | 13.5 | | | | | | 16.8 | |
| M3 | 8.9 | 10.0 | 14.0 | 15.3 | 9.3 | 9.3 | 14.5 | 15.3 | 16.2 | 14.3 | 15.4 | 15.4 |
| M4 | | | 12.7 | 12.6 | 10.9 | 8.8 | 16.6 | 17.1 | 18.2 | | 13.4 | 15.3 |
| M5 | | | | | | 12.5 | | | | | | |
| *Gapdh* | 0.0 | 0.0 | 0.0 | 0.0 | 0.0 | 0.0 | 0.0 | 0.0 | 0.0 | 0.0 | 0.0 | 0.0 |
| *B actin* | -6.3 | -6.1 | -3.8 | -3.7 | -5.8 | -6.4 | -1.6 | -1.8 | -1.4 | -2.3 | -3.9 | -3.7 |

| Macrophages | Tfebfl/fl BMDMs | | TfebΔLysMCre BMDMs | | Wild type RAW264.7 | | Tfeb-/- Tfe3-/- RAW 264.7 | |
|---|---|---|---|---|---|---|---|---|
| α3 | 18.1 | 16.9 | 17.3 | 15.4 | 12.9 | 10.6 | 11.5 | 9.2 |
| α4 | 14.5 | 14.6 | 14.4 | 13.9 | 13.7 | 13.7 | 12.9 | 11.8 |
| α5 | 16.7 | 14.6 | 16.1 | 14.3 | 15.8 | 13.2 | 18.6 | 16.0 |
| α6 | 14.3 | 12.7 | 14.5 | 13.0 | 18.1 | 12.0 | 18.2 | 13.5 |
| α7 | **Oligo pair 1** | **Oligo pair 1** | **Oligo pair 1** | **Oligo pair 1** | **Oligo pair 1** | **Oligo pair 1** | **Oligo pair 1** | **Oligo pair 1** |
| | 15.4 | 13.2 | 14.6 | 13.1 | 17.6 | 20.6 | | 18.4 |
| α9 | 17.7 | 13.9 | | | 13.7 | 13.7 | 14.9 | 14.0 |
| α10 | 14.6 | 13.3 | 14.5 | 12.9 | 9.3 | 8.6 | 9.3 | 7.4 |
| β2 | 11.1 | 9.2 | 9.9 | 8.9 | 8.9 | 11.2 | 11.5 | 9.7 |
| β3 | 13.2 | 12.3 | 12.9 | 11.6 | 15.9 | 14.5 | 16.2 | 13.3 |
| β4 | 14.4 | 14.3 | 14.7 | 13.0 | 14.8 | 13.5 | 13.9 | 12.2 |
| M1 | 18.4 | 17.7 | 15.9 | 16.5 | 17.1 | 14.0 | 15.4 | 12.7 |
| M2 | | | | | | | | 14.0 |
| M3 | 16.8 | | 13.6 | 12.6 | 17.5 | | 20.8 | |
| M4 | 12.3 | 13.1 | 14.9 | 11.5 | 15.0 | 14.8 | 14.3 | 12.7 |
| M5 | | | | | | 19.2 | | 17.2 |
| *Gapdh* | 0.0 | 0.0 | 0.0 | 0.0 | 0.0 | 0.0 | 0.0 | 0.0 |
| *B actin* | -4.0 | -5.5 | -4.5 | -5.6 | -0.4 | -0.9 | -0.6 | -2.4 |

| Brain Lysate | PBS | PBS | PBS | LPS | LPS | LPS |
|---|---|---|---|---|---|---|
| α7 (pair 1) | 6.8 | 6.6 | 7.0 | 7.1 | 6.7 | 6.7 |
| β2 | 6.0 | 5.8 | 6.2 | 5.7 | 6.0 | 6.2 |
| *Gapdh* | 0.0 | 0.0 | 0.0 | 0.0 | 0.0 | 0.0 |
