## Supplementary material for "TFEB-Mediated Pro-inflammatory Response in Murine Macrophages Induced by Acute Alpha7 Nicotinic Receptor Activation": Table_S7

**Table S7. Differential gene expression in Chrna7-/- BMDMs (PNU-282987 v. DMSO).**

| ID | B1_Chrna7KO_DMSO_1 | B2_Chrna7KO_DMSO_2 | B3_Chrna7KO_DMSO_3 | B4_Chrna7KO_DMSO_4 | B5_Chrna7KO_PNU_1 | B6_Chrna7KO_PNU_2 | B7_Chrna7KO_PNU_3 | B8_Chrna7KO_PNU_4 | padj | log2FoldChange | pvalue | stat | foldChange | log10padj |
|---|---|---|---|---|---|---|---|---|---|---|---|---|---|---|
| *Gng4* | 22.85719798 | 29.82131072 | 26.10324011 | 22.5853571 | 77.64106368 | 80.56423075 | 75.3096266 | 86.54885782 | 2.17E-11 | 1.662111819 | 7.36E-14 | 55.96837932 | 3.164794489 | 10.66271883 |
| *Mt2* | 276.2739582 | 356.7906819 | 268.7666944 | 233.7093474 | 522.3126102 | 539.5929874 | 495.7883751 | 500.3916722 | 2.08E-13 | 0.860613402 | 4.81E-16 | 65.87185512 | 1.815810188 | 12.68271914 |
| *Tnf* | 92.42258313 | 120.3502897 | 87.97758702 | 111.9448135 | 180.4902649 | 176.1171556 | 168.4006928 | 205.9265928 | 7.32E-07 | 0.827256905 | 5.94E-09 | 33.85519642 | 1.774308539 | 6.135556691 |
| *Hyal1* | 49.68956082 | 78.81346406 | 74.44257364 | 57.93635082 | 109.9074798 | 110.5416189 | 95.18300029 | 114.4036626 | 0.002060677 | 0.723825115 | 6.03E-05 | 16.09472745 | 1.651555114 | 2.685990171 |
| *Usp27x* | 60.6212642 | 68.16299594 | 55.10684022 | 64.81015516 | 93.77427172 | 105.857652 | 89.95316511 | 104.4555181 | 0.003426053 | 0.666952158 | 0.000117215 | 14.8369672 | 1.58771521 | 2.465205871 |
| *Plcxd2* | 121.2425284 | 96.91925985 | 115.0476138 | 120.7825619 | 189.5651944 | 145.202974 | 196.6418028 | 191.0043759 | 0.00015595 | 0.666580174 | 2.75E-06 | 21.9817489 | 1.587305887 | 3.807013669 |
| *Hspa1a* | 48.69576961 | 61.77271507 | 60.90756025 | 66.77409925 | 85.7076677 | 111.4784123 | 82.63139586 | 95.50218794 | 0.008232066 | 0.658666385 | 0.000387401 | 12.59199995 | 1.578622684 | 2.084491133 |
| *Impact* | 768.2006103 | 790.2647342 | 913.6134037 | 884.7568151 | 1334.01464 | 1290.901279 | 1437.158708 | 1212.678824 | 1.78E-17 | 0.651028848 | 3.01E-20 | 84.98161708 | 1.570287637 | 16.75005246 |
| *Rragd* | 534.6596744 | 571.9301378 | 618.7434692 | 616.6784461 | 899.4263481 | 921.8046868 | 926.726794 | 828.6804433 | 1.09E-15 | 0.610374062 | 1.94E-18 | 76.74528407 | 1.526654989 | 14.96221247 |
| *Ddit3* | 429.3178055 | 423.888631 | 304.5378012 | 260.2225927 | 531.3875397 | 590.1798299 | 538.6730236 | 494.4227855 | 0.000140211 | 0.605196112 | 2.41E-06 | 22.23507595 | 1.521185521 | 3.853216941 |
| *Fnip2* | 4438.271573 | 4627.628396 | 5168.441541 | 5119.020286 | 7495.891784 | 7057.801331 | 7386.619209 | 7338.746255 | 1.83E-29 | 0.597049311 | 3.26E-33 | 144.1739617 | 1.512619698 | 28.73855769 |
| *Gadd45a* | 355.7772555 | 339.7499329 | 395.4157483 | 453.6710861 | 602.9786504 | 575.1911359 | 550.178661 | 605.8420048 | 8.74E-09 | 0.59454735 | 4.13E-11 | 43.55094701 | 1.509998744 | 8.058342822 |
| *Chst3* | 518.759015 | 618.7921975 | 469.8583219 | 426.1758688 | 369.0471339 | 336.3088237 | 312.7441438 | 326.2991421 | 2.96E-06 | -0.595557236 | 2.88E-08 | 30.78747265 | 0.661788792 | 5.528283899 |
| *Rnd3* | 175.9010453 | 164.017209 | 220.4273609 | 258.2586486 | 133.0989663 | 143.3293873 | 134.9297477 | 129.3258795 | 0.001683172 | -0.599635244 | 4.62E-05 | 16.5974563 | 0.659920782 | 2.773871482 |
| *Met* | 126.2114845 | 99.04935347 | 147.9183606 | 136.4941147 | 85.7076677 | 74.94347047 | 83.67736289 | 91.52293011 | 0.00557363 | -0.605214782 | 0.000223482 | 13.62257737 | 0.657373506 | 2.253861868 |
| *F630028O10Rik* | 307.0814859 | 322.7091839 | 399.282895 | 424.2119247 | 245.0230971 | 251.9974195 | 240.5724183 | 214.8799229 | 1.02E-05 | -0.610317109 | 1.19E-07 | 28.04492852 | 0.655052704 | 4.993522701 |
| *Tifab* | 1302.860285 | 1243.974676 | 1769.219607 | 1824.504065 | 991.1839688 | 962.0868021 | 1017.725926 | 1043.560366 | 0.000931877 | -0.613417598 | 2.19E-05 | 18.0156557 | 0.653646445 | 3.030641461 |
| *Bmf* | 1149.816437 | 1126.819527 | 1523.655793 | 1615.344019 | 898.4180226 | 885.2697449 | 866.0607059 | 877.4263517 | 3.50E-08 | -0.619109267 | 1.81E-10 | 40.66429021 | 0.651072782 | 7.456409077 |
| *Id1* | 198.7582433 | 244.9607667 | 170.154454 | 177.7369407 | 107.8908288 | 144.2661807 | 146.4353851 | 115.3984771 | 0.00127108 | -0.620777072 | 3.28E-05 | 17.24886545 | 0.650320554 | 2.895827047 |
| *Cav1* | 604.2250596 | 597.4912613 | 669.9831627 | 854.3156816 | 454.7548016 | 427.1777817 | 451.8577596 | 428.7650313 | 1.07E-07 | -0.630026848 | 6.31E-10 | 38.22290134 | 0.64616439 | 6.969548178 |
| *Scin* | 117.2673635 | 132.0658046 | 123.7486938 | 137.4760867 | 92.76594622 | 73.06988371 | 70.07979142 | 86.54885782 | 0.000985787 | -0.662813434 | 2.39E-05 | 17.84998338 | 0.631645309 | 3.006216761 |
| *Hspb6* | 102.3604953 | 117.1551493 | 91.84473371 | 100.1611489 | 59.49120464 | 59.01798299 | 72.17172549 | 68.64219758 | 0.003304044 | -0.666212738 | 0.000111458 | 14.93197905 | 0.630158767 | 2.48095417 |
| *Matn2* | 94.41016556 | 95.85421304 | 129.5494139 | 133.5481985 | 74.61608717 | 61.82836314 | 76.35559364 | 72.62145542 | 0.004271402 | -0.671347561 | 0.000157255 | 14.28328364 | 0.6279199 | 2.369429543 |
| *Igfbp7* | 120.2487372 | 105.4396343 | 156.6194406 | 136.4941147 | 73.60776167 | 76.81705723 | 76.35559364 | 98.48663132 | 0.00230862 | -0.675514754 | 7.01E-05 | 15.80863189 | 0.626108783 | 2.636647469 |
| *Tmem176a* | 118.2611548 | 119.2852429 | 164.353734 | 198.3583537 | 67.55780866 | 107.7312388 | 108.7805718 | 91.52293011 | 0.009166103 | -0.677444785 | 0.000442862 | 12.34207628 | 0.625271738 | 2.037815248 |
| *Angptl4* | 278.2615406 | 331.2295584 | 340.308908 | 331.9065522 | 163.3487314 | 207.9681306 | 212.3313083 | 214.8799229 | 2.26E-07 | -0.682389415 | 1.47E-09 | 36.57585849 | 0.623132377 | 6.646565828 |
| *Gpr84* | 208.6961555 | 234.3102985 | 217.5270009 | 337.7983845 | 179.4819394 | 137.708627 | 126.5620114 | 178.0717879 | 0.000944135 | -0.68326603 | 2.26E-05 | 17.96034323 | 0.622753863 | 3.024966115 |
| *Ccr2* | 1424.102813 | 1635.911903 | 1747.9503 | 1818.612233 | 1007.317177 | 1117.594503 | 1006.220289 | 914.2344867 | 1.50E-17 | -0.711740079 | 2.27E-20 | 85.5437758 | 0.610583251 | 16.82521893 |
| *Tgfb3* | 80.49708853 | 101.1794471 | 111.1804671 | 98.19720479 | 66.54948315 | 61.82836314 | 44.97658255 | 63.6681253 | 0.003426053 | -0.720377995 | 0.000117276 | 14.83598627 | 0.6069384 | 2.465205871 |
| *Nr4a1* | 194.7830784 | 234.3102985 | 224.2945076 | 206.2141301 | 120.9990603 | 147.0765608 | 122.3781432 | 128.331065 | 7.82E-07 | -0.725880347 | 6.48E-09 | 33.68390201 | 0.604627982 | 6.106757912 |
| *Fos* | 768.2006103 | 924.4606324 | 887.5101636 | 841.550045 | 488.0295431 | 611.7260777 | 448.7198585 | 510.3398168 | 1.94E-13 | -0.731054905 | 4.33E-16 | 66.07920413 | 0.602463229 | 12.71137425 |
| *P2ry13* | 115.2797811 | 127.8056174 | 157.5862273 | 171.8451084 | 89.74096971 | 57.14439623 | 94.13703325 | 98.48663132 | 0.001480892 | -0.758038767 | 3.96E-05 | 16.89031731 | 0.591299611 | 2.829476758 |
| *Ccr1* | 185.8389575 | 193.8385197 | 304.5378012 | 295.5735864 | 142.1738958 | 130.2142799 | 153.7571543 | 149.2221687 | 0.000100287 | -0.770345401 | 1.62E-06 | 23.00225634 | 0.586277095 | 3.998754093 |
| *Cd24a* | 759.2564894 | 934.0460537 | 1344.800259 | 1473.940044 | 669.5281336 | 724.1412834 | 617.1205513 | 618.7745927 | 0.009613099 | -0.778998821 | 0.000469292 | 12.23389129 | 0.582771075 | 2.017136601 |
| *Mblac1* | 78.5095061 | 73.48823 | 53.17326688 | 58.91832287 | 38.31636909 | 31.85097495 | 37.6548133 | 43.77183614 | 0.008315581 | -0.800621928 | 0.000392132 | 12.56931399 | 0.574101636 | 2.080107401 |
| *Fstl1* | 163.9755507 | 195.9686133 | 311.3053079 | 317.1769715 | 156.2904529 | 133.9614535 | 138.0676488 | 134.2999518 | 0.000336016 | -0.81446632 | 6.65E-06 | 20.29147493 | 0.568618791 | 3.473640646 |
| *Nuak1* | 286.2118703 | 298.2131072 | 422.485775 | 412.4282601 | 191.5818454 | 177.9907424 | 224.8829128 | 211.8954795 | 2.46E-07 | -0.818024596 | 1.64E-09 | 36.36412204 | 0.567218072 | 6.608245693 |
| *Ccr5* | 1443.978637 | 1488.935443 | 1321.597379 | 1226.483088 | 745.1525462 | 815.9470347 | 807.4865519 | 723.2301108 | 3.36E-28 | -0.825353683 | 1.20E-31 | 137.0117688 | 0.564343834 | 27.47342492 |
| *H2-Ob* | 64.59642907 | 57.51252782 | 61.87434692 | 64.81015516 | 27.22478256 | 34.66135509 | 39.74674737 | 33.82369156 | 0.002628214 | -0.878789408 | 8.41E-05 | 15.46362367 | 0.543823572 | 2.580339279 |
| *Epha2* | 132.1742318 | 123.5454301 | 141.1508539 | 153.1876395 | 85.7076677 | 68.3859168 | 74.26365957 | 70.6318265 | 1.69E-06 | -0.881893297 | 1.52E-08 | 32.03019827 | 0.54265482 | 5.773149939 |
| *Gpr34* | 69.56538515 | 63.90280869 | 88.94437369 | 128.6383383 | 42.3496711 | 57.14439623 | 49.1604507 | 38.79776385 | 0.004139074 | -0.905691443 | 0.000148694 | 14.3886766 | 0.533776819 | 2.383096842 |
| *Ptgir* | 114.2859899 | 121.4153365 | 119.8815472 | 180.6828568 | 64.53283215 | 65.57553666 | 81.58542882 | 74.61108433 | 4.58E-05 | -0.907663559 | 6.37E-07 | 24.79596492 | 0.533047662 | 4.338860187 |
| *Ptgs2* | 113.2921987 | 117.1551493 | 214.6266409 | 258.2586486 | 83.69101669 | 90.86895794 | 82.63139586 | 104.4555181 | 0.000931877 | -0.960747267 | 2.19E-05 | 18.02054505 | 0.513790718 | 3.030641461 |
| *C630043F03Rik* | 166.9569244 | 129.935711 | 130.5162005 | 127.6563662 | 76.63273818 | 76.81705723 | 61.71205513 | 59.68886746 | 1.80E-07 | -1.012394352 | 1.14E-09 | 37.06709508 | 0.495722844 | 6.743929567 |
| *Inhba* | 81.49087975 | 74.55327681 | 132.4497739 | 107.0349532 | 53.44125162 | 47.77646242 | 49.1604507 | 45.76146506 | 0.000100287 | -1.014970792 | 1.60E-06 | 23.02152249 | 0.494838346 | 3.998754093 |
| *Gpr85* | 56.64609934 | 52.18729377 | 49.3061202 | 53.02649058 | 26.21646306 | 14.05190071 | 18.82740665 | 42.77702168 | 0.00753809 | -1.053646959 | 0.000342902 | 12.82017178 | 0.481748821 | 2.122738672 |
| *Gpr183* | 397.5164866 | 481.4011588 | 488.2272686 | 535.1747661 | 228.889889 | 235.1351386 | 246.8482205 | 204.9317783 | 7.52E-21 | -1.055070435 | 6.71E-24 | 101.6259897 | 0.481273724 | 20.12362327 |
| *Lamb1* | 74.53434123 | 85.20374492 | 122.7819072 | 150.2417233 | 37.30804359 | 56.20760285 | 51.25238477 | 55.70960963 | 0.000103803 | -1.110603442 | 1.68E-06 | 22.92543087 | 0.463100288 | 3.983790685 |
| *Dcstamp* | 37.76406622 | 39.40673203 | 53.17326688 | 78.55776383 | 22.18316105 | 29.97738819 | 27.19514294 | 14.92221687 | 0.004807096 | -1.148516976 | 0.000181262 | 14.01599211 | 0.451088692 | 2.318117164 |
| *Emr4* | 59.62747299 | 55.3824342 | 161.453374 | 140.4220028 | 44.3663221 | 50.58684257 | 48.11448366 | 40.78739277 | 0.002251594 | -1.183047994 | 6.74E-05 | 15.88136685 | 0.440420035 | 2.647509989 |
| *Tnfaip8l3* | 32.79511014 | 25.56112348 | 41.57182684 | 33.38704963 | 18.14985904 | 8.431140428 | 7.321769253 | 15.91703132 | 0.002333741 | -1.423485503 | 7.14E-05 | 15.77454444 | 0.372810525 | 2.631947346 |
