## Supplementary material for "TFEB-Mediated Pro-inflammatory Response in Murine Macrophages Induced by Acute Alpha7 Nicotinic Receptor Activation": Table_S9

**Table S9. Differential gene expression in PNU-282987- treated BMDMs (Chrna7-/- v. WT).**

| ID | A5_WT_PNU_1 | A6_WT_PNU_2 | A7_WT_PNU_3 | A8_WT_PNU_4 | B5_Chrna7KO_PNU_1 | B6_Chrna7KO_PNU_2 | B7_Chrna7KO_PNU_3 | B8_Chrna7KO_PNU_4 | padj | log2FoldChange | pvalue | stat | foldChange | log10padj |
|---|---|---|---|---|---|---|---|---|---|---|---|---|---|---|
| *Mpo* | 0 | 0 | 0 | 0 | 22.02465671 | 10.64030104 | 27.42418901 | 18.25309501 | 3.42E-10 | 6.54188376 | 3.01E-12 | 48.68009017 | 93.1758235 | 9.465434724 |
| *Elane* | 0 | 0 | 0 | 0 | 6.166903878 | 9.003331649 | 10.9696756 | 4.345975003 | 6.88E-05 | 5.182726319 | 2.84E-06 | 21.92206565 | 36.3208565 | 4.162462894 |
| *Ngp* | 0 | 0 | 1.141169275 | 0.976206612 | 8.809862684 | 27.00999495 | 22.85349084 | 19.99148501 | 6.21E-08 | 5.11847567 | 8.55E-10 | 37.63059182 | 34.7387917 | 7.206652661 |
| *Flt3* | 1.179932618 | 2.570596965 | 0 | 0 | 11.45282149 | 18.0066633 | 12.79795487 | 14.77631501 | 4.60E-07 | 4.058342362 | 8.16E-09 | 33.23629014 | 16.66029871 | 6.337648208 |
| *S100a9* | 1.179932618 | 0 | 2.28233855 | 0 | 12.33380776 | 11.45878574 | 15.54037377 | 16.51470501 | 4.37E-07 | 4.03176621 | 7.69E-09 | 33.35116142 | 16.35620574 | 6.359371682 |
| *Chil3* | 2.359865237 | 16.70888027 | 2.28233855 | 2.928619837 | 81.93172296 | 87.57786241 | 82.27256703 | 75.61996505 | 1.11E-11 | 3.809363475 | 7.37E-14 | 55.96657073 | 14.0195047 | 10.95458449 |
| *Gpr128* | 1.179932618 | 0 | 0 | 1.952413225 | 13.21479403 | 9.003331649 | 8.227256703 | 12.16873001 | 3.64E-05 | 3.657772739 | 1.33E-06 | 23.38499035 | 12.62116114 | 4.439446346 |
| *Marveld2* | 1.179932618 | 1.285298482 | 1.141169275 | 0 | 14.97676656 | 5.729392868 | 7.313117069 | 5.215170004 | 0.002645708 | 3.269811676 | 0.0002491 | 13.41891369 | 9.64520349 | 2.577458135 |
| *Cdh1* | 4.719730474 | 1.285298482 | 1.141169275 | 0.976206612 | 20.26268417 | 16.36969391 | 18.28279267 | 19.99148501 | 6.95E-08 | 3.232484146 | 9.83E-10 | 37.35751985 | 9.398849349 | 7.158294044 |
| *Amica1* | 2.359865237 | 0 | 1.141169275 | 1.952413225 | 13.21479403 | 13.91423982 | 10.05553597 | 13.90712001 | 1.12E-05 | 3.179331374 | 3.32E-07 | 26.05492849 | 9.058871719 | 4.952668548 |
| *Slc36a2* | 28.31838284 | 12.85298482 | 0 | 14.64309919 | 107.4803247 | 150.601184 | 124.3229902 | 119.0797151 | 6.51E-05 | 3.168823127 | 2.63E-06 | 22.06600966 | 8.993128774 | 4.186535044 |
| *Galnt9* | 3.539797855 | 0 | 0 | 1.952413225 | 14.09578029 | 10.64030104 | 14.62623414 | 11.29953501 | 3.24E-05 | 3.162321661 | 1.16E-06 | 23.6507436 | 8.952692662 | 4.488828493 |
| *Cyb561* | 3.539797855 | 0 | 0 | 6.833446286 | 24.66761551 | 21.28060208 | 22.85349084 | 20.86068002 | 2.48E-06 | 3.013587866 | 5.76E-08 | 29.44413368 | 8.075703069 | 5.60641363 |
| *Slc27a6* | 3.539797855 | 3.855895447 | 0 | 4.881033062 | 22.90564298 | 12.27727043 | 40.22214388 | 23.46826502 | 6.87E-05 | 2.99335882 | 2.82E-06 | 21.93412849 | 7.963258112 | 4.162755667 |
| *Eef1a2* | 2.359865237 | 5.14119393 | 1.141169275 | 5.857239674 | 23.78662925 | 22.91757147 | 31.99488718 | 35.63699503 | 8.23E-09 | 2.959180332 | 9.41E-11 | 41.94025829 | 7.776819917 | 8.084778353 |
| *A430078G23Rik* | 5.899663092 | 2.570596965 | 0 | 3.904826449 | 20.26268417 | 22.09908678 | 19.19693231 | 30.42182502 | 5.64E-07 | 2.885502839 | 1.04E-08 | 32.77056185 | 7.389633632 | 6.248989643 |
| *Dpp4* | 3.539797855 | 0 | 1.141169275 | 1.952413225 | 9.690848952 | 16.36969391 | 13.71209451 | 6.953560005 | 0.000810643 | 2.788377854 | 5.53E-05 | 16.25669281 | 6.908525634 | 3.091170347 |
| *Scel* | 1.179932618 | 1.285298482 | 1.141169275 | 2.928619837 | 12.33380776 | 10.64030104 | 8.227256703 | 14.77631501 | 0.000214902 | 2.76640469 | 1.11E-05 | 19.31416656 | 6.80410163 | 3.667759308 |
| *Atp8b5* | 2.359865237 | 1.285298482 | 0 | 1.952413225 | 7.047890147 | 8.184846954 | 6.398977436 | 13.90712001 | 0.003319955 | 2.653128135 | 0.000333562 | 12.87184586 | 6.290296995 | 2.478867741 |
| *Htra3* | 29.49831546 | 24.42067117 | 2.28233855 | 25.38137192 | 125.9810364 | 121.1357349 | 115.1815938 | 112.9953501 | 9.27E-06 | 2.542735172 | 2.67E-07 | 26.47562438 | 5.82692674 | 5.033034217 |
| *Xlr3a* | 1.179932618 | 2.570596965 | 2.28233855 | 0.976206612 | 10.57183522 | 13.09575513 | 4.570698168 | 11.29953501 | 0.002943821 | 2.534491323 | 0.000285881 | 13.16070656 | 5.793725461 | 2.531088537 |
| *Cdo1* | 4.719730474 | 5.14119393 | 5.705846376 | 5.857239674 | 36.120437 | 34.37635721 | 27.42418901 | 22.59907002 | 1.34E-08 | 2.486433315 | 1.59E-10 | 40.91234184 | 5.603908173 | 7.871339787 |
| *Igsf11* | 5.899663092 | 5.14119393 | 1.141169275 | 9.762066123 | 29.07254686 | 32.73938782 | 42.05042315 | 19.99148501 | 1.46E-05 | 2.467669988 | 4.57E-07 | 25.43920334 | 5.531497058 | 4.834635766 |
| *Ly6c2* | 5.899663092 | 1.285298482 | 10.27052348 | 0 | 22.90564298 | 23.73605617 | 15.54037377 | 31.29102002 | 0.006232153 | 2.434329695 | 0.000757889 | 11.34162689 | 5.405131418 | 2.205361887 |
| *Skint3* | 0 | 2.570596965 | 3.423507825 | 1.952413225 | 6.166903878 | 11.45878574 | 11.88381524 | 13.03792501 | 0.002287206 | 2.422212789 | 0.000206399 | 13.77191447 | 5.359924909 | 2.640694677 |
| *Kcne3* | 5.899663092 | 5.14119393 | 0 | 5.857239674 | 36.120437 | 13.91423982 | 18.28279267 | 20.86068002 | 0.000925037 | 2.394085116 | 6.62E-05 | 15.91780777 | 5.256436618 | 3.033840842 |
| *Tspan7* | 2.359865237 | 11.56768634 | 5.705846376 | 7.809652899 | 33.4774782 | 30.28393373 | 33.82316645 | 29.55263002 | 4.97E-08 | 2.220754703 | 6.68E-10 | 38.1122337 | 4.661372165 | 7.303652754 |
| *Psg22* | 1.179932618 | 2.570596965 | 4.564677101 | 2.928619837 | 5.28591761 | 13.09575513 | 16.45451341 | 15.64551001 | 0.005406655 | 2.157193718 | 0.000628852 | 11.68857101 | 4.460463765 | 2.267071365 |
| *Gm1966* | 133.3323859 | 39.84425295 | 36.5174168 | 93.71583478 | 340.9416859 | 338.0341792 | 330.0044078 | 318.9945652 | 7.20E-06 | 2.127533096 | 1.96E-07 | 27.06774213 | 4.369696549 | 5.142560084 |
| *Gm14461* | 9.439460947 | 2.570596965 | 4.564677101 | 8.785859511 | 40.52536834 | 24.55454086 | 25.59590974 | 21.72987502 | 4.35E-05 | 2.120914386 | 1.65E-06 | 22.97020724 | 4.349695432 | 4.361667142 |
| *Ptger3* | 5.899663092 | 7.711790894 | 2.28233855 | 19.52413225 | 43.16832715 | 42.56120416 | 37.47972498 | 33.02941002 | 0.00040048 | 2.108692064 | 2.35E-05 | 17.8837089 | 4.31300104 | 3.39741921 |
| *Il1r2* | 12.9792588 | 2.570596965 | 7.988184926 | 7.809652899 | 38.76339581 | 36.0133266 | 32.90902681 | 26.07585002 | 1.28E-06 | 2.07961993 | 2.65E-08 | 30.94929426 | 4.226958447 | 5.893341297 |
| *Erdr1* | 94.39460947 | 97.68268466 | 109.5522504 | 111.2875538 | 435.2072166 | 428.8859804 | 404.0497181 | 477.1880553 | 4.27E-68 | 2.075076937 | 6.83E-72 | 321.4986624 | 4.213668848 | 67.36962277 |
| *Fabp7* | 20.05885451 | 5.14119393 | 9.129354201 | 11.71447935 | 48.45424476 | 60.56786746 | 42.05042315 | 42.59055503 | 3.17E-07 | 2.064926876 | 5.47E-09 | 34.01440447 | 4.184127677 | 6.499477371 |
| *Gfra1* | 2.359865237 | 5.14119393 | 0 | 3.904826449 | 14.09578029 | 11.45878574 | 8.227256703 | 12.16873001 | 0.005300504 | 2.01908372 | 0.000611417 | 11.74089186 | 4.053262807 | 2.275682872 |
| *Notch4* | 126.2527902 | 80.97380439 | 57.05846376 | 92.73962817 | 357.680425 | 385.5062915 | 327.2619889 | 318.1253702 | 2.44E-15 | 1.959416277 | 1.08E-17 | 73.36921725 | 3.889045941 | 14.6118879 |
| *P2ry10* | 8.259528329 | 5.14119393 | 5.705846376 | 1.952413225 | 15.85775283 | 21.28060208 | 24.68177011 | 17.38390001 | 0.000280241 | 1.941966077 | 1.53E-05 | 18.69560047 | 3.842289108 | 3.552467586 |
| *Dio2* | 11.79932618 | 10.28238786 | 23.96455478 | 5.857239674 | 54.62114864 | 46.65362764 | 42.05042315 | 47.80572503 | 1.21E-05 | 1.891753751 | 3.68E-07 | 25.85647765 | 3.710860458 | 4.916120003 |
| *Ccr2* | 315.0420091 | 262.2008904 | 99.28172694 | 278.2188845 | 880.1052821 | 976.4522416 | 879.4023276 | 798.7902056 | 4.88E-07 | 1.88843168 | 8.82E-09 | 33.08549179 | 3.702325347 | 6.311722905 |
| *Spint1* | 151.0313752 | 115.6768634 | 74.17600288 | 137.6451323 | 429.921299 | 405.9684089 | 475.3526095 | 414.6060153 | 7.56E-16 | 1.849126662 | 3.21E-18 | 75.75759372 | 3.602820218 | 15.12121138 |
| *Impg2* | 5.899663092 | 1.285298482 | 4.564677101 | 0.976206612 | 12.33380776 | 13.09575513 | 10.9696756 | 8.691950006 | 0.009455015 | 1.84563282 | 0.001316033 | 10.31990186 | 3.594105655 | 2.024337756 |
| *Pianp* | 16.51905666 | 16.70888027 | 1.141169275 | 15.6193058 | 65.19298386 | 38.46878068 | 32.90902681 | 39.98297003 | 0.006841434 | 1.821899285 | 0.000861952 | 11.10284957 | 3.535463304 | 2.164852839 |
| *Thrb* | 2.359865237 | 5.14119393 | 1.141169275 | 8.785859511 | 18.50071164 | 13.09575513 | 19.19693231 | 12.16873001 | 0.006666777 | 1.815353143 | 0.000832481 | 11.16738114 | 3.51945769 | 2.176084052 |
| *Abca6* | 8.259528329 | 6.426492412 | 4.564677101 | 9.762066123 | 29.07254686 | 20.46211738 | 17.36865304 | 30.42182502 | 0.000331358 | 1.730764049 | 1.86E-05 | 18.32403826 | 3.319035473 | 3.47970212 |
| *Atp6v0d2* | 293.803222 | 226.2125329 | 101.5640655 | 277.2426779 | 730.3376165 | 757.0983432 | 740.4531033 | 725.7778255 | 8.81E-07 | 1.715924828 | 1.74E-08 | 31.76336373 | 3.285071628 | 6.054836518 |
| *Fam171a1* | 23.59865237 | 41.12955144 | 6.847015651 | 38.07205788 | 89.86059937 | 94.12573997 | 78.6160085 | 88.65789006 | 0.001239665 | 1.679780794 | 9.48E-05 | 15.23750204 | 3.203792682 | 2.906695532 |
| *Rbm44* | 15.33912404 | 14.13828331 | 5.705846376 | 20.50033886 | 46.69227222 | 40.92423477 | 49.36354022 | 43.45975003 | 2.81E-06 | 1.679293537 | 6.73E-08 | 29.14069557 | 3.202710813 | 5.550658394 |
| *Grap2* | 81.41535067 | 71.97671501 | 10.27052348 | 63.4534298 | 208.7937456 | 172.7002707 | 197.4541609 | 146.8939551 | 0.006722064 | 1.677092517 | 0.000841466 | 11.14746431 | 3.197828384 | 2.172497356 |
| *Rnf122* | 4.719730474 | 7.711790894 | 7.988184926 | 4.881033062 | 21.14367044 | 18.82514799 | 17.36865304 | 21.72987502 | 0.000350491 | 1.659777567 | 1.99E-05 | 18.19823654 | 3.159678055 | 3.455322905 |
| *Cryab* | 17.69898928 | 8.997089377 | 10.27052348 | 7.809652899 | 22.02465671 | 46.65362764 | 49.36354022 | 20.86068002 | 0.004244111 | 1.639760018 | 0.00045663 | 12.28493293 | 3.116139929 | 2.372213269 |
| *Efna1* | 9.439460947 | 3.855895447 | 4.564677101 | 9.762066123 | 19.3816979 | 25.37302556 | 22.85349084 | 19.99148501 | 0.000418164 | 1.638543373 | 2.47E-05 | 17.78597308 | 3.113513151 | 3.378653023 |
| *Rnf144b* | 337.4607289 | 230.0684283 | 52.49378666 | 296.7668102 | 667.7875914 | 739.0916799 | 726.7410088 | 681.4488805 | 0.005045824 | 1.61828145 | 0.00057316 | 11.8611877 | 3.070091066 | 2.297067874 |
| *Padi4* | 70.7959571 | 52.69723778 | 30.81157043 | 82.00135544 | 180.602185 | 169.4263319 | 196.5400212 | 171.2314151 | 2.85E-07 | 1.596886497 | 4.86E-09 | 34.24519861 | 3.024898003 | 6.545428109 |
| *Xist* | 4614.716471 | 4435.565063 | 4655.970643 | 4309.952193 | 14718.63759 | 12908.32213 | 13931.48802 | 12887.55427 | 5.44E-116 | 1.595609894 | 4.35E-120 | 542.9273446 | 3.022222534 | 115.2644572 |
| *St8sia1* | 8.259528329 | 2.570596965 | 2.28233855 | 7.809652899 | 11.45282149 | 19.64363269 | 18.28279267 | 14.77631501 | 0.008831449 | 1.589890445 | 0.00120522 | 10.48232518 | 3.010264894 | 2.053968017 |
| *Cd109* | 612.385029 | 467.8486476 | 403.9739234 | 605.2480997 | 1543.487942 | 1537.114258 | 1501.931418 | 1528.044811 | 3.90E-22 | 1.547095013 | 9.98E-25 | 105.4003364 | 2.922281207 | 21.4091054 |
| *BC147527* | 35.39797855 | 19.27947724 | 12.55286203 | 31.23861159 | 74.00284654 | 66.29726033 | 70.38875179 | 77.35835506 | 3.37E-06 | 1.53928561 | 8.27E-08 | 28.74090135 | 2.906505442 | 5.47249007 |
| *Pvrl1* | 69.61602449 | 20.56477572 | 29.67040115 | 50.76274384 | 125.1000501 | 135.0499747 | 101.4694993 | 135.5944201 | 0.000236142 | 1.538088466 | 1.25E-05 | 19.08747432 | 2.904094633 | 3.626827514 |
| *Ica1* | 57.8166983 | 55.26783474 | 27.3880626 | 63.4534298 | 154.172597 | 144.8717911 | 140.7775036 | 149.5015401 | 2.53E-09 | 1.527911878 | 2.57E-11 | 44.48110974 | 2.883681598 | 8.597224494 |
| *Plbd1* | 12.9792588 | 12.85298482 | 15.97636985 | 16.59551241 | 43.16832715 | 42.56120416 | 44.79284205 | 39.11377503 | 2.17E-07 | 1.526648369 | 3.52E-09 | 34.87026335 | 2.881157181 | 6.663487624 |
| *Clec2i* | 29.49831546 | 10.28238786 | 9.129354201 | 34.16723143 | 40.52536834 | 63.02332154 | 70.38875179 | 64.32043005 | 0.00683393 | 1.506700051 | 0.00086046 | 11.10606285 | 2.841593233 | 2.165329473 |
| *Mamdc2* | 66.07622663 | 52.69723778 | 50.21144811 | 84.92997527 | 172.6733086 | 175.9742095 | 190.1410438 | 186.8769251 | 1.06E-14 | 1.502638142 | 5.08E-17 | 70.30411445 | 2.833603977 | 13.97507301 |
| *Rapgef3* | 31.8581807 | 14.13828331 | 4.564677101 | 21.47654547 | 59.90706625 | 46.65362764 | 51.19181949 | 44.32894503 | 0.007839051 | 1.484750198 | 0.001034045 | 10.76559193 | 2.798687098 | 2.10573651 |
| *Kit* | 4.719730474 | 11.56768634 | 13.6940313 | 17.57171902 | 40.52536834 | 30.28393373 | 25.59590974 | 33.02941002 | 0.001397363 | 1.425127341 | 0.000109097 | 14.97236231 | 2.685382024 | 2.85469069 |
| *Pacrg* | 14.15919142 | 8.997089377 | 4.564677101 | 5.857239674 | 26.42958805 | 21.28060208 | 16.45451341 | 24.33746002 | 0.005002017 | 1.417299259 | 0.000565466 | 11.88635686 | 2.670850565 | 2.300854847 |
| *Cd300lg* | 73.15582234 | 56.55313323 | 20.54104695 | 92.73962817 | 142.7197755 | 178.4296636 | 134.3785261 | 191.2229001 | 0.003999857 | 1.409108079 | 0.000421071 | 12.43629039 | 2.655729262 | 2.3979555 |
| *Pcyt1b* | 64.89629401 | 56.55313323 | 13.6940313 | 48.81033062 | 134.7908991 | 107.2214951 | 131.6361072 | 113.8645451 | 0.003526477 | 1.407789717 | 0.000360518 | 12.72646052 | 2.653303515 | 2.452658904 |
| *Ptgfrn* | 194.688882 | 186.3682799 | 118.6816046 | 193.2889092 | 448.4220106 | 439.5262814 | 436.9587449 | 502.3947104 | 4.13E-14 | 1.39843308 | 2.15E-16 | 67.46044281 | 2.636151126 | 13.38361717 |
| *Elovl7* | 25.9585176 | 37.27365599 | 6.847015651 | 28.30999176 | 68.71692893 | 54.83847459 | 72.21703106 | 61.71284504 | 0.004275598 | 1.391589176 | 0.000460702 | 12.26836575 | 2.623675281 | 2.369003172 |
| *Mroh6* | 10.61939357 | 15.42358179 | 4.564677101 | 13.66689257 | 31.71550566 | 33.55787251 | 21.93935121 | 27.81424002 | 0.001812944 | 1.381413389 | 0.000154151 | 14.32080987 | 2.605234775 | 2.741615526 |
| *Sirpb1b* | 48.37723735 | 43.7001484 | 36.5174168 | 35.14343804 | 96.02750325 | 90.85180119 | 115.1815938 | 112.1261551 | 1.42E-10 | 1.34559758 | 1.13E-12 | 50.61075564 | 2.541354404 | 9.847051288 |
| *Xlr* | 17.69898928 | 8.997089377 | 7.988184926 | 15.6193058 | 29.07254686 | 30.28393373 | 32.90902681 | 36.50619003 | 0.0003616 | 1.341377247 | 2.07E-05 | 18.12013332 | 2.533931013 | 3.441772143 |
| *Gapt* | 17.69898928 | 23.13537268 | 26.24689333 | 41.97688433 | 88.09862684 | 63.84180624 | 63.07563472 | 59.97445504 | 0.000198517 | 1.317380733 | 1.01E-05 | 19.49848191 | 2.492132428 | 3.702202743 |
| *Nim1k* | 23.59865237 | 12.85298482 | 7.988184926 | 29.28619837 | 45.81128595 | 40.92423477 | 47.53526095 | 50.41331004 | 0.003142612 | 1.306918121 | 0.000309962 | 13.00921165 | 2.474124537 | 2.502709297 |
| *Itgb7* | 1319.164667 | 1244.168931 | 1097.804843 | 1409.642348 | 3036.759667 | 3179.813042 | 3037.686003 | 3197.768407 | 5.41E-43 | 1.295438445 | 4.33E-46 | 203.1309123 | 2.454515785 | 42.26643265 |
| *Olfml3* | 1092.617605 | 664.4993154 | 313.8215507 | 940.0869677 | 1891.477518 | 1704.085136 | 1878.556947 | 1860.946496 | 0.00221739 | 1.28437244 | 0.000198523 | 13.84501143 | 2.435760761 | 2.654157872 |
| *Treml2* | 164.010634 | 133.6710422 | 69.61132578 | 154.2406448 | 306.5832214 | 310.2056996 | 326.3478492 | 318.1253702 | 1.07E-05 | 1.273212766 | 3.17E-07 | 26.14270454 | 2.416992114 | 4.970070473 |
| *Colec12* | 548.6686676 | 372.7365599 | 217.9633316 | 476.3888268 | 911.8207877 | 974.8152722 | 1024.750529 | 971.7600107 | 5.70E-05 | 1.26465171 | 2.26E-06 | 22.35617824 | 2.40269198 | 4.244375511 |
| *Amz1* | 639.5234792 | 561.6754368 | 304.6921965 | 612.0815459 | 1296.811787 | 1222.816135 | 1228.603668 | 1338.560301 | 1.81E-06 | 1.263944399 | 3.93E-08 | 30.18393689 | 2.401514299 | 5.743286227 |
| *Mfge8* | 1452.497053 | 1262.16311 | 1307.779989 | 1468.214745 | 3288.72174 | 3359.879675 | 3162.008993 | 3376.822577 | 1.17E-65 | 1.2631143 | 2.81E-69 | 309.4962448 | 2.400132912 | 64.93121185 |
| *Chst1* | 27.13845022 | 11.56768634 | 14.83520058 | 18.54792563 | 39.64438208 | 36.83181129 | 52.10595912 | 45.19814003 | 0.000374947 | 1.26119084 | 2.17E-05 | 18.03532336 | 2.396935089 | 3.426030676 |
| *Klhl3* | 12.9792588 | 7.711790894 | 10.27052348 | 7.809652899 | 29.95353312 | 19.64363269 | 20.11107194 | 22.59907002 | 0.004298753 | 1.255180816 | 0.0004643 | 12.25384739 | 2.386970631 | 2.36665755 |
| *Cd207* | 307.9624134 | 187.6535784 | 78.74067998 | 252.8375126 | 472.2086398 | 469.8102152 | 539.3423839 | 484.1416154 | 0.005370763 | 1.248131833 | 0.000622341 | 11.70793694 | 2.375336379 | 2.269963985 |
| *Haao* | 38.93777641 | 24.42067117 | 10.27052348 | 32.21481821 | 74.00284654 | 54.0199899 | 67.64633289 | 54.75928504 | 0.004530706 | 1.238741729 | 0.000497613 | 12.12458924 | 2.359926176 | 2.343834142 |
| *Ssbp2* | 24.77858499 | 38.55895447 | 20.54104695 | 25.38137192 | 51.97818983 | 72.02665319 | 74.95944996 | 56.49767504 | 5.52E-05 | 1.236160623 | 2.17E-06 | 22.44122765 | 2.355707837 | 4.257994472 |
| *Angptl4* | 101.4742052 | 64.26492412 | 38.79975535 | 95.66824801 | 142.7197755 | 181.7036024 | 185.5703456 | 187.7461201 | 0.000608539 | 1.21385196 | 3.91E-05 | 16.91273633 | 2.319561279 | 3.2157116 |
| *Abcb1a* | 103.8340704 | 128.5298482 | 131.2344666 | 92.73962817 | 239.628265 | 284.0141893 | 248.6459804 | 280.7499852 | 1.16E-13 | 1.212878773 | 6.42E-16 | 65.30235886 | 2.317997119 | 12.93406906 |
| *Cxcl14* | 679.6411882 | 494.8399157 | 311.5392121 | 589.6287939 | 1215.76105 | 1234.274921 | 1110.679655 | 1216.873001 | 6.94E-06 | 1.202511331 | 1.88E-07 | 27.15699187 | 2.301399322 | 5.15877256 |
| *Rab19* | 42.47757426 | 41.12955144 | 12.55286203 | 45.88171078 | 81.05073669 | 93.30725527 | 79.53014813 | 73.01238005 | 0.004514801 | 1.201231572 | 0.000495348 | 12.13309758 | 2.299358745 | 2.345361395 |
| *Lipn* | 23.59865237 | 8.997089377 | 17.11753913 | 12.69068596 | 26.42958805 | 33.55787251 | 35.65144571 | 46.06733503 | 0.005659871 | 1.182459047 | 0.000665548 | 11.58307613 | 2.269633023 | 2.247193498 |
| *Entpd1* | 930.9668359 | 794.3144621 | 298.9863501 | 930.3249016 | 1709.113361 | 1553.483952 | 1724.981489 | 1717.529321 | 0.003476038 | 1.18218657 | 0.000353971 | 12.76073681 | 2.269204406 | 2.458915432 |
| *Hhat* | 8.259528329 | 11.56768634 | 9.129354201 | 15.6193058 | 34.35846447 | 20.46211738 | 23.76763048 | 23.46826502 | 0.007542559 | 1.17800086 | 0.000986488 | 10.85275689 | 2.262630275 | 2.12248127 |
| *Trp53i11* | 38.93777641 | 47.55604385 | 29.67040115 | 41.00067772 | 89.86059937 | 89.2148318 | 81.3584274 | 86.05030506 | 3.22E-08 | 1.14535491 | 4.07E-10 | 39.0801882 | 2.212005404 | 7.492383993 |
| *Gpnmb* | 54767.75242 | 46293.88074 | 39508.42147 | 53656.22024 | 106177.346 | 104401.8153 | 107106.9985 | 107101.3388 | 6.19E-19 | 1.12899199 | 2.03E-21 | 90.31548323 | 2.187058772 | 18.20822042 |
| *Cytip* | 461.3536538 | 465.2780506 | 268.1747797 | 416.8402235 | 832.5320236 | 977.2707263 | 872.0892105 | 828.3428356 | 1.41E-07 | 1.123692313 | 2.18E-09 | 35.80281728 | 2.179039446 | 6.849495261 |
| *Htr2b* | 261.9450413 | 147.8093255 | 163.1872063 | 190.3602894 | 391.1579031 | 437.889312 | 442.4435827 | 386.7917753 | 2.19E-07 | 1.118482252 | 3.59E-09 | 34.83523091 | 2.171184388 | 6.659931013 |
| *Ly6a* | 1802.937041 | 1236.45714 | 768.0069222 | 1414.523381 | 2791.845484 | 2854.874618 | 2875.883288 | 2809.238242 | 9.17E-05 | 1.11769565 | 4.01E-06 | 21.25867703 | 2.170000913 | 4.037584582 |
| *Tnfsf13* | 276.1042327 | 272.4832783 | 122.1051124 | 322.1481821 | 528.591761 | 491.9093019 | 554.8827576 | 571.9303104 | 0.000909436 | 1.112236253 | 6.42E-05 | 15.97343925 | 2.161804788 | 3.041227717 |
| *Pde7b* | 148.6715099 | 173.5152951 | 231.6573629 | 108.358934 | 410.5396011 | 361.7702354 | 330.9185474 | 324.2097352 | 5.52E-05 | 1.110588493 | 2.17E-06 | 22.43498318 | 2.159337114 | 4.257994472 |
| *Tnfrsf21* | 346.9001898 | 379.1630523 | 252.1984098 | 322.1481821 | 695.0981657 | 729.2698636 | 671.8926307 | 692.7484155 | 1.00E-14 | 1.102670258 | 4.64E-17 | 70.48443058 | 2.147518051 | 14.00004635 |
| *F13a1* | 237.1664563 | 194.0800708 | 166.6107142 | 155.2168514 | 424.6353813 | 352.7669037 | 424.16079 | 411.9984303 | 3.97E-10 | 1.102298593 | 3.65E-12 | 48.30329034 | 2.146964882 | 9.401294845 |
| *Inpp5j* | 16.51905666 | 11.56768634 | 11.41169275 | 23.4289587 | 29.07254686 | 43.37968886 | 30.16660791 | 33.89860502 | 0.005947443 | 1.098608958 | 0.000708403 | 11.46707002 | 2.14148112 | 2.225669676 |
| *5530601H04Rik* | 74.33575496 | 34.70305902 | 30.81157043 | 67.35825625 | 115.4092012 | 108.8584645 | 107.8684768 | 110.3877651 | 0.002359221 | 1.088417016 | 0.000214766 | 13.69727103 | 2.126405905 | 2.627231293 |
| *I830012O16Rik* | 286.7236263 | 191.5094739 | 320.6685663 | 312.3861159 | 540.9255688 | 605.6786746 | 583.2210863 | 628.4279855 | 1.97E-08 | 1.082705678 | 2.40E-10 | 40.10933048 | 2.118004534 | 7.704435693 |
| *Stra6l* | 81.41535067 | 71.97671501 | 74.17600288 | 71.2630827 | 164.7444322 | 175.9742095 | 146.2623414 | 144.2863701 | 9.09E-12 | 1.08149198 | 5.96E-14 | 56.38339785 | 2.116223468 | 11.04139155 |
| *Tnfsf8* | 100.2942726 | 104.1091771 | 47.92910956 | 94.6920414 | 166.5064047 | 197.2548116 | 195.6258816 | 170.3622201 | 0.000149103 | 1.07375041 | 7.14E-06 | 20.15416873 | 2.104898125 | 3.826514058 |
| *LOC100038947* | 158.1109709 | 116.9621619 | 123.2462817 | 100.5492811 | 255.4860178 | 234.9051076 | 287.039845 | 269.4504502 | 2.48E-09 | 1.072479799 | 2.49E-11 | 44.53775288 | 2.103045114 | 8.606356348 |
| *Oxld1* | 27.13845022 | 24.42067117 | 10.27052348 | 31.23861159 | 44.04931342 | 48.29059703 | 56.67665729 | 46.93653003 | 0.004513226 | 1.06666661 | 0.000494249 | 12.137239 | 2.094588163 | 2.345512945 |
| *Zfp174* | 50.73710259 | 30.84716358 | 28.52923188 | 36.11964466 | 81.05073669 | 63.84180624 | 62.16149509 | 96.48064507 | 0.000883828 | 1.052938952 | 6.21E-05 | 16.03826572 | 2.074752078 | 3.053632116 |
| *Gpx3* | 204.128343 | 191.5094739 | 122.1051124 | 219.6464878 | 320.6790017 | 365.0441741 | 436.9587449 | 405.0448703 | 1.07E-05 | 1.049661254 | 3.16E-07 | 26.14866473 | 2.070043743 | 4.970235642 |
| *Arhgef18* | 110.9136661 | 137.5269376 | 89.01120346 | 124.9544464 | 271.3437707 | 229.1757147 | 207.5096968 | 247.7205752 | 2.62E-08 | 1.048729465 | 3.27E-10 | 39.50460478 | 2.068707202 | 7.581251414 |
| *Ldhb* | 56.63676568 | 77.11790894 | 27.3880626 | 62.47722319 | 103.0753934 | 122.7727043 | 105.1260579 | 127.7716651 | 0.002866338 | 1.039591515 | 0.000276572 | 13.22274813 | 2.055645534 | 2.542672649 |
| *Atp1a3* | 1992.906192 | 1586.058327 | 1055.58158 | 1979.74701 | 3594.423975 | 3197.00122 | 3328.382406 | 3402.898427 | 2.02E-05 | 1.031554483 | 6.71E-07 | 24.69654607 | 2.044225689 | 4.694352826 |
| *Cyp4f18* | 129.792588 | 98.96798314 | 65.04664868 | 89.81100834 | 189.4120477 | 173.5187554 | 204.7672779 | 212.0835802 | 2.83E-05 | 1.025428583 | 9.75E-07 | 23.97701358 | 2.035564001 | 4.548560282 |
| *Fam84b* | 341.0005267 | 209.5036526 | 144.9284979 | 249.9088928 | 481.0185025 | 492.7277866 | 485.4081455 | 459.8041553 | 0.000478859 | 1.021198883 | 2.91E-05 | 17.47742398 | 2.029604865 | 3.319792609 |
| *Fam117a* | 62.53642878 | 51.4119393 | 20.54104695 | 50.76274384 | 94.26553071 | 106.4030104 | 82.27256703 | 89.52708506 | 0.007100568 | 1.009224905 | 0.000908232 | 11.005882 | 2.012829408 | 2.148706891 |
| *Dgkg* | 108.5538009 | 133.6710422 | 58.19963303 | 139.5975456 | 242.2712238 | 208.7135973 | 235.8480255 | 196.4380701 | 0.001768785 | 1.004375551 | 0.000149415 | 14.37956542 | 2.00607501 | 2.752324982 |
| *Kdr* | 982.8838711 | 718.4818517 | 449.6206944 | 798.5370089 | 1347.028004 | 1470.816998 | 1614.370593 | 1482.846671 | 0.000234969 | 1.00385104 | 1.24E-05 | 19.10232002 | 2.005345806 | 3.62898998 |
| *Cd97* | 1417.099075 | 1273.730796 | 782.8421227 | 1590.240572 | 2464.999579 | 2516.840438 | 2584.272744 | 2569.340422 | 5.95E-05 | 1.000837403 | 2.38E-06 | 22.25771203 | 2.001161224 | 4.225587894 |
| *Napsa* | 706.7796384 | 665.7846139 | 641.3371326 | 697.9877278 | 1336.456169 | 1384.05762 | 1336.472144 | 1362.897761 | 9.36E-64 | 0.998289994 | 2.99E-67 | 300.1905361 | 1.997630832 | 63.02885522 |
| *Fry* | 420.0560122 | 322.6099191 | 175.7400684 | 402.1971243 | 684.5263305 | 627.7777614 | 627.0997887 | 695.3560005 | 0.001664398 | 0.995985408 | 0.000137801 | 14.53196951 | 1.99444233 | 2.778742855 |
| *Tst* | 20.05885451 | 10.28238786 | 15.97636985 | 20.50033886 | 32.59649193 | 36.0133266 | 36.56558535 | 28.68343502 | 0.00479682 | 0.984720985 | 0.00053605 | 11.98588703 | 1.978930553 | 2.319046595 |
| *Tlr8* | 946.30596 | 990.9651299 | 758.877568 | 865.8952651 | 1660.659116 | 1800.66633 | 1815.481312 | 1689.715081 | 2.60E-19 | 0.968672129 | 8.32E-22 | 92.08169139 | 1.957038488 | 18.58514178 |
| *Kif5c* | 71.97588972 | 68.12081957 | 29.67040115 | 80.04894221 | 127.7430089 | 126.8651278 | 126.1512694 | 107.7801801 | 0.005300504 | 0.965519301 | 0.000611149 | 11.74170952 | 1.952766297 | 2.275682872 |
| *Tbc1d2* | 319.7617396 | 262.2008904 | 123.2462817 | 284.0761242 | 471.3276536 | 485.3614244 | 481.7515869 | 492.8335654 | 0.003679136 | 0.965003477 | 0.000381422 | 12.62107329 | 1.952068226 | 2.434254203 |
| *Mllt3* | 238.3463889 | 177.3711906 | 100.4228962 | 173.764777 | 347.1085897 | 353.5853884 | 332.7468267 | 309.4334202 | 0.001003078 | 0.96154442 | 7.25E-05 | 15.74349809 | 1.947393484 | 2.99866516 |

| | | | | | | | | | | | | | | |
|---|---|---|---|---|---|---|---|---|---|---|---|---|---|---|
| *Igip* | 44.8374395 | 39.84425295 | 19.39987768 | 30.26240498 | 53.74016237 | 72.84513789 | 73.13117069 | 57.36687004 | 0.005773595 | 0.943649479 | 0.000684925 | 11.52971668 | 1.923387542 | 2.238553684 |
| *Mcoln3* | 218.2875344 | 204.3624587 | 119.8227739 | 177.6696034 | 330.3698506 | 366.6811435 | 347.3730608 | 337.2476602 | 1.36E-05 | 0.941871135 | 4.19E-07 | 25.60585523 | 1.92101813 | 4.866543771 |
| *Fabp5* | 2031.843969 | 2462.631892 | 2370.208584 | 2158.39282 | 4434.884875 | 4172.634977 | 4422.607548 | 4292.084913 | 3.62E-30 | 0.941323946 | 5.79E-33 | 143.0289374 | 1.92028966 | 29.44118134 |
| *Zfp467* | 141.5919142 | 89.97089377 | 57.05846376 | 124.9544464 | 188.5310614 | 190.706934 | 214.8228139 | 196.4380701 | 0.004441684 | 0.932546697 | 0.000485704 | 12.16976722 | 1.908642228 | 2.352452371 |
| *Arhgap39* | 358.699516 | 294.3333525 | 139.2226516 | 341.6723143 | 588.4988273 | 545.1108071 | 520.1454516 | 510.2174654 | 0.006776834 | 0.932052118 | 0.000848934 | 11.13107494 | 1.907988027 | 2.168973181 |
| *Tcp11l2* | 292.6232894 | 264.7714874 | 140.3638208 | 259.6709589 | 419.3494637 | 441.1632508 | 479.009168 | 473.7112753 | 0.000591281 | 0.921567842 | 3.76E-05 | 16.98926363 | 1.894172658 | 3.228205903 |
| *Adam33* | 36.57791117 | 17.99417875 | 23.96455478 | 30.26240498 | 55.50213491 | 37.65029599 | 53.93423839 | 59.97445504 | 0.007810543 | 0.918714213 | 0.001029035 | 10.77458141 | 1.890429717 | 2.10731876 |
| *Mid1* | 116.8133292 | 92.54149073 | 110.6934197 | 134.7165125 | 223.7705122 | 196.4363269 | 214.8228139 | 228.5982852 | 4.88E-09 | 0.916611454 | 5.31E-11 | 43.06107509 | 1.887676382 | 8.311848029 |
| *Gprc5b* | 66.07622663 | 47.55604385 | 73.03483361 | 53.69136368 | 109.2422973 | 114.5878574 | 114.2674542 | 115.6029351 | 1.12E-06 | 0.91572052 | 2.28E-08 | 31.23891443 | 1.886511009 | 5.951381497 |
| *Zfp608* | 201.7684777 | 134.9563407 | 74.17600288 | 166.9313307 | 275.748702 | 267.6444954 | 275.1560297 | 272.0580352 | 0.007417403 | 0.915716712 | 0.000964185 | 10.89510871 | 1.88650603 | 2.129748103 |
| *Xk* | 63.71636139 | 47.55604385 | 27.3880626 | 52.71515707 | 86.3366543 | 74.48210728 | 102.383639 | 96.48064507 | 0.004230176 | 0.908698172 | 0.000454454 | 12.29384807 | 1.877350691 | 2.373641545 |
| *Bdh1* | 75.51568758 | 69.40611805 | 79.88184926 | 55.6437769 | 147.1247068 | 118.6802808 | 127.9795487 | 129.5100551 | 1.56E-06 | 0.905615181 | 3.34E-08 | 30.50027278 | 1.873343138 | 5.807678251 |
| *Trim2* | 116.8133292 | 77.11790894 | 54.77612521 | 122.0258265 | 172.6733086 | 178.4296636 | 180.0855078 | 164.2778551 | 0.003431934 | 0.90354663 | 0.000347284 | 12.79641659 | 1.870659045 | 2.464461103 |
| *Fam120c* | 215.9276692 | 109.250371 | 104.9875733 | 168.8837439 | 293.3684274 | 277.4663117 | 298.9236602 | 251.1973552 | 0.002824285 | 0.902192671 | 0.000271787 | 13.25545783 | 1.868904268 | 2.549091557 |
| *Havcr2* | 895.5688574 | 766.0378955 | 398.268077 | 803.418042 | 1363.766743 | 1347.225809 | 1331.901446 | 1283.801016 | 0.00212098 | 0.895562818 | 0.000187649 | 13.95088471 | 1.860335489 | 2.673463367 |
| *Tbc1d32* | 208.8480735 | 177.3711906 | 117.5404353 | 188.4078762 | 319.7980154 | 301.2023679 | 359.256876 | 303.3490552 | 3.74E-05 | 0.890481371 | 1.38E-06 | 23.30851376 | 1.853794559 | 4.42691531 |
| *Lrrc27* | 280.8239632 | 179.9417875 | 114.1169275 | 243.0754465 | 372.6571915 | 391.2356844 | 334.5751059 | 418.0827953 | 0.007260187 | 0.889938424 | 0.00093678 | 10.94852884 | 1.85309703 | 2.13905217 |
| *Zfp473* | 34.21804593 | 28.27656661 | 27.3880626 | 55.6437769 | 75.76481908 | 54.0199899 | 72.21703106 | 69.53560005 | 0.009964407 | 0.883956789 | 0.001402876 | 10.20201045 | 1.84542971 | 2.001548537 |
| *P2ry14* | 198.2286799 | 182.5123845 | 150.6343443 | 147.4071985 | 327.7268918 | 339.6711486 | 273.3277505 | 301.6106652 | 4.78E-08 | 0.876652966 | 6.27E-10 | 38.23502231 | 1.836110603 | 7.320516452 |
| *Ide* | 1393.500422 | 1026.953487 | 740.6188596 | 1152.900009 | 1937.288804 | 1982.369932 | 2009.278915 | 1973.072651 | 8.06E-05 | 0.873131753 | 3.44E-06 | 21.55308181 | 1.831634638 | 4.093824521 |
| *Bambi* | 76.6956202 | 61.69432715 | 37.65858608 | 89.81100834 | 144.481748 | 108.0399798 | 123.4088505 | 111.2569601 | 0.008485787 | 0.868686394 | 0.001147865 | 10.57243322 | 1.82599953 | 2.071307894 |
| *Cd300lf* | 787.0150565 | 535.9694672 | 362.8918295 | 647.224984 | 1094.184945 | 1099.224946 | 1040.290903 | 1022.173321 | 0.00159465 | 0.867018361 | 0.000128965 | 14.65685152 | 1.823889543 | 2.797334567 |
| *Smpdl3a* | 3602.334284 | 3422.749859 | 2647.512718 | 3273.220771 | 6000.397474 | 5955.294644 | 5908.998592 | 5697.573229 | 1.72E-13 | 0.864101446 | 9.91E-16 | 64.44888857 | 1.820205636 | 12.76444899 |
| *Arhgap24* | 388.1978315 | 299.4745464 | 245.3513942 | 350.4581738 | 574.403047 | 570.4838327 | 576.8221088 | 613.6516704 | 4.20E-07 | 0.862714828 | 7.34E-09 | 33.44336942 | 1.818457021 | 6.377242023 |
| *Fcgr1* | 2814.139295 | 1935.659514 | 1913.740874 | 2279.44244 | 4012.892452 | 4141.532559 | 4070.663789 | 4027.849633 | 4.86E-08 | 0.861769315 | 6.42E-10 | 38.18922997 | 1.817265631 | 7.312964761 |
| *Plag1* | 37.75784379 | 26.99126813 | 21.68221623 | 38.07205788 | 48.45424476 | 59.74938276 | 60.33321582 | 58.23606504 | 0.002749885 | 0.856451372 | 0.000262868 | 13.31801737 | 1.810579308 | 2.560685469 |
| *Gramd4* | 192.3290168 | 142.6681315 | 117.5404353 | 239.17062 | 319.7980154 | 320.0275159 | 306.2367773 | 309.4334202 | 0.001403219 | 0.856438101 | 0.000109796 | 14.96032658 | 1.810562653 | 2.852874648 |
| *Pbx1* | 61.35649616 | 59.12373019 | 42.22326318 | 46.85791739 | 93.38454445 | 95.76270936 | 100.5553597 | 86.91950006 | 2.93E-05 | 0.853607315 | 1.02E-06 | 23.88964132 | 1.807013538 | 4.532862408 |
| *Scarb2* | 3109.122449 | 2425.358236 | 1352.285591 | 2840.761242 | 4314.189756 | 4376.437666 | 4421.693408 | 4431.156113 | 0.004704446 | 0.850744457 | 0.000523845 | 12.02880974 | 1.803431288 | 2.327491534 |
| *Fam188b* | 36.57791117 | 29.5618651 | 29.67040115 | 25.38137192 | 48.45424476 | 48.29059703 | 62.16149509 | 58.23606504 | 0.00176115 | 0.84627019 | 0.000148629 | 14.38949526 | 1.797846925 | 2.754203749 |
| *Klf2* | 115.6333966 | 80.97380439 | 130.0932974 | 88.83480172 | 164.7444322 | 184.9775412 | 189.2269042 | 205.9992151 | 6.24E-05 | 0.8420742 | 2.52E-06 | 22.15049274 | 1.792625595 | 4.204788836 |
| *Slc29a3* | 1985.826597 | 1499.943329 | 1053.299241 | 1817.696712 | 2823.56099 | 2741.92373 | 2826.519747 | 2950.917027 | 0.00037738 | 0.835286017 | 2.19E-05 | 18.02030874 | 1.784210727 | 3.423221454 |
| *Aatk* | 225.3671301 | 205.6477572 | 148.3520058 | 219.6464878 | 362.9663426 | 327.3938782 | 369.312412 | 365.9310953 | 6.89E-07 | 0.834310878 | 1.29E-08 | 32.33921808 | 1.783005159 | 6.161941811 |
| *Pdk2* | 132.1524533 | 122.1033558 | 67.32898723 | 165.9551241 | 228.1754435 | 202.9842045 | 225.7924895 | 213.8219702 | 0.007596928 | 0.833963187 | 0.000997853 | 10.83154636 | 1.782575506 | 2.119361978 |
| *Gng4* | 21.23878713 | 51.4119393 | 43.36443246 | 41.97688433 | 67.83594266 | 70.3896838 | 65.81805362 | 75.61996505 | 0.006659466 | 0.825880155 | 0.000831035 | 11.1706059 | 1.772616141 | 2.176560617 |
| *Cetn4* | 35.39797855 | 41.12955144 | 22.8233855 | 31.23861159 | 45.81128595 | 61.38635215 | 58.50493656 | 64.32043005 | 0.004732397 | 0.825292772 | 0.000528472 | 12.01242043 | 1.77189458 | 2.324918838 |
| *Gm5431* | 363.4192465 | 277.6244722 | 349.1977982 | 348.5057606 | 572.6410744 | 601.5862511 | 624.3573698 | 571.0611154 | 2.49E-13 | 0.821053087 | 1.47E-15 | 63.66783057 | 1.766695111 | 12.60410096 |
| *Kif9* | 33.03811332 | 24.42067117 | 17.11753913 | 29.28619837 | 49.33523103 | 49.10908172 | 38.39386461 | 46.93653003 | 0.008230156 | 0.821003173 | 0.001106703 | 10.6399564 | 1.766633989 | 2.084591939 |
| *9930111J21Rik1* | 715.0391668 | 479.4163339 | 524.9378666 | 713.6070336 | 1084.494096 | 1040.294048 | 1105.194817 | 1069.109851 | 4.17E-06 | 0.819909685 | 1.05E-07 | 28.2865102 | 1.765295478 | 5.380359601 |
| *Zfp442* | 49.55716997 | 37.27365599 | 30.81157043 | 47.834124 | 70.47890147 | 72.02665319 | 74.95944996 | 74.75077005 | 0.00045737 | 0.813598129 | 2.75E-05 | 17.58233766 | 1.757589467 | 3.339732158 |
| *Adssl1* | 1230.669721 | 936.9825937 | 562.5964526 | 1206.591373 | 1686.207718 | 1773.656335 | 1661.905854 | 1794.018481 | 0.006389137 | 0.812641754 | 0.000783501 | 11.27991576 | 1.756424732 | 2.194557779 |
| *Ly6e* | 17002.82903 | 12283.5976 | 11954.88933 | 15353.7776 | 24459.70275 | 24296.71818 | 25226.59733 | 25312.69681 | 7.53E-08 | 0.8110149 | 1.08E-09 | 37.17698364 | 1.754445217 | 7.122969013 |
| *Adam10* | 5702.614345 | 4701.621849 | 2980.734147 | 5392.565327 | 8246.912458 | 8143.922719 | 8421.968445 | 8093.943846 | 0.000861504 | 0.80934228 | 6.02E-05 | 16.09535562 | 1.752412342 | 3.064742498 |
| *Runx2* | 263.1249739 | 186.3682799 | 179.1635762 | 311.4099093 | 403.4917109 | 402.6944701 | 405.8779974 | 433.7283053 | 0.000586018 | 0.804715441 | 3.71E-05 | 17.01287838 | 1.746801216 | 3.232088903 |
| *Golm1* | 1133.915246 | 880.4294604 | 502.1144811 | 1007.445224 | 1474.771013 | 1473.272452 | 1567.749472 | 1588.888461 | 0.007134449 | 0.792634697 | 0.000913728 | 10.99469933 | 1.73223504 | 2.146639555 |
| *Fbxo31* | 663.1221315 | 535.9694672 | 342.3507825 | 641.3677443 | 940.0123483 | 946.9867926 | 875.7457691 | 1011.742981 | 0.001981941 | 0.78976407 | 0.000172454 | 14.10968451 | 1.728791723 | 2.702909205 |
| *Acot2* | 108.5538009 | 70.69141653 | 46.78794028 | 82.00135544 | 132.1479403 | 120.3172502 | 149.0047603 | 131.2484451 | 0.009521341 | 0.78925511 | 0.001328311 | 10.30276378 | 1.72818194 | 2.021301892 |
| *Hpse* | 1236.569384 | 929.2708028 | 553.4670984 | 1153.876216 | 1636.872487 | 1583.767886 | 1684.759345 | 1787.934116 | 0.008475406 | 0.789017272 | 0.001144427 | 10.57797858 | 1.727897061 | 2.071839487 |
| *Kdm6a* | 653.6826706 | 516.6899899 | 425.6561396 | 662.8442898 | 971.727854 | 935.5280068 | 1036.634345 | 960.4604757 | 1.80E-05 | 0.788480178 | 5.85E-07 | 24.96032807 | 1.72725391 | 4.745329108 |
| *Plekhg5* | 46.01737212 | 35.98835751 | 33.09390898 | 50.76274384 | 73.12186027 | 71.2081685 | 82.27256703 | 61.71284504 | 0.001800106 | 0.787365772 | 0.000152637 | 14.33939856 | 1.725920212 | 2.744701947 |
| *Tiam1* | 431.8553383 | 307.1863373 | 208.8339774 | 431.4833227 | 592.0227723 | 580.305649 | 598.7614601 | 610.1748904 | 0.006056097 | 0.786763484 | 0.000729096 | 11.41357052 | 1.725199836 | 2.217807186 |
| *2610301B20Rik* | 208.8480735 | 156.8064149 | 102.7052348 | 192.3127026 | 324.2029468 | 269.2814648 | 263.2722145 | 279.0115952 | 0.004536324 | 0.780554772 | 0.000498956 | 12.11956378 | 1.717791303 | 2.343295917 |
| *Tifab* | 422.4158774 | 534.6841687 | 588.843346 | 499.8177855 | 866.0095018 | 840.5837822 | 889.4578636 | 911.7855557 | 5.62E-10 | 0.778047051 | 5.26E-12 | 47.58902521 | 1.714807997 | 9.250569905 |
| *Tnfrsf26* | 156.9310382 | 170.9446982 | 188.2929304 | 167.9075373 | 273.9867295 | 311.0241842 | 299.8377998 | 287.7035452 | 1.36E-11 | 0.777841957 | 9.24E-14 | 55.5226443 | 1.714564238 | 10.86685576 |
| *Cd24a* | 298.5229525 | 290.477457 | 453.0442022 | 299.69543 | 584.9748822 | 632.6886695 | 539.3423839 | 540.6392904 | 0.000137545 | 0.776028379 | 6.48E-06 | 20.34073443 | 1.712410254 | 3.861554418 |
| *Fkbp1b* | 103.8340704 | 87.4002968 | 84.44652636 | 96.64445462 | 181.4831713 | 141.5978523 | 166.3734133 | 148.6323451 | 5.95E-06 | 0.774322988 | 1.57E-07 | 27.50120076 | 1.710387231 | 5.225677821 |
| *Slfn9* | 1404.119816 | 1106.641993 | 717.7954741 | 1329.593406 | 1801.616919 | 2002.013565 | 1980.940586 | 2004.363671 | 0.002537859 | 0.772813836 | 0.000235292 | 13.5259161 | 1.708598991 | 2.595532432 |
| *Arsb* | 2364.584967 | 1844.403322 | 1251.862695 | 2216.965217 | 3323.96119 | 3214.189399 | 3321.069289 | 3254.266082 | 0.001243375 | 0.772166391 | 9.56E-05 | 15.22200681 | 1.707832387 | 2.905397766 |
| *Sash3* | 810.6137088 | 632.3668533 | 383.4328764 | 768.2746039 | 1054.540563 | 1088.584645 | 1156.386637 | 1109.962016 | 0.006608462 | 0.764758484 | 0.000820441 | 11.19441034 | 1.699085537 | 2.179899578 |
| *Tnrc6b* | 1218.870395 | 888.1412513 | 625.3607628 | 1191.948274 | 1839.499328 | 1626.32909 | 1493.704161 | 1692.322666 | 0.005199095 | 0.761013055 | 0.000594313 | 11.79370624 | 1.694680205 | 2.284072249 |
| *Arsg* | 60.17656354 | 50.12664081 | 50.21144811 | 47.834124 | 91.62257191 | 94.12573997 | 77.70186886 | 86.91950006 | 0.000170337 | 0.754602348 | 8.37E-06 | 19.85227548 | 1.687166492 | 3.768689744 |
| *Ctss* | 31035.76766 | 29374.21152 | 26440.89211 | 30188.21328 | 49462.97404 | 49821.16341 | 49427.52999 | 48668.83567 | 4.79E-30 | 0.753982333 | 8.05E-33 | 142.3749734 | 1.686441569 | 29.31938253 |
| *H2-DMa* | 220.6473996 | 201.7918617 | 178.0224069 | 209.8844217 | 340.0606996 | 329.0308475 | 331.832687 | 362.4543153 | 1.41E-11 | 0.749949279 | 9.72E-14 | 55.42331082 | 1.681733705 | 10.84998936 |
| *L1cam* | 1072.55875 | 915.1325195 | 523.7966973 | 1033.802802 | 1517.058354 | 1497.826993 | 1447.08304 | 1499.361376 | 0.005958332 | 0.749609668 | 0.000710201 | 11.46235827 | 1.681337871 | 2.224875299 |
| *Pvrl4* | 335.1008636 | 244.2067117 | 167.7518834 | 319.2195622 | 461.6368046 | 408.423863 | 462.5546546 | 461.5425453 | 0.005486955 | 0.749603093 | 0.000639508 | 11.65730763 | 1.681330208 | 2.260668633 |
| *Card6* | 139.232049 | 114.3915649 | 109.5522504 | 151.3120249 | 214.9606495 | 237.3605617 | 212.080395 | 201.6532401 | 4.97E-06 | 0.746396813 | 1.27E-07 | 27.9072868 | 1.677597728 | 5.303480637 |
| *Dennd1b* | 779.9354608 | 611.8020776 | 459.8912179 | 747.7742651 | 1104.756781 | 1117.231609 | 1108.851376 | 1027.388491 | 0.000325979 | 0.745078024 | 1.83E-05 | 18.35793084 | 1.676064912 | 3.486809788 |
| *Vamp1* | 109.7337335 | 98.96798314 | 68.47015651 | 116.1685869 | 180.602185 | 168.6078472 | 157.232017 | 153.8475151 | 0.000503989 | 0.744667754 | 3.09E-05 | 17.36027057 | 1.675588345 | 3.297579289 |
| *Lrrc1* | 66.07622663 | 39.84425295 | 39.94092463 | 66.38204964 | 80.16975042 | 88.3963471 | 95.98466154 | 93.00386507 | 0.006132882 | 0.743412384 | 0.000739812 | 11.38646531 | 1.674130956 | 2.212335384 |
| *Zfp273* | 40.11770903 | 41.12955144 | 35.37624753 | 46.85791739 | 52.8591761 | 81.02998484 | 63.07563472 | 76.48916006 | 0.006372753 | 0.740096944 | 0.000779963 | 11.28831944 | 1.670288072 | 2.195672943 |
| *Bivm* | 204.128343 | 192.7947724 | 114.1169275 | 206.9558018 | 301.2973038 | 315.1166077 | 269.6711919 | 312.9102002 | 0.002188925 | 0.73950005 | 0.000195399 | 13.87481897 | 1.669597158 | 2.659769103 |
| *Zranb3* | 516.8104869 | 516.6899899 | 332.0802591 | 522.2705376 | 804.340463 | 793.1116698 | 776.104549 | 768.3683806 | 8.00E-05 | 0.734897432 | 3.39E-06 | 21.58133185 | 1.664279144 | 4.096956241 |
| *Slfn3* | 358.699516 | 290.477457 | 279.5864724 | 328.0054217 | 448.4220106 | 524.6486897 | 574.0796899 | 538.9009004 | 4.49E-07 | 0.730044136 | 7.93E-09 | 33.29137493 | 1.658689834 | 6.347989675 |
| *Pira7* | 116.8133292 | 138.8122361 | 178.0224069 | 137.6451323 | 168.2683773 | 287.2881281 | 212.9945346 | 273.7964252 | 0.00900569 | 0.722413956 | 0.001234066 | 10.43863186 | 1.649940451 | 2.045483017 |
| *Cnnm2* | 248.9657825 | 280.1950692 | 179.1635762 | 283.0999176 | 403.4917109 | 412.5162865 | 383.9386461 | 427.6439403 | 0.000109903 | 0.714968161 | 4.99E-06 | 20.83973584 | 1.641446992 | 3.958988841 |
| *Cd300a* | 1928.009898 | 1426.681315 | 1103.510689 | 1640.027109 | 2477.333387 | 2539.75801 | 2378.591327 | 2592.808687 | 0.000592196 | 0.71179718 | 3.77E-05 | 16.98340161 | 1.637843125 | 3.227534433 |
| *Siglec1* | 5054.831337 | 3646.391795 | 3530.777737 | 4264.070483 | 6662.899148 | 6255.678527 | 7144.915377 | 6943.129665 | 4.97E-06 | 0.711103254 | 1.27E-07 | 27.91353909 | 1.637055523 | 5.303524668 |
| *Dock9* | 57.8166983 | 35.98835751 | 38.79975535 | 47.834124 | 81.93172296 | 62.20483685 | 71.30289143 | 80.83513506 | 0.006249706 | 0.710009698 | 0.000760903 | 11.33425563 | 1.635815113 | 2.204140442 |
| *1600014C10Rik* | 530.9696783 | 589.9520034 | 699.5367657 | 461.7457276 | 864.2475293 | 952.7161854 | 923.28103 | 977.8443757 | 1.58E-05 | 0.705302489 | 5.01E-07 | 25.26186252 | 1.630486492 | 4.800241411 |
| *Trpv4* | 60.17656354 | 57.83843171 | 41.08209391 | 67.35825625 | 96.02750325 | 99.85513284 | 77.70186886 | 95.61145007 | 0.002983145 | 0.702218569 | 0.000291008 | 13.12739974 | 1.62700487 | 2.525325685 |
| *4632428N05Rik* | 2936.852287 | 2814.803676 | 2033.563648 | 3036.978771 | 4335.333427 | 4406.7216 | 4382.385404 | 4482.438618 | 1.02E-05 | 0.702064923 | 2.97E-07 | 26.26807243 | 1.626831604 | 4.992356417 |
| *Epas1* | 93.21467685 | 104.1091771 | 55.91729448 | 109.3351406 | 154.172597 | 139.1423982 | 142.6057829 | 153.8475151 | 0.006461142 | 0.700537278 | 0.000794915 | 11.25306755 | 1.625109891 | 2.189690688 |
| *Zfp958* | 112.0935987 | 134.9563407 | 123.2462817 | 144.4785786 | 195.5789516 | 203.8026892 | 226.7066292 | 210.3451902 | 1.01E-06 | 0.696476451 | 2.02E-08 | 31.47899041 | 1.620542044 | 5.997099746 |
| *Tmem119* | 55.45683307 | 65.5502226 | 81.02301854 | 54.66757029 | 90.74158564 | 114.5878574 | 99.64122007 | 109.5185701 | 0.001729619 | 0.695974879 | 0.000145415 | 14.43066893 | 1.619978739 | 2.762049447 |
| *Tob1* | 233.6266584 | 208.2183541 | 163.1872063 | 178.6458101 | 309.2261802 | 304.4763067 | 325.4337096 | 326.8173202 | 3.33E-06 | 0.694938541 | 8.15E-08 | 28.76914004 | 1.61881547 | 5.477405565 |
| *Gpr141* | 307.9624134 | 300.7598449 | 368.5976759 | 230.3847605 | 491.5903377 | 465.7177917 | 485.4081455 | 508.4790754 | 6.87E-05 | 0.694205336 | 2.83E-06 | 21.92864661 | 1.617992966 | 4.162755667 |
| *Ctla2b* | 1156.333966 | 1116.924381 | 722.3601512 | 1179.257588 | 1676.516869 | 1762.197549 | 1680.188647 | 1614.964311 | 0.000563987 | 0.689628155 | 3.53E-05 | 17.1073505 | 1.612867759 | 3.248730879 |
| *BC051226* | 44.8374395 | 42.41484992 | 46.78794028 | 45.88171078 | 70.47890147 | 80.21150015 | 77.70186886 | 61.71284504 | 0.001992909 | 0.687427175 | 0.000174175 | 14.09100479 | 1.610409039 | 2.700512559 |
| *Vps13c* | 2040.103497 | 1551.355268 | 1045.311056 | 1850.887737 | 2678.198256 | 2416.166821 | 2726.878527 | 2624.968902 | 0.006799151 | 0.687082327 | 0.000854449 | 11.11906395 | 1.610024148 | 2.167545294 |
| *F11r* | 960.4651514 | 903.5648331 | 801.1008312 | 872.7287114 | 1429.840714 | 1438.07761 | 1355.669077 | 1468.070356 | 1.08E-17 | 0.686720768 | 3.90E-20 | 84.47279986 | 1.609620705 | 16.96571885 |
| *Ctsd* | 117372.6173 | 107575.6271 | 96313.54565 | 114188.8399 | 173215.9961 | 173900.9878 | 175721.4052 | 176725.5967 | 2.96E-17 | 0.683942624 | 1.14E-19 | 82.35705384 | 1.606524101 | 16.52893591 |
| *Mgat5* | 461.3536538 | 380.4483508 | 314.9627199 | 487.1270996 | 692.4552069 | 594.2198889 | 682.8623064 | 664.0649805 | 0.000240327 | 0.678385954 | 1.28E-05 | 19.03956404 | 1.600348329 | 3.619197176 |
| *Apoc2* | 143.9517794 | 140.0975346 | 106.1287426 | 145.4547852 | 240.5092513 | 198.891781 | 195.6258816 | 222.5139202 | 8.15E-05 | 0.678161851 | 3.51E-06 | 21.51767491 | 1.600099756 | 4.088700223 |
| *Got1* | 1689.66351 | 1461.384374 | 1192.521893 | 1605.859877 | 2331.970652 | 2448.087724 | 2333.798485 | 2390.286252 | 8.02E-07 | 0.675649939 | 1.54E-08 | 32.0065855 | 1.597316207 | 6.09594819 |
| *Fam117b* | 650.1428727 | 547.5371535 | 522.655528 | 566.1998352 | 872.1764057 | 961.7195171 | 865.6902331 | 945.6841607 | 9.17E-11 | 0.673211582 | 7.04E-13 | 51.53275363 | 1.594618799 | 10.03767657 |
| *Nfam1* | 2914.433567 | 2312.25197 | 1686.648189 | 2894.452606 | 3782.955036 | 3703.643247 | 4046.896158 | 4005.250563 | 0.00309568 | 0.663719405 | 0.000304095 | 13.04499952 | 1.584161483 | 2.509243993 |
| *Zc3h6* | 105.014003 | 120.8180573 | 89.01120346 | 103.4779009 | 167.387391 | 151.4196686 | 181.9137871 | 159.0626851 | 5.07E-05 | 0.659855986 | 1.95E-06 | 22.64299214 | 1.579924904 | 4.295008937 |
| *4931414P19Rik* | 120.3531271 | 84.82969984 | 93.57588056 | 96.64445462 | 155.9345695 | 165.3339085 | 144.4340621 | 159.0626851 | 7.73E-05 | 0.659756491 | 3.27E-06 | 21.64937162 | 1.579815948 | 4.111542062 |
| *Ogfrl1* | 863.7106767 | 812.3086409 | 680.136888 | 803.418042 | 1258.048391 | 1278.473094 | 1182.896686 | 1255.986776 | 3.44E-11 | 0.655488957 | 2.48E-13 | 53.58497546 | 1.575149711 | 10.46346627 |
| *Ipp* | 90.85481162 | 111.820968 | 81.02301854 | 122.0258265 | 140.9578029 | 153.8751227 | 159.0602963 | 185.1385351 | 0.002167034 | 0.65261362 | 0.000193111 | 13.89695947 | 1.572013515 | 2.664134289 |
| *Rasal2* | 169.9102971 | 102.8238786 | 110.6934197 | 137.6451323 | 184.1261301 | 217.716929 | 203.8531383 | 212.9527752 | 0.002690328 | 0.649725748 | 0.000255238 | 13.37325557 | 1.56886993 | 2.570194743 |
| *Gm13212* | 69.61602449 | 78.40320743 | 47.92910956 | 62.47722319 | 99.55144832 | 112.1324033 | 104.2119182 | 87.78869506 | 0.004458581 | 0.649602727 | 0.000487908 | 12.16132114 | 1.568736156 | 2.350803359 |
| *Cd84* | 3815.902088 | 3087.286955 | 2338.255845 | 3555.344482 | 5040.122441 | 5016.492698 | 5172.202047 | 4834.462594 | 0.000642523 | 0.648696304 | 4.20E-05 | 16.77916159 | 1.567750852 | 3.192111619 |
| *Tctn3* | 384.6580336 | 353.4570827 | 212.2574852 | 359.2440333 | 497.7572416 | 521.374751 | 499.12024 | 532.8165354 | 0.005163514 | 0.647339561 | 0.000589833 | 11.80779439 | 1.566277197 | 2.287054634 |
| *Mblac2* | 167.5504318 | 124.6739528 | 89.01120346 | 134.7165125 | 192.9359928 | 208.7135973 | 219.3935121 | 186.8769251 | 0.006707173 | 0.646833601 | 0.000838598 | 11.15379857 | 1.565727993 | 2.17346049 |
| *D10Bwg1379e* | 436.5750688 | 285.3362631 | 332.0802591 | 472.4840004 | 648.4058935 | 535.2889908 | 607.9028564 | 601.4829404 | 0.003158523 | 0.646419321 | 0.00031229 | 12.99520427 | 1.565278447 | 2.500515943 |
| *Armc9* | 94.39460947 | 74.54731198 | 58.19963303 | 87.85859511 | 130.3859677 | 118.6802808 | 109.696756 | 134.7252251 | 0.00343362 | 0.645306988 | 0.000348143 | 12.79179518 | 1.564072067 | 2.464247762 |
| *Zbtb38* | 455.4539907 | 437.001484 | 262.4689333 | 476.3888268 | 636.0720858 | 640.8735165 | 626.1856491 | 643.2043005 | 0.005998978 | 0.642071377 | 0.00071934 | 11.43859935 | 1.560568167 | 2.221922701 |
| *Sorl1* | 1559.870922 | 1281.442587 | 869.5709877 | 1496.524737 | 2121.414934 | 1909.524794 | 2001.965798 | 2092.152367 | 0.005531799 | 0.641639287 | 0.000646948 | 11.63579314 | 1.560100844 | 2.257133571 |
| *Axl* | 8386.961052 | 7169.394935 | 6510.370715 | 7914.107006 | 11696.85468 | 11719.88235 | 11764.06295 | 11572.46224 | 3.24E-10 | 0.640960994 | 2.83E-12 | 48.80577868 | 1.559367524 | 9.489300468 |
| *Kif13b* | 1198.81154 | 1123.350874 | 724.6424897 | 1201.71034 | 1659.77813 | 1549.391528 | 1711.269394 | 1702.753006 | 0.002405685 | 0.640430932 | 0.000219572 | 13.65571163 | 1.558794701 | 2.618761247 |
| *Ankrd44* | 1466.656245 | 1413.828331 | 845.6064329 | 1495.54853 | 2088.818442 | 1941.445697 | 2115.319112 | 1987.848966 | 0.006132882 | 0.639279131 | 0.000739716 | 11.38670599 | 1.557550706 | 2.212335384 |
| *Elmod3* | 160.4708361 | 118.2474604 | 103.846404 | 142.5261654 | 193.816979 | 170.2448166 | 230.3631877 | 224.2523102 | 0.004298753 | 0.638313266 | 0.000464572 | 12.25275392 | 1.556508296 | 2.36665755 |
| *Emp1* | 3681.389769 | 3330.208368 | 2015.30494 | 3739.847532 | 4895.640693 | 5011.58179 | 4973.833747 | 4990.048499 | 0.009426523 | 0.638233474 | 0.001306789 | 10.33291223 | 1.556422212 | 2.025648488 |
| *Cd200r4* | 408.256686 | 320.0393221 | 330.9390898 | 412.935397 | 635.1910995 | 572.9392868 | 533.8575461 | 550.2004354 | 9.37E-06 | 0.636658653 | 2.71E-07 | 26.44385139 | 1.554724175 | 5.02829688 |
| *Ctsh* | 2599.391558 | 2552.602786 | 2438.678741 | 2608.424068 | 3936.246647 | 3918.086237 | 3968.28015 | 4004.381368 | 1.91E-53 | 0.633646305 | 7.66E-57 | 252.4328664 | 1.551481296 | 52.71796233 |
| *Foxred2* | 306.7824808 | 249.3479056 | 225.9515165 | 306.5288763 | 443.136093 | 388.7802303 | 433.3021864 | 424.1671603 | 2.71E-05 | 0.631459208 | 9.29E-07 | 24.06922957 | 1.549131064 | 4.567340114 |
| *Frmd4a* | 1405.299749 | 1066.79774 | 818.2183703 | 1376.451323 | 1720.566182 | 1736.824524 | 1917.864951 | 1847.908571 | 0.005277756 | 0.629856126 | 0.000607104 | 11.75406662 | 1.547410669 | 2.277550692 |
| *Lyst* | 2199.394401 | 1916.380037 | 1579.378277 | 2202.322117 | 2968.923724 | 3074.228516 | 3199.488718 | 2933.533127 | 1.45E-05 | 0.624350646 | 4.50E-07 | 25.46705588 | 1.541516836 | 4.839790689 |
| *Fgfr1* | 95.57454209 | 78.40320743 | 85.58769564 | 98.59686785 | 136.5528716 | 118.6802808 | 151.7471792 | 146.8939551 | 0.000554852 | 0.622201997 | 3.45E-05 | 17.15297633 | 1.539222717 | 3.255822601 |
| *Prkd2* | 193.5089494 | 183.797683 | 136.940313 | 188.4078762 | 262.533908 | 233.2681382 | 309.8933358 | 276.4040102 | 0.00085296 | 0.621948535 | 5.94E-05 | 16.12074376 | 1.53895232 | 3.069071219 |
| *Hdgfrp3* | 107.3738683 | 133.6710422 | 83.30535709 | 107.3827274 | 141.8387892 | 195.6178422 | 169.1158322 | 154.7167101 | 0.006232153 | 0.619105415 | 0.000755333 | 11.34790167 | 1.535922492 | 2.205361887 |
| *Trim16* | 173.4500949 | 133.6710422 | 110.6934197 | 169.8599505 | 247.5571414 | 197.2548116 | 228.5349084 | 229.4674802 | 0.003713144 | 0.616362501 | 0.000386369 | 12.59698859 | 1.533005105 | 2.430258242 |
| *Phyh* | 486.1322388 | 375.3071569 | 318.3862278 | 510.5560583 | 628.1432093 | 660.5171492 | 680.1198875 | 617.1284504 | 0.002410414 | 0.611877832 | 0.000220205 | 13.6503058 | 1.528247102 | 2.617908319 |
| *Pard6a* | 112.0935987 | 88.68559529 | 63.90547941 | 90.78721495 | 141.8387892 | 126.8651278 | 131.6361072 | 142.5479801 | 0.00509416 | 0.611388157 | 0.00058028 | 11.83819554 | 1.527728478 | 2.292927404 |
| *Fut8* | 444.8345971 | 435.7161855 | 335.5037669 | 380.7205788 | 570.8791019 | 648.2398787 | 638.0694643 | 572.7995054 | 9.27E-06 | 0.607494422 | 2.66E-07 | 26.4793391 | 1.523610803 | 5.033034217 |
| *Lbh* | 257.2253108 | 251.9185025 | 213.3986545 | 210.8606283 | 351.5135211 | 329.8493322 | 383.9386461 | 353.7623653 | 1.69E-06 | 0.607472791 | 3.65E-08 | 30.32665641 | 1.523587958 | 5.772037934 |
| *Dennd2d* | 90.85481162 | 70.69141653 | 65.04664868 | 79.0727356 | 114.5282149 | 104.766041 | 123.4088505 | 123.4256901 | 0.001403219 | 0.606041383 | 0.000109891 | 14.95868114 | 1.52207704 | 2.852874648 |

| | | | | | | | | | | | | | |
|---|---|---|---|---|---|---|---|---|---|---|---|---|---|
| *Sat1* | 3562.216575 | 3219.672698 | 2304.020767 | 3544.606209 | 4784.636423 | 4951.013922 | 4762.667491 | 4651.062448 | 0.000910382 | 0.600325262 | 6.45E-05 | 15.96644879 | 1.51605833 | 3.040776231 |
| *Ndrg4* | 66.07622663 | 53.98253626 | 60.48197158 | 67.35825625 | 87.21764057 | 114.5878574 | 85.01498593 | 89.52708506 | 0.006368867 | 0.59973773 | 0.000778978 | 11.29066552 | 1.515441047 | 2.195937803 |
| *Pld4* | 7018.239214 | 5668.166307 | 4341.007923 | 6782.683543 | 8843.340162 | 8946.037721 | 8923.831104 | 9235.196882 | 0.002232146 | 0.5942979 | 0.000200341 | 13.82787877 | 1.50973768 | 2.651277367 |
| *Mtmr4* | 648.9629401 | 582.2402125 | 440.4913402 | 541.7946698 | 870.4144331 | 827.488027 | 865.6902331 | 776.1911356 | 0.000118538 | 0.594042652 | 5.49E-06 | 20.65816992 | 1.509470594 | 3.926140774 |
| *Acad12* | 81.41535067 | 75.83261046 | 61.62314086 | 102.5016943 | 130.3859677 | 121.9542196 | 122.4947109 | 112.1261551 | 0.006557117 | 0.592108164 | 0.000810919 | 11.21607244 | 1.507447924 | 2.183287077 |
| *Scai* | 240.7062542 | 217.2154435 | 165.4695449 | 248.9326861 | 333.0128094 | 350.3114496 | 338.2316645 | 294.6571052 | 0.001035835 | 0.592060255 | 7.64E-05 | 15.64542696 | 1.507397866 | 2.984709314 |
| *9330182L06Rik* | 100.2942726 | 78.40320743 | 66.18781796 | 87.85859511 | 123.3380776 | 118.6802808 | 132.5502469 | 127.7716651 | 0.001718694 | 0.591928387 | 0.000144084 | 14.44798665 | 1.50726009 | 2.764801384 |
| *Ift122* | 174.6300275 | 124.6739528 | 119.8227739 | 157.1692646 | 215.8416357 | 196.4363269 | 226.7066292 | 231.2058702 | 0.001402226 | 0.591293988 | 0.000109589 | 14.96387464 | 1.506597445 | 2.853181894 |
| *Osbpl7* | 206.4882082 | 145.2387285 | 136.940313 | 214.7654547 | 251.0810865 | 265.1890413 | 270.5853316 | 271.1888402 | 0.006044113 | 0.585172966 | 0.000726608 | 11.4199217 | 1.500218841 | 2.218667415 |
| *Gadd45a* | 645.4231423 | 754.4702092 | 919.7824358 | 742.893232 | 526.8297885 | 502.549603 | 480.8374473 | 529.3397554 | 5.86E-05 | -0.586521035 | 2.34E-06 | 22.29419292 | 0.66594686 | 4.232101597 |
| *Rapgef5* | 809.4337762 | 1050.08886 | 1003.087793 | 871.7525048 | 583.2129097 | 640.8735165 | 627.0997887 | 627.5587905 | 1.49E-06 | -0.590302147 | 3.16E-08 | 30.6044914 | 0.664203787 | 5.827754566 |
| *Psmd14* | 1379.341231 | 1760.858921 | 2167.080453 | 1366.689257 | 1064.231412 | 1153.244936 | 1108.851376 | 1091.708921 | 0.003703164 | -0.594857109 | 0.000384802 | 12.6045834 | 0.66211003 | 2.431427077 |
| *Jak3* | 496.7516323 | 557.8195414 | 804.524339 | 487.1270996 | 413.1825599 | 374.0475058 | 379.367948 | 384.1841903 | 0.007932489 | -0.597092827 | 0.001054619 | 10.72913145 | 0.661084765 | 2.100590522 |
| *Abhd17c* | 692.620447 | 794.3144621 | 991.6761001 | 645.2725708 | 510.9720356 | 519.7377816 | 518.3171723 | 513.6942454 | 0.000852582 | -0.598190177 | 5.92E-05 | 16.12809162 | 0.660582119 | 3.069263962 |
| *Slc16a10* | 2635.96947 | 2963.8983 | 3585.553863 | 2760.7123 | 1998.076857 | 1969.274177 | 1971.79919 | 1937.435656 | 2.60E-06 | -0.600833896 | 6.14E-08 | 29.31886559 | 0.659372719 | 5.584743908 |
| *Pde1b* | 722.1187625 | 922.8443104 | 1159.427984 | 846.3711329 | 608.7615114 | 612.2265521 | 574.0796899 | 606.6981104 | 0.001128123 | -0.603929073 | 8.45E-05 | 15.45558301 | 0.657959609 | 2.947643736 |
| *Ciapin1* | 463.713519 | 659.3581215 | 634.490117 | 574.0094881 | 408.7776285 | 383.8693221 | 338.2316645 | 402.4372853 | 0.000261628 | -0.603968515 | 1.40E-05 | 18.86326886 | 0.657941621 | 3.582315304 |
| *Zfand5* | 2235.972312 | 2654.141366 | 3521.648383 | 2036.366993 | 1665.064047 | 1842.409049 | 1684.759345 | 1680.153936 | 0.006945943 | -0.604183107 | 0.000883453 | 11.05716086 | 0.657843764 | 2.158268812 |
| *Txnl1* | 1434.798064 | 1799.417875 | 2090.622112 | 1544.358861 | 1147.925108 | 1086.129191 | 1147.24524 | 1127.345916 | 0.000104034 | -0.607285917 | 4.69E-06 | 20.96180802 | 0.656430457 | 3.982825276 |
| *C5ar1* | 3286.112342 | 4029.410742 | 4918.439576 | 3495.795879 | 2601.55245 | 2616.695571 | 2491.944641 | 2604.977417 | 0.00023917 | -0.608635646 | 1.27E-05 | 19.05164168 | 0.655816613 | 3.621293251 |
| *Cklf* | 201.7684777 | 249.3479056 | 321.8097356 | 213.7892481 | 163.8634459 | 161.241485 | 152.6613188 | 168.6238301 | 0.003879358 | -0.60916969 | 0.000406524 | 12.50196515 | 0.655573894 | 2.41124016 |
| *Eif1ax* | 984.0638037 | 1269.874901 | 1725.447944 | 1051.374521 | 793.7686278 | 841.4022669 | 837.3519044 | 810.0897406 | 0.008037711 | -0.615751186 | 0.001077167 | 10.68999121 | 0.652590013 | 2.094867594 |
| *Ets1* | 313.8620765 | 295.618651 | 417.6679547 | 272.3616448 | 222.0085396 | 194.7993575 | 216.6510932 | 213.8219702 | 0.001328586 | -0.616837843 | 0.000102877 | 15.08315067 | 0.652098658 | 2.876610321 |
| *Dot1l* | 1674.324386 | 1259.592513 | 1364.838453 | 1694.694679 | 939.1313621 | 982.1816345 | 991.8415025 | 996.9666657 | 9.34E-06 | -0.616876705 | 2.70E-07 | 26.4550034 | 0.652081093 | 5.02960281 |
| *Tubb2a* | 1289.666352 | 1458.813778 | 1493.790581 | 1333.498232 | 817.555257 | 1032.927686 | 855.6346971 | 919.6083107 | 1.23E-07 | -0.620201051 | 1.86E-09 | 36.11125241 | 0.650580258 | 6.9114385 |
| *Pmp22* | 4330.35271 | 4598.79797 | 5539.235662 | 4288.475648 | 3057.903337 | 3201.912128 | 2946.272039 | 2973.516097 | 9.56E-08 | -0.622839938 | 1.42E-09 | 36.64786552 | 0.649391345 | 7.019427923 |
| *Cct3* | 2749.243001 | 3343.061353 | 4552.124239 | 2703.11611 | 2094.10436 | 2195.994438 | 2197.591679 | 2165.164747 | 0.004970213 | -0.625237953 | 0.000560655 | 11.90227391 | 0.648312838 | 2.303624985 |
| *Sparc* | 624.1843551 | 749.3290152 | 604.8197158 | 673.5825625 | 445.7790518 | 448.5296131 | 387.5952047 | 434.5975003 | 6.07E-08 | -0.62630765 | 8.30E-10 | 37.68811489 | 0.64783232 | 7.216927338 |
| *Herpud1* | 1277.867026 | 1168.33632 | 1140.028106 | 1455.524059 | 865.1285155 | 761.1907667 | 824.5539496 | 816.1741056 | 1.01E-07 | -0.627135699 | 1.50E-09 | 36.53166382 | 0.647460597 | 6.995880022 |
| *Col3a1* | 243.0661194 | 277.6244722 | 237.3632092 | 221.598901 | 193.816979 | 134.23149 | 142.6057829 | 161.6702701 | 0.000661701 | -0.629910928 | 4.35E-05 | 16.71408891 | 0.646216311 | 3.179337968 |
| *Tagap* | 566.3676568 | 721.0524486 | 900.3825581 | 788.7749428 | 451.9459557 | 519.7377816 | 467.1253528 | 480.6648353 | 0.000638214 | -0.633249361 | 4.16E-05 | 16.79658019 | 0.64472268 | 3.195033725 |
| *Nckap1* | 99.11433995 | 143.95343 | 122.1051124 | 102.5016943 | 68.71692893 | 81.02998484 | 71.30289143 | 79.09674506 | 0.002663082 | -0.633375687 | 0.000251801 | 13.39868378 | 0.644666228 | 2.574615394 |
| *Morf4l2* | 3352.188569 | 4474.124017 | 5879.304106 | 3431.366242 | 2748.677157 | 2846.689771 | 2748.817878 | 2633.660852 | 0.00717983 | -0.642438895 | 0.000924688 | 10.97260234 | 0.640629041 | 2.143885824 |
| *Slc7a1* | 2811.77943 | 3006.31315 | 3458.884073 | 2770.474366 | 1924.07401 | 1850.593896 | 1919.693231 | 2012.186426 | 2.12E-10 | -0.644455855 | 1.76E-12 | 49.73465458 | 0.639734035 | 9.674552617 |
| *Pcdh7* | 2363.405035 | 2429.214132 | 3590.11854 | 2589.876143 | 1788.402125 | 1766.289973 | 1697.5573 | 1744.474366 | 0.00029943 | -0.649217325 | 1.66E-05 | 18.54992832 | 0.637626138 | 3.523705242 |
| *Fdps* | 1106.776796 | 1344.422213 | 1605.62517 | 1303.235827 | 855.4376666 | 877.4155935 | 907.7406562 | 777.0603306 | 2.02E-05 | -0.649308019 | 6.70E-07 | 24.70052634 | 0.637586055 | 4.694352826 |
| *Txn1* | 4572.238896 | 5537.065862 | 7579.646326 | 4436.859053 | 3570.637346 | 3586.599935 | 3453.619536 | 3483.733563 | 0.003695647 | -0.650529996 | 0.00038343 | 12.61126159 | 0.637046242 | 2.432309553 |
| *Stip1* | 3011.188042 | 3855.895447 | 4666.241166 | 2966.691895 | 2331.089666 | 2370.331678 | 2186.622004 | 2345.957307 | 0.001023225 | -0.650878543 | 7.51E-05 | 15.67887666 | 0.636892354 | 2.990028973 |
| *Gm5148* | 90.85481162 | 91.25619225 | 79.88184926 | 82.97756205 | 56.38312117 | 57.29392868 | 55.76251765 | 49.54411504 | 0.000676015 | -0.652349737 | 4.48E-05 | 16.65509643 | 0.636243211 | 3.170043949 |
| *Phyhd1* | 69.61602449 | 91.25619225 | 71.89366433 | 76.14411576 | 46.69227222 | 49.10908172 | 56.67665729 | 43.45975003 | 0.003182232 | -0.654558209 | 0.000315907 | 12.97364289 | 0.635269998 | 2.497268109 |
| *Vasn* | 211.2079387 | 185.0829815 | 215.680993 | 224.5275208 | 154.172597 | 114.5878574 | 147.176481 | 116.4721301 | 0.000179579 | -0.655953102 | 8.89E-06 | 19.73579225 | 0.634656074 | 3.745745048 |
| *Psmd7* | 1647.185935 | 2255.698837 | 2944.21673 | 1666.384687 | 1372.576606 | 1292.387334 | 1352.926658 | 1380.281661 | 0.008674885 | -0.657140559 | 0.001181078 | 10.51971363 | 0.634133914 | 2.061736274 |
| *Ezr* | 2628.889874 | 3086.001656 | 3697.388451 | 2879.809506 | 1909.097244 | 1938.171759 | 1927.006348 | 1994.802526 | 7.95E-07 | -0.661859473 | 1.51E-08 | 32.03628585 | 0.632063113 | 6.099737125 |
| *Lpcat2* | 2254.851234 | 2890.636287 | 3909.645937 | 2107.630076 | 1833.332424 | 1715.543922 | 1742.350142 | 1756.643096 | 0.00868458 | -0.663317731 | 0.001183093 | 10.51656406 | 0.631424554 | 2.061251178 |
| *Zfp653* | 171.0902297 | 263.4861889 | 273.880626 | 198.1699423 | 138.3148441 | 157.9675462 | 147.176481 | 127.7716651 | 0.003777178 | -0.664433839 | 0.000394608 | 12.55755341 | 0.630936256 | 2.422832494 |
| *Slc39a1* | 2658.388189 | 3636.109407 | 4935.557115 | 3034.050151 | 2254.443861 | 2138.700509 | 2320.08639 | 2259.907002 | 0.005752765 | -0.66867344 | 0.000680152 | 11.54271485 | 0.629084866 | 2.240123395 |
| *Cat* | 4061.328073 | 5080.784901 | 6688.393122 | 3948.755747 | 3112.524486 | 3071.773062 | 3003.862836 | 3247.312522 | 0.00258119 | -0.669453833 | 0.000240548 | 13.48445741 | 0.628744669 | 2.588179985 |
| *Rbm12* | 624.1843551 | 570.6725262 | 701.8191042 | 526.1753641 | 375.3001503 | 373.2290211 | 393.0800425 | 379.8382153 | 8.57E-08 | -0.670654688 | 1.26E-09 | 36.87282175 | 0.628221539 | 7.067187438 |
| *Smurf1* | 1047.780165 | 1097.644904 | 1348.862083 | 1156.804836 | 739.1474791 | 724.3589554 | 706.6299368 | 750.1152855 | 7.50E-10 | -0.671802973 | 7.14E-12 | 46.98812302 | 0.627721717 | 9.124793378 |
| *Myo1e* | 3461.922302 | 3866.177835 | 4683.358705 | 3529.96311 | 2494.953112 | 2660.07526 | 2274.379409 | 2312.927897 | 2.28E-06 | -0.673639105 | 5.18E-08 | 29.64876054 | 0.626923317 | 5.642163468 |
| *Rapgef1* | 2152.197096 | 2898.348078 | 3971.269078 | 2410.254126 | 1799.854946 | 1765.471488 | 1762.461214 | 1820.094331 | 0.004934183 | -0.67743571 | 0.000555742 | 11.91866823 | 0.625275671 | 2.306784708 |
| *Spi1* | 4235.9581 | 5169.470496 | 6843.592143 | 4729.721037 | 3282.554836 | 3187.997889 | 3183.948344 | 3459.396103 | 0.000323267 | -0.677812676 | 1.81E-05 | 18.37656808 | 0.625112313 | 3.490438826 |
| *Traf2* | 625.3642878 | 826.4469242 | 1208.498262 | 740.9408188 | 535.6396512 | 474.7211233 | 564.024154 | 550.2004354 | 0.009751949 | -0.678944349 | 0.001369064 | 10.24700887 | 0.624622157 | 2.010908582 |
| *Gtpbp6* | 141.5919142 | 179.9417875 | 247.6337327 | 161.074091 | 108.361311 | 99.85513284 | 129.807828 | 117.3413251 | 0.007556534 | -0.681809271 | 0.000989525 | 10.84706533 | 0.623383005 | 2.121677364 |
| *Spata5l1* | 189.9691516 | 192.7947724 | 230.5161936 | 247.9564795 | 139.1958304 | 136.6869441 | 122.4947109 | 137.3328101 | 1.17E-05 | -0.688749966 | 3.51E-07 | 25.94754635 | 0.62039116 | 4.930841199 |
| *Hsp90aa1* | 9689.606662 | 13265.56564 | 17065.04534 | 9176.342156 | 7489.264267 | 7942.575484 | 7611.12659 | 7417.710135 | 0.007134449 | -0.691574518 | 0.000914849 | 10.99242851 | 0.619177727 | 2.146639555 |
| *Rela* | 1869.013268 | 2541.0351 | 3393.837424 | 2142.773514 | 1495.914684 | 1523.200018 | 1524.784909 | 1609.749141 | 0.002805799 | -0.692699104 | 0.000269447 | 13.27166746 | 0.618695264 | 2.551943419 |
| *Cdc42bpg* | 261.9450413 | 314.8981282 | 449.6206944 | 303.6002564 | 216.722622 | 204.6211738 | 202.9389987 | 197.3072651 | 0.001697025 | -0.695061572 | 0.000141587 | 14.48090936 | 0.617682956 | 2.77031165 |
| *Cyfip1* | 9053.622981 | 12065.09685 | 16344.96753 | 9252.486272 | 7269.898686 | 7165.833508 | 7198.849615 | 7180.4199 | 0.004724393 | -0.697077806 | 0.0005272 | 12.01691088 | 0.616820318 | 2.32565398 |
| *Slc20a1* | 1818.276165 | 2686.273828 | 2876.887743 | 1885.054968 | 1309.145595 | 1464.26912 | 1447.08304 | 1475.893111 | 0.001085888 | -0.701726265 | 8.10E-05 | 15.53580225 | 0.614836081 | 2.964214817 |
| *Adam17* | 9205.834289 | 10788.79546 | 13628.98465 | 9396.96485 | 6519.298386 | 6744.31389 | 6611.97197 | 6571.114205 | 1.74E-05 | -0.701907574 | 5.62E-07 | 25.03844784 | 0.614758817 | 4.758636004 |
| *Cyp1b1* | 92.03474424 | 95.1120877 | 89.01120346 | 95.66824801 | 45.81128595 | 71.2081685 | 47.53526095 | 62.58204005 | 0.002089954 | -0.707352797 | 0.000184236 | 13.98540119 | 0.612442881 | 2.679863312 |
| *Ppapdc1b* | 233.6266584 | 311.0422327 | 421.0914625 | 239.17062 | 188.5310614 | 180.8851177 | 171.8582511 | 194.6996801 | 0.006722064 | -0.710289395 | 0.000841535 | 11.14731232 | 0.611197525 | 2.172497356 |
| *Lppr3* | 63.71636139 | 65.5502226 | 61.62314086 | 72.23928931 | 40.52536834 | 36.83181129 | 46.62112132 | 37.37538503 | 0.001611913 | -0.711753711 | 0.000131135 | 14.62540586 | 0.610577482 | 2.792658462 |
| *1810037I17Rik* | 528.609813 | 566.8166307 | 740.6188596 | 528.1277773 | 379.7050817 | 391.2356844 | 320.8630114 | 350.2855853 | 3.51E-05 | -0.712525113 | 1.27E-06 | 23.4757156 | 0.610251097 | 4.454370403 |
| *Tgif1* | 1808.836704 | 2343.099133 | 3246.626588 | 1945.579778 | 1406.935071 | 1372.598834 | 1473.593089 | 1441.994506 | 0.002491807 | -0.714274902 | 0.000230623 | 13.56352804 | 0.609511395 | 2.603485608 |
| *Vav1* | 4900.260164 | 5711.866456 | 7052.42612 | 5005.987508 | 3400.606996 | 3589.873874 | 3376.831807 | 3448.096568 | 3.04E-06 | -0.714462908 | 7.35E-08 | 28.96936397 | 0.609431971 | 5.516585406 |
| *Gm1943* | 86.13508114 | 102.8238786 | 92.43471129 | 113.239967 | 64.31199759 | 68.75271441 | 63.07563472 | 44.32894503 | 0.002159883 | -0.716575286 | 0.000192301 | 13.90485992 | 0.608540301 | 2.665569783 |
| *Pofut2* | 1000.58286 | 1467.810867 | 1817.882655 | 1130.447257 | 814.9122982 | 831.5804505 | 827.2963685 | 820.5200806 | 0.002539657 | -0.717223111 | 0.000235661 | 13.52296848 | 0.608267104 | 2.595224863 |
| *Cald1* | 309.142346 | 339.3187994 | 301.2686886 | 278.2188845 | 204.3888143 | 169.4263319 | 169.1158322 | 202.5224351 | 1.76E-07 | -0.717586744 | 2.78E-09 | 35.33514148 | 0.608113809 | 6.754136199 |
| *Atf5* | 623.0044225 | 660.64342 | 769.1480914 | 613.0577526 | 407.8966422 | 410.8793171 | 413.1911144 | 387.6609703 | 6.48E-11 | -0.718379279 | 4.82E-13 | 52.27795958 | 0.607779838 | 10.18871538 |
| *Serpinh1* | 351.6199203 | 286.6215616 | 348.0566289 | 329.957835 | 181.4831713 | 220.1723831 | 153.5754585 | 244.2437952 | 0.000414599 | -0.718899167 | 2.45E-05 | 17.80484189 | 0.607560858 | 3.382371878 |
| *Actg1* | 33592.68165 | 51640.72243 | 63022.21439 | 38303.41885 | 27657.68291 | 28657.60464 | 28104.3089 | 28616.50701 | 0.003699021 | -0.722841178 | 0.000384076 | 12.6081151 | 0.605903028 | 2.431913202 |
| *Irf1* | 1143.354707 | 1493.516837 | 2073.504573 | 1459.428885 | 971.727854 | 887.2374098 | 936.9931245 | 941.3381857 | 0.001035522 | -0.723309673 | 7.63E-05 | 15.64805205 | 0.605706302 | 2.984840865 |
| *M6pr* | 8063.659514 | 10030.46936 | 13584.47905 | 8232.350362 | 5821.557261 | 6061.697654 | 6127.477965 | 6139.124289 | 0.000858384 | -0.724744226 | 5.99E-05 | 16.10656642 | 0.605104313 | 3.066318463 |
| *Zwint* | 1417.099075 | 1974.218469 | 2345.10286 | 1483.834051 | 1096.827904 | 1125.416456 | 1076.856488 | 1047.379976 | 0.000622305 | -0.731949471 | 4.05E-05 | 16.84914854 | 0.602089778 | 3.20599668 |
| *Cyr61* | 188.7892189 | 230.0684283 | 196.2811153 | 185.4792563 | 122.4570913 | 99.03664814 | 112.4391749 | 146.8939551 | 9.73E-05 | -0.733978465 | 4.29E-06 | 21.13043191 | 0.601243599 | 4.011688767 |
| *Ncoa5* | 566.3676568 | 872.7176695 | 1149.15746 | 636.4867112 | 458.9938458 | 505.8235417 | 468.0394924 | 490.2259804 | 0.009037172 | -0.745380527 | 0.001242692 | 10.42576685 | 0.596510515 | 2.043967451 |
| *Mapk6* | 2487.29796 | 3294.22001 | 4306.772844 | 2756.807473 | 1854.476095 | 1857.141774 | 1934.319465 | 2012.186426 | 0.000565757 | -0.746097896 | 3.55E-05 | 17.09893882 | 0.596213978 | 3.247370063 |
| *Cycs* | 1614.147822 | 2159.30145 | 2665.771427 | 1561.93058 | 1249.238529 | 1229.364012 | 1161.871474 | 1127.345916 | 0.001084473 | -0.74664549 | 8.08E-05 | 15.54029726 | 0.59598772 | 2.964781211 |
| *Acp2* | 3288.472207 | 3555.135602 | 5527.823969 | 3287.86387 | 2319.636845 | 2321.222596 | 2311.859134 | 2371.163962 | 0.000943859 | -0.747996037 | 6.79E-05 | 15.86912984 | 0.59543006 | 3.025092904 |
| *Traf3* | 1602.348496 | 1551.355268 | 1676.377665 | 1770.838795 | 967.3229227 | 955.1716395 | 1029.321228 | 969.1524257 | 9.82E-27 | -0.752889148 | 2.04E-29 | 126.8129172 | 0.593413993 | 26.00810503 |
| *Glipr1* | 4621.796066 | 4827.5811 | 4879.639821 | 4610.62383 | 2762.772938 | 3046.400036 | 2740.590622 | 2681.466577 | 1.47E-32 | -0.753552739 | 1.89E-35 | 154.4072935 | 0.593141105 | 31.83145534 |
| *Cxcl12* | 81.41535067 | 52.69723778 | 62.76431013 | 71.2630827 | 29.07254686 | 41.74271946 | 46.62112132 | 41.72136003 | 0.007134449 | -0.756471985 | 0.000914338 | 10.99346381 | 0.591942117 | 2.146639555 |
| *Nfe2l2* | 5070.170461 | 7597.399329 | 9412.364181 | 5887.502079 | 4123.896722 | 4121.070441 | 4109.057653 | 4157.359688 | 0.001448076 | -0.760247275 | 0.000114099 | 14.88778714 | 0.59039513 | 2.839208542 |
| *Nedd4* | 297.3430198 | 257.0596965 | 219.1045008 | 266.5044052 | 158.5775283 | 144.8717911 | 163.6309944 | 146.8939551 | 1.85E-07 | -0.76127211 | 2.94E-09 | 35.22491143 | 0.589975885 | 6.733942355 |
| *Mtmr14* | 1515.033482 | 1667.032132 | 2065.516388 | 1473.095778 | 981.4187029 | 947.8052773 | 1002.811178 | 1016.088956 | 7.97E-08 | -0.76728128 | 1.16E-09 | 37.03464707 | 0.587523604 | 7.098486222 |
| *Ccrn4l* | 474.3329126 | 625.9403609 | 538.6318979 | 550.5805294 | 333.8937957 | 331.4863016 | 272.4136108 | 342.4628302 | 5.39E-08 | -0.772939685 | 7.29E-10 | 37.94086874 | 0.585223787 | 7.268085945 |
| *Sfn* | 81.41535067 | 100.2532816 | 103.846404 | 86.8823885 | 53.74016237 | 49.10908172 | 50.27767985 | 63.45123505 | 0.000149103 | -0.779823523 | 7.14E-06 | 20.15547921 | 0.582438035 | 3.826514058 |
| *Gca* | 90.85481162 | 138.8122361 | 168.8930527 | 105.4303141 | 61.66903878 | 76.11907667 | 76.78772923 | 77.35835506 | 0.005739759 | -0.785787348 | 0.000677237 | 11.55070057 | 0.580035318 | 2.241106373 |
| *Pgs1* | 917.9875771 | 1255.736617 | 1900.046843 | 981.0876454 | 726.8136714 | 741.547134 | 704.8016576 | 755.3304555 | 0.007391783 | -0.787355018 | 0.000959689 | 10.90376785 | 0.579405378 | 2.13125081 |
| *H2-Q5* | 173.4500949 | 172.2299966 | 228.233855 | 222.5751076 | 125.9810364 | 93.30725527 | 107.8684768 | 134.7252251 | 0.000183152 | -0.789635224 | 9.11E-06 | 19.68890203 | 0.578490341 | 3.737188718 |
| *Gpr176* | 147.4915773 | 154.2358179 | 158.6225292 | 100.5492811 | 81.05073669 | 75.30059198 | 84.1008463 | 82.57352506 | 0.000285575 | -0.79141048 | 1.57E-05 | 18.65408831 | 0.577778938 | 3.54428022 |
| *Rcn1* | 140.4119816 | 192.7947724 | 203.128131 | 120.0734133 | 104.8373659 | 99.03664814 | 84.1008463 | 88.65789006 | 0.001861442 | -0.797473658 | 0.000159847 | 14.25251535 | 0.575355817 | 2.730150446 |
| *Itpripl2* | 3746.286063 | 5531.924668 | 7707.457284 | 4107.877425 | 3175.074511 | 2938.360056 | 2986.494183 | 3027.406187 | 0.005223226 | -0.798498981 | 0.000599161 | 11.77858363 | 0.574947057 | 2.282061184 |
| *Eif1* | 3616.493475 | 4498.544688 | 6116.667315 | 3401.103837 | 2430.641114 | 2534.028617 | 2537.651623 | 2611.930977 | 0.000648842 | -0.801794957 | 4.25E-05 | 16.75827009 | 0.573635034 | 3.187861222 |
| *Lrrc59* | 4283.155405 | 6181.000402 | 8532.52267 | 4493.479037 | 3375.93938 | 3333.688164 | 3402.427717 | 3360.307872 | 0.004123993 | -0.802010673 | 0.000439088 | 12.35805624 | 0.573549269 | 2.384682127 |
| *AA414768* | 125.0728576 | 149.094624 | 162.0460371 | 181.5744299 | 76.64580535 | 85.94089302 | 100.5553597 | 92.13467007 | 4.48E-05 | -0.802832646 | 1.71E-06 | 22.89883498 | 0.573222583 | 4.349199387 |
| *Micall1* | 769.3160672 | 918.9884149 | 1113.781213 | 818.0611411 | 521.5438709 | 505.0050571 | 493.6354022 | 548.4620454 | 1.85E-07 | -0.807047167 | 2.92E-09 | 35.23324348 | 0.571550481 | 6.733942355 |
| *Zfp438* | 56.63676568 | 65.5502226 | 90.15237274 | 57.59619013 | 34.35846447 | 49.92756642 | 32.90902681 | 36.50619003 | 0.008349589 | -0.807558605 | 0.001124767 | 10.6100168 | 0.571347901 | 2.07833489 |
| *Cttn* | 76.6956202 | 91.25619225 | 60.48197158 | 84.92997527 | 36.120437 | 42.56120416 | 51.19181949 | 48.67492004 | 0.001242509 | -0.811890115 | 9.54E-05 | 15.22528862 | 0.569635074 | 2.905700308 |
| *Tuba1a* | 851.9113505 | 1237.742439 | 1311.203497 | 974.2541991 | 579.6889646 | 676.8868431 | 617.0442527 | 612.7824754 | 1.21E-05 | -0.81473844 | 3.65E-07 | 25.8702573 | 0.568511548 | 4.918072916 |
| *11-Sep* | 1366.361972 | 2092.465929 | 2738.80626 | 1396.951662 | 1144.401163 | 1019.83193 | 1106.108957 | 1044.772391 | 0.005740581 | -0.815434894 | 0.000677794 | 11.54917409 | 0.568237168 | 2.241044145 |
| *Sbno2* | 2651.308594 | 4110.384547 | 5728.669761 | 3025.264292 | 2227.133286 | 2147.703841 | 2164.682653 | 2255.561027 | 0.006044113 | -0.818900171 | 0.000726686 | 11.41972111 | 0.566873931 | 2.218667415 |
| *Uck2* | 304.4226155 | 433.1455886 | 591.1256845 | 371.9347193 | 221.1275534 | 229.1757147 | 270.5853316 | 242.5054052 | 0.001812944 | -0.819862718 | 0.000154041 | 14.32215887 | 0.566495846 | 2.741615526 |
| *Fam96b* | 409.4366186 | 592.5226004 | 693.8309193 | 384.6254053 | 291.6064548 | 297.1099444 | 301.6660791 | 286.8343502 | 0.001240286 | -0.820589098 | 9.49E-05 | 15.23458167 | 0.566210694 | 2.906478027 |
| *Arid5a* | 475.5128452 | 619.5138685 | 777.1362764 | 460.769521 | 322.4409742 | 380.5953834 | 298.0095206 | 318.9945652 | 0.000591281 | -0.820597377 | 3.76E-05 | 16.9887238 | 0.566207445 | 3.228205903 |
| *Nab1* | 1844.234683 | 2418.931744 | 3699.67079 | 1800.124993 | 1350.551949 | 1442.170033 | 1394.062941 | 1326.391571 | 0.005531788 | -0.824337894 | 0.000646504 | 11.63706977 | 0.564741324 | 2.257134479 |
| *Nadk* | 3819.441886 | 4736.324908 | 7034.167412 | 4073.710193 | 2788.321539 | 2717.369189 | 2875.883288 | 2709.280817 | 0.00070105 | -0.826141987 | 4.68E-05 | 16.57472049 | 0.564035555 | 3.154251037 |
| *Scn1b* | 181.7096232 | 231.3537268 | 216.8221623 | 228.4323473 | 125.1000501 | 118.6802808 | 108.7826164 | 131.2484451 | 1.97E-08 | -0.827223943 | 2.40E-10 | 40.10963869 | 0.563612712 | 7.704435693 |
| *Ywhag* | 5269.579074 | 8061.392082 | 11485.86875 | 5454.066343 | 4283.355237 | 4238.113753 | 4219.668549 | 4316.422373 | 0.009273962 | -0.827486667 | 0.001280446 | 10.37050356 | 0.563510084 | 2.032734708 |
| *Clic4* | 9817.039385 | 16310.43774 | 21343.28895 | 11184.39916 | 8173.790598 | 8434.484786 | 8155.039672 | 8219.107926 | 0.00625415 | -0.830540846 | 0.000761945 | 11.3317157 | 0.562318397 | 2.203831687 |
| *Mdm2* | 2818.859025 | 3489.58538 | 4642.276611 | 3331.793168 | 1919.669079 | 2003.650534 | 2054.985896 | 2026.093546 | 5.57E-06 | -0.835391342 | 1.47E-07 | 27.63452957 | 0.560430997 | 5.254296809 |
| *Gmfg* | 591.1462418 | 953.6914739 | 1055.58158 | 648.2011906 | 449.3029969 | 439.5262814 | 473.5243302 | 457.1965703 | 0.000795296 | -0.83578386 | 5.38E-05 | 16.30852234 | 0.56027854 | 3.099471022 |
| *Plekhm3* | 1661.345127 | 2429.214132 | 3519.366045 | 1667.360894 | 1327.646306 | 1215.449773 | 1332.815586 | 1309.876866 | 0.007552648 | -0.839053364 | 0.000988412 | 10.84914927 | 0.559010248 | 2.121900732 |
| *Mxra8* | 49.55716997 | 44.98544688 | 30.81157043 | 48.81033062 | 25.54860178 | 24.55454086 | 19.19693231 | 27.81424002 | 0.007830394 | -0.842332689 | 0.001032277 | 10.76875984 | 0.55774103 | 2.106216398 |
| *Ptk7* | 38.93777641 | 47.55604385 | 45.64677101 | 41.00067772 | 22.90564298 | 24.55454086 | 18.28279267 | 30.42182502 | 0.004552295 | -0.842394923 | 0.000501505 | 12.11006228 | 0.557716971 | 2.341769585 |
| *Il2rg* | 2962.810805 | 3643.821198 | 4853.392927 | 2911.048118 | 1885.310614 | 2027.38659 | 2097.03632 | 1999.148501 | 7.25E-05 | -0.84341299 | 3.04E-06 | 21.78940574 | 0.557323545 | 4.139947361 |
| *Chpf2* | 1684.943779 | 2237.704658 | 3177.015262 | 1571.692646 | 1229.856831 | 1255.555523 | 1135.361425 | 1209.919441 | 0.002805799 | -0.843864991 | 0.00026956 | 13.27088711 | 0.557148962 | 2.551943419 |
| *D8Ertd82e* | 299.7028851 | 430.5749916 | 627.6431013 | 353.3867937 | 230.8184023 | 242.2714698 | 232.191467 | 247.7205752 | 0.003293319 | -0.844184499 | 0.000330623 | 12.88841252 | 0.557025586 | 2.482366238 |
| *Rasgef1b* | 1277.867026 | 1327.713332 | 1361.414945 | 1648.812968 | 769.1010123 | 815.2107566 | 811.7559947 | 733.6005805 | 2.49E-13 | -0.84444092 | 1.47E-15 | 63.66750473 | 0.55692659 | 12.60410096 |
| *Fn1* | 2110.899454 | 1975.503767 | 2144.257068 | 2064.676985 | 1167.306806 | 1059.937681 | 1166.442173 | 1221.218976 | 2.14E-34 | -0.846573883 | 2.40E-37 | 163.0808864 | 0.556103806 | 33.66860168 |
| *Loxl1* | 114.453464 | 92.54149073 | 61.62314086 | 82.97756205 | 52.8591761 | 45.01665825 | 53.93423839 | 43.45975003 | 0.002429129 | -0.847351118 | 0.000223226 | 13.62472417 | 0.555804293 | 2.614549366 |
| *Chac1* | 151.0313752 | 160.6623103 | 179.1635762 | 202.0747688 | 103.0753934 | 104.766041 | 87.75740483 | 90.39628007 | 6.50E-07 | -0.847383005 | 1.22E-08 | 32.46059108 | 0.555792008 | 6.187215226 |
| *Asns* | 1680.224049 | 1700.449892 | 2143.115899 | 1482.857844 | 912.701774 | 1014.102538 | 1005.553597 | 957.8528907 | 4.44E-09 | -0.848584729 | 4.69E-11 | 43.30447511 | 0.555329242 | 8.352909601 |
| *Gp49a* | 2982.869659 | 4934.260874 | 6451.029912 | 3385.484532 | 2422.712238 | 2545.487403 | 2459.035615 | 2425.054052 | 0.005002017 | -0.849518812 | 0.000566183 | 11.88399717 | 0.554969807 | 2.300854847 |
| *Pou2f2* | 564.0077916 | 750.6143137 | 1097.804843 | 647.224984 | 398.2057933 | 430.5229498 | 425.9890693 | 442.4202553 | 0.001033624 | -0.850101292 | 7.60E-05 | 15.65415904 | 0.554745786 | 2.985637251 |
| *Enah* | 90.85481162 | 104.1091771 | 82.16418781 | 121.0496199 | 37.88240954 | 63.02332154 | 58.50493656 | 61.71284504 | 0.001733525 | -0.850144804 | 0.000145882 | 14.42463008 | 0.554729055 | 2.761069886 |

| | | | | | | | | | | | | | | |
|---|---|---|---|---|---|---|---|---|---|---|---|---|---|---|
| *Trib1* | 247.7858499 | 323.8952176 | 387.9975535 | 287.004744 | 172.6733086 | 192.3439034 | 155.4037377 | 169.4930251 | 1.10E-05 | -0.852325633 | 3.27E-07 | 26.08087159 | 0.55389114 | 4.957335029 |
| *Trmt61a* | 122.7129923 | 151.6652209 | 162.0460371 | 129.8354794 | 59.02607998 | 93.30725527 | 83.18670666 | 77.35835506 | 0.000100949 | -0.852993633 | 4.51E-06 | 21.03314768 | 0.553634735 | 3.995899994 |
| *Tead1* | 49.55716997 | 56.55313323 | 43.36443246 | 48.81033062 | 28.19156059 | 28.64696434 | 23.76763048 | 28.68343502 | 0.001055781 | -0.853497269 | 7.80E-05 | 15.60526324 | 0.553441499 | 2.97642626 |
| *Zdhhc18* | 945.1260273 | 1174.762813 | 1688.930527 | 999.635571 | 648.4058935 | 671.9759349 | 663.665374 | 676.2337105 | 0.000257182 | -0.853846272 | 1.37E-05 | 18.90451237 | 0.553307632 | 3.589758952 |
| *Xylt2* | 2437.74079 | 3161.834267 | 4293.078813 | 2611.352688 | 1755.805633 | 1785.933605 | 1682.931066 | 1690.584276 | 0.000130243 | -0.854449948 | 6.11E-06 | 20.45490613 | 0.553076156 | 3.88524691 |
| *Rrs1* | 453.0941255 | 785.3173727 | 980.2644073 | 492.0081326 | 394.6818482 | 364.2256894 | 356.5144571 | 382.4458003 | 0.007932489 | -0.855286221 | 0.001054393 | 10.72952932 | 0.552755653 | 2.100590522 |
| *Fyb* | 5995.237634 | 8278.607525 | 12049.60638 | 6635.276344 | 4552.056049 | 4531.949758 | 4594.465799 | 4532.851928 | 0.00151903 | -0.855804753 | 0.000121513 | 14.76906427 | 0.552557018 | 2.818433558 |
| *Arl8a* | 1767.539062 | 2556.458681 | 3478.283951 | 1727.885704 | 1348.789977 | 1235.093405 | 1378.522568 | 1278.585846 | 0.003108302 | -0.862561084 | 0.000306081 | 13.03280916 | 0.54997537 | 2.507476812 |
| *Ifnar1* | 3596.434621 | 5034.514156 | 7658.387005 | 3864.801978 | 2694.936995 | 2804.947051 | 2740.590622 | 2814.453412 | 0.003724452 | -0.866348147 | 0.000387908 | 12.5895577 | 0.548533582 | 2.42893757 |
| *Gm21188* | 453.0941255 | 609.2314807 | 900.3825581 | 412.935397 | 340.9416859 | 338.8526639 | 357.4285968 | 264.2352802 | 0.008511065 | -0.867829255 | 0.001155585 | 10.56004385 | 0.547970732 | 2.070016075 |
| *Plekho2* | 2408.242474 | 2548.746891 | 3877.693197 | 2599.638209 | 1552.297805 | 1608.322426 | 1554.037377 | 1548.905491 | 8.66E-06 | -0.868331146 | 2.44E-07 | 26.65109274 | 0.547780135 | 5.062719361 |
| *Amotl2* | 46.01737212 | 73.2620135 | 90.15237274 | 62.47722319 | 29.07254686 | 35.1948419 | 40.22214388 | 44.32894503 | 0.008039601 | -0.869076562 | 0.00107915 | 10.68658955 | 0.54749718 | 2.094765513 |
| *Col1a1* | 1502.054223 | 1334.139825 | 1346.579745 | 1459.428885 | 824.6031472 | 773.4680371 | 700.2309594 | 790.9674506 | 7.73E-24 | -0.869200586 | 1.67E-26 | 113.5088459 | 0.547450115 | 23.11187178 |
| *Lst1* | 258.4052434 | 344.4599933 | 484.9969419 | 296.7668102 | 204.3888143 | 150.601184 | 188.3127645 | 212.9527752 | 0.001697025 | -0.872693622 | 0.000141588 | 14.48089263 | 0.546126239 | 2.77031165 |
| *Slc12a4* | 1369.90177 | 1699.164594 | 2215.009563 | 1532.644381 | 929.4405131 | 892.148318 | 890.3720032 | 1003.051031 | 3.01E-06 | -0.875639786 | 7.24E-08 | 28.99931033 | 0.545012118 | 5.521858752 |
| *Eif6* | 1740.400612 | 2475.484877 | 3384.70807 | 1857.721183 | 1292.406856 | 1379.146712 | 1203.007758 | 1252.509996 | 0.001010784 | -0.883304997 | 7.33E-05 | 15.72275377 | 0.542124083 | 2.995341448 |
| *Serpinf1* | 108.5538009 | 116.9621619 | 123.2462817 | 124.9544464 | 60.78805252 | 63.02332154 | 69.47461216 | 63.45123505 | 4.97E-08 | -0.887192772 | 6.64E-10 | 38.12442451 | 0.540665134 | 7.303773352 |
| *Slc4a7* | 1630.666879 | 2569.311666 | 3029.804426 | 1931.912886 | 1218.404009 | 1316.941875 | 1283.452046 | 1133.430281 | 0.000269561 | -0.887395325 | 1.46E-05 | 18.79159194 | 0.54058923 | 3.569342429 |
| *Slc25a33* | 261.9450413 | 296.9039494 | 378.8681993 | 201.0985621 | 148.8866794 | 177.6111789 | 156.3178774 | 131.2484451 | 0.000669304 | -0.889190301 | 4.42E-05 | 16.68114663 | 0.539917057 | 3.174376604 |
| *Naip6* | 540.4091392 | 451.1397673 | 600.2550387 | 427.5784962 | 273.9867295 | 302.0208526 | 255.0449578 | 258.1509152 | 3.84E-08 | -0.889585424 | 4.89E-10 | 38.72224488 | 0.539769205 | 7.415491122 |
| *Nr1h3* | 490.8519693 | 499.9811097 | 725.783659 | 510.5560583 | 276.6296883 | 292.1990363 | 309.8933358 | 321.6021502 | 1.03E-06 | -0.892025365 | 2.10E-08 | 31.39945954 | 0.538857098 | 5.985588268 |
| *Hsd17b7* | 391.7376293 | 525.6870793 | 639.0547941 | 406.1019507 | 302.17829 | 252.9117709 | 253.2166785 | 247.7205752 | 5.69E-05 | -0.893499931 | 2.26E-06 | 22.36102106 | 0.538306619 | 4.244595951 |
| *Syn1* | 97.93440733 | 92.54149073 | 118.6816046 | 59.54860335 | 44.93029969 | 52.3830205 | 59.41907619 | 40.85216503 | 0.004538273 | -0.896451495 | 0.000499533 | 12.11740735 | 0.53720644 | 2.34310943 |
| *Cxcl16* | 7843.012115 | 7944.42992 | 9282.270884 | 8505.688213 | 4484.220106 | 4438.642503 | 4524.077047 | 4591.087993 | 7.43E-34 | -0.89649103 | 8.92E-37 | 160.4735048 | 0.537191719 | 33.12895234 |
| *Eef1e1* | 348.0801224 | 494.8399157 | 572.8669761 | 348.5057606 | 222.8895259 | 246.3638933 | 250.4742596 | 226.8598952 | 9.80E-05 | -0.897259377 | 4.35E-06 | 21.10355253 | 0.536905699 | 4.008738472 |
| *Gm16907* | 92.03474424 | 134.9563407 | 112.9757582 | 104.4541075 | 74.00284654 | 47.47211233 | 43.87870242 | 72.14318505 | 0.001722727 | -0.902071397 | 0.000144698 | 14.43998153 | 0.535117866 | 2.763783495 |
| *Fam219aos* | 27.13845022 | 33.41776054 | 37.65858608 | 35.14343804 | 15.85775283 | 21.28060208 | 18.28279267 | 15.64551001 | 0.005888387 | -0.90816683 | 0.000699955 | 11.48936519 | 0.532861746 | 2.230003689 |
| *3110043O21Rik* | 1191.731945 | 1879.106381 | 2622.406994 | 1357.903398 | 888.9151448 | 946.9867926 | 1001.897039 | 912.6547507 | 0.002974907 | -0.910685527 | 0.000289852 | 13.134861 | 0.531932272 | 2.526526564 |
| *Tuba4a* | 941.5862295 | 1257.021916 | 1744.847822 | 909.8245627 | 620.2143329 | 643.3289706 | 640.8118832 | 675.3645155 | 0.000618874 | -0.911495582 | 4.01E-05 | 16.86665925 | 0.531633683 | 3.208398004 |
| *Myd88* | 1469.01611 | 2169.583838 | 3376.719885 | 1516.048869 | 1132.948341 | 1164.703722 | 1158.214916 | 1076.932606 | 0.006876143 | -0.91228138 | 0.000869625 | 11.08641384 | 0.531344195 | 2.162655126 |
| *Col4a1* | 572.2673199 | 654.2169275 | 855.8769564 | 558.3901823 | 366.4902876 | 365.8626588 | 310.8074754 | 358.9775353 | 1.57E-06 | -0.912643752 | 3.37E-08 | 30.48203711 | 0.53121075 | 5.805213425 |
| *Bgn* | 614.7448942 | 580.954914 | 608.2432237 | 592.5574137 | 321.5599879 | 296.2914597 | 308.9791962 | 345.0704153 | 4.15E-28 | -0.914508668 | 7.63E-31 | 133.3357818 | 0.530524518 | 27.38201175 |
| *Cfp* | 5017.073493 | 5179.752884 | 5970.597648 | 4564.742119 | 2645.601764 | 2664.986168 | 2818.292491 | 2867.474307 | 4.40E-18 | -0.914736891 | 1.55E-20 | 86.29478103 | 0.5304406 | 17.35612833 |
| *Cdr2* | 258.4052434 | 397.1572311 | 448.4795251 | 231.3609671 | 163.8634459 | 179.2481483 | 178.2572286 | 186.0077301 | 0.001448076 | -0.915249765 | 0.000114011 | 14.88924493 | 0.530252063 | 2.839208542 |
| *Acot9* | 646.6030749 | 1016.6711 | 1318.050513 | 732.1549593 | 485.4234339 | 516.4638428 | 504.6050778 | 459.8041553 | 0.001131913 | -0.916988929 | 8.48E-05 | 15.44722447 | 0.529613232 | 2.946186762 |
| *Nfe2l1* | 4854.242792 | 7182.24792 | 10373.22871 | 5099.703343 | 3745.953613 | 3608.699022 | 3500.240657 | 3692.340363 | 0.002474179 | -0.919145405 | 0.000228398 | 13.58172747 | 0.528822181 | 2.606568851 |
| *Ifi27* | 540.4091392 | 688.9199866 | 896.9590503 | 574.9856947 | 380.5860679 | 367.4996282 | 303.4943584 | 375.4922403 | 3.08E-05 | -0.920043391 | 1.08E-06 | 23.77273837 | 0.528493125 | 4.511450333 |
| *Fkbp10* | 94.39460947 | 87.4002968 | 93.57588056 | 81.02514882 | 41.40635461 | 44.19817355 | 49.36354022 | 53.02089504 | 2.45E-06 | -0.92126349 | 5.61E-08 | 29.49367843 | 0.528046363 | 5.610463199 |
| *Cyb5a* | 1273.147295 | 1706.876385 | 2305.161936 | 1265.16377 | 845.7468176 | 840.5837822 | 920.5386111 | 848.3343206 | 0.000231819 | -0.922656835 | 1.22E-05 | 19.13141581 | 0.527536625 | 3.634850848 |
| *Glrx* | 1956.328281 | 2174.725032 | 3557.024631 | 2034.41458 | 1166.425819 | 1370.961865 | 1301.734838 | 1288.146991 | 0.00025797 | -0.923094847 | 1.38E-05 | 18.89582757 | 0.527376486 | 3.588431456 |
| *Ptpn23* | 1668.424722 | 2696.556216 | 3881.116705 | 1953.389431 | 1289.763897 | 1303.027635 | 1401.376058 | 1382.889246 | 0.004078463 | -0.923546893 | 0.000433261 | 12.3829966 | 0.527211267 | 2.389503467 |
| *Lilrb4* | 9678.987269 | 15824.59492 | 20706.5165 | 10956.94302 | 7315.709972 | 7525.966774 | 7507.828811 | 7546.350995 | 0.001482112 | -0.935218514 | 0.000117967 | 14.82491435 | 0.522963254 | 2.829118855 |
| *Mina* | 608.8452311 | 1052.659457 | 1270.121403 | 684.3208353 | 437.8501754 | 437.0708273 | 488.1505644 | 523.2553904 | 0.002159883 | -0.938506986 | 0.000192014 | 13.90766796 | 0.521772572 | 2.665569783 |
| *1810022K09Rik* | 206.4882082 | 249.3479056 | 370.8800144 | 212.8130415 | 127.7430089 | 124.4096737 | 145.3482018 | 144.2863701 | 0.000317531 | -0.940000712 | 1.78E-05 | 18.41340071 | 0.521232623 | 3.498213842 |
| *Col1a2* | 1451.317121 | 1456.243181 | 1337.45039 | 1557.049547 | 759.4101633 | 675.2498737 | 743.1955222 | 846.5959306 | 2.36E-20 | -0.940491572 | 6.80E-23 | 97.03900289 | 0.52105531 | 19.62700822 |
| *Fermt2* | 62.53642878 | 83.54440136 | 76.45834143 | 61.50101658 | 41.40635461 | 33.55787251 | 36.56558535 | 35.63699503 | 5.52E-05 | -0.943653668 | 2.18E-06 | 22.43192564 | 0.519914513 | 4.257994472 |
| *Itsn1* | 1894.971785 | 2913.77166 | 4060.280281 | 2042.224233 | 1400.768167 | 1366.050957 | 1391.320522 | 1501.968961 | 0.001736998 | -0.946828836 | 0.000146313 | 14.41907147 | 0.518771513 | 2.760200649 |
| *Hvcn1* | 2107.359656 | 3409.896874 | 5148.955769 | 2264.799341 | 1733.780976 | 1630.421513 | 1650.022039 | 1692.322666 | 0.007264804 | -0.947132269 | 0.000938538 | 10.94505502 | 0.518662414 | 2.138776098 |
| *A630033H20Rik* | 339.8205941 | 476.845737 | 641.3371326 | 303.6002564 | 236.1043199 | 264.3705566 | 202.024859 | 208.6068002 | 0.002571273 | -0.950166613 | 0.000239212 | 13.49490501 | 0.517572685 | 2.589851856 |
| *Nab2* | 755.1568758 | 1471.666762 | 1816.741486 | 1022.088323 | 681.8833717 | 644.1474553 | 653.6098381 | 637.9891305 | 0.003402395 | -0.952327835 | 0.000342662 | 12.82148403 | 0.516797919 | 2.468215252 |
| *Plk3* | 401.1770903 | 615.6579731 | 706.3837813 | 400.2447111 | 259.8909492 | 274.192373 | 269.6711919 | 291.1803252 | 0.000213938 | -0.954697832 | 1.10E-05 | 19.32571793 | 0.515949642 | 3.669712769 |
| *Cdkn1a* | 5196.423251 | 7301.780678 | 9851.714352 | 6177.435443 | 3668.426821 | 3666.811435 | 3710.492773 | 3639.319468 | 3.71E-05 | -0.957978361 | 1.36E-06 | 23.33537738 | 0.514777763 | 4.430145445 |
| *Zfp951* | 63.71636139 | 59.12373019 | 46.78794028 | 40.02447111 | 32.59649193 | 22.09908678 | 31.08074754 | 21.72987502 | 0.00285118 | -0.959527836 | 0.000274832 | 13.23457754 | 0.514225181 | 2.544975293 |
| *H1fx* | 180.5296906 | 150.3799224 | 156.3401907 | 145.4547852 | 88.9796131 | 59.74938276 | 70.38875179 | 105.1725951 | 5.47E-05 | -0.964465912 | 2.13E-06 | 22.47219494 | 0.512468093 | 4.261781563 |
| *Cdh2* | 46.01737212 | 34.70305902 | 47.92910956 | 49.78653723 | 21.14367044 | 30.28393373 | 15.54037377 | 24.33746002 | 0.003472331 | -0.966296266 | 0.000353316 | 12.76420272 | 0.511818334 | 2.459378858 |
| *Odc1* | 2422.401666 | 4175.934769 | 4908.169052 | 2533.256159 | 1802.497905 | 1935.716305 | 1685.673484 | 1754.904706 | 0.001343475 | -0.967574716 | 0.000104353 | 15.05627449 | 0.511364986 | 2.871770458 |
| *Pfkp* | 2487.29796 | 3830.189478 | 5970.597648 | 2530.327539 | 1833.332424 | 1911.980248 | 1907.809415 | 1912.229001 | 0.006232153 | -0.969854748 | 0.00075745 | 11.34270215 | 0.510557464 | 2.205361887 |
| *Lcp1* | 31865.26029 | 57049.25844 | 79784.84987 | 35592.49309 | 25554.76869 | 25738.06973 | 26470.74137 | 26199.27571 | 0.006408786 | -0.97455839 | 0.000786423 | 11.2730049 | 0.508895596 | 2.193224246 |
| *Col5a1* | 300.8828177 | 311.0422327 | 275.0217953 | 317.267149 | 165.6254185 | 129.3205819 | 163.6309944 | 153.8475151 | 3.35E-14 | -0.977899061 | 1.72E-16 | 67.90529309 | 0.507718572 | 13.47486966 |
| *Ddr2* | 48.37723735 | 42.41484992 | 57.05846376 | 73.21549593 | 28.19156059 | 20.46211738 | 30.16660791 | 33.89860502 | 0.002752325 | -0.98399261 | 0.000263321 | 13.31478573 | 0.50557863 | 2.560300242 |
| *Zfp263* | 1209.430934 | 1749.291235 | 2543.666314 | 1328.617199 | 868.6524606 | 865.9568077 | 852.8922782 | 852.6802956 | 0.000575989 | -0.989495118 | 3.63E-05 | 17.05768152 | 0.503654002 | 3.239585443 |
| *Myo1b* | 135.6922511 | 209.5036526 | 252.1984098 | 129.8354794 | 79.28876415 | 109.6769492 | 91.41396337 | 84.31191506 | 0.001866845 | -0.993187675 | 0.000160536 | 14.24442304 | 0.502366555 | 2.728891771 |
| *Znrf1* | 887.309329 | 1525.649299 | 2161.374607 | 981.0876454 | 747.0763556 | 647.4213941 | 606.9887168 | 779.6679156 | 0.005864656 | -0.998048207 | 0.000696665 | 11.4981222 | 0.500676898 | 2.231757483 |
| *Rhoc* | 559.2880611 | 849.5822969 | 1316.909344 | 599.39086 | 386.7529718 | 437.0708273 | 415.9335333 | 420.6903803 | 0.003423134 | -1.001599673 | 0.00034612 | 12.80269903 | 0.499445903 | 2.465576065 |
| *Dbn1* | 48.37723735 | 38.55895447 | 47.92910956 | 35.14343804 | 22.02465671 | 31.10241842 | 16.45451341 | 14.77631501 | 0.007140098 | -1.002622013 | 0.000916144 | 10.98980574 | 0.499092105 | 2.146295807 |
| *Ube2f* | 614.7448942 | 917.7031164 | 1256.427372 | 690.1780749 | 423.7543951 | 406.7868936 | 448.8425601 | 454.5889853 | 0.000310943 | -1.004435532 | 1.73E-05 | 18.46975674 | 0.498465122 | 3.507318681 |
| *BC024978* | 75.51568758 | 68.12081957 | 107.2699119 | 59.54860335 | 34.35846447 | 38.46878068 | 49.36354022 | 31.29102002 | 0.001424128 | -1.016217203 | 0.000111757 | 14.92692379 | 0.494411017 | 2.846450985 |
| *Mag* | 35.39797855 | 47.55604385 | 69.61132578 | 56.61998352 | 24.66761551 | 23.73605617 | 26.51004938 | 28.68343502 | 0.001664398 | -1.018971031 | 0.000137791 | 14.53210441 | 0.493468182 | 2.778742855 |
| *Aff1* | 2466.059172 | 4668.204088 | 6444.182897 | 2679.687151 | 2010.410664 | 2020.020228 | 1960.829514 | 2030.439521 | 0.007498544 | -1.019116814 | 0.000977732 | 10.86926745 | 0.49341832 | 2.125023038 |
| *Ltbp2* | 346.9001898 | 320.0393221 | 306.974535 | 404.1495375 | 182.3641575 | 144.8717911 | 162.7168548 | 191.2229001 | 2.12E-10 | -1.019353701 | 1.76E-12 | 49.73493941 | 0.493337308 | 9.674552617 |
| *Scd1* | 50.73710259 | 38.55895447 | 57.05846376 | 76.14411576 | 17.61972537 | 31.10241842 | 31.99488718 | 29.55263002 | 0.005946182 | -1.02083796 | 0.000707777 | 11.46871288 | 0.492830019 | 2.225761832 |
| *Pxdn* | 55.45683307 | 47.55604385 | 70.75249506 | 44.90550417 | 26.42958805 | 18.0066633 | 41.13628352 | 21.72987502 | 0.008017714 | -1.031544517 | 0.001069763 | 10.70275192 | 0.489186157 | 2.095949434 |
| *Tubb6* | 1950.428618 | 2160.586749 | 2608.712963 | 2310.681051 | 1113.566643 | 1140.967665 | 1060.401975 | 1102.139261 | 2.67E-20 | -1.031834039 | 7.90E-23 | 96.74261876 | 0.489087996 | 19.57389725 |
| *Dyrk2* | 305.6025482 | 511.548796 | 788.5479691 | 350.4581738 | 260.7719354 | 233.2681382 | 220.3076517 | 239.8978202 | 0.005370763 | -1.035232527 | 0.000622875 | 11.7063391 | 0.48793723 | 2.269963985 |
| *Eif4ebp1* | 1188.192147 | 1827.694442 | 2690.877151 | 1208.543786 | 870.4144331 | 788.2007617 | 890.3720032 | 821.3892756 | 0.001902756 | -1.036785863 | 0.000164081 | 14.20332221 | 0.487412156 | 2.720616855 |
| *2700081O15Rik* | 100.2942726 | 142.6681315 | 195.139946 | 123.0020332 | 79.28876415 | 83.48543893 | 66.73219326 | 43.45975003 | 0.003155973 | -1.038455141 | 0.000311785 | 12.99823232 | 0.486848519 | 2.500866747 |
| *Sema7a* | 160.4708361 | 137.5269376 | 78.74067998 | 155.2168514 | 59.02607998 | 75.30059198 | 79.53014813 | 45.19814003 | 0.00343362 | -1.038630642 | 0.000348115 | 12.7919462 | 0.486789299 | 2.464247762 |
| *Gm14420* | 184.0694885 | 140.0975346 | 122.1051124 | 164.9789175 | 66.07397013 | 60.56786746 | 90.49982373 | 80.83513506 | 1.04E-05 | -1.039749127 | 3.05E-07 | 26.2186579 | 0.48641205 | 4.984798736 |
| *Hspb1* | 36.57791117 | 30.84716358 | 36.5174168 | 34.16723143 | 22.90564298 | 9.821816345 | 19.19693231 | 15.64551001 | 0.004806166 | -1.039873926 | 0.000537479 | 11.98092609 | 0.486369975 | 2.31820119 |
| *Polr3d* | 232.4467258 | 395.8719326 | 515.8085124 | 238.1944134 | 170.9113361 | 172.7002707 | 169.1158322 | 159.0626851 | 0.002265996 | -1.040129335 | 0.000203923 | 13.79458362 | 0.486283877 | 2.644740949 |
| *Mvp* | 2908.533904 | 4398.291407 | 7102.637568 | 3208.791135 | 2162.821289 | 2099.413244 | 2040.359662 | 2222.531617 | 0.002317971 | -1.047240851 | 0.000210363 | 13.73617587 | 0.483892723 | 2.634892086 |
| *Sertad1* | 503.8312281 | 694.0611805 | 996.2407772 | 442.2215954 | 336.5367545 | 327.3938782 | 319.9488718 | 289.4419352 | 0.000924354 | -1.049489535 | 6.60E-05 | 15.92132426 | 0.483139082 | 3.034161644 |
| *Mif* | 2204.114131 | 3411.182172 | 5890.715798 | 2208.179357 | 1713.518292 | 1595.226671 | 1578.719147 | 1715.790931 | 0.008035576 | -1.054372442 | 0.001074865 | 10.69394899 | 0.481506627 | 2.094982974 |
| *Nrp2* | 11947.99769 | 16600.9152 | 24138.01251 | 13258.83821 | 8075.120136 | 7658.561295 | 7883.540201 | 8109.589356 | 6.75E-05 | -1.055568285 | 2.76E-06 | 21.97632223 | 0.481107673 | 4.170512095 |
| *Egr2* | 96.75447471 | 86.11499832 | 43.36443246 | 115.1923803 | 43.16832715 | 49.92756642 | 31.08074754 | 39.98297003 | 0.008987629 | -1.056964628 | 0.001230128 | 10.44453576 | 0.480642248 | 2.046354877 |
| *Ccdc88b* | 1646.006003 | 2907.345167 | 4791.769786 | 2076.391464 | 1382.267455 | 1328.400661 | 1369.381171 | 1393.319586 | 0.006557117 | -1.061214749 | 0.000810444 | 11.2171612 | 0.479228379 | 2.183287077 |
| *Acot7* | 1635.386609 | 2039.768692 | 2994.428178 | 1666.384687 | 970.8468677 | 1073.85192 | 950.705219 | 998.7050557 | 1.59E-05 | -1.061331894 | 5.10E-07 | 25.22676299 | 0.479189468 | 4.797800826 |
| *Ptpn14* | 48.37723735 | 41.12955144 | 44.50560173 | 38.07205788 | 11.45282149 | 27.00999495 | 18.28279267 | 25.20665502 | 0.003252499 | -1.063190506 | 0.000323923 | 12.92673092 | 0.47857253 | 2.487782828 |
| *Hcls1* | 7419.416305 | 11473.85955 | 18939.98646 | 7791.104973 | 5597.786749 | 5543.596842 | 5315.72197 | 5335.988109 | 0.00383172 | -1.065916866 | 0.000400613 | 12.52933414 | 0.477668993 | 2.416606285 |
| *Ccl6* | 669.0217946 | 1007.67401 | 1637.57791 | 749.7266783 | 496.8762554 | 486.1799091 | 499.12024 | 454.5889853 | 0.001973845 | -1.069129841 | 0.000170842 | 14.12734208 | 0.476606377 | 2.704686854 |
| *Riok3* | 1858.393874 | 3479.302992 | 5117.00303 | 1939.722539 | 1441.293535 | 1510.104263 | 1554.951517 | 1380.281661 | 0.008303705 | -1.07410559 | 0.001117921 | 10.62130473 | 0.474965428 | 2.080728084 |
| *Rcn3* | 58.99663092 | 47.55604385 | 45.64677101 | 88.83480172 | 18.50071164 | 33.55787251 | 30.16660791 | 32.16021502 | 0.00547115 | -1.081332227 | 0.000637229 | 11.66395133 | 0.472592216 | 2.26192141 |
| *Mtdh* | 2762.22226 | 4737.610206 | 6500.100191 | 2795.855738 | 1992.790939 | 1942.264182 | 2108.005995 | 1879.199591 | 0.002038376 | -1.084057955 | 0.000179007 | 14.03953259 | 0.471700176 | 2.690715741 |
| *Nid1* | 108.5538009 | 96.39738618 | 92.43471129 | 95.66824801 | 48.45424476 | 55.65695929 | 31.08074754 | 49.54411504 | 2.61E-06 | -1.084140036 | 6.18E-08 | 29.30741775 | 0.47167334 | 5.58364792 |
| *Arap1* | 4395.249004 | 7686.084925 | 12353.1574 | 4672.124847 | 3471.966884 | 3375.430884 | 3337.523803 | 3515.024583 | 0.00717983 | -1.087150444 | 0.000923556 | 10.974873 | 0.470690146 | 2.143885824 |
| *Tagln* | 265.4848391 | 308.4716358 | 219.1045008 | 258.6947523 | 124.2190638 | 112.1324033 | 119.752292 | 137.3328101 | 5.92E-12 | -1.090313183 | 3.83E-14 | 57.25213397 | 0.469659409 | 11.22791884 |
| *Itpkb* | 1556.331124 | 2619.438307 | 4264.549581 | 2067.605605 | 1233.380776 | 1228.545528 | 1199.351199 | 1265.547921 | 0.002311748 | -1.092711288 | 0.00020952 | 13.74372081 | 0.468879371 | 2.636059569 |
| *Cpeb4* | 2523.875871 | 4974.105127 | 6447.606405 | 2911.048118 | 1978.695159 | 2074.858703 | 1943.460861 | 1900.929466 | 0.002719077 | -1.09369726 | 0.000258835 | 13.34700709 | 0.468559037 | 2.56557856 |
| *AI504432* | 1065.479154 | 2041.05399 | 2713.700536 | 1099.208645 | 825.4841334 | 811.1183331 | 814.4984136 | 788.3598656 | 0.00435266 | -1.094741825 | 0.000472835 | 12.21985844 | 0.468219906 | 2.361245229 |
| *Cd5l* | 532.1496109 | 381.7336493 | 249.9160713 | 512.5084715 | 199.1028966 | 213.6245055 | 199.2824401 | 172.9698051 | 0.000223315 | -1.095031297 | 1.17E-05 | 19.21439265 | 0.468125968 | 3.651081469 |
| *Vgll3* | 35.39797855 | 30.84716358 | 26.24689333 | 28.30999176 | 8.809862684 | 15.55120921 | 11.88381524 | 19.99148501 | 0.005281847 | -1.096440859 | 0.000608366 | 11.75020299 | 0.467668817 | 2.27721419 |
| *Spred2* | 154.571173 | 242.9214132 | 422.2326318 | 174.7409836 | 117.1711737 | 117.8617961 | 97.8129408 | 130.3792501 | 0.007162937 | -1.101535991 | 0.00092022 | 10.98157789 | 0.466020075 | 2.144908897 |
| *Hspg2* | 318.581807 | 269.9126813 | 271.5982875 | 325.0768019 | 164.7444322 | 108.0399798 | 138.9492243 | 139.0712001 | 2.48E-10 | -1.108347426 | 2.08E-12 | 49.40501962 | 0.463825028 | 9.605738366 |
| *Cdkn2a* | 54.27690045 | 35.98835751 | 30.81157043 | 37.09585127 | 21.14367044 | 15.55120921 | 21.02521157 | 15.64551001 | 0.001611072 | -1.110218077 | 0.000130937 | 14.62824339 | 0.463224005 | 2.792885091 |
| *Scd2* | 1106.776796 | 1173.477514 | 1364.838453 | 1399.880282 | 553.2593765 | 567.2098939 | 596.0190412 | 617.1284504 | 4.66E-23 | -1.113288162 | 1.08E-25 | 109.8056157 | 0.462239303 | 22.33168082 |
| *Rhoq* | 2042.463362 | 3846.898358 | 5686.446498 | 1978.770803 | 1464.199178 | 1526.473957 | 1603.400917 | 1649.732111 | 0.008511065 | -1.118215388 | 0.001153855 | 10.56281276 | 0.460663313 | 2.070016075 |
| *Mrpl52* | 297.3430198 | 471.704543 | 742.9011981 | 374.8633391 | 211.4367044 | 237.3605617 | 212.080395 | 206.8684102 | 0.00101246 | -1.120269135 | 7.36E-05 | 15.71545486 | 0.460008003 | 2.994622254 |
| *Ninj1* | 3110.302382 | 4244.055589 | 6180.572794 | 3581.702061 | 1978.695159 | 2036.389922 | 1996.48096 | 1857.469716 | 9.27E-06 | -1.121106792 | 2.65E-07 | 26.48627588 | 0.459740991 | 5.033034217 |
| *Ube2e2* | 194.688882 | 190.2241754 | 314.9627199 | 166.9313307 | 93.38454445 | 113.7693727 | 91.41396337 | 98.21903507 | 3.87E-05 | -1.125975845 | 1.43E-06 | 23.2411755 | 0.458191994 | 4.412650845 |
| *Bcat1* | 30.67824808 | 43.7001484 | 35.37624753 | 31.23861159 | 15.85775283 | 13.91423982 | 11.88381524 | 22.59907002 | 0.001815086 | -1.126183415 | 0.000154633 | 14.31493758 | 0.458126075 | 2.741102788 |
| *Bcl2l1* | 613.5649616 | 1367.557585 | 1807.612132 | 837.5852734 | 515.376967 | 574.5762562 | 490.8929833 | 537.1625104 | 0.004712989 | -1.126932339 | 0.000525257 | 12.02379229 | 0.457888317 | 2.32670353 |
| *Hspa1b* | 129.792588 | 169.6593997 | 279.5864724 | 140.5737522 | 81.93172296 | 82.66695423 | 83.18670666 | 80.83513506 | 0.000420272 | -1.130270525 | 2.49E-05 | 17.76869912 | 0.456830055 | 3.376469413 |
| *Tnc* | 697.3401775 | 907.4207286 | 1210.780601 | 732.1549593 | 414.9445324 | 390.4171997 | 404.0497181 | 409.3908453 | 2.29E-07 | -1.131721955 | 3.77E-09 | 34.74010379 | 0.45637069 | 6.640824047 |
| *Spryd7* | 648.9629401 | 1082.221322 | 1747.13016 | 710.6784138 | 497.7572416 | 523.830205 | 482.6657266 | 405.9140653 | 0.004048751 | -1.132719465 | 0.000428485 | 12.40369354 | 0.456055255 | 2.392678976 |
| *Tpm2* | 71.97588972 | 116.9621619 | 75.31717216 | 108.358934 | 28.19156059 | 48.29059703 | 34.73730608 | 58.23606504 | 0.00157683 | -1.134927773 | 0.000127272 | 14.68176638 | 0.455357713 | 2.802215033 |
| *Ncam1* | 30.67824808 | 43.7001484 | 31.9527397 | 23.4289587 | 12.33380776 | 9.003331649 | 19.19693231 | 18.25309501 | 0.008478476 | -1.13834621 | 0.001145558 | 10.57615374 | 0.454280029 | 2.071682202 |
| *Susd6* | 2294.968943 | 4494.688793 | 7018.191042 | 2560.589944 | 1843.023273 | 1830.950264 | 1897.75388 | 1856.600521 | 0.007904249 | -1.139763377 | 0.0010496 | 10.73795912 | 0.453834007 | 2.102139397 |
| *Osbpl3* | 667.841862 | 1334.139825 | 2034.704818 | 803.418042 | 574.403047 | 543.4738377 | 529.2868479 | 546.7236554 | 0.006471022 | -1.141399031 | 0.000797166 | 11.24781948 | 0.453319765 | 2.189027149 |
| *Actg2* | 49.55716997 | 56.55313323 | 45.64677101 | 37.09585127 | 20.26268417 | 27.00999495 | 19.19693231 | 18.25309501 | 0.000172564 | -1.142815318 | 8.52E-06 | 19.8181333 | 0.452874961 | 3.763049515 |
| *Lrrc32* | 27.13845022 | 35.98835751 | 39.94092463 | 39.04826449 | 21.14367044 | 14.73272452 | 10.05553597 | 18.25309501 | 0.001751979 | -1.14808713 | 0.000147715 | 14.40111186 | 0.451223112 | 2.756471058 |
| *Itga11* | 53.09696783 | 46.27074537 | 50.21144811 | 61.50101658 | 22.90564298 | 26.19151025 | 21.93935121 | 24.33746002 | 7.77E-06 | -1.151252606 | 2.15E-07 | 26.8932505 | 0.45023415 | 5.109675835 |
| *Sulf1* | 31.8581807 | 29.5618651 | 27.3880626 | 31.23861159 | 15.85775283 | 9.003331649 | 9.141396337 | 19.99148501 | 0.004388623 | -1.154813035 | 0.000477444 | 12.20176218 | 0.449124387 | 2.357671758 |
| *Col5a2* | 507.3710259 | 490.9840203 | 520.3731895 | 535.9374302 | 223.7705122 | 199.7102657 | 255.0449578 | 243.3746002 | 1.63E-27 | -1.158969279 | 3.14E-30 | 130.5314811 | 0.447832371 | 26.78700072 |
| *Slc6a12* | 273.7443675 | 429.2896931 | 770.2892607 | 268.4568184 | 159.4585146 | 219.3538984 | 192.8834627 | 206.8684102 | 0.008511065 | -1.161207715 | 0.001154754 | 10.56137263 | 0.447138069 | 2.070016075 |
| *Ifrd1* | 1238.929249 | 2431.784729 | 3627.777126 | 1293.473761 | 883.6292272 | 1015.739507 | 977.2152684 | 963.0680607 | 0.006871256 | -1.161900805 | 0.000867908 | 11.09007961 | 0.446923309 | 2.162963892 |
| *Psmd10* | 768.1361346 | 1087.362516 | 1797.341608 | 754.6077113 | 482.7804751 | 506.6420264 | 514.6606138 | 455.4581803 | 0.000842133 | -1.169270803 | 5.82E-05 | 16.16022345 | 0.444646026 | 3.074619391 |
| *Pgk1* | 13825.27049 | 22612.2562 | 36577.89878 | 13476.53228 | 9397.480525 | 9817.723921 | 9765.753707 | 9203.905862 | 0.002760474 | -1.179554682 | 0.000264322 | 13.30767544 | 0.441487752 | 2.559016319 |
| *Blvrb* | 1336.863657 | 2173.439734 | 3552.459954 | 1503.358183 | 922.392623 | 969.904364 | 915.0537733 | 972.6292057 | 0.001461045 | -1.180178942 | 0.000115822 | 14.85951348 | 0.441296759 | 2.835336314 |

| | | | | | | | | | | | | | | |
|---|---|---|---|---|---|---|---|---|---|---|---|---|---|---|
| *Nfkbid* | 256.0453782 | 375.3071569 | 512.3850045 | 310.4337027 | 156.8155558 | 152.2381533 | 154.4895981 | 177.3157801 | 7.92E-06 | -1.181785335 | 2.20E-07 | 26.85038927 | 0.440805663 | 5.101297628 |
| *Tank* | 979.3440733 | 1640.040864 | 2398.737816 | 1230.996538 | 675.7164678 | 753.8244045 | 647.2108606 | 676.2337105 | 0.000284194 | -1.182506215 | 1.56E-05 | 18.66610787 | 0.440585458 | 3.546384509 |
| *Abca1* | 6552.16583 | 12810.56997 | 20910.7858 | 7674.936386 | 5152.888684 | 5036.136331 | 5701.488895 | 5225.600344 | 0.006616341 | -1.18312908 | 0.000824029 | 11.18631427 | 0.440395283 | 2.179382104 |
| *Col4a2* | 853.0912831 | 1159.339231 | 1751.694837 | 801.4656287 | 481.0185025 | 495.1832407 | 481.7515869 | 545.8544604 | 0.000110312 | -1.187829881 | 5.02E-06 | 20.82925292 | 0.438962657 | 3.957375773 |
| *Tmem171* | 90.85481162 | 186.3682799 | 280.7276417 | 130.8116861 | 70.47890147 | 73.66362258 | 92.328103 | 64.32043005 | 0.005973352 | -1.195053138 | 0.000714834 | 11.45027672 | 0.43677036 | 2.223781918 |
| *Fbn1* | 277.2841653 | 311.0422327 | 373.162353 | 292.8619837 | 137.4338579 | 144.0533064 | 133.4643865 | 129.5100551 | 1.75E-17 | -1.203053082 | 6.57E-20 | 83.44037295 | 0.434355111 | 16.75780534 |
| *Col6a2* | 119.1731945 | 113.1062665 | 122.1051124 | 118.1210001 | 44.93029969 | 52.3830205 | 63.07563472 | 45.19814003 | 1.63E-11 | -1.203122728 | 1.14E-13 | 55.11535731 | 0.434334143 | 10.78696926 |
| *Tpm4* | 6323.258902 | 12779.72281 | 16965.76361 | 7151.689642 | 4552.056049 | 4694.828213 | 4805.632054 | 4711.036903 | 0.001635558 | -1.203756603 | 0.000134059 | 14.5838414 | 0.434143352 | 2.78633398 |
| *5730508B09Rik* | 154.571173 | 289.1921585 | 478.1499263 | 199.1461489 | 117.1711737 | 126.0466431 | 106.0401975 | 135.5944201 | 0.004904628 | -1.208638694 | 0.000550844 | 11.93516058 | 0.43267669 | 2.309393889 |
| *Cnn3* | 87.31501376 | 118.2474604 | 141.5049901 | 80.04894221 | 51.97818983 | 51.56453581 | 29.25246828 | 51.28250504 | 0.000271619 | -1.209786642 | 1.48E-05 | 18.76637774 | 0.432332548 | 3.566039518 |
| *Ptprj* | 2973.430198 | 5513.930489 | 9799.220566 | 3472.36692 | 2458.832675 | 2352.325015 | 2291.748062 | 2299.889972 | 0.006598968 | -1.210421493 | 0.000818207 | 11.19947106 | 0.432142344 | 2.180524008 |
| *Gdpd1* | 476.6927778 | 694.0611805 | 1066.993272 | 536.9136368 | 295.1303999 | 330.6678169 | 267.8429127 | 302.4798602 | 0.000104161 | -1.213559962 | 4.71E-06 | 20.95003986 | 0.431203274 | 3.982296304 |
| *Tpi1* | 7447.734687 | 13302.83929 | 22696.71571 | 7299.09684 | 5561.666312 | 5458.474434 | 5466.555009 | 5376.840274 | 0.006559779 | -1.214763344 | 0.000812298 | 11.21291988 | 0.430843748 | 2.183110768 |
| *Mitf* | 558.1081285 | 924.1296088 | 1467.543688 | 687.2494551 | 410.5396011 | 387.1432609 | 380.2820876 | 388.5301653 | 0.000596071 | -1.21508263 | 3.81E-05 | 16.96150343 | 0.430748408 | 3.224702127 |
| *Cd302* | 589.9663092 | 1165.765724 | 1863.529426 | 640.3915377 | 439.6121479 | 499.2756642 | 454.3273979 | 440.6818653 | 0.006923761 | -1.215619039 | 0.000878209 | 11.06819955 | 0.430588281 | 2.159657923 |
| *Ptafr* | 1827.715626 | 2913.77166 | 4040.880403 | 2188.655225 | 1159.377929 | 1140.967665 | 1160.957335 | 1235.126096 | 2.14E-05 | -1.223997612 | 7.15E-07 | 24.57531131 | 0.428094851 | 4.66911941 |
| *Gclm* | 2480.218364 | 3940.725147 | 5999.126879 | 2315.562084 | 1555.82175 | 1568.216676 | 1574.148449 | 1604.533971 | 0.000868047 | -1.225195785 | 6.08E-05 | 16.07830721 | 0.427739461 | 3.061456687 |
| *Fam49a* | 795.2745848 | 1406.11654 | 2118.010175 | 990.8497115 | 575.2840332 | 584.3980725 | 536.599965 | 568.4535304 | 0.000565757 | -1.229292503 | 3.55E-05 | 17.09655455 | 0.426526563 | 3.247370063 |
| *Timp3* | 346.9001898 | 543.6812581 | 728.0659975 | 369.0060995 | 192.9359928 | 190.706934 | 224.8783499 | 238.1594302 | 6.89E-05 | -1.23084616 | 2.85E-06 | 21.91535125 | 0.426067478 | 4.161784219 |
| *Acta2* | 625.3642878 | 650.3610321 | 578.5728225 | 630.6294716 | 266.9388393 | 277.4663117 | 249.56012 | 261.6276952 | 3.61E-49 | -1.233772124 | 2.31E-52 | 231.8904645 | 0.425204236 | 48.44307426 |
| *Lmo4* | 1098.517268 | 1935.659514 | 3189.568124 | 1223.186885 | 787.6017239 | 766.1016749 | 771.5338508 | 828.3428356 | 0.002287206 | -1.239570244 | 0.000206631 | 13.76980454 | 0.423498791 | 2.640694677 |
| *Osgin2* | 422.4158774 | 802.026253 | 1327.179867 | 432.4595293 | 276.6296883 | 329.8493322 | 352.8578986 | 303.3490552 | 0.007541633 | -1.240570733 | 0.00098516 | 10.85525102 | 0.423205202 | 2.122534625 |
| *Smpdl3b* | 490.8519693 | 714.6259562 | 1087.534319 | 459.7933144 | 281.9156059 | 288.1066128 | 287.039845 | 306.8258352 | 0.000237636 | -1.241597099 | 1.26E-05 | 19.07254891 | 0.422904232 | 3.62408754 |
| *Rnf19b* | 3058.385347 | 3057.72509 | 4537.289038 | 3324.959722 | 1532.035121 | 1468.361544 | 1442.512342 | 1465.462771 | 1.69E-14 | -1.242432526 | 8.51E-17 | 69.28677454 | 0.42265941 | 13.77226937 |
| *Spata13* | 1760.459467 | 2864.930317 | 4492.783436 | 2076.391464 | 1256.286419 | 1140.967665 | 1121.649331 | 1204.704271 | 0.000316026 | -1.244785555 | 1.76E-05 | 18.42790614 | 0.421970616 | 3.500277141 |
| *Lamb1* | 96.75447471 | 106.679774 | 106.1287426 | 106.4065207 | 32.59649193 | 49.10908172 | 44.79284205 | 48.67492004 | 1.65E-10 | -1.246139774 | 1.34E-12 | 50.26528812 | 0.421574709 | 9.783565478 |
| *Tnn* | 73.15582234 | 109.250371 | 117.5404353 | 60.52480997 | 35.23945073 | 31.92090312 | 37.47972498 | 46.93653003 | 0.00019465 | -1.24617202 | 9.83E-06 | 19.54511631 | 0.421565287 | 3.710745033 |
| *Mocs1* | 536.8693414 | 1047.518263 | 1958.246476 | 654.0584303 | 466.9227222 | 455.8959753 | 405.8779974 | 436.3358903 | 0.009571256 | -1.24942566 | 0.00133604 | 10.29205731 | 0.420615622 | 2.019031063 |
| *Clec5a* | 1689.66351 | 2240.275255 | 3134.791999 | 1872.364282 | 989.3475794 | 951.079216 | 877.5740483 | 937.8614057 | 6.87E-08 | -1.250558984 | 9.68E-10 | 37.38867909 | 0.420285333 | 7.162772033 |
| *St3gal6* | 36.57791117 | 38.55895447 | 23.96455478 | 45.88171078 | 7.928876415 | 15.55120921 | 14.62623414 | 22.59907002 | 0.005559784 | -1.257646114 | 0.000651415 | 11.62299267 | 0.418225776 | 2.254942076 |
| *Zswim4* | 568.7275221 | 1002.532816 | 1904.61152 | 628.6770583 | 433.445244 | 439.5262814 | 421.4183711 | 418.9519903 | 0.006609245 | -1.260282011 | 0.000821067 | 11.19299532 | 0.417462348 | 2.179848127 |
| *St18* | 240.7062542 | 463.9927521 | 694.9720886 | 276.2664713 | 182.3641575 | 187.4329952 | 166.3734133 | 162.5394651 | 0.002287206 | -1.261655355 | 0.000206747 | 13.76874948 | 0.417065143 | 2.640694677 |
| *Slit2* | 23.59865237 | 24.42067117 | 17.11753913 | 21.47654547 | 6.166903878 | 7.366362258 | 15.54037377 | 6.953560005 | 0.009171717 | -1.271348429 | 0.001263395 | 10.39525627 | 0.414272388 | 2.03754936 |
| *Kcna3* | 336.2807962 | 694.0611805 | 1133.18109 | 430.507116 | 315.3930841 | 224.2648065 | 269.6711919 | 265.1044752 | 0.005584785 | -1.271524803 | 0.000655378 | 11.61171204 | 0.414221745 | 2.252993529 |
| *Col12a1* | 414.1563491 | 383.0189478 | 445.0560173 | 401.2209177 | 177.0782399 | 139.1423982 | 187.3986249 | 178.1849751 | 9.11E-23 | -1.271624977 | 2.19E-25 | 108.4091222 | 0.414192985 | 22.04042876 |
| *Slc25a37* | 1308.545274 | 2535.893906 | 4358.125462 | 1477.976811 | 968.2039089 | 1023.105869 | 1028.407088 | 977.8443757 | 0.00509416 | -1.275903774 | 0.000579984 | 11.83914749 | 0.412966376 | 2.292927404 |
| *Slc2a6* | 2307.948202 | 3916.304476 | 5497.012398 | 2814.403663 | 1518.820327 | 1493.734569 | 1424.229549 | 1540.213541 | 4.20E-05 | -1.282045772 | 1.57E-06 | 23.05555039 | 0.411211988 | 4.376330879 |
| *Mafg* | 462.5335864 | 963.9738618 | 1660.401295 | 572.0570748 | 355.9184524 | 368.3181129 | 398.5648803 | 377.2306303 | 0.006557117 | -1.286318101 | 0.000810264 | 11.21757238 | 0.409996045 | 2.183287077 |
| *Rnf113a1* | 37.75784379 | 50.12664081 | 75.31717216 | 28.30999176 | 15.85775283 | 19.64363269 | 21.02521157 | 21.72987502 | 0.003580318 | -1.286760679 | 0.000366882 | 12.69374062 | 0.409870289 | 2.44607843 |
| *Cnn2* | 493.2118345 | 975.5415481 | 1499.496428 | 532.0326037 | 331.2508369 | 368.3181129 | 355.6003175 | 377.2306303 | 0.003180787 | -1.288790374 | 0.000315286 | 12.97732825 | 0.409294058 | 2.497465441 |
| *Itga5* | 6796.411882 | 9782.406749 | 14774.71861 | 8396.353073 | 4197.018582 | 4049.043788 | 4065.178951 | 3944.406913 | 2.13E-06 | -1.290002611 | 4.81E-08 | 29.79328072 | 0.408950289 | 5.671470907 |
| *1700047I17Rik2* | 1308.545274 | 3079.575164 | 4275.961274 | 1537.525414 | 984.942648 | 1076.307374 | 1046.689881 | 1063.025486 | 0.004390086 | -1.290251153 | 0.000477954 | 12.19976921 | 0.408879843 | 2.357527011 |
| *Slc31a1* | 2109.719522 | 4529.391852 | 7601.328542 | 2262.846927 | 1696.779553 | 1657.431508 | 1628.082688 | 1764.465851 | 0.009200608 | -1.290443871 | 0.001268846 | 10.38730584 | 0.408825228 | 2.036183488 |
| *Nupr1* | 283.1838284 | 407.4396189 | 608.2432237 | 371.9347193 | 162.9824596 | 174.3372401 | 165.4592737 | 175.5773901 | 2.01E-06 | -1.300270305 | 4.47E-08 | 29.93496573 | 0.406050113 | 5.697003685 |
| *Cemip* | 43.65750688 | 30.84716358 | 51.35261738 | 39.04826449 | 19.3816979 | 18.82514799 | 20.11107194 | 8.691950006 | 0.000715332 | -1.302357934 | 4.79E-05 | 16.52765282 | 0.40546297 | 3.145492458 |
| *Adam19* | 138.0521164 | 176.0858921 | 191.7164382 | 154.2406448 | 68.71692893 | 58.93089807 | 71.30289143 | 68.66640505 | 1.51E-13 | -1.302889978 | 8.56E-16 | 64.73788849 | 0.405313469 | 12.82207393 |
| *Pf4* | 464.8934517 | 975.5415481 | 1451.567318 | 560.3425955 | 360.3233838 | 342.1266027 | 324.51957 | 368.5386803 | 0.002308451 | -1.306555726 | 0.000209037 | 13.74806032 | 0.404284914 | 2.636679414 |
| *Ddr1* | 17.69898928 | 25.70596965 | 17.11753913 | 25.38137192 | 6.166903878 | 13.91423982 | 9.141396337 | 5.215170004 | 0.00964995 | -1.312798605 | 0.001349222 | 10.2739421 | 0.402539258 | 2.015474922 |
| *Hbegf* | 61.35649616 | 71.97671501 | 91.29354201 | 57.59619013 | 35.23945073 | 26.19151025 | 28.33832864 | 23.46826502 | 3.26E-06 | -1.315832293 | 7.96E-08 | 28.81695304 | 0.401693691 | 5.486705481 |
| *Acsl4* | 1743.94041 | 3828.904179 | 6134.926023 | 1784.505687 | 1384.029428 | 1403.701253 | 1348.35596 | 1258.594361 | 0.007805318 | -1.322464257 | 0.001027722 | 10.77694426 | 0.399851372 | 2.107609396 |
| *Fat4* | 21.23878713 | 11.56768634 | 17.11753913 | 25.38137192 | 7.928876415 | 9.003331649 | 5.484837802 | 7.822755006 | 0.007994673 | -1.32623337 | 0.001064166 | 10.71245768 | 0.398808104 | 2.097199271 |
| *Ralgapa2* | 1740.400612 | 2557.74398 | 3085.72172 | 2119.344555 | 926.7975543 | 906.8810425 | 990.9273629 | 961.3296707 | 8.21E-11 | -1.327723282 | 6.17E-13 | 51.79111614 | 0.398396456 | 10.08568339 |
| *Grap* | 147.4915773 | 169.6593997 | 245.3513942 | 144.4785786 | 85.45566803 | 68.75271441 | 58.50493656 | 68.66640505 | 5.74E-07 | -1.327998952 | 1.06E-08 | 32.72642718 | 0.398320337 | 6.241013045 |
| *Egfr* | 47.19730474 | 43.7001484 | 96.99938839 | 31.23861159 | 22.02465671 | 18.0066633 | 21.02521157 | 26.07585002 | 0.006789534 | -1.32951712 | 0.000851584 | 11.12529302 | 0.3979014 | 2.168160004 |
| *Ift57* | 318.581807 | 544.9665565 | 1029.334686 | 334.838868 | 222.8895259 | 208.7135973 | 198.3683005 | 248.5897702 | 0.004147115 | -1.342111482 | 0.000441881 | 12.34621603 | 0.394442939 | 2.382253927 |
| *Gpr132* | 243.0661194 | 398.4425295 | 527.2202051 | 358.2678267 | 147.1247068 | 140.7793676 | 163.6309944 | 149.5015401 | 8.13E-07 | -1.345434224 | 1.57E-08 | 31.95894133 | 0.393535524 | 6.089684101 |
| *Snx18* | 1834.795222 | 4069.254995 | 7034.167412 | 2085.177324 | 1519.701313 | 1467.543059 | 1406.860896 | 1474.154721 | 0.007318647 | -1.356162348 | 0.00094725 | 10.92793559 | 0.390619981 | 2.135569201 |
| *Icosl* | 969.9046123 | 1624.617282 | 2119.151344 | 1312.021687 | 573.5220607 | 536.9259602 | 650.8674192 | 590.1834054 | 8.54E-07 | -1.357563401 | 1.68E-08 | 31.83246187 | 0.39024082 | 6.068525155 |
| *Psma6* | 2171.076018 | 5118.058557 | 8173.054349 | 2318.490704 | 1816.593685 | 1758.105126 | 1692.986602 | 1654.078086 | 0.008037711 | -1.361055248 | 0.00107761 | 10.68923063 | 0.389297437 | 2.094867594 |
| *Slc11a1* | 2704.405561 | 4864.854756 | 9329.058824 | 2501.041341 | 1836.85637 | 1868.60056 | 1939.804303 | 1900.060271 | 0.006616341 | -1.362324838 | 0.0008244 | 11.18547934 | 0.388955001 | 2.179382104 |
| *Col16a1* | 24.77858499 | 25.70596965 | 25.10572405 | 27.33378515 | 9.690848952 | 6.547877563 | 6.398977436 | 17.38390001 | 0.003080438 | -1.364645037 | 0.000302352 | 13.05576917 | 0.388329971 | 2.511387475 |
| *Nfkbib* | 521.5302173 | 1161.909828 | 1857.82358 | 677.487389 | 414.0635461 | 374.8659905 | 452.4991187 | 387.6609703 | 0.003131173 | -1.372671295 | 0.000308584 | 13.0175595 | 0.386175543 | 2.504292958 |
| *Pik3r5* | 5766.330706 | 9763.127272 | 15113.64588 | 7181.952047 | 3786.478981 | 3464.645716 | 3674.841327 | 3677.564048 | 5.81E-05 | -1.37301435 | 2.32E-06 | 22.3137646 | 0.386083726 | 4.235656714 |
| *H2-Q8* | 238.3463889 | 204.3624587 | 237.3632092 | 297.7430168 | 88.9796131 | 89.2148318 | 99.64122007 | 99.95742507 | 2.73E-16 | -1.376927314 | 1.09E-18 | 77.88337067 | 0.385037986 | 15.56337424 |
| *Myo10* | 998.2229952 | 1314.860347 | 2229.844764 | 1140.209323 | 524.1868278 | 546.7477765 | 552.1403387 | 558.8923854 | 3.59E-06 | -1.38101449 | 8.94E-08 | 28.59048829 | 0.383948711 | 5.444379247 |
| *Aebp1* | 458.9937886 | 421.5779022 | 513.5261738 | 378.7681656 | 143.6007617 | 162.0599697 | 164.5451341 | 206.8684102 | 6.10E-16 | -1.387945141 | 2.54E-18 | 76.22031526 | 0.38210866 | 15.21470714 |
| *Rhbdf2* | 1584.649507 | 3039.730911 | 5040.544688 | 1719.099844 | 1169.949764 | 994.4589049 | 1107.937236 | 1076.932606 | 0.001788779 | -1.388118812 | 0.000151247 | 14.35661851 | 0.382062664 | 2.747443233 |
| *Cers6* | 733.9180886 | 1902.241754 | 2789.017708 | 909.8245627 | 619.3333467 | 572.1208021 | 622.5290905 | 603.2213304 | 0.005277756 | -1.389903913 | 0.000606712 | 11.75527059 | 0.381590216 | 2.277550692 |
| *Mrc1* | 4168.701941 | 3958.719326 | 1526.88449 | 5157.299533 | 1319.71743 | 1258.829462 | 1559.522215 | 1506.314936 | 0.000621682 | -1.391907558 | 4.04E-05 | 16.85338669 | 0.381060624 | 3.206431966 |
| *Mapk11* | 56.63676568 | 38.55895447 | 60.48197158 | 64.42963641 | 31.71550566 | 11.45878574 | 15.54037377 | 25.20665502 | 0.00115697 | -1.397600499 | 8.72E-05 | 15.39579017 | 0.379559904 | 2.936677785 |
| *Mrc2* | 76.6956202 | 59.12373019 | 70.75249506 | 52.71515707 | 24.66761551 | 31.10241842 | 27.42418901 | 14.77631501 | 8.11E-06 | -1.40008203 | 2.26E-07 | 26.79945907 | 0.378907597 | 5.091101955 |
| *Uty* | 292.6232894 | 350.8864857 | 308.1157043 | 328.0054217 | 110.1232835 | 132.5945207 | 119.752292 | 121.6873001 | 2.51E-31 | -1.40026421 | 3.62E-34 | 148.5379858 | 0.378859752 | 30.59979639 |
| *Myadm* | 2463.699307 | 5759.4225 | 8610.122181 | 2813.427457 | 1808.664809 | 1858.778743 | 1895.9256 | 1868.769251 | 0.003105713 | -1.40239533 | 0.000305578 | 13.03589153 | 0.378300521 | 2.507838667 |
| *Fstl1* | 270.2045696 | 383.0189478 | 385.715215 | 262.5995787 | 136.5528716 | 117.0433114 | 120.6664316 | 117.3413251 | 2.10E-12 | -1.402988 | 1.33E-14 | 59.34109009 | 0.378145145 | 11.67820726 |
| *Ust* | 293.803222 | 318.7540236 | 332.0802591 | 287.004744 | 112.7662423 | 103.1290716 | 121.5805713 | 126.9024701 | 1.08E-30 | -1.407228023 | 1.64E-33 | 145.5348825 | 0.377035423 | 29.96678506 |
| *Loxl2* | 292.6232894 | 403.5837235 | 408.5386005 | 300.6716366 | 132.1479403 | 121.9542196 | 114.2674542 | 160.8010751 | 2.99E-12 | -1.407698844 | 1.91E-14 | 58.61897929 | 0.376912398 | 11.52429049 |
| *Pirb* | 4056.608342 | 8973.954004 | 16251.39165 | 4222.093598 | 3154.811827 | 3189.634858 | 3005.691116 | 3199.506797 | 0.008039601 | -1.416667813 | 0.00107883 | 10.68713749 | 0.374576472 | 2.094765513 |
| *Col6a1* | 231.2667932 | 227.4978314 | 247.6337327 | 256.742339 | 92.50355818 | 93.30725527 | 83.18670666 | 89.52708506 | 7.66E-32 | -1.427278892 | 1.04E-34 | 151.0118936 | 0.371831554 | 31.1157155 |
| *Lphn2* | 456.6339233 | 1221.033558 | 1878.364627 | 533.0088103 | 386.7529718 | 351.1299343 | 387.5952047 | 394.6145303 | 0.007962674 | -1.427456441 | 0.001059269 | 10.72099072 | 0.371785796 | 2.098941067 |
| *Wisp1* | 138.0521164 | 105.3944756 | 58.19963303 | 110.3113472 | 44.04931342 | 39.28726538 | 35.65144571 | 33.89860502 | 2.25E-05 | -1.429528586 | 7.55E-07 | 24.47011789 | 0.371252183 | 4.648522599 |
| *Fam20c* | 6390.515061 | 8853.135947 | 13687.18429 | 6642.10979 | 3415.583762 | 3290.308475 | 3100.761637 | 3334.232022 | 8.39E-07 | -1.436692971 | 1.64E-08 | 31.87439069 | 0.369413125 | 6.076131052 |
| *Hivep2* | 296.1630872 | 575.8137201 | 870.7121569 | 331.9102482 | 206.1507868 | 172.7002707 | 184.656206 | 195.5688751 | 0.000540895 | -1.450222887 | 3.34E-05 | 17.21363696 | 0.36596488 | 3.266887114 |
| *Kdm5d* | 423.59581 | 419.0073053 | 422.2326318 | 404.1495375 | 162.1014734 | 139.9608829 | 144.4340621 | 163.4086601 | 5.65E-50 | -1.452157642 | 3.16E-53 | 235.8474308 | 0.365474425 | 49.24796999 |
| *Abracl* | 971.0845449 | 2163.157346 | 3132.50966 | 950.8252404 | 619.3333467 | 676.8868431 | 678.2916082 | 652.7654455 | 0.002192528 | -1.457816347 | 0.000195909 | 13.8699238 | 0.364043726 | 2.659054831 |
| *Gyk* | 684.3609187 | 1344.422213 | 2455.79628 | 716.5356535 | 464.2797634 | 486.9983938 | 462.5546546 | 471.1036903 | 0.003051952 | -1.464178824 | 0.000298335 | 13.08081946 | 0.36244178 | 2.515422371 |
| *Esd* | 5109.108238 | 10823.49852 | 19154.52628 | 5205.133657 | 3570.637346 | 3868.977155 | 3586.169783 | 3567.176283 | 0.004332937 | -1.465210724 | 0.000470052 | 12.23087208 | 0.362182633 | 2.363217651 |
| *Jdp2* | 739.8177517 | 1525.649299 | 2079.210419 | 975.2304057 | 480.1375163 | 476.3580927 | 453.4132583 | 504.1331004 | 6.54E-05 | -1.474617082 | 2.65E-06 | 22.05333893 | 0.359828888 | 4.184525704 |
| *Cd82* | 1759.279534 | 2818.659572 | 4343.290261 | 2017.819068 | 946.1792522 | 1019.013446 | 1002.811178 | 950.8993307 | 7.38E-06 | -1.480898215 | 2.02E-07 | 27.01602774 | 0.358265688 | 5.132213103 |
| *Cdh11* | 142.7718468 | 158.0917133 | 144.9284979 | 180.5982233 | 67.83594266 | 65.47877563 | 44.79284205 | 46.06733503 | 2.03E-12 | -1.482052974 | 1.27E-14 | 59.42824043 | 0.357979041 | 11.69190797 |
| *Agpat4* | 372.8587074 | 768.6084925 | 1049.875733 | 453.9360747 | 216.722622 | 220.1723831 | 243.1611426 | 265.9736702 | 0.000159323 | -1.483201998 | 7.74E-06 | 20.0020033 | 0.357694044 | 3.797721445 |
| *Oaf* | 41.29764164 | 98.96798314 | 140.3638208 | 50.76274384 | 26.42958805 | 31.92090312 | 27.42418901 | 32.16021502 | 0.003619805 | -1.487888761 | 0.000372665 | 12.66449451 | 0.356533919 | 2.441314868 |
| *Lacc1* | 1094.97747 | 2318.678462 | 3541.048261 | 1287.616522 | 680.1213992 | 718.6295625 | 761.4783148 | 759.6764306 | 0.000607968 | -1.497046825 | 3.90E-05 | 16.91869603 | 0.35427785 | 3.216119458 |
| *Fgr* | 315.0420091 | 384.3042462 | 794.2538155 | 295.7906035 | 157.696542 | 160.4230003 | 136.2068054 | 179.0541701 | 0.000200213 | -1.497988976 | 1.02E-05 | 19.47621183 | 0.354046565 | 3.698507194 |
| *Csf1* | 749.2572127 | 999.9622193 | 1061.287426 | 1103.113472 | 341.8226721 | 311.0241842 | 374.7972498 | 356.3699503 | 4.98E-21 | -1.500561564 | 1.35E-23 | 100.2329513 | 0.353415798 | 20.30263795 |
| *Cybb* | 18098.98643 | 38889.27618 | 77842.57977 | 20226.0248 | 13744.26677 | 14532.19577 | 14149.05325 | 12239.1348 | 0.006044113 | -1.504112376 | 0.000725324 | 11.4232077 | 0.352547028 | 2.218667415 |
| *Mapkapk2* | 4374.010216 | 9435.376159 | 15041.75222 | 5208.062277 | 2936.327232 | 2908.076123 | 2978.266927 | 3182.992092 | 0.000910382 | -1.504309639 | 6.45E-05 | 15.96505055 | 0.352498827 | 3.040776231 |
| *P3h4* | 25.9585176 | 26.99126813 | 34.23507825 | 22.45275208 | 7.047890147 | 9.003331649 | 11.88381524 | 10.43034001 | 0.00011444 | -1.513494791 | 5.24E-06 | 20.74883852 | 0.350261715 | 3.941424027 |
| *Calcrl* | 607.6652985 | 1510.225717 | 2334.832337 | 705.7973807 | 462.5177909 | 472.2656692 | 446.1001412 | 412.8676253 | 0.00274123 | -1.52382561 | 0.000261383 | 13.32864188 | 0.347762527 | 2.562054556 |
| *Pfkl* | 2199.394401 | 5094.923184 | 8969.590503 | 2213.06039 | 1598.990077 | 1605.866972 | 1609.799895 | 1605.403166 | 0.005308778 | -1.525033499 | 0.000612796 | 11.73669846 | 0.347471487 | 2.275005409 |
| *Col5a3* | 109.7337335 | 150.3799224 | 214.5398237 | 98.59686785 | 47.57325849 | 58.11241337 | 35.65144571 | 55.62848004 | 1.86E-05 | -1.538007206 | 6.08E-07 | 24.88751401 | 0.344360792 | 4.731055168 |
| *Rbpms* | 96.75447471 | 181.227086 | 313.8215507 | 95.66824801 | 62.55002505 | 63.02332154 | 51.19181949 | 59.10526004 | 0.001916786 | -1.542074587 | 0.000165597 | 14.18601077 | 0.343391305 | 2.717426292 |
| *Alas1* | 1188.192147 | 3436.888142 | 5929.515554 | 1503.358183 | 993.7525107 | 1056.663742 | 1006.467737 | 1052.595146 | 0.007777552 | -1.552905134 | 0.001022822 | 10.78579119 | 0.340823062 | 2.109157095 |
| *Emr4* | 100.2942726 | 122.1033558 | 206.5516388 | 43.92929756 | 38.76339581 | 44.19817355 | 42.05042315 | 35.63699503 | 0.003468309 | -1.555526106 | 0.000352352 | 12.769314 | 0.340204444 | 2.459882184 |
| *Abcc1* | 4173.421671 | 8596.07625 | 13757.93678 | 4396.834582 | 2622.696121 | 2583.956183 | 2682.085685 | 2606.715807 | 0.000642326 | -1.558951281 | 4.19E-05 | 16.78206673 | 0.339397706 | 3.192244626 |
| *Ttc39c* | 192.3290168 | 665.7846139 | 954.017514 | 245.0278597 | 171.7923223 | 166.9708779 | 192.8834627 | 165.1470501 | 0.009964851 | -1.561575516 | 0.001404533 | 10.1998339 | 0.338780909 | 2.001529188 |
| *C920009B18Rik* | 179.349758 | 402.298425 | 616.2314086 | 186.455463 | 109.2422973 | 117.0433114 | 119.752292 | 121.6873001 | 0.001621007 | -1.564909853 | 0.000132523 | 14.6055611 | 0.337998827 | 2.790215233 |
| *Ptprn* | 50.73710259 | 88.68559529 | 170.034222 | 50.76274384 | 27.31057432 | 31.10241842 | 35.65144571 | 26.94504502 | 0.00326114 | -1.572834559 | 0.000326081 | 12.91429985 | 0.336147294 | 2.486630492 |
| *Dcstamp* | 61.35649616 | 55.26783474 | 67.32898723 | 62.47722319 | 19.3816979 | 26.19151025 | 23.76763048 | 13.03792501 | 9.40E-09 | -1.580872327 | 1.10E-10 | 41.63885459 | 0.334279705 | 8.026847315 |
| *Zhx2* | 192.3290168 | 600.2343913 | 788.5479691 | 245.0278597 | 135.6718853 | 171.0633013 | 175.5148097 | 127.7716651 | 0.004230176 | -1.581430178 | 0.000454222 | 12.29480121 | 0.334150473 | 2.373641545 |
| *Nfkb1* | 2920.333231 | 8381.431404 | 12282.40491 | 3950.70816 | 2194.536794 | 2340.047744 | 2341.111602 | 2316.404677 | 0.002123772 | -1.582766468 | 0.000188236 | 13.9450153 | 0.333841111 | 2.672892032 |
| *Tubb3* | 27.13845022 | 30.84716358 | 11.41169275 | 27.33378515 | 7.047890147 | 6.547877563 | 9.141396337 | 9.561145007 | 0.001829546 | -1.583532664 | 0.00015645 | 14.29294366 | 0.33366386 | 2.737656671 |
| *Chpf* | 48.37723735 | 62.97962564 | 107.2699119 | 41.97688433 | 21.14367044 | 23.73605617 | 14.62623414 | 26.94504502 | 0.000479757 | -1.587669194 | 2.92E-05 | 17.46885979 | 0.332708541 | 3.318979042 |
| *Hck* | 3201.157194 | 7804.332385 | 14241.79255 | 3622.702738 | 2399.806595 | 2457.90954 | 2386.818584 | 2342.480527 | 0.004201186 | -1.590394793 | 0.000450331 | 12.3108571 | 0.332080567 | 2.376628068 |
| *Htra1* | 49.55716997 | 122.1033558 | 151.7755136 | 43.92929756 | 28.19156059 | 33.55787251 | 36.56558535 | 23.46826502 | 0.00443183 | -1.590592136 | 0.000483917 | 12.17664072 | 0.332035146 | 2.353416879 |
| *Sqstm1* | 17655.33177 | 34627.22641 | 47888.02746 | 21434.56859 | 10112.84137 | 10209.77809 | 9986.061358 | 10011.38802 | 8.96E-06 | -1.592626009 | 2.54E-07 | 26.56858733 | 0.331567381 | 5.047859018 |
| *Mllt6* | 1931.549696 | 3237.666877 | 4600.053348 | 2494.207895 | 1132.948341 | 933.8910374 | 973.5587099 | 1009.135396 | 1.49E-07 | -1.598515502 | 2.33E-09 | 35.6766537 | 0.330216588 | 6.82584783 |
| *Clec2d* | 331.5610658 | 661.9287184 | 1221.051124 | 372.9109259 | 227.2944572 | 246.3638933 | 189.2269042 | 190.3537051 | 0.001467327 | -1.60018693 | 0.000116555 | 14.84761885 | 0.329834238 | 2.833473128 |
| *Igfbp5* | 25.9585176 | 34.70305902 | 37.65858608 | 18.54792563 | 10.57183522 | 9.821816345 | 6.398977436 | 11.29953501 | 0.000293394 | -1.604240136 | 1.61E-05 | 18.5998147 | 0.32890888 | 3.53254937 |
| *Ptpn13* | 15.33912404 | 15.42358179 | 26.24689333 | 14.64309919 | 7.928876415 | 5.729392868 | 1.828279267 | 7.822755006 | 0.005967096 | -1.613567556 | 0.000712653 | 11.45595392 | 0.326789253 | 2.224236954 |
| *Plk2* | 449.5543276 | 1197.898186 | 1949.117122 | 573.0332814 | 309.2261802 | 386.3247762 | 335.4892456 | 329.4249052 | 0.002699677 | -1.61561483 | 0.000256341 | 13.36516912 | 0.326325848 | 2.568688236 |
| *Pcsk1* | 28.31838284 | 35.98835751 | 21.68221623 | 42.95309094 | 8.809862684 | 5.729392868 | 17.36865304 | 10.43034001 | 0.001068245 | -1.619608835 | 7.93E-05 | 15.57490339 | 0.325423686 | 2.97132896 |
| *Bmp1* | 192.3290168 | 237.7802192 | 338.9272747 | 176.6933968 | 73.12186027 | 76.11907667 | 84.1008463 | 69.53560005 | 1.21E-09 | -1.641976035 | 1.18E-11 | 46.00685739 | 0.320417304 | 8.918123107 |
| *Orai2* | 1797.037378 | 3393.187994 | 6350.607016 | 2094.93939 | 1103.875794 | 1062.393135 | 1139.017984 | 1046.510781 | 0.000356189 | -1.647685688 | 2.03E-05 | 18.15824168 | 0.319151716 | 3.44831999 |
| *Ext1* | 407.0767533 | 1154.198037 | 1941.128937 | 455.888488 | 292.4874411 | 286.4696434 | 355.6003175 | 328.5557102 | 0.005754663 | -1.647789643 | 0.000680837 | 11.54084399 | 0.319128721 | 2.239980107 |
| *Rtn2* | 28.31838284 | 42.41484992 | 69.61132578 | 20.50033886 | 16.7387391 | 15.55120921 | 10.05553597 | 8.691950006 | 0.004332937 | -1.650215777 | 0.000470346 | 12.22970642 | 0.318592503 | 2.363217651 |
| *Cebpb* | 2845.997476 | 6396.930547 | 9818.620443 | 2378.039308 | 1436.888604 | 1885.788738 | 1620.769571 | 1886.153151 | 0.001978249 | -1.650352611 | 0.000171808 | 14.11673563 | 0.318562287 | 2.703718961 |
| *Ccr3* | 24.77858499 | 37.27365599 | 37.65858608 | 31.23861159 | 7.928876415 | 13.91423982 | 10.05553597 | 9.561145007 | 8.99E-06 | -1.650716276 | 2.56E-07 | 26.55511316 | 0.318481996 | 5.046051689 |
| *Zmynd15* | 251.3256477 | 399.727828 | 617.3725779 | 308.4812895 | 118.9331462 | 142.416337 | 137.1209451 | 101.6958151 | 1.81E-06 | -1.656236119 | 3.96E-08 | 30.16852937 | 0.317265792 | 5.743017674 |

| | | | | | | | | | | | | | | |
|---|---|---|---|---|---|---|---|---|---|---|---|---|---|---|
| *Fnbp1l* | 1332.143926 | 2110.460108 | 2833.52331 | 1849.91153 | 647.5249072 | 631.0517001 | 657.2663966 | 634.5123505 | 2.94E-11 | -1.660581326 | 2.09E-13 | 53.91408325 | 0.316311667 | 10.5313628 |
| *Rffl* | 476.6927778 | 1461.384374 | 2147.680576 | 551.556736 | 348.8705623 | 355.2223578 | 389.4234839 | 370.2770703 | 0.003713144 | -1.663437263 | 0.000386433 | 12.59667744 | 0.315686121 | 2.430258242 |
| *Ptprg* | 20.05885451 | 25.70596965 | 17.11753913 | 23.4289587 | 7.928876415 | 4.092423477 | 10.05553597 | 5.215170004 | 0.000388088 | -1.671945803 | 2.27E-05 | 17.95133531 | 0.313829787 | 3.411069567 |
| *Junb* | 4799.965892 | 7513.854928 | 11529.23319 | 6104.219947 | 2508.167906 | 2303.215933 | 2248.783499 | 2301.628362 | 2.49E-08 | -1.67755396 | 3.07E-10 | 39.62819453 | 0.312612211 | 7.603132381 |
| *Efna2* | 25.9585176 | 17.99417875 | 19.39987768 | 39.04826449 | 10.57183522 | 6.547877563 | 9.141396337 | 6.084365004 | 0.000762935 | -1.680241447 | 5.14E-05 | 16.39623456 | 0.312030412 | 3.117512284 |
| *Rassf4* | 7300.24311 | 16931.23691 | 26220.64643 | 8707.762982 | 4678.037085 | 4437.005534 | 4595.379938 | 4732.766778 | 0.00031333 | -1.681518963 | 1.75E-05 | 18.44696493 | 0.311754229 | 3.503997994 |
| *Slc39a4* | 309.142346 | 348.3158887 | 535.20839 | 226.4799341 | 121.576105 | 86.75937771 | 144.4340621 | 87.78869506 | 2.47E-06 | -1.687936225 | 5.70E-08 | 29.46406856 | 0.310370593 | 5.606900358 |
| *Pdgfb* | 2831.838284 | 3052.583896 | 2986.439993 | 3714.46616 | 1021.944071 | 965.8119406 | 956.1900568 | 945.6841607 | 3.55E-53 | -1.694470642 | 1.70E-56 | 250.8413167 | 0.308968006 | 52.45018023 |
| *Ell2* | 2382.283957 | 4193.928948 | 6949.720886 | 2756.807473 | 1251.000501 | 1384.05762 | 1242.315762 | 1129.084306 | 1.42E-05 | -1.701428954 | 4.41E-07 | 25.5080445 | 0.3074814 | 4.847899413 |
| *Clec4d* | 3703.808489 | 8650.058787 | 15411.49106 | 3892.135763 | 2387.472787 | 2501.289229 | 2491.944641 | 2334.657772 | 0.001658776 | -1.704185837 | 0.000136805 | 14.54563944 | 0.306894387 | 2.780212238 |
| *Enpp2* | 53.09696783 | 44.98544688 | 77.59951071 | 37.09585127 | 14.97676656 | 20.46211738 | 16.45451341 | 13.03792501 | 6.87E-06 | -1.707442084 | 1.85E-07 | 27.18749126 | 0.306202489 | 5.163046418 |
| *Gm14005* | 25.9585176 | 84.82969984 | 140.3638208 | 46.85791739 | 23.78662925 | 26.19151025 | 21.02521157 | 19.99148501 | 0.006232153 | -1.70962495 | 0.000755355 | 11.3478472 | 0.305739541 | 2.205361887 |
| *Pvr* | 488.492104 | 1381.695869 | 2191.045008 | 551.556736 | 329.4888644 | 336.3972098 | 388.5093443 | 338.1168552 | 0.002465191 | -1.727835485 | 0.000226779 | 13.59507735 | 0.301904573 | 2.608149441 |
| *Dram1* | 908.5481162 | 2044.909885 | 3471.436935 | 918.6104222 | 573.5220607 | 567.2098939 | 525.6302894 | 550.2004354 | 0.000899668 | -1.728039595 | 6.34E-05 | 15.99817549 | 0.301861863 | 3.045917531 |
| *Gata2* | 17.69898928 | 16.70888027 | 10.27052348 | 18.54792563 | 4.404931342 | 5.729392868 | 1.828279267 | 6.953560005 | 0.002713406 | -1.732940325 | 0.000258079 | 13.35249841 | 0.3008382 | 2.566485154 |
| *Chst7* | 108.5538009 | 137.5269376 | 180.3047455 | 124.9544464 | 43.16832715 | 33.55787251 | 41.13628352 | 47.80572503 | 1.42E-12 | -1.735797082 | 8.62E-15 | 60.18728766 | 0.300243084 | 11.84812264 |
| *Mtmr7* | 55.45683307 | 96.39738618 | 212.2574852 | 62.47722319 | 32.59649193 | 39.28726538 | 25.59590974 | 30.42182502 | 0.002123397 | -1.736057758 | 0.000188033 | 13.947045 | 0.300188839 | 2.672968802 |
| *Stk40* | 581.7067809 | 1943.371305 | 3125.662645 | 676.5111824 | 493.3523103 | 459.1699141 | 496.3778211 | 442.4202553 | 0.005701048 | -1.74209839 | 0.000672214 | 11.5645437 | 0.298934561 | 2.244045274 |
| *Samsn1* | 418.8760795 | 1302.007363 | 2177.350977 | 411.9591904 | 300.4163175 | 373.2290211 | 314.464034 | 299.0030802 | 0.007386082 | -1.743425398 | 0.000957751 | 10.9075124 | 0.298659724 | 2.131585857 |
| *Ppp1r15a* | 267.8447044 | 841.870506 | 1435.590948 | 374.8633391 | 199.1028966 | 225.0832912 | 212.080395 | 224.2523102 | 0.003205116 | -1.762564572 | 0.000318435 | 12.95871969 | 0.294723791 | 2.494156218 |
| *Prg4* | 15.33912404 | 21.8500742 | 46.78794028 | 16.59551241 | 4.404931342 | 8.184846954 | 10.05553597 | 6.953560005 | 0.005353868 | -1.763944859 | 0.000618858 | 11.71838137 | 0.294441951 | 2.27133237 |
| *Pim3* | 252.5055803 | 706.9141653 | 1331.744544 | 206.9558018 | 194.6979653 | 184.1590565 | 176.4289493 | 178.1849751 | 0.009953758 | -1.767798812 | 0.001400581 | 10.20503055 | 0.293656441 | 2.00201294 |
| *Arhgef3* | 520.3502847 | 1728.726459 | 2556.219176 | 695.059108 | 408.7776285 | 450.1665825 | 395.8224614 | 359.8467303 | 0.002331103 | -1.768185795 | 0.000211834 | 13.72309341 | 0.293577682 | 2.632438543 |
| *Fos* | 1601.168563 | 1136.203858 | 1618.178032 | 1791.339134 | 426.3973539 | 534.4705061 | 392.1659028 | 445.8970353 | 8.53E-21 | -1.772807586 | 2.39E-23 | 99.11006871 | 0.292638687 | 20.06899664 |
| *Gpr126* | 243.0661194 | 763.4672985 | 938.0411442 | 296.7668102 | 187.6500752 | 132.5945207 | 162.7168548 | 171.2314151 | 0.000923452 | -1.77631861 | 6.58E-05 | 15.92664116 | 0.291927371 | 3.034585478 |
| *Spred1* | 1247.188778 | 2128.454287 | 3326.508437 | 1438.928547 | 647.5249072 | 631.0517001 | 547.5696406 | 538.0317054 | 8.13E-07 | -1.783764302 | 1.57E-08 | 31.95977992 | 0.290424627 | 6.089684101 |
| *Rtn1* | 125.0728576 | 263.4861889 | 385.715215 | 152.2882315 | 66.95495639 | 72.84513789 | 62.16149509 | 66.92801505 | 3.59E-05 | -1.783792168 | 1.30E-06 | 23.42433793 | 0.290419017 | 4.445462936 |
| *Rapgef2* | 671.3816599 | 1987.071454 | 3609.518417 | 784.8701163 | 495.9952691 | 484.5429397 | 527.4585686 | 538.0317054 | 0.004052288 | -1.785312318 | 0.000429508 | 12.39924213 | 0.290113167 | 2.392299732 |
| *Pdgfra* | 29.49831546 | 29.5618651 | 36.5174168 | 24.40516531 | 4.404931342 | 9.003331649 | 4.570698168 | 16.51470501 | 0.000804755 | -1.790820393 | 5.47E-05 | 16.27931546 | 0.289007654 | 3.094336158 |
| *Timp1* | 251.3256477 | 383.0189478 | 532.9260515 | 244.0516531 | 101.3134209 | 98.21816345 | 90.49982373 | 117.3413251 | 3.91E-08 | -1.79175173 | 5.04E-10 | 38.66355855 | 0.288821145 | 7.407862112 |
| *Cnn1* | 31.8581807 | 17.99417875 | 18.2587084 | 19.52413225 | 9.690848952 | 4.092423477 | 4.570698168 | 6.953560005 | 0.00049334 | -1.793774479 | 3.01E-05 | 17.41327207 | 0.288416483 | 3.306853939 |
| *Col27a1* | 51.91703521 | 173.5152951 | 188.2929304 | 90.78721495 | 53.74016237 | 18.82514799 | 33.82316645 | 39.11377503 | 0.002286398 | -1.793987347 | 0.000205942 | 13.77607513 | 0.28837393 | 2.640848191 |
| *Ass1* | 2453.079914 | 4303.179319 | 8556.487225 | 1866.507043 | 1161.139902 | 1226.090074 | 1268.825812 | 1233.387706 | 0.000715332 | -1.812889309 | 4.80E-05 | 16.5274037 | 0.284620344 | 3.145492458 |
| *Rbpj* | 3027.707099 | 6650.134348 | 11104.71822 | 3939.969887 | 1669.468979 | 1722.910284 | 1867.587272 | 1728.828856 | 6.13E-05 | -1.822692477 | 2.46E-06 | 22.19372757 | 0.282692894 | 4.212845568 |
| *Slco2a1* | 11.79932618 | 15.42358179 | 12.55286203 | 15.6193058 | 2.642958805 | 2.455454086 | 4.570698168 | 6.084365004 | 0.001635558 | -1.823147953 | 0.000134105 | 14.58319662 | 0.282603658 | 2.78633398 |
| *Cx3cl1* | 129.792588 | 525.6870793 | 657.3135025 | 243.0754465 | 119.8141325 | 117.0433114 | 101.4694993 | 100.8266201 | 0.001327998 | -1.824422253 | 0.000102726 | 15.08593719 | 0.282354151 | 2.876802535 |
| *Tmem202* | 21.23878713 | 19.27947724 | 38.79975535 | 10.73827274 | 6.166903878 | 6.547877563 | 7.313117069 | 5.215170004 | 0.002631577 | -1.830411062 | 0.000247094 | 13.43408471 | 0.281184493 | 2.57978391 |
| *Grem1* | 88.49494638 | 100.2532816 | 111.834589 | 95.66824801 | 25.54860178 | 26.19151025 | 20.11107194 | 39.11377503 | 6.10E-14 | -1.833792786 | 3.27E-16 | 66.63255674 | 0.280526159 | 13.21439868 |
| *Vasp* | 4344.511901 | 8207.916109 | 12668.12012 | 5021.606814 | 2200.703698 | 2038.845376 | 2121.71809 | 2118.228217 | 2.10E-06 | -1.834489759 | 4.71E-08 | 29.83337476 | 0.280390668 | 5.677361239 |
| *Tubb2b* | 70.7959571 | 88.68559529 | 131.2344666 | 62.47722319 | 23.78662925 | 22.91757147 | 16.45451341 | 35.63699503 | 3.94E-06 | -1.834839751 | 9.84E-08 | 28.40599591 | 0.280322655 | 5.404390421 |
| *Fam162a* | 250.1457151 | 667.0699124 | 1456.131995 | 273.3378515 | 199.9838829 | 188.2514799 | 170.0299719 | 183.4001451 | 0.005775239 | -1.8351647 | 0.000685582 | 11.52793429 | 0.280259523 | 2.238430044 |
| *Igfbp7* | 180.5296906 | 263.4861889 | 366.3153373 | 205.0033886 | 64.31199759 | 67.11574502 | 66.73219326 | 86.05030506 | 3.75E-10 | -1.836229694 | 3.33E-12 | 48.4815472 | 0.280052712 | 9.425397765 |
| *Map1b* | 24.77858499 | 35.98835751 | 61.62314086 | 31.23861159 | 8.809862684 | 7.366362258 | 11.88381524 | 14.77631501 | 0.000159847 | -1.845296068 | 7.79E-06 | 19.98943943 | 0.278298287 | 3.796296505 |
| *Lrrc25* | 554.5683307 | 2309.681373 | 4245.149704 | 785.8463229 | 548.8544452 | 534.4705061 | 551.2261991 | 558.8923854 | 0.009682477 | -1.847729138 | 0.001357762 | 10.26230137 | 0.277829339 | 2.014013511 |
| *St3gal3* | 208.8480735 | 789.1732682 | 1195.9454 | 304.5764631 | 149.7676656 | 172.7002707 | 171.8582511 | 199.0456551 | 0.00327039 | -1.849133944 | 0.000328002 | 12.90330634 | 0.277558938 | 2.485400396 |
| *Eif2s3y* | 538.049274 | 676.0670017 | 716.6543048 | 601.3432732 | 185.0071164 | 162.8784544 | 199.2824401 | 154.7167101 | 6.66E-36 | -1.851678019 | 6.39E-39 | 170.2916135 | 0.277069917 | 35.176761 |
| *Srxn1* | 2547.474523 | 6763.240614 | 10920.98996 | 2898.357432 | 1621.89572 | 1622.236666 | 1597.91608 | 1539.344346 | 0.000584509 | -1.857794613 | 3.69E-05 | 17.02498102 | 0.27589771 | 3.233208861 |
| *Col11a2* | 30.67824808 | 133.6710422 | 144.9284979 | 43.92929756 | 28.19156059 | 29.46544903 | 20.11107194 | 19.12229001 | 0.005017495 | -1.863087775 | 0.000569119 | 11.87436454 | 0.274887312 | 2.299513049 |
| *Nr4a1* | 149.8514425 | 488.4134233 | 826.2065552 | 189.3840828 | 105.7183522 | 128.5020972 | 106.9543371 | 112.1261551 | 0.003232147 | -1.866830225 | 0.000321379 | 12.94149098 | 0.27417516 | 2.490508956 |
| *Rcor2* | 14.15919142 | 25.70596965 | 47.92910956 | 17.57171902 | 3.523945073 | 7.366362258 | 4.570698168 | 13.03792501 | 0.009036413 | -1.881424516 | 0.001241865 | 10.42699663 | 0.271415588 | 2.04400392 |
| *Sh3bp4* | 192.3290168 | 516.6899899 | 716.6543048 | 258.6947523 | 106.5993385 | 108.8584645 | 140.7775036 | 100.8266201 | 0.000138979 | -1.881565658 | 6.56E-06 | 20.31764329 | 0.271389036 | 3.857050114 |
| *Gch1* | 552.2084654 | 1556.496462 | 2769.617831 | 725.321513 | 390.2769169 | 387.9617456 | 351.0296193 | 391.1377503 | 0.001058205 | -1.88179686 | 7.83E-05 | 15.59888206 | 0.271345547 | 2.975430394 |
| *Pcdhga11* | 36.57791117 | 15.42358179 | 34.23507825 | 19.52413225 | 8.809862684 | 6.547877563 | 5.484837802 | 7.822755006 | 0.000240608 | -1.88254696 | 1.28E-05 | 19.03446957 | 0.271204503 | 3.618689346 |
| *Slc11a2* | 1369.90177 | 3066.722179 | 4980.062717 | 1559.00196 | 727.6946577 | 769.3756137 | 759.6500356 | 717.9550705 | 7.34E-05 | -1.883440946 | 3.09E-06 | 21.76114537 | 0.2710365 | 4.134376696 |
| *Ngf* | 10.61939357 | 23.13537268 | 19.39987768 | 14.64309919 | 2.642958805 | 4.910908172 | 5.484837802 | 5.215170004 | 0.00082271 | -1.885029974 | 5.65E-05 | 16.21785667 | 0.270738136 | 3.084753196 |
| *Gsr* | 1202.351338 | 3854.610149 | 7104.919907 | 1258.330323 | 939.1313621 | 861.8643842 | 920.5386111 | 899.6168257 | 0.004849057 | -1.889850383 | 0.000543439 | 11.96037656 | 0.269835041 | 2.314342683 |
| *Plod2* | 140.4119816 | 417.7220068 | 927.7706207 | 155.2168514 | 125.1000501 | 94.12573997 | 118.8381524 | 103.4342051 | 0.008298785 | -1.894101286 | 0.001116595 | 10.6234998 | 0.269041142 | 2.08098549 |
| *Gsap* | 1258.988104 | 2269.83712 | 4031.751049 | 1312.021687 | 567.3551568 | 598.3123123 | 644.4684417 | 558.0231904 | 1.59E-05 | -1.905535271 | 5.06E-07 | 25.24151508 | 0.266917302 | 4.797864665 |
| *Msantd3* | 55.45683307 | 203.0771602 | 292.1393344 | 78.09652899 | 48.45424476 | 37.65029599 | 42.05042315 | 39.11377503 | 0.002570746 | -1.909643361 | 0.000238957 | 13.49690343 | 0.266158333 | 2.589940788 |
| *Serpine1* | 1820.63603 | 5076.929005 | 7911.726585 | 1843.078084 | 1083.61311 | 1077.944344 | 1063.144394 | 1195.143126 | 0.000801043 | -1.913613359 | 5.43E-05 | 16.29264033 | 0.265426927 | 3.096344101 |
| *Stx11* | 312.6821439 | 732.620135 | 1206.215924 | 451.0074549 | 179.7211987 | 171.0633013 | 191.9693231 | 173.8390001 | 4.06E-05 | -1.914984279 | 1.51E-06 | 23.12990211 | 0.265174825 | 4.391264899 |
| *Relb* | 879.0498007 | 1402.260644 | 2087.198604 | 1033.802802 | 363.8473288 | 339.6711486 | 343.7165023 | 370.2770703 | 3.86E-10 | -1.930111185 | 3.49E-12 | 48.39414056 | 0.262408947 | 9.413795997 |
| *Mybpc3* | 7.07959571 | 29.5618651 | 41.08209391 | 36.11964466 | 10.57183522 | 6.547877563 | 6.398977436 | 6.084365004 | 0.004951899 | -1.947275273 | 0.000558133 | 11.91066926 | 0.259305503 | 2.305228194 |
| *Rgs11* | 29.49831546 | 39.84425295 | 54.77612521 | 15.6193058 | 8.809862684 | 4.910908172 | 10.9696756 | 11.29953501 | 0.000868047 | -1.956449763 | 6.08E-05 | 16.07669956 | 0.257661741 | 3.061456687 |
| *Cd300e* | 218.2875344 | 705.6288668 | 1336.309221 | 204.027182 | 177.9592262 | 153.056638 | 136.2068054 | 166.0162451 | 0.006232153 | -1.960096828 | 0.000756107 | 11.34599848 | 0.257011206 | 2.205361887 |
| *Stap1* | 370.4988422 | 930.5561013 | 1345.438575 | 558.3901823 | 207.0317731 | 211.1690514 | 212.080395 | 192.0920951 | 8.96E-06 | -1.962223879 | 2.54E-07 | 26.57062153 | 0.256632559 | 5.047859018 |
| *Antxr1* | 71.97588972 | 100.2532816 | 360.6094909 | 75.16790915 | 37.88240954 | 45.83514294 | 37.47972498 | 34.76780003 | 0.00405864 | -1.962383719 | 0.000430506 | 12.39490838 | 0.256604127 | 2.391619443 |
| *Denr* | 952.2056231 | 3463.87941 | 5196.884879 | 1107.018298 | 709.193946 | 702.2598686 | 651.7815588 | 664.9341755 | 0.001861442 | -1.974226807 | 0.000159922 | 14.25162911 | 0.254506286 | 2.730150446 |
| *Arrdc4* | 431.8553383 | 1277.586691 | 2284.620889 | 476.3888268 | 259.0099629 | 306.1132761 | 287.039845 | 285.0959602 | 0.001613681 | -1.974709887 | 0.000131475 | 14.62052847 | 0.25442108 | 2.792182269 |
| *Jak2* | 1526.832808 | 6345.518608 | 10669.93272 | 2116.415936 | 1251.881487 | 1357.86611 | 1333.729726 | 1307.269281 | 0.003992251 | -1.976125424 | 0.000419104 | 12.4450352 | 0.25417157 | 2.398782152 |
| *Rgs16* | 108.5538009 | 100.2532816 | 102.7052348 | 118.1210001 | 33.4774782 | 30.28393373 | 21.02521157 | 24.33746002 | 5.93E-22 | -1.976780617 | 1.57E-24 | 104.5084229 | 0.254056166 | 21.22698104 |
| *Pla2g4a* | 509.7308912 | 1355.989899 | 2451.231603 | 516.4132979 | 311.869139 | 284.0141893 | 326.3478492 | 298.1338852 | 0.00101246 | -1.985630077 | 7.36E-05 | 15.71558257 | 0.252502561 | 2.994622254 |
| *Osmr* | 43.65750688 | 62.97962564 | 92.43471129 | 32.21481821 | 13.21479403 | 16.36969391 | 15.54037377 | 13.03792501 | 1.53E-05 | -1.986994857 | 4.78E-07 | 25.34969164 | 0.252263808 | 4.816705559 |
| *Pdgfrb* | 126.2527902 | 183.797683 | 293.2805037 | 124.9544464 | 48.45424476 | 42.56120416 | 31.08074754 | 60.84365004 | 1.31E-06 | -1.991747246 | 2.72E-08 | 30.89509651 | 0.251434191 | 5.882886764 |
| *Cacng8* | 35.39797855 | 37.27365599 | 84.44652636 | 33.19102482 | 8.809862684 | 13.09575513 | 11.88381524 | 13.90712001 | 6.02E-05 | -1.99453593 | 2.42E-06 | 22.23204496 | 0.250948646 | 4.220648461 |
| *Chst11* | 464.8934517 | 1230.030648 | 2405.584832 | 579.8667277 | 315.3930841 | 278.2847964 | 279.7267279 | 300.7414702 | 0.000758978 | -1.994922657 | 5.10E-05 | 16.41059892 | 0.250881387 | 3.119770628 |
| *Sod3* | 81.41535067 | 158.0917133 | 197.4222846 | 97.62066123 | 37.00142327 | 32.73938782 | 29.25246828 | 33.89860502 | 7.97E-08 | -2.006095849 | 1.15E-09 | 37.04485943 | 0.248945898 | 7.098486222 |
| *Ankrd37* | 40.11770903 | 110.5356695 | 181.4459147 | 32.21481821 | 23.78662925 | 22.91757147 | 21.93935121 | 21.72987502 | 0.003483071 | -2.009454237 | 0.000354966 | 12.75548763 | 0.248367062 | 2.458037675 |
| *Ddx3y* | 866.0705419 | 1091.218412 | 1199.368908 | 843.4425131 | 229.0564298 | 252.9117709 | 266.0146334 | 239.8978202 | 1.62E-36 | -2.017109131 | 1.42E-39 | 173.2808598 | 0.247052724 | 35.7918149 |
| *St3gal1* | 2142.757635 | 4944.543262 | 7528.293708 | 2738.259548 | 1064.231412 | 1111.502216 | 1086.912024 | 1023.911711 | 5.11E-06 | -2.017321036 | 1.32E-07 | 27.84080358 | 0.247016439 | 5.291263791 |
| *Acsl1* | 2272.550223 | 4439.420958 | 8947.908286 | 2429.778258 | 1095.946918 | 1064.030104 | 1131.704866 | 1174.282446 | 7.54E-05 | -2.018099462 | 3.19E-06 | 21.70154936 | 0.246883194 | 4.122533495 |
| *Ptgir* | 140.4119816 | 267.3420843 | 470.1617414 | 142.5261654 | 56.38312117 | 57.29392868 | 71.30289143 | 65.18962505 | 3.61E-05 | -2.027970753 | 1.31E-06 | 23.40156781 | 0.245199722 | 4.442235544 |
| *Egr3* | 61.35649616 | 47.55604385 | 38.79975535 | 30.26240498 | 9.690848952 | 13.09575513 | 10.05553597 | 10.43034001 | 1.25E-07 | -2.028968256 | 1.91E-09 | 36.06302976 | 0.245030245 | 6.902970899 |
| *Foxp4* | 322.1216048 | 798.1703576 | 1295.227127 | 540.8184632 | 162.9824596 | 194.7993575 | 167.287553 | 191.2229001 | 1.03E-05 | -2.044953556 | 3.02E-07 | 26.23806194 | 0.242330255 | 4.986795781 |
| *Ppfibp2* | 323.3015374 | 854.7234908 | 1450.426149 | 394.3874714 | 165.6254185 | 188.2514799 | 181.9137871 | 193.8304851 | 0.00018814 | -2.050463618 | 9.44E-06 | 19.62225627 | 0.241406492 | 3.725518447 |
| *Aldh1l2* | 66.07622663 | 182.5123845 | 382.2917072 | 86.8823885 | 44.93029969 | 50.74605111 | 53.93423839 | 23.46826502 | 0.003663441 | -2.051713137 | 0.000379209 | 12.63195138 | 0.241197501 | 2.436110741 |
| *Gas7* | 3242.454835 | 7747.779252 | 13045.84715 | 4235.760491 | 1710.875333 | 1705.722105 | 1605.229197 | 1754.904706 | 2.14E-05 | -2.060679832 | 7.11E-07 | 24.58572287 | 0.239703049 | 4.670423207 |
| *Scube3* | 14.15919142 | 12.85298482 | 28.52923188 | 13.66689257 | 4.404931342 | 3.273938782 | 5.484837802 | 3.476780003 | 0.000610104 | -2.063801966 | 3.94E-05 | 16.90079042 | 0.239184869 | 3.214595957 |
| *Flnb* | 1595.2689 | 5330.132806 | 9589.245419 | 1963.151497 | 1136.472286 | 1064.030104 | 1130.790727 | 1076.932606 | 0.001374929 | -2.067496686 | 0.000107126 | 15.00677451 | 0.238573104 | 2.861719635 |
| *Igsf6* | 2461.339442 | 5808.263842 | 11269.04659 | 2850.523308 | 1311.788554 | 1379.146712 | 1295.335861 | 1340.298691 | 0.000169192 | -2.071494679 | 8.30E-06 | 19.86828754 | 0.237912886 | 3.77161956 |
| *Casp4* | 335.1008636 | 913.847221 | 1766.530038 | 488.1033062 | 204.3888143 | 248.8193474 | 170.9441115 | 205.9992151 | 0.000383127 | -2.077127828 | 2.23E-05 | 17.98104809 | 0.236985743 | 3.416657339 |
| *Serping1* | 58.99663092 | 61.69432715 | 67.32898723 | 48.81033062 | 13.21479403 | 8.184846954 | 18.28279267 | 16.51470501 | 1.31E-11 | -2.080334479 | 8.81E-14 | 55.61525229 | 0.236459584 | 10.88217559 |
| *Tnfrsf1b* | 10115.56234 | 44637.131 | 82103.70584 | 14291.6648 | 8806.338738 | 8870.737129 | 9061.866189 | 8981.391942 | 0.004272291 | -2.081138919 | 0.000460004 | 12.27119529 | 0.236327772 | 2.36933913 |
| *Slc28a2* | 220.6473996 | 854.7234908 | 1661.542465 | 257.7185457 | 176.1972537 | 170.2448166 | 156.3178774 | 191.2229001 | 0.004569713 | -2.109233056 | 0.00050409 | 12.1004729 | 0.231770193 | 2.340111115 |
| *Aqp9* | 117.9932618 | 404.869022 | 663.0193489 | 100.5492811 | 72.24087401 | 72.02665319 | 68.56047253 | 85.18111006 | 0.00327039 | -2.109440939 | 0.000328059 | 12.90298041 | 0.231736799 | 2.485400396 |
| *Bcl3* | 582.8867135 | 1242.883632 | 1941.128937 | 761.4411576 | 262.533908 | 280.7402505 | 255.0449578 | 245.1129902 | 5.23E-07 | -2.117459228 | 9.53E-09 | 32.93417584 | 0.230452412 | 6.281748851 |
| *Txnrd1* | 6412.933781 | 18197.25591 | 30477.20783 | 7774.509461 | 3694.856409 | 3521.939644 | 3662.957512 | 3462.003688 | 0.00014038 | -2.131953888 | 6.66E-06 | 20.288747 | 0.228148664 | 3.852693836 |
| *Pilrb2* | 56.63676568 | 120.8180573 | 193.9987768 | 45.88171078 | 32.59649193 | 18.0066633 | 21.93935121 | 22.59907002 | 0.000420272 | -2.13233288 | 2.49E-05 | 17.77284647 | 0.228088738 | 3.376469413 |
| *Prdx5* | 2279.629819 | 7328.771947 | 13828.68928 | 2533.256159 | 1566.393585 | 1402.882768 | 1433.370946 | 1440.256116 | 0.001188547 | -2.152087872 | 8.98E-05 | 15.34090653 | 0.224986779 | 2.924983692 |
| *Ppap2a* | 66.07622663 | 242.9214132 | 537.4907286 | 66.38204964 | 44.93029969 | 54.0199899 | 44.79284205 | 60.84365004 | 0.008396197 | -2.15711169 | 0.001131717 | 10.59862676 | 0.224204682 | 2.075917369 |
| *Ebi3* | 619.4646247 | 2786.52711 | 4698.193906 | 801.4656287 | 507.4480906 | 449.3480978 | 500.0343796 | 529.3397554 | 0.002926469 | -2.164682852 | 0.000283962 | 13.1733288 | 0.223031153 | 2.533656036 |
| *Dst* | 2420.0418 | 5918.799511 | 11701.54975 | 3198.052862 | 1273.906144 | 1249.007645 | 1314.532793 | 1302.054111 | 6.87E-05 | -2.176797475 | 2.83E-06 | 21.9270821 | 0.221166154 | 4.162755667 |
| *Tnfaip2* | 6215.885034 | 18609.83673 | 28347.78596 | 9908.497115 | 3475.490829 | 3311.589078 | 3496.584099 | 3491.556318 | 1.37E-05 | -2.195144476 | 4.24E-07 | 25.58133072 | 0.218371355 | 4.862150601 |
| *Cd83* | 1137.455044 | 2726.118081 | 3608.377248 | 2143.749721 | 523.3058434 | 559.0250469 | 503.6909382 | 498.9179304 | 2.04E-09 | -2.205321869 | 2.04E-11 | 44.93322741 | 0.21683629 | 8.69061413 |
| *Tnip1* | 2425.941463 | 7673.23194 | 14008.99402 | 3292.744903 | 1498.557642 | 1469.180028 | 1422.40127 | 1543.690321 | 0.000299163 | -2.20716746 | 1.65E-05 | 18.55438711 | 0.216559076 | 3.524092169 |
| *Mir22hg* | 166.3704992 | 569.3872277 | 1043.028717 | 176.6933968 | 103.0753934 | 112.1324033 | 94.15638227 | 112.1261551 | 0.001611072 | -2.21359289 | 0.000130828 | 14.62981249 | 0.215596716 | 2.792885091 |
| *Cxcl10* | 2304.408404 | 4166.93768 | 9602.93945 | 3380.603499 | 996.3954695 | 1174.525538 | 988.184944 | 1029.126881 | 5.22E-06 | -2.215692907 | 1.35E-07 | 27.79544491 | 0.215283118 | 5.282428661 |
| *Ikbke* | 1463.116447 | 2713.265096 | 5009.733118 | 1613.66953 | 546.2114864 | 567.2098939 | 615.2159735 | 581.4914554 | 8.26E-07 | -2.224922231 | 1.61E-08 | 31.92062132 | 0.213910287 | 6.082907619 |
| *Dcbld2* | 382.2981684 | 1115.639083 | 1780.224069 | 525.1991574 | 162.1014734 | 212.8060208 | 200.1965798 | 235.5518452 | 5.52E-05 | -2.229914161 | 2.17E-06 | 22.43480469 | 0.213171406 | 4.257994472 |
| *Psd* | 61.35649616 | 115.6768634 | 172.3165605 | 41.00067772 | 20.26268417 | 13.09575513 | 21.02521157 | 28.68343502 | 0.000181339 | -2.231904805 | 9.01E-06 | 19.71098113 | 0.212877473 | 3.741507663 |
| *Gfpt2* | 11.79932618 | 44.98544688 | 61.62314086 | 29.28619837 | 6.166903878 | 4.910908172 | 7.313117069 | 13.03792501 | 0.001610289 | -2.23283261 | 0.000130501 | 14.63453093 | 0.212740614 | 2.793096219 |
| *Rasip1* | 15.33912404 | 35.98835751 | 29.67040115 | 9.762066123 | 5.28591761 | 3.273938782 | 3.656558535 | 6.953560005 | 0.001198054 | -2.233878027 | 9.07E-05 | 15.32160506 | 0.212586512 | 2.921523492 |
| *Igf2bp2* | 214.7477365 | 809.7380439 | 1395.650024 | 262.5995787 | 137.4338579 | 163.6969391 | 132.5502469 | 132.9868351 | 0.001145715 | -2.242833087 | 8.62E-05 | 15.41827917 | 0.211271038 | 2.940923252 |
| *Plek* | 6808.211208 | 29777.79524 | 47163.38497 | 9518.01447 | 4773.183602 | 5039.410269 | 5072.560827 | 4767.534578 | 0.000954966 | -2.246637339 | 6.88E-05 | 15.84488684 | 0.21071467 | 3.020012059 |
| *Gys1* | 362.2393139 | 1634.89967 | 3733.905868 | 430.507116 | 310.1071665 | 309.3872149 | 338.2316645 | 338.1168552 | 0.00708752 | -2.249353057 | 0.000905429 | 11.01161124 | 0.210318395 | 2.14950573 |
| *Ero1l* | 1579.929776 | 7195.100904 | 15823.45317 | 1777.672241 | 1354.075894 | 1415.978523 | 1344.699401 | 1364.636151 | 0.006353227 | -2.267127506 | 0.000776556 | 11.296445 | 0.207743104 | 2.197005602 |
| *Tfec* | 271.3845022 | 794.3144621 | 1224.474632 | 383.6491987 | 133.9099128 | 134.23149 | 142.6057829 | 138.2020051 | 1.57E-05 | -2.283970619 | 4.96E-07 | 25.27872549 | 0.205331857 | 4.802937499 |
| *Mfsd7a* | 90.85481162 | 384.3042462 | 748.6070445 | 98.59686785 | 74.88383281 | 54.83847459 | 65.81805362 | 73.01238005 | 0.004214478 | -2.299650272 | 0.000452093 | 12.30356884 | 0.20311233 | 2.375256224 |
| *Bnip3* | 613.5649616 | 2493.479056 | 5474.189013 | 617.9387856 | 452.8269419 | 506.6420264 | 438.7870242 | 468.4961053 | 0.004704319 | -2.300913043 | 0.000522702 | 12.03288155 | 0.202934627 | 2.327503269 |
| *E230013L22Rik* | 86.13508114 | 200.5065633 | 317.2450585 | 122.0258265 | 36.120437 | 36.83181129 | 37.47972498 | 36.50619003 | 1.38E-06 | -2.303949895 | 2.91E-08 | 30.76443407 | 0.202507901 | 5.858627679 |
| *P4ha2* | 264.3049065 | 1092.50371 | 2171.645131 | 289.9333639 | 194.6979653 | 206.2581432 | 187.3986249 | 184.2693401 | 0.003074781 | -2.304982805 | 0.00030155 | 13.06073929 | 0.202362966 | 2.512185826 |
| *Osm* | 135.6922511 | 924.1296088 | 1190.239554 | 160.0978844 | 136.5528716 | 107.2214951 | 113.3533146 | 130.3792501 | 0.005559966 | -2.305410232 | 0.000652021 | 11.62126439 | 0.202303021 | 2.254927884 |
| *Exoc3l4* | 8.259528329 | 74.54731198 | 83.30535709 | 23.4289587 | 11.45282149 | 9.821816345 | 8.227256703 | 8.691950006 | 0.006874574 | -2.308109519 | 0.000868877 | 11.0880102 | 0.201924865 | 2.162754205 |
| *Slamf7* | 1176.392821 | 5943.220183 | 11141.23563 | 1404.761315 | 922.392623 | 967.4489099 | 1045.775741 | 1035.211246 | 0.004321854 | -2.308139485 | 0.000468106 | 12.23861446 | 0.201920671 | 2.364329921 |
| *Icam1* | 4126.224367 | 6664.272631 | 8278.041922 | 6330.699881 | 1406.054084 | 1170.433114 | 1270.654091 | 1272.501481 | 1.59E-23 | -2.310712749 | 3.57E-26 | 112.0031845 | 0.201560836 | 22.79786545 |
| *P2ry2* | 213.5678039 | 889.4265498 | 1612.472186 | 230.3847605 | 174.4352811 | 130.1390666 | 138.0350847 | 149.5015401 | 0.002429129 | -2.31461327 | 0.000223268 | 13.62437578 | 0.201016625 | 2.614549366 |
| *Lpar1* | 57.8166983 | 141.3828331 | 179.1635762 | 83.95376866 | 23.78662925 | 26.19151025 | 21.02521157 | 21.72987502 | 2.34E-07 | -2.315235408 | 3.91E-09 | 34.66567726 | 0.200929959 | 6.630499511 |
| *Bcar1* | 47.19730474 | 74.54731198 | 205.4104695 | 51.73895045 | 9.690848952 | 22.91757147 | 27.42418901 | 15.64551001 | 0.000899668 | -2.323715105 | 6.34E-05 | 15.99896682 | 0.199752421 | 3.045917531 |

| | | | | | | | | | | | | | |
|---|---|---|---|---|---|---|---|---|---|---|---|---|---|
| *Birc3* | 1435.977997 | 3665.671272 | 6044.77365 | 1693.718472 | 649.2868798 | 634.3256389 | 627.0997887 | 651.0270555 | 7.42E-06 | -2.32541348 | 2.03E-07 | 26.99987753 | 0.199517406 | 5.129852495 |
| *Itga9* | 331.5610658 | 1023.097592 | 2042.693003 | 305.5526697 | 158.5775283 | 199.7102657 | 170.9441115 | 205.1300201 | 0.00104868 | -2.333903338 | 7.74E-05 | 15.62007126 | 0.19834675 | 2.979356975 |
| *Src* | 191.1490842 | 674.7817033 | 1381.955992 | 304.5764631 | 129.5049814 | 134.23149 | 113.3533146 | 125.1640801 | 0.00052623 | -2.345184562 | 3.24E-05 | 17.27078994 | 0.196801816 | 3.278824792 |
| *Gstt1* | 21.23878713 | 70.69141653 | 101.5640655 | 25.38137192 | 2.642958805 | 10.64030104 | 15.54037377 | 13.90712001 | 0.004704446 | -2.355798971 | 0.000523544 | 12.02988282 | 0.19535919 | 2.327491534 |
| *Myc* | 120.3531271 | 113.1062665 | 152.9166829 | 159.1216778 | 27.31057432 | 30.28393373 | 22.85349084 | 26.07585002 | 9.82E-27 | -2.357968809 | 1.99E-29 | 126.862233 | 0.195065587 | 26.00810503 |
| *Clec4n* | 179.349758 | 683.7787926 | 1752.836007 | 191.336496 | 148.8866794 | 120.3172502 | 146.2623414 | 132.1176401 | 0.005223226 | -2.357984149 | 0.000598273 | 11.7813438 | 0.195063513 | 2.282061184 |
| *Ccl8* | 5.899663092 | 8.997089377 | 7.988184926 | 12.69068596 | 0.880986268 | 0.818484695 | 3.656558535 | 1.738390001 | 0.004659963 | -2.358651104 | 0.000516655 | 12.05457097 | 0.194973357 | 2.331617524 |
| *Wnt6* | 10.61939357 | 44.98544688 | 81.02301854 | 11.71447935 | 7.047890147 | 4.910908172 | 9.141396337 | 7.822755006 | 0.00900569 | -2.358687556 | 0.001234319 | 10.43825418 | 0.19496843 | 2.045483017 |
| *Fscn1* | 88.49494638 | 413.8661113 | 726.9248283 | 116.1685869 | 73.12186027 | 42.56120416 | 51.19181949 | 94.74225507 | 0.003679136 | -2.362377306 | 0.00038121 | 12.62211234 | 0.194470428 | 2.434254203 |
| *Mcoln2* | 601.7656354 | 2269.83712 | 3517.083706 | 742.893232 | 353.2754936 | 340.4896333 | 339.1458041 | 348.5471953 | 0.000226048 | -2.367940784 | 1.19E-05 | 19.18244245 | 0.193721934 | 3.645800237 |
| *Slc13a3* | 213.5678039 | 222.3566375 | 430.2208167 | 109.3351406 | 53.74016237 | 48.29059703 | 41.13628352 | 44.32894503 | 3.31E-07 | -2.378211686 | 5.74E-09 | 33.92177235 | 0.192347677 | 6.480806992 |
| *Ccl9* | 10298.45189 | 36039.76945 | 60180.70289 | 13184.64651 | 5634.788172 | 6173.830057 | 5581.736603 | 5586.316269 | 0.000144145 | -2.38121587 | 6.86E-06 | 20.23166901 | 0.191947561 | 3.841199287 |
| *H2-Q7* | 210.0280061 | 560.3901383 | 863.8651413 | 286.0285374 | 107.4803247 | 93.30725527 | 85.01498593 | 80.83513506 | 3.06E-06 | -2.388557451 | 7.43E-08 | 28.94987154 | 0.19097326 | 5.513650894 |
| *Cflar* | 2182.875344 | 9398.102503 | 15950.12296 | 3346.436267 | 1475.651999 | 1442.170033 | 1400.461919 | 1513.268496 | 0.000393922 | -2.404586028 | 2.30E-05 | 17.92033296 | 0.188863258 | 3.404590071 |
| *Wnt11* | 12.9792588 | 7.711790894 | 18.2587084 | 20.50033886 | 5.28591761 | 4.092423477 | 1.828279267 | 0 | 0.003633867 | -2.410605878 | 0.000374585 | 12.65489052 | 0.188076842 | 2.43963096 |
| *Pla1a* | 11.79932618 | 21.8500742 | 52.49378666 | 11.71447935 | 4.404931342 | 4.092423477 | 5.484837802 | 4.345975003 | 0.001916786 | -2.416521287 | 0.000165525 | 14.18682811 | 0.187307258 | 2.717426292 |
| *Slc31a2* | 962.8250166 | 2754.394648 | 5647.646743 | 1327.640993 | 511.8530219 | 468.1732458 | 463.4687943 | 531.0781454 | 6.75E-05 | -2.436950185 | 2.76E-06 | 21.97901862 | 0.184673634 | 4.170512095 |
| *AW112010* | 84.95514853 | 384.3042462 | 849.0299407 | 91.76342156 | 56.38312117 | 71.2081685 | 66.73219326 | 64.32043005 | 0.004514801 | -2.446384228 | 0.000495076 | 12.13412251 | 0.18346996 | 2.345361395 |
| *Ankrd33b* | 319.7617396 | 1147.771545 | 1842.988379 | 410.9829838 | 167.387391 | 155.5120921 | 174.60067 | 179.9233651 | 0.000100949 | -2.457642331 | 4.40E-06 | 21.03491042 | 0.18204382 | 3.995899994 |
| *Trem3* | 17.69898928 | 78.40320743 | 196.2811153 | 33.19102482 | 22.02465671 | 14.73272452 | 11.88381524 | 9.561145007 | 0.005968918 | -2.48308115 | 0.000713348 | 11.45414256 | 0.178862004 | 2.224104414 |
| *AA467197* | 210.0280061 | 1197.898186 | 2089.480943 | 255.7661324 | 154.172597 | 159.6045156 | 177.3430889 | 176.4465851 | 0.002793322 | -2.491021284 | 0.000267691 | 13.28393108 | 0.177880308 | 2.553878925 |
| *Wisp2* | 29.49831546 | 16.70888027 | 9.129354201 | 22.45275208 | 2.642958805 | 5.729392868 | 2.742418901 | 2.607585002 | 0.000180027 | -2.493538511 | 8.93E-06 | 19.72793909 | 0.177570212 | 3.744661487 |
| *Sdc4* | 2350.425776 | 6462.480769 | 10159.83006 | 3329.840755 | 1068.636344 | 1001.825267 | 936.0789849 | 942.2073807 | 6.19E-07 | -2.497710796 | 1.15E-08 | 32.56700743 | 0.177057419 | 6.208239792 |
| *Rgl1* | 176.9898928 | 523.1164823 | 1332.885713 | 195.2413225 | 102.1944071 | 81.84846954 | 102.383639 | 106.9109851 | 0.00091632 | -2.502035264 | 6.51E-05 | 15.948475 | 0.176527486 | 3.037952628 |
| *Aoah* | 771.6759324 | 1731.297056 | 2984.157655 | 786.8225295 | 237.8662925 | 286.4696434 | 290.6964035 | 283.3575702 | 1.19E-06 | -2.513872084 | 2.45E-08 | 31.10445196 | 0.175085063 | 5.924691827 |
| *Pim1* | 631.2639508 | 3435.602843 | 6086.996914 | 1027.945563 | 510.9720356 | 466.5362764 | 456.1556772 | 505.8714904 | 0.000992713 | -2.5273275 | 7.17E-05 | 15.76524231 | 0.173459709 | 3.003176281 |
| *Tlr1* | 593.5061071 | 1483.234449 | 2402.161324 | 670.6539427 | 227.2944572 | 211.1690514 | 242.2470029 | 210.3451902 | 1.02E-06 | -2.530792367 | 2.05E-08 | 31.44295782 | 0.173043617 | 5.992070608 |
| *Rab20* | 191.1490842 | 525.6870793 | 861.5828027 | 297.7430168 | 67.83594266 | 93.30725527 | 72.21703106 | 89.52708506 | 1.52E-06 | -2.538096797 | 3.24E-08 | 30.5604174 | 0.172169704 | 5.819518444 |
| *Gdf15* | 161.6507687 | 921.5590119 | 1221.051124 | 283.0999176 | 127.7430089 | 90.85180119 | 126.1512694 | 99.95742507 | 0.000374655 | -2.540513071 | 2.16E-05 | 18.04200673 | 0.17188159 | 3.426367963 |
| *Cdkn2b* | 17.69898928 | 68.12081957 | 67.32898723 | 19.52413225 | 10.57183522 | 6.547877563 | 4.570698168 | 7.822755006 | 0.000464213 | -2.543493025 | 2.80E-05 | 17.54904642 | 0.171526927 | 3.333282267 |
| *Nfkb2* | 2420.0418 | 6841.643822 | 10927.83698 | 2919.833978 | 956.7510874 | 1009.191629 | 959.8466154 | 1035.211246 | 2.59E-06 | -2.544505016 | 6.10E-08 | 29.33161014 | 0.17140665 | 5.586125486 |
| *Mt2* | 444.8345971 | 2978.036584 | 6530.911762 | 570.1046616 | 456.350887 | 471.4471845 | 433.3021864 | 437.2050853 | 0.006440385 | -2.548915085 | 0.000791331 | 11.26145597 | 0.170883489 | 2.191088182 |
| *Eid3* | 4.719730474 | 29.5618651 | 44.50560173 | 8.785859511 | 2.642958805 | 3.273938782 | 3.656558535 | 5.215170004 | 0.008448004 | -2.562185634 | 0.001140051 | 10.58506084 | 0.169318834 | 2.073245877 |
| *Gadd45b* | 238.3463889 | 1453.672584 | 2346.24403 | 379.7443722 | 217.6036083 | 193.1623881 | 160.8885755 | 172.1006101 | 0.00121587 | -2.570352517 | 9.24E-05 | 15.28600071 | 0.168363053 | 2.915112742 |
| *Slc39a14* | 606.4853659 | 2949.760017 | 4533.86553 | 789.7511494 | 375.3001503 | 360.133266 | 390.3376236 | 362.4543153 | 0.000365299 | -2.576888634 | 2.10E-05 | 18.09280104 | 0.167602011 | 3.437351512 |
| *Pilrb1* | 83.77521591 | 156.8064149 | 276.1629646 | 48.81033062 | 15.85775283 | 27.82847964 | 29.25246828 | 21.72987502 | 7.02E-05 | -2.577482081 | 2.93E-06 | 21.86350932 | 0.167533083 | 4.153398614 |
| *Irak3* | 534.5094761 | 2753.109349 | 5428.542242 | 624.7722319 | 404.3726972 | 341.308118 | 386.681065 | 425.9055503 | 0.001992909 | -2.583590401 | 0.000173984 | 14.09307069 | 0.166825253 | 2.700512559 |
| *Tlr2* | 10821.16204 | 20288.43654 | 30961.0636 | 13746.94152 | 3219.123825 | 3166.717286 | 3050.483958 | 3165.608192 | 4.00E-13 | -2.588876975 | 2.40E-15 | 62.70526263 | 0.166215062 | 12.39777261 |
| *Slc16a3* | 1840.694885 | 7803.047087 | 16817.41161 | 2007.080795 | 1205.189215 | 1196.624625 | 1119.821051 | 1165.590496 | 0.001198054 | -2.602527263 | 9.08E-05 | 15.31986014 | 0.164649808 | 2.921523492 |
| *Rnd1* | 76.6956202 | 413.8661113 | 626.5019321 | 98.59686785 | 49.33523103 | 56.47544398 | 40.22214388 | 53.89009004 | 0.000986818 | -2.603483873 | 7.12E-05 | 15.77860693 | 0.16454067 | 3.005763001 |
| *Ampd3* | 692.620447 | 2897.062779 | 4544.136054 | 973.2779925 | 363.8473288 | 369.9550823 | 373.8831102 | 384.1841903 | 7.53E-05 | -2.609804971 | 3.17E-06 | 21.70932737 | 0.16382132 | 4.123471058 |
| *Trem1* | 192.3290168 | 1350.848705 | 2524.266437 | 259.6709589 | 170.9113361 | 172.7002707 | 165.4592737 | 184.2693401 | 0.003434436 | -2.641664286 | 0.000348636 | 12.78914692 | 0.160243275 | 2.464144555 |
| *Fbln2* | 297.3430198 | 347.0305902 | 285.2923188 | 352.4105871 | 40.52536834 | 45.01665825 | 46.62112132 | 73.01238005 | 2.14E-34 | -2.643793487 | 2.40E-37 | 163.0807444 | 0.160006954 | 33.66860168 |
| *Tmem176b* | 1317.984735 | 1951.083096 | 2618.983486 | 1252.473084 | 282.7965921 | 258.6411637 | 323.6054303 | 265.9736702 | 3.39E-20 | -2.65845216 | 1.04E-22 | 96.20252712 | 0.158389416 | 19.46917234 |
| *Pstpip2* | 337.4607289 | 1501.228627 | 2790.158878 | 505.6750252 | 188.5310614 | 202.1657198 | 201.1107194 | 204.2608251 | 0.000211899 | -2.689172861 | 1.09E-05 | 19.3469678 | 0.155052333 | 3.67387089 |
| *Txlnb* | 53.09696783 | 83.54440136 | 127.8109588 | 60.52480997 | 15.85775283 | 7.366362258 | 14.62623414 | 12.16873001 | 1.74E-09 | -2.703780746 | 1.73E-11 | 45.25684839 | 0.153490285 | 8.758895057 |
| *Mefv* | 44.8374395 | 282.7656661 | 508.9614967 | 56.61998352 | 39.64438208 | 32.73938782 | 34.73730608 | 29.55263002 | 0.002650003 | -2.707862617 | 0.000249928 | 13.41268581 | 0.153056623 | 2.576753603 |
| *Plaur* | 562.827859 | 2975.465987 | 5562.059047 | 771.2032238 | 357.680425 | 377.3214446 | 377.5396587 | 390.2685553 | 0.000682405 | -2.715565938 | 4.53E-05 | 16.63496289 | 0.15224155 | 3.165957742 |
| *Gpr85* | 48.37723735 | 194.0800708 | 310.3980428 | 45.88171078 | 22.90564298 | 12.27727043 | 16.45451341 | 37.37538503 | 0.001212403 | -2.749485854 | 9.20E-05 | 15.29338521 | 0.148703875 | 2.916353141 |
| *Zc3h12c* | 462.5335864 | 1302.007363 | 2111.163159 | 736.0597857 | 172.6733086 | 162.0599697 | 190.1410438 | 160.8010751 | 6.86E-10 | -2.749711971 | 9.60E-10 | 37.40386727 | 0.14868057 | 7.163679493 |
| *Wfdc17* | 372.8587074 | 2321.249059 | 4923.004253 | 453.9360747 | 325.9649193 | 300.3838832 | 285.2115657 | 270.3196452 | 0.002410742 | -2.771620789 | 0.000220806 | 13.64518915 | 0.146439759 | 2.617849185 |
| *Gm16712* | 35.39797855 | 127.2445498 | 201.9869617 | 30.26240498 | 19.3816979 | 16.36969391 | 9.141396337 | 12.16873001 | 0.000542472 | -2.788378629 | 3.35E-05 | 17.20564709 | 0.144748607 | 3.265622792 |
| *Cxcl11* | 16.51905666 | 11.56768634 | 43.36443246 | 24.40516531 | 5.28591761 | 2.455454086 | 1.828279267 | 4.345975003 | 0.000103278 | -2.789361399 | 4.63E-06 | 20.98259318 | 0.144650037 | 3.985991553 |
| *St7* | 152.2113078 | 856.0087893 | 1448.14381 | 169.859505 | 88.09862684 | 90.85180119 | 85.92912557 | 93.87306007 | 0.000608668 | -2.871713683 | 3.92E-05 | 16.90803834 | 0.136624329 | 3.21561962 |
| *Jag1* | 343.360392 | 1096.359605 | 2002.752078 | 582.7953476 | 155.0535832 | 121.9542196 | 133.4643865 | 138.2020051 | 1.02E-06 | -2.87503797 | 2.06E-08 | 31.43609973 | 0.136309879 | 5.992070608 |
| *Fas* | 176.9898928 | 424.1484992 | 765.7245836 | 230.3847605 | 40.52536834 | 46.65362764 | 51.19181949 | 79.09674506 | 5.05E-07 | -2.876511579 | 9.17E-09 | 33.00873727 | 0.136170719 | 6.296495663 |
| *Hilpda* | 93.21467685 | 786.6026712 | 2073.504573 | 121.0496199 | 120.6951188 | 93.30725527 | 95.0705219 | 107.7801801 | 0.007086081 | -2.882612202 | 0.000903091 | 11.01640387 | 0.13559612 | 2.149593874 |
| *Slamf8* | 93.21467685 | 128.5298482 | 292.1393344 | 80.04894221 | 14.97676656 | 14.73272452 | 28.33832864 | 20.86068002 | 3.15E-07 | -2.913833316 | 5.42E-09 | 34.03388553 | 0.132693231 | 6.50181025 |
| *Slc2a1* | 976.984208 | 5231.164823 | 10971.20141 | 1227.091712 | 604.3565801 | 601.5862511 | 592.3624826 | 624.0820105 | 0.000586018 | -2.925683777 | 3.71E-05 | 17.01562149 | 0.131607738 | 3.232088903 |
| *Lcp2* | 554.5683307 | 3913.733879 | 6751.157432 | 829.7756205 | 407.015656 | 401.0575007 | 390.3376236 | 385.9225803 | 0.000666558 | -2.926950999 | 4.39E-05 | 16.69329452 | 0.131492188 | 3.176161734 |
| *Gem* | 82.59528329 | 200.5065633 | 330.9390898 | 129.8354794 | 22.90564298 | 23.73605617 | 20.11107194 | 29.55263002 | 8.85E-09 | -2.9483538 | 1.03E-10 | 41.76942114 | 0.129555862 | 8.052865208 |
| *Marcksl1* | 3097.323123 | 6400.786442 | 10276.22932 | 4385.120103 | 810.5073669 | 793.9301545 | 809.9277154 | 707.5247305 | 1.46E-13 | -2.952081974 | 8.16E-16 | 64.8315887 | 0.129221499 | 12.83656541 |
| *Mmp9* | 101.4742052 | 248.0626071 | 610.5255622 | 146.4309919 | 39.64438208 | 30.28393373 | 42.05042315 | 30.42182502 | 6.68E-07 | -2.958389033 | 1.19E-08 | 27.2471517 | 0.128657812 | 5.175153173 |
| *Gja1* | 74.33575496 | 524.4017808 | 1168.557338 | 116.1685869 | 59.02607998 | 54.0199899 | 58.50493656 | 70.40479505 | 0.001794828 | -2.960446638 | 0.000151902 | 14.3484822 | 0.128474448 | 2.745977124 |
| *Tmem176a* | 342.1804593 | 677.3523002 | 1154.863306 | 397.3160912 | 59.02607998 | 94.12573997 | 95.0705219 | 79.96594006 | 3.76E-10 | -2.969829777 | 3.37E-12 | 48.46134634 | 0.127641575 | 9.424819148 |
| *Cd55* | 22.41871975 | 41.12955144 | 70.75249506 | 21.47654547 | 4.404931342 | 8.184846954 | 2.742418901 | 4.345975003 | 1.03E-05 | -2.971982433 | 3.03E-07 | 26.23005937 | 0.127451262 | 4.986181073 |
| *Treml4* | 73.15582234 | 200.5065633 | 495.2674654 | 100.5492811 | 28.19156059 | 20.46211738 | 22.85349084 | 37.37538503 | 3.90E-05 | -2.997330828 | 1.45E-06 | 23.2193923 | 0.12523148 | 4.408670702 |
| *Dcn* | 44.8374395 | 53.98253626 | 89.01120346 | 65.40584303 | 10.57183522 | 5.729392868 | 5.484837802 | 9.561145007 | 5.39E-14 | -3.017720948 | 2.85E-16 | 66.90558726 | 0.123473988 | 13.26801879 |
| *Tmem45a* | 8.259528329 | 26.99126813 | 21.68221623 | 20.50033886 | 5.28591761 | 2.455454086 | 0.914139634 | 0.869195001 | 0.000100215 | -3.018062683 | 4.47E-06 | 21.05398621 | 0.123444744 | 3.999066664 |
| *Nod2* | 156.9310382 | 710.7700608 | 1378.532484 | 225.5037275 | 80.16975042 | 90.85180119 | 72.21703106 | 60.84365004 | 8.82E-05 | -3.022674708 | 3.83E-06 | 21.34769278 | 0.123050745 | 4.054563885 |
| *Dusp4* | 453.0941255 | 1322.572138 | 2338.255845 | 745.8218518 | 151.5296382 | 175.1557248 | 134.3785261 | 131.2484451 | 4.07E-08 | -3.036209278 | 5.27E-10 | 38.57508075 | 0.121901748 | 7.390863995 |
| *Fzd1* | 25.9585176 | 347.0305902 | 700.6779349 | 62.47722319 | 37.00142327 | 36.0133266 | 28.33832864 | 36.50619003 | 0.004414242 | -3.042462409 | 0.00048129 | 12.18679357 | 0.121374528 | 2.355143894 |
| *Bcam* | 10.61939357 | 29.5618651 | 36.5174168 | 23.4289587 | 1.761972537 | 4.092423477 | 0.914139634 | 5.215170004 | 1.94E-05 | -3.048835426 | 6.40E-07 | 24.78864161 | 0.120839546 | 4.711953467 |
| *Adamts7* | 3.539797855 | 11.56768634 | 21.68221623 | 13.66689257 | 1.761972537 | 1.636969391 | 0.914139634 | 1.738390001 | 0.000804755 | -3.058375103 | 5.47E-05 | 16.27667923 | 0.120043143 | 3.094336158 |
| *Bcl2a1b* | 684.3609187 | 4480.55051 | 6521.782407 | 1106.042092 | 377.0621229 | 393.6911385 | 406.792137 | 356.3699503 | 8.20E-05 | -3.059991162 | 3.54E-06 | 21.49746291 | 0.119908749 | 4.08608141 |
| *Arl5c* | 796.4545174 | 1898.385858 | 2443.243418 | 1087.494166 | 193.816979 | 167.7893626 | 194.711742 | 189.4845101 | 1.13E-14 | -3.061337446 | 5.52E-17 | 70.14274619 | 0.119796906 | 13.94672523 |
| *Hivep3* | 1024.181513 | 3490.870678 | 5267.637374 | 1450.643026 | 361.20437 | 276.647827 | 400.3931595 | 303.3490552 | 1.43E-07 | -3.065801253 | 2.23E-09 | 35.76528437 | 0.119426818 | 6.843372998 |
| *Nfkbia* | 3060.745212 | 10480.32383 | 19387.32482 | 4222.093598 | 1184.926531 | 1074.670405 | 1012.866714 | 1138.645451 | 1.47E-06 | -3.07415065 | 3.11E-08 | 30.63902963 | 0.118737648 | 5.833849733 |
| *Ddit4* | 62.53642878 | 687.6346881 | 1304.356481 | 93.71583478 | 75.76481908 | 60.56786746 | 63.98977436 | 49.54411504 | 0.003177765 | -3.103812778 | 0.000314701 | 12.98080509 | 0.116321302 | 2.49787829 |
| *Spint2* | 20.05885451 | 134.9563407 | 287.5746573 | 32.21481821 | 10.57183522 | 17.1881786 | 8.227256703 | 17.38390001 | 0.001647654 | -3.151042225 | 0.000135492 | 14.563806 | 0.112574953 | 2.783134038 |
| *Pgf* | 12.9792588 | 20.56477572 | 35.37624753 | 24.40516531 | 5.28591761 | 2.455454086 | 0.914139634 | 1.738390001 | 2.47E-06 | -3.165865991 | 5.72E-08 | 29.45755497 | 0.111424161 | 5.606900358 |
| *Malt1* | 866.0705419 | 7759.346938 | 13962.20608 | 1494.572323 | 659.858715 | 688.3456288 | 655.4381173 | 660.5882005 | 0.000613866 | -3.176152133 | 3.97E-05 | 16.884426 | 0.110632554 | 3.21192675 |
| *Cd200* | 22.41871975 | 160.6623103 | 386.8563843 | 24.40516531 | 15.85775283 | 22.09908678 | 9.141396337 | 17.38390001 | 0.003658687 | -3.202639862 | 0.000378214 | 12.63686173 | 0.108619884 | 2.436747743 |
| *Clec4a1* | 564.0077916 | 1980.644961 | 3828.622918 | 544.7232897 | 177.0782399 | 195.6178422 | 172.7723908 | 197.3072651 | 1.02E-05 | -3.219244242 | 2.96E-07 | 26.27776634 | 0.107376914 | 4.99334488 |
| *Zc3h12a* | 507.3710259 | 3404.75568 | 6136.067192 | 912.7531825 | 314.5120978 | 248.8193474 | 267.8429127 | 320.7329552 | 9.41E-05 | -3.25025002 | 4.13E-06 | 21.20199294 | 0.105093838 | 4.026325941 |
| *6530402F18Rik* | 54.27690045 | 825.1616257 | 1655.836618 | 100.5492811 | 58.14509371 | 68.75271441 | 66.73219326 | 78.22755006 | 0.003619805 | -3.277188861 | 0.000372575 | 12.66494782 | 0.103149672 | 2.441314868 |
| *Peg10* | 20.05885451 | 26.99126813 | 39.94092463 | 15.6193058 | 4.404931342 | 0.818484695 | 3.656558535 | 1.738390001 | 6.19E-07 | -3.285072046 | 1.15E-08 | 32.56299755 | 0.102587578 | 6.208239792 |
| *Cd14* | 10437.68394 | 59527.31391 | 112559.2315 | 15141.94076 | 4980.215375 | 5343.886576 | 4931.783324 | 4967.449429 | 5.55E-05 | -3.288965791 | 2.16E-06 | 22.41669898 | 0.102311074 | 4.255431534 |
| *Egln3* | 40.11770903 | 420.2926037 | 1066.993272 | 42.95309094 | 43.16832715 | 35.1948419 | 37.47972498 | 43.45975003 | 0.004712989 | -3.301140945 | 0.000525551 | 12.02275173 | 0.101451286 | 2.32670353 |
| *Pde4b* | 613.5649616 | 3449.741127 | 6301.536737 | 870.7762982 | 285.4395509 | 263.5520719 | 276.984309 | 309.4334202 | 4.42E-05 | -3.30676111 | 1.68E-06 | 22.92920894 | 0.101056841 | 4.35423717 |
| *C3* | 5684.915355 | 16504.51781 | 33515.00044 | 5436.494624 | 1512.653423 | 1524.836988 | 1516.557652 | 1537.605956 | 7.44E-07 | -3.327221316 | 1.41E-08 | 32.18032196 | 0.099633774 | 6.128270401 |
| *Arg2* | 34.21804593 | 472.9898415 | 930.0529592 | 43.92929756 | 36.120437 | 43.37968886 | 25.59590974 | 41.72136003 | 0.00430907 | -3.33414964 | 0.000466233 | 12.24609622 | 0.099156445 | 2.3656165 |
| *Flrt3* | 51.91703521 | 519.2605869 | 1095.522504 | 94.6920414 | 41.40635461 | 47.47211233 | 45.70698168 | 39.98297003 | 0.001126412 | -3.334632332 | 8.42E-05 | 15.46047104 | 0.099123275 | 2.9483027 |
| *Tpbg* | 11.79932618 | 35.98835751 | 90.15237274 | 10.73827274 | 4.404931342 | 5.729392868 | 3.656558535 | 0.869195001 | 0.001688528 | -3.337335802 | 0.000140474 | 14.4957745 | 0.098937702 | 2.772491745 |
| *Ehd1* | 3276.672881 | 11214.22926 | 19498.01824 | 5046.011979 | 947.0602385 | 937.1649762 | 962.5890343 | 990.0131057 | 2.47E-08 | -3.34676728 | 3.03E-10 | 39.65613468 | 0.098293016 | 7.606516369 |
| *Ch25h* | 130.9725206 | 377.8777538 | 494.1262961 | 243.0754465 | 47.57325849 | 34.37635721 | 24.68177011 | 15.64551001 | 3.19E-09 | -3.348433689 | 3.3E-11 | 43.97878411 | 0.098179546 | 8.495919961 |
| *Sntb1* | 12.9792588 | 65.5502226 | 190.5752689 | 26.35757853 | 5.28591761 | 3.273938782 | 10.05553597 | 10.43034001 | 0.00144217 | -3.349152543 | 0.000113288 | 14.9012519 | 0.098130638 | 2.840983666 |
| *Nlrp3* | 1463.116447 | 9251.578476 | 16536.68397 | 2992.073267 | 757.6481908 | 723.5404707 | 705.7157972 | 750.1152855 | 1.76E-05 | -3.364180126 | 5.68E-07 | 25.01820022 | 0.097113783 | 4.755151782 |
| *Dusp1* | 502.6512954 | 3007.598449 | 5403.436518 | 760.464951 | 265.1768668 | 235.7235923 | 227.6207688 | 175.5773901 | 3.51E-05 | -3.419535723 | 1.27E-06 | 23.4723079 | 0.093458149 | 4.454370403 |
| *Tma16* | 264.3049065 | 1496.087434 | 2672.618442 | 260.6471655 | 124.2190638 | 95.76270936 | 93.24224263 | 125.1640801 | 0.000101398 | -3.420341497 | 4.54E-06 | 21.02121584 | 0.093405966 | 3.993971648 |
| *Ralgds* | 535.6894088 | 3137.413596 | 5627.105696 | 760.464951 | 231.6993886 | 227.5387453 | 255.9590974 | 219.0371402 | 2.93E-05 | -3.428772366 | 1.02E-06 | 23.89554097 | 0.092861708 | 4.532862408 |
| *Pilra* | 316.2219417 | 1409.972435 | 2622.406994 | 337.7674879 | 112.7662423 | 109.6769492 | 101.4694993 | 107.7801801 | 1.28E-05 | -3.440262805 | 3.90E-07 | 25.74507972 | 0.092125043 | 4.893340041 |
| *Slco3a1* | 312.6821439 | 2859.789123 | 5345.236885 | 464.6743475 | 211.4367044 | 200.5287504 | 201.1107194 | 212.9527754 | 0.000401023 | -3.442789099 | 2.35E-05 | 17.87853487 | 0.091963864 | 3.396830542 |
| *Batf* | 99.11433995 | 600.2343913 | 1038.46404 | 81.02514882 | 42.28734088 | 43.37968886 | 39.30800425 | 41.72136003 | 0.000223969 | -3.447394925 | 1.17E-05 | 19.20589808 | 0.091670737 | 3.649812507 |
| *Hmox1* | 4100.265849 | 28273.99602 | 67626.83241 | 5298.849492 | 2382.18687 | 2406.345004 | 2307.288435 | 2464.167827 | 0.000356189 | -3.461342783 | 2.03E-05 | 18.15682838 | 0.090788742 | 3.44831999 |
| *Maff* | 129.792588 | 694.0611805 | 1055.58158 | 270.4092316 | 47.57325849 | 49.10908172 | 42.05042315 | 54.75928504 | 8.02E-07 | -3.473504301 | 1.54E-08 | 32.00308119 | 0.090026635 | 6.09594819 |
| *Olr1* | 44.8374395 | 341.8893963 | 548.9024213 | 119.0972067 | 22.02465671 | 33.55787251 | 20.11107194 | 17.38390001 | 3.74E-05 | -3.500921789 | 1.38E-06 | 23.31518698 | 0.088331891 | 4.427478999 |
| *Siglece* | 130.9725206 | 543.6812581 | 1062.428595 | 112.2637604 | 49.33523103 | 42.56120416 | 37.47972498 | 31.29102002 | 3.08E-05 | -3.524559433 | 1.08E-06 | 23.7748782 | 0.086896421 | 4.511450333 |
| *Adhfe1* | 3.539797855 | 65.5502226 | 94.71704984 | 6.833446286 | 4.404931342 | 4.092423477 | 0.914139634 | 5.215170004 | 0.008035576 | -3.539172761 | 0.001075032 | 10.69366136 | 0.086020674 | 2.094982974 |
| *Cd248* | 11.79932618 | 10.28238786 | 6.847015651 | 11.71447935 | 0 | 0 | 2.742418901 | 0.869195001 | 0.000104161 | -3.540098379 | 4.72E-06 | 20.94929354 | 0.085965501 | 3.982296304 |
| *Vcam1* | 47.19730474 | 124.6739528 | 220.2456701 | 55.6437769 | 6.166903878 | 9.003331649 | 8.227256703 | 14.77631501 | 2.85E-07 | -3.550361439 | 4.86E-09 | 34.24621925 | 0.085356129 | 6.545428109 |
| *Rab11fip1* | 527.4298804 | 6689.978601 | 13259.24581 | 891.2766371 | 469.565681 | 397.783562 | 447.9284205 | 463.2809353 | 0.004915713 | -3.58665318 | 0.000552482 | 11.92962839 | 0.083235733 | 2.308413466 |
| *Klf7* | 21.23878713 | 179.9417875 | 335.5037669 | 47.834124 | 7.928876415 | 7.366362258 | 12.79795487 | 19.12229001 | 0.000316222 | -3.630365346 | 1.77E-05 | 18.42399937 | 0.0807516 | 3.50000852 |
| *Cd274* | 495.5716997 | 7101.274115 | 14468.88524 | 835.6328602 | 412.3015736 | 511.5529346 | 438.7870242 | 465.8885203 | 0.006132882 | -3.646650419 | 0.000739271 | 11.38782384 | 0.079845205 | 2.212335384 |
| *Mcemp1* | 10.61939357 | 28.27656661 | 96.99938839 | 7.809652899 | 0.880986268 | 1.636969391 | 2.742418901 | 6.084365004 | 0.002704898 | -3.663574422 | 0.000257053 | 13.35996647 | 0.078914027 | 2.567849173 |
| *Met* | 80.23541805 | 951.120877 | 2689.735982 | 122.0258265 | 74.88383281 | 65.47877563 | 73.13117069 | 79.96594006 | 0.00765815 | -3.711020637 | 0.001006507 | 10.81555813 | 0.076360977 | 2.115876105 |
| *Gbp5* | 297.3430198 | 3678.524257 | 6516.076561 | 608.1767195 | 188.5310614 | 222.6278371 | 210.2521157 | 219.0371402 | 0.000226048 | -3.723221509 | 1.19E-05 | 19.18455753 | 0.075717915 | 3.645800237 |
| *Inhba* | 82.59528329 | 655.502226 | 1423.038086 | 109.3351406 | 46.69227222 | 41.74271946 | 42.96456278 | 39.98297003 | 0.000218396 | -3.727590759 | 1.13E-05 | 19.27451072 | 0.075488947 | 3.660754558 |
| *Abtb2* | 28.31838284 | 266.0567859 | 556.8906063 | 52.71515707 | 17.61972537 | 15.55120921 | 16.45451341 | 17.38390001 | 0.000266123 | -3.753642242 | 1.45E-05 | 18.82510003 | 0.074138039 | 3.574917357 |
| *Ccrl2* | 580.5268483 | 6720.825764 | 13045.84715 | 1040.636249 | 398.2057933 | 407.6053783 | 372.0548309 | 395.4837253 | 0.002143396 | -3.764854637 | 0.000190147 | 13.92602991 | 0.073564083 | 2.668897543 |
| *Htra4* | 23.59865237 | 176.0858921 | 329.7979205 | 14.64309919 | 6.166903878 | 10.64030104 | 7.313117069 | 15.64551001 | 0.001229168 | -3.772811358 | 9.35E-05 | 15.26346756 | 0.073159481 | 2.910388737 |
| *A430093F15Rik* | 2.359865237 | 8.997089377 | 15.97636985 | 7.809652899 | 0 | 2.455454086 | 0 | 0 | 0.008017714 | -3.781316844 | 0.001069798 | 10.70269098 | 0.072729434 | 2.095949434 |
| *Lif* | 93.21467685 | 574.5284216 | 804.524339 | 117.1447935 | 32.59649193 | 23.73605617 | 24.68177011 | 31.29102002 | 4.33E-06 | -3.822774403 | 1.09E-07 | 28.20550398 | 0.07066921 | 5.363840855 |
| *Fcgr2b* | 1135.095179 | 9400.6731 | 17571.7245 | 1662.479861 | 504.8051318 | 522.1932357 | 530.2009875 | 527.6013654 | 4.77E-05 | -3.835858617 | 1.83E-06 | 22.76327764 | 0.070031189 | 4.32129101 |
| *Sod2* | 2928.592759 | 16382.41446 | 32354.43129 | 3712.513747 | 945.2982659 | 955.9901242 | 941.5638227 | 1023.911711 | 6.42E-06 | -3.840100368 | 1.72E-07 | 27.3300729 | 0.069825588 | 5.192479481 |
| *Susd2* | 9.439460947 | 33.41776054 | 93.57588056 | 11.71447935 | 0.880986268 | 3.273938782 | 0.914139634 | 5.215170004 | 0.001064299 | -3.840231481 | 7.88E-05 | 15.58598048 | 0.069819243 | 2.972936327 |
| *Nfkbie* | 429.4954731 | 1980.644961 | 3838.893442 | 721.4166865 | 145.3627343 | 119.4987655 | 117.0098731 | 94.74225507 | 2.62E-07 | -3.870316722 | 4.42E-09 | 34.42935024 | 0.068378343 | 6.581908146 |
| *Lox* | 650.1428727 | 4983.102216 | 12252.73451 | 818.0611411 | 332.1318232 | 302.8393373 | 278.8125883 | 348.5471953 | 0.000149103 | -3.889161667 | 7.13E-06 | 20.15824906 | 0.067490972 | 3.826514058 |

| | | | | | | | | | | | | | | |
|---|---|---|---|---|---|---|---|---|---|---|---|---|---|---|
| *Tnip3* | 702.059908 | 5426.530193 | 8802.979788 | 1034.779009 | 269.5817981 | 279.9217658 | 234.9338859 | 263.3660852 | 9.27E-06 | -3.929536636 | 2.66E-07 | 26.47911885 | 0.065628368 | 5.033034217 |
| *Alpk2* | 3.539797855 | 16.70888027 | 30.81157043 | 2.928619837 | 0 | 0.818484695 | 1.828279267 | 0.869195001 | 0.004659963 | -3.945544306 | 0.000516321 | 12.05577938 | 0.064904202 | 2.331617524 |
| *Vegfc* | 113.2735314 | 266.0567859 | 362.8918295 | 97.62066123 | 10.57183522 | 12.27727043 | 10.05553597 | 20.86068002 | 4.61E-11 | -3.963570443 | 3.36E-13 | 52.98784898 | 0.064098285 | 10.33625119 |
| *Tnnt2* | 3.539797855 | 15.42358179 | 19.39987768 | 2.928619837 | 0.880986268 | 0 | 0.914139634 | 0.869195001 | 0.002631577 | -3.966706646 | 0.000247348 | 13.43215013 | 0.063959096 | 2.57978391 |
| *Tnfsf9* | 151.0313752 | 767.323194 | 1243.87451 | 366.0774796 | 40.52536834 | 30.28393373 | 38.39386461 | 51.28250504 | 1.53E-08 | -3.977822925 | 1.83E-10 | 40.63623945 | 0.063468171 | 7.815808634 |
| *Tmc3* | 23.59865237 | 176.0858921 | 454.1853715 | 31.23861159 | 7.047890147 | 19.64363269 | 11.88381524 | 4.345975003 | 0.000792446 | -3.994497981 | 5.35E-05 | 16.31981161 | 0.062738812 | 3.101030367 |
| *Dusp2* | 99.11433995 | 1071.938934 | 1606.766339 | 138.621339 | 45.81128595 | 40.92423477 | 41.13628352 | 46.06733503 | 5.21E-05 | -4.067414633 | 2.01E-06 | 22.58315101 | 0.059646668 | 4.283284233 |
| *Dusp16* | 217.1076018 | 2818.659572 | 4741.558338 | 380.7205788 | 117.1711737 | 109.6769492 | 136.2068054 | 117.3413251 | 9.11E-05 | -4.085925737 | 3.97E-06 | 21.2811575 | 0.058886236 | 4.040289484 |
| *Nhs* | 2.359865237 | 21.8500742 | 29.67040115 | 5.857239674 | 1.761972537 | 0 | 0 | 1.738390001 | 0.004582126 | -4.094049205 | 0.000506193 | 12.09271253 | 0.058555593 | 2.338932973 |
| *Cd69* | 338.6406615 | 1706.876385 | 2617.842317 | 650.1536038 | 80.16975042 | 91.67028588 | 60.33321582 | 75.61996505 | 4.46E-09 | -4.109243059 | 4.75E-11 | 43.27858719 | 0.057942146 | 8.350441371 |
| *Art2a-ps* | 5.899663092 | 14.13828331 | 34.23507825 | 6.833446286 | 2.642958805 | 0 | 0.914139634 | 0 | 0.001527239 | -4.120368125 | 0.000122291 | 14.75701801 | 0.057497055 | 2.816093061 |
| *Ets2* | 880.2297333 | 5938.078989 | 10681.34442 | 1300.307208 | 258.1289766 | 278.2847964 | 276.0701694 | 265.9736702 | 2.11E-06 | -4.123657654 | 4.75E-08 | 29.81604026 | 0.057366104 | 5.675026494 |
| *Ppfia3* | 38.93777641 | 410.0102159 | 803.3831697 | 92.73962817 | 14.97676656 | 33.55787251 | 14.62623414 | 13.90712001 | 9.77E-05 | -4.123678073 | 4.32E-06 | 21.11720845 | 0.057365292 | 4.010265501 |
| *Phlda1* | 92.03474424 | 615.6579731 | 1450.426149 | 109.3351406 | 29.07254686 | 46.65362764 | 23.76763048 | 30.42182502 | 5.87E-05 | -4.124624736 | 2.35E-06 | 22.28592634 | 0.057327662 | 4.231101447 |
| *Ereg* | 4.719730474 | 20.56477572 | 34.23507825 | 2.928619837 | 0 | 0.818484695 | 1.828279267 | 0.869195001 | 0.002494405 | -4.155581842 | 0.000231063 | 13.55995042 | 0.056110639 | 2.603033018 |
| *Kdm6b* | 53.09696783 | 1068.083039 | 1898.905674 | 144.4785786 | 44.04931342 | 37.65029599 | 53.02009875 | 41.72136003 | 0.000269561 | -4.164835111 | 1.46E-05 | 18.79211124 | 0.055751904 | 3.569342429 |
| *Nox1* | 74.33575496 | 308.4716358 | 597.9727002 | 137.6451323 | 20.26268417 | 10.64030104 | 15.54037377 | 13.03792501 | 3.13E-08 | -4.233936783 | 3.93E-10 | 39.14860038 | 0.053144464 | 7.504844554 |
| *Gpr31b* | 3.539797855 | 51.4119393 | 99.28172694 | 11.71447935 | 1.761972537 | 1.636969391 | 2.742418901 | 2.607585002 | 0.000692256 | -4.247367204 | 4.61E-05 | 16.6009358 | 0.052652024 | 3.159733216 |
| *Fcrlb* | 4.719730474 | 59.12373019 | 123.2462817 | 11.71447935 | 3.523945073 | 1.636969391 | 1.828279267 | 3.476780003 | 0.00076189 | -4.247727193 | 5.13E-05 | 16.40108564 | 0.052638887 | 3.118107679 |
| *Plat* | 22.41871975 | 181.227086 | 255.6219176 | 18.54792563 | 11.45282149 | 3.273938782 | 4.570698168 | 5.215170004 | 0.000171605 | -4.285716873 | 8.44E-06 | 19.83499613 | 0.051270868 | 3.765470729 |
| *Ccl12* | 29.49831546 | 155.5211164 | 199.7046232 | 31.23861159 | 7.047890147 | 5.729392868 | 5.484837802 | 2.607585002 | 2.21E-06 | -4.315834427 | 5.00E-08 | 29.71586937 | 0.050211637 | 5.655666438 |
| *Nfkbiz* | 1093.797537 | 7638.528881 | 13793.31303 | 1915.317373 | 327.7268918 | 310.2056996 | 283.3832864 | 292.0495202 | 4.72E-07 | -4.332184385 | 8.48E-09 | 33.16279899 | 0.049645805 | 6.325711904 |
| *Lrrc63* | 2.359865237 | 15.42358179 | 29.67040115 | 4.881033062 | 0 | 0.818484695 | 0 | 1.738390001 | 0.003606645 | -4.333917595 | 0.000369967 | 12.67808302 | 0.049586197 | 2.442896571 |
| *Ptgs2os2* | 8.259528329 | 73.2620135 | 186.0105918 | 9.762066123 | 2.642958805 | 2.455454086 | 4.570698168 | 3.476780003 | 0.000583294 | -4.400959568 | 3.68E-05 | 17.03134043 | 0.047334649 | 3.234112572 |
| *Mmp14* | 1402.939883 | 6539.598678 | 13854.93617 | 1638.074696 | 264.2958805 | 270.0999495 | 273.3277505 | 295.5263002 | 1.32E-07 | -4.408846591 | 2.02E-09 | 35.95072308 | 0.047076583 | 6.880207637 |
| *Arnt2* | 7.07959571 | 19.27947724 | 57.05846376 | 9.762066123 | 2.642958805 | 0.818484695 | 0.914139634 | 0 | 0.000338486 | -4.416753091 | 1.92E-05 | 18.26999619 | 0.046819291 | 3.470459759 |
| *Gpr18* | 20.05885451 | 159.3770118 | 293.2805037 | 14.64309919 | 2.642958805 | 4.092423477 | 5.484837802 | 10.43034001 | 0.00031333 | -4.427202559 | 1.74E-05 | 18.44960515 | 0.046481403 | 3.503997994 |
| *Ifi205* | 101.4742052 | 808.4527454 | 1740.283145 | 87.85859511 | 19.3816979 | 43.37968886 | 19.19693231 | 42.59055503 | 9.16E-05 | -4.4575675 | 4.00E-06 | 21.26520833 | 0.045513316 | 4.038269695 |
| *Cish* | 27.13845022 | 359.8835751 | 462.1735564 | 36.11964466 | 8.809862684 | 9.003331649 | 10.9696756 | 11.29953501 | 5.21E-05 | -4.465044032 | 2.01E-06 | 22.5858366 | 0.04527806 | 4.283284233 |
| *F3* | 2.359865237 | 77.11790894 | 165.4695449 | 4.881033062 | 3.523945073 | 4.910908172 | 1.828279267 | 0.869195001 | 0.005027547 | -4.48281072 | 0.000570681 | 11.86925802 | 0.044723884 | 2.298643892 |
| *Adora2b* | 149.8514425 | 1106.641993 | 1986.775708 | 232.3371737 | 39.64438208 | 42.56120416 | 39.30800425 | 31.29102002 | 1.01E-06 | -4.507304572 | 2.02E-08 | 31.47399478 | 0.043970978 | 5.997099746 |
| *Ntng2* | 16.51905666 | 222.3566375 | 529.5025437 | 15.6193058 | 6.166903878 | 10.64030104 | 10.05553597 | 6.953560005 | 0.000619636 | -4.53440355 | 4.02E-05 | 16.86198056 | 0.043152754 | 3.207863267 |
| *Mycl* | 51.91703521 | 47.55604385 | 101.5640655 | 65.40584303 | 3.523945073 | 3.273938782 | 2.742418901 | 1.738390001 | 3.39E-20 | -4.562218441 | 1.06E-22 | 96.16096675 | 0.042328746 | 19.46917234 |
| *Ier3* | 248.9657825 | 3857.180746 | 7773.645102 | 439.2929756 | 133.9099128 | 99.85513284 | 127.9795487 | 153.8475151 | 0.000933921 | -4.578527401 | 6.70E-05 | 15.89337664 | 0.041852935 | 3.029689769 |
| *Hrc* | 2.359865237 | 20.56477572 | 53.63495593 | 5.857239674 | 0 | 1.636969391 | 0 | 1.738390001 | 0.0041583 | -4.586681517 | 0.000444006 | 12.33725895 | 0.041617049 | 2.381084178 |
| *Gm6377* | 164.010634 | 2205.572196 | 3639.188818 | 252.8375126 | 75.76481908 | 63.84180624 | 63.07563472 | 54.75928504 | 2.31E-05 | -4.604163722 | 7.80E-07 | 24.40539946 | 0.041115788 | 4.635996825 |
| *Adamts4* | 29.49831546 | 177.3711906 | 262.4689333 | 37.09585127 | 7.047890147 | 6.547877563 | 2.742418901 | 4.345975003 | 1.79E-06 | -4.609555169 | 3.89E-08 | 30.20470287 | 0.040962422 | 5.746336669 |
| *Socs1* | 46.01737212 | 1033.37998 | 1911.458536 | 95.66824801 | 26.42958805 | 31.10241842 | 38.39386461 | 28.68343502 | 0.001640589 | -4.63051333 | 0.000134722 | 14.57455321 | 0.040371659 | 2.785000264 |
| *Spic* | 22.41871975 | 134.9563407 | 395.9857385 | 31.23861159 | 1.761972537 | 7.366362258 | 7.313117069 | 6.953560005 | 8.06E-05 | -4.642082276 | 3.45E-06 | 21.54807632 | 0.040049214 | 4.093503539 |
| *Slpi* | 17.69898928 | 259.6302934 | 612.8079008 | 21.47654547 | 8.809862684 | 7.366362258 | 5.484837802 | 13.90712001 | 0.00042342 | -4.679144133 | 2.52E-05 | 17.74938289 | 0.039033479 | 3.373228979 |
| *Tnfrsf9* | 41.29764164 | 215.930145 | 475.8675877 | 51.73895045 | 2.642958805 | 10.64030104 | 9.141396337 | 6.953560005 | 5.07E-06 | -4.737745107 | 1.30E-07 | 27.86229758 | 0.037479744 | 5.294737587 |
| *Cfb* | 152.2113078 | 1079.650725 | 3509.095521 | 130.8116861 | 45.81128595 | 52.3830205 | 42.05042315 | 41.72136003 | 0.000710866 | -4.742574269 | 4.75E-05 | 16.54531385 | 0.037354497 | 3.148212182 |
| *Hp* | 27.13845022 | 316.1834267 | 813.6536932 | 39.04826449 | 1.761972537 | 19.64363269 | 6.398977436 | 15.64551001 | 0.000655958 | -4.780324172 | 4.30E-05 | 16.73525631 | 0.036389747 | 3.183124103 |
| *F10* | 34.21804593 | 952.4061755 | 2286.903227 | 78.09652899 | 28.19156059 | 28.64696434 | 32.90902681 | 30.42182502 | 0.002749415 | -4.801717375 | 0.000262552 | 13.32027273 | 0.035854118 | 2.560759699 |
| *Hdc* | 7.07959571 | 152.9505194 | 445.0560173 | 16.59551241 | 5.28591761 | 2.455454086 | 6.398977436 | 7.822755006 | 0.000840076 | -4.824813332 | 5.79E-05 | 16.16924455 | 0.035284703 | 3.075681278 |
| *Traf1* | 364.5991791 | 2759.535842 | 5020.003641 | 655.0346369 | 89.86059937 | 64.66029094 | 68.56047253 | 86.91950006 | 1.13E-07 | -4.827026978 | 1.69E-09 | 36.29787045 | 0.035230604 | 6.948430953 |
| *Il27* | 3.539797855 | 50.12664081 | 92.43471129 | 3.904826449 | 1.761972537 | 1.636969391 | 0 | 1.738390001 | 0.002069421 | -4.857571514 | 0.000182094 | 14.00738028 | 0.034492547 | 2.684151123 |
| *Cd40* | 540.4091392 | 5956.073167 | 8150.230963 | 1148.018976 | 126.8620226 | 121.1357349 | 112.4391749 | 171.2314151 | 1.85E-07 | -4.892720391 | 2.97E-09 | 35.20667677 | 0.033662348 | 6.732052858 |
| *Procr* | 119.1731945 | 4145.087606 | 9861.984876 | 300.6716366 | 127.7430089 | 108.8584645 | 125.2371298 | 121.6873001 | 0.00305567 | -4.899018284 | 0.000299187 | 13.0754752 | 0.03351572 | 2.514893496 |
| *Ak4* | 83.77521591 | 764.752597 | 1951.39946 | 98.59686785 | 28.19156059 | 27.00999495 | 19.19693231 | 20.86068002 | 1.97E-05 | -4.926936895 | 6.52E-07 | 24.75313937 | 0.032873368 | 4.705007035 |
| *Pde10a* | 10.61939357 | 89.97089377 | 160.9048678 | 13.66689257 | 3.523945073 | 0.818484695 | 3.656558535 | 0.869195001 | 0.000111919 | -4.96442993 | 5.10E-06 | 20.7981931 | 0.032030055 | 3.951095282 |
| *Clec4e* | 5617.659196 | 33425.47233 | 60931.59228 | 8735.096767 | 777.910875 | 861.0458995 | 973.5587099 | 852.6802956 | 3.03E-09 | -4.971397754 | 3.10E-11 | 44.10957632 | 0.031875731 | 8.518206161 |
| *Upp1* | 11.79932618 | 276.3391737 | 572.8669761 | 15.6193058 | 3.523945073 | 4.092423477 | 9.141396337 | 10.43034001 | 0.000586018 | -5.012083163 | 3.71E-05 | 17.01306629 | 0.030989361 | 3.232088903 |
| *Lix1* | 17.69898928 | 124.6739528 | 222.5280087 | 8.785859511 | 2.642958805 | 4.092423477 | 0 | 4.345975003 | 0.000399111 | -5.067601779 | 2.34E-05 | 17.89282194 | 0.029819466 | 3.398905947 |
| *Robo2* | 5.899663092 | 14.13828331 | 86.72886491 | 11.71447935 | 0 | 0 | 0 | 3.476780003 | 0.00488297 | -5.090145284 | 0.000548021 | 11.94473357 | 0.029357129 | 2.311315976 |
| *Il4i1* | 17.69898928 | 1149.056843 | 2460.360957 | 121.0496199 | 23.78662925 | 24.55454086 | 37.47972498 | 23.46826502 | 0.002673374 | -5.100108042 | 0.000253202 | 13.38827808 | 0.029155098 | 2.572940272 |
| *Ednrb* | 83.77521591 | 460.1368567 | 738.336521 | 142.5261654 | 12.33380776 | 13.91423982 | 8.227256703 | 5.215170004 | 3.17E-09 | -5.163583888 | 3.27E-11 | 44.00843104 | 0.027900139 | 8.499144401 |
| *Ppm1n* | 23.59865237 | 68.12081957 | 77.59951071 | 20.50033886 | 0.880986268 | 1.636969391 | 0 | 2.607585002 | 8.49E-09 | -5.190460082 | 9.78E-11 | 41.86461169 | 0.027385194 | 8.071003143 |
| *Gpr84* | 1772.258793 | 5691.30168 | 9824.32629 | 2849.547101 | 156.8155558 | 120.3172502 | 110.6108957 | 155.5859051 | 1.47E-17 | -5.211875725 | 5.43E-20 | 83.81716023 | 0.026981685 | 16.83123987 |
| *Tnfaip3* | 841.2919569 | 9813.253913 | 18671.81168 | 1744.481216 | 192.9359928 | 202.1657198 | 218.4793724 | 221.6447252 | 5.24E-07 | -5.217241995 | 9.60E-09 | 32.92116501 | 0.02688151 | 6.280742978 |
| *Col19a1* | 4.719730474 | 33.41776054 | 27.3880626 | 3.904826449 | 0 | 0.818484695 | 0 | 0.869195001 | 0.000331798 | -5.317825658 | 1.87E-05 | 18.31880237 | 0.025071191 | 3.479125865 |
| *H2-M2* | 44.8374395 | 284.0509646 | 758.877568 | 33.19102482 | 3.523945073 | 8.184846954 | 10.05553597 | 5.215170004 | 1.18E-05 | -5.376849588 | 3.55E-07 | 25.92217133 | 0.024066171 | 4.926644836 |
| *Tnfsf15* | 51.91703521 | 1190.186395 | 2074.645742 | 126.9068596 | 20.26268417 | 26.19151025 | 11.88381524 | 23.46826502 | 0.000299667 | -5.394963198 | 1.66E-05 | 18.54566442 | 0.023765899 | 3.523361799 |
| *Cdc42ep2* | 60.17656354 | 704.3435684 | 1117.20472 | 137.6451323 | 11.45282149 | 12.27727043 | 8.227256703 | 15.64551001 | 2.29E-07 | -5.405737291 | 3.79E-09 | 34.72786371 | 0.023589076 | 6.64019663 |
| *Mmp13* | 234.8065911 | 9527.91765 | 21521.31136 | 666.7491162 | 177.9592262 | 180.8851177 | 205.6814176 | 181.6617551 | 0.001384227 | -5.42015678 | 0.000107961 | 14.9921193 | 0.023354481 | 2.85879258 |
| *Mgarp* | 3.539797855 | 92.54149073 | 235.0808707 | 11.71447935 | 1.761972537 | 0.818484695 | 2.742418901 | 2.607585002 | 0.000374947 | -5.439500332 | 2.17E-05 | 18.03525446 | 0.023043436 | 3.426030676 |
| *Ccl5* | 164.010634 | 4628.359835 | 9986.372327 | 345.5771408 | 78.40777788 | 79.39301545 | 77.70186886 | 100.8266201 | 0.000682896 | -5.490821323 | 4.54E-05 | 16.63131558 | 0.022238121 | 3.165645557 |
| *Sh2d4a* | 1.179932618 | 17.99417875 | 7.988184926 | 1.952413225 | 0 | 0 | 0 | 0 | 0.00647624 | -5.49746973 | 0.000798327 | 11.24511839 | 0.022135876 | 2.188677061 |
| *Ccl4* | 297.3430198 | 5605.186682 | 8239.242167 | 808.299075 | 80.16975042 | 91.67028588 | 65.81805362 | 93.00386507 | 3.07E-05 | -5.498544058 | 1.08E-06 | 23.78886193 | 0.022119398 | 4.513109815 |
| *Rasgrp1* | 16.51905666 | 255.774398 | 514.6673431 | 33.19102482 | 3.523945073 | 2.455454086 | 3.656558535 | 7.822755006 | 2.24E-05 | -5.554647382 | 7.52E-07 | 24.47766341 | 0.021275733 | 4.649188168 |
| *Arg1* | 1.179932618 | 133.6710422 | 502.1144811 | 12.69068596 | 0.880986268 | 3.273938782 | 5.484837802 | 3.476780003 | 0.008147099 | -5.631202954 | 0.001094882 | 10.65981661 | 0.020176183 | 2.088997018 |
| *Bcl2a1d* | 123.8929249 | 2512.758533 | 3988.386617 | 268.4568184 | 36.120437 | 39.28726538 | 40.22214388 | 22.59907002 | 8.20E-05 | -5.640066032 | 3.53E-06 | 21.50230992 | 0.020052612 | 4.086374132 |
| *Slc7a11* | 1139.814909 | 26088.9886 | 49330.46543 | 2095.915597 | 382.3480405 | 378.958414 | 377.5396687 | 388.5301653 | 0.00019942 | -5.686397413 | 1.01E-05 | 19.48680002 | 0.019418864 | 3.700231204 |
| *Ptx3* | 4.719730474 | 118.2474604 | 188.2929304 | 13.66689257 | 0 | 3.273938782 | 0.914139634 | 1.738390001 | 0.000299163 | -5.762605556 | 1.65E-05 | 18.55620743 | 0.018419714 | 3.524092169 |
| *Mmp3* | 3.539797855 | 33.41776054 | 53.63495593 | 5.857239674 | 1.761972537 | 0 | 0 | 0 | 0.000813806 | -5.781634289 | 5.57E-05 | 16.24266222 | 0.018178358 | 3.089478954 |
| *Ccl20* | 1.179932618 | 17.99417875 | 18.2587084 | 1.952413225 | 0 | 0 | 0 | 0 | 0.004017802 | -5.938018845 | 0.000423559 | 12.42529091 | 0.016310911 | 2.396011511 |
| *Serpinb2* | 31.8581807 | 787.8879697 | 1807.612132 | 42.95309094 | 3.523945073 | 18.0066633 | 10.9696756 | 8.691950006 | 0.000906083 | -6.017405553 | 6.39E-05 | 15.98257755 | 0.015437623 | 3.042832017 |
| *Irg1* | 5604.679937 | 58828.11154 | 115822.9756 | 10754.86825 | 679.2404129 | 751.3689504 | 703.8875179 | 659.7190055 | 6.45E-09 | -6.095052579 | 7.19E-11 | 42.46529016 | 0.014628721 | 8.190696355 |
| *Cp* | 75.51568758 | 992.2504284 | 2281.197381 | 65.40584303 | 11.45282149 | 17.1881786 | 10.05553597 | 10.43034001 | 0.000116024 | -6.118164185 | 5.33E-06 | 20.71581561 | 0.014396239 | 3.935450503 |
| *Tnf* | 1402.939883 | 17744.83085 | 26397.52767 | 4553.02764 | 157.696542 | 153.8751227 | 147.176481 | 179.9233651 | 8.54E-11 | -6.293521285 | 6.49E-13 | 51.69331649 | 0.012748565 | 10.0686463 |
| *Slc1a2* | 2.359865237 | 131.1004452 | 269.3159489 | 7.809652899 | 0 | 2.455454086 | 1.828279267 | 0.869195001 | 0.000710866 | -6.310237235 | 4.75E-05 | 16.5438096 | 0.012601705 | 3.148212182 |
| *Ccl22* | 35.39797855 | 3083.431059 | 6401.959634 | 183.5268431 | 38.76339581 | 36.83181129 | 20.11107194 | 23.46826502 | 0.000837977 | -6.347262108 | 5.77E-05 | 16.17618262 | 0.012282414 | 3.076767892 |
| *Gpr110* | 1.179932618 | 14.13828331 | 37.65858608 | 0.976206612 | 0 | 0 | 0 | 0 | 0.006795725 | -6.395571823 | 0.000853475 | 11.12117985 | 0.011877938 | 2.167764204 |
| *Ccl2* | 266.6647718 | 6977.885461 | 11361.4813 | 816.1087279 | 69.5979152 | 67.93422972 | 44.79284205 | 40.85216503 | 1.91E-05 | -6.443235366 | 6.25E-07 | 24.83413556 | 0.011491928 | 4.720090111 |
| *Socs3* | 850.7314179 | 12684.61072 | 23569.71021 | 2159.369027 | 125.1000501 | 99.85513284 | 110.6108957 | 104.3034001 | 1.38E-06 | -6.480012305 | 2.90E-08 | 30.77099264 | 0.01120268 | 5.858627679 |
| *Dnmt3l* | 14.15919142 | 64.26492412 | 138.0814823 | 18.54792563 | 0 | 1.636969391 | 0 | 0.869195001 | 1.15E-06 | -6.511586591 | 2.36E-08 | 31.17438087 | 0.010960166 | 5.938643964 |
| *Mmp10* | 2.359865237 | 25.70596965 | 59.34080231 | 9.762066123 | 0 | 0.818484695 | 0 | 0 | 0.000184369 | -6.523566232 | 9.20E-06 | 19.67010609 | 0.010869533 | 3.734311146 |
| *Adm* | 3.539797855 | 66.83552108 | 172.3165605 | 4.881033062 | 0 | 0.818484695 | 0 | 1.738390001 | 0.000669304 | -6.58233341 | 4.42E-05 | 16.680898 | 0.010435667 | 3.174376604 |
| *Il1rn* | 665.4819968 | 28343.40213 | 56358.92699 | 1530.691968 | 214.0796632 | 244.7269239 | 207.5096968 | 237.2902352 | 0.000191394 | -6.587461682 | 9.61E-06 | 19.58645543 | 0.010398637 | 3.718072762 |
| *Zfp811* | 16.51905666 | 255.774398 | 529.5025437 | 34.16723143 | 3.523945073 | 0 | 0.914139634 | 3.476780003 | 1.77E-05 | -6.727213573 | 5.73E-07 | 24.99993732 | 0.009438586 | 4.752111826 |
| *Hcar2* | 77.87555281 | 2708.123902 | 4974.35687 | 216.7178679 | 21.14367044 | 22.91757147 | 17.36865304 | 11.29953501 | 6.17E-05 | -6.77692417 | 2.49E-06 | 22.17717851 | 0.009118902 | 4.209964651 |
| *Ccl3* | 755.1568758 | 18770.49904 | 29283.54477 | 2105.677663 | 106.5993385 | 126.8651278 | 99.64122007 | 112.9953501 | 4.33E-06 | -6.834513076 | 1.09E-07 | 28.20059714 | 0.008762067 | 5.363840855 |
| *Marco* | 246.6059172 | 2484.481966 | 5442.236273 | 231.3609671 | 21.14367044 | 18.0066633 | 13.71209451 | 12.16873001 | 2.61E-06 | -7.013682444 | 6.21E-08 | 29.2970806 | 0.007738757 | 5.582795879 |
| *Ccl7* | 99.11433995 | 2222.281076 | 4026.045203 | 222.5751076 | 15.85775283 | 15.55120921 | 6.398977436 | 12.16873001 | 1.18E-05 | -7.037666001 | 3.57E-07 | 25.91347076 | 0.00761117 | 4.926644836 |
| *Adora2a* | 27.13845022 | 800.7409545 | 1531.449167 | 64.42963641 | 4.404931342 | 4.910908172 | 3.656558535 | 3.476780003 | 3.18E-05 | -7.202117378 | 1.13E-06 | 23.70058358 | 0.006791202 | 4.498026621 |
| *Ptges* | 274.9243001 | 5380.259447 | 13796.73654 | 523.2467442 | 32.59649193 | 28.64696434 | 27.42418901 | 44.32894503 | 1.34E-05 | -7.230640889 | 4.11E-07 | 25.6409025 | 0.006658252 | 4.872169619 |
| *Cxcl5* | 1.179932618 | 37.27365599 | 66.18781796 | 0.976206612 | 0 | 0 | 0 | 0 | 0.003468761 | -7.365144199 | 0.000352675 | 12.76759835 | 0.006065557 | 2.459825684 |
| *Lad1* | 11.79932618 | 417.7220068 | 560.3141141 | 32.21481821 | 1.761972537 | 1.636969391 | 1.828279267 | 0.869195001 | 1.46E-06 | -7.390054704 | 3.09E-08 | 30.64818086 | 0.005961724 | 5.834256128 |
| *Il1f6* | 3.539797855 | 25.70596965 | 83.30535709 | 0 | 0 | 0 | 0 | 0 | 0.009410291 | -7.457459613 | 0.001303033 | 10.33822449 | 0.00568959 | 2.026396925 |
| *Slamf1* | 1.179932618 | 159.3770118 | 309.2568736 | 1.952413225 | 0.880986268 | 0.818484695 | 0.914139634 | 0 | 0.005223226 | -7.496897203 | 0.000598856 | 11.77952863 | 0.005536166 | 2.282061184 |
| *Il10* | 14.15919142 | 379.1630523 | 752.0305523 | 26.35757853 | 0 | 0.818484695 | 3.656558535 | 1.738390001 | 0.000503989 | -7.566346998 | 3.09E-05 | 17.36080846 | 0.005275973 | 3.297579289 |
| *Edn1* | 5.899663092 | 461.4221552 | 707.5249506 | 10.73827274 | 4.404931342 | 0.818484695 | 0.914139634 | 0 | 0.002974239 | -7.595880793 | 0.000289549 | 13.13682115 | 0.005169066 | 2.526624207 |
| *Epha4* | 3.539797855 | 165.8035042 | 353.7624753 | 18.54792563 | 0.880986268 | 0 | 1.828279267 | 0 | 0.00167688 | -7.664469123 | 0.000139371 | 14.51062705 | 0.004929069 | 2.775498069 |
| *Cd247* | 7.07959571 | 33.41776054 | 84.44652636 | 7.809652899 | 0 | 0 | 0 | 0 | 1.59E-05 | -7.695130349 | 5.07E-07 | 25.23671408 | 0.004825418 | 4.797864665 |
| *Fpr2* | 115.6333966 | 1485.805046 | 3051.486642 | 138.621339 | 7.928876415 | 7.366362258 | 4.570698168 | 1.738390001 | 3.58E-06 | -7.791921884 | 8.88E-08 | 28.60501886 | 0.004512298 | 5.446239526 |
| *Vgf* | 2.359865237 | 65.5502226 | 74.17600288 | 0.976206612 | 0 | 0 | 0 | 0 | 0.001539571 | -7.802851711 | 0.000123648 | 14.73621234 | 0.004478242 | 2.812600324 |
| *Cd38* | 18.87892189 | 588.6667049 | 1556.554891 | 48.81033062 | 0.880986268 | 4.092423477 | 3.656558535 | 0.869195001 | 0.000129888 | -7.862235588 | 6.08E-06 | 20.4633974 | 0.004297652 | 3.886431943 |
| *Steap4* | 3.539797855 | 46.27074537 | 96.99938839 | 2.928619837 | 0 | 0 | 0 | 0 | 0.000360987 | -7.868955886 | 2.07E-05 | 18.12601992 | 0.00427768 | 3.442508577 |
| *Bcl2a1a* | 34.21804593 | 647.7904351 | 1145.733952 | 80.04894221 | 1.761972537 | 2.455454086 | 0 | 3.476780003 | 2.93E-05 | -7.947582547 | 1.02E-06 | 23.88966173 | 0.004050786 | 4.532862408 |
| *Lcn2* | 22.41871975 | 1906.097649 | 4520.171499 | 99.57307446 | 7.047890147 | 6.547877563 | 5.484837802 | 4.345975003 | 9.20E-05 | -8.126549712 | 4.03E-06 | 21.2488967 | 0.003578201 | 4.03616185 |
| *Il1f9* | 1.179932618 | 266.0567859 | 633.3489477 | 7.809652899 | 0 | 0 | 0.914139634 | 1.738390001 | 0.006184856 | -8.426872088 | 0.000747566 | 11.36709865 | 0.002905753 | 2.208670394 |
| *Ptgs2* | 507.3710259 | 52819.34113 | 92572.79277 | 2255.037275 | 73.12186027 | 79.39301545 | 72.21703106 | 91.26547507 | 1.59E-05 | -8.872932494 | 5.10E-07 | 25.22729089 | 0.002132952 | 4.797800826 |
| *Tarm1* | 15.33912404 | 710.7700608 | 1653.55428 | 36.11964466 | 1.761972537 | 0 | 2.742418901 | 0 | 0.001088734 | -9.079094849 | 8.13E-05 | 15.52882828 | 0.001848929 | 2.963078413 |
| *Fpr1* | 100.2942726 | 1068.083039 | 2343.961691 | 72.23928931 | 0.880986268 | 0 | 2.742418901 | 0 | 4.29E-05 | -9.970551709 | 1.62E-06 | 23.00439543 | 0.000996701 | 4.367550333 |
| *Saa3* | 456.6339233 | 18737.08128 | 47678.05232 | 1090.422786 | 13.21479403 | 13.91423982 | 10.9696756 | 22.59907002 | 9.90E-07 | -10.12868923 | 1.97E-08 | 31.52114042 | 0.000893225 | 6.004169985 |
| *Cxcl2* | 431.8553383 | 26315.20113 | 40390.54532 | 2221.84625 | 15.85775283 | 14.73272452 | 10.9696756 | 10.43034001 | 7.10E-08 | -10.3813655 | 1.01E-09 | 37.30375824 | 0.000749717 | 7.148768738 |
| *Nos2* | 79.05548543 | 5913.658317 | 12654.42609 | 420.7450499 | 1.761972537 | 3.273938782 | 0.914139634 | 5.215170004 | 2.88E-06 | -10.73503003 | 6.90E-08 | 29.09190758 | 0.000586724 | 5.541171958 |
| *Il1b* | 405.8968207 | 71520.43405 | 140414.0323 | 2933.50087 | 25.54860178 | 35.1948419 | 15.54037377 | 22.59907002 | 2.48E-06 | -11.0878143 | 5.78E-08 | 29.43536857 | 0.000459447 | 5.60641363 |
| *Il12b* | 555.7482633 | 13642.15809 | 21458.54705 | 859.0618189 | 4.404931342 | 4.910908172 | 3.656558535 | 1.738390001 | 6.61E-09 | -11.27635655 | 7.45E-11 | 42.39697774 | 0.000403161 | 8.180088469 |
| *Cxcl1* | 66.07622663 | 5985.635033 | 9796.938227 | 452.9598681 | 0 | 4.910908172 | 0.914139634 | 0 | 0.000216854 | -11.43915333 | 1.12E-05 | 19.29099415 | 0.00036014 | 3.663833216 |
| *Cxcl3* | 71.97588972 | 9985.48391 | 15462.84368 | 427.5784962 | 0 | 6.547877563 | 1.828279267 | 0 | 0.000503989 | -11.59072063 | 3.09E-05 | 17.36049038 | 0.000324224 | 3.297579289 |
| *Shisa3* | 23.59865237 | 983.253339 | 1555.413722 | 55.6437769 | 0 | 0 | 0 | 0 | 0.000350491 | -11.99768137 | 1.99E-05 | 18.19932526 | 0.000244533 | 3.455322905 |
| *Il1a* | 134.5123185 | 29194.26973 | 56160.36354 | 1172.424141 | 3.523945073 | 5.729392868 | 3.656558535 | 0.869195001 | 1.81E-06 | -12.61717839 | 3.96E-08 | 30.17045671 | 0.000159166 | 5.743017674 |
| *Csf3* | 7.07959571 | 1474.237359 | 3138.215507 | 71.2630827 | 0 | 0 | 0 | 0 | 0.001662486 | -12.8390942 | 0.000137377 | 14.53777919 | 0.000136473 | 2.779242041 |
| *Il6* | 29.49831546 | 9188.598851 | 16981.73998 | 307.5050829 | 0 | 0.818484695 | 0.914139634 | 0 | 4.03E-05 | -13.9059346 | 1.50E-06 | 23.14766022 | 6.51E-05 | 4.394342701 |
