## Supplementary material for "TFEB-Mediated Pro-inflammatory Response in Murine Macrophages Induced by Acute Alpha7 Nicotinic Receptor Activation": Table_S10

**Table S10. GO and KEGG Pathway analysis of differentially expressed genes from Table S9.**

| **Enrichr-KG_PNU induced DE genes UP in WT and not up in Chrna7KO** | **Library** | **p-value** | **q-value** | **z-score** | **combined score** | **converted _log(Padj)** |
|---|---|---|---|---|---|---|
| **Term** | GO_Biological_Process_2021 | 8.98E-36 | 3.17E-32 | 5.478 | 442.1 | 31.4985573 |
| cytokine-mediated signaling pathway (GO:0019221) | GO_Biological_Process_2021 | 7.52E-32 | 1.33E-28 | 5.825 | 417.4 | 27.87693276 |
| cellular response to cytokine stimulus (GO:0071345) | KEGG_2021_Human | 5.69E-31 | 1.55E-28 | 16.11 | 1122 | 27.81000466 |
| TNF signaling pathway | GO_Biological_Process_2021 | 3.96E-25 | 3.77E-22 | 6.525 | 366.6 | 21.42369321 |
| extracellular matrix organization (GO:0030198) | GO_Biological_Process_2021 | 4.27E-25 | 3.77E-22 | 7.705 | 432.3 | 21.42369321 |
| inflammatory response (GO:0006954) | GO_Biological_Process_2021 | 3.79E-21 | 2.68E-18 | 8.668 | 407.6 | 17.57253012 |
| response to lipopolysaccharide (GO:0032496) | KEGG_2021_Human | 1.65E-19 | 2.24E-17 | 6.687 | 289.2 | 16.64911264 |
| Lipid and atherosclerosis | KEGG_2021_Human | 4.80E-19 | 4.36E-17 | 11.77 | 496.5 | 16.36101184 |
| IL-17 signaling pathway | KEGG_2021_Human | 1.07E-16 | 7.26E-15 | 9.7 | 356.7 | 14.13905142 |
| NF-kappa B signaling pathway | KEGG_2021_Human | 5.92E-15 | 3.22E-13 | 4.676 | 153.2 | 12.49245445 |
| Cytokine-cytokine receptor interaction | | | | | | |

| **Enrichr-KG_PNU induced DE genes UP in Chrna7KO and not UP in WT** | **Library** | **p-value** | **q-value** | **z-score** | **combined score** | **converted -log(Padj)** |
|---|---|---|---|---|---|---|
| **Term** | KEGG_2021_Human | 9.4603E-06 | 0.001712 | 5.194 | 60.09 | 2.76649624 |
| Transcriptional misregulation in cancer | GO_Biological_Process_2021 | 0.00003559 | 0.02283 | 27.7 | 283.7 | 1.641494089 |
| negative regulation of leukocyte activation (GO:0002695) | GO_Biological_Process_2021 | 0.000043664 | 0.02283 | 75.9 | 762 | 1.641494089 |
| negative regulation of mast cell activation (GO:0033004) | GO_Biological_Process_2021 | 0.000043664 | 0.02283 | 75.9 | 762 | 1.641494089 |
| negative regulation of myeloid leukocyte mediated immunity (GO:0002887) | GO_Biological_Process_2021 | 0.000057005 | 0.02283 | 14.15 | 138.3 | 1.641494089 |
| hematopoietic progenitor cell differentiation (GO:0002244) | GO_Biological_Process_2021 | 0.0001781 | 0.05706 | 37.94 | 327.6 | 1.243668233 |
| negative regulation of leukocyte degranulation (GO:0043301) | KEGG_2021_Human | 0.0002953 | 0.02672 | 5.129 | 41.68 | 1.573163546 |
| Lysosome | KEGG_2021_Human | 0.001908 | 0.1151 | 3.444 | 21.56 | 0.938924676 |
| Rap1 signaling pathway | KEGG_2021_Human | 0.004188 | 0.1895 | 6.611 | 36.2 | 0.722390786 |
| Cysteine and methionine metabolism | KEGG_2021_Human | 0.01193 | 0.3535 | 2.733 | 12.1 | 0.451610582 |
| Ras signaling pathway | | | | | | |
