## Supplementary material for "TFEB-Mediated Pro-inflammatory Response in Murine Macrophages Induced by Acute Alpha7 Nicotinic Receptor Activation": Table_S11

**Table S11. Row-normalized expression of macrophage marker genes in WT and Chrna7-/- BMDMs.**

| | WT | | | | *Chrna7*$^{-/-}$ | | | |
|---|---|---|---|---|---|---|---|---|
| **(Cd11a) *Itgal* | 3.47 | 0.90 | 0.00 | 6.28 | 55.65 | 46.26 | 93.68 | 100.00 |
| (Cd11b)*Itgam* | 16.72 | 4.44 | 0.00 | 15.60 | 14.35 | 6.28 | 100.00 | 88.89 |
| ***(Cd11c) *Itgax* | 1.63 | 2.13 | 0.00 | 8.15 | 52.38 | 58.40 | 93.98 | 100.00 |
| (CD15) *Fut4* | 88.60 | 38.60 | 0.00 | 86.84 | 50.00 | 76.32 | 100.00 | 80.70 |
| (CD16) *Fcgr3* | 11.45 | 0.00 | 23.99 | 12.06 | 35.82 | 25.10 | 93.80 | 100.00 |
| (CD32) *Fcgr2b* | 32.85 | 38.16 | 51.91 | 17.21 | 0.00 | 2.49 | 59.46 | 100.00 |
| (CD64) *Fcgr1* | 14.89 | 6.11 | 2.71 | 8.07 | 100.00 | 98.04 | 14.87 | 0.00 |
| (F4/80)*Emr1* | 98.51 | 89.26 | 87.43 | 100.00 | 21.26 | 13.22 | 9.05 | 0.00 |
| *(MHC Class II)*H2-Ab1* | 4.24 | 0.00 | 6.86 | 3.88 | 32.84 | 34.43 | 100.00 | 89.12 |
| *Ccr5* | 100.00 | 74.32 | 76.70 | 79.98 | 84.05 | 97.51 | 33.37 | 0.00 |
| *Cd14* | 100.00 | 91.53 | 96.50 | 90.38 | 31.59 | 39.01 | 0.07 | 0.00 |
| *Cd163* | 54.17 | 100.00 | 87.50 | 60.42 | 0.00 | 75.00 | 41.67 | 66.67 |
| *Cd33* | 28.92 | 20.49 | 19.62 | 3.15 | 0.00 | 9.77 | 100.00 | 83.37 |
| *Cd68* | 6.34 | 0.00 | 18.59 | 16.63 | 68.28 | 78.39 | 100.00 | 72.48 |
| *Cd80* | 37.21 | 52.09 | 42.33 | 0.00 | 59.53 | 14.88 | 58.60 | 100.00 |
| *Cd86* | 25.88 | 35.64 | 2.85 | 0.00 | 22.48 | 41.34 | 100.00 | 69.19 |
| *Csf1r* | 72.50 | 61.45 | 58.76 | 100.00 | 17.28 | 0.00 | 37.04 | 24.24 |
| *Gal3st4* | 67.69 | 20.00 | 100.00 | 73.85 | 44.62 | 0.00 | 32.31 | 24.62 |
| *Lamp1* | 0.00 | 13.44 | 15.10 | 15.07 | 62.93 | 53.09 | 100.00 | 87.80 |
| *Lamp2* | 26.63 | 0.00 | 4.57 | 16.62 | 66.08 | 89.06 | 65.65 | 100.00 |
| *Lilrb4* | 38.00 | 99.66 | 78.14 | 45.44 | 59.93 | 100.00 | 0.00 | 82.66 |
| *Tlr2* | 45.40 | 40.59 | 46.26 | 54.84 | 6.11 | 0.00 | 51.53 | 100.00 |
| *Tlr4* | 25.52 | 0.00 | 37.42 | 46.78 | 100.00 | 52.25 | 7.05 | 7.90 |
