## Supplementary material for "TFEB-Mediated Pro-inflammatory Response in Murine Macrophages Induced by Acute Alpha7 Nicotinic Receptor Activation": Table_S12

**Table S12. Differential gene expression in baseline BMDMs (Chrna7-/- v. WT).**

| ID | A1_WT_DMSO_1 | A2_WT_DMSO_2 | A3_WT_DMSO_3 | A4_WT_DMSO_4 | B1_Chrna7KO_DMSO_1 | B2_Chrna7KO_DMSO_2 | B3_Chrna7KO_DMSO_3 | B4_Chrna7KO_DMSO_4 | padj | log2FoldChange | pvalue | stat | foldChange | log10padj |
|---|---|---|---|---|---|---|---|---|---|---|---|---|---|---|
| *Ngp* | 0 | 0 | 0 | 0 | 13.64888317 | 26.32992526 | 11.46885134 | 15.22765942 | 1.11E-09 | 6.375409535 | 2.55E-11 | 44.49843662 | 83.02129525 | 8.953580766 |
| *Ly6c2* | 1.014503025 | 0 | 0 | 0 | 20.01836199 | 23.40437801 | 21.17326401 | 36.72553154 | 1.32E-13 | 6.262474799 | 1.68E-15 | 63.41299513 | 76.77021583 | 12.88041396 |
| *Elane* | 0 | 0 | 0 | 0 | 13.64888317 | 4.875912085 | 7.939974003 | 8.061702046 | 2.89E-05 | 5.432288881 | 1.61E-06 | 23.01602408 | 43.17992647 | 4.539003471 |
| *Myo18b* | 0 | 0 | 0 | 0 | 4.549627724 | 2.925547251 | 13.23329001 | 8.957446718 | 0.00024203 | 5.216286241 | 1.77E-05 | 18.41768267 | 37.1756546 | 3.616131135 |
| *Ly6i* | 0 | 0 | 0 | 0 | 2.729776635 | 0.975182417 | 14.99772867 | 7.165957374 | 0.001951315 | 5.019541172 | 0.000188137 | 13.94600318 | 32.4363858 | 2.709672679 |
| *Mpo* | 0 | 2.251921267 | 0 | 0 | 21.83821308 | 15.60291867 | 22.93770268 | 0.895744672 | 5.77E-05 | 4.808002943 | 3.45E-06 | 21.54881118 | 28.0125797 | 4.238718104 |
| *Orm1* | 0 | 0 | 0 | 1.040896809 | 2.729776635 | 3.900729668 | 13.23329001 | 6.270212703 | 0.003296689 | 4.311550045 | 0.000355051 | 12.75503947 | 19.8566459 | 2.481922057 |
| *Siglech* | 0 | 1.125960634 | 0 | 0 | 7.279404359 | 3.900729668 | 8.822193337 | 4.478723359 | 0.003139963 | 4.215863347 | 0.000334356 | 12.86739843 | 18.58237949 | 2.503075481 |
| *Amica1* | 0 | 0 | 1.180411119 | 2.081793619 | 9.099255448 | 8.776641753 | 15.87994801 | 19.70638278 | 1.56E-05 | 4.038815928 | 8.11E-07 | 24.33209598 | 16.4363258 | 4.806716321 |
| *Btla* | 1.014503025 | 1.125960634 | 1.180411119 | 0 | 7.279404359 | 5.851094502 | 19.40882534 | 18.81063811 | 5.88E-05 | 3.977985303 | 3.54E-06 | 21.50126554 | 15.75770256 | 4.230778882 |
| *Ltf* | 0 | 1.125960634 | 1.180411119 | 0 | 2.729776635 | 10.72700659 | 16.76216734 | 2.687234015 | 0.003754324 | 3.910021801 | 0.00041792 | 12.45032111 | 15.03259115 | 2.425468201 |
| *Klra17* | 1.014503025 | 0 | 1.180411119 | 0 | 8.189329904 | 2.925547251 | 7.05775467 | 10.74893606 | 0.003361341 | 3.738551254 | 0.000363886 | 12.70907286 | 13.347996 | 2.473487445 |
| *H2-M2* | 1.014503025 | 1.125960634 | 0 | 1.040896809 | 3.639702179 | 10.72700659 | 14.11550934 | 13.43617008 | 0.000380649 | 3.691858455 | 2.95E-05 | 17.44809655 | 12.9229045 | 3.419475264 |
| *Slc36a2* | 22.31906654 | 15.76344887 | 22.42781126 | 22.89972981 | 163.7865981 | 168.7065581 | 316.7167408 | 367.2553154 | 3.98E-18 | 3.606293116 | 2.86E-20 | 85.08628833 | 12.17874118 | 17.39964799 |
| *Chil3* | 7.101521172 | 12.38556697 | 3.541233356 | 4.163587238 | 78.25359686 | 125.7985318 | 70.5775467 | 50.16170162 | 3.94E-19 | 3.576721148 | 2.60E-21 | 89.82552409 | 11.93164578 | 18.40469883 |
| *Ccr7* | 2.029006049 | 1.125960634 | 2.360822237 | 3.122690428 | 23.65806417 | 31.20583735 | 22.93770268 | 24.18510614 | 8.37E-10 | 3.55205553 | 1.88E-11 | 45.08987168 | 11.72938552 | 9.077046558 |
| *S100a9* | 0 | 0 | 2.360822237 | 5.204484047 | 11.82903208 | 23.40437801 | 16.76216734 | 19.70638278 | 1.28E-05 | 3.236847443 | 6.41E-07 | 24.78432137 | 9.427318323 | 4.894265468 |
| *Gsta3* | 3.043509074 | 3.377881901 | 5.902055594 | 5.204484047 | 25.47791526 | 33.15620218 | 46.75762469 | 57.327659 | 1.45E-11 | 3.225507338 | 2.48E-13 | 53.58104925 | 9.353506662 | 10.83973346 |
| *F10* | 3.043509074 | 5.629803168 | 5.902055594 | 2.081793619 | 13.64888317 | 34.1313846 | 48.52206335 | 53.74468031 | 1.31E-08 | 3.193676979 | 3.65E-10 | 39.2917968 | 9.149398994 | 7.881613497 |
| *Cd2* | 1.014503025 | 0 | 2.360822237 | 3.122690428 | 13.64888317 | 4.875912085 | 22.05548334 | 9.85319139 | 0.001162628 | 2.957992536 | 0.000103667 | 15.06871535 | 7.770419762 | 2.934559375 |
| *Wasf1* | 2.029006049 | 0 | 1.180411119 | 1.040896809 | 5.459553269 | 4.875912085 | 11.46885134 | 9.85319139 | 0.008200504 | 2.876913892 | 0.001029976 | 10.77288855 | 7.34577085 | 2.086159431 |
| *Apol7c* | 7.101521172 | 4.503842535 | 1.180411119 | 2.081793619 | 20.01836199 | 32.18101976 | 24.70214134 | 25.08085081 | 5.19E-07 | 2.751846407 | 1.91E-08 | 31.58100674 | 6.735786482 | 6.285223246 |
| *Mreg* | 4.058012098 | 5.629803168 | 1.180411119 | 5.204484047 | 20.01836199 | 20.47883076 | 28.23101868 | 34.03829753 | 7.89E-07 | 2.663976523 | 3.06E-08 | 30.67057722 | 6.337775353 | 6.102808266 |
| *Gbgt1* | 1.014503025 | 5.629803168 | 2.360822237 | 1.040896809 | 11.82903208 | 17.55328351 | 14.99772867 | 17.01914876 | 0.000220066 | 2.639188234 | 1.58E-05 | 18.63495186 | 6.229810301 | 3.657446509 |
| *Dio2* | 15.21754537 | 11.25960634 | 12.98452231 | 6.245380857 | 40.94664952 | 50.70948569 | 76.75308203 | 101.2191479 | 2.49E-11 | 2.563725142 | 4.38E-13 | 52.46490145 | 5.91232322 | 10.60302887 |
| *Tchh* | 2.029006049 | 0 | 4.721644475 | 2.081793619 | 12.73895763 | 14.62773626 | 7.939974003 | 15.22765942 | 0.001678592 | 2.531533267 | 0.000158733 | 14.26568131 | 5.78185836 | 2.775054823 |
| *F7* | 5.072515123 | 2.251921267 | 1.180411119 | 4.163587238 | 14.55880872 | 13.65255384 | 23.81992201 | 20.60212745 | 9.74E-05 | 2.492400469 | 6.13E-06 | 20.44614688 | 5.627134585 | 4.011566457 |
| *Il1r2* | 11.15953327 | 13.5115276 | 7.082466712 | 8.327174476 | 37.30694734 | 45.8335736 | 90.86859137 | 49.26595695 | 6.91E-10 | 2.477170754 | 1.51E-11 | 45.5236235 | 5.568044548 | 9.160595549 |
| *Cdo1* | 11.15953327 | 10.1336457 | 16.52575566 | 3.122690428 | 26.3878408 | 28.28029009 | 87.33971404 | 84.19999915 | 0.001697046 | 2.473717567 | 0.000161025 | 14.23870151 | 5.554732998 | 2.770306306 |
| *Ltb4r1* | 3.043509074 | 6.755763802 | 5.902055594 | 1.040896809 | 25.47791526 | 6.826276919 | 31.75989601 | 25.08085081 | 0.00028379 | 2.44011761 | 2.12E-05 | 18.08128865 | 5.426859693 | 3.547002152 |
| *Tspan7* | 7.101521172 | 9.007685069 | 4.721644475 | 6.245380857 | 33.66724516 | 19.50364834 | 38.81765068 | 47.47446761 | 3.25E-07 | 2.365118883 | 1.14E-08 | 32.58572376 | 5.151951039 | 6.487449464 |
| *Erdr1* | 90.29076919 | 96.8326145 | 74.36590048 | 96.80340328 | 463.1521023 | 428.1050811 | 449.9318602 | 405.7723363 | 2.58E-84 | 2.282211473 | 1.04E-87 | 394.1369003 | 4.864230088 | 83.58774221 |
| *Plekha6* | 3.043509074 | 4.503842535 | 2.360822237 | 1.040896809 | 12.73895763 | 7.801459336 | 20.29104467 | 11.64468073 | 0.003230692 | 2.272665113 | 0.00034688 | 12.79859499 | 4.832149576 | 2.49070446 |
| *Ifitm1* | 21.30456352 | 27.02305521 | 17.70616678 | 17.69524576 | 82.80322458 | 58.51094502 | 147.3306287 | 110.1765946 | 8.77E-13 | 2.254916195 | 1.22E-14 | 59.51272151 | 4.773065702 | 12.05698356 |
| *Plbd1* | 11.15953327 | 9.007685069 | 10.62370007 | 9.368071285 | 47.31612833 | 27.30510768 | 60.87313402 | 55.53616965 | 5.97E-09 | 2.252610112 | 1.57E-10 | 40.93530897 | 4.765442263 | 8.223727634 |
| *Acpp* | 29.42058771 | 18.01537014 | 15.34534454 | 22.89972981 | 90.08262894 | 96.54305929 | 103.219662 | 120.029786 | 6.26E-20 | 2.245124953 | 3.88E-22 | 93.59038757 | 4.740781669 | 19.2036365 |
| *Met* | 25.36257561 | 14.63748824 | 31.87110021 | 28.10421386 | 115.5605442 | 90.69196478 | 134.9795581 | 124.5085094 | 3.82E-20 | 2.22194912 | 2.31E-22 | 94.61910421 | 4.665232946 | 19.41791585 |
| *Flt3* | 5.072515123 | 1.125960634 | 4.721644475 | 7.286277666 | 11.82903208 | 18.52846592 | 29.11323801 | 25.97659548 | 0.000246556 | 2.219494225 | 1.81E-05 | 18.37613331 | 4.657301321 | 3.608084843 |
| *S100a8* | 2.029006049 | 1.125960634 | 7.082466712 | 3.122690428 | 16.37865981 | 18.52846592 | 11.46885134 | 14.33191475 | 0.001544335 | 2.2058818 | 0.00014442 | 14.44359834 | 4.613564415 | 2.811258456 |
| *Kynu* | 18.26105444 | 14.63748824 | 7.082466712 | 11.4498649 | 38.21687288 | 29.25547251 | 97.04412671 | 71.65957374 | 6.95E-07 | 2.192715899 | 2.67E-08 | 30.93452872 | 4.571652987 | 6.158253839 |
| *Spint1* | 138.9869144 | 140.7450792 | 121.5823452 | 151.9709342 | 452.2329958 | 520.7474107 | 755.1797496 | 778.4021198 | 1.96E-20 | 2.179055455 | 1.15E-22 | 95.99620273 | 4.528569676 | 19.70828557 |
| *Ciita* | 38.55111493 | 32.65285838 | 51.93808922 | 34.34959471 | 107.3712143 | 117.02189 | 236.4347814 | 249.9127634 | 3.25E-07 | 2.176609826 | 1.13E-08 | 32.59552105 | 4.520899434 | 6.488639168 |
| *B3gnt5* | 7.101521172 | 18.01537014 | 3.541233356 | 21.858833 | 26.3878408 | 51.6846681 | 67.93088869 | 82.40850981 | 4.91E-06 | 2.170615895 | 2.25E-07 | 26.80176983 | 4.502155526 | 5.308914905 |
| *Ptger3* | 18.26105444 | 13.5115276 | 17.70616678 | 14.57255533 | 45.49627724 | 37.05693185 | 82.92861737 | 97.63616923 | 5.39E-08 | 2.040700001 | 1.66E-09 | 36.33484501 | 4.11445117 | 7.268709143 |
| *Cdh1* | 5.072515123 | 4.503842535 | 3.541233356 | 11.4498649 | 13.64888317 | 24.37956043 | 36.17099268 | 25.97659548 | 0.00029848 | 2.014284147 | 2.24E-05 | 17.97495783 | 4.039800773 | 3.525084162 |
| *Dpp4* | 3.043509074 | 6.755763802 | 0 | 3.122690428 | 14.55880872 | 10.72700659 | 14.99772867 | 11.64468073 | 0.008397658 | 1.999424039 | 0.001056556 | 10.72573672 | 3.998403417 | 2.075841809 |
| *Tnfaip8l3* | 7.101521172 | 10.1336457 | 8.262877831 | 6.245380857 | 30.02754298 | 23.40437801 | 37.93543135 | 30.45531884 | 9.07E-06 | 1.950002926 | 4.41E-07 | 25.50684875 | 3.863753152 | 5.04227907 |
| *Cfb* | 11.15953327 | 19.14133077 | 23.60822237 | 8.327174476 | 50.95583051 | 76.06422853 | 50.28650202 | 54.64042498 | 2.53E-08 | 1.914555551 | 7.31E-10 | 37.93675429 | 3.769976553 | 7.597375834 |
| *Fabp4* | 50.72515123 | 54.04611042 | 70.82466712 | 46.84035643 | 171.975928 | 219.4160438 | 194.0882534 | 230.2063807 | 1.94E-27 | 1.881263241 | 6.10E-30 | 129.2096287 | 3.683974924 | 26.71163812 |
| *H2-Ab1* | 364.2065858 | 308.5132136 | 401.3397804 | 361.1911929 | 801.644405 | 834.756149 | 1821.782924 | 1637.42126 | 2.92E-07 | 1.828115313 | 1.02E-08 | 32.81093956 | 3.550729143 | 6.534764622 |
| *G530011O06Rik* | 3.043509074 | 5.629803168 | 10.62370007 | 6.245380857 | 30.93746852 | 28.28029009 | 15.87994801 | 14.33191475 | 0.001587019 | 1.823419679 | 0.000149562 | 14.37771318 | 3.539191141 | 2.799417744 |
| *Mefv* | 13.18853932 | 15.76344887 | 8.262877831 | 6.245380857 | 32.75731961 | 38.03211426 | 29.11323801 | 51.95319096 | 1.37E-05 | 1.804770073 | 7.00E-07 | 24.61521692 | 3.493734731 | 4.862378703 |
| *H2-Eb1* | 310.4379255 | 308.5132136 | 290.3811352 | 282.0830354 | 675.1647543 | 773.3196567 | 1407.139837 | 1223.587222 | 1.09E-09 | 1.775748428 | 2.48E-11 | 44.55124524 | 3.424155994 | 8.963763205 |
| *Gm14461* | 10.14503025 | 10.1336457 | 5.902055594 | 6.245380857 | 24.56798971 | 34.1313846 | 29.99545735 | 22.3936168 | 0.000114844 | 1.769801428 | 7.39E-06 | 20.08925534 | 3.410070174 | 3.939892489 |
| *H2-Aa* | 353.0470526 | 327.6545444 | 296.2831908 | 339.3323599 | 701.5525951 | 792.823305 | 1567.703756 | 1401.840411 | 1.59E-07 | 1.761727105 | 5.35E-09 | 34.05668314 | 3.39103836 | 6.799672187 |
| *Nfe2* | 14.20304234 | 10.1336457 | 11.80411119 | 3.122690428 | 20.01836199 | 31.20583735 | 40.58208935 | 40.30851023 | 0.000202885 | 1.754972762 | 1.44E-05 | 18.8193286 | 3.375199461 | 3.692749763 |
| *Hp* | 5.072515123 | 6.755763802 | 5.902055594 | 3.122690428 | 10.00918099 | 12.67737142 | 22.93770268 | 23.28936147 | 0.006798288 | 1.73678343 | 0.000829083 | 11.17496959 | 3.332912471 | 2.167600416 |
| *Mmp8* | 77.10222987 | 99.08453576 | 114.4998785 | 114.498649 | 200.1836199 | 215.5153142 | 464.0473695 | 468.4744634 | 4.73E-05 | 1.735372811 | 2.79E-06 | 21.95566014 | 3.329655254 | 4.32492471 |
| *Cd5* | 12.17403629 | 4.503842535 | 9.44328895 | 12.49076171 | 23.65806417 | 40.95766152 | 37.93543135 | 21.49787212 | 0.000349778 | 1.671473312 | 2.69E-05 | 17.62677124 | 3.185397272 | 3.456207907 |
| *Gpr171* | 4.058012098 | 14.63748824 | 9.44328895 | 9.368071285 | 12.73895763 | 26.32992526 | 32.64211535 | 46.57872293 | 0.002386336 | 1.668847011 | 0.000240075 | 13.48814664 | 3.179603809 | 2.622268408 |
| *Gm8979* | 9.130527221 | 9.007685069 | 3.541233356 | 11.4498649 | 39.12679843 | 36.08174943 | 14.11550934 | 14.33191475 | 0.005174626 | 1.630180134 | 0.0006052 | 11.75991482 | 3.095516467 | 2.286121004 |
| *Ccr1* | 78.11673289 | 75.43936246 | 87.35042279 | 56.20842771 | 170.1560769 | 177.4831999 | 277.8990901 | 269.6191462 | 3.23E-15 | 1.595380197 | 3.41E-17 | 71.09346728 | 3.021741392 | 14.49095717 |
| *Xist* | 5089.761674 | 4992.50945 | 5245.747012 | 5024.408899 | 13973.72659 | 13100.60059 | 16044.0408 | 16658.16366 | 2.16E-122 | 1.554500454 | 3.48E-126 | 570.9534075 | 2.937320016 | 121.665158 |
| *Pdgfrl* | 26.37707864 | 22.51921267 | 24.78863349 | 15.61345214 | 50.95583051 | 65.33722194 | 75.8708627 | 68.07659506 | 6.72E-08 | 1.545787544 | 2.11E-09 | 35.87378217 | 2.919634034 | 7.172704198 |
| *Mfge8* | 1594.798755 | 1586.478533 | 1591.194188 | 1590.490325 | 3607.854785 | 3397.535541 | 5827.058699 | 5701.414836 | 7.68E-12 | 1.54244672 | 1.24E-13 | 54.94803394 | 2.912880912 | 11.11450606 |
| *Grap2* | 74.05872079 | 87.82492943 | 118.0411119 | 73.90367347 | 238.4004928 | 235.0189625 | 286.7212835 | 255.2872315 | 1.50E-22 | 1.529611939 | 6.40E-25 | 106.2802749 | 2.88708171 | 21.82407826 |
| *Atp9a* | 9.130527221 | 5.629803168 | 12.98452231 | 26.02242024 | 40.03672397 | 43.88320877 | 35.28877335 | 36.72553154 | 0.000351798 | 1.527436874 | 2.71E-05 | 17.61382283 | 2.882732308 | 3.453706026 |
| *Stra6l* | 64.92819357 | 51.79418915 | 68.46384489 | 65.576499 | 170.1560769 | 137.5007208 | 200.2637887 | 207.8127639 | 1.47E-16 | 1.514382428 | 1.35E-18 | 77.46776078 | 2.856765126 | 15.83292486 |
| *Flt1* | 62.89918752 | 48.41630725 | 64.92261153 | 55.1675309 | 161.0568214 | 148.2277274 | 178.2083054 | 171.982977 | 3.94E-19 | 1.512043559 | 2.60E-21 | 89.82703297 | 2.852137548 | 18.40469883 |
| *Clec4n* | 97.39229036 | 96.8326145 | 107.4174118 | 95.76250647 | 171.0660024 | 170.656923 | 345.8299788 | 437.1233998 | 0.000425514 | 1.50154265 | 3.37E-05 | 17.19729685 | 2.831453132 | 3.371085909 |
| *Lrrc14b* | 26.37707864 | 24.77113394 | 37.7731558 | 41.63587238 | 69.15434141 | 63.38685711 | 114.6885134 | 120.9255307 | 3.95E-07 | 1.498726963 | 1.41E-08 | 32.17102739 | 2.825932415 | 6.403661785 |
| *Arg2* | 11.15953327 | 6.755763802 | 7.082466712 | 9.368071285 | 20.92828753 | 25.35474284 | 31.75989601 | 19.70638278 | 0.00205407 | 1.498306781 | 0.000200195 | 13.8292496 | 2.825109489 | 2.687384753 |
| *I830012O16Rik* | 189.7120656 | 203.7988747 | 195.9482457 | 222.7519172 | 742.4992446 | 629.9678414 | 531.0960389 | 338.5914859 | 1.50E-07 | 1.464703829 | 5.01E-09 | 34.18446424 | 2.760068035 | 6.825070146 |
| *Fabp5* | 1645.523906 | 1741.8611 | 1600.637477 | 1513.463961 | 3876.282821 | 3826.615804 | 5077.172265 | 5152.323352 | 2.51E-28 | 1.463864753 | 7.08E-31 | 133.4846772 | 2.758463237 | 27.59992106 |
| *St8sia1* | 4.058012098 | 11.25960634 | 10.62370007 | 7.286277666 | 20.01836199 | 21.45401317 | 28.23101868 | 20.60212745 | 0.004525257 | 1.460082251 | 0.000519776 | 12.04334616 | 2.751240485 | 2.344356727 |
| *Cd24a* | 355.0760586 | 370.4410485 | 400.1593693 | 374.7228514 | 695.1831163 | 855.2349797 | 1227.167093 | 1344.512752 | 8.65E-09 | 1.458368672 | 2.33E-10 | 40.16445365 | 2.747974605 | 8.062889724 |
| *Dach1* | 25.36257561 | 21.39325204 | 20.06698902 | 30.18600748 | 60.9650115 | 57.5357626 | 71.45976603 | 77.03404178 | 9.81E-08 | 1.45537158 | 3.20E-09 | 35.05721908 | 2.742271818 | 7.008144164 |
| *Trp53i11* | 44.63813308 | 36.03074028 | 33.05151132 | 36.43138833 | 90.08262894 | 110.1956131 | 98.80856537 | 109.28085 | 1.50E-11 | 1.437869581 | 2.61E-13 | 53.47853353 | 2.709205032 | 10.823145 |
| *Ctsk* | 42.60912703 | 48.41630725 | 48.39685587 | 31.22690428 | 121.930023 | 117.02189 | 112.9240747 | 108.3851053 | 1.13E-12 | 1.436973225 | 1.61E-14 | 58.96070486 | 2.707522309 | 11.94759909 |
| *Cd200* | 11.15953327 | 9.007685069 | 9.44328895 | 7.286277666 | 19.10843644 | 27.30510768 | 30.87767668 | 22.3936168 | 0.002541502 | 1.432818388 | 0.000258552 | 13.34906494 | 2.699736096 | 2.594909549 |
| *Pla2g7* | 490.0049609 | 484.1630725 | 453.2778696 | 446.5447313 | 990.9089183 | 997.6116126 | 1483.892919 | 1541.57658 | 9.55E-13 | 1.419979038 | 1.34E-14 | 59.32153194 | 2.67581623 | 12.0198111 |
| *Itgb7* | 1598.856767 | 1571.841045 | 1725.761056 | 1528.036516 | 3401.301687 | 3528.209985 | 5149.514251 | 5032.293566 | 1.60E-15 | 1.413437238 | 1.58E-17 | 72.60788948 | 2.663710396 | 14.79690129 |
| *Cav1* | 216.0891442 | 203.7988747 | 253.7883905 | 270.6331705 | 553.2347313 | 547.077336 | 611.3779983 | 779.2978645 | 6.01E-28 | 1.399986737 | 1.84E-30 | 131.5902182 | 2.63899156 | 27.22120132 |
| *Cd274* | 144.0594295 | 111.4701027 | 128.6648119 | 140.5210693 | 309.3746852 | 370.5693185 | 332.5966888 | 375.3170175 | 7.76E-30 | 1.39979255 | 2.00E-32 | 140.5682455 | 2.638636376 | 29.11024993 |
| *Cyp2s1* | 13.18853932 | 10.1336457 | 15.34534454 | 5.204484047 | 21.83821308 | 35.10656701 | 22.93770268 | 34.9340422 | 0.003089611 | 1.393819338 | 0.000327501 | 12.90616724 | 2.627734174 | 2.510096226 |
| *Gm14446* | 73.04421777 | 57.42399232 | 48.39685587 | 62.45380857 | 169.2461513 | 155.0540043 | 176.4438667 | 131.6744668 | 2.82E-14 | 1.383735534 | 3.46E-16 | 66.52431781 | 2.609431506 | 13.54931858 |
| *Lipn* | 13.18853932 | 14.63748824 | 10.62370007 | 16.65434895 | 35.48709625 | 33.15620218 | 43.22874735 | 32.24680818 | 0.000232124 | 1.383281751 | 1.68E-05 | 18.52050335 | 2.608610869 | 3.634280511 |
| *Mgll* | 27.39158166 | 10.1336457 | 17.70616678 | 5.204484047 | 31.84739407 | 33.15620218 | 44.11096668 | 47.47446761 | 0.002615325 | 1.370847805 | 0.000267536 | 13.28501227 | 2.58622502 | 2.582474313 |
| *Gpr141* | 271.8868106 | 258.9709457 | 237.2626349 | 285.2057258 | 550.5049546 | 537.3255118 | 838.108367 | 797.2127579 | 3.90E-22 | 1.369900833 | 1.79E-24 | 104.2430578 | 2.584528002 | 21.40917473 |
| *Adgb* | 44.63813308 | 42.78650408 | 44.85562251 | 60.37201495 | 79.1635224 | 106.2948835 | 132.3329001 | 180.044679 | 2.27E-07 | 1.368708353 | 7.77E-09 | 33.33246222 | 2.582392606 | 6.6441601 |
| *Gpr97* | 43.62363006 | 50.66822852 | 60.20096706 | 43.717666 | 83.71315013 | 93.61751204 | 162.3283574 | 166.608509 | 3.17E-07 | 1.358781034 | 1.10E-08 | 32.64730345 | 2.564683919 | 6.499207966 |
| *Ms4a4b* | 8.116024197 | 13.5115276 | 4.721644475 | 19.77703938 | 34.5771707 | 21.45401317 | 34.40655401 | 27.76808483 | 0.005202155 | 1.350032916 | 0.000608839 | 11.7487576 | 2.549179415 | 2.283816672 |
| *Bst1* | 72.02971475 | 70.93551992 | 90.89165614 | 64.53560219 | 187.4446622 | 175.5328351 | 200.2637887 | 193.4808491 | 4.19E-18 | 1.348309497 | 3.04E-20 | 84.96235043 | 2.546136027 | 17.37727824 |
| *Gm1966* | 183.6250474 | 126.107591 | 182.9637234 | 189.4432193 | 498.6391986 | 468.0875602 | 400.5275775 | 367.2553154 | 1.59E-22 | 1.344191975 | 7.04E-25 | 106.0919153 | 2.538879591 | 21.79888506 |
| *Fabp3* | 26.37707864 | 41.66054345 | 36.59274468 | 47.88125324 | 104.6414377 | 112.145978 | 88.22193337 | 81.51276513 | 1.61E-08 | 1.341972864 | 4.54E-10 | 38.86704584 | 2.534977363 | 7.793376476 |
| *Ptgfrn* | 273.9158166 | 247.7113394 | 247.8863349 | 237.3244726 | 554.1446568 | 494.4174854 | 719.008757 | 777.5063751 | 1.24E-23 | 1.338247542 | 4.80E-26 | 111.4136721 | 2.528439998 | 22.90580941 |
| *Vcan* | 93.33427826 | 94.58069323 | 82.62877831 | 84.31264157 | 187.4446622 | 219.4160438 | 260.2547034 | 226.623402 | 1.51E-18 | 1.332349279 | 1.04E-20 | 87.09145585 | 2.518123916 | 17.82004027 |
| *Ppap2b* | 34.49310284 | 27.02305521 | 22.42781126 | 26.02242024 | 60.05508596 | 57.5357626 | 73.2242047 | 85.99148849 | 1.44E-06 | 1.327512947 | 5.97E-08 | 29.37232774 | 2.509696567 | 5.841708659 |
| *Slfn1* | 19.27555747 | 12.38556697 | 12.98452231 | 11.4498649 | 52.7756816 | 42.90802635 | 22.93770268 | 21.49787212 | 0.004691937 | 1.313290657 | 0.000541188 | 11.96810894 | 2.485077178 | 2.328647839 |
| *Pnp2* | 36.52210888 | 24.77113394 | 16.52575566 | 16.65434895 | 80.98337349 | 41.93284393 | 65.28423069 | 46.57872293 | 0.000253831 | 1.307347143 | 1.87E-05 | 18.31653362 | 2.47486039 | 3.595455136 |
| *Gpnmb* | 50473.55448 | 53050.76122 | 52011.27471 | 51717.99888 | 107849.8351 | 107033.0965 | 144959.2232 | 145694.6624 | 1.45E-21 | 1.286423722 | 7.47E-24 | 101.412178 | 2.439226488 | 20.83889459 |
| *Gdf15* | 40.58012098 | 34.90477964 | 53.11850034 | 37.47228514 | 84.62307567 | 72.16349886 | 125.2751454 | 120.9255307 | 5.02E-07 | 1.285018343 | 1.84E-08 | 31.66050363 | 2.43685151 | 6.299359647 |
| *F13a1* | 197.8280898 | 198.1690715 | 214.8348236 | 226.9155045 | 525.0270394 | 388.122602 | 551.3870836 | 561.6319092 | 6.94E-25 | 1.275396085 | 2.57E-27 | 117.2182669 | 2.420652665 | 24.15862107 |
| *Klk1b11* | 13.18853932 | 28.14901584 | 12.98452231 | 16.65434895 | 30.93746852 | 31.20583735 | 53.81537936 | 54.64042498 | 0.001736973 | 1.27107062 | 0.000165093 | 14.19174867 | 2.413405972 | 2.760206938 |
| *Ms4a4c* | 264.7852894 | 247.7113394 | 239.6234571 | 225.8746077 | 697.9128929 | 665.0744084 | 548.7404256 | 446.9765912 | 3.91E-22 | 1.269109 | 1.83E-24 | 104.2016786 | 2.410126716 | 21.40765784 |
| *Ablim1* | 64.92819357 | 57.42399232 | 80.26795607 | 79.10815752 | 151.957566 | 125.7985318 | 194.9704727 | 198.8553171 | 1.97E-10 | 1.255915848 | 3.86E-12 | 48.1936034 | 2.388187067 | 9.704908049 |
| *Irg1* | 230.2921866 | 298.3795679 | 227.8193459 | 200.8930842 | 549.5950291 | 516.846681 | 541.6826709 | 655.6850998 | 2.04E-25 | 1.243663117 | 7.23E-28 | 119.7349666 | 2.367990211 | 24.69034837 |
| *Tppp3* | 15.21754537 | 10.1336457 | 14.16493342 | 13.53165852 | 28.20769189 | 29.25547251 | 49.40428269 | 17.91489344 | 0.009095999 | 1.233058577 | 0.001153282 | 10.56373118 | 2.350648098 | 2.041149612 |
| *Dennd3* | 11.15953327 | 13.5115276 | 10.62370007 | 31.22690428 | 30.02754298 | 39.00729668 | 51.16872135 | 36.72553154 | 0.004603335 | 1.230472807 | 0.000529486 | 12.00884883 | 2.346438758 | 2.336927396 |
| *Fcgr4* | 187.6830595 | 182.4056227 | 141.6493342 | 209.2202587 | 415.835974 | 485.6408437 | 424.3474995 | 359.1936134 | 2.80E-21 | 1.22033025 | 1.51E-23 | 100.0164834 | 2.330000477 | 20.55276592 |
| *Ccdc125* | 27.39158166 | 29.27497648 | 25.96904461 | 20.81793619 | 60.05508596 | 64.36203952 | 47.63984402 | 67.18085039 | 2.12E-05 | 1.210254758 | 1.13E-06 | 23.68873652 | 2.313784912 | 4.673554009 |
| *LOC100038947* | 123.769369 | 135.115276 | 127.4844008 | 135.3165852 | 298.4555787 | 268.1751647 | 282.3101868 | 340.3829753 | 3.20E-21 | 1.189879892 | 1.75E-23 | 99.72146836 | 2.281337496 | 20.49450912 |
| *Pdpn* | 132.8998962 | 104.7143389 | 128.6648119 | 163.4207991 | 344.8617815 | 338.3882987 | 232.0236848 | 294.699997 | 1.42E-13 | 1.18832779 | 1.82E-15 | 63.24629393 | 2.278884473 | 12.84640047 |
| *Napsa* | 697.9780809 | 816.3214594 | 797.9579163 | 846.2491061 | 1351.239434 | 1441.319612 | 2205.548334 | 2122.914872 | 3.13E-08 | 1.173068793 | 9.36E-10 | 37.45402581 | 2.254908344 | 7.50418237 |
| *Gprc5b* | 56.81216938 | 48.41630725 | 66.10302265 | 44.75856281 | 124.6597996 | 145.3021801 | 119.09961 | 95.84467988 | 2.87E-08 | 1.167674768 | 8.43E-10 | 37.65882266 | 2.246493314 | 7.542864605 |
| *Ifit3* | 1033.778582 | 1046.017429 | 1071.813296 | 1029.446945 | 3119.224768 | 2927.497616 | 1890.596032 | 1409.902113 | 9.55E-05 | 1.160622876 | 6.00E-06 | 20.48889819 | 2.235539249 | 4.020154983 |
| *Itga1* | 29.42058771 | 19.14133077 | 24.78863349 | 19.77703938 | 37.30694734 | 24.37956043 | 74.98864336 | 70.76382907 | 0.006308037 | 1.157401336 | 0.000759098 | 11.33866572 | 2.230552857 | 2.200105773 |
| *Dmxl2* | 906.965704 | 916.5319558 | 998.6278065 | 908.7029147 | 1771.625036 | 1654.884562 | 2276.125881 | 2567.204229 | 6.26E-12 | 1.148670649 | 9.94E-14 | 55.37967171 | 2.217095091 | 11.20331369 |
| *Htra3* | 61.8846845 | 64.17975612 | 68.46384489 | 61.41291176 | 150.1377149 | 120.9226197 | 136.7439967 | 157.6510622 | 8.64E-11 | 1.146797514 | 1.58E-12 | 49.94770542 | 2.214218375 | 10.06370081 |
| *Fam171a1* | 63.91369055 | 45.03842535 | 50.75767811 | 52.04484047 | 106.4612887 | 90.69196478 | 143.8017514 | 128.0914881 | 1.53E-07 | 1.145785287 | 5.13E-09 | 34.13848211 | 2.212665375 | 6.815850727 |
| *Pianp* | 30.43509074 | 28.14901584 | 15.34534454 | 19.77703938 | 37.30694734 | 40.95766152 | 56.46203736 | 72.55531842 | 0.001598741 | 1.141680328 | 0.000150796 | 14.36224837 | 2.206378538 | 2.796221767 |
| *Slfn4* | 26.37707864 | 34.90477964 | 41.31438916 | 33.3086979 | 117.3803953 | 90.69196478 | 44.11096668 | 46.57872293 | 0.002027738 | 1.138636946 | 0.000196638 | 13.86294217 | 2.201729062 | 2.692988084 |
| *Cd207* | 262.7562834 | 265.7267096 | 326.9738799 | 369.5183674 | 565.9736889 | 573.4072612 | 808.995129 | 746.1553116 | 8.94E-17 | 1.138034721 | 7.71E-19 | 78.57364725 | 2.200810185 | 16.04854767 |
| *Acot1* | 24.34807259 | 28.14901584 | 28.32986685 | 21.858833 | 43.67642615 | 40.95766152 | 78.5175207 | 60.91063768 | 0.000587941 | 1.132528891 | 4.85E-05 | 16.50513533 | 2.192427128 | 3.230666084 |
| *Tmem119* | 79.13123592 | 68.68359865 | 55.47932258 | 83.27174476 | 151.957566 | 139.4510856 | 173.7972087 | 163.9212749 | 9.92E-11 | 1.129419267 | 1.83E-12 | 49.65747916 | 2.1877066 | 10.00326952 |
| *Mmp14* | 196.8135868 | 206.050796 | 219.5564681 | 223.792814 | 309.3746852 | 263.2992526 | 614.9068756 | 661.9553125 | 0.00744413 | 1.128459938 | 0.000924595 | 10.97278869 | 2.186252355 | 2.128186026 |

| | | | | | | | | | | | | | | |
|---|---|---|---|---|---|---|---|---|---|---|---|---|---|---|
| *Slc7a11* | 167.3929991 | 152.0046855 | 174.7008456 | 188.4023225 | 291.1761744 | 283.7780834 | 416.4075255 | 497.1382929 | 9.48E-11 | 1.12553464 | 1.74E-12 | 49.75583837 | 2.181823864 | 10.02314124 |
| *Clec2i* | 49.7106482 | 43.91246471 | 41.31438916 | 39.55407876 | 72.79404359 | 75.08904611 | 103.219662 | 128.0914881 | 1.16E-05 | 1.120164327 | 5.79E-07 | 24.98256917 | 2.173717304 | 4.934731139 |
| *Cbfa2t3* | 65.9426966 | 58.54995295 | 63.74220041 | 56.20842771 | 98.27195884 | 97.5182417 | 182.6194021 | 147.7978708 | 3.84E-06 | 1.108274037 | 1.73E-07 | 27.31575708 | 2.155875755 | 5.415149943 |
| *Tspan33* | 20.29006049 | 21.39325204 | 25.96904461 | 28.10421386 | 53.68560715 | 54.61021535 | 57.34425669 | 40.30851023 | 0.000335849 | 1.105599876 | 2.57E-05 | 17.71205527 | 2.151883354 | 3.473856006 |
| *Ccr2* | 730.4421777 | 641.7975612 | 769.6280494 | 675.5420294 | 1303.923306 | 1497.880193 | 1595.052555 | 1658.919132 | 1.23E-40 | 1.104424087 | 1.99E-43 | 190.9367283 | 2.150130294 | 39.90921721 |
| *Arhgef18* | 141.0159204 | 132.8633548 | 172.3400233 | 135.3165852 | 242.9501205 | 235.9941449 | 343.1833208 | 409.355315 | 2.27E-09 | 1.085256758 | 5.47E-11 | 43.00122279 | 2.121753059 | 8.643692185 |
| *Cd109* | 756.8192563 | 760.0234277 | 750.7414715 | 758.8137741 | 1567.801714 | 1674.38821 | 1528.886105 | 1649.961685 | 1.27E-79 | 1.084792443 | 6.12E-83 | 372.2286723 | 2.121070307 | 78.89725758 |
| *Pkp2* | 29.42058771 | 33.77881901 | 23.60822237 | 37.47228514 | 47.31612833 | 55.58539777 | 75.8708627 | 85.09574382 | 0.000329936 | 1.084393196 | 2.51E-05 | 17.75387284 | 2.12048341 | 3.481570186 |
| *Papss2* | 107.5373206 | 106.9662602 | 75.5463116 | 113.4577522 | 176.5255557 | 169.6817406 | 245.2569748 | 258.8702102 | 9.23E-09 | 1.073249721 | 2.50E-10 | 40.03136916 | 2.104167744 | 8.034596695 |
| *Rnd3* | 76.08772684 | 84.44704753 | 82.62877831 | 112.4168554 | 161.0568214 | 150.1780922 | 201.1460081 | 235.5808487 | 2.08E-08 | 1.072959125 | 5.94E-10 | 38.34153978 | 2.103743953 | 7.681379895 |
| *Tst* | 35.50760586 | 11.25960634 | 18.8865779 | 18.73614257 | 36.39702179 | 38.03211426 | 51.16872135 | 51.95319096 | 0.006891719 | 1.063779884 | 0.000844326 | 11.14117104 | 2.090401237 | 2.161672468 |
| *Cd34* | 1047.981624 | 1034.757822 | 1072.993707 | 1255.321552 | 1677.902705 | 1473.500632 | 3114.234248 | 2918.336141 | 0.001308565 | 1.058042221 | 0.000117945 | 14.82526569 | 2.082104127 | 2.883204796 |
| *C77080* | 667.5429902 | 766.7791915 | 759.0043493 | 700.5235528 | 1178.353581 | 1298.942979 | 1818.254047 | 1723.412749 | 1.02E-09 | 1.056989044 | 2.31E-11 | 44.69203678 | 2.080584733 | 8.991912993 |
| *Abcb1a* | 138.9869144 | 138.4931579 | 159.355501 | 150.9300374 | 287.5364722 | 264.274435 | 345.8299788 | 322.4680818 | 1.38E-16 | 1.056267467 | 1.25E-18 | 77.62337726 | 2.07954437 | 15.85945491 |
| *Tnfsf10* | 65.9426966 | 68.68359865 | 55.47932258 | 81.18995114 | 202.9133965 | 166.7561933 | 99.69078471 | 94.94893521 | 8.40E-05 | 1.053047367 | 5.23E-06 | 20.75115389 | 2.074907996 | 4.075730616 |
| *Cdkn1c* | 84.20375104 | 72.06148056 | 74.36590048 | 82.23084795 | 144.6781616 | 143.3518153 | 170.2683314 | 182.731913 | 6.96E-10 | 1.033464681 | 1.53E-11 | 45.50087824 | 2.046934135 | 9.157328034 |
| *Tnfsf13* | 382.4676403 | 382.8266155 | 279.7574351 | 297.6964875 | 604.1905618 | 575.357626 | 752.5330916 | 803.4829706 | 2.68E-15 | 1.026213478 | 2.79E-17 | 71.48959937 | 2.036671747 | 14.57146947 |
| *Rsad2* | 988.1259459 | 960.4444205 | 952.5917728 | 953.4614775 | 2474.997482 | 2422.353124 | 1662.983444 | 1221.795732 | 0.000313743 | 1.013470689 | 2.37E-05 | 17.86336156 | 2.018761795 | 3.503426028 |
| *Rapgef3* | 12.17403629 | 20.26729141 | 20.06698902 | 19.77703938 | 27.29776635 | 34.1313846 | 38.81765068 | 44.78723359 | 0.009733574 | 1.011906563 | 0.001251292 | 10.41303162 | 2.0165743 | 2.011727645 |
| *Acp5* | 105.5083146 | 154.2566068 | 106.2370007 | 113.4577522 | 218.3821308 | 216.4904966 | 247.0214134 | 279.4723376 | 1.25E-10 | 1.006137434 | 2.40E-12 | 49.12360005 | 2.008526414 | 9.901820021 |
| *Amz1* | 695.9490749 | 675.5763802 | 662.2106376 | 621.4153953 | 1067.342664 | 1080.502118 | 1637.399083 | 1528.14041 | 4.58E-08 | 1.001057038 | 1.39E-09 | 36.67976214 | 2.001465903 | 7.338701626 |
| *Notch1* | 1146.388418 | 1247.564382 | 1336.225386 | 1102.309721 | 1856.248111 | 1842.119586 | 2961.610303 | 2947.895715 | 1.96E-05 | 0.991692161 | 1.04E-06 | 23.85597834 | 1.988515987 | 4.707303947 |
| *Ccl5* | 43.62363006 | 49.54226788 | 44.85562251 | 34.34959471 | 80.07344795 | 109.2204307 | 91.7508107 | 60.91063768 | 0.000150214 | 0.988965046 | 1.00E-05 | 19.50793466 | 1.984760661 | 3.823290993 |
| *Igsf3* | 19.27555747 | 22.51921267 | 16.52575566 | 13.53165852 | 39.12679843 | 30.23065493 | 37.05321202 | 35.82978687 | 0.009085782 | 0.988363608 | 0.001150186 | 10.56869904 | 1.983933417 | 2.041637669 |
| *Alox5* | 454.497355 | 408.72371 | 389.5356692 | 406.9906525 | 696.0930418 | 636.7941183 | 958.9724157 | 997.8595644 | 3.55E-07 | 0.986987535 | 1.25E-08 | 32.40634537 | 1.982042002 | 6.450348577 |
| *Mgst1* | 240.4372168 | 225.1921267 | 230.1801682 | 234.2017821 | 379.4389522 | 429.0802635 | 461.4007115 | 562.5276539 | 1.96E-15 | 0.97842252 | 1.98E-17 | 72.16784416 | 1.970309844 | 14.7070657 |
| *Angptl4* | 143.0449265 | 138.4931579 | 152.2730343 | 161.3390055 | 254.7791526 | 303.2817317 | 310.5412055 | 302.7616991 | 2.05E-15 | 0.976586927 | 2.08E-17 | 72.07003758 | 1.967804541 | 14.68900097 |
| *Sirpb1b* | 76.08772684 | 49.54226788 | 53.11850034 | 57.24932452 | 117.3803953 | 125.7985318 | 104.1018814 | 119.1340413 | 1.11E-06 | 0.9759633 | 4.47E-08 | 29.93378546 | 1.96695411 | 5.953793107 |
| *Sema4a* | 214.0601382 | 252.2151819 | 249.0667461 | 224.8337109 | 377.6191011 | 350.0904877 | 539.9182322 | 573.27659 | 4.06E-10 | 0.971708902 | 8.61E-12 | 46.62151835 | 1.961162257 | 9.390958101 |
| *Pvrl1* | 82.17474499 | 79.94320499 | 106.2370007 | 103.0487841 | 141.948385 | 148.2277274 | 201.1460081 | 235.5808487 | 1.37E-06 | 0.971429148 | 5.62E-08 | 29.49047088 | 1.960782004 | 5.864797351 |
| *Ifit2* | 1801.757372 | 1754.246667 | 1770.616678 | 1828.855694 | 4383.11135 | 4262.522345 | 3003.074612 | 2378.202104 | 3.06E-05 | 0.971000856 | 1.72E-06 | 22.87978936 | 1.960199993 | 4.513852968 |
| *Mrgpre* | 108.5518236 | 103.5883783 | 93.25247838 | 108.2532682 | 157.4171193 | 182.359112 | 225.8481494 | 243.6425507 | 1.56E-08 | 0.967366399 | 4.39E-10 | 38.92918754 | 1.955268046 | 7.805958764 |
| *Ly75* | 24.34807259 | 34.90477964 | 38.95356692 | 19.77703938 | 43.67642615 | 53.63503294 | 73.2242047 | 58.22340367 | 0.003088674 | 0.966420742 | 0.000327153 | 12.90815758 | 1.953986828 | 2.510227905 |
| *Perm1* | 53.7686603 | 30.40093711 | 41.31438916 | 49.96304686 | 92.81240557 | 91.6671472 | 82.92861737 | 77.03404178 | 5.34E-05 | 0.965651239 | 3.18E-06 | 21.70469188 | 1.95294489 | 4.272253484 |
| *Gbp5* | 84.20375104 | 78.81724436 | 112.1390563 | 112.4168554 | 230.2111628 | 198.9372131 | 176.4438667 | 148.6936155 | 2.33E-07 | 0.961109079 | 7.98E-09 | 33.27894382 | 1.946805937 | 6.63322882 |
| *Pde2a* | 218.1181503 | 217.3104023 | 241.9842793 | 191.5250129 | 331.2128983 | 341.313846 | 473.7517822 | 541.9255264 | 1.33E-09 | 0.960299679 | 3.09E-11 | 44.11883353 | 1.94571402 | 8.875440591 |
| *Stac2* | 91.30527221 | 88.95089006 | 131.0256342 | 81.18995114 | 177.4354812 | 145.3021801 | 176.4438667 | 258.8702102 | 6.74E-06 | 0.954917988 | 3.20E-07 | 26.12360633 | 1.938469436 | 5.171423287 |
| *Il18bp* | 31.44959376 | 45.03842535 | 37.7731558 | 29.14511067 | 75.52382022 | 73.13868128 | 74.98864336 | 51.95319096 | 0.000547093 | 0.94732309 | 4.49E-05 | 16.65007816 | 1.928291405 | 3.261938667 |
| *Flt4* | 28.40608469 | 40.53458281 | 23.60822237 | 22.89972981 | 49.13597942 | 57.5357626 | 58.22647602 | 57.327659 | 0.001689589 | 0.94648255 | 0.000159909 | 14.25178869 | 1.927168277 | 2.772218888 |
| *Nim1k* | 36.52210888 | 38.28266155 | 25.96904461 | 33.3086979 | 51.86575606 | 62.41167469 | 69.69532736 | 74.34680776 | 0.000600832 | 0.943202479 | 4.98E-05 | 16.45574226 | 1.922791697 | 3.221247047 |
| *Gfod1* | 134.9289023 | 110.3441421 | 122.7627563 | 78.06726071 | 191.99429 | 157.0043691 | 256.7258261 | 249.0170188 | 3.46E-06 | 0.940325888 | 1.54E-07 | 27.53418842 | 1.918961661 | 5.461078991 |
| *Nod2* | 53.7686603 | 34.90477964 | 50.75767811 | 36.43138833 | 74.61389468 | 90.69196478 | 98.80856537 | 73.45106309 | 0.00015226 | 0.940075113 | 1.02E-05 | 19.47058519 | 1.918628129 | 3.817413779 |
| *Itgal* | 3512.209471 | 3484.848161 | 3404.305666 | 3675.406634 | 5938.174106 | 5554.639047 | 7636.490552 | 7876.282899 | 1.48E-11 | 0.939967279 | 2.57E-13 | 53.51576033 | 1.918484726 | 10.82935933 |
| *Apoc2* | 161.3059809 | 152.0046855 | 147.5513898 | 176.9524576 | 280.2570678 | 253.5474284 | 359.9454881 | 329.6340392 | 1.16E-11 | 0.939259424 | 1.96E-13 | 54.0411184 | 1.917543656 | 10.93526912 |
| *H2-DMa* | 262.7562834 | 215.058481 | 276.2162018 | 231.0790917 | 394.9076865 | 384.2218723 | 589.3225149 | 518.636165 | 1.01E-10 | 0.939130312 | 1.87E-12 | 49.61900522 | 1.917372056 | 9.996645242 |
| *Pde7b* | 145.0739325 | 147.500843 | 184.1441345 | 156.1345214 | 316.6540896 | 312.0583735 | 300.8367928 | 280.3680823 | 4.81E-15 | 0.937670237 | 5.27E-17 | 70.23223882 | 1.915432568 | 14.31763491 |
| *Bhlhe40* | 572.1797059 | 712.7330811 | 692.9013267 | 625.5789825 | 1067.342664 | 1127.310874 | 1415.079811 | 1366.010625 | 6.62E-21 | 0.936185499 | 3.73E-23 | 98.22647807 | 1.91346233 | 20.17924835 |
| *AW112010* | 39.56561796 | 64.17975612 | 51.93808922 | 37.47228514 | 87.35285231 | 78.01459336 | 95.27968804 | 103.9063819 | 0.000126896 | 0.924593446 | 8.27E-06 | 19.87465244 | 1.898149265 | 3.896551622 |
| *Gbp2* | 451.4538459 | 390.7083399 | 374.1903246 | 413.2360334 | 857.1498632 | 850.3590677 | 767.5308203 | 618.0638235 | 1.17E-17 | 0.922578395 | 9.17E-20 | 82.78107209 | 1.895499919 | 16.93062845 |
| *Mamdc2* | 106.5228176 | 137.3671973 | 109.778234 | 107.2123714 | 234.7607906 | 233.0685977 | 184.3838407 | 216.7702106 | 8.12E-10 | 0.915802574 | 1.82E-11 | 45.15851518 | 1.88661831 | 9.090709984 |
| *Irf4* | 43.62363006 | 25.89709457 | 30.69068909 | 37.47228514 | 47.31612833 | 62.41167469 | 81.1641787 | 68.97233973 | 0.002274606 | 0.91088517 | 0.000227369 | 13.5902035 | 1.880198747 | 2.643093843 |
| *Mx2* | 121.7403629 | 88.95089006 | 97.97412286 | 117.6213395 | 261.1486314 | 248.6715163 | 174.6794281 | 115.5510627 | 0.000193668 | 0.904853249 | 1.36E-05 | 18.92184854 | 1.87235403 | 3.712941345 |
| *Ugt1a6a* | 24.34807259 | 40.53458281 | 38.95356692 | 41.63587238 | 90.08262894 | 48.75912085 | 62.63757269 | 68.97233973 | 0.003083411 | 0.900420566 | 0.000326347 | 12.91277389 | 1.866610046 | 2.51096862 |
| *Mak* | 31.44959376 | 30.40093711 | 15.34534454 | 27.06331705 | 49.13597942 | 49.73430327 | 52.05094069 | 44.78723359 | 0.00543808 | 0.900193106 | 0.000643459 | 11.64585179 | 1.866315774 | 2.26455444 |
| *Colec12* | 539.7156091 | 531.4534191 | 534.7262368 | 576.6568325 | 1099.190058 | 1002.487525 | 951.0324417 | 1007.712756 | 2.54E-36 | 0.894908878 | 4.70E-39 | 170.9005776 | 1.859492435 | 35.59530678 |
| *Ptgir* | 50.72515123 | 82.19512626 | 64.92261153 | 67.65829262 | 104.6414377 | 111.1707955 | 109.3951974 | 164.8170196 | 0.000106539 | 0.887936798 | 6.81E-06 | 20.24725475 | 1.850527785 | 3.972491682 |
| *Ly6a* | 1737.843681 | 1899.495589 | 1715.137356 | 1878.818741 | 3223.866205 | 3244.431901 | 3523.584019 | 3330.37869 | 5.95E-64 | 0.881537179 | 3.35E-67 | 299.9649013 | 1.84233725 | 63.22573313 |
| *Padi4* | 92.31977524 | 93.4547326 | 62.56178929 | 105.1305778 | 172.8858535 | 167.7313757 | 154.3883834 | 154.0680836 | 5.50E-07 | 0.870806116 | 2.04E-08 | 31.45332626 | 1.828684405 | 6.259502114 |
| *Cd300lg* | 107.5373206 | 111.4701027 | 120.4019341 | 86.39443519 | 199.2736943 | 132.6248087 | 212.6148594 | 230.2063807 | 6.61E-06 | 0.867844468 | 3.14E-07 | 26.16384811 | 1.824934226 | 5.179737497 |
| *Kdr* | 627.9773722 | 603.5148997 | 656.308582 | 561.0433803 | 1230.219337 | 1288.215973 | 1045.42991 | 893.0574378 | 2.01E-16 | 0.863894416 | 1.86E-18 | 76.83277424 | 1.81994445 | 15.69708862 |
| *Slfn5* | 2590.026222 | 2477.113394 | 2737.373384 | 2711.536189 | 5511.419025 | 5256.233228 | 4322.874735 | 3977.106343 | 9.08E-12 | 0.858476662 | 1.47E-13 | 54.61072851 | 1.813122828 | 11.04213496 |
| *Rhov* | 58.84117543 | 106.9662602 | 77.90713384 | 85.35353838 | 146.4980127 | 136.5255384 | 173.7972087 | 137.9446795 | 1.41E-05 | 0.858346199 | 7.22E-07 | 24.55550666 | 1.812958874 | 4.850974904 |
| *Ifit1* | 500.1499911 | 587.7514508 | 540.6282924 | 596.4338718 | 1219.30023 | 1249.208676 | 853.988315 | 711.2212694 | 0.000450512 | 0.858172527 | 3.61E-05 | 17.06773057 | 1.812740643 | 3.346293288 |
| *Rnf144b* | 383.4821433 | 406.4717888 | 368.288269 | 398.663478 | 624.2089238 | 671.9006853 | 723.4198536 | 800.7957366 | 1.30E-20 | 0.857254582 | 7.52E-23 | 96.840083 | 1.811587616 | 19.88740451 |
| *Ctse* | 39.56561796 | 25.89709457 | 43.67521139 | 42.67676919 | 95.54218221 | 68.26276919 | 58.22647602 | 52.84893564 | 0.00529201 | 0.854712963 | 0.000622765 | 11.70666907 | 1.808398922 | 2.276379318 |
| *Rilpl2* | 427.1057733 | 401.9679462 | 401.3397804 | 370.5592642 | 687.9037119 | 668.9751381 | 817.8173223 | 710.3255247 | 8.03E-22 | 0.850178808 | 3.88E-24 | 102.7096117 | 1.802724341 | 21.09530889 |
| *Cyp4f18* | 109.5663267 | 105.8402996 | 109.778234 | 145.7255533 | 229.3012373 | 194.061301 | 202.9104467 | 223.0404233 | 1.32E-08 | 0.848988006 | 3.68E-10 | 39.27692401 | 1.801236985 | 7.879562439 |
| *Nlrc4* | 110.5808297 | 110.3441421 | 94.4328895 | 124.9076171 | 176.5255557 | 157.9795516 | 222.3192721 | 236.4765934 | 9.79E-07 | 0.848729754 | 3.86E-08 | 30.21667918 | 1.800914581 | 6.009243262 |
| *C1qtnf6* | 41.59462401 | 50.66822852 | 50.75767811 | 57.24932452 | 98.27195884 | 69.23795161 | 91.7508107 | 100.3234032 | 0.000281313 | 0.84770257 | 2.10E-05 | 18.10003909 | 1.799632803 | 3.550809681 |
| *Slc24a5* | 60.87018147 | 87.82492943 | 99.15453397 | 100.9669905 | 125.5697252 | 123.848167 | 165.8572347 | 206.9170192 | 0.000234493 | 0.839158121 | 1.70E-05 | 18.49903075 | 1.78900587 | 3.629870556 |
| *Glyctk* | 35.50760586 | 27.02305521 | 20.06698902 | 45.79945962 | 67.33449032 | 52.65985052 | 48.52206335 | 62.70212703 | 0.009853741 | 0.838797736 | 0.001270708 | 10.38459871 | 1.788559032 | 2.006398836 |
| *Treml2* | 181.5960414 | 189.1613865 | 199.4894791 | 186.3205289 | 376.7091756 | 343.2642108 | 319.3633988 | 309.0319118 | 1.70E-13 | 0.834571429 | 2.20E-15 | 62.88166716 | 1.7833272 | 12.7687271 |
| *Il15* | 59.85567845 | 67.55763802 | 61.38137817 | 80.14905433 | 124.6597996 | 112.145978 | 128.8040227 | 112.8638286 | 1.19E-05 | 0.830398221 | 5.96E-07 | 24.92634589 | 1.778176117 | 4.924169359 |
| *Gm12250* | 405.8012098 | 413.2275526 | 368.288269 | 371.600161 | 839.8612779 | 794.7736699 | 630.7868236 | 502.5127609 | 1.99E-09 | 0.827474274 | 4.77E-11 | 43.26867534 | 1.774575892 | 8.700144212 |
| *Arhgap26* | 240.4372168 | 251.0892213 | 286.8399019 | 252.9379247 | 379.4389522 | 380.3211426 | 539.9182322 | 526.697867 | 1.43E-09 | 0.826875662 | 3.36E-11 | 43.95675204 | 1.773839726 | 8.843979081 |
| *P2ry14* | 214.0601382 | 156.5085281 | 208.932768 | 149.8891406 | 341.2220793 | 311.083191 | 319.3633988 | 319.7808478 | 3.21E-10 | 0.824802608 | 6.65E-12 | 47.12666127 | 1.77129267 | 9.492875395 |
| *Gbp7* | 295.2203802 | 270.2305521 | 321.0718243 | 287.2875194 | 601.4607851 | 521.7225931 | 477.2806595 | 477.4319101 | 2.86E-15 | 0.824340961 | 3.00E-17 | 71.34669205 | 1.770725967 | 14.54336806 |
| *Naaa* | 607.6873117 | 571.9880019 | 636.241593 | 617.251808 | 921.7545769 | 961.5298632 | 1209.522706 | 1196.714882 | 9.82E-18 | 0.818627963 | 7.60E-20 | 83.15220628 | 1.763727847 | 17.00766796 |
| *Npy* | 646.2384267 | 716.110963 | 629.1591263 | 662.0103708 | 1275.715614 | 1463.748808 | 944.8569064 | 983.5276496 | 5.19E-06 | 0.814536568 | 2.40E-07 | 26.68114362 | 1.758733109 | 5.28484117 |
| *Ctss* | 32414.38614 | 33409.50392 | 33351.33575 | 33593.90363 | 53295.24909 | 53036.27093 | 63911.49741 | 62319.64405 | 2.40E-42 | 0.808725134 | 3.29E-45 | 199.0985816 | 1.751662867 | 41.61995846 |
| *Tiam2* | 31.44959376 | 27.02305521 | 38.95356692 | 38.51318195 | 44.5863517 | 64.36203952 | 67.04866936 | 60.91063768 | 0.006829798 | 0.801936019 | 0.000834539 | 11.16279913 | 1.743439159 | 2.165592155 |
| *Csf2rb* | 5075.558632 | 5236.842907 | 5527.865269 | 5079.57643 | 8301.250746 | 8797.120584 | 9901.147582 | 9457.272245 | 4.51E-46 | 0.801601544 | 5.08E-49 | 216.5624321 | 1.743035007 | 45.34595019 |
| *Nxpe4* | 37.53661191 | 42.78650408 | 35.41233356 | 35.39049152 | 62.78486259 | 77.03941095 | 58.22647602 | 65.38936104 | 0.002412257 | 0.800235721 | 0.00024346 | 13.46187619 | 1.741385627 | 2.617576415 |
| *Tifab* | 830.8779771 | 788.1724436 | 822.7465498 | 790.0406784 | 1192.912389 | 1139.013063 | 1614.461381 | 1664.2936 | 6.00E-07 | 0.796073419 | 2.27E-08 | 31.24711706 | 1.736368817 | 6.221772223 |
| *Atp1a3* | 2198.428054 | 2284.574126 | 2373.80676 | 2053.689405 | 3596.935679 | 3462.872763 | 4265.530478 | 4136.548894 | 7.24E-29 | 0.795782903 | 1.98E-31 | 136.013561 | 1.736019199 | 28.1404891 |
| *Emb* | 2590.026222 | 2560.434481 | 2555.590072 | 2520.011176 | 3537.790518 | 3568.192464 | 5253.616132 | 5369.093563 | 3.20E-05 | 0.793899067 | 1.81E-06 | 22.78758326 | 1.733753827 | 4.494877626 |
| *Tnfsf8* | 133.9143992 | 92.32877196 | 145.1905676 | 117.6213395 | 222.0218329 | 209.6642197 | 225.8481494 | 189.8978704 | 3.96E-07 | 0.79351148 | 1.42E-08 | 32.16198757 | 1.733288109 | 6.402618139 |
| *Isg15* | 698.9925839 | 721.7407662 | 740.1177714 | 707.8098305 | 1596.919331 | 1564.192597 | 951.0324417 | 857.2276509 | 0.004721139 | 0.792580087 | 0.000545731 | 11.95253558 | 1.732169471 | 2.325953218 |
| *Gbp3* | 359.1340707 | 347.9218358 | 395.4377248 | 388.2545099 | 716.1114038 | 689.4539688 | 642.2556749 | 523.1148883 | 9.20E-14 | 0.78613262 | 1.15E-15 | 64.15794488 | 1.724445603 | 13.03631453 |
| *Dgkg* | 188.6975626 | 139.6191186 | 162.8967344 | 224.8337109 | 262.9684825 | 290.6043603 | 328.1855921 | 354.71489 | 3.93E-07 | 0.785350157 | 1.40E-08 | 32.18651042 | 1.723510582 | 6.406143454 |
| *Nat8l* | 62.89918752 | 57.42399232 | 73.18548936 | 100.9669905 | 106.4612887 | 83.86568786 | 151.7417254 | 163.9212749 | 0.004068111 | 0.781520753 | 0.0004575 | 12.28137817 | 1.718941864 | 2.390607248 |
| *Notch4* | 242.4662229 | 239.829615 | 289.2007241 | 304.9827652 | 509.5583051 | 475.8890195 | 436.6985702 | 427.2702085 | 9.66E-13 | 0.780804588 | 1.36E-14 | 59.28809871 | 1.71808878 | 12.01492136 |
| *Matn2* | 66.95719962 | 63.05379549 | 67.28343377 | 44.75856281 | 86.44292676 | 87.76641753 | 118.2173907 | 121.8212754 | 0.000821882 | 0.779202227 | 7.05E-05 | 15.79728126 | 1.716181606 | 3.085190701 |
| *Hpse* | 1109.866309 | 1070.788563 | 1028.138084 | 1034.651429 | 1608.748363 | 1577.845151 | 2028.222248 | 2062.899979 | 1.73E-18 | 0.778442906 | 1.20E-20 | 86.79955019 | 1.715278582 | 17.76102013 |
| *Pyhin1* | 1012.474019 | 1154.10965 | 1198.117286 | 1179.336085 | 2404.933215 | 2460.385238 | 1542.119395 | 1380.342539 | 0.003169408 | 0.777208636 | 0.000338397 | 12.84492089 | 1.713811736 | 2.499021911 |
| *Tnfrsf21* | 483.9179427 | 525.8236159 | 504.0355477 | 454.8719057 | 796.1848517 | 714.8087117 | 877.808237 | 980.8404156 | 3.88E-15 | 0.776945835 | 4.12E-17 | 70.71666702 | 1.713499576 | 14.41131518 |
| *Ndrg4* | 58.84117543 | 55.17207105 | 47.21644475 | 72.86277666 | 106.4612887 | 130.6744439 | 73.2242047 | 92.2617012 | 0.002155703 | 0.775539167 | 0.000212358 | 13.71845027 | 1.711829681 | 2.666411063 |
| *Fam84b* | 314.4959376 | 382.8266155 | 357.664569 | 307.0645588 | 574.1630188 | 570.481714 | 579.6181022 | 601.0446748 | 8.41E-18 | 0.774499998 | 6.23E-20 | 83.54395588 | 1.710597099 | 17.07524939 |
| *Entpd1* | 1257.98375 | 1158.613492 | 1168.607008 | 1081.491785 | 1598.739182 | 1570.043691 | 2466.685257 | 2317.291466 | 0.000192482 | 0.769169065 | 1.35E-05 | 18.93794651 | 1.704287899 | 3.715609821 |
| *Adamtsl5* | 46.66713913 | 64.17975612 | 48.39685587 | 49.96304686 | 90.99255448 | 86.79123512 | 84.69305603 | 93.15744587 | 0.00051157 | 0.767737636 | 4.15E-05 | 16.80178683 | 1.702597759 | 3.29109521 |
| *Mcoln3* | 268.8433015 | 197.0431109 | 231.3605793 | 232.1199885 | 351.2312603 | 370.5693185 | 425.2297188 | 436.2276552 | 2.14E-10 | 0.767100076 | 4.28E-12 | 47.99280857 | 1.701845507 | 9.669279086 |
| *Gstm1* | 684.7895416 | 808.439735 | 826.2877831 | 730.7095603 | 1083.721324 | 1139.988246 | 1434.488637 | 1496.789347 | 6.49E-13 | 0.758437077 | 8.78E-15 | 60.15276823 | 1.691657 | 12.18799982 |
| *Mid1* | 112.6098357 | 141.8710398 | 145.1905676 | 123.8667203 | 277.5272912 | 245.7459691 | 182.6194021 | 177.357445 | 3.02E-05 | 0.756175909 | 1.69E-06 | 22.91475061 | 1.689007704 | 4.520033324 |
| *Lilra6* | 76.08772684 | 61.92783485 | 50.75767811 | 71.82187986 | 91.90248003 | 106.2948835 | 119.9818294 | 121.8212754 | 0.000409104 | 0.750410951 | 3.22E-05 | 17.28364912 | 1.682271956 | 3.388165791 |
| *Neo1* | 114.6388418 | 113.722024 | 108.5978229 | 109.294165 | 142.8583105 | 156.0291867 | 255.8436068 | 195.2723385 | 0.000168647 | 0.750358757 | 1.15E-05 | 19.23906044 | 1.682211096 | 3.773021159 |
| *Gapt* | 45.65263611 | 58.54995295 | 35.41233356 | 53.08573728 | 81.89329904 | 65.33722194 | 83.8108367 | 93.15744587 | 0.003124836 | 0.749837574 | 0.000331738 | 12.88210734 | 1.681603496 | 2.505172828 |
| *Tnfaip2* | 2126.39834 | 2352.131764 | 2187.301803 | 2017.258017 | 2820.769189 | 2803.649449 | 4243.474995 | 4700.868038 | 0.001916226 | 0.746698248 | 0.000184137 | 13.98641044 | 1.67794828 | 2.717553318 |
| *Oplah* | 36.52210888 | 50.66822852 | 38.95356692 | 51.00394367 | 75.52382022 | 63.38685711 | 73.2242047 | 84.19999915 | 0.003550693 | 0.743843249 | 0.000390676 | 12.5762668 | 1.674631014 | 2.449686865 |
| *Abcd2* | 370.293604 | 317.5208987 | 360.0253912 | 345.5777408 | 519.5674861 | 522.6977755 | 628.1401656 | 661.0595678 | 1.19E-12 | 0.74292291 | 1.71E-14 | 58.83508725 | 1.673563057 | 11.92475274 |
| *Pira2* | 1059.141158 | 1119.20487 | 1065.91124 | 945.134303 | 1586.000225 | 1665.611568 | 1804.138537 | 1941.078704 | 5.46E-23 | 0.740749539 | 2.29E-25 | 108.3198465 | 1.671043788 | 22.26274718 |
| *Smpdl3a* | 3840.908451 | 3943.114139 | 3993.330815 | 4151.096476 | 6218.431173 | 6447.906141 | 6996.881536 | 6846.176527 | 1.27E-52 | 0.735020918 | 1.33E-55 | 246.7527231 | 1.664421603 | 51.89760311 |
| *Fam46a* | 1038.851097 | 1084.30009 | 984.462873 | 989.8928658 | 1703.38062 | 1710.469959 | 1714.152165 | 1674.146792 | 6.92E-42 | 0.731118473 | 1.00E-44 | 196.8767806 | 1.659925478 | 41.15991127 |
| *Ms4a6c* | 2516.982004 | 2582.953694 | 2612.249806 | 2642.836999 | 4155.629963 | 4253.745703 | 4555.780639 | 4180.440383 | 6.65E-58 | 0.72772613 | 5.36E-61 | 271.4947379 | 1.656026925 | 57.17691236 |
| *Cgnl1* | 1208.273102 | 1272.335516 | 1279.565653 | 1171.008911 | 2071.900466 | 2046.907893 | 1935.589218 | 2104.999979 | 3.31E-41 | 0.727318642 | 5.07E-44 | 193.6548504 | 1.655559249 | 40.48021184 |
| *Vipr1* | 102.4648055 | 140.7450792 | 131.0256342 | 103.0487841 | 155.5972682 | 154.0788219 | 255.8436068 | 218.5616999 | 0.000515155 | 0.720865962 | 4.19E-05 | 16.78477027 | 1.648171035 | 3.288062425 |
| *S100a11* | 1166.678478 | 1368.04217 | 1317.338809 | 1311.52998 | 1835.319824 | 2110.29475 | 2222.310502 | 2320.874445 | 5.39E-21 | 0.717859262 | 3.00E-23 | 98.66036977 | 1.644739682 | 20.26815634 |
| *Ly6e* | 15130.29811 | 16246.48598 | 16029.98299 | 15310.55117 | 26660.81846 | 26170.97053 | 26269.8451 | 24027.45508 | 1.53E-61 | 0.717599225 | 9.87E-65 | 288.6333649 | 1.644443254 | 60.81480715 |
| *Pbx1* | 66.95719962 | 77.69128372 | 57.84014482 | 95.76250647 | 108.2811398 | 107.2700659 | 137.6262161 | 137.0489348 | 0.000850946 | 0.714945633 | 7.35E-05 | 15.71763607 | 1.64142136 | 3.070097811 |
| *Slfn3* | 252.6112531 | 280.3641978 | 266.7729128 | 256.0606151 | 555.0545824 | 456.3853712 | 370.5321202 | 347.5489327 | 8.20E-07 | 0.712085208 | 3.19E-08 | 30.58915583 | 1.638170146 | 6.086385346 |
| *Usp18* | 567.1071907 | 582.1216476 | 596.107615 | 613.0882208 | 1287.544646 | 1096.105037 | 820.4639803 | 659.2680784 | 0.006033901 | 0.711905743 | 0.000722707 | 11.42992193 | 1.637966378 | 2.219401848 |
| *Msrb1* | 1350.303526 | 1420.96232 | 1428.297454 | 1380.229169 | 2159.253318 | 2104.443656 | 2402.283246 | 2467.776571 | 2.42E-30 | 0.71179169 | 6.04E-33 | 142.9467174 | 1.637836892 | 29.61653405 |
| *Cdk5rap1* | 95.36328431 | 142.9970005 | 131.0256342 | 129.0712044 | 213.832503 | 194.061301 | 214.3792981 | 189.8978704 | 4.57E-06 | 0.708488957 | 2.09E-07 | 26.95251887 | 1.634091713 | 5.340498547 |
| *Ctsh* | 3002.928953 | 3161.697459 | 2979.357664 | 3135.18119 | 4392.210605 | 4329.809932 | 5623.266033 | 5700.519091 | 5.97E-09 | 0.707166675 | 1.57E-10 | 40.9386428 | 1.632594695 | 8.223727634 |
| *Fgfr1* | 87.24726011 | 79.94320499 | 86.17001167 | 121.7849267 | 134.6689806 | 137.5007208 | 172.9149894 | 166.608509 | 0.00022565 | 0.703596987 | 1.63E-05 | 18.58292323 | 1.628560129 | 3.646565524 |
| *Cd97* | 1952.918322 | 2032.358944 | 1939.415468 | 1876.736948 | 2762.533954 | 2770.493247 | 3788.249819 | 3346.502094 | 2.63E-15 | 0.699686535 | 2.71E-17 | 71.54409855 | 1.624151863 | 14.58008521 |
| *Emilin2* | 5039.036523 | 4896.802796 | 4912.871076 | 4910.951147 | 7838.098643 | 8029.652022 | 8182.58432 | 7990.042472 | 1.13E-90 | 0.6972207 | 3.63E-94 | 423.7998162 | 1.621378255 | 89.94773318 |
| *Gm5431* | 362.1775798 | 413.2275526 | 357.664569 | 355.9867088 | 703.3724462 | 679.7021447 | 528.4493809 | 500.7212715 | 9.35E-09 | 0.695573562 | 2.54E-10 | 40.00178961 | 1.619528169 | 8.029310859 |
| *Thbs1* | 3360.034017 | 3186.468593 | 3501.099378 | 2555.401667 | 4737.072386 | 4670.148595 | 5677.963632 | 5304.599946 | 1.03E-07 | 0.69425629 | 3.37E-09 | 34.95911954 | 1.618050113 | 6.988403647 |

| | | | | | | | | | | | | | | |
|---|---|---|---|---|---|---|---|---|---|---|---|---|---|---|
| *Cd276* | 188.6975626 | 195.9171503 | 200.6698902 | 164.4616959 | 303.0052064 | 301.3313669 | 324.6567148 | 282.1595716 | 3.02E-09 | 0.694154711 | 7.49E-11 | 42.38613301 | 1.61793619 | 8.520010004 |
| *B3galt5* | 89.27626616 | 55.17207105 | 55.47932258 | 57.24932452 | 100.0918099 | 106.2948835 | 117.3351714 | 94.05319054 | 0.002571669 | 0.694104977 | 0.000261828 | 13.32545323 | 1.617880417 | 2.589784898 |
| *Htr2b* | 163.334987 | 190.2873471 | 167.6183789 | 235.2426789 | 288.4463977 | 303.2817317 | 310.5412055 | 318.8851032 | 1.59E-07 | 0.689560042 | 5.34E-09 | 34.06060614 | 1.612791613 | 6.799672187 |
| *Lyst* | 1933.642765 | 1903.999432 | 2070.441102 | 2002.685461 | 2970.906904 | 3065.973519 | 3403.602189 | 3271.259541 | 3.49E-35 | 0.684729499 | 7.03E-38 | 165.5237542 | 1.60740057 | 34.4570952 |
| *Sorl1* | 1548.131616 | 1517.794934 | 1728.121878 | 1657.107721 | 2148.334211 | 2110.29475 | 3221.865007 | 2886.089333 | 0.000247595 | 0.684483914 | 1.82E-05 | 18.36600734 | 1.607126971 | 3.606257867 |
| *Nod1* | 127.8273811 | 100.2104964 | 116.8607008 | 107.2123714 | 187.4446622 | 165.7810109 | 218.7903948 | 154.0680836 | 6.13E-05 | 0.683248212 | 3.70E-06 | 21.41387458 | 1.605751019 | 4.212814943 |
| *Slamf7* | 583.3392391 | 623.7821911 | 599.6488483 | 645.3560219 | 905.3759171 | 891.3167292 | 1119.536334 | 1017.565947 | 1.18E-15 | 0.682542142 | 1.16E-17 | 73.22562325 | 1.60496534 | 14.92928751 |
| *Ecm1* | 736.5291958 | 852.3521997 | 806.2207941 | 804.6132337 | 1228.399486 | 1067.824747 | 1478.599603 | 1348.095731 | 3.23E-13 | 0.680201297 | 4.24E-15 | 61.58639895 | 1.602363315 | 12.49114598 |
| *Fgr* | 116.6678478 | 136.2412367 | 139.288512 | 154.0527278 | 190.1744389 | 161.8802812 | 245.2569748 | 275.8893589 | 0.000331394 | 0.679102698 | 2.53E-05 | 17.74347364 | 1.601143594 | 3.47965506 |
| *Mansc1* | 54.78316333 | 51.79418915 | 51.93808922 | 59.33111814 | 82.80322458 | 78.01459336 | 84.69305603 | 103.0106373 | 0.002967721 | 0.677314254 | 0.000310995 | 13.00298259 | 1.599159959 | 2.527576861 |
| *Aph1c* | 657.3979599 | 647.4273644 | 725.952838 | 638.0697442 | 1030.945642 | 1002.487525 | 1123.947431 | 1100.870202 | 9.71E-22 | 0.675702685 | 4.77E-24 | 102.3000857 | 1.597374607 | 21.01270864 |
| *Pira6* | 94.34878129 | 99.08453576 | 110.9586452 | 68.69918943 | 181.985109 | 146.2773626 | 108.512978 | 157.6510622 | 0.00204578 | 0.675005528 | 0.000198728 | 13.84307032 | 1.59660289 | 2.689140998 |
| *Sema4b* | 238.4082108 | 251.0892213 | 216.0152347 | 198.8112906 | 346.6816326 | 332.5372042 | 395.2342615 | 367.2553154 | 3.44E-09 | 0.673977905 | 8.60E-11 | 42.1163033 | 1.595466045 | 8.462900422 |
| *Fmnl2* | 300.2928953 | 297.2536073 | 319.8914132 | 310.1872492 | 443.1337403 | 486.6160261 | 500.2183622 | 526.697867 | 5.76E-13 | 0.673305112 | 7.75E-15 | 60.39823744 | 1.594722181 | 12.23956713 |
| *Gpr56* | 110.5808297 | 161.0123706 | 132.2060453 | 144.6846565 | 190.1744389 | 187.2350241 | 262.0191421 | 230.2063807 | 9.13E-05 | 0.668084082 | 5.70E-06 | 20.58557933 | 1.588961404 | 4.039296974 |
| *Nqo1* | 59.85567845 | 100.2104964 | 72.00507824 | 79.10815752 | 116.4704697 | 88.74159995 | 117.3351714 | 170.1914876 | 0.009878103 | 0.666942769 | 0.001274801 | 10.37866146 | 1.587704877 | 2.005326464 |
| *Plekhg1* | 70.0007087 | 99.08453576 | 60.20096706 | 73.90367347 | 105.5513632 | 115.0715252 | 126.1573647 | 134.3617008 | 0.001477353 | 0.666929017 | 0.000136786 | 14.54590252 | 1.587689743 | 2.830515649 |
| *Ifi204* | 2469.300362 | 2415.185559 | 2469.42006 | 2549.156286 | 4647.899683 | 4676.974872 | 3395.662215 | 2972.976566 | 0.000471901 | 0.664134619 | 3.81E-05 | 16.96443752 | 1.584617477 | 3.326149087 |
| *Elovl7* | 51.73965425 | 50.66822852 | 66.10302265 | 44.75856281 | 69.15434141 | 77.03941095 | 93.51524937 | 95.84467988 | 0.008298952 | 0.659735301 | 0.001043469 | 10.74880146 | 1.579792744 | 2.080976729 |
| *Sgms2* | 90.29076919 | 78.81724436 | 89.71124502 | 98.8851969 | 118.2903208 | 137.5007208 | 146.4484094 | 163.0255303 | 0.000298729 | 0.659242737 | 2.24E-05 | 17.97133258 | 1.579253464 | 3.524723272 |
| *Clec4e* | 733.4856868 | 764.5272703 | 668.1126932 | 739.0367347 | 975.4401841 | 943.9765797 | 1207.758268 | 1428.712752 | 2.87E-08 | 0.649305302 | 8.45E-10 | 37.65422274 | 1.56841278 | 7.542864605 |
| *Olfml3* | 1235.664684 | 1495.275722 | 1342.127442 | 1569.672389 | 2289.372671 | 2285.827586 | 2138.499665 | 2135.455298 | 1.27E-19 | 0.649100898 | 8.10E-22 | 92.1349885 | 1.568190579 | 18.89540329 |
| *Tcp11l2* | 308.4089195 | 289.3718829 | 311.6285353 | 289.369313 | 439.4940382 | 402.7503382 | 503.7472395 | 532.0723351 | 2.80E-09 | 0.649039987 | 6.92E-11 | 42.54046033 | 1.568124371 | 8.552865189 |
| *Ldhb* | 77.10222987 | 90.07685069 | 113.3194674 | 96.80340328 | 117.3803953 | 102.3941538 | 189.6771567 | 179.1489344 | 0.009182631 | 0.646422863 | 0.001168631 | 10.53929288 | 1.565282291 | 2.037032876 |
| *Cd300a* | 1768.278772 | 1750.868785 | 1696.250778 | 1526.99562 | 2363.07664 | 2399.923928 | 2956.316987 | 2830.553163 | 2.96E-16 | 0.646281016 | 2.77E-18 | 76.04716934 | 1.565128398 | 15.52808001 |
| *Clec4a2* | 392.6126705 | 462.7698204 | 417.865536 | 384.0909227 | 554.1446568 | 592.9109096 | 727.8309503 | 712.1170141 | 2.66E-09 | 0.644175564 | 6.51E-11 | 42.66076827 | 1.562845934 | 8.575312285 |
| *Kcnj10* | 101.4503025 | 91.20281133 | 114.4998785 | 110.3350618 | 151.957566 | 135.550356 | 171.1505507 | 191.6893598 | 0.000262134 | 0.641629461 | 1.94E-05 | 18.24276867 | 1.560090218 | 3.581476339 |
| *Sh2d3c* | 490.0049609 | 508.9342064 | 505.2159588 | 499.6304686 | 667.8853499 | 707.9824348 | 910.4503524 | 836.6255235 | 2.06E-10 | 0.64113823 | 4.07E-12 | 48.08928013 | 1.559559105 | 9.687130593 |
| *Irs2* | 116.6678478 | 118.2258665 | 106.2370007 | 104.0896809 | 163.7865981 | 164.8058285 | 174.6794281 | 189.8978704 | 2.80E-05 | 0.638972088 | 1.54E-06 | 23.10114482 | 1.557219254 | 4.553166159 |
| *Rnf128* | 1286.389835 | 1243.06054 | 1349.209909 | 1258.444243 | 1990.007167 | 1913.307902 | 1928.531463 | 2160.536148 | 3.56E-28 | 0.638238286 | 1.03E-30 | 132.7382161 | 1.556427402 | 27.44886347 |
| *Fgl2* | 447.3958338 | 363.6852847 | 428.4892361 | 422.6041047 | 644.2272858 | 616.3152876 | 758.708627 | 567.9021219 | 1.47E-09 | 0.637936135 | 3.48E-11 | 43.88632057 | 1.556101467 | 8.831325803 |
| *Axl* | 8467.042243 | 8513.388351 | 8540.274444 | 8868.440817 | 12697.10105 | 12771.96412 | 14206.37793 | 13824.92326 | 7.53E-45 | 0.637641647 | 9.70E-48 | 210.6925294 | 1.555783862 | 44.12339658 |
| *Gdap10* | 55.79766635 | 68.68359865 | 63.74220041 | 63.49470538 | 96.45210775 | 93.61751204 | 88.22193337 | 111.968084 | 0.003020855 | 0.635057399 | 0.000318023 | 12.96114186 | 1.552999536 | 2.519870121 |
| *Ahrr* | 198.8425928 | 221.8142448 | 216.0152347 | 208.1793619 | 285.7166211 | 248.6715163 | 351.1232948 | 424.5829744 | 0.000103724 | 0.635004683 | 6.59E-06 | 20.30815216 | 1.552942791 | 3.984118856 |
| *Psap* | 73452.04799 | 73510.59189 | 74230.1532 | 71909.31519 | 104544.0756 | 103778.9128 | 121795.6723 | 124920.5519 | 6.21E-27 | 0.634600354 | 2.05E-29 | 126.8024328 | 1.552507625 | 26.20661922 |
| *Abi3* | 760.8772684 | 722.8667268 | 751.9218826 | 749.4457028 | 1062.793036 | 993.710883 | 1260.691428 | 1308.682966 | 1.51E-12 | 0.632466291 | 2.20E-14 | 58.34829682 | 1.550212823 | 11.82212588 |
| *Itgax* | 424.0622643 | 435.7467652 | 423.7675916 | 448.6265249 | 596.0012319 | 618.2656524 | 726.948731 | 741.6765883 | 1.31E-11 | 0.631709269 | 2.25E-13 | 53.7776356 | 1.549399596 | 10.88114596 |
| *Nr1h3* | 221.1616594 | 228.5700086 | 197.1286568 | 202.9748779 | 358.5106647 | 274.0262592 | 344.9477595 | 338.5914859 | 2.83E-07 | 0.631686399 | 9.83E-09 | 32.87459382 | 1.549375035 | 6.547978942 |
| *Gramd4* | 228.2631805 | 201.5469534 | 232.5409904 | 232.1199885 | 321.2037173 | 301.3313669 | 404.9386742 | 357.402124 | 4.02E-07 | 0.631515989 | 1.45E-08 | 32.11609579 | 1.549192035 | 6.395262348 |
| *Klf2* | 136.9579083 | 168.8940951 | 126.3039897 | 144.6846565 | 239.3104183 | 220.3912262 | 217.9081754 | 215.8744659 | 5.24E-06 | 0.631077367 | 2.43E-07 | 26.65899763 | 1.548721107 | 5.280619384 |
| *F11r* | 1162.620466 | 1222.793248 | 1244.153319 | 1317.775361 | 1749.786823 | 1761.179445 | 2143.792981 | 2000.197852 | 2.88E-17 | 0.630211821 | 2.32E-19 | 80.94452872 | 1.547792229 | 16.54029697 |
| *Ica1* | 127.8273811 | 104.7143389 | 108.5978229 | 84.31264157 | 167.4263003 | 139.4510856 | 179.9727441 | 171.0872323 | 0.000395756 | 0.629835712 | 3.08E-05 | 17.36623933 | 1.547388773 | 3.402572356 |
| *Ptger2* | 189.7120656 | 188.0354258 | 152.2730343 | 132.1938948 | 290.2662488 | 260.3737053 | 211.7326401 | 261.5574442 | 7.11E-05 | 0.627877506 | 4.32E-06 | 21.11915149 | 1.545289889 | 4.148350097 |
| *Sdc1* | 596.5277785 | 591.1293327 | 631.5199485 | 611.0064272 | 1042.774674 | 1076.601388 | 837.2261477 | 796.3170132 | 5.41E-10 | 0.626471327 | 1.16E-11 | 46.03133967 | 1.543784445 | 9.266699803 |
| *Dusp18* | 85.21825406 | 75.43936246 | 81.44836719 | 94.72160966 | 130.1193529 | 128.724079 | 126.1573647 | 135.2574454 | 0.000406258 | 0.624911879 | 3.19E-05 | 17.30080891 | 1.542116628 | 3.391197772 |
| *Ass1* | 854.2115467 | 894.0127431 | 795.597094 | 841.0446221 | 1319.39204 | 1296.992615 | 1268.631402 | 1326.597859 | 1.63E-25 | 0.622153496 | 5.65E-28 | 120.225415 | 1.539170972 | 24.78773711 |
| *Slc22a4* | 84.20375104 | 81.06916563 | 76.72672272 | 80.14905433 | 102.8215866 | 133.5999911 | 112.0418554 | 146.0063815 | 0.001941246 | 0.616406264 | 0.00018701 | 13.95730007 | 1.533051608 | 2.711919512 |
| *Pira7* | 143.0449265 | 129.4854729 | 128.6648119 | 129.0712044 | 163.7865981 | 252.572246 | 226.7303688 | 171.0872323 | 0.000532288 | 0.616388628 | 4.34E-05 | 16.71518471 | 1.533032867 | 3.273853131 |
| *1600014C10Rik* | 662.470475 | 656.4350494 | 672.8343377 | 638.0697442 | 1122.848122 | 1003.462707 | 997.7900664 | 901.1191398 | 2.14E-15 | 0.614283285 | 2.18E-17 | 71.97039068 | 1.530797325 | 14.67050347 |
| *Akap2* | 87.24726011 | 83.32108689 | 105.0565896 | 83.27174476 | 159.2369703 | 151.1532746 | 104.1018814 | 132.5702114 | 0.002726876 | 0.609082689 | 0.000281363 | 13.19055557 | 1.525289074 | 2.564334689 |
| *Igsf6* | 963.7778733 | 1021.246295 | 1094.241107 | 1056.510262 | 1328.491295 | 1395.486039 | 1704.447753 | 1871.210619 | 1.28E-09 | 0.607851785 | 2.93E-11 | 44.22305551 | 1.523988256 | 8.894015904 |
| *Tns1* | 697.9780809 | 779.1647585 | 797.9579163 | 589.1475942 | 922.6645025 | 993.710883 | 1203.347171 | 1238.814881 | 9.29E-08 | 0.607218385 | 3.02E-09 | 35.16857078 | 1.523319312 | 7.031904275 |
| *Sla* | 544.7881242 | 475.1553874 | 499.3139032 | 551.675309 | 687.9037119 | 724.5605359 | 921.9192037 | 818.71063 | 2.30E-09 | 0.606037628 | 5.55E-11 | 42.97273356 | 1.522073079 | 8.638818009 |
| *H2-T24* | 373.337113 | 397.4641037 | 355.3037467 | 378.8864387 | 658.7860945 | 673.8510502 | 514.3338715 | 438.9148892 | 6.45E-06 | 0.601924901 | 3.04E-07 | 26.2236959 | 1.517740245 | 5.190237822 |
| *Zhx2* | 124.783872 | 141.8710398 | 134.5668675 | 117.6213395 | 191.99429 | 190.1605713 | 190.5593761 | 213.1872319 | 2.80E-05 | 0.601710148 | 1.53E-06 | 23.10512234 | 1.517514339 | 4.55342729 |
| *Nckap5l* | 194.7845807 | 171.1460163 | 165.2575566 | 143.6437597 | 222.0218329 | 229.167868 | 298.1901348 | 273.2021249 | 7.59E-05 | 0.600072017 | 4.63E-06 | 20.98402596 | 1.515792231 | 4.120020721 |
| *Procr* | 72.02971475 | 92.32877196 | 87.35042279 | 111.3759586 | 154.6873426 | 144.3269977 | 142.9195321 | 108.3851053 | 0.003511604 | 0.599685166 | 0.000385646 | 12.60048936 | 1.515385834 | 2.454494454 |
| *Il7r* | 1811.902402 | 1776.76588 | 1891.018612 | 1954.804208 | 2721.587305 | 3115.707822 | 2649.304659 | 2776.808483 | 5.29E-23 | 0.598929499 | 2.17E-25 | 108.4200044 | 1.5145923 | 22.27626016 |
| *Trem1* | 135.9434053 | 120.4777878 | 139.288512 | 104.0896809 | 171.0660024 | 178.4583823 | 208.2037628 | 197.9595725 | 0.000152285 | 0.598797516 | 1.02E-05 | 19.46798748 | 1.514453745 | 3.817344182 |
| *Zfp503* | 120.7258599 | 113.722024 | 134.5668675 | 110.3350618 | 151.0476404 | 178.4583823 | 189.6771567 | 205.1255298 | 0.000239294 | 0.597812818 | 1.74E-05 | 18.45195051 | 1.513420422 | 3.621067858 |
| *Itga4* | 3514.238477 | 3523.130823 | 3851.68148 | 3808.641426 | 5131.980073 | 4933.447848 | 5831.469796 | 6324.853128 | 3.20E-16 | 0.596633506 | 3.02E-18 | 75.8761938 | 1.512183802 | 15.49420435 |
| *Irgm1* | 1245.809714 | 1248.690343 | 1278.385242 | 1286.548457 | 2322.12999 | 2250.721019 | 1599.463652 | 1449.314879 | 0.004118861 | 0.590994305 | 0.000465134 | 12.2504965 | 1.506284521 | 2.38522285 |
| *Cd22* | 59.85567845 | 65.30571675 | 80.26795607 | 65.576499 | 100.0918099 | 93.61751204 | 114.6885134 | 97.63616923 | 0.005713096 | 0.588582454 | 0.000679681 | 11.54400351 | 1.503768466 | 2.243128514 |
| *Il12rb2* | 262.7562834 | 276.9863159 | 302.1852464 | 267.51048 | 342.1320049 | 374.4700481 | 454.3429569 | 493.5553142 | 7.24E-06 | 0.587416743 | 3.47E-07 | 25.96884441 | 1.502553898 | 5.140275932 |
| *Slfn2* | 3137.857855 | 3321.583869 | 3321.676888 | 3267.375085 | 5636.078825 | 5450.294529 | 4411.978888 | 4097.136129 | 4.97E-06 | 0.58660909 | 2.28E-07 | 26.77638856 | 1.50171297 | 5.303972035 |
| *Tmem154* | 450.4393429 | 395.2121824 | 404.8810137 | 393.458994 | 588.7218275 | 607.5386458 | 659.9000616 | 612.6893555 | 4.23E-12 | 0.585868316 | 6.54E-14 | 56.20262768 | 1.500942091 | 11.37394872 |
| *Nme4* | 170.4365081 | 176.7758195 | 171.1596122 | 146.7664501 | 115.5605442 | 110.1956131 | 104.1018814 | 112.8638286 | 0.000167391 | -0.58629525 | 1.14E-05 | 19.25783871 | 0.66605109 | 3.776268576 |
| *Tubb6* | 2206.544078 | 2117.931952 | 2040.930824 | 2046.403127 | 1474.079383 | 1470.575085 | 1280.100253 | 1365.11488 | 4.72E-23 | -0.590373409 | 1.90E-25 | 108.6881148 | 0.664170979 | 22.3264063 |
| *Igf1* | 4582.510162 | 4772.947126 | 4596.520896 | 4382.175568 | 3242.974642 | 3387.783717 | 2515.20732 | 3012.389331 | 9.03E-15 | -0.592821271 | 1.03E-16 | 68.90513317 | 0.663045017 | 14.04417588 |
| *Ccnd2* | 461.5988762 | 510.0601671 | 440.2933473 | 552.7162058 | 334.8526005 | 400.7999734 | 298.1901348 | 264.2446782 | 4.50E-06 | -0.600160864 | 2.05E-07 | 26.98551439 | 0.659680395 | 5.347145676 |
| *Lonrf1* | 136.9579083 | 119.3518272 | 128.6648119 | 139.4801725 | 86.44292676 | 91.6671472 | 77.63530137 | 89.57446718 | 0.000601496 | -0.605585173 | 4.99E-05 | 16.45180602 | 0.657204756 | 3.220767162 |
| *Gm11710* | 447.3958338 | 468.3996236 | 508.7571922 | 401.7861685 | 353.9610369 | 302.3065493 | 236.4347814 | 305.4489331 | 1.83E-06 | -0.607506236 | 7.77E-08 | 28.86335768 | 0.656330219 | 5.737619474 |
| *Lamb2* | 127.8273811 | 137.3671973 | 118.0411119 | 129.0712044 | 69.15434141 | 79.9649582 | 74.98864336 | 111.968084 | 0.003139822 | -0.607987954 | 0.00033365 | 12.8713534 | 0.656111106 | 2.503095008 |
| *Ier5l* | 298.2638892 | 240.9555756 | 259.6904461 | 242.5289566 | 171.0660024 | 182.359112 | 187.9127181 | 142.4234028 | 1.12E-05 | -0.608650587 | 5.58E-07 | 25.05170714 | 0.655809822 | 4.94960059 |
| *Ralgapa2* | 1325.955453 | 1433.347887 | 1464.890198 | 1455.17374 | 1012.747131 | 993.710883 | 803.701813 | 913.6595652 | 1.64E-15 | -0.609229335 | 1.63E-17 | 72.54330405 | 0.655546791 | 14.78620507 |
| *Ust* | 180.5815384 | 150.8787249 | 154.6338566 | 165.5025927 | 109.1910654 | 121.8978021 | 106.7485394 | 88.67872251 | 0.00021385 | -0.615226529 | 1.53E-05 | 18.7002916 | 0.65282738 | 3.669890938 |
| *Col6a2* | 118.6968539 | 110.3441421 | 126.3039897 | 108.2532682 | 57.32530933 | 85.8160527 | 79.39974003 | 79.72127579 | 0.002320167 | -0.617231439 | 0.000232858 | 13.54543041 | 0.651920777 | 2.634480771 |
| *Spred2* | 267.8287985 | 217.3104023 | 185.3245456 | 156.1345214 | 165.6064492 | 131.6496263 | 129.6862421 | 110.1765946 | 0.001822261 | -0.623382565 | 0.000174227 | 14.09044455 | 0.649147142 | 2.739389453 |
| *Hpgd* | 2566.692652 | 2587.457536 | 2652.383784 | 2810.421386 | 1579.630746 | 1451.071437 | 1812.078511 | 2044.089341 | 1.16E-12 | -0.624008746 | 1.66E-14 | 58.89915447 | 0.64886545 | 11.93646064 |
| *Tnfrsf12a* | 286.0898529 | 313.0170562 | 304.5460686 | 300.8191779 | 168.3362258 | 225.2671383 | 212.6148594 | 175.5659557 | 7.80E-07 | -0.624177337 | 3.02E-08 | 30.69746515 | 0.648789629 | 6.107922591 |
| *Micall1* | 856.2405527 | 851.2262391 | 886.4887502 | 837.9219316 | 584.1721998 | 565.6058019 | 582.2647602 | 491.7638248 | 9.92E-17 | -0.625790902 | 8.71E-19 | 78.33142455 | 0.648064404 | 16.00333805 |
| *Utp14b* | 149.1319446 | 137.3671973 | 107.4174118 | 111.3759586 | 62.78486259 | 81.91532303 | 73.2242047 | 109.28085 | 0.005912724 | -0.628321468 | 0.000705978 | 11.47344156 | 0.646928659 | 2.228212429 |
| *Zfp579* | 86.23275709 | 92.32877196 | 69.64425601 | 79.10815752 | 49.13597942 | 46.80875602 | 64.40201136 | 51.05744629 | 0.009695812 | -0.629786437 | 0.001244094 | 10.42368494 | 0.646272076 | 2.013415813 |
| *Fbn1* | 310.4379255 | 238.7036543 | 350.5821023 | 295.6146939 | 144.6781616 | 141.4014505 | 250.5502908 | 234.685104 | 0.002171779 | -0.629872858 | 0.000214466 | 13.69989494 | 0.646233364 | 2.663184314 |
| *Dennd2c* | 282.0318408 | 244.3334575 | 296.2831908 | 271.6740673 | 166.5163747 | 175.5328351 | 187.9127181 | 174.670211 | 1.67E-07 | -0.634582712 | 5.68E-09 | 33.94124755 | 0.644127096 | 6.777004908 |
| *Pmp22* | 4103.664734 | 3999.412171 | 4137.340971 | 3882.545099 | 3078.278118 | 3092.303444 | 2048.513293 | 2164.119127 | 0.000781718 | -0.634972872 | 6.66E-05 | 15.90636028 | 0.643952923 | 3.106949639 |
| *Cacna1a* | 553.9186514 | 576.4918444 | 517.02007 | 549.5935154 | 441.3138893 | 407.6262503 | 292.0145995 | 274.9936142 | 6.54E-06 | -0.635182115 | 3.09E-07 | 26.19379679 | 0.643859533 | 5.184255393 |
| *Pdlim5* | 4619.032271 | 4545.503078 | 4868.015454 | 5305.451038 | 3360.355037 | 3548.688816 | 2727.82218 | 2816.221248 | 5.68E-15 | -0.635265522 | 6.36E-17 | 69.86307746 | 0.643822311 | 14.24583459 |
| *Dmwd* | 229.2776836 | 231.9478905 | 199.4894791 | 226.9155045 | 149.2277894 | 162.8554636 | 128.8040227 | 131.6744668 | 5.00E-06 | -0.635921216 | 2.30E-07 | 26.76159916 | 0.643529765 | 5.301408142 |
| *Pla2g4a* | 495.077476 | 498.8005607 | 530.0045923 | 536.0618569 | 369.4297712 | 364.718224 | 283.1924061 | 293.8042524 | 6.96E-10 | -0.653063157 | 1.53E-11 | 45.4944849 | 0.635928664 | 9.157328034 |
| *Malat1* | 16722.05335 | 16766.6798 | 16644.97719 | 15307.42848 | 10721.65269 | 10612.91024 | 10123.46685 | 10075.33607 | 4.76E-54 | -0.655927325 | 4.61E-57 | 253.4456008 | 0.634667413 | 53.32194677 |
| *Fos* | 1104.793794 | 1311.744138 | 1314.977986 | 1217.849267 | 703.3724462 | 846.458338 | 809.8773483 | 767.6531837 | 3.51E-17 | -0.661729319 | 2.86E-19 | 80.53350062 | 0.632120138 | 16.45428509 |
| *Chst11* | 519.4255486 | 477.4073087 | 524.1025367 | 511.0803335 | 343.0419304 | 397.8744261 | 281.4279674 | 262.4531888 | 7.41E-08 | -0.663436201 | 2.35E-09 | 35.6636129 | 0.631372706 | 7.130296567 |
| *Gatm* | 2801.042851 | 2776.618923 | 2999.424653 | 3315.256338 | 2306.661256 | 2301.430504 | 1428.313101 | 1462.751049 | 0.007340058 | -0.665364703 | 0.00090939 | 11.0035199 | 0.630529291 | 2.134300513 |
| *Fsd1l* | 377.3951251 | 353.551639 | 381.2727914 | 337.2505663 | 235.6707161 | 252.572246 | 202.9104467 | 223.0404233 | 1.00E-09 | -0.666122331 | 2.27E-11 | 44.72600747 | 0.630198257 | 8.997899669 |
| *Cdk5r1* | 219.1326533 | 198.1690715 | 198.3090679 | 176.9524576 | 130.1193529 | 123.848167 | 116.452952 | 128.9872327 | 2.90E-06 | -0.666821785 | 1.28E-07 | 27.89829055 | 0.629892795 | 5.538090027 |
| *AA414768* | 179.5670354 | 127.2335516 | 151.0926232 | 185.2796321 | 136.4888317 | 112.145978 | 80.28195937 | 77.03404178 | 0.002947689 | -0.667653499 | 0.000308184 | 13.01998854 | 0.629529767 | 2.530518301 |
| *Cd14* | 8832.263332 | 8649.629588 | 8712.614467 | 8489.554378 | 6042.815543 | 6430.352858 | 4660.76474 | 4615.772294 | 1.75E-06 | -0.673356974 | 7.39E-08 | 28.96066052 | 0.62704593 | 5.758135782 |
| *Ccl2* | 87.24726011 | 94.58069323 | 106.2370007 | 94.72160966 | 69.15434141 | 76.06422853 | 52.93316002 | 41.2042549 | 0.00547122 | -0.679422585 | 0.000648261 | 11.63201941 | 0.624415136 | 2.261915809 |
| *Itsn1* | 2136.54337 | 2114.55407 | 2241.600714 | 2050.566715 | 1560.522309 | 1444.24516 | 1137.180721 | 1163.572329 | 2.66E-14 | -0.687595952 | 3.23E-16 | 66.65714759 | 0.62088761 | 13.57571658 |
| *Frmd6* | 409.8592219 | 433.494844 | 385.9944358 | 367.4365738 | 242.0401949 | 265.2496174 | 240.8458781 | 239.1638274 | 7.65E-12 | -0.694340611 | 1.23E-13 | 54.96527436 | 0.617991709 | 11.11613869 |
| *Ddah2* | 125.798375 | 128.3595122 | 136.9276898 | 104.0896809 | 99.18188439 | 74.11386369 | 64.40201136 | 67.18085039 | 0.000735635 | -0.698535965 | 6.23E-05 | 16.03215929 | 0.616197201 | 3.133337715 |
| *Etv5* | 530.5850819 | 549.4687892 | 532.3654145 | 508.9985398 | 351.2312603 | 383.2466899 | 313.1878635 | 259.7659548 | 3.76E-10 | -0.699654438 | 7.90E-12 | 46.78984906 | 0.615719669 | 9.424947625 |
| *B3galnt1* | 412.902731 | 347.9218358 | 335.2367577 | 353.9049152 | 239.3104183 | 254.5226108 | 208.2037628 | 190.7936151 | 9.69E-09 | -0.702790774 | 2.64E-10 | 39.92489201 | 0.614382584 | 8.013500411 |
| *Col6a1* | 233.3356957 | 202.6729141 | 198.3090679 | 230.0381949 | 110.1009909 | 107.2700659 | 163.2105767 | 149.5893602 | 2.36E-05 | -0.704161393 | 1.28E-06 | 23.45499371 | 0.613799173 | 4.62665261 |
| *Hcn2* | 122.754866 | 139.6191186 | 139.288512 | 121.7849267 | 73.70396913 | 91.6671472 | 77.63530137 | 77.92978645 | 7.25E-05 | -0.705561594 | 4.42E-06 | 21.07284703 | 0.613203742 | 4.139580233 |
| *Timp1* | 211.0166291 | 239.829615 | 247.8863349 | 219.6292268 | 90.08262894 | 124.8233494 | 169.3861121 | 177.357445 | 0.000305585 | -0.706803669 | 2.30E-05 | 17.9260955 | 0.612676037 | 3.514867288 |
| *Gas6* | 6799.199271 | 6707.347495 | 6219.586185 | 5709.319 | 4169.278846 | 3828.566169 | 3624.157023 | 3819.455281 | 1.11E-31 | -0.720072415 | 2.60E-34 | 149.1919846 | 0.60706697 | 30.95287552 |
| *Fads2* | 1020.590043 | 1048.26935 | 1010.431918 | 994.0564531 | 540.4957736 | 551.953248 | 736.6531436 | 641.353185 | 4.27E-15 | -0.720337463 | 4.61E-17 | 70.49896561 | 0.606955452 | 14.36992101 |
| *Gas7* | 3154.089903 | 2767.611238 | 2947.486563 | 2666.777626 | 1945.420815 | 2055.684535 | 1473.306287 | 1486.936155 | 2.34E-06 | -0.728929785 | 1.02E-07 | 28.33839647 | 0.603351324 | 5.630462527 |
| *Azin2* | 95.36328431 | 94.58069323 | 77.90713384 | 57.24932452 | 61.87493705 | 41.93284393 | 48.52206335 | 42.99574425 | 0.006576595 | -0.733564722 | 0.000798303 | 11.2451749 | 0.601416051 | 2.181998929 |
| *Col5a3* | 104.4938115 | 112.5960634 | 99.15453397 | 114.498649 | 64.60471368 | 51.6846681 | 68.81310803 | 73.45106309 | 0.000259699 | -0.73413311 | 1.92E-05 | 18.26676494 | 0.601179154 | 3.585530518 |
| *Stk38l* | 1245.809714 | 1065.158759 | 1188.673997 | 1195.990434 | 778.8962664 | 804.5254941 | 622.8468496 | 614.4808449 | 4.32E-15 | -0.736538413 | 4.70E-17 | 70.46017412 | 0.600177686 | 14.36461017 |
| *Neat1* | 7400.799564 | 7351.396977 | 7974.857518 | 6768.951952 | 5678.845325 | 5424.939786 | 3311.851379 | 3190.642521 | 0.007469262 | -0.744462208 | 0.000928318 | 10.96534179 | 0.596890332 | 2.126722324 |
| *Igf2bp2* | 244.4952289 | 231.9478905 | 301.0048353 | 299.7782811 | 217.4722052 | 178.4583823 | 121.746268 | 125.4042541 | 9.35E-05 | -0.74530751 | 5.87E-06 | 20.53070408 | 0.596540705 | 4.029083207 |
| *Ell2* | 2672.200967 | 2743.966064 | 2831.806274 | 2931.165416 | 1849.878633 | 2069.337089 | 1318.917904 | 1426.025518 | 3.02E-05 | -0.746526175 | 1.69E-06 | 22.91915549 | 0.596037012 | 4.520255003 |
| *Stap1* | 425.0767673 | 379.4487336 | 396.6181359 | 377.8455418 | 229.3012373 | 290.6043603 | 189.6771567 | 232.8936147 | 2.30E-09 | -0.74726362 | 5.57E-11 | 42.96435727 | 0.595732421 | 8.638403805 |
| *Map3k9* | 1269.143284 | 1201.399996 | 1405.869642 | 1202.235815 | 766.1573088 | 800.6247644 | 698.7177123 | 738.9893542 | 4.23E-26 | -0.757306972 | 1.43E-28 | 122.9482038 | 0.591599619 | 25.37355007 |
| *Dennd5b* | 301.3073983 | 332.1583869 | 352.9429245 | 408.0315493 | 204.7332476 | 246.7211515 | 186.1482794 | 187.2106364 | 9.77E-09 | -0.759447908 | 2.68E-10 | 39.89763395 | 0.590722346 | 8.010001951 |
| *Ankrd33b* | 231.3066896 | 209.4286779 | 236.0822237 | 176.9524576 | 158.3270448 | 147.252545 | 89.98637204 | 109.28085 | 6.11E-05 | -0.759465353 | 3.69E-06 | 21.42083316 | 0.590715203 | 4.213811661 |
| *Smagp* | 354.0615556 | 363.6852847 | 358.8449801 | 306.023662 | 206.5530987 | 224.2919559 | 197.6171307 | 188.1063811 | 9.97E-12 | -0.760116152 | 1.64E-13 | 54.39765201 | 0.590448791 | 11.00147981 |
| *Loxl2* | 309.4234225 | 296.1276467 | 301.0048353 | 307.0645588 | 123.7498741 | 143.3518153 | 215.2615174 | 229.310636 | 1.43E-05 | -0.76831186 | 7.33E-07 | 24.52524948 | 0.587104061 | 4.845518825 |
| *Plxdc2* | 1453.782834 | 1594.360257 | 1604.17871 | 1407.292486 | 926.3042047 | 961.5298632 | 845.1661217 | 808.8574386 | 3.98E-27 | -0.774670117 | 1.28E-29 | 127.7374859 | 0.584522264 | 26.40051081 |
| *Cfh* | 1299.578374 | 1308.366256 | 1345.668675 | 1388.556344 | 739.769468 | 746.014549 | 826.6395157 | 799.8999919 | 1.21E-36 | -0.778805644 | 2.11E-39 | 172.4941312 | 0.582849113 | 35.91586438 |
| *Scd2* | 1042.909109 | 1183.384626 | 1030.498907 | 835.840138 | 486.8101665 | 519.7722283 | 679.3088869 | 691.5148866 | 2.76E-05 | -0.783101186 | 1.51E-06 | 23.13310141 | 0.581116295 | 4.559108513 |
| *Ltbp2* | 354.0615556 | 323.1507019 | 362.3862135 | 362.2320897 | 186.5347367 | 174.5576526 | 213.4970788 | 232.8936147 | 9.82E-12 | -0.794154648 | 1.61E-13 | 54.43674194 | 0.576680982 | 11.00798467 |
| *Nxpe5* | 188.6975626 | 188.0354258 | 153.4534454 | 128.0303076 | 104.6414377 | 59.48612744 | 101.4552234 | 112.8638286 | 0.000240464 | -0.795392432 | 1.76E-05 | 18.43515624 | 0.576186422 | 3.618950192 |
| *Bmp1* | 158.2624718 | 225.1921267 | 146.3709787 | 187.3614257 | 91.90248003 | 63.38685711 | 142.0373127 | 114.655318 | 0.000771414 | -0.796525943 | 6.56E-05 | 15.9350599 | 0.575733896 | 3.112712252 |

| | | | | | | | | | | | | | | |
|---|---|---|---|---|---|---|---|---|---|---|---|---|---|---|
| *Rasgrp3* | 1268.128781 | 1210.407681 | 1250.055375 | 1341.715987 | 782.5359686 | 783.0714809 | 681.0733256 | 672.7042485 | 2.28E-29 | -0.79762436 | 6.05E-32 | 138.3694139 | 0.575295719 | 28.64276755 |
| *Pxdc1* | 82.17474499 | 69.80955929 | 108.5978229 | 72.86277666 | 56.41538378 | 43.88320877 | 44.99318602 | 45.68297826 | 0.001257038 | -0.800381795 | 0.000112896 | 14.90779523 | 0.574197202 | 2.900651434 |
| *Spred1* | 1104.793794 | 1116.952949 | 1186.313174 | 1185.581466 | 781.626043 | 705.0568875 | 564.6203736 | 584.025526 | 1.77E-17 | -0.802234193 | 1.41E-19 | 81.92978325 | 0.573460415 | 16.75244374 |
| *Kitl* | 457.5408641 | 447.0063716 | 453.2778696 | 499.6304686 | 318.4739407 | 342.2890284 | 190.5593761 | 214.0829766 | 6.05E-07 | -0.804121861 | 2.29E-08 | 31.22882753 | 0.572710571 | 6.218603689 |
| *Rgs16* | 74.05872079 | 88.95089006 | 67.28343377 | 107.2123714 | 35.48709625 | 41.93284393 | 47.63984402 | 67.18085039 | 0.003385643 | -0.811897226 | 0.000367425 | 12.69097498 | 0.569632267 | 2.470358898 |
| *Fnbp1l* | 1345.231011 | 1304.988374 | 1409.410876 | 1515.545755 | 873.5285231 | 876.6889929 | 675.7800096 | 745.2595669 | 2.61E-21 | -0.814912292 | 1.39E-23 | 100.1862496 | 0.568443044 | 20.58346058 |
| *Abcg3* | 57.8266724 | 68.68359865 | 64.92261153 | 59.33111814 | 30.02754298 | 30.23065493 | 37.05321202 | 42.99574425 | 0.002155703 | -0.833154462 | 0.000212197 | 13.7198706 | 0.561300612 | 2.666411063 |
| *Myh10* | 94.34878129 | 76.56532309 | 67.28343377 | 64.53560219 | 24.56798971 | 41.93284393 | 44.11096668 | 59.11914834 | 0.00526907 | -0.835616037 | 0.000618553 | 11.71929678 | 0.560343719 | 2.278266013 |
| *Myc* | 72.02971475 | 96.8326145 | 94.4328895 | 78.06726071 | 37.30694734 | 59.48612744 | 42.34652802 | 51.95319096 | 0.00061861 | -0.836623067 | 5.16E-05 | 16.38944559 | 0.559952724 | 3.208583044 |
| *Cdh11* | 162.3204839 | 128.3595122 | 125.1235786 | 147.8073469 | 59.14516042 | 57.5357626 | 113.806294 | 81.51276513 | 0.00023524 | -0.851573897 | 1.71E-05 | 18.48874005 | 0.554179828 | 3.628489647 |
| *Rtn1* | 111.5953327 | 132.8633548 | 152.2730343 | 133.2347916 | 80.07344795 | 59.48612744 | 86.4574947 | 66.28510571 | 9.43E-06 | -0.853147594 | 4.60E-07 | 25.42294024 | 0.553575656 | 5.025380697 |
| *Trib1* | 380.4386342 | 442.502529 | 434.3912917 | 392.4180972 | 224.7516096 | 280.8525361 | 209.9682014 | 197.0638278 | 1.08E-12 | -0.855303497 | 1.53E-14 | 59.06522384 | 0.552749034 | 11.96820704 |
| *Lama3* | 64.92819357 | 93.4547326 | 82.62877831 | 79.10815752 | 42.76650061 | 43.88320877 | 42.34652802 | 47.47446761 | 0.000284333 | -0.85608001 | 2.12E-05 | 18.07559496 | 0.552451603 | 3.546172189 |
| *Jag2* | 176.5235263 | 197.0431109 | 207.7523569 | 194.6477034 | 123.7498741 | 119.9474373 | 82.04639803 | 103.0106373 | 7.65E-08 | -0.856802273 | 2.45E-09 | 35.5752778 | 0.552175096 | 7.116093543 |
| *Ccrn4l* | 575.2232149 | 685.7100259 | 708.2466712 | 575.6159357 | 346.6816326 | 393.9736965 | 353.7699528 | 309.0319118 | 5.21E-17 | -0.857915991 | 4.37E-19 | 79.69524999 | 0.551748998 | 16.28275465 |
| *Cmah* | 321.5974588 | 297.2536073 | 310.4481242 | 332.0460822 | 140.1285339 | 120.9226197 | 221.4370528 | 211.3957425 | 6.41E-07 | -0.859929335 | 2.45E-08 | 31.09923614 | 0.550979545 | 6.193283361 |
| *Lifr* | 393.6271735 | 378.3227729 | 411.9634804 | 373.6819546 | 225.6615351 | 231.1182328 | 193.2060341 | 206.0212745 | 4.46E-17 | -0.864485413 | 3.70E-19 | 80.02468065 | 0.549242278 | 16.3509681 |
| *A230028O05Rik* | 55.79766635 | 57.42399232 | 49.57726699 | 44.75856281 | 37.30694734 | 30.23065493 | 22.93770268 | 23.28936147 | 0.005303101 | -0.869602175 | 0.000624656 | 11.70102711 | 0.547297748 | 2.275470133 |
| *Shc4* | 47.68164216 | 56.29803168 | 56.6597337 | 52.04484047 | 36.39702179 | 30.23065493 | 22.05548334 | 26.87234015 | 0.003371096 | -0.880035678 | 0.000365213 | 12.7022625 | 0.543353994 | 2.472228818 |
| *Pdgfb* | 2501.764459 | 2361.139449 | 2418.662382 | 2592.873952 | 1258.427029 | 1496.90501 | 1253.633673 | 1303.308497 | 8.68E-46 | -0.8954046 | 1.05E-48 | 215.1202216 | 0.537596407 | 45.06130359 |
| *Prkg1* | 61.8846845 | 46.16438598 | 48.39685587 | 42.67676919 | 33.66724516 | 29.25547251 | 27.34879934 | 17.01914876 | 0.007043186 | -0.895886362 | 0.000866286 | 11.09354702 | 0.537416916 | 2.152230863 |
| *S1pr1* | 3305.250854 | 3022.078341 | 3156.419331 | 3112.28146 | 1924.492527 | 1965.967753 | 1353.324458 | 1425.129773 | 5.01E-09 | -0.917683067 | 1.29E-10 | 41.32056267 | 0.529358475 | 8.299932056 |
| *Hes1* | 123.769369 | 149.7527643 | 134.5668675 | 135.3165852 | 74.61389468 | 63.38685711 | 88.22193337 | 59.11914834 | 5.56E-07 | -0.926257052 | 2.08E-08 | 31.41667047 | 0.526221809 | 6.255055243 |
| *Slco2b1* | 672.6155053 | 539.3351435 | 651.5869375 | 538.1436505 | 405.826793 | 321.8101976 | 290.2501608 | 245.4340401 | 9.45E-12 | -0.927314472 | 1.54E-13 | 54.52176673 | 0.525836258 | 11.02463037 |
| *Fcrl1* | 204.929611 | 200.4209928 | 171.1596122 | 185.2796321 | 101.911661 | 114.0963428 | 95.27968804 | 87.78297784 | 3.64E-10 | -0.936305459 | 7.63E-12 | 46.85931571 | 0.522569396 | 9.438674623 |
| *C1qa* | 14262.89802 | 14312.08561 | 13316.21783 | 13851.21384 | 9230.284727 | 8541.622791 | 5880.874078 | 5453.293562 | 1.58E-05 | -0.937487664 | 8.23E-07 | 24.30293421 | 0.522141356 | 4.800814312 |
| *Mrc2* | 52.75415728 | 63.05379549 | 70.82466712 | 66.61739581 | 24.56798971 | 34.1313846 | 31.75989601 | 41.2042549 | 0.000689737 | -0.94118818 | 5.79E-05 | 16.16863507 | 0.520803778 | 3.161316611 |
| *Ets1* | 480.8744337 | 478.5332693 | 498.1334921 | 530.8573728 | 282.9868444 | 325.7109273 | 221.4370528 | 202.4382958 | 3.92E-13 | -0.947752112 | 5.21E-15 | 61.17874478 | 0.518439623 | 12.40652278 |
| *Zbtb16* | 71.01521172 | 55.17207105 | 77.90713384 | 54.12663409 | 23.65806417 | 47.78393843 | 37.93543135 | 23.28936147 | 0.002249158 | -0.963912699 | 0.000223986 | 13.6183452 | 0.512664641 | 2.647980069 |
| *Wtip* | 51.73965425 | 52.92014978 | 44.85562251 | 57.24932452 | 24.56798971 | 27.30510768 | 32.64211535 | 21.49787212 | 0.001333112 | -0.965434284 | 0.000120694 | 14.78180898 | 0.512124227 | 2.875133215 |
| *Chst7* | 82.17474499 | 109.2181815 | 96.79371174 | 102.0078873 | 51.86575606 | 56.56058019 | 45.87540535 | 44.78723359 | 4.50E-06 | -0.971512084 | 2.05E-07 | 26.98769621 | 0.509971283 | 5.347145676 |
| *Plau* | 2838.579463 | 2979.291837 | 3058.445209 | 3187.226031 | 1575.081118 | 2063.485994 | 1355.971116 | 1154.614882 | 4.86E-06 | -0.972389425 | 2.23E-07 | 26.82676171 | 0.509661251 | 5.313768763 |
| *Ak4* | 40.58012098 | 38.28266155 | 37.7731558 | 49.96304686 | 24.56798971 | 20.47883076 | 21.17326401 | 18.81063811 | 0.003845347 | -0.97271435 | 0.000428672 | 12.40288186 | 0.509546477 | 2.415064479 |
| *Tmem176a* | 290.147865 | 280.3641978 | 254.9688016 | 253.9788215 | 108.2811398 | 109.2204307 | 149.9772867 | 180.9404237 | 1.91E-09 | -0.975007443 | 4.55E-11 | 43.36084924 | 0.508737222 | 8.719137653 |
| *Ptchd1* | 269.8578045 | 278.1122765 | 301.0048353 | 350.7822248 | 151.957566 | 161.8802812 | 111.159636 | 173.7744663 | 1.09E-11 | -1.004165356 | 1.81E-13 | 54.19604887 | 0.49855848 | 10.96116032 |
| *Sncaip* | 57.8266724 | 54.04611042 | 75.5463116 | 54.12663409 | 39.12679843 | 30.23065493 | 24.70214134 | 25.97659548 | 0.000491379 | -1.007116986 | 3.97E-05 | 16.8857657 | 0.497539514 | 3.308583722 |
| *Fcrls* | 9741.258042 | 9418.660701 | 8842.45969 | 9487.774419 | 4980.932433 | 4428.303356 | 4786.922105 | 4451.851019 | 1.22E-93 | -1.007525665 | 2.94E-97 | 438.0062439 | 0.497398594 | 92.91480794 |
| *Naip6* | 577.252221 | 587.7514508 | 645.6848819 | 546.470825 | 359.4205902 | 274.0262592 | 294.6612575 | 237.372338 | 5.63E-19 | -1.015165619 | 3.76E-21 | 89.09400898 | 0.494771526 | 18.2493786 |
| *Grap* | 122.754866 | 139.6191186 | 93.25247838 | 93.68071285 | 76.43374577 | 65.33722194 | 43.22874735 | 36.72553154 | 9.33E-05 | -1.022117988 | 5.83E-06 | 20.54342726 | 0.492392951 | 4.03029409 |
| *Itga9* | 315.5104406 | 373.8189304 | 335.2367577 | 342.4550503 | 192.9042155 | 200.8875779 | 160.5639187 | 119.1340413 | 2.28E-12 | -1.022897157 | 3.36E-14 | 57.50968173 | 0.492127091 | 11.64175203 |
| *Enpp2* | 33.47859981 | 34.90477964 | 44.85562251 | 41.63587238 | 20.92828753 | 22.42919559 | 18.52660601 | 14.33191475 | 0.004139897 | -1.023927379 | 0.000468145 | 12.23845991 | 0.491775791 | 2.383010424 |
| *Fosb* | 52.75415728 | 58.54995295 | 89.71124502 | 56.20842771 | 31.84739407 | 48.75912085 | 19.40882534 | 25.08085081 | 0.002232179 | -1.04274912 | 0.00022169 | 13.63769303 | 0.485401637 | 2.651270917 |
| *Lrig1* | 52.75415728 | 56.29803168 | 67.28343377 | 59.33111814 | 37.30694734 | 23.40437801 | 31.75989601 | 20.60212745 | 0.000313011 | -1.055356618 | 2.36E-05 | 17.87022862 | 0.481178265 | 3.50443977 |
| *Serping1* | 39.56561796 | 41.66054345 | 40.13397804 | 41.63587238 | 13.64888317 | 16.57810109 | 22.93770268 | 24.18510614 | 0.002065459 | -1.071628568 | 0.000202137 | 13.81111274 | 0.475781616 | 2.684983525 |
| *Unc5a* | 33.47859981 | 38.28266155 | 51.93808922 | 32.26780109 | 22.74813862 | 11.702189 | 21.17326401 | 17.01914876 | 0.003935007 | -1.091940435 | 0.000440569 | 12.35176965 | 0.469129967 | 2.405054482 |
| *Pcp4l1* | 1340.158495 | 1445.733454 | 1410.591287 | 1394.801725 | 725.2106592 | 825.9795072 | 572.5603476 | 490.8680801 | 2.70E-09 | -1.097022903 | 6.64E-11 | 42.62304731 | 0.467480178 | 8.568363552 |
| *Plxna3* | 61.8846845 | 93.4547326 | 74.36590048 | 80.14905433 | 46.40620279 | 32.18101976 | 42.34652802 | 23.28936147 | 5.88E-05 | -1.10048645 | 3.54E-06 | 21.50087435 | 0.466359221 | 4.230778882 |
| *Nr4a1* | 443.3378217 | 512.3120883 | 492.2314365 | 317.4735269 | 178.3454068 | 214.5401317 | 204.6748854 | 188.1063811 | 1.23E-16 | -1.166965606 | 1.09E-18 | 77.88239727 | 0.445357068 | 15.90858094 |
| *Dpy19l3* | 29.42058771 | 38.28266155 | 37.7731558 | 48.92215005 | 20.92828753 | 14.62773626 | 14.11550934 | 18.81063811 | 0.001450147 | -1.171367554 | 0.000133509 | 14.59158634 | 0.444000266 | 2.838587958 |
| *Grem1* | 104.4938115 | 120.4777878 | 81.44836719 | 96.80340328 | 38.21687288 | 39.9824791 | 48.52206335 | 50.16170162 | 5.15E-08 | -1.187904659 | 1.58E-09 | 36.42919601 | 0.438939905 | 7.288597535 |
| *Btbd11* | 112.6098357 | 140.7450792 | 144.0101565 | 124.9076171 | 55.50545824 | 64.36203952 | 62.63757269 | 43.89148892 | 2.82E-10 | -1.204996539 | 5.76E-12 | 47.4082395 | 0.433770384 | 9.550163798 |
| *Sod3* | 87.24726011 | 81.06916563 | 86.17001167 | 60.37201495 | 31.84739407 | 19.50364834 | 39.69987002 | 41.2042549 | 4.78E-06 | -1.244070375 | 2.19E-07 | 26.85984957 | 0.422179849 | 5.320440438 |
| *Nanos1* | 649.2819357 | 655.3090888 | 670.4735154 | 641.1924346 | 346.6816326 | 338.3882987 | 215.2615174 | 204.2297852 | 2.18E-08 | -1.244797647 | 6.24E-10 | 38.24421124 | 0.421967079 | 7.662165735 |
| *Tmem176b* | 981.0244248 | 984.0895938 | 924.261906 | 887.8849785 | 296.6357276 | 306.2072789 | 504.6294589 | 472.057442 | 2.03E-08 | -1.257263093 | 5.76E-10 | 38.40073235 | 0.418336825 | 7.693324281 |
| *Gm14403* | 53.7686603 | 67.55763802 | 59.02055594 | 73.90367347 | 27.29776635 | 25.35474284 | 27.34879934 | 25.08085081 | 1.75E-06 | -1.274173532 | 7.39E-08 | 28.95854118 | 0.413461949 | 5.758135782 |
| *Gm4980* | 175.5090233 | 170.0200557 | 169.9792011 | 142.6028629 | 64.60471368 | 90.69196478 | 68.81310803 | 47.47446761 | 1.15E-11 | -1.280593632 | 1.94E-13 | 54.06823179 | 0.4116261 | 10.93918892 |
| *Gm14420* | 144.0594295 | 157.6344887 | 193.5874235 | 148.8482438 | 61.87493705 | 50.70948569 | 74.10642403 | 73.45106309 | 4.78E-13 | -1.301687179 | 6.39E-15 | 60.77741385 | 0.405651526 | 12.32061328 |
| *Ctnnd2* | 265.7997924 | 308.5132136 | 309.2677131 | 273.7558609 | 126.4796507 | 157.9795516 | 90.86859137 | 93.15744587 | 6.79E-16 | -1.307714681 | 6.57E-18 | 74.34248636 | 0.403960271 | 15.16783608 |
| *Nox1* | 38.55111493 | 47.29034661 | 48.39685587 | 34.34959471 | 19.10843644 | 22.42919559 | 15.87994801 | 10.74893606 | 0.000234869 | -1.308192738 | 1.70E-05 | 18.49385557 | 0.403826436 | 3.629173578 |
| *Bmp2* | 28.40608469 | 23.64517331 | 34.23192244 | 40.59497557 | 10.00918099 | 17.55328351 | 7.939974003 | 15.22765942 | 0.002027738 | -1.329804846 | 0.000196649 | 13.86283547 | 0.397822052 | 2.692988084 |
| *Cdkn2a* | 48.69614518 | 58.54995295 | 41.31438916 | 53.08573728 | 25.47791526 | 29.25547251 | 14.11550934 | 11.64468073 | 0.00013881 | -1.333726005 | 9.15E-06 | 19.68199044 | 0.396742263 | 3.857578881 |
| *Mapk11* | 45.65263611 | 37.15670091 | 50.75767811 | 48.92215005 | 19.10843644 | 30.23065493 | 17.64438667 | 5.374468031 | 0.0011727 | -1.342995395 | 0.000104849 | 15.04732544 | 0.394201345 | 2.930813095 |
| *Il1b* | 119.7113569 | 144.1229611 | 135.7472787 | 147.8073469 | 38.21687288 | 46.80875602 | 87.33971404 | 37.62127622 | 2.95E-08 | -1.379074181 | 8.78E-10 | 37.57892698 | 0.384465439 | 7.529645487 |
| *Scd1* | 74.05872079 | 73.18744119 | 72.00507824 | 59.33111814 | 22.74813862 | 22.42919559 | 42.34652802 | 18.81063811 | 2.04E-06 | -1.384718074 | 8.75E-08 | 28.63190234 | 0.382964329 | 5.689816897 |
| *Stx1a* | 23.33356957 | 21.39325204 | 27.14945573 | 36.43138833 | 10.00918099 | 9.75182417 | 14.99772867 | 6.270212703 | 0.002703844 | -1.399740817 | 0.000278334 | 13.21084602 | 0.378997223 | 2.56801833 |
| *Chd5* | 30.43509074 | 34.90477964 | 18.8865779 | 34.34959471 | 13.64888317 | 14.62773626 | 7.939974003 | 8.957446718 | 0.001519619 | -1.402413051 | 0.000141374 | 14.48374611 | 0.378295875 | 2.818265252 |
| *Serpinb2* | 25.36257561 | 56.29803168 | 17.70616678 | 36.43138833 | 9.099255448 | 8.776641753 | 17.64438667 | 14.33191475 | 0.002976231 | -1.440585934 | 0.000312366 | 12.99474474 | 0.368417645 | 2.526333337 |
| *St3gal6* | 27.39158166 | 31.52689774 | 47.21644475 | 32.26780109 | 16.37865981 | 14.62773626 | 9.704412671 | 9.85319139 | 0.000314266 | -1.45149352 | 2.39E-05 | 17.85250259 | 0.365642704 | 3.502703208 |
| *Egr1* | 186.6685565 | 242.0815362 | 232.5409904 | 180.075148 | 63.69478814 | 84.84087028 | 80.28195937 | 76.1382971 | 1.46E-19 | -1.462425074 | 9.43E-22 | 91.83329943 | 0.362882635 | 18.83465774 |
| *Nes* | 1229.577666 | 1154.10965 | 1338.586209 | 1186.622363 | 630.5784026 | 624.1167469 | 222.3192721 | 235.5808487 | 0.002430689 | -1.519462256 | 0.000245712 | 13.44460269 | 0.348815909 | 2.614270541 |
| *Tubb2b* | 100.4357994 | 85.57300816 | 75.5463116 | 72.86277666 | 25.47791526 | 29.25547251 | 25.58436068 | 35.82978687 | 3.59E-10 | -1.527627234 | 7.50E-12 | 46.89291939 | 0.34684735 | 9.444447622 |
| *Sulf1* | 28.40608469 | 30.40093711 | 20.06698902 | 35.39049152 | 10.00918099 | 5.851094502 | 11.46885134 | 10.74893606 | 0.000347711 | -1.582212544 | 2.67E-05 | 17.64203336 | 0.333969314 | 3.458781585 |
| *Cxcl2* | 42.60912703 | 46.16438598 | 46.03603363 | 38.51318195 | 15.46873426 | 13.65255384 | 13.23329001 | 14.33191475 | 1.11E-06 | -1.610399247 | 4.46E-08 | 29.93740443 | 0.327507705 | 5.953793107 |
| *Fat3* | 673.6300083 | 614.774506 | 747.2002382 | 700.5235528 | 321.2037173 | 337.4131163 | 87.33971404 | 104.8021266 | 0.006453673 | -1.685755171 | 0.000780263 | 11.28760477 | 0.310840163 | 2.190193054 |
| *Tox2* | 46.66713913 | 40.53458281 | 38.95356692 | 33.3086979 | 21.83821308 | 15.60291867 | 0.882219334 | 9.85319139 | 0.000600005 | -1.734432722 | 4.96E-05 | 16.46204116 | 0.300527159 | 3.221845082 |
| *Adamts4* | 22.31906654 | 18.01537014 | 12.98452231 | 13.53165852 | 5.459553269 | 1.950364834 | 7.05775467 | 5.374468031 | 0.004692125 | -1.744199233 | 0.000541588 | 11.96673362 | 0.298499572 | 2.328630459 |
| *Mrc1* | 5370.779012 | 4760.561559 | 4673.247619 | 4404.034401 | 1372.167722 | 1329.173634 | 1498.890648 | 1340.929774 | 4.86E-172 | -1.793362527 | 3.92E-176 | 800.6402557 | 0.28849885 | 171.3131952 |
| *Cd5l* | 568.1216938 | 524.6976553 | 578.4014482 | 526.6937856 | 183.8049601 | 209.6642197 | 97.04412671 | 121.8212754 | 1.80E-11 | -1.845159305 | 3.14E-13 | 53.11760808 | 0.27832467 | 10.74535715 |
| *Mmp9* | 95.36328431 | 104.7143389 | 89.71124502 | 88.47622881 | 30.02754298 | 40.95766152 | 16.76216734 | 14.33191475 | 1.07E-11 | -1.899722218 | 1.76E-13 | 54.25202087 | 0.267994961 | 10.97141768 |
| *Uty* | 370.293604 | 364.8112453 | 330.5151132 | 361.1911929 | 149.2277894 | 132.6248087 | 44.11096668 | 38.51702089 | 0.000831759 | -1.96984179 | 7.15E-05 | 15.77113082 | 0.255281025 | 3.08000274 |
| *Pcsk1* | 21.30456352 | 33.77881901 | 41.31438916 | 36.43138833 | 14.55880872 | 10.72700659 | 3.528877335 | 2.687234015 | 3.57E-05 | -2.082376314 | 2.04E-06 | 22.5575483 | 0.236125161 | 4.447805729 |
| *Kdm5d* | 411.888228 | 427.8650408 | 457.9995141 | 471.5262547 | 170.1560769 | 148.2277274 | 52.93316002 | 46.57872293 | 0.000176572 | -2.082750527 | 1.21E-05 | 19.14466646 | 0.236063922 | 3.75307758 |
| *Egr3* | 71.01521172 | 69.80955929 | 86.17001167 | 92.63981604 | 14.55880872 | 22.42919559 | 25.58436068 | 8.957446718 | 1.22E-13 | -2.161985392 | 1.54E-15 | 63.58259037 | 0.223448553 | 12.91504667 |
| *Fbln2* | 298.2638892 | 287.1199616 | 319.8914132 | 283.1239322 | 58.23523487 | 55.58539777 | 68.81310803 | 72.55531842 | 2.70E-60 | -2.215592945 | 1.96E-63 | 282.6775899 | 0.215298035 | 59.56818314 |
| *Ddx3y* | 942.4733098 | 916.5319558 | 936.0660172 | 987.8110722 | 224.7516096 | 322.78538 | 119.9818294 | 117.342552 | 5.00E-08 | -2.269841334 | 1.53E-09 | 36.49342048 | 0.20735269 | 7.300627856 |
| *Wisp2* | 42.60912703 | 42.78650408 | 23.60822237 | 20.81793619 | 5.459553269 | 7.801459336 | 5.293316002 | 8.061702046 | 8.81E-07 | -2.291178279 | 3.47E-08 | 30.42456679 | 0.204308583 | 6.054902044 |
| *Cd248* | 11.15953327 | 7.881724436 | 17.70616678 | 9.368071285 | 0.909925545 | 1.950364834 | 3.528877335 | 2.687234015 | 0.003629142 | -2.332796911 | 0.000401646 | 12.52451804 | 0.198498924 | 2.440196033 |
| *Il1a* | 39.56561796 | 42.78650408 | 49.57726699 | 45.79945962 | 4.549627724 | 9.75182417 | 6.175535336 | 14.33191475 | 5.16E-10 | -2.352243806 | 1.10E-11 | 46.13320041 | 0.195841198 | 9.287650068 |
| *Eif2s3y* | 585.3682452 | 605.7668209 | 599.6488483 | 581.8613165 | 173.7957791 | 182.359112 | 37.05321202 | 59.11914834 | 0.000184996 | -2.39183374 | 1.28E-05 | 19.03566914 | 0.190540061 | 3.732837701 |
| *1500017E21Rik* | 10.14503025 | 11.25960634 | 20.06698902 | 11.4498649 | 0 | 4.875912085 | 2.646658001 | 1.791489344 | 0.001377985 | -2.518009188 | 0.000125534 | 14.70768426 | 0.174583705 | 2.860755522 |
| *Slc39a12* | 14.20304234 | 6.755763802 | 10.62370007 | 10.40896809 | 0.909925545 | 2.925547251 | 3.528877335 | 0 | 0.004309771 | -2.521983748 | 0.000490512 | 12.15139374 | 0.174103396 | 2.365545842 |
| *Tmem45a* | 10.14503025 | 10.1336457 | 12.98452231 | 11.4498649 | 0.909925545 | 2.925547251 | 2.646658001 | 0 | 0.000900428 | -2.797906205 | 7.83E-05 | 15.59993971 | 0.143795835 | 3.04555106 |
| *Il6* | 23.33356957 | 7.881724436 | 20.06698902 | 16.65434895 | 0 | 2.925547251 | 4.411096668 | 0.895744672 | 4.59E-05 | -3.047847201 | 2.70E-06 | 22.02092246 | 0.120922348 | 4.337908715 |
| *Dcn* | 75.07322382 | 56.29803168 | 46.03603363 | 79.10815752 | 7.279404359 | 10.72700659 | 3.528877335 | 0.895744672 | 6.72E-19 | -3.542050481 | 4.55E-21 | 88.71971122 | 0.085849261 | 18.17240737 |
