## Supplementary material for "TFEB-Mediated Pro-inflammatory Response in Murine Macrophages Induced by Acute Alpha7 Nicotinic Receptor Activation": Table_S13

**Table S13. GO and KEGG Pathway analysis of differentially expressed genes from Table S12.**

**Enrichr-KG Differentially Expressed Genes UP in WT (v. Chrna7KO)**

| Term | Library | p-value | q-value | z-score | combined score | converted to -log (Padj) |
|---|---|---|---|---|---|---|
| extracellular matrix organization (GO:0030198) | GO_Biological_Process_2021 | 1.77E-09 | 2.6801E-06 | 7.871 | 158.6 | 5.571849001 |
| Pertussis | KEGG_2021_Human | 1.38E-07 | 0.000027395 | 15.72 | 248.4 | 4.562328695 |
| IL-17 signaling pathway | KEGG_2021_Human | 7.22E-07 | 0.000071831 | 12.42 | 175.6 | 4.143688087 |
| extracellular structure organization (GO:0043062) | GO_Biological_Process_2021 | 1.0696E-06 | 0.0003837 | 7.268 | 99.92 | 3.416008201 |
| external encapsulating structure organization (GO:0045229) | GO_Biological_Process_2021 | 1.1192E-06 | 0.0003837 | 7.232 | 99.1 | 3.416008201 |
| cellular response to cytokine stimulus (GO:0071345) | GO_Biological_Process_2021 | 1.2127E-06 | 0.0003837 | 4.752 | 64.74 | 3.416008201 |
| positive regulation of vascular endothelial growth factor production (GO:0010575) | GO_Biological_Process_2021 | 0.000001268 | 0.0003837 | 32.82 | 445.6 | 3.416008201 |
| Transcriptional misregulation in cancer | KEGG_2021_Human | 2.7405E-06 | 0.0001677 | 7.4 | 94.77 | 3.775466937 |
| MAPK signaling pathway | KEGG_2021_Human | 3.4807E-06 | 0.0001677 | 5.781 | 72.65 | 3.775466937 |
| Pathways in cancer | KEGG_2021_Human | 4.2132E-06 | 0.0001677 | 4.289 | 53.09 | 3.775466937 |

**Enrichr-KG UP in Chrna7KO**

| Term | Library | p-value | q-value | z-score | combined score | converted -log10(Padj) |
|---|---|---|---|---|---|---|
| regulation of immune response (GO:0050776) | GO_Biological_Process_2021 | 8.55E-11 | 1.59E-07 | 7.701 | 178.5 | 6.799313624 |
| inflammatory response (GO:0006954) | GO_Biological_Process_2021 | 1.48E-10 | 1.59E-07 | 6.543 | 148.1 | 6.799313624 |
| regulation of inflammatory response (GO:0050727) | GO_Biological_Process_2021 | 6.30E-09 | 3.49E-06 | 6.18 | 116.7 | 5.457249243 |
| cytokine-mediated signaling pathway (GO:0019221) | GO_Biological_Process_2021 | 6.51E-09 | 3.49E-06 | 3.601 | 67.88 | 5.457249243 |
| positive regulation of cell motility (GO:2000147) | GO_Biological_Process_2021 | 1.90E-08 | 8.13E-06 | 5.719 | 101.7 | 5.090075084 |
| Cell adhesion molecules | KEGG_2021_Human | 7.78E-07 | 0.0001557 | 6.131 | 86.23 | 3.807711387 |
| Hematopoietic cell lineage | KEGG_2021_Human | 0.000030088 | 0.003009 | 6.299 | 65.58 | 2.521577812 |
| Fluid shear stress and atherosclerosis | KEGG_2021_Human | 0.00008239 | 0.005493 | 4.889 | 45.98 | 2.260190401 |
| Cytokine-cytokine receptor interaction | KEGG_2021_Human | 0.0003065 | 0.01533 | 3.158 | 25.55 | 1.814457845 |
| Rheumatoid arthritis | KEGG_2021_Human | 0.0007348 | 0.02939 | 5.095 | 36.77 | 1.531800414 |
