## Supplementary material for "TFEB-Mediated Pro-inflammatory Response in Murine Macrophages Induced by Acute Alpha7 Nicotinic Receptor Activation": Table_S14

**Table S14. Row-normalized expression of macrophage marker genes in Tfebfl/fl and TfebΔLysM BMDMs.**

| | *Tfeb* $^{fl/fl}$ | | | | *Tfeb* $^{-/-}$ | | | |
|---|---|---|---|---|---|---|---|---|
| (Cd11a) *Itgal* | 3.10 | 26.40 | 0.00 | 30.66 | 60.52 | 100.00 | 71.23 | 94.22 |
| (Cd11b)*Itgam* | 7.19 | 19.17 | 0.00 | 13.24 | 72.64 | 84.03 | 62.91 | 100.00 |
| (Cd11c) *Itgax* | 30.15 | 2.71 | 30.80 | 0.00 | 99.23 | 74.23 | 63.53 | 100.00 |
| (CD15) *Fut4* | 63.23 | 0.00 | 32.26 | 36.13 | 89.68 | 100.00 | 65.81 | 44.52 |
| (CD16) *Fcgr3* | 5.33 | 0.00 | 5.07 | 3.83 | 49.78 | 62.18 | 72.08 | 100.00 |
| (CD32) *Fcgr2b* | 12.94 | 17.87 | 0.00 | 32.45 | 86.14 | 68.78 | 100.00 | 99.68 |
| (CD64) *Fcgr1* | 6.79 | 8.87 | 11.29 | 0.00 | 97.38 | 100.00 | 62.81 | 92.53 |
| (F4/80)*Emr1* | 0.00 | 20.16 | 16.43 | 12.01 | 88.41 | 91.76 | 75.25 | 100.00 |
| ****(MHC Class II)*H2-Ab1* | 8.35 | 0.30 | 0.00 | 14.63 | 85.47 | 80.07 | 94.91 | 100.00 |
| *Ccr5* | 65.80 | 100.00 | 63.70 | 37.60 | 57.05 | 26.90 | 0.00 | 16.05 |
| *Cd14* | 100.00 | 94.46 | 74.96 | 72.05 | 68.69 | 0.00 | 29.48 | 36.46 |
| *Cd163* | 20.00 | 100.00 | 17.50 | 60.00 | 7.50 | 60.00 | 0.00 | 92.50 |
| *Cd33* | 60.04 | 100.00 | 65.66 | 92.01 | 1.51 | 82.07 | 14.04 | 0.00 |
| *Cd68* | 54.20 | 0.00 | 2.43 | 13.42 | 57.64 | 51.54 | 74.26 | 100.00 |
| *Cd80* | 96.30 | 100.00 | 40.12 | 46.91 | 25.31 | 19.14 | 0.00 | 20.99 |
| *Cd86* | 15.15 | 15.82 | 64.31 | 0.00 | 24.58 | 96.30 | 68.01 | 100.00 |
| *Csf1r* | 12.55 | 0.00 | 3.09 | 6.98 | 69.85 | 82.11 | 80.36 | 100.00 |
| *Gal3st4* | 0.00 | 0.00 | 56.67 | 33.33 | 56.67 | 100.00 | 26.67 | 0.00 |
| *Lamp1* | 0.00 | 40.13 | 19.71 | 24.08 | 70.73 | 74.95 | 100.00 | 71.79 |
| *Lamp2* | 0.00 | 36.10 | 18.79 | 9.98 | 90.29 | 100.00 | 40.27 | 79.07 |
| *Lilrb4* | 76.57 | 82.69 | 100.00 | 93.01 | 1.85 | 0.00 | 19.45 | 11.03 |
| *Tlr2* | 90.66 | 94.08 | 82.04 | 100.00 | 7.58 | 2.66 | 0.00 | 9.16 |
| *Tlr4* | 61.05 | 100.00 | 92.48 | 85.42 | 25.74 | 25.51 | 0.00 | 14.58 |
