## Supplementary material for "TFEB-Mediated Pro-inflammatory Response in Murine Macrophages Induced by Acute Alpha7 Nicotinic Receptor Activation": Table_S15

**Table S15. Differential gene expression in baseline BMDMs (Tfebfl/fl v. TfebΔLysM).**

| ID | C1_Tfeb_fl_DMSO_1 | C2_Tfeb_fl_DMSO_2 | C3_Tfeb_fl_DMSO_3 | C4_Tfeb_fl_DMSO_4 | D1_Tfeb_LysM_Cre_DMSO_1 | D2_Tfeb_LysM_Cre_DMSO_2 | D3_Tfeb_LysM_Cre_DMSO_3 | D4_Tfeb_LysM_Cre_DMSO_4 | padj | log2FoldChange | pvalue | stat | foldChange | log10padj |
|---|---|---|---|---|---|---|---|---|---|---|---|---|---|---|
| *Syce1* | 1.995172633 | 0 | 0 | 0 | 10.56557338 | 8.463343688 | 8.822935746 | 4.728463616 | 0.000440474 | 4.040654244 | 5.32E-05 | 16.33022384 | 16.4572827 | 3.356080057 |
| *Lyz1* | 22.94448528 | 35.53125688 | 34.53326832 | 32.19081562 | 193.3499929 | 161.861448 | 194.1045864 | 178.7359247 | 3.19E-46 | 2.539328201 | 2.82E-48 | 213.1495825 | 5.813182496 | 45.49680533 |
| *Slco2b1* | 74.81897375 | 85.86720412 | 86.82650321 | 90.7195713 | 389.8696578 | 392.4875635 | 376.4452585 | 409.4849491 | 2.70E-84 | 2.212811438 | 9.91E-87 | 389.6399575 | 4.635777868 | 83.56872105 |
| *Apoc2* | 74.81897375 | 53.29688531 | 51.30657008 | 46.82300454 | 223.9901557 | 262.3636543 | 251.9438318 | 226.9662536 | 1.48E-43 | 2.093914263 | 1.40E-45 | 200.7989701 | 4.269047641 | 42.82862383 |
| *Bco2* | 7.980690533 | 9.869793577 | 2.959994428 | 11.70575114 | 29.58360547 | 37.02712863 | 32.3507644 | 24.5880108 | 7.51E-06 | 1.922543042 | 6.33E-07 | 24.8105061 | 3.790906941 | 5.124401984 |
| *C4b* | 8.97827685 | 11.84375229 | 9.866648093 | 8.779313352 | 33.80983482 | 40.20088252 | 36.27206918 | 32.15355259 | 6.99E-07 | 1.850439355 | 5.06E-08 | 29.69350419 | 3.606099875 | 6.155643787 |
| *Derl3* | 4.987931583 | 2.960938073 | 4.933324046 | 5.852875568 | 13.7352454 | 26.44794902 | 14.70489291 | 12.2940054 | 0.002224321 | 1.837208816 | 0.000329437 | 12.89513559 | 3.573180556 | 2.652802475 |
| *Atp9a* | 10.97344948 | 3.947917431 | 5.919988856 | 7.80383409 | 29.58360547 | 22.21627718 | 29.40978582 | 17.96816174 | 0.000181177 | 1.790882695 | 2.00E-05 | 18.19291798 | 3.460265398 | 3.741897864 |
| *Cd5l* | 120.7079443 | 118.4375229 | 153.9197102 | 130.714221 | 388.8131004 | 421.0513485 | 388.2091728 | 451.095429 | 2.95E-51 | 1.655135193 | 2.17E-53 | 236.5986322 | 3.14952703 | 50.52963291 |
| *Col14a1* | 106.7417359 | 113.5026261 | 108.533129 | 89.74409204 | 290.553268 | 346.9970912 | 329.3896012 | 325.3182968 | 3.34E-42 | 1.62714014 | 3.28E-44 | 194.5183613 | 3.089000575 | 41.47599775 |
| *Cfb* | 25.93724423 | 24.67448394 | 26.63994985 | 13.65670966 | 52.82786691 | 60.30132377 | 60.78022402 | 78.49249602 | 1.42E-07 | 1.47718759 | 9.20E-09 | 33.00352082 | 2.784054764 | 6.84642011 |
| *Apoe* | 58492.47609 | 52526.05444 | 54503.36406 | 55453.06957 | 150136.7977 | 150470.8453 | 156815.919 | 154299.2247 | 0 | 1.469005126 | 0 | 1851.444631 | 2.768309268 | Inf |
| *Ctse* | 17.9565537 | 27.63542202 | 27.62661466 | 8.779313352 | 50.71475223 | 48.6642262 | 58.81957164 | 60.52433428 | 7.39E-06 | 1.41825032 | 6.21E-07 | 24.84638446 | 2.672611837 | 5.13124864 |
| *Flt1* | 65.8406969 | 31.58333945 | 62.15988298 | 39.01917045 | 141.5786833 | 139.6451708 | 110.7768599 | 124.8314395 | 4.75E-11 | 1.379470951 | 2.05E-12 | 49.43857262 | 2.601729459 | 10.32353777 |
| *Pnp2* | 8.97827685 | 9.869793577 | 10.8533129 | 17.5586267 | 35.9229495 | 37.02712863 | 33.33109059 | 17.02246902 | 0.001350258 | 1.377113127 | 0.000184123 | 13.98654715 | 2.597480875 | 2.8695834 |
| *C5ar2* | 43.89379793 | 44.4140711 | 52.29323489 | 44.87204602 | 102.4860618 | 114.2551398 | 113.7178385 | 146.5823721 | 2.05E-12 | 1.365395019 | 7.82E-14 | 55.84925469 | 2.576468616 | 11.68818581 |
| *Rtn4rl1* | 123.7007033 | 108.5677293 | 120.3731067 | 97.54792613 | 250.4040891 | 272.9428339 | 288.215901 | 321.5355259 | 9.99E-26 | 1.333874407 | 1.72E-27 | 118.0130168 | 2.520787328 | 25.00050707 |
| *Xlr3b* | 11.9710358 | 7.895834861 | 7.893318474 | 16.58314744 | 27.47049079 | 30.67962087 | 30.39011201 | 19.85954719 | 0.003470482 | 1.2860672 | 0.000543085 | 11.96158888 | 2.438623775 | 2.459610207 |
| *Abca9* | 172.5824328 | 187.526078 | 130.2397548 | 109.2536773 | 384.5868711 | 333.2441577 | 329.3896012 | 387.7340165 | 1.00E-20 | 1.259771947 | 2.21E-22 | 94.7023676 | 2.394578858 | 19.99820024 |
| *Haao* | 10.97344948 | 16.77864908 | 20.71996099 | 6.828354829 | 27.47049079 | 32.79545679 | 36.27206918 | 34.99063076 | 0.001777289 | 1.252444016 | 0.000253432 | 13.3865722 | 2.382446827 | 2.750242052 |
| *Stag3* | 48.88172951 | 29.60938073 | 32.55993871 | 38.04369119 | 76.07212834 | 95.21261649 | 87.24903126 | 87.00373053 | 7.82E-08 | 1.213390915 | 4.85E-09 | 34.24968414 | 2.31882013 | 7.106562482 |
| *Stxbp6* | 10.97344948 | 18.7526078 | 9.866648093 | 21.46054375 | 32.75327748 | 37.02712863 | 34.31141679 | 35.93632348 | 0.00143717 | 1.196246927 | 0.000197245 | 13.85714651 | 2.291427956 | 2.842491728 |
| *Adcy2* | 72.82380111 | 72.04949311 | 58.21322375 | 74.13642386 | 146.86147 | 170.3247917 | 131.36371 | 182.5186956 | 1.83E-12 | 1.187233476 | 6.93E-14 | 56.08793022 | 2.277156548 | 11.73725954 |
| *Pydc3* | 22.94448528 | 12.83073165 | 13.81330733 | 11.70575114 | 23.24426144 | 43.3746364 | 36.27206918 | 34.99063076 | 0.00268272 | 1.171744926 | 0.00040696 | 12.4999656 | 2.252840111 | 2.571424663 |
| *P2ry12* | 78.80931901 | 64.15365825 | 81.89317917 | 87.79313352 | 195.4631076 | 168.2089558 | 169.5964316 | 167.387612 | 3.10E-14 | 1.163013274 | 9.83E-16 | 64.46417418 | 2.239246381 | 13.50844325 |
| *Il18* | 22.94448528 | 11.84375229 | 20.71996099 | 26.33794005 | 50.71475223 | 40.20088252 | 39.21304776 | 52.01309977 | 0.000414625 | 1.154344397 | 4.99E-05 | 16.45112818 | 2.225831523 | 3.382344984 |
| *Nxpe4* | 29.9275895 | 18.7526078 | 29.59994428 | 23.41150227 | 59.16721093 | 50.78006213 | 64.7015288 | 51.06740705 | 4.28E-05 | 1.150934049 | 4.08E-06 | 21.22923827 | 2.220576155 | 4.368370371 |
| *Gm4980* | 52.87207478 | 59.21876146 | 55.25322932 | 63.40615198 | 140.522126 | 116.3709757 | 123.5211004 | 125.7771322 | 1.26E-10 | 1.132632567 | 5.76E-12 | 47.40904444 | 2.192584688 | 9.900589298 |
| *Zbtb16* | 53.8696611 | 42.44011238 | 29.59994428 | 25.36246079 | 76.07212834 | 65.59091358 | 94.11131462 | 92.67788687 | 1.15E-05 | 1.122474423 | 1.01E-06 | 23.91545532 | 2.177200727 | 4.938381004 |
| *Treml2* | 23.9420716 | 18.7526078 | 14.79997214 | 27.31341932 | 54.94098158 | 47.60630824 | 36.27206918 | 43.50186527 | 0.000687265 | 1.101308033 | 8.71E-05 | 15.39679687 | 2.145491274 | 3.162875772 |
| *Hdac11* | 17.9565537 | 16.77864908 | 12.82664252 | 17.5586267 | 34.86639216 | 38.08504659 | 23.52782865 | 41.61047982 | 0.003915074 | 1.084297863 | 0.000625264 | 11.69921612 | 2.120343294 | 2.407259981 |
| *Gdpd3* | 53.8696611 | 70.0755344 | 57.22655894 | 41.94560824 | 122.5606512 | 106.8497141 | 99.99327178 | 138.0711376 | 9.19E-08 | 1.06855199 | 5.79E-09 | 33.90461921 | 2.097327256 | 7.036762378 |
| *2610035D17Rik* | 33.91793477 | 38.49219495 | 41.43992199 | 31.21533636 | 73.95901367 | 70.88050338 | 72.54413835 | 85.11234509 | 4.77E-06 | 1.061856505 | 3.91E-07 | 25.73848354 | 2.087616203 | 5.321317284 |
| *Ang* | 143.6524296 | 131.2682546 | 126.2930956 | 130.714221 | 236.6688437 | 285.6378495 | 284.2945962 | 298.8389005 | 3.53E-18 | 1.056817119 | 8.54E-20 | 82.92071783 | 2.080336806 | 17.45214968 |
| *Slamf9* | 600.5469626 | 514.2162454 | 541.6789803 | 560.9005752 | 1108.328648 | 1031.470012 | 1254.817528 | 1171.713284 | 2.08E-48 | 1.043091228 | 1.61E-50 | 223.4316643 | 2.060638206 | 47.68111821 |
| *Chst1* | 21.94689897 | 36.51823623 | 22.69329061 | 21.46054375 | 47.54508022 | 64.53299562 | 38.23272156 | 60.52433428 | 0.000874778 | 1.038369026 | 0.000113843 | 14.89202267 | 2.053904391 | 3.058102059 |
| *Slfn4* | 25.93724423 | 40.46615367 | 27.62661466 | 30.2398571 | 66.5631123 | 69.82258542 | 66.66628119 | 50.12171433 | 0.000124672 | 1.023947171 | 1.31E-05 | 18.99166141 | 2.033474882 | 3.904232232 |
| *Fgf13* | 32.92034845 | 34.54427752 | 40.45325718 | 28.28889858 | 65.50655496 | 83.57551892 | 77.44576932 | 48.23032888 | 0.000140723 | 1.009882413 | 1.50E-05 | 18.7319347 | 2.013746962 | 3.851634321 |
| *Nrap* | 20.94931265 | 19.73958715 | 16.77330176 | 10.73027187 | 36.97950683 | 30.67962087 | 34.31141679 | 34.04493803 | 0.007447342 | 0.99753142 | 0.001312873 | 10.32433886 | 1.996580748 | 2.127998679 |
| *Fgl2* | 133.6765664 | 222.0703555 | 188.4529786 | 181.4391426 | 394.0958871 | 409.4142509 | 252.924158 | 388.6797092 | 5.13E-05 | 0.99340371 | 4.96E-06 | 20.85378075 | 1.990876474 | 4.289777501 |
| *Pde1c* | 36.91069371 | 38.49219495 | 41.43992199 | 34.14177415 | 62.33688295 | 69.82258542 | 72.54413835 | 89.8408087 | 5.51E-05 | 0.967293028 | 5.36E-06 | 20.70382283 | 1.955168609 | 4.258632917 |
| *Ly86* | 1194.110821 | 1275.17733 | 1340.877476 | 1231.054828 | 2397.3286 | 2357.041217 | 2428.267982 | 2651.722396 | 1.95E-76 | 0.964519178 | 8.56E-79 | 353.1891675 | 1.951413046 | 75.70992289 |
| *Plxdc2* | 1109.315984 | 1076.794479 | 992.5847981 | 1075.953625 | 2107.83189 | 1986.769931 | 2020.452286 | 2185.495883 | 7.97E-77 | 0.964510307 | 3.44E-79 | 355.0093052 | 1.951401047 | 76.09830179 |
| *Pik3ip1* | 20.94931265 | 9.869793577 | 28.61327947 | 22.43602301 | 48.60163755 | 39.14296456 | 36.27206918 | 34.99063076 | 0.009038532 | 0.955200807 | 0.001643173 | 9.910737469 | 1.938849482 | 2.043902107 |
| *A930005H10Rik* | 28.93000318 | 32.5703188 | 15.78663695 | 23.41150227 | 26.41393345 | 53.95381601 | 58.81957164 | 52.9587925 | 0.007575539 | 0.935457586 | 0.001340225 | 10.28628732 | 1.912497134 | 2.120586489 |
| *Fcrls* | 6188.027922 | 5986.029804 | 6350.174712 | 6482.059691 | 11528.09712 | 11740.77353 | 11856.06499 | 12443.42485 | 1.03E-146 | 0.92781422 | 1.40E-149 | 678.528725 | 1.902391561 | 145.9877935 |
| *Hap1* | 31.92276213 | 33.55729816 | 43.41325161 | 44.87204602 | 73.95901367 | 83.57551892 | 70.58348596 | 64.30710518 | 0.000129029 | 0.924506564 | 1.36E-05 | 18.91890635 | 1.898034958 | 3.889314297 |
| *Tmem119* | 147.6427749 | 103.6328326 | 95.7064865 | 126.812304 | 219.7639263 | 240.1473771 | 221.5537198 | 214.6722482 | 9.32E-11 | 0.919158091 | 4.21E-12 | 48.02385117 | 1.89101144 | 10.03068873 |
| *Mroh6* | 39.90345266 | 19.73958715 | 30.58660909 | 31.21533636 | 69.73278432 | 51.83798009 | 50.97696209 | 56.74156339 | 0.001543785 | 0.916256282 | 0.00021385 | 13.70530441 | 1.887211719 | 2.811413239 |
| *Arrdc3* | 1051.455978 | 1206.088775 | 1331.010828 | 1214.47168 | 2225.109754 | 2313.666581 | 2220.438829 | 2266.825457 | 4.26E-56 | 0.909970823 | 2.96E-58 | 258.9144363 | 1.879007497 | 55.37059654 |
| *Rnase6* | 60.85276531 | 52.30990596 | 68.07987184 | 59.50423494 | 111.9950778 | 111.0813859 | 93.13098842 | 134.2883667 | 3.11E-06 | 0.905060125 | 2.49E-07 | 26.6127793 | 1.872622536 | 5.507165712 |
| *Tspan13* | 137.6669117 | 138.1771101 | 95.7064865 | 97.54792613 | 204.9721236 | 229.5681975 | 207.8291531 | 234.5317953 | 5.99E-10 | 0.903513484 | 2.96E-11 | 44.20037518 | 1.870616066 | 9.222699906 |
| *Plekhh2* | 69.83104216 | 90.80210091 | 60.18655336 | 47.7984838 | 126.7868806 | 111.0813859 | 140.1866457 | 123.8857467 | 4.80E-06 | 0.90336194 | 3.94E-07 | 25.72373142 | 1.870419583 | 5.319193458 |
| *Fat3* | 520.7400573 | 526.0599976 | 455.8391419 | 459.4507321 | 950.9016043 | 873.8402357 | 980.3261939 | 864.363149 | 3.54E-35 | 0.903256291 | 4.26E-37 | 161.9396792 | 1.870282617 | 34.45133196 |
| *Hpgd* | 1640.031905 | 1714.383144 | 1653.65022 | 1591.006675 | 2995.340054 | 3182.217227 | 2890.981946 | 3237.106191 | 1.33E-80 | 0.899091009 | 5.32E-83 | 372.5086876 | 1.864890611 | 79.87521229 |
| *Tspan33* | 33.91793477 | 24.67448394 | 24.66662023 | 20.48506449 | 52.82786691 | 38.08504659 | 48.0359835 | 53.90448522 | 0.003718915 | 0.897248621 | 0.000589682 | 11.80827027 | 1.86251058 | 2.429583697 |
| *Itga6* | 4136.990455 | 4176.896642 | 4326.525189 | 4137.983026 | 7670.606275 | 7961.890574 | 7528.90517 | 7839.792675 | 6.13E-150 | 0.88567709 | 7.83E-153 | 693.4783689 | 1.847631555 | 149.2124571 |
| *Fzd7* | 1074.400463 | 1029.41947 | 987.6514741 | 1114.972796 | 1909.19911 | 1948.684884 | 1893.990207 | 2003.92288 | 6.82E-67 | 0.882780451 | 3.76E-69 | 308.9181065 | 1.843925606 | 66.16616214 |
| *H2-Eb1* | 414.9959077 | 396.7657018 | 378.8792868 | 413.6032068 | 710.0065312 | 655.9091358 | 708.7758382 | 871.9286908 | 1.14E-23 | 0.878547554 | 2.17E-25 | 108.4195792 | 1.838523419 | 22.94134276 |
| *Mfng* | 238.4231297 | 238.8490046 | 221.0129173 | 247.7717324 | 435.3016233 | 406.240497 | 446.0484182 | 441.6385017 | 2.53E-21 | 0.87066832 | 5.33E-23 | 97.52024451 | 1.828509751 | 20.59703015 |
| *Sulf2* | 2018.117119 | 1876.247759 | 1841.116534 | 1952.909481 | 3357.739221 | 3399.090409 | 3661.518334 | 3616.328973 | 1.05E-81 | 0.86866212 | 4.01E-84 | 377.6651715 | 1.825968807 | 80.98016933 |
| *Pde3b* | 810.0400891 | 941.5783072 | 969.8915075 | 913.0485886 | 1764.450755 | 1704.305835 | 1629.302134 | 1499.868659 | 7.23E-41 | 0.859422259 | 7.22E-43 | 188.3689365 | 1.814311606 | 40.14076742 |
| *Cbx7* | 120.7079443 | 100.6718945 | 112.4797883 | 139.4935344 | 179.6147475 | 247.5528029 | 194.1045864 | 236.4231808 | 9.88E-09 | 0.857680559 | 5.47E-10 | 38.50022198 | 1.812122592 | 8.005217782 |
| *Slc38a6* | 430.9572888 | 503.3594724 | 497.2790639 | 485.7886721 | 802.983577 | 921.446544 | 859.7460721 | 868.1459199 | 6.89E-32 | 0.848274356 | 9.30E-34 | 146.6623204 | 1.800346197 | 31.16155875 |
| *Jag2* | 145.6476022 | 134.2291926 | 139.1197381 | 145.3464099 | 264.1393345 | 238.0315412 | 278.4126391 | 232.6404099 | 4.46E-12 | 0.844115979 | 1.78E-13 | 54.23477757 | 1.795164409 | 11.35099178 |
| *Rnase4* | 1924.344005 | 2048.969147 | 2044.369485 | 2000.707965 | 3355.626106 | 3575.762708 | 3706.613339 | 3721.300866 | 6.18E-80 | 0.840944555 | 2.52E-82 | 369.409869 | 1.7912225 | 79.20910826 |
| *Myom1* | 30.92517582 | 28.62240137 | 29.59994428 | 23.41150227 | 42.26229352 | 41.25880048 | 49.0163097 | 65.2527979 | 0.007927952 | 0.818371422 | 0.001415321 | 10.18572399 | 1.763414249 | 2.100838993 |
| *Dgkg* | 88.78518218 | 93.76303898 | 74.9865255 | 89.74409204 | 124.6737659 | 132.2397451 | 171.5570839 | 181.5730029 | 2.75E-06 | 0.816180175 | 2.18E-07 | 26.86267312 | 1.760737908 | 5.560281978 |
| *Pyhin1* | 279.3241687 | 393.8047637 | 280.2128058 | 290.6928199 | 550.4663732 | 646.3878741 | 431.3435253 | 553.2302431 | 2.64E-05 | 0.809883096 | 2.42E-06 | 22.22591659 | 1.753069383 | 4.579178082 |
| *Gng7* | 47.8841432 | 71.06251375 | 51.30657008 | 74.13642386 | 102.4860618 | 123.7764014 | 100.973598 | 99.29773593 | 9.93E-05 | 0.801273772 | 1.03E-05 | 19.45874047 | 1.742639044 | 4.003184556 |
| *Cmah* | 34.91552108 | 59.21876146 | 32.55993871 | 23.41150227 | 67.61966964 | 81.45968299 | 50.97696209 | 61.47002701 | 0.008793224 | 0.799167713 | 0.001590882 | 9.970269969 | 1.740096979 | 2.055851845 |
| *Vill* | 108.7369085 | 95.7369977 | 116.4264475 | 79.98929943 | 179.6147475 | 174.5564636 | 161.753822 | 179.6816174 | 1.89E-07 | 0.795419037 | 1.25E-08 | 32.4100654 | 1.735581408 | 6.723380343 |
| *Ltc4s* | 45.88897056 | 77.97136926 | 59.19988856 | 59.50423494 | 102.4860618 | 101.5601243 | 132.3440362 | 84.16665236 | 0.000299623 | 0.793298608 | 3.50E-05 | 17.12763359 | 1.733032377 | 3.523424294 |
| *Trpv4* | 69.83104216 | 77.97136926 | 88.79983283 | 80.96477869 | 145.8049127 | 133.2976631 | 134.3046886 | 136.1797521 | 1.59E-06 | 0.790631909 | 1.21E-07 | 28.00674761 | 1.729831974 | 5.798913396 |
| *Gm5617* | 43.89379793 | 55.27084403 | 46.37324603 | 39.99464971 | 90.86393108 | 83.57551892 | 82.34740029 | 64.30710518 | 0.000680118 | 0.78886808 | 8.60E-05 | 15.42250907 | 1.727718386 | 3.167415424 |
| *Tlr9* | 566.6290278 | 544.8126054 | 574.238919 | 559.925096 | 931.8835722 | 1046.280863 | 903.8607508 | 954.2039577 | 4.81E-31 | 0.772084018 | 6.76E-33 | 142.7237041 | 1.707734879 | 30.31809285 |
| *Mtus1* | 592.5662721 | 723.4558692 | 630.4788131 | 606.7481005 | 953.014719 | 1085.423828 | 1127.375123 | 1185.898675 | 1.52E-23 | 0.770083567 | 2.89E-25 | 107.8552633 | 1.705368563 | 22.81951094 |
| *H2-Aa* | 861.9145776 | 779.7136926 | 832.745099 | 823.3044965 | 1343.940934 | 1346.729564 | 1456.764724 | 1457.312486 | 7.38E-41 | 0.765967511 | 7.42E-43 | 188.3139675 | 1.700510025 | 40.13222887 |
| *Nat8l* | 81.80207796 | 75.01043118 | 91.75982726 | 89.74409204 | 140.522126 | 137.5293349 | 160.7734958 | 136.1797521 | 2.61E-06 | 0.765426004 | 2.06E-07 | 26.97579241 | 1.699871868 | 5.583070778 |
| *Tmc8* | 87.78759586 | 74.02345183 | 82.87984398 | 86.81765426 | 143.691798 | 152.3401864 | 127.4424052 | 136.1797521 | 4.32E-06 | 0.754154633 | 3.52E-07 | 25.94163753 | 1.686642991 | 5.364833055 |
| *Dach1* | 35.9131074 | 39.47917431 | 38.47992756 | 43.89656676 | 66.5631123 | 68.76466746 | 62.74087641 | 68.08987607 | 0.002120667 | 0.753784036 | 0.000311171 | 13.00192437 | 1.686209785 | 2.673527612 |
| *Shisa9* | 45.88897056 | 40.46615367 | 44.39991642 | 30.2398571 | 75.01557101 | 61.35924174 | 71.56381216 | 63.36141245 | 0.002835545 | 0.75325329 | 0.000433312 | 12.38277765 | 1.685589567 | 2.54736342 |
| *Dos* | 43.89379793 | 40.46615367 | 32.55993871 | 52.67588011 | 77.12868568 | 58.18548785 | 61.76055022 | 87.00373053 | 0.004128253 | 0.745671177 | 0.000663816 | 11.5879192 | 1.676754155 | 2.384233707 |
| *F13a1* | 353.1455561 | 346.4297545 | 412.4258903 | 381.4123912 | 573.7106346 | 715.1525416 | 559.7662567 | 655.3650572 | 7.72E-16 | 0.745262692 | 2.18E-17 | 71.97328449 | 1.676279465 | 15.11250888 |
| *Gatm* | 2572.775111 | 2467.448394 | 2415.355453 | 2420.164047 | 3878.621988 | 4071.926232 | 4421.271135 | 4171.450602 | 4.71E-62 | 0.744641043 | 2.82E-64 | 286.5423306 | 1.675557322 | 61.32731701 |
| *Hgf* | 593.5638584 | 683.9766949 | 636.398802 | 669.1787732 | 1096.706517 | 1034.643766 | 1052.870332 | 1136.722653 | 5.41E-31 | 0.74244233 | 7.65E-33 | 142.4775289 | 1.673005664 | 30.26672739 |
| *Pltp* | 4948.02813 | 4922.066057 | 4908.657426 | 4999.331214 | 8245.373467 | 7867.735876 | 7979.855219 | 8920.719458 | 1.30E-75 | 0.739351741 | 5.90E-78 | 349.338917 | 1.669425532 | 74.88699694 |
| *Cytip* | 621.4962753 | 594.1615733 | 648.2387797 | 585.2875568 | 1123.12045 | 914.0411183 | 1057.771963 | 979.7376612 | 1.26E-24 | 0.734360947 | 2.34E-26 | 112.8408483 | 1.663660374 | 23.89884921 |
| *Etohd2* | 29.9275895 | 38.49219495 | 37.49326275 | 27.31341932 | 49.65819489 | 63.47507766 | 54.89826686 | 52.9587925 | 0.009826207 | 0.729548555 | 0.001811884 | 9.730974665 | 1.658120155 | 2.0076141 |
| *Siglec1* | 5731.133389 | 6571.308563 | 6286.0415 | 7005.892055 | 9787.94718 | 11217.10414 | 10366.9495 | 10994.6236 | 3.33E-42 | 0.72708778 | 3.25E-44 | 194.5402121 | 1.655294345 | 41.47722147 |
| *Ppfia4* | 186.5486412 | 159.8906559 | 200.2929563 | 177.5372256 | 332.8155615 | 312.0857985 | 270.5700295 | 275.1965824 | 1.49E-09 | 0.715494275 | 7.66E-11 | 42.3433545 | 1.642045694 | 8.827014947 |
| *Ifi204* | 718.262148 | 854.7241238 | 796.2385011 | 669.1787732 | 1206.58848 | 1303.354928 | 1150.902952 | 1305.055958 | 1.96E-22 | 0.708823309 | 3.94E-24 | 102.6804688 | 1.634470466 | 21.7083384 |
| *Bbc3* | 306.2589992 | 230.9531697 | 221.9995821 | 253.6246079 | 450.093426 | 397.7771533 | 404.8747181 | 394.3538656 | 5.77E-11 | 0.701249751 | 2.53E-12 | 49.02091131 | 1.625912648 | 10.23903864 |
| *Cmklr1* | 205.5027812 | 226.0182729 | 251.5995264 | 213.6299582 | 341.2680202 | 392.4875635 | 339.1928631 | 384.8969383 | 2.67E-11 | 0.701103636 | 1.13E-12 | 50.6105373 | 1.625747986 | 10.57399047 |
| *Cadm1* | 4831.310531 | 4939.831685 | 4888.92413 | 4604.262113 | 8131.265274 | 7730.206541 | 7527.924843 | 7913.556708 | 2.36E-81 | 0.700335828 | 9.25E-84 | 375.9988141 | 1.624882986 | 80.62632407 |
| *Cracr2b* | 69.83104216 | 77.97136926 | 113.4664531 | 75.11190312 | 133.1262246 | 125.8922374 | 151.9505601 | 135.2340594 | 0.000154254 | 0.700097419 | 1.66E-05 | 18.5388169 | 1.624614493 | 3.811764615 |
| *Cd40* | 161.6089833 | 113.5026261 | 109.5197938 | 132.6651795 | 220.8204837 | 209.4677563 | 201.947196 | 207.1067064 | 1.12E-06 | 0.698147217 | 8.29E-08 | 28.73711595 | 1.622419859 | 5.952619518 |
| *Nr4a1* | 450.9090151 | 381.9610114 | 447.9458234 | 436.0392298 | 778.6827582 | 648.5037101 | 699.9529025 | 648.7452081 | 9.20E-17 | 0.69261961 | 2.38E-18 | 76.346556 | 1.61621554 | 16.03617915 |
| *H2-Ab1* | 828.9942291 | 748.1303531 | 760.7185679 | 839.887644 | 1253.077003 | 1192.273542 | 1335.204276 | 1349.503516 | 1.63E-28 | 0.691728535 | 2.55E-30 | 130.9451611 | 1.615217599 | 27.78649755 |
| *Cyp27a1* | 167.5945012 | 177.6562844 | 140.1064029 | 181.4391426 | 253.5737611 | 297.274947 | 253.9044842 | 272.3595043 | 1.63E-08 | 0.691321961 | 9.28E-10 | 37.47105721 | 1.614762469 | 7.787859342 |
| *Metap1d* | 124.6982896 | 138.1771101 | 109.5197938 | 128.7632625 | 200.7458942 | 207.3519203 | 197.045565 | 196.7040864 | 5.21E-07 | 0.677533814 | 3.65E-08 | 30.32514267 | 1.599403349 | 6.282996782 |
| *Mxi1* | 1409.589465 | 1444.93778 | 1520.450471 | 1495.409708 | 2610.753182 | 2312.608663 | 2176.324151 | 2246.96591 | 1.42E-32 | 0.670409077 | 1.86E-34 | 149.864287 | 1.591524181 | 31.84844265 |
| *Rhobtb1* | 78.80931901 | 82.90626605 | 92.74649207 | 79.98929943 | 127.8434379 | 151.2822684 | 133.3243624 | 119.1572831 | 0.000101791 | 0.666980123 | 1.06E-05 | 19.40788918 | 1.587745986 | 3.992290661 |
| *Nr4a2* | 199.5172633 | 216.1484793 | 197.3329619 | 194.120373 | 375.077855 | 286.6957674 | 300.9601415 | 317.752755 | 5.80E-09 | 0.665293859 | 3.15E-10 | 39.57712895 | 1.585891267 | 8.236569668 |
| *Abcd2* | 447.9162562 | 482.6329059 | 400.5859126 | 362.8782852 | 658.2352216 | 682.3570848 | 629.3694165 | 711.1609278 | 2.87E-14 | 0.662859232 | 9.06E-16 | 64.6241685 | 1.583217246 | 13.54261301 |
| *Acsf2* | 162.6065696 | 182.5911812 | 160.8263639 | 184.3655804 | 283.1573666 | 303.6224548 | 263.7077462 | 240.2059517 | 5.66E-08 | 0.657609164 | 3.45E-09 | 34.91104179 | 1.577466278 | 7.247347867 |
| *Tmem37* | 886.8542355 | 845.8413095 | 882.0783395 | 841.8386025 | 1267.868806 | 1376.351267 | 1395.9845 | 1390.168303 | 2.95E-31 | 0.652251854 | 4.12E-33 | 143.7075717 | 1.57161937 | 30.53073401 |
| *Fam129a* | 1976.218493 | 1918.687871 | 1907.223076 | 2002.658923 | 3085.147427 | 2963.228209 | 2962.545758 | 3235.214806 | 1.22E-49 | 0.650152168 | 9.34E-52 | 229.1055641 | 1.569333712 | 48.91408774 |
| *Trf* | 4272.662194 | 4292.373227 | 4320.6052 | 4524.272814 | 6495.714515 | 6772.790786 | 6707.391819 | 7235.495025 | 4.92E-60 | 0.644493597 | 3.10E-62 | 277.1721805 | 1.563190492 | 59.30801136 |
| *Dmpk* | 704.2959395 | 658.3152316 | 654.1587685 | 688.6883585 | 1129.459794 | 1098.118843 | 1012.676958 | 987.303203 | 3.78E-23 | 0.642951413 | 7.34E-25 | 106.0090698 | 1.561520396 | 22.42214138 |
| *F11r* | 745.1969785 | 656.3412729 | 598.9055392 | 657.4730221 | 969.9196364 | 975.40036 | 1138.158711 | 1060.121543 | 8.88E-18 | 0.641508073 | 2.20E-19 | 81.05249147 | 1.559958958 | 17.05155416 |
| *Disc1* | 51.87448846 | 52.30990596 | 67.09320703 | 42.9210875 | 105.6557338 | 84.63343688 | 74.50479074 | 69.98126152 | 0.008029453 | 0.641331986 | 0.00143857 | 10.15568388 | 1.5597 6857 | 2.095314056 |
| *Slc37a1* | 62.84793795 | 60.20574082 | 75.97319031 | 69.25902755 | 105.6557338 | 105.7917961 | 104.8949028 | 102.1348141 | 0.000851017 | 0.640822728 | 0.000110254 | 14.95247055 | 1.559218083 | 3.070061825 |
| *Mnda* | 335.1890024 | 463.8802981 | 381.8392812 | 385.3143082 | 633.9344029 | 649.561628 | 514.6712518 | 640.2339736 | 2.45E-10 | 0.638043206 | 1.15E-11 | 46.05508302 | 1.556216958 | 9.610998231 |
| *Zfp791* | 162.6065696 | 153.9687798 | 157.8663695 | 145.3464099 | 248.2909745 | 235.9157053 | 232.337308 | 246.8258007 | 1.23E-07 | 0.636556091 | 7.82E-09 | 33.31878502 | 1.554613653 | 6.91088865 |
| *Gpr65* | 2139.822649 | 2198.990009 | 2218.022491 | 2322.616121 | 3416.906431 | 3522.86681 | 3252.722312 | 3520.814008 | 6.61E-48 | 0.6268908 | 5.28E-50 | 221.0716202 | 1.544233376 | 47.17958385 |
| *Pla2g16* | 400.032113 | 312.8724564 | 371.9726331 | 354.0989718 | 554.6926025 | 568.101945 | 561.7269091 | 537.1534668 | 3.27E-13 | 0.626491052 | 1.15E-14 | 59.62919019 | 1.543805553 | 12.48480042 |
| *Gas6* | 7156.684235 | 6893.063834 | 7188.8398 | 7532.650856 | 11209.0168 | 11074.28522 | 10540.46724 | 11584.73586 | 1.32E-61 | 0.626210862 | 8.00E-64 | 284.4623006 | 1.543505755 | 60.87982542 |
| *Hexb* | 10402.83011 | 10396.84055 | 10260.32735 | 10105.96515 | 15390.87074 | 15551.39403 | 16602.80442 | 15973.69579 | 4.04E-84 | 0.625842114 | 1.52E-86 | 388.7909733 | 1.543111291 | 83.39323524 |
| *Eml6* | 72.82380111 | 57.24480275 | 69.06653665 | 61.45519346 | 78.18524302 | 105.7917961 | 99.01294559 | 118.2115904 | 0.003081829 | 0.625245782 | 0.000476361 | 12.2059989 | 1.542473584 | 2.51119152 |
| *Hgsnat* | 2557.811316 | 2570.094247 | 2774.501444 | 2711.832346 | 3883.904775 | 4127.995884 | 4064.4324 | 4235.757707 | 2.23E-48 | 0.620113629 | 1.75E-50 | 223.2741982 | 1.536996232 | 47.65122748 |
| *Enc1* | 4135.992869 | 4183.805497 | 4437.031647 | 4338.931754 | 6586.578446 | 6519.948393 | 6561.323216 | 6566.89027 | 1.46E-72 | 0.617809553 | 6.86E-75 | 335.2633808 | 1.534543511 | 71.83663127 |
| *Crem* | 228.4472665 | 253.6536949 | 228.9062357 | 230.2131057 | 396.2090018 | 296.2170291 | 360.7600394 | 388.6797092 | 3.05E-08 | 0.616284447 | 1.80E-09 | 36.17722827 | 1.532922167 | 7.51637358 |
| *Runx3* | 263.3627876 | 268.4583853 | 286.1327947 | 267.2813176 | 452.2065407 | 467.5997387 | 398.0124347 | 339.5036876 | 2.16E-08 | 0.608581126 | 1.25E-09 | 36.89179073 | 1.524758889 | 7.665500578 |
| *Pdk1* | 1046.468046 | 1094.560108 | 965.9448483 | 1043.76281 | 1582.722892 | 1598.514039 | 1486.17451 | 1660.636422 | 1.97E-27 | 0.608477158 | 3.17E-29 | 125.9423062 | 1.524649011 | 26.70497085 |
| *Slc28a2* | 69.83104216 | 78.95834861 | 72.02653108 | 69.25902755 | 115.1647499 | 94.15469853 | 99.99327178 | 128.6142104 | 0.002311398 | 0.596046084 | 0.000343803 | 12.81526279 | 1.511568212 | 2.636125313 |
| *Chn2* | 198.519677 | 242.796922 | 214.1062636 | 224.3602301 | 340.2114629 | 326.8966499 | 317.6256868 | 345.177844 | 1.55E-08 | 0.596018395 | 8.79E-10 | 37.57610867 | 1.511539201 | 7.809418955 |
| *Fam214a* | 838.9700923 | 936.6434104 | 899.838306 | 894.5144826 | 1304.848313 | 1363.656252 | 1383.24026 | 1342.883667 | 6.74E-27 | 0.595672548 | 1.11E-28 | 123.4561596 | 1.511176893 | 26.1715195 |
| *Ccl8* | 66.83828321 | 67.11459632 | 69.06653665 | 73.1609446 | 87.69425906 | 105.7917961 | 105.8752289 | 117.2658977 | 0.002516708 | 0.595098147 | 0.00037836 | 12.63614108 | 1.510575347 | 2.599167148 |
| *Arsb* | 1505.357752 | 1663.060218 | 1577.67703 | 1573.448048 | 2337.104832 | 2397.2421 | 2405.72048 | 2401.113824 | 8.95E-40 | 0.594414206 | 9.57E-42 | 183.2261842 | 1.509859395 | 39.0482799 |
| *Gpr19* | 100.756218 | 111.5286674 | 79.91984955 | 81.94025795 | 165.8795021 | 129.0659912 | 151.9505601 | 118.2115904 | 0.001284891 | 0.593859156 | 0.000174089 | 14.091935 | 1.509278617 | 2.891133704 |
| *Sepp1* | 10051.67973 | 10926.84847 | 10674.72657 | 10265.94375 | 15763.83548 | 16574.40069 | 14429.42125 | 16495.71817 | 3.27E-42 | 0.593729137 | 3.16E-44 | 194.5915544 | 1.509142603 | 41.48485258 |
| *Zfp773* | 61.85035163 | 54.28386467 | 61.17321817 | 76.08738238 | 101.4295045 | 96.27053445 | 97.0522932 | 87.94942326 | 0.004371121 | 0.593273162 | 0.000707057 | 11.47060327 | 1.508665703 | 2.359407128 |
| *AI839979* | 53.8696611 | 54.28386467 | 57.22655894 | 45.84752528 | 83.46802971 | 74.05425727 | 77.44576932 | 83.22095964 | 0.008937341 | 0.591875877 | 0.001621922 | 9.934696041 | 1.50720523 | 2.048791694 |
| *Nek8* | 114.7224264 | 109.5547087 | 146.0263918 | 119.0084699 | 189.1237635 | 186.1935611 | 167.6357792 | 192.9213155 | 6.80E-05 | 0.588468544 | 6.77E-06 | 20.25622214 | 1.503649739 | 4.167637719 |
| *Cdk5r1* | 144.6500159 | 113.5026261 | 124.319766 | 130.714221 | 179.6147475 | 205.2360844 | 175.4783887 | 210.8894773 | 4.58E-05 | 0.588302399 | 4.37E-06 | 21.09579865 | 1.503476584 | 4.339223095 |
| *Rab6b* | 91.77794113 | 95.7369977 | 94.71982169 | 56.57779715 | 128.8999953 | 128.0080733 | 115.6784909 | 136.1797521 | 0.002555022 | 0.587340244 | 0.000384528 | 12.60591384 | 1.502474227 | 2.592605428 |
| *Galnt12* | 323.2179666 | 295.1068279 | 325.5993871 | 336.5403451 | 514.5434237 | 510.9743751 | 460.7533112 | 438.8014236 | 1.22E-10 | 0.586648101 | 5.57E-12 | 47.47552019 | 1.501753578 | 9.913041155 |
| *Pxylp1* | 143.6524296 | 177.6562844 | 186.4796489 | 187.2920182 | 103.5426191 | 100.5022063 | 122.5407742 | 135.2340594 | 0.000254418 | -0.586542 | 2.92E-05 | 17.47143497 | 0.665937182 | 3.59445186 |
| *Nptxr* | 185.5510549 | 222.0703555 | 200.2929563 | 236.0659812 | 145.8049127 | 159.7456121 | 149.9899077 | 106.8632777 | 9.30E-05 | -0.588972774 | 9.55E-06 | 19.59864416 | 0.6648161 | 4.031629341 |
| *Rasgrp4* | 211.4882991 | 243.7839013 | 228.9062357 | 250.6981702 | 142.6352406 | 139.6451708 | 157.8325172 | 177.790232 | 5.50E-06 | -0.594645961 | 4.55E-07 | 25.44371704 | 0.662206942 | 5.259951634 |
| *Adamtsl5* | 113.7248401 | 101.6588738 | 109.5197938 | 119.0084699 | 73.95901367 | 71.93842134 | 84.30805268 | 63.36141245 | 0.001696998 | -0.59737125 | 0.000239409 | 13.49336188 | 0.660957196 | 2.770318563 |
| *Pira2* | 429.9597025 | 391.830805 | 453.8658123 | 437.0147091 | 295.8360547 | 287.7536854 | 257.825789 | 290.327666 | 1.94E-10 | -0.598333 | 9.01E-12 | 46.53207437 | 0.660516726 | 9.712271053 |
| *Slc15a2* | 136.6693254 | 182.5911812 | 151.9463806 | 198.02229 | 113.0516352 | 124.8343194 | 101.9539242 | 102.1348141 | 0.000305795 | -0.601152717 | 3.57E-05 | 17.08761322 | 0.659227021 | 3.51457013 |
| *Fam20c* | 6059.339287 | 5183.615587 | 5193.803556 | 5359.283062 | 3685.271995 | 3337.731167 | 3766.413237 | 3523.651087 | 1.55E-31 | -0.606596949 | 2.12E-33 | 145.0237504 | 0.656744014 | 30.81098218 |
| *Large* | 552.6628194 | 550.7344816 | 636.398802 | 533.5871559 | 396.2090018 | 398.8350713 | 354.8780822 | 342.3407658 | 4.36E-12 | -0.609080382 | 1.74E-13 | 54.28277778 | 0.655614478 | 11.36073103 |
| *Dhcr24* | 1186.13013 | 1064.950727 | 1120.851223 | 1081.806501 | 697.3278432 | 720.4421314 | 755.8314955 | 743.3144804 | 2.02E-23 | -0.609785888 | 3.86E-25 | 107.2810183 | 0.655293948 | 22.69548994 |
| *Spred1* | 1557.23224 | 1676.877929 | 1642.796907 | 1508.090938 | 1142.138483 | 1092.829254 | 1040.126092 | 908.810707 | 8.26E-21 | -0.611135962 | 1.81E-22 | 95.10288513 | 0.65468101 | 20.08291106 |
| *St3gal1* | 1605.116383 | 1506.1305 | 1556.957069 | 1634.903242 | 1108.328648 | 1080.134238 | 990.1294559 | 947.5841086 | 9.04E-26 | -0.612599142 | 1.55E-27 | 118.2208739 | 0.65401737 | 25.0440033 |
| *Scd2* | 514.7545394 | 552.7084403 | 467.6791196 | 466.2790869 | 329.6458895 | 314.2016344 | 305.8617725 | 355.5804639 | 1.74E-11 | -0.615764983 | 7.22E-13 | 51.48283462 | 0.652583772 | 10.76036733 |
| *Trem1* | 134.6741527 | 149.033883 | 146.0263918 | 187.2920182 | 113.0516352 | 78.28592911 | 102.9342504 | 107.8089704 | 0.000401798 | -0.616759272 | 4.81E-05 | 16.52205356 | 0.652134174 | 3.395991706 |
| *Gpr162* | 292.2927908 | 227.0052523 | 265.4128337 | 280.9380273 | 163.7663874 | 167.1510378 | 192.143934 | 171.1703829 | 6.29E-07 | -0.617164521 | 4.52E-08 | 29.91324453 | 0.651951017 | 6.201020389 |
| *Ikbke* | 963.6683819 | 984.0184196 | 968.9048427 | 943.2884457 | 588.5024373 | 607.2449096 | 630.3497427 | 687.5186098 | 2.67E-21 | -0.617242989 | 5.70E-23 | 97.38659348 | 0.651915558 | 20.57424708 |
| *Errfi1* | 517.7472983 | 509.2813486 | 552.5322932 | 461.4016906 | 327.5327748 | 318.4333062 | 338.2125369 | 345.177844 | 1.38E-12 | -0.617494836 | 5.16E-14 | 56.66664094 | 0.651801765 | 11.85861969 |
| *Nr1h3* | 406.0176309 | 388.8698669 | 333.4927055 | 379.4614326 | 223.9901557 | 249.6686388 | 251.9438318 | 255.3370353 | 1.25E-09 | -0.619115479 | 6.37E-11 | 42.70426799 | 0.651069978 | 8.903766369 |
| *Alox5* | 106.7417359 | 107.58075 | 108.533129 | 121.9349077 | 72.90245633 | 82.51760095 | 51.95728828 | 81.32957419 | 0.001736216 | -0.624894026 | 0.000245773 | 13.44413461 | 0.648467409 | 2.760396183 |
| *9430020K01Rik* | 111.7296675 | 147.0599243 | 133.1997492 | 126.812304 | 77.12868568 | 86.7492728 | 80.3867479 | 91.73219415 | 0.000379175 | -0.625694511 | 4.52E-05 | 16.64083758 | 0.648107704 | 3.421160589 |
| *Chac1* | 186.5486412 | 167.7864908 | 227.9195709 | 254.6000872 | 127.8434379 | 163.9772839 | 145.0882767 | 103.0805068 | 0.000203114 | -0.634808143 | 2.28E-05 | 17.93977067 | 0.644026455 | 3.69226033 |
| *Myo10* | 1626.065696 | 1526.857066 | 1490.850527 | 1553.938463 | 1034.369634 | 992.3270474 | 944.0541248 | 1015.673985 | 1.76E-32 | -0.636831931 | 2.32E-34 | 149.4195399 | 0.643123659 | 31.75387013 |
| *Kdr* | 567.6266142 | 646.4714793 | 551.5456284 | 584.3120775 | 395.1524445 | 404.1246611 | 356.8387346 | 350.8520003 | 3.12E-14 | -0.642707153 | 9.90E-16 | 64.45018996 | 0.640509932 | 13.50645343 |
| *Fam83f* | 213.4834718 | 232.9271284 | 250.6128615 | 280.9380273 | 185.9540915 | 149.1664325 | 119.5997957 | 172.1160756 | 1.10E-05 | -0.642880377 | 9.52E-07 | 24.02381068 | 0.640433031 | 4.960035262 |
| *Pira11* | 775.124568 | 811.297032 | 855.4383896 | 788.1872431 | 540.9573571 | 497.2214416 | 505.8483161 | 522.9680759 | 6.77E-21 | -0.644070695 | 1.48E-22 | 95.50479644 | 0.63990485 | 20.16949454 |
| *Emb* | 2115.880578 | 2148.654062 | 2138.102642 | 2222.141757 | 1433.748308 | 1444.058017 | 1329.322319 | 1312.6215 | 1.32E-40 | -0.644884366 | 1.35E-42 | 187.117179 | 0.639544049 | 39.87781892 |
| *Vdr* | 81.80207796 | 70.0755344 | 88.79983283 | 92.67052982 | 53.88442424 | 63.47507766 | 48.0359835 | 48.23032888 | 0.004482461 | -0.645415143 | 0.000729362 | 11.41289298 | 0.6393088 | 2.348483503 |
| *Fkbp9* | 621.4962753 | 676.08086 | 680.7987184 | 650.6446673 | 396.2090018 | 410.4721689 | 423.5009158 | 444.4755799 | 8.06E-18 | -0.649488436 | 1.99E-19 | 81.25035781 | 0.637506326 | 17.09363422 |
| *Acsbg1* | 101.7538043 | 72.04949311 | 106.5597994 | 67.30806903 | 57.05409626 | 50.78006213 | 56.85891925 | 56.74156339 | 0.006035963 | -0.649533722 | 0.001023102 | 10.78528452 | 0.637486316 | 2.219253423 |

| | | | | | | | | | | | | | | |
|---|---|---|---|---|---|---|---|---|---|---|---|---|---|---|
| *Afap1* | 126.6934622 | 135.216172 | 117.4131123 | 128.7632625 | 79.24180036 | 97.32845241 | 72.54413835 | 74.70972513 | 0.000222921 | -0.652815367 | 2.52E-05 | 17.74921733 | 0.636037897 | 3.651848523 |
| *Phlda3* | 278.3265823 | 219.1094174 | 219.0395877 | 282.8889858 | 157.4270434 | 125.8922374 | 165.6751268 | 184.410081 | 5.08E-06 | -0.655010568 | 4.19E-07 | 25.60405993 | 0.635070839 | 5.294366538 |
| *Mmp12* | 2533.869244 | 2639.182802 | 2619.595069 | 2313.836808 | 1786.638459 | 1460.984704 | 1616.557894 | 1526.348055 | 2.44E-28 | -0.661495404 | 3.86E-30 | 130.1176917 | 0.632222636 | 27.61208919 |
| *Ceacam1* | 341.1745203 | 312.8724564 | 357.1726609 | 353.1234926 | 240.8950731 | 238.0315412 | 198.0258912 | 185.3557737 | 2.62E-09 | -0.665257082 | 1.38E-10 | 41.19792246 | 0.630576328 | 8.581268051 |
| *Tnfaip3* | 541.6893699 | 507.3073899 | 470.639114 | 579.4346812 | 348.6639216 | 335.3599936 | 295.0781844 | 344.2321512 | 4.27E-13 | -0.666172775 | 1.50E-14 | 59.09856887 | 0.630176222 | 12.36967554 |
| *Trim2* | 144.6500159 | 161.8646147 | 140.1064029 | 155.1012025 | 87.69425906 | 104.7338781 | 93.13098842 | 93.62357959 | 2.33E-05 | -0.666820948 | 2.13E-06 | 22.47695524 | 0.629893161 | 4.63178155 |
| *Tuba1a* | 1155.204955 | 1054.093954 | 1068.557988 | 1071.076229 | 694.1581711 | 595.607812 | 691.1299667 | 748.042944 | 2.38E-22 | -0.670857261 | 4.80E-24 | 102.2872512 | 0.628133334 | 21.6238546 |
| *Myo1d* | 321.222794 | 345.4427752 | 310.7994149 | 309.2269258 | 208.1417956 | 193.5989869 | 220.5733936 | 185.3557737 | 3.75E-10 | -0.672130145 | 1.81E-11 | 45.16132911 | 0.62757938 | 9.425930829 |
| *Vim* | 29237.25977 | 27171.54172 | 28111.06708 | 29430.20931 | 18466.50916 | 16284.53117 | 18589.92562 | 17988.96698 | 4.69E-59 | -0.675768704 | 3.04E-61 | 272.6274439 | 0.625998583 | 58.3284279 |
| *Syn1* | 119.710358 | 152.9818004 | 165.759688 | 171.68435 | 87.69425906 | 128.0080733 | 99.01294559 | 67.14418335 | 0.000551884 | -0.680302146 | 6.86E-05 | 15.85014109 | 0.624034568 | 3.258152026 |
| *Src* | 178.5679507 | 188.5130573 | 227.9195709 | 214.6054375 | 121.5040939 | 118.4868116 | 118.6194695 | 145.6366794 | 1.91E-06 | -0.681247374 | 1.48E-07 | 27.61631785 | 0.623625846 | 5.718502798 |
| *Ier3* | 300.2734813 | 294.1198486 | 363.0926498 | 358.0008889 | 206.0286809 | 234.8577873 | 181.3603459 | 196.7040864 | 5.40E-09 | -0.685842904 | 2.92E-10 | 39.72650807 | 0.621642522 | 8.26787072 |
| *Nqo1* | 63.84552426 | 102.6458532 | 73.01319588 | 90.7195713 | 45.43196554 | 56.06965193 | 58.81957164 | 44.44755799 | 0.004382163 | -0.690445466 | 0.000709193 | 11.4649977 | 0.619662485 | 2.358311459 |
| *Zc3h12c* | 366.1141782 | 406.6354954 | 420.3192087 | 443.8430639 | 238.7819584 | 260.2478184 | 276.4519867 | 232.6404099 | 2.59E-12 | -0.699745992 | 1.00E-13 | 55.36689534 | 0.615680597 | 11.58614079 |
| *Clec4d* | 4822.332255 | 4774.019153 | 4655.08457 | 4911.538081 | 2985.831038 | 2761.165878 | 3099.791425 | 2948.669911 | 3.86E-66 | -0.699798399 | 2.22E-68 | 305.3775019 | 0.615658232 | 65.41332591 |
| *Adra1a* | 226.4520939 | 222.0703555 | 235.8128894 | 190.218456 | 128.8999953 | 142.8189247 | 117.6391433 | 147.5280648 | 1.97E-07 | -0.703427301 | 1.31E-08 | 32.31680034 | 0.614111574 | 6.704631593 |
| *Lphn1* | 95.7682864 | 98.69793577 | 108.533129 | 112.180115 | 54.94098158 | 77.22801115 | 56.85891925 | 65.2527979 | 0.000365443 | -0.708484974 | 4.34E-05 | 16.71845442 | 0.611962445 | 3.437179966 |
| *Slco3a1* | 222.4617486 | 243.7839013 | 190.4263082 | 222.4092716 | 140.522126 | 116.3709757 | 159.7931696 | 121.0486686 | 5.34E-07 | -0.708587519 | 3.75E-08 | 30.27265584 | 0.611918949 | 6.272232905 |
| *Fam198b* | 307.2565855 | 247.7318188 | 299.946102 | 327.7610318 | 159.5401581 | 207.3519203 | 198.0258912 | 158.8763775 | 2.76E-08 | -0.709432149 | 1.62E-09 | 36.38252673 | 0.611560804 | 7.558591564 |
| *Slc5a3* | 444.9234972 | 475.7240504 | 427.2258624 | 430.1863542 | 230.3294997 | 283.5220135 | 266.6487248 | 303.5673641 | 2.81E-13 | -0.711994906 | 9.77E-15 | 59.94113157 | 0.610475412 | 12.55065072 |
| *Cd109* | 405.0200445 | 406.6354954 | 471.6257788 | 414.578686 | 236.6688437 | 272.9428339 | 262.72742 | 260.0654989 | 6.99E-14 | -0.717396717 | 2.32E-15 | 62.77213041 | 0.608193913 | 13.15571644 |
| *Epn2* | 121.7055306 | 127.3203371 | 102.6131402 | 118.0329906 | 69.73278432 | 74.05425727 | 90.19000984 | 51.06740705 | 0.000285225 | -0.722261628 | 3.30E-05 | 17.23554121 | 0.606146477 | 3.54481203 |
| *Ryk* | 140.6596706 | 171.7344082 | 159.8396991 | 179.4881841 | 94.03360309 | 87.80719076 | 106.8555551 | 104.9718923 | 3.29E-06 | -0.725196532 | 2.64E-07 | 26.49942931 | 0.604914634 | 5.482984248 |
| *Tnfsf9* | 88.78518218 | 78.95834861 | 70.05320146 | 85.84217499 | 42.26229352 | 51.83798009 | 61.76055022 | 39.71909437 | 0.001931698 | -0.726780744 | 0.000278843 | 13.20741942 | 0.604250747 | 2.714060873 |
| *Plaur* | 606.5324805 | 473.7500917 | 522.9323489 | 477.0093588 | 302.1753987 | 272.9428339 | 349.9764512 | 330.0467604 | 8.72E-13 | -0.726878397 | 3.20E-14 | 57.61040151 | 0.604209848 | 12.05952142 |
| *Ehd1* | 2051.037467 | 1834.794626 | 1950.636328 | 1988.026735 | 1187.570448 | 1153.130577 | 1176.391433 | 1204.812529 | 6.74E-48 | -0.728345879 | 5.44E-50 | 221.0128987 | 0.603595569 | 47.17109682 |
| *Slco4a1* | 863.9097502 | 843.8673508 | 824.8517805 | 844.7650403 | 464.8852288 | 457.0205591 | 546.04169 | 566.4699412 | 4.18E-22 | -0.728886883 | 8.50E-24 | 101.1563318 | 0.603369266 | 21.37932931 |
| *Tnfrsf12a* | 595.559031 | 601.0704288 | 598.9055392 | 575.5327642 | 292.6663827 | 404.1246611 | 408.7960229 | 324.372604 | 3.13E-14 | -0.72948627 | 9.99E-16 | 64.43145274 | 0.60311864 | 13.5045003 |
| *Maged1* | 1993.177461 | 1987.776426 | 1865.783154 | 1821.219781 | 1116.781106 | 1178.520609 | 1208.742197 | 1114.026028 | 5.84E-45 | -0.731514346 | 5.22E-47 | 207.3415628 | 0.602271399 | 44.23356163 |
| *Oaf* | 72.82380111 | 68.10157568 | 67.09320703 | 52.67588011 | 47.54508022 | 40.20088252 | 31.37043821 | 37.82770893 | 0.004974751 | -0.73505582 | 0.000822174 | 11.1904941 | 0.600794778 | 2.303228614 |
| *Stac2* | 133.6765664 | 123.3724197 | 99.65314573 | 102.4253224 | 97.20327511 | 59.24340581 | 41.17370015 | 78.49249602 | 0.00183712 | -0.735253445 | 0.00026299 | 13.31714376 | 0.600712485 | 2.735862532 |
| *Cldn12* | 73.82138743 | 80.93230733 | 73.99986069 | 64.38163125 | 55.99753892 | 48.6642262 | 23.52782865 | 48.23032888 | 0.006300111 | -0.736218351 | 0.001075923 | 10.69212897 | 0.60031085 | 2.200651791 |
| *Ecm1* | 1880.450207 | 1911.779016 | 1920.049719 | 1929.497979 | 1086.140944 | 1161.593921 | 1219.525785 | 1111.18895 | 7.83E-50 | -0.738859454 | 5.88E-52 | 230.0282663 | 0.599212883 | 49.10617194 |
| *Emp2* | 1269.927381 | 1238.659094 | 1324.104174 | 1274.951395 | 819.8884944 | 704.573362 | 779.3593242 | 742.3687877 | 4.65E-36 | -0.745783867 | 5.46E-38 | 166.026846 | 0.596343769 | 35.33253778 |
| *Tnk2* | 235.4303707 | 173.708367 | 232.852895 | 251.6736494 | 142.6352406 | 129.0659912 | 138.2259933 | 122.940054 | 1.55E-07 | -0.746921766 | 1.01E-08 | 32.82840177 | 0.595873599 | 6.811041273 |
| *Tpm1* | 2576.765456 | 2263.143667 | 2434.102084 | 2481.619241 | 1488.689289 | 1411.26256 | 1506.76136 | 1405.299387 | 3.87E-50 | -0.747252311 | 2.87E-52 | 231.4523945 | 0.59573709 | 49.41210141 |
| *P4ha2* | 443.9259109 | 420.4532064 | 407.4925662 | 390.1917045 | 255.6868758 | 242.2632131 | 238.2192651 | 253.4456498 | 1.13E-15 | -0.747859949 | 3.27E-17 | 71.17531796 | 0.595486229 | 14.945379 |
| *Myo6* | 84.79483691 | 96.72397705 | 87.81316802 | 83.89121647 | 53.88442424 | 62.4171597 | 49.99663589 | 44.44755799 | 0.000469141 | -0.749011874 | 5.71E-05 | 16.19703158 | 0.595010951 | 3.328696675 |
| *Rap1gap2* | 104.7465632 | 119.4245023 | 90.77316245 | 126.812304 | 85.58114439 | 65.59091358 | 42.15402634 | 69.98126152 | 0.000676081 | -0.749627837 | 8.54E-05 | 15.43495666 | 0.594756963 | 3.170001544 |
| *Fkbp7* | 65.8406969 | 56.25782339 | 56.23989413 | 46.82300454 | 33.80983482 | 31.73753883 | 28.42945962 | 39.71909437 | 0.007575539 | -0.749698076 | 0.001341521 | 10.28450435 | 0.594728008 | 2.120586489 |
| *Prkca* | 115.7200127 | 137.1901307 | 152.9330454 | 121.9349077 | 89.80737374 | 72.99633931 | 84.30805268 | 67.14418335 | 2.81E-05 | -0.749843925 | 2.60E-06 | 22.09176802 | 0.594667887 | 4.55183724 |
| *Cxcl10* | 103.7489769 | 82.90626605 | 86.82650321 | 89.74409204 | 44.3754082 | 48.6642262 | 67.64250738 | 53.90448522 | 0.000514272 | -0.755541136 | 6.33E-05 | 16.00085229 | 0.592324171 | 3.288807061 |
| *Procr* | 88.78518218 | 112.5156468 | 116.4264475 | 104.376281 | 57.05409626 | 67.7067495 | 67.64250738 | 57.68725611 | 0.000114902 | -0.755819006 | 1.20E-05 | 19.16338611 | 0.592210097 | 3.939673963 |
| *Pdia5* | 89.7827685 | 75.99741054 | 109.5197938 | 101.4498432 | 59.16721093 | 64.53299562 | 45.09500492 | 53.90448522 | 0.000451269 | -0.761766041 | 5.48E-05 | 16.27559537 | 0.589773931 | 3.345564607 |
| *Cd274* | 201.512436 | 236.8750458 | 235.8128894 | 275.0851517 | 145.8049127 | 159.7456121 | 124.5014266 | 130.5055958 | 2.32E-08 | -0.762864723 | 1.35E-09 | 36.74156932 | 0.58932496 | 7.634429855 |
| *S100a11* | 1510.345683 | 1332.422133 | 1472.103895 | 1302.264814 | 770.2302995 | 842.1026969 | 872.4903126 | 819.915591 | 1.23E-33 | -0.764795364 | 1.55E-35 | 154.7969782 | 0.588536843 | 32.91031891 |
| *Zeb1* | 123.7007033 | 103.6328326 | 150.9597158 | 116.0820321 | 60.22376827 | 77.22801115 | 72.54413835 | 80.38388147 | 5.17E-05 | -0.765636022 | 4.99E-06 | 20.83931568 | 0.588194002 | 4.2868564 |
| *Efemp2* | 291.2952045 | 320.7682912 | 358.1593258 | 294.5947369 | 187.0106488 | 197.8306587 | 198.0258912 | 161.7134557 | 2.94E-11 | -0.765994082 | 1.25E-12 | 50.40110586 | 0.588048037 | 10.53091667 |
| *Ccl3* | 192.5341591 | 164.8255527 | 161.8130287 | 241.9188568 | 105.6557338 | 124.8343194 | 116.6588171 | 100.2434287 | 1.93E-06 | -0.768047022 | 1.50E-07 | 27.59511336 | 0.587211846 | 5.714637297 |
| *Fabp5* | 3427.706584 | 3033.974546 | 3109.967479 | 3020.083793 | 1855.314686 | 1792.113026 | 1742.039647 | 2000.14011 | 2.97E-48 | -0.768478981 | 2.35E-50 | 222.6848736 | 0.587036055 | 47.5270975 |
| *Pbx1* | 151.6331201 | 173.708367 | 154.9063751 | 155.1012025 | 84.52458705 | 97.32845241 | 96.07196701 | 94.56927232 | 4.19E-07 | -0.769606749 | 2.91E-08 | 30.76876855 | 0.586577343 | 6.377328762 |
| *Smo* | 103.7489769 | 118.4375229 | 115.4397827 | 106.3272395 | 71.84589899 | 62.4171597 | 66.66218119 | 59.57864156 | 3.17E-05 | -0.770574921 | 2.96E-06 | 21.84040413 | 0.586183831 | 4.499426055 |
| *Vwf* | 590.5710994 | 636.6016857 | 607.7855225 | 602.8461835 | 353.9467083 | 369.2133684 | 369.5829751 | 331.9381458 | 2.11E-23 | -0.775775772 | 4.07E-25 | 107.1793769 | 0.584074469 | 22.67502244 |
| *Fkbp14* | 70.82862848 | 63.16667889 | 73.99986069 | 109.2536773 | 48.60163755 | 42.31671844 | 51.95728828 | 41.61047982 | 0.001746277 | -0.782935058 | 0.000247616 | 13.43012408 | 0.581183215 | 2.757886986 |
| *Igsf6* | 1619.082592 | 1719.318041 | 1648.716896 | 1773.421297 | 912.8655401 | 981.7478678 | 1041.106418 | 988.2488957 | 2.67E-46 | -0.784236968 | 2.34E-48 | 213.5208215 | 0.580658983 | 45.57386297 |
| *Dcn* | 188.5438138 | 203.3177477 | 215.0929284 | 216.556396 | 102.4860618 | 131.1818272 | 107.8358813 | 136.1797521 | 2.01E-08 | -0.784530295 | 1.16E-09 | 37.03709304 | 0.580540936 | 7.696663843 |
| *Ifitm6* | 82.79966428 | 79.94532797 | 73.99986069 | 72.18546534 | 49.65819489 | 33.85337475 | 55.87859305 | 39.71909437 | 0.000945268 | -0.785102149 | 0.000124263 | 14.7268621 | 0.580310867 | 3.024444971 |
| *Apobec1* | 11542.07368 | 11704.5882 | 12322.4568 | 11824.75961 | 6870.79237 | 7003.416902 | 6890.712817 | 6694.558787 | 2.14E-130 | -0.787534516 | 3.42E-133 | 603.1738225 | 0.579333294 | 129.6697819 |
| *Spred3* | 64.84311058 | 64.15365825 | 57.22655894 | 52.67588011 | 38.03606417 | 30.67962087 | 39.21304776 | 30.26216714 | 0.003052965 | -0.79008136 | 0.000471168 | 12.22644806 | 0.578311477 | 2.515278231 |
| *Cc2d2a* | 102.7513906 | 94.75001834 | 87.81316802 | 90.7195713 | 52.82786691 | 67.7067495 | 51.95728828 | 45.39325071 | 0.000157561 | -0.791167064 | 1.70E-05 | 18.49523918 | 0.577876431 | 3.802551723 |
| *Clec4n* | 337.184175 | 392.8177844 | 319.6793982 | 366.7802022 | 201.8024516 | 180.9039713 | 182.3406721 | 251.5542644 | 2.50E-11 | -0.792230537 | 1.05E-12 | 50.74236187 | 0.577450611 | 10.60151525 |
| *Pilrb1* | 100.756218 | 58.2317821 | 65.11987741 | 82.91573721 | 43.31885086 | 33.85337475 | 47.05565731 | 52.01309977 | 0.0020788 | -0.796507409 | 0.000303729 | 13.04725727 | 0.575741292 | 2.682187279 |
| *Armcx2* | 85.79242323 | 86.85418348 | 70.05320146 | 62.43067272 | 53.88442424 | 31.73753883 | 41.17370015 | 48.23032888 | 0.001268004 | -0.800203046 | 0.000171294 | 14.12237163 | 0.574268349 | 2.896879466 |
| *Bhlhe40* | 722.2524932 | 688.9115917 | 730.1319588 | 787.2117639 | 415.2270339 | 481.3526722 | 422.5205896 | 363.1460057 | 5.11E-22 | -0.801810078 | 1.05E-23 | 100.7311157 | 0.573629022 | 21.2911769 |
| *Tiam2* | 77.8117327 | 63.16667889 | 67.09320703 | 55.60231789 | 57.05409626 | 38.08504659 | 30.39011201 | 26.47939625 | 0.00485707 | -0.801886267 | 0.000800011 | 11.24120775 | 0.573598729 | 2.313625635 |
| *Car5b* | 260.3700286 | 192.4609747 | 215.0929284 | 207.7770827 | 135.2393393 | 151.2822684 | 114.6981647 | 102.1348141 | 8.40E-08 | -0.802642999 | 5.25E-09 | 34.09509991 | 0.57329794 | 7.075963201 |
| *Cd207* | 571.6169594 | 504.3464518 | 521.9456841 | 497.4944233 | 303.231956 | 308.9120446 | 315.6650345 | 264.7939625 | 4.44E-20 | -0.81421418 | 9.93E-22 | 91.73139838 | 0.568718177 | 19.35253114 |
| *Aldh1l2* | 222.4617486 | 237.8620252 | 227.9195709 | 198.9977693 | 108.8254058 | 134.355581 | 136.265341 | 122.940054 | 5.64E-10 | -0.819684373 | 2.79E-11 | 44.32227033 | 0.56656588 | 9.248339517 |
| *Dusp9* | 88.78518218 | 87.84116283 | 82.87984398 | 78.0383409 | 52.82786691 | 55.01173397 | 46.07533112 | 37.82770893 | 0.00019364 | -0.820661268 | 2.16E-05 | 18.04701717 | 0.56618237 | 3.713005451 |
| *Hoxa1* | 56.86242005 | 51.3229266 | 57.22655894 | 42.9210875 | 28.52704813 | 31.73753883 | 25.48848104 | 32.15355259 | 0.004471043 | -0.820860445 | 0.000727147 | 11.41854339 | 0.566104209 | 2.349591186 |
| *C3* | 4037.231823 | 4225.25863 | 3967.379198 | 4310.642856 | 2350.840077 | 2359.157053 | 2200.832305 | 2451.235538 | 1.52E-78 | -0.821083435 | 6.45E-81 | 362.9398678 | 0.566016716 | 77.81705481 |
| *Jrk* | 72.82380111 | 45.40105045 | 63.14654779 | 55.60231789 | 26.41393345 | 33.85337475 | 43.13435253 | 30.26216714 | 0.003914402 | -0.823776394 | 0.000624742 | 11.70077221 | 0.564961165 | 2.407334594 |
| *Tnfsf8* | 43.89379793 | 63.16667889 | 47.35991084 | 44.87204602 | 20.07458942 | 28.56378495 | 29.40978582 | 34.04493803 | 0.007434921 | -0.825260343 | 0.001309496 | 10.32909223 | 0.564380347 | 2.128723613 |
| *Bst1* | 105.7441496 | 108.5677293 | 80.90651436 | 117.0575114 | 65.50655496 | 70.88050338 | 43.13435253 | 53.90448522 | 0.000122172 | -0.825398719 | 1.28E-05 | 19.03320926 | 0.564326218 | 3.913029475 |
| *Ak1* | 228.4472665 | 223.0573348 | 242.7195431 | 205.8261241 | 113.0516352 | 151.2822684 | 110.7768599 | 131.4512885 | 8.91E-10 | -0.830045749 | 4.47E-11 | 43.39776149 | 0.562511404 | 9.050167067 |
| *Plxnd1* | 7212.549069 | 6981.891976 | 7541.079137 | 7666.291514 | 4196.645747 | 4287.741496 | 4030.120983 | 4022.976844 | 1.30E-99 | -0.830482291 | 3.94E-102 | 460.3976992 | 0.562341221 | 98.88666074 |
| *Hspa1a* | 105.7441496 | 70.0755344 | 79.91984955 | 107.3027187 | 48.60163755 | 33.85337475 | 56.85891925 | 63.36141245 | 0.000575407 | -0.835353347 | 7.17E-05 | 15.76570209 | 0.560445757 | 3.240025039 |
| *Anpep* | 7009.041461 | 6933.529988 | 7365.452801 | 7594.106049 | 4104.725259 | 4154.443833 | 4037.963593 | 3896.254019 | 1.34E-102 | -0.836073608 | 3.65E-105 | 474.3353124 | 0.560166026 | 101.871319 |
| *Ankrd13b* | 172.5824328 | 157.9166972 | 147.0130566 | 163.8805159 | 77.12868568 | 80.40176503 | 99.01294559 | 100.2434287 | 8.14E-08 | -0.84211025 | 5.06E-09 | 34.16586195 | 0.557827031 | 7.089531707 |
| *Cd24a* | 67.83586953 | 62.17969953 | 41.43992199 | 44.87204602 | 36.97950683 | 15.86876941 | 34.31141679 | 33.09924531 | 0.009087673 | -0.843052336 | 0.001653558 | 9.899143097 | 0.557462886 | 2.041547289 |
| *Csrp2* | 283.3145139 | 265.4974472 | 259.4928448 | 251.6736494 | 150.031142 | 151.2822684 | 157.8325172 | 130.5055958 | 8.72E-13 | -0.847282277 | 3.19E-14 | 57.61535634 | 0.555830814 | 12.05966822 |
| *Arhgap29* | 132.6789801 | 169.7604495 | 138.1330733 | 136.5670966 | 100.3729471 | 63.47507766 | 76.46544313 | 80.38388147 | 1.42E-06 | -0.848213762 | 1.07E-07 | 28.23925883 | 0.555472055 | 5.847873998 |
| *Gpr97* | 48.88172951 | 40.46615367 | 47.35991084 | 54.62683863 | 32.75327748 | 27.50586698 | 23.52782865 | 22.69662536 | 0.005345281 | -0.850434076 | 0.000894508 | 11.03410663 | 0.554617838 | 2.272029449 |
| *Creb3l1* | 226.4520939 | 225.0312936 | 240.7462135 | 236.0659812 | 142.6352406 | 114.2551398 | 144.1079505 | 113.4831268 | 4.98E-11 | -0.852208199 | 2.16E-12 | 49.33122031 | 0.553936228 | 10.3023844 |
| *Layn* | 46.88655688 | 49.34896788 | 75.97319031 | 55.60231789 | 31.69672014 | 29.62170291 | 23.52782865 | 40.6647871 | 0.003869527 | -0.857825857 | 0.000616344 | 11.72595567 | 0.55178347 | 2.412342065 |
| *Cdc42bpa* | 348.1576245 | 360.2474656 | 408.479231 | 350.1970548 | 198.6327796 | 225.3365257 | 190.1832816 | 194.812701 | 1.71E-16 | -0.860271172 | 4.53E-18 | 75.07403252 | 0.55084901 | 15.76819416 |
| *Whrn* | 151.6331201 | 168.7734702 | 129.25309 | 131.6897003 | 98.25983244 | 59.24340581 | 90.19000984 | 71.87264696 | 2.10E-06 | -0.863297871 | 1.63E-07 | 27.42698952 | 0.549694568 | 5.678224862 |
| *Col27a1* | 65.8406969 | 46.38802981 | 66.10654222 | 33.16629488 | 30.64016281 | 29.62170291 | 33.33109059 | 22.69662536 | 0.006999142 | -0.864355007 | 0.001221567 | 10.45743652 | 0.549291927 | 2.1549552 |
| *Cacnb3* | 45.88897056 | 38.49219495 | 54.26656451 | 42.9210875 | 28.52704813 | 17.98460534 | 24.50815485 | 28.3707817 | 0.006546314 | -0.866425498 | 0.001125286 | 10.60916248 | 0.548504173 | 2.184003194 |
| *Nptx1* | 824.0062975 | 658.3152316 | 776.5052049 | 705.2715059 | 381.4171991 | 418.9355125 | 459.772985 | 362.200313 | 1.27E-22 | -0.869614781 | 2.50E-24 | 103.5767834 | 0.547292966 | 21.89787412 |
| *Il2rg* | 2717.425126 | 2720.11511 | 2610.715085 | 2753.777955 | 1497.141748 | 1437.710509 | 1484.213858 | 1483.791883 | 1.27E-92 | -0.871688757 | 4.25E-95 | 428.0833767 | 0.546506759 | 91.89742847 |
| *Crip2* | 44.89138425 | 42.44011238 | 64.1332126 | 38.04369119 | 21.13114676 | 32.79545679 | 27.44913343 | 21.75093263 | 0.006699213 | -0.87991204 | 0.001156445 | 10.5586672 | 0.543400561 | 2.173976205 |
| *Eno2* | 71.8262148 | 67.11459632 | 76.95985512 | 81.94025795 | 36.97950683 | 50.78006213 | 47.05565731 | 27.42508897 | 0.000387043 | -0.880874609 | 4.62E-05 | 16.59807469 | 0.543038124 | 3.412241265 |
| *Pls3* | 1053.45115 | 1030.406449 | 1120.851223 | 1143.261694 | 611.7466988 | 599.8394839 | 589.1760426 | 555.1216285 | 3.64E-44 | -0.885124368 | 3.35E-46 | 203.6446939 | 0.541440846 | 43.43840705 |
| *Pik3r3* | 89.7827685 | 56.25782339 | 77.94651993 | 82.91573721 | 35.9229495 | 45.49047232 | 41.17370015 | 42.55617254 | 0.000216578 | -0.893404902 | 2.45E-05 | 17.80548273 | 0.538342077 | 3.664385251 |
| *Ttc30b* | 193.5317454 | 197.3958715 | 204.2396155 | 170.7088707 | 92.97704575 | 106.8497141 | 110.7768599 | 99.29773593 | 1.60E-10 | -0.901181281 | 7.41E-12 | 46.91545167 | 0.535448126 | 9.794988512 |
| *Vasn* | 592.5662721 | 592.1876146 | 523.9190137 | 530.6607181 | 313.7975294 | 337.4758295 | 286.2552486 | 254.3913425 | 1.10E-22 | -0.912440095 | 2.16E-24 | 103.8719128 | 0.531285744 | 21.95905055 |
| *Ddit4* | 197.5220907 | 193.4479541 | 150.9597158 | 113.1555943 | 68.67622698 | 88.86510872 | 87.24903126 | 101.1891214 | 1.61E-06 | -0.917946735 | 1.23E-07 | 27.97578098 | 0.529261738 | 5.792601122 |
| *Adamtsl4* | 73.82138743 | 72.04949311 | 58.21322375 | 67.30806903 | 57.05409626 | 37.02712863 | 27.44913343 | 22.69662536 | 0.001356326 | -0.921132922 | 0.000185283 | 13.97473961 | 0.528094155 | 2.867635984 |
| *Rhov* | 111.7296675 | 113.5026261 | 83.86650879 | 132.6651795 | 62.33688295 | 56.06965193 | 63.72120261 | 51.06740705 | 4.77E-06 | -0.923147535 | 3.92E-07 | 25.73504579 | 0.527357226 | 5.321317284 |
| *Slc24a5* | 146.6451885 | 154.9557592 | 173.6530064 | 137.5425758 | 82.41147237 | 50.78006213 | 88.22935746 | 100.2434287 | 4.83E-07 | -0.924539606 | 3.36E-08 | 30.48736583 | 0.52684862 | 6.316337568 |
| *Socs3* | 182.5582959 | 166.7995114 | 190.4263082 | 185.3410596 | 89.80737374 | 101.5601243 | 92.15066223 | 97.40635049 | 6.91E-11 | -0.928746783 | 3.09E-12 | 48.63149416 | 0.525314465 | 10.16063583 |
| *Hmga2* | 915.7842387 | 902.0991329 | 1040.931374 | 969.6263857 | 551.5229305 | 471.8314106 | 506.8286423 | 476.6291325 | 1.94E-38 | -0.932560039 | 2.13E-40 | 177.0589384 | 0.523927815 | 37.71134077 |
| *Pilra* | 331.1986571 | 359.2604862 | 322.6393926 | 367.7556815 | 183.8409768 | 173.4985456 | 178.4193673 | 187.2471592 | 2.87E-19 | -0.93306508 | 6.60E-21 | 87.98214789 | 0.523744437 | 18.54175583 |
| *Tjp1* | 118.7127717 | 151.0078417 | 98.66648093 | 110.2291565 | 65.50655496 | 75.11217523 | 53.91794067 | 56.74156339 | 2.49E-06 | -0.933074638 | 1.96E-07 | 27.0769283 | 0.523740967 | 5.604472745 |
| *Nckap1* | 305.2614129 | 234.9010871 | 306.8527557 | 262.4039213 | 155.3139287 | 137.5293349 | 134.3046886 | 149.4194503 | 5.70E-14 | -0.944227409 | 1.87E-15 | 63.19275693 | 0.519707791 | 13.24426566 |
| *Gpc1* | 743.2018059 | 648.445438 | 705.4653386 | 731.609446 | 404.6614605 | 366.0396145 | 348.0157989 | 352.7433857 | 2.64E-33 | -0.944240695 | 3.39E-35 | 153.2417643 | 0.519703005 | 32.5786122 |
| *Gpr124* | 61.85035163 | 86.85418348 | 75.97319031 | 67.30806903 | 43.31885086 | 40.20088252 | 32.3507644 | 35.93632348 | 9.53E-05 | -0.946961477 | 9.81E-06 | 19.54776198 | 0.518723819 | 4.020737064 |
| *Map1a* | 58.85759268 | 62.17969953 | 58.21322375 | 63.40615198 | 31.69672014 | 34.91129271 | 25.48848104 | 33.09924531 | 0.000232695 | -0.956373378 | 2.65E-05 | 17.6568804 | 0.515350766 | 3.633213806 |
| *Fzd1* | 92.77552745 | 69.08855504 | 66.10654222 | 52.67588011 | 41.20573619 | 39.14296456 | 33.33109059 | 31.20785986 | 0.00025419 | -0.956828975 | 2.91E-05 | 17.47555807 | 0.515188046 | 3.59484075 |
| *Tnf* | 336.1865887 | 343.4688165 | 270.3461577 | 359.9518474 | 154.2573714 | 171.3827097 | 193.1242602 | 155.0936066 | 1.67E-15 | -0.958746557 | 4.88E-17 | 70.38420036 | 0.514503731 | 14.77720095 |
| *Gpr141* | 127.6910485 | 105.6067913 | 91.75982726 | 101.4498432 | 53.88442424 | 64.53299562 | 54.89826686 | 46.33894344 | 1.62E-06 | -0.960196778 | 1.24E-07 | 27.957522 | 0.513986803 | 5.789684624 |
| *Ppap2a* | 159.6138107 | 140.1510688 | 144.0530622 | 127.7877832 | 66.5631123 | 83.57551892 | 80.3867479 | 63.36141245 | 1.24E-08 | -0.960994578 | 6.96E-10 | 38.03228197 | 0.51370265 | 7.906669815 |
| *Palld* | 498.7931583 | 508.2943692 | 616.6655058 | 624.3067272 | 289.4967106 | 302.5645368 | 314.6847083 | 243.0430299 | 1.27E-21 | -0.968779437 | 2.67E-23 | 98.88730859 | 0.510938149 | 20.89520041 |
| *Rnd3* | 112.7272538 | 142.1250275 | 154.9063751 | 142.4199721 | 83.46802971 | 61.35924174 | 74.50479074 | 62.41571973 | 4.29E-08 | -0.972436423 | 2.58E-09 | 35.47623322 | 0.509644648 | 7.36766823 |
| *Akap12* | 140.6596706 | 142.1250275 | 116.4264475 | 161.9295574 | 83.46802971 | 68.76466746 | 59.79989783 | 73.76403241 | 2.84E-08 | -0.975087236 | 1.68E-09 | 36.31983651 | 0.508709085 | 7.546146204 |
| *Tfec* | 147.6427749 | 103.6328326 | 94.71982169 | 117.0575114 | 71.84589899 | 52.89589805 | 57.83924544 | 52.9587925 | 1.46E-06 | -0.976958661 | 1.11E-07 | 28.1806759 | 0.508049629 | 5.835659179 |
| *Pdlim1* | 111.7296675 | 99.68491513 | 106.5597994 | 76.08738238 | 65.50655496 | 51.83798009 | 41.17370015 | 41.61047982 | 8.48E-06 | -0.982470587 | 7.22E-07 | 24.55514544 | 0.50611229 | 5.071362568 |
| *Egr3* | 114.7224264 | 143.1120069 | 159.8396991 | 150.2238062 | 61.28032561 | 79.34384707 | 78.42609552 | 68.08987607 | 1.60E-08 | -0.984007494 | 9.12E-10 | 37.50536415 | 0.505573414 | 7.794888771 |
| *Col18a1* | 2416.154059 | 2106.213949 | 2368.982207 | 2350.90502 | 1259.416347 | 1178.520609 | 1137.178385 | 1077.144012 | 4.07E-68 | -0.991260806 | 2.15E-70 | 314.6262351 | 0.503037966 | 67.39031204 |
| *Serpine1* | 8318.872294 | 7851.42079 | 8084.731447 | 7712.13904 | 4385.769511 | 3722.813305 | 4113.44871 | 3836.675378 | 8.47E-102 | -0.99334498 | 2.37E-104 | 470.6057669 | 0.502311782 | 101.0723407 |
| *Acpp* | 26.93483055 | 29.60938073 | 38.47992756 | 50.72492159 | 13.7352454 | 17.98460534 | 15.6852191 | 25.53370353 | 0.009874826 | -0.993608024 | 0.001822802 | 9.71993236 | 0.502220205 | 2.005470539 |
| *Btg3* | 123.7007033 | 112.5156468 | 140.1064029 | 88.76861278 | 72.90245633 | 48.6642262 | 50.97696209 | 59.57864156 | 9.72E-07 | -1.003255311 | 7.16E-08 | 29.02015025 | 0.498873067 | 6.012333043 |
| *Fam101b* | 63.84552426 | 45.40105045 | 50.31990527 | 44.87204602 | 23.24426144 | 32.79545679 | 18.62619768 | 27.42508897 | 0.000945268 | -1.003606238 | 0.0001243 | 14.72630465 | 0.498751734 | 3.024444971 |
| *Col4a2* | 938.7287239 | 847.8152683 | 886.0249987 | 850.6179158 | 463.8286714 | 466.5418208 | 423.5009158 | 388.6797092 | 1.44E-43 | -1.017700414 | 1.34E-45 | 200.88704 | 0.493902983 | 42.84041826 |
| *Adamts1* | 45.88897056 | 55.27084403 | 31.5732739 | 42.9210875 | 24.30081878 | 17.98460534 | 27.44913343 | 17.02246902 | 0.002427016 | -1.018274783 | 0.000362938 | 12.7139488 | 0.493706388 | 2.614927309 |
| *Ndrg2* | 60.85276531 | 72.04949311 | 43.41325161 | 50.72492159 | 26.41393345 | 27.50586698 | 33.33109059 | 24.5880108 | 0.00038059 | -1.021080577 | 4.54E-05 | 16.63249993 | 0.492747147 | 3.41954254 |
| *Mmp14* | 939.7263103 | 949.4741421 | 1008.371435 | 1029.130621 | 469.1114581 | 469.7155747 | 510.749947 | 477.5748252 | 1.78E-54 | -1.026368294 | 1.27E-56 | 251.4294646 | 0.490944452 | 53.74880529 |
| *Sgce* | 48.88172951 | 34.54427752 | 34.53326832 | 48.77396306 | 25.35737611 | 17.98460534 | 20.58685007 | 17.96816174 | 0.002202983 | -1.02817901 | 0.000325215 | 12.91928032 | 0.490328658 | 2.656988776 |
| *B4galt2* | 33.91793477 | 46.38802981 | 33.54660351 | 44.87204602 | 19.01803209 | 23.27419514 | 22.54750246 | 13.23969812 | 0.003091582 | -1.028556107 | 0.000478115 | 12.19914071 | 0.490200511 | 2.509819212 |
| *Nfkbie* | 331.1986571 | 292.1458899 | 295.9994428 | 295.5702162 | 147.9180273 | 139.6451708 | 135.2850148 | 170.2246902 | 3.15E-19 | -1.032627559 | 7.30E-21 | 87.78407193 | 0.488819059 | 18.50127082 |
| *Pdgfrl* | 37.90828003 | 49.34896788 | 39.46659237 | 39.01917045 | 13.7352454 | 33.85337475 | 15.6852191 | 17.96816174 | 0.003823214 | -1.034142809 | 0.000607746 | 11.75210057 | 0.488305926 | 2.417571342 |
| *Loxl3* | 689.3321448 | 680.0287774 | 648.2387797 | 632.1105613 | 337.0417909 | 330.0704038 | 298.9994892 | 328.1553749 | 1.72E-40 | -1.034320118 | 1.77E-42 | 186.5816796 | 0.488245916 | 39.7643006 |
| *Tmem26* | 210.4907128 | 180.6172225 | 228.9062357 | 165.8314744 | 89.80737374 | 105.7917961 | 97.0522932 | 90.78650143 | 4.47E-12 | -1.035850875 | 1.79E-13 | 54.22526013 | 0.487728142 | 11.349756 |
| *Ak4* | 60.85276531 | 50.33594724 | 47.35991084 | 72.18546534 | 26.41393345 | 22.21627718 | 32.3507644 | 31.20785986 | 0.000249796 | -1.036044101 | 2.85E-05 | 17.51269379 | 0.487662823 | 3.602414241 |
| *Tnfaip2* | 3007.722745 | 2897.771394 | 2879.087913 | 3039.593378 | 1359.789294 | 1452.52136 | 1468.528639 | 1481.900497 | 4.57E-120 | -1.036352392 | 9.48E-123 | 555.1633593 | 0.487558625 | 119.3404365 |
| *Irx3* | 50.87690215 | 40.46615367 | 44.39991642 | 42.9210875 | 16.90491741 | 13.75293349 | 22.54750246 | 33.09924531 | 0.002034494 | -1.038109708 | 0.000295956 | 13.09581717 | 0.486965102 | 2.69154368 |
| *Efnb2* | 58.85759268 | 49.34896788 | 54.26656451 | 40.97012897 | 20.07458942 | 33.85337475 | 19.60552388 | 25.53370353 | 0.000633665 | -1.040027024 | 7.98E-05 | 15.56221558 | 0.486318364 | 3.198140558 |
| *Rgl1* | 120.7079443 | 120.4114816 | 119.3864419 | 119.9839491 | 59.16721093 | 61.35924174 | 49.0163097 | 63.36141245 | 4.84E-09 | -1.045184311 | 2.59E-10 | 39.96094567 | 0.484582997 | 8.315491405 |
| *Uchl1* | 904.8107892 | 855.7111031 | 1029.091396 | 987.1850124 | 475.4508022 | 384.0242198 | 533.2974495 | 435.9643454 | 1.90E-33 | -1.045557895 | 2.42E-35 | 153.9141784 | 0.484457531 | 32.72013506 |
| *Pvrl1* | 177.5703644 | 164.8255527 | 166.7463528 | 140.4690136 | 71.84589899 | 81.45968299 | 95.09164081 | 66.19849062 | 1.24E-10 | -1.046128609 | 5.66E-12 | 47.44558628 | 0.484265923 | 9.907169295 |
| *Ccl9* | 9149.861696 | 9665.48885 | 10149.82089 | 9432.884457 | 4966.876046 | 4739.472465 | 4290.887751 | 4583.772629 | 8.43E-122 | -1.047509604 | 1.61E-124 | 563.293917 | 0.483802589 | 121.0743485 |
| *Vcam1* | 36.91069371 | 40.46615367 | 39.46659237 | 36.09273267 | 20.07458942 | 22.21627718 | 14.70489291 | 17.02246902 | 0.002035146 | -1.051941528 | 0.000296213 | 13.09418854 | 0.48231864 | 2.691404398 |
| *Hcar2* | 79.80690533 | 58.2317821 | 76.95985512 | 48.77396306 | 29.58360547 | 25.39003106 | 44.11467873 | 27.42508897 | 0.000161697 | -1.056856324 | 1.76E-05 | 18.43743498 | 0.480678332 | 3.791299295 |
| *Cdh2* | 217.473817 | 183.5781605 | 200.2929563 | 222.4092716 | 106.7122911 | 89.92302668 | 94.11131462 | 104.0261995 | 2.16E-14 | -1.060733953 | 6.76E-16 | 65.20110187 | 0.479388115 | 13.66646451 |
| *Mxra8* | 103.7489769 | 103.6328326 | 79.91984955 | 85.84217499 | 57.05409626 | 34.91129271 | 49.99663589 | 36.8820162 | 1.76E-06 | -1.062625074 | 1.35E-07 | 27.78633324 | 0.478760133 | 5.75352285 |
| *Fads3* | 513.7569531 | 487.5678027 | 474.5857732 | 473.1074417 | 210.2549103 | 203.1202485 | 269.5897033 | 244.9344153 | 2.31E-28 | -1.067606247 | 3.63E-30 | 130.241713 | 0.477109975 | 27.63702494 |
| *Sfmbt2* | 33.91793477 | 41.45313302 | 39.46659237 | 61.45519346 | 17.96147475 | 27.50586698 | 16.6655453 | 21.75093263 | 0.001620352 | -1.074452366 | 0.000227173 | 13.59182465 | 0.474851276 | 2.790390619 |
| *Ston1* | 48.88172951 | 51.3229266 | 59.19988856 | 34.14177415 | 39.09262151 | 10.57917961 | 15.6852191 | 26.47939625 | 0.003730409 | -1.07614033 | 0.000591802 | 11.80158785 | 0.474296021 | 2.428243518 |
| *Eya1* | 101.7538043 | 125.3463784 | 116.4264475 | 157.0521611 | 51.77130957 | 67.7067495 | 55.87859305 | 61.47002701 | 1.96E-08 | -1.080337468 | 1.13E-09 | 37.08791956 | 0.472918188 | 7.70738229 |
| *Flrt2* | 354.1431424 | 366.1693417 | 364.0793146 | 306.300488 | 137.352454 | 177.7302174 | 176.4587149 | 165.4962266 | 7.54E-22 | -1.080513866 | 1.56E-23 | 99.94789692 | 0.472860368 | 21.12279397 |
| *Cyr61* | 931.7456197 | 851.7631857 | 957.064865 | 894.5144826 | 477.5639168 | 431.6305281 | 430.3631991 | 380.1684747 | 7.89E-48 | -1.081656243 | 6.55E-50 | 220.6415269 | 0.472486088 | 47.10280447 |
| *Npy* | 1654.995699 | 1389.666936 | 1407.970683 | 1496.385187 | 736.4204647 | 660.1408076 | 697.9922501 | 697.9212297 | 3.29E-64 | -1.091024187 | 1.92E-66 | 296.4889506 | 0.469428004 | 63.48281956 |
| *Gpr176* | 195.5269181 | 222.0703555 | 200.2929563 | 202.8996863 | 97.20327511 | 101.5601243 | 98.03261939 | 86.05803781 | 6.90E-16 | -1.102134759 | 1.94E-17 | 72.20553521 | 0.465826701 | 15.16116229 |
| *Tm4sf1* | 104.7465632 | 82.90626605 | 95.7064865 | 105.3517602 | 54.94098158 | 49.72214416 | 28.42945962 | 48.23032888 | 5.52E-07 | -1.103890608 | 3.89E-08 | 30.20405155 | 0.465260106 | 6.258349256 |
| *Enpp2* | 103.7489769 | 101.6588738 | 103.599805 | 93.64600908 | 35.9229495 | 61.35924174 | 52.93761447 | 36.8820162 | 1.59E-07 | -1.108163129 | 1.04E-08 | 32.7643626 | 0.463884283 | 6.798327689 |
| *Ctsk* | 125.6958759 | 105.6067913 | 121.3597715 | 102.4253224 | 53.88442424 | 38.08504659 | 61.76055022 | 55.79587067 | 1.22E-08 | -1.114829251 | 6.87E-10 | 38.05839509 | 0.461745802 | 7.911864069 |
| *Atp6v0d2* | 404.0224582 | 460.91936 | 446.9591586 | 450.6714187 | 185.9540915 | 226.3944436 | 204.8881745 | 196.7040864 | 1.07E-30 | -1.115416376 | 1.52E-32 | 141.112482 | 0.461557926 | 29.97069598 |
| *Tns1* | 490.8124678 | 476.7110298 | 529.8390026 | 565.7779715 | 229.2729424 | 248.6107208 | 223.5143722 | 250.6085716 | 4.37E-34 | -1.115758018 | 5.34E-36 | 156.915961 | 0.461448638 | 33.35940011 |
| *Bcar1* | 101.7538043 | 114.4896055 | 86.82650321 | 106.3272395 | 50.71475223 | 47.60630824 | 51.95728828 | 38.77340165 | 3.30E-08 | -1.117212496 | 1.97E-09 | 36.00531631 | 0.460983655 | 7.481562112 |
| *Syde1* | 60.85276531 | 56.25782339 | 61.17321817 | 60.4797142 | 28.52704813 | 26.44794902 | 27.44913343 | 27.42508897 | 1.69E-05 | -1.120080455 | 1.51E-06 | 23.13095319 | 0.460068168 | 4.77150142 |
| *Pvrl3* | 149.6379475 | 165.8125321 | 142.0797325 | 127.7877832 | 84.52458705 | 70.88050338 | 49.99663589 | 64.30710518 | 2.92E-10 | -1.121949534 | 1.39E-11 | 45.68441694 | 0.459472514 | 9.534692599 |
| *Gja1* | 184.5534686 | 185.5521192 | 198.3196267 | 193.1448937 | 85.58114439 | 94.15469853 | 75.48511693 | 94.56927232 | 3.15E-15 | -1.122809688 | 9.39E-17 | 69.09370652 | 0.459198652 | 14.50112878 |
| *Hspa1b* | 185.5510549 | 156.9297179 | 140.1064029 | 174.6107878 | 54.94098158 | 60.30132377 | 92.15066223 | 92.67788687 | 5.01E-10 | -1.124228845 | 2.45E-11 | 44.5737362 | 0.458747167 | 9.299886184 |
| *Bok* | 37.90828003 | 27.63542202 | 62.15988298 | 66.33258977 | 32.75327748 | 16.92668738 | 18.62619768 | 20.80523991 | 0.002045488 | -1.126235361 | 0.000298208 | 13.08161368 | 0.45810958 | 2.689203101 |
| *Rgs16* | 250.3941655 | 227.0052523 | 230.8795654 | 228.2621471 | 101.4295045 | 83.57551892 | 115.6784909 | 126.7228249 | 1.55E-16 | -1.126784896 | 4.10E-18 | 75.27125814 | 0.457935116 | 15.81027049 |
| *Gpc6* | 69.83104216 | 95.7369977 | 64.1332126 | 68.28354829 | 32.75327748 | 37.02712863 | 38.23272156 | 28.3707817 | 4.76E-06 | -1.129352999 | 3.90E-07 | 25.74369685 | 0.457120683 | 5.321994769 |
| *Cpe* | 260.3700286 | 230.9531697 | 270.3461577 | 203.8751656 | 96.14671777 | 133.2976631 | 108.8162075 | 103.0805068 | 7.99E-16 | -1.130621958 | 2.27E-17 | 71.89883719 | 0.456718787 | 15.09734908 |

| | | | | | | | | | | | | | |
|---|---|---|---|---|---|---|---|---|---|---|---|---|---|
| *Dusp4* | 461.8824646 | 410.5834128 | 403.545907 | 490.6660684 | 210.2549103 | 235.9157053 | 178.4193673 | 183.4643883 | 4.03E-26 | -1.131545313 | 6.76E-28 | 119.8686905 | 0.456426571 | 25.39456027 |
| *Kdelr3* | 111.7296675 | 81.91928669 | 73.01319588 | 86.81765426 | 47.54508022 | 32.79545679 | 38.23272156 | 42.55617254 | 6.89E-07 | -1.132555369 | 4.98E-08 | 29.72307315 | 0.456107131 | 6.161615244 |
| *Gstm2* | 59.855179 | 59.21876146 | 75.97319031 | 68.28354829 | 35.9229495 | 33.85337475 | 23.52782865 | 26.47939625 | 9.96E-06 | -1.142362383 | 8.58E-07 | 24.22388991 | 0.453017164 | 5.001949531 |
| *Gata2* | 26.93483055 | 20.72656651 | 40.45325718 | 32.19081562 | 14.79180273 | 14.81085145 | 10.78358813 | 14.18539085 | 0.00552114 | -1.142958399 | 0.000927463 | 10.9670492 | 0.452830049 | 2.257971278 |
| *Cdh11* | 568.6242005 | 505.3334311 | 459.7858011 | 478.9603173 | 246.1778598 | 199.9464946 | 237.2389389 | 226.9662536 | 8.03E-32 | -1.143992941 | 1.09E-33 | 146.3475255 | 0.452505445 | 31.09530368 |
| *Unc5b* | 726.2428385 | 671.1459632 | 720.2653108 | 777.4569713 | 313.7975294 | 305.7382907 | 350.9567774 | 337.6123022 | 7.25E-48 | -1.144684161 | 5.96E-50 | 220.8301455 | 0.452288694 | 47.13975029 |
| *Igsf3* | 51.87448846 | 47.37500917 | 47.35991084 | 52.67588011 | 32.75327748 | 22.21627718 | 19.60652388 | 16.07677629 | 0.000179223 | -1.144741024 | 1.96E-05 | 18.22325529 | 0.452270868 | 3.74660697 |
| *Csgalnact1* | 28.93000318 | 33.55729816 | 39.46659237 | 31.21533636 | 15.84836007 | 4.231671844 | 23.52782865 | 16.07677629 | 0.006346981 | -1.148732658 | 0.001084434 | 10.67755286 | 0.451021259 | 2.19743281 |
| *Dysf* | 42.89621161 | 33.55729816 | 27.62661466 | 34.14177415 | 11.62213072 | 14.81085145 | 12.74424052 | 22.69662536 | 0.002498606 | -1.151744471 | 0.00037504 | 12.65261699 | 0.450080676 | 2.602302144 |
| *Glis2* | 35.9131074 | 48.36198853 | 58.21322375 | 53.65135937 | 23.24426144 | 29.62170291 | 12.74424052 | 22.69662536 | 0.000352002 | -1.157056979 | 4.17E-05 | 16.7946753 | 0.448426369 | 3.453455381 |
| *Lilra6* | 166.5969149 | 151.9948211 | 164.7730231 | 162.9050366 | 68.67622698 | 75.11217523 | 76.46544313 | 69.03556879 | 3.83E-14 | -1.159758671 | 1.23E-15 | 64.01467276 | 0.4475874 | 13.41695067 |
| *Sema7a* | 365.1165919 | 458.9454013 | 386.7726052 | 327.7610318 | 182.7844195 | 157.6297762 | 191.1636078 | 156.984992 | 1.12E-22 | -1.160054353 | 2.22E-24 | 103.8175665 | 0.447495676 | 21.94889971 |
| *Myh10* | 122.7031169 | 117.4505436 | 153.9197102 | 128.7632625 | 62.33688295 | 59.24340581 | 52.93761447 | 59.57864156 | 6.70E-11 | -1.160332339 | 2.99E-12 | 48.69585684 | 0.447409459 | 10.17411225 |
| *Fabp4* | 223.4593349 | 201.343789 | 143.0663973 | 180.4636633 | 91.92048842 | 72.99633931 | 95.09164081 | 74.70972513 | 3.00E-12 | -1.16073032 | 1.17E-13 | 55.05662878 | 0.447286053 | 11.52294937 |
| *Phlda1* | 137.6669117 | 143.1120069 | 136.1597437 | 150.2238062 | 59.16721093 | 61.35924174 | 65.68185499 | 67.14418335 | 1.53E-12 | -1.16079594 | 5.73E-14 | 56.46345831 | 0.447265709 | 11.81652553 |
| *Elk1* | 343.1696929 | 384.9219495 | 369.0126387 | 381.4123912 | 180.6713048 | 148.1085145 | 163.7144744 | 168.3333047 | 7.09E-29 | -1.161643987 | 1.09E-30 | 132.6337707 | 0.447002874 | 28.1491827 |
| *Gpc4* | 95.7682864 | 130.2812752 | 131.2264196 | 94.62148835 | 70.78934165 | 33.85337475 | 57.83924544 | 39.71909437 | 3.49E-07 | -1.162084521 | 2.39E-08 | 31.1527596 | 0.446866401 | 6.45669988 |
| *Tagln* | 1439.517055 | 1334.396092 | 1373.437414 | 1369.572883 | 724.7983339 | 594.5498941 | 598.9793045 | 544.7190086 | 1.42E-61 | -1.165122705 | 8.74E-64 | 284.2861572 | 0.445926331 | 60.84711991 |
| *Tmem47* | 87.78759586 | 94.75001834 | 103.599805 | 85.84217499 | 46.48852288 | 46.54839028 | 33.33109059 | 39.71909437 | 3.48E-08 | -1.167202804 | 2.08E-09 | 35.89868545 | 0.445283851 | 7.458376131 |
| *Grem2* | 40.90103898 | 28.62240137 | 38.47992756 | 48.77396306 | 12.67868806 | 21.15835922 | 24.50815485 | 11.34831268 | 0.001493533 | -1.171457421 | 0.000205815 | 13.77723218 | 0.44397261 | 2.825785097 |
| *Mylk* | 83.7972506 | 71.06251375 | 51.30657008 | 77.06286164 | 35.9229495 | 39.14296456 | 23.52782865 | 27.42508897 | 8.55E-06 | -1.17435681 | 7.29E-07 | 24.53765182 | 0.443081252 | 5.068233777 |
| *Ccbe1* | 152.6307064 | 123.3724197 | 152.9330454 | 119.0084699 | 55.99753892 | 52.89589805 | 84.30805268 | 49.17602161 | 1.11E-09 | -1.175064697 | 5.64E-11 | 42.94217685 | 0.442863899 | 8.953856176 |
| *Ppap2b* | 143.6524296 | 89.81512155 | 109.5197938 | 126.812304 | 46.48852288 | 60.30132377 | 52.93761447 | 48.23032888 | 5.39E-09 | -1.176817438 | 2.91E-10 | 39.73336055 | 0.442326187 | 8.26875346 |
| *Ptrf* | 196.5245044 | 207.2656651 | 264.4261689 | 237.0414605 | 105.6557338 | 108.96555 | 74.50479074 | 111.5917413 | 4.67E-15 | -1.177017671 | 1.41E-16 | 68.29230559 | 0.4422648 | 14.33058744 |
| *F2r* | 32.92034845 | 28.62240137 | 31.5732739 | 27.31341932 | 14.79180273 | 10.57917961 | 5.881957164 | 21.75093263 | 0.005931038 | -1.178236028 | 0.001002949 | 10.82211371 | 0.441891465 | 2.226869273 |
| *Pcdh19* | 88.78518218 | 90.80210091 | 102.6131402 | 90.7195713 | 26.41393345 | 46.54839028 | 40.19337395 | 51.06740705 | 9.15E-08 | -1.178473544 | 5.75E-09 | 33.9186443 | 0.441818721 | 7.038790774 |
| *Col8a1* | 282.3169276 | 268.4583853 | 251.5995264 | 288.7418613 | 106.7122911 | 121.6605655 | 122.5407742 | 129.5599031 | 1.28E-22 | -1.181324062 | 2.54E-24 | 103.5454118 | 0.440946625 | 21.89274497 |
| *Amotl2* | 286.3072729 | 271.4193234 | 276.2661466 | 289.7173406 | 132.0696673 | 114.2551398 | 131.36371 | 116.320205 | 3.74E-24 | -1.185999495 | 7.01E-26 | 110.6637854 | 0.439519934 | 23.42752802 |
| *Mmp13* | 279.3241687 | 319.7813119 | 257.5195152 | 224.3602301 | 113.0516352 | 122.7184835 | 122.5407742 | 116.320205 | 2.86E-19 | -1.187192214 | 6.55E-21 | 87.99714687 | 0.43915672 | 18.54353844 |
| *Adcy6* | 88.78518218 | 111.5286674 | 112.4797883 | 93.64600908 | 46.48852288 | 37.02712863 | 55.87859305 | 38.77340165 | 1.26E-08 | -1.188923991 | 7.10E-10 | 37.99207317 | 0.438629883 | 7.898336969 |
| *Cacna1c* | 53.8696611 | 39.47917431 | 44.39991642 | 58.52875568 | 23.24426144 | 30.67962087 | 10.78358813 | 21.75093263 | 0.000307191 | -1.189696096 | 3.59E-05 | 17.07506708 | 0.438395199 | 3.512591297 |
| *Igfbp5* | 46.88655688 | 39.47917431 | 60.18655336 | 39.01917045 | 28.52704813 | 26.44794902 | 13.72456672 | 13.23969812 | 0.000531894 | -1.190657401 | 6.58E-05 | 15.92727854 | 0.438103182 | 3.274175101 |
| *Ctgf* | 2455.059925 | 2512.849445 | 2507.11528 | 2347.978582 | 1113.611434 | 1130.9143 | 1098.945663 | 962.7151922 | 1.29E-111 | -1.19091345 | 3.31E-114 | 515.8962885 | 0.438025435 | 110.8880068 |
| *Traf1* | 142.6548433 | 143.1120069 | 147.0130566 | 135.5916173 | 63.39344029 | 58.18548785 | 71.56381216 | 55.79587067 | 7.85E-13 | -1.191008742 | 2.83E-14 | 57.85039488 | 0.437996504 | 12.10487303 |
| *Fgr* | 359.131074 | 382.9479908 | 383.8126108 | 337.5158244 | 184.8975342 | 124.8343194 | 178.4193673 | 151.3108357 | 7.55E-25 | -1.19383553 | 1.38E-26 | 113.8844806 | 0.437139142 | 24.12178388 |
| *Ghr* | 76.81414638 | 60.20574082 | 55.25322932 | 60.4797142 | 33.80983482 | 26.44794902 | 33.33109059 | 17.02246902 | 1.56E-05 | -1.197018694 | 1.39E-06 | 23.29716786 | 0.436175701 | 4.806704713 |
| *Ccdc80* | 294.2879634 | 304.9766215 | 314.7460742 | 294.5947369 | 153.200814 | 133.2976631 | 123.5211004 | 118.2115904 | 4.95E-25 | -1.197272449 | 8.97E-27 | 114.7405168 | 0.436098989 | 24.30545899 |
| *Col4a1* | 1020.530802 | 1090.61219 | 1078.424637 | 1046.689247 | 449.0368687 | 480.2947543 | 506.8286423 | 408.5392564 | 4.11E-67 | -1.200240431 | 2.23E-69 | 309.9586679 | 0.435202747 | 66.38650257 |
| *Pla1a* | 19.95172633 | 23.68750458 | 20.71996099 | 35.11725341 | 12.67868806 | 7.405425727 | 8.822935746 | 14.18539085 | 0.009524439 | -1.203440424 | 0.001747478 | 9.797515554 | 0.434238509 | 2.021160576 |
| *Pdpn* | 533.7086794 | 544.8126054 | 598.9055392 | 556.9986582 | 228.216385 | 213.6994281 | 273.5110081 | 251.5542644 | 1.84E-40 | -1.205908375 | 1.91E-42 | 186.4327229 | 0.433496312 | 39.73513708 |
| *Gcnt4* | 23.9420716 | 26.64844266 | 23.67995542 | 25.36246079 | 7.395901367 | 8.463343688 | 17.64587149 | 9.456927232 | 0.008020184 | -1.206576364 | 0.001435629 | 10.15945716 | 0.433295643 | 2.095815691 |
| *Nfib* | 40.90103898 | 39.47917431 | 42.4265868 | 46.82300454 | 15.84836007 | 20.10044126 | 14.70489291 | 22.69662536 | 0.000200736 | -1.207349015 | 2.25E-05 | 17.96761132 | 0.433063649 | 3.697375329 |
| *Myo1b* | 173.5800191 | 228.979211 | 191.412973 | 242.8943361 | 87.69425906 | 99.44428833 | 88.22935746 | 87.00373053 | 5.09E-16 | -1.209058876 | 1.41E-17 | 72.83540924 | 0.432550693 | 15.29356179 |
| *Sdc2* | 158.6162243 | 174.6953463 | 160.8263639 | 167.7824329 | 79.24180036 | 57.12756989 | 95.09164081 | 54.85017794 | 2.09E-12 | -1.20931021 | 7.99E-14 | 55.80869741 | 0.432475344 | 11.68013398 |
| *Ngf* | 106.7417359 | 62.17969953 | 68.07987184 | 47.7984838 | 27.47049079 | 35.96921067 | 31.37043821 | 28.3707817 | 2.22E-05 | -1.209539509 | 2.01E-06 | 22.58405131 | 0.432406613 | 4.654085973 |
| *Amotl1* | 189.5414002 | 213.1875413 | 203.2529507 | 174.6107878 | 90.86393108 | 71.93842134 | 90.19000984 | 84.16665236 | 9.49E-17 | -1.210008666 | 2.46E-18 | 76.280055 | 0.432266019 | 16.02289243 |
| *Arhgef17* | 153.6282928 | 124.3593991 | 117.4131123 | 111.2046358 | 58.1106536 | 46.54839028 | 62.74087641 | 51.06740705 | 1.01E-10 | -1.212240174 | 4.57E-12 | 47.8649334 | 0.431597923 | 9.996254339 |
| *Ccr1* | 62.84793795 | 43.42709174 | 65.11987741 | 42.9210875 | 17.96147475 | 21.15835922 | 16.6655453 | 35.93632348 | 0.000153795 | -1.21656477 | 1.66E-05 | 18.54698494 | 0.43030611 | 3.813057363 |
| *Lama4* | 110.7320811 | 121.398461 | 158.8530343 | 115.1065528 | 45.43196554 | 55.01173397 | 60.78022402 | 55.79587067 | 1.57E-10 | -1.219364514 | 7.21E-12 | 46.96915463 | 0.429471853 | 9.804632698 |
| *Dner* | 45.88897056 | 33.55729816 | 43.41325161 | 32.19081562 | 13.7352454 | 16.92668738 | 24.50815485 | 11.34831268 | 0.000743423 | -1.220408291 | 9.50E-05 | 15.23419473 | 0.429161246 | 3.128764113 |
| *Samd4* | 85.79242323 | 98.69793577 | 99.65314573 | 108.278198 | 47.54508022 | 39.14296456 | 38.23272156 | 43.50186527 | 2.38E-09 | -1.221283438 | 1.25E-10 | 41.39054348 | 0.428900993 | 8.623230255 |
| *B3galt5* | 33.91793477 | 55.27084403 | 53.2798997 | 63.40615198 | 29.58360547 | 27.50586698 | 15.6852191 | 16.07677629 | 0.000181048 | -1.22239171 | 1.99E-05 | 18.19564858 | 0.42857164 | 3.74220571 |
| *Fam171a1* | 36.91069371 | 35.53125688 | 22.69329061 | 42.9210875 | 7.395901367 | 14.81085145 | 17.64587149 | 18.91385446 | 0.002052352 | -1.223662619 | 0.000299537 | 13.07328774 | 0.428194266 | 2.687748112 |
| *Gdf15* | 31.92276213 | 39.47917431 | 61.17321817 | 30.2398571 | 11.62213072 | 6.347507766 | 25.48848104 | 25.53370353 | 0.003676538 | -1.225286083 | 0.000581495 | 11.83430237 | 0.42771269 | 2.434560903 |
| *Peg10* | 41.8986253 | 35.53125688 | 33.54660351 | 51.70040085 | 23.24426144 | 10.57917961 | 15.6852191 | 19.85954719 | 0.000490925 | -1.228164157 | 6.02E-05 | 16.09744315 | 0.426860284 | 3.308985194 |
| *Col7a1* | 34.91552108 | 20.72656651 | 30.58660909 | 22.43602301 | 13.7352454 | 7.405425727 | 17.64587149 | 7.565541785 | 0.006780229 | -1.228388693 | 0.001174698 | 10.52972265 | 0.426793854 | 2.168755621 |
| *Zfp507* | 38.90586635 | 38.49219495 | 34.53326832 | 37.06821193 | 14.79180273 | 15.86876941 | 23.52782865 | 9.456927232 | 0.000703534 | -1.228948797 | 8.94E-05 | 15.3490241 | 0.42662819 | 3.152714892 |
| *Foxc1* | 31.92276213 | 27.63542202 | 25.65328504 | 29.26437784 | 21.13114676 | 11.63709757 | 9.803261939 | 6.619849062 | 0.004587138 | -1.231357565 | 0.000748592 | 11.36455124 | 0.425916473 | 2.338458195 |
| *Dlg5* | 132.6789801 | 88.82814219 | 111.4931234 | 111.2046358 | 40.14917885 | 69.82258542 | 34.31141679 | 45.39325071 | 3.26E-08 | -1.231773676 | 1.94E-09 | 36.03157812 | 0.425793645 | 7.486251018 |
| *Spint1* | 163.6041559 | 135.216172 | 130.2397548 | 157.0521611 | 51.77130957 | 58.18548785 | 79.40642171 | 59.57864156 | 2.59E-12 | -1.232350653 | 9.99E-14 | 55.36991243 | 0.425623392 | 11.58614079 |
| *Gprc5b* | 98.76104535 | 110.5416881 | 100.6398105 | 108.278198 | 44.3754082 | 45.49047232 | 44.11467873 | 43.50186527 | 1.68E-10 | -1.237280094 | 7.80E-12 | 46.81550252 | 0.424171591 | 9.773589086 |
| *Gm2115* | 29.9275895 | 43.42709174 | 33.54660351 | 35.11725341 | 15.84836007 | 28.56378495 | 8.822935746 | 7.565541785 | 0.003174077 | -1.239200148 | 0.000491633 | 12.14713435 | 0.423607446 | 2.49838256 |
| *Ly75* | 90.78035481 | 125.3463784 | 109.5197938 | 119.9839491 | 63.39344029 | 45.49047232 | 45.09500492 | 34.99063076 | 2.70E-09 | -1.243808502 | 1.42E-10 | 41.13363547 | 0.422256489 | 8.568304011 |
| *Fermt2* | 274.3362371 | 294.1198486 | 268.3728281 | 252.6491287 | 129.9565526 | 133.2976631 | 92.15066223 | 105.917585 | 1.04E-21 | -1.244043159 | 2.17E-23 | 99.29928173 | 0.422187814 | 20.98389019 |
| *Fscn1* | 336.1865887 | 327.6771468 | 293.0394483 | 302.398571 | 139.4655686 | 141.7610068 | 123.5211004 | 126.7228249 | 5.11E-28 | -1.2464698 | 8.12E-30 | 128.6417564 | 0.421478282 | 27.29134151 |
| *Snai2* | 42.89621161 | 32.5703188 | 48.34657565 | 36.09273267 | 13.7352454 | 14.81085145 | 23.52782865 | 15.13108357 | 0.000342498 | -1.246539336 | 4.04E-05 | 16.85174772 | 0.421457968 | 3.465342353 |
| *Tnfrsf10b* | 101.7538043 | 112.5156468 | 136.1597437 | 119.9839491 | 62.33688295 | 48.6642262 | 48.0359835 | 39.71909437 | 1.81E-10 | -1.247850169 | 8.39E-12 | 46.672942 | 0.421075205 | 9.742744051 |
| *Rhoj* | 113.7248401 | 120.4114816 | 118.3997771 | 126.812304 | 55.99753892 | 48.6642262 | 49.0163097 | 48.23032888 | 5.12E-12 | -1.249021987 | 2.07E-13 | 53.93981913 | 0.420733329 | 11.29097891 |
| *Lpar1* | 54.86724741 | 55.27084403 | 97.67981612 | 60.4797142 | 28.52704813 | 22.21627718 | 41.17370015 | 20.80523991 | 2.77E-05 | -1.250192409 | 2.56E-06 | 22.11765776 | 0.420392137 | 4.557319856 |
| *Cd55* | 37.90828003 | 46.38802981 | 25.65328504 | 36.09273267 | 16.90491741 | 10.57917961 | 10.78358813 | 22.69662536 | 0.001119499 | -1.254123563 | 0.000150071 | 14.3713177 | 0.419248183 | 2.950976454 |
| *Npr2* | 33.91793477 | 27.63542202 | 36.50659794 | 43.89656676 | 15.84836007 | 15.86876941 | 12.74424052 | 15.13108357 | 0.000575407 | -1.254567439 | 7.17E-05 | 15.76509675 | 0.419119212 | 3.240025039 |
| *Edil3* | 303.2662403 | 305.9636009 | 299.946102 | 255.5755665 | 126.7868806 | 122.7184835 | 104.8949028 | 133.342674 | 6.85E-25 | -1.255640591 | 1.25E-26 | 114.0865502 | 0.418807566 | 24.16414056 |
| *Nexn* | 17.9565537 | 33.55729816 | 26.63994985 | 23.41150227 | 15.84836007 | 9.521261649 | 8.822935746 | 8.511234509 | 0.006111454 | -1.257312672 | 0.001038825 | 10.75705559 | 0.41832245 | 2.213855446 |
| *Pdgfra* | 90.78035481 | 121.398461 | 132.2130844 | 127.7877832 | 45.43196554 | 66.64883154 | 43.13435253 | 42.55617254 | 8.06E-10 | -1.26040213 | 4.02E-11 | 43.60347786 | 0.417427592 | 9.093742536 |
| *Myc* | 241.4158886 | 210.2266032 | 231.8662302 | 250.6981702 | 108.8254058 | 93.09678056 | 106.8555551 | 81.32957419 | 1.52E-20 | -1.262062665 | 3.37E-22 | 93.86887609 | 0.416947411 | 19.81689965 |
| *Il1rn* | 540.6917836 | 530.9948944 | 466.6924548 | 475.0584002 | 221.877041 | 173.4985456 | 219.5930674 | 221.2920972 | 6.03E-38 | -1.265731282 | 6.69E-40 | 174.7780985 | 0.415888506 | 37.21957262 |
| *Osbpl5* | 87.78759586 | 92.77605962 | 93.73315688 | 97.54792613 | 42.26229352 | 37.02712863 | 39.21304776 | 35.93632348 | 9.02E-10 | -1.269200051 | 4.53E-11 | 43.3702443 | 0.414889758 | 9.044750778 |
| *Csf1* | 2023.10505 | 2037.125394 | 2101.596044 | 1975.345504 | 909.6958681 | 869.6085639 | 836.2182434 | 756.5541785 | 1.24E-115 | -1.272462258 | 2.88E-118 | 534.5559307 | 0.413952674 | 114.9048309 |
| *Gpr84* | 1386.64498 | 1248.528887 | 1250.104313 | 1321.774399 | 470.1680155 | 503.5689494 | 600.9399569 | 576.8725611 | 1.38E-70 | -1.272910443 | 6.95E-73 | 326.056219 | 0.413824096 | 69.85979925 |
| *Bmpr1a* | 120.7079443 | 142.1250275 | 122.3464363 | 128.7632625 | 41.20573619 | 55.01173397 | 53.91794067 | 61.47002701 | 1.98E-12 | -1.276433472 | 7.54E-14 | 55.92143804 | 0.41281478 | 11.70322194 |
| *Nid1* | 326.2107255 | 336.559961 | 375.9192923 | 409.7012897 | 153.200814 | 158.6876941 | 140.1866457 | 145.6366794 | 2.92E-30 | -1.278566685 | 4.20E-32 | 139.0935439 | 0.41220483 | 29.53405558 |
| *Cd28* | 475.848673 | 580.3438623 | 584.1055671 | 563.827013 | 220.8204837 | 285.6378495 | 185.2816507 | 217.5093263 | 1.04E-31 | -1.280097902 | 1.42E-33 | 145.8161801 | 0.411767565 | 30.98169132 |
| *Cxcl12* | 338.1817613 | 320.7682912 | 298.9594372 | 314.1043221 | 128.8999953 | 126.9501553 | 149.9899077 | 117.2658977 | 4.88E-29 | -1.281925033 | 7.40E-31 | 133.3967247 | 0.411246403 | 28.3115359 |
| *Tpm2* | 540.6917836 | 459.9323807 | 457.8124715 | 496.518944 | 184.8975342 | 224.2786077 | 192.143934 | 202.3782428 | 7.42E-40 | -1.282776744 | 7.82E-42 | 183.6275648 | 0.411003691 | 39.12937785 |
| *Prrx2* | 52.87207478 | 39.47917431 | 45.38658123 | 37.06821193 | 16.90491741 | 22.21627718 | 23.52782865 | 9.456927232 | 0.000182967 | -1.282937847 | 2.02E-05 | 18.17143515 | 0.410957798 | 3.737627898 |
| *Tgfb2* | 58.85759268 | 89.81512155 | 78.93318474 | 77.06286164 | 30.64016281 | 37.02712863 | 26.46880724 | 31.20785986 | 1.48E-07 | -1.28409986 | 9.58E-09 | 32.92505752 | 0.410626927 | 6.830500103 |
| *Rcn1* | 325.2131392 | 318.7943325 | 345.3326832 | 352.1480133 | 148.9745847 | 154.4560223 | 132.3440362 | 116.320205 | 1.11E-29 | -1.284860038 | 1.62E-31 | 136.4150263 | 0.410410618 | 28.95543827 |
| *Msrb3* | 154.6258791 | 164.8255527 | 153.9197102 | 145.3464099 | 68.67622698 | 63.47507766 | 63.72120261 | 57.68725611 | 7.42E-16 | -1.288877791 | 2.09E-17 | 72.05668348 | 0.409269258 | 15.12963512 |
| *Ass1* | 1341.753596 | 1314.656504 | 1320.157515 | 1332.504671 | 470.1680155 | 558.5806834 | 637.2120261 | 503.1085287 | 2.70E-71 | -1.290741747 | 1.32E-73 | 329.3731494 | 0.408740825 | 70.56823545 |
| *Hic1* | 47.8841432 | 36.51823623 | 54.26656451 | 34.14177415 | 12.67868806 | 17.98460534 | 25.48848104 | 14.18539085 | 0.000184866 | -1.294092316 | 2.04E-05 | 18.14901755 | 0.407792652 | 3.733143589 |
| *Scube3* | 29.9275895 | 27.63542202 | 24.66662023 | 27.31341932 | 7.395901367 | 14.81085145 | 14.70489291 | 7.565541785 | 0.002426633 | -1.302667622 | 0.000362687 | 12.71524321 | 0.405375942 | 2.614995978 |
| *Frk* | 46.88655688 | 44.4140711 | 51.30657008 | 39.01917045 | 22.1877041 | 17.98460534 | 11.76391433 | 21.75093263 | 5.11E-05 | -1.302672477 | 4.94E-06 | 20.86153296 | 0.405374578 | 4.291176165 |
| *Mast4* | 145.6476022 | 115.4765848 | 127.2797604 | 126.812304 | 50.71475223 | 55.01173397 | 57.83924544 | 45.39325071 | 4.93E-13 | -1.303110527 | 1.73E-14 | 58.81190088 | 0.405251512 | 12.30738953 |
| *Il17rc* | 20.94931265 | 45.40105045 | 32.55993871 | 30.2398571 | 9.509016043 | 12.69501553 | 20.58685007 | 9.456927232 | 0.002034494 | -1.303597173 | 0.000295869 | 13.09636661 | 0.405114837 | 2.69154368 |
| *Dnaja4* | 45.88897056 | 29.60938073 | 42.4265868 | 28.28889858 | 13.7352454 | 11.63709757 | 10.78358813 | 22.69662536 | 0.00068377 | -1.304585963 | 8.65E-05 | 15.41000508 | 0.404837275 | 3.165090143 |
| *Fhl2* | 81.80207796 | 50.33594724 | 85.8398384 | 69.25902755 | 27.47049079 | 34.91129271 | 29.40978582 | 24.5880108 | 6.01E-07 | -1.306362843 | 4.28E-08 | 30.01736548 | 0.404338969 | 6.220946104 |
| *Ssc5d* | 37.90828003 | 19.73958715 | 52.29323489 | 30.2398571 | 15.84836007 | 16.92668738 | 12.74424052 | 11.34831268 | 0.001354254 | -1.306939271 | 0.000184892 | 13.9787125 | 0.404177448 | 2.868299811 |
| *Fstl3* | 104.7465632 | 99.68491513 | 94.71982169 | 91.69505056 | 45.43196554 | 32.79545679 | 46.07533112 | 33.09924531 | 3.33E-10 | -1.312914916 | 1.60E-11 | 45.40282692 | 0.402506806 | 9.478047547 |
| *Parva* | 218.4714033 | 242.796922 | 263.4395041 | 230.2131057 | 101.4295045 | 88.86510872 | 90.19000984 | 103.0805068 | 2.15E-23 | -1.315180698 | 4.14E-25 | 107.1416265 | 0.401875157 | 22.66854825 |
| *Egr2* | 120.7079443 | 139.1640894 | 116.4264475 | 175.586267 | 58.1106536 | 49.72214416 | 58.81957164 | 54.85017794 | 1.90E-12 | -1.316825168 | 7.20E-14 | 56.0137971 | 0.401417337 | 11.72179924 |
| *Lyz2* | 320211.2414 | 306469.9213 | 310288.3225 | 303716.4435 | 121613.9758 | 122608.46 | 127257.1236 | 126484.5103 | 0 | -1.317010283 | 0 | 2115.411276 | 0.401365834 | Inf |
| *Loxl4* | 45.88897056 | 73.03647247 | 62.15988298 | 43.89656676 | 17.96147475 | 30.67962087 | 20.58685007 | 20.80523991 | 1.21E-05 | -1.324111869 | 1.06E-06 | 23.82504246 | 0.399394987 | 4.918776576 |
| *Cdc42ep1* | 40.90103898 | 66.12761696 | 55.25322932 | 62.43067272 | 17.96147475 | 19.0425233 | 32.3507644 | 19.85954719 | 1.00E-05 | -1.329042645 | 8.62E-07 | 24.21401709 | 0.398032284 | 5.000125602 |
| *Ddah1* | 87.78759586 | 87.84116283 | 82.87984398 | 127.7877832 | 31.69672014 | 45.49047232 | 33.33109059 | 42.55617254 | 4.50E-09 | -1.335500697 | 2.40E-10 | 40.10834228 | 0.396254522 | 8.346315891 |
| *Tgfb1i1* | 97.76345903 | 102.6458532 | 108.533129 | 116.0820321 | 42.26229352 | 42.31671844 | 49.0163097 | 34.99063076 | 1.80E-11 | -1.335694494 | 7.51E-13 | 51.40699782 | 0.396201296 | 10.74442363 |
| *Tead3* | 31.92276213 | 27.63542202 | 18.74663138 | 26.33794005 | 8.452458705 | 13.75293349 | 13.72456672 | 5.674156339 | 0.003283538 | -1.335823383 | 0.000510423 | 12.0771961 | 0.396165902 | 2.483657951 |
| *Cttn* | 290.2976181 | 282.2760963 | 276.2661466 | 309.2269258 | 103.5426191 | 122.7184835 | 125.4817528 | 106.8632777 | 4.24E-29 | -1.336592488 | 6.39E-31 | 133.6887142 | 0.395954761 | 28.37311676 |
| *Prss23* | 132.6789801 | 130.2812752 | 138.1330733 | 159.0031196 | 62.33688295 | 50.78006213 | 51.95728828 | 56.74156339 | 9.68E-15 | -1.336995021 | 2.97E-16 | 66.82359998 | 0.395844299 | 14.01391096 |
| *Fam132b* | 61.85035163 | 60.20574082 | 27.62661466 | 41.94560824 | 24.30081878 | 16.92668738 | 20.58685007 | 14.18539085 | 0.000126893 | -1.33764146 | 1.34E-05 | 18.95507636 | 0.39566697 | 3.896562792 |
| *Ppic* | 181.5607096 | 151.0078417 | 197.3329619 | 181.4391426 | 77.12868568 | 70.88050338 | 59.79989783 | 73.76403241 | 1.52E-17 | -1.338208446 | 3.81E-19 | 79.96377831 | 0.395511502 | 16.81784332 |
| *Arhgap42* | 31.92276213 | 38.49219495 | 39.46659237 | 33.16629488 | 9.509016043 | 15.86876941 | 22.54750246 | 8.511234509 | 0.000475378 | -1.341720662 | 5.80E-05 | 16.1658108 | 0.394549807 | 3.322961233 |
| *Aqp1* | 32.92034845 | 39.47917431 | 59.19988856 | 49.74944233 | 14.79180273 | 16.92668738 | 20.58685007 | 18.91385446 | 5.04E-05 | -1.34519899 | 4.86E-06 | 20.89179673 | 0.393599696 | 4.29731889 |
| *Nipal1* | 22.94448528 | 18.7526078 | 23.67995542 | 19.50958523 | 11.62213072 | 6.347507766 | 9.803261939 | 5.674156339 | 0.006354475 | -1.348228299 | 0.001088251 | 10.67105321 | 0.392774099 | 2.196920341 |
| *Tln2* | 176.572778 | 186.5390986 | 219.0395877 | 172.6598292 | 72.90245633 | 79.34384707 | 70.58348596 | 73.76403241 | 3.05E-19 | -1.348452784 | 7.04E-21 | 87.85494434 | 0.392712988 | 18.51533188 |
| *Smoc2* | 142.6548433 | 105.6067913 | 134.1864141 | 147.2973685 | 51.77130957 | 40.20088252 | 54.89826686 | 60.52433428 | 8.03E-13 | -1.348549603 | 2.90E-14 | 57.79891322 | 0.392686634 | 12.09542843 |
| *Fat4* | 62.84793795 | 78.95834861 | 66.10654222 | 64.38163125 | 31.69672014 | 23.27419514 | 26.46880724 | 25.53370353 | 8.85E-08 | -1.349128412 | 5.54E-09 | 33.99082925 | 0.392529119 | 7.053244629 |
| *Lynx1* | 28.93000318 | 31.58333945 | 35.51993313 | 27.31341932 | 7.395901367 | 16.92668738 | 10.78358813 | 13.23969812 | 0.000749422 | -1.350752576 | 9.58E-05 | 15.21783587 | 0.392087465 | 3.125273557 |
| *Rhobtb3* | 56.86242005 | 50.33594724 | 76.95985512 | 49.74944233 | 16.90491741 | 28.56378495 | 23.52782865 | 22.69662536 | 3.00E-06 | -1.350788264 | 2.39E-07 | 26.69132572 | 0.392077766 | 5.52351739 |
| *Gpr153* | 32.92034845 | 44.4140711 | 34.53326832 | 32.19081562 | 17.96147475 | 9.521261649 | 18.62619768 | 10.40261995 | 0.000295713 | -1.351021945 | 3.44E-05 | 17.15649295 | 0.392014264 | 3.529130158 |
| *Wnt9a* | 27.93241687 | 22.70052523 | 28.61327947 | 29.26437784 | 17.96147475 | 9.521261649 | 4.90163097 | 10.40261995 | 0.002524549 | -1.351395572 | 0.000379741 | 12.62933349 | 0.391912754 | 2.597816193 |
| *Fgf7* | 42.89621161 | 59.21876146 | 38.47992756 | 55.60231789 | 13.7352454 | 11.63709757 | 14.70489291 | 35.93632348 | 0.000180913 | -1.354618041 | 1.99E-05 | 18.19983963 | 0.391038337 | 3.742531071 |
| *Fn1* | 6833.466269 | 7065.785222 | 7260.866331 | 7524.847022 | 2685.768753 | 3035.16663 | 2948.821191 | 2528.782342 | 8.05E-158 | -1.357362435 | 9.64E-161 | 729.854359 | 0.390295184 | 157.0944449 |
| *Dbn1* | 152.6307064 | 151.0078417 | 126.2930956 | 161.9295574 | 82.41147237 | 51.83798009 | 57.83924544 | 39.71909437 | 1.44E-12 | -1.358352593 | 5.38E-14 | 56.56688398 | 0.390027406 | 11.84193377 |
| *Tnc* | 2049.042294 | 2072.656651 | 1972.342954 | 1909.012914 | 816.7188224 | 822.0022557 | 795.0445433 | 689.4099952 | 5.13E-121 | -1.35901278 | 1.02E-123 | 559.6064385 | 0.389848968 | 120.2899303 |
| *Myl9* | 345.1648656 | 288.1979724 | 332.5060407 | 273.1341932 | 132.0696673 | 125.8922374 | 97.0522932 | 127.6685176 | 7.72E-27 | -1.361241444 | 1.28E-28 | 123.1758327 | 0.389247197 | 26.11226449 |
| *Pdgfrb* | 679.3562816 | 581.3308417 | 634.4254723 | 610.6500176 | 234.5557291 | 270.826998 | 276.4519867 | 188.1928519 | 1.57E-42 | -1.371132383 | 1.50E-44 | 196.0726177 | 0.386587693 | 41.80448729 |
| *Eml1* | 78.80931901 | 115.4765848 | 91.75982726 | 80.96477869 | 36.97950683 | 28.56378495 | 36.27206918 | 39.71909437 | 1.26E-09 | -1.37158684 | 6.46E-11 | 42.67577985 | 0.386465935 | 8.898120563 |
| *Erbb2* | 32.92034845 | 45.40105045 | 29.59994428 | 22.43602301 | 8.452458705 | 12.69501553 | 19.60652388 | 9.456927232 | 0.001006095 | -1.373182441 | 0.000133343 | 14.59393952 | 0.386038746 | 2.997361134 |
| *Foxc2* | 51.87448846 | 29.60938073 | 56.23989413 | 32.19081562 | 19.01803209 | 13.75293349 | 18.62619768 | 14.18539085 | 0.000120712 | -1.373748425 | 1.27E-05 | 19.05904302 | 0.385887328 | 3.918248758 |
| *Khdrbs3* | 25.93724423 | 43.42709174 | 53.2798997 | 34.14177415 | 10.56557338 | 20.10044126 | 10.78358813 | 18.91385446 | 0.000290258 | -1.376100712 | 3.37E-05 | 17.19577076 | 0.385258659 | 3.53721549 |
| *Adra2a* | 29.9275895 | 45.40105045 | 38.47992756 | 39.99464971 | 13.7352454 | 12.69501553 | 11.76391433 | 20.80523991 | 0.000117626 | -1.376342599 | 1.23E-05 | 19.11136841 | 0.385194071 | 3.929495031 |
| *Sdc1* | 1248.978068 | 1152.79189 | 1210.637721 | 1135.45786 | 510.3171943 | 429.5146921 | 452.9107016 | 435.9643454 | 2.05E-93 | -1.377295848 | 6.71E-96 | 431.765651 | 0.384939641 | 92.68841005 |
| *Map1b* | 82.79966428 | 80.93230733 | 83.86650879 | 96.57244687 | 38.03606417 | 39.14296456 | 25.48848104 | 30.26216714 | 1.05E-09 | -1.378429501 | 5.31E-11 | 43.0579184 | 0.384637279 | 8.977492501 |
| *Mmp2* | 372.0996961 | 320.7682912 | 372.9592979 | 401.8974557 | 139.4655686 | 142.8189247 | 146.0686029 | 136.1797521 | 4.21E-36 | -1.378805229 | 4.90E-38 | 166.2403638 | 0.384537119 | 35.37621391 |
| *Adamts5* | 18.95414002 | 17.76562844 | 29.59994428 | 25.36246079 | 7.395901367 | 7.405425727 | 13.72456672 | 6.619849062 | 0.00459787 | -1.38061704 | 0.000751077 | 11.35839465 | 0.3840545 | 2.33744334 |
| *C77080* | 500.7883309 | 511.2553073 | 583.1189023 | 586.263036 | 237.7254011 | 221.1048538 | 200.9668698 | 177.790232 | 2.37E-44 | -1.384380387 | 2.16E-46 | 204.5185688 | 0.383053979 | 43.62529451 |
| *Htra1* | 148.6403612 | 145.0859656 | 121.3597715 | 151.1992855 | 55.99753892 | 75.11217523 | 44.11467873 | 42.55617254 | 2.10E-13 | -1.385815015 | 7.24E-15 | 60.53179506 | 0.382673256 | 12.67696353 |
| *Mmp9* | 150.6355338 | 116.4635642 | 126.2930956 | 86.81765426 | 42.26229352 | 55.01173397 | 40.19337395 | 46.33894344 | 3.61E-11 | -1.386459412 | 1.55E-12 | 49.98728482 | 0.382502369 | 10.4425785 |
| *Tubb3* | 130.6838075 | 131.2682546 | 99.65314573 | 140.4690136 | 35.9229495 | 44.43255436 | 48.0359835 | 62.41571973 | 3.17E-12 | -1.38942621 | 1.25E-13 | 54.93371904 | 0.381716589 | 11.49844703 |
| *Gjc1* | 23.9420716 | 24.67448394 | 24.66662023 | 22.43602301 | 11.62213072 | 10.57917961 | 9.803261939 | 4.728463616 | 0.002507394 | -1.392206975 | 0.00037676 | 12.64406565 | 0.380981546 | 2.600777417 |
| *Gpx3* | 461.8824646 | 450.0625871 | 518.9856897 | 511.1511329 | 165.8795021 | 189.367315 | 186.2619768 | 196.7040864 | 3.37E-47 | -1.394114631 | 2.91E-49 | 217.6752448 | 0.380478113 | 46.47216205 |
| *Nbl1* | 85.79242323 | 59.21876146 | 92.74649207 | 72.18546534 | 29.58360547 | 26.44794902 | 32.3507644 | 29.31647442 | 1.37E-08 | -1.395242703 | 7.72E-10 | 37.83001221 | 0.380180726 | 7.863493191 |
| *Faxc* | 19.95172633 | 30.59636009 | 36.50659794 | 26.33794005 | 10.56557338 | 4.231671844 | 17.64587149 | 10.40261995 | 0.002113897 | -1.395555933 | 0.000309383 | 13.01271252 | 0.380098192 | 2.674916228 |
| *Epb4.1l3* | 235.4303707 | 201.343789 | 193.3863026 | 208.7525619 | 98.25983244 | 103.6759602 | 66.66218119 | 52.01309977 | 5.05E-16 | -1.395583023 | 1.39E-17 | 72.856429 | 0.380091055 | 15.29693384 |
| *Grem1* | 293.2903771 | 268.4583853 | 253.572856 | 301.4230917 | 113.0516352 | 112.1393039 | 95.09164081 | 104.0261995 | 1.89E-29 | -1.398150909 | 2.79E-31 | 135.3374227 | 0.379415124 | 28.72446258 |
| *Lhfp* | 239.420716 | 250.6927569 | 221.0129173 | 243.8698153 | 92.97704575 | 96.27053445 | 76.46544313 | 96.46065776 | 4.22E-26 | -1.399752058 | 7.11E-28 | 119.7685608 | 0.37899427 | 25.37470198 |
| *Ptprk* | 67.83586953 | 72.04949311 | 61.17321817 | 82.91573721 | 25.35737611 | 26.44794902 | 20.58685007 | 34.99063076 | 4.16E-08 | -1.399955723 | 2.50E-09 | 35.53898109 | 0.378940771 | 7.381082046 |
| *Wisp1* | 455.8969467 | 491.5157201 | 530.8256674 | 519.9304463 | 200.7458942 | 235.9157053 | 159.7931696 | 162.6591484 | 1.50E-37 | -1.400678475 | 1.68E-39 | 172.9425917 | 0.37875098 | 36.82493483 |
| *Scd1* | 80.80449165 | 82.90626605 | 89.78649764 | 82.91573721 | 21.13114676 | 46.54839028 | 35.29174298 | 24.5880108 | 7.91E-09 | -1.402568933 | 4.33E-10 | 38.95810546 | 0.378255002 | 8.102059751 |
| *Ncam1* | 194.5293317 | 198.3828509 | 189.4396434 | 163.8805159 | 87.69425906 | 80.40176503 | 70.58348596 | 44.44755799 | 2.07E-16 | -1.405159379 | 5.52E-18 | 74.68695467 | 0.377576432 | 15.68434917 |
| *Serpine2* | 231.4400255 | 301.0287041 | 264.4261689 | 251.6736494 | 103.5426191 | 90.98094464 | 113.7178385 | 87.00373053 | 1.33E-25 | -1.408029251 | 2.34E-27 | 117.4065571 | 0.376826087 | 24.87568443 |
| *Inhba* | 417.9886667 | 372.0912178 | 323.6260574 | 361.9028059 | 153.200814 | 112.1393039 | 145.0882767 | 144.6909866 | 6.42E-33 | -1.408824908 | 8.36E-35 | 151.449879 | 0.376618322 | 32.19234935 |
| *Phldb2* | 157.618638 | 152.9818004 | 169.7063472 | 187.2920182 | 65.50655496 | 72.99633931 | 57.83924544 | 55.79587067 | 2.48E-18 | -1.408909323 | 5.90E-20 | 83.65155903 | 0.376596286 | 17.60548573 |
| *Cnn3* | 431.9548751 | 332.6120435 | 325.5993871 | 345.3196585 | 137.352454 | 156.5718582 | 120.5801219 | 126.7228249 | 1.83E-30 | -1.409579411 | 2.62E-32 | 140.0329234 | 0.376421409 | 29.73706113 |
| *Yes1* | 35.9131074 | 38.49219495 | 34.53326832 | 28.28889858 | 16.90491741 | 12.69501553 | 10.78358813 | 11.34831268 | 0.000150823 | -1.412747998 | 1.62E-05 | 18.58984029 | 0.375595583 | 3.821532384 |
| *Cdr2l* | 50.87690215 | 55.27084403 | 69.06653665 | 41.94560824 | 13.7352454 | 22.21627718 | 16.6655453 | 28.3707817 | 4.77E-06 | -1.416736953 | 3.91E-07 | 25.73659833 | 0.374558521 | 5.321317284 |
| *Timp3* | 1418.567742 | 1442.963821 | 1469.143901 | 1426.15068 | 487.0729329 | 603.0132377 | 512.7105994 | 552.2845503 | 1.21E-112 | -1.417692717 | 3.00E-115 | 520.684384 | 0.374310464 | 111.9159387 |
| *Cald1* | 1369.686013 | 1395.588812 | 1496.770516 | 1399.81274 | 544.1270291 | 552.2331756 | 496.0450541 | 527.6965395 | 4.61E-123 | -1.418074682 | 8.47E-126 | 569.1799434 | 0.374211375 | 122.3362488 |
| *Tead1* | 231.4400255 | 275.3672408 | 257.5195152 | 236.0659812 | 104.5991765 | 99.44428833 | 113.7178385 | 57.68725611 | 9.84E-20 | -1.418093302 | 2.21E-21 | 90.15108656 | 0.374206545 | 19.00722432 |
| *Dnm3os* | 47.8841432 | 46.38802981 | 49.33324046 | 44.87204602 | 21.13114676 | 30.67962087 | 8.822935746 | 10.40261995 | 5.83E-05 | -1.423159733 | 5.71E-06 | 20.58454978 | 0.372894718 | 4.234066786 |
| *Plcb1* | 19.95172633 | 32.5703188 | 19.73329619 | 11.70575114 | 7.395901367 | 7.405425727 | 10.78358813 | 5.674156339 | 0.007499331 | -1.425303069 | 0.001324433 | 10.30816001 | 0.372341139 | 2.124977498 |
| *Acta2* | 4039.226996 | 4008.123172 | 4047.299048 | 3822.903225 | 1518.272895 | 1445.115935 | 1534.210494 | 1421.376163 | 2.73E-247 | -1.427362834 | 1.53E-250 | 1142.950025 | 0.37180992 | 246.5632175 |
| *Yap1* | 138.664498 | 108.5677293 | 156.8797047 | 169.7333915 | 43.31885086 | 55.01173397 | 50.97696209 | 63.36141245 | 5.70E-14 | -1.429003501 | 1.87E-15 | 63.19780588 | 0.371387329 | 13.24432347 |
| *Arhgef5* | 65.8406969 | 87.84116283 | 54.26656451 | 46.82300454 | 30.64016281 | 12.69501553 | 28.42945962 | 22.69662536 | 2.74E-06 | -1.429051486 | 2.17E-07 | 26.87384315 | 0.371374977 | 5.562354933 |
| *F5* | 16.95896738 | 22.70052523 | 19.73329619 | 16.58314744 | 4.226229352 | 6.347507766 | 12.74424052 | 4.728463616 | 0.008786751 | -1.43246702 | 0.00158828 | 9.973283763 | 0.370496798 | 2.056171697 |
| *Wnt5a* | 148.6403612 | 169.7604495 | 224.9595765 | 137.5425758 | 65.50655496 | 78.28592911 | 65.68185499 | 43.50186527 | 1.44E-13 | -1.433335468 | 4.88E-15 | 61.30652726 | 0.37027384 | 12.84076065 |
| *Bmp1* | 724.2476659 | 730.3647247 | 764.6652272 | 767.7021786 | 273.6483506 | 302.5645368 | 282.3339439 | 248.7171862 | 1.04E-74 | -1.433621262 | 4.83E-77 | 345.1475186 | 0.370200497 | 73.98179325 |
| *Vgll3* | 107.7393222 | 156.9297179 | 122.3464363 | 144.3709307 | 49.65819489 | 65.59091358 | 39.21304776 | 42.55617254 | 5.07E-13 | -1.437329529 | 1.79E-14 | 58.75072589 | 0.369250165 | 12.29487035 |

| | | | | | | | | | | | | | | |
|---|---|---|---|---|---|---|---|---|---|---|---|---|---|---|
| *Farp1* | 140.6596706 | 130.2812752 | 117.4131123 | 155.1012025 | 57.05409626 | 45.49047232 | 51.95728828 | 46.33894344 | 1.56E-15 | -1.437454421 | 4.54E-17 | 70.52782716 | 0.369218201 | 14.80644031 |
| *Ddr2* | 205.5027812 | 182.5911812 | 205.2262803 | 207.7770827 | 73.95901367 | 68.76466746 | 75.48511693 | 76.60111058 | 5.78E-24 | -1.44087435 | 1.09E-25 | 109.78302 | 0.368344 | 23.23824768 |
| *Thsd4* | 21.94689897 | 36.51823623 | 30.58660909 | 29.26437784 | 11.62213072 | 12.69501553 | 11.76391433 | 7.565541785 | 0.000466338 | -1.445111999 | 5.67E-05 | 16.21087285 | 0.367263646 | 3.331299524 |
| *Ltbp2* | 1421.560501 | 1387.692977 | 1355.677448 | 1522.723127 | 491.2991622 | 557.5227654 | 528.3958185 | 511.6197632 | 2.36E-119 | -1.445435999 | 5.08E-122 | 551.8121217 | 0.367181175 | 118.6278048 |
| *Smoc1* | 40.90103898 | 61.19272018 | 74.9865255 | 71.20998607 | 28.52704813 | 25.39003106 | 18.62619768 | 18.91385446 | 6.86E-07 | -1.447825717 | 4.95E-08 | 29.73601648 | 0.36657347 | 6.163728964 |
| *Chst2* | 137.6669117 | 174.6953463 | 150.9597158 | 146.3218892 | 68.67622698 | 59.24340581 | 52.93761447 | 43.50186527 | 1.34E-16 | -1.447993747 | 3.52E-18 | 75.57101805 | 0.366530778 | 15.87226189 |
| *Flrt3* | 62.84793795 | 58.2317821 | 33.54660351 | 39.01917045 | 26.41393345 | 13.75293349 | 12.74424052 | 17.96816174 | 2.65E-05 | -1.45299333 | 2.44E-06 | 22.2122506 | 0.365262783 | 4.576840279 |
| *Fam115c* | 22.94448528 | 21.71354587 | 37.49326275 | 18.53410596 | 17.96147475 | 10.57917961 | 4.90163097 | 3.782770893 | 0.005258959 | -1.453654095 | 0.000876963 | 11.07083286 | 0.365095529 | 2.279100251 |
| *Tnfrsf11b* | 77.8117327 | 108.5677293 | 68.07987184 | 81.94025795 | 25.35737611 | 30.67962087 | 35.29174298 | 31.20785986 | 1.44E-09 | -1.454334635 | 7.37E-11 | 42.41713727 | 0.364923348 | 8.842723786 |
| *Col16a1* | 129.6862212 | 116.4635642 | 111.4931234 | 104.376281 | 43.31885086 | 47.60630824 | 34.31141679 | 43.50186527 | 6.39E-14 | -1.455120276 | 2.11E-15 | 62.95880272 | 0.364724678 | 13.19478457 |
| *Itgb7* | 522.7352299 | 415.5183096 | 494.3190694 | 498.4699025 | 172.2188461 | 171.3827097 | 189.2029554 | 171.1703829 | 1.99E-47 | -1.45525631 | 1.70E-49 | 218.7416151 | 0.364690289 | 46.70073312 |
| *Foxs1* | 82.79966428 | 57.24480275 | 67.09320703 | 52.67588011 | 19.01803209 | 23.27419514 | 34.31141679 | 17.96816174 | 3.16E-07 | -1.45607674 | 2.15E-08 | 31.35860328 | 0.364482957 | 6.500201617 |
| *Fbln1* | 16.95896738 | 19.73958715 | 28.61327947 | 17.5586267 | 11.62213072 | 2.115835922 | 7.842609552 | 8.511234509 | 0.006351345 | -1.457750312 | 0.001086691 | 10.67370706 | 0.36406039 | 2.197134292 |
| *Ntn4* | 12.96862212 | 32.5703188 | 24.66662023 | 21.46054375 | 13.7352454 | 5.289589805 | 7.842609552 | 6.619849062 | 0.004465706 | -1.458623336 | 0.000725209 | 11.42350079 | 0.363840151 | 2.350109911 |
| *Has2* | 157.618638 | 178.6432637 | 153.9197102 | 169.7333915 | 54.94098158 | 69.82258542 | 45.09500492 | 69.98126152 | 2.70E-18 | -1.46033905 | 6.47E-20 | 83.47017646 | 0.363407714 | 17.5685433 |
| *Ltbp1* | 205.5027812 | 246.7448394 | 221.9995821 | 160.9540781 | 57.05409626 | 86.7492728 | 89.20968365 | 69.98126152 | 2.74E-18 | -1.461850567 | 6.61E-20 | 83.42785939 | 0.36302717 | 17.56213233 |
| *Slco2a1* | 35.9131074 | 24.67448394 | 37.49326275 | 55.60231789 | 13.7352454 | 21.15835922 | 12.74424052 | 8.511234509 | 0.00019694 | -1.462847354 | 2.20E-05 | 18.00666658 | 0.362776434 | 3.705665719 |
| *Podnl1* | 72.82380111 | 85.86720412 | 54.26656451 | 67.30806903 | 17.96147475 | 29.62170291 | 26.46880724 | 27.42508897 | 2.97E-08 | -1.463107832 | 1.75E-09 | 36.230916 | 0.362710941 | 7.527164232 |
| *Epdr1* | 117.7151854 | 104.6198119 | 74.9865255 | 115.1065528 | 45.43196554 | 20.10044126 | 35.29174298 | 48.23032888 | 1.38E-09 | -1.463348782 | 7.06E-11 | 42.50115731 | 0.362650368 | 8.860027064 |
| *Slc2a10* | 18.95414002 | 24.67448394 | 19.73329619 | 19.50958523 | 4.226229352 | 4.231671844 | 13.72456672 | 7.565541785 | 0.005258959 | -1.464595869 | 0.000877409 | 11.06988909 | 0.362337023 | 2.279100251 |
| *Enah* | 419.9838393 | 502.3724931 | 436.1058457 | 445.7940224 | 177.5016328 | 163.9772839 | 154.8915386 | 157.9306848 | 7.20E-48 | -1.464950938 | 5.86E-50 | 220.8631228 | 0.362247857 | 47.14270629 |
| *Plod2* | 478.841432 | 519.1511421 | 519.9723545 | 421.4070409 | 199.6893369 | 183.0198072 | 166.655453 | 154.1479139 | 1.52E-44 | -1.465875592 | 1.38E-46 | 205.4140272 | 0.36201576 | 43.81685406 |
| *Chpf* | 158.6162243 | 144.0989862 | 141.0930677 | 158.0276403 | 63.39344029 | 49.72214416 | 51.95728828 | 52.9587925 | 1.75E-18 | -1.466233987 | 4.16E-20 | 84.34254041 | 0.361925839 | 17.75581673 |
| *Tbx18* | 16.95896738 | 20.72656651 | 24.66662023 | 18.53410596 | 8.452458705 | 7.405425727 | 3.921304776 | 9.456927232 | 0.004225297 | -1.468686035 | 0.00068077 | 11.54102638 | 0.361311221 | 2.374142721 |
| *Fgfr1* | 321.222794 | 332.6120435 | 306.8527557 | 346.2951378 | 110.9385205 | 111.0813859 | 134.3046886 | 115.3745122 | 1.38E-37 | -1.468747349 | 1.54E-39 | 173.1163892 | 0.361295866 | 36.85979905 |
| *P3h4* | 123.7007033 | 119.4245023 | 131.2264196 | 159.0031196 | 47.54508022 | 46.54839028 | 49.99663589 | 48.23032888 | 6.73E-16 | -1.471236237 | 1.89E-17 | 72.26021164 | 0.360673108 | 15.17195968 |
| *Ptprg* | 92.77552745 | 92.77605962 | 77.94651993 | 98.52340539 | 36.97950683 | 31.73753883 | 36.27206918 | 25.53370353 | 2.81E-11 | -1.47521607 | 1.19E-12 | 50.50044886 | 0.359679522 | 10.55126815 |
| *Vegfc* | 21.94689897 | 13.81771101 | 25.65328504 | 23.41150227 | 6.339344029 | 12.69501553 | 10.78358813 | 0.945692723 | 0.008793224 | -1.476168693 | 0.001590332 | 9.970906758 | 0.359442101 | 2.055851845 |
| *Ndrg4* | 79.80690533 | 83.8932454 | 65.11987741 | 67.30806903 | 22.1877041 | 23.27419514 | 21.56717627 | 38.77340165 | 1.06E-08 | -1.477027633 | 5.90E-10 | 38.35317526 | 0.359228163 | 7.975612239 |
| *Kctd17* | 47.8841432 | 56.25782339 | 48.34657565 | 47.7984838 | 23.24426144 | 20.10044126 | 15.6852191 | 13.23969812 | 1.17E-06 | -1.479198646 | 8.71E-08 | 28.64230469 | 0.358687992 | 5.933218848 |
| *Olfml2b* | 229.4448528 | 191.4739954 | 217.066258 | 198.9977693 | 50.71475223 | 92.0388626 | 78.42609552 | 78.49249602 | 2.58E-21 | -1.48009007 | 5.45E-23 | 97.4758445 | 0.358466432 | 20.58893381 |
| *Igfbp7* | 668.3828321 | 665.2240871 | 638.3721316 | 734.5358838 | 238.7819584 | 230.6261155 | 267.6290509 | 230.7490245 | 2.43E-69 | -1.483249308 | 1.24E-71 | 320.305277 | 0.357682315 | 68.61399684 |
| *Kirrel* | 278.3265823 | 259.5755711 | 300.9327668 | 304.3495295 | 97.20327511 | 100.5022063 | 112.7375123 | 96.46065776 | 1.22E-33 | -1.489932132 | 1.53E-35 | 154.8248772 | 0.356029297 | 32.9136583 |
| *Col5a3* | 308.2541718 | 267.4714059 | 257.5195152 | 290.6928199 | 96.14671777 | 101.5601243 | 105.8752289 | 95.51496504 | 2.61E-33 | -1.493713638 | 3.33E-35 | 153.2792481 | 0.355097316 | 32.58409824 |
| *Nsg1* | 74.81897375 | 67.11459632 | 81.89317917 | 79.98929943 | 28.52704813 | 23.27419514 | 33.33109059 | 22.69662536 | 7.17E-10 | -1.494487408 | 3.57E-11 | 43.83704579 | 0.354906916 | 9.144181421 |
| 5-Sep | 68.83345585 | 57.24480275 | 55.25322932 | 74.13642386 | 21.13114676 | 17.98460534 | 37.25239537 | 14.18539085 | 4.21E-07 | -1.495049226 | 2.92E-08 | 30.76037743 | 0.354768734 | 6.375950819 |
| *Gnao1* | 16.95896738 | 19.73958715 | 16.77330176 | 23.41150227 | 9.509016043 | 4.231671844 | 6.862283358 | 6.619849062 | 0.004388342 | -1.498757388 | 0.000710894 | 11.46054642 | 0.353858042 | 2.3576995 |
| *Timp1* | 1235.01186 | 1154.765848 | 1183.997771 | 1021.326787 | 405.7180178 | 395.6613174 | 458.7926588 | 364.0916984 | 2.70E-84 | -1.500408063 | 9.90E-87 | 389.6420018 | 0.353453403 | 83.56872105 |
| *Sytl2* | 37.90828003 | 41.45313302 | 50.31990527 | 27.31341932 | 10.56557338 | 19.0425233 | 4.90163097 | 20.80523991 | 0.000180035 | -1.50339632 | 1.97E-05 | 18.21325767 | 0.352722052 | 3.744643519 |
| *Gpr116* | 77.8117327 | 74.02345183 | 111.4931234 | 79.98929943 | 30.64016281 | 30.67962087 | 30.39011201 | 29.31647442 | 1.59E-10 | -1.504901814 | 7.35E-12 | 46.9324351 | 0.352354168 | 9.798001246 |
| *Cd69* | 132.6789801 | 130.2812752 | 107.5464642 | 184.3655804 | 39.09262151 | 69.82258542 | 49.0163097 | 37.82770893 | 3.11E-12 | -1.507497541 | 1.22E-13 | 54.98261975 | 0.351720775 | 11.50748341 |
| *Atp1b1* | 19.95172633 | 36.51823623 | 19.73329619 | 21.46054375 | 11.62213072 | 7.405425727 | 5.881957164 | 9.456927232 | 0.001618595 | -1.509426317 | 0.000226668 | 13.59600048 | 0.351250865 | 2.790861773 |
| *Fat1* | 891.8421671 | 965.2658118 | 913.6516134 | 1037.909934 | 359.229495 | 378.73463 | 291.1568796 | 311.1329059 | 1.20E-75 | -1.509589162 | 5.36E-78 | 349.5315726 | 0.35121122 | 74.9212638 |
| *Pdgfc* | 114.7224264 | 103.6328326 | 105.5731346 | 105.3517602 | 34.86639216 | 55.01173397 | 30.39011201 | 31.20785986 | 1.03E-12 | -1.509772956 | 3.82E-14 | 57.25693621 | 0.351166479 | 11.98617924 |
| *Vldlr* | 23.9420716 | 44.4140711 | 20.71996099 | 23.41150227 | 15.84836007 | 7.405425727 | 7.842609552 | 8.511234509 | 0.001072079 | -1.511915223 | 0.000143201 | 14.45956723 | 0.350645417 | 2.969773341 |
| *Lamb1* | 406.0176309 | 431.3099793 | 455.8391419 | 384.3388289 | 142.6352406 | 141.7610068 | 141.1669719 | 161.7134557 | 1.07E-47 | -1.512469729 | 9.04E-50 | 220.0016519 | 0.350510671 | 46.97150486 |
| *Fbln5* | 100.756218 | 130.2812752 | 88.79983283 | 141.4444929 | 43.31885086 | 42.31671844 | 41.17370015 | 34.99063076 | 7.73E-13 | -1.514462532 | 2.77E-14 | 57.89024036 | 0.350026843 | 12.1117399 |
| *Mrc2* | 276.3314097 | 258.5885917 | 263.4395041 | 263.3794005 | 91.92048842 | 98.38637037 | 113.7178385 | 68.08987607 | 9.26E-29 | -1.515066014 | 1.43E-30 | 132.0951959 | 0.349880456 | 28.03361807 |
| *Adamts14* | 23.9420716 | 21.71354587 | 24.66662023 | 13.65670966 | 14.79180273 | 2.115835922 | 7.842609552 | 4.728463616 | 0.006465596 | -1.516378034 | 0.001110379 | 10.63382349 | 0.349562412 | 2.189391427 |
| *Serpinf1* | 703.2983532 | 611.9272018 | 604.8255281 | 703.3205474 | 235.6122864 | 223.2206898 | 243.1208961 | 213.7265554 | 2.82E-66 | -1.518994787 | 1.60E-68 | 306.0302312 | 0.348928952 | 65.54945078 |
| *Ptprm* | 45.88897056 | 56.25782339 | 53.2798997 | 67.30806903 | 22.1877041 | 16.92668738 | 18.62619768 | 19.85954719 | 1.38E-07 | -1.521763705 | 8.89E-09 | 33.07085934 | 0.348259906 | 6.86038665 |
| *Unc5c* | 38.90586635 | 19.73958715 | 24.66662023 | 21.46054375 | 10.56557338 | 11.63709757 | 7.842609552 | 6.619849062 | 0.000954526 | -1.522706493 | 0.000125822 | 14.70335803 | 0.348032396 | 3.02021233 |
| *Schip1* | 28.93000318 | 62.17969953 | 48.34657565 | 53.65135937 | 8.452458705 | 16.92668738 | 23.52782865 | 17.96816174 | 1.62E-05 | -1.52326208 | 1.45E-06 | 23.21707063 | 0.347898393 | 4.789785608 |
| *S100a16* | 24.93965792 | 14.80469037 | 42.4265868 | 37.06821193 | 8.452458705 | 12.69501553 | 15.6852191 | 4.728463616 | 0.001456042 | -1.525025462 | 0.000200068 | 13.83044742 | 0.347473422 | 2.836826095 |
| *Gsta3* | 19.95172633 | 30.59636009 | 27.62661466 | 29.26437784 | 9.509016043 | 10.57917961 | 8.822935746 | 8.511234509 | 0.000402229 | -1.525240721 | 4.82E-05 | 16.51875843 | 0.347421581 | 3.395526609 |
| *Slit2* | 92.77552745 | 127.3203371 | 117.4131123 | 115.1065528 | 33.80983482 | 45.49047232 | 36.27206918 | 41.61047982 | 3.46E-14 | -1.525966775 | 1.11E-15 | 64.22431223 | 0.347246781 | 13.46100736 |
| *Dlc1* | 119.710358 | 141.1380481 | 114.4531179 | 119.9839491 | 51.77130957 | 48.6642262 | 38.23272156 | 34.04493803 | 2.57E-15 | -1.52657873 | 7.61E-17 | 69.50708231 | 0.347099519 | 14.58997976 |
| *Nedd4* | 1319.806697 | 1307.747649 | 1390.210716 | 1268.12304 | 526.1655544 | 481.3526722 | 424.481242 | 404.7564855 | 3.88E-101 | -1.527673661 | 1.12E-103 | 467.5104183 | 0.346836188 | 100.4110018 |
| *Hspg2* | 1165.180818 | 1211.023672 | 1197.811078 | 1268.12304 | 425.7926073 | 484.5264261 | 415.6583062 | 352.7433857 | 1.56E-93 | -1.531038286 | 4.97E-96 | 432.3639193 | 0.346028246 | 92.8078978 |
| *Slc7a11* | 418.986253 | 534.9428119 | 437.0925105 | 499.4453818 | 177.5016328 | 170.3247917 | 158.8128434 | 148.4737575 | 8.92E-47 | -1.53133074 | 7.76E-49 | 215.7205622 | 0.345958108 | 46.04976978 |
| *Clip3* | 31.92276213 | 33.55729816 | 54.26656451 | 51.70040085 | 12.67868806 | 10.57917961 | 20.58685007 | 15.13108357 | 1.73E-05 | -1.5335536 | 1.55E-06 | 23.08912286 | 0.345425477 | 4.762828219 |
| *Cacna1b* | 24.93965792 | 24.67448394 | 11.83997771 | 26.33794005 | 6.339344029 | 6.347507766 | 12.74424052 | 4.728463616 | 0.003278948 | -1.539433739 | 0.000509448 | 12.08076171 | 0.344020457 | 2.484265414 |
| *Hspb1* | 181.5607096 | 172.7213876 | 163.7863583 | 224.3602301 | 72.90245633 | 62.4171597 | 51.95728828 | 68.08987607 | 4.71E-21 | -1.541213152 | 1.02E-22 | 96.23926033 | 0.343596405 | 20.32741852 |
| *C1s1* | 29.9275895 | 30.59636009 | 27.62661466 | 31.21533636 | 4.226229352 | 9.521261649 | 9.803261939 | 17.02246902 | 0.000289586 | -1.541662745 | 3.36E-05 | 17.20409329 | 0.343489346 | 3.538221724 |
| *Rpl39l* | 31.92276213 | 25.6614633 | 23.67995542 | 21.46054375 | 12.67868806 | 4.231671844 | 7.842609552 | 10.40261995 | 0.000737203 | -1.543633062 | 9.41E-05 | 15.2512421 | 0.343020556 | 3.132412615 |
| *Serpinh1* | 1654.995699 | 1489.351851 | 1514.530482 | 1482.728477 | 574.7671919 | 487.70018 | 564.6678877 | 479.4662107 | 1.20E-118 | -1.544250369 | 2.68E-121 | 548.4940225 | 0.342873814 | 117.9217781 |
| *Rbpms2* | 17.9565537 | 13.81771101 | 24.66662023 | 11.70575114 | 6.339344029 | 8.463343688 | 3.921304776 | 4.728463616 | 0.008209765 | -1.548467973 | 0.001475464 | 10.10900805 | 0.341872913 | 2.085669269 |
| *Ggt7* | 16.95896738 | 26.64844266 | 16.77330176 | 13.65670966 | 4.226229352 | 11.63709757 | 3.921304776 | 5.674156339 | 0.006678544 | -1.548557389 | 0.001152282 | 10.56533388 | 0.341851725 | 2.175318178 |
| *Rhou* | 72.82380111 | 60.20574082 | 72.02653108 | 69.25902755 | 31.69672014 | 14.81085145 | 25.48848104 | 21.75093263 | 4.69E-09 | -1.549132822 | 2.51E-10 | 40.02669041 | 0.341715401 | 8.328812299 |
| *Stk39* | 37.90828003 | 36.51823623 | 41.43992199 | 46.82300454 | 20.07458942 | 12.69501553 | 10.78358813 | 12.2940054 | 7.42E-06 | -1.549465106 | 6.24E-07 | 24.83780381 | 0.341636706 | 5.129728093 |
| *Gpx8* | 103.7489769 | 105.6067913 | 70.05320146 | 87.79313352 | 25.35737611 | 19.0425233 | 54.89826686 | 25.53370353 | 1.55E-08 | -1.55048321 | 8.81E-10 | 37.57298791 | 0.3413957 | 7.80933536 |
| *Kif26b* | 92.77552745 | 98.69793577 | 98.66648093 | 112.180115 | 43.31885086 | 33.85337475 | 29.40978582 | 31.20785986 | 1.97E-13 | -1.55075486 | 6.76E-15 | 60.66666399 | 0.341331423 | 12.70469848 |
| *Anxa8* | 37.90828003 | 47.37500917 | 48.34657565 | 49.74944233 | 22.1877041 | 10.57917961 | 18.62619768 | 11.34831268 | 2.79E-06 | -1.551111453 | 2.22E-07 | 26.83271781 | 0.341247066 | 5.55398706 |
| *Pard3* | 34.91552108 | 46.38802981 | 40.45325718 | 31.21533636 | 6.339344029 | 14.81085145 | 12.74424052 | 17.96816174 | 2.86E-05 | -1.551763776 | 2.66E-06 | 22.04655745 | 0.341092804 | 4.543476339 |
| *Chsy3* | 49.87931583 | 39.47917431 | 43.41325161 | 31.21533636 | 22.1877041 | 15.86876941 | 8.822935746 | 9.456927232 | 2.32E-05 | -1.55395655 | 2.11E-06 | 22.49071054 | 0.340574766 | 4.634510723 |
| *Sspn* | 20.94931265 | 17.76562844 | 22.69329061 | 12.6812304 | 5.282786691 | 7.405425727 | 6.862283358 | 5.674156339 | 0.004052673 | -1.554861242 | 0.000650693 | 11.62505653 | 0.340361263 | 2.392258449 |
| *Steap2* | 77.8117327 | 111.5286674 | 68.07987184 | 93.64600908 | 29.58360547 | 35.96921067 | 24.50815485 | 29.31647442 | 1.10E-10 | -1.559522213 | 5.00E-12 | 47.68842145 | 0.339263419 | 9.957918803 |
| *Slc6a12* | 243.4110613 | 194.4349335 | 214.1062636 | 187.2920182 | 87.69425906 | 57.12756989 | 81.3670741 | 58.63294884 | 1.40E-22 | -1.560349854 | 2.79E-24 | 103.3639942 | 0.339068848 | 21.85471753 |
| *Glipr2* | 108.7369085 | 89.81512155 | 118.3997771 | 106.3272395 | 40.14917885 | 32.79545679 | 49.0163097 | 21.75093263 | 3.42E-12 | -1.561552817 | 1.35E-13 | 54.78132865 | 0.338786239 | 11.46653472 |
| *Loxl2* | 1364.698081 | 1311.695566 | 1391.197381 | 1497.360666 | 492.3557196 | 459.1363951 | 495.0647279 | 437.8557308 | 1.83E-128 | -1.562994489 | 3.07E-131 | 594.1892226 | 0.338447863 | 127.736742 |
| *Mpdz* | 100.756218 | 106.5937706 | 99.65314573 | 103.4008017 | 44.3754082 | 19.0425233 | 39.21304776 | 34.99063076 | 7.79E-13 | -1.573747752 | 2.80E-14 | 57.87042994 | 0.335934588 | 12.10833235 |
| *Ptprf* | 41.8986253 | 46.38802981 | 52.29323489 | 35.11725341 | 9.509016043 | 29.62170291 | 14.70489291 | 5.674156339 | 6.75E-05 | -1.574124811 | 6.72E-06 | 20.2718436 | 0.3358468 | 4.170487252 |
| *Pgf* | 31.92276213 | 15.79166972 | 29.59994428 | 15.60766818 | 8.452458705 | 6.347507766 | 6.862283358 | 9.456927232 | 0.001649779 | -1.574156853 | 0.000231957 | 13.55270484 | 0.335839341 | 2.782574319 |
| *Adamts12* | 34.91552108 | 15.79166972 | 27.62661466 | 30.2398571 | 12.67868806 | 8.463343688 | 8.822935746 | 6.619849062 | 0.000529382 | -1.576126747 | 6.54E-05 | 15.93868332 | 0.33538109 | 3.276231091 |
| *Prrx1* | 165.5993286 | 182.5911812 | 208.1862748 | 146.3218892 | 73.95901367 | 72.99633931 | 49.99663589 | 39.71909437 | 2.67E-17 | -1.578165857 | 6.78E-19 | 78.82580446 | 0.334907396 | 16.57319434 |
| *Il1r1* | 30.92517582 | 25.6614633 | 38.47992756 | 53.65135937 | 17.96147475 | 14.81085145 | 6.862283358 | 10.40261995 | 7.95E-05 | -1.582502872 | 8.04E-06 | 19.92727884 | 0.333902113 | 4.099846319 |
| *Gjb3* | 54.86724741 | 33.55729816 | 34.53326832 | 34.14177415 | 13.7352454 | 10.57917961 | 16.6655453 | 11.34831268 | 1.39E-05 | -1.58408188 | 1.23E-06 | 23.5362612 | 0.333536862 | 4.857159268 |
| *Ptgs2* | 753.177669 | 827.0887017 | 670.9320703 | 737.4623215 | 274.7049079 | 224.2786077 | 262.72742 | 234.5317953 | 1.66E-72 | -1.585284238 | 7.95E-75 | 334.9695315 | 0.333259004 | 71.77993266 |
| *Nfatc4* | 61.85035163 | 59.21876146 | 44.39991642 | 59.50423494 | 11.62213072 | 34.91129271 | 12.74424052 | 16.07677629 | 1.39E-06 | -1.5856671 | 1.05E-07 | 28.27835779 | 0.333170576 | 5.855739902 |
| *Bgn* | 2786.258582 | 2722.089068 | 2811.994706 | 2703.053033 | 867.4335746 | 925.6782158 | 932.2902104 | 943.8013377 | 6.89E-255 | -1.58628584 | 3.30E-258 | 1178.221719 | 0.333027717 | 254.1620104 |
| *Ptk7* | 302.2686539 | 253.6536949 | 222.9862469 | 281.9135065 | 77.12868568 | 74.05425727 | 105.8752289 | 94.56927232 | 1.77E-28 | -1.588647192 | 2.78E-30 | 130.7724326 | 0.332483075 | 27.7509291 |
| *Mgll* | 34.91552108 | 31.58333945 | 29.59994428 | 32.19081562 | 17.96147475 | 11.63709757 | 6.862283358 | 6.619849062 | 9.89E-05 | -1.589185325 | 1.02E-05 | 19.46883945 | 0.33235908 | 4.004810605 |
| *Megf10* | 17.9565537 | 11.84375229 | 21.7066258 | 21.46054375 | 7.395901367 | 6.347507766 | 5.881957164 | 4.728463616 | 0.003836922 | -1.589487464 | 0.000610231 | 11.74450489 | 0.332289483 | 2.41601705 |
| *Evc* | 31.92276213 | 40.46615367 | 33.54660351 | 42.9210875 | 14.79180273 | 12.69501553 | 10.78358813 | 11.34831268 | 9.66E-06 | -1.589703199 | 8.31E-07 | 24.28505115 | 0.332239797 | 5.014934438 |
| *Alpl* | 99.75863166 | 125.3463784 | 108.533129 | 91.69505056 | 32.75327748 | 52.89589805 | 30.39011201 | 25.53370353 | 1.71E-12 | -1.595049755 | 6.41E-14 | 56.2407882 | 0.331010812 | 11.7682664 |
| *Tgfbr3* | 21.94689897 | 34.54427752 | 31.5732739 | 39.99464971 | 11.62213072 | 9.521261649 | 15.6852191 | 5.674156339 | 0.000132361 | -1.595192225 | 1.40E-05 | 18.8645133 | 0.330978125 | 3.878241207 |
| *Armcx4* | 52.87207478 | 31.58333945 | 43.41325161 | 43.89656676 | 21.13114676 | 16.92668738 | 9.803261939 | 9.456927232 | 7.75E-06 | -1.595788734 | 6.54E-07 | 24.7464168 | 0.330841304 | 5.110439757 |
| *Adam12* | 213.4834718 | 193.4479541 | 200.2929563 | 202.8996863 | 72.90245633 | 55.01173397 | 77.44576932 | 62.41571973 | 4.00E-27 | -1.596101708 | 6.55E-29 | 124.499163 | 0.33076954 | 26.39768995 |
| *Dzip1l* | 66.83828321 | 57.24480275 | 53.2798997 | 70.23450681 | 29.58360547 | 16.92668738 | 16.6655453 | 18.91385446 | 1.39E-08 | -1.597606195 | 7.87E-10 | 37.79214417 | 0.330424783 | 7.855677711 |
| *Ptges* | 79.80690533 | 66.12761696 | 59.19988856 | 79.01382016 | 22.1877041 | 25.39003106 | 13.72456672 | 32.15355259 | 3.19E-09 | -1.60209496 | 1.68E-10 | 40.80248009 | 0.329398306 | 8.496688617 |
| *Col5a2* | 2904.971354 | 2891.849518 | 2795.221405 | 2954.726682 | 965.693407 | 951.0682469 | 986.2081511 | 900.2994725 | 4.68E-259 | -1.602679344 | 1.87E-262 | 1197.763019 | 0.329264905 | 258.3297344 |
| *Gfpt2* | 42.89621161 | 28.62240137 | 65.11987741 | 32.19081562 | 6.339344029 | 22.21627718 | 14.70489291 | 12.2940054 | 6.18E-05 | -1.604971491 | 6.09E-06 | 20.460239 | 0.328742186 | 4.208689494 |
| *Ddr1* | 151.6331201 | 124.3593991 | 119.3864419 | 103.4008017 | 55.99753892 | 37.02712863 | 33.33109059 | 37.82770893 | 5.43E-15 | -1.60708592 | 1.64E-16 | 67.99172708 | 0.328260731 | 14.26553345 |
| *Lox* | 2496.95855 | 2892.836497 | 2982.687718 | 2873.761904 | 982.5983244 | 887.5931692 | 927.3885795 | 885.1683889 | 1.41E-166 | -1.61106252 | 1.46E-169 | 770.4164718 | 0.327357169 | 165.8519615 |
| *Pcolce* | 262.3652013 | 255.6276536 | 239.7595486 | 270.2077554 | 86.63770173 | 90.98094464 | 85.28837887 | 73.76403241 | 2.29E-35 | -1.613203033 | 2.74E-37 | 162.8158591 | 0.326871833 | 34.63986403 |
| *Sparc* | 3678.100749 | 3305.393869 | 3485.886771 | 3499.04411 | 1213.984381 | 1189.099788 | 1121.493166 | 1033.642146 | 6.77E-211 | -1.616899083 | 5.40E-214 | 974.7971003 | 0.326035489 | 210.1696845 |
| *Lrrc32* | 122.7031169 | 123.3724197 | 153.9197102 | 111.2046358 | 34.86639216 | 30.67962087 | 49.99663589 | 50.12171433 | 9.47E-16 | -1.617808656 | 2.71E-17 | 71.54650171 | 0.325829999 | 15.0234562 |
| *Shroom3* | 6.983104216 | 23.68750458 | 13.81330733 | 26.33794005 | 4.226229352 | 5.289589805 | 5.881957164 | 7.565541785 | 0.008281926 | -1.618683837 | 0.001489755 | 10.09124417 | 0.325632401 | 2.081868643 |
| *Antxr1* | 219.4689897 | 183.5781605 | 216.0795932 | 197.0468108 | 59.16721093 | 80.40176503 | 74.50479074 | 52.01309977 | 1.79E-25 | -1.619970485 | 3.16E-27 | 116.8082479 | 0.32534212 | 24.7466416 |
| *Osmr* | 73.82138743 | 100.6718945 | 69.06653665 | 79.98929943 | 24.30081878 | 27.50586698 | 33.33109059 | 19.85954719 | 6.08E-11 | -1.625156087 | 2.69E-12 | 48.90406875 | 0.324174814 | 10.21631476 |
| *Tbx15* | 69.83104216 | 70.0755344 | 57.22655894 | 49.74944233 | 13.7352454 | 32.79545679 | 20.58685007 | 13.23969812 | 7.99E-08 | -1.626173199 | 4.96E-09 | 34.2040175 | 0.323946349 | 7.097489127 |
| *Wwtr1* | 87.78759586 | 85.86720412 | 107.5464642 | 101.4498432 | 28.52704813 | 30.67962087 | 31.37043821 | 33.09924531 | 5.27E-14 | -1.627672213 | 1.72E-15 | 63.35921978 | 0.323609931 | 13.27779318 |
| *Rcn3* | 304.2638266 | 314.8464151 | 293.0394483 | 314.1043221 | 110.9385205 | 83.57551892 | 80.3867479 | 121.0486686 | 1.21E-36 | -1.628797857 | 1.39E-38 | 168.7434702 | 0.323357537 | 35.91697381 |
| *Srpx2* | 374.0948687 | 318.7943325 | 345.3326832 | 309.2269258 | 115.1647499 | 113.1972218 | 119.5997957 | 87.94942326 | 9.03E-41 | -1.630692501 | 9.16E-43 | 187.8946743 | 0.322933161 | 40.04413467 |
| *Fstl1* | 1147.224264 | 1188.323147 | 1226.424358 | 1179.354427 | 390.9262151 | 382.9663019 | 428.4025468 | 325.3182968 | 4.84E-115 | -1.63523941 | 1.16E-117 | 531.7776339 | 0.321916982 | 114.3151274 |
| *Bcat1* | 113.7248401 | 129.2942959 | 123.3331012 | 130.714221 | 47.54508022 | 44.43255436 | 31.37043821 | 36.8820162 | 1.05E-17 | -1.63897459 | 2.62E-19 | 80.70689472 | 0.321084607 | 16.97839545 |
| *Pappa* | 88.78518218 | 92.77605962 | 125.3064308 | 111.2046358 | 33.80983482 | 30.67962087 | 40.19337395 | 29.31647442 | 3.12E-14 | -1.641543485 | 9.95E-16 | 64.44000189 | 0.320513386 | 13.50529761 |
| *Serpina3n* | 50.87690215 | 79.94532797 | 58.21322375 | 48.77396306 | 9.509016043 | 16.92668738 | 32.3507644 | 17.02246902 | 4.87E-07 | -1.641704253 | 3.39E-08 | 30.46804819 | 0.320477671 | 6.312510745 |
| *Pcdh1* | 44.89138425 | 69.08855504 | 54.26656451 | 64.38163125 | 16.90491741 | 21.15835922 | 13.72456672 | 22.69662536 | 2.33E-08 | -1.642474625 | 1.36E-09 | 36.72919433 | 0.320306588 | 7.632269188 |
| *Ptx3* | 22.94448528 | 38.49219495 | 31.5732739 | 17.5586267 | 8.452458705 | 12.69501553 | 6.862283358 | 7.565541785 | 0.000319377 | -1.642918302 | 3.75E-05 | 16.99343454 | 0.320208098 | 3.495695998 |
| *Srgap1* | 22.94448528 | 16.77864908 | 26.63994985 | 18.53410596 | 7.395901367 | 5.289589805 | 3.921304776 | 10.40261995 | 0.001350258 | -1.645629521 | 0.000184131 | 13.98646962 | 0.319606904 | 2.8695834 |
| *Fosl1* | 37.90828003 | 41.45313302 | 33.54660351 | 30.2398571 | 17.96147475 | 11.63709757 | 13.72456672 | 2.83707817 | 5.69E-05 | -1.646492169 | 5.55E-06 | 20.63585379 | 0.319415855 | 4.2449953 |
| *Col12a1* | 2498.953723 | 2745.776573 | 2796.208069 | 2717.685222 | 877.999148 | 855.8556304 | 853.8641149 | 845.4492945 | 6.25E-234 | -1.648246969 | 3.99E-237 | 1081.2162 | 0.319027575 | 233.2038438 |
| *Ehd2* | 271.3434781 | 277.3411995 | 285.1461299 | 291.6682991 | 109.8819632 | 94.15469853 | 62.74087641 | 92.67788687 | 6.46E-34 | -1.650473044 | 8.00E-36 | 156.1130176 | 0.318535695 | 33.18958322 |
| *Sulf1* | 126.6934622 | 135.216172 | 120.3731067 | 117.0575114 | 49.65819489 | 23.27419514 | 45.09500492 | 40.6647871 | 2.13E-16 | -1.650936848 | 5.69E-18 | 74.62435492 | 0.318433307 | 15.67187665 |
| *Sod3* | 458.8897056 | 466.8412362 | 482.4790917 | 487.7396306 | 148.9745847 | 155.5139403 | 149.0095815 | 150.365143 | 1.67E-67 | -1.651024801 | 8.95E-70 | 311.7776305 | 0.318413895 | 66.77631371 |
| *Cyp1b1* | 579.59765 | 655.3542935 | 655.1454333 | 671.1297318 | 154.2573714 | 249.6686388 | 214.6914365 | 196.7040864 | 4.81E-57 | -1.651526261 | 3.26E-59 | 263.3078364 | 0.318303238 | 56.31813723 |
| *Ptprn* | 104.7465632 | 81.91928669 | 81.89317917 | 79.01382016 | 28.52704813 | 30.67962087 | 28.42945962 | 22.69662536 | 8.85E-13 | -1.658922933 | 3.25E-14 | 57.57694978 | 0.316675479 | 12.05308054 |
| *Fam20a* | 107.7393222 | 108.5677293 | 106.5597994 | 107.3027187 | 40.14917885 | 31.73753883 | 25.48848104 | 38.77340165 | 5.54E-16 | -1.660435182 | 1.54E-17 | 72.66010601 | 0.31634371 | 15.25623517 |
| *Itga1* | 50.87690215 | 38.49219495 | 42.4265868 | 34.14177415 | 8.452458705 | 16.92668738 | 17.64587149 | 9.456927232 | 3.10E-06 | -1.661439223 | 2.48E-07 | 26.62067639 | 0.316123628 | 5.508506773 |
| *Adamts2* | 310.2493445 | 348.4037133 | 297.9727724 | 298.496654 | 94.03360309 | 106.8497141 | 95.09164081 | 100.2434287 | 1.48E-43 | -1.663784196 | 1.39E-45 | 200.8089769 | 0.315610215 | 42.82862383 |
| *Postn* | 1070.410118 | 1263.333578 | 1199.784408 | 1152.041008 | 384.5868711 | 415.7617587 | 354.8780822 | 311.1329059 | 8.65E-100 | -1.678854441 | 2.56E-102 | 461.2609693 | 0.312330542 | 99.06294062 |
| *Mid2* | 15.96138107 | 27.63542202 | 18.74663138 | 32.19081562 | 6.339344029 | 15.86876941 | 0.980326194 | 6.619849062 | 0.002637413 | -1.680433851 | 0.000399455 | 12.53474165 | 0.311988801 | 2.578821909 |
| *Aebp1* | 2131.841959 | 2141.745206 | 2234.795793 | 2298.22914 | 686.7622698 | 715.1525416 | 677.4054 | 667.6590626 | 1.30E-229 | -1.681519262 | 9.34E-233 | 1061.115507 | 0.311754165 | 228.886119 |
| *Fbn1* | 1406.596706 | 1420.263296 | 1407.970683 | 1365.670966 | 426.8491646 | 423.1671844 | 428.4025468 | 464.3351271 | 1.57E-173 | -1.682968032 | 1.50E-176 | 802.5544357 | 0.311441255 | 172.8047216 |
| *Rbms3* | 52.87207478 | 66.12761696 | 51.30657008 | 56.57779715 | 19.01803209 | 19.0425233 | 13.72456672 | 18.91385446 | 4.90E-09 | -1.684318038 | 2.63E-10 | 39.93104706 | 0.311149959 | 8.309490499 |
| *Fkbp11* | 30.92517582 | 23.68750458 | 15.78663695 | 39.99464971 | 6.339344029 | 10.57917961 | 10.78358813 | 6.619849062 | 0.000296176 | -1.687194195 | 3.45E-05 | 17.15221704 | 0.310530268 | 3.52845029 |
| *Akap2* | 340.176934 | 344.4557958 | 321.6527278 | 340.4422622 | 119.3909792 | 98.38637037 | 90.19000984 | 109.7003559 | 8.34E-48 | -1.689996695 | 6.99E-50 | 220.5130772 | 0.309927635 | 47.07894261 |
| *Lrrc15* | 30.92517582 | 27.63542202 | 33.54660351 | 50.72492159 | 11.62213072 | 7.405425727 | 12.74424052 | 12.2940054 | 1.44E-05 | -1.691928547 | 1.28E-06 | 23.45835517 | 0.309512902 | 4.840750921 |
| *Fkbp10* | 433.9500477 | 436.2448761 | 416.3725495 | 398.9710179 | 147.9180273 | 128.0080733 | 116.6588171 | 128.6142104 | 4.69E-59 | -1.695164579 | 3.03E-61 | 272.6286896 | 0.308819428 | 58.3284279 |
| *Fbln2* | 1334.770492 | 1316.630463 | 1374.424079 | 1258.368247 | 419.4532632 | 387.1979737 | 451.9303754 | 369.7658548 | 3.23E-141 | -1.698542976 | 4.64E-144 | 653.1389986 | 0.308097103 | 140.4910514 |
| *Ccl2* | 194.5293317 | 154.9557592 | 189.4396434 | 216.556396 | 62.33688295 | 52.89589805 | 58.81957164 | 58.63294884 | 4.77E-26 | -1.698780844 | 8.07E-28 | 119.5170651 | 0.308046309 | 25.32169465 |
| *Loxl1* | 392.0514224 | 373.0781972 | 381.8392812 | 424.3334787 | 117.2778645 | 132.2397451 | 114.6981647 | 120.1029758 | 3.17E-56 | -1.699104409 | 2.18E-58 | 259.5259748 | 0.307977229 | 55.49887804 |
| *Pdk4* | 32.92034845 | 15.79166972 | 24.66662023 | 18.53410596 | 6.339344029 | 9.521261649 | 5.881957164 | 6.619849062 | 0.000644478 | -1.700002156 | 8.13E-05 | 15.52783103 | 0.307785643 | 3.190791775 |
| *Cnn1* | 105.7441496 | 79.94532797 | 115.4397827 | 123.8858662 | 28.52704813 | 28.56378495 | 34.31141679 | 38.77340165 | 1.01E-14 | -1.701998939 | 3.09E-16 | 66.74425985 | 0.307359943 | 13.9975612 |
| *Ptpn14* | 214.4810581 | 219.1094174 | 214.1062636 | 234.1150227 | 66.5631123 | 63.47507766 | 73.52446455 | 67.14418335 | 5.22E-34 | -1.702581448 | 6.42E-36 | 156.5502094 | 0.307235867 | 33.28230699 |
| *Prkg2* | 13.96620843 | 13.81771101 | 17.75996657 | 13.65670966 | 7.395901367 | 3.173753883 | 2.940978582 | 4.728463616 | 0.006202032 | -1.703609302 | 0.0010564 | 10.72600943 | 0.307017053 | 2.207466004 |
| *Cd248* | 27.93241687 | 25.6614633 | 25.65328504 | 26.33794005 | 11.62213072 | 6.347507766 | 7.842609552 | 6.619849062 | 0.000100678 | -1.708062483 | 1.04E-05 | 19.43035551 | 0.306070842 | 3.997064974 |
| *Rgs4* | 23.9420716 | 46.38802981 | 26.63994985 | 28.28889858 | 11.62213072 | 7.405425727 | 12.74424052 | 6.619849062 | 6.66E-05 | -1.708235832 | 6.60E-06 | 20.30700781 | 0.306034068 | 4.176722877 |
| *Sphk1* | 59.855179 | 43.42709174 | 62.15988298 | 67.30806903 | 24.30081878 | 21.15835922 | 11.76391433 | 14.18539085 | 1.50E-08 | -1.715050548 | 8.49E-10 | 37.64335493 | 0.304591894 | 7.823166129 |
| *Galnt18* | 40.90103898 | 36.51823623 | 50.31990527 | 38.04369119 | 8.452458705 | 14.81085145 | 21.56717627 | 5.674156339 | 5.47E-06 | -1.716425674 | 4.53E-07 | 25.4562574 | 0.304301706 | 5.262356029 |
| *Col6a2* | 686.3393858 | 658.3152316 | 599.892204 | 613.5764554 | 211.3114676 | 191.4831509 | 188.2226292 | 180.6273101 | 5.30E-85 | -1.730742868 | 1.86E-87 | 392.9752049 | 0.301296774 | 84.27549716 |
| *Cemip* | 214.4810581 | 214.1745206 | 212.132934 | 192.1694145 | 58.1106536 | 56.06965193 | 64.7015288 | 70.92695424 | 3.11E-32 | -1.733278435 | 4.17E-34 | 148.2569253 | 0.300767704 | 31.50756277 |
| *Svep1* | 66.83828321 | 88.82814219 | 82.87984398 | 91.69505056 | 29.58360547 | 28.56378495 | 21.56717627 | 19.85954719 | 9.67E-13 | -1.737247652 | 3.57E-14 | 57.39055195 | 0.299941353 | 12.0147464 |
| *Cxcl2* | 73.82138743 | 70.0755344 | 60.18655336 | 84.86669573 | 12.67868806 | 26.44794902 | 32.3507644 | 15.13108357 | 6.32E-10 | -1.737755961 | 3.14E-11 | 44.08871766 | 0.299835693 | 9.199325691 |
| *Tpbg* | 47.8841432 | 33.55729816 | 16.77330176 | 37.06821193 | 12.67868806 | 10.57917961 | 6.862283358 | 10.40261995 | 4.93E-05 | -1.74312327 | 4.73E-06 | 20.94456683 | 0.298722276 | 4.307534443 |
| *Ugt1a6a* | 56.86242005 | 106.5937706 | 95.7064865 | 87.79313352 | 25.35737611 | 28.56378495 | 25.48848104 | 23.64231808 | 4.21E-12 | -1.753800765 | 1.67E-13 | 54.35957619 | 0.296519571 | 11.37596005 |
| *Tbx3* | 16.95896738 | 26.64844266 | 41.43992199 | 31.21533636 | 2.113114676 | 17.98460534 | 9.803261939 | 4.728463616 | 0.000626765 | -1.75532308 | 7.88E-05 | 15.58771534 | 0.296206852 | 3.202895571 |
| *Pth1r* | 83.7972506 | 43.42709174 | 43.41325161 | 51.70040085 | 11.62213072 | 26.44794902 | 13.72456672 | 14.18539085 | 2.56E-07 | -1.757579015 | 1.72E-08 | 31.78962623 | 0.295744037 | 6.591458327 |
| *H2-M2* | 18.95414002 | 15.79166972 | 18.74663138 | 17.5586267 | 2.113114676 | 3.173753883 | 8.822935746 | 6.619849062 | 0.002120667 | -1.75828149 | 0.000310879 | 13.00368335 | 0.295600069 | 2.673527612 |
| *Robo1* | 37.90828003 | 36.51823623 | 32.55993871 | 30.2398571 | 15.84836007 | 10.57917961 | 8.822935746 | 5.674156339 | 7.51E-06 | -1.758703002 | 6.32E-07 | 24.81305145 | 0.295513716 | 5.124563614 |
| *Mmp23* | 64.84311058 | 55.27084403 | 53.2798997 | 58.52875568 | 11.62213072 | 20.10044126 | 18.62619768 | 17.96816174 | 8.89E-10 | -1.760362372 | 4.45E-11 | 43.40576974 | 0.295174015 | 9.051252336 |
| *Slit3* | 64.84311058 | 35.53125688 | 50.31990527 | 61.45519346 | 19.01803209 | 11.63709757 | 19.60652388 | 12.2940054 | 5.68E-08 | -1.762742521 | 3.47E-09 | 34.90227071 | 0.29468744 | 7.245959519 |
| *Pxdn* | 205.5027812 | 260.5625504 | 200.2929563 | 215.5809167 | 72.90245633 | 66.64883154 | 73.52446455 | 47.28463616 | 3.32E-29 | -1.764305137 | 4.96E-31 | 134.1917107 | 0.29436843 | 28.47852458 |
| *Col6a3* | 1326.789801 | 1350.187761 | 1415.864001 | 1375.425758 | 419.4532632 | 436.9201179 | 414.67798 | 328.1553749 | 7.86E-136 | -1.776485334 | 1.19E-138 | 628.2640311 | 0.291893637 | 135.1045964 |
| *Col1a2* | 7362.187017 | 7239.493589 | 7322.039549 | 7524.847022 | 2323.369587 | 2160.268476 | 2163.57991 | 1942.452853 | 0 | -1.778360683 | 0 | 1575.925917 | 0.291514453 | Inf |
| *Cav1* | 846.9507828 | 799.4532797 | 759.7319031 | 788.1872431 | 201.8024516 | 212.6415102 | 242.1405699 | 270.4681188 | 1.39E-95 | -1.780707316 | 4.33E-98 | 441.8280936 | 0.291040672 | 94.85670242 |
| *Pdzrn3* | 69.83104216 | 51.3229266 | 44.39991642 | 58.52875568 | 11.62213072 | 9.521261649 | 20.58685007 | 22.69662536 | 1.92E-08 | -1.784234107 | 1.10E-09 | 37.13668729 | 0.290330067 | 7.717037401 |

| | | | | | | | | | | | | | | |
|---|---|---|---|---|---|---|---|---|---|---|---|---|---|---|
| *Fmod* | 104.7465632 | 127.3203371 | 140.1064029 | 140.4690136 | 33.80983482 | 43.3746364 | 41.17370015 | 30.26216714 | 1.26E-19 | -1.789411768 | 2.85E-21 | 89.64384577 | 0.289289974 | 18.90049101 |
| *Bicc1* | 79.80690533 | 59.21876146 | 80.90651436 | 70.23450681 | 24.30081878 | 17.98460534 | 21.56717627 | 19.85954719 | 3.73E-12 | -1.793938143 | 1.48E-13 | 54.59815638 | 0.288383766 | 11.42781125 |
| *Pak3* | 11.9710358 | 14.80469037 | 8.879983283 | 16.58314744 | 2.113114676 | 4.231671844 | 2.940978582 | 5.674156339 | 0.008445519 | -1.794998039 | 0.001521205 | 10.05275247 | 0.288171978 | 2.073373648 |
| *Sfrp2* | 33.91793477 | 23.68750458 | 23.67995542 | 33.16629488 | 5.282786691 | 4.231671844 | 6.862283358 | 16.07677629 | 0.000102591 | -1.797783044 | 1.06E-05 | 19.39147577 | 0.287616222 | 3.988892347 |
| *Adamts4* | 84.79483691 | 87.84116283 | 91.75982726 | 73.1609446 | 24.30081878 | 32.79545679 | 20.58685007 | 19.85954719 | 7.95E-14 | -1.798207158 | 2.65E-15 | 62.50896405 | 0.287531683 | 13.0997683 |
| *Col6a1* | 1045.47046 | 1067.911665 | 1070.531318 | 1016.44939 | 316.9672014 | 287.7536854 | 255.8651366 | 338.5579949 | 5.04E-126 | -1.807862974 | 8.84E-129 | 582.8843471 | 0.285613688 | 125.2979657 |
| *Wnt11* | 38.90586635 | 41.45313302 | 27.62661466 | 25.36246079 | 6.339344029 | 3.173753883 | 19.60652388 | 8.511234509 | 6.31E-05 | -1.810945115 | 6.24E-06 | 20.41458364 | 0.28500416 | 4.200086821 |
| *Adora2a* | 21.94689897 | 13.81771101 | 16.77330176 | 18.53410596 | 3.169672014 | 9.521261649 | 2.940978582 | 4.728463616 | 0.001756843 | -1.812492505 | 0.000250096 | 13.4114283 | 0.284698638 | 2.755267018 |
| *Fgfr2* | 14.96379475 | 22.70052523 | 22.69329061 | 10.73027187 | 3.169672014 | 8.463343688 | 5.881957164 | 2.83707817 | 0.002290632 | -1.812556706 | 0.000340531 | 12.83315299 | 0.284685969 | 2.640044727 |
| *Tmem45a* | 48.88172951 | 46.38802981 | 41.43992199 | 44.87204602 | 12.67868806 | 21.15835922 | 7.842609552 | 10.40261995 | 1.08E-07 | -1.813489226 | 6.88E-09 | 33.56984819 | 0.284502015 | 6.965315229 |
| *Shox2* | 28.93000318 | 38.49219495 | 20.71996099 | 15.60766818 | 11.62213072 | 7.405425727 | 8.822935746 | 1.891385446 | 0.000402975 | -1.815858699 | 4.83E-05 | 16.51398368 | 0.284035134 | 3.394722416 |
| *Grb10* | 165.5993286 | 126.3333578 | 178.5863305 | 181.4391426 | 25.35737611 | 64.53299562 | 47.05565731 | 47.28463616 | 2.66E-19 | -1.822636909 | 6.08E-21 | 88.1472959 | 0.282703782 | 18.57498926 |
| *Dpysl3* | 478.841432 | 428.3490412 | 449.919153 | 440.9166261 | 134.1827819 | 124.8343194 | 134.3046886 | 115.3745122 | 8.45E-71 | -1.822776599 | 4.18E-73 | 327.0671647 | 0.282676411 | 70.07304264 |
| *Plac8* | 19.95172633 | 46.38802981 | 26.63994985 | 35.11725341 | 7.395901367 | 6.347507766 | 10.78358813 | 11.34831268 | 2.46E-05 | -1.827152924 | 2.25E-06 | 22.37246949 | 0.281820229 | 4.609296218 |
| *Csmd1* | 33.91793477 | 43.42709174 | 36.50659794 | 53.65135937 | 11.62213072 | 4.231671844 | 20.58685007 | 10.40261995 | 1.89E-06 | -1.829585236 | 1.46E-07 | 27.63592518 | 0.281345494 | 5.72245663 |
| *Evc2* | 60.85276531 | 41.45313302 | 32.55993871 | 37.06821193 | 9.509016043 | 10.57917961 | 14.70489291 | 13.23969812 | 3.51E-07 | -1.832936109 | 2.40E-08 | 31.14281941 | 0.280692786 | 6.454983613 |
| *Clcf1* | 55.86483373 | 48.36198853 | 52.29323489 | 52.67588011 | 15.84836007 | 16.92668738 | 7.842609552 | 17.96816174 | 3.23E-09 | -1.838381428 | 1.71E-10 | 40.77372098 | 0.279635334 | 8.490952812 |
| *Thbs3* | 16.95896738 | 13.81771101 | 15.78663695 | 26.33794005 | 8.452458705 | 7.405425727 | 2.940978582 | 1.891385446 | 0.001890769 | -1.840385288 | 0.00027218 | 13.25275054 | 0.279247198 | 2.723361459 |
| *Six4* | 27.93241687 | 20.72656651 | 29.59994428 | 34.14177415 | 9.509016043 | 7.405425727 | 7.842609552 | 6.619849062 | 2.21E-05 | -1.845230478 | 2.00E-06 | 22.59083965 | 0.278310939 | 4.655238034 |
| *Lurap1l* | 41.8986253 | 37.50521559 | 22.69329061 | 36.09273267 | 13.7352454 | 9.521261649 | 7.842609552 | 7.565541785 | 4.12E-06 | -1.845927968 | 3.35E-07 | 26.03784738 | 0.278176419 | 5.385198166 |
| *Gria3* | 26.93483055 | 29.60938073 | 11.83997771 | 25.36246079 | 4.226229352 | 4.231671844 | 5.881957164 | 11.34831268 | 0.000405858 | -1.850839217 | 4.88E-05 | 16.49541692 | 0.277231056 | 3.391625476 |
| *Tead2* | 23.9420716 | 32.5703188 | 27.62661466 | 24.38698153 | 9.509016043 | 2.115835922 | 6.862283358 | 11.34831268 | 5.90E-05 | -1.851179656 | 5.78E-06 | 20.56042926 | 0.277165644 | 4.228949109 |
| *Lif* | 197.5220907 | 214.1745206 | 216.0795932 | 199.9732486 | 55.99753892 | 55.01173397 | 61.76055022 | 55.79587067 | 2.50E-36 | -1.855788685 | 2.89E-38 | 167.2892378 | 0.276281586 | 35.602339 |
| *Lmcd1* | 12.96862212 | 11.84375229 | 12.82664252 | 9.754792613 | 1.056557338 | 3.173753883 | 4.90163097 | 3.782770893 | 0.009472783 | -1.861699465 | 0.001732706 | 9.813127161 | 0.275151965 | 2.023522396 |
| *Scx* | 16.95896738 | 15.79166972 | 9.866648093 | 16.58314744 | 7.395901367 | 3.173753883 | 5.881957164 | 0 | 0.006089149 | -1.864512588 | 0.001033178 | 10.76714347 | 0.274615966 | 2.21544337 |
| *Ptger3* | 11.9710358 | 26.64844266 | 33.54660351 | 26.33794005 | 3.169672014 | 5.289589805 | 11.76391433 | 6.619849062 | 0.000331331 | -1.864854114 | 3.90E-05 | 16.91980983 | 0.274550965 | 3.479738373 |
| *Il11* | 290.2976181 | 258.5885917 | 226.9329061 | 253.6246079 | 51.77130957 | 70.88050338 | 87.24903126 | 71.87264696 | 7.11E-37 | -1.865055994 | 8.11E-39 | 169.816704 | 0.274512549 | 36.14835802 |
| *Sp7* | 42.89621161 | 38.49219495 | 34.53326832 | 48.77396306 | 11.62213072 | 9.521261649 | 5.881957164 | 17.96816174 | 3.81E-07 | -1.865365632 | 2.62E-08 | 30.97119988 | 0.274453638 | 6.419130588 |
| *Egfr* | 104.7465632 | 113.5026261 | 102.6131402 | 119.9839491 | 20.07458942 | 37.02712863 | 31.37043821 | 32.15355259 | 9.87E-19 | -1.867634464 | 2.30E-20 | 85.51369555 | 0.274022362 | 18.00568404 |
| *Spry4* | 28.93000318 | 19.73958715 | 28.61327947 | 22.43602301 | 1.056557338 | 9.521261649 | 8.822935746 | 7.565541785 | 0.000130848 | -1.878417203 | 1.39E-05 | 18.88787901 | 0.271981946 | 3.883233705 |
| *Lpin3* | 13.96620843 | 16.77864908 | 18.74663138 | 17.5586267 | 3.169672014 | 9.521261649 | 0.980326194 | 4.728463616 | 0.002230047 | -1.878668421 | 0.000330634 | 12.88834619 | 0.27193459 | 2.651686047 |
| *Ebf1* | 38.90586635 | 52.30990596 | 34.53326832 | 52.67588011 | 15.84836007 | 12.69501553 | 10.78358813 | 9.456927232 | 5.50E-08 | -1.878886881 | 3.35E-09 | 34.97027394 | 0.271893416 | 7.259421129 |
| *Casc4* | 35.9131074 | 50.33594724 | 21.7066258 | 30.2398571 | 16.90491741 | 4.231671844 | 10.78358813 | 5.674156339 | 2.46E-05 | -1.884357213 | 2.25E-06 | 22.36900814 | 0.270864417 | 4.60889295 |
| *Flnc* | 160.611397 | 155.9427385 | 144.0530622 | 147.2973685 | 47.54508022 | 42.31671844 | 38.23272156 | 36.8820162 | 2.87E-27 | -1.88551094 | 4.65E-29 | 125.1812731 | 0.270647893 | 26.54271861 |
| *Tmem200a* | 11.9710358 | 13.81771101 | 20.71996099 | 12.6812304 | 2.113114676 | 2.115835922 | 5.881957164 | 5.674156339 | 0.003486174 | -1.887180743 | 0.000546098 | 11.95128281 | 0.270334821 | 2.457650932 |
| *Col5a1* | 1774.706057 | 1812.094101 | 1769.090003 | 1943.154688 | 435.3016233 | 580.7969606 | 533.2974495 | 421.7789545 | 1.30E-55 | -1.889211921 | 9.14E-58 | 256.6684226 | 0.269954483 | 54.88596678 |
| *Thbs2* | 1319.806697 | 1247.541908 | 1362.584102 | 1358.842611 | 359.229495 | 329.0124859 | 358.799387 | 377.3313965 | 1.01E-182 | -1.891211023 | 8.89E-186 | 844.9994018 | 0.269580674 | 181.9949042 |
| *Kif1a* | 38.90586635 | 31.58333945 | 31.5732739 | 37.06821193 | 14.79180273 | 11.63709757 | 0.980326194 | 10.40261995 | 8.88E-06 | -1.891324087 | 7.59E-07 | 24.45867165 | 0.269559547 | 5.051649666 |
| *Snai1* | 103.7489769 | 77.97136926 | 101.6264754 | 76.08738238 | 29.58360547 | 21.15835922 | 23.52782865 | 22.69662536 | 2.49E-15 | -1.89184509 | 7.35E-17 | 69.576173 | 0.269462218 | 14.60402037 |
| *Pcolce2* | 60.85276531 | 55.27084403 | 65.11987741 | 80.96477869 | 19.01803209 | 16.92668738 | 19.60652388 | 15.13108357 | 8.66E-12 | -1.893476233 | 3.53E-13 | 52.89100263 | 0.269157731 | 11.06253359 |
| *Serping1* | 339.1793477 | 362.2214243 | 351.2526721 | 396.0445801 | 100.3729471 | 103.6759602 | 99.01294559 | 86.05803781 | 4.72E-60 | -1.899227772 | 2.94E-62 | 277.2799435 | 0.268086825 | 59.32596314 |
| *Dclk1* | 86.79000955 | 99.68491513 | 141.0930677 | 101.4498432 | 34.86639216 | 20.10044126 | 42.15402634 | 17.02246902 | 5.93E-14 | -1.911886987 | 1.96E-15 | 63.10935718 | 0.265744735 | 13.22693013 |
| *Tcf7l1* | 24.93965792 | 18.7526078 | 27.62661466 | 11.70575114 | 3.169672014 | 2.115835922 | 9.803261939 | 6.619849062 | 0.000712848 | -1.916296864 | 9.08E-05 | 15.31943242 | 0.264933675 | 3.147002893 |
| *Lgr6* | 77.8117327 | 62.17969953 | 67.09320703 | 90.7195713 | 22.1877041 | 13.75293349 | 16.6655453 | 25.53370353 | 7.90E-13 | -1.925233052 | 2.85E-14 | 57.83549617 | 0.263297723 | 12.10254556 |
| *Tspan6* | 53.8696611 | 58.2317821 | 50.31990527 | 33.16629488 | 11.62213072 | 16.92668738 | 8.822935746 | 14.18539085 | 1.21E-08 | -1.925913573 | 6.76E-10 | 38.08930914 | 0.263173554 | 7.918125952 |
| *C1qtnf5* | 37.90828003 | 24.67448394 | 15.78663695 | 22.43602301 | 9.509016043 | 3.173753883 | 12.74424052 | 0.945692723 | 0.000487843 | -1.938646076 | 5.97E-05 | 16.11060271 | 0.260861135 | 3.311719955 |
| *Actg2* | 374.0948687 | 319.7813119 | 368.0259739 | 357.0254096 | 76.07212834 | 107.907632 | 95.09164081 | 88.89511598 | 6.01E-58 | -1.947416617 | 3.99E-60 | 267.496923 | 0.259280099 | 57.22084714 |
| *Kalm* | 68.83345585 | 49.34896788 | 59.19988856 | 51.70040085 | 14.79180273 | 12.69501553 | 13.72456672 | 17.96816174 | 6.26E-11 | -1.947946962 | 2.77E-12 | 48.84154898 | 0.259184804 | 10.20325455 |
| *Crlf1* | 225.4545076 | 197.3958715 | 212.132934 | 232.1640642 | 62.33688295 | 50.78006213 | 63.72120261 | 47.28463616 | 3.31E-38 | -1.953614667 | 3.64E-40 | 175.9876811 | 0.25816858 | 37.48057547 |
| *Col1a1* | 7613.578768 | 7593.819178 | 7898.251798 | 8369.612062 | 2090.926972 | 2358.099135 | 2034.176852 | 1638.885489 | 5.86E-64 | -1.954622661 | 3.46E-66 | 295.3096603 | 0.257988263 | 63.2317891 |
| *Osbp2* | 36.91069371 | 29.60938073 | 36.50659794 | 37.06821193 | 8.452458705 | 4.231671844 | 12.74424052 | 10.40261995 | 6.36E-07 | -1.95542387 | 4.57E-08 | 29.89036971 | 0.257845028 | 6.196379491 |
| *P4ha3* | 83.7972506 | 48.36198853 | 85.8398384 | 59.50423494 | 17.96147475 | 13.75293349 | 10.78358813 | 28.3707817 | 2.51E-10 | -1.96155461 | 1.18E-11 | 45.99789701 | 0.256751639 | 9.599795957 |
| *Eda2r* | 26.93483055 | 32.5703188 | 31.5732739 | 41.94560824 | 9.509016043 | 4.231671844 | 8.822935746 | 11.34831268 | 1.49E-06 | -1.962821009 | 1.13E-07 | 28.13307529 | 0.256526361 | 5.825435979 |
| *Itga11* | 260.3700286 | 316.8203738 | 302.9060964 | 361.9028059 | 84.52458705 | 86.7492728 | 90.19000984 | 57.68725611 | 1.26E-42 | -1.964661496 | 1.19E-44 | 196.5301985 | 0.256199312 | 41.90071657 |
| *Setbp1* | 9.975863166 | 13.81771101 | 16.77330176 | 18.53410596 | 3.169672014 | 4.231671844 | 3.921304776 | 3.782770893 | 0.001916486 | -1.9682006 | 0.000276341 | 13.2243118 | 0.255571594 | 2.71749442 |
| *Mdga1* | 35.9131074 | 41.45313302 | 53.2798997 | 39.99464971 | 11.62213072 | 11.63709757 | 12.74424052 | 7.565541785 | 2.02E-08 | -1.974753686 | 1.17E-09 | 37.0218766 | 0.254413356 | 7.693875777 |
| *Atoh8* | 59.855179 | 51.3229266 | 53.2798997 | 76.08738238 | 11.62213072 | 17.98460534 | 16.6655453 | 14.18539085 | 1.89E-11 | -1.99251927 | 7.93E-13 | 51.30045721 | 0.251299678 | 10.72251496 |
| *Wisp2* | 273.3386508 | 258.5885917 | 241.7328783 | 200.9487278 | 59.16721093 | 60.30132377 | 58.81957164 | 66.19849062 | 4.34E-41 | -1.993334766 | 4.29E-43 | 189.4022472 | 0.251157669 | 40.36283534 |
| *Sorcs2* | 65.8406969 | 61.19272018 | 77.94651993 | 55.60231789 | 6.339344029 | 17.98460534 | 21.56717627 | 18.91385446 | 1.83E-11 | -1.997914202 | 7.64E-13 | 51.37367172 | 0.250361703 | 10.73788235 |
| *Prg4* | 121.7055306 | 141.1380481 | 110.5064586 | 129.7387418 | 24.30081878 | 34.91129271 | 41.17370015 | 24.5880108 | 1.91E-22 | -2.009271166 | 3.82E-24 | 102.7416214 | 0.24839858 | 21.72001757 |
| *Gli3* | 30.92517582 | 37.50521559 | 30.58660909 | 39.01917045 | 5.282786691 | 10.57917961 | 7.842609552 | 10.40261995 | 3.10E-07 | -2.012841049 | 2.10E-08 | 31.39826857 | 0.247784689 | 6.508562867 |
| *Dkk3* | 236.427957 | 246.7448394 | 246.6662023 | 201.9242071 | 79.24180036 | 57.12756989 | 54.89826686 | 40.6647871 | 6.42E-36 | -2.01370622 | 7.59E-38 | 165.3718885 | 0.24763614 | 35.19241168 |
| *Hbegf* | 321.222794 | 297.0807867 | 310.7994149 | 289.7173406 | 76.07212834 | 70.88050338 | 80.3867479 | 71.87264696 | 7.17E-59 | -2.025809966 | 4.70E-61 | 271.7584606 | 0.245567243 | 58.14437116 |
| *Tnn* | 135.6717391 | 181.6042018 | 133.1997492 | 163.8805159 | 42.26229352 | 47.60630824 | 36.27206918 | 25.53370353 | 4.03E-25 | -2.026719519 | 7.21E-27 | 115.1728891 | 0.245412473 | 24.39437271 |
| *Gpx7* | 29.9275895 | 23.68750458 | 30.58660909 | 18.53410596 | 4.226229352 | 9.521261649 | 0.980326194 | 10.40261995 | 6.66E-05 | -2.028961107 | 6.61E-06 | 20.30398284 | 0.24503146 | 4.176386079 |
| *Igsf10* | 42.89621161 | 32.5703188 | 51.30657008 | 45.84752528 | 14.79180273 | 3.173753883 | 12.74424052 | 11.34831268 | 4.50E-08 | -2.031530786 | 2.72E-09 | 35.37402117 | 0.244595406 | 7.346602373 |
| *Jam2* | 13.96620843 | 16.77864908 | 8.879983283 | 5.852875568 | 1.056557338 | 2.115835922 | 4.90163097 | 2.83707817 | 0.009169417 | -2.043660258 | 0.00167136 | 9.879438137 | 0.242547589 | 2.037658257 |
| *Cacna2d1* | 50.87690215 | 60.20574082 | 52.29323489 | 45.84752528 | 16.90491741 | 16.92668738 | 5.881957164 | 10.40261995 | 2.58E-10 | -2.075311988 | 1.22E-11 | 45.94164254 | 0.237284211 | 9.588060668 |
| *Kif21a* | 21.94689897 | 21.71354587 | 23.67995542 | 17.5586267 | 6.339344029 | 3.173753883 | 4.90163097 | 5.674156339 | 5.61E-05 | -2.075827043 | 5.47E-06 | 20.66394475 | 0.237199514 | 4.250656363 |
| *Sox9* | 31.92276213 | 17.76562844 | 27.62661466 | 24.38698153 | 6.339344029 | 3.173753883 | 7.842609552 | 6.619849062 | 1.47E-05 | -2.075962377 | 1.30E-06 | 23.42354257 | 0.237177264 | 4.833283743 |
| *Masp1* | 24.93965792 | 21.71354587 | 25.65328504 | 16.58314744 | 2.113114676 | 3.173753883 | 9.803261939 | 5.674156339 | 8.28E-05 | -2.079524062 | 8.40E-06 | 19.84482861 | 0.236592449 | 4.082142079 |
| *Clec4e* | 1213.064961 | 1478.495078 | 1331.010828 | 1387.13151 | 355.0032656 | 289.8695213 | 323.507644 | 311.1329059 | 3.72E-165 | -2.080390556 | 4.16E-168 | 763.7272035 | 0.236450393 | 164.4297012 |
| *Casp12* | 18.95414002 | 22.70052523 | 14.79997214 | 24.38698153 | 3.169672014 | 3.173753883 | 0.980326194 | 11.34831268 | 0.000403613 | -2.092636267 | 4.84E-05 | 16.50971799 | 0.234451877 | 3.394034613 |
| *Steap1* | 27.93241687 | 15.79166972 | 12.82664252 | 25.36246079 | 4.226229352 | 3.173753883 | 7.842609552 | 3.782770893 | 0.00018079 | -2.099986392 | 1.99E-05 | 18.20251984 | 0.233260448 | 3.742826728 |
| *Gspt2* | 28.93000318 | 46.38802981 | 29.59994428 | 30.2398571 | 3.169672014 | 4.231671844 | 11.76391433 | 11.34831268 | 9.77E-07 | -2.125844527 | 7.21E-08 | 29.00856218 | 0.229116851 | 6.010205416 |
| *Slc6a17* | 46.88655688 | 39.47917431 | 49.33324046 | 43.89656676 | 13.7352454 | 12.69501553 | 7.842609552 | 5.674156339 | 5.44E-10 | -2.1843166 | 2.67E-11 | 44.4049015 | 0.220016465 | 9.264557281 |
| *Vcan* | 364.1190056 | 369.1302798 | 369.0126387 | 369.70664 | 79.24180036 | 102.6180422 | 59.79989783 | 81.32957419 | 2.71E-69 | -2.192462504 | 1.41E-71 | 320.0600817 | 0.218777685 | 68.56732148 |
| *Hhipl1* | 11.9710358 | 15.79166972 | 17.75996657 | 25.36246079 | 5.282786691 | 1.057917961 | 3.921304776 | 4.728463616 | 0.000228226 | -2.234257238 | 2.59E-05 | 17.69643643 | 0.212530642 | 3.641635059 |
| *Ror2* | 27.93241687 | 35.53125688 | 12.82664252 | 18.53410596 | 1.056557338 | 6.347507766 | 5.881957164 | 6.619849062 | 4.67E-05 | -2.24042506 | 4.46E-06 | 21.05599996 | 0.211623968 | 4.330929069 |
| *A4galt* | 23.9420716 | 13.81771101 | 6.906653665 | 31.21533636 | 2.113114676 | 1.057917961 | 8.822935746 | 3.782770893 | 0.000888183 | -2.248011223 | 0.000115872 | 14.85871314 | 0.210514101 | 3.051497455 |
| *Col3a1* | 1673.949839 | 1638.385734 | 1608.263639 | 1577.349966 | 361.3426096 | 340.6495834 | 324.4879702 | 328.1553749 | 7.72E-297 | -2.263454807 | 2.47E-300 | 1372.068155 | 0.208272634 | 296.1121603 |
| *Twist2* | 9.975863166 | 6.908855504 | 21.7066258 | 6.828354829 | 3.169672014 | 5.289589805 | 0 | 0.945692723 | 0.007720053 | -2.304183673 | 0.001367728 | 10.24880945 | 0.202475089 | 2.112379692 |
| *Olfm2* | 19.95172633 | 9.869793577 | 3.946659237 | 11.70575114 | 5.282786691 | 2.115835922 | 1.960652388 | 0 | 0.007498965 | -2.308016872 | 0.001323769 | 10.30908454 | 0.201937833 | 2.124998692 |
| *Ebf3* | 7.980690533 | 13.81771101 | 9.866648093 | 13.65670966 | 0 | 3.173753883 | 0.980326194 | 4.728463616 | 0.003593746 | -2.331406938 | 0.000566105 | 11.88425506 | 0.198690261 | 2.444452645 |
| *Met* | 84.79483691 | 71.06251375 | 62.15988298 | 59.50423494 | 29.58360547 | 9.521261649 | 7.842609552 | 8.511234509 | 1.90E-12 | -2.338556354 | 7.22E-14 | 56.00793583 | 0.197708067 | 11.72141785 |
| *Marco* | 15.96138107 | 19.73958715 | 19.73329619 | 21.46054375 | 4.226229352 | 2.115835922 | 4.90163097 | 3.782770893 | 2.76E-05 | -2.349542634 | 2.55E-06 | 22.12573081 | 0.196208217 | 4.558771335 |
| *Rgs17* | 26.93483055 | 17.76562844 | 8.879983283 | 18.53410596 | 2.113114676 | 4.231671844 | 2.940978582 | 4.728463616 | 0.000143432 | -2.356403897 | 1.54E-05 | 18.69083738 | 0.195277293 | 3.843353983 |
| *Lama5* | 23.9420716 | 26.64844266 | 40.45325718 | 30.2398571 | 9.509016043 | 5.289589805 | 6.862283358 | 1.891385446 | 2.54E-07 | -2.38017629 | 1.70E-08 | 31.81144608 | 0.192085924 | 6.595819953 |
| *Gzme* | 63.84552426 | 45.40105045 | 66.10654222 | 83.89121647 | 15.84836007 | 8.463343688 | 5.881957164 | 18.91385446 | 3.36E-13 | -2.395768005 | 1.18E-14 | 59.5735148 | 0.190021161 | 12.47350453 |
| *Fam26e* | 19.95172633 | 9.869793577 | 8.879983283 | 4.877396306 | 2.113114676 | 5.289589805 | 0.980326194 | 0 | 0.006710302 | -2.412704148 | 0.001159369 | 10.55400079 | 0.1878035 | 2.173257954 |
| *Ccl7* | 98.76104535 | 89.81512155 | 112.4797883 | 115.1065528 | 15.84836007 | 17.98460534 | 21.56717627 | 21.75093263 | 1.05E-25 | -2.425733021 | 1.82E-27 | 117.9047526 | 0.186115095 | 24.98080448 |
| *Ak5* | 11.9710358 | 8.882814219 | 11.83997771 | 11.70575114 | 1.056557338 | 2.115835922 | 1.960652388 | 2.83707817 | 0.001524343 | -2.465735543 | 0.000210791 | 13.7323582 | 0.181025452 | 2.816917364 |
| *Tfeb* | 702.3007669 | 579.356883 | 590.0255559 | 663.3258977 | 92.97704575 | 135.413499 | 128.4227314 | 93.62357959 | 5.84E-110 | -2.494279511 | 1.54E-112 | 508.2310119 | 0.177479031 | 109.2336391 |
| *Plbd1* | 45.88897056 | 50.33594724 | 37.49326275 | 61.45519346 | 6.339344029 | 14.81085145 | 5.881957164 | 7.565541785 | 4.93E-12 | -2.506008062 | 1.98E-13 | 54.0204433 | 0.176042045 | 11.30707954 |
| *Dio2* | 35.9131074 | 35.53125688 | 47.35991084 | 43.89656676 | 7.395901367 | 16.92668738 | 1.960652388 | 2.83707817 | 7.24E-09 | -2.510986081 | 3.95E-10 | 39.13673349 | 0.175435658 | 8.140528329 |
| *S1pr3* | 14.96379475 | 11.84375229 | 13.81330733 | 11.70575114 | 1.056557338 | 5.289589805 | 0.980326194 | 1.891385446 | 0.000545935 | -2.521641531 | 6.78E-05 | 15.87186994 | 0.1741447 | 3.262859439 |
| *Irg1* | 265.3579602 | 279.3151582 | 262.4528393 | 308.2514466 | 61.28032561 | 49.72214416 | 64.7015288 | 17.96816174 | 1.11E-10 | -2.528469153 | 5.06E-12 | 47.66548498 | 0.173322499 | 9.95360166 |
| *Nbea* | 28.93000318 | 27.63542202 | 21.7066258 | 33.16629488 | 1.056557338 | 11.63709757 | 4.90163097 | 1.891385446 | 7.41E-07 | -2.53270451 | 5.37E-08 | 29.57868747 | 0.172814418 | 6.130397964 |
| *Ildr2* | 9.975863166 | 9.869793577 | 20.71996099 | 6.828354829 | 1.056557338 | 3.173753883 | 1.960652388 | 1.891385446 | 0.001395909 | -2.554108245 | 0.000191248 | 13.91517812 | 0.170269479 | 2.855143019 |
| *Tspan18* | 43.89379793 | 58.2317821 | 61.17321817 | 35.11725341 | 4.226229352 | 13.75293349 | 9.803261939 | 5.674156339 | 5.07E-12 | -2.573776011 | 2.04E-13 | 53.96313204 | 0.167964004 | 11.29527274 |
| *Ifitm1* | 56.86242005 | 58.2317821 | 58.21322375 | 38.04369119 | 13.7352454 | 5.289589805 | 3.921304776 | 12.2940054 | 6.04E-13 | -2.583216441 | 2.14E-14 | 58.39497924 | 0.166868502 | 12.21928143 |
| *Syt13* | 33.91793477 | 49.34896788 | 46.37324603 | 36.09273267 | 6.339344029 | 9.521261649 | 4.90163097 | 6.619849062 | 1.60E-11 | -2.604267403 | 6.60E-13 | 51.65883509 | 0.164451332 | 10.79595295 |
| *Rasd2* | 10.97344948 | 15.79166972 | 15.78663695 | 14.63218892 | 1.056557338 | 4.231671844 | 0.980326194 | 2.83707817 | 0.000140655 | -2.655411022 | 1.50E-05 | 18.73428003 | 0.158723646 | 3.851843991 |
| *Nnmt* | 23.9420716 | 14.80469037 | 5.919988856 | 2.926437784 | 2.113114676 | 2.115835922 | 0.980326194 | 1.891385446 | 0.002157391 | -2.749483387 | 0.000317106 | 12.96654698 | 0.148704129 | 2.666071139 |
| *Col11a1* | 40.90103898 | 37.50521559 | 51.30657008 | 14.63218892 | 8.452458705 | 6.347507766 | 2.940978582 | 3.782770893 | 1.28E-08 | -2.760831138 | 7.23E-10 | 37.95720093 | 0.147539061 | 7.891190944 |
| *Cthrc1* | 12.96862212 | 17.76562844 | 30.58660909 | 7.80383409 | 2.113114676 | 3.173753883 | 1.960652388 | 2.83707817 | 5.20E-05 | -2.777481657 | 5.03E-06 | 20.82544135 | 0.145846063 | 4.284068862 |
| *Ogn* | 19.95172633 | 36.51823623 | 18.74663138 | 43.89656676 | 4.226229352 | 2.115835922 | 4.90163097 | 5.674156339 | 3.63E-08 | -2.803787046 | 2.17E-09 | 35.81523156 | 0.143210875 | 7.440354082 |
| *Ank3* | 20.94931265 | 17.76562844 | 8.879983283 | 16.58314744 | 3.169672014 | 2.115835922 | 1.960652388 | 1.891385446 | 2.86E-05 | -2.819933814 | 2.66E-06 | 22.04770996 | 0.141616982 | 4.543476339 |
| *Cyp7b1* | 16.95896738 | 5.921876146 | 12.82664252 | 7.80383409 | 0 | 5.289589805 | 0 | 0.945692723 | 0.002113897 | -2.833081545 | 0.000309532 | 13.01181449 | 0.140332246 | 2.674916228 |
| *Cpa6* | 16.95896738 | 4.934896788 | 6.906653665 | 7.80383409 | 0 | 1.057917961 | 1.960652388 | 1.891385446 | 0.002990335 | -2.869406574 | 0.000459592 | 12.27286373 | 0.136842989 | 2.524280103 |
| *Mgst2* | 3.990345266 | 4.934896788 | 12.82664252 | 7.80383409 | 2.113114676 | 0 | 0.980326194 | 0.945692723 | 0.007927952 | -2.874079519 | 0.001415063 | 10.18606038 | 0.136400466 | 2.100838993 |
| *Dpep1* | 14.96379475 | 15.79166972 | 16.77330176 | 11.70575114 | 2.113114676 | 1.057917961 | 2.940978582 | 1.891385446 | 3.05E-05 | -2.87923366 | 2.85E-06 | 21.91578801 | 0.135914034 | 4.51537405 |
| *Fam19a5* | 17.9565537 | 6.908855504 | 11.83997771 | 7.80383409 | 0 | 2.115835922 | 1.960652388 | 1.891385446 | 0.000723018 | -2.883623351 | 9.22E-05 | 15.29030476 | 0.135501117 | 3.140850635 |
| *Epha3* | 12.96862212 | 9.869793577 | 14.79997214 | 15.60766818 | 0 | 4.231671844 | 1.960652388 | 0.945692723 | 0.000126809 | -2.91220303 | 1.34E-05 | 18.95778891 | 0.132843263 | 3.896851357 |
| *Tshz2* | 14.96379475 | 6.908855504 | 12.82664252 | 10.73027187 | 0 | 1.057917961 | 3.921304776 | 0.945692723 | 0.000559648 | -2.916039848 | 6.96E-05 | 15.82127484 | 0.132490438 | 3.252084772 |
| *Rasl12* | 1.995172633 | 5.921876146 | 12.82664252 | 10.73027187 | 1.056557338 | 0 | 1.960652388 | 0.945692723 | 0.00640681 | -2.972354569 | 0.001098749 | 10.65329674 | 0.127418391 | 2.193358124 |
| *Hoxc9* | 13.96620843 | 12.83073165 | 3.946659237 | 10.73027187 | 0 | 4.231671844 | 0.980326194 | 0 | 0.00147714 | -3.027553817 | 0.000203321 | 13.80014475 | 0.122635297 | 2.830578221 |
| *Elfn1* | 10.97344948 | 9.869793577 | 14.79997214 | 6.828354829 | 2.113114676 | 1.057917961 | 0.980326194 | 0.945692723 | 0.000395675 | -3.071332847 | 4.73E-05 | 16.55370723 | 0.118969788 | 3.402661507 |
| *Smarcd3* | 8.97827685 | 16.77864908 | 13.81330733 | 14.63218892 | 0 | 5.289589805 | 0 | 0.945692723 | 9.21E-05 | -3.153994139 | 9.44E-06 | 19.62216585 | 0.112344847 | 4.035962398 |
| *Pawr* | 40.90103898 | 27.63542202 | 25.65328504 | 14.63218892 | 6.339344029 | 1.057917961 | 1.960652388 | 2.83707817 | 1.65E-08 | -3.165691324 | 9.38E-10 | 37.4493339 | 0.111437652 | 7.783630911 |
| *Col2a1* | 17.9565537 | 17.76562844 | 20.71996099 | 17.5586267 | 2.113114676 | 2.115835922 | 0.980326194 | 2.83707817 | 3.82E-07 | -3.198916683 | 2.63E-08 | 30.96355754 | 0.108900563 | 6.418411018 |
| *Col10a1* | 12.96862212 | 6.908855504 | 4.933324046 | 8.779313352 | 1.056557338 | 0 | 0.980326194 | 0.945692723 | 0.000931146 | -3.47673037 | 0.000122146 | 14.75926915 | 0.089825547 | 3.030982074 |
| *Ifi202b* | 215.4786444 | 228.979211 | 256.5328504 | 231.1885849 | 20.07458942 | 15.86876941 | 15.6852191 | 26.47939625 | 4.32E-86 | -3.569732454 | 1.48E-88 | 398.0239262 | 0.084217715 | 85.36458702 |
| *Cyp4b1* | 0.997586317 | 2.960938073 | 12.82664252 | 22.43602301 | 0 | 1.057917961 | 0.980326194 | 0.945692723 | 0.001670547 | -3.703787534 | 0.00023541 | 13.52496795 | 0.076744782 | 2.777141312 |
| *C1qtnf3* | 19.95172633 | 2.960938073 | 12.82664252 | 7.80383409 | 2.113114676 | 0 | 0.980326194 | 0 | 0.000195291 | -3.840814369 | 2.18E-05 | 18.02403728 | 0.06979104 | 3.709318207 |
| *Gp1bb* | 8.97827685 | 2.960938073 | 9.866648093 | 7.80383409 | 2.113114676 | 0 | 0 | 0 | 0.00153199 | -3.869954728 | 0.000211971 | 13.72187473 | 0.068395503 | 2.814744002 |
| *Rspo3* | 10.97344948 | 3.947917431 | 4.933324046 | 1.950958523 | 1.056557338 | 0 | 0 | 0 | 0.004990474 | -4.153433253 | 0.000825531 | 11.18293361 | 0.056194266 | 2.301858193 |
| *Pcsk9* | 0.997586317 | 0.986979358 | 10.8533129 | 3.901917045 | 0 | 0 | 0 | 0 | 0.008769121 | -4.495646224 | 0.001584393 | 9.977795311 | 0.044327745 | 2.057043962 |
| *Unc5a* | 10.97344948 | 5.921876146 | 1.973329619 | 10.73027187 | 0 | 0 | 0 | 0.945692723 | 0.000475378 | -4.595955306 | 5.80E-05 | 16.1667467 | 0.041350388 | 3.322961233 |
| *8430408G22Rik* | 37.90828003 | 30.59636009 | 21.7066258 | 38.04369119 | 4.226229352 | 0 | 0 | 0.945692723 | 4.34E-15 | -4.665895071 | 1.30E-16 | 68.4481313 | 0.039393596 | 14.362602 |
| *Ptn* | 5.9855179 | 18.7526078 | 5.919988856 | 4.877396306 | 0 | 0 | 0.980326194 | 0 | 8.75E-05 | -4.859525131 | 8.92E-06 | 19.72898767 | 0.034445871 | 4.057866004 |
| *Phyhip* | 6.983104216 | 1.973958715 | 11.83997771 | 6.828354829 | 0 | 0 | 0 | 0 | 0.000150765 | -5.21776307 | 1.62E-05 | 18.59199413 | 0.026871803 | 3.821700292 |
| *Gm5886* | 11.9710358 | 18.7526078 | 20.71996099 | 7.80383409 | 0 | 0 | 0 | 0 | 1.75E-09 | -6.318230003 | 9.05E-11 | 42.01651332 | 0.012532083 | 8.75712049 |
