## Supplementary material for "TFEB-Mediated Pro-inflammatory Response in Murine Macrophages Induced by Acute Alpha7 Nicotinic Receptor Activation": Table_S16

**Table S16. GO and KEGG Pathway analysis of differentially expressed genes from Table S15.**

**Enrichr-KG_DMSO (Tfebfl vs TfebLysM) DE UP genes in Tfeb$^{fl/fl}$ only**

| Term | Library | p-value | q-value | z-score | combined score | converted -Log10(Padj) |
|---|---|---|---|---|---|---|
| extracellular matrix organization (GO:0030198) | GO_Biological_Process_2021 | 5.39E-54 | 1.73E-50 | 12.8 | 1570 | 49.76210455 |
| extracellular structure organization (GO:0043062) | GO_Biological_Process_2021 | 9.68E-42 | 1.42E-38 | 13.12 | 1239 | 37.84909013 |
| external encapsulating structure organization (GO:0045229) | GO_Biological_Process_2021 | 1.32E-41 | 1.42E-38 | 13.03 | 1227 | 37.84909013 |
| collagen fibril organization (GO:0030199) | GO_Biological_Process_2021 | 4.18E-34 | 3.36E-31 | 24.03 | 1847 | 30.47426864 |
| regulation of cell migration (GO:0030334) | GO_Biological_Process_2021 | 1.06E-24 | 6.78E-22 | 5.76 | 318 | 21.16853977 |
| Focal adhesion | KEGG_2021_Human | 5.10E-21 | 1.32E-18 | 7.874 | 367.9 | 17.87932738 |
| Proteoglycans in cancer | KEGG_2021_Human | 1.36E-15 | 1.76E-13 | 6.263 | 214.4 | 12.7532799 |
| ECM-receptor interaction | KEGG_2021_Human | 7.87E-15 | 6.80E-13 | 10.62 | 344.9 | 12.16775302 |
| PI3K-Akt signaling pathway | KEGG_2021_Human | 4.87E-14 | 3.15E-12 | 4.348 | 133.3 | 11.50127603 |
| Protein digestion and absorption | KEGG_2021_Human | 2.10E-11 | 1.09E-09 | 7.656 | 188.2 | 8.963690556 |

**Enrichr-KG_DMSO (TfebflvsTfebLysM) DE UP genes in Tfeb$^{\Delta LysM}$ only**

| Term | Library | p-value | q-value | z-score | combined score | converted -log10(Padj) |
|---|---|---|---|---|---|---|
| positive regulation of multicellular organismal process (GO:0051240) | GO_Biological_Process_2021 | 0.0001437 | 0.09216 | 4.554 | 40.29 | 1.035457534 |
| regulation of neuron projection development (GO:0010975) | GO_Biological_Process_2021 | 0.0001501 | 0.09216 | 6.664 | 58.67 | 1.035457534 |
| high-density lipoprotein particle remodeling (GO:0034375) | GO_Biological_Process_2021 | 0.0002431 | 0.09216 | 29.4 | 244.7 | 1.035457534 |
| Glycosaminoglycan degradation | KEGG_2021_Human | 0.0002872 | 0.02123 | 27.56 | 224.8 | 1.673050006 |
| positive regulation of phosphorylation (GO:0042327) | GO_Biological_Process_2021 | 0.0003787 | 0.09216 | 4.927 | 38.82 | 1.035457534 |
| neuron migration (GO:0001764) | GO_Biological_Process_2021 | 0.0003907 | 0.09216 | 12.86 | 100.9 | 1.035457534 |
| Malaria | KEGG_2021_Human | 0.0003907 | 0.02123 | 12.86 | 100.9 | 1.673050006 |
| Cholesterol metabolism | KEGG_2021_Human | 0.0003907 | 0.02123 | 12.86 | 100.9 | 1.673050006 |
| Morphine addiction | KEGG_2021_Human | 0.003647 | 0.1192 | 6.785 | 38.09 | 0.923723745 |
| Cell adhesion molecules | KEGG_2021_Human | 0.003656 | 0.1192 | 5.184 | 29.09 | 0.923723745 |
