## Supplementary material for "TFEB-Mediated Pro-inflammatory Response in Murine Macrophages Induced by Acute Alpha7 Nicotinic Receptor Activation": Table_S17

**Table S17. Differential gene expression in PNU-282987-treated BMDMs (Tfebfl/fl v. TfebΔLysM).**

| ID | C5_Tfeb_fl_PNU_1 | C6_Tfeb_fl_PNU_2 | C7_Tfeb_fl_PNU_3 | C8_Tfeb_fl_PNU_4 | D5_Tfeb_LysMCre_PNU_1 | D6_Tfeb_LysMCre_PNU_2 | D7_Tfeb_LysMCre_PNU_3 | D8_Tfeb_LysMCre_PNU_4 | padj | log2FoldChange | pvalue | stat | foldChange | log10padj |
|---|---|---|---|---|---|---|---|---|---|---|---|---|---|---|
| *Syce1* | 0 | 0 | 0 | 0 | 15.58288114 | 0 | 7.992785046 | 8.088763935 | 0.00377378 | 5.369381185 | 0.00024323 | 13.4636665 | 41.3375559 | 2.42322314 |
| *Dscam* | 0 | 0 | 1.020331787 | 0 | 9.739300714 | 7.531052841 | 2.997294392 | 5.39250929 | 0.00645932 | 4.346543714 | 0.00046919 | 12.2342975 | 20.3441728 | 2.1898132 |
| *Egfem1* | 0 | 2.13154045 | 0 | 0 | 8.765370643 | 3.765526421 | 10.99007944 | 11.68377013 | 0.00265576 | 4.078916494 | 0.00015829 | 14.2708873 | 16.8995918 | 2.57581174 |
| *Chrm3* | 2.08919105 | 0 | 0 | 0 | 8.765370643 | 2.824144815 | 7.992785046 | 12.58252168 | 0.00636465 | 3.949902013 | 0.0004613 | 12.2659365 | 15.4539316 | 2.19622563 |
| *Gm11992* | 1.044595525 | 0 | 3.060995361 | 4.121847259 | 8.765370643 | 9.413816051 | 18.98286448 | 18.87378252 | 0.00262289 | 2.757768756 | 0.00015592 | 14.2993565 | 6.76349412 | 2.58122068 |
| *Slco2b1* | 82.52304646 | 73.53814551 | 65.30123437 | 55.64493799 | 343.7973152 | 175.0969786 | 324.7068925 | 369.3868864 | 6.22E-20 | 2.131164342 | 1.98E-22 | 94.924269 | 4.38070887 | 19.206028 |
| *Apoc2* | 56.40815834 | 50.09120056 | 49.99625757 | 50.49262892 | 248.3521682 | 62.13118594 | 257.7673177 | 220.1941293 | 0.00050353 | 1.929262573 | 2.15E-05 | 18.0525609 | 3.80860474 | 3.29797509 |
| *Lyz1* | 43.87301204 | 42.63080899 | 42.85393506 | 52.55355255 | 197.7078045 | 82.84158125 | 181.8358598 | 196.8265891 | 8.56E-11 | 1.857109099 | 7.28E-13 | 51.4670486 | 3.62280988 | 10.0674148 |
| *Cd5l* | 137.8866093 | 127.892427 | 159.1717588 | 153.5388104 | 491.8346861 | 539.4116597 | 441.6013738 | 418.8182215 | 3.82E-41 | 1.70829709 | 4.25E-44 | 194.002098 | 3.26774881 | 40.4175378 |
| *Col14a1* | 96.10278828 | 100.1824011 | 89.78919726 | 78.31509791 | 268.8046997 | 165.6831625 | 371.6645046 | 319.9555512 | 2.17E-12 | 1.627799012 | 1.43E-14 | 59.190013 | 3.09041163 | 11.6634766 |
| *Zbtb16* | 32.38246127 | 9.591932023 | 27.54895825 | 17.51785085 | 83.75798614 | 37.65526421 | 73.93326167 | 71.00137232 | 0.00014724 | 1.611027866 | 5.16E-06 | 20.7768077 | 3.054694 | 3.83197 |
| *Atp9a* | 6.267573149 | 7.460391573 | 8.162654296 | 12.36554178 | 21.42646157 | 20.71039531 | 26.97564953 | 26.96254645 | 0.0060707 | 1.486479171 | 0.00043517 | 12.3747795 | 2.80204315 | 2.2167612 |
| *Abca9* | 107.5933391 | 114.0374141 | 133.6634641 | 143.2341922 | 387.6241684 | 378.4354053 | 275.7510841 | 337.0318306 | 2.04E-21 | 1.467386879 | 6.15E-24 | 101.797528 | 2.76520584 | 20.6911466 |
| *Rell2* | 19.84731497 | 13.85501292 | 6.121990722 | 13.39600359 | 39.93113293 | 30.12421136 | 37.96572897 | 24.26629181 | 0.00901084 | 1.313433811 | 0.00071057 | 11.4613983 | 2.48532378 | 2.0452349 |
| *P2ry12* | 80.43385541 | 76.73545618 | 82.64687475 | 73.16278884 | 147.0634408 | 162.8590177 | 189.8286448 | 193.2315829 | 1.08E-10 | 1.147054549 | 9.62E-13 | 50.9200453 | 2.2146129 | 9.96623143 |
| *Flt1* | 69.98790016 | 50.09120056 | 41.83360327 | 56.67539981 | 120.7673289 | 85.66572607 | 141.8719346 | 127.6227199 | 1.34E-05 | 1.122344102 | 3.20E-07 | 26.1279535 | 2.17700407 | 4.87427806 |
| *Plxdc2* | 1020.569828 | 1023.139416 | 985.6405063 | 1119.081531 | 2287.761738 | 2275.31934 | 2150.059177 | 2130.939921 | 7.99E-70 | 1.092071619 | 1.90E-73 | 328.636657 | 2.1317993 | 69.0976099 |
| *Nr4a2* | 135.7974182 | 98.05086068 | 124.480478 | 112.3203378 | 283.4136508 | 202.3970451 | 229.7925701 | 282.2079862 | 4.57E-10 | 1.084009645 | 4.50E-12 | 47.8915753 | 2.11991974 | 9.33998796 |
| *Ang* | 122.2176764 | 153.4709124 | 133.6634641 | 129.8381886 | 316.5272732 | 188.276321 | 288.7393598 | 342.4243399 | 4.88E-08 | 1.075566303 | 6.63E-10 | 38.1279936 | 2.10754919 | 7.31200796 |
| *Arrdc3* | 940.1359723 | 937.8777978 | 984.6201745 | 921.2328623 | 2008.243807 | 1570.224517 | 2113.092547 | 2167.788735 | 4.96E-28 | 1.054495937 | 9.46E-31 | 132.91023 | 2.0769924 | 27.3045641 |
| *Hpgd* | 1230.533528 | 1322.620849 | 1175.422219 | 1069.619364 | 2321.84929 | 2551.14415 | 2607.646121 | 2453.591727 | 6.29E-47 | 1.050128991 | 3.00E-50 | 222.197635 | 2.07071498 | 46.2013332 |
| *Ly86* | 1228.444337 | 1192.596882 | 1215.215158 | 1207.701247 | 2458.1995 | 2072.922295 | 2475.765168 | 2538.973124 | 6.59E-42 | 0.978652954 | 6.28E-45 | 197.807827 | 1.97062458 | 41.181171 |
| *Cmah* | 25.0702926 | 24.51271517 | 25.50829468 | 22.67015992 | 43.82685321 | 37.65526421 | 48.95580841 | 60.21635374 | 0.00831899 | 0.965054974 | 0.00064328 | 11.6463718 | 1.95213791 | 2.07992941 |
| *Sncaip* | 54.31896729 | 63.94621349 | 71.42322509 | 76.25417428 | 141.2198604 | 140.2658592 | 118.8926776 | 106.9514343 | 1.90E-05 | 0.930441823 | 4.82E-07 | 25.3333173 | 1.90585957 | 4.72200472 |
| *Fcrls* | 6273.840722 | 7063.92505 | 5749.56962 | 6279.634299 | 12451.69596 | 11161.96169 | 12167.01704 | 12426.13891 | 1.16E-44 | 0.926353874 | 8.29E-48 | 211.005135 | 1.90046687 | 43.9358821 |
| *Gatm* | 1764.321841 | 1686.048496 | 1606.002233 | 1602.368122 | 3017.235361 | 2248.019273 | 3699.660378 | 3648.032535 | 4.78E-07 | 0.921590038 | 7.60E-09 | 33.374789 | 1.8942018 | 6.32041592 |
| *Pde3b* | 686.2992598 | 633.0675135 | 613.219404 | 687.3180304 | 1413.172534 | 1138.130361 | 1278.845607 | 1124.338187 | 1.06E-23 | 0.918782556 | 2.78E-26 | 112.494383 | 1.89051927 | 22.9739782 |
| *Itga6* | 3753.231721 | 3351.847357 | 3573.201918 | 3648.865286 | 6904.190276 | 4721.02875 | 7732.020434 | 7650.17318 | 2.35E-07 | 0.914589227 | 3.52E-09 | 34.8703107 | 1.88503228 | 6.62962452 |
| *Rtn4rl1* | 100.2811704 | 107.6427927 | 94.8908562 | 104.0766433 | 198.6817346 | 146.8555304 | 181.8358598 | 236.3716572 | 2.51E-06 | 0.909668011 | 4.99E-08 | 29.7201578 | 1.87861315 | 5.59994131 |
| *Rnase4* | 1963.839587 | 2095.304262 | 1740.686029 | 1944.481444 | 3430.181711 | 3253.414827 | 3720.641439 | 4078.534526 | 1.49E-28 | 0.903359277 | 2.48E-31 | 135.565834 | 1.87041613 | 27.8274635 |
| *Rnase6* | 47.00679862 | 54.35428146 | 60.19957544 | 58.73632344 | 109.080168 | 49.89322507 | 120.8908738 | 131.2177261 | 0.00967799 | 0.899654937 | 0.00078221 | 11.282984 | 1.86561971 | 2.01421491 |
| *Fgl2* | 152.5109466 | 151.3393719 | 172.436072 | 188.5745121 | 284.3875808 | 338.8973779 | 321.7095981 | 292.0942532 | 2.92E-11 | 0.894995094 | 2.32E-13 | 53.7130655 | 1.85960356 | 10.5346994 |
| *Cytip* | 423.0611876 | 401.7953747 | 472.4136174 | 446.1899657 | 971.0082812 | 646.7291627 | 813.2658784 | 791.8001141 | 7.75E-13 | 0.885795473 | 4.80E-15 | 61.3389715 | 1.84778317 | 12.1106395 |
| *Disc1* | 49.09598967 | 25.57848539 | 59.17924365 | 58.73632344 | 89.60156657 | 82.84158125 | 84.92334111 | 97.96391877 | 0.00384486 | 0.881902884 | 0.00024903 | 13.4194298 | 1.84280432 | 2.41511933 |
| *Fat3* | 615.2667641 | 403.9269152 | 571.3858008 | 565.7235362 | 968.086491 | 858.5400239 | 1068.035902 | 1040.754293 | 7.51E-12 | 0.867519923 | 5.43E-14 | 56.5665642 | 1.82452374 | 11.124235 |
| *Gpr65* | 1290.075473 | 1398.290535 | 1222.357481 | 1206.670785 | 2348.145402 | 1724.611101 | 2593.658747 | 2637.835794 | 4.82E-13 | 0.862557929 | 2.91E-15 | 62.3244762 | 1.81825927 | 12.3167154 |
| *Abcd2* | 268.4610499 | 276.0344882 | 248.960956 | 234.9452937 | 438.2685321 | 429.2700119 | 493.5544766 | 503.3008671 | 7.51E-15 | 0.858927201 | 3.82E-17 | 70.8679188 | 1.81368914 | 14.1243793 |
| *Gdpd3* | 45.96220309 | 63.94621349 | 84.68753833 | 54.61447618 | 116.8716086 | 111.0830294 | 117.8935794 | 105.1539312 | 0.00052219 | 0.854289719 | 2.25E-05 | 17.9639014 | 1.80786847 | 3.28217129 |
| *Tmem119* | 101.3257659 | 136.4185888 | 85.70787011 | 97.89387239 | 176.2813429 | 143.090004 | 226.7952757 | 213.004117 | 0.00014084 | 0.850326987 | 4.90E-06 | 20.8750417 | 1.80290951 | 3.85128041 |
| *Eml6* | 68.94330464 | 57.55159214 | 49.99625757 | 73.16278884 | 112.9758883 | 112.024411 | 90.9179299 | 130.3189745 | 0.0004783 | 0.839681491 | 2.03E-05 | 18.1646638 | 1.78965499 | 3.32029642 |
| *Cbx7* | 106.5487435 | 136.4185888 | 123.4601462 | 131.8991123 | 279.5179305 | 159.0934913 | 247.7763364 | 200.4215953 | 8.66E-05 | 0.830528166 | 2.67E-06 | 22.039889 | 1.77833629 | 4.06246688 |
| *Gm4980* | 52.22977624 | 53.28851124 | 52.03692114 | 57.70586162 | 118.8194687 | 73.4277652 | 82.92514485 | 106.0526827 | 0.00199459 | 0.824913172 | 0.00011144 | 14.932351 | 1.77142842 | 2.70014716 |
| *Adcy2* | 85.65683304 | 103.3797118 | 77.54521582 | 78.31509791 | 121.7412589 | 144.0313856 | 168.8475841 | 172.5602973 | 9.38E-05 | 0.817516926 | 2.97E-06 | 21.8340236 | 1.7623701 | 4.02802071 |
| *Ralgps2* | 65.80951806 | 66.07775394 | 44.89459863 | 56.67539981 | 114.9237484 | 96.96230533 | 105.9044019 | 90.77390639 | 0.00086199 | 0.807539693 | 4.04E-05 | 16.8516818 | 1.75022414 | 3.06449838 |
| *Rhobtb1* | 41.78382099 | 45.82811967 | 58.15891186 | 42.2489344 | 107.1323079 | 78.13467323 | 76.93055607 | 65.60886303 | 0.00710685 | 0.799850483 | 0.00053147 | 12.0018615 | 1.74092069 | 2.1483231 |
| *Nr4a1* | 161.9123063 | 205.6936534 | 194.8833713 | 224.6406756 | 387.6241684 | 369.9629708 | 336.6960701 | 257.0429428 | 4.63E-06 | 0.778802375 | 9.61E-08 | 28.4509474 | 1.71570602 | 5.33418639 |
| *F11r* | 438.7301204 | 452.9523455 | 400.9903923 | 392.6059514 | 730.4475535 | 452.8045521 | 848.234313 | 816.9651575 | 2.31E-05 | 0.757200934 | 5.95E-07 | 24.9275858 | 1.69020816 | 4.63606461 |
| *Vill* | 99.23657486 | 103.3797118 | 135.7041277 | 99.95479602 | 200.6295947 | 153.4452016 | 187.8304486 | 197.7253406 | 6.46E-05 | 0.754188321 | 1.92E-06 | 22.6776874 | 1.68668238 | 4.18957533 |
| *Sulf2* | 2010.846385 | 1779.836275 | 1766.194323 | 2022.796542 | 3495.435026 | 2326.153946 | 3458.877729 | 3482.66225 | 2.47E-05 | 0.751748187 | 6.51E-07 | 24.7538836 | 1.68383198 | 4.60752331 |
| *H2-Aa* | 789.7142168 | 882.4577461 | 801.9807846 | 768.7245137 | 1313.831666 | 884.8987088 | 1552.598495 | 1655.500352 | 0.00401986 | 0.737554464 | 0.00026323 | 13.3154217 | 1.66734709 | 2.39578955 |
| *Zfp791* | 139.9758003 | 117.2347247 | 125.5008098 | 136.0209595 | 202.5774549 | 155.3279648 | 248.7754346 | 239.0679119 | 0.00037142 | 0.7048185 | 1.52E-05 | 18.7153705 | 1.6299396 | 3.43014037 |
| *Hist3h2a* | 102.3703614 | 94.85355001 | 81.62654296 | 126.7468032 | 181.1509933 | 164.7417809 | 135.8773458 | 176.1553035 | 0.00083042 | 0.698697929 | 3.85E-05 | 16.9451499 | 1.62303929 | 3.08070018 |
| *Dgkg* | 84.61223751 | 101.2481714 | 84.68753833 | 77.2846361 | 142.1937904 | 100.7278318 | 163.8520934 | 157.281521 | 0.00305756 | 0.697716582 | 0.00018929 | 13.9345196 | 1.62193565 | 2.51462546 |
| *Fam129a* | 1845.800292 | 1516.59103 | 1790.682286 | 1819.795565 | 2988.991389 | 2119.991375 | 3101.200598 | 3078.224053 | 4.52E-09 | 0.694895041 | 5.14E-11 | 43.124413 | 1.61876666 | 8.34486521 |
| *Fcrl1* | 90.87981066 | 73.53814551 | 73.46388867 | 77.2846361 | 127.5848394 | 136.5003327 | 112.8980888 | 128.5214714 | 0.0008788 | 0.682469271 | 4.15E-05 | 16.8022054 | 1.60488427 | 3.0561105 |
| *Qpct* | 552.5910326 | 524.3589506 | 466.2916267 | 482.2561293 | 841.4755817 | 721.0983095 | 903.1847102 | 780.116344 | 1.27E-11 | 0.680062886 | 9.52E-14 | 55.4629593 | 1.60220959 | 10.894539 |
| *Ccl8* | 58.49734939 | 71.40660506 | 59.17924365 | 75.22371247 | 99.34086728 | 94.13816051 | 99.90981307 | 124.0277137 | 0.00642928 | 0.660957488 | 0.0004665 | 12.245039 | 1.58113164 | 2.19183794 |
| *Gpr155* | 369.7868158 | 333.5860804 | 283.6522368 | 294.712079 | 455.7992734 | 469.749421 | 591.4660934 | 506.8958733 | 5.79E-07 | 0.659069027 | 9.44E-09 | 32.9542894 | 1.57906332 | 6.23722155 |
| *Chst3* | 271.5948365 | 249.3902326 | 242.8389653 | 224.6406756 | 410.0245601 | 378.4354053 | 438.6040794 | 331.6393213 | 6.93E-07 | 0.656086506 | 1.16E-08 | 32.5587231 | 1.57580226 | 6.15930856 |
| *Nanos1* | 587.062685 | 630.9359731 | 547.9181697 | 535.8401436 | 887.250295 | 726.7465992 | 995.1017382 | 1014.690498 | 3.37E-08 | 0.655009425 | 4.42E-10 | 38.9195232 | 1.57462624 | 7.47262977 |
| *Arg2* | 75.21087779 | 102.3139416 | 70.40289331 | 83.46740699 | 159.7245317 | 108.2588846 | 124.8872663 | 125.8252168 | 0.00685291 | 0.646759463 | 0.00050599 | 12.0934447 | 1.56564753 | 2.16412489 |
| *Cadm1* | 4863.636764 | 5014.448908 | 5002.686752 | 4878.206231 | 8388.459705 | 5236.905869 | 8961.910233 | 8341.31312 | 0.00092579 | 0.646416557 | 4.44E-05 | 16.6748359 | 1.56527545 | 3.03348658 |
| *Slc38a6* | 658.0951806 | 663.97485 | 628.5243808 | 638.8863251 | 969.060421 | 1182.375296 | 927.1630653 | 973.3479269 | 4.50E-12 | 0.646309745 | 3.08E-14 | 57.6855021 | 1.56515957 | 11.3467824 |
| *Hgsnat* | 3124.385215 | 3331.597723 | 3065.076688 | 3169.700542 | 4974.834805 | 4673.018288 | 5062.430228 | 5131.871341 | 5.67E-36 | 0.644886218 | 8.11E-39 | 169.816782 | 1.56361596 | 35.246305 |
| *S1pr1* | 2312.734492 | 2112.356586 | 1966.179354 | 2054.740858 | 3200.334215 | 2668.816851 | 3724.637831 | 3586.91743 | 2.53E-10 | 0.642090386 | 2.39E-12 | 49.1317347 | 1.56058873 | 9.59675359 |
| *Sepp1* | 10480.4269 | 11599.84313 | 10835.92358 | 10995.02756 | 17297.972 | 19487.54061 | 15611.90739 | 16124.50153 | 1.68E-19 | 0.641998934 | 5.73E-22 | 92.8205156 | 1.56048981 | 18.7758441 |
| *Hexb* | 10122.13064 | 10724.84577 | 9420.72339 | 9781.143545 | 15639.36909 | 12670.99641 | 17258.42111 | 16905.51662 | 1.12E-12 | 0.641511223 | 7.18E-15 | 60.547086 | 1.55996236 | 11.9523384 |
| *Acsf2* | 196.3839587 | 197.1674916 | 191.822376 | 191.6658975 | 352.5626858 | 386.9078397 | 225.7961775 | 242.6629181 | 0.00194943 | 0.636560889 | 0.00010845 | 14.9836296 | 1.55461882 | 2.71009269 |
| *Fzd7* | 1265.005181 | 1210.714975 | 1176.44255 | 1252.011105 | 1949.808003 | 1842.283801 | 1957.233238 | 1862.213208 | 4.96E-28 | 0.63398468 | 9.26E-31 | 132.951549 | 1.55184523 | 27.3045641 |
| *Chn2* | 227.7218244 | 212.0882747 | 183.6597217 | 206.0923629 | 279.5179305 | 297.4765872 | 375.6608972 | 332.5380729 | 1.77E-05 | 0.631609926 | 4.46E-07 | 25.4828911 | 1.54929291 | 4.7516255 |
| *Tbxas1* | 1747.608313 | 1691.377347 | 1621.30721 | 1797.125405 | 2589.68006 | 2344.040197 | 2768.50092 | 2892.182483 | 7.13E-17 | 0.627641912 | 3.17E-19 | 80.3266538 | 1.54503756 | 16.1468527 |
| *Hgf* | 690.4776419 | 674.6325523 | 689.7442881 | 690.4094158 | 801.5444488 | 1083.530228 | 1099.007944 | 1250.163404 | 6.13E-07 | 0.625548351 | 1.00E-08 | 32.8349878 | 1.54279711 | 6.21268326 |
| *Mxi1* | 1355.884991 | 1428.132101 | 1454.993128 | 1366.392366 | 2477.678102 | 1831.928604 | 2122.08443 | 2192.055026 | 3.96E-12 | 0.62140288 | 2.68E-14 | 57.959937 | 1.53837037 | 11.4022934 |
| *Mtus1* | 543.1896729 | 669.3037012 | 574.4467961 | 568.8149217 | 877.5109943 | 692.8568614 | 1015.083701 | 1035.361784 | 5.91E-06 | 0.620308895 | 1.27E-07 | 27.9082364 | 1.53720428 | 5.22867364 |
| *H2-Eb1* | 436.6409294 | 454.0181158 | 434.6613413 | 465.7687402 | 715.8386025 | 441.5079728 | 763.3109719 | 830.4464307 | 0.00057331 | 0.619222419 | 2.52E-05 | 17.7496959 | 1.53604706 | 3.24161318 |
| *Tspan13* | 120.1284854 | 138.5501292 | 173.4564038 | 146.3255777 | 229.8474968 | 266.4109943 | 216.8042944 | 174.3578004 | 0.00207881 | 0.616191551 | 0.00011713 | 14.8382985 | 1.53282346 | 2.68218455 |
| *Fam214a* | 619.4451462 | 581.9105427 | 620.3617265 | 678.043874 | 918.4160573 | 882.074564 | 949.1432242 | 1081.198113 | 2.23E-11 | 0.616102217 | 1.76E-13 | 54.2583389 | 1.53272855 | 10.6508579 |
| *F13a1* | 352.0286919 | 313.3364461 | 400.9903923 | 396.7277986 | 663.2463786 | 362.431918 | 603.455271 | 612.0498044 | 0.00138732 | 0.614788304 | 7.27E-05 | 15.7403707 | 1.53133328 | 2.85782472 |
| *Cmklr1* | 247.5691394 | 227.0090579 | 248.960956 | 282.3465372 | 431.4510216 | 305.9490217 | 427.614 | 365.7918802 | 6.84E-05 | 0.604696346 | 2.04E-06 | 22.5554757 | 1.52065866 | 4.16502218 |
| *Siglec1* | 6758.533046 | 5957.655557 | 6469.923862 | 7415.203218 | 10874.90318 | 9662.340795 | 9725.221205 | 10171.17127 | 2.56E-17 | 0.604017813 | 1.06E-19 | 82.4987042 | 1.51994362 | 16.5919822 |
| *P2rx7* | 1608.677108 | 1453.710587 | 1544.782326 | 1560.119187 | 2445.538409 | 2333.684999 | 2217.99785 | 2349.336547 | 2.04E-24 | 0.599780798 | 5.20E-27 | 115.82331 | 1.51548629 | 23.6897079 |
| *Crem* | 258.0150946 | 322.9283781 | 248.960956 | 277.1942281 | 474.3039448 | 384.0836949 | 408.6311355 | 408.033203 | 4.55E-06 | 0.597719775 | 9.40E-08 | 28.4936299 | 1.51332282 | 5.3420277 |
| *Trf* | 4418.63907 | 4691.520529 | 4297.637487 | 4579.372304 | 6980.156822 | 7110.255264 | 6283.328144 | 6818.827997 | 2.78E-27 | 0.596370587 | 5.75E-30 | 129.327255 | 1.51190824 | 26.5554462 |
| *Pltp* | 5311.768244 | 5670.963366 | 5357.762214 | 5517.092556 | 8580.323929 | 8108.119765 | 7975.800378 | 8308.958065 | 1.51E-39 | 0.593222397 | 1.92E-42 | 186.426874 | 1.50861262 | 38.8220304 |
| *Cyp27a1* | 184.8934079 | 195.0359511 | 186.720717 | 220.5188283 | 312.6315529 | 310.6559297 | 268.7573972 | 294.7905079 | 9.37E-06 | 0.592642096 | 2.12E-07 | 26.9178781 | 1.50800592 | 5.02813914 |
| *Bmyc* | 157.7339242 | 182.2467084 | 102.0331787 | 164.8738903 | 208.4210353 | 235.3454013 | 232.7898645 | 233.6754026 | 0.00366284 | 0.585999252 | 0.00023433 | 13.5335938 | 1.50107832 | 2.43618214 |
| *Tbc1d2* | 647.6492254 | 606.4232579 | 639.7480305 | 659.4955614 | 474.3039448 | 467.8666578 | 354.6798364 | 400.8431906 | 1.56E-07 | -0.588613691 | 2.29E-09 | 35.7130633 | 0.66498159 | 6.80590839 |
| *Emb* | 2120.528915 | 2082.515019 | 2275.339885 | 2139.238727 | 1317.727387 | 1777.328471 | 1253.868154 | 1364.30485 | 1.65E-08 | -0.592794494 | 2.05E-10 | 40.4156044 | 0.66305732 | 7.7837373 |
| *Gda* | 582.8843029 | 646.9225264 | 669.3376523 | 700.714034 | 403.2070496 | 472.5735658 | 402.6365467 | 442.1857618 | 3.95E-09 | -0.594949774 | 4.45E-11 | 43.4035886 | 0.6620675 | 8.4037826 |
| *Rgs3* | 235.0339931 | 271.7714073 | 264.2659328 | 274.1028427 | 202.5774549 | 186.3935578 | 152.862014 | 150.0915086 | 0.00016257 | -0.595631487 | 5.77E-06 | 20.5613625 | 0.66175473 | 3.78897143 |
| *Gdpd1* | 441.863907 | 466.8073585 | 491.7999214 | 460.6164312 | 317.5012033 | 274.8834287 | 326.7050888 | 309.1705326 | 1.06E-08 | -0.600238452 | 1.27E-10 | 41.3559608 | 0.65964492 | 7.97290035 |
| *Bsn* | 320.6908261 | 218.4828961 | 237.7373064 | 344.1742461 | 201.6035248 | 213.6936244 | 166.8493878 | 157.281521 | 0.00357093 | -0.600915832 | 0.00022732 | 13.5906432 | 0.65933527 | 2.44721883 |
| *Gpnmb* | 98763.37309 | 94327.05951 | 99318.07582 | 97616.67816 | 63179.81766 | 69096.46844 | 62061.97769 | 62083.06071 | 1.63E-37 | -0.605050456 | 2.20E-40 | 176.986803 | 0.65744839 | 36.7873111 |
| *Pira6* | 527.52074 | 545.6743551 | 727.4965642 | 565.7235362 | 373.0152173 | 433.0355384 | 355.6789345 | 390.058172 | 3.52E-06 | -0.608648629 | 7.18E-08 | 29.0155863 | 0.65581071 | 5.45403483 |
| *Ikbke* | 1059.219862 | 868.6027332 | 1115.222643 | 1065.497516 | 696.360001 | 814.2950885 | 553.5003644 | 629.1260839 | 2.48E-06 | -0.609159505 | 4.89E-08 | 29.7611835 | 0.65557852 | 5.60564374 |
| *Gpr176* | 247.5691394 | 206.7594236 | 194.8833713 | 219.4883665 | 123.6891191 | 181.6866498 | 127.8845607 | 134.8127323 | 0.00110254 | -0.611833343 | 5.55E-05 | 16.2515459 | 0.65436462 | 2.9576058 |
| *Nkain1* | 498.2720653 | 451.8865753 | 499.9625757 | 523.4746018 | 293.1529515 | 325.7180354 | 317.7132056 | 350.5131039 | 4.64E-09 | -0.616029279 | 5.31E-11 | 43.0599045 | 0.65246423 | 8.33357234 |
| *Tpm1* | 2426.595404 | 2510.95465 | 2636.537338 | 2573.063151 | 1468.686548 | 2124.698283 | 1458.683271 | 1562.928943 | 6.13E-08 | -0.617101576 | 8.43E-10 | 37.6578511 | 0.65197946 | 7.21239997 |
| *Tnk2* | 241.3015662 | 239.7983006 | 251.0016196 | 263.7982246 | 160.6984618 | 224.9902036 | 158.8566028 | 105.1539312 | 0.00725152 | -0.617283366 | 0.00054402 | 11.9583759 | 0.65189731 | 2.13957078 |
| *Myo1d* | 352.0286919 | 287.7579607 | 294.8758865 | 283.376999 | 158.7506016 | 249.4661254 | 194.8241355 | 183.3453159 | 0.00023796 | -0.630862486 | 9.15E-06 | 19.6805307 | 0.64579023 | 3.62350171 |
| *Dhcr24* | 1389.312048 | 1125.453357 | 1227.45914 | 1193.274781 | 784.9876375 | 861.3641687 | 739.3326167 | 787.3063564 | 2.37E-13 | -0.637131412 | 1.36E-15 | 63.8316259 | 0.6429017 | 12.6255435 |
| *Cpne8* | 102.3703614 | 109.7743332 | 87.74853369 | 111.289876 | 71.09689521 | 56.48289631 | 72.93416354 | 63.81135993 | 0.00727376 | -0.638569842 | 0.00054743 | 11.9467595 | 0.6423494 | 2.13824108 |
| *Pbx1* | 158.7785198 | 151.3393719 | 211.2086799 | 170.0261994 | 94.47121693 | 125.2037535 | 121.8899719 | 102.4576765 | 0.00149107 | -0.639971846 | 7.90E-05 | 15.5811548 | 0.64172547 | 2.82650332 |
| *Ralgds* | 426.1949741 | 408.1899961 | 468.3322903 | 539.9619909 | 327.240504 | 350.1939571 | 240.7826495 | 262.4354521 | 3.97E-05 | -0.642314337 | 1.12E-06 | 23.7108901 | 0.64068435 | 4.4014306 |
| *Fam20c* | 8628.359035 | 8696.685034 | 8190.203255 | 8774.382352 | 5615.680792 | 6231.004844 | 5054.437443 | 5037.502429 | 1.56E-19 | -0.644284683 | 5.14E-22 | 93.0337871 | 0.63980994 | 18.8062601 |
| *Nr1h3* | 420.9719965 | 410.3215365 | 467.3119585 | 363.7530206 | 285.3615109 | 322.8938906 | 239.7835514 | 215.7003716 | 3.07E-05 | -0.644564264 | 8.37E-07 | 24.270482 | 0.63968597 | 4.51272146 |
| *Emp2* | 1329.770103 | 1141.439911 | 1289.699379 | 1236.554178 | 808.3619593 | 899.0194329 | 766.3082663 | 714.507481 | 3.35E-13 | -0.648757502 | 1.95E-15 | 63.1195409 | 0.6378294 | 12.4745419 |
| *Slco3a1* | 209.9637005 | 215.2855854 | 214.2696753 | 256.5849918 | 174.3334828 | 141.2072408 | 119.8917757 | 134.8127323 | 0.00016461 | -0.652302204 | 5.88E-06 | 20.527982 | 0.63626417 | 3.78355111 |
| *Mrgpre* | 337.4043545 | 409.2557663 | 315.2825222 | 388.4841041 | 205.4992451 | 291.8282976 | 228.7934719 | 195.9278375 | 7.86E-05 | -0.653498059 | 2.39E-06 | 22.2567786 | 0.63573699 | 4.10476297 |
| *Cd109* | 462.7558175 | 430.5711708 | 461.1899678 | 479.1647438 | 247.3782381 | 324.7766538 | 295.7330467 | 297.4867625 | 9.86E-09 | -0.653531662 | 1.17E-10 | 41.5193235 | 0.63572218 | 8.00628813 |
| *S100a11* | 1429.006678 | 1298.108134 | 1464.176114 | 1283.955421 | 839.5277215 | 1076.940556 | 789.2875233 | 773.8250831 | 3.53E-09 | -0.653990289 | 3.87E-11 | 43.6777701 | 0.63552012 | 8.45222189 |
| *Fam198b* | 403.2138726 | 412.453077 | 409.1530466 | 412.1847259 | 215.2385458 | 337.9559962 | 240.7826495 | 245.3591727 | 1.21E-05 | -0.654759716 | 2.86E-07 | 26.3399097 | 0.63518127 | 4.91759435 |
| *Plxna1* | 6947.604836 | 5545.20248 | 7048.451985 | 7130.795757 | 4419.694664 | 4967.67073 | 3866.509766 | 3652.526293 | 6.40E-10 | -0.65781927 | 6.36E-12 | 47.2163674 | 0.63383566 | 9.1938913 |
| *Cd274* | 474.2463683 | 440.1631028 | 486.6982624 | 435.8853476 | 268.8046997 | 310.6559297 | 257.7673177 | 325.3480605 | 1.18E-08 | -0.658509343 | 1.42E-10 | 41.1303545 | 0.63353255 | 7.9285267 |
| *Rnd3* | 122.2176764 | 121.4978056 | 140.8057866 | 130.8686505 | 81.810126 | 102.610595 | 75.93145794 | 65.60886303 | 0.00292898 | -0.661993461 | 0.000179 | 14.0395848 | 0.63200441 | 2.53328315 |
| *Sema4b* | 241.3015662 | 285.6264202 | 272.4285871 | 286.4683845 | 186.0206436 | 206.1625715 | 140.8728364 | 152.7877632 | 5.49E-05 | -0.662526019 | 1.60E-06 | 23.0189955 | 0.63177116 | 4.26055858 |
| *Bmpr1a* | 121.1730809 | 134.2870483 | 102.0331787 | 120.5640323 | 74.9926155 | 92.2553973 | 61.94408411 | 71.00137232 | 0.00372677 | -0.669606728 | 0.00023975 | 13.4907173 | 0.62867804 | 2.42866775 |
| *Rhov* | 156.6893287 | 168.3916955 | 165.2937495 | 165.9043522 | 103.2365876 | 121.4382271 | 92.91612616 | 93.47016103 | 0.0001551 | -0.674726305 | 5.48E-06 | 20.6604013 | 0.62645105 | 3.80938287 |
| *Nckap5l* | 236.0785886 | 204.6278832 | 214.2696753 | 314.2908535 | 138.2980701 | 176.9797418 | 151.8629159 | 139.30649 | 0.00041423 | -0.6768902 | 1.72E-05 | 18.4783909 | 0.62551214 | 3.38275782 |
| *Igsf6* | 1911.60981 | 1952.491052 | 2245.750263 | 2076.380557 | 1308.962016 | 1464.789778 | 1190.924972 | 1146.806976 | 2.68E-14 | -0.679611037 | 1.41E-16 | 68.294732 | 0.62433358 | 13.5711675 |
| *Slco4a1* | 759.4209465 | 782.275345 | 713.2119192 | 725.4451175 | 441.1903223 | 548.8254758 | 424.6167056 | 434.9957494 | 1.53E-11 | -0.687769165 | 1.16E-13 | 55.0685419 | 0.62081307 | 10.8149113 |
| *P2ry14* | 91.92440618 | 101.2481714 | 124.480478 | 116.4421851 | 71.09689521 | 72.4863836 | 60.94498597 | 64.71011148 | 0.00276029 | -0.689506679 | 0.0001654 | 14.188223 | 0.62006584 | 2.55904505 |
| *Pvrl3* | 135.7974182 | 125.7608865 | 125.5008098 | 118.5031087 | 80.83619593 | 90.37263409 | 75.93145794 | 66.50761458 | 0.00081517 | -0.689568394 | 3.77E-05 | 16.9836178 | 0.62003932 | 3.08875435 |
| *Vim* | 28396.28475 | 30006.76068 | 28874.36924 | 28909.60621 | 17191.81362 | 19528.02002 | 17564.14514 | 17473.5276 | 2.91E-43 | -0.695229272 | 2.32E-46 | 204.377658 | 0.61761116 | 42.5356366 |
| *Large* | 604.8208089 | 645.8567562 | 654.0326755 | 624.4598597 | 355.4844761 | 480.1046186 | 366.669014 | 359.5006193 | 1.46E-09 | -0.695246044 | 1.53E-11 | 45.493897 | 0.61760398 | 8.83577148 |
| *Csrp2* | 292.486747 | 296.2841225 | 238.7576382 | 250.402221 | 151.9330911 | 219.341914 | 145.8683271 | 146.4965024 | 0.00012187 | -0.698987563 | 4.08E-06 | 21.2280846 | 0.61600435 | 3.91410834 |
| *Adam19* | 296.6651291 | 272.8371775 | 310.1808633 | 297.8034644 | 186.9945737 | 240.0523093 | 145.8683271 | 151.8890117 | 9.36E-05 | -0.699949204 | 2.96E-06 | 21.8417084 | 0.61559388 | 4.02867027 |
| *Scd2* | 677.9424956 | 604.2917174 | 550.979165 | 556.4493799 | 280.4918606 | 484.8115266 | 342.6906588 | 362.196874 | 1.51E-05 | -0.700117456 | 3.66E-07 | 25.8647476 | 0.61552209 | 4.82222346 |
| *Fabp5* | 5587.541462 | 5318.193422 | 5554.686249 | 5351.188204 | 3105.862998 | 3961.333794 | 3091.209616 | 3243.594338 | 4.64E-19 | -0.702544454 | 1.66E-21 | 90.7161842 | 0.61448749 | 18.3337619 |
| *Adra1a* | 590.1964715 | 432.7027113 | 502.0032392 | 559.5407654 | 317.5012033 | 340.7801411 | 318.7123037 | 299.2842656 | 5.00E-09 | -0.7081985 | 5.84E-11 | 42.8732473 | 0.61208397 | 8.30109125 |
| *Flrt2* | 503.495043 | 366.6249573 | 454.0476452 | 485.3475147 | 193.8120842 | 353.9594835 | 283.7438691 | 273.2204707 | 0.00012231 | -0.711657494 | 4.10E-06 | 21.2166483 | 0.6106182 | 3.9125474 |
| *P4ha2* | 418.8828055 | 468.9388989 | 442.8239956 | 475.0428966 | 221.0821262 | 367.138826 | 273.7528878 | 239.9666634 | 1.05E-05 | -0.712393734 | 2.41E-07 | 26.6718323 | 0.61030667 | 4.97889676 |
| *Anpep* | 9069.178347 | 9261.543253 | 9164.620111 | 9687.37152 | 5286.492428 | 7619.542712 | 4907.570018 | 4775.965728 | 0.00066233 | -0.718968508 | 2.98E-05 | 17.4276656 | 0.60753166 | 3.17892678 |

| | | | | | | | | | | | | | | |
|---|---|---|---|---|---|---|---|---|---|---|---|---|---|---|
| *Vwf* | 738.5290361 | 621.344041 | 678.5206384 | 730.5974266 | 414.8942104 | 573.3013975 | 365.6699158 | 328.0443152 | 1.10E-05 | -0.719399426 | 2.58E-07 | 26.5399641 | 0.60735022 | 4.95670403 |
| *Pira11* | 836.7210154 | 772.683413 | 1044.81975 | 877.9534661 | 582.4101827 | 566.7117263 | 485.5616915 | 505.0983702 | 8.97E-11 | -0.72339691 | 7.84E-13 | 51.3213418 | 0.60566968 | 10.0471902 |
| *Hmga2* | 1276.495731 | 1270.398108 | 1227.45914 | 1201.518476 | 708.0471619 | 787.9364035 | 763.3109719 | 745.0650336 | 1.15E-27 | -0.727724081 | 2.28E-30 | 131.166481 | 0.60385577 | 26.9408167 |
| *Pvrl1* | 141.0203959 | 108.7085629 | 110.195833 | 153.5388104 | 71.09689521 | 98.84506854 | 76.93055607 | 62.91260839 | 0.00302905 | -0.72982008 | 0.0001868 | 13.9593791 | 0.60297911 | 2.51869287 |
| *Pstpip2* | 161.9123063 | 170.523236 | 225.4933249 | 216.3969811 | 124.6630491 | 121.4382271 | 111.8989906 | 108.7489374 | 6.44E-05 | -0.731377052 | 1.90E-06 | 22.6908571 | 0.60232872 | 4.19138602 |
| *Clec4d* | 5361.908829 | 5843.618142 | 6236.267883 | 5224.4414 | 3603.541264 | 3456.753254 | 3237.077944 | 3325.380729 | 4.51E-28 | -0.73453919 | 7.88E-31 | 133.271831 | 0.60100996 | 27.3458777 |
| *Dusp9* | 120.1284854 | 105.5112523 | 107.1348376 | 102.0157197 | 60.38366443 | 63.07256754 | 73.93326167 | 63.81135993 | 0.00069708 | -0.736114977 | 3.16E-05 | 17.3204457 | 0.60035387 | 3.15671837 |
| *Trim2* | 175.4920482 | 171.5890062 | 224.4729932 | 196.8182066 | 88.6276365 | 151.5624384 | 128.8836589 | 87.17890019 | 0.00109637 | -0.753311922 | 5.50E-05 | 16.2681771 | 0.59324012 | 2.9600436 |
| *Tfec* | 138.9312048 | 118.300495 | 158.151427 | 115.4117232 | 89.60156657 | 96.96230533 | 57.94769158 | 70.10262077 | 0.00203284 | -0.754290891 | 0.00011406 | 14.8884824 | 0.5928377 | 2.69189763 |
| *Hspa1b* | 137.8866093 | 167.3259253 | 134.6837959 | 110.2594142 | 75.96654557 | 78.13467323 | 70.93596728 | 97.96391877 | 0.00061515 | -0.765236255 | 2.74E-05 | 17.5917756 | 0.58835701 | 3.21101831 |
| *Pls3* | 1091.602323 | 1092.41448 | 1120.324302 | 1107.746451 | 539.5572596 | 807.7054172 | 617.4426448 | 626.4298292 | 5.47E-12 | -0.76752908 | 3.78E-14 | 57.2775536 | 0.5874227 | 11.2617267 |
| *Kdr* | 1202.329449 | 1076.427927 | 1289.699379 | 1264.376647 | 738.2389941 | 738.98456 | 724.3461448 | 635.4173447 | 1.56E-19 | -0.76931192 | 5.21E-22 | 93.0056949 | 0.58669723 | 18.8062601 |
| *Tmem26* | 174.4474526 | 175.8520871 | 217.3306706 | 187.5440503 | 114.9237484 | 116.731319 | 101.9080093 | 108.7489374 | 2.92E-06 | -0.771899254 | 5.85E-08 | 29.4109701 | 0.58564599 | 5.53412431 |
| *Cdc42bpa* | 382.3219621 | 366.6249573 | 407.112383 | 416.3065731 | 191.8642241 | 354.9008651 | 200.8187243 | 170.7627942 | 0.00044165 | -0.775845528 | 1.85E-05 | 18.338076 | 0.58404623 | 3.35491798 |
| *Samd4* | 115.9501033 | 94.85355001 | 96.93151977 | 104.0766433 | 49.67043364 | 87.54848928 | 47.95671028 | 54.82384445 | 0.00531403 | -0.777348057 | 0.00037122 | 12.6717718 | 0.58343828 | 2.27457614 |
| *Nptx1* | 915.0656797 | 717.2633613 | 781.5741489 | 896.5017788 | 490.860756 | 508.3460668 | 491.5562803 | 435.894501 | 2.96E-14 | -0.781542566 | 1.58E-16 | 68.0693086 | 0.58174444 | 13.52805 |
| *Nckap1* | 267.4164544 | 236.6009899 | 242.8389653 | 292.6511554 | 121.7412589 | 206.1625715 | 136.8764439 | 138.4077384 | 3.96E-05 | -0.78460874 | 1.11E-06 | 23.7193404 | 0.58050937 | 4.40211239 |
| *Pde2a* | 310.2448709 | 321.8626079 | 353.0347983 | 291.6206935 | 235.6910773 | 214.635006 | 144.869229 | 145.5977508 | 2.32E-05 | -0.785852259 | 5.99E-07 | 24.9166173 | 0.58000922 | 4.63528427 |
| *Tnfaip2* | 4229.56728 | 3943.349832 | 4256.824216 | 4354.731629 | 2528.322465 | 3094.321336 | 1959.231434 | 2136.33243 | 1.09E-05 | -0.788342036 | 2.52E-07 | 26.5840645 | 0.57900911 | 4.96365326 |
| *Slc24a5* | 195.3393631 | 208.8909641 | 164.2734177 | 184.4526648 | 90.57549664 | 144.0313856 | 123.8881682 | 76.39388161 | 0.00054096 | -0.793035007 | 2.35E-05 | 17.8844371 | 0.5771287 | 3.26683462 |
| *Bst1* | 84.61223751 | 105.5112523 | 115.2974919 | 110.2594142 | 51.61829378 | 55.5415147 | 74.93235981 | 57.5200991 | 0.00110543 | -0.796810041 | 5.58E-05 | 16.2406106 | 0.57562053 | 2.95646919 |
| *Alox5* | 92.96900171 | 91.65623933 | 91.82986084 | 111.289876 | 48.69650357 | 75.31052841 | 48.95580841 | 48.53258361 | 0.0020279 | -0.807285635 | 0.00011362 | 14.895741 | 0.57145602 | 2.69295315 |
| *Gpc1* | 736.439845 | 686.3560248 | 694.845947 | 747.0848156 | 422.685651 | 569.5358711 | 331.7005794 | 310.0692842 | 2.10E-06 | -0.81004403 | 4.05E-08 | 30.1239981 | 0.57036445 | 5.67806176 |
| *Ccdc80* | 297.7097246 | 335.7176208 | 336.7094897 | 268.9505336 | 123.6891191 | 268.2937575 | 149.8647196 | 160.8765272 | 0.00044549 | -0.817368708 | 1.87E-05 | 18.3143812 | 0.567476 | 3.35116157 |
| *Col4a2* | 908.7981066 | 804.6565197 | 911.1562858 | 860.4356152 | 448.9817629 | 755.9294289 | 388.6491729 | 382.8681596 | 0.00927382 | -0.818227429 | 0.00074143 | 11.3823972 | 0.56713833 | 2.03274147 |
| *Src* | 217.2758692 | 203.5621129 | 217.3306706 | 239.067141 | 136.35021 | 157.2107281 | 115.8953832 | 87.17890019 | 2.75E-05 | -0.822233109 | 7.44E-07 | 24.4973242 | 0.56556584 | 4.56004529 |
| *Il2rg* | 2684.610499 | 2829.619947 | 2871.213649 | 2850.257379 | 1497.90445 | 1720.845574 | 1521.626453 | 1605.170265 | 1.16E-42 | -0.823980011 | 1.01E-45 | 201.436303 | 0.56488143 | 41.9352045 |
| *Creb3l1* | 238.1677797 | 221.6802068 | 242.8389653 | 221.5492902 | 151.9330911 | 152.50382 | 100.9089112 | 115.0401982 | 1.58E-06 | -0.828516971 | 2.94E-08 | 30.7443539 | 0.5631078 | 5.8005542 |
| *Bhlhe40* | 1524.064871 | 1720.153143 | 1864.146175 | 1992.91315 | 1000.226183 | 1314.168721 | 853.2298036 | 825.0539214 | 0.00014101 | -0.830734617 | 4.92E-06 | 20.8683603 | 0.56224288 | 3.85075594 |
| *Amotl1* | 213.0974871 | 172.6547764 | 214.2696753 | 209.1837484 | 68.175105 | 194.8659923 | 91.91702803 | 98.86267032 | 0.00796252 | -0.833541703 | 0.00060496 | 11.7606645 | 0.56114997 | 2.09894971 |
| *Whrn* | 182.8042168 | 106.5770225 | 119.3788191 | 145.2951159 | 81.810126 | 105.4347398 | 67.93867289 | 54.82384445 | 0.0031987 | -0.838528634 | 0.00020082 | 13.8233753 | 0.55921361 | 2.49502646 |
| *Myo6* | 97.14738381 | 66.07775394 | 123.4601462 | 98.92433421 | 63.30545464 | 38.59664581 | 65.94047663 | 47.63383206 | 0.00654427 | -0.843261465 | 0.00047744 | 12.2017739 | 0.55738208 | 2.18418881 |
| *Procr* | 103.414957 | 115.1031843 | 114.2771602 | 90.68063969 | 52.59222386 | 59.30704112 | 55.94949532 | 67.40636613 | 0.00017727 | -0.845757938 | 6.46E-06 | 20.3469317 | 0.55641841 | 3.75135949 |
| *Acsbg1* | 86.70142856 | 66.07775394 | 85.70787011 | 93.77202513 | 30.19183221 | 45.18631705 | 62.94318224 | 45.83632897 | 0.00544997 | -0.853263012 | 0.00038288 | 12.6139463 | 0.55353137 | 2.26360616 |
| *Gpr162* | 591.2410671 | 595.7655557 | 601.9957544 | 638.8863251 | 376.9109376 | 412.3251431 | 298.7303411 | 256.1441913 | 1.47E-09 | -0.853620359 | 1.55E-11 | 45.4717381 | 0.55339428 | 8.83413587 |
| *Tagln* | 1393.49043 | 1431.329412 | 1235.621794 | 1279.833574 | 607.7323646 | 1152.251085 | 629.4318224 | 551.8334507 | 0.00999785 | -0.860442184 | 0.00081362 | 11.2098996 | 0.55078372 | 2.0000935 |
| *Rilpl2* | 1156.367246 | 1331.147011 | 1210.113499 | 1188.122472 | 721.6821829 | 694.7396246 | 668.3966495 | 603.062289 | 1.19E-26 | -0.86256699 | 2.65E-29 | 126.291598 | 0.54997312 | 25.9233663 |
| *C3* | 4117.795559 | 3782.418528 | 4876.16561 | 4842.140067 | 2471.834521 | 2620.806389 | 2242.975304 | 2305.297722 | 6.01E-23 | -0.869909572 | 1.67E-25 | 108.941388 | 0.54718115 | 22.221161 |
| *Erdr1* | 988.1873665 | 1120.124506 | 975.4371884 | 1089.198138 | 713.8907423 | 481.9873818 | 539.5129906 | 537.4534259 | 6.60E-14 | -0.876920615 | 3.57E-16 | 66.4617934 | 0.54452847 | 13.180371 |
| *Traf1* | 149.3771601 | 140.6816697 | 159.1717588 | 141.1732686 | 91.54942671 | 103.5519766 | 53.95129906 | 71.00137232 | 0.00012306 | -0.882479995 | 4.17E-06 | 21.1868179 | 0.54242418 | 3.90988513 |
| *Loxl3* | 747.9303958 | 609.6205686 | 636.6870351 | 760.4808192 | 312.6315529 | 581.773832 | 313.7168131 | 284.9042408 | 0.00863171 | -0.883616802 | 0.00067569 | 11.5549482 | 0.54200693 | 2.06390332 |
| *Mylk* | 82.52304646 | 76.73545618 | 83.66720654 | 75.22371247 | 46.74864343 | 64.95533075 | 29.97294392 | 30.55755264 | 0.00728936 | -0.885191531 | 0.00054918 | 11.9408009 | 0.54141564 | 2.13731048 |
| *Steap2* | 69.98790016 | 87.39315843 | 89.78919726 | 78.31509791 | 34.0875525 | 60.24842273 | 45.95851401 | 35.95006193 | 0.00248879 | -0.885511788 | 0.00014636 | 14.4184119 | 0.54129547 | 2.60401244 |
| *Ccl9* | 8847.724095 | 9290.319049 | 10441.05518 | 9262.821252 | 5077.097462 | 5823.386609 | 4878.596172 | 4692.381834 | 3.56E-32 | -0.88640012 | 5.38E-35 | 152.32418 | 0.54096227 | 31.447995 |
| *Tubb3* | 135.7974182 | 124.6951163 | 117.3381555 | 145.2951159 | 56.48794414 | 115.7899374 | 63.94228037 | 46.73508052 | 0.00386007 | -0.886816892 | 0.00025124 | 13.4028397 | 0.54080602 | 2.41340461 |
| *Rgs16* | 266.3718588 | 220.6144365 | 204.0663574 | 234.9452937 | 99.34086728 | 212.7522428 | 93.91522429 | 92.57140948 | 0.00189037 | -0.892592552 | 0.00010471 | 15.049797 | 0.53864529 | 2.72345413 |
| *Btg3* | 115.9501033 | 150.2736017 | 115.2974919 | 97.89387239 | 52.59222386 | 108.2588846 | 41.96212149 | 54.82384445 | 0.0085576 | -0.893929853 | 0.00066853 | 11.574759 | 0.53814623 | 2.06764778 |
| *Hspb1* | 144.1541824 | 175.8520871 | 185.7003852 | 195.7877448 | 81.810126 | 156.2693465 | 69.93686915 | 69.20386922 | 0.00226082 | -0.894610658 | 0.00013008 | 14.6405858 | 0.53789234 | 2.64573459 |
| *Chac1* | 213.0974871 | 204.6278832 | 211.2086799 | 274.1028427 | 139.2720002 | 141.2072408 | 98.91071494 | 104.2551796 | 1.22E-06 | -0.901132768 | 2.16E-08 | 31.3493175 | 0.53546613 | 5.91305208 |
| *Pilra* | 348.8949053 | 378.3484298 | 378.543093 | 334.9000898 | 174.3334828 | 211.8108612 | 171.8448785 | 211.2066139 | 4.81E-13 | -0.903421196 | 2.86E-15 | 62.3571235 | 0.53461744 | 12.3181627 |
| *Pik3r3* | 87.74602409 | 77.80122641 | 64.28090258 | 88.61971676 | 45.77471336 | 51.77598828 | 24.97745327 | 46.73508052 | 0.00342565 | -0.90814109 | 0.00021698 | 13.6780274 | 0.53287125 | 2.4652576 |
| *Phospho1* | 152.5109466 | 159.8655337 | 202.0256938 | 213.3055956 | 123.6891191 | 85.66572607 | 96.91251868 | 79.9888878 | 1.23E-05 | -0.916260578 | 2.93E-07 | 26.2932094 | 0.52988068 | 4.90877215 |
| *Myc* | 149.3771601 | 151.3393719 | 175.4970674 | 143.2341922 | 74.01868543 | 119.5554639 | 71.93506541 | 61.11510529 | 0.00012005 | -0.923641885 | 4.01E-06 | 21.2614568 | 0.52717655 | 3.92063693 |
| *Cdh2* | 208.919105 | 221.6802068 | 214.2696753 | 237.0062174 | 76.94047564 | 197.6901371 | 76.93055607 | 110.5464404 | 0.00174313 | -0.930779016 | 9.42E-05 | 15.2495234 | 0.52457501 | 2.7586714 |
| *Pdpn* | 548.4126505 | 556.3320573 | 633.6260398 | 563.6626126 | 253.2218186 | 361.4905364 | 285.7420654 | 301.9805203 | 3.45E-15 | -0.936317614 | 1.73E-17 | 72.4347477 | 0.52256499 | 14.462397 |
| *Ifitm6* | 85.65683304 | 59.68313259 | 106.1145059 | 81.40648336 | 45.77471336 | 58.36565952 | 32.97023831 | 36.84881348 | 0.00305756 | -0.936437748 | 0.00018916 | 13.9358434 | 0.52252148 | 2.51462546 |
| *Rhoj* | 129.5298451 | 90.59046911 | 118.3584873 | 110.2594142 | 38.95720286 | 78.13467323 | 54.95039719 | 62.01385684 | 0.00046839 | -0.937314311 | 1.98E-05 | 18.2081169 | 0.5222041 | 3.32938974 |
| *Ak1* | 275.7732186 | 241.929841 | 231.6153157 | 258.6459155 | 111.0280281 | 175.0969786 | 128.8836589 | 110.5464404 | 1.16E-07 | -0.939567369 | 1.68E-09 | 36.3187485 | 0.52138921 | 6.93376462 |
| *Cd28* | 1134.43074 | 1062.572914 | 1090.73468 | 1079.923982 | 413.9202803 | 856.6572607 | 501.5472616 | 504.1996186 | 0.00111112 | -0.940024865 | 5.63E-05 | 16.2219528 | 0.5212239 | 2.95423912 |
| *Fads3* | 512.8964027 | 481.7281416 | 537.7148518 | 480.1952056 | 219.1342661 | 385.9664581 | 223.7979813 | 218.3966263 | 2.30E-07 | -0.942045452 | 3.44E-09 | 34.9188491 | 0.5204944 | 6.63814734 |
| *Fkbp14* | 108.6379346 | 88.45892866 | 87.74853369 | 72.13232703 | 37.00934271 | 67.77947557 | 45.95851401 | 35.05131039 | 0.00213799 | -0.942309853 | 0.00012132 | 14.7721092 | 0.52039902 | 2.66999365 |
| *Cd207* | 571.3937521 | 512.6354781 | 556.0808239 | 528.6269109 | 279.5179305 | 237.2281645 | 289.7384579 | 315.4617935 | 1.19E-18 | -0.950506912 | 4.55E-21 | 88.7186262 | 0.51745062 | 17.9232936 |
| *Ddit4* | 130.5744406 | 160.9313039 | 153.0497681 | 153.5388104 | 53.56615393 | 135.5589511 | 63.94228037 | 55.722596 | 0.00298097 | -0.953157534 | 0.0001836 | 13.9918855 | 0.51650079 | 2.52564177 |
| *Lhfp* | 227.7218244 | 225.9432877 | 260.1846057 | 222.579752 | 94.47121693 | 200.5142819 | 106.9035 | 81.7863909 | 0.00040683 | -0.953237201 | 1.68E-05 | 18.523733 | 0.51647227 | 3.39058658 |
| *Ddr2* | 201.6069363 | 158.7997635 | 207.1273528 | 226.7015992 | 68.175105 | 176.9797418 | 90.9179299 | 71.90012227 | 0.00212844 | -0.961076833 | 0.00012061 | 14.7831948 | 0.51367336 | 2.67193814 |
| *Csf1* | 1996.222048 | 1949.293741 | 2021.27727 | 2099.050716 | 877.5109943 | 1702.959324 | 783.2929345 | 779.2175924 | 0.00916303 | -0.961120488 | 0.00073134 | 11.4078522 | 0.51365782 | 2.03796092 |
| *Ptgir* | 85.65683304 | 73.53814551 | 77.54521582 | 91.7111015 | 42.85292314 | 50.83460668 | 33.96933645 | 40.44381968 | 0.00022133 | -0.965161211 | 8.39E-06 | 19.8468249 | 0.51222117 | 3.65496198 |
| *Fpr2* | 44.91760757 | 58.61736236 | 56.11824829 | 49.4621671 | 32.13969236 | 16.94486889 | 33.96933645 | 24.26629181 | 0.00831627 | -0.966313018 | 0.00064241 | 11.6488927 | 0.51181239 | 2.08007138 |
| *Slc4a3* | 56.40815834 | 60.74890281 | 69.38256152 | 89.65017788 | 44.80078328 | 36.7138826 | 20.98106075 | 38.64631658 | 0.00491878 | -0.966580703 | 0.0003354 | 12.8615777 | 0.51171744 | 2.308143 |
| *Fam101b* | 56.40815834 | 59.68313259 | 71.42322509 | 64.91909432 | 26.29611193 | 41.42079063 | 37.96572897 | 23.36754026 | 0.00280791 | -0.971003128 | 0.00016937 | 14.1436046 | 0.51015122 | 2.55161622 |
| *Msrb3* | 141.0203959 | 151.3393719 | 138.765123 | 174.1480467 | 69.14903507 | 127.0865167 | 52.95220093 | 59.31760219 | 0.00089457 | -0.971358193 | 4.25E-05 | 16.7570728 | 0.51002568 | 3.04838462 |
| *Egr2* | 150.4217556 | 172.6547764 | 152.0294363 | 140.1428068 | 88.6276365 | 119.5554639 | 51.9531028 | 53.02634135 | 0.00070272 | -0.973978603 | 3.19E-05 | 17.3017903 | 0.50910015 | 3.15321609 |
| *Chst2* | 147.287969 | 164.1286146 | 152.0294363 | 176.2089703 | 62.33152457 | 151.5624384 | 50.95490467 | 60.21635374 | 0.00511292 | -0.975483141 | 0.00035392 | 12.7610175 | 0.5085695 | 2.29133092 |
| *Dusp4* | 644.5154388 | 557.3978276 | 583.6297822 | 597.6678525 | 339.9015949 | 323.8352722 | 296.7321448 | 251.6504335 | 1.04E-18 | -0.976536153 | 3.90E-21 | 89.0232644 | 0.50819844 | 17.9810301 |
| *Cttn* | 270.5502409 | 306.9418247 | 307.1198679 | 322.534548 | 134.4023499 | 233.4626381 | 110.8998925 | 133.0152292 | 6.86E-06 | -0.979593218 | 1.52E-07 | 27.568289 | 0.50712271 | 5.16343348 |
| *Phlda1* | 141.0203959 | 116.1689545 | 111.2161648 | 126.7468032 | 46.74864343 | 90.37263409 | 61.94408411 | 51.22883826 | 0.0001227 | -0.984079904 | 4.14E-06 | 21.1969981 | 0.50554804 | 3.91117128 |
| *Rhou* | 83.56764199 | 71.40660506 | 73.46388867 | 64.91909432 | 29.21790214 | 56.48289631 | 32.97023831 | 28.76004955 | 0.00219626 | -0.992095527 | 0.0001255 | 14.708241 | 0.502747 | 2.6583126 |
| *Edil3* | 334.2705679 | 296.2841225 | 307.1198679 | 272.0419191 | 116.8716086 | 197.6901371 | 138.8746402 | 153.6865148 | 3.65E-09 | -0.99324596 | 4.03E-11 | 43.5986922 | 0.50234626 | 8.43780882 |
| *Ppic* | 177.5812392 | 174.7863169 | 182.6393899 | 149.4169631 | 63.30545464 | 146.8555304 | 55.94949532 | 77.29263316 | 0.00100341 | -0.993255547 | 4.90E-05 | 16.4878724 | 0.50234292 | 2.99852274 |
| *Ccbe1* | 173.4028571 | 134.2870483 | 136.7244595 | 176.2089703 | 61.3575945 | 126.1451351 | 62.94318224 | 60.21635374 | 0.00045115 | -0.997784523 | 1.90E-05 | 18.2867135 | 0.50076842 | 3.34567424 |
| *Spint1* | 113.8609122 | 154.5366826 | 125.5008098 | 140.1428068 | 68.175105 | 53.65875149 | 62.94318224 | 81.7863909 | 3.95E-06 | -1.000286842 | 8.10E-08 | 28.7819595 | 0.4999006 | 5.40333908 |
| *Ppap2b* | 168.1798795 | 163.0628444 | 186.720717 | 161.7825049 | 86.67977635 | 126.1451351 | 49.95490654 | 76.39388161 | 0.00010847 | -1.001432562 | 3.56E-06 | 21.4881064 | 0.49950376 | 3.96467923 |
| *Egfr* | 91.92440618 | 92.72200956 | 91.82986084 | 105.1071051 | 33.11362243 | 77.19329162 | 48.95580841 | 30.55755264 | 0.00240121 | -1.008226828 | 0.00013995 | 14.502882 | 0.49715691 | 2.6195691 |
| *Lilra6* | 157.7339242 | 151.3393719 | 160.1920906 | 191.6658975 | 84.73191621 | 78.13467323 | 84.92334111 | 79.09013626 | 4.91E-09 | -1.016910801 | 5.70E-11 | 42.9205018 | 0.49417338 | 8.30861714 |
| *Parva* | 238.1677797 | 249.3902326 | 234.676311 | 260.7068391 | 83.75798614 | 206.1625715 | 90.9179299 | 104.2551796 | 0.00016904 | -1.017729882 | 6.11E-06 | 20.4526779 | 0.49389289 | 3.77201562 |
| *Eya1* | 195.3393631 | 219.5486663 | 261.2049375 | 261.7373009 | 84.73191621 | 155.3279648 | 128.8836589 | 94.36891258 | 1.95E-06 | -1.018150067 | 3.71E-08 | 30.2953803 | 0.49374907 | 5.7110432 |
| *Ncam1* | 169.224475 | 143.8789803 | 162.2327541 | 203.0009775 | 55.51401407 | 151.5624384 | 66.93957476 | 60.21635374 | 0.00223518 | -1.02068007 | 0.00012843 | 14.6646893 | 0.49288396 | 2.65068818 |
| *Amotl2* | 273.6840275 | 280.2975691 | 272.4285871 | 277.1942281 | 110.0540981 | 228.7557301 | 109.9007944 | 93.47016103 | 8.27E-05 | -1.025154241 | 2.53E-06 | 22.1444212 | 0.49135776 | 4.08274526 |
| *Npy* | 1632.702805 | 1723.350453 | 1529.477349 | 1524.053024 | 787.9094278 | 849.1262078 | 779.296542 | 732.4825119 | 1.45E-45 | -1.025476578 | 8.07E-49 | 215.643299 | 0.49124799 | 44.8385802 |
| *Gpr84* | 1004.900895 | 1157.426464 | 994.8234924 | 1068.588902 | 542.4790498 | 534.7047517 | 429.6121962 | 567.112227 | 3.63E-26 | -1.025694964 | 8.66E-29 | 123.945779 | 0.49117364 | 25.4399328 |
| *Gdf15* | 55.36356282 | 73.53814551 | 64.28090258 | 46.37078166 | 37.98327278 | 32.00697457 | 19.98196261 | 26.96254645 | 0.00287441 | -1.033323419 | 0.00017521 | 14.0798552 | 0.48858334 | 2.54145153 |
| *Gem* | 50.14058519 | 56.48582191 | 40.81327148 | 47.40124347 | 19.47860143 | 31.06559297 | 22.97925701 | 21.57003716 | 0.00386791 | -1.033392781 | 0.00025206 | 13.3967462 | 0.48855985 | 2.4125237 |
| *Pilrb1* | 96.10278828 | 74.60391573 | 82.64687475 | 97.89387239 | 40.905063 | 46.12769865 | 31.97114018 | 52.12758981 | 9.13E-05 | -1.033621321 | 2.87E-06 | 21.9045474 | 0.48847881 | 4.0396063 |
| *Atp6v0d2* | 671.6749225 | 621.344041 | 761.1675131 | 715.1404994 | 254.1957486 | 428.3286303 | 323.7077944 | 339.7280853 | 1.72E-13 | -1.04039128 | 9.70E-16 | 64.4895072 | 0.48619559 | 12.7644885 |
| *Epb4.1l3* | 190.1163855 | 177.9836275 | 197.9443667 | 206.0923629 | 63.30545464 | 150.6210568 | 68.93777102 | 91.67265793 | 8.99E-05 | -1.041960803 | 2.81E-06 | 21.9391351 | 0.48566694 | 4.04633284 |
| *Arhgef17* | 145.198778 | 139.6158994 | 129.582137 | 144.2646541 | 53.56615393 | 118.6140822 | 50.95400467 | 47.63383206 | 0.00083623 | -1.044190478 | 3.88E-05 | 16.928667 | 0.48491693 | 3.07767363 |
| *Col18a1* | 2463.156248 | 2165.645097 | 2544.707477 | 2541.118835 | 1199.881848 | 1249.21339 | 1144.966458 | 1114.45192 | 8.45E-50 | -1.045122521 | 3.36E-53 | 235.728832 | 0.48460375 | 49.0731407 |
| *Ptpm* | 108.6379346 | 86.32738821 | 89.78919726 | 87.58925425 | 39.93113293 | 70.60362039 | 38.9648271 | 30.55755264 | 0.00066727 | -1.048495672 | 3.01E-05 | 17.4101721 | 0.48347203 | 3.17569521 |
| *Fstl3* | 82.52304646 | 83.13007753 | 111.2161648 | 87.58925425 | 37.00934271 | 69.66223878 | 26.97564953 | 42.24132277 | 0.00138581 | -1.048621318 | 7.25E-05 | 15.7453016 | 0.48342992 | 2.85829723 |
| *Nqo1* | 68.94330464 | 87.39315843 | 96.93151977 | 91.7111015 | 30.19183221 | 53.65875149 | 33.96933645 | 48.53258361 | 0.00025372 | -1.048713528 | 9.84E-06 | 19.5422234 | 0.48339903 | 3.59563838 |
| *Ctsk* | 145.198778 | 154.5366826 | 134.6837959 | 151.4778868 | 52.59222386 | 127.0865167 | 53.95129906 | 50.33008671 | 0.0009879 | -1.05455766 | 4.80E-05 | 16.5266838 | 0.48144481 | 3.00528533 |
| *Has2* | 168.1798795 | 184.9447506 | 180.5987263 | 164.8738903 | 61.3575945 | 142.1486224 | 56.94859345 | 55.722596 | 0.00089457 | -1.058112658 | 4.25E-05 | 16.7585603 | 0.48025993 | 3.04838462 |
| *Bcar1* | 94.01359723 | 108.7085629 | 128.5618052 | 110.2594142 | 49.67043364 | 91.3140157 | 46.95761214 | 24.26629181 | 0.00452163 | -1.058430483 | 0.00030293 | 13.052207 | 0.48015414 | 2.34470505 |
| *Gpr97* | 58.49734939 | 85.26161798 | 57.13858008 | 62.85817069 | 23.37432171 | 41.42079063 | 32.97023831 | 28.76004955 | 0.00110728 | -1.058523884 | 5.61E-05 | 16.2314793 | 0.48012305 | 2.95574169 |
| *Dlg5* | 128.4852496 | 96.98509045 | 106.1145059 | 121.5944941 | 37.00934271 | 102.610595 | 50.95400467 | 26.0637949 | 0.00916303 | -1.065475324 | 0.00072917 | 11.4133765 | 0.47781521 | 2.03796092 |
| *Gpx8* | 78.34466436 | 93.78777978 | 105.0941741 | 74.19325066 | 25.32218186 | 75.31052841 | 32.97023831 | 33.25380729 | 0.00508318 | -1.073136957 | 0.00035105 | 12.7762301 | 0.47528443 | 2.29386451 |
| *Uchl1* | 975.6522202 | 1050.849442 | 1018.291123 | 925.3547096 | 451.9035531 | 466.9252762 | 466.5788271 | 491.617097 | 3.70E-45 | -1.080141907 | 2.35E-48 | 213.514648 | 0.4729823 | 44.4322071 |
| *Osmr* | 120.1284854 | 96.98509045 | 123.4601462 | 75.22371247 | 37.00934271 | 84.72434446 | 39.96392523 | 34.15255884 | 0.00243354 | -1.085750971 | 0.00014254 | 14.4683378 | 0.47114696 | 2.6137607 |
| *Il1m* | 577.6613252 | 577.6474618 | 625.4633855 | 531.7182964 | 223.0299863 | 322.8938906 | 297.731243 | 244.4604212 | 2.47E-18 | -1.088041257 | 9.62E-21 | 87.2384402 | 0.4703996 | 17.6072554 |
| *Myo1b* | 215.1866781 | 206.7594236 | 196.9240349 | 252.4631446 | 87.65370643 | 139.3244776 | 84.92334111 | 97.06516722 | 1.43E-08 | -1.089892473 | 1.76E-10 | 40.7177005 | 0.46979639 | 7.84531441 |
| *Ak4* | 59.54194492 | 50.09120056 | 73.46388867 | 69.04094158 | 21.42646157 | 26.35868494 | 36.96663084 | 33.25380729 | 0.00070483 | -1.096282623 | 3.20E-05 | 17.2927815 | 0.46772011 | 3.15191704 |
| *Mmp14* | 1076.977986 | 1077.493697 | 1082.572026 | 1130.416611 | 477.225735 | 663.6740316 | 454.5896495 | 445.780768 | 8.74E-21 | -1.097171914 | 2.71E-23 | 98.8613623 | 0.4674319 | 20.0586128 |
| *Gpc6* | 61.63113596 | 68.20929439 | 109.1755012 | 98.92433421 | 26.29611193 | 67.77947557 | 28.97384579 | 34.15255884 | 0.00506321 | -1.102927648 | 0.00034927 | 12.7857462 | 0.46557076 | 2.29557373 |
| *Nbl1* | 78.34466436 | 69.27506461 | 89.78919726 | 103.0461815 | 35.06148257 | 68.72085718 | 34.96843458 | 19.77253406 | 0.00459806 | -1.104267738 | 0.00030892 | 13.0155172 | 0.4651385 | 2.33742572 |
| *Elk1* | 393.8125129 | 424.1765495 | 446.9053227 | 403.9410313 | 171.4116926 | 195.8073739 | 216.8042944 | 191.4340798 | 1.69E-22 | -1.106340887 | 4.84E-25 | 106.834081 | 0.46447058 | 21.7716332 |
| *Grb10* | 128.4852496 | 155.6024528 | 167.3344131 | 173.1175849 | 44.80078328 | 134.6175695 | 57.94769158 | 52.12758981 | 0.00134577 | -1.108610961 | 7.03E-05 | 15.8036468 | 0.46374031 | 2.87103011 |
| *Loxl2* | 1365.286351 | 1241.622312 | 1387.65123 | 1436.46377 | 453.8514133 | 1083.530228 | 488.5589859 | 480.8320784 | 0.00658623 | -1.115330808 | 0.00048155 | 12.1857916 | 0.4615853 | 2.18136315 |
| *Adam12* | 222.4988468 | 163.0628444 | 180.5987263 | 218.4570947 | 75.96654557 | 158.1521097 | 67.93867289 | 59.31760219 | 0.00024096 | -1.1183066 | 9.29E-06 | 19.6526311 | 0.46063419 | 3.61805713 |
| *Smoc2* | 124.3068675 | 134.2870483 | 121.4194827 | 138.0818832 | 48.69650357 | 109.2002662 | 39.96392523 | 39.54506813 | 0.0009986 | -1.12500729 | 4.86E-05 | 16.5031569 | 0.45849957 | 3.00060682 |
| *Gpr116* | 69.98790016 | 75.66968596 | 104.0738423 | 73.16278884 | 23.37432171 | 67.77947557 | 26.97564953 | 29.6588011 | 0.00469408 | -1.126052499 | 0.00031672 | 12.9688342 | 0.45816765 | 2.32844914 |
| *Adcy6* | 96.10278828 | 100.1824011 | 116.3178237 | 99.95479602 | 18.50467136 | 90.37263409 | 32.97023831 | 44.93757742 | 0.00829388 | -1.140910892 | 0.00063941 | 11.6575912 | 0.45347317 | 2.08124231 |
| *Cemip* | 192.2055766 | 187.5755596 | 156.1107634 | 181.3612794 | 42.85292314 | 139.3244776 | 74.93235981 | 65.60886303 | 0.00022024 | -1.151313129 | 8.31E-06 | 19.864318 | 0.45021526 | 3.65711145 |
| *Mpdz* | 99.23657486 | 76.73545618 | 98.97218334 | 126.7468032 | 32.13969236 | 85.66572607 | 23.97835514 | 38.64631658 | 0.00480886 | -1.15264723 | 0.00032545 | 12.9179299 | 0.44979913 | 2.31868111 |
| *Cnn1* | 130.5744406 | 131.0897376 | 107.1348376 | 139.112345 | 28.24397207 | 114.8485558 | 45.95851401 | 38.64631658 | 0.00493127 | -1.156526929 | 0.00033719 | 12.8515832 | 0.44859115 | 2.30704106 |
| *Ndrg4* | 75.21087779 | 87.39315843 | 96.93151977 | 69.04094158 | 22.40039164 | 58.36565952 | 30.97204205 | 35.05131039 | 0.00054543 | -1.160095434 | 2.38E-05 | 17.8583475 | 0.44748293 | 3.26326033 |
| *Bcat1* | 168.1798795 | 130.0239674 | 177.5377309 | 120.5640323 | 45.77471336 | 122.3796087 | 52.95220093 | 44.93757742 | 0.00084105 | -1.164008382 | 3.91E-05 | 16.9145169 | 0.44627089 | 3.07517934 |
| *Ccl2* | 104.4595525 | 155.6024528 | 168.3547449 | 168.9957376 | 79.86226585 | 96.02092372 | 45.95851401 | 43.1400732 | 9.65E-05 | -1.173371164 | 3.10E-06 | 21.7556051 | 0.44338407 | 4.01567871 |
| *Syde1* | 77.30006884 | 67.14352416 | 60.19957544 | 68.01047977 | 22.40039164 | 40.47940902 | 28.97384579 | 28.76004955 | 9.18E-05 | -1.174946821 | 2.90E-06 | 21.8845679 | 0.44290008 | 4.03727804 |
| *Abtb2* | 49.09598967 | 26.64425562 | 40.81327148 | 49.4621671 | 14.60895107 | 24.47592173 | 21.98015888 | 12.58252168 | 0.00760965 | -1.176318676 | 0.00057573 | 11.8528606 | 0.44247913 | 2.1186351 |
| *Cdr2l* | 57.45275387 | 51.15697079 | 52.03692114 | 48.43170529 | 24.34825178 | 39.53802742 | 13.98737383 | 14.38002477 | 0.00504755 | -1.17974794 | 0.00034699 | 12.7980204 | 0.44142862 | 2.29691955 |
| *Gpr141* | 145.198778 | 101.2481714 | 123.4601462 | 103.0461815 | 53.56615393 | 48.01046186 | 49.95490654 | 55.722596 | 9.00E-08 | -1.18980439 | 1.27E-09 | 36.8540652 | 0.43836229 | 7.04577207 |
| *Mxra8* | 98.19197933 | 115.1031843 | 141.8261184 | 126.7468032 | 41.87899307 | 98.84506854 | 32.97023831 | 36.84881348 | 0.00107621 | -1.193572733 | 5.37E-05 | 16.3123716 | 0.43721878 | 2.96810303 |
| *Fgfr1* | 400.080086 | 349.5726337 | 364.258448 | 357.5702497 | 134.4023499 | 284.2972448 | 121.8899719 | 101.558925 | 0.00666548 | -1.196224456 | 0.0004884 | 12.1594313 | 0.43641589 | 2.1761687 |
| *Cxcl2* | 59.54194492 | 76.73545618 | 63.2605708 | 95.83294876 | 27.270042 | 54.6001331 | 20.98106075 | 25.16504335 | 0.0014501 | -1.204425206 | 7.65E-05 | 15.6423599 | 0.4339422 | 2.83860269 |
| *Ptges* | 65.80951806 | 59.68313259 | 85.70787011 | 84.4978688 | 22.40039164 | 44.24493544 | 26.97564953 | 34.15255884 | 0.00012832 | -1.20810394 | 4.39E-06 | 21.0886621 | 0.4328371 | 3.89169976 |

| | | | | | | | | | | | | | | |
|---|---|---|---|---|---|---|---|---|---|---|---|---|---|---|
| *Fgr* | 433.5071428 | 373.0195787 | 489.7592578 | 431.7635003 | 220.1081961 | 210.8694796 | 147.8665233 | 168.9652911 | 1.20E-16 | -1.208511539 | 5.55E-19 | 79.223541 | 0.43271483 | 15.919623 |
| *Pdzrn3* | 66.85411359 | 46.89388989 | 51.01658935 | 56.67539981 | 25.32218186 | 39.53802742 | 8.991883177 | 21.57003716 | 0.00605135 | -1.211047563 | 0.00043234 | 12.3869621 | 0.43195485 | 2.21814799 |
| *Gpc4* | 121.1730809 | 106.5770225 | 108.1551694 | 104.0766433 | 32.13969236 | 90.37263409 | 38.9648271 | 27.861298 | 0.00108688 | -1.216226957 | 5.43E-05 | 16.2906585 | 0.43040688 | 2.96381768 |
| *Ugt1a6a* | 133.7082272 | 102.3139416 | 164.2734177 | 70.0714034 | 44.80078328 | 83.78296286 | 36.96663084 | 36.84881348 | 0.00104504 | -1.216254162 | 5.15E-05 | 16.3923653 | 0.43039876 | 2.9808659 |
| *Alpl* | 91.92440618 | 100.1824011 | 108.1551694 | 108.1984905 | 25.32218186 | 80.95881804 | 34.96843458 | 34.15255884 | 0.00071265 | -1.218179417 | 3.26E-05 | 17.2618715 | 0.42982479 | 3.147124 |
| *Sdc1* | 1342.305249 | 1294.910823 | 1466.216778 | 1387.001603 | 530.7918889 | 821.8261413 | 490.5571822 | 512.2883826 | 2.56E-18 | -1.220691353 | 1.02E-20 | 87.1267766 | 0.42907705 | 17.5915098 |
| *Nfatc4* | 67.89870911 | 66.07775394 | 58.15891186 | 54.61447618 | 18.50467136 | 46.12769865 | 25.9765514 | 15.27877632 | 0.00219628 | -1.221187922 | 0.00012537 | 14.7100989 | 0.42892939 | 2.6583126 |
| *Pappa* | 91.92440618 | 79.93276686 | 122.4398144 | 110.2594142 | 30.19183221 | 81.90019965 | 29.97294392 | 30.55755264 | 0.00141669 | -1.22791578 | 7.44E-05 | 15.6950118 | 0.42693378 | 2.84872656 |
| *Rcn3* | 315.4678485 | 344.2437826 | 327.5265036 | 319.4431625 | 132.4544897 | 238.1695461 | 93.91522429 | 92.57140948 | 0.00296675 | -1.229705063 | 0.00018249 | 14.0033038 | 0.42640461 | 2.52771841 |
| *Fkbp10* | 427.2395697 | 454.0181158 | 477.5152763 | 410.1238022 | 133.4284198 | 342.6629043 | 146.8674252 | 130.3189745 | 0.00720658 | -1.231454017 | 0.00054008 | 11.9719344 | 0.425888 | 2.14227065 |
| *Sparc* | 3504.617986 | 3789.878919 | 3648.706471 | 3633.408358 | 1035.287666 | 2744.127379 | 1180.933991 | 1233.087124 | 0.00411737 | -1.234807642 | 0.00027136 | 13.2583985 | 0.42489915 | 2.38538048 |
| *Rgl1* | 175.4920482 | 155.6024528 | 204.0663574 | 178.2698939 | 83.75798614 | 105.4347398 | 57.94769158 | 55.722596 | 4.74E-08 | -1.236644547 | 6.41E-10 | 38.1926439 | 0.42435849 | 7.32385012 |
| *Fgf7* | 36.56084337 | 52.22274101 | 53.05725293 | 46.37078166 | 13.635021 | 36.7138826 | 12.9882757 | 15.27877632 | 0.00613185 | -1.256868912 | 0.00044199 | 12.3457477 | 0.41845114 | 2.21240859 |
| *Vcam1* | 41.78382099 | 50.09120056 | 46.9352622 | 55.64493799 | 21.42646157 | 33.88973779 | 11.98917757 | 13.48127323 | 0.00274986 | -1.26632354 | 0.00016456 | 14.1978477 | 0.41571781 | 2.56069012 |
| *Lrrc32* | 124.3068675 | 119.3662652 | 108.1551694 | 103.0461815 | 25.32218186 | 77.19329162 | 39.96392523 | 44.93757742 | 4.42E-05 | -1.27672111 | 1.27E-06 | 23.4726042 | 0.41273248 | 4.35492851 |
| *Map6* | 80.43385541 | 66.07775394 | 71.42322509 | 69.04094158 | 25.32218186 | 39.53802742 | 25.9765514 | 26.96254645 | 6.74E-06 | -1.28332694 | 1.48E-07 | 27.6164013 | 0.41084698 | 5.17110037 |
| *Timp1* | 1166.813201 | 1394.027454 | 1163.178237 | 1235.523716 | 366.1977068 | 931.0264075 | 351.682542 | 384.6656627 | 0.00571524 | -1.286085421 | 0.0004047 | 12.5103899 | 0.41006218 | 2.24296551 |
| *Ass1* | 1375.732306 | 1419.605939 | 1404.996871 | 1298.381886 | 555.1401407 | 412.3251431 | 636.4255093 | 628.2273323 | 1.78E-29 | -1.300794417 | 2.82E-32 | 139.883348 | 0.40590263 | 28.7505228 |
| *Lyz2* | 317391.9935 | 325518.1998 | 335273.8829 | 316411.5439 | 130265.0949 | 139389.4329 | 124331.7678 | 131348.045 | 1.86E-180 | -1.301189073 | 1.48E-184 | 839.38621 | 0.40579161 | 179.731172 |
| *Pdgfra* | 122.2176764 | 115.1031843 | 129.582137 | 131.8991123 | 38.95720286 | 95.07954212 | 39.96392523 | 27.861298 | 0.00021628 | -1.30523323 | 8.10E-06 | 19.9150946 | 0.40465569 | 3.66497377 |
| *Slc7a11* | 751.0641824 | 647.9882967 | 942.7865712 | 760.4808192 | 298.9965319 | 422.6803407 | 260.7646121 | 260.637949 | 2.13E-16 | -1.319436492 | 1.01E-18 | 78.0321581 | 0.40069142 | 15.6724293 |
| *Actg2* | 322.7800172 | 377.2826596 | 336.7094897 | 385.3927187 | 121.7412589 | 252.2902702 | 104.9053037 | 90.77390639 | 0.00244631 | -1.319676882 | 0.00014348 | 14.455929 | 0.40062466 | 2.61148841 |
| *Lgr6* | 104.4595525 | 93.78777978 | 68.36222973 | 62.85817069 | 19.47860143 | 70.60362039 | 20.98106075 | 20.67128561 | 0.0059319 | -1.320738475 | 0.00042192 | 12.4325203 | 0.40032997 | 2.22680652 |
| *Evc2* | 40.73922547 | 45.82811967 | 41.83360327 | 39.15754896 | 18.50467136 | 24.47592173 | 13.98737383 | 9.886267032 | 0.00110655 | -1.327468845 | 5.59E-05 | 16.2357032 | 0.39846672 | 2.95602765 |
| *Svep1* | 84.61223751 | 71.40660506 | 91.82986084 | 82.43694517 | 27.270042 | 64.01394915 | 17.98376635 | 21.57003716 | 0.00104304 | -1.334571802 | 5.11E-05 | 16.4052022 | 0.39650974 | 2.98170058 |
| *Slc6a12* | 199.5177452 | 166.2601551 | 210.1883481 | 172.087123 | 72.07082528 | 103.5519766 | 60.94498597 | 56.62134755 | 1.77E-10 | -1.351541587 | 1.63E-12 | 49.8801292 | 0.39187309 | 9.75133825 |
| *Wisp2* | 186.9825989 | 203.5621129 | 191.822376 | 218.4579047 | 52.59222386 | 148.7382936 | 61.94408411 | 50.33008671 | 0.00654427 | -1.352299682 | 0.00047716 | 12.202866 | 0.39166723 | 2.18413881 |
| *Ccl7* | 83.56764199 | 60.74890281 | 65.30123437 | 81.40648336 | 29.21790214 | 48.01046186 | 19.98196261 | 16.17752787 | 0.00027287 | -1.360401312 | 1.07E-05 | 19.3764229 | 0.38947394 | 3.56404229 |
| *Srpx2* | 367.6976247 | 286.6921905 | 321.4045129 | 382.3013332 | 71.09689521 | 237.2281645 | 98.91071494 | 117.7364528 | 0.00544524 | -1.370818456 | 0.00038211 | 12.6176848 | 0.38667182 | 2.26398321 |
| *Ntn1* | 30.29327022 | 27.71002584 | 41.83360327 | 32.97477807 | 14.60895107 | 19.76901371 | 3.996392523 | 12.58252168 | 0.00842217 | -1.377503418 | 0.00065327 | 11.6177172 | 0.38488426 | 2.07457603 |
| *Fam20a* | 112.8163167 | 114.0374141 | 121.4194827 | 149.4169631 | 45.77471336 | 86.60710767 | 30.97204205 | 27.861298 | 7.06E-05 | -1.379903402 | 2.12E-06 | 22.4813552 | 0.38424452 | 4.15097604 |
| *Ndn* | 31.33786574 | 28.77579607 | 27.54895825 | 34.00523988 | 7.791440571 | 20.71039531 | 9.990981307 | 8.088763935 | 0.00670699 | -1.384226588 | 0.00049304 | 12.141792 | 0.38309482 | 2.17347204 |
| *Itga11* | 321.7354216 | 269.6398669 | 316.302854 | 294.712079 | 74.9926155 | 219.341914 | 91.91702803 | 71.90012387 | 0.00675617 | -1.391893893 | 0.0004972 | 12.1261541 | 0.38106423 | 2.1702996 |
| *Sphk1* | 43.87301204 | 56.48582191 | 60.19957544 | 42.2489344 | 12.66109093 | 40.47940902 | 11.98917757 | 11.68377013 | 0.00528128 | -1.398871022 | 0.00036772 | 12.6894566 | 0.37922579 | 2.27726093 |
| *Zfp462* | 33.42705679 | 30.90733652 | 41.83360327 | 42.2489344 | 15.58288114 | 21.65177692 | 9.990981307 | 8.987515484 | 0.00185418 | -1.401987608 | 0.00010182 | 15.1025947 | 0.37840745 | 2.73184695 |
| *Fabp4* | 527.52074 | 580.8447725 | 546.8978379 | 543.0533763 | 170.4377625 | 301.2421136 | 159.8557009 | 198.6240922 | 1.04E-16 | -1.403686817 | 4.71E-19 | 79.5470743 | 0.37796202 | 15.9831882 |
| *Rgs4* | 25.0702926 | 33.03887697 | 33.67094897 | 26.79200718 | 13.635021 | 15.06210568 | 6.993686915 | 8.987515484 | 0.00380735 | -1.407033278 | 0.00024569 | 13.444738 | 0.37708632 | 2.4193767 |
| *Pcolce2* | 62.67573149 | 49.02543034 | 68.36222973 | 59.76678525 | 10.71323079 | 30.12421136 | 28.97384579 | 20.67128561 | 0.00011743 | -1.407284705 | 3.91E-06 | 21.3082859 | 0.37702061 | 3.93021128 |
| *Kdelr3* | 80.43385541 | 121.4978056 | 113.2568284 | 90.68063969 | 26.29611193 | 71.54500199 | 27.97474766 | 26.0637949 | 0.00013377 | -1.416855117 | 4.59E-06 | 21.0000335 | 0.37452784 | 3.87362642 |
| *Gstm2* | 87.74602409 | 65.01198371 | 63.2605708 | 73.16278884 | 18.50467136 | 48.01046186 | 18.98286448 | 22.46878871 | 0.0001227 | -1.418731339 | 4.14E-06 | 21.1998692 | 0.37404109 | 3.91117128 |
| *Ddr1* | 159.8231153 | 130.0239674 | 133.6634641 | 112.3203378 | 47.7225735 | 86.60710767 | 28.97384579 | 36.84881348 | 8.65E-06 | -1.41926596 | 1.95E-07 | 27.0864013 | 0.3739025 | 5.0629447 |
| *Thbs2* | 1320.368743 | 1195.794192 | 1258.069093 | 1356.087748 | 296.0747417 | 903.7263409 | 373.6627009 | 302.8792718 | 0.00505054 | -1.451083979 | 0.00034759 | 12.7947507 | 0.36574651 | 2.29666231 |
| *Dkk3* | 207.8745094 | 242.9956112 | 257.1236103 | 237.0062174 | 58.43580428 | 170.3900705 | 66.93957476 | 48.53258361 | 0.00852264 | -1.456559339 | 0.00066445 | 11.5861586 | 0.36436105 | 2.06942561 |
| *Ptprf* | 54.31896729 | 43.69657922 | 45.91493042 | 30.91385444 | 10.71323079 | 26.35868494 | 11.98917757 | 14.38002477 | 0.00085349 | -1.459056589 | 4.00E-05 | 16.8737013 | 0.3637309 | 3.06880016 |
| *Mmp23* | 47.00679862 | 76.73545618 | 63.2605708 | 58.73632344 | 18.50467136 | 40.47940902 | 11.98917757 | 17.07627942 | 0.00039777 | -1.477651653 | 1.63E-05 | 18.5740219 | 0.35907282 | 3.40036761 |
| *Cacnb3* | 56.40815834 | 60.74890281 | 42.85393506 | 57.70586162 | 19.47860143 | 32.94835618 | 13.98737383 | 11.68377013 | 0.00019873 | -1.478810657 | 7.33E-06 | 20.1056123 | 0.35878447 | 3.70174049 |
| *Mreg* | 32.38246127 | 35.17041742 | 26.52862646 | 31.94431625 | 16.55681121 | 13.17934247 | 11.98917757 | 3.595006193 | 0.00401986 | -1.483335495 | 0.00026324 | 13.3153587 | 0.35766095 | 2.39578955 |
| *Tbx3* | 37.60543889 | 34.10464719 | 33.67094897 | 43.27939622 | 8.765370643 | 28.24144815 | 6.993686915 | 8.987515484 | 0.00602244 | -1.48503936 | 0.0004298 | 12.397981 | 0.35723879 | 2.22022745 |
| *Csgalnact1* | 31.33786574 | 44.76234944 | 42.85393506 | 43.27939622 | 9.739300714 | 29.18282976 | 7.992785046 | 10.78501858 | 0.0031987 | -1.487867198 | 0.00020023 | 13.8288794 | 0.35653925 | 2.49502646 |
| *Itgb7* | 479.4693459 | 551.0032062 | 569.3451372 | 527.5964491 | 219.1342661 | 181.6866498 | 177.8394673 | 170.7627942 | 1.11E-40 | -1.506275691 | 1.32E-43 | 191.75244 | 0.35201878 | 39.9564869 |
| *Flrt3* | 85.65683304 | 55.42005169 | 55.0979165 | 50.49262892 | 24.34825178 | 24.47592173 | 18.98286448 | 18.87378252 | 6.60E-06 | -1.508987544 | 1.44E-07 | 27.6709259 | 0.35135771 | 5.18018351 |
| *Akap2* | 509.7626161 | 444.4261837 | 504.0439028 | 523.4746018 | 128.5587694 | 330.4249434 | 117.8935794 | 114.1414466 | 0.00209767 | -1.519789737 | 0.00011836 | 14.818614 | 0.34873674 | 2.67826385 |
| *Sp7* | 57.45275387 | 34.10464719 | 45.91493042 | 27.822469 | 8.765370643 | 26.35868494 | 6.993686915 | 15.27877632 | 0.00381416 | -1.520073593 | 0.00024644 | 13.4390794 | 0.34866813 | 2.41860127 |
| *Adamts4* | 79.38925989 | 75.66968596 | 69.38256152 | 86.55879243 | 16.55681121 | 36.7138826 | 24.97745327 | 26.96254645 | 9.81E-08 | -1.560715424 | 1.40E-09 | 36.6744759 | 0.33898294 | 7.00819815 |
| *Pxdn* | 214.1420826 | 177.9836275 | 220.391666 | 243.1889883 | 50.64436371 | 135.5589511 | 47.95671028 | 53.02634135 | 0.00101615 | -1.574594391 | 4.97E-05 | 16.4608538 | 0.3357375 | 2.99304028 |
| *G530011O06Rik* | 44.91760757 | 52.22274101 | 47.95559399 | 39.15754896 | 14.60895107 | 22.59315852 | 6.993686915 | 17.07627942 | 9.14E-05 | -1.581350891 | 2.88E-06 | 21.8981115 | 0.33416884 | 4.03924786 |
| *Prrx2* | 48.05139414 | 52.22274101 | 42.85393506 | 38.12708714 | 7.791440571 | 27.30006655 | 17.98376635 | 7.190012387 | 0.00089975 | -1.59084921 | 4.28E-05 | 16.7415875 | 0.33197599 | 3.04587678 |
| *Foxc2* | 50.14058519 | 56.48582191 | 31.6302854 | 37.09662533 | 10.71323079 | 32.94835618 | 8.991883177 | 5.39250929 | 0.00676845 | -1.593973209 | 0.00049864 | 12.120755 | 0.33125791 | 2.16951094 |
| *Pdk4* | 13.57974182 | 25.57848539 | 23.4676311 | 21.63969811 | 5.843580428 | 11.29657926 | 5.994588784 | 4.493757742 | 0.00949686 | -1.609236783 | 0.00076304 | 11.3290503 | 0.3277717 | 2.02242014 |
| *Dner* | 37.60543889 | 42.63080899 | 48.97592578 | 39.15754896 | 15.58288114 | 12.23796087 | 10.99007944 | 15.27877632 | 1.58E-05 | -1.636073847 | 3.88E-07 | 25.7516119 | 0.32173084 | 4.80100402 |
| *Syt13* | 36.56084337 | 52.22274101 | 46.9352622 | 39.15754896 | 4.869650357 | 30.12421136 | 10.99007944 | 9.886267032 | 0.00220281 | -1.643293884 | 0.00012605 | 14.7000207 | 0.32012475 | 2.65702244 |
| *Ifitm1* | 60.58654044 | 51.15697079 | 44.89459863 | 40.18801077 | 13.635021 | 22.59315852 | 16.98466822 | 9.886267032 | 1.74E-05 | -1.643812582 | 4.38E-07 | 25.5205248 | 0.32000967 | 4.75872457 |
| *Cav1* | 663.3181583 | 579.7790023 | 706.0695966 | 632.7035542 | 182.1249234 | 209.9280979 | 205.8142149 | 223.7891355 | 1.68E-55 | -1.650939179 | 5.35E-59 | 262.322818 | 0.31843279 | 54.7740699 |
| *Gzme* | 53.27437177 | 55.42005169 | 72.44355688 | 53.58401436 | 12.66109093 | 40.47940902 | 6.993686915 | 13.48127323 | 0.00085034 | -1.669292545 | 3.96E-05 | 16.890417 | 0.31440748 | 3.07040717 |
| *Rapgef3* | 90.87981066 | 115.1031843 | 96.93151977 | 116.4421851 | 41.87899307 | 35.772501 | 29.97294392 | 23.36754026 | 1.68E-11 | -1.681622686 | 1.30E-13 | 54.8588644 | 0.31173182 | 10.7747065 |
| *Saa3* | 57.45275387 | 89.52469888 | 80.60621118 | 75.22371247 | 25.32218186 | 10.35519766 | 34.96843458 | 22.46878871 | 1.24E-06 | -1.706170532 | 2.23E-08 | 31.2865787 | 0.30647249 | 5.90484722 |
| *Gprc5b* | 123.2622719 | 120.4320354 | 159.1717588 | 107.1680287 | 39.93113293 | 62.13118594 | 32.97023831 | 20.67128561 | 1.39E-08 | -1.713551256 | 1.70E-10 | 40.7855839 | 0.3049086 | 7.85758994 |
| *Kctd15* | 19.84731497 | 18.11809382 | 26.52862646 | 23.70062174 | 6.8175105 | 11.29657926 | 1.998196261 | 6.291260839 | 0.00512851 | -1.733756284 | 0.0003554 | 12.7531778 | 0.3006681 | 2.29000896 |
| *Il11* | 262.1934767 | 284.56065 | 183.6597217 | 247.3108355 | 37.98327278 | 128.0278983 | 61.94408411 | 52.12758981 | 0.00020809 | -1.803122947 | 7.72E-06 | 20.0052471 | 0.28655363 | 3.68174693 |
| *Sfrp2* | 30.29327022 | 27.71002584 | 33.67094897 | 45.34031984 | 12.66109093 | 17.8862505 | 4.995490654 | 3.595006193 | 0.00112124 | -1.809768401 | 5.71E-05 | 16.1958873 | 0.28523671 | 2.95030095 |
| *Dio2* | 62.67573149 | 46.89388989 | 49.99625757 | 55.64493799 | 13.635021 | 26.35868494 | 9.990981307 | 10.78501858 | 2.51E-06 | -1.822625916 | 4.96E-08 | 29.7314774 | 0.28270594 | 5.60073655 |
| *Osbp2* | 74.16628226 | 72.47237528 | 75.50455224 | 98.92433421 | 17.53074129 | 24.47592173 | 29.97294392 | 18.87378252 | 2.15E-10 | -1.825209725 | 2.02E-12 | 49.466416 | 0.28220007 | 9.66718341 |
| *Zfp532* | 19.84731497 | 20.24963427 | 23.4676311 | 27.822469 | 5.843580428 | 8.472434446 | 4.995490654 | 6.291260839 | 0.00090242 | -1.833109756 | 4.31E-05 | 16.7296431 | 0.280659 | 3.04459035 |
| *Gjb3* | 30.29327022 | 53.28851124 | 38.77260791 | 37.09662533 | 9.739300714 | 13.17934247 | 7.992785046 | 13.48127323 | 1.44E-05 | -1.837910706 | 3.48E-07 | 25.961013 | 0.27972659 | 4.84103329 |
| *Dock3* | 34.47165232 | 40.49926854 | 32.65061719 | 42.2489344 | 11.68716086 | 11.29657926 | 3.996392523 | 14.38002477 | 3.45E-05 | -1.847500823 | 9.48E-07 | 24.0306798 | 0.27787331 | 4.4624247 |
| *Prkcdbp* | 31.33786574 | 28.77579607 | 30.60995361 | 36.06616351 | 7.791440571 | 20.71039531 | 3.996392523 | 2.696254645 | 0.00319722 | -1.848536416 | 0.00019946 | 13.8361541 | 0.27767392 | 2.49522738 |
| *Tspan18* | 40.73922547 | 43.69657922 | 58.15891186 | 64.91909432 | 3.895720286 | 31.06559297 | 6.993686915 | 14.38002477 | 0.00071265 | -1.875231724 | 3.25E-05 | 17.2647844 | 0.27258315 | 3.147124 |
| *Col7a1* | 36.56084337 | 43.69657922 | 35.71161255 | 31.94431625 | 9.739300714 | 9.413816051 | 4.995490654 | 15.27877632 | 2.06E-05 | -1.895981785 | 5.26E-07 | 25.1644245 | 0.26869069 | 4.68536855 |
| *Dpep1* | 20.8919105 | 11.72347247 | 28.56929004 | 17.51785085 | 5.843580428 | 5.648289631 | 3.996392523 | 5.39250929 | 0.00296396 | -1.913613891 | 0.00018208 | 14.0075077 | 0.26542683 | 2.52812792 |
| *Grb14* | 22.98110155 | 20.24963427 | 31.6302854 | 34.00523988 | 2.921790214 | 16.00348729 | 5.994588784 | 3.595006193 | 0.00242366 | -1.932534083 | 0.00014176 | 14.4785654 | 0.26196862 | 2.61552915 |
| *Mgll* | 40.73922547 | 23.44694495 | 24.48796289 | 27.822469 | 10.71323079 | 6.589671236 | 4.995490654 | 8.088763935 | 0.0001551 | -1.936820643 | 5.49E-06 | 20.6599267 | 0.26119141 | 3.80938287 |
| *Clec4e* | 1643.148761 | 1709.495441 | 2179.428697 | 1853.800805 | 485.0171756 | 647.6705443 | 355.6789345 | 421.5144762 | 4.79E-17 | -1.951152392 | 2.05E-19 | 81.1865022 | 0.25860958 | 16.3200302 |
| *Rasd2* | 47.00679862 | 39.43349832 | 57.13858008 | 52.55355255 | 11.68716086 | 19.76901371 | 7.992785046 | 7.190012387 | 3.47E-07 | -2.072783764 | 5.38E-09 | 34.0486309 | 0.2377004 | 6.45985808 |
| *H2-M2* | 10.44595525 | 8.526161798 | 16.32530859 | 21.63969811 | 2.921790214 | 3.765526421 | 2.997294392 | 3.595006193 | 0.00811647 | -2.0990457 | 0.00062181 | 11.7095147 | 0.23341259 | 2.09063279 |
| *Plbd1* | 52.22977624 | 46.89388989 | 53.05725293 | 56.67539981 | 11.68716086 | 8.472434446 | 14.98647196 | 12.58252168 | 9.22E-10 | -2.132634208 | 9.38E-12 | 46.453713 | 0.2280411 | 9.0351839 |
| *Met* | 54.31896729 | 61.81467304 | 84.68753833 | 81.40648336 | 13.635021 | 18.8276321 | 22.97925701 | 8.088763935 | 9.52E-10 | -2.160048165 | 9.76E-12 | 46.3765249 | 0.2237488 | 9.02145545 |
| *Gsta3* | 41.78382099 | 50.09120056 | 39.7929397 | 39.15754896 | 6.8175105 | 4.706908026 | 12.9882757 | 13.48127323 | 3.24E-07 | -2.166611204 | 4.99E-09 | 34.1922866 | 0.22273324 | 6.48968681 |
| *Irg1* | 500.3612564 | 553.1347467 | 684.6426291 | 549.2361472 | 118.8194687 | 191.1004658 | 66.93957476 | 112.3439435 | 4.09E-10 | -2.224487957 | 3.99E-12 | 48.1274016 | 0.21397469 | 9.38870666 |
| *Marco* | 26.11488812 | 24.51271517 | 38.77260791 | 27.822469 | 3.895720286 | 3.765526421 | 7.992785046 | 8.987515484 | 1.73E-05 | -2.247519172 | 4.32E-07 | 25.5473862 | 0.21058591 | 4.76214613 |
| *Gcm2* | 17.75812392 | 7.460391573 | 19.38630395 | 14.42646541 | 3.895720286 | 0.941381605 | 3.996392523 | 0.898751548 | 0.00196929 | -2.622137373 | 0.00010971 | 14.9618063 | 0.16242692 | 2.70568984 |
| *Tfeb* | 768.8223063 | 835.5638562 | 745.8625363 | 761.511281 | 123.6891191 | 159.0934913 | 102.9071075 | 99.76142187 | 1.57E-99 | -2.680182924 | 2.49E-103 | 465.906959 | 0.15602153 | 98.8047932 |
| *Cxcl3* | 9.401359723 | 4.263080899 | 11.22364966 | 13.39600359 | 1.947860143 | 0.941381605 | 1.998196261 | 0.898751548 | 0.00947719 | -2.740852735 | 0.00076071 | 11.3347372 | 0.14959639 | 2.02332034 |
| *Emx2* | 9.401359723 | 8.526161798 | 7.142322509 | 18.54831266 | 1.947860143 | 0.941381605 | 0.999098131 | 1.797503097 | 0.00311808 | -2.935608502 | 0.00019428 | 13.8856509 | 0.13070548 | 2.5061126 |
| *Gm5886* | 14.62433735 | 8.526161798 | 13.26431323 | 15.45692722 | 0.973930071 | 1.88276321 | 0 | 0 | 3.49E-05 | -4.183677164 | 9.63E-07 | 24.0002859 | 0.0550285 | 4.45682249 |
