## Supplementary material for "TFEB-Mediated Pro-inflammatory Response in Murine Macrophages Induced by Acute Alpha7 Nicotinic Receptor Activation": Table_S18

**Table S18. GO and KEGG Pathway analysis of differentially expressed genes from Table S17.**

**Enrichr-KG_PNU (Tfebfl vs TfebΔLysM) DE genes only UP in Tfebfl**

| Term | Library | p-value | q-value | z-score | combined score |
|---|---|---|---|---|---|
| extracellular matrix organization (GO:0030198) | GO_Biological_Process_2021 | 7.08E-13 | 1.50E-09 | 7.044 | 197.1 |
| regulation of cell migration (GO:0030334) | GO_Biological_Process_2021 | 3.23E-12 | 3.41E-09 | 5.747 | 152.1 |
| positive regulation of cell migration (GO:0030335) | GO_Biological_Process_2021 | 2.59E-11 | 1.83E-08 | 6.83 | 166.5 |
| transmembrane receptor protein tyrosine kinase signaling pathway | GO_Biological_Process_2021 | 8.27E-11 | 4.37E-08 | 5.323 | 123.6 |
| positive regulation of cell motility (GO:2000147) | GO_Biological_Process_2021 | 2.62E-10 | 1.11E-07 | 7.145 | 157.6 |
| Focal adhesion | KEGG_2021_Human | 2.10E-08 | 4.9E-06 | 6.499 | 114.9 |
| Fluid shear stress and atherosclerosis | KEGG_2021_Human | 6.87E-08 | 8.01E-06 | 7.688 | 126.8 |
| PI3K-Akt signaling pathway | KEGG_2021_Human | 2.3E-06 | 0.000179 | 4.026 | 52.26 |
| Rap1 signaling pathway | KEGG_2021_Human | 7.52E-06 | 0.000438 | 4.9 | 57.81 |
| TNF signaling pathway | KEGG_2021_Human | 2.53E-05 | 0.001141 | 6.421 | 67.96 |

**Enrichr-KG_UP in PNU-282987 treated TfebΔLysM cells**

| Term | Library | p-value | q-value | z-score | combined score |
|---|---|---|---|---|---|
| positive regulation of phosphate metabolic process (GO:0045937) | GO_Biological_Process_2021 | 8.02E-06 | 0.005301 | 37.31 | 437.8 |
| positive regulation of phosphorylation (GO:0042327) | GO_Biological_Process_2021 | 1.33E-05 | 0.005301 | 8.234 | 92.45 |
| organic substance transport (GO:0071702) | GO_Biological_Process_2021 | 0.000303 | 0.08036 | 9.322 | 75.54 |
| negative regulation of bone resorption (GO:0045779) | GO_Biological_Process_2021 | 0.000645 | 0.09526 | 67.71 | 497.4 |
| positive regulation of ossification (GO:0045778) | GO_Biological_Process_2021 | 0.000676 | 0.09526 | 19.42 | 141.7 |
| Cholesterol metabolism | KEGG_2021_Human | 0.0013 | 0.1137 | 15.28 | 101.5 |
| Glycosaminoglycan degradation | KEGG_2021_Human | 0.00298 | 0.1137 | 27.87 | 162.1 |
| Calcium signaling pathway | KEGG_2021_Human | 0.003777 | 0.1137 | 5.169 | 28.84 |
| Cell adhesion molecules | KEGG_2021_Human | 0.003823 | 0.1137 | 6.697 | 37.28 |
| Aldosterone synthesis and secretion | KEGG_2021_Human | 0.008704 | 0.2071 | 7.541 | 35.77 |
