## Supplementary material for "TFEB-Mediated Pro-inflammatory Response in Murine Macrophages Induced by Acute Alpha7 Nicotinic Receptor Activation": Table_S19

**Table S19. Genes that were induced by PNU-282987 in Tfebfl/fl but not TfebΔLysM BMDMs.**

*6330416G13Rik*
*Adi1*
*Angpt2*
*Ankrd12*
*Blnk*
*Bsn*
*C77080*
*Chst10*
*Ctns*
*Fam20c*
*Fam219a*
*Flcn*
*Fn3k*
*Gng4*
*Gpr137b*
*Hilpda*
*Htr2b*
*Id2*
*Irg1*
*Klhl21*
*Mfsd7b*
*Mreg*
*Nat6*
*Nat8l*
*Osbp2*
*Osgin1*
*P2ry2*
*Pcyt1a*
*Pfkfb3*
*Phospho1*
*Plxna1*
*Ppargc1b*
*Ptpre*
*Rapgef3*
*Rasd2*
*Rnf144b*
*Sh2d3c*
*Sik1*
*Slc2a4rg-ps*
*Smad6*
*Snx24*
*Sorbs3*
*Tm4sf19*
*Tmem104*
*Tmem144*
*Tmem189*
*Tnfsf15*
*Tns1*
*Tuft1*
*Usp35*
*Vash2*
*Zfp667*
