## Supplementary material for "TFEB-Mediated Pro-inflammatory Response in Murine Macrophages Induced by Acute Alpha7 Nicotinic Receptor Activation": Table_S20

**Table S20. Differential gene expression in PNU-282987-treated RAW264.7 cells (WT v. Tfeb Tfe3 dKO).**

| ID | Gene | counts_WT_PN1 | counts_WT_PN2 | counts_WT_PN3 | counts_KO_PN1 | counts_KO_PN2 | counts_KO_PN3 | padj | log2FoldChange | pvalue | stat | foldChange | log10padj |
|---|---|---|---|---|---|---|---|---|---|---|---|---|---|
| ENSMUSG00000039007 | Cpq | 0 | 0 | 0 | 131.3806293 | 152.4397261 | 103.9205409 | 8.87E-43 | 9.498850088 | 3.83E-44 | 194.2094403 | 723.500442 | 42.05220916 |
| ENSMUSG00000041444 | Arhgap32 | 0.850611819 | 1.099678033 | 0 | 416.2324362 | 322.0289213 | 402.2109823 | 2.04E-112 | 9.204490555 | 3.04E-114 | 516.0618885 | 589.9670433 | 111.6895446 |
| ENSMUSG00000019846 | Lama4 | 0 | 0 | 0 | 46.50641745 | 52.40115583 | 40.41354368 | 2.34E-17 | 8.022330771 | 2.46E-18 | 76.28495667 | 259.9933245 | 16.6315448 |
| ENSMUSG00000030043 | Tacr1 | 6.804894552 | 4.39871213 | 1.958179327 | 817.3502867 | 683.1205224 | 777.4796021 | 8.88E-203 | 7.397707627 | 6.16E-205 | 933.1324685 | 168.6288564 | 202.0513794 |
| ENSMUSG00000042340 | Ctf1 | 0 | 0 | 0 | 17.43990655 | 20.96046233 | 23.09345353 | 4.17E-08 | 6.84445081 | 8.00E-09 | 33.27451018 | 114.9171899 | 7.379889487 |
| ENSMUSG00000027200 | Sema6d | 5.103670914 | 6.598068195 | 17.62361394 | 1118.47934 | 867.000942 | 1120.994723 | 6.18E-47 | 6.734945088 | 2.40E-48 | 213.4704791 | 106.5173837 | 46.20926986 |
| ENSMUSG00000022994 | Adcy6 | 0 | 0 | 0 | 22.09054829 | 19.05496576 | 14.43340846 | 2.68E-07 | 6.693849068 | 5.51E-08 | 29.52735962 | 103.5259813 | 6.572276268 |
| ENSMUSG00000074259 | Gramd2 | 0 | 0 | 0.979089664 | 29.06651091 | 36.20443494 | 34.64018029 | 1.12E-11 | 6.582674718 | 1.61E-12 | 49.9068907 | 95.84788669 | 10.95143513 |
| ENSMUSG00000093456 | Gm20662 | 0 | 0 | 0 | 9.301283491 | 25.72420377 | 13.47118123 | 5.37E-06 | 6.505127366 | 1.27E-06 | 23.46514081 | 90.83191341 | 5.269635138 |
| ENSMUSG00000089837 | Npcd | 0 | 0 | 0 | 13.95192524 | 13.33847603 | 11.54672676 | 2.16E-05 | 6.178291024 | 5.49E-06 | 20.65879545 | 72.41873265 | 4.665525392 |
| ENSMUSG00000042351 | Grap2 | 6.804894552 | 7.697746228 | 1.958179327 | 368.5633583 | 380.1465669 | 370.4574837 | 1.47E-109 | 6.091241443 | 2.33E-111 | 502.8104059 | 68.17833408 | 108.831663 |
| ENSMUSG00000032068 | Plet1 | 7.655506371 | 4.39871213 | 3.916358654 | 370.8886792 | 342.0366354 | 376.2308471 | 1.09E-105 | 6.055081766 | 1.88E-107 | 484.8525963 | 66.49075091 | 104.9622545 |
| ENSMUSG00000040612 | Ildr2 | 3.402447276 | 7.697746228 | 4.895448318 | 327.870243 | 305.8322004 | 326.1950311 | 8.15E-94 | 5.952070108 | 1.57E-95 | 430.0751186 | 61.9086935 | 93.08904078 |
| ENSMUSG00000022338 | Eny2 | 0 | 0 | 0 | 11.62660436 | 5.716489727 | 12.50895399 | 0.000372985 | 5.797293463 | 0.000113835 | 14.89215411 | 55.61081064 | 3.428308462 |
| ENSMUSG00000082706 | Gm11663 | 0 | 0 | 0 | 6.975962618 | 3.810993152 | 16.35786292 | 0.001279363 | 5.664883205 | 0.00042979 | 12.39801662 | 50.73407735 | 2.893006228 |
| ENSMUSG00000032502 | Stac | 0 | 0 | 0 | 11.62660436 | 8.574734591 | 6.735590613 | 0.00066994 | 5.647472802 | 0.000213844 | 13.70535512 | 50.12549946 | 3.173964164 |
| ENSMUSG00000032036 | Kirrel3 | 34.87508458 | 37.38905311 | 31.33086923 | 1604.471402 | 1623.483083 | 1635.786292 | 0 | 5.556331706 | 0 | 1815.857813 | 47.05681289 | Inf |
| ENSMUSG00000015342 | Xk | 0 | 0 | 0 | 12.7892648 | 4.763741439 | 7.697817843 | 0.001358247 | 5.551320687 | 0.000458527 | 12.27719369 | 46.89365051 | 2.867021088 |
| ENSMUSG00000050157 | Gm867 | 0 | 0 | 0 | 10.46394393 | 3.810993152 | 10.58449953 | 0.001602701 | 5.532086665 | 0.00054754 | 11.94636889 | 46.27261302 | 2.795147613 |
| ENSMUSG00000042401 | Crtac1 | 4.253059095 | 3.299034098 | 2.937268991 | 165.097782 | 178.1639298 | 142.4096301 | 1.57E-45 | 5.516748258 | 6.33E-47 | 206.9595041 | 45.78325902 | 44.80529814 |
| ENSMUSG00000037846 | Rtkn2 | 0.850611819 | 0 | 0.979089664 | 30.22917134 | 16.19672089 | 41.37577091 | 2.23E-08 | 5.515466413 | 4.19E-09 | 34.53336626 | 45.74259834 | 7.651316191 |
| ENSMUSG00000037855 | Zfp365 | 9.356730009 | 9.897102293 | 14.68634495 | 466.226835 | 452.5554367 | 508.0559776 | 8.33E-124 | 5.39871114 | 1.09E-125 | 568.6840273 | 42.18654812 | 123.0795981 |
| ENSMUSG00000038048 | Cntnap5c | 2.551835457 | 1.099678033 | 4.895448318 | 105.8020997 | 99.08582194 | 146.258539 | 2.04E-29 | 5.340267646 | 1.32E-30 | 132.2504916 | 40.51172597 | 28.6898228 |
| ENSMUSG00000089996 | Tmsb15b2 | 0.850611819 | 0 | 0.979089664 | 17.43990655 | 37.15718323 | 19.24454461 | 2.79E-07 | 5.271242259 | 5.74E-08 | 29.44767611 | 38.61908984 | 6.554999249 |
| ENSMUSG00000028348 | Cavin4 | 0 | 0 | 0 | 2.325320873 | 2.858244864 | 15.39563569 | 0.008949475 | 5.269536739 | 0.003497136 | 8.528053141 | 38.57346224 | 2.048202458 |
| ENSMUSG00000034336 | Ina | 5.103670914 | 10.99678033 | 8.811806972 | 344.1474892 | 265.8167723 | 304.0638048 | 2.79E-72 | 5.233246565 | 6.71E-74 | 330.7173083 | 37.61527062 | 71.55517119 |
| ENSMUSG00000031410 | Nxf7 | 45.08242641 | 49.48551147 | 50.9126625 | 1533.549116 | 1697.797449 | 1614.617293 | 0 | 5.063198397 | 0 | 1540.768636 | 33.43294173 | Inf |
| ENSMUSG00000022505 | Emp2 | 41.67997913 | 46.18647737 | 36.22631755 | 1312.643633 | 1280.493699 | 1293.233398 | 1.52E-298 | 4.973404025 | 5.75E-301 | 1374.977636 | 31.41548669 | 297.816907 |
| ENSMUSG00000024140 | Epas1 | 0 | 3.299034098 | 0.979089664 | 55.80770094 | 30.48794521 | 34.64018029 | 4.24E-10 | 4.934967274 | 6.86E-11 | 42.55764578 | 30.58955627 | 9.373006529 |
| ENSMUSG00000034557 | Zfyve9 | 25.51835457 | 20.89388262 | 16.64452428 | 665.0417696 | 606.9006594 | 609.0898368 | 6.71E-147 | 4.885578914 | 7.23E-149 | 675.2451657 | 29.56009322 | 146.1733771 |
| ENSMUSG00000020838 | Slc6a4 | 53.5885446 | 48.38583343 | 50.9126625 | 1475.416094 | 1499.625805 | 1547.261386 | 0 | 4.881737096 | 0 | 1535.515645 | 29.48148103 | Inf |
| ENSMUSG00000040875 | Osbpl10 | 8.50611819 | 6.598068195 | 13.70725529 | 309.2676761 | 253.4310446 | 268.4613973 | 3.37E-65 | 4.841867962 | 9.04E-67 | 297.9876595 | 28.67790961 | 64.47240056 |
| ENSMUSG00000113409 | Gm8288 | 0 | 4.39871213 | 6.853627645 | 106.9647601 | 95.27482879 | 101.0338592 | 8.05E-24 | 4.81662346 | 6.25E-25 | 106.3277789 | 28.18046398 | 23.09442744 |
| ENSMUSG00000068551 | Zfp467 | 5.103670914 | 6.598068195 | 1.958179327 | 127.892648 | 107.6605565 | 118.3539493 | 4.91E-30 | 4.706804782 | 3.11E-31 | 135.1173393 | 26.11496349 | 29.30855264 |
| ENSMUSG00000031289 | Il13ra2 | 2.551835457 | 1.099678033 | 0 | 32.55449222 | 33.34619008 | 30.79127137 | 4.03E-09 | 4.660911658 | 7.07E-10 | 38.00058198 | 25.29730263 | 8.394757834 |
| ENSMUSG00000015962 | 1700016C15Rik | 0.850611819 | 0 | 0 | 5.813302182 | 13.33847603 | 6.735590613 | 0.003902693 | 4.637389799 | 0.001423007 | 10.17573797 | 24.88819674 | 2.408635651 |
| ENSMUSG00000074415 | Mir100hg | 3.402447276 | 5.498390163 | 0.979089664 | 77.89824923 | 72.40886988 | 90.44935965 | 3.01E-20 | 4.634527658 | 2.72E-21 | 89.73816243 | 24.83887035 | 19.52133165 |
| ENSMUSG00000050953 | Gja1 | 19.56407184 | 6.598068195 | 13.70725529 | 324.3822617 | 311.5486901 | 360.8352114 | 6.08E-70 | 4.59902985 | 1.53E-71 | 319.8937536 | 24.23516249 | 69.2158583 |
| ENSMUSG00000066456 | Hmgn3 | 0 | 0 | 0.979089664 | 6.975962618 | 9.527482879 | 7.697817843 | 0.004919038 | 4.537435426 | 0.001825991 | 9.716719935 | 23.22224308 | 2.308119834 |
| ENSMUSG00000026475 | Rgs16 | 17.01223638 | 16.49517049 | 19.58179327 | 448.7869284 | 412.5400087 | 369.4952565 | 4.16E-91 | 4.533136082 | 8.17E-93 | 417.5882443 | 23.15314199 | 90.38039029 |
| ENSMUSG00000095028 | Sirpb1b | 0 | 0 | 0.979089664 | 8.138623054 | 11.43297945 | 3.848908921 | 0.008084791 | 4.487618596 | 0.003131511 | 8.72921604 | 22.43405635 | 2.092331221 |
| ENSMUSG00000006221 | Hspb7 | 62.09466279 | 48.38583343 | 55.80811082 | 1285.902443 | 1199.510094 | 1249.933172 | 1.72E-251 | 4.477760793 | 8.71E-254 | 1157.876388 | 22.28128902 | 250.7655812 |
| ENSMUSG00000043300 | B3galnt1 | 10.20734183 | 7.697746228 | 6.853627645 | 174.3990655 | 189.5969093 | 191.4832188 | 3.57E-46 | 4.472541243 | 1.42E-47 | 209.934672 | 22.20082283 | 45.44698127 |
| ENSMUSG00000094942 | Gm3604 | 0.850611819 | 2.199356065 | 0 | 20.92788785 | 20.00771405 | 21.16899907 | 5.11E-06 | 4.407815754 | 1.21E-06 | 23.56540911 | 21.22681121 | 5.291385887 |
| ENSMUSG00000001131 | Timp1 | 6.804894552 | 8.797424261 | 11.74907596 | 189.5136511 | 211.5101199 | 171.276447 | 6.77E-44 | 4.399570221 | 2.84E-45 | 199.3881238 | 21.10583821 | 43.16942846 |
| ENSMUSG00000009394 | Syn2 | 1.701223638 | 1.099678033 | 0.979089664 | 31.39183178 | 23.8187072 | 24.05568076 | 1.65E-07 | 4.362312375 | 3.34E-08 | 30.50102253 | 20.56775429 | 6.781719955 |
| ENSMUSG00000056313 | Tcim | 3.402447276 | 8.797424261 | 3.916358654 | 102.3141184 | 99.08582194 | 107.7694498 | 1.42E-24 | 4.309125387 | 1.07E-25 | 109.8250321 | 19.82330197 | 23.84631687 |
| ENSMUSG00000041540 | Sox5 | 1.701223638 | 0 | 0 | 12.7892648 | 12.38572774 | 10.58449953 | 0.000969886 | 4.2499049 | 0.000319901 | 12.95011831 | 19.02605964 | 3.013279385 |
| ENSMUSG00000038668 | Lpar1 | 68.04894552 | 48.38583343 | 51.89175217 | 1102.202094 | 1001.338451 | 1082.505634 | 2.07E-188 | 4.226724911 | 1.56E-190 | 866.8686493 | 18.72280791 | 187.6834855 |
| ENSMUSG00000027412 | Lpin3 | 8.50611819 | 3.299034098 | 3.916358654 | 89.5248536 | 115.2825428 | 93.33604135 | 4.32E-22 | 4.200068401 | 3.58E-23 | 98.30737305 | 18.38004509 | 21.36432661 |
| ENSMUSG00000002459 | Rgs20 | 2.551835457 | 10.99678033 | 3.916358654 | 88.36219316 | 98.13307365 | 114.5050404 | 3.33E-20 | 4.172530785 | 3.02E-21 | 89.53026734 | 18.03254083 | 19.47734175 |
| ENSMUSG00000028347 | Tmeff1 | 15.31101274 | 10.99678033 | 20.56088293 | 274.387863 | 261.0530309 | 292.517078 | 2.01E-59 | 4.132108271 | 6.07E-61 | 271.2458601 | 17.53430413 | 58.69649216 |
| ENSMUSG00000047443 | Erfe | 2.551835457 | 3.299034098 | 3.916358654 | 49.99439876 | 56.21214899 | 54.84695213 | 1.76E-13 | 4.056797046 | 2.26E-14 | 58.2892027 | 16.64246289 | 12.75523434 |
| ENSMUSG00000044912 | Syt16 | 0.850611819 | 1.099678033 | 0.979089664 | 15.11458567 | 18.10221747 | 13.47118123 | 0.000125326 | 4.007202339 | 3.55E-05 | 17.09763658 | 16.08007621 | 3.90195871 |
| ENSMUSG00000015854 | Cd5l | 31.4726373 | 38.48873114 | 35.24722789 | 526.6851777 | 580.2237073 | 522.4893861 | 4.24E-109 | 3.962358414 | 6.84E-111 | 500.6558987 | 15.58794038 | 108.3722289 |
| ENSMUSG00000021488 | Nsd1 | 55.28976823 | 56.08357966 | 47.97539351 | 772.0065297 | 686.9315156 | 864.0800529 | 5.41E-120 | 3.865876263 | 7.43E-122 | 551.0528529 | 14.57956998 | 119.266499 |
| ENSMUSG00000063760 | Rnf217 | 7.655506371 | 9.897102293 | 8.811806972 | 122.0793458 | 107.6605565 | 139.5229484 | 3.62E-26 | 3.819399037 | 2.57E-27 | 117.2165701 | 14.11736604 | 25.44079999 |
| ENSMUSG00000021250 | Fos | 164.1680811 | 248.5272354 | 178.1943188 | 2907.813751 | 2548.60167 | 2753.894333 | 6.47E-184 | 3.805833087 | 5.20E-186 | 846.0707572 | 13.98523981 | 183.1888128 |
| ENSMUSG00000026589 | Sec16b | 3.402447276 | 0 | 1.958179327 | 22.09054829 | 19.05496576 | 34.64018029 | 1.11E-05 | 3.745164924 | 2.72E-06 | 22.00429863 | 13.40932697 | 4.953969001 |
| ENSMUSG00000020712 | Tcam1 | 48.48487368 | 51.68486753 | 48.95448318 | 672.0177322 | 662.1600601 | 654.3145166 | 2.32E-132 | 3.739603619 | 2.84E-134 | 608.142321 | 13.35773617 | 131.6340421 |
| ENSMUSG00000024727 | Trpm6 | 0 | 0 | 3.916358654 | 16.27724611 | 14.29122432 | 21.16899907 | 0.000671723 | 3.733592866 | 0.000214505 | 13.69955716 | 13.30219911 | 3.172810072 |
| ENSMUSG00000003849 | Nqo1 | 12.75917728 | 10.99678033 | 13.70725529 | 147.6578754 | 157.2034675 | 184.7476282 | 3.95E-33 | 3.703884613 | 2.22E-34 | 149.5089377 | 13.03107874 | 32.40393968 |
| ENSMUSG00000089768 | Tmsb15b1 | 1.701223638 | 0 | 0.979089664 | 6.975962618 | 13.33847603 | 16.35786292 | 0.002160437 | 3.700903349 | 0.000754537 | 11.34985832 | 13.00417841 | 2.665458333 |
| ENSMUSG00000028755 | Cda | 28.92080185 | 38.48873114 | 27.41451058 | 423.2083988 | 423.0202398 | 359.8729842 | 9.79E-71 | 3.680049044 | 2.43E-72 | 323.5582455 | 12.81755377 | 70.00927831 |
| ENSMUSG00000002997 | Prkar2b | 43.38120277 | 48.38583343 | 31.33086923 | 565.0529721 | 500.1928511 | 507.0937504 | 5.44E-89 | 3.677849128 | 1.09E-90 | 407.8240119 | 12.79802361 | 88.26401785 |
| ENSMUSG00000054932 | Afp | 215.2047902 | 260.6236937 | 242.8142366 | 2603.196717 | 3595.672038 | 2898.228418 | 4.23E-162 | 3.66612272 | 4.12E-164 | 745.3478744 | 12.69442127 | 161.3737642 |
| ENSMUSG00000030854 | Ptpn5 | 85.91179372 | 124.2636177 | 100.8462353 | 1304.50501 | 1292.879427 | 1222.028583 | 7.11E-172 | 3.63102678 | 6.34E-174 | 790.4784361 | 12.38933442 | 171.1481286 |
| ENSMUSG00000036225 | Kctd1 | 3.402447276 | 7.697746228 | 5.874537981 | 62.78366356 | 59.07039385 | 76.97817843 | 1.34E-13 | 3.583620086 | 1.70E-14 | 58.84880935 | 11.9888393 | 12.87435342 |
| ENSMUSG00000074923 | Pak6 | 1.701223638 | 0 | 1.958179327 | 12.7892648 | 16.19672089 | 16.35786292 | 0.000523582 | 3.577474026 | 0.00016357 | 14.20918847 | 11.937874 | 3.281015635 |
| ENSMUSG00000040478 | Prdm13 | 4.253059095 | 10.99678033 | 2.937268991 | 69.75962618 | 69.55062502 | 69.28036059 | 2.95E-13 | 3.563357168 | 3.87E-14 | 57.23311461 | 11.82163083 | 12.53008787 |
| ENSMUSG00000045620 | Odf3l1 | 18.71346002 | 15.39549246 | 21.5399726 | 226.7187851 | 215.3211131 | 218.4255813 | 2.71E-44 | 3.561731593 | 1.12E-45 | 201.2442221 | 11.80831816 | 43.56769884 |
| ENSMUSG00000063275 | Hacd1 | 43.38120277 | 17.59484852 | 18.60270361 | 286.0144673 | 370.619084 | 285.7814874 | 2.94E-32 | 3.537888281 | 1.71E-33 | 145.453328 | 11.61476678 | 31.53196385 |
| ENSMUSG00000022010 | Tsc22d1 | 171.8235874 | 179.2475193 | 145.8843599 | 1920.715041 | 1947.4175 | 1876.343099 | 9.21E-267 | 3.532111955 | 4.11E-269 | 1228.397322 | 11.56835604 | 266.035506 |
| ENSMUSG00000029309 | Sparcl1 | 0.850611819 | 1.099678033 | 0.979089664 | 10.46394393 | 10.48023117 | 12.50895399 | 0.002558453 | 3.525925087 | 0.000905131 | 11.01222057 | 11.51885241 | 2.59202248 |
| ENSMUSG00000044927 | H1f10 | 28.92080185 | 17.59484852 | 26.43542092 | 272.0625421 | 307.737697 | 254.0279888 | 1.83E-47 | 3.49723095 | 7.03E-49 | 215.9167296 | 11.29201425 | 46.73674687 |
| ENSMUSG00000037624 | Kcnk2 | 10.20734183 | 15.39549246 | 7.832717308 | 116.2660436 | 121.9517808 | 132.7873578 | 7.79E-24 | 3.490910838 | 6.04E-25 | 106.3964956 | 11.24265475 | 23.10871885 |
| ENSMUSG00000114456 | H2bc9 | 0.850611819 | 1.099678033 | 0.979089664 | 3.487981309 | 11.43297945 | 17.32009015 | 0.007520935 | 3.480283942 | 0.002894533 | 8.872784458 | 11.16014559 | 2.123728155 |
| ENSMUSG00000041609 | Bicdl1 | 27.21957821 | 21.99356065 | 34.26813822 | 317.4062991 | 298.2102141 | 317.534986 | 1.04E-56 | 3.475698699 | 3.31E-58 | 258.6918196 | 11.12473218 | 55.98420325 |
| ENSMUSG00000068323 | Slc4a5 | 0 | 1.099678033 | 1.958179327 | 11.62660436 | 10.48023117 | 9.622272304 | 0.005884275 | 3.424662421 | 0.00221377 | 9.363268951 | 10.73806711 | 2.230307039 |
| ENSMUSG00000037370 | Enpp1 | 6.804894552 | 1.099678033 | 3.916358654 | 43.01843614 | 34.29893836 | 47.14913429 | 6.75E-08 | 3.340095624 | 1.32E-08 | 32.29626771 | 10.12672395 | 7.170550624 |
| ENSMUSG00000013419 | Zfp651 | 138.6497265 | 120.9645836 | 144.9052702 | 1402.168486 | 1372.910283 | 1322.100215 | 1.33E-198 | 3.335600695 | 9.30E-201 | 913.9073361 | 10.09522175 | 197.8764629 |
| ENSMUSG00000031750 | Il34 | 2.551835457 | 5.498390163 | 0.979089664 | 32.55449222 | 35.25168665 | 19.24454461 | 1.34E-05 | 3.302730677 | 3.33E-06 | 21.61794894 | 9.867815057 | 4.871483819 |
| ENSMUSG00000015243 | Abca1 | 46.78365004 | 49.48551147 | 37.20540721 | 474.365458 | 375.3828254 | 453.2090255 | 3.03E-62 | 3.287521007 | 8.66E-64 | 284.3040717 | 9.764329685 | 61.5188679 |
| ENSMUSG00000032911 | Cspg4 | 12.75917728 | 23.09323868 | 20.56088293 | 223.2308038 | 151.4869778 | 163.5786292 | 4.36E-25 | 3.270900212 | 3.21E-26 | 112.2119952 | 9.652483691 | 24.36081789 |
| ENSMUSG00000031844 | Hsd17b2 | 1.701223638 | 3.299034098 | 0.979089664 | 10.46394393 | 25.72420377 | 18.28231738 | 0.000602257 | 3.223150423 | 0.000190545 | 13.92209567 | 9.338238421 | 3.220218123 |
| ENSMUSG00000036957 | Lrfn3 | 13.6097891 | 13.19613639 | 10.7699863 | 129.0553084 | 121.9517808 | 97.18495027 | 2.01E-20 | 3.208405128 | 1.80E-21 | 90.55619787 | 9.243281549 | 19.69724944 |
| ENSMUSG00000034107 | Ano7 | 3.402447276 | 7.697746228 | 0 | 37.20513396 | 32.39344179 | 29.82904414 | 1.51E-05 | 3.199196981 | 3.76E-06 | 21.38221876 | 9.184473242 | 4.820857767 |
| ENSMUSG00000040148 | Hmx3 | 6.804894552 | 10.99678033 | 7.832717308 | 72.08494705 | 71.45612159 | 83.71376904 | 3.95E-14 | 3.168258165 | 4.91E-15 | 61.29671033 | 8.989607738 | 13.40286689 |
| ENSMUSG00000023873 | 1700010I14Rik | 4.253059095 | 1.099678033 | 0.979089664 | 24.41586916 | 14.29122432 | 20.20677184 | 0.000401793 | 3.155886649 | 0.000123234 | 14.74253809 | 8.912848908 | 3.395997233 |
| ENSMUSG00000024039 | Cbs | 12.75917728 | 8.797424261 | 4.895448318 | 75.57292836 | 80.03085618 | 74.09149674 | 4.08E-13 | 3.099762802 | 5.40E-14 | 56.57709001 | 8.57277811 | 12.38883082 |
| ENSMUSG00000040841 | Six5 | 17.8628482 | 15.39549246 | 21.5399726 | 175.5617259 | 150.5342295 | 143.3718573 | 3.76E-26 | 3.092496069 | 2.68E-27 | 117.138096 | 8.529706312 | 25.42487353 |
| ENSMUSG00000110195 | Pde2a | 456.7785468 | 270.520796 | 225.1906226 | 2514.834524 | 2717.238117 | 2781.798923 | 1.86E-19 | 3.071135228 | 1.75E-20 | 86.05372576 | 8.404344077 | 18.73045719 |
| ENSMUSG00000063455 | D630045J12Rik | 19.56407184 | 15.39549246 | 7.832717308 | 133.7059502 | 95.27482879 | 132.7873578 | 1.56E-16 | 3.066957399 | 1.70E-17 | 72.46035531 | 8.380041563 | 15.80738556 |
| ENSMUSG00000044165 | Bcl2l15 | 5.954282733 | 8.797424261 | 4.895448318 | 43.01843614 | 62.881387 | 54.84695213 | 1.51E-09 | 3.051212865 | 2.54E-10 | 39.99599268 | 8.289085045 | 8.822400769 |
| ENSMUSG00000100720 | Gm10075 | 7.655506371 | 0 | 8.811806972 | 51.1570592 | 60.97589042 | 27.90458968 | 9.21E-06 | 3.049431499 | 2.23E-06 | 22.38446418 | 8.278856423 | 5.035802235 |
| ENSMUSG00000049811 | Fam161a | 42.53059095 | 26.39227278 | 32.3099589 | 286.0144673 | 265.8167723 | 282.8948057 | 1.24E-41 | 3.025917078 | 5.52E-43 | 188.9010385 | 8.145013433 | 40.90578207 |
| ENSMUSG00000009378 | Slc16a12 | 152.2595156 | 215.5368944 | 163.5079738 | 1433.560318 | 1340.516841 | 1469.320981 | 7.87E-122 | 3.006324169 | 1.05E-123 | 559.5610044 | 8.035145614 | 121.1041076 |
| ENSMUSG00000030616 | Sytl2 | 1.701223638 | 3.299034098 | 1.958179327 | 16.27724611 | 12.38572774 | 25.98013522 | 0.00076845 | 3.000338585 | 0.000248557 | 13.42300371 | 8.001877733 | 3.114384142 |
| ENSMUSG00000039911 | Spsb1 | 21.26529547 | 14.29581442 | 9.790896635 | 138.3565919 | 111.4715497 | 109.6939043 | 2.92E-17 | 2.967631414 | 3.08E-18 | 75.83728527 | 7.822509004 | 16.53478528 |
| ENSMUSG00000038235 | F11r | 25.51835457 | 31.89066294 | 41.12176587 | 219.7428225 | 284.8717381 | 257.8768977 | 2.79E-33 | 2.961113401 | 1.56E-34 | 150.2061441 | 7.787247077 | 32.55420965 |
| ENSMUSG00000042821 | Snai1 | 1.701223638 | 3.299034098 | 0 | 8.138623054 | 13.33847603 | 16.35786292 | 0.008549319 | 2.956157091 | 0.003331383 | 8.616461694 | 7.760540261 | 2.068068473 |
| ENSMUSG00000039814 | Xkr5 | 9.356730009 | 5.498390163 | 7.832717308 | 54.64504051 | 58.11764556 | 64.46922443 | 1.48E-10 | 2.947618182 | 2.32E-11 | 44.68371345 | 7.714743455 | 9.829091465 |
| ENSMUSG00000039693 | Msantd3 | 113.9819837 | 92.37295474 | 109.6580423 | 773.1691902 | 812.6942896 | 813.0820097 | 1.53E-108 | 2.917964637 | 2.51E-110 | 498.0612751 | 7.557791073 | 107.8150198 |
| ENSMUSG00000022235 | Cmbl | 2.551835457 | 3.299034098 | 3.916358654 | 33.71715265 | 27.62970035 | 11.54672676 | 0.000304138 | 2.903039357 | 9.14E-05 | 15.30687798 | 7.480005633 | 3.516929917 |
| ENSMUSG00000111890 | Gm48836 | 2.551835457 | 10.99678033 | 2.937268991 | 49.99439876 | 34.29893836 | 33.67795306 | 1.05E-05 | 2.884493525 | 2.56E-06 | 22.12223613 | 7.384465628 | 4.979633645 |
| ENSMUSG00000034783 | Cd207 | 4.253059095 | 8.797424261 | 8.811806972 | 60.45834269 | 43.82642124 | 53.8847249 | 1.12E-08 | 2.876136166 | 2.03E-09 | 35.94567521 | 7.341811969 | 7.951572951 |
| ENSMUSG00000063873 | Slc24a3 | 30.62202548 | 36.28937507 | 24.47724159 | 223.2308038 | 220.0848545 | 207.8410818 | 1.52E-32 | 2.838443023 | 8.74E-34 | 146.7868114 | 7.152477347 | 31.81797019 |
| ENSMUSG00000046598 | Bdh1 | 10.20734183 | 6.598068195 | 7.832717308 | 44.18109658 | 71.45612159 | 61.58254274 | 2.60E-09 | 2.834993139 | 4.48E-10 | 38.89348996 | 7.135394221 | 8.585579213 |
| ENSMUSG00000070305 | Mpzl3 | 11.05795365 | 13.19613639 | 12.72816563 | 81.38623054 | 100.0385702 | 78.90263289 | 4.80E-14 | 2.823135024 | 6.00E-15 | 60.90045989 | 7.076985796 | 13.31856643 |
| ENSMUSG00000034416 | Pkd1l2 | 394.683884 | 445.3696032 | 383.8031481 | 3048.495664 | 2703.899641 | 2803.930149 | 3.09E-208 | 2.807685542 | 2.03E-210 | 958.3471433 | 7.001604361 | 207.5104039 |
| ENSMUSG00000038042 | Ptpdc1 | 16.16162456 | 6.598068195 | 7.832717308 | 76.7355888 | 72.40886988 | 69.28036059 | 1.10E-10 | 2.802628398 | 1.70E-11 | 45.2946195 | 6.97710429 | 9.960169081 |
| ENSMUSG00000082082 | Gm13230 | 6.804894552 | 1.099678033 | 4.895448318 | 26.74119004 | 30.48794521 | 34.64018029 | 3.36E-05 | 2.800012646 | 8.80E-06 | 19.75511176 | 6.964465554 | 4.473532715 |
| ENSMUSG00000026435 | Slc45a3 | 51.88732096 | 51.68486753 | 53.84993149 | 355.7740935 | 366.8080908 | 373.3441654 | 1.16E-55 | 2.799846583 | 3.79E-57 | 253.8355835 | 6.963663947 | 54.93392677 |
| ENSMUSG00000046668 | Cxxc5 | 24.66774275 | 18.69452655 | 22.51906226 | 146.495215 | 150.5342295 | 162.6164019 | 6.41E-24 | 2.793456202 | 4.94E-25 | 106.7919131 | 6.93288679 | 23.19344783 |
| ENSMUSG00000002565 | Scin | 8.50611819 | 9.897102293 | 20.56088293 | 87.19953273 | 82.88910105 | 92.37381411 | 7.45E-12 | 2.757321492 | 1.06E-12 | 50.72174207 | 6.761397628 | 11.12782172 |
| ENSMUSG00000079481 | Nhsl2 | 15.31101274 | 18.69452655 | 12.72816563 | 119.7540249 | 79.07810789 | 115.4672676 | 3.50E-14 | 2.753987996 | 4.32E-15 | 61.54628122 | 6.745792755 | 13.45599136 |
| ENSMUSG00000026204 | Ptprn | 12.75917728 | 9.897102293 | 9.790896635 | 87.19953273 | 66.69238015 | 65.43145166 | 2.26E-11 | 2.743833726 | 3.31E-12 | 48.49392987 | 6.698479839 | 10.64619071 |
| ENSMUSG00000003032 | Klf4 | 69.75016916 | 112.1671593 | 85.18080073 | 606.9087478 | 564.0269864 | 579.2607927 | 2.16E-57 | 2.724687944 | 6.78E-59 | 261.8497615 | 6.610172605 | 56.66501974 |
| ENSMUSG00000031434 | Morc4 | 51.03670914 | 58.28293573 | 66.57809712 | 383.677944 | 394.4377912 | 379.1175288 | 6.49E-53 | 2.72271846 | 2.21E-54 | 241.1519938 | 6.601154936 | 52.18750451 |
| ENSMUSG00000048216 | Gpr85 | 30.62202548 | 16.49517049 | 19.58179327 | 169.7484237 | 114.3297945 | 161.6541747 | 1.02E-16 | 2.719774031 | 1.11E-17 | 73.31213037 | 6.587696229 | 15.99043724 |
| ENSMUSG00000107523 | Gm44451 | 2.551835457 | 9.897102293 | 2.937268991 | 32.55449222 | 29.53519692 | 35.60240752 | 4.98E-05 | 2.713187278 | 1.34E-05 | 18.95747076 | 6.557688068 | 4.302913825 |
| ENSMUSG00000073433 | Arhgdig | 18.71346002 | 10.99678033 | 20.56088293 | 94.17549534 | 141.0067466 | 92.37381411 | 3.77E-13 | 2.692586345 | 4.98E-14 | 56.73849868 | 6.464713096 | 12.42335101 |
| ENSMUSG00000049097 | Ankrd34a | 53.5885446 | 41.78776524 | 50.9126625 | 319.73162 | 313.4541867 | 292.517078 | 6.60E-42 | 2.653049694 | 2.90E-43 | 190.1805558 | 6.289954991 | 41.18035955 |
| ENSMUSG00000045326 | Fndc7 | 5.103670914 | 2.199356065 | 2.937268991 | 16.27724611 | 23.8187072 | 25.01790799 | 0.000546172 | 2.642819234 | 0.000171377 | 14.12146915 | 6.245509343 | 3.262670924 |
| ENSMUSG00000004105 | Angptl2 | 455.927935 | 506.951573 | 461.1512315 | 3110.116667 | 2930.653734 | 2802.967922 | 2.09E-217 | 2.636574685 | 1.35E-219 | 1000.570029 | 6.218534755 | 216.6791869 |
| ENSMUSG00000042734 | Ttc9 | 1.701223638 | 8.797424261 | 1.958179327 | 15.11458567 | 32.39344179 | 26.94236245 | 0.00142591 | 2.634136082 | 0.000483424 | 12.17854245 | 6.208032379 | 2.845907846 |
| ENSMUSG00000031520 | Vegfc | 11.05795365 | 15.39549246 | 18.60270361 | 98.82613709 | 82.88910105 | 95.26049581 | 2.98E-13 | 2.629566521 | 3.92E-14 | 57.20963701 | 6.188400297 | 12.52513064 |
| ENSMUSG00000002489 | Tiam1 | 96.11913554 | 81.37617441 | 93.01351804 | 601.0954456 | 496.381858 | 581.1852471 | 1.22E-61 | 2.627742876 | 3.55E-63 | 281.4911916 | 6.180582767 | 60.91314441 |
| ENSMUSG00000042485 | Mustn1 | 11.90856547 | 8.797424261 | 17.62361394 | 69.75962618 | 97.18032536 | 70.24258782 | 1.63E-10 | 2.624723577 | 2.55E-11 | 44.49544982 | 6.167661453 | 9.788663924 |
| ENSMUSG00000029762 | Akr1b8 | 1100.691694 | 1143.665154 | 1125.953113 | 6659.718979 | 7246.603478 | 6672.083615 | 0 | 2.610982879 | 0 | 1589.453235 | 6.10919749 | Inf |
| ENSMUSG00000068699 | Flnc | 128.4423847 | 153.9549246 | 153.7170772 | 935.9416513 | 837.4657451 | 874.6645524 | 2.49E-93 | 2.60666255 | 4.81E-95 | 427.8330694 | 6.09093011 | 92.6041473 |
| ENSMUSG00000040093 | Bmf | 39.12814367 | 49.48551147 | 43.0799452 | 263.923919 | 268.6750172 | 257.8768977 | 3.46E-36 | 2.593645156 | 1.76E-37 | 163.6976805 | 6.036219029 | 35.46101968 |
| ENSMUSG00000042659 | Arrdc4 | 23.81713093 | 26.39227278 | 30.35177957 | 175.5617259 | 155.2979709 | 149.1452207 | 5.11E-22 | 2.576983321 | 4.25E-23 | 97.97039166 | 5.966907121 | 21.29185657 |
| ENSMUSG00000017832 | Hspb9 | 3.402447276 | 2.199356065 | 0.979089664 | 12.7892648 | 13.33847603 | 13.47118123 | 0.008417538 | 2.568127781 | 0.003274257 | 8.647971481 | 5.930393278 | 2.074814935 |
| ENSMUSG00000008658 | Rbfox1 | 35.7256964 | 23.09323868 | 50.9126625 | 238.3453894 | 174.3529367 | 236.7078987 | 7.70E-19 | 2.55634864 | 7.58E-20 | 83.15717904 | 5.882170652 | 18.11340635 |
| ENSMUSG00000029648 | Flt1 | 32.32324912 | 40.68808721 | 43.0799452 | 226.7187851 | 226.7540925 | 223.2367174 | 3.30E-30 | 2.550966729 | 2.07E-31 | 135.9231458 | 5.86026835 | 29.48102347 |
| ENSMUSG00000025141 | Myadml2 | 5.103670914 | 7.697746228 | 3.916358654 | 25.5785296 | 40.01542809 | 30.79127137 | 3.91E-05 | 2.545995264 | 1.04E-05 | 19.44544847 | 5.840108871 | 4.407783426 |
| ENSMUSG00000081738 | Hmgb1-ps2 | 4.253059095 | 0 | 4.895448318 | 30.22917134 | 15.24397261 | 9.622272304 | 0.009876135 | 2.545372042 | 0.003897868 | 8.330791128 | 5.837586579 | 2.005412993 |
| ENSMUSG00000075410 | Prcd | 2.551835457 | 4.39871213 | 0.979089664 | 13.95192524 | 20.00771405 | 11.54672676 | 0.006704612 | 2.54407224 | 0.002552298 | 9.102719199 | 5.832329552 | 2.173626349 |
| ENSMUSG00000025268 | Maged2 | 48.48487368 | 57.18325769 | 46.01721419 | 261.5985982 | 329.6509076 | 286.7437146 | 2.89E-34 | 2.539855298 | 1.56E-35 | 154.7831505 | 5.815306765 | 33.5393652 |
| ENSMUSG00000036766 | Dner | 46.78365004 | 50.5851895 | 52.87084183 | 294.1530904 | 278.2025001 | 296.365987 | 4.89E-39 | 2.534130242 | 2.30E-40 | 176.8986866 | 5.792275573 | 38.31078322 |
| ENSMUSG00000040616 | Tmem51 | 119.0856547 | 119.8649056 | 133.1561942 | 761.5425858 | 746.0019094 | 642.7677899 | 3.21E-73 | 2.530360234 | 7.68E-75 | 335.038301 | 5.777159131 | 72.49378323 |
| ENSMUSG00000100230 | Gm28650 | 4.253059095 | 3.299034098 | 5.874537981 | 23.25320873 | 30.48794521 | 24.05568076 | 0.000168107 | 2.530297643 | 4.84E-05 | 16.50842644 | 5.776908497 | 3.774413697 |
| ENSMUSG00000075610 | Tmem92 | 22.96651911 | 19.79420459 | 16.64452428 | 108.1274206 | 101.9440668 | 133.749585 | 4.44E-15 | 2.528031671 | 5.22E-16 | 65.71306972 | 5.767842105 | 14.35285079 |
| ENSMUSG00000036862 | Dchs1 | 3.402447276 | 10.99678033 | 15.66543462 | 66.27164487 | 60.02314214 | 44.2624526 | 2.57E-06 | 2.524474154 | 5.89E-07 | 24.94918315 | 5.753636803 | 5.589816216 |

| | | | | | | | | | | | | | |
|---|---|---|---|---|---|---|---|---|---|---|---|---|---|
| ENSMUSG00000027670 | *Ocstamp* | 96.11913554 | 112.1671593 | 81.26444207 | 553.4263677 | 562.1214899 | 542.6961579 | 3.03E-61 | 2.522350696 | 8.84E-63 | 279.6758292 | 5.745174434 | 60.51857146 |
| ENSMUSG00000022114 | *Spry2* | 40.82936731 | 46.18647737 | 44.05903486 | 246.4840125 | 265.8167723 | 229.9723081 | 3.23E-32 | 2.505834829 | 1.89E-33 | 145.2584648 | 5.67977913 | 31.4903857 |
| ENSMUSG00000060594 | *Layn* | 50.18609732 | 87.97424261 | 101.825325 | 410.419134 | 495.4291097 | 444.5489804 | 2.17E-28 | 2.502667316 | 1.44E-29 | 127.5003702 | 5.667322556 | 27.66280775 |
| ENSMUSG00000043079 | *Synpo* | 50.18609732 | 52.78454556 | 47.97539351 | 297.6410717 | 281.0607449 | 267.49917 | 7.41E-37 | 2.487599028 | 3.71E-38 | 166.7923022 | 5.608438017 | 36.12994615 |
| ENSMUSG00000102918 | *Pcdhgc3* | 535.0348341 | 451.9676714 | 469.9630385 | 2719.462761 | 2642.923751 | 2554.713297 | 1.48E-174 | 2.43816241 | 1.27E-176 | 802.8949447 | 5.41950997 | 173.8306818 |
| ENSMUSG00000053395 | *Cacng8* | 153.9607392 | 156.1542806 | 196.7970224 | 974.3094456 | 838.4184933 | 930.4737318 | 1.26E-73 | 2.435968158 | 3.01E-75 | 336.9075274 | 5.411273491 | 72.89838568 |
| ENSMUSG00000022421 | *Nptxr* | 14.46040092 | 19.79420459 | 15.66543462 | 93.01283491 | 92.41658392 | 81.78931458 | 3.06E-12 | 2.430296252 | 4.27E-13 | 52.51407748 | 5.390041017 | 11.51416606 |
| ENSMUSG00000045349 | *Sh2d5* | 53.5885446 | 50.5851895 | 46.01721419 | 302.2917134 | 245.8090583 | 260.7635794 | 2.52E-31 | 2.425013497 | 1.52E-32 | 141.1105097 | 5.370340253 | 30.59803255 |
| ENSMUSG00000055452 | *Gm7353* | 84.21057008 | 92.37295474 | 83.2226214 | 508.0826107 | 404.9180224 | 471.4913429 | 8.02E-47 | 2.414991114 | 3.13E-48 | 212.9428423 | 5.333161859 | 46.09569173 |
| ENSMUSG00000067199 | *Frat1* | 1.701223638 | 2.199356065 | 10.7699863 | 29.06651091 | 20.96046233 | 27.90458968 | 0.002356805 | 2.413330721 | 0.000828617 | 11.17601105 | 5.327027472 | 2.627676366 |
| ENSMUSG00000032420 | *Nt5e* | 11.05795365 | 7.697746228 | 9.790896635 | 58.13302182 | 38.10993152 | 56.77140659 | 5.76E-07 | 2.408428797 | 1.23E-07 | 27.97186069 | 5.308958252 | 6.239647831 |
| ENSMUSG00000020014 | *Cfap54* | 5.954282733 | 2.199356065 | 6.853627645 | 32.55449222 | 24.77145549 | 23.09345353 | 0.000395484 | 2.395122517 | 0.000121109 | 14.77533777 | 5.260217732 | 3.402871341 |
| ENSMUSG00000030465 | *Psd3* | 18.71346002 | 12.09645836 | 6.853627645 | 74.41026793 | 53.35390412 | 71.20481505 | 1.54E-07 | 2.383232994 | 3.10E-08 | 30.64261168 | 5.21704542 | 6.812675577 |
| ENSMUSG00000039611 | *Pgap4* | 15.31101274 | 9.897102293 | 15.66543462 | 63.946324 | 75.26711474 | 74.09149674 | 1.51E-09 | 2.372933638 | 2.55E-10 | 39.98890613 | 5.179933721 | 8.821001406 |
| ENSMUSG00000058317 | *Ube2e2* | 105.4758656 | 122.0642616 | 102.8044147 | 555.7516886 | 577.3654625 | 563.865157 | 3.48E-63 | 2.365695861 | 9.83E-65 | 288.6412138 | 5.154011873 | 62.45869256 |
| ENSMUSG00000055069 | *Rab39* | 44.23181459 | 40.68808721 | 56.78720049 | 234.8574081 | 221.9903511 | 262.6880339 | 3.98E-26 | 2.342734551 | 2.84E-27 | 117.0216373 | 5.072632171 | 25.40020837 |
| ENSMUSG00000028565 | *Nfia* | 10.20734183 | 12.09645836 | 4.895448318 | 40.69311527 | 47.63741439 | 48.11136152 | 4.31E-06 | 2.333657735 | 1.01E-06 | 23.90565461 | 5.040817542 | 5.365377793 |
| ENSMUSG00000046442 | *Ppm1e* | 85.91179372 | 91.2732767 | 87.13898005 | 447.624268 | 373.4773289 | 483.0380696 | 2.31E-40 | 2.303854938 | 1.07E-41 | 183.0052865 | 4.937753915 | 39.63580749 |
| ENSMUSG00000034686 | *Prr7* | 16.16162456 | 19.79420459 | 14.68634495 | 82.54889098 | 74.31436646 | 87.56267796 | 1.88E-10 | 2.277305356 | 2.98E-11 | 44.19179792 | 4.847716583 | 9.724709556 |
| ENSMUSG00000074794 | *Arrdc3* | 108.8783128 | 103.3697351 | 109.6580423 | 513.8959129 | 512.5785789 | 534.0361129 | 1.50E-58 | 2.275952607 | 4.61E-60 | 267.2071808 | 4.843173235 | 57.82312522 |
| ENSMUSG00000043648 | *Pld6* | 6.804894552 | 6.598068195 | 10.7699863 | 38.3677944 | 44.77916953 | 31.7534986 | 3.14E-05 | 2.249265547 | 8.17E-06 | 19.89740279 | 4.754407451 | 4.503465547 |
| ENSMUSG00000039741 | *Bahcc1* | 193.0888829 | 197.9420459 | 221.274264 | 1092.90081 | 825.0800173 | 971.8495027 | 1.32E-58 | 2.237992444 | 4.02E-60 | 267.4775741 | 4.717401662 | 57.88011086 |
| ENSMUSG00000003452 | *Bicd1* | 59.54282733 | 68.18003802 | 60.70355914 | 304.6170343 | 255.3365412 | 324.2705766 | 4.11E-29 | 2.233651643 | 2.67E-30 | 130.8498953 | 4.703229211 | 28.38662783 |
| ENSMUSG00000106964 | *None* | 15.31101274 | 23.09323868 | 23.49815192 | 94.17549534 | 97.18032536 | 96.22272304 | 1.98E-11 | 2.228595352 | 2.90E-12 | 48.75605126 | 4.686774401 | 10.70240435 |
| ENSMUSG00000000942 | *Hoxa4* | 18.71346002 | 12.09645836 | 21.5399726 | 101.151458 | 74.31436646 | 72.16704228 | 4.93E-09 | 2.227966275 | 8.71E-10 | 37.59381231 | 4.684731212 | 8.307078555 |
| ENSMUSG00000078653 | *Cntd1* | 16.16162456 | 17.59484852 | 0 | 49.99439876 | 46.68466611 | 60.62031551 | 0.002103444 | 2.221564854 | 0.000733189 | 11.4031701 | 4.663990512 | 2.677069095 |
| ENSMUSG00000075611 | *Gm11545* | 28.07019003 | 27.49195081 | 27.41451058 | 122.0793458 | 126.7155223 | 134.7118123 | 1.35E-15 | 2.208608576 | 1.54E-16 | 68.12072956 | 4.622292562 | 14.86961415 |
| ENSMUSG00000013033 | *Adgrl1* | 122.4881019 | 111.0674813 | 112.5953113 | 573.1915951 | 489.71262 | 538.847249 | 6.00E-51 | 2.206157391 | 2.12E-52 | 232.0564276 | 4.614445805 | 50.22219243 |
| ENSMUSG00000038545 | *Cul7* | 65.49711006 | 57.18325769 | 60.70355914 | 317.4062991 | 265.8167723 | 261.7258067 | 1.83E-28 | 2.197942999 | 1.21E-29 | 127.8524579 | 4.588246811 | 27.73805407 |
| ENSMUSG00000030688 | *Stard10* | 32.32324912 | 50.5851895 | 48.95448318 | 208.1162181 | 220.0848545 | 162.6164019 | 9.36E-17 | 2.173806634 | 1.01E-17 | 73.49310771 | 4.512123743 | 16.0288794 |
| ENSMUSG00000113637 | *Gm7049* | 62.09466279 | 85.77488654 | 68.53627645 | 322.0569409 | 336.3201456 | 309.8371682 | 3.49E-31 | 2.170895099 | 2.11E-32 | 140.4596561 | 4.50302691 | 30.45768323 |
| ENSMUSG00000069755 | *Zfp125* | 15.31101274 | 9.897102293 | 17.62361394 | 65.10898443 | 51.44840755 | 76.97817843 | 3.26E-07 | 2.165123927 | 6.77E-08 | 29.12957527 | 4.485049559 | 6.487016346 |
| ENSMUSG00000050010 | *Shisa3* | 9.356730009 | 12.09645836 | 9.790896635 | 51.1570592 | 41.92092467 | 46.18690706 | 3.99E-06 | 2.162487616 | 9.33E-07 | 24.06124127 | 4.476861282 | 5.398691762 |
| ENSMUSG00000032012 | *Nectin1* | 563.1050242 | 569.6332209 | 599.2028741 | 2811.312935 | 2439.035617 | 2466.188391 | 4.75E-132 | 2.155042713 | 5.90E-134 | 606.6798841 | 4.453818332 | 131.3232154 |
| ENSMUSG00000040722 | *Scamp5* | 247.5280393 | 265.0224059 | 256.5214918 | 1161.497776 | 1161.400163 | 1068.072226 | 6.40E-96 | 2.141934724 | 1.20E-97 | 439.7885851 | 4.413535263 | 95.19371353 |
| ENSMUSG00000031647 | *Mfap3l* | 80.8081228 | 61.58196982 | 71.47354544 | 359.2620748 | 302.0212073 | 278.0836696 | 1.69E-26 | 2.12599212 | 1.19E-27 | 118.7472381 | 4.365031665 | 25.77258183 |
| ENSMUSG00000002504 | *Nherf2* | 27.21957821 | 38.48873114 | 29.37268991 | 129.0553084 | 145.770488 | 135.6740395 | 1.15E-14 | 2.121787143 | 1.38E-15 | 63.79225499 | 4.352327572 | 13.93852688 |
| ENSMUSG00000020886 | *Dlg4* | 121.6374901 | 76.97746228 | 124.3443873 | 524.3598568 | 460.177423 | 426.2666631 | 2.18E-28 | 2.118039856 | 1.45E-29 | 127.4960262 | 4.341037412 | 27.66230558 |
| ENSMUSG00000019990 | *Pde7b* | 7.655506371 | 7.697746228 | 8.811806972 | 41.85577571 | 26.67695206 | 36.56463475 | 0.000132502 | 2.117352537 | 3.76E-05 | 16.98823189 | 4.338969777 | 3.877777766 |
| ENSMUSG00000032418 | *Me1* | 1098.99047 | 1211.845192 | 1153.367624 | 4902.93906 | 5093.392347 | 4995.88378 | 1.53E-235 | 2.115365907 | 8.92E-238 | 1084.212388 | 4.33299901 | 234.8155855 |
| ENSMUSG00000051682 | *Treml4* | 77.40567553 | 72.57875015 | 76.36899376 | 316.2436387 | 341.0838871 | 318.4972133 | 4.38E-34 | 2.106666444 | 2.38E-35 | 153.9423622 | 4.306949607 | 33.3589167 |
| ENSMUSG00000024883 | *Rin1* | 104.6252537 | 80.27649638 | 101.825325 | 448.7869284 | 387.7685532 | 404.1354368 | 1.04E-35 | 2.105519334 | 5.39E-37 | 161.4726818 | 4.303526445 | 34.98132218 |
| ENSMUSG00000019732 | *Calr3* | 49.3354855 | 21.99356065 | 25.45633125 | 130.2179689 | 161.0144607 | 128.9384489 | 3.46E-10 | 2.100224603 | 5.57E-11 | 42.96496827 | 4.287761329 | 9.460853993 |
| ENSMUSG00000033209 | *Ttc28* | 17.8628482 | 20.89388262 | 13.70725529 | 83.71155142 | 78.12535961 | 61.58254274 | 3.63E-08 | 2.092801466 | 6.92E-09 | 33.55856438 | 4.265756059 | 7.440359875 |
| ENSMUSG00000029392 | *Rilpl1* | 67.1983337 | 68.18003802 | 73.43172477 | 292.99043 | 305.8322004 | 284.8192602 | 1.10E-30 | 2.081993667 | 6.81E-32 | 138.1349283 | 4.233918994 | 29.95902826 |
| ENSMUSG00000031373 | *Car5b* | 295.1623012 | 246.3278793 | 271.2078368 | 1126.617963 | 1098.518776 | 1186.426175 | 3.99E-83 | 2.065474421 | 8.46E-85 | 380.7694931 | 4.185715978 | 82.39894876 |
| ENSMUSG00000043153 | *Crppa* | 4.253059095 | 5.498390163 | 7.832717308 | 16.27724611 | 20.00771405 | 36.56463475 | 0.004066342 | 2.062638075 | 0.001486584 | 10.09517086 | 4.177494924 | 2.390796064 |
| ENSMUSG00000079598 | *Clec2l* | 11.05795365 | 8.797424261 | 5.874537981 | 41.85577571 | 34.29893836 | 30.79127137 | 0.00019757 | 2.043312315 | 5.75E-05 | 16.18351025 | 4.121908028 | 3.704279009 |
| ENSMUSG00000100774 | *Gm7329* | 66.34772188 | 72.57875015 | 82.24353174 | 349.9607913 | 281.0607449 | 279.0458968 | 4.70E-25 | 2.040973641 | 3.47E-26 | 112.0582742 | 4.115231643 | 24.32795321 |
| ENSMUSG00000060206 | *Zfp462* | 8.50611819 | 8.797424261 | 3.916358654 | 26.74119004 | 22.86595891 | 36.56463475 | 0.001374523 | 2.022828333 | 0.000464767 | 12.25197135 | 4.063796989 | 2.861848127 |
| ENSMUSG00000040466 | *Blvrb* | 752.7914598 | 769.7746228 | 829.288945 | 2968.272094 | 3725.245806 | 2856.852647 | 6.38E-77 | 2.022288117 | 1.43E-78 | 352.1640302 | 4.062275588 | 76.19507353 |
| ENSMUSG00000001014 | *Icam4* | 17.8628482 | 7.697746228 | 17.62361394 | 58.13302182 | 68.59787673 | 50.03581598 | 7.00E-06 | 2.01521656 | 1.68E-06 | 22.93530985 | 4.042412538 | 5.154898969 |
| ENSMUSG00000078532 | *Nkain1* | 45.08242641 | 52.78454556 | 49.93357284 | 211.6041994 | 186.7386644 | 188.5965372 | 2.95E-19 | 1.992707537 | 2.82E-20 | 85.1128026 | 3.979832014 | 18.53008884 |
| ENSMUSG00000028047 | *Thbs3* | 10.20734183 | 10.99678033 | 10.7699863 | 54.64504051 | 34.29893836 | 38.48908921 | 6.75E-05 | 1.991585808 | 1.85E-05 | 18.33674057 | 3.976738806 | 4.170757955 |
| ENSMUSG00000110632 | *Gm45880* | 26.36896639 | 15.39549246 | 33.28904856 | 96.50081622 | 97.18032536 | 106.8072226 | 4.75E-09 | 1.991059629 | 8.37E-10 | 37.67262184 | 3.975288676 | 8.323611973 |
| ENSMUSG00000039787 | *Cercam* | 3.402447276 | 5.498390163 | 8.811806972 | 32.55449222 | 14.29122432 | 23.09345353 | 0.007364097 | 1.985464149 | 0.002829624 | 8.914195493 | 3.959900414 | 2.132880473 |
| ENSMUSG00000061013 | *Mkx* | 64.64649824 | 69.27971605 | 59.72446948 | 259.2732773 | 249.6200514 | 253.0657616 | 5.83E-25 | 1.977793618 | 4.31E-26 | 111.6301735 | 3.93890226 | 24.23457882 |
| ENSMUSG00000097214 | *Gm19791* | 7.655506371 | 3.299034098 | 5.874537981 | 27.90385047 | 15.24397261 | 24.05568076 | 0.006111098 | 1.970951576 | 0.00230833 | 9.286625845 | 3.920266075 | 2.213880753 |
| ENSMUSG00000027187 | *Cat* | 2495.695077 | 2549.05368 | 2424.226007 | 9967.487921 | 9234.036406 | 9973.485243 | 6.78E-244 | 1.965822235 | 3.67E-246 | 1122.792667 | 3.906352757 | 243.1689738 |
| ENSMUSG00000023830 | *Igf2r* | 1718.235874 | 1851.857807 | 1655.640621 | 7036.420961 | 6129.982484 | 7114.708141 | 1.12E-134 | 1.956848794 | 1.34E-136 | 618.8308436 | 3.882130981 | 133.9489257 |
| ENSMUSG00000089876 | *Tmem102* | 14.46040092 | 21.99356065 | 21.5399726 | 83.71155142 | 63.83413529 | 74.09149674 | 2.21E-07 | 1.943199234 | 4.53E-08 | 29.90949749 | 3.845574745 | 6.654690882 |
| ENSMUSG00000027784 | *Ppm1l* | 651.5686533 | 719.1894333 | 630.5337433 | 2698.534873 | 2317.083836 | 2630.729248 | 1.36E-98 | 1.934788895 | 2.51E-100 | 452.1041158 | 3.823221788 | 97.86758088 |
| ENSMUSG00000043154 | *Ppp2r3a* | 86.76240554 | 112.1671593 | 76.36899376 | 373.2140001 | 315.3596833 | 358.9107569 | 1.85E-24 | 1.933024057 | 1.40E-25 | 109.2963859 | 3.818547729 | 23.73365474 |
| ENSMUSG00000038860 | *Garnl3* | 4.253059095 | 12.09645836 | 10.7699863 | 22.09054829 | 49.54291097 | 29.82904414 | 0.00262327 | 1.929965576 | 0.000929322 | 10.96333829 | 3.81046107 | 2.581156929 |
| ENSMUSG00000042207 | *Kdm5b* | 159.0644101 | 184.7459095 | 183.0897671 | 701.0842431 | 606.9006594 | 672.596834 | 5.47E-47 | 1.912997315 | 2.12E-48 | 213.7172586 | 3.76590685 | 46.26233931 |
| ENSMUSG00000027624 | *Epb41l1* | 44.23181459 | 43.9871213 | 42.10085553 | 155.7964985 | 178.1639298 | 155.8808113 | 1.13E-15 | 1.910960466 | 1.28E-16 | 68.47509634 | 3.760593758 | 14.94661203 |
| ENSMUSG00000101309 | *Gm29397* | 11.05795365 | 8.797424261 | 11.74907596 | 34.87981309 | 51.44840755 | 31.7534986 | 0.000260865 | 1.898133767 | 7.75E-05 | 15.61953417 | 3.727307297 | 3.583584807 |
| ENSMUSG00000000562 | *Adora3* | 16.16162456 | 15.39549246 | 20.56088293 | 72.08494705 | 80.03085618 | 42.33799814 | 1.18E-05 | 1.897337901 | 2.90E-06 | 21.88169171 | 3.725251687 | 4.927925038 |
| ENSMUSG00000032174 | *Icam5* | 37.42692004 | 28.59162885 | 34.26813822 | 117.4287041 | 137.1957535 | 118.3539493 | 1.12E-11 | 1.888586926 | 1.61E-12 | 49.91343928 | 3.702723772 | 10.95267783 |
| ENSMUSG00000035539 | *Ccdc180* | 16.16162456 | 3.299034098 | 8.811806972 | 37.20513396 | 34.29893836 | 35.60240752 | 0.001846819 | 1.888037278 | 0.000638163 | 11.66122661 | 3.70131335 | 2.733575675 |
| ENSMUSG00000074738 | *Fndc10* | 31.4726373 | 34.09001901 | 30.35177957 | 120.9166854 | 124.8100257 | 108.731677 | 1.28E-11 | 1.887692251 | 1.85E-12 | 49.63894517 | 3.70042827 | 10.89295141 |
| ENSMUSG00000038569 | *Rad9b* | 13.6097891 | 17.59484852 | 28.39360024 | 70.92228662 | 61.92863871 | 86.60045073 | 3.25E-06 | 1.886804196 | 7.51E-07 | 24.4795695 | 3.698151163 | 5.488657351 |
| ENSMUSG00000047880 | *Cxcr5* | 28.07019003 | 32.99034098 | 33.28904856 | 124.4046667 | 108.6133048 | 113.5428132 | 3.19E-11 | 1.880263167 | 4.72E-12 | 47.79926503 | 3.681422081 | 10.49677751 |
| ENSMUSG00000034488 | *Edil3* | 60.39343915 | 56.08357966 | 53.84993149 | 216.2548412 | 195.313399 | 216.5011268 | 3.38E-19 | 1.879598172 | 3.24E-20 | 84.83721479 | 3.679725561 | 18.4711143 |
| ENSMUSG00000031257 | *Nox1* | 14.46040092 | 14.29581442 | 17.62361394 | 45.34375702 | 58.11764556 | 66.3936789 | 8.40E-06 | 1.87648807 | 2.02E-06 | 22.57199823 | 3.671801505 | 5.075612751 |
| ENSMUSG00000001663 | *Gstt1* | 15.31101274 | 5.498390163 | 5.874537981 | 40.69311527 | 32.39344179 | 26.94236245 | 0.002329571 | 1.874953129 | 0.000818083 | 11.19975078 | 3.667897006 | 2.632724049 |
| ENSMUSG00000037568 | *Vash2* | 35.7256964 | 25.29259475 | 23.49815192 | 106.9647601 | 103.8495634 | 100.071632 | 1.59E-09 | 1.867783554 | 2.69E-10 | 39.88487054 | 3.649714351 | 8.798398245 |
| ENSMUSG00000037868 | *Egr2* | 69.75016916 | 74.77810622 | 67.55718678 | 272.0625421 | 223.8958477 | 277.1214423 | 7.88E-21 | 1.866721463 | 6.93E-22 | 92.44286667 | 3.647028474 | 20.10359657 |
| ENSMUSG00000016529 | *Il10* | 53.5885446 | 39.58840917 | 50.9126625 | 190.6763116 | 161.0144607 | 177.0498104 | 4.46E-15 | 1.86630847 | 5.24E-16 | 65.70307826 | 3.645984605 | 14.35090254 |
| ENSMUSG00000015568 | *Lpl* | 15904.73979 | 15179.95556 | 15238.55152 | 57185.45356 | 51224.5117 | 58986.45368 | 1.05E-185 | 1.853375139 | 8.32E-188 | 854.3302814 | 3.613445511 | 184.9769622 |
| ENSMUSG00000024538 | *Ppic* | 414.2479558 | 352.9966485 | 384.7822378 | 1407.981788 | 1321.461875 | 1440.454164 | 1.00E-79 | 1.852372966 | 2.20E-81 | 365.0857015 | 3.61093629 | 78.99936289 |
| ENSMUSG00000049709 | *Nlrp10* | 222.0096848 | 235.331099 | 230.0860709 | 801.0730406 | 792.6865755 | 878.5134613 | 1.88E-59 | 1.848416422 | 5.64E-61 | 271.392987 | 3.601046982 | 58.7265787 |
| ENSMUSG00000025002 | *Cyp2c55* | 45.93303823 | 34.09001901 | 37.20540721 | 125.5673271 | 157.2034675 | 139.5229484 | 5.01E-12 | 1.842608335 | 7.10E-13 | 51.51655605 | 3.586578816 | 11.30010828 |
| ENSMUSG00000102543 | *Pcdhgc5* | 11.90856547 | 24.19291672 | 19.58179327 | 69.75962618 | 62.881387 | 63.5069972 | 5.31E-06 | 1.833480495 | 1.26E-06 | 23.48908157 | 3.563958394 | 5.27466156 |
| ENSMUSG00000034957 | *Cebpa* | 763.8494134 | 701.5945848 | 776.4181032 | 2779.921103 | 2649.592989 | 2541.242115 | 4.23E-121 | 1.828192763 | 5.72E-123 | 556.1729822 | 3.550919768 | 120.3737583 |
| ENSMUSG00000021485 | *Mxd3* | 41.67997913 | 40.68808721 | 54.82902116 | 165.097782 | 157.2034675 | 161.6541747 | 7.44E-14 | 1.818064548 | 9.35E-15 | 60.0285242 | 3.526078389 | 13.12856957 |
| ENSMUSG00000001435 | *Col18a1* | 119.0856547 | 101.170379 | 109.6580423 | 403.4431714 | 383.95756 | 373.3441654 | 6.55E-31 | 1.809718647 | 4.02E-32 | 139.1825056 | 3.505739132 | 30.18377427 |
| ENSMUSG00000048897 | *Zfp710* | 1600.000832 | 1719.896443 | 1770.194112 | 6293.480942 | 5791.756842 | 5718.51643 | 3.89E-149 | 1.806833471 | 4.08E-151 | 685.582599 | 3.49873518 | 148.4102035 |
| ENSMUSG00000037224 | *Zfyve28* | 16.16162456 | 13.19613639 | 5.874537981 | 39.53045484 | 37.15718323 | 46.18690706 | 0.000633981 | 1.795126189 | 0.000201365 | 13.81830127 | 3.470458289 | 3.197923973 |
| ENSMUSG00000041911 | *Dlx1* | 26.36896639 | 26.39227278 | 23.49815192 | 87.19953273 | 76.21986303 | 101.0338592 | 4.66E-08 | 1.793938344 | 8.99E-09 | 33.0492681 | 3.467602057 | 7.331595619 |
| ENSMUSG00000046841 | *Ckap4* | 1007.975005 | 1134.86773 | 1093.643154 | 3779.809079 | 3831.000866 | 3484.224801 | 1.89E-121 | 1.778814661 | 2.52E-123 | 557.8058455 | 3.431441266 | 120.7245328 |
| ENSMUSG00000027254 | *Map1a* | 34.02447276 | 34.09001901 | 33.28904856 | 130.2179689 | 111.4715497 | 105.8449953 | 2.73E-10 | 1.774632431 | 4.35E-11 | 43.44999299 | 3.421508261 | 9.56460838 |
| ENSMUSG00000024985 | *Tcf7l2* | 40.82936731 | 46.18647737 | 38.18449688 | 140.6819128 | 122.9045291 | 163.5786292 | 1.92E-11 | 1.773846334 | 2.80E-12 | 48.82341688 | 3.419644455 | 10.71670884 |
| ENSMUSG00000111817 | *Gm47154* | 11.90856547 | 9.897102293 | 6.853627645 | 29.06651091 | 24.77145549 | 44.2624526 | 0.002300903 | 1.770797929 | 0.000807226 | 11.22454338 | 3.412426397 | 2.638101711 |
| ENSMUSG00000031788 | *Kifc3* | 518.8732096 | 506.951573 | 536.5411356 | 1860.256698 | 1754.962346 | 1716.613379 | 5.41E-101 | 1.769852618 | 9.73E-103 | 463.1878398 | 3.410191174 | 100.2665551 |
| ENSMUSG00000042073 | *Abhd14b* | 22.11590729 | 17.59484852 | 20.56088293 | 73.24760749 | 67.64512844 | 64.46922443 | 1.26E-06 | 1.760721388 | 2.79E-07 | 26.38686484 | 3.388675258 | 5.898536559 |
| ENSMUSG00000022947 | *Cbr3* | 203.2962247 | 173.7491291 | 187.0061257 | 618.5353521 | 735.5216782 | 557.1295664 | 8.67E-32 | 1.758037495 | 5.15E-33 | 143.2625715 | 3.382377056 | 31.06211548 |
| ENSMUSG00000039193 | *Nlrc4* | 66.34772188 | 53.8842236 | 61.6826488 | 195.3269533 | 200.0771405 | 220.3500358 | 1.32E-16 | 1.754406129 | 1.44E-17 | 72.7987547 | 3.373874078 | 15.87966855 |
| ENSMUSG00000035919 | *Bbs9* | 199.0431656 | 205.6397921 | 162.5288841 | 567.3782929 | 698.364495 | 629.2966087 | 5.63E-33 | 1.742026044 | 3.18E-34 | 148.7919724 | 3.345045985 | 32.24986246 |
| ENSMUSG00000037217 | *Syn1* | 20.41468366 | 15.39549246 | 14.68634495 | 48.83173833 | 60.97589042 | 59.65808828 | 2.19E-05 | 1.740775367 | 5.56E-06 | 20.63353286 | 3.342147411 | 4.660029554 |
| ENSMUSG00000040488 | *Ltbp4* | 7.655506371 | 7.697746228 | 11.74907596 | 22.09054829 | 21.91321062 | 46.18690706 | 0.006297394 | 1.739283602 | 0.002386909 | 9.225320827 | 3.338693374 | 2.200839106 |
| ENSMUSG00000017493 | *Igfbp4* | 757.8951307 | 773.0736569 | 787.1880895 | 2640.401851 | 2636.254513 | 2448.868301 | 1.12E-122 | 1.736772626 | 1.47E-124 | 563.4801924 | 3.332887512 | 121.9521582 |
| ENSMUSG00000000223 | *Drp2* | 9.356730009 | 4.39871213 | 9.790896635 | 23.25320873 | 25.72420377 | 29.82904414 | 0.005807321 | 1.728207195 | 0.00218163 | 9.390077106 | 3.313158436 | 2.236024202 |
| ENSMUSG00000031066 | *Usp11* | 18.71346002 | 31.89066294 | 17.62361394 | 70.92228662 | 80.98360447 | 71.20481505 | 4.43E-06 | 1.724583024 | 1.04E-06 | 23.85167889 | 3.30484595 | 5.35383518 |
| ENSMUSG00000045934 | *Mtmr11* | 60.39343915 | 59.38261376 | 76.36899376 | 231.3694268 | 217.2266096 | 200.1432639 | 4.08E-16 | 1.724474443 | 4.55E-17 | 70.52455166 | 3.304597228 | 15.38967018 |
| ENSMUSG00000071757 | *Zhx2* | 56.14038005 | 62.68164786 | 71.47354544 | 223.2308038 | 182.9276713 | 221.312263 | 3.66E-15 | 1.722944235 | 4.27E-16 | 66.1085208 | 3.301094034 | 14.43667947 |
| ENSMUSG00000038236 | *Hoxa7* | 102.9240301 | 111.0674813 | 112.5953113 | 359.2620748 | 362.0443494 | 336.7795306 | 1.03E-26 | 1.6976976 | 7.19E-28 | 119.7469354 | 3.243828619 | 25.9880582 |
| ENSMUSG00000027848 | *Olfml3* | 39.97875549 | 39.58840917 | 46.99630385 | 138.3565919 | 146.7232363 | 125.0895399 | 3.87E-11 | 1.696221585 | 5.79E-12 | 47.40108616 | 3.240511569 | 10.41217107 |
| ENSMUSG00000079037 | *Prnp* | 165.0186929 | 152.8552465 | 164.4870635 | 499.9439876 | 507.8148374 | 555.2051119 | 3.89E-36 | 1.694905521 | 1.99E-37 | 163.4595617 | 3.237556838 | 35.40958541 |
| ENSMUSG00000027111 | *Itga6* | 364.0618585 | 325.5046976 | 333.8695753 | 1116.154019 | 1036.590137 | 1160.44604 | 1.81E-60 | 1.692209102 | 5.34E-62 | 276.0903769 | 3.231511448 | 59.7433067 |
| ENSMUSG00000070526 | *Peg12* | 79.10689917 | 107.7684472 | 101.825325 | 349.9607913 | 282.0134932 | 292.517078 | 4.26E-18 | 1.683705091 | 4.32E-19 | 79.71592043 | 3.212519232 | 17.37030721 |
| ENSMUSG00000002908 | *Kcnn1* | 12.75917728 | 8.797424261 | 6.853627645 | 26.74119004 | 33.34619008 | 31.7534986 | 0.003085022 | 1.683105002 | 0.001106239 | 10.64073185 | 3.211183263 | 2.51074172 |
| ENSMUSG00000006310 | *Zbtb32* | 1253.801821 | 1293.221366 | 1236.590245 | 4244.873253 | 3976.771354 | 3894.133601 | 7.56E-141 | 1.679041733 | 8.51E-143 | 647.3299169 | 3.20215187 | 140.1214616 |
| ENSMUSG00000113924 | *Gm48493* | 190.5370475 | 129.7620078 | 111.6162216 | 452.2749097 | 292.4937244 | 638.918881 | 2.49E-09 | 1.676270472 | 4.28E-10 | 38.98140591 | 3.196006788 | 8.604272462 |
| ENSMUSG00000020453 | *Patz1* | 657.5229361 | 693.8968386 | 677.5300472 | 2102.090069 | 2170.3606 | 2112.088771 | 1.99E-111 | 1.655142314 | 3.04E-113 | 511.4701795 | 3.149542577 | 110.7004276 |
| ENSMUSG00000026069 | *Il1rl1* | 791.0689917 | 833.5559487 | 787.1880895 | 2447.400218 | 2400.925685 | 2742.347607 | 2.59E-89 | 1.655130013 | 5.18E-91 | 409.3115517 | 3.149515723 | 88.58632995 |
| ENSMUSG00000053128 | *Rnf26* | 803.8281689 | 852.2504752 | 830.2680347 | 2712.486798 | 2616.246799 | 2487.35739 | 5.48E-111 | 1.652991709 | 8.47E-113 | 509.4234067 | 3.144851096 | 110.2609388 |
| ENSMUSG00000072623 | *Zfp9* | 17.8628482 | 30.79098491 | 20.56088293 | 75.57292836 | 66.69238015 | 71.20481505 | 1.05E-05 | 1.638040772 | 2.56E-06 | 22.12204407 | 3.112428664 | 4.979633645 |
| ENSMUSG00000089762 | *Ier5l* | 43.38120277 | 56.08357966 | 57.76629015 | 172.0737446 | 173.4001884 | 140.4851756 | 4.10E-11 | 1.633131188 | 6.15E-12 | 47.28131032 | 3.101854871 | 10.38682852 |
| ENSMUSG00000029192 | *Tbc1d14* | 1497.076801 | 1462.571783 | 1420.659102 | 4509.959833 | 4484.586191 | 4559.994845 | 4.60E-170 | 1.629287865 | 4.17E-172 | 782.1170664 | 3.093602563 | 169.3368076 |
| ENSMUSG00000037235 | *Mxd4* | 222.0096848 | 205.6397921 | 233.0233399 | 703.409564 | 682.1677741 | 654.3145166 | 1.43E-42 | 1.624229445 | 6.22E-44 | 193.2448182 | 3.082774676 | 41.84442247 |
| ENSMUSG00000067925 | *Rtl8a* | 42.53059095 | 38.48873114 | 32.3099589 | 113.9407228 | 139.10125 | 96.22272304 | 4.80E-08 | 1.621457164 | 9.29E-09 | 32.98512336 | 3.076856509 | 7.318501932 |
| ENSMUSG00000006241 | *Ccdc159* | 51.88732096 | 45.08679934 | 44.05903486 | 162.7724611 | 162.9199572 | 109.6939043 | 4.75E-09 | 1.621146265 | 8.37E-10 | 37.67134576 | 3.076193521 | 8.323496886 |
| ENSMUSG00000018821 | *Avpi1* | 243.2749802 | 250.7265914 | 299.601437 | 791.7717571 | 913.6856081 | 731.2926951 | 1.04E-32 | 1.619401642 | 5.92E-34 | 147.5594883 | 3.072475789 | 31.98254993 |
| ENSMUSG00000050821 | *Fam131a* | 24.66774275 | 19.79420459 | 20.56088293 | 66.27164487 | 65.73963186 | 68.31813336 | 7.37E-06 | 1.617052257 | 1.77E-06 | 22.83480071 | 3.067476427 | 5.132652545 |
| ENSMUSG00000037583 | *Nr0b2* | 70.60078098 | 78.07714031 | 77.34808342 | 246.4840125 | 234.3760788 | 209.7655362 | 2.73E-16 | 1.612759112 | 3.02E-17 | 71.32854199 | 3.058361857 | 15.563432 |
| ENSMUSG00000050947 | *Amigo1* | 120.7868783 | 118.7652275 | 129.2398356 | 411.5817945 | 364.9025943 | 350.2507119 | 3.32E-24 | 1.609281329 | 2.54E-25 | 108.1159888 | 3.050998201 | 23.47893801 |
| ENSMUSG00000050199 | *Lgr4* | 32.32324912 | 35.18969704 | 33.28904856 | 101.151458 | 97.18032536 | 107.7694498 | 2.41E-08 | 1.604983857 | 4.54E-09 | 34.37662218 | 3.04192347 | 7.617451937 |
| ENSMUSG00000029334 | *Prkg2* | 20.41468366 | 14.29581442 | 21.5399726 | 47.66907789 | 52.40115583 | 71.20481505 | 0.000136708 | 1.601382385 | 3.89E-05 | 16.92294917 | 3.034339232 | 3.864206183 |
| ENSMUSG00000049932 | *H2ax* | 881.2338445 | 868.7456457 | 953.6333323 | 2932.22962 | 2652.451233 | 2573.957841 | 6.98E-84 | 1.592647638 | 1.47E-85 | 384.2607124 | 3.016023433 | 83.156209 |
| ENSMUSG00000066357 | *Wdr6* | 871.0265026 | 960.0189224 | 896.8461318 | 2808.987614 | 2764.875531 | 2630.729248 | 1.28E-103 | 1.58993307 | 2.25E-105 | 475.3037528 | 3.010353834 | 102.8930143 |
| ENSMUSG00000030410 | *Dmwd* | 476.3426186 | 456.3663835 | 472.9003075 | 1459.138848 | 1391.965249 | 1362.513758 | 1.09E-74 | 1.582371137 | 2.54E-76 | 341.8380668 | 2.994616247 | 73.96332467 |
| ENSMUSG00000038816 | *Ctnnal1* | 102.9240301 | 82.47585244 | 96.92987669 | 290.6651091 | 257.2420377 | 296.365987 | 9.06E-18 | 1.574478108 | 9.36E-19 | 78.1895934 | 2.978277347 | 17.0426337 |
| ENSMUSG00000045095 | *Magi1* | 157.3631865 | 138.5594321 | 121.4071183 | 467.3894954 | 399.2015326 | 380.079756 | 3.97E-21 | 1.57411664 | 3.45E-22 | 93.82167869 | 2.97753123 | 20.40069677 |
| ENSMUSG00000089704 | *Galnt2* | 108.027701 | 81.37617441 | 134.1352839 | 277.8758443 | 362.0443494 | 322.3461222 | 1.07E-13 | 1.570118645 | 1.35E-14 | 59.29825919 | 2.969291321 | 12.97049001 |
| ENSMUSG00000074604 | *Mgst2* | 284.1043475 | 345.2989022 | 302.538706 | 876.645969 | 982.2834848 | 898.7202332 | 4.20E-42 | 1.569477472 | 1.84E-43 | 191.0916548 | 2.96797198 | 41.37719203 |
| ENSMUSG00000053080 | *Zfta* | 53.5885446 | 57.18325769 | 62.66173847 | 208.1162181 | 166.7309504 | 140.4851756 | 3.70E-10 | 1.569330095 | 5.97E-11 | 42.82868626 | 2.967668806 | 9.431524034 |
| ENSMUSG00000000881 | *Dlg3* | 65.49711006 | 42.88744327 | 69.51536611 | 181.3750281 | 189.5969093 | 159.7297202 | 2.44E-10 | 1.568414179 | 3.88E-11 | 43.67523969 | 2.965785335 | 9.61328745 |
| ENSMUSG00000020331 | *Hcn2* | 38.27753185 | 60.48229179 | 50.9126625 | 141.8445732 | 172.4474401 | 126.0517672 | 1.41E-08 | 1.56718536 | 2.59E-09 | 35.47277503 | 2.963260296 | 7.850073375 |
| ENSMUSG00000042724 | *Map3k9* | 127.5917728 | 122.0642616 | 125.3234769 | 373.2140001 | 345.8476285 | 392.58871 | 3.51E-24 | 1.566509455 | 2.68E-25 | 108.0035738 | 2.961872328 | 23.45508519 |
| ENSMUSG00000032324 | *Tspan3* | 1178.097369 | 1119.472237 | 1133.78583 | 3318.232885 | 3508.971944 | 3324.495081 | 1.75E-124 | 1.564265982 | 2.26E-126 | 571.8196607 | 2.957270029 | 123.7570864 |
| ENSMUSG00000056367 | *Actr3b* | 17.01223638 | 23.09323868 | 23.49815192 | 74.41026793 | 55.2594007 | 57.73363382 | 6.13E-05 | 1.562899946 | 1.67E-05 | 18.53376074 | 2.954471222 | 4.212262208 |
| ENSMUSG00000013523 | *Bcas1* | 14.46040092 | 19.79420459 | 14.68634495 | 56.97036138 | 44.77916953 | 42.33799814 | 0.000406744 | 1.562576169 | 0.000124864 | 14.71776237 | 2.953808237 | 3.390678523 |
| ENSMUSG00000050195 | *Scd4* | 17.8628482 | 10.99678033 | 10.7699863 | 25.5785296 | 46.68466611 | 45.22467983 | 0.003363663 | 1.561305626 | 0.00121331 | 10.46996588 | 2.951208043 | 2.473187579 |
| ENSMUSG00000036867 | *Smad6* | 129.2929965 | 139.6591101 | 127.2816563 | 389.4912462 | 403.9652741 | 371.4197109 | 2.04E-25 | 1.557665092 | 1.48E-26 | 113.7513758 | 2.943770277 | 24.69106935 |
| ENSMUSG00000032491 | *Nradd* | 273.0463939 | 278.2185422 | 280.0196438 | 776.6571715 | 859.3789557 | 803.4597374 | 2.17E-46 | 1.55440763 | 8.58E-48 | 210.9362135 | 2.937131034 | 45.66323104 |
| ENSMUSG00000034457 | *Eda2r* | 39.97875549 | 36.28937507 | 28.39360024 | 87.19953273 | 117.1880394 | 102.9583136 | 4.17E-07 | 1.553936216 | 8.77E-08 | 28.62886596 | 2.936171457 | 6.379739233 |

| | | | | | | | | | | | | | |
|---|---|---|---|---|---|---|---|---|---|---|---|---|---|
| ENSMUSG00000083846 | *Gm13797* | 13.6097891 | 9.897102293 | 12.72816563 | 36.04247353 | 25.72420377 | 45.22467983 | 0.003541887 | 1.553887331 | 0.001282701 | 10.36725575 | 2.936071967 | 2.450765277 |
| ENSMUSG00000023262 | *Acy1* | 310.4733139 | 284.8166104 | 302.538706 | 769.6812089 | 982.2834848 | 858.3066895 | 1.23E-34 | 1.539620095 | 6.57E-36 | 156.5032687 | 2.907179386 | 33.91026532 |
| ENSMUSG00000028108 | *Ecm1* | 94.41791191 | 126.4629737 | 111.6162216 | 334.8462057 | 307.737697 | 313.6860771 | 2.26E-18 | 1.529918426 | 2.27E-19 | 80.98881718 | 2.887695108 | 17.64504827 |
| ENSMUSG00000032194 | *Kank2* | 220.3084611 | 243.0288452 | 232.0442503 | 739.4520375 | 622.144632 | 642.7677899 | 5.37E-33 | 1.52770038 | 3.03E-34 | 148.8883742 | 2.883258883 | 32.27040794 |
| ENSMUSG00000003545 | *Fosb* | 56.14038005 | 49.48551147 | 44.05903486 | 149.9831963 | 148.6287329 | 133.749585 | 5.61E-10 | 1.525833997 | 9.15E-11 | 41.99442092 | 2.879531286 | 9.251049676 |
| ENSMUSG00000021453 | *Gadd45g* | 98.670971 | 133.0610419 | 120.4280286 | 358.0994144 | 350.6113699 | 298.2904414 | 3.70E-17 | 1.52098704 | 3.93E-18 | 75.35561025 | 2.869873289 | 16.43137566 |
| ENSMUSG00000027261 | *Hao1* | 83.35995826 | 67.08035999 | 82.24353174 | 226.7187851 | 237.2343237 | 205.9166273 | 4.53E-14 | 1.520243351 | 5.64E-15 | 61.02170468 | 2.86839429 | 13.34411971 |
| ENSMUSG00000031398 | *Plxna3* | 43.38120277 | 47.2861554 | 65.59900746 | 168.5857633 | 135.2902569 | 143.3718573 | 9.27E-09 | 1.517013101 | 1.67E-09 | 36.32326128 | 2.861979029 | 8.032922447 |
| ENSMUSG00000027408 | *Cpxm1* | 11.05795365 | 12.09645836 | 12.72816563 | 30.22917134 | 40.96817638 | 30.79127137 | 0.003569484 | 1.512066678 | 0.001293185 | 10.35222819 | 2.852183252 | 2.447394516 |
| ENSMUSG00000074796 | *Slc4a11* | 13.6097891 | 7.697746228 | 9.790896635 | 37.20513396 | 27.62970035 | 25.01790799 | 0.008785487 | 1.511178642 | 0.003430041 | 8.563315781 | 2.850428162 | 2.056234141 |
| ENSMUSG00000010154 | *Spire2* | 188.8358238 | 185.8455875 | 218.336995 | 588.3061808 | 580.2237073 | 518.6404772 | 6.15E-29 | 1.507613315 | 4.02E-30 | 130.0391515 | 2.843392606 | 28.21107597 |
| ENSMUSG00000053580 | *Tanc2* | 181.1803174 | 200.1414019 | 219.3160846 | 583.655539 | 463.9884162 | 653.3522894 | 4.91E-20 | 1.502969336 | 4.49E-21 | 88.74702121 | 2.834254552 | 19.30866979 |
| ENSMUSG00000008153 | *Clstn3* | 84.21057008 | 107.7684472 | 96.92987669 | 274.387863 | 267.7222689 | 272.3103062 | 1.04E-16 | 1.501018421 | 1.13E-17 | 73.27433541 | 2.830424461 | 15.98239619 |
| ENSMUSG00000070337 | *Gpr179* | 370.8667531 | 422.2763645 | 350.5140995 | 1199.86557 | 906.0636218 | 1122.919178 | 9.55E-30 | 1.497999973 | 6.12E-31 | 133.7736219 | 2.824508755 | 29.01977414 |
| ENSMUSG00000095098 | *Ccdc85b* | 188.8358238 | 207.8391482 | 200.713381 | 544.1250842 | 562.1214899 | 577.3363382 | 4.50E-33 | 1.497698875 | 2.54E-34 | 149.2434524 | 2.823919327 | 32.34643865 |
| ENSMUSG00000097739 | *Gm2539* | 35.7256964 | 31.89066294 | 27.41451058 | 98.82613709 | 88.60559077 | 79.86486012 | 2.36E-06 | 1.486385541 | 5.38E-07 | 25.12081744 | 2.801861302 | 5.626517341 |
| ENSMUSG00000035504 | *Reep6* | 159.915022 | 177.0481632 | 173.2988704 | 487.1547228 | 492.5708648 | 446.4734349 | 3.13E-27 | 1.484902493 | 2.16E-28 | 122.1320712 | 2.798982551 | 26.50505372 |
| ENSMUSG00000034255 | *Arhgap27* | 1160.234521 | 1214.044548 | 1179.803045 | 3422.872325 | 3221.241961 | 3232.121267 | 4.01E-111 | 1.474753678 | 6.16E-113 | 510.0586709 | 2.779361857 | 110.3972197 |
| ENSMUSG00000031137 | *Fgf13* | 139.5003383 | 129.7620078 | 120.4280286 | 401.1178505 | 373.4773289 | 310.7993954 | 7.62E-18 | 1.474525099 | 7.84E-19 | 78.54053099 | 2.778921534 | 17.11776321 |
| ENSMUSG00000032281 | *Acsbg1* | 626.9009106 | 677.4016681 | 696.1327508 | 1888.160549 | 1833.087706 | 1802.251602 | 2.38E-74 | 1.466372361 | 5.60E-76 | 340.2615125 | 2.763262006 | 73.62379747 |
| ENSMUSG00000032601 | *Prkar2a* | 2324.722101 | 2426.989418 | 2526.051332 | 6923.642898 | 6261.461748 | 6824.115518 | 4.04E-109 | 1.459242878 | 6.48E-111 | 500.7636723 | 2.749640254 | 108.3938262 |
| ENSMUSG00000031803 | *B3gnt3* | 134.3966674 | 124.2636177 | 146.8634495 | 387.1659253 | 371.5718323 | 356.9863025 | 1.95E-21 | 1.457718268 | 1.67E-22 | 95.26117483 | 2.746736027 | 20.70890592 |
| ENSMUSG00000020658 | *Efr3b* | 119.0856547 | 105.5690911 | 125.3234769 | 370.8886792 | 289.6354795 | 303.1015776 | 6.70E-16 | 1.45642644 | 7.56E-17 | 69.52109699 | 2.744277626 | 15.17379249 |
| ENSMUSG00000096472 | *Cdkn2d* | 417.6504031 | 413.4789402 | 403.3849414 | 1134.756586 | 1158.541918 | 1094.052361 | 5.72E-57 | 1.455894219 | 1.81E-58 | 259.8979753 | 2.743265429 | 56.24242395 |
| ENSMUSG00000007872 | *Id3* | 380.2234831 | 411.2795842 | 434.7158106 | 1152.196492 | 1162.352911 | 1043.054318 | 7.35E-46 | 1.454337443 | 2.94E-47 | 208.4819988 | 2.740306837 | 45.13377785 |
| ENSMUSG00000042507 | *Mideas* | 893.1424099 | 942.4240739 | 858.6616349 | 2621.799284 | 2286.595891 | 2427.699302 | 8.18E-69 | 1.44515797 | 2.08E-70 | 314.6919126 | 2.722926368 | 68.08747553 |
| ENSMUSG00000059994 | *Fcrl1* | 1558.320852 | 1429.581442 | 1468.634495 | 3793.761004 | 4218.769419 | 4105.823592 | 1.10E-91 | 1.442699358 | 2.15E-93 | 420.2517787 | 2.718289962 | 90.9586224 |
| ENSMUSG00000004698 | *Hdac9* | 282.4031239 | 291.4146786 | 280.0196438 | 795.2597385 | 705.9864813 | 804.4219646 | 2.58E-36 | 1.433185935 | 1.31E-37 | 164.2884738 | 2.700423979 | 35.58832239 |
| ENSMUSG00000001036 | *Epn2* | 339.3941158 | 350.7972924 | 303.5177957 | 953.3815578 | 860.331704 | 859.2689167 | 4.15E-38 | 1.427085447 | 2.00E-39 | 172.6008854 | 2.689029247 | 37.38206364 |
| ENSMUSG00000008398 | *Elk3* | 1480.915177 | 1513.156973 | 1608.644317 | 4377.416543 | 3955.810891 | 4025.958732 | 4.62E-91 | 1.424816807 | 9.11E-93 | 417.3727518 | 2.684804069 | 90.33500101 |
| ENSMUSG00000021508 | *Cxcl14* | 347.900234 | 329.9034098 | 319.1832303 | 896.4111964 | 925.1185875 | 857.3444623 | 2.04E-43 | 1.424699977 | 8.69E-45 | 197.1625398 | 2.684586661 | 42.69116077 |
| ENSMUSG00000046295 | *Ankle1* | 149.7076801 | 147.3568564 | 140.0098219 | 420.883078 | 336.3201456 | 414.7199363 | 1.27E-18 | 1.421205983 | 1.26E-19 | 82.15228656 | 2.678092856 | 17.89653638 |
| ENSMUSG00000036882 | *Arhgap33* | 22.96651911 | 30.79098491 | 33.28904856 | 70.92228662 | 60.02314214 | 100.071632 | 0.000160479 | 1.4152959 | 4.61E-05 | 16.60101635 | 2.667144336 | 3.79458301 |
| ENSMUSG00000046785 | *Epm2aip1* | 510.3670914 | 486.0576904 | 459.1930522 | 1392.867203 | 1078.511062 | 1408.700665 | 1.01E-30 | 1.413188414 | 6.24E-32 | 138.3071213 | 2.663251021 | 29.99572326 |
| ENSMUSG00000078503 | *Zfp990* | 66.34772188 | 27.49195081 | 39.16358654 | 105.8020997 | 193.4079024 | 56.77140659 | 0.003607439 | 1.412917975 | 0.001307183 | 10.33235501 | 2.66275183 | 2.44280097 |
| ENSMUSG00000047888 | *Tnrc6b* | 271.3451703 | 305.7104931 | 277.0823748 | 809.2116637 | 532.5862929 | 927.5870501 | 7.38E-14 | 1.41031664 | 9.27E-15 | 60.04477883 | 2.657954927 | 13.13191908 |
| ENSMUSG00000030409 | *Dmpk* | 558.8519651 | 640.012615 | 563.9556462 | 1660.279103 | 1422.453194 | 1598.25943 | 9.53E-46 | 1.410168123 | 3.83E-47 | 207.9586742 | 2.657681321 | 45.02108348 |
| ENSMUSG00000025171 | *Ubtd1* | 700.053527 | 697.1958727 | 760.7526686 | 1918.38972 | 2028.401105 | 1780.120376 | 2.25E-59 | 1.408035497 | 6.82E-61 | 271.0147968 | 2.653755576 | 58.64711961 |
| ENSMUSG00000039765 | *Cc2d2a* | 34.87508458 | 69.27971605 | 46.99630385 | 143.0072337 | 113.3770463 | 141.4474029 | 4.46E-06 | 1.407596749 | 1.05E-06 | 23.83498328 | 2.652948646 | 5.350321936 |
| ENSMUSG00000022265 | *Ank* | 5483.894397 | 5638.049273 | 5555.354751 | 15316.88859 | 13973.00639 | 14923.18212 | 6.69E-146 | 1.406593039 | 7.30E-148 | 670.6270668 | 2.651103582 | 145.1745813 |
| ENSMUSG00000026321 | *Tnfrsf11a* | 793.6208271 | 778.5720471 | 764.6690272 | 2210.217489 | 1760.678836 | 2225.631584 | 1.41E-41 | 1.406173657 | 6.27E-43 | 188.6478696 | 2.650333036 | 40.85118861 |
| ENSMUSG00000091337 | *Eid1* | 1182.350428 | 1148.063866 | 1205.259376 | 3101.978044 | 3088.809949 | 3117.616226 | 3.98E-109 | 1.3960843 | 6.37E-111 | 500.798951 | 2.631862831 | 108.3996422 |
| ENSMUSG00000022707 | *Gbe1* | 599.6813324 | 593.8261376 | 615.8473984 | 1599.82076 | 1610.144607 | 1547.261386 | 2.46E-69 | 1.394353979 | 6.20E-71 | 317.1014831 | 2.628708154 | 68.60881747 |
| ENSMUSG00000036995 | *Asap3* | 28.92080185 | 31.89066294 | 28.39360024 | 89.5248536 | 71.45612159 | 73.12926951 | 2.59E-05 | 1.391329879 | 6.67E-06 | 20.28679021 | 2.623203767 | 4.586708026 |
| ENSMUSG00000025477 | *Inpp5a* | 144.6040092 | 142.9581442 | 157.6334358 | 376.7019814 | 383.95756 | 404.1354368 | 2.31E-21 | 1.387983768 | 1.98E-22 | 94.92072652 | 2.617126697 | 20.63630625 |
| ENSMUSG00000021384 | *Susd3* | 545.242176 | 612.5206641 | 619.763757 | 1673.068368 | 1576.798416 | 1389.456121 | 1.62E-41 | 1.384827477 | 7.23E-43 | 188.3664623 | 2.611407273 | 40.79176388 |
| ENSMUSG00000059495 | *Arhgef12* | 617.5441806 | 613.6203422 | 563.9556462 | 1576.567552 | 1433.886173 | 1653.106382 | 1.34E-49 | 1.376990734 | 4.91E-51 | 225.8013611 | 2.597260524 | 48.87293078 |
| ENSMUSG00000027168 | *Pax6* | 217.7566257 | 208.9388262 | 256.5214918 | 616.2100313 | 581.1764556 | 577.3363382 | 2.80E-26 | 1.375905336 | 1.98E-27 | 117.7341795 | 2.595307235 | 25.55329855 |
| ENSMUSG00000040990 | *Sh3kbp1* | 221.1590729 | 215.5368944 | 241.8351469 | 613.8847104 | 523.05881 | 619.6743364 | 8.60E-25 | 1.371115893 | 6.41E-26 | 110.8409676 | 2.586705647 | 24.06570468 |
| ENSMUSG00000020057 | *Dram1* | 660.9253833 | 723.5881454 | 709.8400061 | 1898.624493 | 1732.096387 | 1733.933469 | 2.03E-58 | 1.357695954 | 6.27E-60 | 266.5941451 | 2.562755694 | 57.69241609 |
| ENSMUSG00000074731 | *Zfp345* | 39.12814367 | 38.48873114 | 15.66543462 | 84.87421185 | 73.36161817 | 80.82708735 | 0.000630524 | 1.355725917 | 0.000200094 | 13.83019905 | 2.559258573 | 3.200298527 |
| ENSMUSG00000037419 | *Endod1* | 1444.338869 | 1340.507522 | 1414.784564 | 3658.892393 | 3356.532218 | 3720.9327 | 2.08E-80 | 1.353117729 | 4.52E-82 | 368.2425191 | 2.554635977 | 79.68262588 |
| ENSMUSG00000074480 | *Mex3a* | 92.71668827 | 86.87456457 | 86.15989039 | 268.5745608 | 196.2661473 | 215.5388996 | 9.76E-11 | 1.351677375 | 1.51E-11 | 45.52653768 | 2.552086759 | 10.01043668 |
| ENSMUSG00000033565 | *Rbfox2* | 220.3084611 | 201.24108 | 231.0651606 | 530.173159 | 499.2401029 | 635.069972 | 1.48E-20 | 1.349822324 | 1.32E-21 | 91.17404772 | 2.548807336 | 19.83018898 |
| ENSMUSG00000030259 | *Rassf8* | 1158.533297 | 1110.674813 | 1174.907596 | 2993.850624 | 2673.411696 | 3093.560546 | 1.54E-64 | 1.346238711 | 4.26E-66 | 294.8976598 | 2.542484029 | 63.81229549 |
| ENSMUSG00000044229 | *Nxpe4* | 48.48487368 | 49.48551147 | 66.57809712 | 151.1458567 | 125.762774 | 141.4474029 | 1.66E-07 | 1.345404305 | 3.35E-08 | 30.4958615 | 2.541013967 | 6.780859731 |
| ENSMUSG00000003420 | *Fcgrt* | 60.39343915 | 41.78776524 | 33.28904856 | 111.6154019 | 129.5737672 | 104.8827681 | 1.25E-05 | 1.345076065 | 3.07E-06 | 21.77085109 | 2.540435904 | 4.904290323 |
| ENSMUSG00000026463 | *Atp2b4* | 976.5023682 | 974.3147369 | 940.9051667 | 2510.183882 | 2289.454136 | 2514.299753 | 1.29E-71 | 1.338306941 | 3.16E-73 | 327.6277334 | 2.5285441 | 70.88839546 |
| ENSMUSG00000052911 | *Lamb2* | 112.2807601 | 116.5658715 | 122.3862079 | 313.9183178 | 300.1157107 | 271.348079 | 2.75E-15 | 1.333619567 | 3.18E-16 | 66.68653159 | 2.520342091 | 14.56095371 |
| ENSMUSG00000023232 | *Serinc2* | 278.1500648 | 277.1188642 | 264.3542092 | 666.20443 | 720.2777056 | 678.3701974 | 1.59E-32 | 1.333482403 | 9.14E-34 | 146.6963544 | 2.520102481 | 31.79959039 |
| ENSMUSG00000028698 | *Pik3r3* | 54.43915641 | 19.79420459 | 52.87084183 | 91.85017447 | 107.6605565 | 121.240631 | 0.000498361 | 1.325085758 | 0.000155041 | 14.30997487 | 2.505477805 | 3.302455776 |
| ENSMUSG00000015944 | *Castor2* | 717.0657634 | 677.4016681 | 656.9691642 | 1792.822393 | 1590.136892 | 1747.40465 | 6.23E-52 | 1.320890162 | 2.16E-53 | 236.6078027 | 2.498202046 | 51.20533514 |
| ENSMUSG00000025203 | *Scd2* | 29428.6171 | 30163.06876 | 29428.49802 | 76075.19767 | 71213.17078 | 74985.40583 | 4.37E-175 | 1.320141067 | 3.72E-177 | 805.3438182 | 2.496905234 | 174.3596193 |
| ENSMUSG00000100349 | *Gm28188* | 23.81713093 | 17.59484852 | 20.56088293 | 39.53045484 | 53.35390412 | 61.58254274 | 0.001997058 | 1.314994604 | 0.000693544 | 11.50646991 | 2.488014001 | 2.69960934 |
| ENSMUSG00000061393 | *Acvr2b* | 109.7289246 | 131.9613639 | 118.4698493 | 295.3157508 | 293.4464727 | 303.1015776 | 3.27E-15 | 1.312693236 | 3.80E-16 | 66.33720141 | 2.484048317 | 14.48604251 |
| ENSMUSG00000042210 | *Abhd14a* | 71.45139279 | 91.2732767 | 62.66173847 | 170.9110841 | 181.974923 | 203.9921728 | 9.35E-09 | 1.311635973 | 1.69E-09 | 36.3047092 | 2.482228577 | 8.029283872 |
| ENSMUSG00000032431 | *Crtap* | 602.2331678 | 517.9483533 | 634.450102 | 1497.506642 | 1466.279615 | 1396.191711 | 3.55E-41 | 1.310802479 | 1.60E-42 | 186.7800203 | 2.480794924 | 40.44943319 |
| ENSMUSG00000045730 | *Adrb2* | 260.2872166 | 278.2185422 | 242.8142366 | 701.0842431 | 593.5621834 | 640.8433354 | 3.36E-25 | 1.308578912 | 2.45E-26 | 112.744751 | 2.476974321 | 24.47386067 |
| ENSMUSG00000049848 | *Ceacam19* | 107.1770892 | 95.67198883 | 92.03442837 | 289.5024486 | 201.0298887 | 242.4812621 | 5.00E-10 | 1.308473551 | 8.14E-11 | 42.22473664 | 2.476793432 | 9.300918801 |
| ENSMUSG00000004558 | *Ndrg2* | 1359.277687 | 1376.796897 | 1419.680012 | 3470.541402 | 3381.303674 | 3439.962349 | 6.35E-102 | 1.308390322 | 1.13E-103 | 467.4806509 | 2.476650551 | 101.197394 |
| ENSMUSG00000050288 | *Fzd2* | 36.57630822 | 28.59162885 | 29.37268991 | 75.57292836 | 81.93635276 | 76.97817843 | 5.87E-05 | 1.304721902 | 1.59E-05 | 18.62460034 | 2.470361035 | 4.231421894 |
| ENSMUSG00000040563 | *Plppr2* | 68.89955734 | 60.48229179 | 65.59900746 | 162.7724611 | 147.6759846 | 172.2386742 | 6.69E-09 | 1.304276592 | 1.19E-09 | 36.98089252 | 2.469598637 | 8.174447019 |
| ENSMUSG00000036667 | *Tcaf1* | 126.741161 | 124.2636177 | 116.51167 | 326.7075826 | 292.4937244 | 289.6303963 | 2.88E-15 | 1.30406964 | 3.34E-16 | 66.58889374 | 2.469244403 | 14.54047057 |
| ENSMUSG00000024646 | *Cyb5a* | 2337.481279 | 2322.520005 | 2357.64791 | 5222.67068 | 6458.680644 | 5630.953752 | 9.93E-57 | 1.303078952 | 3.16E-58 | 258.7830267 | 2.467549373 | 56.00314962 |
| ENSMUSG00000059588 | *Calcrl* | 3438.172972 | 3083.497203 | 3154.626896 | 8246.750475 | 6972.211971 | 8646.573892 | 5.24E-51 | 1.301903476 | 1.84E-52 | 232.3363694 | 2.465539688 | 50.28070828 |
| ENSMUSG00000035877 | *Zhx3* | 356.4063522 | 303.511137 | 358.3468169 | 810.3743241 | 846.9932279 | 846.7599627 | 4.67E-32 | 1.295723446 | 2.73E-33 | 144.5198361 | 2.455000718 | 31.3304324 |
| ENSMUSG00000032177 | *Pde4a* | 446.571205 | 506.951573 | 479.7539351 | 1331.2462 | 1076.605565 | 1104.63686 | 1.32E-29 | 1.293467772 | 8.48E-31 | 133.1281817 | 2.451165288 | 28.88043949 |
| ENSMUSG00000041040 | *Fam117b* | 381.0740949 | 289.2153226 | 346.5977409 | 887.1099129 | 724.0886988 | 883.3245975 | 8.83E-21 | 1.290416052 | 7.78E-22 | 92.21263709 | 2.445985841 | 20.05408762 |
| ENSMUSG00000074994 | *Qser1* | 569.9099187 | 461.8647737 | 327.9950373 | 1549.826362 | 650.7270806 | 1124.843632 | 0.006276314 | 1.289616104 | 0.002378057 | 9.232123289 | 2.444629961 | 2.202295339 |
| ENSMUSG00000041762 | *Gpr155* | 62.09466279 | 60.48229179 | 46.01721419 | 145.3325545 | 128.6210189 | 138.5607212 | 2.70E-07 | 1.288707128 | 5.55E-08 | 29.5138305 | 2.443090197 | 6.569389452 |
| ENSMUSG00000050232 | *Cxcr3* | 55.28976823 | 59.38261376 | 62.66173847 | 165.097782 | 121.9517808 | 146.258539 | 2.35E-07 | 1.28778724 | 4.81E-08 | 29.78997515 | 2.441532936 | 6.629373212 |
| ENSMUSG00000049823 | *Zbtb12* | 159.915022 | 174.8488072 | 183.0897671 | 446.4616076 | 414.4455052 | 402.2109823 | 2.26E-19 | 1.287297306 | 2.15E-20 | 85.65045933 | 2.440703941 | 18.64504542 |
| ENSMUSG00000074652 | *Myh7b* | 69.75016916 | 57.18325769 | 70.49445577 | 174.3990655 | 151.4869778 | 156.8430385 | 1.78E-08 | 1.283759602 | 3.30E-09 | 35.00091911 | 2.434726303 | 7.750185612 |
| ENSMUSG00000030148 | *Clec4a2* | 73.15261643 | 59.38261376 | 61.6826488 | 147.6578754 | 186.7386644 | 138.5607212 | 1.93E-07 | 1.280684812 | 3.92E-08 | 30.18783268 | 2.42954274 | 6.714687342 |
| ENSMUSG00000034342 | *Cbl* | 2479.533452 | 2632.62921 | 2434.016904 | 6339.987359 | 5222.966114 | 6764.457429 | 1.36E-42 | 1.280298454 | 5.91E-44 | 193.3483767 | 2.428892189 | 41.86633953 |
| ENSMUSG00000028655 | *Mfsd2a* | 171.8235874 | 171.5497731 | 179.1734084 | 472.0401372 | 399.2015326 | 396.4376189 | 5.47E-18 | 1.276516846 | 5.58E-19 | 79.21243932 | 2.422533886 | 17.26197885 |
| ENSMUSG00000051238 | *Swsap1* | 152.2595156 | 119.8649056 | 140.0098219 | 323.2196013 | 357.280608 | 317.534986 | 1.85E-14 | 1.271564868 | 2.26E-15 | 62.82473185 | 2.414232915 | 13.73210827 |
| ENSMUSG00000026389 | *Steap3* | 360.6594112 | 366.1927848 | 361.2840858 | 901.0618382 | 895.5833906 | 828.4776453 | 4.20E-36 | 1.270349779 | 2.15E-37 | 163.3056808 | 2.412200419 | 35.37713598 |
| ENSMUSG00000032501 | *Trib1* | 281.5525121 | 288.1156445 | 294.7059887 | 695.2709409 | 669.7820464 | 716.8592866 | 9.06E-31 | 1.268664676 | 5.59E-32 | 138.526904 | 2.409384555 | 30.04282337 |
| ENSMUSG00000043290 | *Zfp784* | 20.41468366 | 32.99034098 | 42.10085553 | 75.57292836 | 83.84184933 | 69.28036059 | 0.000674385 | 1.268568646 | 0.00021554 | 13.69051403 | 2.409224184 | 3.171091911 |
| ENSMUSG00000028124 | *Gclm* | 2285.593958 | 2186.159929 | 2409.539662 | 5672.620269 | 5309.666208 | 5514.524257 | 1.44E-89 | 1.260840467 | 2.84E-91 | 410.5076997 | 2.396353039 | 88.84220026 |
| ENSMUSG00000034460 | *Six4* | 23.81713093 | 28.59162885 | 41.12176587 | 91.85017447 | 65.73963186 | 66.3936789 | 0.000749209 | 1.259550916 | 0.00024182 | 13.47456134 | 2.39421202 | 3.125396835 |
| ENSMUSG00000029844 | *Hoxa1* | 129.2929965 | 119.8649056 | 105.7416837 | 297.6410717 | 275.3442552 | 278.0836696 | 2.53E-13 | 1.258882179 | 3.30E-14 | 57.54978394 | 2.393102479 | 12.59773401 |
| ENSMUSG00000027002 | *Nckap1* | 55.28976823 | 61.58196982 | 56.78720049 | 138.3565919 | 131.4792637 | 144.3340846 | 1.21E-07 | 1.256560399 | 2.43E-08 | 31.11617576 | 2.389254273 | 6.915535263 |
| ENSMUSG00000000686 | *Abhd15* | 24.66774275 | 38.48873114 | 25.45633125 | 72.08494705 | 71.45612159 | 66.3936789 | 0.00045604 | 1.254520017 | 0.00014078 | 14.49168195 | 2.385877575 | 3.340996857 |
| ENSMUSG00000053897 | *Slc39a8* | 48.48487368 | 78.07714031 | 58.74537981 | 143.0072337 | 160.0617124 | 135.6740395 | 1.64E-06 | 1.253629457 | 3.68E-07 | 25.85476413 | 2.384405253 | 5.784876172 |
| ENSMUSG00000031840 | *Rab3a* | 45.93303823 | 46.18647737 | 65.59900746 | 120.9166854 | 132.432012 | 121.240631 | 3.28E-06 | 1.249143879 | 7.61E-07 | 24.45507444 | 2.377003254 | 5.483522076 |
| ENSMUSG00000051212 | *Gpr183* | 503.5621968 | 476.1605881 | 482.6912041 | 1175.449701 | 1124.24298 | 1170.068312 | 1.07E-44 | 1.24502836 | 4.40E-46 | 203.1011155 | 2.370232134 | 43.96927113 |
| ENSMUSG00000018381 | *Abi3* | 105.4758656 | 107.7684472 | 103.7835043 | 252.2973147 | 282.0134932 | 216.5011268 | 6.85E-11 | 1.244243212 | 1.05E-11 | 46.23805972 | 2.36894255 | 10.16410018 |
| ENSMUSG00000044345 | *Marveld1* | 290.9092421 | 276.0191862 | 279.0405541 | 699.9215827 | 630.7193666 | 674.5212885 | 1.36E-27 | 1.243005899 | 9.26E-29 | 123.81166 | 2.366911721 | 26.86748494 |
| ENSMUSG00000030380 | *Mzf1* | 30.62202548 | 20.89388262 | 32.3099589 | 59.29568225 | 76.21986303 | 63.5069972 | 0.000767278 | 1.241952574 | 0.000248125 | 13.42626708 | 2.365184249 | 3.115047508 |
| ENSMUSG00000020262 | *Adarb1* | 139.5003383 | 130.8616859 | 130.2189253 | 342.9848287 | 311.5486901 | 294.4415325 | 1.95E-14 | 1.241124223 | 2.38E-15 | 62.72524456 | 2.363826623 | 13.71065944 |
| ENSMUSG00000072437 | *Nanos1* | 70.60078098 | 63.78132589 | 69.51536611 | 149.9831963 | 161.9672089 | 168.3897653 | 2.19E-08 | 1.235147195 | 4.12E-09 | 34.56802457 | 2.354053642 | 7.658572415 |
| ENSMUSG00000029033 | *Acap3* | 569.9099187 | 539.941914 | 597.2446948 | 1385.89124 | 1272.871713 | 1356.740395 | 1.77E-43 | 1.23263117 | 7.56E-45 | 197.4406974 | 2.349951807 | 42.75116623 |
| ENSMUSG00000103906 | *Tigd5* | 55.28976823 | 57.18325769 | 67.55718678 | 155.7964985 | 143.8649915 | 123.1650855 | 6.72E-07 | 1.230753013 | 1.44E-07 | 27.66245903 | 2.346894538 | 6.172697879 |
| ENSMUSG00000030653 | *Gm45837* | 1768.421972 | 2020.108546 | 2040.422859 | 4841.318057 | 4561.758802 | 4247.270995 | 1.15E-52 | 1.228205407 | 3.92E-54 | 240.0055618 | 2.342753894 | 51.93928482 |
| ENSMUSG00000040364 | *Sec1* | 17.01223638 | 20.89388262 | 17.62361394 | 38.3677944 | 38.10993152 | 52.92249767 | 0.00679654 | 1.227684506 | 0.002591024 | 9.075177973 | 2.341908169 | 2.167712106 |
| ENSMUSG00000027654 | *Fam83d* | 166.7199165 | 185.8455875 | 160.5707048 | 389.4912462 | 443.0279539 | 363.7218931 | 2.58E-15 | 1.223706072 | 2.98E-16 | 66.81506875 | 2.335458925 | 14.5890143 |
| ENSMUSG00000032086 | *Bace1* | 141.201562 | 168.250739 | 153.7170772 | 380.1899627 | 297.2574658 | 401.2487551 | 5.45E-12 | 1.22178796 | 7.74E-13 | 51.34704001 | 2.332355916 | 11.26344853 |
| ENSMUSG00000034135 | *Sik3* | 1928.336994 | 1982.719493 | 1914.120292 | 4663.43101 | 4462.67298 | 4434.905305 | 1.32E-97 | 1.219119384 | 2.45E-99 | 447.5590885 | 2.328045707 | 96.88006188 |
| ENSMUSG00000048240 | *Gng7* | 39.97875549 | 38.48873114 | 24.47724159 | 84.87421185 | 79.07810789 | 76.0159512 | 0.000308153 | 1.218314528 | 9.27E-05 | 15.27994876 | 2.326747291 | 3.511234221 |
| ENSMUSG00000032403 | *2300009A05Rik* | 168.4211402 | 144.0578223 | 150.7798082 | 411.5817945 | 354.4223631 | 315.6105316 | 5.84E-13 | 1.218139235 | 7.81E-14 | 55.85322021 | 2.326464599 | 12.23342185 |
| ENSMUSG00000036777 | *Anln* | 2864.860606 | 2766.78993 | 2715.994727 | 6336.499378 | 6142.368212 | 6940.545013 | 6.47E-76 | 1.217901548 | 1.47E-77 | 347.514116 | 2.326081341 | 75.18907782 |
| ENSMUSG00000070705 | *Eid2b* | 75.70445189 | 91.2732767 | 92.03442837 | 208.1162181 | 210.5573716 | 180.8987193 | 4.17E-09 | 1.214342611 | 7.33E-10 | 37.93065288 | 2.32035028 | 8.379700727 |
| ENSMUSG00000038175 | *Mylip* | 99.52158282 | 131.9613639 | 129.2398356 | 305.7796948 | 243.9035617 | 283.857033 | 3.58E-10 | 1.21188364 | 5.78E-11 | 42.89494383 | 2.316398777 | 9.445678964 |
| ENSMUSG00000021892 | *Sh3bp5* | 472.9401714 | 454.1670275 | 428.8412726 | 1131.268605 | 986.094478 | 1026.696455 | 1.52E-32 | 1.211409796 | 8.70E-34 | 146.7953535 | 2.315638095 | 31.81915933 |
| ENSMUSG00000029804 | *Herc3* | 478.8944541 | 500.3535048 | 488.5657421 | 1170.799059 | 1100.424273 | 1121.956951 | 1.15E-41 | 1.209236874 | 5.10E-43 | 189.0593036 | 2.312153011 | 40.93965816 |
| ENSMUSG00000044066 | *Cep68* | 398.0863313 | 380.4885993 | 384.7822378 | 923.1523865 | 821.2690242 | 938.1715496 | 3.72E-30 | 1.203925386 | 2.34E-31 | 135.6818253 | 2.30365614 | 29.42966259 |
| ENSMUSG00000110003 | *Gm45403* | 24.66774275 | 27.49195081 | 33.28904856 | 56.97036138 | 58.11764556 | 80.82708735 | 0.001358764 | 1.201973619 | 0.000458795 | 12.27610535 | 2.300541719 | 2.866856079 |
| ENSMUSG00000041552 | *Ptchd1* | 523.1262687 | 577.3309671 | 589.4119774 | 1387.053901 | 1171.880394 | 1320.17576 | 7.30E-32 | 1.199345073 | 4.31E-33 | 143.6160721 | 2.296354019 | 31.13638804 |
| ENSMUSG00000113996 | *Gm48455* | 57.84160369 | 41.78776524 | 46.01721419 | 122.0793458 | 115.2825428 | 99.10940473 | 1.88E-05 | 1.198853727 | 4.74E-06 | 20.94087106 | 2.295572071 | 4.725602529 |
| ENSMUSG00000028035 | *Dnajb4* | 535.0348341 | 476.1605881 | 518.9175217 | 1189.401626 | 1046.11762 | 1274.95108 | 1.91E-28 | 1.196208735 | 1.27E-29 | 127.754703 | 2.291367296 | 27.71801272 |
| ENSMUSG00000028551 | *Cdkn2c* | 511.2177032 | 472.861554 | 474.8584868 | 1135.919246 | 1144.250694 | 1062.298862 | 1.57E-37 | 1.194198078 | 7.71E-39 | 169.9178887 | 2.288176084 | 36.80483996 |
| ENSMUSG00000050100 | *Hmx2* | 130.9942201 | 125.3632957 | 134.1352839 | 303.4543739 | 280.1079966 | 310.7993954 | 1.79E-13 | 1.194181565 | 2.31E-14 | 58.24952652 | 2.288149895 | 12.74670692 |
| ENSMUSG00000025648 | *Pfkfb4* | 669.4315015 | 669.7039218 | 619.763757 | 1428.909676 | 1584.420403 | 1446.227527 | 1.56E-41 | 1.18735685 | 6.99E-43 | 188.4333032 | 2.27735129 | 40.80568783 |
| ENSMUSG00000021575 | *Ahrr* | 23.81713093 | 21.99356065 | 25.45633125 | 56.97036138 | 42.87367295 | 62.54476997 | 0.003214179 | 1.185173954 | 0.001156522 | 10.55854436 | 2.273908108 | 2.492929886 |
| ENSMUSG00000026525 | *Opn3* | 54.43915641 | 43.9871213 | 31.33086923 | 105.8020997 | 93.36933221 | 96.22272304 | 0.000157153 | 1.179984328 | 4.51E-05 | 16.64209275 | 2.265743158 | 3.803677344 |
| ENSMUSG00000063160 | *Numbl* | 39.12814367 | 27.49195081 | 42.10085553 | 72.08494705 | 83.84184933 | 90.44935965 | 0.000363813 | 1.175759198 | 0.000110886 | 14.94167544 | 2.259117326 | 3.43912219 |
| ENSMUSG00000051339 | *2900026A02Rik* | 448.2724286 | 524.5464215 | 492.4821008 | 1148.708511 | 1024.204409 | 1119.070269 | 4.43E-30 | 1.169597948 | 2.79E-31 | 135.3319004 | 2.249489992 | 29.35406975 |
| ENSMUSG00000054723 | *Vmac* | 45.93303823 | 37.38905311 | 49.93357284 | 119.7540249 | 80.03085618 | 101.0338592 | 0.000155861 | 1.167211394 | 4.47E-05 | 16.65866335 | 2.245771886 | 3.80726374 |
| ENSMUSG00000080316 | *Spaca6* | 31.4726373 | 40.68808721 | 28.39360024 | 74.41026793 | 67.64512844 | 82.75154181 | 0.000565491 | 1.166861646 | 0.000177982 | 14.05033974 | 2.245227518 | 3.247574682 |
| ENSMUSG00000031438 | *Rnf128* | 476.3426186 | 449.7683153 | 443.5276176 | 1085.924848 | 871.7646834 | 1120.032496 | 4.01E-22 | 1.166649882 | 3.31E-23 | 98.46142727 | 2.244897979 | 21.39739287 |
| ENSMUSG00000002732 | *Fkbp7* | 27.21957821 | 19.79420459 | 19.58179327 | 46.50641745 | 58.11764556 | 45.22467983 | 0.005555687 | 1.163573431 | 0.002079538 | 9.477971599 | 2.240115984 | 2.255262228 |
| ENSMUSG00000025006 | *Sorbs1* | 177.7778702 | 184.7459095 | 209.525188 | 423.2083988 | 429.6894778 | 425.3044358 | 7.69E-17 | 1.161000755 | 8.26E-18 | 73.88890727 | 2.236124871 | 16.11402584 |
| ENSMUSG00000024388 | *Myo7b* | 23.81713093 | 34.09001901 | 31.33086923 | 66.27164487 | 73.36161817 | 58.69586105 | 0.001302904 | 1.160391141 | 0.000438056 | 12.3624472 | 2.235180193 | 2.885087562 |
| ENSMUSG00000040061 | *Plcb2* | 784.2640971 | 798.3662516 | 734.3172477 | 1720.737446 | 1673.025994 | 1782.044831 | 1.11E-48 | 1.159824044 | 4.16E-50 | 221.5473385 | 2.234301756 | 47.95394288 |
| ENSMUSG00000033124 | *Atg9a* | 1026.688466 | 1063.388657 | 957.5496909 | 2367.176648 | 2227.525497 | 2207.349266 | 1.67E-52 | 1.158231256 | 5.73E-54 | 239.2490565 | 2.231836367 | 51.77694547 |
| ENSMUSG00000110185 | *Igip* | 42.53059095 | 46.18647737 | 37.20540721 | 104.6394393 | 65.73963186 | 110.6561315 | 0.00054834 | 1.157571272 | 0.000172132 | 14.11319169 | 2.230815611 | 3.260949784 |
| ENSMUSG00000027605 | *Acss2* | 545.242176 | 609.22163 | 618.7846674 | 1277.76382 | 1363.3828 | 1296.120079 | 1.06E-36 | 1.153027628 | 5.33E-38 | 166.0725504 | 2.223800901 | 35.97450703 |
| ENSMUSG00000014547 | *Wdfy2* | 1149.176567 | 1218.44326 | 1140.639458 | 2707.836156 | 2424.744393 | 2663.444974 | 3.17E-52 | 1.152247112 | 1.10E-53 | 237.9591025 | 2.222598123 | 51.49826919 |
| ENSMUSG00000019590 | *Cyb561* | 46.78365004 | 50.5851895 | 42.10085553 | 104.6394393 | 100.0385702 | 104.8827681 | 2.56E-05 | 1.151743102 | 6.57E-06 | 20.31514776 | 2.221821786 | 4.592565438 |
| ENSMUSG00000040430 | *Pitpnc1* | 95.26852373 | 120.9645836 | 115.5325803 | 254.6226356 | 206.7463785 | 272.3103062 | 1.49E-08 | 1.148626192 | 2.73E-09 | 35.36852675 | 2.217026773 | 7.827966424 |
| ENSMUSG00000039145 | *Camk1d* | 1261.457328 | 1242.636177 | 1192.53121 | 2822.939539 | 2465.712569 | 2894.379509 | 7.62E-44 | 1.145947129 | 3.21E-45 | 199.1465704 | 2.212913608 | 43.11781043 |
| ENSMUSG00000033594 | *Spata2l* | 56.99099187 | 28.59162885 | 39.16358654 | 91.85017447 | 95.27482879 | 91.41158688 | 0.000619811 | 1.145899877 | 0.000196525 | 13.86402527 | 2.21284113 | 3.207740487 |
| ENSMUSG00000074203 | *G430095P16Rik* | 27.21957821 | 32.99034098 | 36.22631755 | 62.78366356 | 73.36161817 | 76.0159512 | 0.000828945 | 1.144427167 | 0.00026949 | 13.27137472 | 2.210583404 | 3.081474418 |
| ENSMUSG00000040857 | *Erf* | 669.4315015 | 676.30199 | 767.6062962 | 1625.39929 | 1584.420403 | 1452.000891 | 1.30E-34 | 1.141075151 | 6.96E-36 | 156.3904533 | 2.205453207 | 33.88617028 |
| ENSMUSG00000041607 | *Mbp* | 1600.851443 | 1532.951177 | 1558.710744 | 3594.946069 | 3306.036559 | 3446.697939 | 1.01E-68 | 1.13989462 | 2.59E-70 | 314.2514099 | 2.203649263 | 67.99386053 |

| | | | | | | | | | | | | | |
|---|---|---|---|---|---|---|---|---|---|---|---|---|---|
| ENSMUSG00000024909 | *Efemp2* | 135.2472792 | 94.5723108 | 116.51167 | 273.2252025 | 260.1002826 | 231.8967625 | 8.29E-09 | 1.136003674 | 1.49E-09 | 36.54494275 | 2.197714031 | 8.08132433 |
| ENSMUSG00000032288 | *Imp3* | 1181.499817 | 1124.970627 | 1288.481997 | 2726.438723 | 2670.553451 | 2502.753026 | 1.23E-47 | 1.13503401 | 4.72E-49 | 216.7111153 | 2.196237401 | 46.90925816 |
| ENSMUSG00000042216 | *Sgsm1* | 1026.688466 | 1051.292199 | 1092.664065 | 2466.002785 | 2184.651824 | 2305.496444 | 1.79E-48 | 1.133138575 | 6.75E-50 | 220.5834857 | 2.193353845 | 47.74767269 |
| ENSMUSG00000022070 | *Bora* | 339.3941158 | 325.5046976 | 351.4931892 | 716.1988288 | 745.0491611 | 763.0461937 | 1.57E-26 | 1.12972287 | 1.10E-27 | 118.8964848 | 2.188167032 | 25.80483558 |
| ENSMUSG00000025574 | *Tk1* | 1185.752876 | 1199.748734 | 1249.318411 | 2565.991583 | 2972.574658 | 2409.416985 | 2.04E-35 | 1.1290641 | 1.06E-36 | 160.1213626 | 2.187168091 | 34.69124408 |
| ENSMUSG00000053846 | *Lipg* | 49.3354855 | 34.09001901 | 36.22631755 | 91.85017447 | 91.46383564 | 79.86486012 | 0.000334353 | 1.12764275 | 0.000101265 | 15.11297604 | 2.185014343 | 3.475794914 |
| ENSMUSG00000005447 | *Pafah1b3* | 392.9826604 | 387.0866675 | 383.8031481 | 775.494511 | 956.559281 | 802.4975101 | 1.54E-21 | 1.123693248 | 1.31E-22 | 95.74412691 | 2.179040859 | 20.81170044 |
| ENSMUSG00000025764 | *Jade1* | 933.9717772 | 875.3437139 | 905.6579388 | 2071.860898 | 1868.339393 | 1980.26364 | 1.03E-46 | 1.123181977 | 4.05E-48 | 212.4297738 | 2.178268773 | 45.98680366 |
| ENSMUSG00000056069 | *Otulinl* | 63.79588642 | 61.58196982 | 43.0799452 | 74.41026793 | 173.4001884 | 117.3917221 | 0.002113704 | 1.119951279 | 0.0007372 | 11.39303428 | 2.173396326 | 2.674955921 |
| ENSMUSG00000078881 | *Gm14434* | 57.84160369 | 40.68808721 | 74.41081443 | 97.66347665 | 108.6133048 | 169.3519925 | 0.000926736 | 1.117587667 | 0.000304715 | 13.04118455 | 2.169838499 | 3.033043776 |
| ENSMUSG00000032481 | *Smarcc1* | 3662.734493 | 3705.91497 | 3511.015533 | 8092.116637 | 7465.735584 | 7981.674876 | 2.04E-82 | 1.113340826 | 4.34E-84 | 377.5091802 | 2.163460568 | 81.69052644 |
| ENSMUSG00000020101 | *Vsir* | 522.2756569 | 539.941914 | 507.1684457 | 1110.340717 | 1214.754067 | 1068.072226 | 3.68E-31 | 1.113284263 | 2.23E-32 | 140.348222 | 2.163375748 | 30.43380722 |
| ENSMUSG00000049580 | *Tsku* | 21.26529547 | 23.09323868 | 24.47724159 | 58.13302182 | 48.59016268 | 42.33799814 | 0.006905019 | 1.112973613 | 0.002637117 | 9.042936844 | 2.162909966 | 2.1608351 |
| ENSMUSG00000039958 | *Etfbkmt* | 62.09466279 | 50.5851895 | 37.20540721 | 104.6394393 | 91.46383564 | 128.9384489 | 0.000360461 | 1.112589162 | 0.000109692 | 14.96211327 | 2.162333669 | 3.443141537 |
| ENSMUSG00000037979 | *Ccdc92* | 175.2260347 | 155.0546026 | 180.1524981 | 399.9551901 | 365.8553425 | 339.6662123 | 5.30E-13 | 1.111008161 | 7.06E-14 | 56.05091158 | 2.159965338 | 12.27552855 |
| ENSMUSG00000025355 | *Mmp19* | 69.75016916 | 96.77166687 | 72.4526351 | 176.7243863 | 155.2979709 | 181.8609465 | 1.50E-06 | 1.110719334 | 3.35E-07 | 26.03726404 | 2.159532957 | 5.82313416 |
| ENSMUSG00000063382 | *Bcl9l* | 193.0888829 | 188.0449436 | 181.1315878 | 440.6483054 | 401.1070292 | 374.3063926 | 1.36E-14 | 1.110650618 | 1.65E-15 | 63.44429043 | 2.1594301 | 13.8655231 |
| ENSMUSG00000054545 | *Ugt1a6a* | 48.48487368 | 79.17681835 | 34.26813822 | 139.5192524 | 112.424298 | 96.22272304 | 0.00166408 | 1.109770472 | 0.000570222 | 11.87075814 | 2.158113097 | 2.778825838 |
| ENSMUSG00000068566 | *Myadm* | 2176.715645 | 2226.848016 | 2166.725425 | 4833.179434 | 4342.626696 | 4993.959326 | 3.92E-56 | 1.108793615 | 1.27E-57 | 256.0125817 | 2.156652321 | 55.40665971 |
| ENSMUSG00000028158 | *Mttp* | 91.01546463 | 92.37295474 | 106.7207733 | 184.8630094 | 214.3683648 | 225.1611719 | 3.50E-08 | 1.108504697 | 6.66E-09 | 33.63296613 | 2.156220468 | 7.456189431 |
| ENSMUSG00000010205 | *Raver1* | 3277.407339 | 3299.034098 | 3210.435007 | 7354.98992 | 7059.864813 | 6630.707844 | 2.68E-76 | 1.10436325 | 6.07E-78 | 349.2822713 | 2.150039627 | 75.57149953 |
| ENSMUSG00000089951 | *Zfp968-ps* | 47.63426186 | 54.98390163 | 41.12176587 | 94.17549534 | 131.4792637 | 81.78931458 | 0.000530605 | 1.101138399 | 0.00016591 | 14.18246409 | 2.145239019 | 3.275228933 |
| ENSMUSG00000057982 | *Zfp809* | 571.6111424 | 547.6396602 | 527.7293286 | 1225.4441 | 1051.83411 | 1255.706536 | 2.19E-26 | 1.099874064 | 1.55E-27 | 118.2287396 | 2.143359818 | 25.66032326 |
| ENSMUSG00000029330 | *Cds1* | 130.9942201 | 147.3568564 | 130.2189253 | 319.73162 | 283.9189898 | 271.348079 | 8.19E-11 | 1.099321365 | 1.26E-11 | 45.87975999 | 2.14253885 | 10.08661937 |
| ENSMUSG00000052673 | *Gm9887* | 32.32324912 | 36.28937507 | 42.10085553 | 73.24760749 | 88.60559077 | 74.09149674 | 0.000713522 | 1.095945175 | 0.000228832 | 13.57816202 | 2.137530751 | 3.14659283 |
| ENSMUSG00000031378 | *Abcd1* | 912.7064818 | 927.0285815 | 908.5952078 | 2068.372916 | 1883.583365 | 1921.567779 | 3.78E-47 | 1.095138997 | 1.46E-48 | 214.4657527 | 2.136336632 | 46.42254486 |
| ENSMUSG00000037703 | *Lzts3* | 85.0611819 | 70.37939408 | 81.26444207 | 168.5857633 | 182.9276713 | 154.9185841 | 5.82E-07 | 1.093293114 | 1.25E-07 | 27.94876771 | 2.133605005 | 6.234743651 |
| ENSMUSG00000038009 | *Dnajc22* | 143.7533974 | 164.9517049 | 143.9261805 | 323.2196013 | 316.3124316 | 323.3083494 | 2.26E-12 | 1.091870172 | 3.12E-13 | 53.12806207 | 2.13150165 | 11.64649784 |
| ENSMUSG00000058099 | *Nfam1* | 982.4566509 | 1013.903146 | 969.2987669 | 2093.951446 | 2105.573716 | 2108.239862 | 1.79E-56 | 1.089360367 | 5.77E-58 | 257.5836491 | 2.127796776 | 55.74636185 |
| ENSMUSG00000038648 | *Creb3l2* | 251.7810984 | 288.1156445 | 254.5633125 | 577.8422369 | 558.3104967 | 551.356203 | 4.71E-19 | 1.089099648 | 4.58E-20 | 84.15388473 | 2.127412282 | 18.32657429 |
| ENSMUSG00000035759 | *Bbs10* | 39.12814367 | 34.09001901 | 54.82902116 | 83.71155142 | 81.93635276 | 106.8072226 | 0.000851112 | 1.088710531 | 0.000277806 | 13.21440611 | 2.126838564 | 3.070013527 |
| ENSMUSG00000081534 | *Slc48a1* | 992.6639928 | 1133.768052 | 1032.939595 | 2198.590885 | 2441.893862 | 2070.713 | 3.72E-33 | 1.088398197 | 2.09E-34 | 149.6295962 | 2.126378168 | 32.42925272 |
| ENSMUSG00000113308 | *Gm48445* | 102.9240301 | 102.270057 | 115.5325803 | 186.0256698 | 252.4782963 | 240.5568076 | 1.17E-07 | 1.085295062 | 2.33E-08 | 31.19846595 | 2.121809393 | 6.933498836 |
| ENSMUSG00000039735 | *Fnbp1l* | 3731.63405 | 3694.918189 | 3576.614541 | 7671.233559 | 7128.46269 | 8505.126489 | 1.15E-49 | 1.082690998 | 4.22E-51 | 226.1013218 | 2.117982984 | 48.93753815 |
| ENSMUSG00000016526 | *Dyrk3* | 456.7785468 | 455.2667055 | 457.2348729 | 917.3390843 | 961.3230225 | 1016.111955 | 2.46E-29 | 1.080804446 | 1.59E-30 | 131.8803845 | 2.115215196 | 28.60977748 |
| ENSMUSG00000035024 | *Ncapd3* | 1215.524289 | 1270.128128 | 1203.301196 | 2641.564511 | 2444.752107 | 2712.518562 | 1.66E-49 | 1.080387843 | 6.09E-51 | 225.3710846 | 2.114604478 | 48.78071037 |
| ENSMUSG00000030583 | *Sipa1l3* | 781.7122616 | 834.6556267 | 719.6309027 | 1748.641296 | 1532.971995 | 1656.955291 | 1.14E-31 | 1.080159833 | 6.82E-33 | 142.7036764 | 2.114270304 | 30.94191939 |
| ENSMUSG00000035847 | *Ids* | 512.9189268 | 476.1605881 | 482.6912041 | 1104.527415 | 907.0163701 | 1102.712406 | 5.74E-22 | 1.079499046 | 4.79E-23 | 97.73246089 | 2.113302141 | 21.24074649 |
| ENSMUSG00000033033 | *Calhm2* | 323.2324912 | 328.8037317 | 324.0786786 | 709.2228662 | 675.4985361 | 677.4079702 | 5.90E-24 | 1.07873559 | 4.54E-25 | 106.9590505 | 2.112184105 | 23.22930199 |
| ENSMUSG00000025558 | *Dock9* | 121.6374901 | 112.1671593 | 121.4071183 | 270.8998817 | 211.5101199 | 268.4613973 | 1.48E-08 | 1.076748402 | 2.72E-09 | 35.37292149 | 2.109276757 | 7.828621658 |
| ENSMUSG00000097829 | *None* | 307.0708667 | 284.8166104 | 242.8142366 | 684.806997 | 556.4050001 | 522.4893861 | 1.72E-12 | 1.076235166 | 2.35E-13 | 53.68631569 | 2.108526519 | 11.76493464 |
| ENSMUSG00000032187 | *Smarca4* | 5331.634881 | 5347.734272 | 5358.557729 | 11624.27904 | 11164.30444 | 11024.23738 | 6.69E-109 | 1.075933634 | 1.09E-110 | 499.7307603 | 2.108085869 | 108.1746181 |
| ENSMUSG00000020034 | *Tcp11l2* | 158.2137983 | 138.5594321 | 139.0307322 | 302.2917134 | 326.7926627 | 290.5926236 | 3.73E-11 | 1.074813153 | 5.56E-12 | 47.47886582 | 2.106449242 | 10.42880449 |
| ENSMUSG00000027488 | *Snta1* | 240.7231448 | 259.5240157 | 271.2078368 | 613.8847104 | 499.2401029 | 511.9048866 | 8.41E-15 | 1.074075449 | 1.00E-15 | 64.42772479 | 2.105372411 | 14.07536728 |
| ENSMUSG00000042684 | *Npl* | 137.7991147 | 116.5658715 | 141.9680012 | 237.182729 | 302.9739555 | 294.4415325 | 7.19E-09 | 1.073987111 | 1.28E-09 | 36.83849079 | 2.105243501 | 8.143562353 |
| ENSMUSG00000046380 | *Jrk* | 183.7321529 | 190.2442996 | 187.0061257 | 416.2324362 | 417.3037501 | 347.3640302 | 7.66E-13 | 1.073581797 | 1.03E-13 | 55.30885934 | 2.104652132 | 12.11560455 |
| ENSMUSG00000005718 | *Tfap4* | 244.125592 | 244.1285232 | 225.1906226 | 511.570592 | 542.1137758 | 446.4734349 | 5.52E-15 | 1.071861748 | 6.51E-16 | 65.27650907 | 2.102144363 | 14.25791238 |
| ENSMUSG00000052056 | *Zfp217* | 707.7090334 | 755.4788084 | 730.400889 | 1655.628461 | 1438.649915 | 1514.545661 | 1.12E-33 | 1.07104701 | 6.18E-35 | 152.0480302 | 2.100957546 | 32.94980816 |
| ENSMUSG00000037239 | *Spred3* | 92.71668827 | 103.3697351 | 96.92987669 | 205.7908972 | 221.0376028 | 184.7476282 | 7.28E-08 | 1.063842872 | 1.43E-08 | 32.14365119 | 2.090492507 | 7.137954968 |
| ENSMUSG00000058515 | *Or8d1b* | 28.07019003 | 35.18969704 | 21.5399726 | 69.75962618 | 47.63741439 | 59.65808828 | 0.00793948 | 1.063310904 | 0.003071414 | 8.764552685 | 2.089721817 | 2.100207953 |
| ENSMUSG00000039046 | *Usp6nl* | 512.9189268 | 586.1283914 | 542.4156736 | 1181.263003 | 1140.439701 | 1102.712406 | 1.21E-29 | 1.062605877 | 7.78E-31 | 133.2979074 | 2.088700846 | 28.91690232 |
| ENSMUSG00000032305 | *Fam219b* | 400.6381667 | 409.0802281 | 436.6739899 | 918.5017447 | 783.1590926 | 900.6446876 | 7.21E-22 | 1.061508469 | 6.03E-23 | 97.27639519 | 2.087112647 | 21.14178825 |
| ENSMUSG00000041801 | *Phlda3* | 134.3966674 | 115.4661934 | 155.6752565 | 286.0144673 | 319.1706764 | 242.4812621 | 4.54E-08 | 1.0613739 | 8.73E-09 | 33.10429 | 2.086917979 | 7.343421663 |
| ENSMUSG00000062121 | *Or10d1b* | 28.92080185 | 21.99356065 | 31.33086923 | 55.80770094 | 64.78688358 | 50.99804321 | 0.006146619 | 1.056143928 | 0.002323434 | 9.274679109 | 2.079366304 | 2.211363687 |
| ENSMUSG00000044042 | *Fmn1* | 916.108929 | 991.9095854 | 926.2188217 | 2025.35448 | 1705.419435 | 2154.426769 | 1.29E-25 | 1.054549934 | 9.34E-27 | 114.6607034 | 2.077070138 | 24.88816802 |
| ENSMUSG00000049288 | *Lix1l* | 509.5164796 | 580.6300012 | 605.0774121 | 1189.401626 | 1224.28155 | 1101.750179 | 3.43E-25 | 1.053922735 | 2.51E-26 | 112.7004978 | 2.076167346 | 24.46498222 |
| ENSMUSG00000048895 | *Cdk5r1* | 66.34772188 | 79.17681835 | 68.53627645 | 176.7243863 | 127.6682706 | 138.5607212 | 2.49E-05 | 1.048958152 | 6.39E-06 | 20.36678409 | 2.069035146 | 4.60347281 |
| ENSMUSG00000024579 | *Pcyox1l* | 318.1288203 | 371.691175 | 334.8486649 | 724.3374518 | 684.0732707 | 706.2747871 | 1.41E-20 | 1.047724922 | 1.25E-21 | 91.26873848 | 2.06726727 | 19.85030133 |
| ENSMUSG00000060044 | *Tmem26* | 1503.881696 | 1406.488204 | 1370.725529 | 3067.098231 | 2756.300797 | 3028.129094 | 4.47E-44 | 1.04684818 | 1.86E-45 | 200.2293506 | 2.06601135 | 43.34982733 |
| ENSMUSG00000032180 | *Tmed1* | 494.2054668 | 528.9451337 | 548.2902116 | 1028.954486 | 1191.888108 | 1020.923091 | 2.01E-23 | 1.04635326 | 1.58E-24 | 104.4852379 | 2.065302722 | 22.69782655 |
| ENSMUSG00000021996 | *Esd* | 8703.460132 | 8376.247574 | 8493.602831 | 16741.14762 | 19231.22419 | 16762.96058 | 5.32E-56 | 1.044137539 | 1.73E-57 | 255.3996573 | 2.062133221 | 55.2739675 |
| ENSMUSG00000034880 | *Mrpl34* | 422.754074 | 461.8647737 | 441.5694383 | 921.989726 | 999.432954 | 810.195328 | 6.51E-20 | 1.04365298 | 5.98E-21 | 88.17941198 | 2.061440728 | 19.18632963 |
| ENSMUSG00000030223 | *Ptpro* | 552.0470705 | 519.0480314 | 498.3566387 | 1061.508978 | 1079.46381 | 1095.976815 | 2.91E-30 | 1.04341802 | 1.82E-31 | 136.1822029 | 2.061105024 | 29.53625835 |
| ENSMUSG00000031274 | *Col4a5* | 1796.492162 | 1733.092579 | 1650.745173 | 3817.014212 | 3107.864915 | 3750.761744 | 4.68E-31 | 1.042573889 | 2.85E-32 | 139.8648554 | 2.05989941 | 30.329578 |
| ENSMUSG00000024013 | *Fgd2* | 524.8274923 | 558.6364405 | 512.063894 | 1085.924848 | 1083.274803 | 1112.334678 | 2.74E-31 | 1.041604368 | 1.66E-32 | 140.9389285 | 2.058515581 | 30.56150063 |
| ENSMUSG00000089715 | *Cbx6* | 1265.710387 | 1250.333923 | 1315.896508 | 2772.945141 | 2545.743425 | 2564.335569 | 5.36E-49 | 1.039909886 | 1.99E-50 | 223.0123929 | 2.05609922 | 48.27109682 |
| ENSMUSG00000032320 | *Rcn2* | 1423.924185 | 1393.292067 | 1402.056398 | 2814.800916 | 2962.094427 | 2888.606146 | 8.01E-61 | 1.038310259 | 2.35E-62 | 277.7281049 | 2.053820729 | 60.09616059 |
| ENSMUSG00000062867 | *Impdh2* | 6103.139801 | 5936.06202 | 6091.895887 | 11792.86481 | 13687.1819 | 11672.77853 | 1.66E-48 | 1.035048403 | 6.24E-50 | 220.7399907 | 2.049182396 | 47.78101962 |
| ENSMUSG00000043487 | *Acot6* | 42.53059095 | 32.99034098 | 63.64082813 | 94.17549534 | 92.41658392 | 99.10940473 | 0.00159296 | 1.03457541 | 0.000543994 | 11.95847557 | 2.048510674 | 2.797795089 |
| ENSMUSG00000027544 | *Nfatc2* | 612.4405097 | 753.2794523 | 677.5300472 | 1453.325545 | 1291.926678 | 1430.831892 | 7.98E-24 | 1.033233967 | 6.20E-25 | 106.3446685 | 2.046606818 | 23.09774501 |
| ENSMUSG00000021240 | *Abcd4* | 777.4592025 | 853.3501533 | 790.1253585 | 1718.412125 | 1611.097355 | 1614.617293 | 5.22E-37 | 1.030981648 | 2.60E-38 | 167.4979337 | 2.043414171 | 36.2822897 |
| ENSMUSG00000036854 | *Hspb6* | 45.93303823 | 47.2861554 | 52.87084183 | 102.3141184 | 100.9913185 | 94.29826858 | 0.000200022 | 1.026669202 | 5.83E-05 | 16.15702604 | 2.0373152 | 3.698923029 |
| ENSMUSG00000046329 | *Slc25a23* | 502.711585 | 453.0673494 | 516.9593423 | 1003.375957 | 962.2757708 | 1037.280954 | 1.87E-25 | 1.026130936 | 1.36E-26 | 113.9204644 | 2.036555225 | 24.72769251 |
| ENSMUSG00000048429 | *Timm29* | 1185.752876 | 1201.94809 | 1283.586549 | 2549.714337 | 2430.460882 | 2496.017436 | 2.29E-49 | 1.025945843 | 8.44E-51 | 224.7226682 | 2.036293958 | 48.64009781 |
| ENSMUSG00000021054 | *Sgpp1* | 528.2299396 | 532.2441678 | 499.3357284 | 1113.828698 | 991.8109677 | 1070.958907 | 8.41E-27 | 1.025347596 | 5.86E-28 | 120.15196 | 2.035449737 | 26.07544917 |
| ENSMUSG00000019066 | *Rab3d* | 385.327154 | 410.1799062 | 407.3013 | 833.6275329 | 817.458031 | 791.9130106 | 1.63E-24 | 1.023078115 | 1.23E-25 | 109.5512965 | 2.03225032 | 23.78673993 |
| ENSMUSG00000029162 | *Khk* | 233.9182502 | 271.620474 | 248.6887745 | 477.8534393 | 600.2314214 | 449.3601166 | 4.49E-11 | 1.021062388 | 6.74E-12 | 47.10185497 | 2.029412851 | 10.34766247 |
| ENSMUSG00000041912 | *Tdrkh* | 1508.134755 | 1419.68434 | 1364.850991 | 2884.560543 | 2859.197612 | 2969.433233 | 1.16E-50 | 1.020594587 | 4.18E-52 | 230.7069442 | 2.028754911 | 49.93375207 |
| ENSMUSG00000005583 | *Mef2c* | 1493.674354 | 1410.886916 | 1383.453695 | 2982.224019 | 2649.592989 | 3061.807047 | 1.33E-37 | 1.018830273 | 6.54E-39 | 170.2465426 | 2.026275403 | 36.8748001 |
| ENSMUSG00000038119 | *Cdon* | 28.92080185 | 34.09001901 | 35.24722789 | 74.41026793 | 60.97589042 | 63.5069972 | 0.003455464 | 1.018359221 | 0.00124832 | 10.41742232 | 2.025613915 | 2.461493649 |
| ENSMUSG00000041064 | *Pif1* | 871.0265026 | 942.4240739 | 845.9334693 | 1854.443396 | 1676.836987 | 1848.43851 | 2.03E-33 | 1.017030883 | 1.13E-34 | 150.8496062 | 2.023749723 | 32.69325235 |
| ENSMUSG00000028961 | *Pgd* | 16586.93047 | 16448.98401 | 16605.36069 | 33356.72792 | 35804.28066 | 31200.21794 | 3.71E-65 | 1.015585244 | 1.00E-66 | 297.7873016 | 2.021722859 | 64.4309641 |
| ENSMUSG00000064307 | *Lrrc51* | 42.53059095 | 37.38905311 | 42.10085553 | 74.41026793 | 79.07810789 | 93.33604135 | 0.001077836 | 1.015573091 | 0.000357726 | 12.74100444 | 2.021705829 | 2.967447389 |
| ENSMUSG00000020649 | *Rrm2* | 1989.581045 | 2033.304682 | 1958.179327 | 4136.745832 | 4080.620917 | 3872.002375 | 2.23E-63 | 1.015247396 | 6.25E-65 | 289.5441562 | 2.021249471 | 62.65226373 |
| ENSMUSG00000054752 | *Fsd1l* | 109.7289246 | 89.07392064 | 78.32717308 | 218.580162 | 146.7232363 | 197.2565822 | 3.86E-05 | 1.01517018 | 1.02E-05 | 19.4735855 | 2.021141292 | 4.413730631 |
| ENSMUSG00000026383 | *Epb41l5* | 217.7566257 | 184.7459095 | 208.5460983 | 431.3470219 | 362.0443494 | 443.5867532 | 3.64E-11 | 1.013796234 | 5.43E-12 | 47.52607884 | 2.019217382 | 10.43846479 |
| ENSMUSG00000034066 | *Farp2* | 100.3721946 | 117.6655495 | 140.0098219 | 218.580162 | 216.2738614 | 285.7814874 | 2.53E-06 | 1.012857312 | 5.80E-07 | 24.97916666 | 2.01790368 | 5.596309793 |
| ENSMUSG00000070047 | *Fat1* | 847.2093717 | 861.0478995 | 890.9715938 | 1824.214225 | 1513.917029 | 1906.172143 | 1.89E-23 | 1.012425153 | 1.49E-24 | 104.6012102 | 2.017299307 | 22.72286863 |
| ENSMUSG00000060244 | *Alyref2* | 51.03670914 | 47.2861554 | 54.82902116 | 102.3141184 | 100.0385702 | 106.8072226 | 0.000174059 | 1.011922126 | 5.03E-05 | 16.43795582 | 2.016596055 | 3.759303862 |
| ENSMUSG00000032997 | *Chpf* | 691.5474088 | 699.3952287 | 710.8190957 | 1513.783888 | 1431.980677 | 1293.233398 | 1.51E-28 | 1.011404983 | 9.95E-30 | 128.238433 | 2.015873322 | 27.82070643 |
| ENSMUSG00000045691 | *Thtpa* | 396.3851076 | 382.6879553 | 338.7650236 | 765.0305671 | 752.6711474 | 734.1793768 | 3.59E-20 | 1.009312902 | 3.26E-21 | 89.37681254 | 2.012952183 | 19.44496791 |
| ENSMUSG00000083307 | *AA414768* | 335.9916685 | 316.7072734 | 280.0196438 | 609.2340686 | 613.5698974 | 653.3522894 | 8.96E-17 | 1.007433913 | 9.66E-18 | 73.57986511 | 2.010332189 | 16.047692 |
| ENSMUSG00000024042 | *Sik1* | 36.57630822 | 39.58840917 | 37.20540721 | 79.06090967 | 69.55062502 | 78.90263289 | 0.001523378 | 1.005714797 | 0.000518559 | 12.047715 | 2.007938103 | 2.817192245 |
| ENSMUSG00000029471 | *Camkk2* | 1193.408382 | 1178.854851 | 1126.932203 | 2357.875365 | 2342.80804 | 2323.778761 | 6.71E-51 | 1.005040581 | 2.39E-52 | 231.8197171 | 2.00699995 | 50.1730893 |
| ENSMUSG00000075271 | *Ttc30a1* | 38.27753185 | 31.89066294 | 51.89175217 | 88.36219316 | 65.73963186 | 91.41158688 | 0.003690573 | 1.00395338 | 0.00133908 | 10.28786442 | 2.005488065 | 2.432906219 |
| ENSMUSG00000038400 | *Pmepa1* | 214.3541784 | 193.5433337 | 187.0061257 | 423.2083988 | 394.4377912 | 378.1553015 | 2.43E-12 | 1.003845007 | 3.37E-13 | 52.97961414 | 2.005337422 | 11.61432135 |
| ENSMUSG00000007035 | *Msh5* | 130.1436083 | 133.0610419 | 117.4907596 | 247.6466729 | 257.2420377 | 257.8768977 | 4.28E-09 | 1.003561118 | 7.52E-10 | 37.88015124 | 2.004942856 | 8.368628245 |
| ENSMUSG00000000579 | *Dynlt1c* | 51.88732096 | 64.88100392 | 75.38990409 | 137.1939315 | 87.65284249 | 159.7297202 | 0.001494267 | 1.003016372 | 0.000508137 | 12.08556591 | 2.004185955 | 2.825571921 |
| ENSMUSG00000004500 | *Zfp324* | 240.7231448 | 238.6301331 | 265.3332988 | 465.0641745 | 504.0038443 | 521.5271589 | 9.05E-15 | 1.002315425 | 1.08E-15 | 64.27883743 | 2.003212437 | 14.04354846 |
| ENSMUSG00000021697 | *Depdc1b* | 132.6954438 | 135.260398 | 151.7588978 | 284.8518069 | 283.9189898 | 271.348079 | 1.12E-09 | 1.001386878 | 1.88E-10 | 40.59092517 | 2.001923546 | 8.949867424 |
| ENSMUSG00000028737 | *Aldh4a1* | 423.6046859 | 414.5786183 | 440.5903486 | 801.0730406 | 901.2998803 | 855.4200078 | 1.67E-22 | 1.000858523 | 1.36E-23 | 100.2216122 | 2.00119052 | 21.7768551 |
| ENSMUSG00000023495 | *Pcbp4* | 44.23181459 | 45.08679934 | 35.24722789 | 90.68751403 | 84.79459762 | 74.09149674 | 0.001349148 | 1.000837164 | 0.000454345 | 12.29429605 | 2.001160893 | 2.869940319 |
| ENSMUSG00000021611 | *Tert* | 32.32324912 | 39.58840917 | 32.3099589 | 76.7355888 | 66.69238015 | 64.46922443 | 0.003255509 | 0.998500749 | 0.001173134 | 10.53218545 | 1.997922677 | 2.487381072 |
| ENSMUSG00000071176 | *Arhgef10* | 370.8667531 | 413.4789402 | 387.7195068 | 785.958455 | 760.2931337 | 791.9130106 | 5.51E-22 | 0.998259684 | 4.59E-23 | 97.81484119 | 1.997588864 | 21.25845695 |
| ENSMUSG00000050965 | *Prkca* | 313.0251494 | 399.1831258 | 322.1204993 | 731.3134145 | 617.3808906 | 712.0481505 | 2.32E-13 | 0.996947195 | 3.02E-14 | 57.72207203 | 1.995772388 | 12.63394367 |
| ENSMUSG00000031628 | *Casp3* | 1087.081905 | 1153.562256 | 1188.614852 | 2248.585284 | 2248.485959 | 2324.740989 | 2.26E-44 | 0.993252861 | 9.30E-46 | 201.6090375 | 1.990668317 | 43.64658494 |
| ENSMUSG00000043252 | *Tmem64* | 640.5106997 | 601.5238838 | 596.2656051 | 1240.558686 | 1125.195728 | 1291.308943 | 1.25E-25 | 0.991039953 | 9.02E-27 | 114.7289976 | 1.98761723 | 24.90229861 |
| ENSMUSG00000055675 | *Kbtbd11* | 736.6298352 | 780.7714031 | 856.7034556 | 1704.4602 | 1392.917997 | 1619.428429 | 2.19E-21 | 0.990249335 | 1.88E-22 | 95.02841547 | 1.986528285 | 20.65923647 |
| ENSMUSG00000073542 | *Cep76* | 618.3947924 | 596.0254937 | 606.0565017 | 1221.956119 | 1227.139795 | 1161.408267 | 3.81E-31 | 0.986957029 | 2.31E-32 | 140.2785994 | 1.982000091 | 30.41907359 |
| ENSMUSG00000021514 | *Zfp369* | 188.8358238 | 197.9420459 | 209.525188 | 419.7204175 | 350.6113699 | 409.9088001 | 3.97E-11 | 0.984619385 | 5.94E-12 | 47.34925686 | 1.978791194 | 10.40108665 |
| ENSMUSG00000040204 | *Pclaf* | 2955.025459 | 2847.066426 | 3034.198867 | 5459.853409 | 6180.478144 | 5825.323653 | 5.22E-50 | 0.983141304 | 1.89E-51 | 227.7038614 | 1.976764907 | 49.28212902 |
| ENSMUSG00000027890 | *Gstm4* | 30.62202548 | 26.39227278 | 33.28904856 | 54.64504051 | 54.30665241 | 69.28036059 | 0.008646775 | 0.979582101 | 0.003372325 | 8.594215169 | 1.971894136 | 2.06314586 |
| ENSMUSG00000044813 | *Shb* | 202.4456129 | 215.5368944 | 192.8806637 | 406.9311527 | 413.4927569 | 382.9664377 | 1.56E-12 | 0.979198836 | 2.13E-13 | 53.88070579 | 1.971370354 | 11.80659585 |
| ENSMUSG00000110949 | *Nudt8* | 130.9942201 | 141.8584662 | 139.0307322 | 256.9479564 | 288.6827312 | 264.6124883 | 4.80E-09 | 0.978967117 | 8.47E-10 | 37.64903799 | 1.971053748 | 8.318868099 |
| ENSMUSG00000023050 | *Map3k12* | 120.7868783 | 133.0610419 | 161.5497945 | 290.6651091 | 275.3442552 | 251.1413071 | 1.11E-07 | 0.976808434 | 2.21E-08 | 31.29960153 | 1.968106695 | 6.955676333 |
| ENSMUSG00000032411 | *Tfdp2* | 266.2414993 | 257.3246596 | 262.3960298 | 494.1306854 | 522.1060618 | 530.1872039 | 7.11E-16 | 0.976539792 | 8.03E-17 | 69.40186973 | 1.96774025 | 15.14833171 |
| ENSMUSG00000046574 | *Prr12* | 567.3580833 | 619.1187323 | 610.95195 | 1244.046667 | 1083.274803 | 1204.708492 | 6.87E-24 | 0.975043594 | 5.32E-25 | 106.6469294 | 1.965700595 | 23.16283186 |
| ENSMUSG00000058290 | *Espl1* | 1023.286018 | 665.3052097 | 901.7415801 | 1367.288673 | 1854.048168 | 1861.909691 | 1.26E-09 | 0.971603554 | 2.12E-10 | 40.35375348 | 1.961019055 | 8.898395684 |
| ENSMUSG00000032050 | *Rdx* | 4073.580001 | 3835.676978 | 3854.676005 | 7910.741609 | 7117.029711 | 8039.40851 | 4.32E-49 | 0.971026474 | 1.60E-50 | 223.4544383 | 1.960234801 | 48.36429496 |
| ENSMUSG00000008855 | *Hdac5* | 1422.222961 | 1457.073393 | 1459.822688 | 3122.905932 | 2862.055857 | 2521.035344 | 2.19E-28 | 0.970595823 | 1.46E-29 | 127.4808007 | 1.95964975 | 27.65942182 |
| ENSMUSG00000006930 | *Hap1* | 249.229263 | 290.3150006 | 253.5842229 | 516.2212337 | 518.2950686 | 514.7915682 | 1.33E-14 | 0.969029573 | 1.61E-15 | 63.49530848 | 1.957523427 | 13.87627829 |
| ENSMUSG00000050493 | *Fam167b* | 1038.597031 | 1051.292199 | 1073.082271 | 2088.138144 | 2101.762723 | 1998.545957 | 1.69E-43 | 0.968339569 | 7.21E-45 | 197.535071 | 1.956587417 | 42.77106204 |
| ENSMUSG00000026566 | *Mpzl1* | 366.613694 | 347.4982583 | 345.6186512 | 688.2949783 | 707.8919779 | 678.3701974 | 7.26E-20 | 0.968082347 | 6.67E-21 | 87.96249343 | 1.956238603 | 19.13902751 |
| ENSMUSG00000027010 | *Slc25a12* | 2329.825772 | 2251.040933 | 2376.250613 | 4575.068817 | 4425.515797 | 4597.521707 | 4.40E-64 | 0.966387246 | 1.22E-65 | 292.7988728 | 1.953941461 | 63.35608541 |
| ENSMUSG00000037112 | *Sik2* | 420.2022386 | 450.8679934 | 386.7404171 | 825.4889098 | 768.8678683 | 861.1933712 | 1.15E-18 | 0.966079831 | 1.14E-19 | 82.34771246 | 1.953525153 | 17.93857534 |
| ENSMUSG00000032604 | *Qars* | 4648.593591 | 4834.184631 | 4833.765669 | 9017.594344 | 9842.842562 | 9084.387282 | 3.09E-63 | 0.965268791 | 8.70E-65 | 288.8854833 | 1.952427248 | 62.5098041 |
| ENSMUSG00000032109 | *Nlrx1* | 530.781775 | 581.7296792 | 535.562046 | 1052.207695 | 1140.439701 | 1018.03641 | 2.35E-23 | 0.963863027 | 1.87E-24 | 104.1610298 | 1.950525727 | 22.62826985 |
| ENSMUSG00000075595 | *Zfp652* | 868.4746672 | 819.2601343 | 869.4316212 | 1818.400922 | 1484.381833 | 1689.671017 | 1.00E-23 | 0.963651836 | 7.81E-25 | 105.8871776 | 1.950240217 | 22.99939855 |
| ENSMUSG00000094649 | *Gm7102* | 39.97875549 | 32.99034098 | 27.41451058 | 65.10898443 | 61.92863871 | 69.28036059 | 0.00671742 | 0.962764304 | 0.002558556 | 9.098239644 | 1.949040817 | 2.172797495 |
| ENSMUSG00000068854 | *H2bc21* | 101.2228065 | 108.8681252 | 130.2189253 | 225.5561246 | 194.3606507 | 241.5190348 | 2.15E-06 | 0.959703079 | 4.89E-07 | 25.30757663 | 1.944909572 | 5.667000123 |
| ENSMUSG00000044469 | *Tnfaip8l1* | 69.75016916 | 80.27649638 | 81.26444207 | 136.0312711 | 141.0067466 | 171.276447 | 4.48E-05 | 0.959471154 | 1.20E-05 | 19.17083384 | 1.944596937 | 4.348685963 |
| ENSMUSG00000044551 | *9930012K11Rik* | 76.55506371 | 71.47907212 | 70.49445577 | 148.8205359 | 146.7232363 | 129.9006761 | 3.07E-05 | 0.958454679 | 7.98E-06 | 19.94243508 | 1.943227321 | 4.512893394 |
| ENSMUSG00000055485 | *Soga1* | 764.7000253 | 847.8517631 | 756.8363099 | 1527.735813 | 1468.185112 | 1603.070566 | 1.14E-28 | 0.958271654 | 7.50E-30 | 128.7989891 | 1.942980812 | 27.94154922 |
| ENSMUSG00000036278 | *Macrod1* | 76.55506371 | 78.07714031 | 83.2226214 | 167.4231028 | 164.8254538 | 129.9006761 | 3.69E-05 | 0.957085031 | 9.74E-06 | 19.56191421 | 1.941383359 | 4.432570741 |
| ENSMUSG00000041115 | *Iqsec2* | 209.2505075 | 223.2346406 | 213.4415467 | 440.6483054 | 405.8707706 | 407.0221184 | 1.22E-12 | 0.956781058 | 1.65E-13 | 54.38093405 | 1.940974356 | 11.91474914 |
| ENSMUSG00000054920 | *Klhl5* | 1309.942201 | 1303.118469 | 1270.858383 | 2481.117371 | 2549.554418 | 2506.601935 | 2.99E-50 | 0.956617465 | 1.08E-51 | 228.8171761 | 1.940754274 | 49.52410741 |
| ENSMUSG00000079260 | *Tmppe* | 315.5769848 | 316.7072734 | 288.8314507 | 605.7460873 | 571.6489727 | 610.0520641 | 1.61E-16 | 0.956076913 | 1.76E-17 | 72.39915205 | 1.940027245 | 15.79446127 |
| ENSMUSG00000035473 | *Galm* | 110.5795365 | 126.4629737 | 109.6580423 | 201.1402555 | 233.4233305 | 235.7456714 | 4.32E-07 | 0.955971574 | 9.11E-08 | 28.55490346 | 1.939885598 | 6.364141905 |
| ENSMUSG00000028063 | *Lmna* | 16954.39478 | 17847.77447 | 18358.91028 | 35407.66093 | 36140.6008 | 31571.63766 | 4.49E-49 | 0.955913594 | 1.66E-50 | 223.3705049 | 1.939807639 | 48.34759968 |
| ENSMUSG00000012187 | *Mogat1* | 101.2228065 | 112.1671593 | 100.8462353 | 166.2604424 | 248.6673031 | 192.4454461 | 2.17E-05 | 0.955472053 | 5.53E-06 | 20.64580524 | 1.939214046 | 4.662696112 |
| ENSMUSG00000037966 | *Ninj1* | 2553.536681 | 2687.613112 | 2756.137403 | 5154.073714 | 5594.537946 | 4750.515836 | 3.59E-37 | 0.954931446 | 1.78E-38 | 168.2553758 | 1.938487519 | 36.44473737 |
| ENSMUSG00000034487 | *Poglut3* | 591.1752142 | 592.7264596 | 572.7674532 | 1164.985757 | 998.4802057 | 1236.461991 | 3.77E-19 | 0.952288513 | 3.63E-20 | 84.61181948 | 1.934939574 | 18.42408546 |
| ENSMUSG00000032254 | *Kif23* | 2222.648683 | 2123.478281 | 2214.700819 | 4363.464618 | 4066.329693 | 4271.326676 | 1.43E-55 | 0.952244644 | 4.68E-57 | 253.4123352 | 1.934880738 | 54.84348861 |
| ENSMUSG00000032375 | *Aph1b* | 109.7289246 | 70.37939408 | 82.24353174 | 188.3509907 | 159.1089641 | 163.5786292 | 8.58E-05 | 0.952220562 | 2.38E-05 | 17.85958681 | 1.93484844 | 4.066342563 |
| ENSMUSG00000030589 | *Rasgrp4* | 127.5917728 | 141.8584662 | 135.1143736 | 262.7612586 | 252.4782963 | 264.6124883 | 1.45E-08 | 0.948928147 | 2.67E-09 | 35.41369175 | 1.930437902 | 7.837550031 |
| ENSMUSG00000003123 | *Lipe* | 541.8397287 | 635.6139028 | 609.9728604 | 1213.817496 | 1185.21887 | 1046.903227 | 2.07E-19 | 0.948404292 | 1.96E-20 | 85.82743684 | 1.929737071 | 18.68328497 |
| ENSMUSG00000010607 | *Pigyl* | 157.3631865 | 186.9452655 | 191.9015741 | 292.99043 | 411.5872604 | 325.2328039 | 3.20E-07 | 0.946063677 | 6.63E-08 | 29.16966333 | 1.926608822 | 6.49542825 |

| | | | | | | | | | | | | | |
|---|---|---|---|---|---|---|---|---|---|---|---|---|---|
| ENSMUSG00000039936 | *Pik3cd* | 1984.477374 | 2176.262826 | 2222.533536 | 4220.457384 | 4133.022073 | 3937.433827 | 2.88E-43 | 0.945779621 | 1.23E-44 | 196.4702376 | 1.926229524 | 42.54077067 |
| ENSMUSG00000074026 | *Gm6091* | 75.70445189 | 75.87778425 | 68.53627645 | 146.495215 | 150.5342295 | 127.0139944 | 4.94E-05 | 0.945150152 | 1.33E-05 | 18.9716075 | 1.925389265 | 4.30590965 |
| ENSMUSG00000097530 | *Kansl2-ps* | 368.3149176 | 356.2956826 | 405.3431207 | 720.8494705 | 712.6557193 | 737.0660585 | 2.15E-18 | 0.94136458 | 2.16E-19 | 81.09066267 | 1.920343746 | 17.66683536 |
| ENSMUSG00000031988 | *Vps26b* | 1338.863003 | 1284.423942 | 1376.600067 | 2490.418655 | 2665.78971 | 2522.959798 | 4.20E-42 | 0.94083271 | 1.84E-43 | 191.086777 | 1.919635915 | 41.37680753 |
| ENSMUSG00000083610 | *Oaz2-ps* | 43.38120277 | 48.38583343 | 51.89175217 | 74.41026793 | 111.4715497 | 88.52490519 | 0.002597084 | 0.939724558 | 0.000919333 | 10.98336658 | 1.918161985 | 2.585514037 |
| ENSMUSG00000028980 | *H6pd* | 757.0445189 | 801.6652857 | 815.5816897 | 1655.628461 | 1431.980677 | 1467.396526 | 3.81E-25 | 0.939603179 | 2.80E-26 | 112.4858046 | 1.918000609 | 24.41958025 |
| ENSMUSG00000032239 | *Rp9* | 567.3580833 | 582.8293573 | 573.7465428 | 1073.135583 | 1134.723211 | 1095.014588 | 3.53E-27 | 0.938999012 | 2.44E-28 | 121.8897236 | 1.917197563 | 26.45243516 |
| ENSMUSG00000005442 | *Cic* | 3307.178752 | 3553.059723 | 3500.245547 | 7050.372886 | 6299.57168 | 6493.10935 | 4.27E-47 | 0.937613539 | 1.65E-48 | 214.2142188 | 1.915357293 | 46.36921527 |
| ENSMUSG00000029442 | *Wdr66* | 100.3721946 | 134.16072 | 105.7416837 | 218.580162 | 236.2815754 | 194.3699005 | 5.36E-06 | 0.937348044 | 1.27E-06 | 23.47245745 | 1.915004847 | 5.271160867 |
| ENSMUSG00000067924 | *Rtl8b* | 84.21057008 | 120.9645836 | 123.3652976 | 213.9295203 | 237.2343237 | 175.1253559 | 6.05E-05 | 0.93578574 | 1.65E-05 | 18.56075281 | 1.912932199 | 4.217973874 |
| ENSMUSG00000032245 | *Cln6* | 646.4649824 | 702.6942628 | 665.7809712 | 1309.155651 | 1273.824461 | 1267.253262 | 4.80E-29 | 0.935317881 | 3.13E-30 | 130.5375031 | 1.912311945 | 28.31919661 |
| ENSMUSG00000022641 | *Bbx* | 1239.34142 | 1261.330703 | 1232.673886 | 2452.05086 | 2129.392423 | 2552.788842 | 9.59E-28 | 0.93414685 | 6.51E-29 | 124.5111907 | 1.910760356 | 27.01796377 |
| ENSMUSG00000066894 | *Vsig10* | 141.201562 | 153.9549246 | 163.5079738 | 313.9183178 | 275.3442552 | 286.7437146 | 1.41E-08 | 0.933761906 | 2.58E-09 | 35.47773612 | 1.91025059 | 7.85101677 |
| ENSMUSG00000022992 | *Kansl2* | 756.1939071 | 776.372691 | 655.0109849 | 1357.98739 | 1514.869778 | 1300.931215 | 8.81E-20 | 0.932554693 | 8.14E-21 | 87.56850047 | 1.908652806 | 19.05510628 |
| ENSMUSG00000006676 | *Usp19* | 3877.088671 | 4043.516126 | 3982.936751 | 7872.373814 | 7386.657476 | 7420.696401 | 3.39E-65 | 0.93004455 | 9.11E-67 | 297.9729058 | 1.905334831 | 64.47029546 |
| ENSMUSG00000000876 | *Pxmp4* | 131.8448319 | 130.8616859 | 101.825325 | 224.3934642 | 241.9980651 | 228.0478536 | 1.02E-06 | 0.930041942 | 2.23E-07 | 26.82028867 | 1.905331387 | 5.991360462 |
| ENSMUSG00000061046 | *Haghl* | 259.4366048 | 240.8294891 | 239.8769676 | 495.2933459 | 445.8861987 | 471.4913429 | 6.06E-13 | 0.929917953 | 8.11E-14 | 55.77945889 | 1.905167645 | 12.21757295 |
| ENSMUSG00000019132 | *BC005537* | 10958.43206 | 11097.9507 | 10551.6493 | 21909.17326 | 18320.39683 | 21883.9339 | 1.87E-35 | 0.92961872 | 9.73E-37 | 160.2993528 | 1.904772531 | 34.7289911 |
| ENSMUSG00000032415 | *Ube2cbp* | 45.08242641 | 37.38905311 | 49.93357284 | 88.36219316 | 83.84184933 | 80.82708735 | 0.00227232 | 0.92960891 | 0.000796419 | 11.24955954 | 1.904759578 | 2.643530443 |
| ENSMUSG00000063856 | *Gpx1* | 6325.149486 | 6040.531433 | 6295.546537 | 11318.49935 | 12725.85888 | 11499.57763 | 5.68E-51 | 0.929546838 | 2.01E-52 | 232.1686866 | 1.904677627 | 50.24583083 |
| ENSMUSG00000020941 | *Map3k14* | 391.2814367 | 424.4757206 | 442.5485279 | 818.5129472 | 808.8832964 | 766.8951026 | 6.55E-19 | 0.929180503 | 6.42E-20 | 83.48374669 | 1.904194046 | 18.18363448 |
| ENSMUSG00000078870 | *Gm14410* | 134.3966674 | 115.4661934 | 121.4071183 | 259.2732773 | 250.5727997 | 197.2565822 | 2.80E-06 | 0.924945425 | 6.43E-07 | 24.77887669 | 1.898612419 | 5.55209855 |
| ENSMUSG00000002105 | *Slc39a13* | 974.8011446 | 1019.401536 | 970.2778566 | 1934.666966 | 1920.740548 | 1766.649195 | 6.08E-33 | 0.92352809 | 3.45E-34 | 148.6330867 | 1.896748096 | 32.21618346 |
| ENSMUSG00000068267 | *Cenpb* | 5076.451336 | 5302.647473 | 5223.443355 | 10300.00881 | 10041.96695 | 9180.610005 | 1.60E-51 | 0.920054149 | 5.56E-53 | 234.7248229 | 1.892186312 | 50.79644724 |
| ENSMUSG00000026202 | *Tuba4a* | 3023.925016 | 3025.214268 | 2911.812659 | 5527.287714 | 5986.117493 | 5428.886034 | 5.85E-50 | 0.919058979 | 2.12E-51 | 227.4709551 | 1.890881533 | 49.23297881 |
| ENSMUSG00000030802 | *Bckdk* | 2034.663471 | 1975.021747 | 2178.474501 | 3939.093558 | 4189.234222 | 3568.900797 | 2.16E-31 | 0.918368707 | 1.30E-32 | 141.4258895 | 1.889977038 | 30.66600527 |
| ENSMUSG00000038451 | *Spsb2* | 292.6104657 | 283.7169324 | 306.4550647 | 569.7036138 | 605.9479111 | 489.7736603 | 4.85E-12 | 0.91496598 | 6.85E-13 | 51.58557153 | 1.885524609 | 11.31453313 |
| ENSMUSG00000050148 | *Ubqln2* | 704.3065861 | 776.372691 | 729.4217993 | 1484.717377 | 1279.540951 | 1401.002847 | 8.55E-23 | 0.914675775 | 6.91E-24 | 101.5674203 | 1.885145365 | 22.06826421 |
| ENSMUSG00000043964 | *Orai3* | 619.2454042 | 614.7200202 | 569.8301842 | 1114.991358 | 1154.730925 | 1129.654768 | 2.25E-25 | 0.914327155 | 1.63E-26 | 113.5541852 | 1.884689885 | 24.64829024 |
| ENSMUSG00000084883 | *Ccdc85c* | 315.5769848 | 309.0095272 | 308.413244 | 583.655539 | 603.0896662 | 571.5629748 | 4.99E-16 | 0.913902569 | 5.58E-17 | 70.118997 | 1.8841353 | 15.30225289 |
| ENSMUSG00000029108 | *Pcdh7* | 5995.1121 | 6084.518554 | 5829.499857 | 11775.4249 | 10345.89366 | 11600.61149 | 3.57E-46 | 0.912883376 | 1.42E-47 | 209.9302944 | 1.882804721 | 45.44677562 |
| ENSMUSG00000032329 | *Hmg20a* | 942.4778954 | 910.533411 | 1006.504174 | 1820.726243 | 1759.726088 | 1803.21383 | 5.86E-33 | 0.912154391 | 3.32E-34 | 148.7068288 | 1.881853592 | 32.23177718 |
| ENSMUSG00000001504 | *Irx2* | 616.6935688 | 614.7200202 | 651.0946263 | 1266.137215 | 1077.558314 | 1198.935129 | 6.55E-21 | 0.911132132 | 5.73E-22 | 92.8207512 | 1.880520628 | 20.18380223 |
| ENSMUSG00000046417 | *Lrrc75a* | 612.4405097 | 614.7200202 | 620.7428467 | 1145.22053 | 1210.943074 | 1114.259133 | 1.56E-25 | 0.909505214 | 1.13E-26 | 114.2876646 | 1.878401173 | 24.80729278 |
| ENSMUSG00000025786 | *Zdhhc3* | 3598.087994 | 3582.75103 | 3566.823644 | 6929.456201 | 6644.46656 | 6618.19889 | 3.78E-70 | 0.909488598 | 9.47E-72 | 320.8482469 | 1.878379538 | 69.42257864 |
| ENSMUSG00000083282 | *Ctsf* | 589.4739906 | 657.6074635 | 599.2028741 | 1190.564287 | 1175.691387 | 1095.976815 | 1.15E-22 | 0.908331956 | 9.27E-24 | 100.9839394 | 1.876874202 | 21.94107217 |
| ENSMUSG00000041774 | *Ydjc* | 592.025826 | 534.4435238 | 563.9556462 | 1054.533016 | 1111.857252 | 1007.45191 | 1.32E-21 | 0.907272369 | 1.12E-22 | 96.05196775 | 1.875496238 | 20.8781742 |
| ENSMUSG00000109946 | *Brd3os* | 110.5795365 | 80.27649638 | 89.09715938 | 199.977595 | 141.9594949 | 185.7098555 | 0.000187719 | 0.9066111 | 5.44E-05 | 16.2875704 | 1.874636788 | 3.726491389 |
| ENSMUSG00000034908 | *Sidt2* | 1091.334964 | 1099.678033 | 1118.120396 | 2205.566848 | 1968.377963 | 2027.412774 | 1.70E-32 | 0.905397126 | 9.79E-34 | 146.5601046 | 1.873060015 | 31.77032173 |
| ENSMUSG00000026856 | *Dolpp1* | 338.543504 | 348.5979363 | 306.4550647 | 616.2100313 | 634.5303597 | 609.0898368 | 2.33E-15 | 0.905054518 | 2.69E-16 | 67.01494616 | 1.872615257 | 14.63201671 |
| ENSMUSG00000037960 | *Card19* | 1687.613849 | 1735.291935 | 1775.08956 | 3177.550973 | 3419.413605 | 3133.011862 | 7.99E-42 | 0.905020507 | 3.52E-43 | 189.7952707 | 1.872571112 | 41.09761307 |
| ENSMUSG00000000530 | *Acvrl1* | 415.9491795 | 383.7876334 | 385.7613274 | 770.8438693 | 719.3249574 | 733.2171495 | 6.88E-18 | 0.904926397 | 7.03E-19 | 78.75411444 | 1.872448965 | 17.16268368 |
| ENSMUSG00000024975 | *Pdcd4* | 105.4758656 | 67.08035999 | 112.5953113 | 170.9110841 | 167.6836987 | 197.2565822 | 0.000269579 | 0.904567536 | 8.03E-05 | 15.55216534 | 1.871983263 | 3.569313512 |
| ENSMUSG00000032480 | *Dhx30* | 2820.628792 | 3039.510082 | 2950.976246 | 5564.492848 | 5569.766491 | 5340.361129 | 2.96E-55 | 0.903391028 | 9.72E-57 | 251.9580123 | 1.870457295 | 54.52826727 |
| ENSMUSG00000036611 | *Eepd1* | 228.8145793 | 218.8359285 | 211.4833673 | 437.1603241 | 417.3037501 | 380.079756 | 8.34E-11 | 0.903177724 | 1.28E-11 | 45.84343989 | 1.870180767 | 10.07876148 |
| ENSMUSG00000032392 | *Parp16* | 115.6832074 | 150.6558905 | 113.574401 | 237.182729 | 254.3837929 | 216.5011268 | 4.44E-06 | 0.90307691 | 1.04E-06 | 23.8451298 | 1.870050085 | 5.352484404 |
| ENSMUSG00000061143 | *Maml3* | 415.0985677 | 457.4660616 | 422.9667346 | 908.0378008 | 714.5612159 | 800.5730557 | 1.35E-13 | 0.903014824 | 1.73E-14 | 58.81875125 | 1.86996961 | 12.86840832 |
| ENSMUSG00000062861 | *Zfp28* | 97.82035918 | 102.270057 | 99.86714568 | 180.2123676 | 186.7386644 | 192.4454461 | 3.87E-06 | 0.900893902 | 9.03E-07 | 24.12377194 | 1.867222567 | 5.412028743 |
| ENSMUSG00000048612 | *Myof* | 1910.474145 | 2031.105326 | 1912.162113 | 3714.700094 | 3470.862013 | 3732.479427 | 1.21E-43 | 0.899685527 | 5.11E-45 | 198.2209529 | 1.865659269 | 42.91863813 |
| ENSMUSG00000054836 | *Elp6* | 342.796563 | 396.9837698 | 354.4304582 | 661.5537883 | 709.7974745 | 663.936789 | 2.97E-15 | 0.898411352 | 3.45E-16 | 66.52818054 | 1.864012263 | 14.52735094 |
| ENSMUSG00000046057 | *Gm15428* | 747.6877889 | 711.4916871 | 762.7108479 | 1311.480972 | 1492.956567 | 1336.533623 | 2.29E-23 | 0.898236395 | 1.81E-24 | 104.2174069 | 1.863786228 | 22.64025115 |
| ENSMUSG00000042978 | *Sbk1* | 170.9729756 | 163.8520269 | 134.1352839 | 336.0088661 | 280.1079966 | 259.8013522 | 1.33E-06 | 0.898009791 | 2.95E-07 | 26.28480382 | 1.863493505 | 5.876529003 |
| ENSMUSG00000015961 | *Adss* | 2914.196092 | 3019.715877 | 2982.307115 | 5749.355858 | 5369.689351 | 5494.317485 | 4.82E-57 | 0.897834491 | 1.52E-58 | 260.2448144 | 1.863267088 | 56.31708138 |
| ENSMUSG00000054517 | *Trim65* | 213.5035666 | 216.6365724 | 206.587919 | 432.5096823 | 369.6663357 | 383.9286649 | 3.22E-10 | 0.895846095 | 5.17E-11 | 43.1127264 | 1.860700807 | 9.492170703 |
| ENSMUSG00000029822 | *Osbpl3* | 348.7508458 | 366.1927848 | 380.8658791 | 733.6387353 | 652.6325772 | 652.3900622 | 3.55E-15 | 0.895113552 | 4.13E-16 | 66.17295053 | 1.859756258 | 14.45036454 |
| ENSMUSG00000041220 | *Elovl6* | 1398.40583 | 1330.610419 | 1350.164646 | 2632.263228 | 2153.211131 | 2797.194559 | 6.29E-19 | 0.893832059 | 6.16E-20 | 83.56584592 | 1.858105037 | 18.20136669 |
| ENSMUSG00000074865 | *Zfp934* | 68.89955734 | 47.2861554 | 43.0799452 | 102.3141184 | 91.46383564 | 103.9205409 | 0.003503206 | 0.893828725 | 0.001266769 | 10.39033062 | 1.858100743 | 2.45553429 |
| ENSMUSG00000038518 | *Jarid2* | 909.3040345 | 895.1379185 | 912.5115664 | 1798.635695 | 1508.20054 | 1744.517969 | 8.29E-23 | 0.893734617 | 6.69E-24 | 101.6303801 | 1.857979543 | 22.08169898 |
| ENSMUSG00000004099 | *Dnmt1* | 5816.483618 | 6158.196982 | 5787.399001 | 11446.392 | 10592.65546 | 10949.18365 | 8.11E-58 | 0.893237262 | 2.53E-59 | 263.8160998 | 1.857339133 | 57.09075675 |
| ENSMUSG00000011267 | *Zfp296* | 107.1770892 | 144.0578223 | 110.637132 | 232.5320873 | 237.2343237 | 200.1432639 | 1.18E-05 | 0.893190057 | 2.91E-06 | 21.87541945 | 1.857278362 | 4.926626777 |
| ENSMUSG00000042557 | *Sin3a* | 1472.409059 | 1514.256651 | 1534.233503 | 2881.072561 | 2718.190865 | 2797.194559 | 8.63E-44 | 0.893167279 | 3.63E-45 | 198.8972071 | 1.857249038 | 43.06410036 |
| ENSMUSG00000041995 | *Zbed3* | 522.2756569 | 547.6396602 | 491.5030111 | 991.7493522 | 948.9372947 | 956.453867 | 3.90E-21 | 0.892087531 | 3.39E-22 | 93.85945019 | 1.855859548 | 20.4086412 |
| ENSMUSG00000070565 | *Rasal2* | 647.3155942 | 569.6332209 | 578.6419911 | 1155.684474 | 930.8350773 | 1248.008718 | 1.50E-12 | 0.89141074 | 2.05E-13 | 53.95420045 | 1.854989139 | 11.82262252 |
| ENSMUSG00000048911 | *Rnf24* | 627.7515224 | 641.112293 | 627.5964743 | 1206.841533 | 1095.660531 | 1211.444083 | 2.82E-23 | 0.889590119 | 2.24E-24 | 103.8001179 | 1.852649696 | 22.55028554 |
| ENSMUSG00000037111 | *Setd7* | 1288.676906 | 1250.333923 | 1344.290108 | 2656.679097 | 2093.187988 | 2448.868301 | 2.54E-20 | 0.889516329 | 2.29E-21 | 90.0799261 | 1.85255494 | 19.5946995 |
| ENSMUSG00000074918 | *Inafm2* | 333.439833 | 289.2153226 | 328.9741269 | 620.860673 | 561.1687416 | 583.1097016 | 2.51E-13 | 0.887840772 | 3.28E-14 | 57.55972939 | 1.850404618 | 12.59970156 |
| ENSMUSG00000031584 | *Gsr* | 2570.548917 | 2385.201653 | 2495.699552 | 4563.442213 | 4647.506148 | 4580.201617 | 1.87E-53 | 0.887474161 | 6.31E-55 | 243.6452603 | 1.849934462 | 52.72760226 |
| ENSMUSG00000031983 | *2310022B05Rik* | 207.5492838 | 186.9452655 | 196.7970224 | 361.5873957 | 374.4300771 | 357.9485297 | 4.23E-10 | 0.885619239 | 6.86E-11 | 42.55970147 | 1.847557467 | 9.37327898 |
| ENSMUSG00000019471 | *Cdc37* | 9953.859506 | 10135.73243 | 10534.02569 | 18823.47246 | 20357.37267 | 17392.25719 | 1.52E-40 | 0.885517531 | 7.01E-42 | 183.8453948 | 1.847427221 | 39.8172813 |
| ENSMUSG00000055334 | *Snupn* | 369.1655294 | 324.4050196 | 339.7441132 | 602.258106 | 652.6325772 | 655.2767439 | 1.82E-14 | 0.885033209 | 2.22E-15 | 62.86321454 | 1.846807132 | 13.74010465 |
| ENSMUSG00000040883 | *Tmem205* | 632.8551933 | 711.4916871 | 653.0528056 | 1225.4441 | 1376.721276 | 1082.505634 | 2.49E-15 | 0.884707877 | 2.88E-16 | 66.88417051 | 1.846390718 | 14.60372258 |
| ENSMUSG00000022021 | *Diaph3* | 351.3026812 | 380.4885993 | 347.5768306 | 691.7829596 | 677.4040327 | 622.561018 | 4.36E-15 | 0.884550185 | 5.12E-16 | 65.74999603 | 1.846188912 | 14.36022651 |
| ENSMUSG00000038836 | *Agbl3* | 104.6252537 | 116.5658715 | 89.09715938 | 198.8149346 | 181.974923 | 191.4832188 | 1.56E-05 | 0.884334991 | 3.89E-06 | 21.31735833 | 1.845913553 | 4.806762461 |
| ENSMUSG00000042215 | *Bag2* | 569.9099187 | 556.4370845 | 605.0774121 | 974.3094456 | 1128.053973 | 1088.278998 | 5.64E-19 | 0.882820984 | 5.51E-20 | 83.78870984 | 1.843977413 | 18.24849765 |
| ENSMUSG00000009108 | *Gnat2* | 50.18609732 | 45.08679934 | 40.1426762 | 81.38623054 | 79.07810789 | 89.48713242 | 0.003751459 | 0.882009063 | 0.001363231 | 10.25488545 | 1.84293995 | 2.425799765 |
| ENSMUSG00000023982 | *Guca1a* | 258.585993 | 268.3214399 | 237.9187882 | 451.1122493 | 539.2555309 | 417.606618 | 3.26E-09 | 0.881622022 | 5.68E-10 | 38.42909435 | 1.842445599 | 8.486378407 |
| ENSMUSG00000038178 | *Slc43a2* | 1062.414162 | 1107.375779 | 1123.994934 | 2152.084468 | 1886.44161 | 2030.299456 | 2.49E-28 | 0.881417964 | 1.66E-29 | 127.2244786 | 1.842185017 | 27.60467285 |
| ENSMUSG00000054409 | *Tmem74* | 92.71668827 | 114.3665154 | 108.6789527 | 193.0016324 | 168.636447 | 218.4255813 | 3.58E-05 | 0.881201529 | 9.42E-06 | 19.62570746 | 1.841908671 | 4.445940162 |
| ENSMUSG00000052364 | *B630019K06Rik* | 282.4031239 | 259.5240157 | 261.4169402 | 540.6371029 | 476.3741439 | 465.7179795 | 1.98E-11 | 0.880841719 | 2.90E-12 | 48.75676034 | 1.841449354 | 10.70240435 |
| ENSMUSG00000062995 | *Ica1* | 527.3793278 | 539.941914 | 556.1229289 | 867.3446855 | 1131.864966 | 979.5473205 | 9.95E-14 | 0.877379962 | 1.26E-14 | 59.44463815 | 1.83703608 | 13.00232346 |
| ENSMUSG00000042498 | *Radx* | 273.0463939 | 243.0288452 | 260.4378505 | 501.1066481 | 356.3278597 | 570.6007476 | 2.23E-06 | 0.877143962 | 5.07E-07 | 25.2351982 | 1.836735598 | 5.651489777 |
| ENSMUSG00000058360 | *Gm10040* | 176.0766465 | 152.8552465 | 159.5916152 | 374.3766605 | 238.187072 | 287.7059419 | 2.05E-05 | 0.876766246 | 5.19E-06 | 20.76725604 | 1.83625478 | 4.689185201 |
| ENSMUSG00000001281 | *Itgb7* | 735.7792234 | 691.6974825 | 685.3627645 | 1317.294274 | 1357.66631 | 1206.632947 | 1.84E-22 | 0.876301364 | 1.50E-23 | 100.0312064 | 1.835663176 | 21.73583446 |
| ENSMUSG00000024208 | *Uqcc2* | 953.5358491 | 910.533411 | 1049.584119 | 1523.085172 | 2305.650857 | 1516.470115 | 1.00E-08 | 0.875827901 | 1.81E-09 | 36.16891101 | 1.835060848 | 7.999351642 |
| ENSMUSG00000017390 | *Aldoc* | 1695.269355 | 1657.214795 | 1625.288841 | 3054.308966 | 3115.486901 | 2961.735415 | 9.09E-45 | 0.875019358 | 3.72E-46 | 203.4361095 | 1.834032696 | 44.04164052 |
| ENSMUSG00000003779 | *Kif20a* | 3067.306219 | 2951.535839 | 3035.177957 | 5555.191565 | 5549.758777 | 5503.939758 | 9.08E-64 | 0.874905662 | 2.53E-65 | 291.3463607 | 1.833888164 | 63.04174846 |
| ENSMUSG00000044982 | *Sft2d3* | 88.46362917 | 85.77488654 | 126.3025666 | 195.3269533 | 177.2111815 | 178.9742648 | 0.000134484 | 0.874572933 | 3.82E-05 | 16.95683241 | 1.833465264 | 3.871329341 |
| ENSMUSG00000018983 | *E2f2* | 432.110804 | 470.6621979 | 486.6075628 | 861.5313833 | 843.1822348 | 839.0621449 | 2.21E-18 | 0.873798384 | 2.22E-19 | 81.0356355 | 1.832481183 | 17.65503995 |
| ENSMUSG00000001288 | *Rarg* | 214.3541784 | 211.1381823 | 210.5042777 | 405.7684923 | 391.5795463 | 368.5330292 | 2.09E-10 | 0.873070955 | 3.31E-11 | 43.98382496 | 1.83155745 | 9.680448904 |
| ENSMUSG00000043432 | *Leng9* | 56.14038005 | 50.5851895 | 50.9126625 | 104.6394393 | 82.88910105 | 101.9960864 | 0.002251647 | 0.872725096 | 0.000788401 | 11.26834217 | 1.831118422 | 2.6474996 |
| ENSMUSG00000046410 | *Kcnk6* | 674.5351725 | 741.182994 | 729.4217993 | 1391.704542 | 1193.793605 | 1340.382532 | 8.81E-20 | 0.872326771 | 8.15E-21 | 87.56611029 | 1.830612923 | 19.05490375 |
| ENSMUSG00000040936 | *Ulk4* | 54.43915641 | 59.38261376 | 66.57809712 | 102.3141184 | 119.093536 | 107.7694498 | 0.000987144 | 0.871229957 | 0.000326108 | 12.91414758 | 1.829221723 | 3.005619463 |
| ENSMUSG00000036430 | *Tbcc* | 284.9549594 | 288.1156445 | 306.4550647 | 552.2637073 | 603.0896662 | 453.2090255 | 1.81E-09 | 0.870839381 | 3.08E-10 | 39.62542829 | 1.828726571 | 8.742639019 |
| ENSMUSG00000027087 | *Itgav* | 1767.57136 | 1744.08936 | 1683.055132 | 3190.340237 | 2811.560198 | 3489.035937 | 2.73E-23 | 0.869326519 | 2.17E-24 | 103.8624184 | 1.826809908 | 22.56356661 |
| ENSMUSG00000050860 | *Phospho1* | 222.0096848 | 246.3278793 | 224.2115329 | 460.4135328 | 395.3905395 | 408.9465729 | 6.28E-10 | 0.869023529 | 1.03E-10 | 41.76537145 | 1.826426287 | 9.202361569 |
| ENSMUSG00000047821 | *Trim16* | 106.3264774 | 115.4661934 | 90.07624905 | 211.6041994 | 176.2584333 | 181.8609465 | 3.63E-05 | 0.868547712 | 9.55E-06 | 19.59983294 | 1.825824011 | 4.440356702 |
| ENSMUSG00000025578 | *Cbx8* | 223.7109084 | 207.8391482 | 216.3788156 | 410.419134 | 421.1147432 | 353.1373935 | 2.50E-09 | 0.868542513 | 4.30E-10 | 38.97146705 | 1.825817431 | 8.602407754 |
| ENSMUSG00000001517 | *Foxm1* | 1859.437436 | 1929.934947 | 1845.584016 | 3556.578275 | 3242.202424 | 3485.187028 | 1.64E-39 | 0.867908601 | 7.65E-41 | 179.0925919 | 1.825015354 | 38.78602132 |
| ENSMUSG00000045767 | *B230219D22Rik* | 1040.298255 | 978.713449 | 930.1351804 | 1809.099639 | 1644.443545 | 1930.227824 | 8.69E-22 | 0.867505537 | 7.28E-23 | 96.9037623 | 1.824505546 | 21.06112335 |
| ENSMUSG00000022474 | *Pmm1* | 313.8757612 | 285.9162885 | 324.0786786 | 606.9087478 | 559.263245 | 522.4893861 | 4.72E-12 | 0.867236725 | 6.67E-13 | 51.6395914 | 1.824165625 | 11.32627185 |
| ENSMUSG00000043987 | *Cep164* | 795.3220507 | 813.7617441 | 785.2299102 | 1506.807925 | 1404.350976 | 1455.8498 | 6.33E-28 | 0.866799686 | 4.27E-29 | 125.3493302 | 1.82361311 | 27.19875505 |
| ENSMUSG00000058173 | *Smco4* | 463.5834413 | 461.8647737 | 502.2729974 | 884.7845921 | 879.3866697 | 840.0243721 | 8.06E-19 | 0.866791465 | 7.95E-20 | 83.06197082 | 1.823602718 | 18.09339666 |
| ENSMUSG00000018401 | *Mtmr4* | 641.3613115 | 667.5045658 | 719.6309027 | 1202.190891 | 1259.533237 | 1233.575309 | 5.77E-23 | 0.866354639 | 4.64E-24 | 102.3573679 | 1.823050643 | 22.23849059 |
| ENSMUSG00000032398 | *Snapc5* | 245.8268157 | 237.530455 | 252.6051332 | 420.883078 | 442.0752056 | 478.2269335 | 5.31E-11 | 0.866329158 | 8.02E-12 | 46.75963602 | 1.823018444 | 10.27479689 |
| ENSMUSG00000032290 | *Ptpn9* | 1711.43098 | 1778.179379 | 1613.539766 | 3242.659957 | 2912.551516 | 3145.520816 | 2.37E-32 | 0.865871118 | 1.37E-33 | 145.8904352 | 1.822439748 | 31.62598427 |
| ENSMUSG00000047412 | *Zbtb44* | 1122.807601 | 1100.777711 | 1065.249554 | 2033.493103 | 1854.048168 | 2103.428726 | 1.80E-27 | 0.864700533 | 1.23E-28 | 123.2425829 | 1.820961642 | 26.74510355 |
| ENSMUSG00000039852 | *Rere* | 1064.115386 | 1151.3629 | 1201.343017 | 2193.940243 | 1895.969093 | 2130.371088 | 6.47E-23 | 0.864528443 | 5.20E-24 | 102.1314652 | 1.820744444 | 22.18933666 |
| ENSMUSG00000022679 | *Mpv17l* | 113.1313719 | 115.4661934 | 136.0934632 | 218.580162 | 226.7540925 | 217.4633541 | 3.26E-06 | 0.862736778 | 7.55E-07 | 24.46868492 | 1.818484688 | 5.486461369 |
| ENSMUSG00000060032 | *H2aj* | 649.8674297 | 642.211971 | 738.2336063 | 1204.516212 | 1323.367372 | 1163.332722 | 3.65E-18 | 0.862619188 | 3.69E-19 | 80.03001505 | 1.818336475 | 17.43728399 |
| ENSMUSG00000033720 | *Sfxn5* | 102.0734183 | 81.37617441 | 88.11806972 | 170.9110841 | 161.0144607 | 163.5786292 | 6.11E-05 | 0.861928717 | 1.66E-05 | 18.5419325 | 1.817466431 | 4.214014594 |
| ENSMUSG00000024960 | *Plcb3* | 915.2583172 | 949.0221421 | 913.4906561 | 1738.177352 | 1678.742483 | 1625.201792 | 3.66E-30 | 0.860134683 | 2.30E-31 | 135.7172644 | 1.815207762 | 29.43694065 |
| ENSMUSG00000005802 | *Slc30a4* | 1509.835979 | 1481.26631 | 1412.826384 | 2899.675128 | 2280.879401 | 2816.439103 | 1.82E-18 | 0.859997636 | 1.81E-19 | 81.43221106 | 1.815035336 | 17.74040225 |
| ENSMUSG00000032038 | *St3gal4* | 639.6600879 | 673.0029559 | 651.0946263 | 1178.937682 | 1225.234298 | 1157.559358 | 3.57E-24 | 0.859996668 | 2.73E-25 | 107.9669678 | 1.815034119 | 23.44745332 |
| ENSMUSG00000032849 | *Abcc4* | 871.8771145 | 855.5495093 | 1070.145002 | 1670.743047 | 1539.641233 | 1866.720827 | 2.66E-14 | 0.85981098 | 3.27E-15 | 62.09880473 | 1.814800522 | 13.57494942 |
| ENSMUSG00000061536 | *Sec22c* | 240.7231448 | 313.4082393 | 264.3542092 | 532.4984798 | 473.5158991 | 476.302479 | 2.51E-09 | 0.859693267 | 4.33E-10 | 38.95961684 | 1.814652455 | 8.599944557 |
| ENSMUSG00000026873 | *Phf19* | 586.9221551 | 566.3341868 | 569.8301842 | 1101.039433 | 1034.684641 | 992.0562745 | 1.32E-20 | 0.858578138 | 1.17E-21 | 91.40097416 | 1.813250364 | 19.87865255 |
| ENSMUSG00000020493 | *Prr11* | 1127.06066 | 1127.169983 | 1030.981416 | 1949.781552 | 1966.472466 | 2036.072819 | 1.33E-30 | 0.857643697 | 8.24E-32 | 137.7559295 | 1.812076293 | 29.87758191 |
| ENSMUSG00000030867 | *Plk1* | 1714.833427 | 1716.597409 | 1644.870635 | 3176.388312 | 3213.619975 | 2806.816831 | 7.61E-30 | 0.857075976 | 4.85E-31 | 134.2360662 | 1.811363354 | 29.11860007 |
| ENSMUSG00000032185 | *Carm1* | 2274.536004 | 2359.909058 | 2338.066117 | 4348.350032 | 4045.36923 | 4210.70636 | 1.80E-46 | 0.85420217 | 7.09E-48 | 211.3151077 | 1.807758764 | 45.74513832 |
| ENSMUSG00000003308 | *Keap1* | 2362.149021 | 2310.423546 | 2473.18049 | 4451.826811 | 4485.538939 | 3981.696279 | 6.43E-34 | 0.85398564 | 3.51E-35 | 153.1720017 | 1.807487462 | 33.19165198 |
| ENSMUSG00000051098 | *Mblac2* | 96.11913554 | 73.67842818 | 101.825325 | 179.0497072 | 118.1407877 | 195.3321278 | 0.001836627 | 0.853787769 | 0.000634515 | 11.67189174 | 1.807239576 | 2.735979096 |
| ENSMUSG00000030852 | *Tacc2* | 575.8642015 | 633.4145468 | 616.826488 | 1188.238966 | 1020.393416 | 1089.241225 | 8.30E-18 | 0.85295101 | 8.56E-19 | 78.36676611 | 1.806191685 | 17.08071936 |
| ENSMUSG00000062397 | *Zfp706* | 1323.55199 | 1281.124908 | 1276.732921 | 2417.171047 | 2252.296953 | 2343.985533 | 1.37E-35 | 0.852612243 | 7.12E-37 | 160.9220994 | 1.805767614 | 34.86333289 |
| ENSMUSG00000049807 | *Arhgap23* | 234.768862 | 261.7233718 | 190.9224844 | 434.8350032 | 382.0520634 | 423.3799814 | 8.16E-08 | 0.852199086 | 1.61E-08 | 31.91261216 | 1.805250555 | 7.088118319 |
| ENSMUSG00000073557 | *Ppp1r12b* | 458.4797704 | 442.0705691 | 432.7576313 | 804.5610219 | 794.5920721 | 806.346419 | 2.84E-18 | 0.850494533 | 2.86E-19 | 80.53467263 | 1.8031189 | 17.54642688 |
| ENSMUSG00000063851 | *Rnf183* | 147.1558447 | 140.7587882 | 141.9680012 | 254.6226356 | 269.6277655 | 251.1413071 | 2.84E-07 | 0.850374302 | 5.86E-08 | 29.40835831 | 1.802968638 | 6.547054664 |
| ENSMUSG00000005057 | *Sh2b2* | 554.598906 | 586.1283914 | 584.5165291 | 1089.412829 | 1034.684641 | 984.3584567 | 6.50E-20 | 0.849477552 | 5.96E-21 | 88.18468353 | 1.801848298 | 19.18716233 |
| ENSMUSG00000024891 | *Slc29a2* | 209.2505075 | 158.3536367 | 178.1943188 | 341.8221683 | 334.414649 | 310.7993954 | 2.79E-07 | 0.8491434 | 5.75E-08 | 29.44486648 | 1.801431009 | 6.554514141 |
| ENSMUSG00000009772 | *Nuak2* | 319.8300439 | 317.8069514 | 327.9950373 | 569.7036138 | 546.8775172 | 622.561018 | 7.16E-13 | 0.849010201 | 9.62E-14 | 55.44357374 | 1.801264696 | 12.1449245 |
| ENSMUSG00000053559 | *Smagp* | 108.027701 | 104.4694131 | 113.574401 | 199.977595 | 204.8408819 | 182.8231738 | 1.08E-05 | 0.848695743 | 2.63E-06 | 22.06617739 | 1.800872125 | 4.967360441 |
| ENSMUSG00000058135 | *Gstm1* | 223.7109084 | 227.6333527 | 204.6297397 | 399.9551901 | 403.9652741 | 377.1930743 | 7.73E-10 | 0.848403138 | 1.28E-10 | 41.3393821 | 1.800506912 | 9.111708514 |
| ENSMUSG00000035413 | *Tmem98* | 1001.170111 | 1023.800248 | 1045.667761 | 1821.888904 | 1927.409786 | 1777.233694 | 9.05E-30 | 0.848362876 | 5.79E-31 | 133.8832153 | 1.800456666 | 29.04328209 |
| ENSMUSG00000079003 | *Samd1* | 802.9775571 | 765.3759107 | 807.7489724 | 1518.43453 | 1434.838922 | 1327.873578 | 3.19E-22 | 0.847898124 | 2.64E-23 | 98.91482931 | 1.799876757 | 21.49574457 |
| ENSMUSG00000010277 | *Bltp2* | 10833.39213 | 11133.1404 | 10564.37747 | 19944.27712 | 19080.68996 | 19513.96823 | 3.58E-71 | 0.847604396 | 8.85E-73 | 325.5738529 | 1.799510345 | 70.44589616 |
| ENSMUSG00000040524 | *Zfp609* | 1080.27701 | 1122.771271 | 1013.357802 | 1957.920175 | 1766.395326 | 2057.241819 | 2.95E-22 | 0.846192589 | 2.43E-23 | 99.0783548 | 1.797750223 | 21.53052108 |
| ENSMUSG00000015357 | *Clpx* | 883.7856799 | 921.5301913 | 884.1179662 | 1644.001857 | 1592.995137 | 1594.410521 | 8.67E-30 | 0.845470416 | 5.54E-31 | 133.9732408 | 1.796850545 | 29.06204277 |
| ENSMUSG00000026198 | *Abcb6* | 988.4109337 | 1020.501214 | 1043.709581 | 1741.665334 | 1916.929555 | 1817.647238 | 4.66E-28 | 0.844250246 | 3.14E-29 | 125.9605923 | 1.795331488 | 27.33164458 |
| ENSMUSG00000003131 | *Pafah1b2* | 3651.676539 | 3623.439117 | 3489.475561 | 6287.66764 | 6432.95644 | 6597.992119 | 3.24E-56 | 0.843804598 | 1.05E-57 | 256.3980095 | 1.794776996 | 55.48975738 |
| ENSMUSG00000021559 | *Dapk1* | 85.91179372 | 90.17359867 | 65.59900746 | 179.0497072 | 121.9517808 | 133.749585 | 0.001593042 | 0.84374186 | 0.000544131 | 11.95800594 | 1.794698949 | 2.797772878 |
| ENSMUSG00000034485 | *Uaca* | 603.9343915 | 648.8100392 | 592.3492464 | 1203.353552 | 968.9450088 | 1139.277041 | 1.28E-14 | 0.843568881 | 1.54E-15 | 63.57363534 | 1.794483778 | 13.89280656 |
| ENSMUSG00000042213 | *Zfand4* | 105.4758656 | 91.2732767 | 83.2226214 | 170.9110841 | 161.0144607 | 171.276447 | 8.63E-05 | 0.842075708 | 2.39E-05 | 17.84979049 | 1.792627468 | 4.064214359 |
| ENSMUSG00000039202 | *Abhd2* | 1768.421972 | 1734.192257 | 1668.368787 | 3132.207215 | 2791.552484 | 3346.626307 | 8.85E-25 | 0.841904977 | 6.61E-26 | 110.7817177 | 1.792415338 | 24.05312397 |
| ENSMUSG00000025154 | *Arhgap19* | 64.64649824 | 56.08357966 | 79.30626275 | 131.3806293 | 92.41658392 | 135.6740395 | 0.003707219 | 0.841796546 | 0.001346392 | 10.27781673 | 1.792280628 | 2.430951702 |
| ENSMUSG00000005774 | *Rfx5* | 355.5557403 | 407.9805501 | 349.5350099 | 660.3911278 | 642.152346 | 688.9546969 | 2.86E-13 | 0.84173174 | 3.75E-14 | 57.29433866 | 1.79220012 | 12.54315267 |
| ENSMUSG00000025510 | *Cd151* | 819.9897935 | 898.4369526 | 889.9925042 | 1514.946549 | 1696.844701 | 1456.812027 | 5.66E-20 | 0.841221212 | 5.18E-21 | 88.46143637 | 1.791566025 | 19.24694868 |
| ENSMUSG00000043391 | *2510009E07Rik* | 3161.724131 | 3284.738283 | 3205.539558 | 6031.882344 | 5219.155121 | 6035.089189 | 1.76E-31 | 0.840665439 | 1.05E-32 | 141.8425961 | 1.790875988 | 30.75563396 |
| ENSMUSG00000071041 | *Impdh2-ps* | 673.6845606 | 662.0061756 | 507.1684457 | 955.7068787 | 1242.383767 | 1099.825724 | 1.45E-09 | 0.840376016 | 2.45E-10 | 40.06682211 | 1.790516752 | 8.83726606 |

| | | | | | | | | | | | | | |
|---|---|---|---|---|---|---|---|---|---|---|---|---|---|
| ENSMUSG00000029461 | *Fam168a* | 604.7850033 | 646.6106832 | 635.4291916 | 1139.407228 | 1029.920899 | 1204.708492 | 9.02E-18 | 0.839222566 | 9.31E-19 | 78.19964214 | 1.789085786 | 17.04455433 |
| ENSMUSG00000040158 | *Tax1bp3* | 461.8822177 | 461.8647737 | 508.1475354 | 854.5554207 | 881.2921663 | 823.6665092 | 5.60E-17 | 0.838020725 | 5.99E-18 | 74.52356273 | 1.787596004 | 16.25142542 |
| ENSMUSG00000041731 | *Pgm5* | 142.9027856 | 136.360076 | 167.4243325 | 274.387863 | 282.9662415 | 241.5190348 | 2.42E-06 | 0.837523019 | 5.52E-07 | 25.07337429 | 1.786979419 | 5.617007667 |
| ENSMUSG00000034413 | *Neurl1b* | 224.5615202 | 204.5401141 | 215.399726 | 417.3950966 | 337.2728939 | 398.3620734 | 3.49E-08 | 0.835910989 | 6.63E-09 | 33.64041359 | 1.784983811 | 7.457695553 |
| ENSMUSG00000047514 | *Tspyl1* | 843.8069244 | 835.7553048 | 788.1671791 | 1467.277471 | 1527.255505 | 1410.62512 | 1.10E-24 | 0.83586133 | 8.25E-26 | 110.3419313 | 1.78492237 | 23.95876804 |
| ENSMUSG00000030539 | *Sema4b* | 537.5866696 | 563.0351527 | 516.9593423 | 1044.069072 | 923.213091 | 919.8892322 | 1.51E-16 | 0.835181161 | 1.64E-17 | 72.53236327 | 1.784081054 | 15.82241385 |
| ENSMUSG00000032598 | *Nckipsd* | 372.5679767 | 390.3857016 | 371.0749825 | 699.9215827 | 621.1918837 | 701.4636509 | 1.14E-13 | 0.834799649 | 1.44E-14 | 59.18003584 | 1.783609327 | 12.94486883 |
| ENSMUSG00000032044 | *Rpusd4* | 1360.97891 | 1450.475325 | 1415.763653 | 2517.159845 | 2535.263194 | 2470.999528 | 4.97E-37 | 0.832512358 | 2.48E-38 | 167.5987026 | 1.780783781 | 36.30370269 |
| ENSMUSG00000032312 | *Csk* | 4458.907155 | 4506.480577 | 4789.706634 | 8368.829821 | 8420.389368 | 7675.686617 | 3.06E-41 | 0.83069516 | 1.38E-42 | 187.0826986 | 1.778542144 | 40.51484635 |
| ENSMUSG00000032078 | *Zpr1* | 1872.196614 | 1771.58131 | 1895.517589 | 3222.89473 | 3400.358639 | 3229.234585 | 1.74E-38 | 0.830417903 | 8.31E-40 | 174.3474402 | 1.778200376 | 37.7586956 |
| ENSMUSG00000039206 | *Daglb* | 986.70971 | 1110.674813 | 1062.312285 | 1900.949813 | 1905.496576 | 1799.364921 | 9.13E-26 | 0.828446056 | 6.55E-27 | 115.3640832 | 1.775771627 | 25.03931142 |
| ENSMUSG00000074476 | *Spc24* | 717.9163752 | 680.7007022 | 694.1745714 | 1224.281439 | 1333.847603 | 1159.483813 | 2.18E-19 | 0.828405951 | 2.07E-20 | 85.72438103 | 1.775722264 | 18.66096614 |
| ENSMUSG00000040188 | *Scamp2* | 2208.188282 | 2403.896179 | 2414.43511 | 4213.481421 | 4236.871636 | 4021.147596 | 5.24E-38 | 0.828390459 | 2.53E-39 | 172.1317739 | 1.775703196 | 37.28041154 |
| ENSMUSG00000031627 | *Irf2* | 705.1571979 | 668.6042438 | 678.5091368 | 1268.462536 | 1152.825428 | 1222.028583 | 2.36E-21 | 0.826410132 | 2.02E-22 | 94.88071595 | 1.773267434 | 20.62787565 |
| ENSMUSG00000043067 | *Dpy19l1* | 1730.995052 | 1615.42703 | 1640.954276 | 3061.284929 | 2791.552484 | 2990.602232 | 1.79E-32 | 0.825177022 | 1.04E-33 | 146.4461957 | 1.771752423 | 31.74593622 |
| ENSMUSG00000105383 | *None* | 158.2137983 | 222.1349626 | 215.399726 | 324.3822617 | 365.8553425 | 360.8352114 | 2.64E-06 | 0.825022113 | 6.05E-07 | 24.89658978 | 1.771562192 | 5.578228771 |
| ENSMUSG00000014778 | *Fhod1* | 331.7386094 | 373.8905311 | 395.5522241 | 708.0602057 | 641.1995977 | 600.4297917 | 5.03E-11 | 0.824639573 | 7.58E-12 | 46.87226533 | 1.771092513 | 10.29817551 |
| ENSMUSG00000042790 | *Rnf214* | 634.556417 | 629.0158346 | 596.2656051 | 1121.967321 | 1024.204409 | 1146.974859 | 9.90E-19 | 0.823717509 | 9.79E-20 | 82.6510197 | 1.769960923 | 18.00431452 |
| ENSMUSG00000039981 | *Zc3h12d* | 111.4301483 | 91.2732767 | 103.7835043 | 170.9110841 | 182.9276713 | 189.5587644 | 5.66E-05 | 0.823329204 | 1.53E-05 | 18.69790821 | 1.769484598 | 4.247239995 |
| ENSMUSG00000047434 | *Xxylt1* | 603.9343915 | 703.7939408 | 626.6173847 | 1173.12438 | 1064.219838 | 1181.615039 | 1.39E-16 | 0.823325725 | 1.52E-17 | 72.68677294 | 1.76948033 | 15.85557367 |
| ENSMUSG00000049086 | *Bmyc* | 201.5950011 | 237.530455 | 214.4206363 | 429.021701 | 368.7135874 | 356.9863025 | 7.10E-08 | 0.821974952 | 1.39E-08 | 32.1947798 | 1.767824369 | 7.148777465 |
| ENSMUSG00000032249 | *Anp32a* | 3361.617909 | 3414.500291 | 3393.524774 | 5849.344655 | 5838.441508 | 6264.09927 | 2.94E-48 | 0.820149892 | 1.11E-49 | 219.58399 | 1.765589423 | 47.53202 |
| ENSMUSG00000020592 | *Sdc1* | 339.3941158 | 301.3117809 | 347.5768306 | 619.6980126 | 564.0269864 | 564.8273842 | 1.40E-11 | 0.820052784 | 2.02E-12 | 49.46084662 | 1.765470585 | 10.85435051 |
| ENSMUSG00000055633 | *Zfp580* | 64.64649824 | 75.87778425 | 70.49445577 | 134.8686106 | 136.2430052 | 101.0338592 | 0.001793638 | 0.819955491 | 0.000618678 | 11.71892075 | 1.765351528 | 2.746265274 |
| ENSMUSG00000028678 | *Kif2c* | 1578.735536 | 1606.629606 | 1508.777172 | 2603.196717 | 2824.898674 | 2853.003738 | 1.79E-31 | 0.819749332 | 1.08E-32 | 141.8001051 | 1.76509928 | 30.74683961 |
| ENSMUSG00000039153 | *Runx2* | 66.34772188 | 58.28293573 | 68.53627645 | 117.4287041 | 96.22757708 | 127.9762216 | 0.002274272 | 0.819354956 | 0.000797415 | 11.24723928 | 1.764616838 | 2.643157562 |
| ENSMUSG00000040943 | *Tet2* | 980.7554273 | 1012.803468 | 1008.462353 | 1759.10524 | 1454.846636 | 2079.373045 | 2.19E-11 | 0.818402848 | 3.20E-12 | 48.55905336 | 1.763452661 | 10.66000276 |
| ENSMUSG00000028159 | *Dapp1* | 358.1075758 | 355.1960045 | 390.6567758 | 598.7701247 | 691.695257 | 653.3522894 | 2.62E-12 | 0.81749753 | 3.64E-13 | 52.82827874 | 1.762346409 | 11.5815304 |
| ENSMUSG00000039952 | *Dag1* | 2736.418222 | 2741.497335 | 2699.350202 | 5052.922256 | 4643.695155 | 4713.951202 | 1.03E-42 | 0.817001618 | 4.45E-44 | 193.9107667 | 1.761740724 | 41.98771098 |
| ENSMUSG00000027811 | *4930579G24Rik* | 385.327154 | 391.4853796 | 367.1586238 | 658.065807 | 669.7820464 | 686.0680152 | 1.16E-14 | 0.816611482 | 1.40E-15 | 63.76958161 | 1.761264376 | 13.93377671 |
| ENSMUSG00000092060 | *Bend4* | 1481.765789 | 1506.558905 | 1391.286412 | 2695.046891 | 2257.060694 | 2762.554378 | 5.89E-19 | 0.816542468 | 5.76E-20 | 83.69959032 | 1.761180125 | 18.23013965 |
| ENSMUSG00000048497 | *Mmgt2* | 529.0805514 | 489.3567245 | 539.4784046 | 903.387159 | 926.0713358 | 914.1158688 | 7.22E-18 | 0.815354554 | 7.39E-19 | 78.65573913 | 1.759730567 | 17.14134802 |
| ENSMUSG00000064264 | *Zfp428* | 71.45139279 | 79.17681835 | 90.07624905 | 144.1698941 | 119.093536 | 159.7297202 | 0.001041004 | 0.81476698 | 0.000344716 | 12.81030365 | 1.759014017 | 2.982547591 |
| ENSMUSG00000051517 | *Arhgef39* | 427.8577449 | 506.951573 | 493.4611904 | 870.8326668 | 838.4184933 | 799.6108284 | 7.46E-14 | 0.81468817 | 9.38E-15 | 60.02127973 | 1.758917931 | 13.12720805 |
| ENSMUSG00000060600 | *Eno3* | 393.8332722 | 426.6750766 | 411.2176587 | 710.3855266 | 804.119555 | 646.6166988 | 2.27E-11 | 0.812520414 | 3.33E-12 | 48.48223273 | 1.756277012 | 10.64380338 |
| ENSMUSG00000029570 | *Lfng* | 781.7122616 | 795.0672176 | 775.4390135 | 1403.331147 | 1396.72899 | 1330.76026 | 1.06E-24 | 0.812365103 | 7.90E-26 | 110.4258749 | 1.756087952 | 23.97636302 |
| ENSMUSG00000039713 | *Plekhg5* | 91.01546463 | 90.17359867 | 84.20171106 | 163.9351215 | 149.5814812 | 152.9941296 | 0.000153609 | 0.812198056 | 4.41E-05 | 16.687626 | 1.755884631 | 3.813582984 |
| ENSMUSG00000029270 | *Dipk1a* | 184.5827647 | 167.151061 | 161.5497945 | 288.3397882 | 306.7849487 | 305.9882593 | 2.43E-07 | 0.810657406 | 5.00E-08 | 29.71836226 | 1.754010526 | 6.6142013 |
| ENSMUSG00000032508 | *Myd88* | 2597.768495 | 2626.031142 | 2726.764713 | 4633.201839 | 4755.166705 | 4552.297027 | 8.14E-47 | 0.810330357 | 3.18E-48 | 212.9096116 | 1.75361295 | 46.08920479 |
| ENSMUSG00000026739 | *Bmi1* | 2020.20307 | 2013.510478 | 2046.297397 | 3617.036617 | 3354.626722 | 3683.405838 | 3.42E-36 | 0.809160724 | 1.74E-37 | 163.7255626 | 1.752191823 | 35.46652506 |
| ENSMUSG00000021607 | *Mrpl36* | 588.6233787 | 565.2345087 | 552.2065702 | 959.19486 | 1020.393416 | 1008.414137 | 7.61E-19 | 0.808437393 | 7.48E-20 | 83.18261703 | 1.751313538 | 18.11869208 |
| ENSMUSG00000037071 | *Scd1* | 7325.468985 | 7626.267156 | 7201.204475 | 13343.85383 | 12366.67278 | 13080.51697 | 1.22E-50 | 0.808255955 | 4.40E-52 | 230.6042077 | 1.751093301 | 49.91217682 |
| ENSMUSG00000000594 | *Gm2a* | 969.6974736 | 902.8356647 | 945.800615 | 1687.020293 | 1643.490797 | 1608.843929 | 1.76E-26 | 0.808013089 | 1.24E-27 | 118.6658958 | 1.750798544 | 25.75519484 |
| ENSMUSG00000042599 | *Kdm7a* | 995.2158282 | 987.5108733 | 959.5078703 | 1820.726243 | 1438.649915 | 1892.700962 | 2.25E-13 | 0.80769345 | 2.92E-14 | 57.78984231 | 1.750410685 | 12.64844761 |
| ENSMUSG00000063849 | *Ppcdc* | 1157.682686 | 1132.668374 | 1082.873168 | 1986.986686 | 1912.165814 | 2005.281548 | 2.45E-29 | 0.807161513 | 1.59E-30 | 131.8847553 | 1.749765409 | 28.6102729 |
| ENSMUSG00000040212 | *Emp3* | 4428.28513 | 4629.644517 | 4587.035074 | 7806.10217 | 8572.829094 | 7476.50558 | 1.35E-34 | 0.806236004 | 7.26E-36 | 156.3064285 | 1.74864327 | 33.86836571 |
| ENSMUSG00000031824 | *6430548M08Rik* | 1156.832074 | 1190.951309 | 1268.900204 | 2181.150979 | 2059.841798 | 2080.335272 | 6.66E-28 | 0.805500551 | 4.49E-29 | 125.2465565 | 1.747752079 | 27.17670424 |
| ENSMUSG00000032892 | *Rangrf* | 273.0463939 | 234.2314209 | 189.9433947 | 386.0032649 | 434.4532193 | 399.3243006 | 1.35E-06 | 0.805214838 | 2.99E-07 | 26.25466714 | 1.747405986 | 5.870423461 |
| ENSMUSG00000025868 | *Higd2a* | 512.9189268 | 517.9483533 | 573.7465428 | 935.9416513 | 1048.975865 | 817.8931458 | 2.22E-11 | 0.804845966 | 3.26E-12 | 48.52571213 | 1.746959262 | 10.65302589 |
| ENSMUSG00000036768 | *Kif15* | 1536.204945 | 1615.42703 | 1518.568068 | 2674.119004 | 2511.444487 | 2969.433233 | 5.55E-23 | 0.804713233 | 4.45E-24 | 102.4392561 | 1.746798543 | 22.25607145 |
| ENSMUSG00000003575 | *Crtc1* | 346.1990103 | 329.9034098 | 351.4931892 | 635.9752587 | 607.8534077 | 553.2806575 | 9.02E-12 | 0.804473519 | 1.29E-12 | 50.33958539 | 1.746508324 | 11.04471227 |
| ENSMUSG00000030677 | *Kif22* | 1217.225513 | 1254.732635 | 1211.133914 | 2070.698237 | 2310.414598 | 2035.110592 | 2.30E-24 | 0.801531161 | 1.75E-25 | 108.8499352 | 1.742949973 | 23.63740811 |
| ENSMUSG00000021238 | *Aldh6a1* | 62.09466279 | 64.88100392 | 48.95448318 | 103.4767788 | 119.093536 | 83.71376904 | 0.007109839 | 0.80043678 | 0.002723145 | 8.984263386 | 1.74162833 | 2.148140234 |
| ENSMUSG00000028013 | *Ppa2* | 474.641395 | 466.2634858 | 460.1721419 | 837.1155142 | 819.3635276 | 785.17742 | 2.12E-16 | 0.800418676 | 2.34E-17 | 71.83803174 | 1.741606474 | 15.67368257 |
| ENSMUSG00000032126 | *Hmbs* | 1295.4818 | 1287.722976 | 1315.896508 | 2191.614922 | 2446.657603 | 2148.653405 | 6.85E-25 | 0.799934209 | 5.09E-26 | 111.3003186 | 1.741021729 | 24.16433175 |
| ENSMUSG00000032485 | *Scap* | 2328.124549 | 2496.269134 | 2374.292434 | 4221.620044 | 4268.31233 | 4033.65655 | 2.56E-39 | 0.799405309 | 1.20E-40 | 178.1937409 | 1.740383578 | 38.59230109 |
| ENSMUSG00000078619 | *Smarcd2* | 1541.308616 | 1559.34345 | 1552.836206 | 2779.921103 | 2852.528374 | 2467.150619 | 2.38E-25 | 0.799402204 | 1.73E-26 | 113.4381226 | 1.740379832 | 24.62368944 |
| ENSMUSG00000004100 | *Ppan* | 2481.234676 | 2447.883301 | 2593.608519 | 4423.92296 | 4582.719265 | 4083.692366 | 1.07E-32 | 0.799003033 | 6.09E-34 | 147.5038473 | 1.739898363 | 31.97143333 |
| ENSMUSG00000038502 | *Ptov1* | 731.5261643 | 726.8871795 | 800.8953448 | 1332.40886 | 1377.674024 | 1220.104128 | 3.16E-18 | 0.798489684 | 3.18E-19 | 80.32047509 | 1.739279371 | 17.49964607 |
| ENSMUSG00000073434 | *Wdr90* | 335.1410567 | 387.0866675 | 346.5977409 | 696.4336014 | 608.806156 | 553.2806575 | 1.93E-09 | 0.798386328 | 3.29E-10 | 39.49334507 | 1.739154773 | 8.71414028 |
| ENSMUSG00000037411 | *Serpine1* | 1933.440665 | 1956.32722 | 1821.106774 | 3442.637552 | 3239.344179 | 3243.667994 | 1.92E-34 | 0.797125076 | 1.03E-35 | 155.6060505 | 1.73763501 | 33.71698253 |
| ENSMUSG00000026970 | *Rbms1* | 2368.103304 | 2303.825478 | 2327.29613 | 4239.059951 | 3549.940121 | 4372.360535 | 9.36E-21 | 0.79664247 | 8.26E-22 | 92.09564144 | 1.737053839 | 20.02875017 |
| ENSMUSG00000090115 | *Usp49* | 189.6864356 | 213.3375383 | 190.9224844 | 359.2620748 | 345.8476285 | 325.2328039 | 8.30E-08 | 0.796093654 | 1.64E-08 | 31.8774298 | 1.736393172 | 7.080704441 |
| ENSMUSG00000015488 | *Cacfd1* | 435.5132513 | 427.7747547 | 437.6530796 | 862.6940438 | 647.8688358 | 751.4994669 | 6.31E-10 | 0.79603385 | 1.04E-10 | 41.75403915 | 1.736321194 | 9.200026072 |
| ENSMUSG00000022656 | *Nectin3* | 406.5924495 | 530.0448117 | 454.2976039 | 881.2966107 | 725.0414471 | 804.4219646 | 1.73E-09 | 0.79524429 | 2.94E-10 | 39.71101736 | 1.735371197 | 8.761145991 |
| ENSMUSG00000024867 | *Pip5k1b* | 1677.406507 | 1594.533147 | 1631.163379 | 2948.506867 | 2773.450266 | 2790.458968 | 2.71E-34 | 0.79477758 | 1.46E-35 | 154.9146864 | 1.734809898 | 33.56645343 |
| ENSMUSG00000062661 | *Ncs1* | 634.556417 | 699.3952287 | 599.2028741 | 1163.823097 | 1118.52649 | 1066.147771 | 2.45E-16 | 0.793309713 | 2.71E-17 | 71.54266499 | 1.733045717 | 15.61002321 |
| ENSMUSG00000000811 | *Txnrd3* | 80.8081228 | 119.8649056 | 80.28535241 | 182.5376885 | 142.9122432 | 159.7297202 | 0.001785394 | 0.79247926 | 0.00061559 | 11.7282343 | 1.732048418 | 2.748265952 |
| ENSMUSG00000026285 | *Pdcd1* | 883.7856799 | 943.5237519 | 915.4488354 | 1617.260667 | 1662.545762 | 1467.396526 | 9.47E-21 | 0.792040515 | 8.36E-22 | 92.07144123 | 1.731521757 | 20.02377685 |
| ENSMUSG00000036138 | *Acaa1a* | 2981.394426 | 2982.326824 | 2940.20626 | 5141.284449 | 5459.24769 | 4813.060606 | 3.05E-34 | 0.791704068 | 1.65E-35 | 154.6723589 | 1.731118 | 33.51625444 |
| ENSMUSG00000026509 | *Capn2* | 4139.927723 | 4105.098096 | 4158.193801 | 7282.904973 | 7122.7462 | 7060.823416 | 5.07E-59 | 0.791152081 | 1.54E-60 | 269.3912012 | 1.730455787 | 58.29523011 |
| ENSMUSG00000072381 | *Gm10355* | 61.24405097 | 61.58196982 | 89.09715938 | 132.5432897 | 131.4792637 | 102.9583136 | 0.005081455 | 0.791132325 | 0.001889769 | 9.653638896 | 1.73043209 | 2.294011887 |
| ENSMUSG00000032815 | *Fanca* | 608.1874506 | 638.9129369 | 567.8720049 | 1006.863938 | 1081.369307 | 1047.865454 | 2.79E-17 | 0.79039604 | 2.94E-18 | 75.92874075 | 1.729549183 | 16.55461734 |
| ENSMUSG00000074886 | *Grk6* | 1429.027856 | 1447.176291 | 1521.505337 | 2463.677465 | 2719.143614 | 2420.001484 | 1.25E-25 | 0.790243443 | 8.98E-27 | 114.7375395 | 1.729366254 | 24.90375553 |
| ENSMUSG00000078429 | *Ctdsp2* | 1044.551314 | 1050.192521 | 1087.768616 | 1923.040362 | 1746.387612 | 1834.005101 | 2.70E-25 | 0.789560572 | 1.97E-26 | 113.1796343 | 1.728547887 | 24.56789109 |
| ENSMUSG00000025239 | *Limd1* | 3422.86196 | 3451.889344 | 3311.281242 | 6021.4184 | 5825.103032 | 5755.081065 | 4.43E-49 | 0.788953035 | 1.64E-50 | 223.3995874 | 1.727820127 | 48.35313789 |
| ENSMUSG00000057337 | *Chst3* | 227.1133557 | 215.5368944 | 229.1069813 | 404.6058318 | 383.95756 | 373.3441654 | 7.89E-09 | 0.788488164 | 1.41E-09 | 36.65034385 | 1.727263472 | 8.103147494 |
| ENSMUSG00000042105 | *Inpp5f* | 261.1378284 | 215.5368944 | 264.3542092 | 416.2324362 | 411.5872604 | 454.1712527 | 3.98E-08 | 0.788137 | 7.63E-09 | 33.3675634 | 1.726843092 | 7.399737629 |
| ENSMUSG00000027843 | *Ptpn22* | 1565.976359 | 1540.648924 | 1529.338054 | 2645.052493 | 2532.404949 | 2823.174694 | 3.57E-29 | 0.787120523 | 2.31E-30 | 131.1363952 | 1.725626843 | 28.4479319 |
| ENSMUSG00000024948 | *Map4k2* | 387.8789895 | 340.9001901 | 368.1377135 | 662.7164487 | 620.2391354 | 612.9387457 | 5.75E-12 | 0.786232646 | 8.17E-13 | 51.2411948 | 1.724565168 | 11.24024228 |
| ENSMUSG00000070808 | *Bicra* | 227.9639675 | 234.2314209 | 240.8560572 | 476.6907789 | 375.3828254 | 361.7974386 | 7.41E-07 | 0.785540827 | 1.60E-07 | 27.46614226 | 1.723738381 | 6.130415972 |
| ENSMUSG00000035900 | *Gramd4* | 861.6697726 | 858.8485434 | 836.1425727 | 1575.404891 | 1424.35869 | 1410.62512 | 7.47E-21 | 0.785540703 | 6.55E-22 | 92.55368746 | 1.723738232 | 20.12689834 |
| ENSMUSG00000097327 | *E030030I06Rik* | 102.0734183 | 103.3697351 | 122.3862079 | 208.1162181 | 181.974923 | 175.1253559 | 0.000142073 | 0.784072591 | 4.05E-05 | 16.84619464 | 1.721985018 | 3.847488877 |
| ENSMUSG00000026879 | *Gsn* | 6599.897103 | 6468.306188 | 6483.531752 | 11466.15722 | 11750.24463 | 10427.6565 | 1.01E-38 | 0.782910653 | 4.80E-40 | 175.4379027 | 1.7205987 | 37.99431863 |
| ENSMUSG00000039234 | *Sec24d* | 1514.089038 | 1476.867598 | 1485.27902 | 2567.154243 | 2550.507167 | 2584.542341 | 2.43E-36 | 0.782620312 | 1.23E-37 | 164.4108432 | 1.720252466 | 35.61446626 |
| ENSMUSG00000078931 | *Pdf* | 409.9948967 | 380.4885993 | 415.1340173 | 673.1803926 | 708.8447262 | 692.8036059 | 1.44E-13 | 0.782436134 | 1.84E-14 | 58.69629085 | 1.720032869 | 12.84235841 |
| ENSMUSG00000028944 | *Prkag2* | 370.8667531 | 407.9805501 | 359.3259065 | 649.9271839 | 648.8215841 | 656.2389711 | 1.27E-12 | 0.782218647 | 1.72E-13 | 54.29916623 | 1.719773592 | 11.89711668 |
| ENSMUSG00000097487 | *Ptges3l* | 105.4758656 | 105.5690911 | 130.2189253 | 206.9535577 | 213.4156165 | 166.4653109 | 0.00027058 | 0.781274794 | 8.06E-05 | 15.54472564 | 1.718648835 | 3.567704551 |
| ENSMUSG00000044134 | *Pheta1* | 406.5924495 | 413.4789402 | 462.1303212 | 742.9400188 | 732.6634334 | 727.4437862 | 1.96E-13 | 0.781041974 | 2.53E-14 | 58.07212477 | 1.718371503 | 12.70846734 |
| ENSMUSG00000043415 | *Otud1* | 129.2929965 | 155.0546026 | 121.4071183 | 220.9054829 | 233.4233305 | 240.5568076 | 2.70E-05 | 0.78063712 | 6.96E-06 | 20.20404867 | 1.717889356 | 4.569081242 |
| ENSMUSG00000095762 | *Gm16378* | 227.1133557 | 230.9323868 | 290.7896301 | 448.7869284 | 407.7762672 | 430.115572 | 1.98E-07 | 0.780550881 | 4.03E-08 | 30.13362359 | 1.71778667 | 6.702841146 |
| ENSMUSG00000032123 | *Dpagt1* | 970.5480855 | 1047.993165 | 1056.437747 | 1776.545147 | 1742.576619 | 1758.951377 | 2.25E-25 | 0.780407868 | 1.64E-26 | 113.5481714 | 1.717616396 | 24.64738333 |
| ENSMUSG00000039157 | *Eeig1* | 1515.790261 | 1622.025098 | 1595.916152 | 2800.848991 | 2657.214975 | 2669.218337 | 8.49E-32 | 0.780192162 | 5.02E-33 | 143.3120041 | 1.717359603 | 31.0709153 |
| ENSMUSG00000032336 | *Nptn* | 3685.701012 | 3547.561333 | 3617.736307 | 6228.371957 | 6182.38364 | 6226.572408 | 3.93E-54 | 0.7799939 | 1.31E-55 | 246.7774129 | 1.717123612 | 53.40604687 |
| ENSMUSG00000050737 | *Ptges* | 159.915022 | 159.4533147 | 191.9015741 | 259.2732773 | 331.5564042 | 285.7814874 | 1.34E-05 | 0.779973132 | 3.31E-06 | 21.6299739 | 1.717098894 | 4.873965677 |
| ENSMUSG00000032058 | *Ppp2r1b* | 1605.104502 | 1697.902882 | 1684.034221 | 2862.469994 | 2887.780061 | 2808.741285 | 2.22E-35 | 0.779940527 | 1.16E-36 | 159.9475701 | 1.717060088 | 34.65384212 |
| ENSMUSG00000029463 | *Fam216a* | 347.900234 | 347.4982583 | 359.3259065 | 548.775726 | 646.9160875 | 612.9387457 | 6.20E-11 | 0.779693387 | 9.44E-12 | 46.44083435 | 1.716765973 | 10.20748225 |
| ENSMUSG00000066839 | *Ecsit* | 634.556417 | 609.22163 | 655.0109849 | 1073.135583 | 1189.029863 | 996.8674107 | 1.46E-14 | 0.779087233 | 1.76E-15 | 63.3126753 | 1.716044819 | 13.83699764 |
| ENSMUSG00000033166 | *Dis3* | 125.0399374 | 141.8584662 | 177.2152291 | 296.4784113 | 201.0298887 | 264.6124883 | 0.000496482 | 0.778398265 | 0.000154423 | 14.31750101 | 1.715225506 | 3.304096272 |
| ENSMUSG00000033857 | *Engase* | 170.1223638 | 162.7523488 | 169.3825118 | 252.2973147 | 320.1234247 | 287.7059419 | 4.72E-06 | 0.777741479 | 1.11E-06 | 23.72109892 | 1.714444828 | 5.32563811 |
| ENSMUSG00000042293 | *Gm5617* | 85.91179372 | 74.77810622 | 89.09715938 | 148.8205359 | 161.0144607 | 119.3161766 | 0.001459669 | 0.777688801 | 0.000495771 | 12.13150652 | 1.71438223 | 2.835745722 |
| ENSMUSG00000032231 | *Anxa2* | 16716.22347 | 16652.42445 | 17060.63739 | 26750.49132 | 31656.01461 | 27886.30736 | 3.09E-30 | 0.77505481 | 1.94E-31 | 136.0587696 | 1.711255063 | 29.50973697 |
| ENSMUSG00000024799 | *Tm7sf2* | 214.3541784 | 190.2442996 | 256.5214918 | 399.9551901 | 425.8784847 | 306.9504865 | 3.18E-05 | 0.775037031 | 8.27E-06 | 19.87337167 | 1.711233975 | 4.498235054 |
| ENSMUSG00000045658 | *Pid1* | 2676.024783 | 2738.198301 | 2576.963994 | 4654.129727 | 4260.690343 | 4751.478064 | 1.56E-31 | 0.774122729 | 9.34E-33 | 142.0798296 | 1.710149831 | 30.8070093 |
| ENSMUSG00000048534 | *Jaml* | 73.15261643 | 82.47585244 | 56.78720049 | 95.33815578 | 124.8100257 | 141.4474029 | 0.007333833 | 0.773108153 | 0.002815983 | 8.923021466 | 1.70894759 | 2.134668984 |
| ENSMUSG00000038384 | *Setd1b* | 1006.273782 | 1056.790589 | 964.4033186 | 1870.720642 | 1546.310471 | 1757.026923 | 1.96E-16 | 0.772782063 | 2.16E-17 | 71.99254456 | 1.708561363 | 15.70714782 |
| ENSMUSG00000070284 | *Gmppb* | 660.0747715 | 601.5238838 | 678.5091368 | 1119.642 | 1129.006721 | 1068.072226 | 3.50E-17 | 0.772077439 | 3.71E-18 | 75.46833121 | 1.707727091 | 16.45560736 |
| ENSMUSG00000027203 | *Dut* | 1721.638322 | 1717.697087 | 1672.285145 | 2811.312935 | 3127.872629 | 2782.76115 | 1.40E-26 | 0.771186778 | 9.83E-28 | 119.1247758 | 1.706673136 | 25.85397076 |
| ENSMUSG00000022749 | *Tbc1d23* | 1070.92028 | 859.9482215 | 941.8842563 | 1646.327178 | 1606.333613 | 1651.181927 | 4.94E-16 | 0.769185082 | 5.53E-17 | 70.13787489 | 1.70430682 | 15.30587652 |
| ENSMUSG00000020091 | *Eif4ebp2* | 2494.844465 | 2531.458831 | 2341.982475 | 4200.692156 | 4179.706739 | 4142.388227 | 2.88E-38 | 0.765215661 | 1.38E-39 | 173.3328868 | 1.699624048 | 37.54024824 |
| ENSMUSG00000057842 | *Zfp595* | 102.9240301 | 137.4597541 | 126.3025666 | 187.1883303 | 194.3606507 | 238.6323531 | 0.000355375 | 0.764486788 | 0.000108022 | 14.99105464 | 1.698765587 | 3.449312655 |
| ENSMUSG00000046179 | *E2f8* | 459.3303823 | 466.2634858 | 468.0048592 | 777.8198319 | 748.8601543 | 838.0999176 | 8.47E-14 | 0.763356285 | 1.07E-14 | 59.76847228 | 1.697434946 | 13.07237008 |
| ENSMUSG00000047067 | *Dusp28* | 57.84160369 | 76.97746228 | 67.55718678 | 116.2660436 | 116.2352911 | 109.6939043 | 0.003433041 | 0.76328082 | 0.00123985 | 10.42999552 | 1.69734616 | 2.464320949 |
| ENSMUSG00000043881 | *Kbtbd7* | 198.1925538 | 179.2475193 | 182.1106774 | 313.9183178 | 291.5409761 | 345.4395757 | 1.40E-06 | 0.763189255 | 3.12E-07 | 26.17345628 | 1.697238435 | 5.853098053 |
| ENSMUSG00000040713 | *Creg1* | 4735.355996 | 4952.949859 | 4938.528263 | 8054.911503 | 8547.104891 | 8213.571638 | 4.49E-47 | 0.763078899 | 1.74E-48 | 214.1117915 | 1.697108613 | 46.3476399 |
| ENSMUSG00000037936 | *Scarb1* | 3957.046182 | 4024.821599 | 4029.933055 | 7067.812792 | 6924.574556 | 6391.113264 | 6.02E-38 | 0.762819849 | 2.91E-39 | 171.8530041 | 1.696803908 | 37.22076151 |
| ENSMUSG00000030515 | *Tarsl2* | 253.4823221 | 241.9291672 | 219.3160846 | 456.9255515 | 378.2410703 | 380.079756 | 3.98E-07 | 0.762658429 | 8.35E-08 | 28.72405952 | 1.696614067 | 6.399809896 |
| ENSMUSG00000039568 | *Ubald1* | 478.8944541 | 489.3567245 | 511.0848044 | 889.4352338 | 799.3558135 | 821.7420547 | 4.12E-14 | 0.7622953 | 5.12E-15 | 61.21271171 | 1.69618708 | 13.3852973 |
| ENSMUSG00000040390 | *Map3k10* | 554.598906 | 555.3374064 | 536.5411356 | 983.6107291 | 944.1735533 | 865.0422801 | 8.67E-15 | 0.761409019 | 1.03E-15 | 64.36575192 | 1.695145393 | 14.06195721 |
| ENSMUSG00000064360 | *mt-Nd3* | 1144.923508 | 869.8453238 | 1090.705885 | 1820.726243 | 1864.528399 | 1584.788248 | 4.82E-11 | 0.761098565 | 7.25E-12 | 46.9599799 | 1.694780654 | 10.31682025 |
| ENSMUSG00000032279 | *Idh3a* | 4161.193018 | 3890.660879 | 3865.445992 | 6337.662038 | 7111.313221 | 6729.817249 | 1.81E-31 | 0.759731013 | 1.09E-32 | 141.7798618 | 1.693174906 | 30.7429093 |
| ENSMUSG00000096210 | *H1f0* | 2065.285496 | 2010.211444 | 2055.109204 | 3353.112698 | 3588.050052 | 3436.11344 | 1.27E-35 | 0.759394233 | 6.57E-37 | 161.0801571 | 1.692779702 | 34.89671637 |
| ENSMUSG00000001911 | *Nfix* | 284.1043475 | 284.8166104 | 281.9778231 | 529.0104985 | 447.7916953 | 464.7557523 | 6.75E-09 | 0.758647381 | 1.20E-09 | 36.96409764 | 1.691903613 | 8.17087328 |
| ENSMUSG00000060019 | *Gm10073* | 64.64649824 | 53.8842236 | 76.36899376 | 116.2660436 | 110.5188014 | 103.9205409 | 0.005213142 | 0.758593256 | 0.001941246 | 9.604277645 | 1.69184014 | 2.282900426 |
| ENSMUSG00000028613 | *Lrp8* | 1702.924862 | 1750.687428 | 1747.675049 | 3079.887496 | 2611.483057 | 3106.0695 | 7.80E-20 | 0.758019514 | 7.19E-21 | 87.81513671 | 1.691167449 | 19.10764512 |
| ENSMUSG00000074748 | *Atxn7l3b* | 1405.210725 | 1421.883696 | 1437.303626 | 2366.013988 | 2387.587209 | 2455.603892 | 4.22E-32 | 0.757894175 | 2.47E-33 | 144.7241042 | 1.69102053 | 31.37458183 |
| ENSMUSG00000057469 | *E2f6* | 505.2634205 | 503.6525389 | 564.9347359 | 855.7180811 | 919.4020978 | 882.3623702 | 4.53E-14 | 0.756395366 | 5.65E-15 | 61.01957464 | 1.689264649 | 13.34388878 |
| ENSMUSG00000040652 | *Oaz2* | 1124.508825 | 1095.27932 | 1137.702189 | 1753.291938 | 2007.440643 | 1908.096598 | 4.14E-21 | 0.756272211 | 3.60E-22 | 93.73940444 | 1.689120451 | 20.38298741 |
| ENSMUSG00000033629 | *Hacd3* | 1758.21463 | 1734.192257 | 1626.267931 | 2876.421919 | 2865.86685 | 2903.039554 | 5.53E-32 | 0.755970602 | 3.25E-33 | 144.1779163 | 1.688767362 | 31.25720274 |
| ENSMUSG00000024186 | *Rgs11* | 70.60078098 | 95.67198883 | 77.34808342 | 160.4471402 | 107.6605565 | 142.4096301 | 0.005441748 | 0.754683776 | 0.002033128 | 9.519383854 | 1.687261721 | 2.264261581 |
| ENSMUSG00000051705 | *Senp8* | 222.8602966 | 244.1285232 | 225.1906226 | 392.9792275 | 312.5014384 | 460.9068433 | 2.12E-05 | 0.753718259 | 5.39E-06 | 20.69465789 | 1.686132906 | 4.673190249 |
| ENSMUSG00000028337 | *Coro2a* | 102.9240301 | 101.170379 | 128.2607459 | 189.5136511 | 163.8727055 | 206.8788545 | 0.000518419 | 0.75290276 | 0.000161744 | 14.23031782 | 1.685180071 | 3.285319227 |
| ENSMUSG00000027001 | *Dusp19* | 390.4308249 | 344.1992242 | 428.8412726 | 585.9808599 | 707.8919779 | 666.8234706 | 1.09E-08 | 0.75289636 | 1.97E-09 | 36.00018749 | 1.685172595 | 7.963558805 |
| ENSMUSG00000060510 | *Zfp266* | 747.6877889 | 697.1958727 | 710.8190957 | 1213.817496 | 1153.778177 | 1262.442126 | 7.40E-18 | 0.750645275 | 7.58E-19 | 78.60547107 | 1.682545215 | 17.13087872 |
| ENSMUSG00000059713 | *Rcan3* | 756.1939071 | 798.3662516 | 841.038021 | 1313.806293 | 1331.942106 | 1379.833848 | 4.49E-19 | 0.749971005 | 4.34E-20 | 84.25723081 | 1.68175903 | 18.3480467 |
| ENSMUSG00000074305 | *Peak1* | 8122.492259 | 8175.006494 | 7898.316316 | 14043.77541 | 12261.87047 | 14384.33487 | 1.02E-27 | 0.749840655 | 6.93E-29 | 124.3879934 | 1.681607088 | 26.99187754 |
| ENSMUSG00000056124 | *B4galt6* | 1100.691694 | 1109.575135 | 1118.120396 | 1988.149346 | 1708.27768 | 1899.436553 | 1.65E-19 | 0.748833468 | 1.55E-20 | 86.29207553 | 1.680433518 | 18.78121601 |
| ENSMUSG00000043885 | *Slc36a4* | 1417.969902 | 1376.796897 | 1304.147432 | 2369.501969 | 2254.202449 | 2262.196219 | 2.19E-26 | 0.747571875 | 1.55E-27 | 118.2214886 | 1.678964673 | 25.65915705 |
| ENSMUSG00000027189 | *Trim44* | 1730.14444 | 1788.076481 | 1726.135077 | 2988.037321 | 2553.365412 | 3261.950311 | 1.37E-15 | 0.747335696 | 1.57E-16 | 68.0847004 | 1.678689838 | 14.86246295 |
| ENSMUSG00000006398 | *Cdc20* | 2624.988073 | 2853.664495 | 2643.542092 | 4550.652948 | 4752.30846 | 4323.286946 | 2.01E-29 | 0.747047889 | 1.29E-30 | 132.2874042 | 1.678354985 | 28.69743623 |
| ENSMUSG00000030774 | *Pak1* | 1693.568132 | 1615.42703 | 1598.853421 | 2775.270462 | 2737.245831 | 2727.914198 | 2.38E-32 | 0.746860364 | 1.38E-33 | 145.8761482 | 1.678136842 | 31.62337456 |
| ENSMUSG00000017754 | *Pltp* | 130.1436083 | 149.5562124 | 141.9680012 | 256.9479564 | 236.2815754 | 213.6144451 | 4.39E-05 | 0.746101259 | 1.17E-05 | 19.21279626 | 1.677254087 | 4.357450178 |
| ENSMUSG00000004846 | *Plod3* | 2730.463939 | 2796.481237 | 2778.656465 | 4827.366132 | 4719.915018 | 4379.096125 | 1.66E-33 | 0.745541502 | 9.24E-35 | 151.2508457 | 1.676603449 | 32.7793443 |
| ENSMUSG00000087598 | *Zfp111* | 420.2022386 | 394.7844137 | 395.5522241 | 667.3670905 | 596.4204282 | 765.9328754 | 6.16E-09 | 0.744886781 | 1.10E-09 | 37.14637811 | 1.675842748 | 8.210133179 |
| ENSMUSG00000032293 | *Ireb2* | 3393.090546 | 3457.387734 | 3307.364883 | 5805.163559 | 5162.942972 | 6046.635916 | 3.48E-25 | 0.744144289 | 2.55E-26 | 112.6656171 | 1.674980487 | 24.45815519 |
| ENSMUSG00000000823 | *Zfp512b* | 368.3149176 | 368.3921409 | 347.5768306 | 627.8366356 | 581.1764556 | 608.1276096 | 2.69E-11 | 0.744103226 | 3.96E-12 | 48.14212515 | 1.674932813 | 10.57090848 |
| ENSMUSG00000043683 | *Fem1a* | 1844.977035 | 1950.82883 | 1823.064954 | 3132.207215 | 3117.392398 | 3144.558589 | 2.03E-33 | 0.742126987 | 1.13E-34 | 150.8453164 | 1.67264002 | 32.69284861 |

| | | | | | | | | | | | | | |
|---|---|---|---|---|---|---|---|---|---|---|---|---|---|
| ENSMUSG00000029385 | *Ccng2* | 323.2324912 | 284.8166104 | 281.9778231 | 487.1547228 | 499.2401029 | 504.2070687 | 3.43E-09 | 0.741915189 | 5.97E-10 | 38.32969868 | 1.672394483 | 8.465284916 |
| ENSMUSG00000075312 | *Gm13597* | 145.454621 | 168.250739 | 151.7588978 | 256.9479564 | 284.8717381 | 234.7834442 | 2.78E-05 | 0.741461905 | 7.18E-06 | 20.1447215 | 1.671869112 | 4.55664956 |
| ENSMUSG00000047180 | *Neurl3* | 635.4070288 | 656.5077854 | 628.575564 | 1046.394393 | 1127.101225 | 1032.469818 | 1.87E-16 | 0.74020645 | 2.05E-17 | 72.094214 | 1.670414859 | 15.7287105 |
| ENSMUSG00000049881 | *2810025M15Rik* | 901.6485281 | 895.1379185 | 922.3024631 | 1435.885639 | 1682.553476 | 1417.36071 | 3.53E-15 | 0.738783099 | 4.12E-16 | 66.18002547 | 1.668767654 | 14.45166756 |
| ENSMUSG00000024769 | *Cdc42bpg* | 512.9189268 | 513.5496412 | 493.4611904 | 874.3206481 | 824.127269 | 839.0621449 | 1.06E-14 | 0.738719549 | 1.26E-15 | 63.96861727 | 1.668694147 | 13.97665995 |
| ENSMUSG00000040446 | *Rprd1a* | 882.9350681 | 834.6556267 | 833.2053037 | 1356.824729 | 1400.539983 | 1499.150025 | 9.93E-19 | 0.738538741 | 9.82E-20 | 82.6440144 | 1.668485029 | 18.00307663 |
| ENSMUSG00000025104 | *Hdgfl3* | 84.21057008 | 82.47585244 | 93.01351804 | 122.0793458 | 154.3452226 | 155.8808113 | 0.001756531 | 0.738194459 | 0.000605156 | 11.76005013 | 1.668086912 | 2.755344139 |
| ENSMUSG00000050796 | *B3galt6* | 307.0708667 | 262.8230498 | 329.9532166 | 476.6907789 | 526.8698032 | 498.4337053 | 5.02E-08 | 0.737725634 | 9.72E-09 | 32.89639417 | 1.667544931 | 7.299142893 |
| ENSMUSG00000060671 | *Atp8b2* | 2088.252016 | 2216.950914 | 2141.269094 | 3669.356337 | 3560.420352 | 3515.016073 | 1.41E-34 | 0.737427669 | 7.54E-36 | 156.2303785 | 1.667200562 | 33.85230301 |
| ENSMUSG00000025651 | *Uqcrc1* | 6656.037484 | 6600.267552 | 7007.344722 | 10855.76049 | 12687.74895 | 10224.62655 | 1.19E-19 | 0.736835161 | 1.11E-20 | 86.95362697 | 1.666515991 | 18.92394316 |
| ENSMUSG00000047649 | *Polr1g* | 658.3735479 | 656.5077854 | 655.9900746 | 1142.895209 | 1125.195728 | 1018.03641 | 8.31E-16 | 0.736828498 | 9.41E-17 | 69.08964881 | 1.666508295 | 15.08037148 |
| ENSMUSG00000071537 | *Klrg2* | 609.8886742 | 587.2280694 | 609.9728604 | 1002.213296 | 1002.291199 | 1005.527456 | 3.72E-17 | 0.73546296 | 3.96E-18 | 75.34369104 | 1.664931659 | 16.42903459 |
| ENSMUSG00000096255 | *Dynlt1b* | 2952.473624 | 2935.040669 | 2946.080798 | 4472.754699 | 5479.255404 | 4747.629155 | 9.39E-20 | 0.735099026 | 8.71E-21 | 87.43567203 | 1.664511717 | 19.02722821 |
| ENSMUSG00000031310 | *Zmym3* | 290.0586303 | 316.7072734 | 348.5559202 | 568.5409534 | 500.1928511 | 519.6027044 | 2.08E-08 | 0.733602393 | 3.89E-09 | 34.67846546 | 1.66278587 | 7.682416529 |
| ENSMUSG00000050332 | *Amer1* | 201.5950011 | 201.24108 | 171.3406911 | 319.73162 | 320.1234247 | 314.6483043 | 1.97E-06 | 0.732976333 | 4.45E-07 | 25.48917008 | 1.662064457 | 5.706435729 |
| ENSMUSG00000026335 | *Pam* | 1924.934546 | 1940.931727 | 1981.677479 | 3144.99648 | 3212.667227 | 3352.399671 | 1.10E-32 | 0.73208225 | 6.31E-34 | 147.4333826 | 1.661034743 | 31.95707369 |
| ENSMUSG00000113640 | *Adat3* | 107.1770892 | 145.1575003 | 123.3652976 | 208.1162181 | 218.1793579 | 195.3321278 | 0.000287762 | 0.731947317 | 8.61E-05 | 15.41923151 | 1.660879396 | 3.540966337 |
| ENSMUSG00000047714 | *Ppp1r2* | 2628.390521 | 2508.365592 | 2814.882783 | 4584.3701 | 4264.501337 | 4362.738262 | 1.10E-27 | 0.731871358 | 7.47E-29 | 124.2381108 | 1.660791952 | 26.96038592 |
| ENSMUSG00000032436 | *Cmtm7* | 1691.016296 | 1700.102238 | 1702.636925 | 2847.355409 | 2816.323939 | 2792.383423 | 2.98E-34 | 0.731186255 | 1.61E-35 | 154.7187703 | 1.660003467 | 33.5258468 |
| ENSMUSG00000031938 | *4931406C07Rik* | 421.0528504 | 427.7747547 | 464.0885005 | 733.6387353 | 754.576644 | 690.8791514 | 7.34E-12 | 0.731139539 | 1.05E-12 | 50.75095139 | 1.659949715 | 11.13407631 |
| ENSMUSG00000006494 | *Pdk1* | 930.56933 | 833.5559487 | 793.0626275 | 1498.669302 | 1365.288297 | 1386.569439 | 4.78E-15 | 0.731006176 | 5.62E-16 | 65.56628198 | 1.659796277 | 14.32101118 |
| ENSMUSG00000040829 | *Zmynd15* | 143.7533974 | 147.3568564 | 121.4071183 | 226.7187851 | 225.8013442 | 231.8967625 | 5.51E-05 | 0.730896972 | 1.49E-05 | 18.75203049 | 1.659670644 | 4.25868439 |
| ENSMUSG00000009418 | *Nav1* | 1558.320852 | 1739.690648 | 1584.167076 | 2807.824954 | 2473.334555 | 2802.967922 | 5.70E-19 | 0.728072717 | 5.57E-20 | 83.7669618 | 1.656424811 | 18.24402533 |
| ENSMUSG00000007379 | *Dennd2c* | 801.2763335 | 850.0511192 | 808.7280621 | 1366.126013 | 1348.138827 | 1355.778168 | 2.50E-20 | 0.727192267 | 2.25E-21 | 90.11525166 | 1.655414233 | 19.60179065 |
| ENSMUSG00000051444 | *Bbs12* | 250.9304866 | 263.9227278 | 260.4378505 | 395.3045484 | 406.8235189 | 479.1891607 | 2.99E-07 | 0.726986574 | 6.18E-08 | 29.30506803 | 1.655178228 | 6.524482894 |
| ENSMUSG00000091537 | *Tma7* | 1760.766465 | 1601.131215 | 1747.675049 | 2704.348175 | 3133.589119 | 2621.106976 | 1.06E-16 | 0.726985387 | 1.15E-17 | 73.23142634 | 1.655176867 | 15.97350441 |
| ENSMUSG00000019055 | *Plod1* | 2309.411089 | 2462.179115 | 2536.821318 | 3979.786674 | 4228.296902 | 3882.586875 | 2.35E-27 | 0.726886677 | 1.62E-28 | 122.7069546 | 1.655063623 | 26.62916326 |
| ENSMUSG00000026281 | *Dtymk* | 1506.433531 | 1504.359549 | 1563.606193 | 2286.953078 | 2959.236182 | 2319.929852 | 3.91E-12 | 0.726510837 | 5.49E-13 | 52.0230856 | 1.654632514 | 11.40771014 |
| ENSMUSG00000041625 | *Ggact* | 270.4945584 | 267.2217619 | 195.8179327 | 411.5817945 | 406.8235189 | 395.4753917 | 7.58E-06 | 0.726078876 | 1.82E-06 | 22.77932365 | 1.65413717 | 5.120613371 |
| ENSMUSG00000032396 | *Dis3l* | 849.7612072 | 917.1314792 | 947.7587943 | 1463.789489 | 1560.601696 | 1459.698708 | 4.00E-18 | 0.725409639 | 4.05E-19 | 79.8438574 | 1.653370028 | 17.39754595 |
| ENSMUSG00000032258 | *Lca5* | 237.3206975 | 243.0288452 | 222.2533536 | 388.3285857 | 372.5245806 | 400.2865278 | 1.12E-07 | 0.725059659 | 2.24E-08 | 31.27125826 | 1.652968989 | 6.949485097 |
| ENSMUSG00000029468 | *P2rx7* | 641.3613115 | 593.8261376 | 626.6173847 | 1038.25577 | 925.1185875 | 1115.22136 | 4.29E-12 | 0.724378657 | 6.04E-13 | 51.83416833 | 1.652188915 | 11.3678332 |
| ENSMUSG00000037946 | *Fgd3* | 2651.35704 | 2654.622771 | 2615.148491 | 4347.187371 | 4288.320044 | 4451.263168 | 2.16E-40 | 0.72436807 | 9.99E-42 | 183.1421064 | 1.65217679 | 39.66503328 |
| ENSMUSG00000030341 | *Tnfrsf1a* | 1656.141212 | 1705.600629 | 1760.403215 | 2850.84339 | 2853.481122 | 2749.083197 | 7.07E-30 | 0.723017732 | 4.50E-31 | 134.3862993 | 1.650631105 | 29.15052618 |
| ENSMUSG00000039616 | *Mocos* | 546.9433996 | 522.3470655 | 537.5202253 | 867.3446855 | 979.4252399 | 805.3841918 | 3.98E-11 | 0.722688417 | 5.96E-12 | 47.34222105 | 1.650254368 | 10.39972698 |
| ENSMUSG00000037905 | *Bri3bp* | 4198.619938 | 4145.786183 | 4408.840755 | 7194.54278 | 7221.832022 | 6625.896708 | 5.30E-33 | 0.722230861 | 2.99E-34 | 148.9163373 | 1.649731067 | 32.27599291 |
| ENSMUSG00000002393 | *Nr2f6* | 665.1784424 | 755.4788084 | 715.714544 | 1170.799059 | 1200.462843 | 1147.937086 | 6.87E-16 | 0.72203487 | 7.76E-17 | 69.46874544 | 1.649506965 | 15.16279333 |
| ENSMUSG00000032479 | *Map4* | 4360.236184 | 4362.422755 | 4158.193801 | 7332.899372 | 7065.581303 | 6851.05788 | 5.37E-39 | 0.721965865 | 2.53E-40 | 176.7104528 | 1.649428071 | 38.27031182 |
| ENSMUSG00000023909 | *Paqr4* | 175.2260347 | 204.5401141 | 218.336995 | 318.5689596 | 344.8948802 | 320.4216677 | 5.33E-06 | 0.721448547 | 1.26E-06 | 23.48386132 | 1.648836729 | 5.273609177 |
| ENSMUSG00000024913 | *Lrp5* | 1732.696275 | 1745.189038 | 1739.842332 | 3008.965209 | 2831.567912 | 2764.478833 | 2.29E-28 | 0.721251943 | 1.52E-29 | 127.3919166 | 1.648612048 | 27.64041922 |
| ENSMUSG00000002820 | *Atg4d* | 926.3162709 | 911.633089 | 934.051539 | 1490.530679 | 1668.262252 | 1410.62512 | 1.15E-15 | 0.721217392 | 1.31E-16 | 68.43857012 | 1.648572566 | 14.93882994 |
| ENSMUSG00000034201 | *Gas2l1* | 1275.917728 | 1378.996253 | 1296.314715 | 2340.435458 | 2117.959444 | 2053.39291 | 2.88E-19 | 0.720708317 | 2.75E-20 | 85.16061826 | 1.647990946 | 18.53996668 |
| ENSMUSG00000060261 | *Gtf2i* | 2692.186407 | 2622.732108 | 2725.785623 | 4475.080019 | 4469.342218 | 4298.269038 | 8.10E-38 | 0.719415946 | 3.93E-39 | 171.2583248 | 1.646515332 | 37.09149906 |
| ENSMUSG00000042992 | *Borcs5* | 122.4881019 | 117.6655495 | 112.5953113 | 173.236405 | 229.6123374 | 177.0498104 | 0.000720317 | 0.718524129 | 0.000231456 | 13.55675967 | 1.645497835 | 3.142476321 |
| ENSMUSG00000006941 | *Eif1b* | 1099.841082 | 1141.465798 | 1114.204037 | 1724.225427 | 2122.723185 | 1671.388699 | 3.30E-12 | 0.71850497 | 4.62E-13 | 52.3615265 | 1.645475983 | 11.48085721 |
| ENSMUSG00000022102 | *Dok2* | 3105.583751 | 3102.19173 | 3277.992194 | 5215.694717 | 5514.50709 | 4873.680922 | 4.20E-27 | 0.718072791 | 2.91E-28 | 121.5419854 | 1.644983133 | 26.37674482 |
| ENSMUSG00000021022 | *Ppp2r3c* | 755.3432953 | 810.46271 | 837.1216623 | 1240.558686 | 1298.595916 | 1406.776211 | 3.87E-15 | 0.717251055 | 4.53E-16 | 65.99120762 | 1.644046444 | 14.4118511 |
| ENSMUSG00000025026 | *Add3* | 681.340067 | 622.4177664 | 621.7219363 | 1052.207695 | 1004.196695 | 1110.410224 | 3.64E-14 | 0.716434073 | 4.51E-15 | 61.46443318 | 1.643115702 | 13.43842004 |
| ENSMUSG00000007880 | *Arid1a* | 3095.376409 | 3181.368548 | 3020.491612 | 5483.106618 | 4352.154179 | 5417.339307 | 3.99E-15 | 0.713978872 | 4.67E-16 | 65.93200946 | 1.6403218 | 14.39957094 |
| ENSMUSG00000004849 | *Ap1s1* | 5538.333553 | 5708.428667 | 5841.248933 | 9048.986176 | 10230.61112 | 8727.400979 | 3.93E-24 | 0.713007057 | 3.01E-25 | 107.7739073 | 1.639217234 | 23.40592664 |
| ENSMUSG00000026994 | *Galnt3* | 590.3246024 | 638.9129369 | 554.1647496 | 1061.508978 | 843.1822348 | 1018.998637 | 2.04E-09 | 0.712933768 | 3.49E-10 | 39.38102441 | 1.639133963 | 8.690207451 |
| ENSMUSG00000051586 | *Mical3* | 552.0470705 | 552.0383724 | 512.063894 | 932.4536699 | 825.0800173 | 892.9468698 | 1.72E-12 | 0.712666513 | 2.36E-13 | 53.68209067 | 1.638830346 | 11.76421833 |
| ENSMUSG00000050912 | *Tmem123* | 3805.637278 | 3546.461655 | 3843.906019 | 6162.100313 | 5957.535044 | 6226.572408 | 7.92E-35 | 0.711971804 | 4.21E-36 | 157.3893373 | 1.638041381 | 34.10108699 |
| ENSMUSG00000039987 | *Phtf2* | 614.9923451 | 590.5271035 | 627.5964743 | 973.1467852 | 922.2603427 | 1107.523542 | 1.03E-11 | 0.711901225 | 1.48E-12 | 50.07140136 | 1.637961248 | 10.98701907 |
| ENSMUSG00000042826 | *Fgf11* | 245.8268157 | 287.0159665 | 258.4796712 | 481.3414206 | 348.7058734 | 465.7179795 | 2.20E-05 | 0.711781148 | 5.59E-06 | 20.62268333 | 1.637824924 | 4.657685661 |
| ENSMUSG00000032020 | *Ubash3b* | 1246.146315 | 1258.031669 | 1241.485693 | 1955.594854 | 2062.700043 | 2113.050998 | 4.99E-24 | 0.711732734 | 3.85E-25 | 107.2902096 | 1.637769964 | 23.30148466 |
| ENSMUSG00000001062 | *Vps9d1* | 242.4243684 | 287.0159665 | 274.1451058 | 455.762891 | 392.5322946 | 465.7179795 | 8.51E-07 | 0.711364971 | 1.85E-07 | 27.1878594 | 1.637352525 | 6.06984291 |
| ENSMUSG00000042303 | *Sgsm3* | 716.2151516 | 702.6942628 | 728.4427097 | 1152.196492 | 1250.005754 | 1113.296906 | 1.74E-15 | 0.711222861 | 2.00E-16 | 67.6026346 | 1.63719125 | 14.75836739 |
| ENSMUSG00000021900 | *Btd* | 461.8822177 | 473.961232 | 483.6702938 | 716.1988288 | 833.6547519 | 770.7440115 | 3.23E-11 | 0.710851181 | 4.79E-12 | 47.77046097 | 1.636769516 | 10.49099964 |
| ENSMUSG00000028524 | *Sgip1* | 1044.551314 | 1146.964188 | 1043.709581 | 1699.809558 | 1770.206319 | 1814.760556 | 1.49E-19 | 0.709646282 | 1.39E-20 | 86.50576761 | 1.635403102 | 18.82782614 |
| ENSMUSG00000037151 | *Lrrc20* | 384.4765422 | 400.2828039 | 406.3222104 | 639.46324 | 664.0655567 | 642.7677899 | 2.30E-11 | 0.709572918 | 3.38E-12 | 48.45736172 | 1.635319941 | 10.63870135 |
| ENSMUSG00000026836 | *Acvr1* | 643.0625352 | 571.8325769 | 552.2065702 | 998.7253148 | 956.559281 | 939.1337768 | 1.45E-12 | 0.709526369 | 1.98E-13 | 54.02246226 | 1.635267178 | 11.83749209 |
| ENSMUSG00000073062 | *Zxdb* | 168.4211402 | 200.1414019 | 162.5288841 | 292.99043 | 282.9662415 | 290.5926236 | 1.62E-05 | 0.709277011 | 4.05E-06 | 21.24257771 | 1.634984559 | 4.79125131 |
| ENSMUSG00000064210 | *Ano6* | 1551.515958 | 1596.732503 | 1556.752565 | 2740.390648 | 2388.539958 | 2565.297796 | 2.67E-21 | 0.709133954 | 2.30E-22 | 94.62756798 | 1.634822443 | 20.5740691 |
| ENSMUSG00000031556 | *Tm2d2* | 787.6665444 | 812.6620661 | 789.1462688 | 1290.553084 | 1310.981644 | 1301.893443 | 1.81E-19 | 0.708729416 | 1.70E-20 | 86.11584901 | 1.634364096 | 18.74346679 |
| ENSMUSG00000040495 | *Chrm4* | 134.3966674 | 133.0610419 | 122.3862079 | 206.9535577 | 240.0925685 | 189.5587644 | 0.000290292 | 0.707866027 | 8.69E-05 | 15.40095719 | 1.633386295 | 3.537164659 |
| ENSMUSG00000033405 | *Nudt15* | 113.9819837 | 120.9645836 | 108.6789527 | 173.236405 | 183.8804196 | 203.0299456 | 0.000377987 | 0.707720013 | 0.000115388 | 14.86660395 | 1.633220988 | 3.422523607 |
| ENSMUSG00000028521 | *Slc35d1* | 461.0316059 | 511.3502851 | 473.8793972 | 784.7957945 | 794.5920721 | 779.4040566 | 1.59E-12 | 0.707590088 | 2.17E-13 | 53.84328776 | 1.633073912 | 11.79897947 |
| ENSMUSG00000047260 | *Emc6* | 1114.301483 | 1112.874169 | 1126.932203 | 1775.382486 | 1935.984521 | 1763.762513 | 1.18E-20 | 0.707372085 | 1.05E-21 | 91.62609994 | 1.63282716 | 19.92738881 |
| ENSMUSG00000026918 | *Brd3* | 1175.545534 | 1220.642616 | 1182.740314 | 2020.703838 | 1824.512971 | 1996.621503 | 1.60E-20 | 0.70682667 | 1.43E-21 | 91.01006559 | 1.632209982 | 19.79486713 |
| ENSMUSG00000000568 | *Hnrnpd* | 2384.264929 | 2356.610024 | 2385.06242 | 4070.474188 | 3732.867792 | 3830.626604 | 1.26E-30 | 0.706579918 | 7.83E-32 | 137.8562457 | 1.63193084 | 29.89856138 |
| ENSMUSG00000050627 | *Gpd1l* | 2017.651235 | 1936.533015 | 2000.280183 | 3203.129502 | 3343.193742 | 3170.538724 | 8.83E-31 | 0.706210712 | 5.44E-32 | 138.5798509 | 1.631513259 | 30.0539191 |
| ENSMUSG00000018809 | *Smyd4* | 418.5010149 | 450.8679934 | 466.0466798 | 704.5722244 | 718.3722091 | 752.4616941 | 2.30E-11 | 0.70583231 | 3.37E-12 | 48.45830035 | 1.631085389 | 10.63870632 |
| ENSMUSG00000030541 | *Idh2* | 2893.781408 | 2845.966748 | 2868.732714 | 4600.647347 | 4947.621859 | 4491.676711 | 1.38E-30 | 0.705734902 | 8.59E-32 | 137.6734864 | 1.630975264 | 29.8605091 |
| ENSMUSG00000020167 | *Tcf3* | 1610.208173 | 1725.394833 | 1753.549587 | 2914.789714 | 2675.317192 | 2703.858517 | 7.14E-23 | 0.704751161 | 5.76E-24 | 101.9279864 | 1.629863518 | 22.1462078 |
| ENSMUSG00000032026 | *Rexo2* | 3668.688775 | 3641.033966 | 3735.227066 | 5404.045708 | 6875.984394 | 5717.554203 | 1.82E-15 | 0.704721646 | 2.10E-16 | 67.51096071 | 1.629830175 | 14.73895313 |
| ENSMUSG00000011148 | *Adssl1* | 939.0754482 | 975.4144149 | 889.9925042 | 1542.850399 | 1581.562158 | 1443.340846 | 1.68E-17 | 0.704069626 | 1.75E-18 | 76.94920013 | 1.629093746 | 16.77591559 |
| ENSMUSG00000050538 | *B230217C12Rik* | 84.21057008 | 85.77488654 | 80.28535241 | 123.2420063 | 136.2430052 | 147.2207662 | 0.002536591 | 0.702930988 | 0.000896179 | 11.03064726 | 1.627808502 | 2.595749597 |
| ENSMUSG00000042066 | *Tmcc2* | 449.1230404 | 434.3728229 | 414.1549277 | 733.6387353 | 719.3249574 | 660.08788 | 8.36E-11 | 0.702050251 | 1.28E-11 | 45.83825084 | 1.62681506 | 10.0778051 |
| ENSMUSG00000022892 | *App* | 301.9671957 | 439.871213 | 282.9569128 | 496.4560063 | 528.7752998 | 636.9944265 | 0.000184995 | 0.701762147 | 5.36E-05 | 16.31661979 | 1.62649022 | 3.73284065 |
| ENSMUSG00000026365 | *Cfh* | 480.5956777 | 429.9741107 | 436.6739899 | 774.3318506 | 596.4204282 | 822.704282 | 8.93E-07 | 0.701355555 | 1.94E-07 | 27.08926937 | 1.626031893 | 6.048930267 |
| ENSMUSG00000047045 | *Tmem164* | 1523.445768 | 1601.131215 | 1494.090827 | 2597.383415 | 2451.421345 | 2460.415028 | 6.88E-25 | 0.701282596 | 5.11E-26 | 111.2904175 | 1.625949665 | 24.16256396 |
| ENSMUSG00000058756 | *Thra* | 182.8815411 | 163.8520269 | 163.5079738 | 268.5745608 | 296.3047175 | 265.5747156 | 1.99E-05 | 0.701194645 | 5.05E-06 | 20.8199998 | 1.625850545 | 4.700204452 |
| ENSMUSG00000029475 | *Kdm2b* | 970.5480855 | 968.8163467 | 973.2151256 | 1619.585988 | 1560.601696 | 1555.921431 | 1.59E-21 | 0.700924874 | 1.35E-22 | 95.67732786 | 1.625546554 | 20.79739818 |
| ENSMUSG00000030095 | *Tmem43* | 2006.593281 | 2041.002428 | 1856.354002 | 3220.569409 | 3240.296927 | 3133.011862 | 1.11E-26 | 0.700512169 | 7.77E-28 | 119.5932042 | 1.625081608 | 25.95525297 |
| ENSMUSG00000032440 | *Tgfbr2* | 3469.64561 | 3544.262299 | 3463.04014 | 5878.411166 | 5562.144505 | 5579.955709 | 4.15E-38 | 0.699913045 | 2.00E-39 | 172.6000559 | 1.624406883 | 37.38206364 |
| ENSMUSG00000059851 | *Kmt5c* | 362.3606349 | 363.9934288 | 379.8867895 | 637.1379191 | 544.0192724 | 616.7876547 | 4.78E-09 | 0.699579091 | 8.43E-10 | 37.65726653 | 1.62403091 | 8.32053123 |
| ENSMUSG00000022476 | *Polr3h* | 312.1745376 | 289.2153226 | 315.2668717 | 468.5521558 | 566.8852313 | 452.2467983 | 7.18E-07 | 0.698516237 | 1.55E-07 | 27.5296377 | 1.622834903 | 6.143983284 |
| ENSMUSG00000032405 | *Pias1* | 1097.289246 | 985.3115172 | 944.8215253 | 1671.905707 | 1561.554444 | 1681.973199 | 1.49E-15 | 0.697684651 | 1.70E-16 | 67.9206544 | 1.621899752 | 14.82737471 |
| ENSMUSG00000014837 | *Matcap1* | 163.3174692 | 155.0546026 | 183.0897671 | 284.8518069 | 267.7222689 | 261.7258067 | 2.72E-05 | 0.697517473 | 7.02E-06 | 20.18713098 | 1.621711819 | 4.565932683 |
| ENSMUSG00000062762 | *Ei24* | 1765.870136 | 1749.58775 | 1810.336788 | 2784.571745 | 2967.810917 | 2879.9461 | 2.88E-28 | 0.697080344 | 1.93E-29 | 126.923477 | 1.621220524 | 27.54014161 |
| ENSMUSG00000022253 | *Nadk2* | 397.2357195 | 339.8005121 | 318.2041406 | 531.3358194 | 551.6412587 | 629.2966087 | 4.18E-07 | 0.697038428 | 8.78E-08 | 28.62573407 | 1.621173422 | 6.379320303 |
| ENSMUSG00000059058 | *Tma7-ps* | 351.3026812 | 339.8005121 | 336.8068443 | 509.2452711 | 746.9546577 | 408.9465729 | 0.000740689 | 0.696384762 | 0.000238765 | 13.49841745 | 1.620439056 | 3.130364068 |
| ENSMUSG00000026458 | *Ppfia4* | 482.2969014 | 449.7683153 | 487.5866524 | 769.6812089 | 838.4184933 | 692.8036059 | 1.28E-09 | 0.695736496 | 2.15E-10 | 40.32536297 | 1.619711085 | 8.892617483 |
| ENSMUSG00000050382 | *Kif7* | 80.8081228 | 91.2732767 | 91.05533871 | 158.1218193 | 140.0539983 | 127.9762216 | 0.002839816 | 0.695488423 | 0.001010908 | 10.80748105 | 1.619432597 | 2.546709761 |
| ENSMUSG00000050705 | *2310061I04Rik* | 447.4218168 | 448.6686373 | 399.4685827 | 723.1747914 | 708.8447262 | 666.8234706 | 2.21E-10 | 0.695388395 | 3.51E-11 | 43.87134921 | 1.619320319 | 9.655867461 |
| ENSMUSG00000032171 | *Pin1* | 2259.224991 | 2314.822259 | 2407.581483 | 3508.909197 | 4346.437689 | 3446.697939 | 2.22E-13 | 0.695383549 | 2.88E-14 | 57.81474133 | 1.619314881 | 12.65325575 |
| ENSMUSG00000001576 | *Ergic1* | 2943.116894 | 2895.45226 | 2869.711804 | 4862.245945 | 4800.898623 | 4438.754214 | 5.55E-30 | 0.695024795 | 3.52E-31 | 134.874199 | 1.618912256 | 29.25584017 |
| ENSMUSG00000051065 | *Mb21d2* | 316.4275967 | 314.5079173 | 284.9150921 | 491.8053646 | 512.5785789 | 478.2269335 | 1.97E-08 | 0.69488788 | 3.69E-09 | 34.78199558 | 1.618758625 | 7.704875043 |
| ENSMUSG00000035004 | *Igsf6* | 715.3645398 | 660.9064976 | 632.4919226 | 1064.99696 | 994.6692126 | 1193.161766 | 5.46E-11 | 0.69457225 | 8.27E-12 | 46.69992249 | 1.618404516 | 10.26254858 |
| ENSMUSG00000030315 | *Vgll4* | 651.5686533 | 587.2280694 | 610.95195 | 1037.093109 | 963.2285191 | 997.8296379 | 1.27E-13 | 0.694418746 | 1.62E-14 | 58.94759936 | 1.618232325 | 12.89521093 |
| ENSMUSG00000066829 | *Zfp810* | 91.01546463 | 96.77166687 | 88.11806972 | 141.8445732 | 131.4792637 | 172.2386742 | 0.003006065 | 0.693413129 | 0.001074626 | 10.69436083 | 1.617104744 | 2.522001635 |
| ENSMUSG00000039016 | *Timm8b* | 621.7972397 | 601.5238838 | 688.3000335 | 883.6219316 | 1267.155223 | 938.1715496 | 1.69E-06 | 0.693212777 | 3.80E-07 | 25.79309681 | 1.616880187 | 5.772194801 |
| ENSMUSG00000031145 | *Prickle3* | 80.8081228 | 83.57553048 | 81.26444207 | 138.3565919 | 109.5660531 | 149.1452207 | 0.005555687 | 0.692299775 | 0.002079856 | 9.477690359 | 1.615857276 | 2.255262228 |
| ENSMUSG00000033065 | *Pfkm* | 286.656183 | 170.450095 | 278.0614644 | 403.4431714 | 443.9807022 | 343.5151212 | 0.001185997 | 0.691800512 | 0.00039639 | 12.54913291 | 1.615298185 | 2.925916466 |
| ENSMUSG00000027947 | *Il6ra* | 1949.602289 | 2047.600497 | 1954.262968 | 3315.907564 | 2948.755951 | 3347.588534 | 5.12E-21 | 0.691697817 | 4.46E-22 | 93.31550407 | 1.615183207 | 20.29065944 |
| ENSMUSG00000025236 | *Adpgk* | 1611.909397 | 1602.230893 | 1555.773475 | 2655.516437 | 2522.877466 | 2528.733161 | 1.16E-26 | 0.691673389 | 8.15E-28 | 119.4983451 | 1.615155859 | 25.9349102 |
| ENSMUSG00000023004 | *Tuba1b* | 36070.19418 | 36436.73193 | 36870.55855 | 58276.02905 | 63311.07648 | 55069.22662 | 6.51E-31 | 0.691645952 | 3.99E-32 | 139.1953298 | 1.615125143 | 30.18609268 |
| ENSMUSG00000033228 | *Scaf11* | 359.8087994 | 435.4725009 | 425.9040036 | 752.2413023 | 514.4840755 | 705.3125599 | 2.45E-05 | 0.691637897 | 6.26E-06 | 20.40617313 | 1.615116126 | 4.611598142 |
| ENSMUSG00000012429 | *Mplkip* | 253.4823221 | 229.8327088 | 270.2287471 | 395.3045484 | 406.8235189 | 415.6821635 | 4.47E-07 | 0.691406586 | 9.42E-08 | 28.48969425 | 1.61485719 | 6.35008085 |
| ENSMUSG00000035769 | *Xylb* | 303.6684194 | 314.5079173 | 291.7687197 | 531.3358194 | 481.1378854 | 458.0201617 | 6.76E-08 | 0.691400115 | 1.33E-08 | 32.29165355 | 1.614849947 | 7.169823237 |
| ENSMUSG00000032041 | *Tirap* | 807.2306162 | 822.5591684 | 812.6444207 | 1339.384823 | 1170.927646 | 1433.718573 | 2.53E-12 | 0.69127797 | 3.51E-13 | 52.8979029 | 1.614713233 | 11.59646926 |
| ENSMUSG00000030008 | *Pradc1* | 174.3754229 | 161.6526708 | 160.5707048 | 281.3638256 | 260.1002826 | 261.7258067 | 2.26E-05 | 0.690867988 | 5.75E-06 | 20.5691767 | 1.614254432 | 4.646484009 |
| ENSMUSG00000063146 | *Clip2* | 1111.749647 | 1241.536499 | 1215.050272 | 2076.511539 | 1881.677869 | 1800.327148 | 1.90E-15 | 0.690674343 | 2.18E-16 | 67.42860624 | 1.614037774 | 14.72158997 |
| ENSMUSG00000031367 | *Ap1s2* | 675.3857843 | 619.1187323 | 624.6592053 | 1067.322281 | 953.7010362 | 1078.656725 | 2.07E-12 | 0.689837485 | 2.86E-13 | 53.30418292 | 1.613101798 | 11.6839268 |
| ENSMUSG00000049091 | *Sephs2* | 2955.876071 | 3181.368548 | 3126.233296 | 4980.837309 | 5033.369205 | 4921.792283 | 6.69E-34 | 0.68979545 | 3.66E-35 | 153.0903907 | 1.613054798 | 33.17436138 |
| ENSMUSG00000033392 | *Clasp2* | 1475.811506 | 1382.295287 | 1233.652976 | 2275.326474 | 2007.440643 | 2315.118716 | 2.23E-13 | 0.68845365 | 2.90E-14 | 57.80323107 | 1.61155525 | 12.65109847 |
| ENSMUSG00000060166 | *Zdhhc8* | 256.0341575 | 260.6236937 | 256.5214918 | 451.1122493 | 395.3905395 | 400.2865278 | 3.66E-07 | 0.687863254 | 7.63E-08 | 28.89661244 | 1.610895886 | 6.436649556 |
| ENSMUSG00000024654 | *Asrgl1* | 371.7173649 | 394.7844137 | 362.2631755 | 626.6739752 | 640.2468495 | 550.3939758 | 1.04E-08 | 0.687459529 | 1.89E-09 | 36.08533819 | 1.610445154 | 7.981879365 |
| ENSMUSG00000032307 | *Ube2q2* | 1454.54621 | 1405.388526 | 1388.349143 | 2463.677465 | 1922.646045 | 2453.679437 | 3.21E-11 | 0.686372445 | 4.76E-12 | 47.78352979 | 1.609232127 | 10.49369381 |
| ENSMUSG00000019718 | *L3hypdh* | 126.741161 | 96.77166687 | 126.3025666 | 180.2123676 | 206.7463785 | 177.0498104 | 0.001296683 | 0.685562633 | 0.000435787 | 12.37214522 | 1.608329088 | 2.887166133 |
| ENSMUSG00000055932 | *Fto* | 1106.645976 | 1089.78093 | 1138.681279 | 1919.55238 | 1764.489829 | 1681.973199 | 2.69E-17 | 0.685032954 | 2.83E-18 | 76.00534917 | 1.607738705 | 16.57090073 |
| ENSMUSG00000028799 | *Zfp362* | 216.9060138 | 227.6333527 | 243.7933262 | 376.7019814 | 386.8158049 | 342.552894 | 2.13E-06 | 0.684822387 | 4.83E-07 | 25.33072003 | 1.607504066 | 5.671816969 |
| ENSMUSG00000014592 | *Camta1* | 132.6954438 | 139.6591101 | 143.9261805 | 208.1162181 | 274.3915069 | 185.7098555 | 0.001438169 | 0.684786884 | 0.000487975 | 12.16106611 | 1.607464508 | 2.842190164 |
| ENSMUSG00000028431 | *Elp1* | 1377.991147 | 1409.787238 | 1335.478301 | 2289.278399 | 2109.384709 | 2228.518266 | 4.59E-22 | 0.68437141 | 3.81E-23 | 98.18456871 | 1.60700165 | 21.33775319 |
| ENSMUSG00000033855 | *Ston1* | 96.96974736 | 130.8616859 | 125.3234769 | 199.977595 | 159.1089641 | 206.8788545 | 0.002552365 | 0.683918535 | 0.000902802 | 11.01699619 | 1.606497276 | 2.593057152 |
| ENSMUSG00000047921 | *Trappc9* | 543.5409523 | 526.7457776 | 489.5448318 | 865.0193646 | 875.5756766 | 766.8951026 | 1.56E-10 | 0.683812143 | 2.45E-11 | 44.57356262 | 1.606378809 | 9.805610867 |
| ENSMUSG00000075419 | *Dolk* | 405.7418377 | 393.6847357 | 380.8658791 | 658.065807 | 640.2468495 | 598.5053373 | 6.41E-10 | 0.68305452 | 1.05E-10 | 41.71995882 | 1.60553545 | 9.193181879 |
| ENSMUSG00000074916 | *Chst14* | 289.2080185 | 293.6140347 | 286.8732714 | 489.4800437 | 460.177423 | 446.4734349 | 4.35E-08 | 0.682009704 | 8.36E-09 | 33.18908145 | 1.604373124 | 7.361275642 |
| ENSMUSG00000036737 | *Oxsr1* | 2545.881174 | 2455.581047 | 2480.034118 | 4057.684923 | 3913.889967 | 4034.618777 | 1.35E-33 | 0.68182534 | 7.44E-35 | 151.6814504 | 1.604168112 | 32.87076923 |
| ENSMUSG00000024818 | *Slc25a45* | 904.2003636 | 1013.903146 | 1007.483264 | 1569.591589 | 1628.246824 | 1490.48998 | 1.46E-15 | 0.681703774 | 1.67E-16 | 67.96401036 | 1.604032946 | 14.83640273 |
| ENSMUSG00000051185 | *Fam174a* | 1188.304711 | 1123.870949 | 1217.987541 | 1896.299172 | 1952.181242 | 1813.798329 | 1.18E-19 | 0.68093071 | 1.10E-20 | 86.97326204 | 1.60317366 | 18.92793514 |
| ENSMUSG00000014498 | *Ankrd52* | 3365.020356 | 3529.966485 | 3298.553076 | 5762.145122 | 5120.069299 | 5460.639532 | 3.93E-25 | 0.680914517 | 2.89E-26 | 112.4205925 | 1.603155665 | 24.40610608 |
| ENSMUSG00000013822 | *Elof1* | 1354.174016 | 1282.224586 | 1392.265502 | 2004.426592 | 2399.020189 | 2049.544001 | 5.69E-14 | 0.679867123 | 7.13E-15 | 60.56071615 | 1.601992199 | 13.24456669 |
| ENSMUSG00000020150 | *Gamt* | 134.3966674 | 137.4597541 | 120.4280286 | 217.4175016 | 209.6046233 | 201.1054911 | 0.000249306 | 0.678642145 | 7.38E-05 | 15.71050021 | 1.600632541 | 3.603267578 |
| ENSMUSG00000026177 | *Slc11a1* | 1943.648006 | 2031.105326 | 2034.548321 | 3337.998113 | 3231.722193 | 3047.373639 | 8.01E-25 | 0.678416903 | 5.97E-26 | 110.9837564 | 1.60038266 | 24.09618579 |
| ENSMUSG00000070282 | *3000002C10Rik* | 147.1558447 | 147.3568564 | 152.7379875 | 226.7187851 | 230.5650857 | 257.8768977 | 9.68E-05 | 0.678308533 | 2.71E-05 | 17.61411705 | 1.600262451 | 4.013945573 |
| ENSMUSG00000032733 | *Snx33* | 277.299453 | 267.2217619 | 257.5005815 | 398.7925297 | 444.9334504 | 438.775617 | 3.04E-07 | 0.677925225 | 6.29E-08 | 29.27117257 | 1.599837334 | 6.517605705 |
| ENSMUSG00000031918 | *Mtmr2* | 2463.371828 | 2322.520005 | 2236.240792 | 3612.385976 | 3861.488811 | 3761.346243 | 5.99E-25 | 0.677847235 | 4.43E-26 | 111.5722734 | 1.599750852 | 24.22229832 |
| ENSMUSG00000005161 | *Prdx2* | 4093.144073 | 3821.381163 | 3886.006875 | 6124.895179 | 6788.331551 | 5966.771055 | 2.97E-23 | 0.677825866 | 2.36E-24 | 103.6965166 | 1.599727157 | 22.52795082 |
| ENSMUSG00000034055 | *Phka1* | 413.397344 | 420.0770084 | 398.4894931 | 659.2284674 | 654.5380738 | 656.2389711 | 8.93E-11 | 0.677424506 | 1.37E-11 | 45.70464236 | 1.599282172 | 10.04895878 |
| ENSMUSG00000035382 | *Pcsk7* | 1122.807601 | 1217.343582 | 1170.991238 | 1954.432193 | 1893.110848 | 1761.838059 | 7.78E-18 | 0.676294089 | 8.00E-19 | 78.50043366 | 1.598029553 | 17.10923778 |
| ENSMUSG00000039835 | *Nhsl1* | 122.4881019 | 160.5529928 | 135.1143736 | 231.3694268 | 182.9276713 | 252.1035344 | 0.001663164 | 0.675600837 | 0.00056968 | 11.87252969 | 1.597261843 | 2.779064983 |
| ENSMUSG00000030431 | *Tmem238* | 230.5158029 | 210.0385042 | 238.8978779 | 389.4912462 | 370.619084 | 327.1572583 | 7.46E-06 | 0.675334109 | 1.79E-06 | 22.80807253 | 1.596966566 | 5.126985761 |
| ENSMUSG00000040181 | *Fmo1* | 644.7637588 | 600.4242058 | 612.9101294 | 982.4480687 | 967.0395122 | 1018.998637 | 8.63E-14 | 0.674797108 | 1.09E-14 | 59.72894999 | 1.596372252 | 13.06388509 |
| ENSMUSG00000032376 | *Usp3* | 849.7612072 | 880.8421041 | 947.7587943 | 1337.059502 | 1523.444512 | 1408.700665 | 8.79E-14 | 0.673883278 | 1.11E-14 | 59.69078996 | 1.595361401 | 13.05617255 |
| ENSMUSG00000038256 | *Bcl9* | 857.4167135 | 845.652407 | 797.9580758 | 1447.512243 | 1260.485985 | 1282.648898 | 2.23E-13 | 0.672694788 | 2.90E-14 | 57.80288878 | 1.594047686 | 12.65109847 |

| | | | | | | | | | | | | | |
|---|---|---|---|---|---|---|---|---|---|---|---|---|---|
| ENSMUSG00000014496 | *Ankrd28* | 325.7843267 | 433.2731448 | 384.7822378 | 652.2525048 | 670.7347947 | 497.4714781 | 3.45E-05 | 0.672366342 | 9.07E-06 | 19.69839813 | 1.593684824 | 4.461553889 |
| ENSMUSG00000040043 | *Rbms2* | 533.3336105 | 514.6493192 | 443.5276176 | 778.9824923 | 851.7569694 | 746.6883308 | 7.23E-09 | 0.672342024 | 1.29E-09 | 36.8251086 | 1.593657961 | 8.140948351 |
| ENSMUSG00000029575 | *Mmab* | 317.2782085 | 371.691175 | 342.6813822 | 567.3782929 | 527.8225515 | 545.5828396 | 3.81E-08 | 0.671407894 | 7.27E-09 | 33.46150113 | 1.592626418 | 7.419311678 |
| ENSMUSG00000039577 | *Nphp4* | 72.30200461 | 68.18003802 | 83.2226214 | 115.1033832 | 132.432012 | 108.731677 | 0.009096089 | 0.671241715 | 0.00355942 | 8.495928724 | 1.592442979 | 2.041145319 |
| ENSMUSG00000025192 | *Entpd7* | 687.2943497 | 708.192653 | 744.1081443 | 1153.359153 | 1080.416558 | 1171.992767 | 4.25E-14 | 0.670826805 | 5.29E-15 | 61.1504118 | 1.591985069 | 13.37203345 |
| ENSMUSG00000025584 | *Pde8a* | 540.1385051 | 596.0254937 | 657.9482539 | 1008.026598 | 888.9141526 | 957.4160942 | 1.22E-09 | 0.670284001 | 2.04E-10 | 40.4259611 | 1.591386208 | 8.914090837 |
| ENSMUSG00000032294 | *Pkm* | 168501.9483 | 172098.5124 | 173897.0942 | 273549.5848 | 290370.0484 | 254673.6433 | 4.40E-31 | 0.669985967 | 2.67E-32 | 139.9916615 | 1.591057491 | 30.35681777 |
| ENSMUSG00000040102 | *Klhl42* | 380.2234831 | 396.9837698 | 358.3468169 | 579.0048973 | 628.81387 | 596.5808828 | 3.81E-09 | 0.669623282 | 6.68E-10 | 38.11246312 | 1.590657559 | 8.418641197 |
| ENSMUSG00000018474 | *Chd3* | 1435.83275 | 1523.054075 | 1406.951847 | 2496.231957 | 2092.23524 | 2352.645578 | 7.33E-15 | 0.668667428 | 8.69E-16 | 64.70697164 | 1.589604022 | 14.13516232 |
| ENSMUSG00000044030 | *Irf2bp1* | 832.7489708 | 836.8549828 | 887.0552352 | 1375.427296 | 1420.547697 | 1266.291035 | 1.30E-14 | 0.667928245 | 1.57E-15 | 63.54240372 | 1.588789777 | 13.88616795 |
| ENSMUSG00000035443 | *Thyn1* | 1835.620305 | 1997.015307 | 1984.614748 | 2807.824954 | 3570.900583 | 2855.89042 | 3.02E-11 | 0.66755378 | 4.47E-12 | 47.90793044 | 1.588377445 | 10.51984179 |
| ENSMUSG00000032381 | *Ciao2a* | 3139.608224 | 3074.699779 | 3087.069709 | 4812.251546 | 5080.053871 | 4874.643149 | 4.91E-34 | 0.666893212 | 2.68E-35 | 153.7111886 | 1.58765034 | 33.30894239 |
| ENSMUSG00000071369 | *Map3k5* | 742.584118 | 782.9707592 | 785.2299102 | 1212.654835 | 1149.967183 | 1301.893443 | 1.22E-13 | 0.666049477 | 1.55E-14 | 59.03645962 | 1.586722101 | 12.91412022 |
| ENSMUSG00000045038 | *Prkce* | 282.4031239 | 295.8133908 | 258.4796712 | 449.9495889 | 417.3037501 | 459.9446161 | 5.27E-07 | 0.665987013 | 1.12E-07 | 28.15264536 | 1.586653404 | 6.278121814 |
| ENSMUSG00000032047 | *Acat1* | 3696.758965 | 3840.07569 | 3757.746129 | 5649.36706 | 6342.445352 | 5901.339604 | 3.07E-26 | 0.664337759 | 2.18E-27 | 117.5468273 | 1.584840616 | 25.51269559 |
| ENSMUSG00000032066 | *Bco2* | 78.25628735 | 90.17359867 | 84.20171106 | 130.2179689 | 125.762774 | 143.3718573 | 0.004774264 | 0.664005943 | 0.001767008 | 9.777078829 | 1.584476149 | 2.321093604 |
| ENSMUSG00000089847 | *Timm10b* | 419.3516268 | 433.2731448 | 475.8375765 | 697.5962618 | 711.7029711 | 693.7658331 | 5.51E-10 | 0.663467414 | 8.99E-11 | 42.03063265 | 1.583884807 | 9.258543879 |
| ENSMUSG00000060935 | *Tmem263* | 329.1867739 | 307.9098491 | 269.2496575 | 502.2693085 | 397.296036 | 536.9227945 | 3.81E-05 | 0.662773783 | 1.01E-05 | 19.49750148 | 1.583123476 | 4.418830084 |
| ENSMUSG00000026811 | *St6galnac6* | 964.5938027 | 1021.600892 | 1062.312285 | 1735.852031 | 1619.672089 | 1471.245435 | 3.73E-13 | 0.662692509 | 4.92E-14 | 56.76323256 | 1.583034294 | 12.42858757 |
| ENSMUSG00000038893 | *Fam117a* | 119.9362665 | 100.070701 | 110.637132 | 193.0016324 | 199.1243922 | 132.7873578 | 0.006716956 | 0.662521248 | 0.002557919 | 9.098695391 | 1.582846384 | 2.172827466 |
| ENSMUSG00000032198 | *Dock6* | 213.5035666 | 268.3214399 | 285.8941818 | 416.2324362 | 403.0125258 | 393.5509372 | 1.97E-05 | 0.662505374 | 4.98E-06 | 20.84403639 | 1.582828968 | 4.705301251 |
| ENSMUSG00000052005 | *Gm9864* | 766.4012489 | 764.2762326 | 753.8990409 | 1170.799059 | 1163.30566 | 1279.762216 | 1.87E-14 | 0.661999407 | 2.28E-15 | 62.80579732 | 1.582273952 | 13.7281775 |
| ENSMUSG00000001403 | *Ube2c* | 2045.721425 | 2067.394701 | 2115.812763 | 3243.822617 | 3322.23328 | 3277.345947 | 6.86E-30 | 0.660465795 | 4.36E-31 | 134.4474363 | 1.580592858 | 29.16343051 |
| ENSMUSG00000034926 | *Dhcr24* | 6049.551257 | 5994.344956 | 5937.19972 | 9804.71546 | 9555.112579 | 9053.59601 | 1.10E-35 | 0.659857459 | 5.67E-37 | 161.3739456 | 1.579926516 | 34.96032703 |
| ENSMUSG00000036752 | *Tubb4b* | 6517.387757 | 6857.592211 | 6662.70516 | 10312.79807 | 11271.96499 | 10057.19901 | 2.97E-27 | 0.659392978 | 2.05E-28 | 122.2398887 | 1.579417935 | 26.52779118 |
| ENSMUSG00000052698 | *Tln2* | 425.3059095 | 435.4725009 | 447.4439762 | 696.4336014 | 681.2150258 | 687.9924697 | 1.19E-10 | 0.659203722 | 1.84E-11 | 45.12965959 | 1.579210757 | 9.924742438 |
| ENSMUSG00000028212 | *Ccne2* | 593.7270496 | 615.8196982 | 584.5165291 | 921.989726 | 970.8505054 | 938.1715496 | 3.84E-13 | 0.65916914 | 5.07E-14 | 56.70091334 | 1.579172903 | 12.41527602 |
| ENSMUSG00000026162 | *Nhej1* | 96.11913554 | 95.67198883 | 93.9926077 | 140.6819128 | 169.5891952 | 140.4851756 | 0.003311102 | 0.658881395 | 0.001193896 | 10.49976668 | 1.578857969 | 2.480027387 |
| ENSMUSG00000063894 | *Zkscan8* | 500.1597496 | 456.3663835 | 483.6702938 | 744.1026793 | 657.3963186 | 873.7023252 | 4.78E-07 | 0.658875769 | 1.01E-07 | 28.34921224 | 1.578851812 | 6.320118352 |
| ENSMUSG00000004936 | *Map2k1* | 4349.17823 | 4573.560937 | 4516.540618 | 7373.592487 | 7060.817562 | 6782.739747 | 8.48E-31 | 0.658808748 | 5.21E-32 | 138.6664532 | 1.578778467 | 30.07140503 |
| ENSMUSG00000021460 | *Auh* | 366.613694 | 351.8969704 | 353.4513685 | 537.1491216 | 602.1369179 | 552.3184302 | 1.79E-08 | 0.658396768 | 3.33E-09 | 34.98217976 | 1.578327692 | 7.746648428 |
| ENSMUSG00000039477 | *Tnrc18* | 1554.067793 | 1662.713185 | 1611.581586 | 2713.649458 | 2316.131088 | 2590.315704 | 3.74E-16 | 0.658197067 | 4.17E-17 | 70.69697517 | 1.578109231 | 15.42709282 |
| ENSMUSG00000039357 | *Fut11* | 357.256964 | 380.4885993 | 376.9495205 | 603.4207665 | 544.9720207 | 610.0520641 | 1.56E-08 | 0.658121179 | 2.88E-09 | 35.26509064 | 1.578026222 | 7.806197745 |
| ENSMUSG00000032558 | *Nphp3* | 294.3116894 | 274.9195081 | 281.9778231 | 477.8534393 | 393.4850429 | 473.4157973 | 2.86E-06 | 0.657376366 | 6.56E-07 | 24.73906318 | 1.577211752 | 5.543517525 |
| ENSMUSG00000050373 | *Snx21* | 106.3264774 | 124.2636177 | 136.0934632 | 187.1883303 | 192.4551542 | 197.2565822 | 0.001048636 | 0.657328461 | 0.000347459 | 12.79547447 | 1.577159382 | 2.979375062 |
| ENSMUSG00000021643 | *Serf1* | 186.2839884 | 168.250739 | 174.2779601 | 230.2067664 | 340.1311388 | 262.6880339 | 0.000923444 | 0.656912744 | 0.000303506 | 13.04863111 | 1.576704984 | 3.034589243 |
| ENSMUSG00000046711 | *Hmga1* | 3316.535482 | 3064.802677 | 3104.693323 | 4740.166599 | 5199.147407 | 5014.166097 | 1.89E-24 | 0.656429466 | 1.43E-25 | 109.2448866 | 1.576176904 | 23.72276482 |
| ENSMUSG00000040687 | *Madd* | 1301.436083 | 1185.452919 | 1297.293804 | 2175.337676 | 1912.165814 | 1884.040917 | 4.08E-14 | 0.656364711 | 5.07E-15 | 61.23373823 | 1.576106159 | 13.38921624 |
| ENSMUSG00000028807 | *Zbtb8a* | 124.1893256 | 147.3568564 | 105.7416837 | 205.7908972 | 181.0221747 | 206.8788545 | 0.001850844 | 0.656189065 | 0.000640124 | 11.65551849 | 1.575914282 | 2.732630228 |
| ENSMUSG00000051451 | *Crebzf* | 690.696797 | 673.0029559 | 692.2163921 | 1195.214929 | 972.7560019 | 1074.807816 | 2.51E-10 | 0.655716941 | 3.99E-11 | 43.61714332 | 1.575398647 | 9.60058212 |
| ENSMUSG00000066233 | *Tmem42* | 152.2595156 | 155.0546026 | 141.9680012 | 246.4840125 | 260.1002826 | 201.1054911 | 0.000544207 | 0.654987156 | 0.000170685 | 14.12907158 | 1.574601934 | 3.264236079 |
| ENSMUSG00000073755 | *5730409E04Rik* | 253.4823221 | 272.7201521 | 285.8941818 | 426.6963801 | 436.3587159 | 413.7577091 | 6.59E-07 | 0.654216111 | 1.41E-07 | 27.70488757 | 1.573760617 | 6.181251925 |
| ENSMUSG00000108608 | *Gm6916* | 162.4668574 | 171.5497731 | 187.9852154 | 274.387863 | 320.1234247 | 226.1233991 | 0.000716732 | 0.653606654 | 0.000230058 | 13.56812992 | 1.573095933 | 3.144643222 |
| ENSMUSG00000038717 | *Atp5l* | 1543.00984 | 1574.738943 | 1626.267931 | 2075.348879 | 2963.047175 | 2419.039257 | 3.14E-08 | 0.653412589 | 5.95E-09 | 33.85095203 | 1.572884341 | 7.503128173 |
| ENSMUSG00000029478 | *Ncor2* | 3453.483985 | 3664.127205 | 3436.604719 | 5933.056207 | 5229.635352 | 5430.810488 | 5.51E-22 | 0.652690653 | 4.59E-23 | 97.81508951 | 1.572097455 | 21.25845695 |
| ENSMUSG00000033961 | *Zfp446* | 222.8602966 | 252.9259475 | 245.7515055 | 426.6963801 | 322.9816696 | 384.8908921 | 5.34E-05 | 0.652668776 | 1.44E-05 | 18.81736266 | 1.572073616 | 4.272567405 |
| ENSMUSG00000013236 | *Ptprs* | 2049.974484 | 2060.796633 | 2132.457287 | 3491.46929 | 2949.708699 | 3376.455351 | 1.52E-16 | 0.652617094 | 1.66E-17 | 72.51078862 | 1.5720173 | 15.81821165 |
| ENSMUSG00000028152 | *Tspan5* | 631.1539697 | 629.0158346 | 660.8855229 | 970.8214643 | 1031.826396 | 1015.149728 | 2.59E-13 | 0.652249959 | 3.39E-14 | 57.4966779 | 1.571617305 | 12.58646457 |
| ENSMUSG00000047090 | *Tmem198b* | 146.3052329 | 140.7587882 | 140.9889115 | 226.7187851 | 223.8958477 | 222.2744902 | 0.000183111 | 0.651477886 | 5.30E-05 | 16.33736465 | 1.570776463 | 3.737286051 |
| ENSMUSG00000035047 | *Kri1* | 1466.454776 | 1501.060514 | 1452.969061 | 2490.418655 | 2216.092518 | 2239.102765 | 4.34E-18 | 0.651267017 | 4.40E-19 | 79.67929509 | 1.57054689 | 17.36284397 |
| ENSMUSG00000075307 | *Klhl41* | 306.2202548 | 346.3985803 | 271.2078368 | 422.0457384 | 553.5467553 | 470.5291156 | 4.02E-05 | 0.649709807 | 1.07E-05 | 19.39004018 | 1.568852595 | 4.395970996 |
| ENSMUSG00000074211 | *Sdhaf1* | 233.0676384 | 221.0352845 | 235.9606089 | 354.6114331 | 402.0597775 | 326.1950311 | 1.94E-05 | 0.649630845 | 4.89E-06 | 20.87793911 | 1.568766731 | 4.712162307 |
| ENSMUSG00000028795 | *Ccdc28b* | 112.2807601 | 123.1639396 | 98.88805602 | 154.633838 | 202.9353853 | 165.5030836 | 0.003788272 | 0.649617901 | 0.001377128 | 10.2361755 | 1.568752656 | 2.421558808 |
| ENSMUSG00000023348 | *Trip6* | 1110.048424 | 1064.488336 | 1142.597637 | 1687.020293 | 1835.945951 | 1678.12429 | 1.81E-16 | 0.648791054 | 1.99E-17 | 72.15816695 | 1.567853818 | 15.74251389 |
| ENSMUSG00000061119 | *Prcp* | 946.7309545 | 908.3340549 | 891.9506835 | 1370.776654 | 1424.35869 | 1509.734524 | 2.87E-15 | 0.648175618 | 3.33E-16 | 66.59549945 | 1.567185135 | 14.54166917 |
| ENSMUSG00000004897 | *Hdgf* | 14895.91417 | 15225.04236 | 15223.86518 | 24003.12471 | 24782.88846 | 22259.20252 | 7.37E-32 | 0.647819061 | 4.35E-33 | 143.5961408 | 1.566797858 | 31.13253499 |
| ENSMUSG00000032673 | *Prorsd1* | 208.3998957 | 205.6397921 | 207.5670087 | 308.1050156 | 320.1234247 | 344.4773485 | 1.26E-05 | 0.646836123 | 3.10E-06 | 21.75246302 | 1.56573073 | 4.900384981 |
| ENSMUSG00000039347 | *Atp6v0e2* | 649.8674297 | 693.8968386 | 675.5718678 | 1015.002561 | 1129.006721 | 1012.263046 | 7.00E-12 | 0.64591494 | 9.98E-13 | 50.84860863 | 1.564731305 | 11.1550582 |
| ENSMUSG00000054612 | *Mgmt* | 204.9974484 | 174.8488072 | 214.4206363 | 308.1050156 | 317.2651799 | 305.9882593 | 4.30E-05 | 0.645710773 | 1.14E-05 | 19.25396771 | 1.564509884 | 4.366368361 |
| ENSMUSG00000032437 | *Stt3b* | 4839.98125 | 4831.985275 | 4991.399105 | 7859.58455 | 7447.633366 | 7626.613028 | 4.10E-36 | 0.645051146 | 2.10E-37 | 163.3524793 | 1.563794725 | 35.38677628 |
| ENSMUSG00000037570 | *Mcrs1* | 669.4315015 | 935.8260057 | 926.2188217 | 1423.096374 | 1416.736704 | 1115.22136 | 1.34E-05 | 0.644941402 | 3.31E-06 | 21.62876601 | 1.563675773 | 4.873812637 |
| ENSMUSG00000020156 | *Pwwp3a* | 888.038739 | 834.6556267 | 891.9506835 | 1435.885639 | 1352.902569 | 1302.85567 | 1.82E-14 | 0.644416797 | 2.21E-15 | 62.86855195 | 1.563107279 | 13.7410368 |
| ENSMUSG00000032397 | *Tipin* | 2004.041446 | 1843.060383 | 1857.333092 | 2876.421919 | 2980.196645 | 3055.071456 | 1.32E-21 | 0.643104067 | 1.12E-22 | 96.05907949 | 1.561685631 | 20.8793816 |
| ENSMUSG00000029056 | *Pank4* | 589.4739906 | 559.7361186 | 584.5165291 | 883.6219316 | 946.0790499 | 878.5134613 | 9.76E-12 | 0.643056997 | 1.40E-12 | 50.17936783 | 1.56163468 | 11.01049998 |
| ENSMUSG00000021180 | *Rps6ka5* | 284.9549594 | 252.9259475 | 253.5842229 | 396.4672088 | 423.0202398 | 417.606618 | 1.93E-06 | 0.642891744 | 4.36E-07 | 25.52904492 | 1.561455813 | 5.715016129 |
| ENSMUSG00000037408 | *Cnnm4* | 346.1990103 | 377.1895652 | 384.7822378 | 632.4872774 | 542.1137758 | 556.1673392 | 2.13E-07 | 0.642862452 | 4.34E-08 | 29.99225283 | 1.561424111 | 6.672496777 |
| ENSMUSG00000020520 | *Galnt10* | 406.5924495 | 390.3857016 | 394.5731344 | 667.3670905 | 656.4435704 | 538.847249 | 4.39E-07 | 0.642841055 | 9.26E-08 | 28.5221866 | 1.561400953 | 6.357086417 |
| ENSMUSG00000008226 | *Scrn3* | 392.9826604 | 350.7972924 | 374.0122515 | 573.1915951 | 576.4127142 | 597.5431101 | 1.71E-08 | 0.642622038 | 3.16E-09 | 35.08184431 | 1.561163933 | 7.767107984 |
| ENSMUSG00000024228 | *Nudt12* | 158.2137983 | 144.0578223 | 136.0934632 | 237.182729 | 203.8881336 | 244.4057165 | 0.00057405 | 0.642208878 | 0.000180873 | 14.0200397 | 1.560716909 | 3.241050179 |
| ENSMUSG00000078249 | *Hmga1b* | 17134.72448 | 18002.82907 | 18061.26702 | 28388.67987 | 28406.1902 | 26223.57871 | 1.52E-32 | 0.642077962 | 8.74E-34 | 146.7851668 | 1.56057529 | 31.81797019 |
| ENSMUSG00000038034 | *Igsf8* | 3448.380314 | 3588.24942 | 3666.69079 | 5705.174761 | 5786.040352 | 5208.535998 | 1.88E-23 | 0.641862245 | 1.48E-24 | 104.6135973 | 1.560341964 | 22.72482773 |
| ENSMUSG00000032235 | *Ice2* | 540.9891169 | 448.6686373 | 458.2139625 | 698.7589222 | 768.8678683 | 793.837465 | 7.46E-08 | 0.64185742 | 1.47E-08 | 32.09464102 | 1.560336746 | 7.127453404 |
| ENSMUSG00000036114 | *Rpp25l* | 585.2209315 | 651.0093953 | 655.0109849 | 980.1227478 | 1074.700069 | 891.9846425 | 3.88E-09 | 0.641174486 | 6.80E-10 | 38.0760823 | 1.559598298 | 8.41071365 |
| ENSMUSG00000000168 | *Dlat* | 2670.921112 | 2644.725668 | 2613.190312 | 4227.433347 | 3927.228443 | 4213.593042 | 1.22E-26 | 0.641100029 | 8.59E-28 | 119.3942393 | 1.55951781 | 25.91254242 |
| ENSMUSG00000032040 | *Dcps* | 1338.863003 | 1295.420722 | 1277.712011 | 1910.251097 | 2344.713536 | 1841.702919 | 3.45E-10 | 0.640363436 | 5.56E-11 | 42.97109208 | 1.558721775 | 9.461843171 |
| ENSMUSG00000070319 | *Eif3g* | 4213.930951 | 4447.097964 | 4379.468065 | 6557.404861 | 7298.051885 | 6463.280306 | 2.50E-22 | 0.640233442 | 2.06E-23 | 99.40716134 | 1.558581333 | 21.60153904 |
| ENSMUSG00000042496 | *Prdm10* | 461.8822177 | 433.2731448 | 468.0048592 | 716.1988288 | 660.2545635 | 749.5750125 | 4.58E-09 | 0.639968415 | 8.06E-10 | 37.74669072 | 1.558295043 | 8.339594797 |
| ENSMUSG00000014301 | *Pam16* | 233.9182502 | 223.2346406 | 303.5177957 | 413.9071153 | 435.4059676 | 335.8173034 | 0.000372985 | 0.638796462 | 0.00011382 | 14.89241032 | 1.557029698 | 3.428308462 |
| ENSMUSG00000038387 | *Rras* | 148.0064565 | 170.450095 | 190.9224844 | 272.0625421 | 263.9112757 | 255.9524433 | 0.000276541 | 0.638522199 | 8.25E-05 | 15.49917935 | 1.556733728 | 3.558240339 |
| ENSMUSG00000057963 | *Itpk1* | 606.4862269 | 626.8164786 | 621.7219363 | 961.5201809 | 1005.149444 | 919.8892322 | 3.86E-12 | 0.638512735 | 5.41E-13 | 52.0509137 | 1.556723515 | 11.41343982 |
| ENSMUSG00000039740 | *Alg2* | 645.6143706 | 597.1251717 | 616.826488 | 988.2613709 | 969.8977571 | 939.1337768 | 2.70E-12 | 0.638003934 | 3.75E-13 | 52.7713581 | 1.556174597 | 11.56934665 |
| ENSMUSG00000032434 | *Cmtm6* | 1940.245559 | 2064.095667 | 1999.301093 | 3254.286561 | 2900.165788 | 3185.93436 | 3.53E-19 | 0.637726464 | 3.39E-20 | 84.74572789 | 1.55587533 | 18.45226084 |
| ENSMUSG00000004730 | *Adgre1* | 2247.316426 | 2011.311122 | 2052.171935 | 3238.009315 | 3289.839838 | 3295.628264 | 2.78E-21 | 0.637499969 | 2.40E-22 | 94.53944413 | 1.555631086 | 20.55542601 |
| ENSMUSG00000033751 | *Gadd45gip1* | 712.8127043 | 698.2955507 | 775.4390135 | 1048.719714 | 1319.556379 | 1029.583136 | 1.02E-07 | 0.636593736 | 2.04E-08 | 31.45789776 | 1.554654218 | 6.989889057 |
| ENSMUSG00000002031 | *Ift46* | 879.5326208 | 842.353373 | 832.226214 | 1309.155651 | 1380.532269 | 1280.724444 | 9.75E-15 | 0.636008997 | 1.17E-15 | 64.12524763 | 1.554024228 | 14.01093862 |
| ENSMUSG00000042489 | *Clspn* | 706.8584216 | 719.1894333 | 739.212696 | 1134.756586 | 1099.471524 | 1129.654768 | 3.08E-14 | 0.63565747 | 3.79E-15 | 61.80710144 | 1.553645621 | 13.51206915 |
| ENSMUSG00000113890 | *Gm48368* | 2015.950011 | 2145.471842 | 2119.729122 | 3232.196013 | 3177.41554 | 3323.532854 | 1.84E-24 | 0.632571611 | 1.39E-25 | 109.3099114 | 1.550325996 | 23.73582998 |
| ENSMUSG00000043008 | *Klhl6* | 483.1475132 | 424.4757206 | 451.3603349 | 749.9159814 | 687.8842639 | 673.5590613 | 1.35E-08 | 0.632449424 | 2.47E-09 | 35.55904363 | 1.550194699 | 7.868167081 |
| ENSMUSG00000035585 | *Tsen34* | 659.2241597 | 602.6235619 | 701.0281991 | 1052.207695 | 1048.023117 | 945.8693674 | 4.71E-10 | 0.632424486 | 7.65E-11 | 42.34482469 | 1.550167903 | 9.326852774 |
| ENSMUSG00000020441 | *2310033P09Rik* | 364.9124703 | 369.4918189 | 374.9913411 | 539.4744425 | 606.9006594 | 571.5629748 | 4.20E-08 | 0.632366615 | 8.06E-09 | 33.26050409 | 1.550105722 | 7.376916739 |
| ENSMUSG00000003355 | *Fkbp11* | 280.7019003 | 250.7265914 | 278.0614644 | 370.8886792 | 522.1060618 | 360.8352114 | 0.000481036 | 0.631543993 | 0.000149222 | 14.38199612 | 1.549222107 | 3.317822186 |
| ENSMUSG00000030811 | *Fbxl19* | 546.9433996 | 560.8357966 | 579.6210808 | 944.0802743 | 853.6624659 | 817.8931458 | 7.08E-10 | 0.631370302 | 1.17E-10 | 41.52012495 | 1.549035602 | 9.149696578 |
| ENSMUSG00000006611 | *Hfe* | 288.3574066 | 267.2217619 | 275.1241955 | 420.883078 | 435.4059676 | 431.0777992 | 7.90E-07 | 0.630959932 | 1.71E-07 | 27.33953423 | 1.548595046 | 6.102534521 |
| ENSMUSG00000029998 | *Pcyox1* | 1206.167559 | 1200.848412 | 1230.715707 | 1814.912941 | 1918.835052 | 1892.700962 | 3.14E-19 | 0.629662369 | 3.00E-20 | 84.9859252 | 1.547202863 | 18.5025329 |
| ENSMUSG00000024847 | *Aip* | 1382.244206 | 1332.809775 | 1409.889115 | 2116.041994 | 2352.335523 | 1913.869961 | 4.19E-12 | 0.629507311 | 5.88E-13 | 51.8850883 | 1.547036581 | 11.37826327 |
| ENSMUSG00000035342 | *Lzts2* | 688.9955734 | 698.2955507 | 704.9445577 | 1177.775022 | 1044.212124 | 1016.111955 | 5.52E-11 | 0.628893377 | 8.38E-12 | 46.67571292 | 1.546378385 | 10.25777325 |
| ENSMUSG00000039470 | *Zdhhc2* | 813.1848989 | 835.7553048 | 847.8916486 | 1320.782256 | 1237.620026 | 1300.931215 | 1.09E-14 | 0.628197248 | 1.30E-15 | 63.90788104 | 1.545632407 | 13.96351963 |
| ENSMUSG00000019838 | *Slc16a10* | 3698.460189 | 3674.024307 | 3845.864198 | 5899.339054 | 5730.780952 | 5709.856385 | 2.12E-32 | 0.628015601 | 1.23E-33 | 146.1149258 | 1.545437812 | 31.67454697 |
| ENSMUSG00000032526 | *Ss18l2* | 259.4366048 | 241.9291672 | 235.9606089 | 388.3285857 | 376.3355737 | 376.2308471 | 4.13E-06 | 0.627862495 | 9.68E-07 | 23.9907874 | 1.54527381 | 5.383564961 |
| ENSMUSG00000057191 | *AB124611* | 1740.351782 | 1742.989682 | 1629.2052 | 2612.498 | 2822.993177 | 2454.641665 | 6.78E-16 | 0.626096206 | 7.65E-17 | 69.49837054 | 1.543383092 | 15.1690525 |
| ENSMUSG00000021910 | *Nisch* | 3764.807911 | 3992.930936 | 3941.814985 | 6329.523415 | 5764.127142 | 5960.997692 | 6.55E-25 | 0.62592524 | 4.85E-26 | 111.3940824 | 1.543200205 | 24.1840685 |
| ENSMUSG00000032477 | *Cdc25a* | 1469.006611 | 1466.970495 | 1442.199074 | 2289.278399 | 2183.699076 | 2284.327445 | 2.28E-21 | 0.625825566 | 1.95E-22 | 94.95100984 | 1.543093591 | 20.6426025 |
| ENSMUSG00000025212 | *Sfxn3* | 449.9736522 | 511.3502851 | 542.4156736 | 781.3078132 | 695.5062502 | 840.0243721 | 4.09E-07 | 0.624895412 | 8.58E-08 | 28.67088363 | 1.542099027 | 6.388311048 |
| ENSMUSG00000060862 | *Zbtb40* | 571.6111424 | 663.1058536 | 556.1229289 | 954.5442182 | 867.9536903 | 936.2470951 | 6.21E-09 | 0.624718922 | 1.10E-09 | 37.132238 | 1.541910388 | 8.207151552 |
| ENSMUSG00000026938 | *Fcna* | 207.5492838 | 207.8391482 | 192.8806637 | 259.2732773 | 363.949846 | 312.7238499 | 0.000492892 | 0.62455815 | 0.00015317 | 14.33282739 | 1.541738569 | 3.307248541 |
| ENSMUSG00000019659 | *Ccdc12* | 949.28279 | 1012.803468 | 1021.190519 | 1569.591589 | 1646.349041 | 1380.796076 | 2.48E-11 | 0.624144648 | 3.65E-12 | 48.30177441 | 1.541296743 | 10.60525697 |
| ENSMUSG00000024213 | *Nudt3* | 2007.443893 | 2061.896311 | 2055.109204 | 3196.153539 | 3185.990275 | 3055.071456 | 3.36E-25 | 0.6238973 | 2.46E-26 | 112.7396073 | 1.541032513 | 24.47314159 |
| ENSMUSG00000093930 | *Hmgcs1* | 6525.043263 | 6241.772513 | 6440.451807 | 10260.47835 | 9006.329565 | 10334.32045 | 1.41E-20 | 0.623724271 | 1.26E-21 | 91.2647764 | 1.5408477 | 19.84976698 |
| ENSMUSG00000030122 | *Ptms* | 2489.740794 | 2604.037581 | 2571.089456 | 4006.527864 | 4139.691311 | 3660.312384 | 8.56E-20 | 0.623456977 | 7.90E-21 | 87.62852623 | 1.540562248 | 19.06731796 |
| ENSMUSG00000050549 | *Fam241a* | 234.768862 | 266.1220839 | 296.6641681 | 431.3470219 | 436.3587159 | 359.8729842 | 7.79E-05 | 0.623122724 | 2.15E-05 | 18.05205898 | 1.540205362 | 4.10864178 |
| ENSMUSG00000001942 | *Siae* | 602.2331678 | 641.112293 | 582.5583498 | 878.9712899 | 951.7955396 | 976.6606388 | 1.88E-10 | 0.622590592 | 2.97E-11 | 44.19432432 | 1.539637368 | 9.725081361 |
| ENSMUSG00000021594 | *Srd5a1* | 102.0734183 | 96.77166687 | 112.5953113 | 186.0256698 | 165.7782021 | 128.9384489 | 0.009509222 | 0.622562292 | 0.003741799 | 8.405050786 | 1.539607167 | 2.021855014 |
| ENSMUSG00000029674 | *Limk1* | 276.4488412 | 285.9162885 | 237.9187882 | 451.1122493 | 422.0674915 | 359.8729842 | 5.71E-05 | 0.622510688 | 1.55E-05 | 18.6779224 | 1.539552097 | 4.243128129 |
| ENSMUSG00000030847 | *Bag3* | 1491.122519 | 1543.947958 | 1426.53364 | 2348.574081 | 2288.501388 | 2231.404947 | 1.89E-19 | 0.622485461 | 1.78E-20 | 86.01754253 | 1.539525177 | 18.72282721 |
| ENSMUSG00000027304 | *Rtf1* | 1485.168236 | 1391.092711 | 1390.307322 | 2263.69987 | 1959.803228 | 2346.872215 | 1.92E-12 | 0.622032763 | 2.64E-13 | 53.46065803 | 1.53904217 | 11.71743842 |
| ENSMUSG00000030019 | *Fbxl14* | 1344.817286 | 1481.26631 | 1413.805474 | 2352.062063 | 2109.384709 | 2060.1285 | 3.91E-14 | 0.621246252 | 4.85E-15 | 61.32124868 | 1.538203363 | 13.40755879 |
| ENSMUSG00000028756 | *Pink1* | 275.5982294 | 258.4243377 | 320.16232 | 445.2989471 | 452.5554367 | 416.6443907 | 1.01E-05 | 0.620783262 | 2.45E-06 | 22.20830862 | 1.537709802 | 4.997680515 |
| ENSMUSG00000009927 | *Rps25* | 16056.1487 | 16058.59831 | 17058.67921 | 22482.36486 | 30130.6646 | 22952.96835 | 9.28E-10 | 0.61992584 | 1.54E-10 | 40.97473888 | 1.536796182 | 9.032304641 |
| ENSMUSG00000058192 | *Zfp846* | 188.8358238 | 159.4533147 | 185.0479464 | 260.4359377 | 261.0530309 | 299.2526686 | 0.00029431 | 0.619912491 | 8.82E-05 | 15.37368836 | 1.536781963 | 3.5311956 |
| ENSMUSG00000025810 | *Nrp1* | 668.5808897 | 683.9997363 | 620.7428467 | 1018.490542 | 969.8977571 | 1042.09209 | 2.36E-11 | 0.619048465 | 3.47E-12 | 48.40055897 | 1.535861864 | 10.62632446 |
| ENSMUSG00000038773 | *Kdm3b* | 1868.794166 | 2130.076349 | 1918.036651 | 3106.628686 | 2863.008605 | 3112.80509 | 6.41E-16 | 0.618993077 | 7.22E-17 | 69.61258355 | 1.5358029 | 15.1931432 |
| ENSMUSG00000030844 | *Rgs10* | 888.8893508 | 924.8292254 | 948.737884 | 1326.595558 | 1545.357723 | 1364.438213 | 1.48E-11 | 0.618162934 | 2.14E-12 | 49.34752817 | 1.534919435 | 10.83008694 |
| ENSMUSG00000029196 | *Tada2b* | 518.0225978 | 465.1638078 | 472.9003075 | 782.4704737 | 706.9392296 | 748.6127852 | 7.92E-09 | 0.6174662 | 1.42E-09 | 36.64006187 | 1.534178342 | 8.101189219 |
| ENSMUSG00000051950 | *B3glct* | 282.4031239 | 258.4243377 | 267.2914781 | 410.419134 | 418.2564984 | 411.8332546 | 2.20E-06 | 0.616728797 | 5.00E-07 | 25.26547012 | 1.53339438 | 5.658044055 |
| ENSMUSG00000054892 | *Txk* | 113.1313719 | 119.8649056 | 111.6162216 | 146.495215 | 189.5969093 | 190.5209916 | 0.004472393 | 0.616260383 | 0.001650066 | 9.903033513 | 1.532896599 | 2.349460079 |
| ENSMUSG00000025037 | *Maoa* | 565.6568596 | 574.031933 | 545.3529426 | 794.097078 | 940.3625601 | 845.7977355 | 5.65E-09 | 0.616191924 | 1.00E-09 | 37.31877121 | 1.53282386 | 8.247854398 |
| ENSMUSG00000032115 | *Hyou1* | 15170.66179 | 15420.78505 | 15034.90087 | 23976.38352 | 22875.48639 | 23062.66226 | 2.37E-42 | 0.615682137 | 1.04E-43 | 192.2312247 | 1.532282321 | 41.62524323 |
| ENSMUSG00000018599 | *Mief2* | 253.4823221 | 255.1253036 | 302.538706 | 432.5096823 | 425.8784847 | 384.8908921 | 2.01E-05 | 0.61556674 | 5.10E-06 | 20.7997413 | 1.532159764 | 4.695846148 |
| ENSMUSG00000039461 | *Tcta* | 199.8937775 | 168.250739 | 185.0479464 | 266.2492399 | 309.6431936 | 272.3103062 | 0.000265209 | 0.614862918 | 7.88E-05 | 15.58655285 | 1.531412478 | 3.576410981 |
| ENSMUSG00000074527 | *Gm14296* | 144.6040092 | 123.1639396 | 127.2816563 | 190.6763116 | 181.0221747 | 233.821217 | 0.003042467 | 0.61429828 | 0.001089101 | 10.66961039 | 1.530813236 | 2.516774095 |
| ENSMUSG00000031400 | *G6pdx* | 9943.652164 | 9859.71324 | 9570.601461 | 14682.07599 | 15449.76624 | 14826.95939 | 3.62E-37 | 0.61411758 | 1.80E-38 | 168.2351483 | 1.530621511 | 36.44092032 |
| ENSMUSG00000022272 | *Myo10* | 4208.82728 | 4407.509555 | 4463.669776 | 7007.35445 | 6037.5659 | 6975.185193 | 2.95E-17 | 0.614041059 | 3.11E-18 | 75.81839835 | 1.530540328 | 16.53091358 |
| ENSMUSG00000083327 | *Vcp-rs* | 1963.212078 | 1929.934947 | 1496.049006 | 2634.588549 | 2769.639273 | 2840.494784 | 4.88E-09 | 0.613489897 | 8.63E-10 | 37.61279328 | 1.529955718 | 8.311135962 |
| ENSMUSG00000082691 | *Dynlt1-ps1* | 2479.533452 | 2432.487808 | 2382.125151 | 3404.269758 | 4200.667201 | 3550.61848 | 1.59E-12 | 0.613326715 | 2.17E-13 | 53.8437242 | 1.529782676 | 11.79897947 |
| ENSMUSG00000044456 | *Rin3* | 617.5441806 | 583.9290353 | 568.8510945 | 931.2910095 | 942.2680567 | 837.1376904 | 1.07E-09 | 0.613308938 | 1.79E-10 | 40.68247366 | 1.529763826 | 8.969144001 |
| ENSMUSG00000022426 | *Josd1* | 1600.851443 | 1653.915761 | 1590.041614 | 2556.6903 | 2477.145549 | 2377.663486 | 4.27E-20 | 0.61324833 | 3.90E-21 | 89.02527787 | 1.529699561 | 19.36910466 |
| ENSMUSG00000062619 | *2310039H08Rik* | 115.6832074 | 104.4694131 | 125.3234769 | 163.9351215 | 206.7463785 | 157.8052658 | 0.005931091 | 0.613230782 | 0.002233418 | 9.347073959 | 1.529680955 | 2.226865396 |
| ENSMUSG00000032306 | *Mpi* | 611.5898978 | 657.6074635 | 597.2446948 | 946.4055952 | 979.4252399 | 926.6248228 | 5.31E-11 | 0.613166288 | 8.02E-12 | 46.76158874 | 1.529612574 | 10.27503236 |
| ENSMUSG00000046806 | *Cyren* | 272.1957821 | 272.7201521 | 209.525188 | 372.0513396 | 404.9180224 | 376.2308471 | 0.000105628 | 0.612617573 | 2.96E-05 | 17.44103954 | 1.529030912 | 3.976221789 |
| ENSMUSG00000032741 | *Tpcn1* | 1784.583596 | 1860.655231 | 1814.253147 | 2860.144673 | 2842.048143 | 2642.275975 | 4.20E-20 | 0.612176897 | 3.83E-21 | 89.0607378 | 1.528563935 | 19.37656241 |
| ENSMUSG00000078183 | *Gm15610* | 375.970424 | 363.9934288 | 415.1340173 | 560.4023303 | 633.5776114 | 570.6007476 | 4.58E-07 | 0.611807381 | 9.68E-08 | 28.43652311 | 1.528172476 | 6.338718471 |
| ENSMUSG00000031295 | *Phka2* | 1081.978234 | 1130.469017 | 1149.451265 | 1754.454598 | 1740.671122 | 1641.559655 | 4.95E-16 | 0.611806423 | 5.54E-17 | 70.13332195 | 1.528171461 | 15.3051405 |
| ENSMUSG00000067719 | *Gm10221* | 3051.144595 | 2873.458699 | 3137.003282 | 4027.455751 | 5760.316149 | 4055.787776 | 1.15E-06 | 0.611552866 | 2.54E-07 | 26.56770266 | 1.527902905 | 5.93757332 |
| ENSMUSG00000010751 | *Tnfrsf22* | 693.2486325 | 696.0961946 | 680.4673162 | 1085.924848 | 945.1263016 | 1131.579223 | 1.63E-09 | 0.611178784 | 2.76E-10 | 39.83742874 | 1.52750678 | 8.788201329 |
| ENSMUSG00000033186 | *Mzt1* | 492.5042432 | 413.4789402 | 491.5030111 | 737.1267166 | 686.9315156 | 714.9348322 | 1.02E-07 | 0.611110589 | 2.03E-08 | 31.46967537 | 1.527434578 | 6.99237364 |
| ENSMUSG00000038884 | *Shfl* | 1028.389689 | 1009.504434 | 1009.441443 | 1687.020293 | 1341.469589 | 1629.050701 | 2.17E-09 | 0.610997904 | 3.71E-10 | 39.26088857 | 1.527315279 | 8.664184054 |
| ENSMUSG00000046152 | *Fut10* | 175.2260347 | 180.3471973 | 150.7798082 | 256.9479564 | 253.4310446 | 262.6880339 | 0.000319466 | 0.610929788 | 9.65E-05 | 15.20417496 | 1.527243169 | 3.495575651 |

| | | | | | | | | | | | | | |
|---|---|---|---|---|---|---|---|---|---|---|---|---|---|
| ENSMUSG00000020069 | *Hnmph3* | 342.796563 | 274.9195081 | 284.9150921 | 479.0160998 | 392.5322946 | 509.9804321 | 0.00014057 | 0.610649522 | 4.01E-05 | 16.86733006 | 1.526946508 | 3.852108008 |
| ENSMUSG00000032777 | *Gtf3c1* | 1868.794166 | 1910.140743 | 1857.333092 | 2988.037321 | 2874.441585 | 2741.385379 | 1.03E-20 | 0.609980483 | 9.11E-22 | 91.90159039 | 1.526238561 | 19.98683885 |
| ENSMUSG00000019866 | *Crybg1* | 2289.847017 | 2331.317429 | 2253.864405 | 3565.879558 | 3257.446396 | 3668.972429 | 5.81E-19 | 0.609831343 | 5.68E-20 | 83.72831792 | 1.526080793 | 18.23614594 |
| ENSMUSG00000033543 | *Gtf2a2* | 818.2885699 | 727.9868576 | 847.8916486 | 1178.937682 | 1344.327834 | 1131.579223 | 7.36E-09 | 0.609593589 | 1.32E-09 | 36.7896747 | 1.525829319 | 8.133354857 |
| ENSMUSG00000026657 | *Frmd4a* | 666.8796661 | 681.8003802 | 673.6136885 | 1074.298243 | 943.220805 | 1069.034453 | 2.02E-10 | 0.609551083 | 3.20E-11 | 44.0512689 | 1.525784363 | 9.694661219 |
| ENSMUSG00000043991 | *Pura* | 634.556417 | 605.9225959 | 644.2409986 | 989.4240313 | 866.0481937 | 1021.885319 | 3.69E-09 | 0.609277475 | 6.45E-10 | 38.18104941 | 1.525495025 | 8.433395903 |
| ENSMUSG00000021608 | *Lpcat1* | 1853.483154 | 1707.799985 | 1843.625836 | 2793.873029 | 2723.907355 | 2731.763107 | 2.44E-20 | 0.60897399 | 2.19E-21 | 90.16774545 | 1.525174156 | 19.61298142 |
| ENSMUSG00000037523 | *Mavs* | 1085.380681 | 1116.173203 | 1046.64685 | 1760.267901 | 1621.577586 | 1572.279294 | 6.36E-14 | 0.608406936 | 7.98E-15 | 60.34048714 | 1.524574802 | 13.19669065 |
| ENSMUSG00000041498 | *Kif14* | 637.1082524 | 625.7168005 | 647.1782676 | 960.3575204 | 913.6856081 | 1036.318727 | 2.07E-10 | 0.607492405 | 3.28E-11 | 44.00144911 | 1.523608672 | 9.684171479 |
| ENSMUSG00000020895 | *Tmem107* | 152.2595156 | 177.0481632 | 158.6125255 | 260.4359377 | 283.9189898 | 198.2188095 | 0.002942293 | 0.607336468 | 0.001050011 | 10.73723438 | 1.523443999 | 2.531314065 |
| ENSMUSG00000033083 | *Tbc1d4* | 263.6896639 | 244.1285232 | 236.9396986 | 370.8886792 | 359.1861045 | 405.097664 | 1.94E-05 | 0.606943037 | 4.89E-06 | 20.88027847 | 1.523028604 | 4.712574585 |
| ENSMUSG00000001768 | *Rin2* | 654.9711006 | 636.7135809 | 595.2865154 | 1011.51458 | 893.677894 | 970.8872754 | 9.00E-10 | 0.606635426 | 1.49E-10 | 41.03706548 | 1.522703899 | 9.045615823 |
| ENSMUSG00000063952 | *Brpf3* | 332.5892212 | 347.4982583 | 330.9323063 | 560.4023303 | 449.6971919 | 530.1872039 | 6.48E-06 | 0.606385556 | 1.54E-06 | 23.0916112 | 1.522440194 | 5.188426193 |
| ENSMUSG00000050592 | *Fam78a* | 613.2911215 | 699.3952287 | 595.2865154 | 955.7068787 | 951.7955396 | 992.0562745 | 1.46E-09 | 0.605775388 | 2.47E-10 | 40.05336364 | 1.521796436 | 8.834626978 |
| ENSMUSG00000038055 | *Dexi* | 176.9272583 | 184.7459095 | 161.5497945 | 282.526486 | 276.2970035 | 237.6701259 | 0.000455015 | 0.605672124 | 0.00014037 | 14.49717648 | 1.521687513 | 3.341973919 |
| ENSMUSG00000030337 | *Vamp1* | 268.7933348 | 273.8198301 | 268.2705678 | 454.6002306 | 359.1861045 | 420.4932997 | 3.75E-05 | 0.604392331 | 9.91E-06 | 19.52920537 | 1.520338246 | 4.425699541 |
| ENSMUSG00000028556 | *Dock7* | 1604.253891 | 1545.047636 | 1535.212592 | 2428.797652 | 2102.715471 | 2591.277931 | 1.92E-11 | 0.604052885 | 2.80E-12 | 48.82099651 | 1.519980574 | 10.71637713 |
| ENSMUSG00000050954 | *Zfp169* | 142.0521738 | 136.360076 | 158.6125255 | 222.0681433 | 208.651875 | 233.821217 | 0.000956568 | 0.60389179 | 0.000315312 | 12.97717308 | 1.519810858 | 3.019284173 |
| ENSMUSG00000042476 | *Abcb4* | 558.0013533 | 591.6267815 | 561.0183772 | 904.5498195 | 808.8832964 | 886.2112792 | 1.27E-09 | 0.603843885 | 2.13E-10 | 40.34179122 | 1.519760394 | 8.896091838 |
| ENSMUSG00000070056 | *Mfhas1* | 494.2054668 | 504.752217 | 522.8338803 | 824.3262494 | 739.3326714 | 750.5372397 | 5.14E-09 | 0.603808091 | 9.10E-10 | 37.50885549 | 1.519722688 | 8.289091203 |
| ENSMUSG00000027510 | *Rbm38* | 943.3285073 | 1033.697351 | 978.1105739 | 1559.127645 | 1556.790702 | 1371.173803 | 1.58E-11 | 0.603074398 | 2.29E-12 | 49.22038666 | 1.518950019 | 10.80214453 |
| ENSMUSG00000056394 | *Lig1* | 2366.40208 | 2415.992638 | 2460.452324 | 3637.964505 | 3814.804145 | 3540.033981 | 8.25E-23 | 0.602229841 | 6.66E-24 | 101.6410176 | 1.518061082 | 22.08366203 |
| ENSMUSG00000003031 | *Cdkn1b* | 649.8674297 | 586.1283914 | 588.4328878 | 963.8455017 | 899.3943838 | 910.2669599 | 6.04E-10 | 0.601887203 | 9.89E-11 | 41.84380907 | 1.517700587 | 9.219056552 |
| ENSMUSG00000032612 | *Usp4* | 3789.475654 | 3781.792754 | 3748.934322 | 5515.66111 | 6057.573614 | 5596.313572 | 4.24E-24 | 0.601213417 | 3.25E-25 | 107.6211616 | 1.516991937 | 23.3728441 |
| ENSMUSG00000054792 | *Klhl18* | 515.4707623 | 506.951573 | 537.5202253 | 770.8438693 | 792.6865755 | 801.5352829 | 6.12E-10 | 0.600594444 | 1.00E-10 | 41.8155535 | 1.516341227 | 9.213326259 |
| ENSMUSG00000031971 | *Ccsap* | 134.3966674 | 122.0642616 | 110.637132 | 177.8870468 | 194.3606507 | 184.7476282 | 0.002760187 | 0.600399006 | 0.000981046 | 10.86300035 | 1.516135826 | 2.55906152 |
| ENSMUSG00000028906 | *Epb41* | 5243.171252 | 5221.271299 | 5356.599549 | 8117.695166 | 7931.629497 | 7938.37465 | 4.35E-36 | 0.600305633 | 2.23E-37 | 163.2306925 | 1.516037703 | 35.36133647 |
| ENSMUSG00000021254 | *Gpatch2l* | 626.0502988 | 625.7168005 | 566.8929152 | 935.9416513 | 898.4416355 | 920.8514595 | 3.02E-10 | 0.598859016 | 4.83E-11 | 43.2433974 | 1.514518306 | 9.519873622 |
| ENSMUSG00000102914 | *Gm35106* | 358.9581876 | 281.5175763 | 411.2176587 | 488.3173833 | 694.5535019 | 409.9088001 | 0.006069197 | 0.598165531 | 0.002291253 | 9.300228199 | 1.513790472 | 2.216868744 |
| ENSMUSG00000016933 | *Plcg1* | 318.1288203 | 294.7137127 | 274.1451058 | 481.3414206 | 419.2092467 | 444.5489804 | 1.13E-05 | 0.597880884 | 2.76E-06 | 21.97381588 | 1.513491827 | 4.94731488 |
| ENSMUSG00000022817 | *Itgb5* | 788.5171562 | 654.3084294 | 798.9371654 | 1203.353552 | 1185.21887 | 1009.376365 | 2.80E-07 | 0.597852017 | 5.77E-08 | 29.43764691 | 1.513461544 | 6.553040754 |
| ENSMUSG00000068101 | *Cenpm* | 201.5950011 | 201.24108 | 214.4206363 | 334.8462057 | 327.745411 | 272.3103062 | 0.000243168 | 0.597554614 | 7.19E-05 | 15.76028797 | 1.513149584 | 3.614093681 |
| ENSMUSG00000041193 | *Pla2g5* | 2744.073728 | 2600.738547 | 2555.424022 | 4071.636848 | 3948.188905 | 3939.358281 | 1.25E-23 | 0.597418725 | 9.77E-25 | 105.4414122 | 1.513007066 | 22.90322882 |
| ENSMUSG00000020564 | *Atxn7l1* | 559.7025769 | 558.6364405 | 546.3320323 | 840.6034955 | 805.0723033 | 872.7400979 | 4.71E-10 | 0.597163217 | 7.65E-11 | 42.34487913 | 1.512739129 | 9.326852774 |
| ENSMUSG00000018750 | *Zbtb4* | 366.613694 | 417.8776524 | 335.8277546 | 591.7941621 | 491.6181165 | 610.0520641 | 3.14E-05 | 0.596921061 | 8.16E-06 | 19.89920335 | 1.512485238 | 4.503760153 |
| ENSMUSG00000043445 | *Pgp* | 983.3072627 | 974.3147369 | 1062.312285 | 1461.464168 | 1693.033708 | 1411.587347 | 7.63E-10 | 0.596853974 | 1.26E-10 | 41.36610448 | 1.512414907 | 9.117465348 |
| ENSMUSG00000040964 | *Arhgef10l* | 1884.105179 | 1857.356197 | 1803.48316 | 2927.578979 | 2828.709667 | 2628.804793 | 1.18E-17 | 0.596001734 | 1.22E-18 | 77.66500361 | 1.511521745 | 16.92988337 |
| ENSMUSG00000015127 | *Unkl* | 307.9214785 | 392.5850576 | 310.3714233 | 477.8534393 | 524.9643066 | 520.5649316 | 2.85E-05 | 0.595945045 | 7.38E-06 | 20.09173313 | 1.511462352 | 4.545538277 |
| ENSMUSG00000021185 | *Dglucy* | 801.2763335 | 789.5688274 | 759.7735789 | 1190.564287 | 1203.321088 | 1157.559358 | 5.48E-13 | 0.595002533 | 7.32E-14 | 55.98082754 | 1.510475238 | 12.26094131 |
| ENSMUSG00000006442 | *Srm* | 4335.568441 | 4391.014384 | 4458.774328 | 6652.743017 | 7024.613127 | 6231.383544 | 9.86E-22 | 0.594534941 | 8.27E-23 | 96.64996489 | 1.509985757 | 21.00616447 |
| ENSMUSG00000039910 | *Cited2* | 6676.452167 | 6649.753063 | 7090.567343 | 11047.59947 | 9992.424043 | 9765.644161 | 5.86E-20 | 0.593178499 | 5.37E-21 | 88.39050941 | 1.508566714 | 19.23202728 |
| ENSMUSG00000038252 | *Ncapd2* | 3042.638476 | 3216.558245 | 3212.393186 | 5034.319689 | 4746.59197 | 4503.223438 | 1.07E-19 | 0.592689108 | 9.98E-21 | 87.16585761 | 1.508055065 | 18.96926614 |
| ENSMUSG00000006526 | *Stimate* | 230.5158029 | 227.6333527 | 237.9187882 | 362.7500561 | 335.3673973 | 352.1751663 | 2.68E-05 | 0.592411153 | 6.91E-06 | 20.21916344 | 1.507764545 | 4.5721661 |
| ENSMUSG00000026939 | *Tmem141* | 608.1874506 | 620.2184104 | 664.8018815 | 899.8991777 | 1078.511062 | 873.7023252 | 2.24E-07 | 0.592173628 | 4.59E-08 | 29.88332378 | 1.507516328 | 6.649119692 |
| ENSMUSG00000025646 | *Atrip* | 630.3033579 | 559.7361186 | 572.7674532 | 916.1764238 | 988.9527228 | 754.3861486 | 2.94E-06 | 0.591802011 | 6.76E-07 | 24.68369347 | 1.507128064 | 5.531948911 |
| ENSMUSG00000025870 | *Arl10* | 174.3754229 | 183.6462314 | 210.5042777 | 281.3638256 | 292.4937244 | 281.9325785 | 0.000291154 | 0.59139646 | 8.72E-05 | 15.39492555 | 1.50670446 | 3.535877886 |
| ENSMUSG00000020897 | *Aurkb* | 1316.747096 | 1444.976935 | 1347.227377 | 2104.41539 | 2131.29792 | 1951.396823 | 1.46E-14 | 0.591267443 | 1.77E-15 | 63.30697537 | 1.506569725 | 13.83598679 |
| ENSMUSG00000037221 | *Mospd3* | 457.6291586 | 490.4564025 | 466.0466798 | 719.6868101 | 758.3876372 | 650.4656077 | 1.22E-07 | 0.590804773 | 2.43E-08 | 31.11297883 | 1.506086647 | 6.915117762 |
| ENSMUSG00000003299 | *Mrpl4* | 2294.100076 | 2428.089096 | 2394.853317 | 3489.143969 | 3790.032689 | 3432.264531 | 1.46E-18 | 0.59049481 | 1.46E-19 | 81.86478982 | 1.5057631 | 17.834263 |
| ENSMUSG00000005886 | *Ncoa2* | 839.5538653 | 843.453051 | 838.100752 | 1287.065103 | 1190.93536 | 1318.251306 | 2.81E-12 | 0.59025418 | 3.90E-13 | 52.69119432 | 1.505511971 | 11.55204917 |
| ENSMUSG00000002274 | *Metrn* | 793.6208271 | 763.1765546 | 869.4316212 | 1249.859969 | 1334.800351 | 1069.034453 | 8.63E-08 | 0.590107884 | 1.71E-08 | 31.80096854 | 1.505359314 | 7.064059996 |
| ENSMUSG00000042308 | *Setd1a* | 2117.172817 | 2183.960573 | 2058.046473 | 3367.064624 | 2988.771379 | 3218.650086 | 5.02E-17 | 0.590071133 | 5.36E-18 | 74.7424758 | 1.505320966 | 16.29930385 |
| ENSMUSG00000032536 | *Trak1* | 1817.757457 | 1881.549114 | 1781.943188 | 2829.915502 | 2719.143614 | 2701.934063 | 1.91E-20 | 0.590068274 | 1.71E-21 | 90.66049805 | 1.505317984 | 19.71980974 |
| ENSMUSG00000032218 | *Ccnb2* | 2094.206298 | 2019.008868 | 2108.959135 | 3120.580611 | 3265.068383 | 2979.055505 | 3.26E-19 | 0.589502234 | 3.12E-20 | 84.90942594 | 1.504727489 | 18.4866644 |
| ENSMUSG00000022571 | *Pycrl* | 1023.286018 | 991.9095854 | 1026.085967 | 1503.319944 | 1698.750197 | 1372.13603 | 1.60E-09 | 0.588762677 | 2.72E-10 | 39.86693294 | 1.50395633 | 8.794585775 |
| ENSMUSG00000026126 | *Ptpn18* | 931.4199418 | 902.8356647 | 1086.789527 | 1387.053901 | 1641.5853 | 1363.475985 | 1.07E-07 | 0.588740999 | 2.12E-08 | 31.37684979 | 1.503933733 | 6.972358759 |
| ENSMUSG00000033272 | *Slc35a4* | 3083.467844 | 3102.19173 | 2965.662591 | 4802.950263 | 4551.278571 | 4409.887397 | 3.01E-22 | 0.588496041 | 2.48E-23 | 99.03615179 | 1.503678398 | 21.52162784 |
| ENSMUSG00000002222 | *Rmnd5a* | 1145.77412 | 1164.559036 | 1134.76492 | 1821.888904 | 1532.971995 | 1826.307283 | 1.47E-10 | 0.588261047 | 2.30E-11 | 44.6950333 | 1.503433491 | 9.831220688 |
| ENSMUSG00000024642 | *Tle4* | 874.4289499 | 834.6556267 | 816.5607794 | 1390.541882 | 1097.566028 | 1312.477942 | 5.07E-08 | 0.588053889 | 9.83E-09 | 32.87527605 | 1.503217626 | 7.294886732 |
| ENSMUSG00000007987 | *Ift22* | 643.913147 | 622.4177664 | 654.0318952 | 888.2725734 | 1094.707783 | 901.6069149 | 1.87E-07 | 0.587658145 | 3.79E-08 | 30.25534639 | 1.502805337 | 6.729218379 |
| ENSMUSG00000048922 | *Cdca2* | 1108.3472 | 1134.86773 | 1125.953113 | 1717.249464 | 1695.891952 | 1649.257473 | 2.02E-16 | 0.587487625 | 2.23E-17 | 71.93003329 | 1.502627723 | 15.69366062 |
| ENSMUSG00000037649 | *H2-DMa* | 105.4758656 | 118.7652275 | 136.0934632 | 175.5617259 | 202.9353853 | 161.6541747 | 0.007686075 | 0.586682365 | 0.00296409 | 8.829441739 | 1.501789245 | 2.114295393 |
| ENSMUSG00000040274 | *Cdk6* | 4986.286483 | 5374.126545 | 4877.824704 | 8199.081397 | 6396.752005 | 8273.229727 | 2.19E-09 | 0.585673163 | 3.74E-10 | 39.24228389 | 1.500739072 | 8.660393984 |
| ENSMUSG00000055053 | *Nfic* | 1440.936421 | 1550.546026 | 1492.132647 | 2418.333708 | 2164.64411 | 2145.766724 | 8.41E-14 | 0.585523991 | 1.06E-14 | 59.78153561 | 1.500583907 | 13.07501625 |
| ENSMUSG00000031730 | *Dhodh* | 335.9916685 | 347.4982583 | 391.6358654 | 568.5409534 | 564.0269864 | 481.1136152 | 1.19E-05 | 0.585502034 | 2.94E-06 | 21.85779414 | 1.500561069 | 4.923001368 |
| ENSMUSG00000025351 | *Cd63* | 3617.652066 | 3448.59031 | 3664.732611 | 4999.439876 | 6013.747193 | 5086.33314 | 6.64E-13 | 0.585366008 | 8.89E-14 | 55.59732029 | 1.500419594 | 12.17800294 |
| ENSMUSG00000025521 | *Tmem192* | 1172.993698 | 1127.169983 | 1140.639458 | 699.9215827 | 865.0954454 | 726.4815589 | 1.91E-08 | -0.585227504 | 3.56E-09 | 34.85120814 | 0.66654422 | 7.718876926 |
| ENSMUSG00000029720 | *Gm20605* | 456.7785468 | 448.6686373 | 387.7195068 | 316.2436387 | 271.533262 | 275.1969879 | 4.62E-05 | -0.586273393 | 1.23E-05 | 19.10879257 | 0.666061181 | 4.335015972 |
| ENSMUSG00000004207 | *Psap* | 19160.88183 | 19495.09216 | 19398.7035 | 12956.6879 | 13104.09995 | 12605.17672 | 1.10E-40 | -0.586332116 | 5.06E-42 | 184.4964169 | 0.66603407 | 39.95811202 |
| ENSMUSG00000036931 | *Nfkbid* | 940.7766718 | 1022.70057 | 1026.085967 | 697.5962618 | 630.7193666 | 662.9745617 | 7.24E-10 | -0.58656557 | 1.19E-10 | 41.47601338 | 0.665926303 | 9.140258791 |
| ENSMUSG00000078942 | *Naip6* | 322.3818794 | 331.0030878 | 280.0196438 | 188.3509907 | 189.5969093 | 242.4812621 | 0.000874919 | -0.587028271 | 0.000286177 | 13.15876939 | 0.665712761 | 3.058032011 |
| ENSMUSG00000026187 | *Xrcc5* | 974.8011446 | 932.5269716 | 866.4943522 | 562.7276512 | 653.5853255 | 628.3343814 | 1.94E-08 | -0.587172789 | 3.62E-09 | 34.8182391 | 0.665646079 | 7.712002281 |
| ENSMUSG00000070738 | *Dgkd* | 12203.72777 | 12869.53202 | 12388.42151 | 8465.330637 | 8200.304514 | 8260.720773 | 8.08E-34 | -0.587721204 | 4.42E-35 | 152.7142728 | 0.665393093 | 33.0927056 |
| ENSMUSG00000029826 | *Zc3hav1* | 7721.854093 | 7840.704372 | 7463.600505 | 5256.387833 | 4725.631508 | 5328.814402 | 2.96E-20 | -0.588903992 | 2.67E-21 | 89.77068487 | 0.664847796 | 19.52814069 |
| ENSMUSG00000022150 | *Dab2* | 6964.809574 | 7066.531037 | 6835.024941 | 4806.438244 | 4307.37501 | 4759.175881 | 1.03E-21 | -0.58919118 | 8.67E-23 | 96.55779773 | 0.664715463 | 20.98665717 |
| ENSMUSG00000028048 | *Gba* | 4265.818272 | 4220.564289 | 4125.883842 | 2806.662293 | 2877.299829 | 2698.085154 | 1.78E-24 | -0.589572244 | 1.34E-25 | 109.3797414 | 0.664539912 | 23.75073497 |
| ENSMUSG00000054720 | *Lrrc8c* | 646.4649824 | 620.2184104 | 623.6801157 | 413.9071153 | 385.8630566 | 455.13348 | 2.98E-07 | -0.591414401 | 6.16E-08 | 29.31085425 | 0.663691912 | 6.525635636 |
| ENSMUSG00000007867 | *Ift43* | 227.1133557 | 246.3278793 | 226.1697123 | 145.3325545 | 176.2584333 | 141.4474029 | 0.001381657 | -0.592527929 | 0.00046742 | 12.24135002 | 0.663179847 | 2.859599747 |
| ENSMUSG00000001128 | *Cfp* | 7160.450292 | 7044.537477 | 7168.894516 | 4669.244312 | 5007.645001 | 4495.52562 | 9.65E-24 | -0.592699294 | 7.51E-25 | 105.9630916 | 0.663101078 | 23.01565339 |
| ENSMUSG00000052821 | *Cysltr1* | 384.4765422 | 437.671857 | 384.7822378 | 254.6226356 | 236.2815754 | 307.9127137 | 0.000162427 | -0.593225712 | 4.67E-05 | 16.57676716 | 0.662859167 | 3.789340948 |
| ENSMUSG00000022394 | *L3mbtl2* | 1352.472792 | 1339.407844 | 1321.771046 | 895.248536 | 878.4339214 | 887.1735064 | 1.25E-14 | -0.593398684 | 1.51E-15 | 63.61468058 | 0.662779698 | 13.9013614 |
| ENSMUSG00000044350 | *Lacc1* | 329.1867739 | 321.1059855 | 326.036858 | 208.1162181 | 216.2738614 | 222.2744902 | 4.25E-05 | -0.593645222 | 1.13E-05 | 19.27850424 | 0.662666447 | 4.371389285 |
| ENSMUSG00000024299 | *Adamts10* | 444.0193695 | 497.0544707 | 498.3566387 | 338.334187 | 304.8794521 | 310.7993954 | 4.29E-06 | -0.593760782 | 1.01E-06 | 23.91672989 | 0.662613369 | 5.367622241 |
| ENSMUSG00000018774 | *Cd68* | 17057.31881 | 16567.74924 | 16635.71247 | 11094.10588 | 11338.65737 | 10853.92316 | 1.38E-38 | -0.594554443 | 6.58E-40 | 174.8131742 | 0.662248951 | 37.85914972 |
| ENSMUSG00000108218 | *Or2y7* | 546.0927878 | 513.5496412 | 525.7711493 | 344.1474892 | 310.5959419 | 395.4753917 | 8.73E-06 | -0.594561735 | 2.11E-06 | 22.49103538 | 0.662245603 | 5.058790927 |
| ENSMUSG00000038217 | *Tlcd2* | 176.0766465 | 171.5497731 | 161.5497945 | 90.68751403 | 118.1407877 | 127.0139944 | 0.007725218 | -0.595499042 | 0.002980045 | 8.819645821 | 0.661815487 | 2.112089251 |
| ENSMUSG00000024349 | *Sting1* | 1241.893256 | 1150.263222 | 1214.071183 | 777.8198319 | 823.1745207 | 785.17742 | 2.23E-12 | -0.596055655 | 3.09E-13 | 53.1494069 | 0.661560198 | 11.65078639 |
| ENSMUSG00000027164 | *Traf6* | 1825.412964 | 1870.552333 | 1819.148595 | 1217.305477 | 1142.345197 | 1288.422261 | 9.69E-15 | -0.596127508 | 1.16E-15 | 64.13960679 | 0.66152725 | 14.01367017 |
| ENSMUSG00000044906 | *4930503L19Rik* | 714.5139279 | 751.0800962 | 706.9027371 | 529.0104985 | 387.7685532 | 520.5649316 | 1.52E-05 | -0.597420251 | 3.79E-06 | 21.36741049 | 0.660934747 | 4.817862113 |
| ENSMUSG00000037965 | *Zc3h7a* | 2325.572713 | 2160.867334 | 2025.736514 | 1570.754249 | 1174.738639 | 1559.77034 | 3.26E-07 | -0.598060603 | 6.78E-08 | 29.12603412 | 0.660641451 | 6.486365959 |
| ENSMUSG00000031433 | *Rbm41* | 539.2878932 | 514.6493192 | 532.624777 | 344.1474892 | 313.4541867 | 390.6642555 | 4.27E-06 | -0.598172496 | 1.00E-06 | 23.92516882 | 0.660590214 | 5.369271776 |
| ENSMUSG00000049985 | *Ankrd55* | 313.0251494 | 315.6075953 | 313.3086923 | 201.1402555 | 213.4156165 | 206.8788545 | 4.76E-05 | -0.598957825 | 1.28E-05 | 19.04750935 | 0.660230721 | 4.322274399 |
| ENSMUSG00000020070 | *Rufy2* | 603.0837797 | 579.5303232 | 606.0565017 | 399.9551901 | 385.8630566 | 395.4753917 | 3.92E-08 | -0.59953841 | 7.49E-09 | 33.40244615 | 0.659965077 | 7.406904432 |
| ENSMUSG00000033088 | *Triobp* | 909.3040345 | 921.5301913 | 972.2360359 | 637.1379191 | 633.5776114 | 579.2607927 | 3.45E-10 | -0.600100584 | 5.55E-11 | 42.97202705 | 0.659707959 | 9.461865483 |
| ENSMUSG00000035929 | *H2-Q4* | 3368.422803 | 3370.51317 | 3429.751091 | 2364.851328 | 2241.816721 | 2098.617589 | 2.05E-19 | -0.601551206 | 1.94E-20 | 85.85435913 | 0.65904496 | 18.6885678 |
| ENSMUSG00000021929 | *Kpna3* | 5466.031549 | 5360.930409 | 5244.983328 | 3443.800212 | 3303.178314 | 3842.173331 | 1.01E-17 | -0.601856784 | 1.04E-18 | 77.97710372 | 0.658905382 | 16.99706821 |
| ENSMUSG00000031207 | *Msn* | 25391.61341 | 25736.86467 | 25090.15172 | 16801.60597 | 16471.1124 | 16940.97262 | 1.41E-45 | -0.602058653 | 5.66E-47 | 207.1804412 | 0.658813191 | 44.85202633 |
| ENSMUSG00000021282 | *Eif5* | 13300.1664 | 13480.953 | 13203.02411 | 8766.45969 | 8569.018101 | 9002.597967 | 3.60E-38 | -0.602235671 | 1.73E-39 | 172.8886847 | 0.65873236 | 37.44386001 |
| ENSMUSG00000023022 | *Lima1* | 783.4134853 | 870.9450018 | 904.6788491 | 573.1915951 | 564.0269864 | 547.5072941 | 5.43E-09 | -0.602437296 | 9.63E-10 | 37.39864433 | 0.658640305 | 8.265136911 |
| ENSMUSG00000098374 | *Gm28043* | 1007.124394 | 1062.288979 | 863.5570832 | 666.20443 | 527.8225515 | 738.0282857 | 2.08E-05 | -0.60246746 | 5.27E-06 | 20.73597752 | 0.658626534 | 4.682326701 |
| ENSMUSG00000034041 | *Lyl1* | 1412.01562 | 1528.552465 | 1496.049006 | 1001.050636 | 994.6692126 | 925.6625956 | 1.53E-13 | -0.602616984 | 1.96E-14 | 58.57220362 | 0.658558276 | 12.81608302 |
| ENSMUSG00000045886 | *Pam16l* | 519.7238214 | 506.951573 | 497.3775491 | 262.7612586 | 369.6663357 | 368.5330292 | 0.000109806 | -0.602785834 | 3.09E-05 | 17.36405348 | 0.658481204 | 3.9593743 |
| ENSMUSG00000083902 | *Tent2-ps1* | 310.4733139 | 240.8294891 | 313.3086923 | 220.9054829 | 179.1166781 | 171.276447 | 0.001623812 | -0.603692796 | 0.000555644 | 11.91899734 | 0.658067375 | 2.789464137 |
| ENSMUSG00000042097 | *Zfp239* | 274.7476175 | 256.2249816 | 228.1278916 | 153.4711776 | 176.2584333 | 169.3519925 | 0.000649483 | -0.603808211 | 0.000206646 | 13.76967027 | 0.658014732 | 3.187432193 |
| ENSMUSG00000034083 | *Ccdc174* | 527.3793278 | 577.3309671 | 564.9347359 | 410.419134 | 335.3673973 | 354.0996208 | 2.39E-06 | -0.604121759 | 5.47E-07 | 25.09122439 | 0.657871738 | 5.620766989 |
| ENSMUSG00000021819 | *Zswim8* | 2107.816087 | 2128.976671 | 2076.649176 | 1424.259035 | 1298.595916 | 1431.794119 | 9.40E-17 | -0.604205425 | 1.02E-17 | 73.48188394 | 0.657833587 | 16.02696171 |
| ENSMUSG00000059839 | *Zfp874b* | 189.6864356 | 195.7426898 | 173.2988704 | 131.3806293 | 107.6605565 | 128.9384489 | 0.003035519 | -0.60439471 | 0.001086405 | 10.6741941 | 0.657747284 | 2.517767024 |
| ENSMUSG00000051978 | *Erich1* | 428.7083568 | 434.3728229 | 491.5030111 | 301.129053 | 276.2970035 | 313.6860771 | 8.53E-06 | -0.60440595 | 2.06E-06 | 22.53998784 | 0.657742159 | 5.068870807 |
| ENSMUSG00000019845 | *Tube1* | 187.1346002 | 205.6397921 | 220.2951743 | 143.0072337 | 135.2902569 | 125.0895399 | 0.001696464 | -0.604416523 | 0.000582599 | 11.83076893 | 0.657737339 | 2.770455353 |
| ENSMUSG00000032497 | *Lrrfip2* | 2610.527672 | 2580.944342 | 2488.845925 | 1775.382486 | 1602.52262 | 1677.162063 | 4.14E-18 | -0.604461604 | 4.19E-19 | 79.7756553 | 0.657716786 | 17.38314335 |
| ENSMUSG00000054381 | *Zfp747* | 400.6381667 | 358.4950386 | 390.6567758 | 245.3213521 | 262.0057792 | 249.2168527 | 1.26E-05 | -0.604524457 | 3.10E-06 | 21.7519973 | 0.657688132 | 4.900384981 |
| ENSMUSG00000049606 | *Zfp644* | 1829.666023 | 1795.774227 | 1752.570498 | 1183.588324 | 983.2362331 | 1368.287122 | 1.47E-07 | -0.605511433 | 2.95E-08 | 30.73816153 | 0.657238349 | 6.833025477 |
| ENSMUSG00000021661 | *Ankra2* | 413.397344 | 405.781194 | 419.050376 | 265.0865795 | 246.7618066 | 301.1771231 | 1.47E-05 | -0.606661788 | 3.65E-06 | 21.43799858 | 0.656714499 | 4.833013598 |
| ENSMUSG00000027944 | *Hax1* | 11704.41863 | 11329.98277 | 11610.04523 | 6959.685372 | 8548.057639 | 7238.835454 | 1.32E-14 | -0.606747337 | 1.60E-15 | 63.5037649 | 0.656675558 | 13.8778961 |
| ENSMUSG00000063019 | *Manbal* | 645.6143706 | 703.7939408 | 647.1782676 | 447.624268 | 473.5158991 | 388.7398011 | 4.43E-07 | -0.60769503 | 9.34E-08 | 28.50705233 | 0.656244335 | 6.353833093 |
| ENSMUSG00000043259 | *Fam13c* | 383.6259304 | 437.671857 | 385.7613274 | 289.5024486 | 248.6673031 | 254.0279888 | 2.48E-05 | -0.608547791 | 6.35E-06 | 20.38087228 | 0.655856551 | 4.606205282 |
| ENSMUSG00000022844 | *Pdia5* | 632.0045815 | 632.3148687 | 714.7354544 | 429.021701 | 445.8861987 | 422.4177541 | 6.40E-08 | -0.608972919 | 1.25E-08 | 32.40296435 | 0.655663315 | 7.193484028 |
| ENSMUSG00000061981 | *Flot2* | 609.8886742 | 623.5174445 | 676.5509575 | 437.1603241 | 429.6894778 | 385.8531194 | 1.38E-07 | -0.609340549 | 2.77E-08 | 30.86407539 | 0.655496259 | 6.8606117 |
| ENSMUSG00000027829 | *Ccnl1* | 2115.471594 | 2152.06991 | 1990.489286 | 1489.368019 | 1118.52649 | 1496.263343 | 8.70E-08 | -0.60936986 | 1.72E-08 | 31.78325811 | 0.655482941 | 7.060250398 |
| ENSMUSG00000042408 | *Zmym6* | 315.5769848 | 257.3246596 | 255.5424022 | 176.7243863 | 167.6836987 | 199.1810367 | 0.000701146 | -0.609861509 | 0.000224623 | 13.61301667 | 0.6552596 | 3.154191542 |
| ENSMUSG00000027365 | *Trpm7* | 4583.096481 | 4551.567377 | 4385.342603 | 2951.994848 | 2533.357697 | 3371.644215 | 2.88E-10 | -0.610258588 | 4.60E-11 | 43.34088896 | 0.655079275 | 9.541139674 |
| ENSMUSG00000079659 | *Tmem243* | 501.8609732 | 534.4435238 | 474.8584868 | 334.8462057 | 348.7058734 | 305.9882593 | 1.56E-06 | -0.610400967 | 3.48E-07 | 25.96051776 | 0.655014629 | 5.807471327 |
| ENSMUSG00000099693 | *Gm29284* | 587.7727669 | 590.5271035 | 617.8055777 | 342.9848287 | 427.7839813 | 403.1732095 | 9.45E-07 | -0.610482804 | 2.06E-07 | 26.97558938 | 0.654977474 | 6.0244844 |
| ENSMUSG00000029401 | *Rilpl2* | 1833.06847 | 1903.542674 | 2018.882886 | 1252.18529 | 1379.579521 | 1134.465905 | 3.14E-11 | -0.611497553 | 4.64E-12 | 47.83175246 | 0.654516944 | 10.50337038 |
| ENSMUSG00000039105 | *Atp6v1g1* | 3761.405464 | 3692.718833 | 3724.45708 | 2390.429857 | 2567.656636 | 2353.607805 | 4.88E-23 | -0.612260207 | 3.91E-24 | 102.696235 | 0.654171038 | 22.31129168 |
| ENSMUSG00000025059 | *Gk* | 2639.448474 | 2527.060119 | 2554.444932 | 1704.4602 | 1611.097355 | 1736.820151 | 4.17E-20 | -0.612424186 | 3.80E-21 | 89.07748755 | 0.654096688 | 19.37991239 |
| ENSMUSG00000020077 | *Srgn* | 5386.92465 | 5391.721394 | 5499.54664 | 3600.759371 | 3501.349958 | 3545.807344 | 6.18E-33 | -0.612578112 | 3.51E-34 | 148.5989364 | 0.654026904 | 32.20924339 |
| ENSMUSG00000027009 | *Itga4* | 15305.90907 | 15132.66941 | 14593.33143 | 10102.35653 | 8450.877314 | 10895.29893 | 1.40E-12 | -0.612832226 | 1.90E-13 | 54.10518372 | 0.653911715 | 11.85511959 |
| ENSMUSG00000041236 | *Vps41* | 2274.536004 | 2363.208092 | 2259.738943 | 1447.512243 | 1530.11375 | 1528.979069 | 9.24E-20 | -0.612963682 | 8.56E-21 | 87.46842658 | 0.653852134 | 19.03409876 |
| ENSMUSG00000042508 | *Dmtf1* | 1554.067793 | 1470.26953 | 1409.889115 | 930.1283491 | 871.7646834 | 1094.052361 | 2.79E-09 | -0.614562737 | 4.82E-10 | 38.74763656 | 0.653127819 | 8.554165244 |
| ENSMUSG00000091780 | *Sco2* | 1018.182347 | 1063.388657 | 1115.183127 | 702.2469035 | 733.6161817 | 650.4656077 | 7.86E-11 | -0.615349197 | 1.20E-11 | 45.96486623 | 0.652771874 | 10.10470925 |
| ENSMUSG00000009905 | *Kdsr* | 581.8184842 | 575.131611 | 540.4574943 | 356.936754 | 348.7058734 | 401.2487551 | 3.03E-07 | -0.616217812 | 6.28E-08 | 29.2750992 | 0.652378973 | 6.518341945 |
| ENSMUSG00000028709 | *Mob3c* | 242.4243684 | 226.5336747 | 266.3123885 | 181.3750281 | 156.2507192 | 143.3718573 | 0.000833525 | -0.616581666 | 0.000271436 | 13.25788325 | 0.652214461 | 3.079081563 |
| ENSMUSG00000026014 | *Raph1* | 441.4675341 | 439.871213 | 474.8584868 | 310.4303365 | 268.6750172 | 305.9882593 | 3.15E-06 | -0.617207322 | 7.28E-07 | 24.53910462 | 0.651931675 | 5.501044928 |
| ENSMUSG00000002108 | *Nr1h3* | 261.1378284 | 280.4178983 | 308.413244 | 154.633838 | 199.1243922 | 198.2188095 | 0.000568309 | -0.617910931 | 0.000178986 | 14.03976174 | 0.651613803 | 3.245415784 |
| ENSMUSG00000041671 | *Pyroxd1* | 415.0985677 | 374.9902091 | 431.7785416 | 258.1106169 | 191.5024059 | 346.4018029 | 0.003666542 | -0.618584133 | 0.001330109 | 10.30026819 | 0.651309812 | 2.43574339 |
| ENSMUSG00000027722 | *Spata5* | 1167.890027 | 1118.372559 | 1080.914989 | 724.3374518 | 701.2227399 | 766.8951026 | 3.78E-12 | -0.619321859 | 5.29E-13 | 52.0928484 | 0.650976848 | 11.42250182 |
| ENSMUSG00000023249 | *Parp3* | 551.1964587 | 504.752217 | 514.0220734 | 344.1474892 | 350.6113699 | 328.1194856 | 2.12E-07 | -0.619514565 | 4.33E-08 | 29.99594863 | 0.650889901 | 6.67317847 |
| ENSMUSG00000032661 | *Oas3* | 8078.260445 | 8530.202499 | 8047.137945 | 5655.180362 | 4955.243845 | 5426.961579 | 2.03E-21 | -0.620713772 | 1.74E-22 | 95.18341716 | 0.650349088 | 20.69219625 |
| ENSMUSG00000022329 | *Stk3* | 1463.90294 | 1369.099151 | 1124.974023 | 775.494511 | 822.2217724 | 973.7739571 | 1.06E-06 | -0.621525047 | 2.33E-07 | 26.74010218 | 0.649983478 | 5.974019539 |
| ENSMUSG00000023961 | *Enpp4* | 873.5783381 | 845.652407 | 853.7661866 | 605.7460873 | 477.3268922 | 590.8075194 | 9.28E-08 | -0.622257257 | 1.84E-08 | 31.65583003 | 0.649653677 | 7.032356809 |
| ENSMUSG00000025255 | *Zfhx4* | 969.6974736 | 938.0253618 | 889.9925042 | 642.9512213 | 497.3346063 | 678.3701974 | 1.37E-06 | -0.622600583 | 3.04E-07 | 26.22491331 | 0.649499093 | 5.864134648 |
| ENSMUSG00000021143 | *Pacs2* | 1815.205622 | 1884.848148 | 1783.901367 | 1238.233365 | 1170.927646 | 1153.710449 | 2.34E-17 | -0.622736617 | 2.45E-18 | 76.28617948 | 0.649437854 | 16.6315448 |
| ENSMUSG00000079555 | *Haus3* | 887.1881272 | 910.533411 | 906.6370284 | 595.2821434 | 483.9961302 | 675.4835157 | 1.36E-06 | -0.624230404 | 3.02E-07 | 26.23612834 | 0.648765765 | 5.866522686 |
| ENSMUSG00000073131 | *Vma21* | 1671.452224 | 1572.539587 | 1718.30236 | 1042.906411 | 1041.353879 | 1134.465905 | 1.17E-14 | -0.624502451 | 1.41E-15 | 63.75594843 | 0.64864344 | 13.93101933 |
| ENSMUSG00000101249 | *Gm29216* | 49271.68961 | 52254.50075 | 52047.42742 | 31767.3711 | 35739.49378 | 32083.54254 | 1.08E-24 | -0.62477099 | 8.10E-26 | 110.3775978 | 0.648522714 | 23.96618436 |
| ENSMUSG00000007038 | *Neu1* | 2357.895962 | 2321.420327 | 2530.94678 | 1568.428929 | 1674.93149 | 1432.756346 | 2.49E-14 | -0.624848731 | 3.06E-15 | 62.22785276 | 0.648487769 | 13.60316634 |

| | | | | | | | | | | | | |
|---|---|---|---|---|---|---|---|---|---|---|---|---|
| ENSMUSG00000068036 | *Afdn* | 4007.232279 | 4038.017736 | 3958.45951 | 2721.788081 | 2375.201482 | 2689.425109 | 5.31E-19 | -0.624986002 | 5.17E-20 | 83.9145727 | 0.648426069 | 18.27522982 |
| ENSMUSG00000038280 | *Ostm1* | 1255.503045 | 1249.234245 | 1149.451265 | 825.4889098 | 746.9546577 | 797.686374 | 2.62E-12 | -0.625687997 | 3.64E-13 | 52.8274287 | 0.64811063 | 11.5815304 |
| ENSMUSG00000066595 | *Flvcr1* | 1469.006611 | 1460.372427 | 1491.153558 | 1028.954486 | 846.0404796 | 990.13182 | 2.39E-11 | -0.627031125 | 3.52E-12 | 48.37786333 | 0.64750753 | 10.62150086 |
| ENSMUSG00000049988 | *Lrrc25* | 1405.210725 | 1336.10881 | 1329.603763 | 881.2966107 | 959.4175259 | 795.7619195 | 3.31E-11 | -0.627117348 | 4.91E-12 | 47.72163751 | 0.647468833 | 10.48078128 |
| ENSMUSG00000030105 | *Arl8b* | 6737.696218 | 6842.196719 | 6856.564914 | 4441.362867 | 4285.461799 | 4505.147893 | 6.34E-35 | -0.627136801 | 3.36E-36 | 157.8392433 | 0.647460102 | 34.19770846 |
| ENSMUSG00000025532 | *Crcp* | 1807.550115 | 1693.50417 | 1731.030525 | 1044.069072 | 1228.092543 | 1113.296906 | 2.72E-13 | -0.627202318 | 3.56E-14 | 57.39928948 | 0.6474307 | 12.56518868 |
| ENSMUSG00000022263 | *Trio* | 2207.33767 | 2227.947694 | 2197.077205 | 1485.880038 | 1273.824461 | 1535.71466 | 2.39E-13 | -0.627244185 | 3.10E-14 | 57.66722567 | 0.647411912 | 12.62206335 |
| ENSMUSG00000114277 | *Gm48583* | 2212.441341 | 2009.111766 | 2013.987438 | 1275.438499 | 1422.453194 | 1337.49585 | 4.41E-15 | -0.627626824 | 5.18E-16 | 65.72482902 | 0.647240225 | 14.35518844 |
| ENSMUSG00000000804 | *Usp32* | 1293.780577 | 1299.819435 | 1153.367624 | 828.9768911 | 736.4744265 | 857.3444623 | 1.96E-10 | -0.629583461 | 3.11E-11 | 44.10800601 | 0.646363008 | 9.706873082 |
| ENSMUSG00000047123 | *Ticam1* | 1441.787033 | 1587.935079 | 1499.965365 | 970.8214643 | 941.3153084 | 1013.225274 | 1.34E-14 | -0.629846909 | 1.62E-15 | 63.48100858 | 0.646244988 | 13.87337204 |
| ENSMUSG00000039497 | *Dse* | 940.7766718 | 1008.404756 | 899.7834008 | 616.2100313 | 594.5149316 | 629.2966087 | 4.86E-11 | -0.630115436 | 7.31E-12 | 46.94187777 | 0.646124714 | 10.31313315 |
| ENSMUSG00000026097 | *Ormdl1* | 997.7676637 | 923.7295474 | 963.4242289 | 563.8903116 | 660.2545635 | 637.9566537 | 4.04E-10 | -0.630294439 | 6.53E-11 | 42.65466247 | 0.646044551 | 9.393441942 |
| ENSMUSG00000020385 | *Clk4* | 1222.329184 | 1101.877389 | 1096.580423 | 774.3318506 | 604.0424145 | 833.2887815 | 8.00E-07 | -0.630466613 | 1.73E-07 | 27.31367408 | 0.645967455 | 6.097139832 |
| ENSMUSG00000028293 | *Slc35a1* | 654.9711006 | 605.9225959 | 690.2582128 | 437.1603241 | 412.5400087 | 411.8332546 | 1.84E-08 | -0.631054245 | 3.43E-09 | 34.92265059 | 0.645704396 | 7.734332534 |
| ENSMUSG00000058486 | *Wdr91* | 2125.678936 | 2233.446084 | 2173.579053 | 1407.981788 | 1418.642201 | 1388.493893 | 2.09E-21 | -0.63166863 | 1.79E-22 | 95.12687442 | 0.645429476 | 20.68014039 |
| ENSMUSG00000071793 | *2610005L07Rik* | 1285.274458 | 1305.317825 | 1312.959239 | 885.9472525 | 720.2777056 | 911.2291872 | 1.64E-09 | -0.633671652 | 2.77E-10 | 39.82721734 | 0.64453399 | 8.786457944 |
| ENSMUSG00000062232 | *Rapgef2* | 3771.612805 | 3698.217224 | 3617.736307 | 2502.045259 | 2169.407852 | 2477.735118 | 5.00E-18 | -0.633773567 | 5.09E-19 | 79.39398135 | 0.644488461 | 17.30130057 |
| ENSMUSG00000018848 | *Rars* | 10249.02181 | 10032.36269 | 10253.02696 | 6482.994593 | 6311.957407 | 6883.773606 | 4.82E-32 | -0.63379098 | 2.82E-33 | 144.4556668 | 0.644480682 | 31.3169119 |
| ENSMUSG00000045210 | *Vcpip1* | 3353.962402 | 3569.554894 | 3350.444829 | 2274.163813 | 2035.070343 | 2312.232035 | 3.38E-18 | -0.633861577 | 3.41E-19 | 80.18514013 | 0.644449146 | 17.47049296 |
| ENSMUSG00000048234 | *Rnf149* | 3512.176201 | 3558.558113 | 3760.683398 | 2475.304069 | 2183.699076 | 2324.740989 | 1.95E-19 | -0.633867988 | 1.84E-20 | 85.95999909 | 0.644446282 | 18.71082185 |
| ENSMUSG00000089782 | *Btf3-ps1* | 559.7025769 | 635.6139028 | 594.3074258 | 301.129053 | 426.831233 | 421.4555269 | 3.90E-05 | -0.634328241 | 1.03E-05 | 19.4498867 | 0.644240721 | 4.408680019 |
| ENSMUSG00000049775 | *Tmsb4x* | 20756.62961 | 20638.75732 | 21423.46093 | 12442.79199 | 14799.9919 | 13209.45542 | 3.21E-20 | -0.634801472 | 2.90E-21 | 89.60864142 | 0.644029433 | 19.49355904 |
| ENSMUSG00000024843 | *Chka* | 998.6182755 | 910.533411 | 925.239732 | 625.5113147 | 566.8852313 | 634.1077448 | 1.38E-10 | -0.635629206 | 2.16E-11 | 44.82383549 | 0.643660032 | 9.8592163 |
| ENSMUSG00000030823 | *9130019O22Rik* | 149.7076801 | 203.440436 | 170.3616015 | 108.1274206 | 127.6682706 | 100.071632 | 0.005999413 | -0.635674827 | 0.002262026 | 9.323749494 | 0.643639679 | 2.22189126 |
| ENSMUSG00000039286 | *Fndc3b* | 5983.203535 | 6218.679274 | 5900.973402 | 4125.119228 | 3488.96423 | 4036.543231 | 1.23E-17 | -0.636144609 | 1.28E-18 | 77.57101661 | 0.643430126 | 16.90950454 |
| ENSMUSG00000024019 | *Cmtr1* | 4782.139646 | 4855.078514 | 4960.068235 | 3097.327402 | 3315.564042 | 2971.357687 | 2.96E-24 | -0.637171273 | 2.25E-25 | 108.3495365 | 0.642972405 | 23.5293314 |
| ENSMUSG00000033730 | *Egr3* | 200.7443893 | 167.151061 | 209.525188 | 141.8445732 | 125.762774 | 104.8827681 | 0.003338205 | -0.638085401 | 0.001203898 | 10.48435292 | 0.642565131 | 2.476486987 |
| ENSMUSG00000026433 | *Rab29* | 740.8828943 | 696.0961946 | 715.714544 | 458.0882119 | 507.8148374 | 417.606618 | 2.52E-08 | -0.638185848 | 4.74E-09 | 34.29135466 | 0.642520394 | 7.598582967 |
| ENSMUSG00000034765 | *Dusp5* | 1840.723976 | 1904.642352 | 1822.085864 | 1261.486573 | 1144.250694 | 1171.992767 | 1.83E-17 | -0.638871499 | 1.92E-18 | 76.77081722 | 0.642215104 | 16.73754154 |
| ENSMUSG00000022948 | *Setd4* | 200.7443893 | 234.2314209 | 211.4833673 | 132.5432897 | 123.8572774 | 157.8052658 | 0.00114222 | -0.639282991 | 0.000380662 | 12.62480508 | 0.642031955 | 2.942250302 |
| ENSMUSG00000030474 | *Siglece* | 162.4668574 | 173.7491291 | 148.8216289 | 96.50081622 | 128.6210189 | 85.6382235 | 0.008587902 | -0.639515334 | 0.003347597 | 8.607618857 | 0.641928565 | 2.066112895 |
| ENSMUSG00000030276 | *Ttll3* | 297.7141366 | 331.0030878 | 331.9113959 | 229.044106 | 174.3529367 | 213.6144451 | 0.000138435 | -0.640511232 | 3.94E-05 | 16.89774202 | 0.641485592 | 3.858752881 |
| ENSMUSG00000089739 | *Gm20431* | 1486.86946 | 1729.793545 | 1637.037917 | 976.6347665 | 1334.800351 | 799.6108284 | 6.35E-05 | -0.640917783 | 1.73E-05 | 18.46323219 | 0.641304848 | 4.197503136 |
| ENSMUSG00000025340 | *Rabgef1* | 1168.740639 | 1186.552597 | 1127.911292 | 741.7773584 | 762.1986303 | 729.3682406 | 1.67E-14 | -0.640926665 | 2.02E-15 | 63.04050034 | 0.641300899 | 13.77821546 |
| ENSMUSG00000021027 | *Ralgapa1* | 1167.039416 | 1260.231025 | 1184.698493 | 831.302212 | 714.5612159 | 771.7062388 | 6.69E-12 | -0.641105689 | 9.54E-13 | 50.93640308 | 0.641221325 | 11.17427462 |
| ENSMUSG00000025969 | *Nrp2* | 20347.48532 | 21361.24578 | 20436.53855 | 13817.05663 | 12400.01897 | 13578.95067 | 1.76E-28 | -0.643098396 | 1.16E-29 | 127.9299613 | 0.640336256 | 27.75456259 |
| ENSMUSG00000079190 | *ENSMUSG00000079190* | 1451.994375 | 1342.706878 | 1292.398356 | 847.5794581 | 904.1581252 | 865.0422801 | 1.02E-13 | -0.643421671 | 1.30E-14 | 59.38691202 | 0.640192788 | 12.98981957 |
| ENSMUSG00000026991 | *Pkp4* | 1762.467689 | 1768.282276 | 1732.009615 | 1167.311078 | 1048.975865 | 1154.672676 | 7.54E-17 | -0.643484212 | 8.10E-18 | 73.92901914 | 0.640165036 | 16.12257346 |
| ENSMUSG00000056708 | *Ier5* | 3260.395102 | 3410.101579 | 3392.545684 | 2312.531608 | 2105.573716 | 2026.450547 | 1.03E-19 | -0.643655664 | 9.60E-21 | 87.24229936 | 0.640088962 | 18.98540997 |
| ENSMUSG00000095193 | *Gm20939* | 429.5589686 | 420.0770084 | 407.3013 | 279.0385047 | 216.2738614 | 308.8749409 | 7.02E-05 | -0.645572228 | 1.93E-05 | 18.25779296 | 0.639239194 | 4.153395818 |
| ENSMUSG00000056501 | *Cebpb* | 4035.302469 | 3985.23319 | 4957.130966 | 2954.320169 | 2818.229436 | 2524.884252 | 6.36E-11 | -0.645666603 | 9.70E-12 | 46.38807993 | 0.639197379 | 10.19676708 |
| ENSMUSG00000002043 | *Trappc6a* | 800.4257217 | 829.1572366 | 916.4279251 | 588.3061808 | 568.7907279 | 471.4913429 | 7.78E-08 | -0.645788326 | 1.53E-08 | 32.01143963 | 0.63914345 | 7.109156003 |
| ENSMUSG00000033644 | *Piwil2* | 347.0496221 | 345.2989022 | 332.8904856 | 187.1883303 | 215.3211131 | 251.1413071 | 6.41E-05 | -0.645919636 | 1.75E-05 | 18.44354698 | 0.63908528 | 4.19323534 |
| ENSMUSG00000039967 | *Zfp292* | 1957.257795 | 1877.150402 | 1869.082168 | 1202.190891 | 931.7878256 | 1508.772297 | 8.30E-06 | -0.646926606 | 2.00E-06 | 22.59684471 | 0.638639368 | 5.080981963 |
| ENSMUSG00000020859 | *Spag9* | 4777.886587 | 4808.892036 | 4692.776757 | 3162.436387 | 2602.908323 | 3343.739626 | 7.12E-13 | -0.648837057 | 9.55E-14 | 55.4575941 | 0.637794227 | 12.14780009 |
| ENSMUSG00000031107 | *Rbmx2* | 581.8184842 | 550.9386943 | 547.3111219 | 381.3526231 | 302.9739555 | 388.7398011 | 1.61E-06 | -0.648919588 | 3.62E-07 | 25.88771986 | 0.637757742 | 5.792158027 |
| ENSMUSG00000027312 | *Atrn* | 3251.038372 | 3184.667582 | 3024.407971 | 2106.740711 | 1772.111815 | 2151.540087 | 4.74E-14 | -0.650199335 | 5.92E-15 | 60.92720021 | 0.637192268 | 13.32398846 |
| ENSMUSG00000066643 | *Wdr35* | 452.5254877 | 437.671857 | 424.924914 | 284.8518069 | 270.5805138 | 282.8948057 | 3.03E-07 | -0.650748988 | 6.27E-08 | 29.27673848 | 0.63694955 | 6.518565475 |
| ENSMUSG00000056952 | *Tatdn2* | 3661.883881 | 3856.57086 | 3846.843288 | 2461.352144 | 2446.657603 | 2329.552125 | 5.88E-27 | -0.651018073 | 4.09E-28 | 120.8656418 | 0.63683076 | 26.23040443 |
| ENSMUSG00000019715 | *Gle1* | 1698.671803 | 1783.677769 | 1675.222414 | 1058.020997 | 1133.770463 | 1086.354543 | 4.67E-18 | -0.652631481 | 4.75E-19 | 79.5315635 | 0.636118971 | 17.33066437 |
| ENSMUSG00000003573 | *Homer3* | 518.8732096 | 433.2731448 | 516.9593423 | 302.2917134 | 322.0289213 | 309.8371682 | 1.01E-06 | -0.654362702 | 2.21E-07 | 26.84322957 | 0.635356092 | 5.996243511 |
| ENSMUSG00000060550 | *H2-Q7* | 825.9440762 | 833.5559487 | 854.7452763 | 546.4504051 | 526.8698032 | 524.4138405 | 3.89E-12 | -0.65471256 | 5.45E-13 | 52.03635831 | 0.635202035 | 11.41043319 |
| ENSMUSG00000037876 | *Jmjd1c* | 7257.42004 | 6818.003802 | 6646.060636 | 4511.122493 | 3586.144556 | 5066.126368 | 1.63E-09 | -0.6547736 | 2.77E-10 | 39.83295083 | 0.635175161 | 8.78755716 |
| ENSMUSG00000079442 | *St6galnac4* | 2465.923663 | 2496.269134 | 2403.665124 | 1627.724611 | 1637.774307 | 1414.474029 | 1.04E-16 | -0.654864191 | 1.13E-17 | 73.27292488 | 0.635135277 | 15.98236079 |
| ENSMUSG00000002307 | *Daxx* | 2790.006766 | 2790.982847 | 2685.642947 | 1862.582019 | 1684.458973 | 1706.991107 | 2.62E-21 | -0.654937512 | 2.25E-22 | 94.66524508 | 0.635102999 | 20.58198937 |
| ENSMUSG00000003283 | *Hck* | 4230.943188 | 4481.187983 | 4473.460673 | 2860.144673 | 2943.039461 | 2568.184478 | 1.84E-20 | -0.655411116 | 1.64E-21 | 90.73336436 | 0.634894544 | 19.73513666 |
| ENSMUSG00000041187 | *Prkd2* | 403.1900022 | 451.9676714 | 462.1303212 | 308.1050156 | 271.533262 | 256.9146705 | 3.40E-06 | -0.656429181 | 7.88E-07 | 24.38701168 | 0.634446676 | 5.468821818 |
| ENSMUSG00000005514 | *Por* | 10590.96776 | 11347.57762 | 11361.35646 | 7222.446631 | 7176.100104 | 6726.930567 | 1.28E-30 | -0.656514026 | 7.94E-32 | 137.8289129 | 0.634409366 | 29.89306383 |
| ENSMUSG00000030469 | *Zfp719* | 652.4192652 | 632.3148687 | 626.6173847 | 375.5393209 | 427.7839813 | 407.0221184 | 3.22E-09 | -0.657617386 | 5.60E-10 | 38.45602042 | 0.63392436 | 8.49185632 |
| ENSMUSG00000031995 | *St14* | 1623.817962 | 1623.124776 | 1597.874331 | 1055.695676 | 1068.030831 | 947.7938219 | 3.45E-16 | -0.65808349 | 3.84E-17 | 70.85807181 | 0.633719586 | 15.46202018 |
| ENSMUSG00000058291 | *Zfp68* | 1059.011715 | 987.5108733 | 973.2151256 | 653.4151652 | 548.7830138 | 712.0481505 | 1.31E-08 | -0.65866851 | 2.39E-09 | 35.62608941 | 0.633462662 | 7.882624924 |
| ENSMUSG00000068250 | *Amn1* | 257.7353812 | 299.1124249 | 251.6260435 | 190.6763116 | 152.4397261 | 169.3519925 | 0.000225221 | -0.659168235 | 6.62E-05 | 15.91593475 | 0.633243279 | 3.647390369 |
| ENSMUSG00000027782 | *Kpna4* | 6126.10632 | 5961.354615 | 5960.697872 | 3829.803477 | 3640.451208 | 3958.602826 | 2.56E-31 | -0.659381411 | 1.55E-32 | 141.0775958 | 0.633149717 | 30.59132924 |
| ENSMUSG00000104434 | *Gm37421* | 238.1713093 | 225.4339967 | 203.65065 | 145.3325545 | 159.1089641 | 118.3539493 | 0.000912076 | -0.659675607 | 0.000299388 | 13.07421809 | 0.633020617 | 3.039968941 |
| ENSMUSG00000043822 | *Adamtsl5* | 220.3084611 | 240.8294891 | 248.6887745 | 149.9831963 | 135.2902569 | 163.5786292 | 0.000267553 | -0.660228651 | 7.95E-05 | 15.56904701 | 0.632778001 | 3.572590441 |
| ENSMUSG00000027550 | *Lrrcc1* | 1048.804373 | 961.1186005 | 1010.420533 | 659.2284674 | 507.8148374 | 744.7638763 | 1.79E-06 | -0.660742871 | 4.04E-07 | 25.67589337 | 0.6325525 | 5.7471434 |
| ENSMUSG00000035529 | *Prdm4* | 1423.924185 | 1447.176291 | 1387.370053 | 921.989726 | 864.1426971 | 908.3425055 | 3.59E-17 | -0.660850938 | 3.81E-18 | 75.41730186 | 0.63250512 | 16.44466427 |
| ENSMUSG00000030641 | *Ddias* | 600.5319442 | 561.9354746 | 551.2274806 | 372.0513396 | 324.8871662 | 387.7775738 | 9.58E-08 | -0.661414356 | 1.90E-08 | 31.58978916 | 0.632258154 | 7.018488414 |
| ENSMUSG00000024197 | *Plin3* | 1317.597708 | 1410.886916 | 1384.432784 | 895.248536 | 872.7174317 | 832.3265543 | 5.58E-16 | -0.661535642 | 6.26E-17 | 69.89201909 | 0.632205003 | 15.25334317 |
| ENSMUSG00000056758 | *Hmga2* | 10652.21181 | 10509.62296 | 10788.589 | 6842.256668 | 6285.280455 | 7064.672325 | 3.69E-29 | -0.662183111 | 2.40E-30 | 131.0657167 | 0.631921339 | 28.43292825 |
| ENSMUSG00000025911 | *Adhfe1* | 304.5190312 | 313.4082393 | 295.6850784 | 211.6041994 | 190.5496576 | 176.0875832 | 2.64E-05 | -0.66243301 | 6.81E-06 | 20.24457481 | 0.631811889 | 4.577703666 |
| ENSMUSG00000006850 | *Tmco6* | 468.6871123 | 521.2473874 | 474.8584868 | 295.3157508 | 353.4696148 | 275.1969879 | 3.41E-06 | -0.662602133 | 7.91E-07 | 24.38007183 | 0.631737828 | 5.467393979 |
| ENSMUSG00000033159 | *Cnppd1* | 2406.380836 | 2496.269134 | 2320.442503 | 1630.049932 | 1516.775274 | 1418.322938 | 9.63E-18 | -0.662800002 | 9.96E-19 | 78.06651147 | 0.63165119 | 17.01643779 |
| ENSMUSG00000048440 | *Cyp4f16* | 436.3638631 | 415.6782963 | 480.7330248 | 309.2676761 | 261.0530309 | 273.2725334 | 2.07E-06 | -0.662941302 | 4.69E-07 | 25.38665069 | 0.631589328 | 5.684146357 |
| ENSMUSG00000034189 | *Hsdl1* | 1365.231969 | 1354.803336 | 1376.600067 | 865.0193646 | 852.7097177 | 869.8534162 | 4.77E-18 | -0.663008667 | 4.85E-19 | 79.48667364 | 0.631559837 | 17.32109011 |
| ENSMUSG00000096592 | *Gm15801* | 549.4952351 | 579.5303232 | 530.6665976 | 367.4006979 | 302.0212073 | 379.1175288 | 5.63E-07 | -0.663330701 | 1.20E-07 | 28.0200351 | 0.631418878 | 6.249622224 |
| ENSMUSG00000031138 | *F9* | 199.0431656 | 181.4468754 | 201.6924707 | 117.4287041 | 126.7155223 | 123.1650855 | 0.000520221 | -0.664428855 | 0.000162377 | 14.22295813 | 0.630938436 | 3.283811763 |
| ENSMUSG00000033319 | *Fem1c* | 1494.524966 | 1470.26953 | 1513.67262 | 952.2188974 | 843.1822348 | 1030.545364 | 1.08E-12 | -0.664651666 | 1.46E-13 | 54.62480106 | 0.630841 | 11.96710337 |
| ENSMUSG00000034792 | *Gna15* | 391.2814367 | 391.4853796 | 420.0294657 | 242.9960312 | 266.7695206 | 248.2546254 | 8.64E-07 | -0.664817307 | 1.87E-07 | 27.15824703 | 0.630768575 | 6.063465675 |
| ENSMUSG00000027953 | *Slc50a1* | 866.7734435 | 925.9289034 | 944.8215253 | 609.2340686 | 564.0269864 | 554.2428847 | 9.66E-12 | -0.664900125 | 1.39E-12 | 50.20178697 | 0.630732367 | 11.0150469 |
| ENSMUSG00000040659 | *Efhd2* | 12636.68918 | 12548.42603 | 13029.72524 | 8087.465995 | 8203.162759 | 7799.813929 | 8.63E-43 | -0.665733772 | 3.73E-44 | 194.2658303 | 0.63036801 | 42.0638266 |
| ENSMUSG00000050565 | *Tor1aip2* | 6644.128918 | 6616.762722 | 6480.594483 | 4215.806742 | 3920.559205 | 4306.929083 | 9.98E-31 | -0.666068671 | 6.16E-32 | 138.3335435 | 0.630221697 | 30.0010194 |
| ENSMUSG00000039178 | *Tbc1d19* | 580.9678724 | 560.8357966 | 519.8966113 | 329.0329035 | 351.5641182 | 365.6463475 | 2.95E-08 | -0.666391683 | 5.58E-09 | 33.9753048 | 0.63008061 | 7.530256633 |
| ENSMUSG00000039634 | *Zfp189* | 269.6439466 | 281.5175763 | 270.2287471 | 183.7003489 | 154.3452226 | 179.9364921 | 5.67E-05 | -0.66664882 | 1.54E-05 | 18.69351055 | 0.629968318 | 4.246348474 |
| ENSMUSG00000029462 | *Vps29* | 2994.153603 | 2770.088964 | 3072.383364 | 1640.513876 | 2060.794547 | 1857.098555 | 4.37E-13 | -0.668023776 | 5.78E-14 | 56.44453258 | 0.629368215 | 12.36000386 |
| ENSMUSG00000044098 | *Rsbn1* | 3496.865188 | 3489.278397 | 3490.454651 | 2342.760779 | 1870.244889 | 2382.474622 | 9.45E-13 | -0.668164268 | 1.28E-13 | 54.88868875 | 0.629306929 | 12.02475568 |
| ENSMUSG00000043017 | *Ptgir* | 2249.868261 | 2241.14383 | 2174.558143 | 1446.349583 | 1304.312406 | 1444.303073 | 8.21E-20 | -0.668780478 | 7.57E-21 | 87.71240115 | 0.629038194 | 19.08541116 |
| ENSMUSG00000040528 | *Milr1* | 2662.414993 | 2543.555289 | 2646.479361 | 1534.711776 | 1748.293108 | 1652.144155 | 4.98E-20 | -0.669260264 | 4.55E-21 | 88.72062274 | 0.628829034 | 19.30320039 |
| ENSMUSG00000004317 | *Clcn5* | 904.2003636 | 967.7166687 | 831.2471243 | 551.1010468 | 522.1060618 | 625.4476997 | 3.28E-09 | -0.66926344 | 5.71E-10 | 38.41704608 | 0.62882765 | 8.483868439 |
| ENSMUSG00000027291 | *Vps39* | 2587.561153 | 2522.661407 | 2585.775801 | 1674.231028 | 1607.286362 | 1560.732568 | 8.05E-25 | -0.669369763 | 6.00E-26 | 110.9718971 | 0.628781308 | 24.09398786 |
| ENSMUSG00000001054 | *Rmnd5b* | 288.3574066 | 299.1124249 | 314.287782 | 177.8870468 | 198.1716439 | 189.5587644 | 1.52E-05 | -0.670556553 | 3.79E-06 | 21.368727 | 0.628264273 | 4.81804078 |
| ENSMUSG00000026142 | *Rhbdd1* | 2065.285496 | 2036.603716 | 1932.722996 | 1276.601159 | 1244.289264 | 1270.139944 | 1.16E-21 | -0.671056658 | 9.79E-23 | 96.31608266 | 0.628046525 | 20.93469816 |
| ENSMUSG00000016427 | *Ndufa1* | 1122.807601 | 1122.771271 | 1173.928507 | 723.1747914 | 746.0019094 | 676.4457429 | 1.84E-14 | -0.672592882 | 2.24E-15 | 62.84395267 | 0.627378119 | 13.73610201 |
| ENSMUSG00000030595 | *Nfkbib* | 1947.050454 | 2091.587618 | 2059.025562 | 1274.275838 | 1359.571807 | 1189.312857 | 3.34E-17 | -0.672923084 | 3.53E-18 | 75.56697791 | 0.627234542 | 16.47674226 |
| ENSMUSG00000038650 | *Rnh1* | 13160.66606 | 13603.01726 | 13742.50252 | 8232.79855 | 9431.255302 | 7740.155841 | 3.75E-19 | -0.67293774 | 3.62E-20 | 84.61982247 | 0.62722817 | 18.42553415 |
| ENSMUSG00000029456 | *Acad10* | 215.2047902 | 200.1414019 | 206.587919 | 134.8686106 | 121.9517808 | 133.749585 | 0.000268892 | -0.673514775 | 8.00E-05 | 15.55742998 | 0.626977347 | 3.570422773 |
| ENSMUSG00000027244 | *Atg13* | 3108.986198 | 3181.368548 | 3112.52604 | 2000.938611 | 1995.054915 | 1897.512098 | 1.41E-28 | -0.674123333 | 9.28E-30 | 128.3773089 | 0.626712932 | 27.85064182 |
| ENSMUSG00000046688 | *Tifa* | 330.0373858 | 357.3953606 | 350.5140995 | 232.5320873 | 205.7936302 | 212.6522179 | 3.92E-06 | -0.674176226 | 9.16E-07 | 24.09754491 | 0.626689955 | 5.406496879 |
| ENSMUSG00000083563 | *Gm13340* | 233.9182502 | 268.3214399 | 248.6887745 | 155.7964985 | 163.8727055 | 150.1074479 | 8.39E-05 | -0.674537515 | 2.32E-05 | 17.90311973 | 0.626533035 | 4.07606195 |
| ENSMUSG00000040040 | *Ift88* | 278.1500648 | 238.6301331 | 210.5042777 | 145.3325545 | 121.9517808 | 188.5965372 | 0.002640514 | -0.674653696 | 0.000936337 | 10.9494047 | 0.626482582 | 2.578311587 |
| ENSMUSG00000024300 | *Myo1f* | 2710.899867 | 2789.883169 | 2706.20383 | 1712.598823 | 1750.198605 | 1676.199835 | 2.12E-27 | -0.675095466 | 1.45E-28 | 122.9159832 | 0.626290775 | 26.67448381 |
| ENSMUSG00000029994 | *Anxa4* | 5526.424988 | 5522.58308 | 5394.784046 | 3510.071857 | 3360.343211 | 3430.340076 | 8.66E-38 | -0.675130045 | 4.21E-39 | 171.1209921 | 0.626275764 | 37.06273224 |
| ENSMUSG00000025402 | *Nab2* | 3769.06097 | 3811.484061 | 3876.215978 | 2550.876997 | 2385.681713 | 2239.102765 | 1.22E-23 | -0.675788947 | 9.53E-25 | 105.4919462 | 0.625989799 | 22.91392317 |
| ENSMUSG00000059108 | *Ifitm6* | 557.1507414 | 590.5271035 | 594.3074258 | 355.7740935 | 381.0993152 | 352.1751663 | 3.03E-09 | -0.676260885 | 5.25E-10 | 38.582978 | 0.625785057 | 8.51890558 |
| ENSMUSG00000055024 | *Ep300* | 2930.357716 | 3117.587222 | 2923.561735 | 2043.957047 | 1584.420403 | 1984.112549 | 6.29E-12 | -0.677296681 | 8.95E-13 | 51.06124376 | 0.62533593 | 11.20168679 |
| ENSMUSG00000020448 | *Rnf185* | 1157.682686 | 1201.94809 | 1155.325803 | 702.2469035 | 759.3403854 | 734.1793768 | 1.36E-15 | -0.677300196 | 1.55E-16 | 68.10449915 | 0.625334407 | 14.86656236 |
| ENSMUSG00000023952 | *Gtpbp2* | 1896.864356 | 1951.928508 | 1894.538499 | 1228.932081 | 1172.833142 | 1190.275084 | 1.92E-22 | -0.677426025 | 1.57E-23 | 99.93725097 | 0.625279869 | 21.71596142 |
| ENSMUSG00000015839 | *Nfe2l2* | 5396.28138 | 5236.666791 | 4967.900953 | 3311.256923 | 3129.778126 | 3306.212764 | 4.15E-29 | -0.678947882 | 2.70E-30 | 130.8245359 | 0.624620627 | 28.3815375 |
| ENSMUSG00000027534 | *Snx16* | 505.2634205 | 472.861554 | 476.8166661 | 312.7556574 | 292.4937244 | 304.0638048 | 2.73E-08 | -0.680034742 | 5.16E-09 | 34.12815625 | 0.624150244 | 7.563268681 |
| ENSMUSG00000038843 | *Gcnt1* | 749.3890125 | 733.4852477 | 702.9863784 | 420.883078 | 449.6971919 | 491.6981147 | 4.38E-10 | -0.680626261 | 7.10E-11 | 42.49034416 | 0.623894389 | 9.358246938 |
| ENSMUSG00000018286 | *Psmb6* | 7573.847636 | 7438.222212 | 7692.707486 | 4482.055982 | 5476.397159 | 4183.763998 | 1.65E-13 | -0.682809891 | 2.12E-14 | 58.41640652 | 0.622950791 | 12.78285168 |
| ENSMUSG00000052031 | *Tagap1* | 934.8223891 | 902.8356647 | 906.6370284 | 595.2821434 | 515.4368237 | 600.4297917 | 1.80E-11 | -0.682944776 | 2.62E-12 | 48.95767492 | 0.622892551 | 10.74582343 |
| ENSMUSG00000014771 | *Pdcd2* | 874.4289499 | 804.9643198 | 975.1733049 | 559.2396699 | 532.5862929 | 561.9407025 | 3.07E-10 | -0.683733013 | 4.91E-11 | 43.21341529 | 0.622552317 | 9.513404949 |
| ENSMUSG00000026470 | *Stx6* | 1799.894609 | 1832.063602 | 1788.796815 | 1134.756586 | 1202.368339 | 1037.280954 | 3.35E-17 | -0.6838249 | 3.55E-18 | 75.55866537 | 0.622512667 | 16.47519529 |
| ENSMUSG00000033793 | *Atp6v1h* | 5202.341885 | 5368.628155 | 5582.769261 | 3334.510131 | 3348.910232 | 3369.719761 | 6.35E-35 | -0.683982831 | 3.36E-36 | 157.8338983 | 0.622444525 | 34.19710347 |
| ENSMUSG00000021720 | *Rnf180* | 147.1558447 | 127.5626518 | 172.3197808 | 98.82613709 | 82.88910105 | 97.18495027 | 0.004356222 | -0.684132169 | 0.001604515 | 9.954559739 | 0.622380097 | 2.360889979 |
| ENSMUSG00000093445 | *Lrch4* | 2588.411765 | 2586.442733 | 2727.743803 | 1750.966617 | 1583.467654 | 1587.67493 | 1.45E-21 | -0.684148207 | 1.23E-22 | 95.87077048 | 0.622373179 | 20.83913028 |
| ENSMUSG00000062352 | *Itgb1bp1* | 962.0419673 | 973.2150588 | 1032.939595 | 560.4023303 | 653.5853255 | 630.2588359 | 5.84E-12 | -0.684387446 | 8.31E-13 | 51.20824317 | 0.62226998 | 11.23337163 |
| ENSMUSG00000055401 | *Fbxo6* | 808.081228 | 738.9836379 | 808.7280621 | 482.5040811 | 532.5862929 | 450.3223438 | 5.84E-10 | -0.685387631 | 9.55E-11 | 41.91136205 | 0.621838725 | 9.233332074 |
| ENSMUSG00000033955 | *Tnks1bp1* | 461.0316059 | 524.5464215 | 426.8830933 | 295.3157508 | 277.2497518 | 305.026032 | 7.19E-07 | -0.685459768 | 1.55E-07 | 27.52491408 | 0.621807633 | 6.143060861 |
| ENSMUSG00000078651 | *Aoc2* | 330.8879976 | 332.1027658 | 302.538706 | 209.2788785 | 192.4551542 | 199.1810367 | 5.01E-06 | -0.685587645 | 1.18E-06 | 23.60688848 | 0.62175252 | 5.300495401 |
| ENSMUSG00000035845 | *Alg12* | 464.4340532 | 489.3567245 | 516.9593423 | 316.2436387 | 306.7849487 | 291.5548508 | 3.95E-08 | -0.685724774 | 7.55E-09 | 33.38847296 | 0.621693425 | 7.403939904 |
| ENSMUSG00000073987 | *Ggh* | 418.5010149 | 433.2731448 | 442.5485279 | 248.8093334 | 278.2025001 | 276.1592151 | 1.66E-07 | -0.685793669 | 3.34E-08 | 30.49756082 | 0.621663737 | 6.781092602 |
| ENSMUSG00000012640 | *Zfp715* | 1030.090913 | 990.8099073 | 1026.085967 | 660.3911278 | 580.2237073 | 653.3522894 | 4.85E-13 | -0.68754489 | 6.45E-14 | 56.23022797 | 0.620909586 | 12.31446567 |
| ENSMUSG00000036533 | *Cdc42ep3* | 967.99625 | 973.2150588 | 1030.981416 | 591.7941621 | 610.7116525 | 639.8811082 | 6.28E-14 | -0.688941338 | 7.88E-15 | 60.36533067 | 0.620308871 | 13.20193448 |
| ENSMUSG00000008348 | *Ubc* | 46276.6854 | 48102.1165 | 46230.65573 | 30150.11044 | 27433.4342 | 29545.18711 | 1.81E-38 | -0.690516776 | 8.63E-40 | 174.2722039 | 0.619631857 | 37.74351354 |
| ENSMUSG00000012519 | *Mlkl* | 771.5049198 | 835.7553048 | 855.7243659 | 465.0641745 | 545.924769 | 511.9048866 | 1.82E-10 | -0.690793157 | 2.87E-11 | 44.26288473 | 0.619513164 | 9.73896873 |
| ENSMUSG00000029060 | *Mib2* | 873.5783381 | 1000.70701 | 929.1560907 | 585.9808599 | 579.270959 | 569.6385204 | 2.05E-12 | -0.691237172 | 2.82E-13 | 53.32682807 | 0.619322527 | 11.68828482 |
| ENSMUSG00000005410 | *Mcm5* | 7591.710484 | 7891.289562 | 7896.358136 | 4861.083284 | 5117.211054 | 4494.563393 | 8.04E-28 | -0.691808136 | 5.45E-29 | 124.8640916 | 0.619077471 | 27.09476099 |
| ENSMUSG00000019808 | *Adat2* | 330.0373858 | 341.9998681 | 377.9286101 | 223.2308038 | 228.6595891 | 198.2188095 | 4.87E-06 | -0.691866107 | 1.15E-06 | 23.66298275 | 0.619052595 | 5.312776098 |
| ENSMUSG00000035168 | *Tanc1* | 2196.279717 | 2238.944474 | 2172.599963 | 1480.066735 | 1280.493699 | 1332.684714 | 4.94E-19 | -0.692246479 | 4.81E-20 | 84.05688155 | 0.618889401 | 18.30618529 |
| ENSMUSG00000035310 | *None* | 1365.231969 | 1260.231025 | 1300.231073 | 838.2781746 | 724.0886988 | 866.9667346 | 8.09E-13 | -0.693730249 | 1.09E-13 | 55.19811954 | 0.618253219 | 12.09180155 |
| ENSMUSG00000014470 | *Rnf166* | 710.2608688 | 665.3052097 | 705.9236474 | 458.0882119 | 441.1224573 | 389.7020283 | 7.39E-10 | -0.693779971 | 1.22E-10 | 41.43560777 | 0.618231911 | 9.131643799 |
| ENSMUSG00000052748 | *Swt1* | 942.4778954 | 984.2118392 | 928.177001 | 608.0714082 | 531.6335446 | 625.4476997 | 8.82E-12 | -0.694071387 | 1.26E-12 | 50.38466083 | 0.618107045 | 11.05447964 |
| ENSMUSG00000031662 | *Snx20* | 2093.355687 | 2110.282145 | 2196.098115 | 1305.66767 | 1450.082894 | 1197.972902 | 8.42E-16 | -0.69454835 | 9.54E-17 | 69.06178674 | 0.617902729 | 15.07449922 |
| ENSMUSG00000028708 | *Mknk1* | 1511.537202 | 1562.642484 | 1492.132647 | 945.2429347 | 931.7878256 | 943.944913 | 1.77E-20 | -0.69462061 | 1.58E-21 | 90.80839008 | 0.617871781 | 19.75127033 |
| ENSMUSG00000055413 | *H2-Q5* | 1098.139858 | 1152.462578 | 1295.335625 | 722.012131 | 765.0568752 | 702.4258782 | 2.05E-12 | -0.694904849 | 2.83E-13 | 53.32489228 | 0.61775006 | 11.68807309 |
| ENSMUSG00000026104 | *Stat1* | 4305.797028 | 4176.577168 | 4274.705471 | 2664.81772 | 2630.538023 | 2584.542341 | 8.21E-37 | -0.695503641 | 4.12E-38 | 166.5862807 | 0.617493716 | 36.08553841 |
| ENSMUSG00000022237 | *Ankrd33b* | 919.5113763 | 1027.099282 | 910.5533871 | 574.3542556 | 565.932483 | 621.5987908 | 5.28E-12 | -0.695918245 | 7.49E-13 | 51.41152674 | 0.617316285 | 11.27750453 |
| ENSMUSG00000026600 | *Soat1* | 5893.889294 | 5602.859576 | 5498.56755 | 3434.498929 | 3387.020163 | 3663.199066 | 1.59E-30 | -0.696907393 | 9.91E-32 | 137.3904052 | 0.616893182 | 29.79907614 |
| ENSMUSG00000069892 | *9930111J21Rik2* | 1511.537202 | 1414.18595 | 1314.917418 | 873.1579877 | 703.1282365 | 1038.243182 | 2.32E-07 | -0.698098867 | 4.76E-08 | 29.81230497 | 0.616383921 | 6.63423002 |
| ENSMUSG00000038712 | *Mindy1* | 1954.70596 | 1928.835269 | 2044.339217 | 1240.558686 | 1213.801319 | 1199.897356 | 2.09E-23 | -0.698456395 | 1.65E-24 | 104.4023256 | 0.616231188 | 22.6800303 |
| ENSMUSG00000032413 | *Rasa2* | 1612.760009 | 1488.964056 | 1563.606193 | 983.6107291 | 874.6229283 | 1018.03641 | 2.98E-15 | -0.698931734 | 3.46E-16 | 66.52080146 | 0.616028185 | 14.52598163 |
| ENSMUSG00000025949 | *Pikfyve* | 2693.887631 | 2663.420195 | 2556.403111 | 1656.791122 | 1491.051071 | 1728.160106 | 8.66E-20 | -0.699063798 | 8.00E-21 | 87.60260132 | 0.615971797 | 19.06227136 |
| ENSMUSG00000033713 | *Foxn3* | 1129.612496 | 1165.658715 | 1055.458657 | 753.4039627 | 600.2314214 | 712.0481505 | 1.25E-10 | -0.699360694 | 1.95E-11 | 45.0242154 | 0.615845048 | 9.902509415 |
| ENSMUSG00000035521 | *Gnptg* | 550.3458469 | 530.0448117 | 535.562046 | 322.0569409 | 372.5245806 | 300.2148959 | 5.75E-08 | -0.699457048 | 1.12E-08 | 32.62330237 | 0.615803918 | 7.24027818 |
| ENSMUSG00000042726 | *Trafd1* | 3546.200673 | 3412.300935 | 3516.890071 | 2189.289602 | 2190.368314 | 2072.637454 | 5.67E-31 | -0.699677111 | 3.46E-32 | 139.4771729 | 0.615709993 | 30.24675351 |

| | | | | | | | | | | | | | |
|---|---|---|---|---|---|---|---|---|---|---|---|---|---|
| ENSMUSG00000007617 | *Homer1* | 1103.243529 | 1061.189301 | 1093.643154 | 733.6387353 | 539.2555309 | 735.141604 | 3.68E-08 | -0.699841806 | 7.01E-09 | 33.5318625 | 0.615639709 | 7.434710377 |
| ENSMUSG00000027132 | *Katnbl1* | 1674.854672 | 1606.629606 | 1677.180594 | 992.9120126 | 1032.779144 | 1025.734228 | 1.82E-21 | -0.700303657 | 1.55E-22 | 95.41238585 | 0.615442655 | 20.74102991 |
| ENSMUSG00000000275 | *Trim25* | 13509.41691 | 13665.69891 | 13332.26395 | 8608.337871 | 7538.144454 | 8775.512341 | 6.93E-26 | -0.700902799 | 4.96E-27 | 115.9148938 | 0.615187119 | 25.15951977 |
| ENSMUSG00000038437 | *Mllt6* | 2947.369953 | 3109.889476 | 2769.844658 | 1917.22706 | 1692.080959 | 1818.609465 | 1.17E-19 | -0.702063099 | 1.09E-20 | 86.99286044 | 0.614692548 | 18.93191882 |
| ENSMUSG00000021009 | *Ptpn21* | 333.439833 | 409.0802281 | 383.8031481 | 247.6466729 | 222.9430994 | 221.312263 | 4.02E-06 | -0.702610421 | 9.39E-07 | 24.04950938 | 0.614459394 | 5.396173185 |
| ENSMUSG00000035314 | *Gdpd5* | 1908.772922 | 1958.526576 | 1967.970224 | 1290.553084 | 1138.534204 | 1159.483813 | 1.84E-19 | -0.702691579 | 1.73E-20 | 86.08021671 | 0.614424829 | 18.73595833 |
| ENSMUSG00000110669 | *Gm10060* | 140.3509501 | 172.6494511 | 162.5288841 | 102.3141184 | 95.27482879 | 94.29826858 | 0.001390774 | -0.702858401 | 0.000470748 | 12.22811063 | 0.614353785 | 2.856743457 |
| ENSMUSG00000018567 | *Gabarap* | 6681.555838 | 6792.711207 | 6938.808445 | 3906.539066 | 4609.396217 | 4007.676414 | 3.04E-22 | -0.704362891 | 2.51E-23 | 99.0113783 | 0.613713451 | 21.51655652 |
| ENSMUSG00000033863 | *Klf9* | 628.6021342 | 612.5206641 | 646.1991779 | 423.2083988 | 309.6431936 | 426.2666631 | 9.93E-07 | -0.705368648 | 2.17E-07 | 26.87503402 | 0.613285757 | 6.003118031 |
| ENSMUSG00000023106 | *Denr* | 3893.250295 | 3785.091788 | 3761.662487 | 2357.875365 | 2319.942081 | 2336.287715 | 2.96E-36 | -0.706133382 | 1.51E-37 | 164.0099783 | 0.612960757 | 35.52807109 |
| ENSMUSG00000029924 | *Slc37a3* | 1135.566778 | 1143.665154 | 1135.74401 | 723.1747914 | 656.4435704 | 713.0103777 | 3.84E-16 | -0.70747918 | 4.27E-17 | 70.64586676 | 0.612389232 | 15.4161094 |
| ENSMUSG00000025348 | *Itga7* | 327.4855503 | 345.2989022 | 385.7613274 | 232.5320873 | 247.7145549 | 168.3897653 | 0.000122316 | -0.707501137 | 3.46E-05 | 17.14750053 | 0.612379912 | 3.912518199 |
| ENSMUSG00000057143 | *Trim12c* | 1463.052329 | 1417.484984 | 1461.780868 | 924.3150469 | 828.8910105 | 905.4558238 | 5.10E-18 | -0.709094025 | 5.20E-19 | 79.35142702 | 0.611704153 | 17.29223891 |
| ENSMUSG00000026578 | *Ccdc181* | 474.641395 | 494.8551147 | 478.7748455 | 251.1346542 | 286.7772347 | 345.4395757 | 1.52E-06 | -0.709687547 | 3.39E-07 | 26.01610288 | 0.61145255 | 5.818908546 |
| ENSMUSG00000022228 | *Zscan26* | 1138.969226 | 1142.565476 | 1173.928507 | 690.6202992 | 656.4435704 | 764.9706481 | 2.34E-14 | -0.709861783 | 2.87E-15 | 62.35183028 | 0.611378709 | 13.63002098 |
| ENSMUSG00000026088 | *Mitd1* | 828.4959117 | 796.1668956 | 855.7243659 | 520.8718755 | 444.9334504 | 551.356203 | 3.72E-10 | -0.710285842 | 6.01E-11 | 42.8174635 | 0.61119903 | 9.429217136 |
| ENSMUSG00000028256 | *Odf2l* | 483.998125 | 495.9547927 | 468.0048592 | 261.5985982 | 275.3442552 | 346.4018029 | 1.08E-06 | -0.710316963 | 2.37E-07 | 26.70633781 | 0.611185845 | 5.967109938 |
| ENSMUSG00000026213 | *Stk11ip* | 1833.06847 | 2019.008868 | 1864.186719 | 1194.052268 | 1149.967183 | 1148.899313 | 7.39E-21 | -0.710345345 | 6.48E-22 | 92.57522309 | 0.611173822 | 20.13128453 |
| ENSMUSG00000037325 | *Bbs7* | 366.613694 | 367.2924629 | 369.1168032 | 238.3453894 | 239.1398203 | 197.2565822 | 1.54E-06 | -0.710618872 | 3.44E-07 | 25.98764658 | 0.611057957 | 5.813041061 |
| ENSMUSG00000052825 | *Gm9892* | 2705.796196 | 2777.78671 | 3007.763446 | 1670.743047 | 1652.065531 | 1862.871918 | 1.42E-19 | -0.710911944 | 1.33E-20 | 86.59383176 | 0.610933839 | 18.8462103 |
| ENSMUSG00000032698 | *Lmo2* | 1355.024628 | 1320.713317 | 1371.704619 | 955.7068787 | 800.3085618 | 720.7081955 | 1.07E-10 | -0.71095521 | 1.65E-11 | 45.34514328 | 0.610915517 | 9.971179876 |
| ENSMUSG00000055371 | *Stam2* | 2619.033791 | 2726.101843 | 2602.420326 | 1550.989022 | 1653.018279 | 1643.484109 | 2.98E-27 | -0.712178968 | 2.06E-28 | 122.229021 | 0.610397531 | 26.52584422 |
| ENSMUSG00000004043 | *Stat5a* | 1178.947981 | 1373.497863 | 1220.92481 | 811.5369846 | 741.238168 | 749.5750125 | 1.39E-13 | -0.712642217 | 1.77E-14 | 58.76989783 | 0.610201564 | 12.85828246 |
| ENSMUSG00000024451 | *Arap3* | 2746.625563 | 2814.076085 | 2703.266561 | 1762.593221 | 1594.900634 | 1684.85988 | 6.77E-26 | -0.713452482 | 4.84E-27 | 115.9631479 | 0.609858951 | 25.16967071 |
| ENSMUSG00000073664 | *Nbeal1* | 868.4746672 | 831.3565926 | 831.2471243 | 524.3598568 | 403.0125258 | 616.7876547 | 1.58E-06 | -0.713537295 | 3.54E-07 | 25.9316381 | 0.6098231 | 5.801240646 |
| ENSMUSG00000039531 | *Zup1* | 595.4282733 | 548.7393383 | 563.9556462 | 355.7740935 | 261.0530309 | 425.3044358 | 4.03E-05 | -0.713862858 | 1.07E-05 | 19.38422041 | 0.609685501 | 4.394759859 |
| ENSMUSG00000029098 | *Acox3* | 615.8429569 | 607.022274 | 597.2446948 | 382.5152836 | 351.5641182 | 376.2308471 | 1.05E-10 | -0.714324568 | 1.63E-11 | 45.37668456 | 0.609490413 | 9.977981141 |
| ENSMUSG00000053470 | *Kdm3a* | 1593.195937 | 1597.832181 | 1605.707048 | 880.1339503 | 994.6692126 | 1042.09209 | 1.10E-16 | -0.715812123 | 1.20E-17 | 73.15797003 | 0.608862294 | 15.95789028 |
| ENSMUSG00000102525 | *Gm37784* | 148.8570683 | 147.3568564 | 147.8425392 | 88.36219316 | 95.27482879 | 86.60045073 | 0.001013381 | -0.715936414 | 0.000335151 | 12.86295304 | 0.608809842 | 2.994227371 |
| ENSMUSG00000025366 | *Esyt1* | 4122.915487 | 4384.416316 | 4262.956395 | 2675.281664 | 2573.373126 | 2524.884252 | 2.11E-33 | -0.716258623 | 1.18E-34 | 150.7619782 | 0.608673886 | 32.67516669 |
| ENSMUSG00000028527 | *Ak4* | 1192.55777 | 1138.166764 | 1142.597637 | 725.5001123 | 711.7029711 | 675.4835157 | 4.43E-17 | -0.718526689 | 4.72E-18 | 74.99592901 | 0.60771774 | 16.35394002 |
| ENSMUSG00000024205 | *Rpl36-ps2* | 699.2029152 | 732.3855697 | 1073.082271 | 488.3173833 | 644.0578426 | 388.7398011 | 0.000537016 | -0.718995005 | 0.000168172 | 14.15697927 | 0.6075205 | 3.270012975 |
| ENSMUSG00000026080 | *Chst10* | 2240.511531 | 2262.037713 | 2184.349039 | 1373.101975 | 1361.477303 | 1326.911351 | 1.09E-27 | -0.719555836 | 7.42E-29 | 124.2515924 | 0.607284379 | 26.96289948 |
| ENSMUSG00000033653 | *Vps8* | 945.0297309 | 985.3115172 | 1016.295071 | 608.0714082 | 538.3027827 | 642.7677899 | 2.37E-12 | -0.719935344 | 3.28E-13 | 53.03219077 | 0.607124651 | 11.62573121 |
| ENSMUSG00000022946 | *Dop1b* | 371.7173649 | 366.1927848 | 374.0122515 | 187.1883303 | 276.2970035 | 209.7655362 | 3.53E-05 | -0.720224401 | 9.28E-06 | 19.65410325 | 0.60700302 | 4.452054202 |
| ENSMUSG00000004707 | *Ly9* | 2653.908875 | 2497.368812 | 2578.922174 | 1590.519477 | 1532.971995 | 1570.35484 | 1.85E-28 | -0.720592052 | 1.22E-29 | 127.826998 | 0.606848353 | 27.73293322 |
| ENSMUSG00000009569 | *Mrtfb* | 1426.47602 | 1583.536367 | 1377.579157 | 948.7309161 | 790.7810789 | 923.7381411 | 1.14E-12 | -0.720713835 | 1.54E-13 | 54.51849109 | 0.606797129 | 11.94470999 |
| ENSMUSG00000057561 | *Eif1a* | 8269.648104 | 8014.453501 | 7875.797253 | 5113.380599 | 4415.035566 | 5132.520047 | 1.84E-24 | -0.721000013 | 1.39E-25 | 109.303867 | 0.606676774 | 23.73489993 |
| ENSMUSG00000025612 | *Bach1* | 1472.409059 | 1616.526708 | 1476.467213 | 1013.8399 | 771.7261132 | 983.3962294 | 1.71E-10 | -0.722222144 | 2.68E-11 | 44.39784579 | 0.606163066 | 9.76739213 |
| ENSMUSG00000025579 | *Gaa* | 2630.942356 | 2805.278661 | 2732.639251 | 1711.436162 | 1666.356756 | 1572.279294 | 3.67E-26 | -0.722860859 | 2.61E-27 | 117.1918499 | 0.605894763 | 25.43580642 |
| ENSMUSG00000028173 | *Wls* | 1974.270032 | 1876.050724 | 1858.312181 | 1135.919246 | 1114.715497 | 1208.557401 | 4.23E-22 | -0.722896413 | 3.50E-23 | 98.35213249 | 0.605879832 | 21.37378327 |
| ENSMUSG00000022035 | *Ccdc25* | 1261.457328 | 1238.237465 | 1200.363927 | 723.1747914 | 725.9941954 | 786.1396472 | 1.43E-17 | -0.726600974 | 1.50E-18 | 77.26013682 | 0.604326045 | 16.84314906 |
| ENSMUSG00000038507 | *Parp12* | 5427.754017 | 5455.50272 | 5344.850473 | 3325.208848 | 3256.493648 | 3226.347903 | 4.44E-45 | -0.726691053 | 1.81E-46 | 204.8695778 | 0.604288314 | 44.35223277 |
| ENSMUSG00000003198 | *Zfp959* | 330.0373858 | 285.9162885 | 317.225051 | 213.9295203 | 123.8572774 | 227.0856264 | 0.001759244 | -0.727423655 | 0.000606211 | 11.75680675 | 0.603981533 | 2.75467386 |
| ENSMUSG00000074220 | *Zfp382* | 336.8422803 | 316.7072734 | 280.0196438 | 197.6522742 | 161.9672089 | 204.9544001 | 2.33E-05 | -0.72782819 | 5.96E-06 | 20.5013549 | 0.603812199 | 4.631797671 |
| ENSMUSG00000021504 | *B4galt7* | 1759.065242 | 1774.880345 | 1960.137506 | 1139.407228 | 1125.195728 | 1050.752136 | 4.56E-19 | -0.729173774 | 4.42E-20 | 84.22122706 | 0.603249293 | 18.34075301 |
| ENSMUSG00000053398 | *Phgdh* | 14810.00238 | 15096.38003 | 15334.50231 | 8838.544637 | 10096.27361 | 8342.510087 | 1.89E-23 | -0.729807305 | 1.49E-24 | 104.6035641 | 0.602984447 | 22.72300676 |
| ENSMUSG00000020744 | *Slc25a19* | 893.9930217 | 803.8646418 | 894.8879525 | 506.9199502 | 570.6962244 | 485.9247513 | 2.21E-11 | -0.72991234 | 3.24E-12 | 48.539323 | 0.602940548 | 10.6558368 |
| ENSMUSG00000031447 | *Lamp1* | 25198.52453 | 24948.39553 | 25893.00524 | 15033.19944 | 16112.87904 | 14634.51395 | 2.08E-42 | -0.73197668 | 9.09E-44 | 192.4921083 | 0.602078423 | 41.68150256 |
| ENSMUSG00000031792 | *Usb1* | 1952.154125 | 1980.520137 | 2031.611052 | 1189.401626 | 1218.56506 | 1180.652812 | 1.43E-26 | -0.732615131 | 1.01E-27 | 119.079339 | 0.601812038 | 25.84444668 |
| ENSMUSG00000038633 | *Degs1* | 5740.779166 | 5415.91431 | 5720.820904 | 3328.696829 | 3396.547646 | 3424.566713 | 1.15E-40 | -0.733735677 | 5.27E-42 | 184.4125713 | 0.60134479 | 39.94045619 |
| ENSMUSG00000008435 | *Rdh13* | 1382.244206 | 1371.298507 | 1474.509033 | 862.6940438 | 833.6547519 | 845.7977355 | 4.88E-20 | -0.734456565 | 4.45E-21 | 88.76322103 | 0.601044384 | 19.31189991 |
| ENSMUSG00000024661 | *Fth1* | 140185.9314 | 140030.8013 | 142866.8055 | 84105.6933 | 88150.17709 | 81977.91112 | 2.50E-56 | -0.734779834 | 8.05E-58 | 256.9195737 | 0.600909721 | 55.60252846 |
| ENSMUSG00000035367 | *Rmi1* | 1050.505596 | 927.0285815 | 997.6923671 | 624.3486543 | 566.8852313 | 599.4675645 | 7.82E-14 | -0.73482744 | 9.84E-15 | 59.92770916 | 0.600889893 | 13.10703474 |
| ENSMUSG00000020315 | *Sptbn1* | 2072.090391 | 2030.005648 | 2031.611052 | 1283.577122 | 1125.195728 | 1274.95108 | 6.52E-21 | -0.736624108 | 5.69E-22 | 92.83218751 | 0.600142037 | 20.18597077 |
| ENSMUSG00000075705 | *Msrb1* | 1024.13663 | 1016.102502 | 1030.002326 | 618.5353521 | 658.3490669 | 565.7896115 | 1.23E-14 | -0.736709113 | 1.48E-15 | 63.65216361 | 0.600106677 | 13.90937766 |
| ENSMUSG00000033355 | *Rtp4* | 3031.580523 | 2867.960309 | 2824.673679 | 1740.502673 | 1735.907381 | 1757.98915 | 8.55E-31 | -0.737401968 | 5.25E-32 | 138.6495319 | 0.599818545 | 30.06818945 |
| ENSMUSG00000082838 | *Gm14519* | 477.1932304 | 499.2538268 | 425.9040036 | 311.5929969 | 234.3760788 | 296.365987 | 1.48E-06 | -0.73752936 | 3.31E-07 | 26.06261297 | 0.599765582 | 5.828702417 |
| ENSMUSG00000059142 | *Zfp945* | 150.558292 | 129.7620078 | 168.4034221 | 95.33815578 | 80.98360447 | 93.33604135 | 0.001591511 | -0.73838347 | 0.00054339 | 11.96054529 | 0.599410612 | 2.798190237 |
| ENSMUSG00000034714 | *Ttyh2* | 1752.260347 | 1722.095799 | 1734.946884 | 1056.858337 | 1001.338451 | 1063.26109 | 2.64E-24 | -0.739465091 | 2.01E-25 | 108.5787564 | 0.598961388 | 23.57916561 |
| ENSMUSG00000038963 | *Slco4a1* | 12038.70907 | 12569.31991 | 12235.68353 | 7744.481166 | 7097.974745 | 7217.666455 | 1.00E-41 | -0.740215946 | 4.42E-43 | 189.344347 | 0.598649738 | 40.9998599 |
| ENSMUSG00000010358 | *Ifi35* | 2966.934025 | 2930.641957 | 3004.826177 | 1640.513876 | 1930.268031 | 1753.178014 | 1.30E-22 | -0.740691727 | 1.05E-23 | 100.7305273 | 0.598452345 | 21.88587705 |
| ENSMUSG00000029376 | *Mthfd2l* | 179.4790938 | 173.7491291 | 135.1143736 | 105.8020997 | 73.36161817 | 113.5428132 | 0.004183184 | -0.74113954 | 0.001535614 | 10.03538587 | 0.598266614 | 2.378493023 |
| ENSMUSG00000052760 | *A630001G21Rik* | 1351.62218 | 1310.816215 | 1310.02197 | 795.2597385 | 755.5293923 | 826.5531909 | 1.98E-19 | -0.741155918 | 1.87E-20 | 85.92102092 | 0.598259822 | 18.70289277 |
| ENSMUSG00000028381 | *Ugcg* | 2280.490287 | 2090.48794 | 2103.084597 | 1297.529047 | 1197.604598 | 1377.909394 | 1.38E-19 | -0.741902997 | 1.29E-20 | 86.65780672 | 0.597950102 | 18.8599404 |
| ENSMUSG00000039770 | *Ypel5* | 498.4585259 | 413.4789402 | 396.5313137 | 268.5745608 | 243.9035617 | 271.348079 | 6.69E-07 | -0.742054733 | 1.44E-07 | 27.67149702 | 0.597887215 | 6.174449673 |
| ENSMUSG00000036478 | *Btg1* | 4952.26201 | 5038.724745 | 4989.440925 | 2979.898698 | 2863.008605 | 3104.145045 | 1.54E-38 | -0.743515266 | 7.35E-40 | 174.5931019 | 0.597282242 | 37.81171736 |
| ENSMUSG00000001542 | *Ell2* | 5945.776615 | 5751.31611 | 5890.203416 | 3383.34187 | 3434.657578 | 3672.821338 | 1.99E-38 | -0.745189798 | 9.55E-40 | 174.071867 | 0.59658938 | 37.7003866 |
| ENSMUSG00000020143 | *Dock2* | 9053.912201 | 9325.269716 | 8954.754063 | 5416.834973 | 5347.77614 | 5534.731029 | 2.29E-52 | -0.74573872 | 7.87E-54 | 238.6185152 | 0.596362431 | 51.64032453 |
| ENSMUSG00000034854 | *Mfsd12* | 4734.505384 | 5139.895124 | 5095.182609 | 3099.652723 | 2922.078999 | 2904.964008 | 1.07E-32 | -0.745953268 | 6.13E-34 | 147.4905688 | 0.59627375 | 31.96905295 |
| ENSMUSG00000072825 | *Cep170b* | 280.7019003 | 298.0127468 | 309.3923337 | 187.1883303 | 177.2111815 | 165.5030836 | 2.36E-06 | -0.746010228 | 5.39E-07 | 25.11948461 | 0.596250209 | 5.626347926 |
| ENSMUSG00000028497 | *Hacd4* | 101.2228065 | 92.37295474 | 85.18080073 | 59.29568225 | 49.54291097 | 57.73363382 | 0.009495197 | -0.746838424 | 0.003735142 | 8.408288082 | 0.595908023 | 2.022496041 |
| ENSMUSG00000107928 | *Gm45140* | 370.8667531 | 340.9001901 | 407.3013 | 270.8998817 | 206.7463785 | 191.4832188 | 2.04E-05 | -0.747204737 | 5.18E-06 | 20.77108103 | 0.595756736 | 4.689935072 |
| ENSMUSG00000021242 | *Npc2* | 9145.778278 | 9018.459545 | 9030.143967 | 5348.238007 | 5808.906311 | 5040.146233 | 6.43E-32 | -0.74750797 | 3.78E-33 | 143.8751797 | 0.59563153 | 31.19152106 |
| ENSMUSG00000042644 | *Itpr3* | 4103.351415 | 4331.63177 | 4164.068339 | 2613.660661 | 2359.004761 | 2530.657616 | 3.16E-31 | -0.748055493 | 1.91E-32 | 140.6556929 | 0.595405522 | 30.50005865 |
| ENSMUSG00000074580 | *4931440P22Rik* | 108.8783128 | 93.47263277 | 126.3025666 | 61.62100313 | 68.59787673 | 65.43145166 | 0.005945199 | -0.748307511 | 0.00223936 | 9.342204418 | 0.595301523 | 2.225833578 |
| ENSMUSG00000110126 | *Gm9347* | 3247.635925 | 3334.223795 | 3410.169298 | 1899.787153 | 2239.911225 | 1806.100511 | 9.20E-18 | -0.74837009 | 9.51E-19 | 78.15782365 | 0.595275701 | 17.03622562 |
| ENSMUSG00000021891 | *Mettl6* | 856.5661017 | 843.453051 | 845.9334693 | 502.2693085 | 499.2401029 | 513.829341 | 1.32E-15 | -0.748618762 | 1.50E-16 | 68.17032065 | 0.595173104 | 14.88027471 |
| ENSMUSG00000020888 | *Dvl2* | 1944.498618 | 1940.931727 | 2071.753728 | 1184.750985 | 1219.517808 | 1141.201495 | 1.74E-24 | -0.748669101 | 1.31E-25 | 109.4174602 | 0.595152338 | 23.75820872 |
| ENSMUSG00000042766 | *Trim46* | 232.2170266 | 276.0191862 | 272.1869265 | 159.2844798 | 147.6759846 | 156.8430385 | 1.58E-05 | -0.749462549 | 3.93E-06 | 21.29694471 | 0.594825108 | 4.802615365 |
| ENSMUSG00000018736 | *Ndel1* | 3040.937253 | 3079.098491 | 3083.15335 | 1760.267901 | 1857.859161 | 1852.287418 | 9.77E-35 | -0.749693991 | 5.21E-36 | 156.9654068 | 0.594729692 | 34.01013325 |
| ENSMUSG00000031458 | *Coprs* | 413.397344 | 490.4564025 | 469.9630385 | 263.923919 | 276.2970035 | 274.2347607 | 2.41E-08 | -0.751466805 | 4.54E-09 | 34.37898869 | 0.593999324 | 7.617821644 |
| ENSMUSG00000082729 | *Gm14845* | 402.3393904 | 336.501478 | 379.8867895 | 195.3269533 | 289.6354795 | 178.9742648 | 0.000221004 | -0.751742559 | 6.49E-05 | 15.95436561 | 0.593885799 | 3.655599416 |
| ENSMUSG00000110277 | *Gm45871* | 386.1777658 | 426.6750766 | 363.2422652 | 216.2548412 | 212.4628682 | 268.4613973 | 1.85E-06 | -0.751808342 | 4.19E-07 | 25.60618641 | 0.59385872 | 5.7321153 |
| ENSMUSG00000021094 | *Dhrs7* | 687.2943497 | 620.2184104 | 656.9691642 | 376.7019814 | 383.0048117 | 406.0598912 | 1.03E-11 | -0.753272131 | 1.49E-12 | 50.06637737 | 0.593256484 | 10.98611419 |
| ENSMUSG00000025417 | *Pip4k2c* | 3154.919237 | 3146.178851 | 3247.640414 | 1921.877701 | 1887.394358 | 1856.136327 | 1.27E-36 | -0.753490418 | 6.40E-38 | 165.7111748 | 0.593166728 | 35.89675121 |
| ENSMUSG00000003518 | *Dusp3* | 5973.846805 | 5850.287133 | 5919.576106 | 3461.240119 | 3550.892869 | 3510.204936 | 1.53E-50 | -0.75377194 | 5.51E-52 | 230.1561701 | 0.593050991 | 49.81529601 |
| ENSMUSG00000027950 | *Chrnb2* | 444.0193695 | 494.8551147 | 452.3394246 | 297.6410717 | 264.864024 | 262.6880339 | 1.14E-08 | -0.754579667 | 2.08E-09 | 35.90167872 | 0.592719051 | 7.942094637 |
| ENSMUSG00000045294 | *Insig1* | 6029.136573 | 5915.168137 | 6085.042259 | 3550.764973 | 3479.436747 | 3649.727885 | 2.88E-47 | -0.755524679 | 1.11E-48 | 215.0122125 | 0.592330928 | 46.5409812 |
| ENSMUSG00000024413 | *Npc1* | 3220.416347 | 3257.246332 | 3146.794179 | 1925.365683 | 1775.922809 | 1995.659276 | 1.41E-28 | -0.756709942 | 9.26E-30 | 128.3827427 | 0.591844491 | 27.8513777 |
| ENSMUSG00000028793 | *Rnf19b* | 4886.7649 | 5197.078382 | 5091.26625 | 3065.935571 | 2998.298862 | 2893.417282 | 1.80E-39 | -0.760544304 | 8.42E-41 | 178.9004965 | 0.590273589 | 38.7447151 |
| ENSMUSG00000020611 | *Gna13* | 6397.451491 | 6358.338384 | 6265.194757 | 3757.71853 | 3511.830189 | 3955.716144 | 2.64E-35 | -0.760936924 | 1.38E-36 | 159.6032839 | 0.590112972 | 34.57918792 |
| ENSMUSG00000041143 | *Tmco4* | 1150.027179 | 1134.86773 | 1173.928507 | 665.0417696 | 662.1600601 | 713.0103777 | 6.83E-19 | -0.761155854 | 6.71E-20 | 83.39785163 | 0.590023428 | 18.16567492 |
| ENSMUSG00000021431 | *Snrnp48* | 1360.128299 | 1287.722976 | 1316.875597 | 777.8198319 | 656.4435704 | 904.4935965 | 1.04E-10 | -0.761830916 | 1.60E-11 | 45.41025257 | 0.58974741 | 9.98503865 |
| ENSMUSG00000033157 | *Abhd10* | 523.1262687 | 477.2602661 | 534.5829563 | 269.7372212 | 314.406935 | 319.4594405 | 5.60E-09 | -0.762633578 | 9.93E-10 | 37.33793919 | 0.589419388 | 8.251954966 |
| ENSMUSG00000067869 | *Tcea1-ps1* | 178.628482 | 204.5401141 | 177.2152291 | 87.19953273 | 120.0462843 | 121.240631 | 0.000505744 | -0.763301189 | 0.000157477 | 14.28063328 | 0.589146696 | 3.296069139 |
| ENSMUSG00000045251 | *Zfp688* | 172.6741993 | 175.9484852 | 193.8597534 | 122.0793458 | 103.8495634 | 94.29826858 | 0.000277363 | -0.764575545 | 8.28E-05 | 15.49313616 | 0.588626523 | 3.556951695 |
| ENSMUSG00000029313 | *Aff1* | 6124.405097 | 6272.563498 | 5865.726174 | 3787.947702 | 3224.100206 | 3737.290563 | 8.91E-25 | -0.765023459 | 6.67E-26 | 110.7639199 | 0.5884438 | 24.05002254 |
| ENSMUSG00000049659 | *Aftph* | 2980.543814 | 2940.539059 | 2867.753624 | 1804.448997 | 1534.877492 | 1834.005101 | 1.04E-20 | -0.765257334 | 9.16E-22 | 91.89093875 | 0.588348415 | 19.98483828 |
| ENSMUSG00000002668 | *Dennd1c* | 370.8667531 | 393.6847357 | 372.0540721 | 219.7428225 | 223.8958477 | 223.2367174 | 1.84E-08 | -0.768024336 | 3.43E-09 | 34.92562524 | 0.58722108 | 7.734835866 |
| ENSMUSG00000091549 | *Gm6548* | 619.2454042 | 673.0029559 | 625.638295 | 369.7260188 | 331.5564042 | 423.3799814 | 1.59E-09 | -0.769069544 | 2.68E-10 | 39.89269478 | 0.586795802 | 8.799961937 |
| ENSMUSG00000030423 | *Pop4* | 483.1475132 | 515.7489973 | 478.7748455 | 246.4840125 | 331.5564042 | 285.7814874 | 8.76E-08 | -0.770402562 | 1.74E-08 | 31.76912879 | 0.586253867 | 7.057391857 |
| ENSMUSG00000090213 | *Peds1* | 5523.022541 | 5728.222872 | 5631.723745 | 3427.522966 | 3234.580437 | 3237.89463 | 3.54E-44 | -0.770403372 | 1.46E-45 | 200.7080237 | 0.586253537 | 43.45141193 |
| ENSMUSG00000017760 | *Ctsa* | 21999.37347 | 22074.93683 | 22509.27136 | 13032.26083 | 13329.9013 | 12666.75926 | 6.70E-65 | -0.770618257 | 1.83E-66 | 296.5832149 | 0.586166223 | 64.17411655 |
| ENSMUSG00000032737 | *Inppl1* | 6276.664612 | 6526.589123 | 6409.120938 | 3879.797876 | 3659.506174 | 3717.083791 | 8.35E-47 | -0.771538785 | 3.27E-48 | 212.8559176 | 0.585792332 | 46.07825227 |
| ENSMUSG00000031897 | *Psmb10* | 2392.771047 | 2424.790062 | 2439.891442 | 1417.283072 | 1572.034675 | 1257.63099 | 3.74E-18 | -0.77291987 | 3.78E-19 | 79.98100887 | 0.585231825 | 17.42710263 |
| ENSMUSG00000028232 | *Tmem68* | 1825.412964 | 1673.709966 | 1893.559409 | 1017.327882 | 1073.74732 | 1063.26109 | 9.97E-22 | -0.773662465 | 8.37E-23 | 96.62625623 | 0.584930667 | 21.00131894 |
| ENSMUSG00000031402 | *Mpp1* | 2186.072375 | 2231.246728 | 2263.655302 | 1169.636399 | 1462.468622 | 1271.102171 | 1.47E-17 | -0.773923581 | 1.54E-18 | 77.20502357 | 0.584824809 | 16.83131544 |
| ENSMUSG00000101431 | *Gm7901* | 2865.711218 | 2812.976407 | 2799.217348 | 1613.772686 | 1766.395326 | 1575.165976 | 1.56E-28 | -0.774415412 | 1.03E-29 | 128.167294 | 0.58462547 | 27.80559225 |
| ENSMUSG00000029714 | *Gigyf1* | 1077.725175 | 1201.94809 | 1058.395926 | 716.1988288 | 596.4204282 | 639.8811082 | 1.83E-13 | -0.774830356 | 2.35E-14 | 58.21091176 | 0.584457345 | 12.73841359 |
| ENSMUSG00000052688 | *Rab7b* | 2701.543137 | 2849.265782 | 2701.308382 | 1530.061134 | 1618.719341 | 1668.502017 | 4.48E-30 | -0.775320278 | 2.83E-31 | 135.3052063 | 0.584258905 | 29.34870278 |
| ENSMUSG00000025139 | *Tollip* | 4283.68112 | 4263.451732 | 4254.144588 | 2534.599751 | 2498.106011 | 2446.943847 | 3.75E-46 | -0.775573572 | 1.49E-47 | 209.8332353 | 0.584156335 | 45.42634817 |
| ENSMUSG00000001962 | *Fam50a* | 1207.018171 | 1152.462578 | 1163.15852 | 667.3670905 | 665.9710532 | 723.5948772 | 4.10E-19 | -0.77602492 | 3.96E-20 | 84.43782239 | 0.58397361 | 18.3867918 |
| ENSMUSG00000024293 | *Esco1* | 1767.57136 | 1851.857807 | 1659.55698 | 1005.701277 | 893.677894 | 1178.728357 | 6.76E-13 | -0.777854336 | 9.06E-14 | 55.56037373 | 0.583233569 | 12.17006285 |
| ENSMUSG00000039813 | *Tbc1d2* | 569.9099187 | 631.2151907 | 553.1856599 | 370.8886792 | 339.1783905 | 313.6860771 | 5.36E-10 | -0.778153653 | 8.73E-11 | 42.08645678 | 0.583112578 | 9.270575966 |
| ENSMUSG00000037857 | *Nufip2* | 4753.218844 | 4658.236146 | 4499.896094 | 2761.318536 | 2402.831182 | 2948.264234 | 7.95E-22 | -0.778330145 | 6.66E-23 | 97.08084755 | 0.583041247 | 21.09960977 |
| ENSMUSG00000071267 | *Zfp942* | 418.5010149 | 425.5753986 | 408.2803897 | 249.9719938 | 191.5024059 | 288.6681691 | 4.61E-06 | -0.779027151 | 1.08E-06 | 23.7722174 | 0.582759632 | 5.336666815 |
| ENSMUSG00000023951 | *Vegfa* | 551.1964587 | 543.2409481 | 611.9310397 | 365.075377 | 303.9267038 | 327.1572583 | 9.82E-10 | -0.779190914 | 1.63E-10 | 40.8604756 | 0.582693485 | 9.007808856 |
| ENSMUSG00000038550 | *Ciart* | 189.6864356 | 163.8520269 | 172.3197808 | 88.36219316 | 101.9440668 | 115.4672676 | 0.000285342 | -0.780151349 | 8.53E-05 | 15.43606053 | 0.582305702 | 3.544634576 |
| ENSMUSG00000029007 | *Agtrap* | 787.6665444 | 763.1765546 | 739.212696 | 437.1603241 | 439.2169607 | 457.0579344 | 8.30E-15 | -0.780198388 | 9.88E-16 | 64.45469203 | 0.582286716 | 14.08106115 |
| ENSMUSG00000069011 | *Gm10254* | 466.1352768 | 555.3374064 | 541.4365839 | 315.0809782 | 344.8948802 | 249.2168527 | 5.74E-07 | -0.780863377 | 1.23E-07 | 27.98101765 | 0.582018382 | 6.241424029 |
| ENSMUSG00000041235 | *Chd7* | 3094.525797 | 3258.346011 | 3196.727751 | 1983.498704 | 1585.373151 | 1986.037003 | 4.17E-17 | -0.782187232 | 4.44E-18 | 75.11614722 | 0.581484551 | 16.37954442 |
| ENSMUSG00000013495 | *Tmem175* | 440.6169222 | 442.0705691 | 405.3431207 | 273.2252025 | 217.2266096 | 259.8013522 | 6.12E-08 | -0.782311515 | 1.20E-08 | 32.49456687 | 0.58143446 | 7.213040322 |
| ENSMUSG00000021715 | *Cwc27* | 837.8526417 | 867.6459677 | 837.1216623 | 481.3414206 | 447.7916953 | 548.4695213 | 8.89E-13 | -0.782318915 | 1.20E-13 | 55.01045425 | 0.581431478 | 12.05099926 |
| ENSMUSG00000037860 | *Aim2* | 2246.465814 | 2267.536103 | 2009.09199 | 1166.148418 | 1319.556379 | 1303.817897 | 3.80E-20 | -0.782333531 | 3.46E-21 | 89.26106346 | 0.581425588 | 19.41988554 |
| ENSMUSG00000020300 | *Cpeb4* | 1453.695599 | 1552.745382 | 1454.92724 | 959.19486 | 661.2073118 | 971.8495027 | 1.06E-08 | -0.784140288 | 1.92E-09 | 36.04992038 | 0.580697896 | 7.974314591 |
| ENSMUSG00000033949 | *Trim36* | 171.8235874 | 169.350417 | 187.9852154 | 99.98879752 | 90.51108735 | 116.4294949 | 0.000214464 | -0.785413247 | 6.28E-05 | 16.0169949 | 0.580185744 | 3.668645437 |
| ENSMUSG00000022043 | *Trim35* | 6835.516577 | 7034.640374 | 6858.523093 | 4253.011876 | 3835.764607 | 3940.320508 | 2.41E-40 | -0.785602456 | 1.11E-41 | 182.9231818 | 0.580109658 | 39.61852613 |
| ENSMUSG00000022390 | *Zc3h7b* | 1771.824419 | 1957.426898 | 1879.852154 | 1134.756586 | 1041.353879 | 1073.845589 | 1.88E-23 | -0.787530493 | 1.48E-24 | 104.6217473 | 0.57933491 | 22.72623563 |
| ENSMUSG00000024397 | *Aif1* | 141.201562 | 146.2571783 | 154.6961668 | 91.85017447 | 93.36933221 | 71.20481505 | 0.000721355 | -0.787681146 | 0.000231889 | 13.55325581 | 0.579274416 | 3.141851014 |
| ENSMUSG00000069874 | *Irgm2* | 1361.829522 | 1350.404624 | 1316.875597 | 791.7717571 | 714.5612159 | 827.5154181 | 4.56E-19 | -0.788310528 | 4.42E-20 | 84.22477094 | 0.57902176 | 18.34122402 |
| ENSMUSG00000030872 | *Gga2* | 1830.516634 | 1834.262958 | 1902.371216 | 1075.460904 | 1086.133048 | 1061.336635 | 3.96E-29 | -0.788544945 | 2.57E-30 | 130.9229472 | 0.578927685 | 28.4021519 |
| ENSMUSG00000054434 | *Tmem120b* | 392.1320486 | 417.8776524 | 448.4230659 | 231.3694268 | 255.3365412 | 240.5568076 | 9.31E-09 | -0.78884878 | 1.68E-09 | 36.3134358 | 0.578805774 | 8.031063375 |
| ENSMUSG00000091144 | *Phf11c* | 117.384431 | 128.6623298 | 121.4071183 | 86.03687229 | 60.97589042 | 66.3936789 | 0.002362239 | -0.789400027 | 0.000831014 | 11.17065256 | 0.578584658 | 2.626676248 |
| ENSMUSG00000030108 | *Slc6a13* | 566.5074714 | 614.7200202 | 629.5546537 | 379.0273022 | 321.076173 | 348.3262574 | 7.98E-11 | -0.789790377 | 1.22E-11 | 45.933435 | 0.578428131 | 10.09793581 |
| ENSMUSG00000049891 | *Gm7984* | 428.7083568 | 437.671857 | 345.6186512 | 241.8333708 | 282.9662415 | 176.0875832 | 6.22E-05 | -0.790435008 | 1.70E-05 | 18.50310213 | 0.578169733 | 4.205932974 |
| ENSMUSG00000049313 | *Sorl1* | 3802.234831 | 3807.085349 | 3843.906019 | 2285.790418 | 2038.881336 | 2295.874172 | 1.55E-31 | -0.791248606 | 9.27E-33 | 142.0948087 | 0.577843771 | 30.8097868 |
| ENSMUSG00000030659 | *Nucb2* | 1064.115386 | 1078.78415 | 1003.566905 | 583.655539 | 574.5072176 | 656.2389711 | 2.82E-16 | -0.79337055 | 3.12E-17 | 71.26464597 | 0.576994491 | 15.54963682 |
| ENSMUSG00000024528 | *Srfbp1* | 1242.743868 | 1140.36612 | 1214.071183 | 699.9215827 | 679.3095293 | 696.6525148 | 2.11E-20 | -0.794362135 | 1.89E-21 | 90.45531574 | 0.576598051 | 19.67543886 |
| ENSMUSG00000040327 | *Cul9* | 239.872533 | 255.1253036 | 261.4169402 | 186.0256698 | 120.0462843 | 131.8251306 | 0.000137337 | -0.794525854 | 3.91E-05 | 16.91378303 | 0.576532621 | 3.862213748 |
| ENSMUSG00000030089 | *Slc41a3* | 125.0399374 | 120.9645836 | 121.4071183 | 79.06090967 | 60.02314214 | 73.12926951 | 0.001501392 | -0.796427377 | 0.000510766 | 12.07594401 | 0.575773232 | 2.823505824 |
| ENSMUSG00000056260 | *Lrif1* | 1414.567455 | 1374.597541 | 1353.101915 | 883.6219316 | 664.0655567 | 838.0999176 | 1.18E-12 | -0.797770668 | 1.59E-13 | 54.45053729 | 0.57523738 | 11.92991205 |
| ENSMUSG00000032840 | *Spring1* | 1299.734859 | 1420.784018 | 1363.871901 | 746.4280001 | 814.5997861 | 783.2529655 | 6.14E-21 | -0.79904366 | 5.36E-22 | 92.95231762 | 0.574730031 | 20.21199048 |
| ENSMUSG00000034247 | *Plekhm1* | 4344.925171 | 4593.355142 | 4320.722685 | 2662.492399 | 2484.767535 | 2473.886209 | 9.41E-38 | -0.799270272 | 4.59E-39 | 170.9491184 | 0.574639762 | 37.02641798 |
| ENSMUSG00000034116 | *Vav1* | 6904.416135 | 7006.048746 | 6855.585824 | 4163.487023 | 3814.804145 | 3949.942781 | 7.21E-47 | -0.800322898 | 2.81E-48 | 213.1584683 | 0.574220643 | 46.14196814 |
| ENSMUSG00000022391 | *Rangap1* | 6623.714234 | 6593.669483 | 6674.454236 | 3826.315496 | 3931.039436 | 3645.878976 | 8.06E-50 | -0.802812907 | 2.94E-51 | 226.8246489 | 0.573230426 | 49.09366153 |
| ENSMUSG00000044252 | *Osbpl1a* | 522.2756569 | 470.6621979 | 501.2939077 | 283.6891465 | 302.0212073 | 271.348079 | 1.86E-10 | -0.802931734 | 2.94E-11 | 44.21880475 | 0.573183214 | 9.729829427 |

| | | | | | | | | | | | | | |
|---|---|---|---|---|---|---|---|---|---|---|---|---|---|
| ENSMUSG00000052477 | *C130026I21Rik* | 2450.61265 | 2412.693603 | 2565.214918 | 1270.787857 | 1535.83024 | 1447.189754 | 1.86E-21 | -0.802942622 | 1.59E-22 | 95.36188868 | 0.573178889 | 20.73065128 |
| ENSMUSG00000078546 | *Zfp995* | 182.0309293 | 167.151061 | 161.5497945 | 111.6154019 | 72.40886988 | 109.6939043 | 0.000868678 | -0.803051563 | 0.000283837 | 13.17415163 | 0.573135608 | 3.061141312 |
| ENSMUSG00000034917 | *Tjp3* | 176.0766465 | 194.6430118 | 171.3406911 | 80.22357011 | 112.424298 | 116.4294949 | 0.000349793 | -0.803130967 | 0.000106205 | 15.02306431 | 0.573104064 | 3.456188637 |
| ENSMUSG00000035399 | *Oser1* | 3450.081538 | 3358.416711 | 3347.50756 | 1762.593221 | 2158.92762 | 1885.003144 | 1.50E-22 | -0.805705709 | 1.22E-23 | 100.441954 | 0.572082172 | 21.82480381 |
| ENSMUSG00000024835 | *Coro1b* | 3951.091899 | 3976.435766 | 4025.037607 | 2247.422623 | 2386.634461 | 2200.613676 | 1.01E-40 | -0.806081302 | 4.61E-42 | 184.6812658 | 0.571933255 | 39.99716634 |
| ENSMUSG00000084817 | *Gm5526* | 354.7051285 | 215.5368944 | 242.8142366 | 162.7724611 | 201.0298887 | 101.9960864 | 0.004802575 | -0.806549452 | 0.001779464 | 9.764164484 | 0.571747694 | 2.31852587 |
| ENSMUSG00000049969 | *Plekhf2* | 1377.991147 | 1395.491423 | 1356.039184 | 746.4280001 | 823.1745207 | 789.0263289 | 5.28E-23 | -0.806594676 | 4.23E-24 | 102.5405236 | 0.571729772 | 22.27752831 |
| ENSMUSG00000080242 | *Atp6v0c-ps2* | 6417.866174 | 6358.338384 | 6585.357077 | 3908.864387 | 3826.237124 | 3328.34399 | 1.20E-29 | -0.80784277 | 7.73E-31 | 133.3115943 | 0.571235375 | 28.91917042 |
| ENSMUSG00000010095 | *Slc3a2* | 11332.70126 | 11434.45218 | 11279.11292 | 6727.153285 | 6679.718246 | 6042.787007 | 6.77E-44 | -0.807957555 | 2.84E-45 | 199.386354 | 0.571189928 | 43.16942846 |
| ENSMUSG00000009292 | *Trpm2* | 118.2350428 | 104.4694131 | 108.6789527 | 73.24760749 | 65.73963186 | 50.99804321 | 0.003150775 | -0.809077302 | 0.001131546 | 10.59890538 | 0.570746771 | 2.501582618 |
| ENSMUSG00000026170 | *Cyp27a1* | 347.900234 | 266.1220839 | 338.7650236 | 176.7243863 | 171.4946918 | 196.294355 | 3.51E-06 | -0.809235756 | 8.16E-07 | 24.3199701 | 0.570684088 | 5.454734099 |
| ENSMUSG00000028229 | *Rmdn1* | 214.3541784 | 227.6333527 | 189.9433947 | 113.9407228 | 120.9990326 | 125.0895399 | 2.23E-05 | -0.809272358 | 5.68E-06 | 20.59315226 | 0.57066961 | 4.651338025 |
| ENSMUSG00000025531 | *Chm* | 1249.548762 | 1240.436821 | 1209.175734 | 623.1859939 | 682.1677741 | 802.4975101 | 2.92E-14 | -0.809574814 | 3.58E-15 | 61.91578411 | 0.570549984 | 13.53531291 |
| ENSMUSG00000021930 | *Spryd7* | 3227.221241 | 3004.320385 | 3272.117656 | 1766.081203 | 1803.552509 | 1849.400737 | 1.17E-34 | -0.810526955 | 6.27E-36 | 156.5967468 | 0.57017356 | 33.93013404 |
| ENSMUSG00000004151 | *Etv1* | 1679.958342 | 1726.494511 | 1716.34418 | 980.1227478 | 888.9141526 | 1051.714363 | 2.54E-21 | -0.810594256 | 2.19E-22 | 94.72695954 | 0.570146962 | 20.59518298 |
| ENSMUSG00000053870 | *Fpgt* | 679.6388434 | 720.2891113 | 738.2336063 | 424.3710593 | 339.1783905 | 455.13348 | 7.41E-10 | -0.811623502 | 1.22E-10 | 41.42488826 | 0.569740354 | 9.129984354 |
| ENSMUSG00000030720 | *Cln3* | 2263.47805 | 2342.314209 | 2291.069813 | 1334.734181 | 1270.966216 | 1324.024669 | 3.95E-34 | -0.811659137 | 2.15E-35 | 154.1520087 | 0.569726281 | 33.40363735 |
| ENSMUSG00000052684 | *Jun* | 3706.966307 | 3601.445557 | 3721.519811 | 2197.428225 | 2100.809975 | 1988.923685 | 4.37E-37 | -0.811867105 | 2.17E-38 | 167.8617629 | 0.56964416 | 36.35996407 |
| ENSMUSG00000014850 | *Msh3* | 865.9228317 | 886.3404943 | 811.6653311 | 502.2693085 | 461.1301713 | 497.4714781 | 5.57E-16 | -0.812203093 | 6.25E-17 | 69.89779348 | 0.569511511 | 15.25434898 |
| ENSMUSG00000024121 | *Atp6v0c* | 12771.93646 | 12307.59654 | 12677.25296 | 7200.356082 | 7486.696046 | 6814.493245 | 5.23E-50 | -0.812397179 | 1.89E-51 | 227.6975069 | 0.569434899 | 49.28156646 |
| ENSMUSG00000030830 | *Itgal* | 901.6485281 | 912.732767 | 952.6542426 | 515.0585733 | 519.2478169 | 538.847249 | 2.11E-18 | -0.81402418 | 2.11E-19 | 81.13123989 | 0.568793081 | 17.67545492 |
| ENSMUSG00000025981 | *Coq10b* | 604.7850033 | 607.022274 | 559.0601979 | 347.6354705 | 311.5486901 | 346.4018029 | 4.06E-12 | -0.817560318 | 5.70E-13 | 51.94900147 | 0.56740064 | 11.39196137 |
| ENSMUSG00000021598 | *Med10* | 1601.702055 | 1644.018659 | 1700.678746 | 901.0618382 | 994.6692126 | 908.3425055 | 2.30E-24 | -0.817764591 | 1.75E-25 | 108.851426 | 0.567320306 | 23.63740811 |
| ENSMUSG00000096727 | *Psmb9* | 1309.091589 | 1298.719756 | 1225.820259 | 728.9880936 | 820.3162759 | 622.561018 | 1.48E-13 | -0.819842776 | 1.89E-14 | 58.6416553 | 0.566503677 | 12.83118117 |
| ENSMUSG00000020078 | *Vps26a* | 5376.717308 | 5242.165181 | 5266.5233 | 2978.736038 | 2923.984496 | 3096.447227 | 6.33E-51 | -0.819898835 | 2.25E-52 | 231.9444977 | 0.566481664 | 50.19862412 |
| ENSMUSG00000021707 | *Dhfr* | 2439.554697 | 2426.989418 | 2417.372379 | 1227.769421 | 1428.169684 | 1461.623163 | 8.17E-24 | -0.82137347 | 6.36E-25 | 106.2945811 | 0.565902937 | 23.08753551 |
| ENSMUSG00000045763 | *Basp1* | 834.4501944 | 825.8582025 | 924.2606424 | 517.3838942 | 467.7994094 | 477.2647063 | 2.78E-15 | -0.823182033 | 3.22E-16 | 66.66251894 | 0.565193965 | 14.55617767 |
| ENSMUSG00000021684 | *Pde8b* | 674.5351725 | 715.8903992 | 714.7354544 | 355.7740935 | 393.4850429 | 437.8133898 | 1.88E-12 | -0.823449202 | 2.59E-13 | 53.49814081 | 0.565089308 | 11.72529146 |
| ENSMUSG00000049232 | *Tigd2* | 639.6600879 | 592.7264596 | 620.7428467 | 349.9607913 | 338.2256422 | 358.9107569 | 1.31E-13 | -0.824655383 | 1.67E-14 | 58.88487635 | 0.564617057 | 12.88206905 |
| ENSMUSG00000049502 | *Dtx3l* | 2824.881851 | 2974.629078 | 2813.903693 | 1639.351215 | 1466.279615 | 1753.178014 | 1.77E-24 | -0.825949128 | 1.34E-25 | 109.3864511 | 0.56411096 | 23.75181006 |
| ENSMUSG00000040423 | *Rc3h1* | 5648.062478 | 5584.165049 | 5429.052184 | 3278.70243 | 2712.474376 | 3400.511032 | 3.35E-22 | -0.82739876 | 2.77E-23 | 98.81537892 | 0.563544421 | 21.47465639 |
| ENSMUSG00000033220 | *Rac2* | 6814.251282 | 6854.293177 | 6620.604305 | 3787.947702 | 3904.362484 | 3735.366108 | 1.40E-57 | -0.828063057 | 4.37E-59 | 262.7256692 | 0.563284994 | 56.85403561 |
| ENSMUSG00000016534 | *Lamp2* | 12464.01498 | 11896.31696 | 11927.27028 | 6595.772655 | 6975.070216 | 6854.906789 | 5.36E-60 | -0.828971435 | 1.60E-61 | 273.9092825 | 0.562930439 | 59.27099057 |
| ENSMUSG00000050552 | *Lamtor4* | 822.541629 | 891.8388844 | 921.3233734 | 501.1066481 | 534.4917895 | 446.4734349 | 4.75E-14 | -0.830074438 | 5.94E-15 | 60.92131535 | 0.562500219 | 13.3229288 |
| ENSMUSG00000000555 | *Itga5* | 10156.30512 | 10416.15032 | 10260.85967 | 5876.085845 | 5734.591945 | 5726.214248 | 2.87E-71 | -0.830727843 | 7.06E-73 | 326.0229316 | 0.562245517 | 70.54250182 |
| ENSMUSG00000044583 | *Tlr7* | 5737.376719 | 5645.747019 | 5539.689316 | 3161.273726 | 2894.449299 | 3453.43353 | 4.27E-30 | -0.831642794 | 2.69E-31 | 135.404303 | 0.561889056 | 29.36943395 |
| ENSMUSG00000063268 | *Parp10* | 1808.400727 | 1947.529796 | 1921.95301 | 1110.340717 | 1053.739606 | 1023.809773 | 8.04E-28 | -0.833087237 | 5.45E-29 | 124.8645113 | 0.561326768 | 27.09476099 |
| ENSMUSG00000069793 | *Slfn9* | 1690.165684 | 1666.012219 | 1673.264235 | 960.3575204 | 789.8283307 | 1072.883362 | 3.68E-14 | -0.833545588 | 4.56E-15 | 61.44201975 | 0.56114846 | 13.43371702 |
| ENSMUSG00000029752 | *Asns* | 12932.7021 | 12806.85037 | 12991.54075 | 6879.461802 | 7921.149265 | 6927.073831 | 4.69E-37 | -0.833730201 | 2.33E-38 | 167.7177291 | 0.561076658 | 36.32910272 |
| ENSMUSG00000035469 | *Rcbtb1* | 1396.704607 | 1380.095931 | 1328.624673 | 791.7717571 | 689.7897604 | 822.704282 | 4.26E-19 | -0.834043875 | 4.12E-20 | 84.36321778 | 0.56095468 | 18.37102043 |
| ENSMUSG00000037816 | *Fbxw17* | 811.4836753 | 831.3565926 | 879.2225179 | 467.3894954 | 476.3741439 | 469.5668884 | 1.27E-17 | -0.834949136 | 1.32E-18 | 77.50417647 | 0.560602803 | 16.89567227 |
| ENSMUSG00000023074 | *Mospd1* | 279.8512884 | 296.9130688 | 337.7859339 | 158.1218193 | 161.0144607 | 192.4454461 | 1.12E-06 | -0.83549158 | 2.46E-07 | 26.63520968 | 0.56039206 | 5.951800579 |
| ENSMUSG00000021285 | *Ppp1r13b* | 1116.853318 | 1123.870949 | 1216.029362 | 680.1563553 | 625.9556251 | 632.1832903 | 3.41E-20 | -0.835871646 | 3.09E-21 | 89.4824489 | 0.560244449 | 19.46717319 |
| ENSMUSG00000028007 | *Snx7* | 718.766987 | 632.3148687 | 713.7563647 | 345.3101496 | 418.2564984 | 391.6264828 | 6.25E-12 | -0.836541176 | 8.90E-13 | 51.07200889 | 0.559984509 | 11.2038593 |
| ENSMUSG00000038375 | *Trp53inp2* | 481.4462895 | 461.8647737 | 460.1721419 | 283.6891465 | 253.4310446 | 250.1790799 | 7.26E-11 | -0.837085889 | 1.11E-11 | 46.12427504 | 0.559773118 | 10.13926797 |
| ENSMUSG00000020234 | *Tektip1* | 461.8822177 | 470.6621979 | 465.0675902 | 302.2917134 | 257.2420377 | 224.1989447 | 7.92E-09 | -0.838252725 | 1.42E-09 | 36.64231855 | 0.559320563 | 8.101525804 |
| ENSMUSG00000030615 | *Tmem126a* | 1179.798593 | 1054.591233 | 1195.468479 | 584.8181995 | 665.9710532 | 659.1256528 | 2.46E-17 | -0.843771282 | 2.59E-18 | 76.17850638 | 0.557185152 | 16.60870026 |
| ENSMUSG00000057835 | *Zfp119a* | 125.0399374 | 159.4533147 | 150.7798082 | 76.7355888 | 66.69238015 | 98.1471775 | 0.000972186 | -0.845694484 | 0.000320832 | 12.94468079 | 0.556442884 | 3.01225068 |
| ENSMUSG00000083240 | *Gm13453* | 1061.56355 | 1107.375779 | 1099.517692 | 556.914349 | 692.6480053 | 563.865157 | 5.72E-15 | -0.847955038 | 6.75E-16 | 65.20440121 | 0.555571679 | 14.24277892 |
| ENSMUSG00000027322 | *Siglec1* | 1523.445768 | 1514.256651 | 1423.596371 | 834.7901933 | 836.5129968 | 807.3086463 | 8.61E-27 | -0.848259455 | 6.01E-28 | 120.10175 | 0.555454462 | 26.06488415 |
| ENSMUSG00000038642 | *Ctss* | 11938.33688 | 11678.58071 | 11753.97141 | 6522.525048 | 6533.947758 | 6586.445392 | 1.54E-80 | -0.848598046 | 3.35E-82 | 368.8402887 | 0.555324116 | 79.81140806 |
| ENSMUSG00000033871 | *Ppargc1b* | 375.1198122 | 447.5689593 | 437.6530796 | 231.3694268 | 221.9903511 | 245.3679437 | 3.66E-09 | -0.848899443 | 6.39E-10 | 38.19912808 | 0.555208114 | 8.436737592 |
| ENSMUSG00000026637 | *Traf5* | 266.2414993 | 263.9227278 | 259.4587608 | 169.7484237 | 130.5265154 | 139.5229484 | 1.57E-06 | -0.849318575 | 3.51E-07 | 25.94435178 | 0.555046838 | 5.803967706 |
| ENSMUSG00000025525 | *Apool* | 677.0870079 | 632.3148687 | 732.3590683 | 358.0994144 | 364.9025943 | 408.9465729 | 6.83E-13 | -0.850395347 | 9.16E-14 | 55.53915811 | 0.554632727 | 12.16559795 |
| ENSMUSG00000027078 | *Ube2l6* | 3695.908353 | 3604.744591 | 3779.286101 | 2027.679801 | 2125.58143 | 1990.84814 | 4.70E-44 | -0.85062679 | 1.96E-45 | 200.1224534 | 0.554543758 | 43.32792867 |
| ENSMUSG00000028233 | *Tgs1* | 2323.871489 | 2223.548982 | 2149.101811 | 1318.456935 | 1048.023117 | 1348.08035 | 5.84E-17 | -0.851187499 | 6.25E-18 | 74.43985085 | 0.554328274 | 16.23356685 |
| ENSMUSG00000047407 | *Tgif1* | 2748.326787 | 2743.696691 | 2856.983638 | 1535.874436 | 1543.452226 | 1547.261386 | 1.34E-42 | -0.851569535 | 5.81E-44 | 193.3805149 | 0.554181503 | 41.87266735 |
| ENSMUSG00000019528 | *Gyg* | 1399.256442 | 1479.066954 | 1376.600067 | 781.3078132 | 762.1986303 | 813.0820097 | 1.86E-25 | -0.85179448 | 1.35E-26 | 113.9369621 | 0.554095102 | 24.73089439 |
| ENSMUSG00000031553 | *Adam3* | 170.9729756 | 181.4468754 | 169.3825118 | 103.4767788 | 82.88910105 | 102.9583136 | 5.61E-05 | -0.852589083 | 1.52E-05 | 18.71445234 | 0.553790004 | 4.250787755 |
| ENSMUSG00000032905 | *Atg12* | 1506.433531 | 1479.066954 | 1486.258109 | 799.9103802 | 782.2063444 | 891.0224153 | 4.39E-24 | -0.854057208 | 3.37E-25 | 107.5514838 | 0.553226739 | 23.35796355 |
| ENSMUSG00000056116 | *H2-T22* | 1635.726528 | 1708.899663 | 1652.703352 | 824.3262494 | 1088.991293 | 844.8355083 | 1.76E-14 | -0.855900551 | 2.14E-15 | 62.93514104 | 0.552520328 | 13.75547465 |
| ENSMUSG00000044827 | *Tlr1* | 812.3342871 | 724.6878235 | 742.149965 | 418.5577571 | 404.9180224 | 436.8511626 | 1.36E-15 | -0.855944892 | 1.55E-16 | 68.11008906 | 0.552503347 | 14.86753214 |
| ENSMUSG00000024079 | *Eif2ak2* | 5344.394059 | 5279.554234 | 5217.568817 | 2770.61982 | 2755.348049 | 3219.612313 | 1.35E-31 | -0.856714516 | 8.08E-33 | 142.3679515 | 0.552208685 | 30.86901202 |
| ENSMUSG00000031770 | *Herpud1* | 4089.741626 | 4315.1366 | 4277.64274 | 2360.200686 | 2399.020189 | 2240.064992 | 3.30E-45 | -0.857407779 | 1.34E-46 | 205.4676267 | 0.551943394 | 44.48199139 |
| ENSMUSG00000028744 | *Slc66a1* | 256.0341575 | 267.2217619 | 288.8314507 | 148.8205359 | 154.3452226 | 144.3340846 | 2.11E-07 | -0.858990274 | 4.31E-08 | 30.00405064 | 0.551338298 | 6.674846957 |
| ENSMUSG00000022898 | *Vps26c* | 1991.282268 | 1956.32722 | 1944.472072 | 1055.695676 | 1085.1803 | 1106.561315 | 4.27E-35 | -0.859045912 | 2.25E-36 | 158.6325073 | 0.551317036 | 34.36934566 |
| ENSMUSG00000028680 | *Plk3* | 916.9595409 | 975.4144149 | 977.1314842 | 527.8478381 | 514.4840755 | 538.847249 | 2.53E-20 | -0.859157483 | 2.28E-21 | 90.08818927 | 0.551274402 | 19.59618193 |
| ENSMUSG00000034248 | *Slc25a37* | 3011.165839 | 3029.61298 | 2833.485486 | 1734.689371 | 1452.941139 | 1703.142198 | 6.19E-25 | -0.860377343 | 4.59E-26 | 111.5051614 | 0.550808473 | 24.20799939 |
| ENSMUSG00000058325 | *Dock1* | 1034.343972 | 1008.404756 | 960.4869599 | 585.9808599 | 536.3972861 | 533.0738856 | 1.02E-19 | -0.86124768 | 9.46E-21 | 87.27097843 | 0.550476286 | 18.99138626 |
| ENSMUSG00000038459 | *Abhd17c* | 1694.418743 | 1670.410931 | 1724.176897 | 1028.954486 | 872.7174317 | 899.6824604 | 1.82E-23 | -0.863358552 | 1.43E-24 | 104.6847776 | 0.549671448 | 22.73967207 |
| ENSMUSG00000052949 | *Rnf157* | 796.1726626 | 842.353373 | 744.1081443 | 447.624268 | 441.1224573 | 420.4932997 | 1.93E-16 | -0.863907689 | 2.12E-17 | 72.03026068 | 0.549462265 | 15.71490669 |
| ENSMUSG00000021532 | *Fastkd3* | 1192.55777 | 1151.3629 | 1116.162216 | 630.1619565 | 662.1600601 | 609.0898368 | 1.32E-22 | -0.863920782 | 1.07E-23 | 100.6967848 | 0.549457279 | 21.87884617 |
| ENSMUSG00000035868 | *Zfp983* | 122.4881019 | 137.4597541 | 103.7835043 | 59.29568225 | 71.45612159 | 68.31813336 | 0.000941819 | -0.864634121 | 0.000310127 | 13.00821916 | 0.549185667 | 3.026032775 |
| ENSMUSG00000034947 | *Tmem106a* | 2391.920435 | 2453.381691 | 2373.313344 | 1359.15005 | 1254.769495 | 1350.967031 | 4.69E-36 | -0.864882079 | 2.41E-37 | 163.0776936 | 0.549091286 | 35.32912304 |
| ENSMUSG00000053835 | *H2-T24* | 3334.39833 | 3467.284837 | 3511.994623 | 1880.021926 | 1764.489829 | 2015.866048 | 2.81E-34 | -0.865375072 | 1.52E-35 | 154.842426 | 0.548903684 | 33.55176733 |
| ENSMUSG00000028224 | *Nbn* | 1352.472792 | 1305.317825 | 1396.18186 | 730.150754 | 670.7347947 | 822.704282 | 8.97E-19 | -0.866493592 | 8.86E-20 | 82.84875102 | 0.548478285 | 18.04715517 |
| ENSMUSG00000016356 | *Col20a1* | 123.3387138 | 127.5626518 | 117.4907596 | 62.78366356 | 54.30665241 | 84.67599627 | 0.001391862 | -0.866512165 | 0.000471212 | 12.22627272 | 0.548471224 | 2.856403678 |
| ENSMUSG00000091512 | *Lamtor3* | 1655.2906 | 1497.76148 | 1570.45982 | 827.8142307 | 909.8746149 | 851.5710989 | 4.41E-25 | -0.867223259 | 3.25E-26 | 112.1850378 | 0.548200953 | 24.35531707 |
| ENSMUSG00000032812 | *Arap1* | 8325.788484 | 8754.536817 | 8708.023467 | 4811.088886 | 4765.646936 | 4543.636982 | 1.30E-59 | -0.868974673 | 3.90E-61 | 272.1281702 | 0.547535847 | 58.88481728 |
| ENSMUSG00000040165 | *Cd209c* | 294.3116894 | 296.9130688 | 281.9778231 | 166.2604424 | 118.1407877 | 193.4076733 | 1.85E-05 | -0.87105048 | 4.66E-06 | 20.97215045 | 0.546748598 | 4.732456759 |
| ENSMUSG00000023827 | *Agpat4* | 2277.938451 | 2202.655099 | 2305.756158 | 1246.371988 | 1289.068434 | 1175.841676 | 2.78E-34 | -0.871088908 | 1.50E-35 | 154.8622791 | 0.546734034 | 33.55555367 |
| ENSMUSG00000026024 | *Als2* | 1260.606716 | 1132.668374 | 1180.782134 | 608.0714082 | 639.2941012 | 704.3503326 | 1.79E-19 | -0.872331837 | 1.68E-20 | 86.13117956 | 0.546263208 | 18.74619911 |
| ENSMUSG00000029552 | *Tes* | 7411.380779 | 7377.739921 | 7711.31019 | 4047.220979 | 4155.888032 | 4075.994548 | 3.91E-66 | -0.873613035 | 1.03E-67 | 302.3255641 | 0.54577831 | 65.40736892 |
| ENSMUSG00000046962 | *Zbtb21* | 544.3915641 | 561.9354746 | 619.763757 | 310.4303365 | 302.9739555 | 328.1194856 | 5.31E-13 | -0.873777538 | 7.08E-14 | 56.04681242 | 0.545716081 | 12.27484676 |
| ENSMUSG00000069516 | *Lyz2* | 119556.043 | 119906.6933 | 121296.4811 | 59850.27128 | 74328.65768 | 62464.90511 | 1.21E-29 | -0.875417307 | 7.78E-31 | 133.2969838 | 0.545096172 | 28.91690232 |
| ENSMUSG00000069607 | *Cd300ld3* | 285.8055712 | 271.620474 | 266.3123885 | 161.6098007 | 146.7232363 | 141.4474029 | 9.96E-08 | -0.876283189 | 1.98E-08 | 31.51274565 | 0.544769112 | 7.001557465 |
| ENSMUSG00000080859 | *Rpl10-ps1* | 4431.687577 | 3610.242981 | 4740.752151 | 2392.755178 | 2127.486927 | 2437.321575 | 1.25E-17 | -0.878122755 | 1.30E-18 | 77.54605766 | 0.544074925 | 16.90430316 |
| ENSMUSG00000025464 | *Paox* | 112.2807601 | 118.7652275 | 129.2398356 | 60.45834269 | 67.64512844 | 67.35590613 | 0.000432739 | -0.878888749 | 0.00013323 | 14.59552726 | 0.543786127 | 3.363774003 |
| ENSMUSG00000021624 | *Cd180* | 4009.784115 | 3892.860235 | 3822.366046 | 2139.295203 | 1990.291173 | 2240.064992 | 1.61E-39 | -0.88061068 | 7.51E-41 | 179.1287528 | 0.543137477 | 38.79327872 |
| ENSMUSG00000034858 | *Atosa* | 128.4423847 | 131.9613639 | 127.2816563 | 60.45834269 | 74.31436646 | 74.09149674 | 0.000242063 | -0.887371251 | 7.16E-05 | 15.76934141 | 0.540598251 | 3.616071203 |
| ENSMUSG00000047648 | *Fbxo30* | 1162.786357 | 1102.977067 | 1146.513996 | 660.3911278 | 537.3500344 | 647.578926 | 1.26E-17 | -0.888866588 | 1.31E-18 | 77.52769193 | 0.540038217 | 16.90055129 |
| ENSMUSG00000040938 | *Slc16a11* | 67.1983337 | 64.88100392 | 75.38990409 | 38.3677944 | 34.29893836 | 39.45131644 | 0.007774113 | -0.88937681 | 0.00300104 | 8.806835803 | 0.539847262 | 2.109349132 |
| ENSMUSG00000078153 | *Psme2b* | 132.6954438 | 120.9645836 | 165.4661531 | 69.75962618 | 55.2594007 | 101.0338592 | 0.00272852 | -0.890000705 | 0.000968481 | 10.88687564 | 0.539613855 | 2.564072805 |
| ENSMUSG00000018986 | *Slfn3* | 605.6356151 | 670.8035999 | 584.5165291 | 319.73162 | 324.8871662 | 357.9485297 | 1.59E-13 | -0.89011971 | 2.03E-14 | 58.49813432 | 0.539569345 | 12.79996515 |
| ENSMUSG00000075256 | *Cerkl* | 64.64649824 | 61.58196982 | 64.61991779 | 36.04247353 | 34.29893836 | 32.71572583 | 0.009717484 | -0.891480264 | 0.003829252 | 8.363060201 | 0.539060736 | 2.012446149 |
| ENSMUSG00000069727 | *Zfp975* | 242.4243684 | 201.24108 | 219.3160846 | 134.8686106 | 95.27482879 | 127.9762216 | 1.93E-05 | -0.8942728 | 4.86E-06 | 20.89260058 | 0.538018319 | 4.715250185 |
| ENSMUSG00000024824 | *Rad9a* | 1110.048424 | 1073.28576 | 1115.183127 | 613.8847104 | 547.8302655 | 613.900973 | 3.33E-22 | -0.894876535 | 2.75E-23 | 98.82834917 | 0.537793217 | 21.47714023 |
| ENSMUSG00000036052 | *Dnajb5* | 182.0309293 | 133.0610419 | 181.1315878 | 96.50081622 | 100.9913185 | 70.24258782 | 0.000400298 | -0.894988214 | 0.000122721 | 14.7504138 | 0.537751588 | 3.397616963 |
| ENSMUSG00000056019 | *Zfp709* | 200.7443893 | 212.2378603 | 222.2533536 | 116.2660436 | 106.7078082 | 118.3539493 | 1.60E-06 | -0.896170868 | 3.59E-07 | 25.90498834 | 0.537310945 | 5.795910235 |
| ENSMUSG00000056429 | *Tgoln1* | 7667.414936 | 7540.492269 | 7292.259814 | 4060.010244 | 3739.53703 | 4256.893267 | 5.34E-45 | -0.900296087 | 2.18E-46 | 204.4997962 | 0.535776761 | 44.27227892 |
| ENSMUSG00000018395 | *Kif3a* | 970.5480855 | 936.9256838 | 973.2151256 | 508.0826107 | 477.3268922 | 556.1673392 | 1.07E-19 | -0.902154982 | 9.96E-21 | 87.16902401 | 0.535086864 | 18.9696408 |
| ENSMUSG00000019876 | *Pkib* | 176.9272583 | 217.7362504 | 188.9643051 | 105.8020997 | 90.51108735 | 115.4672676 | 1.83E-05 | -0.902730009 | 4.61E-06 | 20.99440733 | 0.534873633 | 4.737383675 |
| ENSMUSG00000029449 | *Rhof* | 3520.682319 | 3641.033966 | 3727.394349 | 1952.106873 | 1921.693297 | 1945.62346 | 2.90E-52 | -0.90382236 | 9.98E-54 | 238.1459443 | 0.534468801 | 51.53814613 |
| ENSMUSG00000032596 | *Uba7* | 2202.233999 | 2155.368944 | 2136.373646 | 1268.462536 | 1025.157158 | 1178.728357 | 2.48E-24 | -0.904806983 | 1.89E-25 | 108.7022813 | 0.534104157 | 23.60544803 |
| ENSMUSG00000037822 | *Smim14* | 745.9865652 | 821.4594903 | 744.1081443 | 391.816567 | 420.161995 | 420.4932997 | 1.52E-17 | -0.904923149 | 1.59E-18 | 77.14476583 | 0.534061152 | 16.81835029 |
| ENSMUSG00000006717 | *Acot13* | 702.6053625 | 727.9868576 | 703.9654681 | 365.075377 | 382.0520634 | 390.6642555 | 3.57E-18 | -0.906097207 | 3.60E-19 | 80.07902159 | 0.533626713 | 17.44776009 |
| ENSMUSG00000035725 | *Prkx* | 1502.180472 | 1580.237333 | 1664.452428 | 859.2060625 | 777.4426029 | 888.1357336 | 6.76E-25 | -0.910921115 | 5.01E-26 | 111.329089 | 0.531845417 | 24.17023275 |
| ENSMUSG00000069893 | *9930111J21Rik1* | 392.1320486 | 431.0737888 | 420.0294657 | 222.0681433 | 186.7386644 | 252.1035344 | 4.29E-09 | -0.911029979 | 7.54E-10 | 37.87530256 | 0.531805286 | 8.367718356 |
| ENSMUSG00000027709 | *Mccc1* | 734.9286116 | 693.8968386 | 748.0245029 | 402.280511 | 385.8630566 | 370.4574837 | 7.50E-18 | -0.911740214 | 7.70E-19 | 78.5747037 | 0.531543544 | 17.1246957 |
| ENSMUSG00000032691 | *Nlrp3* | 9153.433784 | 9111.932178 | 8896.987773 | 4827.366132 | 4927.614145 | 4681.235476 | 1.10E-74 | -0.911959049 | 2.58E-76 | 341.8086101 | 0.531462922 | 73.95818509 |
| ENSMUSG00000042997 | *Nhlrc3* | 1665.497942 | 1669.311253 | 1613.539766 | 868.5073459 | 806.9777998 | 953.5671853 | 6.38E-26 | -0.912294017 | 4.56E-27 | 116.0810294 | 0.531339541 | 25.19506873 |
| ENSMUSG00000033792 | *Atp7a* | 1299.734859 | 1256.931991 | 1177.844865 | 726.6627727 | 503.051096 | 754.3861486 | 1.50E-10 | -0.913736889 | 2.34E-11 | 44.66272491 | 0.530808402 | 9.824817261 |
| ENSMUSG00000094685 | *Gm5900* | 501.8609732 | 446.4692812 | 549.2693012 | 232.5320873 | 289.6354795 | 271.348079 | 3.46E-10 | -0.914294959 | 5.58E-11 | 42.96402204 | 0.530603112 | 9.460828944 |
| ENSMUSG00000028211 | *Trp53inp1* | 217.7566257 | 238.6301331 | 192.8806637 | 105.8020997 | 107.6605565 | 129.9006761 | 4.68E-06 | -0.915841138 | 1.10E-06 | 23.74042794 | 0.530034753 | 5.329746808 |
| ENSMUSG00000068245 | *Phf11d* | 4236.046859 | 4217.265255 | 4164.068339 | 1954.432193 | 2418.075155 | 2309.345353 | 1.23E-28 | -0.91596456 | 8.11E-30 | 128.645202 | 0.529989411 | 27.90835327 |
| ENSMUSG00000004567 | *Mcoln1* | 1200.213277 | 1307.517181 | 1265.962935 | 677.8310344 | 630.7193666 | 689.9169242 | 3.46E-25 | -0.916761405 | 2.54E-26 | 112.6793927 | 0.529696762 | 24.46076617 |
| ENSMUSG00000064289 | *Tank* | 3074.961726 | 2858.063207 | 2793.34281 | 1588.194156 | 1366.241045 | 1656.955291 | 5.77E-26 | -0.920939015 | 4.12E-27 | 116.2819311 | 0.528165139 | 25.23864629 |
| ENSMUSG00000032261 | *Sh3bgrl2* | 3428.816242 | 3506.873246 | 3373.942981 | 1749.803957 | 1825.46572 | 1864.796372 | 4.76E-51 | -0.921469461 | 1.67E-52 | 232.5297814 | 0.52797098 | 50.32203891 |
| ENSMUSG00000034613 | *Ppm1h* | 1412.866231 | 1472.468886 | 1500.944454 | 851.0674394 | 691.695257 | 775.5551477 | 6.53E-22 | -0.921628614 | 5.44E-23 | 97.47848509 | 0.527912739 | 21.18539964 |
| ENSMUSG00000096433 | *Zfp994* | 844.6575362 | 880.8421041 | 871.3898005 | 449.9495889 | 346.8003768 | 573.4874293 | 1.18E-08 | -0.922142167 | 2.14E-09 | 35.84353499 | 0.527724853 | 7.929298303 |
| ENSMUSG00000032251 | *Irak1bp1* | 365.7630822 | 340.9001901 | 327.0159476 | 160.4471402 | 192.4551542 | 191.4832188 | 2.97E-09 | -0.922430596 | 5.14E-10 | 38.62350874 | 0.527619359 | 8.527580194 |
| ENSMUSG00000095742 | *ENSMUSG00000095742* | 741.7335062 | 769.7746228 | 910.5533871 | 409.2564736 | 436.3587159 | 431.0777992 | 5.70E-15 | -0.922590832 | 6.73E-16 | 65.21018525 | 0.527560761 | 14.24380119 |
| ENSMUSG00000035778 | *Ggta1* | 3006.062168 | 2930.641957 | 3007.763446 | 1497.506642 | 1542.499478 | 1675.237608 | 5.07E-41 | -0.923008752 | 2.30E-42 | 186.0636148 | 0.527407959 | 40.29501751 |
| ENSMUSG00000026632 | *Tatdn3* | 116.5338192 | 106.6687692 | 146.8634495 | 48.83173833 | 78.12535961 | 67.35590613 | 0.001348541 | -0.923240037 | 0.000454048 | 12.29551609 | 0.527323415 | 2.870135717 |
| ENSMUSG00000016495 | *Plgrkt* | 837.0020299 | 807.1636759 | 810.6862414 | 440.6483054 | 410.6345121 | 443.5867532 | 1.56E-20 | -0.923964846 | 1.39E-21 | 91.06113719 | 0.527058554 | 19.80574201 |
| ENSMUSG00000005413 | *Hmox1* | 7692.082679 | 7632.865224 | 7543.885858 | 4010.015845 | 4066.329693 | 3974.960689 | 1.51E-81 | -0.924201307 | 3.22E-83 | 373.5085668 | 0.526972175 | 80.82229586 |
| ENSMUSG00000057531 | *Dtnbp1* | 1430.72908 | 1504.359549 | 1536.191682 | 809.2116637 | 819.3635276 | 727.4437862 | 7.32E-27 | -0.924536852 | 5.09E-28 | 120.4310312 | 0.526849625 | 26.13568869 |
| ENSMUSG00000058298 | *Mcm9* | 731.5261643 | 719.1894333 | 763.6899376 | 405.7684923 | 359.1861045 | 402.2109823 | 1.06E-17 | -0.925403943 | 1.09E-18 | 77.88022325 | 0.526533072 | 16.97605516 |
| ENSMUSG00000052435 | *Cebpe* | 170.9729756 | 182.5465534 | 202.6715604 | 97.66347665 | 84.79459762 | 109.6939043 | 1.30E-05 | -0.92830831 | 3.21E-06 | 21.68465076 | 0.525474147 | 4.88562164 |
| ENSMUSG00000038774 | *Ascc3* | 4585.648316 | 4380.017604 | 4307.99452 | 2339.272798 | 2035.070343 | 2597.051295 | 1.04E-25 | -0.929265766 | 7.48E-27 | 115.1016866 | 0.525125527 | 24.98267687 |
| ENSMUSG00000031129 | *Slc9a9* | 77.40567553 | 68.18003802 | 63.64082813 | 37.20513396 | 27.62970035 | 45.22467983 | 0.009527621 | -0.92938192 | 0.003750512 | 8.400822786 | 0.52508325 | 2.021015506 |
| ENSMUSG00000041028 | *Ghitm* | 17856.89392 | 18019.32424 | 18321.70487 | 9026.895628 | 9812.354617 | 9604.952213 | 1.72E-73 | -0.929814449 | 4.10E-75 | 336.289853 | 0.52492585 | 72.76511299 |
| ENSMUSG00000073647 | *Gm10557* | 890.5905745 | 848.9514411 | 809.7071517 | 445.2989471 | 419.2092467 | 473.4157973 | 3.69E-19 | -0.930652335 | 3.55E-20 | 84.65867362 | 0.524621073 | 18.43344893 |
| ENSMUSG00000024104 | *Washc2* | 5304.415303 | 5133.297056 | 4974.75458 | 2714.812119 | 2585.758853 | 2778.912241 | 1.06E-54 | -0.932069089 | 3.51E-56 | 249.4021875 | 0.524106138 | 53.97559956 |
| ENSMUSG00000026946 | *Nmi* | 1305.689142 | 1289.922332 | 1278.691101 | 590.6315017 | 734.56893 | 700.5014237 | 7.12E-21 | -0.933048552 | 6.24E-22 | 92.65153804 | 0.523750437 | 20.1473512 |
| ENSMUSG00000055945 | *Prr18* | 102.9240301 | 101.170379 | 94.97169736 | 52.31971964 | 54.30665241 | 50.03581598 | 0.00060368 | -0.933108607 | 0.000191078 | 13.916844 | 0.523728635 | 3.219193064 |
| ENSMUSG00000038855 | *Itpkb* | 2400.426553 | 2348.912278 | 2396.811496 | 1313.806293 | 1235.714529 | 1192.199538 | 8.53E-41 | -0.934776369 | 3.89E-42 | 185.0171869 | 0.523123553 | 40.06919693 |
| ENSMUSG00000078151 | *Gm12226* | 199.0431656 | 232.0320649 | 288.8314507 | 117.4287041 | 151.4869778 | 106.8072226 | 3.69E-05 | -0.935296496 | 9.72E-06 | 19.56609682 | 0.522934988 | 4.433295482 |
| ENSMUSG00000051498 | *Tlr6* | 704.3065861 | 660.9064976 | 657.9482539 | 362.7500561 | 373.4773289 | 321.3838949 | 3.29E-16 | -0.937239163 | 3.65E-17 | 70.95536054 | 0.522231301 | 15.48290007 |
| ENSMUSG00000045005 | *Fzd5* | 405.7418377 | 420.0770084 | 389.6776861 | 244.1586916 | 186.7386644 | 204.9544001 | 5.15E-10 | -0.938780909 | 8.39E-11 | 42.16552376 | 0.521673513 | 9.287952272 |
| ENSMUSG00000028893 | *Sesn2* | 1213.823066 | 1300.919113 | 1326.666494 | 712.7108475 | 652.6325772 | 638.918881 | 2.39E-25 | -0.939551032 | 1.74E-26 | 113.4306062 | 0.521395114 | 24.62245264 |
| ENSMUSG00000029787 | *Avl9* | 2266.029886 | 2189.458963 | 2115.812763 | 1176.612362 | 1031.826396 | 1219.141901 | 6.81E-29 | -0.939771316 | 4.46E-30 | 129.8301608 | 0.521315509 | 28.16671655 |
| ENSMUSG00000099875 | *Rbm3-ps* | 1552.36657 | 1675.909322 | 1497.028096 | 741.7773584 | 974.6614985 | 743.8016491 | 1.09E-15 | -0.9399988 | 1.24E-16 | 68.54028876 | 0.521233314 | 14.96070706 |
| ENSMUSG00000038172 | *Ttc39b* | 1242.743868 | 1245.935211 | 1262.046576 | 661.5537883 | 617.3808906 | 676.4457429 | 1.63E-28 | -0.940165465 | 1.08E-29 | 128.0833635 | 0.521173103 | 27.78767855 |
| ENSMUSG00000079293 | *Clec7a* | 95.26852373 | 80.27649638 | 87.13898005 | 44.18109658 | 55.2594007 | 37.52686198 | 0.002550627 | -0.940347455 | 0.00090174 | 11.01917955 | 0.521107363 | 2.593353097 |
| ENSMUSG00000078920 | *Ifi47* | 559.7025769 | 372.790853 | 424.924914 | 254.6226356 | 238.187072 | 216.5011268 | 1.66E-07 | -0.940581968 | 3.35E-08 | 30.49408279 | 0.521022663 | 6.780609036 |
| ENSMUSG00000044641 | *Pard6b* | 438.0650868 | 489.3567245 | 450.3812452 | 244.1586916 | 233.4233305 | 239.5945804 | 5.14E-13 | -0.941276813 | 6.84E-14 | 56.1149815 | 0.520771783 | 12.28945727 |

| | | | | | | | | | | | | | |
|---|---|---|---|---|---|---|---|---|---|---|---|---|---|
| ENSMUSG00000014905 | *Dnajb9* | 814.8861226 | 744.4820281 | 760.7526686 | 363.9127166 | 429.6894778 | 412.7954818 | 2.98E-17 | -0.941860125 | 3.14E-18 | 75.79699155 | 0.520561267 | 16.52648745 |
| ENSMUSG00000034744 | *Nagk* | 909.3040345 | 825.8582025 | 898.8043111 | 466.226835 | 475.4213957 | 430.115572 | 3.23E-20 | -0.942653889 | 2.93E-21 | 89.59309539 | 0.520274935 | 19.49047577 |
| ENSMUSG00000060802 | *B2m* | 30556.52837 | 30916.34821 | 30744.39452 | 16194.69722 | 15561.23779 | 16160.60633 | 1.80E-106 | -0.94456484 | 3.03E-108 | 488.4929606 | 0.519586251 | 105.7456286 |
| ENSMUSG00000016239 | *Lonrf3* | 2027.007965 | 2161.967012 | 2122.666391 | 1147.545851 | 1018.48792 | 1113.296906 | 4.28E-34 | -0.945197795 | 2.33E-35 | 153.9890725 | 0.519358342 | 33.36857746 |
| ENSMUSG00000024610 | *Cd74* | 7180.014364 | 7129.212685 | 7202.183565 | 3775.158437 | 3797.654676 | 3593.918705 | 7.55E-75 | -0.946102924 | 1.76E-76 | 342.5727307 | 0.519032605 | 74.12203918 |
| ENSMUSG00000082419 | *Gm11425* | 247.5280393 | 211.1381823 | 260.4378505 | 123.2420063 | 110.5188014 | 139.5229484 | 8.91E-07 | -0.947500956 | 1.94E-07 | 27.09432738 | 0.518529885 | 6.049929522 |
| ENSMUSG00000025492 | *Ifitm3* | 26500.81122 | 25192.52405 | 26234.70753 | 13641.4949 | 14251.20889 | 12507.02954 | 3.36E-55 | -0.947871334 | 1.11E-56 | 251.6972193 | 0.518396781 | 54.473227 |
| ENSMUSG00000022969 | *Il10rb* | 3006.91278 | 2897.651616 | 3004.826177 | 1417.283072 | 1633.010565 | 1561.694795 | 2.13E-38 | -0.94884629 | 1.02E-39 | 173.9372762 | 0.518046573 | 37.67161624 |
| ENSMUSG00000041842 | *Fhdc1* | 71.45139279 | 94.5723108 | 90.07624905 | 44.18109658 | 37.15718323 | 50.99804321 | 0.002967941 | -0.949267962 | 0.001060183 | 10.71939598 | 0.51789518 | 2.527544664 |
| ENSMUSG00000020220 | *Vps13d* | 2899.735691 | 3070.301067 | 3003.847088 | 1670.743047 | 1338.611344 | 1634.824064 | 1.33E-25 | -0.951016159 | 9.63E-27 | 114.5987927 | 0.517267997 | 24.87502159 |
| ENSMUSG00000110519 | *Or7g23* | 115.6832074 | 108.8681252 | 117.4907596 | 55.80770094 | 45.73191782 | 75.05372397 | 0.000832457 | -0.9532127 | 0.000270935 | 13.26134675 | 0.516481042 | 3.079638056 |
| ENSMUSG00000114367 | *Gm48216* | 113.9819837 | 102.270057 | 71.47354544 | 51.1570592 | 58.11764556 | 39.45131644 | 0.003855594 | -0.954114137 | 0.001404247 | 10.20020969 | 0.516158431 | 2.41390869 |
| ENSMUSG00000026627 | *Pacc1* | 1333.759332 | 1321.812995 | 1397.16095 | 705.7348849 | 682.1677741 | 702.4258782 | 6.85E-32 | -0.95562151 | 4.04E-33 | 143.7472811 | 0.515619414 | 31.16457004 |
| ENSMUSG00000018899 | *Irf1* | 1503.031084 | 1361.401404 | 1481.362661 | 706.8975453 | 795.5448204 | 736.1038312 | 5.02E-27 | -0.956615698 | 3.48E-28 | 121.1833194 | 0.515264213 | 26.29951927 |
| ENSMUSG00000068129 | *Cst7* | 227.1133557 | 202.340758 | 204.6297397 | 111.6154019 | 115.2825428 | 100.071632 | 5.21E-07 | -0.957821276 | 1.11E-07 | 28.17635957 | 0.514833817 | 6.2830225 |
| ENSMUSG00000028164 | *Manba* | 3438.172972 | 3262.744723 | 3274.075835 | 1730.038729 | 1712.088673 | 1695.44438 | 3.03E-55 | -0.957828185 | 9.97E-57 | 251.9062881 | 0.514831351 | 54.51789888 |
| ENSMUSG00000073643 | *Wdfy1* | 1447.741316 | 1400.989813 | 1434.366357 | 713.8735079 | 729.8051885 | 760.159512 | 1.76E-33 | -0.958217301 | 9.80E-35 | 151.1326446 | 0.514692512 | 32.75404552 |
| ENSMUSG00000073725 | *Lmbrd1* | 1446.890704 | 1348.205268 | 1464.718137 | 731.3134145 | 735.5216782 | 724.5571045 | 9.41E-32 | -0.959635733 | 5.61E-33 | 143.0932925 | 0.514186724 | 31.02656982 |
| ENSMUSG00000026121 | *Sema4c* | 704.3065861 | 721.3887894 | 658.9273436 | 366.2380374 | 349.6586217 | 356.0240752 | 6.62E-19 | -0.960121396 | 6.50E-20 | 83.46092922 | 0.51401366 | 18.17922912 |
| ENSMUSG00000041328 | *Pcf11* | 2725.360268 | 2711.806028 | 2651.374809 | 1424.259035 | 1211.895822 | 1519.356797 | 4.59E-27 | -0.961299302 | 3.18E-28 | 121.3621293 | 0.513594158 | 26.3378033 |
| ENSMUSG00000040560 | *Wdr7* | 1247.847538 | 1206.346802 | 1061.333195 | 692.9456201 | 564.0269864 | 551.356203 | 2.45E-16 | -0.961819595 | 2.70E-17 | 71.5503184 | 0.513408969 | 15.61143808 |
| ENSMUSG00000005615 | *Pcyt1a* | 3564.914133 | 3415.599969 | 3476.747395 | 1773.057165 | 1766.395326 | 1826.307283 | 1.92E-58 | -0.962803691 | 5.92E-60 | 266.7069588 | 0.51305888 | 57.7160383 |
| ENSMUSG00000030966 | *Trim21* | 870.1758908 | 983.1121611 | 932.0933597 | 433.6723428 | 451.6026885 | 540.7717035 | 1.22E-16 | -0.96317275 | 1.33E-17 | 72.95180299 | 0.51292765 | 15.91307295 |
| ENSMUSG00000017830 | *Dhx58* | 2476.131005 | 2407.195213 | 2455.556876 | 1273.113178 | 1282.399195 | 1206.632947 | 3.95E-47 | -0.964562213 | 1.53E-48 | 214.3749918 | 0.512433886 | 46.4035174 |
| ENSMUSG00000031530 | *Dusp4* | 10203.08877 | 10375.46224 | 10506.61118 | 5620.300549 | 5245.832073 | 5067.088595 | 6.13E-65 | -0.964608985 | 1.67E-66 | 296.7648099 | 0.512417274 | 64.21258886 |
| ENSMUSG00000033880 | *Lgals3bp* | 14320.90058 | 14404.68255 | 13925.59228 | 7500.322475 | 7319.012348 | 7038.69219 | 9.65E-85 | -0.964630346 | 2.01E-86 | 388.2271636 | 0.512409687 | 84.01546669 |
| ENSMUSG00000068739 | *Sars* | 19604.05059 | 19583.0664 | 20365.065 | 9637.292357 | 11545.40375 | 9327.830771 | 9.39E-34 | -0.964719473 | 5.15E-35 | 152.4108647 | 0.512378032 | 33.02744366 |
| ENSMUSG00000079192 | *ENSMUSG00000079192* | 903.3497518 | 906.1346988 | 875.3061592 | 437.1603241 | 451.6026885 | 481.1136152 | 1.64E-23 | -0.969506828 | 1.29E-24 | 104.8934344 | 0.510680604 | 22.78502669 |
| ENSMUSG00000031812 | *Map1lc3b* | 2454.015098 | 2397.298111 | 2516.260435 | 1235.908044 | 1310.028896 | 1213.368537 | 9.08E-45 | -0.97055276 | 3.71E-46 | 203.439912 | 0.510310503 | 44.04174095 |
| ENSMUSG00000063234 | *Gpr84* | 5282.299396 | 5338.936848 | 5502.483909 | 2732.252025 | 2946.850454 | 2540.279888 | 1.54E-45 | -0.971975176 | 6.23E-47 | 206.9904255 | 0.509807613 | 44.8113068 |
| ENSMUSG00000052459 | *Atp6v1a* | 15641.05013 | 15572.54062 | 15303.17144 | 8083.978014 | 7408.570687 | 8220.307229 | 1.60E-67 | -0.972231819 | 4.15E-69 | 308.7188899 | 0.509716931 | 66.79675271 |
| ENSMUSG00000050335 | *Lgals3* | 29750.99898 | 30676.6184 | 30726.77091 | 14658.82278 | 17336.20785 | 14450.72855 | 3.29E-40 | -0.972649451 | 1.53E-41 | 182.2948646 | 0.509569399 | 39.4826345 |
| ENSMUSG00000066442 | *Mthfs* | 196.4913302 | 196.8423678 | 178.1943188 | 99.98879752 | 101.9440668 | 89.48713242 | 1.08E-06 | -0.972669704 | 2.36E-07 | 26.71206462 | 0.509562246 | 5.968261352 |
| ENSMUSG00000007613 | *Tgfbr1* | 1557.470241 | 1462.571783 | 1539.128951 | 767.355888 | 647.8688358 | 905.4558238 | 1.51E-16 | -0.974493758 | 1.65E-17 | 72.52071905 | 0.508918394 | 15.82012427 |
| ENSMUSG00000031827 | *Cotl1* | 4218.18401 | 4243.657528 | 4386.321693 | 2264.86253 | 2321.847578 | 1952.35905 | 7.26E-37 | -0.974758426 | 3.63E-38 | 166.8366193 | 0.50882504 | 36.13903117 |
| ENSMUSG00000018476 | *Kdm6b* | 2294.950688 | 2345.613243 | 2265.613481 | 1292.878405 | 1023.251661 | 1200.859583 | 3.70E-26 | -0.974927224 | 2.63E-27 | 117.1716811 | 0.50876551 | 25.43180906 |
| ENSMUSG00000022906 | *Parp9* | 4815.313507 | 4828.686241 | 4755.438496 | 2325.320873 | 2466.665317 | 2525.84648 | 1.21E-62 | -0.975767707 | 3.46E-64 | 286.134306 | 0.5084692 | 61.91559408 |
| ENSMUSG00000021752 | *Kctd6* | 137.7991147 | 136.360076 | 151.7588978 | 80.22357011 | 73.36161817 | 63.5069972 | 3.31E-05 | -0.976013948 | 8.64E-06 | 19.79024098 | 0.508382422 | 4.480603836 |
| ENSMUSG00000000682 | *Cd52* | 4218.18401 | 4320.63499 | 4345.199927 | 1943.96825 | 2514.302732 | 2086.108635 | 2.48E-26 | -0.976323146 | 1.75E-27 | 117.9776083 | 0.508273477 | 25.60617903 |
| ENSMUSG00000053113 | *Socs3* | 1514.93965 | 1622.025098 | 1572.418 | 810.3743241 | 809.8360447 | 769.7817843 | 2.99E-35 | -0.978294497 | 1.57E-36 | 159.3489153 | 0.507579428 | 34.52417842 |
| ENSMUSG00000020806 | *Rhbdf2* | 6185.649148 | 6260.467039 | 6303.379254 | 3339.160773 | 3054.511011 | 3126.276271 | 1.37E-65 | -0.978463279 | 3.66E-67 | 299.7913969 | 0.507520049 | 64.86203394 |
| ENSMUSG00000079017 | *Ifi27l2a* | 5245.723088 | 5453.303364 | 5761.94267 | 2606.684698 | 3276.501362 | 2464.263937 | 2.20E-22 | -0.979219985 | 1.80E-23 | 99.66986308 | 0.50725392 | 21.65841987 |
| ENSMUSG00000030717 | *Nupr1* | 1038.597031 | 965.5173126 | 950.6960633 | 525.5225172 | 469.7049059 | 505.1692959 | 2.48E-23 | -0.97990398 | 1.97E-24 | 104.0556897 | 0.507013484 | 22.60555569 |
| ENSMUSG00000058715 | *Fcer1g* | 9248.702308 | 8701.752272 | 9192.672851 | 4576.231477 | 4754.213957 | 4428.169714 | 3.21E-69 | -0.980403207 | 8.12E-71 | 316.5633491 | 0.506838068 | 68.49395694 |
| ENSMUSG00000002233 | *Rhoc* | 5080.704395 | 5158.589651 | 5095.182609 | 2464.840125 | 2776.308511 | 2522.959798 | 8.43E-53 | -0.981183038 | 2.87E-54 | 240.6292519 | 0.506564177 | 52.07440165 |
| ENSMUSG00000027901 | *Dennd2d* | 1705.476697 | 1535.150533 | 1510.735351 | 740.6146979 | 800.3085618 | 864.0800529 | 9.93E-26 | -0.981609091 | 7.13E-27 | 115.1971344 | 0.506414602 | 25.0031651 |
| ENSMUSG00000025993 | *Slc40a1* | 1689.315072 | 1680.308034 | 1595.916152 | 881.2966107 | 817.458031 | 817.8931458 | 1.50E-35 | -0.981782002 | 7.81E-37 | 160.7380892 | 0.50635391 | 34.82370344 |
| ENSMUSG00000035235 | *Trim13* | 314.726373 | 304.610815 | 296.6641681 | 145.3325545 | 169.5891952 | 148.1829935 | 7.41E-10 | -0.982379225 | 1.22E-10 | 41.42615265 | 0.506144342 | 9.130084862 |
| ENSMUSG00000044934 | *Zfp367* | 1065.816609 | 1034.797029 | 1056.437747 | 562.7276512 | 457.3191782 | 578.2985654 | 3.74E-19 | -0.9835167 | 3.60E-20 | 84.62780856 | 0.505745436 | 18.42697891 |
| ENSMUSG00000048647 | *Exd1* | 59.54282733 | 56.08357966 | 53.84993149 | 31.39183178 | 27.62970035 | 26.94236245 | 0.008186192 | -0.983803741 | 0.003173596 | 8.704878459 | 0.505644822 | 2.086918077 |
| ENSMUSG00000027956 | *Tmem144* | 975.6517564 | 889.6395284 | 892.9297731 | 479.0160998 | 477.3268922 | 439.7378443 | 1.42E-22 | -0.984092648 | 1.16E-23 | 100.5487536 | 0.505543574 | 21.8478554 |
| ENSMUSG00000037110 | *Ralgapa2* | 780.8616498 | 864.3469336 | 752.9199513 | 381.3526231 | 389.6740497 | 438.775617 | 4.33E-18 | -0.984879591 | 4.40E-19 | 79.68252741 | 0.505267892 | 17.36326092 |
| ENSMUSG00000034445 | *Cyb561a3* | 242.4243684 | 243.0288452 | 242.8142366 | 109.290081 | 118.1407877 | 139.5229484 | 6.20E-08 | -0.98545705 | 1.21E-08 | 32.46599447 | 0.505065692 | 7.207265337 |
| ENSMUSG00000066861 | *Oas1g* | 5557.897625 | 5387.322682 | 5350.725011 | 2650.865795 | 2955.425189 | 2622.069203 | 1.15E-50 | -0.985537924 | 4.13E-52 | 230.7290061 | 0.50503738 | 49.93773215 |
| ENSMUSG00000049323 | *Smcr8* | 9294.635346 | 9402.247179 | 8901.883221 | 4706.449446 | 4291.178289 | 4938.150146 | 2.99E-51 | -0.985901959 | 1.04E-52 | 233.4754149 | 0.50490996 | 50.52484825 |
| ENSMUSG00000051650 | *B3gnt2* | 2865.711218 | 2804.178983 | 2737.534699 | 1275.438499 | 1513.917029 | 1450.076436 | 1.72E-35 | -0.986404696 | 8.97E-37 | 160.4626462 | 0.504734044 | 34.76409642 |
| ENSMUSG00000046675 | *Tmem251* | 1411.165008 | 1422.983374 | 1525.421696 | 776.6571715 | 668.8292981 | 755.3483758 | 2.39E-27 | -0.987403623 | 1.65E-28 | 122.6680734 | 0.504384685 | 26.62108534 |
| ENSMUSG00000041515 | *Irf8* | 3720.576096 | 3934.648001 | 3909.505026 | 1919.55238 | 2118.912192 | 1783.007058 | 5.68E-38 | -0.989873619 | 2.75E-39 | 171.9695869 | 0.503521882 | 37.24560715 |
| ENSMUSG00000024589 | *Nedd4l* | 146.3052329 | 159.4533147 | 156.6543462 | 73.24760749 | 80.98360447 | 77.94040566 | 6.21E-06 | -0.990765029 | 1.48E-06 | 23.17895757 | 0.503210863 | 5.207152709 |
| ENSMUSG00000025498 | *Irf7* | 10233.71079 | 10595.39784 | 10775.86084 | 5428.461577 | 5648.844599 | 4825.56956 | 1.06E-47 | -0.990916673 | 4.05E-49 | 217.0141756 | 0.503157972 | 46.97458858 |
| ENSMUSG00000090215 | *Trim34b* | 240.7231448 | 230.9323868 | 194.838843 | 93.01283491 | 109.5660531 | 131.8251306 | 2.39E-06 | -0.990984169 | 5.46E-07 | 25.0926504 | 0.503134433 | 5.620957637 |
| ENSMUSG00000032462 | *Pik3cb* | 2452.313874 | 2610.635649 | 2583.817622 | 1317.294274 | 1247.147509 | 1279.762216 | 4.87E-48 | -0.992308202 | 1.85E-49 | 218.5724495 | 0.502672893 | 47.31215774 |
| ENSMUSG00000039886 | *Tmem120a* | 1732.696275 | 1624.224454 | 1796.629533 | 810.3743241 | 970.8505054 | 803.4597374 | 1.74E-24 | -0.994854161 | 1.31E-25 | 109.4206531 | 0.501786596 | 23.75851257 |
| ENSMUSG00000074785 | *Plxnc1* | 1628.921633 | 1548.34667 | 1512.69353 | 759.2172649 | 765.0568752 | 828.4776453 | 1.59E-33 | -0.994959505 | 8.83E-35 | 151.3403488 | 0.501749958 | 32.79836998 |
| ENSMUSG00000037580 | *Gch1* | 1791.388491 | 1843.060383 | 2067.837369 | 916.1764238 | 1072.794572 | 866.0045073 | 1.89E-22 | -0.997327463 | 1.54E-23 | 99.97506681 | 0.500927089 | 21.72388912 |
| ENSMUSG00000040751 | *Lat2* | 9798.197543 | 9451.73269 | 10013.14999 | 4478.568001 | 5346.823392 | 4824.607333 | 4.37E-44 | -0.997762176 | 1.81E-45 | 200.2794645 | 0.500776173 | 43.35932991 |
| ENSMUSG00000052776 | *Oas1a* | 8637.963022 | 8778.729734 | 9107.49205 | 4146.047116 | 4984.779042 | 4140.463772 | 1.24E-36 | -0.998626873 | 6.26E-38 | 165.7548826 | 0.500476116 | 35.90570754 |
| ENSMUSG00000057554 | *Lgals8* | 3268.050609 | 3074.699779 | 3217.288634 | 1623.073969 | 1539.641233 | 1617.503974 | 1.65E-54 | -1.000754547 | 5.50E-56 | 248.504004 | 0.499738562 | 53.78157882 |
| ENSMUSG00000016256 | *Ctsz* | 24668.59336 | 25321.18638 | 25568.92656 | 12699.73995 | 13090.76148 | 11949.89997 | 1.60E-81 | -1.001484858 | 3.44E-83 | 373.3790397 | 0.499485652 | 80.79548425 |
| ENSMUSG00000037997 | *Parp11* | 1118.554542 | 1087.581574 | 1128.890382 | 560.4023303 | 547.8302655 | 558.0917936 | 4.85E-31 | -1.00162925 | 2.96E-32 | 139.7920326 | 0.499435664 | 30.31463135 |
| ENSMUSG00000034707 | *Gns* | 6745.351725 | 6834.498972 | 6934.892087 | 3557.740935 | 3335.571756 | 3352.399671 | 1.92E-76 | -1.002145238 | 4.33E-78 | 349.9547874 | 0.49925707 | 75.71663184 |
| ENSMUSG00000030530 | *Furin* | 5235.515746 | 5347.734272 | 5326.24777 | 2791.547708 | 2579.089615 | 2570.108932 | 5.15E-65 | -1.003239339 | 1.40E-66 | 297.1184468 | 0.49887859 | 64.2885417 |
| ENSMUSG00000037434 | *Slc30a1* | 3113.239257 | 3256.146654 | 3197.706841 | 1704.4602 | 1472.948853 | 1598.25943 | 6.73E-43 | -1.003294932 | 2.89E-44 | 194.772329 | 0.498859367 | 42.17229483 |
| ENSMUSG00000062753 | *Smim29* | 77.40567553 | 83.57553048 | 86.15989039 | 41.85577571 | 40.96817638 | 40.41354368 | 0.000912076 | -1.003437885 | 0.000299394 | 13.07418001 | 0.498809938 | 3.039968941 |
| ENSMUSG00000040296 | *Rigi* | 5170.869248 | 4943.052756 | 4989.440925 | 2506.695901 | 2463.807072 | 2561.448887 | 8.14E-73 | -1.004028018 | 1.95E-74 | 333.1758958 | 0.498605942 | 72.0894064 |
| ENSMUSG00000090641 | *Zfp712* | 69.75016916 | 84.67520851 | 79.30626275 | 49.99439876 | 27.62970035 | 39.45131644 | 0.004174194 | -1.004252489 | 0.001530595 | 10.04141601 | 0.49852837 | 2.379427389 |
| ENSMUSG00000040415 | *Dtx3* | 612.4405097 | 542.1412701 | 622.701026 | 302.2917134 | 267.7222689 | 316.5727588 | 4.25E-15 | -1.005239024 | 4.98E-16 | 65.80609116 | 0.498187586 | 14.37207911 |
| ENSMUSG00000010609 | *Psen2* | 1265.710387 | 1450.475325 | 1337.43648 | 725.5001123 | 731.7106851 | 560.9784753 | 1.38E-17 | -1.00655409 | 1.44E-18 | 77.33546603 | 0.497733677 | 16.85942762 |
| ENSMUSG00000028459 | *Cd72* | 7795.857321 | 8049.643198 | 7856.21546 | 3993.738599 | 3951.999898 | 3821.004332 | 2.28E-88 | -1.010369621 | 4.63E-90 | 404.938 | 0.496419048 | 87.64170256 |
| ENSMUSG00000030560 | *Ctsc* | 2655.610099 | 2640.326956 | 2446.745069 | 1304.50501 | 1207.132081 | 1329.798032 | 5.55E-42 | -1.011637423 | 2.44E-43 | 190.5276507 | 0.495983 | 41.2554441 |
| ENSMUSG00000094568 | *Smarce1-ps1* | 187.1346002 | 222.1349626 | 221.274264 | 90.68751403 | 145.770488 | 75.05372397 | 0.000145237 | -1.013245618 | 4.15E-05 | 16.80032389 | 0.495430428 | 3.837922393 |
| ENSMUSG00000018143 | *Mafk* | 3019.671957 | 3173.670802 | 3215.330455 | 1571.91691 | 1545.357723 | 1528.016842 | 5.29E-58 | -1.018170333 | 1.64E-59 | 264.6756741 | 0.493742133 | 57.27619465 |
| ENSMUSG00000031808 | *Slc27a1* | 8578.420194 | 8863.404943 | 8862.719634 | 4600.647347 | 4074.904427 | 4276.137812 | 1.62E-63 | -1.022661026 | 4.51E-65 | 290.1938089 | 0.492207646 | 62.79168848 |
| ENSMUSG00000043832 | *Clec4a3* | 124.1893256 | 135.260398 | 146.8634495 | 59.29568225 | 60.97589042 | 78.90263289 | 4.40E-05 | -1.024136062 | 1.17E-05 | 19.20921668 | 0.491704662 | 4.356747713 |
| ENSMUSG00000025195 | *Dnmbp* | 632.0045815 | 656.5077854 | 622.701026 | 352.2861122 | 264.864024 | 323.3083494 | 7.83E-15 | -1.025905121 | 9.31E-16 | 64.5717426 | 0.491102094 | 14.10610939 |
| ENSMUSG00000107283 | *Mpv17* | 931.4199418 | 984.2118392 | 963.4242289 | 462.7388537 | 460.177423 | 488.811433 | 1.04E-27 | -1.027042895 | 7.06E-29 | 124.3497282 | 0.490714941 | 26.9839406 |
| ENSMUSG00000026956 | *Uap1l1* | 4253.909707 | 4301.940463 | 4294.287264 | 2040.469066 | 2320.894829 | 1929.265597 | 7.84E-41 | -1.030119098 | 3.57E-42 | 185.1874837 | 0.489669724 | 40.10572053 |
| ENSMUSG00000015947 | *Fcgr1* | 2001.48961 | 1854.057163 | 1879.852154 | 921.989726 | 1002.291199 | 880.4379158 | 4.58E-36 | -1.032240054 | 2.35E-37 | 163.1278966 | 0.488950372 | 35.33946125 |
| ENSMUSG00000028979 | *Masp2* | 124.1893256 | 96.77166687 | 97.90896635 | 47.66907789 | 43.82642124 | 64.46922443 | 0.00052138 | -1.032483575 | 0.000162775 | 14.21835938 | 0.488867846 | 3.282845917 |
| ENSMUSG00000071713 | *Csf2rb* | 734.0779998 | 768.6749448 | 690.2582128 | 359.2620748 | 352.5168665 | 359.8729842 | 7.20E-22 | -1.032761478 | 6.01E-23 | 97.28308166 | 0.488773686 | 21.14289812 |
| ENSMUSG00000050103 | *Agmo* | 642.2119233 | 658.7071415 | 613.889219 | 298.8037321 | 269.6277655 | 366.6085748 | 5.58E-14 | -1.033127278 | 6.98E-15 | 60.60305254 | 0.488649772 | 13.25343109 |
| ENSMUSG00000021213 | *Akr1c13* | 182.0309293 | 210.0385042 | 215.399726 | 93.01283491 | 89.55833906 | 113.5428132 | 5.95E-07 | -1.033295374 | 1.27E-07 | 27.90540366 | 0.48859284 | 6.225150089 |
| ENSMUSG00000015766 | *Eps8* | 3524.084766 | 3407.902223 | 3568.781824 | 1711.436162 | 1615.861096 | 1799.364921 | 6.62E-54 | -1.03466225 | 2.22E-55 | 245.7254716 | 0.488130143 | 53.17937309 |
| ENSMUSG00000020097 | *Sgpl1* | 10924.40759 | 11176.02785 | 10788.589 | 5355.21397 | 5394.460806 | 5296.098676 | 2.18E-108 | -1.035339653 | 3.60E-110 | 497.3401208 | 0.487901 | 107.6617289 |
| ENSMUSG00000112226 | *Gm48786* | 106.3264774 | 94.5723108 | 83.2226214 | 39.53045484 | 35.25168665 | 63.5069972 | 0.002003053 | -1.036523755 | 0.00069586 | 11.500273 | 0.487500717 | 2.698307484 |
| ENSMUSG00000072621 | *Slfn10-ps* | 1797.342774 | 1784.777447 | 1796.629533 | 881.2966107 | 769.8206166 | 970.8872754 | 9.65E-28 | -1.036879408 | 6.56E-29 | 124.497087 | 0.487380553 | 27.01531587 |
| ENSMUSG00000033450 | *Tagap* | 429.5589686 | 427.7747547 | 388.6985964 | 173.236405 | 192.4551542 | 239.5945804 | 9.30E-11 | -1.038174896 | 1.43E-11 | 45.62358992 | 0.486943099 | 10.0315688 |
| ENSMUSG00000058402 | *Zfp420* | 245.8268157 | 219.9356065 | 239.8769676 | 106.9647601 | 101.9440668 | 133.749585 | 6.75E-08 | -1.041637391 | 1.32E-08 | 32.29672666 | 0.485775828 | 7.170550624 |
| ENSMUSG00000002227 | *Mov10* | 3103.882527 | 3145.079173 | 3033.219778 | 1502.157284 | 1516.775274 | 1484.716616 | 1.60E-65 | -1.043272766 | 4.27E-67 | 299.4828247 | 0.485225486 | 64.79592408 |
| ENSMUSG00000042628 | *Zfyve1* | 768.9530844 | 696.0961946 | 720.6099924 | 379.0273022 | 363.949846 | 318.4972133 | 2.42E-19 | -1.044864715 | 2.30E-20 | 85.51802084 | 0.484690357 | 18.61627231 |
| ENSMUSG00000071266 | *Zfp946* | 335.9916685 | 324.4050196 | 306.4550647 | 152.3085172 | 128.6210189 | 187.6343099 | 7.11E-09 | -1.044996 | 1.27E-09 | 36.85901814 | 0.484646252 | 8.147968083 |
| ENSMUSG00000028917 | *Plekhm2* | 2691.335795 | 2803.079305 | 2792.36372 | 1362.638031 | 1345.280582 | 1307.666806 | 3.22E-59 | -1.045275728 | 9.76E-61 | 270.2999673 | 0.484552292 | 58.49231069 |
| ENSMUSG00000027580 | *Helz2* | 7617.228839 | 7867.096645 | 7537.03223 | 3944.90686 | 3434.657578 | 3776.741879 | 5.61E-58 | -1.045579719 | 1.74E-59 | 264.5544285 | 0.484450202 | 57.25072751 |
| ENSMUSG00000023045 | *Soat2* | 270.4945584 | 281.5175763 | 288.8314507 | 134.8686106 | 143.8649915 | 127.9762216 | 1.94E-10 | -1.046995198 | 3.06E-11 | 44.13779706 | 0.483975124 | 9.712917379 |
| ENSMUSG00000028420 | *Tmem38b* | 1901.968027 | 1903.542674 | 1930.764816 | 891.7605547 | 925.1185875 | 957.4160942 | 2.36E-47 | -1.047056829 | 9.09E-49 | 215.4064983 | 0.48395445 | 46.62621867 |
| ENSMUSG00000022246 | *Rai14* | 1492.823742 | 1425.18273 | 1425.55455 | 769.6812089 | 528.7752998 | 805.3841918 | 8.77E-14 | -1.047121438 | 1.11E-14 | 59.69710521 | 0.483932777 | 13.05717637 |
| ENSMUSG00000068959 | *Zfp619* | 326.6349385 | 298.0127468 | 293.7268991 | 130.2179689 | 142.9122432 | 170.3142198 | 9.84E-10 | -1.047886746 | 1.64E-10 | 40.85519114 | 0.483676133 | 9.006813475 |
| ENSMUSG00000090272 | *Mndal* | 2431.89919 | 2411.593925 | 2320.442503 | 1173.12438 | 1068.983579 | 1222.028583 | 8.48E-43 | -1.048698585 | 3.65E-44 | 194.3052337 | 0.483404033 | 42.07173553 |
| ENSMUSG00000022270 | *Retreg1* | 2159.703408 | 2155.368944 | 2365.480627 | 1109.178056 | 1122.337483 | 998.7918651 | 8.20E-38 | -1.048802656 | 3.98E-39 | 171.2316769 | 0.483369163 | 37.08629299 |
| ENSMUSG00000046079 | *Lrrc8d* | 6492.720014 | 6428.717778 | 6268.132026 | 3175.225652 | 2962.094427 | 3139.747453 | 4.33E-79 | -1.049026532 | 9.57E-81 | 362.153111 | 0.48329416 | 78.36352304 |
| ENSMUSG00000003746 | *Man1a* | 923.7644354 | 982.0124831 | 893.9088628 | 494.1306854 | 437.3114641 | 420.4932997 | 2.11E-23 | -1.052193482 | 1.67E-24 | 104.3809923 | 0.482234415 | 22.6757311 |
| ENSMUSG00000031292 | *Cdkl5* | 118.2350428 | 133.0610419 | 140.0098219 | 65.10898443 | 56.21214899 | 67.35590613 | 2.14E-05 | -1.052200972 | 5.42E-06 | 20.68157608 | 0.482231911 | 4.670457882 |
| ENSMUSG00000030203 | *Dusp16* | 2192.877269 | 2159.767656 | 1961.116596 | 990.5866918 | 980.3779882 | 1069.99668 | 1.80E-38 | -1.053686946 | 8.62E-40 | 174.275593 | 0.481735469 | 37.7436301 |
| ENSMUSG00000069910 | *Spdl1* | 2243.913978 | 2198.256387 | 2170.641784 | 1088.250168 | 995.6219608 | 1098.863497 | 9.09E-47 | -1.055806946 | 3.57E-48 | 212.6834491 | 0.481028092 | 46.04138672 |
| ENSMUSG00000052837 | *Junb* | 9871.350159 | 10202.81279 | 10623.12285 | 5216.857378 | 4962.865832 | 4586.937207 | 9.30E-61 | -1.056096101 | 2.73E-62 | 277.4275305 | 0.48093169 | 60.0316741 |
| ENSMUSG00000004267 | *Eno2* | 206.698672 | 230.9323868 | 230.0860709 | 117.4287041 | 97.18032536 | 106.8072226 | 2.60E-08 | -1.056760364 | 4.90E-09 | 34.22940169 | 0.480710305 | 7.585548165 |
| ENSMUSG00000003541 | *Ier3* | 3752.899345 | 3556.358757 | 3849.780557 | 1711.436162 | 1844.520685 | 1797.440466 | 1.36E-58 | -1.059256276 | 4.17E-60 | 267.4077378 | 0.479879378 | 57.86586466 |
| ENSMUSG00000085028 | *Slc2a4rg-ps* | 92.71668827 | 106.6687692 | 104.762594 | 66.27164487 | 36.20443494 | 44.2624526 | 0.000823151 | -1.060822313 | 0.000267437 | 13.28571075 | 0.479358755 | 3.084520337 |
| ENSMUSG00000027309 | *Dnaaf9* | 1622.967351 | 1733.092579 | 1574.376179 | 832.4648724 | 732.6634334 | 799.6108284 | 1.85E-34 | -1.060842221 | 9.92E-36 | 155.6843778 | 0.47935214 | 33.73354469 |
| ENSMUSG00000006585 | *Cdt1* | 4102.500803 | 4072.107755 | 4078.887538 | 2121.855296 | 1915.976807 | 1837.85401 | 1.23E-52 | -1.06158163 | 4.19E-54 | 239.8721501 | 0.479106526 | 51.91106804 |
| ENSMUSG00000006599 | *Gtf2h1* | 4961.61874 | 4800.094612 | 4788.727544 | 2247.422623 | 2259.918939 | 2462.339483 | 2.72E-65 | -1.061603571 | 7.27E-67 | 298.423064 | 0.47909924 | 64.56615026 |
| ENSMUSG00000047945 | *Marcksl1* | 10408.08622 | 10366.66481 | 10642.70464 | 5119.193901 | 5130.54953 | 4803.438334 | 7.26E-93 | -1.061697197 | 1.41E-94 | 425.6818895 | 0.479068149 | 92.13900423 |
| ENSMUSG00000027397 | *Slc20a1* | 4859.545322 | 4777.001374 | 4762.292123 | 2392.755178 | 2187.510069 | 2318.005398 | 6.35E-71 | -1.062424794 | 1.57E-72 | 324.4254531 | 0.4788266 | 70.19696636 |
| ENSMUSG00000033545 | *Znrf1* | 2246.465814 | 2361.008736 | 2272.467109 | 1087.087508 | 1117.573742 | 1086.354543 | 1.03E-54 | -1.063184426 | 3.42E-56 | 249.4517048 | 0.478574547 | 53.98549479 |
| ENSMUSG00000037922 | *Bank1* | 341.0953394 | 325.5046976 | 332.8904856 | 148.8205359 | 171.4946918 | 156.8430385 | 2.85E-12 | -1.065009693 | 3.97E-13 | 52.6588163 | 0.477969448 | 11.54510364 |
| ENSMUSG00000063889 | *Crem* | 459.3303823 | 405.781194 | 422.9667346 | 202.3029159 | 205.7936302 | 207.8410818 | 6.10E-15 | -1.065440034 | 7.22E-16 | 65.07330343 | 0.477826895 | 14.21464325 |
| ENSMUSG00000103847 | *Gm20056* | 711.9620925 | 762.0768766 | 680.4673162 | 345.3101496 | 321.076173 | 362.7596658 | 5.20E-21 | -1.065486055 | 4.53E-22 | 93.28302159 | 0.477811653 | 20.28387399 |
| ENSMUSG00000071637 | *Cebpd* | 2457.417545 | 2617.233718 | 2715.015637 | 1226.60676 | 1246.194761 | 1247.046491 | 3.32E-51 | -1.065637708 | 1.16E-52 | 233.2578589 | 0.477761429 | 50.47911229 |
| ENSMUSG00000000058 | *Cav2* | 110.5795365 | 122.0642616 | 132.1771046 | 59.29568225 | 54.30665241 | 60.62031551 | 2.71E-05 | -1.065746997 | 6.99E-06 | 20.19601832 | 0.477725239 | 4.567719765 |
| ENSMUSG00000028145 | *Them4* | 426.1565213 | 373.8905311 | 385.7613274 | 172.0737446 | 192.4551542 | 201.1054911 | 4.42E-13 | -1.066730098 | 5.86E-14 | 56.41741632 | 0.477399812 | 12.35446452 |
| ENSMUSG00000055013 | *Agap1* | 1329.506273 | 1342.706878 | 1329.603763 | 687.1323179 | 595.4676799 | 629.2966087 | 1.05E-33 | -1.067649483 | 5.79E-35 | 152.1781356 | 0.477095676 | 32.97770237 |
| ENSMUSG00000028587 | *Orc1* | 661.7759952 | 640.012615 | 691.2373025 | 313.9183178 | 303.9267038 | 332.9306217 | 1.62E-21 | -1.067908509 | 1.37E-22 | 95.64957522 | 0.477010025 | 20.79166126 |
| ENSMUSG00000030123 | *Plxnd1* | 2966.083413 | 3110.989154 | 2931.394453 | 1476.578754 | 1386.248759 | 1426.982983 | 1.94E-59 | -1.070739609 | 5.86E-61 | 271.3180809 | 0.476074873 | 58.71124394 |
| ENSMUSG00000051435 | *Fhad1* | 57.84160369 | 60.48229179 | 77.34808342 | 26.74119004 | 40.96817638 | 25.01790799 | 0.005441748 | -1.071994607 | 0.002033468 | 9.519077505 | 0.475660916 | 2.264261581 |
| ENSMUSG00000010461 | *Eya4* | 162.4668574 | 155.0546026 | 158.6125255 | 95.33815578 | 58.11764556 | 74.09149674 | 1.43E-05 | -1.07369068 | 3.54E-06 | 21.4992023 | 0.475102044 | 4.845675987 |
| ENSMUSG00000017631 | *Abr* | 15492.19306 | 15968.42471 | 15962.09878 | 7697.974749 | 7278.996919 | 7539.05035 | 1.71E-110 | -1.074762943 | 2.66E-112 | 507.1413777 | 0.474749063 | 109.7663573 |
| ENSMUSG00000048388 | *Fam171b* | 127.5917728 | 140.7587882 | 121.4071183 | 66.27164487 | 51.44840755 | 67.35590613 | 2.14E-05 | -1.076011786 | 5.43E-06 | 20.67932497 | 0.474338283 | 4.670064623 |
| ENSMUSG00000071551 | *Akr1c19* | 153.1101274 | 150.6558905 | 163.5079738 | 81.38623054 | 67.64512844 | 73.12926951 | 1.33E-06 | -1.076532647 | 2.95E-07 | 26.28527487 | 0.474167061 | 5.876529003 |
| ENSMUSG00000028378 | *Ptgr1* | 529.0805514 | 575.131611 | 539.4784046 | 302.2917134 | 232.4705822 | 244.4057165 | 6.88E-15 | -1.080584026 | 8.15E-16 | 64.83416535 | 0.472837372 | 14.1626936 |
| ENSMUSG00000031765 | *Mt1* | 21011.81315 | 20265.96646 | 22256.66623 | 10246.52643 | 10731.75671 | 9050.709329 | 9.96E-51 | -1.081263672 | 3.56E-52 | 231.0253535 | 0.472614674 | 50.00152676 |
| ENSMUSG00000007039 | *Ddah2* | 55.28976823 | 73.67842818 | 65.59900746 | 31.39183178 | 20.96046233 | 39.45131644 | 0.005840216 | -1.08166907 | 0.00219519 | 9.378717781 | 0.472481887 | 2.233571061 |
| ENSMUSG00000097715 | *Gpr137b-ps* | 3866.881329 | 3863.168928 | 3696.06348 | 1821.888904 | 1766.395326 | 1808.024966 | 4.34E-74 | -1.082582451 | 1.03E-75 | 339.0514299 | 0.47218285 | 73.3627872 |
| ENSMUSG00000079808 | *ENSMUSG00000079808* | 2167.358915 | 2110.282145 | 2188.265398 | 971.9841248 | 1077.558314 | 1000.71632 | 1.60E-48 | -1.083050651 | 6.00E-50 | 220.8158737 | 0.472029637 | 47.79677959 |
| ENSMUSG00000074151 | *Nlrc5* | 1863.690495 | 1989.317561 | 1943.492982 | 939.4296326 | 776.4898546 | 1019.960864 | 1.26E-25 | -1.083402318 | 9.09E-27 | 114.7144858 | 0.47191459 | 24.89953382 |
| ENSMUSG00000049832 | *Rbx1-ps* | 848.0599835 | 826.9578805 | 874.3270695 | 359.2620748 | 438.2642124 | 403.1732095 | 4.58E-23 | -1.08363645 | 3.66E-24 | 102.827133 | 0.471838011 | 22.3392417 |
| ENSMUSG00000041135 | *Ripk2* | 1592.345325 | 1594.533147 | 1628.22611 | 785.958455 | 725.0414471 | 761.1217392 | 5.45E-44 | -1.084590412 | 2.28E-45 | 199.8231473 | 0.471526118 | 43.26332586 |
| ENSMUSG00000001156 | *Mxd1* | 416.7997913 | 438.771535 | 424.924914 | 227.8814455 | 201.0298887 | 176.0875832 | 1.34E-13 | -1.08501637 | 1.70E-14 | 58.84780173 | 0.471386919 | 12.87435342 |

| | | | | | | | | | | | | |
|---|---|---|---|---|---|---|---|---|---|---|---|---|
| ENSMUSG00000021796 | *Bmpr1a* | 5298.46102 | 5227.869367 | 5061.89356 | 2455.538842 | 2139.872655 | 2752.932106 | 8.89E-35 | -1.085059714 | 4.73E-36 | 157.157986 | 0.471372757 | 34.05109493 |
| ENSMUSG00000031639 | *Tlr3* | 762.9988016 | 752.1797743 | 745.087234 | 352.2861122 | 308.6904453 | 404.1354368 | 1.15E-18 | -1.085582898 | 1.14E-19 | 82.34534182 | 0.471201848 | 17.9383551 |
| ENSMUSG00000030591 | *Psmd8* | 11456.89059 | 11580.70936 | 11903.77213 | 5257.550493 | 5960.393289 | 5232.591679 | 2.03E-63 | -1.086510076 | 5.68E-65 | 289.733702 | 0.470899118 | 62.69250108 |
| ENSMUSG00000046311 | *Zfp62* | 1681.659566 | 1592.333791 | 1572.418 | 775.494511 | 647.8688358 | 859.2689167 | 7.61E-24 | -1.086898316 | 5.90E-25 | 106.4430351 | 0.470772412 | 23.11853353 |
| ENSMUSG00000063972 | *Nr6a1* | 330.8879976 | 340.9001901 | 302.538706 | 163.9351215 | 144.8177398 | 150.1074479 | 4.85E-12 | -1.088506937 | 6.86E-13 | 51.58427343 | 0.470247788 | 11.31445665 |
| ENSMUSG00000037172 | *Dennd11* | 1187.454099 | 1142.565476 | 1020.211429 | 577.8422369 | 460.177423 | 538.847249 | 1.64E-21 | -1.089520804 | 1.40E-22 | 95.61476541 | 0.469917433 | 20.78437588 |
| ENSMUSG00000029860 | *Zyx* | 945.8803427 | 1108.475457 | 1010.420533 | 520.8718755 | 433.500471 | 485.9247513 | 1.38E-22 | -1.090195116 | 1.12E-23 | 100.6067325 | 0.469697846 | 21.86020184 |
| ENSMUSG00000025044 | *Msr1* | 11723.13209 | 11548.8187 | 11499.4081 | 5223.83334 | 5250.595815 | 5848.417106 | 9.65E-74 | -1.090774561 | 2.29E-75 | 337.4512355 | 0.469509235 | 73.01554512 |
| ENSMUSG00000021665 | *Hexb* | 3813.292784 | 3878.564421 | 4073.99209 | 1862.582019 | 1879.772372 | 1773.384786 | 7.68E-68 | -1.093084461 | 1.99E-69 | 310.1823239 | 0.468758104 | 67.11440633 |
| ENSMUSG00000093954 | *Gm16867* | 130.9942201 | 114.3665154 | 151.7588978 | 52.31971964 | 49.54291097 | 83.71376904 | 0.000195881 | -1.09485649 | 5.69E-05 | 16.20156615 | 0.468182693 | 3.708008723 |
| ENSMUSG00000026065 | *Slc9a4* | 1681.659566 | 1540.648924 | 1610.602497 | 678.9936948 | 779.3480995 | 800.5730557 | 2.87E-33 | -1.095806751 | 1.61E-34 | 150.1500505 | 0.467874416 | 32.54248092 |
| ENSMUSG00000026307 | *Scly* | 890.5905745 | 960.0189224 | 966.3614979 | 445.2989471 | 489.71262 | 382.0042105 | 3.16E-21 | -1.096493552 | 2.74E-22 | 94.27998191 | 0.467651736 | 20.49987846 |
| ENSMUSG00000053693 | *Mast1* | 104.6252537 | 116.5658715 | 94.97169736 | 63.946324 | 22.86595891 | 61.58254274 | 0.00500724 | -1.097024323 | 0.001860794 | 9.682025258 | 0.467479717 | 2.300401576 |
| ENSMUSG00000027368 | *Dusp2* | 1656.991823 | 1709.999341 | 1670.326966 | 878.9712899 | 740.2854197 | 738.0282857 | 1.97E-33 | -1.097751626 | 1.10E-34 | 150.9124681 | 0.467244107 | 32.70645744 |
| ENSMUSG00000021306 | *Gpr137b* | 3149.815566 | 3104.391086 | 3064.550647 | 1485.880038 | 1372.910283 | 1497.22557 | 8.23E-62 | -1.097757123 | 2.37E-63 | 282.2980857 | 0.467242327 | 61.0848599 |
| ENSMUSG00000056888 | *Glipr1* | 5354.6014 | 5153.091261 | 5341.913204 | 2367.176648 | 2551.459915 | 2481.584027 | 1.08E-79 | -1.09833975 | 2.37E-81 | 364.9357421 | 0.467053671 | 78.96806594 |
| ENSMUSG00000023030 | *Slc11a2* | 5249.976147 | 5155.290617 | 5129.450747 | 2477.62939 | 2346.619033 | 2430.585984 | 2.51E-88 | -1.099045794 | 5.11E-90 | 404.7421497 | 0.466825154 | 87.6005344 |
| ENSMUSG00000037617 | *Spag1* | 80.8081228 | 109.9678033 | 104.762594 | 46.50641745 | 34.29893836 | 56.77140659 | 0.000602775 | -1.100661573 | 0.00019075 | 13.92007321 | 0.466302616 | 3.219845015 |
| ENSMUSG00000041420 | *Meis3* | 173.5248111 | 234.2314209 | 179.1734084 | 93.01283491 | 78.12535961 | 101.9960864 | 1.38E-06 | -1.10126785 | 3.06E-07 | 26.21137599 | 0.466106698 | 5.861358283 |
| ENSMUSG00000026872 | *Zeb2* | 5117.280703 | 5188.280958 | 5085.391712 | 2425.30967 | 2250.391456 | 2498.904117 | 1.27E-71 | -1.101295289 | 3.10E-73 | 327.6680239 | 0.466097833 | 70.89594934 |
| ENSMUSG00000044701 | *Il27* | 95.26852373 | 89.07392064 | 90.07624905 | 40.69311527 | 41.92092467 | 45.22467983 | 0.00013644 | -1.10139208 | 3.88E-05 | 16.92759329 | 0.466066564 | 3.865059318 |
| ENSMUSG00000024644 | *Cndp2* | 8047.638419 | 7974.865092 | 7865.027267 | 3503.095895 | 4032.030754 | 3593.918705 | 7.25E-63 | -1.101448563 | 2.06E-64 | 287.1671336 | 0.466048317 | 62.13959902 |
| ENSMUSG00000020844 | *Nxn* | 45.08242641 | 52.78454556 | 47.97539351 | 17.43990655 | 23.8187072 | 25.98013522 | 0.008208891 | -1.106708019 | 0.003183522 | 8.699185854 | 0.464352395 | 2.085715522 |
| ENSMUSG00000019818 | *Cd164* | 2524.615879 | 2517.163017 | 2405.623303 | 1238.233365 | 1012.77143 | 1208.557401 | 7.53E-36 | -1.107342345 | 3.88E-37 | 162.126321 | 0.464148273 | 35.12296979 |
| ENSMUSG00000005054 | *Cstb* | 7117.069089 | 6946.666132 | 7321.632504 | 3004.314567 | 3766.213982 | 3140.70968 | 3.91E-39 | -1.108959724 | 1.84E-40 | 177.3478462 | 0.463628216 | 38.40822898 |
| ENSMUSG00000020184 | *Mdm2* | 7940.46133 | 7855.000187 | 7902.232674 | 3736.790642 | 3333.666259 | 3909.529237 | 1.04E-59 | -1.110098855 | 3.10E-61 | 272.5845236 | 0.463262287 | 58.98327715 |
| ENSMUSG00000021038 | *Vipas39* | 2086.550792 | 2077.291804 | 2060.983742 | 891.7605547 | 927.0240841 | 1062.298862 | 2.25E-39 | -1.110131443 | 1.05E-40 | 178.4572732 | 0.463251822 | 38.64857392 |
| ENSMUSG00000013160 | *Atp6v0d1* | 3692.505906 | 3565.156182 | 3862.508723 | 1696.321577 | 1814.03274 | 1639.635201 | 1.51E-58 | -1.110471267 | 4.65E-60 | 267.1886498 | 0.463142717 | 57.8200568 |
| ENSMUSG00000028646 | *Rragc* | 5607.233111 | 5774.409349 | 5709.071828 | 2564.828923 | 2836.331653 | 2511.413071 | 5.48E-66 | -1.110506753 | 1.45E-67 | 301.6373509 | 0.463131326 | 65.26083101 |
| ENSMUSG00000060791 | *Gmfg* | 175.2260347 | 178.1478413 | 133.1561942 | 77.89824923 | 91.46383564 | 55.80917936 | 2.80E-05 | -1.112132379 | 7.24E-06 | 20.12886677 | 0.462609764 | 4.553279951 |
| ENSMUSG00000030041 | *M1ap* | 192.2382711 | 191.3439777 | 253.5842229 | 103.4767788 | 119.093536 | 72.16704228 | 6.07E-06 | -1.11320217 | 1.44E-06 | 23.22630012 | 0.462266855 | 5.217091766 |
| ENSMUSG00000074170 | *Plekhf1* | 75.70445189 | 76.97746228 | 79.30626275 | 30.22917134 | 47.63741439 | 28.86681691 | 0.001279746 | -1.114337373 | 0.000430007 | 12.39707593 | 0.461903258 | 2.892876139 |
| ENSMUSG00000036086 | *Zranb3* | 529.9311632 | 542.1412701 | 565.9138255 | 247.6466729 | 254.3837929 | 254.0279888 | 1.41E-20 | -1.114437017 | 1.25E-21 | 91.27097993 | 0.461871356 | 19.85045778 |
| ENSMUSG00000026131 | *Dst* | 7326.319597 | 7872.595035 | 7248.200779 | 3662.380374 | 2691.513913 | 3993.243006 | 2.99E-21 | -1.117450404 | 2.58E-22 | 94.39576575 | 0.460907643 | 20.52424811 |
| ENSMUSG00000039997 | *Ifi203* | 3628.71002 | 3579.451996 | 3515.910982 | 1721.900106 | 1460.563125 | 1760.875832 | 3.05E-43 | -1.117976082 | 1.31E-44 | 196.3500361 | 0.460739731 | 42.51523422 |
| ENSMUSG00000096100 | *ENSMUSG00000096100* | 211.8023429 | 235.331099 | 196.7970224 | 112.7780623 | 78.12535961 | 105.8449953 | 1.71E-07 | -1.12099801 | 3.47E-08 | 30.42661941 | 0.459775657 | 6.765948158 |
| ENSMUSG00000017713 | *Tha1* | 80.8081228 | 100.070701 | 101.825325 | 38.3677944 | 47.63741439 | 43.30022537 | 0.000171893 | -1.12171755 | 4.96E-05 | 16.46394911 | 0.459546403 | 3.764742376 |
| ENSMUSG00000020275 | *Rel* | 3299.523246 | 3238.551806 | 3234.912248 | 1633.537913 | 1249.053005 | 1609.806156 | 1.70E-29 | -1.122283597 | 1.09E-30 | 132.620286 | 0.459366133 | 28.76979612 |
| ENSMUSG00000026207 | *Speg* | 1772.675031 | 1943.131084 | 1856.354002 | 937.1043117 | 789.8283307 | 833.2887815 | 2.39E-36 | -1.123347116 | 1.21E-37 | 164.4497647 | 0.459027625 | 35.62237987 |
| ENSMUSG00000028037 | *Ifi44* | 11742.69616 | 11277.19822 | 10931.53609 | 5265.689116 | 4578.908272 | 5736.798747 | 1.64E-43 | -1.123777208 | 6.96E-45 | 197.6037953 | 0.458890801 | 42.78535909 |
| ENSMUSG00000079491 | *H2-T10* | 230.5158029 | 236.430777 | 298.6223474 | 102.3141184 | 141.9594949 | 104.8827681 | 1.01E-07 | -1.128985132 | 2.00E-08 | 31.4914123 | 0.457237257 | 6.997085674 |
| ENSMUSG00000015149 | *Sirt2* | 1804.99828 | 1913.439777 | 1844.604926 | 875.4833086 | 887.008656 | 778.4418294 | 1.48E-42 | -1.130777918 | 6.46E-44 | 193.1701251 | 0.456669417 | 41.82880444 |
| ENSMUSG00000071203 | *Naip5* | 403.1900022 | 368.3921409 | 384.7822378 | 159.2844798 | 157.2034675 | 210.7277634 | 6.06E-12 | -1.131577181 | 8.62E-13 | 51.13519343 | 0.456416489 | 11.21741972 |
| ENSMUSG00000041827 | *Oasl1* | 9249.55292 | 9248.292254 | 9103.575692 | 4293.704991 | 4301.65852 | 4003.827506 | 4.37E-100 | -1.131649458 | 7.91E-102 | 459.0068336 | 0.456393624 | 99.36000322 |
| ENSMUSG00000041308 | *Sntb2* | 2935.461387 | 2936.140347 | 2896.147225 | 1401.005826 | 1261.438733 | 1340.382532 | 1.40E-62 | -1.132287956 | 4.00E-64 | 285.8443241 | 0.456191681 | 61.85345189 |
| ENSMUSG00000022304 | *Dpys* | 91.01546463 | 107.7684472 | 91.05533871 | 40.69311527 | 45.73191782 | 45.22467983 | 8.07E-05 | -1.133462511 | 2.23E-05 | 17.98189443 | 0.455820429 | 4.093174504 |
| ENSMUSG00000023905 | *Tnfrsf12a* | 1308.240978 | 1425.18273 | 1402.056398 | 599.9327851 | 705.9864813 | 576.374111 | 5.90E-29 | -1.133989381 | 3.85E-30 | 130.1251059 | 0.455653994 | 28.22942566 |
| ENSMUSG00000028073 | *Pear1* | 186.2839884 | 167.151061 | 189.9433947 | 83.71155142 | 80.03085618 | 83.71376904 | 3.53E-08 | -1.136779503 | 6.72E-09 | 33.61466993 | 0.454773627 | 7.452574058 |
| ENSMUSG00000007891 | *Ctsd* | 57780.35964 | 58440.18969 | 59926.16195 | 26645.85188 | 28222.30978 | 25132.41303 | 2.86E-93 | -1.138683531 | 5.55E-95 | 427.549101 | 0.454173825 | 92.54387817 |
| ENSMUSG00000089976 | *Gm2389* | 58.69221551 | 48.38583343 | 46.99630385 | 22.09054829 | 23.8187072 | 24.05568076 | 0.004413052 | -1.139820742 | 0.001626961 | 9.928986502 | 0.453815962 | 2.355260952 |
| ENSMUSG00000023393 | *Slc17a9* | 271.3451703 | 305.7104931 | 283.9360024 | 143.0072337 | 119.093536 | 128.9384489 | 2.04E-11 | -1.140639077 | 2.98E-12 | 48.70095395 | 0.453558618 | 10.69081531 |
| ENSMUSG00000022558 | *Mroh1* | 3126.849047 | 3054.905574 | 3139.940551 | 1488.205359 | 1397.681738 | 1342.306986 | 9.62E-65 | -1.141835629 | 2.65E-66 | 295.8467366 | 0.453182599 | 64.01691731 |
| ENSMUSG00000032515 | *Csrnp1* | 1831.367246 | 2050.899531 | 2048.255576 | 945.2429347 | 887.008656 | 854.4577806 | 1.15E-41 | -1.142706451 | 5.09E-43 | 189.0653591 | 0.452909137 | 40.94030822 |
| ENSMUSG00000081953 | *Gm12435* | 1237.640197 | 1525.253431 | 1362.892812 | 718.5241497 | 520.2005652 | 629.2966087 | 8.03E-19 | -1.144445695 | 7.90E-20 | 83.07426741 | 0.45236346 | 18.09549338 |
| ENSMUSG00000107169 | *None* | 116.5338192 | 126.4629737 | 133.1561942 | 40.69311527 | 54.30665241 | 74.09149674 | 6.92E-05 | -1.144578772 | 1.90E-05 | 18.28921073 | 0.452321735 | 4.160124532 |
| ENSMUSG00000041390 | *Mdfic* | 3475.599892 | 3346.320253 | 3416.043836 | 1530.061134 | 1546.310471 | 1543.412478 | 3.79E-82 | -1.148167787 | 8.09E-84 | 376.2669655 | 0.451197886 | 81.42146968 |
| ENSMUSG00000035107 | *Dcbld2* | 2310.2617 | 2402.796501 | 2239.178061 | 1120.804661 | 988.9527228 | 1027.658682 | 1.01E-49 | -1.149169424 | 3.67E-51 | 226.379977 | 0.450884736 | 48.99749691 |
| ENSMUSG00000037321 | *Tap1* | 2368.953916 | 2388.500687 | 2362.543358 | 1073.135583 | 1115.668245 | 1020.923091 | 1.02E-60 | -1.149382805 | 3.01E-62 | 277.2314145 | 0.450818053 | 59.99068241 |
| ENSMUSG00000070034 | *Sp110* | 2584.158706 | 2447.883301 | 2428.142366 | 1047.557053 | 1107.093511 | 1196.048447 | 5.43E-51 | -1.153896044 | 1.91E-52 | 232.2625777 | 0.449409945 | 50.26546198 |
| ENSMUSG00000025779 | *Ly96* | 326.6349385 | 317.8069514 | 326.036858 | 143.0072337 | 153.3924743 | 139.5229484 | 5.98E-14 | -1.154310296 | 7.49E-15 | 60.46409467 | 0.449280921 | 13.22348727 |
| ENSMUSG00000075122 | *Cd80* | 58.69221551 | 98.97102293 | 76.36899376 | 33.71715265 | 40.96817638 | 29.82904414 | 0.001459175 | -1.156100155 | 0.000495503 | 12.13251348 | 0.448723873 | 2.835892478 |
| ENSMUSG00000110040 | *Gm49369* | 11128.55443 | 11406.96023 | 10593.75016 | 4723.889353 | 5243.926577 | 4880.416512 | 6.24E-81 | -1.157403895 | 1.35E-82 | 370.6541236 | 0.448318552 | 80.20496034 |
| ENSMUSG00000027253 | *Lrp4* | 206.698672 | 210.0385042 | 188.9643051 | 105.8020997 | 80.03085618 | 86.60045073 | 1.95E-08 | -1.158295589 | 3.64E-09 | 34.80797308 | 0.448041543 | 7.710191337 |
| ENSMUSG00000059456 | *Ptk2b* | 9024.991399 | 9077.842159 | 9060.495746 | 4050.70896 | 4087.290155 | 4012.487551 | 4.15E-135 | -1.160615144 | 4.92E-137 | 620.8350238 | 0.447321763 | 134.3823246 |
| ENSMUSG00000037280 | *Galnt6* | 1533.65311 | 1538.449568 | 1514.651709 | 723.1747914 | 654.5380738 | 669.7101523 | 1.50E-46 | -1.165340073 | 5.93E-48 | 211.6736957 | 0.445859149 | 45.82261354 |
| ENSMUSG00000066760 | *Psg16* | 675.3857843 | 741.182994 | 658.9273436 | 306.9423552 | 268.6750172 | 347.3640302 | 2.52E-19 | -1.16841235 | 2.40E-20 | 85.43324311 | 0.444910684 | 18.59828004 |
| ENSMUSG00000020392 | *Cdkn2aipnl* | 5477.089502 | 5267.457776 | 5399.679494 | 2257.886567 | 2621.01054 | 2299.723081 | 1.92E-60 | -1.168695257 | 5.68E-62 | 275.9676062 | 0.444823448 | 59.71755795 |
| ENSMUSG00000000489 | *Pdgfb* | 12858.69887 | 12809.04972 | 12853.4891 | 5988.863908 | 5650.750095 | 5476.997395 | 3.40E-109 | -1.170691321 | 5.41E-111 | 501.1240899 | 0.44420843 | 108.4685176 |
| ENSMUSG00000017418 | *Arl5b* | 1634.025304 | 1623.124776 | 1632.142469 | 767.355888 | 652.6325772 | 753.4239214 | 1.82E-40 | -1.171176821 | 8.37E-42 | 183.4925447 | 0.444058969 | 39.74089489 |
| ENSMUSG00000027737 | *Slc7a11* | 9831.371404 | 9794.832236 | 9551.998757 | 4448.338829 | 3783.363451 | 4726.460156 | 2.65E-48 | -1.171214419 | 1.00E-49 | 219.7936082 | 0.444047397 | 47.57695732 |
| ENSMUSG00000001123 | *Lgals9* | 4160.342407 | 4127.091656 | 4187.566491 | 1830.027527 | 1798.788768 | 1910.021052 | 2.20E-89 | -1.171282363 | 4.37E-91 | 409.6470733 | 0.444026485 | 88.65786961 |
| ENSMUSG00000053178 | *Mterf1b* | 108.8783128 | 98.97102293 | 102.8044147 | 56.97036138 | 22.86595891 | 58.69586105 | 0.001610233 | -1.171863053 | 0.000550666 | 11.93576275 | 0.443847799 | 2.793111177 |
| ENSMUSG00000025283 | *Sat1* | 6284.320119 | 6179.090865 | 6606.89705 | 2481.117371 | 3098.337432 | 2869.361601 | 9.06E-46 | -1.173655808 | 3.64E-47 | 208.0622299 | 0.443296597 | 45.04293565 |
| ENSMUSG00000062081 | *Gm6055* | 976.5023682 | 1011.70379 | 989.8596498 | 453.4375702 | 502.0983477 | 364.6841203 | 2.52E-21 | -1.173835311 | 2.17E-22 | 94.74371157 | 0.443241444 | 20.5985118 |
| ENSMUSG00000079484 | *Phyhd1* | 165.0186929 | 182.5465534 | 146.8634495 | 74.41026793 | 77.17261132 | 67.35590613 | 1.80E-07 | -1.174547979 | 3.64E-08 | 30.33078161 | 0.443022544 | 6.745668213 |
| ENSMUSG00000031389 | *Arhgap4* | 100.3721946 | 144.0578223 | 96.92987669 | 43.01843614 | 40.96817638 | 66.3936789 | 0.000274892 | -1.175665088 | 8.20E-05 | 15.51178892 | 0.442679635 | 3.56083778 |
| ENSMUSG00000110206 | *Flt3l* | 117.384431 | 111.0674813 | 117.4907596 | 55.80770094 | 39.0626798 | 58.69586105 | 2.22E-05 | -1.17580781 | 5.66E-06 | 20.60134932 | 0.442635844 | 4.653080441 |
| ENSMUSG00000063458 | *Lrmda* | 155.6619629 | 175.9484852 | 180.1524981 | 61.62100313 | 84.79459762 | 78.90263289 | 2.39E-07 | -1.175875271 | 4.90E-08 | 29.75596953 | 0.442615147 | 6.622191037 |
| ENSMUSG00000086414 | *Gm15645* | 769.8036962 | 740.0833159 | 724.526351 | 316.2436387 | 354.4223631 | 317.534986 | 8.33E-27 | -1.176109095 | 5.80E-28 | 120.1723388 | 0.442543417 | 26.07948294 |
| ENSMUSG00000078495 | *Zfp984* | 343.6471749 | 354.0963265 | 302.538706 | 153.4711776 | 128.6210189 | 160.6919475 | 2.13E-12 | -1.176877553 | 2.94E-13 | 53.24581965 | 0.442307756 | 11.67145538 |
| ENSMUSG00000040146 | *Rgl3* | 67.1983337 | 76.97746228 | 67.55718678 | 29.06651091 | 29.53519692 | 34.64018029 | 0.000495963 | -1.178941827 | 0.000154211 | 14.32007928 | 0.441675335 | 3.304550473 |
| ENSMUSG00000029821 | *Gsdme* | 437.214475 | 413.4789402 | 476.8166661 | 194.1642929 | 181.974923 | 209.7655362 | 9.37E-17 | -1.180461731 | 1.01E-17 | 73.48947589 | 0.441210267 | 16.02835631 |
| ENSMUSG00000056592 | *Zfp658* | 162.4668574 | 160.5529928 | 140.0098219 | 56.97036138 | 77.17261132 | 68.31813336 | 5.11E-07 | -1.188104941 | 1.09E-07 | 28.2152277 | 0.438878974 | 6.291322836 |
| ENSMUSG00000018927 | *Ccl6* | 47.63426186 | 51.68486753 | 50.9126625 | 19.76522742 | 26.67695206 | 19.24454461 | 0.003821846 | -1.188310514 | 0.00139012 | 10.21885698 | 0.438816441 | 2.417726823 |
| ENSMUSG00000039096 | *Rsad1* | 134.3966674 | 137.4597541 | 106.7207733 | 65.10898443 | 54.30665241 | 47.14913429 | 1.04E-05 | -1.190080805 | 2.53E-06 | 22.14360273 | 0.438278312 | 4.983900005 |
| ENSMUSG00000082516 | *Gm13342* | 68.04894552 | 73.67842818 | 95.95078703 | 27.90385047 | 40.96817638 | 34.64018029 | 0.000555982 | -1.19154518 | 0.000174684 | 14.08551411 | 0.437833673 | 3.254938907 |
| ENSMUSG00000056919 | *Cep162* | 1210.420618 | 1164.559036 | 1108.329499 | 524.3598568 | 404.9180224 | 593.6942011 | 3.50E-19 | -1.194400003 | 3.36E-20 | 84.76376311 | 0.436968139 | 18.45591221 |
| ENSMUSG00000040033 | *Stat2* | 2596.917883 | 2667.818907 | 2679.768409 | 1226.60676 | 1115.668245 | 1126.768087 | 4.18E-65 | -1.196616021 | 1.13E-66 | 297.535802 | 0.43629746 | 64.37837395 |
| ENSMUSG00000080950 | *Gm7278* | 553.7482942 | 532.2441678 | 628.575564 | 248.8093334 | 244.85631 | 254.0279888 | 8.06E-21 | -1.197641717 | 7.10E-22 | 92.3934625 | 0.435987381 | 20.0934319 |
| ENSMUSG00000047843 | *Bri3* | 1945.34923 | 1815.568432 | 2163.788156 | 874.3206481 | 946.0790499 | 763.0461937 | 1.03E-30 | -1.197865641 | 6.35E-32 | 138.2727475 | 0.435919715 | 29.98868761 |
| ENSMUSG00000030789 | *Itgax* | 84.21057008 | 109.9678033 | 76.36899376 | 44.18109658 | 40.96817638 | 32.71572583 | 0.000209534 | -1.199566635 | 6.13E-05 | 16.06235797 | 0.435406052 | 3.678744713 |
| ENSMUSG00000028886 | *Eya3* | 5489.84868 | 5446.705295 | 5301.770528 | 2384.616555 | 2309.46185 | 2374.776805 | 4.34E-108 | -1.200144622 | 7.20E-110 | 495.958609 | 0.43523165 | 107.3629341 |
| ENSMUSG00000021262 | *Evl* | 677.9376197 | 619.1187323 | 659.9064332 | 289.5024486 | 302.0212073 | 260.7635794 | 6.85E-24 | -1.200643064 | 5.29E-25 | 106.656397 | 0.435081306 | 23.16452149 |
| ENSMUSG00000011179 | *Odc1* | 71522.84419 | 69951.61933 | 71654.67703 | 29683.8836 | 32869.81593 | 30124.4479 | 5.56E-110 | -1.201370748 | 8.70E-112 | 504.7735402 | 0.43486191 | 109.255002 |
| ENSMUSG00000074342 | *I830077J02Rik* | 295.1623012 | 273.8198301 | 265.3332988 | 130.2179689 | 107.6605565 | 125.0895399 | 4.34E-12 | -1.204529246 | 6.11E-13 | 51.81102647 | 0.433910906 | 11.36292595 |
| ENSMUSG00000022489 | *Pde1b* | 711.1114807 | 690.5978045 | 677.5300472 | 277.8758443 | 261.0530309 | 361.7974386 | 6.75E-18 | -1.205597977 | 6.90E-19 | 78.79209733 | 0.43358959 | 17.17045082 |
| ENSMUSG00000019810 | *Fuca2* | 2824.881851 | 2915.246464 | 2764.94921 | 1237.070704 | 1210.943074 | 1237.424218 | 2.33E-77 | -1.206433595 | 5.21E-79 | 354.1820726 | 0.433338524 | 76.63185535 |
| ENSMUSG00000107092 | *Gm7993* | 7499.844408 | 7621.868444 | 7495.910464 | 3161.273726 | 3323.186028 | 3308.137218 | 3.18E-121 | -1.207125659 | 4.28E-123 | 556.7524741 | 0.433130701 | 120.4976086 |
| ENSMUSG00000033540 | *Idua* | 454.2267113 | 443.1702471 | 470.9421282 | 195.3269533 | 180.0694264 | 215.5388996 | 1.57E-18 | -1.211479874 | 1.57E-19 | 81.72066097 | 0.431825435 | 17.80289142 |
| ENSMUSG00000048249 | *Crebrf* | 797.0232744 | 797.2665736 | 784.2508205 | 333.6835452 | 340.1311388 | 352.1751663 | 1.40E-32 | -1.212248517 | 8.04E-34 | 146.952447 | 0.431595427 | 31.85246073 |
| ENSMUSG00000022295 | *Atp6v1c1* | 5026.265238 | 4891.367889 | 5006.08545 | 1993.962648 | 2300.887115 | 2131.333315 | 6.86E-72 | -1.214702698 | 1.66E-73 | 328.9084632 | 0.43086186 | 71.16366923 |
| ENSMUSG00000022564 | *Grina* | 1793.940326 | 1637.420591 | 1599.83251 | 805.7236824 | 683.1205224 | 680.2946519 | 4.69E-36 | -1.216590354 | 2.41E-37 | 163.0752587 | 0.430298479 | 35.32912304 |
| ENSMUSG00000010051 | *Hyal1* | 211.8023429 | 258.4243377 | 212.462457 | 89.5248536 | 102.8968151 | 100.071632 | 6.65E-10 | -1.21763745 | 1.09E-10 | 41.6449503 | 0.429986285 | 9.177063713 |
| ENSMUSG00000029254 | *Stap1* | 11480.70772 | 11493.8348 | 11411.29003 | 4691.334861 | 4831.386568 | 5230.667224 | 2.88E-101 | -1.220386178 | 5.16E-103 | 464.4540403 | 0.429167824 | 100.5404365 |
| ENSMUSG00000022895 | *Ets2* | 14269.86388 | 14251.8273 | 13861.95146 | 6248.137185 | 5887.031671 | 6042.787007 | 6.91E-137 | -1.221621447 | 8.06E-139 | 629.045876 | 0.428800518 | 136.1605353 |
| ENSMUSG00000024235 | *Map3k8* | 430.4095804 | 484.9580124 | 495.4193697 | 197.6522742 | 191.5024059 | 214.5766724 | 1.26E-18 | -1.222715795 | 1.25E-19 | 82.17061848 | 0.428475377 | 17.90026421 |
| ENSMUSG00000020593 | *Lpin1* | 1406.911949 | 1432.880476 | 1388.349143 | 640.6259004 | 615.475394 | 556.1673392 | 1.06E-43 | -1.223655267 | 4.46E-45 | 198.4897469 | 0.428196448 | 42.97659291 |
| ENSMUSG00000003812 | *Dnase2a* | 3527.487213 | 3414.500291 | 3713.687094 | 1567.266268 | 1534.877492 | 1462.58539 | 1.84E-74 | -1.223750357 | 4.32E-76 | 340.7796495 | 0.428168226 | 73.73536995 |
| ENSMUSG00000036390 | *Gadd45a* | 154.8113511 | 207.8391482 | 171.3406911 | 94.17549534 | 58.11764556 | 76.97817843 | 1.38E-06 | -1.223827343 | 3.06E-07 | 26.20916811 | 0.428145378 | 5.86099572 |
| ENSMUSG00000053746 | *Ptrh1* | 427.8577449 | 477.2602661 | 425.9040036 | 198.8149346 | 198.1716439 | 172.2386742 | 7.45E-18 | -1.22571972 | 7.64E-19 | 78.58962845 | 0.42758415 | 17.12768642 |
| ENSMUSG00000083373 | *Gm13392* | 373.4185885 | 357.3953606 | 373.0331618 | 153.4711776 | 188.644161 | 128.9384489 | 1.10E-12 | -1.228062965 | 1.50E-13 | 54.57533307 | 0.426890226 | 11.95683194 |
| ENSMUSG00000002250 | *Ppard* | 1343.116062 | 1443.877257 | 1367.78826 | 618.5353521 | 588.7984419 | 566.7518387 | 6.89E-46 | -1.228322904 | 2.75E-47 | 208.6166528 | 0.426813317 | 45.16166663 |
| ENSMUSG00000092029 | *Gm9027* | 164.1680811 | 129.7620078 | 76.36899376 | 56.97036138 | 59.07039385 | 42.33799814 | 0.00041278 | -1.22910557 | 0.000126859 | 14.68788709 | 0.426581833 | 3.384281071 |
| ENSMUSG00000029798 | *Herc6* | 840.4044772 | 798.3662516 | 808.7280621 | 366.2380374 | 304.8794521 | 373.3441654 | 1.09E-26 | -1.230607184 | 7.62E-28 | 119.6299403 | 0.42613806 | 25.96287017 |
| ENSMUSG00000020120 | *Plek* | 26876.78164 | 26374.67793 | 25899.85887 | 11160.37753 | 10794.6381 | 11770.92571 | 1.19E-125 | -1.230754746 | 1.52E-127 | 577.2026107 | 0.426094476 | 124.9256905 |
| ENSMUSG00000024395 | *Lims2* | 246.6774275 | 247.4275573 | 225.1906226 | 111.6154019 | 89.55833906 | 105.8449953 | 5.42E-11 | -1.231647063 | 8.21E-12 | 46.71557701 | 0.425831015 | 10.26562412 |
| ENSMUSG00000001751 | *Naglu* | 1135.566778 | 1106.276101 | 1130.848561 | 480.1787602 | 509.720334 | 446.4734349 | 2.52E-39 | -1.231711044 | 1.18E-40 | 178.2218486 | 0.425812131 | 38.597804 |
| ENSMUSG00000069792 | *Wfdc17* | 184.5827647 | 200.1414019 | 199.7342914 | 60.45834269 | 94.3220805 | 92.37381411 | 6.62E-08 | -1.23257845 | 1.30E-08 | 32.33720317 | 0.425556193 | 7.179395719 |
| ENSMUSG00000020903 | *Stx8* | 599.6813324 | 558.6364405 | 572.7674532 | 227.8814455 | 296.3047175 | 211.6899907 | 1.13E-16 | -1.232738714 | 1.23E-17 | 73.09868191 | 0.425508922 | 15.94511875 |
| ENSMUSG00000052539 | *Magi3* | 78.25628735 | 69.27971605 | 53.84993149 | 31.39183178 | 22.86595891 | 31.7534986 | 0.000944418 | -1.232768345 | 0.000311048 | 13.00266777 | 0.425500182 | 3.024835742 |
| ENSMUSG00000033192 | *Lpcat2* | 1295.4818 | 1291.02201 | 1333.520122 | 595.2821434 | 501.1455994 | 572.5252021 | 5.90E-39 | -1.233844301 | 2.79E-40 | 176.5174549 | 0.425182964 | 38.22879898 |
| ENSMUSG00000031144 | *Syp* | 147.1558447 | 145.1575003 | 126.3025666 | 55.80770094 | 67.64512844 | 53.8847249 | 4.96E-07 | -1.237115348 | 1.05E-07 | 28.2777716 | 0.424220032 | 6.304512651 |
| ENSMUSG00000032369 | *Plscr1* | 534.1842223 | 580.6300012 | 553.1856599 | 212.7668598 | 269.6277655 | 223.2367174 | 8.88E-20 | -1.237362184 | 8.22E-21 | 87.55057353 | 0.424147457 | 19.05181435 |
| ENSMUSG00000038608 | *Dock10* | 8197.3461 | 7946.273463 | 7642.773914 | 3378.691228 | 2937.322972 | 3732.479427 | 1.81E-48 | -1.243292267 | 6.85E-50 | 220.552875 | 0.42240761 | 47.74178471 |
| ENSMUSG00000039934 | *Gsap* | 1764.168913 | 1708.899663 | 1686.97149 | 713.8735079 | 717.4194608 | 745.7261035 | 1.70E-59 | -1.24489128 | 5.09E-61 | 271.5982737 | 0.421939694 | 58.77032647 |
| ENSMUSG00000045664 | *Cdc42ep2* | 677.0870079 | 679.6010241 | 749.0035926 | 260.4359377 | 346.8003768 | 279.0458968 | 2.70E-20 | -1.245359824 | 2.43E-21 | 89.95693966 | 0.421802683 | 19.56836495 |
| ENSMUSG00000050957 | *Insl6* | 65.49711006 | 63.78132589 | 77.34808342 | 24.41586916 | 30.48794521 | 31.7534986 | 0.000366582 | -1.247970743 | 0.000111755 | 14.92694546 | 0.421040015 | 3.435829268 |
| ENSMUSG00000033721 | *Vav3* | 1204.466336 | 1163.459358 | 1180.782134 | 511.570592 | 449.6971919 | 534.0361129 | 1.36E-37 | -1.247981343 | 6.67E-39 | 170.2076239 | 0.421036922 | 36.86690776 |
| ENSMUSG00000026471 | *Mr1* | 130.1436083 | 128.6623298 | 105.7416837 | 58.13302182 | 37.15718323 | 58.69586105 | 1.47E-05 | -1.248156242 | 3.65E-06 | 21.44275633 | 0.420985882 | 4.833851603 |
| ENSMUSG00000024696 | *Lpxn* | 6846.574531 | 7055.534257 | 7137.563647 | 2795.035689 | 3053.558263 | 2968.471006 | 3.29E-104 | -1.253944057 | 5.73E-106 | 478.0310357 | 0.419300351 | 103.4830672 |
| ENSMUSG00000061132 | *Blnk* | 652.4192652 | 690.5978045 | 697.1118404 | 301.129053 | 288.6827312 | 265.5747156 | 7.79E-28 | -1.255183135 | 5.27E-29 | 124.9297374 | 0.418940384 | 27.10824814 |
| ENSMUSG00000059810 | *Rgs3* | 1899.416192 | 2006.912409 | 1970.907493 | 817.3502867 | 858.4262074 | 772.668466 | 8.01E-59 | -1.262806238 | 2.44E-60 | 268.4736576 | 0.416732569 | 58.09623185 |
| ENSMUSG00000022747 | *St3gal6* | 4972.676694 | 4940.8534 | 4994.336374 | 2132.31924 | 1975.999949 | 2102.466498 | 5.43E-104 | -1.263846657 | 9.50E-106 | 477.022095 | 0.416432145 | 103.2652281 |
| ENSMUSG00000032633 | *Flcn* | 2463.371828 | 2458.880081 | 2527.030422 | 1064.99696 | 1020.393416 | 1014.187501 | 1.63E-78 | -1.26586306 | 3.61E-80 | 359.5017109 | 0.41585052 | 77.78753856 |
| ENSMUSG00000062901 | *Klhl24* | 1309.091589 | 1195.350021 | 1264.004756 | 484.829402 | 524.0115583 | 556.1673392 | 6.51E-41 | -1.266679561 | 2.96E-42 | 185.5634361 | 0.415615233 | 40.18648322 |
| ENSMUSG00000025856 | *Pdgfa* | 409.1442849 | 409.0802281 | 391.6358654 | 179.0497072 | 141.9594949 | 181.8609465 | 1.42E-16 | -1.269006518 | 1.55E-17 | 72.65239301 | 0.414945418 | 15.84828148 |
| ENSMUSG00000002083 | *Bbc3* | 1342.26545 | 1433.980154 | 1549.898937 | 634.8125982 | 636.4358563 | 519.6027044 | 3.29E-32 | -1.273197664 | 1.92E-33 | 145.2215462 | 0.413741717 | 31.4828251 |
| ENSMUSG00000079625 | *Tm4sf19* | 1336.311168 | 1417.484984 | 1292.398356 | 548.775726 | 556.4050001 | 567.7140659 | 1.18E-48 | -1.27320837 | 4.42E-50 | 221.4246496 | 0.413738647 | 47.92797824 |
| ENSMUSG00000026222 | *Sp100* | 3295.270187 | 3361.715746 | 3095.881516 | 1364.963352 | 1313.839889 | 1348.08035 | 1.85E-81 | -1.276425578 | 3.98E-83 | 373.0863129 | 0.412817038 | 80.73313533 |
| ENSMUSG00000001750 | *Tcirg1* | 13794.37187 | 14098.97206 | 14068.53938 | 6150.473708 | 5372.547595 | 5800.305745 | 1.58E-97 | -1.276761579 | 2.95E-99 | 447.1882298 | 0.412720905 | 96.80095135 |
| ENSMUSG00000019806 | *Aig1* | 1639.979587 | 1740.790326 | 1648.786993 | 670.8550718 | 686.9315156 | 713.9726049 | 3.12E-58 | -1.27824054 | 9.65E-60 | 265.7353637 | 0.412298026 | 57.50620266 |
| ENSMUSG00000067212 | *H2-T23* | 3629.560632 | 3974.23641 | 4020.142158 | 1589.356816 | 1682.553476 | 1510.696752 | 2.83E-68 | -1.28074373 | 7.29E-70 | 312.1882375 | 0.411583277 | 67.54791531 |
| ENSMUSG00000042406 | *Atf4* | 14301.33651 | 14296.9141 | 14179.17651 | 5799.350256 | 6027.085669 | 5777.212291 | 3.83E-167 | -1.280840612 | 3.60E-169 | 768.6112236 | 0.411555639 | 166.4164249 |
| ENSMUSG00000014444 | *Piezo1* | 11165.98135 | 12084.3619 | 11477.86813 | 4965.722724 | 4561.758802 | 4754.364745 | 4.52E-107 | -1.282122339 | 7.60E-109 | 491.2542515 | 0.411190164 | 106.3446853 |
| ENSMUSG00000029560 | *Snx8* | 3993.62249 | 4163.381031 | 4345.199927 | 1767.243863 | 1817.843733 | 1553.996977 | 4.11E-62 | -1.282817112 | 1.18E-63 | 283.6911381 | 0.410992191 | 61.38634586 |
| ENSMUSG00000027438 | *Napb* | 435.5132513 | 473.961232 | 398.4894931 | 194.1642929 | 149.5814812 | 194.3699005 | 1.03E-15 | -1.283122693 | 1.17E-16 | 68.65956614 | 0.410905147 | 14.9867126 |
| ENSMUSG00000030894 | *Tpp1* | 5054.335428 | 4961.747283 | 4648.717722 | 1985.824025 | 1871.197637 | 2165.011268 | 8.23E-69 | -1.284015336 | 2.10E-70 | 314.6731658 | 0.410650985 | 68.08456404 |

| | | | | | | | | | | | | |
|---|---|---|---|---|---|---|---|---|---|---|---|---|
| ENSMUSG00000021281 | *Tnfaip2* | 9201.068046 | 9301.0768 | 9482.483391 | 3792.598343 | 3923.41745 | 3736.328336 | 4.20E-144 | -1.288842135 | 4.64E-146 | 662.3371844 | 0.409279373 | 143.377192 |
| ENSMUSG00000010307 | *Tmem86a* | 167.5705283 | 179.2475193 | 215.399726 | 75.57292836 | 80.98360447 | 73.12926951 | 3.65E-09 | -1.290089354 | 6.36E-10 | 38.20631687 | 0.408925702 | 8.438166898 |
| ENSMUSG00000022359 | *Wdyhv1* | 194.7901065 | 149.5562124 | 184.0688567 | 80.22357011 | 68.59787673 | 67.35590613 | 1.38E-08 | -1.296308253 | 2.52E-09 | 35.52507426 | 0.407166775 | 7.861082656 |
| ENSMUSG00000029605 | *Oas1b* | 523.9768805 | 577.3309671 | 596.2656051 | 241.8333708 | 236.2815754 | 212.6522179 | 1.17E-23 | -1.297743699 | 9.16E-25 | 105.5691694 | 0.406761856 | 22.93008474 |
| ENSMUSG00000042712 | *Tceal9* | 1086.231293 | 931.4272936 | 996.7132775 | 417.3950966 | 370.619084 | 437.8133898 | 2.30E-31 | -1.299940085 | 1.38E-32 | 141.3009791 | 0.406143065 | 30.639187 |
| ENSMUSG00000024236 | *Svil* | 153.9607392 | 186.9452655 | 166.4452428 | 72.08494705 | 60.97589042 | 72.16704228 | 7.45E-09 | -1.30550366 | 1.33E-09 | 36.76420928 | 0.404579841 | 8.127848553 |
| ENSMUSG00000022125 | *Cln5* | 3365.020356 | 3190.165972 | 3229.03771 | 1311.480972 | 1377.674024 | 1267.253262 | 1.06E-84 | -1.306444401 | 2.21E-86 | 388.0407563 | 0.404316111 | 83.97631137 |
| ENSMUSG00000001418 | *Glmp* | 6969.913245 | 6893.881586 | 6938.808445 | 2813.638256 | 2969.716413 | 2625.918112 | 3.46E-99 | -1.306759263 | 6.36E-101 | 454.8448489 | 0.404227881 | 98.46080124 |
| ENSMUSG00000015837 | *Sqstm1* | 75801.42164 | 75848.09294 | 77880.7082 | 30536.1137 | 32317.22193 | 29656.80547 | 7.76E-152 | -1.310978415 | 7.94E-154 | 698.0500165 | 0.403047446 | 151.1098658 |
| ENSMUSG00000034959 | *Rubcnl* | 120.7868783 | 142.9581442 | 119.448939 | 46.50641745 | 45.73191782 | 61.58254274 | 8.62E-07 | -1.311721012 | 1.87E-07 | 27.16285206 | 0.402840039 | 6.064362779 |
| ENSMUSG00000028965 | *Tnfrsf9* | 50.18609732 | 53.8842236 | 72.4526351 | 22.09054829 | 23.8187072 | 25.01790799 | 0.000819665 | -1.312033314 | 0.000266079 | 13.29525252 | 0.402752845 | 3.086363574 |
| ENSMUSG00000111928 | *Gm48082* | 74.85384007 | 76.97746228 | 78.32717308 | 27.90385047 | 22.86595891 | 41.37577091 | 0.000208775 | -1.31759225 | 6.11E-05 | 16.06967923 | 0.40120396 | 3.680321948 |
| ENSMUSG00000094915 | *ENSMUSG00000094915* | 229.6651911 | 225.4339967 | 227.1488019 | 101.151458 | 89.55833906 | 83.71376904 | 3.05E-12 | -1.318147613 | 4.26E-13 | 52.52088248 | 0.401049547 | 11.51545733 |
| ENSMUSG00000027313 | *Chac1* | 488.2511841 | 510.2506071 | 481.7121145 | 194.1642929 | 212.4628682 | 185.7098555 | 1.55E-23 | -1.320089199 | 1.22E-24 | 105.0050976 | 0.400510175 | 22.80874164 |
| ENSMUSG00000039623 | *Ap5z1* | 1140.670449 | 1204.147446 | 1128.890382 | 479.0160998 | 425.8784847 | 480.151388 | 1.43E-43 | -1.327255538 | 6.07E-45 | 197.8754745 | 0.398525643 | 42.8439467 |
| ENSMUSG00000028300 | *C9orf72* | 2866.56183 | 2746.995725 | 2933.352632 | 1063.834299 | 1166.163904 | 1164.294949 | 7.54E-77 | -1.331221808 | 1.70E-78 | 351.8265183 | 0.397431518 | 76.1229028 |
| ENSMUSG00000100838 | *Gm29094* | 426.1565213 | 341.9998681 | 393.5940447 | 158.1218193 | 142.9122432 | 161.6541747 | 7.21E-17 | -1.331317735 | 7.73E-18 | 74.02032763 | 0.397405092 | 16.14182876 |
| ENSMUSG00000022305 | *Lrp12* | 1478.363341 | 1477.967276 | 1441.219985 | 596.4448038 | 545.924769 | 600.4297917 | 1.64E-56 | -1.336181495 | 5.25E-58 | 257.7708553 | 0.396067576 | 55.78624274 |
| ENSMUSG00000045322 | *Tlr9* | 676.2363961 | 699.3952287 | 624.6592053 | 286.0144673 | 244.85631 | 261.7258067 | 3.51E-28 | -1.337949786 | 2.36E-29 | 126.525078 | 0.395582419 | 27.45429092 |
| ENSMUSG00000005824 | *Tnfsf14* | 63.79588642 | 68.18003802 | 87.13898005 | 20.92788785 | 31.4406935 | 33.67795306 | 0.000249602 | -1.338123292 | 7.39E-05 | 15.70738077 | 0.395534847 | 3.602752599 |
| ENSMUSG00000025880 | *Smad7* | 254.3329339 | 300.2121029 | 255.5424022 | 101.151458 | 103.8495634 | 113.5428132 | 1.19E-13 | -1.342997476 | 1.51E-14 | 59.080464 | 0.394200777 | 12.92359769 |
| ENSMUSG00000000600 | *Krit1* | 3307.178752 | 3267.143435 | 3138.961461 | 1277.76382 | 1080.416558 | 1465.472072 | 1.53E-39 | -1.345145363 | 7.14E-41 | 179.2296383 | 0.393614326 | 38.81466753 |
| ENSMUSG00000025978 | *Rftn2* | 102.9240301 | 87.97424261 | 104.762594 | 52.31971964 | 33.34619008 | 31.7534986 | 2.34E-05 | -1.345578482 | 5.97E-06 | 20.49749034 | 0.393496175 | 4.631037551 |
| ENSMUSG00000020818 | *Mfsd11* | 1503.881696 | 1501.060514 | 1442.199074 | 575.516916 | 614.5226457 | 558.0917936 | 2.97E-57 | -1.346652019 | 9.32E-59 | 261.2153146 | 0.393203476 | 56.52767781 |
| ENSMUSG00000030683 | *Sez6l2* | 90.16485281 | 101.170379 | 97.90896635 | 49.99439876 | 25.72420377 | 38.48908921 | 3.84E-05 | -1.350305297 | 1.01E-05 | 19.48389911 | 0.392209042 | 4.415850181 |
| ENSMUSG00000000078 | *Klf6* | 5157.259458 | 5231.168401 | 5056.019022 | 2062.559614 | 1879.772372 | 2115.93768 | 1.28E-95 | -1.35043117 | 2.43E-97 | 438.3899239 | 0.392174824 | 94.89246804 |
| ENSMUSG00000009145 | *Dqx1* | 62.9452746 | 57.18325769 | 43.0799452 | 20.92788785 | 21.91321062 | 21.16899907 | 0.000884707 | -1.351084553 | 0.000289803 | 13.13517599 | 0.391997252 | 3.053200693 |
| ENSMUSG00000038037 | *Socs1* | 107.1770892 | 108.8681252 | 109.6580423 | 80.22357011 | 26.67695206 | 22.1312263 | 0.003198247 | -1.351151683 | 0.001150351 | 10.56843527 | 0.391979013 | 2.495088038 |
| ENSMUSG00000056204 | *Pgpep1* | 769.8036962 | 737.8839599 | 784.2508205 | 320.8942804 | 288.6827312 | 289.6303963 | 5.95E-35 | -1.352702938 | 3.14E-36 | 157.9728544 | 0.391557765 | 34.22577519 |
| ENSMUSG00000035969 | *Rusc2* | 4930.996715 | 5252.062284 | 5103.994416 | 2098.602088 | 1894.063596 | 1994.697049 | 1.31E-98 | -1.352898062 | 2.43E-100 | 452.174587 | 0.39150481 | 97.88130572 |
| ENSMUSG00000022014 | *Epsti1* | 1659.543659 | 1733.092579 | 1867.123988 | 733.6387353 | 686.9315156 | 639.8811082 | 3.84E-51 | -1.353293324 | 1.35E-52 | 232.9621408 | 0.391397562 | 50.4154755 |
| ENSMUSG00000056144 | *Trim34a* | 271.3451703 | 263.9227278 | 277.0823748 | 105.8020997 | 94.3220805 | 117.3917221 | 2.21E-14 | -1.355781381 | 2.71E-15 | 62.46782826 | 0.390723144 | 13.65511461 |
| ENSMUSG00000028838 | *Extl1* | 73.15261643 | 53.8842236 | 74.41081443 | 22.09054829 | 34.29893836 | 22.1312263 | 0.000423178 | -1.356497466 | 0.0001302 | 14.63889054 | 0.390529256 | 3.373476426 |
| ENSMUSG00000020865 | *Abcc3* | 222.0096848 | 248.5272354 | 222.2533536 | 94.17549534 | 80.03085618 | 96.22272304 | 2.07E-12 | -1.357640654 | 2.85E-13 | 53.31018599 | 0.390219924 | 11.68503794 |
| ENSMUSG00000007812 | *Zfp655* | 2281.340899 | 2284.031274 | 2202.951743 | 851.0674394 | 826.0327656 | 944.9071402 | 2.98E-67 | -1.367582962 | 7.77E-69 | 307.469234 | 0.387539975 | 66.52566263 |
| ENSMUSG00000029209 | *Gnpda2* | 717.0657634 | 673.0029559 | 675.5718678 | 239.5080499 | 273.4387586 | 282.8948057 | 4.96E-31 | -1.373508135 | 3.03E-32 | 139.743336 | 0.385951606 | 30.30447065 |
| ENSMUSG00000056832 | *Ttc26* | 254.3329339 | 337.601156 | 242.8142366 | 97.66347665 | 100.9913185 | 122.2028583 | 3.31E-11 | -1.374536141 | 4.93E-12 | 47.71521966 | 0.38567669 | 10.47956485 |
| ENSMUSG00000021939 | *Ctsb* | 122212.5037 | 120023.2592 | 122100.3138 | 46451.77241 | 47895.60918 | 46046.42188 | 1.24E-238 | -1.375783449 | 7.06E-241 | 1098.480783 | 0.385343391 | 237.9064211 |
| ENSMUSG00000071714 | *Csf2rb2* | 318.9794321 | 396.9837698 | 353.4513685 | 129.0553084 | 124.8100257 | 156.8430385 | 1.80E-15 | -1.376942223 | 2.06E-16 | 67.54082176 | 0.385034007 | 14.74527124 |
| ENSMUSG00000092224 | *Gm8815* | 362.3606349 | 469.5625199 | 422.9667346 | 180.2123676 | 193.4079024 | 109.6939043 | 1.29E-10 | -1.377222216 | 2.01E-11 | 44.96387623 | 0.384959288 | 9.889510573 |
| ENSMUSG00000049037 | *Clec4a1* | 56.99099187 | 50.5851895 | 50.9126625 | 19.76522742 | 11.43297945 | 29.82904414 | 0.002768049 | -1.377929399 | 0.000984221 | 10.85701772 | 0.384770634 | 2.557826162 |
| ENSMUSG00000078853 | *Igtp* | 1546.412287 | 1507.658583 | 1450.031792 | 527.8478381 | 596.4204282 | 606.2031551 | 1.91E-53 | -1.377982239 | 6.45E-55 | 243.6002957 | 0.384756542 | 52.71868217 |
| ENSMUSG00000079415 | *Cntf* | 59.54282733 | 58.28293573 | 36.22631755 | 22.09054829 | 17.14946918 | 20.20677184 | 0.001731986 | -1.378592294 | 0.000595511 | 11.78995584 | 0.384593879 | 2.761455502 |
| ENSMUSG00000021458 | *Aopep* | 5352.049565 | 5264.158742 | 5344.850473 | 1999.77595 | 2136.061661 | 1999.508185 | 7.89E-126 | -1.378968965 | 1.01E-127 | 578.0270644 | 0.384493479 | 125.1026985 |
| ENSMUSG00000030921 | *Trim30a* | 6698.568074 | 6444.113271 | 6298.483806 | 2507.858561 | 2321.847578 | 2645.162656 | 1.01E-95 | -1.379241281 | 1.91E-97 | 438.8694152 | 0.384420911 | 94.99524818 |
| ENSMUSG00000035954 | *Dock4* | 794.4714389 | 798.3662516 | 760.7526686 | 306.9423552 | 263.9112757 | 333.8928489 | 8.64E-31 | -1.379939135 | 5.32E-32 | 138.6251823 | 0.384235005 | 30.06334883 |
| ENSMUSG00000020134 | *Peli1* | 2169.060138 | 2037.703394 | 2161.829977 | 865.0193646 | 746.0019094 | 838.0999176 | 1.17E-61 | -1.380531073 | 3.40E-63 | 281.5790223 | 0.384077386 | 60.93125838 |
| ENSMUSG00000063383 | *Zfp947* | 118.2350428 | 98.97102293 | 100.8462353 | 47.66907789 | 22.86595891 | 51.96027044 | 6.92E-05 | -1.383428788 | 1.90E-05 | 18.2878697 | 0.383306724 | 4.15992739 |
| ENSMUSG00000028967 | *Errfi1* | 684.7425143 | 652.1090733 | 679.4882265 | 247.6466729 | 271.533262 | 253.0657616 | 9.39E-34 | -1.383718043 | 5.15E-35 | 152.4106889 | 0.38322988 | 33.02744366 |
| ENSMUSG00000034993 | *Vat1* | 8381.078252 | 8612.678351 | 8620.884487 | 3344.974075 | 3407.027877 | 3054.109229 | 6.23E-117 | -1.385374341 | 8.93E-119 | 536.8943938 | 0.382790162 | 116.2055134 |
| ENSMUSG00000020661 | *Dnmt3a* | 8419.355784 | 8453.225036 | 8385.902968 | 3279.865091 | 3134.541867 | 3253.290266 | 2.42E-167 | -1.385811367 | 2.24E-169 | 769.5619093 | 0.382674224 | 166.6167446 |
| ENSMUSG00000038987 | *Cfap157* | 31.4726373 | 37.38905311 | 32.3099589 | 12.7892648 | 14.29122432 | 11.54672676 | 0.006638175 | -1.386192127 | 0.002526096 | 9.121595314 | 0.382573241 | 2.17795129 |
| ENSMUSG00000066152 | *Slc31a2* | 2141.84056 | 2165.266046 | 2095.25188 | 852.2300998 | 745.0491611 | 840.0243721 | 7.17E-68 | -1.394524358 | 1.86E-69 | 310.3252074 | 0.380370072 | 67.14438201 |
| ENSMUSG00000070327 | *Rnf213* | 38297.94654 | 39659.88824 | 36063.78867 | 14748.34763 | 12399.06622 | 16116.34388 | 9.46E-57 | -1.398103361 | 3.01E-58 | 258.8830458 | 0.379427628 | 56.02401476 |
| ENSMUSG00000039232 | *Stx11* | 698.3523034 | 696.0961946 | 653.0528056 | 263.923919 | 239.1398203 | 271.348079 | 3.73E-33 | -1.403639667 | 2.10E-34 | 149.6206102 | 0.377974375 | 32.42781914 |
| ENSMUSG00000023206 | *Il15ra* | 203.2962247 | 265.0224059 | 212.462457 | 95.33815578 | 86.7000942 | 75.05372397 | 3.72E-11 | -1.405827008 | 5.55E-12 | 47.48300912 | 0.377401743 | 10.4295227 |
| ENSMUSG00000002799 | *Jag2* | 90.16485281 | 76.97746228 | 82.24353174 | 37.20513396 | 28.58244864 | 28.86681691 | 1.28E-05 | -1.406900066 | 3.17E-06 | 21.71095671 | 0.377121141 | 4.891335735 |
| ENSMUSG00000054051 | *Ercc6* | 1180.649205 | 1054.591233 | 1090.705885 | 446.4616076 | 345.8476285 | 463.793525 | 7.65E-30 | -1.406968995 | 4.88E-31 | 134.2231739 | 0.377103123 | 29.11624721 |
| ENSMUSG00000021709 | *Erbin* | 6547.159171 | 6378.132589 | 6347.438289 | 2528.786449 | 1935.031773 | 2792.383423 | 3.11E-37 | -1.409567317 | 1.54E-38 | 168.5456899 | 0.376424565 | 36.5069436 |
| ENSMUSG00000040483 | *Xaf1* | 2550.134233 | 2429.188774 | 2572.068546 | 920.8270656 | 1009.913185 | 909.3047327 | 2.32E-78 | -1.41042569 | 5.15E-80 | 358.7949127 | 0.376200667 | 77.63497679 |
| ENSMUSG00000086513 | *Gvin-ps1* | 244.9762039 | 311.2088832 | 270.2287471 | 110.4527415 | 72.40886988 | 127.9762216 | 9.64E-10 | -1.411003313 | 1.60E-10 | 40.89859908 | 0.376050074 | 9.016101844 |
| ENSMUSG00000033538 | *Casp4* | 846.3587599 | 910.533411 | 878.2434282 | 286.0144673 | 323.9344179 | 378.1553015 | 4.75E-31 | -1.411235956 | 2.90E-32 | 139.8330841 | 0.375989439 | 30.32311966 |
| ENSMUSG00000020399 | *Havcr2* | 1051.356208 | 1047.993165 | 1059.375016 | 365.075377 | 350.6113699 | 469.5668884 | 6.14E-31 | -1.412628131 | 3.76E-32 | 139.3151666 | 0.375626791 | 30.21181258 |
| ENSMUSG00000025432 | *Avil* | 69.75016916 | 79.17681835 | 60.70355914 | 29.06651091 | 28.58244864 | 21.16899907 | 8.81E-05 | -1.413544636 | 2.45E-05 | 17.80634404 | 0.375388241 | 4.0549429 |
| ENSMUSG00000039146 | *Ifi44l* | 4838.280026 | 4834.184631 | 4867.054717 | 1716.086804 | 1891.205351 | 1838.816237 | 2.87E-118 | -1.415620149 | 4.06E-120 | 543.0671321 | 0.374848582 | 117.5421006 |
| ENSMUSG00000036587 | *Fut7* | 102.0734183 | 92.37295474 | 86.15989039 | 29.06651091 | 45.73191782 | 29.82904414 | 1.10E-05 | -1.418033686 | 2.68E-06 | 22.03097909 | 0.374222009 | 4.959779178 |
| ENSMUSG00000049409 | *Prokr1* | 61.24405097 | 75.87778425 | 78.32717308 | 24.41586916 | 27.62970035 | 27.90458968 | 4.47E-05 | -1.424075722 | 1.19E-05 | 19.17710273 | 0.372658037 | 4.350000533 |
| ENSMUSG00000029561 | *Oasl2* | 2551.835457 | 2470.976539 | 2356.66882 | 935.9416513 | 885.1031594 | 921.8136867 | 1.77E-85 | -1.428736464 | 3.67E-87 | 391.6251131 | 0.371456078 | 84.75233814 |
| ENSMUSG00000046879 | *Irgm1* | 5956.834568 | 5992.145599 | 5773.691746 | 2134.644561 | 2114.148451 | 2328.589897 | 6.99E-118 | -1.429608695 | 9.93E-120 | 541.280912 | 0.371231569 | 117.1556182 |
| ENSMUSG00000040613 | *Apobec1* | 7488.786454 | 7178.698197 | 7291.280724 | 2653.191116 | 2491.436773 | 2984.828869 | 2.54E-84 | -1.433506334 | 5.34E-86 | 386.2799262 | 0.370229989 | 83.594391 |
| ENSMUSG00000095788 | *Sirpb1a* | 116.5338192 | 139.6591101 | 125.3234769 | 44.18109658 | 60.02314214 | 36.56463475 | 4.15E-07 | -1.434022379 | 8.72E-08 | 28.63980539 | 0.370097583 | 6.382050867 |
| ENSMUSG00000019970 | *Sgk1* | 290.0586303 | 287.0159665 | 245.7515055 | 113.9407228 | 95.27482879 | 96.22272304 | 7.78E-15 | -1.434221835 | 9.24E-16 | 64.58578883 | 0.37004642 | 14.10895454 |
| ENSMUSG00000030064 | *Frmd4b* | 529.0805514 | 493.7554366 | 490.5239214 | 201.1402555 | 190.5496576 | 169.3519925 | 3.97E-26 | -1.434956196 | 2.83E-27 | 117.0294481 | 0.369858107 | 25.40150092 |
| ENSMUSG00000036499 | *Eea1* | 7202.980883 | 6969.75937 | 6797.819534 | 2636.91387 | 2139.872655 | 2947.302007 | 2.42E-46 | -1.441144521 | 9.58E-48 | 210.717298 | 0.368275028 | 45.61697612 |
| ENSMUSG00000015659 | *Serac1* | 453.3760995 | 400.2828039 | 396.5313137 | 148.8205359 | 129.5737672 | 181.8609465 | 8.39E-18 | -1.442109242 | 8.65E-19 | 78.34530756 | 0.368028847 | 17.0762907 |
| ENSMUSG00000112328 | *Gm47325* | 48.48487368 | 38.48873114 | 49.93357284 | 29.06651091 | 13.33847603 | 8.660045073 | 0.007151316 | -1.446601095 | 0.002740994 | 8.972325894 | 0.366884766 | 2.145614007 |
| ENSMUSG00000095649 | *Gvin-ps3* | 81.65873462 | 53.8842236 | 103.7835043 | 39.53045484 | 17.14946918 | 31.7534986 | 0.000677901 | -1.448614455 | 0.000216711 | 13.68034609 | 0.366373115 | 3.168833662 |
| ENSMUSG00000053040 | *Aph1c* | 322.3818794 | 305.7104931 | 313.3086923 | 111.6154019 | 115.2825428 | 117.3917221 | 1.94E-19 | -1.450781391 | 1.83E-20 | 85.97073543 | 0.365823234 | 18.71286378 |
| ENSMUSG00000000318 | *Clec10a* | 209.2505075 | 237.530455 | 216.3788156 | 62.78366356 | 95.27482879 | 82.75154181 | 1.43E-11 | -1.452581258 | 2.07E-12 | 49.41656135 | 0.365367127 | 10.8449584 |
| ENSMUSG00000060131 | *Atp8b4* | 5992.560265 | 5863.48327 | 5815.792601 | 2111.391352 | 1975.999949 | 2366.116759 | 2.08E-85 | -1.453194894 | 4.32E-87 | 391.2984691 | 0.365211755 | 84.68266371 |
| ENSMUSG00000001313 | *Rnd2* | 422.754074 | 417.8776524 | 449.4021556 | 146.495215 | 181.974923 | 141.4474029 | 3.08E-21 | -1.454621861 | 2.66E-22 | 94.33454321 | 0.364850703 | 20.51116058 |
| ENSMUSG00000066800 | *Rnasel* | 216.9060138 | 218.8359285 | 238.8978779 | 105.8020997 | 65.73963186 | 76.0159512 | 6.17E-11 | -1.455431606 | 9.39E-12 | 46.45163876 | 0.36464598 | 10.20948533 |
| ENSMUSG00000060519 | *Tor3a* | 11923.87648 | 12074.4648 | 11940.97754 | 4519.261116 | 4281.650806 | 4303.080174 | 1.66E-191 | -1.455960165 | 1.23E-193 | 881.1516959 | 0.364512409 | 190.7805814 |
| ENSMUSG00000052373 | *Mpp3* | 51.88732096 | 61.58196982 | 47.97539351 | 26.74119004 | 16.19672089 | 16.35786292 | 0.000616762 | -1.456636323 | 0.000195473 | 13.87410832 | 0.364341611 | 3.209882618 |
| ENSMUSG00000052397 | *Ezr* | 3237.428583 | 3261.645045 | 3194.769572 | 1282.414461 | 1150.919932 | 1094.052361 | 1.70E-83 | -1.460195591 | 3.59E-85 | 382.4786314 | 0.363443853 | 82.76964315 |
| ENSMUSG00000020108 | *Ddit4* | 314.726373 | 377.1895652 | 376.9495205 | 132.5432897 | 125.762774 | 128.9384489 | 3.43E-19 | -1.463815174 | 3.29E-20 | 84.8053001 | 0.36253315 | 18.46472555 |
| ENSMUSG00000015745 | *Plekho1* | 11054.5512 | 10941.79642 | 11546.4044 | 3953.045484 | 4473.153212 | 3716.121564 | 8.20E-88 | -1.465793038 | 1.68E-89 | 402.3736686 | 0.362036475 | 87.0864169 |
| ENSMUSG00000057948 | *Unc13d* | 1171.292475 | 987.5108733 | 1107.350409 | 375.5393209 | 545.924769 | 255.9524433 | 2.42E-05 | -1.471891214 | 6.18E-06 | 20.43169462 | 0.360509401 | 4.617040509 |
| ENSMUSG00000022350 | *Washc5* | 4701.331524 | 4529.573816 | 4660.466798 | 1652.14048 | 1673.025994 | 1676.199835 | 3.11E-147 | -1.473814616 | 3.33E-149 | 676.7900189 | 0.36002909 | 146.5065574 |
| ENSMUSG00000021806 | *Nid2* | 165.0186929 | 174.8488072 | 172.3197808 | 74.41026793 | 47.63741439 | 62.54476997 | 6.32E-10 | -1.479554679 | 1.04E-10 | 41.7508684 | 0.358599485 | 9.19950337 |
| ENSMUSG00000042770 | *Hebp1* | 124.1893256 | 112.1671593 | 107.699863 | 43.01843614 | 40.01542809 | 40.41354368 | 4.04E-08 | -1.482081662 | 7.73E-09 | 33.34068526 | 0.357971922 | 7.394046422 |
| ENSMUSG00000046718 | *Bst2* | 12760.87851 | 13204.93382 | 13959.86042 | 4866.896587 | 4792.323888 | 4618.690706 | 5.07E-152 | -1.483655142 | 5.15E-154 | 698.9147102 | 0.357581712 | 151.2949751 |
| ENSMUSG00000070031 | *Sp140* | 1417.11929 | 1444.976935 | 1405.972757 | 511.570592 | 510.6730823 | 502.2826142 | 2.93E-72 | -1.485246537 | 7.09E-74 | 330.6075464 | 0.357187491 | 71.53250059 |
| ENSMUSG00000032589 | *Bsn* | 97.82035918 | 100.070701 | 92.03442837 | 31.39183178 | 35.25168665 | 36.56463475 | 4.12E-07 | -1.4862017 | 8.66E-08 | 28.6534699 | 0.356951087 | 6.38468973 |
| ENSMUSG00000051413 | *Plagl2* | 4650.294814 | 4995.837302 | 4577.244177 | 1777.707807 | 1601.569872 | 1697.368834 | 3.45E-103 | -1.486883284 | 6.11E-105 | 473.3059407 | 0.35678249 | 102.4616638 |
| ENSMUSG00000028385 | *Snx30* | 3821.798903 | 3945.644781 | 3731.310708 | 1452.162885 | 1305.265154 | 1343.269214 | 4.67E-106 | -1.488671433 | 7.92E-108 | 486.5766219 | 0.356340549 | 105.3304127 |
| ENSMUSG00000044811 | *Cd300c2* | 817.437958 | 754.3791303 | 707.8818267 | 262.7612586 | 256.2892894 | 291.5548508 | 1.35E-35 | -1.491782126 | 7.01E-37 | 160.9523236 | 0.355573047 | 34.86936219 |
| ENSMUSG00000073412 | *Lst1* | 1585.540431 | 1518.655363 | 1612.560676 | 549.9383864 | 598.3259248 | 525.3760678 | 3.93E-65 | -1.494630098 | 1.06E-66 | 297.6633197 | 0.354871815 | 64.40505562 |
| ENSMUSG00000078937 | *Cpt1b* | 91.86607645 | 50.5851895 | 84.20171106 | 29.06651091 | 35.25168665 | 16.35786292 | 0.000414517 | -1.496598911 | 0.000127421 | 14.67955337 | 0.35438786 | 3.382458009 |
| ENSMUSG00000020577 | *Tspan13* | 92.71668827 | 52.78454556 | 78.32717308 | 43.01843614 | 23.8187072 | 13.47118123 | 0.001393145 | -1.49783798 | 0.000471838 | 12.22379803 | 0.354083622 | 2.856003643 |
| ENSMUSG00000028249 | *Sdcbp* | 27404.16097 | 26581.4174 | 27359.68156 | 9059.45012 | 9971.463581 | 9741.58848 | 8.36E-172 | -1.499116184 | 7.52E-174 | 790.1385233 | 0.353770049 | 171.0775523 |
| ENSMUSG00000034422 | *Parp14* | 9269.116991 | 8934.884015 | 8467.16741 | 3226.382711 | 2666.742458 | 3536.185072 | 1.15E-56 | -1.500269107 | 3.69E-58 | 258.4723045 | 0.353487448 | 55.93821712 |
| ENSMUSG00000083594 | *Gm13722* | 155.6619629 | 144.0578223 | 143.9261805 | 49.99439876 | 53.35390412 | 52.92249767 | 1.79E-10 | -1.504719356 | 2.81E-11 | 44.30316586 | 0.352398733 | 9.747336812 |
| ENSMUSG00000039081 | *Zfp503* | 193.0888829 | 219.9356065 | 283.9360024 | 102.3141184 | 68.59787673 | 75.05372397 | 4.72E-10 | -1.508679376 | 7.67E-11 | 42.34122119 | 0.351432769 | 9.326236029 |
| ENSMUSG00000005949 | *Ctns* | 902.4991399 | 869.8453238 | 862.5779936 | 312.7556574 | 288.6827312 | 314.6483043 | 9.84E-49 | -1.525567667 | 3.67E-50 | 221.7950595 | 0.347342857 | 48.00717774 |
| ENSMUSG00000048752 | *Prss50* | 71.45139279 | 83.57553048 | 88.11806972 | 17.43990655 | 34.29893836 | 31.7534986 | 2.39E-05 | -1.526895269 | 6.10E-06 | 20.4565179 | 0.347023371 | 4.622207297 |
| ENSMUSG00000074802 | *Gas2l3* | 6297.929908 | 6052.627891 | 5988.112382 | 2120.692636 | 1918.835052 | 2320.89208 | 6.99E-89 | -1.527852444 | 1.40E-90 | 407.3196534 | 0.34679321 | 88.15571515 |
| ENSMUSG00000079138 | *Gm8818* | 59.54282733 | 86.87456457 | 87.13898005 | 43.01843614 | 14.29122432 | 24.05568076 | 0.000508272 | -1.535299312 | 0.000158438 | 14.2691779 | 0.345007753 | 3.29390423 |
| ENSMUSG00000042286 | *Stab1* | 196.4913302 | 195.7426898 | 177.2152291 | 75.57292836 | 72.40886988 | 49.07358875 | 5.49E-11 | -1.536070964 | 8.32E-12 | 46.6883414 | 0.344823269 | 10.26017875 |
| ENSMUSG00000021703 | *Serinc5* | 25.51835457 | 41.78776524 | 30.35177957 | 10.46394393 | 13.33847603 | 9.622272304 | 0.00555044 | -1.537863312 | 0.00207713 | 9.480096431 | 0.34439514 | 2.255672587 |
| ENSMUSG00000062980 | *Cped1* | 335.9916685 | 314.5079173 | 321.1414096 | 119.7540249 | 87.65284249 | 127.9762216 | 7.28E-17 | -1.537908361 | 7.80E-18 | 74.00171813 | 0.344384386 | 16.1380125 |
| ENSMUSG00000069633 | *Pex11g* | 45.08242641 | 39.58840917 | 41.12176587 | 10.46394393 | 18.10221747 | 14.43340846 | 0.001098721 | -1.538884106 | 0.00036526 | 12.70202238 | 0.344151546 | 2.959112505 |
| ENSMUSG00000045827 | *Serpinb9* | 1391.600936 | 1436.179511 | 1371.704619 | 454.6002306 | 472.5631508 | 514.7915682 | 4.05E-66 | -1.540266163 | 1.06E-67 | 302.2515069 | 0.343822017 | 65.39237016 |
| ENSMUSG00000052749 | *Trim30b* | 28.92080185 | 32.99034098 | 27.41451058 | 11.62660436 | 10.48023117 | 8.660045073 | 0.005525474 | -1.541521604 | 0.002066271 | 9.489713399 | 0.343522951 | 2.257630466 |
| ENSMUSG00000007659 | *Bcl2l1* | 7746.521835 | 7985.861873 | 7989.371654 | 2796.198349 | 2659.120471 | 2668.25611 | 6.27E-185 | -1.546459812 | 4.99E-187 | 850.7531563 | 0.342349115 | 184.2030506 |
| ENSMUSG00000028789 | *Azin2* | 48.48487368 | 46.18647737 | 53.84993149 | 23.25320873 | 11.43297945 | 16.35786292 | 0.000532823 | -1.556249582 | 0.00016664 | 14.17419967 | 0.340033883 | 3.273416642 |
| ENSMUSG00000045312 | *Lhfpl2* | 4474.218168 | 4387.71535 | 4314.848147 | 1503.319944 | 1448.177398 | 1523.205706 | 4.22E-150 | -1.558476105 | 4.40E-152 | 690.0306151 | 0.339509511 | 149.3746303 |
| ENSMUSG00000034708 | *Grn* | 26652.22012 | 27246.72261 | 26492.20812 | 9305.934132 | 9123.517605 | 8820.737021 | 2.32E-247 | -1.560905416 | 1.24E-249 | 1138.765179 | 0.338938302 | 246.6348902 |
| ENSMUSG00000090222 | *Ifi203-ps* | 268.7933348 | 258.4243377 | 246.7305952 | 102.3141184 | 81.93635276 | 78.90263289 | 2.21E-16 | -1.563215039 | 2.44E-17 | 71.75216111 | 0.338396127 | 15.65505339 |
| ENSMUSG00000017417 | *Plxdc1* | 470.3883359 | 478.3599442 | 523.81297 | 172.0737446 | 169.5891952 | 154.9185841 | 6.82E-30 | -1.569424371 | 4.33E-31 | 134.4637557 | 0.336942806 | 29.16653095 |
| ENSMUSG00000026822 | *Lcn2* | 52.73793278 | 39.58840917 | 40.1426762 | 13.95192524 | 8.574734591 | 22.1312263 | 0.002007689 | -1.570341014 | 0.000697745 | 11.49524266 | 0.336728791 | 2.69730365 |
| ENSMUSG00000112557 | *Gm47626* | 101.2228065 | 129.7620078 | 86.15989039 | 61.62100313 | 14.29122432 | 31.7534986 | 0.000651874 | -1.572108261 | 0.000207496 | 13.76195872 | 0.336316564 | 3.185836644 |
| ENSMUSG00000038085 | *Cnbd2* | 92.71668827 | 80.27649638 | 97.90896635 | 26.74119004 | 29.53519692 | 34.64018029 | 6.14E-07 | -1.572118545 | 1.32E-07 | 27.84312488 | 0.336314166 | 6.211588731 |
| ENSMUSG00000031883 | *Car7* | 33.17386094 | 32.99034098 | 30.35177957 | 9.301283491 | 7.621986303 | 15.39563569 | 0.004679947 | -1.573626726 | 0.001730174 | 9.815817179 | 0.33596277 | 2.329759082 |
| ENSMUSG00000053687 | *Dpep2* | 1099.841082 | 1182.153885 | 1103.434051 | 347.6354705 | 409.6817638 | 375.2686198 | 5.25E-54 | -1.576553741 | 1.76E-55 | 246.189134 | 0.335281842 | 53.27957315 |
| ENSMUSG00000027398 | *Il1b* | 96.11913554 | 76.97746228 | 103.7835043 | 34.87981309 | 30.48794521 | 27.90458968 | 7.39E-07 | -1.576976367 | 1.60E-07 | 27.47036498 | 0.335183639 | 6.131226158 |
| ENSMUSG00000062585 | *Cnr2* | 393.8332722 | 391.4853796 | 355.4095479 | 134.8686106 | 114.3297945 | 133.749585 | 1.28E-23 | -1.577149273 | 1.00E-24 | 105.3899943 | 0.335143469 | 22.89234036 |
| ENSMUSG00000042268 | *Slc26a9* | 49.3354855 | 41.78776524 | 41.12176587 | 10.46394393 | 18.10221747 | 15.39563569 | 0.000670847 | -1.579040484 | 0.000214179 | 13.70240974 | 0.334704422 | 3.1733764 |
| ENSMUSG00000072620 | *Slfn2* | 13564.70668 | 13222.52866 | 13718.02528 | 4162.324362 | 5005.739505 | 4357.927126 | 2.90E-104 | -1.581933186 | 5.03E-106 | 478.292917 | 0.334033989 | 103.5383393 |
| ENSMUSG00000022848 | *Slc49a4* | 281.5525121 | 332.1027658 | 302.538706 | 95.33815578 | 105.75506 | 103.9205409 | 8.62E-20 | -1.58243543 | 7.96E-21 | 87.61323338 | 0.333917722 | 19.06428309 |
| ENSMUSG00000021792 | *Prxl2a* | 1412.01562 | 1386.693999 | 1263.025666 | 409.2564736 | 447.7916953 | 494.5847964 | 8.55E-54 | -1.585163409 | 2.88E-55 | 245.2096256 | 0.333286917 | 53.06779308 |
| ENSMUSG00000082593 | *Gm11331* | 91.86607645 | 72.57875015 | 101.825325 | 32.55449222 | 27.62970035 | 28.86681691 | 1.18E-06 | -1.586608074 | 2.61E-07 | 26.52008798 | 0.332953342 | 5.927274813 |
| ENSMUSG00000043157 | *Arl11* | 1776.077478 | 1655.015439 | 1740.821422 | 513.8959129 | 671.687543 | 534.0361129 | 5.30E-48 | -1.586805456 | 2.02E-49 | 218.4004163 | 0.332907792 | 47.27541496 |
| ENSMUSG00000026821 | *Ralgds* | 3113.239257 | 3174.77048 | 3058.676109 | 1069.647601 | 1003.243947 | 1038.243182 | 1.71E-127 | -1.587769595 | 2.14E-129 | 585.7154553 | 0.332685387 | 126.768009 |
| ENSMUSG00000005045 | *Chd5* | 165.0186929 | 178.1478413 | 193.8597534 | 59.29568225 | 58.11764556 | 60.62031551 | 5.38E-13 | -1.591878127 | 7.18E-14 | 56.01811628 | 0.331739308 | 12.26895495 |
| ENSMUSG00000041695 | *Kcnj2* | 3719.725484 | 3487.079041 | 3360.235725 | 1228.932081 | 984.1889814 | 1286.497807 | 7.76E-57 | -1.595181423 | 2.45E-58 | 259.2879528 | 0.330980603 | 56.11039593 |
| ENSMUSG00000112105 | *Rps7-ps2* | 60.39343915 | 70.37939408 | 63.64082813 | 12.7892648 | 15.24397261 | 35.60240752 | 0.000465164 | -1.598169142 | 0.000143884 | 14.4506045 | 0.330295875 | 3.332393639 |
| ENSMUSG00000084408 | *Gm11870* | 39.97875549 | 39.58840917 | 40.1426762 | 11.62660436 | 11.43297945 | 16.35786292 | 0.000842545 | -1.598177523 | 0.000274605 | 13.23612817 | 0.330293956 | 3.07440712 |
| ENSMUSG00000035112 | *Wnk4* | 41.67997913 | 42.88744327 | 40.1426762 | 18.60256698 | 10.48023117 | 12.50895399 | 0.000845222 | -1.598523375 | 0.000275593 | 13.22939231 | 0.330214786 | 3.073029252 |
| ENSMUSG00000031762 | *Mt2* | 23802.67053 | 22855.70823 | 25505.28574 | 7884.000419 | 8584.262074 | 7284.060134 | 2.03E-108 | -1.603202681 | 3.35E-110 | 497.4886151 | 0.329145486 | 107.6922328 |
| ENSMUSG00000052485 | *Tmem171* | 1499.628637 | 1475.76792 | 1463.739047 | 498.7813272 | 515.4368237 | 440.7000715 | 2.54E-69 | -1.610224751 | 6.41E-71 | 317.0345361 | 0.32754732 | 68.5954163 |
| ENSMUSG00000041845 | *Rhod* | 159.0644101 | 126.4629737 | 147.8425392 | 43.01843614 | 60.97589042 | 37.52686198 | 5.14E-09 | -1.613210082 | 9.10E-10 | 37.50848749 | 0.326870235 | 8.289091203 |
| ENSMUSG00000062007 | *Hsh2d* | 250.0798748 | 182.5465534 | 219.3160846 | 60.45834269 | 70.5033733 | 81.78931458 | 1.12E-12 | -1.613545842 | 1.52E-13 | 54.53955525 | 0.326794171 | 11.94914523 |
| ENSMUSG00000024423 | *Impact* | 3305.477529 | 3029.61298 | 3006.784357 | 1059.183658 | 855.5679625 | 1137.352586 | 9.51E-55 | -1.614760961 | 3.14E-56 | 249.6229441 | 0.326519043 | 54.0219248 |
| ENSMUSG00000042182 | *Bend6* | 232.2170266 | 263.9227278 | 259.4587608 | 74.41026793 | 90.51108735 | 80.82708735 | 3.05E-17 | -1.615180964 | 3.23E-18 | 75.74385938 | 0.326423999 | 16.51508338 |
| ENSMUSG00000026437 | *Cdk18* | 6574.378749 | 6741.02634 | 6697.952388 | 2228.820056 | 2109.384709 | 2193.878085 | 2.04E-192 | -1.615722399 | 1.48E-194 | 885.3725001 | 0.326301517 | 191.6900348 |
| ENSMUSG00000029298 | *Gbp9* | 386.1777658 | 405.781194 | 358.3468169 | 111.6154019 | 113.3770463 | 149.1452207 | 2.52E-21 | -1.61659514 | 2.17E-22 | 94.74467319 | 0.326104185 | 20.5985118 |

| | | | | | | | | | | | | | |
|---|---|---|---|---|---|---|---|---|---|---|---|---|---|
| ENSMUSG00000025357 | Dgka | 60.39343915 | 80.27649638 | 67.55718678 | 27.90385047 | 20.00771405 | 20.20677184 | 1.40E-05 | -1.618059224 | 3.48E-06 | 21.53240083 | 0.325773414 | 4.852713667 |
| ENSMUSG00000042349 | Ikbke | 1261.457328 | 1256.931991 | 1275.753832 | 418.5577571 | 399.2015326 | 415.6821635 | 2.70E-75 | -1.62181856 | 6.22E-77 | 344.6425138 | 0.324925627 | 74.56892698 |
| ENSMUSG00000029158 | Yipf7 | 341.0953394 | 381.5882773 | 377.9286101 | 117.4287041 | 119.093536 | 120.2784038 | 3.13E-25 | -1.623293503 | 2.28E-26 | 112.8891448 | 0.324593608 | 24.50508063 |
| ENSMUSG00000027583 | Zbtb46 | 54.43915641 | 70.37939408 | 70.49445577 | 27.90385047 | 17.14946918 | 18.28231738 | 3.43E-05 | -1.634824158 | 8.99E-06 | 19.71384214 | 0.322009653 | 4.464836532 |
| ENSMUSG00000046798 | Cldn12 | 451.6748759 | 511.3502851 | 512.063894 | 153.4711776 | 158.1562158 | 160.6919475 | 8.65E-32 | -1.640671983 | 5.12E-33 | 143.2723402 | 0.32070706 | 31.06274661 |
| ENSMUSG00000027068 | Dhrs9 | 5087.509289 | 4839.683021 | 4805.372069 | 1560.290306 | 1464.374118 | 1687.746562 | 1.58E-118 | -1.644609175 | 2.23E-120 | 544.264256 | 0.319833026 | 117.8004158 |
| ENSMUSG00000031712 | Il15 | 155.6619629 | 148.4565344 | 132.1771046 | 54.64504051 | 41.92092467 | 43.30022537 | 7.63E-11 | -1.649005066 | 1.17E-11 | 46.02271632 | 0.318859978 | 10.11733907 |
| ENSMUSG00000033066 | Gas7 | 8109.733082 | 8055.141589 | 7675.083872 | 2619.473963 | 2403.78393 | 2580.693432 | 1.61E-167 | -1.649108821 | 1.47E-169 | 770.4075501 | 0.318837048 | 166.7941289 |
| ENSMUSG00000025466 | Fuom | 299.4153603 | 280.4178983 | 302.538706 | 79.06090967 | 113.3770463 | 87.56267796 | 3.80E-18 | -1.65079276 | 3.85E-19 | 79.94718305 | 0.318465112 | 17.4199625 |
| ENSMUSG00000009739 | Pou6f1 | 48.48487368 | 39.58840917 | 37.20540721 | 15.11458567 | 12.38572774 | 12.50895399 | 0.000461996 | -1.655420949 | 0.000142809 | 14.46473096 | 0.317445108 | 3.335361791 |
| ENSMUSG00000058818 | Pirb | 222.8602966 | 180.3471973 | 158.6125255 | 69.75962618 | 49.54291097 | 59.65808828 | 2.41E-11 | -1.658669414 | 3.54E-12 | 48.3612933 | 0.316731132 | 10.61803374 |
| ENSMUSG00000079222 | ENSMUSG00000079222 | 74.00322825 | 87.97424261 | 73.43172477 | 33.71715265 | 15.24397261 | 25.98013522 | 1.20E-05 | -1.662969787 | 2.94E-06 | 21.85360673 | 0.315788429 | 4.922174784 |
| ENSMUSG00000045136 | Tubb2b | 247.5280393 | 234.2314209 | 269.2496575 | 87.19953273 | 79.07810789 | 71.20481505 | 6.20E-18 | -1.665597052 | 6.33E-19 | 78.96366225 | 0.315213876 | 17.20787493 |
| ENSMUSG00000045374 | Wdr81 | 3518.130483 | 3669.625595 | 3662.774431 | 1221.956119 | 1099.471524 | 1097.90127 | 4.58E-124 | -1.667399122 | 5.94E-126 | 569.887586 | 0.314820388 | 123.3391133 |
| ENSMUSG00000035891 | Cerk | 1963.212078 | 1954.127864 | 1996.363824 | 620.860673 | 627.8611217 | 611.9765185 | 2.02E-110 | -1.668266757 | 3.15E-112 | 506.7992596 | 0.314631112 | 109.6938387 |
| ENSMUSG00000063286 | Gvin-ps7 | 5469.433996 | 5167.387075 | 4959.089146 | 1648.652499 | 1233.809033 | 1996.621503 | 1.99E-15 | -1.676561405 | 2.30E-16 | 67.32974589 | 0.312827358 | 14.70084759 |
| ENSMUSG00000094728 | ENSMUSG00000094728 | 517.1719859 | 539.941914 | 565.9138255 | 163.9351215 | 192.4551542 | 148.1829935 | 9.41E-32 | -1.684027294 | 5.61E-33 | 143.0931178 | 0.311212671 | 31.02656982 |
| ENSMUSG00000055994 | Nod2 | 532.4829987 | 525.6460996 | 544.3738529 | 186.0256698 | 163.8727055 | 149.1452207 | 8.91E-35 | -1.687640368 | 4.74E-36 | 157.151162 | 0.310434248 | 34.05016384 |
| ENSMUSG00000036526 | Card11 | 685.5931261 | 608.121952 | 718.651813 | 230.2067664 | 206.7463785 | 189.5587644 | 1.40E-36 | -1.687747419 | 7.08E-38 | 165.5093015 | 0.310411214 | 35.85324385 |
| ENSMUSG00000019302 | Atp6v0a1 | 3150.666178 | 3238.551806 | 3218.267724 | 1005.701277 | 1034.684641 | 942.0204585 | 4.69E-134 | -1.68775487 | 5.67E-136 | 615.9557328 | 0.31040961 | 133.3285587 |
| ENSMUSG00000028927 | Padi2 | 609.0380624 | 602.6235619 | 565.9138255 | 186.0256698 | 201.982637 | 163.5786292 | 9.89E-37 | -1.688619434 | 4.97E-38 | 166.2135281 | 0.310223647 | 36.00470917 |
| ENSMUSG00000022372 | Sla | 313.8757612 | 284.8166104 | 302.538706 | 76.7355888 | 98.13307365 | 102.9583136 | 1.87E-20 | -1.692156062 | 1.67E-21 | 90.70156055 | 0.309464095 | 19.72848945 |
| ENSMUSG00000036636 | Clcn7 | 2796.811661 | 3003.220707 | 2696.412933 | 921.989726 | 811.7415413 | 895.8335515 | 3.97E-96 | -1.692883556 | 7.43E-98 | 440.7517731 | 0.309308084 | 95.40175215 |
| ENSMUSG00000040852 | Plekhh2 | 85.91179372 | 80.27649638 | 93.01351804 | 32.55449222 | 15.24397261 | 32.71572583 | 4.18E-06 | -1.695683846 | 9.78E-07 | 23.97130888 | 0.308708295 | 5.379298473 |
| ENSMUSG00000098188 | Sowahc | 740.8828943 | 888.5398503 | 823.414407 | 245.3213521 | 232.4705822 | 278.0836696 | 1.10E-40 | -1.696181427 | 5.03E-42 | 184.5080141 | 0.308601841 | 39.95999416 |
| ENSMUSG00000041992 | Rapgef5 | 2018.501846 | 2014.610156 | 1834.814029 | 619.6980126 | 566.8852313 | 623.5232453 | 7.54E-87 | -1.697632675 | 1.55E-88 | 397.931838 | 0.308291566 | 86.12239208 |
| ENSMUSG00000050106 | Tmc8 | 384.4765422 | 437.671857 | 390.6567758 | 126.7299876 | 131.4792637 | 115.4672676 | 7.39E-28 | -1.697640832 | 5.00E-29 | 125.0367618 | 0.308289823 | 27.13123058 |
| ENSMUSG00000020605 | Hs1bp3 | 749.3890125 | 806.0639979 | 784.2508205 | 244.1586916 | 239.1398203 | 235.7456714 | 4.15E-53 | -1.702108557 | 1.41E-54 | 242.0494601 | 0.30733659 | 52.38230868 |
| ENSMUSG00000052142 | Rasal3 | 1001.170111 | 1064.488336 | 983.0060222 | 326.7075826 | 326.7926627 | 283.857033 | 1.94E-57 | -1.70241408 | 6.07E-59 | 262.070653 | 0.307271512 | 56.7122161 |
| ENSMUSG00000041754 | Trem3 | 479.7450659 | 453.0673494 | 478.7748455 | 148.8205359 | 140.0539983 | 145.2963118 | 7.06E-35 | -1.70286232 | 3.75E-36 | 157.6199489 | 0.307176058 | 34.15091622 |
| ENSMUSG00000026031 | Cflar | 10605.42816 | 10637.18561 | 10019.02453 | 3254.286561 | 2965.90542 | 3372.606442 | 2.46E-151 | -1.704578398 | 2.53E-153 | 695.7376019 | 0.306810891 | 150.6099178 |
| ENSMUSG00000039783 | Kmo | 121.6374901 | 115.4661934 | 99.86714568 | 29.06651091 | 40.96817638 | 32.71572583 | 5.54E-09 | -1.707671856 | 9.83E-10 | 37.35870652 | 0.306153726 | 8.256411567 |
| ENSMUSG00000061175 | Fnip2 | 10975.4443 | 11343.17891 | 10701.45002 | 3514.722499 | 3061.180249 | 3508.280482 | 1.08E-138 | -1.711625998 | 1.23E-140 | 637.403313 | 0.305315767 | 137.9678459 |
| ENSMUSG00000055632 | Hmcn2 | 41.67997913 | 49.48551147 | 47.97539351 | 20.92788785 | 9.527482879 | 12.50895399 | 0.000347975 | -1.712584432 | 0.000105629 | 15.03332621 | 0.305113003 | 3.458451897 |
| ENSMUSG00000059089 | Fcgr4 | 144.6040092 | 146.2571783 | 146.8634495 | 36.04247353 | 41.92092467 | 54.84695213 | 3.31E-11 | -1.713244237 | 4.91E-12 | 47.72161575 | 0.304973494 | 10.48078128 |
| ENSMUSG00000042677 | Zc3h12a | 2516.960372 | 2590.841445 | 2551.507663 | 824.3262494 | 784.1118409 | 728.4060134 | 1.09E-114 | -1.714051524 | 1.59E-116 | 526.5471005 | 0.304802888 | 113.9626504 |
| ENSMUSG00000026958 | Dpp7 | 3176.184532 | 3193.465007 | 3203.581379 | 974.3094456 | 1028.015403 | 905.4558238 | 3.58E-127 | -1.71910856 | 4.52E-129 | 584.224598 | 0.303736341 | 126.4460887 |
| ENSMUSG00000079800 | ENSMUSG00000079800 | 497.6079141 | 499.2538268 | 504.2311767 | 161.6098007 | 150.5342295 | 144.3340846 | 3.08E-37 | -1.719905787 | 1.52E-38 | 168.5696101 | 0.303568545 | 36.51156467 |
| ENSMUSG00000074874 | Ctla2b | 39.97875549 | 37.38905311 | 29.37268991 | 9.301283491 | 11.43297945 | 11.54672676 | 0.000914851 | -1.720404612 | 0.000300556 | 13.06692363 | 0.303463601 | 3.038649439 |
| ENSMUSG00000004837 | Grap | 4125.467322 | 4139.188115 | 4460.732507 | 1234.745383 | 1397.681738 | 1226.839719 | 1.17E-113 | -1.720515238 | 1.73E-115 | 521.7899629 | 0.303440332 | 112.9317467 |
| ENSMUSG00000090231 | Cfb | 114.8325956 | 102.270057 | 93.9926077 | 39.53045484 | 28.58244864 | 26.94236245 | 2.32E-08 | -1.722166356 | 4.35E-09 | 34.45973713 | 0.303093253 | 7.635363927 |
| ENSMUSG00000024330 | Col11a2 | 228.8145793 | 273.8198301 | 213.4415467 | 72.08494705 | 70.5033733 | 74.09149674 | 4.52E-17 | -1.722559727 | 4.82E-18 | 74.95505672 | 0.303010622 | 16.34522887 |
| ENSMUSG00000034731 | Dgkh | 133.5460556 | 188.0449436 | 146.8634495 | 44.18109658 | 50.49565926 | 46.18690706 | 3.14E-11 | -1.727643199 | 4.64E-12 | 47.83366493 | 0.301944814 | 10.50359264 |
| ENSMUSG00000001627 | Ifrd1 | 11983.41931 | 11803.944 | 12028.11652 | 3790.273022 | 3365.106953 | 3655.501248 | 1.04E-185 | -1.728712235 | 8.13E-188 | 854.3771871 | 0.301721156 | 184.9833666 |
| ENSMUSG00000006219 | Fblim1 | 312.1745376 | 303.511137 | 318.2041406 | 90.68751403 | 87.65284249 | 102.9583136 | 4.39E-24 | -1.730247542 | 3.38E-25 | 107.5456518 | 0.301400237 | 23.35707318 |
| ENSMUSG00000079243 | Xirp1 | 51.03670914 | 56.08357966 | 42.10085553 | 6.975962618 | 25.72420377 | 11.54672676 | 0.001068438 | -1.734733263 | 0.000354313 | 12.75893079 | 0.30046456 | 2.971250487 |
| ENSMUSG00000028238 | Atp6v0d2 | 49.3354855 | 47.2861554 | 45.03812452 | 12.7892648 | 20.96046233 | 8.660045073 | 0.000313442 | -1.735765159 | 9.45E-05 | 15.24436158 | 0.300249727 | 3.503843312 |
| ENSMUSG00000020696 | Rffl | 1164.48758 | 1309.716537 | 1125.953113 | 358.0994144 | 341.0838871 | 379.1175288 | 1.02E-60 | -1.738386407 | 3.01E-62 | 277.2301395 | 0.299704696 | 59.99068241 |
| ENSMUSG00000073902 | Gvin3 | 153.1101274 | 151.7555685 | 141.9680012 | 49.99439876 | 39.0626798 | 45.22467983 | 1.76E-12 | -1.739878388 | 2.42E-13 | 53.63180668 | 0.299394913 | 11.75332029 |
| ENSMUSG00000020261 | Slc36a1 | 1005.42317 | 991.9095854 | 1101.475871 | 318.5689596 | 280.1079966 | 328.1194856 | 8.10E-57 | -1.742618925 | 2.57E-58 | 259.1960823 | 0.298826723 | 56.09131119 |
| ENSMUSG00000049307 | Fut4 | 102.0734183 | 78.07714031 | 54.82902116 | 32.55449222 | 17.14946918 | 21.16899907 | 2.54E-05 | -1.744854946 | 6.52E-06 | 20.3299911 | 0.298363933 | 4.595586544 |
| ENSMUSG00000031387 | Renbp | 1929.187605 | 1907.941387 | 1918.036651 | 567.3782929 | 627.8611217 | 521.5271589 | 3.24E-86 | -1.744872222 | 6.68E-88 | 395.020502 | 0.29836036 | 85.49006217 |
| ENSMUSG00000043740 | B430306N03Rik | 327.4855503 | 403.581838 | 301.5596164 | 98.82613709 | 81.93635276 | 126.0517672 | 4.40E-17 | -1.74872022 | 4.69E-18 | 75.00767166 | 0.297565625 | 16.35624332 |
| ENSMUSG00000055172 | C1ra | 550.3458469 | 494.8551147 | 520.875701 | 162.7724611 | 156.2507192 | 147.2207662 | 8.88E-38 | -1.751005863 | 4.33E-39 | 171.0671364 | 0.297094569 | 37.05158216 |
| ENSMUSG00000020189 | Osbpl8 | 10145.24716 | 10203.91246 | 9903.491947 | 3107.791346 | 2584.806105 | 3251.365811 | 1.49E-99 | -1.758398153 | 2.72E-101 | 456.5404093 | 0.295576166 | 98.82816843 |
| ENSMUSG00000021114 | Atp6v1d | 2348.539232 | 2328.018395 | 2388.978779 | 655.7404861 | 703.1282365 | 723.5948772 | 1.66E-122 | -1.76086131 | 2.20E-124 | 562.6753587 | 0.29507195 | 121.7793376 |
| ENSMUSG00000039158 | Akna | 3575.121475 | 3709.214004 | 3546.262761 | 1154.521813 | 962.2757708 | 1078.656725 | 1.84E-109 | -1.762569946 | 2.91E-111 | 502.3611974 | 0.294722693 | 108.7358094 |
| ENSMUSG00000028525 | Pde4b | 1227.432855 | 1175.555817 | 1099.517692 | 358.0994144 | 318.2179282 | 356.9863025 | 7.59E-66 | -1.762922796 | 2.01E-67 | 300.9849594 | 0.294650619 | 65.1198255 |
| ENSMUSG00000028494 | Plin2 | 9542.163385 | 8886.498181 | 9220.087362 | 2578.780848 | 2832.52066 | 2725.989744 | 3.38E-186 | -1.764105055 | 2.62E-188 | 856.6367808 | 0.294409258 | 185.4707769 |
| ENSMUSG00000096292 | None | 80.8081228 | 67.08035999 | 70.49445577 | 22.09054829 | 7.621986303 | 34.64018029 | 0.000277873 | -1.765758807 | 8.30E-05 | 15.48922854 | 0.294071972 | 3.556153678 |
| ENSMUSG00000026896 | Ifih1 | 3319.087318 | 3393.606409 | 3288.76218 | 1012.67724 | 885.1031594 | 1044.016545 | 2.21E-112 | -1.76612882 | 3.30E-114 | 515.8999515 | 0.29399656 | 111.6563093 |
| ENSMUSG00000014686 | Ceacam16 | 70.60078098 | 80.27649638 | 65.59900746 | 24.41586916 | 19.05496576 | 20.20677184 | 8.85E-07 | -1.770513447 | 1.92E-07 | 27.10791377 | 0.293104405 | 6.052844666 |
| ENSMUSG00000105867 | Gm42517 | 53.5885446 | 58.28293573 | 69.51536611 | 16.27724611 | 12.38572774 | 24.05568076 | 2.49E-05 | -1.779424569 | 6.38E-06 | 20.36958728 | 0.291299561 | 4.603992837 |
| ENSMUSG00000036534 | Slc38a7 | 2807.869614 | 2939.439381 | 2827.610948 | 906.8751403 | 834.6075002 | 759.1972848 | 5.99E-106 | -1.779571221 | 1.02E-107 | 486.0651307 | 0.291269951 | 105.2226188 |
| ENSMUSG00000020099 | Unc5b | 157.3631865 | 158.3536367 | 161.5497945 | 52.31971964 | 50.49565926 | 36.56463475 | 5.62E-13 | -1.780700786 | 7.50E-14 | 55.93260797 | 0.291041989 | 12.25051283 |
| ENSMUSG00000026749 | Nek6 | 2171.611974 | 2080.590838 | 2068.816459 | 609.2340686 | 617.3808906 | 608.1276096 | 1.09E-123 | -1.784805111 | 1.43E-125 | 568.1308906 | 0.29021518 | 122.9615552 |
| ENSMUSG00000036820 | Amdhd2 | 1666.348553 | 1655.015439 | 1728.093256 | 537.1491216 | 519.2478169 | 407.9843457 | 4.96E-61 | -1.788021172 | 1.45E-62 | 278.6904276 | 0.289568952 | 60.30485384 |
| ENSMUSG00000017929 | B4galt5 | 11075.81649 | 11453.14671 | 10984.40694 | 3378.691228 | 3078.329718 | 3232.121267 | 1.80E-218 | -1.790829372 | 1.15E-220 | 1005.488134 | 0.289005855 | 217.7435576 |
| ENSMUSG00000032860 | P2ry2 | 865.9228317 | 885.2408162 | 938.9469873 | 260.4359377 | 287.7299829 | 228.0478536 | 3.01E-51 | -1.792977153 | 1.05E-52 | 233.4578203 | 0.288575924 | 50.52186549 |
| ENSMUSG00000032487 | Ptgs2 | 14326.85487 | 13902.12969 | 13331.28486 | 4070.474188 | 3536.601645 | 4343.493718 | 4.07E-117 | -1.798316036 | 5.81E-119 | 537.7548595 | 0.287509985 | 116.3906246 |
| ENSMUSG00000035640 | Cbarp | 138.6497265 | 131.9613639 | 144.9052702 | 41.85577571 | 41.92092467 | 35.60240752 | 1.78E-12 | -1.802072822 | 2.44E-13 | 53.61132401 | 0.286762281 | 11.74922716 |
| ENSMUSG00000095403 | Gm21092 | 89.31424099 | 34.09001901 | 74.41081443 | 18.60256698 | 30.48794521 | 7.697817843 | 0.001397936 | -1.803830904 | 0.000473556 | 12.21701485 | 0.286413043 | 2.854512645 |
| ENSMUSG00000055301 | Adh7 | 423.6046859 | 415.6782963 | 405.3431207 | 105.8020997 | 137.1957535 | 111.6183587 | 4.62E-30 | -1.807246908 | 2.92E-31 | 135.2411992 | 0.285735678 | 29.33517383 |
| ENSMUSG00000072812 | None | 882.0844563 | 1017.20218 | 949.7169736 | 308.1050156 | 222.9430994 | 283.857033 | 1.54E-41 | -1.808611219 | 6.86E-43 | 188.4702675 | 0.285465594 | 40.81308906 |
| ENSMUSG00000030055 | Rab43 | 1355.024628 | 1328.411063 | 1404.014578 | 431.3470219 | 361.0916011 | 374.3063926 | 1.99E-75 | -1.812399156 | 4.56E-77 | 345.2617586 | 0.28471706 | 74.70118905 |
| ENSMUSG00000078606 | Gvin2 | 7347.584892 | 7077.527818 | 6694.03603 | 2048.607689 | 1559.648947 | 2388.247986 | 1.51E-49 | -1.816560213 | 5.54E-51 | 225.5593349 | 0.283897055 | 48.82095712 |
| ENSMUSG00000005125 | Ndrg1 | 10044.87497 | 10285.28864 | 9770.335752 | 2800.848991 | 2902.071285 | 2838.57033 | 4.40E-264 | -1.816863141 | 2.02E-266 | 1216.010102 | 0.28383745 | 263.3566195 |
| ENSMUSG00000105338 | Gm43802 | 49.3354855 | 28.59162885 | 41.12176587 | 13.95192524 | 10.48023117 | 9.622272304 | 0.00046694 | -1.82120589 | 0.000144529 | 14.44217632 | 0.282984337 | 3.330739118 |
| ENSMUSG00000045362 | Tnfrsf26 | 1538.756781 | 1475.76792 | 1416.742743 | 395.3045484 | 392.5322946 | 464.7557523 | 4.55E-75 | -1.821641075 | 1.05E-76 | 343.5958958 | 0.282898989 | 74.34229074 |
| ENSMUSG00000055561 | Spink5 | 3307.178752 | 3302.333132 | 3229.03771 | 974.3094456 | 817.458031 | 979.5473205 | 2.89E-107 | -1.829094569 | 4.84E-109 | 492.1535688 | 0.281441197 | 106.5385889 |
| ENSMUSG00000074361 | C5ar2 | 42.53059095 | 35.18969704 | 27.41451058 | 10.46394393 | 2.858244864 | 16.35786292 | 0.004823732 | -1.829659249 | 0.001787965 | 9.755402387 | 0.281331061 | 2.316616796 |
| ENSMUSG00000049719 | Prss46 | 87.61301736 | 84.67520851 | 78.32717308 | 16.27724611 | 27.62970035 | 25.98013522 | 1.29E-07 | -1.829821557 | 2.58E-08 | 31.0035643 | 0.281299412 | 6.89093407 |
| ENSMUSG00000020788 | Atp2a3 | 1846.678259 | 1931.034625 | 1815.232236 | 541.7997633 | 526.8698032 | 505.1692959 | 5.96E-112 | -1.8300491 | 9.00E-114 | 513.8971478 | 0.281255049 | 111.2245308 |
| ENSMUSG00000048285 | Frmd6 | 228.8145793 | 203.440436 | 218.336995 | 70.92228662 | 62.881387 | 50.03581598 | 3.09E-17 | -1.830836001 | 3.26E-18 | 75.7223647 | 0.281101684 | 16.5106375 |
| ENSMUSG00000037887 | Dusp8 | 52.73793278 | 65.98068195 | 34.26813822 | 16.27724611 | 13.33847603 | 13.47118123 | 0.000110242 | -1.830979541 | 3.10E-05 | 17.35558988 | 0.281073717 | 3.957652265 |
| ENSMUSG00000025804 | Ccr1 | 72.30200461 | 90.17359867 | 74.41081443 | 25.5785296 | 21.91321062 | 19.24454461 | 1.71E-07 | -1.831507727 | 3.46E-08 | 30.43256407 | 0.280970832 | 6.767131686 |
| ENSMUSG00000013698 | Pea15a | 7474.326053 | 7603.173917 | 7288.343455 | 2146.271165 | 2172.266096 | 1960.056868 | 4.31E-192 | -1.833113316 | 3.16E-194 | 883.8625222 | 0.28065831 | 191.3658554 |
| ENSMUSG00000030523 | Trpm1 | 124.1893256 | 105.5690911 | 101.825325 | 25.5785296 | 37.15718323 | 29.82904414 | 1.08E-09 | -1.835244175 | 1.81E-10 | 40.66695321 | 0.280244084 | 8.965873181 |
| ENSMUSG00000024399 | Ltb | 3484.956622 | 3632.236542 | 3574.656362 | 942.9176139 | 1079.46381 | 953.5671853 | 3.89E-134 | -1.84367455 | 4.67E-136 | 616.3413643 | 0.278611256 | 133.4099584 |
| ENSMUSG00000023341 | Mx2 | 1275.067117 | 1199.748734 | 1161.200341 | 339.4968474 | 295.3519692 | 375.2686198 | 1.07E-61 | -1.848680078 | 3.11E-63 | 281.7579823 | 0.277646271 | 60.96922815 |
| ENSMUSG00000026360 | Rgs2 | 4279.428061 | 4282.146259 | 4174.838325 | 1073.135583 | 1267.155223 | 1186.426175 | 7.42E-138 | -1.85076243 | 8.61E-140 | 633.5127545 | 0.277245811 | 137.1294752 |
| ENSMUSG00000044562 | Rasip1 | 66.34772188 | 59.38261376 | 65.59900746 | 9.301283491 | 19.05496576 | 24.05568076 | 1.24E-05 | -1.850821904 | 3.05E-06 | 21.78269196 | 0.277234383 | 4.906849729 |
| ENSMUSG00000047735 | Samd9l | 7333.975103 | 7042.338121 | 6970.139315 | 2052.09567 | 1532.019247 | 2329.552125 | 3.89E-53 | -1.852164784 | 1.32E-54 | 242.1817621 | 0.276976449 | 52.41027536 |
| ENSMUSG00000035992 | Fnip1 | 2624.988073 | 2458.880081 | 2434.016904 | 661.5537883 | 678.356781 | 740.9149674 | 1.53E-118 | -1.852354452 | 2.15E-120 | 544.337487 | 0.276940038 | 117.8142231 |
| ENSMUSG00000017009 | Sdc4 | 13080.70855 | 13151.04959 | 12879.92452 | 3537.975708 | 3655.695181 | 3615.087704 | 0 | -1.855086616 | 0 | 1517.714009 | 0.276416067 | Inf |
| ENSMUSG00000073490 | Ifi207 | 6595.644044 | 6610.164654 | 6301.421075 | 1899.787153 | 1456.752132 | 2031.261683 | 1.24E-68 | -1.85670315 | 3.16E-70 | 313.8526982 | 0.276106518 | 67.90817332 |
| ENSMUSG00000033777 | Tlr13 | 2411.484507 | 2558.950782 | 2443.8078 | 698.7589222 | 691.695257 | 651.427835 | 1.25E-135 | -1.860771037 | 1.47E-137 | 623.2439942 | 0.275329092 | 134.9037465 |
| ENSMUSG00000053007 | Creb5 | 348.7508458 | 399.1831258 | 349.5350099 | 123.2420063 | 66.69238015 | 111.6183587 | 8.17E-18 | -1.869442278 | 8.42E-19 | 78.39942707 | 0.273679205 | 17.08761066 |
| ENSMUSG00000027298 | Tyro3 | 36.57630822 | 40.68808721 | 46.99630385 | 13.95192524 | 13.33847603 | 6.735590613 | 0.000229794 | -1.876874255 | 6.77E-05 | 15.87392705 | 0.272272984 | 3.63866213 |
| ENSMUSG00000032470 | Mras | 23.81713093 | 37.38905311 | 43.0799452 | 6.975962618 | 14.29122432 | 6.735590613 | 0.001545742 | -1.881885308 | 0.000526596 | 12.01904821 | 0.271328912 | 2.810863042 |
| ENSMUSG00000021822 | Plau | 35593.00095 | 34241.77458 | 34137.9193 | 9424.525497 | 9006.329565 | 9721.381708 | 1.65E-296 | -1.884958021 | 6.46E-299 | 1365.540809 | 0.270751639 | 295.7817448 |
| ENSMUSG00000032265 | Tent5a | 652.4192652 | 635.6139028 | 647.1782676 | 181.3750281 | 141.9594949 | 198.2188095 | 5.71E-40 | -1.893559228 | 2.66E-41 | 181.1965671 | 0.269142247 | 39.24348508 |
| ENSMUSG00000022178 | Ajuba | 79.10689917 | 43.9871213 | 59.72446948 | 34.87981309 | 9.527482879 | 5.773363382 | 0.002404925 | -1.89411838 | 0.000847351 | 11.13453715 | 0.269037954 | 2.618898484 |
| ENSMUSG00000024670 | Cd6 | 32.32324912 | 24.19291672 | 35.24722789 | 11.62660436 | 8.574734591 | 4.811136152 | 0.001670443 | -1.895015787 | 0.000572631 | 11.86290515 | 0.268870655 | 2.777168458 |
| ENSMUSG00000000552 | Zfp385a | 562.2544123 | 710.392009 | 629.5546537 | 106.9647601 | 223.8958477 | 178.0120376 | 3.29E-18 | -1.895051237 | 3.32E-19 | 80.23997366 | 0.268864049 | 17.48224892 |
| ENSMUSG00000079057 | Cyp4v3 | 807.2306162 | 691.6974825 | 734.3172477 | 208.1162181 | 220.0848545 | 172.2386742 | 1.64E-44 | -1.897343983 | 6.73E-46 | 202.2529583 | 0.268437107 | 43.7863731 |
| ENSMUSG00000072115 | Ang | 37.42692004 | 53.8842236 | 64.61991779 | 13.95192524 | 15.24397261 | 12.50895399 | 3.73E-05 | -1.898965179 | 9.85E-06 | 19.53988535 | 0.268135626 | 4.427901753 |
| ENSMUSG00000029101 | Rgs12 | 32.32324912 | 36.28937507 | 27.41451058 | 11.62660436 | 5.716489727 | 8.660045073 | 0.000938294 | -1.899319414 | 0.000308709 | 13.01680199 | 0.268069797 | 3.027661172 |
| ENSMUSG00000098470 | C1rb | 28.92080185 | 37.38905311 | 20.56088293 | 5.813302182 | 4.763741439 | 12.50895399 | 0.003821932 | -1.900159009 | 0.00139055 | 10.21828645 | 0.267913835 | 2.417717021 |
| ENSMUSG00000095672 | ENSMUSG00000095672 | 171.8235874 | 172.6494511 | 145.8843599 | 43.01843614 | 50.49565926 | 37.52686198 | 2.26E-14 | -1.902847893 | 2.76E-15 | 62.42872719 | 0.267414965 | 13.64673537 |
| ENSMUSG00000039908 | Slc26a11 | 210.1011193 | 177.0481632 | 178.1943188 | 54.64504051 | 43.82642124 | 52.92249767 | 1.76E-16 | -1.906554075 | 1.93E-17 | 72.21852225 | 0.266728876 | 15.7555252 |
| ENSMUSG00000027570 | Col9a3 | 46.78365004 | 43.9871213 | 35.24722789 | 20.92788785 | 7.621986303 | 5.773363382 | 0.000995629 | -1.907539699 | 0.000329007 | 12.89758106 | 0.266546714 | 3.001902483 |
| ENSMUSG00000099974 | Bcl2a1d | 3571.719028 | 3465.085481 | 3516.890071 | 939.4296326 | 962.2757708 | 907.3802782 | 4.47E-188 | -1.909832848 | 3.40E-190 | 865.3170328 | 0.266123377 | 187.3500985 |
| ENSMUSG00000000392 | Fap | 46.78365004 | 58.28293573 | 37.20540721 | 13.95192524 | 8.574734591 | 15.39563569 | 8.12E-05 | -1.910062575 | 2.25E-05 | 17.9686559 | 0.266081004 | 4.090261881 |
| ENSMUSG00000032715 | Trib3 | 2023.605517 | 2010.211444 | 1990.489286 | 549.9383864 | 558.3104967 | 494.5847964 | 8.10E-120 | -1.911100848 | 1.12E-121 | 550.2386697 | 0.265889581 | 119.0915475 |
| ENSMUSG00000021451 | Sema4d | 9143.226442 | 9530.909508 | 9157.425623 | 2502.045259 | 2385.681713 | 2499.866344 | 7.23E-273 | -1.913715151 | 3.13E-275 | 1256.55243 | 0.2654082 | 272.1406503 |
| ENSMUSG00000037921 | Ddx60 | 545.242176 | 545.4403042 | 531.6456873 | 133.7059502 | 123.8572774 | 172.2386742 | 5.52E-36 | -1.914238542 | 2.84E-37 | 162.7464535 | 0.265311931 | 35.25787029 |
| ENSMUSG00000004508 | Gab2 | 2897.183855 | 3067.002033 | 2930.415363 | 808.0490033 | 755.5293923 | 796.7241467 | 5.42E-159 | -1.914447297 | 5.35E-161 | 731.0287971 | 0.265273543 | 158.2662893 |
| ENSMUSG00000049526 | Tmem202 | 42.53059095 | 56.08357966 | 53.84993149 | 8.138623054 | 17.14946918 | 14.43340846 | 3.43E-05 | -1.920666023 | 9.01E-06 | 19.71012446 | 0.264132545 | 4.464105252 |
| ENSMUSG00000028121 | Bcar3 | 6562.470183 | 6689.341472 | 6554.026208 | 1774.219826 | 1713.99417 | 1704.104425 | 2.11E-271 | -1.931990501 | 9.28E-274 | 1249.7764 | 0.262067344 | 270.6749212 |
| ENSMUSG00000061852 | Gm53056 | 56.14038005 | 56.08357966 | 60.70355914 | 19.76522742 | 8.574734591 | 17.32009015 | 1.08E-05 | -1.934629043 | 2.63E-06 | 22.06562772 | 0.261588488 | 4.96735802 |
| ENSMUSG00000071669 | Snx29 | 529.9311632 | 534.4435238 | 585.4956188 | 152.3085172 | 130.5265154 | 147.2207662 | 3.84E-44 | -1.942338026 | 1.59E-45 | 200.5414842 | 0.260194428 | 43.41578947 |
| ENSMUSG00000021003 | Galc | 574.1629778 | 559.7361186 | 503.2520871 | 134.8686106 | 140.0539983 | 149.1452207 | 4.39E-44 | -1.947299435 | 1.83E-45 | 200.2666166 | 0.25930116 | 43.35724348 |
| ENSMUSG00000047786 | Lix1 | 129.2929965 | 69.27971605 | 101.825325 | 24.41586916 | 24.77145549 | 28.86681691 | 5.61E-08 | -1.947372315 | 1.09E-08 | 32.67323553 | 0.259288061 | 7.251128868 |
| ENSMUSG00000034640 | Tiparp | 3080.065397 | 3196.764041 | 3072.383364 | 767.355888 | 780.3008478 | 871.7778707 | 3.05E-146 | -1.948733912 | 3.31E-148 | 672.2051619 | 0.259043464 | 145.5150291 |
| ENSMUSG00000051457 | Spn | 508.6658678 | 548.7393383 | 519.8966113 | 139.5192524 | 132.432012 | 136.6362667 | 2.40E-46 | -1.949011735 | 9.48E-48 | 210.7373238 | 0.258993584 | 45.62059308 |
| ENSMUSG00000020902 | Ntn1 | 21.26529547 | 24.19291672 | 25.45633125 | 5.813302182 | 4.763741439 | 7.697817843 | 0.00307679 | -1.95298791 | 0.001102865 | 10.64638116 | 0.258280761 | 2.511902088 |
| ENSMUSG00000072872 | Rybp | 7297.398795 | 7319.456985 | 7201.204475 | 1899.787153 | 1789.261285 | 1935.03896 | 1.01E-255 | -1.956143693 | 4.94E-258 | 1177.415796 | 0.25771641 | 254.994164 |
| ENSMUSG00000030084 | Plxna1 | 10231.15896 | 10711.96372 | 10489.96666 | 2774.107801 | 2597.191833 | 2688.462882 | 4.61E-285 | -1.963895495 | 1.93E-287 | 1312.735279 | 0.256335377 | 284.3360738 |
| ENSMUSG00000019726 | Lyst | 3149.815566 | 3183.567904 | 3043.989764 | 840.6034955 | 726.9469437 | 832.3265543 | 1.69E-138 | -1.967560747 | 1.94E-140 | 636.4887889 | 0.255684968 | 137.7715571 |
| ENSMUSG00000019850 | Tnfaip3 | 1793.089714 | 1827.66489 | 1685.992401 | 454.6002306 | 434.4532193 | 463.793525 | 6.42E-117 | -1.972037459 | 9.25E-119 | 536.8249833 | 0.254892802 | 116.1924861 |
| ENSMUSG00000035692 | Isg15 | 3598.938606 | 3590.448776 | 3709.770735 | 833.6275329 | 1046.11762 | 893.909097 | 5.21E-112 | -1.972452902 | 7.82E-114 | 514.1772593 | 0.254819413 | 111.2834955 |
| ENSMUSG00000017724 | Etv4 | 172.6741993 | 185.8455875 | 193.8597534 | 48.83173833 | 46.68466611 | 45.22467983 | 2.62E-18 | -1.973592183 | 2.63E-19 | 80.69460172 | 0.254618265 | 17.58098246 |
| ENSMUSG00000020205 | Phlda1 | 1635.726528 | 1672.610288 | 1752.570498 | 449.9495889 | 400.1542809 | 439.7378443 | 1.50E-108 | -1.973716924 | 2.44E-110 | 498.1165082 | 0.25459625 | 107.8252166 |
| ENSMUSG00000021298 | Gpr132 | 300.2659721 | 351.8969704 | 330.9323063 | 80.22357011 | 92.41658392 | 76.0159512 | 4.01E-28 | -1.979972388 | 2.70E-29 | 126.2581579 | 0.253494722 | 27.39632176 |
| ENSMUSG00000030657 | Xylt1 | 1830.516634 | 2020.108546 | 1790.754995 | 501.1066481 | 423.0202398 | 502.2826142 | 9.75E-89 | -1.984690108 | 1.97E-90 | 406.6472822 | 0.252667129 | 88.01083431 |
| ENSMUSG00000030344 | Akap3 | 49.3354855 | 48.38583343 | 44.05903486 | 13.95192524 | 9.527482879 | 12.50895399 | 1.42E-05 | -1.986481347 | 3.52E-06 | 21.51233275 | 0.252353614 | 4.848409375 |
| ENSMUSG00000037447 | Arid5a | 438.0650868 | 429.9741107 | 431.7785416 | 113.9407228 | 102.8968151 | 108.731677 | 3.84E-41 | -1.999842612 | 1.73E-42 | 186.6242041 | 0.250027275 | 40.41607831 |
| ENSMUSG00000089901 | Gm8113 | 717.9163752 | 621.3180884 | 686.3418541 | 153.4711776 | 148.6287329 | 202.0677184 | 4.90E-41 | -2.004763302 | 2.22E-42 | 186.1349594 | 0.249175944 | 40.30993465 |
| ENSMUSG00000029490 | Mfsd7a | 46.78365004 | 58.28293573 | 48.95448318 | 19.76522742 | 11.43297945 | 7.697817843 | 3.08E-05 | -2.005392114 | 8.00E-06 | 19.93674878 | 0.249067362 | 4.511830937 |
| ENSMUSG00000035208 | Slfn8 | 1569.378806 | 1568.140874 | 1423.596371 | 374.3766605 | 290.5882278 | 466.6802067 | 4.28E-43 | -2.011036889 | 1.83E-44 | 195.6768428 | 0.24809475 | 42.36900923 |
| ENSMUSG00000026655 | Fam107b | 839.5538653 | 764.2762326 | 697.1118404 | 195.3269533 | 193.4079024 | 180.8987193 | 2.68E-55 | -2.015977424 | 8.77E-57 | 252.1635073 | 0.247246598 | 54.57215607 |
| ENSMUSG00000022500 | Litaf | 10123.13126 | 10312.78059 | 10385.20406 | 2617.148642 | 2565.751139 | 2423.850393 | 4.27E-297 | -2.019048198 | 1.64E-299 | 1368.280746 | 0.246720894 | 296.3694624 |
| ENSMUSG00000017652 | Cd40 | 1649.336317 | 1558.243772 | 1578.292538 | 355.7740935 | 423.0202398 | 396.4376189 | 2.91E-102 | -2.023063802 | 5.17E-104 | 469.0451095 | 0.246035124 | 101.5361607 |
| ENSMUSG00000016496 | Cd274 | 571.6111424 | 589.4274255 | 553.1856599 | 144.1698941 | 128.6210189 | 148.1829935 | 6.52E-51 | -2.02679508 | 2.32E-52 | 231.8802552 | 0.24539962 | 50.18545361 |
| ENSMUSG00000053581 | Zfand2a | 3217.864511 | 3256.146654 | 3198.685931 | 826.6515702 | 743.1436646 | 795.7619195 | 8.67E-182 | -2.033209878 | 7.02E-184 | 836.2700483 | 0.244310898 | 181.0617879 |
| ENSMUSG00000021118 | Plek2 | 31.4726373 | 45.08679934 | 30.35177957 | 13.95192524 | 9.527482879 | 2.886681691 | 0.001140503 | -2.038976779 | 0.000380011 | 12.62800226 | 0.24333626 | 2.942903664 |

| | | | | | | | | | | | | | |
|---|---|---|---|---|---|---|---|---|---|---|---|---|---|
| ENSMUSG00000062743 | *Zfp677* | 86.76240554 | 82.47585244 | 78.32717308 | 20.92788785 | 11.43297945 | 27.90458968 | 1.20E-07 | -2.041497233 | 2.40E-08 | 31.13847402 | 0.242911512 | 6.920375178 |
| ENSMUSG00000050390 | *C77080* | 1461.351105 | 1570.340231 | 1532.275323 | 361.5873957 | 374.4300771 | 369.4952565 | 2.02E-116 | -2.044200193 | 2.94E-118 | 534.5173044 | 0.242456831 | 115.6945646 |
| ENSMUSG00000024677 | *Ms4a6b* | 979.0542036 | 973.2150588 | 950.6960633 | 198.8149346 | 262.9585275 | 236.7078987 | 7.83E-66 | -2.050014888 | 2.08E-67 | 300.9163319 | 0.24148159 | 65.10599769 |
| ENSMUSG00000052102 | *Gnpda1* | 2682.829677 | 2626.031142 | 2594.587608 | 670.8550718 | 629.7666183 | 606.2031551 | 9.56E-166 | -2.053292966 | 9.05E-168 | 762.1723575 | 0.240933521 | 165.0195794 |
| ENSMUSG00000094054 | *ENSMUSG00000094054* | 41.67997913 | 46.18647737 | 51.89175217 | 11.62660436 | 16.19672089 | 5.773363382 | 5.06E-05 | -2.056129258 | 1.36E-05 | 18.9244885 | 0.240460319 | 4.295738717 |
| ENSMUSG00000029470 | *P2rx4* | 1004.572558 | 922.6298693 | 1027.065057 | 253.4599751 | 218.1793579 | 233.821217 | 5.48E-75 | -2.069915088 | 1.27E-76 | 343.2162264 | 0.238173517 | 74.26089466 |
| ENSMUSG00000091649 | *Phf11b* | 1258.05488 | 1099.678033 | 1256.172038 | 267.4119004 | 323.9344179 | 268.4613973 | 2.38E-70 | -2.070035882 | 5.94E-72 | 321.7776339 | 0.238153576 | 69.62382524 |
| ENSMUSG00000054364 | *Rhob* | 364.9124703 | 400.2828039 | 352.4722789 | 77.89824923 | 87.65284249 | 99.10940473 | 1.33E-33 | -2.072439269 | 7.35E-35 | 151.703797 | 0.237757166 | 32.87511347 |
| ENSMUSG00000039982 | *Dtx4* | 6136.313662 | 6374.833555 | 6402.26731 | 1538.199757 | 1431.980677 | 1525.13016 | 1.72E-257 | -2.073410116 | 8.15E-260 | 1185.616908 | 0.237597224 | 256.7641083 |
| ENSMUSG00000017400 | *Stac2* | 90.16485281 | 97.8713449 | 84.20171106 | 24.41586916 | 20.96046233 | 19.24454461 | 2.95E-10 | -2.080195085 | 4.72E-11 | 43.28907145 | 0.236482432 | 9.52982503 |
| ENSMUSG00000018752 | *Tnfsfm13* | 227.9639675 | 236.430777 | 249.6678642 | 63.946324 | 55.2594007 | 50.03581598 | 6.46E-24 | -2.082626358 | 4.99E-25 | 106.7730413 | 0.23608424 | 23.18969805 |
| ENSMUSG00000040253 | *Gbp7* | 438.0650868 | 471.761876 | 340.7232029 | 111.6154019 | 64.78688358 | 119.3161766 | 1.98E-19 | -2.083612029 | 1.87E-20 | 85.92853696 | 0.235922999 | 18.70422793 |
| ENSMUSG00000031304 | *Il2rg* | 2044.020201 | 2153.169588 | 2006.154721 | 477.8534393 | 455.4136816 | 526.338295 | 5.83E-126 | -2.086673614 | 7.41E-128 | 578.641446 | 0.235422871 | 125.2339985 |
| ENSMUSG00000036564 | *Ndrg4* | 560.5531887 | 576.2312891 | 588.4328878 | 124.4046667 | 142.9122432 | 136.6362667 | 1.64E-54 | -2.090745285 | 5.45E-56 | 248.5225705 | 0.234759381 | 53.78473021 |
| ENSMUSG00000028751 | *Pla2g2e* | 115.6832074 | 140.7587882 | 130.2189253 | 40.69311527 | 24.77145549 | 25.98013522 | 2.91E-12 | -2.09277726 | 4.06E-13 | 52.61451402 | 0.234428965 | 11.53594878 |
| ENSMUSG00000005803 | *Sqor* | 382.7753185 | 423.3760425 | 392.6149551 | 86.03687229 | 108.6133048 | 84.67599627 | 1.97E-35 | -2.097051325 | 1.03E-36 | 160.1936017 | 0.233735484 | 34.70645703 |
| ENSMUSG00000055202 | *Zfp811* | 59.54282733 | 54.98390163 | 57.76629015 | 16.27724611 | 5.716489727 | 18.28231738 | 9.28E-06 | -2.107454149 | 2.25E-06 | 22.36998379 | 0.232056152 | 5.032650699 |
| ENSMUSG00000019960 | *Dusp6* | 1920.681487 | 1796.873905 | 1981.677479 | 442.9736262 | 432.5477227 | 446.4734349 | 1.06E-133 | -2.10873928 | 1.28E-135 | 614.3217388 | 0.231849532 | 132.9756271 |
| ENSMUSG00000076435 | *Acsf2* | 130.9942201 | 130.8616859 | 150.7798082 | 39.53045484 | 29.53519692 | 26.94236245 | 3.48E-14 | -2.114347126 | 4.30E-15 | 61.5574013 | 0.230950068 | 13.45820289 |
| ENSMUSG00000035299 | *Mid1* | 407.4430613 | 409.0802281 | 461.1512315 | 104.6394393 | 77.17261132 | 113.5428132 | 1.54E-32 | -2.115670682 | 8.89E-34 | 146.7517384 | 0.230738287 | 31.81118044 |
| ENSMUSG00000035561 | *Aldh1b1* | 147.1558447 | 152.8552465 | 129.2398356 | 25.5785296 | 42.87367295 | 29.82904414 | 9.85E-14 | -2.116558611 | 1.25E-14 | 59.46471166 | 0.230596319 | 13.00651769 |
| ENSMUSG00000015968 | *Cacna1d* | 51.88732096 | 73.67842818 | 59.72446948 | 11.62660436 | 16.19672089 | 14.43340846 | 3.98E-07 | -2.120211631 | 8.33E-08 | 28.72834791 | 0.230013169 | 6.400629356 |
| ENSMUSG00000091844 | *Gm8251* | 97.82035918 | 84.67520851 | 105.7416837 | 22.09054829 | 10.48023117 | 33.67795306 | 1.49E-07 | -2.123195832 | 2.99E-08 | 30.71122306 | 0.229537881 | 6.827144424 |
| ENSMUSG00000089929 | *Bcl2a1b* | 2851.250817 | 2687.613112 | 2955.871694 | 652.2525048 | 658.3490669 | 638.918881 | 2.58E-172 | -2.123894355 | 2.28E-174 | 792.5212367 | 0.22942677 | 171.5889226 |
| ENSMUSG00000022876 | *Samsn1* | 714.5139279 | 721.3887894 | 728.4427097 | 144.1698941 | 164.8254538 | 183.785401 | 6.47E-62 | -2.129868669 | 1.86E-63 | 282.7816982 | 0.22847866 | 61.18920886 |
| ENSMUSG00000039384 | *Dusp10* | 182.0309293 | 136.360076 | 130.2189253 | 41.85577571 | 29.53519692 | 31.7534986 | 1.22E-13 | -2.132818087 | 1.55E-14 | 59.02912296 | 0.22801204 | 12.91273551 |
| ENSMUSG00000000134 | *Tfe3* | 3018.821346 | 3040.60976 | 3086.090619 | 724.3374518 | 657.3963186 | 704.3503326 | 3.49E-194 | -2.133662213 | 2.49E-196 | 893.5375515 | 0.227878669 | 193.4567696 |
| ENSMUSG00000039208 | *Metrnl* | 2714.302314 | 2818.474797 | 2871.669983 | 606.9087478 | 673.5930395 | 627.3721542 | 2.12E-173 | -2.137402897 | 1.83E-175 | 797.5584045 | 0.22728858 | 172.6738828 |
| ENSMUSG00000028179 | *Cth* | 1236.789585 | 1280.02523 | 1278.691101 | 297.6410717 | 286.7772347 | 278.0836696 | 1.12E-111 | -2.138834636 | 1.70E-113 | 512.6243741 | 0.227063129 | 110.9495847 |
| ENSMUSG00000020644 | *Id2* | 1949.602289 | 2032.205004 | 2042.381038 | 398.7925297 | 485.9016268 | 470.5291156 | 5.38E-119 | -2.148298712 | 7.50E-121 | 546.4372905 | 0.225578471 | 118.2688885 |
| ENSMUSG00000053820 | *Bcl2a1c* | 40.82936731 | 50.5851895 | 43.0799452 | 17.43990655 | 6.669238015 | 6.735590613 | 5.76E-05 | -2.152958952 | 1.56E-05 | 18.66139397 | 0.224850975 | 4.239583293 |
| ENSMUSG00000026875 | *Traf1* | 7970.232744 | 8104.6271 | 8058.887021 | 1813.750281 | 1872.150386 | 1738.744605 | 4.59481050632359e-321 | -2.153415328 | 1.48219693752374e-323 | 1478.849386 | 0.224779858 | 320.3377324 |
| ENSMUSG00000040428 | *Plekha4* | 27.21957821 | 29.69130688 | 24.47724159 | 5.813302182 | 10.48023117 | 1.924454461 | 0.002065276 | -2.155201943 | 0.00071946 | 11.43829 | 0.224501666 | 2.685021916 |
| ENSMUSG00000030785 | *Cox6a2* | 129.2929965 | 98.97102293 | 133.1561942 | 34.87981309 | 23.8187072 | 23.09345353 | 5.79E-12 | -2.158322495 | 8.23E-13 | 51.2261423 | 0.224016593 | 11.2371219 |
| ENSMUSG00000111860 | *None* | 70.60078098 | 87.97424261 | 90.07624905 | 15.11458567 | 30.48794521 | 9.622272304 | 6.72E-07 | -2.161585875 | 1.44E-07 | 27.66275036 | 0.22351044 | 6.172697879 |
| ENSMUSG00000058755 | *Osm* | 1023.286018 | 1045.793809 | 986.9223808 | 224.3934642 | 237.2343237 | 220.3500358 | 1.43E-93 | -2.163008534 | 2.75E-95 | 428.9474125 | 0.223290142 | 92.84514329 |
| ENSMUSG00000041153 | *Osgin2* | 4538.864666 | 4466.892168 | 4471.502493 | 961.5201809 | 980.3779882 | 1046.903227 | 9.33E-249 | -2.171993474 | 4.86E-251 | 1145.237212 | 0.221903838 | 248.0302217 |
| ENSMUSG00000081665 | *Pira1* | 86.76240554 | 81.37617441 | 78.32717308 | 13.95192524 | 14.29122432 | 25.98013522 | 7.23E-09 | -2.175536343 | 1.29E-09 | 36.82451047 | 0.221359571 | 8.140948351 |
| ENSMUSG00000025743 | *Sdc3* | 4544.818949 | 4653.837434 | 4397.091679 | 1098.714112 | 910.8273632 | 991.0940473 | 2.84E-167 | -2.181845356 | 2.65E-169 | 769.2233924 | 0.220393662 | 166.5463464 |
| ENSMUSG00000033948 | *Zswim5* | 21.26529547 | 17.59484852 | 16.64452428 | 4.650641745 | 3.810993152 | 3.848901921 | 0.004392082 | -2.182150512 | 0.001618326 | 9.938781973 | 0.22034705 | 2.35732952 |
| ENSMUSG00000027546 | *Atp9a* | 55.28976823 | 41.78776524 | 59.72446948 | 11.62660436 | 13.33847603 | 9.622272304 | 1.74E-06 | -2.18347237 | 3.91E-07 | 25.73757985 | 0.220145251 | 5.760364259 |
| ENSMUSG00000026193 | *Fn1* | 26.36986639 | 13.19613639 | 31.33086923 | 9.301283491 | 4.763741439 | 1.924454461 | 0.005878587 | -2.184042545 | 0.002211227 | 9.365375833 | 0.220058264 | 2.230727027 |
| ENSMUSG00000028575 | *Eqtn* | 16.16162456 | 20.89388262 | 18.60270361 | 3.487981309 | 2.858244864 | 5.773363382 | 0.005044069 | -2.190229744 | 0.001875172 | 9.667883172 | 0.219116534 | 2.297219016 |
| ENSMUSG00000018217 | *Pmp22* | 6465.500436 | 6433.116491 | 6401.28822 | 1399.843165 | 1432.933425 | 1385.607212 | 0 | -2.193707567 | 0 | 1567.732154 | 0.218588958 | Inf |
| ENSMUSG00000079794 | *ENSMUSG00000079794* | 119.9362665 | 85.77488654 | 87.13898005 | 19.76522742 | 14.29122432 | 29.82904414 | 3.72E-09 | -2.197443372 | 6.51E-10 | 38.16201754 | 0.218023663 | 8.429330309 |
| ENSMUSG00000016206 | *H2-M3* | 2617.332567 | 2694.21118 | 2610.253043 | 592.9568225 | 591.6566868 | 537.8850218 | 1.07E-181 | -2.20219577 | 8.77E-184 | 835.8266437 | 0.21730665 | 180.9690539 |
| ENSMUSG00000091199 | *Gm2619* | 440.6169222 | 450.8679934 | 418.0712863 | 101.151458 | 103.8495634 | 79.86486012 | 1.05E-41 | -2.203118013 | 4.67E-43 | 189.2365284 | 0.217167781 | 40.97699883 |
| ENSMUSG00000036896 | *Tnfsf9* | 3777.567088 | 3917.053152 | 3772.432474 | 860.3687229 | 764.1041269 | 849.6466444 | 3.00E-205 | -2.213638524 | 2.00E-207 | 944.5803491 | 0.215589897 | 204.5223476 |
| ENSMUSG00000026536 | *Ifi211* | 858.2673254 | 828.0575585 | 801.8744344 | 155.7964985 | 165.7782021 | 212.6522179 | 2.39E-60 | -2.215563966 | 7.10E-62 | 275.523931 | 0.215302359 | 59.62187237 |
| ENSMUSG00000074384 | *AI429214* | 36.57630822 | 35.18969704 | 32.3099589 | 6.975962618 | 4.763741439 | 10.58449953 | 9.63E-05 | -2.219370962 | 2.69E-05 | 17.62635035 | 0.214734967 | 4.016525787 |
| ENSMUSG00000024401 | *Tnf* | 30922.29146 | 30600.74061 | 30456.54216 | 6508.573123 | 6377.697039 | 6841.435608 | 0 | -2.221027369 | 0 | 2045.74794 | 0.214488564 | Inf |
| ENSMUSG00000010142 | *Tnfrsf13b* | 76.55506371 | 80.27649638 | 66.57809712 | 13.95192524 | 17.14946918 | 16.35786292 | 2.14E-09 | -2.228420316 | 3.67E-10 | 39.28271222 | 0.21339225 | 8.668689776 |
| ENSMUSG00000024164 | *C3* | 462.7328295 | 478.3599442 | 481.7121145 | 77.89824923 | 109.5660531 | 113.5428132 | 2.06E-41 | -2.231459945 | 9.24E-43 | 187.8772126 | 0.212943125 | 40.68629593 |
| ENSMUSG00000027555 | *Car13* | 1544.711063 | 1592.333791 | 1438.282716 | 320.8942804 | 352.5168665 | 300.2148959 | 1.21E-109 | -2.231691843 | 1.91E-111 | 503.2043945 | 0.212908899 | 108.9154936 |
| ENSMUSG00000024679 | *Ms4a6d* | 1314.19526 | 1320.713317 | 1308.06379 | 223.2308038 | 324.8871662 | 286.7437146 | 2.15E-71 | -2.233897435 | 5.27E-73 | 326.606043 | 0.212583653 | 70.66829713 |
| ENSMUSG00000027698 | *Nceh1* | 6180.545477 | 6070.22274 | 6352.333737 | 1269.625196 | 1310.028896 | 1359.627077 | 1.13E-300 | -2.238768783 | 4.17E-303 | 1384.82335 | 0.211867061 | 299.9484548 |
| ENSMUSG00000030748 | *Il4ra* | 177.7778702 | 192.4436557 | 173.2988704 | 27.90385047 | 44.7788953 | 41.37577091 | 4.79E-19 | -2.239477444 | 4.66E-20 | 84.1176863 | 0.211763017 | 18.31923598 |
| ENSMUSG00000043263 | *Ifi209* | 289.2080185 | 246.3278793 | 253.5842229 | 59.29568225 | 56.21214899 | 51.96027044 | 1.90E-28 | -2.240970764 | 1.26E-29 | 127.7664063 | 0.211543936 | 27.72012431 |
| ENSMUSG00000029596 | *Sdsl* | 24.66774275 | 20.89388262 | 27.41451058 | 6.975962618 | 5.716489727 | 2.886681691 | 0.001084321 | -2.24605031 | 0.000360175 | 12.72823978 | 0.210800426 | 2.964842299 |
| ENSMUSG00000062593 | *Gm49339* | 8794.475596 | 9400.047822 | 7896.358136 | 1780.033128 | 1741.62387 | 1971.603595 | 1.51E-163 | -2.247422049 | 1.47E-165 | 752.0117906 | 0.210600089 | 162.8196737 |
| ENSMUSG00000074141 | *Il4i1* | 155.6619629 | 169.350417 | 217.3579053 | 46.50641745 | 36.20443494 | 31.7534986 | 5.79E-17 | -2.2527875 | 6.19E-18 | 74.45782134 | 0.209818312 | 16.2372417 |
| ENSMUSG00000025877 | *Hk3* | 7228.499238 | 7525.096777 | 7367.649718 | 1595.170119 | 1528.208254 | 1519.356797 | 0 | -2.252992433 | 0 | 1578.151811 | 0.20978851 | Inf |
| ENSMUSG00000040435 | *Ppp1r15a* | 2977.141366 | 2930.641957 | 2942.164439 | 592.9568225 | 620.2391354 | 636.0321993 | 2.88E-221 | -2.257474385 | 1.80E-223 | 1018.401125 | 0.209137781 | 220.5409018 |
| ENSMUSG00000022041 | *Chrna2* | 48.48487368 | 31.89066294 | 28.39360024 | 11.62660436 | 4.763741439 | 6.735590613 | 0.000152912 | -2.261933222 | 4.38E-05 | 16.69716274 | 0.208492411 | 3.815558424 |
| ENSMUSG00000060477 | *Irak2* | 3520.682319 | 3593.74781 | 3505.140995 | 746.4280001 | 741.238168 | 720.7081955 | 2.03E-255 | -2.266011021 | 1.00E-257 | 1176.001678 | 0.207903936 | 254.692906 |
| ENSMUSG00000034855 | *Cxcl10* | 3012.867063 | 2968.03101 | 2878.523611 | 670.8550718 | 589.7511902 | 583.1097016 | 5.21E-178 | -2.267535489 | 4.29E-180 | 818.8570793 | 0.207684364 | 177.2833507 |
| ENSMUSG00000057137 | *Tmem140* | 395.5344958 | 429.9741107 | 400.4476724 | 65.10898443 | 87.65284249 | 99.10940473 | 5.32E-37 | -2.273215738 | 2.66E-38 | 167.456124 | 0.206868267 | 36.27375424 |
| ENSMUSG00000026535 | *Ifi202b* | 4277.726838 | 4198.570728 | 4267.851843 | 856.8807416 | 786.0173375 | 989.1695928 | 3.76E-192 | -2.275265761 | 3.79E-154 | 699.5242009 | 0.206574523 | 151.4245687 |
| ENSMUSG00000030107 | *Usp18* | 8508.670025 | 8523.60443 | 8257.642222 | 1637.025894 | 1838.804196 | 1721.424515 | 7.68E-293 | -2.28162495 | 3.06E-295 | 1348.63089 | 0.205665976 | 292.1146539 |
| ENSMUSG00000050533 | *Gm9845* | 299.4153603 | 299.1124249 | 304.4968854 | 51.15705592 | 72.40886988 | 60.62031551 | 3.01E-32 | -2.285572689 | 1.76E-33 | 145.4003632 | 0.205103968 | 31.52089651 |
| ENSMUSG00000026004 | *Kansl1l* | 1414.567455 | 1393.292067 | 1288.481997 | 283.6891465 | 253.4310446 | 303.1015776 | 7.43E-105 | -2.286271126 | 1.29E-106 | 481.0148012 | 0.205004698 | 104.1288961 |
| ENSMUSG00000022799 | *Arhgap31* | 128.4423847 | 134.16072 | 115.5325803 | 25.5785296 | 25.72420377 | 25.98013522 | 1.00E-15 | -2.289857883 | 1.14E-16 | 68.71542132 | 0.204495658 | 14.99875052 |
| ENSMUSG00000058163 | *Gm5431* | 328.3361621 | 298.0127468 | 312.3296027 | 34.87981309 | 64.78688358 | 90.44935965 | 7.23E-18 | -2.291675522 | 7.41E-19 | 78.65135986 | 0.204238178 | 17.14067625 |
| ENSMUSG00000031101 | *Sash3* | 48.48487368 | 61.58196982 | 45.03812452 | 11.62660436 | 14.29122432 | 5.773363382 | 2.97E-06 | -2.292356469 | 6.82E-07 | 24.66449518 | 0.204141801 | 5.527881754 |
| ENSMUSG00000020638 | *Cmpk2* | 4541.416502 | 4780.300408 | 4478.356121 | 932.4536699 | 884.1504112 | 989.1695928 | 2.83E-231 | -2.29797592 | 1.69E-233 | 1064.529489 | 0.203348194 | 230.5476278 |
| ENSMUSG00000040552 | *C3ar1* | 8487.40473 | 8286.073975 | 8019.723434 | 1710.273502 | 1524.397261 | 1788.780421 | 6.13E-226 | -2.303649905 | 3.75E-228 | 1039.937342 | 0.202550015 | 225.2123161 |
| ENSMUSG00000036944 | *Tmem71* | 32.32324912 | 35.18969704 | 56.78720049 | 4.650641745 | 11.43297945 | 8.660045073 | 4.65E-05 | -2.307882382 | 1.24E-05 | 19.09878917 | 0.201956659 | 4.332962783 |
| ENSMUSG00000000204 | *Slfn4* | 13538.33771 | 12611.10768 | 12489.26775 | 2719.462761 | 1974.094453 | 3108.956181 | 1.64E-30 | -2.308131732 | 1.02E-31 | 137.3231347 | 0.201921756 | 29.7848412 |
| ENSMUSG00000028773 | *Fabp3* | 39.97875549 | 50.5851895 | 45.03812452 | 5.813302182 | 3.810993152 | 17.32009015 | 9.13E-05 | -2.316847731 | 2.54E-05 | 17.73503383 | 0.200705529 | 4.039737181 |
| ENSMUSG00000020387 | *Jade2* | 3341.203225 | 3339.722185 | 3286.804 | 680.1563553 | 674.5457878 | 642.7677899 | 5.32E-250 | -2.319845753 | 2.74E-252 | 1150.988448 | 0.200288882 | 249.2744204 |
| ENSMUSG00000046245 | *Pilra* | 90.16485281 | 92.37295474 | 80.28535241 | 13.95192524 | 13.33847603 | 25.01790799 | 4.18E-10 | -2.320228655 | 6.77E-11 | 42.58464447 | 0.200235731 | 9.378633073 |
| ENSMUSG00000029923 | *Rab19* | 225.412132 | 205.6397921 | 198.7552017 | 41.85577571 | 34.29893836 | 50.03581598 | 3.46E-23 | -2.320504677 | 2.76E-24 | 103.3869126 | 0.200197425 | 22.46083264 |
| ENSMUSG00000024066 | *Xdh* | 358.1075758 | 329.9034098 | 320.16232 | 59.29568225 | 72.40886988 | 69.28036059 | 1.78E-37 | -2.321910057 | 8.76E-39 | 169.6651442 | 0.200002501 | 36.75024294 |
| ENSMUSG00000025408 | *Ddit3* | 1163.636968 | 1145.86451 | 1121.057665 | 237.182729 | 201.0298887 | 244.4057165 | 1.08E-100 | -2.331366408 | 1.94E-102 | 461.8109292 | 0.198695843 | 99.96857352 |
| ENSMUSG00000070501 | *Ifi214* | 40.82936731 | 48.38583343 | 39.16358654 | 8.138623054 | 5.716489727 | 11.54672676 | 6.69E-06 | -2.334421712 | 1.60E-06 | 23.02873955 | 0.198275495 | 5.1747243 |
| ENSMUSG00000022667 | *Cd200r1* | 205.8480602 | 236.430777 | 194.838843 | 47.66907789 | 32.39344179 | 46.18690706 | 2.00E-22 | -2.340440103 | 1.64E-23 | 99.85577768 | 0.197450085 | 21.69846019 |
| ENSMUSG00000021062 | *Rab15* | 112.2807601 | 111.0674813 | 119.448939 | 25.5785296 | 28.58244864 | 13.47118123 | 1.69E-12 | -2.347036686 | 2.32E-13 | 53.71565723 | 0.196549325 | 11.77120293 |
| ENSMUSG00000031266 | *Gla* | 2812.973285 | 2861.362241 | 2760.053762 | 535.9864612 | 566.8852313 | 551.356203 | 5.53E-227 | -2.348901853 | 3.34E-229 | 1044.768356 | 0.196295383 | 226.2574546 |
| ENSMUSG00000028270 | *Gbp2* | 148.0064565 | 160.5529928 | 144.9052702 | 22.09054829 | 37.15718323 | 28.86681691 | 2.13E-17 | -2.350524083 | 2.24E-18 | 76.46725086 | 0.196074784 | 16.67106767 |
| ENSMUSG00000031227 | *Magee1* | 16.16162456 | 12.09645836 | 18.60270361 | 3.487981309 | 2.858244864 | 2.886681691 | 0.006412832 | -2.353644374 | 0.002434623 | 9.189086676 | 0.195651168 | 2.19295011 |
| ENSMUSG00000039236 | *Isg20* | 1139.819837 | 1221.742294 | 1211.133914 | 227.8814455 | 247.7145549 | 220.3500358 | 9.74E-113 | -2.358483062 | 1.44E-114 | 517.5496237 | 0.194996068 | 112.0112189 |
| ENSMUSG00000004665 | *Cnn2* | 123.3387138 | 116.5658715 | 111.6162216 | 24.41586916 | 29.53519692 | 14.43340846 | 3.99E-13 | -2.364966198 | 5.27E-14 | 56.62811311 | 0.194121767 | 12.39942341 |
| ENSMUSG00000095307 | *Gm12573* | 22.96651911 | 49.48551147 | 39.16358654 | 9.301283491 | 4.763741439 | 7.697817843 | 9.41E-05 | -2.36539664 | 2.69E-05 | 17.62548116 | 0.194063858 | 4.016433998 |
| ENSMUSG00000034664 | *Itga2b* | 23.81713093 | 15.39549246 | 14.68634495 | 6.975962618 | 1.905496576 | 1.924454461 | 0.006876032 | -2.368775498 | 0.002625574 | 9.050956656 | 0.193609883 | 2.162662138 |
| ENSMUSG00000039304 | *Tnfsf10* | 92.71668827 | 90.17359867 | 80.28535241 | 16.27724611 | 15.24397261 | 19.24454461 | 9.71E-12 | -2.372829708 | 1.39E-12 | 50.19151445 | 0.193066571 | 11.01298088 |
| ENSMUSG00000030047 | *Arhgap25* | 2945.668729 | 3001.021351 | 2961.746232 | 581.3302182 | 583.0819522 | 546.5450668 | 9.67E-239 | -2.380920046 | 5.44E-241 | 1099.001903 | 0.191986924 | 238.0144196 |
| ENSMUSG00000023349 | *Clec4n* | 82.50934664 | 76.97746228 | 81.26444207 | 8.138623054 | 17.14946918 | 20.20677184 | 1.14E-09 | -2.382758545 | 1.90E-10 | 40.56652236 | 0.191742421 | 8.944621768 |
| ENSMUSG00000047798 | *Cd300lf* | 1207.868783 | 1198.649056 | 1232.673886 | 238.3453894 | 227.7068408 | 231.8967625 | 3.38E-128 | -2.38364931 | 4.23E-130 | 588.9560262 | 0.191624069 | 127.4705044 |
| ENSMUSG00000035211 | *Xrra1* | 31.4726373 | 31.89066294 | 37.20540721 | 4.650641745 | 7.621986303 | 6.735590613 | 3.40E-05 | -2.389124785 | 8.90E-06 | 19.73477849 | 0.190898175 | 4.469139675 |
| ENSMUSG00000040264 | *Gbp2b* | 166.7199165 | 128.6623298 | 112.5953113 | 32.55449222 | 18.10221747 | 27.90458968 | 1.44E-13 | -2.389817923 | 1.84E-14 | 58.69545318 | 0.190806481 | 12.84235841 |
| ENSMUSG00000067297 | *Ifit1bl2* | 54.43915641 | 83.57553048 | 51.89175217 | 16.27724611 | 5.716489727 | 14.43340846 | 8.67E-07 | -2.393949902 | 1.88E-07 | 27.15016798 | 0.190260779 | 6.061925396 |
| ENSMUSG00000041592 | *Sdk2* | 21.26529547 | 26.39227278 | 16.64452428 | 3.487981309 | 1.905496576 | 6.735590613 | 0.002047917 | -2.398730425 | 0.000712851 | 11.45543884 | 0.189631373 | 2.688687621 |
| ENSMUSG00000096802 | *None* | 291.7598539 | 224.3343186 | 302.538706 | 49.99439876 | 63.83413529 | 40.41354368 | 2.57E-24 | -2.408220993 | 1.96E-25 | 108.6270758 | 0.188388004 | 23.58936151 |
| ENSMUSG00000037820 | *Tgm2* | 413.397344 | 414.5786183 | 464.0885005 | 81.38623054 | 74.31436646 | 86.60045073 | 9.71E-49 | -2.415295697 | 3.62E-50 | 221.8240196 | 0.187466446 | 48.01269537 |
| ENSMUSG00000029602 | *Rasal1* | 264.5402757 | 304.610815 | 271.2078368 | 59.29568225 | 49.54291097 | 49.07358875 | 5.10E-33 | -2.41666762 | 2.88E-34 | 148.994696 | 0.187288261 | 32.29259303 |
| ENSMUSG00000021895 | *Arhgef3* | 419.3516268 | 370.591497 | 429.8203623 | 68.59696574 | 82.88910105 | 73.12926951 | 1.77E-45 | -2.430844818 | 7.18E-47 | 206.7086656 | 0.184533568 | 44.75130511 |
| ENSMUSG00000079419 | *Ms4a6c* | 946.7309545 | 922.6298693 | 826.351676 | 139.5192524 | 177.2111815 | 178.0120376 | 4.33E-76 | -2.440075318 | 9.84E-78 | 348.319826 | 0.184274032 | 75.36322546 |
| ENSMUSG00000032135 | *Mcam* | 18.71346002 | 23.09323868 | 20.56088293 | 2.325320873 | 0.952748288 | 3.848901921 | 0.00258716 | -2.442306824 | 0.000915642 | 10.99082096 | 0.183989224 | 2.587176668 |
| ENSMUSG00000032688 | *Malt1* | 6428.073516 | 6304.454161 | 6289.671999 | 1212.654835 | 1050.881362 | 1234.537537 | 2.67E-242 | -2.444028438 | 1.47E-244 | 1115.426249 | 0.183769795 | 241.5734125 |
| ENSMUSG00000034647 | *Ankrd12* | 1431.579691 | 1350.404624 | 1266.942025 | 244.1586916 | 204.8408819 | 291.5548508 | 2.17E-80 | -2.451084705 | 4.74E-82 | 368.1461249 | 0.182873165 | 79.66300548 |
| ENSMUSG00000027463 | *Slc52a3* | 31.4726373 | 32.99034098 | 35.24722789 | 3.487981309 | 4.763741439 | 9.622272304 | 5.25E-05 | -2.459398914 | 1.41E-05 | 18.85239991 | 0.181822304 | 4.279770214 |
| ENSMUSG00000062488 | *Ifit3b* | 261.9884402 | 216.6365724 | 248.6887745 | 36.04247353 | 52.40115583 | 42.33799814 | 1.53E-27 | -2.466662839 | 1.05E-28 | 123.5694821 | 0.180658515 | 26.81578558 |
| ENSMUSG00000029810 | *Tmem176b* | 745.1359534 | 830.2569146 | 781.3135515 | 131.3806293 | 161.0144607 | 131.8251306 | 4.88E-77 | -2.469690532 | 1.09E-78 | 352.7066138 | 0.18052987 | 76.31189634 |
| ENSMUSG00000025375 | *Aatk* | 155.6619629 | 160.5529928 | 171.3406911 | 32.55449222 | 24.77145549 | 30.79127137 | 2.12E-21 | -2.473638049 | 1.81E-22 | 95.09523126 | 0.180036578 | 20.67354677 |
| ENSMUSG00000024014 | *Pim1* | 4138.226499 | 4355.824687 | 4286.454587 | 806.8863428 | 759.1972848 | 795.9197833 | 3.71E-278 | -2.476515989 | 1.51E-280 | 1279.55462 | 0.179677792 | 277.1324965 |
| ENSMUSG00000009731 | *Kcnd1* | 148.8570683 | 163.8520269 | 152.7379875 | 29.06651091 | 20.00771405 | 34.64018029 | 9.88E-19 | -2.476828344 | 9.76E-20 | 82.65745259 | 0.179638895 | 18.00542643 |
| ENSMUSG00000062310 | *Glrp1* | 479.7450659 | 424.4757206 | 424.924914 | 86.03687229 | 60.02314214 | 93.33604135 | 9.95E-41 | -2.477639462 | 4.55E-42 | 184.7067051 | 0.179537926 | 40.00206838 |
| ENSMUSG00000027470 | *Mylk2* | 91.01546463 | 87.97424261 | 104.762594 | 17.43990655 | 17.14946918 | 16.35786292 | 2.78E-13 | -2.479226819 | 3.63E-14 | 57.35763737 | 0.179340494 | 12.55644719 |
| ENSMUSG00000004891 | *Nes* | 217.7566257 | 208.9388262 | 191.9015741 | 41.85577571 | 25.72420377 | 43.30022537 | 4.78E-23 | -2.487206112 | 3.82E-24 | 102.7401883 | 0.178351331 | 22.32055457 |
| ENSMUSG00000044703 | *Phf11a* | 340.2447276 | 335.4017999 | 331.9113959 | 51.1570592 | 58.11764556 | 68.31813336 | 1.99E-41 | -2.497756033 | 8.90E-43 | 187.9509509 | 0.177051868 | 40.70172777 |
| ENSMUSG00000020669 | *Sh3yl1* | 11.90856547 | 20.89388262 | 24.47724159 | 2.325320873 | 2.858244864 | 4.811136152 | 0.002615898 | -2.498926319 | 0.000926531 | 10.96891223 | 0.176908305 | 2.582379133 |
| ENSMUSG00000027562 | *Car2* | 10721.11137 | 9771.738997 | 10157.07617 | 1713.761483 | 1882.630617 | 1818.609465 | 3.1323761946335e-321 | -2.500153824 | 9.88131291682493e-324 | 1480.163245 | 0.176757848 | 320.5041261 |
| ENSMUSG00000079049 | *Serpinb1c* | 229.6651911 | 210.0385042 | 239.8769676 | 38.3677944 | 41.92092467 | 39.45131644 | 7.47E-30 | -2.503545474 | 4.76E-31 | 134.2745419 | 0.176342794 | 29.12654856 |
| ENSMUSG00000079339 | *Ifit1bl1* | 112.2807601 | 111.0674813 | 126.3025666 | 25.5785296 | 16.19672089 | 20.20677184 | 1.68E-15 | -2.508677749 | 1.92E-16 | 67.67942181 | 0.175716583 | 14.77502125 |
| ENSMUSG00000055523 | *Gucy2g* | 127.5917728 | 115.4661934 | 116.51167 | 16.27724611 | 20.00771405 | 25.98013522 | 4.44E-16 | -2.520105394 | 4.95E-17 | 70.35538651 | 0.174330222 | 15.35296299 |
| ENSMUSG00000029484 | *Anxa3* | 2488.890182 | 2395.098755 | 2379.187882 | 432.5096823 | 401.1070292 | 426.2866631 | 2.21E-219 | -2.528887369 | 1.39E-221 | 1009.706786 | 0.173272262 | 218.6558385 |
| ENSMUSG00000048965 | *Mrgpre* | 61.24405097 | 89.07392064 | 73.43172477 | 12.7892648 | 10.48023117 | 15.39563569 | 4.46E-10 | -2.529050973 | 7.24E-11 | 42.45311202 | 0.173252614 | 9.350347362 |
| ENSMUSG00000024663 | *Rab3il1* | 1309.942201 | 1372.398185 | 1333.520122 | 210.441539 | 252.4782963 | 226.1233991 | 3.85E-132 | -2.538895475 | 4.75E-134 | 607.1109773 | 0.172074417 | 131.4145738 |
| ENSMUSG00000024334 | *H2-Oa* | 15.31101274 | 20.89388262 | 23.49815192 | 2.325320873 | 1.905496576 | 5.773363382 | 0.001682427 | -2.559931938 | 0.000577202 | 11.84810145 | 0.169583541 | 2.774063695 |
| ENSMUSG00000037606 | *Osbpl5* | 18.71346002 | 19.79420459 | 27.41451058 | 3.487981309 | 4.763741439 | 2.886681691 | 0.000593589 | -2.560500226 | 0.000187599 | 13.95138578 | 0.169516754 | 3.226513921 |
| ENSMUSG00000040723 | *Rcsd1* | 1446.040092 | 1418.584662 | 1477.446302 | 266.2492399 | 240.0925685 | 230.9345353 | 4.02E-149 | -2.561965497 | 4.25E-151 | 685.5011259 | 0.169344672 | 148.3953154 |
| ENSMUSG00000030157 | *Clec2d* | 91.01546463 | 83.57553048 | 74.41081443 | 15.11458567 | 20.96046233 | 5.773363382 | 2.78E-09 | -2.576139618 | 4.81E-10 | 38.75328912 | 0.167689049 | 8.555250392 |
| ENSMUSG00000056290 | *Ms4a4b* | 18.71346002 | 24.19291672 | 18.60270361 | 4.650641745 | 1.905496576 | 3.848901921 | 0.000908098 | -2.580051546 | 0.000297777 | 13.08432621 | 0.167234969 | 3.041867224 |
| ENSMUSG00000054203 | *Ifi205* | 50.18609732 | 45.08679934 | 56.78720049 | 8.574734591 | 8.660045073 | | 4.85E-08 | -2.582280653 | 9.38E-09 | 32.96536643 | 0.166976774 | 7.31424266 |
| ENSMUSG00000018849 | *Wwc1* | 277.299453 | 283.7169324 | 262.3960298 | 45.34375702 | 56.21214899 | 35.60240752 | 1.05E-32 | -2.584613006 | 5.98E-34 | 147.5401718 | 0.166707047 | 31.97885092 |
| ENSMUSG00000041488 | *Stx3* | 3618.502678 | 3558.558113 | 3639.276279 | 596.4448038 | 587.8456936 | 611.9765185 | 1.19063985544718e-313 | -2.590129396 | 4.00311190592232e-316 | 1444.730992 | 0.166070831 | 312.9242196 |
| ENSMUSG00000090877 | *Hspa1b* | 140.3509501 | 158.3536367 | 177.2152291 | 20.92788785 | 25.72420377 | 31.7534986 | 2.75E-20 | -2.590788729 | 2.48E-21 | 89.91866574 | 0.165994952 | 19.56029405 |
| ENSMUSG00000074417 | *Pira12* | 42.53059095 | 30.79098491 | 41.12176587 | 0 | 8.574734591 | 9.622272304 | 0.00012468 | -2.612892896 | 3.53E-05 | 17.10791002 | 0.163471054 | 3.904202739 |
| ENSMUSG00000027360 | *Hdc* | 205.8480602 | 194.6430118 | 219.3160846 | 39.53045484 | 27.62970035 | 34.64018029 | 1.43E-27 | -2.615522766 | 9.77E-29 | 123.7058003 | 0.163128536 | 26.84475119 |
| ENSMUSG00000079554 | *Aox2* | 87.61301736 | 74.77810622 | 58.74537981 | 13.95192524 | 2.858244864 | 19.24454461 | 5.20E-07 | -2.625319734 | 1.10E-07 | 28.18177634 | 0.162069022 | 6.28398474 |
| ENSMUSG00000037754 | *Ppp1r16b* | 147.1558447 | 180.3471973 | 151.7588978 | 29.06651091 | 26.67695206 | 22.1312263 | 1.28E-21 | -2.625534424 | 1.08E-22 | 96.12268609 | 0.162044906 | 20.89298155 |
| ENSMUSG00000079173 | *Zan* | 91.01546463 | 103.3697351 | 60.70355914 | 17.43990655 | 9.527482879 | 14.43340846 | 2.05E-10 | -2.635207042 | 3.25E-11 | 44.02240554 | 0.160962102 | 9.688445255 |
| ENSMUSG00000044770 | *Scml4* | 26.36986639 | 28.59162885 | 26.43542092 | 2.325320873 | 1.905496576 | 8.660045073 | 0.000309974 | -2.637046572 | 9.33E-05 | 15.26752413 | 0.160756996 | 3.508674577 |
| ENSMUSG00000032554 | *Trf* | 153.1101274 | 123.1639396 | 114.5534906 | 29.06651091 | 18.10221747 | 16.35786292 | 5.97E-16 | -2.641293448 | 6.71E-17 | 69.75557323 | 0.16028447 | 15.22409701 |
| ENSMUSG00000063506 | *Arhgap22* | 1206.167559 | 1251.433601 | 1221.9039 | 210.441539 | 200.0771405 | 178.0120376 | 7.69E-136 | -2.647306223 | 9.02E-138 | 624.2235888 | 0.159617836 | 135.1142704 |
| ENSMUSG00000079457 | *Gm7609* | 1558.320852 | 1638.520269 | 1651.724262 | 245.3213521 | 253.4310446 | 269.4236245 | 3.75E-173 | -2.655796389 | 3.27E-175 | 796.403748 | 0.158681254 | 172.4262717 |
| ENSMUSG00000010021 | *Kif19a* | 23.81713093 | 12.09645836 | 16.64452428 | 5.813302182 | 1.905496576 | 0.96222723 | 0.004244359 | -2.655857656 | 0.001559818 | 10.00658126 | 0.158674516 | 2.37218786 |

| | | | | | | | | | | | | | |
|---|---|---|---|---|---|---|---|---|---|---|---|---|---|
| ENSMUSG00000039103 | *Nexn* | 25.51835457 | 21.99356065 | 16.64452428 | 3.487981309 | 2.858244864 | 3.848908921 | 0.000477335 | -2.657639904 | 0.000147943 | 14.39820949 | 0.158478617 | 3.32117716 |
| ENSMUSG00000076441 | *Ass1* | 499.3091377 | 532.2441678 | 469.9630385 | 81.38623054 | 80.03085618 | 76.0159512 | 7.25E-65 | -2.661612713 | 1.99E-66 | 296.4188203 | 0.158042808 | 64.13938894 |
| ENSMUSG00000034171 | *Faah* | 72.30200461 | 58.28293573 | 62.66173847 | 10.46394393 | 8.574734591 | 11.54672676 | 4.12E-10 | -2.66385008 | 6.67E-11 | 42.61316867 | 0.157797901 | 9.384597926 |
| ENSMUSG00000021356 | *Irf4* | 156.5125747 | 171.5497731 | 145.8843599 | 25.5785296 | 24.77145549 | 24.05568076 | 3.92E-23 | -2.671919376 | 3.13E-24 | 103.1356055 | 0.156917768 | 22.40649078 |
| ENSMUSG00000092556 | *Or11a3-ps1* | 30.62202548 | 26.39227278 | 35.24722789 | 8.138623054 | 2.858244864 | 3.848908921 | 4.31E-05 | -2.672279004 | 1.15E-05 | 19.24921059 | 0.156878657 | 4.365398219 |
| ENSMUSG00000072572 | *Slc39a2* | 95.26852373 | 80.27649638 | 91.05533871 | 13.95192524 | 16.19672089 | 11.54672676 | 1.62E-13 | -2.6782909 | 2.08E-14 | 58.45515488 | 0.156226284 | 12.79117353 |
| ENSMUSG00000053338 | *Tarm1* | 75.70445189 | 73.67842818 | 93.01351804 | 12.7892648 | 13.33847603 | 11.54672676 | 2.17E-12 | -2.686703138 | 3.00E-13 | 53.21124052 | 0.155317992 | 11.6640261 |
| ENSMUSG00000024737 | *Slc15a3* | 7158.749069 | 7550.389372 | 7444.018712 | 1210.329514 | 1116.620993 | 1084.430089 | 0 | -2.700569293 | 0 | 1696.23939 | 0.153832337 | Inf |
| ENSMUSG00000097328 | *Tnfsf12* | 797.8738862 | 886.3404943 | 857.6825453 | 123.2420063 | 135.2902569 | 128.9384489 | 2.90E-103 | -2.710477613 | 5.12E-105 | 473.6598082 | 0.152779448 | 102.5369802 |
| ENSMUSG00000032204 | *Aqp9* | 568.2086951 | 546.5399822 | 589.4119774 | 95.33815578 | 96.22757708 | 69.28036059 | 7.48E-65 | -2.712309282 | 2.05E-66 | 296.3538646 | 0.1525856 | 64.1263238 |
| ENSMUSG00000061972 | *Or1o4* | 133.5460556 | 138.5594321 | 142.9470909 | 19.76522742 | 20.96046233 | 22.1312263 | 1.82E-21 | -2.719602839 | 1.55E-22 | 95.40566817 | 0.151816148 | 20.73990602 |
| ENSMUSG00000030427 | *Lilra6* | 22.11590729 | 21.99356065 | 29.37268991 | 2.325320873 | 3.810993152 | 4.811136152 | 0.000143123 | -2.726855184 | 4.09E-05 | 16.8295231 | 0.151054892 | 3.84429186 |
| ENSMUSG00000040133 | *Gpr176* | 115.6832074 | 91.2732767 | 76.36899376 | 13.95192524 | 12.38572774 | 16.35786292 | 3.33E-13 | -2.732205079 | 4.38E-14 | 56.99201681 | 0.150495778 | 12.47820779 |
| ENSMUSG00000027204 | *Fbn1* | 11.90856547 | 15.39549246 | 14.68634495 | 3.487981309 | 1.905496576 | 0.96222723 | 0.004798962 | -2.757716306 | 0.001777466 | 9.766229063 | 0.147857947 | 2.318852693 |
| ENSMUSG00000023990 | *Tfeb* | 397.2357195 | 404.681516 | 491.5030111 | 74.41026793 | 57.16489727 | 60.62031551 | 4.85E-49 | -2.758088692 | 1.80E-50 | 223.2131248 | 0.147819787 | 48.31407683 |
| ENSMUSG00000074577 | *Ripor3* | 35.7256964 | 32.99034098 | 33.28904856 | 2.325320873 | 4.763741439 | 7.697817843 | 7.81E-06 | -2.759069478 | 1.88E-06 | 22.7184749 | 0.147719329 | 5.107606584 |
| ENSMUSG00000040855 | *Reps2* | 20.41468366 | 12.09645836 | 22.51906226 | 2.325320873 | 1.905496576 | 3.848908921 | 0.001265647 | -2.766014745 | 0.000425009 | 12.4189083 | 0.147009904 | 2.897687465 |
| ENSMUSG00000024675 | *Ms4a4c* | 49.3354855 | 50.5851895 | 53.84993149 | 8.138623054 | 9.527482879 | 4.811136152 | 1.50E-08 | -2.777887819 | 2.76E-09 | 35.34416519 | 0.145805008 | 7.82302083 |
| ENSMUSG00000022180 | *Slc7a8* | 1051.356208 | 999.6073316 | 983.9851119 | 161.6098007 | 124.8100257 | 153.9563569 | 3.87E-111 | -2.790187114 | 5.92E-113 | 510.1386718 | 0.144567272 | 110.4126824 |
| ENSMUSG00000073489 | *Ifi204* | 2697.290078 | 2634.828566 | 2592.629429 | 377.8646418 | 401.1070292 | 364.6841203 | 1.27E-264 | -2.792383828 | 5.73E-267 | 1218.530305 | 0.144347314 | 263.8977927 |
| ENSMUSG00000045932 | *Ifit2* | 3126.849047 | 3105.490764 | 2947.059887 | 432.5096823 | 392.5322946 | 491.6981147 | 7.18E-204 | -2.801170545 | 4.88E-206 | 938.1976366 | 0.143470841 | 203.1437535 |
| ENSMUSG00000040026 | *Saa3* | 120.7868783 | 109.9678033 | 123.3652976 | 16.27724611 | 17.14946918 | 17.32009015 | 4.60E-19 | -2.801701083 | 4.47E-20 | 84.20258386 | 0.14341809 | 18.33696494 |
| ENSMUSG00000030630 | *Fah* | 63.79588642 | 54.98390163 | 44.05903486 | 5.813302182 | 7.621986303 | 9.622272304 | 8.20E-09 | -2.808847335 | 1.47E-09 | 36.57008242 | 0.142709439 | 8.086428244 |
| ENSMUSG00000019558 | *Slc6a8* | 2216.6944 | 2322.520005 | 2256.801674 | 320.8942804 | 333.4619008 | 314.6483043 | 4.89E-248 | -2.809401901 | 2.59E-250 | 1141.898935 | 0.142654593 | 247.3103684 |
| ENSMUSG00000076617 | *Ighm* | 313.8757612 | 340.9001901 | 338.7650236 | 49.99439876 | 38.10993152 | 53.8847249 | 2.83E-45 | -2.809462917 | 1.15E-46 | 205.7775137 | 0.142648559 | 44.54887242 |
| ENSMUSG00000081176 | *H3f3a-ps2* | 27.21957821 | 21.99356065 | 19.58179327 | 0 | 9.527482879 | 0 | 0.009397251 | -2.813911207 | 0.003692744 | 8.429044016 | 0.142209406 | 2.026999172 |
| ENSMUSG00000089844 | *A530032D15Rik* | 210.1011193 | 203.440436 | 194.838843 | 16.27724611 | 31.4406935 | 37.52686198 | 8.89E-25 | -2.815135458 | 6.64E-26 | 110.7703804 | 0.14208878 | 24.05103925 |
| ENSMUSG00000035356 | *Nfkbiz* | 3086.019679 | 3039.510082 | 3000.909819 | 475.5281185 | 389.6740497 | 432.0400264 | 3.47E-239 | -2.818462738 | 1.93E-241 | 1101.072685 | 0.141761459 | 238.4591634 |
| ENSMUSG00000026411 | *Tmem9* | 403.1900022 | 345.2989022 | 385.7613274 | 43.01843614 | 68.59787673 | 48.11136152 | 1.01E-43 | -2.819629833 | 4.25E-45 | 198.5867184 | 0.141646825 | 42.99704943 |
| ENSMUSG00000073555 | *Gm4951* | 16.16162456 | 18.69452655 | 17.62361394 | 5.813302182 | 0 | 1.924454461 | 0.003655408 | -2.827680482 | 0.001325066 | 10.3072771 | 0.140858596 | 2.437064175 |
| ENSMUSG00000034459 | *Ifit1* | 3692.505906 | 3533.265519 | 3684.314404 | 483.6667415 | 495.4291097 | 552.3184302 | 5.01E-290 | -2.831087047 | 2.03E-292 | 1335.644421 | 0.140526386 | 289.3000041 |
| ENSMUSG00000026548 | *Slamf9* | 71.45139279 | 45.08679934 | 61.6826488 | 3.487981309 | 15.24397261 | 5.773363382 | 1.72E-07 | -2.837841284 | 3.49E-08 | 30.41580494 | 0.139870024 | 6.763821565 |
| ENSMUSG00000000184 | *Ccnd2* | 2586.710542 | 2519.362373 | 2353.731551 | 381.3526231 | 311.5486901 | 352.1751663 | 5.99E-196 | -2.839290123 | 4.23E-198 | 901.6812487 | 0.139729629 | 195.2229206 |
| ENSMUSG00000029491 | *Pde6b* | 69.75016916 | 71.47907212 | 56.78720049 | 10.46394393 | 6.669238015 | 10.58449953 | 8.04E-11 | -2.844683527 | 1.23E-11 | 45.91841457 | 0.139208236 | 10.09480031 |
| ENSMUSG00000108900 | *Ccdc194* | 19.56407184 | 21.99356065 | 24.47724159 | 3.487981309 | 2.858244864 | 2.886681691 | 0.000167186 | -2.844908804 | 4.81E-05 | 16.52019514 | 0.1391865 | 3.776798866 |
| ENSMUSG00000043421 | *Hilpda* | 2306.859253 | 2255.439645 | 2320.442503 | 322.0569409 | 304.8794521 | 329.0817128 | 1.11E-256 | -2.848624634 | 5.34E-259 | 1181.861608 | 0.13882847 | 255.9542165 |
| ENSMUSG00000034220 | *Gpc1* | 32.32324912 | 31.89066294 | 31.33086923 | 4.650641745 | 4.763741439 | 3.848908921 | 4.02E-06 | -2.851657894 | 9.40E-07 | 24.04637265 | 0.138536891 | 5.395593157 |
| ENSMUSG00000001827 | *Folr1* | 38.27753185 | 21.99356065 | 27.41451058 | 3.487981309 | 2.858244864 | 5.773363382 | 3.24E-05 | -2.854131789 | 8.45E-06 | 19.83281013 | 0.138299535 | 4.489591274 |
| ENSMUSG00000000317 | *Bcl6b* | 1222.329184 | 1206.346802 | 1237.569335 | 182.5376885 | 170.5419435 | 152.0319024 | 5.41E-151 | -2.863809273 | 5.60E-153 | 694.1466704 | 0.137374937 | 150.2668415 |
| ENSMUSG00000049723 | *Mmp12* | 366.613694 | 338.700834 | 336.8068443 | 54.64504051 | 42.87367295 | 46.18690706 | 7.90E-51 | -2.867021939 | 2.82E-52 | 231.4926008 | 0.137069364 | 50.10258887 |
| ENSMUSG00000021876 | *Rnase4* | 395.5344958 | 373.8905311 | 392.6149551 | 48.83173833 | 48.59016268 | 60.62031551 | 1.12E-56 | -2.875126173 | 3.57E-58 | 258.5385836 | 0.136301546 | 55.95173373 |
| ENSMUSG00000021125 | *Arg2* | 126.741161 | 115.4661934 | 122.3862079 | 12.7892648 | 21.91321062 | 14.43340846 | 8.06E-19 | -2.879536856 | 7.95E-20 | 83.06198401 | 0.135885474 | 18.09339666 |
| ENSMUSG00000050022 | *Amz1* | 329.1867739 | 341.9998681 | 331.9113959 | 37.20513396 | 48.59016268 | 49.07358875 | 5.86E-50 | -2.884895917 | 2.13E-51 | 227.4638867 | 0.135381647 | 49.23225742 |
| ENSMUSG00000038781 | *Stap2* | 32.32324912 | 26.39227278 | 40.1426762 | 6.975962618 | 2.858244864 | 3.848908921 | 7.69E-06 | -2.885112591 | 1.85E-06 | 22.74805113 | 0.135361316 | 5.11404273 |
| ENSMUSG00000048058 | *Ldlrad3* | 91.01546463 | 58.28293573 | 81.26444207 | 5.813302182 | 10.48023117 | 14.43340846 | 6.27E-11 | -2.889496187 | 9.56E-12 | 46.4163512 | 0.134950648 | 10.20264251 |
| ENSMUSG00000000730 | *Dnmt3l* | 176.9272583 | 201.24108 | 203.65065 | 16.27724611 | 40.96817638 | 20.20677184 | 1.22E-20 | -2.89116292 | 1.09E-21 | 91.55559497 | 0.134794831 | 19.91225144 |
| ENSMUSG00000062783 | *Csprs* | 4312.601922 | 4009.426107 | 3940.835896 | 516.2212337 | 541.1610275 | 587.9208378 | 1.24E-285 | -2.896265032 | 5.10E-288 | 1315.396988 | 0.134318969 | 284.907317 |
| ENSMUSG00000036067 | *Slc2a6* | 5678.684504 | 5857.98488 | 6111.47768 | 808.0490033 | 759.3403854 | 803.4597374 | 0 | -2.896706324 | 0 | 1844.711974 | 0.13427789 | Inf |
| ENSMUSG00000020303 | *Stc2* | 753.6420716 | 811.562388 | 761.7317582 | 99.98879752 | 122.9045291 | 87.56267796 | 1.47E-89 | -2.90339724 | 2.92E-91 | 410.4512472 | 0.133656579 | 88.8314174 |
| ENSMUSG00000023367 | *Tmem176a* | 214.3541784 | 234.2314209 | 206.587919 | 26.74119004 | 26.67695206 | 32.71572583 | 3.46E-34 | -2.922726525 | 1.88E-35 | 154.415541 | 0.131877786 | 33.46068022 |
| ENSMUSG00000068859 | *Sp9* | 66.34772188 | 23.09323868 | 49.93357284 | 6.975962618 | 11.43297945 | 0 | 0.000169347 | -2.927957904 | 4.88E-05 | 16.4940427 | 0.131400448 | 3.771222458 |
| ENSMUSG00000074896 | *Ifit3* | 1780.330537 | 1705.600629 | 1638.996097 | 218.580162 | 202.9353853 | 250.1790799 | 1.72E-172 | -2.931124996 | 1.51E-174 | 793.3386425 | 0.131112306 | 171.7632635 |
| ENSMUSG00000017309 | *Cd300lg* | 50.18609732 | 50.5851895 | 55.80811082 | 9.301283491 | 5.716489727 | 5.773363382 | 2.48E-09 | -2.932975827 | 4.26E-10 | 38.9911336 | 0.13094421 | 8.60626324 |
| ENSMUSG00000038418 | *Egr1* | 1525.997603 | 1488.964056 | 1507.798082 | 206.9535577 | 162.9199572 | 223.2367174 | 2.70E-140 | -2.933237277 | 3.06E-142 | 644.7727985 | 0.130920482 | 139.5679737 |
| ENSMUSG00000023046 | *Igfbp6* | 31.4726373 | 32.99034098 | 21.5399726 | 4.650641745 | 6.669238015 | 0 | 0.000190934 | -2.934598306 | 5.54E-05 | 16.25315215 | 0.130797031 | 3.719116088 |
| ENSMUSG00000027579 | *Srms* | 21.26529547 | 21.99356065 | 19.58179327 | 3.487981309 | 0.952748288 | 3.848908921 | 0.00028056 | -2.938984384 | 8.38E-05 | 15.47017384 | 0.130399986 | 3.551975019 |
| ENSMUSG00000045502 | *Hcar2* | 923.7644354 | 848.9514411 | 850.8289176 | 120.9166854 | 93.36933221 | 127.0139944 | 4.56E-100 | -2.946372884 | 8.30E-102 | 458.9101014 | 0.129733873 | 99.34059448 |
| ENSMUSG00000089809 | *Rasgef1b* | 3117.492317 | 2939.439381 | 2982.307115 | 409.2564736 | 385.8630566 | 379.1175288 | 1.52E-303 | -2.946826588 | 5.53E-306 | 1398.067072 | 0.12969308 | 302.8182386 |
| ENSMUSG00000079455 | *Gm16026* | 183.7321529 | 181.4468754 | 192.8806637 | 30.22917134 | 24.77145549 | 17.32009015 | 2.33E-28 | -2.96185739 | 1.55E-29 | 127.3562589 | 0.12834888 | 27.63306341 |
| ENSMUSG00000037681 | *Esyt3* | 17.01223638 | 8.797424261 | 21.5399726 | 1.162660436 | 1.905496576 | 2.886681691 | 0.002217777 | -2.973522788 | 0.000775476 | 11.29902923 | 0.127315256 | 2.654082143 |
| ENSMUSG00000047963 | *Stbd1* | 25.51835457 | 23.09323868 | 24.47724159 | 4.650641745 | 2.858244864 | 1.924454461 | 5.27E-05 | -2.983366307 | 1.42E-05 | 18.84339289 | 0.12644954 | 4.277940986 |
| ENSMUSG00000021403 | *Serpinb9b* | 290.0586303 | 288.1156445 | 266.3123885 | 34.87981309 | 36.20443494 | 35.60240752 | 7.14E-46 | -2.983652562 | 2.86E-47 | 208.5430036 | 0.126424453 | 45.14634352 |
| ENSMUSG00000038352 | *Arl5c* | 1373.738088 | 1227.240684 | 1323.729225 | 165.097782 | 202.9353853 | 127.9762216 | 4.89E-94 | -2.984154942 | 9.33E-96 | 431.1068748 | 0.126380436 | 93.31048847 |
| ENSMUSG00000039062 | *Anpep* | 1155.981462 | 1173.356461 | 1158.263072 | 130.2179689 | 139.10125 | 165.5030836 | 3.35E-145 | -2.998752141 | 3.67E-147 | 667.4002767 | 0.125108166 | 144.4755713 |
| ENSMUSG00000031024 | *Dennd2b* | 27.21957821 | 19.79420459 | 44.05903486 | 5.813302182 | 0.952748288 | 4.811136152 | 6.41E-05 | -3.005594555 | 1.75E-05 | 18.44244562 | 0.124516207 | 4.193156159 |
| ENSMUSG00000024736 | *Tmem132a* | 1365.231969 | 1338.308166 | 1292.398356 | 184.8630094 | 151.4869778 | 160.6919475 | 6.81E-165 | -3.0131471 | 6.54E-167 | 758.223278 | 0.123866065 | 164.1671715 |
| ENSMUSG00000027514 | *Zbp1* | 1234.237749 | 1321.812995 | 1275.753832 | 144.1698941 | 158.1562158 | 169.3519925 | 5.90E-165 | -3.017441471 | 5.62E-167 | 758.5235618 | 0.12349791 | 164.2293498 |
| ENSMUSG00000048621 | *Gm6377* | 41.67997913 | 21.99356065 | 33.28904856 | 1.162660436 | 1.905496576 | 8.660045073 | 6.67E-05 | -3.019466687 | 1.83E-05 | 18.36215488 | 0.123324668 | 4.176191734 |
| ENSMUSG00000030103 | *Bhlhe40* | 3023.925016 | 3065.902355 | 3044.968854 | 362.7500561 | 394.4377912 | 365.6463475 | 0 | -3.022615876 | 0 | 1537.110205 | 0.123055763 | Inf |
| ENSMUSG00000020598 | *Nrcam* | 24.66774275 | 27.49195081 | 31.33086923 | 4.650641745 | 4.763741439 | 0.96222723 | 2.69E-05 | -3.025566156 | 6.93E-06 | 20.21302696 | 0.122804373 | 4.570888534 |
| ENSMUSG00000028716 | *Pdzk1ip1* | 29.77141366 | 21.99356065 | 22.51906226 | 2.325320873 | 5.716489727 | 0.96222723 | 0.000107783 | -3.032444456 | 3.03E-05 | 17.40078446 | 0.122220276 | 3.967449097 |
| ENSMUSG00000061666 | *Gdpd1* | 473.7907832 | 460.7650956 | 472.9003075 | 56.97036138 | 60.97589042 | 53.8847249 | 1.78E-75 | -3.033805543 | 4.06E-77 | 345.4943483 | 0.122105023 | 74.75053844 |
| ENSMUSG00000079652 | *Garin1a* | 54.43915641 | 50.5851895 | 53.84993149 | 4.650641745 | 4.763741439 | 9.622272304 | 1.51E-09 | -3.047306392 | 2.56E-10 | 39.98422197 | 0.120967685 | 8.820136104 |
| ENSMUSG00000018819 | *Lsp1* | 3150.666178 | 3248.448908 | 3299.532166 | 361.5873957 | 418.2564984 | 388.7398011 | 1.53582299142597e-310 | -3.049712815 | 5.26905102039929e-313 | 1430.375897 | 0.120766079 | 309.8136588 |
| ENSMUSG00000023935 | *Spats1* | 16.16162456 | 8.797424261 | 17.62361394 | 2.325320873 | 1.905496576 | 0.96222723 | 0.002624743 | -3.060677814 | 0.000930204 | 10.96158083 | 0.119851692 | 2.580913235 |
| ENSMUSG00000000982 | *Ccl3* | 61136.87388 | 57503.264 | 58044.35161 | 6835.280705 | 7054.148324 | 7117.594823 | 0 | -3.072077826 | 0 | 3599.493636 | 0.11890837 | Inf |
| ENSMUSG00000015355 | *Cd48* | 83.35995826 | 60.48229179 | 71.47354544 | 10.46394393 | 5.716489727 | 9.622272304 | 2.89E-12 | -3.077295763 | 4.02E-13 | 52.63142088 | 0.118479079 | 11.53947361 |
| ENSMUSG00000052631 | *Sh2d6* | 182.8815411 | 206.7394701 | 179.1734084 | 22.09054829 | 28.58244864 | 16.35786292 | 3.24E-29 | -3.082446951 | 2.10E-30 | 131.3303311 | 0.1180568 | 28.48990091 |
| ENSMUSG00000006720 | *Zfp184* | 14.46040092 | 19.79420459 | 17.62361394 | 1.162660436 | 1.905496576 | 2.886681691 | 0.000565457 | -3.095179681 | 0.000177932 | 14.05086077 | 0.117019456 | 3.247600315 |
| ENSMUSG00000001930 | *Vwf* | 109.7289246 | 82.47585244 | 113.574401 | 15.11458567 | 7.621986303 | 13.47118123 | 5.77E-16 | -3.095340329 | 6.48E-17 | 69.82504826 | 0.117006426 | 15.23912864 |
| ENSMUSG00000098557 | *Kctd12* | 1355.024628 | 1305.317825 | 1305.126521 | 148.8205359 | 136.2430052 | 177.0498104 | 1.42E-157 | -3.100103403 | 1.41E-159 | 724.4882269 | 0.116620765 | 156.8470818 |
| ENSMUSG00000040522 | *Tlr8* | 8.50611819 | 24.19291672 | 11.74907596 | 2.325320873 | 0 | 2.886681691 | 0.005103374 | -3.104327512 | 0.001898971 | 9.644715922 | 0.116279807 | 2.29214259 |
| ENSMUSG00000042622 | *Maff* | 1564.275135 | 1583.536367 | 1703.616015 | 197.6522742 | 223.8958477 | 140.4851756 | 6.17E-108 | -3.111086428 | 1.03E-109 | 495.246532 | 0.115736319 | 107.2097887 |
| ENSMUSG00000043051 | *Disc1* | 33.17386094 | 35.18969704 | 29.37268991 | 4.650641745 | 4.763741439 | 1.924454461 | 1.59E-06 | -3.119847345 | 3.57E-07 | 25.91430595 | 0.115035628 | 5.797740561 |
| ENSMUSG00000057596 | *Trim30d* | 119.9362665 | 129.7620078 | 131.1980149 | 24.41586916 | 11.43297945 | 8.660045073 | 6.12E-18 | -3.131067602 | 6.25E-19 | 78.98825842 | 0.114144433 | 17.21298997 |
| ENSMUSG00000031504 | *Rab20* | 1428.177244 | 1520.854719 | 1467.655406 | 168.5857633 | 186.7386644 | 148.1829935 | 7.50E-177 | -3.132386672 | 6.28E-179 | 813.4946554 | 0.114040117 | 176.1246705 |
| ENSMUSG00000071068 | *Treml2* | 831.898359 | 831.3565926 | 801.8744344 | 116.2660436 | 76.21986303 | 89.48713242 | 1.05E-99 | -3.140040662 | 1.92E-101 | 457.2394166 | 0.113436697 | 98.97865708 |
| ENSMUSG00000015312 | *Gadd45b* | 3612.548395 | 3675.123985 | 3762.641577 | 426.6963801 | 412.5400087 | 395.4753917 | 0 | -3.163379899 | 0 | 1811.247233 | 0.111616336 | Inf |
| ENSMUSG00000090176 | *Cd200r2* | 8.50611819 | 12.09645836 | 16.64452428 | 2.325320873 | 0.952748288 | 0.96222723 | 0.004183184 | -3.171722881 | 0.001535435 | 10.03560009 | 0.110972731 | 2.378493023 |
| ENSMUSG00000015340 | *Cybb* | 31264.23741 | 30368.70855 | 30311.63689 | 3413.571041 | 3176.462792 | 3549.656253 | 0 | -3.180936575 | 0 | 2915.569729 | 0.110266268 | Inf |
| ENSMUSG00000006457 | *Actn3* | 28.07019003 | 47.2861554 | 38.18449688 | 5.813302182 | 5.716489727 | 0.96222723 | 1.40E-06 | -3.200608217 | 3.12E-07 | 26.17277314 | 0.108772954 | 5.853078306 |
| ENSMUSG00000018930 | *Ccl4* | 54135.488 | 52667.97969 | 53283.03858 | 5620.300549 | 5646.939102 | 6090.898368 | 0 | -3.204943806 | 0 | 3541.280168 | 0.10844656 | Inf |
| ENSMUSG00000044734 | *Serpinb1a* | 22.96651911 | 19.79420459 | 23.49815192 | 3.487981309 | 0.952748288 | 2.886681691 | 7.06E-05 | -3.206261633 | 1.94E-05 | 18.24589251 | 0.108347545 | 4.150899438 |
| ENSMUSG00000074973 | *Gm11382* | 34.02447276 | 35.18969704 | 46.01721419 | 4.650641745 | 2.858244864 | 4.811136152 | 9.33E-08 | -3.234009101 | 1.85E-08 | 31.64544279 | 0.1062836 | 7.030184154 |
| ENSMUSG00000055541 | *Lair1* | 684.7425143 | 519.0480314 | 578.6419911 | 54.64504051 | 80.03085618 | 51.96027044 | 8.39E-62 | -3.25102788 | 2.42E-63 | 282.2527137 | 0.105037189 | 61.07600563 |
| ENSMUSG00000056366 | *Fabp3-ps1* | 24.66774275 | 15.39549246 | 29.37268991 | 4.650641745 | 2.858244864 | 0 | 0.000205582 | -3.259968883 | 6.00E-05 | 16.10287044 | 0.104388241 | 3.68701578 |
| ENSMUSG00000035373 | *Ccl7* | 338.543504 | 291.4146786 | 357.3677272 | 23.25320873 | 44.77916953 | 33.67795306 | 1.39E-44 | -3.263119292 | 5.71E-46 | 202.5802507 | 0.104160538 | 43.85706704 |
| ENSMUSG00000045659 | *Plekha7* | 17.8628482 | 31.89066294 | 29.37268991 | 2.325320873 | 1.905496576 | 3.848908921 | 1.71E-05 | -3.279268763 | 4.29E-06 | 21.12941017 | 0.103001071 | 4.766677793 |
| ENSMUSG00000021367 | *Edn1* | 136.097891 | 133.0610419 | 106.7207733 | 6.975962618 | 19.05496576 | 11.54672676 | 1.46E-19 | -3.296954769 | 1.37E-20 | 86.54522673 | 0.101746088 | 18.8361734 |
| ENSMUSG00000066621 | *Tecpr1* | 3624.456961 | 3913.754118 | 3828.240584 | 425.5337197 | 345.8476285 | 387.7775738 | 1.35E-305 | -3.2973433 | 4.71E-308 | 1407.589642 | 0.101718691 | 304.8708046 |
| ENSMUSG00000046971 | *Pla2g4f* | 13.6097891 | 14.29581442 | 12.72816563 | 1.162660436 | 0 | 2.886681691 | 0.002452669 | -3.321376276 | 0.000865014 | 11.09627183 | 0.100038257 | 2.610361017 |
| ENSMUSG00000032841 | *Prr5l* | 39.97875549 | 27.49195081 | 13.70725529 | 2.325320873 | 4.763741439 | 0.96222723 | 7.38E-05 | -3.328068527 | 2.03E-05 | 18.16065034 | 0.099575282 | 4.132093797 |
| ENSMUSG00000046733 | *Gprc5a* | 449.1230404 | 406.8808721 | 442.5485279 | 45.34375702 | 43.82642124 | 39.45131644 | 2.23E-75 | -3.33996238 | 5.12E-77 | 345.0296825 | 0.098757739 | 74.65194752 |
| ENSMUSG00000038305 | *Spats2l* | 127.5917728 | 104.4694131 | 99.86714568 | 15.11458567 | 9.527482879 | 8.660045073 | 1.16E-19 | -3.341093295 | 1.08E-20 | 87.00767121 | 0.098680354 | 18.93485094 |
| ENSMUSG00000030577 | *Cd22* | 7.655506371 | 15.39549246 | 8.811806972 | 2.325320873 | 0 | 0.96222723 | 0.008949242 | -3.341957616 | 0.003496431 | 8.528419971 | 0.098621252 | 2.048213726 |
| ENSMUSG00000030142 | *Clec4e* | 24340.2572 | 24783.44382 | 24347.02266 | 2184.63896 | 2503.822501 | 2485.432936 | 0 | -3.355091623 | 0 | 2620.251902 | 0.097727499 | Inf |
| ENSMUSG00000082292 | *Gm12250* | 180.3297056 | 164.9517049 | 191.9015741 | 16.27724611 | 16.19672089 | 19.24454461 | 1.39E-33 | -3.374283518 | 7.68E-35 | 151.6168909 | 0.096436058 | 32.85719821 |
| ENSMUSG00000045991 | *Onecut2* | 12.75917728 | 19.79420459 | 10.7699863 | 2.325320873 | 0.952748288 | 0.96222723 | 0.00122544 | -3.389633488 | 0.000410582 | 12.48341026 | 0.095415437 | 2.911708085 |
| ENSMUSG00000055782 | *Abcd2* | 34.02447276 | 35.18969704 | 26.43542092 | 2.325320873 | 1.905496576 | 4.811136152 | 7.93E-07 | -3.390012256 | 1.71E-07 | 27.33207032 | 0.09539039 | 6.100996017 |
| ENSMUSG00000017144 | *Rnd3* | 9.356730009 | 15.39549246 | 6.853627645 | 0 | 0.952748288 | 1.924454461 | 0.007686075 | -3.392416233 | 0.00296314 | 8.830027287 | 0.095231573 | 2.114295393 |
| ENSMUSG00000050350 | *Gpr18* | 35.7256964 | 41.78776524 | 40.1426762 | 3.487981309 | 3.810993152 | 3.848908921 | 1.34E-08 | -3.394065809 | 2.44E-09 | 35.58424252 | 0.095122747 | 7.873621965 |
| ENSMUSG00000112148 | *Lilrb4a* | 31582.36623 | 29351.50637 | 29698.72676 | 2950.832187 | 2213.234273 | 3443.811257 | 3.98E-63 | -3.396360952 | 1.13E-64 | 288.3686971 | 0.094971539 | 62.40036544 |
| ENSMUSG00000039115 | *Itga9* | 43.38120277 | 32.99034098 | 38.18449688 | 0 | 9.527482879 | 0.96222723 | 1.91E-05 | -3.398840601 | 4.80E-06 | 20.91502918 | 0.094808446 | 4.719980529 |
| ENSMUSG00000028128 | *F3* | 16.16162456 | 9.897102293 | 6.853627645 | 2.325320873 | 0.952748288 | 0 | 0.0075212 | -3.402855414 | 0.002895821 | 8.871972302 | 0.094544975 | 2.123712853 |
| ENSMUSG00000105096 | *Gbp10* | 56.14038005 | 57.18325769 | 95.95078703 | 11.62660436 | 7.621986303 | 0.96222723 | 3.06E-09 | -3.407614597 | 5.32E-10 | 38.55780544 | 0.094233602 | 8.513647979 |
| ENSMUSG00000028268 | *Gbp3* | 476.3426186 | 477.2602661 | 492.4821008 | 44.18109658 | 41.92092467 | 48.11136152 | 1.85E-88 | -3.42897148 | 3.74E-90 | 405.3639926 | 0.092848893 | 87.73296319 |
| ENSMUSG00000054404 | *Slfn5* | 4248.806036 | 4134.789402 | 4059.305745 | 416.2324362 | 363.949846 | 374.3063926 | 0 | -3.433872479 | 0 | 1945.106041 | 0.09253401 | Inf |
| ENSMUSG00000007594 | *Hapln4* | 97.82035918 | 89.07392064 | 97.90896635 | 3.487981309 | 10.48023117 | 11.54672676 | 1.71E-17 | -3.44576943 | 1.79E-18 | 76.90469069 | 0.091774081 | 16.76641242 |
| ENSMUSG00000022758 | *P2rx6* | 8.50611819 | 14.29581442 | 10.7699863 | 1.162660436 | 1.905496576 | 0 | 0.004853582 | -3.452168022 | 0.001800361 | 9.74270129 | 0.091367949 | 2.313937606 |
| ENSMUSG00000031444 | *F10* | 25.51835457 | 17.59484852 | 23.49815192 | 1.162660436 | 1.905496576 | 2.886681691 | 2.85E-05 | -3.461602896 | 7.39E-06 | 20.08985926 | 0.090772374 | 4.545227699 |
| ENSMUSG00000072919 | *Noxred1* | 96.96974736 | 118.7652275 | 99.86714568 | 9.301283491 | 7.621986303 | 11.54672676 | 1.77E-20 | -3.467994236 | 1.58E-21 | 90.81422863 | 0.090371129 | 19.75221788 |
| ENSMUSG00000022123 | *Scel* | 15.31101274 | 20.89388262 | 9.790896635 | 2.325320873 | 0.952748288 | 0.96222723 | 0.000803944 | -3.47767025 | 0.0002607 | 13.33354775 | 0.089767047 | 3.094774218 |
| ENSMUSG00000112023 | *Lilrb4b* | 21586.82674 | 20401.22686 | 19784.46483 | 1913.739078 | 1419.594949 | 2163.086814 | 1.35E-71 | -3.490537596 | 3.31E-73 | 327.5326622 | 0.088969978 | 70.8689066 |
| ENSMUSG00000028051 | *Hcn3* | 56.14038005 | 61.58196982 | 55.80811082 | 4.650641745 | 2.858244864 | 7.697817843 | 7.22E-12 | -3.507826917 | 1.03E-12 | 50.7850376 | 0.087910121 | 11.14140984 |
| ENSMUSG00000036634 | *Mag* | 51.88732096 | 38.48873114 | 43.0799452 | 6.975962618 | 0.952748288 | 3.848908921 | 1.04E-08 | -3.555138716 | 1.88E-09 | 36.08978093 | 0.085073952 | 7.982704981 |
| ENSMUSG00000028339 | *Col15a1* | 34.87508458 | 31.89066294 | 53.84993149 | 1.162660436 | 2.858244864 | 5.773363382 | 6.41E-08 | -3.585042313 | 1.26E-08 | 32.39951491 | 0.083328723 | 7.193017675 |
| ENSMUSG00000044071 | *Tafa2* | 568.2086951 | 549.8390163 | 547.3111219 | 48.83173833 | 45.73191782 | 44.2624526 | 5.45E-106 | -3.58830382 | 9.27E-108 | 486.2632331 | 0.083140554 | 105.2639766 |
| ENSMUSG00000026832 | *Cytip* | 30.62202548 | 39.58840917 | 42.10085553 | 4.650641745 | 0.952748288 | 3.848908921 | 5.47E-08 | -3.600924502 | 1.06E-08 | 32.72421777 | 0.082416414 | 7.26236666 |
| ENSMUSG00000026981 | *Il1rn* | 3795.429936 | 3755.400481 | 3745.017963 | 302.2917134 | 293.4464727 | 330.04394 | 0 | -3.608142608 | 0 | 2120.539386 | 0.082005097 | Inf |
| ENSMUSG00000073758 | *Sh3d21* | 16.16162456 | 14.29581442 | 6.853627645 | 1.162660436 | 0.952748288 | 0.96222723 | 0.002009573 | -3.610277519 | 0.000698538 | 11.49313231 | 0.081883835 | 2.696896287 |
| ENSMUSG00000066677 | *Ifi208* | 95.26852373 | 103.3697351 | 110.637132 | 5.813302182 | 11.43297945 | 7.697817843 | 6.67E-21 | -3.6111103 | 5.84E-22 | 92.78267126 | 0.081836582 | 20.17578675 |
| ENSMUSG00000031779 | *Ccl22* | 32.32324912 | 28.59162885 | 41.12176587 | 4.650641745 | 0.952748288 | 2.886681691 | 2.00E-07 | -3.631596264 | 4.07E-08 | 30.11458542 | 0.080682731 | 6.699016909 |
| ENSMUSG00000079363 | *Gbp4* | 39.12814367 | 30.79098491 | 44.05903486 | 3.487981309 | 4.763741439 | 0.96222723 | 3.90E-08 | -3.637414597 | 7.45E-09 | 33.41436275 | 0.080357996 | 7.409253712 |
| ENSMUSG00000049130 | *C5ar1* | 2451.463262 | 2455.581047 | 2467.305952 | 201.1402555 | 168.636447 | 216.5011268 | 1.50E-303 | -3.654696211 | 5.34E-306 | 1398.134474 | 0.079401154 | 302.8246126 |
| ENSMUSG00000090186 | *Gm52951* | 198.1925538 | 155.0546026 | 145.8843599 | 11.62660436 | 15.24397261 | 12.50895399 | 1.34E-30 | -3.658669749 | 8.34E-32 | 137.7327689 | 0.079182765 | 29.87299574 |
| ENSMUSG00000042807 | *Hecw2* | 71.45139279 | 82.47585244 | 68.53627645 | 4.650641745 | 7.621986303 | 4.811136152 | 5.65E-16 | -3.688387901 | 6.34E-17 | 69.86721144 | 0.077568359 | 15.24814661 |
| ENSMUSG00000026656 | *Fcgr2b* | 3368.422803 | 3301.233454 | 3257.431311 | 273.2252025 | 247.7145549 | 248.2546254 | 0 | -3.693639536 | 0 | 2027.56581 | 0.077286512 | Inf |
| ENSMUSG00000021457 | *Syk* | 3631.261855 | 3738.905311 | 3728.373439 | 301.129053 | 276.2970035 | 273.2725334 | 0 | -3.708709914 | 0 | 2193.38243 | 0.07648338 | Inf |
| ENSMUSG00000032221 | *Mns1* | 54.43915641 | 54.98390163 | 49.93357284 | 2.325320873 | 4.763741439 | 4.811136152 | 4.49E-12 | -3.715463803 | 6.34E-13 | 51.73814866 | 0.076126164 | 11.34743974 |
| ENSMUSG00000001761 | *Smo* | 37.42692004 | 50.5851895 | 33.28904856 | 3.487981309 | 2.858244864 | 2.886681691 | 3.90E-09 | -3.721466386 | 6.84E-10 | 38.06654701 | 0.075810086 | 8.408761321 |
| ENSMUSG00000035385 | *Ccl2* | 6354.9209 | 6170.293441 | 6117.352218 | 452.2749097 | 458.2719265 | 492.6603419 | 0 | -3.730478964 | 0 | 2878.336941 | 0.075337973 | Inf |
| ENSMUSG00000028356 | *Ambp* | 256.8847693 | 230.9323868 | 217.3579053 | 18.60256698 | 9.527482879 | 25.01790799 | 1.10E-38 | -3.73511558 | 5.23E-40 | 175.2700387 | 0.075096237 | 37.95828991 |
| ENSMUSG00000053819 | *Camk2d* | 540.1385051 | 445.3696032 | 481.7121145 | 34.87981309 | 24.77145549 | 50.03581598 | 5.74E-67 | -3.741203621 | 1.50E-68 | 306.1587015 | 0.074780005 | 66.2413039 |
| ENSMUSG00000019823 | *Mical1* | 29.77141366 | 38.48873114 | 41.12176587 | 2.325320873 | 1.905496576 | 3.848908921 | 1.77E-08 | -3.748563951 | 3.29E-09 | 35.00583581 | 0.074399465 | 7.751121466 |
| ENSMUSG00000017466 | *Timp2* | 22.96651911 | 25.29259475 | 36.22631755 | 4.650641745 | 0.952748288 | 0.96222723 | 3.11E-06 | -3.759770953 | 7.16E-07 | 24.57075278 | 0.073823761 | 5.507790462 |
| ENSMUSG00000004415 | *Col26a1* | 107.1770892 | 104.4694131 | 88.11806972 | 2.325320873 | 7.621986303 | 11.54672676 | 6.56E-19 | -3.760805897 | 6.44E-20 | 83.47902854 | 0.073770821 | 18.18290176 |
| ENSMUSG00000032359 | *Ctsh* | 73.15261643 | 78.07714031 | 73.43172477 | 9.301283491 | 1.905496576 | 5.773363382 | 3.66E-15 | -3.769018041 | 4.28E-16 | 66.10445784 | 0.073352094 | 14.43603963 |

| | | | | | | | | | | | | | |
|---|---|---|---|---|---|---|---|---|---|---|---|---|---|
| ENSMUSG00000031448 | *Adprhl1* | 9.356730009 | 16.49517049 | 15.66543462 | 0 | 1.905496576 | 0.96222723 | 0.000862573 | -3.783234786 | 0.000281665 | 13.1885491 | 0.07263281 | 3.064204238 |
| ENSMUSG00000032531 | *Amotl2* | 173.5248111 | 172.6494511 | 148.8216289 | 17.43990655 | 4.763741439 | 14.43340846 | 3.07E-28 | -3.783882435 | 2.06E-29 | 126.7986258 | 0.072600212 | 27.5132648 |
| ENSMUSG00000002602 | *Axl* | 590.3246024 | 600.4242058 | 599.2028741 | 46.50641745 | 37.15718323 | 46.18690706 | 8.08E-117 | -3.790104096 | 1.17E-118 | 536.3567384 | 0.072287795 | 116.0926824 |
| ENSMUSG00000034227 | *Foxj1* | 9.356730009 | 12.09645836 | 7.832717308 | 2.325320873 | 0 | 0 | 0.007847924 | -3.79078072 | 0.003033303 | 8.787327504 | 0.0722539 | 2.105245193 |
| ENSMUSG00000059498 | *Fcgr3* | 973.9505327 | 906.1346988 | 958.5287806 | 65.10898443 | 69.55062502 | 68.31813336 | 2.43E-175 | -3.803938757 | 2.05E-177 | 806.5309235 | 0.071597908 | 174.6141984 |
| ENSMUSG00000031026 | *Trim66* | 11.90856547 | 8.797424261 | 8.811806972 | 2.325320873 | 0 | 0 | 0.007173992 | -3.808200951 | 0.002751655 | 8.965233985 | 0.071386696 | 2.144239131 |
| ENSMUSG00000050014 | *Apol10b* | 37.42692004 | 32.99034098 | 29.37268991 | 1.162660436 | 2.858244864 | 2.886681691 | 4.16E-08 | -3.820541646 | 7.98E-09 | 33.2788411 | 0.070778665 | 7.380546053 |
| ENSMUSG00000018925 | *Heatr9* | 12.75917728 | 6.598068195 | 9.790896635 | 1.162660436 | 0.952748288 | 0 | 0.005087482 | -3.833339901 | 0.001892359 | 9.651122691 | 0.070153559 | 2.293497128 |
| ENSMUSG00000039431 | *Mtmr7* | 333.439833 | 282.6172544 | 393.5940447 | 29.06651091 | 22.86595891 | 18.28231738 | 5.20E-54 | -3.861214139 | 1.74E-55 | 246.2123854 | 0.068811135 | 53.28375159 |
| ENSMUSG00000029275 | *Gfi1* | 15.31101274 | 17.59484852 | 10.7699863 | 0 | 2.858244864 | 0 | 0.000986724 | -3.863429199 | 0.000325861 | 12.91556388 | 0.068705567 | 3.005804413 |
| ENSMUSG00000029919 | *Hpgds* | 3690.804683 | 3459.58709 | 3550.17912 | 252.2973147 | 205.7936302 | 265.5747156 | 0 | -3.888938356 | 0 | 1650.988952 | 0.067501419 | Inf |
| ENSMUSG00000031385 | *Plxnb3* | 28.07019003 | 19.79420459 | 12.72816563 | 1.162660436 | 1.905496576 | 0.96222723 | 4.05E-05 | -3.903803845 | 1.07E-05 | 19.37403611 | 0.066809458 | 4.392556103 |
| ENSMUSG00000058046 | *4933430I17Rik* | 180.3297056 | 139.6591101 | 159.5916152 | 13.95192524 | 3.810993152 | 14.43340846 | 1.44E-27 | -3.918960933 | 9.84E-29 | 123.6923715 | 0.066111225 | 26.84224779 |
| ENSMUSG00000027669 | *Gnb4* | 71.45139279 | 48.38583343 | 65.59900746 | 3.487981309 | 0.952748288 | 7.697817843 | 5.06E-12 | -3.930103677 | 7.18E-13 | 51.49495145 | 0.065602578 | 11.29553931 |
| ENSMUSG00000002007 | *Srpk3* | 28.07019003 | 21.99356065 | 26.43542092 | 0 | 1.905496576 | 2.886681691 | 2.11E-06 | -3.932932648 | 4.79E-07 | 25.3471072 | 0.065474065 | 5.675375078 |
| ENSMUSG00000030256 | *Bhlhe41* | 893.1424099 | 909.4337329 | 856.7034556 | 54.64504051 | 44.77916953 | 74.09149674 | 3.17E-138 | -3.935280462 | 3.66E-140 | 635.2224331 | 0.0653676 | 137.4987331 |
| ENSMUSG00000105504 | *Gbp5* | 153.1101274 | 161.6526708 | 184.0688567 | 9.301283491 | 6.669238015 | 16.35786292 | 2.77E-32 | -3.939832657 | 1.61E-33 | 145.5731196 | 0.065161668 | 31.55764023 |
| ENSMUSG00000013483 | *Card14* | 5.103670914 | 13.19613639 | 12.72816563 | 0 | 0.952748288 | 0.96222723 | 0.003979097 | -3.943559843 | 0.001452231 | 10.13826083 | 0.064993541 | 2.400215477 |
| ENSMUSG00000068735 | *Trp53i11* | 61.24405097 | 52.78454556 | 59.72446948 | 5.813302182 | 2.858244864 | 2.886681691 | 1.47E-13 | -3.947278893 | 1.89E-14 | 58.64447999 | 0.064826213 | 12.83157224 |
| ENSMUSG00000026358 | *Rgs1* | 10449.7662 | 10038.96076 | 10479.19667 | 609.2340686 | 652.6325772 | 728.4060134 | 0 | -3.957151503 | 0 | 2826.927127 | 0.064384111 | Inf |
| ENSMUSG00000020689 | *Itgb3* | 131.8448319 | 93.47263277 | 122.3862079 | 2.325320873 | 14.29122432 | 4.811136152 | 8.80E-19 | -3.980440619 | 8.68E-20 | 82.88922123 | 0.063353116 | 18.05574489 |
| ENSMUSG00000025481 | *Urah* | 36.57630822 | 37.38905311 | 41.12176587 | 4.650641745 | 1.905496576 | 0.96222723 | 3.22E-09 | -3.995883113 | 5.60E-10 | 38.45758579 | 0.062678605 | 8.492033176 |
| ENSMUSG00000029361 | *Nos1* | 76.55506371 | 103.3697351 | 96.92987669 | 5.813302182 | 8.574734591 | 2.886681691 | 3.23E-19 | -3.998746805 | 3.09E-20 | 84.92821144 | 0.062554314 | 18.49016794 |
| ENSMUSG00000021886 | *Gpr65* | 273.8970057 | 300.2121029 | 291.7687197 | 6.975962618 | 15.24397261 | 29.82904414 | 7.53E-42 | -4.022743514 | 3.32E-43 | 189.9155074 | 0.061522438 | 41.12318229 |
| ENSMUSG00000055407 | *Map6* | 17.8628482 | 29.69130688 | 19.58179327 | 2.325320873 | 0.952748288 | 0.96222723 | 8.16E-06 | -4.025563878 | 1.97E-06 | 22.6287261 | 0.061402284 | 5.088064794 |
| ENSMUSG00000062515 | *Fabp4* | 498.4585259 | 436.5721789 | 459.1930522 | 31.39183178 | 24.77145549 | 29.82904414 | 7.07E-94 | -4.026996939 | 1.35E-95 | 430.3655535 | 0.061341322 | 93.15070001 |
| ENSMUSG00000022440 | *C1qtnf6* | 11.05795365 | 64.88100392 | 72.4526351 | 1.162660436 | 3.810993152 | 3.848908921 | 1.73E-06 | -4.036707266 | 3.88E-07 | 25.7512412 | 0.060929839 | 5.763173778 |
| ENSMUSG00000028327 | *Stra6l* | 43.38120277 | 49.48551147 | 42.10085553 | 3.487981309 | 1.905496576 | 2.886681691 | 3.32E-11 | -4.044993367 | 4.94E-12 | 47.7118417 | 0.060580892 | 10.47901691 |
| ENSMUSG00000020364 | *Zfp354a* | 33.17386094 | 21.99356065 | 28.39360024 | 0 | 1.905496576 | 2.886681691 | 6.09E-07 | -4.061145928 | 1.30E-07 | 27.86113878 | 0.059906405 | 6.215493033 |
| ENSMUSG00000024481 | *Lvrn* | 42.53059095 | 35.18969704 | 41.12176587 | 1.162660436 | 0.952748288 | 4.811136152 | 2.33E-09 | -4.071531242 | 4.00E-10 | 39.11374806 | 0.059476714 | 8.632845524 |
| ENSMUSG00000090639 | *None* | 234.768862 | 282.6172544 | 344.6395616 | 4.650641745 | 14.29122432 | 31.7534986 | 1.17E-07 | -4.076067863 | 2.33E-08 | 31.19952066 | 0.059289981 | 6.933585682 |
| ENSMUSG00000041548 | *Hspb8* | 11.05795365 | 10.99678033 | 13.70725529 | 2.325320873 | 0 | 0 | 0.001764844 | -4.089183501 | 0.000608383 | 11.75015053 | 0.058753414 | 2.75329367 |
| ENSMUSG00000056529 | *Ptafr* | 2509.304866 | 2350.011956 | 2336.107937 | 138.3565919 | 139.10125 | 143.3718573 | 0 | -4.095394802 | 0 | 1748.241822 | 0.058501004 | Inf |
| ENSMUSG00000045672 | *Col27a1* | 128.4423847 | 98.97102293 | 86.15989039 | 4.650641745 | 8.574734591 | 4.811136152 | 1.28E-21 | -4.105872222 | 1.08E-22 | 96.12560014 | 0.058077686 | 20.89326755 |
| ENSMUSG00000047959 | *Kcna3* | 39.97875549 | 31.89066294 | 35.24722789 | 1.162660436 | 0 | 4.811136152 | 3.88E-08 | -4.142518417 | 7.41E-09 | 33.42478972 | 0.056621021 | 7.411426128 |
| ENSMUSG00000004872 | *Pax3* | 88.46362917 | 101.170379 | 79.30626275 | 2.325320873 | 3.810993152 | 8.660045073 | 1.80E-19 | -4.148635477 | 1.70E-20 | 86.11767245 | 0.056381455 | 18.74355013 |
| ENSMUSG00000066026 | *Dhrs3* | 5457.525431 | 5231.168401 | 5430.031274 | 312.7556574 | 279.1552484 | 307.9127137 | 0 | -4.165555999 | 0 | 2943.12072 | 0.055724053 | Inf |
| ENSMUSG00000047881 | *Rell1* | 151.4089038 | 136.360076 | 117.4907596 | 9.301283491 | 7.621986303 | 5.773363382 | 3.52E-30 | -4.175081482 | 2.21E-31 | 135.7968011 | 0.055357343 | 29.45386338 |
| ENSMUSG00000026009 | *Icos* | 11.05795365 | 8.797424261 | 16.64452428 | 0 | 1.905496576 | 0 | 0.001374523 | -4.187339619 | 0.00046487 | 12.25155567 | 0.054888982 | 2.861848127 |
| ENSMUSG00000037902 | *Sirpa* | 10489.74495 | 10884.61317 | 10610.39468 | 612.7220499 | 538.3027827 | 601.392019 | 0 | -4.192016115 | 0 | 3702.972164 | 0.054711347 | Inf |
| ENSMUSG00000066682 | *Pilrb2* | 17.01223638 | 9.897102293 | 10.7699863 | 1.162660436 | 0 | 0.96222723 | 0.000747273 | -4.203453418 | 0.000241041 | 13.48061425 | 0.054279325 | 3.126520782 |
| ENSMUSG00000092077 | *Or12d12* | 15.31101274 | 9.897102293 | 11.74907596 | 0 | 0 | 1.924454461 | 0.001083191 | -4.203754342 | 0.000359726 | 12.73057459 | 0.054268004 | 2.965294759 |
| ENSMUSG00000030134 | *Rasgef1a* | 23.81713093 | 12.09645836 | 22.51906226 | 3.487981309 | 0 | 0 | 0.000116624 | -4.212037529 | 3.29E-05 | 17.24179568 | 0.053957319 | 3.933211451 |
| ENSMUSG00000069515 | *Lyz1* | 1146.624732 | 1161.260002 | 1194.48939 | 54.64504051 | 71.45612159 | 60.62031551 | 1.03E-225 | -4.220776779 | 6.35E-228 | 1038.882718 | 0.053631456 | 224.9879399 |
| ENSMUSG00000040086 | *Tnni3k* | 20.41468366 | 4.39871213 | 12.72816563 | 0 | 0 | 1.924454461 | 0.002550955 | -4.2351234 | 0.000902129 | 11.01838046 | 0.05310077 | 2.593297219 |
| ENSMUSG00000020017 | *Hal* | 170.1223638 | 136.360076 | 145.8843599 | 16.27724611 | 5.716489727 | 2.886681691 | 2.09E-26 | -4.248422001 | 1.47E-27 | 118.322508 | 0.052613542 | 25.68043232 |
| ENSMUSG00000046805 | *Mpeg1* | 29826.70343 | 29027.10135 | 28996.71948 | 1496.343982 | 1548.215968 | 1551.110295 | 0 | -4.256038485 | 0 | 6392.381086 | 0.052336509 | Inf |
| ENSMUSG00000039114 | *Nrn1* | 14.46040092 | 10.99678033 | 13.70725529 | 1.162660436 | 0.952748288 | 0 | 0.000436598 | -4.258631766 | 0.000134478 | 14.57795707 | 0.052242518 | 3.359918118 |
| ENSMUSG00000020641 | *Rsad2* | 13237.22113 | 12718.87612 | 12345.34157 | 681.3190157 | 617.3808906 | 691.8413786 | 0 | -4.267355134 | 0 | 3716.471991 | 0.051927582 | Inf |
| ENSMUSG00000095366 | *Gm21860* | 68.89955734 | 57.18325769 | 34.26813822 | 0 | 3.810993152 | 3.848908921 | 4.01E-11 | -4.323503117 | 6.00E-12 | 47.32822445 | 0.049945444 | 10.396825 |
| ENSMUSG00000030742 | *Lat* | 81.65873462 | 102.270057 | 87.13898005 | 9.301283491 | 3.810993152 | 0.96222723 | 6.11E-19 | -4.334002227 | 5.98E-20 | 83.62502453 | 0.049583289 | 18.21406318 |
| ENSMUSG00000073889 | *Il11ra1* | 163.3174692 | 163.8520269 | 145.8843599 | 9.301283491 | 8.574734591 | 5.773363382 | 1.14E-37 | -4.334554014 | 5.58E-39 | 170.559876 | 0.049564328 | 36.94201609 |
| ENSMUSG00000004098 | *Col5a3* | 13.6097891 | 16.49517049 | 12.72816563 | 2.325320873 | 0 | 0 | 0.00036328 | -4.353523643 | 0.000110699 | 14.94486622 | 0.048916885 | 3.439758867 |
| ENSMUSG00000042189 | *Tekt3* | 10.20734183 | 6.598068195 | 5.874537981 | 0 | 0.952748288 | 0 | 0.008445699 | -4.37327728 | 0.003288688 | 8.639959129 | 0.048251671 | 2.073364416 |
| ENSMUSG00000038067 | *Csf3* | 1577.884924 | 1580.237333 | 1610.602497 | 69.75962618 | 78.12535961 | 79.86486012 | 3.80634102343846e-316 | -4.382537058 | 1.25363228844466e-318 | 1456.255455 | 0.047942965 | 315.4194923 |
| ENSMUSG00000033213 | *AA467197* | 3.402447276 | 7.697746228 | 12.72816563 | 0 | 0.952748288 | 0 | 0.009281647 | -4.43491361 | 0.003643495 | 8.453460468 | 0.046233628 | 2.032374967 |
| ENSMUSG00000028859 | *Csf3r* | 614.9923451 | 668.6042438 | 596.2656051 | 33.71715265 | 19.05496576 | 34.64018029 | 2.49E-120 | -4.437811344 | 3.39E-122 | 552.6212798 | 0.046140858 | 119.6033287 |
| ENSMUSG00000004552 | *Ctse* | 747.6877889 | 752.1797743 | 747.0454133 | 30.22917134 | 28.58244864 | 44.2624526 | 1.73E-155 | -4.43859804 | 1.73E-157 | 714.885732 | 0.046115705 | 154.762022 |
| ENSMUSG00000013974 | *Mcemp1* | 75.70445189 | 68.18003802 | 78.32717308 | 4.650641745 | 2.858244864 | 2.886681691 | 4.08E-19 | -4.444515415 | 3.94E-20 | 84.45004499 | 0.045926944 | 18.38916807 |
| ENSMUSG00000028583 | *Pdpn* | 16.16162456 | 12.09645836 | 16.64452428 | 1.162660436 | 0.952748288 | 0 | 0.000116423 | -4.456806248 | 3.28E-05 | 17.24590925 | 0.045537337 | 3.933960451 |
| ENSMUSG00000099871 | *Gm21742* | 16.16162456 | 15.39549246 | 14.68634495 | 2.325320873 | 0 | 0 | 0.000153278 | -4.469074281 | 4.40E-05 | 16.69217229 | 0.04515175 | 3.814519883 |
| ENSMUSG00000055333 | *Fat2* | 11.90856547 | 5.498390163 | 6.853627645 | 0 | 0 | 0.96222723 | 0.006364856 | -4.473174666 | 0.002414662 | 9.204156488 | 0.045023603 | 2.196211416 |
| ENSMUSG00000059901 | *Adamts14* | 46.78365004 | 42.88744327 | 46.99630385 | 3.487981309 | 2.858244864 | 0 | 1.19E-11 | -4.475111851 | 1.72E-12 | 49.77821587 | 0.044963188 | 10.92316033 |
| ENSMUSG00000028480 | *Glipr2* | 101.2228065 | 107.7684472 | 108.6789527 | 3.487981309 | 3.810993152 | 6.735590613 | 8.86E-27 | -4.482328004 | 6.19E-28 | 120.0433439 | 0.04473885 | 26.05252431 |
| ENSMUSG00000055027 | *Smyd1* | 106.3264774 | 104.4694131 | 113.574401 | 9.301283491 | 0 | 5.773363382 | 1.37E-22 | -4.490537986 | 1.11E-23 | 100.6285384 | 0.044484977 | 21.86461644 |
| ENSMUSG00000031383 | *Dusp9* | 11.90856547 | 8.797424261 | 3.916358654 | 1.162660436 | 0 | 0 | 0.007464923 | -4.490743063 | 0.002872464 | 8.886757574 | 0.044478654 | 2.126974654 |
| ENSMUSG00000014599 | *Csf1* | 441.4675341 | 488.2570465 | 515.9802527 | 23.25320873 | 20.00771405 | 21.16899907 | 1.00E-105 | -4.493122932 | 1.72E-107 | 485.0294356 | 0.044405342 | 104.9989998 |
| ENSMUSG00000034394 | *Lif* | 2391.069823 | 2478.674285 | 2243.094419 | 109.290081 | 92.41658392 | 110.6561315 | 0 | -4.51241213 | 0 | 1724.971615 | 0.043815583 | Inf |
| ENSMUSG00000002992 | *Apoc2* | 29.77141366 | 16.49517049 | 22.51906226 | 0 | 0.952748288 | 1.924454461 | 2.29E-06 | -4.514776983 | 5.22E-07 | 25.18019518 | 0.04374382 | 5.639496804 |
| ENSMUSG00000025701 | *Alox5* | 1116.002707 | 1068.887048 | 1076.01954 | 49.99439876 | 43.82642124 | 49.07358875 | 9.50E-235 | -4.516364338 | 5.61E-237 | 1080.539085 | 0.043695716 | 234.0222819 |
| ENSMUSG00000026604 | *Ptpn14* | 220.3084611 | 203.440436 | 213.4415467 | 15.11458567 | 5.716489727 | 7.697817843 | 1.32E-48 | -4.525782777 | 4.97E-50 | 221.1908907 | 0.043411384 | 47.87778549 |
| ENSMUSG00000061603 | *Akap6* | 10.20734183 | 7.697746228 | 7.832717308 | 0 | 0 | 0.96222723 | 0.003747337 | -4.553955249 | 0.001361476 | 10.25726194 | 0.042571884 | 2.426277215 |
| ENSMUSG00000027646 | *Src* | 215.2047902 | 197.9420459 | 234.0024296 | 9.301283491 | 7.621986303 | 10.58449953 | 4.43E-52 | -4.558583363 | 1.53E-53 | 237.2909318 | 0.042435534 | 51.35343476 |
| ENSMUSG00000058427 | *Cxcl2* | 21364.81706 | 21364.54482 | 21168.89762 | 868.5073459 | 832.7020036 | 957.4160942 | 0 | -4.586471493 | 0 | 4810.727245 | 0.041623108 | Inf |
| ENSMUSG00000020432 | *Tcn2* | 114.8325956 | 119.8649056 | 133.1561942 | 4.650641745 | 8.574734591 | 1.924454461 | 6.32E-29 | -4.592991995 | 4.13E-30 | 129.9838064 | 0.04143541 | 28.19942283 |
| ENSMUSG00000057135 | *Scimp* | 886.3375154 | 907.2343769 | 881.1806972 | 47.66907789 | 31.4406935 | 31.7534986 | 7.59E-188 | -4.613257868 | 5.84E-190 | 864.2387858 | 0.040857426 | 187.119585 |
| ENSMUSG00000000732 | *Icosl* | 682.1906788 | 784.0704372 | 685.3627645 | 27.90385047 | 26.67695206 | 32.71572583 | 7.69E-149 | -4.619814687 | 8.18E-151 | 684.195337 | 0.040672157 | 148.1141651 |
| ENSMUSG00000034438 | *Gbp8* | 58.69221551 | 40.68808721 | 50.9126625 | 2.325320873 | 2.858244864 | 0.96222723 | 4.00E-13 | -4.623570111 | 5.28E-14 | 56.62203657 | 0.040566423 | 12.39833217 |
| ENSMUSG00000035095 | *Fam167a* | 13.6097891 | 9.897102293 | 27.41451058 | 1.162660436 | 0.952748288 | 0 | 8.76E-05 | -4.636438121 | 2.43E-05 | 17.81806826 | 0.040206202 | 4.057404135 |
| ENSMUSG00000038179 | *Slamf7* | 13.6097891 | 23.09323868 | 13.70725529 | 0 | 1.905496576 | 0 | 6.58E-05 | -4.642483709 | 1.80E-05 | 18.3875608 | 0.040038071 | 4.181456401 |
| ENSMUSG00000035352 | *Ccl12* | 21.26529547 | 48.38583343 | 36.22631755 | 3.487981309 | 0.952748288 | 0 | 3.46E-08 | -4.668996072 | 6.57E-09 | 33.65859185 | 0.039309012 | 7.461440649 |
| ENSMUSG00000027219 | *Slc28a2* | 50.18609732 | 62.68164786 | 47.97539351 | 4.650641745 | 0 | 1.924454461 | 2.54E-13 | -4.698215964 | 3.31E-14 | 57.54089677 | 0.038520868 | 12.596 |
| ENSMUSG00000013367 | *Iglon5* | 17.01223638 | 21.99356065 | 13.70725529 | 0 | 0.952748288 | 0.96222723 | 1.86E-05 | -4.705799473 | 4.68E-06 | 20.9637497 | 0.038318915 | 4.730670754 |
| ENSMUSG00000025746 | *Il6* | 71.45139279 | 75.87778425 | 66.57809712 | 1.162660436 | 3.810993152 | 2.886681691 | 3.43E-19 | -4.723146976 | 3.29E-20 | 84.80680812 | 0.037860913 | 18.46474624 |
| ENSMUSG00000045875 | *Adra1a* | 13.6097891 | 6.598068195 | 8.811806972 | 0 | 0 | 0.96222723 | 0.002003634 | -4.730881055 | 0.000696199 | 11.49936706 | 0.03765849 | 2.69818158 |
| ENSMUSG00000053318 | *Slamf8* | 31.4726373 | 29.69130688 | 21.5399726 | 2.325320873 | 0 | 0.96222723 | 9.49E-08 | -4.743075073 | 1.88E-08 | 31.60989956 | 0.037341532 | 7.022535787 |
| ENSMUSG00000054072 | *Iigp1* | 90.16485281 | 74.77810622 | 84.20171106 | 3.487981309 | 2.858244864 | 2.886681691 | 2.21E-22 | -4.76610551 | 1.81E-23 | 99.65806379 | 0.036750164 | 21.65619575 |
| ENSMUSG00000024486 | *Hbegf* | 307.0708667 | 306.8101711 | 305.475975 | 11.62660436 | 5.716489727 | 16.35786292 | 7.67E-72 | -4.774292661 | 1.86E-73 | 328.6796751 | 0.036542201 | 71.1150669 |
| ENSMUSG00000026628 | *Atf3* | 18337.48959 | 18656.03782 | 18504.79464 | 634.8125982 | 702.1754882 | 668.7479251 | 0 | -4.7879198 | 0 | 5993.9595 | 0.036198662 | Inf |
| ENSMUSG00000026768 | *Itga8* | 33.17386094 | 39.58840917 | 41.12176587 | 1.162660436 | 0.952748288 | 1.924454461 | 6.23E-11 | -4.806404 | 9.49E-12 | 46.43067135 | 0.035737834 | 10.20542538 |
| ENSMUSG00000025422 | *Agap2* | 472.0895595 | 504.752217 | 448.4230659 | 18.60256698 | 16.19672089 | 16.35786292 | 2.02E-114 | -4.80646501 | 2.97E-116 | 525.3047632 | 0.035736323 | 113.6944035 |
| ENSMUSG00000028599 | *Tnfrsf1b* | 8610.743444 | 8801.822973 | 8673.755329 | 287.1771278 | 312.5014384 | 303.1015776 | 0 | -4.84996191 | 0 | 4792.390382 | 0.034674961 | Inf |
| ENSMUSG00000042265 | *Trem1* | 317.2782085 | 277.1188642 | 295.6850784 | 8.138623054 | 12.38572774 | 9.622272304 | 3.89E-74 | -4.870082628 | 9.18E-76 | 339.2751349 | 0.03419472 | 73.41024588 |
| ENSMUSG00000035042 | *Ccl5* | 1041.999478 | 1035.896707 | 1039.793223 | 43.01843614 | 33.34619008 | 29.82904414 | 3.26E-236 | -4.891683918 | 1.88E-238 | 1087.326875 | 0.03368654 | 235.4873699 |
| ENSMUSG00000028358 | *Zfp618* | 22.96651911 | 24.19291672 | 14.68634495 | 2.325320873 | 0 | 0 | 6.26E-06 | -4.893667072 | 1.49E-06 | 23.16234332 | 0.033640266 | 5.203525756 |
| ENSMUSG00000028602 | *Tnfrsf8* | 129.2929965 | 97.8713449 | 81.26444207 | 4.650641745 | 1.905496576 | 3.848908921 | 3.34E-24 | -4.918857499 | 2.55E-25 | 108.1021533 | 0.033057983 | 23.47629639 |
| ENSMUSG00000021768 | *Dusp13* | 10.20734183 | 15.39549246 | 7.832717308 | 0 | 0.952748288 | 0 | 0.000706414 | -4.924406207 | 0.000226456 | 13.59775806 | 0.032931084 | 3.150940598 |
| ENSMUSG00000037759 | *Ptger2* | 123.3387138 | 100.070701 | 90.07624905 | 6.975962618 | 0.952748288 | 2.886681691 | 7.63E-25 | -4.932142625 | 5.68E-26 | 111.0818257 | 0.032754964 | 24.11726929 |
| ENSMUSG00000090675 | *Or5v1b* | 128.4423847 | 149.5562124 | 128.2607459 | 3.487981309 | 2.858244864 | 6.735590613 | 3.70E-35 | -4.941911774 | 1.95E-36 | 158.9217864 | 0.032533914 | 34.43198675 |
| ENSMUSG00000024669 | *Cd5* | 545.242176 | 535.5432019 | 552.2065702 | 18.60256698 | 12.38572774 | 22.1312263 | 3.28E-132 | -4.947269958 | 4.03E-134 | 607.4418895 | 0.032413307 | 131.484129 |
| ENSMUSG00000022441 | *Efcab6* | 9.356730009 | 10.99678033 | 13.70725529 | 1.162660436 | 0 | 0 | 0.000546477 | -4.95379316 | 0.00017151 | 14.12000614 | 0.03226708 | 3.26242812 |
| ENSMUSG00000000386 | *Mx1* | 1030.941525 | 1058.989945 | 983.9851119 | 20.92788785 | 33.34619008 | 42.33799814 | 9.89E-209 | -4.966906335 | 6.45E-211 | 960.6424299 | 0.031975122 | 208.0047898 |
| ENSMUSG00000010122 | *Slc47a1* | 12.75917728 | 10.99678033 | 10.7699863 | 0 | 0.952748288 | 0 | 0.000406584 | -4.976362074 | 0.000124787 | 14.71892872 | 0.031766235 | 3.390850125 |
| ENSMUSG00000038508 | *Gdf15* | 192.2382711 | 162.7523488 | 209.525188 | 10.46394393 | 2.858244864 | 4.811136152 | 4.55E-44 | -5.018444588 | 1.90E-45 | 200.1888181 | 0.030853018 | 43.34169716 |
| ENSMUSG00000060913 | *Trim55* | 7.655506371 | 7.697746228 | 2.937268991 | 0 | 0 | 0 | 0.00899099 | -5.02020609 | 0.003514592 | 8.518991297 | 0.03081537 | 2.046192487 |
| ENSMUSG00000022900 | *Ildr1* | 11.05795365 | 13.19613639 | 11.74907596 | 1.162660436 | 0 | 0 | 0.0003108 | -5.033573293 | 9.36E-05 | 15.26163665 | 0.03053117 | 3.507518805 |
| ENSMUSG00000004668 | *Abca13* | 69.75016916 | 51.68486753 | 78.32717308 | 0 | 2.858244864 | 2.886681691 | 2.70E-17 | -5.04715447 | 2.85E-18 | 75.99264334 | 0.030245106 | 16.56838912 |
| ENSMUSG00000006529 | *Itih1* | 9.356730009 | 18.69452655 | 8.811806972 | 0 | 0 | 0.96222723 | 0.000387114 | -5.062777962 | 0.000118413 | 14.81778759 | 0.029919337 | 3.412160683 |
| ENSMUSG00000048806 | *Ifnb1* | 148.8570683 | 147.3568564 | 145.8843599 | 3.487981309 | 3.810993152 | 5.773363382 | 7.71E-41 | -5.06520635 | 3.51E-42 | 185.2239879 | 0.029869019 | 40.11303512 |
| ENSMUSG00000038729 | *Pakap* | 67.1983337 | 63.78132589 | 75.38990409 | 1.162660436 | 1.905496576 | 2.886681691 | 1.37E-19 | -5.084179876 | 1.28E-20 | 86.67774151 | 0.02947877 | 18.86399906 |
| ENSMUSG00000031709 | *Tbc1d9* | 27.21957821 | 30.79098491 | 10.7699863 | 0 | 0.952748288 | 0.96222723 | 2.36E-06 | -5.091192287 | 5.38E-07 | 25.12162924 | 0.029335832 | 5.626569377 |
| ENSMUSG00000054675 | *Tmem119* | 7.655506371 | 5.498390163 | 6.853627645 | 0 | 0 | 0 | 0.00460526 | -5.152305123 | 0.001700666 | 9.847457814 | 0.028119113 | 2.336745858 |
| ENSMUSG00000066007 | *Zfp600* | 5.103670914 | 4.39871213 | 10.7699863 | 0 | 0 | 0 | 0.005386367 | -5.169090165 | 0.002008708 | 9.541561742 | 0.027793856 | 2.268704077 |
| ENSMUSG00000036853 | *Mcoln3* | 5.954282733 | 6.598068195 | 7.832717308 | 0 | 0 | 0 | 0.004143461 | -5.175579197 | 0.001518757 | 10.05571899 | 0.027669124 | 2.382636732 |
| ENSMUSG00000022126 | *Acod1* | 29279.76003 | 29188.75402 | 28702.99258 | 754.5666232 | 807.9305481 | 784.2151927 | 0 | -5.213571387 | 0 | 7927.957799 | 0.026949991 | Inf |
| ENSMUSG00000020826 | *Nos2* | 1886.657015 | 1895.844928 | 1882.789423 | 40.69311527 | 58.11764556 | 51.96027044 | 0 | -5.217237369 | 0 | 1859.263797 | 0.026881596 | Inf |
| ENSMUSG00000043727 | *F830045P16Rik* | 11.90856547 | 8.797424261 | 20.56088293 | 0 | 0 | 0.96222723 | 0.000140451 | -5.233884071 | 4.00E-05 | 16.8693958 | 0.026573203 | 3.852476171 |
| ENSMUSG00000022901 | *Cd86* | 72.30200461 | 81.37617441 | 78.32717308 | 2.325320873 | 1.905496576 | 1.924454461 | 1.34E-22 | -5.245788089 | 1.08E-23 | 100.6740473 | 0.026354843 | 21.87422794 |
| ENSMUSG00000102037 | *Bcl2a1a* | 193.9394947 | 168.250739 | 140.0098219 | 0 | 3.810993152 | 8.660045073 | 1.56E-37 | -5.261122466 | 7.64E-39 | 169.9358077 | 0.026076201 | 36.80814752 |
| ENSMUSG00000032579 | *Hemk1* | 290.9092421 | 295.8133908 | 281.9778231 | 5.813302182 | 7.621986303 | 8.660045073 | 7.60E-80 | -5.280505754 | 1.66E-81 | 365.6411337 | 0.025728198 | 79.11894248 |
| ENSMUSG00000104713 | *Gbp6* | 179.4790938 | 114.3665154 | 185.0479464 | 5.813302182 | 0.952748288 | 5.773363382 | 1.42E-33 | -5.289481965 | 7.87E-35 | 151.5686943 | 0.025568619 | 32.84720278 |
| ENSMUSG00000046186 | *Cd109* | 793.6208271 | 824.7585244 | 792.0835378 | 18.60256698 | 23.8187072 | 17.32009015 | 5.06E-203 | -5.328266844 | 3.47E-205 | 934.276445 | 0.024890399 | 202.2957348 |
| ENSMUSG00000015396 | *Cd83* | 6759.812125 | 6944.466776 | 6952.515701 | 162.7724611 | 179.1166781 | 163.5786292 | 0 | -5.350301351 | 0 | 4547.737244 | 0.024513132 | Inf |
| ENSMUSG00000037977 | *6430571L13Rik* | 6.804894552 | 7.697746228 | 8.811806972 | 0 | 0 | 0 | 0.001850844 | -5.369120282 | 0.000640188 | 11.65533062 | 0.024195452 | 2.732630228 |
| ENSMUSG00000026167 | *Wnt10a* | 6.804894552 | 10.99678033 | 5.874537981 | 0 | 0 | 0 | 0.001933997 | -5.38799431 | 0.000670941 | 11.56806848 | 0.023880977 | 2.713544116 |
| ENSMUSG00000066755 | *Tnfsf18* | 10.20734183 | 8.797424261 | 4.895448318 | 0 | 0 | 0 | 0.001868965 | -5.407622453 | 0.000646969 | 11.63573064 | 0.023558272 | 2.728398765 |
| ENSMUSG00000045092 | *S1pr1* | 143.7533974 | 145.1575003 | 157.6334358 | 4.650641745 | 2.858244864 | 2.886681691 | 8.03E-43 | -5.452621256 | 3.45E-44 | 194.4170121 | 0.022834812 | 42.09543943 |
| ENSMUSG00000024810 | *Il33* | 9.356730009 | 5.498390163 | 9.790896635 | 0 | 0 | 0 | 0.001434133 | -5.454215333 | 0.000486507 | 12.16668476 | 0.022809595 | 2.843410594 |
| ENSMUSG00000023802 | *Nox3* | 39.12814367 | 30.79098491 | 19.58179327 | 0 | 0 | 1.924454461 | 1.49E-08 | -5.470587549 | 2.74E-09 | 35.35954643 | 0.022552207 | 7.826126244 |
| ENSMUSG00000028341 | *Nr4a3* | 126.741161 | 134.16072 | 109.6580423 | 5.813302182 | 1.905496576 | 0.96222723 | 1.46E-33 | -5.49776616 | 8.07E-35 | 151.518873 | 0.022131328 | 32.83685157 |
| ENSMUSG00000090523 | *Gypc* | 8.50611819 | 4.39871213 | 12.72816563 | 0 | 0 | 0 | 0.001429339 | -5.510592418 | 0.000484783 | 12.17330699 | 0.021935442 | 2.844864635 |
| ENSMUSG00000056498 | *Tmem154* | 2081.447121 | 2152.06991 | 2049.234666 | 45.34375702 | 44.77916953 | 47.14913429 | 0 | -5.51548725 | 0 | 2136.304167 | 0.021861145 | Inf |
| ENSMUSG00000020053 | *Igf1* | 1054.758656 | 994.1089414 | 1018.25325 | 18.60256698 | 24.77145549 | 23.09345353 | 2.74E-255 | -5.515730992 | 1.37E-257 | 1175.370359 | 0.021857452 | 254.5616907 |
| ENSMUSG00000001700 | *Gramd3* | 10.20734183 | 7.697746228 | 8.811806972 | 0 | 0 | 0 | 0.000733322 | -5.569184809 | 0.000236088 | 13.51957282 | 0.021062423 | 3.134705047 |
| ENSMUSG00000072663 | *Spef2* | 10.20734183 | 12.09645836 | 4.895448318 | 0 | 0 | 0 | 0.000899671 | -5.591017502 | 0.000294828 | 13.1029671 | 0.020746079 | 3.045916177 |
| ENSMUSG00000103793 | *Pcdhga6* | 15.31101274 | 15.39549246 | 22.51906226 | 1.162660436 | 0 | 0 | 5.05E-06 | -5.59912072 | 1.19E-06 | 23.59024346 | 0.020629881 | 5.296864903 |
| ENSMUSG00000023809 | *Rps6ka2* | 3193.196768 | 3132.982715 | 3091.965157 | 68.59696574 | 70.5033733 | 54.84695213 | 0 | -5.605936118 | 0 | 2833.058049 | 0.020532653 | Inf |
| ENSMUSG00000035273 | *Hpse* | 7.655506371 | 13.19613639 | 7.832717308 | 0 | 0 | 0 | 0.000519483 | -5.664372877 | 0.000162111 | 14.22604548 | 0.019717591 | 3.284428778 |
| ENSMUSG00000090626 | *Tex9* | 53.5885446 | 58.28293573 | 43.0799452 | 0 | 0 | 2.886681691 | 5.83E-15 | -5.672875235 | 6.89E-16 | 65.16564151 | 0.01960173 | 14.2344892 |
| ENSMUSG00000028634 | *Hivep3* | 3368.422803 | 3586.050064 | 3164.417793 | 52.31971964 | 60.02314214 | 80.82708735 | 0 | -5.698342716 | 0 | 2234.456624 | 0.019258742 | Inf |
| ENSMUSG00000029178 | *Klf3* | 347.0496221 | 320.0063075 | 337.7859339 | 6.975962618 | 4.763741439 | 7.697817843 | 9.23E-95 | -5.699986612 | 1.75E-96 | 434.4421779 | 0.01923681 | 94.03484119 |
| ENSMUSG00000087207 | *Gm13147* | 8.50611819 | 13.19613639 | 7.832717308 | 0 | 0 | 0 | 0.000399922 | -5.707393411 | 0.00012255 | 14.75303045 | 0.019138301 | 3.398025227 |
| ENSMUSG00000054640 | *Slc8a1* | 17.8628482 | 15.39549246 | 26.43542092 | 0 | 0.952748288 | 0 | 1.13E-06 | -5.765701029 | 2.49E-07 | 26.60922244 | 0.018380234 | 5.946365434 |
| ENSMUSG00000054385 | *Ceacam2* | 13.6097891 | 6.598068195 | 10.7699863 | 0 | 0 | 0 | 0.00029619 | -5.785808177 | 8.88E-05 | 15.36036032 | 0.018125842 | 3.528430272 |
| ENSMUSG00000049866 | *Arl4c* | 124.1893256 | 138.5594321 | 131.1980149 | 2.325320873 | 3.810993152 | 0.96222723 | 7.16E-39 | -5.789181944 | 3.39E-40 | 176.1309922 | 0.018083504 | 38.14503886 |
| ENSMUSG00000066684 | *Pilrb1* | 19.56407184 | 18.69452655 | 22.51906226 | 0 | 0 | 0.96222723 | 4.95E-07 | -5.791061966 | 1.05E-07 | 28.28334027 | 0.018059954 | 6.305621373 |
| ENSMUSG00000064267 | *Hvcn1* | 108.8783128 | 156.1542806 | 131.1980149 | 3.487981309 | 1.905496576 | 1.924454461 | 1.00E-35 | -5.791339589 | 5.17E-37 | 161.558564 | 0.018056479 | 34.99950821 |

| | | | | | | | | | | | | | |
|---|---|---|---|---|---|---|---|---|---|---|---|---|---|
| ENSMUSG00000025854 | *Fam20c* | 9151.73256 | 9611.186005 | 9569.622371 | 160.4471402 | 162.9199572 | 182.8231738 | 0 | -5.803425886 | 0 | 5231.270643 | 0.017905841 | Inf |
| ENSMUSG00000028553 | *Angptl3* | 10.20734183 | 16.49517049 | 4.895448318 | 0 | 0 | 0 | 0.000406352 | -5.804437039 | 0.00012466 | 14.7208446 | 0.017893296 | 3.391097241 |
| ENSMUSG00000031214 | *Ophn1* | 58.69221551 | 65.98068195 | 46.99630385 | 1.162660436 | 1.905496576 | 0 | 4.18E-17 | -5.812126675 | 4.45E-18 | 75.11227625 | 0.017798177 | 16.37897304 |
| ENSMUSG00000024987 | *Cyp26a1* | 12.75917728 | 12.09645836 | 6.853627645 | 0 | 0 | 0 | 0.000238625 | -5.814539863 | 7.05E-05 | 15.79816445 | 0.017768431 | 3.622284038 |
| ENSMUSG00000107758 | *Gm19253* | 17.8628482 | 21.99356065 | 22.51906226 | 1.162660436 | 0 | 0 | 3.88E-07 | -5.826243543 | 8.13E-08 | 28.77495472 | 0.01762487 | 6.410653869 |
| ENSMUSG00000003617 | *Cp* | 398.0863313 | 369.4918189 | 396.5313137 | 4.650641745 | 9.527482879 | 5.773363382 | 5.07E-109 | -5.841444426 | 8.21E-111 | 500.291662 | 0.017440142 | 108.2948227 |
| ENSMUSG00000082289 | *Gm15596* | 11.90856547 | 13.19613639 | 7.832717308 | 0 | 0 | 0 | 0.000158669 | -5.867808415 | 4.56E-05 | 16.62297418 | 0.017124333 | 3.799507255 |
| ENSMUSG00000031125 | *3830403N18Rik* | 11.05795365 | 14.29581442 | 8.811806972 | 0 | 0 | 0 | 0.000110929 | -5.919148565 | 3.12E-05 | 17.34285198 | 0.016525657 | 3.954953315 |
| ENSMUSG00000031586 | *Rbpms* | 147.1558447 | 161.6526708 | 130.2189253 | 3.487981309 | 0 | 3.848908921 | 2.60E-41 | -5.941482326 | 1.17E-42 | 187.4070475 | 0.0162718 | 40.58498852 |
| ENSMUSG00000043953 | *Ccrl2* | 1691.016296 | 1585.735723 | 1670.326966 | 29.06651091 | 20.96046233 | 29.82904414 | 0 | -5.961578069 | 0 | 1781.323191 | 0.016046716 | Inf |
| ENSMUSG00000102418 | *Sh2d1b1* | 12.75917728 | 10.99678033 | 11.74907596 | 0 | 0 | 0 | 6.48E-05 | -5.978310428 | 1.77E-05 | 18.42004235 | 0.015861682 | 4.188641731 |
| ENSMUSG00000047139 | *Cd24a* | 23.81713093 | 19.79420459 | 27.41451058 | 1.162660436 | 0 | 0 | 4.54E-08 | -6.01700384 | 8.75E-09 | 33.10025892 | 0.015441922 | 7.34267597 |
| ENSMUSG00000020340 | *Cyfip2* | 74.00322825 | 54.98390163 | 68.53627645 | 1.162660436 | 0.952748288 | 0.96222723 | 2.55E-20 | -6.017812882 | 2.30E-21 | 90.07045897 | 0.015433265 | 19.59295254 |
| ENSMUSG00000062082 | *Cd200r4* | 227.9639675 | 227.6333527 | 230.0860709 | 4.650641745 | 0.952748288 | 4.811136152 | 5.12E-68 | -6.072182351 | 1.32E-69 | 311.0039615 | 0.014862469 | 67.29108952 |
| ENSMUSG00000063660 | *Or1o3* | 32.32324912 | 23.09323868 | 18.60270361 | 1.162660436 | 0 | 0 | 3.97E-08 | -6.078406811 | 7.60E-09 | 33.37495488 | 0.014798484 | 7.401232606 |
| ENSMUSG00000102433 | *Gm38050* | 23.81713093 | 7.697746228 | 8.811806972 | 0 | 0 | 0 | 6.66E-05 | -6.169241099 | 1.82E-05 | 18.36442348 | 0.013895474 | 4.176509139 |
| ENSMUSG00000051379 | *Flrt3* | 14.46040092 | 16.49517049 | 9.790896635 | 0 | 0 | 0 | 2.04E-05 | -6.174476687 | 5.17E-06 | 20.77362383 | 0.013845139 | 4.69027652 |
| ENSMUSG00000113749 | *Mrto4-ps1* | 9.356730009 | 52.78454556 | 152.7379875 | 0 | 2.858244864 | 0 | 0.005324062 | -6.193841953 | 0.001985107 | 9.563255938 | 0.013660537 | 2.273756936 |
| ENSMUSG00000013584 | *Aldh1a2* | 17.01223638 | 16.49517049 | 7.832717308 | 0 | 0 | 0 | 2.56E-05 | -6.196975448 | 6.57E-06 | 20.31539062 | 0.013630899 | 4.592565438 |
| ENSMUSG00000021365 | *Nedd9* | 140.3509501 | 111.0674813 | 124.3443873 | 2.325320873 | 0.952748288 | 1.924454461 | 1.21E-37 | -6.206737686 | 5.91E-39 | 170.4465417 | 0.013538975 | 36.9178727 |
| ENSMUSG00000040584 | *Abcb1a* | 21.26529547 | 7.697746228 | 12.72816563 | 0 | 0 | 0 | 3.03E-05 | -6.216163919 | 7.86E-06 | 19.97070866 | 0.013450803 | 4.51920085 |
| ENSMUSG00000102805 | *Gm37240* | 3.402447276 | 16.49517049 | 24.47724159 | 0 | 0 | 0 | 8.71E-05 | -6.293323709 | 2.41E-05 | 17.83108683 | 0.012750311 | 4.060160602 |
| ENSMUSG00000032578 | *Cish* | 145.454621 | 179.2475193 | 176.2361394 | 4.650641745 | 0 | 1.924454461 | 7.86E-48 | -6.349529664 | 3.00E-49 | 217.613067 | 0.012263124 | 47.10444884 |
| ENSMUSG00000110768 | *Gm18541* | 34.87508458 | 149.5562124 | 83.2226214 | 3.487981309 | 0 | 0 | 5.71E-05 | -6.362589056 | 1.55E-05 | 18.68103688 | 0.012152618 | 4.2437275 |
| ENSMUSG00000036353 | *P2ry12* | 13.6097891 | 18.69452655 | 14.68634495 | 0 | 0 | 0 | 3.22E-06 | -6.378510857 | 7.45E-07 | 24.49418857 | 0.012019237 | 5.491566022 |
| ENSMUSG00000024544 | *Ldlrad4* | 18.71346002 | 13.19613639 | 15.66543462 | 0 | 0 | 0 | 2.72E-06 | -6.402208556 | 6.24E-07 | 24.83692297 | 0.011823422 | 5.565046707 |
| ENSMUSG00000041544 | *Disp3* | 404.040614 | 429.9741107 | 415.1340173 | 4.650641745 | 0.952748288 | 8.660045073 | 2.02E-119 | -6.453930068 | 2.80E-121 | 548.4068213 | 0.011407053 | 118.6952005 |
| ENSMUSG00000030739 | *Myh14* | 476.3426186 | 489.3567245 | 468.0048592 | 6.975962618 | 4.763741439 | 4.811136152 | 5.47E-143 | -6.459108372 | 6.08E-145 | 657.198632 | 0.011366183 | 142.2623725 |
| ENSMUSG00000004609 | *Cd33* | 3210.209005 | 3075.799457 | 3039.094316 | 31.39183178 | 38.10993152 | 34.64018029 | 0 | -6.476738766 | 0 | 3060.940382 | 0.011228128 | Inf |
| ENSMUSG00000015709 | *Amt2* | 17.8628482 | 20.89388262 | 11.74907596 | 0 | 0 | 0 | 1.73E-06 | -6.4840233 | 3.90E-07 | 25.74525401 | 0.011171577 | 5.761958853 |
| ENSMUSG00000090894 | *Or5v1* | 41.67997913 | 31.89066294 | 25.45633125 | 0 | 0.952748288 | 0 | 6.76E-11 | -6.49762249 | 1.03E-11 | 46.26471318 | 0.011066766 | 10.16981308 |
| ENSMUSG00000070691 | *Runx3* | 656.6723243 | 651.0093953 | 638.3664606 | 6.975962618 | 5.716489727 | 8.660045073 | 1.06E-190 | -6.509361946 | 7.92E-193 | 877.4290299 | 0.010977079 | 189.9753002 |
| ENSMUSG00000063531 | *Sema3e* | 468.6871123 | 469.5625199 | 457.2348729 | 4.650641745 | 4.763741439 | 5.773363382 | 6.42E-141 | -6.515265386 | 7.18E-143 | 647.6698923 | 0.010932253 | 140.1927438 |
| ENSMUSG00000001588 | *Acap1* | 1271.664669 | 1346.005912 | 1300.231073 | 16.27724611 | 11.43297945 | 14.43340846 | 0 | -6.552117816 | 0 | 1595.097248 | 0.010656535 | Inf |
| ENSMUSG00000021200 | *Asb2* | 36.57630822 | 28.59162885 | 42.10085553 | 0 | 0 | 0.96222723 | 5.13E-12 | -6.612217368 | 7.27E-13 | 51.46946544 | 0.010221726 | 11.29011158 |
| ENSMUSG00000021725 | *Parp8* | 111.4301483 | 87.97424261 | 107.699863 | 2.325320873 | 0 | 0.96222723 | 8.98E-32 | -6.648965727 | 5.34E-33 | 143.1890084 | 0.009964646 | 31.04653502 |
| ENSMUSG00000022893 | *Adamts1* | 23.81713093 | 15.39549246 | 19.58179327 | 0 | 0 | 0 | 1.53E-07 | -6.708329446 | 3.09E-08 | 30.64916766 | 0.009562944 | 6.813995119 |
| ENSMUSG00000017002 | *Slpi* | 642.2119233 | 698.2955507 | 694.1745714 | 9.301283491 | 5.716489727 | 4.811136152 | 3.04E-193 | -6.715166518 | 2.19E-195 | 889.1960611 | 0.009517732 | 192.5171276 |
| ENSMUSG00000094796 | *BC147527* | 33.17386094 | 36.28937507 | 47.97539351 | 0 | 0.952748288 | 0 | 4.39E-13 | -6.740211552 | 5.82E-14 | 56.43099082 | 0.009353931 | 12.35723787 |
| ENSMUSG00000038094 | *Atp13a4* | 80.8081228 | 82.47585244 | 64.61991779 | 0 | 1.905496576 | 0 | 3.11E-24 | -6.811704612 | 2.37E-25 | 108.2453595 | 0.008901693 | 23.5068952 |
| ENSMUSG00000029816 | *Gpnmb* | 37378.43516 | 38420.5511 | 37405.14151 | 315.0809782 | 307.737697 | 373.3441654 | 0 | -6.825801604 | 0 | 8028.679112 | 0.008815135 | Inf |
| ENSMUSG00000063354 | *Slc39a4* | 6.804894552 | 48.38583343 | 9.790896635 | 0 | 0 | 0 | 9.03E-06 | -6.840307649 | 2.18E-06 | 22.42691702 | 0.008726945 | 5.044540152 |
| ENSMUSG00000057315 | *Arhgap24* | 280.7019003 | 277.1188642 | 267.2914781 | 2.325320873 | 2.858244864 | 1.924454461 | 2.02E-87 | -6.856579318 | 4.15E-89 | 400.5627983 | 0.008629069 | 86.69367166 |
| ENSMUSG00000017667 | *Zfp334* | 34.87508458 | 41.78776524 | 52.87084183 | 0 | 0.952748288 | 0 | 2.76E-14 | -6.880956796 | 3.38E-15 | 62.02912414 | 0.008484487 | 13.55982465 |
| ENSMUSG00000031503 | *Col4a2* | 23.81713093 | 24.19291672 | 19.58179327 | 0 | 0 | 0 | 1.08E-08 | -6.905724533 | 1.96E-09 | 36.01382814 | 0.008340071 | 7.966434772 |
| ENSMUSG00000031451 | *Gas6* | 187.1346002 | 225.4339967 | 213.4415467 | 1.162660436 | 0.952748288 | 2.886681691 | 4.85E-64 | -6.942849341 | 1.35E-65 | 292.6017258 | 0.008128195 | 63.31420386 |
| ENSMUSG00000061126 | *Cyp4f39* | 34.87508458 | 52.78454556 | 48.95448318 | 1.162660436 | 0 | 0 | 6.23E-15 | -6.955821455 | 7.38E-16 | 65.030093 | 0.008055437 | 14.20537195 |
| ENSMUSG00000032238 | *Rora* | 39.97875549 | 49.48551147 | 50.9126625 | 0 | 0 | 0.96222723 | 6.27E-16 | -6.996538133 | 7.06E-17 | 69.65594703 | 0.007831269 | 15.20242632 |
| ENSMUSG00000020435 | *Osbp2* | 516.3213741 | 534.4435238 | 511.0848044 | 4.650641745 | 4.763741439 | 2.886681691 | 4.51E-160 | -6.998572183 | 4.43E-162 | 736.0064763 | 0.007820236 | 159.3456218 |
| ENSMUSG00000089945 | *Pakap* | 37.42692004 | 30.79098491 | 7.832717308 | 0 | 0 | 0 | 1.56E-07 | -7.077565085 | 3.15E-08 | 30.61221676 | 0.00740356 | 6.806167747 |
| ENSMUSG00000006800 | *Sulf2* | 22.96651911 | 32.99034098 | 20.56088293 | 0 | 0 | 0 | 1.93E-09 | -7.08152758 | 3.29E-10 | 39.49609187 | 0.007383254 | 8.714576287 |
| ENSMUSG00000020674 | *Pxdn* | 27.21957821 | 21.99356065 | 27.41451058 | 0 | 0 | 0 | 9.38E-10 | -7.088608807 | 1.56E-10 | 40.95438596 | 0.007347103 | 9.027961242 |
| ENSMUSG00000044468 | *Tent5c* | 1225.731631 | 1248.134567 | 1290.440177 | 8.138623054 | 5.716489727 | 13.47118123 | 0 | -7.098726638 | 0 | 1579.5173 | 0.007295757 | Inf |
| ENSMUSG00000049604 | *Hoxb13* | 46.78365004 | 58.28293573 | 46.01721419 | 0 | 0.952748288 | 0 | 4.03E-17 | -7.102443752 | 4.28E-18 | 75.18685042 | 0.007276983 | 16.39481633 |
| ENSMUSG00000073176 | *Zfp449* | 53.5885446 | 54.98390163 | 46.99630385 | 0 | 0.952748288 | 0 | 6.81E-18 | -7.146448178 | 6.96E-19 | 78.77543315 | 0.007058375 | 17.16707898 |
| ENSMUSG00000036106 | *Prr5* | 93.56730009 | 95.67198883 | 100.8462353 | 1.162660436 | 0.952748288 | 0 | 1.87E-32 | -7.153399629 | 1.08E-33 | 146.3667182 | 0.007024447 | 31.72907705 |
| ENSMUSG00000034266 | *Batf* | 22.96651911 | 28.59162885 | 29.37268991 | 0 | 0 | 0 | 3.16E-10 | -7.163769858 | 5.07E-11 | 43.15033525 | 0.006974136 | 9.499961416 |
| ENSMUSG00000070729 | *Gm12966* | 22.11590729 | 16.49517049 | 43.0799452 | 0 | 0 | 0 | 4.95E-09 | -7.180385196 | 8.74E-10 | 37.58735465 | 0.006894276 | 8.305809377 |
| ENSMUSG00000022676 | *Snai2* | 108.8783128 | 87.97424261 | 97.90896635 | 0 | 0 | 1.924454461 | 6.82E-32 | -7.18324652 | 4.01E-33 | 143.7575442 | 0.006880616 | 31.16630808 |
| ENSMUSG00000014773 | *Dll1* | 30.62202548 | 26.39227278 | 25.45633125 | 0 | 0 | 0 | 1.78E-10 | -7.194249511 | 2.80E-11 | 44.31160797 | 0.006828339 | 9.749020074 |
| ENSMUSG00000007682 | *Dio2* | 24.66774275 | 39.58840917 | 22.51906226 | 0 | 0 | 0 | 2.44E-10 | -7.262179175 | 3.88E-11 | 43.67534153 | 0.006514277 | 9.61328745 |
| ENSMUSG00000004113 | *Cacna1b* | 127.5917728 | 83.57553048 | 114.5534906 | 1.162660436 | 0 | 0.96222723 | 1.43E-32 | -7.323289394 | 8.20E-34 | 146.9129645 | 0.006244105 | 31.84435005 |
| ENSMUSG00000021902 | *Phf7* | 20.41468366 | 21.99356065 | 53.84993149 | 0 | 0 | 0 | 6.09E-10 | -7.415372552 | 9.97E-11 | 41.82641172 | 0.005858014 | 9.215556221 |
| ENSMUSG00000021322 | *Aoah* | 204.9974484 | 188.0449436 | 143.9261805 | 2.325320873 | 0.952748288 | 0 | 1.31E-51 | -7.455765024 | 4.54E-53 | 235.1299311 | 0.005696277 | 50.88392926 |
| ENSMUSG00000060568 | *Fam78b* | 114.8325956 | 140.7587882 | 115.5325803 | 0 | 0.952748288 | 0.96222723 | 3.16E-40 | -7.511712422 | 1.47E-41 | 182.3784433 | 0.005479605 | 39.50023987 |
| ENSMUSG00000026012 | *Cd28* | 72.30200461 | 63.78132589 | 68.53627645 | 0 | 0 | 0.96222723 | 1.21E-23 | -7.543177523 | 9.43E-25 | 105.512066 | 0.005361389 | 22.91795133 |
| ENSMUSG00000049511 | *Htr1b* | 557.1507414 | 516.8486753 | 575.7047222 | 4.650641745 | 1.905496576 | 1.924454461 | 2.41E-167 | -7.661991482 | 2.22E-169 | 769.5796158 | 0.004937541 | 166.6173655 |
| ENSMUSG00000028214 | *Gem* | 55.28976823 | 30.79098491 | 42.10085553 | 0 | 0 | 0 | 9.69E-15 | -7.833537006 | 1.16E-15 | 64.13876866 | 0.004383998 | 14.01367017 |
| ENSMUSG00000036528 | *Ppfibp2* | 153.1101274 | 166.0513829 | 149.8007185 | 1.162660436 | 0 | 0.96222723 | 1.41E-52 | -7.846391996 | 4.83E-54 | 239.589187 | 0.004345109 | 51.85024037 |
| ENSMUSG00000046058 | *Eid2* | 92.71668827 | 64.88100392 | 96.92987669 | 0 | 0 | 0.96222723 | 4.89E-27 | -7.859434206 | 3.39E-28 | 121.235644 | 0.004306005 | 26.3105443 |
| ENSMUSG00000052595 | *A1cf* | 53.5885446 | 38.48873114 | 42.10085553 | 0 | 0 | 0 | 3.37E-16 | -7.897939257 | 3.75E-17 | 70.90533502 | 0.0041926 | 15.47215623 |
| ENSMUSG00000020439 | *Smtn* | 41.67997913 | 50.5851895 | 46.01721419 | 0 | 0 | 0 | 4.89E-17 | -7.936790124 | 5.22E-18 | 74.79702931 | 0.004081202 | 16.31102543 |
| ENSMUSG00000047250 | *Ptgs1* | 47.63426186 | 38.48873114 | 52.87084183 | 0 | 0 | 0 | 7.89E-17 | -7.947550415 | 8.49E-18 | 73.83666746 | 0.004050876 | 16.10280967 |
| ENSMUSG00000041707 | *Tmem273* | 637.1082524 | 560.8357966 | 581.5792601 | 3.487981309 | 0.952748288 | 2.886681691 | 1.12E-177 | -7.965049158 | 9.30E-180 | 817.3104916 | 0.004002039 | 176.9507081 |
| ENSMUSG00000079362 | *Gm43302* | 174.3754229 | 193.5433337 | 153.7170772 | 0 | 0 | 1.924454461 | 1.21E-55 | -8.002988412 | 3.93E-57 | 253.7614169 | 0.003898167 | 54.918674 |
| ENSMUSG00000025586 | *Cpeb1* | 57.84160369 | 50.5851895 | 42.10085553 | 0 | 0 | 0 | 4.07E-18 | -8.06222701 | 4.12E-19 | 79.80994349 | 0.003741346 | 17.3903859 |
| ENSMUSG00000047409 | *Ctdspl* | 57.84160369 | 51.68486753 | 43.0799452 | 0 | 0 | 0 | 1.99E-18 | -8.081817033 | 1.99E-19 | 81.25185305 | 0.003690886 | 17.70106629 |
| ENSMUSG00000045382 | *Cxcr4* | 208.3998957 | 221.0352845 | 174.2779601 | 2.325320873 | 0 | 0 | 3.27E-63 | -8.208347047 | 9.22E-65 | 288.7681775 | 0.00338097 | 62.48530183 |
| ENSMUSG00000075394 | *Hoxc4* | 135.2472792 | 150.6558905 | 117.4907596 | 0 | 0 | 0.96222723 | 1.18E-44 | -8.520385958 | 4.86E-46 | 202.9015448 | 0.00272338 | 43.92645034 |
| ENSMUSG00000038068 | *Rnf144b* | 882.0844563 | 828.0575585 | 920.3442837 | 1.162660436 | 1.905496576 | 3.848908921 | 1.13E-259 | -8.530157831 | 5.29E-262 | 1195.68462 | 0.002704996 | 258.9457772 |
| ENSMUSG00000034612 | *Chst11* | 439.7663104 | 404.681516 | 356.3886375 | 1.162660436 | 1.905496576 | 0 | 2.86E-120 | -8.620645434 | 3.91E-122 | 552.3364558 | 0.002540546 | 119.5435563 |
| ENSMUSG00000032352 | *Lrrc1* | 75.70445189 | 83.57553048 | 99.86714568 | 0 | 0 | 0 | 2.77E-30 | -8.843835877 | 1.73E-31 | 136.2786054 | 0.002176407 | 29.55686691 |
| ENSMUSG00000011008 | *Mcoln2* | 639.6600879 | 627.9161566 | 616.826488 | 2.325320873 | 0.952748288 | 0.96222723 | 5.35E-204 | -8.854523359 | 3.60E-206 | 938.8074717 | 0.002160344 | 203.2719092 |
| ENSMUSG00000030156 | *Cd69* | 177.7778702 | 212.2378603 | 188.9643051 | 0 | 0 | 0.96222723 | 1.04E-64 | -9.041126159 | 2.88E-66 | 295.6809685 | 0.001898234 | 63.98188201 |
| ENSMUSG00000022885 | *St6gal1* | 226.2627438 | 190.2442996 | 186.0270361 | 0 | 0.952748288 | 0 | 4.73E-66 | -9.101611202 | 1.25E-67 | 301.9357635 | 0.001820296 | 65.32471431 |
| ENSMUSG00000021375 | *Kif13a* | 199.8937775 | 213.3375383 | 198.7552017 | 0 | 0.952748288 | 0 | 1.31E-70 | -9.122028552 | 3.27E-72 | 322.9650864 | 0.001794716 | 69.88127748 |
| ENSMUSG00000049744 | *Arhgap15* | 234.768862 | 184.7459095 | 214.4206363 | 0 | 0 | 0.96222723 | 1.88E-68 | -9.175280782 | 4.83E-70 | 313.007351 | 0.001729678 | 67.72518928 |
| ENSMUSG00000044716 | *Dok7* | 102.9240301 | 124.2636177 | 134.1352839 | 0 | 0 | 0 | 2.31E-41 | -9.322684794 | 1.04E-42 | 187.6416735 | 0.001561681 | 40.63554323 |
| ENSMUSG00000003882 | *Il7r* | 1662.946106 | 1514.256651 | 1675.222414 | 0 | 0 | 0 | 0 | -13.07204132 | 0 | 2043.760944 | 0.000116124 | Inf |
