## Supplementary material for "TFEB-Mediated Pro-inflammatory Response in Murine Macrophages Induced by Acute Alpha7 Nicotinic Receptor Activation": Table_S21

**Table S21. GO and KEGG Pathway analysis of differentially expressed genes from Table S20.**

| Term | Library | p-value | q-value | z-score | combined score |
|---|---|---|---|---|---|
| Immune System R-HSA-168256 | Reactome_2022 | 1.33E-25 | 1.82E-22 | 2.173 | 124.5 |
| cytokine-mediated signaling pathway (GO:0019221) | GO_Biological_Process_2021 | 1.09E-21 | 4.80E-18 | 2.987 | 144.2 |
| Cytokine Signaling In Immune System R-HSA-1280215 | Reactome_2022 | 5.14E-21 | 3.50E-18 | 2.801 | 130.9 |
| abnormal macrophage physiology MP:0002451 | MGI_Mammalian_Phenotype_Level_4_2021 | 1.92E-20 | 6.56E-17 | 5.686 | 258.1 |
| Innate Immune System R-HSA-168249 | Reactome_2022 | 1.92E-17 | 8.70E-15 | 2.26 | 87.02 |
| Lysosome | KEGG_2021_Human | 1.31E-15 | 3.88E-13 | 5.612 | 192.3 |
| positive regulation of cytokine production (GO:0001819) | GO_Biological_Process_2021 | 5.39E-15 | 1.10E-11 | 3.234 | 106.2 |
| cellular response to cytokine stimulus (GO:0071345) | GO_Biological_Process_2021 | 9.35E-15 | 1.10E-11 | 2.752 | 88.9 |
| positive regulation of intracellular signal transduction (GO:1902533) | GO_Biological_Process_2021 | 1.00E-14 | 1.10E-11 | 2.621 | 84.49 |
| abnormal immune system physiology MP:0001790 | MGI_Mammalian_Phenotype_Level_4_2021 | 3.66E-14 | 6.25E-11 | 5.731 | 177.3 |
| Signaling By Interleukins R-HSA-449147 | Reactome_2022 | 2.68E-13 | 9.12E-11 | 2.686 | 77.75 |
| defense response to symbiont (GO:0140546) | GO_Biological_Process_2021 | 3.39E-13 | 2.99E-10 | 5.064 | 145.4 |
| Interleukin-10 Signaling R-HSA-6783783 | Reactome_2022 | 6.95E-12 | 1.90E-09 | 9.957 | 255.8 |
| decreased tumor necrosis factor secretion MP:0008561 | MGI_Mammalian_Phenotype_Level_4_2021 | 3.86E-10 | 4.39E-07 | 5.069 | 109.9 |
| Tuberculosis | KEGG_2021_Human | 4.21E-09 | 4.93E-07 | 3.269 | 63.05 |
| NOD-like receptor signaling pathway | KEGG_2021_Human | 4.98E-09 | 4.93E-07 | 3.246 | 62.05 |
| decreased IgG2a level MP:0008496 | MGI_Mammalian_Phenotype_Level_4_2021 | 1.39E-08 | 1.05E-05 | 5.981 | 108.2 |
| decreased IgG1 level MP:0008495 | MGI_Mammalian_Phenotype_Level_4_2021 | 1.54E-08 | 1.05E-05 | 4.419 | 79.5 |
| Rheumatoid arthritis | KEGG_2021_Human | 1.54E-08 | 9.29E-07 | 4.419 | 79.5 |
| TNF signaling pathway | KEGG_2021_Human | 1.56E-08 | 9.29E-07 | 3.983 | 71.59 |

**Enrichr-KG_UP in PNUtreated dKO RAW cells_DE UP in dKO from PNU(WTvdKORAW) padj0.01Fc1.5**

| Term | Library | p-value | q-value | z-score | combined score | converted -log(Padj) |
|---|---|---|---|---|---|---|
| Hepatocellular carcinoma | KEGG_2021_Human | 3.99E-05 | 0.01193 | 2.683 | 27.18 | 1.92336 |
| peptidyl-threonine modification (GO:0018210) | GO_Biological_Process_2021 | 0.000203 | 0.2933 | 3.677 | 31.26 | 0.532688 |
| regulation of ubiquitin-protein transferase activity (GO:0051438) | GO_Biological_Process_2021 | 0.00028 | 0.2933 | 5.804 | 47.48 | 0.532688 |
| phosphorylation (GO:0016310) | GO_Biological_Process_2021 | 0.000304 | 0.2933 | 1.855 | 15.02 | 0.532688 |
| regulation of striated muscle cell differentiation (GO:0051153) | GO_Biological_Process_2021 | 0.00044 | 0.2933 | 20.27 | 156.7 | 0.532688 |
| bleb assembly (GO:0032060) | GO_Biological_Process_2021 | 0.00044 | 0.2933 | 20.27 | 156.7 | 0.532688 |
| Glutathione metabolism | KEGG_2021_Human | 0.000647 | 0.09672 | 3.648 | 26.79 | 1.014484 |
| Fc gamma R-mediated phagocytosis | KEGG_2021_Human | 0.002535 | 0.1831 | 2.574 | 15.39 | 0.737312 |
| AGE-RAGE signaling pathway in diabetic complications | KEGG_2021_Human | 0.003374 | 0.1831 | 2.484 | 14.14 | 0.737312 |
| Fructose and mannose metabolism | KEGG_2021_Human | 0.00353 | 0.1831 | 4.098 | 23.14 | 0.737312 |
