## Supplementary material for "TFEB-Mediated Pro-inflammatory Response in Murine Macrophages Induced by Acute Alpha7 Nicotinic Receptor Activation": Table_S22

**Table S22. Differential gene expression in baseline RAW264.7 cells (WT v. Tfeb Tfe3 dKO).**

| ID | Gene | counts_WT_CO1 | counts_WT_CO2 | counts_WT_CO3 | counts_KO_CO1 | counts_KO_CO2 | counts_KO_CO3 | padj | log2FoldChange | pvalue | stat | foldChange | log10padj |
|---|---|---|---|---|---|---|---|---|---|---|---|---|---|
| ENSMUSG00000030043 | *Tacr1* | 1.506846974 | 2.877574023 | 2.521136538 | 839.636646 | 1071.712006 | 919.2269967 | 1.07E-194 | 8.733795526 | 1.40E-197 | 899.2944953 | 425.7301711 | 193.9700454 |
| ENSMUSG00000069622 | *Gm10273* | 0 | 0 | 0 | 38.41474851 | 57.90192569 | 41.32389096 | 5.28E-16 | 8.006921816 | 4.95E-17 | 70.35472665 | 257.2311975 | 15.27757586 |
| ENSMUSG00000037846 | *Rtkn2* | 0 | 0 | 0 | 35.12205578 | 42.11049141 | 39.48727358 | 1.08E-14 | 7.769899157 | 1.13E-15 | 64.18460591 | 218.2592757 | 13.9655572 |
| ENSMUSG00000041444 | *Arhgap32* | 1.506846974 | 0 | 1.260568269 | 175.6102789 | 228.4494159 | 177.2335768 | 1.35E-55 | 7.676722807 | 2.12E-57 | 254.9876494 | 204.6085762 | 54.87107955 |
| ENSMUSG00000042351 | *Grap2* | 3.013693948 | 4.795956706 | 0 | 556.4650713 | 516.906282 | 562.9232257 | 3.56E-158 | 7.586684433 | 8.31E-161 | 730.1489554 | 192.2292989 | 157.4481494 |
| ENSMUSG00000038048 | *Cntnap5c* | 0.753423487 | 0 | 0 | 54.87821216 | 58.95468797 | 61.5266821 | 1.08E-20 | 7.395644744 | 7.51E-22 | 92.28430548 | 168.3879094 | 19.96700987 |
| ENSMUSG00000022994 | *Adcy6* | 0 | 0 | 0 | 48.2928267 | 16.84419656 | 21.12109982 | 1.56E-08 | 7.330186841 | 2.85E-09 | 35.28145638 | 160.9185509 | 7.805515878 |
| ENSMUSG00000019846 | *Lama4* | 0 | 0 | 0 | 31.82936305 | 20.00248342 | 33.97742146 | 2.91E-10 | 7.326878685 | 4.46E-11 | 43.39979472 | 160.5499811 | 9.535416827 |
| ENSMUSG00000032036 | *Kirrel3* | 11.3013523 | 12.46948743 | 8.823977885 | 1558.541225 | 1973.929285 | 1759.479446 | 3.54E-265 | 7.319291796 | 2.19E-268 | 1225.056865 | 159.707892 | 264.4509335 |
| ENSMUSG00000072693 | *None* | 0 | 0 | 0 | 35.12205578 | 10.52762285 | 37.65065621 | 6.37E-08 | 7.284298501 | 1.24E-08 | 32.42628838 | 155.8806996 | 7.195545429 |
| ENSMUSG00000020838 | *Slc6a4* | 10.54792882 | 20.14301816 | 7.563409615 | 1672.687907 | 1817.067704 | 1731.930186 | 0 | 7.069787297 | 0 | 1609.387572 | 134.3439272 | Inf |
| ENSMUSG00000032068 | *Plet1* | 1.506846974 | 3.836765364 | 1.260568269 | 331.4644014 | 223.1856045 | 327.8362016 | 9.41E-63 | 7.053670156 | 1.29E-64 | 288.0966938 | 132.8514493 | 62.0262367 |
| ENSMUSG00000028464 | *Tpm2* | 0 | 0 | 0 | 9.878078189 | 29.47734399 | 12.85632163 | 7.50E-06 | 6.608034177 | 1.88E-06 | 22.71168266 | 97.54757996 | 5.125177358 |
| ENSMUSG00000037855 | *Zfp365* | 4.520540922 | 4.795956706 | 2.521136538 | 331.4644014 | 414.7883404 | 381.0981055 | 2.54E-96 | 6.51935058 | 1.54E-98 | 443.8954795 | 91.73183408 | 95.59431537 |
| ENSMUSG00000089768 | *Tmsb15b1* | 0 | 0 | 0 | 16.46346365 | 8.422098282 | 16.52955638 | 2.33E-05 | 6.275763154 | 6.29E-06 | 20.3964839 | 77.48059641 | 4.633564256 |
| ENSMUSG00000041202 | *Pla2g2d* | 0 | 0 | 0 | 19.75615638 | 9.474860567 | 11.93801294 | 2.64E-05 | 6.263755663 | 7.18E-06 | 20.14373679 | 76.83840481 | 4.579153469 |
| ENSMUSG00000028348 | *Cavin4* | 0 | 0 | 0 | 7.682949702 | 15.79143428 | 11.93801294 | 9.69E-05 | 6.049799017 | 2.87E-05 | 17.50331416 | 66.24772545 | 4.013770784 |
| ENSMUSG00000042401 | *Crtac1* | 3.013693948 | 0.959191341 | 6.302841346 | 221.7079771 | 181.0751131 | 224.9856286 | 5.81E-54 | 6.038739877 | 9.68E-56 | 247.3779107 | 65.74183755 | 53.23599649 |
| ENSMUSG00000001131 | *Timp1* | 3.013693948 | 3.836765364 | 7.563409615 | 280.9764463 | 256.8739976 | 277.3292238 | 3.72E-76 | 5.928822571 | 4.03E-78 | 350.0994072 | 60.91909359 | 75.42964563 |
| ENSMUSG00000027200 | *Sema6d* | 12.05477579 | 5.755148047 | 2.521136538 | 502.6844234 | 416.8938649 | 406.8107488 | 9.18E-88 | 5.889325414 | 7.18E-90 | 404.0635 | 59.27391374 | 87.03701287 |
| ENSMUSG00000085738 | *Gm12335* | 0 | 0 | 0 | 10.97564243 | 5.263811426 | 14.69293901 | 0.000434943 | 5.85678867 | 0.000143359 | 14.45748234 | 57.95208532 | 3.361567826 |
| ENSMUSG00000006221 | *Hspb7* | 30.13693948 | 39.32684499 | 27.73250192 | 1515.73622 | 1901.288687 | 1722.747099 | 2.56E-220 | 5.7181452 | 2.63E-223 | 1017.638364 | 52.64210229 | 219.5922175 |
| ENSMUSG00000052572 | *Dlg2* | 0 | 0 | 0 | 4.390256973 | 11.58038514 | 11.01970426 | 0.001085056 | 5.660060187 | 0.000384967 | 12.60378227 | 50.56475339 | 2.96454792 |
| ENSMUSG00000040612 | *Ildr2* | 4.520540922 | 17.26544414 | 0 | 315.0009378 | 382.1527095 | 329.672819 | 5.37E-05 | 5.543656553 | 1.53E-05 | 18.69601678 | 46.64519448 | 4.269920817 |
| ENSMUSG00000047443 | *Erfe* | 5.273964409 | 0 | 0 | 91.09783218 | 105.2762285 | 90.91256011 | 4.79E-22 | 5.487599675 | 3.06E-23 | 98.61932336 | 44.86752427 | 21.31921603 |
| ENSMUSG00000051190 | *Or7c70* | 0 | 0 | 0 | 12.07320667 | 2.10552457 | 8.264778192 | 0.005542712 | 5.390107148 | 0.002275357 | 9.312983026 | 41.93570336 | 2.256277657 |
| ENSMUSG00000040875 | *Osbpl10* | 15.06846974 | 14.38787012 | 2.521136538 | 374.2694069 | 546.383626 | 492.2134568 | 1.31E-61 | 5.36865147 | 1.82E-63 | 282.8258628 | 41.31665263 | 60.88417435 |
| ENSMUSG00000111409 | *Gm49380* | 0 | 0 | 0 | 3.29269273 | 11.58038514 | 6.428160816 | 0.004932128 | 5.31603765 | 0.002003043 | 9.546745504 | 39.83701515 | 2.306965629 |
| ENSMUSG00000050439 | *Enthd1* | 4.520540922 | 1.918382682 | 3.781704808 | 160.2443795 | 104.2234662 | 134.0730684 | 1.55E-30 | 5.262094373 | 6.52E-32 | 138.2206605 | 38.37498745 | 29.80960993 |
| ENSMUSG00000027849 | *Syt6* | 0.753423487 | 0 | 0 | 13.17077092 | 6.316573711 | 19.28448245 | 0.000291698 | 5.221762294 | 9.32E-05 | 15.26923617 | 37.31703078 | 3.535066431 |
| ENSMUSG00000026726 | *Cubn* | 0 | 0 | 0 | 8.780513945 | 7.369335996 | 2.754926064 | 0.008251725 | 5.137622992 | 0.003512589 | 8.520028787 | 35.20291501 | 2.083455241 |
| ENSMUSG00000038668 | *Lpar1* | 5.273964409 | 23.97978353 | 15.12681923 | 515.8551943 | 535.8560032 | 444.461405 | 7.12E-54 | 5.124807438 | 1.20E-55 | 246.9485902 | 34.89159002 | 53.14772536 |
| ENSMUSG00000034336 | *Ina* | 15.82189323 | 4.795956706 | 6.302841346 | 317.1960663 | 332.6728821 | 339.7742146 | 3.77E-65 | 5.102046404 | 5.02E-67 | 299.1616566 | 34.34543387 | 64.42386357 |
| ENSMUSG00000113409 | *Gm8288* | 0 | 1.918382682 | 6.302841346 | 69.14654732 | 75.79888453 | 64.28160816 | 4.62E-15 | 4.907683536 | 4.70E-16 | 65.91800749 | 30.01649326 | 14.33515992 |
| ENSMUSG00000066554 | *Gm10167* | 0.753423487 | 0 | 0 | 15.3658994 | 10.52762285 | 4.59154344 | 0.001805716 | 4.864549656 | 0.000671987 | 11.56517131 | 29.13233946 | 2.743350468 |
| ENSMUSG00000042340 | *Ctf1* | 0 | 1.918382682 | 1.260568269 | 26.34154184 | 24.21353256 | 30.3041867 | 7.37E-08 | 4.798351958 | 1.44E-08 | 32.13575047 | 27.82581344 | 7.132750368 |
| ENSMUSG00000066456 | *Hmgn3* | 0.753423487 | 0 | 0 | 15.3658994 | 5.263811426 | 7.346469504 | 0.002816369 | 4.74306659 | 0.001086579 | 10.67389833 | 26.7796758 | 2.550310445 |
| ENSMUSG00000027670 | *Ocstamp* | 20.34243415 | 21.1022095 | 13.86625096 | 456.5867252 | 543.2253392 | 499.5599263 | 5.56E-102 | 4.728417544 | 2.86E-104 | 470.2246887 | 26.5091323 | 101.2546673 |
| ENSMUSG00000002565 | *Scin* | 3.767117435 | 5.755148047 | 0 | 99.87834613 | 80.00993367 | 91.8308688 | 1.45E-21 | 4.724382856 | 9.54E-23 | 96.36690064 | 26.43509958 | 20.83807217 |
| ENSMUSG00000054901 | *Arhgef33* | 0.753423487 | 0 | 0 | 10.97564243 | 5.263811426 | 11.01970426 | 0.002371693 | 4.710850361 | 0.000903138 | 11.01630812 | 26.18829737 | 2.6249416 |
| ENSMUSG00000071757 | *Zhx2* | 5.273964409 | 10.55110475 | 5.042273077 | 172.3175862 | 191.6027359 | 179.9885028 | 7.17E-44 | 4.701701962 | 1.73E-45 | 200.3783718 | 26.02275789 | 43.14452487 |
| ENSMUSG00000030854 | *Ptpn5* | 67.05469034 | 77.69449863 | 51.68329904 | 1668.29765 | 1609.673534 | 1543.676905 | 1.24E-234 | 4.59795519 | 9.38E-238 | 1084.110824 | 24.2171165 | 233.9055442 |
| ENSMUSG00000037370 | *Enpp1* | 0.753423487 | 2.877574023 | 1.260568269 | 40.609877 | 41.05772912 | 30.3041867 | 5.17E-10 | 4.546688083 | 8.14E-11 | 42.2228967 | 23.37165645 | 9.286420943 |
| ENSMUSG00000037868 | *Egr2* | 15.06846974 | 18.22463548 | 0 | 197.5615638 | 301.0900136 | 261.7179761 | 0.000661157 | 4.495373506 | 0.000225861 | 13.60269877 | 22.55497069 | 3.17969526 |
| ENSMUSG00000044165 | *Bcl2l15* | 4.520540922 | 0.959191341 | 1.260568269 | 39.51231275 | 48.42706512 | 66.11822554 | 4.00E-11 | 4.392310116 | 5.68E-12 | 47.4372904 | 20.99989364 | 10.39795438 |
| ENSMUSG00000034557 | *Zfyve9* | 25.61639856 | 19.18382682 | 27.73250192 | 466.4648033 | 507.4314215 | 463.7458874 | 4.21E-99 | 4.316701943 | 2.31E-101 | 456.8651997 | 19.92768122 | 98.37548784 |
| ENSMUSG00000042671 | *Rgs8* | 0 | 0.959191341 | 1.260568269 | 5.487821216 | 15.79143428 | 17.44786507 | 0.001234869 | 4.294094418 | 0.000443543 | 12.33920919 | 19.61784162 | 2.908378947 |
| ENSMUSG00000029309 | *Sparcl1* | 0 | 0 | 2.521136538 | 5.487821216 | 25.26629484 | 10.10139557 | 0.004366875 | 4.287871288 | 0.001756699 | 9.787837755 | 19.53340143 | 2.35982927 |
| ENSMUSG00000024727 | *Trpm6* | 0 | 2.877574023 | 1.260568269 | 18.65859213 | 30.53010627 | 25.71264326 | 4.00E-06 | 4.264854493 | 9.65E-07 | 23.99644622 | 19.22423778 | 5.39769373 |
| ENSMUSG00000090291 | *Lrrc10b* | 1.506846974 | 0 | 1.260568269 | 16.46346365 | 22.10800799 | 15.6112477 | 2.67E-05 | 4.263625953 | 7.29E-06 | 20.11477316 | 19.20787417 | 4.573014081 |
| ENSMUSG00000050953 | *Gja1* | 5.273964409 | 16.3062528 | 17.64795577 | 276.5861893 | 221.0800799 | 206.6194548 | 1.55E-30 | 4.235309868 | 6.50E-32 | 138.2277853 | 18.83455273 | 29.81045905 |
| ENSMUSG00000056313 | *Tcim* | 3.767117435 | 0 | 0 | 28.53667032 | 22.10800799 | 26.63095195 | 1.27E-05 | 4.126151901 | 3.30E-06 | 21.63335329 | 17.46206052 | 4.896295142 |
| ENSMUSG00000074259 | *Gramd2* | 0 | 0.959191341 | 1.260568269 | 12.07320667 | 7.369335996 | 14.69293901 | 0.001567882 | 4.11400383 | 0.000574883 | 11.85559698 | 17.31564028 | 2.804686517 |
| ENSMUSG00000021488 | *Nsd1* | 36.91775086 | 30.69412292 | 20.16909231 | 524.6357082 | 493.7455118 | 506.9063958 | 2.65E-89 | 4.072043143 | 1.98E-91 | 411.2249304 | 16.8192695 | 88.57641579 |
| ENSMUSG00000024140 | *Epas1* | 3.013693948 | 0 | 0 | 25.24397759 | 16.84419656 | 17.44786507 | 0.000108234 | 4.067520037 | 3.23E-05 | 17.27598723 | 16.76662067 | 3.965636464 |
| ENSMUSG00000040478 | *Prdm13* | 7.534234869 | 2.877574023 | 6.302841346 | 88.9027037 | 109.4872777 | 85.40270798 | 3.53E-20 | 4.065490538 | 2.54E-21 | 89.8699497 | 16.74305096 | 19.45266726 |
| ENSMUSG00000036225 | *Kctd1* | 6.780811383 | 4.795956706 | 0 | 63.6587261 | 81.06269596 | 67.95484291 | 7.09E-14 | 4.057421439 | 7.93E-15 | 60.35243748 | 16.64966724 | 13.14936081 |
| ENSMUSG00000070524 | *Fcrlb* | 1.506846974 | 0 | 0 | 13.17077092 | 8.422098282 | 8.264778192 | 0.003637741 | 4.050645343 | 0.001437173 | 10.15747593 | 16.57164992 | 2.439168166 |
| ENSMUSG00000026475 | *Rgs16* | 59.52045547 | 29.73493157 | 23.95079712 | 582.8066131 | 604.2855517 | 617.1034383 | 8.63E-45 | 3.949188903 | 2.01E-46 | 204.6619092 | 15.44629477 | 44.06412459 |
| ENSMUSG00000023140 | *Reg2* | 0 | 1.918382682 | 0 | 5.487821216 | 13.68590971 | 10.10139557 | 0.007623483 | 3.931576657 | 0.003213247 | 8.682246416 | 15.2588746 | 2.117846565 |
| ENSMUSG00000039814 | *Xkr5* | 3.767117435 | 12.46948743 | 3.781704808 | 105.3661673 | 98.95965481 | 98.25902962 | 7.85E-18 | 3.91636563 | 6.57E-19 | 78.88775652 | 15.09883811 | 17.10500486 |
| ENSMUSG00000038463 | *Olfml2b* | 0.753423487 | 1.918382682 | 0 | 10.97564243 | 9.474860567 | 21.12109982 | 0.000887351 | 3.892623308 | 0.000309038 | 13.01480185 | 14.85239113 | 3.051904344 |
| ENSMUSG00000079243 | *Xirp1* | 2.260270461 | 1.918382682 | 1.260568269 | 35.12205578 | 26.31905713 | 21.12109982 | 6.33E-07 | 3.871564015 | 1.38E-07 | 27.74444181 | 14.63716265 | 6.198720633 |
| ENSMUSG00000045620 | *Odf3l1* | 27.87666902 | 17.26544414 | 7.563409615 | 217.3177201 | 293.7206776 | 284.6756933 | 3.06E-27 | 3.849188904 | 1.50E-28 | 122.8576691 | 14.41190262 | 26.51400106 |
| ENSMUSG00000063760 | *Rnf217* | 11.3013523 | 8.63272207 | 3.781704808 | 116.3418098 | 124.2259497 | 116.6252034 | 2.37E-24 | 3.832680644 | 1.35E-25 | 109.370252 | 14.24793213 | 23.62587733 |
| ENSMUSG00000068551 | *Zfp467* | 9.79450533 | 14.38787012 | 13.86625096 | 153.658994 | 176.8640639 | 197.4363679 | 9.20E-34 | 3.823256069 | 3.37E-35 | 153.254012 | 14.15515919 | 33.03626285 |
| ENSMUSG00000034783 | *Cd207* | 5.273964409 | 6.714339388 | 6.302841346 | 90.00026794 | 74.74612225 | 81.72947323 | 1.22E-18 | 3.767044047 | 9.76E-20 | 82.65675929 | 13.61423532 | 17.91217734 |
| ENSMUSG00000074415 | *Mir100hg* | 3.013693948 | 2.877574023 | 0 | 31.82936305 | 27.37181941 | 28.46756933 | 4.57E-07 | 3.765397938 | 9.84E-08 | 28.40605973 | 13.5987104 | 6.33977473 |
| ENSMUSG00000026204 | *Ptprn* | 5.273964409 | 4.795956706 | 5.042273077 | 65.85385459 | 57.90192569 | 72.54638635 | 6.55E-15 | 3.696436532 | 6.75E-16 | 65.20503687 | 12.96397764 | 14.18386767 |
| ENSMUSG00000029828 | *4921507P07Rik* | 0 | 0.959191341 | 1.260568269 | 8.780513945 | 7.369335996 | 9.18308688 | 0.008521831 | 3.674980785 | 0.003634585 | 8.457913568 | 12.77260406 | 2.069467081 |
| ENSMUSG00000042485 | *Mustn1* | 10.54792882 | 9.591913411 | 7.563409615 | 127.3174522 | 86.32650739 | 141.419538 | 1.27E-19 | 3.656561369 | 9.56E-21 | 87.25110516 | 12.61056814 | 18.89512269 |
| ENSMUSG00000002459 | *Rgs20* | 6.027387896 | 2.877574023 | 7.563409615 | 43.90256973 | 87.37926967 | 70.70976898 | 5.42E-11 | 3.640009729 | 7.77E-12 | 46.82388453 | 12.46671735 | 10.26600766 |
| ENSMUSG00000032911 | *Cspg4* | 23.3561281 | 18.22463548 | 15.12681923 | 185.4883571 | 280.0347679 | 224.0673199 | 1.53E-31 | 3.574481604 | 6.14E-33 | 142.914442 | 11.91313825 | 30.81644065 |
| ENSMUSG00000009394 | *Syn2* | 2.260270461 | 4.795956706 | 3.781704808 | 51.58551943 | 31.58286856 | 39.48727358 | 1.82E-08 | 3.534002766 | 3.32E-09 | 34.98387476 | 11.58352759 | 7.74078131 |
| ENSMUSG00000031410 | *Nxf7* | 158.2189323 | 137.1643618 | 157.5710337 | 1724.273426 | 1741.26882 | 1678.668282 | 3.65E-223 | 3.505367858 | 3.51E-226 | 1030.867656 | 11.355882 | 222.4377579 |
| ENSMUSG00000043300 | *B3galnt1* | 19.58901066 | 8.63272207 | 20.16909231 | 188.7810498 | 185.2861622 | 171.7237247 | 7.29E-29 | 3.498839747 | 3.31E-30 | 130.4259959 | 11.30461338 | 28.13733776 |
| ENSMUSG00000028347 | *Tmeff1* | 23.3561281 | 23.02059219 | 36.55647981 | 261.2202899 | 326.3563084 | 309.4700279 | 6.69E-43 | 3.470779566 | 1.70E-44 | 195.8236207 | 11.08686496 | 42.17459633 |
| ENSMUSG00000015854 | *Cd5l* | 39.93144481 | 65.2250112 | 68.07068654 | 631.0994398 | 568.491634 | 636.3879208 | 1.31E-60 | 3.433466082 | 1.85E-62 | 278.2024074 | 10.80379363 | 59.88302092 |
| ENSMUSG00000044433 | *Camsap3* | 0.753423487 | 0.959191341 | 1.260568269 | 10.97564243 | 12.63314742 | 7.346469504 | 0.003980427 | 3.421612701 | 0.001588947 | 9.972509944 | 10.71539183 | 2.400070387 |
| ENSMUSG00000101355 | *H3c10* | 0 | 3.836765364 | 6.302841346 | 55.9757764 | 32.63563084 | 12.85632163 | 0.000955677 | 3.417058113 | 0.000334802 | 12.86490676 | 10.68161668 | 3.019689089 |
| ENSMUSG00000040829 | *Zmynd15* | 27.12324553 | 15.34706146 | 13.86625096 | 204.1469492 | 217.921793 | 179.0701942 | 4.91E-28 | 3.367193083 | 2.29E-29 | 126.5815315 | 10.31872692 | 27.30906906 |
| ENSMUSG00000040148 | *Hmx3* | 15.06846974 | 13.42867878 | 8.823977885 | 115.2442455 | 143.1756708 | 127.6449076 | 4.97E-23 | 3.331780695 | 3.05E-24 | 103.184878 | 10.06852675 | 22.30359489 |
| ENSMUSG00000041609 | *Bicdl1* | 22.60270461 | 42.20441901 | 21.42966058 | 239.269005 | 269.507145 | 303.9601757 | 1.37E-28 | 3.23309613 | 6.28E-30 | 129.1510691 | 9.402837139 | 27.86231389 |
| ENSMUSG00000027315 | *Spint1* | 2.260270461 | 2.877574023 | 0 | 18.65859213 | 21.0552457 | 11.93801294 | 0.000695288 | 3.225648896 | 0.000238284 | 13.50220251 | 9.354424477 | 3.157835205 |
| ENSMUSG00000002997 | *Prkar2b* | 34.6574804 | 44.12280169 | 34.03534327 | 341.3424796 | 389.5220455 | 325.9995842 | 7.63E-52 | 3.225502191 | 1.40E-53 | 237.4658565 | 9.353473292 | 51.11734171 |
| ENSMUSG00000054932 | *Afp* | 256.917409 | 308.8596118 | 434.8960529 | 3150.009378 | 2928.784677 | 3028.582053 | 1.57E-69 | 3.195415117 | 1.89E-71 | 319.4686921 | 9.160428702 | 68.80536161 |
| ENSMUSG00000054304 | *D130007C19Rik* | 17.3287402 | 10.55110475 | 6.302841346 | 114.1466813 | 98.95965481 | 109.2787339 | 3.05E-16 | 3.170816213 | 2.82E-17 | 71.46546448 | 9.005561385 | 15.51632886 |
| ENSMUSG00000041064 | *Pif1* | 192.1229892 | 189.9198855 | 104.6271663 | 1365.369918 | 1491.764158 | 1478.476988 | 7.26E-52 | 3.14121407 | 1.33E-53 | 237.5715704 | 8.822662333 | 51.13876951 |
| ENSMUSG00000023873 | *1700010I14Rik* | 1.506846974 | 0 | 2.521136538 | 14.26833516 | 11.58038514 | 8.264778192 | 0.007021668 | 3.140589821 | 0.002939348 | 8.844739977 | 8.818845625 | 2.153559708 |
| ENSMUSG00000008658 | *Rbfox1* | 22.60270461 | 20.14301816 | 16.3873875 | 154.7565583 | 207.3941702 | 166.2138725 | 2.06E-26 | 3.136909472 | 1.05E-27 | 118.9912405 | 8.79637721 | 25.68548278 |
| ENSMUSG00000038042 | *Ptpdc1* | 12.80819928 | 12.46948743 | 3.781704808 | 86.70757521 | 95.80136795 | 83.56609061 | 8.36E-14 | 3.128485244 | 9.42E-15 | 60.01323977 | 8.745162812 | 13.07796871 |
| ENSMUSG00000044927 | *H1f10* | 76.09577218 | 53.7147151 | 74.37352789 | 541.0991719 | 600.0745026 | 575.7795474 | 2.96E-68 | 3.07127645 | 3.64E-70 | 313.572226 | 8.405166801 | 67.52828044 |
| ENSMUSG00000027848 | *Olfml3* | 24.86297507 | 41.24522767 | 28.99307019 | 185.4883571 | 335.831169 | 254.3715066 | 3.43E-20 | 3.032953849 | 2.47E-21 | 89.92905635 | 8.184837923 | 19.46481463 |
| ENSMUSG00000044912 | *Syt16* | 1.506846974 | 1.918382682 | 7.563409615 | 39.51231275 | 24.21353256 | 19.28448245 | 0.000282486 | 3.025586371 | 9.01E-05 | 15.33350683 | 8.143146576 | 3.549002392 |
| ENSMUSG00000032340 | *Neo1* | 2.260270461 | 0.959191341 | 5.042273077 | 19.75615638 | 28.4245817 | 14.69293901 | 0.000514169 | 3.001579476 | 0.000171801 | 14.11681288 | 8.008763272 | 3.288894103 |
| ENSMUSG00000032595 | *Cdhr4* | 3.013693948 | 0 | 3.781704808 | 25.24397759 | 16.84419656 | 10.10139557 | 0.003817949 | 2.962191029 | 0.001518848 | 10.05560912 | 7.793065968 | 2.418169849 |
| ENSMUSG00000042821 | *Snai1* | 0 | 10.55110475 | 3.781704808 | 31.82936305 | 37.89944227 | 39.48727358 | 0.001192526 | 2.961163866 | 0.000426615 | 12.41186182 | 7.787519475 | 2.923532115 |
| ENSMUSG00000034416 | *Pkd1l2* | 414.3829178 | 404.778746 | 289.9307019 | 3089.643345 | 2652.960959 | 2891.754059 | 4.02E-87 | 2.953182439 | 3.29E-89 | 401.0306851 | 7.744555508 | 86.39545962 |
| ENSMUSG00000053846 | *Lipg* | 11.3013523 | 3.836765364 | 2.521136538 | 60.36603337 | 26.31905713 | 56.01682997 | 2.41E-05 | 2.940149861 | 6.52E-06 | 20.32823911 | 7.674910152 | 4.618735699 |
| ENSMUSG00000074671 | *Tspyl3* | 0.753423487 | 4.795956706 | 1.260568269 | 10.97564243 | 17.89695885 | 22.03940851 | 0.003085322 | 2.917644439 | 0.001201141 | 10.48858775 | 7.556113848 | 2.510699458 |
| ENSMUSG00000089996 | *Tmsb15b2* | 0 | 3.836765364 | 2.521136538 | 20.85372062 | 13.68590971 | 11.01970426 | 0.007534451 | 2.914386204 | 0.003171067 | 8.706331449 | 7.539068099 | 2.122948359 |
| ENSMUSG00000003452 | *Bicd1* | 21.09585763 | 24.93897487 | 12.60568269 | 170.1224577 | 128.4369988 | 147.8476988 | 1.80E-19 | 2.898987553 | 1.37E-20 | 86.53817299 | 7.459027537 | 18.74448288 |
| ENSMUSG00000015243 | *Abca1* | 51.23279711 | 50.83714108 | 36.55647981 | 354.5132505 | 335.831169 | 345.2840667 | 2.94E-48 | 2.879051536 | 6.05E-50 | 220.8001101 | 7.356663159 | 47.5317415 |
| ENSMUSG00000020712 | *Tcam1* | 119.0409109 | 121.8173003 | 118.4934173 | 875.856266 | 863.2650739 | 835.6609061 | 2.45E-115 | 2.840291049 | 1.01E-117 | 532.0536241 | 7.161645209 | 114.6104587 |
| ENSMUSG00000021250 | *Fos* | 485.9581491 | 533.3103857 | 312.6209308 | 3002.935769 | 3387.789034 | 3147.962182 | 2.08E-53 | 2.835087698 | 3.59E-55 | 244.765752 | 7.135861917 | 52.68228094 |
| ENSMUSG00000050010 | *Shisa3* | 7.534234869 | 14.38787012 | 5.042273077 | 64.75629035 | 64.2184994 | 66.11822554 | 8.45E-10 | 2.83437818 | 1.36E-10 | 41.218547 | 7.132353356 | 9.073241861 |
| ENSMUSG00000026589 | *Sec16b* | 3.767117435 | 7.673530729 | 0 | 36.21962002 | 23.16077027 | 24.79433458 | 0.000488954 | 2.816292705 | 0.000162403 | 14.22265872 | 7.043501004 | 3.310731693 |
| ENSMUSG00000078653 | *Cntd1* | 12.05477579 | 2.877574023 | 3.781704808 | 46.09769821 | 48.42706512 | 44.07881702 | 4.25E-06 | 2.813024815 | 1.03E-06 | 23.87091165 | 7.027564622 | 5.371593982 |
| ENSMUSG00000039911 | *Spsb1* | 29.38351599 | 21.1022095 | 18.90852404 | 164.6346365 | 163.1781542 | 161.6223291 | 7.86E-24 | 2.786881693 | 4.62E-25 | 106.926666 | 6.901364794 | 23.10435882 |
| ENSMUSG00000015568 | *Lpl* | 6285.812152 | 5539.329995 | 4572.081113 | 36407.30351 | 38021.56269 | 36794.79251 | 3.84E-112 | 2.761192931 | 1.69E-114 | 517.2396773 | 6.779566046 | 111.4155711 |
| ENSMUSG00000043648 | *Pld6* | 9.041081843 | 12.46948743 | 10.08454615 | 62.56116186 | 61.06021254 | 86.32101667 | 1.51E-10 | 2.741449592 | 2.25E-11 | 44.73996847 | 6.687419371 | 9.821725087 |
| ENSMUSG00000023830 | *Igf2r* | 846.8479993 | 877.6600771 | 589.94595 | 4708.550603 | 5637.542037 | 5153.548357 | 5.98E-66 | 2.740114661 | 7.71E-68 | 302.8943186 | 6.681234339 | 65.22343193 |
| ENSMUSG00000022421 | *Nptxr* | 18.08216369 | 25.89816621 | 10.08454615 | 122.9271952 | 126.3314742 | 113.8702773 | 7.69E-15 | 2.718311068 | 7.96E-16 | 64.88119638 | 6.58101937 | 14.1142278 |
| ENSMUSG00000110195 | *Pde2a* | 393.2870602 | 559.2085519 | 255.8953587 | 2629.763927 | 2031.83121 | 3255.404299 | 2.79E-11 | 2.710660274 | 3.91E-12 | 48.16799902 | 6.546211767 | 10.55368854 |
| ENSMUSG00000016529 | *Il10* | 33.15063343 | 28.77574023 | 35.29591154 | 221.7079771 | 198.9720719 | 210.2926896 | 5.59E-31 | 2.703079263 | 2.30E-32 | 140.2929515 | 6.511903242 | 30.25275519 |
| ENSMUSG00000049134 | *Nrap* | 0.753423487 | 3.836765364 | 1.260568269 | 10.97564243 | 14.73867199 | 11.93801294 | 0.009063073 | 2.701770627 | 0.003899634 | 8.32996819 | 6.505999117 | 2.042724523 |
| ENSMUSG00000031520 | *Vegfc* | 6.780811383 | 14.38787012 | 5.042273077 | 58.17090489 | 51.58535197 | 61.5266821 | 5.92E-08 | 2.693748344 | 1.15E-08 | 32.57539995 | 6.469922111 | 7.22748395 |
| ENSMUSG00000003526 | *Prodh* | 7.534234869 | 8.63272207 | 1.260568269 | 26.34154184 | 29.47734399 | 60.60837341 | 0.000205246 | 2.6881887 | 6.42E-05 | 15.9745691 | 6.445037266 | 3.68772613 |
| ENSMUSG00000102543 | *Pcdhgc5* | 21.09585763 | 13.42867878 | 8.823977885 | 41.70744124 | 160.0198673 | 78.97454717 | 1.19E-05 | 2.667673069 | 3.08E-06 | 21.76840192 | 6.354035143 | 4.92480054 |
| ENSMUSG00000038235 | *F11r* | 40.6848683 | 36.44927096 | 47.90159423 | 243.659262 | 261.0850467 | 279.1658412 | 9.80E-35 | 2.661441678 | 3.48E-36 | 157.7646386 | 6.326649527 | 34.00874184 |
| ENSMUSG00000031137 | *Fgf13* | 50.47937363 | 57.55148047 | 32.774775 | 327.0741445 | 323.1980216 | 250.6982718 | 7.66E-26 | 2.657074225 | 4.02E-27 | 116.3337526 | 6.307525897 | 25.11569656 |
| ENSMUSG00000046295 | *Ankle1* | 132.6025337 | 104.5518562 | 40.33818462 | 518.0503228 | 689.5592968 | 559.249991 | 5.61E-14 | 2.654300792 | 6.19E-15 | 60.83919083 | 6.295411975 | 13.25138721 |
| ENSMUSG00000003032 | *Klf4* | 89.65739495 | 95.91913411 | 64.28898173 | 488.4160882 | 582.1775437 | 516.0894827 | 9.77E-46 | 2.650974328 | 2.20E-47 | 209.0637194 | 6.280913183 | 45.01021687 |
| ENSMUSG00000039007 | *Cpq* | 14.31504625 | 7.673530729 | 16.3873875 | 68.04898308 | 103.1707039 | 66.11822554 | 3.03E-09 | 2.638776069 | 5.16E-10 | 38.61466746 | 6.228030755 | 8.518734759 |
| ENSMUSG00000024039 | *Cbs* | 22.60270461 | 9.591913411 | 10.08454615 | 104.2686031 | 88.43203196 | 78.97454717 | 1.03E-09 | 2.635752495 | 1.68E-10 | 40.81025645 | 6.214991831 | 8.98638244 |
| ENSMUSG00000074652 | *Myh7b* | 41.43829178 | 40.28603633 | 27.73250192 | 210.7323347 | 260.0322844 | 220.3940851 | 1.29E-26 | 2.635405164 | 6.51E-28 | 119.9422168 | 6.213495745 | 25.88839264 |
| ENSMUSG00000004902 | *Slc25a18* | 2.260270461 | 0.959191341 | 7.563409615 | 18.65859213 | 23.16077027 | 20.20279114 | 0.002278161 | 2.623136285 | 0.000864394 | 11.09760161 | 6.160879359 | 2.642415502 |
| ENSMUSG00000042607 | *Asb4* | 5.273964409 | 0.959191341 | 2.521136538 | 23.04884911 | 14.73867199 | 18.36617376 | 0.002323749 | 2.62142268 | 0.000882808 | 11.05851611 | 6.153565929 | 2.633810812 |
| ENSMUSG00000063275 | *Hacd1* | 45.20540922 | 28.77574023 | 81.9369375 | 334.7570942 | 302.1427758 | 301.2052497 | 5.96E-15 | 2.61214859 | 6.13E-16 | 65.39639419 | 6.114135773 | 14.22459044 |
| ENSMUSG00000060044 | *Tmem26* | 391.0267897 | 426.8401468 | 298.7546798 | 2051.34757 | 2464.51651 | 2274.65062 | 1.00E-67 | 2.59881857 | 1.26E-69 | 311.0934841 | 6.057903387 | 66.99791406 |
| ENSMUSG00000029648 | *Flt1* | 33.90405691 | 34.53088828 | 23.95079712 | 188.7810498 | 174.7585393 | 194.6814419 | 1.28E-24 | 2.573706469 | 7.15E-26 | 110.6260578 | 5.953369625 | 23.89361112 |
| ENSMUSG00000102918 | *Pcdhgc3* | 613.2867184 | 648.4133466 | 514.3118539 | 3345.375813 | 3658.348941 | 3566.710944 | 2.46E-123 | 2.569031494 | 9.10E-126 | 569.0351959 | 5.934109274 | 122.6096868 |
| ENSMUSG00000003849 | *Nqo1* | 35.41090389 | 24.93897487 | 46.64102596 | 207.439642 | 206.3414079 | 209.3743809 | 7.29E-23 | 2.562151799 | 4.53E-24 | 102.401736 | 5.905879002 | 22.13721928 |
| ENSMUSG00000022010 | *Tsc22d1* | 140.8901921 | 157.3073799 | 138.6625096 | 819.8804896 | 865.3705984 | 879.7397231 | 1.69E-91 | 2.55287414 | 1.18E-93 | 421.4467899 | 5.868021432 | 90.77254439 |
| ENSMUSG00000029581 | *Fscn1* | 2.260270461 | 3.836765364 | 2.521136538 | 16.46346365 | 12.63314742 | 21.12109982 | 0.002307258 | 2.551763304 | 0.000876385 | 11.07205605 | 5.863504955 | 2.63690377 |
| ENSMUSG00000000942 | *Hoxa4* | 21.84928112 | 27.81654889 | 31.51420673 | 131.7077092 | 149.4922445 | 185.498355 | 2.56E-17 | 2.546663683 | 2.21E-18 | 76.49637882 | 5.842815293 | 16.5911391 |
| ENSMUSG00000020889 | *Nr1d1* | 73.08207823 | 50.83714108 | 37.81704808 | 270.0008038 | 346.3587918 | 344.365758 | 1.46E-21 | 2.544976374 | 9.64E-23 | 96.34726237 | 5.835985806 | 20.83421749 |
| ENSMUSG00000026435 | *Slc45a3* | 82.12316008 | 122.7764917 | 78.15523269 | 540.0016076 | 532.6977163 | 547.3119781 | 2.15E-38 | 2.513802288 | 6.47E-40 | 174.8462351 | 5.711233179 | 37.6681603 |
| ENSMUSG00000018920 | *Cxcl16* | 844.5877289 | 733.781376 | 704.6576625 | 4150.987968 | 4591.096326 | 4292.174808 | 1.10E-135 | 2.509375551 | 3.31E-138 | 626.2224972 | 5.693735799 | 134.959536 |
| ENSMUSG00000039693 | *Msantd3* | 112.2600996 | 96.87832545 | 138.6625096 | 658.5385459 | 622.1825105 | 669.4470336 | 5.93E-55 | 2.499585855 | 9.64E-57 | 251.9742859 | 5.655230608 | 54.227195 |
| ENSMUSG00000027186 | *Elf5* | 3.013693948 | 4.795956706 | 8.823977885 | 28.53667032 | 38.95220455 | 21.12109982 | 0.000324781 | 2.477011068 | 0.000104419 | 15.05507162 | 5.567428279 | 3.488409646 |
| ENSMUSG00000045326 | *Fndc7* | 7.534234869 | 2.877574023 | 2.521136538 | 26.34154184 | 26.31905713 | 22.9577172 | 0.000453602 | 2.473549912 | 0.000149852 | 14.37406992 | 5.554087524 | 3.343325219 |
| ENSMUSG00000073433 | *Arhgdig* | 34.6574804 | 27.81654889 | 22.69022885 | 170.1224577 | 162.1253919 | 148.7660075 | 8.31E-20 | 2.470067649 | 6.16E-21 | 88.11972415 | 5.540697674 | 19.08025196 |
| ENSMUSG00000074026 | *Gm6091* | 54.99991455 | 70.0209679 | 61.76784519 | 330.3668372 | 346.3587918 | 346.2023754 | 1.46E-40 | 2.459989854 | 4.05E-42 | 184.9356575 | 5.502128579 | 39.83501767 |
| ENSMUSG00000043079 | *Synpo* | 55.75333803 | 59.46986315 | 41.59875288 | 261.2202899 | 277.9292433 | 293.8587802 | 3.54E-30 | 2.393356581 | 1.52E-31 | 136.5430554 | 5.253782883 | 29.45043233 |
| ENSMUSG00000028565 | *Nfia* | 8.287658356 | 4.795956706 | 7.563409615 | 29.63423457 | 43.16325369 | 35.81403883 | 1.85E-05 | 2.389771005 | 4.93E-06 | 20.86335199 | 5.240741702 | 4.73191668 |
| ENSMUSG00000070305 | *Mpzl3* | 9.041081843 | 23.02059219 | 20.16909231 | 70.24411156 | 108.4345154 | 87.23932536 | 8.27E-08 | 2.377914821 | 1.62E-08 | 31.89784974 | 5.197849353 | 7.082608668 |
| ENSMUSG00000110003 | *Gm45403* | 13.56162277 | 0.959191341 | 6.302841346 | 31.82936305 | 36.84667998 | 42.24219965 | 0.001860639 | 2.374374182 | 0.000694228 | 11.50463588 | 5.185108514 | 2.730337869 |
| ENSMUSG00000079710 | *Dynlt2a2* | 3.767117435 | 1.918382682 | 7.563409615 | 16.46346365 | 16.84419656 | 31.22249539 | 0.003817147 | 2.342803704 | 0.001518005 | 10.05663191 | 5.072875324 | 2.4182611 |
| ENSMUSG00000022346 | *Myc* | 133.3559572 | 134.2867878 | 80.67636923 | 491.7087809 | 655.8709037 | 627.2048339 | 3.55E-25 | 2.336143772 | 1.94E-26 | 113.2152653 | 5.049511312 | 24.44945104 |
| ENSMUSG00000036862 | *Dchs1* | 27.12324553 | 22.06140085 | 15.12681923 | 88.9027037 | 132.6480479 | 104.6871904 | 9.40E-11 | 2.313026387 | 1.38E-11 | 45.69691683 | 4.969244015 | 10.02709295 |
| ENSMUSG00000110632 | *Gm45880* | 8.287658356 | 8.63272207 | 12.60568269 | 42.80500548 | 44.21601598 | 55.09852128 | 1.30E-06 | 2.299252047 | 2.95E-07 | 26.28331104 | 4.92202521 | 5.88753048 |
| ENSMUSG00000061013 | *Mkx* | 28.6300925 | 41.24522767 | 36.55647981 | 189.8786141 | 160.0198673 | 168.0504899 | 1.29E-18 | 2.294995172 | 1.03E-19 | 82.55137725 | 4.907523485 | 17.88977008 |
| ENSMUSG00000063873 | *Slc24a3* | 28.6300925 | 41.24522767 | 60.50727692 | 222.8055414 | 197.9193096 | 197.4363679 | 1.19E-14 | 2.27711496 | 1.25E-15 | 63.98721898 | 4.847076861 | 13.92431808 |
| ENSMUSG00000015468 | *Notch4* | 6.027387896 | 6.714339388 | 3.781704808 | 26.34154184 | 32.63563084 | 20.20279114 | 0.000543925 | 2.23072813 | 0.00018264 | 14.00176017 | 4.693708118 | 3.264461156 |
| ENSMUSG00000029762 | *Akr1b8* | 1643.970049 | 1563.481886 | 1948.838544 | 7778.437791 | 8587.38196 | 7723.894375 | 3.52E-95 | 2.225745848 | 2.24E-97 | 438.5457702 | 4.677526568 | 94.45401995 |
| ENSMUSG00000063455 | *D630045J12Rik* | 51.9862206 | 25.89816621 | 27.73250192 | 122.9271952 | 210.552457 | 168.0504899 | 2.46E-09 | 2.225636845 | 4.15E-10 | 39.04296586 | 4.67717317 | 8.609630824 |
| ENSMUSG00000079481 | *Nhsl2* | 26.36982204 | 28.77574023 | 11.34511442 | 97.68321764 | 91.59031881 | 125.8082903 | 1.18E-08 | 2.213689194 | 2.12E-09 | 35.86239188 | 4.638599198 | 7.929937949 |
| ENSMUSG00000053080 | *Zfta* | 46.71225619 | 62.34743717 | 26.47193365 | 214.0250274 | 215.8162685 | 207.5377635 | 4.34E-14 | 2.213535537 | 4.75E-15 | 61.36295174 | 4.638105179 | 13.36204258 |
| ENSMUSG00000027784 | *Ppm1l* | 515.3416651 | 488.2283926 | 383.2127538 | 1935.005761 | 2335.026749 | 2173.636665 | 3.14E-56 | 2.210626064 | 4.89E-58 | 257.9116189 | 4.628760971 | 55.5027664 |

| | | | | | | | | | | | | | |
|---|---|---|---|---|---|---|---|---|---|---|---|---|---|
| ENSMUSG00000025574 | *Tk1* | 1461.641565 | 1279.561249 | 1167.286217 | 5830.26126 | 6026.01132 | 5759.632091 | 5.34E-103 | 2.169184514 | 2.64E-105 | 474.9847752 | 4.497690893 | 102.2727561 |
| ENSMUSG00000045349 | *Sh2d5* | 42.19171527 | 57.55148047 | 40.33818462 | 183.2932286 | 230.5549405 | 215.8025417 | 1.54E-18 | 2.165877818 | 1.24E-19 | 82.18172402 | 4.487393867 | 17.81226382 |
| ENSMUSG00000046668 | *Cxxc5* | 45.9588327 | 56.59228913 | 49.1621625 | 193.1713068 | 234.7659896 | 245.1884197 | 5.73E-21 | 2.153952494 | 3.92E-22 | 93.5695253 | 4.450453932 | 20.24212627 |
| ENSMUSG00000068699 | *Flnc* | 345.8213805 | 401.9011719 | 301.2758163 | 1423.540823 | 1706.527664 | 1539.085361 | 3.67E-54 | 2.150514142 | 6.07E-56 | 248.3097107 | 4.439859867 | 53.435555 |
| ENSMUSG00000058290 | *Espl1* | 777.5330385 | 575.5148047 | 524.3964 | 2569.397893 | 2963.525833 | 2805.433042 | 1.54E-42 | 2.146478164 | 3.97E-44 | 194.1383432 | 4.427456597 | 41.81144223 |
| ENSMUSG00000033209 | *Ttc28* | 20.34243415 | 7.673530729 | 16.3873875 | 70.24411156 | 73.69335996 | 53.2619039 | 3.94E-06 | 2.137737935 | 9.50E-07 | 24.02630519 | 4.400714977 | 5.404181445 |
| ENSMUSG00000070526 | *Peg12* | 104.7258647 | 102.6334735 | 47.90159423 | 346.8303008 | 367.4140375 | 422.4219965 | 3.20E-14 | 2.137621411 | 3.47E-15 | 61.97870098 | 4.400359552 | 13.49456775 |
| ENSMUSG00000039741 | *Bahcc1* | 287.807772 | 244.593792 | 170.1767163 | 908.7831933 | 1113.822498 | 1038.607126 | 3.26E-27 | 2.113634377 | 1.60E-28 | 122.7266853 | 4.327801642 | 26.48715378 |
| ENSMUSG00000032254 | *Kif23* | 885.2725972 | 825.8637447 | 574.8191308 | 3344.278249 | 3284.61833 | 3282.95356 | 8.37E-42 | 2.11221282 | 2.23E-43 | 190.7069044 | 4.323539351 | 41.07728201 |
| ENSMUSG00000034686 | *Prr7* | 31.64378645 | 33.57169694 | 30.25363846 | 132.8052734 | 132.6480479 | 146.9293901 | 6.23E-16 | 2.108955821 | 5.89E-17 | 70.01242314 | 4.313789626 | 15.20569292 |
| ENSMUSG00000025268 | *Maged2* | 80.6163131 | 79.61288131 | 137.4019413 | 423.6597979 | 410.5772912 | 427.9318486 | 1.22E-19 | 2.10353923 | 9.13E-21 | 87.34104356 | 4.297623892 | 18.91367886 |
| ENSMUSG00000070583 | *Fv1* | 7.534234869 | 8.63272207 | 6.302841346 | 31.82936305 | 30.53010627 | 34.89573014 | 0.000100552 | 2.103525308 | 2.99E-05 | 17.42596944 | 4.29758242 | 3.997608819 |
| ENSMUSG00000021697 | *Depdc1b* | 94.17793587 | 84.40883802 | 73.11295962 | 329.269273 | 372.677849 | 370.0784013 | 8.65E-31 | 2.077859744 | 3.58E-32 | 139.4090901 | 4.221804414 | 30.06308381 |
| ENSMUSG00000058317 | *Ube2e2* | 97.9450533 | 94.00075143 | 136.1413731 | 463.1721106 | 456.8988318 | 445.3797137 | 1.29E-30 | 2.0731167 | 5.42E-32 | 138.5886705 | 4.207947484 | 29.88796099 |
| ENSMUSG00000003779 | *Kif20a* | 2056.092696 | 1786.973469 | 1352.589753 | 6844.41062 | 7675.689821 | 7267.494957 | 5.17E-43 | 2.066186282 | 1.30E-44 | 196.3565138 | 4.187781823 | 42.28618899 |
| ENSMUSG00000045934 | *Mtmr11* | 64.04099639 | 55.63309779 | 23.95079712 | 231.5860553 | 193.7082605 | 185.498355 | 1.48E-09 | 2.061809033 | 2.45E-10 | 40.07284612 | 4.175095024 | 8.828840883 |
| ENSMUSG00000028755 | *Cda* | 70.06838429 | 98.79670814 | 112.190576 | 425.8549263 | 341.0949804 | 389.3628837 | 7.07E-21 | 2.056788676 | 4.85E-22 | 93.14766429 | 4.160591583 | 20.1508626 |
| ENSMUSG00000009876 | *Cox4i2* | 6.027387896 | 7.673530729 | 12.60568269 | 36.21962002 | 27.37181941 | 42.24219965 | 0.000300486 | 2.055260322 | 9.62E-05 | 15.21074427 | 4.156186294 | 3.522175354 |
| ENSMUSG00000067199 | *Frat1* | 9.041081843 | 3.836765364 | 3.781704808 | 20.85372062 | 22.10800799 | 28.46756933 | 0.003080141 | 2.049969108 | 0.001198701 | 10.49234494 | 4.140971025 | 2.511429467 |
| ENSMUSG00000049811 | *Fam161a* | 39.17802132 | 38.36765364 | 61.76784519 | 189.8786141 | 191.6027359 | 183.6617376 | 1.88E-15 | 2.045638374 | 1.85E-16 | 67.75207391 | 4.128559148 | 14.72521904 |
| ENSMUSG00000006310 | *Zbtb32* | 1223.559743 | 1106.906808 | 837.0173308 | 4464.891341 | 4229.998862 | 4351.864872 | 4.70E-48 | 2.038991605 | 9.80E-50 | 219.8397488 | 4.109581844 | 47.32801589 |
| ENSMUSG00000053580 | *Tanc2* | 80.6163131 | 85.36802936 | 61.76784519 | 324.879016 | 351.6226033 | 263.5545935 | 9.48E-21 | 2.032710627 | 6.55E-22 | 92.5543694 | 4.091729086 | 20.02326651 |
| ENSMUSG00000051682 | *Treml4* | 96.43820633 | 125.6540657 | 148.7470558 | 443.4159542 | 570.5971586 | 481.1937525 | 6.75E-23 | 2.025101096 | 4.18E-24 | 102.5627418 | 4.070203976 | 22.1705944 |
| ENSMUSG00000049932 | *H2ax* | 1637.942661 | 1465.644369 | 1109.300077 | 5138.795787 | 6124.970975 | 5839.524947 | 7.20E-41 | 2.019219891 | 1.97E-42 | 186.3743081 | 4.053645398 | 40.14258342 |
| ENSMUSG00000034391 | *Fbxo15* | 31.64378645 | 33.57169694 | 20.16909231 | 124.0247595 | 105.2762285 | 122.1350555 | 3.88E-11 | 2.018534388 | 5.50E-12 | 47.50167209 | 4.051719747 | 10.41116713 |
| ENSMUSG00000023232 | *Serinc2* | 317.191288 | 297.3493157 | 220.5994471 | 1038.295774 | 1251.734357 | 1107.480278 | 6.56E-36 | 2.01606252 | 2.18E-37 | 163.2704571 | 4.044783601 | 35.18308603 |
| ENSMUSG00000020886 | *Dlg4* | 138.6299216 | 149.6338492 | 143.7047827 | 523.538144 | 602.1800271 | 618.021747 | 2.14E-43 | 2.015623662 | 5.32E-45 | 198.1377729 | 4.043553394 | 42.66896858 |
| ENSMUSG00000020493 | *Prr11* | 486.7115726 | 419.1666161 | 302.5363846 | 1603.541359 | 1697.052804 | 1572.144474 | 1.28E-32 | 2.004411148 | 5.01E-34 | 147.89301 | 4.012249016 | 31.89207442 |
| ENSMUSG00000030688 | *Stard10* | 77.60261916 | 76.73530729 | 71.85239135 | 276.5861893 | 308.4593496 | 322.3263495 | 7.49E-28 | 2.000852312 | 3.53E-29 | 125.7236131 | 4.002363809 | 27.12576157 |
| ENSMUSG00000029392 | *Rilpl1* | 106.9861351 | 83.44964668 | 100.8454615 | 346.8303008 | 440.0546352 | 380.1797968 | 4.90E-26 | 2.000308242 | 2.54E-27 | 117.2411964 | 4.000854719 | 25.30983344 |
| ENSMUSG00000018476 | *Kdm6b* | 225.2736226 | 211.9812864 | 115.9722808 | 694.7581659 | 751.6722716 | 774.134224 | 7.32E-18 | 1.994164964 | 6.09E-19 | 79.03963784 | 3.983854518 | 17.13553596 |
| ENSMUSG00000028068 | *Iqgap3* | 1417.189579 | 1239.275213 | 665.5800462 | 4057.695007 | 4814.28193 | 4327.070538 | 1.92E-08 | 1.989284877 | 3.52E-09 | 34.8702839 | 3.970401424 | 7.716745549 |
| ENSMUSG00000039883 | *Lrrc17* | 3.767117435 | 4.795956706 | 8.823977885 | 23.04884911 | 21.0552457 | 22.03940851 | 0.003744921 | 1.983142983 | 0.001485596 | 10.09639592 | 3.953534416 | 2.426557364 |
| ENSMUSG00000013419 | *Zfp651* | 819.7247538 | 798.0471958 | 629.0235664 | 2889.886652 | 3061.432725 | 2933.077949 | 7.03E-64 | 1.97895499 | 9.51E-66 | 293.2969859 | 3.942074358 | 63.1527604 |
| ENSMUSG00000049097 | *Ankrd34a* | 119.7943344 | 134.2867878 | 74.37352789 | 417.0744124 | 456.8988318 | 430.6867747 | 1.53E-19 | 1.976005937 | 1.16E-20 | 86.87118327 | 3.934024486 | 18.81486099 |
| ENSMUSG00000080999 | *H2ac5-ps* | 10.54792882 | 9.591913411 | 6.302841346 | 19.75615638 | 26.31905713 | 58.77175603 | 0.003538595 | 1.972905662 | 0.001394787 | 10.21267555 | 3.925579554 | 2.451169106 |
| ENSMUSG00000002489 | *Tiam1* | 131.0956867 | 147.7154665 | 129.8385317 | 557.5626355 | 508.4841837 | 535.3739651 | 1.19E-43 | 1.969447048 | 2.91E-45 | 199.3418952 | 3.916179921 | 42.92326293 |
| ENSMUSG00000026605 | *Cenpf* | 326.2323698 | 196.6342249 | 93.28205192 | 779.2706126 | 837.998779 | 800.7651759 | 0.000312697 | 1.968669586 | 0.000100253 | 15.13193722 | 3.914070076 | 3.504876791 |
| ENSMUSG00000026069 | *Il1rl1* | 397.8076011 | 386.5541105 | 374.388776 | 1475.126343 | 1569.668567 | 1496.843161 | 8.50E-99 | 1.967908101 | 4.73E-101 | 455.4389444 | 3.912004693 | 98.07049947 |
| ENSMUSG00000060594 | *Layn* | 78.35604264 | 111.2661956 | 102.1060298 | 407.1963342 | 343.200505 | 382.9347229 | 6.53E-24 | 1.967629606 | 3.82E-25 | 107.3029079 | 3.9112496 | 23.18477365 |
| ENSMUSG00000026285 | *Pdcd1* | 734.5878998 | 696.3729137 | 591.2065183 | 2775.739971 | 2523.471198 | 2633.709317 | 2.69E-74 | 1.96720541 | 3.01E-76 | 341.4953243 | 3.910099743 | 73.5694508 |
| ENSMUSG00000031434 | *Morc4* | 100.2053238 | 97.83751679 | 54.20443558 | 309.5131166 | 335.831169 | 347.1206841 | 6.50E-16 | 1.957843093 | 6.16E-17 | 69.92661903 | 3.884807448 | 15.18711644 |
| ENSMUSG00000037224 | *Zfyve28* | 8.287658356 | 13.42867878 | 11.34511442 | 34.02449154 | 52.63811426 | 40.40558227 | 9.69E-05 | 1.957753385 | 2.87E-05 | 17.50381815 | 3.884565896 | 4.013785226 |
| ENSMUSG00000089876 | *Tmem102* | 27.12324553 | 27.81654889 | 31.51420673 | 129.5125807 | 101.0651794 | 102.8505731 | 6.74E-11 | 1.954780157 | 9.73E-12 | 46.3828134 | 3.876568497 | 10.17136387 |
| ENSMUSG00000024883 | *Rin1* | 187.6024482 | 187.0423115 | 123.5356904 | 624.5140544 | 654.8181414 | 666.6921075 | 2.60E-28 | 1.953447146 | 1.20E-29 | 127.8637074 | 3.872988307 | 27.58451333 |
| ENSMUSG00000030708 | *Dnajb13* | 10.54792882 | 16.3062528 | 8.823977885 | 57.07334064 | 47.37430283 | 33.05911277 | 0.000145729 | 1.933328687 | 4.45E-05 | 16.67107964 | 3.819354114 | 3.836454086 |
| ENSMUSG00000075611 | *Gm11545* | 36.16432737 | 49.87794974 | 31.51420673 | 140.4882231 | 155.8088182 | 151.5209335 | 9.36E-13 | 1.922499019 | 1.15E-13 | 55.09269747 | 3.790791267 | 12.0287218 |
| ENSMUSG00000078503 | *Zfp990* | 49.72595014 | 57.55148047 | 64.28898173 | 229.3909268 | 209.4996948 | 200.191294 | 1.55E-18 | 1.910367307 | 1.25E-19 | 82.17185997 | 3.75904792 | 17.81083563 |
| ENSMUSG00000028678 | *Kif2c* | 1113.559914 | 915.0685394 | 823.1510798 | 3492.449422 | 3673.087613 | 3561.201092 | 1.21E-54 | 1.907577995 | 1.98E-56 | 250.5431561 | 3.751787184 | 53.91885149 |
| ENSMUSG00000030867 | *Plk1* | 1732.120596 | 1437.82782 | 889.9611981 | 4585.623408 | 5494.366366 | 5075.492119 | 4.51E-19 | 1.898376009 | 3.50E-20 | 84.68357872 | 3.727933202 | 18.34607746 |
| ENSMUSG00000032501 | *Trib1* | 228.2873165 | 255.1448967 | 160.0921702 | 805.6121545 | 829.5766807 | 778.7257674 | 1.63E-27 | 1.898342954 | 7.80E-29 | 124.1519002 | 3.727847787 | 26.78866752 |
| ENSMUSG00000069755 | *Zfp125* | 6.027387896 | 15.34706146 | 8.823977885 | 31.82936305 | 37.89944227 | 41.32389096 | 0.000719523 | 1.893099412 | 0.000247034 | 13.43453866 | 3.714323343 | 3.14295509 |
| ENSMUSG00000024013 | *Fgd2* | 416.6431883 | 408.6155113 | 322.7054769 | 1345.613762 | 1427.545659 | 1495.006544 | 5.14E-50 | 1.888044804 | 1.00E-51 | 228.9655625 | 3.701332659 | 49.28886086 |
| ENSMUSG00000039193 | *Nlrc4* | 27.87666902 | 38.36765364 | 37.81704808 | 150.3663013 | 124.2259497 | 106.5238078 | 4.98E-10 | 1.884401642 | 7.82E-11 | 42.30159178 | 3.691997668 | 9.302379984 |
| ENSMUSG00000036957 | *Lrfn3* | 37.67117435 | 43.16361035 | 20.16909231 | 130.6101449 | 136.8590971 | 111.1153512 | 1.05E-08 | 1.881234966 | 1.88E-09 | 36.09603495 | 3.683902724 | 7.97869343 |
| ENSMUSG00000032187 | *Smarca4* | 3850.747442 | 3730.295126 | 2790.898148 | 11535.4002 | 13605.89977 | 12734.18658 | 1.54E-42 | 1.867673481 | 3.95E-44 | 194.1513212 | 3.6494359 | 41.81311507 |
| ENSMUSG00000036777 | *Anln* | 1671.846718 | 1483.869005 | 1360.153163 | 5311.113373 | 5662.808332 | 5508.015511 | 2.66E-82 | 1.865307635 | 2.39E-84 | 378.6968235 | 3.64345617 | 81.57506763 |
| ENSMUSG00000024985 | *Tcf7l2* | 60.27387896 | 35.49007962 | 23.95079712 | 114.1466813 | 164.2309165 | 164.3772552 | 9.72E-07 | 1.861274872 | 2.18E-07 | 26.86393588 | 3.633285842 | 6.012383936 |
| ENSMUSG00000034295 | *Fhod3* | 48.21910316 | 43.16361035 | 50.42273077 | 133.9028377 | 185.2861622 | 193.7631332 | 2.70E-12 | 1.859469442 | 3.46E-13 | 52.9273471 | 3.628741888 | 11.56886738 |
| ENSMUSG00000006930 | *Hap1* | 319.4515585 | 290.6349764 | 162.6133067 | 962.5638413 | 879.0565081 | 955.9593442 | 6.73E-16 | 1.847796491 | 6.39E-17 | 69.85360504 | 3.599499931 | 15.17198396 |
| ENSMUSG00000043415 | *Otud1* | 45.9588327 | 61.38824583 | 49.1621625 | 167.9273292 | 193.7082605 | 200.191294 | 6.04E-15 | 1.847488832 | 6.21E-16 | 65.36912997 | 3.59873241 | 14.21887213 |
| ENSMUSG00000074738 | *Fndc10* | 69.3149608 | 88.24560338 | 68.07068654 | 263.4154184 | 274.7709564 | 273.655989 | 1.68E-21 | 1.845855768 | 1.11E-22 | 96.06047252 | 3.594661116 | 20.7749152 |
| ENSMUSG00000026321 | *Tnfrsf11a* | 808.4234015 | 722.2710799 | 457.5862817 | 2166.591816 | 2662.435819 | 2335.258994 | 2.54E-20 | 1.845500058 | 1.81E-21 | 90.54699598 | 3.593774927 | 19.59503852 |
| ENSMUSG00000074476 | *Spc24* | 644.1770813 | 571.6780393 | 451.2834404 | 1935.005761 | 2060.255792 | 1992.729853 | 1.02E-40 | 1.838923662 | 2.81E-42 | 185.6680627 | 3.577430307 | 39.99058533 |
| ENSMUSG00000033952 | *Aspm* | 339.0405691 | 254.1857054 | 204.2120596 | 935.1247352 | 988.5437858 | 935.7565531 | 2.83E-23 | 1.83263978 | 1.72E-24 | 104.3272236 | 3.561882141 | 22.54862811 |
| ENSMUSG00000010205 | *Raver1* | 3762.596894 | 3579.702085 | 2537.523926 | 11171.00887 | 11950.95746 | 11955.46081 | 9.92E-36 | 1.826980837 | 3.34E-37 | 162.4272403 | 3.547938094 | 35.00332719 |
| ENSMUSG00000053128 | *Rnf26* | 1307.18975 | 1273.806101 | 921.4754048 | 3876.596907 | 4343.697189 | 4210.445335 | 9.57E-38 | 1.824852638 | 2.98E-39 | 171.806958 | 3.542708194 | 37.01901594 |
| ENSMUSG00000038545 | *Cul7* | 88.15054797 | 117.0213436 | 122.2751221 | 389.6353063 | 382.1527095 | 375.5882534 | 4.87E-24 | 1.822441238 | 2.82E-25 | 107.9055407 | 3.536791661 | 23.31284103 |
| ENSMUSG00000033581 | *Igf2bp2* | 110.7532526 | 239.7978353 | 114.7117125 | 547.6845573 | 511.6424706 | 583.1260169 | 8.10E-10 | 1.81902752 | 1.30E-10 | 41.30724856 | 3.528432766 | 9.091652988 |
| ENSMUSG00000050100 | *Hmx2* | 131.0956867 | 108.3886215 | 141.1836462 | 439.0256973 | 444.2656844 | 453.6444919 | 8.56E-31 | 1.81704161 | 3.54E-32 | 139.4319103 | 3.523579123 | 30.06735457 |
| ENSMUSG00000040943 | *Tet2* | 229.7941635 | 258.9816621 | 191.6063769 | 807.807283 | 779.044091 | 813.6214976 | 2.96E-39 | 1.812259555 | 8.63E-41 | 178.8533898 | 3.511918963 | 38.52807053 |
| ENSMUSG00000032218 | *Ccnb2* | 1295.888398 | 1035.926648 | 864.7498327 | 3672.449958 | 3797.313563 | 3759.555769 | 1.15E-35 | 1.809468807 | 3.88E-37 | 162.125816 | 3.505132075 | 34.93836063 |
| ENSMUSG00000032012 | *Nectin1* | 1283.080198 | 1245.030361 | 863.4892644 | 3583.547254 | 4213.154665 | 4085.555353 | 3.75E-30 | 1.80632363 | 1.61E-31 | 136.4289912 | 3.497498963 | 29.42618013 |
| ENSMUSG00000074604 | *Mgst2* | 273.4927258 | 296.3901244 | 305.0575212 | 1019.637182 | 1027.49599 | 1000.038161 | 1.45E-65 | 1.805021706 | 1.90E-67 | 301.0931853 | 3.494344158 | 64.83790968 |
| ENSMUSG00000038569 | *Rad9b* | 12.80819928 | 12.46948743 | 32.774775 | 65.85385459 | 55.79640112 | 73.46469504 | 0.000265349 | 1.792415473 | 8.43E-05 | 15.45869017 | 3.463943678 | 3.576182245 |
| ENSMUSG00000052336 | *Cx3cr1* | 1252.189835 | 1106.906808 | 611.3756106 | 2977.691792 | 3989.969061 | 3301.319733 | 7.71E-14 | 1.787284869 | 8.65E-15 | 60.18056272 | 3.45164686 | 13.11276858 |
| ENSMUSG00000044229 | *Nxpe4* | 45.20540922 | 20.14301816 | 23.95079712 | 91.09783218 | 109.4872777 | 112.0336599 | 3.88E-06 | 1.783115965 | 9.35E-07 | 24.05784024 | 3.441687157 | 5.41092301 |
| ENSMUSG00000096001 | *Gpr15lg* | 15.82189323 | 14.38787012 | 26.47193365 | 65.85385459 | 78.95717139 | 46.83374309 | 6.51E-05 | 1.781916057 | 1.88E-05 | 18.3082168 | 3.438825852 | 4.186538153 |
| ENSMUSG00000034413 | *Neurl1b* | 104.7258647 | 102.6334735 | 57.98614039 | 267.8056753 | 369.5195621 | 274.5742977 | 8.18E-11 | 1.76626017 | 1.19E-11 | 45.98007694 | 3.401710037 | 10.0870196 |
| ENSMUSG00000046598 | *Bdh1* | 10.54792882 | 19.18382682 | 28.99307019 | 63.6587261 | 77.9044091 | 52.34359522 | 0.000225421 | 1.76061976 | 7.08E-05 | 15.78925116 | 3.388436557 | 3.647006328 |
| ENSMUSG00000006398 | *Cdc20* | 2665.612297 | 2214.772807 | 1361.413731 | 6700.629705 | 7309.328546 | 7078.323367 | 3.71E-17 | 1.755075574 | 3.23E-18 | 75.74503036 | 3.375439999 | 16.43071714 |
| ENSMUSG00000084353 | *Gm15452* | 233.561281 | 208.144521 | 186.5641038 | 756.2217635 | 694.8231082 | 683.2216639 | 9.28E-38 | 1.754879472 | 2.88E-39 | 171.8733174 | 3.374981215 | 37.03255096 |
| ENSMUSG00000028885 | *Smpdl3b* | 1482.737422 | 1313.132946 | 1081.567575 | 4294.768884 | 4400.546352 | 4395.025381 | 5.02E-47 | 1.752427946 | 1.08E-48 | 215.054865 | 3.369251085 | 46.29969307 |
| ENSMUSG00000038754 | *Elovl3* | 13.56162277 | 23.02059219 | 16.3873875 | 46.09769821 | 72.64059768 | 58.77175603 | 5.77E-05 | 1.752033896 | 1.65E-05 | 18.55507725 | 3.368330953 | 4.239170212 |
| ENSMUSG00000020057 | *Dram1* | 523.6293234 | 530.4328116 | 524.3964 | 1683.663549 | 1833.911901 | 1788.865324 | 1.48E-87 | 1.749460446 | 1.16E-89 | 403.099177 | 3.362327947 | 86.83088948 |
| ENSMUSG00000073434 | *Wdr90* | 311.9173236 | 318.4515253 | 143.7047827 | 850.6122885 | 923.2725241 | 832.90598 | 7.41E-11 | 1.744739032 | 1.07E-11 | 46.18953664 | 3.351342263 | 10.13037777 |
| ENSMUSG00000075610 | *Tmem92* | 41.43829178 | 30.69412292 | 39.07761635 | 117.439374 | 134.7535725 | 121.2167468 | 1.71E-10 | 1.744573771 | 2.56E-11 | 44.48553159 | 3.350958388 | 9.767477425 |
| ENSMUSG00000025141 | *Myadml2* | 17.3287402 | 12.46948743 | 3.781704808 | 24.14641335 | 49.4798274 | 42.24219965 | 0.007518311 | 1.743283605 | 0.003163758 | 8.71053818 | 3.347963049 | 2.123879736 |
| ENSMUSG00000023015 | *Racgap1* | 1741.915102 | 1435.909438 | 1100.476099 | 4429.769285 | 5070.103165 | 4834.895242 | 3.23E-27 | 1.742198102 | 1.58E-28 | 122.7469488 | 3.345444944 | 26.49098224 |
| ENSMUSG00000022021 | *Diaph3* | 119.7943344 | 133.3275964 | 181.5218308 | 480.7331385 | 471.6375038 | 485.785296 | 2.53E-21 | 1.741332535 | 1.69E-22 | 95.24028792 | 3.343438395 | 20.5972292 |
| ENSMUSG00000021485 | *Mxd3* | 103.2190177 | 85.36802936 | 75.63409615 | 275.488625 | 303.1955381 | 310.3883365 | 2.71E-19 | 1.736888247 | 2.08E-20 | 85.71146005 | 3.333154629 | 18.56721232 |
| ENSMUSG00000029385 | *Ccng2* | 88.15054797 | 52.75552376 | 50.42273077 | 184.3907929 | 225.291129 | 233.2504068 | 9.17E-10 | 1.730817699 | 1.48E-10 | 41.05164277 | 3.319158903 | 9.037817886 |
| ENSMUSG00000004105 | *Angptl2* | 1329.039031 | 1339.031112 | 1227.793494 | 4351.842224 | 4153.147215 | 4429.002802 | 1.17E-114 | 1.729420733 | 4.88E-117 | 528.9083608 | 3.315946505 | 113.9333694 |
| ENSMUSG00000028047 | *Thbs3* | 10.54792882 | 22.06140085 | 25.21136538 | 69.14654732 | 58.95468797 | 58.77175603 | 0.000105522 | 1.726069261 | 3.14E-05 | 17.32858519 | 3.308252293 | 3.976658528 |
| ENSMUSG00000001420 | *Tmem79* | 15.82189323 | 16.3062528 | 8.823977885 | 46.09769821 | 32.63563084 | 58.77175603 | 0.000611346 | 1.725230874 | 0.000207544 | 13.76152229 | 3.306330342 | 3.213712886 |
| ENSMUSG00000031750 | *Il34* | 18.08216369 | 6.714339388 | 10.08454615 | 41.70744124 | 42.11049141 | 33.97742146 | 0.001666457 | 1.723198586 | 0.000615144 | 11.72958299 | 3.301676077 | 2.778205767 |
| ENSMUSG00000071041 | *Impdh2-ps* | 537.9443697 | 245.5529833 | 659.2772048 | 1167.808355 | 1728.635672 | 1837.535685 | 0.000573484 | 1.714878362 | 0.000193667 | 13.89155181 | 3.282689651 | 3.241478499 |
| ENSMUSG00000062248 | *Cks2* | 288.5611955 | 258.9816621 | 209.2543327 | 825.3683109 | 871.6871721 | 802.6017933 | 2.49E-32 | 1.713772066 | 9.83E-34 | 146.5526474 | 3.280173362 | 31.60362296 |
| ENSMUSG00000039145 | *Camk1d* | 506.3005832 | 556.3309779 | 486.5793519 | 1535.492376 | 1877.075154 | 1664.893651 | 1.21E-47 | 1.711487812 | 2.56E-49 | 217.9306987 | 3.274983896 | 46.91727108 |
| ENSMUSG00000019838 | *Slc16a10* | 2291.914247 | 2195.58898 | 1516.463628 | 5930.139606 | 7066.140458 | 6682.532323 | 1.15E-26 | 1.711223826 | 5.77E-28 | 120.1834822 | 3.274384689 | 25.94002715 |
| ENSMUSG00000030653 | *Gm45837* | 1847.39439 | 1799.442956 | 1801.352057 | 5502.089551 | 6858.746288 | 5429.959272 | 8.19E-53 | 1.70701229 | 1.44E-54 | 241.9946305 | 3.264839998 | 52.08661957 |
| ENSMUSG00000035759 | *Bbs10* | 24.10955158 | 26.85735755 | 34.03534327 | 95.48808916 | 92.6430811 | 85.40270798 | 5.49E-08 | 1.705181551 | 1.06E-08 | 32.72766003 | 3.260699636 | 7.26058051 |
| ENSMUSG00000058057 | *Mettl7a3* | 8.287658356 | 8.63272207 | 8.823977885 | 19.75615638 | 23.16077027 | 40.40558227 | 0.006284496 | 1.704351632 | 0.002605746 | 9.064817231 | 3.258824438 | 2.201729513 |
| ENSMUSG00000046442 | *Ppm1e* | 96.43820633 | 75.77611595 | 55.46500385 | 246.9519547 | 240.029801 | 264.4729021 | 7.09E-13 | 1.702864915 | 8.61E-14 | 55.66212006 | 3.255467905 | 12.14964923 |
| ENSMUSG00000026463 | *Atp2b4* | 691.642761 | 689.6585743 | 543.304924 | 1917.444733 | 2246.594717 | 2098.335352 | 1.16E-40 | 1.698196002 | 3.20E-42 | 185.4079035 | 3.244949446 | 39.93487556 |
| ENSMUSG00000091996 | *Smim35* | 7.534234869 | 7.673530729 | 6.302841346 | 21.95128486 | 26.31905713 | 22.03940851 | 0.005370636 | 1.697309983 | 0.002197346 | 9.376918353 | 3.242957199 | 2.269974298 |
| ENSMUSG00000001014 | *Icam4* | 30.13693948 | 25.89816621 | 22.69022885 | 88.9027037 | 78.95717139 | 89.99425142 | 1.03E-07 | 1.696095405 | 2.06E-08 | 31.44165822 | 3.240228163 | 6.985867172 |
| ENSMUSG00000035004 | *Igsf6* | 278.7666902 | 288.7165937 | 300.0152481 | 960.3687128 | 907.4810898 | 928.4100836 | 2.64E-57 | 1.691859371 | 4.06E-59 | 262.8703448 | 3.230728175 | 56.5778664 |
| ENSMUSG00000027654 | *Fam83d* | 188.3558717 | 180.3279721 | 133.6202365 | 476.3428815 | 569.5443963 | 576.6978561 | 4.95E-21 | 1.681959615 | 3.38E-22 | 93.86105612 | 3.208634845 | 20.30522198 |
| ENSMUSG00000034342 | *Cbl* | 1720.065821 | 1575.951373 | 1038.708254 | 4395.744794 | 5011.148478 | 4461.143606 | 3.05E-21 | 1.675740184 | 2.06E-22 | 94.84721523 | 3.194832255 | 20.5154397 |
| ENSMUSG00000025154 | *Arhgap19* | 45.20540922 | 38.36765364 | 16.3873875 | 103.1710389 | 103.1707039 | 119.3801294 | 1.35E-05 | 1.67477701 | 3.52E-06 | 21.51118984 | 3.192700029 | 4.870462506 |
| ENSMUSG00000047880 | *Cxcr5* | 46.71225619 | 48.9187584 | 63.02841346 | 176.7078431 | 156.8615805 | 167.1321812 | 2.11E-12 | 1.674314214 | 2.67E-13 | 53.44032831 | 3.19167602 | 11.67666917 |
| ENSMUSG00000024538 | *Ppic* | 337.5337222 | 289.675785 | 308.839226 | 1044.881159 | 964.3302532 | 967.8973572 | 8.24E-48 | 1.665687668 | 1.74E-49 | 218.701626 | 3.172648445 | 47.08402342 |
| ENSMUSG00000039611 | *Pgap4* | 34.6574804 | 23.02059219 | 26.47193365 | 85.61001097 | 94.74860567 | 89.07594274 | 2.32E-07 | 1.664162853 | 4.82E-08 | 29.78618556 | 3.169296978 | 6.635216675 |
| ENSMUSG00000052909 | *Gm9894* | 13.56162277 | 7.673530729 | 8.823977885 | 43.90256973 | 27.37181941 | 25.71264326 | 0.004226554 | 1.663595704 | 0.00169532 | 9.853249482 | 3.168051315 | 2.374013569 |
| ENSMUSG00000074480 | *Mex3a* | 112.2600996 | 86.3272207 | 44.11988942 | 262.3178541 | 281.0875301 | 235.0870241 | 1.85E-07 | 1.662649685 | 3.81E-08 | 30.24301559 | 3.165974608 | 6.731913775 |
| ENSMUSG00000063522 | *Ly6m* | 12.05477579 | 15.34706146 | 10.08454615 | 49.39039094 | 21.0552457 | 48.67036046 | 0.003802192 | 1.661577287 | 0.001511536 | 10.06450021 | 3.163622121 | 2.419965975 |
| ENSMUSG00000004099 | *Dnmt1* | 5053.96475 | 5100.020361 | 3416.14001 | 13070.89257 | 15410.33433 | 14238.37621 | 1.13E-25 | 1.653834784 | 6.03E-27 | 115.526857 | 3.146689406 | 24.94576341 |
| ENSMUSG00000013033 | *Adgrl1* | 240.3420923 | 258.9816621 | 170.1767163 | 589.3919986 | 795.8882876 | 727.3004809 | 6.90E-17 | 1.651424368 | 6.12E-18 | 74.48247177 | 3.14143639 | 16.16114054 |
| ENSMUSG00000068101 | *Cenpm* | 213.9722703 | 136.2051704 | 172.6978529 | 509.2698088 | 547.4363883 | 589.5541777 | 7.33E-18 | 1.649674549 | 6.10E-19 | 79.03502876 | 3.137628508 | 17.13488093 |
| ENSMUSG00000046785 | *Epm2aip1* | 279.5201137 | 245.5529833 | 209.2543327 | 812.1975399 | 740.0918865 | 758.5229763 | 4.80E-31 | 1.644258934 | 1.97E-32 | 140.6018997 | 3.125872501 | 30.31886104 |
| ENSMUSG00000017493 | *Igfbp4* | 1330.545878 | 1498.256875 | 1246.702018 | 3883.182292 | 4497.400482 | 4300.439586 | 2.11E-55 | 1.636783937 | 3.36E-57 | 254.0718705 | 3.109718387 | 54.67520557 |
| ENSMUSG00000031955 | *Bcar1* | 36.16432737 | 39.32684499 | 17.64795577 | 87.80513945 | 106.3289908 | 100.095647 | 5.51E-06 | 1.634416337 | 1.36E-06 | 23.3386394 | 3.104619228 | 5.258899295 |
| ENSMUSG00000031844 | *Hsd17b2* | 8.287658356 | 21.1022095 | 15.12681923 | 49.39039094 | 40.00496684 | 46.83374309 | 0.001233977 | 1.633428282 | 0.000442987 | 12.34154857 | 3.102493703 | 2.908692771 |
| ENSMUSG00000034135 | *Sik3* | 1818.764297 | 1737.095519 | 1310.991 | 4780.989843 | 5159.58796 | 5111.306157 | 2.07E-35 | 1.626811877 | 7.10E-37 | 160.9266678 | 3.088297801 | 34.68339526 |
| ENSMUSG00000102049 | *None* | 36.91775086 | 24.93897487 | 51.68329904 | 105.3661673 | 101.0651794 | 138.6646119 | 3.84E-06 | 1.621880606 | 9.25E-07 | 24.07885833 | 3.077759722 | 5.41541584 |
| ENSMUSG00000031274 | *Col4a5* | 917.6698071 | 990.8446554 | 650.4532269 | 2475.007368 | 2826.666736 | 2578.610796 | 2.27E-24 | 1.619824729 | 1.29E-25 | 109.4550943 | 3.07337696 | 23.64341701 |
| ENSMUSG00000001036 | *Epn2* | 224.5201991 | 248.4305574 | 200.4303548 | 700.2459871 | 704.2979688 | 669.4470336 | 6.40E-36 | 1.617660107 | 2.13E-37 | 163.3238457 | 3.068769117 | 35.19385299 |
| ENSMUSG00000059994 | *Fcrl1* | 1540.751031 | 1411.929654 | 1602.18227 | 4714.038424 | 4506.875343 | 4653.070122 | 5.94E-93 | 1.607827223 | 4.12E-95 | 428.1450082 | 3.04792462 | 92.22617335 |
| ENSMUSG00000041431 | *Ccnb1* | 2885.611955 | 2347.141212 | 2071.113666 | 7730.144965 | 7212.474416 | 7290.452674 | 1.98E-36 | 1.604437731 | 6.47E-38 | 165.6895401 | 3.040772179 | 35.70340448 |
| ENSMUSG00000062867 | *Impdh2* | 5942.251042 | 5860.659094 | 5366.239122 | 17647.73547 | 17463.22079 | 17042.89094 | 3.25E-122 | 1.602228584 | 1.25E-124 | 563.8048536 | 3.036119518 | 121.4877285 |
| ENSMUSG00000022272 | *Myo10* | 1506.846974 | 1485.787387 | 902.5668808 | 3535.254427 | 4556.35517 | 3743.944521 | 5.81E-16 | 1.601536541 | 5.47E-17 | 70.15802429 | 3.034663479 | 15.23585515 |
| ENSMUSG00000037406 | *Htra4* | 29.38351599 | 20.14301816 | 10.08454615 | 76.82949702 | 54.74363883 | 54.18021259 | 0.00081255 | 1.601294379 | 0.000281426 | 13.19013862 | 3.034154141 | 3.0901498 |
| ENSMUSG00000113640 | *Adat3* | 85.13685402 | 112.2253869 | 90.76091539 | 266.7081111 | 278.9820056 | 325.0812756 | 1.67E-15 | 1.598728682 | 1.63E-16 | 68.00672897 | 3.028762978 | 14.77827189 |
| ENSMUSG00000032185 | *Carm1* | 1983.764041 | 2021.975347 | 1464.780329 | 5075.13706 | 5794.403618 | 5698.105409 | 2.22E-31 | 1.597061088 | 9.02E-33 | 142.1498496 | 3.025264091 | 30.65295233 |
| ENSMUSG00000036768 | *Kif15* | 705.2043838 | 557.2901692 | 531.9598096 | 1860.371392 | 1863.389245 | 1720.910481 | 1.62E-34 | 1.596351686 | 5.80E-36 | 156.7524461 | 3.023776874 | 33.79173853 |
| ENSMUSG00000001403 | *Ube2c* | 1843.627273 | 1604.727114 | 1124.426896 | 4492.330447 | 4651.103776 | 4678.782765 | 1.17E-21 | 1.593767256 | 7.68E-23 | 96.79807105 | 3.018364957 | 20.93037732 |
| ENSMUSG00000022505 | *Emp2* | 475.4102203 | 509.3306021 | 588.6853817 | 1612.321873 | 1560.193707 | 1539.085361 | 6.58E-48 | 1.588026162 | 1.38E-49 | 219.1556629 | 3.006377473 | 47.18164564 |
| ENSMUSG00000091269 | *Gm6682* | 14.31504625 | 17.26544414 | 6.302841346 | 26.34154184 | 45.26877826 | 44.07881702 | 0.00512472 | 1.584285728 | 0.002087024 | 9.471379017 | 2.998593021 | 2.290329845 |
| ENSMUSG00000031004 | *Mki67* | 2490.818048 | 1758.197728 | 965.5952942 | 5046.60039 | 5343.82136 | 5234.359522 | 8.17E-05 | 1.582569348 | 2.40E-05 | 17.84476186 | 2.995027704 | 4.087788558 |
| ENSMUSG00000040722 | *Scamp5* | 675.8208678 | 697.332105 | 569.7768577 | 1945.981403 | 1944.451941 | 1945.89611 | 6.63E-54 | 1.582560282 | 1.11E-55 | 247.0996094 | 2.995008883 | 53.17823738 |
| ENSMUSG00000091775 | *Gm16494* | 86.643701 | 59.46986315 | 104.6271663 | 276.5861893 | 215.8162685 | 253.4531979 | 4.74E-10 | 1.580635199 | 7.44E-11 | 42.40064774 | 2.99101511 | 9.323805977 |
| ENSMUSG00000092097 | *Gm5819* | 13.56162277 | 12.46948743 | 6.302841346 | 30.73179881 | 36.84667998 | 31.22249539 | 0.003397233 | 1.574994454 | 0.001334694 | 10.29391802 | 2.979343474 | 2.468874649 |
| ENSMUSG00000026482 | *Rgl1* | 984.7244974 | 918.9053048 | 842.0596039 | 2682.44701 | 2741.392991 | 2773.292238 | 2.65E-71 | 1.574402176 | 3.10E-73 | 327.6680024 | 2.978120596 | 70.57622484 |
| ENSMUSG00000024190 | *Dusp1* | 360.8898502 | 350.1048395 | 310.0997942 | 963.6614055 | 1042.234662 | 1045.035287 | 1.46E-44 | 1.573894962 | 3.42E-46 | 203.6018137 | 2.977073751 | 43.83663728 |
| ENSMUSG00000035539 | *Ccdc180* | 12.05477579 | 11.51029609 | 21.42966058 | 39.51231275 | 52.63811426 | 37.65065621 | 0.001488455 | 1.559293038 | 0.000542491 | 11.96362815 | 2.947093921 | 2.827264322 |
| ENSMUSG00000021508 | *Cxcl14* | 602.7387896 | 575.5148047 | 530.6992414 | 1423.540823 | 2030.778448 | 1583.164178 | 2.59E-24 | 1.557869909 | 1.48E-25 | 109.1782115 | 2.944188228 | 23.58589178 |
| ENSMUSG00000040093 | *Bmf* | 70.06838429 | 94.00075143 | 98.324325 | 234.878748 | 267.4016204 | 262.6362848 | 1.18E-13 | 1.557571556 | 1.35E-14 | 59.30758072 | 2.943579426 | 12.92724148 |
| ENSMUSG00000026955 | *Sapcd2* | 512.3279711 | 415.3298507 | 286.1489971 | 1220.491438 | 1132.772219 | 1231.451951 | 3.89E-15 | 1.555950712 | 3.93E-16 | 66.27044882 | 2.940274221 | 14.40958132 |
| ENSMUSG00000074203 | *G430095P16Rik* | 31.64378645 | 28.77574023 | 37.81704808 | 110.8539886 | 100.0124171 | 76.2196211 | 1.72E-06 | 1.553720768 | 3.98E-07 | 25.70182915 | 2.93573301 | 5.763866609 |
| ENSMUSG00000035024 | *Ncapd3* | 793.3549318 | 806.6799179 | 675.6645923 | 2146.83566 | 2284.494159 | 2264.549225 | 5.47E-55 | 1.553422323 | 8.86E-57 | 252.1414644 | 2.935125768 | 54.2618039 |
| ENSMUSG00000061393 | *Acvr2b* | 101.7121707 | 115.1029609 | 84.45807404 | 277.6837535 | 307.4065873 | 300.286941 | 9.63E-17 | 1.547255912 | 8.62E-18 | 73.80516523 | 2.922607138 | 16.01647985 |
| ENSMUSG00000026921 | *Egfl7* | 27.87666902 | 27.81654889 | 21.42966058 | 62.56116186 | 68.42954854 | 95.50410355 | 2.78E-05 | 1.54583141 | 7.61E-06 | 20.03340546 | 2.91972281 | 4.555409504 |
| ENSMUSG00000032324 | *Tspan3* | 1029.929907 | 1062.784006 | 1051.313937 | 3095.131166 | 3117.229126 | 2924.813171 | 5.44E-96 | 1.53938272 | 3.32E-98 | 442.3569282 | 2.90670109 | 95.26437997 |
| ENSMUSG00000074794 | *Arrdc3* | 114.52037 | 98.79670814 | 117.232849 | 334.7570942 | 342.1477427 | 283.7573846 | 1.73E-17 | 1.53931648 | 1.47E-18 | 77.29683618 | 2.906567635 | 16.7619351 |
| ENSMUSG00000023072 | *Cep89* | 467.1225619 | 473.8405225 | 388.2550269 | 1182.07669 | 1365.432684 | 1325.119437 | 5.89E-36 | 1.538339882 | 1.95E-37 | 163.4936651 | 2.90460077 | 35.23005355 |
| ENSMUSG00000040061 | *Plcb2* | 666.7797859 | 732.8221846 | 547.0866289 | 1784.639459 | 2006.564916 | 1873.349724 | 1.81E-33 | 1.537615796 | 6.75E-35 | 151.8730746 | 2.903143322 | 32.74171411 |
| ENSMUSG00000063894 | *Zkscan8* | 156.7120853 | 123.735683 | 128.5779635 | 385.2450494 | 446.3712089 | 361.8136231 | 1.08E-17 | 1.536925389 | 9.11E-19 | 78.24418827 | 2.901754344 | 16.96847501 |
| ENSMUSG00000059588 | *Calcrl* | 1422.463543 | 1404.256123 | 1324.857251 | 4318.915297 | 3599.394253 | 4127.797553 | 4.99E-56 | 1.535876983 | 7.81E-58 | 256.9798202 | 2.899646407 | 55.30155792 |
| ENSMUSG00000039206 | *Daglb* | 1054.039458 | 1046.477753 | 839.5384673 | 2811.959591 | 2960.367546 | 2758.599299 | 1.99E-43 | 1.533480488 | 4.92E-45 | 198.2929317 | 2.894833732 | 42.70042312 |
| ENSMUSG00000056531 | *Ccdc18* | 18.08216369 | 11.51029609 | 6.302841346 | 36.21962002 | 27.37181941 | 43.16050834 | 0.007894185 | 1.532139424 | 0.003341431 | 8.610976585 | 2.892144076 | 2.102692715 |
| ENSMUSG00000072082 | *Ccnf* | 1527.942832 | 1361.092513 | 995.8489327 | 3553.913019 | 4061.556896 | 3626.401009 | 2.33E-22 | 1.530528802 | 1.47E-23 | 100.0655086 | 2.888917094 | 21.63307044 |

| | | | | | | | | | | | | |
|---|---|---|---|---|---|---|---|---|---|---|---|---|
| ENSMUSG00000039981 | *Zc3h12d* | 38.42459783 | 26.85735755 | 36.55647981 | 90.00026794 | 98.95965481 | 104.6871904 | 4.75E-07 | 1.527130379 | 1.02E-07 | 28.32707829 | 2.882119948 | 6.323439667 |
| ENSMUSG00000021108 | *Prkch* | 1054.792882 | 1006.191717 | 806.7636923 | 2702.203167 | 2739.287466 | 2810.024585 | 4.23E-39 | 1.521199297 | 1.24E-40 | 178.1327601 | 2.870295551 | 38.37378267 |
| ENSMUSG00000039568 | *Ubald1* | 393.2870602 | 381.7581538 | 358.0013885 | 1054.759238 | 1133.824981 | 1069.829622 | 4.17E-50 | 1.520154204 | 8.09E-52 | 229.3921796 | 2.868217052 | 49.38036845 |
| ENSMUSG00000063856 | *Gpx1* | 8390.877374 | 7972.798427 | 7801.657018 | 21428.84428 | 23833.48537 | 23432.48279 | 2.92E-94 | 1.507002953 | 1.94E-96 | 434.2385275 | 2.842189903 | 93.53486929 |
| ENSMUSG00000034485 | *Uaca* | 317.191288 | 315.5739512 | 172.6978529 | 744.1485569 | 814.8380087 | 741.0751112 | 2.51E-11 | 1.506212242 | 3.49E-12 | 48.39254659 | 2.840632586 | 10.60107219 |
| ENSMUSG00000029177 | *Cenpa* | 1915.202504 | 1626.788515 | 1410.575893 | 4656.965084 | 4816.387455 | 4607.154688 | 3.26E-36 | 1.505138747 | 1.07E-37 | 164.686823 | 2.838519686 | 35.48709317 |
| ENSMUSG00000001021 | *S100a3* | 11.3013523 | 12.46948743 | 7.563409615 | 29.63423457 | 35.7939177 | 24.79433458 | 0.006270758 | 1.503294152 | 0.002598758 | 9.069727132 | 2.834892742 | 2.202679951 |
| ENSMUSG00000042606 | *Hirip3* | 803.1494371 | 803.8023439 | 502.9667394 | 1852.688442 | 2155.004398 | 1982.628457 | 1.73E-16 | 1.501938807 | 1.58E-17 | 72.61149878 | 2.832230742 | 15.76206965 |
| ENSMUSG00000038518 | *Jarid2* | 543.9717576 | 507.4122195 | 363.0436615 | 1288.540421 | 1395.96279 | 1336.139141 | 1.67E-21 | 1.501296959 | 1.11E-22 | 96.07632168 | 2.830970978 | 20.77794257 |
| ENSMUSG00000044201 | *Cdc25c* | 343.56111 | 284.8798283 | 180.2612625 | 724.3924005 | 786.413427 | 788.827163 | 1.07E-11 | 1.499405139 | 1.46E-12 | 50.10732158 | 2.827261131 | 10.96943694 |
| ENSMUSG00000038252 | *Ncapd2* | 2743.968339 | 2447.856303 | 1737.063075 | 6206.725795 | 6957.705943 | 6433.670668 | 1.02E-20 | 1.498631237 | 7.09E-22 | 92.39889056 | 2.825744915 | 19.99043903 |
| ENSMUSG00000032400 | *Zwilch* | 456.5746331 | 392.3092585 | 414.7269606 | 1218.29631 | 1229.626349 | 1126.76476 | 6.14E-43 | 1.496861626 | 1.56E-44 | 195.9987591 | 2.822280978 | 42.21164715 |
| ENSMUSG00000051378 | *Kif18b* | 790.3412378 | 797.0880045 | 495.4033298 | 1861.468956 | 2077.099989 | 1954.160888 | 1.07E-16 | 1.496770665 | 9.64E-18 | 73.58568287 | 2.822103042 | 15.97051294 |
| ENSMUSG00000037217 | *Syn1* | 32.39720994 | 21.1022095 | 20.16909231 | 72.43924005 | 66.32402397 | 72.54638635 | 4.24E-05 | 1.496571631 | 1.19E-05 | 19.17546881 | 2.821713731 | 4.372645803 |
| ENSMUSG00000078532 | *Nkain1* | 82.87658356 | 118.9397263 | 103.3665981 | 248.049519 | 260.0322844 | 346.2023754 | 2.50E-10 | 1.492343326 | 3.81E-11 | 43.71143197 | 2.813455854 | 9.602626362 |
| ENSMUSG00000020381 | *Mrnip* | 35.41090389 | 56.59228913 | 44.11988942 | 141.5857874 | 121.0676628 | 117.5435121 | 2.85E-07 | 1.488004007 | 5.98E-08 | 29.36860184 | 2.805006292 | 6.54549918 |
| ENSMUSG00000039958 | *Etfbkmt* | 30.13693948 | 22.06140085 | 25.21136538 | 73.53680429 | 71.58783539 | 73.46469504 | 9.64E-06 | 1.487881198 | 2.46E-06 | 22.2010931 | 2.804767526 | 5.016094208 |
| ENSMUSG00000001517 | *Foxm1* | 1983.010618 | 1803.279721 | 1125.687464 | 4256.354135 | 4885.869766 | 4635.622257 | 8.57E-15 | 1.486306785 | 8.90E-16 | 64.66002988 | 2.801708354 | 14.06720647 |
| ENSMUSG00000033016 | *Nfatc1* | 870.9575509 | 843.1291888 | 508.0090125 | 1814.273694 | 2351.870945 | 2056.093152 | 5.39E-13 | 1.482494046 | 6.51E-14 | 56.20943993 | 2.794313804 | 12.26809244 |
| ENSMUSG00000028175 | *Depdc1a* | 241.0955158 | 108.3886215 | 100.8454615 | 403.9036415 | 451.6350203 | 410.4839835 | 7.16E-06 | 1.480938871 | 1.79E-06 | 22.80590273 | 2.791303254 | 5.14492453 |
| ENSMUSG00000055069 | *Rab39* | 39.93144481 | 56.59228913 | 51.68329904 | 138.2930946 | 123.1731874 | 146.0110814 | 2.85E-08 | 1.470638348 | 5.30E-09 | 34.07767395 | 2.771444944 | 7.545700238 |
| ENSMUSG00000070047 | *Fat1* | 550.752569 | 613.8824583 | 360.522525 | 1217.198746 | 1621.253919 | 1397.665823 | 6.87E-13 | 1.470309398 | 8.33E-14 | 55.72735227 | 2.770813096 | 12.16307759 |
| ENSMUSG00000024186 | *Rgs11* | 33.90405691 | 49.87794974 | 27.73250192 | 110.8539886 | 104.2234662 | 95.50410355 | 4.87E-06 | 1.467919666 | 1.19E-06 | 23.59591737 | 2.766227221 | 5.31223785 |
| ENSMUSG00000036766 | *Dner* | 62.53414942 | 69.06177656 | 61.76784519 | 181.0981001 | 177.9168262 | 175.3969594 | 2.56E-12 | 1.465377062 | 3.26E-13 | 53.04347797 | 2.761356319 | 11.59244799 |
| ENSMUSG00000030677 | *Kif22* | 1137.669465 | 1096.355703 | 887.4400615 | 2718.66663 | 2990.897652 | 2922.058245 | 6.80E-37 | 1.464348504 | 2.17E-38 | 167.8611948 | 2.759388333 | 36.16761398 |
| ENSMUSG00000022114 | *Spry2* | 43.69856224 | 59.46986315 | 56.72557212 | 177.8054074 | 131.5952856 | 128.5632163 | 1.73E-07 | 1.461271607 | 3.56E-08 | 30.37561019 | 2.75350954 | 6.760731379 |
| ENSMUSG00000040703 | *Cyp2s1* | 17.3287402 | 16.3062528 | 20.16909231 | 50.48795519 | 41.05772912 | 55.09852128 | 0.000463736 | 1.457264969 | 0.000153359 | 14.33051366 | 2.745873129 | 3.333729004 |
| ENSMUSG00000054793 | *Cadm4* | 9.79450533 | 14.38787012 | 16.3873875 | 28.53667032 | 33.68839313 | 46.83374309 | 0.005536282 | 1.454851809 | 0.002270818 | 9.316642007 | 2.741284015 | 2.256781797 |
| ENSMUSG00000051517 | *Arhgef39* | 539.4512167 | 457.5342697 | 403.3818462 | 1322.564913 | 1327.533242 | 1204.820999 | 6.05E-28 | 1.454583148 | 2.84E-29 | 126.1590306 | 2.740773576 | 27.21851822 |
| ENSMUSG00000029330 | *Cds1* | 105.4792882 | 110.3070042 | 113.4511442 | 302.9277311 | 332.6728821 | 261.7179761 | 3.31E-15 | 1.448198762 | 3.32E-16 | 66.60306759 | 2.728671576 | 14.48049688 |
| ENSMUSG00000032086 | *Bace1* | 77.60261916 | 75.77611595 | 44.11988942 | 172.3175862 | 201.0775965 | 171.7237247 | 8.90E-08 | 1.446038935 | 1.76E-08 | 31.74804067 | 2.724589599 | 7.050625527 |
| ENSMUSG00000113637 | *Gm7049* | 94.17793587 | 89.20479472 | 95.80318846 | 279.878882 | 229.5021782 | 248.8616544 | 2.33E-14 | 1.441817828 | 2.50E-15 | 62.62752987 | 2.716629512 | 13.63351045 |
| ENSMUSG00000045328 | *Cenpe* | 510.0677007 | 379.8397711 | 184.0429673 | 971.3443552 | 1004.33522 | 942.1847139 | 2.15E-06 | 1.436552797 | 5.02E-07 | 25.25608779 | 2.706733402 | 5.667631566 |
| ENSMUSG00000040860 | *Crocc* | 174.0408255 | 161.1441453 | 83.19750577 | 338.0497869 | 409.5245289 | 392.1178098 | 1.49E-07 | 1.434237723 | 3.03E-08 | 30.6885845 | 2.702393426 | 6.826559635 |
| ENSMUSG00000097739 | *Gm2539* | 15.82189323 | 18.22463548 | 15.12681923 | 38.41474851 | 54.74363883 | 39.48727358 | 0.001381272 | 1.428006294 | 0.000499351 | 12.11808751 | 2.690746155 | 2.859720762 |
| ENSMUSG00000032481 | *Smarcc1* | 2383.078489 | 2460.32579 | 2271.544021 | 6270.384521 | 6759.786633 | 6120.527406 | 1.58E-78 | 1.427709623 | 1.54E-80 | 361.2086724 | 2.690192897 | 77.80236896 |
| ENSMUSG00000025420 | *Katnal2* | 24.10955158 | 12.46948743 | 25.21136538 | 45.00013397 | 61.06021254 | 58.77175603 | 0.001334483 | 1.421073584 | 0.000481245 | 12.18696843 | 2.677847092 | 2.874687062 |
| ENSMUSG00000030802 | *Bckdk* | 2589.516525 | 2473.754469 | 2155.57174 | 6469.043649 | 6502.912636 | 6362.04259 | 1.22E-57 | 1.419557543 | 1.86E-59 | 264.4226513 | 2.675034583 | 56.91427131 |
| ENSMUSG00000053395 | *Cacng8* | 442.2595868 | 451.7791217 | 400.8607096 | 1019.637182 | 1275.94789 | 1171.761886 | 4.52E-29 | 1.418792985 | 2.02E-30 | 131.4030537 | 2.67361732 | 28.34514608 |
| ENSMUSG00000079598 | *Clec2l* | 20.34243415 | 11.51029609 | 25.21136538 | 51.58551943 | 53.69087654 | 45.9154344 | 0.001522017 | 1.418251885 | 0.000556081 | 11.91753066 | 2.672614738 | 2.817580488 |
| ENSMUSG00000054409 | *Tmem74* | 27.12324553 | 41.24522767 | 40.33818462 | 100.9759104 | 91.59031881 | 94.58579486 | 5.36E-06 | 1.418003839 | 1.32E-06 | 23.39843524 | 2.672155267 | 5.270892857 |
| ENSMUSG00000027469 | *Tpx2* | 2205.270546 | 1825.341122 | 1776.140691 | 5031.234491 | 5530.160284 | 4951.520446 | 3.12E-38 | 1.415988877 | 9.50E-40 | 174.0824457 | 2.668425766 | 37.50530468 |
| ENSMUSG00000026622 | *Nek2* | 1489.518234 | 1326.561625 | 990.8066596 | 3264.156059 | 3635.188171 | 3275.60709 | 7.43E-21 | 1.415794062 | 5.11E-22 | 93.04520871 | 2.668065459 | 20.1288142 |
| ENSMUSG00000042659 | *Arrdc4* | 61.02730244 | 61.38824583 | 100.8454615 | 214.0250274 | 196.8665473 | 176.3152681 | 1.46E-07 | 1.412529378 | 2.96E-08 | 30.73066254 | 2.662034701 | 6.835011707 |
| ENSMUSG00000007033 | *Hspa1l* | 17.3287402 | 11.51029609 | 15.12681923 | 40.609877 | 31.58286856 | 44.99712571 | 0.003230753 | 1.40956723 | 0.001262636 | 10.39636487 | 2.656574608 | 2.490696277 |
| ENSMUSG00000027168 | *Pax6* | 202.670918 | 162.1033366 | 192.8669452 | 512.5625016 | 480.059602 | 486.7036046 | 9.88E-22 | 1.405696708 | 6.42E-23 | 97.15365919 | 2.64945699 | 21.00517114 |
| ENSMUSG00000039046 | *Usp6nl* | 366.1638147 | 356.8191789 | 289.9307019 | 849.5147242 | 954.8553927 | 888.92281 | 1.17E-26 | 1.40428153 | 5.89E-28 | 120.1400605 | 2.646859341 | 25.93111393 |
| ENSMUSG00000040204 | *Pclaf* | 1955.887372 | 1656.523446 | 1927.408884 | 4947.819608 | 4891.133577 | 4781.633338 | 1.70E-55 | 1.400047426 | 2.69E-57 | 254.5135714 | 2.639102576 | 54.76961796 |
| ENSMUSG00000021990 | *Spata13* | 577.122391 | 631.1479025 | 482.7976471 | 1427.93108 | 1495.975207 | 1533.575509 | 7.79E-31 | 1.394676341 | 3.21E-32 | 139.6275977 | 2.62929559 | 30.10870505 |
| ENSMUSG00000098549 | *Gm5269* | 130.3422632 | 90.16398607 | 83.19750577 | 241.4641335 | 245.2936124 | 317.734806 | 1.97E-08 | 1.394340381 | 3.62E-09 | 34.81859955 | 2.628683377 | 7.705865551 |
| ENSMUSG00000027254 | *Map1a* | 55.75333803 | 59.46986315 | 35.29591154 | 111.9515528 | 136.8590971 | 150.6026248 | 2.86E-06 | 1.393437909 | 6.78E-07 | 24.67564974 | 2.627039529 | 5.543834907 |
| ENSMUSG00000030259 | *Rassf8* | 742.8755581 | 729.9446106 | 709.6999356 | 1893.298319 | 1951.821277 | 1894.470823 | 5.10E-70 | 1.393276465 | 6.10E-72 | 321.7258559 | 2.626745569 | 69.29205451 |
| ENSMUSG00000040003 | *Magi2* | 20.34243415 | 11.51029609 | 12.60568269 | 43.90256973 | 47.37430283 | 27.54926064 | 0.008031686 | 1.392615743 | 0.003409553 | 8.574223455 | 2.625542853 | 2.095193255 |
| ENSMUSG00000034255 | *Arhgap27* | 2291.914247 | 2089.118741 | 1396.709642 | 4645.989441 | 5365.929368 | 5147.120196 | 1.57E-15 | 1.390105656 | 1.53E-16 | 68.13488345 | 2.620978749 | 14.80466541 |
| ENSMUSG00000031373 | *Car5b* | 487.4649961 | 556.3309779 | 520.6146952 | 1275.369651 | 1491.764158 | 1327.874363 | 4.42E-35 | 1.389600316 | 1.53E-36 | 159.3998843 | 2.620060847 | 34.35500434 |
| ENSMUSG00000032783 | *Troap* | 521.369053 | 531.392003 | 296.2335433 | 1146.954634 | 1196.990718 | 1200.229455 | 4.09E-11 | 1.388703603 | 5.81E-12 | 47.39363335 | 2.618432843 | 10.38849196 |
| ENSMUSG00000040540 | *Ctdsp2-ps* | 44.45198573 | 74.81692461 | 52.94386731 | 159.1468153 | 102.1179417 | 188.253281 | 6.65E-05 | 1.388501034 | 1.92E-05 | 18.26230432 | 2.618065213 | 4.177102006 |
| ENSMUSG00000098404 | *Mrip-ps* | 141.6436155 | 118.9397263 | 112.190576 | 319.3911948 | 340.0422181 | 323.2446582 | 3.43E-16 | 1.388120307 | 3.18E-17 | 71.2281615 | 2.617374397 | 15.46505991 |
| ENSMUSG00000006676 | *Usp19* | 3527.528766 | 3787.846606 | 2853.926562 | 8269.049008 | 9259.044298 | 9097.684172 | 1.92E-31 | 1.387809203 | 7.73E-33 | 142.455105 | 2.616810046 | 30.71748702 |
| ENSMUSG00000029022 | *Miip* | 324.7255229 | 259.9408534 | 209.2543327 | 682.6849592 | 660.0819528 | 743.8300373 | 6.51E-15 | 1.385449645 | 6.71E-16 | 65.21652609 | 2.612533696 | 14.18611045 |
| ENSMUSG00000027364 | *Usp50* | 24.10955158 | 35.49007962 | 39.07761635 | 91.09783218 | 92.6430811 | 70.70976898 | 6.80E-05 | 1.38422529 | 1.97E-05 | 18.21567399 | 2.610317489 | 4.167395822 |
| ENSMUSG00000032601 | *Prkar2a* | 2273.07866 | 2249.303695 | 1863.119902 | 5255.137596 | 6026.01132 | 5396.900159 | 2.13E-37 | 1.383615083 | 6.71E-39 | 170.1944148 | 2.609213654 | 36.67158902 |
| ENSMUSG00000030795 | *Fus* | 4320.883698 | 3905.827141 | 2450.544715 | 8736.611376 | 9987.5558 | 9137.171446 | 2.57E-13 | 1.382920642 | 3.03E-14 | 57.71643284 | 2.607958012 | 12.59065411 |
| ENSMUSG00000046434 | *Hnrnpa1* | 3653.350488 | 2802.757099 | 2153.050604 | 7419.534284 | 7711.483739 | 7331.776565 | 9.96E-16 | 1.382449383 | 9.55E-17 | 69.06067444 | 2.607106257 | 15.00176322 |
| ENSMUSG00000024925 | *Rnaseh2c* | 485.9581491 | 474.7997139 | 504.2273077 | 1262.19888 | 1208.571103 | 1341.648993 | 1.53E-45 | 1.381735993 | 3.46E-47 | 208.1612016 | 2.605817401 | 44.81594707 |
| ENSMUSG00000025855 | *Prkar1b* | 26.36982204 | 21.1022095 | 23.95079712 | 52.68308367 | 78.95717139 | 55.09852128 | 0.000399212 | 1.381398333 | 0.000130568 | 14.63356284 | 2.605207586 | 3.398796799 |
| ENSMUSG00000040841 | *Six5* | 100.9587473 | 104.5518562 | 70.59182308 | 235.9763123 | 231.6077027 | 257.1264326 | 9.83E-11 | 1.379461653 | 1.45E-11 | 45.6035929 | 2.601712691 | 10.00761698 |
| ENSMUSG00000015127 | *Unkl* | 166.5065906 | 160.184954 | 137.4019413 | 397.318256 | 414.7883404 | 402.2192053 | 3.44E-20 | 1.379026993 | 2.48E-21 | 89.91814109 | 2.600928956 | 19.46283242 |
| ENSMUSG00000020897 | *Aurkb* | 1233.354248 | 1225.846534 | 859.7075596 | 2710.983681 | 3040.37748 | 2894.508985 | 2.02E-20 | 1.378707652 | 1.43E-21 | 91.00914301 | 2.600353304 | 19.69479479 |
| ENSMUSG00000032815 | *Fanca* | 681.8482557 | 640.7398159 | 485.3187837 | 1453.175058 | 1702.316615 | 1558.369844 | 8.56E-21 | 1.378095954 | 5.91E-22 | 92.75859392 | 2.599250997 | 20.06765062 |
| ENSMUSG00000003545 | *Fosb* | 64.04099639 | 69.06177656 | 42.85932115 | 156.9516868 | 157.9143428 | 146.9293901 | 8.21E-08 | 1.375302559 | 1.61E-08 | 31.9134885 | 2.594223107 | 7.085654712 |
| ENSMUSG00000061046 | *Haghl* | 311.9173236 | 282.9614456 | 238.2474029 | 725.4899647 | 714.8255916 | 732.810333 | 1.94E-24 | 1.374583906 | 1.10E-25 | 109.7757823 | 2.592931163 | 23.71157203 |
| ENSMUSG00000035919 | *Bbs9* | 276.5064197 | 248.4305574 | 265.9799048 | 718.9045793 | 650.6070922 | 683.2216639 | 5.92E-30 | 1.374359541 | 2.56E-31 | 135.506784 | 2.592527946 | 29.22793401 |
| ENSMUSG00000096361 | *Gm5814* | 67.80811383 | 36.44927096 | 37.81704808 | 96.5856534 | 115.8038514 | 158.867403 | 0.000243253 | 1.371616206 | 7.67E-05 | 15.63699377 | 2.587602849 | 3.613941798 |
| ENSMUSG00000039735 | *Fnbp1l* | 1115.820184 | 955.3545758 | 1014.757457 | 2532.080709 | 2788.767293 | 2668.605047 | 3.08E-47 | 1.37085898 | 6.57E-49 | 216.0511515 | 2.586245052 | 46.5114783 |
| ENSMUSG00000040084 | *Bub1b* | 2988.077549 | 2472.795277 | 2037.078323 | 6319.774912 | 6699.779183 | 6372.143986 | 2.95E-23 | 1.369409867 | 1.79E-24 | 104.2399238 | 2.583648606 | 22.52998375 |
| ENSMUSG00000041774 | *Ydjc* | 610.2730244 | 520.8408982 | 417.2480971 | 1344.516198 | 1390.698979 | 1275.530768 | 4.60E-20 | 1.367216583 | 3.34E-21 | 89.33221924 | 2.57972375 | 19.33750792 |
| ENSMUSG00000038384 | *Setd1b* | 780.5467325 | 739.536524 | 412.205824 | 1443.29698 | 1808.645606 | 1737.440038 | 1.92E-09 | 1.365280511 | 3.20E-10 | 39.54890682 | 2.576264128 | 8.717189819 |
| ENSMUSG00000070705 | *Eid2b* | 80.6163131 | 58.51067181 | 59.24670865 | 189.8786141 | 164.2309165 | 161.6223291 | 1.95E-08 | 1.364255316 | 3.58E-09 | 34.84214886 | 2.574434056 | 7.710794267 |
| ENSMUSG00000030409 | *Dmpk* | 1228.080284 | 1154.866375 | 645.4109539 | 2378.421715 | 2759.289949 | 2670.441665 | 3.85E-10 | 1.364117627 | 5.96E-11 | 42.83266085 | 2.574188366 | 9.414560254 |
| ENSMUSG00000020422 | *Tns3* | 3956.22673 | 4265.523894 | 3085.871123 | 8890.27037 | 10778.18028 | 9452.151326 | 2.50E-23 | 1.364084937 | 1.50E-24 | 104.5862771 | 2.574130038 | 22.60245177 |
| ENSMUSG00000059495 | *Arhgef12* | 419.6568822 | 400.9419806 | 368.0859346 | 897.8075509 | 1156.985751 | 1009.221248 | 1.24E-23 | 1.363147027 | 7.38E-25 | 105.996957 | 2.572457115 | 22.90565694 |
| ENSMUSG00000113242 | *Gm48016* | 12.05477579 | 9.591913411 | 21.42966058 | 39.51231275 | 33.68839313 | 34.89573014 | 0.009427832 | 1.359294068 | 0.004064345 | 8.254841621 | 2.565596105 | 2.025588168 |
| ENSMUSG00000041498 | *Kif14* | 379.7254374 | 358.7375616 | 233.2051298 | 711.2216296 | 969.5940647 | 820.9679671 | 1.07E-10 | 1.358559902 | 1.58E-11 | 45.43642107 | 2.564290842 | 9.971560535 |
| ENSMUSG00000002028 | *Kmt2a* | 895.0671025 | 949.5994277 | 448.7623038 | 1858.176264 | 2181.323455 | 1847.63708 | 4.73E-08 | 1.357488167 | 9.05E-09 | 33.0359714 | 2.562386614 | 7.325583009 |
| ENSMUSG00000032782 | *Cntrob* | 208.6983059 | 217.7364344 | 128.5779635 | 429.1476191 | 499.0093232 | 501.3965437 | 4.07E-10 | 1.356053983 | 6.32E-11 | 42.71886617 | 2.559840611 | 9.390257729 |
| ENSMUSG00000028066 | *Pmf1* | 1714.038433 | 1627.747706 | 1187.45531 | 3815.133309 | 3991.021823 | 3795.369808 | 5.98E-22 | 1.354754185 | 3.84E-23 | 98.16821867 | 2.557535357 | 21.22307652 |
| ENSMUSG00000037624 | *Kcnk2* | 30.89036296 | 23.97978353 | 15.12681923 | 72.43924005 | 49.4798274 | 60.60837341 | 0.001424 | 1.354072143 | 0.000516418 | 12.055429 | 2.556326554 | 2.846489895 |
| ENSMUSG00000001151 | *Pcnt* | 765.4782627 | 703.087253 | 402.1212779 | 1384.028511 | 1736.005008 | 1671.321812 | 2.28E-09 | 1.353426831 | 3.84E-10 | 39.19161399 | 2.555183374 | 8.641458843 |
| ENSMUSG00000061143 | *Maml3* | 294.5885834 | 296.3901244 | 202.9514913 | 667.3190598 | 706.4034934 | 662.1005641 | 1.07E-15 | 1.350101746 | 1.03E-16 | 68.90565079 | 2.549301038 | 14.9691771 |
| ENSMUSG00000032657 | *Entrep3* | 16.57531671 | 21.1022095 | 10.08454615 | 42.80500548 | 42.11049141 | 38.5689649 | 0.004231679 | 1.348060761 | 0.001697666 | 9.850705601 | 2.545697083 | 2.373487317 |
| ENSMUSG00000027715 | *Ccna2* | 2490.064624 | 1970.179015 | 1663.950115 | 4906.112167 | 5493.313604 | 5196.708865 | 5.88E-20 | 1.346964847 | 4.31E-21 | 88.82678598 | 2.543764032 | 19.23063421 |
| ENSMUSG00000035279 | *Ssc5d* | 40.6848683 | 39.32684499 | 28.99307019 | 88.9027037 | 111.5928022 | 79.89285586 | 2.90E-05 | 1.346785053 | 7.96E-06 | 19.94794405 | 2.543447038 | 4.53763114 |
| ENSMUSG00000037465 | *Klf10* | 296.0954304 | 226.3691565 | 320.1843404 | 569.6358422 | 677.9789117 | 889.8411187 | 2.69E-10 | 1.345584001 | 4.12E-11 | 43.5580573 | 2.541330484 | 9.569761405 |
| ENSMUSG00000056394 | *Lig1* | 2456.913991 | 2503.4894 | 2020.690936 | 5553.67507 | 6401.847456 | 5792.691204 | 6.04E-34 | 1.3448883 | 2.20E-35 | 154.105088 | 2.54010529 | 33.21899366 |
| ENSMUSG00000002055 | *Spag5* | 1333.559572 | 1101.15166 | 864.7498327 | 2734.03253 | 2780.345195 | 2882.570972 | 1.66E-18 | 1.344692637 | 1.34E-19 | 82.02776334 | 2.539760817 | 17.78064777 |
| ENSMUSG00000012443 | *Kif11* | 954.587558 | 751.0468201 | 742.4747106 | 2077.689112 | 2077.099989 | 2067.112857 | 1.03E-28 | 1.342050226 | 4.68E-30 | 129.7362833 | 2.5351133 | 27.98774634 |
| ENSMUSG00000017716 | *Birc5* | 1326.778761 | 1073.335111 | 932.8205192 | 2747.203301 | 2915.098768 | 2777.883781 | 6.23E-23 | 1.33741635 | 3.85E-24 | 102.725753 | 2.526983685 | 22.20584851 |
| ENSMUSG00000040990 | *Sh3kbp1* | 226.0270461 | 217.7364344 | 199.1697865 | 519.147887 | 502.16761 | 607.0020428 | 7.64E-20 | 1.337343558 | 5.65E-21 | 88.29138447 | 2.526856187 | 19.11713605 |
| ENSMUSG00000108312 | *Gm44521* | 21.09585763 | 17.26544414 | 10.08454615 | 34.02449154 | 54.74363883 | 35.81403883 | 0.009404816 | 1.332838323 | 0.004053778 | 8.259567809 | 2.518977652 | 2.026649695 |
| ENSMUSG00000037313 | *Tacc3* | 1676.367258 | 1498.256875 | 1179.8919 | 3403.546718 | 3793.102514 | 3777.003634 | 2.49E-22 | 1.331233115 | 1.58E-23 | 99.93063125 | 2.516176482 | 21.60443494 |
| ENSMUSG00000030091 | *Nup210* | 2150.270632 | 2072.812488 | 1264.349974 | 4194.890537 | 4946.929978 | 4669.599679 | 2.63E-12 | 1.330243226 | 3.36E-13 | 52.98629611 | 2.514450628 | 11.58073795 |
| ENSMUSG00000066357 | *Wdr6* | 1712.531586 | 1928.933787 | 1448.392941 | 4063.182828 | 4421.601598 | 4312.377599 | 7.99E-30 | 1.328795056 | 3.49E-31 | 134.8882553 | 2.511927901 | 29.09744181 |
| ENSMUSG00000045282 | *Tmem86b* | 30.13693948 | 27.81654889 | 21.42966058 | 70.24411156 | 65.27126168 | 66.11822554 | 0.000120707 | 1.327007259 | 3.63E-05 | 17.05531538 | 2.508817033 | 3.918266642 |
| ENSMUSG00000046057 | *Gm15428* | 468.6294089 | 375.0438144 | 470.1919644 | 1078.905651 | 1094.872777 | 1117.581673 | 2.29E-27 | 1.325614593 | 1.11E-28 | 123.4465104 | 2.506396384 | 26.6398321 |
| ENSMUSG00000079037 | *Prnp* | 134.8628042 | 181.2871635 | 151.2681923 | 358.9035075 | 404.2607175 | 404.9741314 | 2.73E-14 | 1.325086221 | 2.95E-15 | 62.29900432 | 2.505478609 | 13.56327827 |
| ENSMUSG00000001435 | *Col18a1* | 212.4654233 | 252.2673227 | 209.2543327 | 481.8307028 | 595.8634534 | 610.6752775 | 2.99E-16 | 1.324914139 | 2.77E-17 | 71.50123606 | 2.505179779 | 15.52388056 |
| ENSMUSG00000039765 | *Cc2d2a* | 51.23279711 | 74.81692461 | 44.11988942 | 151.4638656 | 118.9621382 | 157.0307856 | 7.82E-06 | 1.323844043 | 1.97E-06 | 22.62512825 | 2.50332229 | 5.106562467 |
| ENSMUSG00000038236 | *Hoxa7* | 161.2326262 | 150.5930406 | 155.0498971 | 396.2206918 | 386.3637587 | 386.6079577 | 1.31E-20 | 1.322384584 | 9.19E-22 | 91.88417304 | 2.50079116 | 19.88133434 |
| ENSMUSG00000015961 | *Adss* | 1848.147813 | 1842.606566 | 1873.204448 | 4442.940056 | 4959.563126 | 4507.977349 | 1.16E-66 | 1.322165967 | 1.48E-68 | 306.178846 | 2.500412235 | 65.93666864 |
| ENSMUSG00000052056 | *Zfp217* | 525.1361704 | 564.9636999 | 354.2196837 | 1148.052198 | 1265.420267 | 1199.311147 | 6.41E-14 | 1.3181106 | 7.12E-15 | 60.56435799 | 2.49339353 | 13.19317585 |
| ENSMUSG00000041912 | *Tdrkh* | 877.7383623 | 796.1288131 | 951.7290433 | 2117.201425 | 2304.496642 | 2115.783217 | 1.81E-39 | 1.317906665 | 5.24E-41 | 179.8448307 | 2.493041096 | 38.74146277 |
| ENSMUSG00000027261 | *Hao1* | 78.35604264 | 66.18420254 | 86.97921058 | 207.439642 | 173.7057771 | 193.7631332 | 1.57E-09 | 1.317649587 | 2.60E-10 | 39.95385656 | 2.492596894 | 8.802927364 |
| ENSMUSG00000028873 | *Cdca8* | 2015.407828 | 1776.422364 | 1531.590447 | 4186.110023 | 4676.370071 | 4416.146481 | 1.56E-29 | 1.316688305 | 6.84E-31 | 133.5536185 | 2.490936604 | 28.80821951 |
| ENSMUSG00000020898 | *Ctc1* | 603.492213 | 595.6578228 | 422.2903702 | 1320.369785 | 1379.118594 | 1349.913771 | 1.38E-18 | 1.315326321 | 1.11E-19 | 82.39979858 | 2.488586132 | 17.85869812 |
| ENSMUSG00000021338 | *Carmil1* | 34.6574804 | 38.36765364 | 45.38045769 | 84.51244672 | 95.80136795 | 111.1153512 | 1.26E-05 | 1.315001749 | 3.27E-06 | 21.64884423 | 2.488026322 | 4.899573185 |
| ENSMUSG00000024590 | *Lmnb1* | 3670.679228 | 3502.007586 | 2458.108125 | 7265.87529 | 8650.547697 | 8036.119329 | 3.35E-17 | 1.31353302 | 2.90E-18 | 75.95517417 | 2.485494688 | 16.47556381 |
| ENSMUSG00000034906 | *Ncaph* | 821.2316008 | 752.0060114 | 615.1573159 | 1770.371124 | 1893.919351 | 1791.62025 | 3.20E-26 | 1.313466946 | 1.64E-27 | 118.1107792 | 2.485380858 | 25.49503823 |
| ENSMUSG00000037568 | *Vash2* | 21.09585763 | 22.06140085 | 18.90852404 | 40.609877 | 65.27126168 | 48.67036046 | 0.001828264 | 1.311055896 | 0.00068127 | 11.53966131 | 2.481230727 | 2.737961084 |
| ENSMUSG00000034023 | *Fancd2* | 525.1361704 | 474.7997139 | 380.6916173 | 1032.807953 | 1284.369988 | 1117.581673 | 2.37E-17 | 1.309813002 | 2.03E-18 | 76.65921569 | 2.479094046 | 16.62590885 |
| ENSMUSG00000111497 | *Gm38431* | 143.903886 | 117.980535 | 108.4088712 | 339.1473511 | 368.4667998 | 213.0476156 | 7.25E-07 | 1.306330486 | 1.60E-07 | 27.46397799 | 2.473116988 | 6.139546885 |
| ENSMUSG00000027412 | *Lpin3* | 34.6574804 | 50.83714108 | 60.50727692 | 113.049117 | 132.6480479 | 109.2787339 | 2.27E-05 | 1.301620162 | 6.15E-06 | 20.44114741 | 2.465055556 | 4.64325922 |
| ENSMUSG00000027330 | *Cdc25b* | 1414.929308 | 1276.683675 | 920.2148365 | 2897.569602 | 2942.470587 | 3071.742561 | 4.74E-17 | 1.300384368 | 4.15E-18 | 75.25031462 | 2.462944925 | 16.32463491 |
| ENSMUSG00000034303 | *Ccdc15* | 29.38351599 | 15.34706146 | 22.69022885 | 54.87821216 | 69.48231082 | 43.16050834 | 0.003744921 | 1.30008821 | 0.001485249 | 10.09682675 | 2.462439382 | 2.426557364 |
| ENSMUSG00000034832 | *Tet3* | 1373.491017 | 1485.787387 | 665.5800462 | 2657.203033 | 3145.653708 | 2871.551267 | 0.000817402 | 1.298451148 | 0.000283331 | 13.17749714 | 2.459646775 | 3.087564232 |
| ENSMUSG00000022303 | *Dcstamp* | 388.7665193 | 268.5735755 | 344.1351375 | 844.026903 | 687.4537722 | 933.001627 | 2.57E-12 | 1.29801262 | 3.29E-13 | 53.02793986 | 2.458899244 | 11.58924594 |
| ENSMUSG00000045658 | *Pid1* | 1859.449166 | 1810.953252 | 1419.399871 | 3907.328706 | 4478.450761 | 4134.225713 | 1.01E-25 | 1.296695376 | 5.33E-27 | 115.7734455 | 2.456655185 | 24.99695739 |
| ENSMUSG00000032174 | *Icam5* | 51.9862206 | 32.6125056 | 36.55647981 | 96.5856534 | 109.4872777 | 94.58579486 | 3.46E-05 | 1.295212883 | 9.60E-06 | 19.58979717 | 2.454132057 | 4.461216865 |
| ENSMUSG00000024579 | *Pcyox1l* | 415.1363413 | 359.6967529 | 258.4164952 | 758.416892 | 894.8479424 | 887.0861926 | 4.47E-12 | 1.291236742 | 5.85E-13 | 51.89738482 | 2.447377658 | 11.34922868 |
| ENSMUSG00000027496 | *Aurka* | 2516.434446 | 2345.222829 | 1916.063769 | 5516.357886 | 5673.335955 | 5391.390307 | 1.99E-30 | 1.288924345 | 8.41E-32 | 137.7167166 | 2.443458066 | 29.70222255 |
| ENSMUSG00000021775 | *Nr1d2* | 169.5202846 | 112.2253869 | 114.7117125 | 320.488759 | 324.2507838 | 330.5911277 | 1.01E-09 | 1.287362707 | 1.64E-10 | 40.85909812 | 2.440814587 | 8.996686287 |
| ENSMUSG00000030583 | *Sipa1l3* | 824.9987182 | 892.0479472 | 504.2273077 | 1632.07803 | 1928.660506 | 1869.676489 | 2.20E-10 | 1.286893032 | 3.34E-11 | 43.96736038 | 2.440020099 | 9.657259747 |
| ENSMUSG00000052911 | *Lamb2* | 174.794249 | 140.0419358 | 158.8316019 | 362.1962002 | 398.9969061 | 395.7910445 | 8.37E-16 | 1.285332669 | 7.97E-17 | 69.41674116 | 2.437382495 | 15.07736174 |
| ENSMUSG00000059273 | *Zc3h4* | 1550.545536 | 1564.441077 | 1131.990306 | 3254.277981 | 3656.243416 | 3444.575889 | 1.41E-20 | 1.283752721 | 9.90E-22 | 91.73675097 | 2.434714689 | 19.84983213 |
| ENSMUSG00000040274 | *Cdk6* | 2025.955756 | 1751.483389 | 1304.688159 | 4092.817063 | 4387.913205 | 3901.893615 | 2.77E-16 | 1.282854376 | 2.56E-17 | 71.65666886 | 2.4331991 | 15.55680189 |
| ENSMUSG00000030330 | *Ing4* | 395.5473306 | 343.3905001 | 344.1351375 | 891.2221655 | 878.0037458 | 872.3932536 | 2.33E-30 | 1.281280877 | 9.87E-32 | 137.3974767 | 2.430546739 | 29.63310693 |
| ENSMUSG00000078762 | *Haus5* | 469.3828324 | 416.289042 | 378.1704808 | 1016.344489 | 1046.445711 | 1022.07757 | 2.12E-28 | 1.281098817 | 9.77E-30 | 128.2752563 | 2.430240037 | 27.67392156 |
| ENSMUSG00000025786 | *Zdhhc3* | 3280.405862 | 3475.150229 | 2976.201684 | 7340.509658 | 8461.050486 | 7829.499874 | 2.53E-42 | 1.279273533 | 6.56E-44 | 193.1395225 | 2.427167264 | 41.59690279 |
| ENSMUSG00000037845 | *Fdxacb1* | 120.5477579 | 125.6540657 | 121.0145538 | 312.8058093 | 278.9820056 | 298.4503236 | 5.87E-15 | 1.277467564 | 6.03E-16 | 65.42787553 | 2.424130832 | 14.23123785 |
| ENSMUSG00000059540 | *Tcea2* | 30.13693948 | 18.22463548 | 28.99307019 | 46.09769821 | 76.85164682 | 64.28160816 | 0.002359913 | 1.275997685 | 0.000898004 | 11.02687426 | 2.421662282 | 2.627103933 |
| ENSMUSG00000028551 | *Cdkn2c* | 579.3826615 | 616.7600323 | 547.0866289 | 1301.711192 | 1507.555592 | 1412.358762 | 2.24E-33 | 1.274575522 | 8.41E-35 | 151.4373328 | 2.419276261 | 32.65045696 |
| ENSMUSG00000070713 | *Hmgn2-ps* | 537.1909462 | 480.5548619 | 467.6708279 | 1099.759372 | 1226.468062 | 1273.69415 | 6.44E-29 | 1.274140467 | 2.91E-30 | 130.678216 | 2.418546821 | 28.19120394 |
| ENSMUSG00000059791 | *Nrm* | 482.9444551 | 443.1463996 | 273.5433144 | 934.0271709 | 999.0714086 | 977.9987527 | 3.76E-10 | 1.273027102 | 5.81E-11 | 42.88226813 | 2.416681085 | 9.424609958 |
| ENSMUSG00000061331 | *Gm17132* | 14.31504625 | 19.18382682 | 15.12681923 | 43.90256973 | 27.37181941 | 45.9154344 | 0.009716871 | 1.272416842 | 0.004196285 | 8.196864315 | 2.415659046 | 2.012473575 |
| ENSMUSG00000038178 | *Slc43a2* | 1814.99718 | 1735.177136 | 1314.772705 | 3417.815053 | 4539.510974 | 3800.87966 | 3.06E-16 | 1.271683212 | 2.84E-17 | 71.45505625 | 2.414430962 | 15.51435952 |
| ENSMUSG00000029192 | *Tbc1d14* | 2394.379842 | 2558.163307 | 2572.819838 | 5785.261126 | 6213.403007 | 6149.913284 | 5.96E-71 | 1.271037398 | 7.03E-73 | 326.0313332 | 2.413350398 | 70.22482536 |
| ENSMUSG00000079260 | *Tmppe* | 253.1502916 | 257.0632794 | 157.5710337 | 489.5136525 | 563.2278226 | 567.5147692 | 8.41E-10 | 1.270695043 | 1.36E-10 | 41.22738781 | 2.412777772 | 9.07502116 |
| ENSMUSG00000050796 | *B3galt6* | 255.4105621 | 262.8184275 | 235.7262663 | 579.5139204 | 667.4512888 | 573.0246213 | 3.32E-21 | 1.267957951 | 2.24E-22 | 94.67472538 | 2.408204572 | 20.47892141 |
| ENSMUSG00000022614 | *Lmf2* | 283.2872311 | 410.533894 | 347.9168423 | 881.3440873 | 992.7548349 | 631.7963773 | 2.78E-09 | 1.267444668 | 4.72E-10 | 38.78965712 | 2.407347931 | 8.555566781 |
| ENSMUSG00000050555 | *Hyls1* | 145.410733 | 138.1235531 | 134.8808048 | 336.9522227 | 371.6250867 | 300.286941 | 6.39E-14 | 1.265424519 | 7.10E-15 | 60.57008712 | 2.403979375 | 13.19417149 |
| ENSMUSG00000039621 | *Prex1* | 2444.859215 | 2415.243797 | 1584.534314 | 4682.209061 | 5584.903923 | 5238.951065 | 7.32E-14 | 1.265411952 | 8.19E-15 | 60.28805292 | 2.403958436 | 13.13568777 |
| ENSMUSG00000016756 | *Cmah* | 443.0130103 | 366.4110923 | 311.3603625 | 858.2952382 | 911.692139 | 933.9199357 | 4.42E-17 | 1.263647265 | 3.86E-18 | 75.39275568 | 2.40101974 | 16.35494373 |
| ENSMUSG00000106964 | *None* | 33.15063343 | 46.04118437 | 74.37352789 | 140.4882231 | 127.3842365 | 95.50410355 | 0.00078009 | 1.262760014 | 0.000269541 | 13.27101659 | 2.399543578 | 3.107855474 |
| ENSMUSG00000039852 | *Rere* | 922.9437715 | 1027.293926 | 697.0942529 | 1839.517672 | 2356.081994 | 2156.188799 | 1.48E-14 | 1.260545102 | 1.57E-15 | 63.54732037 | 2.395862481 | 13.83044612 |
| ENSMUSG00000029516 | *Cit* | 337.5337222 | 287.7574023 | 147.4864875 | 651.9531604 | 688.5065345 | 518.8444087 | 1.09E-05 | 1.259419561 | 2.81E-06 | 21.94115586 | 2.393994041 | 4.960778765 |
| ENSMUSG00000036864 | *Proser3* | 57.26018501 | 48.9187584 | 35.29591154 | 121.829631 | 101.0651794 | 120.2984381 | 1.81E-05 | 1.259371859 | 4.80E-06 | 20.91553032 | 2.393914886 | 4.742625248 |
| ENSMUSG00000032180 | *Tmed1* | 692.3961845 | 699.2504877 | 661.7983414 | 1627.687773 | 1721.266336 | 1563.879696 | 2.70E-45 | 1.256733519 | 6.23E-47 | 206.9892959 | 2.389540996 | 44.56808191 |
| ENSMUSG00000004897 | *Hdgf* | 13629.43088 | 12998.96105 | 11520.33341 | 29228.1358 | 31902.90829 | 29884.51963 | 5.36E-47 | 1.254160682 | 1.16E-48 | 214.9150413 | 2.385283394 | 46.2705673 |
| ENSMUSG00000053641 | *Dennd4a* | 504.7937363 | 577.4331874 | 519.3541269 | 1278.662343 | 1308.58352 | 1231.451951 | 7.36E-36 | 1.253830177 | 2.47E-37 | 163.0282241 | 2.384737015 | 35.13284532 |
| ENSMUSG00000066306 | *Numa1* | 3293.967485 | 3443.496915 | 2148.008331 | 6224.286823 | 7833.604164 | 7074.650132 | 2.94E-12 | 1.249141084 | 3.78E-13 | 52.75552076 | 2.376998651 | 11.53157071 |
| ENSMUSG00000038034 | *Igsf8* | 3765.610588 | 3642.049522 | 2958.553728 | 7823.437925 | 8315.769291 | 8473.234264 | 8.51E-31 | 1.246462831 | 3.52E-32 | 139.4465262 | 2.372590027 | 30.06982999 |
| ENSMUSG00000047888 | *Tnrc6b* | 210.2051529 | 181.2871635 | 119.7539856 | 454.3915967 | 424.2632009 | 342.5291406 | 1.67E-07 | 1.245172044 | 3.41E-08 | 30.45722222 | 2.370468208 | 6.777401969 |

| | | | | | | | | | | | | | |
|---|---|---|---|---|---|---|---|---|---|---|---|---|---|
| ENSMUSG00000072915 | *Gm12258* | 21.84928112 | 23.02059219 | 20.16909231 | 59.26846913 | 50.53258969 | 44.99712571 | 0.001529027 | 1.243982236 | 0.000559062 | 11.90757182 | 2.36851406 | 2.815584862 |
| ENSMUSG00000027306 | *Nusap1* | 519.862206 | 361.6151356 | 354.2196837 | 1016.344489 | 999.0714086 | 921.0636141 | 4.63E-13 | 1.242887474 | 5.56E-14 | 56.52252792 | 2.366717439 | 12.33477074 |
| ENSMUSG00000008398 | *Elk3* | 1897.12034 | 1828.218696 | 1534.111584 | 4018.182694 | 4263.687255 | 4180.141148 | 6.02E-35 | 1.242473698 | 2.11E-36 | 158.7645008 | 2.366038744 | 34.22045464 |
| ENSMUSG00000051212 | *Gpr183* | 667.5332094 | 658.00526 | 555.9106067 | 1384.028511 | 1586.512764 | 1484.905149 | 2.24E-27 | 1.23999004 | 1.08E-28 | 123.5004162 | 2.361969017 | 26.64978679 |
| ENSMUSG00000044042 | *Fmn1* | 683.3551027 | 775.985795 | 736.1718692 | 1562.931482 | 1917.080121 | 1693.361221 | 5.81E-27 | 1.237730892 | 2.89E-28 | 121.5571724 | 2.358273251 | 26.23597634 |
| ENSMUSG00000021453 | *Gadd45g* | 229.7941635 | 236.9202613 | 200.4303548 | 479.6355743 | 591.6524043 | 504.1514697 | 7.34E-16 | 1.235083576 | 6.97E-17 | 69.68078773 | 2.353949836 | 15.13456128 |
| ENSMUSG00000002897 | *Il17ra* | 1291.367857 | 1321.765668 | 939.1233606 | 2687.934832 | 2916.15153 | 2770.537312 | 1.10E-18 | 1.234824765 | 8.70E-20 | 82.88418547 | 2.353527588 | 17.95989704 |
| ENSMUSG00000032288 | *Imp3* | 1423.216967 | 1344.78626 | 1211.406107 | 3105.009244 | 3191.975249 | 3085.517192 | 2.49E-45 | 1.234641356 | 5.70E-47 | 207.1660948 | 2.353228405 | 44.60406413 |
| ENSMUSG00000006241 | *Ccdc159* | 88.15054797 | 74.81692461 | 78.15523269 | 206.3420777 | 178.9695885 | 184.5800463 | 2.79E-09 | 1.234608654 | 4.74E-10 | 38.78256171 | 2.353175064 | 8.554297224 |
| ENSMUSG00000035545 | *Leng8* | 853.6288107 | 790.3736651 | 506.7484442 | 1632.07803 | 1703.369377 | 1736.521729 | 2.65E-11 | 1.234014915 | 3.70E-12 | 48.27628173 | 2.352206818 | 10.5770319 |
| ENSMUSG00000034173 | *Zbed5* | 54.99991455 | 60.42905449 | 78.15523269 | 140.4882231 | 158.9671051 | 150.6026248 | 5.11E-07 | 1.233588127 | 1.11E-07 | 28.18036411 | 2.351511075 | 6.291763423 |
| ENSMUSG00000040249 | *Lrp1* | 10521.559 | 9906.528171 | 5404.05617 | 16987.00179 | 23833.48537 | 19875.87324 | 0.000218203 | 1.232313411 | 6.84E-05 | 15.85538238 | 2.349434278 | 3.66113978 |
| ENSMUSG00000037411 | *Serpine1* | 2086.229635 | 2113.098524 | 2008.085253 | 4598.794179 | 5083.789075 | 4885.40222 | 2.22E-62 | 1.230200248 | 3.08E-64 | 286.3667883 | 2.345995504 | 61.65361499 |
| ENSMUSG00000020415 | *Pttg1* | 900.3410669 | 770.2306469 | 684.4885702 | 1882.322677 | 1814.96218 | 1839.372302 | 4.57E-25 | 1.228946203 | 2.50E-26 | 112.7085739 | 2.343957162 | 24.34010474 |
| ENSMUSG00000033752 | *Mnd1* | 84.38343054 | 74.81692461 | 78.15523269 | 201.9518207 | 187.3916868 | 168.9687986 | 4.42E-09 | 1.22888593 | 7.65E-10 | 37.84613787 | 2.343859237 | 8.354798284 |
| ENSMUSG00000047534 | *Mis18bp1* | 299.1091243 | 238.8386439 | 207.9937644 | 542.1967361 | 609.5493631 | 606.0837341 | 3.63E-13 | 1.228277219 | 4.31E-14 | 57.02122097 | 2.34287051 | 12.44041593 |
| ENSMUSG00000022070 | *Bora* | 407.6021064 | 379.8397711 | 346.656274 | 839.636646 | 909.5866144 | 914.6354533 | 1.54E-26 | 1.227101131 | 7.82E-28 | 119.5799832 | 2.340961375 | 25.81144868 |
| ENSMUSG00000049904 | *Tmem17* | 42.19171527 | 47.00037571 | 45.38045769 | 97.68321764 | 108.4345154 | 107.4421165 | 4.30E-06 | 1.22464158 | 1.04E-06 | 23.84617751 | 2.336973832 | 5.366382443 |
| ENSMUSG00000100774 | *Gm7329* | 101.7121707 | 107.4294302 | 139.9230779 | 282.0740105 | 258.9795222 | 267.2278282 | 2.99E-10 | 1.22419624 | 4.59E-11 | 43.34553932 | 2.336252552 | 9.524153025 |
| ENSMUSG00000109645 | *Gm45246* | 148.4244269 | 147.7154665 | 171.4372846 | 384.1474851 | 369.5195621 | 334.2643624 | 1.56E-14 | 1.223533697 | 1.66E-15 | 63.43564813 | 2.335179898 | 13.80638825 |
| ENSMUSG00000034379 | *Wdr5b* | 34.6574804 | 42.20441901 | 35.29591154 | 85.61001097 | 82.11545824 | 93.66748618 | 3.96E-05 | 1.22225758 | 1.11E-05 | 19.31449929 | 2.333115259 | 4.40221158 |
| ENSMUSG00000042507 | *Mideas* | 1007.327202 | 1081.008641 | 673.1434558 | 2037.079235 | 2226.592233 | 2190.166221 | 1.30E-12 | 1.222123495 | 1.61E-13 | 54.42715859 | 2.332898427 | 11.88719606 |
| ENSMUSG00000032418 | *Me1* | 1429.244355 | 1785.055086 | 2008.085253 | 3962.206918 | 4072.084519 | 4118.614466 | 3.87E-21 | 1.220601099 | 2.63E-22 | 94.36253294 | 2.330437947 | 20.41262585 |
| ENSMUSG00000026478 | *Lamc1* | 3367.802987 | 3243.025924 | 1940.014566 | 5909.285885 | 7417.763061 | 6570.498663 | 9.03E-10 | 1.217576821 | 1.46E-10 | 41.08333482 | 2.325557839 | 9.044491315 |
| ENSMUSG00000040721 | *Zfhx2* | 157.4655088 | 169.7768674 | 84.45807404 | 253.5373402 | 356.8864147 | 351.7122275 | 2.93E-05 | 1.216166231 | 8.04E-06 | 19.9275065 | 2.323285144 | 4.533205901 |
| ENSMUSG00000057110 | *Cntrl* | 627.6017646 | 503.5754541 | 434.8960529 | 1265.491572 | 1167.513374 | 1218.595629 | 3.52E-16 | 1.215801123 | 3.27E-17 | 71.17257196 | 2.322697255 | 15.45346522 |
| ENSMUSG00000032403 | *2300009A05Rik* | 152.9449679 | 150.5930406 | 132.3596683 | 294.1472172 | 343.200505 | 377.4248708 | 7.33E-12 | 1.215388419 | 9.79E-13 | 50.88485525 | 2.322032909 | 11.13508124 |
| ENSMUSG00000007880 | *Arid1a* | 2652.050674 | 2466.080938 | 1379.061687 | 4518.671989 | 5457.519686 | 5074.57381 | 2.98E-08 | 1.210832614 | 5.55E-09 | 33.98703771 | 2.314711859 | 7.526430483 |
| ENSMUSG00000086699 | *Gm16510* | 178.5613664 | 151.5522319 | 157.5710337 | 362.1962002 | 390.5748078 | 377.4248708 | 2.08E-15 | 1.207483619 | 2.06E-16 | 67.54107589 | 2.309344842 | 14.68175872 |
| ENSMUSG00000033589 | *Reep4* | 664.5195155 | 579.35157 | 438.6777577 | 1162.320534 | 1478.078248 | 1254.409668 | 4.75E-12 | 1.206569583 | 6.23E-13 | 51.77270887 | 2.307882194 | 11.32362455 |
| ENSMUSG00000080316 | *Spaca6* | 18.08216369 | 23.97978353 | 21.42966058 | 50.48795519 | 48.42706512 | 46.83374309 | 0.002441378 | 1.205824077 | 0.000931852 | 10.95830191 | 2.306689914 | 2.612365008 |
| ENSMUSG00000042524 | *Sun2* | 1725.339785 | 1670.911316 | 1195.018719 | 3202.692462 | 3781.522128 | 3608.953144 | 1.94E-15 | 1.204343288 | 1.91E-16 | 67.69097611 | 2.304323531 | 14.71236618 |
| ENSMUSG00000061119 | *Prcp* | 585.4100494 | 560.1677432 | 468.9313962 | 1213.906053 | 1263.314742 | 1255.327977 | 4.37E-26 | 1.203808144 | 2.26E-27 | 117.4752217 | 2.303468938 | 25.35992687 |
| ENSMUSG00000096210 | *H1f0* | 2300.201906 | 2088.15955 | 1699.246027 | 4984.039228 | 4531.088875 | 4512.568893 | 2.84E-22 | 1.202506841 | 1.81E-23 | 99.66452018 | 2.301392159 | 21.54655094 |
| ENSMUSG00000034908 | *Sidt2* | 1213.765237 | 1242.152787 | 978.2009769 | 2554.031994 | 2760.342712 | 2595.140352 | 3.00E-26 | 1.201001887 | 1.54E-27 | 118.2386693 | 2.298992702 | 25.5224586 |
| ENSMUSG00000032485 | *Scap* | 2675.406802 | 2726.021791 | 2197.170493 | 5529.528657 | 6137.604123 | 5811.975686 | 7.98E-30 | 1.200574365 | 3.48E-31 | 134.8928789 | 2.29831153 | 29.09777083 |
| ENSMUSG00000048537 | *Phldb1* | 1278.559657 | 1306.418607 | 987.0249548 | 2410.251078 | 3025.638808 | 2765.02746 | 2.52E-17 | 1.197154441 | 2.17E-18 | 76.53172396 | 2.292869811 | 16.59856558 |
| ENSMUSG00000031295 | *Phka2* | 923.697195 | 830.6597014 | 781.5523269 | 1859.273828 | 2082.3638 | 1879.777884 | 1.97E-30 | 1.195695737 | 8.34E-32 | 137.731083 | 2.29055267 | 29.70466007 |
| ENSMUSG00000021559 | *Dapk1* | 76.84919567 | 77.69449863 | 63.02841346 | 127.3174522 | 188.444449 | 183.6617376 | 5.46E-06 | 1.193586651 | 1.35E-06 | 23.35645104 | 2.28720654 | 5.262679045 |
| ENSMUSG00000048897 | *Zfp710* | 3442.391912 | 3576.824511 | 2490.8829 | 6663.31252 | 7534.619675 | 7543.905872 | 1.30E-16 | 1.192080302 | 1.17E-17 | 73.20118827 | 2.284819665 | 15.88756442 |
| ENSMUSG00000008153 | *Clstn3* | 121.3011814 | 112.2253869 | 112.190576 | 256.8300329 | 276.876481 | 257.1264326 | 2.67E-12 | 1.189848505 | 3.43E-13 | 52.94747575 | 2.281287865 | 11.57308554 |
| ENSMUSG00000026683 | *Nuf2* | 805.4097075 | 647.4541553 | 692.0519798 | 1650.736622 | 1634.939829 | 1611.631747 | 2.26E-27 | 1.188099921 | 1.10E-28 | 123.4740021 | 2.278524559 | 26.64523561 |
| ENSMUSG00000023505 | *Cdca3* | 1426.984084 | 1140.478505 | 858.4469914 | 2532.080709 | 2668.752393 | 2606.160057 | 5.54E-12 | 1.185574377 | 7.32E-13 | 51.45684573 | 2.274539323 | 11.25646975 |
| ENSMUSG00000078429 | *Ctdsp2* | 1009.587473 | 754.8835855 | 572.2979942 | 1627.687773 | 1962.3489 | 1734.685112 | 2.28E-09 | 1.184810711 | 3.84E-10 | 39.19412777 | 2.273335654 | 8.641590597 |
| ENSMUSG00000094627 | *Gm10182* | 872.4643979 | 888.2111819 | 772.728349 | 1941.591146 | 1957.085088 | 1872.431415 | 7.83E-39 | 1.184571143 | 2.31E-40 | 176.8899396 | 2.272958184 | 38.10645334 |
| ENSMUSG00000050821 | *Fam131a* | 80.6163131 | 80.57207265 | 55.46500385 | 174.5127147 | 153.7032936 | 168.9687986 | 8.64E-07 | 1.183911464 | 1.92E-07 | 27.10890106 | 2.271919101 | 6.063291013 |
| ENSMUSG00000035413 | *Tmem98* | 791.8480848 | 716.5159318 | 626.5024298 | 1823.054208 | 1378.065831 | 1658.465491 | 2.70E-16 | 1.183829805 | 2.49E-17 | 71.70844333 | 2.27179051 | 15.5678743 |
| ENSMUSG00000032178 | *Ilf3* | 2225.61298 | 2148.588604 | 1929.93002 | 4481.354805 | 5010.095715 | 4819.283995 | 1.02E-39 | 1.181305677 | 2.90E-41 | 181.0202637 | 2.267819279 | 38.99291541 |
| ENSMUSG00000048922 | *Cdca2* | 1001.299814 | 1010.028482 | 811.8059654 | 2057.932956 | 2298.180069 | 2044.155139 | 6.01E-24 | 1.177814239 | 3.51E-25 | 107.472434 | 2.262337609 | 23.22090179 |
| ENSMUSG00000024660 | *Incenp* | 2804.242218 | 2959.105287 | 2014.388094 | 5334.162222 | 6368.159063 | 5887.276999 | 1.12E-14 | 1.176309751 | 1.18E-15 | 64.10801196 | 2.259979602 | 13.95009486 |
| ENSMUSG00000020808 | *Pimreg* | 1253.696682 | 1104.029234 | 741.2141423 | 2236.835928 | 2333.973986 | 2436.272949 | 8.38E-11 | 1.17432285 | 1.23E-11 | 45.92987131 | 2.256869266 | 10.07670567 |
| ENSMUSG00000052512 | *Nav2* | 817.4644833 | 732.8221846 | 436.1566212 | 1246.83298 | 1724.424623 | 1517.045953 | 3.96E-07 | 1.172880454 | 8.45E-08 | 28.70072275 | 2.254613993 | 6.402527421 |
| ENSMUSG00000030539 | *Sema4b* | 318.698135 | 302.1452725 | 269.7616096 | 714.5143223 | 623.2352728 | 677.7118117 | 1.47E-19 | 1.17260134 | 1.11E-20 | 86.95961877 | 2.254177841 | 18.83349173 |
| ENSMUSG00000027947 | *Il6ra* | 1426.984084 | 1412.888845 | 1062.659051 | 2907.44768 | 3039.324717 | 2863.286489 | 4.29E-20 | 1.172203077 | 3.11E-21 | 89.47146302 | 2.253555651 | 19.36725361 |
| ENSMUSG00000048316 | *Gm9823* | 105.4792882 | 89.20479472 | 41.59875288 | 167.9273292 | 189.4972113 | 180.9068115 | 0.000610231 | 1.171227262 | 0.000207124 | 13.76533053 | 2.252031899 | 3.214505519 |
| ENSMUSG00000040616 | *Tmem51* | 342.0542631 | 417.2482334 | 467.6708279 | 878.0513945 | 1006.440745 | 859.536932 | 1.03E-14 | 1.166763756 | 1.08E-15 | 64.27580908 | 2.245075179 | 13.98537625 |
| ENSMUSG00000034317 | *Trim59* | 527.3964409 | 410.533894 | 339.0928644 | 969.1492267 | 921.1669995 | 986.2635309 | 6.98E-12 | 1.165308656 | 9.31E-13 | 50.9848609 | 2.24281194 | 11.15631555 |
| ENSMUSG00000022033 | *Pbk* | 1251.436412 | 1091.559746 | 878.6160837 | 2409.153514 | 2467.674796 | 2361.889946 | 6.18E-18 | 1.164693559 | 5.12E-19 | 79.38068471 | 2.241855915 | 17.20907037 |
| ENSMUSG00000043290 | *Zfp784* | 36.16432737 | 32.6125056 | 31.51420673 | 74.63436854 | 85.2737451 | 66.11822554 | 0.000324409 | 1.163751078 | 0.000104255 | 15.05803739 | 2.240391838 | 3.48890661 |
| ENSMUSG00000039840 | *Epg5* | 455.8212096 | 476.7180965 | 264.7193365 | 839.636646 | 988.5437858 | 863.2101667 | 1.22E-07 | 1.16363653 | 2.44E-08 | 31.10477723 | 2.240213962 | 6.914085886 |
| ENSMUSG00000051444 | *Bbs12* | 134.8628042 | 117.980535 | 98.324325 | 271.0983681 | 270.5599073 | 251.6165805 | 9.75E-10 | 1.162750875 | 1.58E-10 | 40.92669201 | 2.238839141 | 9.010972474 |
| ENSMUSG00000048895 | *Cdk5r1* | 61.78072593 | 87.28641204 | 49.1621625 | 156.9516868 | 141.0701462 | 147.8476988 | 2.94E-05 | 1.161312225 | 8.08E-06 | 19.91830891 | 2.236607691 | 4.53122514 |
| ENSMUSG00000035683 | *Melk* | 801.6425901 | 673.3523215 | 664.3194779 | 1591.468153 | 1648.625739 | 1548.268448 | 1.56E-26 | 1.158482391 | 7.90E-28 | 119.5595728 | 2.232224904 | 25.8075682 |
| ENSMUSG00000044052 | *Ccr10* | 42.19171527 | 39.32684499 | 22.69022885 | 59.26846913 | 92.6430811 | 83.56609061 | 0.002904548 | 1.157550326 | 0.001124386 | 10.61064242 | 2.230783223 | 2.53692144 |
| ENSMUSG00000042496 | *Prdm10* | 199.657224 | 258.0224708 | 196.64865 | 474.147753 | 468.4792169 | 515.171174 | 6.91E-13 | 1.155159054 | 8.38E-14 | 55.71416059 | 2.227088754 | 12.16040919 |
| ENSMUSG00000040857 | *Erf* | 1529.449679 | 1613.359836 | 1239.138609 | 3119.277579 | 3234.08574 | 3392.232293 | 2.60E-23 | 1.15133787 | 1.57E-24 | 104.5045334 | 2.221197797 | 22.58502905 |
| ENSMUSG00000074886 | *Grk6* | 1903.147728 | 1888.647751 | 1542.935562 | 3888.670114 | 4017.34088 | 3956.992137 | 1.47E-29 | 1.150960758 | 6.44E-31 | 133.6734636 | 2.220617264 | 28.83307832 |
| ENSMUSG00000071054 | *Safb* | 1570.88797 | 1578.828947 | 1009.715184 | 2754.88625 | 3429.899525 | 3061.641166 | 1.66E-10 | 1.150754377 | 2.49E-11 | 44.54161828 | 2.220299623 | 9.779123584 |
| ENSMUSG00000029505 | *Ep400* | 1515.134632 | 1531.828572 | 781.5523269 | 2499.153782 | 3134.073323 | 2866.041415 | 1.09E-06 | 1.148926216 | 2.47E-07 | 26.62490678 | 2.217487875 | 5.961307993 |
| ENSMUSG00000035504 | *Reep6* | 394.0404837 | 371.207049 | 330.2688865 | 794.6365121 | 807.4686727 | 833.8242887 | 3.24E-23 | 1.147242031 | 1.97E-24 | 104.0520608 | 2.214900716 | 22.48978565 |
| ENSMUSG00000055633 | *Zfp580* | 86.643701 | 101.6742822 | 79.41580096 | 208.5372062 | 176.8640639 | 207.5377635 | 6.73E-08 | 1.143490736 | 1.31E-08 | 32.31461164 | 2.209149014 | 7.171812349 |
| ENSMUSG00000020235 | *Fzr1* | 2015.407828 | 1915.505108 | 1624.872499 | 3952.32884 | 4270.003829 | 4064.434253 | 3.92E-29 | 1.143063839 | 1.75E-30 | 131.6883803 | 2.208495418 | 28.40623381 |
| ENSMUSG00000020330 | *Hmmr* | 1130.13523 | 857.517059 | 727.3478914 | 2083.176934 | 1899.183162 | 2021.197422 | 2.26E-12 | 1.141528675 | 2.87E-13 | 53.29692941 | 2.206146619 | 11.64637713 |
| ENSMUSG00000028874 | *Fgr* | 285.5475016 | 249.3897487 | 257.1559269 | 569.6358422 | 599.0217403 | 582.2077082 | 5.84E-20 | 1.140368132 | 4.28E-21 | 88.84134672 | 2.20437265 | 19.23342403 |
| ENSMUSG00000022322 | *Shcbp1* | 322.4652524 | 260.9000448 | 242.0291077 | 582.8066131 | 624.2880351 | 618.021747 | 2.79E-14 | 1.137162269 | 3.02E-15 | 62.25267463 | 2.199479673 | 13.553854 |
| ENSMUSG00000030102 | *Itpr1* | 583.1497789 | 584.1475267 | 466.4102596 | 1141.466813 | 1220.151489 | 1239.716729 | 1.31E-20 | 1.135895218 | 9.13E-22 | 91.89817192 | 2.197548823 | 19.88398015 |
| ENSMUSG00000033031 | *Cip2a* | 606.505907 | 508.3714108 | 437.4171894 | 1201.832846 | 1111.716973 | 1110.235204 | 6.66E-16 | 1.135426942 | 6.31E-17 | 69.87671648 | 2.196835648 | 15.17675832 |
| ENSMUSG00000039936 | *Pik3cd* | 2434.311286 | 2503.4894 | 1929.93002 | 4806.233821 | 5059.575543 | 5235.27783 | 6.79E-23 | 1.13542685 | 4.21E-24 | 102.5505274 | 2.196835509 | 22.1683979 |
| ENSMUSG00000025241 | *Fyco1* | 206.4380354 | 197.5934163 | 141.1836462 | 378.6596639 | 416.8938649 | 410.4839835 | 6.21E-10 | 1.1352756 | 9.86E-11 | 41.84970082 | 2.196605209 | 9.206929985 |
| ENSMUSG00000038400 | *Pmepa1* | 678.0811383 | 714.5975491 | 587.4248135 | 1125.003349 | 1809.698368 | 1408.685527 | 7.92E-10 | 1.132089476 | 1.27E-10 | 41.35483941 | 2.191759462 | 9.101483695 |
| ENSMUSG00000103906 | *Tigd5* | 71.57523126 | 95.91913411 | 55.46500385 | 162.439508 | 153.7032936 | 174.4786507 | 1.93E-05 | 1.129981662 | 5.16E-06 | 20.77589296 | 2.188559584 | 4.713761845 |
| ENSMUSG00000032316 | *Clk3* | 877.7383623 | 861.3538243 | 806.7636923 | 1839.517672 | 1818.120467 | 1921.101775 | 6.97E-42 | 1.129673623 | 1.85E-43 | 191.0767654 | 2.18809234 | 41.15689197 |
| ENSMUSG00000040945 | *Rcc2* | 13841.8963 | 13450.74018 | 9861.42557 | 25420.68544 | 28188.76295 | 27690.68018 | 5.68E-17 | 1.129469532 | 5.00E-18 | 74.88162899 | 2.187782824 | 16.24591199 |
| ENSMUSG00000032480 | *Dhx30* | 3299.241449 | 3524.068987 | 2949.72975 | 6774.166509 | 7413.552012 | 7195.866879 | 2.80E-35 | 1.12886862 | 9.68E-37 | 160.3104621 | 2.186871756 | 34.55223231 |
| ENSMUSG00000041126 | *H2az2* | 370.6843556 | 301.1860811 | 284.8884288 | 657.4409817 | 721.1421654 | 720.8723201 | 4.46E-15 | 1.127369398 | 4.53E-16 | 65.99135851 | 2.184600381 | 14.35073608 |
| ENSMUSG00000042207 | *Kdm5b* | 163.4928967 | 175.5320154 | 173.9584212 | 329.269273 | 413.7355781 | 375.5882534 | 4.14E-12 | 1.127365153 | 5.39E-13 | 52.05765294 | 2.184593952 | 11.38300155 |
| ENSMUSG00000073633 | *Fbxo36* | 24.86297507 | 23.97978353 | 26.47193365 | 57.07334064 | 54.74363883 | 52.34359522 | 0.001949881 | 1.126708309 | 0.000730603 | 11.40973313 | 2.183599556 | 2.709991876 |
| ENSMUSG00000003038 | *Hmgn2* | 3382.118033 | 3008.024046 | 2920.73668 | 7122.094374 | 6666.09079 | 6531.011389 | 1.90E-38 | 1.124669943 | 5.68E-40 | 175.103759 | 2.180516554 | 37.7214189 |
| ENSMUSG00000020069 | *Hnmph3* | 178.5613664 | 165.940102 | 118.4934173 | 327.0741445 | 353.7281278 | 337.0192885 | 6.52E-09 | 1.124620719 | 1.15E-09 | 37.05686753 | 2.180442157 | 8.185538158 |
| ENSMUSG00000058922 | *Gm10052* | 1775.065735 | 1547.175633 | 1154.680535 | 3338.790428 | 3122.492938 | 3311.421129 | 2.40E-13 | 1.124054163 | 2.81E-14 | 57.86133161 | 2.179586051 | 12.62061968 |
| ENSMUSG00000011148 | *Adssl1* | 882.2589032 | 926.5788355 | 993.3277962 | 2043.664621 | 2088.680374 | 1962.425666 | 2.16E-38 | 1.123678715 | 6.51E-40 | 174.8335003 | 2.179018908 | 37.66636749 |
| ENSMUSG00000040524 | *Zfp609* | 961.3683693 | 1014.824439 | 565.9951529 | 1695.736756 | 1926.554982 | 1922.938393 | 5.83E-08 | 1.12265652 | 1.13E-08 | 32.60842488 | 2.177475551 | 7.234555446 |
| ENSMUSG00000040195 | *Nemp1* | 433.218505 | 380.7989624 | 282.3672923 | 774.8803557 | 855.8957379 | 766.7877545 | 7.34E-11 | 1.121779911 | 1.06E-11 | 46.21077705 | 2.176152877 | 10.13446865 |
| ENSMUSG00000048911 | *Rnf24* | 310.4104766 | 427.7993381 | 273.5433144 | 713.4167581 | 733.7753128 | 757.6046676 | 4.54E-10 | 1.121473652 | 7.10E-11 | 42.49246839 | 2.175690966 | 9.342670572 |
| ENSMUSG00000053293 | *Pom121* | 2672.393108 | 2555.285733 | 1706.809437 | 4497.818268 | 5539.635145 | 5047.024549 | 8.62E-11 | 1.120041954 | 1.26E-11 | 45.87336153 | 2.173532931 | 10.06458535 |
| ENSMUSG00000029471 | *Camkk2* | 1292.12128 | 1411.929654 | 1188.715878 | 2670.373804 | 2946.681636 | 2841.247081 | 2.22E-31 | 1.118305888 | 8.99E-33 | 142.1565268 | 2.170918985 | 30.65367686 |
| ENSMUSG00000019732 | *Calr3* | 54.24649106 | 64.26581986 | 50.42273077 | 107.5612958 | 129.4897611 | 129.481525 | 1.74E-05 | 1.11517761 | 4.62E-06 | 20.98831995 | 2.166216757 | 4.758452261 |
| ENSMUSG00000012114 | *Med15* | 1298.902091 | 1142.396887 | 1041.22939 | 2252.201827 | 2748.762327 | 2550.143227 | 9.51E-20 | 1.114436606 | 7.06E-21 | 87.84924673 | 2.165104421 | 19.02175205 |
| ENSMUSG00000015165 | *Hnmpl* | 4952.25258 | 5134.551249 | 4319.967459 | 10261.12811 | 10449.71844 | 10496.2683 | 3.08E-39 | 1.114408589 | 8.97E-41 | 178.7755322 | 2.165062374 | 38.51209351 |
| ENSMUSG00000025236 | *Adpgk* | 1342.600654 | 1533.746954 | 1113.081782 | 2640.739569 | 3169.867241 | 2831.145685 | 6.01E-16 | 1.113725314 | 5.67E-17 | 70.08980714 | 2.164037222 | 15.22146854 |
| ENSMUSG00000001020 | *S100a4* | 15658.40033 | 14331.27783 | 17930.32306 | 36793.64612 | 32776.70099 | 34085.78188 | 1.44E-27 | 1.113271535 | 6.89E-29 | 124.3990444 | 2.163356662 | 26.84026699 |
| ENSMUSG00000031706 | *Rfx1* | 401.5747185 | 417.2482334 | 281.106724 | 780.3681769 | 841.1570659 | 764.9511371 | 3.78E-11 | 1.111310807 | 5.35E-12 | 47.55567601 | 2.160418499 | 10.42228876 |
| ENSMUSG00000042557 | *Sin3a* | 1343.354077 | 1387.949871 | 1166.025649 | 2663.788418 | 2912.993243 | 2850.430168 | 3.44E-30 | 1.110601807 | 1.47E-31 | 136.6033952 | 2.15935704 | 29.462933 |
| ENSMUSG00000113996 | *Gm48455* | 87.39712449 | 87.28641204 | 74.37352789 | 171.2200219 | 176.8640639 | 191.9265158 | 8.21E-08 | 1.11019575 | 1.61E-08 | 31.91283054 | 2.15874936 | 7.085654712 |
| ENSMUSG00000034163 | *Zfc3h1* | 397.0541776 | 328.0434387 | 173.9584212 | 665.1239314 | 636.9211825 | 647.407625 | 4.00E-05 | 1.109900303 | 1.12E-05 | 19.29292537 | 2.158307319 | 4.397666803 |
| ENSMUSG00000024795 | *Kif20b* | 403.834989 | 246.5121747 | 151.2681923 | 543.2943004 | 585.3358306 | 608.8386601 | 0.000400449 | 1.109173497 | 0.000131028 | 14.62693843 | 2.157220272 | 3.397452511 |
| ENSMUSG00000045767 | *B230219D22Rik* | 587.6703198 | 537.147151 | 496.6638981 | 1149.149763 | 1102.242113 | 1251.654742 | 1.34E-22 | 1.107722203 | 8.40E-24 | 101.1789732 | 2.155051286 | 21.87438123 |
| ENSMUSG00000021575 | *Ahrr* | 18.83558717 | 35.49007962 | 22.69022885 | 62.56116186 | 51.58535197 | 51.42528653 | 0.008974302 | 1.107346538 | 0.003854663 | 8.351041373 | 2.154490202 | 2.046999325 |
| ENSMUSG00000026657 | *Frmd4a* | 483.6978786 | 463.2894178 | 313.881499 | 878.0513945 | 919.061475 | 930.2467009 | 1.59E-10 | 1.107207866 | 2.38E-11 | 44.63282613 | 2.154283123 | 9.798159462 |
| ENSMUSG00000032245 | *Cln6* | 936.5053943 | 983.1711247 | 784.0734635 | 1793.419973 | 2108.682857 | 1926.611627 | 5.00E-20 | 1.105804646 | 3.65E-21 | 89.15662289 | 2.152188804 | 19.30059833 |
| ENSMUSG00000074994 | *Qser1* | 190.6161422 | 156.3481886 | 137.4019413 | 397.318256 | 484.2706512 | 163.4589465 | 0.003838384 | 1.105341404 | 0.001528031 | 10.04450489 | 2.151497858 | 2.415851635 |
| ENSMUSG00000010048 | *Ifrd2* | 1801.435557 | 1800.402147 | 1395.449074 | 3470.498137 | 3681.509711 | 3610.789761 | 3.96E-21 | 1.104850865 | 2.70E-22 | 94.31182746 | 2.150766439 | 20.40281498 |
| ENSMUSG00000005682 | *Pan2* | 230.547587 | 262.8184275 | 179.0006942 | 463.1721106 | 503.2203723 | 485.785296 | 1.12E-10 | 1.104347468 | 1.66E-11 | 45.33264628 | 2.150016108 | 9.949959775 |
| ENSMUSG00000032741 | *Tpcn1* | 2104.311799 | 2063.220575 | 1473.604307 | 3754.767276 | 4364.752434 | 4009.335732 | 8.49E-14 | 1.102822919 | 9.58E-15 | 59.9798332 | 2.147745306 | 13.07086163 |
| ENSMUSG00000022678 | *Nde1* | 1477.463458 | 1357.255748 | 1135.772011 | 2880.008574 | 2904.571145 | 2754.007755 | 1.83E-22 | 1.101828732 | 1.16E-23 | 100.5472823 | 2.146265766 | 21.73729555 |
| ENSMUSG00000036430 | *Tbcc* | 391.0267897 | 400.9419806 | 340.3534327 | 820.9780539 | 841.1570659 | 775.9708414 | 3.95E-21 | 1.101439008 | 2.69E-22 | 94.31691079 | 2.145686061 | 20.40349271 |
| ENSMUSG00000025402 | *Nab2* | 1179.107757 | 1046.477753 | 816.8482385 | 2124.884375 | 2201.325938 | 2214.960555 | 3.31E-15 | 1.10107788 | 3.32E-16 | 66.60572368 | 2.14514903 | 14.48065348 |
| ENSMUSG00000058173 | *Smco4* | 535.6840992 | 451.7791217 | 518.0935587 | 1081.10078 | 1079.081342 | 1067.993004 | 1.00E-24 | 1.099701369 | 5.57E-26 | 111.1201748 | 2.143103267 | 23.9991857 |
| ENSMUSG00000021240 | *Abcd4* | 761.7111453 | 703.087253 | 683.2280019 | 1537.687505 | 1546.507797 | 1522.555805 | 1.47E-35 | 1.097478962 | 4.97E-37 | 161.6343174 | 2.139804454 | 34.83362712 |
| ENSMUSG00000079553 | *Kifc1* | 1243.148753 | 1195.152411 | 753.819825 | 2088.664755 | 2442.408502 | 2306.791424 | 3.99E-09 | 1.096643011 | 6.90E-10 | 38.04933444 | 2.138564931 | 8.398824473 |
| ENSMUSG00000026873 | *Phf19* | 918.4232306 | 905.476626 | 652.9743635 | 1671.590342 | 1780.221024 | 1856.820167 | 6.57E-14 | 1.096550682 | 7.31E-15 | 60.51304227 | 2.138428073 | 13.18265835 |
| ENSMUSG00000025332 | *Kdm5c* | 2060.613237 | 1980.730119 | 1559.322949 | 3777.816125 | 4313.167082 | 3872.507737 | 2.90E-18 | 1.093230064 | 2.36E-19 | 80.91201252 | 2.133511761 | 17.53729686 |
| ENSMUSG00000027187 | *Cat* | 3302.255143 | 3858.826765 | 4385.517009 | 7452.461211 | 9230.619717 | 7917.657508 | 7.97E-17 | 1.092012184 | 7.10E-18 | 74.18860465 | 2.131711475 | 16.09850353 |
| ENSMUSG00000046380 | *Jrk* | 294.5885834 | 327.0842473 | 259.6770635 | 664.0263671 | 619.0242237 | 599.6555733 | 9.56E-16 | 1.090299775 | 9.15E-17 | 69.14361064 | 2.129182739 | 15.01971517 |
| ENSMUSG00000030774 | *Pak1* | 1799.92871 | 1703.523822 | 1578.231473 | 3483.668908 | 3761.519645 | 3585.995427 | 6.60E-39 | 1.089907156 | 1.95E-40 | 177.2340648 | 2.128603376 | 38.18058777 |
| ENSMUSG00000054150 | *Syne3* | 3116.159542 | 2754.797532 | 1783.704101 | 4899.526781 | 5918.629567 | 5486.894411 | 1.20E-08 | 1.089839245 | 2.17E-09 | 35.81685688 | 2.12850318 | 7.920943999 |
| ENSMUSG00000030735 | *Gm9755* | 327.7392168 | 257.0632794 | 537.0020827 | 628.9043113 | 799.0465745 | 951.3678008 | 9.77E-05 | 1.089328022 | 2.90E-05 | 17.48506583 | 2.127749071 | 4.009927485 |
| ENSMUSG00000051238 | *Swsap1* | 216.9859642 | 207.1853297 | 196.64865 | 398.4158203 | 466.3736923 | 458.2360353 | 6.71E-14 | 1.088664912 | 7.48E-15 | 60.46810504 | 2.126771314 | 13.17301165 |
| ENSMUSG00000050107 | *Haspin* | 469.3828324 | 434.5136775 | 305.0575212 | 819.8804896 | 921.1669995 | 838.4158321 | 4.68E-10 | 1.087105444 | 7.33E-11 | 42.42795529 | 2.124473642 | 9.329679428 |
| ENSMUSG00000066406 | *Akap13* | 1698.969963 | 1715.993309 | 1043.750527 | 2918.423323 | 3427.794001 | 3131.432626 | 6.29E-09 | 1.086219217 | 1.11E-09 | 37.12953074 | 2.123169008 | 8.201213172 |
| ENSMUSG00000063445 | *Nmral1* | 1096.231174 | 887.2519905 | 689.5308433 | 1884.517806 | 1889.708302 | 1911.00038 | 3.20E-11 | 1.085263052 | 4.50E-12 | 47.89197065 | 2.121762316 | 10.49424884 |
| ENSMUSG00000105175 | *Gm43097* | 201.164071 | 213.8996691 | 138.6625096 | 390.7328706 | 387.416521 | 403.137514 | 1.55E-08 | 1.08399894 | 2.83E-09 | 35.29528748 | 2.11990401 | 7.808436634 |
| ENSMUSG00000070337 | *Gpr179* | 590.6840138 | 690.6177656 | 506.7484442 | 1201.832846 | 1350.694012 | 1243.389964 | 1.95E-15 | 1.083415007 | 1.93E-16 | 67.67425469 | 2.119046148 | 14.70898512 |
| ENSMUSG00000023004 | *Tuba1b* | 48646.29428 | 47122.19302 | 41629.00652 | 92628.9343 | 100889.3681 | 97628.15155 | 3.52E-38 | 1.083303692 | 1.08E-39 | 173.8311618 | 2.118882654 | 37.45336005 |
| ENSMUSG00000079083 | *Jrkl* | 51.9862206 | 40.28603633 | 28.99307019 | 107.5612958 | 68.42954854 | 84.4843993 | 0.003376982 | 1.082986911 | 0.001326042 | 10.30591835 | 2.118417449 | 2.471471257 |
| ENSMUSG00000028063 | *Lmna* | 26308.79474 | 26950.39911 | 24365.52408 | 51809.42253 | 56850.21616 | 55183.92399 | 2.15E-49 | 1.077632548 | 4.24E-51 | 226.0955852 | 2.110569809 | 48.66750129 |
| ENSMUSG00000061607 | *Mdc1* | 1130.888654 | 1137.600931 | 742.4747106 | 1908.664219 | 2315.024265 | 2138.740934 | 8.13E-10 | 1.076945981 | 1.31E-10 | 41.29856579 | 2.109565644 | 9.08990936 |
| ENSMUSG00000044982 | *Sft2d3* | 76.09577218 | 104.5518562 | 71.85239135 | 169.0248934 | 174.7585393 | 188.253281 | 5.22E-06 | 1.073215477 | 1.28E-06 | 23.45401702 | 2.104117801 | 5.282644962 |
| ENSMUSG00000050737 | *Ptges* | 103.2190177 | 104.5518562 | 71.85239135 | 193.1713068 | 216.8690307 | 183.6617376 | 2.81E-06 | 1.07315037 | 6.65E-07 | 24.71471441 | 2.104022846 | 5.551895117 |
| ENSMUSG00000100622 | *Gm20379* | 375.2048965 | 322.2882906 | 344.1351375 | 752.9290708 | 723.2476899 | 719.9540114 | 2.82E-20 | 1.07299467 | 2.02E-21 | 90.32910359 | 2.103795785 | 19.55013615 |
| ENSMUSG00000094484 | *Gm21244* | 47.46567968 | 42.20441901 | 35.29591154 | 105.3661673 | 88.43203196 | 71.62807766 | 0.000960424 | 1.071618184 | 0.000336596 | 12.85490127 | 2.101789496 | 3.017537144 |
| ENSMUSG00000022314 | *Rad21* | 2679.927343 | 2614.755596 | 2444.241874 | 5119.03963 | 5616.486791 | 5537.401389 | 1.83E-43 | 1.071335613 | 4.51E-45 | 198.4699554 | 2.101377873 | 42.73663573 |
| ENSMUSG00000028158 | *Mttp* | 90.41081843 | 76.73530729 | 107.1483029 | 189.8786141 | 205.2886456 | 178.1518855 | 9.63E-07 | 1.070802424 | 2.16E-07 | 26.8837084 | 2.100601393 | 6.016428865 |
| ENSMUSG00000040466 | *Blvrb* | 1649.244013 | 1880.015029 | 2432.89676 | 4161.96361 | 4237.368198 | 4084.637044 | 8.86E-14 | 1.067977869 | 1.00E-14 | 59.89431768 | 2.096492791 | 13.05251994 |
| ENSMUSG00000036611 | *Eepd1* | 206.4380354 | 184.1647375 | 147.4864875 | 356.708379 | 398.9969061 | 380.1797968 | 1.07E-09 | 1.067781838 | 1.75E-10 | 40.73314445 | 2.096207942 | 8.969794755 |
| ENSMUSG00000041598 | *Cdc42ep4* | 702.9441133 | 626.3519458 | 466.4102596 | 1231.467081 | 1283.317226 | 1261.756137 | 9.56E-12 | 1.067607607 | 1.29E-12 | 50.34066802 | 2.095954804 | 11.01953925 |
| ENSMUSG00000048327 | *Ckap2l* | 601.2319426 | 472.8813312 | 563.4740164 | 1107.442321 | 1231.731874 | 1092.787339 | 7.70E-17 | 1.066310382 | 6.85E-18 | 74.25860399 | 2.094071035 | 16.11356859 |
| ENSMUSG00000021068 | *Nin* | 1103.011985 | 1175.968584 | 848.3624452 | 2099.640397 | 2275.019298 | 2185.574677 | 1.58E-15 | 1.066276187 | 1.54E-16 | 68.11976553 | 2.094021401 | 14.80194741 |
| ENSMUSG00000041731 | *Pgm5* | 94.93135936 | 72.89854193 | 83.19750577 | 180.0005359 | 176.8640639 | 170.805416 | 6.60E-07 | 1.065670063 | 1.45E-07 | 27.65661617 | 2.093141818 | 6.180635272 |
| ENSMUSG00000030978 | *Rrm1* | 5672.525433 | 5156.61265 | 5043.533645 | 10737.47099 | 11546.69674 | 10885.63119 | 1.55E-43 | 1.062668888 | 3.80E-45 | 198.8078996 | 2.088792074 | 42.80917036 |
| ENSMUSG00000002346 | *Slc25a42* | 31.64378645 | 32.6125056 | 32.774775 | 52.68308367 | 86.32650739 | 63.36329947 | 0.004020598 | 1.062054137 | 0.001607191 | 9.951492375 | 2.087902202 | 2.395709394 |
| ENSMUSG00000036941 | *Elac1* | 129.5888398 | 119.8989176 | 117.232849 | 263.4154184 | 249.5046616 | 255.2898153 | 3.32E-10 | 1.061972853 | 5.10E-11 | 43.13768714 | 2.08778457 | 9.479182532 |
| ENSMUSG00000036278 | *Macrod1* | 82.87658356 | 114.1437696 | 129.8385317 | 226.0982341 | 214.7635062 | 234.1687154 | 2.99E-06 | 1.060373005 | 7.10E-07 | 24.58761992 | 2.085470645 | 5.525011576 |
| ENSMUSG00000018678 | *Sp2* | 414.3829178 | 397.1052152 | 226.9022885 | 678.2947023 | 786.413427 | 708.9343071 | 3.02E-06 | 1.059800621 | 7.19E-07 | 24.56230227 | 2.084643406 | 5.520056685 |
| ENSMUSG00000034883 | *Lrr1* | 188.3558717 | 166.8992934 | 175.2189894 | 395.1231275 | 373.7306112 | 339.7742146 | 6.07E-12 | 1.059434139 | 8.05E-13 | 51.27010087 | 2.08411392 | 11.21673363 |
| ENSMUSG00000042548 | *Asxl1* | 1417.189579 | 1261.336614 | 800.460851 | 2387.202229 | 2672.963442 | 2194.757764 | 1.32E-07 | 1.057973896 | 2.66E-08 | 30.94093812 | 2.082005523 | 6.879649059 |
| ENSMUSG00000083282 | *Ctsf* | 750.409793 | 789.4144737 | 705.9182308 | 1531.102119 | 1600.198673 | 1550.105065 | 3.54E-33 | 1.057842077 | 1.35E-34 | 150.502784 | 2.0818153 | 32.45092631 |
| ENSMUSG00000025006 | *Sorbs1* | 180.0682134 | 173.6136327 | 133.6202365 | 330.3668372 | 348.4643164 | 342.5291406 | 8.39E-10 | 1.057465633 | 1.35E-10 | 41.23338053 | 2.08127216 | 9.07616757 |
| ENSMUSG00000055452 | *Gm7353* | 398.5610246 | 331.880204 | 268.5010413 | 661.8312386 | 689.5592968 | 737.4018765 | 2.28E-10 | 1.057128799 | 3.46E-11 | 43.89985924 | 2.08078629 | 9.642674977 |
| ENSMUSG00000042724 | *Map3k9* | 309.6570531 | 269.5327669 | 238.2474029 | 534.5137864 | 644.2905185 | 529.864113 | 5.57E-11 | 1.056931413 | 7.99E-12 | 46.76817543 | 2.080501622 | 10.25449329 |
| ENSMUSG00000021144 | *Mta1* | 1483.490846 | 1448.378925 | 1024.842003 | 2591.349178 | 2915.098768 | 2734.723273 | 2.06E-12 | 1.056321429 | 2.61E-13 | 53.4820371 | 2.079622154 | 11.68518297 |
| ENSMUSG00000026281 | *Dtymk* | 1685.40834 | 1538.542911 | 1922.366611 | 3627.449824 | 3420.424665 | 3640.175639 | 7.05E-25 | 1.055961522 | 3.89E-26 | 111.8326269 | 2.079103419 | 24.15202768 |

| | | | | | | | | | | | | | |
|---|---|---|---|---|---|---|---|---|---|---|---|---|---|
| ENSMUSG00000026565 | *Pou2f1* | 388.7665193 | 372.1662404 | 220.5994471 | 612.4408477 | 766.4109436 | 669.4470336 | 2.81E-06 | 1.055308092 | 6.65E-07 | 24.71324698 | 2.078161957 | 5.551690245 |
| ENSMUSG00000054920 | *Klhl5* | 1092.464056 | 1187.47888 | 1104.257804 | 2282.933626 | 2335.026749 | 2400.45891 | 1.50E-40 | 1.052638281 | 4.17E-42 | 184.8804404 | 2.074319724 | 39.82403694 |
| ENSMUSG00000020770 | *Unk* | 318.698135 | 314.6147599 | 269.7616096 | 614.6359762 | 608.4966008 | 656.5907119 | 9.89E-17 | 1.052389315 | 8.87E-18 | 73.74814091 | 2.073961791 | 16.00459868 |
| ENSMUSG00000068267 | *Cenpb* | 7684.919567 | 7885.512015 | 6662.103303 | 14825.8978 | 15743.00721 | 15509.31543 | 6.07E-34 | 1.050947796 | 2.21E-35 | 154.0908808 | 2.071890554 | 33.2167075 |
| ENSMUSG00000056493 | *Foxk1* | 999.0395437 | 1028.253118 | 617.6784519 | 1654.029314 | 2028.672924 | 1806.313189 | 8.33E-08 | 1.05075504 | 1.64E-08 | 31.88192256 | 2.071613752 | 7.079199214 |
| ENSMUSG00000033287 | *Kctd17* | 876.2315153 | 907.3950087 | 670.6223192 | 1581.590074 | 1851.80886 | 1661.220417 | 7.10E-14 | 1.05068806 | 7.95E-15 | 60.34765209 | 2.071517575 | 13.14857135 |
| ENSMUSG00000024042 | *Sik1* | 76.09577218 | 58.51067181 | 37.81704808 | 120.7320667 | 122.1204251 | 119.3801294 | 0.000865803 | 1.049650566 | 0.00030088 | 13.06490665 | 2.070028407 | 3.062580927 |
| ENSMUSG00000058503 | *Fam133b* | 447.5335512 | 461.3710351 | 531.9598096 | 1077.808087 | 923.2725241 | 969.7339745 | 1.78E-17 | 1.047905427 | 1.52E-18 | 77.23463801 | 2.067525935 | 16.74895887 |
| ENSMUSG00000051705 | *Senp8* | 115.2737935 | 133.3275964 | 124.7962587 | 267.8056753 | 229.5021782 | 272.7376803 | 1.01E-08 | 1.04767947 | 1.80E-09 | 36.17683977 | 2.067202143 | 7.996201533 |
| ENSMUSG00000060600 | *Eno3* | 536.4375227 | 596.6170142 | 647.9320904 | 1210.61336 | 1274.895127 | 1181.863281 | 1.03E-21 | 1.046402508 | 6.73E-23 | 97.05801925 | 2.065373224 | 20.98511071 |
| ENSMUSG00000040455 | *Usp45* | 533.4238288 | 546.7390644 | 489.1004885 | 1173.296176 | 1053.815047 | 1018.404335 | 2.78E-21 | 1.045270738 | 1.86E-22 | 95.04951433 | 2.063753609 | 20.5567149 |
| ENSMUSG00000050860 | *Phospho1* | 276.5064197 | 300.2268898 | 332.7900231 | 554.2699428 | 720.0894031 | 596.9006472 | 2.59E-11 | 1.045224727 | 3.62E-12 | 48.32196317 | 2.063687791 | 10.58629549 |
| ENSMUSG00000035047 | *Kri1* | 1494.792198 | 1344.78626 | 1183.673605 | 2557.324687 | 2985.633841 | 2757.68099 | 2.96E-19 | 1.042629995 | 2.28E-20 | 85.53088967 | 2.059979521 | 18.52871587 |
| ENSMUSG00000005442 | *Cic* | 4310.335769 | 4186.870204 | 2725.348598 | 6926.727939 | 8354.721495 | 7841.437887 | 2.02E-09 | 1.042198821 | 3.37E-10 | 39.44551447 | 2.059363952 | 8.695266292 |
| ENSMUSG00000051639 | *Fbl-ps2* | 149.9312739 | 104.5518562 | 119.7539856 | 252.4397759 | 216.8690307 | 303.041867 | 7.82E-06 | 1.04087235 | 1.97E-06 | 22.626219 | 2.057471362 | 5.10669057 |
| ENSMUSG00000034271 | *Jdp2* | 1011.847743 | 1119.376295 | 1157.201671 | 2124.884375 | 2368.715142 | 2255.366138 | 2.95E-28 | 1.039673932 | 1.37E-29 | 127.6105327 | 2.055762971 | 27.53040195 |
| ENSMUSG00000038943 | *Prc1* | 1637.189237 | 1363.010896 | 1297.124749 | 2853.667032 | 3099.332168 | 2892.672367 | 1.27E-20 | 1.039225285 | 8.88E-22 | 91.9521577 | 2.055123773 | 19.89540108 |
| ENSMUSG00000082500 | *Gm15195* | 55.75333803 | 51.79633242 | 57.98614039 | 106.4637316 | 110.5400399 | 122.1350555 | 4.10E-05 | 1.038563252 | 1.15E-05 | 19.24432747 | 2.054180922 | 4.387250084 |
| ENSMUSG00000053877 | *Srcap* | 2579.722019 | 2460.32579 | 1309.730432 | 3912.816527 | 4725.849898 | 4412.473246 | 5.58E-06 | 1.038314119 | 1.38E-06 | 23.31396311 | 2.053826224 | 5.253690099 |
| ENSMUSG00000044030 | *Irf2bp1* | 1214.518661 | 1189.397263 | 980.7221135 | 2243.421313 | 2466.622034 | 2253.52952 | 9.52E-21 | 1.03802672 | 6.58E-22 | 92.54410225 | 2.053417122 | 20.0214445 |
| ENSMUSG00000039477 | *Tnrc18* | 1522.668867 | 1604.727114 | 925.2571096 | 2483.787882 | 3069.854824 | 2773.292238 | 2.41E-07 | 1.037287419 | 5.03E-08 | 29.70495092 | 2.052365129 | 6.618161475 |
| ENSMUSG00000048429 | *Timm29* | 1407.395074 | 1372.602809 | 1314.772705 | 2703.300731 | 2805.61149 | 2901.855454 | 8.07E-43 | 1.037259657 | 2.06E-44 | 195.4407195 | 2.052325636 | 42.09336453 |
| ENSMUSG00000028832 | *Stmn1* | 2808.009336 | 2538.020289 | 2519.87597 | 5353.918378 | 5553.321054 | 5242.6243 | 1.81E-41 | 1.036587485 | 4.88E-43 | 189.1460646 | 2.051369651 | 40.74324822 |
| ENSMUSG00000062944 | *9130023H24Rik* | 55.75333803 | 74.81692461 | 52.94386731 | 109.7564243 | 133.7008102 | 133.1547598 | 0.000131495 | 1.035683211 | 3.98E-05 | 16.88028576 | 2.050084267 | 3.881092006 |
| ENSMUSG00000019132 | *BC005537* | 11113.74986 | 10863.80113 | 7318.859371 | 19131.64232 | 20588.87201 | 20306.56002 | 2.50E-11 | 1.034568793 | 3.48E-12 | 48.39852337 | 2.048501278 | 10.60218174 |
| ENSMUSG00000024665 | *Fads2* | 4648.622914 | 4331.708097 | 3114.864193 | 8181.243869 | 8272.606037 | 8324.468257 | 1.06E-12 | 1.033790053 | 1.31E-13 | 54.83237202 | 2.047395833 | 11.97313541 |
| ENSMUSG00000037993 | *Dhx38* | 1992.805123 | 1778.340746 | 1292.082476 | 3175.253355 | 3673.087613 | 3529.060288 | 1.12E-10 | 1.033548931 | 1.65E-11 | 45.34502809 | 2.047053675 | 9.952302889 |
| ENSMUSG00000046574 | *Prr12* | 967.3957572 | 900.6806693 | 639.1081125 | 1447.687237 | 1890.761064 | 1800.803337 | 8.23E-09 | 1.032863176 | 1.46E-09 | 36.58749704 | 2.046080882 | 8.084356667 |
| ENSMUSG00000022265 | *Ank* | 7980.261574 | 8649.028323 | 7013.80185 | 14946.62986 | 17130.5479 | 16299.0609 | 5.04E-24 | 1.032590492 | 2.93E-25 | 107.8297806 | 2.045694188 | 23.29767113 |
| ENSMUSG00000010277 | *Bltp2* | 11425.66718 | 11870.95204 | 9919.411711 | 21372.86851 | 24268.2762 | 22257.04767 | 5.70E-27 | 1.031229223 | 2.82E-28 | 121.600807 | 2.043764864 | 26.24432681 |
| ENSMUSG00000028248 | *Pnisr* | 385.7528253 | 317.4923339 | 284.8884288 | 682.6849592 | 697.9813951 | 645.5710077 | 3.60E-12 | 1.028438594 | 4.65E-13 | 52.34601784 | 2.039815397 | 11.44402066 |
| ENSMUSG00000035569 | *Ankrd11* | 1869.243671 | 1770.667216 | 1101.736667 | 3060.00911 | 3440.427148 | 3182.857913 | 4.33E-08 | 1.028390437 | 8.25E-09 | 33.21493586 | 2.039747309 | 7.363683841 |
| ENSMUSG00000028851 | *Nudc* | 4297.52757 | 4148.50255 | 3673.295937 | 8154.902327 | 8594.751296 | 7979.18419 | 2.80E-33 | 1.027882736 | 1.06E-34 | 150.9786927 | 2.039029625 | 32.55258715 |
| ENSMUSG00000028690 | *Mmachc* | 357.1227328 | 377.9213884 | 334.0505913 | 696.9532944 | 714.8255916 | 767.7060632 | 9.13E-19 | 1.025853119 | 7.20E-20 | 83.25787942 | 2.036163087 | 18.03973828 |
| ENSMUSG00000070808 | *Bicra* | 256.1639856 | 235.9610699 | 156.3104654 | 398.4158203 | 477.9540775 | 451.8078745 | 1.22E-06 | 1.025742693 | 2.77E-07 | 26.40165905 | 2.036007242 | 5.913093024 |
| ENSMUSG00000050567 | *Maml1* | 526.6430174 | 584.1475267 | 359.2619567 | 898.9051152 | 1016.968367 | 1082.685943 | 6.34E-08 | 1.024803579 | 1.23E-08 | 32.43813386 | 2.034682345 | 7.197885651 |
| ENSMUSG00000111527 | *None* | 64.04099639 | 81.531264 | 30.25363846 | 132.8052734 | 126.3314742 | 101.9322644 | 0.008730578 | 1.02307142 | 0.003736135 | 8.407805091 | 2.032240888 | 2.058957005 |
| ENSMUSG00000026389 | *Steap3* | 612.5332949 | 651.2909206 | 660.5377731 | 1185.369383 | 1398.068315 | 1320.527893 | 6.76E-22 | 1.022585919 | 4.36E-23 | 97.91814878 | 2.031557106 | 21.17008344 |
| ENSMUSG00000003308 | *Keap1* | 2671.639685 | 2707.797156 | 2286.67084 | 5138.795787 | 5243.808943 | 5204.973644 | 6.35E-31 | 1.022423979 | 2.62E-32 | 140.0352935 | 2.031329081 | 30.19713622 |
| ENSMUSG00000028433 | *Ubap2* | 2248.215685 | 2329.875768 | 1642.520455 | 3718.547656 | 4686.897694 | 4235.239669 | 2.95E-11 | 1.021799209 | 4.14E-12 | 48.05772514 | 2.030449589 | 10.52990219 |
| ENSMUSG00000041995 | *Zbed3* | 617.8072593 | 580.3107614 | 383.2127538 | 979.0273049 | 1140.141555 | 1100.133808 | 4.79E-08 | 1.021121485 | 9.17E-09 | 33.00950162 | 2.029495984 | 7.319981551 |
| ENSMUSG00000032119 | *Hinfp* | 704.4509603 | 624.4335631 | 568.5162894 | 1222.686567 | 1343.324676 | 1292.060324 | 6.39E-19 | 1.019562793 | 5.00E-20 | 83.97943032 | 2.027304494 | 18.19445952 |
| ENSMUSG00000022325 | *Pop1* | 739.8618642 | 858.4762503 | 797.9397144 | 1463.053136 | 1644.414689 | 1742.031581 | 1.11E-19 | 1.018646799 | 8.26E-21 | 87.53949884 | 2.026017729 | 18.95565983 |
| ENSMUSG00000008855 | *Hdac5* | 2651.297251 | 2247.385312 | 1660.168411 | 4136.719632 | 4628.995768 | 4530.935067 | 4.63E-10 | 1.018118379 | 7.25E-11 | 42.45001663 | 2.02527579 | 9.33400702 |
| ENSMUSG00000037583 | *Nr0b2* | 178.5613664 | 172.6544414 | 138.6625096 | 321.5863232 | 349.5170787 | 326.9178929 | 1.33E-09 | 1.017353614 | 2.18E-10 | 40.29557619 | 2.024202487 | 8.875641 |
| ENSMUSG00000029822 | *Osbpl3* | 484.4513021 | 485.3508186 | 388.2550269 | 830.8561321 | 942.2222452 | 980.7536788 | 1.27E-13 | 1.015902676 | 1.46E-14 | 59.15612529 | 2.022167741 | 12.89459866 |
| ENSMUSG00000028034 | *Fubp1* | 1769.791771 | 1851.239288 | 1797.570352 | 3769.035611 | 3537.281278 | 3644.767183 | 4.49E-49 | 1.015242607 | 8.91E-51 | 224.6137492 | 2.021242761 | 48.34732515 |
| ENSMUSG00000051451 | *Crebzf* | 399.3144481 | 409.5747027 | 269.7616096 | 665.1239314 | 731.6697882 | 790.6637804 | 3.30E-08 | 1.014767726 | 6.20E-09 | 33.76978025 | 2.020577553 | 7.481435643 |
| ENSMUSG00000029478 | *Ncor2* | 4207.870175 | 4014.215763 | 2371.128914 | 6520.629169 | 7784.124337 | 7095.771232 | 4.05E-07 | 1.01377386 | 8.65E-08 | 28.65432235 | 2.019186066 | 6.392540157 |
| ENSMUSG00000111619 | *Gm48348* | 163.4928967 | 146.7562752 | 78.15523269 | 277.6837535 | 218.9745553 | 292.0221628 | 0.000729611 | 1.011109202 | 0.000250597 | 13.40767221 | 2.015460072 | 3.136908698 |
| ENSMUSG00000031378 | *Abcd1* | 1496.299045 | 1578.828947 | 1249.223155 | 2629.763927 | 3083.540733 | 3007.460953 | 2.92E-17 | 1.01049293 | 2.52E-18 | 76.23030389 | 2.014599316 | 16.53435187 |
| ENSMUSG00000000568 | *Hnrnpd* | 2414.722276 | 2378.794526 | 1609.74568 | 3960.011789 | 4648.998251 | 4289.419882 | 4.42E-10 | 1.009007075 | 6.90E-11 | 42.54639884 | 2.012525517 | 9.354263397 |
| ENSMUSG00000031647 | *Mfap3l* | 58.01360849 | 40.28603633 | 76.89466442 | 108.6588601 | 113.6983268 | 126.7265989 | 0.001350231 | 1.007606534 | 0.00048711 | 12.16437451 | 2.010572742 | 2.86959179 |
| ENSMUSG00000028687 | *Mutyh* | 114.52037 | 116.0621523 | 80.67636923 | 204.1469492 | 240.029801 | 186.4166637 | 1.61E-05 | 1.006876415 | 4.26E-06 | 21.14382749 | 2.009555489 | 4.792233556 |
| ENSMUSG00000038884 | *Shfl* | 928.2177359 | 987.00789 | 825.6722164 | 1880.127549 | 1797.065221 | 1838.453993 | 2.71E-25 | 1.006349629 | 1.47E-26 | 113.7644551 | 2.008821854 | 24.56754572 |
| ENSMUSG00000067925 | *Rtl8a* | 80.6163131 | 102.6334735 | 56.72557212 | 190.9761783 | 170.5474902 | 123.9716729 | 0.001243226 | 1.006219303 | 0.000446886 | 12.32519373 | 2.008640395 | 2.905450015 |
| ENSMUSG00000002983 | *Relb* | 389.5199428 | 334.7577781 | 245.8108125 | 659.6361101 | 614.8131746 | 683.2216639 | 4.08E-08 | 1.005660226 | 7.74E-09 | 33.33957796 | 2.007862153 | 7.389796968 |
| ENSMUSG00000096472 | *Cdkn2d* | 599.7250956 | 589.9026748 | 612.6361789 | 1220.491438 | 1216.993202 | 1179.108355 | 1.22E-30 | 1.005373274 | 5.10E-32 | 138.7094796 | 2.007462829 | 29.91366837 |
| ENSMUSG00000047044 | *D030056L22Rik* | 512.3279711 | 465.2078004 | 307.5786577 | 807.807283 | 885.3730818 | 896.2692795 | 1.91E-07 | 1.005241613 | 3.94E-08 | 30.17791581 | 2.007279635 | 6.718205307 |
| ENSMUSG00000005161 | *Prdx2* | 4375.883612 | 4376.79009 | 4336.354846 | 8708.074705 | 8774.773647 | 8786.377527 | 6.23E-81 | 1.004902034 | 5.77E-83 | 372.3479838 | 2.00680722 | 80.20584504 |
| ENSMUSG00000093505 | *Gm20691* | 70.06838429 | 77.69449863 | 63.02841346 | 151.4638656 | 142.1229085 | 131.3181424 | 1.50E-05 | 1.004887766 | 3.95E-06 | 21.29123229 | 2.006787373 | 4.823937543 |
| ENSMUSG00000090115 | *Usp49* | 168.7668611 | 262.8184275 | 171.4372846 | 368.7815857 | 380.047185 | 460.9909614 | 8.43E-06 | 1.004797484 | 2.13E-06 | 22.47394056 | 2.006661796 | 5.074392495 |
| ENSMUSG00000032536 | *Trak1* | 1619.860497 | 1653.645872 | 1174.849627 | 2899.76473 | 3051.957865 | 2979.911693 | 5.13E-13 | 1.003624199 | 6.19E-14 | 56.31127903 | 2.005030523 | 12.28984581 |
| ENSMUSG00000022508 | *Bcl6* | 865.6835865 | 999.4773774 | 589.94595 | 1532.199683 | 1771.798926 | 1625.406378 | 8.91E-08 | 1.003199352 | 1.76E-08 | 31.74462186 | 2.004440165 | 7.050012127 |
| ENSMUSG00000037428 | *Vgf* | 67.05469034 | 84.40883802 | 65.54955 | 138.2930946 | 161.0726296 | 135.9096858 | 3.37E-05 | 1.001912533 | 9.35E-06 | 19.64002198 | 2.002653092 | 4.471880859 |
| ENSMUSG00000071064 | *Zfp827* | 392.5336367 | 398.0644066 | 312.6209308 | 694.7581659 | 779.044091 | 742.9117286 | 3.27E-13 | 1.001200189 | 3.88E-14 | 57.22861218 | 2.001664507 | 12.48546036 |
| ENSMUSG00000024301 | *Kifc5b* | 546.9854515 | 587.0251008 | 466.4102596 | 1081.10078 | 1086.450678 | 1043.19867 | 1.04E-17 | 1.000353805 | 8.77E-19 | 78.31861027 | 2.000490538 | 16.98307374 |
| ENSMUSG00000043800 | *Kifc5c-ps* | 24.86297507 | 36.44927096 | 32.774775 | 50.48795519 | 65.27126168 | 70.70976898 | 0.008099987 | 0.999606624 | 0.003441883 | 8.557041461 | 1.999454739 | 2.091515664 |
| ENSMUSG00000045039 | *Megf8* | 1417.943002 | 1450.297308 | 1023.581435 | 2267.567726 | 2887.726948 | 2631.8727 | 3.62E-10 | 0.998909742 | 5.58E-11 | 42.96314069 | 1.998489152 | 9.44179111 |
| ENSMUSG00000040936 | *Ulk4* | 76.09577218 | 52.75552376 | 66.81011827 | 148.1711728 | 124.2259497 | 120.2984381 | 0.000183045 | 0.998859349 | 5.68E-05 | 16.20595413 | 1.998419347 | 3.737442411 |
| ENSMUSG00000032249 | *Anp32a* | 1869.997095 | 1762.993685 | 2185.825379 | 3963.304482 | 3883.64007 | 3751.290991 | 5.02E-23 | 0.996625467 | 3.09E-24 | 103.1611451 | 1.995327371 | 22.29887731 |
| ENSMUSG00000047248 | *C2cd3* | 612.5332949 | 651.2909206 | 467.6708279 | 999.8810255 | 1314.900094 | 1144.212625 | 2.13E-09 | 0.995328565 | 3.57E-10 | 39.33505327 | 1.993534489 | 8.671410846 |
| ENSMUSG00000026355 | *Mcm6* | 5648.415882 | 5691.841418 | 4932.603638 | 10088.81052 | 11432.99842 | 10914.09876 | 6.16E-28 | 0.994593482 | 2.90E-29 | 126.1194225 | 1.992519001 | 27.21048475 |
| ENSMUSG00000044786 | *Zfp36* | 1295.888398 | 1197.070794 | 704.6576625 | 2088.664755 | 2078.152751 | 2211.287321 | 1.68E-06 | 0.99378228 | 3.89E-07 | 25.75038459 | 1.991398958 | 5.773890378 |
| ENSMUSG00000037725 | *Ckap2* | 529.6567113 | 455.615887 | 385.7338904 | 915.3685788 | 947.4860567 | 878.8214144 | 1.84E-12 | 0.993514026 | 2.32E-13 | 53.71180776 | 1.991028712 | 11.73408577 |
| ENSMUSG00000031386 | *Hcfc1* | 5584.374885 | 5542.207569 | 3562.365929 | 8929.782682 | 10483.40684 | 9843.350827 | 6.75E-09 | 0.993466155 | 1.19E-09 | 36.98686045 | 1.990962648 | 8.170791775 |
| ENSMUSG00000046841 | *Ckap4* | 3077.734944 | 3149.025173 | 2634.587683 | 5499.894423 | 6127.0765 | 6016.758524 | 2.65E-24 | 0.99260806 | 1.52E-25 | 109.133748 | 1.989778803 | 23.57667081 |
| ENSMUSG00000009575 | *Cbx5* | 2548.831656 | 2419.080562 | 1945.056839 | 4412.208258 | 4904.819487 | 4443.695741 | 9.77E-17 | 0.991654407 | 8.75E-18 | 73.77567122 | 1.988463949 | 16.01032339 |
| ENSMUSG00000032044 | *Rpusd4* | 1376.504711 | 1352.459791 | 1424.442144 | 2720.861759 | 2749.815089 | 2776.047164 | 1.06E-46 | 0.990585786 | 2.33E-48 | 213.5288149 | 1.986991617 | 45.97362263 |
| ENSMUSG00000035847 | *Ids* | 347.3282275 | 338.5945434 | 277.3250192 | 587.1968701 | 689.5592968 | 644.652699 | 4.79E-12 | 0.990542934 | 6.30E-13 | 51.75229437 | 1.986932599 | 11.31956279 |
| ENSMUSG00000026708 | *Cenpl* | 488.971843 | 433.5544862 | 395.8184365 | 923.0515285 | 832.7349676 | 873.3115623 | 1.04E-15 | 0.990298022 | 1.00E-16 | 68.9650636 | 1.986595325 | 14.98133021 |
| ENSMUSG00000099703 | *None* | 54.24649106 | 76.73530729 | 90.76091539 | 143.7809159 | 168.4419656 | 123.9716729 | 0.00068244 | 0.989591717 | 0.000233646 | 13.53908462 | 1.98562298 | 3.16593539 |
| ENSMUSG00000020850 | *Prpf8* | 10462.03854 | 10443.67532 | 7418.444264 | 17478.71057 | 20019.32762 | 18749.10848 | 3.18E-12 | 0.989418013 | 4.10E-13 | 52.59478062 | 1.98538392 | 11.49718528 |
| ENSMUSG00000083218 | *Gm16425* | 79.86288962 | 70.98015924 | 34.03534327 | 105.3661673 | 135.8063348 | 129.481525 | 0.006724971 | 0.986147443 | 0.002805677 | 8.929717893 | 1.98088818 | 2.172309559 |
| ENSMUSG00000028868 | *Wasf2* | 3277.392168 | 3065.575526 | 1894.634109 | 5044.405262 | 5840.725158 | 5437.305742 | 2.40E-07 | 0.985569795 | 5.00E-08 | 29.71679885 | 1.9800952 | 6.620530245 |
| ENSMUSG00000032312 | *Csk* | 5750.128052 | 5868.332625 | 5163.287631 | 10613.44623 | 11390.88793 | 11237.34342 | 3.92E-36 | 0.985555407 | 1.29E-37 | 164.3144482 | 1.980075453 | 35.40665297 |
| ENSMUSG00000027699 | *Ect2* | 834.7932235 | 814.3534486 | 790.3763048 | 1623.297516 | 1566.51028 | 1645.609169 | 1.58E-33 | 0.985543185 | 5.84E-35 | 152.1618658 | 1.980058678 | 32.80242551 |
| ENSMUSG00000031788 | *Kifc3* | 1398.353992 | 1453.174882 | 1082.828143 | 2510.129424 | 2638.222287 | 2652.075491 | 3.78E-15 | 0.985233137 | 3.81E-16 | 66.33421848 | 1.979633193 | 14.42304083 |
| ENSMUSG00000013236 | *Ptprs* | 1811.983486 | 1844.524949 | 1289.561339 | 2910.740373 | 3576.233483 | 3312.339438 | 2.67E-10 | 0.984894874 | 4.07E-11 | 43.57919626 | 1.97916909 | 9.574257611 |
| ENSMUSG00000031971 | *Ccsap* | 85.89027751 | 98.79670814 | 61.76784519 | 155.8541225 | 148.4394822 | 186.4166637 | 0.000155979 | 0.984815879 | 4.78E-05 | 16.53155955 | 1.979060723 | 3.806933391 |
| ENSMUSG00000024913 | *Lrp5* | 2235.407486 | 2366.325039 | 1594.618861 | 3691.10855 | 4610.046047 | 3958.828754 | 1.37E-09 | 0.983309423 | 2.25E-10 | 40.23424486 | 1.976995275 | 8.862927758 |
| ENSMUSG00000032777 | *Gtf3c1* | 2029.722874 | 2097.751463 | 1497.555104 | 3465.010316 | 3806.788423 | 3847.713403 | 1.58E-12 | 0.981695391 | 1.97E-13 | 54.03190836 | 1.974784725 | 11.80199007 |
| ENSMUSG00000052698 | *Tln2* | 364.6569677 | 341.4721174 | 271.0221779 | 619.0262331 | 669.5568134 | 650.1625511 | 2.13E-11 | 0.981309543 | 2.94E-12 | 48.72777148 | 1.974256641 | 10.67251666 |
| ENSMUSG00000049881 | *2810025M15Rik* | 1033.697024 | 1090.600555 | 1184.934173 | 2305.982475 | 2107.630095 | 2104.763513 | 1.16E-25 | 0.980243389 | 6.20E-27 | 115.4733399 | 1.972798201 | 24.93460267 |
| ENSMUSG00000032077 | *Bud13* | 759.4508748 | 692.5361483 | 552.1289019 | 1326.95517 | 1349.64125 | 1289.305398 | 2.84E-13 | 0.979897622 | 3.35E-14 | 57.51501122 | 1.972325442 | 12.54719054 |
| ENSMUSG00000084883 | *Ccdc85c* | 604.2456365 | 529.4736203 | 369.3465029 | 945.0028134 | 1035.918089 | 990.8550744 | 9.12E-08 | 0.97820301 | 1.80E-08 | 31.69928582 | 1.970010078 | 7.040176244 |
| ENSMUSG00000068854 | *H2bc21* | 83.63000705 | 90.16398607 | 89.50034712 | 190.9761783 | 173.7057771 | 153.3575509 | 5.11E-06 | 0.977065283 | 1.25E-06 | 23.49975686 | 1.968457116 | 5.291875181 |
| ENSMUSG00000074846 | *Gm10774* | 371.4377791 | 422.0441901 | 381.9521856 | 795.7340763 | 788.5189516 | 727.3004809 | 3.00E-17 | 0.975553057 | 2.59E-18 | 76.1775762 | 1.96639487 | 16.52310047 |
| ENSMUSG00000022556 | *Hsf1* | 597.4648251 | 691.5769569 | 490.3610567 | 1133.783863 | 1162.249563 | 1208.494233 | 4.75E-12 | 0.974886089 | 6.23E-13 | 51.77404858 | 1.965486002 | 11.3236938 |
| ENSMUSG00000019214 | *Chtf18* | 855.1356577 | 908.3542 | 661.7983414 | 1527.809426 | 1706.527664 | 1532.6572 | 2.39E-12 | 0.971835722 | 3.04E-13 | 53.18181269 | 1.96133466 | 11.621864 |
| ENSMUSG00000007080 | *Pole* | 1295.134974 | 1136.641739 | 926.5176779 | 2145.738095 | 2328.710175 | 2113.028291 | 7.60E-13 | 0.968979373 | 9.27E-14 | 55.51624297 | 1.957455313 | 12.11913562 |
| ENSMUSG00000004849 | *Ap1s1* | 5905.333291 | 6067.844424 | 7018.844123 | 12199.42656 | 12431.01706 | 12472.4686 | 5.38E-29 | 0.966846407 | 2.43E-30 | 131.0395358 | 1.954563435 | 28.26893423 |
| ENSMUSG00000071042 | *Rasgrp3* | 1826.298532 | 1898.239664 | 2151.790036 | 3658.181622 | 3951.016856 | 3856.89649 | 3.94E-26 | 0.966311695 | 2.03E-27 | 117.6858851 | 1.95383914 | 25.40490105 |
| ENSMUSG00000036854 | *Hspb6* | 79.86288962 | 96.87832545 | 76.89466442 | 148.1711728 | 185.2861622 | 162.5406378 | 2.54E-05 | 0.965776052 | 6.90E-06 | 20.22186415 | 1.953113855 | 4.595796513 |
| ENSMUSG00000045095 | *Magi1* | 140.1367686 | 160.184954 | 123.5356904 | 259.0251614 | 250.5574239 | 319.5714234 | 1.27E-06 | 0.965588174 | 2.88E-07 | 26.32882473 | 1.952859523 | 5.897371994 |
| ENSMUSG00000026434 | *Nucks1* | 2734.927258 | 2701.082817 | 2566.516996 | 5109.161552 | 5392.248425 | 5132.427257 | 7.99E-47 | 0.9651771 | 1.75E-48 | 214.1018014 | 1.952303164 | 46.09726237 |
| ENSMUSG00000024773 | *Atg2a* | 846.0945758 | 789.4144737 | 596.2487914 | 1407.07736 | 1469.65615 | 1491.333309 | 1.53E-11 | 0.964832514 | 2.09E-12 | 49.39466719 | 1.951836914 | 10.81519013 |
| ENSMUSG00000005774 | *Rfx5* | 308.1502062 | 336.6761607 | 221.8600154 | 531.2210937 | 619.0242237 | 548.2302867 | 1.39E-07 | 0.964548147 | 2.82E-08 | 30.82919854 | 1.951452229 | 6.855529673 |
| ENSMUSG00000038759 | *Nup205* | 4610.95174 | 4533.138278 | 3843.472653 | 8187.829254 | 8877.944351 | 8256.513414 | 3.61E-24 | 0.962406815 | 2.08E-25 | 108.5093956 | 1.948557919 | 23.44246638 |
| ENSMUSG00000022463 | *Srebf2* | 4006.706104 | 3596.008338 | 2749.299395 | 6462.458264 | 6969.286328 | 6733.0393 | 4.29E-12 | 0.960900404 | 5.59E-13 | 51.98681551 | 1.946524367 | 11.36779175 |
| ENSMUSG00000027331 | *Knstrn* | 1286.847316 | 1067.579963 | 1066.440756 | 2169.884509 | 2344.501609 | 2152.515565 | 8.93E-19 | 0.960156012 | 7.04E-20 | 83.30316481 | 1.945520271 | 18.04930947 |
| ENSMUSG00000075271 | *Ttc30a1* | 32.39720994 | 35.49007962 | 34.03534327 | 61.46359762 | 57.90192569 | 78.05623848 | 0.005397822 | 0.959040627 | 0.002210321 | 9.366126364 | 1.94401672 | 2.267781464 |
| ENSMUSG00000024393 | *Prrc2a* | 12437.51492 | 11611.97038 | 7776.445653 | 18939.56858 | 22342.77398 | 20595.82725 | 1.49E-08 | 0.958947318 | 2.71E-09 | 35.37958031 | 1.943890991 | 7.826250512 |
| ENSMUSG00000000881 | *Dlg3* | 57.26018501 | 93.04156009 | 56.72557212 | 117.439374 | 154.7560559 | 130.3998337 | 0.001735211 | 0.958079729 | 0.0006435 | 11.64573251 | 1.942722351 | 2.760647699 |
| ENSMUSG00000020901 | *Pik3r5* | 2652.804098 | 2821.940926 | 2236.24811 | 4545.013531 | 5564.90144 | 4878.974059 | 3.47E-15 | 0.958052959 | 3.49E-16 | 66.50568404 | 1.942686302 | 14.45993112 |
| ENSMUSG00000060727 | *Gm12571* | 91.91766541 | 74.81692461 | 107.1483029 | 167.9273292 | 182.1278753 | 179.0701942 | 1.84E-05 | 0.957738525 | 4.87E-06 | 20.88582869 | 1.942262942 | 4.736115809 |
| ENSMUSG00000024056 | *Ndc80* | 381.2322844 | 325.1658646 | 375.6493442 | 721.0997078 | 688.5065345 | 692.4047508 | 1.13E-15 | 0.957668663 | 1.09E-16 | 68.79572298 | 1.942168891 | 14.9462066 |
| ENSMUSG00000038773 | *Kdm3b* | 1601.778333 | 1668.033742 | 1235.356904 | 2668.178675 | 3181.447626 | 2908.283615 | 2.46E-12 | 0.95733174 | 3.13E-13 | 53.12355086 | 1.941715375 | 11.60945079 |
| ENSMUSG00000027189 | *Trim44* | 1400.614262 | 1324.643242 | 1005.933479 | 2195.128486 | 2583.478648 | 2473.005297 | 2.44E-11 | 0.956413988 | 3.40E-12 | 48.44601795 | 1.94048057 | 10.61205711 |
| ENSMUSG00000005417 | *Mprip* | 2941.365293 | 3069.412292 | 2449.284147 | 5183.79592 | 5794.403618 | 5446.488829 | 5.27E-19 | 0.956089376 | 4.10E-20 | 84.37241421 | 1.940044004 | 18.2784941 |
| ENSMUSG00000043262 | *Uevld* | 345.067957 | 366.4110923 | 385.7338904 | 665.1239314 | 708.5090179 | 745.6666547 | 4.29E-16 | 0.954283884 | 4.01E-17 | 70.77318605 | 1.937617612 | 15.36746878 |
| ENSMUSG00000072980 | *Oip5* | 169.5202846 | 125.6540657 | 131.0991 | 266.7081111 | 267.4016204 | 295.6953975 | 8.48E-07 | 0.953417149 | 1.88E-07 | 27.14782726 | 1.93645389 | 6.071499543 |
| ENSMUSG00000062031 | *Pgghg* | 461.8485975 | 402.8603633 | 306.3180894 | 760.6120205 | 789.5717139 | 727.3004809 | 1.15E-08 | 0.952767346 | 2.08E-09 | 35.90165541 | 1.93558189 | 7.938027791 |
| ENSMUSG00000032290 | *Ptpn9* | 1455.614177 | 1549.094016 | 1239.138609 | 2572.690586 | 2828.77226 | 2819.207672 | 1.91E-18 | 0.95223326 | 1.55E-19 | 81.74866055 | 1.93486547 | 17.72005379 |
| ENSMUSG00000028961 | *Pgd* | 19924.28411 | 21430.25294 | 21903.63425 | 38778.04228 | 41895.7279 | 41511.22593 | 1.12E-41 | 0.949915795 | 2.99E-43 | 190.1240145 | 1.931759905 | 40.95228403 |
| ENSMUSG00000032892 | *Rangrf* | 350.3419214 | 338.5945434 | 332.7900231 | 593.7822556 | 655.8709037 | 722.7089375 | 1.99E-13 | 0.948628366 | 2.31E-14 | 58.24467174 | 1.930036814 | 12.70116891 |
| ENSMUSG00000039087 | *Rreb1* | 810.683672 | 844.0883802 | 466.4102596 | 1191.954768 | 1447.548142 | 1460.110814 | 1.91E-05 | 0.947904668 | 5.09E-06 | 20.80437059 | 1.929068894 | 4.719884794 |
| ENSMUSG00000043987 | *Cep164* | 858.1493516 | 905.476626 | 683.2280019 | 1424.638388 | 1748.638156 | 1554.696609 | 1.96E-11 | 0.947760793 | 2.71E-12 | 48.8845712 | 1.928876524 | 10.70680583 |
| ENSMUSG00000038002 | *Cramp1* | 345.067957 | 394.2276412 | 226.9022885 | 544.3918646 | 674.8206248 | 650.1625511 | 1.32E-05 | 0.947484002 | 3.45E-06 | 21.55076241 | 1.928506491 | 4.878738583 |
| ENSMUSG00000046329 | *Slc25a23* | 611.7798714 | 633.0662851 | 558.4317433 | 1081.10078 | 1195.937956 | 1203.90269 | 1.70E-19 | 0.946855064 | 1.29E-20 | 86.65902086 | 1.927665949 | 18.76984465 |
| ENSMUSG00000052551 | *Adarb2* | 58.01360849 | 55.63309779 | 68.07068654 | 100.9759104 | 106.3289908 | 140.5012293 | 0.000746995 | 0.946564924 | 0.000257029 | 13.3601382 | 1.927278315 | 3.126682423 |
| ENSMUSG00000045775 | *Slc16a5* | 26.36982204 | 31.65331426 | 40.33818462 | 68.04898308 | 64.2184994 | 55.09852128 | 0.009654427 | 0.946409205 | 0.004166006 | 8.210004083 | 1.927070303 | 2.015273507 |
| ENSMUSG00000036138 | *Acaa1a* | 3496.638403 | 3587.375616 | 3349.329891 | 6495.385191 | 6797.686075 | 6800.994143 | 2.27E-48 | 0.94508978 | 4.62E-50 | 221.3377446 | 1.925308696 | 47.64464598 |
| ENSMUSG00000028841 | *Cnksr1* | 85.13685402 | 89.20479472 | 79.41580096 | 140.4882231 | 186.3389245 | 162.5406378 | 3.36E-05 | 0.945067702 | 9.31E-06 | 19.64836637 | 1.925279233 | 4.473454664 |
| ENSMUSG00000078695 | *Cisd3* | 341.3008396 | 366.4110923 | 368.0859346 | 715.6118865 | 729.5642636 | 623.5315992 | 3.48E-14 | 0.944483934 | 3.78E-15 | 61.80893665 | 1.924500351 | 13.45822424 |
| ENSMUSG00000022568 | *Scrib* | 2228.626674 | 2250.262886 | 1684.119208 | 3730.620863 | 4168.938649 | 3962.501989 | 2.31E-13 | 0.943163963 | 2.71E-14 | 57.93408184 | 1.922740365 | 12.6361734 |
| ENSMUSG00000024312 | *Wdr46* | 1231.847401 | 1171.172628 | 882.3977885 | 1944.883839 | 2297.127306 | 2081.805796 | 9.57E-11 | 0.942132626 | 1.41E-11 | 45.65686151 | 1.92136635 | 10.0190228 |
| ENSMUSG00000041219 | *Arhgap11a* | 769.9988037 | 747.2100547 | 592.4670865 | 1291.833114 | 1463.339576 | 1308.58988 | 7.68E-13 | 0.941860042 | 9.37E-14 | 55.49491949 | 1.921003359 | 12.11466896 |
| ENSMUSG00000033685 | *Ucp2* | 15167.16822 | 14236.31788 | 11272.00146 | 24557.99994 | 27131.78961 | 26442.69867 | 5.33E-15 | 0.941493783 | 5.44E-16 | 65.63173689 | 1.920515733 | 14.27354149 |
| ENSMUSG00000022962 | *Gart* | 947.0533231 | 846.9659542 | 668.1011827 | 1442.199416 | 1626.517731 | 1668.566886 | 1.60E-10 | 0.940870566 | 2.39E-11 | 44.6210371 | 1.919686287 | 9.795744004 |
| ENSMUSG00000042423 | *Fbrs* | 1248.422718 | 1176.927776 | 809.2848289 | 1876.834856 | 2287.652446 | 2055.174844 | 5.37E-08 | 0.940810606 | 1.03E-08 | 32.77701603 | 1.919606504 | 7.270217055 |
| ENSMUSG00000034659 | *Tmem109* | 3721.912026 | 3670.825262 | 2666.101889 | 6046.481416 | 6682.934986 | 6581.518367 | 8.69E-12 | 0.940021702 | 1.17E-12 | 50.53595127 | 1.918557099 | 11.06120883 |
| ENSMUSG00000046341 | *Gm11223* | 989.9984618 | 879.5784598 | 926.5176779 | 1724.273426 | 1886.550015 | 1754.887903 | 3.24E-24 | 0.939018746 | 1.86E-25 | 108.7297744 | 1.917223789 | 23.48919814 |
| ENSMUSG00000052488 | *Cherp* | 2771.091585 | 2852.635048 | 1850.514219 | 4367.208124 | 5034.309248 | 4925.807802 | 1.72E-08 | 0.937733062 | 3.14E-09 | 35.09621259 | 1.915515981 | 7.765021351 |
| ENSMUSG00000034311 | *Kif4* | 696.9167254 | 579.35157 | 477.755374 | 1117.3204 | 1153.827465 | 1096.460573 | 5.06E-10 | 0.935875617 | 7.94E-11 | 42.27135213 | 1.913051374 | 9.29604447 |
| ENSMUSG00000034022 | *Cpsf1* | 968.1491807 | 1193.234028 | 661.7983414 | 1554.150968 | 1392.804503 | 2456.47574 | 0.000387836 | 0.935765235 | 0.000126502 | 14.6932009 | 1.91290501 | 3.411351653 |
| ENSMUSG00000032336 | *Nptn* | 2740.201222 | 2879.492406 | 2875.356222 | 5223.308233 | 5615.434029 | 5377.615677 | 2.30E-43 | 0.933380857 | 5.72E-45 | 197.9957702 | 1.909746116 | 42.639177 |
| ENSMUSG00000003123 | *Lipe* | 696.9167254 | 782.7001344 | 608.854474 | 1275.369651 | 1348.588487 | 1371.034871 | 3.98E-15 | 0.933375582 | 4.02E-16 | 66.22644082 | 1.909739132 | 14.40018005 |
| ENSMUSG00000029547 | *Ints1* | 2456.913991 | 2319.324663 | 1696.72489 | 3665.864572 | 4410.021213 | 4288.501573 | 1.07E-09 | 0.932420218 | 1.73E-10 | 40.7459101 | 1.908474907 | 8.972448702 |
| ENSMUSG00000040354 | *Mars1* | 4272.664594 | 3836.765364 | 3186.716585 | 7192.338485 | 7286.167776 | 7082.914911 | 5.51E-16 | 0.931537462 | 5.19E-17 | 70.26373946 | 1.907307507 | 15.25849607 |
| ENSMUSG00000074622 | *Mafb* | 693.149608 | 661.8420254 | 479.0159423 | 1037.19821 | 1268.578554 | 1201.147764 | 2.08E-08 | 0.931443341 | 3.82E-09 | 34.7118757 | 1.907183079 | 7.682867713 |
| ENSMUSG00000031015 | *Swap70* | 2162.325408 | 2099.669846 | 1819.000013 | 3842.572415 | 3854.162726 | 3906.485159 | 2.38E-25 | 0.930279771 | 1.29E-26 | 114.0241425 | 1.905645507 | 24.62332204 |
| ENSMUSG00000052593 | *Adam17* | 1631.161849 | 1565.400269 | 1332.420661 | 2816.349848 | 2928.784677 | 2898.182219 | 6.33E-21 | 0.93001562 | 4.35E-22 | 93.36597102 | 1.905296625 | 20.19833119 |
| ENSMUSG00000027381 | *Bcl2l11* | 400.8212951 | 331.880204 | 249.5925173 | 613.5384119 | 642.184994 | 625.3682165 | 4.78E-07 | 0.929774419 | 1.03E-07 | 28.3124796 | 1.904978109 | 6.32030256 |
| ENSMUSG00000034066 | *Farp2* | 106.9861351 | 123.735683 | 93.28205192 | 167.9273292 | 233.7132273 | 216.7208504 | 6.20E-05 | 0.929031387 | 1.78E-05 | 18.40738643 | 1.90399724 | 4.207281014 |
| ENSMUSG00000027630 | *Tbl1xr1* | 644.1770813 | 579.35157 | 586.1642452 | 1190.857204 | 1147.510891 | 1113.908439 | 2.86E-21 | 0.928753458 | 1.92E-22 | 94.98697803 | 1.903630478 | 20.54388521 |
| ENSMUSG00000027203 | *Dut* | 1935.544938 | 1877.137455 | 1970.268205 | 3514.400707 | 3760.466883 | 3727.414965 | 6.86E-40 | 0.928484779 | 1.95E-41 | 181.8159316 | 1.90327599 | 39.16349137 |
| ENSMUSG00000027544 | *Nfatc2* | 1151.984512 | 1086.763789 | 598.7699279 | 1549.760711 | 2022.35635 | 1834.780759 | 8.28E-05 | 0.927964053 | 2.43E-05 | 17.81822036 | 1.902589146 | 4.081933961 |
| ENSMUSG00000029480 | *Dhx37* | 1744.175372 | 1842.606566 | 1429.484417 | 2990.862563 | 3363.575501 | 3193.877617 | 2.02E-15 | 0.927055789 | 2.00E-16 | 67.60024039 | 1.901391727 | 14.6941895 |
| ENSMUSG00000059851 | *Kmt5c* | 374.451473 | 401.9011719 | 282.3672923 | 656.3434174 | 679.0316739 | 685.0582813 | 2.88E-09 | 0.926679822 | 4.90E-10 | 38.71599022 | 1.900896288 | 8.540053951 |
| ENSMUSG00000039515 | *Ptpa* | 7776.837232 | 8068.717562 | 7158.767201 | 14333.09145 | 14857.63413 | 14549.68285 | 3.23E-39 | 0.926651258 | 9.44E-41 | 178.6733027 | 1.900858654 | 38.49079248 |
| ENSMUSG00000020899 | *Pfas* | 2230.886945 | 2349.059594 | 1583.273746 | 3631.840081 | 4182.624559 | 3907.403467 | 1.85E-09 | 0.926166393 | 3.08E-10 | 39.62234925 | 1.900219915 | 8.732807153 |

| | | | | | | | | | | | | |
|---|---|---|---|---|---|---|---|---|---|---|---|---|
| ENSMUSG00000037628 | *Cdkn3* | 107.7395586 | 102.6334735 | 80.67636923 | 175.6102789 | 189.4972113 | 191.9265158 | 1.23E-05 | 0.925805241 | 3.19E-06 | 21.69801146 | 1.89974429 | 4.909901102 |
| ENSMUSG00000032434 | *Cmtm6* | 1110.54622 | 1181.723732 | 993.3277962 | 2027.201157 | 2102.366284 | 2120.374761 | 9.58E-23 | 0.925709235 | 5.99E-24 | 101.8491979 | 1.899617873 | 22.01847445 |
| ENSMUSG00000029033 | *Acap3* | 1052.532611 | 1068.539154 | 936.602224 | 1891.103191 | 2037.095022 | 1887.124354 | 2.76E-23 | 0.92494891 | 1.67E-24 | 104.3829321 | 1.898617005 | 22.55936249 |
| ENSMUSG00000030067 | *Foxp1* | 872.4643979 | 839.2924235 | 670.6223192 | 1499.272756 | 1471.761675 | 1563.879696 | 4.77E-14 | 0.924927574 | 5.23E-15 | 61.17309906 | 1.898588926 | 13.32179778 |
| ENSMUSG00000048216 | *Gpr85* | 32.39720994 | 37.4084623 | 36.55647981 | 70.24411156 | 52.63811426 | 78.05623848 | 0.008963517 | 0.924717339 | 0.003849416 | 8.353516556 | 1.898312277 | 2.047521541 |
| ENSMUSG00000008450 | *Nutf2* | 198.9038006 | 299.2676984 | 122.2751221 | 444.5135185 | 320.0397347 | 415.9938357 | 0.003863657 | 0.923877984 | 0.001538887 | 10.03146326 | 1.897208167 | 2.4130014 |
| ENSMUSG00000023908 | *Pkmyt1* | 705.9578073 | 697.332105 | 499.1850346 | 1104.149629 | 1284.369988 | 1226.860407 | 4.64E-09 | 0.922442085 | 8.05E-10 | 37.74711522 | 1.895320836 | 8.333784398 |
| ENSMUSG00000091228 | *Gm20390* | 1139.929736 | 1725.585223 | 1508.900218 | 3333.302606 | 2302.391118 | 2645.64733 | 9.48E-07 | 0.921473533 | 2.13E-07 | 26.91590639 | 1.894048841 | 6.023041987 |
| ENSMUSG00000095098 | *Ccdc85b* | 446.7801278 | 419.1666161 | 414.7269606 | 716.7094508 | 841.1570659 | 868.7200189 | 1.69E-13 | 0.921046599 | 1.95E-14 | 58.58216464 | 1.89348842 | 12.77232577 |
| ENSMUSG00000018921 | *Pelp1* | 1892.599799 | 1900.158047 | 1392.927938 | 3073.179881 | 3481.484877 | 3271.933855 | 1.89E-11 | 0.920453376 | 2.61E-12 | 48.96034718 | 1.892709996 | 10.7231527 |
| ENSMUSG00000007035 | *Msh5* | 157.4655088 | 128.5316397 | 131.0991 | 275.488625 | 251.6101862 | 265.3912108 | 2.51E-07 | 0.918824444 | 5.24E-08 | 29.62422866 | 1.890574163 | 6.600789382 |
| ENSMUSG00000027379 | *Bub1* | 1162.53244 | 1018.661204 | 1065.180188 | 2049.152442 | 1982.351383 | 2110.273365 | 7.80E-25 | 0.918389854 | 4.32E-26 | 111.6231494 | 1.890004742 | 24.10775758 |
| ENSMUSG00000032547 | *Ryk* | 1515.888056 | 1405.215315 | 1273.173952 | 2675.861625 | 2816.139113 | 2446.374345 | 7.74E-19 | 0.918082021 | 6.07E-20 | 83.59657828 | 1.889601508 | 18.1114958 |
| ENSMUSG00000057278 | *Snrpg* | 784.3138499 | 706.9240184 | 630.2841346 | 1342.321069 | 1251.734357 | 1422.460158 | 9.18E-15 | 0.917310608 | 9.57E-16 | 64.51769374 | 1.888591402 | 14.03697972 |
| ENSMUSG00000003316 | *Glg1* | 4176.979812 | 4164.808803 | 2846.363152 | 6478.921727 | 7850.448361 | 6802.830761 | 1.06E-08 | 0.91672231 | 1.90E-09 | 36.07744161 | 1.887821435 | 7.975215634 |
| ENSMUSG00000068566 | *Myadm* | 2414.722276 | 2336.590107 | 1611.006248 | 3936.96294 | 3889.956644 | 4192.079161 | 3.25E-09 | 0.916285891 | 5.55E-10 | 38.47295662 | 1.887250451 | 8.488423296 |
| ENSMUSG00000025898 | *Cwf19l2* | 208.6983059 | 166.8992934 | 175.2189894 | 354.5132505 | 366.3612752 | 323.2446582 | 3.83E-08 | 0.915938206 | 7.25E-09 | 33.46613166 | 1.886795685 | 7.416327348 |
| ENSMUSG00000029730 | *Mcm7* | 4999.718259 | 5049.18322 | 4705.701349 | 8818.928694 | 9681.201975 | 9338.281048 | 3.34E-38 | 0.915499658 | 1.02E-39 | 173.9380605 | 1.886222227 | 37.47572995 |
| ENSMUSG00000044348 | *Slc25a53* | 51.9862206 | 37.4084623 | 49.1621625 | 84.51244672 | 84.22098282 | 92.74917749 | 0.002263614 | 0.915490419 | 0.000858253 | 11.11082407 | 1.886210148 | 2.645197543 |
| ENSMUSG00000041911 | *Dlx1* | 64.79441988 | 40.28603633 | 36.55647981 | 92.19539643 | 90.53755653 | 88.15763405 | 0.007098007 | 0.915251976 | 0.002974714 | 8.822913195 | 1.885898428 | 2.148863569 |
| ENSMUSG00000037905 | *Bri3bp* | 4839.99248 | 4959.019234 | 4424.594625 | 8505.02532 | 9100.077193 | 9221.655845 | 2.03E-32 | 0.914871536 | 8.01E-34 | 146.9591231 | 1.885401179 | 31.69172561 |
| ENSMUSG00000040725 | *Hnrnpul1* | 3983.349975 | 3820.459112 | 2833.757469 | 6306.604141 | 7149.308679 | 6611.822554 | 3.36E-11 | 0.914785443 | 4.74E-12 | 47.79365258 | 1.885288671 | 10.47352905 |
| ENSMUSG00000024388 | *Myo7b* | 56.50676152 | 59.46986315 | 52.94386731 | 107.5612958 | 103.1707039 | 108.3604252 | 0.000312022 | 0.914574856 | 1.00E-04 | 15.13718118 | 1.8850135 | 3.505814196 |
| ENSMUSG00000026811 | *St6galnac6* | 1209.99812 | 1277.642866 | 1201.321561 | 2203.909 | 2349.765421 | 2394.949058 | 3.99E-30 | 0.913571539 | 1.72E-31 | 136.297659 | 1.88370303 | 29.39871389 |
| ENSMUSG00000035365 | *Parpbp* | 290.0680425 | 253.2265141 | 242.0291077 | 511.4649373 | 485.3234135 | 488.540222 | 1.89E-11 | 0.912845994 | 2.61E-12 | 48.95806038 | 1.882755936 | 10.72286216 |
| ENSMUSG00000020167 | *Tcf3* | 2054.585849 | 1990.322033 | 1503.857945 | 3310.253757 | 3688.879047 | 3455.595593 | 6.07E-12 | 0.912175218 | 8.06E-13 | 51.2679055 | 1.881880759 | 11.21669729 |
| ENSMUSG00000033124 | *Atg9a* | 1376.504711 | 1434.950246 | 1100.476099 | 2292.811704 | 2519.260148 | 2549.224918 | 1.02E-13 | 0.91012685 | 1.15E-14 | 59.61836649 | 1.879210722 | 12.99346921 |
| ENSMUSG00000005198 | *Polr2a* | 4778.965178 | 4490.933859 | 2855.18713 | 6677.580855 | 8278.922611 | 7829.499874 | 1.36E-06 | 0.909514013 | 3.11E-07 | 26.18188013 | 1.878412629 | 5.866288583 |
| ENSMUSG00000042462 | *Dctpp1* | 1404.38138 | 1448.378925 | 1703.027732 | 2900.862295 | 2825.613973 | 2812.779511 | 2.79E-20 | 0.908810155 | 1.99E-21 | 90.35050992 | 1.877496417 | 19.55441826 |
| ENSMUSG00000108608 | *Gm6916* | 134.8628042 | 156.3481886 | 194.1275135 | 352.3181221 | 257.9267599 | 294.7770888 | 3.32E-05 | 0.907357116 | 9.19E-06 | 19.67312402 | 1.875606412 | 4.478758571 |
| ENSMUSG00000039952 | *Dag1* | 2545.817962 | 2812.349012 | 2111.451851 | 4417.696079 | 4965.879699 | 4635.622257 | 5.32E-14 | 0.907315446 | 5.85E-15 | 60.95074365 | 1.875552238 | 13.27437477 |
| ENSMUSG00000035632 | *Cnot3* | 1491.025081 | 1455.093264 | 1105.518372 | 2385.0071 | 2681.38554 | 2540.96014 | 8.46E-12 | 0.906646667 | 1.14E-12 | 50.59102848 | 1.874683004 | 11.07273246 |
| ENSMUSG00000001855 | *Nup214* | 2360.475785 | 2502.530209 | 1546.717266 | 3553.913019 | 4545.827547 | 3924.851333 | 1.05E-06 | 0.906596795 | 2.37E-07 | 26.70463172 | 1.874618201 | 5.978303661 |
| ENSMUSG00000038664 | *Herc1* | 1289.107586 | 1266.13257 | 843.3201721 | 1990.981537 | 2342.396085 | 2041.400213 | 1.60E-07 | 0.905114804 | 3.26E-08 | 30.54723741 | 1.872693511 | 6.796823385 |
| ENSMUSG00000046591 | *Ticrr* | 611.0264479 | 584.1475267 | 352.9591154 | 895.6124224 | 1107.505924 | 902.6974403 | 3.20E-05 | 0.904207721 | 8.83E-06 | 19.74874021 | 1.87151644 | 4.49513229 |
| ENSMUSG00000026365 | *Cfh* | 142.397039 | 150.5930406 | 166.3950115 | 248.049519 | 297.9317267 | 309.4700279 | 5.89E-07 | 0.904076515 | 1.28E-07 | 27.8905534 | 1.871346242 | 6.229744126 |
| ENSMUSG00000029844 | *Hoxa1* | 127.3285693 | 162.1033366 | 136.1413731 | 241.4641335 | 286.3513416 | 267.2278282 | 1.53E-06 | 0.903667478 | 3.53E-07 | 25.93792853 | 1.870815747 | 5.814405895 |
| ENSMUSG00000048277 | *Syngr2* | 2439.585251 | 2448.815494 | 2090.02219 | 4098.304884 | 4660.578637 | 4295.848042 | 5.70E-20 | 0.902355141 | 4.17E-21 | 88.8923811 | 1.869114747 | 19.24422084 |
| ENSMUSG00000033392 | *Clasp2* | 881.5054797 | 948.6402364 | 729.8690279 | 1572.80956 | 1687.577943 | 1533.575509 | 1.40E-13 | 0.902168204 | 1.60E-14 | 58.97001328 | 1.868872573 | 12.85456422 |
| ENSMUSG00000026785 | *Pkn3* | 177.0545194 | 188.0015029 | 138.6625096 | 299.6350384 | 323.1980216 | 323.2446582 | 3.38E-07 | 0.901701719 | 7.16E-08 | 29.02198326 | 1.868268384 | 6.470915216 |
| ENSMUSG00000060244 | *Alyref2* | 61.78072593 | 57.55148047 | 56.72557212 | 91.09783218 | 125.2787119 | 112.9519686 | 0.000951815 | 0.901170476 | 0.000333383 | 12.87285075 | 1.86758056 | 3.021447559 |
| ENSMUSG00000039396 | *Neil3* | 140.1367686 | 157.3073799 | 132.3596683 | 252.4397759 | 285.2985793 | 266.3095195 | 1.72E-07 | 0.901158357 | 3.53E-08 | 30.39340326 | 1.867564872 | 6.764279039 |
| ENSMUSG00000098134 | *Rnf113a2* | 186.0956013 | 199.511799 | 207.9937644 | 390.7328706 | 363.2029884 | 351.7122275 | 7.35E-10 | 0.900866611 | 1.17E-10 | 41.50790267 | 1.867187245 | 9.133626121 |
| ENSMUSG00000022565 | *Plec* | 14882.37414 | 13306.86148 | 8189.912045 | 19436.76518 | 25408.41775 | 23038.52836 | 1.33E-05 | 0.899746118 | 3.47E-06 | 21.53540201 | 1.865737626 | 4.875488769 |
| ENSMUSG00000055809 | *Dnaaf3* | 104.7258647 | 126.613257 | 78.15523269 | 160.2443795 | 200.0248342 | 219.4757764 | 0.000572803 | 0.899488029 | 0.000193398 | 13.89416698 | 1.865403887 | 3.24199453 |
| ENSMUSG00000001995 | *Sipa1l2* | 457.3280566 | 407.65632 | 286.1489971 | 707.9289368 | 689.5592968 | 758.5229763 | 1.15E-06 | 0.899142146 | 2.60E-07 | 26.52940374 | 1.864956714 | 5.940498355 |
| ENSMUSG00000037331 | *Larp1* | 5933.963383 | 5622.779642 | 3505.640357 | 8925.392425 | 9950.70912 | 9209.717832 | 1.00E-06 | 0.898318218 | 2.25E-07 | 26.80679479 | 1.863891936 | 6.00007333 |
| ENSMUSG00000036114 | *Rpp25l* | 1056.299729 | 929.4564095 | 945.4262019 | 1806.590744 | 1804.434557 | 1858.656785 | 1.50E-23 | 0.897807572 | 8.94E-25 | 105.6190021 | 1.863232322 | 22.82482479 |
| ENSMUSG00000024294 | *Mib1* | 1140.683159 | 1169.254245 | 786.5946 | 1894.395884 | 1953.926801 | 1930.284862 | 9.96E-09 | 0.897401173 | 1.78E-09 | 36.20266522 | 1.862707535 | 8.001736608 |
| ENSMUSG00000064264 | *Zfp428* | 134.1093807 | 135.2459791 | 117.232849 | 225.0006698 | 241.0825633 | 256.208124 | 5.25E-07 | 0.897338062 | 1.14E-07 | 28.12238656 | 1.862626052 | 6.27944195 |
| ENSMUSG00000023118 | *Sympk* | 7151.495738 | 6746.951893 | 5401.535034 | 11491.49763 | 12376.27342 | 12085.86064 | 8.19E-15 | 0.89695087 | 8.49E-16 | 64.75226448 | 1.862126225 | 14.08696151 |
| ENSMUSG00000044700 | *Tmem201* | 1203.217309 | 1257.499848 | 843.3201721 | 1863.664085 | 2166.584783 | 2129.557847 | 3.50E-08 | 0.89652623 | 6.59E-09 | 33.65241564 | 1.861578211 | 7.456500223 |
| ENSMUSG00000052533 | *Nup188* | 3706.090132 | 3722.621595 | 2508.530856 | 5620.626489 | 6607.136102 | 6278.4765 | 2.08E-08 | 0.896268456 | 3.83E-09 | 34.70910479 | 1.861245623 | 7.682411341 |
| ENSMUSG00000029162 | *Khk* | 339.0405691 | 305.0228465 | 310.0997942 | 578.4163561 | 642.184994 | 556.4950649 | 4.07E-12 | 0.893860512 | 5.30E-13 | 52.09051339 | 1.858141684 | 11.39004031 |
| ENSMUSG00000022521 | *Crebbp* | 620.8209532 | 667.5971734 | 402.1212779 | 1092.076422 | 1062.237146 | 994.5283091 | 2.83E-06 | 0.893303557 | 6.71E-07 | 24.69690986 | 1.857424484 | 5.54826035 |
| ENSMUSG00000025278 | *Flnb* | 2047.805038 | 1993.199607 | 1203.842697 | 2892.081781 | 3627.818835 | 3229.691656 | 8.27E-06 | 0.893066535 | 2.09E-06 | 22.51189592 | 1.857119351 | 5.082383943 |
| ENSMUSG00000036667 | *Tcaf1* | 113.7669465 | 126.613257 | 124.7962587 | 229.3909268 | 228.4494159 | 218.5574677 | 4.55E-07 | 0.892625555 | 9.78E-08 | 28.41708255 | 1.856551783 | 6.341970219 |
| ENSMUSG00000028655 | *Mfsd2a* | 236.5749749 | 161.1441453 | 129.8385317 | 363.2937645 | 275.8237187 | 346.2023754 | 0.000638233 | 0.891877364 | 0.000217548 | 13.67310699 | 1.855589212 | 3.195020515 |
| ENSMUSG00000000171 | *Sdhd* | 3507.186332 | 3696.723429 | 3832.127538 | 6868.557034 | 6841.902091 | 6726.61114 | 3.93E-42 | 0.889883354 | 1.04E-43 | 192.2310625 | 1.853026296 | 41.40521362 |
| ENSMUSG00000027597 | *Ahcy* | 539.4512167 | 443.1463996 | 460.1074183 | 863.7830594 | 893.7951801 | 921.9819228 | 1.35E-13 | 0.88971513 | 1.54E-14 | 59.04376195 | 1.852810237 | 12.87058115 |
| ENSMUSG00000022557 | *Bop1* | 4179.240082 | 3983.52164 | 3298.907161 | 6815.87395 | 7033.504827 | 7398.813099 | 5.76E-17 | 0.889591469 | 5.08E-18 | 74.8489048 | 1.85265143 | 16.23938996 |
| ENSMUSG00000097829 | *None* | 354.1090389 | 312.6963772 | 170.1767163 | 535.6113507 | 596.9162157 | 425.1769225 | 0.001521374 | 0.889513767 | 0.000555742 | 11.91866766 | 1.852551651 | 2.817763979 |
| ENSMUSG00000055491 | *Pprc1* | 2374.037407 | 2270.405904 | 1546.717266 | 3497.937243 | 4133.144732 | 3848.631711 | 8.75E-08 | 0.889449334 | 1.72E-08 | 31.78470654 | 1.852468916 | 7.058219446 |
| ENSMUSG00000044066 | *Cep68* | 604.99906 | 576.473996 | 552.1289019 | 1042.686031 | 1176.988235 | 996.3649265 | 2.50E-15 | 0.888876401 | 2.50E-16 | 67.16362355 | 1.851733396 | 14.60130933 |
| ENSMUSG00000049848 | *Ceacam19* | 125.8217223 | 126.613257 | 109.6694394 | 197.5615638 | 200.0248342 | 273.655989 | 6.91E-05 | 0.888779325 | 2.01E-05 | 18.18362009 | 1.851608801 | 4.160497467 |
| ENSMUSG00000039231 | *Suv39h1* | 1851.914931 | 1824.381931 | 1755.971599 | 3267.448752 | 3420.424665 | 3371.111194 | 8.54E-39 | 0.887850618 | 2.53E-40 | 176.7109799 | 1.850417248 | 38.06838185 |
| ENSMUSG00000034957 | *Cebpa* | 2174.380183 | 2263.691565 | 2173.219696 | 3735.011119 | 4296.322886 | 4200.343939 | 6.12E-26 | 0.887757517 | 3.20E-27 | 116.7832682 | 1.850297839 | 25.21299291 |
| ENSMUSG00000004100 | *Ppan* | 3625.473819 | 3897.194419 | 3480.428991 | 6724.776118 | 6646.088306 | 6992.920659 | 7.05E-34 | 0.887735499 | 2.57E-35 | 153.7914376 | 1.8502696 | 33.15208427 |
| ENSMUSG00000041936 | *Agrn* | 521.369053 | 430.6769122 | 329.0083183 | 736.4656072 | 855.8957379 | 786.9905456 | 1.20E-06 | 0.887427243 | 2.72E-07 | 26.4407902 | 1.849874301 | 5.921234225 |
| ENSMUSG00000020331 | *Hcn2* | 126.5751458 | 160.184954 | 156.3104654 | 230.4884911 | 301.0900136 | 283.7573846 | 1.16E-05 | 0.886544536 | 2.98E-06 | 21.82957712 | 1.84874281 | 4.937378759 |
| ENSMUSG00000032294 | *Pkm* | 191288.9494 | 191906.3708 | 177078.3276 | 346453.8363 | 340999.179 | 348304.384 | 5.12E-49 | 0.886461854 | 1.02E-50 | 224.3456656 | 1.848636859 | 48.29035464 |
| ENSMUSG00000074220 | *Zfp382* | 89.65739495 | 76.73530729 | 46.64102596 | 147.0736086 | 122.1204251 | 129.481525 | 0.004005486 | 0.886312077 | 0.00160005 | 9.959689455 | 1.848444949 | 2.39734477 |
| ENSMUSG00000022560 | *Slc52a2* | 188.3558717 | 233.0834959 | 197.9092183 | 379.7572281 | 353.7281278 | 409.5656749 | 3.92E-08 | 0.885892649 | 7.43E-09 | 33.41869091 | 1.847907636 | 7.406521018 |
| ENSMUSG00000013076 | *Amotl1* | 2041.77765 | 1946.199231 | 1500.07624 | 3156.594763 | 3573.075196 | 3420.699863 | 1.83E-11 | 0.885445682 | 2.53E-12 | 49.02460009 | 1.847335219 | 10.73716333 |
| ENSMUSG00000032109 | *Nlrx1* | 775.2727681 | 826.822936 | 732.3901644 | 1416.955438 | 1481.236535 | 1417.868614 | 2.36E-22 | 0.884850951 | 1.49E-23 | 100.0387311 | 1.846573837 | 21.62766904 |
| ENSMUSG00000032479 | *Map4* | 3765.610588 | 3996.950318 | 3005.194754 | 6313.189527 | 6780.841879 | 6799.157526 | 7.95E-14 | 0.884754578 | 8.94E-15 | 60.11672785 | 1.846450488 | 13.09947979 |
| ENSMUSG00000009073 | *Nf2* | 3655.610759 | 3559.559067 | 2794.679853 | 5654.650981 | 6580.817045 | 6250.927239 | 1.11E-12 | 0.884122216 | 1.37E-13 | 54.75359018 | 1.845641329 | 11.95620923 |
| ENSMUSG00000028430 | *Nol6* | 2743.214916 | 2819.063352 | 2058.507984 | 4357.330045 | 5020.623338 | 4686.129235 | 7.01E-11 | 0.882880587 | 1.01E-11 | 46.30065856 | 1.844053596 | 10.15397954 |
| ENSMUSG00000031145 | *Prickle3* | 79.10946613 | 70.0209679 | 73.11295962 | 139.3906589 | 122.1204251 | 148.7660075 | 0.000168379 | 0.882414958 | 5.19E-05 | 16.37681407 | 1.843458524 | 3.773712383 |
| ENSMUSG00000030216 | *Wbp11* | 2159.311714 | 2146.670221 | 1720.675688 | 3415.619925 | 3979.441438 | 3724.660039 | 9.38E-14 | 0.882262276 | 1.06E-14 | 59.77924171 | 1.843263439 | 13.02791765 |
| ENSMUSG00000052915 | *Msl1* | 1434.518319 | 1443.582968 | 1212.666675 | 2427.812106 | 2594.006271 | 2524.430583 | 2.80E-19 | 0.88113911 | 2.16E-20 | 85.64182266 | 1.841828982 | 18.55230575 |
| ENSMUSG00000006307 | *Kmt2b* | 1459.381294 | 1428.235907 | 866.010401 | 2124.884375 | 2453.988887 | 2342.605463 | 6.06E-06 | 0.880822154 | 1.50E-06 | 23.14293139 | 1.841424382 | 5.217595829 |
| ENSMUSG00000014498 | *Ankrd52* | 4206.363328 | 4136.992254 | 2764.426214 | 5963.066533 | 7638.843141 | 6838.6448 | 3.81E-07 | 0.879179533 | 8.11E-08 | 28.77902444 | 1.839328969 | 6.418968028 |
| ENSMUSG00000037438 | *Uqcrh-ps1* | 78.35604264 | 50.83714108 | 90.76091539 | 138.2930946 | 100.0124171 | 164.3772552 | 0.009310079 | 0.879135504 | 0.004011026 | 8.278816583 | 1.839272837 | 2.031046645 |
| ENSMUSG00000034235 | *Usp54* | 245.6160567 | 259.9408534 | 195.3880817 | 392.9279991 | 468.4792169 | 432.5233921 | 1.64E-07 | 0.878455193 | 3.34E-08 | 30.49842799 | 1.838405721 | 6.786480764 |
| ENSMUSG00000058099 | *Nfam1* | 1756.983572 | 1779.299938 | 1570.668063 | 3067.69206 | 3273.037945 | 3054.294696 | 6.40E-25 | 0.877704197 | 3.52E-26 | 112.0322175 | 1.837448987 | 24.19358275 |
| ENSMUSG00000074305 | *Peak1* | 4328.417933 | 4240.584919 | 3128.730444 | 6892.703447 | 7431.448971 | 7182.092249 | 3.40E-11 | 0.877660525 | 4.79E-12 | 47.76990353 | 1.837393366 | 10.46869106 |
| ENSMUSG00000002319 | *Ipo4* | 2958.694033 | 3094.351266 | 2379.952892 | 4911.599988 | 5329.082688 | 5257.317239 | 2.12E-14 | 0.87691077 | 2.27E-15 | 62.81753422 | 1.836438738 | 13.67373947 |
| ENSMUSG00000032320 | *Rcn2* | 1065.340811 | 1196.111602 | 1434.52669 | 2210.494386 | 2230.803282 | 2330.66745 | 1.76E-12 | 0.87665001 | 2.21E-13 | 53.80598635 | 1.836106841 | 11.75395558 |
| ENSMUSG00000092118 | *Fancf* | 165.7531671 | 197.5934163 | 166.3950115 | 317.1960663 | 326.3563084 | 328.7545103 | 3.13E-08 | 0.876044025 | 5.87E-09 | 33.8771286 | 1.835335771 | 7.503939511 |
| ENSMUSG00000020453 | *Patz1* | 1764.517806 | 1688.17676 | 1411.836462 | 2734.03253 | 3196.186298 | 3005.624336 | 4.40E-14 | 0.875531113 | 4.81E-15 | 61.33485045 | 1.834683381 | 13.35611671 |
| ENSMUSG00000037111 | *Setd7* | 1096.231174 | 996.5998034 | 734.911301 | 1697.931884 | 1750.74368 | 1749.378051 | 9.19E-09 | 0.875040808 | 1.63E-09 | 36.36834429 | 1.834059964 | 8.036869987 |
| ENSMUSG00000008373 | *Prpf31* | 3086.776026 | 2992.676984 | 2915.694407 | 5324.284144 | 5681.758053 | 5494.24088 | 5.67E-40 | 0.874560279 | 1.60E-41 | 182.2059379 | 1.833449182 | 39.24653224 |
| ENSMUSG00000021701 | *Plk2* | 413.6294943 | 433.5544862 | 383.2127538 | 824.2707466 | 696.9286328 | 738.3201852 | 1.68E-12 | 0.874185683 | 2.11E-13 | 53.89942369 | 1.832973189 | 11.77342093 |
| ENSMUSG00000042506 | *Usp22* | 950.067017 | 813.3942573 | 659.2772048 | 1457.565315 | 1551.771608 | 1441.74464 | 1.71E-09 | 0.87355791 | 2.83E-10 | 39.78741561 | 1.832175763 | 8.768079091 |
| ENSMUSG00000051586 | *Mical3* | 657.7387041 | 641.6990072 | 461.3679865 | 905.4905006 | 1231.731874 | 1095.542265 | 1.49E-06 | 0.873336119 | 3.42E-07 | 25.99851256 | 1.831894118 | 5.827516346 |
| ENSMUSG00000044469 | *Tnfaip8l1* | 120.5477579 | 143.8787012 | 152.5287606 | 266.7081111 | 246.3463747 | 246.1067284 | 1.72E-06 | 0.872618493 | 3.97E-07 | 25.70687387 | 1.830983123 | 5.764486228 |
| ENSMUSG00000035696 | *Rnf38* | 399.3144481 | 361.6151356 | 279.8461558 | 588.2944343 | 679.0316739 | 646.4893164 | 1.17E-07 | 0.872291334 | 2.35E-08 | 31.18067777 | 1.830567959 | 6.930472452 |
| ENSMUSG00000007850 | *Hnrnph1* | 2132.188468 | 1930.85217 | 1750.929326 | 3578.059433 | 3613.080163 | 3465.696989 | 3.57E-20 | 0.872101812 | 2.58E-21 | 89.84525189 | 1.8303275 | 19.44765879 |
| ENSMUSG00000025731 | *Mettl26* | 242.6023628 | 298.3085071 | 236.9868346 | 455.4891609 | 507.4314215 | 461.9092701 | 1.03E-08 | 0.871798031 | 1.85E-09 | 36.12591377 | 1.829942138 | 7.985185723 |
| ENSMUSG00000079429 | *Mroh2a* | 106.2327117 | 113.1845783 | 63.02841346 | 181.0981001 | 170.5474902 | 169.8871073 | 0.001284615 | 0.871277414 | 0.000462557 | 12.26086683 | 1.829281895 | 2.891227028 |
| ENSMUSG00000027115 | *Kif18a* | 250.1365977 | 145.7970839 | 160.0921702 | 344.6351724 | 347.4115541 | 330.5911277 | 0.000265171 | 0.870758779 | 8.42E-05 | 15.4607776 | 1.828624404 | 3.576474306 |
| ENSMUSG00000058006 | *Mdn1* | 2233.900639 | 2168.731622 | 1318.55441 | 3320.131836 | 3699.40667 | 3448.249123 | 4.90E-06 | 0.870168916 | 1.19E-06 | 23.58529387 | 1.827876902 | 5.310206572 |
| ENSMUSG00000032381 | *Ciao2a* | 2488.557777 | 2599.408534 | 2953.511455 | 4705.25791 | 5013.254002 | 4962.54015 | 6.77E-22 | 0.869944444 | 4.37E-23 | 97.91406745 | 1.827592521 | 21.16965008 |
| ENSMUSG00000027133 | *Nop10* | 627.6017646 | 729.9446106 | 831.9750577 | 1479.5166 | 1226.468062 | 1285.632163 | 6.65E-10 | 0.869629278 | 1.06E-10 | 41.71204354 | 1.827193315 | 9.176916309 |
| ENSMUSG00000028737 | *Aldh4a1* | 556.0265334 | 585.1067181 | 481.5370788 | 1004.271282 | 974.8578761 | 992.6916917 | 5.42E-15 | 0.869117634 | 5.53E-16 | 65.59749785 | 1.826545427 | 14.26628845 |
| ENSMUSG00000032598 | *Nckipsd* | 450.5472452 | 485.3508186 | 335.3111596 | 724.3924005 | 755.8833208 | 847.598919 | 1.72E-07 | 0.868758229 | 3.52E-08 | 30.3971841 | 1.826090453 | 6.764980089 |
| ENSMUSG00000029063 | *Nadk* | 6397.318828 | 6729.686449 | 6518.39852 | 11997.47474 | 12240.46709 | 11629.46122 | 6.64E-49 | 0.868521001 | 1.33E-50 | 223.8214287 | 1.825790206 | 48.17750945 |
| ENSMUSG00000024613 | *Tcof1* | 6076.360422 | 5692.80061 | 3731.282077 | 9212.954257 | 9566.450886 | 9528.370947 | 3.29E-07 | 0.868291872 | 6.96E-08 | 29.07695867 | 1.825500257 | 6.482675899 |
| ENSMUSG00000032604 | *Qars* | 5962.593476 | 6230.906952 | 6016.692349 | 11326.86299 | 10662.37642 | 11231.83356 | 6.86E-46 | 0.867401657 | 1.53E-47 | 209.7860304 | 1.82437418 | 45.16381617 |
| ENSMUSG00000032431 | *Crtap* | 976.4368391 | 1037.845031 | 959.2924529 | 1646.346365 | 1864.442007 | 1913.755306 | 1.24E-18 | 0.867360625 | 9.90E-20 | 82.62947863 | 1.824322293 | 17.90655696 |
| ENSMUSG00000049643 | *2310022A10Rik* | 1555.819501 | 1439.746203 | 1317.293841 | 2470.617111 | 2741.392991 | 2662.176887 | 3.26E-19 | 0.866440612 | 2.51E-20 | 85.33847287 | 1.823159285 | 18.48722762 |
| ENSMUSG00000031310 | *Zmym3* | 385.7528253 | 344.3496915 | 320.1843404 | 627.8067471 | 660.0819528 | 634.5513034 | 3.60E-12 | 0.866289403 | 4.66E-13 | 52.34434442 | 1.822968211 | 11.44388092 |
| ENSMUSG00000020914 | *Top2a* | 5744.854088 | 5193.061921 | 5251.52741 | 9721.126502 | 10214.95245 | 9582.551159 | 1.47E-32 | 0.866008561 | 5.76E-34 | 147.6140899 | 1.822613377 | 31.83262698 |
| ENSMUSG00000040481 | *Bptf* | 1185.135145 | 1282.438823 | 766.4255077 | 1929.517939 | 2178.165168 | 1795.293485 | 6.74E-06 | 0.865978778 | 1.68E-06 | 22.92878834 | 1.822575751 | 5.171617428 |
| ENSMUSG00000022707 | *Gbe1* | 498.0129249 | 575.5148047 | 742.4747106 | 1060.247059 | 1196.990718 | 1038.607126 | 1.96E-07 | 0.864356429 | 4.03E-08 | 30.13408023 | 1.820527368 | 6.708822787 |
| ENSMUSG00000000628 | *Hk2* | 1158.011899 | 1142.396887 | 771.4677808 | 1736.346633 | 1978.140334 | 1887.124354 | 1.97E-07 | 0.864149119 | 4.06E-08 | 30.1222192 | 1.820265784 | 6.706455571 |
| ENSMUSG00000038256 | *Bcl9* | 977.1902626 | 978.3751679 | 597.5093596 | 1415.857874 | 1682.314132 | 1557.451535 | 9.98E-06 | 0.864018217 | 2.55E-06 | 22.12949794 | 1.82010063 | 5.000833241 |
| ENSMUSG00000052557 | *Gan* | 771.5056506 | 703.087253 | 496.6638981 | 1202.930411 | 1216.993202 | 1178.190047 | 2.59E-07 | 0.863743456 | 5.42E-08 | 29.55989383 | 1.819754025 | 6.587228777 |
| ENSMUSG00000020185 | *E2f7* | 357.1227328 | 379.8397711 | 233.2051298 | 509.2698088 | 646.3960431 | 615.266821 | 4.58E-05 | 0.863185379 | 1.29E-05 | 19.02015676 | 1.819050227 | 4.339417967 |
| ENSMUSG00000041552 | *Ptchd1* | 344.3145335 | 330.9210127 | 263.4587683 | 545.4894289 | 595.8634534 | 573.94293 | 3.67E-09 | 0.862909989 | 6.31E-10 | 38.22379058 | 1.81870303 | 8.435406159 |
| ENSMUSG00000041406 | *BC055324* | 720.2728535 | 724.1894625 | 750.0381202 | 1370.85774 | 1360.168872 | 1255.327977 | 1.51E-21 | 0.861757689 | 9.95E-23 | 96.28468731 | 1.817250987 | 20.82139597 |
| ENSMUSG00000036223 | *Ska1* | 321.7118289 | 289.675785 | 300.0152481 | 521.3430155 | 514.8007575 | 619.8583644 | 1.02E-09 | 0.861000938 | 1.66E-10 | 40.82676455 | 1.816298016 | 8.989867733 |
| ENSMUSG00000097327 | *E030030I06Rik* | 106.9861351 | 105.5110475 | 155.0498971 | 206.3420777 | 217.921793 | 236.0053328 | 0.000156385 | 0.858242932 | 4.80E-05 | 16.52621271 | 1.812829109 | 3.805805922 |
| ENSMUSG00000042178 | *Armc5* | 858.1493516 | 845.0475715 | 681.9674337 | 1404.882231 | 1451.759191 | 1476.64037 | 1.80E-13 | 0.857732203 | 2.08E-14 | 58.4540633 | 1.812187463 | 12.74533767 |
| ENSMUSG00000039844 | *Rapgef1* | 3966.774659 | 3864.581913 | 2807.285536 | 6002.578846 | 6887.17087 | 6396.938321 | 1.68E-09 | 0.857410731 | 2.79E-10 | 39.81321741 | 1.811783703 | 8.773457115 |
| ENSMUSG00000029463 | *Fam216a* | 379.7254374 | 431.6361035 | 544.5654923 | 830.8561321 | 744.3029356 | 869.6383275 | 1.54E-07 | 0.856458043 | 3.14E-08 | 30.61988827 | 1.810587681 | 6.812355138 |
| ENSMUSG00000078183 | *Gm15610* | 320.204982 | 292.553359 | 361.7830933 | 639.8799538 | 572.7026831 | 548.2302867 | 1.44E-09 | 0.856441617 | 2.38E-10 | 40.12846143 | 1.810567065 | 8.840560465 |
| ENSMUSG00000023942 | *Slc29a1* | 1995.065393 | 1983.607693 | 1692.943186 | 3257.570674 | 3513.067746 | 3500.592719 | 2.60E-19 | 0.855155827 | 1.99E-20 | 85.79601129 | 1.808954132 | 18.58524677 |
| ENSMUSG00000032177 | *Pde4a* | 763.2179923 | 793.2512391 | 543.304924 | 1224.881695 | 1347.535725 | 1234.206877 | 6.63E-08 | 0.854677185 | 1.29E-08 | 32.34627997 | 1.808354076 | 7.1787372 |
| ENSMUSG00000045482 | *Trrap* | 2746.22861 | 2696.28686 | 1583.273746 | 3962.206918 | 4572.146605 | 4175.549604 | 1.76E-05 | 0.854127508 | 4.67E-06 | 20.9668573 | 1.807665211 | 4.753699389 |
| ENSMUSG00000091337 | *Eid1* | 1408.901921 | 1293.949119 | 1300.906454 | 2494.763525 | 2397.139723 | 2351.78855 | 9.92E-27 | 0.85358437 | 4.97E-28 | 120.4787412 | 1.806984799 | 26.00347793 |
| ENSMUSG00000014592 | *Camta1* | 118.2874875 | 117.0213436 | 129.8385317 | 231.5860553 | 209.4996948 | 216.7208504 | 2.30E-06 | 0.853067603 | 5.38E-07 | 25.12141949 | 1.806337661 | 5.638961518 |
| ENSMUSG00000020739 | *Nup85* | 2332.599116 | 2174.48677 | 2059.768552 | 3936.96294 | 4024.710216 | 3913.831628 | 1.36E-28 | 0.853010853 | 6.20E-30 | 129.1776149 | 1.806266608 | 27.86747059 |
| ENSMUSG00000005371 | *Fbxo11* | 373.6980495 | 405.7379373 | 209.2543327 | 620.1237974 | 581.1247814 | 591.3907951 | 0.000408343 | 0.852977585 | 0.000133863 | 14.5865963 | 1.806224956 | 3.388974482 |
| ENSMUSG00000030752 | *Kdm8* | 190.6161422 | 184.1647375 | 153.7893288 | 334.7570942 | 293.7206776 | 330.5911277 | 4.06E-07 | 0.852263287 | 8.68E-08 | 28.64882627 | 1.805330891 | 6.391447032 |
| ENSMUSG00000113308 | *Gm48445* | 263.6982204 | 211.022095 | 194.1275135 | 368.7815857 | 414.7883404 | 429.768466 | 1.20E-06 | 0.852063003 | 2.71E-07 | 26.44570077 | 1.805080281 | 5.922206769 |
| ENSMUSG00000029703 | *Lrwd1* | 1454.10733 | 1549.094016 | 1195.018719 | 2461.836597 | 2591.900746 | 2528.103818 | 8.21E-14 | 0.850782296 | 9.24E-15 | 60.05146467 | 1.803478589 | 13.08587416 |
| ENSMUSG00000064215 | *Ifi27* | 1238.628213 | 1405.215315 | 1676.555798 | 2448.665827 | 2836.141596 | 2493.208088 | 1.86E-10 | 0.85041539 | 2.80E-11 | 44.31470455 | 1.803019987 | 9.73136569 |
| ENSMUSG00000026970 | *Rbms1* | 1134.655771 | 1104.988425 | 907.6091539 | 1958.05461 | 1799.170745 | 1928.448245 | 6.29E-14 | 0.850283852 | 6.98E-15 | 60.60459496 | 1.802855605 | 13.20151617 |
| ENSMUSG00000032235 | *Ice2* | 259.931103 | 242.6754093 | 331.5294548 | 501.5868591 | 489.5344626 | 505.0697784 | 8.18E-08 | 0.85014991 | 1.60E-08 | 31.92206816 | 1.802688232 | 7.087416454 |
| ENSMUSG00000032293 | *Ireb2* | 1430.751202 | 1473.3179 | 1329.899524 | 2550.739301 | 2527.682247 | 2562.08124 | 1.05E-28 | 0.850077176 | 4.79E-30 | 129.6886403 | 1.802597352 | 27.97797678 |
| ENSMUSG00000020150 | *Gamt* | 180.8216369 | 188.9606942 | 160.0921702 | 329.269273 | 311.6176364 | 317.734806 | 3.95E-08 | 0.850037406 | 7.49E-09 | 33.40400982 | 1.80254766 | 7.403557128 |
| ENSMUSG00000036737 | *Oxsr1* | 1915.202504 | 1998.954755 | 1943.796271 | 3467.205444 | 3527.806418 | 3553.854623 | 1.69E-39 | 0.848955681 | 4.87E-41 | 179.9892409 | 1.801196626 | 38.7709254 |
| ENSMUSG00000021809 | *Nudt13* | 269.7256083 | 293.5125504 | 305.0575212 | 495.0014737 | 535.8560032 | 527.1091869 | 2.47E-11 | 0.848218861 | 3.43E-12 | 48.42374786 | 1.800276946 | 10.60733955 |
| ENSMUSG00000042390 | *Gatad2b* | 1199.450191 | 1164.458288 | 831.9750577 | 1849.39575 | 2060.255792 | 1852.228624 | 2.99E-08 | 0.847576713 | 5.57E-09 | 33.97863503 | 1.799475815 | 7.524714761 |
| ENSMUSG00000005057 | *Sh2b2* | 1010.340896 | 941.925897 | 709.6999356 | 1493.784935 | 1643.361927 | 1658.465491 | 8.50E-09 | 0.84604116 | 1.51E-09 | 36.52382597 | 1.797561537 | 8.070338875 |
| ENSMUSG00000003868 | *Ruvbl2* | 4083.555299 | 3800.316094 | 3733.803213 | 6958.557302 | 7102.987138 | 6829.461713 | 1.26E-35 | 0.845648919 | 4.25E-37 | 161.9479872 | 1.797072881 | 34.90039376 |
| ENSMUSG00000056665 | *Them6* | 1653.01113 | 1900.158047 | 1679.076935 | 2858.057289 | 3316.201198 | 3224.181804 | 6.62E-17 | 0.845169386 | 5.86E-18 | 74.56912882 | 1.796475657 | 16.17919442 |
| ENSMUSG00000025646 | *Atrip* | 630.6154586 | 695.4137223 | 547.0866289 | 1099.759372 | 1155.932989 | 1115.745056 | 7.24E-13 | 0.844859639 | 8.81E-14 | 55.61688712 | 1.796089994 | 12.14014678 |
| ENSMUSG00000042303 | *Sgsm3* | 756.4371809 | 860.394633 | 762.6438029 | 1402.687103 | 1445.442618 | 1425.215084 | 3.73E-19 | 0.844669544 | 2.89E-20 | 85.0625126 | 1.79585335 | 18.42815732 |
| ENSMUSG00000047547 | *Cltb* | 2036.503685 | 2098.710654 | 2240.029814 | 3571.474047 | 4049.976511 | 3817.409216 | 2.67E-23 | 0.844597777 | 1.61E-24 | 104.4472587 | 1.795764017 | 22.57296893 |
| ENSMUSG00000020649 | *Rrm2* | 4541.636779 | 4390.218768 | 3401.01319 | 7059.533212 | 7718.853075 | 7380.446925 | 4.81E-12 | 0.844515242 | 6.32E-13 | 51.74552524 | 1.795661287 | 11.31829222 |
| ENSMUSG00000040121 | *Rep15* | 79.86288962 | 63.30662851 | 44.11988942 | 106.4637316 | 101.0651794 | 132.2364511 | 0.008564972 | 0.844082279 | 0.003654161 | 8.44814453 | 1.795122477 | 2.067274073 |
| ENSMUSG00000020721 | *Helz* | 812.1905189 | 834.4964668 | 568.5162894 | 1177.686433 | 1490.711396 | 1315.93635 | 1.21E-06 | 0.844004521 | 2.73E-07 | 26.42970056 | 1.795025726 | 5.918872372 |
| ENSMUSG00000021549 | *Rasa1* | 1546.778419 | 1298.745076 | 949.2079067 | 2435.495056 | 2050.780932 | 2326.075907 | 2.24E-06 | 0.841888489 | 5.25E-07 | 25.16969872 | 1.792394853 | 5.649070663 |
| ENSMUSG00000054717 | *Hmgb2* | 2749.995727 | 2063.220575 | 2368.607778 | 4642.696749 | 4041.554413 | 4192.079161 | 1.42E-11 | 0.841556923 | 1.94E-12 | 49.54424052 | 1.791982965 | 10.84699249 |
| ENSMUSG00000024050 | *Wiz* | 1202.463885 | 1277.642866 | 818.1088067 | 1699.029448 | 2212.906323 | 2005.586175 | 5.29E-06 | 0.841477503 | 1.30E-06 | 23.42364242 | 1.791884319 | 5.276395008 |
| ENSMUSG00000035877 | *Zhx3* | 474.6567968 | 453.6975044 | 424.8115067 | 729.8802217 | 903.2700407 | 795.2553238 | 6.68E-11 | 0.841161666 | 9.63E-12 | 46.40245326 | 1.791492081 | 10.17551039 |
| ENSMUSG00000041949 | *Tango6* | 185.3421778 | 196.6342249 | 151.2681923 | 270.0008038 | 345.3060295 | 342.5291406 | 1.28E-05 | 0.840449171 | 3.32E-06 | 21.62231575 | 1.790607545 | 4.894025243 |
| ENSMUSG00000030847 | *Bag3* | 1588.970134 | 1567.318651 | 1235.356904 | 2304.884911 | 2806.664252 | 2760.435916 | 4.94E-10 | 0.840408745 | 7.75E-11 | 42.3198328 | 1.790557372 | 9.30624067 |
| ENSMUSG00000031353 | *Rbbp7* | 3539.583542 | 3529.824135 | 3409.837168 | 6122.213348 | 6534.495504 | 6111.344319 | 8.60E-38 | 0.84023319 | 2.67E-39 | 172.0294654 | 1.7903395 | 37.06569326 |

| | | | | | | | | | | | | |
|---|---|---|---|---|---|---|---|---|---|---|---|---|
| ENSMUSG00000032508 | *Myd88* | 2942.87214 | 2991.717793 | 2909.391565 | 5188.186177 | 5309.080204 | 5335.373477 | 3.70E-46 | 0.839993183 | 8.22E-48 | 211.0230063 | 1.790041684 | 45.4323485 |
| ENSMUSG00000029394 | *Cdk2ap1* | 4628.28048 | 4412.280169 | 4304.840639 | 7714.779065 | 8236.812119 | 7939.696916 | 1.14E-36 | 0.839524626 | 3.67E-38 | 166.8161555 | 1.78946041 | 35.94491417 |
| ENSMUSG00000031627 | *Irf2* | 544.7251811 | 608.1273103 | 447.5017356 | 939.5149922 | 988.5437858 | 943.1030226 | 7.04E-10 | 0.839505893 | 1.12E-10 | 41.59825886 | 1.789437175 | 9.152577266 |
| ENSMUSG00000075470 | *Alg10b* | 1172.326946 | 1155.825566 | 700.8759577 | 1523.41917 | 1917.080121 | 1986.301692 | 6.01E-05 | 0.839278453 | 1.72E-05 | 18.47184455 | 1.789155093 | 4.221140745 |
| ENSMUSG00000038838 | *Vars2* | 553.7662629 | 591.8210575 | 392.0367317 | 858.2952382 | 999.0714086 | 898.1058969 | 1.37E-06 | 0.837476024 | 3.14E-07 | 26.16325755 | 1.786921211 | 5.862361176 |
| ENSMUSG00000027351 | *Spred1* | 3862.802218 | 3807.989624 | 2925.778953 | 6321.970041 | 6236.563777 | 6384.081999 | 1.79E-12 | 0.837018415 | 2.25E-13 | 53.77651714 | 1.786354506 | 11.74791589 |
| ENSMUSG00000044791 | *Setd2* | 802.3960136 | 755.8427768 | 549.6077654 | 1234.759774 | 1262.26198 | 1278.285694 | 4.68E-08 | 0.836739092 | 8.95E-09 | 33.05748345 | 1.78600868 | 7.330076725 |
| ENSMUSG00000028820 | *Sfpq* | 5782.525262 | 5700.47414 | 4053.987554 | 8723.440605 | 9711.732081 | 9318.996566 | 5.05E-09 | 0.836387798 | 8.79E-10 | 37.57612901 | 1.785573843 | 8.296915325 |
| ENSMUSG00000020766 | *Galk1* | 2746.982033 | 2489.10153 | 2429.115055 | 4802.941128 | 4203.679805 | 4682.456 | 8.98E-20 | 0.835657983 | 6.66E-21 | 87.96623939 | 1.784670804 | 19.0469547 |
| ENSMUSG00000024251 | *Thada* | 695.4098785 | 605.2497362 | 432.3749163 | 973.5394837 | 1132.772219 | 995.4466178 | 5.77E-06 | 0.835273383 | 1.43E-06 | 23.24433611 | 1.784195102 | 5.238691897 |
| ENSMUSG00000001525 | *Tubb5* | 33393.23579 | 31427.90429 | 26771.9489 | 51470.27518 | 57685.05665 | 54217.86325 | 2.95E-16 | 0.834735005 | 2.73E-17 | 71.53396395 | 1.783529409 | 15.53076204 |
| ENSMUSG00000019471 | *Cdc37* | 13015.39074 | 13469.924 | 13959.53301 | 23661.28995 | 24034.56297 | 24417.82801 | 4.23E-47 | 0.834615706 | 9.08E-49 | 215.4069615 | 1.783381932 | 46.37373562 |
| ENSMUSG00000039994 | *Timeless* | 1251.436412 | 1198.989176 | 782.8128952 | 1900.981269 | 1993.931768 | 1881.614502 | 2.10E-06 | 0.834603535 | 4.89E-07 | 25.304998 | 1.783366886 | 5.678005742 |
| ENSMUSG00000040658 | *Dnph1* | 843.0808819 | 867.1089724 | 744.9958471 | 1457.565315 | 1497.02797 | 1423.378466 | 8.84E-18 | 0.831355412 | 7.43E-19 | 78.64487703 | 1.779356282 | 17.05338694 |
| ENSMUSG00000046982 | *Tshz1* | 429.4513876 | 425.8809555 | 278.5855875 | 627.8067471 | 725.3532145 | 672.2019596 | 1.22E-05 | 0.83119029 | 3.17E-06 | 21.7091294 | 1.77915264 | 4.912188202 |
| ENSMUSG00000004846 | *Plod3* | 3495.884979 | 3642.049522 | 3021.582141 | 5829.163695 | 6035.486181 | 6220.623053 | 3.16E-19 | 0.831118589 | 2.44E-20 | 85.40133418 | 1.779064219 | 18.50064792 |
| ENSMUSG00000002308 | *Cd320* | 751.91664 | 753.9243941 | 723.5661865 | 1365.369918 | 1274.895127 | 1328.792672 | 8.20E-22 | 0.830860827 | 5.31E-23 | 97.52873495 | 1.778746387 | 21.0860587 |
| ENSMUSG00000030122 | *Ptms* | 3532.049307 | 3621.906504 | 3194.279994 | 5667.821752 | 6390.267071 | 6351.022886 | 1.07E-20 | 0.830457204 | 7.44E-22 | 92.30113771 | 1.778248818 | 19.97027522 |
| ENSMUSG00000020737 | *Jpt1* | 3965.267812 | 3643.008714 | 3684.641051 | 6512.946219 | 6834.532755 | 6724.774522 | 1.10E-33 | 0.829142732 | 4.06E-35 | 152.8842292 | 1.776629353 | 32.95707111 |
| ENSMUSG00000032397 | *Tipin* | 1869.997095 | 1673.78889 | 1766.056145 | 3169.765534 | 3164.603429 | 3102.965057 | 1.60E-27 | 0.828775625 | 7.66E-29 | 124.187859 | 1.776177331 | 26.79591604 |
| ENSMUSG00000027087 | *Itgav* | 1407.395074 | 1406.174506 | 1244.180882 | 2156.713738 | 2641.380574 | 2415.151849 | 5.50E-14 | 0.828707029 | 6.07E-15 | 60.87859967 | 1.77609288 | 13.25927107 |
| ENSMUSG00000032305 | *Fam219b* | 491.985537 | 527.5552376 | 462.6285548 | 831.9536963 | 875.8982213 | 925.6551575 | 6.03E-14 | 0.828008301 | 6.68E-15 | 60.69076465 | 1.775232889 | 13.2197203 |
| ENSMUSG00000020747 | *Tmem94* | 580.8895084 | 553.4534038 | 453.8045769 | 815.4902327 | 1020.126654 | 989.9367657 | 2.80E-08 | 0.827829403 | 5.20E-09 | 34.11390822 | 1.775012769 | 7.553146024 |
| ENSMUSG00000025409 | *Mbd6* | 641.9168109 | 644.5765812 | 500.4456029 | 929.636914 | 1121.191834 | 1127.683069 | 1.98E-08 | 0.827794892 | 3.64E-09 | 34.80834081 | 1.77497031 | 7.703739223 |
| ENSMUSG00000048087 | *Ahcyl* | 5738.073277 | 5553.717865 | 5046.054782 | 9581.735843 | 9840.16908 | 9591.734246 | 5.12E-29 | 0.827728276 | 2.30E-30 | 131.1437659 | 1.774888353 | 28.29041354 |
| ENSMUSG00000031388 | *Naa10* | 1833.079344 | 1697.768674 | 1738.323643 | 3117.082451 | 2980.370029 | 3255.404299 | 2.37E-26 | 0.827266558 | 1.21E-27 | 118.7109678 | 1.774320411 | 25.62470266 |
| ENSMUSG00000054874 | *Pcnx3* | 1711.778162 | 1551.012399 | 1135.772011 | 2451.958519 | 2836.141596 | 2527.185509 | 1.90E-07 | 0.827198325 | 3.90E-08 | 30.19772597 | 1.774236495 | 6.722351923 |
| ENSMUSG00000024002 | *Brd4* | 1986.777735 | 2049.791896 | 1154.680535 | 2799.886384 | 3324.623297 | 3093.78197 | 6.61E-05 | 0.827094179 | 1.91E-05 | 18.2746504 | 1.77410842 | 4.179607737 |
| ENSMUSG00000040188 | *Scamp2* | 2524.722105 | 2752.879149 | 2316.924479 | 4368.305688 | 4571.093842 | 4541.036462 | 3.73E-21 | 0.827004147 | 2.53E-22 | 94.43718306 | 1.77399771 | 20.42856527 |
| ENSMUSG00000041607 | *Mbp* | 2526.228952 | 2529.387567 | 1956.401954 | 3916.10922 | 4346.855476 | 4187.487617 | 7.52E-12 | 0.82695229 | 1.01E-12 | 50.83025146 | 1.773933945 | 11.12362236 |
| ENSMUSG00000040651 | *Tasor* | 199.657224 | 147.7154665 | 119.7539856 | 289.7569602 | 244.2408502 | 300.286941 | 0.000541031 | 0.826453307 | 0.000181482 | 14.01371124 | 1.773320503 | 3.26677747 |
| ENSMUSG00000034297 | *Med13* | 1212.25839 | 1013.865248 | 663.0589096 | 1713.297784 | 1722.319099 | 1695.197838 | 4.45E-05 | 0.825666477 | 1.26E-05 | 19.07651976 | 1.772353618 | 4.351400605 |
| ENSMUSG00000032329 | *Hmg20a* | 617.8072593 | 702.1280617 | 579.8614039 | 1075.612958 | 1093.820014 | 1199.311147 | 1.04E-12 | 0.82534502 | 1.28E-13 | 54.88753044 | 1.77195875 | 11.98484056 |
| ENSMUSG00000027111 | *Itga6* | 595.9579782 | 571.6780393 | 508.0090125 | 973.5394837 | 962.2247287 | 1039.525435 | 4.09E-14 | 0.824605536 | 4.47E-15 | 61.48254966 | 1.771050729 | 13.38787705 |
| ENSMUSG00000033272 | *Slc35a4* | 4086.568993 | 4080.399965 | 3426.224556 | 6544.775582 | 7147.203154 | 6850.582813 | 8.13E-18 | 0.824442636 | 6.82E-19 | 78.81469558 | 1.770850763 | 17.08965486 |
| ENSMUSG00000015944 | *Castor2* | 946.2998996 | 953.4361931 | 709.6999356 | 1386.223639 | 1603.35696 | 1638.262699 | 1.55E-08 | 0.824022732 | 2.83E-09 | 35.29658633 | 1.770335423 | 7.808562723 |
| ENSMUSG00000042684 | *Npl* | 170.273708 | 148.6746579 | 144.965351 | 309.5131166 | 238.9770387 | 275.4926064 | 1.26E-05 | 0.823142472 | 3.29E-06 | 21.64142075 | 1.769255583 | 4.898007095 |
| ENSMUSG00000030512 | *Snrpa1* | 1415.682732 | 1327.520816 | 1268.131679 | 2286.226319 | 2465.569272 | 2352.706859 | 1.06E-22 | 0.822666483 | 6.66E-24 | 101.6404809 | 1.768671948 | 21.97366743 |
| ENSMUSG00000025728 | *Pigq* | 1851.161507 | 1786.014277 | 1382.843391 | 2883.301267 | 3047.746816 | 2959.708901 | 3.95E-11 | 0.822572449 | 5.61E-12 | 47.46109403 | 1.768556671 | 10.40301808 |
| ENSMUSG00000015289 | *Lage3* | 723.2865475 | 680.0666609 | 746.2564154 | 1303.906321 | 1279.106177 | 1214.922394 | 5.49E-19 | 0.821754105 | 4.29E-20 | 84.28390867 | 1.767553772 | 18.26019998 |
| ENSMUSG00000036639 | *Nudt1* | 333.7666047 | 331.880204 | 335.3111596 | 576.2212277 | 594.8106911 | 596.9006472 | 1.80E-13 | 0.821181711 | 2.09E-14 | 58.44490877 | 1.766852629 | 12.74383142 |
| ENSMUSG00000010362 | *Rdm1* | 79.86288962 | 78.65368997 | 93.28205192 | 164.6346365 | 156.8615805 | 122.1350555 | 0.000847312 | 0.820525585 | 0.000294047 | 13.10793785 | 1.76604926 | 3.071956765 |
| ENSMUSG00000037405 | *Icam1* | 3671.432652 | 3585.457233 | 2971.159411 | 5925.749349 | 6051.277615 | 6099.406306 | 1.37E-16 | 0.820452617 | 1.25E-17 | 73.07713431 | 1.765959941 | 15.86191325 |
| ENSMUSG00000047832 | *Cdca4* | 1821.024568 | 1801.361339 | 1545.456698 | 3009.521155 | 3079.329684 | 3049.703153 | 1.81E-19 | 0.820345518 | 1.38E-20 | 86.52277306 | 1.765828849 | 18.74149246 |
| ENSMUSG00000031683 | *Lsm6* | 730.0673589 | 689.6585743 | 758.8620981 | 1227.076824 | 1327.533242 | 1288.387089 | 9.55E-19 | 0.820290338 | 7.55E-20 | 83.16392307 | 1.76576131 | 18.01985094 |
| ENSMUSG00000041040 | *Fam117b* | 413.6294943 | 400.9419806 | 329.0083183 | 666.2214956 | 729.5642636 | 631.7963773 | 4.69E-09 | 0.820285989 | 8.15E-10 | 37.72451396 | 1.765755988 | 8.32892422 |
| ENSMUSG00000039512 | *Uhrf1bp1* | 1025.409366 | 939.048323 | 621.4601567 | 1434.516466 | 1627.570493 | 1510.617792 | 1.04E-05 | 0.819286649 | 2.67E-06 | 22.04068266 | 1.764533291 | 4.982137751 |
| ENSMUSG00000074211 | *Sdhaf1* | 302.1228183 | 262.8184275 | 276.064451 | 511.4649373 | 479.0068398 | 496.8050002 | 2.83E-10 | 0.819269848 | 4.33E-11 | 43.45776347 | 1.764512742 | 9.547892899 |
| ENSMUSG00000028809 | *Srrm1* | 3799.514645 | 3898.15361 | 2619.460863 | 5555.870199 | 6530.284455 | 6117.772479 | 2.40E-07 | 0.818424022 | 5.01E-08 | 29.71400795 | 1.763478543 | 6.620047791 |
| ENSMUSG00000031060 | *Rbm10* | 1802.188981 | 1568.277843 | 1285.779635 | 2717.569066 | 2774.028621 | 2731.050038 | 3.67E-10 | 0.818106996 | 5.66E-11 | 42.93374562 | 1.763091069 | 9.43565187 |
| ENSMUSG00000032306 | *Mpi* | 760.9577218 | 760.6387335 | 709.6999356 | 1315.979528 | 1325.427717 | 1298.488485 | 1.60E-21 | 0.817938971 | 1.06E-22 | 96.16695215 | 1.76288574 | 20.79647258 |
| ENSMUSG00000042535 | *Gtpbp1* | 1675.613835 | 1671.870508 | 1255.525996 | 2645.129826 | 2790.872818 | 2689.726147 | 1.83E-10 | 0.817817508 | 2.75E-11 | 44.34865571 | 1.762737326 | 9.738501724 |
| ENSMUSG00000059552 | *Trp53* | 4786.499413 | 4799.793471 | 3888.853111 | 7641.242261 | 8133.641415 | 7991.122203 | 1.49E-15 | 0.817804117 | 1.45E-16 | 68.23718327 | 1.762720964 | 14.82658386 |
| ENSMUSG00000024392 | *Bag6* | 6631.633532 | 6390.132715 | 5196.062406 | 10326.98196 | 10793.97171 | 11003.1747 | 2.36E-14 | 0.817691978 | 2.54E-15 | 62.59321968 | 1.762583956 | 13.62622204 |
| ENSMUSG00000028614 | *Ndc1* | 2578.968596 | 2352.89636 | 2015.648663 | 3922.694605 | 4285.795263 | 4047.904697 | 8.10E-14 | 0.817363135 | 9.11E-15 | 60.07908021 | 1.762182244 | 13.09170282 |
| ENSMUSG00000030254 | *Rad18* | 802.3960136 | 670.4747474 | 762.6438029 | 1315.979528 | 1352.799536 | 1269.102607 | 1.22E-14 | 0.81576203 | 1.29E-15 | 63.93307931 | 1.760227657 | 13.91266654 |
| ENSMUSG00000026335 | *Pam* | 1507.600397 | 1514.563128 | 1507.63965 | 2499.153782 | 2763.500999 | 2703.500777 | 6.86E-26 | 0.814659876 | 3.59E-27 | 116.5566549 | 1.758883435 | 25.16393919 |
| ENSMUSG00000038342 | *Mlxip* | 910.8889957 | 991.8038467 | 611.3756106 | 1376.345561 | 1583.354477 | 1469.293901 | 8.30E-06 | 0.814338775 | 2.10E-06 | 22.50323467 | 1.758492004 | 5.080661809 |
| ENSMUSG00000028013 | *Ppa2* | 525.8895939 | 559.2085519 | 504.2273077 | 863.7830594 | 939.0639584 | 992.6916917 | 1.62E-13 | 0.814219608 | 1.87E-14 | 58.66742295 | 1.758346758 | 12.79011119 |
| ENSMUSG00000046152 | *Fut10* | 149.9312739 | 141.0011271 | 142.4442144 | 233.7811838 | 263.1905713 | 265.3912108 | 1.68E-06 | 0.813848274 | 3.89E-07 | 25.74885448 | 1.757894237 | 5.773675146 |
| ENSMUSG00000013629 | *Cad* | 7490.536307 | 7585.285126 | 5493.556517 | 11351.0094 | 12635.25295 | 12167.59012 | 1.37E-09 | 0.813193476 | 2.25E-10 | 40.2335007 | 1.75709656 | 8.862927758 |
| ENSMUSG00000030871 | *Ears2* | 360.1364268 | 313.6555685 | 297.4941115 | 564.148021 | 588.4941174 | 560.1682997 | 5.14E-10 | 0.812297216 | 8.09E-11 | 42.23483386 | 1.75600532 | 9.288693199 |
| ENSMUSG00000026965 | *Anapc2* | 2579.722019 | 2653.12325 | 2395.079712 | 4213.54913 | 4705.847415 | 4482.264706 | 3.82E-23 | 0.812294683 | 2.33E-24 | 103.7180314 | 1.756002237 | 22.41754485 |
| ENSMUSG00000032405 | *Pias1* | 731.5742058 | 823.945362 | 775.2494856 | 1280.857472 | 1447.548142 | 1360.933476 | 8.55E-16 | 0.812244577 | 8.15E-17 | 69.37372673 | 1.755941251 | 15.06820309 |
| ENSMUSG00000021366 | *Hivep1* | 357.8761563 | 322.2882906 | 190.3458087 | 491.7087809 | 562.1750603 | 482.1120612 | 0.000763107 | 0.812214548 | 0.00026293 | 13.31757368 | 1.755904703 | 3.117414758 |
| ENSMUSG00000032342 | *Mto1* | 539.4512167 | 560.1677432 | 508.0090125 | 935.1247352 | 963.2774909 | 928.4100836 | 1.05E-16 | 0.811981776 | 9.49E-18 | 73.6158863 | 1.755621418 | 15.97682722 |
| ENSMUSG00000034765 | *Dusp5* | 487.4649961 | 487.2692013 | 426.072075 | 868.1733163 | 730.6170259 | 865.0467841 | 2.78E-10 | 0.811758226 | 4.24E-11 | 43.49756564 | 1.755349401 | 9.556531499 |
| ENSMUSG00000006019 | *Dhx34* | 650.2044692 | 645.5357726 | 543.304924 | 1117.3204 | 1003.282458 | 1114.826747 | 5.04E-12 | 0.811356473 | 6.64E-13 | 51.64790445 | 1.754860649 | 11.29737807 |
| ENSMUSG00000020329 | *Polrmt* | 913.9026897 | 946.7218537 | 705.9182308 | 1424.638388 | 1611.779059 | 1476.64037 | 2.73E-09 | 0.811282907 | 4.63E-10 | 38.8285811 | 1.754771168 | 8.563699427 |
| ENSMUSG00000036057 | *Ptpn23* | 1909.928539 | 1902.076429 | 1590.837156 | 3030.374875 | 3311.990149 | 3148.880491 | 1.05E-15 | 0.81108926 | 1.01E-16 | 68.94772662 | 1.754535648 | 14.97782274 |
| ENSMUSG00000041926 | *Rnpep* | 3789.720139 | 3707.274533 | 2778.292465 | 5642.577774 | 6298.676752 | 6097.569688 | 7.59E-10 | 0.810989143 | 1.21E-10 | 41.44284579 | 1.754413894 | 9.119919214 |
| ENSMUSG00000021048 | *Mthfd1* | 2827.598347 | 2974.452349 | 3272.435227 | 5151.966557 | 5523.84371 | 5225.176435 | 5.04E-22 | 0.810468794 | 3.22E-23 | 98.51780742 | 1.753781229 | 21.29788577 |
| ENSMUSG00000024853 | *Sf3b2* | 6558.551454 | 6217.478273 | 5623.395049 | 10535.51917 | 11192.96862 | 10549.53021 | 8.51E-23 | 0.810208116 | 5.31E-24 | 102.0900418 | 1.75346437 | 22.07032183 |
| ENSMUSG00000049807 | *Arhgap23* | 234.3147044 | 283.920637 | 146.2259192 | 342.4400439 | 407.4190044 | 419.6670704 | 0.001548877 | 0.810194316 | 0.00056717 | 11.8807523 | 1.753447598 | 2.809983195 |
| ENSMUSG00000012076 | *Brms1l* | 608.7661775 | 622.5151804 | 632.8052712 | 1069.027573 | 1053.815047 | 1141.457699 | 4.62E-18 | 0.810101451 | 3.81E-19 | 79.96593709 | 1.753334733 | 17.33518974 |
| ENSMUSG00000024429 | *Gnl1* | 2217.325322 | 2280.957009 | 2037.078323 | 3756.962404 | 3948.911332 | 3759.555769 | 1.30E-25 | 0.809694847 | 6.96E-27 | 115.2435713 | 1.75284065 | 24.8848423 |
| ENSMUSG00000030541 | *Idh2* | 3226.912795 | 3351.414546 | 3310.252275 | 5834.651517 | 5792.298093 | 5700.860335 | 1.89E-43 | 0.809518851 | 4.65E-45 | 198.4059548 | 1.752626832 | 42.72388025 |
| ENSMUSG00000063015 | *Ccni* | 2330.338845 | 2266.569139 | 1955.141386 | 3469.400573 | 4183.677321 | 3837.612007 | 2.58E-13 | 0.809304276 | 3.05E-14 | 57.70058488 | 1.752366179 | 12.58766003 |
| ENSMUSG00000042271 | *Nxt2* | 692.3961845 | 624.4335631 | 689.5308433 | 1227.076824 | 1174.88271 | 1114.826747 | 5.28E-16 | 0.809237434 | 4.96E-17 | 70.35167706 | 1.752284992 | 15.27722184 |
| ENSMUSG00000032047 | *Acat1* | 2821.570959 | 2727.940174 | 2983.765093 | 5134.40553 | 4930.085782 | 4879.892368 | 9.77E-31 | 0.809105459 | 4.08E-32 | 139.1535016 | 1.752124704 | 30.01005715 |
| ENSMUSG00000055485 | *Soga1* | 1097.73802 | 933.2931749 | 583.6431087 | 1403.784667 | 1603.35696 | 1582.245869 | 0.000183039 | 0.808868944 | 5.68E-05 | 16.20643008 | 1.751837484 | 3.737455494 |
| ENSMUSG00000091537 | *Tma7* | 1432.258049 | 1472.358709 | 1811.436603 | 2915.13063 | 2665.594106 | 2663.095195 | 3.56E-12 | 0.807766123 | 4.61E-13 | 52.36512203 | 1.750498861 | 11.44801509 |
| ENSMUSG00000035164 | *Zc3h12c* | 595.2045547 | 560.1677432 | 552.1289019 | 1065.73488 | 923.2725241 | 1003.711396 | 5.02E-14 | 0.807476553 | 5.52E-15 | 61.06584401 | 1.750147546 | 13.29922388 |
| ENSMUSG00000007050 | *Lsm2* | 1552.805807 | 1500.175258 | 1310.991 | 2483.787882 | 2551.895779 | 2611.669909 | 4.27E-18 | 0.807068301 | 3.51E-19 | 80.13002508 | 1.749652361 | 17.36980678 |
| ENSMUSG00000033732 | *Sf3b3* | 7368.481702 | 7427.977746 | 5943.579389 | 11426.74134 | 12478.39137 | 12392.57574 | 4.96E-14 | 0.806981913 | 5.45E-15 | 61.09036435 | 1.749547596 | 13.30436175 |
| ENSMUSG00000031803 | *B3gnt3* | 284.7940781 | 287.7574023 | 234.4656981 | 471.9526246 | 486.3761758 | 460.0726527 | 1.06E-08 | 0.806935589 | 1.91E-09 | 36.06680114 | 1.74949142 | 7.973010536 |
| ENSMUSG00000010154 | *Spire2* | 360.8898502 | 347.2272655 | 365.5647981 | 655.2458532 | 585.3358306 | 636.3879208 | 4.76E-12 | 0.80644494 | 6.26E-13 | 51.76460973 | 1.748896533 | 11.32206004 |
| ENSMUSG00000046179 | *E2f8* | 921.4369245 | 902.599052 | 557.171175 | 1318.174656 | 1527.558076 | 1325.119437 | 4.50E-05 | 0.80558256 | 1.27E-05 | 19.05584372 | 1.747851431 | 4.346906598 |
| ENSMUSG00000074918 | *Inafm2* | 403.0815655 | 339.5537348 | 243.289676 | 550.9772501 | 566.3861094 | 613.4302036 | 5.12E-05 | 0.805122068 | 1.46E-05 | 18.79183022 | 1.747293626 | 4.290381514 |
| ENSMUSG00000021384 | *Susd3* | 1335.819842 | 1303.541033 | 1218.969516 | 2270.860419 | 2266.5972 | 2213.123938 | 2.08E-25 | 0.80481979 | 1.12E-26 | 114.3036119 | 1.746927566 | 24.68232052 |
| ENSMUSG00000059409 | *Ppp2r5d* | 2015.407828 | 1970.179015 | 1904.718655 | 3441.961467 | 3382.525222 | 3472.125149 | 4.80E-34 | 0.804693241 | 1.74E-35 | 154.5651045 | 1.746774337 | 33.3187039 |
| ENSMUSG00000021025 | *Nfkbia* | 2024.448909 | 2007.587477 | 2062.289688 | 3670.254829 | 3450.954771 | 3518.040584 | 4.03E-33 | 0.804121691 | 1.54E-34 | 150.2373474 | 1.746082458 | 32.39447414 |
| ENSMUSG00000032491 | *Nradd* | 489.7252665 | 476.7180965 | 500.4456029 | 844.026903 | 779.044091 | 934.8382444 | 6.50E-12 | 0.803750419 | 8.65E-13 | 51.12798393 | 1.745633168 | 11.18686279 |
| ENSMUSG00000053774 | *Ubxn7* | 123.5614519 | 111.2661956 | 93.28205192 | 185.4883571 | 204.2358833 | 187.3349724 | 0.000101594 | 0.803696914 | 3.02E-05 | 17.40417208 | 1.74556843 | 3.993129888 |
| ENSMUSG00000050271 | *Prag1* | 1310.956867 | 1329.439199 | 1047.532232 | 1922.932554 | 2313.971503 | 2206.695777 | 6.29E-10 | 0.803211983 | 9.99E-11 | 41.82287507 | 1.744981793 | 9.201159311 |
| ENSMUSG00000032050 | *Rdx* | 2584.995984 | 2803.71629 | 2836.278606 | 4779.892279 | 4854.286897 | 4706.332026 | 6.81E-30 | 0.803070297 | 2.95E-31 | 135.2206488 | 1.744810428 | 29.16672459 |
| ENSMUSG00000040549 | *Ckap5* | 1555.819501 | 1487.70577 | 1401.751915 | 2490.373268 | 2746.656802 | 2528.103818 | 1.06E-20 | 0.803012893 | 7.38E-22 | 92.31794086 | 1.744741004 | 19.97353376 |
| ENSMUSG00000037979 | *Ccdc92* | 269.7256083 | 294.4717417 | 255.8953587 | 432.4403118 | 521.1173312 | 478.4388265 | 2.76E-08 | 0.80265527 | 5.12E-09 | 34.14279819 | 1.744308561 | 7.559433291 |
| ENSMUSG00000040225 | *Prrc2c* | 4809.855541 | 4745.119565 | 2775.771329 | 6820.264207 | 7682.006395 | 7010.368524 | 4.54E-05 | 0.802278847 | 1.28E-05 | 19.03722771 | 1.743853503 | 4.343092188 |
| ENSMUSG00000038481 | *Cdk19* | 388.0130958 | 368.329475 | 389.5155952 | 610.2457192 | 740.0918865 | 647.407625 | 3.87E-10 | 0.802179775 | 6.00E-11 | 42.82210088 | 1.743733754 | 9.412408272 |
| ENSMUSG00000096006 | *Gm21596* | 840.0671879 | 843.1291888 | 814.3271019 | 1528.906991 | 1327.533242 | 1499.598088 | 1.80E-16 | 0.801908906 | 1.65E-17 | 72.52982007 | 1.743406394 | 15.74442167 |
| ENSMUSG00000019866 | *Crybg1* | 1958.901066 | 2018.138582 | 1536.63272 | 3191.716819 | 3276.196232 | 3153.472035 | 1.48E-11 | 0.801456401 | 2.02E-12 | 49.46534864 | 1.742859657 | 10.83040142 |
| ENSMUSG00000026918 | *Brd3* | 1112.80649 | 1171.172628 | 888.7006298 | 1797.81023 | 1964.454424 | 1776.009003 | 2.55E-10 | 0.801167863 | 3.89E-11 | 43.66741409 | 1.74251112 | 9.593249154 |
| ENSMUSG00000055302 | *Mrfap1* | 4787.252836 | 4416.116935 | 3240.92102 | 6961.849994 | 7614.629609 | 7119.647258 | 3.82E-08 | 0.801127521 | 7.23E-09 | 33.4719909 | 1.742462395 | 7.417478078 |
| ENSMUSG00000043964 | *Orai3* | 899.5876434 | 952.4770017 | 853.4047183 | 1477.321471 | 1689.683468 | 1550.105065 | 5.51E-16 | 0.800877159 | 5.18E-17 | 70.26544746 | 1.742160038 | 15.25855521 |
| ENSMUSG00000011267 | *Zfp296* | 270.4790318 | 221.5731998 | 166.3950115 | 376.4645354 | 368.4667998 | 408.6473662 | 0.000108297 | 0.799965393 | 3.24E-05 | 17.27443949 | 1.741059363 | 3.965382487 |
| ENSMUSG00000027010 | *Slc25a12* | 2701.023201 | 2603.2453 | 2606.855181 | 4416.598515 | 4745.852382 | 4606.236379 | 8.46E-33 | 0.79901911 | 3.27E-34 | 148.7362457 | 1.739917752 | 32.07256603 |
| ENSMUSG00000070282 | *3000002C10Rik* | 150.6846974 | 177.4503981 | 150.007624 | 278.7813178 | 267.4016204 | 285.594002 | 1.87E-06 | 0.798506378 | 4.34E-07 | 25.53556978 | 1.739299498 | 5.728375938 |
| ENSMUSG00000032300 | *1700017B05Rik* | 3762.596894 | 3627.661652 | 2831.236333 | 5542.699428 | 6299.729515 | 5945.130446 | 1.65E-10 | 0.798295073 | 2.47E-11 | 44.55942269 | 1.739044769 | 9.782873947 |
| ENSMUSG00000042744 | *Hectd4* | 708.2180777 | 736.65895 | 368.0859346 | 946.1003776 | 1127.508407 | 1085.440869 | 0.001424 | 0.798068572 | 0.000516459 | 12.05527929 | 1.738771763 | 2.846489895 |
| ENSMUSG00000028991 | *Mtor* | 1627.394732 | 1601.84954 | 1216.44838 | 2496.958653 | 2716.126696 | 2527.185509 | 8.02E-10 | 0.797806593 | 1.29E-10 | 41.32795195 | 1.738456048 | 9.095695852 |
| ENSMUSG00000020458 | *Rtn4* | 7190.673759 | 7312.874785 | 5879.290408 | 11450.88775 | 12088.86932 | 11902.19891 | 3.99E-15 | 0.797577509 | 4.03E-16 | 66.22167553 | 1.738180022 | 14.39942504 |
| ENSMUSG00000003346 | *Abhd17a* | 1875.271059 | 1864.667967 | 1844.211378 | 3061.106674 | 3306.726338 | 3338.052081 | 1.08E-27 | 0.797508343 | 5.13E-29 | 124.9841002 | 1.738096692 | 26.96580662 |
| ENSMUSG00000014633 | *Cmc2* | 509.3142772 | 474.7997139 | 470.1919644 | 810.0024115 | 842.2098282 | 878.8214144 | 2.05E-14 | 0.797480049 | 2.19E-15 | 62.88796352 | 1.738062605 | 13.68871163 |
| ENSMUSG00000004880 | *Lbr* | 2322.80461 | 2258.895608 | 1796.309784 | 3628.547388 | 3783.627653 | 3684.254456 | 2.78E-12 | 0.797217156 | 3.57E-13 | 52.86759975 | 1.737745917 | 11.55588895 |
| ENSMUSG00000027878 | *Notch2* | 1740.408255 | 1859.87201 | 1113.081782 | 2427.812106 | 2931.942964 | 2834.81892 | 3.17E-05 | 0.796534151 | 8.76E-06 | 19.7637668 | 1.736923424 | 4.498278936 |
| ENSMUSG00000020692 | *Nle1* | 1864.72313 | 1850.280097 | 1742.105348 | 3099.521423 | 3128.809512 | 3248.057829 | 7.42E-29 | 0.79510869 | 3.37E-30 | 130.3876088 | 1.735208097 | 28.12959566 |
| ENSMUSG00000032512 | *Wdr48* | 1518.148326 | 1609.52307 | 1836.647968 | 2960.130764 | 2821.402924 | 2818.289363 | 2.67E-16 | 0.794875109 | 2.46E-17 | 71.73705695 | 1.734927178 | 15.57348878 |
| ENSMUSG00000105171 | *Gm5288* | 65.54784336 | 70.98015924 | 83.19750577 | 124.0247595 | 140.0173839 | 114.788586 | 0.001502591 | 0.794576781 | 0.000548159 | 11.94426286 | 1.734568459 | 2.823159147 |
| ENSMUSG00000022540 | *Rogdi* | 405.341836 | 425.8809555 | 392.0367317 | 648.6604677 | 727.4587391 | 745.6666547 | 1.80E-11 | 0.794239602 | 2.49E-12 | 49.05502559 | 1.734163112 | 10.74368385 |
| ENSMUSG00000039670 | *Oxld1* | 88.15054797 | 80.57207265 | 103.3665981 | 160.2443795 | 147.3867199 | 161.6223291 | 0.000316861 | 0.794129275 | 0.000101677 | 15.10529907 | 1.734030501 | 3.499130591 |
| ENSMUSG00000039148 | *Sart1* | 1622.120767 | 1518.399893 | 1234.096336 | 2271.957983 | 2768.76481 | 2551.061535 | 2.80E-09 | 0.793434009 | 4.76E-10 | 38.77404559 | 1.733195035 | 8.552795559 |
| ENSMUSG00000020994 | *Pnn* | 929.7245829 | 770.2306469 | 659.2772048 | 1315.979528 | 1341.219151 | 1441.74464 | 3.14E-08 | 0.793306106 | 5.89E-09 | 33.87026473 | 1.733041385 | 7.502596932 |
| ENSMUSG00000057982 | *Zfp809* | 360.8898502 | 355.8599876 | 434.8960529 | 657.4409817 | 671.662338 | 658.4273293 | 5.09E-10 | 0.792985612 | 8.01E-11 | 42.25633133 | 1.732656433 | 9.292898542 |
| ENSMUSG00000008690 | *Ncaph2* | 4789.513107 | 4726.894929 | 4371.650758 | 8055.023981 | 7982.043646 | 8038.874255 | 1.93E-33 | 0.792926899 | 7.22E-35 | 151.7394327 | 1.732585921 | 32.71410207 |
| ENSMUSG00000015880 | *Ncapg* | 867.1904335 | 782.7001344 | 741.2141423 | 1424.638388 | 1342.271914 | 1385.72781 | 1.59E-15 | 0.792855271 | 1.55E-16 | 68.10407528 | 1.732499902 | 14.79879719 |
| ENSMUSG00000024769 | *Cdc42bpg* | 457.3280566 | 453.6975044 | 329.0083183 | 660.7336744 | 751.6722716 | 742.9117286 | 1.08E-06 | 0.792374476 | 2.43E-07 | 26.65716731 | 1.731922622 | 5.967899501 |
| ENSMUSG00000057337 | *Chst3* | 540.9580636 | 539.0655337 | 272.2827462 | 717.807015 | 919.061475 | 711.6892332 | 0.003053968 | 0.792291746 | 0.001187678 | 10.5094164 | 1.73182331 | 2.515135488 |
| ENSMUSG00000024891 | *Slc29a2* | 298.3557008 | 299.2676984 | 249.5925173 | 497.1966022 | 481.1123643 | 494.9683828 | 4.53E-09 | 0.79222673 | 7.85E-10 | 37.79698686 | 1.731745265 | 8.344200202 |
| ENSMUSG00000041193 | *Pla2g5* | 1647.737166 | 1797.524573 | 1694.203754 | 3020.496797 | 2868.777227 | 3003.787718 | 9.22E-26 | 0.791452846 | 4.86E-27 | 115.9540243 | 1.73081658 | 25.03537438 |
| ENSMUSG00000061118 | *Dnajc30* | 216.9859642 | 226.3691565 | 229.423425 | 376.4645354 | 401.1024307 | 384.7713403 | 5.35E-09 | 0.791133656 | 9.34E-10 | 37.45802702 | 1.730433687 | 8.271814757 |
| ENSMUSG00000015749 | *Anp32e* | 4304.308381 | 3963.378622 | 4341.397119 | 7068.313726 | 7485.139848 | 7257.393561 | 2.78E-29 | 0.790759888 | 1.23E-30 | 132.3862766 | 1.729985431 | 28.55623024 |
| ENSMUSG00000025239 | *Limd1* | 4001.432139 | 3952.827517 | 3337.984777 | 6246.238108 | 6713.465093 | 6587.946528 | 7.55E-17 | 0.790715663 | 6.71E-18 | 74.30150893 | 1.729932399 | 16.12200183 |
| ENSMUSG00000074748 | *Atxn7l3b* | 1225.06659 | 1244.071169 | 1178.631332 | 2034.884107 | 2278.177585 | 1994.56647 | 3.29E-18 | 0.789115233 | 2.69E-19 | 80.65348865 | 1.728014392 | 17.48267383 |
| ENSMUSG00000089715 | *Cbx6* | 2483.283813 | 2594.612578 | 1820.260581 | 3744.889198 | 4312.11432 | 3871.589429 | 6.49E-08 | 0.788936769 | 1.26E-08 | 32.38916564 | 1.727800646 | 7.187709131 |
| ENSMUSG00000032078 | *Zpr1* | 2007.120169 | 2022.934538 | 2255.156634 | 3601.108282 | 3717.303629 | 3529.060288 | 8.95E-23 | 0.788848743 | 5.59E-24 | 101.985967 | 1.727695228 | 22.04798245 |
| ENSMUSG00000030374 | *Strn4* | 2284.380012 | 2369.202613 | 1923.627179 | 3623.059567 | 3955.227905 | 3788.941647 | 2.00E-14 | 0.787981301 | 2.14E-15 | 62.93490946 | 1.726656737 | 13.69878405 |
| ENSMUSG00000049354 | *Dcaf7* | 3356.501634 | 3150.943556 | 2441.720738 | 4737.087274 | 5575.429062 | 5150.793431 | 6.09E-09 | 0.787971433 | 1.07E-09 | 37.1955288 | 1.726644928 | 8.215572273 |
| ENSMUSG00000050947 | *Amigo1* | 395.5473306 | 441.2280169 | 310.0997942 | 633.2945683 | 659.0291905 | 692.4047508 | 5.88E-07 | 0.786878647 | 1.28E-07 | 27.89454847 | 1.725337555 | 6.23050403 |
| ENSMUSG00000030498 | *Gas2* | 83.63000705 | 119.8989176 | 97.06375673 | 200.8542565 | 155.8088182 | 160.7040204 | 0.001620605 | 0.78543265 | 0.00059655 | 11.78671236 | 1.723609136 | 2.790322799 |
| ENSMUSG00000073423 | *Zfp414* | 495.7526544 | 470.0037571 | 467.6708279 | 758.416892 | 841.1570659 | 872.3932536 | 2.00E-12 | 0.785393861 | 2.52E-13 | 53.5503666 | 1.723562794 | 11.69910781 |
| ENSMUSG00000024498 | *Tcerg1* | 1668.0796 | 1570.196225 | 1160.983376 | 2377.324151 | 2686.649352 | 2527.185509 | 9.30E-08 | 0.784977678 | 1.84E-08 | 31.65740233 | 1.72306566 | 7.031564159 |
| ENSMUSG00000045691 | *Thtpa* | 449.0403982 | 533.3103857 | 529.4386731 | 861.5879309 | 891.6896556 | 844.843993 | 1.66E-11 | 0.784922637 | 2.28E-12 | 49.23029488 | 1.722999923 | 10.78097373 |
| ENSMUSG00000026790 | *Odf2* | 1396.847145 | 1316.01052 | 1091.652121 | 2045.859749 | 2404.509059 | 2113.028291 | 3.93E-10 | 0.784426702 | 6.10E-11 | 42.78975466 | 1.722407733 | 9.405419486 |
| ENSMUSG00000043885 | *Slc36a4* | 868.6972804 | 966.8648719 | 997.109501 | 1615.614566 | 1565.457518 | 1686.014751 | 1.92E-16 | 0.783890653 | 1.76E-17 | 72.40205227 | 1.721767873 | 15.71695638 |
| ENSMUSG00000104913 | *Gm6560* | 139.3833451 | 110.3070042 | 81.9369375 | 195.3664353 | 195.813785 | 185.498355 | 0.00168448 | 0.783642511 | 0.000622952 | 11.70610901 | 1.721471756 | 2.773534133 |
| ENSMUSG00000007029 | *Vars* | 10523.06584 | 9387.605656 | 6565.039546 | 14887.36139 | 15495.60808 | 15187.90739 | 1.28E-06 | 0.783062081 | 2.91E-07 | 26.30536464 | 1.720779306 | 5.892227695 |
| ENSMUSG00000035960 | *Apex1* | 3966.774659 | 3661.233349 | 3877.507996 | 6811.483693 | 6413.427841 | 6559.478958 | 1.77E-30 | 0.781831519 | 7.47E-32 | 137.9505471 | 1.719312176 | 29.75124462 |
| ENSMUSG00000074282 | *Zfp94* | 99.45190028 | 103.5926648 | 115.9722808 | 159.1468153 | 184.2333999 | 202.0279114 | 0.000243903 | 0.781809298 | 7.70E-05 | 15.63030513 | 1.719285695 | 3.612783437 |
| ENSMUSG00000029512 | *Ulk1* | 379.7254374 | 403.8195546 | 257.1559269 | 579.5139204 | 594.8106911 | 621.6949818 | 3.06E-05 | 0.781661537 | 8.42E-06 | 19.84054955 | 1.719109614 | 4.514643847 |
| ENSMUSG00000052833 | *Sae1* | 3969.034929 | 3942.276412 | 3325.379094 | 6038.798466 | 6835.585518 | 6447.445298 | 8.81E-15 | 0.781179981 | 9.17E-16 | 64.60090463 | 1.71853589 | 14.05503436 |
| ENSMUSG00000027323 | *Rad51* | 1179.107757 | 1099.233277 | 976.9404087 | 1702.322141 | 1963.401662 | 1936.713023 | 3.22E-12 | 0.78107363 | 4.16E-13 | 52.56663138 | 1.71840921 | 11.49165413 |
| ENSMUSG00000027995 | *Tlr2* | 3268.351086 | 3577.783702 | 3956.923797 | 5810.505103 | 6465.013193 | 6272.048339 | 9.19E-16 | 0.780876363 | 8.79E-17 | 69.22395122 | 1.718174258 | 15.03647196 |
| ENSMUSG00000002052 | *Supt6* | 3262.323698 | 3380.190286 | 2645.932797 | 4772.209329 | 5904.943658 | 5285.784808 | 9.01E-10 | 0.780451387 | 1.45E-10 | 41.08869355 | 1.717668209 | 9.045497582 |
| ENSMUSG00000078812 | *Eif5a* | 39343.77449 | 39164.74165 | 36895.57267 | 65514.70724 | 66333.49883 | 66304.6422 | 2.40E-44 | 0.77982334 | 5.66E-46 | 202.5984208 | 1.71692062 | 43.62022314 |
| ENSMUSG00000032298 | *Neil1* | 201.9174945 | 241.716218 | 231.9445615 | 387.4401778 | 411.6300535 | 358.1403883 | 4.42E-07 | 0.779742754 | 9.48E-08 | 28.47786584 | 1.71682472 | 6.35463337 |
| ENSMUSG00000036103 | *Colec12* | 3177.186844 | 3327.434762 | 3530.851722 | 5423.064925 | 6099.70468 | 5695.350483 | 2.48E-21 | 0.779687893 | 1.65E-22 | 95.28015583 | 1.716759436 | 20.60508095 |
| ENSMUSG00000094111 | *Nutf2-ps2* | 452.8075157 | 498.7794974 | 442.4594625 | 709.0265011 | 869.5816476 | 813.6214976 | 1.51E-09 | 0.778764972 | 2.50E-10 | 40.03244144 | 1.715661541 | 8.820038459 |
| ENSMUSG00000027134 | *Lpcat4* | 956.8478284 | 914.1093481 | 670.6223192 | 1337.930812 | 1514.924928 | 1515.209335 | 4.05E-07 | 0.778192422 | 8.64E-08 | 28.65749853 | 1.714980796 | 6.39297337 |
| ENSMUSG00000020091 | *Eif4ebp2* | 2832.118887 | 2863.186153 | 2108.930714 | 4197.085666 | 4849.023086 | 4344.518403 | 1.33E-08 | 0.777749467 | 2.41E-09 | 35.60682667 | 1.714454322 | 7.875604532 |
| ENSMUSG00000025261 | *Huwe1* | 6337.044949 | 6236.6621 | 4074.156646 | 8622.464694 | 10473.93198 | 9453.987943 | 7.00E-06 | 0.777681377 | 1.75E-06 | 22.85341679 | 1.714373407 | 5.155184352 |
| ENSMUSG00000032612 | *Usp4* | 4056.432054 | 4262.64632 | 3987.177436 | 6893.801011 | 7204.052318 | 6988.329116 | 1.67E-34 | 0.77665982 | 5.99E-36 | 156.6878418 | 1.713159908 | 33.77845108 |

| | | | | | | | | | | | | | |
|---|---|---|---|---|---|---|---|---|---|---|---|---|---|
| ENSMUSG00000047454 | *Gphn* | 509.3142772 | 548.6574471 | 504.2273077 | 883.5392157 | 951.6971058 | 840.2524495 | 8.61E-13 | 0.775206235 | 1.05E-13 | 55.26349408 | 1.711434686 | 12.06524474 |
| ENSMUSG00000037221 | *Mospd3* | 543.9717576 | 649.3725379 | 569.7768577 | 985.6126904 | 989.5965481 | 1039.525435 | 3.48E-12 | 0.774941903 | 4.49E-13 | 52.41576742 | 1.711121143 | 11.45898459 |
| ENSMUSG00000044783 | *Hjurp* | 3426.570019 | 3251.658646 | 2250.114361 | 4820.502156 | 5320.660589 | 5145.283579 | 9.56E-07 | 0.774755347 | 2.14E-07 | 26.8987328 | 1.710899892 | 6.019672171 |
| ENSMUSG00000005470 | *Asf1b* | 2593.283642 | 2562.000072 | 2546.347904 | 4235.500414 | 4476.345237 | 4463.898532 | 1.29E-33 | 0.774506311 | 4.77E-35 | 152.5627322 | 1.710604584 | 32.8892316 |
| ENSMUSG00000015143 | *Actn1* | 2629.447969 | 2718.348261 | 2354.741527 | 4088.426806 | 4610.046047 | 4485.019632 | 4.51E-17 | 0.774479078 | 3.95E-18 | 75.34717993 | 1.710572294 | 16.34560082 |
| ENSMUSG00000038170 | *Pde4dip* | 1591.230404 | 1625.829323 | 1560.583517 | 2648.422519 | 2870.882752 | 2652.9938 | 2.42E-24 | 0.773914175 | 1.38E-25 | 109.3225362 | 1.709902631 | 23.61646937 |
| ENSMUSG00000028901 | *Gmeb1* | 457.3280566 | 464.2486091 | 339.0928644 | 754.0266351 | 712.7200671 | 696.9962942 | 4.13E-07 | 0.773514072 | 8.84E-08 | 28.61214264 | 1.709428489 | 6.383777265 |
| ENSMUSG00000038685 | *Rtel1* | 1099.998291 | 987.9670814 | 819.369375 | 1576.102253 | 1729.688435 | 1674.995047 | 1.76E-09 | 0.773433683 | 2.93E-10 | 39.71901472 | 1.709333241 | 8.753586732 |
| ENSMUSG00000037679 | *Inf2* | 2655.064368 | 2606.122874 | 1984.134456 | 3719.64522 | 4597.412899 | 4071.780723 | 3.95E-08 | 0.772796029 | 7.48E-09 | 33.40486904 | 1.708577902 | 7.403591787 |
| ENSMUSG00000001729 | *Akt1* | 9579.779637 | 10062.87636 | 9010.541989 | 15638.09534 | 17095.80675 | 16212.73989 | 4.25E-25 | 0.772288233 | 2.32E-26 | 112.853061 | 1.707976627 | 24.37120791 |
| ENSMUSG00000066878 | *Gm10184* | 3659.377876 | 3366.761607 | 3411.097737 | 5950.993326 | 6258.671785 | 5609.029466 | 6.07E-23 | 0.770950919 | 3.75E-24 | 102.7772717 | 1.706394143 | 22.21666023 |
| ENSMUSG00000039715 | *Dync2i2* | 560.5470743 | 506.4530281 | 439.938326 | 845.1244672 | 876.9509836 | 858.6186233 | 5.63E-10 | 0.770665432 | 8.90E-11 | 42.05011705 | 1.706056508 | 9.249372511 |
| ENSMUSG00000022177 | *Haus4* | 347.3282275 | 287.7574023 | 301.2758163 | 491.7087809 | 579.0192569 | 529.864113 | 9.97E-08 | 0.769893153 | 1.98E-08 | 31.51179499 | 1.705143495 | 7.00110462 |
| ENSMUSG00000007836 | *Hnrnpa0* | 5246.841163 | 5288.981055 | 3741.366623 | 7399.778127 | 8513.6886 | 8435.583608 | 1.41E-07 | 0.769566445 | 2.86E-08 | 30.79961154 | 1.704757398 | 6.849497856 |
| ENSMUSG00000022952 | *Runx1* | 681.8482557 | 801.8839612 | 547.0866289 | 1146.954634 | 1150.669178 | 1170.843577 | 3.07E-07 | 0.76951874 | 6.48E-08 | 29.2154011 | 1.704701027 | 6.512437844 |
| ENSMUSG00000028431 | *Elp1* | 1488.011387 | 1512.644745 | 1283.258498 | 2288.421447 | 2602.428369 | 2420.661702 | 7.36E-14 | 0.769473521 | 8.25E-15 | 60.27455818 | 1.704647598 | 13.13297634 |
| ENSMUSG00000091105 | *Mrto4-ps2* | 128.8354163 | 164.0217193 | 204.2120596 | 296.3423457 | 251.6101862 | 292.9404715 | 0.000564198 | 0.769385326 | 0.000190028 | 13.92720719 | 1.704543391 | 3.248568144 |
| ENSMUSG00000032307 | *Ube2q2* | 887.5328676 | 808.5983006 | 610.1150423 | 1308.296578 | 1326.480479 | 1306.753263 | 2.58E-07 | 0.769198061 | 5.40E-08 | 29.56593634 | 1.704322153 | 6.588440456 |
| ENSMUSG00000061436 | *Hipk2* | 1130.13523 | 1111.702764 | 636.586976 | 1516.833784 | 1844.439524 | 1551.941683 | 0.000447189 | 0.769019331 | 0.00014758 | 14.40283808 | 1.704111024 | 3.349508929 |
| ENSMUSG00000112808 | *Gm4739* | 2556.365891 | 1780.259129 | 1918.584906 | 3606.596103 | 3530.964705 | 3530.896905 | 7.28E-08 | 0.768970005 | 1.42E-08 | 32.15904404 | 1.70405276 | 7.137804572 |
| ENSMUSG00000018932 | *Map2k3* | 5438.210729 | 5280.348333 | 4377.953599 | 8188.926818 | 8897.946834 | 8649.549532 | 6.93E-14 | 0.76886448 | 7.74E-15 | 60.40040255 | 1.703928124 | 13.15941031 |
| ENSMUSG00000037992 | *Rara* | 422.6705762 | 514.1265588 | 371.8676394 | 762.807149 | 757.9888453 | 711.6892332 | 2.32E-07 | 0.767504667 | 4.84E-08 | 29.77858332 | 1.702322842 | 6.63394273 |
| ENSMUSG00000050965 | *Prkca* | 240.3420923 | 265.6960015 | 236.9868346 | 418.1719766 | 403.2079552 | 443.5430963 | 2.42E-08 | 0.767493964 | 4.48E-09 | 34.40397601 | 1.702310212 | 7.615787016 |
| ENSMUSG00000098923 | *Tmem185b* | 1543.011301 | 1583.624904 | 1166.025649 | 2383.909536 | 2480.307944 | 2453.720814 | 4.13E-09 | 0.767462773 | 7.14E-10 | 37.9811522 | 1.70227341 | 8.383820083 |
| ENSMUSG00000080775 | *Gm6368* | 190.6161422 | 183.2055462 | 148.7470558 | 305.1228596 | 306.353825 | 282.8390759 | 8.48E-06 | 0.765922904 | 2.15E-06 | 22.45950228 | 1.700457447 | 5.07136373 |
| ENSMUSG00000054702 | *Ap1s3* | 519.1087825 | 517.9633242 | 426.072075 | 867.0757521 | 791.6772385 | 836.5792148 | 6.94E-10 | 0.765393934 | 1.10E-10 | 41.62712749 | 1.699834082 | 9.158428448 |
| ENSMUSG00000035900 | *Gramd4* | 590.6840138 | 678.1482782 | 531.9598096 | 972.4419194 | 1043.287425 | 1048.708522 | 5.54E-10 | 0.76497424 | 8.74E-11 | 42.08490989 | 1.699339655 | 9.256722123 |
| ENSMUSG00000023348 | *Trip6* | 1406.64165 | 1421.521568 | 1546.717266 | 2532.080709 | 2486.624518 | 2404.132145 | 5.50E-22 | 0.764396097 | 3.53E-23 | 98.33894837 | 1.698658802 | 21.25959103 |
| ENSMUSG00000037126 | *Psd* | 156.7120853 | 129.4908311 | 105.8877346 | 239.269005 | 213.7107439 | 218.5574677 | 0.000385702 | 0.763987905 | 0.000125753 | 14.70439818 | 1.698178256 | 3.413748483 |
| ENSMUSG00000067150 | *Xpo5* | 2496.092012 | 2585.020664 | 2015.648663 | 3738.303812 | 4224.73505 | 4094.73844 | 7.85E-11 | 0.763568653 | 1.14E-11 | 46.06884342 | 1.697684832 | 10.10526569 |
| ENSMUSG00000021714 | *Cenpk* | 114.52037 | 91.12317741 | 102.1060298 | 171.2200219 | 182.1278753 | 170.805416 | 0.000243327 | 0.763543075 | 7.68E-05 | 15.63600745 | 1.697654733 | 3.613809749 |
| ENSMUSG00000042129 | *Rassf4* | 2164.585678 | 2227.242294 | 2121.536397 | 3535.254427 | 3899.431504 | 3625.4827 | 2.43E-24 | 0.763540529 | 1.39E-25 | 109.3130953 | 1.697651737 | 23.61492318 |
| ENSMUSG00000024208 | *Uqcc2* | 1378.011558 | 1514.563128 | 1940.014566 | 2765.861893 | 2740.340228 | 2685.134604 | 2.12E-08 | 0.763509257 | 3.91E-09 | 34.6679764 | 1.697614938 | 7.673397748 |
| ENSMUSG00000021466 | *Ptch1* | 673.5605973 | 694.454531 | 422.2903702 | 930.7344782 | 1128.56117 | 986.2635309 | 0.000163841 | 0.762938082 | 5.04E-05 | 16.43281826 | 1.696942972 | 3.785576273 |
| ENSMUSG00000049339 | *Retreg2* | 2616.63977 | 2864.145345 | 2589.207225 | 4630.623542 | 4575.304891 | 4489.611176 | 1.64E-25 | 0.762727173 | 8.81E-27 | 114.7764385 | 1.696694912 | 24.78420988 |
| ENSMUSG00000109324 | *Prmt1* | 5774.991027 | 5593.04471 | 5922.149729 | 10025.1518 | 9563.292599 | 9735.90871 | 6.21E-37 | 0.762370303 | 1.98E-38 | 168.0440777 | 1.696275264 | 36.20662523 |
| ENSMUSG00000033624 | *Pdpr* | 407.6021064 | 379.8397711 | 303.7969529 | 542.1967361 | 686.4010099 | 628.1231426 | 4.88E-06 | 0.761795389 | 1.19E-06 | 23.59022297 | 1.695599433 | 5.31119696 |
| ENSMUSG00000035049 | *Rrp12* | 2248.215685 | 2046.914322 | 1692.943186 | 3295.985422 | 3562.547573 | 3307.747894 | 2.04E-10 | 0.761783691 | 3.08E-11 | 44.1283106 | 1.695585684 | 9.691395631 |
| ENSMUSG00000076437 | *Selenoh* | 1371.230746 | 1327.520816 | 1309.730432 | 2219.2749 | 2242.383667 | 2336.177302 | 5.55E-25 | 0.761261157 | 3.04E-26 | 112.3180213 | 1.694971666 | 24.25564485 |
| ENSMUSG00000029366 | *Dck* | 1549.792113 | 1484.828196 | 1374.019413 | 2524.397759 | 2436.091928 | 2522.593966 | 7.71E-21 | 0.761169515 | 5.31E-22 | 92.96907328 | 1.694864002 | 20.11297346 |
| ENSMUSG00000047702 | *Or2y16* | 178.5613664 | 142.9195098 | 110.9300077 | 250.2446474 | 233.7132273 | 254.3715066 | 0.000714318 | 0.760061963 | 0.000245198 | 13.44853532 | 1.693563361 | 3.146108427 |
| ENSMUSG00000049823 | *Zbtb12* | 328.4926403 | 301.1860811 | 284.8884288 | 468.6599318 | 561.122298 | 522.5176435 | 6.52E-08 | 0.759501866 | 1.27E-08 | 32.38083655 | 1.692905997 | 7.186000928 |
| ENSMUSG00000025651 | *Uqcrc1* | 9987.381743 | 9850.895073 | 10136.22945 | 16455.7807 | 16929.47031 | 17335.83141 | 4.48E-43 | 0.759013328 | 1.13E-44 | 196.648717 | 1.692332827 | 42.34877484 |
| ENSMUSG00000059058 | *Tma7-ps* | 277.2598432 | 260.9000448 | 390.7761635 | 536.7089149 | 477.9540775 | 549.1485954 | 6.94E-05 | 0.758310289 | 2.02E-05 | 18.17448922 | 1.691508338 | 4.158517917 |
| ENSMUSG00000038644 | *Pold1* | 2748.48888 | 2749.042384 | 2009.345821 | 3885.377421 | 4485.820097 | 4336.253625 | 6.59E-08 | 0.758257932 | 1.28E-08 | 32.35843907 | 1.691446953 | 7.181301612 |
| ENSMUSG00000003660 | *Snrnp200* | 4994.444295 | 4984.9174 | 3635.478888 | 7194.533614 | 8109.427883 | 7729.404227 | 2.83E-08 | 0.75785793 | 5.27E-09 | 34.08809888 | 1.690978047 | 7.54754612 |
| ENSMUSG00000033253 | *Szt2* | 601.2319426 | 514.1265588 | 351.6985471 | 774.8803557 | 931.6946224 | 782.3990022 | 0.00026472 | 0.757753105 | 8.41E-05 | 15.46439892 | 1.690855187 | 3.577212592 |
| ENSMUSG00000037408 | *Cnnm4* | 601.9853661 | 588.9434834 | 403.3818462 | 838.5390818 | 908.5338521 | 955.9593442 | 1.36E-05 | 0.757386169 | 3.54E-06 | 21.49844681 | 1.690425187 | 4.867690731 |
| ENSMUSG00000099083 | *Atf7* | 830.2726826 | 799.9655785 | 569.7768577 | 1089.881293 | 1373.854782 | 1260.837829 | 1.16E-05 | 0.75649468 | 2.99E-06 | 21.8214582 | 1.689380941 | 4.935772102 |
| ENSMUSG00000005899 | *Smpd4* | 1005.066932 | 983.1711247 | 718.5239135 | 1363.17479 | 1660.206124 | 1557.451535 | 1.47E-06 | 0.756382258 | 3.38E-07 | 26.02181882 | 1.6892493 | 5.832230265 |
| ENSMUSG00000017421 | *Zfp207* | 3205.063513 | 2934.166312 | 3034.187824 | 5315.50363 | 4850.075848 | 5329.863625 | 4.83E-22 | 0.75591749 | 3.09E-23 | 98.60222386 | 1.688705192 | 21.31593273 |
| ENSMUSG00000054836 | *Elp6* | 581.6429319 | 603.3313536 | 539.5232192 | 941.7101206 | 1009.599031 | 965.1424311 | 5.95E-14 | 0.755709873 | 6.57E-15 | 60.72184946 | 1.68846219 | 13.22577009 |
| ENSMUSG00000036046 | *5031439G07Rik* | 5606.224166 | 5618.942876 | 4504.010426 | 8230.63426 | 9472.755042 | 8859.842222 | 2.32E-11 | 0.755392442 | 3.23E-12 | 48.54579688 | 1.688090725 | 10.63372469 |
| ENSMUSG00000021277 | *Traf3* | 1329.792454 | 1425.358333 | 939.1233606 | 2030.49385 | 2167.637545 | 2045.073448 | 2.19E-06 | 0.754888531 | 5.12E-07 | 25.21912161 | 1.687501204 | 5.659690387 |
| ENSMUSG00000041341 | *Atg2b* | 920.683501 | 1017.702013 | 729.8690279 | 1369.760175 | 1601.251436 | 1536.330435 | 2.18E-07 | 0.75418817 | 4.53E-08 | 29.90714463 | 1.6866822 | 6.660878346 |
| ENSMUSG00000037316 | *Bag4* | 365.4103912 | 406.6971286 | 390.7761635 | 662.9288029 | 608.4966008 | 685.9765899 | 3.34E-10 | 0.75361318 | 5.14E-11 | 43.12129225 | 1.686010101 | 9.47573708 |
| ENSMUSG00000053286 | *Trmt1l* | 546.9854515 | 554.4125952 | 506.7484442 | 847.3195957 | 953.8026304 | 912.7988359 | 1.74E-12 | 0.753013901 | 2.19E-13 | 53.83063186 | 1.685309898 | 11.75889776 |
| ENSMUSG00000052364 | *B630019K06Rik* | 371.4377791 | 309.8188032 | 196.64865 | 475.2453173 | 522.1700935 | 489.4585307 | 0.001423864 | 0.752859538 | 0.000516214 | 12.05616279 | 1.685129586 | 2.846531342 |
| ENSMUSG00000022892 | *App* | 249.3831742 | 232.1243046 | 148.7470558 | 339.1473511 | 368.4667998 | 360.8953144 | 0.000661413 | 0.752580814 | 0.000225993 | 13.60159541 | 1.684804055 | 3.179527242 |
| ENSMUSG00000038902 | *Pogz* | 944.7930526 | 992.7630381 | 659.2772048 | 1343.418634 | 1586.512764 | 1452.764344 | 9.08E-06 | 0.752401829 | 2.31E-06 | 22.32163357 | 1.684595046 | 5.042118032 |
| ENSMUSG00000026385 | *Dbi* | 1179.861181 | 1096.355703 | 1152.159398 | 1907.566655 | 1859.178196 | 2008.341101 | 6.86E-20 | 0.752260308 | 5.05E-21 | 88.51111871 | 1.684429804 | 19.16376443 |
| ENSMUSG00000020841 | *Cpd* | 3546.364353 | 3348.536972 | 2308.100501 | 4851.233955 | 5443.833777 | 5205.891952 | 3.15E-06 | 0.751191786 | 7.53E-07 | 24.47388661 | 1.683182706 | 5.501377177 |
| ENSMUSG00000041837 | *Pdcd7* | 779.0398855 | 736.65895 | 630.2841346 | 1131.588735 | 1223.309775 | 1265.429372 | 1.88E-10 | 0.751078907 | 2.83E-11 | 44.289525 | 1.683051016 | 9.725977444 |
| ENSMUSG00000097487 | *Ptges3l* | 96.43820633 | 93.04156009 | 131.0991 | 169.0248934 | 193.7082605 | 172.6420333 | 0.001152119 | 0.751068082 | 0.000410579 | 12.48342444 | 1.683038387 | 2.938502581 |
| ENSMUSG00000041215 | *Yeats2* | 427.9445406 | 543.8614904 | 383.2127538 | 805.6121545 | 784.3079025 | 694.2413681 | 4.21E-06 | 0.750761981 | 1.02E-06 | 23.88845212 | 1.682681329 | 5.37542818 |
| ENSMUSG00000087598 | *Zfp111* | 423.4239997 | 462.3302264 | 476.4948058 | 761.7095848 | 806.4159105 | 719.0357027 | 6.59E-11 | 0.750340865 | 9.50E-12 | 46.42922635 | 1.682190234 | 10.18123779 |
| ENSMUSG00000042790 | *Rnf214* | 430.9582345 | 467.1261831 | 417.2480971 | 737.5631714 | 759.0416076 | 718.117394 | 5.55E-12 | 0.75007544 | 7.34E-13 | 51.45062739 | 1.681880776 | 11.25531969 |
| ENSMUSG00000042185 | *Nfrkb* | 880.7520562 | 963.9872978 | 700.8759577 | 1397.199282 | 1473.867199 | 1417.868614 | 2.34E-08 | 0.749786923 | 4.33E-09 | 34.47073633 | 1.681544459 | 7.630199403 |
| ENSMUSG00000030357 | *Fkbp4* | 11280.25645 | 11452.74461 | 11190.06453 | 18491.76237 | 19240.28352 | 19280.80921 | 1.51E-44 | 0.749006127 | 3.55E-46 | 203.5271446 | 1.680634642 | 43.82161221 |
| ENSMUSG00000025225 | *Nfkb2* | 1724.586362 | 1598.971966 | 1288.300771 | 2439.885313 | 2666.646868 | 2654.830417 | 2.43E-09 | 0.748860656 | 4.10E-10 | 39.06323925 | 1.680465187 | 8.613787341 |
| ENSMUSG00000029554 | *Mad1l1* | 1154.244782 | 1186.519689 | 989.5460914 | 1804.395616 | 2039.200546 | 1760.397755 | 3.66E-11 | 0.748572722 | 5.16E-12 | 47.62346265 | 1.680129833 | 10.43688408 |
| ENSMUSG00000045817 | *Zfp36l2* | 2593.283642 | 2602.286108 | 1589.576588 | 3275.131702 | 4095.245289 | 4018.518819 | 0.000143119 | 0.746187489 | 4.36E-05 | 16.7079558 | 1.677354339 | 3.844301574 |
| ENSMUSG00000047221 | *Fam185a* | 128.0819928 | 152.5114232 | 150.007624 | 230.4884911 | 277.9292433 | 211.2109982 | 0.000251705 | 0.744267085 | 7.96E-05 | 15.56666324 | 1.67512306 | 3.599108913 |
| ENSMUSG00000027959 | *Sass6* | 753.4234869 | 711.7199751 | 645.4109539 | 1318.174656 | 1043.287425 | 1180.026664 | 7.24E-09 | 0.744104468 | 1.28E-09 | 36.845317 | 1.674934256 | 8.140445612 |
| ENSMUSG00000061028 | *Clasrp* | 445.2732808 | 475.7589052 | 364.3042298 | 696.9532944 | 725.3532145 | 736.4835678 | 4.00E-08 | 0.743670553 | 7.59E-09 | 33.37735472 | 1.674430567 | 7.397761956 |
| ENSMUSG00000050232 | *Cxcr3* | 187.6024482 | 164.0217193 | 122.2751221 | 228.2933626 | 286.3513416 | 283.7573846 | 0.001002319 | 0.743613658 | 0.000352586 | 12.76807159 | 1.674364534 | 2.998994204 |
| ENSMUSG00000042608 | *Stk40* | 1470.682647 | 1455.093264 | 1056.35621 | 2219.2749 | 2152.898873 | 2301.281572 | 1.51E-07 | 0.742658773 | 3.08E-08 | 30.65762234 | 1.67325668 | 6.819776139 |
| ENSMUSG00000022992 | *Kansl2* | 993.0121558 | 1081.008641 | 978.2009769 | 1820.859079 | 1565.457518 | 1716.318938 | 1.52E-13 | 0.740880741 | 1.74E-14 | 58.80292202 | 1.671195764 | 12.81872361 |
| ENSMUSG00000015488 | *Cacfd1* | 394.0404837 | 376.9621971 | 335.3111596 | 604.757898 | 639.0267071 | 611.5935862 | 1.08E-09 | 0.740728699 | 1.76E-10 | 40.71690017 | 1.671019651 | 8.966733477 |
| ENSMUSG00000017210 | *Med24* | 1557.326348 | 1543.338868 | 1244.180882 | 2267.567726 | 2535.051583 | 2467.495445 | 1.81E-10 | 0.740670225 | 2.73E-11 | 44.36500219 | 1.670951924 | 9.741930002 |
| ENSMUSG00000025227 | *Mfsd13a* | 760.2042983 | 768.3122642 | 729.8690279 | 1154.637584 | 1388.593454 | 1232.370259 | 7.72E-12 | 0.740584099 | 1.04E-12 | 50.77608434 | 1.670852175 | 11.11234958 |
| ENSMUSG00000022191 | *Drosha* | 1351.641736 | 1467.562752 | 1108.039509 | 2076.591548 | 2340.29056 | 2150.678947 | 4.15E-09 | 0.740003891 | 7.17E-10 | 37.97324017 | 1.670180343 | 8.382403564 |
| ENSMUSG00000003131 | *Pafah1b2* | 3524.515072 | 3545.171197 | 3953.142092 | 6128.798734 | 6132.340311 | 6135.220345 | 2.25E-23 | 0.74000275 | 1.36E-24 | 104.794232 | 1.670179023 | 22.64753715 |
| ENSMUSG00000052798 | *Nup107* | 1721.572668 | 1596.094392 | 1637.478182 | 2696.715345 | 2849.827506 | 2730.131729 | 4.12E-23 | 0.739239877 | 2.53E-24 | 103.5596084 | 1.669296094 | 22.38476763 |
| ENSMUSG00000034032 | *Rpap1* | 1029.929907 | 1130.886591 | 766.4255077 | 1554.150968 | 1714.949763 | 1623.56976 | 1.56E-06 | 0.738683298 | 3.58E-07 | 25.90815769 | 1.668652218 | 5.807967301 |
| ENSMUSG00000030056 | *Isy1* | 1411.915615 | 1410.970463 | 1476.125443 | 2413.543771 | 2476.096895 | 2279.242164 | 1.26E-21 | 0.738354468 | 8.30E-23 | 96.64408601 | 1.66827193 | 20.89796598 |
| ENSMUSG00000016757 | *Ttll12* | 2069.654319 | 2092.955506 | 1726.978529 | 3125.862965 | 3444.638197 | 3265.505695 | 9.63E-13 | 0.738335582 | 1.19E-13 | 55.03316827 | 1.668250091 | 12.01629553 |
| ENSMUSG00000026879 | *Gsn* | 9614.437117 | 9520.933252 | 8090.327152 | 14279.3108 | 15930.3989 | 15216.37496 | 7.88E-15 | 0.738174225 | 8.16E-16 | 64.83005225 | 1.668063517 | 14.10353128 |
| ENSMUSG00000106864 | *Gtf3c2* | 2728.146446 | 2834.410413 | 2115.233556 | 4154.28066 | 4426.865409 | 4234.32136 | 2.32E-09 | 0.737894149 | 3.91E-10 | 39.1582959 | 1.667739721 | 8.634578668 |
| ENSMUSG00000028108 | *Ecm1* | 308.9036296 | 294.4717417 | 240.7685394 | 453.2940324 | 499.0093232 | 461.9092701 | 9.47E-07 | 0.737380173 | 2.12E-07 | 26.91995912 | 1.667145676 | 6.023777073 |
| ENSMUSG00000007817 | *Zmiz1* | 3133.488282 | 3152.861938 | 1757.232167 | 3842.572415 | 5011.148478 | 4563.075871 | 0.000816568 | 0.737322797 | 0.00028293 | 13.18015303 | 1.667079374 | 3.088007744 |
| ENSMUSG00000021947 | *Cryl1* | 745.8892521 | 811.4758746 | 821.8905115 | 1274.272086 | 1326.480479 | 1356.341932 | 1.06E-15 | 0.736703538 | 1.02E-16 | 68.93407159 | 1.666363955 | 14.97512584 |
| ENSMUSG00000030822 | *Prr14* | 611.7798714 | 604.2905449 | 531.9598096 | 929.636914 | 999.0714086 | 990.8550744 | 5.31E-12 | 0.736595173 | 7.01E-13 | 51.54173738 | 1.666238795 | 11.27457162 |
| ENSMUSG00000005583 | *Mef2c* | 1064.587387 | 1073.335111 | 1048.7928 | 1683.663549 | 1728.635672 | 1896.307441 | 5.56E-17 | 0.736557089 | 4.89E-18 | 74.9232925 | 1.66619481 | 16.25473823 |
| ENSMUSG00000053553 | *3110082I17Rik* | 607.2593305 | 591.8210575 | 669.361751 | 1055.856802 | 1042.234662 | 1009.221248 | 1.16E-13 | 0.736529245 | 1.32E-14 | 59.34754483 | 1.666162653 | 12.9357999 |
| ENSMUSG00000021391 | *Cenpp* | 152.1915444 | 126.613257 | 131.0991 | 225.0006698 | 234.7659896 | 225.9039372 | 4.88E-05 | 0.736300439 | 1.38E-05 | 18.89046319 | 1.665898426 | 4.311685187 |
| ENSMUSG00000039187 | *Fanci* | 494.9992309 | 489.187584 | 502.9667394 | 756.2217635 | 891.6896556 | 827.3961279 | 7.36E-11 | 0.735843688 | 1.07E-11 | 46.20351848 | 1.665371094 | 10.13327135 |
| ENSMUSG00000025364 | *Pa2g4* | 11968.13209 | 11502.62256 | 12338.44222 | 20374.08505 | 19681.39092 | 19556.30182 | 7.09E-35 | 0.735391038 | 2.50E-36 | 158.4229262 | 1.66484866 | 34.14921363 |
| ENSMUSG00000034187 | *Nsf* | 2407.941464 | 2479.509617 | 2140.444921 | 3593.425332 | 4223.682288 | 3887.200676 | 2.44E-13 | 0.73497651 | 2.86E-14 | 57.82771381 | 1.66437037 | 12.61345125 |
| ENSMUSG00000027160 | *Ccdc34* | 305.8899357 | 282.0022543 | 247.0713808 | 531.2210937 | 374.7833735 | 486.7036046 | 8.61E-05 | 0.733990946 | 2.53E-05 | 17.7396682 | 1.663233759 | 4.065118683 |
| ENSMUSG00000020895 | *Tmem107* | 229.04074 | 175.5320154 | 197.9092183 | 347.9278651 | 338.9894558 | 318.6531147 | 1.45E-05 | 0.733944074 | 3.82E-06 | 21.35405796 | 1.663179723 | 4.837149458 |
| ENSMUSG00000002111 | *Spi1* | 8581.493516 | 8535.843745 | 8047.467831 | 13671.26021 | 14087.01214 | 14104.30314 | 9.25E-35 | 0.733893197 | 3.28E-36 | 157.8837695 | 1.663121071 | 34.03393347 |
| ENSMUSG00000091876 | *Gm3851* | 320.204982 | 237.8794526 | 306.3180894 | 440.1232615 | 459.0043563 | 537.2105825 | 2.83E-05 | 0.733223102 | 7.74E-06 | 20.00100263 | 1.662348773 | 4.548703969 |
| ENSMUSG00000095562 | *Gm55594* | 255.4105621 | 317.4923339 | 305.0575212 | 433.5378761 | 516.906282 | 504.1514697 | 2.83E-06 | 0.733104589 | 6.70E-07 | 24.69892215 | 1.662212221 | 5.548588081 |
| ENSMUSG00000059237 | *Ppp1r2-ps4* | 199.657224 | 187.0423115 | 186.5641038 | 327.0741445 | 270.5599073 | 355.3854623 | 3.21E-05 | 0.732488568 | 8.87E-06 | 19.74138016 | 1.66150262 | 4.493623576 |
| ENSMUSG00000043432 | *Leng9* | 64.04099639 | 71.93935058 | 74.37352789 | 99.87834613 | 120.0149005 | 127.6449076 | 0.004533036 | 0.732482009 | 0.001827586 | 9.715115052 | 1.661495067 | 2.343610795 |
| ENSMUSG00000021733 | *Slc4a7* | 1018.628554 | 1157.743949 | 897.5246077 | 1689.15137 | 1658.100599 | 1764.989298 | 2.94E-10 | 0.73223096 | 4.50E-11 | 43.38198223 | 1.661205968 | 9.531658073 |
| ENSMUSG00000026833 | *Olfm1* | 8210.055737 | 7757.939567 | 5923.410297 | 11516.7416 | 12447.86126 | 12409.1053 | 1.62E-08 | 0.732052576 | 2.95E-09 | 35.21658112 | 1.661000579 | 7.791375516 |
| ENSMUSG00000038215 | *Cep44* | 240.3420923 | 198.5526076 | 173.9584212 | 357.8059433 | 352.6753655 | 314.0615713 | 5.94E-05 | 0.73176112 | 1.70E-05 | 18.4948947 | 1.660665054 | 4.226185507 |
| ENSMUSG00000037936 | *Scarb1* | 4113.692239 | 4311.565078 | 3311.512843 | 6364.775046 | 6480.804628 | 6654.064753 | 6.95E-11 | 0.731611512 | 1.00E-11 | 46.31947296 | 1.660492851 | 10.15794363 |
| ENSMUSG00000013833 | *Med16* | 924.4506185 | 898.7622866 | 693.3125481 | 1253.418366 | 1518.083215 | 1415.113688 | 4.90E-07 | 0.731499397 | 1.06E-07 | 28.2648862 | 1.660363816 | 6.309900713 |
| ENSMUSG00000038489 | *Polr2l* | 626.8483411 | 634.9846678 | 658.0166365 | 1072.320266 | 1082.239629 | 1030.342348 | 8.06E-16 | 0.73148451 | 7.67E-17 | 69.4914604 | 1.660346683 | 15.09350113 |
| ENSMUSG00000027185 | *Nat10* | 2225.61298 | 2228.201485 | 1750.929326 | 3322.326964 | 3518.331557 | 3465.696989 | 1.75E-10 | 0.730487687 | 2.63E-11 | 44.43823277 | 1.659199871 | 9.757977218 |
| ENSMUSG00000040463 | *Mybbp1a* | 17069.56252 | 17386.30225 | 15049.92457 | 26387.63953 | 28222.45134 | 27522.62969 | 2.07E-19 | 0.730148171 | 1.58E-20 | 86.2540148 | 1.65880945 | 18.68364378 |
| ENSMUSG00000048163 | *Selplg* | 852.1219637 | 960.1505325 | 775.2494856 | 1374.150432 | 1445.442618 | 1476.64037 | 3.68E-11 | 0.729941185 | 5.19E-12 | 47.61199288 | 1.658571474 | 10.43455412 |
| ENSMUSG00000039197 | *Adk* | 1090.203786 | 1189.397263 | 1251.744291 | 1938.298453 | 1917.080121 | 1989.974927 | 1.37E-16 | 0.729833096 | 1.25E-17 | 73.075382 | 1.658447216 | 15.86185606 |
| ENSMUSG00000028412 | *Slc44a1* | 1564.860582 | 1600.890348 | 1479.907148 | 2427.812106 | 2805.61149 | 2473.923605 | 4.93E-15 | 0.729465984 | 5.02E-16 | 65.79000571 | 1.658025258 | 14.3072475 |
| ENSMUSG00000041879 | *Ipo9* | 3702.323015 | 3678.498793 | 3065.702031 | 5606.358154 | 6096.546394 | 5628.313949 | 3.27E-13 | 0.729316076 | 3.88E-14 | 57.22919693 | 1.657852984 | 12.48546036 |
| ENSMUSG00000001419 | *Mef2d* | 3455.200111 | 3317.842849 | 2453.065852 | 4872.087675 | 5374.351466 | 5052.534401 | 1.46E-07 | 0.728559782 | 2.96E-08 | 30.73615891 | 1.656984126 | 6.835885602 |
| ENSMUSG00000071708 | *Sms* | 1869.243671 | 1917.423491 | 1839.169105 | 3174.155791 | 3008.794611 | 3140.615713 | 2.51E-26 | 0.728459046 | 1.29E-27 | 118.5932411 | 1.656868432 | 25.59950999 |
| ENSMUSG00000035478 | *Mbd3* | 3853.761136 | 4065.052904 | 3853.557199 | 6326.360298 | 6844.007616 | 6334.49333 | 2.31E-25 | 0.728320982 | 1.24E-26 | 114.093678 | 1.656709879 | 24.63689508 |
| ENSMUSG00000036672 | *Cenpt* | 439.9993164 | 464.2486091 | 398.3395731 | 712.3191938 | 724.3004522 | 724.5455548 | 9.52E-11 | 0.727222891 | 1.40E-11 | 45.66775616 | 1.655449373 | 10.02123578 |
| ENSMUSG00000038175 | *Mylip* | 192.8764127 | 249.3897487 | 211.7754692 | 320.488759 | 396.8913815 | 363.6502404 | 4.41E-05 | 0.727193325 | 1.24E-05 | 19.0979332 | 1.655415447 | 4.355639009 |
| ENSMUSG00000002825 | *Qtrt1* | 895.0671025 | 840.2516148 | 810.5453971 | 1395.004153 | 1324.374955 | 1498.679779 | 1.23E-13 | 0.726821731 | 1.41E-14 | 59.22096992 | 1.654989118 | 12.90838797 |
| ENSMUSG00000047434 | *Xxylt1* | 729.3139354 | 695.4137223 | 647.9320904 | 1058.05193 | 1191.726907 | 1185.536516 | 5.84E-12 | 0.726724474 | 7.74E-13 | 51.34689889 | 1.654877553 | 11.23349869 |
| ENSMUSG00000024867 | *Pip5k1b* | 930.4780064 | 1012.906056 | 1013.496888 | 1640.858544 | 1571.774092 | 1672.240121 | 4.77E-17 | 0.726345849 | 4.18E-18 | 75.23583129 | 1.6544433 | 16.32178936 |
| ENSMUSG00000066036 | *Ubr4* | 5642.388494 | 5147.020736 | 3113.603625 | 6996.97205 | 8370.51293 | 7640.328284 | 0.000383648 | 0.726153492 | 0.000125004 | 14.71565122 | 1.654222724 | 3.416066523 |
| ENSMUSG00000003437 | *Paf1* | 1234.861095 | 1245.030361 | 1092.912689 | 1910.859347 | 2020.250825 | 1985.383383 | 6.01E-16 | 0.725477706 | 5.67E-17 | 70.08707109 | 1.653448036 | 15.2211819 |
| ENSMUSG00000028962 | *Slc4a2* | 2785.406631 | 2856.471814 | 2377.431756 | 4271.720034 | 4583.72699 | 4408.800011 | 3.01E-14 | 0.724789654 | 3.26E-15 | 62.1051475 | 1.65265966 | 13.5216292 |
| ENSMUSG00000024201 | *Kdm4b* | 504.7937363 | 606.2089276 | 477.755374 | 919.7588358 | 813.7852465 | 894.4326621 | 5.45E-08 | 0.724680776 | 1.05E-08 | 32.74264299 | 1.652534941 | 7.26361943 |
| ENSMUSG00000056211 | *R3hdm1* | 940.2725117 | 805.7207265 | 592.4670865 | 1238.052466 | 1312.79457 | 1320.527893 | 2.58E-05 | 0.723647835 | 7.01E-06 | 20.19040632 | 1.651352182 | 4.589202481 |
| ENSMUSG00000002105 | *Slc39a13* | 1219.792625 | 1374.521192 | 1157.201671 | 1984.396152 | 2123.421529 | 2087.315648 | 1.09E-13 | 0.722792046 | 1.24E-14 | 59.47164209 | 1.65037291 | 12.96240164 |
| ENSMUSG00000035293 | *G2e3* | 299.8625478 | 254.1857054 | 298.7546798 | 502.6844234 | 437.9491106 | 466.5008135 | 4.98E-07 | 0.722449247 | 1.08E-07 | 28.23009809 | 1.649980812 | 6.302647053 |
| ENSMUSG00000063576 | *Klhdc3* | 2484.037236 | 2518.836462 | 2395.079712 | 3932.572683 | 4181.571797 | 4096.575057 | 6.78E-27 | 0.722428558 | 3.38E-28 | 121.243228 | 1.64995715 | 26.16904737 |
| ENSMUSG00000059208 | *Hnrnpm* | 3422.049478 | 3402.251687 | 2325.748457 | 4864.404726 | 5160.640722 | 5078.247045 | 2.34E-06 | 0.722009406 | 5.49E-07 | 25.08485962 | 1.649477851 | 5.631489802 |
| ENSMUSG00000041740 | *Rnf10* | 5788.55265 | 6107.171269 | 5196.062406 | 8771.733431 | 9760.159146 | 9667.035559 | 2.13E-15 | 0.721949922 | 2.11E-16 | 67.495355 | 1.649409843 | 14.67228826 |
| ENSMUSG00000035842 | *Ddx11* | 768.4919567 | 714.5975491 | 679.4462971 | 1116.222835 | 1272.789603 | 1182.78159 | 7.43E-12 | 0.721331341 | 9.94E-13 | 50.85528033 | 1.64870278 | 11.12876031 |
| ENSMUSG00000071497 | *Nutf2-ps1* | 1603.28518 | 1430.15429 | 1674.034662 | 2622.080977 | 2655.066483 | 2480.351766 | 4.07E-15 | 0.721232908 | 4.12E-16 | 66.17643037 | 1.648590296 | 14.39033955 |
| ENSMUSG00000006378 | *Gcat* | 1432.258049 | 1427.276716 | 1329.899524 | 2178.665023 | 2422.406018 | 2310.464659 | 1.31E-17 | 0.721022778 | 1.11E-18 | 77.84966959 | 1.648350195 | 16.88243849 |
| ENSMUSG00000020496 | *Rnf187* | 3858.281677 | 3677.539602 | 3237.139315 | 5564.650713 | 6408.16403 | 5794.527821 | 6.96E-13 | 0.720916292 | 8.45E-14 | 55.69891428 | 1.648228534 | 12.15728674 |
| ENSMUSG00000049482 | *Ctu2* | 1320.751373 | 1386.031488 | 1307.209295 | 2154.518609 | 2270.808249 | 2192.002838 | 1.08E-21 | 0.720875398 | 7.02E-23 | 96.97398859 | 1.648181815 | 20.96759456 |
| ENSMUSG00000039849 | *Pcif1* | 1396.093721 | 1347.663834 | 1062.659051 | 2015.12795 | 2062.361317 | 2205.777469 | 8.69E-09 | 0.720754014 | 1.54E-09 | 36.47965622 | 1.648043147 | 8.061001437 |
| ENSMUSG00000032436 | *Cmtm7* | 2384.585336 | 2537.061097 | 2328.269593 | 3771.23074 | 4164.7276 | 4010.254041 | 4.73E-20 | 0.720240882 | 3.44E-21 | 89.27329368 | 1.647457082 | 19.32539236 |
| ENSMUSG00000062075 | *Lmnb2* | 1608.559145 | 1722.707649 | 1502.597377 | 2513.422117 | 2762.448236 | 2688.807838 | 8.79E-16 | 0.719520422 | 8.39E-17 | 69.31621243 | 1.646634573 | 15.05616365 |
| ENSMUSG00000078249 | *Hmga1b* | 22788.80021 | 23810.96585 | 21911.19766 | 37016.45167 | 37459.38763 | 38265.00472 | 1.10E-32 | 0.71852061 | 4.27E-34 | 148.2113072 | 1.645493822 | 31.9593567 |
| ENSMUSG00000073838 | *Tufm* | 2887.872225 | 3053.106039 | 2724.08803 | 4671.233419 | 4925.874732 | 4666.844752 | 2.91E-21 | 0.718340459 | 1.95E-22 | 94.94931578 | 1.64528836 | 20.53651069 |
| ENSMUSG00000021185 | *Dglucy* | 896.5739495 | 863.272207 | 945.4262019 | 1491.589806 | 1540.191223 | 1415.113688 | 1.12E-15 | 0.718072184 | 1.08E-16 | 68.81428021 | 1.64498244 | 14.94998423 |
| ENSMUSG00000029290 | *Zfp326* | 271.9858788 | 283.920637 | 233.2051298 | 378.6596639 | 477.9540775 | 444.461405 | 1.09E-05 | 0.717889384 | 2.80E-06 | 21.9463308 | 1.644774023 | 4.961833796 |
| ENSMUSG00000026786 | *Apbb1ip* | 1934.038091 | 1883.851794 | 1368.97714 | 2608.910206 | 3035.113668 | 2891.754059 | 1.40E-06 | 0.717029476 | 3.20E-07 | 26.12666605 | 1.643793959 | 5.854913068 |
| ENSMUSG00000014355 | *Anapc1* | 5243.827469 | 5463.553879 | 4268.28416 | 7793.803691 | 8914.791031 | 7917.657508 | 8.04E-10 | 0.716978495 | 1.29E-10 | 41.32302903 | 1.643735873 | 9.094973459 |
| ENSMUSG00000037544 | *Dlgap5* | 1043.491529 | 903.5582433 | 987.0249548 | 1571.711996 | 1627.570493 | 1625.406378 | 1.11E-14 | 0.716688583 | 1.16E-15 | 64.13567951 | 1.643405594 | 13.95562569 |
| ENSMUSG00000107968 | *Gm44419* | 100.9587473 | 85.36802936 | 95.80318846 | 160.2443795 | 177.9168262 | 126.7265989 | 0.003470173 | 0.716639005 | 0.001365732 | 10.2515044 | 1.64334912 | 2.459648843 |
| ENSMUSG00000070834 | *Gm13015* | 72.32865475 | 83.44964668 | 104.6271663 | 135.0004019 | 145.2811954 | 143.2561553 | 0.003668389 | 0.715965079 | 0.001450791 | 10.1400899 | 1.642581642 | 2.435524641 |
| ENSMUSG00000078619 | *Smarcd2* | 2247.462262 | 2368.243421 | 2205.994471 | 3480.376215 | 3838.371292 | 3886.282368 | 9.26E-19 | 0.715897856 | 7.32E-20 | 83.22631619 | 1.642505106 | 18.03318143 |
| ENSMUSG00000033047 | *Eif3l* | 5658.96381 | 5521.10536 | 5201.104679 | 8654.294057 | 9222.197618 | 9034.320873 | 5.08E-26 | 0.71558579 | 2.64E-27 | 117.1673205 | 1.642149858 | 25.29423096 |
| ENSMUSG00000055932 | *Fto* | 1277.806234 | 1382.194723 | 1201.321561 | 1918.542297 | 2306.602167 | 2118.538143 | 3.38E-11 | 0.715493242 | 4.76E-12 | 47.78363445 | 1.64204452 | 10.47152134 |
| ENSMUSG00000027395 | *Polr1b* | 1653.01113 | 1468.521943 | 1423.181576 | 2244.518877 | 2711.915647 | 2511.574262 | 9.29E-11 | 0.715224649 | 1.36E-11 | 45.7218292 | 1.641738841 | 10.03200729 |
| ENSMUSG00000027245 | *Hypk* | 141.6436155 | 177.4503981 | 119.7539856 | 227.1957983 | 256.8739976 | 238.7602589 | 0.000696504 | 0.714909889 | 0.000238748 | 13.4985483 | 1.641380694 | 3.157076437 |
| ENSMUSG00000030315 | *Vgll4* | 576.3689675 | 636.9030505 | 491.621625 | 876.9538303 | 931.6946224 | 993.6100004 | 6.81E-08 | 0.714380105 | 1.33E-08 | 32.29072011 | 1.640778059 | 7.166625447 |
| ENSMUSG00000058301 | *Upf1* | 2977.52962 | 2962.942053 | 2169.437991 | 4032.451029 | 4734.271997 | 4549.30124 | 5.37E-07 | 0.714376843 | 1.17E-07 | 28.07682695 | 1.640774349 | 6.269630591 |
| ENSMUSG00000042992 | *Borcs5* | 130.3422632 | 189.9198855 | 184.0429673 | 244.7568262 | 277.9292433 | 299.3686323 | 0.001073115 | 0.713336581 | 0.000380362 | 12.62627508 | 1.639591688 | 2.969353648 |
| ENSMUSG00000028163 | *Nfkb1* | 3522.254801 | 3656.437392 | 2829.975764 | 5416.47954 | 5443.833777 | 5560.359106 | 1.31E-10 | 0.713283414 | 1.95E-11 | 45.02034428 | 1.639531265 | 9.88271014 |

| | | | | | | | | | | | | | |
|---|---|---|---|---|---|---|---|---|---|---|---|---|---|
| ENSMUSG00000026797 | *Stxbp1* | 2395.886688 | 2460.32579 | 1943.796271 | 3538.54712 | 3864.690349 | 3754.045917 | 1.75E-10 | 0.713039585 | 2.62E-11 | 44.43920023 | 1.639254193 | 9.757993571 |
| ENSMUSG00000019942 | *Cdk1* | 3070.200709 | 2722.185026 | 2894.264746 | 4965.380636 | 4707.952939 | 4564.912488 | 1.04E-18 | 0.71225055 | 8.25E-20 | 82.98835566 | 1.638357901 | 17.9821013 |
| ENSMUSG00000039202 | *Abhd2* | 2092.257023 | 2017.17939 | 1550.498971 | 2932.691658 | 3249.877174 | 3101.128439 | 4.45E-08 | 0.712202981 | 8.48E-09 | 33.16106827 | 1.638303882 | 7.352121539 |
| ENSMUSG00000049086 | *Bmyc* | 343.56111 | 351.0640309 | 250.8530856 | 485.1233955 | 532.6977163 | 537.2105825 | 2.57E-05 | 0.711850034 | 7.01E-06 | 20.19141173 | 1.637903129 | 4.589321252 |
| ENSMUSG00000043140 | *Tmem186* | 364.6569677 | 420.1258074 | 385.7338904 | 660.7336744 | 639.0267071 | 615.266821 | 2.45E-09 | 0.711082686 | 4.13E-10 | 39.05149786 | 1.637032184 | 8.611352155 |
| ENSMUSG00000047676 | *Rpsa-ps10* | 11473.13286 | 11722.27738 | 10963.16224 | 18830.90972 | 18421.23447 | 18656.35931 | 2.16E-35 | 0.710579238 | 7.44E-37 | 160.8338535 | 1.636461019 | 34.66485429 |
| ENSMUSG00000013787 | *Ehmt2* | 5182.046743 | 5235.26634 | 4028.776188 | 7528.193144 | 8125.219317 | 7997.550364 | 1.41E-09 | 0.710530355 | 2.33E-10 | 40.16985713 | 1.636405573 | 8.849322921 |
| ENSMUSG00000024181 | *Mrpl28* | 1805.956098 | 1857.953628 | 2057.247415 | 3042.448082 | 3146.70647 | 3158.063578 | 5.64E-18 | 0.7102044 | 4.66E-19 | 79.56748238 | 1.636035894 | 17.24905137 |
| ENSMUSG00000014778 | *Fhod1* | 817.4644833 | 658.9644514 | 397.0790048 | 864.8806236 | 1148.563653 | 1057.891609 | 0.005581222 | 0.709864275 | 0.002292698 | 9.299073503 | 1.635650232 | 2.253270741 |
| ENSMUSG00000035671 | *Zswim4* | 626.8483411 | 589.9026748 | 499.1850346 | 892.3197297 | 950.6443435 | 969.7339745 | 9.19E-09 | 0.708621481 | 1.63E-09 | 36.36900202 | 1.634241825 | 8.036869987 |
| ENSMUSG00000025580 | *Eif4a3* | 4352.527484 | 4286.626103 | 4205.255746 | 6785.142151 | 7201.946793 | 7007.613598 | 2.59E-31 | 0.70854571 | 1.06E-32 | 141.833354 | 1.634155997 | 30.58594773 |
| ENSMUSG00000037966 | *Ninj1* | 3122.940353 | 3484.742142 | 4196.431768 | 5915.871271 | 5655.438996 | 6068.18381 | 3.58E-09 | 0.708358048 | 6.14E-10 | 38.27633958 | 1.633943444 | 8.44658105 |
| ENSMUSG00000028010 | *Gar1* | 1178.354334 | 1230.642491 | 997.109501 | 1725.37099 | 1966.559949 | 1880.696193 | 5.82E-10 | 0.708258382 | 9.22E-11 | 41.98031225 | 1.633830569 | 9.235187857 |
| ENSMUSG00000057406 | *Nsd2* | 2948.146104 | 2625.306701 | 2387.516302 | 4138.914761 | 4536.352687 | 4336.253625 | 3.05E-12 | 0.707408559 | 3.93E-13 | 52.67985165 | 1.632868441 | 11.51530226 |
| ENSMUSG00000019718 | *L3hypdh* | 105.4792882 | 88.24560338 | 158.8316019 | 185.4883571 | 190.5499736 | 194.6814419 | 0.009198877 | 0.706643422 | 0.003961224 | 8.301505067 | 1.632002675 | 2.036265198 |
| ENSMUSG00000018362 | *Kpna2* | 846.8479993 | 851.7619109 | 855.9258548 | 1348.906455 | 1407.543175 | 1410.522145 | 4.29E-18 | 0.706534476 | 3.53E-19 | 80.11737795 | 1.631879438 | 17.36775206 |
| ENSMUSG00000030096 | *Slc6a6* | 6760.468948 | 7854.817892 | 5939.797685 | 10575.03148 | 11981.48757 | 10985.72683 | 2.25E-09 | 0.706197657 | 3.77E-10 | 39.22678834 | 1.631498495 | 8.64821878 |
| ENSMUSG00000037112 | *Sik2* | 497.2595014 | 549.6166385 | 450.0228721 | 775.9779199 | 830.629443 | 839.3341408 | 3.89E-09 | 0.706116362 | 6.72E-10 | 38.10138585 | 1.631406564 | 8.409892544 |
| ENSMUSG00000021514 | *Zfp369* | 125.0682988 | 142.9195098 | 110.9300077 | 192.0737426 | 213.7107439 | 213.9659243 | 0.0003599 | 0.70590705 | 0.000116758 | 14.84433826 | 1.631169891 | 3.443817594 |
| ENSMUSG00000031928 | *Mre11a* | 1159.518746 | 1020.579587 | 1133.250874 | 1861.468956 | 1849.703335 | 1694.279529 | 1.72E-13 | 0.705480889 | 1.99E-14 | 58.54318303 | 1.630688127 | 12.76397917 |
| ENSMUSG00000020827 | *Mink1* | 1458.627871 | 1568.277843 | 1186.194741 | 2178.665023 | 2442.408502 | 2257.202755 | 1.77E-08 | 0.705368276 | 3.24E-09 | 35.03483662 | 1.630560844 | 7.751821917 |
| ENSMUSG00000094483 | *Purb* | 1989.038006 | 1794.646999 | 1421.921008 | 2775.739971 | 2894.043522 | 2829.309068 | 9.28E-08 | 0.705197721 | 1.83E-08 | 31.66189363 | 1.630368091 | 7.032417823 |
| ENSMUSG00000018861 | *Fdxr* | 356.3693093 | 320.3699079 | 223.1205837 | 482.928267 | 494.798274 | 496.8050002 | 0.000285441 | 0.704939104 | 9.11E-05 | 15.31181905 | 1.630075858 | 3.544483568 |
| ENSMUSG00000061536 | *Sec22c* | 244.1092098 | 342.4313088 | 200.4303548 | 414.8792839 | 423.2104386 | 447.2163311 | 0.001246878 | 0.704676861 | 0.000448284 | 12.31936237 | 1.629779582 | 2.904176183 |
| ENSMUSG00000024277 | *Mapre2* | 1188.902262 | 1147.192844 | 902.5668808 | 1679.273292 | 1890.761064 | 1716.318938 | 7.46E-08 | 0.704077447 | 1.46E-08 | 32.11069567 | 1.629102578 | 7.127454922 |
| ENSMUSG00000030243 | *Recql* | 525.8895939 | 525.6368549 | 497.9244664 | 824.2707466 | 882.214795 | 820.9679671 | 3.76E-12 | 0.703868623 | 4.87E-13 | 52.25612686 | 1.628866788 | 11.42529139 |
| ENSMUSG00000005621 | *Zfp592* | 904.1081843 | 916.0277308 | 588.6853817 | 1262.19888 | 1369.643733 | 1300.325102 | 8.03E-05 | 0.70386443 | 2.35E-05 | 17.87938931 | 1.628862055 | 4.095071773 |
| ENSMUSG00000018042 | *Cyb5r3* | 2193.215771 | 2183.119492 | 1933.711725 | 3170.863099 | 3726.77849 | 3386.722441 | 1.52E-12 | 0.703635586 | 1.90E-13 | 54.10981151 | 1.628603701 | 11.81825426 |
| ENSMUSG00000025785 | *Exosc7* | 800.1357431 | 844.0883802 | 785.3340317 | 1365.369918 | 1293.844848 | 1298.488485 | 9.42E-16 | 0.703030825 | 9.02E-17 | 69.17333245 | 1.627921153 | 15.02594877 |
| ENSMUSG00000028920 | *Fbxo42* | 945.5464761 | 921.7828788 | 731.1295962 | 1336.833248 | 1457.023003 | 1444.499566 | 2.87E-08 | 0.70262512 | 5.34E-09 | 34.06006942 | 1.627463423 | 7.54193121 |
| ENSMUSG00000004996 | *Mri1* | 893.5602555 | 867.1089724 | 731.1295962 | 1312.686835 | 1366.485446 | 1383.891193 | 1.47E-10 | 0.701948324 | 2.20E-11 | 44.78684012 | 1.626700129 | 9.831721795 |
| ENSMUSG00000089762 | *Ier5l* | 115.2737935 | 163.062528 | 142.4442144 | 207.439642 | 238.9770387 | 235.0870241 | 0.000938083 | 0.701584024 | 0.000328246 | 12.90191343 | 1.626289416 | 3.027758828 |
| ENSMUSG00000049124 | *Gm8186* | 1067.601081 | 1007.150908 | 1172.32849 | 1980.005895 | 1639.150878 | 1655.710564 | 3.12E-09 | 0.700890043 | 5.32E-10 | 38.55642168 | 1.625507309 | 8.506298384 |
| ENSMUSG00000048921 | *Zfp689* | 116.027217 | 112.2253869 | 76.89466442 | 165.7322007 | 175.8113016 | 158.867403 | 0.003842941 | 0.700459082 | 0.001530372 | 10.04168415 | 1.625021811 | 2.415336329 |
| ENSMUSG00000040236 | *Trappc5* | 1264.244611 | 1246.948743 | 1331.160092 | 2039.274364 | 2049.728169 | 2147.924021 | 1.26E-19 | 0.700435338 | 9.43E-21 | 87.27852394 | 1.624995067 | 18.90074619 |
| ENSMUSG00000004980 | *Hnrnpa2b1* | 9387.656647 | 9112.317741 | 8386.560695 | 14595.40931 | 14622.86814 | 14487.23786 | 1.24E-24 | 0.700433741 | 6.91E-26 | 110.6927767 | 1.624993268 | 23.90769441 |
| ENSMUSG00000036606 | *Plxnb2* | 14773.88116 | 14460.76866 | 10619.0271 | 20259.93836 | 22870.20788 | 21628.92453 | 2.80E-07 | 0.700121658 | 5.89E-08 | 29.39926891 | 1.624641788 | 6.552088012 |
| ENSMUSG00000046417 | *Lrrc75a* | 996.7792732 | 1140.478505 | 958.0318846 | 1536.58994 | 1771.798926 | 1720.910481 | 3.80E-10 | 0.699777929 | 5.88E-11 | 42.86023728 | 1.624254756 | 9.420297276 |
| ENSMUSG00000064210 | *Ano6* | 1940.818902 | 1873.300689 | 1783.704101 | 2786.715613 | 3276.196232 | 3029.500362 | 4.80E-14 | 0.698946333 | 5.27E-15 | 61.15703208 | 1.623318775 | 13.31852526 |
| ENSMUSG00000010097 | *Nxf1* | 2199.996582 | 2212.854424 | 1829.084559 | 3375.010048 | 3290.934904 | 3475.798384 | 2.60E-12 | 0.698670922 | 3.32E-13 | 53.00938725 | 1.623008912 | 11.58537724 |
| ENSMUSG00000028790 | *Khdrbs1* | 3173.419727 | 3256.454603 | 2444.241874 | 4597.696615 | 4942.718929 | 4867.954355 | 1.68E-08 | 0.698178266 | 3.07E-09 | 35.13810715 | 1.622454777 | 7.774201514 |
| ENSMUSG00000027201 | *Myef2* | 1067.601081 | 946.7218537 | 1125.687464 | 1789.029716 | 1572.826854 | 1728.256951 | 8.17E-11 | 0.697990108 | 1.19E-11 | 45.98382723 | 1.622243188 | 10.08764675 |
| ENSMUSG00000064302 | *Clasp1* | 1164.039287 | 1222.009769 | 1018.539162 | 1657.322007 | 2081.311038 | 1787.028707 | 3.38E-08 | 0.697236869 | 6.36E-09 | 33.72037056 | 1.621396426 | 7.470734741 |
| ENSMUSG00000004929 | *Thop1* | 2415.475699 | 2570.632794 | 2240.029814 | 3728.425734 | 3943.64752 | 4049.741314 | 1.85E-16 | 0.697209949 | 1.70E-17 | 72.47092843 | 1.621366173 | 15.73178785 |
| ENSMUSG00000020918 | *Kat2a* | 2552.598774 | 2610.918831 | 2071.113666 | 3704.279321 | 4065.767945 | 3966.175223 | 4.16E-10 | 0.696725459 | 6.46E-11 | 42.67462692 | 1.620821773 | 9.380819608 |
| ENSMUSG00000078923 | *Ube2v1* | 3049.104852 | 3186.433635 | 3161.505219 | 5132.210401 | 5273.286287 | 4819.283995 | 4.11E-23 | 0.696308752 | 2.52E-24 | 103.566655 | 1.620353683 | 22.38582566 |
| ENSMUSG00000067547 | *Gm7666* | 1961.91476 | 2019.097773 | 2194.649357 | 3439.766338 | 3338.309206 | 3217.753643 | 2.92E-18 | 0.696094519 | 2.37E-19 | 80.90076466 | 1.620113086 | 17.53519127 |
| ENSMUSG00000062397 | *Zfp706* | 1882.805294 | 1887.688559 | 1613.527385 | 2843.788954 | 3043.535766 | 2839.410463 | 2.20E-13 | 0.694923124 | 2.56E-14 | 58.0455882 | 1.618798171 | 12.65849781 |
| ENSMUSG00000005566 | *Trim28* | 11506.28349 | 11470.96925 | 9899.242618 | 16942.00166 | 18250.68698 | 18033.74602 | 8.59E-16 | 0.694763504 | 8.19E-17 | 69.36310229 | 1.618619077 | 15.06617621 |
| ENSMUSG00000033712 | *Ccar2* | 1774.312312 | 1779.299938 | 1410.575893 | 2779.032664 | 2656.119246 | 2611.669909 | 1.08E-09 | 0.694762675 | 1.75E-10 | 40.72257085 | 1.618618147 | 8.967810861 |
| ENSMUSG00000021608 | *Lpcat1* | 1638.696084 | 1620.074175 | 1264.349974 | 2358.665559 | 2564.528927 | 2406.887071 | 1.47E-08 | 0.694369403 | 2.68E-09 | 35.40334193 | 1.618176979 | 7.831220912 |
| ENSMUSG00000024966 | *Stip1* | 6538.962443 | 6787.23793 | 7820.565542 | 11234.66759 | 11764.61854 | 11203.36599 | 9.72E-15 | 0.694265246 | 1.01E-15 | 64.4049196 | 1.618060157 | 14.0124082 |
| ENSMUSG00000061273 | *Mmgt1* | 729.3139354 | 737.6181413 | 777.7706221 | 1148.052198 | 1288.581037 | 1191.046368 | 6.63E-13 | 0.694217015 | 8.02E-14 | 55.79984416 | 1.618006064 | 12.17859985 |
| ENSMUSG00000020893 | *Per1* | 641.9168109 | 666.6379821 | 468.9313962 | 920.8564 | 993.8075972 | 968.8156658 | 7.44E-06 | 0.693954149 | 1.87E-06 | 22.72600289 | 1.617711283 | 5.128176364 |
| ENSMUSG00000019432 | *Ddx39b* | 7329.303681 | 6648.155185 | 5821.304267 | 10612.34867 | 10843.45154 | 10581.67101 | 1.96E-12 | 0.693684945 | 2.46E-13 | 53.59548979 | 1.617409449 | 11.70860973 |
| ENSMUSG00000014907 | *Naf1* | 555.2731099 | 562.0861259 | 383.2127538 | 729.8802217 | 865.3705984 | 837.4975235 | 9.87E-05 | 0.692952634 | 2.93E-05 | 17.46587248 | 1.616588662 | 4.00582128 |
| ENSMUSG00000028333 | *Anp32b* | 8267.315922 | 7739.714932 | 7149.943223 | 12178.57284 | 12858.43855 | 12404.51376 | 1.18E-18 | 0.692645074 | 9.38E-20 | 82.73612495 | 1.616244067 | 17.92886687 |
| ENSMUSG00000092060 | *Bend4* | 1163.285864 | 1016.742822 | 645.4109539 | 1407.07736 | 1640.20364 | 1527.147348 | 0.000946533 | 0.692446171 | 0.000331403 | 12.88399821 | 1.616021252 | 3.023864215 |
| ENSMUSG00000038013 | *Wipf2* | 626.8483411 | 637.8622418 | 426.072075 | 860.4903666 | 982.2272121 | 897.1875882 | 8.59E-05 | 0.692308792 | 2.53E-05 | 17.74413949 | 1.615867376 | 4.066038129 |
| ENSMUSG00000081752 | *Sms-ps* | 1136.162618 | 1047.436945 | 1171.067922 | 1940.493582 | 1796.012459 | 1682.341516 | 2.32E-12 | 0.691958399 | 2.95E-13 | 53.24071231 | 1.615474971 | 11.63441747 |
| ENSMUSG00000051403 | *Ppp1r37* | 977.9436861 | 1011.946865 | 857.1864231 | 1463.053136 | 1512.819404 | 1629.079613 | 4.30E-11 | 0.691808141 | 6.12E-12 | 47.29141041 | 1.615306727 | 10.3664726 |
| ENSMUSG00000026202 | *Tuba4a* | 5901.566173 | 6085.109868 | 5949.882231 | 9215.149386 | 10153.89224 | 9594.489172 | 3.15E-25 | 0.691387628 | 1.71E-26 | 113.4570601 | 1.61483597 | 24.50131623 |
| ENSMUSG00000047767 | *Atg16l2* | 428.6979641 | 412.4522767 | 329.0083183 | 640.977518 | 654.8181414 | 601.4921906 | 1.42E-06 | 0.690806236 | 3.24E-07 | 26.09882981 | 1.614185338 | 5.849042451 |
| ENSMUSG00000027478 | *Dnmt3b* | 94.93135936 | 112.2253869 | 78.15523269 | 162.439508 | 153.7032936 | 146.9293901 | 0.003304979 | 0.690730709 | 0.001295047 | 10.34957173 | 1.614100836 | 2.480831324 |
| ENSMUSG00000003824 | *Syce2* | 1817.257451 | 1638.298811 | 1842.95081 | 2935.98435 | 2756.131663 | 2855.94002 | 1.02E-16 | 0.690350897 | 9.21E-18 | 73.67415531 | 1.613675954 | 15.98931622 |
| ENSMUSG00000002129 | *Sf3a1* | 4400.746587 | 4345.136775 | 3424.963988 | 6147.457326 | 6920.859263 | 6581.518367 | 4.92E-09 | 0.690278988 | 8.56E-10 | 37.6280269 | 1.613595525 | 8.307956703 |
| ENSMUSG00000025224 | *Gbf1* | 2294.927941 | 2203.262511 | 1563.104654 | 3088.54578 | 3496.223549 | 3205.81563 | 7.76E-06 | 0.69026301 | 1.95E-06 | 22.64335649 | 1.613577654 | 5.110208727 |
| ENSMUSG00000078348 | *Sf3b5* | 1805.202675 | 1815.749209 | 1950.099113 | 2937.081915 | 3030.902619 | 3008.379262 | 1.18E-21 | 0.689727828 | 7.73E-23 | 96.78537401 | 1.612979193 | 20.92804774 |
| ENSMUSG00000027327 | *1700037H04Rik* | 806.163131 | 844.0883802 | 834.4961942 | 1323.662477 | 1302.266947 | 1377.463032 | 4.56E-16 | 0.689413039 | 4.27E-17 | 70.64956164 | 1.612627287 | 15.34089222 |
| ENSMUSG00000030079 | *Ruvbl1* | 2163.078831 | 2216.691189 | 2062.289688 | 3404.644282 | 3507.803934 | 3475.798384 | 5.77E-24 | 0.688510806 | 3.36E-25 | 107.5554036 | 1.611619097 | 23.23857179 |
| ENSMUSG00000114827 | *Gm46432* | 223.0133521 | 171.6952501 | 194.1275135 | 318.2936305 | 314.7759233 | 318.6531147 | 4.30E-05 | 0.688394198 | 1.21E-05 | 19.14535335 | 1.611488841 | 4.366111605 |
| ENSMUSG00000023944 | *Hsp90ab1* | 52118.06971 | 52311.41817 | 43377.41471 | 76299.37349 | 83240.86113 | 78546.61532 | 8.92E-13 | 0.687698307 | 1.09E-13 | 55.1897381 | 1.610711718 | 12.04943482 |
| ENSMUSG00000026496 | *Parp1* | 5545.196864 | 5396.410485 | 4438.460876 | 7853.07216 | 8767.404311 | 8148.152989 | 1.84E-10 | 0.686754172 | 2.76E-11 | 44.33805698 | 1.609657974 | 9.736348443 |
| ENSMUSG00000001056 | *Nhp2* | 4241.020808 | 4134.11468 | 4654.01805 | 7258.19234 | 6723.992716 | 6980.064338 | 1.43E-18 | 0.686720639 | 1.15E-19 | 82.34079233 | 1.60962056 | 17.84610412 |
| ENSMUSG00000019952 | *Poc1b* | 426.4376936 | 416.289042 | 415.9875288 | 606.9530265 | 744.3029356 | 674.9568857 | 5.97E-08 | 0.686511244 | 1.16E-08 | 32.55926944 | 1.609386955 | 7.224033396 |
| ENSMUSG00000031985 | *Gnpat* | 747.3960991 | 785.5777084 | 656.7560683 | 1098.661807 | 1261.209218 | 1169.925269 | 2.43E-09 | 0.686407464 | 4.10E-10 | 39.06401863 | 1.609271188 | 8.613787341 |
| ENSMUSG00000019947 | *Arid5b* | 152.1915444 | 203.3485643 | 153.7893288 | 245.8543905 | 301.0900136 | 272.7376803 | 0.000699016 | 0.686309635 | 0.000239705 | 13.49104185 | 1.609162068 | 3.155512972 |
| ENSMUSG00000021699 | *Pde4d* | 121.3011814 | 113.1845783 | 99.58489327 | 183.2932286 | 170.5474902 | 186.4166637 | 0.000658803 | 0.685984949 | 0.000224921 | 13.61052663 | 1.608799958 | 3.181244329 |
| ENSMUSG00000068917 | *Clk2* | 558.2868038 | 610.045693 | 456.3257135 | 828.6610036 | 924.3252864 | 866.8834015 | 9.00E-07 | 0.685796274 | 2.01E-07 | 27.02418283 | 1.608589574 | 6.045995841 |
| ENSMUSG00000034349 | *Smc4* | 1238.628213 | 895.8847126 | 587.4248135 | 1520.126477 | 1421.229085 | 1444.499566 | 0.004620902 | 0.685448951 | 0.001866499 | 9.676400456 | 1.608202359 | 2.335273233 |
| ENSMUSG00000060261 | *Gtf2i* | 2795.95456 | 3138.474068 | 2735.433144 | 4461.598648 | 4874.28938 | 4608.072996 | 3.10E-15 | 0.685277514 | 3.10E-16 | 66.73679114 | 1.608011266 | 14.5089522 |
| ENSMUSG00000032000 | *Birc3* | 595.9579782 | 563.0453172 | 548.3471971 | 973.5394837 | 904.322803 | 873.3115623 | 5.37E-11 | 0.685264733 | 7.69E-12 | 46.84303516 | 1.60799702 | 10.26983533 |
| ENSMUSG00000029326 | *Enoph1* | 1085.683245 | 1023.457161 | 985.7643865 | 1577.199817 | 1757.060254 | 1648.364095 | 8.32E-14 | 0.685132927 | 9.37E-15 | 60.02361317 | 1.607850119 | 13.07999323 |
| ENSMUSG00000003778 | *Brd8* | 794.8617787 | 945.7626623 | 831.9750577 | 1432.321337 | 1292.792086 | 1409.603836 | 3.50E-10 | 0.685109874 | 5.39E-11 | 43.02974281 | 1.607824427 | 9.456382622 |
| ENSMUSG00000031400 | *G6pdx* | 11079.09238 | 11666.64428 | 11669.08047 | 17967.12666 | 19228.70314 | 18124.65858 | 1.30E-27 | 0.684962631 | 6.19E-29 | 124.6111628 | 1.60766034 | 26.88481344 |
| ENSMUSG00000029135 | *Fosl2* | 909.3821487 | 962.0689151 | 615.1573154 | 1283.0526 | 1384.382405 | 1337.975758 | 8.69E-05 | 0.684885336 | 2.56E-05 | 17.72012321 | 1.607574207 | 4.060757939 |
| ENSMUSG00000032258 | *Lca5* | 185.3421778 | 180.3279721 | 202.9514913 | 334.7570942 | 288.4568661 | 289.2672367 | 2.75E-05 | 0.684773323 | 7.50E-06 | 20.06030398 | 1.607449398 | 4.561409117 |
| ENSMUSG00000036246 | *Gmip* | 2416.229123 | 2562.959263 | 2150.529467 | 3759.157533 | 3834.160243 | 3871.589429 | 1.43E-14 | 0.684171046 | 1.52E-15 | 63.6103405 | 1.606778482 | 13.84377679 |
| ENSMUSG00000008958 | *Vps72* | 1708.764468 | 1707.360587 | 1728.239097 | 2659.398161 | 2799.294916 | 2803.596424 | 3.88E-23 | 0.684023382 | 2.37E-24 | 103.685624 | 1.606614032 | 22.41092931 |
| ENSMUSG00000036054 | *Sugp2* | 410.6158004 | 439.3096342 | 302.5363846 | 609.148155 | 650.6070922 | 598.7372646 | 4.52E-05 | 0.683969946 | 1.28E-05 | 19.04706001 | 1.606554525 | 4.345224338 |
| ENSMUSG00000032058 | *Ppp2r1b* | 1769.791771 | 1920.301065 | 1911.021496 | 2881.106138 | 3214.083257 | 2897.263911 | 1.97E-15 | 0.683826449 | 1.95E-16 | 67.65534163 | 1.606394738 | 14.70542278 |
| ENSMUSG00000037275 | *Gemin5* | 2108.83234 | 2175.445962 | 1821.521149 | 3204.88759 | 3388.841796 | 3224.181804 | 4.34E-13 | 0.683687605 | 5.20E-14 | 56.65386873 | 1.606240147 | 12.36203796 |
| ENSMUSG00000061410 | *Zcchc14* | 955.3409814 | 917.9461135 | 559.6923115 | 1184.271818 | 1328.586004 | 1401.339058 | 0.000948504 | 0.683268739 | 0.000332159 | 12.87973821 | 1.605773866 | 3.022960775 |
| ENSMUSG00000046865 | *Fbl* | 3602.117691 | 3756.193292 | 3416.14001 | 5888.432165 | 5871.255265 | 5550.25771 | 1.61E-21 | 0.683267412 | 1.06E-22 | 96.15037467 | 1.605772389 | 20.79328688 |
| ENSMUSG00000014547 | *Wdfy2* | 1524.175714 | 1645.972341 | 1483.688853 | 2347.689916 | 2725.601556 | 2400.45891 | 2.55E-12 | 0.682896731 | 3.26E-13 | 53.04612102 | 1.605359861 | 11.59279881 |
| ENSMUSG00000019961 | *Tmpo* | 4985.403213 | 4704.833528 | 4346.439392 | 7490.87596 | 7490.403659 | 7565.026972 | 1.58E-19 | 0.682785507 | 1.19E-20 | 86.81116891 | 1.605236101 | 18.80214378 |
| ENSMUSG00000027488 | *Snta1* | 458.8349036 | 520.8408982 | 531.9598096 | 799.026769 | 848.5264019 | 773.2159153 | 3.63E-09 | 0.6827137 | 6.24E-10 | 38.24341844 | 1.605156206 | 8.439601177 |
| ENSMUSG00000029404 | *Arl6ip4* | 1312.463714 | 1382.194723 | 1280.737362 | 2077.689112 | 2247.647479 | 2058.848079 | 2.91E-16 | 0.682657197 | 2.68E-17 | 71.56410749 | 1.605093341 | 15.53675188 |
| ENSMUSG00000035382 | *Pcsk7* | 1096.984597 | 1237.35683 | 1005.933479 | 1815.371258 | 1738.110533 | 1812.74135 | 5.91E-11 | 0.682503873 | 8.51E-12 | 46.64532195 | 1.604922767 | 10.22809661 |
| ENSMUSG00000047514 | *Tspyl1* | 708.2180777 | 804.7615352 | 789.1157365 | 1198.540154 | 1278.053414 | 1210.330851 | 5.65E-12 | 0.681735133 | 7.48E-13 | 51.41492873 | 1.604067812 | 11.24809841 |
| ENSMUSG00000049504 | *Proser1* | 633.6291525 | 640.7398159 | 344.1351375 | 864.8806236 | 912.7449013 | 825.5595105 | 0.003836981 | 0.681681744 | 0.001527209 | 10.04549578 | 1.604008453 | 2.41601037 |
| ENSMUSG00000029147 | *Ppm1g* | 6161.497276 | 5911.496235 | 5111.604332 | 8713.562527 | 9701.204458 | 9149.109459 | 1.72E-12 | 0.681027846 | 2.16E-13 | 53.85742611 | 1.603281604 | 11.76437568 |
| ENSMUSG00000060862 | *Zbtb40* | 534.1772522 | 591.8210575 | 481.5370788 | 818.7829254 | 946.4332944 | 816.3764236 | 1.96E-07 | 0.680900338 | 4.04E-08 | 30.12796821 | 1.603139909 | 6.70759856 |
| ENSMUSG00000021910 | *Nisch* | 4708.896793 | 4913.937241 | 3694.725597 | 6877.337548 | 7154.57249 | 7323.511787 | 1.24E-08 | 0.680580825 | 2.24E-09 | 35.75021655 | 1.602784902 | 7.906583348 |
| ENSMUSG00000033961 | *Zfp446* | 278.7666902 | 284.8798283 | 209.2543327 | 419.2695409 | 391.6275701 | 433.4417007 | 6.26E-05 | 0.680329719 | 1.80E-05 | 18.3878771 | 1.602505956 | 4.203145334 |
| ENSMUSG00000029468 | *P2rx7* | 769.2453802 | 808.5983006 | 771.4677808 | 1266.589137 | 1274.895127 | 1224.105481 | 2.05E-15 | 0.680290988 | 2.03E-16 | 67.57039156 | 1.602462935 | 14.68791551 |
| ENSMUSG00000002107 | *Celf2* | 1296.641821 | 1244.071169 | 1105.518372 | 1846.103057 | 2087.627612 | 1917.428541 | 3.15E-11 | 0.68004473 | 4.42E-12 | 47.92910334 | 1.60218943 | 10.50204969 |
| ENSMUSG00000049553 | *Polr1a* | 3278.899015 | 3314.965275 | 2432.89676 | 4496.720704 | 5179.590443 | 4794.48966 | 7.29E-07 | 0.679880709 | 1.61E-07 | 27.45366452 | 1.602007286 | 6.13750086 |
| ENSMUSG00000039509 | *Nup133* | 884.5191737 | 803.8023439 | 729.8690279 | 1287.442857 | 1290.686562 | 1303.998337 | 8.80E-11 | 0.679130657 | 1.29E-11 | 45.82888271 | 1.601174625 | 10.05533496 |
| ENSMUSG00000000876 | *Pxmp4* | 125.8217223 | 131.4092137 | 118.4934173 | 216.2201559 | 204.2358833 | 182.7434289 | 0.000399161 | 0.678641691 | 0.000130525 | 14.63419651 | 1.600632037 | 3.398851685 |
| ENSMUSG00000021957 | *Tkt* | 16769.69997 | 17030.44226 | 16179.39374 | 25927.76012 | 27261.27938 | 26814.61369 | 1.12E-32 | 0.678615186 | 4.34E-34 | 148.1772038 | 1.600602631 | 31.95266853 |
| ENSMUSG00000022100 | *Xpo7* | 1677.874105 | 1763.952876 | 1536.63272 | 2606.715078 | 2823.508449 | 2541.878448 | 1.66E-13 | 0.677876276 | 1.92E-14 | 58.61359213 | 1.599783055 | 12.77900467 |
| ENSMUSG00000031540 | *Kat6a* | 1257.4638 | 1268.050953 | 971.8981356 | 1763.785739 | 1970.770998 | 1869.676489 | 1.55E-07 | 0.677811507 | 3.17E-08 | 30.60316133 | 1.599711235 | 6.808757247 |
| ENSMUSG00000032570 | *Atp2c1* | 1423.216967 | 1567.318651 | 1539.153857 | 2343.299659 | 2496.099378 | 2399.540602 | 5.53E-17 | 0.677658842 | 4.86E-18 | 74.9364863 | 1.599541964 | 16.25730161 |
| ENSMUSG00000029414 | *Kntc1* | 1056.299729 | 931.3747922 | 718.5239135 | 1371.955304 | 1549.666084 | 1416.950306 | 1.36E-05 | 0.677645591 | 3.55E-06 | 21.49629334 | 1.599527272 | 4.867317149 |
| ENSMUSG00000023909 | *Paqr4* | 365.4103912 | 406.6971286 | 355.4802519 | 470.8550603 | 682.1899608 | 650.1625511 | 0.000164483 | 0.677551583 | 5.06E-05 | 16.42456575 | 1.599423048 | 3.783879314 |
| ENSMUSG00000023988 | *Bysl* | 2436.571557 | 2609.959639 | 2420.291077 | 3785.499075 | 4017.34088 | 4138.817257 | 6.77E-19 | 0.677496243 | 5.31E-20 | 83.8611148 | 1.599361698 | 18.16922963 |
| ENSMUSG00000033294 | *Noc4l* | 1967.942148 | 2023.89373 | 1714.372846 | 2999.643077 | 3077.22416 | 3056.131314 | 5.47E-14 | 0.67691626 | 6.03E-15 | 60.89108578 | 1.598718862 | 13.26175523 |
| ENSMUSG00000041133 | *Smc1a* | 1445.819671 | 1348.623026 | 1264.349974 | 2150.128352 | 2233.961569 | 2112.109982 | 8.23E-15 | 0.676105422 | 8.55E-16 | 64.74014432 | 1.597820586 | 14.0845773 |
| ENSMUSG00000000561 | *Wdr77* | 3894.446004 | 4084.23673 | 3530.851722 | 5846.724723 | 6451.327284 | 6096.65138 | 5.30E-15 | 0.675811603 | 5.41E-16 | 65.64254742 | 1.597495207 | 14.2753392 |
| ENSMUSG00000018974 | *Sart3* | 2168.352795 | 2177.364344 | 1733.28137 | 3032.570004 | 3437.268861 | 3245.302903 | 8.19E-09 | 0.674934709 | 1.45E-09 | 36.599467 | 1.596524519 | 8.086855336 |
| ENSMUSG00000040731 | *Eif4h* | 11222.99626 | 11354.9071 | 9393.754742 | 16486.5125 | 17364.26113 | 17196.24849 | 7.09E-13 | 0.674688526 | 8.60E-14 | 55.662495 | 1.596252109 | 12.14964923 |
| ENSMUSG00000020238 | *Ncln* | 4315.609733 | 4370.07575 | 3680.859346 | 6159.530533 | 6859.79905 | 6728.447757 | 3.90E-12 | 0.674542169 | 5.07E-13 | 52.17703581 | 1.596090182 | 11.40871778 |
| ENSMUSG00000042308 | *Setd1a* | 2124.654233 | 2138.037499 | 1424.442144 | 2811.959591 | 3195.133536 | 3076.334105 | 4.50E-05 | 0.674116393 | 1.27E-05 | 19.05268476 | 1.595619205 | 4.346398868 |
| ENSMUSG00000064127 | *Med14* | 823.4918712 | 870.9457377 | 656.7560683 | 1200.735282 | 1298.055898 | 1260.837829 | 2.50E-07 | 0.674114917 | 5.22E-08 | 29.63232596 | 1.595617572 | 6.602461066 |
| ENSMUSG00000035649 | *Zcchc7* | 179.3147899 | 188.9606942 | 147.4864875 | 279.878882 | 257.9267599 | 288.348928 | 0.000147343 | 0.673578576 | 4.50E-05 | 16.64723025 | 1.59502449 | 3.831669161 |
| ENSMUSG00000062762 | *Ei24* | 1873.764212 | 1893.443707 | 2258.938338 | 3020.496797 | 3294.09319 | 3284.790177 | 9.31E-11 | 0.67353931 | 1.37E-11 | 45.71758946 | 1.594981079 | 10.03127039 |
| ENSMUSG00000016253 | *Nelfcd* | 999.0395437 | 987.9670814 | 905.0880173 | 1458.662879 | 1558.088182 | 1598.775426 | 2.57E-13 | 0.672633221 | 3.03E-14 | 57.71194392 | 1.593979661 | 12.58991557 |
| ENSMUSG00000024799 | *Tm7sf2* | 205.6846119 | 211.022095 | 173.9584212 | 310.6106808 | 313.723161 | 320.4897321 | 2.04E-05 | 0.671804861 | 5.46E-06 | 20.6669932 | 1.5930647 | 4.690511765 |
| ENSMUSG00000032498 | *Mlh1* | 731.5742058 | 697.332105 | 744.9958471 | 1119.515528 | 1190.674145 | 1148.804169 | 1.06E-13 | 0.670817086 | 1.20E-14 | 59.53787979 | 1.591974344 | 12.97649498 |
| ENSMUSG00000038845 | *Phb* | 2899.927001 | 3088.596118 | 3480.428991 | 4785.3801 | 4958.510363 | 5314.252377 | 4.37E-12 | 0.670584322 | 5.71E-13 | 51.94444317 | 1.591717516 | 11.35933372 |
| ENSMUSG00000033036 | *Ppihl* | 236.5749749 | 205.266947 | 268.5010413 | 329.269273 | 407.4190044 | 390.2811924 | 0.000146672 | 0.670509405 | 4.48E-05 | 16.65674739 | 1.591634863 | 3.833653015 |
| ENSMUSG00000040446 | *Rprd1a* | 501.7800423 | 489.187584 | 563.4740164 | 834.1488248 | 811.6797219 | 821.8862758 | 5.12E-10 | 0.670433386 | 8.05E-11 | 42.24561125 | 1.591550999 | 9.290707562 |
| ENSMUSG00000030852 | *Tacc2* | 632.875729 | 696.3729137 | 482.7976471 | 913.1734503 | 1023.284941 | 954.1227268 | 1.90E-05 | 0.670296246 | 5.08E-06 | 20.80657372 | 1.591399716 | 4.720272693 |
| ENSMUSG00000037210 | *Fam193a* | 559.7936508 | 586.0659094 | 384.4733221 | 754.0266351 | 849.5791641 | 837.4975235 | 0.000195025 | 0.669607284 | 6.08E-05 | 16.07794393 | 1.59063992 | 3.709910687 |
| ENSMUSG00000027404 | *Snrpb* | 9206.081587 | 9640.83217 | 8685.315375 | 14429.6771 | 14617.60433 | 14749.87415 | 3.25E-24 | 0.669442877 | 1.87E-25 | 108.7224631 | 1.590458665 | 23.48811535 |
| ENSMUSG00000029771 | *Irf5* | 6906.633105 | 6975.239433 | 5523.810156 | 9918.688065 | 10526.57009 | 10428.31346 | 5.70E-10 | 0.669352508 | 9.01E-11 | 42.02502015 | 1.590359042 | 9.244364299 |
| ENSMUSG00000001504 | *Irx2* | 708.9715012 | 670.4747474 | 529.4386731 | 969.1492267 | 1043.287425 | 1032.178965 | 1.06E-06 | 0.669217981 | 2.39E-07 | 26.69170634 | 1.590210753 | 5.975530675 |
| ENSMUSG00000000168 | *Dlat* | 2248.215685 | 2259.8548 | 2200.952198 | 3512.205578 | 3667.823802 | 3489.573014 | 1.81E-24 | 0.668825074 | 1.02E-25 | 109.9168444 | 1.589777731 | 23.74194927 |
| ENSMUSG00000056167 | *Cnot10* | 1784.106817 | 1749.565006 | 1689.161481 | 2775.739971 | 2762.448236 | 2771.45562 | 5.24E-23 | 0.66875675 | 3.23E-24 | 103.0749812 | 1.589702442 | 22.2809573 |
| ENSMUSG00000028741 | *Mrto4* | 2463.694802 | 2574.46956 | 2803.503831 | 4227.817465 | 4064.715183 | 4158.101739 | 1.93E-17 | 0.668435227 | 1.65E-18 | 77.06492061 | 1.589348196 | 16.71373383 |
| ENSMUSG00000035769 | *Xylb* | 497.2595014 | 474.7997139 | 441.1988942 | 630.0018756 | 789.5717139 | 828.3144366 | 5.51E-06 | 0.668377544 | 1.36E-06 | 23.34016589 | 1.589284651 | 5.259123067 |
| ENSMUSG00000001440 | *Kpnb1* | 15137.7847 | 15650.16592 | 13789.3563 | 22885.31203 | 24690.43387 | 23263.51399 | 4.29E-18 | 0.668030001 | 3.53E-19 | 80.11869227 | 1.588901841 | 17.36775206 |
| ENSMUSG00000029165 | *Agbl5* | 178.5613664 | 150.5930406 | 114.7117125 | 211.8298989 | 276.876481 | 221.3123938 | 0.005707869 | 0.667651298 | 0.00235099 | 9.253085313 | 1.588484815 | 2.243526037 |
| ENSMUSG00000025047 | *Pdcd11* | 2263.284155 | 2422.917328 | 1792.528079 | 3150.009378 | 3702.564957 | 3446.412506 | 5.81E-07 | 0.667597229 | 1.27E-07 | 27.91729262 | 1.588425282 | 6.235472124 |
| ENSMUSG00000036752 | *Tubb4b* | 12686.8981 | 11700.21598 | 11353.9384 | 18228.34695 | 19817.19726 | 18732.57893 | 5.27E-19 | 0.667461804 | 4.10E-20 | 84.37352041 | 1.588276185 | 18.2784941 |
| ENSMUSG00000039164 | *Naif1* | 485.9581491 | 493.9835407 | 485.3187837 | 772.6852272 | 819.0490579 | 736.4835678 | 2.39E-10 | 0.66737946 | 3.64E-11 | 43.80011562 | 1.588185534 | 9.621326941 |
| ENSMUSG00000018821 | *Avpi1* | 660.7523981 | 698.2912963 | 862.2286962 | 1221.589003 | 1164.355087 | 1132.274612 | 2.96E-07 | 0.667376007 | 6.23E-08 | 29.28902077 | 1.588181733 | 6.528372399 |
| ENSMUSG00000032281 | *Acsbg1* | 1326.778761 | 1427.276716 | 1550.498971 | 2205.006565 | 2304.496642 | 2314.137894 | 1.74E-13 | 0.667214669 | 2.01E-14 | 58.51981947 | 1.588004134 | 12.75907986 |
| ENSMUSG00000029538 | *Srsf9* | 2371.777137 | 2220.527955 | 1986.655592 | 3198.302205 | 3740.464399 | 3517.122275 | 2.05E-10 | 0.667129093 | 3.10E-11 | 44.11367951 | 1.587909943 | 9.68834665 |
| ENSMUSG00000022965 | *Ifngr2* | 2486.297507 | 2628.184275 | 2987.546798 | 4300.256705 | 4271.056591 | 4269.217091 | 2.51E-13 | 0.665884038 | 2.96E-14 | 57.75784462 | 1.586540157 | 12.59954461 |
| ENSMUSG00000070319 | *Eif3g* | 5875.196351 | 5908.618661 | 5862.90302 | 9350.149788 | 9352.740142 | 9289.610688 | 3.62E-38 | 0.665589127 | 1.11E-39 | 173.770154 | 1.586215874 | 37.44100966 |
| ENSMUSG00000025314 | *Ptprj* | 1280.819928 | 1300.663459 | 1019.79973 | 1868.054342 | 1933.924318 | 1918.346849 | 1.30E-08 | 0.665048947 | 2.36E-09 | 35.65421649 | 1.585622069 | 7.885840716 |
| ENSMUSG00000045962 | *Wnk1* | 4254.582431 | 4238.666536 | 2768.207919 | 5442.821082 | 6626.085823 | 5793.609513 | 0.000145943 | 0.664825951 | 4.45E-05 | 16.66704841 | 1.585377 | 3.83581619 |
| ENSMUSG00000071652 | *Ints5* | 2129.174774 | 2041.159174 | 1658.907842 | 2908.545244 | 3203.555634 | 3136.942478 | 1.17E-08 | 0.664320207 | 2.11E-09 | 35.86875117 | 1.584821335 | 7.931190043 |
| ENSMUSG00000050721 | *Plekho2* | 5530.881818 | 5441.492478 | 4036.339598 | 8101.121679 | 7607.260273 | 8085.707998 | 4.21E-07 | 0.664119668 | 9.02E-08 | 28.57370745 | 1.584601056 | 6.375574483 |
| ENSMUSG00000047067 | *Dusp28* | 102.4655942 | 79.61288131 | 103.3665981 | 158.049251 | 163.1781542 | 131.3181424 | 0.006055806 | 0.663930184 | 0.002502197 | 9.138986456 | 1.584392947 | 2.21782801 |
| ENSMUSG00000074649 | *BC029722* | 850.6151168 | 935.2115576 | 883.6583567 | 1352.199148 | 1477.025486 | 1397.665823 | 5.29E-13 | 0.663780818 | 6.39E-14 | 56.24744202 | 1.584228919 | 12.27623961 |
| ENSMUSG00000029684 | *Wasl* | 260.6845265 | 255.1448967 | 160.0921702 | 332.5619657 | 374.7833735 | 369.1600926 | 0.002851415 | 0.663576513 | 0.001101274 | 10.64905083 | 1.584004587 | 2.544939504 |
| ENSMUSG00000025857 | *Dnaaf5* | 969.6560277 | 796.1288131 | 583.6431087 | 1112.930143 | 1352.799536 | 1263.592755 | 0.000598422 | 0.663389734 | 0.000202746 | 13.80546492 | 1.583799526 | 3.222992441 |
| ENSMUSG00000031728 | *Zfp821* | 133.3559572 | 143.8787012 | 114.7117125 | 208.5372062 | 174.7585393 | 238.7602589 | 0.00260151 | 0.663384203 | 0.000997793 | 10.83165699 | 1.583793454 | 2.584774474 |
| ENSMUSG00000038886 | *Man2a2* | 2602.324724 | 2606.122874 | 1811.436603 | 3416.717489 | 4003.654971 | 3701.702321 | 2.12E-05 | 0.662718525 | 5.70E-06 | 20.58682024 | 1.58306284 | 4.673657913 |

| | | | | | | | | | | | | | |
|---|---|---|---|---|---|---|---|---|---|---|---|---|---|
| ENSMUSG00000032220 | *Myo1e* | 4420.335598 | 4439.137527 | 3268.653522 | 5919.163963 | 6862.957337 | 6427.242507 | 1.33E-06 | 0.662718145 | 3.02E-07 | 26.23442456 | 1.583062423 | 5.877452627 |
| ENSMUSG00000038717 | *Atp5l* | 1231.847401 | 1141.437696 | 1575.710337 | 2252.201827 | 1873.916868 | 2116.701526 | 6.91E-06 | 0.662592637 | 1.73E-06 | 22.87816868 | 1.58292471 | 5.16041922 |
| ENSMUSG00000021258 | *Ccnk* | 1119.587302 | 1309.296181 | 895.0034712 | 1647.443929 | 1791.801409 | 1826.51598 | 1.01E-05 | 0.662142882 | 2.59E-06 | 22.10033523 | 1.582431316 | 4.994702333 |
| ENSMUSG00000041642 | *Kif21b* | 996.0258497 | 1004.273334 | 611.3756106 | 1270.979394 | 1484.394822 | 1383.891193 | 0.000863504 | 0.661751913 | 0.000300022 | 13.07025618 | 1.582002537 | 3.063735522 |
| ENSMUSG00000019832 | *Rab32* | 668.2866329 | 677.1890868 | 766.4255077 | 1041.588467 | 1240.153972 | 1053.300065 | 6.12E-08 | 0.661504362 | 1.19E-08 | 32.50906517 | 1.581731105 | 7.213276031 |
| ENSMUSG00000047090 | *Tmem198b* | 133.3559572 | 202.389373 | 142.4442144 | 273.2934965 | 212.6579816 | 269.9827543 | 0.006351904 | 0.661394459 | 0.002635003 | 9.044402992 | 1.581610615 | 2.197096092 |
| ENSMUSG00000024989 | *Cep55* | 626.0949177 | 463.2894178 | 592.4670865 | 931.8320424 | 842.2098282 | 887.0861926 | 5.46E-06 | 0.661378204 | 1.35E-06 | 23.35756269 | 1.581592795 | 5.262809075 |
| ENSMUSG00000028106 | *Rprd2* | 906.3684548 | 921.7828788 | 694.5731164 | 1239.150031 | 1351.746774 | 1405.930601 | 1.82E-06 | 0.661089469 | 4.23E-07 | 25.58748938 | 1.581276294 | 5.739420211 |
| ENSMUSG00000021671 | *Poc5* | 362.3966972 | 328.0434387 | 340.3534327 | 561.9528925 | 537.9615277 | 532.619039 | 3.38E-08 | 0.660818827 | 6.37E-09 | 33.7197178 | 1.580979682 | 7.470734741 |
| ENSMUSG00000022982 | *Sod1* | 6656.496507 | 6185.824959 | 7820.565542 | 11000.88641 | 11101.3783 | 10554.12175 | 1.20E-10 | 0.660773477 | 1.79E-11 | 45.19200663 | 1.580929986 | 9.919977766 |
| ENSMUSG00000069895 | *Atxn1l* | 1280.066504 | 1318.888094 | 992.0672279 | 1933.908196 | 1910.763548 | 1842.127228 | 2.18E-07 | 0.660639771 | 4.52E-08 | 29.91079014 | 1.580783476 | 6.661551305 |
| ENSMUSG00000067719 | *Gm10221* | 2416.982546 | 2335.630916 | 2981.243957 | 4273.915163 | 3787.838702 | 4151.673578 | 2.46E-08 | 0.660433585 | 4.54E-09 | 34.37544166 | 1.580557571 | 7.609742196 |
| ENSMUSG00000021244 | *Ylpm1* | 962.1217928 | 940.9667056 | 531.9598096 | 1097.564243 | 1401.226602 | 1354.505315 | 0.004594107 | 0.659722579 | 0.001854415 | 9.688334968 | 1.579778814 | 2.33779892 |
| ENSMUSG00000025764 | *Jade1* | 1303.422632 | 1315.051329 | 1169.807354 | 2006.347437 | 2108.682857 | 1877.941267 | 2.97E-12 | 0.659600482 | 3.82E-13 | 52.73582821 | 1.579645121 | 11.5274482 |
| ENSMUSG00000001847 | *Rac1* | 5823.210131 | 5740.760177 | 5174.632745 | 8138.438863 | 9269.571921 | 9028.81102 | 6.53E-14 | 0.658928196 | 7.26E-15 | 60.52673349 | 1.578909188 | 13.18541114 |
| ENSMUSG00000028035 | *Dnajb4* | 205.6846119 | 254.1857054 | 301.2758163 | 397.318256 | 403.2079552 | 393.9544272 | 0.000275617 | 0.658750494 | 8.78E-05 | 15.3832727 | 1.57871472 | 3.559693474 |
| ENSMUSG00000021901 | *Bap1* | 1943.079173 | 2121.731247 | 2044.641733 | 3060.00911 | 3284.61833 | 3293.973264 | 2.77E-17 | 0.658608914 | 2.39E-18 | 76.33714294 | 1.578559799 | 16.55715785 |
| ENSMUSG00000030431 | *Tmem238* | 393.2870602 | 488.2283926 | 436.1566212 | 616.8311047 | 763.2526568 | 696.9962942 | 7.20E-06 | 0.65846447 | 1.80E-06 | 22.79306003 | 1.57840176 | 5.142379579 |
| ENSMUSG00000048497 | *Mmgt2* | 678.0811383 | 756.8019681 | 664.3194779 | 1127.198478 | 1122.244596 | 1066.156387 | 6.45E-11 | 0.658274676 | 9.29E-12 | 46.47173466 | 1.578194127 | 10.1902457 |
| ENSMUSG00000013822 | *Elof1* | 1841.367002 | 1917.423491 | 2179.522538 | 3272.936573 | 2999.319751 | 3088.272118 | 5.43E-12 | 0.658219983 | 7.16E-13 | 51.49939965 | 1.578134298 | 11.26543189 |
| ENSMUSG00000069565 | *Dazap1* | 2746.22861 | 2687.654138 | 2408.945963 | 4005.011923 | 4313.167082 | 4063.515944 | 5.51E-15 | 0.657445217 | 5.63E-16 | 65.5610087 | 1.577287025 | 14.25912225 |
| ENSMUSG00000000594 | *Gm2a* | 1299.655515 | 1397.541784 | 1220.230085 | 2090.859883 | 2091.838661 | 2000.994631 | 4.68E-14 | 0.657065013 | 5.12E-15 | 61.21247369 | 1.576871405 | 13.33021151 |
| ENSMUSG00000002833 | *Hdgfl2* | 1244.6556 | 1333.275964 | 1052.574505 | 1871.347035 | 1981.298621 | 1880.696193 | 4.14E-09 | 0.657045511 | 7.16E-10 | 37.97754715 | 1.57685009 | 8.383189983 |
| ENSMUSG00000046580 | *Gm7862* | 278.7666902 | 308.8596118 | 354.2196837 | 543.2943004 | 442.1601598 | 494.0500741 | 3.82E-05 | 0.656993541 | 1.07E-05 | 19.39007899 | 1.576793289 | 4.418484335 |
| ENSMUSG00000043881 | *Kbtbd7* | 126.5751458 | 113.1845783 | 90.76091539 | 188.7810498 | 160.0198673 | 176.3152681 | 0.003744921 | 0.65665212 | 0.001485336 | 10.0967181 | 1.576420176 | 2.426557364 |
| ENSMUSG00000037419 | *Endod1* | 2221.09244 | 2428.672476 | 2005.564116 | 3563.791098 | 3423.582951 | 3509.775806 | 2.22E-12 | 0.656347619 | 2.82E-13 | 53.32917411 | 1.576087487 | 11.65327111 |
| ENSMUSG00000000555 | *Itga5* | 4875.403384 | 4883.243118 | 3852.296631 | 6519.531604 | 7857.817697 | 7078.323367 | 1.46E-07 | 0.655969771 | 2.95E-08 | 30.7373783 | 1.575674756 | 6.836011372 |
| ENSMUSG00000047714 | *Ppp1r2* | 2344.653891 | 2333.712533 | 2315.663911 | 3624.157131 | 3619.396736 | 3776.085325 | 6.85E-25 | 0.655863393 | 3.77E-26 | 111.8938716 | 1.575558577 | 24.16436197 |
| ENSMUSG00000001924 | *Uba1* | 15502.44167 | 15736.49314 | 13653.21492 | 22972.01961 | 23694.52075 | 24061.52424 | 1.33E-16 | 0.655581391 | 1.21E-17 | 73.139237 | 1.575250635 | 15.87459175 |
| ENSMUSG00000031935 | *Med17* | 1626.641308 | 1608.563879 | 1844.211378 | 2655.007904 | 2614.008754 | 2720.948643 | 4.53E-14 | 0.655554946 | 4.96E-15 | 61.27570086 | 1.575221761 | 13.34361428 |
| ENSMUSG00000037773 | *Pced1a* | 570.3415796 | 586.0659094 | 477.755374 | 882.4416515 | 880.1092704 | 819.1313497 | 7.85E-08 | 0.655271868 | 1.54E-08 | 32.00374118 | 1.574912709 | 7.104916201 |
| ENSMUSG00000027936 | *Crtc2* | 1157.258476 | 1269.969336 | 850.8835817 | 1690.248934 | 1758.113016 | 1720.910481 | 2.26E-05 | 0.654848605 | 6.11E-06 | 20.45281178 | 1.574450724 | 4.645354737 |
| ENSMUSG00000011257 | *Pabpc4* | 5662.730928 | 5876.006156 | 5081.350693 | 8207.58541 | 9146.398734 | 8820.354948 | 6.89E-14 | 0.654771683 | 7.70E-15 | 60.41098968 | 1.574366778 | 13.16147932 |
| ENSMUSG00000042625 | *Safb2* | 1429.244355 | 1412.888845 | 980.7221135 | 1791.224845 | 2241.330905 | 1993.648162 | 0.000115918 | 0.65468685 | 3.48E-05 | 17.13734333 | 1.574274206 | 3.935850002 |
| ENSMUSG00000021978 | *Extl3* | 4008.212951 | 3924.051777 | 3051.83578 | 5179.405664 | 6268.146646 | 5847.789725 | 7.93E-07 | 0.654205528 | 1.76E-07 | 27.28470534 | 1.573749073 | 6.1009025 |
| ENSMUSG00000058440 | *Nrf1* | 1094.724327 | 1074.294302 | 959.2924529 | 1537.687505 | 1746.532631 | 1644.69086 | 9.49E-11 | 0.653745104 | 1.40E-11 | 45.67615393 | 1.573246904 | 10.02289507 |
| ENSMUSG00000025159 | *Mms19* | 1607.805721 | 1519.359084 | 1152.159398 | 2174.274766 | 2322.393601 | 2243.428125 | 2.13E-06 | 0.653101127 | 4.98E-07 | 25.27206101 | 1.57254481 | 5.670972899 |
| ENSMUSG00000093752 | *Gm20716* | 659.9989746 | 629.2295198 | 534.4809462 | 971.3443552 | 1058.026097 | 845.7623017 | 3.41E-06 | 0.652792973 | 8.17E-07 | 24.31837305 | 1.572208956 | 5.467315657 |
| ENSMUSG00000025384 | *Faap100* | 629.1086116 | 548.6574471 | 467.6708279 | 777.0754842 | 995.9131218 | 820.9679671 | 6.33E-05 | 0.652770609 | 1.82E-05 | 18.36635494 | 1.572184585 | 4.198757431 |
| ENSMUSG00000030342 | *Cd9* | 5794.580038 | 5598.799858 | 6108.713833 | 9119.661296 | 8961.112572 | 9413.582361 | 6.02E-23 | 0.652102677 | 3.72E-24 | 102.7951741 | 1.571456871 | 22.22010199 |
| ENSMUSG00000028367 | *Txn1* | 5084.855113 | 4920.65158 | 5245.224568 | 8230.63426 | 7890.453328 | 7837.764652 | 3.01E-25 | 0.651919025 | 1.63E-26 | 113.5540504 | 1.57125684 | 24.52201112 |
| ENSMUSG00000020929 | *Eftud2* | 5909.853832 | 6190.620916 | 5575.493455 | 8963.807174 | 9547.501164 | 9262.061427 | 5.03E-20 | 0.651540871 | 3.67E-21 | 89.14332213 | 1.570845042 | 19.29808715 |
| ENSMUSG00000031756 | *Cenpn* | 573.3552736 | 576.473996 | 593.7276548 | 878.0513945 | 968.5413024 | 888.92281 | 9.40E-11 | 0.650713247 | 1.38E-11 | 45.69462854 | 1.569944161 | 10.02678841 |
| ENSMUSG00000031988 | *Vps26b* | 1662.805636 | 1791.769425 | 1687.900913 | 2460.739033 | 2809.822539 | 2799.92319 | 8.18E-13 | 0.650593928 | 1.00E-13 | 55.36790169 | 1.569814323 | 12.08733849 |
| ENSMUSG00000045038 | *Prkce* | 207.1914589 | 218.6956258 | 167.6555798 | 308.4155523 | 317.9342101 | 309.4700279 | 0.00011462 | 0.649880897 | 3.43E-05 | 17.16054361 | 1.569038657 | 3.940740522 |
| ENSMUSG00000030208 | *Emp1* | 2166.845948 | 2339.467681 | 2333.311866 | 3511.108014 | 3698.353908 | 3512.530732 | 8.55E-19 | 0.649518438 | 6.73E-20 | 83.39172551 | 1.568644505 | 18.06786066 |
| ENSMUSG00000032010 | *Usp2* | 561.3004978 | 553.4534038 | 562.2134481 | 810.0024115 | 949.5915812 | 868.7200189 | 3.33E-09 | 0.648520001 | 5.71E-10 | 38.4191575 | 1.567559279 | 8.477147995 |
| ENSMUSG00000014077 | *Chp1* | 3694.035356 | 3892.398462 | 3713.634121 | 5799.529461 | 5981.795304 | 5931.355816 | 3.93E-26 | 0.648493642 | 2.03E-27 | 117.6899599 | 1.567530639 | 25.40507146 |
| ENSMUSG00000037415 | *Ranbp10* | 1103.765408 | 1117.457912 | 896.2640394 | 1620.004823 | 1642.309165 | 1633.671156 | 7.90E-09 | 0.64828731 | 1.40E-09 | 36.66951391 | 1.567306468 | 8.102124419 |
| ENSMUSG00000034390 | *Cmip* | 1648.490589 | 1460.848413 | 1017.278593 | 2100.737961 | 2241.330905 | 2134.149391 | 0.000204935 | 0.647995053 | 6.41E-05 | 15.9778505 | 1.566989 | 3.688383628 |
| ENSMUSG00000031918 | *Mtmr2* | 2064.380354 | 2160.0989 | 2407.685394 | 3657.084058 | 3346.731305 | 3374.784428 | 2.06E-12 | 0.647456185 | 2.60E-13 | 53.49008525 | 1.566403816 | 11.68649035 |
| ENSMUSG00000111228 | *Gm48789* | 297.6022773 | 285.8390197 | 265.9799048 | 460.9769821 | 437.9491106 | 435.2783181 | 6.75E-07 | 0.646795069 | 1.48E-07 | 27.61016399 | 1.565686175 | 6.170748413 |
| ENSMUSG00000026728 | *Vim* | 33163.44162 | 30907.06339 | 24597.46864 | 43405.37312 | 48636.56481 | 46772.21641 | 1.35E-07 | 0.646547127 | 2.72E-08 | 30.89473348 | 1.565417119 | 6.8696051 |
| ENSMUSG00000034201 | *Gas2l1* | 1839.860155 | 2018.138582 | 1739.584212 | 2758.178943 | 3021.427758 | 2984.503236 | 1.74E-12 | 0.646191255 | 2.19E-13 | 53.83049311 | 1.565031023 | 11.75889776 |
| ENSMUSG00000049734 | *Trex1* | 1518.148326 | 1457.011647 | 1295.864181 | 2471.714676 | 1789.695885 | 2426.171554 | 1.43E-05 | 0.646040472 | 3.75E-06 | 21.38898359 | 1.564867462 | 4.844663297 |
| ENSMUSG00000031833 | *Mast3* | 1747.94249 | 1607.604688 | 1042.489959 | 2146.83566 | 2432.933641 | 2308.628042 | 0.000638142 | 0.645457054 | 0.000217473 | 13.6737539 | 1.564234767 | 3.195082685 |
| ENSMUSG00000037552 | *Plekhg2* | 100.9587473 | 81.531264 | 73.11295962 | 142.6833516 | 126.3314742 | 134.0730684 | 0.009314784 | 0.645088374 | 0.004013693 | 8.277609944 | 1.563835078 | 2.030827206 |
| ENSMUSG00000029415 | *Sdad1* | 1583.69617 | 1543.338868 | 1631.17534 | 2574.885714 | 2411.878395 | 2450.965888 | 3.93E-18 | 0.644947916 | 3.22E-19 | 80.29597862 | 1.563682834 | 17.40591886 |
| ENSMUSG00000032855 | *Pkd1* | 1079.655857 | 937.1299403 | 699.6153894 | 1245.735416 | 1613.884583 | 1394.910897 | 0.00042891 | 0.644603101 | 0.000141253 | 14.48536071 | 1.563309146 | 3.367633834 |
| ENSMUSG00000055900 | *Tmem69* | 317.9447115 | 324.2066733 | 310.0997942 | 504.8795519 | 513.7479952 | 471.0923569 | 1.53E-07 | 0.643809109 | 3.11E-08 | 30.63651353 | 1.56244901 | 6.81563741 |
| ENSMUSG00000030435 | *U2af2* | 10135.80617 | 10569.32939 | 9042.056195 | 14853.3369 | 15820.91162 | 15794.90943 | 1.33E-14 | 0.643227423 | 1.41E-15 | 63.75861513 | 1.561819168 | 13.87561929 |
| ENSMUSG00000005233 | *Spc25* | 439.9993164 | 377.9213884 | 457.5862817 | 668.4166241 | 669.5568134 | 651.0808598 | 1.99E-07 | 0.642356888 | 4.11E-08 | 30.09531734 | 1.560877036 | 6.700864109 |
| ENSMUSG00000053536 | *Cstf2t* | 714.9988891 | 775.985795 | 635.3264077 | 1056.954366 | 1115.928022 | 1150.640786 | 1.32E-08 | 0.642226172 | 2.39E-09 | 35.6237334 | 1.560735619 | 7.879209341 |
| ENSMUSG00000004667 | *Polr2e* | 2861.502403 | 2969.656392 | 2882.919632 | 4481.354805 | 4596.360137 | 4521.75198 | 3.93E-26 | 0.642200647 | 2.03E-27 | 117.6893027 | 1.560708006 | 25.40507146 |
| ENSMUSG00000022702 | *Hira* | 1020.135401 | 1030.1715 | 809.2848289 | 1532.199683 | 1502.291781 | 1438.071405 | 1.29E-07 | 0.64202055 | 2.60E-08 | 30.98153284 | 1.560513189 | 6.888141149 |
| ENSMUSG00000025200 | *Cwf19l1* | 518.355359 | 526.5960463 | 455.0651452 | 738.6607357 | 823.260107 | 784.2356196 | 3.85E-08 | 0.642012142 | 7.28E-09 | 33.45825005 | 1.560504094 | 7.41488246 |
| ENSMUSG00000075528 | *Aarsd1* | 1486.50454 | 1404.256123 | 1217.708948 | 2064.518341 | 2304.496642 | 2051.501609 | 5.48E-09 | 0.641755341 | 9.58E-10 | 37.40827765 | 1.560226347 | 8.260907526 |
| ENSMUSG00000032279 | *Idh3a* | 3330.885236 | 3486.660525 | 4380.474736 | 6027.822823 | 5648.06966 | 5775.243339 | 7.79E-08 | 0.641092453 | 1.53E-08 | 32.02093412 | 1.559509621 | 7.10830328 |
| ENSMUSG00000073557 | *Ppp1r12b* | 364.6569677 | 343.3905001 | 305.0575212 | 459.8794179 | 596.9162157 | 527.1091869 | 4.92E-05 | 0.641068153 | 1.40E-05 | 18.87318175 | 1.559483353 | 4.307855805 |
| ENSMUSG00000033307 | *Mif* | 12944.56893 | 12232.56717 | 12603.16156 | 19826.40049 | 18444.39524 | 20638.06945 | 1.42E-19 | 0.640835136 | 1.07E-20 | 87.02820029 | 1.559231494 | 18.84736543 |
| ENSMUSG00000074634 | *Tmem267* | 176.3010959 | 179.3687808 | 144.965351 | 243.659262 | 280.0347679 | 259.8813587 | 0.00037363 | 0.639786246 | 0.000121432 | 14.77031254 | 1.55809829 | 3.427558346 |
| ENSMUSG00000083708 | *Gm13123* | 85.13685402 | 95.91913411 | 78.15523269 | 125.1223237 | 129.4897611 | 149.6843161 | 0.00715779 | 0.639700161 | 0.003001733 | 8.806414623 | 1.558005321 | 2.145221037 |
| ENSMUSG00000021690 | *Jmy* | 205.6846119 | 205.266947 | 200.4303548 | 307.3179881 | 343.200505 | 303.041867 | 2.47E-05 | 0.639683616 | 6.70E-06 | 20.27622459 | 1.557987454 | 4.607478532 |
| ENSMUSG00000026836 | *Acvr1* | 889.0397146 | 903.5582433 | 766.4255077 | 1238.052466 | 1373.854782 | 1381.136267 | 6.08E-09 | 0.639598815 | 1.07E-09 | 37.1997441 | 1.557895879 | 8.216341237 |
| ENSMUSG00000026596 | *Abl2* | 1804.449251 | 1912.627534 | 1310.991 | 2364.15338 | 2845.616457 | 2629.117774 | 5.53E-05 | 0.639235898 | 1.58E-05 | 18.63849293 | 1.557504032 | 4.25744425 |
| ENSMUSG00000029504 | *Ddx51* | 1313.217138 | 1446.460542 | 1196.279288 | 1952.566789 | 2119.21048 | 2094.662117 | 3.49E-10 | 0.639193956 | 5.38E-11 | 43.03210471 | 1.557458753 | 9.4567136 |
| ENSMUSG00000028792 | *Ak2* | 3885.404922 | 3620.947313 | 4028.776188 | 5950.993326 | 6254.460736 | 5755.958856 | 7.49E-17 | 0.639141671 | 6.65E-18 | 74.31846423 | 1.55740231 | 16.12539617 |
| ENSMUSG00000062203 | *Gspt1* | 5261.909633 | 5195.939495 | 4933.864206 | 7854.169724 | 8234.706595 | 7888.27163 | 6.71E-24 | 0.638915986 | 3.93E-25 | 107.2471889 | 1.5571587 | 23.17358192 |
| ENSMUSG00000020538 | *Srebf1* | 4748.828238 | 4811.303767 | 3390.928644 | 6392.214152 | 6707.148519 | 7076.48675 | 9.05E-06 | 0.638888012 | 2.30E-06 | 22.32805975 | 1.557128507 | 5.043404941 |
| ENSMUSG00000022283 | *Pabpc1* | 28141.12066 | 27577.71025 | 21125.86362 | 38137.06476 | 42097.85826 | 39422.99198 | 2.11E-07 | 0.638757993 | 4.38E-08 | 29.97448121 | 1.556988181 | 6.674810386 |
| ENSMUSG00000068039 | *Tcp1* | 10352.79213 | 9305.1152 | 9155.507339 | 14966.38602 | 15109.24432 | 14785.68819 | 3.75E-18 | 0.638303392 | 3.07E-19 | 80.3930663 | 1.556497643 | 17.42616669 |
| ENSMUSG00000000088 | *Cox5a* | 4280.952253 | 4348.014349 | 4086.762329 | 6489.89737 | 6647.141069 | 6657.737988 | 1.77E-25 | 0.638006225 | 9.50E-27 | 114.6270578 | 1.556177067 | 24.75204992 |
| ENSMUSG00000007670 | *Khsrp* | 4268.144054 | 4327.871331 | 2816.109513 | 5526.235964 | 6094.440869 | 6147.158358 | 0.000135944 | 0.637987811 | 4.13E-05 | 16.81240328 | 1.556157205 | 3.866639872 |
| ENSMUSG00000038379 | *Ttk* | 388.0130958 | 368.329475 | 405.9029827 | 631.0994398 | 594.8106911 | 580.3710908 | 4.64E-08 | 0.637197379 | 8.88E-09 | 33.0717179 | 1.555304843 | 7.333100428 |
| ENSMUSG00000023147 | *Get1* | 800.1357431 | 896.8439039 | 858.4469914 | 1337.930812 | 1328.586004 | 1302.16172 | 2.40E-12 | 0.636452238 | 3.06E-13 | 53.17010187 | 1.554501747 | 11.61950892 |
| ENSMUSG00000029472 | *Anapc5* | 8054.850499 | 7756.980376 | 7245.746412 | 11745.03497 | 12241.51985 | 11868.22148 | 1.98E-20 | 0.636393867 | 1.40E-21 | 91.05501746 | 1.554438854 | 19.70402098 |
| ENSMUSG00000069833 | *Ahnak* | 11999.77588 | 12711.20365 | 10588.77346 | 17316.27106 | 19708.76274 | 17831.7181 | 7.50E-11 | 0.635784692 | 1.09E-11 | 46.16328219 | 1.553782633 | 10.12517453 |
| ENSMUSG00000058192 | *Zfp846* | 107.7395586 | 92.08236875 | 117.232849 | 175.6102789 | 149.4922445 | 166.2138725 | 0.00377604 | 0.635729249 | 0.001499325 | 10.0794444 | 1.553722923 | 2.422963378 |
| ENSMUSG00000035198 | *Tubg1* | 1843.627273 | 1838.769801 | 1714.372846 | 2577.080843 | 2979.317267 | 2832.982302 | 1.55E-12 | 0.635704666 | 1.93E-13 | 54.07303369 | 1.553696447 | 11.81060294 |
| ENSMUSG00000075600 | *Zc3h3* | 449.7938217 | 566.8820826 | 462.6285548 | 735.3680429 | 809.5741973 | 752.0948155 | 2.99E-06 | 0.634900258 | 7.13E-07 | 24.58065051 | 1.55283039 | 5.523691371 |
| ENSMUSG00000023947 | *Nfkbie* | 241.0955158 | 188.9606942 | 187.8246721 | 324.879016 | 327.4090707 | 312.2249539 | 0.000257284 | 0.634683284 | 8.15E-05 | 15.52234973 | 1.55259687 | 3.589587934 |
| ENSMUSG00000075284 | *Wipf1* | 2083.969365 | 1904.954003 | 1513.942491 | 2626.471234 | 3030.902619 | 2894.508985 | 3.42E-06 | 0.63437715 | 8.20E-07 | 24.30983275 | 1.55226745 | 5.465514574 |
| ENSMUSG00000024969 | *Mark2* | 2206.02397 | 2072.812488 | 1592.097724 | 2808.666898 | 3235.138502 | 3078.170722 | 5.39E-06 | 0.634197081 | 1.33E-06 | 23.38512462 | 1.552073717 | 5.268184218 |
| ENSMUSG00000042719 | *Naa25* | 2038.763956 | 2018.138582 | 2113.972988 | 3323.424528 | 3113.018077 | 3136.02417 | 3.89E-19 | 0.633897484 | 3.02E-20 | 84.97468297 | 1.55175144 | 18.40963469 |
| ENSMUSG00000028896 | *Rcc1* | 3568.967058 | 3633.4168 | 3334.203072 | 5288.064524 | 5564.90144 | 5503.423967 | 7.54E-20 | 0.633758279 | 5.57E-21 | 88.31896064 | 1.551601719 | 19.12278743 |
| ENSMUSG00000026274 | *Pask* | 394.7939072 | 434.5136775 | 335.3111596 | 564.148021 | 642.184994 | 605.1654254 | 1.16E-05 | 0.633645683 | 2.98E-06 | 21.82962016 | 1.551480628 | 4.937378759 |
| ENSMUSG00000040212 | *Emp3* | 7320.262599 | 7646.673371 | 8058.812945 | 11987.59666 | 11874.10581 | 11846.18208 | 6.87E-23 | 0.633525543 | 4.26E-24 | 102.5240846 | 1.551351434 | 22.16308164 |
| ENSMUSG00000010651 | *Acaa1b* | 189.1092952 | 184.1647375 | 172.6978529 | 294.1472172 | 267.4016204 | 287.4306193 | 6.05E-05 | 0.633306112 | 1.74E-05 | 18.45651269 | 1.551115494 | 4.217958343 |
| ENSMUSG00000025583 | *Rptor* | 2997.118631 | 2925.53359 | 2445.502442 | 4150.987968 | 4435.287508 | 4398.698616 | 4.60E-10 | 0.63260757 | 7.20E-11 | 42.46446274 | 1.550364638 | 9.337024186 |
| ENSMUSG00000039201 | *Tbc1d25* | 253.9037151 | 286.798211 | 288.6701337 | 391.8304348 | 474.7957906 | 415.9938357 | 2.95E-05 | 0.632537786 | 8.11E-06 | 19.91087236 | 1.550289647 | 4.529644356 |
| ENSMUSG00000044465 | *Fhip1b* | 292.3283129 | 320.3699079 | 262.1982 | 402.8060772 | 474.7957906 | 480.2754438 | 4.12E-05 | 0.632028425 | 1.16E-05 | 19.23414319 | 1.549742396 | 4.385464167 |
| ENSMUSG00000038957 | *Edc3* | 1528.696255 | 1642.135576 | 1561.844086 | 2447.568262 | 2478.202419 | 2405.968763 | 5.47E-18 | 0.631689173 | 4.52E-19 | 79.62738021 | 1.549378014 | 17.26185699 |
| ENSMUSG00000026192 | *Atic* | 5896.292209 | 6292.295198 | 6291.496232 | 9420.393899 | 9716.995892 | 9486.128747 | 1.47E-24 | 0.631682326 | 8.25E-26 | 110.3406207 | 1.549370661 | 23.83267366 |
| ENSMUSG00000022673 | *Mcm4* | 2957.94061 | 2758.634297 | 2461.88983 | 4045.6218 | 4524.772302 | 4108.51307 | 2.29E-10 | 0.631245091 | 3.47E-11 | 43.88938848 | 1.548901167 | 9.640547901 |
| ENSMUSG00000083669 | *Mif-ps4* | 4224.445491 | 4277.03419 | 4432.158035 | 6858.678956 | 6243.933113 | 6920.374273 | 7.55E-18 | 0.630988165 | 6.31E-19 | 78.96822597 | 1.548625351 | 17.12198214 |
| ENSMUSG00000034154 | *Ino80* | 766.9851097 | 786.5368997 | 636.586976 | 1117.3204 | 1164.355087 | 1116.663365 | 4.77E-08 | 0.630204475 | 9.13E-09 | 33.01862811 | 1.547784347 | 7.321864611 |
| ENSMUSG00000064145 | *Arih2* | 1348.628042 | 1370.684426 | 1491.252263 | 2224.762721 | 2182.376217 | 2102.008587 | 1.77E-14 | 0.630154901 | 1.89E-15 | 63.17930082 | 1.547731163 | 13.75127197 |
| ENSMUSG00000024177 | *Nme4* | 297.6022773 | 291.5941677 | 346.656274 | 452.1964682 | 476.9013152 | 513.3345566 | 8.26E-06 | 0.63002012 | 2.08E-06 | 22.51523216 | 1.547586576 | 5.083020141 |
| ENSMUSG00000049288 | *Lix1l* | 856.6425047 | 798.0471958 | 823.1510798 | 1278.662343 | 1292.792086 | 1265.429372 | 1.59E-13 | 0.629688822 | 1.83E-14 | 58.70438141 | 1.547231232 | 12.7977329 |
| ENSMUSG00000001052 | *Sec24b* | 1473.69634 | 1539.502103 | 992.0672279 | 1877.93242 | 2212.906323 | 2113.028291 | 0.0003599 | 0.629519213 | 0.000116773 | 14.84410393 | 1.547049344 | 3.443817594 |
| ENSMUSG00000020212 | *Mdm1* | 336.0268752 | 345.3088828 | 335.3111596 | 533.4162222 | 496.9037986 | 541.8021259 | 1.44E-07 | 0.629139961 | 2.91E-08 | 30.76911217 | 1.546642713 | 6.842966529 |
| ENSMUSG00000047777 | *Phf13* | 366.1638147 | 455.615887 | 363.0436615 | 586.0993059 | 637.9739448 | 608.8386601 | 9.21E-06 | 0.628860162 | 2.34E-06 | 22.29061755 | 1.546342783 | 5.035523441 |
| ENSMUSG00000026116 | *Tmem131* | 1334.312995 | 1300.663459 | 751.2986885 | 1569.516868 | 1961.296137 | 1711.727394 | 0.004467293 | 0.628575981 | 0.001800467 | 9.742592905 | 1.546038216 | 2.349955561 |
| ENSMUSG00000024847 | *Aip* | 1860.202589 | 1971.138206 | 2101.367305 | 3016.10654 | 3149.864757 | 2995.52294 | 6.24E-15 | 0.628523543 | 6.42E-16 | 65.30274081 | 1.545982022 | 14.20482099 |
| ENSMUSG00000034773 | *Hrob* | 568.0813092 | 525.6368549 | 447.5017356 | 792.4413836 | 854.8429756 | 743.8300373 | 3.34E-06 | 0.628464669 | 7.99E-07 | 24.35995723 | 1.545918935 | 5.476566478 |
| ENSMUSG00000023940 | *Sgo1* | 414.3829178 | 361.6151356 | 360.522525 | 610.2457192 | 576.9137323 | 574.8612387 | 4.48E-07 | 0.628044946 | 9.62E-08 | 28.44988552 | 1.545469246 | 6.348773668 |
| ENSMUSG00000023088 | *Abcc1* | 5447.251811 | 5481.778515 | 4220.382565 | 7354.777993 | 8250.498029 | 7807.460465 | 2.32E-07 | 0.627391748 | 4.84E-08 | 29.77922321 | 1.544769674 | 6.633943098 |
| ENSMUSG00000039781 | *Cep131* | 202.670918 | 206.2261383 | 139.9230779 | 252.4397759 | 275.8237187 | 323.2446582 | 0.004833654 | 0.627279028 | 0.001959401 | 9.587184895 | 1.544648984 | 2.315724411 |
| ENSMUSG00000003344 | *Btbd2* | 329.9994873 | 353.9416049 | 283.6278606 | 443.4159542 | 484.2706512 | 567.5147692 | 0.000137662 | 0.625883883 | 4.18E-05 | 16.78728519 | 1.543155968 | 3.861185818 |
| ENSMUSG00000027198 | *Ext2* | 1894.86007 | 2027.730495 | 1876.986153 | 2897.569602 | 3119.334651 | 2929.404715 | 6.81E-16 | 0.625038591 | 6.47E-17 | 69.82808654 | 1.542252079 | 15.16667938 |
| ENSMUSG00000060860 | *Ube2s* | 9244.506185 | 8399.638574 | 7720.980649 | 12874.42857 | 12948.97611 | 13287.92672 | 3.74E-13 | 0.624239631 | 4.45E-14 | 56.96117696 | 1.541398221 | 12.42765737 |
| ENSMUSG00000025025 | *Mxi1* | 702.1906898 | 720.3526972 | 736.1718692 | 1022.929875 | 1168.566137 | 1132.274612 | 5.14E-10 | 0.624237972 | 8.10E-11 | 42.23379275 | 1.541396449 | 9.288651359 |
| ENSMUSG00000029823 | *Luc7l2* | 1654.517977 | 1548.134825 | 1245.44145 | 2295.006832 | 2347.659896 | 2225.061951 | 3.79E-07 | 0.624194346 | 8.07E-08 | 28.79020528 | 1.541349839 | 6.421335195 |
| ENSMUSG00000017167 | *Cntnap1* | 223.7667756 | 229.2467305 | 158.8316019 | 317.1960663 | 321.092497 | 310.3883365 | 0.001169578 | 0.624095542 | 0.000417764 | 12.45101905 | 1.541244281 | 2.931970738 |
| ENSMUSG00000021607 | *Mrpl36* | 809.9302485 | 805.7207265 | 825.6722164 | 1204.027975 | 1242.259497 | 1313.181424 | 1.76E-12 | 0.62406661 | 2.21E-13 | 53.80900885 | 1.541213373 | 11.7543863 |
| ENSMUSG00000006678 | *Pola1* | 563.5607682 | 470.0037571 | 502.9667394 | 803.417026 | 791.6772385 | 777.8074587 | 8.78E-08 | 0.623996234 | 1.73E-08 | 31.77661861 | 1.541138194 | 7.056562237 |
| ENSMUSG00000034724 | *Cnot6l* | 656.9852806 | 620.5967977 | 509.2695808 | 906.5880649 | 919.061475 | 936.6748618 | 9.80E-07 | 0.623794991 | 2.20E-07 | 26.84702136 | 1.540923234 | 6.008715577 |
| ENSMUSG00000109946 | *Brd3os* | 135.6162277 | 138.1235531 | 143.7047827 | 249.1470832 | 193.7082605 | 200.191294 | 0.001803586 | 0.623790489 | 0.000670961 | 11.56801344 | 1.540918425 | 2.743863204 |
| ENSMUSG00000072437 | *Nanos1* | 202.670918 | 164.9809107 | 153.7893288 | 259.0251614 | 287.4041039 | 261.7179761 | 0.000861949 | 0.623442111 | 0.000299422 | 13.07400389 | 1.540546373 | 3.06451868 |
| ENSMUSG00000046330 | *Rpl37a* | 4946.225192 | 5164.286181 | 5406.577307 | 7928.804093 | 7830.445877 | 8130.705124 | 2.93E-21 | 0.62331903 | 1.97E-22 | 94.92876278 | 1.54041495 | 20.53244498 |
| ENSMUSG00000074243 | *Josd1-ps* | 151.4381209 | 121.8173003 | 133.6202365 | 193.1713068 | 222.1328422 | 213.0476156 | 0.001498511 | 0.623268045 | 0.000546465 | 11.9500298 | 1.540360512 | 2.824339984 |
| ENSMUSG00000020196 | *Cabin1* | 2247.462262 | 2466.080938 | 1831.605695 | 3098.423858 | 3598.341491 | 3392.232293 | 1.74E-06 | 0.623254174 | 4.04E-07 | 25.67635076 | 1.540345703 | 5.758647904 |
| ENSMUSG00000040563 | *Plppr2* | 123.5614519 | 130.4500224 | 107.1483029 | 186.5859213 | 176.8640639 | 194.6814419 | 0.001664687 | 0.622935224 | 0.000614262 | 11.73225381 | 1.540005202 | 2.778667523 |
| ENSMUSG00000031848 | *Lsm4* | 2514.927599 | 2619.551553 | 2700.137233 | 3932.572683 | 3972.072102 | 4148.000344 | 2.60E-19 | 0.62260993 | 2.00E-20 | 85.79534168 | 1.539658005 | 18.58524677 |
| ENSMUSG00000030835 | *Nomo1* | 6745.400479 | 6915.769569 | 5811.219721 | 9418.198771 | 10607.63279 | 9960.894339 | 1.02E-10 | 0.622432708 | 1.51E-11 | 45.52155922 | 1.539468884 | 9.990034571 |
| ENSMUSG00000033671 | *Cep350* | 494.9992309 | 582.2291441 | 415.9875288 | 725.4899647 | 828.5239184 | 749.3398894 | 6.33E-05 | 0.622363111 | 1.82E-05 | 18.36733981 | 1.53939462 | 4.198878432 |
| ENSMUSG00000033855 | *Ston1* | 114.52037 | 113.1845783 | 122.2751221 | 156.9516868 | 205.2886456 | 175.3969594 | 0.003546035 | 0.622152772 | 0.001398265 | 10.20808216 | 1.5391702 | 2.450256952 |
| ENSMUSG00000080875 | *Gm7332* | 1945.339443 | 2012.383434 | 2205.994471 | 3038.057825 | 3324.623297 | 3113.984761 | 6.66E-13 | 0.622101405 | 8.06E-14 | 55.79036955 | 1.539115399 | 12.17675294 |
| ENSMUSG00000027694 | *Gm8325* | 516.0950886 | 476.7180965 | 480.2765106 | 739.7582999 | 782.2023779 | 748.4215807 | 3.31E-09 | 0.621790571 | 5.66E-10 | 38.4355464 | 1.538783827 | 8.480621019 |
| ENSMUSG00000029714 | *Gigyf1* | 442.2595868 | 444.1055909 | 272.2827462 | 577.3187919 | 602.1800271 | 609.7569688 | 0.002720902 | 0.621513601 | 0.001046386 | 10.74363511 | 1.538488437 | 2.565287094 |
| ENSMUSG00000019362 | *D8Ertd738e* | 1306.436326 | 1345.745452 | 1458.477488 | 2155.616174 | 2012.881489 | 2146.087404 | 4.33E-13 | 0.621410761 | 5.17E-14 | 56.66488925 | 1.538378773 | 12.36397328 |
| ENSMUSG00000028937 | *Acot7* | 4112.185392 | 4188.788587 | 3767.838557 | 6109.042577 | 6128.129262 | 6328.065169 | 6.13E-18 | 0.620605368 | 5.07E-19 | 79.40030185 | 1.537520204 | 17.21266294 |
| ENSMUSG00000040105 | *Plpp6* | 114.52037 | 94.95994277 | 123.5356904 | 156.9516868 | 176.8640639 | 176.3152681 | 0.004257462 | 0.619772912 | 0.001708887 | 9.838587437 | 1.536633288 | 2.370849175 |
| ENSMUSG00000024870 | *Rab1b* | 5259.649362 | 5573.860883 | 5458.260606 | 8038.560517 | 8604.226157 | 8375.893543 | 6.67E-22 | 0.619161353 | 4.29E-23 | 97.94876868 | 1.535982047 | 21.17587402 |
| ENSMUSG00000040331 | *Nsmce4a* | 1600.271486 | 1386.990679 | 1372.758845 | 2185.250408 | 2198.167651 | 2321.484363 | 1.23E-10 | 0.619097991 | 1.83E-11 | 45.14476552 | 1.535914589 | 9.909902484 |
| ENSMUSG00000020941 | *Map3k14* | 906.3684548 | 920.8236875 | 757.6015298 | 1286.345293 | 1375.960307 | 1314.099733 | 2.63E-08 | 0.618285826 | 4.88E-09 | 34.23486568 | 1.535050189 | 7.579498444 |
| ENSMUSG00000024169 | *Ift140* | 630.6154586 | 612.923267 | 412.205824 | 758.416892 | 887.4786064 | 901.7791316 | 0.001050823 | 0.617900487 | 0.000371884 | 12.6684173 | 1.534640237 | 2.978470243 |
| ENSMUSG00000028438 | *Kif24* | 433.9719285 | 409.5747027 | 336.5717279 | 611.3432834 | 637.9739448 | 569.3513866 | 1.78E-05 | 0.617661772 | 4.72E-06 | 20.9481496 | 1.53438633 | 4.749683251 |
| ENSMUSG00000022708 | *Zbtb20* | 182.3284838 | 182.2463548 | 147.4864875 | 253.5373402 | 300.0372513 | 236.0053328 | 0.001594922 | 0.617543171 | 0.00058633 | 11.81888427 | 1.534260196 | 2.797260482 |
| ENSMUSG00000082127 | *Gm13577* | 505.5471597 | 462.3302264 | 453.8045769 | 769.3925345 | 737.9863619 | 679.5484291 | 1.27E-07 | 0.617460298 | 2.56E-08 | 31.01816102 | 1.534172065 | 6.896040911 |
| ENSMUSG00000040681 | *Hmgn1* | 3597.59715 | 3281.393578 | 3755.232874 | 5451.601596 | 5583.851161 | 5272.928487 | 1.11E-14 | 0.617268716 | 1.16E-15 | 64.14093777 | 1.53396835 | 13.95650048 |
| ENSMUSG00000008604 | *Ubqln4* | 4194.308552 | 4069.84886 | 3129.991013 | 5371.479406 | 6206.033671 | 5908.398099 | 2.52E-06 | 0.617084025 | 5.95E-07 | 24.92863071 | 1.533771986 | 5.59782031 |
| ENSMUSG00000025821 | *Zfp282* | 1076.642163 | 1065.66158 | 920.2148365 | 1466.345829 | 1619.148395 | 1618.978217 | 4.59E-09 | 0.61706053 | 7.96E-10 | 37.76971808 | 1.533747009 | 8.338473101 |
| ENSMUSG00000032518 | *Rpsa* | 81327.54487 | 79449.81879 | 83062.62497 | 125188.1776 | 124788.1247 | 124005.6503 | 2.50E-35 | 0.617054453 | 8.61E-37 | 160.5434264 | 1.533740548 | 34.60226713 |
| ENSMUSG00000052296 | *Ppp6r1* | 5435.197035 | 5279.389142 | 3988.438004 | 7116.606553 | 7838.867976 | 7598.086085 | 2.32E-06 | 0.61655803 | 5.45E-07 | 25.096372 | 1.533212887 | 5.63370171 |
| ENSMUSG00000002326 | *Gmpr2* | 581.6429319 | 564.9636999 | 584.9036769 | 905.4905006 | 938.0111961 | 811.7848802 | 1.33E-08 | 0.61615703 | 2.42E-09 | 35.60523517 | 1.532786788 | 7.875414068 |
| ENSMUSG00000074800 | *Gm4149* | 5142.115298 | 5463.553879 | 5691.465736 | 8548.92789 | 8117.849981 | 8299.673922 | 1.10E-18 | 0.616023094 | 8.74E-20 | 82.87497368 | 1.532644494 | 17.95862387 |
| ENSMUSG00000041459 | *Tardbp* | 3956.22673 | 3590.25319 | 4012.388801 | 6010.261796 | 5954.423485 | 5743.102535 | 2.06E-16 | 0.615504686 | 1.88E-17 | 72.26262284 | 1.532093863 | 15.68692305 |
| ENSMUSG00000022253 | *Nadk2* | 228.2873165 | 169.7768674 | 185.3035356 | 299.6350384 | 303.1955381 | 294.7770888 | 0.000669185 | 0.615451046 | 0.000228787 | 13.57853453 | 1.5320369 | 3.174453632 |
| ENSMUSG00000063382 | *Bcl9l* | 427.9445406 | 468.0853745 | 310.0997942 | 564.148021 | 666.3985265 | 622.6132905 | 0.000815048 | 0.615221449 | 0.000282347 | 13.18401579 | 1.531793104 | 3.088816912 |
| ENSMUSG00000006498 | *Ptbp1* | 15697.57835 | 14849.24115 | 11617.39717 | 20383.96312 | 22426.99496 | 21772.18068 | 6.70E-07 | 0.614881289 | 1.47E-07 | 27.62433029 | 1.531431979 | 6.173793503 |
| ENSMUSG00000038766 | *Gabpb2* | 2108.83234 | 2133.241543 | 1755.971599 | 3005.130898 | 3225.663642 | 2954.199049 | 4.80E-09 | 0.613119644 | 8.34E-10 | 37.67915418 | 1.529563121 | 8.319168437 |

| | | | | | | | | | | | | |
|---|---|---|---|---|---|---|---|---|---|---|---|---|
| ENSMUSG00000084786 | *Ubl5* | 1790.134205 | 1759.15692 | 2018.169799 | 2889.886652 | 2825.613973 | 2788.903485 | 8.03E-13 | 0.612824572 | 9.81E-14 | 55.40444648 | 1.529250314 | 12.09516869 |
| ENSMUSG00000048498 | *Cd300e* | 268.9721848 | 304.0636551 | 306.3180894 | 442.31839 | 487.428938 | 412.3206009 | 2.47E-05 | 0.612753492 | 6.72E-06 | 20.27230317 | 1.529174971 | 4.606698153 |
| ENSMUSG00000038914 | *Dido1* | 1439.792284 | 1493.460918 | 1230.314631 | 2019.518207 | 2146.5823 | 2207.614086 | 5.09E-09 | 0.612592177 | 8.86E-10 | 37.56068801 | 1.529003996 | 8.293648407 |
| ENSMUSG00000033629 | *Hacd3* | 2123.147386 | 2175.445962 | 2277.846863 | 3314.644014 | 3404.63323 | 3325.195759 | 3.26E-19 | 0.612234689 | 2.52E-20 | 85.33252824 | 1.528625168 | 18.48630784 |
| ENSMUSG00000040363 | *Bcor* | 1414.929308 | 1339.031112 | 775.2494856 | 1668.29765 | 1953.926801 | 1777.84562 | 0.005428948 | 0.611605417 | 0.002224185 | 9.35466622 | 1.527958561 | 2.265284319 |
| ENSMUSG00000020362 | *Cnot6* | 1712.531586 | 1679.544038 | 1356.371458 | 2319.153246 | 2528.735009 | 2415.151849 | 1.13E-07 | 0.61106987 | 2.26E-08 | 31.26073534 | 1.527391468 | 6.947488936 |
| ENSMUSG00000066043 | *Phactr4* | 622.3278002 | 680.0666609 | 433.6354846 | 781.4657411 | 907.4810898 | 967.8973572 | 0.001442938 | 0.61105366 | 0.000523724 | 12.0292423 | 1.527374307 | 2.840752245 |
| ENSMUSG00000005447 | *Pafah1b3* | 741.3687112 | 715.5567405 | 820.6299433 | 1175.491304 | 1131.719457 | 1164.415416 | 4.37E-10 | 0.610901119 | 6.81E-11 | 42.57242193 | 1.52721282 | 9.359468075 |
| ENSMUSG00000047710 | *Champ1* | 1529.449679 | 1381.235531 | 1135.772011 | 1935.005761 | 2301.338355 | 1950.487653 | 1.41E-05 | 0.610337437 | 3.70E-06 | 21.41422211 | 1.526616232 | 4.850095884 |
| ENSMUSG00000049091 | *Sephs2* | 3724.925719 | 3810.867198 | 3745.148328 | 5743.553684 | 5874.413551 | 5604.437923 | 1.13E-24 | 0.610326959 | 6.31E-26 | 110.8716183 | 1.526605145 | 23.94580548 |
| ENSMUSG00000015305 | *Sash1* | 1181.368028 | 1316.01052 | 974.4192721 | 1758.297918 | 1750.74368 | 1798.048411 | 1.40E-06 | 0.610261328 | 3.20E-07 | 26.12493733 | 1.526535699 | 5.854654411 |
| ENSMUSG00000039630 | *Hnrnpu* | 9572.998825 | 9104.64421 | 7042.79492 | 12794.30638 | 13255.32993 | 13218.13526 | 7.60E-07 | 0.609992436 | 1.68E-07 | 27.36940902 | 1.526251207 | 6.119118674 |
| ENSMUSG00000003233 | *Dvl3* | 565.8210387 | 471.9221398 | 340.3534327 | 639.8799538 | 768.5164682 | 701.5878376 | 0.002768365 | 0.609491115 | 0.001066919 | 10.7076777 | 1.525720943 | 2.557776641 |
| ENSMUSG00000002835 | *Chaf1a* | 2658.831485 | 2779.736507 | 2009.345821 | 3507.815321 | 4005.760495 | 3856.89649 | 1.14E-05 | 0.609308878 | 2.94E-06 | 21.85315158 | 1.525528231 | 4.941905041 |
| ENSMUSG00000051396 | *Gm45902* | 137.8764981 | 156.3481886 | 163.873875 | 226.0982341 | 250.5574239 | 219.4757764 | 0.000886767 | 0.609222525 | 0.000308774 | 13.01640373 | 1.525436923 | 3.052190264 |
| ENSMUSG00000056091 | *St3gal5* | 3833.418702 | 3478.986994 | 3490.513538 | 5003.795385 | 6043.908279 | 5435.469124 | 6.21E-10 | 0.609136504 | 9.85E-11 | 41.85167564 | 1.52534597 | 9.207180817 |
| ENSMUSG00000029363 | *Rfc5* | 2361.229208 | 2280.957009 | 2189.607084 | 3427.693131 | 3553.072713 | 3447.330815 | 1.54E-18 | 0.608900184 | 1.25E-19 | 82.17483938 | 1.525096133 | 17.81112084 |
| ENSMUSG00000032295 | *Man2c1* | 1593.490675 | 1673.78889 | 1444.611237 | 2374.031458 | 2360.293043 | 2458.312358 | 3.27E-12 | 0.608797989 | 4.22E-13 | 52.53638827 | 1.524988105 | 11.48519726 |
| ENSMUSG00000096768 | *Gm47283* | 882.2589032 | 857.517059 | 670.6223192 | 1135.978992 | 1253.839882 | 1292.978633 | 9.67E-06 | 0.60848158 | 2.47E-06 | 22.19280392 | 1.524653683 | 5.014453065 |
| ENSMUSG00000030530 | *Furin* | 2388.352454 | 2556.244924 | 1922.366611 | 3302.570808 | 3793.102514 | 3381.212589 | 2.20E-06 | 0.608314362 | 5.14E-07 | 25.21038517 | 1.524476977 | 5.657978036 |
| ENSMUSG00000037071 | *Scd1* | 4702.115982 | 4117.808427 | 3014.018732 | 6283.555292 | 5561.743153 | 6195.828718 | 0.000181195 | 0.607544115 | 5.61E-05 | 16.22855364 | 1.523663283 | 3.741854395 |
| ENSMUSG00000026819 | *Slc25a25* | 1178.354334 | 1190.356454 | 980.7221135 | 1625.492644 | 1765.482352 | 1719.992173 | 2.72E-08 | 0.607220676 | 5.05E-09 | 34.16905631 | 1.52332173 | 7.565132995 |
| ENSMUSG00000059183 | *Mtfmt* | 507.8074302 | 475.7589052 | 560.9528798 | 794.6365121 | 745.3556979 | 807.1933368 | 1.49E-07 | 0.607039369 | 3.03E-08 | 30.69125979 | 1.523130303 | 6.827011731 |
| ENSMUSG00000025894 | *Aasdhppt* | 737.6015937 | 640.7398159 | 727.3478914 | 1071.222701 | 1045.392949 | 1090.032413 | 3.35E-09 | 0.606888389 | 5.74E-10 | 38.40666279 | 1.522970913 | 8.474541475 |
| ENSMUSG00000001016 | *Ilf2* | 2622.667158 | 2497.734252 | 2643.411661 | 4041.231543 | 3896.273217 | 3884.44575 | 6.71E-20 | 0.606711066 | 4.94E-21 | 88.55595512 | 1.522783735 | 19.17320359 |
| ENSMUSG00000009418 | *Nav1* | 1858.695742 | 1872.341498 | 1234.096336 | 2321.348374 | 2782.45072 | 2464.740519 | 0.000534526 | 0.606625325 | 0.000179097 | 14.03859118 | 1.522693237 | 3.272031561 |
| ENSMUSG00000036932 | *Aifm1* | 1118.833878 | 1124.172252 | 1150.89883 | 1652.93175 | 1780.221024 | 1731.011877 | 2.45E-14 | 0.606477964 | 2.64E-15 | 62.51924302 | 1.522537713 | 13.61074022 |
| ENSMUSG00000036285 | *Noa1* | 1333.559572 | 1340.949495 | 1217.708948 | 1882.322677 | 2130.790865 | 1916.510232 | 1.61E-10 | 0.60579724 | 2.41E-11 | 44.60618479 | 1.521819486 | 9.792848305 |
| ENSMUSG00000066839 | *Ecsit* | 991.5053088 | 979.3343593 | 952.9896115 | 1405.979795 | 1551.771608 | 1494.088235 | 2.87E-12 | 0.605792033 | 3.68E-13 | 52.80680202 | 1.521813993 | 11.54267788 |
| ENSMUSG00000033222 | *Ttf2* | 1440.545707 | 1566.35946 | 1297.124749 | 1983.298587 | 2353.97647 | 2216.797173 | 9.67E-08 | 0.605708779 | 1.92E-08 | 31.57466379 | 1.521726176 | 7.014565051 |
| ENSMUSG00000005087 | *Cd44* | 8888.890299 | 8435.128654 | 7192.802544 | 12470.52493 | 12471.02203 | 12378.80111 | 2.01E-10 | 0.605652347 | 3.04E-11 | 44.15378392 | 1.521666653 | 9.696652649 |
| ENSMUSG00000046806 | *Cyren* | 317.9447115 | 375.0438144 | 371.8676394 | 508.1722446 | 547.4363883 | 559.249991 | 6.57E-06 | 0.60553551 | 1.64E-06 | 22.98180246 | 1.521543426 | 5.182757732 |
| ENSMUSG00000022221 | *Ripk3* | 561.3004978 | 682.9442349 | 554.6500385 | 895.6124224 | 854.8429756 | 986.2635309 | 5.24E-06 | 0.605334355 | 1.29E-06 | 23.4432167 | 1.521331292 | 5.280571206 |
| ENSMUSG00000047417 | *Rexo1* | 1601.02491 | 1636.380428 | 1234.096336 | 2187.445537 | 2367.662379 | 2256.284446 | 2.40E-06 | 0.605183088 | 5.65E-07 | 25.0287579 | 1.521171789 | 5.619234594 |
| ENSMUSG00000063849 | *Ppcdc* | 1334.312995 | 1402.337741 | 1459.738056 | 2035.981671 | 2221.328422 | 2118.538143 | 4.30E-13 | 0.605174778 | 5.14E-14 | 56.67713723 | 1.521163027 | 12.36617911 |
| ENSMUSG00000041959 | *S100a10* | 6580.400735 | 5705.270097 | 7371.803239 | 10490.51904 | 9610.666902 | 9792.843849 | 3.56E-08 | 0.605094332 | 6.72E-09 | 33.61397887 | 1.521078207 | 7.448234646 |
| ENSMUSG00000031949 | *Adat1* | 292.3283129 | 344.3496915 | 305.0575212 | 496.0990379 | 519.0118066 | 417.830453 | 9.98E-05 | 0.604963432 | 2.96E-05 | 17.4430309 | 1.520940202 | 4.001004919 |
| ENSMUSG00000020740 | *Gga3* | 1212.25839 | 1320.806477 | 942.9050654 | 1543.175326 | 1843.386761 | 1905.490528 | 0.000101475 | 0.604939348 | 3.02E-05 | 17.40728899 | 1.520914811 | 3.993641597 |
| ENSMUSG00000035754 | *Wdr18* | 3382.871456 | 3535.579283 | 3307.731138 | 5116.844502 | 5150.113099 | 5286.703117 | 3.67E-21 | 0.604741808 | 2.49E-22 | 94.47016563 | 1.520706575 | 20.4353623 |
| ENSMUSG00000066551 | *Hmgb1* | 2720.612211 | 2335.630916 | 2527.43938 | 3977.572817 | 3879.429021 | 3676.907987 | 7.79E-12 | 0.604143191 | 1.05E-12 | 50.75613188 | 1.52007572 | 11.10837898 |
| ENSMUSG00000026238 | *Ptma* | 23179.07358 | 21069.597 | 25266.83039 | 35208.76336 | 35776.02074 | 34671.66282 | 1.20E-12 | 0.604085815 | 1.49E-13 | 54.58679716 | 1.520015267 | 11.92127629 |
| ENSMUSG00000048170 | *Mcmbp* | 2384.585336 | 2472.795277 | 2053.465711 | 3337.692863 | 3599.394253 | 3573.139105 | 7.21E-10 | 0.603667519 | 1.15E-10 | 41.54660573 | 1.519574617 | 9.141850593 |
| ENSMUSG00000006611 | *Hfe* | 391.7802132 | 368.329475 | 381.9521856 | 621.2213616 | 562.1750603 | 553.7401389 | 5.39E-07 | 0.603506533 | 1.17E-07 | 28.07095641 | 1.519405062 | 6.268450555 |
| ENSMUSG00000034575 | *Tent4a* | 1353.902006 | 1412.888845 | 1031.144844 | 1887.810498 | 1992.879006 | 1898.144058 | 8.41E-06 | 0.60329257 | 2.13E-06 | 22.47788493 | 1.519179739 | 5.075166394 |
| ENSMUSG00000073435 | *Nme3* | 361.6432737 | 360.6559443 | 418.5086654 | 604.757898 | 580.0720191 | 543.6387433 | 3.02E-06 | 0.603152362 | 7.21E-07 | 24.55933267 | 1.519032105 | 5.519512454 |
| ENSMUSG00000028960 | *Ube4b* | 2625.680852 | 2582.14309 | 1916.063769 | 3252.082853 | 3936.278184 | 3638.339022 | 3.66E-05 | 0.60271174 | 1.02E-05 | 19.4718939 | 1.51856824 | 4.436126894 |
| ENSMUSG00000029056 | *Pank4* | 681.8482557 | 745.2916721 | 729.8690279 | 1088.783729 | 1081.186867 | 1101.052117 | 5.68E-11 | 0.602627282 | 8.15E-12 | 46.72825119 | 1.518479343 | 10.24585314 |
| ENSMUSG00000019849 | *Prep* | 4196.568822 | 4387.341194 | 4241.812226 | 6087.091293 | 6761.892158 | 6623.760567 | 1.09E-16 | 0.602581593 | 9.80E-18 | 73.55252955 | 1.518431255 | 15.96387996 |
| ENSMUSG00000028524 | *Sgip1* | 803.9028606 | 676.2298955 | 728.6084596 | 1183.174254 | 1102.242113 | 1073.502856 | 5.92E-08 | 0.602520507 | 1.15E-08 | 32.57742538 | 1.518366962 | 7.227782565 |
| ENSMUSG00000051786 | *Tubgcp6* | 784.3138499 | 752.9652028 | 586.1642452 | 1062.442187 | 1145.405366 | 1024.832496 | 1.86E-05 | 0.60220178 | 4.96E-06 | 20.85132949 | 1.518031555 | 4.729526624 |
| ENSMUSG00000027883 | *Gpsm2* | 749.6563695 | 672.3931301 | 661.7983414 | 1056.954366 | 1082.239629 | 1031.260657 | 2.22E-09 | 0.602115516 | 3.72E-10 | 39.25696095 | 1.517940789 | 8.654574798 |
| ENSMUSG00000021022 | *Ppp2r3c* | 676.5742913 | 561.1269346 | 772.728349 | 1067.930009 | 941.169483 | 1037.688817 | 3.79E-05 | 0.602074074 | 1.06E-05 | 19.40272182 | 1.517897186 | 4.42114595 |
| ENSMUSG00000031931 | *Ankrd49* | 519.1087825 | 509.3306021 | 495.4033298 | 827.5634393 | 732.7225505 | 754.8497415 | 3.42E-08 | 0.601472694 | 6.45E-09 | 33.69364905 | 1.51726459 | 7.465389681 |
| ENSMUSG00000025404 | *R3hdm2* | 1577.668782 | 1571.155417 | 1070.222461 | 2098.542833 | 2165.532021 | 2145.169095 | 0.000161756 | 0.601176096 | 4.97E-05 | 16.45837398 | 1.516952694 | 3.791140131 |
| ENSMUSG00000020131 | *Pcsk4* | 98.69847679 | 92.08236875 | 93.28205192 | 133.9028377 | 158.9671051 | 138.6646119 | 0.006834122 | 0.600799431 | 0.00285568 | 8.897457525 | 1.516556694 | 2.165317269 |
| ENSMUSG00000048668 | *Rhno1* | 682.6016792 | 763.5163075 | 658.0166365 | 1035.003081 | 1110.664211 | 1046.871904 | 1.19E-08 | 0.600310087 | 2.14E-09 | 35.84053921 | 1.516042384 | 7.925562731 |
| ENSMUSG00000048546 | *Tob2* | 1043.491529 | 1129.9274 | 866.010401 | 1475.126343 | 1609.673534 | 1528.983966 | 1.21E-06 | 0.599843326 | 2.75E-07 | 26.41530224 | 1.515551972 | 5.915897913 |
| ENSMUSG00000021235 | *Coq6* | 650.9578927 | 647.4541553 | 591.2065183 | 964.7589697 | 966.4357778 | 938.5114791 | 7.10E-10 | 0.599646925 | 1.13E-10 | 41.57750252 | 1.515345666 | 9.148527367 |
| ENSMUSG00000029108 | *Pcdh7* | 5826.223825 | 5982.476395 | 4246.854499 | 7906.852808 | 8078.897777 | 8339.161196 | 1.36E-05 | 0.598784599 | 3.57E-06 | 21.48481432 | 1.514440186 | 4.864945572 |
| ENSMUSG00000032198 | *Dock6* | 427.1911171 | 352.0232222 | 310.0997942 | 532.3186579 | 565.3333471 | 558.3316823 | 0.000152116 | 0.597829568 | 4.66E-05 | 16.58208444 | 1.513437993 | 3.817823835 |
| ENSMUSG00000034880 | *Mrpl34* | 702.9441133 | 907.3950087 | 1022.320866 | 1263.296444 | 1289.633799 | 1420.62354 | 0.000137992 | 0.597428321 | 4.19E-05 | 16.78188605 | 1.513017129 | 3.86014632 |
| ENSMUSG00000004936 | *Map2k1* | 3456.706958 | 3773.458736 | 3296.386024 | 5464.772367 | 5085.8946 | 5372.105825 | 3.52E-13 | 0.596614005 | 4.18E-14 | 57.0801269 | 1.512163362 | 12.45317333 |
| ENSMUSG00000044950 | *Pwwp2a* | 257.6708325 | 241.716218 | 175.2189894 | 375.3669712 | 312.6703987 | 337.9375972 | 0.003064327 | 0.596473019 | 0.001191916 | 10.5028335 | 1.512015595 | 2.513664932 |
| ENSMUSG00000001833 | *Septin7* | 2408.694888 | 2191.752214 | 2333.311866 | 3666.962136 | 3236.191265 | 3582.322192 | 6.94E-12 | 0.596467045 | 9.25E-13 | 50.99742171 | 1.512009333 | 11.15887121 |
| ENSMUSG00000057522 | *Spop* | 1646.983742 | 1640.217193 | 1513.942491 | 2383.909536 | 2476.096895 | 2406.887071 | 6.85E-15 | 0.596316275 | 7.06E-16 | 65.11623599 | 1.511851328 | 14.16458638 |
| ENSMUSG00000095865 | *Gm13237* | 756.4371809 | 564.0045086 | 681.9674337 | 1017.442053 | 1027.49599 | 985.3452222 | 1.11E-05 | 0.596231331 | 2.87E-06 | 21.90432055 | 1.511762315 | 4.952906599 |
| ENSMUSG00000034793 | *G6pc3* | 835.546647 | 822.0269793 | 755.0803933 | 1149.149763 | 1289.633799 | 1214.004086 | 2.27E-09 | 0.596193506 | 3.81E-10 | 39.20703731 | 1.51172268 | 8.644003026 |
| ENSMUSG00000024642 | *Tle4* | 693.149608 | 701.1688704 | 535.7415144 | 942.8076849 | 968.5413024 | 1013.812792 | 1.24E-05 | 0.596114034 | 3.21E-06 | 21.68877033 | 1.511639407 | 4.907923859 |
| ENSMUSG00000024960 | *Plcb3* | 1200.957038 | 1196.111602 | 864.7498327 | 1524.516734 | 1707.580427 | 1704.380925 | 7.58E-05 | 0.595420893 | 2.21E-05 | 17.9969044 | 1.510913317 | 4.120366603 |
| ENSMUSG00000041164 | *Zmiz2* | 3525.268495 | 3500.089204 | 2706.440074 | 4272.817599 | 5385.931851 | 5051.616093 | 2.19E-05 | 0.595326561 | 5.89E-06 | 20.52339405 | 1.510814527 | 4.660154451 |
| ENSMUSG00000049489 | *Ccnq* | 467.1225619 | 440.2688256 | 500.4456029 | 664.0263671 | 749.5667471 | 708.9343071 | 4.31E-07 | 0.594753149 | 9.23E-08 | 28.52906172 | 1.510214159 | 6.365838889 |
| ENSMUSG00000041132 | *N4bp2l1* | 284.7940781 | 235.9610699 | 244.5502442 | 437.928133 | 349.5170787 | 371.9150186 | 0.000462226 | 0.594619584 | 0.000152796 | 14.33743517 | 1.51007435 | 3.335145406 |
| ENSMUSG00000041961 | *Znrf3* | 192.1229892 | 206.2261383 | 134.8808048 | 251.3422117 | 277.9292433 | 280.0841498 | 0.005562733 | 0.594313617 | 0.002283958 | 9.306071015 | 1.509754128 | 2.254711821 |
| ENSMUSG00000051149 | *Adnp* | 997.5326967 | 1095.396512 | 704.6576625 | 1261.101315 | 1438.073282 | 1529.902274 | 0.000962651 | 0.594183077 | 0.000337443 | 12.85020088 | 1.509617526 | 3.016531281 |
| ENSMUSG00000041528 | *Rnf123* | 1558.079771 | 1438.787012 | 1274.43452 | 2098.542833 | 2232.908807 | 2124.966304 | 6.64E-09 | 0.593572253 | 1.17E-09 | 37.01924347 | 1.508978502 | 8.177666221 |
| ENSMUSG00000027454 | *Gins1* | 445.2732808 | 335.7169694 | 399.6001413 | 569.6358422 | 620.076986 | 593.2274125 | 6.61E-05 | 0.592794279 | 1.91E-05 | 18.27743337 | 1.508165005 | 4.180036051 |
| ENSMUSG00000029790 | *Cep41* | 270.4790318 | 286.798211 | 229.423425 | 362.1962002 | 439.0018729 | 388.444575 | 0.000307434 | 0.592208711 | 9.84E-05 | 15.16636543 | 1.507552987 | 3.512247974 |
| ENSMUSG00000101330 | *Gm10193* | 432.4650815 | 428.7585295 | 361.7830933 | 576.2212277 | 601.1272648 | 670.3653422 | 1.61E-05 | 0.591968627 | 4.25E-06 | 21.1487338 | 1.507302131 | 4.793232537 |
| ENSMUSG00000045983 | *Eif4g1* | 18600.51905 | 20554.51125 | 17511.8144 | 26595.07918 | 30489.04854 | 28326.14979 | 1.75E-10 | 0.591827074 | 2.62E-11 | 44.44095117 | 1.507154246 | 9.757993571 |
| ENSMUSG00000032637 | *Atxn2l* | 4571.020295 | 4240.584919 | 2841.320879 | 5292.454781 | 6219.719581 | 6059.000723 | 0.000880566 | 0.591805116 | 0.000306192 | 13.03213209 | 1.507131308 | 3.055238245 |
| ENSMUSG00000037103 | *Dcaf15* | 981.7108035 | 910.2725827 | 854.6652865 | 1313.784399 | 1441.231568 | 1391.237662 | 2.31E-09 | 0.59154223 | 3.88E-10 | 39.17085902 | 1.506856705 | 8.63701925 |
| ENSMUSG00000025645 | *Ccdc51* | 399.3144481 | 479.5956706 | 466.4102596 | 698.0508587 | 610.6021254 | 713.5258506 | 1.58E-05 | 0.591461083 | 4.17E-06 | 21.18440545 | 1.506771951 | 4.800977269 |
| ENSMUSG00000066979 | *Bub3* | 5390.745049 | 5286.103481 | 5758.275854 | 8321.732092 | 8236.812119 | 8195.90504 | 3.42E-20 | 0.591455834 | 2.46E-21 | 89.93663057 | 1.506766469 | 19.46606305 |
| ENSMUSG00000029550 | *Sppl3* | 924.4506185 | 966.8648719 | 770.2072125 | 1238.052466 | 1409.6487 | 1368.279945 | 1.68E-06 | 0.591144005 | 3.87E-07 | 25.75966596 | 1.506440827 | 5.775849788 |
| ENSMUSG00000001062 | *Vps9d1* | 348.081651 | 450.8199303 | 267.2404731 | 515.8551943 | 566.3861094 | 527.1091869 | 0.00444411 | 0.590518485 | 0.001790209 | 9.753096558 | 1.50578781 | 2.352215185 |
| ENSMUSG00000023262 | *Acy1* | 669.0400564 | 685.8218089 | 907.6091539 | 1170.003483 | 1061.184383 | 1166.252034 | 6.96E-05 | 0.590472764 | 2.02E-05 | 18.16787507 | 1.505740091 | 4.157214834 |
| ENSMUSG00000029111 | *Nelfa* | 678.0811383 | 632.1070938 | 490.3610567 | 841.8317745 | 938.0111961 | 938.5114791 | 9.36E-05 | 0.590132193 | 2.76E-05 | 17.57354073 | 1.505384678 | 4.028903995 |
| ENSMUSG00000032392 | *Parp16* | 155.9586618 | 127.5724484 | 163.873875 | 262.3178541 | 211.6052193 | 199.2729853 | 0.006519591 | 0.589658807 | 0.002712171 | 8.991642293 | 1.504890803 | 2.185779626 |
| ENSMUSG00000016528 | *Mapkapk2* | 9922.587323 | 9829.792864 | 8029.819875 | 13471.50352 | 13935.41437 | 14386.22391 | 8.07E-09 | 0.588717585 | 1.43E-09 | 36.62825282 | 1.503909324 | 8.093100327 |
| ENSMUSG00000001436 | *Slc19a1* | 2314.516952 | 2206.140085 | 2009.345821 | 3079.765266 | 3396.211132 | 3351.826711 | 7.95E-11 | 0.588490911 | 1.16E-11 | 46.04068124 | 1.503673051 | 10.09963692 |
| ENSMUSG00000047260 | *Emc6* | 1465.408682 | 1337.11273 | 1449.65351 | 2063.420777 | 2140.265726 | 2187.411295 | 5.23E-13 | 0.588076734 | 6.31E-14 | 56.27275551 | 1.50324143 | 12.2815839 |
| ENSMUSG00000057691 | *Zfp746* | 693.9030315 | 793.2512391 | 613.8967471 | 1014.149361 | 1086.450678 | 1062.483152 | 2.65E-06 | 0.588068806 | 6.26E-07 | 24.83171969 | 1.50323317 | 5.576872228 |
| ENSMUSG00000028906 | *Epb41* | 4907.800594 | 4688.527275 | 4277.108138 | 6921.240117 | 7016.660631 | 6928.639051 | 1.74E-14 | 0.587937255 | 1.85E-15 | 63.2190098 | 1.503096105 | 13.75974699 |
| ENSMUSG00000070699 | *Sars2* | 628.3551881 | 587.9842921 | 484.0582154 | 806.7097187 | 894.8479424 | 862.291858 | 1.34E-05 | 0.587931074 | 3.50E-06 | 21.51913728 | 1.503089665 | 4.872148155 |
| ENSMUSG00000037731 | *Themis2* | 1133.902348 | 1059.906432 | 791.6368731 | 1534.394812 | 1458.075765 | 1504.189631 | 7.30E-05 | 0.587774294 | 2.13E-05 | 18.07247114 | 1.50292633 | 4.136480508 |
| ENSMUSG00000024654 | *Asrgl1* | 553.7662629 | 575.5148047 | 560.9528798 | 879.1489588 | 828.5239184 | 831.9876713 | 2.00E-09 | 0.587363328 | 3.35E-10 | 39.46135484 | 1.502498268 | 8.698432658 |
| ENSMUSG00000030990 | *Pgap2* | 1371.230746 | 1367.806852 | 1280.737362 | 1953.664353 | 2177.112406 | 1913.755306 | 1.69E-10 | 0.587147748 | 2.53E-11 | 44.51342003 | 1.502273769 | 9.773067467 |
| ENSMUSG00000049305 | *Ccdc71* | 1133.148924 | 1127.049826 | 1270.652815 | 1728.663683 | 1792.854172 | 1771.417459 | 3.58E-11 | 0.586526817 | 5.05E-12 | 47.6667949 | 1.501627334 | 10.44627244 |
| ENSMUSG00000058799 | *Nap1l1* | 7771.563268 | 7039.505252 | 6863.794226 | 10996.49615 | 10755.01951 | 10808.49326 | 2.01E-15 | 0.586507964 | 1.99E-16 | 67.61533204 | 1.501607711 | 14.69721267 |
| ENSMUSG00000085795 | *Zfp703* | 367.6706616 | 414.3706594 | 296.2335433 | 495.0014737 | 615.8659368 | 512.4162479 | 0.001020843 | 0.58625781 | 0.000359943 | 12.72944567 | 1.501347364 | 2.991040948 |
| ENSMUSG00000041836 | *Ptpre* | 287.807772 | 326.125056 | 282.3672923 | 439.0256973 | 469.5319792 | 437.1149355 | 1.65E-05 | 0.585058095 | 4.36E-06 | 21.09871479 | 1.500099395 | 4.7822354 |
| ENSMUSG00000032393 | *Dpp8* | 2471.982461 | 2483.346382 | 2697.616096 | 3886.474985 | 4017.34088 | 3570.384179 | 4.35E-12 | 0.584972589 | 5.67E-13 | 51.95668951 | 1.500010489 | 11.36181408 |
| ENSMUSG00000062526 | *Mppe1* | 470.8896793 | 459.4526524 | 518.0935587 | 335.8546584 | 310.5648741 | 317.734806 | 6.22E-06 | -0.585026475 | 1.54E-06 | 23.09118683 | 0.666637105 | 5.206508573 |
| ENSMUSG00000075307 | *Klhl41* | 510.8211241 | 522.7592809 | 687.0097067 | 379.7572281 | 395.8386192 | 368.2417839 | 0.000340657 | -0.585061605 | 0.000110108 | 14.95496401 | 0.666620873 | 3.467682878 |
| ENSMUSG00000021262 | *Evl* | 467.1225619 | 463.2894178 | 370.6070712 | 309.5131166 | 272.6654319 | 286.5123107 | 0.000227488 | -0.587469409 | 7.15E-05 | 15.77032388 | 0.665509235 | 3.643040936 |
| ENSMUSG00000063904 | *Dpp3* | 3758.829776 | 3932.684499 | 3564.887065 | 2490.373268 | 2490.835567 | 2511.574262 | 2.59E-16 | -0.587697392 | 2.38E-17 | 71.80247513 | 0.665404076 | 15.5872757 |
| ENSMUSG00000031299 | *Pdha1* | 2102.051529 | 2262.732374 | 2697.616096 | 1654.029314 | 1474.919962 | 1565.716313 | 1.14E-06 | -0.587754233 | 2.58E-07 | 26.53862094 | 0.66537786 | 5.942438734 |
| ENSMUSG00000058407 | *Txndc9* | 2071.161166 | 1919.341874 | 2566.516996 | 1504.760577 | 1444.389855 | 1407.767219 | 5.15E-06 | -0.588583846 | 1.26E-06 | 23.48027869 | 0.664995348 | 5.288208617 |
| ENSMUSG00000037190 | *Cyb561d2* | 649.4510457 | 751.0468201 | 734.911301 | 459.8794179 | 461.1098809 | 495.8866915 | 9.05E-07 | -0.588793303 | 2.02E-07 | 27.01097303 | 0.664898808 | 6.043161304 |
| ENSMUSG00000046671 | *Mtfr1l* | 2201.503429 | 2480.468808 | 2499.706878 | 1550.858276 | 1634.939829 | 1582.245869 | 2.66E-11 | -0.589703787 | 3.71E-12 | 48.27004206 | 0.664479323 | 10.57589449 |
| ENSMUSG00000074151 | *Nlrc5* | 1153.491359 | 1176.927776 | 845.8413087 | 752.9290708 | 644.2905185 | 716.2807766 | 0.000237078 | -0.589998278 | 7.46E-05 | 15.68933134 | 0.6643437 | 3.625109242 |
| ENSMUSG00000050989 | *Selenon* | 2193.969194 | 2156.262135 | 1825.302854 | 1355.49184 | 1409.6487 | 1341.648993 | 1.05E-08 | -0.590291463 | 1.89E-09 | 36.08857695 | 0.664208705 | 7.977197894 |
| ENSMUSG00000028463 | *Car9* | 161.2326262 | 159.2257626 | 168.9161481 | 119.6345025 | 96.85413024 | 108.3604252 | 0.00491949 | -0.590359952 | 0.001997234 | 9.552075674 | 0.664177174 | 2.308079962 |
| ENSMUSG00000021819 | *Zswim8* | 2945.13241 | 2841.124752 | 2159.353445 | 1713.297784 | 1842.333999 | 1725.502025 | 1.27E-05 | -0.590587729 | 3.30E-06 | 21.63201387 | 0.66407232 | 4.896106505 |
| ENSMUSG00000026727 | *Rsu1* | 1873.010789 | 2076.649254 | 2432.89676 | 1385.126075 | 1422.281847 | 1424.296775 | 5.57E-07 | -0.591075487 | 1.21E-07 | 28.0082354 | 0.663847843 | 6.254512156 |
| ENSMUSG00000040760 | *Appl1* | 825.7521417 | 729.9446106 | 857.1864231 | 592.6846913 | 526.3811426 | 483.0303699 | 8.73E-06 | -0.591439402 | 2.21E-06 | 22.40066528 | 0.663680411 | 5.058883944 |
| ENSMUSG00000026150 | *Mff* | 2522.461834 | 2216.691189 | 2853.926562 | 1738.541761 | 1552.824371 | 1743.868199 | 8.90E-07 | -0.591709699 | 1.99E-07 | 27.04689717 | 0.663556079 | 6.050698667 |
| ENSMUSG00000078193 | *Gm2000* | 14140.252 | 13498.69974 | 19063.57394 | 10582.71443 | 10008.61105 | 10391.58111 | 2.02E-05 | -0.591808119 | 5.40E-06 | 20.68908742 | 0.663510813 | 4.695300626 |
| ENSMUSG00000014504 | *Srp19* | 1133.902348 | 989.885464 | 1338.723502 | 775.9779199 | 747.4612225 | 769.5426805 | 1.40E-05 | -0.59273042 | 3.67E-06 | 21.4321752 | 0.663086772 | 4.853820951 |
| ENSMUSG00000030105 | *Arl8b* | 5548.963981 | 5644.841043 | 6325.531575 | 3679.035343 | 4051.029273 | 3878.935898 | 5.46E-12 | -0.593042291 | 7.21E-13 | 51.48696726 | 0.662943446 | 11.26290745 |
| ENSMUSG00000053768 | *Chchd3* | 3530.54246 | 3531.742518 | 4597.292478 | 2705.495859 | 2442.408502 | 2576.774179 | 3.60E-06 | -0.593123145 | 8.65E-07 | 24.20712516 | 0.662906294 | 5.443418015 |
| ENSMUSG00000021468 | *Sptlc1* | 1734.380867 | 1798.483765 | 1922.366611 | 1278.662343 | 1165.40785 | 1169.925269 | 6.04E-11 | -0.593184024 | 8.69E-12 | 46.60312059 | 0.662878321 | 10.219158 |
| ENSMUSG00000038059 | *Smim3* | 973.4231451 | 1029.212309 | 1028.623708 | 749.6363781 | 574.8082077 | 684.1399726 | 3.97E-06 | -0.593286201 | 9.58E-07 | 24.01109729 | 0.662831375 | 5.400874842 |
| ENSMUSG00000062997 | *Rpl35* | 9285.191053 | 8918.56109 | 11492.60091 | 7160.509122 | 5947.054149 | 6567.743737 | 2.42E-06 | -0.593632171 | 5.69E-07 | 25.01343083 | 0.662672442 | 5.615909101 |
| ENSMUSG00000030104 | *Edem1* | 14483.05969 | 13861.27407 | 13286.38956 | 9117.466168 | 9005.328588 | 9461.334413 | 6.59E-20 | -0.593998758 | 4.84E-21 | 88.59624797 | 0.662504079 | 19.18123948 |
| ENSMUSG00000021009 | *Ptpn21* | 323.9720994 | 384.6357278 | 296.2335433 | 215.1225917 | 227.3966536 | 224.0673199 | 0.000469611 | -0.594033445 | 0.000155592 | 14.30330223 | 0.66248815 | 3.328261696 |
| ENSMUSG00000066152 | *Slc31a2* | 523.6293234 | 589.9026748 | 616.4178837 | 428.0500548 | 345.3060295 | 370.99671 | 4.28E-05 | -0.594431133 | 1.20E-05 | 19.15779363 | 0.662305556 | 4.36883622 |
| ENSMUSG00000047264 | *Zfp358* | 284.7940781 | 244.593792 | 243.289676 | 145.9760443 | 166.3364411 | 199.2729853 | 0.002988343 | -0.594748833 | 0.0011599 | 10.5531536 | 0.662159724 | 2.524569613 |
| ENSMUSG00000081855 | *Rpl17-ps5* | 4807.59527 | 4354.728689 | 6674.708986 | 3486.961601 | 3418.31914 | 3576.81234 | 0.000211965 | -0.594784258 | 6.64E-05 | 15.91193507 | 0.662143465 | 3.673736677 |
| ENSMUSG00000027708 | *Dcun1d1* | 682.6016792 | 675.2707042 | 818.1088067 | 477.4404458 | 466.3736923 | 493.1317655 | 3.48E-06 | -0.594985091 | 8.34E-07 | 24.2768024 | 0.662051297 | 5.458440348 |
| ENSMUSG00000024121 | *Atp6v0c* | 14353.47085 | 15611.79827 | 15686.51154 | 9691.492267 | 10251.79913 | 10257.50805 | 8.65E-18 | -0.595815739 | 7.26E-19 | 78.69270728 | 0.661670223 | 17.06319238 |
| ENSMUSG00000021891 | *Mettl6* | 470.8896793 | 466.1669918 | 636.586976 | 387.4401778 | 371.6250867 | 281.0024585 | 0.003458878 | -0.595916565 | 0.001360812 | 10.2581621 | 0.661623982 | 2.461064705 |
| ENSMUSG00000018286 | *Psmb6* | 9530.80711 | 9828.833672 | 11720.76377 | 6943.191402 | 6742.942437 | 6865.275752 | 1.34E-09 | -0.596358326 | 2.20E-10 | 40.27633297 | 0.66142142 | 8.871723509 |
| ENSMUSG00000030760 | *Acer3* | 955.3409814 | 995.6406121 | 1048.7928 | 705.7338084 | 651.6598545 | 624.4499078 | 9.52E-09 | -0.597362548 | 1.69E-09 | 36.29698316 | 0.660961183 | 8.021470946 |
| ENSMUSG00000043421 | *Hilpda* | 320.204982 | 301.1860811 | 417.2480971 | 238.1714408 | 211.6052193 | 234.1687154 | 0.001649298 | -0.597534607 | 0.000608017 | 11.75127035 | 0.66088236 | 2.782700941 |
| ENSMUSG00000046861 | *Hectd3* | 1559.586618 | 1560.604312 | 1396.709642 | 981.2224334 | 1028.548753 | 978.9170614 | 8.77E-11 | -0.597658209 | 1.28E-11 | 45.83833928 | 0.660825742 | 10.05722809 |
| ENSMUSG00000009030 | *Pdcl* | 1036.710718 | 1008.1101 | 1350.068616 | 750.7339423 | 749.5667471 | 739.2384938 | 2.73E-05 | -0.597692146 | 7.46E-06 | 20.07178163 | 0.660810197 | 4.563906531 |
| ENSMUSG00000024854 | *Pold4* | 493.4923839 | 510.2897935 | 526.9175365 | 315.0009378 | 357.939177 | 337.0192885 | 1.75E-06 | -0.597852493 | 4.06E-07 | 25.66561977 | 0.660736756 | 5.756361577 |
| ENSMUSG00000020412 | *Ascc2* | 3028.762418 | 3105.861563 | 2829.975764 | 1892.200755 | 2008.67044 | 2023.952348 | 2.32E-14 | -0.597906856 | 2.49E-15 | 62.63701328 | 0.660711859 | 13.63532406 |
| ENSMUSG00000096592 | *Gm15801* | 247.1229037 | 297.3493157 | 317.6632038 | 171.2200219 | 203.183121 | 192.8448245 | 0.001301838 | -0.598051246 | 0.000469205 | 12.23423795 | 0.660645735 | 2.885442892 |
| ENSMUSG00000031897 | *Psmb10* | 3526.021919 | 3164.372234 | 2795.940421 | 2180.860151 | 1929.713269 | 2160.780343 | 1.71E-07 | -0.598068686 | 3.50E-08 | 30.40836065 | 0.660637749 | 6.767191339 |
| ENSMUSG00000022724 | *Riox2* | 1381.778675 | 1408.092889 | 1530.329879 | 969.1492267 | 925.3780487 | 954.1227268 | 2.15E-11 | -0.599076534 | 2.97E-12 | 48.70599388 | 0.660176398 | 10.66812491 |
| ENSMUSG00000025277 | *Abhd6* | 588.4237433 | 601.4129709 | 705.9182308 | 430.2451833 | 408.4717667 | 409.5656749 | 3.60E-06 | -0.599561959 | 8.65E-07 | 24.20687523 | 0.659954305 | 5.443418015 |
| ENSMUSG00000020423 | *Btg2* | 334.5200282 | 343.3905001 | 293.7124067 | 233.7811838 | 175.8113016 | 232.3320981 | 0.001223795 | -0.59971793 | 0.000438474 | 12.36066913 | 0.65988296 | 2.912291167 |
| ENSMUSG00000028029 | *Aimp1* | 2596.297336 | 2421.958136 | 3034.187824 | 1886.712934 | 1727.58291 | 1692.442912 | 1.16E-07 | -0.600884871 | 2.33E-08 | 31.19722611 | 0.659349422 | 6.934025865 |
| ENSMUSG00000022861 | *Dgkg* | 262.9447969 | 237.8794526 | 244.5502442 | 158.049251 | 160.0198673 | 173.560342 | 0.000370098 | -0.600902935 | 0.000120233 | 14.78902062 | 0.659341166 | 3.43168362 |
| ENSMUSG00000029777 | *Gars* | 13403.40383 | 12278.60836 | 11497.64318 | 8304.171064 | 8010.468228 | 8205.088127 | 5.97E-14 | -0.600925928 | 6.60E-15 | 60.71268353 | 0.659330658 | 13.22401745 |
| ENSMUSG00000001630 | *Stk38l* | 654.7250102 | 685.8218089 | 734.911301 | 429.1476191 | 416.8938649 | 518.8444087 | 1.18E-05 | -0.60100357 | 3.05E-06 | 21.78477063 | 0.659295176 | 4.928044769 |
| ENSMUSG00000097175 | *Gm27018* | 1051.025764 | 933.2931749 | 1207.624402 | 780.3681769 | 710.6145425 | 612.5118949 | 5.68E-05 | -0.601307599 | 1.63E-05 | 18.58474229 | 0.659156253 | 4.245616366 |
| ENSMUSG00000020803 | *Txndc17* | 2007.120169 | 1943.321657 | 2482.058922 | 1472.931214 | 1312.79457 | 1449.09111 | 1.47E-06 | -0.601647136 | 3.38E-07 | 26.02021145 | 0.659001139 | 5.8320081 |
| ENSMUSG00000038736 | *Nudcd1* | 397.0541776 | 389.4316845 | 432.3749163 | 284.269139 | 232.660465 | 284.6756933 | 8.59E-05 | -0.601860624 | 2.53E-05 | 17.74420473 | 0.658903628 | 4.066038129 |
| ENSMUSG00000033157 | *Abhd10* | 381.2322844 | 385.5949191 | 436.1566212 | 242.5616977 | 275.8237187 | 270.901063 | 3.67E-05 | -0.603649624 | 1.02E-05 | 19.47048287 | 0.658087067 | 4.435912994 |
| ENSMUSG00000034111 | *Tmed8* | 1115.066761 | 1221.050577 | 1033.665981 | 691.4654732 | 826.4183939 | 701.5878376 | 1.18E-06 | -0.603680536 | 2.68E-07 | 26.46927822 | 0.658072967 | 5.927376072 |
| ENSMUSG00000049969 | *Plekhf2* | 731.5742058 | 703.087253 | 855.9258548 | 496.0990379 | 514.8007575 | 493.1317655 | 1.21E-06 | -0.604121453 | 2.74E-07 | 26.42506637 | 0.657871877 | 5.917961868 |
| ENSMUSG00000059316 | *Slc27a4* | 1741.161678 | 1808.075678 | 1759.753304 | 1187.564511 | 1118.033547 | 1185.536516 | 1.28E-14 | -0.604456478 | 1.35E-15 | 63.84156076 | 0.657719123 | 13.89334003 |
| ENSMUSG00000037965 | *Zc3h7a* | 1126.368113 | 1119.376295 | 1039.968822 | 752.9290708 | 652.7126168 | 756.6863589 | 1.98E-08 | -0.604460614 | 3.65E-09 | 34.80219248 | 0.657717237 | 7.702529731 |
| ENSMUSG00000038780 | *Smurf1* | 1236.367942 | 1322.724859 | 1034.926549 | 794.6365121 | 799.0465745 | 773.2159153 | 7.80E-07 | -0.604791492 | 1.73E-07 | 27.31587086 | 0.657566409 | 6.107767826 |
| ENSMUSG00000041571 | *Selenow* | 2047.805038 | 2057.465427 | 1750.929326 | 1359.882097 | 1168.566137 | 1325.119437 | 6.23E-08 | -0.604959709 | 1.21E-08 | 32.47250173 | 0.657489742 | 7.205412511 |
| ENSMUSG00000001039 | *B9d1* | 231.3010105 | 215.8180518 | 278.5855875 | 164.6346365 | 146.3339576 | 164.3772552 | 0.001742923 | -0.605019628 | 0.000646838 | 11.63610687 | 0.657462435 | 2.758721723 |
| ENSMUSG00000024754 | *Cemip2* | 577.8758145 | 590.8618661 | 434.8960529 | 345.7327366 | 322.1452593 | 387.5262663 | 0.00049165 | -0.606080556 | 0.000163535 | 14.20959425 | 0.656979129 | 3.308344258 |
| ENSMUSG00000021548 | *Ccnh* | 968.1491807 | 989.885464 | 1019.79973 | 651.9531604 | 673.7678625 | 628.1231426 | 9.75E-11 | -0.607383468 | 1.44E-11 | 45.61996504 | 0.656386072 | 10.01104466 |
| ENSMUSG00000024194 | *Cuta* | 2161.571984 | 2355.773934 | 2587.946657 | 1550.858276 | 1460.18129 | 1646.527478 | 4.77E-09 | -0.607609975 | 8.29E-10 | 37.69033404 | 0.656283026 | 8.321486075 |

| | | | | | | | | | | | | |
|---|---|---|---|---|---|---|---|---|---|---|---|---|
| ENSMUSG00000008668 | *Rps18* | 9656.628832 | 8974.194188 | 10898.87326 | 6832.337414 | 6157.606606 | 6385.000308 | 4.33E-10 | -0.607737758 | 6.74E-11 | 42.59185688 | 0.6562249 | 9.363591966 |
| ENSMUSG00000073702 | *Rpl31* | 8103.069602 | 7744.510888 | 10896.35212 | 6481.116856 | 5585.956685 | 5467.609928 | 4.42E-05 | -0.608746692 | 1.24E-05 | 19.09352179 | 0.655766136 | 4.354740971 |
| ENSMUSG00000024968 | *Rcor2* | 351.8487684 | 310.7779945 | 273.5433144 | 200.8542565 | 196.8665473 | 217.6391591 | 0.000448341 | -0.609692378 | 0.000148021 | 14.39720964 | 0.655336423 | 3.348391434 |
| ENSMUSG00000027078 | *Ube2l6* | 3551.638317 | 3099.147223 | 2579.122679 | 2211.59195 | 1764.42959 | 2076.295944 | 3.39E-05 | -0.609767142 | 9.40E-06 | 19.63017771 | 0.655302462 | 4.469750869 |
| ENSMUSG00000024180 | *Pgap6* | 332.2597577 | 374.084623 | 298.7546798 | 194.268871 | 220.0273176 | 244.270111 | 0.000581604 | -0.610017512 | 0.000196848 | 13.86093401 | 0.655188749 | 3.235372797 |
| ENSMUSG00000038080 | *Kdm1b* | 531.1635583 | 637.8622418 | 642.8898173 | 384.1474851 | 418.9993895 | 381.0981055 | 9.45E-06 | -0.610818652 | 2.41E-06 | 22.23883048 | 0.654825019 | 5.024396077 |
| ENSMUSG00000033318 | *Gstt2* | 281.0269606 | 301.1860811 | 313.881499 | 205.2445135 | 180.0223508 | 200.191294 | 0.000123638 | -0.610909874 | 3.73E-05 | 17.00457029 | 0.654783615 | 3.907847902 |
| ENSMUSG00000013878 | *Rnf170* | 190.6161422 | 255.1448967 | 297.4941115 | 150.3663013 | 157.9143428 | 175.3969594 | 0.00889028 | -0.611652473 | 0.003812473 | 8.371040244 | 0.654446665 | 2.05108455 |
| ENSMUSG00000026615 | *Eprs* | 11090.39373 | 10250.87786 | 10805.5912 | 6981.606151 | 7098.776089 | 6958.024929 | 4.65E-23 | -0.61175803 | 2.85E-24 | 103.3181613 | 0.654398783 | 22.3323266 |
| ENSMUSG00000029254 | *Stap1* | 3959.240424 | 2744.246427 | 2644.672229 | 2089.762319 | 1937.082605 | 2093.743809 | 0.000221969 | -0.611904944 | 6.97E-05 | 15.81968678 | 0.654332147 | 3.653707556 |
| ENSMUSG00000041362 | *Shtn1* | 2230.886945 | 2343.304446 | 2814.848945 | 1534.394812 | 1678.103083 | 1616.223291 | 1.09E-07 | -0.61221545 | 2.17E-08 | 31.33308662 | 0.654191332 | 6.962927835 |
| ENSMUSG00000055093 | *Rps27a-ps3* | 10010.73787 | 8210.67788 | 12744.3452 | 7262.582597 | 6568.183897 | 6422.650964 | 0.000117778 | -0.612324696 | 3.54E-05 | 17.10457077 | 0.654141796 | 3.928935622 |
| ENSMUSG00000037499 | *Nenf* | 546.9854515 | 563.0453172 | 535.7415144 | 339.1473511 | 361.0974638 | 375.5882534 | 2.30E-07 | -0.612695833 | 4.78E-08 | 29.80498641 | 0.653973539 | 6.638568876 |
| ENSMUSG00000029767 | *Calu* | 6443.27766 | 6895.626551 | 8578.167072 | 4886.356011 | 4617.415383 | 4814.692451 | 3.47E-07 | -0.613631455 | 7.35E-08 | 28.97091913 | 0.653549559 | 6.459889348 |
| ENSMUSG00000002257 | *Def6* | 2410.201735 | 2347.141212 | 2163.13515 | 1497.077628 | 1458.075765 | 1567.55293 | 3.45E-13 | -0.614564418 | 4.09E-14 | 57.12316143 | 0.653127058 | 12.46242556 |
| ENSMUSG00000071715 | *Ncf4* | 4541.636779 | 4662.629109 | 4943.948752 | 3083.057959 | 3039.324717 | 3110.311526 | 1.58E-19 | -0.614993563 | 1.19E-20 | 86.80968461 | 0.652932807 | 18.80214378 |
| ENSMUSG00000026078 | *Pdcl3* | 992.2587323 | 1040.722605 | 1453.435214 | 751.8315066 | 739.0391242 | 780.5623848 | 0.000245521 | -0.615431407 | 7.76E-05 | 15.61615942 | 0.652734678 | 3.609912077 |
| ENSMUSG00000025856 | *Pdgfa* | 300.6159713 | 312.6963772 | 258.4164952 | 171.2200219 | 203.183121 | 195.5997505 | 0.000410609 | -0.615492886 | 0.00013469 | 14.57498957 | 0.652706863 | 3.386572063 |
| ENSMUSG00000062116 | *Zfp954* | 418.9034587 | 498.7794974 | 554.6500385 | 311.7082451 | 335.831169 | 310.3883365 | 9.96E-05 | -0.615831921 | 2.95E-05 | 17.44700379 | 0.652553494 | 4.001812123 |
| ENSMUSG00000086583 | *Gm15500* | 41265.7578 | 37511.09578 | 50771.90818 | 30181.91912 | 26484.34081 | 27841.2828 | 9.11E-07 | -0.616282424 | 2.04E-07 | 26.99747225 | 0.652349757 | 6.040261303 |
| ENSMUSG00000022035 | *Ccdc25* | 528.1498643 | 500.6978801 | 576.079699 | 363.2937645 | 343.200505 | 338.8559059 | 1.09E-06 | -0.616939198 | 2.46E-07 | 26.63415584 | 0.652052847 | 5.963255095 |
| ENSMUSG00000112567 | *Gm32899* | 1353.902006 | 1478.113857 | 1902.197518 | 1038.295774 | 925.3780487 | 1118.499982 | 8.33E-05 | -0.616994254 | 2.44E-05 | 17.80688083 | 0.652027964 | 4.079548784 |
| ENSMUSG00000054894 | *Dmac2l* | 144.6573095 | 138.1235531 | 182.782399 | 97.68321764 | 105.2762285 | 99.1773383 | 0.007827462 | -0.617028991 | 0.00331104 | 8.627619232 | 0.652012265 | 2.106379017 |
| ENSMUSG00000009563 | *Tor2a* | 1952.120255 | 1996.077181 | 2008.085253 | 1256.711058 | 1267.525791 | 1351.750389 | 1.18E-15 | -0.618741347 | 1.14E-16 | 68.71452385 | 0.651238842 | 14.92894228 |
| ENSMUSG00000069117 | *Rps18-ps6* | 20300.99586 | 18584.33223 | 23749.10619 | 14221.1399 | 13123.73465 | 13435.77441 | 5.80E-09 | -0.618940758 | 1.01E-09 | 37.29703666 | 0.651148833 | 8.236306433 |
| ENSMUSG00000026426 | *Arl8a* | 2437.32498 | 2761.511871 | 2504.749151 | 1630.980465 | 1666.522697 | 1717.237247 | 4.32E-13 | -0.618942298 | 5.16E-14 | 56.66742426 | 0.651148138 | 12.36428363 |
| ENSMUSG00000034501 | *Pcnx4* | 400.8212951 | 382.7173451 | 408.4241192 | 274.3910608 | 247.399137 | 254.3715066 | 5.22E-06 | -0.618981431 | 1.28E-06 | 23.45141535 | 0.651130476 | 5.282300606 |
| ENSMUSG00000031521 | *Aga* | 671.3003269 | 763.5163075 | 871.052674 | 498.2941664 | 513.7479952 | 485.785296 | 5.93E-06 | -0.619113416 | 1.47E-06 | 23.18455643 | 0.651070909 | 5.226636351 |
| ENSMUSG00000057561 | *Eif1a* | 2893.899613 | 2871.818875 | 3467.823309 | 2074.39642 | 1929.713269 | 2002.831249 | 2.66E-09 | -0.619178493 | 4.51E-10 | 38.87841817 | 0.651041542 | 8.574435226 |
| ENSMUSG00000024248 | *Cox7a2l* | 4448.96569 | 4306.769122 | 5457.000038 | 3298.180551 | 2924.573628 | 3025.827127 | 1.14E-07 | -0.619302247 | 2.27E-08 | 31.24451299 | 0.650985698 | 6.944306889 |
| ENSMUSG00000027822 | *Slc33a1* | 1265.751458 | 1300.663459 | 1245.44145 | 807.807283 | 792.7300007 | 879.7397231 | 1.40E-11 | -0.61936434 | 1.90E-12 | 49.58197666 | 0.65095768 | 10.85512741 |
| ENSMUSG00000045410 | *Akr1e1* | 797.1220492 | 692.5361483 | 993.3277962 | 566.3431495 | 522.1700935 | 525.2725695 | 0.000121667 | -0.619454544 | 3.66E-05 | 17.03789099 | 0.650916981 | 3.914828212 |
| ENSMUSG00000042363 | *Lgalsl* | 504.0403128 | 517.9633242 | 524.3964 | 345.7327366 | 330.5673575 | 329.672819 | 1.37E-07 | -0.619786124 | 2.77E-08 | 30.86516032 | 0.650767395 | 6.863134798 |
| ENSMUSG00000022526 | *Zfp251* | 641.9168109 | 461.3710351 | 514.3118539 | 274.3910608 | 378.9944227 | 399.4642793 | 0.003014807 | -0.619962553 | 0.001171827 | 10.53424497 | 0.650687817 | 2.520740413 |
| ENSMUSG00000025190 | *Got1* | 4448.212267 | 4457.362162 | 4124.579377 | 2836.106004 | 2812.980826 | 2832.063994 | 1.08E-19 | -0.620234179 | 8.03E-21 | 87.59607864 | 0.650565319 | 18.96728344 |
| ENSMUSG00000022415 | *Syngr1* | 910.1355722 | 1079.090259 | 1338.723502 | 658.5385459 | 751.6722716 | 750.2581981 | 0.000180276 | -0.620345782 | 5.58E-05 | 16.23902535 | 0.650514995 | 3.74406244 |
| ENSMUSG00000015013 | *Trappc2l* | 1468.422376 | 1535.665337 | 1866.901607 | 1092.076422 | 1005.387982 | 1066.156387 | 3.88E-07 | -0.620610333 | 8.27E-08 | 28.74150003 | 0.650395719 | 6.411252793 |
| ENSMUSG00000018999 | *Slc35b4* | 2189.448653 | 2369.202613 | 2262.720043 | 1424.638388 | 1597.040387 | 1412.358762 | 4.32E-12 | -0.621445933 | 5.64E-13 | 51.9691512 | 0.650019123 | 11.36434195 |
| ENSMUSG00000016984 | *Etaa1* | 461.8485975 | 431.6361035 | 442.4594625 | 296.3423457 | 274.7709564 | 297.5320149 | 1.38E-06 | -0.621685181 | 3.15E-07 | 26.15788208 | 0.649911337 | 5.861412943 |
| ENSMUSG00000021706 | *Zfyve16* | 433.218505 | 403.8195546 | 479.0159423 | 272.1959323 | 289.5096284 | 291.1038541 | 8.75E-06 | -0.622541539 | 2.22E-06 | 22.39567914 | 0.649525676 | 5.057874041 |
| ENSMUSG00000029217 | *Tec* | 1008.834049 | 984.130316 | 1007.194047 | 666.2214956 | 682.1899608 | 601.4921906 | 4.91E-10 | -0.622837427 | 7.69E-11 | 42.33444421 | 0.649392476 | 9.309295253 |
| ENSMUSG00000023572 | *Ccndbp1* | 1035.203871 | 1079.090259 | 1148.377693 | 673.9044453 | 688.5065345 | 752.0948155 | 1.22E-09 | -0.622898758 | 1.99E-10 | 40.47645359 | 0.649364869 | 8.914752187 |
| ENSMUSG00000047694 | *Yipf6* | 691.642761 | 784.618517 | 984.5038183 | 559.757764 | 535.8560032 | 498.6416176 | 0.000124132 | -0.623418004 | 3.75E-05 | 16.99570526 | 0.649131196 | 3.906116519 |
| ENSMUSG00000028461 | *Ccdc107* | 1245.409024 | 1277.642866 | 1510.160787 | 871.4660091 | 845.368115 | 896.2692795 | 2.97E-08 | -0.623578485 | 5.54E-09 | 33.99159818 | 0.649058993 | 7.527288534 |
| ENSMUSG00000041328 | *Pcf11* | 992.2587323 | 889.1703732 | 794.1580096 | 552.0748143 | 607.4438386 | 580.3710908 | 7.97E-07 | -0.623720025 | 1.77E-07 | 27.27103748 | 0.648995318 | 6.098370397 |
| ENSMUSG00000052712 | *BC004004* | 1827.805379 | 1689.135952 | 2005.564116 | 1193.052332 | 1155.932989 | 1227.778716 | 2.25E-10 | -0.625247281 | 3.41E-11 | 43.92381484 | 0.648308646 | 9.647794084 |
| ENSMUSG00000021785 | *Ngly1* | 252.3968681 | 263.7776188 | 297.4941115 | 152.5614298 | 190.5499736 | 181.8251202 | 0.00056316 | -0.626916859 | 0.000189639 | 13.93105345 | 0.647558817 | 3.249368288 |
| ENSMUSG00000003559 | *As3mt* | 834.7932235 | 877.6600771 | 879.8766519 | 559.757764 | 524.275618 | 592.3091038 | 2.03E-09 | -0.626974679 | 3.39E-10 | 39.43315436 | 0.647532864 | 8.692695525 |
| ENSMUSG00000028466 | *Creb3* | 2523.968681 | 2540.897863 | 2461.88983 | 1666.102521 | 1610.726296 | 1598.775426 | 1.42E-19 | -0.62715337 | 1.07E-20 | 87.02395943 | 0.647452666 | 18.84682989 |
| ENSMUSG00000084235 | *Gm15421* | 481.4376082 | 496.8611147 | 700.8759577 | 447.8062112 | 336.8839313 | 301.2052497 | 0.005637043 | -0.62728873 | 0.002318336 | 9.278702044 | 0.647391922 | 2.248948644 |
| ENSMUSG00000009905 | *Kdsr* | 327.7392168 | 368.329475 | 379.431049 | 195.3664353 | 234.7659896 | 263.5545935 | 0.000379459 | -0.627380198 | 0.000123535 | 14.73794329 | 0.647350879 | 3.420835604 |
| ENSMUSG00000029125 | *Stx18* | 930.4780064 | 856.5578676 | 882.3977885 | 588.2944343 | 565.3333471 | 575.7795474 | 2.06E-10 | -0.62747384 | 3.12E-11 | 44.10149514 | 0.647308862 | 9.685840411 |
| ENSMUSG00000027835 | *Pdcd10* | 1157.258476 | 1146.233653 | 1684.119208 | 907.6856291 | 788.5189516 | 880.6580318 | 0.000380386 | -0.62784041 | 0.000123863 | 14.73294238 | 0.64714441 | 3.419775186 |
| ENSMUSG00000099875 | *Rbm3-ps* | 1163.285864 | 1113.621147 | 1174.849627 | 852.8074169 | 700.0869197 | 682.3033552 | 2.09E-07 | -0.627941242 | 4.33E-08 | 29.99667976 | 0.647099182 | 6.67963825 |
| ENSMUSG00000031534 | *Smim19* | 456.5746331 | 417.2482334 | 395.8184365 | 254.6349044 | 286.3513416 | 281.9207672 | 9.98E-06 | -0.628049117 | 2.55E-06 | 22.13014274 | 0.647050798 | 5.000862383 |
| ENSMUSG00000044408 | *Sptssa* | 1964.17503 | 1915.505108 | 2715.264052 | 1540.980197 | 1380.171356 | 1342.567302 | 6.58E-05 | -0.62816846 | 1.90E-05 | 18.28539637 | 0.646997275 | 4.181748368 |
| ENSMUSG00000024521 | *Pmaip1* | 281.0269606 | 282.0022543 | 288.6701337 | 201.9518207 | 154.7560559 | 193.7631332 | 0.000359896 | -0.628592753 | 0.000116722 | 14.84492695 | 0.646807022 | 3.443823353 |
| ENSMUSG00000059482 | *Cfap418* | 509.3142772 | 505.4938368 | 741.2141423 | 413.7817197 | 324.2507838 | 394.8727358 | 0.002192775 | -0.628660509 | 0.00082989 | 11.17316409 | 0.646776646 | 2.659005954 |
| ENSMUSG00000073131 | *Vma21* | 1349.381465 | 1331.357581 | 1324.857251 | 871.4660091 | 851.6846887 | 866.8834015 | 6.05E-15 | -0.629421267 | 6.22E-16 | 65.36459535 | 0.646435679 | 14.21816263 |
| ENSMUSG00000040659 | *Efhd2* | 14855.25089 | 16267.88515 | 16270.15465 | 9961.493071 | 10371.81403 | 10291.48547 | 1.53E-20 | -0.629739095 | 1.07E-21 | 91.57473927 | 0.646293284 | 19.81639748 |
| ENSMUSG00000026005 | *Rpe* | 969.6560277 | 940.9667056 | 1351.329185 | 702.4411156 | 717.9838785 | 684.1399726 | 0.000132029 | -0.629810214 | 4.00E-05 | 16.87215825 | 0.646261425 | 3.879330763 |
| ENSMUSG00000021477 | *Ctsl* | 16881.96007 | 16886.56356 | 20736.34803 | 12080.88962 | 11327.72219 | 11808.53142 | 8.48E-10 | -0.629890084 | 1.37E-10 | 41.20998537 | 0.646225648 | 9.071524503 |
| ENSMUSG00000028277 | *Ube2j1* | 3484.583627 | 3660.274158 | 4348.960529 | 2485.983011 | 2529.787771 | 2406.887071 | 2.78E-09 | -0.629925658 | 4.71E-10 | 38.79530756 | 0.646209713 | 8.556648214 |
| ENSMUSG00000022685 | *Pam* | 944.7930526 | 961.1097238 | 1097.954963 | 668.4166241 | 603.2327894 | 665.7737988 | 3.16E-08 | -0.630061638 | 5.94E-09 | 33.85484999 | 0.646148808 | 7.499632749 |
| ENSMUSG00000030314 | *Atg7* | 2416.229123 | 2427.713284 | 2195.909925 | 1457.565315 | 1577.037903 | 1515.209335 | 9.65E-14 | -0.630743155 | 1.09E-14 | 59.71845234 | 0.645843645 | 13.0152927 |
| ENSMUSG00000027502 | *Rtf2* | 3999.171869 | 4147.543359 | 4451.066559 | 2719.764195 | 2764.553761 | 2643.810713 | 3.83E-17 | -0.631474109 | 3.33E-18 | 75.68086205 | 0.645516506 | 16.41694607 |
| ENSMUSG00000039982 | *Dtx4* | 1583.69617 | 1633.502854 | 1304.688159 | 878.0513945 | 1028.548753 | 1013.812792 | 8.88E-07 | -0.631845616 | 1.98E-07 | 27.05126821 | 0.645350301 | 6.051413527 |
| ENSMUSG00000007617 | *Homer1* | 376.7117435 | 414.3706594 | 375.6493442 | 272.1959323 | 240.029801 | 241.5151849 | 1.18E-05 | -0.63185363 | 3.06E-06 | 21.77862875 | 0.645346716 | 4.926769795 |
| ENSMUSG00000028931 | *Kcnab2* | 13335.59572 | 14145.19471 | 12569.12621 | 8266.85388 | 8901.105121 | 8675.262176 | 2.89E-17 | -0.63213992 | 2.49E-18 | 76.25673079 | 0.645218666 | 16.53981872 |
| ENSMUSG00000016552 | *Foxred2* | 1695.956269 | 1946.199231 | 1385.364528 | 1050.368981 | 1100.136588 | 1095.542265 | 6.20E-06 | -0.632325956 | 1.54E-06 | 23.09538953 | 0.64513547 | 5.207337923 |
| ENSMUSG00000027286 | *Lrrc57* | 400.8212951 | 465.2078004 | 523.1358317 | 343.5376081 | 272.6654319 | 278.2475325 | 0.000320469 | -0.63266503 | 0.000102923 | 15.08231672 | 0.644983862 | 3.494214541 |
| ENSMUSG00000075033 | *Nxpe3* | 671.3003269 | 558.2493605 | 471.4525327 | 357.8059433 | 363.2029884 | 378.3431795 | 0.000155841 | -0.633744225 | 4.78E-05 | 16.53366796 | 0.644501568 | 3.807319121 |
| ENSMUSG00000081642 | *Gm13532* | 4541.636779 | 3956.664282 | 5542.71868 | 3276.229266 | 2907.729432 | 2861.449872 | 4.73E-06 | -0.634007866 | 1.15E-06 | 23.65684497 | 0.644383802 | 5.32525463 |
| ENSMUSG00000005483 | *Dnajb1* | 2183.421265 | 2157.221326 | 2159.353445 | 1305.003885 | 1443.337093 | 1435.316479 | 2.24E-15 | -0.635285635 | 2.23E-16 | 67.3854384 | 0.643813336 | 14.64897553 |
| ENSMUSG00000028967 | *Errfi1* | 272.7393023 | 283.920637 | 265.9799048 | 155.8541225 | 190.5499736 | 182.7434289 | 0.000162997 | -0.635923985 | 5.01E-05 | 16.44346403 | 0.64352853 | 3.78782138 |
| ENSMUSG00000019792 | *Trmt11* | 344.3145335 | 307.9004205 | 384.4733221 | 230.4884911 | 190.5499736 | 244.270111 | 0.000394253 | -0.636153151 | 0.000128784 | 14.65949278 | 0.643426317 | 3.404224534 |
| ENSMUSG00000021504 | *B4galt7* | 1075.888739 | 1131.845783 | 1004.672911 | 681.587395 | 664.293002 | 721.7906288 | 3.05E-10 | -0.636279597 | 4.67E-11 | 43.30931286 | 0.643369925 | 9.516306812 |
| ENSMUSG00000083364 | *Llph-ps2* | 707.4646542 | 651.2909206 | 913.9119952 | 501.5868591 | 481.1123643 | 476.6022091 | 6.69E-05 | -0.63628137 | 1.94E-05 | 18.25037076 | 0.643369135 | 4.17448415 |
| ENSMUSG00000036299 | *BC031181* | 1544.518148 | 1562.522695 | 2076.155939 | 1114.027707 | 1119.086309 | 1096.460573 | 4.84E-06 | -0.63667445 | 1.18E-06 | 23.60764344 | 0.643193864 | 5.314762158 |
| ENSMUSG00000028138 | *Adh5* | 4064.719712 | 4090.95107 | 5412.880148 | 3080.862831 | 2861.407891 | 2780.638707 | 1.63E-06 | -0.636748056 | 3.75E-07 | 25.8186258 | 0.64316105 | 5.788470407 |
| ENSMUSG00000059182 | *Skap2* | 4623.759939 | 5013.69314 | 5857.860747 | 3348.668506 | 3275.143469 | 3335.297155 | 1.26E-09 | -0.636928807 | 2.06E-10 | 40.41233345 | 0.643080475 | 8.901228021 |
| ENSMUSG00000050786 | *Ccdc126* | 292.3283129 | 292.553359 | 347.9168423 | 198.659128 | 209.4996948 | 190.0898984 | 0.00011099 | -0.637078379 | 3.32E-05 | 17.22428204 | 0.643013807 | 3.954714937 |
| ENSMUSG00000030979 | *Uros* | 437.7390459 | 403.8195546 | 535.7415144 | 318.2936305 | 287.4041039 | 278.2475325 | 0.000124704 | -0.637092447 | 3.77E-05 | 16.9856734 | 0.643007537 | 3.904117983 |
| ENSMUSG00000044252 | *Osbpl1a* | 400.8212951 | 439.3096342 | 364.3042298 | 237.0738765 | 297.9317267 | 240.5968763 | 0.000144001 | -0.637459797 | 4.39E-05 | 16.69501771 | 0.64284383 | 3.841633156 |
| ENSMUSG00000005107 | *Slc2a9* | 977.9436861 | 1074.294302 | 1071.483029 | 677.197138 | 656.923666 | 671.2836509 | 4.45E-11 | -0.637687289 | 6.33E-12 | 47.22398686 | 0.642742471 | 10.35174331 |
| ENSMUSG00000022964 | *Tmem50b* | 1134.655771 | 1257.499848 | 1173.589059 | 756.2217635 | 769.5692305 | 764.9511371 | 5.60E-12 | -0.637865593 | 7.41E-13 | 51.43326131 | 0.642663039 | 11.25170343 |
| ENSMUSG00000031207 | *Msn* | 20481.81749 | 21681.56107 | 20782.98905 | 13277.23465 | 13723.80915 | 13441.28427 | 1.28E-29 | -0.63823034 | 5.61E-31 | 133.9481631 | 0.642500579 | 28.89248775 |
| ENSMUSG00000105796 | *Gm42845* | 2152.530902 | 2194.629788 | 2983.765093 | 1659.517136 | 1531.769125 | 1513.372718 | 1.37E-05 | -0.638739244 | 3.59E-06 | 21.47363574 | 0.64227398 | 4.862755842 |
| ENSMUSG00000021408 | *Ripk1* | 1642.463202 | 1825.341122 | 2005.564116 | 1182.07669 | 1134.877743 | 1191.964677 | 1.08E-09 | -0.639393022 | 1.77E-10 | 40.70747482 | 0.64198299 | 8.965004295 |
| ENSMUSG00000042229 | *Rabif* | 1023.902519 | 1001.39576 | 1193.758151 | 701.3435514 | 642.184994 | 719.0357027 | 5.74E-08 | -0.639589751 | 1.11E-08 | 32.63918463 | 0.641895454 | 7.241120384 |
| ENSMUSG00000054226 | *Tprkb* | 843.8343054 | 731.8629933 | 934.0810875 | 544.3918646 | 525.3283803 | 539.0471999 | 7.35E-07 | -0.639800772 | 1.62E-07 | 27.43664276 | 0.641801572 | 6.133948176 |
| ENSMUSG00000040213 | *Kyat3* | 398.5610246 | 451.7791217 | 579.8614039 | 340.2449154 | 267.4016204 | 307.6334105 | 0.001115132 | -0.639807716 | 0.000396326 | 12.54943393 | 0.641798483 | 2.95267381 |
| ENSMUSG00000042628 | *Zfyve1* | 358.6295798 | 478.6364792 | 418.5086654 | 261.2202899 | 237.9242765 | 304.8784844 | 0.00050302 | -0.639867156 | 0.000167593 | 14.16347008 | 0.641772041 | 3.29841448 |
| ENSMUSG00000081895 | *Rpl17-ps10* | 4098.623769 | 4506.280921 | 5338.50662 | 3384.888126 | 2709.810122 | 2849.511859 | 2.02E-06 | -0.640101677 | 4.69E-07 | 25.38523169 | 0.641667724 | 5.695430318 |
| ENSMUSG00000020231 | *Dip2a* | 1258.217223 | 1315.051329 | 954.2501798 | 718.9045793 | 776.9385665 | 770.4609892 | 1.34E-05 | -0.640241412 | 3.50E-06 | 21.52353521 | 0.641605577 | 4.872915675 |
| ENSMUSG00000068747 | *Sort1* | 313.4241706 | 314.6147599 | 311.3603625 | 187.6834856 | 214.7635062 | 200.191294 | 2.08E-05 | -0.640283542 | 5.57E-06 | 20.6294873 | 0.641586841 | 4.682893109 |
| ENSMUSG00000021103 | *Mnat1* | 649.4510457 | 640.7398159 | 676.9251606 | 445.6110827 | 427.4214878 | 389.3628837 | 3.13E-08 | -0.640503274 | 5.87E-09 | 33.8769917 | 0.641489131 | 7.503939511 |
| ENSMUSG00000040548 | *Tex2* | 872.4643979 | 1000.436569 | 941.6444971 | 580.6114846 | 589.5468797 | 631.7963773 | 3.68E-09 | -0.641379293 | 6.33E-10 | 38.21713125 | 0.64109973 | 8.434097366 |
| ENSMUSG00000054293 | *P2ry10b* | 324.7255229 | 277.2062976 | 414.7269606 | 225.0006698 | 221.0800799 | 203.8645287 | 0.001529807 | -0.641852849 | 0.000559452 | 11.90627223 | 0.640889328 | 2.815363373 |
| ENSMUSG00000062963 | *Ufc1* | 2129.928198 | 2072.812488 | 2553.911313 | 1579.394946 | 1378.065831 | 1367.361636 | 8.89E-08 | -0.642870532 | 1.75E-08 | 31.75074877 | 0.640437401 | 7.051079951 |
| ENSMUSG00000042298 | *Ttc19* | 507.8074302 | 469.0445658 | 535.7415144 | 287.5618317 | 353.7281278 | 325.9995842 | 6.57E-06 | -0.643031336 | 1.64E-06 | 22.9802429 | 0.640366021 | 5.182644625 |
| ENSMUSG00000035051 | *Dhx57* | 448.2869747 | 434.5136775 | 403.3818462 | 279.878882 | 289.5096284 | 256.208124 | 2.68E-06 | -0.643643316 | 6.33E-07 | 24.80858287 | 0.640094441 | 5.571911806 |
| ENSMUSG00000095597 | *Rps7-ps3* | 26836.94461 | 26623.31486 | 36478.32458 | 19899.93729 | 18378.07121 | 19272.54444 | 1.70E-06 | -0.643991447 | 3.92E-07 | 25.73529447 | 0.639940001 | 5.770752813 |
| ENSMUSG00000026095 | *Asnsd1* | 2482.530389 | 2390.304822 | 3116.124762 | 1813.17613 | 1654.942312 | 1640.099317 | 3.59E-07 | -0.644245959 | 7.63E-08 | 28.89850371 | 0.639827116 | 6.444496707 |
| ENSMUSG00000020910 | *Adprm* | 382.7391314 | 380.7989624 | 496.6638981 | 261.2202899 | 287.4041039 | 255.2898153 | 0.000175952 | -0.644318218 | 5.44E-05 | 16.28796387 | 0.639795071 | 3.754605648 |
| ENSMUSG00000053317 | *Sec61b* | 4147.596296 | 4445.851866 | 5542.71868 | 3230.131568 | 2806.664252 | 3001.032792 | 6.32E-07 | -0.644653106 | 1.38E-07 | 27.74710491 | 0.639646574 | 6.199045934 |
| ENSMUSG00000024339 | *Tap2* | 1756.983572 | 1794.646999 | 1472.343738 | 1090.978858 | 1007.493507 | 1116.663365 | 1.26E-08 | -0.645308534 | 2.28E-09 | 35.71619843 | 0.639356044 | 7.899329286 |
| ENSMUSG00000025058 | *Tasl* | 478.4239142 | 394.2276412 | 519.3541269 | 310.6106808 | 268.4543827 | 309.4700279 | 9.39E-05 | -0.645509574 | 2.77E-05 | 17.56675498 | 0.639266955 | 4.027555928 |
| ENSMUSG00000038822 | *Hace1* | 427.1911171 | 385.5949191 | 465.1496913 | 299.6350384 | 256.8739976 | 259.8813587 | 2.34E-05 | -0.645593198 | 6.33E-06 | 20.3856893 | 0.639229902 | 4.631334611 |
| ENSMUSG00000058672 | *Tubb2a* | 1166.299558 | 1173.09101 | 1334.941797 | 702.4411156 | 850.6319264 | 791.5820891 | 5.40E-08 | -0.646078861 | 1.04E-08 | 32.76350284 | 0.639014751 | 7.26766201 |
| ENSMUSG00000035840 | *Lysmd3* | 368.4240851 | 313.6555685 | 332.7900231 | 206.3420777 | 211.6052193 | 230.4954807 | 3.57E-05 | -0.6470766 | 9.93E-06 | 19.52550478 | 0.638572973 | 4.447673632 |
| ENSMUSG00000057657 | *Rps18-ps3* | 14014.43028 | 12932.77685 | 15893.24474 | 9425.88172 | 8684.236091 | 9224.410771 | 2.13E-11 | -0.64804076 | 2.95E-12 | 48.72482082 | 0.638146355 | 10.67207865 |
| ENSMUSG00000020225 | *Tmbim4* | 1213.011814 | 1217.213812 | 1458.477488 | 881.3440873 | 790.6244762 | 806.2750281 | 4.12E-08 | -0.648228533 | 7.83E-09 | 33.31662996 | 0.638063303 | 7.385299584 |
| ENSMUSG00000025939 | *Ube2w* | 650.2044692 | 698.2912963 | 821.8905115 | 477.4404458 | 451.6350203 | 451.8078745 | 1.34E-06 | -0.649035182 | 3.05E-07 | 26.21546077 | 0.637706644 | 5.873579742 |
| ENSMUSG00000023047 | *Amhr2* | 332.2597577 | 376.0030057 | 322.7054769 | 239.269005 | 191.6027359 | 226.8222459 | 0.000102608 | -0.649304828 | 3.05E-05 | 17.38399343 | 0.637587465 | 3.988819741 |
| ENSMUSG00000044328 | *Trp53i13* | 459.588327 | 436.4320602 | 390.7761635 | 316.098502 | 250.5574239 | 256.208124 | 6.02E-05 | -0.64959359 | 1.73E-05 | 18.46761496 | 0.637459862 | 4.220280755 |
| ENSMUSG00000053119 | *Chmp3* | 2101.298105 | 2115.016907 | 2557.693018 | 1365.369918 | 1428.598421 | 1517.045953 | 7.04E-09 | -0.650163259 | 1.24E-09 | 36.90208498 | 0.637208201 | 8.1524157 |
| ENSMUSG00000025290 | *Rps24* | 24577.42757 | 22234.05529 | 32458.37236 | 17387.61274 | 16372.55906 | 16734.33922 | 4.48E-06 | -0.650531628 | 1.09E-06 | 23.76519874 | 0.637045521 | 5.349093165 |
| ENSMUSG00000040997 | *Abhd4* | 990.7518853 | 1074.294302 | 1188.715878 | 684.8800877 | 729.5642636 | 654.7540945 | 1.52E-08 | -0.651067493 | 2.77E-09 | 35.33750009 | 0.636808946 | 7.817195022 |
| ENSMUSG00000031168 | *Ebp* | 2185.681536 | 2265.609948 | 2552.650745 | 1519.028913 | 1397.015552 | 1538.167052 | 3.09E-11 | -0.65143893 | 4.34E-12 | 47.96671708 | 0.636645013 | 10.50995655 |
| ENSMUSG00000020621 | *Rdh14* | 213.9722703 | 199.511799 | 268.5010413 | 137.1955304 | 145.2811954 | 149.6843161 | 0.001390344 | -0.651443384 | 0.000503108 | 12.1041114 | 0.636643048 | 2.856877671 |
| ENSMUSG00000027865 | *Gdap2* | 1026.162789 | 976.4567853 | 1159.722808 | 698.0508587 | 631.6573711 | 680.4667378 | 9.58E-09 | -0.651675541 | 1.71E-09 | 36.28108025 | 0.636540608 | 8.018596327 |
| ENSMUSG00000020644 | *Id2* | 1659.038518 | 1602.808731 | 2020.690936 | 1110.735014 | 1156.985751 | 1090.032413 | 3.38E-08 | -0.652156521 | 6.35E-09 | 33.72485901 | 0.636328427 | 7.471565634 |
| ENSMUSG00000028555 | *Ttc39a* | 261.43795 | 239.7978353 | 297.4941115 | 155.8541225 | 158.9671051 | 191.0082071 | 0.000555104 | -0.653126273 | 0.000186736 | 13.96005487 | 0.635900843 | 3.255625597 |
| ENSMUSG00000047446 | *Arl4a* | 284.0406546 | 251.3081314 | 386.9944587 | 212.9274632 | 185.2861622 | 186.4166637 | 0.003090239 | -0.653459988 | 0.001203268 | 10.48532031 | 0.635753768 | 2.510007962 |
| ENSMUSG00000083624 | *Gm12469* | 1093.21748 | 959.1913411 | 1232.835767 | 709.0265011 | 603.2327894 | 773.2159153 | 1.02E-05 | -0.653704524 | 2.62E-06 | 22.07951672 | 0.635646017 | 4.990108909 |
| ENSMUSG00000033955 | *Tnks1bp1* | 369.1775086 | 365.451901 | 291.1912702 | 240.3665693 | 198.9720719 | 214.884233 | 0.000226959 | -0.654147208 | 7.13E-05 | 15.77514566 | 0.635451002 | 3.644052989 |
| ENSMUSG00000053716 | *Dusp7* | 645.6839283 | 674.3115128 | 453.8045769 | 386.3426136 | 377.9416604 | 365.4868578 | 0.000326665 | -0.654706688 | 0.000105162 | 15.04168706 | 0.63520462 | 3.48589717 |
| ENSMUSG00000025044 | *Msr1* | 5329.717747 | 4670.30264 | 5376.323668 | 3475.985958 | 3157.234093 | 3133.269243 | 1.25E-12 | -0.654903473 | 1.55E-13 | 54.50688196 | 0.635117984 | 11.90409565 |
| ENSMUSG00000029186 | *Pi4k2b* | 724.7933944 | 676.2298955 | 697.0942529 | 463.1721106 | 439.0018729 | 431.6050834 | 1.22E-09 | -0.655198326 | 2.00E-10 | 40.46674683 | 0.634988194 | 8.912958954 |
| ENSMUSG00000027944 | *Hax1* | 14999.15478 | 14132.72522 | 15981.48452 | 9983.444356 | 9199.036848 | 9454.906252 | 1.32E-17 | -0.655539492 | 1.12E-18 | 77.83258389 | 0.634838051 | 16.8790331 |
| ENSMUSG00000028917 | *Plekhm2* | 1995.818817 | 2173.527579 | 1928.669452 | 1254.51593 | 1285.42275 | 1331.547598 | 5.42E-14 | -0.655596114 | 5.97E-15 | 60.9106698 | 0.634813136 | 13.26580505 |
| ENSMUSG00000026342 | *Slc35f5* | 541.7114871 | 573.596422 | 642.8898173 | 351.2205578 | 373.7306112 | 387.5262663 | 4.98E-07 | -0.655857546 | 1.08E-07 | 28.23157215 | 0.634698111 | 6.302702047 |
| ENSMUSG00000035356 | *Nfkbiz* | 462.602021 | 474.7997139 | 359.2619567 | 319.3911948 | 216.8690307 | 288.348928 | 0.001534038 | -0.655895102 | 0.00056121 | 11.90042832 | 0.634681589 | 2.814163826 |
| ENSMUSG00000028032 | *Papss1* | 1947.599714 | 2137.078308 | 2270.283453 | 1279.759908 | 1332.797053 | 1415.113688 | 1.49E-11 | -0.655985021 | 2.04E-12 | 49.44866876 | 0.634642032 | 10.82692697 |
| ENSMUSG00000040528 | *Milr1* | 2311.503258 | 2152.425369 | 2282.889136 | 1580.49251 | 1318.058381 | 1383.891193 | 4.70E-11 | -0.656266742 | 6.70E-12 | 47.1127701 | 0.634518115 | 10.3279374 |
| ENSMUSG00000028692 | *Akr1a1* | 28652.69521 | 32465.74932 | 36391.34536 | 21137.98976 | 20037.22457 | 20679.39335 | 7.77E-11 | -0.656534507 | 1.13E-11 | 46.08912003 | 0.634400359 | 10.10935059 |
| ENSMUSG00000005686 | *Ampd3* | 525.1361704 | 655.127686 | 489.1004885 | 354.5132505 | 354.7808901 | 350.7939188 | 2.63E-05 | -0.656580043 | 7.16E-06 | 20.15101346 | 0.634380336 | 4.580696308 |
| ENSMUSG00000021824 | *Ap3m1* | 2658.831485 | 2552.408159 | 3404.794895 | 1865.859213 | 1786.537598 | 1807.231498 | 2.90E-07 | -0.657132174 | 6.10E-08 | 29.32988307 | 0.6341376 | 6.537105445 |
| ENSMUSG00000020078 | *Vps26a* | 2571.434361 | 2691.490903 | 3273.695795 | 1784.639459 | 1766.535115 | 1855.901858 | 8.63E-09 | -0.657363302 | 1.53E-09 | 36.49559979 | 0.634036016 | 8.064217938 |
| ENSMUSG00000006476 | *Nsmf* | 898.0807964 | 1036.88584 | 900.0457442 | 600.367641 | 601.1272648 | 595.9823385 | 1.80E-09 | -0.657681974 | 3.00E-10 | 39.67521482 | 0.633895981 | 8.744026095 |
| ENSMUSG00000021716 | *Srek1ip1* | 455.8212096 | 452.738313 | 550.8683337 | 289.7569602 | 320.0397347 | 312.2249539 | 7.28E-06 | -0.658057732 | 1.82E-06 | 22.77344058 | 0.633730901 | 5.13806493 |
| ENSMUSG00000029993 | *Nfu1* | 965.8889103 | 929.4564095 | 1090.391553 | 656.3434174 | 603.2327894 | 629.0414513 | 3.09E-09 | -0.658800803 | 5.26E-10 | 38.57694722 | 0.633404577 | 8.510516286 |
| ENSMUSG00000061232 | *H2-K1* | 13498.33519 | 13662.72146 | 13191.84694 | 8503.927756 | 8485.264019 | 8570.574985 | 1.18E-36 | -0.658897215 | 3.83E-38 | 166.733349 | 0.633362249 | 35.9277475 |
| ENSMUSG00000026670 | *Uap1* | 1585.95644 | 1504.971214 | 1556.801813 | 971.3443552 | 983.2799744 | 988.1001483 | 8.81E-17 | -0.659498649 | 7.87E-18 | 73.98578174 | 0.633098267 | 16.05488429 |
| ENSMUSG00000034422 | *Parp14* | 1500.066163 | 1325.602433 | 1350.068616 | 1006.466411 | 690.6120591 | 945.8579486 | 3.40E-05 | -0.660320387 | 9.44E-06 | 19.62202001 | 0.632737766 | 4.468004162 |
| ENSMUSG00000004994 | *Ccdc130* | 299.8625478 | 291.5941677 | 326.4871817 | 158.049251 | 222.1328422 | 199.2729853 | 0.000426726 | -0.660547753 | 0.000140446 | 14.49615667 | 0.632638055 | 3.369851279 |
| ENSMUSG00000091705 | *H2-Q2* | 2067.394048 | 2047.873513 | 1923.627179 | 1263.296444 | 1227.520825 | 1327.874363 | 5.66E-16 | -0.661735153 | 5.33E-17 | 70.20997416 | 0.632117582 | 15.24697595 |
| ENSMUSG00000024480 | *Ap3s1* | 803.9028606 | 823.945362 | 1100.476099 | 588.2944343 | 552.7001997 | 578.5344734 | 1.91E-05 | -0.662887722 | 5.09E-06 | 20.80277206 | 0.631612784 | 4.719633963 |
| ENSMUSG00000036249 | *Rbm43* | 714.2454656 | 813.3942573 | 964.334726 | 509.2698088 | 586.3885929 | 474.7655917 | 3.01E-05 | -0.663893386 | 8.29E-06 | 19.86979037 | 0.631172658 | 4.521178758 |
| ENSMUSG00000100865 | *Gm9320* | 1249.929565 | 1058.947241 | 1598.400565 | 813.2951042 | 760.0943699 | 888.92281 | 9.06E-05 | -0.664177883 | 2.67E-05 | 17.63646419 | 0.631048204 | 4.042667939 |
| ENSMUSG00000060126 | *Tpt1* | 24172.08573 | 21954.93061 | 29429.22681 | 16549.07366 | 15340.85202 | 15782.97142 | 1.90E-08 | -0.664265927 | 3.49E-09 | 34.88948931 | 0.631009693 | 7.72070465 |
| ENSMUSG00000021000 | *Mia2* | 4641.08868 | 4794.997514 | 5586.838569 | 3241.10721 | 3048.799578 | 3180.102987 | 4.57E-12 | -0.664939605 | 5.98E-13 | 51.85280998 | 0.630715107 | 11.33997785 |
| ENSMUSG00000025007 | *Aldh18a1* | 5648.415882 | 5270.756419 | 4475.017356 | 3232.326696 | 3247.77165 | 3234.283199 | 1.18E-10 | -0.664993113 | 1.74E-11 | 45.23881074 | 0.630691715 | 9.929352785 |
| ENSMUSG00000040483 | *Xaf1* | 1098.491444 | 983.1711247 | 920.2148365 | 752.9290708 | 493.7455118 | 648.3259337 | 0.000146579 | -0.665000529 | 4.48E-05 | 16.65837247 | 0.630688473 | 3.833927474 |
| ENSMUSG00000041650 | *Pcca* | 797.1220492 | 956.3137671 | 1072.743597 | 564.148021 | 632.7101334 | 581.2893995 | 4.12E-06 | -0.6655986 | 9.97E-07 | 23.93453582 | 0.630427075 | 5.3850857 |
| ENSMUSG00000057531 | *Dtnbp1* | 1722.326091 | 1761.075302 | 1895.894677 | 1071.222701 | 1172.777186 | 1143.294317 | 5.54E-14 | -0.665629594 | 6.11E-15 | 60.86481368 | 0.630413531 | 13.25676996 |
| ENSMUSG00000006782 | *Cnp* | 1240.888483 | 1271.887718 | 1442.0901 | 795.7340763 | 827.4711562 | 865.0467841 | 1.63E-10 | -0.665772865 | 2.45E-11 | 44.57701055 | 0.630350929 | 9.786576148 |
| ENSMUSG00000022313 | *Utp23* | 540.2046401 | 511.2489848 | 618.9390202 | 367.6840215 | 347.4115541 | 336.1009798 | 8.74E-07 | -0.666015961 | 1.95E-07 | 27.08604425 | 0.630244723 | 6.058498663 |
| ENSMUSG00000040918 | *Slc19a2* | 1433.764896 | 1386.031488 | 1158.462239 | 863.7830594 | 873.7926967 | 774.134224 | 5.34E-08 | -0.666221175 | 1.03E-08 | 32.79034167 | 0.630155081 | 7.272729668 |
| ENSMUSG00000045312 | *Lhfpl2* | 1595.750945 | 1923.178639 | 1773.619555 | 999.8810255 | 1193.832431 | 1138.702773 | 1.53E-08 | -0.666357491 | 2.78E-09 | 35.33171622 | 0.630095542 | 7.816232698 |
| ENSMUSG00000020424 | *Castor1* | 157.4655088 | 161.1441453 | 134.8808048 | 108.6588601 | 84.22098282 | 93.66748618 | 0.003836363 | -0.667793713 | 0.0015267 | 10.04610988 | 0.629468586 | 2.41608026 |

| | | | | | | | | | | | | | |
|---|---|---|---|---|---|---|---|---|---|---|---|---|---|
| ENSMUSG00000057143 | *Trim12c* | 406.848683 | 420.1258074 | 486.5793519 | 313.9033735 | 233.7132273 | 277.3292238 | 9.05E-05 | -0.668566204 | 2.67E-05 | 17.63885089 | 0.629131628 | 4.043111904 |
| ENSMUSG00000030474 | *Siglece* | 182.3284838 | 164.9809107 | 137.4019413 | 94.39052491 | 104.2234662 | 107.4421165 | 0.00324493 | -0.668999516 | 0.001269291 | 10.3866595 | 0.628942697 | 2.488794648 |
| ENSMUSG00000022337 | *Emc2* | 756.4371809 | 643.6173899 | 881.1372202 | 470.8550603 | 479.0068398 | 482.1120612 | 5.93E-06 | -0.669384791 | 1.47E-06 | 23.18896457 | 0.628774759 | 5.227200006 |
| ENSMUSG00000020149 | *Rab1a* | 4868.622573 | 4986.835782 | 5773.402673 | 3364.034405 | 3283.565568 | 3168.164974 | 4.05E-13 | -0.670423854 | 4.82E-14 | 56.80009834 | 0.628322063 | 12.39258521 |
| ENSMUSG00000026142 | *Rhbdd1* | 1197.943344 | 1367.806852 | 1268.131679 | 803.417026 | 832.7349676 | 772.2976066 | 1.39E-11 | -0.670497243 | 1.89E-12 | 49.59572291 | 0.628290102 | 10.85795162 |
| ENSMUSG00000015846 | *Rxra* | 199.657224 | 188.9606942 | 176.4795577 | 127.3174522 | 105.2762285 | 123.0533642 | 0.000779728 | -0.671206543 | 0.000269362 | 13.27225923 | 0.62798128 | 3.10805708 |
| ENSMUSG00000019139 | *Isyna1* | 9830.669658 | 10427.36907 | 10532.04789 | 6059.652187 | 6613.452676 | 6651.309827 | 1.45E-20 | -0.671670341 | 1.02E-21 | 91.68225961 | 0.627779428 | 19.83829829 |
| ENSMUSG00000042460 | *C1galt1* | 824.9987182 | 911.2317741 | 785.3340317 | 512.5625016 | 493.7455118 | 574.8612387 | 3.42E-08 | -0.673034928 | 6.45E-09 | 33.69527571 | 0.627185918 | 7.465594602 |
| ENSMUSG00000064284 | *Cdpf1* | 413.6294943 | 437.3912516 | 495.4033298 | 287.5618317 | 276.876481 | 277.3292238 | 1.62E-06 | -0.673750287 | 3.73E-07 | 25.82887285 | 0.626875005 | 5.790517295 |
| ENSMUSG00000063145 | *Bbs5* | 172.5339785 | 191.8382682 | 235.7262663 | 117.439374 | 129.4897611 | 126.7265989 | 0.001729689 | -0.675443672 | 0.000641333 | 11.65200653 | 0.626139633 | 2.762032073 |
| ENSMUSG00000032456 | *Nmnat3* | 201.164071 | 285.8390197 | 244.5502442 | 143.7809159 | 148.4394822 | 164.3772552 | 0.001476934 | -0.675655253 | 0.000537887 | 11.97951162 | 0.626047812 | 2.830639046 |
| ENSMUSG00000064351 | *mt-Co1* | 228574.3709 | 231120.0312 | 281365.1405 | 151571.4269 | 158292.2844 | 153970.0628 | 8.22E-12 | -0.675964043 | 1.10E-12 | 50.64903924 | 0.625913829 | 11.08512582 |
| ENSMUSG00000040374 | *Pex2* | 1186.641992 | 1063.743197 | 1319.814978 | 749.6363781 | 695.8758705 | 786.0722369 | 2.30E-08 | -0.676010449 | 4.24E-09 | 34.51039708 | 0.625893696 | 7.63888717 |
| ENSMUSG00000045251 | *Zfp688* | 160.4792027 | 139.0827445 | 161.3527385 | 100.9759104 | 85.2737451 | 101.9322644 | 0.00234348 | -0.676035188 | 0.000890947 | 11.04150125 | 0.625882964 | 2.630138805 |
| ENSMUSG00000058569 | *Tmed9* | 8701.287851 | 9377.054551 | 10428.68129 | 6095.871807 | 5890.204986 | 5849.626343 | 1.63E-14 | -0.676139309 | 1.73E-15 | 63.35339691 | 0.625837795 | 13.78853497 |
| ENSMUSG00000034786 | *Gpsm3* | 1422.463543 | 1549.094016 | 1381.582823 | 837.4415175 | 958.0136795 | 926.5734662 | 2.55E-11 | -0.677399245 | 3.55E-12 | 48.35989876 | 0.625291476 | 10.59426964 |
| ENSMUSG00000032487 | *Ptgs2* | 5464.580551 | 5161.408607 | 4601.074183 | 3128.058093 | 3155.128569 | 3239.793051 | 1.53E-14 | -0.677865151 | 1.62E-15 | 63.48133471 | 0.625089576 | 13.81617971 |
| ENSMUSG00000084319 | *Tpt1-ps3* | 26570.23269 | 24394.15419 | 32005.82836 | 18001.15115 | 16735.76205 | 17118.19225 | 1.70E-09 | -0.678001008 | 2.82E-10 | 39.79341571 | 0.625030715 | 8.769233699 |
| ENSMUSG00000040006 | *Ginm1* | 651.7113162 | 648.4133466 | 787.8551683 | 408.2938985 | 424.2632009 | 467.4191222 | 7.97E-07 | -0.67935848 | 1.77E-07 | 27.27301018 | 0.624442882 | 6.098544657 |
| ENSMUSG00000068220 | *Lgals1* | 14694.77169 | 14418.56424 | 20061.944 | 11471.74147 | 9381.164723 | 9843.350827 | 5.77E-06 | -0.679721524 | 1.43E-06 | 23.24444923 | 0.624285765 | 5.238691897 |
| ENSMUSG00000034265 | *Zdhhc14* | 368.4240851 | 409.5747027 | 303.7969529 | 219.5128486 | 222.1328422 | 235.0870241 | 8.33E-05 | -0.679779353 | 2.45E-05 | 17.80490047 | 0.624260742 | 4.079198248 |
| ENSMUSG00000020076 | *Ddx50* | 867.943857 | 859.4354416 | 1200.060992 | 620.1237974 | 609.5493631 | 594.1457211 | 2.31E-05 | -0.680022356 | 6.25E-06 | 20.40857121 | 0.624155603 | 4.636197228 |
| ENSMUSG00000000804 | *Usp32* | 512.3279711 | 690.6177656 | 712.2210721 | 374.2694069 | 368.4667998 | 449.0529484 | 0.000148857 | -0.680196099 | 4.55E-05 | 16.6252866 | 0.62408044 | 3.827229732 |
| ENSMUSG00000005043 | *Sgsh* | 851.3685402 | 764.4754989 | 693.3125481 | 435.7330045 | 553.752962 | 453.6444919 | 5.11E-06 | -0.680754998 | 1.25E-06 | 23.49752566 | 0.623838719 | 5.291493335 |
| ENSMUSG00000003762 | *Coq8b* | 886.0260206 | 837.3740408 | 859.7075596 | 550.9772501 | 551.6474374 | 510.5796305 | 1.43E-11 | -0.680775479 | 1.95E-12 | 49.53600891 | 0.623829863 | 10.84538878 |
| ENSMUSG00000062691 | *Cebpzos* | 251.6434446 | 273.3695322 | 397.0790048 | 195.3664353 | 169.4947279 | 207.5377635 | 0.003885765 | -0.680991566 | 0.00154796 | 10.02063655 | 0.623736432 | 2.410523439 |
| ENSMUSG00000038876 | *Rnf146* | 441.5061634 | 435.4728689 | 484.0582154 | 317.1960663 | 253.7157107 | 277.3292238 | 4.13E-06 | -0.6810607 | 1.00E-06 | 23.928652 | 0.623706544 | 5.383881583 |
| ENSMUSG00000070476 | *Fam217b* | 293.0817364 | 283.920637 | 269.7616096 | 184.3907929 | 202.1303588 | 143.2561553 | 0.000404256 | -0.681387808 | 0.000132385 | 14.60752824 | 0.623565144 | 3.393343703 |
| ENSMUSG00000058325 | *Dock1* | 606.505907 | 660.882834 | 559.6923115 | 369.8791499 | 366.3612752 | 403.137514 | 9.64E-08 | -0.681977957 | 1.91E-08 | 31.58083588 | 0.623310121 | 7.01579521 |
| ENSMUSG00000026031 | *Cflar* | 1578.422205 | 1751.483389 | 1719.415119 | 1028.417696 | 989.5965481 | 1124.928143 | 2.34E-12 | -0.682286866 | 2.98E-13 | 53.22403746 | 0.623176672 | 11.63096521 |
| ENSMUSG00000019975 | *Ikbip* | 190.6161422 | 222.5323911 | 283.6278606 | 164.6346365 | 126.3314742 | 141.419538 | 0.003887849 | -0.682506949 | 0.001549057 | 10.01933178 | 0.623081614 | 2.410290588 |
| ENSMUSG00000033610 | *Pank1* | 496.5060779 | 540.0247251 | 475.2342375 | 317.1960663 | 321.092497 | 303.9601757 | 7.59E-08 | -0.683992858 | 1.48E-08 | 32.07489602 | 0.622440199 | 7.119909618 |
| ENSMUSG00000055148 | *Klf2* | 394.7939072 | 419.1666161 | 350.4379788 | 259.0251614 | 248.4518993 | 219.4757764 | 1.13E-05 | -0.684294751 | 2.91E-06 | 21.87240003 | 0.622309963 | 4.946145754 |
| ENSMUSG00000069520 | *Tmem19* | 782.8070029 | 816.2718313 | 850.8835817 | 476.3428815 | 551.6474374 | 494.9683828 | 1.22E-09 | -0.684391102 | 2.00E-10 | 40.46698372 | 0.622268403 | 8.912958954 |
| ENSMUSG00000078453 | *Abracl* | 755.6837574 | 732.8221846 | 1068.961892 | 496.0990379 | 504.2731346 | 586.7992516 | 0.000182291 | -0.684736098 | 5.65E-05 | 16.21586802 | 0.622119616 | 3.739234799 |
| ENSMUSG00000067889 | *Sptbn2* | 176.3010959 | 162.1033366 | 137.4019413 | 105.3661673 | 95.80136795 | 96.42241224 | 0.002086976 | -0.684765372 | 0.000786555 | 11.27269491 | 0.622106993 | 2.680482486 |
| ENSMUSG00000035929 | *H2-Q4* | 3648.076524 | 3814.703964 | 3394.710349 | 2287.323883 | 2164.479258 | 2304.036498 | 1.07E-17 | -0.68499921 | 9.05E-19 | 78.25730414 | 0.622006167 | 16.97065415 |
| ENSMUSG00000034947 | *Tmem106a* | 2248.969109 | 2128.445586 | 1861.859334 | 1307.199014 | 1267.525791 | 1310.426498 | 8.81E-12 | -0.685050176 | 1.19E-12 | 50.50498903 | 0.621984194 | 11.05479946 |
| ENSMUSG00000024953 | *Prdx5* | 3672.186075 | 3926.929351 | 4738.476124 | 2613.300463 | 2531.893296 | 2522.593966 | 1.20E-09 | -0.685273993 | 1.97E-10 | 40.49957793 | 0.621887708 | 8.919709825 |
| ENSMUSG00000050106 | *Tmc8* | 137.8764981 | 131.4092137 | 127.3173952 | 68.04898308 | 88.43203196 | 89.99425142 | 0.00464935 | -0.685274845 | 0.001879391 | 9.663755036 | 0.621887341 | 2.33260772 |
| ENSMUSG00000024516 | *Sec11c* | 2627.941122 | 2566.796029 | 3401.01319 | 1778.054074 | 1751.796443 | 1809.068115 | 8.02E-08 | -0.685837362 | 1.57E-08 | 31.96118421 | 0.62164491 | 7.095640341 |
| ENSMUSG00000096887 | *Gm20594* | 396.3007541 | 407.65632 | 530.6992414 | 232.6836196 | 296.8789644 | 296.6137062 | 0.000310277 | -0.686616797 | 9.94E-05 | 15.14858148 | 0.621309149 | 3.50825048 |
| ENSMUSG00000055994 | *Nod2* | 167.2600141 | 170.7360587 | 126.0568269 | 91.09783218 | 102.1179417 | 96.42241224 | 0.003756203 | -0.686630181 | 0.001490676 | 10.09010568 | 0.621303385 | 2.425250895 |
| ENSMUSG00000030614 | *Tmem126b* | 531.1635583 | 449.860739 | 571.037426 | 351.2205578 | 309.5121118 | 303.041867 | 7.35E-06 | -0.686954613 | 1.84E-06 | 22.75165014 | 0.621163682 | 5.133853833 |
| ENSMUSG00000069682 | *Gm10275* | 37222.13395 | 36146.1665 | 47051.97122 | 26307.51734 | 23394.4835 | 25082.6835 | 7.49E-09 | -0.687179469 | 1.33E-09 | 36.77561881 | 0.621066876 | 8.125256666 |
| ENSMUSG00000032932 | *Hspa13* | 1303.422632 | 1302.581841 | 1728.239097 | 897.8075509 | 912.7449013 | 876.984797 | 8.37E-07 | -0.687467493 | 1.86E-07 | 27.17506818 | 0.620942897 | 6.077350596 |
| ENSMUSG00000092358 | *Gm20430* | 5438.210729 | 5502.880724 | 7139.858677 | 4070.865778 | 3403.580468 | 3748.536064 | 2.76E-07 | -0.687526316 | 5.80E-08 | 29.43011719 | 0.62091758 | 6.558717058 |
| ENSMUSG00000055681 | *Cope* | 5628.073447 | 6019.884857 | 6818.413768 | 3795.377153 | 3690.984572 | 3973.521693 | 4.91E-13 | -0.687530359 | 5.90E-14 | 56.40448099 | 0.62091584 | 12.30894611 |
| ENSMUSG00000067203 | *H2-K2* | 1790.134205 | 1883.851794 | 1766.056145 | 1175.491304 | 1094.872777 | 1104.725352 | 4.23E-17 | -0.689394684 | 3.69E-18 | 75.47913742 | 0.620113978 | 16.37326089 |
| ENSMUSG00000021661 | *Ankra2* | 216.9859642 | 252.2673227 | 293.7124067 | 160.2443795 | 162.1253919 | 148.7660075 | 0.00042716 | -0.689944962 | 0.000140618 | 14.49384582 | 0.619877497 | 3.369409062 |
| ENSMUSG00000027523 | *Gnas* | 5502.251725 | 5570.024118 | 5211.189225 | 3359.644148 | 3328.834346 | 3406.006924 | 2.44E-27 | -0.690234511 | 1.19E-28 | 123.315456 | 0.6197531 | 26.61237204 |
| ENSMUSG00000022205 | *Sub1* | 2123.90081 | 2248.344504 | 2886.701337 | 1564.029047 | 1413.859749 | 1516.127644 | 4.19E-07 | -0.69035948 | 8.97E-08 | 28.58478299 | 0.619699419 | 6.377919422 |
| ENSMUSG00000001288 | *Rarg* | 508.5608537 | 519.8817069 | 415.9875288 | 296.3423457 | 267.4016204 | 332.4277451 | 2.19E-05 | -0.690413548 | 5.89E-06 | 20.52482031 | 0.619676195 | 4.660367383 |
| ENSMUSG00000021286 | *Zfyve21* | 264.4516439 | 245.5529833 | 252.1136538 | 165.7322007 | 133.7008102 | 172.6420333 | 0.000165478 | -0.690551667 | 5.10E-05 | 16.41229052 | 0.619616872 | 3.781260571 |
| ENSMUSG00000060477 | *Irak2* | 748.902946 | 794.2104304 | 811.8059654 | 443.4159542 | 491.6399872 | 520.6810261 | 2.09E-09 | -0.690973793 | 3.50E-10 | 39.37394928 | 0.619435601 | 8.679883941 |
| ENSMUSG00000041231 | *Ublcp1* | 502.5334658 | 498.7794974 | 665.5800462 | 342.4400439 | 343.200505 | 343.4474493 | 1.99E-05 | -0.691502237 | 5.31E-06 | 20.72219389 | 0.61920875 | 4.7021412 |
| ENSMUSG00000066643 | *Wdr35* | 400.8212951 | 335.7169694 | 438.6777577 | 238.1714408 | 250.5574239 | 236.9236415 | 2.10E-05 | -0.693514656 | 5.64E-06 | 20.60562106 | 0.618345616 | 4.677590182 |
| ENSMUSG00000038301 | *Snx10* | 1981.503771 | 2121.731247 | 2664.841321 | 1474.028779 | 1311.741807 | 1394.910897 | 2.30E-07 | -0.693590834 | 4.78E-08 | 29.80353596 | 0.618312967 | 6.638530624 |
| ENSMUSG00000069255 | *Dusp22* | 479.9307612 | 511.2489848 | 594.9882231 | 315.0009378 | 355.8336524 | 307.6334105 | 3.00E-06 | -0.693750583 | 7.15E-07 | 24.57336565 | 0.618244505 | 5.522299912 |
| ENSMUSG00000021764 | *Ndufs4* | 1914.44908 | 1734.217945 | 2343.396413 | 1318.174656 | 1186.463095 | 1196.55622 | 2.32E-07 | -0.69402239 | 4.83E-08 | 29.78308245 | 0.618128037 | 6.634664603 |
| ENSMUSG00000082035 | *Rpl17-ps8* | 14431.07347 | 13639.70087 | 18842.97449 | 9958.200378 | 9044.280792 | 9969.159117 | 1.79E-07 | -0.695146327 | 3.68E-08 | 30.31342888 | 0.61764667 | 6.747389915 |
| ENSMUSG00000027997 | *Casp6* | 376.7117435 | 344.3496915 | 481.5370788 | 244.7568262 | 241.0825633 | 254.3715066 | 0.000105725 | -0.695215566 | 3.15E-05 | 17.32352797 | 0.617617028 | 3.975820782 |
| ENSMUSG00000030272 | *Camk1* | 5741.086971 | 6195.416872 | 7216.753341 | 4145.500146 | 3716.250867 | 3959.747063 | 9.94E-11 | -0.695603384 | 1.47E-11 | 45.5792271 | 0.617451026 | 10.0024168 |
| ENSMUSG00000068250 | *Amn1* | 105.4792882 | 99.75589948 | 124.7962587 | 77.92706126 | 61.06021254 | 64.28160816 | 0.009951479 | -0.696085109 | 0.00430648 | 8.149846736 | 0.617244889 | 2.002112364 |
| ENSMUSG00000070871 | *Ccnyl1* | 659.9989746 | 607.1681189 | 652.9743635 | 467.5623676 | 364.2557507 | 354.4671536 | 3.69E-06 | -0.696606339 | 8.87E-07 | 24.15916414 | 0.617021926 | 5.4330311 |
| ENSMUSG00000034574 | *Daam1* | 327.7392168 | 380.7989624 | 413.4663923 | 266.7081111 | 198.9720719 | 224.9856286 | 0.000152347 | -0.697311703 | 4.67E-05 | 16.57879252 | 0.616720324 | 3.817167278 |
| ENSMUSG00000026553 | *Copa* | 15019.49721 | 15589.73687 | 15275.56629 | 9236.003106 | 9643.302532 | 9389.706335 | 5.90E-37 | -0.69873349 | 1.87E-38 | 168.1510964 | 0.616112841 | 36.2290637 |
| ENSMUSG00000026755 | *Arpc5l* | 3299.994873 | 3027.207873 | 3237.139315 | 2052.445135 | 1891.813826 | 1938.54964 | 3.38E-19 | -0.701475544 | 2.62E-20 | 85.26001302 | 0.61494294 | 18.47115156 |
| ENSMUSG00000044952 | *Kctd21* | 125.8217223 | 170.7360587 | 189.0852404 | 105.3661673 | 83.16822053 | 108.3604252 | 0.008725463 | -0.701580992 | 0.003732812 | 8.409422495 | 0.614897995 | 2.059211497 |
| ENSMUSG00000030946 | *Lhpp* | 787.3275439 | 812.4350659 | 837.0173308 | 482.928267 | 528.4866672 | 485.785296 | 1.72E-11 | -0.701644943 | 2.36E-12 | 49.15600528 | 0.614870739 | 10.76539259 |
| ENSMUSG00000029735 | *Tpk1* | 185.3421778 | 206.2261383 | 204.2120596 | 121.829631 | 127.3842365 | 116.6252034 | 0.00018646 | -0.701849978 | 5.80E-05 | 16.16759318 | 0.61478336 | 3.729415045 |
| ENSMUSG00000019790 | *Stxbp5* | 1530.203102 | 1645.01315 | 1510.160787 | 986.7102546 | 963.2774909 | 932.0833183 | 4.28E-16 | -0.701907485 | 4.00E-17 | 70.77878686 | 0.614758855 | 15.36838249 |
| ENSMUSG00000030720 | *Cln3* | 2166.845948 | 2445.93792 | 2258.938338 | 1441.101851 | 1379.118594 | 1403.175675 | 1.17E-16 | -0.70204082 | 1.06E-17 | 73.40521038 | 0.614702041 | 15.93212628 |
| ENSMUSG00000017760 | *Ctsa* | 26406.73979 | 28628.02477 | 28062.77081 | 16066.14539 | 17504.27852 | 17502.04528 | 3.09E-23 | -0.702137422 | 1.88E-24 | 104.1451405 | 0.614660882 | 22.50969834 |
| ENSMUSG00000003863 | *Ppfia3* | 171.0271315 | 152.5114232 | 124.7962587 | 102.0734746 | 83.16822053 | 91.8308688 | 0.004059308 | -0.702523753 | 0.001623779 | 9.932589843 | 0.614496308 | 2.391547953 |
| ENSMUSG00000038871 | *Bpgm* | 381.2322844 | 499.7386887 | 495.4033298 | 305.1228596 | 248.4518993 | 290.1855454 | 7.19E-05 | -0.702601437 | 2.09E-05 | 18.10410836 | 0.61446322 | 4.1433895 |
| ENSMUSG00000031792 | *Usb1* | 1619.107073 | 1765.871259 | 1598.400565 | 1021.83231 | 1008.546269 | 1030.342348 | 2.59E-16 | -0.70348047 | 2.39E-17 | 71.79649683 | 0.614088942 | 15.58628391 |
| ENSMUSG00000042541 | *Sem1* | 2816.296994 | 2673.266268 | 3600.182977 | 1863.664085 | 1873.916868 | 1839.372302 | 4.18E-08 | -0.703727955 | 7.95E-09 | 33.28722027 | 0.613983608 | 7.379201603 |
| ENSMUSG00000079659 | *Tmem243* | 391.7802132 | 385.5949191 | 405.9029827 | 249.1470832 | 209.4996948 | 266.3095195 | 4.37E-06 | -0.704388595 | 1.06E-06 | 23.81479622 | 0.613702517 | 5.359793375 |
| ENSMUSG00000066270 | *Gm10157* | 1869.243671 | 1941.403274 | 1863.119902 | 1234.759774 | 1067.500957 | 1180.026664 | 7.00E-15 | -0.704404442 | 7.22E-16 | 65.07118339 | 0.613695776 | 14.15494619 |
| ENSMUSG00000025531 | *Chm* | 462.602021 | 427.7993381 | 460.1074183 | 270.0008038 | 250.5574239 | 306.7151018 | 8.92E-07 | -0.705002491 | 1.99E-07 | 27.04075911 | 0.613441429 | 6.04958664 |
| ENSMUSG00000006010 | *Odr4* | 642.6702344 | 587.0251008 | 726.0873231 | 422.5622336 | 385.3109964 | 390.2811924 | 1.32E-07 | -0.705065778 | 2.65E-08 | 30.94537625 | 0.61341452 | 6.880346442 |
| ENSMUSG00000047557 | *Lxn* | 505.5471597 | 500.6978801 | 537.0020827 | 306.2204238 | 322.1452593 | 316.8164974 | 4.59E-09 | -0.705276784 | 7.97E-10 | 37.76687094 | 0.61332481 | 8.338010946 |
| ENSMUSG00000031490 | *Eif4ebp1* | 5973.894828 | 5934.516828 | 6634.370801 | 3986.353331 | 3467.798967 | 3905.56685 | 8.16E-15 | -0.706474688 | 8.47E-16 | 64.75852235 | 0.612815763 | 14.08805344 |
| ENSMUSG00000030083 | *Abtb1* | 331.5063343 | 307.9004205 | 356.7408202 | 199.7566923 | 200.0248342 | 209.3743809 | 3.07E-06 | -0.706495327 | 7.32E-07 | 24.52938311 | 0.612806996 | 5.51351183 |
| ENSMUSG00000022684 | *Bfar* | 945.5464761 | 943.8442797 | 983.24325 | 582.8066131 | 567.4388717 | 605.1654254 | 6.46E-14 | -0.708998839 | 7.18E-15 | 60.54819765 | 0.611744513 | 13.18987866 |
| ENSMUSG00000030934 | *Oat* | 6642.934884 | 6867.810002 | 7685.684738 | 4326.598247 | 4277.373165 | 4354.619799 | 1.49E-18 | -0.709277154 | 1.20E-19 | 82.24778639 | 0.611626511 | 17.82640948 |
| ENSMUSG00000018750 | *Zbtb4* | 659.2455511 | 813.3942573 | 636.586976 | 454.3915967 | 427.4214878 | 409.5656749 | 7.97E-07 | -0.709319882 | 1.77E-07 | 27.27167137 | 0.611608396 | 6.098378382 |
| ENSMUSG00000068134 | *Zfp120* | 354.1090389 | 290.6349764 | 341.614001 | 181.0981001 | 200.0248342 | 221.3123938 | 4.06E-05 | -0.70954123 | 1.14E-05 | 19.26497965 | 0.611514567 | 4.39162917 |
| ENSMUSG00000026353 | *Ubxn4* | 2523.968681 | 2619.551553 | 3043.011802 | 1710.005091 | 1680.208607 | 1608.876821 | 1.56E-12 | -0.710489229 | 1.95E-13 | 54.05779975 | 0.61111287 | 11.80747438 |
| ENSMUSG00000085791 | *Rpl30-ps9* | 6425.195497 | 6576.215835 | 8429.420016 | 4580.135587 | 4343.697189 | 4167.284826 | 7.05E-09 | -0.71067396 | 1.24E-09 | 36.89817709 | 0.611034625 | 8.151714108 |
| ENSMUSG00000075706 | *Gpx4* | 7119.098528 | 7685.041025 | 9655.952942 | 5103.673731 | 4775.329726 | 5059.880871 | 1.19E-08 | -0.710867116 | 2.15E-09 | 35.82928308 | 0.610952822 | 7.923218812 |
| ENSMUSG00000041654 | *Slc39a11* | 2630.954816 | 2723.144217 | 2995.110208 | 1724.273426 | 1665.469935 | 1706.217542 | 2.19E-17 | -0.711005433 | 1.87E-18 | 76.81875717 | 0.61089425 | 16.65995051 |
| ENSMUSG00000043372 | *Hexim2* | 197.3969536 | 215.8180518 | 200.4303548 | 115.2442455 | 120.0149005 | 138.6646119 | 0.000204684 | -0.711507747 | 6.40E-05 | 15.98058735 | 0.610681588 | 3.688916173 |
| ENSMUSG00000025880 | *Smad7* | 188.3558717 | 227.3283478 | 172.6978529 | 129.5125807 | 140.0173839 | 90.91256011 | 0.005072523 | -0.712415876 | 0.002064585 | 9.491211037 | 0.610297305 | 2.294775993 |
| ENSMUSG00000069516 | *Lyz2* | 93878.07332 | 91610.44661 | 109147.5642 | 60672.2538 | 61356.03874 | 57632.13495 | 1.11E-15 | -0.713630414 | 1.07E-16 | 68.83137043 | 0.60978374 | 14.95312913 |
| ENSMUSG00000037860 | *Aim2* | 779.793309 | 772.1490296 | 1129.469169 | 559.757764 | 537.9615277 | 533.5373477 | 5.34E-05 | -0.714211755 | 1.52E-05 | 18.70976834 | 0.609538074 | 4.272843962 |
| ENSMUSG00000038506 | *Dcun1d2* | 1329.792454 | 1495.379301 | 1687.900913 | 881.3440873 | 892.7424178 | 970.6522832 | 3.12E-09 | -0.7148114 | 5.33E-10 | 38.55274463 | 0.609284777 | 8.505654759 |
| ENSMUSG00000030641 | *Ddias* | 308.9036296 | 283.920637 | 248.331949 | 195.3664353 | 162.1253919 | 157.0307856 | 0.000133032 | -0.715828314 | 4.03E-05 | 16.8560681 | 0.608855461 | 3.876042503 |
| ENSMUSG00000058838 | *Rps27a-ps2* | 6224.784849 | 5495.207193 | 7773.924516 | 4217.939386 | 3546.756139 | 4096.575057 | 5.81E-07 | -0.716364383 | 1.27E-07 | 27.91802122 | 0.608629268 | 6.235498763 |
| ENSMUSG00000037638 | *Zbtb42* | 447.5335512 | 434.5136775 | 380.6916173 | 264.5129826 | 253.7157107 | 252.5348892 | 3.87E-07 | -0.716580176 | 8.24E-08 | 28.74972129 | 0.608538238 | 6.412676865 |
| ENSMUSG00000015247 | *Nipsnap3b* | 1097.73802 | 1276.683675 | 1432.005554 | 763.9047132 | 736.9335996 | 810.8665715 | 2.90E-08 | -0.716661242 | 5.41E-09 | 34.03593212 | 0.608504045 | 7.536864021 |
| ENSMUSG00000037172 | *Dennd11* | 488.2184195 | 518.9225155 | 526.9175365 | 296.3423457 | 313.723161 | 321.4080408 | 5.81E-09 | -0.716881071 | 1.02E-09 | 37.2942235 | 0.608411332 | 8.235850442 |
| ENSMUSG00000024592 | *C330018D20Rik* | 319.4515585 | 268.5735755 | 381.9521856 | 194.268871 | 194.7610228 | 199.2729853 | 0.000147811 | -0.717200904 | 4.52E-05 | 16.63994282 | 0.608276467 | 3.83029318 |
| ENSMUSG00000020134 | *Peli1* | 697.6701489 | 640.7398159 | 645.4109539 | 397.318256 | 407.4190044 | 403.137514 | 9.33E-11 | -0.717508191 | 1.37E-11 | 45.7120868 | 0.608146921 | 10.03025339 |
| ENSMUSG00000053012 | *Krcc1* | 834.0398 | 762.5571162 | 910.1302904 | 522.4405797 | 485.3234135 | 514.2528653 | 9.07E-10 | -0.717992717 | 1.47E-10 | 41.07245697 | 0.607942711 | 9.042258602 |
| ENSMUSG00000027130 | *Slc12a6* | 2151.777479 | 2412.366223 | 2021.951504 | 1384.028511 | 1322.26943 | 1299.406794 | 2.86E-13 | -0.718326913 | 3.39E-14 | 57.49460543 | 0.607801899 | 12.54293688 |
| ENSMUSG00000022130 | *Tgds* | 664.5195155 | 571.6780393 | 855.9258548 | 450.0013397 | 429.5270124 | 390.2811924 | 6.43E-05 | -0.718371988 | 1.86E-05 | 18.33099346 | 0.607782909 | 4.191523889 |
| ENSMUSG00000022620 | *Arsa* | 889.0397146 | 932.3339836 | 942.9050654 | 526.8308367 | 559.0167734 | 591.3907951 | 2.11E-12 | -0.718452372 | 2.68E-13 | 53.43229736 | 0.607749046 | 11.67512908 |
| ENSMUSG00000041801 | *Phlda3* | 486.7115726 | 587.9842921 | 640.3686808 | 344.6351724 | 331.6201198 | 362.7319318 | 2.34E-06 | -0.718493394 | 5.49E-07 | 25.0849333 | 0.607731765 | 5.631489802 |
| ENSMUSG00000027663 | *Zmat3* | 1701.230234 | 1993.199607 | 1899.676382 | 1069.027573 | 1186.463095 | 1141.457699 | 1.16E-12 | -0.718508532 | 1.43E-13 | 54.65992554 | 0.607725388 | 11.93671562 |
| ENSMUSG00000020376 | *Rnf130* | 2178.147301 | 2110.22095 | 2390.037438 | 1376.345561 | 1364.379922 | 1315.018041 | 7.67E-17 | -0.718672475 | 6.81E-18 | 74.27017594 | 0.607656332 | 16.11545796 |
| ENSMUSG00000028098 | *Rnf115* | 1990.544853 | 2062.261383 | 2410.206531 | 1310.491706 | 1298.055898 | 1312.263115 | 2.08E-12 | -0.719340365 | 2.64E-13 | 53.46287525 | 0.607375085 | 11.68141817 |
| ENSMUSG00000016427 | *Ndufa1* | 886.7794441 | 862.3130157 | 1135.772011 | 658.5385459 | 546.383626 | 543.6387433 | 4.33E-06 | -0.720799822 | 1.05E-06 | 23.83111574 | 0.606760964 | 5.363107102 |
| ENSMUSG00000004285 | *Atp6v1f* | 4193.555128 | 4290.462869 | 5818.783131 | 3016.10654 | 2857.196842 | 2799.004881 | 2.56E-07 | -0.721154483 | 5.35E-08 | 29.58462034 | 0.606611821 | 6.592342211 |
| ENSMUSG00000061111 | *Mcrip1* | 1558.833194 | 1550.053207 | 1656.386706 | 1017.442053 | 967.4885401 | 904.5340577 | 1.79E-15 | -0.72175169 | 1.76E-16 | 67.8598959 | 0.606360765 | 14.74745221 |
| ENSMUSG00000023905 | *Tnfrsf12a* | 1631.915273 | 1671.870508 | 2020.690936 | 1064.637316 | 1036.970851 | 1120.336599 | 3.66E-10 | -0.722541947 | 5.64E-11 | 42.9403381 | 0.606028713 | 9.436922288 |
| ENSMUSG00000043017 | *Ptgir* | 1701.983657 | 1903.994812 | 1603.442838 | 1040.490903 | 1030.654277 | 1086.359178 | 4.93E-13 | -0.72280451 | 5.93E-14 | 56.39477986 | 0.605918429 | 12.30705136 |
| ENSMUSG00000019578 | *Ubxn6* | 2995.611784 | 3154.780321 | 3685.901619 | 1960.249738 | 1920.238408 | 2073.541018 | 4.53E-12 | -0.722890065 | 5.92E-13 | 51.87294916 | 0.605882497 | 11.34420457 |
| ENSMUSG00000025201 | *Bloc1s2-ps* | 278.0132667 | 298.3085071 | 442.4594625 | 208.5372062 | 218.9745553 | 187.3349724 | 0.001739112 | -0.723589117 | 0.000645066 | 11.64121101 | 0.605588991 | 2.759672428 |
| ENSMUSG00000041695 | *Kcnj2* | 452.8075157 | 511.2489848 | 476.4948058 | 273.2934965 | 261.0850467 | 336.1009798 | 3.47E-06 | -0.723598546 | 8.31E-07 | 24.28463228 | 0.605585033 | 5.460081449 |
| ENSMUSG00000068290 | *Ddrgk1* | 1844.380696 | 1897.280473 | 2392.558575 | 1231.467081 | 1252.787119 | 1225.942098 | 5.13E-09 | -0.723712238 | 8.94E-10 | 37.54320907 | 0.605537311 | 8.289927627 |
| ENSMUSG00000021624 | *Cd180* | 2798.968254 | 2990.758602 | 2828.715196 | 1694.639191 | 1773.904451 | 1745.704816 | 1.18E-23 | -0.72479477 | 7.02E-25 | 106.0965437 | 0.605083114 | 22.92647803 |
| ENSMUSG00000037824 | *Tspan14* | 2234.654062 | 2589.816621 | 2516.094265 | 1427.93108 | 1501.239019 | 1507.862866 | 2.98E-15 | -0.725107429 | 2.98E-16 | 66.81822278 | 0.604951996 | 14.52611323 |
| ENSMUSG00000060450 | *Rnf14* | 1521.16202 | 1563.481886 | 1797.570352 | 1043.783595 | 934.8529092 | 971.5705919 | 1.27E-11 | -0.72519349 | 1.72E-12 | 49.77966355 | 0.60491591 | 10.89779214 |
| ENSMUSG00000002668 | *Dennd1c* | 415.1363413 | 386.5541105 | 297.4941115 | 237.0738765 | 214.7635062 | 215.8025417 | 8.06E-05 | -0.725354014 | 2.36E-05 | 17.8729217 | 0.604848606 | 4.093808661 |
| ENSMUSG00000039234 | *Sec24d* | 3394.926232 | 3815.663155 | 3111.082488 | 1960.249738 | 2193.956602 | 2090.070574 | 3.72E-12 | -0.725478988 | 4.81E-13 | 52.27923988 | 0.604796213 | 11.42994316 |
| ENSMUSG00000027230 | *Creb3l1* | 147.6710034 | 133.3275964 | 137.4019413 | 71.34167581 | 102.1179417 | 79.89285586 | 0.003409033 | -0.725543626 | 0.001339798 | 10.28687563 | 0.604769117 | 2.46736883 |
| ENSMUSG00000017747 | *Ghdc* | 458.8349036 | 467.1261831 | 465.1496913 | 308.4155523 | 283.1930547 | 250.6982718 | 1.71E-07 | -0.72600465 | 3.49E-08 | 30.41230456 | 0.604575889 | 6.767783127 |
| ENSMUSG00000028057 | *Rit1* | 1388.559486 | 1509.767171 | 1839.169105 | 974.6370479 | 887.4786064 | 996.3649265 | 5.17E-08 | -0.726643132 | 9.93E-09 | 32.85534063 | 0.604308386 | 7.286786047 |
| ENSMUSG00000021615 | *Xrcc4* | 326.2323698 | 397.1052152 | 443.7200308 | 233.7811838 | 224.2383667 | 244.270111 | 2.96E-05 | -0.726947238 | 8.14E-06 | 19.90501607 | 0.604181017 | 4.528747772 |
| ENSMUSG00000027775 | *Mfsd1* | 7105.536905 | 7493.202757 | 8288.23637 | 4590.013665 | 4724.797136 | 4505.222423 | 1.43E-18 | -0.727272112 | 1.15E-19 | 82.33024288 | 0.60404498 | 17.84415687 |
| ENSMUSG00000024955 | *Esrra* | 2526.982375 | 2651.204867 | 2222.381859 | 1475.126343 | 1494.922445 | 1502.353014 | 1.68E-14 | -0.727655473 | 1.79E-15 | 63.28657676 | 0.603884491 | 13.77436383 |
| ENSMUSG00000039623 | *Ap5z1* | 1038.217565 | 1118.417104 | 878.6160837 | 593.7822556 | 672.7151002 | 569.3513866 | 9.38E-08 | -0.727695848 | 1.86E-08 | 31.63828702 | 0.603867591 | 7.027590661 |
| ENSMUSG00000019986 | *Ahi1* | 540.9580636 | 573.596422 | 641.629249 | 399.5133845 | 345.3060295 | 314.97988 | 1.00E-06 | -0.727861333 | 2.26E-07 | 26.79761557 | 0.603798328 | 5.998275423 |
| ENSMUSG00000021631 | *Mrps36-ps1* | 190.6161422 | 169.7768674 | 221.8600154 | 117.439374 | 110.5400399 | 122.1350555 | 0.000473665 | -0.728447665 | 0.000157065 | 14.28556177 | 0.603552986 | 3.324528317 |
| ENSMUSG00000031532 | *Saraf* | 2325.064881 | 2294.385688 | 2379.952892 | 1402.687103 | 1441.231568 | 1379.299649 | 1.32E-24 | -0.728722979 | 7.36E-26 | 110.566138 | 0.603437819 | 23.88101665 |
| ENSMUSG00000021124 | *Vti1b* | 1977.736653 | 2057.465427 | 2698.876664 | 1339.028377 | 1295.950373 | 1421.541849 | 1.70E-07 | -0.729549223 | 3.48E-08 | 30.41985739 | 0.603092324 | 6.769328388 |
| ENSMUSG00000034792 | *Gna15* | 709.7249247 | 753.9243941 | 777.7706221 | 430.2451833 | 437.9491106 | 480.2754438 | 8.68E-11 | -0.729873404 | 1.27E-11 | 45.85770984 | 0.602956821 | 10.06131901 |
| ENSMUSG00000041187 | *Prkd2* | 527.3964409 | 493.9835407 | 389.5155952 | 275.488625 | 302.1427758 | 275.4926064 | 1.15E-05 | -0.730658086 | 2.95E-06 | 21.84889826 | 0.602628962 | 4.941058107 |
| ENSMUSG00000037361 | *Sf3b6* | 2178.900724 | 1956.750336 | 2590.467793 | 1381.833382 | 1323.322192 | 1344.403919 | 2.14E-09 | -0.730892776 | 3.58E-10 | 39.32945607 | 0.602530937 | 8.670343811 |
| ENSMUSG00000040370 | *Etfrf1* | 216.2325408 | 189.9198855 | 242.0291077 | 119.6345025 | 153.7032936 | 116.6252034 | 0.000763107 | -0.73151086 | 0.000262949 | 13.3174407 | 0.602272854 | 3.117414758 |
| ENSMUSG00000020590 | *Snx13* | 467.1225619 | 515.0857502 | 524.3964 | 305.1228596 | 291.615153 | 308.5517192 | 9.74E-09 | -0.731788028 | 1.74E-09 | 36.24784133 | 0.602157158 | 8.011523469 |
| ENSMUSG00000024425 | *Ndfip1* | 1715.54528 | 1839.728992 | 2153.050604 | 1219.393874 | 1113.822498 | 1100.133808 | 3.77E-10 | -0.732098979 | 5.83E-11 | 42.8760923 | 0.602027386 | 9.423431735 |
| ENSMUSG00000020227 | *Irak3* | 416.6431883 | 348.1864568 | 398.3395731 | 234.878748 | 224.2383667 | 240.5968763 | 9.65E-07 | -0.733441495 | 2.17E-07 | 26.87792379 | 0.601467423 | 6.01526178 |
| ENSMUSG00000006445 | *Epha2* | 598.9716721 | 519.8817069 | 398.3395731 | 272.1959323 | 329.5145953 | 313.1432626 | 0.000145729 | -0.733518506 | 4.45E-05 | 16.6706878 | 0.601435318 | 3.836454086 |
| ENSMUSG00000019877 | *Serinc1* | 3257.803158 | 3299.618213 | 4611.158729 | 2295.006832 | 2273.966536 | 2143.332478 | 7.79E-07 | -0.733766752 | 1.72E-07 | 27.32128624 | 0.601331837 | 6.108714866 |
| ENSMUSG00000028037 | *Ifi44* | 4101.637463 | 3139.433259 | 2971.159411 | 2565.007636 | 1464.392339 | 2109.355056 | 0.000541431 | -0.7347561 | 0.000181691 | 14.0115543 | 0.600919607 | 3.266456954 |
| ENSMUSG00000024413 | *Npc1* | 3296.227755 | 3224.801289 | 2592.98893 | 1715.492912 | 1894.972113 | 1869.676489 | 4.31E-10 | -0.73479167 | 6.71E-11 | 42.6024805 | 0.600904791 | 9.365567919 |
| ENSMUSG00000051412 | *None* | 597.4648251 | 587.9842921 | 687.0097067 | 386.3426136 | 313.723161 | 422.4219965 | 3.16E-06 | -0.735014009 | 7.55E-07 | 24.47054178 | 0.600812191 | 5.500747924 |
| ENSMUSG00000027615 | *Hps3* | 836.3000705 | 765.4346902 | 781.5523269 | 485.1233955 | 460.0571186 | 487.6219133 | 3.22E-12 | -0.735228084 | 4.15E-13 | 52.57127808 | 0.600723046 | 11.4924506 |
| ENSMUSG00000035735 | *Dagla* | 142.397039 | 128.5316397 | 112.190576 | 57.07334064 | 81.06269596 | 91.8308688 | 0.009925036 | -0.736627357 | 0.004292392 | 8.155789044 | 0.600140685 | 2.003267915 |
| ENSMUSG00000083563 | *Gm13340* | 184.5887543 | 181.2871635 | 288.6701337 | 126.219888 | 127.3842365 | 136.8279945 | 0.004217964 | -0.737196724 | 0.001691296 | 9.857621999 | 0.599903883 | 2.374897158 |
| ENSMUSG00000017386 | *Traf4* | 762.4645688 | 753.9243941 | 787.8551683 | 446.708647 | 482.1651266 | 452.7261832 | 1.03E-12 | -0.737248549 | 1.26E-13 | 54.90680569 | 0.599882334 | 11.98885794 |
| ENSMUSG00000029924 | *Slc37a3* | 694.656455 | 799.9655785 | 779.0311904 | 445.6110827 | 456.8988318 | 459.154344 | 1.08E-10 | -0.737326633 | 1.59E-11 | 45.41689376 | 0.599849867 | 9.967432371 |
| ENSMUSG00000035530 | *Eif1* | 15027.03145 | 14772.50584 | 14665.45124 | 9182.222458 | 8441.048003 | 9048.095503 | 5.15E-32 | -0.737433251 | 2.05E-33 | 145.0963806 | 0.599805538 | 31.2882696 |
| ENSMUSG00000020553 | *Pctp* | 275.7529962 | 316.5331426 | 308.839226 | 156.9516868 | 205.2886456 | 177.2335768 | 3.70E-05 | -0.737535591 | 1.03E-05 | 19.44933466 | 0.599762992 | 4.431425009 |
| ENSMUSG00000036977 | *Anapc10* | 399.3144481 | 383.6765364 | 535.7415144 | 274.3910608 | 274.7709564 | 239.6785676 | 7.34E-05 | -0.738060795 | 2.14E-05 | 18.06287465 | 0.599544692 | 4.134393868 |
| ENSMUSG00000039531 | *Zup1* | 183.0819073 | 188.0015029 | 247.0713808 | 142.6833516 | 125.2787119 | 101.9322644 | 0.002095949 | -0.738308613 | 0.000790512 | 11.26337962 | 0.599441714 | 2.678619343 |
| ENSMUSG00000015312 | *Gadd45b* | 468.6294089 | 392.3092585 | 447.5017356 | 295.2447814 | 229.5021782 | 259.8813587 | 5.46E-06 | -0.739023077 | 1.34E-06 | 23.35835017 | 0.599144927 | 5.262845524 |
| ENSMUSG00000030304 | *Ergic2* | 1678.627529 | 1666.11536 | 2164.395718 | 1114.027707 | 1101.18935 | 1080.849326 | 8.67E-09 | -0.739419274 | 1.54E-09 | 36.48455151 | 0.598980411 | 8.06192425 |
| ENSMUSG00000035914 | *Cd276* | 1661.298789 | 1669.952125 | 1239.138609 | 830.8561321 | 980.1216875 | 929.3283923 | 6.00E-07 | -0.739445431 | 1.31E-07 | 27.85314649 | 0.598969551 | 6.221621283 |

| | | | | | | | | | | | | |
|---|---|---|---|---|---|---|---|---|---|---|---|---|
| ENSMUSG00000052760 | *A630001G21Rik* | 1063.833964 | 1149.111227 | 1186.194741 | 695.8557302 | 672.7151002 | 664.8554901 | 1.66E-14 | -0.739737559 | 1.77E-15 | 63.30721278 | 0.598848279 | 13.77863314 |
| ENSMUSG00000037251 | *Pomk* | 913.1492662 | 1100.192468 | 1230.314631 | 626.7091828 | 672.7151002 | 639.1428469 | 9.53E-08 | -0.739919919 | 1.89E-08 | 31.60469099 | 0.598772588 | 7.020679306 |
| ENSMUSG00000003518 | *Dusp3* | 4638.828409 | 5075.081386 | 4851.927268 | 2892.081781 | 2864.566178 | 2961.545519 | 1.40E-26 | -0.740065824 | 7.06E-28 | 119.7835786 | 0.598712035 | 25.85484378 |
| ENSMUSG00000028295 | *Smim8* | 228.2873165 | 289.675785 | 320.1843404 | 182.1956644 | 168.4419656 | 149.6843161 | 0.000342134 | -0.740584191 | 0.000110609 | 14.94639815 | 0.598496954 | 3.465803421 |
| ENSMUSG00000022024 | *Sugt1* | 488.971843 | 471.9221398 | 574.8191308 | 347.9278651 | 260.0322844 | 309.4700279 | 8.98E-06 | -0.741080675 | 2.28E-06 | 22.3431566 | 0.598291024 | 5.046584303 |
| ENSMUSG00000054099 | *Slc25a40* | 329.2460638 | 300.2268898 | 345.3957058 | 192.0737426 | 193.7082605 | 196.5180592 | 1.03E-06 | -0.741477644 | 2.32E-07 | 26.74374332 | 0.598126423 | 5.986697508 |
| ENSMUSG00000020377 | *Ltc4s* | 116.027217 | 108.3886215 | 122.2751221 | 53.78064792 | 73.69335996 | 78.97454717 | 0.00627933 | -0.741832784 | 0.002603173 | 9.066623783 | 0.597979203 | 2.2020867 |
| ENSMUSG00000091512 | *Lamtor3* | 944.7930526 | 1071.416728 | 1255.525996 | 670.6117526 | 621.1297483 | 660.2639467 | 4.23E-08 | -0.742262045 | 8.06E-09 | 33.26090595 | 0.597801307 | 7.373481366 |
| ENSMUSG00000047866 | *Lonp2* | 1112.80649 | 1186.519689 | 1259.307701 | 667.3190598 | 752.7250339 | 702.5061463 | 7.94E-13 | -0.743533982 | 9.70E-14 | 55.42787808 | 0.597274493 | 12.10010151 |
| ENSMUSG00000032840 | *Spring1* | 1203.970732 | 1484.828196 | 1312.251568 | 782.5633054 | 821.1545824 | 783.3173109 | 5.21E-11 | -0.74455964 | 7.44E-12 | 46.90821648 | 0.596850023 | 10.28344595 |
| ENSMUSG00000022822 | *Abcc5* | 1117.327031 | 1401.378549 | 1372.758845 | 780.3681769 | 771.674755 | 765.8694458 | 7.57E-10 | -0.745740287 | 1.21E-10 | 41.44724147 | 0.596361783 | 9.120709404 |
| ENSMUSG00000061244 | *Exoc5* | 1160.27217 | 1104.988425 | 1149.638262 | 703.5386799 | 639.0267071 | 693.3230594 | 1.92E-15 | -0.746124159 | 1.89E-16 | 67.71498036 | 0.596203124 | 14.71735117 |
| ENSMUSG00000009621 | *Vav2* | 953.080711 | 1009.069291 | 886.1794933 | 555.367507 | 594.8106911 | 550.0669041 | 2.13E-12 | -0.74615089 | 2.70E-13 | 53.41733415 | 0.596192078 | 11.67205633 |
| ENSMUSG00000004446 | *Bid* | 4286.226217 | 4581.097845 | 4879.65977 | 2843.788954 | 2669.805155 | 2675.951517 | 4.25E-20 | -0.746647439 | 3.08E-21 | 89.49295224 | 0.595986915 | 19.37114767 |
| ENSMUSG00000044434 | *Gm9791* | 259.1776795 | 241.716218 | 250.8530856 | 163.5370722 | 137.9118594 | 146.9293901 | 1.37E-05 | -0.747348664 | 3.59E-06 | 21.47228398 | 0.595697305 | 4.862563632 |
| ENSMUSG00000018507 | *Trpv2* | 7944.097246 | 7748.347654 | 6952.034005 | 4572.452637 | 4372.12177 | 4547.464623 | 2.52E-21 | -0.747562401 | 1.68E-22 | 95.25158154 | 0.595609058 | 20.59925978 |
| ENSMUSG00000039096 | *Rsad1* | 135.6162277 | 145.7970839 | 141.1836462 | 82.31731824 | 88.43203196 | 80.81116454 | 0.0006244 | -0.748006665 | 0.000212361 | 13.71842173 | 0.595425674 | 3.204536866 |
| ENSMUSG00000044092 | *C130050O18Rik* | 1345.614348 | 1284.357206 | 1211.406107 | 740.8558641 | 815.890771 | 732.810333 | 4.43E-14 | -0.748527928 | 4.85E-15 | 61.32087626 | 0.595210578 | 13.35330681 |
| ENSMUSG00000074166 | *AW146154* | 113.7669465 | 129.4908311 | 152.5287606 | 65.85385459 | 70.53507311 | 97.34072093 | 0.008113478 | -0.748647721 | 0.003449045 | 8.553257351 | 0.595161158 | 2.09079293 |
| ENSMUSG00000110841 | *Gpx4-ps2* | 3887.665193 | 4010.378997 | 5270.435934 | 2603.422385 | 2689.807639 | 2535.450288 | 1.16E-08 | -0.74947901 | 2.09E-09 | 35.88576561 | 0.594818321 | 7.934651572 |
| ENSMUSG00000027207 | *Galk2* | 1267.258305 | 1266.13257 | 1720.675688 | 824.2707466 | 844.3153527 | 856.7820059 | 3.22E-07 | -0.75006088 | 6.80E-08 | 29.12207781 | 0.594578467 | 6.49250805 |
| ENSMUSG00000039089 | *L3mbtl3* | 176.3010959 | 178.4095894 | 115.9722808 | 82.31731824 | 104.2234662 | 94.58579486 | 0.005884763 | -0.750257048 | 0.002427889 | 9.194157079 | 0.594497625 | 2.23027104 |
| ENSMUSG00000037509 | *Arhgef4* | 88.15054797 | 109.3478129 | 104.6271663 | 58.17090489 | 61.06021254 | 59.69006472 | 0.005361252 | -0.750975824 | 0.002193139 | 9.380431464 | 0.59420151 | 2.270733748 |
| ENSMUSG00000079316 | *Rab9* | 274.2461492 | 267.6143842 | 302.5363846 | 181.0981001 | 172.6530148 | 147.8476988 | 1.39E-05 | -0.751539965 | 3.65E-06 | 21.44095853 | 0.593969203 | 4.855696509 |
| ENSMUSG00000068396 | *Rpl34-ps1* | 3154.58414 | 3072.289866 | 4214.079724 | 1974.518073 | 1900.235925 | 2321.484363 | 5.99E-07 | -0.751695111 | 1.31E-07 | 27.85780534 | 0.593905331 | 6.222530302 |
| ENSMUSG00000025492 | *Ifitm3* | 25499.61792 | 23830.14968 | 21994.39516 | 16043.09654 | 11171.91337 | 15120.87086 | 1.32E-07 | -0.752558954 | 2.66E-08 | 30.94422593 | 0.593549825 | 6.88023693 |
| ENSMUSG00000041396 | *Mettl18* | 299.1091243 | 300.2268898 | 388.2550269 | 203.049385 | 183.1806376 | 197.4363679 | 3.13E-05 | -0.753341031 | 8.64E-06 | 19.79173288 | 0.593228152 | 4.504263576 |
| ENSMUSG00000025791 | *Pgm1* | 2484.037236 | 2890.043511 | 3301.428297 | 1887.810498 | 1531.769125 | 1721.82879 | 2.84E-08 | -0.753862075 | 5.27E-09 | 34.08686817 | 0.59301394 | 7.547431785 |
| ENSMUSG00000036086 | *Zranb3* | 339.0405691 | 321.3290993 | 389.5155952 | 205.2445135 | 195.813785 | 219.4757764 | 2.25E-06 | -0.75390948 | 5.27E-07 | 25.16209795 | 0.592994455 | 5.647486334 |
| ENSMUSG00000029449 | *Rhof* | 2551.091927 | 2461.284981 | 2343.396413 | 1477.321471 | 1361.221635 | 1522.555805 | 8.90E-19 | -0.754702038 | 7.01E-20 | 83.31064463 | 0.592668778 | 18.0505746 |
| ENSMUSG00000086784 | *Isoc2a* | 879.2452093 | 950.5586191 | 1090.391553 | 611.3432834 | 520.0645689 | 595.9823385 | 1.07E-08 | -0.75479359 | 1.92E-09 | 36.05349383 | 0.592631169 | 7.970377222 |
| ENSMUSG00000019210 | *Atp6v1e1* | 2798.214831 | 2987.881028 | 3885.071406 | 1893.298319 | 1860.230958 | 1971.608753 | 3.17E-08 | -0.755164988 | 5.95E-09 | 33.85189421 | 0.592478625 | 7.499131755 |
| ENSMUSG00000021036 | *Sptlc2* | 2273.832084 | 2176.405153 | 2280.367999 | 1346.711326 | 1250.681595 | 1388.482736 | 1.49E-20 | -0.755308368 | 1.05E-21 | 91.62390558 | 0.592419746 | 19.82634054 |
| ENSMUSG00000012640 | *Zfp715* | 493.4923839 | 427.7993381 | 462.6285548 | 308.4155523 | 244.2408502 | 268.1461369 | 7.08E-07 | -0.755625279 | 1.56E-07 | 27.51404012 | 0.592289625 | 6.150248096 |
| ENSMUSG00000039699 | *Batf2* | 161.2326262 | 166.8992934 | 142.4442144 | 94.39052491 | 88.43203196 | 96.42241224 | 0.000390228 | -0.755974952 | 0.000127389 | 14.68002567 | 0.592146087 | 3.408681564 |
| ENSMUSG00000039031 | *Arhgap18* | 467.8759854 | 403.8195546 | 477.755374 | 276.5861893 | 245.2936124 | 276.4109151 | 2.28E-07 | -0.756172733 | 4.74E-08 | 29.8187226 | 0.592064914 | 6.641502502 |
| ENSMUSG00000071657 | *Bscl2* | 1649.997436 | 1824.381931 | 1850.514219 | 1043.783595 | 1038.023613 | 1067.074695 | 1.08E-17 | -0.756221846 | 9.10E-19 | 78.24618634 | 0.592044759 | 16.96856221 |
| ENSMUSG00000112449 | *Srp54b* | 385.7528253 | 383.6765364 | 470.1919644 | 296.3423457 | 194.7610228 | 241.5151849 | 0.000184424 | -0.756975021 | 5.73E-05 | 16.19006522 | 0.591735756 | 3.734183661 |
| ENSMUSG00000001158 | *Snrnp27* | 1264.244611 | 1151.029609 | 1544.19613 | 846.2220315 | 754.8305585 | 739.2384938 | 7.51E-08 | -0.757398473 | 1.47E-08 | 32.09566719 | 0.591562099 | 7.124400491 |
| ENSMUSG00000037697 | *Ddhd1* | 360.1364268 | 399.0235979 | 298.7546798 | 205.2445135 | 230.5549405 | 191.9265158 | 2.82E-05 | -0.757536672 | 7.71E-06 | 20.00766584 | 0.591505434 | 4.549999565 |
| ENSMUSG00000029276 | *Glmn* | 263.6982204 | 228.2875392 | 345.3957058 | 187.6834856 | 164.2309165 | 142.3378466 | 0.001016049 | -0.758235741 | 0.000358183 | 12.73861415 | 0.591218885 | 2.993085432 |
| ENSMUSG00000018986 | *Slfn3* | 456.5746331 | 449.860739 | 491.621625 | 264.5129826 | 302.1427758 | 258.96305 | 3.15E-08 | -0.758908731 | 5.91E-09 | 33.86279771 | 0.590943157 | 7.501089088 |
| ENSMUSG00000052298 | *Cdc42se2* | 4214.650986 | 4492.852242 | 4206.516314 | 2466.226854 | 2530.840534 | 2630.954391 | 5.94E-27 | -0.759309165 | 2.95E-28 | 121.5114092 | 0.590779158 | 26.22655796 |
| ENSMUSG00000028893 | *Sesn2* | 917.6698071 | 885.3336079 | 630.2841346 | 479.6355743 | 469.5319792 | 490.3768394 | 6.46E-06 | -0.76028962 | 1.61E-06 | 23.01489924 | 0.590377801 | 5.189875398 |
| ENSMUSG00000098188 | *Sowahc* | 195.8901066 | 188.9606942 | 147.4864875 | 100.9759104 | 103.1707039 | 111.1153512 | 0.000564982 | -0.761265085 | 0.000190331 | 13.92421411 | 0.589978757 | 3.247965092 |
| ENSMUSG00000030062 | *Rpn1* | 20767.36499 | 20684.00208 | 23761.71188 | 12744.91599 | 12957.39821 | 12765.40907 | 5.47E-22 | -0.761319009 | 3.50E-23 | 98.35385204 | 0.589956706 | 21.26239436 |
| ENSMUSG00000028048 | *Gba* | 6468.894059 | 6726.808875 | 7220.535046 | 4045.6218 | 3966.808291 | 4008.417423 | 4.38E-27 | -0.763590765 | 2.16E-28 | 122.1329466 | 0.589028455 | 26.35900438 |
| ENSMUSG00000052825 | *Gm9892* | 1032.943601 | 750.0876288 | 1293.343044 | 707.9289368 | 541.1198146 | 562.0049171 | 0.000432129 | -0.763600174 | 0.000142343 | 14.47088319 | 0.589024614 | 3.364386359 |
| ENSMUSG00000024293 | *Esco1* | 504.7937363 | 412.4522767 | 462.6285548 | 267.8056753 | 280.0347679 | 265.3912108 | 1.14E-07 | -0.764685468 | 2.27E-08 | 31.2482834 | 0.588581675 | 6.944852395 |
| ENSMUSG00000033416 | *Gucd1* | 687.8756436 | 850.8027196 | 828.1933529 | 487.318524 | 437.9491106 | 465.5825048 | 8.87E-09 | -0.764843429 | 1.58E-09 | 36.43813677 | 0.588517235 | 8.052085944 |
| ENSMUSG00000041992 | *Rapgef5* | 711.2317717 | 710.7607838 | 660.5377731 | 358.9035075 | 427.4214878 | 438.0332442 | 1.91E-09 | -0.766028557 | 3.18E-10 | 39.56006703 | 0.588033985 | 8.719493046 |
| ENSMUSG00000014763 | *Fam120b* | 504.7937363 | 556.3309779 | 567.2557212 | 339.1473511 | 318.9869724 | 298.4503236 | 3.24E-09 | -0.766539293 | 5.53E-10 | 38.48076451 | 0.587825849 | 8.489986401 |
| ENSMUSG00000030469 | *Zfp719* | 283.2872311 | 292.553359 | 346.656274 | 164.6346365 | 192.6554982 | 182.7434289 | 9.66E-06 | -0.766990998 | 2.46E-06 | 22.19572088 | 0.587641831 | 5.014995965 |
| ENSMUSG00000004317 | *Clcn5* | 565.8210387 | 679.1074695 | 601.2910644 | 331.4644014 | 369.5195621 | 382.0164142 | 1.80E-08 | -0.767612257 | 3.29E-09 | 35.00425852 | 0.587388833 | 7.745164834 |
| ENSMUSG00000021038 | *Vipas39* | 1970.955842 | 1907.831577 | 2006.824685 | 1174.39374 | 1186.463095 | 1096.460573 | 9.31E-22 | -0.767888509 | 6.04E-23 | 97.27328487 | 0.587276368 | 21.03095017 |
| ENSMUSG00000025759 | *Mfsd8* | 259.1776795 | 252.2673227 | 320.1843404 | 177.8054074 | 137.9118594 | 170.805416 | 0.000111469 | -0.768664875 | 3.34E-05 | 17.21479542 | 0.586960419 | 3.952844972 |
| ENSMUSG00000044600 | *Smim7* | 1928.764127 | 2109.261759 | 2591.728362 | 1334.63812 | 1258.050931 | 1291.142015 | 1.95E-09 | -0.769919516 | 3.26E-10 | 39.51088591 | 0.58645019 | 8.70909141 |
| ENSMUSG00000022051 | *Bnip3l* | 1636.435814 | 1536.624528 | 1960.183659 | 1104.149629 | 940.1167207 | 963.3058137 | 2.91E-09 | -0.770316811 | 4.95E-10 | 38.6950764 | 0.586288713 | 8.535751163 |
| ENSMUSG00000101316 | *Gm12663* | 91.91766541 | 119.8989176 | 104.6271663 | 52.68308367 | 73.69335996 | 58.77175603 | 0.007790272 | -0.770747646 | 0.00329317 | 8.637477903 | 0.586113655 | 2.10844738 |
| ENSMUSG00000010911 | *Apip* | 565.8210387 | 611.0048843 | 751.2986885 | 346.8303008 | 396.8913815 | 382.0164142 | 6.42E-07 | -0.772056003 | 1.40E-07 | 27.7167544 | 0.585582359 | 6.192777334 |
| ENSMUSG00000028771 | *Ptpn12* | 1266.504882 | 1291.071545 | 1551.759539 | 819.8804896 | 757.9888453 | 823.7228931 | 5.94E-11 | -0.772544847 | 8.55E-12 | 46.63610748 | 0.585383973 | 10.22626163 |
| ENSMUSG00000096727 | *Psmb9* | 1623.627614 | 1544.298059 | 1371.498277 | 1004.271282 | 784.3079025 | 871.4749449 | 1.89E-09 | -0.772755077 | 3.14E-10 | 39.58424554 | 0.585298677 | 8.724512441 |
| ENSMUSG00000033720 | *Sfxn5* | 302.8762418 | 329.9618213 | 302.5363846 | 188.7810498 | 178.9695885 | 179.9885028 | 3.48E-07 | -0.772863476 | 7.37E-08 | 28.96603557 | 0.585254701 | 6.458935126 |
| ENSMUSG00000002768 | *Mea1* | 2875.064026 | 2935.125504 | 3737.584918 | 1938.298453 | 1741.26882 | 1903.65391 | 8.71E-10 | -0.772968467 | 1.41E-10 | 41.15648676 | 0.585212112 | 9.060191926 |
| ENSMUSG00000056234 | *Ncoa4* | 2383.831913 | 2617.63317 | 3277.4775 | 1675.980599 | 1509.661117 | 1653.873947 | 1.18E-08 | -0.773289466 | 2.12E-09 | 35.85727984 | 0.585081917 | 7.928963805 |
| ENSMUSG00000030560 | *Ctsc* | 2047.805038 | 2047.873513 | 1861.859334 | 1119.515528 | 1232.784636 | 1135.947847 | 7.50E-18 | -0.773478368 | 6.26E-19 | 78.98464337 | 0.585005313 | 17.12487749 |
| ENSMUSG00000032251 | *Irak1bp1* | 310.4104766 | 287.7574023 | 332.7900231 | 159.1468153 | 184.2333999 | 199.2729853 | 5.56E-06 | -0.774235635 | 1.37E-06 | 23.31970168 | 0.584698325 | 5.254865043 |
| ENSMUSG00000022292 | *Rrm2b* | 316.4378645 | 327.0842473 | 374.388776 | 193.1713068 | 196.8665473 | 202.94622 | 6.01E-07 | -0.774475041 | 1.31E-07 | 27.8492587 | 0.584601307 | 6.22088529 |
| ENSMUSG00000003721 | *Insig2* | 473.9033733 | 537.147151 | 608.854474 | 311.7082451 | 293.7206776 | 337.9375972 | 3.50E-07 | -0.774979954 | 7.42E-08 | 28.95255327 | 0.584396744 | 6.456193586 |
| ENSMUSG00000027742 | *Cog6* | 1073.628469 | 1151.029609 | 1202.582129 | 682.6849592 | 646.3960431 | 670.3653422 | 1.45E-15 | -0.775486999 | 1.41E-16 | 68.28838258 | 0.58419139 | 14.83755346 |
| ENSMUSG00000029128 | *Rab28* | 1737.394561 | 1770.667216 | 1986.655592 | 1129.393606 | 1022.232179 | 1054.218374 | 1.62E-15 | -0.77610078 | 1.58E-16 | 68.06300229 | 0.583942904 | 14.7900562 |
| ENSMUSG00000026922 | *Agpat2* | 1411.162191 | 1538.542911 | 1674.034662 | 915.3685788 | 846.4208773 | 933.9199357 | 1.80E-13 | -0.776181579 | 2.09E-14 | 58.44992929 | 0.583910201 | 12.74468246 |
| ENSMUSG00000021610 | *Clptm1l* | 9313.821146 | 10062.87636 | 10369.43458 | 5545.992121 | 6021.800271 | 5796.364439 | 3.71E-25 | -0.776205816 | 2.02E-26 | 113.1266016 | 0.583900392 | 24.43057734 |
| ENSMUSG00000036503 | *Rnf13* | 1374.997864 | 1361.092513 | 1893.37354 | 949.3930703 | 890.6368933 | 859.536932 | 4.41E-07 | -0.776513235 | 9.45E-08 | 28.48255293 | 0.583775983 | 6.355545743 |
| ENSMUSG00000019590 | *Cyb561* | 248.6297507 | 221.5731998 | 239.5079712 | 130.6101449 | 152.6505314 | 131.3181424 | 1.67E-05 | -0.777092895 | 4.42E-06 | 21.07223148 | 0.583541476 | 4.77645868 |
| ENSMUSG00000083022 | *Rps15a-ps6* | 1548.285266 | 1594.176009 | 1810.176035 | 886.8319085 | 906.4283275 | 1091.86903 | 3.40E-10 | -0.777256994 | 5.23E-11 | 43.08897805 | 0.583475104 | 9.4689514 |
| ENSMUSG00000028803 | *Nipal3* | 704.4509603 | 831.6188928 | 641.629249 | 372.0742784 | 484.2706512 | 415.075527 | 1.40E-06 | -0.777349385 | 3.20E-07 | 26.12224214 | 0.58343774 | 5.854178337 |
| ENSMUSG00000050538 | *B230217C12Rik* | 194.3832596 | 186.0831202 | 215.557174 | 138.2930946 | 90.53755653 | 118.4618208 | 0.00074729 | -0.777535177 | 0.000257182 | 13.35902401 | 0.583362609 | 3.126511012 |
| ENSMUSG00000039233 | *Tbce* | 1147.463971 | 1010.987674 | 1251.744291 | 662.9288029 | 672.7151002 | 651.9991685 | 1.55E-11 | -0.777576346 | 2.12E-12 | 49.36607065 | 0.583345962 | 10.80929496 |
| ENSMUSG00000032802 | *Srxn1* | 7907.179496 | 9241.808572 | 9961.010464 | 4990.624614 | 5669.124906 | 5148.956814 | 9.11E-13 | -0.777726715 | 1.12E-13 | 55.14723872 | 0.583285164 | 12.04028746 |
| ENSMUSG00000035762 | *Tmem161b* | 302.8762418 | 255.1448967 | 344.1351375 | 196.4639995 | 157.9143428 | 170.805416 | 6.41E-05 | -0.778616517 | 1.85E-05 | 18.34045781 | 0.582925526 | 4.193371382 |
| ENSMUSG00000030203 | *Dusp16* | 792.6015083 | 683.9034262 | 723.5661865 | 469.7574961 | 404.2607175 | 410.4839835 | 8.17E-10 | -0.7787288 | 1.31E-10 | 41.2885207 | 0.582880159 | 9.088003408 |
| ENSMUSG00000029390 | *Tmed2* | 12934.77442 | 13814.27369 | 15733.15257 | 7986.974998 | 8458.944962 | 8292.327453 | 2.71E-17 | -0.779784671 | 2.34E-18 | 76.38138336 | 0.582453721 | 16.56654161 |
| ENSMUSG00000025188 | *Hps1* | 1618.35365 | 1568.277843 | 1365.195436 | 908.7831933 | 896.953467 | 849.4355364 | 6.89E-14 | -0.780106176 | 7.68E-15 | 60.41484603 | 0.582323935 | 13.16206314 |
| ENSMUSG00000047205 | *Dusp18* | 233.561281 | 312.6963772 | 230.6839933 | 139.3906589 | 156.8615805 | 156.112477 | 0.000160286 | -0.780186198 | 4.92E-05 | 16.47737224 | 0.582291637 | 3.795104241 |
| ENSMUSG00000054720 | *Lrrc8c* | 879.9986328 | 808.5983006 | 607.5939058 | 407.1963342 | 467.4264546 | 463.7458874 | 4.56E-06 | -0.781474132 | 1.11E-06 | 23.72880158 | 0.58177204 | 5.341125347 |
| ENSMUSG00000003812 | *Dnase2a* | 3241.981264 | 3480.905377 | 3281.259205 | 1916.347169 | 1963.401662 | 1936.713023 | 4.81E-29 | -0.782306979 | 2.16E-30 | 131.2756329 | 0.581436289 | 28.317935 |
| ENSMUSG00000021120 | *Pigh* | 245.6160567 | 272.4103409 | 262.1982 | 167.9273292 | 152.6505314 | 133.1547598 | 1.23E-05 | -0.783733948 | 3.18E-06 | 21.70614284 | 0.580861474 | 4.911627044 |
| ENSMUSG00000066442 | *Mthfs* | 117.534064 | 128.5316397 | 158.8316019 | 65.85385459 | 83.16822053 | 84.4843993 | 0.002700726 | -0.784099952 | 0.001038071 | 10.7584004 | 0.580714132 | 2.568519502 |
| ENSMUSG00000025287 | *Acot9* | 1851.161507 | 1586.502478 | 1849.253651 | 1146.954634 | 901.1645161 | 1022.07757 | 4.59E-10 | -0.784230295 | 7.18E-11 | 42.46960469 | 0.580661669 | 9.33797532 |
| ENSMUSG00000021796 | *Bmpr1a* | 1844.380696 | 1717.911692 | 2091.282759 | 1156.832712 | 1080.134105 | 1043.19867 | 4.56E-13 | -0.784631428 | 5.47E-14 | 56.55142212 | 0.580500241 | 12.34090416 |
| ENSMUSG00000044350 | *Lacc1* | 235.8215514 | 232.1243046 | 268.5010413 | 155.8541225 | 127.3842365 | 143.2561553 | 1.96E-05 | -0.784992202 | 5.22E-06 | 20.75343114 | 0.580355094 | 4.708779533 |
| ENSMUSG00000053907 | *Mat2a* | 14788.1962 | 14934.60918 | 16418.90171 | 9556.491865 | 8500.002691 | 8709.239597 | 8.40E-22 | -0.785563191 | 5.44E-23 | 97.47936539 | 0.580125447 | 21.07569074 |
| ENSMUSG00000025417 | *Pip4k2c* | 3263.830545 | 3577.783702 | 3464.041604 | 1886.712934 | 2108.682857 | 1980.79184 | 1.54E-21 | -0.785652426 | 1.02E-22 | 96.23938795 | 0.580089566 | 20.81191086 |
| ENSMUSG00000021285 | *Ppp1r13b* | 1122.600996 | 1084.845407 | 936.602224 | 619.0262331 | 602.1800271 | 606.0837341 | 2.17E-12 | -0.785715084 | 2.75E-13 | 53.37853511 | 0.580064372 | 11.66394888 |
| ENSMUSG00000020829 | *Slc46a1* | 278.0132667 | 304.0636551 | 262.1982 | 153.658994 | 165.2836788 | 170.805416 | 2.05E-06 | -0.786066456 | 4.78E-07 | 25.35054682 | 0.579923113 | 5.687877399 |
| ENSMUSG00000041278 | *Ttc1* | 1546.024995 | 1513.603936 | 2107.670146 | 1087.686165 | 920.1142373 | 981.6719875 | 4.75E-07 | -0.788105372 | 1.02E-07 | 28.32733698 | 0.579104105 | 6.323439667 |
| ENSMUSG00000002233 | *Rhoc* | 5872.936081 | 5767.617534 | 5717.937669 | 3426.595567 | 3337.256444 | 3288.463412 | 1.31E-40 | -0.788555191 | 3.62E-42 | 185.1592463 | 0.578923574 | 39.88167367 |
| ENSMUSG00000029456 | *Acad10* | 377.465167 | 305.9820378 | 265.9799048 | 173.4151504 | 193.7082605 | 184.5800463 | 5.11E-05 | -0.78871835 | 1.45E-05 | 18.79735116 | 0.578858105 | 4.291429143 |
| ENSMUSG00000034473 | *Sec22a* | 756.4371809 | 723.2302712 | 825.6722164 | 474.147753 | 439.0018729 | 420.5853791 | 1.63E-11 | -0.788840731 | 2.24E-12 | 49.26473932 | 0.578809004 | 10.78794855 |
| ENSMUSG00000038286 | *Bphl* | 422.6705762 | 411.4930853 | 413.4663923 | 248.049519 | 203.183121 | 269.9827543 | 5.81E-07 | -0.789266218 | 1.26E-07 | 27.91914667 | 0.578638324 | 6.23561443 |
| ENSMUSG00000048787 | *Dcun1d3* | 281.0269606 | 292.553359 | 320.1843404 | 193.1713068 | 169.4947279 | 154.2758596 | 4.56E-06 | -0.789501601 | 1.11E-06 | 23.72784716 | 0.578543923 | 5.341032473 |
| ENSMUSG00000049625 | *Tifab* | 510.8211241 | 477.6772879 | 434.8960529 | 248.049519 | 285.2985793 | 291.1038541 | 4.61E-08 | -0.78954739 | 8.81E-09 | 33.08849018 | 0.578525562 | 7.336534987 |
| ENSMUSG00000039745 | *Htatip2* | 2342.393621 | 2542.816245 | 3243.442157 | 1524.516734 | 1627.570493 | 1544.595213 | 6.01E-09 | -0.790079889 | 1.05E-09 | 37.22278344 | 0.578312067 | 8.220792288 |
| ENSMUSG00000073402 | *Gm8909* | 1592.737251 | 1605.686305 | 1547.977835 | 979.0273049 | 866.4233607 | 900.8608229 | 5.48E-19 | -0.790297042 | 4.27E-20 | 84.29203045 | 0.578225027 | 18.26160192 |
| ENSMUSG00000028563 | *Tm2d1* | 690.8893375 | 678.1482782 | 985.7643865 | 482.928267 | 428.4742501 | 445.3797137 | 1.03E-05 | -0.792540289 | 2.62E-06 | 22.07427659 | 0.577326643 | 4.989039894 |
| ENSMUSG00000042348 | *Arl15* | 413.6294943 | 462.3302264 | 466.4102596 | 252.4397759 | 260.0322844 | 260.7996674 | 3.46E-09 | -0.792541706 | 5.94E-10 | 38.34217912 | 0.577326076 | 8.460538912 |
| ENSMUSG00000008435 | *Rdh13* | 1313.217138 | 1432.072672 | 1452.174646 | 804.5145902 | 871.6871721 | 744.748346 | 2.71E-14 | -0.793572647 | 2.92E-15 | 62.31866198 | 0.57691367 | 13.5670608 |
| ENSMUSG00000001467 | *Cyp51* | 3646.569677 | 3507.762734 | 3876.247428 | 2344.397223 | 1833.911901 | 2180.064825 | 1.14E-12 | -0.794513655 | 1.41E-13 | 54.68682825 | 0.576537496 | 11.94242006 |
| ENSMUSG00000074063 | *Osgin1* | 742.8755581 | 859.4354416 | 765.1649394 | 454.3915967 | 452.6877826 | 457.3177266 | 2.66E-12 | -0.794708941 | 3.41E-13 | 52.95447417 | 0.57645946 | 11.57440038 |
| ENSMUSG00000028643 | *Svbp* | 499.5197718 | 558.2493605 | 673.1434558 | 346.8303008 | 341.0949804 | 307.6334105 | 6.52E-07 | -0.794830158 | 1.43E-07 | 27.68355992 | 0.576411027 | 6.185869334 |
| ENSMUSG00000032679 | *Cd59a* | 220.7530817 | 208.144521 | 323.9660452 | 138.2930946 | 147.3867199 | 146.0110814 | 0.000770616 | -0.795461433 | 0.000265646 | 13.29830849 | 0.576158865 | 3.113162175 |
| ENSMUSG00000049988 | *Lrrc25* | 1506.09355 | 1529.910189 | 1520.245333 | 850.6122885 | 759.0416076 | 1012.894483 | 2.32E-10 | -0.795632933 | 3.53E-11 | 43.86054174 | 0.576090378 | 9.634343214 |
| ENSMUSG00000069874 | *Irgm2* | 632.1223055 | 699.2504877 | 675.6645923 | 460.9769821 | 309.5121118 | 385.689649 | 2.51E-06 | -0.795677319 | 5.91E-07 | 24.94142907 | 0.576072654 | 5.600323784 |
| ENSMUSG00000028199 | *Cryz* | 796.3686257 | 835.4556581 | 1131.990306 | 530.1235294 | 512.6952329 | 544.557052 | 9.51E-07 | -0.796741188 | 2.13E-07 | 26.9096812 | 0.575648005 | 6.021866566 |
| ENSMUSG00000041891 | *Lman1* | 5650.676152 | 6432.337134 | 7693.248147 | 3714.157399 | 3873.112447 | 3788.023338 | 5.05E-11 | -0.797248484 | 7.21E-12 | 46.96968658 | 0.575445625 | 10.29685874 |
| ENSMUSG00000024963 | *Dnajc4* | 332.2597577 | 340.5129261 | 388.2550269 | 209.6347704 | 201.0775965 | 198.3546766 | 1.46E-07 | -0.79737992 | 2.96E-08 | 30.73218448 | 0.575393201 | 6.835143163 |
| ENSMUSG00000025138 | *Sirt7* | 1747.189066 | 1866.58635 | 1844.211378 | 1031.710389 | 1005.387982 | 1100.133808 | 5.12E-21 | -0.797472398 | 3.50E-22 | 93.79284586 | 0.575356319 | 20.29069147 |
| ENSMUSG00000010025 | *Aldh3a2* | 1619.860497 | 1648.849915 | 1750.929326 | 961.466277 | 966.4357778 | 955.9593442 | 1.08E-22 | -0.798288627 | 6.81E-24 | 101.5968194 | 0.575030894 | 21.96457033 |
| ENSMUSG00000025355 | *Mmp19* | 366.1638147 | 374.084623 | 500.4456029 | 206.3420777 | 276.876481 | 227.7405546 | 7.56E-05 | -0.799160119 | 2.21E-05 | 18.00231262 | 0.574683639 | 4.121498334 |
| ENSMUSG00000008892 | *Vdac3* | 7169.577902 | 7183.383954 | 8681.53367 | 4504.403654 | 4208.943616 | 4511.650584 | 1.96E-15 | -0.799997411 | 1.94E-16 | 67.66747268 | 0.574350208 | 14.70779317 |
| ENSMUSG00000034591 | *Slc41a2* | 1325.271914 | 1405.215315 | 1353.850321 | 779.2706126 | 716.9311162 | 846.6806103 | 2.15E-15 | -0.800598123 | 2.14E-16 | 67.47021813 | 0.574111109 | 14.66705124 |
| ENSMUSG00000031447 | *Lamp1* | 26827.90352 | 29084.59985 | 31686.90458 | 15842.24229 | 17370.57771 | 17064.01204 | 3.37E-20 | -0.800861794 | 2.42E-21 | 89.96590213 | 0.574006193 | 19.47207386 |
| ENSMUSG00000036499 | *Eea1* | 1519.655173 | 1272.84691 | 1197.539856 | 822.0756181 | 721.1421654 | 750.2581981 | 4.01E-10 | -0.801256239 | 6.22E-11 | 42.75107282 | 0.573849276 | 9.397023913 |
| ENSMUSG00000021916 | *Glt8d1* | 302.8762418 | 362.5743269 | 423.5509385 | 234.878748 | 192.6554982 | 195.5997505 | 3.55E-05 | -0.801693334 | 9.88E-06 | 19.53418193 | 0.573675442 | 4.449412652 |
| ENSMUSG00000029082 | *Bst1* | 878.4917858 | 950.5586191 | 1137.032579 | 580.6114846 | 559.0167734 | 556.4950649 | 9.29E-10 | -0.803512383 | 1.50E-10 | 41.02371125 | 0.572952569 | 9.031979649 |
| ENSMUSG00000073664 | *Nbeal1* | 254.6571386 | 239.7978353 | 185.3035356 | 122.9271952 | 133.7008102 | 134.0730684 | 0.000108156 | -0.804749331 | 3.23E-05 | 17.2782288 | 0.572461537 | 3.965949174 |
| ENSMUSG00000020661 | *Dnmt3a* | 4733.759768 | 4904.345327 | 3868.684018 | 2517.812374 | 2681.38554 | 2535.450288 | 2.20E-13 | -0.80502072 | 2.56E-14 | 58.04324874 | 0.57235386 | 12.65823672 |
| ENSMUSG00000028459 | *Cd72* | 9095.328334 | 8741.110692 | 8135.70761 | 5218.917976 | 4776.382488 | 4867.036046 | 2.35E-25 | -0.805811623 | 1.27E-26 | 114.0525312 | 0.572040175 | 24.62843604 |
| ENSMUSG00000037594 | *Clba1* | 232.8078575 | 237.8794526 | 325.2266135 | 153.658994 | 146.3339576 | 152.4392422 | 0.000100304 | -0.808214505 | 2.98E-05 | 17.43197577 | 0.571088206 | 3.998680025 |
| ENSMUSG00000037235 | *Mxd4* | 755.6837574 | 714.5975491 | 651.7137952 | 437.928133 | 383.2054718 | 393.0361185 | 7.51E-11 | -0.808931248 | 1.09E-11 | 46.15860385 | 0.570804555 | 10.12434279 |
| ENSMUSG00000031903 | *Pla2g15* | 1645.476895 | 1730.381179 | 1628.654204 | 949.3930703 | 922.2197618 | 984.4269135 | 1.05E-22 | -0.808981029 | 6.58E-24 | 101.6625499 | 0.570784859 | 21.97802998 |
| ENSMUSG00000078878 | *Gm14305* | 135.6162277 | 142.9195098 | 124.7962587 | 102.0734746 | 60.00745026 | 68.8731516 | 0.004836349 | -0.809568825 | 0.001960826 | 9.585850796 | 0.570552352 | 2.315482364 |
| ENSMUSG00000033793 | *Atp6v1h* | 3746.021577 | 4165.767994 | 4841.842722 | 2469.519547 | 2443.461264 | 2353.625167 | 2.49E-13 | -0.810731797 | 2.93E-14 | 57.77959571 | 0.570092609 | 12.60384107 |
| ENSMUSG00000007812 | *Zfp655* | 644.1770813 | 608.1273103 | 756.3409615 | 385.2450494 | 369.5195621 | 387.5262663 | 1.14E-09 | -0.811089393 | 1.86E-10 | 40.61248266 | 0.56995132 | 8.944257298 |
| ENSMUSG00000006599 | *Gtf2h1* | 2053.079002 | 2135.159925 | 3054.356916 | 1468.540957 | 1245.417783 | 1409.603836 | 1.61E-06 | -0.811382355 | 3.71E-07 | 25.83865183 | 0.569835594 | 5.792588149 |
| ENSMUSG00000001891 | *Ugp2* | 1543.011301 | 1511.685554 | 1638.73875 | 853.9049812 | 930.6418601 | 887.0861926 | 2.99E-20 | -0.811758679 | 2.14E-21 | 90.2073367 | 0.569686973 | 19.52382423 |
| ENSMUSG00000052151 | *Plpp2* | 1711.778162 | 1854.116862 | 2008.085253 | 1056.954366 | 981.1744498 | 1128.601378 | 1.01E-14 | -0.813683348 | 1.06E-15 | 64.31508618 | 0.568927472 | 13.99374892 |
| ENSMUSG00000027287 | *Snap23* | 950.067017 | 876.7008858 | 835.7567625 | 498.2941664 | 522.1700935 | 496.8050002 | 2.74E-14 | -0.813937288 | 2.96E-15 | 62.28996239 | 0.56882734 | 13.56156026 |
| ENSMUSG00000057375 | *Yipf1* | 1084.176398 | 1266.13257 | 1622.351363 | 735.3680429 | 736.9335996 | 783.3173109 | 4.75E-07 | -0.813999146 | 1.02E-07 | 28.32862319 | 0.568802951 | 6.323509904 |
| ENSMUSG00000002014 | *Ssr4* | 6001.771497 | 6786.278738 | 9354.677126 | 4430.86685 | 3924.697799 | 4234.32136 | 5.77E-07 | -0.814115395 | 1.26E-07 | 27.93361682 | 0.56875712 | 6.238588193 |
| ENSMUSG00000026600 | *Soat1* | 5516.566771 | 5760.903195 | 5812.480289 | 2993.057691 | 3353.047878 | 3363.764724 | 1.55E-25 | -0.815054055 | 8.29E-27 | 114.8968354 | 0.56838719 | 24.81002116 |
| ENSMUSG00000035469 | *Rcbtb1* | 1032.190177 | 1078.131067 | 1013.496888 | 587.1968701 | 606.3910763 | 581.2893995 | 1.30E-18 | -0.81635935 | 1.04E-19 | 82.53509195 | 0.567873167 | 17.88656485 |
| ENSMUSG00000039178 | *Tbc1d19* | 409.1089534 | 386.5541105 | 423.5509385 | 253.5373402 | 232.660465 | 206.6194548 | 3.91E-08 | -0.816998856 | 7.40E-09 | 33.42698287 | 0.567621501 | 7.408215318 |
| ENSMUSG00000030556 | *Lrrc28* | 201.164071 | 181.2871635 | 254.6347904 | 115.2442455 | 109.4872777 | 134.9913771 | 0.00030509 | -0.817030879 | 9.77E-05 | 15.18122058 | 0.567608902 | 3.515571595 |
| ENSMUSG00000029148 | *Nrbp1* | 3287.940097 | 3305.373362 | 3595.140704 | 2000.859615 | 1851.80886 | 1925.693319 | 2.38E-25 | -0.817465774 | 1.29E-26 | 114.0253687 | 0.567437824 | 24.62332204 |
| ENSMUSG00000026123 | *Plekhb2* | 5122.526288 | 5614.14692 | 5661.212097 | 3085.253088 | 3059.327201 | 3151.635417 | 2.01E-29 | -0.81809327 | 8.86E-31 | 133.0410844 | 0.567191072 | 28.69745423 |
| ENSMUSG00000047789 | *Slc38a9* | 429.4513876 | 440.2688256 | 579.8614039 | 266.7081111 | 277.9292433 | 274.5742977 | 1.31E-06 | -0.818439353 | 2.99E-07 | 26.2555944 | 0.567055027 | 5.881821158 |
| ENSMUSG00000025040 | *Fundc1* | 694.656455 | 728.0262279 | 1041.22939 | 496.0990379 | 446.3712089 | 451.8078745 | 8.21E-06 | -0.818582559 | 2.07E-06 | 22.52872337 | 0.566998743 | 5.085597635 |
| ENSMUSG00000028671 | *Gale* | 1257.4638 | 1328.480007 | 1307.209295 | 773.7827914 | 689.5592968 | 741.9934199 | 1.31E-18 | -0.819461554 | 1.05E-19 | 82.515448 | 0.566653391 | 17.88299365 |
| ENSMUSG00000029798 | *Herc6* | 319.4515585 | 322.2882906 | 274.8038827 | 212.9274632 | 156.8615805 | 151.5209335 | 4.43E-05 | -0.819488472 | 1.25E-05 | 19.08827142 | 0.566642818 | 4.35375774 |
| ENSMUSG00000020696 | *Rffl* | 291.5748894 | 282.0022543 | 248.331949 | 190.9761783 | 115.8038514 | 159.7857117 | 0.000339844 | -0.819816031 | 0.000109799 | 14.96027425 | 0.566514178 | 3.468720687 |
| ENSMUSG00000040040 | *Ift88* | 223.0133521 | 171.6952501 | 228.1628567 | 118.5369383 | 120.0149005 | 113.8702773 | 9.46E-05 | -0.821140855 | 2.80E-05 | 17.55142382 | 0.565994188 | 4.024155204 |
| ENSMUSG00000022765 | *Snap29* | 1424.723814 | 1582.665713 | 1762.27444 | 870.3684448 | 905.3755653 | 917.3903793 | 8.76E-14 | -0.822228028 | 9.89E-15 | 59.91850185 | 0.565567833 | 13.0575925 |
| ENSMUSG00000030876 | *Mettl9* | 1370.477323 | 1287.23478 | 1580.75261 | 775.9779199 | 819.0490579 | 798.9285586 | 1.65E-13 | -0.822272624 | 1.90E-14 | 58.63361033 | 0.56555035 | 12.78316495 |
| ENSMUSG00000024787 | *Snx15* | 933.4917003 | 1100.192468 | 1062.659051 | 557.5626355 | 633.7628957 | 558.3316823 | 5.65E-12 | -0.822321039 | 7.49E-13 | 51.41203753 | 0.565531371 | 11.2476839 |
| ENSMUSG00000030921 | *Trim30a* | 1591.983828 | 1591.298435 | 1594.618861 | 1056.954366 | 714.8255916 | 928.4100836 | 3.55E-08 | -0.82343356 | 6.70E-09 | 33.61886716 | 0.565095435 | 7.449168042 |
| ENSMUSG00000035142 | *Nubpl* | 301.3693948 | 248.4305574 | 267.2404731 | 158.049251 | 134.7535725 | 168.9687986 | 9.71E-06 | -0.824042164 | 2.48E-06 | 22.18554955 | 0.564857098 | 5.01292899 |
| ENSMUSG00000051316 | *Taf7* | 177.8079429 | 174.5728241 | 255.8953587 | 122.9271952 | 88.43203196 | 130.3998337 | 0.002263986 | -0.824362906 | 0.000858705 | 11.10984865 | 0.564731533 | 2.645126316 |
| ENSMUSG00000021822 | *Plau* | 17651.20545 | 16763.78707 | 13754.06039 | 9366.613251 | 8838.992146 | 8993.91529 | 1.18E-14 | -0.824795861 | 1.24E-15 | 64.00175981 | 0.564562081 | 13.92695652 |
| ENSMUSG00000051498 | *Tlr6* | 312.6707471 | 272.4103409 | 342.8745692 | 159.1468153 | 195.813785 | 168.0504899 | 5.22E-06 | -0.82503217 | 1.28E-06 | 23.4527306 | 0.564469615 | 5.282476056 |
| ENSMUSG00000080115 | *Eef1akmt3* | 135.6162277 | 163.062528 | 104.6271663 | 79.02462551 | 69.48231082 | 79.89285586 | 0.002561451 | -0.825214601 | 0.000981022 | 10.86304559 | 0.564398242 | 2.591513995 |

| | | | | | | | | | | | | | |
|---|---|---|---|---|---|---|---|---|---|---|---|---|---|
| ENSMUSG00000025498 | *Irf7* | 11655.46134 | 9817.323376 | 6833.540588 | 6314.287091 | 4096.298052 | 5565.868958 | 0.000117309 | -0.825421045 | 3.52E-05 | 17.1125892 | 0.564317484 | 3.93066993 |
| ENSMUSG00000025650 | *Col7a1* | 67.80811383 | 85.36802936 | 86.97921058 | 49.39039094 | 46.32154055 | 39.48727358 | 0.008818421 | -0.825861184 | 0.003778632 | 8.387245247 | 0.564145348 | 2.054609187 |
| ENSMUSG00000055675 | *Kbtbd11* | 1942.325749 | 1898.239664 | 1343.765775 | 1058.05193 | 903.2700407 | 966.0607398 | 3.02E-07 | -0.826257601 | 6.37E-08 | 29.24841336 | 0.563990356 | 6.51969569 |
| ENSMUSG00000040913 | *Fbxw4* | 855.1356577 | 857.517059 | 790.3763048 | 492.8063452 | 450.5822581 | 470.1740483 | 5.32E-15 | -0.82633046 | 5.43E-16 | 65.63457904 | 0.563961874 | 14.27387564 |
| ENSMUSG00000039126 | *Prune2* | 738.3550172 | 771.1898383 | 627.7629981 | 399.5133845 | 427.4214878 | 381.0981055 | 4.42E-10 | -0.826440322 | 6.90E-11 | 42.54735968 | 0.56391893 | 9.354285711 |
| ENSMUSG00000008822 | *Acyp1* | 192.1229892 | 239.7978353 | 378.1704808 | 166.829765 | 141.0701462 | 146.9293901 | 0.004451355 | -0.826658441 | 0.001793433 | 9.749789225 | 0.563833678 | 2.351507816 |
| ENSMUSG00000040327 | *Cul9* | 367.6706616 | 360.6559443 | 225.6417202 | 164.6346365 | 181.0751131 | 193.7631332 | 0.000504079 | -0.826683404 | 0.000168084 | 14.15796684 | 0.563823922 | 3.297501809 |
| ENSMUSG00000002428 | *Hltf* | 608.012754 | 577.4331874 | 702.136526 | 431.3427476 | 301.0900136 | 331.5094364 | 2.73E-06 | -0.826686127 | 6.46E-07 | 24.76869454 | 0.563822858 | 5.563554729 |
| ENSMUSG00000029007 | *Agtrap* | 876.2315153 | 965.9056805 | 907.6091539 | 487.318524 | 525.3283803 | 535.3739651 | 3.31E-15 | -0.827473416 | 3.32E-16 | 66.60512624 | 0.56351526 | 14.48065348 |
| ENSMUSG00000024235 | *Map3k8* | 195.1366831 | 146.7562752 | 150.007624 | 102.0734746 | 75.79888453 | 100.095647 | 0.001074169 | -0.827856952 | 0.00038081 | 12.62407806 | 0.563365471 | 2.968927296 |
| ENSMUSG00000022400 | *Rbx1* | 406.848683 | 423.0033814 | 404.6424144 | 215.1225917 | 250.5574239 | 229.577172 | 2.28E-09 | -0.828258002 | 3.83E-10 | 39.19814284 | 0.563208885 | 8.642202036 |
| ENSMUSG00000027165 | *Iftap* | 313.4241706 | 286.798211 | 413.4663923 | 197.5615638 | 202.1303588 | 169.8871073 | 4.80E-05 | -0.828471409 | 1.36E-05 | 18.92237822 | 0.56312558 | 4.318425293 |
| ENSMUSG00000027737 | *Slc7a11* | 1652.257707 | 1794.646999 | 1866.901607 | 999.8810255 | 996.9658841 | 989.018457 | 4.00E-21 | -0.830051799 | 2.73E-22 | 94.28943365 | 0.562509046 | 20.39833884 |
| ENSMUSG00000039637 | *Coro7* | 2748.48888 | 2856.471814 | 2361.044368 | 1449.882365 | 1513.872166 | 1518.88257 | 9.01E-17 | -0.830461613 | 8.06E-18 | 73.93825647 | 0.562349281 | 16.04542861 |
| ENSMUSG00000028212 | *Ccne2* | 1784.860241 | 1436.868629 | 1764.795577 | 986.7102546 | 881.1620327 | 933.001627 | 4.73E-12 | -0.832205451 | 6.20E-13 | 51.78332991 | 0.56166996 | 11.32529224 |
| ENSMUSG00000047037 | *Nipa1* | 290.0680425 | 339.5537348 | 364.3042298 | 196.4639995 | 188.444449 | 171.7237247 | 1.10E-06 | -0.833430352 | 2.49E-07 | 26.60752496 | 0.561193284 | 5.957532703 |
| ENSMUSG00000034714 | *Ttyh2* | 1698.969963 | 1875.219072 | 1617.309089 | 943.9052491 | 1007.493507 | 961.4691963 | 3.75E-18 | -0.834437358 | 3.07E-19 | 80.39010193 | 0.560801705 | 17.42587904 |
| ENSMUSG00000018809 | *Smyd4* | 1124.107843 | 1259.418231 | 1317.293841 | 759.5144563 | 666.3985265 | 640.0611555 | 1.29E-12 | -0.84035651 | 1.60E-13 | 54.43874924 | 0.558505537 | 11.889518 |
| ENSMUSG00000032698 | *Lmo2* | 1538.49076 | 1460.848413 | 1338.723502 | 780.3681769 | 820.1018202 | 823.7228931 | 3.21E-18 | -0.841128455 | 2.62E-19 | 80.70171763 | 0.558206777 | 17.4929086 |
| ENSMUSG00000021748 | *Pdhb* | 5383.210814 | 5599.759049 | 6936.907186 | 3564.888662 | 3145.653708 | 3284.790177 | 1.47E-12 | -0.841710843 | 1.83E-13 | 54.17779219 | 0.557981485 | 11.83280143 |
| ENSMUSG00000014470 | *Rnf166* | 691.642761 | 759.6795422 | 699.6153894 | 354.5132505 | 371.6250867 | 472.0106656 | 1.51E-08 | -0.84175165 | 2.75E-09 | 35.35305161 | 0.557965703 | 7.820498972 |
| ENSMUSG00000021451 | *Sema4d* | 5472.114786 | 5587.289562 | 4028.776188 | 2571.593022 | 2944.576112 | 2901.855454 | 2.28E-09 | -0.842336862 | 3.84E-10 | 39.19241658 | 0.557739417 | 8.641460213 |
| ENSMUSG00000032383 | *Ppib* | 16175.24884 | 18642.84291 | 24516.79227 | 11614.42482 | 10451.82397 | 11019.70426 | 2.00E-08 | -0.842484856 | 3.68E-09 | 34.78379053 | 0.557682206 | 7.69874862 |
| ENSMUSG00000019897 | *Ccdc59* | 892.806832 | 790.3736651 | 1183.673605 | 576.2212277 | 495.8510363 | 524.3542609 | 1.37E-06 | -0.842618373 | 3.12E-07 | 26.17494643 | 0.557630597 | 5.864859643 |
| ENSMUSG00000029552 | *Tes* | 6632.386956 | 6521.541928 | 6929.343776 | 3912.816527 | 3643.610269 | 3637.420713 | 6.23E-35 | -0.843224236 | 2.19E-36 | 158.6912796 | 0.557396467 | 34.20530662 |
| ENSMUSG00000073079 | *Srp54a* | 2481.776966 | 2397.978353 | 3162.765788 | 1521.224041 | 1350.694012 | 1605.203587 | 5.65E-10 | -0.843785303 | 8.93E-11 | 42.04194299 | 0.557179737 | 9.247745745 |
| ENSMUSG00000022159 | *Rab2b* | 304.3830887 | 279.1246803 | 286.1489971 | 163.5370722 | 146.3339576 | 174.4786507 | 3.02E-07 | -0.844246321 | 6.36E-08 | 29.25114759 | 0.557001717 | 6.520167035 |
| ENSMUSG00000006736 | *Tspan31* | 4733.006345 | 5038.632115 | 5492.295949 | 2797.691256 | 2883.515899 | 2814.616129 | 7.73E-25 | -0.844502302 | 4.28E-26 | 111.6441164 | 0.556902895 | 24.1118121 |
| ENSMUSG00000044715 | *Gskip* | 537.9443697 | 579.35157 | 637.8475442 | 311.7082451 | 327.4090707 | 335.1826711 | 6.56E-11 | -0.844898747 | 9.45E-12 | 46.4390327 | 0.556749882 | 10.18320452 |
| ENSMUSG00000003363 | *Pld3* | 3886.158346 | 4387.341194 | 4239.291089 | 2253.299391 | 2313.971503 | 2394.03075 | 5.10E-26 | -0.845194777 | 2.65E-27 | 117.1557028 | 0.556635653 | 25.29226035 |
| ENSMUSG00000030284 | *Creld1* | 1989.791429 | 2027.730495 | 2211.036744 | 1133.783863 | 1148.563653 | 1176.353429 | 4.87E-24 | -0.846951141 | 2.82E-25 | 107.9021474 | 0.555958407 | 23.31284103 |
| ENSMUSG00000033808 | *Tmem87a* | 1162.53244 | 1068.539154 | 1440.829532 | 683.7825235 | 719.0366408 | 636.3879208 | 2.32E-09 | -0.847035273 | 3.91E-10 | 39.15848903 | 0.555925987 | 8.634578668 |
| ENSMUSG00000112368 | *Gm49376* | 156.7120853 | 155.3889973 | 163.873875 | 102.0734746 | 100.0124171 | 63.36329947 | 0.001272025 | -0.847102402 | 0.000457936 | 12.27960281 | 0.55590012 | 2.895504506 |
| ENSMUSG00000024104 | *Washc2* | 3191.501891 | 3178.760104 | 3491.774106 | 1717.688041 | 1857.072671 | 1902.735602 | 6.70E-24 | -0.847147186 | 3.92E-25 | 107.2513945 | 0.555882864 | 23.17399468 |
| ENSMUSG00000023186 | *Vwa5a* | 4702.115982 | 5044.387263 | 6310.404756 | 3049.033468 | 2912.993243 | 2952.362432 | 3.83E-12 | -0.848290313 | 4.97E-13 | 52.21783119 | 0.555442582 | 11.4170517 |
| ENSMUSG00000082064 | *Rpl5-ps2* | 78.35604264 | 62.34743717 | 85.71864231 | 45.00013397 | 41.05772912 | 39.48727358 | 0.008673406 | -0.849241926 | 0.003708756 | 8.421177041 | 0.555076328 | 2.061810302 |
| ENSMUSG00000014867 | *Surf4* | 16966.3435 | 19064.8871 | 19361.06805 | 10141.49361 | 10426.55767 | 10157.4124 | 3.62E-29 | -0.850047817 | 1.61E-30 | 131.856593 | 0.554766348 | 28.44169554 |
| ENSMUSG00000030826 | *Bcat2* | 3293.214061 | 3606.559443 | 4135.924491 | 1982.201023 | 2004.459391 | 2129.557847 | 1.38E-15 | -0.850201255 | 1.34E-16 | 68.39768789 | 0.554707349 | 14.86039806 |
| ENSMUSG00000031696 | *Vps35* | 6183.346557 | 5831.883354 | 8057.552377 | 3979.767946 | 3494.118025 | 3654.868578 | 4.07E-10 | -0.850541903 | 6.32E-11 | 42.71983439 | 0.554576388 | 9.39028065 |
| ENSMUSG00000039232 | *Stx11* | 140.8901921 | 179.3687808 | 186.5641038 | 100.9759104 | 75.79888453 | 102.8505731 | 0.00061232 | -0.851285055 | 0.000207959 | 13.75777163 | 0.554290791 | 3.213021297 |
| ENSMUSG00000051319 | *Mtln* | 1151.984512 | 1273.806101 | 1822.781717 | 853.9049812 | 694.8231082 | 800.7651759 | 6.95E-06 | -0.852755501 | 1.74E-06 | 22.86638467 | 0.553726127 | 5.157875948 |
| ENSMUSG00000054855 | *Rnd1* | 334.5200282 | 336.6761607 | 379.431049 | 192.0737426 | 202.1303588 | 186.4166637 | 1.48E-08 | -0.852899398 | 2.69E-09 | 35.39826522 | 0.553670899 | 7.830252906 |
| ENSMUSG00000024661 | *Fth1* | 183026.9074 | 195020.8651 | 207339.5295 | 106259.5846 | 107711.2677 | 110013.3808 | 1.38E-34 | -0.853439765 | 4.93E-36 | 157.0752573 | 0.553463559 | 33.86144612 |
| ENSMUSG00000020038 | *Cry1* | 1049.518917 | 1045.518562 | 883.6583567 | 556.4650713 | 530.5921917 | 563.8415344 | 1.56E-13 | -0.853693364 | 1.79E-14 | 58.74676488 | 0.553366279 | 12.8065878 |
| ENSMUSG00000039050 | *Osbpl2* | 1439.03886 | 1376.439575 | 1421.921008 | 808.9048472 | 788.5189516 | 747.503272 | 3.34E-23 | -0.854650611 | 2.04E-24 | 103.9860751 | 0.552999235 | 22.47581116 |
| ENSMUSG00000032596 | *Uba7* | 2179.654148 | 1835.892227 | 1401.751915 | 1048.173852 | 908.5338521 | 1040.443744 | 4.68E-07 | -0.855504018 | 1.01E-07 | 28.35672189 | 0.552672212 | 6.329398832 |
| ENSMUSG00000037049 | *Smpd1* | 1115.066761 | 1126.090634 | 1036.187117 | 594.8798198 | 617.9714614 | 599.6555733 | 1.00E-19 | -0.856103873 | 7.46E-21 | 87.74242872 | 0.552442465 | 18.99901716 |
| ENSMUSG00000016481 | *Cr1l* | 3907.254203 | 3967.215387 | 4670.405438 | 2431.104799 | 2238.172618 | 2252.611212 | 7.17E-18 | -0.857175305 | 5.96E-19 | 79.0813133 | 0.552032341 | 17.14433894 |
| ENSMUSG00000027309 | *Dnaaf9* | 791.0946613 | 743.3732894 | 605.0727692 | 446.708647 | 332.6728821 | 403.137514 | 3.58E-07 | -0.858209328 | 7.60E-08 | 28.90553024 | 0.551636824 | 6.445931711 |
| ENSMUSG00000028470 | *Hint2* | 606.505907 | 613.8824583 | 805.503124 | 384.1474851 | 349.5170787 | 380.1797968 | 5.07E-08 | -0.859235635 | 9.73E-09 | 32.89529752 | 0.55124454 | 7.295402132 |
| ENSMUSG00000068523 | *Gng5* | 3012.187101 | 3090.514501 | 3845.993789 | 1834.02985 | 1726.530148 | 1914.673614 | 5.00E-13 | -0.860355794 | 6.01E-14 | 56.36773231 | 0.5508167 | 12.30132511 |
| ENSMUSG00000041390 | *Mdfic* | 1299.655515 | 1429.195098 | 1676.555798 | 822.0756181 | 786.413427 | 813.6214976 | 1.67E-12 | -0.860712112 | 2.09E-13 | 53.91945705 | 0.550680676 | 11.77761103 |
| ENSMUSG00000058715 | *Fcer1g* | 10012.99814 | 10249.91867 | 11270.7409 | 6065.140008 | 5490.155317 | 5784.426426 | 6.72E-27 | -0.862517978 | 3.35E-28 | 121.262906 | 0.549991803 | 26.17275728 |
| ENSMUSG00000079884 | *Tmed2b* | 2443.352368 | 2467.999321 | 3037.969529 | 1477.321471 | 1550.718846 | 1340.730684 | 7.30E-13 | -0.862828539 | 8.88E-14 | 55.60095155 | 0.549873422 | 12.13687127 |
| ENSMUSG00000030591 | *Psmd8* | 11549.22863 | 10648.94227 | 12193.47687 | 6597.458666 | 5938.632051 | 6372.143986 | 3.44E-25 | -0.862894388 | 1.87E-26 | 113.2787806 | 0.549848325 | 24.46281562 |
| ENSMUSG00000055725 | *Paqr3* | 377.465167 | 409.5747027 | 421.0298019 | 216.2201559 | 225.291129 | 221.3123938 | 4.27E-10 | -0.863044808 | 6.65E-11 | 42.61952404 | 0.549790999 | 9.369160535 |
| ENSMUSG00000050668 | *Gpatch11* | 499.5197718 | 483.4324359 | 704.6576625 | 311.7082451 | 308.4593496 | 303.9601757 | 2.70E-06 | -0.864915938 | 6.37E-07 | 24.79610192 | 0.549078399 | 5.569313193 |
| ENSMUSG00000004661 | *Arid3b* | 137.1230746 | 147.7154665 | 108.4088712 | 85.61001097 | 65.27126168 | 66.11822554 | 0.001232536 | -0.865948056 | 0.000442197 | 12.34488063 | 0.548685724 | 2.909200211 |
| ENSMUSG00000058298 | *Mcm9* | 743.6289816 | 815.31264 | 804.2425558 | 433.5378761 | 440.0546352 | 421.5036878 | 4.38E-16 | -0.866571462 | 4.10E-17 | 70.72920289 | 0.548448681 | 15.35810552 |
| ENSMUSG00000026571 | *Dcaf6* | 516.0950886 | 566.8820826 | 645.4109539 | 311.7082451 | 325.3035461 | 306.7151018 | 4.58E-10 | -0.86931857 | 7.16E-11 | 42.47370048 | 0.547405346 | 9.338694075 |
| ENSMUSG00000078622 | *Ccdc47* | 2300.955329 | 2131.32316 | 2517.354834 | 1276.467215 | 1245.417783 | 1275.530768 | 7.52E-21 | -0.870876739 | 5.17E-22 | 93.02104942 | 0.546814445 | 20.12394605 |
| ENSMUSG00000021257 | *Angel1* | 263.6982204 | 260.9000448 | 207.9937644 | 133.9028377 | 136.8590971 | 131.3181424 | 4.12E-06 | -0.871497431 | 9.95E-07 | 23.93738451 | 0.546579239 | 5.385605097 |
| ENSMUSG00000035725 | *Prkx* | 769.9988037 | 841.2108062 | 773.9889173 | 437.928133 | 408.4717667 | 456.3994179 | 4.36E-15 | -0.871734923 | 4.42E-16 | 66.03775679 | 0.546489271 | 14.36037237 |
| ENSMUSG00000038774 | *Ascc3* | 1457.874447 | 1267.091762 | 1598.400565 | 857.1976739 | 733.7753128 | 770.4609892 | 3.89E-12 | -0.871889858 | 5.05E-13 | 52.18384512 | 0.546430585 | 11.40999439 |
| ENSMUSG00000036339 | *Tmem260* | 415.1363413 | 418.2074247 | 359.2619567 | 256.8300329 | 210.552457 | 186.4166637 | 8.61E-07 | -0.872460202 | 1.91E-07 | 27.11815507 | 0.546214606 | 6.065101944 |
| ENSMUSG00000020733 | *Nherf1* | 1439.792284 | 1558.685929 | 1455.956351 | 847.3195957 | 724.3004522 | 859.536932 | 1.55E-16 | -0.873092654 | 1.41E-17 | 72.83010286 | 0.545975208 | 15.80952129 |
| ENSMUSG00000025702 | *Marchf8* | 3524.515072 | 4032.440398 | 3552.281383 | 1898.786141 | 2141.318488 | 2024.870657 | 1.22E-20 | -0.873117738 | 8.50E-22 | 92.03792995 | 0.545965715 | 19.91379732 |
| ENSMUSG00000002885 | *Adgre5* | 91.91766541 | 107.4294302 | 90.76091539 | 65.85385459 | 54.74363883 | 38.5689649 | 0.006216222 | -0.873436372 | 0.002573171 | 9.087821998 | 0.545845146 | 2.206473479 |
| ENSMUSG00000054708 | *Ankrd24* | 155.2052383 | 178.4095894 | 229.423425 | 104.2686031 | 98.95965481 | 101.9322644 | 0.000212962 | -0.875603725 | 6.67E-05 | 15.9026325 | 0.545025742 | 3.671697298 |
| ENSMUSG00000037752 | *Xkr8* | 331.5063343 | 405.7379373 | 384.4733221 | 195.3664353 | 224.2383667 | 191.0082071 | 1.07E-07 | -0.875655482 | 2.14E-08 | 31.36233411 | 0.545006189 | 6.969171412 |
| ENSMUSG00000078974 | *Sec61g* | 186.0956013 | 192.7974596 | 233.2051298 | 97.68321764 | 108.4345154 | 124.8899816 | 4.22E-05 | -0.87832362 | 1.19E-05 | 19.18507066 | 0.543999179 | 4.374724376 |
| ENSMUSG00000021556 | *Golm1* | 1377.258134 | 1560.604312 | 1774.880123 | 823.1731824 | 928.5363355 | 807.1933368 | 7.09E-12 | -0.87945738 | 9.47E-13 | 50.95079606 | 0.543571839 | 11.14944823 |
| ENSMUSG00000073411 | *H2-D1* | 29758.72089 | 31190.02484 | 28824.15404 | 15959.68166 | 16265.17731 | 16565.37042 | 2.69E-48 | -0.879712635 | 5.52E-50 | 220.9822979 | 0.543475673 | 47.57003053 |
| ENSMUSG00000052949 | *Rnf157* | 917.6698071 | 810.5166832 | 684.4885702 | 435.7330045 | 426.3687255 | 451.8078745 | 7.68E-10 | -0.879833937 | 1.23E-10 | 41.41484843 | 0.54342998 | 9.114443883 |
| ENSMUSG00000038225 | *Primpol* | 413.6294943 | 410.533894 | 465.1496913 | 244.7568262 | 197.9193096 | 256.208124 | 4.57E-08 | -0.879870019 | 8.73E-09 | 33.1055471 | 0.543416389 | 7.34018887 |
| ENSMUSG00000035151 | *Elmod2* | 506.3005832 | 526.5960463 | 606.3333375 | 268.9032396 | 300.0372513 | 318.6531147 | 9.46E-10 | -0.880099799 | 1.53E-10 | 40.9868677 | 0.543329845 | 9.024160638 |
| ENSMUSG00000022722 | *Arl6* | 307.3967827 | 361.6151356 | 448.7623038 | 242.5616977 | 186.3389245 | 177.2335768 | 5.32E-05 | -0.880455036 | 1.52E-05 | 18.71503277 | 0.543196077 | 4.273938471 |
| ENSMUSG00000031765 | *Mt1* | 9557.176932 | 11125.66037 | 11908.58844 | 6090.383985 | 5500.68294 | 6101.242923 | 2.34E-17 | -0.881001137 | 2.01E-18 | 76.68113794 | 0.5429905 | 16.63038248 |
| ENSMUSG00000062007 | *Hsh2d* | 111.5066761 | 115.1029609 | 104.6271663 | 72.43924005 | 44.21601598 | 63.36329947 | 0.002356407 | -0.881511173 | 0.000896347 | 11.03029961 | 0.542798571 | 2.627749741 |
| ENSMUSG00000027074 | *Slc43a3* | 1772.805465 | 1804.238913 | 1822.781717 | 972.4419194 | 1008.546269 | 949.5311834 | 7.38E-30 | -0.881688065 | 3.22E-31 | 135.0524782 | 0.542732021 | 29.13199708 |
| ENSMUSG00000062352 | *Itgb1bp1* | 1073.628469 | 1158.70314 | 1479.907148 | 705.7338084 | 636.9211825 | 666.6921075 | 1.89E-09 | -0.883264175 | 3.15E-10 | 39.57827827 | 0.542139423 | 8.723364237 |
| ENSMUSG00000025464 | *Paox* | 180.0682134 | 165.940102 | 187.8246721 | 104.2686031 | 88.43203196 | 96.42241224 | 1.49E-05 | -0.883808458 | 3.91E-06 | 21.30611498 | 0.54193493 | 4.826855838 |
| ENSMUSG00000030861 | *Acadsb* | 690.135914 | 674.3115128 | 970.6375673 | 451.0989039 | 391.6275701 | 419.6670704 | 6.32E-07 | -0.884425577 | 1.38E-07 | 27.74602753 | 0.541703164 | 6.198940363 |
| ENSMUSG00000067235 | *H2-Q10* | 864.1767395 | 874.7825031 | 804.2425558 | 452.1964682 | 453.7405449 | 472.0106656 | 6.94E-18 | -0.885146113 | 5.76E-19 | 79.1475319 | 0.541432685 | 17.15853677 |
| ENSMUSG00000081251 | *Gm12164* | 192.8764127 | 192.7974596 | 219.3388788 | 103.1710389 | 115.8038514 | 107.4421165 | 4.19E-06 | -0.886312228 | 1.01E-06 | 23.89997857 | 0.540995227 | 5.377782461 |
| ENSMUSG00000023791 | *Pigx* | 590.6840138 | 557.2901692 | 717.2633452 | 323.7814517 | 326.3563084 | 355.3854623 | 1.68E-09 | -0.88725791 | 2.79E-10 | 39.81641097 | 0.540640723 | 8.773919337 |
| ENSMUSG00000034563 | *Ccpg1* | 648.6976223 | 728.0262279 | 751.2986885 | 336.9522227 | 400.0496684 | 410.4839835 | 5.21E-11 | -0.887408212 | 7.45E-12 | 46.90477968 | 0.540584401 | 10.28289291 |
| ENSMUSG00000031105 | *Slc25a14* | 191.3695657 | 155.3889973 | 205.4726279 | 127.3174522 | 93.69584338 | 78.05623848 | 0.000881149 | -0.887589297 | 0.000306455 | 13.03052307 | 0.540516552 | 3.054950814 |
| ENSMUSG00000015363 | *Trabd* | 4063.212865 | 4143.706594 | 3798.092195 | 2186.347972 | 2118.157718 | 2186.492986 | 1.51E-35 | -0.887716956 | 5.13E-37 | 161.5723452 | 0.540468726 | 34.82096601 |
| ENSMUSG00000079108 | *Srp54c* | 1271.778846 | 1384.113105 | 1824.042286 | 823.1731824 | 866.4233607 | 729.1370983 | 3.09E-08 | -0.887788717 | 5.77E-09 | 33.91014792 | 0.540441843 | 7.510225706 |
| ENSMUSG00000042444 | *Mindy2* | 743.6289816 | 775.985795 | 750.0381202 | 428.0500548 | 374.7833735 | 423.3403052 | 2.11E-15 | -0.88783845 | 2.09E-16 | 67.51510461 | 0.540423213 | 14.67633837 |
| ENSMUSG00000095595 | *Fam177a* | 1267.258305 | 1486.746579 | 1554.280676 | 769.3925345 | 720.0894031 | 833.8242887 | 1.86E-13 | -0.888503212 | 2.16E-14 | 58.38252522 | 0.540174256 | 12.73083114 |
| ENSMUSG00000026222 | *Sp100* | 1723.832938 | 1645.972341 | 1237.87804 | 971.3443552 | 679.0316739 | 839.3341408 | 6.44E-07 | -0.88953691 | 1.41E-07 | 27.70955104 | 0.539787357 | 6.19129631 |
| ENSMUSG00000000600 | *Krit1* | 860.4096221 | 843.1291888 | 1034.926549 | 502.6844234 | 442.1601598 | 529.864113 | 7.98E-11 | -0.88989272 | 1.16E-11 | 46.03228651 | 0.539654246 | 10.09798051 |
| ENSMUSG00000029119 | *Man2b2* | 4084.308723 | 4542.730192 | 4191.389495 | 2165.494252 | 2441.355739 | 2302.199881 | 1.11E-25 | -0.891468675 | 5.90E-27 | 115.5718642 | 0.539065066 | 24.95505974 |
| ENSMUSG00000028405 | *Aco1* | 3013.693948 | 3322.638806 | 3165.286924 | 1789.029716 | 1639.150878 | 1684.178134 | 1.79E-28 | -0.894064222 | 8.25E-30 | 128.6101665 | 0.538096108 | 27.74655733 |
| ENSMUSG00000036636 | *Clcn7* | 1688.422034 | 1803.279721 | 1515.20306 | 857.1976739 | 894.8479424 | 942.1847139 | 7.44E-18 | -0.894652599 | 6.20E-19 | 79.00403492 | 0.5378767 | 17.12842545 |
| ENSMUSG00000100838 | *Gm29094* | 189.8627187 | 150.5930406 | 271.0221779 | 117.439374 | 97.90689252 | 112.0336599 | 0.001395158 | -0.895445694 | 0.000505233 | 12.09625285 | 0.537581094 | 2.855376604 |
| ENSMUSG00000021831 | *Ero1a* | 3380.611186 | 3399.374113 | 3868.684018 | 2067.811034 | 1644.414689 | 2009.259409 | 2.16E-14 | -0.895559514 | 2.31E-15 | 62.7783187 | 0.537538683 | 13.66537082 |
| ENSMUSG00000035235 | *Trim13* | 141.6436155 | 138.1235531 | 166.3950115 | 92.19539643 | 75.79888453 | 71.62807766 | 0.000165385 | -0.895609902 | 5.09E-05 | 16.41377956 | 0.53751991 | 3.781504929 |
| ENSMUSG00000027088 | *Phospho2* | 659.2455511 | 625.3927544 | 889.9611981 | 396.2206918 | 368.4667998 | 401.3008967 | 1.25E-07 | -0.895729946 | 2.50E-08 | 31.05880384 | 0.537475185 | 6.904541918 |
| ENSMUSG00000029314 | *Gpat3* | 230.547587 | 262.8184275 | 166.3950115 | 117.439374 | 100.0124171 | 137.7463032 | 0.000468495 | -0.895801748 | 0.000155126 | 14.3089509 | 0.537448436 | 3.329294946 |
| ENSMUSG00000028636 | *Ppcs* | 619.3141063 | 600.4537795 | 625.2418615 | 325.9765802 | 350.569841 | 314.97988 | 1.01E-14 | -0.896474294 | 1.05E-15 | 64.32974817 | 0.53719795 | 13.99641011 |
| ENSMUSG00000028862 | *Map3k6* | 192.8764127 | 208.144521 | 115.9722808 | 94.39052491 | 109.4872777 | 75.30131242 | 0.003833864 | -0.897402927 | 0.001525442 | 10.04762837 | 0.536852279 | 2.416363349 |
| ENSMUSG00000028756 | *Pink1* | 785.8206969 | 860.394633 | 848.3624452 | 433.5378761 | 421.1049141 | 481.1937525 | 2.24E-15 | -0.898340261 | 2.23E-16 | 67.38884882 | 0.536503593 | 14.64942716 |
| ENSMUSG00000042742 | *Bmt2* | 817.4644833 | 716.5159318 | 897.5246077 | 448.9037755 | 375.8361358 | 477.5205178 | 1.38E-09 | -0.898643503 | 2.28E-10 | 40.21354126 | 0.536390837 | 8.858671956 |
| ENSMUSG00000036002 | *Atosb* | 855.8890812 | 952.4770017 | 884.918925 | 466.4648033 | 481.1123643 | 495.8866915 | 9.30E-18 | -0.898673794 | 7.82E-19 | 78.54414439 | 0.536379575 | 17.03159586 |
| ENSMUSG00000047843 | *Bri3* | 1708.764468 | 1851.239288 | 2204.733903 | 1045.978724 | 1010.651794 | 1031.260657 | 3.34E-14 | -0.89880074 | 3.62E-15 | 61.89609648 | 0.53633238 | 13.47662427 |
| ENSMUSG00000025950 | *Idh1* | 2926.296823 | 3002.268898 | 3859.86004 | 1817.566387 | 1711.791476 | 1716.318938 | 9.69E-13 | -0.898966884 | 1.19E-13 | 55.01988692 | 0.536270618 | 12.01360301 |
| ENSMUSG00000002580 | *Mien1* | 1686.161764 | 1840.688184 | 2335.833003 | 1097.564243 | 986.4382612 | 1054.218374 | 1.75E-10 | -0.89997122 | 2.62E-11 | 44.43975027 | 0.535897422 | 9.757993571 |
| ENSMUSG00000025521 | *Tmem192* | 1633.42212 | 1836.851418 | 2226.163563 | 1011.954232 | 992.7548349 | 1043.19867 | 8.70E-12 | -0.900256525 | 1.17E-12 | 50.53228378 | 0.535791454 | 11.06061844 |
| ENSMUSG00000041957 | *Pkp2* | 978.6971095 | 1039.763414 | 1068.961892 | 554.2699428 | 532.6977163 | 563.8415344 | 2.65E-20 | -0.901142134 | 1.89E-21 | 90.45299686 | 0.535462655 | 19.57607964 |
| ENSMUSG00000066595 | *Flvcr1* | 844.5877289 | 1052.232901 | 848.3624452 | 481.8307028 | 504.2731346 | 483.9486786 | 6.64E-12 | -0.901363264 | 8.84E-13 | 51.08578521 | 0.535380588 | 11.17775049 |
| ENSMUSG00000024614 | *Tmx3* | 2267.804696 | 2216.691189 | 2901.828156 | 1378.540689 | 1202.25453 | 1370.116563 | 1.46E-11 | -0.901378596 | 1.99E-12 | 49.49333003 | 0.535374899 | 10.8363776 |
| ENSMUSG00000032353 | *Tmed3* | 3046.091158 | 3195.066357 | 4074.156646 | 1883.420241 | 1836.017425 | 1796.211794 | 8.54E-13 | -0.902217303 | 1.05E-13 | 55.28042331 | 0.53506375 | 12.06849829 |
| ENSMUSG00000029605 | *Oas1b* | 242.6023628 | 216.7772431 | 213.0360375 | 142.6833516 | 105.2762285 | 112.9519686 | 1.08E-05 | -0.902793428 | 2.78E-06 | 21.96472564 | 0.534850121 | 4.965648094 |
| ENSMUSG00000030717 | *Nupr1* | 284.0406546 | 227.3283478 | 240.7685394 | 147.0736086 | 122.1204251 | 134.0730684 | 2.84E-06 | -0.903378912 | 6.73E-07 | 24.6903077 | 0.534633108 | 5.546898293 |
| ENSMUSG00000031155 | *Pim2* | 481.4376082 | 472.8813312 | 437.4171894 | 260.1227256 | 261.0850467 | 224.9856286 | 7.86E-11 | -0.903564268 | 1.14E-11 | 46.06533707 | 0.534564424 | 10.10469323 |
| ENSMUSG00000006095 | *Tbcb* | 3809.30915 | 4246.340067 | 5327.161506 | 2382.811972 | 2351.870945 | 2414.233541 | 5.64E-12 | -0.903698867 | 7.46E-13 | 51.41936535 | 0.534514553 | 11.24885469 |
| ENSMUSG00000079317 | *Trappc2* | 221.5065052 | 164.0217193 | 255.8953587 | 109.7564243 | 112.6455645 | 119.3801294 | 0.000140819 | -0.903856928 | 4.28E-05 | 16.74169787 | 0.534455995 | 3.851340014 |
| ENSMUSG00000028251 | *Tstd3* | 497.2595014 | 511.2489848 | 641.629249 | 327.0741445 | 278.9820056 | 273.655989 | 3.29E-08 | -0.90507664 | 6.18E-09 | 33.77822134 | 0.534004335 | 7.483161465 |
| ENSMUSG00000007659 | *Bcl2l1* | 4552.184708 | 4704.833528 | 4303.580071 | 2559.519815 | 2205.536987 | 2474.841914 | 7.67E-25 | -0.905678406 | 4.24E-26 | 111.6627083 | 0.533781642 | 24.11534564 |
| ENSMUSG00000025512 | *Chid1* | 3681.227157 | 3632.457609 | 4195.1712 | 2138.055146 | 1950.768514 | 2044.155139 | 2.78E-23 | -0.907338902 | 1.68E-24 | 104.3683766 | 0.53316763 | 22.55666476 |
| ENSMUSG00000067567 | *Hdac8* | 405.341836 | 381.7581538 | 404.6424144 | 195.3664353 | 226.3438913 | 213.0476156 | 1.49E-10 | -0.908031654 | 2.22E-11 | 44.76546484 | 0.532911675 | 9.827180548 |
| ENSMUSG00000055401 | *Fbxo6* | 1022.395672 | 1045.518562 | 1140.814284 | 616.8311047 | 547.4363883 | 544.557052 | 5.20E-17 | -0.908228373 | 4.56E-18 | 75.06135888 | 0.532839015 | 16.28409126 |
| ENSMUSG00000038160 | *Atg5* | 1233.354248 | 1321.765668 | 1634.957045 | 773.7827914 | 709.5617802 | 744.748346 | 1.10E-11 | -0.908975542 | 1.50E-12 | 50.04873118 | 0.53256313 | 10.95668902 |
| ENSMUSG00000022014 | *Epsti1* | 1130.13523 | 960.1505325 | 881.1372202 | 646.4653392 | 425.3159632 | 511.4979392 | 5.76E-07 | -0.910299139 | 1.25E-07 | 27.93895041 | 0.532074755 | 6.239648216 |
| ENSMUSG00000060147 | *Serpinb6a* | 4099.377192 | 3980.644066 | 5454.478901 | 2388.299793 | 2405.561822 | 2400.45891 | 6.99E-12 | -0.910945552 | 9.33E-13 | 50.98058734 | 0.531836408 | 11.15559335 |
| ENSMUSG00000056383 | *AI987944* | 195.8901066 | 160.184954 | 168.9161481 | 94.39052491 | 94.74860567 | 90.91256011 | 1.52E-05 | -0.911142361 | 4.00E-06 | 21.26559696 | 0.531763861 | 4.818356066 |
| ENSMUSG00000101249 | *Gm29216* | 40532.67675 | 37974.3852 | 49279.39535 | 21556.16174 | 22912.31837 | 23408.60677 | 1.54E-16 | -0.912640775 | 1.41E-17 | 72.83976889 | 0.531211846 | 15.81132116 |
| ENSMUSG00000022175 | *Lrp10* | 3283.419556 | 3393.618965 | 3011.497595 | 1669.395214 | 1698.105566 | 1775.090694 | 7.05E-29 | -0.914218558 | 3.19E-30 | 130.4954027 | 0.530631211 | 28.15186555 |
| ENSMUSG00000022898 | *Vps26c* | 2034.243415 | 2315.487897 | 2792.158716 | 1321.467349 | 1208.571103 | 1255.327977 | 5.30E-12 | -0.914501608 | 6.98E-13 | 51.54959914 | 0.530527114 | 11.2760847 |
| ENSMUSG00000041153 | *Osgin2* | 739.8618642 | 730.9038019 | 984.5038183 | 463.1721106 | 389.5220455 | 447.2163311 | 2.00E-08 | -0.914529206 | 3.68E-09 | 34.78651984 | 0.530516966 | 7.699195559 |
| ENSMUSG00000113948 | *Rpl17-ps3* | 3526.775342 | 3177.800913 | 5564.14834 | 1960.249738 | 2320.288077 | 2224.143642 | 6.59E-06 | -0.914742512 | 1.64E-06 | 22.971193 | 0.530438533 | 5.180839214 |
| ENSMUSG00000015536 | *Mocs2* | 798.6288962 | 760.6387335 | 935.3416558 | 458.7818536 | 457.9515941 | 404.9741314 | 4.29E-12 | -0.915185999 | 5.60E-13 | 51.98335989 | 0.530275501 | 11.36725608 |
| ENSMUSG00000038172 | *Ttc39b* | 871.7109744 | 901.6398607 | 843.3201721 | 490.6112167 | 456.8988318 | 441.7064789 | 1.62E-18 | -0.915407421 | 1.31E-19 | 82.07045198 | 0.530194121 | 17.78965967 |
| ENSMUSG00000022681 | *Ntan1* | 1093.21748 | 1163.499097 | 1539.153857 | 649.758032 | 708.5090179 | 649.2442424 | 2.54E-09 | -0.916785238 | 4.30E-10 | 38.97200376 | 0.529688012 | 8.594550506 |
| ENSMUSG00000020311 | *Erlec1* | 554.5196864 | 549.6166385 | 613.8967471 | 307.3179881 | 250.5574239 | 349.8756101 | 1.22E-08 | -0.917065616 | 2.20E-09 | 35.79126397 | 0.529585081 | 7.915403922 |
| ENSMUSG00000025232 | *Hexa* | 12646.96665 | 13964.86674 | 15410.44709 | 7225.265413 | 7699.903354 | 7295.962526 | 2.12E-23 | -0.918929113 | 1.27E-24 | 104.9219244 | 0.528901469 | 22.67353948 |
| ENSMUSG00000038524 | *Fchsd1* | 397.8076011 | 388.4724932 | 389.5155952 | 196.4639995 | 195.813785 | 228.6588633 | 2.16E-10 | -0.91904463 | 3.27E-11 | 44.01094759 | 0.528859121 | 9.666340987 |
| ENSMUSG00000073409 | *H2-Q6* | 3935.884296 | 4384.46362 | 4163.656993 | 2293.909268 | 2002.353866 | 2303.11819 | 1.28E-23 | -0.919271935 | 7.64E-25 | 105.9288406 | 0.528775803 | 22.89173041 |
| ENSMUSG00000036181 | *H1f2* | 947.8067466 | 1090.600555 | 1390.406801 | 572.9285349 | 569.5443963 | 665.7737988 | 3.65E-08 | -0.919997066 | 6.89E-09 | 33.56538959 | 0.528510095 | 7.437859595 |
| ENSMUSG00000056758 | *Hmga2* | 9532.313957 | 10184.69366 | 10711.04858 | 5577.821484 | 5242.75618 | 5246.297535 | 5.92E-35 | -0.921050064 | 2.07E-36 | 158.8027818 | 0.528124486 | 34.22796658 |
| ENSMUSG00000037348 | *Paqr7* | 2069.654319 | 2373.998569 | 2511.051992 | 1269.881829 | 1206.465579 | 1190.12806 | 2.80E-19 | -0.922455026 | 2.16E-20 | 85.63988245 | 0.527610424 | 18.55226687 |
| ENSMUSG00000028419 | *Chmp5* | 1663.559059 | 1692.013526 | 2332.051298 | 1037.19821 | 986.4382612 | 971.5705919 | 8.17E-10 | -0.923683962 | 1.31E-10 | 41.28831872 | 0.527161179 | 9.088003408 |
| ENSMUSG00000051285 | *Pcmtd1* | 137.8764981 | 129.4908311 | 167.6555798 | 72.43924005 | 75.79888453 | 79.89285586 | 9.87E-05 | -0.923954437 | 2.93E-05 | 17.46513775 | 0.527062357 | 4.005753949 |
| ENSMUSG00000038302 | *Afg1l* | 286.300925 | 305.0228465 | 397.0790048 | 160.2443795 | 191.6027359 | 167.1321812 | 3.20E-06 | -0.924085138 | 7.66E-07 | 24.44135879 | 0.52701461 | 5.494418925 |
| ENSMUSG00000099974 | *Bcl2a1d* | 1576.915358 | 1267.091762 | 1327.378388 | 770.4900987 | 751.6722716 | 679.5484291 | 9.16E-14 | -0.924126233 | 1.04E-14 | 59.82620433 | 0.526999598 | 13.03801725 |
| ENSMUSG00000026068 | *Il18rap* | 1280.066504 | 1172.131819 | 1295.864181 | 700.2459871 | 629.5518465 | 644.652699 | 2.16E-20 | -0.924926826 | 1.53E-21 | 90.8755978 | 0.526707232 | 19.66590423 |
| ENSMUSG00000091383 | *H2af-ps2* | 81.36973659 | 103.5926648 | 126.0568269 | 46.09769821 | 64.2184994 | 52.34359522 | 0.003559178 | -0.92574088 | 0.001403936 | 10.20061812 | 0.526410116 | 2.448650324 |
| ENSMUSG00000052776 | *Oas1a* | 11056.48967 | 9958.324504 | 9333.247465 | 5985.017818 | 4439.498557 | 5553.012636 | 5.96E-13 | -0.925744979 | 7.20E-14 | 56.01221377 | 0.526408621 | 12.22477381 |
| ENSMUSG00000049686 | *Orai1* | 1226.573437 | 1203.785133 | 1183.673605 | 639.8799538 | 603.2327894 | 659.345638 | 2.18E-24 | -0.925791983 | 1.24E-25 | 109.5394296 | 0.52639147 | 23.66136904 |
| ENSMUSG00000026024 | *Als2* | 1106.779102 | 1210.499472 | 958.0318846 | 488.4160882 | 650.6070922 | 584.9626343 | 7.25E-10 | -0.926591022 | 1.16E-10 | 41.53575091 | 0.526100008 | 9.139625962 |
| ENSMUSG00000021598 | *Med10* | 1264.244611 | 1388.909062 | 1549.238403 | 774.8803557 | 663.2402397 | 768.6243719 | 3.73E-14 | -0.927193011 | 4.06E-15 | 61.67014159 | 0.52588053 | 13.4287071 |
| ENSMUSG00000017929 | *B4galt5* | 3177.940268 | 3698.641811 | 2923.257816 | 1724.273426 | 1619.148395 | 1808.149807 | 1.47E-16 | -0.927976413 | 1.34E-17 | 72.93470001 | 0.525595048 | 15.83155569 |
| ENSMUSG00000019188 | *H13* | 8338.891154 | 8605.864713 | 8627.329235 | 4471.476727 | 4394.229778 | 4564.912488 | 4.18E-59 | -0.928709885 | 6.08E-61 | 271.2425075 | 0.525327901 | 58.37870429 |
| ENSMUSG00000027274 | *Mkks* | 435.4787755 | 423.0033814 | 608.854474 | 265.6105468 | 248.4518993 | 254.3715066 | 6.98E-07 | -0.929007383 | 1.54E-07 | 27.54331746 | 0.525219584 | 6.156416364 |
| ENSMUSG00000004207 | *Psap* | 27924.1347 | 33317.51123 | 32623.50681 | 15523.94866 | 17149.49763 | 16597.51123 | 1.27E-23 | -0.929739226 | 7.55E-25 | 105.953527 | 0.524953221 | 22.89664005 |
| ENSMUSG00000032192 | *Gnb5* | 319.4515585 | 304.0636551 | 307.5786577 | 138.2930946 | 168.4419656 | 180.9068115 | 5.35E-08 | -0.93073825 | 1.03E-08 | 32.78646812 | 0.524589832 | 7.272019163 |
| ENSMUSG00000070291 | *Ddx43* | 150.6846974 | 137.1643618 | 102.1060298 | 63.6587261 | 76.85164682 | 65.19991685 | 0.000658988 | -0.93156417 | 0.000225074 | 13.60924515 | 0.524289599 | 3.181122435 |
| ENSMUSG00000066760 | *Psg16* | 237.3283984 | 248.4305574 | 252.1136538 | 132.8052734 | 112.6455645 | 140.5012293 | 2.52E-07 | -0.931608857 | 5.26E-08 | 29.61764344 | 0.524273359 | 6.599456421 |
| ENSMUSG00000022895 | *Ets2* | 6394.305134 | 6732.564023 | 5109.083195 | 3438.668774 | 2855.091317 | 3260.914151 | 5.02E-13 | -0.932937792 | 6.04E-14 | 56.35689131 | 0.523790649 | 12.29917828 |

| | | | | | | | | | | | | | |
|---|---|---|---|---|---|---|---|---|---|---|---|---|---|
| ENSMUSG00000028293 | *Slc35a1* | 418.1500353 | 533.3103857 | 644.1503856 | 280.9764463 | 282.1402924 | 269.9827543 | 8.07E-07 | -0.933402697 | 1.79E-07 | 27.24559807 | 0.523621885 | 6.092925001 |
| ENSMUSG00000024197 | *Plin3* | 1714.038433 | 1792.728617 | 2008.085253 | 972.4419194 | 975.9106384 | 935.7565531 | 5.73E-22 | -0.933716672 | 3.68E-23 | 98.25620925 | 0.523507941 | 21.24190996 |
| ENSMUSG00000031458 | *Coprs* | 554.5196864 | 613.8824583 | 560.9528798 | 311.7082451 | 321.092497 | 273.655989 | 1.17E-12 | -0.933752281 | 1.45E-13 | 54.63537553 | 0.52349502 | 11.93153092 |
| ENSMUSG00000032860 | *P2ry2* | 519.862206 | 507.4122195 | 453.8045769 | 268.9032396 | 251.6101862 | 256.208124 | 1.78E-12 | -0.934782496 | 2.23E-13 | 53.78685555 | 0.523121331 | 11.74996404 |
| ENSMUSG00000021619 | *Atg10* | 127.3285693 | 121.8173003 | 152.5287606 | 69.14654732 | 68.42954854 | 71.62807766 | 9.87E-05 | -0.934789746 | 2.93E-05 | 17.46369819 | 0.523118702 | 4.005525577 |
| ENSMUSG00000078920 | *Ifi47* | 307.3967827 | 284.8798283 | 323.9660452 | 161.3419437 | 153.7032936 | 163.4589465 | 3.19E-09 | -0.93500615 | 5.44E-10 | 38.51153172 | 0.52304024 | 8.496658489 |
| ENSMUSG00000037636 | *Slc25a43* | 299.1091243 | 328.0434387 | 358.0013885 | 174.5127147 | 150.5450068 | 188.253281 | 6.47E-08 | -0.93505976 | 1.26E-08 | 32.3946711 | 0.523020804 | 7.188786219 |
| ENSMUSG00000041236 | *Vps41* | 1414.175885 | 1392.745827 | 2063.550257 | 893.4172939 | 805.3631482 | 843.9256843 | 4.11E-08 | -0.936044182 | 7.82E-09 | 33.31954192 | 0.522664044 | 7.385793132 |
| ENSMUSG00000021226 | *Acot2* | 240.3420923 | 202.389373 | 160.0921702 | 129.5125807 | 91.59031881 | 95.50410355 | 0.000312697 | -0.937559573 | 0.000100255 | 15.13189337 | 0.522115331 | 3.504876791 |
| ENSMUSG00000026946 | *Nmi* | 922.9437715 | 673.3523215 | 685.7491385 | 452.1964682 | 396.8913815 | 344.365758 | 2.06E-07 | -0.938849167 | 4.25E-08 | 30.03157367 | 0.521648832 | 6.687165178 |
| ENSMUSG00000041096 | *Tspyl2* | 599.7250956 | 623.4743717 | 486.5793519 | 302.9277311 | 282.1402924 | 308.5517192 | 2.59E-10 | -0.938964238 | 3.95E-11 | 43.63886961 | 0.521607226 | 9.587109992 |
| ENSMUSG00000025533 | *Asl* | 2144.243244 | 2165.854048 | 2435.417896 | 1180.979126 | 1086.450678 | 1245.226581 | 2.57E-21 | -0.939718432 | 1.71E-22 | 95.20831 | 0.521334618 | 20.5906604 |
| ENSMUSG00000029787 | *Avl9* | 649.4510457 | 771.1898383 | 784.0734635 | 391.8304348 | 369.5195621 | 385.689649 | 2.51E-13 | -0.9398331 | 2.96E-14 | 57.76135394 | 0.521293184 | 12.60006656 |
| ENSMUSG00000030041 | *M1ap* | 223.7667756 | 271.4511495 | 249.5925173 | 129.5125807 | 120.0149005 | 137.7463032 | 3.69E-07 | -0.939890899 | 7.84E-08 | 28.84519179 | 0.521272299 | 6.432823845 |
| ENSMUSG00000030894 | *Tpp1* | 3490.611015 | 3580.661276 | 2928.300089 | 1655.126879 | 1840.228475 | 1719.992173 | 1.03E-18 | -0.939973405 | 8.15E-20 | 83.01449525 | 0.521242489 | 17.98739829 |
| ENSMUSG00000082241 | *Gng5-ps* | 269.7256083 | 295.4309331 | 345.3957058 | 156.9516868 | 146.3339576 | 168.9687986 | 1.87E-07 | -0.940549023 | 3.84E-08 | 30.22641358 | 0.521034561 | 6.72834122 |
| ENSMUSG00000021772 | *Nkiras1* | 118.2874875 | 97.83751679 | 121.0145538 | 71.34167581 | 48.42706512 | 56.01682997 | 0.000931742 | -0.940886791 | 0.000325713 | 12.91641421 | 0.520912589 | 3.030704389 |
| ENSMUSG00000024587 | *Nars* | 13624.91034 | 13279.04493 | 14834.36739 | 7374.53415 | 7179.838785 | 7183.010558 | 1.72E-41 | -0.940978926 | 4.64E-43 | 189.2481221 | 0.520879323 | 40.76442198 |
| ENSMUSG00000026544 | *Dusp23* | 67.05469034 | 86.3272207 | 69.33125481 | 35.12205578 | 43.16325369 | 37.65065621 | 0.00391208 | -0.94152573 | 0.001559516 | 10.00693766 | 0.520681939 | 2.407592315 |
| ENSMUSG00000027746 | *Ufm1* | 1808.216369 | 1764.912068 | 2267.762316 | 1083.295908 | 955.908155 | 999.1198525 | 2.06E-13 | -0.941628033 | 2.40E-14 | 58.17592519 | 0.520645018 | 12.68624861 |
| ENSMUSG00000024487 | *Yipf5* | 1790.134205 | 1690.095143 | 2042.120596 | 952.6857631 | 942.2222452 | 974.325518 | 5.43E-20 | -0.942976016 | 3.97E-21 | 88.98830802 | 0.520158781 | 19.26487272 |
| ENSMUSG00000040219 | *Ttc12* | 758.6974514 | 861.3538243 | 1109.300077 | 497.1966022 | 483.2178889 | 436.1966268 | 1.97E-08 | -0.94380383 | 3.61E-09 | 34.82207124 | 0.519860401 | 7.706477832 |
| ENSMUSG00000051615 | *Rap2a* | 556.0265334 | 664.7195994 | 544.5654923 | 305.1228596 | 304.2483004 | 308.5517192 | 6.30E-12 | -0.944022649 | 8.38E-13 | 51.19117267 | 0.519781558 | 11.20055744 |
| ENSMUSG00000013495 | *Tmem175* | 543.2183341 | 565.9228913 | 412.205824 | 290.8545244 | 260.0322844 | 242.4334936 | 8.03E-08 | -0.944235561 | 1.58E-08 | 31.95743711 | 0.519704854 | 7.095020252 |
| ENSMUSG00000042997 | *Nhlrc3* | 971.1628747 | 1057.988049 | 955.5107481 | 542.1967361 | 482.1651266 | 527.1091869 | 1.04E-18 | -0.944303924 | 8.26E-20 | 82.98695971 | 0.519680228 | 17.9821013 |
| ENSMUSG00000014177 | *Tvp23b* | 686.3687966 | 646.4949639 | 751.2986885 | 377.5620996 | 341.0949804 | 362.7319318 | 9.75E-15 | -0.944762333 | 1.02E-15 | 64.39659139 | 0.519515129 | 14.0108583 |
| ENSMUSG00000025525 | *Apool* | 452.0540922 | 448.9015476 | 525.6569683 | 240.3665693 | 276.876481 | 223.1490112 | 9.42E-10 | -0.944764684 | 1.53E-10 | 40.995121 | 0.519514282 | 9.025810645 |
| ENSMUSG00000019088 | *Dnase1l1* | 1565.614006 | 1510.726362 | 1793.788647 | 859.3928024 | 849.5791641 | 815.4581149 | 4.69E-20 | -0.946730828 | 3.41E-21 | 89.29157933 | 0.518806756 | 19.32899666 |
| ENSMUSG00000034853 | *Acot11* | 518.355359 | 481.5140532 | 426.072075 | 209.6347704 | 265.2960959 | 265.3912108 | 6.62E-09 | -0.946963827 | 1.17E-09 | 37.02693266 | 0.518722974 | 8.179093255 |
| ENSMUSG00000036632 | *Alg5* | 1692.942575 | 1561.563503 | 1945.056839 | 932.9296067 | 914.8504258 | 847.598919 | 1.36E-16 | -0.947107158 | 1.24E-17 | 73.09467594 | 0.518671442 | 15.86511563 |
| ENSMUSG00000061132 | *Blnk* | 901.8479139 | 1002.354951 | 815.5876702 | 464.2696749 | 473.7430283 | 473.847283 | 2.50E-15 | -0.947403623 | 2.49E-16 | 67.16911597 | 0.518564869 | 14.60222047 |
| ENSMUSG00000029571 | *Tmem106b* | 786.5741204 | 808.5983006 | 1068.961892 | 497.1966022 | 425.3159632 | 456.3994179 | 2.66E-09 | -0.947532085 | 4.50E-10 | 38.88260468 | 0.518518696 | 8.575190549 |
| ENSMUSG00000007613 | *Tgfbr1* | 651.7113162 | 636.9030505 | 644.1503856 | 347.9278651 | 310.5648741 | 343.4474493 | 1.50E-16 | -0.947792192 | 1.36E-17 | 72.90412986 | 0.51842522 | 15.82515642 |
| ENSMUSG00000079017 | *Ifi27l2a* | 4852.800679 | 4155.21689 | 4622.503843 | 2927.203837 | 1829.700852 | 2306.791424 | 1.29E-08 | -0.948552486 | 2.34E-09 | 35.6711496 | 0.518152084 | 7.889451155 |
| ENSMUSG00000064109 | *Hcst* | 220.7530817 | 240.7570266 | 265.9799048 | 149.2687371 | 102.1179417 | 124.8899816 | 8.76E-06 | -0.94871522 | 2.22E-06 | 22.39390274 | 0.518093641 | 5.057707423 |
| ENSMUSG00000042208 | *Sanbr* | 643.4236579 | 629.2295198 | 853.4047183 | 389.6353063 | 349.5170787 | 359.9770057 | 4.30E-09 | -0.948860954 | 7.44E-10 | 37.90115074 | 0.518041308 | 8.366700655 |
| ENSMUSG00000020070 | *Rufy2* | 381.2322844 | 404.778746 | 398.3395731 | 204.1469492 | 200.0248342 | 208.4560722 | 3.21E-12 | -0.94940992 | 4.14E-13 | 52.57546583 | 0.517844223 | 11.49314546 |
| ENSMUSG00000024589 | *Nedd4l* | 103.2190177 | 106.4702389 | 162.6133067 | 62.56116186 | 44.21601598 | 84.4843993 | 0.008250178 | -0.949754772 | 0.003511365 | 8.520663383 | 0.517720456 | 2.083536662 |
| ENSMUSG00000015542 | *Nat9* | 554.5196864 | 576.473996 | 519.3541269 | 285.3667032 | 289.5096284 | 280.0841498 | 5.81E-15 | -0.950933672 | 5.96E-16 | 65.45169526 | 0.517297573 | 14.23619663 |
| ENSMUSG00000035778 | *Ggta1* | 2501.365977 | 2668.470311 | 2898.046451 | 1522.321605 | 1337.008102 | 1312.263115 | 1.53E-20 | -0.951173282 | 1.08E-21 | 91.56163829 | 0.517211665 | 19.8139453 |
| ENSMUSG00000073412 | *Lst1* | 1120.340725 | 1069.498345 | 1125.687464 | 636.587261 | 505.3258969 | 573.0246213 | 4.26E-16 | -0.951363735 | 3.98E-17 | 70.78812039 | 0.517143391 | 15.37011766 |
| ENSMUSG00000028385 | *Snx30* | 1378.764981 | 1558.685929 | 1439.568963 | 714.5143223 | 736.9335996 | 809.0299541 | 1.16E-20 | -0.951873329 | 8.08E-22 | 92.13965721 | 0.516960756 | 19.93569437 |
| ENSMUSG00000025762 | *Larp1b* | 447.5335512 | 482.4732446 | 456.3257135 | 225.0006698 | 250.5574239 | 240.5968763 | 5.12E-13 | -0.952038741 | 6.17E-14 | 56.31777774 | 0.516901487 | 12.29103397 |
| ENSMUSG00000049411 | *Tmem241* | 419.6568822 | 391.3500672 | 443.7200308 | 206.3420777 | 207.3941702 | 233.2504068 | 3.89E-11 | -0.95206836 | 5.52E-12 | 47.49364941 | 0.516890875 | 10.40981038 |
| ENSMUSG00000002504 | *Nherf2* | 168.0134376 | 294.4717417 | 297.4941115 | 149.2687371 | 105.2762285 | 136.8279945 | 0.001146908 | -0.952703708 | 0.000408328 | 12.4936929 | 0.516663292 | 2.940471404 |
| ENSMUSG00000039474 | *Wfs1* | 187.6024482 | 178.4095894 | 104.6271663 | 59.26846913 | 87.37926967 | 96.42241224 | 0.004268234 | -0.956737972 | 0.001713826 | 9.833278556 | 0.515220545 | 2.369751811 |
| ENSMUSG00000038702 | *Dsel* | 213.9722703 | 187.0423115 | 194.1275135 | 96.5856534 | 106.3289908 | 103.7688817 | 7.65E-07 | -0.958290868 | 1.69E-07 | 27.35554951 | 0.514666267 | 6.116140771 |
| ENSMUSG00000020056 | *Washc3* | 1151.231088 | 1014.824439 | 1227.793494 | 583.9041774 | 579.0192569 | 582.2077082 | 7.82E-18 | -0.9584406 | 6.54E-19 | 78.89773872 | 0.514612855 | 17.10684316 |
| ENSMUSG00000004707 | *Ly9* | 2890.132496 | 2950.472565 | 3351.851028 | 1703.419705 | 1449.653667 | 1573.062783 | 1.52E-20 | -0.959051029 | 1.07E-21 | 91.5830029 | 0.514395159 | 19.81778759 |
| ENSMUSG00000090136 | *Gm10177* | 2764.310774 | 2801.797907 | 4177.523244 | 1828.542029 | 1535.980174 | 1644.69086 | 3.58E-08 | -0.959068818 | 6.76E-09 | 33.60383074 | 0.514388817 | 7.446126817 |
| ENSMUSG00000071714 | *Csf2rb2* | 345.067957 | 315.5739512 | 293.7124067 | 158.049251 | 186.3389245 | 147.8476988 | 3.73E-08 | -0.960174137 | 7.04E-09 | 33.52261006 | 0.513994869 | 7.428623188 |
| ENSMUSG00000051978 | *Erich1* | 328.4926403 | 303.1044638 | 452.5440087 | 165.7322007 | 175.8113016 | 213.0476156 | 1.15E-05 | -0.960619893 | 2.96E-06 | 21.84070466 | 0.513836082 | 4.93955062 |
| ENSMUSG00000069808 | *Tlcd3a* | 638.9031169 | 701.1688704 | 736.1718692 | 349.0254293 | 343.200505 | 371.9150186 | 1.65E-15 | -0.960626094 | 1.61E-16 | 68.02954953 | 0.513833874 | 14.78299329 |
| ENSMUSG00000020319 | *Wdpcp* | 365.4103912 | 384.6357278 | 320.1843404 | 217.3177201 | 165.2836788 | 168.9687986 | 1.11E-07 | -0.960908206 | 2.21E-08 | 31.29632724 | 0.513733406 | 6.955152696 |
| ENSMUSG00000020387 | *Jade2* | 1506.09355 | 1374.521192 | 1140.814284 | 653.0507247 | 692.7175837 | 721.7906288 | 2.75E-13 | -0.96153016 | 3.25E-14 | 57.57993581 | 0.51351198 | 12.56127373 |
| ENSMUSG00000023249 | *Parp3* | 452.0540922 | 487.2692013 | 457.5862817 | 219.5128486 | 242.1353256 | 254.3715066 | 1.21E-12 | -0.96214553 | 1.50E-13 | 54.57492652 | 0.513292993 | 11.91889311 |
| ENSMUSG00000056429 | *Tgoln1* | 3764.857164 | 4357.606263 | 5377.584237 | 2296.104397 | 2251.858528 | 2376.582885 | 1.28E-12 | -0.96227368 | 1.59E-13 | 54.45360279 | 0.5132474 | 11.89256091 |
| ENSMUSG00000028542 | *Slc6a9* | 1264.244611 | 1201.86675 | 934.0810875 | 619.0262331 | 576.9137323 | 551.9035215 | 2.63E-11 | -0.963062527 | 3.67E-12 | 48.29137899 | 0.51296684 | 10.58016232 |
| ENSMUSG00000031827 | *Cotl1* | 7258.481873 | 7403.038771 | 7456.261313 | 3443.059031 | 4088.928716 | 3802.716277 | 2.42E-30 | -0.964242801 | 1.03E-31 | 137.3187418 | 0.512547352 | 29.6165893 |
| ENSMUSG00000000409 | *Lck* | 369.1775086 | 349.1456482 | 360.522525 | 222.8055414 | 151.5977691 | 179.0701942 | 1.61E-07 | -0.964698652 | 3.29E-08 | 30.52755776 | 0.512385427 | 6.792564015 |
| ENSMUSG00000039824 | *Myl6b* | 125.8217223 | 144.8378925 | 127.3173952 | 71.34167581 | 68.42954854 | 64.28160816 | 3.95E-05 | -0.965536131 | 1.10E-05 | 19.32322762 | 0.512088075 | 4.403813703 |
| ENSMUSG00000082908 | *Gm13736* | 745.8892521 | 736.65895 | 1043.750527 | 445.6110827 | 417.9466272 | 427.0135399 | 9.96E-09 | -0.966099327 | 1.78E-09 | 36.20242579 | 0.511888207 | 8.001736608 |
| ENSMUSG00000001054 | *Rmnd5b* | 544.7251811 | 519.8817069 | 520.6146952 | 268.9032396 | 263.1905713 | 279.1658412 | 1.26E-15 | -0.966795029 | 1.22E-16 | 68.57625211 | 0.511641421 | 14.89910792 |
| ENSMUSG00000060802 | *B2m* | 28392.76411 | 29291.78518 | 26366.04592 | 15166.14271 | 12962.66202 | 14850.8881 | 9.26E-28 | -0.967654952 | 4.38E-29 | 125.2986512 | 0.511336546 | 27.0333891 |
| ENSMUSG00000061981 | *Flot2* | 980.2039565 | 1043.600179 | 1096.694394 | 492.8063452 | 507.4314215 | 591.3907951 | 2.38E-16 | -0.96805702 | 2.18E-17 | 71.97102854 | 0.511194061 | 15.62404827 |
| ENSMUSG00000038085 | *Cnbd2* | 54.24649106 | 63.30662851 | 69.33125481 | 29.63423457 | 40.00496684 | 25.71264326 | 0.009565573 | -0.968399033 | 0.004126351 | 8.22735988 | 0.511072889 | 2.019289009 |
| ENSMUSG00000025579 | *Gaa* | 3816.089961 | 4349.932732 | 3692.204461 | 1921.83499 | 2160.268209 | 1978.955223 | 5.24E-23 | -0.968508247 | 3.22E-24 | 103.0764205 | 0.511034202 | 22.2809573 |
| ENSMUSG00000029407 | *Uso1* | 5674.03228 | 5858.740712 | 7049.097762 | 3225.741311 | 3115.123602 | 3148.880491 | 3.30E-21 | -0.968828896 | 2.23E-22 | 94.68739802 | 0.510920633 | 20.48126059 |
| ENSMUSG00000094685 | *Gm5900* | 330.7529108 | 345.3088828 | 415.9875288 | 186.5859213 | 180.0223508 | 189.1715897 | 3.83E-09 | -0.969120082 | 6.60E-10 | 38.13418145 | 0.510817522 | 8.416673774 |
| ENSMUSG00000020120 | *Plek* | 10107.9295 | 9198.644961 | 9344.59258 | 5254.040032 | 4506.875343 | 4877.137442 | 2.55E-29 | -0.969125808 | 1.13E-30 | 132.5621714 | 0.510815494 | 28.59336278 |
| ENSMUSG00000045502 | *Hcar2* | 140.8901921 | 99.75589948 | 122.2751221 | 71.34167581 | 44.21601598 | 69.79146029 | 0.001805716 | -0.970392916 | 0.000672001 | 11.56513231 | 0.510367046 | 2.743350468 |
| ENSMUSG00000024835 | *Coro1b* | 4979.375825 | 5100.020361 | 5150.681948 | 2540.861223 | 2616.114279 | 2605.241748 | 4.25E-58 | -0.971934019 | 6.36E-60 | 266.5659861 | 0.509822157 | 57.37158226 |
| ENSMUSG00000041168 | *Lonp1* | 10736.28469 | 10110.83593 | 8116.799086 | 4750.258044 | 4996.409806 | 5022.230215 | 3.65E-17 | -0.972050159 | 3.17E-18 | 75.78131594 | 0.509781117 | 16.43801232 |
| ENSMUSG00000074064 | *Mlycd* | 421.1637292 | 532.3511943 | 489.1004885 | 211.8298989 | 250.5574239 | 270.901063 | 6.82E-09 | -0.972312302 | 1.20E-09 | 36.96421066 | 0.509688497 | 8.165915882 |
| ENSMUSG00000026131 | *Dst* | 2998.625478 | 2881.410789 | 1715.633414 | 1268.784265 | 1312.79457 | 1292.060324 | 1.03E-06 | -0.972504117 | 2.33E-07 | 26.74165776 | 0.509620735 | 5.986361043 |
| ENSMUSG00000040774 | *Cept1* | 1686.161764 | 1662.278594 | 2184.564811 | 974.6370479 | 916.9559504 | 921.9819228 | 1.78E-13 | -0.974141911 | 2.06E-14 | 58.47338067 | 0.509042525 | 12.74934435 |
| ENSMUSG00000024856 | *Cdk2ap2* | 2487.804354 | 2427.713284 | 2449.284147 | 1273.174522 | 1206.465579 | 1267.265989 | 6.96E-42 | -0.975034866 | 1.84E-43 | 191.084574 | 0.508727551 | 41.15747272 |
| ENSMUSG00000089917 | *Uckl1* | 910.8889957 | 968.7832545 | 1008.454615 | 490.6112167 | 473.7430283 | 502.3148523 | 1.03E-21 | -0.975166787 | 6.71E-23 | 97.06405935 | 0.508681035 | 20.98597719 |
| ENSMUSG00000044583 | *Tlr7* | 2222.599286 | 2260.813991 | 3072.004872 | 1353.296712 | 1221.204251 | 1264.511063 | 1.71E-11 | -0.975613813 | 2.35E-12 | 49.16636748 | 0.508523442 | 10.76747035 |
| ENSMUSG00000074513 | *Arfip1* | 1440.545707 | 1527.032615 | 1639.999318 | 793.5389478 | 780.0968533 | 766.7877545 | 1.06E-25 | -0.975675633 | 5.64E-27 | 115.659769 | 0.508501652 | 24.97374967 |
| ENSMUSG00000018442 | *Derl2* | 2912.735201 | 2855.512623 | 3656.908549 | 1614.517002 | 1566.51028 | 1607.958513 | 7.67E-17 | -0.975679726 | 6.82E-18 | 74.26871481 | 0.50850021 | 16.11545796 |
| ENSMUSG00000031021 | *Tmem9b* | 1744.175372 | 1729.421988 | 2113.972988 | 943.9052491 | 890.6368933 | 1001.874779 | 6.00E-17 | -0.976134637 | 5.30E-18 | 74.76693761 | 0.508339894 | 16.2216969 |
| ENSMUSG00000038642 | *Ctss* | 7419.714499 | 7021.280617 | 8823.977885 | 4266.232213 | 3608.869114 | 3948.727358 | 1.76E-17 | -0.976149461 | 1.49E-18 | 77.26500841 | 0.508334671 | 16.75528691 |
| ENSMUSG00000035578 | *Iqcg* | 136.3696511 | 145.7970839 | 184.0429673 | 48.2928267 | 101.0651794 | 86.32101667 | 0.002972183 | -0.97735254 | 0.001152812 | 10.5644841 | 0.507910942 | 2.526924487 |
| ENSMUSG00000021282 | *Eif5* | 7923.754812 | 8015.002846 | 9863.946707 | 4609.769821 | 4042.607175 | 4449.205593 | 2.92E-18 | -0.977373684 | 2.38E-19 | 80.89384543 | 0.507903498 | 17.53403667 |
| ENSMUSG00000071648 | *Rom1* | 51.23279711 | 56.59228913 | 73.11295962 | 29.63423457 | 27.37181941 | 33.97742146 | 0.008667447 | -0.978468416 | 0.003703807 | 8.423604955 | 0.507518242 | 2.062108794 |
| ENSMUSG00000024696 | *Lpxn* | 5909.100408 | 5871.210199 | 6068.375648 | 3151.106942 | 2886.674186 | 3018.480657 | 1.42E-48 | -0.978729313 | 2.87E-50 | 222.2823673 | 0.507426471 | 47.84627568 |
| ENSMUSG00000031808 | *Slc27a1* | 9453.957914 | 9732.914538 | 7941.580096 | 4366.110559 | 4622.679194 | 4765.103782 | 8.50E-23 | -0.980220238 | 5.29E-24 | 102.0942551 | 0.506902351 | 22.0707675 |
| ENSMUSG00000051890 | *Klhdc1* | 107.7395586 | 121.8173003 | 144.965351 | 68.04898308 | 63.16573711 | 57.85344734 | 0.000177093 | -0.980870695 | 5.48E-05 | 16.27445509 | 0.50667386 | 3.751798468 |
| ENSMUSG00000030786 | *Itgam* | 6815.468863 | 7085.546437 | 6173.002814 | 3360.741713 | 3260.404797 | 3542.834918 | 3.30E-32 | -0.982090225 | 1.30E-33 | 145.9901479 | 0.506245742 | 31.48140838 |
| ENSMUSG00000003161 | *Sri* | 3230.679912 | 3515.436265 | 4444.763717 | 1909.761783 | 1832.859139 | 1914.673614 | 4.23E-14 | -0.983107427 | 4.62E-15 | 61.41481872 | 0.505888929 | 13.37321039 |
| ENSMUSG00000089716 | *Gm6264* | 176.3010959 | 197.5934163 | 255.8953587 | 96.5856534 | 124.2259497 | 96.42241224 | 5.27E-05 | -0.983712321 | 1.50E-05 | 18.73617712 | 0.505676864 | 4.278544989 |
| ENSMUSG00000053870 | *Fpgt* | 285.5475016 | 351.0640309 | 442.4594625 | 211.8298989 | 156.8615805 | 175.3969594 | 1.34E-05 | -0.983924748 | 3.50E-06 | 21.52259878 | 0.505602412 | 4.87281785 |
| ENSMUSG00000023048 | *Prr13* | 1327.532184 | 1497.297683 | 1531.590447 | 653.0507247 | 756.9360831 | 784.2356196 | 1.84E-17 | -0.987096548 | 1.57E-18 | 77.16489446 | 0.504492054 | 16.73467145 |
| ENSMUSG00000026380 | *Tfcp2l1* | 836.3000705 | 772.1490296 | 528.1781048 | 370.9767142 | 363.2029884 | 346.2023754 | 2.38E-07 | -0.987527036 | 4.96E-08 | 29.7334096 | 0.504341541 | 6.62410868 |
| ENSMUSG00000024976 | *Shoc2* | 1060.82027 | 1134.723357 | 1413.09703 | 632.1970041 | 563.2278226 | 618.9400557 | 2.05E-12 | -0.989479579 | 2.58E-13 | 53.50101451 | 0.503659427 | 11.68867036 |
| ENSMUSG00000030922 | *Lyrm1* | 124.3148753 | 131.4092137 | 151.2681923 | 74.63436854 | 57.90192569 | 71.62807766 | 6.55E-05 | -0.989501283 | 1.89E-05 | 18.29542251 | 0.503651849 | 4.183827808 |
| ENSMUSG00000031767 | *Nudt7* | 169.5202846 | 156.3481886 | 252.1136538 | 111.9515528 | 93.69584338 | 84.4843993 | 0.000326665 | -0.990106779 | 0.000105182 | 15.04133457 | 0.503440512 | 3.48589717 |
| ENSMUSG00000026915 | *Strbp* | 658.4921276 | 711.7199751 | 845.8413087 | 352.3181221 | 368.4667998 | 390.2811924 | 2.04E-12 | -0.991702403 | 2.58E-13 | 53.50641761 | 0.502884014 | 11.68962858 |
| ENSMUSG00000033066 | *Gas7* | 3637.528595 | 3674.662028 | 2788.377012 | 1792.322409 | 1505.450068 | 1783.355472 | 4.43E-13 | -0.991911652 | 5.31E-14 | 56.61232808 | 0.502811081 | 12.35311252 |
| ENSMUSG00000041736 | *Tspo* | 4483.623171 | 4689.486467 | 5030.927963 | 2532.080709 | 2255.016815 | 2351.78855 | 1.47E-31 | -0.992076907 | 5.89E-33 | 142.9957384 | 0.502753489 | 30.83198669 |
| ENSMUSG00000000204 | *Slfn4* | 2914.242048 | 2195.58898 | 1380.322255 | 1236.954902 | 870.6344099 | 1157.068947 | 0.000101405 | -0.992300539 | 3.01E-05 | 17.40904096 | 0.502675563 | 3.993941631 |
| ENSMUSG00000029462 | *Vps29* | 2981.296738 | 3078.045014 | 4033.818462 | 1828.542029 | 1621.253919 | 1614.386674 | 5.03E-13 | -0.994223341 | 6.06E-14 | 56.3515228 | 0.502006051 | 12.29824005 |
| ENSMUSG00000035227 | *Spcs2* | 7174.098443 | 7531.57041 | 9008.020852 | 4053.30475 | 3916.275701 | 3930.361185 | 4.36E-22 | -0.994273753 | 2.78E-23 | 98.81254826 | 0.50198851 | 21.36065615 |
| ENSMUSG00000020986 | *Sec23a* | 1187.395415 | 1255.581466 | 1432.005554 | 631.0994398 | 623.2352728 | 685.0582813 | 2.28E-18 | -0.99561986 | 1.85E-19 | 81.39605285 | 0.501520348 | 17.64293634 |
| ENSMUSG00000025724 | *Sec11a* | 3097.323955 | 2975.41154 | 3385.886371 | 1640.858544 | 1581.248952 | 1519.800879 | 3.80E-31 | -0.995759608 | 1.55E-32 | 141.0708131 | 0.50147177 | 30.42066928 |
| ENSMUSG00000031176 | *Dynlt3* | 679.5879852 | 550.5758298 | 871.052674 | 320.488759 | 361.0974638 | 369.1600926 | 1.98E-07 | -0.996799128 | 4.09E-08 | 30.1043294 | 0.501110569 | 6.702738021 |
| ENSMUSG00000031007 | *Atp6ap2* | 3151.570446 | 3454.048019 | 3714.894689 | 1776.95651 | 1649.678501 | 1735.60342 | 1.63E-27 | -0.998417775 | 7.82E-29 | 124.1485319 | 0.500548658 | 26.78855118 |
| ENSMUSG00000029452 | *Tmem116* | 97.19162982 | 128.5316397 | 121.0145538 | 41.70744124 | 68.42954854 | 62.44499078 | 0.001083522 | -0.999525051 | 0.000384274 | 12.60714912 | 0.500164632 | 2.965162165 |
| ENSMUSG00000021476 | *Habp4* | 656.9852806 | 784.618517 | 882.3977885 | 395.1231275 | 365.308513 | 398.5459706 | 1.33E-11 | -1.00015227 | 1.81E-12 | 49.68150292 | 0.49994723 | 10.87650218 |
| ENSMUSG00000035954 | *Dock4* | 356.3693093 | 367.3702836 | 286.1489971 | 169.0248934 | 167.3892033 | 169.8871073 | 5.35E-09 | -1.000528509 | 9.34E-10 | 37.45817806 | 0.499816866 | 8.271814757 |
| ENSMUSG00000074657 | *Kif5a* | 1093.970903 | 1167.335862 | 974.4192721 | 503.7819876 | 563.2278226 | 550.0669041 | 6.67E-18 | -1.001804359 | 5.54E-19 | 79.22715609 | 0.499375048 | 17.17568103 |
| ENSMUSG00000030660 | *Pik3c2a* | 1070.614775 | 1047.436945 | 1055.095641 | 594.8798198 | 492.6927495 | 498.6416176 | 8.55E-19 | -1.00184087 | 6.73E-20 | 83.39105 | 0.49936241 | 18.06786066 |
| ENSMUSG00000025911 | *Adhfe1* | 299.1091243 | 231.1651132 | 297.4941115 | 141.5857874 | 133.7008102 | 137.7463032 | 1.24E-07 | -1.002761622 | 2.48E-08 | 31.07316471 | 0.49904381 | 6.907309908 |
| ENSMUSG00000024782 | *Ak3* | 1711.024739 | 1818.626783 | 1950.099113 | 890.1246012 | 899.0589916 | 941.2664052 | 6.74E-28 | -1.002850885 | 3.18E-29 | 125.9342265 | 0.499012934 | 27.1712219 |
| ENSMUSG00000028825 | *Rhd* | 324.7255229 | 338.5945434 | 413.4663923 | 171.2200219 | 180.0223508 | 183.6617376 | 2.95E-09 | -1.003537187 | 5.02E-10 | 38.66891048 | 0.498775606 | 8.530104337 |
| ENSMUSG00000053253 | *Ndfip2* | 1210.751544 | 1447.419734 | 1603.442838 | 717.807015 | 647.4488054 | 755.7680502 | 2.21E-13 | -1.004214767 | 2.58E-14 | 58.0322776 | 0.498541405 | 12.6565799 |
| ENSMUSG00000001123 | *Lgals9* | 3851.500865 | 3357.169694 | 2644.672229 | 1894.395884 | 1289.633799 | 1728.256951 | 3.16E-08 | -1.004869846 | 5.92E-09 | 33.85980818 | 0.498315085 | 7.50058068 |
| ENSMUSG00000037321 | *Tap1* | 2378.557948 | 2257.936417 | 1894.634109 | 1238.052466 | 887.4786064 | 1129.519686 | 3.25E-11 | -1.005542987 | 4.57E-12 | 47.8628399 | 0.498082633 | 10.48821998 |
| ENSMUSG00000028646 | *Rragc* | 3447.665876 | 3779.213884 | 4148.530174 | 1756.102789 | 1933.924318 | 1960.589049 | 8.59E-24 | -1.008220148 | 5.06E-25 | 106.7472852 | 0.497159214 | 23.06606223 |
| ENSMUSG00000036257 | *Pnpla8* | 437.7390459 | 426.8401468 | 549.6077654 | 259.0251614 | 196.8665473 | 245.1884197 | 2.39E-08 | -1.00886158 | 4.42E-09 | 34.42920206 | 0.496938223 | 7.621254573 |
| ENSMUSG00000057948 | *Unc13d* | 984.7244974 | 965.9056805 | 775.2494856 | 447.8062112 | 423.2104386 | 482.1120612 | 2.19E-13 | -1.01231959 | 2.55E-14 | 58.05265566 | 0.495748534 | 12.65974086 |
| ENSMUSG00000021715 | *Cwc27* | 441.5061634 | 470.0037571 | 587.4248135 | 248.049519 | 235.8187519 | 256.208124 | 5.38E-10 | -1.013143382 | 8.48E-11 | 42.14427303 | 0.495465537 | 9.269527877 |
| ENSMUSG00000025949 | *Pikfyve* | 1105.272255 | 1168.295053 | 1050.053368 | 579.5139204 | 563.2278226 | 504.1514697 | 3.40E-21 | -1.015329005 | 2.30E-22 | 94.62801371 | 0.494715496 | 20.46911377 |
| ENSMUSG00000027068 | *Dhrs9* | 1723.832938 | 1700.646248 | 2275.325726 | 910.9783218 | 945.3805321 | 959.632579 | 1.09E-13 | -1.01544054 | 1.23E-14 | 59.48231627 | 0.494677251 | 12.96449525 |
| ENSMUSG00000032557 | *Uba5* | 1778.079429 | 1732.299562 | 2076.155939 | 968.0516625 | 906.4283275 | 885.2495752 | 2.90E-21 | -1.016398595 | 1.95E-22 | 94.95404912 | 0.494348858 | 20.5371053 |
| ENSMUSG00000095500 | *ENSMUSG00000095500* | 92.67108889 | 77.69449863 | 92.02148365 | 48.2928267 | 31.58286856 | 49.58866915 | 0.001667557 | -1.016485986 | 0.000615779 | 11.72766351 | 0.494318914 | 2.77791928 |
| ENSMUSG00000078495 | *Zfp984* | 131.8491102 | 110.3070042 | 115.9722808 | 83.41488248 | 49.4798274 | 44.99712571 | 0.001828264 | -1.017576364 | 0.000681145 | 11.54000348 | 0.493945453 | 2.737961084 |
| ENSMUSG00000073405 | *H2-T-ps* | 195.1366831 | 229.2467305 | 218.0783106 | 120.7320667 | 111.5928022 | 85.40270798 | 2.86E-06 | -1.018396182 | 6.78E-07 | 24.67559849 | 0.493664846 | 5.543834907 |
| ENSMUSG00000040560 | *Wdr7* | 853.6288107 | 940.0075143 | 673.1434558 | 384.1474851 | 414.7883404 | 417.830453 | 4.12E-11 | -1.021503281 | 5.86E-12 | 47.376601 | 0.492602795 | 10.38513777 |
| ENSMUSG00000051154 | *Commd3* | 1616.09338 | 1426.317524 | 2169.437991 | 881.3440873 | 830.629443 | 852.1904625 | 3.73E-10 | -1.021908193 | 5.76E-11 | 42.89891571 | 0.492464559 | 9.428112789 |
| ENSMUSG00000026511 | *Srp9* | 1388.559486 | 1332.316773 | 1957.662522 | 786.9535624 | 716.9311162 | 796.1736325 | 7.05E-10 | -1.022494869 | 1.12E-10 | 41.59305631 | 0.492264337 | 9.151608635 |
| ENSMUSG00000021242 | *Npc2* | 10585.59999 | 10782.26987 | 10885.007 | 5650.260724 | 4917.452634 | 5293.131278 | 2.17E-42 | -1.023944403 | 5.63E-44 | 193.4461479 | 0.491769988 | 41.6626762 |
| ENSMUSG00000022236 | *Ropn1l* | 76.09577218 | 90.16398607 | 83.19750577 | 50.48795519 | 44.21601598 | 28.46756933 | 0.002896943 | -1.024409799 | 0.001121045 | 10.6161459 | 0.491611375 | 2.53806004 |
| ENSMUSG00000047735 | *Samd9l* | 913.9026897 | 681.0258522 | 531.9598096 | 377.5620996 | 263.1905713 | 405.8924401 | 2.96E-05 | -1.024792341 | 8.13E-06 | 19.90749515 | 0.491481037 | 4.529094109 |
| ENSMUSG00000033653 | *Vps8* | 1341.093807 | 1533.746954 | 1463.519761 | 676.0995738 | 753.7777962 | 699.7512203 | 8.77E-24 | -1.025588443 | 5.17E-25 | 106.7025239 | 0.491209905 | 23.05675887 |
| ENSMUSG00000019853 | *Hebp2* | 229.04074 | 255.1448967 | 267.2404731 | 122.9271952 | 108.4345154 | 135.9096858 | 4.90E-08 | -1.026953425 | 9.39E-09 | 32.96376224 | 0.490745374 | 7.310230647 |
| ENSMUSG00000001962 | *Fam50a* | 1326.025337 | 1454.134073 | 1484.949421 | 673.9044453 | 721.1421654 | 694.2413681 | 4.38E-27 | -1.027868124 | 2.16E-28 | 122.1330506 | 0.49043433 | 26.35900438 |
| ENSMUSG00000104217 | *Gm37988* | 219.9996582 | 293.5125504 | 254.6347904 | 128.4150165 | 128.4369988 | 119.3801294 | 1.73E-07 | -1.02847271 | 3.55E-08 | 30.37876798 | 0.490228849 | 6.761293097 |
| ENSMUSG00000052821 | *Cysltr1* | 238.8352454 | 177.4503981 | 243.289676 | 124.0247595 | 91.59031881 | 107.4421165 | 1.03E-05 | -1.029722947 | 2.64E-06 | 22.06431225 | 0.489804201 | 4.987134913 |
| ENSMUSG00000032115 | *Hyou1* | 48155.81559 | 51139.28635 | 44054.33987 | 22538.48173 | 24156.6834 | 23508.70241 | 1.23E-38 | -1.029962391 | 3.66E-40 | 175.9790857 | 0.489722915 | 37.91056772 |
| ENSMUSG00000026433 | *Rab29* | 629.8620351 | 654.1684946 | 826.9327846 | 357.8059433 | 342.1477427 | 330.5911277 | 6.26E-12 | -1.031510453 | 8.32E-13 | 51.20592664 | 0.489197707 | 11.20343434 |
| ENSMUSG00000029482 | *Aacs* | 2856.981863 | 2841.124752 | 2240.029814 | 1367.565047 | 1129.613932 | 1387.564428 | 3.53E-14 | -1.031711737 | 3.84E-15 | 61.78169007 | 0.489129459 | 13.45248894 |
| ENSMUSG00000044072 | *Eml6* | 81.36973659 | 85.36802936 | 55.46500385 | 31.82936305 | 36.84667998 | 40.40558227 | 0.00341989 | -1.032068969 | 0.001344299 | 10.28068621 | 0.489008359 | 2.465987836 |
| ENSMUSG00000020462 | *Cfap36* | 568.8347326 | 468.0853745 | 675.6645923 | 327.0741445 | 221.0800799 | 288.348928 | 6.32E-07 | -1.032144509 | 1.38E-07 | 27.74909662 | 0.488982755 | 6.199356718 |
| ENSMUSG00000078771 | *Evi2a* | 3138.762247 | 2819.063352 | 3332.942504 | 1705.614834 | 1309.636283 | 1524.392422 | 6.59E-17 | -1.033050478 | 5.83E-18 | 74.57902762 | 0.488675785 | 16.18103496 |
| ENSMUSG00000052713 | *Zfp608* | 155.9586618 | 164.0217193 | 109.6694394 | 53.78064792 | 92.6430811 | 64.28160816 | 0.00105412 | -1.03324879 | 0.000373123 | 12.66219805 | 0.488608616 | 2.977109816 |
| ENSMUSG00000074863 | *Platr25* | 56.50676152 | 54.67390644 | 51.68329904 | 23.04884911 | 23.16077027 | 33.05911277 | 0.007031399 | -1.033987405 | 0.002944869 | 8.841315081 | 0.488358528 | 2.152958268 |
| ENSMUSG00000031441 | *Atp11a* | 2243.695144 | 2636.816997 | 1995.47957 | 1104.149629 | 1136.983268 | 1117.581673 | 1.05E-17 | -1.034400732 | 8.86E-19 | 78.29861441 | 0.488218635 | 16.97938359 |
| ENSMUSG00000034707 | *Gns* | 7950.878058 | 8797.702981 | 8202.517728 | 3808.547924 | 4218.418477 | 4143.4088 | 1.05E-39 | -1.035503455 | 3.02E-41 | 180.9431357 | 0.487845608 | 38.97711846 |
| ENSMUSG00000062300 | *Nectin2* | 149.1778504 | 179.3687808 | 168.9161481 | 73.53680429 | 70.53507311 | 97.34072093 | 1.63E-05 | -1.035854783 | 4.30E-06 | 21.12493327 | 0.487726821 | 4.788064539 |
| ENSMUSG00000022940 | *Pigp* | 232.8078575 | 218.6956258 | 358.0013885 | 133.9028377 | 121.0676628 | 137.7463032 | 3.45E-05 | -1.036933571 | 9.56E-06 | 19.59800614 | 0.487362255 | 4.462760647 |
| ENSMUSG00000025494 | *Sigirr* | 493.4923839 | 500.6978801 | 606.3333375 | 274.3910608 | 244.2408502 | 258.96305 | 2.32E-12 | -1.038011137 | 2.95E-13 | 53.24140946 | 0.486998375 | 11.63441747 |
| ENSMUSG00000050552 | *Lamtor4* | 1267.258305 | 1124.172252 | 1457.216919 | 637.6848253 | 594.8106911 | 638.2245382 | 2.33E-16 | -1.039033531 | 2.14E-17 | 72.01143125 | 0.486653376 | 15.63261584 |
| ENSMUSG00000043822 | *Adamtsl5* | 383.4925549 | 460.4118437 | 448.7623038 | 214.0250274 | 176.8640639 | 236.0053328 | 1.66E-09 | -1.040010927 | 2.74E-10 | 39.85012152 | 0.48632379 | 8.780943576 |
| ENSMUSG00000021027 | *Ralgapa1* | 545.4786045 | 533.3103857 | 412.205824 | 128.4150165 | 326.3563084 | 269.9827543 | 0.001150735 | -1.040685941 | 0.000409967 | 12.48621183 | 0.4860963 | 2.939024851 |
| ENSMUSG00000031357 | *Syap1* | 498.7663484 | 552.4942125 | 690.7914115 | 290.8545244 | 253.7157107 | 299.3686323 | 1.23E-09 | -1.041023161 | 2.01E-10 | 40.45728918 | 0.485982692 | 8.911038795 |
| ENSMUSG00000041000 | *Trim62* | 70.82180777 | 68.10258522 | 71.85239135 | 32.9269273 | 43.16325369 | 26.63095195 | 0.002726691 | -1.04156286 | 0.001048986 | 10.73904203 | 0.485800924 | 2.564364141 |
| ENSMUSG00000033021 | *Gmppa* | 2654.310945 | 2863.186153 | 2909.391565 | 1390.613896 | 1320.163906 | 1375.626415 | 5.32E-40 | -1.043196193 | 1.49E-41 | 182.341309 | 0.485251241 | 39.27396873 |
| ENSMUSG00000083373 | *Gm13392* | 288.5611955 | 361.6151356 | 363.0436615 | 182.1956644 | 161.0726296 | 147.8476988 | 1.39E-08 | -1.043346782 | 2.51E-09 | 35.52797402 | 0.485200593 | 7.858353015 |
| ENSMUSG00000040613 | *Apobec1* | 4425.609562 | 4332.667288 | 4617.46157 | 2135.860017 | 2232.908807 | 2117.619835 | 8.03E-53 | -1.043857293 | 1.41E-54 | 242.0430402 | 0.48502893 | 52.09548154 |
| ENSMUSG00000022092 | *Ppp3cc* | 524.3827469 | 414.3706594 | 476.4948058 | 178.9029716 | 248.4518993 | 258.0447413 | 5.37E-08 | -1.044676637 | 1.03E-08 | 32.77747182 | 0.484753548 | 7.270217055 |

| | | | | | | | | | | | | | |
|---|---|---|---|---|---|---|---|---|---|---|---|---|---|
| ENSMUSG00000049832 | *Rbx1-ps* | 811.4370954 | 889.1703732 | 1089.130985 | 445.6110827 | 432.6852992 | 470.1740483 | 4.29E-13 | -1.045242577 | 5.11E-14 | 56.68518866 | 0.484563426 | 12.36770749 |
| ENSMUSG00000033502 | *Cdc14a* | 412.1226474 | 412.4522767 | 394.5578683 | 201.9518207 | 186.3389245 | 202.0279114 | 2.30E-14 | -1.047041897 | 2.47E-15 | 62.64859403 | 0.483959459 | 13.63759997 |
| ENSMUSG00000023022 | *Lima1* | 813.6973659 | 822.0269793 | 918.9542683 | 410.4890269 | 417.9466272 | 405.8924401 | 1.48E-21 | -1.047191272 | 9.77E-23 | 96.31985249 | 0.483909353 | 20.82865757 |
| ENSMUSG00000028383 | *Hsdl2* | 409.8623769 | 393.2684499 | 564.7345846 | 203.049385 | 208.4469325 | 247.0250371 | 1.26E-07 | -1.048890983 | 2.53E-08 | 31.03449015 | 0.483339571 | 6.899250027 |
| ENSMUSG00000064105 | *Cnnm2* | 568.8347326 | 569.7596566 | 510.530149 | 268.9032396 | 250.5574239 | 278.2475325 | 4.00E-16 | -1.049309688 | 3.73E-17 | 70.91453157 | 0.483199314 | 15.39762501 |
| ENSMUSG00000047617 | *Paxx* | 202.670918 | 212.9404777 | 267.2404731 | 104.2686031 | 114.7510891 | 109.2787339 | 3.40E-07 | -1.049942865 | 7.21E-08 | 29.00861363 | 0.482987292 | 6.468199472 |
| ENSMUSG00000029428 | *Stx2* | 1594.244098 | 1829.177888 | 1860.598765 | 878.0513945 | 834.8404922 | 836.5792148 | 9.10E-26 | -1.050170012 | 4.80E-27 | 115.9815203 | 0.482911253 | 25.04083056 |
| ENSMUSG00000020392 | *Cdkn2aipnl* | 5166.978273 | 5346.532535 | 6650.758189 | 2799.886384 | 2645.591623 | 2833.900611 | 5.11E-20 | -1.051087621 | 3.73E-21 | 89.10995295 | 0.482604201 | 19.29116991 |
| ENSMUSG00000037325 | *Bbs7* | 266.7119144 | 262.8184275 | 365.5647981 | 159.1468153 | 132.6480479 | 138.6646119 | 5.71E-07 | -1.051761039 | 1.24E-07 | 27.95826733 | 0.482378985 | 6.243709449 |
| ENSMUSG00000056144 | *Trim34a* | 106.2327117 | 92.08236875 | 110.9300077 | 61.46359762 | 37.89944227 | 49.58866915 | 0.000490196 | -1.054142248 | 0.000162849 | 14.21749847 | 0.481583461 | 3.309630521 |
| ENSMUSG00000030882 | *Dnhd1* | 90.41081843 | 104.5518562 | 84.45807404 | 41.70744124 | 43.16325369 | 49.58866915 | 0.000261095 | -1.054266528 | 8.28E-05 | 15.4929142 | 0.481541977 | 3.583201157 |
| ENSMUSG00000054277 | *Arfgap3* | 634.382576 | 657.0460687 | 753.819825 | 426.9524906 | 248.4518993 | 308.5517192 | 4.65E-07 | -1.055503562 | 1.00E-07 | 28.37290334 | 0.481129258 | 6.332752169 |
| ENSMUSG00000042500 | *Ago4* | 139.3833451 | 146.7562752 | 132.3596683 | 70.24411156 | 64.2184994 | 67.03653422 | 3.09E-06 | -1.056592549 | 7.37E-07 | 24.51611521 | 0.480766225 | 5.510646129 |
| ENSMUSG00000024942 | *Capn1* | 2243.695144 | 2431.55005 | 2265.24118 | 1076.710523 | 1078.02858 | 1179.108355 | 6.36E-35 | -1.057043543 | 2.24E-36 | 158.6462893 | 0.480615958 | 34.19632534 |
| ENSMUSG00000024777 | *Ppp2r5b* | 941.0259352 | 963.0281065 | 985.7643865 | 474.147753 | 414.7883404 | 496.8050002 | 2.65E-21 | -1.058511531 | 1.77E-22 | 95.14280032 | 0.480127165 | 20.57673479 |
| ENSMUSG00000032018 | *Sc5d* | 1524.929138 | 1425.358333 | 1776.140691 | 885.7343442 | 633.7628957 | 748.4215807 | 4.02E-12 | -1.058829199 | 5.22E-13 | 52.11939484 | 0.480021457 | 11.39619885 |
| ENSMUSG00000060224 | *Pyroxd2* | 86.643701 | 134.2867878 | 102.1060298 | 50.48795519 | 43.16325369 | 60.60837341 | 0.000886689 | -1.060697224 | 0.000308625 | 13.01730694 | 0.47940032 | 3.052228514 |
| ENSMUSG00000020873 | *Slc35b1* | 1048.765494 | 1214.336238 | 1150.89883 | 577.3187919 | 639.0267071 | 421.5036878 | 3.19E-10 | -1.060752513 | 4.90E-11 | 43.21675407 | 0.479381948 | 9.495956882 |
| ENSMUSG00000045679 | *Pqlc3* | 856.6425047 | 963.9872978 | 1109.300077 | 445.6110827 | 468.4792169 | 486.7036046 | 1.87E-15 | -1.061074612 | 1.84E-16 | 67.76474525 | 0.479274932 | 14.72740436 |
| ENSMUSG00000056116 | *H2-T22* | 1964.17503 | 1887.688559 | 2001.782412 | 979.0273049 | 851.6846887 | 972.4889006 | 3.38E-30 | -1.061696401 | 1.44E-31 | 136.6416046 | 0.479068413 | 29.47059226 |
| ENSMUSG00000040714 | *Klc3* | 173.287402 | 244.593792 | 191.6063769 | 91.09783218 | 91.59031881 | 108.3604252 | 5.25E-06 | -1.062738597 | 1.29E-06 | 23.43794137 | 0.478722461 | 5.279501691 |
| ENSMUSG00000037997 | *Parp11* | 853.6288107 | 823.945362 | 757.6015298 | 429.1476191 | 362.1502261 | 375.5882534 | 3.45E-18 | -1.064350814 | 2.82E-19 | 80.55974221 | 0.478187786 | 17.46243491 |
| ENSMUSG00000029060 | *Mib2* | 1166.299558 | 1119.376295 | 973.1587039 | 587.1968701 | 449.5294958 | 523.4359522 | 2.25E-14 | -1.064653729 | 2.41E-15 | 62.69993902 | 0.478087394 | 13.64836594 |
| ENSMUSG00000038312 | *Edem2* | 3069.447286 | 3387.863817 | 3353.111596 | 1622.199951 | 1481.236535 | 1583.164178 | 1.57E-37 | -1.065104448 | 4.95E-39 | 170.7996846 | 0.477938056 | 36.80283657 |
| ENSMUSG00000095762 | *Gm16378* | 543.2183341 | 634.0254765 | 482.7976471 | 246.9519547 | 254.768473 | 291.1038541 | 2.81E-11 | -1.066284487 | 3.94E-12 | 48.15470736 | 0.477547291 | 10.55095747 |
| ENSMUSG00000034613 | *Ppm1h* | 1347.121195 | 1480.032239 | 1379.061687 | 677.197138 | 622.1825105 | 707.0976898 | 5.81E-27 | -1.066929803 | 2.88E-28 | 121.5601293 | 0.477333732 | 26.23602336 |
| ENSMUSG00000013539 | *Tango2* | 535.6840992 | 554.4125952 | 723.5661865 | 278.7813178 | 285.2985793 | 298.4503236 | 5.39E-11 | -1.067659755 | 7.72E-12 | 46.83492701 | 0.477092279 | 10.26824668 |
| ENSMUSG00000018931 | *Natd1* | 383.4925549 | 442.1872083 | 392.0367317 | 169.0248934 | 181.0751131 | 228.6588633 | 1.01E-09 | -1.069467141 | 1.65E-10 | 40.84782491 | 0.47649496 | 8.994364566 |
| ENSMUSG00000000127 | *Fer* | 501.0266188 | 439.3096342 | 652.9743635 | 255.7324687 | 254.768473 | 246.1067284 | 5.03E-09 | -1.071226386 | 8.75E-10 | 37.58529366 | 0.475914268 | 8.298784706 |
| ENSMUSG00000019710 | *Mrpl24* | 4584.581918 | 4593.567333 | 5738.106762 | 2522.202631 | 2224.486709 | 2348.115315 | 2.70E-20 | -1.071456809 | 1.93E-21 | 90.42037138 | 0.475838263 | 19.56933581 |
| ENSMUSG00000001105 | *Ift20* | 1153.491359 | 1214.336238 | 1647.562728 | 654.1482889 | 615.8659368 | 637.3062295 | 7.67E-12 | -1.071779344 | 1.03E-12 | 50.79170475 | 0.475731894 | 11.11536116 |
| ENSMUSG00000028527 | *Ak4* | 1112.80649 | 1285.316397 | 1206.363834 | 614.6359762 | 508.4841837 | 590.4724864 | 2.56E-19 | -1.071921554 | 1.96E-20 | 85.8267144 | 0.475685003 | 18.59136 |
| ENSMUSG00000044894 | *Uqcrq* | 5690.607597 | 5537.411612 | 7539.458818 | 3188.424126 | 2720.337745 | 3005.624336 | 7.06E-15 | -1.073551443 | 7.29E-16 | 65.05362605 | 0.4751479 | 14.15136541 |
| ENSMUSG00000040462 | *Os9* | 5358.347839 | 6362.316166 | 6570.081819 | 2747.203301 | 3039.324717 | 2877.979428 | 9.00E-26 | -1.077337219 | 4.73E-27 | 116.0075966 | 0.473902699 | 25.04597511 |
| ENSMUSG00000071266 | *Zfp946* | 130.3422632 | 95.91913411 | 152.5287606 | 48.2928267 | 64.2184994 | 66.11822554 | 0.000379362 | -1.077389732 | 0.000123452 | 14.7392165 | 0.473885449 | 3.420945751 |
| ENSMUSG00000055128 | *Cgrrf1* | 449.7938217 | 500.6978801 | 630.2841346 | 254.6349044 | 234.7659896 | 256.208124 | 3.82E-10 | -1.07938731 | 5.91E-11 | 42.85122121 | 0.473229754 | 9.418488244 |
| ENSMUSG00000092224 | *Gm8815* | 678.0811383 | 737.6181413 | 533.2203779 | 289.7569602 | 290.5623907 | 340.6925232 | 1.09E-10 | -1.082593941 | 1.61E-11 | 45.39341249 | 0.47217909 | 9.962427326 |
| ENSMUSG00000020315 | *Sptbn1* | 1781.093123 | 1886.729368 | 1584.534314 | 729.8802217 | 892.7424178 | 857.7003146 | 1.90E-19 | -1.082778328 | 1.45E-20 | 86.43125376 | 0.472118746 | 18.7217856 |
| ENSMUSG00000020952 | *Scfd1* | 1527.189408 | 1284.357206 | 1769.83785 | 794.6365121 | 659.0291905 | 707.0976898 | 1.86E-13 | -1.083486348 | 2.15E-14 | 58.38647779 | 0.471887105 | 12.73144691 |
| ENSMUSG00000022946 | *Dop1b* | 392.5336367 | 352.0232222 | 306.3180894 | 93.29296067 | 220.0273176 | 181.8251202 | 0.000412138 | -1.08557804 | 0.000135249 | 14.56719642 | 0.471203434 | 3.384957542 |
| ENSMUSG00000004508 | *Gab2* | 915.4095366 | 908.3542 | 734.911301 | 398.4158203 | 427.4214878 | 382.0164142 | 9.16E-16 | -1.086316086 | 8.75E-17 | 69.23321595 | 0.47096244 | 15.03820016 |
| ENSMUSG00000097012 | *Gm16531* | 143.1504625 | 148.6746579 | 117.232849 | 76.82949702 | 74.74612225 | 42.24219965 | 0.000502642 | -1.087409076 | 0.000167363 | 14.16604862 | 0.470605773 | 3.298741243 |
| ENSMUSG00000093954 | *Gm16867* | 50.47937363 | 61.38824583 | 80.67636923 | 27.43910608 | 33.68839313 | 28.46756933 | 0.004157532 | -1.092408449 | 0.001665923 | 9.885433177 | 0.468977805 | 2.381164366 |
| ENSMUSG00000000958 | *Slc7a7* | 302.8762418 | 286.798211 | 233.2051298 | 163.5370722 | 124.2259497 | 100.095647 | 5.13E-06 | -1.092672365 | 1.26E-06 | 23.48791215 | 0.468892021 | 5.289566757 |
| ENSMUSG00000039081 | *Zfp503* | 259.931103 | 276.2471062 | 170.1767163 | 83.41488248 | 114.7510891 | 132.2364511 | 9.75E-05 | -1.097492224 | 2.89E-05 | 17.48973862 | 0.467328127 | 4.010871003 |
| ENSMUSG00000025130 | *P4hb* | 64484.0094 | 72771.92867 | 82417.21401 | 34503.02955 | 33387.30311 | 34758.90215 | 9.72E-28 | -1.097571397 | 4.60E-29 | 125.198997 | 0.467302481 | 27.0122092 |
| ENSMUSG00000029560 | *Snx8* | 3883.898075 | 3936.521264 | 4240.551658 | 1884.517806 | 1862.336483 | 1885.287736 | 1.03E-52 | -1.097738011 | 1.82E-54 | 241.5333604 | 0.467248517 | 51.98773168 |
| ENSMUSG00000026749 | *Nek6* | 1456.3676 | 1518.399893 | 1600.921702 | 675.0020095 | 742.1974111 | 717.1990853 | 1.95E-32 | -1.098401879 | 7.68E-34 | 147.0433472 | 0.467033558 | 31.70938014 |
| ENSMUSG00000035824 | *Tk2* | 620.0675298 | 712.6791665 | 839.5384673 | 313.9033735 | 364.2557507 | 333.3460537 | 1.84E-12 | -1.099181415 | 2.31E-13 | 53.71903942 | 0.466781272 | 11.73544748 |
| ENSMUSG00000030872 | *Gga2* | 2667.119144 | 2893.880276 | 3166.547492 | 1324.760042 | 1424.387372 | 1317.772967 | 3.38E-30 | -1.1006001 | 1.44E-31 | 136.6448106 | 0.466322485 | 29.47059468 |
| ENSMUSG00000027006 | *Dnajc10* | 4788.759683 | 4937.917024 | 6548.652159 | 2618.788284 | 2473.99137 | 2489.534853 | 8.70E-17 | -1.101365736 | 7.76E-18 | 74.01272612 | 0.466075074 | 16.06047831 |
| ENSMUSG00000024423 | *Impact* | 712.7386187 | 790.3736651 | 960.5530212 | 399.5133845 | 345.3060295 | 400.382588 | 9.56E-13 | -1.101781368 | 1.18E-13 | 55.04866509 | 0.46594082 | 12.01947744 |
| ENSMUSG00000018398 | *Septin8* | 1975.476383 | 2136.119117 | 2008.085253 | 878.0513945 | 1025.390466 | 945.8579486 | 2.54E-31 | -1.102546541 | 1.03E-32 | 141.8793265 | 0.465693761 | 30.59526819 |
| ENSMUSG00000001750 | *Tcirg1* | 12130.87156 | 13500.61813 | 13209.49489 | 6094.774242 | 5833.355822 | 6153.586518 | 1.44E-53 | -1.102789512 | 2.45E-55 | 245.5274055 | 0.465615338 | 52.8413923 |
| ENSMUSG00000029994 | *Anxa4* | 5634.100835 | 5953.700654 | 6238.552364 | 2745.008172 | 2838.247121 | 2713.602173 | 3.81E-53 | -1.102875067 | 6.62E-55 | 243.5490467 | 0.465587727 | 52.41872329 |
| ENSMUSG00000032763 | *Ilvbl* | 1133.148924 | 1234.479256 | 1162.243944 | 563.0504567 | 491.6399872 | 586.7992516 | 1.30E-23 | -1.103515001 | 7.76E-25 | 105.8986867 | 0.465381253 | 22.88562192 |
| ENSMUSG00000031513 | *Leprotl1* | 2713.077976 | 2726.021791 | 2763.165646 | 1274.272086 | 1222.257013 | 1318.691276 | 2.56E-52 | -1.10352248 | 4.57E-54 | 239.7029216 | 0.46537884 | 51.59197215 |
| ENSMUSG00000009828 | *Cilk1* | 421.1637292 | 436.4320602 | 439.938326 | 201.9518207 | 204.2358833 | 196.5180592 | 6.75E-17 | -1.105699084 | 5.98E-18 | 74.52858233 | 0.464677249 | 16.17094806 |
| ENSMUSG00000027956 | *Tmem144* | 818.2179068 | 989.885464 | 1179.8919 | 410.4890269 | 509.536946 | 463.7458874 | 4.55E-11 | -1.107756767 | 6.49E-12 | 47.17608217 | 0.464014963 | 10.34154763 |
| ENSMUSG00000027397 | *Slc20a1* | 5004.992224 | 4941.753789 | 4543.088042 | 2083.176934 | 2360.293043 | 2270.977385 | 1.99E-39 | -1.110014139 | 5.78E-41 | 179.6509045 | 0.463289491 | 38.70117879 |
| ENSMUSG00000025935 | *Tram1* | 6958.619325 | 7145.0163 | 8260.503868 | 3395.863768 | 3422.530189 | 3527.223671 | 2.11E-35 | -1.11151347 | 7.23E-37 | 160.8903516 | 0.462808264 | 34.67633034 |
| ENSMUSG00000020143 | *Dock2* | 9830.669658 | 10377.49112 | 9805.960566 | 4622.940592 | 4720.586087 | 4548.382932 | 4.41E-75 | -1.111534851 | 4.84E-77 | 345.1414247 | 0.462801405 | 74.35541673 |
| ENSMUSG00000022500 | *Litaf* | 4307.322075 | 4746.078756 | 3885.071406 | 1921.83499 | 2008.67044 | 2055.174844 | 3.99E-30 | -1.112392306 | 1.72E-31 | 136.2970685 | 0.462526424 | 29.39871389 |
| ENSMUSG00000024349 | *Sting1* | 1989.791429 | 1882.892603 | 1749.668758 | 908.7831933 | 840.1043036 | 854.9453885 | 5.67E-33 | -1.112404988 | 2.17E-34 | 149.5504556 | 0.462522358 | 32.24667226 |
| ENSMUSG00000036775 | *Decr2* | 498.7663484 | 492.065158 | 549.6077654 | 240.3665693 | 245.2936124 | 225.9039372 | 2.25E-17 | -1.113019602 | 1.93E-18 | 76.76272496 | 0.462325357 | 16.64797656 |
| ENSMUSG00000056888 | *Glipr1* | 5266.430174 | 4418.994509 | 4514.094972 | 2315.860553 | 1977.087572 | 2273.732312 | 7.99E-25 | -1.113023106 | 4.43E-26 | 111.5746134 | 0.462324234 | 24.09766311 |
| ENSMUSG00000028343 | *Erp44* | 3770.131129 | 3984.480831 | 4536.785201 | 1894.395884 | 1885.497253 | 1891.715897 | 7.35E-33 | -1.114824353 | 2.83E-34 | 149.0255537 | 0.461747369 | 32.1334887 |
| ENSMUSG00000038543 | *BC028528* | 582.3963554 | 668.5563648 | 963.0741577 | 376.4645354 | 315.8286856 | 326.9178929 | 1.04E-07 | -1.11646491 | 2.07E-08 | 31.4238977 | 0.461222593 | 6.982194135 |
| ENSMUSG00000002043 | *Trappc6a* | 1203.217309 | 1330.39839 | 1530.329879 | 639.8799538 | 606.3910763 | 623.5315992 | 2.56E-20 | -1.117720968 | 1.82E-21 | 90.52727258 | 0.460821212 | 19.59112833 |
| ENSMUSG00000027932 | *Slc27a3* | 72.32865475 | 86.3272207 | 60.50727692 | 40.609877 | 30.53010627 | 30.3041867 | 0.001367158 | -1.119951705 | 0.000493779 | 12.1390121 | 0.460109228 | 2.864181366 |
| ENSMUSG00000074749 | *Kiz* | 556.7799569 | 536.1879597 | 753.819825 | 317.1960663 | 281.0875301 | 249.7799631 | 1.64E-09 | -1.12114345 | 2.72E-10 | 39.86949231 | 0.459729309 | 8.784890573 |
| ENSMUSG00000030047 | *Arhgap25* | 756.4371809 | 926.5788355 | 862.2286962 | 426.9524906 | 343.200505 | 398.5459706 | 1.72E-15 | -1.121733069 | 1.69E-16 | 67.93697786 | 0.459541459 | 14.76321358 |
| ENSMUSG00000020321 | *Mdh1* | 10825.94208 | 12567.32495 | 17355.50393 | 6397.701973 | 6208.139196 | 6115.017553 | 4.69E-12 | -1.121866391 | 6.14E-13 | 51.80031834 | 0.459498994 | 11.3288225 |
| ENSMUSG00000078945 | *Naip2* | 1341.093807 | 1500.175258 | 1733.28137 | 730.9777859 | 697.9813951 | 669.4470336 | 1.75E-19 | -1.122737704 | 1.33E-20 | 86.59564764 | 0.459221564 | 18.75671258 |
| ENSMUSG00000023191 | *P3h3* | 734.5878998 | 875.7416944 | 859.7075596 | 385.2450494 | 426.3687255 | 321.4080408 | 7.59E-14 | -1.124604103 | 8.51E-15 | 60.21388799 | 0.458627859 | 13.11985588 |
| ENSMUSG00000020205 | *Phlda1* | 494.9992309 | 516.0449415 | 274.8038827 | 223.9031056 | 155.8088182 | 211.2109982 | 5.43E-05 | -1.124838229 | 1.55E-05 | 18.67587853 | 0.458553437 | 4.26543838 |
| ENSMUSG00000022305 | *Lrp12* | 932.7382768 | 1159.662331 | 1150.89883 | 509.2698088 | 493.7455118 | 481.1937525 | 8.03E-19 | -1.126102318 | 6.31E-20 | 83.52059709 | 0.458151828 | 18.0951835 |
| ENSMUSG00000029401 | *Rilpl2* | 2044.03792 | 2236.834207 | 2773.250192 | 1164.515662 | 918.0087127 | 1139.621082 | 6.97E-14 | -1.129180609 | 7.79E-15 | 60.38685388 | 0.457175308 | 13.15668772 |
| ENSMUSG00000054484 | *Tmem62* | 464.108868 | 434.5136775 | 521.8752635 | 215.1225917 | 196.8665473 | 235.0870241 | 9.04E-14 | -1.130240813 | 1.02E-14 | 59.85436272 | 0.456839464 | 13.04396718 |
| ENSMUSG00000056671 | *Prelid2* | 256.1639856 | 202.389373 | 225.6417202 | 116.3418098 | 97.90689252 | 99.1773383 | 2.89E-08 | -1.130702834 | 5.37E-09 | 34.04982218 | 0.456693185 | 7.539804188 |
| ENSMUSG00000072680 | *None* | 567.3278857 | 487.2692013 | 603.812201 | 259.0251614 | 253.7157107 | 243.3518023 | 1.75E-15 | -1.132443624 | 1.71E-16 | 67.90750893 | 0.45614246 | 14.75733154 |
| ENSMUSG00000020230 | *Prmt2* | 1250.682988 | 1314.092137 | 1334.941797 | 619.0262331 | 514.8007575 | 641.8977729 | 1.34E-22 | -1.133459926 | 8.46E-24 | 101.1662698 | 0.455821246 | 21.87206933 |
| ENSMUSG00000020184 | *Mdm2* | 4441.431456 | 4404.606638 | 4913.695114 | 2031.591414 | 2242.383667 | 1996.403088 | 3.64E-38 | -1.133460521 | 1.12E-39 | 173.7584287 | 0.455821057 | 37.43941956 |
| ENSMUSG00000033813 | *Tcea1* | 5718.484266 | 5397.369676 | 7790.311904 | 2972.20397 | 2703.493548 | 2933.996258 | 7.65E-15 | -1.134309443 | 7.91E-16 | 64.89138262 | 0.455552918 | 14.11618474 |
| ENSMUSG00000040033 | *Stat2* | 1284.587045 | 1237.35683 | 970.6375673 | 569.6358422 | 491.6399872 | 531.7007304 | 1.88E-15 | -1.135025408 | 1.85E-16 | 67.75682875 | 0.455326898 | 14.72596361 |
| ENSMUSG00000020585 | *Laptm4a* | 6880.263283 | 7174.751232 | 8740.780379 | 3536.351991 | 3389.894558 | 3441.820963 | 3.74E-26 | -1.136112581 | 1.92E-27 | 117.7929062 | 0.454983906 | 25.42659853 |
| ENSMUSG00000022365 | *Derl1* | 9541.355039 | 10242.24514 | 10700.96404 | 4504.403654 | 4541.616498 | 4812.855834 | 3.74E-52 | -1.13672672 | 6.78E-54 | 238.9155101 | 0.454790265 | 51.42690985 |
| ENSMUSG00000052485 | *Tmem171* | 724.7933944 | 825.8637447 | 833.235626 | 407.1963342 | 305.3010627 | 370.0784013 | 7.69E-15 | -1.137554294 | 7.96E-16 | 64.87938973 | 0.454529459 | 14.11411752 |
| ENSMUSG00000030123 | *Plxnd1* | 4378.143883 | 4492.852242 | 3421.182283 | 1661.712264 | 1993.931768 | 1932.12148 | 1.01E-17 | -1.137831705 | 8.51E-19 | 78.37794931 | 0.454442067 | 16.99541223 |
| ENSMUSG00000081254 | *Gm12112* | 633.6291525 | 658.9644514 | 892.4823346 | 366.5864572 | 306.353825 | 317.734806 | 1.10E-10 | -1.138672456 | 1.63E-11 | 45.37054385 | 0.454177312 | 9.957557977 |
| ENSMUSG00000005949 | *Ctns* | 527.3964409 | 610.045693 | 589.94595 | 253.5373402 | 253.7157107 | 275.4926064 | 6.17E-18 | -1.138708622 | 5.11E-19 | 79.38580855 | 0.454165926 | 17.20983702 |
| ENSMUSG00000054676 | *1600014C10Rik* | 781.300156 | 775.985795 | 829.4539212 | 352.3181221 | 351.6226033 | 377.4248708 | 4.17E-26 | -1.139675982 | 2.16E-27 | 117.5669926 | 0.4538615 | 25.37944511 |
| ENSMUSG00000039497 | *Dse* | 655.4784336 | 640.7398159 | 773.9889173 | 331.4644014 | 301.0900136 | 304.8784844 | 1.80E-17 | -1.140790792 | 1.53E-18 | 77.21819297 | 0.453510924 | 16.74569257 |
| ENSMUSG00000048440 | *Cyp4f16* | 750.409793 | 785.5777084 | 761.3832346 | 312.8058093 | 363.2029884 | 362.7319318 | 8.08E-23 | -1.143459311 | 5.03E-24 | 102.1950473 | 0.452672851 | 22.09238594 |
| ENSMUSG00000046949 | *Nqo2* | 1514.381209 | 1577.869756 | 1958.92309 | 806.7097187 | 760.0943699 | 717.1990853 | 1.34E-18 | -1.143787514 | 1.07E-19 | 82.47466523 | 0.452569883 | 17.87440488 |
| ENSMUSG00000114380 | *Gm47942* | 143.1504625 | 157.3073799 | 228.1628567 | 84.51244672 | 71.58783539 | 81.72947323 | 2.49E-05 | -1.143956234 | 6.76E-06 | 20.25965724 | 0.452516959 | 4.604047197 |
| ENSMUSG00000003573 | *Homer3* | 911.6424192 | 1060.865623 | 1210.145538 | 462.0745464 | 490.5872249 | 482.1120612 | 3.71E-17 | -1.146439315 | 3.22E-18 | 75.74695041 | 0.451738783 | 16.4307968 |
| ENSMUSG00000041058 | *Wwp1* | 540.9580636 | 550.5758298 | 726.0873231 | 263.4154184 | 267.4016204 | 287.4306193 | 3.34E-12 | -1.146757805 | 4.32E-13 | 52.49377111 | 0.451639068 | 11.47600381 |
| ENSMUSG00000048138 | *Dmrt2* | 189.8627187 | 182.2463548 | 182.782399 | 80.12218975 | 92.6430811 | 78.05623848 | 1.53E-08 | -1.147957996 | 2.78E-09 | 35.33416507 | 0.451263502 | 7.81661509 |
| ENSMUSG00000033429 | *Mcee* | 424.1774232 | 411.4930853 | 630.2841346 | 242.5616977 | 188.444449 | 227.7405546 | 1.52E-07 | -1.15017638 | 3.10E-08 | 30.64665244 | 0.450570143 | 6.817613884 |
| ENSMUSG00000019726 | *Lyst* | 857.3959281 | 953.4361931 | 923.9965414 | 377.5620996 | 455.8460695 | 397.6276619 | 1.66E-21 | -1.150698494 | 1.10E-22 | 96.08297804 | 0.45040711 | 20.77895292 |
| ENSMUSG00000047554 | *Tmem41b* | 2577.461749 | 2570.632794 | 3616.570364 | 1313.784399 | 1360.168872 | 1270.939224 | 1.12E-14 | -1.150722849 | 1.17E-15 | 64.12099301 | 0.450399506 | 13.9526724 |
| ENSMUSG00000045763 | *Basp1* | 684.8619496 | 757.7611595 | 862.2286962 | 392.9279991 | 242.1353256 | 401.3008967 | 5.51E-08 | -1.151062209 | 1.06E-08 | 32.72047909 | 0.450293573 | 7.259130335 |
| ENSMUSG00000042622 | *Maff* | 158.9723557 | 171.6952501 | 185.3035356 | 74.63436854 | 67.37678625 | 89.07594274 | 3.33E-07 | -1.151228982 | 7.05E-08 | 29.05151181 | 0.450241523 | 6.477112315 |
| ENSMUSG00000078349 | *AW011738* | 56.50676152 | 52.75552376 | 39.07761635 | 25.24397759 | 20.00248342 | 22.03940851 | 0.00512472 | -1.151887233 | 0.002087237 | 9.471192231 | 0.45003614 | 2.290329845 |
| ENSMUSG00000038650 | *Rnh1* | 23255.92277 | 24934.17891 | 27864.86159 | 11195.15528 | 11343.51362 | 11675.37666 | 4.12E-41 | -1.15224024 | 1.12E-42 | 187.4883313 | 0.449926036 | 40.38467854 |
| ENSMUSG00000039166 | *Akap7* | 100.2053238 | 110.3070042 | 100.8454615 | 45.00013397 | 55.79640112 | 39.48727358 | 4.39E-05 | -1.153923081 | 1.24E-05 | 19.10718944 | 0.449401523 | 4.357533605 |
| ENSMUSG00000027828 | *Ssr3* | 9808.066953 | 10549.18637 | 12775.85941 | 4902.819474 | 5004.831904 | 4976.31478 | 7.74E-26 | -1.154120826 | 4.06E-27 | 116.3112405 | 0.44933993 | 25.11133488 |
| ENSMUSG00000052397 | *Ezr* | 3039.310346 | 2945.676609 | 2785.855875 | 1211.710924 | 1404.384888 | 1326.037745 | 9.78E-38 | -1.154133322 | 3.05E-39 | 171.7606267 | 0.449336037 | 37.00985287 |
| ENSMUSG00000020326 | *Ccng1* | 3677.46004 | 3877.051401 | 5874.248135 | 2037.079235 | 1926.554982 | 2065.276239 | 5.42E-11 | -1.154612527 | 7.77E-12 | 46.82249654 | 0.449186811 | 10.26590802 |
| ENSMUSG00000075486 | *Commd6* | 1023.149095 | 1220.091386 | 1797.570352 | 696.9532944 | 560.0695357 | 555.5767562 | 9.25E-08 | -1.15521313 | 1.83E-08 | 31.66840778 | 0.44899985 | 7.033723797 |
| ENSMUSG00000030747 | *Dgat2* | 91.91766541 | 101.6742822 | 103.3665981 | 39.51231275 | 60.00745026 | 33.97742146 | 0.000383931 | -1.155308096 | 0.000125123 | 14.71386689 | 0.448970296 | 3.415746933 |
| ENSMUSG00000027953 | *Slc50a1* | 1524.929138 | 1556.767547 | 1703.027732 | 713.4167581 | 695.8758705 | 734.6469504 | 4.56E-36 | -1.156116378 | 1.51E-37 | 164.0052478 | 0.448718827 | 35.3409126 |
| ENSMUSG00000035521 | *Gnptg* | 843.0808819 | 954.3953844 | 1125.687464 | 465.3672391 | 375.8361358 | 467.4191222 | 1.76E-13 | -1.15667027 | 2.03E-14 | 58.50250987 | 0.448546584 | 12.75551674 |
| ENSMUSG00000026209 | *Dnpep* | 4045.130701 | 4166.727186 | 4564.517703 | 1977.810766 | 1751.796443 | 1991.811544 | 1.96E-36 | -1.158198678 | 6.38E-38 | 165.7159296 | 0.44807164 | 35.70825572 |
| ENSMUSG00000018171 | *Vmp1* | 4250.06189 | 4527.38313 | 5636.000732 | 2288.421447 | 1996.037293 | 2164.453578 | 4.00E-20 | -1.159599909 | 2.89E-21 | 89.61477258 | 0.447636658 | 19.39747751 |
| ENSMUSG00000025059 | *Gk* | 2169.106219 | 2302.059219 | 2605.594613 | 1185.369383 | 948.538819 | 1028.505731 | 1.36E-21 | -1.161439147 | 8.94E-23 | 96.49648864 | 0.447066346 | 20.86604432 |
| ENSMUSG00000057137 | *Tmem140* | 187.6024482 | 171.6952501 | 200.4303548 | 79.02462551 | 84.22098282 | 86.32101667 | 1.23E-08 | -1.161745143 | 2.22E-09 | 35.7736214 | 0.446971533 | 7.911635955 |
| ENSMUSG00000069206 | *Zfp874a* | 53.49306757 | 67.14339388 | 66.81011827 | 24.14641335 | 22.10800799 | 36.73234752 | 0.002891783 | -1.162085979 | 0.001118651 | 10.62009836 | 0.446865949 | 2.538834269 |
| ENSMUSG00000026632 | *Tatdn3* | 193.6298361 | 181.2871635 | 205.4726279 | 115.2442455 | 66.32402397 | 78.05623848 | 8.27E-06 | -1.162832209 | 2.09E-06 | 22.51108049 | 0.446634869 | 5.082317621 |
| ENSMUSG00000007038 | *Neu1* | 3904.240509 | 4412.280169 | 5144.379107 | 1956.957046 | 1981.298621 | 2051.501609 | 3.65E-24 | -1.167208599 | 2.10E-25 | 108.4861255 | 0.445282062 | 23.43788491 |
| ENSMUSG00000020097 | *Sgpl1* | 9442.656562 | 11127.57875 | 12532.56973 | 4707.453039 | 5039.573059 | 4988.252793 | 1.18E-24 | -1.167275967 | 6.60E-26 | 110.7838485 | 0.44526127 | 23.92711203 |
| ENSMUSG00000055413 | *H2-Q5* | 1720.065821 | 1770.667216 | 1777.40126 | 778.1730484 | 773.7802796 | 791.5820891 | 2.87E-50 | -1.167777372 | 5.53E-52 | 230.1481125 | 0.445106547 | 49.54214538 |
| ENSMUSG00000005823 | *Gpr108* | 2202.256852 | 2312.610323 | 2547.608472 | 1047.076288 | 1028.548753 | 1058.809917 | 2.18E-38 | -1.170363633 | 6.60E-40 | 174.8072838 | 0.444309338 | 37.66162836 |
| ENSMUSG00000053398 | *Phgdh* | 24416.19494 | 26160.98464 | 28110.6724 | 11964.54781 | 10921.35595 | 12035.35366 | 3.13E-45 | -1.17186456 | 7.25E-47 | 206.6892262 | 0.443847335 | 44.50390117 |
| ENSMUSG00000007867 | *Ift43* | 330.7529108 | 364.4927096 | 452.5440087 | 166.829765 | 183.1806376 | 157.9490943 | 2.39E-10 | -1.172038478 | 3.63E-11 | 43.80137941 | 0.443793832 | 9.621411351 |
| ENSMUSG00000028744 | *Slc66a1* | 308.1502062 | 308.8596118 | 337.8322962 | 161.3419437 | 137.9118594 | 124.8899816 | 1.28E-11 | -1.172069794 | 1.74E-12 | 49.76232514 | 0.443784199 | 10.89417346 |
| ENSMUSG00000037280 | *Galnt6* | 1566.367429 | 1781.21832 | 1774.880123 | 759.5144563 | 772.7275173 | 737.4018765 | 3.18E-34 | -1.173329174 | 1.15E-35 | 155.3917456 | 0.443396973 | 33.49770543 |
| ENSMUSG00000029156 | *Sgcb* | 712.7386187 | 806.6799179 | 825.6722164 | 368.7815857 | 305.3010627 | 362.7319318 | 1.19E-18 | -1.17509946 | 9.49E-20 | 82.71244638 | 0.442853228 | 17.92403863 |
| ENSMUSG00000008206 | *Cers4* | 431.711658 | 441.2280169 | 347.9168423 | 197.5615638 | 147.3867199 | 196.5180592 | 5.31E-10 | -1.175104327 | 8.36E-11 | 42.17162372 | 0.442851734 | 9.275223737 |
| ENSMUSG00000045374 | *Wdr81* | 2624.927429 | 2700.123625 | 2369.868346 | 1098.661807 | 1194.885194 | 1115.745056 | 2.05E-38 | -1.175469564 | 6.16E-40 | 174.9447292 | 0.442739635 | 37.68867853 |
| ENSMUSG00000031861 | *Lpar2* | 71.57523126 | 63.30662851 | 71.85239135 | 29.63423457 | 30.53010627 | 31.22249539 | 0.00033055 | -1.176325686 | 0.000106524 | 15.01741564 | 0.442476983 | 3.480763175 |
| ENSMUSG00000023495 | *Pcbp4* | 238.0818219 | 268.5735755 | 239.5079712 | 113.049117 | 101.0651794 | 115.7068947 | 8.90E-11 | -1.176862137 | 1.30E-11 | 45.80717763 | 0.442312483 | 10.05072623 |
| ENSMUSG00000068874 | *Selenbp1* | 209.4517294 | 164.0217193 | 163.873875 | 100.9759104 | 61.06021254 | 76.2196211 | 1.16E-05 | -1.17937752 | 3.01E-06 | 21.81302312 | 0.441541969 | 4.934093735 |
| ENSMUSG00000015659 | *Serac1* | 153.6983913 | 166.8992934 | 161.3527385 | 72.43924005 | 69.48231082 | 70.70976898 | 3.04E-08 | -1.17952565 | 5.67E-09 | 33.94566539 | 0.441496636 | 7.517834683 |
| ENSMUSG00000054408 | *Spcs3* | 8398.411609 | 8476.373881 | 11109.38816 | 4510.989039 | 3835.213005 | 3993.724484 | 3.81E-19 | -1.181009022 | 2.96E-20 | 85.01700853 | 0.441042925 | 18.41854713 |
| ENSMUSG00000073471 | *Rsph3a* | 190.6161422 | 248.4305574 | 307.5786577 | 172.3175862 | 70.53507311 | 86.32101667 | 0.001505818 | -1.1814398 | 0.000549439 | 11.93991633 | 0.440911252 | 2.822227642 |
| ENSMUSG00000042349 | *Ikbke* | 653.9715867 | 576.473996 | 465.1496913 | 200.8542565 | 256.8739976 | 290.1855454 | 3.00E-09 | -1.181593209 | 5.12E-10 | 38.63278445 | 0.44086437 | 8.522591099 |
| ENSMUSG00000019820 | *Utrn* | 827.2589887 | 873.8233118 | 721.04505 | 373.1718427 | 370.5723244 | 325.9995842 | 2.21E-19 | -1.182534121 | 1.69E-20 | 86.11870064 | 0.440576936 | 18.65470695 |
| ENSMUSG00000003380 | *Rabac1* | 2914.242048 | 3572.028554 | 4332.573141 | 1629.882901 | 1493.869683 | 1637.344391 | 1.45E-15 | -1.183176951 | 1.40E-16 | 68.29898961 | 0.44038067 | 14.83958274 |
| ENSMUSG00000084817 | *Gm5526* | 297.6022773 | 309.8188032 | 415.9875288 | 188.7810498 | 124.2259497 | 136.8279945 | 5.17E-07 | -1.184032362 | 1.12E-07 | 28.15607741 | 0.440119634 | 6.286589204 |
| ENSMUSG00000023951 | *Vegfa* | 338.2871456 | 379.8397711 | 344.1351375 | 187.6834856 | 131.5952856 | 148.7660075 | 1.61E-10 | -1.184265505 | 2.40E-11 | 44.61421357 | 0.440048515 | 9.794429876 |
| ENSMUSG00000022125 | *Cln5* | 1943.079173 | 2260.813991 | 2797.200989 | 1017.442053 | 1033.812564 | 1025.750805 | 8.83E-17 | -1.184553795 | 7.90E-18 | 73.97840279 | 0.439960591 | 16.05392798 |
| ENSMUSG00000058240 | *Cryzl1* | 1280.066504 | 1168.295053 | 1503.857945 | 653.0507247 | 525.3283803 | 559.249991 | 5.69E-17 | -1.18478886 | 5.01E-18 | 74.87656866 | 0.439888911 | 16.24513722 |
| ENSMUSG00000028927 | *Padi2* | 467.8759854 | 451.7791217 | 296.2335433 | 184.3907929 | 158.9671051 | 192.8448245 | 9.44E-08 | -1.184829559 | 1.87E-08 | 31.62496461 | 0.439876502 | 7.024912385 |
| ENSMUSG00000050002 | *Idnk* | 687.1222201 | 699.2504877 | 813.0665337 | 345.7327366 | 305.3010627 | 314.97988 | 1.19E-19 | -1.18494731 | 8.91E-21 | 87.3896738 | 0.439840601 | 18.92316118 |
| ENSMUSG00000021236 | *Entpd5* | 1871.503942 | 2071.853297 | 2391.298007 | 901.1002436 | 924.3252864 | 954.1227268 | 9.36E-25 | -1.186448639 | 5.19E-26 | 111.2584186 | 0.439383123 | 24.02893378 |
| ENSMUSG00000051650 | *B3gnt2* | 1698.21654 | 2148.588604 | 2339.614708 | 1003.173718 | 934.8529092 | 778.7257674 | 1.79E-14 | -1.186840873 | 1.90E-15 | 63.16405223 | 0.439263682 | 13.74818982 |
| ENSMUSG00000003541 | *Ier3* | 1941.572326 | 2187.915449 | 2067.331962 | 1005.368847 | 733.7753128 | 980.7536788 | 1.09E-17 | -1.187482575 | 9.25E-19 | 78.21358387 | 0.439068343 | 16.96209823 |
| ENSMUSG00000063954 | *H2ac19* | 112.2600996 | 93.04156009 | 105.8877346 | 24.14641335 | 56.8491634 | 55.09852128 | 0.001743974 | -1.188205644 | 0.000647348 | 11.63464185 | 0.43884834 | 2.758459971 |
| ENSMUSG00000031497 | *Tnfsf13b* | 208.6983059 | 260.9000448 | 244.5502442 | 100.9759104 | 101.0651794 | 110.1970426 | 1.11E-09 | -1.188859356 | 1.81E-10 | 40.66262045 | 0.438649535 | 8.955218098 |
| ENSMUSG00000020102 | *Slc16a7* | 244.1092098 | 231.1651132 | 361.7830933 | 138.2930946 | 133.7008102 | 94.58579486 | 7.12E-06 | -1.189561047 | 1.78E-06 | 22.8180467 | 0.438436238 | 5.147549595 |
| ENSMUSG00000073643 | *Wdfy1* | 1362.943088 | 1434.950246 | 1501.336809 | 651.9531604 | 599.0217403 | 630.8780687 | 7.17E-37 | -1.190561037 | 2.30E-38 | 167.7454266 | 0.438132446 | 36.14419109 |
| ENSMUSG00000031402 | *Mpp1* | 3067.187015 | 2860.308579 | 3136.293854 | 1343.418634 | 1282.264463 | 1340.730684 | 5.89E-53 | -1.191852452 | 1.03E-54 | 242.6676159 | 0.437740431 | 52.22996171 |
| ENSMUSG00000026004 | *Kansl1l* | 174.794249 | 208.144521 | 205.4726279 | 75.73193278 | 86.32650739 | 93.66748618 | 1.83E-08 | -1.195536885 | 3.36E-09 | 34.96443498 | 0.436623932 | 7.736608208 |
| ENSMUSG00000090272 | *Mndal* | 1156.505052 | 1010.028482 | 923.9965414 | 530.1235294 | 391.6275701 | 428.8501573 | 1.42E-14 | -1.196891319 | 1.50E-15 | 63.62897685 | 0.436214212 | 13.84760364 |
| ENSMUSG00000044811 | *Cd300c2* | 783.5604264 | 777.9041777 | 776.5100539 | 332.5619657 | 354.7808901 | 332.4277451 | 6.94E-30 | -1.197567611 | 3.02E-31 | 135.1760328 | 0.436009776 | 29.15833789 |
| ENSMUSG00000037722 | *Gnpnat1* | 814.4507894 | 716.5159318 | 936.602224 | 392.9279991 | 303.1955381 | 377.4248708 | 7.97E-14 | -1.198898301 | 8.97E-15 | 60.11007772 | 0.435607802 | 13.09827751 |
| ENSMUSG00000027322 | *Siglec1* | 2206.02397 | 1937.566509 | 1119.384623 | 849.5147242 | 659.0291905 | 784.2356196 | 3.88E-07 | -1.200078948 | 8.27E-08 | 28.74155213 | 0.435251463 | 6.411252793 |
| ENSMUSG00000110126 | *Gm9347* | 5565.539298 | 5929.720871 | 5970.051323 | 2541.958787 | 2405.561822 | 2648.402256 | 6.26E-59 | -1.200518152 | 9.15E-61 | 270.4294 | 0.435118978 | 58.20353677 |
| ENSMUSG00000030588 | *Yif1b* | 3308.282531 | 3739.887039 | 4270.805296 | 1566.224175 | 1610.726296 | 1742.94989 | 2.52E-25 | -1.200752916 | 1.36E-26 | 113.9088544 | 0.435048179 | 24.59862236 |
| ENSMUSG00000040435 | *Ppp1r15a* | 1141.436583 | 1077.171876 | 979.4615452 | 451.0989039 | 451.6350203 | 488.540222 | 2.48E-27 | -1.202077541 | 1.21E-28 | 123.2806296 | 0.434648918 | 26.60536045 |

| | | | | | | | | | | | | |
|---|---|---|---|---|---|---|---|---|---|---|---|---|
| ENSMUSG00000079491 | H2-T10 | 289.314619 | 301.1860811 | 335.3111596 | 128.4150165 | 125.2787119 | 146.9293901 | 2.06E-12 | -1.202411111 | 2.61E-13 | 53.48450771 | 0.434548433 | 11.6854933 |
| ENSMUSG00000018567 | Gabarap | 8967.999765 | 10130.97894 | 13656.99663 | 4659.160212 | 4630.04853 | 4915.706407 | 1.58E-15 | -1.204995686 | 1.54E-16 | 68.12058353 | 0.433770641 | 14.80194741 |
| ENSMUSG00000033540 | Idua | 388.7665193 | 416.289042 | 320.1843404 | 151.4638656 | 191.6027359 | 146.0110814 | 2.63E-10 | -1.206332163 | 4.02E-11 | 43.60538943 | 0.433368992 | 9.579875323 |
| ENSMUSG00000019810 | Fuca2 | 2646.023286 | 2739.45047 | 3320.336821 | 1210.61336 | 1332.797053 | 1225.942098 | 9.78E-26 | -1.206549633 | 5.17E-27 | 115.8344796 | 0.433303672 | 25.00975968 |
| ENSMUSG00000027398 | Il1b | 52.73964409 | 60.42905449 | 60.50727692 | 26.34154184 | 20.00248342 | 28.46756933 | 0.001236938 | -1.208012851 | 0.000444455 | 12.33537217 | 0.432864427 | 2.907652204 |
| ENSMUSG00000021376 | Tpmt | 87.39712449 | 52.75552376 | 97.06375673 | 43.90256973 | 26.31905713 | 32.14080408 | 0.002889773 | -1.212235237 | 0.001117675 | 10.62171241 | 0.4315994 | 2.539136294 |
| ENSMUSG00000027950 | Chmb2 | 577.122391 | 546.7390644 | 471.4525327 | 184.3907929 | 247.399137 | 255.2898153 | 4.63E-12 | -1.215385068 | 6.07E-13 | 51.82477267 | 0.430658118 | 11.33400401 |
| ENSMUSG00000101431 | Gm7901 | 4702.115982 | 4951.345703 | 5572.972318 | 2202.811436 | 2195.009365 | 2150.678947 | 7.28E-43 | -1.216623875 | 1.86E-44 | 195.6509577 | 0.430288481 | 42.13808242 |
| ENSMUSG00000061778 | Mospd2 | 531.9169818 | 599.4945882 | 739.953574 | 289.7569602 | 231.6077027 | 281.0024585 | 6.22E-12 | -1.218055336 | 8.26E-13 | 51.21843397 | 0.429861755 | 11.20597702 |
| ENSMUSG00000028042 | Zbtb7b | 3977.322588 | 4273.197425 | 3516.985471 | 1659.517136 | 1634.939829 | 1763.152681 | 4.04E-35 | -1.218637371 | 1.40E-36 | 159.5818392 | 0.429688368 | 34.39389922 |
| ENSMUSG00000026080 | Chst10 | 1661.298789 | 1729.421988 | 1679.076935 | 724.3924005 | 741.1446488 | 711.6892332 | 1.74E-51 | -1.219662263 | 3.25E-53 | 235.7940735 | 0.429383226 | 50.75923132 |
| ENSMUSG00000027248 | Pdia3 | 34407.3438 | 38537.43051 | 44682.10287 | 16821.26959 | 17128.44238 | 16539.65778 | 3.11E-31 | -1.220062762 | 1.27E-32 | 141.4729783 | 0.429264043 | 30.50787875 |
| ENSMUSG00000031812 | Map1lc3b | 2822.324382 | 3100.106415 | 3581.274453 | 1352.199148 | 1337.008102 | 1384.809502 | 2.89E-29 | -1.220698333 | 1.28E-30 | 132.3037295 | 0.429074975 | 28.53884365 |
| ENSMUSG00000019818 | Cd164 | 1745.682219 | 1619.114984 | 1531.590447 | 678.2947023 | 674.8206248 | 743.8300373 | 1.21E-34 | -1.223992166 | 4.31E-36 | 157.3430877 | 0.428096467 | 33.91830298 |
| ENSMUSG00000042042 | Csgalnact2 | 873.9712449 | 1010.028482 | 1236.617472 | 485.1233955 | 391.6275701 | 455.4811093 | 9.53E-14 | -1.22558786 | 1.08E-14 | 59.74727208 | 0.427623232 | 13.02112626 |
| ENSMUSG00000029752 | Asns | 13120.1166 | 12303.54733 | 12415.33688 | 5746.846377 | 4899.555675 | 5522.70845 | 4.68E-48 | -1.226731341 | 9.72E-50 | 219.8564011 | 0.427284432 | 47.33021724 |
| ENSMUSG00000039908 | Slc26a11 | 94.93135936 | 85.36802936 | 68.07068654 | 36.21962002 | 42.11049141 | 28.46756933 | 0.000272553 | -1.230148997 | 8.68E-05 | 15.40480833 | 0.426273419 | 3.564549485 |
| ENSMUSG00000027955 | Gask1b | 313.4241706 | 321.3290993 | 376.9099125 | 130.6101449 | 144.2284331 | 154.2758596 | 7.21E-13 | -1.230987459 | 8.76E-14 | 55.62753211 | 0.426025751 | 12.14225334 |
| ENSMUSG00000043298 | Smco3 | 57.26018501 | 43.16361035 | 94.54262019 | 19.75615638 | 28.4245817 | 33.97742146 | 0.008945865 | -1.231121992 | 0.003838708 | 8.358578086 | 0.425986025 | 2.048377665 |
| ENSMUSG00000051736 | Fam229b | 90.41081843 | 80.57207265 | 102.1060298 | 36.21962002 | 40.00496684 | 39.48727358 | 2.97E-05 | -1.233631001 | 8.18E-06 | 19.89484914 | 0.425245831 | 4.526654891 |
| ENSMUSG00000026187 | Xrcc5 | 1093.970903 | 1250.785509 | 1512.681923 | 537.8064792 | 582.1775437 | 517.0077913 | 3.07E-17 | -1.23467464 | 2.66E-18 | 76.1285367 | 0.424938322 | 16.51300411 |
| ENSMUSG00000016496 | Cd274 | 131.0956867 | 172.6544414 | 171.4372846 | 77.92706126 | 44.21601598 | 78.97454717 | 4.74E-05 | -1.234948724 | 1.34E-05 | 18.94855587 | 0.4248576 | 4.324068754 |
| ENSMUSG00000028173 | Wls | 2121.640539 | 2341.386064 | 2666.101889 | 983.4175619 | 1015.915605 | 1023.914187 | 1.47E-29 | -1.235908606 | 6.46E-31 | 133.6672865 | 0.424575019 | 28.83240622 |
| ENSMUSG00000038712 | Mindy1 | 2545.817962 | 2974.452349 | 2949.72975 | 1175.491304 | 1176.988235 | 1218.595629 | 1.87E-39 | -1.244772026 | 5.40E-41 | 179.7835177 | 0.421974573 | 38.72910594 |
| ENSMUSG00000027253 | Lrp4 | 235.8215514 | 269.5327669 | 189.0852404 | 83.41488248 | 104.2234662 | 105.6054991 | 4.69E-08 | -1.245311259 | 8.98E-09 | 33.0511728 | 0.421816882 | 7.328822868 |
| ENSMUSG00000002732 | Fkbp7 | 52.73964409 | 70.0209679 | 81.9369375 | 25.24397759 | 25.26629484 | 34.89573014 | 0.001145045 | -1.245322365 | 0.000407586 | 12.49709002 | 0.421813635 | 2.94117741 |
| ENSMUSG00000026896 | Ifih1 | 821.2316008 | 870.9457377 | 781.5523269 | 396.2206918 | 287.4041039 | 359.9770057 | 2.48E-17 | -1.245867206 | 2.13E-18 | 76.56184059 | 0.421654365 | 16.60484191 |
| ENSMUSG00000024150 | Mcfd2 | 3162.871798 | 3870.337061 | 4632.588389 | 1677.078164 | 1592.829337 | 1637.344391 | 1.30E-18 | -1.248409104 | 1.04E-19 | 82.52954078 | 0.420912102 | 17.88571779 |
| ENSMUSG00000091498 | Mpc1-ps | 433.9719285 | 406.6971286 | 518.0935587 | 188.7810498 | 155.8088182 | 224.9856286 | 7.44E-11 | -1.249380993 | 1.08E-11 | 46.17887414 | 0.420628645 | 10.12821986 |
| ENSMUSG00000003814 | Calr | 83688.02066 | 95055.86191 | 105523.4304 | 38047.06449 | 41456.72603 | 40020.81093 | 7.28E-36 | -1.249891622 | 2.43E-37 | 163.0563718 | 0.420479794 | 35.1381037 |
| ENSMUSG00000057554 | Lgals8 | 2163.078831 | 2206.140085 | 2529.960516 | 1018.539618 | 911.692139 | 965.1424311 | 1.05E-34 | -1.251523153 | 3.74E-36 | 157.6210808 | 0.420004546 | 33.978211 |
| ENSMUSG00000003746 | Man1a | 700.6838429 | 681.9850435 | 605.0727692 | 288.659396 | 236.8715142 | 309.4700279 | 3.93E-17 | -1.251788423 | 3.43E-18 | 75.62753452 | 0.419927326 | 16.40555898 |
| ENSMUSG00000032369 | Plscr1 | 602.7387896 | 583.1883354 | 681.9674337 | 308.4155523 | 253.7157107 | 222.2307025 | 1.38E-14 | -1.252977517 | 1.46E-15 | 63.68402444 | 0.419581357 | 13.85945746 |
| ENSMUSG00000020407 | Upp1 | 63.2875729 | 53.7147151 | 76.89466442 | 34.02449154 | 14.73867199 | 32.14080408 | 0.004399269 | -1.254604154 | 0.00177124 | 9.772680492 | 0.419108546 | 2.356619435 |
| ENSMUSG00000036661 | Dennd3 | 472.3965263 | 493.9835407 | 345.3957058 | 201.9518207 | 164.2309165 | 184.5800463 | 1.12E-10 | -1.256124707 | 1.66E-11 | 45.3383592 | 0.418667052 | 9.951025391 |
| ENSMUSG00000066861 | Oas1g | 7862.72751 | 7076.913715 | 6348.221804 | 3302.570808 | 2479.255182 | 3129.596009 | 4.70E-21 | -1.256586556 | 3.21E-22 | 93.96440331 | 0.418533046 | 20.32746141 |
| ENSMUSG00000056131 | Pgm3 | 667.5332094 | 694.454531 | 700.8759577 | 277.6837535 | 276.876481 | 305.7967931 | 1.85E-27 | -1.25890179 | 8.92E-29 | 123.885529 | 0.417861924 | 26.73222654 |
| ENSMUSG00000038280 | Ostm1 | 1099.244867 | 1080.04945 | 1287.040203 | 459.8794179 | 489.5344626 | 495.8866915 | 2.16E-27 | -1.259271633 | 1.04E-28 | 123.5766196 | 0.417754816 | 26.66584955 |
| ENSMUSG00000056220 | Pla2g4a | 6755.948407 | 6437.13309 | 8971.464372 | 3142.326428 | 3074.065873 | 3039.601757 | 3.89E-21 | -1.259372898 | 2.64E-22 | 94.35048843 | 0.417725495 | 20.41042171 |
| ENSMUSG00000073002 | Vamp5 | 88.90397146 | 121.8173003 | 105.8877346 | 37.31718427 | 49.4798274 | 44.99712571 | 2.27E-05 | -1.259418274 | 6.13E-06 | 20.44792912 | 0.417712356 | 4.644467399 |
| ENSMUSG00000035790 | Cep19 | 324.7255229 | 355.8599876 | 444.980599 | 169.0248934 | 150.5450068 | 148.7660075 | 1.58E-11 | -1.261183869 | 2.17E-12 | 49.32419856 | 0.417201466 | 10.80024274 |
| ENSMUSG00000028015 | Ctso | 574.1086971 | 606.2089276 | 794.1580096 | 254.6349044 | 269.507145 | 296.6137062 | 1.59E-13 | -1.261619981 | 1.83E-14 | 58.70429888 | 0.417075369 | 12.7977329 |
| ENSMUSG00000040253 | Gbp7 | 78.35604264 | 52.75552376 | 39.07761635 | 26.34154184 | 25.26629484 | 20.20279114 | 0.004885516 | -1.262255802 | 0.001982436 | 9.565727677 | 0.416891597 | 2.311089535 |
| ENSMUSG00000062081 | Gm6055 | 873.2178214 | 1060.865623 | 1237.87804 | 491.7087809 | 396.8913815 | 429.768466 | 1.28E-14 | -1.264769435 | 1.35E-15 | 63.83339538 | 0.416165872 | 13.89182245 |
| ENSMUSG00000024049 | Myom1 | 38.42459783 | 32.6125056 | 36.55647981 | 18.65859213 | 10.52762285 | 15.6112477 | 0.009745903 | -1.266093457 | 0.004210161 | 8.190875302 | 0.415784114 | 2.011177915 |
| ENSMUSG00000023861 | Mpc1 | 1009.587473 | 987.9670814 | 1503.857945 | 511.4649373 | 448.4767335 | 493.1317655 | 8.26E-12 | -1.266649723 | 1.11E-12 | 50.63736116 | 0.415623829 | 11.08276344 |
| ENSMUSG00000075702 | Selenom | 1219.039202 | 1284.357206 | 1725.717961 | 624.5140544 | 563.2278226 | 567.5147692 | 6.02E-16 | -1.266937782 | 5.69E-17 | 70.08073269 | 0.415540851 | 15.22010193 |
| ENSMUSG00000040415 | Dtx3 | 857.3959281 | 907.3950087 | 756.3409615 | 342.4400439 | 329.5145953 | 374.6699447 | 1.65E-23 | -1.268965706 | 9.87E-25 | 105.4222654 | 0.414957157 | 22.78220533 |
| ENSMUSG00000011263 | Exoc3l2 | 58.01360849 | 83.44964668 | 54.20443558 | 29.63423457 | 33.68839313 | 18.36617376 | 0.002460038 | -1.269584823 | 0.00093965 | 10.9428612 | 0.41477912 | 2.609058145 |
| ENSMUSG00000013707 | Tnfaip8l2 | 2495.338589 | 2739.45047 | 3272.435227 | 1273.174522 | 1054.86781 | 1195.637912 | 4.49E-22 | -1.270576112 | 2.86E-23 | 98.75122392 | 0.41449422 | 21.34767527 |
| ENSMUSG00000024395 | Lims2 | 244.1092098 | 236.9202613 | 196.64865 | 92.19539643 | 86.32650739 | 102.8505731 | 4.26E-10 | -1.270592621 | 6.63E-11 | 42.62666108 | 0.414489476 | 9.370361834 |
| ENSMUSG00000060798 | Intu | 186.0956013 | 197.5934163 | 210.514901 | 88.9027037 | 77.9044091 | 78.97454717 | 2.48E-10 | -1.271946966 | 3.78E-11 | 43.72260502 | 0.414100553 | 9.60471451 |
| ENSMUSG00000034738 | Nostrin | 495.7526544 | 449.860739 | 692.0519798 | 256.8300329 | 195.813785 | 224.0673199 | 1.65E-09 | -1.272330731 | 2.73E-10 | 39.85766541 | 0.413990414 | 8.782440953 |
| ENSMUSG00000022270 | Retreg1 | 1883.558717 | 1593.216818 | 2260.198907 | 830.8561321 | 784.3079025 | 755.7680502 | 6.11E-19 | -1.274095625 | 4.78E-20 | 84.06894994 | 0.413484277 | 18.21374347 |
| ENSMUSG00000019822 | Smpd2 | 375.95832 | 409.5747027 | 404.6424144 | 128.4150165 | 182.1278753 | 179.9885028 | 2.08E-12 | -1.274829151 | 2.63E-13 | 53.46447082 | 0.413274097 | 11.68153532 |
| ENSMUSG00000030683 | Sez6l2 | 103.9724412 | 115.1029609 | 80.67636923 | 28.53667032 | 49.4798274 | 45.9154344 | 0.000168412 | -1.274831196 | 5.19E-05 | 16.37601475 | 0.413273512 | 3.773625877 |
| ENSMUSG00000063851 | Rnf183 | 599.7250956 | 599.4945882 | 767.686076 | 283.1715747 | 267.4016204 | 259.8813587 | 6.63E-17 | -1.27640736 | 5.87E-18 | 74.56481399 | 0.412822251 | 16.17858201 |
| ENSMUSG00000040747 | Cd53 | 13231.62328 | 12139.52561 | 16280.2392 | 6088.188857 | 5472.258358 | 5613.62101 | 5.43E-26 | -1.277959313 | 2.83E-27 | 117.02539 | 0.412378404 | 25.26486997 |
| ENSMUSG00000025920 | Stau2 | 1381.025252 | 1390.827445 | 1719.415119 | 649.758032 | 585.3358306 | 611.5935862 | 1.33E-24 | -1.280471926 | 7.48E-26 | 110.535207 | 0.411660826 | 23.87477226 |
| ENSMUSG00000057069 | Ero1b | 2266.297849 | 2243.548547 | 2359.7838 | 955.9784558 | 969.5940647 | 898.1058969 | 9.93E-59 | -1.282834545 | 1.46E-60 | 269.4998399 | 0.410987225 | 58.00297727 |
| ENSMUSG00000055013 | Agap1 | 1866.229977 | 1809.034869 | 1177.370763 | 628.9043113 | 743.2501733 | 624.4499078 | 9.32E-12 | -1.282900943 | 1.26E-12 | 50.39310478 | 0.41096831 | 11.03070249 |
| ENSMUSG00000024993 | Dennd10 | 628.3551881 | 650.3317293 | 845.8413087 | 304.0252954 | 273.7181941 | 292.9404715 | 1.23E-15 | -1.283523062 | 1.19E-16 | 68.63340452 | 0.410791131 | 14.91138626 |
| ENSMUSG00000021458 | Aopep | 5038.896281 | 5398.328868 | 6249.897479 | 2415.738899 | 2137.107439 | 2296.690029 | 1.85E-33 | -1.284051063 | 6.90E-35 | 151.8306946 | 0.410640816 | 32.73325063 |
| ENSMUSG00000042097 | Zfp239 | 326.9857933 | 382.7173451 | 365.5647981 | 129.5125807 | 180.0223508 | 131.3181424 | 2.32E-11 | -1.286009977 | 3.22E-12 | 48.54911749 | 0.41008362 | 10.63424532 |
| ENSMUSG00000024644 | Cndp2 | 10152.38149 | 10449.43047 | 11052.66258 | 4485.745062 | 4229.998862 | 4261.870621 | 5.77E-80 | -1.286183544 | 5.47E-82 | 367.8614776 | 0.410034287 | 79.2385396 |
| ENSMUSG00000033029 | 1700088E04Rik | 91.16424192 | 87.28641204 | 123.5356904 | 37.31718427 | 35.7939177 | 49.58866915 | 5.83E-05 | -1.286631953 | 1.67E-05 | 18.5331912 | 0.409906862 | 4.234599822 |
| ENSMUSG00000021482 | Prxl2c | 885.2725972 | 859.4354416 | 1308.469863 | 399.5133845 | 409.5245289 | 438.9515529 | 3.83E-12 | -1.287777562 | 4.98E-13 | 52.21455062 | 0.409581494 | 11.41655577 |
| ENSMUSG00000073491 | Ifi213 | 606.505907 | 527.5552376 | 723.5661865 | 280.9764463 | 250.5574239 | 228.6588633 | 3.63E-14 | -1.288360787 | 3.95E-15 | 61.7242109 | 0.40941595 | 13.44035911 |
| ENSMUSG00000043510 | Hscb | 509.3142772 | 575.5148047 | 664.3194779 | 259.0251614 | 246.3463747 | 209.3743809 | 4.07E-15 | -1.289913307 | 4.13E-16 | 66.17497669 | 0.408975604 | 14.39031339 |
| ENSMUSG00000037573 | Tob1 | 237.3283984 | 253.2265141 | 279.8461558 | 114.1466813 | 83.16822053 | 116.6252034 | 5.75E-10 | -1.29025743 | 9.10E-11 | 42.00519155 | 0.408878064 | 9.24033714 |
| ENSMUSG00000049232 | Tigd2 | 261.43795 | 330.9210127 | 408.4241192 | 154.7565583 | 122.1204251 | 130.3998337 | 1.20E-08 | -1.292937165 | 2.16E-09 | 35.82408088 | 0.408119298 | 7.922224292 |
| ENSMUSG00000026637 | Traf5 | 249.3831742 | 260.9000448 | 259.6770635 | 73.53680429 | 120.0149005 | 119.3801294 | 2.14E-08 | -1.294362496 | 3.95E-09 | 34.64867992 | 0.40771629 | 7.669416069 |
| ENSMUSG00000078853 | Igtp | 834.7932235 | 795.1696218 | 864.7498327 | 400.6109488 | 268.4543827 | 346.2023754 | 1.04E-15 | -1.297012073 | 9.95E-17 | 68.97949159 | 0.406968187 | 14.98419686 |
| ENSMUSG00000001418 | Glmp | 8540.055225 | 9158.358925 | 9188.282114 | 3593.425332 | 3709.934293 | 3633.747478 | 3.50E-89 | -1.297319649 | 2.65E-91 | 410.6508325 | 0.406881432 | 88.45541631 |
| ENSMUSG00000052459 | Atp6v1a | 11337.51663 | 13273.28978 | 15399.10198 | 5406.601462 | 5652.280709 | 5211.401804 | 1.79E-27 | -1.297869351 | 8.60E-29 | 123.9584178 | 0.40672643 | 26.74756078 |
| ENSMUSG00000019872 | Smpdl3a | 732.3276293 | 829.7005101 | 1153.419966 | 339.1473511 | 385.3109964 | 377.4248708 | 7.78E-12 | -1.297951715 | 1.04E-12 | 50.76092155 | 0.406703211 | 11.10921671 |
| ENSMUSG00000034445 | Cyb561a3 | 313.4241706 | 302.1452725 | 329.0083183 | 119.6345025 | 132.6480479 | 130.3998337 | 4.89E-16 | -1.301004333 | 4.58E-17 | 70.50816992 | 0.405843572 | 15.31040271 |
| ENSMUSG00000022453 | Naga | 149.9312739 | 189.9198855 | 107.1483029 | 38.41474851 | 57.90192569 | 84.4843993 | 0.00051723 | -1.303053638 | 0.000172931 | 14.10448729 | 0.405267492 | 3.2863159 |
| ENSMUSG00000073725 | Lmbrd1 | 1135.409195 | 1166.376671 | 1303.42759 | 553.1723786 | 452.6877826 | 450.8895658 | 6.66E-25 | -1.307260876 | 3.66E-26 | 111.9525853 | 0.404087358 | 24.1766816 |
| ENSMUSG00000028909 | Ptpru | 51.9862206 | 73.85773327 | 42.85932115 | 17.56102789 | 31.58286856 | 19.28448245 | 0.004273737 | -1.307597392 | 0.001716886 | 9.82999782 | 0.403993114 | 2.369192217 |
| ENSMUSG00000027602 | Map1lc3a | 287.807772 | 326.125056 | 421.0298019 | 132.8052734 | 130.5425234 | 151.5209335 | 2.49E-10 | -1.311697288 | 3.79E-11 | 43.71820315 | 0.402846663 | 9.603933368 |
| ENSMUSG00000034744 | Nagk | 1173.833793 | 1196.111602 | 1139.553715 | 484.0258312 | 437.9491106 | 490.3768394 | 2.08E-40 | -1.31310533 | 5.82E-42 | 184.2170907 | 0.402453685 | 39.68135948 |
| ENSMUSG00000026930 | Gpsm1 | 425.6842701 | 469.0445658 | 395.8184365 | 160.2443795 | 185.2861622 | 173.560342 | 7.51E-18 | -1.315025939 | 6.28E-19 | 78.97966 | 0.401918269 | 17.12413899 |
| ENSMUSG00000020605 | Hs1bp3 | 612.5332949 | 670.4747474 | 582.3825404 | 251.3422117 | 263.1905713 | 236.0053328 | 3.45E-24 | -1.315851579 | 1.98E-25 | 108.6031922 | 0.401688321 | 23.46250042 |
| ENSMUSG00000046027 | Stard5 | 392.5336367 | 431.6361035 | 417.2480971 | 171.2200219 | 157.9143428 | 168.9687986 | 4.53E-20 | -1.315966743 | 3.29E-21 | 89.36158086 | 0.401656257 | 19.34354288 |
| ENSMUSG00000029403 | Cdkl2 | 207.1914589 | 293.5125504 | 351.6985471 | 117.439374 | 118.9621382 | 104.6871904 | 1.10E-07 | -1.316675383 | 2.20E-08 | 31.31031797 | 0.401459015 | 6.958133188 |
| ENSMUSG00000027709 | Mccc1 | 839.3137645 | 838.3332321 | 932.8205192 | 336.9522227 | 356.8864147 | 351.7122275 | 1.80E-32 | -1.317548424 | 7.07E-34 | 147.2070505 | 0.401216148 | 31.74440771 |
| ENSMUSG00000039384 | Dusp10 | 58.01360849 | 54.67390644 | 41.59875288 | 18.65859213 | 25.26629484 | 18.36617376 | 0.00145081 | -1.319292737 | 0.000526979 | 12.01769291 | 0.400731344 | 2.838389484 |
| ENSMUSG00000038831 | Ralgps1 | 296.0954304 | 286.798211 | 206.7331962 | 96.5856534 | 120.0149005 | 101.0139557 | 4.42E-09 | -1.31964353 | 7.66E-10 | 37.84398124 | 0.400633918 | 8.354490148 |
| ENSMUSG00000026097 | Ormdl1 | 982.464227 | 980.2935506 | 1246.702018 | 459.8794179 | 393.7330947 | 429.768466 | 1.82E-20 | -1.320254811 | 1.29E-21 | 91.21861631 | 0.400464202 | 19.73950585 |
| ENSMUSG00000041084 | Ostc | 5954.305817 | 5663.065678 | 7790.311904 | 2670.373804 | 2488.730042 | 2606.160057 | 9.40E-24 | -1.321017559 | 5.56E-25 | 106.5598375 | 0.400252534 | 23.02700366 |
| ENSMUSG00000022246 | Rai14 | 1607.805721 | 1644.053959 | 1745.887053 | 651.9531604 | 765.3581813 | 581.2893995 | 8.47E-27 | -1.32275665 | 4.24E-28 | 120.7950811 | 0.399770342 | 26.07213527 |
| ENSMUSG00000038963 | Slco4a1 | 18652.50527 | 19964.60857 | 16421.42284 | 7525.998015 | 6941.914509 | 7524.62139 | 5.08E-45 | -1.323557676 | 1.18E-46 | 205.7211986 | 0.399548439 | 44.2939737 |
| ENSMUSG00000035299 | Mid1 | 314.9310175 | 335.7169694 | 225.6417202 | 122.9271952 | 112.6455645 | 115.7068947 | 8.61E-10 | -1.324187128 | 1.39E-10 | 41.18044149 | 0.399374153 | 9.065145025 |
| ENSMUSG00000089753 | Cd300ld2 | 131.0956867 | 124.6948743 | 134.8808048 | 63.6587261 | 30.53010627 | 61.5266821 | 5.34E-05 | -1.324518519 | 1.52E-05 | 18.70621594 | 0.399282426 | 4.27213934 |
| ENSMUSG00000038843 | Gcnt1 | 865.6835865 | 874.7825031 | 1019.79973 | 356.708379 | 361.0974638 | 381.0981055 | 3.73E-28 | -1.325332465 | 1.73E-29 | 127.1383775 | 0.399057221 | 27.42836688 |
| ENSMUSG00000024352 | Spata24 | 113.7669465 | 114.1437696 | 109.6694394 | 37.31718427 | 48.42706512 | 48.67036046 | 7.25E-07 | -1.325754531 | 1.60E-07 | 27.46546321 | 0.398940492 | 6.139745344 |
| ENSMUSG00000094127 | None | 109.2464056 | 86.3272207 | 142.4442144 | 58.17090489 | 35.7939177 | 40.40558227 | 0.000115918 | -1.32901515 | 3.48E-05 | 17.13698057 | 0.398039869 | 3.935850002 |
| ENSMUSG00000045664 | Cdc42ep2 | 256.917409 | 307.9004205 | 221.8600154 | 129.5125807 | 77.9044091 | 105.6054991 | 1.10E-07 | -1.332473499 | 2.20E-08 | 31.31252074 | 0.397086853 | 6.958476673 |
| ENSMUSG00000068739 | Sars | 26373.58916 | 24971.58737 | 26882.87891 | 10888.93486 | 9512.760009 | 10634.01461 | 6.74E-64 | -1.333710996 | 9.06E-66 | 293.3927195 | 0.396746391 | 63.17140892 |
| ENSMUSG00000038539 | Atf5 | 3101.844496 | 3181.637679 | 2710.221779 | 1258.906187 | 1030.654277 | 1279.204002 | 3.90E-29 | -1.333799632 | 1.74E-30 | 131.7040629 | 0.396722016 | 28.40899608 |
| ENSMUSG00000034640 | Tiparp | 647.9441988 | 601.4129709 | 738.6930058 | 256.8300329 | 252.6629484 | 276.4109151 | 2.32E-22 | -1.335396624 | 1.46E-23 | 100.0784593 | 0.396283108 | 21.63543945 |
| ENSMUSG00000096100 | ENSMUSG00000096100 | 337.5337222 | 314.6147599 | 369.3465029 | 116.3418098 | 122.1204251 | 164.3772552 | 8.49E-12 | -1.336013368 | 1.14E-12 | 50.5815753 | 0.396113735 | 11.07086213 |
| ENSMUSG00000074364 | Ehd2 | 873.2178214 | 931.3747922 | 610.1150423 | 329.269273 | 277.9292433 | 349.8756101 | 4.23E-12 | -1.336752219 | 5.51E-13 | 52.01324921 | 0.395910924 | 11.3734095 |
| ENSMUSG00000022747 | St3gal6 | 2660.338332 | 2139.955882 | 2004.303548 | 1027.320132 | 735.8808373 | 932.0833183 | 1.17E-16 | -1.337010051 | 1.06E-17 | 73.40534968 | 0.395840175 | 15.93212628 |
| ENSMUSG00000009292 | Trpm2 | 146.91758 | 185.1239288 | 173.9584212 | 64.75629035 | 61.06021254 | 73.46469504 | 9.54E-09 | -1.338074841 | 1.70E-09 | 36.2922041 | 0.395548131 | 8.020573379 |
| ENSMUSG00000073490 | Ifi207 | 1001.299814 | 999.4773774 | 635.3264077 | 403.9036415 | 304.2483004 | 336.1009798 | 2.01E-10 | -1.33870153 | 3.04E-11 | 44.15406753 | 0.395376347 | 9.696652649 |
| ENSMUSG00000073987 | Ggh | 457.3280566 | 474.7997139 | 608.854474 | 231.5860553 | 190.5499736 | 185.498355 | 1.30E-13 | -1.340745365 | 1.49E-14 | 59.11462629 | 0.394816622 | 12.88596101 |
| ENSMUSG00000030825 | Hsd17b14 | 48.97252665 | 90.16398607 | 89.50034712 | 30.73179881 | 30.53010627 | 28.46756933 | 0.000784313 | -1.341167688 | 0.000271162 | 13.25977886 | 0.394701064 | 3.105510757 |
| ENSMUSG00000034959 | Rubcnl | 134.8628042 | 130.4500224 | 139.9230779 | 64.75629035 | 51.58535197 | 44.07881702 | 2.06E-07 | -1.342037225 | 4.25E-08 | 30.03077543 | 0.394463242 | 6.687130471 |
| ENSMUSG00000021585 | Cast | 1271.025422 | 1226.805725 | 1484.949421 | 548.7821216 | 501.1148478 | 517.9261 | 9.16E-31 | -1.343469904 | 3.80E-32 | 139.2924734 | 0.394071712 | 30.03830004 |
| ENSMUSG00000025353 | Ormdl2 | 3217.871713 | 3267.005708 | 4509.052699 | 1537.687505 | 1363.327159 | 1428.888319 | 8.25E-20 | -1.343518258 | 6.11E-21 | 88.13696465 | 0.394058504 | 19.08363494 |
| ENSMUSG00000037921 | Ddx60 | 144.6573095 | 146.7562752 | 110.9300077 | 45.00013397 | 45.26877826 | 67.95484291 | 4.84E-06 | -1.344502783 | 1.18E-06 | 23.60947982 | 0.393789682 | 5.315054366 |
| ENSMUSG00000072872 | Rybp | 2355.955244 | 2295.344879 | 2435.417896 | 913.1734503 | 947.4860567 | 925.6551575 | 3.07E-70 | -1.346188106 | 3.64E-72 | 322.7537813 | 0.393329935 | 69.513451 |
| ENSMUSG00000107283 | Mpv17 | 1100.751714 | 1196.111602 | 1391.667369 | 501.5868591 | 505.3258969 | 442.6247876 | 7.01E-25 | -1.346312332 | 3.86E-26 | 111.845616 | 0.393296068 | 24.15433251 |
| ENSMUSG00000003228 | Grk5 | 397.8076011 | 477.6772879 | 409.6846875 | 166.829765 | 183.1806376 | 155.1941683 | 4.50E-17 | -1.348381696 | 3.94E-18 | 75.35339035 | 0.392732339 | 16.34662609 |
| ENSMUSG00000044339 | Alkbh2 | 681.0948322 | 695.4137223 | 918.9542683 | 302.9277311 | 269.507145 | 325.9995842 | 7.94E-16 | -1.349358662 | 7.55E-17 | 69.52364396 | 0.392466478 | 15.10027431 |
| ENSMUSG00000031712 | Il15 | 55.75333803 | 39.32684499 | 41.59875288 | 16.46346365 | 20.00248342 | 17.44786507 | 0.001988213 | -1.349973395 | 0.000746058 | 11.37084931 | 0.392299283 | 2.701537006 |
| ENSMUSG00000029098 | Acox3 | 1028.42306 | 1053.192093 | 978.2009769 | 368.7815857 | 430.5797746 | 400.382588 | 1.81E-35 | -1.351125084 | 6.17E-37 | 161.2068448 | 0.39198624 | 34.74286395 |
| ENSMUSG00000074093 | Svip | 76.09577218 | 86.3272207 | 88.23977885 | 39.51231275 | 28.4245817 | 30.3041867 | 2.26E-05 | -1.351419306 | 6.10E-06 | 20.4577993 | 0.391906306 | 4.646376126 |
| ENSMUSG00000010609 | Psen2 | 2080.202247 | 2152.425369 | 1963.965363 | 786.9535624 | 865.3705984 | 776.8891501 | 4.15E-50 | -1.352119524 | 8.04E-52 | 229.4042968 | 0.391716139 | 49.38147372 |
| ENSMUSG00000012519 | Mlkl | 1258.970647 | 1281.479632 | 1553.020108 | 538.9040434 | 537.9615277 | 524.3542609 | 1.67E-29 | -1.352177382 | 7.35E-31 | 133.410438 | 0.39170043 | 28.7775757 |
| ENSMUSG00000001751 | Naglu | 1680.887799 | 1662.278594 | 1546.717266 | 645.367775 | 634.815658 | 631.7963773 | 1.98E-54 | -1.35629929 | 3.26E-56 | 249.5458292 | 0.390582904 | 53.70323871 |
| ENSMUSG00000059714 | Flot1 | 1179.861181 | 1259.418231 | 1382.843391 | 498.2941664 | 481.1123643 | 509.6613218 | 5.36E-37 | -1.357397564 | 1.70E-38 | 168.3482143 | 0.39028568 | 36.27117977 |
| ENSMUSG00000026792 | Lrsam1 | 698.4235724 | 728.0262279 | 723.5661865 | 278.7813178 | 270.5599073 | 288.348928 | 4.35E-34 | -1.358299793 | 1.57E-35 | 154.7665068 | 0.390041681 | 33.36189531 |
| ENSMUSG00000000552 | Zfp385a | 913.9026897 | 912.1909654 | 834.4961942 | 443.4159542 | 244.2408502 | 350.7939188 | 5.46E-11 | -1.358317459 | 7.83E-12 | 46.80693573 | 0.390036905 | 10.26287525 |
| ENSMUSG00000015766 | Eps8 | 1853.421778 | 1856.994436 | 2282.889136 | 864.8806236 | 749.5667471 | 719.0357027 | 5.28E-26 | -1.360384597 | 2.75E-27 | 117.0850625 | 0.389478448 | 25.27736474 |
| ENSMUSG00000074500 | Zfp558 | 25.61639856 | 35.49007962 | 36.55647981 | 14.26833516 | 12.63314742 | 11.01970426 | 0.00856813 | -1.360511034 | 0.003656096 | 8.447181475 | 0.389444316 | 2.067113945 |
| ENSMUSG00000027900 | Dram2 | 771.5056506 | 823.945362 | 1128.208601 | 380.8547924 | 332.6728821 | 345.2840667 | 8.74E-15 | -1.360655075 | 9.10E-16 | 64.61694259 | 0.389405435 | 14.05828293 |
| ENSMUSG00000018669 | Cdk5rap3 | 2567.667244 | 2293.426497 | 2561.474723 | 937.3198637 | 996.9658841 | 953.2044181 | 1.77E-53 | -1.36202064 | 3.05E-55 | 245.0933961 | 0.389037022 | 52.75198926 |
| ENSMUSG00000005413 | Hmox1 | 11760.18721 | 14185.48074 | 15699.11723 | 5279.28401 | 5451.203113 | 5444.652211 | 3.90E-33 | -1.364047462 | 1.48E-34 | 150.3087223 | 0.388490854 | 32.40929298 |
| ENSMUSG00000026170 | Cyp27a1 | 479.1773377 | 444.1055909 | 433.6354846 | 175.6102789 | 157.9143428 | 192.8448245 | 4.95E-20 | -1.365960658 | 3.60E-21 | 89.18219168 | 0.387976007 | 19.30580233 |
| ENSMUSG00000000202 | Btbd17 | 42.94513876 | 51.79633242 | 45.38045769 | 24.14641335 | 15.79143428 | 14.69293901 | 0.001830414 | -1.365992599 | 0.000682323 | 11.53679177 | 0.387967417 | 2.737450685 |
| ENSMUSG00000031639 | Tlr3 | 310.4104766 | 324.2066733 | 365.5647981 | 119.6345025 | 122.1204251 | 144.174464 | 3.88E-15 | -1.367134805 | 3.91E-16 | 66.27892245 | 0.387660379 | 14.41115299 |
| ENSMUSG00000038418 | Egr1 | 410.6158004 | 359.6967529 | 181.5218308 | 122.9271952 | 113.6983268 | 133.1547598 | 1.28E-05 | -1.368246426 | 3.32E-06 | 21.62068076 | 0.387361795 | 4.893769617 |
| ENSMUSG00000026958 | Dpp7 | 2187.188383 | 2400.855927 | 2603.073476 | 928.5393497 | 934.8529092 | 919.2269967 | 1.37E-45 | -1.368398304 | 3.09E-47 | 208.384922 | 0.387321018 | 44.86344035 |
| ENSMUSG00000067071 | Hes6 | 529.6567113 | 632.1070938 | 675.6645923 | 223.9031056 | 228.4494159 | 256.208124 | 3.27E-19 | -1.369803101 | 2.53E-20 | 85.3271965 | 0.386944055 | 18.48552233 |
| ENSMUSG00000028573 | Fggy | 60.27387896 | 40.28603633 | 28.99307019 | 17.56102789 | 13.68590971 | 19.28448245 | 0.007139074 | -1.369982308 | 0.002992904 | 8.811789031 | 0.386895993 | 2.146358126 |
| ENSMUSG00000038217 | Tlcd2 | 153.6983913 | 161.1441453 | 185.3035356 | 55.9757764 | 67.37678625 | 68.8731516 | 2.73E-09 | -1.371849082 | 4.62E-10 | 38.83063704 | 0.386395693 | 8.563980904 |
| ENSMUSG00000032261 | Sh3bgrl2 | 2694.995813 | 3121.208624 | 3693.465029 | 1164.515662 | 1286.475513 | 1212.167468 | 5.98E-26 | -1.375162986 | 3.12E-27 | 116.8330345 | 0.385509152 | 25.22332021 |
| ENSMUSG00000078606 | Gvin2 | 1482.737422 | 1145.274461 | 763.9043712 | 515.8551943 | 318.9869724 | 472.9289743 | 1.74E-07 | -1.376511864 | 3.58E-08 | 30.36315278 | 0.38514888 | 6.758232694 |
| ENSMUSG00000105867 | Gm42517 | 45.9588327 | 59.46986315 | 37.81704808 | 18.65859213 | 22.10800799 | 14.69293901 | 0.00205952 | -1.379379409 | 0.000774652 | 11.30100376 | 0.384384107 | 2.686233992 |
| ENSMUSG00000095687 | Rnaset2a | 1208.491273 | 1362.051704 | 1571.928632 | 520.2454513 | 547.4363883 | 520.6810261 | 2.59E-27 | -1.380787721 | 1.26E-28 | 123.1934449 | 0.384009067 | 26.5868877 |
| ENSMUSG00000000982 | Ccl3 | 13383.81482 | 14318.80834 | 13649.43322 | 5807.212411 | 4462.659327 | 5601.682997 | 5.74E-35 | -1.381464305 | 2.00E-36 | 158.8698487 | 0.383829019 | 34.24091071 |
| ENSMUSG00000029442 | Wdr66 | 356.3693093 | 425.8809555 | 554.6500385 | 177.8054074 | 150.5450068 | 182.7434289 | 6.86E-11 | -1.381934981 | 9.91E-12 | 46.34715835 | 0.383703816 | 10.16387385 |
| ENSMUSG00000021665 | Hexb | 4682.526971 | 4561.914018 | 4587.207932 | 1694.639191 | 1821.278753 | 1790.701942 | 8.18E-94 | -1.381986968 | 5.61E-96 | 432.1218311 | 0.38368999 | 93.08740654 |
| ENSMUSG00000024193 | Phf1 | 397.8076011 | 532.3511943 | 410.9452558 | 177.8054074 | 152.6505314 | 183.6617376 | 6.83E-14 | -1.382327057 | 7.62E-15 | 60.43156472 | 0.383599553 | 13.16548462 |
| ENSMUSG00000044816 | D630023F18Rik | 124.3148753 | 137.1643618 | 171.4372846 | 68.04898308 | 44.21601598 | 53.2619039 | 1.09E-06 | -1.383072618 | 2.46E-07 | 26.63556273 | 0.383401366 | 5.963439388 |
| ENSMUSG00000020101 | Vsir | 3649.583371 | 3738.927848 | 3669.514232 | 1340.125941 | 1478.078248 | 1418.786923 | 5.53E-80 | -1.383702455 | 5.20E-82 | 367.9613918 | 0.383234021 | 79.25713583 |
| ENSMUSG00000060550 | H2-Q7 | 1263.491188 | 1342.867878 | 1251.744291 | 489.5136525 | 446.3712089 | 539.0471999 | 6.94E-37 | -1.385959743 | 2.22E-38 | 167.8152713 | 0.382634869 | 36.15851605 |
| ENSMUSG00000021213 | Akr1c13 | 281.0269606 | 245.5529833 | 350.4379788 | 115.2442455 | 101.0651794 | 117.5435121 | 3.89E-11 | -1.388339269 | 5.51E-12 | 47.49579071 | 0.382004286 | 10.41007451 |
| ENSMUSG00000003809 | Gcdh | 628.3551881 | 645.5357726 | 734.911301 | 255.7324687 | 255.8212353 | 253.4531979 | 1.36E-27 | -1.389888926 | 6.47E-29 | 124.5233666 | 0.38159418 | 26.86622519 |
| ENSMUSG00000031442 | Mcf2l | 313.4241706 | 373.1254317 | 286.1489971 | 124.0247595 | 109.4872777 | 136.8279945 | 4.34E-13 | -1.39286776 | 5.19E-14 | 56.65763842 | 0.380807089 | 12.36262128 |
| ENSMUSG00000072825 | Cep170b | 672.0537504 | 745.2916721 | 521.8752635 | 234.878748 | 251.6101862 | 252.5348892 | 1.98E-16 | -1.39399546 | 1.81E-17 | 72.3387269 | 0.380509543 | 15.70334604 |
| ENSMUSG00000028007 | Snx7 | 852.1219637 | 885.3336079 | 1094.173258 | 351.2205578 | 373.7306112 | 349.8756101 | 9.31E-24 | -1.394877431 | 5.50E-25 | 106.5809633 | 0.380276995 | 23.03112921 |
| ENSMUSG00000030064 | Frmd4b | 652.4647397 | 632.1070938 | 645.4109539 | 238.1714408 | 270.5599073 | 225.9039372 | 3.98E-28 | -1.39517061 | 1.85E-29 | 127.0068922 | 0.380199724 | 27.40023496 |
| ENSMUSG00000035845 | Alg12 | 975.6834156 | 1063.743197 | 876.0949471 | 349.0254293 | 373.7306112 | 385.689649 | 6.92E-30 | -1.395801131 | 3.01E-31 | 135.1848744 | 0.380033596 | 29.15958627 |
| ENSMUSG00000038936 | Sccpdh | 556.0265334 | 573.596422 | 705.9182308 | 260.1227256 | 218.9745553 | 217.6391591 | 3.12E-18 | -1.39583138 | 2.54E-19 | 80.76505938 | 0.380025628 | 17.50609895 |
| ENSMUSG00000013698 | Pea15a | 4060.199171 | 4223.319475 | 4483.841334 | 1715.492912 | 1562.299231 | 1571.226165 | 2.76E-65 | -1.39643981 | 3.65E-67 | 299.7948622 | 0.379865393 | 64.55957412 |
| ENSMUSG00000052539 | Magi3 | 51.23279711 | 66.18420254 | 55.46500385 | 20.85372062 | 16.84419656 | 27.54926064 | 0.000503067 | -1.39706255 | 0.000167643 | 14.16290921 | 0.37970146 | 3.298374456 |
| ENSMUSG00000030966 | Trim21 | 514.5882416 | 517.0041329 | 560.9528798 | 229.3909268 | 151.5977691 | 222.2307025 | 3.19E-14 | -1.398744106 | 3.46E-15 | 61.98698516 | 0.379259151 | 13.49611984 |
| ENSMUSG00000040751 | Lat2 | 12701.21314 | 14262.21605 | 18992.98211 | 6125.506041 | 5483.838744 | 5789.936278 | 3.03E-21 | -1.401018454 | 2.04E-22 | 94.86034513 | 0.378661735 | 20.51787779 |

| | | | | | | | | | | | | |
|---|---|---|---|---|---|---|---|---|---|---|---|---|
| ENSMUSG00000052142 | *Rasal3* | 1029.929907 | 1026.334735 | 897.5246077 | 357.8059433 | 364.2557507 | 394.8727358 | 1.10E-33 | -1.403831189 | 4.03E-35 | 152.8982646 | 0.377924201 | 32.95932754 |
| ENSMUSG00000034647 | *Ankrd12* | 221.5065052 | 206.2261383 | 110.9300077 | 93.29296067 | 60.00745026 | 51.42528653 | 0.000135051 | -1.405263096 | 4.10E-05 | 16.82619661 | 0.377549288 | 3.869501772 |
| ENSMUSG00000068129 | *Cst7* | 391.7802132 | 382.7173451 | 405.9029827 | 140.4882231 | 123.1731874 | 179.9885028 | 1.48E-14 | -1.406913011 | 1.56E-15 | 63.55030322 | 0.377117757 | 13.83082237 |
| ENSMUSG00000078942 | *Naip6* | 305.1365122 | 389.4316845 | 376.9099125 | 131.7077092 | 143.1756708 | 126.7265989 | 3.55E-15 | -1.413376835 | 3.58E-16 | 66.45688186 | 0.375431905 | 14.44977101 |
| ENSMUSG00000105835 | *Gm43552* | 151.4381209 | 163.062528 | 166.3950115 | 61.46359762 | 45.26877826 | 72.54638635 | 2.92E-08 | -1.416912097 | 5.45E-09 | 34.02307432 | 0.374513052 | 7.534154163 |
| ENSMUSG00000075705 | *Msrb1* | 1991.298276 | 2204.221702 | 2469.453239 | 788.0511266 | 906.4283275 | 796.1736325 | 1.66E-33 | -1.418935041 | 6.18E-35 | 152.0497727 | 0.373988278 | 32.77953233 |
| ENSMUSG00000039105 | *Atp6v1g1* | 4439.924609 | 4876.528778 | 6585.208639 | 2003.054744 | 1981.298621 | 1955.997505 | 1.12E-21 | -1.419917951 | 7.32E-23 | 96.89148486 | 0.373733567 | 20.94995482 |
| ENSMUSG00000000278 | *Scpep1* | 7311.974941 | 8499.394474 | 9031.971649 | 3187.326562 | 3069.854824 | 3014.807423 | 1.23E-47 | -1.421549539 | 2.61E-49 | 217.8917375 | 0.373311139 | 46.91018005 |
| ENSMUSG00000019791 | *Hint3* | 323.2186759 | 298.3085071 | 384.4733221 | 119.6345025 | 122.1204251 | 132.2364511 | 3.50E-15 | -1.422170983 | 3.52E-16 | 66.48708016 | 0.373150369 | 14.45612843 |
| ENSMUSG00000071669 | *Snx29* | 284.7940781 | 307.9004205 | 272.2827462 | 113.049117 | 115.8038514 | 94.58579486 | 2.54E-15 | -1.423808499 | 2.54E-16 | 67.13053841 | 0.372727069 | 14.59431955 |
| ENSMUSG00000054434 | *Tmem120b* | 447.5335512 | 476.7180965 | 583.6431087 | 183.2932286 | 176.8640639 | 198.3546766 | 4.87E-18 | -1.427482373 | 4.02E-19 | 79.85984775 | 0.371779113 | 17.31259388 |
| ENSMUSG00000039770 | *Ypel5* | 420.4103057 | 403.8195546 | 553.3894702 | 178.9029716 | 172.6530148 | 157.0307856 | 5.48E-15 | -1.435030223 | 5.61E-16 | 65.5707113 | 0.369839129 | 14.26096899 |
| ENSMUSG00000039005 | *Tlr4* | 827.2589887 | 968.7832545 | 1288.300771 | 361.098636 | 388.4692832 | 387.5262663 | 1.33E-15 | -1.436619167 | 1.29E-16 | 68.47469289 | 0.369432023 | 14.87704946 |
| ENSMUSG00000033644 | *Piwil2* | 482.1910316 | 605.2497362 | 534.4809462 | 197.5615638 | 222.1328422 | 179.0701942 | 5.41E-19 | -1.438264324 | 4.22E-20 | 84.3163247 | 0.369010987 | 18.26655605 |
| ENSMUSG00000026932 | *Nacc2* | 75.34234869 | 110.3070042 | 92.02148365 | 41.70744124 | 27.37181941 | 33.05911277 | 2.19E-05 | -1.442029113 | 5.88E-06 | 20.52585315 | 0.368049288 | 4.660491028 |
| ENSMUSG00000018752 | *Tnfsfm13* | 210.9585763 | 201.4301816 | 157.5710337 | 75.73193278 | 58.95468797 | 75.30131242 | 1.66E-09 | -1.445033417 | 2.76E-10 | 39.83897031 | 0.367283651 | 8.778644018 |
| ENSMUSG00000030615 | *Tmem126a* | 1392.326604 | 1351.5006 | 1844.211378 | 604.757898 | 517.9590443 | 558.3316823 | 1.26E-21 | -1.447000532 | 8.26E-23 | 96.6533674 | 0.366783201 | 20.89954733 |
| ENSMUSG00000046879 | *Irgm1* | 2481.776966 | 2408.529458 | 2876.61679 | 1148.052198 | 715.8783539 | 983.5086049 | 6.12E-17 | -1.447385007 | 5.41E-18 | 74.7258823 | 0.366685467 | 16.21300315 |
| ENSMUSG00000036052 | *Dnajb5* | 159.7257792 | 174.5728241 | 161.3527385 | 65.85385459 | 49.4798274 | 66.11822554 | 5.63E-10 | -1.44795101 | 8.89E-11 | 42.05190538 | 0.366541636 | 9.249580981 |
| ENSMUSG00000002058 | *Unc119* | 2459.174261 | 2881.410789 | 2916.954975 | 1020.734746 | 965.3830155 | 1035.8522 | 2.13E-46 | -1.448948043 | 4.71E-48 | 212.1311824 | 0.36628841 | 45.67142358 |
| ENSMUSG00000026121 | *Sema4c* | 647.9441988 | 667.5971734 | 569.7768577 | 250.2446474 | 204.2358833 | 236.9236415 | 4.51E-24 | -1.449277667 | 2.61E-25 | 108.0572135 | 0.366204731 | 23.34545112 |
| ENSMUSG00000035678 | *Tnfsf9* | 1528.696255 | 1644.053959 | 1432.005554 | 636.587261 | 517.9590443 | 533.5373477 | 8.27E-34 | -1.449472663 | 3.02E-35 | 153.4702192 | 0.366155237 | 33.08269986 |
| ENSMUSG00000004460 | *Dnajb11* | 11552.24233 | 12621.03967 | 15062.53025 | 4721.721374 | 4946.929978 | 4693.475704 | 1.60E-39 | -1.449607039 | 4.58E-41 | 180.1108705 | 0.366121134 | 38.7964443 |
| ENSMUSG00000023206 | *Il15ra* | 152.1915444 | 134.2867878 | 121.0145538 | 59.26846913 | 48.42706512 | 42.24219965 | 5.41E-08 | -1.452871641 | 1.04E-08 | 32.75971073 | 0.365293594 | 7.266969352 |
| ENSMUSG00000049357 | *Brd8dc* | 149.1778504 | 152.5114232 | 176.4795577 | 63.6587261 | 53.69087654 | 56.93513866 | 4.27E-10 | -1.452946896 | 6.64E-11 | 42.62258268 | 0.36527454 | 9.36964802 |
| ENSMUSG00000022295 | *Atp6v1c1* | 4649.376338 | 4630.975795 | 6404.947376 | 1886.712934 | 1960.243375 | 1879.777884 | 2.02E-25 | -1.4529998 | 1.09E-26 | 114.3580227 | 0.365261145 | 24.693683 |
| ENSMUSG00000063889 | *Crem* | 246.3694802 | 235.0018786 | 399.6001413 | 106.4637316 | 96.85413024 | 116.6252034 | 4.62E-08 | -1.454552568 | 8.83E-09 | 33.08252711 | 0.364868227 | 7.335358967 |
| ENSMUSG00000034930 | *Rtkn* | 66.30126685 | 71.93935058 | 100.8454615 | 23.04884911 | 38.95220455 | 24.79433458 | 0.000173256 | -1.455099654 | 5.35E-05 | 16.31933099 | 0.364729892 | 3.761312379 |
| ENSMUSG00000042106 | *Inka1* | 558.2868038 | 508.3714108 | 567.2557212 | 216.2201559 | 174.7585393 | 203.8645287 | 4.87E-24 | -1.45727788 | 2.82E-25 | 107.9013515 | 0.364179626 | 23.31284103 |
| ENSMUSG00000017417 | *Plxdc1* | 533.4238288 | 582.2291441 | 670.6223192 | 197.5615638 | 253.7157107 | 198.3546766 | 3.83E-18 | -1.457304473 | 3.14E-19 | 80.34726996 | 0.364172914 | 17.41682877 |
| ENSMUSG00000021266 | *Wars1* | 13284.36292 | 13829.62076 | 13656.99663 | 4905.014603 | 4980.618371 | 4941.41905 | 1.69E-157 | -1.459186512 | 4.17E-160 | 726.9303652 | 0.363698149 | 156.7731082 |
| ENSMUSG00000016495 | *Plgrkt* | 775.2727681 | 784.618517 | 1007.194047 | 309.5131166 | 278.9820056 | 341.6108319 | 2.41E-20 | -1.460810446 | 1.71E-21 | 90.65523485 | 0.363288991 | 19.61837687 |
| ENSMUSG00000084333 | *Gm11599* | 48.97252665 | 42.20441901 | 54.20443558 | 25.24397759 | 6.316573711 | 21.12109982 | 0.008763292 | -1.461269441 | 0.003752604 | 8.399809194 | 0.363173429 | 2.057332694 |
| ENSMUSG00000027624 | *Epb41l1* | 500.2731953 | 543.8614904 | 564.7345846 | 180.0005359 | 178.9695885 | 222.2307025 | 3.53E-22 | -1.463607787 | 2.25E-23 | 99.22971051 | 0.362585268 | 21.45167074 |
| ENSMUSG00000053581 | *Zfand2a* | 1653.764554 | 1835.892227 | 1641.259887 | 625.6116186 | 645.3432808 | 586.7992516 | 1.04E-50 | -1.466722526 | 1.98E-52 | 232.1968501 | 0.3618033 | 49.9827047 |
| ENSMUSG00000026956 | *Uap1l1* | 7646.494969 | 8436.087845 | 9693.76999 | 2948.057557 | 3220.39983 | 3136.942478 | 8.63E-44 | -1.469367253 | 2.09E-45 | 199.9976341 | 0.361140656 | 43.06409395 |
| ENSMUSG00000025366 | *Esyt1* | 9233.204833 | 8956.928743 | 7409.620287 | 3021.594361 | 3043.535766 | 3176.429752 | 1.20E-46 | -1.470270393 | 2.63E-48 | 213.2888208 | 0.360914649 | 45.92262166 |
| ENSMUSG00000031337 | *Mtm1* | 169.5202846 | 165.940102 | 250.8530856 | 61.46359762 | 76.85164682 | 71.62807766 | 3.12E-08 | -1.47374632 | 5.83E-09 | 33.89019581 | 0.360046134 | 7.506250188 |
| ENSMUSG00000029446 | *Psph* | 4784.992566 | 4465.035693 | 4809.067947 | 1755.005225 | 1520.18874 | 1771.417459 | 8.41E-60 | -1.477788984 | 1.21E-61 | 274.4580143 | 0.359038639 | 59.07536679 |
| ENSMUSG00000034071 | *Zfp551* | 36.91775086 | 43.16361035 | 25.21136538 | 13.17077092 | 15.79143428 | 9.18308688 | 0.005707869 | -1.481199989 | 0.002350926 | 9.253135094 | 0.358190756 | 2.243526037 |
| ENSMUSG00000002279 | *Lmf1* | 1650.75086 | 1789.851043 | 1992.958434 | 643.1726465 | 605.338314 | 694.2413681 | 4.35E-39 | -1.481396123 | 1.28E-40 | 178.0714117 | 0.358142064 | 38.36140293 |
| ENSMUSG00000020108 | *Ddit4* | 190.6161422 | 239.7978353 | 204.2120596 | 91.09783218 | 70.53507311 | 66.11822554 | 1.22E-10 | -1.481907296 | 1.82E-11 | 45.15600691 | 0.35801519 | 9.912194959 |
| ENSMUSG00000000078 | *Klf6* | 1669.586447 | 1739.013901 | 2106.409578 | 664.0263671 | 671.662338 | 627.2048339 | 5.78E-35 | -1.489060315 | 2.01E-36 | 158.8543741 | 0.35624451 | 34.2383853 |
| ENSMUSG00000022769 | *Sdf2l1* | 11191.35248 | 11886.2991 | 12303.14631 | 4146.597711 | 4109.983961 | 4339.92686 | 9.26E-103 | -1.489465305 | 4.64E-105 | 473.8583004 | 0.356144519 | 102.0336221 |
| ENSMUSG00000030523 | *Trpm1* | 119.0409109 | 133.3275964 | 136.1413731 | 54.87821216 | 44.21601598 | 39.48727358 | 1.86E-08 | -1.489657523 | 3.40E-09 | 34.94067265 | 0.356097072 | 7.731633535 |
| ENSMUSG00000046718 | *Bst2* | 12283.81653 | 10920.39342 | 9715.199651 | 4508.793911 | 3253.035461 | 3962.501989 | 3.25E-26 | -1.48968016 | 1.67E-27 | 118.0768213 | 0.356091484 | 25.48818393 |
| ENSMUSG00000027646 | *Src* | 39.17802132 | 64.26581986 | 39.07761635 | 19.75615638 | 12.63314742 | 18.36617376 | 0.001866479 | -1.489860468 | 0.000696663 | 11.49812774 | 0.356046983 | 2.728976976 |
| ENSMUSG00000010461 | *Eya4* | 109.9998291 | 119.8989176 | 151.2681923 | 42.80500548 | 52.63811426 | 39.48727358 | 1.72E-07 | -1.493749346 | 3.51E-08 | 30.40117525 | 0.355088527 | 6.765728174 |
| ENSMUSG00000019935 | *Slc17a8* | 23.3561281 | 30.69412292 | 44.11988942 | 9.878078189 | 11.58038514 | 12.85632163 | 0.007333224 | -1.496292507 | 0.003078827 | 8.760156275 | 0.354463134 | 2.134705071 |
| ENSMUSG00000052477 | *C130026I21Rik* | 2882.598261 | 2778.777315 | 3336.724209 | 1208.418232 | 953.8026304 | 1026.669113 | 2.21E-33 | -1.496351209 | 8.28E-35 | 151.4672178 | 0.354448712 | 32.65540259 |
| ENSMUSG00000023045 | *Soat2* | 266.7119144 | 295.4309331 | 197.9092183 | 86.70757521 | 89.48479424 | 92.74917749 | 2.83E-11 | -1.502696346 | 3.96E-12 | 48.14403694 | 0.352893229 | 10.54880685 |
| ENSMUSG00000039963 | *Ccdc40* | 31.64378645 | 35.49007962 | 36.55647981 | 8.780513945 | 13.68590971 | 13.77463032 | 0.002682584 | -1.505887043 | 0.001030361 | 10.77219744 | 0.352113625 | 2.571446695 |
| ENSMUSG00000030824 | *Nucb1* | 9346.971779 | 11289.68209 | 11930.0181 | 3770.133175 | 3780.469366 | 3907.403467 | 3.50E-46 | -1.506626425 | 7.77E-48 | 211.1354524 | 0.351933213 | 45.45553797 |
| ENSMUSG00000004567 | *Mcoln1* | 1820.271144 | 1955.791145 | 2009.345821 | 664.0263671 | 695.8758705 | 672.2019596 | 9.32E-66 | -1.508215769 | 1.21E-67 | 301.9993687 | 0.351545719 | 65.03076162 |
| ENSMUSG00000031570 | *Plpp5* | 2016.914675 | 2247.385312 | 2393.819143 | 766.0998417 | 764.305419 | 805.3567194 | 1.73E-51 | -1.509308249 | 3.22E-53 | 235.8147558 | 0.351279612 | 50.7621417 |
| ENSMUSG00000031292 | *Cdkl5* | 62.53414942 | 64.26581986 | 84.45807404 | 29.63423457 | 25.26629484 | 19.28448245 | 5.27E-05 | -1.509746981 | 1.50E-05 | 18.7366512 | 0.351172802 | 4.278548396 |
| ENSMUSG00000056832 | *Ttc26* | 223.0133521 | 195.6750336 | 223.1205837 | 66.95141883 | 80.00993367 | 78.05623848 | 1.46E-13 | -1.509787746 | 1.67E-14 | 58.88175472 | 0.351162879 | 12.8356042 |
| ENSMUSG00000002083 | *Bbc3* | 816.7110599 | 786.5368997 | 627.7629981 | 273.2934965 | 248.4518993 | 262.6362848 | 4.09E-24 | -1.511377755 | 2.36E-25 | 108.2570324 | 0.350776072 | 23.38871984 |
| ENSMUSG00000023074 | *Mospd1* | 186.0956013 | 174.5728241 | 272.2827462 | 74.63436854 | 74.74612225 | 71.62807766 | 4.28E-09 | -1.511976565 | 7.41E-10 | 37.90922509 | 0.350630508 | 8.368325896 |
| ENSMUSG00000083240 | *Gm13453* | 1383.285522 | 1501.134449 | 2039.59946 | 609.148155 | 547.4363883 | 567.5147692 | 2.01E-20 | -1.512366976 | 1.42E-21 | 91.01638104 | 0.350535636 | 19.69596239 |
| ENSMUSG00000022350 | *Washc5* | 3969.034929 | 3930.766116 | 4284.671547 | 1380.735818 | 1440.178806 | 1446.336184 | 2.29E-89 | -1.512690153 | 1.68E-91 | 411.5536908 | 0.350457122 | 88.63993519 |
| ENSMUSG00000052373 | *Mpp3* | 54.99991455 | 68.10258522 | 52.94386731 | 14.26833516 | 24.21353256 | 22.9577172 | 0.000188114 | -1.514254997 | 5.85E-05 | 16.14960346 | 0.350077198 | 3.725577729 |
| ENSMUSG00000020964 | *Sel1l* | 10986.42129 | 11504.54095 | 10877.4436 | 3707.572013 | 4142.619592 | 3794.451499 | 2.71E-95 | -1.518900214 | 1.69E-97 | 439.1111687 | 0.348951826 | 94.56763162 |
| ENSMUSG00000050103 | *Agmo* | 437.7390459 | 393.2684499 | 456.3257135 | 147.0736086 | 134.7535725 | 166.2138725 | 2.40E-21 | -1.518968343 | 1.60E-22 | 95.34828337 | 0.348935348 | 20.61957881 |
| ENSMUSG00000025150 | *Cbr2* | 81.36973659 | 75.77611595 | 95.80318846 | 29.63423457 | 22.10800799 | 35.81403883 | 8.24E-06 | -1.52037943 | 2.08E-06 | 22.52135412 | 0.348594224 | 5.084286095 |
| ENSMUSG00000029304 | *Spp1* | 218773.0848 | 207205.4727 | 257766.042 | 90993.56358 | 66036.61986 | 81245.52455 | 1.18E-28 | -1.520818992 | 5.37E-30 | 129.463134 | 0.34848803 | 27.92928998 |
| ENSMUSG00000020638 | *Cmpk2* | 1551.29896 | 1481.950622 | 993.3277962 | 575.1236634 | 345.3060295 | 483.0303699 | 2.62E-11 | -1.521953301 | 3.66E-12 | 48.29953121 | 0.348214141 | 10.58175446 |
| ENSMUSG00000024330 | *Col11a2* | 39.93144481 | 37.4084623 | 26.47193365 | 13.17077092 | 7.369335996 | 15.6112477 | 0.005203772 | -1.524006943 | 0.002122648 | 9.440334656 | 0.347718819 | 2.283681742 |
| ENSMUSG00000020571 | *Pdia6* | 30104.54227 | 32709.38392 | 40682.31975 | 12210.40221 | 11782.5155 | 11780.06385 | 2.57E-38 | -1.53252693 | 7.80E-40 | 174.4735472 | 0.345671381 | 37.58973171 |
| ENSMUSG00000039063 | *Echdc3* | 142.397039 | 126.613257 | 99.58489327 | 58.17090489 | 28.4245817 | 41.32389096 | 8.22E-06 | -1.534802931 | 2.07E-06 | 22.52648831 | 0.345126479 | 5.085210533 |
| ENSMUSG00000060803 | *Gstp1* | 1182.874875 | 1464.685178 | 1830.345127 | 512.5625016 | 531.644954 | 498.6416176 | 7.47E-20 | -1.535421582 | 5.52E-21 | 88.33846966 | 0.344978515 | 19.12666726 |
| ENSMUSG00000026223 | *Itm2c* | 10107.9295 | 11632.11339 | 12605.68269 | 4093.914627 | 3735.200588 | 4013.008967 | 1.64E-51 | -1.53585653 | 3.05E-53 | 235.9222303 | 0.344874525 | 50.78397209 |
| ENSMUSG00000075701 | *Selenos* | 3907.254203 | 3841.561321 | 4515.35554 | 1520.126477 | 1366.485446 | 1315.018041 | 5.49E-50 | -1.545302017 | 1.07E-51 | 228.828223 | 0.342623968 | 49.26043531 |
| ENSMUSG00000070034 | *Sp110* | 2668.625991 | 2773.022167 | 2928.300089 | 1071.222701 | 848.5264019 | 942.1847139 | 8.13E-46 | -1.548118801 | 1.82E-47 | 209.4420228 | 0.341955666 | 45.09009707 |
| ENSMUSG00000029821 | *Gsdme* | 494.9992309 | 435.4728689 | 545.8260606 | 163.5370722 | 169.4947279 | 170.805416 | 8.40E-24 | -1.548263947 | 4.94E-25 | 106.7944724 | 0.341921264 | 23.07589659 |
| ENSMUSG00000067212 | *H2-T23* | 5346.293063 | 4858.304143 | 4210.298019 | 1874.639727 | 1348.588487 | 1699.789381 | 1.83E-26 | -1.550417347 | 9.34E-28 | 119.2280723 | 0.341411285 | 25.73675004 |
| ENSMUSG00000076617 | *Ighm* | 134.8628042 | 105.5110475 | 57.98614039 | 54.87821216 | 11.58038514 | 35.81403883 | 0.007769766 | -1.551189056 | 0.003282902 | 8.64316718 | 0.341228711 | 2.109592036 |
| ENSMUSG00000021792 | *Prxl2a* | 849.8616933 | 829.7005101 | 1048.7928 | 322.6838875 | 304.2483004 | 302.1235584 | 4.25E-29 | -1.552115717 | 1.90E-30 | 131.526302 | 0.341009606 | 28.37144309 |
| ENSMUSG00000056204 | *Pgpep1* | 785.0672734 | 845.0475715 | 916.4331317 | 282.0740105 | 283.1930547 | 300.286941 | 5.04E-38 | -1.553306002 | 1.56E-39 | 173.1002846 | 0.340728374 | 37.29762505 |
| ENSMUSG00000021752 | *Kctd6* | 109.2464056 | 94.95994277 | 84.45807404 | 31.82936305 | 32.63563084 | 33.97742146 | 2.15E-07 | -1.557336795 | 4.47E-08 | 29.93587253 | 0.339777731 | 6.666738314 |
| ENSMUSG00000015149 | *Sirt2* | 2716.09167 | 3040.636551 | 3113.603625 | 980.1248691 | 1005.387982 | 1021.159261 | 3.67E-68 | -1.559477769 | 4.53E-70 | 313.136952 | 0.339273871 | 67.43588162 |
| ENSMUSG00000021939 | *Ctsb* | 121114.3324 | 127361.4263 | 132949.6142 | 44153.91194 | 42473.6944 | 42695.84414 | 1.78E-134 | -1.560393614 | 5.76E-137 | 620.522728 | 0.339058563 | 133.7485129 |
| ENSMUSG00000035868 | *Zfp983* | 64.04099639 | 81.531264 | 80.67636923 | 28.53667032 | 23.16077027 | 24.79433458 | 5.46E-06 | -1.560969891 | 1.34E-06 | 23.35826 | 0.338923155 | 5.262845524 |
| ENSMUSG00000027438 | *Napb* | 204.9311884 | 180.3279721 | 196.64865 | 92.19539643 | 63.16573711 | 42.24219965 | 2.77E-07 | -1.566210166 | 5.82E-08 | 29.423239 | 0.337694324 | 6.557317693 |
| ENSMUSG00000019876 | *Pkib* | 197.3969536 | 213.8996691 | 221.8600154 | 70.24411156 | 65.27126168 | 77.13792979 | 1.23E-14 | -1.568593034 | 1.29E-15 | 63.92652912 | 0.337137022 | 13.9115056 |
| ENSMUSG00000042675 | *Ypel3* | 177.0545194 | 174.5728241 | 181.5218308 | 62.56116186 | 62.11297483 | 55.09852128 | 2.21E-13 | -1.570490821 | 2.58E-14 | 58.03339838 | 0.336693828 | 12.6565799 |
| ENSMUSG00000021094 | *Dhrs7* | 1197.189921 | 1233.520065 | 1224.011789 | 369.8791499 | 475.8485529 | 383.8530316 | 1.49E-36 | -1.571845432 | 4.82E-38 | 166.2730218 | 0.336377839 | 35.82811452 |
| ENSMUSG00000016534 | *Lamp2* | 9229.437715 | 9802.935506 | 13203.19205 | 3829.401644 | 3399.369419 | 3593.341896 | 3.12E-28 | -1.574373566 | 1.45E-29 | 127.4982941 | 0.335788897 | 27.50648437 |
| ENSMUSG00000023341 | *Mx2* | 369.9309321 | 342.4313088 | 330.2688865 | 138.2930946 | 84.22098282 | 127.6449076 | 2.31E-13 | -1.574984093 | 2.71E-14 | 57.93438277 | 0.335646826 | 12.6361734 |
| ENSMUSG00000041895 | *Wipi1* | 150.6846974 | 142.9195098 | 139.9230779 | 54.87821216 | 43.16325369 | 47.75205178 | 7.78E-11 | -1.575806133 | 1.13E-11 | 46.08728425 | 0.335455631 | 10.10914865 |
| ENSMUSG00000006717 | *Acot13* | 651.7113162 | 660.882834 | 913.9119952 | 237.0738765 | 253.7157107 | 252.5348892 | 1.24E-19 | -1.579016505 | 9.26E-21 | 87.31447153 | 0.334709985 | 18.90824206 |
| ENSMUSG00000021684 | *Pde8b* | 761.7111453 | 728.0262279 | 1008.454615 | 285.3667032 | 251.6101862 | 296.6137062 | 9.71E-21 | -1.580143064 | 6.73E-22 | 92.50219179 | 0.334448722 | 20.01267805 |
| ENSMUSG00000099693 | *Gm29284* | 867.1904335 | 821.067788 | 1067.701324 | 330.3668372 | 263.1905713 | 323.2446582 | 1.20E-23 | -1.585458138 | 7.13E-25 | 106.0675878 | 0.333218837 | 22.92063447 |
| ENSMUSG00000024737 | *Slc15a3* | 3333.89893 | 3404.17007 | 2739.214849 | 1141.466813 | 932.7473847 | 1077.176091 | 7.63E-37 | -1.589273132 | 2.46E-38 | 167.6138056 | 0.332338852 | 36.11729744 |
| ENSMUSG00000079190 | *ENSMUSG00000079190* | 1920.476468 | 1881.933411 | 2180.783106 | 717.807015 | 552.7001997 | 714.4441593 | 4.94E-33 | -1.590626863 | 1.89E-34 | 149.8320702 | 0.332027154 | 32.3066707 |
| ENSMUSG00000020189 | *Osbpl8* | 3095.063684 | 2984.044262 | 3095.955669 | 1077.808087 | 924.3252864 | 1040.443744 | 4.45E-73 | -1.592323452 | 5.10E-75 | 335.8555632 | 0.331636924 | 72.35172604 |
| ENSMUSG00000021895 | *Arhgef3* | 173.287402 | 150.5930406 | 190.3458087 | 61.46359762 | 54.74363883 | 54.18021259 | 8.87E-12 | -1.592822882 | 1.20E-12 | 50.491836 | 0.331522138 | 11.05210976 |
| ENSMUSG00000102525 | *Gm37784* | 84.38343054 | 70.0209679 | 78.15523269 | 30.73179881 | 17.89695885 | 28.46756933 | 8.31E-06 | -1.59311166 | 2.10E-06 | 22.50132419 | 0.331455785 | 5.080347843 |
| ENSMUSG00000022389 | *Tef* | 947.8067466 | 1015.78363 | 1034.926549 | 311.7082451 | 314.7759233 | 362.7319318 | 4.26E-43 | -1.59633578 | 1.07E-44 | 196.7552422 | 0.330715878 | 42.37083469 |
| ENSMUSG00000010051 | *Hyal1* | 109.2464056 | 176.4912068 | 181.5218308 | 52.68308367 | 42.11049141 | 58.77175603 | 3.69E-07 | -1.596553611 | 7.83E-08 | 28.84732462 | 0.330665948 | 6.433161737 |
| ENSMUSG00000053007 | *Creb5* | 103.9724412 | 124.6948743 | 84.45807404 | 28.53667032 | 31.58286856 | 43.16050834 | 1.31E-06 | -1.596998787 | 2.99E-07 | 26.25639945 | 0.330563929 | 5.881871369 |
| ENSMUSG00000026437 | *Cdk18* | 5958.826358 | 5954.659846 | 4609.898161 | 1791.224845 | 1786.537598 | 1882.53281 | 3.09E-41 | -1.597936397 | 8.39E-43 | 188.0698392 | 0.330349165 | 40.51052153 |
| ENSMUSG00000031169 | *Porcn* | 53.49306757 | 46.04118437 | 42.85932115 | 20.85372062 | 14.73867199 | 11.93801294 | 0.000349339 | -1.598448067 | 0.000113034 | 14.90548031 | 0.330232023 | 3.456752905 |
| ENSMUSG00000046312 | *Myorg* | 83.63000705 | 73.85773327 | 103.3665981 | 24.14641335 | 17.89695885 | 43.16050834 | 0.000125545 | -1.599605818 | 3.79E-05 | 16.97248072 | 0.329967121 | 3.901198872 |
| ENSMUSG00000029376 | *Mthfd2l* | 139.3833451 | 152.5114232 | 180.2612625 | 35.12205578 | 42.11049141 | 77.13792979 | 3.52E-06 | -1.600469884 | 8.45E-07 | 24.25177903 | 0.329769555 | 5.452922432 |
| ENSMUSG00000024055 | *Cyp4f13* | 285.5475016 | 332.8393954 | 248.331949 | 104.2686031 | 73.69335996 | 107.4421165 | 1.48E-12 | -1.602864285 | 1.85E-13 | 54.15656134 | 0.329222699 | 11.82834807 |
| ENSMUSG00000034041 | *Lyl1* | 3505.679485 | 3656.437392 | 3451.435921 | 1194.149897 | 1070.659244 | 1218.595629 | 3.28E-81 | -1.607055215 | 3.01E-83 | 373.6426298 | 0.328267718 | 80.48449459 |
| ENSMUSG00000031553 | *Adam3* | 162.7394732 | 138.1235531 | 194.1275135 | 50.48795519 | 49.4798274 | 61.5266821 | 5.47E-10 | -1.607738174 | 8.63E-11 | 42.11014297 | 0.328112355 | 9.262137154 |
| ENSMUSG00000020399 | *Havcr2* | 849.8616933 | 930.4156009 | 1202.582129 | 277.6837535 | 362.1502261 | 334.2643624 | 6.95E-20 | -1.611518136 | 5.13E-21 | 88.48208269 | 0.327253803 | 19.1577938 |
| ENSMUSG00000020593 | *Lpin1* | 246.3694802 | 249.3897487 | 189.0852404 | 70.24411156 | 57.90192569 | 94.58579486 | 3.41E-10 | -1.618754571 | 5.26E-11 | 43.07976194 | 0.325616436 | 9.467099132 |
| ENSMUSG00000015745 | *Plekho1* | 10222.44987 | 11403.82585 | 12714.09156 | 3889.767678 | 3410.949804 | 3866.997885 | 6.81E-52 | -1.620207092 | 1.24E-53 | 237.7079563 | 0.325288767 | 51.16687931 |
| ENSMUSG00000005045 | *Chd5* | 81.36973659 | 72.89854193 | 30.25363846 | 23.04884911 | 21.0552457 | 16.52955638 | 0.00156046 | -1.621860114 | 0.000571947 | 11.86513112 | 0.324916268 | 2.806747301 |
| ENSMUSG00000040322 | *Slc25a24* | 3272.871627 | 3148.065982 | 3515.724903 | 1069.027573 | 1066.448195 | 1088.195795 | 1.04E-90 | -1.623136311 | 7.48E-93 | 417.7643767 | 0.324628977 | 89.98361097 |
| ENSMUSG00000000058 | *Cav2* | 128.0819928 | 140.0419358 | 151.2681923 | 48.2928267 | 46.32154055 | 41.32389096 | 7.93E-11 | -1.624329601 | 1.16E-11 | 46.04533299 | 0.324360579 | 10.1004635 |
| ENSMUSG00000039813 | *Tbc1d2* | 509.3142772 | 583.1883354 | 557.171175 | 196.4639995 | 155.8088182 | 181.8251202 | 2.40E-27 | -1.625954186 | 1.17E-28 | 123.353195 | 0.32399553 | 26.62002025 |
| ENSMUSG00000020818 | *Mfsd11* | 1656.778248 | 1865.627158 | 2130.360375 | 530.1235294 | 649.55433 | 643.7343903 | 1.33E-32 | -1.630111174 | 5.21E-34 | 147.8120349 | 0.323063311 | 31.87513613 |
| ENSMUSG00000040552 | *C3ar1* | 3089.78972 | 3385.945434 | 3224.533633 | 1066.832444 | 1010.651794 | 1052.381756 | 9.65E-95 | -1.63158159 | 6.36E-97 | 436.4676966 | 0.322734208 | 94.01553352 |
| ENSMUSG00000074607 | *Tox2* | 24.86297507 | 38.36765364 | 27.73250192 | 8.780513945 | 8.422098282 | 11.93801294 | 0.003744921 | -1.633010532 | 0.001485684 | 10.09628658 | 0.322414709 | 2.426557364 |
| ENSMUSG00000026207 | *Speg* | 1967.942148 | 2038.2816 | 1807.654898 | 671.7093168 | 537.9615277 | 664.8554901 | 5.90E-43 | -1.633206245 | 1.49E-44 | 196.085314 | 0.322370974 | 42.22936187 |
| ENSMUSG00000028229 | *Rmdn1* | 250.8900212 | 245.5529833 | 313.881499 | 83.41488248 | 103.1707039 | 74.38300373 | 1.03E-13 | -1.633409161 | 1.17E-14 | 59.59212921 | 0.322325635 | 12.98820438 |
| ENSMUSG00000041028 | *Ghitm* | 17637.64383 | 18025.12368 | 21959.09925 | 6788.434844 | 5643.858611 | 6019.51345 | 8.96E-45 | -1.64275165 | 2.09E-46 | 204.5799595 | 0.320245089 | 44.04752252 |
| ENSMUSG00000030187 | *Klra2* | 95.68478284 | 90.16398607 | 115.9722808 | 28.53667032 | 31.58286856 | 35.81403883 | 6.20E-08 | -1.64380557 | 1.20E-08 | 32.4815966 | 0.320011228 | 7.207291324 |
| ENSMUSG00000033721 | *Vav3* | 796.3686257 | 852.7211023 | 796.6791462 | 298.5374741 | 229.5021782 | 254.3715066 | 2.13E-34 | -1.645697727 | 7.67E-36 | 156.1971243 | 0.319591795 | 33.67204845 |
| ENSMUSG00000015837 | *Sqstm1* | 59495.59249 | 64757.88501 | 68925.35183 | 21062.25783 | 19643.49148 | 20957.64088 | 4.52E-94 | -1.647378101 | 3.07E-96 | 433.323767 | 0.319219768 | 93.34463919 |
| ENSMUSG00000079444 | *Gm21981* | 42.19171527 | 25.89816621 | 23.95079712 | 14.26833516 | 5.263811426 | 10.10139557 | 0.008835675 | -1.649254374 | 0.003787238 | 8.383110279 | 0.318804882 | 2.053760258 |
| ENSMUSG00000035992 | *Fnip1* | 611.0264479 | 581.2699527 | 794.1580096 | 227.1957983 | 210.552457 | 193.7631332 | 8.05E-22 | -1.651801221 | 5.20E-23 | 97.56755248 | 0.318242579 | 21.09411176 |
| ENSMUSG00000079800 | *ENSMUSG00000079800* | 515.3416651 | 435.4728689 | 479.0159423 | 181.0981001 | 117.9093759 | 156.112477 | 4.51E-18 | -1.652791729 | 3.71E-19 | 80.0159461 | 0.318024159 | 17.34581986 |
| ENSMUSG00000027808 | *Serp1* | 12674.84332 | 13279.04493 | 17126.08051 | 4727.209195 | 4370.016246 | 4567.667414 | 6.43E-40 | -1.656288033 | 1.82E-41 | 181.9507485 | 0.317254376 | 39.19187289 |
| ENSMUSG00000028073 | *Pear1* | 284.0406546 | 345.3088828 | 271.0221779 | 107.5612958 | 101.0651794 | 78.05623848 | 4.40E-15 | -1.656780181 | 4.47E-16 | 66.01807519 | 0.317146169 | 14.3563294 |
| ENSMUSG00000039997 | *Ifi203* | 1335.066419 | 1186.519689 | 1236.617472 | 459.8794179 | 331.6201198 | 398.5459706 | 4.75E-32 | -1.66011956 | 1.88E-33 | 145.2606153 | 0.316412925 | 31.32342435 |
| ENSMUSG00000015599 | *Ttbk1* | 168.7668611 | 169.7768674 | 102.1060298 | 52.68308367 | 46.32154055 | 41.32389096 | 9.53E-08 | -1.661227278 | 1.89E-08 | 31.6053652 | 0.316170073 | 7.020679681 |
| ENSMUSG00000079808 | *ENSMUSG00000079808* | 2734.173834 | 2634.898614 | 2987.546798 | 997.685897 | 725.3532145 | 917.3903793 | 8.95E-36 | -1.661766138 | 3.00E-37 | 162.636801 | 0.316052003 | 35.04822215 |
| ENSMUSG00000022178 | *Ajuba* | 76.09577218 | 70.98015924 | 49.1621625 | 20.85372062 | 25.26629484 | 16.52955638 | 3.88E-05 | -1.661873767 | 1.08E-05 | 19.35668038 | 0.316028425 | 4.41120885 |
| ENSMUSG00000030108 | *Slc6a13* | 1053.286035 | 1026.334735 | 1235.356904 | 389.6353063 | 314.7759233 | 340.6925232 | 1.31E-34 | -1.664478163 | 4.69E-36 | 157.1747101 | 0.315458436 | 33.88234499 |
| ENSMUSG00000034449 | *Dhrs11* | 609.5196009 | 665.6787907 | 739.953574 | 221.7079771 | 218.9745553 | 193.7631332 | 1.22E-31 | -1.666387646 | 4.86E-33 | 143.3782622 | 0.315041186 | 30.91487739 |
| ENSMUSG00000031304 | *Il2rg* | 1806.709522 | 1730.381179 | 1404.273052 | 505.9771161 | 544.2781014 | 506.9063958 | 4.73E-39 | -1.667985849 | 1.39E-40 | 177.9003625 | 0.31469238 | 38.32506721 |
| ENSMUSG00000042766 | *Trim46* | 436.2321989 | 446.983165 | 349.1774106 | 137.1955304 | 118.9621382 | 132.2364511 | 2.82E-21 | -1.669474604 | 1.89E-22 | 95.01682498 | 0.314367808 | 20.5499885 |
| ENSMUSG00000045362 | *Tnfrsf26* | 751.1632165 | 851.7619109 | 843.3201721 | 287.5618317 | 229.5021782 | 247.9433458 | 6.41E-35 | -1.67687187 | 2.26E-36 | 158.6278046 | 0.312760046 | 34.19313386 |
| ENSMUSG00000049515 | *Espnl* | 140.8901921 | 144.8378925 | 136.1413731 | 38.41474851 | 56.8491634 | 36.73234752 | 7.63E-10 | -1.680446809 | 1.22E-10 | 41.42884664 | 0.311985999 | 9.117367726 |
| ENSMUSG00000015980 | *Lrrc27* | 320.9584054 | 358.7375616 | 386.9944587 | 115.2442455 | 94.74860567 | 121.2167468 | 2.25E-20 | -1.682019842 | 1.60E-21 | 90.78902602 | 0.311646012 | 19.64732298 |
| ENSMUSG00000018217 | *Pmp22* | 3675.953193 | 3715.907256 | 3355.632733 | 1097.564243 | 1053.815047 | 1197.474529 | 1.51E-77 | -1.682103637 | 1.55E-79 | 356.5976055 | 0.311627912 | 76.82211686 |
| ENSMUSG00000047875 | *Gpr157* | 45.9588327 | 32.6125056 | 31.51420673 | 9.878078189 | 12.63314742 | 11.93801294 | 0.000907635 | -1.683208837 | 0.000316601 | 12.96953367 | 0.311389276 | 3.042088844 |
| ENSMUSG00000004267 | *Eno2* | 235.8215514 | 245.5529833 | 287.4095654 | 92.19539643 | 51.58535197 | 93.66748618 | 1.71E-10 | -1.690339927 | 2.56E-11 | 44.48704255 | 0.309853909 | 9.767613885 |
| ENSMUSG00000039648 | *Kyat1* | 78.35604264 | 80.57207265 | 64.28898173 | 28.53667032 | 18.94972113 | 22.03940851 | 2.20E-06 | -1.691248854 | 5.14E-07 | 25.21195065 | 0.309658756 | 5.658203105 |
| ENSMUSG00000039886 | *Tmem120a* | 3379.104339 | 3778.254693 | 4423.334057 | 1149.149763 | 1218.045964 | 1215.840703 | 6.09E-47 | -1.691316187 | 1.33E-48 | 214.6542322 | 0.309644304 | 46.21504423 |
| ENSMUSG00000035640 | *Cbarp* | 97.9450533 | 117.980535 | 132.3596683 | 27.43910608 | 29.47734399 | 49.58866915 | 6.56E-07 | -1.694205659 | 1.44E-07 | 27.66833428 | 0.30902476 | 6.18285878 |
| ENSMUSG00000027570 | *Col9a3* | 27.87666902 | 23.02059219 | 21.42966058 | 9.878078189 | 5.263811426 | 7.346469504 | 0.006628156 | -1.696063352 | 0.002760974 | 8.959057036 | 0.308627099 | 2.178607268 |
| ENSMUSG00000031504 | *Rab20* | 312.6707471 | 330.9210127 | 385.7338904 | 124.0247595 | 88.43203196 | 104.6871904 | 7.97E-18 | -1.6964138 | 6.68E-19 | 78.85645774 | 0.308552139 | 17.09848014 |
| ENSMUSG00000022564 | *Grina* | 2707.804012 | 3102.983989 | 2956.032591 | 839.636646 | 885.3730818 | 976.1621353 | 7.10E-64 | -1.696920731 | 9.65E-66 | 293.2677561 | 0.308443739 | 63.14859069 |
| ENSMUSG00000018340 | *Anxa6* | 3144.789635 | 3666.029306 | 3815.740151 | 1047.076288 | 1141.194317 | 1081.767634 | 2.96E-60 | -1.699338064 | 4.24E-62 | 276.5510085 | 0.307927354 | 59.5294196 |
| ENSMUSG00000042712 | *Tceal9* | 668.2866329 | 648.4133466 | 1007.194047 | 250.2446474 | 221.0800799 | 242.4334936 | 5.46E-17 | -1.700100914 | 4.80E-18 | 74.96144954 | 0.307764575 | 16.26245366 |
| ENSMUSG00000071203 | *Naip5* | 579.3826615 | 599.4945882 | 577.3402673 | 166.829765 | 187.3916868 | 185.498355 | 9.85E-40 | -1.700873709 | 2.81E-41 | 181.0876806 | 0.307599762 | 39.00658956 |
| ENSMUSG00000048285 | *Frmd6* | 82.12316008 | 123.735683 | 95.80318846 | 28.53667032 | 43.16325369 | 21.12109982 | 7.50E-06 | -1.705045144 | 1.88E-06 | 22.71023189 | 0.306711647 | 5.124968147 |
| ENSMUSG00000040813 | *Tex264* | 3191.501891 | 3672.743645 | 3977.092889 | 1049.271416 | 1156.985751 | 1112.99013 | 2.14E-55 | -1.706497691 | 3.42E-57 | 254.0388401 | 0.306402996 | 54.66987303 |
| ENSMUSG00000055172 | *C1ra* | 417.3966118 | 441.2280169 | 402.1212779 | 126.219888 | 133.7008102 | 126.7265989 | 9.37E-31 | -1.706499772 | 3.90E-32 | 139.2435982 | 0.306402554 | 30.02832904 |
| ENSMUSG00000047821 | *Trim16* | 290.0680425 | 289.675785 | 307.5786577 | 92.19539643 | 92.6430811 | 86.32101667 | 1.76E-23 | -1.710312671 | 1.05E-24 | 105.2959182 | 0.305593832 | 22.75501204 |
| ENSMUSG00000064179 | *Tnnt1* | 61.02730244 | 44.12280169 | 80.67636923 | 15.3658994 | 16.84419656 | 23.87602589 | 0.000115116 | -1.713087089 | 3.45E-05 | 17.15146666 | 0.305006715 | 3.938864051 |
| ENSMUSG00000022304 | *Dpys* | 59.52045547 | 59.46986315 | 69.33125481 | 27.43910608 | 11.58038514 | 18.36617376 | 6.25E-05 | -1.71568645 | 1.80E-05 | 18.39275439 | 0.304457667 | 4.20415338 |
| ENSMUSG00000007783 | *Cpt1c* | 30.89036296 | 24.93897487 | 31.51420673 | 10.97564243 | 7.369335996 | 8.264778192 | 0.002122048 | -1.718220094 | 0.000801583 | 11.23756464 | 0.303923451 | 2.673244862 |
| ENSMUSG00000018459 | *Slc13a3* | 48.21910316 | 85.36802936 | 78.15523269 | 17.56102789 | 25.26629484 | 21.12109982 | 4.00E-05 | -1.718844978 | 1.12E-05 | 19.29362713 | 0.303791839 | 4.39772002 |
| ENSMUSG00000036390 | *Gadd45a* | 70.06838429 | 83.44964668 | 76.89466442 | 16.46346365 | 28.4245817 | 24.79433458 | 1.74E-06 | -1.719015449 | 4.03E-07 | 25.67846083 | 0.303755945 | 5.758994069 |
| ENSMUSG00000035692 | *Isg15* | 2536.023457 | 2094.873889 | 2020.690936 | 955.9784558 | 423.2104386 | 629.95976 | 1.36E-11 | -1.72788176 | 1.86E-12 | 49.62882087 | 0.301894889 | 10.86505945 |

| | | | | | | | | | | | | | |
|---|---|---|---|---|---|---|---|---|---|---|---|---|---|
| ENSMUSG00000028420 | *Tmem38b* | 1612.326262 | 2028.689686 | 2344.656981 | 671.7093168 | 553.752962 | 580.3710908 | 3.13E-27 | -1.727929281 | 1.53E-28 | 122.8117639 | 0.301884945 | 26.50456168 |
| ENSMUSG00000021806 | *Nid2* | 216.2325408 | 222.5323911 | 155.0498971 | 51.58551943 | 57.90192569 | 69.79146029 | 2.58E-11 | -1.72843481 | 3.60E-12 | 48.32909463 | 0.301779182 | 10.58766124 |
| ENSMUSG00000033114 | *Slc35d2* | 163.4928967 | 188.0015029 | 239.5079712 | 53.78064792 | 60.00745026 | 63.36329947 | 6.30E-12 | -1.728938017 | 8.38E-13 | 51.19089608 | 0.301673941 | 11.20055744 |
| ENSMUSG00000035208 | *Slfn8* | 569.5881561 | 479.5956706 | 554.6500385 | 196.4639995 | 132.6480479 | 155.1941683 | 3.46E-21 | -1.729459383 | 2.34E-22 | 94.58795831 | 0.30156494 | 20.460766 |
| ENSMUSG00000036526 | *Card11* | 831.0261061 | 919.8644961 | 813.0665337 | 262.3178541 | 275.8237187 | 236.0053328 | 2.80E-43 | -1.730113691 | 7.00E-45 | 197.5917069 | 0.301428202 | 42.5521913 |
| ENSMUSG00000010307 | *Tmem86a* | 226.7804696 | 304.0636551 | 352.9591154 | 85.61001097 | 100.0124171 | 79.89285586 | 4.55E-13 | -1.731207262 | 5.46E-14 | 56.55725682 | 0.301199804 | 12.34194444 |
| ENSMUSG00000094728 | *ENSMUSG00000094728* | 466.3691384 | 432.5952948 | 444.980599 | 167.9273292 | 90.53755653 | 146.0110814 | 2.72E-14 | -1.732660575 | 2.93E-15 | 62.31326319 | 0.300896541 | 13.56614658 |
| ENSMUSG00000038020 | *Rapgefl1* | 23.3561281 | 27.81654889 | 22.69022885 | 5.487821216 | 7.369335996 | 9.18308688 | 0.004549982 | -1.735247241 | 0.001834731 | 9.707944857 | 0.300357534 | 2.341990277 |
| ENSMUSG00000029810 | *Tmem176b* | 647.9441988 | 547.6982558 | 597.5093596 | 166.829765 | 193.7082605 | 178.1518855 | 1.63E-33 | -1.735840594 | 6.05E-35 | 152.0905132 | 0.300234028 | 32.78763443 |
| ENSMUSG00000029158 | *Yipf7* | 333.0131812 | 238.8386439 | 245.8108125 | 77.92706126 | 88.43203196 | 79.89285586 | 1.01E-14 | -1.73621535 | 1.06E-15 | 64.31886953 | 0.300156049 | 13.99429761 |
| ENSMUSG00000050335 | *Lgals3* | 56655.93937 | 62707.13393 | 76501.36712 | 19453.22865 | 19683.49645 | 19618.74681 | 1.51E-50 | -1.73699317 | 2.90E-52 | 231.4369979 | 0.299994266 | 49.82012359 |
| ENSMUSG00000003623 | *Crot* | 616.3004123 | 636.9030505 | 819.369375 | 233.7811838 | 196.8665473 | 190.0898984 | 7.09E-24 | -1.738167035 | 4.16E-25 | 107.135458 | 0.299750271 | 23.14960581 |
| ENSMUSG00000044377 | *Gm13080* | 24.10955158 | 42.20441901 | 65.54955 | 15.3658994 | 14.73867199 | 9.18308688 | 0.003204411 | -1.739611458 | 0.001251022 | 10.41342908 | 0.299450312 | 2.494251723 |
| ENSMUSG00000030055 | *Rab43* | 959.8615224 | 1041.681796 | 961.8135894 | 273.2934965 | 310.5648741 | 301.2052497 | 6.63E-54 | -1.742494249 | 1.12E-55 | 247.0966553 | 0.298852548 | 53.17823738 |
| ENSMUSG00000000325 | *None* | 83.63000705 | 93.04156009 | 35.29591154 | 23.04884911 | 14.73867199 | 25.71264326 | 0.000619347 | -1.744673532 | 0.000210557 | 13.73444356 | 0.298401453 | 3.208066226 |
| ENSMUSG00000013160 | *Atp6v0d1* | 7200.468265 | 7539.243941 | 8827.759589 | 2349.885045 | 2260.280626 | 2408.723689 | 5.26E-69 | -1.746849498 | 6.42E-71 | 317.0302826 | 0.297951724 | 68.27912156 |
| ENSMUSG00000082088 | *Gm15753* | 80.6163131 | 61.38824583 | 52.94386731 | 24.14641335 | 16.84419656 | 17.44786507 | 1.49E-05 | -1.752148966 | 3.92E-06 | 21.30481308 | 0.296859263 | 4.826674297 |
| ENSMUSG00000020048 | *Hsp90b1* | 52634.1648 | 58221.95521 | 78658.19943 | 19479.57019 | 18278.0588 | 18485.55389 | 3.46E-33 | -1.752503809 | 1.31E-34 | 150.5493624 | 0.296786256 | 32.46032237 |
| ENSMUSG00000035202 | *Lars2* | 14089.77263 | 36682.35446 | 29981.35572 | 7699.413166 | 7756.752517 | 8438.338534 | 9.24E-06 | -1.756813266 | 2.35E-06 | 22.28480648 | 0.295901053 | 5.034326348 |
| ENSMUSG00000005054 | *Cstb* | 8277.863851 | 8899.377263 | 13242.26967 | 3052.32616 | 2949.839923 | 2977.156767 | 5.00E-25 | -1.760000908 | 2.74E-26 | 112.5272464 | 0.29524798 | 24.30093085 |
| ENSMUSG00000070031 | *Sp140* | 1293.628127 | 1269.010144 | 1424.442144 | 426.9524906 | 303.1955381 | 444.461405 | 1.75E-28 | -1.761529359 | 8.03E-30 | 128.6646365 | 0.294935347 | 27.7578282 |
| ENSMUSG00000018930 | *Ccl4* | 11898.81713 | 10656.6158 | 8913.478232 | 3639.52303 | 2219.222897 | 3394.98722 | 5.79E-21 | -1.765930076 | 3.97E-22 | 93.54540383 | 0.294037064 | 20.23726846 |
| ENSMUSG00000046719 | *Nxph3* | 18.08216369 | 24.93897487 | 26.47193365 | 6.585385459 | 6.316573711 | 7.346469504 | 0.005542253 | -1.766139164 | 0.002274788 | 9.313441312 | 0.293994452 | 2.256313681 |
| ENSMUSG00000017652 | *Cd40* | 915.4095366 | 898.7622866 | 1144.595988 | 365.488893 | 229.5021782 | 270.901063 | 1.10E-19 | -1.772719285 | 8.19E-21 | 87.55650835 | 0.292656599 | 18.95899496 |
| ENSMUSG00000028100 | *Nudt17* | 45.20540922 | 51.79633242 | 42.85932115 | 17.56102789 | 11.58038514 | 11.93801294 | 7.24E-05 | -1.776060393 | 2.11E-05 | 18.08886539 | 0.291979626 | 4.140117592 |
| ENSMUSG00000089901 | *Gm8113* | 182.3284838 | 247.471366 | 245.8108125 | 82.31731824 | 49.4798274 | 65.19991685 | 2.66E-11 | -1.77616105 | 3.72E-12 | 48.26922177 | 0.291959255 | 10.57589449 |
| ENSMUSG00000030084 | *Plxna1* | 8434.575936 | 8626.007731 | 5462.042311 | 1996.469358 | 2373.978953 | 2205.777469 | 2.42E-24 | -1.776342616 | 1.38E-25 | 109.3233383 | 0.291922514 | 23.61646937 |
| ENSMUSG00000030030 | *1700003E16Rik* | 28.6300925 | 19.18382682 | 28.99307019 | 7.682949702 | 7.369335996 | 7.346469504 | 0.003521169 | -1.777350131 | 0.001386593 | 10.22354247 | 0.291718719 | 2.453313069 |
| ENSMUSG00000026456 | *Cyb5r1* | 7657.796321 | 7629.407927 | 8589.512187 | 2305.982475 | 2387.664863 | 2268.222459 | 1.45E-109 | -1.777725746 | 6.75E-112 | 505.2798614 | 0.291642778 | 108.8397988 |
| ENSMUSG00000020773 | *Trim47* | 37.67117435 | 42.20441901 | 34.03534327 | 8.780513945 | 9.474860567 | 14.69293901 | 0.000395798 | -1.779894645 | 0.000129371 | 14.6509338 | 0.291204662 | 3.40252605 |
| ENSMUSG00000035064 | *Eef2k* | 852.1219637 | 964.9464892 | 733.6507327 | 265.6105468 | 223.1856045 | 254.3715066 | 2.10E-31 | -1.780725158 | 8.50E-33 | 142.26806 | 0.291037073 | 30.67732726 |
| ENSMUSG00000045294 | *Insig1* | 3035.543229 | 2943.758226 | 2695.09496 | 1030.612824 | 699.0341574 | 792.5003977 | 8.17E-33 | -1.783145594 | 3.15E-34 | 148.813605 | 0.290549204 | 32.08793305 |
| ENSMUSG00000079192 | *ENSMUSG00000079192* | 1004.313508 | 973.5792112 | 1254.265428 | 299.6350384 | 302.1427758 | 334.2643624 | 1.64E-35 | -1.784430965 | 5.60E-37 | 161.3997777 | 0.290290454 | 34.78414068 |
| ENSMUSG00000059108 | *Ifitm6* | 954.587558 | 933.2931749 | 1163.504513 | 367.6840215 | 210.552457 | 306.7151018 | 8.70E-18 | -1.785110628 | 7.31E-19 | 78.67838217 | 0.290153728 | 17.06039809 |
| ENSMUSG00000042286 | *Stab1* | 427.1911171 | 339.5537348 | 255.8953587 | 88.9027037 | 104.2234662 | 103.7688817 | 1.24E-13 | -1.788195743 | 1.41E-14 | 59.21410072 | 0.289533915 | 12.90713301 |
| ENSMUSG00000024236 | *Svil* | 273.4927258 | 325.1658646 | 262.1982 | 81.21975399 | 84.22098282 | 83.56609061 | 1.05E-20 | -1.790431417 | 7.30E-22 | 92.34111326 | 0.289085586 | 19.9781896 |
| ENSMUSG00000019528 | *Gyg* | 2629.447969 | 2756.715914 | 3196.801131 | 831.9536963 | 842.2098282 | 805.3567194 | 1.29E-65 | -1.790433095 | 1.68E-67 | 301.337308 | 0.28908525 | 64.88881531 |
| ENSMUSG00000042682 | *Selenok* | 3221.63883 | 3337.985867 | 4623.764412 | 1127.198478 | 1001.176933 | 1098.297191 | 1.50E-31 | -1.792355395 | 6.00E-33 | 142.9582708 | 0.288700319 | 30.82453888 |
| ENSMUSG00000029185 | *Fam114a1* | 128.0819928 | 155.3889973 | 205.4726279 | 45.00013397 | 38.95220455 | 56.01682997 | 2.52E-09 | -1.792867594 | 4.26E-10 | 38.99139005 | 0.28859784 | 8.598686874 |
| ENSMUSG00000024827 | *Gldc* | 21.09585763 | 33.57169694 | 30.25363846 | 5.487821216 | 9.474860567 | 9.18308688 | 0.002873021 | -1.799681652 | 0.00111021 | 10.63410461 | 0.287237964 | 2.541661136 |
| ENSMUSG00000049037 | *Clec4a1* | 88.90397146 | 110.3070042 | 129.8385317 | 25.24397759 | 26.31905713 | 41.32389096 | 1.47E-07 | -1.808803077 | 2.98E-08 | 30.72266554 | 0.285427634 | 6.833453663 |
| ENSMUSG00000030107 | *Usp18* | 3781.432481 | 3677.539602 | 3774.141398 | 1364.272354 | 688.5065345 | 1140.539391 | 1.71E-17 | -1.814673074 | 1.45E-18 | 77.32548393 | 0.284268653 | 16.76788372 |
| ENSMUSG00000109899 | *Gm6012* | 33.15063343 | 25.89816621 | 31.51420673 | 7.682949702 | 13.68590971 | 4.59154344 | 0.00327447 | -1.815808954 | 0.001282194 | 10.36798586 | 0.284044928 | 2.484859014 |
| ENSMUSG00000006179 | *Prss16* | 384.9994018 | 331.880204 | 316.4026356 | 93.29296067 | 106.3289908 | 94.58579486 | 6.21E-24 | -1.816583349 | 3.63E-25 | 107.4055218 | 0.283892502 | 23.2067498 |
| ENSMUSG00000036186 | *Dipk1b* | 45.9588327 | 32.6125056 | 42.85932115 | 7.682949702 | 12.63314742 | 13.77463032 | 0.000282854 | -1.824327526 | 9.03E-05 | 15.33023811 | 0.282372691 | 3.548437808 |
| ENSMUSG00000037447 | *Arid5a* | 262.9447969 | 272.4103409 | 185.3035356 | 72.43924005 | 38.95220455 | 91.8308688 | 5.03E-08 | -1.824714789 | 9.65E-09 | 32.91013769 | 0.282296904 | 7.298562121 |
| ENSMUSG00000038067 | *Csf3* | 137.1230746 | 179.3687808 | 163.873875 | 60.36603337 | 37.89944227 | 37.65065621 | 3.35E-10 | -1.825374676 | 5.15E-11 | 43.1193217 | 0.282167811 | 9.475493338 |
| ENSMUSG00000028121 | *Bcar3* | 3678.213463 | 4135.073872 | 3457.738763 | 1116.222835 | 972.7523515 | 1085.440869 | 1.89E-67 | -1.828345021 | 2.39E-69 | 309.8199632 | 0.281587457 | 66.72285413 |
| ENSMUSG00000049588 | *Ccdc69* | 290.0680425 | 300.2268898 | 439.938326 | 120.7320667 | 82.11545824 | 86.32101667 | 9.36E-13 | -1.831497099 | 1.15E-13 | 55.09095146 | 0.280972901 | 12.02857856 |
| ENSMUSG00000003355 | *Fkbp11* | 1542.257878 | 1998.954755 | 2809.806672 | 636.587261 | 553.752962 | 592.3091038 | 1.10E-18 | -1.831765898 | 8.71E-20 | 82.8814046 | 0.280920556 | 17.95966159 |
| ENSMUSG00000029860 | *Zyx* | 2794.447713 | 2707.797156 | 2146.747763 | 739.7582999 | 637.9739448 | 766.7877545 | 2.58E-41 | -1.835256057 | 6.99E-43 | 188.4312957 | 0.280241776 | 40.58832533 |
| ENSMUSG00000069607 | *Cd300ld3* | 504.7937363 | 542.9022991 | 506.7484442 | 132.8052734 | 138.9646216 | 162.5406378 | 1.82E-35 | -1.835794825 | 6.22E-37 | 161.1902121 | 0.280137141 | 34.74010301 |
| ENSMUSG00000031762 | *Mt2* | 7308.961247 | 7896.06312 | 9802.178862 | 2554.031994 | 1993.931768 | 2416.070158 | 1.97E-38 | -1.843985585 | 5.91E-40 | 175.0243088 | 0.278551196 | 37.70506401 |
| ENSMUSG00000049719 | *Prss46* | 64.04099639 | 69.06177656 | 71.85239135 | 20.85372062 | 16.84419656 | 19.28448245 | 2.83E-07 | -1.844131425 | 5.94E-08 | 29.38334225 | 0.278523039 | 6.548660724 |
| ENSMUSG00000058755 | *Osm* | 334.5200282 | 423.0033814 | 441.1988942 | 131.7077092 | 100.0124171 | 101.0139557 | 6.61E-20 | -1.84865616 | 4.86E-21 | 88.58815256 | 0.277650874 | 19.17986748 |
| ENSMUSG00000030759 | *Far1* | 2696.50266 | 2517.87727 | 3264.871817 | 828.6610036 | 726.4059768 | 797.0919412 | 1.59E-48 | -1.849020915 | 3.23E-50 | 222.0509415 | 0.277580684 | 47.79727268 |
| ENSMUSG00000032575 | *Manf* | 21960.03437 | 21561.66216 | 24501.66545 | 6292.335806 | 6151.290032 | 6389.591851 | 1.77E-130 | -1.852534961 | 5.95E-133 | 602.0640241 | 0.27690539 | 129.7518183 |
| ENSMUSG00000022848 | *Slc49a4* | 290.821466 | 271.4511495 | 276.064451 | 87.80513945 | 67.37678625 | 77.13792979 | 1.24E-22 | -1.853814969 | 7.80E-24 | 101.3258464 | 0.276659819 | 21.90610954 |
| ENSMUSG00000000134 | *Tfe3* | 2780.132667 | 2741.368853 | 2416.509372 | 721.0997078 | 742.1974111 | 733.7286417 | 3.55E-87 | -1.85434311 | 2.82E-89 | 401.332263 | 0.276558558 | 86.45001988 |
| ENSMUSG00000029209 | *Gnpda2* | 354.8624624 | 316.5331426 | 523.1358317 | 128.4150165 | 84.22098282 | 115.7068947 | 1.53E-12 | -1.859707396 | 1.91E-13 | 54.0910671 | 0.275532156 | 11.8143501 |
| ENSMUSG00000027562 | *Car2* | 6283.551881 | 6929.198248 | 6897.829569 | 1969.030252 | 1690.73623 | 1866.921563 | 6.37E-95 | -1.863227096 | 4.15E-97 | 437.3165184 | 0.274860768 | 94.1957252 |
| ENSMUSG00000022844 | *Pdia5* | 1355.408853 | 1593.216818 | 1864.38047 | 421.4646694 | 429.5270124 | 469.2557396 | 3.32E-38 | -1.86343733 | 1.01E-39 | 173.9570711 | 0.274820717 | 37.47890449 |
| ENSMUSG00000042182 | *Bend6* | 138.6299216 | 145.7970839 | 200.4303548 | 41.70744124 | 51.58535197 | 39.48727358 | 4.10E-11 | -1.863871051 | 5.82E-12 | 47.38911038 | 0.27473811 | 10.38769963 |
| ENSMUSG00000024084 | *Qpct* | 886.0260206 | 1129.9274 | 1558.062381 | 341.3424796 | 281.0875301 | 354.4671536 | 7.44E-18 | -1.868779263 | 6.21E-19 | 79.00149562 | 0.273805008 | 17.12822483 |
| ENSMUSG00000029298 | *Gbp9* | 223.7667756 | 200.4709903 | 181.5218308 | 50.48795519 | 41.05772912 | 73.46469504 | 6.83E-12 | -1.871384282 | 9.10E-13 | 51.02859102 | 0.273311054 | 11.1653202 |
| ENSMUSG00000070960 | *Gm19680* | 142.397039 | 164.9809107 | 147.4864875 | 51.58551943 | 43.16325369 | 30.3041867 | 2.17E-11 | -1.871742744 | 3.01E-12 | 48.68068867 | 0.273243154 | 10.66295153 |
| ENSMUSG00000071552 | *Tigit* | 27.12324553 | 34.53088828 | 42.85932115 | 6.585385459 | 10.52762285 | 11.01970426 | 0.000638071 | -1.874202123 | 0.000217405 | 13.67434128 | 0.27277775 | 3.195131068 |
| ENSMUSG00000072812 | *None* | 394.7939072 | 492.065158 | 316.4026356 | 129.5125807 | 116.8566137 | 82.64778192 | 4.07E-14 | -1.876913839 | 4.44E-15 | 61.49633415 | 0.272265514 | 13.39064429 |
| ENSMUSG00000024610 | *Cd74* | 14220.86832 | 14954.7522 | 13670.86288 | 4048.914493 | 3619.396736 | 3991.887867 | 3.14E-127 | -1.877632144 | 1.12E-129 | 587.0074324 | 0.272129989 | 126.502647 |
| ENSMUSG00000028249 | *Sdcbp* | 13275.32184 | 12396.58889 | 18085.37296 | 4284.890805 | 3627.818835 | 3959.747063 | 1.40E-35 | -1.881730573 | 4.74E-37 | 161.7280009 | 0.271358015 | 34.85321452 |
| ENSMUSG00000032757 | *Bet1* | 784.3138499 | 747.2100547 | 1016.018025 | 245.8543905 | 233.7132273 | 210.2926896 | 6.02E-30 | -1.883735237 | 2.61E-31 | 135.4697961 | 0.270981218 | 29.22053256 |
| ENSMUSG00000032489 | *Kif9* | 22.60270461 | 37.4084623 | 31.51420673 | 9.878078189 | 9.474860567 | 5.509852128 | 0.001570693 | -1.886349977 | 0.000576021 | 11.8519135 | 0.270490536 | 2.803908767 |
| ENSMUSG00000095672 | *ENSMUSG00000095672* | 177.0545194 | 166.8992934 | 119.7539856 | 20.85372062 | 72.64059768 | 32.14080408 | 3.36E-05 | -1.887398318 | 9.32E-06 | 19.64612573 | 0.270294054 | 4.473053014 |
| ENSMUSG00000035891 | *Cerk* | 2451.640027 | 2894.839468 | 2871.574517 | 665.1239314 | 839.0515413 | 713.5258506 | 3.10E-52 | -1.889092813 | 5.58E-54 | 239.3034889 | 0.269976771 | 51.50820519 |
| ENSMUSG00000025283 | *Sat1* | 5336.498558 | 4883.243118 | 6461.672948 | 1718.785605 | 1251.734357 | 1529.902274 | 4.19E-36 | -1.88985059 | 1.38E-37 | 164.1766713 | 0.269835003 | 35.37745894 |
| ENSMUSG00000058427 | *Cxcl2* | 990.7518853 | 1207.621898 | 893.7429029 | 307.3179881 | 201.0775965 | 323.2446582 | 3.32E-20 | -1.894059767 | 2.38E-21 | 90.00019538 | 0.269048885 | 19.47918656 |
| ENSMUSG00000063506 | *Arhgap22* | 843.8343054 | 845.0475715 | 835.7567625 | 235.9763123 | 233.7132273 | 210.2926896 | 1.71E-61 | -1.895021954 | 2.40E-63 | 282.2765664 | 0.268869506 | 60.76661107 |
| ENSMUSG00000018395 | *Kif3a* | 805.4097075 | 819.1494053 | 993.3277962 | 227.1957983 | 282.1402924 | 194.6814419 | 1.38E-27 | -1.895078908 | 6.57E-29 | 124.4943857 | 0.268858892 | 26.86050855 |
| ENSMUSG00000005125 | *Ndrg1* | 6807.934628 | 6882.197873 | 5970.051323 | 2104.030654 | 1380.171356 | 1801.721646 | 7.74E-35 | -1.895316008 | 2.74E-36 | 158.2441469 | 0.26881471 | 34.11099413 |
| ENSMUSG00000059708 | *Akap17b* | 83.63000705 | 99.75589948 | 98.324325 | 31.82936305 | 16.84419656 | 26.63095195 | 4.33E-08 | -1.901005795 | 8.26E-09 | 33.21404487 | 0.26775663 | 7.363641153 |
| ENSMUSG00000020865 | *Abcc3* | 696.1633019 | 756.8019681 | 674.404024 | 166.829765 | 201.0775965 | 200.191294 | 7.33E-44 | -1.903284786 | 1.77E-45 | 200.3298269 | 0.267333995 | 43.13516692 |
| ENSMUSG00000040502 | *Marchf9* | 537.9443697 | 570.718848 | 451.2834404 | 136.0979662 | 133.7008102 | 146.9293901 | 3.86E-32 | -1.905650515 | 1.53E-33 | 145.6735331 | 0.266895981 | 31.41294449 |
| ENSMUSG00000060791 | *Gmfg* | 195.8901066 | 166.8992934 | 250.8530856 | 69.14654732 | 32.63563084 | 60.60837341 | 3.79E-09 | -1.914429612 | 6.52E-10 | 38.1599415 | 0.265276795 | 8.421888324 |
| ENSMUSG00000024617 | *Camk2a* | 16.57531671 | 36.44927096 | 31.51420673 | 5.487821216 | 8.422098282 | 8.264778192 | 0.003538595 | -1.914769914 | 0.001394379 | 10.21321496 | 0.265214229 | 2.451169106 |
| ENSMUSG00000001313 | *Rnd2* | 470.1362559 | 522.7592809 | 684.4885702 | 195.3664353 | 112.6455645 | 135.9096858 | 4.99E-15 | -1.917432938 | 5.08E-16 | 65.76395559 | 0.26472513 | 14.30179997 |
| ENSMUSG00000032265 | *Tent5a* | 204.9311884 | 210.0629037 | 194.1275135 | 68.04898308 | 41.05772912 | 52.34359522 | 3.46E-15 | -1.919613968 | 3.48E-16 | 66.51096904 | 0.264325228 | 14.46050262 |
| ENSMUSG00000000154 | *Slc22a18* | 31.64378645 | 51.79633242 | 39.07761635 | 10.97564243 | 7.369335996 | 13.77463032 | 0.000267588 | -1.920078767 | 8.51E-05 | 15.44158514 | 0.264240083 | 3.572532897 |
| ENSMUSG00000040264 | *Gbp2b* | 23.3561281 | 29.73493157 | 23.95079712 | 3.29269273 | 10.52762285 | 6.428160816 | 0.00356731 | -1.924438746 | 0.001408123 | 10.19512655 | 0.263442727 | 2.447659133 |
| ENSMUSG00000021356 | *Irf4* | 61.78072593 | 62.34743717 | 69.33125481 | 15.3658994 | 16.84419656 | 18.36617376 | 2.37E-07 | -1.928175836 | 4.94E-08 | 29.74061624 | 0.2627612 | 6.625580266 |
| ENSMUSG00000028145 | *Them4* | 489.7252665 | 462.3302264 | 574.8191308 | 163.5370722 | 121.0676628 | 116.6252034 | 1.75E-24 | -1.929691402 | 9.82E-26 | 109.9951032 | 0.262485312 | 23.75803683 |
| ENSMUSG00000061451 | *Tmem151a* | 39.93144481 | 61.38824583 | 45.38045769 | 6.585385459 | 16.84419656 | 14.69293901 | 0.000155758 | -1.934536167 | 4.78E-05 | 16.53509733 | 0.261605329 | 3.8075493 |
| ENSMUSG00000060131 | *Atp8b4* | 5432.936764 | 5514.39102 | 5628.437322 | 1480.614164 | 1354.905061 | 1484.905149 | 1.93E-156 | -1.939333444 | 4.91E-159 | 722.0039281 | 0.260736878 | 155.7137727 |
| ENSMUSG00000044629 | *Cnrip1* | 122.0546049 | 134.2867878 | 141.1836462 | 25.24397759 | 29.47734399 | 47.75205178 | 8.67E-10 | -1.941348758 | 1.40E-10 | 41.16583634 | 0.260372907 | 9.062084649 |
| ENSMUSG00000032507 | *Fbxl2* | 35.41090389 | 39.32684499 | 27.73250192 | 4.390256973 | 6.316573711 | 15.6112477 | 0.002289075 | -1.941355726 | 0.000868692 | 11.08840379 | 0.260371649 | 2.640340019 |
| ENSMUSG00000001827 | *Folr1* | 46.71225619 | 50.83714108 | 50.42273077 | 14.26833516 | 8.422098282 | 15.6112477 | 1.55E-05 | -1.94200027 | 4.09E-06 | 21.22319939 | 0.260255351 | 4.80908934 |
| ENSMUSG00000019806 | *Aig1* | 1677.120682 | 1840.688184 | 1996.740138 | 484.0258312 | 446.3712089 | 497.7233089 | 1.86E-69 | -1.946963932 | 2.26E-71 | 319.1146831 | 0.259361468 | 68.73071085 |
| ENSMUSG00000031389 | *Arhgap4* | 162.7394732 | 196.6342249 | 139.9230779 | 63.6587261 | 27.37181941 | 38.5689649 | 1.84E-08 | -1.952297737 | 3.37E-09 | 34.95585524 | 0.25840435 | 7.734857284 |
| ENSMUSG00000046688 | *Tifa* | 329.2460638 | 373.1254317 | 410.9452558 | 100.9759104 | 102.1179417 | 84.4843993 | 4.76E-26 | -1.952544066 | 2.47E-27 | 117.3016491 | 0.258360233 | 25.32249585 |
| ENSMUSG00000033777 | *Tlr13* | 1268.765152 | 1206.662707 | 1613.527385 | 372.0742784 | 318.9869724 | 361.8136231 | 1.04E-38 | -1.955495457 | 3.10E-40 | 176.3108827 | 0.257832234 | 37.98201921 |
| ENSMUSG00000040133 | *Gpr176* | 32.39720994 | 46.04118437 | 45.38045769 | 3.29269273 | 11.58038514 | 16.52955638 | 0.001470418 | -1.955940626 | 0.00053511 | 11.98915637 | 0.257752687 | 2.832559326 |
| ENSMUSG00000091649 | *Phf11b* | 905.6150313 | 771.1898383 | 656.7560683 | 233.7811838 | 150.5450068 | 216.7208504 | 1.06E-21 | -1.958639199 | 6.91E-23 | 97.00675505 | 0.257271009 | 20.97432423 |
| ENSMUSG00000108578 | *Gm45700* | 32.39720994 | 34.53088828 | 50.42273077 | 6.585385459 | 7.369335996 | 15.6112477 | 0.000629225 | -1.960701059 | 0.000214218 | 13.70207123 | 0.256903587 | 3.201193899 |
| ENSMUSG00000040146 | *Rgl3* | 107.7395586 | 118.9397263 | 133.6202365 | 27.43910608 | 43.16325369 | 22.03940851 | 4.58E-09 | -1.962210724 | 7.94E-10 | 37.77521367 | 0.256634899 | 8.339524802 |
| ENSMUSG00000037031 | *Tspan15* | 26.36982204 | 27.81654889 | 34.03534327 | 8.780513945 | 7.369335996 | 6.428160816 | 0.00057385 | -1.965862516 | 0.000193948 | 13.8888265 | 0.255986118 | 3.241201618 |
| ENSMUSG00000037580 | *Gch1* | 3477.802816 | 3511.5995 | 3907.761635 | 979.0273049 | 820.1018202 | 981.6719875 | 1.71E-76 | -1.96943269 | 1.84E-78 | 351.6656005 | 0.255353424 | 75.76794657 |
| ENSMUSG00000020689 | *Itgb3* | 25.61639856 | 27.81654889 | 17.64795577 | 3.29269273 | 8.422098282 | 6.428160816 | 0.003773624 | -1.971984095 | 0.001498107 | 10.08094243 | 0.254902231 | 2.423241394 |
| ENSMUSG00000025140 | *Pycr1* | 784.3138499 | 775.985795 | 794.1580096 | 225.0006698 | 177.9168262 | 197.4363679 | 2.66E-52 | -1.972566661 | 4.76E-54 | 239.6182378 | 0.254799321 | 51.57517436 |
| ENSMUSG00000041420 | *Meis3* | 417.3966118 | 456.5750784 | 501.7061712 | 128.4150165 | 108.4345154 | 112.9519686 | 3.45E-33 | -1.973950696 | 1.31E-34 | 150.5606122 | 0.254554999 | 32.4619952 |
| ENSMUSG00000079455 | *Gm16026* | 135.6162277 | 119.8989176 | 172.6978529 | 63.6587261 | 17.89695885 | 27.54926064 | 1.83E-05 | -1.976834586 | 4.87E-06 | 20.88808068 | 0.254046662 | 4.736514132 |
| ENSMUSG00000074194 | *Zfp791* | 20.34243415 | 17.26544414 | 22.69022885 | 5.487821216 | 4.211049141 | 5.509852128 | 0.004184088 | -1.981239621 | 0.001677138 | 9.873087848 | 0.253272155 | 2.378399236 |
| ENSMUSG00000113014 | *Selenok-ps8* | 339.0405691 | 235.9610699 | 475.2342375 | 102.0734746 | 80.00993367 | 83.56609061 | 9.88E-12 | -1.981312198 | 1.34E-12 | 50.27570071 | 0.253259414 | 11.00538193 |
| ENSMUSG00000026271 | *Gpr35* | 103.2190177 | 83.44964668 | 76.89466442 | 16.46346365 | 28.4245817 | 22.03940851 | 3.84E-08 | -1.981406787 | 7.26E-09 | 33.46380502 | 0.25324281 | 7.415965371 |
| ENSMUSG00000015947 | *Fcgr1* | 5855.607341 | 5763.780769 | 5864.163589 | 1499.272756 | 1444.389855 | 1480.313605 | 3.56E-219 | -1.982576614 | 3.90E-222 | 1012.250425 | 0.253037548 | 218.4491197 |
| ENSMUSG00000021114 | *Atp6v1d* | 1805.202675 | 1759.15692 | 2395.079712 | 512.5625016 | 507.4314215 | 484.8669873 | 1.62E-42 | -1.984416294 | 4.18E-44 | 194.0349061 | 0.252715088 | 41.79002316 |
| ENSMUSG00000025348 | *Itga7* | 886.7794441 | 914.1093481 | 1019.79973 | 253.5373402 | 216.8690307 | 238.7602589 | 1.08E-55 | -1.990103382 | 1.69E-57 | 255.4435976 | 0.251720849 | 54.96858004 |
| ENSMUSG00000074272 | *Ceacam1* | 659.9989746 | 712.6791665 | 577.3402673 | 184.3907929 | 161.0726296 | 146.0110814 | 2.20E-36 | -1.993256661 | 7.21E-38 | 165.4744713 | 0.251171267 | 35.65733557 |
| ENSMUSG00000040852 | *Plekhh2* | 67.05469034 | 64.26581986 | 71.85239135 | 6.585385459 | 18.94972113 | 24.79433458 | 3.46E-05 | -1.996531392 | 9.59E-06 | 19.59114004 | 0.250601787 | 4.461414739 |
| ENSMUSG00000089929 | *Bcl2a1b* | 1884.312141 | 1608.563879 | 1619.830226 | 441.2208258 | 395.8386192 | 440.7881702 | 5.00E-68 | -2.001303606 | 6.21E-70 | 312.5077847 | 0.249774204 | 67.30123176 |
| ENSMUSG00000007891 | *Ctsd* | 103594.976 | 128521.0886 | 157308.8355 | 31980.82692 | 32839.86672 | 32385.9925 | 4.20E-44 | -2.002187575 | 1.00E-45 | 201.466082 | 0.249621209 | 43.37690463 |
| ENSMUSG00000022372 | *Sla* | 309.6570531 | 315.5739512 | 323.9660452 | 55.9757764 | 91.59031881 | 87.23932536 | 1.34E-20 | -2.009813348 | 9.41E-22 | 91.83835272 | 0.248305247 | 19.87170449 |
| ENSMUSG00000024736 | *Tmem132a* | 446.0267043 | 435.4728689 | 284.8884288 | 105.3661673 | 77.9044091 | 105.6054991 | 2.25E-16 | -2.016333157 | 2.07E-17 | 72.07853225 | 0.247185641 | 15.64705891 |
| ENSMUSG00000015396 | *Cd83* | 195.8901066 | 199.511799 | 200.4303548 | 55.9757764 | 49.4798274 | 42.24219965 | 1.22E-19 | -2.017740485 | 9.08E-21 | 87.35230192 | 0.246944632 | 18.91535409 |
| ENSMUSG00000049858 | *Suox* | 343.56111 | 424.9217641 | 375.6493442 | 84.51244672 | 98.95965481 | 98.25902962 | 6.44E-29 | -2.018624641 | 2.91E-30 | 130.6811374 | 0.246793339 | 28.19120394 |
| ENSMUSG00000022876 | *Samsn1* | 226.7804696 | 240.7570266 | 276.064451 | 45.00013397 | 68.42954854 | 68.8731516 | 1.02E-17 | -2.019568605 | 8.62E-19 | 78.35359529 | 0.246631913 | 16.99041177 |
| ENSMUSG00000050390 | *C77080* | 1124.861266 | 1261.336614 | 1097.954963 | 272.1959323 | 285.2985793 | 301.2052497 | 2.16E-67 | -2.019601245 | 2.74E-69 | 309.545351 | 0.246626333 | 66.66538542 |
| ENSMUSG00000035165 | *Kcne3* | 100.2053238 | 77.69449863 | 80.67636923 | 18.65859213 | 22.10800799 | 22.9577172 | 3.29E-09 | -2.021813323 | 5.63E-10 | 38.44516395 | 0.246248472 | 8.482587087 |
| ENSMUSG00000020903 | *Stx8* | 952.3272875 | 998.5181861 | 1316.033273 | 266.7081111 | 260.0322844 | 275.4926064 | 1.02E-36 | -2.022867254 | 3.28E-38 | 167.0365772 | 0.246068646 | 35.99213849 |
| ENSMUSG00000028494 | *Plin2* | 10560.73702 | 10594.26836 | 10780.37984 | 2695.617781 | 2469.780321 | 2685.134604 | 1.87E-199 | -2.024079676 | 2.31E-202 | 921.293923 | 0.24586194 | 198.728916 |
| ENSMUSG00000022558 | *Mroh1* | 6741.633361 | 7444.283998 | 7049.097762 | 1688.053806 | 1747.585393 | 1776.927311 | 1.04E-164 | -2.025901695 | 2.29E-167 | 760.3220059 | 0.24555163 | 163.9825847 |
| ENSMUSG00000059033 | *Rpl18a-ps1* | 24.10955158 | 16.3062528 | 42.85932115 | 7.682949702 | 4.211049141 | 8.264778192 | 0.004226526 | -2.027760846 | 0.001695019 | 9.853576469 | 0.2452354 | 2.374016454 |
| ENSMUSG00000017724 | *Etv4* | 201.9174945 | 213.8996691 | 200.4303548 | 53.78064792 | 29.47734399 | 67.03653422 | 1.55E-11 | -2.029648531 | 2.12E-12 | 49.37050264 | 0.244914733 | 10.81005838 |
| ENSMUSG00000038515 | *Grtp1* | 144.6573095 | 150.5930406 | 166.3950115 | 35.12205578 | 37.89944227 | 39.48727358 | 2.91E-16 | -2.030726543 | 2.69E-17 | 71.56203192 | 0.244731796 | 15.53661758 |
| ENSMUSG00000051457 | *Spn* | 724.039971 | 678.1482782 | 448.7623038 | 125.1223237 | 164.2309165 | 159.7857117 | 1.39E-19 | -2.04455937 | 1.04E-20 | 87.08228513 | 0.242396476 | 18.85844926 |
| ENSMUSG00000016493 | *Cd46* | 21.84928112 | 23.97978353 | 30.25363846 | 6.585385459 | 5.263811426 | 6.428160816 | 0.001041172 | -2.047328849 | 0.000367969 | 12.68820888 | 0.241931604 | 2.982477339 |
| ENSMUSG00000025466 | *Fuom* | 489.7252665 | 467.1261831 | 626.5024298 | 125.1223237 | 134.7535725 | 122.1350555 | 2.90E-30 | -2.048559977 | 1.23E-31 | 136.9553051 | 0.241725239 | 29.5378037 |
| ENSMUSG00000020788 | *Atp2a3* | 2731.16014 | 2987.881028 | 2391.298007 | 633.2945683 | 664.293002 | 658.4273293 | 5.42E-76 | -2.052476881 | 5.91E-78 | 349.3362439 | 0.241069848 | 75.26619568 |
| ENSMUSG00000049422 | *Chchd10* | 244.8626333 | 327.0842473 | 450.0228721 | 82.31731824 | 73.69335996 | 89.07594274 | 3.36E-14 | -2.053447603 | 3.64E-15 | 61.88328746 | 0.240907698 | 13.47407384 |
| ENSMUSG00000062585 | *Cnr2* | 467.8759854 | 516.0449415 | 505.487876 | 108.6588601 | 115.8038514 | 130.3998337 | 9.20E-42 | -2.064383286 | 2.45E-43 | 190.5146883 | 0.239088512 | 41.03644395 |
| ENSMUSG00000027636 | *Sla2* | 18.08216369 | 20.14301816 | 28.99307019 | 4.390256973 | 2.10552457 | 9.18308688 | 0.00718245 | -2.066448301 | 0.003012568 | 8.799841172 | 0.238746535 | 2.143727367 |
| ENSMUSG00000034993 | *Vat1* | 16328.94723 | 19743.99457 | 20562.38961 | 4411.110693 | 4562.671744 | 4525.425214 | 2.30E-89 | -2.06859969 | 1.70E-91 | 411.5307409 | 0.238390774 | 88.6389796 |
| ENSMUSG00000025068 | *Gsto1* | 3254.789464 | 4115.890045 | 5662.472665 | 1094.27155 | 999.0714086 | 1003.711396 | 4.74E-28 | -2.072637474 | 2.21E-29 | 126.6534337 | 0.237724504 | 27.32416568 |
| ENSMUSG00000034708 | *Grn* | 47571.91239 | 50205.99318 | 41685.7321 | 10746.2515 | 11209.81281 | 11064.70138 | 2.30E-121 | -2.078501211 | 9.14E-124 | 559.8331931 | 0.23676025 | 120.6390037 |
| ENSMUSG00000050957 | *Insl6* | 143.1504625 | 94.00075143 | 171.4372846 | 27.43910608 | 32.63563084 | 35.81403883 | 1.15E-09 | -2.083696781 | 1.87E-10 | 40.59662168 | 0.23590914 | 8.940914564 |
| ENSMUSG00000079222 | *ENSMUSG00000079222* | 64.79441988 | 62.34743717 | 85.71864231 | 29.63423457 | 2.10552457 | 18.36617376 | 0.002736242 | -2.086664154 | 0.001053224 | 10.73158096 | 0.235424414 | 2.562845434 |
| ENSMUSG00000025743 | *Sdc3* | 4197.322246 | 3872.255444 | 2635.848251 | 1029.51526 | 597.968978 | 880.6580318 | 2.61E-20 | -2.094190642 | 1.86E-21 | 90.49088805 | 0.234199412 | 19.58397886 |
| ENSMUSG00000079794 | *ENSMUSG00000079794* | 74.58892521 | 62.34743717 | 44.11988942 | 17.56102789 | 9.474860567 | 15.6112477 | 5.00E-06 | -2.094688204 | 1.22E-06 | 23.54200968 | 0.234118654 | 5.300925065 |
| ENSMUSG00000045932 | *Ifit2* | 737.6015937 | 574.5556133 | 571.037426 | 153.658994 | 118.9621382 | 168.0504899 | 6.40E-28 | -2.095509264 | 3.01E-29 | 126.0395336 | 0.233985452 | 27.19363516 |
| ENSMUSG00000024663 | *Rab3il1* | 2077.188554 | 2228.201485 | 2255.156634 | 541.0991719 | 488.4817003 | 504.1514697 | 3.50E-114 | -2.096540632 | 1.49E-116 | 526.6787027 | 0.233818238 | 113.4553532 |
| ENSMUSG00000110040 | *Gm49369* | 18987.02529 | 24689.58512 | 28613.63914 | 5571.236098 | 5607.011931 | 5703.615261 | 4.94E-48 | -2.098161724 | 1.03E-49 | 219.7354498 | 0.233555654 | 47.30669179 |
| ENSMUSG00000038065 | *Mturn* | 18.83558717 | 11.51029609 | 22.69022885 | 5.487821216 | 4.211049141 | 2.754926064 | 0.008991934 | -2.098812265 | 0.003864705 | 8.34631433 | 0.233450363 | 2.046146908 |
| ENSMUSG00000014905 | *Dnajb9* | 693.149608 | 559.2085519 | 800.460851 | 163.5370722 | 135.8063348 | 178.1518855 | 2.74E-28 | -2.100744043 | 1.27E-29 | 127.7590573 | 0.23313798 | 27.56225856 |
| ENSMUSG00000024486 | *Hbegf* | 38.42459783 | 37.4084623 | 37.81704808 | 8.780513945 | 8.422098282 | 9.18308688 | 2.18E-05 | -2.105510069 | 5.87E-06 | 20.53147037 | 0.232369066 | 4.661654757 |
| ENSMUSG00000030103 | *Bhlhe40* | 728.5605119 | 745.2916721 | 676.9251606 | 159.1468153 | 183.1806376 | 157.9490943 | 1.14E-56 | -2.10646396 | 1.77E-58 | 259.9436277 | 0.232215478 | 55.94378531 |
| ENSMUSG00000028268 | *Gbp3* | 98.69847679 | 63.30662851 | 49.1621625 | 14.26833516 | 12.63314742 | 22.03940851 | 7.82E-06 | -2.10982464 | 1.97E-06 | 22.62657409 | 0.231675174 | 5.10669057 |
| ENSMUSG00000030089 | *Slc41a3* | 371.4377791 | 395.1868325 | 375.6493442 | 82.31731824 | 85.2737451 | 95.50410355 | 9.97E-38 | -2.114640796 | 3.12E-39 | 171.717319 | 0.230903061 | 37.00134803 |
| ENSMUSG00000018849 | *Wwc1* | 91.16424192 | 95.91913411 | 115.9722808 | 26.34154184 | 22.10800799 | 21.12109982 | 3.49E-11 | -2.121546475 | 4.93E-12 | 47.71473713 | 0.22980045 | 10.45668176 |
| ENSMUSG00000028497 | *Hacd4* | 181.5750604 | 225.4099652 | 197.9092183 | 39.51231275 | 49.4798274 | 48.67036046 | 2.00E-19 | -2.131043441 | 1.53E-20 | 86.32431334 | 0.228292688 | 18.69869157 |
| ENSMUSG00000037822 | *Smim14* | 1506.09355 | 1519.359084 | 1527.808742 | 363.2937645 | 294.7734399 | 380.1797968 | 1.83E-71 | -2.131149655 | 2.12E-73 | 328.4216014 | 0.228275882 | 70.73780041 |
| ENSMUSG00000083365 | *Gm12895* | 113.013523 | 362.5743269 | 273.5433144 | 46.09769821 | 56.8491634 | 67.03653422 | 6.53E-07 | -2.134832779 | 1.43E-07 | 27.68009499 | 0.227693849 | 6.185363294 |
| ENSMUSG00000083558 | *Gm12896* | 113.013523 | 362.5743269 | 273.5433144 | 46.09769821 | 56.8491634 | 67.03653422 | 6.53E-07 | -2.134832779 | 1.43E-07 | 27.68009499 | 0.227693849 | 6.185363294 |
| ENSMUSG00000063458 | *Lrmda* | 311.1639001 | 293.5125504 | 355.4802519 | 82.31731824 | 74.74612225 | 61.5266821 | 1.44E-26 | -2.137897021 | 7.28E-28 | 119.7219647 | 0.227210747 | 25.84194421 |
| ENSMUSG00000031266 | *Gla* | 2145.750091 | 2085.281976 | 2610.636886 | 544.3918646 | 463.2154055 | 543.6387433 | 2.63E-60 | -2.13943943 | 3.75E-62 | 276.7946952 | 0.226967962 | 59.58044245 |

| | | | | | | | | | | | | |
|---|---|---|---|---|---|---|---|---|---|---|---|---|
| ENSMUSG00000053040 | *Aph1c* | 449.0403982 | 441.2280169 | 549.6077654 | 94.39052491 | 96.85413024 | 132.2364511 | 1.94E-27 | -2.146196594 | 9.38E-29 | 123.7866889 | 0.225907396 | 26.71121127 |
| ENSMUSG00000048458 | *Inka2* | 109.9998291 | 111.2661956 | 92.02148365 | 28.53667032 | 10.52762285 | 31.22249539 | 1.58E-07 | -2.152517119 | 3.23E-08 | 30.56363216 | 0.224919848 | 6.800201203 |
| ENSMUSG00000031144 | *Syp* | 252.3968681 | 329.00263 | 311.3603625 | 79.02462551 | 60.00745026 | 61.5266821 | 1.20E-22 | -2.154978597 | 7.51E-24 | 101.4006745 | 0.224536424 | 21.92204116 |
| ENSMUSG00000045827 | *Serpinb9* | 1084.929821 | 1081.008641 | 1418.139303 | 285.3667032 | 262.137809 | 255.2898153 | 1.83E-45 | -2.157303671 | 4.16E-47 | 207.793372 | 0.224174849 | 44.73831759 |
| ENSMUSG00000037922 | *Bank1* | 626.8483411 | 583.1883354 | 818.1088067 | 154.7565583 | 155.8088182 | 143.2561553 | 2.41E-33 | -2.158103716 | 9.09E-35 | 151.2822052 | 0.224050567 | 32.6173453 |
| ENSMUSG00000023393 | *Slc17a9* | 696.1633019 | 735.6997586 | 673.1434558 | 143.7809159 | 181.0751131 | 146.9293901 | 1.56E-51 | -2.159229917 | 2.88E-53 | 236.0337761 | 0.223875736 | 50.80668426 |
| ENSMUSG00000041488 | *Stx3* | 1370.477323 | 1468.521943 | 1934.972293 | 368.7815857 | 375.8361358 | 323.2446582 | 1.37E-40 | -2.159738808 | 3.80E-42 | 185.064076 | 0.223796781 | 39.86197622 |
| ENSMUSG00000040904 | *Gm21988* | 119.7943344 | 140.0419358 | 153.7893288 | 29.63423457 | 27.37181941 | 34.89573014 | 1.90E-14 | -2.160038097 | 2.02E-15 | 63.04138602 | 0.223750359 | 13.72142325 |
| ENSMUSG00000052102 | *Gnpda1* | 3056.639087 | 3172.045765 | 2861.489971 | 675.0020095 | 700.0869197 | 652.9174772 | 1.26E-138 | -2.165476331 | 3.73E-141 | 639.7815412 | 0.22290852 | 137.8984978 |
| ENSMUSG00000020261 | *Slc36a1* | 1412.669038 | 1518.399893 | 1213.927243 | 280.9764463 | 330.5673575 | 300.286941 | 1.28E-60 | -2.185680777 | 1.80E-62 | 278.2575439 | 0.219808521 | 59.89292302 |
| ENSMUSG00000028378 | *Ptgr1* | 746.6426756 | 829.7005101 | 1023.581435 | 206.3420777 | 180.0223508 | 181.8251202 | 9.12E-40 | -2.192343411 | 2.59E-41 | 181.2446387 | 0.218795745 | 39.03981092 |
| ENSMUSG00000036534 | *Slc38a7* | 3384.378303 | 3375.394329 | 2919.476112 | 726.587529 | 686.4010099 | 705.2610724 | 8.12E-112 | -2.193152862 | 3.68E-114 | 515.6846492 | 0.21867302 | 111.0906137 |
| ENSMUSG00000027298 | *Tyro3* | 147.6710034 | 163.062528 | 119.7539856 | 39.51231275 | 30.53010627 | 24.79433458 | 3.09E-13 | -2.194795862 | 3.66E-14 | 57.34194821 | 0.218424128 | 12.50998409 |
| ENSMUSG00000033538 | *Casp4* | 659.2455511 | 500.6978801 | 663.0589096 | 143.7809159 | 111.5928022 | 142.3378466 | 1.53E-31 | -2.195284251 | 6.16E-33 | 142.9074729 | 0.218350199 | 30.81565738 |
| ENSMUSG00000060675 | *Plaat3* | 79.86288962 | 87.28641204 | 100.8454615 | 18.65859213 | 25.26629484 | 14.69293901 | 9.83E-10 | -2.195395138 | 1.59E-10 | 40.90899537 | 0.218333417 | 9.007407475 |
| ENSMUSG00000101111 | *Gm28437* | 47611.09041 | 24006.64089 | 86451.03247 | 10800.03215 | 9612.772426 | 14072.16233 | 0.000203609 | -2.196493385 | 6.36E-05 | 15.99096742 | 0.218167274 | 3.691202075 |
| ENSMUSG00000072115 | *Ang* | 53.49306757 | 32.6125056 | 49.1621625 | 8.780513945 | 10.52762285 | 10.10139557 | 1.13E-05 | -2.200621724 | 2.91E-06 | 21.87738155 | 0.217543871 | 4.947097238 |
| ENSMUSG00000026655 | *Fam107b* | 505.5471597 | 555.3717865 | 549.6077654 | 130.6101449 | 113.6983268 | 105.6054991 | 3.42E-47 | -2.204937779 | 7.31E-49 | 215.838412 | 0.216894026 | 46.46646474 |
| ENSMUSG00000027399 | *Il1a* | 15.82189323 | 12.46948743 | 18.90852404 | 2.195128486 | 5.263811426 | 2.754926064 | 0.008857479 | -2.205660487 | 0.003797191 | 8.378339797 | 0.216785401 | 2.05268987 |
| ENSMUSG00000026307 | *Scly* | 2682.187614 | 2967.738009 | 3306.47057 | 647.5629035 | 608.4966008 | 679.5484291 | 2.35E-86 | -2.208425593 | 2.02E-88 | 397.409013 | 0.216370303 | 85.62842637 |
| ENSMUSG00000029470 | *P2rx4* | 821.9850243 | 941.925897 | 947.9473385 | 218.4152844 | 177.9168262 | 190.0898984 | 9.16E-58 | -2.209083411 | 1.40E-59 | 264.9997894 | 0.216271668 | 57.03811331 |
| ENSMUSG00000029419 | *Ajm1* | 40.6848683 | 45.08199303 | 25.21136538 | 5.487821216 | 6.316573711 | 11.93801294 | 0.000192313 | -2.214943196 | 5.99E-05 | 16.1057102 | 0.215395021 | 3.715991403 |
| ENSMUSG00000056185 | *Snx32* | 217.7393877 | 200.4709903 | 325.2266135 | 40.609877 | 66.32402397 | 52.34359522 | 9.55E-14 | -2.216856367 | 1.08E-14 | 59.74141609 | 0.215109573 | 13.02009715 |
| ENSMUSG00000027583 | *Zbtb46* | 82.87658356 | 47.95956706 | 57.98614039 | 10.97564243 | 17.89695885 | 11.93801294 | 1.51E-06 | -2.218319778 | 3.47E-07 | 25.96939394 | 0.214891485 | 5.821225299 |
| ENSMUSG00000025978 | *Rftn2* | 137.8764981 | 117.0213436 | 152.5287606 | 32.9269273 | 23.16077027 | 31.22249539 | 3.54E-14 | -2.218663218 | 3.85E-15 | 61.77608072 | 0.214840335 | 13.45152568 |
| ENSMUSG00000049307 | *Fut4* | 120.5477579 | 125.6540657 | 102.1060298 | 29.63423457 | 24.21353256 | 21.12109982 | 3.02E-13 | -2.226952048 | 3.57E-14 | 57.39008494 | 0.213609535 | 12.52036165 |
| ENSMUSG00000050777 | *Tmem37* | 150.6846974 | 161.1441453 | 210.514901 | 35.12205578 | 40.00496684 | 35.81403883 | 1.63E-16 | -2.22917258 | 1.48E-17 | 72.73402365 | 0.21328101 | 15.78870559 |
| ENSMUSG00000035211 | *Xrra1* | 45.20540922 | 43.16361035 | 35.29591154 | 9.878078189 | 6.316573711 | 10.10139557 | 9.21E-06 | -2.235422883 | 2.34E-06 | 22.29175384 | 0.212358994 | 5.035663226 |
| ENSMUSG00000029923 | *Rab19* | 156.7120853 | 183.2055462 | 228.1628567 | 51.58551943 | 27.37181941 | 41.32389096 | 2.49E-13 | -2.236183351 | 2.93E-14 | 57.78423037 | 0.212247086 | 12.60461118 |
| ENSMUSG00000034245 | *Hdac11* | 13.56162277 | 24.93897487 | 18.90852404 | 3.29269273 | 3.158286856 | 5.509852128 | 0.003961675 | -2.242144978 | 0.001580374 | 9.982471362 | 0.21137183 | 2.402121201 |
| ENSMUSG00000006219 | *Fblim1* | 553.7662629 | 696.3729137 | 666.8406144 | 133.9028377 | 144.2284331 | 126.7265989 | 2.54E-44 | -2.242614605 | 6.02E-46 | 202.4755885 | 0.211303035 | 43.59468067 |
| ENSMUSG00000045136 | *Tubb2b* | 219.9996582 | 168.817676 | 175.2189894 | 41.70744124 | 20.00248342 | 56.93513866 | 8.03E-10 | -2.243658313 | 1.29E-10 | 41.3244753 | 0.211150225 | 9.095109147 |
| ENSMUSG00000032591 | *Mst1* | 686.3687966 | 762.5571162 | 835.7567625 | 182.1956644 | 140.0173839 | 159.7857117 | 4.55E-47 | -2.243660832 | 9.81E-49 | 215.2541518 | 0.211149856 | 46.34178603 |
| ENSMUSG00000079339 | *Ifit1bl1* | 49.72595014 | 33.57169694 | 51.68329904 | 5.487821216 | 13.68590971 | 9.18308688 | 2.72E-05 | -2.246275817 | 7.43E-06 | 20.0788749 | 0.210767479 | 4.565189997 |
| ENSMUSG00000046733 | *Gprc5a* | 103.2190177 | 122.7764917 | 146.2259192 | 21.95128486 | 25.26629484 | 30.3041867 | 9.20E-13 | -2.249899265 | 1.13E-13 | 55.12795948 | 0.210238783 | 12.03627046 |
| ENSMUSG00000004891 | *Nes* | 97.9450533 | 83.44964668 | 54.20443558 | 18.65859213 | 12.63314742 | 18.36617376 | 9.16E-08 | -2.253777407 | 1.81E-08 | 31.68827684 | 0.209674394 | 7.037896191 |
| ENSMUSG00000024164 | *C3* | 374.451473 | 342.4313088 | 288.6701337 | 66.95141883 | 82.11545824 | 62.44499078 | 3.40E-27 | -2.255767897 | 1.67E-28 | 122.639773 | 0.209385305 | 26.46873571 |
| ENSMUSG00000028211 | *Trp53inp1* | 138.6299216 | 111.2661956 | 177.740126 | 25.24397759 | 29.47734399 | 33.97742146 | 1.17E-12 | -2.258993287 | 1.45E-13 | 54.63241979 | 0.208917712 | 11.93111815 |
| ENSMUSG00000002992 | *Apoc2* | 18.08216369 | 11.51029609 | 34.03534327 | 3.29269273 | 3.158286856 | 6.428160816 | 0.006457976 | -2.272506398 | 0.00268388 | 9.010806217 | 0.206970004 | 2.189903602 |
| ENSMUSG00000079419 | *Ms4a6c* | 668.2866329 | 492.065158 | 576.079699 | 124.0247595 | 104.2234662 | 129.481525 | 5.16E-35 | -2.279090475 | 1.79E-36 | 159.0854627 | 0.2060276 | 34.28716417 |
| ENSMUSG00000036815 | *Dpp10* | 27.87666902 | 33.57169694 | 30.25363846 | 12.07320667 | 5.263811426 | 1.836617376 | 0.001474447 | -2.285080738 | 0.00053688 | 11.98300252 | 0.20517392 | 2.83137078 |
| ENSMUSG00000015968 | *Cacna1d* | 83.63000705 | 80.57207265 | 68.07068654 | 23.04884911 | 16.84419656 | 8.264778192 | 1.29E-07 | -2.291247871 | 2.59E-08 | 30.98934883 | 0.204298728 | 6.889742028 |
| ENSMUSG00000022136 | *Dnajc3* | 8082.727168 | 8821.682764 | 11676.64388 | 1948.176532 | 1977.087572 | 1905.490528 | 3.37E-58 | -2.292963999 | 5.02E-60 | 267.0366397 | 0.204055853 | 57.47216687 |
| ENSMUSG00000091401 | *Gm9252* | 20.34243415 | 16.3062528 | 22.69022885 | 2.195128486 | 4.211049141 | 5.509852128 | 0.002263209 | -2.293478414 | 0.000857945 | 11.11149182 | 0.203983107 | 2.645275295 |
| ENSMUSG00000059762 | *Or6c8* | 27.87666902 | 26.85735755 | 40.33818462 | 2.195128486 | 10.52762285 | 6.428160816 | 0.000604498 | -2.297670868 | 0.00020497 | 13.78496436 | 0.203391195 | 3.218605218 |
| ENSMUSG00000071324 | *Armc2* | 207.1914589 | 170.7360587 | 205.4726279 | 35.12205578 | 38.95220455 | 44.07881702 | 3.27E-21 | -2.298603753 | 2.21E-22 | 94.70612012 | 0.20325972 | 20.48492653 |
| ENSMUSG00000095788 | *Sirpb1a* | 152.9449679 | 138.1235531 | 136.1413731 | 36.21962002 | 23.16077027 | 27.54926064 | 3.50E-16 | -2.303651279 | 3.26E-17 | 71.18322289 | 0.202549822 | 15.45548905 |
| ENSMUSG00000111715 | *Or4a73* | 22.60270461 | 13.42867878 | 25.21136538 | 5.487821216 | 4.211049141 | 2.754926064 | 0.002555799 | -2.309396239 | 0.000978507 | 10.86779982 | 0.201744851 | 2.592473226 |
| ENSMUSG00000014686 | *Ceacam16* | 118.2874875 | 142.9195098 | 163.873875 | 28.53667032 | 26.31905713 | 30.3041867 | 1.84E-15 | -2.310887698 | 1.81E-16 | 67.80013738 | 0.201536395 | 14.73489675 |
| ENSMUSG00000020844 | *Nxn* | 174.794249 | 165.940102 | 163.873875 | 30.73179881 | 29.47734399 | 40.40558227 | 6.57E-20 | -2.319694346 | 4.83E-21 | 88.60273791 | 0.200309903 | 19.18225863 |
| ENSMUSG00000016206 | *H2-M3* | 2827.598347 | 2762.471062 | 3121.167035 | 636.587261 | 556.9112489 | 551.9035215 | 9.93E-113 | -2.319782101 | 4.29E-115 | 519.9721457 | 0.200297719 | 112.0032194 |
| ENSMUSG00000064202 | *Spata6l* | 23.3561281 | 16.3062528 | 11.34511442 | 4.390256973 | 4.211049141 | 1.836617376 | 0.006351904 | -2.323646937 | 0.00263468 | 9.044627008 | 0.19976186 | 2.197096092 |
| ENSMUSG00000074489 | *Bglap3* | 24.10955158 | 37.4084623 | 56.72557212 | 6.585385459 | 8.422098282 | 8.264778192 | 0.000109436 | -2.326526703 | 3.27E-05 | 17.25281692 | 0.199363513 | 3.96083909 |
| ENSMUSG00000021003 | *Galc* | 501.7800423 | 544.8206818 | 660.5377731 | 114.1466813 | 121.0676628 | 104.6871904 | 5.36E-40 | -2.326702815 | 1.51E-41 | 182.3222019 | 0.199339177 | 39.27085772 |
| ENSMUSG00000038375 | *Trp53inp2* | 952.3272875 | 1192.274837 | 940.3839289 | 215.1225917 | 211.6052193 | 188.253281 | 8.25E-54 | -2.328828852 | 1.40E-55 | 246.6465672 | 0.199045636 | 53.08364011 |
| ENSMUSG00000029166 | *Mapre3* | 30.89036296 | 35.49007962 | 36.55647981 | 9.878078189 | 3.158286856 | 7.346469504 | 7.65E-05 | -2.334797461 | 2.23E-05 | 17.9780822 | 0.198223861 | 4.116276497 |
| ENSMUSG00000021062 | *Rab15* | 54.24649106 | 72.89854193 | 40.33818462 | 7.682949702 | 10.52762285 | 14.69293901 | 2.43E-06 | -2.339729377 | 5.72E-07 | 25.00532037 | 0.197547381 | 5.614335721 |
| ENSMUSG00000026360 | *Rgs2* | 1967.188724 | 1663.237786 | 1728.239097 | 389.6353063 | 349.5170787 | 314.97988 | 3.10E-77 | -2.348904864 | 3.25E-79 | 355.1205708 | 0.196294974 | 76.50908674 |
| ENSMUSG00000025375 | *Aatk* | 216.9859642 | 226.3691565 | 168.9161481 | 25.24397759 | 46.32154055 | 47.75205178 | 1.77E-15 | -2.354874449 | 1.73E-16 | 67.88540429 | 0.195484422 | 14.75276685 |
| ENSMUSG00000027463 | *Slc52a3* | 58.01360849 | 53.7147151 | 47.90159423 | 5.487821216 | 11.58038514 | 13.77463032 | 7.50E-07 | -2.359363754 | 1.66E-07 | 27.39511896 | 0.194877069 | 6.124757866 |
| ENSMUSG00000024677 | *Ms4a6b* | 1161.025593 | 939.048323 | 1108.039509 | 245.8543905 | 176.8640639 | 199.2729853 | 1.69E-48 | -2.368590852 | 3.43E-50 | 221.9314724 | 0.193634664 | 47.77268415 |
| ENSMUSG00000110206 | *Flt3l* | 255.4105621 | 240.7570266 | 210.514901 | 41.70744124 | 48.42706512 | 46.83374309 | 6.85E-27 | -2.3694366 | 3.42E-28 | 121.2187807 | 0.193521183 | 26.16429258 |
| ENSMUSG00000028965 | *Tnfrsf9* | 63.2875729 | 78.65368997 | 69.33125481 | 18.65859213 | 7.369335996 | 14.69293901 | 3.63E-08 | -2.3750094 | 6.86E-09 | 33.57524823 | 0.192775097 | 7.439902969 |
| ENSMUSG00000035385 | *Ccl2* | 1588.21671 | 1464.685178 | 1603.442838 | 343.5376081 | 257.9267599 | 294.7770888 | 1.06E-76 | -2.378165878 | 1.13E-78 | 352.6353656 | 0.192353785 | 75.97634797 |
| ENSMUSG00000052435 | *Cebpe* | 588.4237433 | 542.9022991 | 568.5162894 | 100.9759104 | 100.0124171 | 124.8899816 | 1.67E-53 | -2.378338532 | 2.87E-55 | 245.2140912 | 0.192330766 | 52.77657022 |
| ENSMUSG00000035042 | *Ccl5* | 350.3419214 | 305.9820378 | 186.5641038 | 46.09769821 | 52.63811426 | 63.36329947 | 3.47E-15 | -2.379502369 | 3.48E-16 | 66.50790265 | 0.192175674 | 14.46012358 |
| ENSMUSG00000038987 | *Cfap157* | 24.10955158 | 33.57169694 | 31.51420673 | 2.195128486 | 6.316573711 | 8.264778192 | 0.000281286 | -2.383292111 | 8.97E-05 | 15.34277378 | 0.19167152 | 3.55085224 |
| ENSMUSG00000058163 | *Gm5431* | 143.1504625 | 162.1033366 | 177.740126 | 34.02449154 | 20.00248342 | 37.65065621 | 1.84E-15 | -2.388102217 | 1.80E-16 | 67.80707662 | 0.19103353 | 14.73612182 |
| ENSMUSG00000036564 | *Ndrg4* | 619.3141063 | 804.7615352 | 748.7775519 | 129.5125807 | 147.3867199 | 136.8279945 | 9.44E-49 | -2.390764267 | 1.89E-50 | 223.115203 | 0.190681361 | 48.02496411 |
| ENSMUSG00000074678 | *Defb25* | 11.3013523 | 24.93897487 | 12.60568269 | 3.29269273 | 4.211049141 | 1.836617376 | 0.007687009 | -2.40164902 | 0.003241605 | 8.666233356 | 0.189248134 | 2.114242593 |
| ENSMUSG00000039691 | *Tspan10* | 18.08216369 | 38.36765364 | 36.55647981 | 2.195128486 | 12.63314742 | 2.754926064 | 0.003702886 | -2.404541106 | 0.001465959 | 10.12091931 | 0.188869139 | 2.431459665 |
| ENSMUSG00000029370 | *Rassf6* | 28.6300925 | 15.34706146 | 25.21136538 | 4.390256973 | 2.10552457 | 6.428160816 | 0.001421093 | -2.405912667 | 0.000515014 | 12.06050124 | 0.188689668 | 2.847377609 |
| ENSMUSG00000073489 | *Ifi204* | 916.9163836 | 755.8427768 | 742.4747106 | 205.2445135 | 105.2762285 | 146.0110814 | 1.02E-24 | -2.406093714 | 5.70E-26 | 111.0746127 | 0.18866599 | 23.98974034 |
| ENSMUSG00000009739 | *Pou6f1* | 113.013523 | 96.87832545 | 74.37352789 | 13.17077092 | 18.94972113 | 21.12109982 | 1.07E-10 | -2.415469585 | 1.58E-11 | 45.43833687 | 0.187443852 | 9.971783479 |
| ENSMUSG00000055177 | *Cstl1* | 16.57531671 | 10.55110475 | 26.47193365 | 1.097564243 | 3.158286856 | 5.509852128 | 0.008071015 | -2.416228512 | 0.003429018 | 8.563858991 | 0.187345274 | 2.093071862 |
| ENSMUSG00000027555 | *Car13* | 1160.27217 | 1139.519313 | 1420.660439 | 246.9519547 | 191.6027359 | 254.3715066 | 2.12E-52 | -2.421985316 | 3.77E-54 | 240.0825728 | 0.186599197 | 51.67307961 |
| ENSMUSG00000019970 | *Sgk1* | 313.4241706 | 338.5945434 | 271.0221779 | 60.36603337 | 55.79640112 | 56.01682997 | 2.24E-33 | -2.426651861 | 8.41E-35 | 151.4373246 | 0.185996598 | 32.65045696 |
| ENSMUSG00000026535 | *Ifi202b* | 1187.395415 | 1008.1101 | 1280.737362 | 261.2202899 | 143.1756708 | 241.5151849 | 4.94E-28 | -2.427259919 | 2.31E-29 | 126.5643773 | 0.185918222 | 27.30595236 |
| ENSMUSG00000019302 | *Atp6v0a1* | 4002.938986 | 4286.626103 | 3378.322962 | 739.7582999 | 697.9813951 | 721.7906288 | 4.27E-100 | -2.434518968 | 2.28E-102 | 461.4854542 | 0.184985107 | 99.36994355 |
| ENSMUSG00000062310 | *Glrp1* | 171.0271315 | 141.0011271 | 146.2259192 | 25.24397759 | 30.53010627 | 28.46756933 | 2.18E-19 | -2.445224899 | 1.67E-20 | 86.14712061 | 0.183617453 | 18.66055867 |
| ENSMUSG00000038204 | *Asb10* | 28.6300925 | 27.81654889 | 15.12681923 | 4.390256973 | 3.158286856 | 5.509852128 | 0.000644565 | -2.457675563 | 0.000219794 | 13.65381352 | 0.182039627 | 3.190733079 |
| ENSMUSG00000039304 | *Tnfsf10* | 57.26018501 | 37.4084623 | 74.37352789 | 12.07320667 | 7.369335996 | 11.01970426 | 1.26E-06 | -2.465267145 | 2.87E-07 | 26.33398533 | 0.181084235 | 5.89827016 |
| ENSMUSG00000021703 | *Serinc5* | 49.72595014 | 59.46986315 | 63.02841346 | 15.3658994 | 11.58038514 | 4.59154344 | 1.08E-06 | -2.467632622 | 2.45E-07 | 26.64187029 | 0.180787568 | 5.964725204 |
| ENSMUSG00000089844 | *A530032D15Rik* | 186.8490248 | 127.5724484 | 160.0921702 | 27.43910608 | 34.74115541 | 23.87602589 | 5.23E-16 | -2.470029533 | 4.91E-17 | 70.37421726 | 0.180487455 | 15.28154916 |
| ENSMUSG00000038022 | *Mindy4* | 50.47937363 | 58.51067181 | 79.41580096 | 13.17077092 | 11.58038514 | 9.18308688 | 3.48E-08 | -2.471779676 | 6.57E-09 | 33.65906755 | 0.180268637 | 7.457827221 |
| ENSMUSG00000040350 | *Trim7* | 24.10955158 | 23.97978353 | 56.72557212 | 9.878078189 | 6.316573711 | 2.754926064 | 0.000772619 | -2.474369118 | 0.000266536 | 13.29203801 | 0.179945369 | 3.112034507 |
| ENSMUSG00000018927 | *Ccl6* | 65.54784336 | 78.65368997 | 93.28205192 | 24.14641335 | 10.52762285 | 8.264778192 | 1.99E-07 | -2.48004184 | 4.12E-08 | 30.09318356 | 0.179239208 | 6.700537949 |
| ENSMUSG00000074342 | *I830077J02Rik* | 412.8760708 | 320.3699079 | 363.0436615 | 63.6587261 | 47.37430283 | 84.4843993 | 2.09E-24 | -2.481695457 | 1.18E-25 | 109.6242094 | 0.179033882 | 23.67941772 |
| ENSMUSG00000028599 | *Tnfrsf1b* | 1264.998035 | 1366.847661 | 1139.553715 | 239.269005 | 215.8162685 | 220.3940851 | 3.68E-87 | -2.483322872 | 2.96E-89 | 401.2400614 | 0.178832038 | 86.43367678 |
| ENSMUSG00000048752 | *Prss50* | 125.8217223 | 121.8173003 | 152.5287606 | 26.34154184 | 22.10800799 | 22.9577172 | 2.62E-17 | -2.484310848 | 2.26E-18 | 76.44985111 | 0.178709614 | 16.58125306 |
| ENSMUSG00000030157 | *Clec2d* | 48.97252665 | 30.69412292 | 50.42273077 | 5.487821216 | 7.369335996 | 10.10139557 | 5.36E-06 | -2.486786031 | 1.32E-06 | 23.39769047 | 0.178403271 | 5.270892857 |
| ENSMUSG00000033276 | *Stk36* | 32.39720994 | 19.18382682 | 28.99307019 | 5.487821216 | 6.316573711 | 2.754926064 | 0.00028724 | -2.487396444 | 9.17E-05 | 15.29954211 | 0.178327803 | 3.541754418 |
| ENSMUSG00000029101 | *Rgs12* | 71.57523126 | 86.3272207 | 52.94386731 | 9.878078189 | 16.84419656 | 11.01970426 | 1.59E-08 | -2.489334602 | 2.90E-09 | 35.24769824 | 0.178088393 | 7.798151338 |
| ENSMUSG00000020099 | *Unc5b* | 282.5338076 | 269.5327669 | 190.3458087 | 40.609877 | 41.05772912 | 50.50697784 | 1.82E-21 | -2.490350086 | 1.21E-22 | 95.90373375 | 0.177963084 | 20.74098113 |
| ENSMUSG00000031883 | *Car7* | 42.19171527 | 38.36765364 | 45.38045769 | 6.585385459 | 8.422098282 | 7.346469504 | 7.03E-07 | -2.491076483 | 1.55E-07 | 27.52807414 | 0.177873503 | 6.153128893 |
| ENSMUSG00000074896 | *Ifit3* | 656.9852806 | 560.1677432 | 553.3894702 | 138.2930946 | 67.37678625 | 109.2787339 | 2.28E-24 | -2.492902944 | 1.30E-25 | 109.4456563 | 0.177648456 | 23.641874 |
| ENSMUSG00000053687 | *Dpep2* | 2062.120084 | 2211.895233 | 2434.157328 | 440.1232615 | 360.0447015 | 391.1995011 | 1.15E-96 | -2.492986534 | 6.84E-99 | 445.5115857 | 0.177638164 | 95.94107059 |
| ENSMUSG00000038352 | *Arl5c* | 858.9027751 | 919.8644961 | 1110.560645 | 195.3664353 | 150.5450068 | 165.2955638 | 4.10E-52 | -2.497994824 | 7.45E-54 | 238.7275476 | 0.177022565 | 51.38756675 |
| ENSMUSG00000076435 | *Acsf2* | 246.3694802 | 246.5121747 | 252.1136538 | 30.73179881 | 62.11297483 | 38.5689649 | 1.75E-20 | -2.503511796 | 1.23E-21 | 91.30217805 | 0.17634691 | 19.75742348 |
| ENSMUSG00000035561 | *Aldh1b1* | 298.3557008 | 298.3085071 | 305.0575212 | 60.36603337 | 46.32154055 | 52.34359522 | 7.85E-38 | -2.504856305 | 2.43E-39 | 172.2149046 | 0.176182642 | 37.10523029 |
| ENSMUSG00000031387 | *Renbp* | 3423.556325 | 3954.745899 | 4986.808073 | 737.5631714 | 712.7200671 | 724.5455548 | 1.59E-64 | -2.506480548 | 2.13E-66 | 296.2795882 | 0.1759844 | 63.79817503 |
| ENSMUSG00000095366 | *Gm21860* | 49.72595014 | 27.81654889 | 23.95079712 | 6.585385459 | 1.052762285 | 10.10139557 | 0.001141402 | -2.50806 | 0.000406133 | 12.50376355 | 0.175791839 | 2.942561275 |
| ENSMUSG00000026981 | *Il1rn* | 1455.614177 | 1205.703516 | 1207.624402 | 255.7324687 | 230.5549405 | 194.6814419 | 4.77E-63 | -2.51078946 | 6.51E-65 | 289.4617427 | 0.17545957 | 62.32149584 |
| ENSMUSG00000059900 | *Tmem40* | 27.87666902 | 29.73493157 | 23.95079712 | 5.487821216 | 5.263811426 | 3.673234752 | 7.36E-05 | -2.514474163 | 2.15E-05 | 18.05484716 | 0.17501201 | 4.13297187 |
| ENSMUSG00000017309 | *Cd300lg* | 42.94513876 | 57.55148047 | 32.774775 | 4.390256973 | 10.52762285 | 8.264778192 | 6.27E-06 | -2.521550018 | 1.56E-06 | 23.07464111 | 0.174155746 | 5.202891194 |
| ENSMUSG00000026864 | *Hspa5* | 136304.8567 | 131964.5855 | 155633.5402 | 25646.78367 | 24126.15329 | 23831.02876 | 6.54E-178 | -2.525875796 | 1.30E-180 | 821.2394045 | 0.17363434 | 177.1845717 |
| ENSMUSG00000028238 | *Atp6v0d2* | 52.73964409 | 42.20441901 | 65.54955 | 6.585385459 | 15.79143428 | 5.509852128 | 3.72E-06 | -2.530334758 | 8.94E-07 | 24.14330386 | 0.173098514 | 5.429578352 |
| ENSMUSG00000035498 | *Cdcp1* | 59.52045547 | 93.04156009 | 134.8808048 | 25.24397759 | 10.52762285 | 13.77463032 | 3.21E-07 | -2.535943273 | 6.78E-08 | 29.12565187 | 0.172426895 | 6.493168099 |
| ENSMUSG00000025823 | *Pdia4* | 23449.55261 | 27394.5047 | 33817.26496 | 5041.112569 | 4778.488012 | 4763.267165 | 3.92E-77 | -2.537326394 | 4.17E-79 | 354.6256653 | 0.172261667 | 76.40695848 |
| ENSMUSG00000036820 | *Amdhd2* | 3632.254631 | 3597.926721 | 3900.198225 | 688.1727805 | 554.8057243 | 673.1202683 | 4.67E-122 | -2.537534407 | 1.83E-124 | 563.046562 | 0.172236832 | 121.3304632 |
| ENSMUSG00000030142 | *Clec4e* | 8138.480506 | 8028.431525 | 11251.83237 | 1655.126879 | 1525.452551 | 1538.167052 | 8.51E-68 | -2.538436582 | 1.06E-69 | 311.4357086 | 0.172129159 | 67.0700847 |
| ENSMUSG00000050157 | *Gm867* | 36.16432737 | 64.26581986 | 112.190576 | 23.04884911 | 5.263811426 | 8.264778192 | 0.000203246 | -2.542058489 | 6.35E-05 | 15.99476476 | 0.171697568 | 3.691977829 |
| ENSMUSG00000048058 | *Ldlrad3* | 97.9450533 | 86.3272207 | 71.85239135 | 17.56102789 | 23.16077027 | 3.673234752 | 4.96E-06 | -2.548035563 | 1.21E-06 | 23.55876758 | 0.170987698 | 5.304585569 |
| ENSMUSG00000030344 | *Akap3* | 67.05469034 | 70.0209679 | 45.38045769 | 7.682949702 | 10.52762285 | 12.85632163 | 3.11E-08 | -2.550303668 | 5.80E-09 | 33.89915101 | 0.170719094 | 7.507930436 |
| ENSMUSG00000024253 | *Dync2li1* | 42.94513876 | 29.73493157 | 51.68329904 | 7.682949702 | 2.10552457 | 11.01970426 | 6.32E-05 | -2.558854373 | 1.82E-05 | 18.37080706 | 0.169710252 | 4.199358129 |
| ENSMUSG00000055561 | *Spink5* | 2421.503087 | 2406.611075 | 3229.575906 | 497.1966022 | 442.1601598 | 427.0135399 | 2.51E-65 | -2.559802294 | 3.30E-67 | 299.995686 | 0.169598781 | 64.60107082 |
| ENSMUSG00000026536 | *Ifi211* | 229.04074 | 257.0632794 | 268.5010413 | 45.00013397 | 26.31905713 | 55.09852128 | 1.49E-20 | -2.568286648 | 1.04E-21 | 91.6325182 | 0.168604314 | 19.82780641 |
| ENSMUSG00000059089 | *Fcgr4* | 292.3283129 | 283.920637 | 389.5155952 | 71.34167581 | 47.37430283 | 44.07881702 | 3.92E-23 | -2.572015396 | 2.40E-24 | 103.6624088 | 0.168169106 | 22.40632816 |
| ENSMUSG00000025993 | *Slc40a1* | 2516.434446 | 2863.186153 | 2751.820532 | 492.8063452 | 426.3687255 | 448.1346397 | 1.89E-135 | -2.572375261 | 5.83E-138 | 625.0955628 | 0.168127164 | 134.7241953 |
| ENSMUSG00000066800 | *Rnasel* | 239.5886688 | 225.4099652 | 244.5502442 | 42.80500548 | 38.95220455 | 37.65065621 | 7.26E-33 | -2.573331544 | 2.79E-34 | 149.0543476 | 0.168015758 | 32.13900781 |
| ENSMUSG00000082292 | *Gm12250* | 71.57523126 | 61.38824583 | 66.81011827 | 10.97564243 | 11.58038514 | 11.01970426 | 1.28E-10 | -2.574762536 | 1.91E-11 | 45.06323683 | 0.167849188 | 9.892022349 |
| ENSMUSG00000036587 | *Fut7* | 278.7666902 | 333.7985867 | 334.0505913 | 60.36603337 | 55.79640112 | 43.16050834 | 8.46E-33 | -2.575392177 | 3.27E-34 | 148.7376947 | 0.167775949 | 32.07256603 |
| ENSMUSG00000044770 | *Scml4* | 15.06846974 | 18.22463548 | 16.3873875 | 3.29269273 | 2.10552457 | 2.754926064 | 0.001585947 | -2.607591296 | 0.000582377 | 11.83147879 | 0.164072881 | 2.799711391 |
| ENSMUSG00000023367 | *Tmem176a* | 175.5476725 | 216.7772431 | 175.2189894 | 21.95128486 | 45.26877826 | 25.71264326 | 9.96E-17 | -2.612965695 | 8.94E-18 | 73.7345478 | 0.163462805 | 16.00194012 |
| ENSMUSG00000080917 | *Pira2-ps1* | 23.3561281 | 10.55110475 | 15.12681923 | 2.195128486 | 2.10552457 | 3.673234752 | 0.003538595 | -2.613779769 | 0.001395089 | 10.2122766 | 0.163370594 | 2.451169106 |
| ENSMUSG00000079173 | *Zan* | 30.13693948 | 28.77574023 | 22.69022885 | 8.780513945 | 1.052762285 | 3.673234752 | 0.000570633 | -2.615584047 | 0.000192469 | 13.90321362 | 0.163166405 | 3.243643336 |
| ENSMUSG00000025357 | *Dgka* | 183.8353308 | 192.7974596 | 213.0360375 | 25.24397759 | 30.53010627 | 39.48727358 | 3.87E-24 | -2.616423407 | 2.23E-25 | 108.3681206 | 0.163071503 | 23.41254502 |
| ENSMUSG00000040723 | *Rcsd1* | 1448.079942 | 1445.501351 | 1521.505901 | 268.9032396 | 203.183121 | 245.1884197 | 4.03E-98 | -2.62164653 | 2.27E-100 | 452.3109955 | 0.162482187 | 97.39511284 |
| ENSMUSG00000022180 | *Slc7a8* | 1338.080113 | 1251.7447 | 864.7498327 | 153.658994 | 245.2936124 | 158.867403 | 1.44E-28 | -2.632840952 | 6.60E-30 | 129.054644 | 0.161226304 | 27.84251595 |
| ENSMUSG00000023990 | *Tfeb* | 823.4918712 | 853.6802936 | 626.5024298 | 119.6345025 | 114.7510891 | 136.8279945 | 1.35E-48 | -2.633551647 | 2.71E-50 | 222.3963013 | 0.161146901 | 47.86964672 |
| ENSMUSG00000089809 | *Rasgef1b* | 1002.053238 | 968.7832545 | 985.7643865 | 147.0736086 | 144.2284331 | 182.7434289 | 5.05E-88 | -2.635998321 | 3.92E-90 | 405.2734787 | 0.160873843 | 87.29657611 |
| ENSMUSG00000074417 | *Pira12* | 49.72595014 | 77.69449863 | 63.02841346 | 8.780513945 | 12.63314742 | 9.18308688 | 6.33E-09 | -2.636543212 | 1.11E-09 | 37.11813194 | 0.160813094 | 8.198844078 |
| ENSMUSG00000047798 | *Cd300lf* | 1605.545451 | 1765.871259 | 1752.189894 | 257.9275971 | 296.8789644 | 266.3095195 | 5.68E-130 | -2.640853555 | 1.95E-132 | 599.6952286 | 0.16033335 | 129.2453616 |
| ENSMUSG00000053746 | *Ptrh1* | 961.3683693 | 1036.88584 | 1448.392941 | 192.0737426 | 167.3892033 | 191.0082071 | 1.31E-43 | -2.643861822 | 3.20E-45 | 199.1495087 | 0.159999375 | 42.88250077 |
| ENSMUSG00000039208 | *Metrnl* | 3287.940097 | 3263.168942 | 3059.399189 | 601.4652053 | 429.5270124 | 506.9063958 | 7.62E-91 | -2.644656155 | 5.44E-93 | 418.400764 | 0.159911306 | 90.11797373 |
| ENSMUSG00000032715 | *Trib3* | 1567.120853 | 1480.991431 | 1285.779635 | 268.9032396 | 178.9695885 | 238.7602589 | 7.29E-57 | -2.65927318 | 1.13E-58 | 260.8409509 | 0.158299304 | 56.13745331 |
| ENSMUSG00000038305 | *Spats2l* | 45.20540922 | 20.14301816 | 42.85932115 | 7.682949702 | 0 | 9.18308688 | 0.001919608 | -2.669593505 | 0.000718074 | 11.44187118 | 0.15717095 | 2.716787558 |
| ENSMUSG00000038527 | *C1rl* | 15.82189323 | 15.34706146 | 13.86625096 | 2.195128486 | 1.052762285 | 3.673234752 | 0.002935156 | -2.680589735 | 0.001137041 | 10.58994971 | 0.155977546 | 2.532368773 |
| ENSMUSG00000035085 | *1700020L24Rik* | 14.31504625 | 18.22463548 | 6.302841346 | 2.195128486 | 2.10552457 | 1.836617376 | 0.008730578 | -2.681268192 | 0.003736631 | 8.407563797 | 0.155904212 | 2.058957005 |
| ENSMUSG00000059430 | *Actg2* | 15.82189323 | 16.3062528 | 20.16909231 | 3.29269273 | 1.052762285 | 3.673234752 | 0.001446256 | -2.686881105 | 0.000525027 | 12.02460982 | 0.155298833 | 2.839754969 |
| ENSMUSG00000002602 | *Axl* | 295.3420069 | 272.4103409 | 148.7470558 | 46.09769821 | 33.68839313 | 32.14080408 | 4.54E-15 | -2.68792353 | 4.62E-16 | 65.95318126 | 0.155186662 | 14.34261723 |
| ENSMUSG00000020641 | *Rsad2* | 1700.47681 | 1586.502478 | 1274.43452 | 280.9764463 | 178.9695885 | 247.9433458 | 2.62E-45 | -2.689097376 | 6.01E-47 | 207.0609188 | 0.155060446 | 44.58241456 |
| ENSMUSG00000038578 | *Susd1* | 20.34243415 | 16.3062528 | 21.42966058 | 1.097564243 | 1.052762285 | 6.428160816 | 0.00279881 | -2.708784183 | 0.001079421 | 10.6861246 | 0.152958885 | 2.553026533 |
| ENSMUSG00000043157 | *Arl11* | 1403.627956 | 1609.52307 | 2116.494124 | 286.4642675 | 218.9745553 | 275.4926064 | 6.14E-47 | -2.713855106 | 1.34E-48 | 214.6337729 | 0.152422194 | 46.21194889 |
| ENSMUSG00000095440 | *Fignl2* | 19.58901066 | 37.4084623 | 23.95079712 | 4.390256973 | 3.158286856 | 4.59154344 | 0.000108162 | -2.730835989 | 3.23E-05 | 17.27769629 | 0.150638664 | 3.965927288 |
| ENSMUSG00000047793 | *Sned1* | 21.09585763 | 23.02059219 | 31.51420673 | 4.390256973 | 5.263811426 | 1.836617376 | 0.000154749 | -2.731625622 | 4.74E-05 | 16.54870211 | 0.150556237 | 3.810373463 |
| ENSMUSG00000074577 | *Ripor3* | 48.97252665 | 53.7147151 | 37.81704808 | 7.682949702 | 3.158286856 | 10.10139557 | 6.58E-07 | -2.735883603 | 1.45E-07 | 27.66118225 | 0.150112539 | 6.181524675 |
| ENSMUSG00000039236 | *Isg20* | 1253.696682 | 1093.478129 | 1010.975752 | 217.3177201 | 127.3842365 | 159.7857117 | 5.64E-43 | -2.73780784 | 1.42E-44 | 196.1807288 | 0.149912455 | 42.24900698 |
| ENSMUSG00000081723 | *Gm15931* | 43.69856224 | 60.42905449 | 30.25363846 | 8.780513945 | 1.052762285 | 10.10139557 | 7.64E-05 | -2.745614775 | 2.23E-05 | 17.98162599 | 0.149103417 | 4.116982964 |
| ENSMUSG00000048388 | *Fam171b* | 244.8626333 | 189.9198855 | 216.8177423 | 23.04884911 | 37.89944227 | 35.81403883 | 1.79E-24 | -2.746705729 | 1.01E-25 | 109.9453105 | 0.148990709 | 23.74765718 |
| ENSMUSG00000074785 | *Plxnc1* | 3584.035527 | 4157.135272 | 3756.493442 | 567.4407137 | 570.5971586 | 573.0246213 | 3.54E-183 | -2.748085277 | 6.80E-186 | 845.532502 | 0.148848308 | 182.450826 |
| ENSMUSG00000100846 | *Gm28845* | 9.79450533 | 33.57169694 | 37.81704808 | 0 | 6.316573711 | 5.509852128 | 0.00582594 | -2.752034755 | 0.002402021 | 9.213765694 | 0.148441382 | 2.234633957 |
| ENSMUSG00000054364 | *Rhob* | 395.5473306 | 440.2688256 | 446.2411673 | 50.48795519 | 58.95468797 | 78.97454717 | 3.93E-42 | -2.75539097 | 1.04E-43 | 192.2299373 | 0.148096457 | 41.40521362 |
| ENSMUSG00000029596 | *Sdsl* | 67.05469034 | 57.55148047 | 94.54262019 | 7.682949702 | 10.52762285 | 13.77463032 | 1.05E-09 | -2.756114574 | 1.70E-10 | 40.77997501 | 0.148022196 | 8.979836087 |
| ENSMUSG00000069792 | *Wfdc17* | 297.6022773 | 343.3905001 | 513.0512856 | 77.92706126 | 36.84667998 | 52.34359522 | 1.05E-17 | -2.787606943 | 8.86E-19 | 78.29861373 | 0.144826053 | 16.97938359 |
| ENSMUSG00000023272 | *Creld2* | 22061.74654 | 24460.33839 | 29040.97179 | 3665.864572 | 3572.022434 | 3688.846 | 7.50E-131 | -2.789626163 | 2.47E-133 | 603.8203745 | 0.144623494 | 130.1248805 |
| ENSMUSG00000043953 | *Ccrl2* | 151.4381209 | 137.1643618 | 150.007624 | 19.75615638 | 23.16077027 | 20.20279114 | 2.20E-23 | -2.797687969 | 1.32E-24 | 104.8453204 | 0.143817589 | 22.65774324 |
| ENSMUSG00000034459 | *Ifit1* | 1319.997949 | 1127.049826 | 1409.315325 | 223.9031056 | 127.3842365 | 201.1096027 | 6.89E-42 | -2.80283453 | 1.82E-43 | 191.1100663 | 0.143305459 | 41.1619104 |
| ENSMUSG00000109032 | *Gm7972* | 15.06846974 | 12.46948743 | 23.95079712 | 4.390256973 | 3.158286856 | 0 | 0.005519837 | -2.803942018 | 0.002263315 | 9.322705822 | 0.143195492 | 2.258073709 |
| ENSMUSG00000026471 | *Mr1* | 131.0956867 | 133.3275964 | 220.5994471 | 15.3658994 | 31.58286856 | 22.03940851 | 1.85E-13 | -2.806119929 | 2.15E-14 | 58.39351941 | 0.142979485 | 12.73274444 |
| ENSMUSG00000041754 | *Trem3* | 929.7245829 | 941.925897 | 1087.870416 | 142.6833516 | 126.3314742 | 152.4392422 | 4.65E-87 | -2.80773569 | 3.83E-89 | 400.7258419 | 0.142819444 | 86.33273374 |
| ENSMUSG00000098470 | *C1rb* | 60.27387896 | 14.38787012 | 18.90852404 | 7.682949702 | 2.10552457 | 3.673234752 | 0.001799582 | -2.820194909 | 0.000669101 | 11.57317498 | 0.141591355 | 2.74482833 |

| | | | | | | | | | | | | | |
|---|---|---|---|---|---|---|---|---|---|---|---|---|---|
| ENSMUSG00000022126 | *Acod1* | 2144.243244 | 2156.262135 | 1679.076935 | 291.9520887 | 218.9745553 | 334.2643624 | 2.22E-55 | -2.821656035 | 3.57E-57 | 253.9526102 | 0.141448028 | 54.65293524 |
| ENSMUSG00000044645 | *Gm7334* | 52.73964409 | 81.531264 | 70.59182308 | 7.682949702 | 16.84419656 | 4.59154344 | 1.15E-07 | -2.825837975 | 2.31E-08 | 31.21339902 | 0.141038606 | 6.937495182 |
| ENSMUSG00000097328 | *Tnfsf12* | 861.9164691 | 1076.212685 | 1119.384623 | 125.1223237 | 147.3867199 | 153.3575509 | 4.11E-67 | -2.839296919 | 5.24E-69 | 308.253912 | 0.139728971 | 66.38639215 |
| ENSMUSG00000004044 | *Cavin1* | 12.80819928 | 14.38787012 | 15.12681923 | 0 | 1.052762285 | 4.59154344 | 0.008620105 | -2.841465257 | 0.003681842 | 8.434420453 | 0.139519119 | 2.064487457 |
| ENSMUSG00000062980 | *Cped1* | 270.4790318 | 249.3897487 | 334.0505913 | 41.70744124 | 33.68839313 | 43.16050834 | 9.63E-33 | -2.842325487 | 3.74E-34 | 148.4750107 | 0.139435954 | 32.01623217 |
| ENSMUSG00000024063 | *Lbh* | 18.08216369 | 19.18382682 | 13.86625096 | 3.29269273 | 2.10552457 | 1.836617376 | 0.00082482 | -2.842574591 | 0.000286015 | 13.15982748 | 0.13941188 | 3.083640996 |
| ENSMUSG00000029275 | *Gfi1* | 15.82189323 | 18.22463548 | 17.64795577 | 1.097564243 | 5.263811426 | 0.918308688 | 0.002216343 | -2.843929572 | 0.000839114 | 11.15265803 | 0.139281005 | 2.654363081 |
| ENSMUSG00000027514 | *Zbp1* | 1072.121622 | 844.0883802 | 620.1995885 | 148.1711728 | 81.06269596 | 123.9716729 | 1.02E-25 | -2.846389091 | 5.40E-27 | 115.7459246 | 0.13904376 | 24.99149346 |
| ENSMUSG00000090231 | *Cfb* | 220.7530817 | 230.2059219 | 216.8177423 | 31.82936305 | 30.53010627 | 30.3041867 | 6.68E-36 | -2.850969085 | 2.23E-37 | 163.2310977 | 0.138603051 | 35.17538205 |
| ENSMUSG00000026628 | *Atf3* | 1693.695999 | 1778.340746 | 1708.070005 | 232.6836196 | 238.9770387 | 245.1884197 | 5.54E-186 | -2.85208786 | 1.03E-188 | 858.5122746 | 0.138495609 | 185.256854 |
| ENSMUSG00000027698 | *Nceh1* | 4076.021064 | 4434.34157 | 5614.571071 | 643.1726465 | 707.4562556 | 604.2471167 | 2.15E-86 | -2.852889001 | 1.83E-88 | 397.6024503 | 0.138418722 | 85.66704756 |
| ENSMUSG00000028716 | *Pdzk1ip1* | 16.57531671 | 17.26544414 | 25.21136538 | 2.195128486 | 2.10552457 | 3.673234752 | 0.000401026 | -2.859825656 | 0.000131244 | 14.62382967 | 0.137754786 | 3.396827207 |
| ENSMUSG00000021298 | *Gpr132* | 365.4103912 | 459.4526524 | 393.2973 | 37.31718427 | 66.32402397 | 63.36329947 | 2.50E-33 | -2.859996491 | 9.42E-35 | 151.2122378 | 0.137738474 | 32.60284324 |
| ENSMUSG00000090675 | *Or5v1b* | 31.64378645 | 11.51029609 | 16.3873875 | 5.487821216 | 1.052762285 | 1.836617376 | 0.002723092 | -2.862009089 | 0.001047415 | 10.74181606 | 0.13754646 | 2.564937688 |
| ENSMUSG00000048120 | *Entpd1* | 11.3013523 | 24.93897487 | 34.03534327 | 7.682949702 | 2.10552457 | 0 | 0.007756994 | -2.871985497 | 0.003275376 | 8.647348741 | 0.136598591 | 2.110306557 |
| ENSMUSG00000021876 | *Rnase4* | 373.6980495 | 327.0842473 | 419.7692337 | 71.34167581 | 46.32154055 | 35.81403883 | 1.51E-27 | -2.877047052 | 7.23E-29 | 124.3034427 | 0.136120187 | 26.82059095 |
| ENSMUSG00000043740 | *B430306N03Rik* | 511.5745476 | 496.8611147 | 533.2203779 | 69.14654732 | 65.27126168 | 74.38300373 | 3.00E-73 | -2.880624644 | 3.41E-75 | 336.6555131 | 0.135783055 | 72.52333814 |
| ENSMUSG00000032470 | *Mras* | 91.16424192 | 140.0419358 | 177.740126 | 15.3658994 | 16.84419656 | 22.03940851 | 1.30E-13 | -2.898977897 | 1.49E-14 | 59.11622965 | 0.134066631 | 12.88605428 |
| ENSMUSG00000025804 | *Ccr1* | 65.54784336 | 79.61288131 | 89.50034712 | 8.780513945 | 8.422098282 | 13.77463032 | 3.72E-12 | -2.900443947 | 4.82E-13 | 52.27771039 | 0.133930463 | 11.42983487 |
| ENSMUSG00000015340 | *Cybb* | 11093.40742 | 9588.076646 | 9391.233606 | 1420.248131 | 1251.734357 | 1355.423623 | 2.52E-177 | -2.900913011 | 5.18E-180 | 818.4796892 | 0.133886926 | 176.5993013 |
| ENSMUSG00000000184 | *Ccnd2* | 1938.558632 | 1864.667967 | 1447.132373 | 260.1227256 | 201.0775965 | 241.5151849 | 4.76E-71 | -2.902815406 | 5.58E-73 | 326.4925565 | 0.133710493 | 70.32275152 |
| ENSMUSG00000074121 | *Ntf5* | 26.36982204 | 18.22463548 | 15.12681923 | 3.29269273 | 0 | 4.59154344 | 0.001597288 | -2.906329082 | 0.000587529 | 11.81508173 | 0.133385238 | 2.796616665 |
| ENSMUSG00000044703 | *Phf11a* | 358.6295798 | 214.8588604 | 247.0713808 | 31.82936305 | 44.21601598 | 33.05911277 | 2.00E-22 | -2.915798758 | 1.27E-23 | 100.3669443 | 0.132512581 | 21.69822504 |
| ENSMUSG00000004665 | *Cnn2* | 212.4654233 | 232.1243046 | 205.4726279 | 35.12205578 | 29.47734399 | 22.03940851 | 1.42E-30 | -2.919865733 | 5.97E-32 | 138.3966229 | 0.132139552 | 29.84667283 |
| ENSMUSG00000054545 | *Ugt1a6a* | 686.3687966 | 845.0475715 | 1138.293147 | 121.829631 | 96.85413024 | 132.2364511 | 4.91E-37 | -2.923293682 | 1.55E-38 | 168.525687 | 0.131825952 | 36.30900258 |
| ENSMUSG00000054404 | *Slfn5* | 1161.779017 | 1232.560873 | 1305.948727 | 194.268871 | 137.9118594 | 155.1941683 | 3.05E-88 | -2.926071255 | 2.34E-90 | 406.300069 | 0.131572396 | 87.51618362 |
| ENSMUSG00000031101 | *Sash3* | 134.1093807 | 147.7154665 | 133.6202365 | 25.24397759 | 20.00248342 | 10.10139557 | 3.02E-17 | -2.927984873 | 2.62E-18 | 76.15874615 | 0.131397991 | 16.51930376 |
| ENSMUSG00000017400 | *Stac2* | 178.5613664 | 195.6750336 | 201.6909231 | 27.43910608 | 27.37181941 | 21.12109982 | 5.31E-30 | -2.928449915 | 2.29E-31 | 135.7262988 | 0.131355643 | 29.27456662 |
| ENSMUSG00000026832 | *Cytip* | 64.04099639 | 64.26581986 | 44.11988942 | 7.682949702 | 12.63314742 | 2.754926064 | 2.15E-07 | -2.928580568 | 4.46E-08 | 29.94019748 | 0.131343748 | 6.66756324 |
| ENSMUSG00000039431 | *Mtmr7* | 35.41090389 | 53.7147151 | 54.20443558 | 9.878078189 | 6.316573711 | 2.754926064 | 5.18E-07 | -2.939902916 | 1.12E-07 | 28.14973435 | 0.130316989 | 6.285441102 |
| ENSMUSG00000069633 | *Pex11g* | 79.86288962 | 100.7150908 | 144.965351 | 20.85372062 | 11.58038514 | 10.10139557 | 1.70E-11 | -2.941879058 | 2.34E-12 | 49.17441997 | 0.130138609 | 10.76903661 |
| ENSMUSG00000029602 | *Rasal1* | 496.5060779 | 526.5960463 | 395.8184365 | 66.95141883 | 68.42954854 | 49.58866915 | 7.13E-44 | -2.948416057 | 1.71E-45 | 200.3994456 | 0.129550271 | 43.14675246 |
| ENSMUSG00000062593 | *Gm49339* | 5246.841163 | 3553.803919 | 6408.729081 | 846.2220315 | 604.2855517 | 520.6810261 | 2.49E-34 | -2.94846161 | 8.97E-36 | 155.8840296 | 0.129546181 | 33.60445694 |
| ENSMUSG00000056366 | *Fabp3-ps1* | 40.6848683 | 4.795956706 | 54.20443558 | 1.097564243 | 4.211049141 | 7.346469504 | 0.005671145 | -2.953578554 | 0.002333918 | 9.266432655 | 0.129087521 | 2.246329213 |
| ENSMUSG00000027360 | *Hdc* | 232.8078575 | 207.1853297 | 88.23977885 | 29.63423457 | 16.84419656 | 22.03940851 | 4.67E-11 | -2.954810494 | 6.65E-12 | 47.12684443 | 0.128977338 | 10.33084691 |
| ENSMUSG00000096802 | *None* | 169.5202846 | 207.1853297 | 236.9868346 | 19.75615638 | 22.10800799 | 35.81403883 | 6.42E-22 | -2.963455912 | 4.13E-23 | 98.02700156 | 0.128206747 | 21.19256883 |
| ENSMUSG00000098934 | *Gvin-ps4* | 21.84928112 | 7.673530729 | 10.08454615 | 1.097564243 | 3.158286856 | 0.918308688 | 0.007789624 | -2.965144989 | 0.003292362 | 8.637925155 | 0.128056733 | 2.108483491 |
| ENSMUSG00000010830 | *Kdelr3* | 41.43829178 | 50.83714108 | 76.89466442 | 9.878078189 | 5.263811426 | 6.428160816 | 3.40E-08 | -2.968689816 | 6.40E-09 | 33.70876819 | 0.127742472 | 7.468448565 |
| ENSMUSG00000045007 | *Tubg2* | 11.3013523 | 17.26544414 | 21.42966058 | 4.390256973 | 2.10552457 | 0 | 0.003919545 | -2.983559445 | 0.001562761 | 10.00310954 | 0.126432613 | 2.406764323 |
| ENSMUSG00000028051 | *Hcn3* | 29.38351599 | 42.20441901 | 47.90159423 | 1.097564243 | 5.263811426 | 8.264778192 | 7.20E-06 | -2.99046251 | 1.80E-06 | 22.7934703 | 0.125829098 | 5.142379579 |
| ENSMUSG00000091304 | *Speer6-ps1* | 11.3013523 | 20.14301816 | 8.823977885 | 2.195128486 | 1.052762285 | 1.836617376 | 0.003792848 | -2.991126854 | 0.001506519 | 10.07062469 | 0.125771169 | 2.421034618 |
| ENSMUSG00000030659 | *Nucb2* | 2225.61298 | 2506.366974 | 3753.972306 | 379.7572281 | 313.723161 | 359.9770057 | 4.07E-48 | -3.008975735 | 8.44E-50 | 220.137628 | 0.124224726 | 47.39012432 |
| ENSMUSG00000043263 | *Ifi209* | 122.8080284 | 108.3886215 | 131.0991 | 21.95128486 | 8.422098282 | 14.69293901 | 2.60E-16 | -3.011594919 | 2.40E-17 | 71.78798308 | 0.123999403 | 15.58473383 |
| ENSMUSG00000037754 | *Ppp1r16b* | 275.7529962 | 274.3287236 | 248.331949 | 31.82936305 | 42.11049141 | 25.71264326 | 2.87E-36 | -3.011973966 | 9.40E-38 | 164.9449619 | 0.123966828 | 35.54257579 |
| ENSMUSG00000055538 | *Zcchc24* | 146.1641565 | 154.4298059 | 167.6555798 | 18.65859213 | 18.94972113 | 19.28448245 | 4.77E-27 | -3.036942054 | 2.35E-28 | 121.9604627 | 0.121839847 | 26.32185153 |
| ENSMUSG00000025746 | *Il6* | 23.3561281 | 30.69412292 | 30.25363846 | 4.390256973 | 3.158286856 | 2.754926064 | 7.18E-06 | -3.039486476 | 1.80E-06 | 22.80102763 | 0.121625153 | 5.143941976 |
| ENSMUSG00000030486 | *Zfp108* | 35.41090389 | 25.89816621 | 13.86625096 | 1.097564243 | 6.316573711 | 1.836617376 | 0.000522404 | -3.04237716 | 0.00017484 | 14.08383715 | 0.1213817 | 3.281993408 |
| ENSMUSG00000030630 | *Fah* | 80.6163131 | 72.89854193 | 88.23977885 | 4.390256973 | 12.63314742 | 11.93801294 | 1.97E-12 | -3.043795433 | 2.49E-13 | 53.57660129 | 0.121262432 | 11.70467083 |
| ENSMUSG00000071068 | *Treml2* | 809.9302485 | 916.0277308 | 874.8343789 | 113.049117 | 102.1179417 | 98.25902962 | 1.50E-105 | -3.054770036 | 7.08E-108 | 486.7994717 | 0.120343486 | 104.8251147 |
| ENSMUSG00000030748 | *Il4ra* | 301.3693948 | 290.6349764 | 300.0152481 | 42.80500548 | 35.7939177 | 29.38587802 | 1.54E-45 | -3.056015805 | 3.49E-47 | 208.1448989 | 0.120239615 | 44.81370418 |
| ENSMUSG00000017607 | *Tns4* | 66.30126685 | 59.46986315 | 61.76784519 | 7.682949702 | 10.52762285 | 4.59154344 | 4.50E-11 | -3.058289761 | 6.40E-12 | 47.20189131 | 0.120050244 | 10.34705696 |
| ENSMUSG00000015355 | *Cd48* | 51.9862206 | 72.89854193 | 109.6694394 | 16.46346365 | 7.369335996 | 4.59154344 | 7.66E-08 | -3.058419672 | 1.50E-08 | 32.05563126 | 0.120039434 | 7.115755374 |
| ENSMUSG00000034171 | *Faah* | 113.7669465 | 124.6948743 | 102.1060298 | 19.75615638 | 9.474860567 | 11.93801294 | 3.42E-17 | -3.060740076 | 2.97E-18 | 75.91107203 | 0.11984652 | 16.46620761 |
| ENSMUSG00000079057 | *Cyp4v3* | 974.9299921 | 1033.049074 | 1181.152468 | 120.7320667 | 138.9646216 | 122.1350555 | 2.83E-99 | -3.060765593 | 1.53E-101 | 457.685536 | 0.1198444 | 98.54854659 |
| ENSMUSG00000032554 | *Trf* | 219.9996582 | 228.2875392 | 161.3527385 | 13.17077092 | 33.68839313 | 25.71264326 | 2.46E-19 | -3.067800334 | 1.89E-20 | 85.90603717 | 0.119261448 | 18.60839191 |
| ENSMUSG00000089942 | *Pira2* | 30.13693948 | 43.16361035 | 21.42966058 | 5.487821216 | 3.158286856 | 2.754926064 | 1.04E-05 | -3.077330748 | 2.65E-06 | 22.05201471 | 0.118476206 | 4.984468974 |
| ENSMUSG00000031024 | *Dennd2b* | 47.46567968 | 63.30662851 | 57.98614039 | 8.780513945 | 11.58038514 | 0 | 3.49E-05 | -3.077417405 | 9.70E-06 | 19.56989817 | 0.11846909 | 4.457015421 |
| ENSMUSG00000055541 | *Lair1* | 568.0813092 | 486.3100099 | 582.3825404 | 71.34167581 | 43.16325369 | 78.05623848 | 8.63E-44 | -3.082358924 | 2.10E-45 | 199.9928593 | 0.118064004 | 43.06409395 |
| ENSMUSG00000058818 | *Pirb* | 335.2734517 | 388.4724932 | 360.522525 | 39.51231275 | 54.74363883 | 33.05911277 | 2.07E-43 | -3.096408246 | 5.12E-45 | 198.2158683 | 0.116919847 | 42.68481014 |
| ENSMUSG00000024011 | *Pi16* | 1503.83328 | 1698.727865 | 1797.570352 | 159.1468153 | 211.6052193 | 209.3743809 | 2.88E-100 | -3.103529394 | 1.52E-102 | 462.2920605 | 0.116344152 | 99.53986957 |
| ENSMUSG00000073913 | *Or52e5* | 14.31504625 | 9.591913411 | 10.08454615 | 0 | 1.052762285 | 2.754926064 | 0.007802667 | -3.104624431 | 0.003300016 | 8.633694604 | 0.116255878 | 2.107756932 |
| ENSMUSG00000050824 | *Sstr5* | 30.13693948 | 19.18382682 | 36.55647981 | 1.097564243 | 2.10552457 | 6.428160816 | 8.53E-05 | -3.107763691 | 2.51E-05 | 17.75954206 | 0.116003184 | 4.069148848 |
| ENSMUSG00000094054 | *ENSMUSG00000094054* | 39.93144481 | 25.89816621 | 63.02841346 | 0 | 5.263811426 | 9.18308688 | 0.000135359 | -3.118410889 | 4.11E-05 | 16.82102212 | 0.115150223 | 3.868514085 |
| ENSMUSG00000047959 | *Kcna3* | 15.82189323 | 12.46948743 | 7.563409615 | 2.195128486 | 2.10552457 | 0 | 0.007413431 | -3.121546915 | 0.003115554 | 8.73853109 | 0.11490019 | 2.129980741 |
| ENSMUSG00000046971 | *Pla2g4f* | 16.57531671 | 22.06140085 | 15.12681923 | 3.29269273 | 2.10552457 | 0.918308688 | 0.000419842 | -3.124494657 | 0.000137921 | 14.53033218 | 0.114665663 | 3.376914091 |
| ENSMUSG00000010142 | *Tnfrsf13b* | 116.027217 | 95.91913411 | 132.3596683 | 9.878078189 | 11.58038514 | 17.44786507 | 1.93E-17 | -3.126352108 | 1.65E-18 | 77.07093059 | 0.114518128 | 16.71470699 |
| ENSMUSG00000070570 | *Slc17a7* | 24.10955158 | 49.87794974 | 47.90159423 | 2.195128486 | 2.10552457 | 9.18308688 | 2.30E-05 | -3.130833234 | 6.23E-06 | 20.41756684 | 0.114162977 | 4.638128509 |
| ENSMUSG00000010021 | *Kif19a* | 48.97252665 | 45.08199303 | 39.07761635 | 2.195128486 | 6.316573711 | 6.428160816 | 2.37E-08 | -3.140786175 | 4.38E-09 | 34.44742023 | 0.113378094 | 7.625158307 |
| ENSMUSG00000048965 | *Mrgpre* | 97.19162982 | 116.0621523 | 113.4511442 | 5.487821216 | 23.16077027 | 8.264778192 | 5.01E-11 | -3.149034794 | 7.15E-12 | 46.98450949 | 0.112731703 | 10.29993462 |
| ENSMUSG00000072919 | *Noxred1* | 154.4518148 | 132.3684051 | 100.8454615 | 19.75615638 | 10.52762285 | 13.77463032 | 1.83E-17 | -3.149577113 | 1.56E-18 | 77.18491287 | 0.112689335 | 16.7387241 |
| ENSMUSG00000041845 | *Rhod* | 464.108868 | 500.6978801 | 521.8752635 | 64.75629035 | 37.89944227 | 64.28160816 | 1.02E-50 | -3.150161547 | 1.93E-52 | 232.2507418 | 0.112643694 | 49.99288643 |
| ENSMUSG00000056529 | *Ptafr* | 470.8896793 | 607.1681189 | 742.4747106 | 65.85385459 | 66.32402397 | 69.79146029 | 2.28E-45 | -3.167621083 | 5.21E-47 | 207.3461707 | 0.111288692 | 44.64205177 |
| ENSMUSG00000023349 | *Clec4n* | 119.7943344 | 142.9195098 | 180.2612625 | 20.85372062 | 12.63314742 | 15.6112477 | 4.29E-20 | -3.172049697 | 3.11E-21 | 89.47439149 | 0.110947595 | 19.36748501 |
| ENSMUSG00000052276 | *Ostn* | 23.3561281 | 21.1022095 | 37.81704808 | 1.097564243 | 3.158286856 | 4.59154344 | 3.19E-05 | -3.1762577 | 8.80E-06 | 19.75516652 | 0.110624459 | 4.496432633 |
| ENSMUSG00000038781 | *Stap2* | 35.41090389 | 38.36765364 | 27.73250192 | 3.29269273 | 5.263811426 | 2.754926064 | 5.62E-07 | -3.181471341 | 1.22E-07 | 27.98980542 | 0.110225403 | 6.250513236 |
| ENSMUSG00000061666 | *Gdpd1* | 326.2323698 | 344.3496915 | 332.7900231 | 38.41474851 | 28.4245817 | 43.16050834 | 3.78E-51 | -3.182437122 | 7.08E-53 | 234.2443421 | 0.11015164 | 50.42288521 |
| ENSMUSG00000079049 | *Serpinb1c* | 146.91758 | 177.4503981 | 234.4656981 | 30.73179881 | 15.79143428 | 14.69293901 | 2.56E-18 | -3.198709776 | 2.08E-19 | 81.16120293 | 0.108916182 | 17.59169341 |
| ENSMUSG00000043051 | *Disc1* | 36.16432737 | 36.44927096 | 39.07761635 | 4.390256973 | 2.10552457 | 5.509852128 | 1.07E-07 | -3.200348941 | 2.12E-08 | 31.37893837 | 0.108792504 | 6.972586545 |
| ENSMUSG00000036523 | *Greb1* | 16.57531671 | 19.18382682 | 20.16909231 | 1.097564243 | 1.052762285 | 3.673234752 | 0.000261194 | -3.214667535 | 8.29E-05 | 15.49178663 | 0.10771809 | 3.583036039 |
| ENSMUSG00000019558 | *Slc6a8* | 3502.665791 | 4327.871331 | 4098.107443 | 378.6596639 | 473.7430283 | 426.0952312 | 2.27E-135 | -3.221008607 | 7.15E-138 | 624.6859097 | 0.107245676 | 134.6446437 |
| ENSMUSG00000025877 | *Hk3* | 16928.67233 | 17599.24273 | 16407.55659 | 1915.249604 | 1769.693401 | 1776.009003 | 0 | -3.221946483 | 0 | 1960.391523 | 0.10717598 | Inf |
| ENSMUSG00000022676 | *Snai2* | 22.60270461 | 24.93897487 | 28.99307019 | 5.487821216 | 0 | 2.754926064 | 9.48E-05 | -3.222527825 | 2.80E-05 | 17.54728562 | 0.107132802 | 4.023310839 |
| ENSMUSG00000018819 | *Lsp1* | 4109.171698 | 3940.358029 | 3958.184365 | 412.6841554 | 470.5847415 | 402.2192053 | 2.65E-264 | -3.224941123 | 1.82E-267 | 1220.822319 | 0.106953743 | 263.5764202 |
| ENSMUSG00000091199 | *Gm2619* | 436.2321989 | 391.3500672 | 394.5578683 | 58.17090489 | 16.84419656 | 54.18021259 | 9.51E-20 | -3.239673965 | 7.07E-21 | 87.84779835 | 0.105867086 | 19.02175205 |
| ENSMUSG00000000673 | *Haao* | 11.3013523 | 8.63272207 | 8.823977885 | 1.097564243 | 1.052762285 | 0.918308688 | 0.006498114 | -3.24024625 | 0.002701453 | 8.998878629 | 0.105825099 | 2.187212664 |
| ENSMUSG00000068859 | *Sp9* | 32.39720994 | 34.53088828 | 18.90852404 | 3.29269273 | 2.10552457 | 3.673234752 | 7.33E-06 | -3.243972518 | 1.84E-06 | 22.75616492 | 0.105552122 | 5.134636497 |
| ENSMUSG00000062783 | *Csprs* | 1979.996924 | 1970.179015 | 2108.930714 | 248.049519 | 135.8063348 | 253.4531979 | 3.71E-52 | -3.247454222 | 6.70E-54 | 238.9377746 | 0.105297696 | 51.43011642 |
| ENSMUSG00000024066 | *Xdh* | 937.2588178 | 910.2725827 | 714.7422087 | 84.51244672 | 94.74860567 | 89.07594274 | 1.90E-78 | -3.257451277 | 1.86E-80 | 360.8244701 | 0.104570565 | 77.72175835 |
| ENSMUSG00000079554 | *Aox2* | 69.3149608 | 76.73530729 | 81.9369375 | 6.585385459 | 1.052762285 | 15.6112477 | 1.09E-07 | -3.262109245 | 2.17E-08 | 31.34049169 | 0.104233487 | 6.964434921 |
| ENSMUSG00000002799 | *Jag2* | 287.0543485 | 340.5129261 | 204.2120596 | 30.73179881 | 30.53010627 | 24.79433458 | 5.04E-29 | -3.281354154 | 2.26E-30 | 131.1805266 | 0.102852292 | 28.29779246 |
| ENSMUSG00000031129 | *Slc9a9* | 190.6161422 | 199.511799 | 192.8669452 | 19.75615638 | 17.89695885 | 22.03940851 | 1.69E-36 | -3.282723597 | 5.51E-38 | 166.0095543 | 0.102754708 | 35.77148016 |
| ENSMUSG00000029491 | *Pde6b* | 77.60261916 | 83.44964668 | 88.23977885 | 13.17077092 | 5.263811426 | 7.346469504 | 1.27E-14 | -3.284120889 | 1.34E-15 | 63.85292623 | 0.102655235 | 13.89556289 |
| ENSMUSG00000108900 | *Ccdc194* | 36.91775086 | 31.65331426 | 21.42966058 | 5.487821216 | 2.10552457 | 1.836617376 | 7.04E-06 | -3.291595034 | 1.76E-06 | 22.84204781 | 0.102124786 | 5.152734657 |
| ENSMUSG00000020303 | *Stc2* | 676.5742913 | 593.7394402 | 559.6923115 | 66.95141883 | 37.89944227 | 80.81116454 | 8.03E-41 | -3.295850322 | 2.20E-42 | 186.1530027 | 0.101824009 | 40.0953621 |
| ENSMUSG00000079457 | *Gm7609* | 1172.326946 | 1128.968209 | 1273.173952 | 150.3663013 | 92.6430811 | 120.2984381 | 6.38E-83 | -3.299804077 | 5.70E-85 | 381.5592099 | 0.101545339 | 82.19492563 |
| ENSMUSG00000017639 | *Rab11fip4* | 12.05477579 | 27.81654889 | 20.16909231 | 1.097564243 | 2.10552457 | 2.754926064 | 0.000249403 | -3.304071036 | 7.89E-05 | 15.58526097 | 0.101245449 | 3.603098136 |
| ENSMUSG00000021403 | *Serpinb9b* | 266.7119144 | 259.9408534 | 389.5155952 | 32.9269273 | 40.00496684 | 20.20279114 | 1.77E-26 | -3.304922346 | 9.00E-28 | 119.3013955 | 0.101185723 | 25.75221764 |
| ENSMUSG00000050022 | *Amz1* | 346.574804 | 524.6776636 | 555.9106067 | 64.75629035 | 38.95220455 | 39.48727358 | 4.90E-31 | -3.320971472 | 2.01E-32 | 140.5565419 | 0.10006633 | 30.30966947 |
| ENSMUSG00000101751 | *Gm2427* | 30.13693948 | 70.98015924 | 27.73250192 | 0 | 5.263811426 | 7.346469504 | 7.39E-05 | -3.324052661 | 2.16E-05 | 18.04689823 | 0.099852845 | 4.131260847 |
| ENSMUSG00000020577 | *Tspan13* | 261.43795 | 289.675785 | 300.0152481 | 28.53667032 | 35.7939177 | 21.12109982 | 3.07E-42 | -3.324500662 | 8.05E-44 | 192.7330551 | 0.099821842 | 41.51275823 |
| ENSMUSG00000090186 | *Gm52951* | 68.56153731 | 56.59228913 | 84.45807404 | 6.585385459 | 0 | 13.77463032 | 2.73E-06 | -3.335391205 | 6.46E-07 | 24.77127052 | 0.099071149 | 5.564009169 |
| ENSMUSG00000022469 | *Rapgef3* | 27.12324553 | 45.08199303 | 26.47193365 | 1.097564243 | 0 | 8.264778192 | 0.00042245 | -3.34386481 | 0.000138836 | 14.51787849 | 0.098490965 | 3.374224637 |
| ENSMUSG00000046245 | *Pilra* | 158.2189323 | 152.5114232 | 210.514901 | 16.46346365 | 18.94972113 | 15.6112477 | 5.52E-27 | -3.350439272 | 2.73E-28 | 121.6649912 | 0.098043156 | 26.25777507 |
| ENSMUSG00000052271 | *Bhlha15* | 37.67117435 | 24.93897487 | 21.42966058 | 5.487821216 | 2.10552457 | 0.918308688 | 2.72E-05 | -3.352760996 | 7.44E-06 | 20.07653917 | 0.097885502 | 4.564768721 |
| ENSMUSG00000028838 | *Extl1* | 207.1914589 | 240.7570266 | 213.0360375 | 29.63423457 | 20.00248342 | 15.6112477 | 5.26E-33 | -3.356173591 | 2.01E-34 | 149.7014634 | 0.097654234 | 32.2789019 |
| ENSMUSG00000004791 | *Pgf* | 14.31504625 | 22.06140085 | 16.3873875 | 3.29269273 | 1.052762285 | 0.918308688 | 0.000337562 | -3.357548148 | 0.000109015 | 14.97378868 | 0.097561236 | 3.471646679 |
| ENSMUSG00000066621 | *Tecpr1* | 3464.994616 | 3700.560194 | 3094.695101 | 308.4155523 | 331.6201198 | 349.8756101 | 6.22E-193 | -3.37259316 | 8.97E-196 | 890.9799714 | 0.096549115 | 192.2060227 |
| ENSMUSG00000071335 | *Mfsd4b3-ps* | 6.780811383 | 15.34706146 | 20.16909231 | 0 | 2.10552457 | 1.836617376 | 0.003286895 | -3.378235984 | 0.001287284 | 10.36067044 | 0.096172219 | 2.483214185 |
| ENSMUSG00000029814 | *Igf2bp3* | 22.60270461 | 34.53088828 | 27.73250192 | 4.390256973 | 0 | 3.673234752 | 1.99E-05 | -3.38578376 | 5.31E-06 | 20.72152581 | 0.095670387 | 4.702101111 |
| ENSMUSG00000067001 | *Serpinb7* | 48.21910316 | 25.89816621 | 56.72557212 | 4.390256973 | 5.263811426 | 2.754926064 | 1.63E-07 | -3.408774142 | 3.34E-08 | 30.50035038 | 0.094157894 | 6.786765256 |
| ENSMUSG00000029650 | *Slc46a3* | 364.6569677 | 313.6555685 | 293.7124067 | 34.02449154 | 28.4245817 | 29.38587802 | 6.94E-51 | -3.410382834 | 1.30E-52 | 233.0258639 | 0.09405296 | 50.15876095 |
| ENSMUSG00000049130 | *C5ar1* | 1498.559316 | 1473.3179 | 1140.814284 | 118.5369383 | 111.5928022 | 154.2758596 | 1.41E-79 | -3.416018322 | 1.35E-81 | 366.0531243 | 0.093686285 | 78.85104528 |
| ENSMUSG00000020826 | *Nos2* | 425.6842701 | 539.0655337 | 369.3465029 | 41.70744124 | 28.4245817 | 54.18021259 | 7.22E-37 | -3.416743503 | 2.32E-38 | 167.727723 | 0.093639205 | 36.14125311 |
| ENSMUSG00000020872 | *Tac4* | 30.13693948 | 20.14301816 | 36.55647981 | 3.29269273 | 1.052762285 | 3.673234752 | 5.43E-06 | -3.418337566 | 1.34E-06 | 23.36969477 | 0.093535798 | 5.265064059 |
| ENSMUSG00000034220 | *Gpc1* | 18.83558717 | 17.26544414 | 17.64795577 | 1.097564243 | 1.052762285 | 2.754926064 | 0.000135052 | -3.418452029 | 4.10E-05 | 16.82575512 | 0.093528378 | 3.869498984 |
| ENSMUSG00000061852 | *Gm53056* | 40.6848683 | 29.73493157 | 26.47193365 | 0 | 5.263811426 | 3.673234752 | 6.52E-06 | -3.423568781 | 1.62E-06 | 22.99565462 | 0.093197252 | 5.185767397 |
| ENSMUSG00000026807 | *Ak8* | 12.05477579 | 15.34706146 | 25.21136538 | 0 | 0 | 4.59154344 | 0.004190371 | -3.428907057 | 0.001679944 | 9.870011734 | 0.092853039 | 2.377747489 |
| ENSMUSG00000006457 | *Actn3* | 56.50676152 | 50.83714108 | 55.46500385 | 1.097564243 | 6.316573711 | 7.346469504 | 5.14E-10 | -3.432518321 | 8.09E-11 | 42.23510546 | 0.092620906 | 9.288693199 |
| ENSMUSG00000021457 | *Syk* | 2723.625905 | 3139.433259 | 2884.1802 | 283.1715747 | 238.9770387 | 285.594002 | 2.47E-191 | -3.435581744 | 3.73E-194 | 883.534676 | 0.092424443 | 190.6076826 |
| ENSMUSG00000074419 | *Pira13* | 46.71225619 | 69.06177656 | 39.07761635 | 4.390256973 | 5.263811426 | 4.59154344 | 1.08E-09 | -3.446328292 | 1.75E-10 | 40.72437106 | 0.091738537 | 8.968027973 |
| ENSMUSG00000076441 | *Ass1* | 1065.340811 | 1111.702764 | 1114.34235 | 102.0734746 | 94.74860567 | 102.8505731 | 1.55E-168 | -3.45561578 | 3.29E-171 | 777.9903886 | 0.091149858 | 167.8104124 |
| ENSMUSG00000024334 | *H2-Oa* | 22.60270461 | 27.81654889 | 17.64795577 | 3.29269273 | 2.10552457 | 0.918308688 | 2.73E-05 | -3.469211637 | 7.46E-06 | 20.07236798 | 0.090294903 | 4.563930605 |
| ENSMUSG00000038742 | *Angptl6* | 1438.285437 | 1413.848037 | 1524.027038 | 151.4638656 | 126.3314742 | 117.5435121 | 3.40E-162 | -3.473054759 | 7.69E-165 | 748.7005713 | 0.090054691 | 161.4690503 |
| ENSMUSG00000014599 | *Csf1* | 338.2871456 | 325.1658646 | 202.9514913 | 17.56102789 | 32.63563084 | 27.54926064 | 1.00E-26 | -3.477436652 | 5.02E-28 | 120.4574174 | 0.089781583 | 25.99940422 |
| ENSMUSG00000057596 | *Trim30d* | 63.2875729 | 41.24522767 | 54.20443558 | 6.585385459 | 3.158286856 | 4.59154344 | 2.42E-10 | -3.478387933 | 3.68E-11 | 43.77963241 | 0.089722403 | 9.616977615 |
| ENSMUSG00000019768 | *Esr1* | 34.6574804 | 23.97978353 | 10.08454615 | 3.29269273 | 2.10552457 | 0.918308688 | 0.000218562 | -3.489369798 | 6.85E-05 | 15.851853 | 0.089042024 | 3.6604249 |
| ENSMUSG00000036067 | *Slc2a6* | 7673.618215 | 7802.062369 | 7142.379814 | 677.197138 | 637.9739448 | 685.9765899 | 0 | -3.49845297 | 0 | 1807.956615 | 0.088483179 | Inf |
| ENSMUSG00000112148 | *Lilrb4a* | 9105.12284 | 9155.481351 | 7971.833735 | 817.6853612 | 550.5946752 | 944.0213313 | 1.58E-77 | -3.503236975 | 1.63E-79 | 356.4928604 | 0.088190253 | 76.80219402 |
| ENSMUSG00000026411 | *Tmem9* | 646.4373518 | 790.3736651 | 860.9681279 | 61.46359762 | 66.32402397 | 73.46469504 | 6.34E-79 | -3.506673905 | 6.14E-81 | 363.0381311 | 0.087980408 | 78.19765268 |
| ENSMUSG00000019960 | *Dusp6* | 1799.175287 | 1880.015029 | 1787.485806 | 206.3420777 | 93.69584338 | 179.9885028 | 1.40E-47 | -3.509166795 | 2.97E-49 | 217.6334199 | 0.087828514 | 46.85523506 |
| ENSMUSG00000028634 | *Hivep3* | 594.4511312 | 600.4537795 | 281.106724 | 48.2928267 | 43.16325369 | 37.65065621 | 8.45E-26 | -3.521113105 | 4.44E-27 | 116.1347898 | 0.087104248 | 25.0732622 |
| ENSMUSG00000070501 | *Ifi214* | 30.13693948 | 23.97978353 | 15.12681923 | 1.097564243 | 1.052762285 | 3.673234752 | 4.71E-05 | -3.538572737 | 1.33E-05 | 18.9637182 | 0.086056457 | 4.327098991 |
| ENSMUSG00000074874 | *Ctla2b* | 57.26018501 | 54.67390644 | 64.28898173 | 0 | 9.474860567 | 5.509852128 | 1.15E-08 | -3.538907626 | 2.07E-09 | 35.9075514 | 0.086036484 | 7.939176325 |
| ENSMUSG00000022667 | *Cd200r1* | 192.8764127 | 220.6140085 | 390.7761635 | 19.75615638 | 13.68590971 | 34.89573014 | 4.25E-17 | -3.540218696 | 3.71E-18 | 75.46849813 | 0.085958332 | 16.37126231 |
| ENSMUSG00000039062 | *Anpep* | 1902.394305 | 2375.916952 | 2033.296618 | 186.5859213 | 152.6505314 | 201.1096027 | 2.51E-123 | -3.543588169 | 9.46E-126 | 568.9586363 | 0.085757807 | 122.6010019 |
| ENSMUSG00000025481 | *Urah* | 42.19171527 | 62.34743717 | 64.28898173 | 8.780513945 | 2.10552457 | 3.673234752 | 1.33E-09 | -3.550213973 | 2.18E-10 | 40.29477292 | 0.085364854 | 8.875641 |
| ENSMUSG00000025408 | *Ddit3* | 1394.586874 | 1480.991431 | 1801.352057 | 154.7565583 | 121.0676628 | 123.0533642 | 3.46E-104 | -3.553031155 | 1.68E-106 | 480.4756488 | 0.085198323 | 103.4614955 |
| ENSMUSG00000045659 | *Plekha7* | 39.17802132 | 49.87794974 | 31.51420673 | 6.585385459 | 2.10552457 | 1.836617376 | 1.06E-07 | -3.554047524 | 2.11E-08 | 31.38673945 | 0.085138323 | 6.974181938 |
| ENSMUSG00000032268 | *Tmprss5* | 9.79450533 | 12.46948743 | 26.47193365 | 0 | 3.158286856 | 0.918308688 | 0.001771867 | -3.564318788 | 0.000658431 | 11.60306813 | 0.084534333 | 2.751568993 |
| ENSMUSG00000029919 | *Hpgds* | 1097.73802 | 1201.86675 | 1594.618861 | 102.0734746 | 126.3314742 | 99.1773383 | 6.18E-74 | -3.57105893 | 7.00E-76 | 339.8157453 | 0.084140317 | 73.20896674 |
| ENSMUSG00000020598 | *Nrcam* | 18.83558717 | 30.69412292 | 36.55647981 | 1.097564243 | 5.263811426 | 0.918308688 | 2.04E-05 | -3.573986147 | 5.47E-06 | 20.66508686 | 0.08396977 | 4.6903405 |
| ENSMUSG00000066026 | *Dhrs3* | 2332.599116 | 2343.304446 | 2330.79073 | 238.1714408 | 135.8063348 | 210.2926896 | 5.12E-86 | -3.583904136 | 4.43E-88 | 395.8417818 | 0.083394489 | 85.29071321 |
| ENSMUSG00000049723 | *Mmp12* | 188.3558717 | 208.144521 | 236.9868346 | 14.26833516 | 12.63314742 | 24.79433458 | 8.20E-32 | -3.592089289 | 3.27E-33 | 144.1648108 | 0.082922689 | 31.08609941 |
| ENSMUSG00000039114 | *Nrn1* | 16.57531671 | 13.42867878 | 7.563409615 | 1.097564243 | 2.10552457 | 0 | 0.002599116 | -3.612643251 | 0.000996696 | 10.83369325 | 0.081749672 | 2.585174372 |
| ENSMUSG00000036944 | *Tmem71* | 36.16432737 | 44.12280169 | 32.774775 | 0 | 7.369335996 | 1.836617376 | 3.44E-06 | -3.623936181 | 8.25E-07 | 24.29914008 | 0.081112261 | 5.463228146 |
| ENSMUSG00000035373 | *Ccl7* | 205.6846119 | 209.1037124 | 226.9022885 | 13.17077092 | 20.00248342 | 18.36617376 | 1.30E-41 | -3.629711918 | 3.49E-43 | 189.8139475 | 0.080788182 | 40.88682452 |
| ENSMUSG00000029484 | *Anxa3* | 3042.32404 | 2969.656392 | 3162.765788 | 284.269139 | 205.2886456 | 250.6982718 | 2.79E-186 | -3.632066965 | 4.97E-189 | 859.9584412 | 0.080656412 | 185.5548593 |
| ENSMUSG00000112023 | *Lilrb4b* | 8106.83672 | 7464.427017 | 7989.48169 | 743.0509926 | 455.8460695 | 690.5681334 | 7.54E-96 | -3.640133114 | 4.66E-98 | 441.6831561 | 0.080206718 | 95.12259519 |
| ENSMUSG00000028751 | *Pla2g2e* | 320.9584054 | 369.2886663 | 452.5440087 | 28.53667032 | 34.74115541 | 27.54926064 | 6.63E-50 | -3.651965849 | 1.30E-51 | 228.4445072 | 0.079551567 | 49.17827234 |
| ENSMUSG00000002007 | *Srpk3* | 48.97252665 | 33.57169694 | 20.16909231 | 4.390256973 | 1.052762285 | 2.754926064 | 2.02E-06 | -3.666938708 | 4.70E-07 | 25.38227647 | 0.078730218 | 5.694892982 |
| ENSMUSG00000025176 | *Hoga1* | 63.2875729 | 83.44964668 | 92.02148365 | 6.585385459 | 8.422098282 | 3.673234752 | 5.76E-15 | -3.689404485 | 5.90E-16 | 65.46976085 | 0.077513721 | 14.23988715 |
| ENSMUSG00000034586 | *Hid1* | 174.794249 | 191.8382682 | 136.1413731 | 29.63423457 | 1.052762285 | 8.264778192 | 0.002585319 | -3.695418512 | 0.000990873 | 10.84454309 | 0.077191269 | 2.587485906 |
| ENSMUSG00000098557 | *Kctd12* | 1352.395159 | 1356.296556 | 1048.7928 | 111.9515528 | 72.64059768 | 103.7688817 | 3.77E-82 | -3.704463542 | 3.42E-84 | 377.985407 | 0.07670883 | 81.42348195 |
| ENSMUSG00000026639 | *Lamb3* | 19.58901066 | 18.22463548 | 15.12681923 | 0 | 3.158286856 | 0.918308688 | 0.000202223 | -3.705278901 | 6.32E-05 | 16.00473484 | 0.076665489 | 3.69416954 |
| ENSMUSG00000047963 | *Stbd1* | 19.58901066 | 14.38787012 | 20.16909231 | 0 | 4.211049141 | 0 | 0.001049599 | -3.713384093 | 0.000371307 | 12.67132331 | 0.076235983 | 2.978976788 |
| ENSMUSG00000104713 | *Gbp6* | 56.50676152 | 78.65368997 | 50.42273077 | 2.195128486 | 6.316573711 | 5.509852128 | 6.85E-12 | -3.713568748 | 9.13E-13 | 51.02227149 | 0.076226226 | 11.16414569 |
| ENSMUSG00000047591 | *Mafa* | 14.31504625 | 20.14301816 | 18.90852404 | 1.097564243 | 1.052762285 | 1.836617376 | 7.11E-05 | -3.715647765 | 2.07E-05 | 18.126226 | 0.076116458 | 4.148024097 |
| ENSMUSG00000061126 | *Cyp4f39* | 24.86297507 | 20.14301816 | 21.42966058 | 0 | 3.158286856 | 1.836617376 | 2.05E-05 | -3.717268973 | 5.51E-06 | 20.64991088 | 0.076030971 | 4.687414866 |
| ENSMUSG00000025885 | *Myo5b* | 4.520540922 | 12.46948743 | 10.08454615 | 0 | 1.052762285 | 0.918308688 | 0.008583341 | -3.735613893 | 0.003663765 | 8.443371001 | 0.075070302 | 2.066343645 |
| ENSMUSG00000044071 | *Tafa2* | 147.6710034 | 131.4092137 | 143.7047827 | 9.878078189 | 12.63314742 | 9.18308688 | 5.47E-30 | -3.743482139 | 2.36E-31 | 135.6651037 | 0.074661995 | 29.26187273 |
| ENSMUSG00000028356 | *Ambp* | 171.0271315 | 174.5728241 | 199.1697865 | 9.878078189 | 11.58038514 | 18.36617376 | 1.12E-33 | -3.750209443 | 4.14E-35 | 152.8477091 | 0.074314655 | 32.94989916 |
| ENSMUSG00000020256 | *Aldh1l2* | 6001.018074 | 7124.873282 | 8028.559307 | 574.0260992 | 428.4742501 | 555.5767562 | 1.34E-124 | -3.762480198 | 4.87E-127 | 574.8801097 | 0.073685257 | 123.8729872 |

| | | | | | | | | | | | | | |
|---|---|---|---|---|---|---|---|---|---|---|---|---|---|
| ENSMUSG00000028179 | *Cth* | 2460.681108 | 2210.936041 | 2180.783106 | 185.4883571 | 117.9093759 | 196.5180592 | 2.55E-97 | -3.774730315 | 1.49E-99 | 448.5584757 | 0.073062234 | 96.59406749 |
| ENSMUSG00000026358 | *Rgs1* | 2924.036553 | 2602.286108 | 2628.284841 | 218.4152844 | 168.4419656 | 202.0279114 | 1.95E-203 | -3.792294283 | 2.28E-206 | 939.7188717 | 0.072178137 | 202.7093099 |
| ENSMUSG00000061972 | *Or1o4* | 138.6299216 | 130.4500224 | 187.8246721 | 14.26833516 | 6.316573711 | 11.93801294 | 2.01E-24 | -3.80730131 | 1.13E-25 | 109.7088055 | 0.071431226 | 23.69742538 |
| ENSMUSG00000013974 | *Mcemp1* | 101.7121707 | 89.20479472 | 80.67636923 | 9.878078189 | 5.263811426 | 4.59154344 | 1.81E-18 | -3.808512756 | 1.47E-19 | 81.85331161 | 0.071371269 | 17.74268229 |
| ENSMUSG00000030745 | *Il21r* | 67.80811383 | 52.75552376 | 37.81704808 | 3.29269273 | 7.369335996 | 0.918308688 | 7.87E-09 | -3.811324109 | 1.39E-09 | 36.67779621 | 0.071232325 | 8.103801206 |
| ENSMUSG00000089937 | *Gm7281* | 26.36982204 | 17.26544414 | 12.60568269 | 0 | 2.10552457 | 1.836617376 | 0.000136396 | -3.814947369 | 4.14E-05 | 16.80566959 | 0.071053653 | 3.865197017 |
| ENSMUSG00000058046 | *4933430I17Rik* | 75.34234869 | 72.89854193 | 113.4511442 | 9.878078189 | 3.158286856 | 5.509852128 | 1.09E-14 | -3.824620072 | 1.14E-15 | 64.17138086 | 0.07057886 | 13.96292666 |
| ENSMUSG00000032531 | *Amotl2* | 96.43820633 | 53.7147151 | 36.55647981 | 5.487821216 | 2.10552457 | 5.509852128 | 3.88E-09 | -3.835841742 | 6.69E-10 | 38.10798134 | 0.070032008 | 8.411187854 |
| ENSMUSG00000020669 | *Sh3yl1* | 21.09585763 | 21.1022095 | 30.25363846 | 1.097564243 | 1.052762285 | 2.754926064 | 4.20E-06 | -3.838011953 | 1.02E-06 | 23.89515245 | 0.069926739 | 5.376816753 |
| ENSMUSG00000052131 | *Akr1b7* | 131.0956867 | 179.3687808 | 359.2619567 | 16.46346365 | 21.0552457 | 9.18308688 | 5.44E-15 | -3.847419006 | 5.56E-16 | 65.58703532 | 0.069472267 | 14.26427465 |
| ENSMUSG00000114576 | *Naip3-ps1* | 12.80819928 | 12.46948743 | 17.64795577 | 0 | 0 | 2.754926064 | 0.001192472 | -3.851921804 | 0.000426514 | 12.41230522 | 0.069255775 | 2.923551914 |
| ENSMUSG00000057135 | *Scimp* | 409.8623769 | 423.0033814 | 395.8184365 | 34.02449154 | 22.10800799 | 28.46756933 | 2.27E-77 | -3.863388813 | 2.37E-79 | 355.7548949 | 0.06870749 | 76.644366 |
| ENSMUSG00000041592 | *Sdk2* | 33.15063343 | 26.85735755 | 28.99307019 | 2.195128486 | 2.10552457 | 1.836617376 | 9.44E-08 | -3.867320182 | 1.87E-08 | 31.62714434 | 0.068520516 | 7.025249339 |
| ENSMUSG00000102433 | *Gm38050* | 12.80819928 | 17.26544414 | 16.3873875 | 3.29269273 | 0 | 0 | 0.000971544 | -3.89079862 | 0.000340761 | 12.83189435 | 0.067414436 | 3.012537365 |
| ENSMUSG00000090176 | *Cd200r2* | 24.86297507 | 12.46948743 | 23.95079712 | 1.097564243 | 3.158286856 | 0 | 0.000125771 | -3.891527801 | 3.80E-05 | 16.96821591 | 0.067380372 | 3.90042056 |
| ENSMUSG00000060913 | *Trim55* | 15.06846974 | 7.673530729 | 7.563409615 | 0 | 2.10552457 | 0 | 0.006961124 | -3.892830101 | 0.002913048 | 8.861145072 | 0.067319576 | 2.157320648 |
| ENSMUSG00000026656 | *Fcgr2b* | 2847.940781 | 2471.836086 | 2091.282759 | 160.2443795 | 165.2836788 | 171.7237247 | 1.45E-142 | -3.897846539 | 3.99E-145 | 658.0371641 | 0.067085904 | 141.8373381 |
| ENSMUSG00000030256 | *Bhlhe41* | 577.122391 | 540.0247251 | 479.0159423 | 37.31718427 | 38.95220455 | 30.3041867 | 3.87E-87 | -3.914411727 | 3.13E-89 | 401.1253317 | 0.066320021 | 86.4123979 |
| ENSMUSG00000001930 | *Vwf* | 128.0819928 | 171.6952501 | 207.9937644 | 12.07320667 | 7.369335996 | 13.77463032 | 1.51E-25 | -3.919382712 | 8.08E-27 | 114.9483505 | 0.0660919 | 24.82074609 |
| ENSMUSG00000000244 | *Tspan32* | 14.31504625 | 8.63272207 | 23.95079712 | 2.195128486 | 0 | 0.918308688 | 0.000702788 | -3.922611497 | 0.000241047 | 13.48056901 | 0.06594415 | 3.153175914 |
| ENSMUSG00000028111 | *Ctsk* | 15.82189323 | 19.18382682 | 11.34511442 | 0 | 2.10552457 | 0.918308688 | 0.000279136 | -3.934372291 | 8.89E-05 | 15.35848742 | 0.065408762 | 3.554183699 |
| ENSMUSG00000021751 | *Acox2* | 7.534234869 | 6.714339388 | 17.64795577 | 0 | 1.052762285 | 0.918308688 | 0.003702602 | -3.963524919 | 0.001465338 | 10.12169985 | 0.064100307 | 2.431492997 |
| ENSMUSG00000089722 | *Cd300ld5* | 18.08216369 | 25.89816621 | 70.59182308 | 0 | 7.369335996 | 0 | 0.001006811 | -3.969991628 | 0.000354719 | 12.75678971 | 0.063813628 | 2.997052029 |
| ENSMUSG00000069515 | *Lyz1* | 978.6971095 | 935.2115576 | 1184.934173 | 61.46359762 | 71.58783539 | 61.5266821 | 4.76E-118 | -3.992735647 | 1.93E-120 | 544.5515069 | 0.062815498 | 117.3220634 |
| ENSMUSG00000046805 | *Mpeg1* | 20979.07699 | 20140.14059 | 21048.96896 | 1251.223237 | 1427.545659 | 1224.105481 | 0 | -3.994030178 | 0 | 2063.607594 | 0.062759159 | Inf |
| ENSMUSG00000079484 | *Phyhd1* | 730.0673589 | 903.5582433 | 1212.666675 | 38.41474851 | 80.00993367 | 59.69006472 | 1.35E-44 | -3.994295607 | 3.16E-46 | 203.7561956 | 0.062747613 | 43.86905218 |
| ENSMUSG00000027546 | *Atp9a* | 216.2325408 | 301.1860811 | 344.1351375 | 17.56102789 | 13.68590971 | 22.03940851 | 3.23E-37 | -4.001256901 | 1.02E-38 | 169.3643903 | 0.062445573 | 36.49124719 |
| ENSMUSG00000037902 | *Sirpa* | 12081.89904 | 12630.63158 | 10641.71733 | 619.0262331 | 834.8404922 | 745.6666547 | 7.52E-194 | -4.006215771 | 1.03E-196 | 895.3003476 | 0.062231301 | 193.1240392 |
| ENSMUSG00000000318 | *Clec10a* | 1035.203871 | 1131.845783 | 1609.74568 | 103.1710389 | 55.79640112 | 76.2196211 | 7.45E-55 | -4.006267314 | 1.22E-56 | 251.5111727 | 0.062229078 | 54.12806349 |
| ENSMUSG00000005803 | *Sqor* | 1206.984426 | 1146.233653 | 1279.476793 | 92.19539643 | 58.95468797 | 73.46469504 | 7.91E-141 | -4.018196426 | 2.28E-143 | 649.9584829 | 0.06171665 | 140.1015898 |
| ENSMUSG00000042770 | *Hebp1* | 384.9994018 | 456.5750784 | 694.5731164 | 28.53667032 | 32.63563084 | 33.05911277 | 2.55E-42 | -4.020936281 | 6.63E-44 | 193.1192303 | 0.061599554 | 41.59362124 |
| ENSMUSG00000043557 | *Mdga1* | 6.780811383 | 17.26544414 | 8.823977885 | 0 | 1.052762285 | 0.918308688 | 0.002641597 | -4.023600671 | 0.001013893 | 10.80202177 | 0.061485896 | 2.57813351 |
| ENSMUSG00000027669 | *Gnb4* | 36.16432737 | 33.57169694 | 32.774775 | 5.487821216 | 1.052762285 | 0 | 6.04E-07 | -4.040332498 | 1.32E-07 | 27.83995216 | 0.060776925 | 6.218932923 |
| ENSMUSG00000067206 | *Lrrc66* | 21.09585763 | 21.1022095 | 23.95079712 | 0 | 1.052762285 | 2.754926064 | 8.12E-06 | -4.043307933 | 2.05E-06 | 22.55189718 | 0.060651707 | 5.090481801 |
| ENSMUSG00000032221 | *Mns1* | 27.87666902 | 37.4084623 | 39.07761635 | 0 | 6.316573711 | 0 | 1.86E-05 | -4.071429638 | 4.95E-06 | 20.85841821 | 0.059480903 | 4.730910007 |
| ENSMUSG00000043832 | *Clec4a3* | 874.7246683 | 990.8446554 | 1316.033273 | 72.43924005 | 62.11297483 | 54.18021259 | 1.24E-77 | -4.078353526 | 1.27E-79 | 357.0020371 | 0.059196122 | 76.90727787 |
| ENSMUSG00000024481 | *Lvrn* | 26.36982204 | 29.73493157 | 31.51420673 | 4.390256973 | 0 | 0.918308688 | 1.05E-06 | -4.087500663 | 2.36E-07 | 26.71176672 | 0.058821987 | 5.979775134 |
| ENSMUSG00000017466 | *Timp2* | 69.3149608 | 49.87794974 | 52.94386731 | 2.195128486 | 4.211049141 | 3.673234752 | 2.20E-13 | -4.089222552 | 2.57E-14 | 58.03643982 | 0.058751824 | 12.65698876 |
| ENSMUSG00000028583 | *Pdpn* | 35.41090389 | 38.36765364 | 47.90159423 | 1.097564243 | 2.10552457 | 3.673234752 | 6.11E-10 | -4.100759373 | 9.69E-11 | 41.88267214 | 0.058283876 | 9.213689641 |
| ENSMUSG00000050014 | *Apol10b* | 74.58892521 | 72.89854193 | 81.9369375 | 3.29269273 | 5.263811426 | 4.59154344 | 8.54E-19 | -4.116649055 | 6.71E-20 | 83.39778726 | 0.057645466 | 18.06858285 |
| ENSMUSG00000029005 | *Draxin* | 4.520540922 | 19.18382682 | 12.60568269 | 1.097564243 | 1.052762285 | 0 | 0.00292357 | -4.125705692 | 0.001132151 | 10.59791725 | 0.057284725 | 2.534086447 |
| ENSMUSG00000047746 | *Fbxo40* | 11.3013523 | 9.591913411 | 15.12681923 | 0 | 2.10552457 | 0 | 0.001645419 | -4.126552655 | 0.000606362 | 11.75634526 | 0.057251105 | 2.783723362 |
| ENSMUSG00000082160 | *Gm11578* | 32.39720994 | 43.16361035 | 60.50727692 | 0 | 0 | 7.346469504 | 1.05E-05 | -4.127339737 | 2.70E-06 | 22.02048102 | 0.057219879 | 4.977799473 |
| ENSMUSG00000024427 | *Spry4* | 21.84928112 | 15.34706146 | 17.64795577 | 2.195128486 | 1.052762285 | 0 | 4.58E-05 | -4.15067215 | 1.30E-05 | 19.01741266 | 0.056301917 | 4.338898871 |
| ENSMUSG00000074973 | *Gm11382* | 12.05477579 | 23.02059219 | 39.07761635 | 3.29269273 | 0 | 0.918308688 | 6.80E-05 | -4.158192622 | 1.97E-05 | 18.21501506 | 0.05600919 | 4.167348327 |
| ENSMUSG00000063594 | *Gng8* | 21.84928112 | 15.34706146 | 39.07761635 | 2.195128486 | 2.10552457 | 0 | 1.49E-05 | -4.194085936 | 3.92E-06 | 21.30357728 | 0.054632909 | 4.826507701 |
| ENSMUSG00000055301 | *Adh7* | 586.9168963 | 596.6170142 | 757.6015298 | 25.24397759 | 41.05772912 | 36.73234752 | 6.77E-81 | -4.227374083 | 6.32E-83 | 372.1659102 | 0.053386764 | 80.16940775 |
| ENSMUSG00000039518 | *Cdsn* | 183.8353308 | 266.6551928 | 327.74775 | 17.56102789 | 12.63314742 | 11.01970426 | 1.14E-32 | -4.245383808 | 4.45E-34 | 148.1277428 | 0.052724459 | 31.94262196 |
| ENSMUSG00000068735 | *Trp53i11* | 36.91775086 | 19.18382682 | 22.69022885 | 1.097564243 | 3.158286856 | 0 | 5.05E-06 | -4.264675997 | 1.24E-06 | 23.52100944 | 0.052024103 | 5.296420185 |
| ENSMUSG00000040026 | *Saa3* | 196.6435301 | 167.8584847 | 239.5079712 | 5.487821216 | 10.52762285 | 14.69293901 | 1.23E-33 | -4.268992936 | 4.53E-35 | 152.6669463 | 0.051868666 | 32.91120131 |
| ENSMUSG00000053819 | *Camk2d* | 400.8212951 | 438.3504429 | 385.7338904 | 25.24397759 | 23.16077027 | 14.69293901 | 5.95E-78 | -4.29737659 | 5.94E-80 | 358.5119125 | 0.050858172 | 77.22576758 |
| ENSMUSG00000040522 | *Tlr8* | 13.56162277 | 8.63272207 | 18.90852404 | 2.195128486 | 0 | 0 | 0.000917188 | -4.302021137 | 0.000320248 | 12.94809089 | 0.050694704 | 3.037541635 |
| ENSMUSG00000075602 | *Ly6a* | 18.83558717 | 15.34706146 | 26.47193365 | 1.097564243 | 0 | 1.836617376 | 1.43E-05 | -4.317981196 | 3.75E-06 | 21.38873262 | 0.050136976 | 4.844663297 |
| ENSMUSG00000000386 | *Mx1* | 488.2184195 | 470.0037571 | 369.3465029 | 16.46346365 | 23.16077027 | 24.79433458 | 5.85E-70 | -4.358219054 | 7.02E-72 | 321.4441243 | 0.048757939 | 69.233177 |
| ENSMUSG00000032579 | *Hemk1* | 378.2185904 | 294.4717417 | 368.0859346 | 14.26833516 | 13.68590971 | 22.03940851 | 2.03E-59 | -4.362820573 | 2.94E-61 | 272.6937654 | 0.048602672 | 58.69293788 |
| ENSMUSG00000072572 | *Slc39a2* | 143.1504625 | 147.7154665 | 143.7047827 | 2.195128486 | 5.263811426 | 12.85632163 | 1.34E-26 | -4.367317581 | 6.74E-28 | 119.875011 | 0.048451409 | 25.87427061 |
| ENSMUSG00000066677 | *Ifi208* | 80.6163131 | 95.91913411 | 52.94386731 | 2.195128486 | 3.158286856 | 5.509852128 | 2.79E-15 | -4.375586589 | 2.79E-16 | 66.94519139 | 0.048174497 | 14.55378667 |
| ENSMUSG00000022799 | *Arhgap31* | 354.8624624 | 402.8603633 | 239.5079712 | 19.75615638 | 14.73867199 | 13.77463032 | 2.81E-42 | -4.380591263 | 7.34E-44 | 192.9169528 | 0.04800767 | 41.55175774 |
| ENSMUSG00000062515 | *Fabp4* | 119.7943344 | 108.3886215 | 112.190576 | 5.487821216 | 6.316573711 | 4.59154344 | 1.38E-28 | -4.385705025 | 6.33E-30 | 129.1378941 | 0.047837804 | 27.86008178 |
| ENSMUSG00000045672 | *Col27a1* | 67.05469034 | 91.12317741 | 31.51420673 | 2.195128486 | 3.158286856 | 3.673234752 | 1.19E-10 | -4.386739549 | 1.77E-11 | 45.20808037 | 0.047803513 | 9.922940054 |
| ENSMUSG00000079362 | *Gm43302* | 50.47937363 | 9.591913411 | 26.47193365 | 0 | 4.211049141 | 0 | 0.00034595 | -4.393314813 | 0.00011189 | 14.92467271 | 0.047586138 | 3.460986821 |
| ENSMUSG00000028773 | *Fabp3* | 87.39712449 | 99.75589948 | 117.232849 | 6.585385459 | 5.263811426 | 2.754926064 | 7.05E-23 | -4.404659687 | 4.38E-24 | 102.4697838 | 0.047213404 | 22.15165763 |
| ENSMUSG00000026650 | *Meig1* | 3.767117435 | 8.63272207 | 11.34511442 | 0 | 1.052762285 | 0 | 0.008939032 | -4.4199479 | 0.003834607 | 8.360520728 | 0.046715726 | 2.048709514 |
| ENSMUSG00000022758 | *P2rx6* | 14.31504625 | 21.1022095 | 8.823977885 | 2.195128486 | 0 | 0 | 0.000548804 | -4.422851464 | 0.000184428 | 13.9834344 | 0.046621801 | 3.260583104 |
| ENSMUSG00000066682 | *Pilrb2* | 15.82189323 | 7.673530729 | 21.42966058 | 2.195128486 | 0 | 0 | 0.00067579 | -4.430405336 | 0.000231184 | 13.55897068 | 0.046378329 | 3.170188521 |
| ENSMUSG00000055435 | *Maf* | 18.08216369 | 19.18382682 | 5.042273077 | 0 | 0 | 1.836617376 | 0.001298029 | -4.431049377 | 0.000467654 | 12.24041694 | 0.04635763 | 2.886715691 |
| ENSMUSG00000000732 | *Icosl* | 416.6431883 | 411.4930853 | 255.8953587 | 19.75615638 | 12.63314742 | 17.44786507 | 7.13E-44 | -4.446944643 | 1.71E-45 | 200.3942578 | 0.045849676 | 43.14675246 |
| ENSMUSG00000024679 | *Ms4a6d* | 3730.199684 | 4126.44115 | 6287.714527 | 226.0982341 | 200.0248342 | 221.3123938 | 3.21E-95 | -4.448614665 | 2.02E-97 | 438.7517868 | 0.045796633 | 94.49416228 |
| ENSMUSG00000044055 | *Otos* | 8.287658356 | 5.755148047 | 10.08454615 | 0 | 1.052762285 | 0 | 0.005578402 | -4.454730157 | 0.002291157 | 9.30030541 | 0.045602915 | 2.253490219 |
| ENSMUSG00000034685 | *Fam171a2* | 57.26018501 | 59.46986315 | 63.02841346 | 3.29269273 | 3.158286856 | 1.836617376 | 5.20E-16 | -4.458656291 | 4.88E-17 | 70.38614317 | 0.04547898 | 15.28385684 |
| ENSMUSG00000092556 | *Or11a3-ps1* | 18.83558717 | 41.24522767 | 31.51420673 | 2.195128486 | 1.052762285 | 0.918308688 | 1.95E-07 | -4.479268009 | 4.02E-08 | 30.13968013 | 0.044833843 | 6.709932273 |
| ENSMUSG00000025854 | *Fam20c* | 4856.567797 | 5061.652707 | 4024.994484 | 189.8786141 | 231.6077027 | 198.3546766 | 3.91E-224 | -4.492379892 | 3.49E-227 | 1035.479763 | 0.044428219 | 223.408047 |
| ENSMUSG00000055407 | *Map6* | 12.80819928 | 15.34706146 | 18.90852404 | 2.195128486 | 0 | 0 | 0.000151108 | -4.502399061 | 4.63E-05 | 16.59555019 | 0.044120744 | 3.820712779 |
| ENSMUSG00000004872 | *Pax3* | 44.45198573 | 59.46986315 | 59.24670865 | 2.195128486 | 4.211049141 | 0.918308688 | 2.93E-13 | -4.50509053 | 3.46E-14 | 57.45211714 | 0.04403851 | 12.53380713 |
| ENSMUSG00000036634 | *Mag* | 141.6436155 | 107.4294302 | 157.5710337 | 5.487821216 | 11.58038514 | 0.918308688 | 1.22E-19 | -4.533763996 | 9.13E-21 | 87.34143044 | 0.043171888 | 18.91367886 |
| ENSMUSG00000031385 | *Plxnb3* | 53.49306757 | 42.20441901 | 46.64102596 | 2.195128486 | 3.158286856 | 0.918308688 | 1.47E-12 | -4.538731871 | 1.83E-13 | 54.1802839 | 0.043023483 | 11.83311293 |
| ENSMUSG00000000730 | *Dnmt3l* | 634.382576 | 676.2298955 | 931.559951 | 36.21962002 | 31.58286856 | 28.46756933 | 1.10E-79 | -4.544715522 | 1.05E-81 | 366.5561684 | 0.04284541 | 78.95746328 |
| ENSMUSG00000028278 | *Rragd* | 8.287658356 | 18.22463548 | 21.42966058 | 0 | 1.052762285 | 0.918308688 | 0.000145765 | -4.554056822 | 4.45E-05 | 16.66978772 | 0.042568887 | 3.836345691 |
| ENSMUSG00000103793 | *Pcdhga6* | 18.83558717 | 34.53088828 | 20.16909231 | 1.097564243 | 1.052762285 | 0.918308688 | 8.85E-07 | -4.589642899 | 1.97E-07 | 27.05966826 | 0.04153171 | 6.053167093 |
| ENSMUSG00000047881 | *Rell1* | 161.9860497 | 181.2871635 | 172.6978529 | 8.780513945 | 4.211049141 | 8.264778192 | 1.27E-41 | -4.595035401 | 3.42E-43 | 189.8554538 | 0.041376763 | 40.89477518 |
| ENSMUSG00000035352 | *Ccl12* | 18.83558717 | 20.14301816 | 10.08454615 | 1.097564243 | 0 | 0.918308688 | 6.00E-05 | -4.60558449 | 1.72E-05 | 18.47655551 | 0.041075317 | 4.222110392 |
| ENSMUSG00000004552 | *Ctse* | 835.546647 | 729.9446106 | 607.5939058 | 27.43910608 | 26.31905713 | 33.05911277 | 4.37E-98 | -4.641443937 | 2.49E-100 | 452.1228952 | 0.040066938 | 97.35944169 |
| ENSMUSG00000047250 | *Ptgs1* | 192.1229892 | 132.3684051 | 128.5779635 | 7.682949702 | 1.052762285 | 9.18308688 | 2.99E-24 | -4.643320769 | 1.72E-25 | 108.8897738 | 0.040014848 | 23.52373546 |
| ENSMUSG00000057207 | *Or5m11* | 26.36982204 | 23.97978353 | 25.21136538 | 0 | 1.052762285 | 1.836617376 | 1.71E-07 | -4.6440083 | 3.50E-08 | 30.40921139 | 0.039995783 | 6.767236239 |
| ENSMUSG00000040270 | *Bach2* | 18.08216369 | 26.85735755 | 32.774775 | 1.097564243 | 2.10552457 | 0 | 7.21E-07 | -4.646668794 | 1.59E-07 | 27.47588134 | 0.039922094 | 6.141949734 |
| ENSMUSG00000022440 | *C1qtnf6* | 113.7669465 | 164.0217193 | 156.3104654 | 6.585385459 | 4.211049141 | 6.428160816 | 1.94E-30 | -4.647470166 | 8.18E-32 | 137.7712502 | 0.039899925 | 29.71273909 |
| ENSMUSG00000044468 | *Tent5c* | 198.1503771 | 240.7570266 | 272.2827462 | 3.29269273 | 12.63314742 | 11.93801294 | 1.75E-40 | -4.647603493 | 4.88E-42 | 184.5652824 | 0.039896238 | 39.75630567 |
| ENSMUSG00000017667 | *Zfp334* | 17.3287402 | 17.26544414 | 17.64795577 | 1.097564243 | 1.052762285 | 0 | 1.37E-05 | -4.669094568 | 3.59E-06 | 21.47132178 | 0.039306329 | 4.862459622 |
| ENSMUSG00000027803 | *Wwtr1* | 14.31504625 | 17.26544414 | 20.16909231 | 0 | 1.052762285 | 0.918308688 | 1.72E-05 | -4.669282562 | 4.56E-06 | 21.01456229 | 0.039301207 | 4.763725423 |
| ENSMUSG00000040950 | *Mgl2* | 35.41090389 | 52.75552376 | 63.02841346 | 0 | 0 | 5.509852128 | 1.06E-08 | -4.669902828 | 1.89E-09 | 36.08168418 | 0.039284314 | 7.975994751 |
| ENSMUSG00000029490 | *Mfsd7a* | 182.3284838 | 231.1651132 | 239.5079712 | 13.17077092 | 5.263811426 | 7.346469504 | 2.63E-42 | -4.674709299 | 6.86E-44 | 193.0506128 | 0.039153652 | 41.57978913 |
| ENSMUSG00000022123 | *Scel* | 7.534234869 | 14.38787012 | 6.302841346 | 0 | 0 | 0.918308688 | 0.00261028 | -4.690081041 | 0.001001515 | 10.82476365 | 0.038738689 | 2.583312967 |
| ENSMUSG00000025196 | *Cpn1* | 9.041081843 | 14.38787012 | 5.042273077 | 1.097564243 | 0 | 0 | 0.00300469 | -4.707459136 | 0.001167483 | 10.54111033 | 0.038274859 | 2.522200279 |
| ENSMUSG00000026220 | *Slc16a14* | 12.80819928 | 28.77574023 | 12.60568269 | 0 | 2.10552457 | 0 | 0.000113683 | -4.724950696 | 3.41E-05 | 17.17656109 | 0.037813608 | 3.944303121 |
| ENSMUSG00000039115 | *Itga9* | 9.79450533 | 10.55110475 | 8.823977885 | 0 | 1.052762285 | 0 | 0.001382563 | -4.736073012 | 0.000499913 | 12.11599131 | 0.037523208 | 2.859314995 |
| ENSMUSG00000006720 | *Zfp184* | 13.56162277 | 15.34706146 | 26.47193365 | 0 | 2.10552457 | 0 | 4.71E-05 | -4.744417369 | 1.33E-05 | 18.96263517 | 0.037306805 | 4.327063086 |
| ENSMUSG00000018672 | *Copz2* | 27.87666902 | 29.73493157 | 54.20443558 | 1.097564243 | 1.052762285 | 1.836617376 | 3.55E-09 | -4.774767366 | 6.10E-10 | 38.29032252 | 0.036530179 | 8.449519419 |
| ENSMUSG00000032936 | *Camkv* | 11.3013523 | 8.63272207 | 10.08454615 | 1.097564243 | 0 | 0 | 0.001179804 | -4.777223121 | 0.000421821 | 12.43296659 | 0.03646805 | 2.928190022 |
| ENSMUSG00000024669 | *Cd5* | 412.1226474 | 370.2478577 | 330.2688865 | 14.26833516 | 11.58038514 | 14.69293901 | 8.98E-78 | -4.777829901 | 9.06E-80 | 357.6699597 | 0.036452715 | 77.04685013 |
| ENSMUSG00000032359 | *Ctsh* | 158.2189323 | 165.940102 | 211.7754692 | 8.780513945 | 7.369335996 | 3.673234752 | 1.34E-37 | -4.781139726 | 4.19E-39 | 171.1303477 | 0.036369182 | 36.87410509 |
| ENSMUSG00000031502 | *Col4a1* | 197.3969536 | 199.511799 | 192.8669452 | 4.390256973 | 8.422098282 | 8.264778192 | 2.57E-50 | -4.790993646 | 4.94E-52 | 230.3758955 | 0.036121619 | 49.59027253 |
| ENSMUSG00000021768 | *Dusp13* | 14.31504625 | 9.591913411 | 6.302841346 | 1.097564243 | 0 | 0 | 0.00165194 | -4.798415857 | 0.000609218 | 11.74759817 | 0.035936262 | 2.782005742 |
| ENSMUSG00000055963 | *Triqk* | 17.3287402 | 24.93897487 | 42.85932115 | 0 | 0 | 2.754926064 | 2.47E-06 | -4.824638367 | 5.82E-07 | 24.97108456 | 0.035288982 | 5.60687739 |
| ENSMUSG00000047786 | *Lix1* | 234.3147044 | 219.6548171 | 300.0152481 | 12.07320667 | 5.263811426 | 9.18308688 | 3.70E-48 | -4.829326119 | 7.64E-50 | 220.3365825 | 0.035174504 | 47.43208072 |
| ENSMUSG00000059498 | *Fcgr3* | 1821.024568 | 1793.687808 | 2257.67777 | 63.6587261 | 73.69335996 | 67.03653422 | 4.02E-191 | -4.84312858 | 6.34E-194 | 882.4718793 | 0.034839589 | 190.3959436 |
| ENSMUSG00000044405 | *Adig* | 18.83558717 | 47.95956706 | 52.94386731 | 3.29269273 | 0 | 0.918308688 | 1.03E-07 | -4.857799038 | 2.04E-08 | 31.4548273 | 0.034487108 | 6.988662869 |
| ENSMUSG00000046186 | *Cd109* | 324.7255229 | 286.798211 | 308.839226 | 4.390256973 | 18.94972113 | 8.264778192 | 4.29E-54 | -4.866709538 | 7.12E-56 | 247.9906274 | 0.034274762 | 53.36778823 |
| ENSMUSG00000042807 | *Hecw2* | 58.76703198 | 62.34743717 | 57.98614039 | 2.195128486 | 2.10552457 | 1.836617376 | 3.24E-17 | -4.874573391 | 2.81E-18 | 76.02185863 | 0.034088445 | 16.48988628 |
| ENSMUSG00000021200 | *Asb2* | 20.34243415 | 25.89816621 | 13.86625096 | 0 | 2.10552457 | 0 | 1.20E-05 | -4.881215346 | 3.10E-06 | 21.75349846 | 0.033931868 | 4.921542042 |
| ENSMUSG00000020682 | *Mmp28* | 12.80819928 | 5.755148047 | 13.86625096 | 0 | 0 | 0.918308688 | 0.001103947 | -4.883697338 | 0.000391915 | 12.57035257 | 0.033873542 | 2.957051811 |
| ENSMUSG00000028859 | *Csf3r* | 934.9985473 | 851.7619109 | 584.9036769 | 20.85372062 | 32.63563084 | 26.63095195 | 1.89E-74 | -4.887541809 | 2.08E-76 | 342.2315967 | 0.033783397 | 73.72443356 |
| ENSMUSG00000033948 | *Zswim5* | 40.6848683 | 36.44927096 | 42.85932115 | 0 | 1.052762285 | 2.754926064 | 3.27E-11 | -4.895946747 | 4.60E-12 | 47.85004038 | 0.033587151 | 10.48559653 |
| ENSMUSG00000094796 | *BC147527* | 39.17802132 | 32.6125056 | 18.90852404 | 2.195128486 | 0 | 0.918308688 | 8.36E-08 | -4.899168985 | 1.64E-08 | 31.87460786 | 0.033512219 | 7.077715303 |
| ENSMUSG00000044734 | *Serpinb1a* | 116.027217 | 139.0827445 | 172.6978529 | 4.390256973 | 4.211049141 | 5.509852128 | 1.51E-31 | -4.90585864 | 6.06E-33 | 142.9384629 | 0.033357185 | 30.82095098 |
| ENSMUSG00000030156 | *Cd69* | 35.41090389 | 39.32684499 | 15.12681923 | 0 | 1.052762285 | 1.836617376 | 3.17E-07 | -4.906418755 | 6.69E-08 | 29.15317571 | 0.033344237 | 6.499055428 |
| ENSMUSG00000081665 | *Pira1* | 151.4381209 | 135.2459791 | 267.2404731 | 0 | 9.474860567 | 8.264778192 | 1.42E-19 | -4.933514331 | 1.07E-20 | 87.02014543 | 0.032723836 | 18.84638774 |
| ENSMUSG00000041548 | *Hspb8* | 21.84928112 | 21.1022095 | 18.90852404 | 0 | 0 | 1.836617376 | 2.29E-06 | -4.951115603 | 5.36E-07 | 25.13045604 | 0.032327021 | 5.640742284 |
| ENSMUSG00000108337 | *Gm4971* | 7.534234869 | 4.795956706 | 5.042273077 | 0 | 0 | 0 | 0.009634237 | -4.954708694 | 0.004156633 | 8.214091307 | 0.03224661 | 2.016182669 |
| ENSMUSG00000055027 | *Smyd1* | 27.87666902 | 23.97978353 | 12.60568269 | 1.097564243 | 0 | 0.918308688 | 3.18E-06 | -5.000715843 | 7.61E-07 | 24.45485385 | 0.031234498 | 5.497336323 |
| ENSMUSG00000027603 | *Ggt7* | 12.05477579 | 18.22463548 | 5.042273077 | 1.097564243 | 0 | 0 | 0.001003435 | -5.02033019 | 0.000353254 | 12.76452962 | 0.030812719 | 2.998510604 |
| ENSMUSG00000028076 | *Cd1d1* | 14.31504625 | 12.46948743 | 8.823977885 | 0 | 1.052762285 | 0 | 0.000336907 | -5.030446006 | 0.00010878 | 14.97785527 | 0.030597424 | 3.472490335 |
| ENSMUSG00000090639 | *None* | 592.9442842 | 554.4125952 | 584.9036769 | 34.02449154 | 14.73867199 | 4.59154344 | 1.41E-09 | -5.031368772 | 2.33E-10 | 40.17147106 | 0.030577859 | 8.849500571 |
| ENSMUSG00000027221 | *Chst1* | 70.06838429 | 94.00075143 | 112.190576 | 0 | 3.158286856 | 4.59154344 | 6.38E-20 | -5.092946121 | 4.68E-21 | 88.6647432 | 0.029300191 | 19.19546549 |
| ENSMUSG00000024155 | *Meiob* | 6.027387896 | 9.591913411 | 3.781704808 | 0 | 0 | 0 | 0.006876662 | -5.113904819 | 0.002874871 | 8.885228177 | 0.028877611 | 2.162622352 |
| ENSMUSG00000049866 | *Arl4c* | 161.9860497 | 189.9198855 | 104.6271663 | 5.487821216 | 2.10552457 | 5.509852128 | 1.50E-27 | -5.117508454 | 7.15E-29 | 124.325012 | 0.028805569 | 26.82468822 |
| ENSMUSG00000054385 | *Ceacam2* | 25.61639856 | 22.06140085 | 23.95079712 | 2.195128486 | 0 | 0 | 3.34E-07 | -5.123537719 | 7.06E-08 | 29.04767773 | 0.028685436 | 6.476393731 |
| ENSMUSG00000032766 | *Gng11* | 7.534234869 | 5.755148047 | 6.302841346 | 0 | 0 | 0 | 0.005043802 | -5.124347181 | 0.002051929 | 9.502492718 | 0.028669346 | 2.297241932 |
| ENSMUSG00000090877 | *Hspa1b* | 235.0681279 | 289.675785 | 221.8600154 | 8.780513945 | 5.263811426 | 7.346469504 | 2.55E-55 | -5.128408425 | 4.12E-57 | 253.6698536 | 0.028588754 | 54.59314737 |
| ENSMUSG00000031026 | *Trim66* | 6.780811383 | 6.714339388 | 6.302841346 | 0 | 0 | 0 | 0.004644936 | -5.137795167 | 0.001876845 | 9.666245514 | 0.028403349 | 2.333020287 |
| ENSMUSG00000028358 | *Zfp618* | 29.38351599 | 28.77574023 | 13.86625096 | 1.097564243 | 1.052762285 | 0 | 8.61E-07 | -5.143876662 | 1.92E-07 | 27.11625831 | 0.02828387 | 6.064809855 |
| ENSMUSG00000087207 | *Gm13147* | 4.520540922 | 6.714339388 | 8.823977885 | 0 | 0 | 0 | 0.005217459 | -5.14459036 | 0.002129305 | 9.434591802 | 0.028269881 | 2.282540947 |
| ENSMUSG00000006800 | *Sulf2* | 36.91775086 | 24.93897487 | 46.64102596 | 1.097564243 | 0 | 1.836617376 | 1.15E-09 | -5.156172232 | 1.88E-10 | 40.5888668 | 0.028043841 | 8.939373398 |
| ENSMUSG00000026768 | *Itga8* | 22.60270461 | 24.93897487 | 27.73250192 | 1.097564243 | 1.052762285 | 0 | 5.47E-08 | -5.195428988 | 1.06E-08 | 32.73314758 | 0.027291037 | 7.261652271 |
| ENSMUSG00000003271 | *Sult2b1* | 12.05477579 | 14.38787012 | 13.86625096 | 0 | 0 | 0.918308688 | 8.04E-05 | -5.197983249 | 2.36E-05 | 17.87701512 | 0.027242761 | 4.094641128 |
| ENSMUSG00000032204 | *Aqp9* | 1194.92965 | 1275.724484 | 1010.975752 | 38.41474851 | 24.21353256 | 32.14080408 | 1.15E-157 | -5.202660851 | 2.76E-160 | 727.7555359 | 0.027154576 | 156.940303 |
| ENSMUSG00000030742 | *Lat* | 203.4243415 | 228.2875392 | 287.4095654 | 5.487821216 | 6.316573711 | 7.346469504 | 9.55E-51 | -5.215617691 | 1.80E-52 | 232.3831884 | 0.026911793 | 50.02019321 |
| ENSMUSG00000018924 | *Alox15* | 51.9862206 | 58.51067181 | 80.67636923 | 1.097564243 | 2.10552457 | 1.836617376 | 1.02E-16 | -5.226601547 | 9.15E-18 | 73.68764835 | 0.02670768 | 15.9919533 |
| ENSMUSG00000026497 | *Mixl1* | 5.273964409 | 11.51029609 | 5.042273077 | 0 | 0 | 0 | 0.003985116 | -5.279125515 | 0.001591093 | 9.970025456 | 0.025752825 | 2.39955901 |
| ENSMUSG00000031503 | *Col4a2* | 9.79450533 | 5.755148047 | 6.302841346 | 0 | 0 | 0 | 0.00309685 | -5.286315442 | 0.001206054 | 10.48104615 | 0.0256248 | 2.50907979 |
| ENSMUSG00000048621 | *Gm6377* | 9.041081843 | 6.714339388 | 6.302841346 | 0 | 0 | 0 | 0.002696202 | -5.298552425 | 0.001035962 | 10.7621641 | 0.025408369 | 2.56924757 |
| ENSMUSG00000037759 | *Ptger2* | 58.01360849 | 71.93935058 | 68.07068654 | 0 | 0 | 4.59154344 | 6.51E-16 | -5.30450194 | 6.17E-17 | 69.92201938 | 0.025303803 | 15.18641833 |
| ENSMUSG00000021886 | *Gpr65* | 183.8353308 | 141.9603185 | 205.4726279 | 2.195128486 | 7.369335996 | 3.673234752 | 4.10E-38 | -5.326016056 | 1.26E-39 | 173.5171684 | 0.024929261 | 37.38770054 |
| ENSMUSG00000020674 | *Pxdn* | 14.31504625 | 13.42867878 | 16.3873875 | 0 | 1.052762285 | 0 | 3.30E-05 | -5.328110269 | 9.13E-06 | 19.68519435 | 0.0248931 | 4.481178557 |
| ENSMUSG00000102412 | *None* | 9.041081843 | 4.795956706 | 8.823977885 | 0 | 0 | 0 | 0.002702123 | -5.330885943 | 0.001038793 | 10.75711305 | 0.024845253 | 2.568294945 |
| ENSMUSG00000036362 | *P2ry13* | 12.80819928 | 4.795956706 | 5.042273077 | 0 | 0 | 0 | 0.003795015 | -5.344474362 | 0.001507641 | 10.06925364 | 0.02461234 | 2.420786469 |
| ENSMUSG00000025422 | *Agap2* | 799.3823197 | 808.5983006 | 641.629249 | 21.95128486 | 23.16077027 | 11.01970426 | 4.31E-112 | -5.349661773 | 1.92E-114 | 516.9788009 | 0.024524002 | 111.3655468 |
| ENSMUSG00000004668 | *Abca13* | 112.2600996 | 115.1029609 | 147.4864875 | 4.390256973 | 1.052762285 | 3.673234752 | 2.31E-31 | -5.35236181 | 9.40E-33 | 142.0669178 | 0.024478148 | 30.63555315 |
| ENSMUSG00000068466 | *Gm5518* | 2.260270461 | 13.42867878 | 7.563409615 | 0 | 0 | 0 | 0.005606013 | -5.361300876 | 0.002303651 | 9.290342052 | 0.024326948 | 2.251345865 |
| ENSMUSG00000025701 | *Alox5* | 2437.32498 | 2450.733877 | 2366.086641 | 38.41474851 | 56.8491634 | 76.2196211 | 8.16E-234 | -5.384976853 | 6.72E-237 | 1080.176836 | 0.023930978 | 233.0882902 |
| ENSMUSG00000066684 | *Pilrb1* | 15.82189323 | 37.4084623 | 32.774775 | 0 | 2.10552457 | 0 | 3.13E-07 | -5.385775993 | 6.60E-08 | 29.17960459 | 0.023917725 | 6.504696966 |
| ENSMUSG00000020053 | *Igf1* | 644.9305048 | 490.1467753 | 515.5724221 | 10.97564243 | 10.52762285 | 17.44786507 | 3.41E-94 | -5.387064637 | 2.29E-96 | 433.9099625 | 0.023896371 | 93.46781302 |
| ENSMUSG00000039543 | *Cfap70* | 45.20540922 | 29.73493157 | 8.823977885 | 0 | 0 | 1.836617376 | 8.23E-06 | -5.407935778 | 2.08E-06 | 22.52283686 | 0.023553156 | 5.084503183 |
| ENSMUSG00000026548 | *Slamf9* | 342.8076866 | 341.4721174 | 400.8607096 | 8.780513945 | 6.316573711 | 10.10139557 | 2.13E-86 | -5.414993301 | 1.80E-88 | 397.6355526 | 0.023438218 | 85.67073705 |
| ENSMUSG00000017943 | *Gdap1l1* | 9.79450533 | 6.714339388 | 7.563409615 | 0 | 0 | 0 | 0.001587777 | -5.422812972 | 0.000583158 | 11.82898346 | 0.023311522 | 2.799210538 |
| ENSMUSG00000029178 | *Klf3* | 149.9312739 | 137.1643618 | 108.4088712 | 4.390256973 | 4.211049141 | 0.918308688 | 5.46E-32 | -5.429500497 | 2.17E-33 | 144.9777908 | 0.023203713 | 31.26309239 |
| ENSMUSG00000024672 | *Ms4a7* | 24.10955158 | 23.97978353 | 40.33818462 | 1.097564243 | 0 | 0.918308688 | 1.18E-08 | -5.434060754 | 2.14E-09 | 35.84549676 | 0.023130484 | 7.926502579 |
| ENSMUSG00000028327 | *Stra6l* | 61.78072593 | 42.20441901 | 73.11295962 | 2.195128486 | 0 | 1.836617376 | 5.58E-15 | -5.442742597 | 5.72E-16 | 65.53064826 | 0.022991707 | 14.25301401 |
| ENSMUSG00000022441 | *Efcab6* | 5.273964409 | 10.55110475 | 8.823977885 | 0 | 0 | 0 | 0.001677949 | -5.446292357 | 0.000619961 | 11.71506499 | 0.022935205 | 2.775221197 |
| ENSMUSG00000094447 | *9430069I07Rik* | 9.041081843 | 6.714339388 | 8.823977885 | 0 | 0 | 0 | 0.001362634 | -5.448561702 | 0.000492052 | 12.14554798 | 0.022899157 | 2.865620837 |
| ENSMUSG00000020787 | *P2rx1* | 7.534234869 | 8.63272207 | 8.823977885 | 0 | 0 | 0 | 0.001167855 | -5.469919242 | 0.000417068 | 12.45413134 | 0.022562657 | 2.932610949 |
| ENSMUSG00000024424 | *Ttc39c* | 16.57531671 | 13.42867878 | 18.90852404 | 0 | 0 | 0.918308688 | 1.07E-05 | -5.476306274 | 2.75E-06 | 21.98027304 | 0.022462989 | 4.968933888 |
| ENSMUSG00000040855 | *Reps2* | 38.42459783 | 33.57169694 | 65.54955 | 1.097564243 | 1.052762285 | 0.918308688 | 7.52E-12 | -5.483894738 | 1.01E-12 | 50.8300464 | 0.022345146 | 11.12362236 |

| | | | | | | | | | | | | | |
|---|---|---|---|---|---|---|---|---|---|---|---|---|---|
| ENSMUSG00000007594 | *Hapln4* | 365.4103912 | 396.1460239 | 340.3534327 | 7.682949702 | 3.158286856 | 12.85632163 | 2.33E-81 | -5.504427225 | 2.12E-83 | 374.3413661 | 0.022029381 | 80.63337365 |
| ENSMUSG00000054675 | *Tmem119* | 130.3422632 | 144.8378925 | 144.965351 | 6.585385459 | 3.158286856 | 0 | 8.68E-35 | -5.506471201 | 3.07E-36 | 158.0129107 | 0.021998193 | 34.06131065 |
| ENSMUSG00000029361 | *Nos1* | 267.4653379 | 298.3085071 | 278.5855875 | 4.390256973 | 9.474860567 | 4.59154344 | 4.52E-73 | -5.525381008 | 5.21E-75 | 335.8124644 | 0.021711737 | 72.34493237 |
| ENSMUSG00000037977 | *6430571L13Rik* | 10.54792882 | 11.51029609 | 3.781704808 | 0 | 0 | 0 | 0.00178603 | -5.532872736 | 0.000663817 | 11.58791814 | 0.021599283 | 2.748111222 |
| ENSMUSG00000047139 | *Cd24a* | 5.273964409 | 8.63272207 | 12.60568269 | 0 | 0 | 0 | 0.001235143 | -5.544511819 | 0.000443726 | 12.33843875 | 0.02142573 | 2.908282677 |
| ENSMUSG00000073889 | *Il11ra1* | 214.7256938 | 237.8794526 | 219.3388788 | 5.487821216 | 2.10552457 | 6.428160816 | 2.21E-60 | -5.563087266 | 3.14E-62 | 277.1486034 | 0.021151631 | 59.65547515 |
| ENSMUSG00000053338 | *Tarm1* | 712.7386187 | 805.7207265 | 917.6937 | 13.17077092 | 24.21353256 | 13.77463032 | 2.74E-127 | -5.577990053 | 9.59E-130 | 587.3210169 | 0.020934263 | 126.5624234 |
| ENSMUSG00000042265 | *Trem1* | 437.7390459 | 437.3912516 | 548.3471971 | 9.878078189 | 15.79143428 | 4.59154344 | 1.97E-83 | -5.582835118 | 1.73E-85 | 383.934315 | 0.020864077 | 82.70528031 |
| ENSMUSG00000020364 | *Zfp354a* | 21.84928112 | 17.26544414 | 13.86625096 | 0 | 0 | 0.918308688 | 4.61E-06 | -5.602982639 | 1.12E-06 | 23.70816175 | 0.020574731 | 5.336712788 |
| ENSMUSG00000026009 | *Icos* | 11.3013523 | 3.836765364 | 12.60568269 | 0 | 0 | 0 | 0.001337206 | -5.619953247 | 0.000482319 | 12.1828107 | 0.020334125 | 2.8738017 |
| ENSMUSG00000030785 | *Cox6a2* | 1007.327202 | 1151.029609 | 1489.991694 | 36.21962002 | 9.474860567 | 28.46756933 | 1.77E-22 | -5.620100006 | 1.12E-23 | 100.616166 | 0.020332057 | 21.75192702 |
| ENSMUSG00000052631 | *Sh2d6* | 1363.696511 | 1351.5006 | 1163.504513 | 46.09769821 | 13.68590971 | 19.28448245 | 2.80E-23 | -5.623585734 | 1.70E-24 | 104.3455579 | 0.020282992 | 22.55215525 |
| ENSMUSG00000095649 | *Gvin-ps3* | 12.80819928 | 4.795956706 | 10.08454615 | 0 | 0 | 0 | 0.001030493 | -5.623738436 | 0.000363699 | 12.71003159 | 0.020280845 | 2.986954916 |
| ENSMUSG00000064294 | *Aox3* | 5.273964409 | 7.673530729 | 15.12681923 | 0 | 0 | 0 | 0.001106185 | -5.624178628 | 0.000392919 | 12.56556933 | 0.020274658 | 2.956172049 |
| ENSMUSG00000069873 | *4930438A08Rik* | 6.780811383 | 13.42867878 | 7.563409615 | 0 | 0 | 0 | 0.000809966 | -5.624583612 | 0.000280442 | 13.19670182 | 0.020268967 | 3.09153306 |
| ENSMUSG00000064267 | *Hvcn1* | 144.6573095 | 248.4305574 | 231.9445615 | 7.682949702 | 5.263811426 | 0 | 9.69E-31 | -5.654639858 | 4.04E-32 | 139.1739773 | 0.019851064 | 30.01382002 |
| ENSMUSG00000101800 | *Rnf170-ps* | 0.753423487 | 8.63272207 | 20.16909231 | 0 | 0 | 0 | 0.006662479 | -5.697775976 | 0.002776185 | 8.949019616 | 0.019266309 | 2.176364139 |
| ENSMUSG00000001700 | *Gramd3* | 6.027387896 | 5.755148047 | 18.90852404 | 0 | 0 | 0 | 0.000988913 | -5.75120583 | 0.000347463 | 12.79544982 | 0.018565837 | 3.004841699 |
| ENSMUSG00000042641 | *Rgsl1* | 12.05477579 | 5.755148047 | 12.60568269 | 0 | 0 | 0 | 0.000465403 | -5.754029749 | 0.000153974 | 14.32297956 | 0.018529532 | 3.332171166 |
| ENSMUSG00000004415 | *Col26a1* | 139.3833451 | 182.2463548 | 170.1767163 | 1.097564243 | 1.052762285 | 6.428160816 | 1.64E-38 | -5.757746629 | 4.91E-40 | 175.3946963 | 0.018481855 | 37.7839544 |
| ENSMUSG00000089945 | *Pakap* | 9.041081843 | 17.26544414 | 34.03534327 | 1.097564243 | 0 | 0 | 1.91E-05 | -5.766333993 | 5.10E-06 | 20.79962139 | 0.018372172 | 4.719031134 |
| ENSMUSG00000025129 | *Ppp1r27* | 3.767117435 | 20.14301816 | 7.563409615 | 0 | 0 | 0 | 0.001262891 | -5.802796169 | 0.000454561 | 12.29340814 | 0.017913659 | 2.898634248 |
| ENSMUSG00000018906 | *P4ha2* | 9.79450533 | 10.55110475 | 11.34511442 | 0 | 0 | 0 | 0.000189489 | -5.812596385 | 5.90E-05 | 16.13539947 | 0.017792384 | 3.722416588 |
| ENSMUSG00000045092 | *S1pr1* | 370.6843556 | 366.4110923 | 296.2335433 | 5.487821216 | 9.474860567 | 3.673234752 | 1.20E-77 | -5.818034188 | 1.22E-79 | 357.0771313 | 0.017725447 | 76.92070523 |
| ENSMUSG00000020017 | *Hal* | 654.7250102 | 776.9449863 | 931.559951 | 19.75615638 | 14.73867199 | 7.346469504 | 9.50E-101 | -5.847729632 | 4.95E-103 | 464.535075 | 0.017364328 | 100.0223034 |
| ENSMUSG00000054072 | *Iigp1* | 27.12324553 | 18.22463548 | 17.64795577 | 0 | 0 | 0.918308688 | 4.69E-07 | -5.85219974 | 1.01E-07 | 28.35371857 | 0.017310609 | 6.328863395 |
| ENSMUSG00000025876 | *Unc5a* | 15.82189323 | 11.51029609 | 5.042273077 | 0 | 0 | 0 | 0.000416483 | -5.860211752 | 0.000136789 | 14.54586149 | 0.017214741 | 3.380403044 |
| ENSMUSG00000074418 | *Pira2-ps2* | 4.520540922 | 15.34706146 | 13.86625096 | 0 | 0 | 0 | 0.000423908 | -5.894757914 | 0.000139402 | 14.51020995 | 0.01680742 | 3.372728768 |
| ENSMUSG00000030666 | *Calcb* | 7.534234869 | 20.14301816 | 6.302841346 | 0 | 0 | 0 | 0.000423618 | -5.918924462 | 0.000139278 | 14.51188961 | 0.016528224 | 3.373025349 |
| ENSMUSG00000024544 | *Ldlrad4* | 12.80819928 | 12.46948743 | 8.823977885 | 0 | 0 | 0 | 0.000112978 | -5.927192586 | 3.38E-05 | 17.18881991 | 0.016433771 | 3.947006254 |
| ENSMUSG00000055782 | *Abcd2* | 78.35604264 | 70.0209679 | 103.3665981 | 3.29269273 | 1.052762285 | 0 | 6.76E-22 | -5.936310094 | 4.35E-23 | 97.92140427 | 0.016330241 | 21.17033518 |
| ENSMUSG00000002104 | *Rapsn* | 9.79450533 | 27.81654889 | 30.25363846 | 0 | 0 | 0.918308688 | 2.87E-06 | -5.940659168 | 6.81E-07 | 24.66843609 | 0.016281087 | 5.542597486 |
| ENSMUSG00000040717 | *Il17rd* | 9.041081843 | 14.38787012 | 11.34511442 | 0 | 0 | 0 | 0.000103821 | -5.946401173 | 3.09E-05 | 17.36034466 | 0.016216416 | 3.983716176 |
| ENSMUSG00000021365 | *Nedd9* | 85.13685402 | 107.4294302 | 123.5356904 | 3.29269273 | 0 | 1.836617376 | 2.25E-27 | -5.951633148 | 1.09E-28 | 123.486922 | 0.016157713 | 26.64744878 |
| ENSMUSG00000043629 | *1700019D03Rik* | 27.12324553 | 19.18382682 | 21.42966058 | 1.097564243 | 0 | 0 | 1.14E-07 | -5.953486021 | 2.27E-08 | 31.24745399 | 0.016136975 | 6.944815858 |
| ENSMUSG00000049551 | *Fzd9* | 9.041081843 | 16.3062528 | 10.08454615 | 0 | 0 | 0 | 0.000108432 | -5.9757085 | 3.24E-05 | 17.27163997 | 0.015890315 | 3.964842373 |
| ENSMUSG00000040065 | *Pfpl* | 7.534234869 | 19.18382682 | 8.823977885 | 0 | 0 | 0 | 0.000197743 | -5.980512988 | 6.17E-05 | 16.05006506 | 0.015837484 | 3.703898968 |
| ENSMUSG00000031722 | *Hp* | 30.13693948 | 11.51029609 | 28.99307019 | 0 | 1.052762285 | 0 | 8.90E-07 | -6.009367009 | 1.99E-07 | 27.04536831 | 0.01552388 | 6.050488705 |
| ENSMUSG00000001761 | *Smo* | 85.13685402 | 84.40883802 | 93.28205192 | 1.097564243 | 0 | 2.754926064 | 1.68E-26 | -6.018384257 | 8.53E-28 | 119.4066473 | 0.015427154 | 25.77467468 |
| ENSMUSG00000028214 | *Gem* | 14.31504625 | 8.63272207 | 13.86625096 | 0 | 0 | 0 | 6.69E-05 | -6.031233766 | 1.94E-05 | 18.24932194 | 0.015290361 | 4.174347966 |
| ENSMUSG00000063660 | *Or1o3* | 9.79450533 | 12.46948743 | 15.12681923 | 0 | 0 | 0 | 5.11E-05 | -6.046460132 | 1.45E-05 | 18.79593731 | 0.015129833 | 4.291211984 |
| ENSMUSG00000102037 | *Bcl2a1a* | 174.0408255 | 174.5728241 | 155.0498971 | 0 | 0 | 7.346469504 | 0.000985736 | -6.063600314 | 0.000346279 | 12.80183647 | 0.014951144 | 3.00623948 |
| ENSMUSG00000034266 | *Batf* | 46.71225619 | 59.46986315 | 102.1060298 | 3.29269273 | 0 | 0 | 2.57E-14 | -6.068927927 | 2.77E-15 | 62.42520423 | 0.014896034 | 13.59027951 |
| ENSMUSG00000022901 | *Cd86* | 216.9859642 | 188.9606942 | 226.9022885 | 4.390256973 | 3.158286856 | 1.836617376 | 1.77E-58 | -6.10728951 | 2.62E-60 | 268.3322168 | 0.014505165 | 57.75253516 |
| ENSMUSG00000079363 | *Gbp4* | 24.10955158 | 22.06140085 | 30.25363846 | 0 | 0 | 0.918308688 | 1.12E-08 | -6.118532584 | 2.01E-09 | 35.9655407 | 0.014392564 | 7.951438413 |
| ENSMUSG00000028445 | *Enho* | 13.56162277 | 5.755148047 | 20.16909231 | 0 | 0 | 0 | 0.000121106 | -6.12446626 | 3.64E-05 | 17.04861334 | 0.01433349 | 3.916832954 |
| ENSMUSG00000102418 | *Sh2d1b1* | 9.041081843 | 22.06140085 | 8.823977885 | 0 | 0 | 0 | 8.53E-05 | -6.150170802 | 2.51E-05 | 17.75892388 | 0.014080371 | 4.069109118 |
| ENSMUSG00000023046 | *Igfbp6* | 87.39712449 | 88.24560338 | 114.7117125 | 1.097564243 | 1.052762285 | 1.836617376 | 5.04E-28 | -6.154833526 | 2.36E-29 | 126.5224859 | 0.014034938 | 27.2974213 |
| ENSMUSG00000023809 | *Rps6ka2* | 2698.009507 | 2928.411164 | 2393.819143 | 28.53667032 | 32.63563084 | 47.75205178 | 1.80E-271 | -6.184635501 | 8.66E-275 | 1254.515986 | 0.01374799 | 270.7439074 |
| ENSMUSG00000020340 | *Cyfip2* | 69.3149608 | 74.81692461 | 83.19750577 | 2.195128486 | 0 | 0.918308688 | 2.23E-23 | -6.214093952 | 1.34E-24 | 104.8178359 | 0.013470116 | 22.6522152 |
| ENSMUSG00000026822 | *Lcn2* | 801.6425901 | 1099.233277 | 1576.970905 | 18.65859213 | 11.58038514 | 16.52955638 | 1.99E-78 | -6.214711786 | 1.97E-80 | 360.7153576 | 0.013464348 | 77.70102601 |
| ENSMUSG00000030134 | *Rasgef1a* | 8.287658356 | 7.673530729 | 26.47193365 | 0 | 0 | 0 | 0.000123058 | -6.219788518 | 3.71E-05 | 17.01393027 | 0.013417052 | 3.90988996 |
| ENSMUSG00000071553 | *Cpa2* | 21.84928112 | 13.42867878 | 6.302841346 | 0 | 0 | 0 | 7.36E-05 | -6.221625382 | 2.14E-05 | 18.05688989 | 0.01339998 | 4.133233318 |
| ENSMUSG00000030762 | *Aqp8* | 18.83558717 | 14.38787012 | 10.08454615 | 0 | 0 | 0 | 1.43E-05 | -6.275480022 | 3.76E-06 | 21.38408142 | 0.01290899 | 4.843723358 |
| ENSMUSG00000026012 | *Cd28* | 57.26018501 | 75.77611595 | 105.8877346 | 0 | 3.158286856 | 0 | 1.56E-18 | -6.277858946 | 1.26E-19 | 82.15031066 | 0.012887722 | 17.80683858 |
| ENSMUSG00000107758 | *Gm19253* | 10.54792882 | 18.22463548 | 15.12681923 | 0 | 0 | 0 | 1.16E-05 | -6.280772601 | 2.99E-06 | 21.82331156 | 0.01286172 | 4.936076111 |
| ENSMUSG00000056498 | *Tmem154* | 1448.833365 | 1526.073424 | 1744.626485 | 30.73179881 | 21.0552457 | 10.10139557 | 7.29E-188 | -6.286083793 | 1.25E-190 | 867.3160453 | 0.012814457 | 187.137355 |
| ENSMUSG00000004609 | *Cd33* | 1256.710376 | 1257.499848 | 1232.835767 | 7.682949702 | 22.10800799 | 17.44786507 | 5.69E-271 | -6.294719629 | 3.12E-274 | 1251.954599 | 0.01273798 | 270.2452582 |
| ENSMUSG00000021725 | *Parp8* | 177.0545194 | 164.9809107 | 139.9230779 | 0 | 4.211049141 | 1.836617376 | 6.05E-44 | -6.307923097 | 1.45E-45 | 200.7328701 | 0.012621935 | 43.21815541 |
| ENSMUSG00000026604 | *Ptpn14* | 242.6023628 | 283.920637 | 279.8461558 | 3.29269273 | 2.10552457 | 4.59154344 | 3.01E-74 | -6.309179462 | 3.38E-76 | 341.2642774 | 0.012610948 | 73.52178973 |
| ENSMUSG00000035273 | *Hpse* | 34.6574804 | 25.89816621 | 31.51420673 | 1.097564243 | 0 | 0 | 2.17E-10 | -6.393509211 | 3.28E-11 | 44.00087242 | 0.011894932 | 9.664302248 |
| ENSMUSG00000048065 | *Cyb5r2* | 10.54792882 | 15.34706146 | 23.95079712 | 0 | 0 | 0 | 5.46E-06 | -6.456183355 | 1.34E-06 | 23.35834768 | 0.011389251 | 5.262845524 |
| ENSMUSG00000017002 | *Slpi* | 721.7797005 | 732.8221846 | 688.270275 | 7.682949702 | 5.263811426 | 9.18308688 | 1.12E-190 | -6.583555387 | 1.85E-193 | 880.3335122 | 0.010426831 | 189.9495609 |
| ENSMUSG00000009596 | *Taf7l* | 20.34243415 | 23.97978353 | 61.76784519 | 1.097564243 | 0 | 0 | 1.75E-08 | -6.583591044 | 3.20E-09 | 35.06089409 | 0.010426574 | 7.757307504 |
| ENSMUSG00000031586 | *Rbpms* | 397.0541776 | 373.1254317 | 335.3111596 | 1.097564243 | 5.263811426 | 4.59154344 | 4.65E-98 | -6.63125539 | 2.68E-100 | 451.974726 | 0.010087725 | 97.33239749 |
| ENSMUSG00000090626 | *Tex9* | 42.94513876 | 26.85735755 | 41.59875288 | 0 | 0 | 0.918308688 | 7.92E-12 | -6.666923025 | 1.06E-12 | 50.72308714 | 0.009841384 | 11.10128906 |
| ENSMUSG00000031383 | *Dusp9* | 15.82189323 | 16.3062528 | 26.47193365 | 0 | 0 | 0 | 3.51E-07 | -6.692761439 | 7.45E-08 | 28.94396433 | 0.009666696 | 6.454408452 |
| ENSMUSG00000030427 | *Lilra6* | 45.20540922 | 35.49007962 | 37.81704808 | 1.097564243 | 0 | 0 | 2.28E-13 | -6.759416705 | 2.66E-14 | 57.96901762 | 0.009230236 | 12.64286867 |
| ENSMUSG00000028972 | *Car6* | 581.6429319 | 801.8839612 | 1411.836462 | 9.878078189 | 8.422098282 | 7.346469504 | 7.00E-58 | -6.774425191 | 1.06E-59 | 265.5454823 | 0.009134711 | 57.1550917 |
| ENSMUSG00000011008 | *Mcoln2* | 206.4380354 | 281.0430629 | 312.6209308 | 3.29269273 | 1.052762285 | 2.754926064 | 5.23E-58 | -6.808836432 | 7.89E-60 | 266.135792 | 0.008919407 | 57.28178485 |
| ENSMUSG00000040086 | *Tnni3k* | 25.61639856 | 17.26544414 | 20.16909231 | 0 | 0 | 0 | 6.34E-08 | -6.811566216 | 1.23E-08 | 32.43681749 | 0.008902547 | 7.197745008 |
| ENSMUSG00000038094 | *Atp13a4* | 41.43829178 | 38.36765364 | 46.64102596 | 0 | 0 | 0.918308688 | 1.99E-14 | -6.847264628 | 2.12E-15 | 62.94744751 | 0.008684963 | 13.70126943 |
| ENSMUSG00000030739 | *Myh14* | 207.1914589 | 174.5728241 | 92.02148365 | 0 | 4.211049141 | 0 | 1.05E-25 | -6.861521512 | 5.55E-27 | 115.6922267 | 0.00859956 | 24.9802962 |
| ENSMUSG00000028480 | *Glipr2* | 470.1362559 | 487.2692013 | 473.9736692 | 4.390256973 | 4.211049141 | 3.673234752 | 5.61E-141 | -6.873427565 | 1.58E-143 | 650.6931119 | 0.008528882 | 140.2508916 |
| ENSMUSG00000038179 | *Slamf7* | 15.82189323 | 21.1022095 | 30.25363846 | 0 | 0 | 0 | 6.00E-08 | -6.888884944 | 1.16E-08 | 32.54870066 | 0.00843799 | 7.22182557 |
| ENSMUSG00000029797 | *Sspo* | 33.15063343 | 59.46986315 | 41.59875288 | 1.097564243 | 0 | 0 | 1.99E-13 | -6.933821277 | 2.31E-14 | 58.24874565 | 0.008179219 | 12.70181195 |
| ENSMUSG00000022893 | *Adamts1* | 17.3287402 | 27.81654889 | 23.95079712 | 0 | 0 | 0 | 1.87E-08 | -6.935507067 | 3.42E-09 | 34.92767824 | 0.008169667 | 7.728897763 |
| ENSMUSG00000003617 | *Cp* | 460.3417505 | 519.8817069 | 670.6223192 | 3.29269273 | 1.052762285 | 8.264778192 | 2.56E-91 | -6.969563331 | 1.81E-93 | 420.595235 | 0.007979072 | 90.59143152 |
| ENSMUSG00000031214 | *Ophn1* | 149.9312739 | 180.3279721 | 186.5641038 | 1.097564243 | 2.10552457 | 0.918308688 | 1.08E-50 | -6.984975034 | 2.06E-52 | 232.1193473 | 0.007894289 | 49.96736833 |
| ENSMUSG00000020432 | *Tcn2* | 533.4238288 | 664.7195994 | 886.1794933 | 6.585385459 | 2.10552457 | 7.346469504 | 1.36E-88 | -7.002049479 | 1.03E-90 | 407.9320746 | 0.00780141 | 87.86754035 |
| ENSMUSG00000054640 | *Slc8a1* | 23.3561281 | 22.06140085 | 27.73250192 | 0 | 0 | 0 | 2.95E-09 | -7.018547609 | 5.02E-10 | 38.66757798 | 0.007712704 | 8.529982935 |
| ENSMUSG00000029816 | *Gpnmb* | 49350.74524 | 55486.34151 | 55742.32887 | 394.0255633 | 388.4692832 | 447.2163311 | 0 | -7.026551217 | 0 | 3947.963719 | 0.007670035 | Inf |
| ENSMUSG00000003882 | *Il7r* | 317.9447115 | 343.3905001 | 450.0228721 | 0 | 0 | 8.264778192 | 0.000740269 | -7.042683823 | 0.00025441 | 13.37934681 | 0.007584744 | 3.13061044 |
| ENSMUSG00000105096 | *Gbp10* | 21.09585763 | 21.1022095 | 32.774775 | 0 | 0 | 0 | 5.16E-09 | -7.04946313 | 9.01E-10 | 37.52878078 | 0.007549186 | 8.287056873 |
| ENSMUSG00000036353 | *P2ry12* | 23.3561281 | 23.02059219 | 28.99307019 | 0 | 0 | 0 | 1.74E-09 | -7.060925744 | 2.89E-10 | 39.74655518 | 0.007489444 | 8.759352188 |
| ENSMUSG00000044716 | *Dok7* | 30.13693948 | 22.06140085 | 23.95079712 | 0 | 0 | 0 | 1.71E-09 | -7.083801352 | 2.85E-10 | 39.77687887 | 0.007371626 | 8.765915633 |
| ENSMUSG00000032352 | *Lrrc1* | 103.2190177 | 89.20479472 | 83.19750577 | 0 | 0 | 1.836617376 | 5.19E-29 | -7.092126414 | 2.34E-30 | 131.1141562 | 0.007329211 | 28.28459811 |
| ENSMUSG00000073176 | *Zfp449* | 20.34243415 | 25.89816621 | 32.774775 | 0 | 0 | 0 | 1.62E-09 | -7.125414284 | 2.67E-10 | 39.90009193 | 0.007162037 | 8.791333917 |
| ENSMUSG00000075394 | *Hoxc4* | 131.8491102 | 139.0827445 | 161.3527385 | 1.097564243 | 0 | 1.836617376 | 3.57E-44 | -7.145642976 | 8.48E-46 | 201.7931749 | 0.007062316 | 43.44702742 |
| ENSMUSG00000004113 | *Cacna1b* | 54.99991455 | 51.79633242 | 54.20443558 | 1.097564243 | 0 | 0 | 1.37E-18 | -7.198514882 | 1.10E-19 | 82.4173526 | 0.006808181 | 17.86218377 |
| ENSMUSG00000020435 | *Osbp2* | 407.6021064 | 460.4118437 | 330.2688865 | 2.195128486 | 2.10552457 | 3.673234752 | 1.37E-89 | -7.206747714 | 9.94E-92 | 412.6036051 | 0.00676944 | 88.86439386 |
| ENSMUSG00000038729 | *Pakap* | 22.60270461 | 42.20441901 | 23.95079712 | 0 | 0 | 0 | 7.08E-10 | -7.301333551 | 1.13E-10 | 41.58499654 | 0.006339859 | 9.150005206 |
| ENSMUSG00000025432 | *Avil* | 4210.130445 | 6335.458808 | 8173.524658 | 47.19526246 | 26.31905713 | 41.32389096 | 1.60E-53 | -7.348547227 | 2.74E-55 | 245.3064722 | 0.006135739 | 52.79497098 |
| ENSMUSG00000001588 | *Acap1* | 1511.367515 | 1429.195098 | 1338.723502 | 10.97564243 | 6.316573711 | 7.346469504 | 2.06E-307 | -7.454250682 | 8.4756928753008e-311 | 1420.221966 | 0.005702259 | 306.6864569 |
| ENSMUSG00000026809 | *Spaca9* | 25.61639856 | 26.85735755 | 55.46500385 | 0 | 0 | 0 | 8.23E-11 | -7.572377843 | 1.20E-11 | 45.96757306 | 0.005253964 | 10.08445182 |
| ENSMUSG00000038319 | *Kcnh2* | 36.16432737 | 38.36765364 | 32.774775 | 0 | 0 | 0 | 2.44E-13 | -7.576699572 | 2.87E-14 | 57.82304646 | 0.005238249 | 12.61267438 |
| ENSMUSG00000030205 | *Gprc5d* | 25.61639856 | 24.93897487 | 57.98614039 | 0 | 0 | 0 | 1.90E-10 | -7.580246396 | 2.87E-11 | 44.26657442 | 0.005225387 | 9.721083545 |
| ENSMUSG00000041544 | *Disp3* | 391.0267897 | 379.8397711 | 265.9799048 | 2.195128486 | 1.052762285 | 1.836617376 | 2.55E-74 | -7.675345589 | 2.83E-76 | 341.6207274 | 0.004892048 | 73.5940884 |
| ENSMUSG00000021186 | *Fbln5* | 33.15063343 | 32.6125056 | 57.98614039 | 0 | 0 | 0 | 3.81E-13 | -7.77156783 | 4.54E-14 | 56.91892393 | 0.00457641 | 12.4185767 |
| ENSMUSG00000048078 | *Tenm4* | 441.5061634 | 619.6376064 | 515.5724221 | 4.390256973 | 3.158286856 | 0 | 8.63E-105 | -7.78501896 | 4.15E-107 | 483.2718444 | 0.00453394 | 104.0637763 |
| ENSMUSG00000062082 | *Cd200r4* | 553.7662629 | 697.332105 | 1002.151774 | 0 | 5.263811426 | 4.59154344 | 2.21E-83 | -7.794708449 | 1.95E-85 | 383.6925902 | 0.004503591 | 82.65603413 |
| ENSMUSG00000036106 | *Prr5* | 164.2463202 | 174.5728241 | 128.5779635 | 1.097564243 | 0 | 0.918308688 | 8.91E-46 | -7.848293616 | 2.00E-47 | 209.2535735 | 0.004339385 | 45.05031234 |
| ENSMUSG00000073198 | *Bnip3l-ps* | 67.80811383 | 26.85735755 | 35.29591154 | 0 | 0 | 0 | 4.92E-12 | -7.858600659 | 6.47E-13 | 51.69953552 | 0.004308494 | 11.30834604 |
| ENSMUSG00000023802 | *Nox3* | 33.15063343 | 53.7147151 | 45.38045769 | 0 | 0 | 0 | 7.08E-15 | -7.87252634 | 7.32E-16 | 65.04464528 | 0.004267106 | 14.14967461 |
| ENSMUSG00000063531 | *Sema3e* | 561.3004978 | 612.923267 | 528.1781048 | 4.390256973 | 1.052762285 | 1.836617376 | 2.34E-153 | -7.90157854 | 6.26E-156 | 707.7220964 | 0.004182037 | 152.6310488 |
| ENSMUSG00000013584 | *Aldh1a2* | 52.73964409 | 37.4084623 | 45.38045769 | 0 | 0 | 0 | 6.13E-16 | -7.913920673 | 5.80E-17 | 70.04436257 | 0.004146412 | 15.21240982 |
| ENSMUSG00000033207 | *Mamdc2* | 74.58892521 | 77.69449863 | 129.8385317 | 0 | 0 | 0.918308688 | 1.57E-24 | -8.000364906 | 8.81E-26 | 110.2119129 | 0.003905262 | 23.80500668 |
| ENSMUSG00000049511 | *Htr1b* | 356.3693093 | 277.2062976 | 211.7754692 | 0 | 1.052762285 | 1.836617376 | 4.52E-58 | -8.122005089 | 6.79E-60 | 266.4337192 | 0.003589491 | 57.34474152 |
| ENSMUSG00000041707 | *Tmem273* | 798.6288962 | 817.2310226 | 1061.398483 | 4.390256973 | 1.052762285 | 3.673234752 | 2.64E-156 | -8.190662698 | 6.88E-159 | 721.3308019 | 0.003422669 | 155.5789851 |
| ENSMUSG00000038068 | *Rnf144b* | 350.3419214 | 401.9011719 | 477.755374 | 0 | 1.052762285 | 2.754926064 | 9.23E-97 | -8.240145324 | 5.45E-99 | 445.9643388 | 0.003307266 | 96.03458347 |
| ENSMUSG00000045382 | *Cxcr4* | 42.94513876 | 58.51067181 | 73.11295962 | 0 | 0 | 0 | 1.55E-18 | -8.269408727 | 1.25E-19 | 82.16938067 | 0.003240857 | 17.81066 |
| ENSMUSG00000020439 | *Smtn* | 74.58892521 | 51.79633242 | 46.64102596 | 0 | 0 | 0 | 1.12E-18 | -8.270819061 | 8.92E-20 | 82.83501383 | 0.003237691 | 17.95021924 |
| ENSMUSG00000031451 | *Gas6* | 314.9310175 | 469.0445658 | 502.9667394 | 2.195128486 | 1.052762285 | 0.918308688 | 5.95E-78 | -8.300781267 | 5.96E-80 | 358.5052372 | 0.003171143 | 77.22576758 |
| ENSMUSG00000036528 | *Ppfibp2* | 274.9995727 | 340.5129261 | 363.0436615 | 2.195128486 | 1.052762285 | 0 | 9.70E-85 | -8.319502857 | 8.45E-87 | 389.9579245 | 0.003130258 | 84.01324741 |
| ENSMUSG00000049744 | *Arhgap15* | 348.081651 | 284.8798283 | 427.3326433 | 0 | 0 | 2.754926064 | 3.14E-77 | -8.443638836 | 3.32E-79 | 355.0818331 | 0.002872179 | 76.50348068 |
| ENSMUSG00000047409 | *Ctdspl* | 145.410733 | 164.0217193 | 109.6694394 | 0 | 0 | 0.918308688 | 4.49E-40 | -8.582324491 | 1.26E-41 | 182.6819458 | 0.002608933 | 39.34727489 |
| ENSMUSG00000057315 | *Arhgap24* | 258.424256 | 272.4103409 | 292.4518385 | 0 | 2.10552457 | 0 | 7.29E-87 | -8.658060507 | 6.06E-89 | 399.8100401 | 0.002475507 | 86.13697964 |
| ENSMUSG00000035860 | *Cdhr3* | 59.52045547 | 75.77611595 | 98.324325 | 0 | 0 | 0 | 5.77E-24 | -8.690295361 | 3.36E-25 | 107.5554877 | 0.002420808 | 23.23857179 |
| ENSMUSG00000027219 | *Slc28a2* | 79.86288962 | 85.36802936 | 71.85239135 | 0 | 0 | 0 | 4.09E-28 | -8.720436681 | 1.91E-29 | 126.9494959 | 0.002370757 | 27.38831493 |
| ENSMUSG00000034612 | *Chst11* | 459.588327 | 483.4324359 | 370.6070712 | 1.097564243 | 1.052762285 | 0.918308688 | 6.61E-110 | -8.752323292 | 3.04E-112 | 506.8717273 | 0.002318933 | 109.1797143 |
| ENSMUSG00000046058 | *Eid2* | 76.09577218 | 89.20479472 | 81.9369375 | 0 | 0 | 0 | 2.76E-29 | -8.777941342 | 1.22E-30 | 132.3991332 | 0.002278119 | 28.55836998 |
| ENSMUSG00000070691 | *Runx3* | 1215.272084 | 1041.681796 | 663.0589096 | 0 | 2.10552457 | 2.754926064 | 4.71E-102 | -9.171282243 | 2.39E-104 | 470.5837827 | 0.001734479 | 101.3269791 |
| ENSMUSG00000049604 | *Hoxb13* | 110.7532526 | 105.5110475 | 124.7962587 | 0 | 0 | 0 | 1.71E-39 | -9.240796639 | 4.94E-41 | 179.961838 | 0.001652886 | 38.7659775 |
| ENSMUSG00000025586 | *Cpeb1* | 128.8354163 | 149.6338492 | 163.873875 | 0 | 0 | 0 | 2.67E-48 | -9.614565528 | 5.46E-50 | 221.0055182 | 0.001275638 | 47.57364044 |
| ENSMUSG00000021322 | *Aoah* | 415.8897648 | 517.9633242 | 665.5800462 | 0 | 1.052762285 | 0.918308688 | 5.58E-95 | -9.616984139 | 3.60E-97 | 437.6013039 | 0.001273501 | 94.25311054 |
| ENSMUSG00000022885 | *St6gal1* | 272.7393023 | 278.1654889 | 336.5717279 | 1.097564243 | 0 | 0 | 1.02E-86 | -9.657914824 | 8.51E-89 | 399.1308978 | 0.001237878 | 85.99271288 |
| ENSMUSG00000060568 | *Fam78b* | 174.0408255 | 195.6750336 | 161.3527385 | 0 | 0 | 0 | 1.08E-58 | -9.883353175 | 1.59E-60 | 269.3289872 | 0.001058801 | 57.96776508 |
| ENSMUSG00000034438 | *Gbp8* | 176.3010959 | 176.4912068 | 200.4303548 | 0 | 0 | 0 | 1.88E-62 | -9.9386571 | 2.59E-64 | 286.7070331 | 0.001018981 | 61.72559899 |
| ENSMUSG00000021375 | *Kif13a* | 283.2872311 | 291.5941677 | 158.8316019 | 0 | 0 | 0 | 2.76E-49 | -10.3521688 | 5.46E-51 | 225.5884488 | 0.000765044 | 48.55840523 |
| ENSMUSG00000074486 | *Bglap2* | 142.397039 | 316.5331426 | 588.6853817 | 0 | 0 | 0 | 2.63E-12 | -10.86124904 | 3.37E-13 | 52.98158256 | 0.000537574 | 11.5799287 |
| ENSMUSG00000072214 | *Septin5* | 227.5338931 | 424.9217641 | 528.1781048 | 0 | 0 | 0 | 5.54E-53 | -11.03151674 | 9.65E-55 | 242.7980705 | 0.00047773 | 52.25669888 |
