## Supplementary material for "TFEB-Mediated Pro-inflammatory Response in Murine Macrophages Induced by Acute Alpha7 Nicotinic Receptor Activation": Table_S23

**Table S23. Row-normalized expression of macrophage marker genes in WT and Tfeb Tfe3 dKO RAW264.7 cells.**

| | WT | | | *Tfeb Tfe3* dKO | | |
|---|---|---|---|---|---|---|
| (Cd11a) *Itgal* | 100.00 | 50.15 | 0.00 | 39.02 | 82.01 | 69.97 |
| ***(Cd11b)*Itgam* | 100.00 | 72.28 | 30.67 | 0.00 | 0.59 | 13.31 |
| (Cd11c) *Itgax* | 100.00 | 98.04 | 7.84 | 0.00 | 56.86 | 45.10 |
| ***(CD15) *Fut4* | 100.00 | 78.83 | 42.34 | 2.92 | 0.00 | 0.00 |
| ***(CD16) *Fcgr3* | 100.00 | 76.83 | 73.46 | 0.00 | 0.50 | 0.66 |
| ***(CD32) *Fcgr2b* | 100.00 | 66.88 | 41.62 | 0.00 | 0.29 | 1.11 |
| ****(CD64) *Fcgr1* | 100.00 | 72.48 | 51.30 | 0.00 | 0.09 | 3.84 |
| (MHC Class II)*H2-Ab1* | 0.00 | 100.00 | 100.00 | 0.00 | 0.00 | 0.00 |
| *Ccr5* | 100.00 | 0.00 | 9.61 | 43.05 | 33.66 | 54.53 |
| *Cd14* | 99.73 | 46.19 | 0.00 | 70.14 | 57.16 | 100.00 |
| *Cd163* | 100.00 | 0.00 | 0.00 | 0.00 | 96.24 | 0.00 |
| *****Cd33* | 100.00 | 78.51 | 58.48 | 0.00 | 0.84 | 0.72 |
| *Cd68* | 100.00 | 64.01 | 16.98 | 0.00 | 19.25 | 43.62 |
| *Cd80* | 89.21 | 100.00 | 87.85 | 0.00 | 10.51 | 49.05 |
| *****Cd86* | 100.00 | 68.18 | 62.24 | 0.70 | 0.35 | 0.00 |
| *Csf1r* | 100.00 | 56.91 | 0.00 | 33.95 | 48.29 | 64.88 |
| *Gal3st4* | 62.16 | 10.81 | 0.00 | 64.86 | 75.68 | 100.00 |
| ***Lamp1* | 100.00 | 75.03 | 50.55 | 0.00 | 9.76 | 19.59 |
| ***Lamp2* | 100.00 | 77.50 | 80.32 | 2.88 | 0.00 | 7.58 |
| *****Lilrb4* | 100.00 | 71.16 | 57.18 | 2.37 | 0.00 | 3.09 |
| ****Tlr2* | 32.48 | 16.01 | 0.00 | 58.39 | 81.33 | 100.00 |
| ***Tlr4* | 100.00 | 88.69 | 90.12 | 0.00 | 5.20 | 12.09 |
