## Supplementary material for "TFEB-Mediated Pro-inflammatory Response in Murine Macrophages Induced by Acute Alpha7 Nicotinic Receptor Activation": Table_S24

**Table S24. Oligonucleotide sequences.**

| Target gene | Oligo Name | Pair Designation | Oligo Sequence (5' - 3') | Reference |
|---|---|---|---|---|
| *Chrna2* | Chrna2__FWD1 | | F - CTGGATGGGCTGCAGAG | This study |
| | Chrna2__REV1 | | R - CAGAAGCAGACACCAGAGAT | |
| *Chrna3* | Chrna3__FWD1 | | F: CTACGACAAGGCAAAGATCG | (Tan et al., 2022) |
| | Chrna3__REV1 | | R: CCTCACAGCAGTTGTACTTG | |
| *Chrna4* | Chrna4__FWD1 | | F: CATCTCGGTGCTGCTTTCTC | (Tan et al., 2022) |
| | Chrna4__REV1 | | R: GGTGGTGTACATTGAGCACG | |
| *Chrna5* | Chrna5__FWD1 | | F: CACTCAGAGAGAAGAAGCCG | (Tan et al., 2022) |
| | Chrna5__REV1 | | R: CATCCGATCGAGAACTTGGG | |
| *Chrna6* | Chrna6__FWD1 | | F: GACTTCCAGGTCGAAGGCAA | (Tan et al., 2022) |
| | Chrna6__REV1 | | R: AGGTCGATTTCTGCCTTGTC | |
| *Chrna7* | Chrna7_FWD1 | Pair 1 | F: TGCAGTGGAACATGTCTGAG | (Tan et al., 2022) |
| | Chrna7_REV1 | | R: AATGCCCAGATGCATTCACC | |
| | Chrna7_FWD2 | Pair 2 | F: CCGTGCCCTTGATAGCACA | This study |
| | Chrna7_REV2 | | R: GGCATTTTGCCACCATCAGG | |
| | Chrna7_FWD3 | Pair 3 | F: CCAGTATCTCCCTCCAGGCAT | This study |
| | Chrna7_REV3 | | R: AGGACCACCCTCCATAGGAC | |
| | Chrna7_FWD4 | Pair 4 | F: CGTGCCCTTGATAGCACAGTA | This study |
| | Chrna7_REV4 | | R: TTCATGCGCAGAAACCATGC | |
| *Chrna9* | Chrna9__FWD1 | | F: GCCGTGAAGAACGTCATCTC | (Tan et al., 2022) |
| | Chrna9__REV1 | | R: GCGGAGCGAGGAACGATATG | |
| *Chrna10* | Chrna10__FWD1 | | F: CAAGTGCCTTGAGACCAGTG | (Tan et al., 2022) |
| | Chrna10__REV1 | | R: GCCACACTAGACTGCTGGGA | |
| *Chrnb2* | Chrnb2__FWD1 | | F: GGTGACTGTACAGCTCATGG | (Tan et al., 2022) |
| | Chrnb2__REV1 | | R: GCTTAGAAGGGAGTCGGACT | |
| *Chrnb3* | Chrnb3__FWD1 | | F: CAATGAAGTTCGGATCCTGG | (Tan et al., 2022) |
| | Chrnb3__REV1 | | R: AAGCCTTCTCTTCTGTTGCC | |
| *Chrnb4* | Chrnb4__FWD1 | | F: GGACCTATGACCACACAGAG | (Tan et al., 2022) |
| | Chrnb4__REV1 | | R: AAGTCATAGGTCACGTCCAC | |
| *M1* | Chrm1__FWD1 | | F: GCAGCAGCTCAGAGAGGTCACAG | (Kawashima et al., 2007) |
| | Chrm1__REV1 | | R: GCCTTTGCCGCCTCGGTCTCG | |
| *M2* | Chrm2__FWD1 | | F: TGTCAGCAATGCCTCCGTTATG | (Kawashima et al., 2007) |
| | Chrm2__REV1 | | R: GCCTTGCCATTCTGGATCTTG | |
| *M3* | Chrm3__FWD1 | | F: GGTGTGATGATTGGTCTGGCTTG | (Kawashima et al., 2007) |
| | Chrm3__REV1 | | R: AGAAGCAGAGTTTTCCAGGGAG | |
| *M4* | Chrm4__FWD1 | | F: TCAAGAGCCCTCTGATGAAGCC | (Kawashima et al., 2007) |
| | Chrm4__REV1 | | R: AGATTGTCCGAGTCACTTTGCG | |
| *M5* | Chrm5__FWD1 | | F: GCTGACCTCCAAGGTTCCGATTC | (Kawashima et al., 2007) |
| | Chrm5__REV1 | | R: CCGTCAGCTTTTACCACCAATC | |
| *β-actin* | B actin_FWD | | F: CTGTATTCCCCTCCATCGTGG | (Tan et al., 2022) |
| | B actin_REV | | R: TCTGGGTCATCTTTTCACGGT | |
| *Gapdh* | Gapdh_FWD | | F: CATCACTGCCACCCAGAAG | (Tan et al., 2022) |
| | Gapdh_REV | | R: CGTTCAGCTCTGGGATGAC | |
